# Expanding the Landscape of BREX Diversity: Uncovering Multi-Layered Functional Frameworks and Identification of Novel BREX-Related Defense Systems

**DOI:** 10.1101/2025.10.09.681413

**Authors:** Siuli Rakesh, Arunkumar Krishnan

## Abstract

Despite extensive scrutiny of BREX systems, several overarching questions persist regarding the functional modalities of individual components and the collective mechanistic framework underlying their defense responses. Using comparative genomics and sequence–structure analyses, we comprehensively map the phyletic distribution and domain-level functional annotations of BREX proteins across all subtypes. Our analysis uncovers numerous previously uncharacterized domains with key functional implications and demonstrates that BrxA- and BrxB-like homologs are universally present across all subtypes, thereby redefining the core machinery of BREX. Our survey strikingly expands the BREX landscape by characterizing three newly defined RM-like systems, which we term BREX-related (BR) systems, and establishes them as new subtypes that share multiple core components with BREX. Furthermore, we identified a novel composite anti-invader system that intriguingly integrates the BREX machinery with components derived from HerA/FtsK-based capture and Dpd defense systems, forming a unique multifaceted defense machinery. Notably, we identify an extensive repertoire of auxiliary effectors recruited alongside the primary effectors across all BREX and BR systems, functioning to reinforce initial restriction responses and counter phage anti-defense mechanisms. Based on these findings, we propose a unified model organized around a multi-modal “sensing-modifying-restricting” functional architecture, defining the fundamental basis of the multi-layered BREX defense systems.

## Introduction

Recent advances in genome sequencing and large-scale computational analyses have uncovered a vast landscape of complex, multi-component Restriction-Modification (RM)-like immune systems (**1–9**). Within this expanding landscape, the BacteRiophage EXclusion (BREX) system has emerged as a widely distributed prokaryotic defense system (**10**), and has drawn significant attention due to its complex proteomic organization featuring multiple functionally diverse proteins that vary across its subtypes. BREX systems are classified into six subtypes based on their protein composition. All subtypes, excluding Type-4 BREX, share three conserved core proteins: (i) BrxC-ATPase (also known as PglY), a AAA+ ATPase; (ii) BrxX-Methyltransferase (PglX), an N6-adenine methyltransferase, and (iii) PglZ (BrxZ), an alkaline-phosphatase (**10,11**). Type-1 BREX systems, in addition to these, harbor three additional conserved components: (i) BrxA (DNA-binding protein); (ii) BrxB, a AAA+ ATPase-like protein; and (iii) BrxL, a AAA+ ATPase fused to a C-terminal Lon-protease (**11**). Recent findings indicate that DNA methylation in Type-1 BREX is mediated by a multifunctional protein complex comprising BrxB, BrxC, BrxX, and PglZ. Within this complex, BrxX plays a central role in recognizing and modifying self-DNA at non-palindromic sites, resulting in the modification of only one DNA strand (**10,12,13**). BrxC-ATPase, homologous to the ORC/Cdc6 clade of AAA+ ATPases, is thought to function as a scaffold for BREX complex (**12,13**), whereas BrxL is essential to disrupt phage replication (**11,14**). Until recently, the nuclease responsible for DNA degradation in BREX systems remained unidentified, leaving a major gap in understanding the effector deployment of BREX immunity. Emerging evidence now implicates PglZ as a metal-dependent nuclease in Type-1 BREX, capable of nicking plasmid and dsDNA, thereby providing protection against invasive elements (**15,16**).

Type-2 BREX, previously identified as phage growth limitation (Pgl) systems, are characterized by the presence of an additional protein, PglW, which contains both kinase and endonuclease domains (**17–20**). Type-3 BREX systems remain less explored and are distinguished by the inclusion of an additional helicase alongside the three core BREX components. Although the methyltransferases of Type-1 and Type-2 BREX are closely related, the Type-3 methyltransferase (BrxXI-MTase) constitutes a distinct variant (**10**). Type-4 BREX differs fundamentally by replacing the methyltransferase with a Phosphoadenylyl Sulfate (PAPS)-reductase involved in DNA PT-modification (**1,4–6**). Notably, these systems are now recognized as a subset of PT-dependent defense systems (**2,21**). Type-5 and Type-6 BREX systems exhibit a limited distribution, retaining the Type-1 core components, while incorporating minor variations in their protein composition (**10**).

The diversity of BREX subtypes, distinguished by their unique protein components, underscores a highly modular architecture whose functional versatility is still far from being fully clarified. Since its discovery, studies have begun to elucidate the molecular basis of self-DNA modification and non-self-DNA degradation (**10,12–15,22**). However, several components across all BREX-subtypes remain uncharacterized, impeding a comprehensive understanding of these systems **(Figure 1A)**. For instance, the Type-2 specific PglW, the Type-3 specific MTase, the helicases found across various subtypes, and even core components such as BrxC and PglZ contain multiple uncharacterized domains with unresolved functional roles **(Figure 1A)**. These gaps at the component-level reflect broader limitations in our understanding of system-level organization and diversity. While canonical Type-1 BREX has been extensively studied, Types 2, 3, and 4 remain largely unexplored. The classification and evolutionary placement of the sparsely distributed Type-5 and Type-6 BREX are also ambiguous, raising the question of whether they represent truly distinct BREX subtypes or merely subsets of canonical Type-1 BREX. Furthermore, a systematic mapping of the complete phyletic distribution of BREX and their subtypes across bacteria and archaea is still lacking. Equally important, despite the coexistence of multiple subtypes, no comparative evolutionary analysis has been conducted to examine how the conserved tripartite core coevolves with type-specific components. Finally, the potential recruitment of BREX-specific components into conflict systems beyond known BREX subtypes remains unexplored, offering an opportunity to expand the known landscape of BREX-associated prokaryotic immunity.

**Figure 1.**
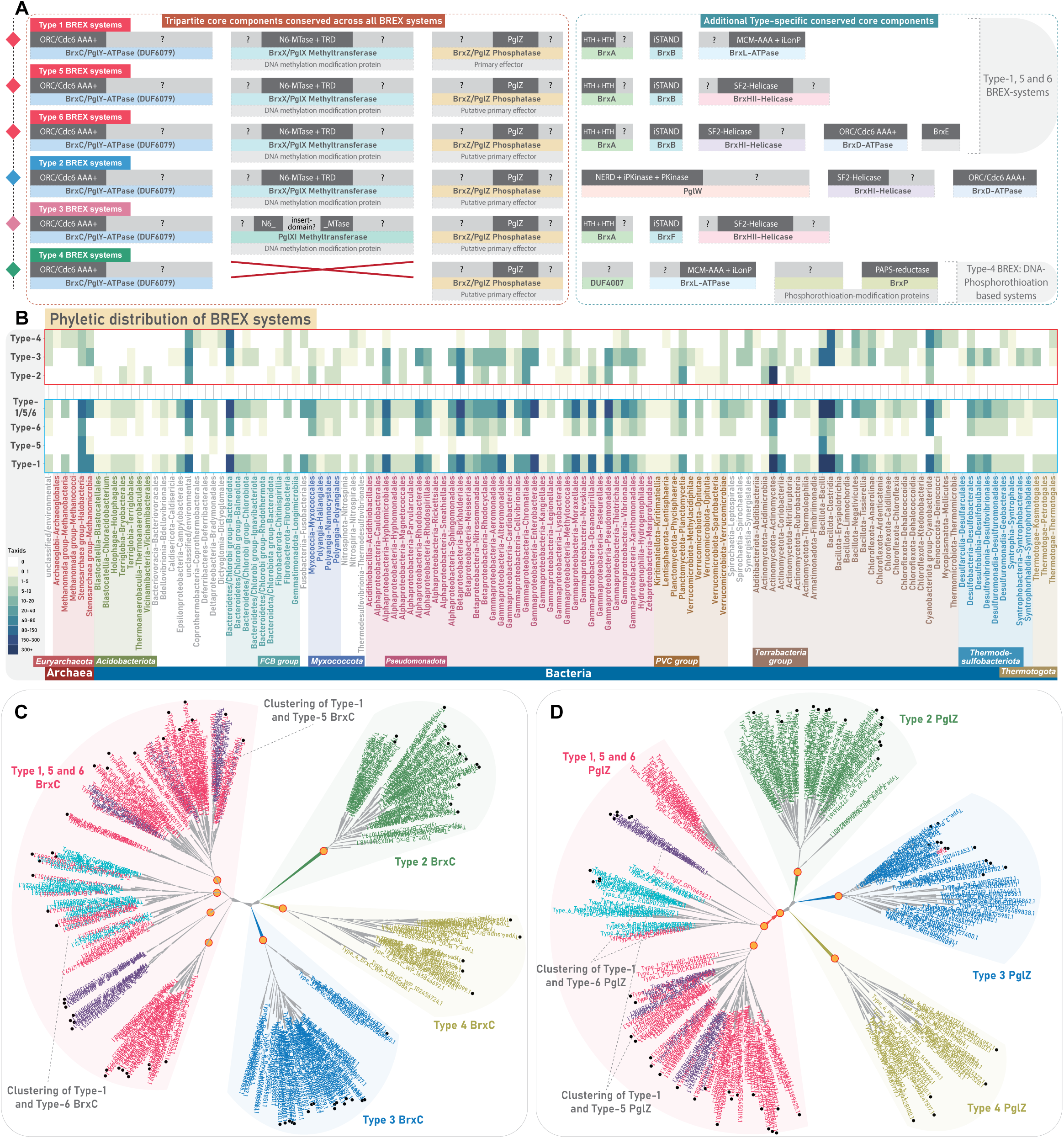
Overview of BREX system diversity, phyletic distribution, and their evolutionary relationships. **(A)** A schematic illustration of various core protein components and their domain architectures across six BREX subtypes. Each protein component is shown as a rectangular box, with its common name and predicted function indicated in dotted, coloured boxes below the corresponding protein. Within each protein, previously annotated domains are shown in dark grey, while uncharacterized regions encoding putative functional domains are highlighted in light grey. **(B)** The heatmap depicts the updated phyletic distribution of all BREX subtypes across major taxonomic groups (Y-axis). The colour intensity of each box represents the number of unique NCBI species-level Tax IDs. Taxonomic groups with a high representation of BREX subtypes are colour-coded for emphasis. Type-1 BREX and its related Type-5 and 6 are grouped within a rectangular box with blue border, while other subtypes are grouped in a rectangular box with red border **(C and D)** Maximum-likelihood phylogenies of BrxC and PglZ, respectively. Both trees resolve into four major clades, each colored separately. Type-5 and Type-6 representatives are clustered with Type-1 sequences and are labelled accordingly. Previously classified sequences from earlier studies are marked with black circular dots. Nodes with high bootstrap support are marked with orange dots.

To address these gaps, we performed a systematic and comprehensive analysis of all BREX systems across prokaryotes. We first extended and refined their phyletic distribution by identifying an exhaustive collection of BREX components and mapped the distribution of each subtype to assess their prevalence across prokaryotes. This was complemented by a detailed examination of their contextual genomic organization. Our neighbourhood-based classification and large-scale phylogenetic reconstructions resolved four major groups of BREX systems—showing that the previously designated Type-5 and Type-6 systems are more appropriately regarded as subtypes of canonical Type-1 BREX. Building on this framework, we conducted comparative sequence-structure analyses of key BREX components—including BrxC, BrxX/PglZ, PglZ/BrxZ, BrxA, BrxB, BrxL, PglW, and BrxH helicases—across all subtypes, uncovering multiple previously unrecognized functional domains that provide new insights into their mechanistic diversity. By leveraging gene-neighborhood reconstructions from ∼21,500 prokaryotic assemblies spanning 8,475 taxa, we uncovered a remarkable diversity of auxiliary nuclease effectors associated with BREX and related systems—many of which were previously unknown. Strikingly, we further expand the landscape of BREX immunity by identifying three newly defined RM-like defense systems—centered on DUF499-ATPases— each sharing multiple homologous components with BREX. Through comprehensive analyses of their domain architectures, sequence-structure synapomorphies, and contextual gene neighbourhoods, we firmly establish these systems as novel subsets of the BREX systems. Furthermore, we uncover a novel composite immune system that integrates components from three distinct conflict operons—including BREX, Dpd and HerA/FtsK anchored capture systems—to form a multi-layered immune apparatus.

## Materials and Methods

### Sequence analysis

Sequence searches were performed against the NCBI non-redundant (nr) database using PSI-BLAST (**23**) (<u>RRID</u>: SCR_001010) and JACKHMMER programs (**24**) (<u>RRID</u>: SCR_005305). For iterative searches, position-specific scoring matrices (PSSMs) and HMM-based profiles were generated and refined at each step. To verify homology, sensitive HMM profile-profile searches were carried out against both the PDB (**25**) (<u>RRID</u>: SCR_012820), and Pfam databases (**26**), using the online HHpred program (**27,28**) (<u>RRID</u>: SCR_010276). Multiple sequence alignments (MSAs) for HHpred were generated using HHblits (**29,30**) against the UniRef30 database, with default parameters. Distant homologs with borderline scores obtained in the initial searches were examined to confirm homology and further used to conduct reciprocal searches against the NCBI nr database using BLASTP (**31**) (<u>RRID</u>: SCR_001010). Additionally, RPS-BLAST searches were conducted locally against the Pfam database and a custom in-house profile database containing a diverse range of protein domains. All retrieved homologous sequences were then clustered using the BLASTCLUST program (ftp://ftp.ncbi.nih.gov/blast/documents/blastclust.html) (<u>RRID</u>: SCR_016641; version 2.2.26) to group them into sets of closely related sequences for downstream analyses. The BLASTCLUST parameters for minimum alignment coverage (L, 0.2–0.5) and bit-score threshold (S, 20–50) were manually optimized to achieve appropriate sequence clustering. To reduce redundancy, nearly identical sequences within each cluster were filtered out using CD-HIT (**32**) (<u>RRID</u>: SCR_007105), with the sequence identity threshold (-c, 0.4–0.9) and word length (-n, 2–5) adjusted as needed. Representative MSAs were subsequently built from the clustered sequences using: (i) the MAFFT program with the local-pair algorithm and *– maxiterate 1000* setting (**33,34**) (<u>RRID</u>:SCR_011811); (ii) Kalign V3 with default parameters (**35,36**) (<u>RRID</u>:SCR_011810); and (iii) GISMO with the *-fast* parameter (**37**). The resulting alignments were manually refined using information from profile-based searches, predicted 3D structural models, structural alignments, and secondary structure predictions.

### Structure and domain architecture analysis

The representative MSAs for all analyzed proteins were used as reference to construct 3D structural model using AlphaFold3 (AF3) (**38**) (<u>RRID</u>: SCR_025885). Each predicted structure was subsequently analyzed for structural similarity using: (i) DALI server (**39**) (<u>RRID</u>: SCR_013433), searched against the PDB clustered at 50–75% sequence identity, and (ii) the FOLDSEEK server (**40**), queried against both AlphaFoldDB UniProt50 and the PDB at 100% identity. A DALI Z-score of 4 and above was used as the minimum threshold to infer potential homology. Homologous structures were further examined through comparative topology assessments and structure-guided sequence analysis to validate the inferred relationships. Secondary structure predictions for all MSAs were carried out using the JPred v4 program (**41**) (<u>RRID</u>: SCR_016504). The predicted secondary structure elements were then cross-referenced with the AF3 structural models to accurately define the boundaries of individual domains and secondary structural elements. Fully annotated 3D structural models, incorporating both domain architecture and secondary structure information, were then visualized and rendered using PyMOL (https://www.pymol.org/) (<u>RRID</u>: SCR_000305).

### Comparative genomics and phyletic distribution analysis

Contextual gene neighbourhood information was retrieved using an in-house Perl script designed to extract upstream and downstream genes flanking the query gene of interest. The script uses GenBank genome files corresponding to unique GenBank assembly IDs (GCA) to extract the neighboring genes. The corresponding protein products of the neighboring genes were then clustered using BLASTCLUST to group sequences based on conserved sequence identity and domain composition. Each cluster was then individually annotated using the above-mentioned sequence-structure analysis pipeline to delineate the complete domain architectures. Sequences forming small, divergent clusters or remaining as unclustered singlets were further examined for shared sequence motifs or structural synapomorphies. Based on these features, they were either merged with existing clusters or retained as distinct groups if no clear relationship could be established. Finally, the protein IDs of the annotated cluster file were mapped back to their corresponding genes in the contextual neighborhood data to reconstruct genomic neighborhoods. Filtering parameters for inclusion of a protein sequence within a gene neighbourhood includes: (i) genomic proximity, with adjacent genes separated by no more than ∼100 nucleotides; (ii) conservation of gene orientation; and (iii) occurrence of conserved gene-neighbourhoods across multiple phyla. The taxonomic distribution of all analyzed protein sequences was determined using taxonomy information from the NCBI Taxonomy Database.

### Phylogenetic analysis

Phylogenetic relationships were inferred using the approximate maximum likelihood (ML) method implemented in FastTree (**42**) (<u>RRID</u>: SCR_015501). Local support values for the tree nodes were estimated accordingly. To improve the accuracy of the resulting tree topologies, the number of minimum-evolution subtree-prune-regraft (SPR) rounds in FastTree was increased to four (*-spr 4*). The options ‘*-mlacc*’ and ‘*-slownni*’ were enabled to allow for a more exhaustive search during ML nearest-neighbor interchanges (NNIs). To complement these analyses, phylogenetic trees were also generated using IQ-TREE (**43,44**) (<u>RRID</u>: SCR_017254), employing the ML methods based on the edge-linked partition model. Branch support values were calculated using the ultrafast bootstrap method with 1000 replicates (**45**). Final trees were visualized and rendered using FigTree (<u>RRID</u>: SCR_008515) (https://tree.bio.ed.ac.uk/software/figtree/).

### Entropy analysis

Position-wise Shannon entropy (H) for each column in a given MSA was calculated using the following equation:

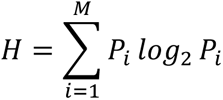

where *P_i_* is the fraction of residues of amino acid type *i*, and *M* is the number of amino acid types. The entropy value for a given alignment position ranges from 0 (indicating complete conservation, with only one residue type present) to 4.32 (indicating maximum variability, where all 20 amino acid residues are equally represented) (**46–48**). The resulting entropy values were analyzed and visualized using the R programming language.

## Results and Discussion

### Multi-pronged sequence, structure, and profile-based search strategies to establish the phyletic spread of all BREX and related systems

To ensure a complete recovery of BREX and related systems, we anchored our initial sequence searches on BrxC-ATPase and PglZ—the only two proteins conserved across all known subtypes. After retrieval, these anchoring components were also used to recover their gene-neighbourhoods for contextual genomic analysis. We first compiled a dataset of all previously characterized BrxC and PglZ from earlier studies (**10**). Using this as a starting point, we performed multiple iterative sequence searches with PSI-BLAST and JACKHMMER against the NCBI *nr* database. Sequences retrieved in the initial rounds were clustered using BLASTCLUST, and representatives from each cluster were used as seeds for subsequent searches. These iterations continued until convergence or until the Position-Specific Scoring Matrices (PSSMs) and HMM profiles began detecting false positives.

*Systematic Curation of BrxC and PglZ Homologs*.

To systematically screen for true homologs of BrxC and PglZ from BREX systems, we employed a multi-step approach:

i. *Homology Confirmation:* Candidates retrieved from iterative searches were clustered and screened using HHpred for sensitive HMM-HMM profile-based alignments against PFAM and PDB databases to establish homology with known profiles and structures;
ii. *Sequence comparison:* Representative sequences from each cluster were aligned with previously classified BREX members to assess shared synapomorphies and membership;
iii. *3D Structure Analysis:* After defining the structural synapomorphies of previously characterized BREX members, 3D structures of representative sequences across each cluster were modeled, superimposed, and compared to establish their structural similarities.
iv. *Genomic Context Screening:* Gene neighborhoods of candidates were examined and compared with BREX-associated genomic contexts (see methods section);
v. *Extended searches using HMM profiles:* Qualified BREX members using the above steps were used to build HMM profiles for HMMSCAN and reverse HMMSEARCH to recover distant homologs.
vi. *Reciprocal Searches:* Distant candidates with higher E-values were used as seeds in new reciprocal searches against the *nr* database, with resulting sequences being screened again using the above steps to confirm their inclusion.

*Concurrent recovery of DUF499-associated systems alongside canonical BREX*.

While performing the sequence searches for BrxC-ATPase, we consistently recovered a substantial number of hits corresponding to a distinct AAA+ ATPase. Further scrutiny using RPS-BLAST and HHpred against the PFAM database classified these proteins as members of the DUF499 family (InterPro entry IPR007555). Earlier studies reported that BrxC-ATPase corresponds to the DUF499 family (**10**). In contrast, our analysis consistently identified DUF6079 (InterPro entry A0A1Q8DLA8) as the top hit for all BrxC-ATPases, with DUF499 appearing only as a secondary match. This discrepancy likely arose because the DUF6079 profile was only released in 2018, three years after BrxC was first annotated as DUF499. Consequently, in earlier studies, DUF499 appeared as the primary match, while DUF6079 was unavailable. Notably, both DUF499 and DUF6079 represent erroneous profile definitions, as each corresponds to the full-length multidomain protein rather than capturing the individual constituent domains—thus obscuring their domain-level resolution. This observation prompted us to compare the characteristics of both protein families, revealing a similar domain architecture anchored on N-terminal AAA+ ATPases. However, in their C-terminal ends, they differ in terms of their domain architecture and sequence features, which allowed us to clearly distinguish DUF499 homologs from the canonical BrxC-ATPase/DUF6079 homologs **(Supplementary Figure S1)**. To further recover all homologs of DUF499-associated systems, we conducted separate searches using the DUF499-ATPases with the same methods described above. In the course of our survey, we also noted instances where homologs of both BrxC and PglZ appeared together in conflict-related contexts outside the known BREX-systems—an observation suggestive of novel, uncharacterized BREX-related variants that we explore in detail in later sections.

All validated sequences from this workflow were used to establish the phylogenetic distribution of BREX and related systems and to perform subsequent analyses. To avoid redundancy caused by multiple genome assemblies for the same taxon, we based our distribution analysis on unique NCBI Taxonomy IDs, each representing a distinct prokaryotic strain, thus ensuring a robust phylogenetic mapping.

### An updated phyletic distribution and broader classification of BREX systems

In the foundational study on BREX systems, PglZ-containing proteins were identified in 126 taxa, including bacteria and archaea, and were categorized into six subtypes (**10**). Type-1 (71 bacteria; 4 archaea) was the canonical and most widespread form. Types 2–4 were moderately distributed and compositionally distinct from Type-1, whereas the sparsely retained Type-5 and 6 closely resembled Type-1 but included additional components, such as helicases in both and BrxE (unknown function) exclusively in Type-6 (**10**). In our current analysis using the greatly expanded NCBI *nr* dataset, we revisited BREX distribution and phylogeny. Leveraging the exponential growth of genomic data and a robust comparative genomics approach, we identified BREX and related systems in 8,475 prokaryotic taxa, including 552 archaea, with overall subtype distribution trends broadly consistent with earlier reports. Excluding the DUF499-associated systems, we recovered genes encoding BREX systems from 5934 prokaryotic taxa, including 227 archaea **(Figure 1B)**. Following the retrieval of all qualified homologs and their corresponding gene neighborhoods, we segregated the various subtypes based on their genomic associations, in accordance with the previous classification (**10**) **(Supplementary Data S1)**. After segregation, a small fraction of protein hits remained unassigned, largely due to incomplete gene neighborhoods from assembly artifacts or low-quality genomes, which were subsequently discarded.

Given the expanded taxonomic coverage of BREX systems, we examined whether this larger dataset aligns with the six previously defined subtypes or shows divergence due to the emergence of new variants and rearrangement among existing subtypes. Once the neighbourhoods were grouped, we constructed multiple phylogenetic trees using various combinations of representatives (BrxC and PglZ) from each subtype **(Supplementary Figure S2-S5)**. To improve topological clarity and clade-level resolution, we included sequences from both the original study and our expanded dataset. The resulting phylogenies were broadly consistent with earlier reports (**10**). All previously defined subtypes clustered congruently with their corresponding groups from our dataset **(Figure 1C)**. However, at a broader level, phylogenetic reconstructions of both PglZ and BrxC-ATPase resolved into four major monophyletic clusters, where Type-5 and Type-6 consistently grouped with the canonical Type-1 **(Figure 1C, Supplementary Figure S2-S5)**. Unlike the more divergent Type-2, 3, and 4, the core components of Type-5 and 6 closely resembled Type-1, indicating a greater degree of functional overlap. Therefore, while conducting contextual neighborhood analysis and protein synapomorphy comparisons, we addressed Type-5 and Type-6 in conjunction with canonical Type-1 BREX.

Our phylogenetic survey identified 3464 taxa harboring Type-1 BREX systems, including 122 archaeal representatives. In contrast, the so-called Type-5 and Type-6 variants were detected in only 160 and 506 taxa, respectively, and consistently clustered with Type-1 in the phylogeny **(Figure 1B–D)**. We also recovered substantial numbers of unique taxa representing Type-2 and Type-3 BREX systems (**10**). Specifically, Type-2 BREX were identified in 1309 taxa, which notably did not include archaeal lineages, whereas Type-3 BREX were found in 919 taxa, including 28 archaea. Although Type-2 BREX systems were more numerous overall, their distribution was relatively restricted, with 1039 representatives confined to the actinomycetes group of bacteria. Type-4 BREX were moderately distributed, identified across 457 prokaryotic taxa, including 20 archaea **(Figure 1B)**. Collectively, our expanded phyletic analysis reinforces earlier findings that BREX systems are among the most widespread innate immune mechanisms in prokaryotes (**10,11**), with canonical Type-1 systems as the most abundant, followed by the more moderately distributed Type-2, Type-3, and Type-4 systems **(Figure 1B)**. The detailed phyletic distribution of individual protein components across various subtypes, inferring their presence-absence within a system, is further discussed in subsequent sections.

### Distribution of the canonical BREX systems

Gene neighbourhood analysis of protein components within our broader dataset of canonical BREX systems (Type-1, Type-5 and Type-6), highlight the remarkable conservation of six core genes: (i) BrxC-ATPase, (ii) BrxX-MTase, (iii) PglZ, (iv) BrxA, (v) BrxB, and (vi) BrxL. Among 3937 taxa (anchored on BrxC), we found BrxB in 3868 taxa (98%), followed by BrxX-MTase in 3769 taxa (96%), BrxA in 3770 taxa (96%), PglZ in 3473 taxa (88%), and BrxL in 2913 taxa (74%) **(Figure 2A and 2B)**. This high degree of conservation spans all major higher-order prokaryotic lineages. Notably, BrxL shows a relatively lower representation, and is frequently replaced by distinct versions of helicases in the previously defined Type-5 and Type-6 BREX-subtypes (**10**) **(Figure 2B, Supplementary Data S1)**. Despite extensive study of Type-1 BREX, the precise domain composition and unique features of their core components remain incompletely defined. Here, through comparative sequence–structure analysis of all encoded components, we place particular emphasis on newly characterized domains and any novel or overlooked features of previously known domains, describing them in detail and highlighting their distinctive structure–function characteristics relative to other subtypes, thereby providing deeper insights into the functional spectrum of these components.

**Figure 2.**
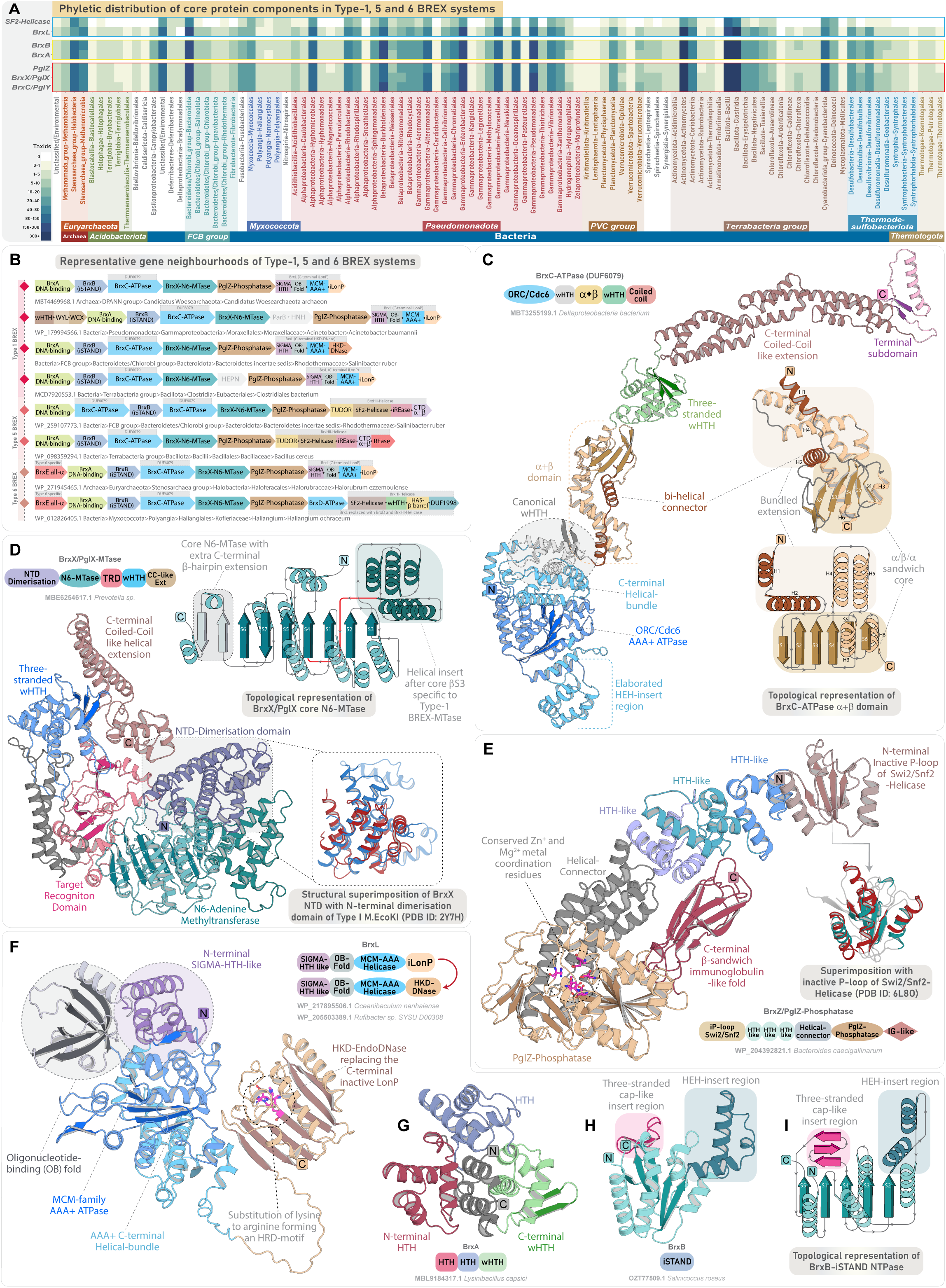
Genomic organization and domain characterization of Type-1 BREX core components. **(A)** Phyletic distribution and presence-absence patterns of Type-1 BREX core components. The heatmap shows the occurrence of core protein components across diverse taxonomic groups. Colour intensity represents the number of unique NCBI species-level Tax IDs, following the scheme used in Figure 1B. **(B)** Representative gene neighbourhoods of Type-1 BREX and the previously classified Type-5 and Type-6 systems. Genes are depicted as box arrows, with the arrow direction indicating gene orientation. Domain names and architectures are illustrated within each box and colour-coded by domain type. All neighbourhoods are labeled with the organism’s name and NCBI accession**. (C)** 3D structure and domain architecture of BrxC-ATPase. Structural topology diagram and the accompanying 3D structure highlight the key features of the characteristic α+β domain present in all BrxC. **(D)** 3D structure and domain architecture of BrxX/PglX. The structural topology diagram illustrates the key features of BrxX core N6-MTase. Insert shows a superimposition of the BrxX NTD with the N-terminal dimerization domain of Type I M.EcoKI (PDB ID: 2Y7H) reveals key structural similarities. **(E)** 3D structure and domain architecture of PglZ. Insert shows a superimposition of PglZ NTD with inactive P-loop of Swi2/Snf2-Helicase (PDB ID: 6L80). **(F-H)** 3D structure and domain architectures of BrxL **(F)**, BrxA **(G)** and BrxB **(H)**. (**I)** Structural topology diagram of BrxB inactive STAND showing key features.

### BrxC ATPase

BrxC-ATPase is a large, highly conserved component (1,200 to 1,300 residues), present across all BREX subtypes. Alongside the previously identified N-terminal ATPase that has been classified within the ORC/Cdc6 clade of AAA+ ATPases (**10,11,13,49**), we have characterized four additional conserved domains. Immediately following the ATPase’s C-terminal helical bundle, we identified a universally conserved winged helix-turn-helix (wHTH) domain across all BREX systems **(Figure 2C)**. Subsequently, we identified a ∼200-residue α+β domain exhibiting a unique structural arrangement, with no significant matches in DALI or FoldSeek searches, potentially pointing to a novel fold **(Supplementary Data S2)**. The domain begins with a bi-helical connector extending from the wHTH, followed by a six-stranded α/β/α-sandwich core, with two long helices after strand-4 forming a bundled extension with the connector. While this bundle is positioned away, the core features a unique six-stranded β-sheet with central parallel strands, flanked by β-hairpins and two other helices on either side of the sheet, forming an α/β/α-sandwich with a distinctive architecture **(Figure 2C)**. At the sequence level, the α+β domain shows rapid sequence evolution, lacking any notable residue conservation that would indicate an enzymatic function. Instead, the overall domain alongside the helical bundle extends into a spatially contiguous exposed surface enriched with positively charged (arginine and lysine), polar (asparagine and threonine), and aromatic (phenylalanine and tryptophan) residues, suggestive of potential DNA-binding functionality **(Supplementary Data S3).** Following this, BrxC contains another wHTH, which is occasionally duplicated into two or even three tandem copies. Given that BrxC is proposed to serve as an oligomeric scaffold for BREX complex assembly (**14**), we suggest that the α+β domain, together with its flanking wHTH modules, forms an extended DNA-binding interface, anchoring the BREX complex to target DNA. Notably, the α+β domain and downstream wHTH modules display high sequence variability across BREX subtypes, suggesting arms-race driven evolution to complement and potentially participate in the recognition of rapidly evolving invasive elements **(Supplementary Data S3)**.

The C-terminus of BrxC comprises an extended coiled-coil of ∼250–500 residues, exhibiting considerable variation across BREX subtypes. At its distal end is a small, distinct subdomain, where two antiparallel strands are separated by one or two intervening helices **(Figure 2C)**. Although belonging to a different P-loop NTPase superfamily, comparable structural features are found in many coiled-coil–containing ABC superfamily members, where a Zn-hook or a hinge domain is positioned at the apex of the coiled-coil to stabilize the long helical arms and facilitate toroidal ring formation around DNA (**50**). By analogy, the terminal subdomain likely stabilizes the elongated coiled-coil tails within the BrxC multimeric assembly through interactions with equivalent regions of other subunits, thereby allowing the domains C-terminal to the ATPase to form a continuous, open, and extended DNA-binding interface. As a AAA+ ATPase bearing a conserved arginine finger, BrxC likely forms a canonical hexameric toroid around DNA, with the C-terminal domains of each subunit extending as long arms to recognize and interact with the DNA.

### BrxX Methyltransferase

BrxX-MTase is the most extensively studied BREX component, functioning as a DNA adenine methyltransferase (DAM), analogous to those in canonical Type I and Type II RM systems (**51–53**). Architecturally, it comprises four distinct regions: an N-terminally located moderately large helical-bundle of unknown function, a central N6-MTase domain, a target recognition domain (TRD) with a helical spacer, and a C-terminal region of ∼250–300 residues (**22,54**). While the core N6-MTase and TRD domains are well characterized, previous studies have described the TRD, the intervening helical spacer, and a distal segment of ∼250 residues collectively as the ‘C-terminal region’ or as a single motif-recognition unit (**14,22**). Although this description is likely due to the presence of the TRD, the precise structural and functional composition of the C-terminal region beyond the helical spacer remains unresolved.

Here, using structural similarity searches and comparative analysis, we identify that the N-terminal helical bundle shares features with the N-terminal dimerization domain of the modification subunit M.EcoKI from Type I RM systems **(Figure 2D)** (**55**). DALI analysis and structure-based alignments reveal significant homology (Z-score ≥ 7.4) to multiple such helical domains located in the N-terminus of DAM from Type I RM systems **(Supplementary Data S2)**. This homology is further supported by a superimposable core structural framework and conserved sequence motifs positioned within the central and terminal helices across these domains **(Supplementary Data S3, Supplementary Figure S6)**. Downstream of the TRD, BrxX-MTase contains a previously described double-helical spacer, proposed to mimic the coiled-coil spacers found in the specificity subunit (HsdS) of M.EcoKI, which mediates dimerization and promotes DNA-clamping within the modification complex (**22,55,56**). Beyond this spacer, we identified a previously uncharacterized three-stranded wHTH that extends the DNA binding interface, followed by a coiled-coil-like helical extension (∼130 residues) at the extreme C-terminus **(Figure 2D)**. This extension, composed of two elongated helices, consistently recovers coiled-coil domains in both structural and sequence profile searches **(Supplementary Data S2)**. Collectively, these features—including the N-terminal dimerization domain and the helical spacer, which also mediates dimerization—suggest that BrxX-MTase likely functions as a dimer during DNA modification. The newly identified wHTH and C-terminal coiled-coil–like extension likely extends the DNA-binding interface and may also contribute to motif recognition alongside the TRD. Across subtypes, Type-1 and Type-2 BrxX-MTases share a conserved domain architecture, with the wHTH module unique to Type-1 enzymes. In contrast, Type-3 BrxXI-MTase shows marked architectural divergence, particularly in the TRD and terminal regions, as described in subsequent sections.

### PglZ

PglZ constitutes the third core and defining component of all BREX systems (**10**). Its central phosphatase domain is homologous to the PglZ domain of the PorX response regulator in PorXY two-component systems, where it displays phosphodiesterase activity against both cyclic and linear oligonucleotides (**16**). More recently, PglZ from Type-1 BREX was shown to act as a metal-dependent nuclease capable of degrading both plasmid and dsDNA, suggesting a role as the principal effector component. Structural studies also revealed an uncharacterized N-terminal domain (∼first 100 residues) (**15**). Yet, beyond the well-defined central phosphatase core (residues ∼430-720), the extended N-terminal (∼430 residues) and the shorter C-terminal (∼80 residues) regions flanking this domain remain poorly characterized.

The N-terminal domain adopts an α/β/α sandwich architecture with a central parallel β-sheet flanked by α-helices **(Figure 2E)**. Structure-based searches with DALI, along with topological comparisons and structural superimposition, identify this as a shortened variant of the inactive second P-loop domain of Swi2/Snf2-type SF2 helicases **(Supplementary Data S2, Supplementary Figure S7)**. In line with its homology, the domain is catalytically inactive owing to the absence of conserved ATP-binding motifs **(Supplementary Data S3)**, but it preserves the core structural architecture associated with DNA binding. This interpretation is reinforced by the downstream extended helical region (residues ∼120 to 260), which comprises two to three rapidly evolving tri-helical units **(Figure 2E)**. These assemble into compact triangular modules with sharp angular turns typical of HTH-like folds **(Figure 2E)** (**57**). Notably, a similar arrangement—an inactive P-loop domain of Swi2/Snf2-helicase followed by a rapidly evolving HTH domain—was also identified by us in the N-terminal region of DndH, a key component of the DndFGH defense system (**1**).

Downstream of these domains, PglZ contains a helical segment (residues ∼260 to 430) comprising five to six elongated helices, resembling the linker region in the PorX–PglZ counterpart (**16**). This is followed by the core phosphatase domain and a C-terminal region distinguished by a rapidly evolving β-sandwich domain **(Figure 2E, Supplementary Data S3)**. Although previously described as a β-barrel (**15**), our topology-based analysis, supported by profile-based and DALI searches, reveals similarity to the β-sandwich immunoglobulin (IG)-like domains of macroglobulin complexes (**58**), and carbohydrate-binding β-sandwich domains of β-galactosidases **(Supplementary Data S2)**. Like the N-terminal iSwi2/Snf2-Helicase domain, the C-terminal β-sandwich domain has a structural counterpart in DndH, where we hypothesized that it may bind to the sugar-phosphate backbone of DNA or selectively bind invasive proteins, akin to the IG-like β-sandwich domains (**1,59**). Consistent with observations in DndH, both the N-terminal inactive P-loop and C-terminal β-sandwich domains of BREX-PglZ are predominantly fast-evolving yet consistently retained **(Supplementary Data S3)**. Given PglZ’s role as the primary effector in Type-1 BREX, these rapidly evolving domains likely act as invader recognition modules, triggering the downstream restriction response of PglZ (**60,61**).

In Type-2 BREX-PglZ, the C-terminal β-sandwich domain is replaced by a three-stranded wHTH, while Type-3 PglZ variants lack C-terminal domains altogether **(Supplementary Figure S8)**. In addition, most Type-4 BREX-PglZ representatives feature an additional inactive STAND-NTPase domain at their N-terminus, followed by the usual core architecture (see BrxB inactive STAND section). This clade-specific architectural diversification of BREX-PglZ—marked by recurrent recruitment of fast-evolving DNA- and macromolecule-binding modules—likely provides multiple functional layers to sense, discriminate, and neutralize invasive elements. Future experimental studies targeting these domains will be key to elucidate the mechanisms of phage restriction mediated by PglZ—a newly characterized principal effector of BREX systems.

### BrxL

BrxL component is specific to Type-1 and Type-4 systems (**10,11**). Structurally, it has been categorized into three regions: (i) an uncharacterized N-terminal region (residues ∼1-180); (ii) a central MCM family AAA+ ATPase domain (residues ∼180-500) known for assembling into hexameric or heptameric helicases that drive DNA-unwinding at replication origin (**62–64**); and (iii) a C-terminal Lon-protease (LonP) domain (residues ∼500-700), notably lacking the catalytic serine–lysine dyad essential for the proteolytic function (**11,49,65,66**).

Extending upon the previous findings, we identified two distinct DNA-binding domains in the N-terminal region of BrxL. First, we observed a SIGMA-HTH-like tri-helical domain (residues ∼1-80) exhibiting significant structural homology to RNA-polymerase-associated SIGMA factors **(Figure 2F, Supplementary Data S2)**. This is immediately followed by an Oligonucleotide-Binding (OB) fold domain (residues ∼80-180) **(Figure 2F, Supplementary Data S2)**, a member of the small β-barrel assemblage of protein folds (**67**). Both SIGMA-HTH and OB-fold are well-established DNA-binding domains (**57,68–71**), likely contributing to BrxL substrate recognition (**11**). These two domains, alongside the AAA+ ATPase domain, are consistently retained and exhibit conserved sequence and structural features across Type-1 and Type-4 BREX systems **(Supplementary Data S3)**. In contrast, the C-terminal region displays significant variation in its domain composition.

A recent study reported that certain BrxL proteins carry a C-terminal HKD-endoDNase (Phospholipase-D) (**72**). In our dataset of 2913 Type-1 BREX taxa harboring BrxL, we identified 1030 instances where the LonP domain is substituted by an HKD-endoDNase **(Figure 2F, Supplementary Data S2-S3)**. These enzymes are well-established restriction factors in Type-I/Type-III-like ATP-dependent RM-systems (**73–77**), and are characterized by a conserved histidine-lysine-aspartate (HKD) catalytic triad (**74–76,78**). However, BrxL-HKD variants exhibit a conservative lysine-to-arginine substitution—forming an HRD motif that likely preserves its enzymatic activity, unlike the inactive LonP found in other BrxL variants **(Supplementary Data S3)**. Consistent with the original proposal that BrxL promotes phage restriction (**11**), recruitment of an HKD-endoDNase likely strengthens this function by directly degrading invasive DNA. Intriguingly, nearly all Type-1 BREX loci carrying BrxL-HKD fusions also retain an active PglZ (identified as nuclease effector in Type-1 BREX), underscoring an evolved multilayered defense strategy in which auxiliary effectors act as fail-safes against phage invasion. Supporting this, we also uncovered—for the first time—a diverse repertoire of endoDNases and endoRNases from distinct superfamilies consistently associated with core BREX loci across multiple subtypes (see backup effector section).

Unlike Type-1 counterparts, Type-4 BrxL retains only the N-terminal DNA-binding domains and the central MCM-family AAA+ ATPase, lacking any additional C-terminal extensions **(Supplementary Data S3)**. Notably, BrxL occurs in 76% of Type-1 BREX systems **(Figure 2A, Supplementary Data S1)** but is entirely absent from Type-2 and Type-3 (**10,11**), implying that it does not constitute a central, indispensable component of anti-phage restriction. Instead, PglZ, together with the auxiliary effectors identified in this study, likely constitutes the core nuclease-driven defense machinery (see following sections). In this context, BrxL most likely functions as a helicase that unwinds invasive DNA to facilitate cleavage, with HKD-fused variants likely acting as an additional nuclease effector. Consistent with this view, Type-2 and Type-3 BREX, which lack BrxL, encode SF2-family helicases—BrxHI and BrxHII, respectively (**10,11**) **(Figure 2B)**—presumably fulfilling the same DNA-unwinding role.

### BrxA Tripartite DNA-binding component

BrxA from Type-1 systems have been experimentally shown to contain two distinct DNA-binding HTH modules (**79**), structurally homologous to the DNA-binding domains of restriction enzymes FokI/BpuJI, and the SspB component of SSP-systems (**14,79–81**). Our analysis confirms the presence of this bipartite two-HTH architecture and further identifies a subset of Type-1 BrxA variants carrying an additional C-terminal wHTH, resulting in a composite tripartite DNA-binding architecture **(Figure 2G, Supplementary Data S3)**. Structural modelling of Type-3 BrxA homologs similarly resolves into three distinct HTH-like modules, with the second and third HTH adopting an antiparallel β-sheet, typical of canonical wHTH **(Supplementary Figure S9)**.

Although BrxA homologs were previously thought to be absent from Type-2 and Type-4 BREX systems (**10,79**), we identified their counterparts in both Type-2 and Type-4 systems, often as fused domains with other core components. In 1047 taxa (80%) of Type-2 BREX, we detected a BrxA-like triad of DNA-binding domains embedded within the PglW component (see PglW section). Likewise, in 442 (97%) taxa of Type-4 BREX, our gene neighbourhood analysis— alongside a recent study on Type-4 BREX (**21**)—identified a conserved component designated as DUF4007 (InterPro entry A0A250KUR8), frequently fused to the PAPS-reductase domain **(Supplementary Data S1)**. Through computational structural modeling, we determined that DUF4007 adopts a BrxA-like tripartite DNA-binding architecture: an N-terminal HTH, followed by two wHTH modules **(Supplementary Figure S9)**. Interestingly, despite their conserved architecture, BrxA-like domains across various BREX-subtypes exhibit substantial sequence divergence. Phylogenetic reconstructions—anchored on individual HTH/wHTH modules— consistently group them into distinct, subtype-specific monophyletic clades, suggesting that these domains have undergone clade-specific adaptations to recognize diverse DNA substrates across various BREX systems **(Supplementary Figure S9)**. Collectively, our findings reveal that all BREX systems encode at least one component with a bipartite or tripartite HTH/wHTH architecture, which likely underpins the DNA-binding and recognition capabilities in BREX-mediated immunity.

### BrxB Inactive STAND NTPase

BrxB is a consistently occurring yet functionally enigmatic component of Type-1 BREX (**10**). Structural analyses have revealed its homology to the STAND/ORC-Cdc6 family of AAA+ ATPases, though it lacks the Walker-A/B motifs required for ATP hydrolysis (**11**). Likewise, in Type-3 BREX, BrxF has been identified as a structural homolog of STAND/ORC-Cdc6 AAA+ ATPase, likely serving as the functional counterpart of BrxB (**11**). Contrary to prior reports of its absence from Type-2 and Type-4 BREX, our analysis reveals that BrxB homologs are not only retained in both systems but have also undergone distinct domain fusions—integrated into PglW component in Type-2 and PglZ in Type-4 BREX. These results now position BrxB-like iSTAND as a consistently retained component across all BREX-subtypes—underscoring its essential and adaptable role within BREX-machinery.

Structurally, Type-1 BrxB adopts a compact inactive-STAND (iSTAND) fold, characterized by loss of Walker motifs and absence of the C-terminal helical bundle typical of AAA+ ATPases. It retains the hallmark five-stranded α/β/α fold, with a characteristic “helix-extension-helix” (HEH) insert after strand-2 (**62,63,82**), while strand-4 extends into an insert region comprising a distinctive three-stranded antiparallel β-sheet that forms a cap-like subdomain before connecting to core Strand-5 **(Figure 2H-I)**. At the sequence level, BrxB evolves rapidly with minimal conservation **(Figure 3A)**. Likewise, the Type-3 BrxF-factor mirrors these features, sharing the cap-like β-sheet and similar sequence signatures following strand-4. In contrast, the newly identified Type-2 BREX iSTAND typically lacks this cap-like β-sheet, though it retains subtle sequence similarities to canonical Type-1 BrxB (See PglW sections), whereas the Type-4 iSTAND exhibits highly divergent sequence profiles, distinct from those of canonical BrxB **(Figure 3A-E, Supplementary Data S3)**.

**Figure 3.**
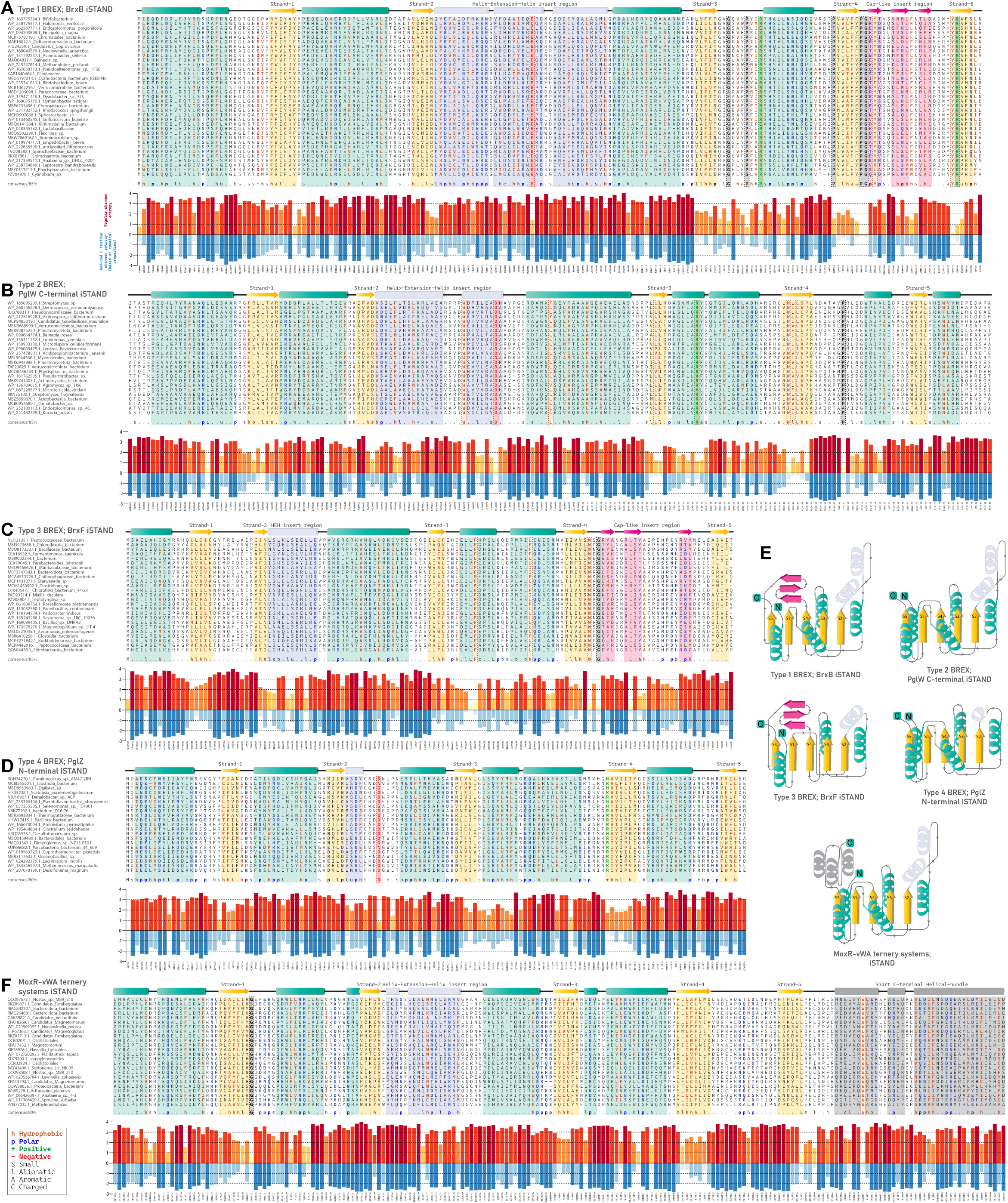
Comparative sequence features and entropy of iSTAND NTPases of all BREX systems. **(A-D)** Representative MSA of iSTAND NTPases from all BREX systems: **(A)** Type-1 BREX BrxB-iSTAND, **(B)** Type-2 BREX PglW C-terminal iSTAND, **(C)** Type-3 BREX BrxF-iSTAND, **(D)** Type-4 BREX PglZ N-terminal iSTAND. In the alignments, a-helices and b-sheet of the P-loop domain are colored in cyan and yellow, respectively, whereas the clade-specific cap-like insert region are colored in magenta. A consensus sequence is displayed at the bottom of each alignment, with highly conserved residues outlined by dotted boxes. Below each alignment, a bar plot shows the entropy plots for the corresponding alignment position. Shannon entropy data computed using regular 20 amino acids are shown above the zero line in yellow to red shades. Shannon entropy data, computed using an 8-residue alphabet (based on chemical properties of amino acids), are shown below the zero line in shades of light blue to dark blue. **(E)** Corresponding structural topology diagrams of iSTAND NTPases. **(F)** Representative MSA of MoxR-vWA-associated iSTAND NTPases.

Interestingly, iSTAND modules have been independently recruited in various MoxR-vWA-centric counter-invader systems (**60,61**). Like BrxB-iSTAND, the MoxR-vWA-centric iSTAND is also rapidly evolving, exhibiting remarkable sequence and structural diversity and a reduced C-terminal helical bundle **(Figure 3F, Supplementary Data S3)**. These systems feature three components: (i) a MoxR AAA+ ATPase; (ii) a vWA component fused to variable peptidase effectors—presumed to remain intrinsically inactive until triggered by invasive elements; and (iii) an iSTAND component proposed to act as a regulatory sensor, detecting invasive elements and inducing conformational changes that activate the effector peptidases, akin to known MoxR-vWA systems (**60,61,83,84**). Given that BrxB-iSTAND is an essential component in the BREX complex, and is closely associated with the primary effector PglZ (**14,15**), we propose that it may serve a similar sensory role—acting as a regulatory switch that directly senses foreign elements and induces conformational changes to trigger PglZ’s nuclease activity. Notably, both BrxB-iSTAND and the iSwi2/Snf2-Helicase of PglZ—predicted to interact during BrxB-PglZ complex formation (**15**)—are rapidly evolving, inactive P-loop NTPases, likely adapted to recognize invasive nucleic acids and favorably positioned to coordinate effector activation.

The iSTAND-mediated sensory function appears to be broadly conserved across all BREX systems, albeit with subtype-specific adaptations **(Figure 3A-F)**. While the iSTAND identified by us in Type-2 BREX is universally retained at the C-terminus of all PglW, in Type-4 BREX, 292 of the 457 identified taxa encode an iSTAND fused to the N-terminus of PglZ. The lower prevalence of iSTAND—present in only 64% of Type-4 taxa compared to over 90% retention in other subtypes—suggests a reduced reliance on iSTAND-mediated sensing for the PT-dependent defense mechanisms unique to Type-4 BREX. Importantly, both these fusion architectures across Type-2 and Type-4 BREX pair the iSTAND module with an effector component—PglW harbors an N-terminal NERD-REase, and PglZ is potentially the metal-dependent nuclease akin to Type-1 BREX PglZ. This modular fusion with various effectors further supports the role of iSTAND as a potential sensor that may also coordinate nuclease response through conformational changes.

### Type-2 BREX systems: Previously characterized Phage growth limitation (Pgl) systems

Type-2 BREX/Pgl-systems were first described in *Streptomyces coelicolor* A3 (**85**), where phage φC31 was observed to infect Pgl+ strains and produce viable progeny during the initial infection cycle, but subsequent infections in other Pgl+ strains were inhibited, thereby limiting phage propagation, though the underlying mechanism remained unclear (**85**). Intriguingly, follow-up studies proposed that PglXI-MTase methylates the phage DNA instead of host DNA. In this context, it was postulated that the resulting methylated phage can lyse the host and reinfect another Pgl^+^ strain, but the methylation mark is then recognized as foreign, triggering a restriction response that blocks further spread (**17–19,85**). Yet, direct experimental evidence confirming phage DNA methylation during primary infection remains lacking. Instead, from a comparative perspective, the conserved genomic organization and shared synapomorphies with Type-1 BREX strongly point toward a BREX-like mechanism incorporating subtype-specific adaptations, as opposed to previously proposed models, for which convincing evidence is still lacking (**10,86**).

In addition to the tripartite core (BrxC/PglY-ATPase, BrxX/PglX Methyltransferase, and BrxZ/PglZ Phosphatase), Type-2 BREX consistently encodes three additional components— PglW, BrxHI-helicase, and the BrxD-ATPase **(Figure 4A and 4B, Supplementary Data S1)** (**10**). In our extended dataset encompassing 1309 taxa harboring Type-2 BREX, PglW was detected in 1035 taxa (∼80%), whereas BrxHI-Helicase and BrxD-ATPase were retained in 1193 (91%) and 1153 (88%) taxa, respectively **(Supplementary Data S1)**. Our structural analysis of the PglW and BrxHI-Helicase revealed several previously uncharacterized domains.

**Figure 4.**
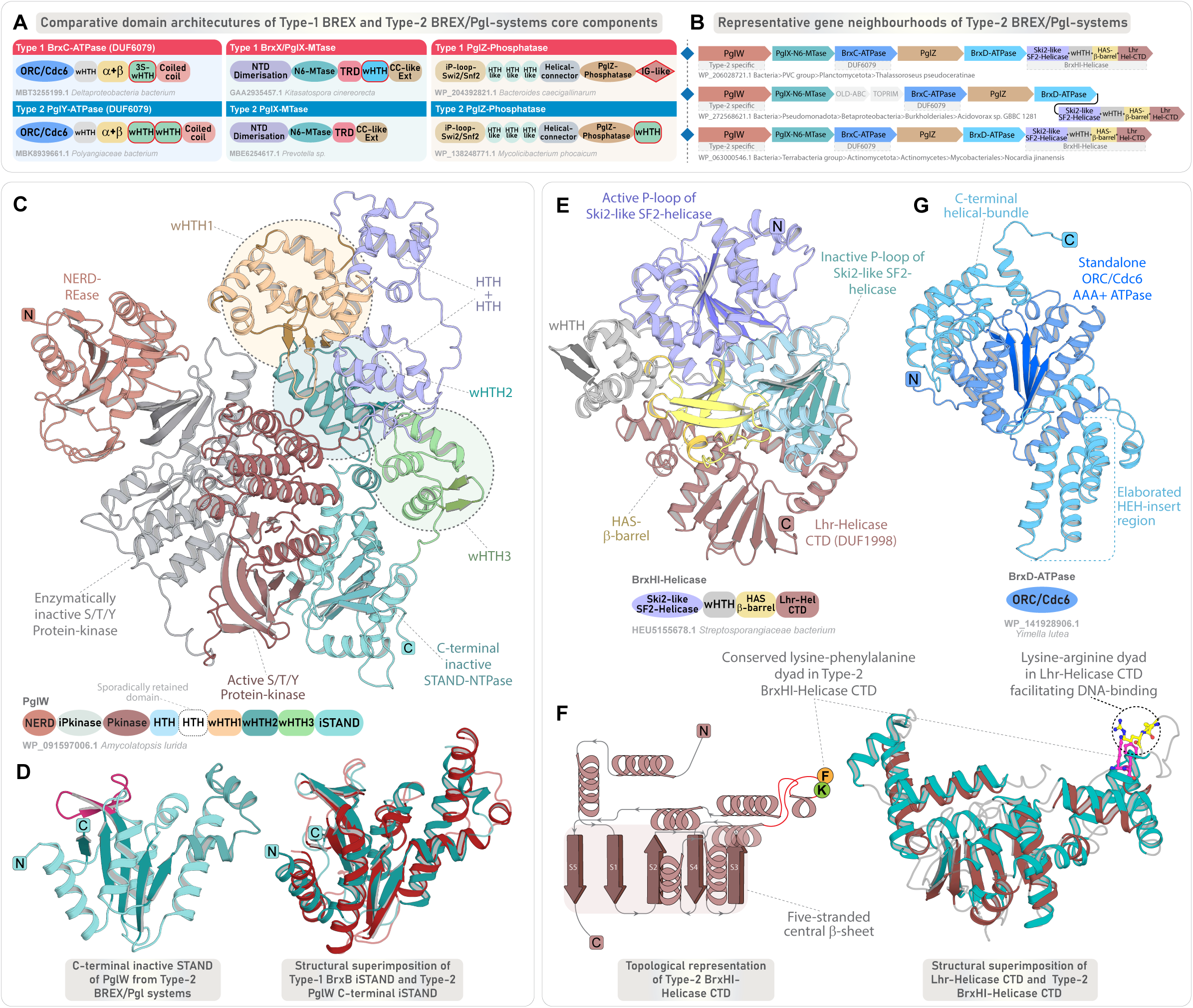
Genomic organization and domain characterization of Type-2 BREX core components. **(A)** Comparison of domain architectures of the tripartite core proteins across Type-1 and Type-2 BREX, highlighting shared and subtype-specific features. Each domain is colored separately, and clade-specific domains are highlighted in red border. **(B)** Representative gene neighbourhoods of Type-2 BREX systems, depicted as in Figure 2B. **(C)** 3D structure and domain architecture of PglW component. **(D)** 3D structure of the PglW C-terminal iSTAND and its structural superimposition with Type-1 BrxB-iSTAND. **(E)** 3D structure and domain architecture of Type-2 specific BrxHI-Helicase. **(F)** Structural topology diagram of BrxHI-Helicase CTD highlighting key features, and its structural superimposition with the CTD of Lhr-Helicase. **(G)** 3D structure and domain architecture of Type-2 specific BrxD-ATPase.

### PglW: A multi-domain enzymatic, DNA-binding, and sensing component specific to Type-2 BREX

PglW is a large, multi-domain protein (1400-1600 residues), organized in two distinct halves: an N-terminal segment with enzymatic domains and a C-terminal segment with multiple uncharacterized domains (residues ∼800-1600). In the N-terminal half, previous studies have identified a NERD-REase, followed by tandem serine/threonine/tyrosine (S/T/Y)-kinase modules—comprising an inactive pseudo-kinase and an active S/T/Y-kinase (**10,20,87**). Interestingly, an identical tripartite arrangement—comprising NERD-REase followed by an inactive and active S/T/Y-kinase—was also identified by us in a subset of DndF proteins from DndFGH systems. In DndF, this arrangement is followed by its core defining domains—a derived P-loop nucleotide-kinase and its associated lid-subdomain, and a unique C-terminal rapidly-evolving α+β domain, both of which were hypothesized to recognize phage-derived modified nucleotides and invader macromolecules (**1**). Strikingly, PglW precisely follows the same architectural “grammar”, where the N-terminal enzymatic modules are followed by a nucleic-acid binding component and a fast-evolving C-terminal sensory module **(Figure 4C)**.

In PglW, the active S/T/Y-kinase is followed by a helical domain structurally similar to the CTD of RNA polymerase α-subunit (**87**). Our analysis reveals that this is succeeded by three distinct wHTH-like modules. The first wHTH adopts an expanded triangular core with three antiparallel β-strands, forming an augmented three-stranded wHTH, while the remaining two display canonical compact wHTH folds **(Figure 4C)**. At the extreme C-terminus, we find that the PglW contains an inactive STAND/ORC-Cdc6-like AAA+ ATPase domain that adopts a compact conformation **(Figure 4C)**. Like BrxB-iSTAND, it is marked by degenerate Walker-A/B motifs and the absence of the characteristic C-terminal helical bundle **(Figure 3B)**. While most PglW-iSTAND lacks the β-sheet cap-like region seen in BrxB-iSTAND, a subset of PglW retains this structural hallmark with conserved sequence elements, indicating a shared origin followed by clade-specific adaptations **(Figure 4D, Supplementary Data S3)**. Likewise, despite sequence divergence, the triple-wHTH configuration parallels the BrxA architecture, supporting the inference that PglW’s C-terminal half encodes structural analogs of both BrxA (triple-wHTH) and BrxB (iSTAND) **(Figure 4C)**. The modular architecture, comprising N-terminal effectors and C-terminal sensory apparatus suggests that, like DndF of DndFGH systems, the PglW in Type-2 BREX may serve as a sensory hub, which can deploy downstream effectors in response to invader recognition. Given that these systems also harbor an active PglZ with putative nuclease-like functionality similar to Type-1 BREX, the enzymatic co-effectors might act as backup modules, targeting self-molecules such as DNA (via NERD-REase) or proteins (via S/T/Y-kinases), through a suicidal mechanism (**1**).

### BrxHI-Helicase and BrxD-ATPase

BrxHI contains an N-terminal SF2-helicase domain that we classify, based on sequence and profile comparisons, as a Ski2-like helicase, within the SF2-superfamily **(Supplementary Figure S10)** (**88**). Ski2-helicases are known participants in antiviral defense, such as the HamB component of Hachiman systems (**89,90**). Following the core helicase, we found two distinct DNA-binding modules: a canonical wHTH and an β-barrel homologous to HAS-barrel domains within the small β-barrel assemblage of DNA-binding proteins **(Figure 4E)**. The C-terminus of BrxHI features an α+β domain homologous to the CTD of the Lhr-helicase (**91**), comprising a five-stranded central β-sheet flanked by α-helices and a distinct helical extension protruding outward **(Figure 4F, Supplementary Data S2)**. The distal end of this extension contains a conserved lysine-arginine dyad, associated with DNA-binding in Lhr-helicase (**91**). A conserved lysine in a comparable position is also conserved in BrxHI, suggesting that BrxHI-CTD likely provides an auxiliary DNA-binding interface, complementing the upstream wHTH and HAS β-barrel domains **(Figure 4F, Supplementary Data S3)**.

BrxD-ATPase has been classified as an ORC-Cdc6 AAA+ ATPase (**62,63,82,92**). Like BrxC, it retains the hallmark features, including the HEH insert after strand-2, the arginine finger at the base of strand-5 essential for oligomerization, and the C-terminal helical bundle **(Supplementary Data S2-S3)**, suggesting that, like the typical ORC-Cdc6 AAA+ ATPase, it may form an oligomeric complex. However, unlike BrxC, BrxD lacks additional DNA-binding or regulatory domains, functioning as a standalone ATPase **(Figure 4G)**. The markedly distinct sequence profile of BrxD compared to BrxC suggests a functionally distinct and non-redundant role specific to Type-2 BREX **(Supplementary Data S3)**. In the genomic locus, BrxD and BrxHI-Helicase co-occur as a tightly linked dyad, pointing to a functional association **(Supplementary Data S1)**. While BrxC likely acts as a scaffolding and recognition unit, BrxD appears to play a distinct, yet uncharacterized role—possibly facilitating the loading or activation of BrxHI-helicase on target DNA substrates, akin to replicative helicase loading mechanisms (**62,93**).

### Type-3 BREX systems

Like the canonical Type-1 and Type-2 BREX, Type-3 BREX comprises six conserved core components **(Figure 5A and 5B)**. In addition to the shared tripartite core, they feature three additional components: (i) a DNA-binding homolog of BrxA; (ii) an iSTAND homolog of BrxB (denoted as BrxF) **(Figure 5C)**; and (iii) BrxHII-helicase—distinct from the BrxHI-helicase found in Type-2 BREX (**10**). Besides being recruited in a smaller subset of the so-called Type-5 BREX systems, BrxHII-helicase is exclusively found in Type-3 BREX as a core component **(Figure 5A)** (**10**). Our gene neighbourhood analyses identify BrxHII-helicase as a well-conserved component, present in 702 taxa (77%) of Type-3 BREX systems. Through detailed sequence-structure analyses of all Type-3-specific components, we identify several clade-specific synapomorphies and define their full-length domain architectures, providing a comparative viewpoint within the broader context of BREX systems.

**Figure 5.**
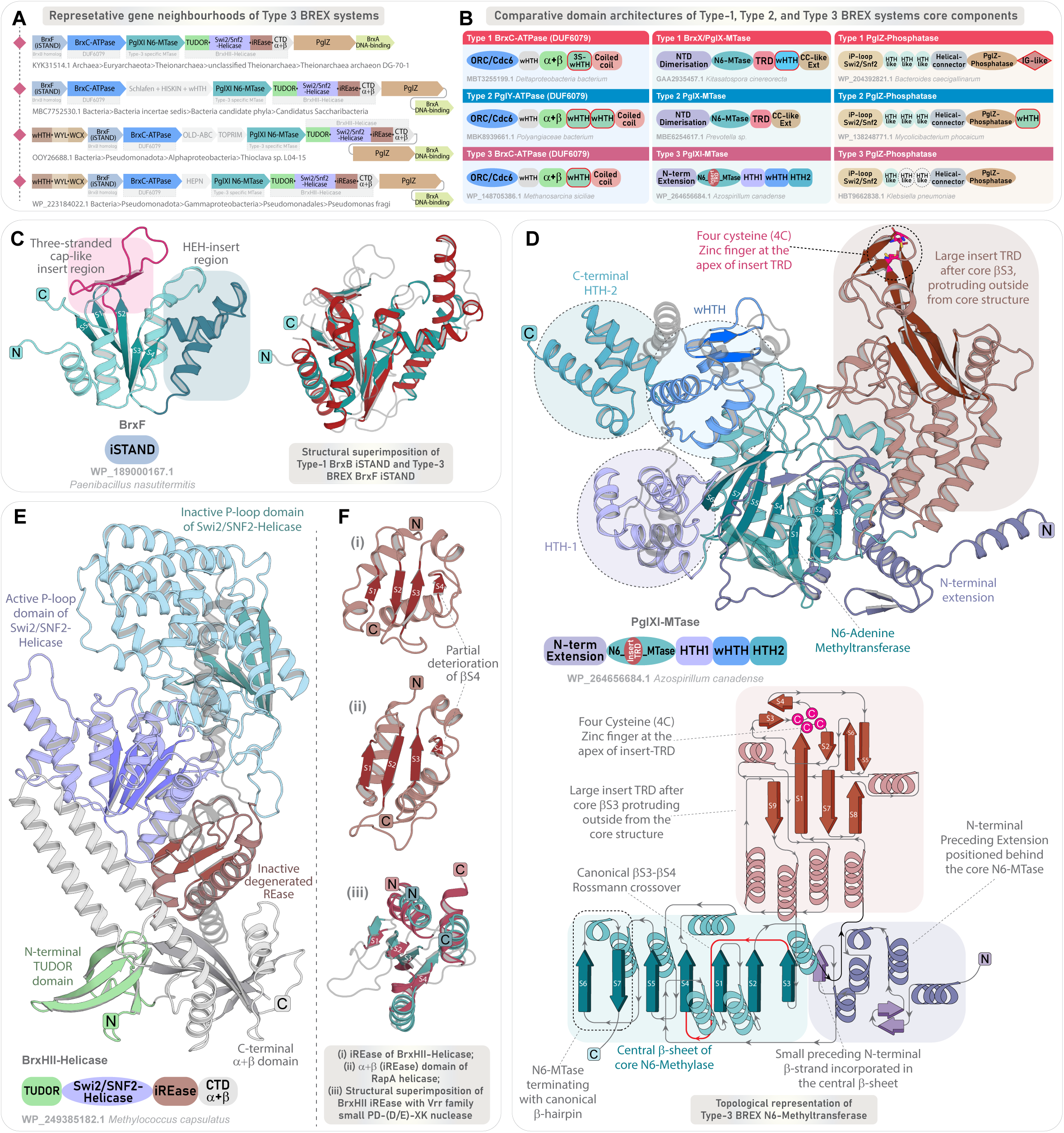
Genomic organization and domain characterization of Type-3 BREX core components. **(A)** Representative gene neighbourhoods of Type-3 BREX systems, depicted as in Figure 2B. **(B)** Comparison of domain architectures of the tripartite core proteins between Type-1, 2, and 3 BREX, highlighting shared and subtype-specific features. Each domain is colored separately, and sporadically retained domains are marked in a white background with dotted borders. **(C)** 3D structure of the BrxF-iSTAND and its structural superimposition with Type-1 BrxB-iSTAND. **(D)** 3D structure and domain architecture of Type-3 BREX specific PglXI-MTase. Structural topology diagram highlighting key features of Type-3 BREX N6-MTase alongside its embedded TRD. **(E)** 3D structure and domain architecture of Type-3 specific BrxHII-Helicase. **(F)** (i-iii) Structural comparison of the inactive REase domain of BrxHII-Helicase with α+β domain (inactive REase) found in RapA helicase. Structural superimposition of BrxHII-Helicase iREase with Vrr family small PD-(D/E)XK nuclease (PDB ID: 4QBN).

### Type-3 specific BrxXI-MTase

The Type-3 BrxXI-MTase differs considerably from the other BREX-MTases, both in terms of its domain architecture and recognition strategies. Notably, its N-terminal region lacks the dimerization domain (∼230 residues) present in other BREX-MTases. Instead, it contains a relatively shorter, distinct N-terminal segment that appears as a simple prelude leading directly into the central MTase domain **(Figure 5D)**. The core MTase, however, adopts a typical Rossmann-fold as observed in all DAM domains, preserving the signature motifs required for adenine methylation **(Supplementary Data S3)**.

The defining hallmark of all Type-3 BREX MTases is a large, approximately 250-residue insert-subdomain positioned between the core strands 2 and 3 of the Rossmann-fold DAM domain **(Figure 5D)**. This insert adopts an α+β module that protrudes from the core MTase and closely parallels the TRD-insert architecture typical of α-class N6-MTases in Type-II RM systems (**51,94**). At the apex of the insert-TRD lies a conserved four-cysteine (4C) Zn-finger motif **(Figure 5D, Supplementary Data S3)**, commonly linked to structural stabilization in nucleic-acid binding proteins (**95,96**), and may further refine TRD-binding specificity (**97**). Although the sequence and structure searches using this insert-TRD yielded no homologs outside BREX systems, it is noteworthy that the TRDs are well-known to undergo rapid evolutionary divergence to accommodate variations in their binding specificities across various host immune systems (**98–102**). Thus, unlike the canonical BREX-MTases that house their TRDs at the C-terminus, the Type-3 MTase internalizes it as a large insert-subdomain— without compromising the structural integrity of the core MTase **(Figure 5D)**.

Beyond the core MTase, the C-terminal region contains a conserved segment of ∼280– 300 residues, consistently predicted to fold into three compact helical domains. Structural homology searches identify these as HTH-like DNA-binding domains, with the first and third adopting simple HTH-like configurations and the second forming a canonical wHTH **(Figure 5D, Supplementary Data S2)**. Given the role of TRDs in directing sequence-specific methylation, these additional DNA-binding HTH domains may act synergistically to enhance target specificity or substrate recognition and binding. Thus, the unique domain organization of Type-3 BrxXI-MTase—featuring a nested TRD and an extended DNA-binding interface with three HTH domains—likely enables a distinct DNA recognition strategy, allowing Type-3 BREX to target unique consensus sites and diversify methylation marking within BREX immunity.

### Type-3 BREX helicase (BrxHII)

The Type-3 BrxHII helicase (∼940 residues), comprises four domains, with only the central helicase unit defined to date (residues ∼75-670), while the flanking regions remain uncharacterized. N-terminal to the helicase unit, we here identify a TUDOR domain (residues ∼1-75) **(Figure 5E, Supplementary Data S2-S3)**, typically involved in protein-protein or nucleic acid interactions (**103,104**). The downstream helicase unit, in contrast to the Type-2 specific BrxHI Ski2-like helicase, has recently been classified as Swi2/Snf2-helicase (**105**). Extending on this classification, our analysis further delineates the C-terminal region of the Type-3 BrxHII helicase, where we identify two additional α+β domains that clearly distinguish it from the Type-2 BrxHI-helicase **(Figure 5E, Supplementary Data S2-S3)**. Notably, we observe that the entire domain architecture parallels the RNA-polymerase (RNAP)-associated Swi2/Snf2-helicase RapA (**106**), except at the N-terminus where RapA carries a tandem TUDOR pair, whereas BrxHII harbors only a single TUDOR domain **(Supplementary Data S2)**.

The first α+β domain following the Swi2/Snf2 helicase of RapA was previously classified as a novel domain, likely due to the limited structural homologs in the PDB at the time or the absence of detailed comparative scrutiny, which may have obscured its similarity to established folds (**106**). Our analysis, however, reveals that it adopts a degenerate REase-like fold **(Figure 5F)**. Structural searches using the first α+β domain of BrxHII retrieved the corresponding α+β domain of RapA as the closest homolog, followed by multiple PD-(D/E)-XK superfamily nucleases—including virus-type replication-repair (VRR) endonucleases and holliday junction resolvases (HJC) **(Figure 5F, Supplementary Data S2)**. A careful examination reveals that, despite retaining the characteristic PD-(D/E)-XK fold, the core β-sheet— particularly β-strand 4—is partially eroded, and its catalytic motif is fully degenerate, lacking conserved residues across all representatives **(Figure 5F, Supplementary Data S2-S3)**. These observations indicate that the REase domain is likely inactive. In prokaryotic immunity, such inactive REases often function as nucleotide sensors when paired with NTPases, particularly in signaling-based immune systems (**107**). The consistent retention of this derived and fast-evolving inactive REase across all BrxHII-helicases points to a similar nucleotide-sensing role in Type-3 BREX immunity. The C-terminal α+β domain adopts a distinct fold, comprising a five-stranded antiparallel β-sheet with two long coiled-coil-like helices forming an insert-like region between strands 4 and 5 **(Figure 5E)**. Structural searches recovered no true homologs beyond the corresponding C-terminal α+β domain of RapA, suggesting that it is unique to these helicases **(Supplementary Data S2)**.

While establishing the phyletic distribution and conserved genomic context of Type-3 BREX, further sequence searches using the ATPase and MTase recovered a distinct group of defense systems centered on the so-called DUF499-ATPases. Strikingly, these searches consistently retrieved both ATPase and MTase homologs together, and our comparative analysis revealed that all three core components of Type-3 BREX—BrxC-ATPase, BrxXI-MTase, and BrxHII-helicase—have homologous counterparts in these DUF499-associated systems **(Figure 6A)**. Previously, these systems were only noted incidentally alongside HEPN-endoRNases and EVE-like cell-death domains (**108,109**). However, their overall diversity and relationship with BREX systems are undetermined as of yet. This unexpected link prompted us to conduct a comprehensive survey, mapping their entire phyletic distribution across prokaryotes. Our in-depth architectural and sequence-structure analysis of DUF499-associated components revealed multiple domain overlaps with core components of Type-3 BREX, indicating a shared functional framework and deep evolutionary linkage, as detailed in the subsequent sections.

**Figure 6.**
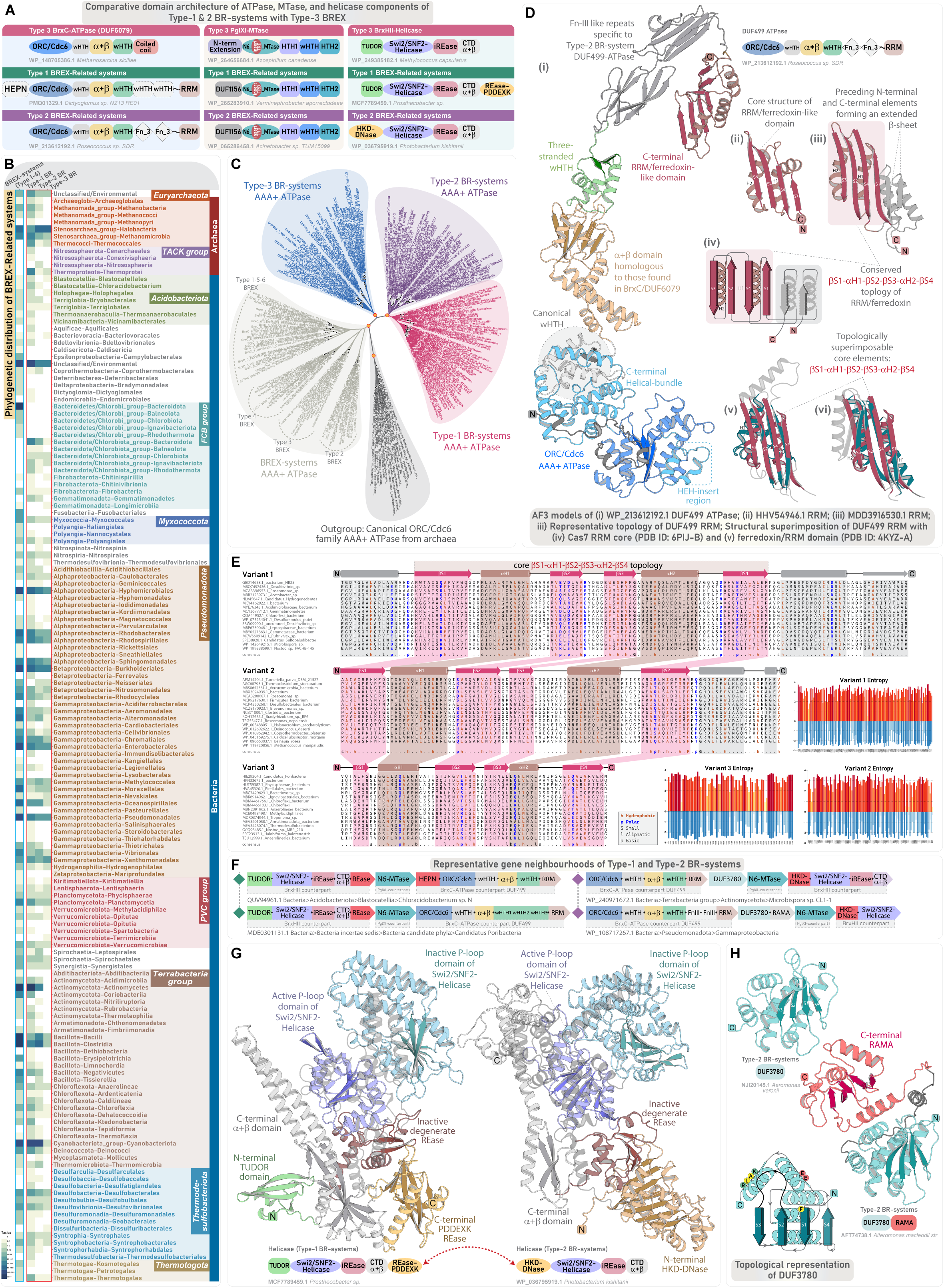
Phyletic distribution, domain characterization, and genomic organization of Type-1 and Type-2 BR systems. **(A)** Comparison of domain architectures of the tripartite core proteins in Type-1 and 2 BR systems, and their homologs from Type-3 BREX, highlighting conserved and subtype-specific features. Domains are labelled as in Figure 4A and 5B. **(B)** Phyletic distribution of all BREX and their related BR systems. The heatmap shows the occurrence of all identified BR subtypes across major taxonomic groups. Colour intensity corresponds to the number of unique NCBI species-level Tax IDs, following the colour scheme in Figure 1B. **(C)** Maximum-likelihood phylogeny of DUF499-ATPases and their BrxC homologs, with canonical ORC/Cdc6 ATPases included as an outgroup. The tree resolves into five major clades, with DUF499-ATPases forming three distinct clades, each representing a specific BR subtype. Nodes with high bootstrap support are marked with orange dots. **(D)** Structural characterization of DUF499-ATPases. (i) Overall 3D structure and domain architecture of DUF499-ATPase. (ii–iii) 3D structures of representative DUF499 C-terminal RRM. (iv) Structural topology diagram highlighting key features and diversity of the DUF499 C-terminal RRM. (v–vi) Structural superimpositions of the DUF499 C-terminal RRM with the Cas7 RRM core (PDB ID: 6PIJ-B) and the RRM/Ferredoxin domain (PDB ID: 4KYZ-A), illustrating shared structural elements. **(E)** Representative MSAs of distinct versions of DUF499 C-terminal RRM. Core α-helices and β-sheets are coloured brown and pink, respectively, with a consensus sequence displayed at the bottom of each alignment. Separate bar plots represent positional entropy for each alignment: yellow-to-red indicates absolute entropy, while light-to-dark blue represents amino acid property-based entropy. **(F)** Representative gene neighbourhoods of Type-1 and Type-2 BR systems, depicted as in Figure 2B. **(G)** Comparative 3D structure and domain architecture of Type-1 and Type-2 BR systems helicases. **(H)** 3D structure of standalone DUF3780 and its fusion with RAMA domain. A corresponding topology diagram highlights conserved residues within DUF3780.

### DUF499-centered BREX-related RM-like defense systems

Unlike the typical six-gene architecture of BREX systems, DUF499-centered systems feature a simpler and conserved tripartite organization, comprising a DUF499 AAA+ ATPase, an N6-MTase, and a helicase-containing protein—mirroring the BrxC-ATPase, BrxXI N6-MTase, and BrxHII-helicase of Type-3 BREX **(Figure 6A)**. Our exhaustive iterative searches, combined with sequence-structure and neighbourhood analysis, recovered a widespread distribution of these systems across 3592 prokaryotic taxa **(Supplementary Data S1)**. Compared to BREX, these are more evenly and abundantly represented in archaea, with 408 taxa spanning both the *Euryarchaeota* and TACK superphylum **(Figure 6B)**. In bacteria, their distribution aligns with that of BREX systems, spanning all major phyla **(Figure 6B)**. Overall, we have delineated three distinct subtypes of these systems—Type-1, Type-2, and Type-3—each characterized by unique architectural and sequence-structure synapomorphies of their tripartite core, along with additional subtype-specific conserved genes in their neighborhoods **(Supplementary Data S1)**. Our phylogenetic analysis shows that DUF499-ATPases form three well-supported monophyletic clades, clearly segregating from BrxC-ATPases, supporting their classification as three evolutionarily distinct lineages **(Figure 6C)**.

Notably, the DUF499-centered gene neighborhood was first noted by L. Aravind and colleagues in 2013 (**108**), during their investigation of HEPN-endoRNase, where they identified a DUF499-ATPase fused to HEPN-endoRNase, along with an associated N6-MTase and helicase. A similar genomic arrangement was later reported by Koonin and coworkers in 2020 (**109**), in their analysis of EVE domains. In both cases, these isolated neighborhoods were identified exclusively in the broader context of their analysis of HEPN- and EVE-anchored genomic associations. Interestingly, these systems were again reported by another group in 2023, but were misclassified as a completely novel defense system, naming it as ‘Hma system’, based on the presence of a Helicase (HmaA), an MTase (HmaB), and an ATPase (HmaC) (**110**). A subsequent experimental study confirmed the phage defense role of so-called HmaA-helicase and annotated HmaC as a DUF499-ATPase (**111**). However, in both instances, the former studies were neither cited nor integrated into these later analyses, and the broader connection to BREX systems remained entirely unrecognized. Thus, to resolve prior nomenclatural inconsistencies and highlight their evolutionary and functional linkage to BREX systems, we refer to them collectively as DUF499-centered BREX-related (BR) systems. The following sections provide a detailed examination of their subtypes, genomic contexts, and comparisons with BREX counterparts.

### Type-1 BREX-related (BR) systems

Type-1, with the simplest genomic organization comprising only the ATPase, MTase, and helicase, is the most widespread BR-subtype identified across 2446 prokaryotic taxa, including 203 archaea **(Figure 6B)**. As previously noted, the central DUF499-ATPase often includes an N-terminal HEPN-endoRNase (**108**). In our dataset, 1370 taxa of Type-1 BR systems featured this HEPN-fused variant, while the remaining taxa retained the canonical DUF499-ATPase **(Supplementary Data S1 and S3)**.

**1. *DUF499-ATPases: Conserved BrxC-Like Core and Hypervariable C-Terminal RRM-fold Domains:*** DUF499-ATPases exhibit a modular architecture closely paralleling that of BrxC-ATPases, with both featuring an N-terminal STAND/ORC-Cdc6 AAA+ ATPase followed by the characteristic triad of domains (wHTH, α+β, and wHTH) **(Figure 6D)**. However, at the C-terminal region, DUF499-ATPases diverge from BrxC by replacing the typical coiled-coil extension with additional wHTH domains and an RRM (RNA recognition motif)/ferredoxin-like domain **(Figure 6D)**. Structurally, the domain preserves all defining hallmarks of RRM domains, featuring a four-stranded antiparallel β-sheet flanked by two α-helices, arranged in the canonical β-α-β-β-α-β topology (**112,113**). Structural similarity searches revealed strong homology to established RRM domains **(Supplementary Data S2)**, including the Cas7-associated RRM of Type III CRISPR complexes (**114,115**). The secondary structural elements closely align with and can be superimposed onto the core unit of Cas7-RRM and ferredoxin/RRM domains **(Figure 6D, Supplementary Data S2)**. Notably, within each BR subtype, we observed substantial sequence and structural variations, with some variants displaying an extended β-sheet of five or six strands (instead of the canonical four), accompanied by additional flanking α-helices at the N- or C-terminal ends **(Figure 6E, Supplementary Data S3)**. Although the RRM fold can occasionally act as an endonuclease (as shown in Cas6-endonuclease) (**114,116–118**), in this context, the lack of conserved catalytic motifs, together with the hypervariable sequence and structural features **(Figure 6E, Supplementary Data S3)**, points to a non-enzymatic role more consistent with molecular interactions involved in recognizing invasive nucleic acids (**60**).
**2. *Helicases:*** Like the Type-3 BrxHII-helicase, the helicase component of all BR systems belongs to the Swi2/Snf2-family of SF2-helicases (**109,111**). These proteins share a common domain architecture, including an N-terminal TUDOR, central Swi2/Snf2-helicase core, an inactive REase, and a C-terminal α+β domain. A notable distinction in Type-1 BR-helicase is the occurrence of an additional C-terminal active REase belonging to the PD-(D/E)XK superfamily (**108,111**). The helicase-nuclease fusion was recovered from 1921 (79%) taxa of Type-1 BR systems **(Supplementary Data S1)**. In the remaining cases, other nucleases—typically belonging to PD-(D/E)XK, HEPN, HNH, or TOPRIM superfamilies—are positioned nearby or, in some instances, within the locus **(Supplementary Data S1)**. These nucleases likely fulfill a compensatory role and are also found alongside the primary helicase-nuclease fusion **(Supplementary Data S4)** (see backup nuclease effector section).
**3. *Adenine Methyltransferases:*** The N6-MTase closely mirrors Type-3 BREX MTases in architecture and sequence-structure features, except for their subtle variability at the N-terminus. While Type-3 MTases possess an extended α+β element preceding the central MTase domain, BR-MTases carry a conserved helical domain (DUF1156, InterPro entry IPR009537). The core MTase remains indistinguishable from that of Type-3 BREX MTases, with the TRD-insert precisely retained after core strand-2, forming the characteristic extended α+β module. Despite subtle variations in the secondary structural elements and sequence profiles, BR-MTases and their Type-3 BREX counterparts retain several shared motifs **(Supplementary Data S3)**. Additionally, all BR-MTases encode a C-terminal triad of HTH-like domains, structurally homologous to those found in Type-3 BREX MTase **(Supplementary Data S3)**. Overall, the MTases across all BR systems exhibit architectural and sequence similarities, indicating a unified mechanism for target recognition and self-DNA modification.

The consistent presence of an ATPase, a DNA-modifying N6-MTase, and a helicase-nuclease effector with known antiphage activity (**111**) positions Type-1 BR systems as an RM-like defense module (**108**). The N6-MTase likely prevents the host genome by selectively modifying the self-DNA. Meanwhile, the DUF499-ATPase, like its BrxC-counterpart, may assemble a scanning and restriction complex, and contribute to invader recognition via its fast-evolving α+β domain, downstream wHTH units, and C-terminal RRM. Upon detection, the helicase-nuclease effector likely mediates the restriction response, causing targeted degradation of foreign DNA. Furthermore, the N-terminal HEPN-endoRNase found in a subset of DUF499-ATPase, may function as a last-resort suicide-effector, triggering self-RNA degradation under overwhelming infection—similar to class 2 CRISPR-Cas effectors—thereby preventing invader proliferation at the population level (**108,119,120**).

### Type-2 BREX-related (BR) systems

We identified Type-2 BR systems in 1120 prokaryotic taxa, including 119 archaea, making them the second most prevalent subtype after Type-1 BR systems **(Figure 6B)**. While both subtypes share a similar genomic architecture, Type-2 BR systems exhibit few notable variations in the domain architecture of their core components **(Figure 6A and 6F)**.

#### Type-2 specific DUF499 and helicase synapomorphies

In 590 (53%) taxa of Type-2 specific DUF499-ATPases, the C-terminal RRM is preceded by two tandem FnIII-like β-sandwich domains **(Figure 6D)**, likely functioning as accessory sensory modules that complement the downstream RRM in invader recognition (**60**). With respect to helicase synapomorphies, the defining feature is the loss of the N-terminal TUDOR domain, which is otherwise retained as the N-terminal-most element in Type-1 and Type-3 BR-helicases. This position is instead occupied by an HKD-endoDNase, which, while architecturally replacing the TUDOR, functionally substitutes for the C-terminal REase found in Type-1 BR-helicases **(Figure 6G)**.

Aside from this replacement—and the occasional loss of the C-terminal inactive REase and α+β domain—Type-2 BR-helicases preserves all other key elements seen in the other BR-helicases **(Figure 6G, Supplementary Data S3)**. Analogous to the helicase-REase fusion effector of Type-1 BR systems, the HKD-helicase fusion effector is retained in 921 (83%) taxa, whereas in the remaining subset, auxiliary nuclease-effectors from distinct superfamilies are encoded alongside the core loci **(Supplementary Data S4)** (see backup nuclease effector section).

#### Type-2 specific DUF3780

Type-2 BR systems consistently feature a DUF3780 (InterPro entry IPR024220), present in 1090 (98%) taxa of Type-2 BR systems. Despite extensive sequence and structural searches, no definitive homologs were identified **(Supplementary Data S2)**. Structural analysis reveals a conserved α+β fold, comprising a four-stranded antiparallel β-sheet, flanked posteriorly by five tightly packed α-helices **(Figure 6H)**. In a subset of their representatives, DUF3780 is fused to designated “reader modules” such as ASCH/PUA domain, and RAMA (restriction enzyme adenine methylase-associated) domain **(Figure 6H, Supplementary Data S3)**, both of which are known to bind and recognize modified bases in DNA and RNA (**108,109,121**). Given that TUDOR domains are known to act as readers of methylation marks (**122–124**), it is plausible that in Type-2 BR systems, DUF3780 substitutes for the missing TUDOR domains, functioning as a reader module that scans self-DNA for modification marks or detects modified bases in phage DNA, including methylated nucleotides.

Overall, Type-2 BR systems mirror the functional architecture of Type-1 BR systems, with few clade-specific variations in their domain organization. Like Type-1 systems, these likely operate as an RM-like defense system, detecting invasive elements and triggering restriction response through their HKD-endoDNase+Swi2/Snf2-helicase effector, while the N6-MTase safeguards self-DNA by selective methylation.

### Type-3 BREX-related (BR) systems

The final subgroup, identified in 569 prokaryotic taxa, including 134 archaea, retains all three core components of canonical BR systems. However, the overall genomic architecture and their domain compositions align closely with typical BREX systems **(Figure 7A)**. Notably, we identified two additional protein components with significant functional implications, previously thought to be unique only to the BREX systems **(Figure 7A and 7B)**. An in-depth comparative analysis of its components reveals the following key features:

1. ***ATPase and MTase Components***: These largely resemble their canonical counterparts, retaining the conserved domain architectures characteristic of Type-1 and Type-2 BR systems. However, a smaller subset of Type-3 BR-system MTases demonstrates notable elaboration within the TRD-insert, incorporating two 4C zinc-finger motifs, instead of the usual singular Zn-finger, which may reflect subtle differences in the target DNA recognition sites **(Supplementary Data S3)**.
2. ***Helicase Component:*** Type-3 BR-system’s helicase demonstrates a key architectural divergence from its canonical BR-counterparts, characterized by loss of its nuclease fusion. Instead, these helicases preserve a domain arrangement identical to the BrxHII-helicases of Type-3 BREX systems **(Figure 7A)**. Furthermore, genomic neighbourhoods of these systems lack conserved standalone nucleases adjacent to core loci **(Figure 7B, Supplementary Data S1)**.
3. ***Recruitment of BREX-like PglZ and iSTAND:*** A defining hallmark of Type-3 BR systems, and a striking finding, is the near-universal presence of an active PglZ across all 569 identified taxa, accompanied by an iSTAND component retained in 466 (82%) taxa **(Supplementary Data S1)**. Sequence and structural comparison of this system’s PglZ to those found in BREX and PorXY two-component systems reveals that it retains all key synapomorphies, including: (i) conserved residues at the active site for coordinating Zn^+^ and Mg^2+^ ions; (ii) the characteristic α/β/α structural-scaffold with a six-stranded central β-sheet; and (iii) a three-stranded β-sheet insert forming a cap-like subdomain (**125**). However, unlike their BREX counterparts, this occurs as a standalone domain without additional fusions **(Figure 7C, Supplementary Data S3)**. Similarly, the iSTAND is characterized by: (i) eroded Walker-A/B motifs; (ii) the hallmark three-stranded antiparallel β-sheet insert region following β-strand-4; and (iii) absence of the C-terminal helical bundle—structurally unifying it with the iSTAND domains found in BREX systems **(Figure 7D and 7E)**. However, as observed in BREX, the iSTAND domain is rapidly evolving at the sequence level **(Figure 7E, Supplementary Data S3)**, consistent with its role as a sensor module adapting to the diverse repertoire and combat strategies of invasive elements.

**Figure 7.**
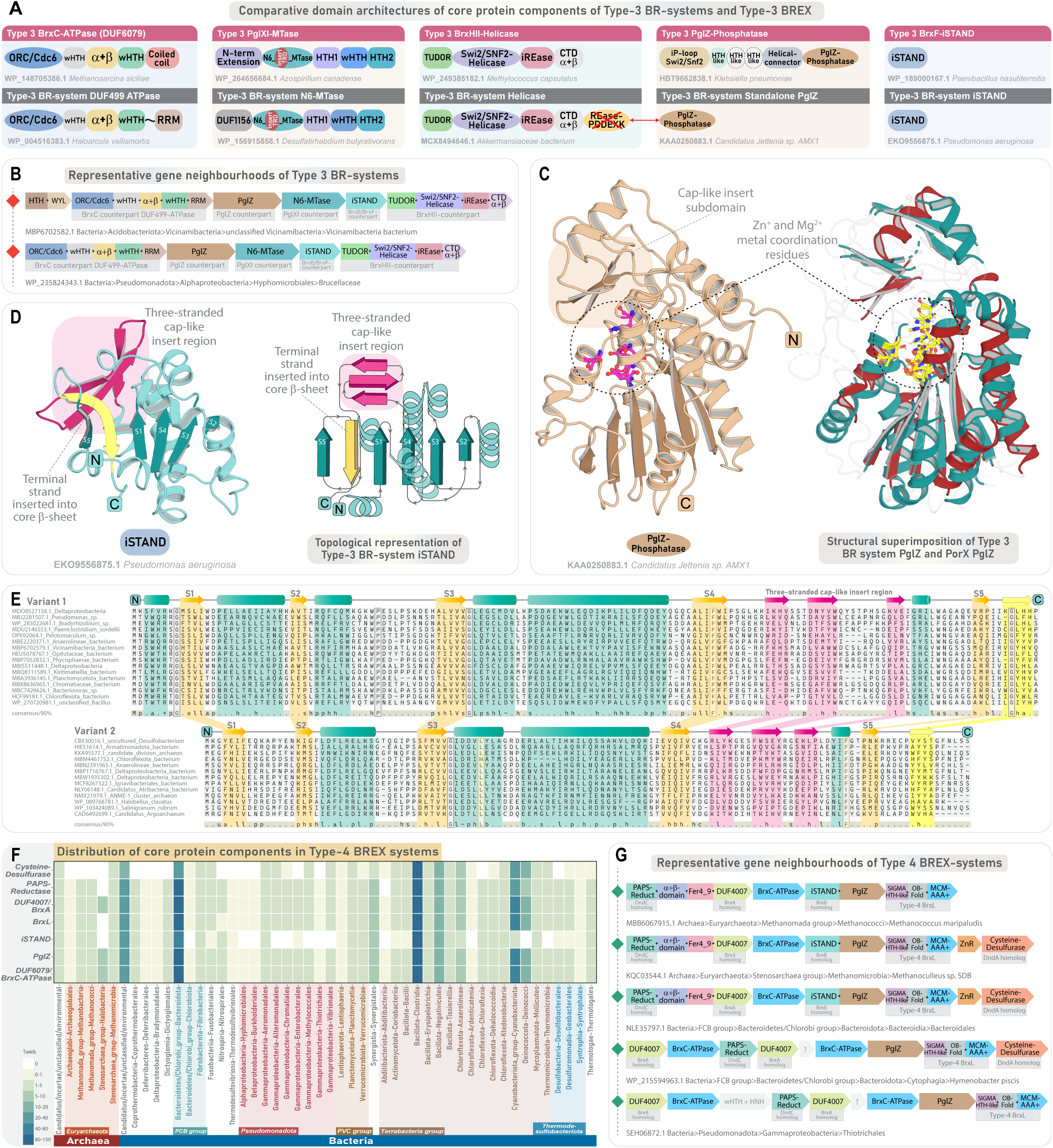
Genomic organization and domain characterization of Type-3 BR and Type-4 BREX systems. Panels **(A-E)** corresponds to Type-3 BR systems, and panels F and G corresponds to phyletic distribution and genomic organization of Type-4 BREX systems. **(A)** Comparison of domain architectures of core protein components in Type-3 BR systems and their homologs from Type-3 BREX. Domains are labelled as in Figure 4A and 5B. **(B)** Representative gene neighbourhoods of Type-3 BR systems, depicted as in Figure 2B. **(C)** 3D structure of standalone PglZ of Type-3 BR system and its structural superimposition with PorX PglZ, highlighting the conservation of key residues for enzymatic activity. **(D)** 3D structure and topology diagram of Type-3 BR-system iSTAND, highlighting key features. **(E)** Representative MSAs of Type-3 BR system iSTAND, labelled as in Figure 3. **(F)** Phyletic distribution heatmap showing the occurrence of core protein components in Type-4 BREX, across diverse taxonomic groups. Colour intensity represents the number of unique NCBI species-level Tax IDs, following the scheme used in Figure 1B. **(G)** Representative gene neighbourhoods of Type-4 BREX systems, depicted as in Figure 2B.

Thus, Type-3 BR systems, while retaining the three core components of BREX, have additionally recruited PglZ and iSTAND, making them the closest in organization and function to BREX machinery. Although it remains unclear which of the three BR-subtype emerged first (potentially from Type-3 BREX), the interchange of the PD-(D/E)XK or HKD-endoDNase of Type-1 and Type-2 BR systems with PglZ, strongly supports its role as the primary nuclease mediating invader restriction—consistent with recent evidence of PglZ’s nuclease activity in Type-1 BREX (**15,16**). The parallel recruitment of an inactive STAND, akin to BrxB/BrxF-iSTAND, further underscores mechanistic convergence with BREX machinery, in terms of recognizing the invasive elements, followed by its restriction via PglZ (see sections below for sensory and effector responses). In summary, our identification and characterization of three new BR systems enabled the delineation and comparison of their functional modalities with those of established BREX systems, thereby refining our understanding of the core components shared across both. Collectively, the relatively streamlined RM-like BR systems described here underscore a deep evolutionary connection and shared ancestry with BREX, offering fresh insights into the emergence of modular complexity in these defense systems.

### Type-4 DNA-Phosphorothioation-based BREX systems

Type-4 BREX is unique among all BREX-subtypes as the typical methylation component is substituted by a PT-modification apparatus, linking it functionally to PT-based Dnd and SSP systems (**1,2,21**). Of the five known major components of Type-4 BREX, three components— BrxC-ATPase (SspC), PAPS-reductase (SspD), and DUF4007 (SspB)—shares homology with counterparts in SSP systems, while PAPS-reductase homolog (DndC) are also found in Dnd-modification systems (**2,21**).

In our gene neighborhood analysis of 457 taxa, PAPS-reductase and DUF4007— frequently occurring as a fusion component—were detected in 406 (89%) and 442 (97%) taxa, respectively **(Figure 7F)**. BrxL was identified in 436 (96%) taxa, while PglZ was retained across all analyzed taxa. Notably, we identified a cysteine-desulfurase that has not been previously associated with BREX systems, present in 255 (56%) taxa, which potentially represents the sixth core component of Type-4 BREX **(Figure 7F)**. This enzyme often co-occurs with a small zinc-finger protein bearing similarity to HTH-type transcriptional regulators **(Figure 7G, Supplementary Data S1)**. In PT-dependent systems, cysteine-desulfurase and PAPS-reductase play critical roles in the initial steps of PT-modification, wherein cysteine-desulfurase mobilizes sulfur from cysteine, serving as a donor, while PAPS-reductase accepts and transfers the sulfur to the DNA backbone (**1,3–6**). The sporadic distribution of cysteine-desulfurase likely reflects its substitution by housekeeping homologs (IscS, NifS, SufS) (**6**), a scenario that has been validated *in vivo* (**126**).

While the cysteine-desulfurase of Type-4 BREX closely resembles its Dnd-system counterpart (DndA), the PAPS-reductase shows notable divergence from DndC. Specifically, it lacks a dyad of cysteines unique to DndC (**1**). Instead, it features an additional α+β domain— structurally homologous to the Mog1p/PsbP-like fold containing DUF1795—followed by a highly conserved cysteine-rich region reminiscent of (4Fe-4S) iron-sulfur cluster-binding ferredoxin-type domain **(Supplementary Data S2-S3)**. The roles of these extra elements remain unclear within the context of BREX-immunity. Interestingly, this same arrangement is also found in a subset of PAPS-reductase (SspD) in Ssp-systems, hinting at a shared evolutionary origin and potentially related biochemical pathways **(Supplementary Data S3)**.

### Identification of a diverse array of auxiliary and backup nuclease effector modules retained across all BREX and their related systems

Evidence from a recent study demonstrating the nuclease activity of PglZ in Type-1 BREX showed that inhibition of PglZ diminishes the system’s defensive capacity but does not completely abolish resistance against phages (**15**). This indicates that residual anti-phage activity is still maintained through yet-uncharacterized compensatory mechanisms (**15**), underscoring the functional plasticity of BREX-mediated immunity. Supporting this, our comprehensive cataloging and gene neighborhood analysis of all BREX and BR systems has, for the first time, revealed a previously unrecognized repertoire of auxiliary nuclease effectors that co-occur alongside the primary effector PglZ. These include endoDNases such as HNH, HKD-endoDNase, PD-(D/E)XK-REase, and TOPRIM/OLD nucleases, as well as endoRNases from the HEPN, PIN, and Schlafen superfamilies. Many of these are encoded within the same operon as the core components—often flanked by BREX-associated genes in both upstream and downstream regions. To ensure functional relevance, we filtered for gene neighborhoods where these nucleases are positioned immediately adjacent to or within the core loci, excluding cases where they were distantly located and their genomic association was uncertain. Even under stringent filtering criteria, we identified numerous conserved neighborhoods where these nucleases are consistently closely linked with core BREX genes, collectively revealing their presence in at least 4,363 taxa across all BREX and related systems—which excludes the known auxiliary effectors already embedded within the core components **(Figure 8A, Supplementary Data S4)**. These associations were not reported in earlier studies, likely due to limited genomic sampling and the absence of large-scale contextual analyses. Thus, through an expansive survey, combined with a thorough reconstruction of gene neighborhoods, we reveal that the retention of these diverse nuclease effectors is not incidental but instead represents a recurring pattern of genomic and potential functional integration with BREX and BR systems across a broad phylogenetic spectrum **(Figure 8A, Supplementary Data S4)**.

**Figure 8.**
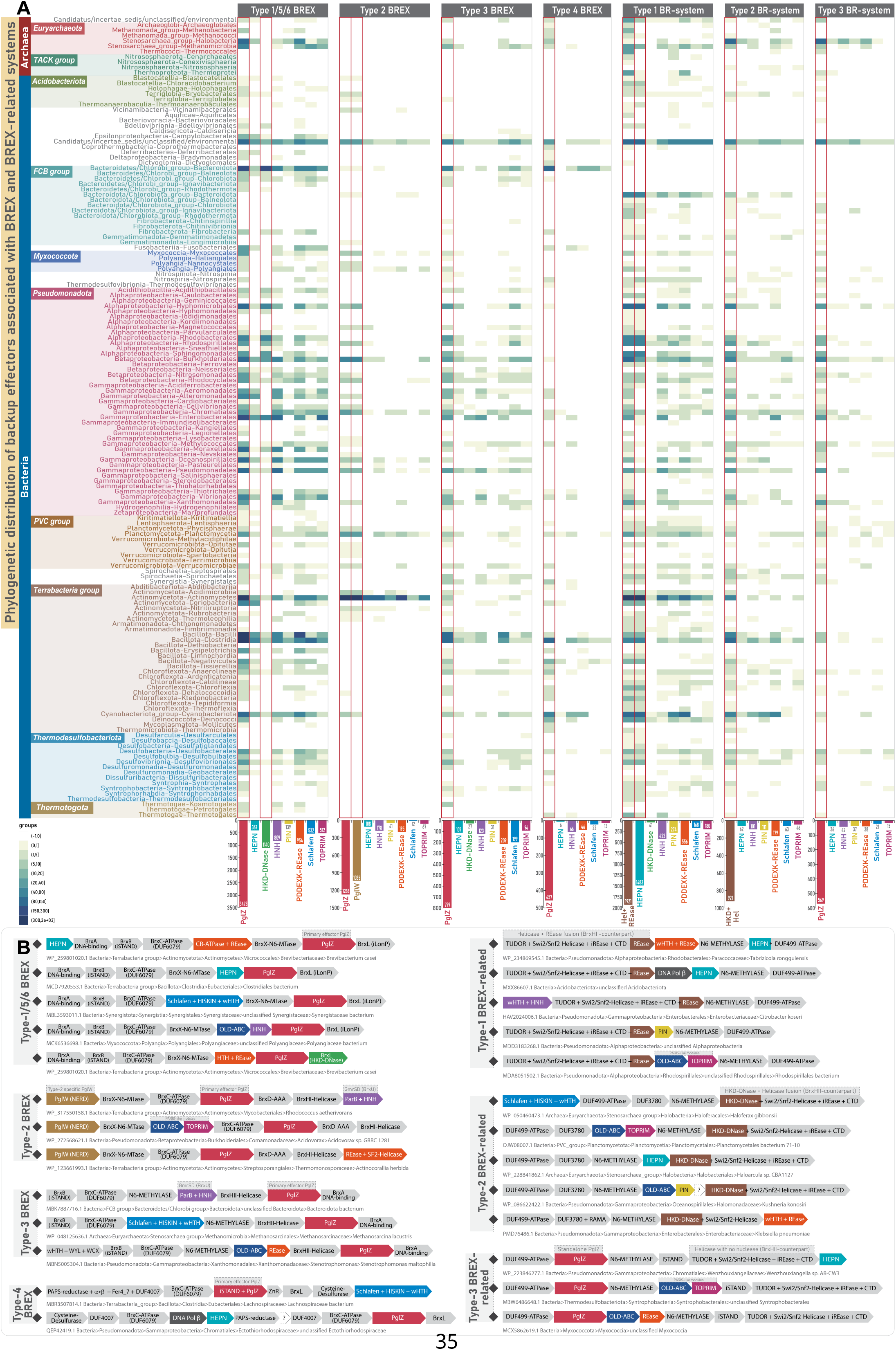
Phyletic distribution and genomic context of nuclease effectors in BREX and BR systems. **(A)** Phyletic distribution of primary and backup nuclease effectors across BREX and BR system subtypes. The panel comprises seven heatmaps, each corresponding to a major subtype of BREX or BR systems. Type-1, Type-5, and Type-6 BREX systems are grouped together in the first heatmap. Within each heatmap, columns represent the phyletic distribution of nuclease effector domains associated with the corresponding system, identified through gene neighbourhood analysis. Colour intensity reflects the number of unique NCBI species-level Tax IDs, following the scheme used in Figure 1B. The total number of unique Tax IDs for each nuclease is summarized in a bar plot positioned below the heatmaps. Primary nuclease effectors, as well as backup effectors embedded within core protein components, are indicated with red borders. **(B)** Representative gene neighbourhoods illustrating the occurrence of distinct nuclease effectors embedded within core loci of BREX and BR systems. The PglZ primary effector in all BREX and Type-3 BR systems are colored in red. The REase and HKD-endoDNase primary nuclease effector in Type-1 and 2 BR system, respectively, are colored in brown. Canonical core components are uniformly colored in grey background, while backup effector modules are color-coded separately to enhance visibility.

Among the identified effectors, members of the PD-(D/E)XK family of REase were the most prevalent across BREX and BR systems. These occur both as standalone REase and as fusions with other conflict-associated modules such as HARE-HTH, CR-ATPases, and SF2-Helicases (**82,127–129**) **(Figure 8B, Supplementary Data S4)**. A subset of these is fused to variable N-terminal sensory modules, forming the basis of *shedu* immune systems, which have been previously linked to BREX (**130**). In some systems, these REases integrate directly into the core machinery—for example, the NERD-REase (PD-(D/E)XK clan) in PglW of Type-2 BREX, and a C-terminal active PD-(D/E)XK-REase in the helicase-nuclease component of Type-1 BR systems. In BR systems, these REase domains act as the primary effector (**111**), whereas in Type-2 BREX, the NERD-REase appears to function as a secondary “backup” nuclease, complementing the putative principal effector PglZ **(see PglW section)**. Notably, in a subset of the so-called Type-5 BREX, we show that the BrxHII-helicase itself carries an active C-terminal REase, thereby establishing a dual-effector configuration with PglZ that had remained unidentified until now **(Figure 2B, Supplementary Data S4)**.

The HNH-endoDNase represents the second-most prominent class of effector, with the most notable version being the GmrSD-family Type IV restriction endonuclease (**131**), characterized by a ParB and HNH domain architecture, which has been shown to complement phage restriction in Type-1 BREX (**132**) **(Figure 8B, Supplementary Data S4)**. Beyond these, our survey uncovered additional variants of HNH-endoDNases, including standalone HNH domains, and versions fused to an N-terminal wHTH and SAD(SRA) domains—previously associated with Dnd-systems and known for recognizing modified DNA (**1,133,134**) **(Figure 8B, Supplementary Data S4)**. Also, we observed a further HNH variant in a subset of our datasets, where HNH is genomically linked to OLD (Overcoming Lysogenization Defect) ABC-ATPases, forming a distinctive dyad architecture.

A notable theme emerging from our survey is the pervasive association of nucleases with OLD-ABC ATPases, spanning at least 1242 taxa across BREX and BR systems **(Figure 8B, Supplementary Data S4)**. The most prominent representatives are the TOPRIM-linked OLD-ABC. While the versions recruited by BREX and BR systems are identified in this study, distinct collections of TOPRIM-linked OLD-ABC and their associated neighborhoods were previously described by L. Aravind and colleagues as part of the broader repertoire of ABC-ATPase– anchored conflict systems (**50**). These operons were later independently re-reported twice, under different nomenclature, as the so-called PARIS (AriA: OLD-ABC and AriB: TOPRIM) and Gabija (GajA: OLD-ABC+TOPRIM and GajB: SF2-helicase) anti-phage systems (**135–140**). Additionally, these ATPases were often misclassified as AAA+ ATPase (**140**), or more recently, as Rad50/SMC-family ABC-ATPases (**135**). However, the sequence-structure synapomorphies unambiguously place their assignment to the OLD-family of ABC ATPases **(Supplementary Data S3)**. In PARIS, the OLD-ABC (AriA) detects the phage-encoded Ocr DNA mimics—potent inhibitors of host restriction systems—and subsequently activates the TOPRIM-nuclease (AriB), which cleaves host tRNAs to abort infection (**135**). Since phage-Ocr proteins suppress BREX activity (**54**), the recruitment of PARIS-like modules symbolizes a direct countermeasure against phage anti-restriction strategies. Beyond the recruitment of canonical OLD-ABC and TOPRIM modules (PARIS), we identified an assemblage of 594 unique taxa, in which the OLD-ABC are paired with alternative nucleases—such as the HNH-endoDNase, PD-(D/E)XK-REase, and PIN-endoRNases—rather than the usual TOPRIM-nuclease **(Figure 8B, Supplementary Data S4)**. The coupling of OLD-ABCs with various such endonucleases has been previously reported in the ABC-ATPase–anchored RM-like systems as well (**50,141**). Collectively, these OLD-ABC-centric modules likely form a versatile backup effector layer within BREX-machinery, reinforcing their capacity to withstand phage counter-defence strategies targeting the core system.

The HKD-endoDNase exhibits a more restricted distribution, and we primarily found it in Type-1 BREX, where it localizes to the C-terminus of the BrxL component **(Figure 2F)**. In Type-2 BR systems, however, the HKD-endoDNase occupies the N-terminus of the helicase component and serves as the primary effector **(Figure 6G, Supplementary Data S3)**.

Among endoRNases, the HEPN and Schlafen family RNases are the most widely represented, whereas PIN-endoRNases are less commonly observed **(Figure 8A)**. HEPN domains are typically found as standalone modules, though in some cases they are fused to minimal nucleotidyltransferases (MNTs) of the DNA polymerase-β superfamily—an architecture reminiscent of Type II toxin–antitoxin (TA) systems (**108,142**) **(Figure 8B, Supplementary Data S4)**. In Type 1 BR systems, HEPN are often fused to the N-terminus of the core DUF499-ATPase, effectively forming a secondary effector alongside the primary helicase-nuclease component. Lastly, the Schlafen-family RNases—fused to a C-terminal minimal histidine kinase and a wHTH—are also widely disseminated across BREX and BR systems, occurring in at least 1017 taxa where they are closely linked with the core loci **(Figure 8A)**. In prokaryotes, several Schlafen-linked systems can operate as an anti-phage defense module, which can sense phage-derived signals and activate the RNase to cleave host and viral tRNAs, thereby inducing abortive infection pathways (**50,143,144**).

The incorporation of diverse effectors as multiple layers of defense is a recurring theme in several prokaryotic conflict systems (**1,60,107,129,145–148**). Here, we present the first comprehensive catalog of various nuclease domains that are consistently retained alongside the primary effectors in BREX and BR systems. These occur both as standalone domains and as part of multidomain architectures fused with NTPases and DNA-binding domains—potentially enabling sensory or self/non-self-discriminatory functions (**1,60,61,107**). We propose two overarching functional themes for these effectors. First, the endoDNases fused with additional domains—such as ParB-or wHTH/SAD(SRA)-linked HNH, BrxL-HKD-endoDNase fusions, PD-(D/E)XK-REase coupled to SF2-helicases or CR-ATPases— likely constitute an auxiliary network of *bona fide* restriction factors, poised to compensate when the primary nuclease is neutralized, and likely complements PglZ by inflicting additional damage on invading DNA (see sections for effector response). Second, the endoRNases and standalone endoDNases, many of which are linked to phage-sensing OLD-ABCs, appear to serve as backup suicidal effectors, unleashing a broader, less discriminating nucleic-acid degradation response during late-stage or overwhelming infections (**1,50,108,149,150**). In a broader context, nearly all identified nuclease superfamilies and their diverse architectures are retained across the entire breadth of BREX and BR systems, suggesting that these effectors are not sporadically acquired. Instead, they reflect a unified functional theme of backup arsenals—underscoring the evolutionary plasticity of BREX systems to mitigate phage counter-defense mechanisms (**151–153**), while expanding their ability to neutralize a broader range of invasive elements.

### Novel HerA/FtsK-centered BREX-related capture systems with potential invader interception mechanisms

Given the recent characterization of PglZ as the primary nuclease effector in Type-1 BREX (**15**), and our identification of a unique Type-3 BR-system where canonical nucleases are interchanged with standalone PglZ, we investigated whether PglZ has also been recruited elsewhere as a central effector in other defense contexts. Strikingly, our systematic searches and neighborhood analyses uncovered a novel system that integrates multiple components associated with diverse immune strategies. Although sparsely distributed—identified in only 45 unique taxa—this system spans a wide phylogenetic range, including representatives from three archaeal and seven bacterial classes, as well as several unclassified groups **(Supplementary Data S1)**. The system exhibits a conserved architecture encompassing seven components: (i) PglZ, (ii) ORC/Cdc6 AAA+ ATPase-containing DUF6079, (iii) STAND-NTPase, (iv) tripartite DNA-binding module as DUF4007, (v) HerA/FtsK-translocase, (vi) GCN5-related N-acetyltransferases (GNAT), and (vii) tRNA-guanine transglycosylase (TGT) **(Figure 9A-9B, Supplementary Data S1)**. Notably, four of these components share homology with known BREX-counterparts, suggesting a potential evolutionary link: (i) the so-called DUF6079 contains all four domains typical of BrxC; (ii) PglZ retains the N-terminal inactive Swi2/Snf2-Helicase, but lacks additional elements (tandem HTH-like units, helical-connector and C-terminal all β-sandwich IG-like or wHTH) commonly found in BREX-PglZ **(Figure 9C)**; (iii) DUF4007 mirrors the tripartite DNA-binding architecture of BrxA **(Figure 9D)**; and (iv) STAND-NTPase represents an enzymatically active homolog of BrxB-iSTAND **(Figure 9E and 9F)**.

**Figure 9.**
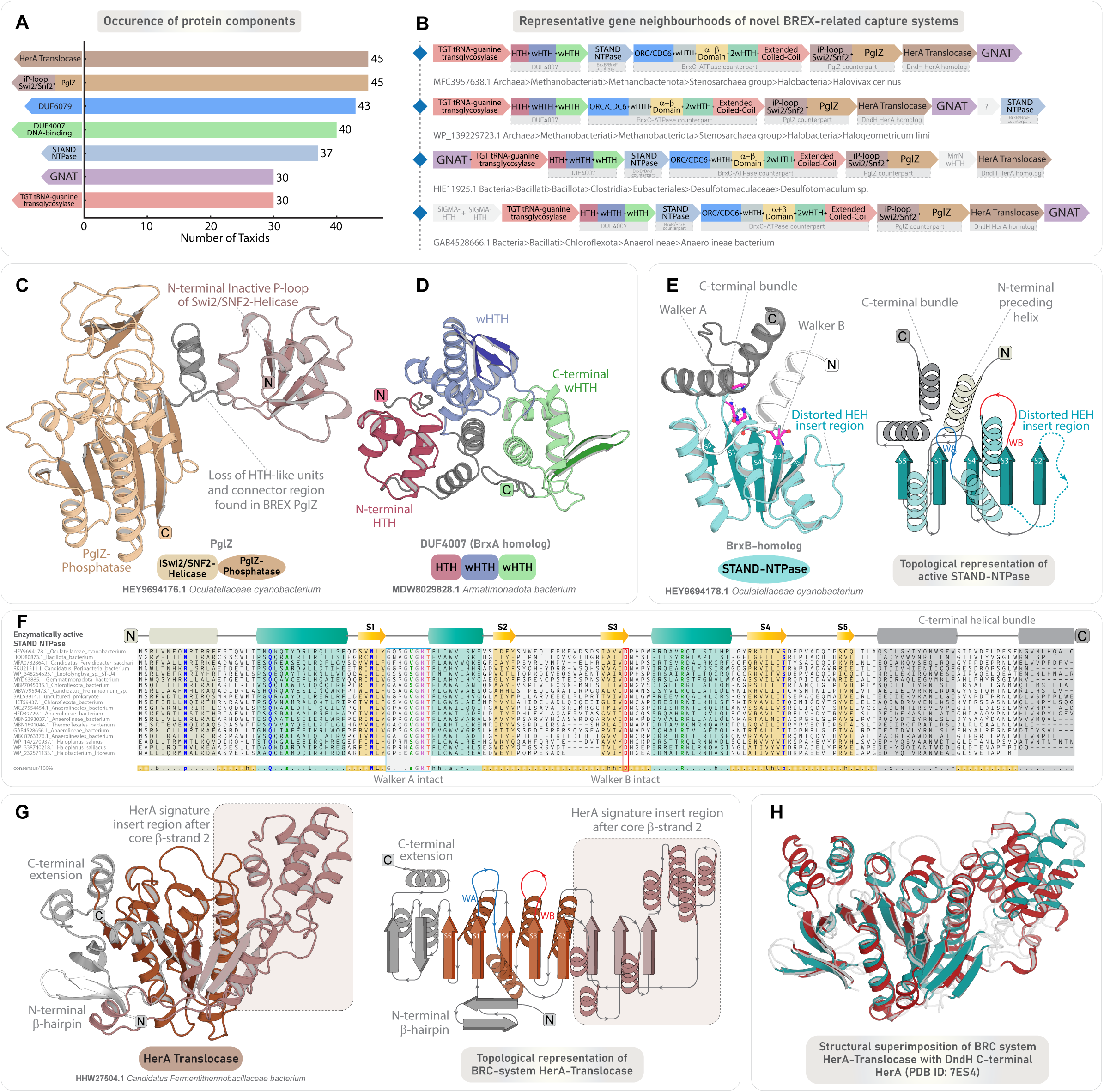
Genomic organization and domain characterization of a novel HerA/FtsK anchored BREX-related capture system. **(A)** A bar-plot depicting the prevalence of all seven protein components associated with the novel BRC system, in which x-axis denotes number of Tax IDs, and y-axis denotes the protein component. **(B)** Representative gene neighbourhoods of BRC system, depicted as in Figure 2B. **(C and D)** 3D structure and domain architecture of PglZ and DUF4007 (BrxA homolog) specific to BRC system. **(E)** 3D structure and topology diagram of enzymatically active STAND NTPase (BrxB homolog) of BRC system, highlighting key features. **(F)** Representative MSAs of BRC system STAND NTPase, labelled as in Figure 3. **(G)** 3D structure and topology diagram of HerA-Translocase of BRC system, highlighting key features. **(H)** Structural superimposition of BRC system HerA-Translocase with DndH C-terminal HerA (PDB ID: 7ES4).

The defining feature distinguishing this system from BREX is the recruitment of HerA/FtsK translocase, GNAT, and TGT domains. While GNAT and TGT appear in a subset of these loci, the HerA/FtsK is universally retained across all identified taxa **(Figure 9A, Supplementary Data S1)**. The consistent recruitment of HerA/FtsK translocases has been documented by us in DndFGH systems and their related HerA/FtsK-anchored anti-invader capture systems (**1**). Structural searches of the HerA/FtsK in the present system also identify DndH-HerA/FtsK as a top hit, indicating potential evolutionary and functional parallels **(Figure 9G-H, Supplementary Data S2)**. Notably, the co-occurrence of HerA/FtsK with ORC/Cdc6-like AAA+ ATPases, rapidly evolving wHTH domains akin to DUF4007, and effector nucleases has been recurrently observed across various anti-invader capture systems (**1**). The fast-evolving DNA-binding and/or sensory domains are hypothesized to facilitate spatial localization of the defense machinery in proximity to invading genetic elements during molecular processes such as plasmid segregation, phage DNA packaging, or conjugative transfer (**1**). Once localized, the HerA/FtsK-mediated DNA translocation then intercepts the foreign DNA, capturing and channeling it toward associated nucleases for targeted degradation (**1,154**).

In contrast, TGTs are central to the sophisticated RM-like Dpd-defense systems, which mark host DNA with modified nucleosides—2′-deoxy-7-cyano-7-deazaguanosine (dPreQ₀) and 2′-deoxy-7-amido-7-deazaguanosine (dADG)—to guide endonuclease-mediated restriction of invaders (**7–9**). In Dpd systems, DpdA (TGT homolog) synthesizes dPreQ₀, which is inserted into DNA via the ATPase activity of DpdB (a ParB-like ATPase). Subsequently, DpdC (DUF328; Pfam PF03883) catalyzes the hydrolysis of dPreQ₀ to its final derivative, dADG. Notably, DpdB has been suggested to be dispensable in certain Dpd clusters (**7,8**). In our system, no homologs of DpdB or DpdC were detected. Instead, the active STAND-NTPase may substitute for DpdB, driving the ATP-dependent dPreQ₀ insertion into DNA, and any potential base modification is perhaps restricted to dPreQ₀ alone. Furthermore, the GNAT— occasionally fused with TGT—are functionally versatile and known to catalyze acetylation across a broad range of substrates, including small molecules, nucleic acids, carbohydrates, and proteins (**155–157**).

Based on the associated protein components, we propose that this newly identified system represents an offshoot of the previously described HerA/FtsK anti-invader capture system, where the homologs of BREX and Dpd system components have been co-opted to form a multi-layered immune apparatus, and hence we refer to it as BREX-related capture (BRC) systems. The BREX-counterparts are likely to oligomerize into a macromolecular sensory and restriction unit, analogous to BREX systems, whereas the HerA/FtsK-translocase may help to mobilize the invader DNA near the PglZ-effector of the restriction apparatus (**1,154**). Additionally, a subset of the BRC-system may also modify host-DNA via GNAT and TGT modules to establish a self/non-self-discrimination signal. Collectively, BRC systems exemplify the modularity of prokaryotic immunity, showcasing how multiple strategies—spanning distinct mechanistic themes—can be co-opted into a unified, multifaceted conflict apparatus to protect against invasive elements.

### Overall functional modalities and organizational features of BREX and their related systems

Through meticulous analysis of all protein components in BREX and related systems, we identified multiple functional domains within various uncharacterized regions that had previously eluded annotation or functional characterization. These include a myriad of potential DNA-binding HTH/wHTH modules in various configurations, rapidly evolving and catalytically inactive enzymatic domains, regulatory elements that may be critical for complex assembly, domains potentially facilitating site-specific DNA-binding, and additional components exhibiting novel folds. By leveraging a thorough comparative analysis of an extensive dataset, we elucidate the modular organization and evolutionary innovations that underpin BREX and related systems—offering fresh insights into their mechanistic basis, spanning invader sensing, self/non-self-discrimination, effector action, and the regulatory controls that fine-tune their deployment, thereby providing opportunities for targeted experimental exploration.

### Modular network of domains associated with invader sensing and self/non-self-recognition apparatus

The coordination and regulation of the multi-protein BREX complex are likely mediated through multiple DNA-anchoring interfaces, as suggested by the widespread identification of nucleic acid-binding domains. Moreover, unlike traditional RM-systems, BREX systems harbor a complex network of functional domains capable of interacting with foreign molecular entities. Several of these domains exhibit dynamic sequence and structural variability, often bearing hallmarks of positive diversifying selection—reflecting adaptive pressures driven by the evolutionary arms race with invasive elements **(Supplementary Data S3)**. Nearly all major components of BREX and BR systems encode one or more such modules, potentially contributing to invader detection and self/non-self-recognition.

The iSwi2/Snf2 helicase of PglZ and the iSTAND NTPases display marked sequence divergence across all BREX subtypes, with substantial variability evident even within individual subtypes **(Supplementary Data S3)**. Despite this variation, they retain the signature P-loop fold, suggesting a conserved ancestral role in nucleic-acid binding—potentially enabling recognition of phage-derived nucleotides and their derivatives. PglZ further harbors additional binding modules, including a variable number of HTH-like elements with predicted DNA-anchoring functions, as well as a variably retained C-terminal β-sandwich IG-like domains, which may recognize foreign proteins akin to classical immunoglobulin domains **(Figure 10)**. Comparable to the inactive P-loop NTPases, the penultimate domain of the BR-system helicase-nuclease fusion effector—a highly derived and inactive REase—likely serves as a non-enzymatic nucleotide sensor by recognizing and sequestering invasive nucleic-acids, allowing for a precise effector response, as also observed in other conflict systems (**60,61,107,128**). Within Type-1 and Type-2 BR-system helicases that possess active endonucleases, the preserved inactive domains (iREases) may function as molecular decoys, intercepting phage-encoded inhibitors that would otherwise target the active nuclease (**60,61,107**). Furthermore, the C-terminus of DUF499-ATPases in BR systems consistently features a hypervariable RRM domain and occasionally includes an FnIII-like repeat. While the RRM is likely to bind foreign nucleic acids, the FnIII-like repeats may facilitate peptide-based sensing, thereby broadening the system’s capabilities to recognize foreign elements. Overall, the differential recruitment of distinct binding modules across BREX and BR systems underscores their potential to recognize and respond to a diverse repertoire of phages and their molecular cues **(Figure 10)**.

**Figure 10.**
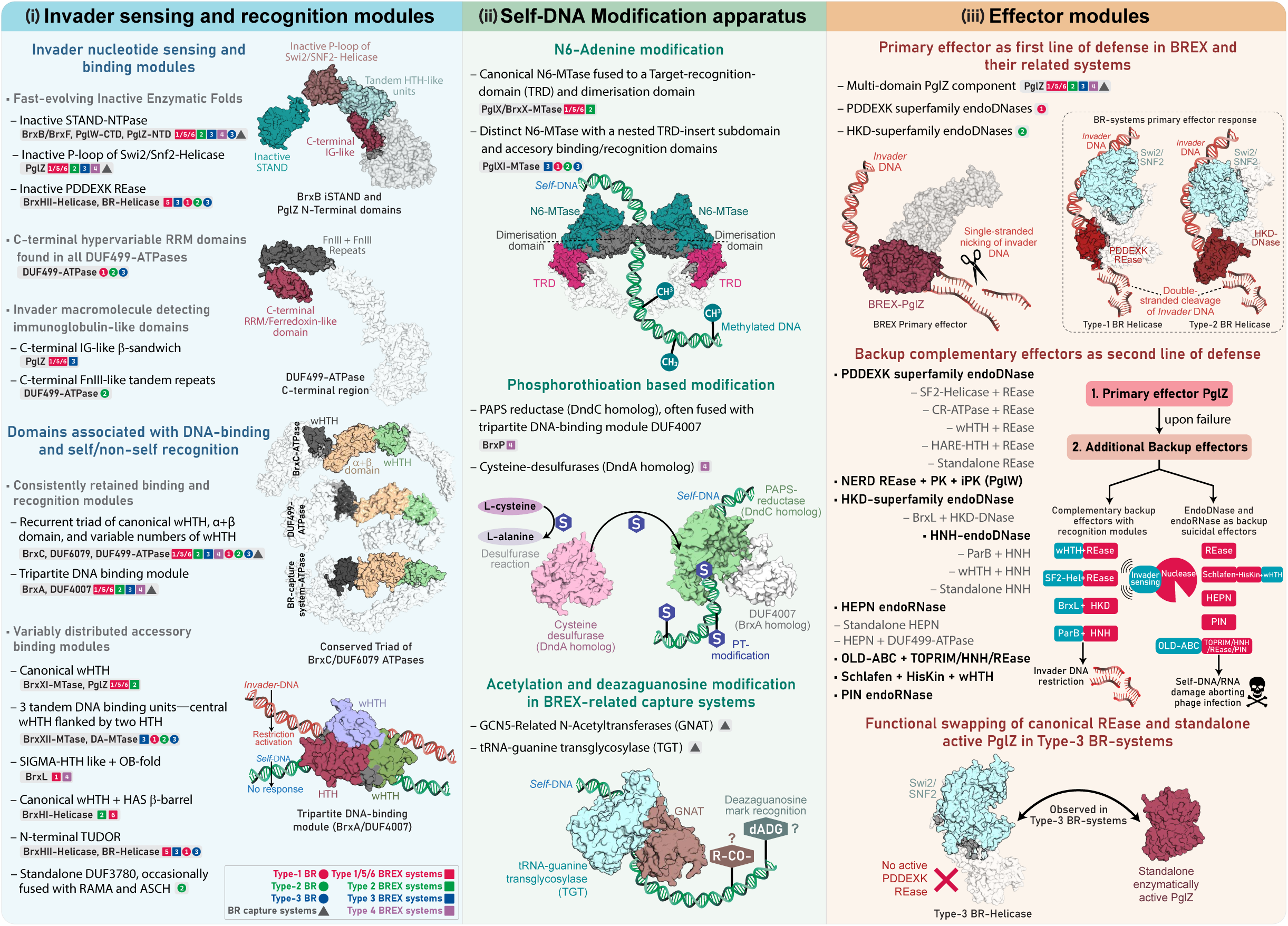
Schematic illustration summarizing the functional organization of BREX and their related systems. The diagram highlights three major groups of functional modules: (i) Invader sensing and recognition: components responsible for detecting foreign genetic elements, potentially through nucleotide-based signals or direct DNA-protein interactions. (ii) Self-DNA modification and tagging enzymatic modules that mark host DNA to distinguish self from non-self and prevent autoimmunity. (iii) Effector modules: primary and backup nuclease effectors that mediate targeted degradation of invading nucleic acids or trigger non-specific suicidal responses to halt invader replication. The distribution of these components across different systems is depicted using colored geometric shapes with numbers, where the shape denotes the system and the number corresponds to the subtype. Squares denote BREX systems, circles denote BREX-related (BR) systems, triangles denote BREX-related capture (BRC) systems, and the color/number denotes the specific subtype.

Besides the repertoire described above, the central ATPases of BREX and related systems consistently encode a recurrent triad—a canonical wHTH, an α+β domain, and one or more additional wHTH modules—that is conserved across all lineages. This triad likely facilitates the scaffolding of ATPase-anchored oligomeric complexes along the DNA substrate and may also contribute to recognition of invasive elements. In parallel, the tripartite DNA-binding module—denoted as BrxA or DUF4007—likely forms a dedicated scanning unit involved in invader recognition and self/non-self-discrimination during the immune response. Beyond these core components, several auxiliary nucleic-acid binding domains are selectively associated with specific BREX components, exhibiting subtype-specific distribution. These include canonical wHTH, SIGMA-HTH-like, OB-fold, HAS b-barrel, and TUDOR domains **(Figure 10)**. Notably, these modules show relatively conserved sequence profiles and retain an invariant structural scaffold **(Supplementary Data S3)**, indicative of archetypal roles as accessory domains that likely enhance the binding specificity or stability of the larger protein assemblies in which they reside.

Collectively, this diverse array of domain configurations—spanning multiple distinct protein folds—underscores the functional versatility of BREX components in recognizing host DNA and, more critically, foreign DNA. Multiple fast-evolving nucleotide-binding modules with putative sensory functions likely constitute a frontline recognition layer for primary surveillance, which likely operates in concert with the DNA-scanning mechanisms of the overall complex to distinguish self from non-self. In agreement with this, the recent experimental evidence suggests that Type-1 BrxX-MTase—functioning in a complex with BrxC, PglZ, and BrxB-iSTAND—can bind non-methylated BREX consensus sites on phage-DNA *in vivo* (**14**). Together, these observations suggest that BREX systems operate as a multivalent immune complex orchestrated around an oligomeric BrxC-ATPase scaffold, which together can sense invasive elements, evaluate their modification status, and coordinate the effector activation accordingly.

### Modification apparatus for tagging self-DNA

BREX systems employ at least two distinct strategies to mark self-DNA, including the well-characterized adenine methylation observed in Type-1 BREX (**10,14,22,54**), and the PT-based sulfur modification specific to Type-4 BREX (**21**). The novel BRC-system identified in this study potentially expands this modification repertoire by utilizing acyl and 7-deazaguanine nucleoside derivatives as alternative self-markers.

The MTase associated with Type-2 BREX/Pgl-systems is closely conserved and structurally congruent to its Type-1 counterpart, indicating a certain degree of functional homology. However, sequence divergence in their TRDs and accessory domains suggests that they methylate distinct consensus sites. The Type-3 BREX MTase, while retaining an N6-MTase core, features a large and distinct TRD embedded within the core MTase, suggesting a varied mode of DNA target recognition and consensus sites compared to Type-1 BREX. Notably, this nested TRD architecture is consistently observed in all BR systems, pointing to a direct evolutionary linkage between Type-3 BREX and BR systems, in addition to their other shared components. Contrastingly, in Type-4 BREX, the MTase is entirely absent and is replaced by a PT-modification apparatus. An inversely analogous configuration is also reported by us in a subset of Dnd-systems, where the typical PT-modification genes are substituted by cytosine methylases (**1**).

Collectively, the DNA modification machinery across BREX and BR systems is predominantly built around an N6-MTase core, exhibiting subtype-specific adaptations. The incorporation of non-canonical strategies—such as the Type-4 PT-based mechanism and the putative GNAT/TGT-driven modification in BRC-systems—highlights the mechanistic plasticity of the BREX-machinery and opens promising avenues for future experimental exploration **(Figure 10)**.

### Effector responses in BREX-mediated immunity

A longstanding puzzle in BREX immunity has been the apparent absence of a nuclease responsible for phage genome degradation. Recent studies show that in Type-1 BREX, PglZ acts as a nicking nuclease on phage dsDNA—introducing non-specific single-strand nicks— rather than the double-strand breaks typical of restriction endonucleases (**15**). This activity likely reflects an evolutionary adaptation, as PglZ’s core domain belongs to the nucleotide pyrophosphatase (NPP) superfamily, whose members typically hydrolyze phosphoric ester bonds in small soluble nucleotide substrates (**15,16,125,158–163**), rather than long DNA polymers. Thus, BREX-PglZ appears to have been repurposed from an ancestral NPP phosphodiesterase to act on DNA, rather than canonical small-molecule substrates. Comparable functional shifts are seen in phospholipase-D (PLD) superfamily enzymes, where several homologs have evolved to cleave nucleic acids alongside their canonical phospholipid substrates (**74,75,164–168**). PglZ’s nuclease activity is further inhibited by ATP (**15**), suggesting that the domain retains its ancestral affinity for substrates such as nucleotide di-and triphosphates or cyclic dinucleotides. Parallel examples also occur in the ParB/Sulfiredoxin (ParB/Srx) NTPase superfamily, which catalyze non-specific single-strand DNA nicking, with ATP binding acting as a negative regulatory switch (**169,170**). Notably, DndB—a ParB superfamily protein in Dnd systems—is proposed to act as a negative regulator by limiting ATP pools required for phosphorothioation (**1**). Moreover, a diverse repertoire of domains including phosphoesterases, have been recently identified in nucleotide-targeting immune systems, where they act as antiviral effectors by depleting NTP pools critical for phage DNA replication (**107**).

Building on these biochemical analogies and our comparative analysis, supported by recent biochemical evidence, we propose a layered model for PglZ-mediated phage inhibition. In the initial stage, PglZ, together with its inactive P-loop domains (iSwi2/Snf2 helicase and associated BrxB) and C-terminal IG-like domains, likely forms a surveillance module that contributes to phage DNA detection in concert with other invader-sensing components of the BREX machinery **(Figure 10)**. Upon recognition of invasive DNA by the multi-module sensing machinery of BREX, PglZ’s nicking activity is triggered, introducing non-specific single strand breaks in phage dsDNA (on one strand) to destabilize the genome and impede its replication. In addition, as highlighted above, PglZ may further impede phage replication by depleting essential nucleotide cofactors. Hydrolysis of ATP, other nucleotide di- and triphosphates, or cyclic dinucleotides could reduce the intracellular pool of energy carriers and signaling molecules, directly impairing phage genome replication.

In the archetypal versions of BREX systems harboring only PglZ as a sole effector, its activity alone is likely sufficient to block phage replication through the steps outlined above. However, as phages evolved diverse anti-restriction strategies, BREX systems appear to have recruited additional auxiliary effectors (endoDNases fused to NTPase or DNA-binding domains), as identified in this study **(Figure 10, Supplementary Data S4)**. The initial single-strand nicking by PglZ may serve as a signal for a secondary defensive response by auxiliary effectors—either through direct recognition of the nicks as a non-self-feature or via conformational changes if these effectors are organized in a multi-protein complex— ultimately leading to more extensive degradation of the phage DNA. Moreover, some of these auxiliary effectors may also compensate for PglZ activity if it is neutralized by phage-encoded inhibiting factors. This second line of defense is strongly supported by our identification of a diverse repertoire of nucleases spanning seven distinct superfamilies across all BREX and BR systems **(Figure 10, Supplementary Data S4)**. These auxiliary effectors are not taxonomically restricted but instead occur in over half of all BREX and BR systems (including both self and non-self-targeting), indicating strong selection pressure to maintain them within their genomic neighborhoods.

The incorporation of PglZ-centric restriction, followed by the subsequent recruitment of canonical restriction endonucleases, potentially represents a layered network of effector deployment, where invasive DNA is inhibited while minimizing the risk of autoimmunity that could arise from uncontrolled double-stranded cleavage by more uncompromising nucleases (**171,172**). Even so, certain BREX systems, including the two novel BREX-related systems identified here (Type-3 BR and BRC), either lack auxiliary nucleases or retain them in limited instances and consequently rely on PglZ as the primary effector to mediate phage inhibition. This central role is further highlighted by the remarkable conservation of PglZ across BREX and related systems, both in phyletic distribution and in core structural and catalytic motifs, encompassing nearly all BREX, BR, and BRC systems, with the exception of Type-1 and Type-2 BR systems, where canonical restriction endonucleases are found instead of PglZ. Collectively, the co-option of an ancestral NPP-like enzyme as a first line of defense (in the majority of cases), together with the subsequent recruitment of diverse auxiliary nucleases— including backup suicidal effectors such as endoRNases and standalone endoDNases, many associated with phage-sensing OLD-ABCs—highlights the evolutionary plasticity and resilience of BREX and BR systems, positioning them among the most sophisticated RM-like defense systems.

### Conclusion

Our comprehensive survey substantially extends the phyletic distribution of BREX systems across prokaryotes, including a broad representation of all major subtypes across multiple archaeal lineages that were previously underrepresented. By resolving the complete domain architectures of all BREX-associated proteins, we uncover a dynamic landscape of domain fusions and modular rearrangements that illuminate the evolutionary and functional framework of BREX defense. We also establish the universal conservation of DNA-binding/sensing HTHs (BrxA homologs) and invader-sensing iSTAND (BrxB homologs) across all subtypes. Together with the tripartite core, these components assemble into an oligomeric immune complex—armed with multiple specialized domains that detect invaders, mediate self/non-self discrimination, and coordinate effector responses.

A striking outcome of this study is the expansion of the known BREX landscape through the identification of three BREX-related (BR) systems anchored on DUF499-type ATPases, which can be unequivocally unified with BREX, with Type-3 BREX emerging as their closest relative. Notably, the Type-3 BR system, in particular, exhibits significant mechanistic parallels to BREX, in which, instead of a canonical restriction endonuclease, an enzymatically active PglZ operates as the primary effector. We also uncovered a novel BRC (BREX-Related Capture) system that integrates BREX machinery with HerA/FtsK-based capture modules and Dpd-derived components, forming a hybrid, multi-layered immune architecture that combines strategies from diverse conflict origins.

By integrating emerging biochemical evidence with comparative genomics, we propose a unified three-layered functional framework underlying BREX immunity. First, we identify a diverse repertoire of conserved and subtype-specific sensory domains positioned to detect invading macromolecules. Second, we revisit the canonical self-DNA modification strategies and suggest an alternative pathway—centered on GNAT/TGT apparatus—unique to BRC systems. Third, we reveal an extensive arsenal of effectors across all BREX and relatives, composed of multiple strategically organized nucleases. While PglZ and canonical restriction endonucleases serve as primary effectors in distinct subtypes, we uncovered numerous secondary backup nucleases that reinforce primary restriction and provide protection against phage countermeasures. Collectively, our expanded framework redefines the evolutionary and functional landscape of BREX systems and opens several promising avenues for experimental exploration of the molecular mechanisms governing the broader biology of complex RM-like immune systems.

## Data availability

The data underlying this article are available in the article and in its online supplementary material, compiled as Supplementary Data S1 to S4, and a separate file containing the supplementary figures. This data is also available in various computer-readable formats at https://doi.org/10.5281/zenodo.17293984.

