## Supplementary Figures S1-S10 for "Expanding the Landscape of BREX Diversity: Uncovering Multi-Layered Functional Frameworks and Identification of Novel BREX-Related Defense Systems"

### Supplementary Figures S1-S10 Index

1. **Supplementary Figure S1: (A and B)** Comparison of structure and domain architecture of DUF6079 (BrxC ATPase) with DUF499 ATPase; **(C and D)** Comparison of MSAs of C-terminal region from DUF6079 (BrxC ATPase) and DUF499 ATPase…………………………………….....2
2. **Supplementary Figure S2-S5:** Mirror trees showing maximum-likelihood phylogeny of BrxC ATPase (including homologs across various subtypes) and the corresponding PglZ, both derived from the same genomic neighborhoods……………………………………………..3-6
3. **Supplementary Figure S6: (A)** Structural superimposition of BrxX-MTase dimerisation NTD with NTD localized at C-terminal of multiple Adenine Methyltransferases from Type-I RM. **(B)** DALI hitlist for BrxX-MTase N-terminal Dimerisation domain. **(C)** Multiple sequence alignment highlighting conserved residues across various N-terminal dimerisation domains from Type-1 RM systems……………………………………………………………………………………...7
4. **Supplementary Figure S7: (A)** Structure based alignment of PglZ N-terminal inactive P-loop domain, with multiple inactive P-loop from Swi2/Snf2 Helicase. **(B)** Structural superimposition of PglZ N-terminal inactive P-loop with inactive P-loop domain of Swi2/Snf2-Helicases……....8
5. **Supplementary Figure S8: (A-F)** 3D structure and domain architecture of PglZ from **(A)** Type-1 BREX, **(B)** Type-2 BREX, **(C)** Type-3 BREX, **(D)** Type-4 BREX, **(E)** BREX-related Capture systems, and **(F)** Type-3 BR-system standalone PglZ……………………………….…9
6. **Supplementary Figure S9: (A-F)** 3D structure and domain architecture of DNA binding HTHs (BrxA/DUF4007) from **(A)** Type-1 BREX (version 1), **(B)** Type-1 BREX (version 2), **(C)** Type-3 BREX, **(D)** Type-2 BREX, **(E)** Type-4 BREX, and **(F)** BRC system. **(G)** Maximum likelihood phylogeny of individual HTH/wHTH modules across BrxA/DUF4007 protein components…...10
7. **Supplementary Figure S10: (A)** Maximum likelihood phylogeny of BrxHI-Helicase core unit with previously classified members of SF1 and SF2 helicase superfamilies. **(B)** MSA of BrxHI-Helicase core unit with Ski2 like SF2-helicase superfamily………………………………..11

***# Titles are internally hyperlinked. Click on title to access material.***

### Supplementary Figure S1

A

B

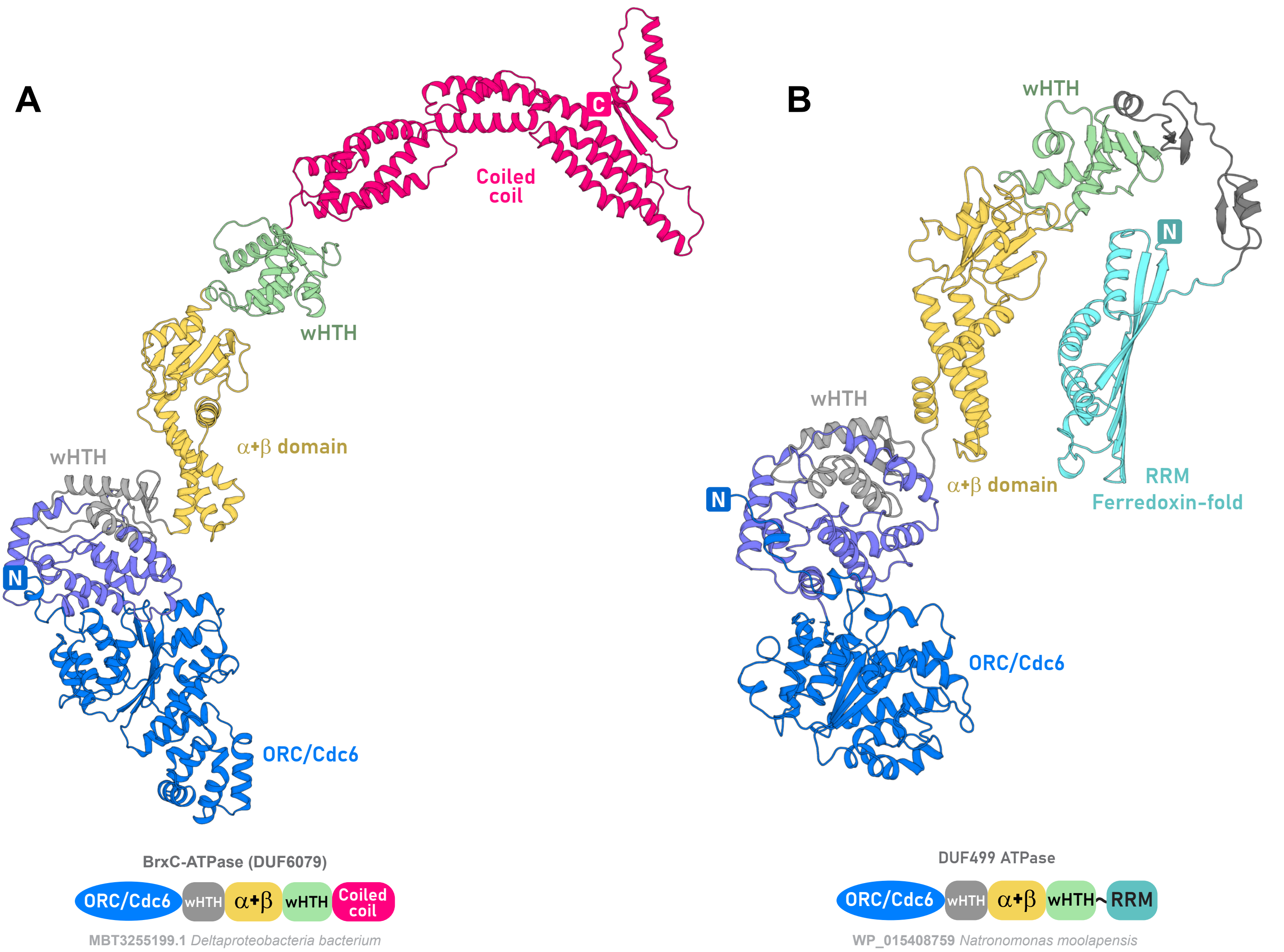


C

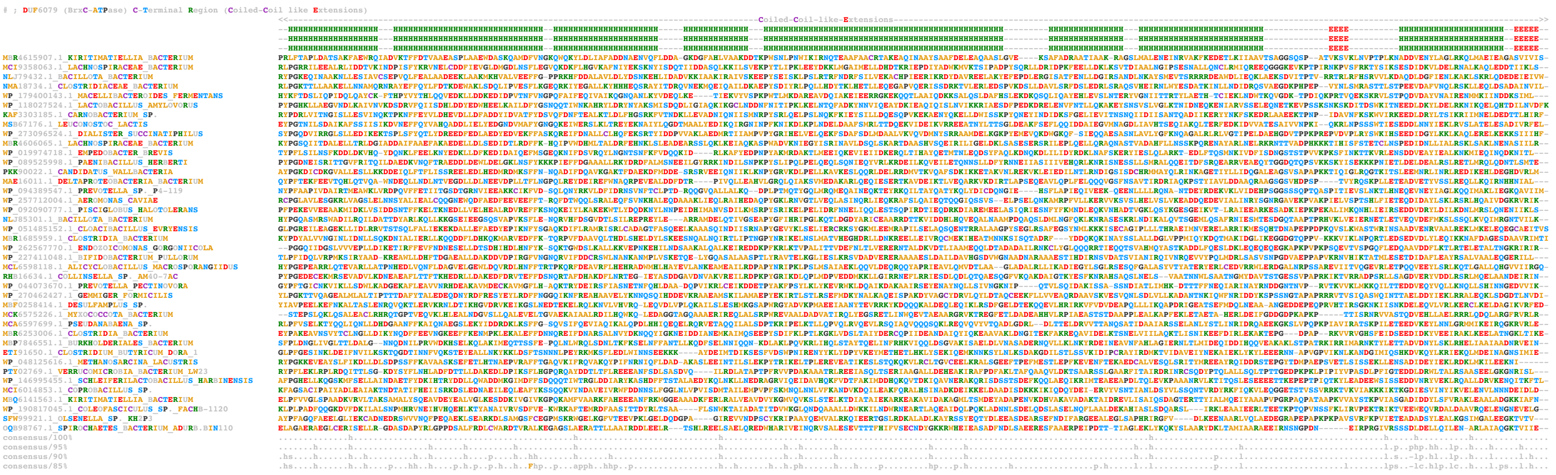


D

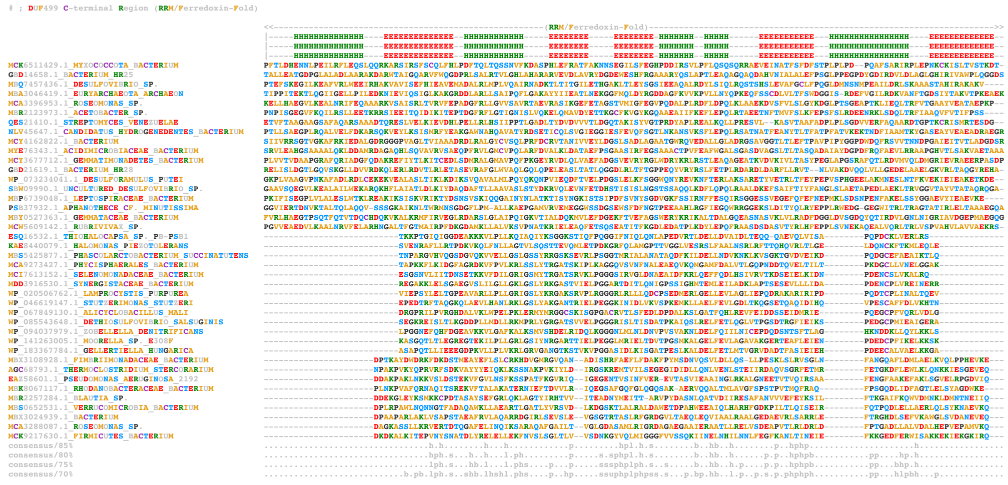

### Supplementary Figure S2

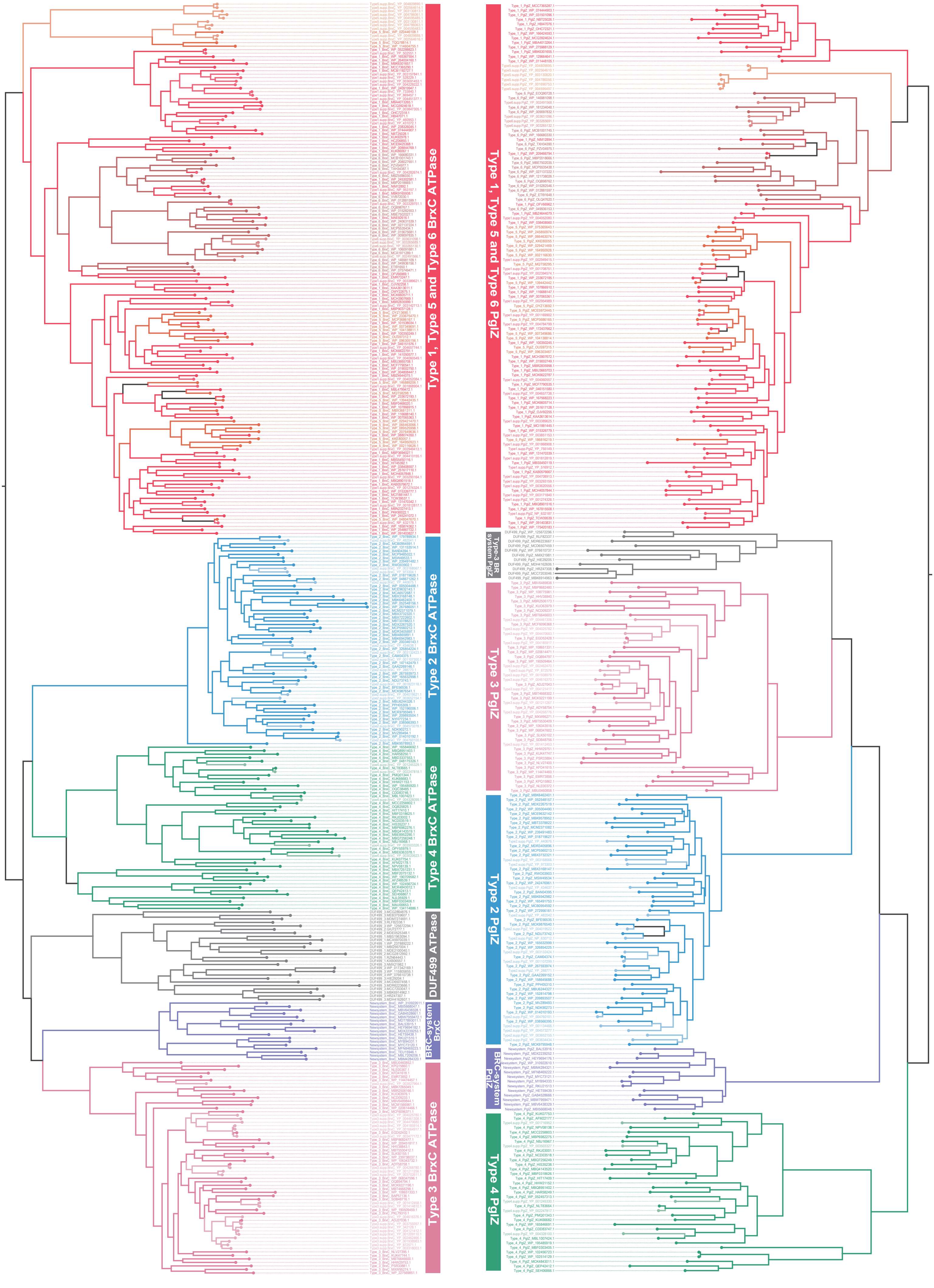

### Supplementary Figure S3

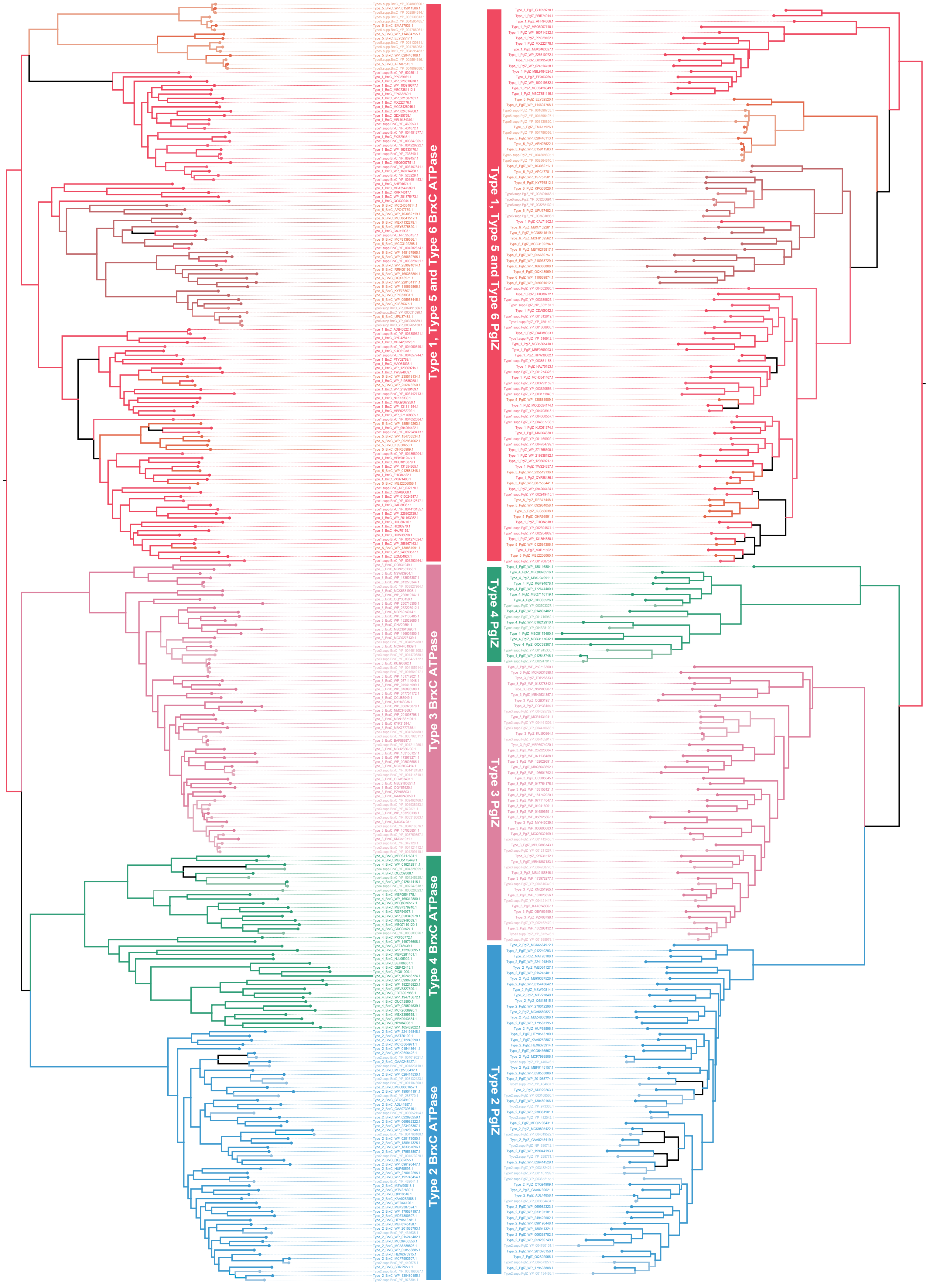

### Supplementary Figure S4

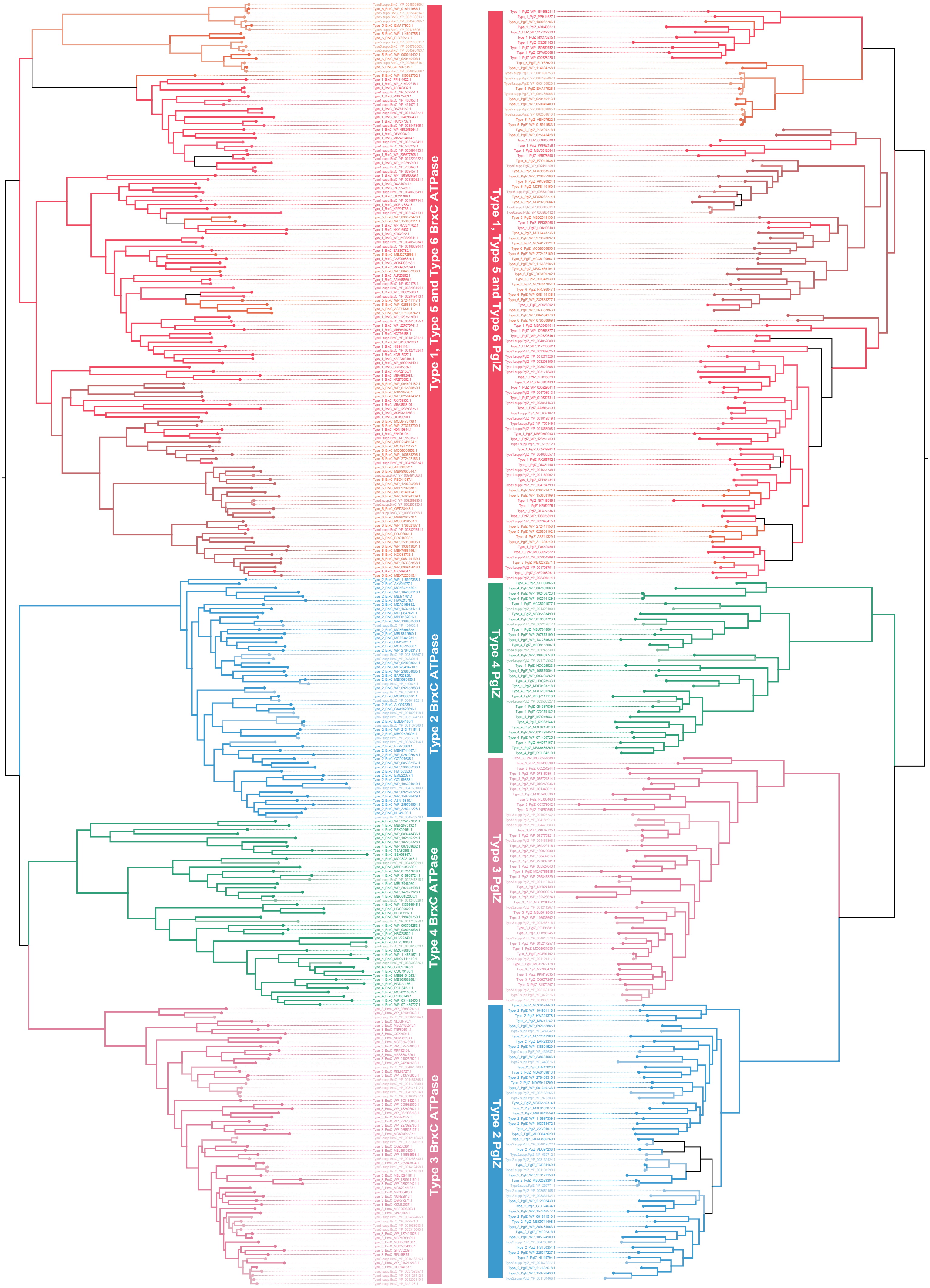

### Supplementary Figure S5

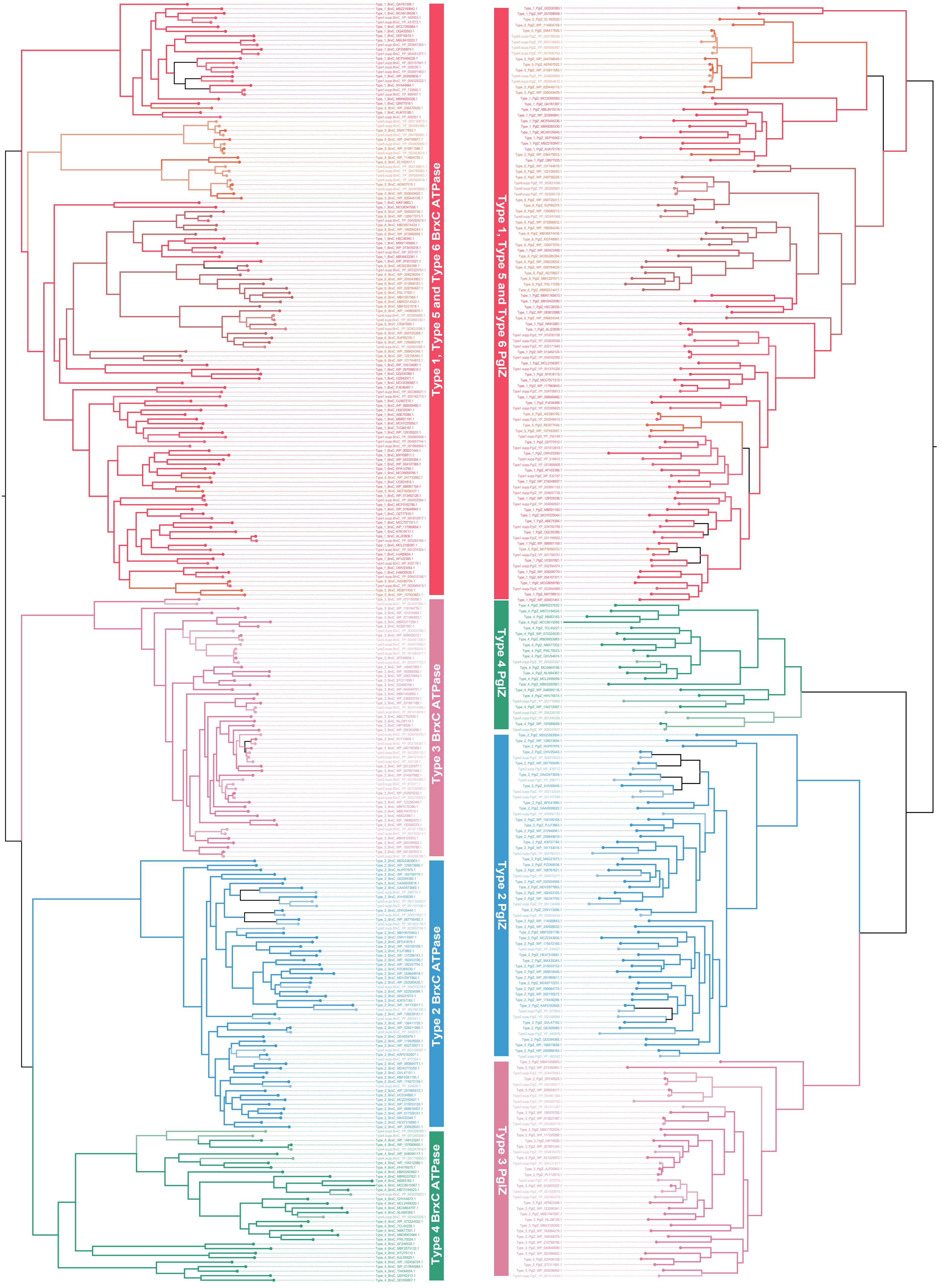

### Supplementary Figure S6

#### Structural superimposition of BrxX-MTase dimerisation NTD with NTD localized at C-terminal of multiple Adenine Methyltransferases from Type-I RM

**A**

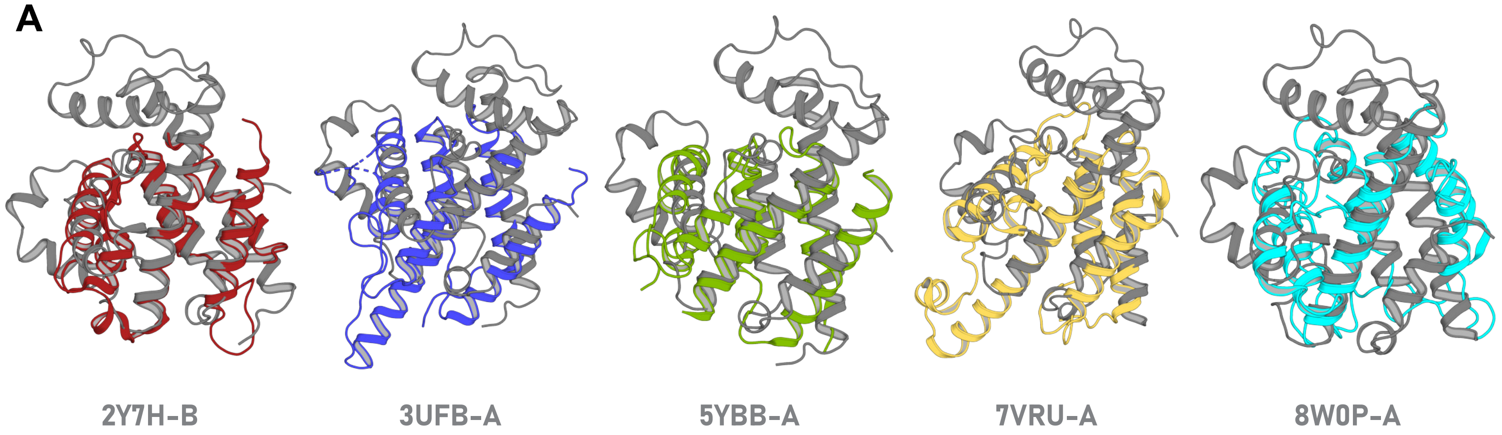


**B**

#### DALI hitlist for BrxX-MTase N-terminal Dimerisation domain

```
# Job: BREX MTase MBZ5596028 NTD Dimerisation domain DALI-hitlist
# Query: s001A
# No:  Chain     Z    rmsd lali nres  %id PDB  Description
   1:  8c45-A 17.8   2.4  192  1219   23   MOLECULE: SITE-SPECIFIC DNA-METHYLTRANSFERASE (ADENINE-SPEC
   2:  3khk-A  7.7   4.0  122   488   12   MOLECULE: TYPE I RESTRICTION-MODIFICATION SYSTEM
   3:  2y7h-B  7.4   3.5  123   530   11   MOLECULE: TYPE-1 RESTRICTION ENZYME ECOKI SPECIFICITY PROTE
   4:  5ybb-A  6.8   4.0  127   488   15   MOLECULE: TYPE I RESTRICTION-MODIFICATION SYSTEM METHYLTRAN
   5:  3ufb-A  6.7   3.8  123   484   10   MOLECULE: TYPE I RESTRICTION-MODIFICATION SYSTEM METHYLTRAN
   6:  2okc-B  6.6   3.5  121   427   15   MOLECULE: TYPE I RESTRICTION ENZYME STYSJI M PROTEIN;
   9:  7vru-A  5.9   3.8  120   498   11   MOLECULE: SITE-SPECIFIC DNA-METHYLTRANSFERASE (ADENINE-SPEC
  10:  3lkd-B  5.9   3.6  118   472   10   MOLECULE: TYPE I RESTRICTION-MODIFICATION SYSTEM
  11:  5hr4-J  5.6   3.5  126   752   13   MOLECULE: MMEI;
  12:  7eew-A  4.3   4.0  111   611   12   MOLECULE: TYPE I RESTRICTION-MODIFICATION SYSTEM METHYLTRAN
  13:  8w0p-A  3.9   4.4  118   917    9   MOLECULE: RM.BSAXI;
  16:  2f8l-A  3.6   3.5   84   324    8   MOLECULE: HYPOTHETICAL PROTEIN LMO1582;
  18:  3s1s-A  3.4   4.0  109   871    8   MOLECULE: RESTRICTION ENDONUCLEASE BPUSI;
```

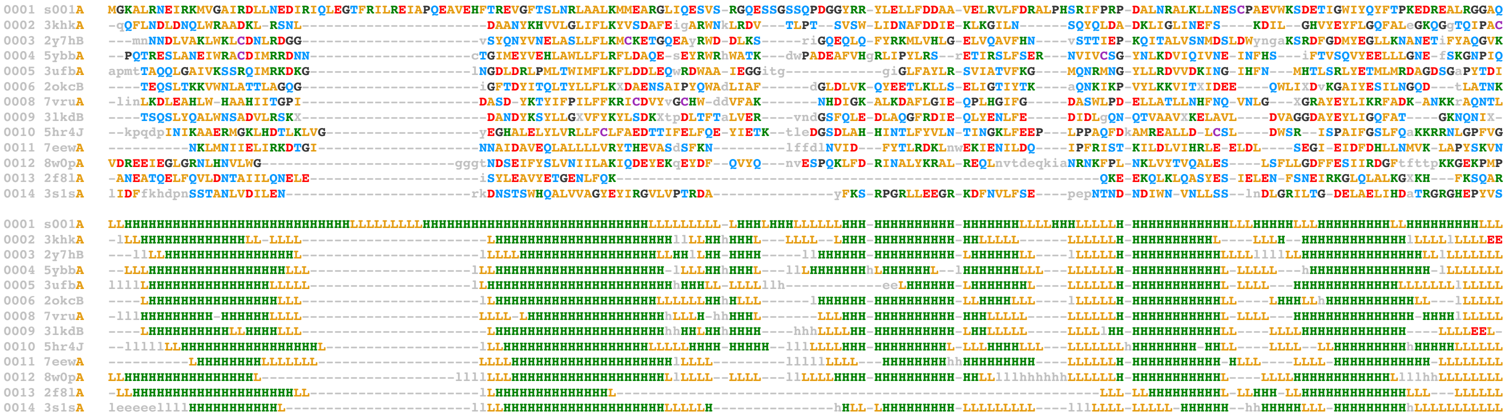

**C**

#### Multiple sequence alignment highlighting conserved residues across various N-terminal dimerisation domains from Type-1 RM systems

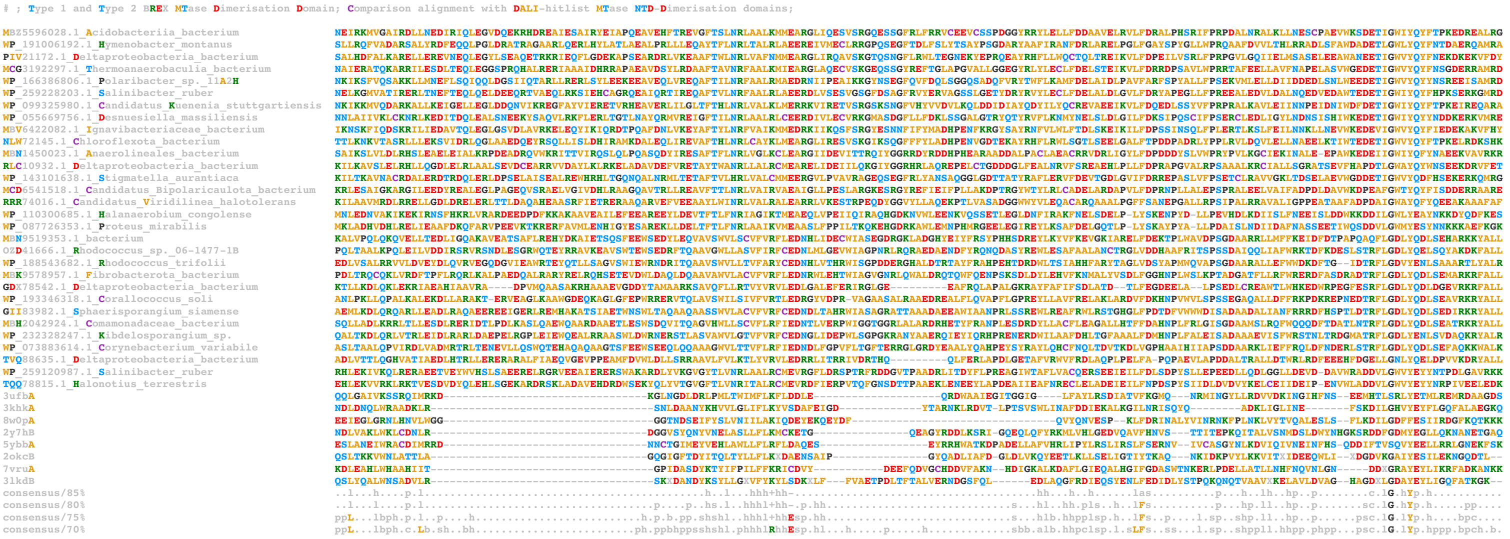

### Supplementary Figure S7

**A**

#### Structure based alignment of PglZ N-terminal inactive P-loop domain

```
# Job: Structure based Alignment of BREX PglZ N-terminal Inactive P-loop Swi2/Snf2-Helicase with various inactive P-loop domain from Swi2/Snf2-Helicases;
# Query: s001A
# No:  Chain     Z    rmsd lali nres  %id PDB
   1:  s001-A  22.6   0.0   95    95  100 Type1.PglZ.NTD
   2:  s002-A   9.0   2.1   82    97   11 Type3.PglZ.NTD
   3:  s022-B   7.2   3.0   90   164    6 6nmi
   4:  s003-A   6.8   2.7   88   139   11 6l80
   5:  s007-A   6.5   2.8   86   155    8 7vdv
   6:  s015-A   6.4   2.9   86   149   10 9ii8
   7:  s021-A   6.4   2.5   83   150   11 8zns
   8:  s016-A   6.3   3.1   81   143    7 6acx
   9:  s010-K   6.3   3.1   88   156    8 9n6k
  10:  s009-A   5.9   3.0   85   149    8 6bog
  11:  s012-G   5.9   2.7   83   142   10 9c57
  12:  s013-A   5.8   3.2   84   160    8 3i5x
  13:  s011-A   5.7   2.7   78   148    8 2d7d
  14:  s008-E   5.7   2.7   85   158   12 7zke
  15:  s018-A   5.5   2.3   74   136    7 4buj
  16:  s005-A   5.3   2.9   83   140    6 8t5s
  17:  s019-A   5.2   2.8   79   144    9 5lst
  18:  s020-A   5.1   2.5   77   124   10 2fzl
  19:  s014-C   5.1   3.1   83   161    7 7smo
  20:  s006-A   4.3   2.8   67   123    3 9el5
  21:  s004-A   4.1   2.9   69   114    6 8qy7
  22:  s017-A   3.5   3.3   70   158    9 7w0b

s001A_Type1.PglZ.NTD  MASSRITEALLNA-F-NKYRIVFWDDDKE-KL-IDEFNSL---ELP-D-VTKIKL-D-N------NE-FSVKY-RILVEETDK--KFLIYRPA-G--EPDVKDNWLLDVQL-AS-YKFVAE----------------
s002A_Type3.PglZ.NTD  SWRDHILKEFT---P-RVSRLTLVAD--P-DG-L--LLEEileGIR-EgFELIPF-E-D----HIAF-RYAYEsRFRS-RWDA--ALVVVLRSgA--LDV----lPYDLLQ-AG-RKLSF-----------------
s003A_6l80            TKIKALLRHLKQI-QeTSPGQIIVFSQFS-SFlDILEIES---HLPdQ-VIIYKF-D-G-RLDMKER-TRILE-QFH--DKDLscIKLLLLSL-T--GGV--GLNL-TCAS----RAFMMDpwwspgmedqaidrih
s005A_8t5s            PMISKTVQYIVEQ-N-LADSRTIIFVRTR-YEaTILNKVLsneELL-MgIKSEWM-SglISASKQKQ-MEKLK-MFAD-----geiRILVSTS-V--AEE--GLDVP---E-CS-LVIKYNyatneiahvqrrgrgr
s007A_7vdv            GKFELLDRILPKL-R-ATNHKVLLFCQMT-SL-MTIMEDY---FyR-G-FKYLRL-D--GTTKAEDR-GMLLK-TFNE-PGSE--YFIFLLST--raGGLG---LNLQSA---D-TVIIFDsdwnphqdlqaqdrah
s008E_7zke            PKLGALRDLLVDC-GigKPHRALIFCQMK-EMlDMVQNTVlkqMLP-S-VSYLRL-D--GSVEANKR-QDIVN-KFNS--DPS--YDVLLLTT--svGGLG---LNLTGA---D-TVIFVEhdwnpqkdlqamdrah
s009A_6bog            wNFDPRVEWLXGY-L-TSHQKVLVICAKA-AT-ALQLEQV-lrERE-G-IRAAVF-H--EGXSIIER-DRAAA-WFA--EEDT-gaQVLLCSE----IGSE-GRNF-QFAS----HXVXFDlpfnpdlleqrigrld
s010K_9n6k            GKMVLLDQLLTRL-K-KDGHRVLIFSQMV-RM-LDILGDY---LiK-G-INFQRL-D--GTVPSAQR-RISID-HFNSPDS-N--DFVFLLS--T--RAGGLGIN-LMTA---D-TVVIFDsdwnpqadlqamarah
s011A_2d7d            GQIDDLIGEIQAR-I-ERNERVLVTTLTK-KMsEDLTDYK---EIG---IKVNYL-H--SEIKTLER-IEIIR-DL-----rlgkyDVLVG-I-N--LL---rEGLD--IP-EVsLVAILDadkegflrsersliqt
s012G_9c57            GKLEALAILLQKL-KsEGRR-VLILSQMI-LM-LDILEMN---FHY---LTYVRI-D--ENASSEQR-QELMR-SFN--RDRR--IFCAILST-H--SR---tTGINLV--eaD-TVVFYDndlnpvmdakaqewcd
s013A_3i5x            NSIFAAVEHIKKQ-I-KSNYKAIIFAPTV-KFtSFLCSIL-knEFK-KdLPILEF-H--GKITQNKR-TSLVK-RFKK---dE--SGILVCTD-G--ARG----MDFPNV---H-EVLQIGvpselanyihrigrta
s014C_7smo            RNESEKWRHIANL-C-RAEAQSIVFTFSR-KRtHELAAY-----SK-G-LKAKPY-H-S-GLPYKQR-KLTEM-EFLAQR-----lDVVVTT--A--ALGA-GVDF----P-AS-QVIFESlamgnkwlsvrefhqm
s015A_9ii8            EYVASIVSYIIEY-T-TTSSKCLVLFTSY-KMmHMVQDML---NFE-D-YVVLTQ-QqN------QN-Y-KIV-QQF-NNFDK--AILLGTST-F--FEG-----FDFQangIK-CVMIAKlpfmntstfkeyvlpd
s016A_6acx            -DDKQVAAALRRE-L-LRDGQAFYIHNRVrTI-DEAAARV-rqLVP-E-ARVVVA-H-G-QMNEETL-EKTVE-GFWN-----reyDILVC-T-T--IV---eTGLD--IS-NAnTLIVERadtfglsqlhqlrgrv
s018A_4buj            -SKKTWPEIVNYL-R-KELLPMVVFVFSK-KRcEEYADWL---EGI-N---IAVHvK-E------LIeILFSK-GF---------iKVLFA-T-E--TF--aMGLNL----ptR-TVIFS----------------
s019A_5lst            DTDQALLTLLQGkrF-QNLDSIIIYCNRR-EDtERIAALL-rtCLH-AktTAEAY-H--AGMCSRER-RRVQR-AFMQ-----gqlRVVVA-T-V--AF---gMGLDR-pD-VR-AVLHLGlppsfesyvqavgrag
s020A_2fzl            NKIRKLREILER--H-RKD-KIIIFTRHN-ELvYRISKVF----------lIPAI-T--HRTSREER-EEILE-GFR---TGR--FRAIVSSQ----VLDE-GIDV---pD-AN-VGVIMSgsgsareyiqrlgril
s021A_8zns            -SWQKIADEIAAR------PCVLAVVNTR-KH-AQKLFAA---LPS-N-GIKLHLhC-S------EViALVRR-YLALySLHK--PLWLVSTQ-L--IEA--GVDL----D-FP-CVYRAMagldsiaqaagrcnre
s022B_6nmi            AVIRNYGNLLLEM-S-AVVPGIVAFFTSYqYM-ESTVASWileNIQ-RnKLLFIE-T-Q---DGAET-SVALE-KYQE-ENGR--GAILLSVA-RgkVSEG--IDFVHHY--gR-AVIMFGvpirendfltfdamrh
s004A_8qy7            ------------------KDKVIIFTELR-DLqRELQHAQ---NFG---frPVII-N-GKSQSQNSR-QRLID-DFQ--AQPG--FGVIIL---S--TVAVGFGVN---VQ-KAnHVIHFTrcwnpakedqatdray
s006A_9el5            ------------------ETRVMIFSSFR-DSvQEIAEMS---QHQpI-IRVMTF-V-GKGFTQKEQ-LEVVK-QFRD-----ggyNTLVSTC---vGEEG--LDIG---E-VD-LIICFDsqkspirlvqrmgrtg
s017A_7w0b            -------------dkdPKDICCLVFVERR-YTcKCIYGlN---YIQnV-LTPQFM-V-gSVLERKWQ-KSAIQ-QFRDGN-----aNLMICSS-V--LEE--GIDV---QA-CN-HVFILDpvktfnmyvqskgrar

s001A_Type1.PglZ.NTD  LHHHHHHHHHHHH-H-HHLLEEEEEELHHH-LL-HHHHHHL---LLL-L-EEEEEL-L-L------LH-HHHHH-HHHLLLLLL--EEEEEELL-L--LLLLLLLLLHHHHH-LL-EEELLL----------------
s002A_Type3.PglZ.NTD  LHHHHHHLLLL---L-LLLLEEEEEL--L-LL-H--HHLHhhhHHH-HlEEEEEL-L-L----HHHH-HHHHHhHLLL-LLLL--EEEEEELLlH--HHH----lLHHHHH-HL-EEEEL-----------------
s003A_6l80            HHHHHHHHHHHHH-HhHLLLEEEEEEELLH-HHhHHHHHHH---HLLlL-EEEEEL-L-L-LLLHHHH-HHHHH-HHL--LLLLllEEEEEEEL-L--LLL--LLLL-LLEE----EEEELLlllhhhhhhhhhlll
s005A_8t5s            HHHHHHHHHHHHH-H-HLLLLEEEELLLH-HHhHHHHHHHllhHHH-HlLLEELL-LllLLLLHHHH-HHHHH-HHHH-----lllLEEEELH-H--HHL--LLLLL---L-LL-EEEEELllllhhhhhhhhhlll
s007A_7vdv            HHHHHHHHHHHHH-H-HHLLLEEEEEELLH-HH-HHHHHHH---HhL-L-LLEEEE-L--LLLLHHHH-HHHHH-HHHL-LLLL--LLEEEEEH--hhHLLL---LLLLLL---L-EEEELLllllhhhhhhhhhllll
s008E_7zke            HHHHHHHHHHHHL-LllLLLLEEEEELLH-HHhHHHHHHHhhlLLL-L-LLEEEE-L--LLLLHHHH-HHHHH-HHHH--LLL--LLEEEEEH--hhHLLL---LLLLLL---L-EEEELLllllhhhhhhhhhlll
s009A_6bog            lLLLHHHHHHHHH-H-HHLLLEEEEEELLH-HH-HHHHHHH-hhHHH-L-LLEEEE-L--LLLLHHHH-HHHHH-HHH--LLLL-llLEEEEEL----LLLL-LLLL-LLLL----EEEELLllllhhhhhhhhhlll
s010K_9n6k            LLHHHHHHHHHHH-H-HLLLLEEEEEELLH-HH-HHHHHHH---HhH-L-LLEEEE-L--LLLLHHHH-HHHHH-HHLLLLL-L--LLEEEEE--L--LLLLLLLL-LLLL---L-EEEELLllllhhhhhhhhhllll
s011A_2d7d            LHHHHHHHHHHHH-H-LLLLEEEEELLLH-HHhHHHHHHH---HLL---LLEEEE-L--LLLLHHHH-HHHHH-HH-----hhlllLEEEE-L-L--LL---lLLLL--LL-LEeEEEELLlllllllllhhhhhhh
s012G_9c57            HHHHHHHHHHHHH-HhHLLL-EEEEELLH-HH-HHHHHHH---HLL---LLEEEE-L--LLLLHHHH-HHHHH-HHH--HLLL--LLEEEEEL-L--LL---lLLLLLL--llL-EEEELLllllhhhhhhhhhhhh
s013A_3i5x            HHHHHHHHHHHHH-H-HLLLEEEEELLLH-HHhHHHHHHH-hhHHL-LlLLEEEE-L--LLLLHHHH-HHHHH-HHHH---lL--LEEEEELH-H--LLL----LLLLLL---L-EEEEELllllllhhhhhhllll
s014C_7smo            LLHHHHHHHHHHH-H-HHHHLEEEELLLH-HHhHHHHHH-----HL-L-LLEEEL-L-L-LLLHHHH-HHHHH-HHHLLL-----lLEEEEL--L--LLLL-LLLL----L-LL-EEEELLlllllllllhhhhhhh
s015A_9ii8            HHHHHHHHHHHHH-H-HLLLLEEEELLLH-HHhHHHHHHH---HHL-L-LEEEEL-LlL------LL-L-HHH-HLL-LLLLL--LEEEEELL-L--LLL-----LLLLlllLE-EEEELLlllllllhhhhlhhhh
s016A_6acx            -LHHHHHHHHHHH-H-HLLLEEEEEELLLHhHH-HHHHHHH-hhHLL-L-LLEEEE-L-L-LLLHHHH-HHHHH-HHHL-----lllLEEEE-L-H--HH---hLLLL--LL-LEeEEEELLhhhllhhhhhhhhhll
s018A_4buj            -LLLHHHHHHHHH-H-HLLLLEEEELLLH-HHhHHHHHLL---LLL-L---LEEEhH-H------HHhHHHHL-LL---------lLEEEE-L-H--HH--hHLLLL----llL-EEEEL----------------
s019A_5lst            LHHHHHHHHHHLllL-LLLLLEEEEELLH-HHhHHHHHHH-hhHLL-LleEEEEE-L--LLLLHHHH-HHHHH-HHHL-----lleEEEEE-E-L--LL---lLLLLL-lL-LL-EEEEELllllhhhhhhhhllll
s020A_2fzl            HHHHHHHHHHHH--L-LLL-LEEEELLLH-HHhHHHHHHL----------lLLEE-L--LLLLLHHH-HHHHH-HHL---LLL--LLEEEELL----LLLL-LLLL---lL-LL-EEEEELlllllhhhhhhhhhl
s021A_8zns            -LHHHHHHHHHLL------LEEEEEELLH-HH-HHHHHHH---LLL-L-LEEEEEhH-H------HHhHHHHH-HHHHhLLLL--LEEEEEEH-H--HLL--LLLL----L-LL-EEEEELllhhhhhhhhlllllll
s022B_6nmi            HHHHHHHHHHHHH-H-HHLLLEEEEEELLHhHH-HHHHHLLlhhHHH-HhLEEEEL-L-L---LLLHH-HHHHH-HHHH-LLLL--LEEEEEEL-LlhHHLL--LLLLLLL--lL-EEEEELllllhhhhhhhhhhhh
s004A_8qy7            ------------------LLEEEEELLLH-HHhHHHHHHH---HHL---llLEEE-L-LLLHHHHHH-HHHHH-HHH--HLLL--LEEEEE---L--LLLLLLLL---LL-LEeEEEELLllllhhhhhhhhhlll
s006A_9el5            ------------------LLLEEEELLLH-HHhHHHHHHH---LLLlL-LLEEEE-L-LLLLLHHHH-HHHHH-HHHL-----lllLEEEELH---hHLLL--LLLL---L-LL-EEEELLlllllhhhhhhhllll
s017A_7w0b            -------------lllHHHLLEEEELLLH-HHhHHHHHhH---HHHlL-LLLLLL-L-lLLLHHHHH-HHHHH-HHHLLL------lLLLEELL-L--HHH--LLLL---LL-LL-EEEELLllllhhhhhhhhhlll
```

#### Structural superimposition of PglZ N-terminal inactive P-loop with inactive P-loop domain of Swi2/Snf2-Helicases

**B**

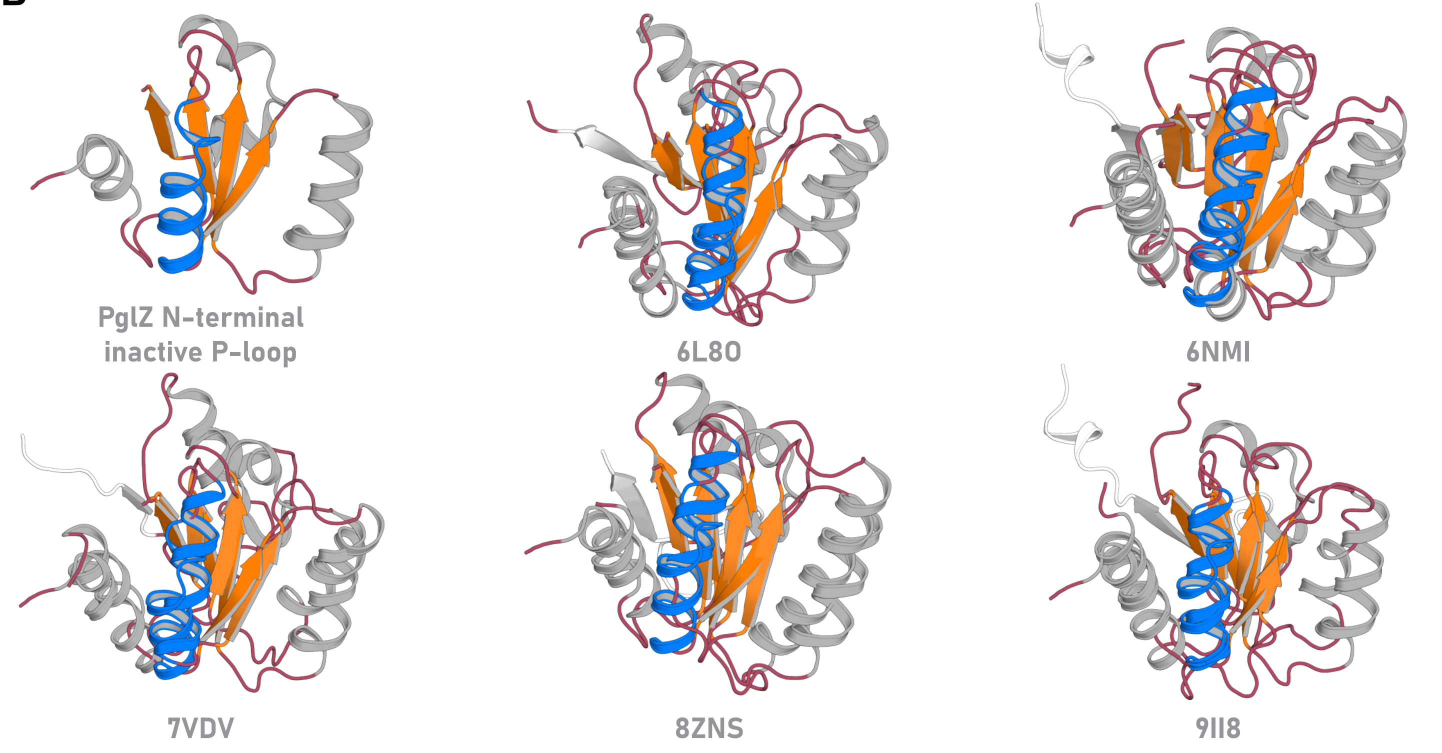

### Supplementary Figure S8

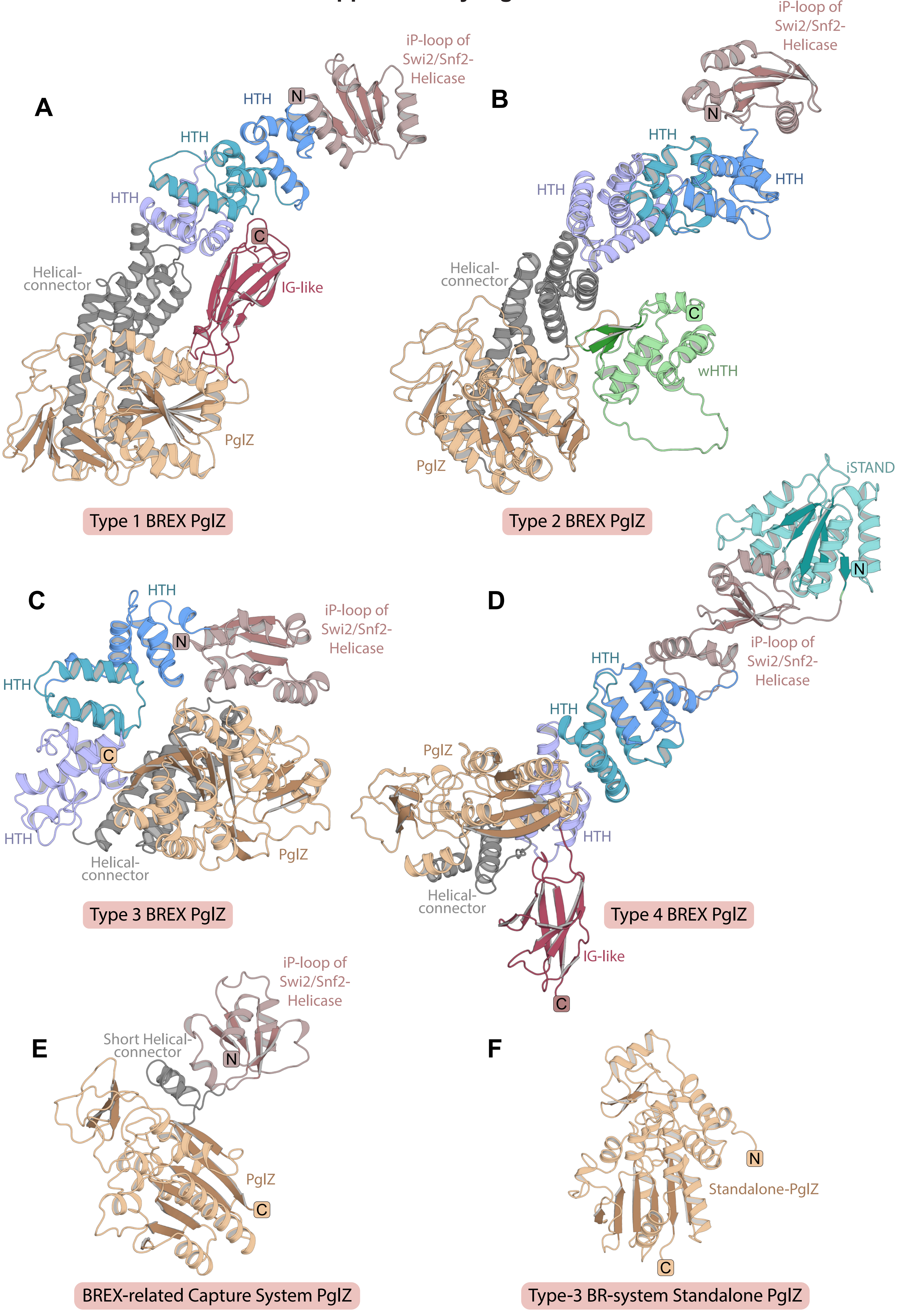

#### Supplementary Figure S9

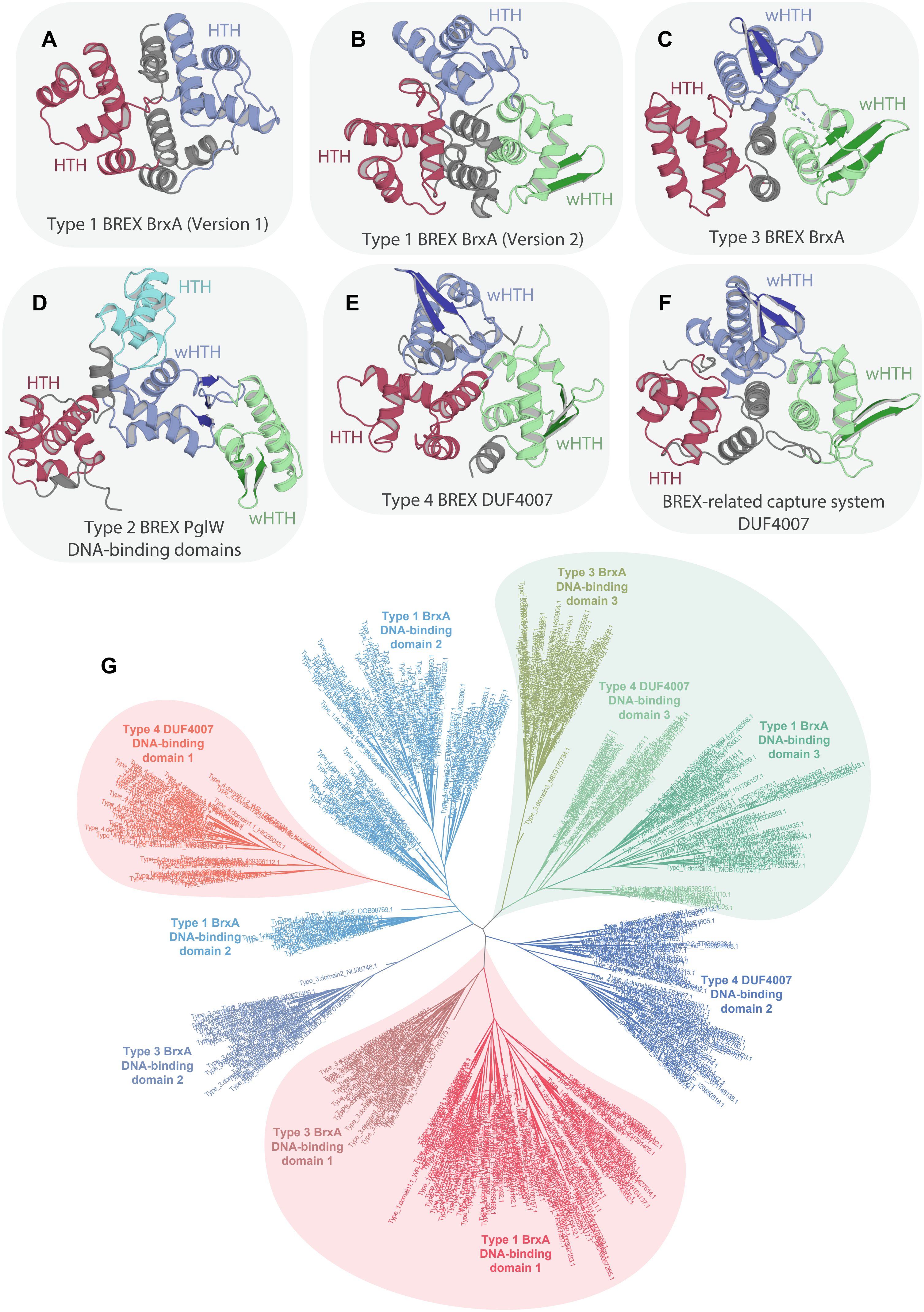

### Supplementary Figure S10

Phylogenetic clustering and multiple sequence alignment comparison of BrxHI-Helicase core unit with previously classified members of SF1 and SF2 helicase superfamilies

A

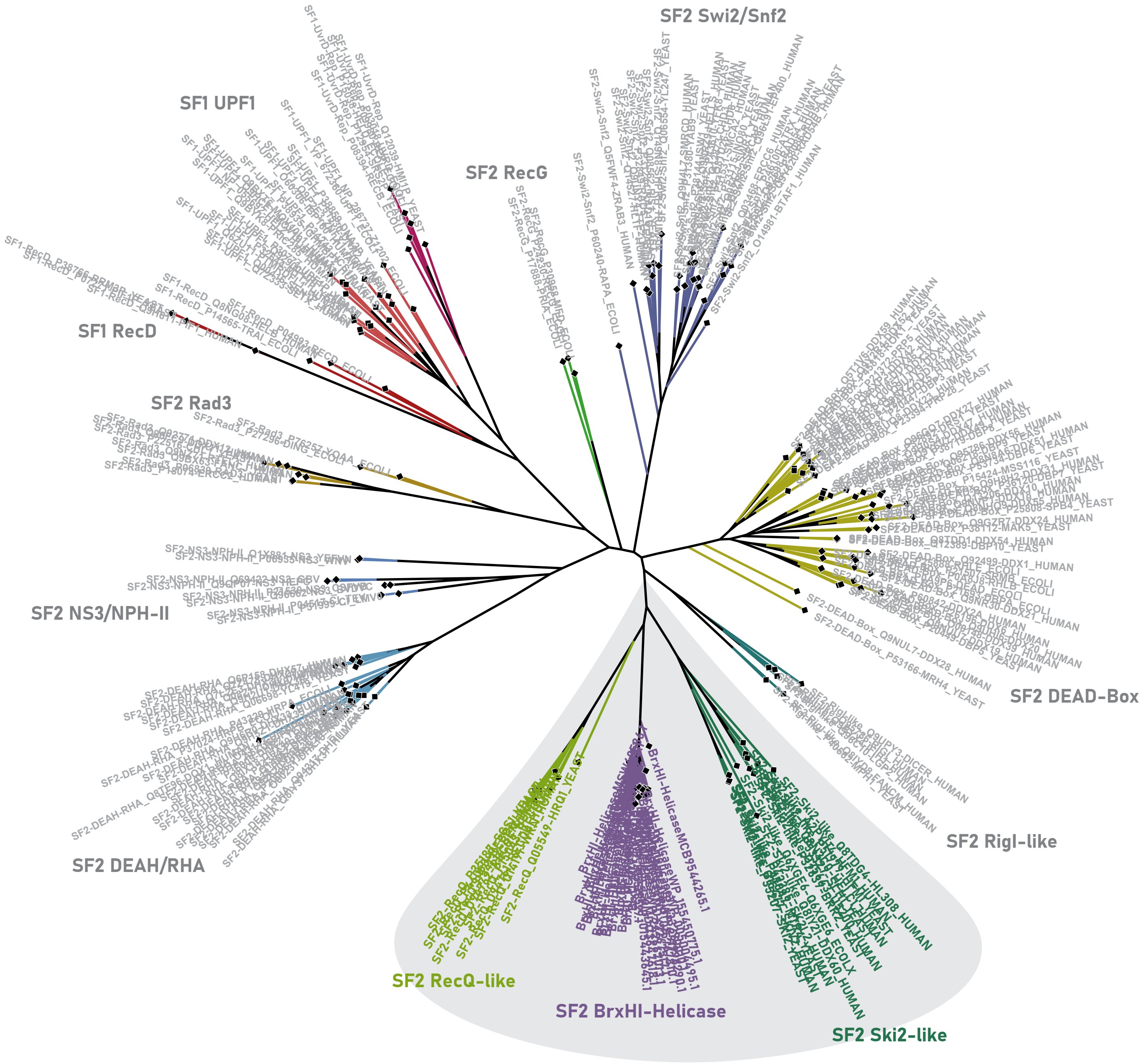


B

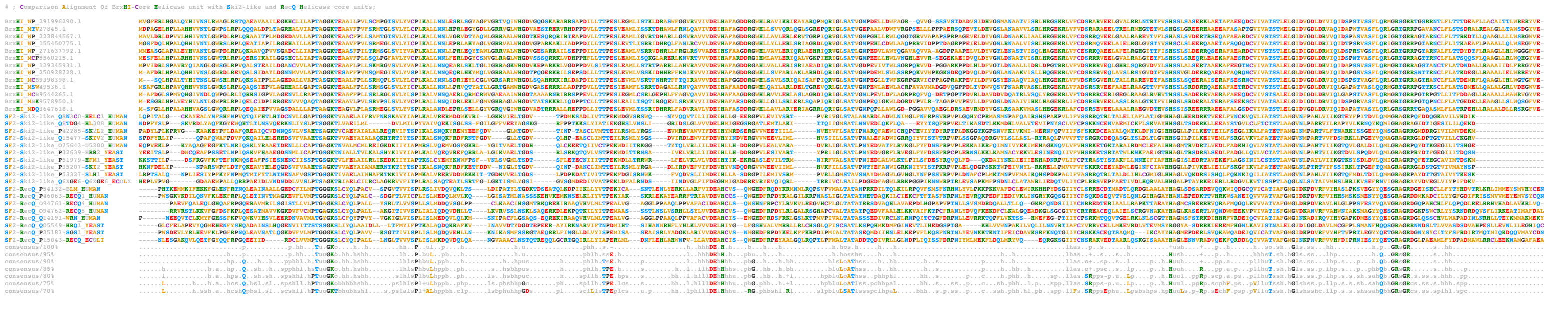
