## Supplementary Data S1 for "Expanding the Landscape of BREX Diversity: Uncovering Multi-Layered Functional Frameworks and Identification of Novel BREX-Related Defense Systems"

This PDF contains the conserved gene neighbourhoods with corresponding NCBI node information parsed for all ten systems examined in this study. Following these, the original treefiles used for the main figures, generated with IQ-TREE bootstrap analyses, are provided in Newick format.


1. Type-1 BREX systems……………………………………………………2-59
2. Type-5 BREX systems……………………………………………………59-61
3. Type-6 BREX systems……………………………………………………61-66
4. Type-2 BREX/Pgl systems………………………………………………67-80
5. Type-3 BREX systems…………………………………………………..81-92
6. Type-4 BREX systems…………………………………………………..93-98
7. Type-1 DUF499 centered BREX-related systems…………………99-140
8. Type-2 DUF499 centered BREX-related systems…………………141-155
9. Type-3 DUF499 centered BREX-related systems…………………156-163
10. BREX related capture systems………………………………………164
11. Newick tree (BrxC-ATPase)…………………………………………..165
12. Newick tree (PglZ)……………………………………………………166
13. Newick tree (DUF499-ATPase)………………………………………167


***Note: Titles are internally hyperlinked. Click on the titles to access the material***

| Protein-ID | System | Gene-Neighbourhoods | Length | Taxonomy | Assembly-ID | Strain | Locus-Tag | Genomic-coordinates | TaxID | Fasta-defline |
|---|---|---|---|---|---|---|---|---|---|---|
| -- | | | | | | | | | | |
| --Type-1-BREX | | | | | | | | | | |
| --Canonical-BREX | | | | | | | | | | |
| -- | | | | | | | | | | |
| MCC7571311.1 | Type1.BREX | nSTAND1+HNH->HNH->TrwI->?->?->VirB4-FtsK->Lysozyme+NLPC->BrxA-DNA-binding->BrxB-BrxF-iSTAND->DUF6079*->N6-METHYLASE->PglZ->SIGMA+OB-Fold+AAA-ATPase+Lon-Protease-BrxL->?->?-> [illegible] | 1210 | Archaea>DPANN_group>Candidatus_Micrarchaeota>unclassified_Candidatus_Micrarchaeota | GCA_020854835.1 | Candidatus_Micrarchaeota_archaeon | K0465_08330 | 10159..13791 | 2250274 | BREX-system-P-loop-protein-BrxC-[Candidatus-Micrarchaeota-archaeon] |
| MBT4111448.1 | Type1.BREX | <-HIT\|\|CopG/RHH->ZnR->?->DnaB->?->BrxA-DNA-binding->BrxB-BrxF-iSTAND->DUF6079-p*-> | 882 | Archaea>DPANN_group>Candidatus_Woesearchaeota | GCA_018650945.1 | Candidatus_Woesearchaeota_archaeon | HOE37_06330 | 27255..29901 | 2026803 | BREX-system-P-loop-protein-BrxC-[Candidatus-Woesearchaeota-archaeon] |
| MBT4336377.1 | Type1.BREX | <-HIT\|\|CopG/RHH->ZnR->?->DnaB->?->BrxA-DNA-binding->BrxB-BrxF-iSTAND->DUF6079*->N6-METHYLASE->PglZ->SIGMA+OB-Fold+AAA-ATPase+Lon-Protease-BrxL->SF1-SF2-Helicase+REase->?-><-?<-?<- [illegible] | 1173 | Archaea>DPANN_group>Candidatus_Woesearchaeota | GCA_018652365.1 | Candidatus_Woesearchaeota_archaeon | HOD66_03370 | 27503..31024 | 2026803 | BREX-system-P-loop-protein-BrxC-[Candidatus-Woesearchaeota-archaeon] |
| MBT3814300.1 | Type1.BREX | <-DUF6079-p*<-BrxB-BrxF-iSTAND<-BrxA-DNA-binding<-?<-DnaB<-?<-ZnR<-CopG/RHH\|\|HIT-> | 353 | Archaea>DPANN_group>Candidatus_Woesearchaeota | GCA_018654495.1 | Candidatus_Woesearchaeota_archaeon | HOG07_02565 | 1..1059 | 2026803 | BREX-system-P-loop-protein-BrxC-[Candidatus-Woesearchaeota-archaeon] |
| MBT4469968.1 | Type1.BREX | Phage_integrase->?->?-><-?<-SF1-SF2-Helicase+REase<-SIGMA+OB-Fold+AAA-ATPase+Lon-Protease-BrxL<-PglZ<-N6-METHYLASE<-DUF6079*<-BrxB-BrxF-iSTAND<-BrxA-DNA-binding<-?<-DnaB<-?<-ZnR<- [illegible] | 1173 | Archaea>DPANN_group>Candidatus_Woesearchaeota | GCA_018656325.1 | Candidatus_Woesearchaeota_archaeon | HOC62_06290 | 58853..62374 | 2026803 | BREX-system-P-loop-protein-BrxC-[Candidatus-Woesearchaeota-archaeon] |
| MBT6744308.1 | Type1.BREX | <-HIT\|\|CopG/RHH->ZnR->?->DnaB->?->BrxA-DNA-binding->BrxB-BrxF-iSTAND->DUF6079*->N6-METHYLASE->PglZ->SIGMA+OB-Fold+AAA-ATPase+Lon-Protease-BrxL->SF1-SF2-Helicase+REase->?-><-?<-?<- [illegible] | 1173 | Archaea>DPANN_group>Candidatus_Woesearchaeota | GCA_018696595.1 | Candidatus_Woesearchaeota_archaeon | HOA70_03205 | 27503..31024 | 2026803 | BREX-system-P-loop-protein-BrxC-[Candidatus-Woesearchaeota-archaeon] |
| MCL2156391.1 | Type1.BREX | BrxA-DNA-binding->BrxB-BrxF-iSTAND->DUF6079*->Phage_integrase+Phage_integrase-><-N6-METHYLASE\|\|N6-METHYLASE->?->PIN->PglZ->SIGMA+OB-Fold+AAA-ATPase+Lon-Protease-BrxL-> | 1194 | Archaea>Euryarchaeota>Methanomada_group>Methanobacteria>Methanobacteriales>Methanobacteriaceae | GCA_009776495.1 | Methanobrevibacter_sp. | FWH54_00015 | 1416..5000 | 66852 | BREX-system-P-loop-protein-BrxC-[Methanobrevibacter-sp.] |
| MBE6500276.1 | Type1.BREX | N6-METHYLASE->BrxA-DNA-binding->BrxB-BrxF-iSTAND->DUF6079*->?->PglZ->SIGMA+OB-Fold+AAA-ATPase+Lon-Protease-BrxL-> | 1195 | Archaea>Euryarchaeota>Methanomada_group>Methanobacteria>Methanobacteriales>Methanobacteriaceae | GCA_015062925.1 | Methanobrevibacter_thaueri | E7Z80_07025 | 4963..8550 | 190975 | BREX-system-P-loop-protein-BrxC-[Methanobrevibacter-thaueri] |
| MBR3112546.1 | Type1.BREX | <-TPR+TPR+TPR+TPR<-?<-OLD-ABC<-SF1-SF2-Helicase+REase<-?<-?<-SIGMA+OB-Fold+AAA-ATPase+Lon-Protease-BrxL<-PglZ<-?<-DUF6079*<-BrxB-BrxF-iSTAND<-BrxA-DNA-binding<-N6-METHYLASE\|\| [illegible] | 1187 | Archaea>Euryarchaeota>Methanomada_group>Methanobacteria>Methanobacteriales>Methanobacteriaceae | GCA_017545145.1 | Methanobrevibacter_sp. | IKH29_02390 | 52350..55913 | 66852 | BREX-system-P-loop-protein-BrxC-[Methanobrevibacter-sp.] |
| MBR5504577.1 | Type1.BREX | <-SbcC<-?\|\|?->McrC-NTD+REase-><-URI-DNase+RAMA\|\|BrxA-DNA-binding->BrxB-BrxF-iSTAND->DUF6079*->N6-METHYLASE-> | 1186 | Archaea>Euryarchaeota>Methanomada_group>Methanobacteria>Methanobacteriales>Methanobacteriaceae | GCA_017558445.1 | Methanobrevibacter_sp. | IKV87_09095 | 7776..11336 | 66852 | BREX-system-P-loop-protein-BrxC-[Methanobrevibacter-sp.] |
| MBS6828080.1 | Type1.BREX | <-N6-METHYLASE<-DUF6079-p*<-DUF6079-p<-BrxB-BrxF-iSTAND<-BrxA-DNA-binding\|\|?->?->\|\|CR-ATPase5+REase-><-Pkinase+STYKIN | 905 | Archaea>Euryarchaeota>Methanomada_group>Methanobacteria>Methanobacteriales>Methanobacteriaceae | GCA_018379875.1 | Methanobrevibacter_smithii | KH189_08115 | 2976..5693 | 2173 | BREX-system-P-loop-protein-BrxC-[Methanobrevibacter-smithii] |
| WP_011954749.1 | Type1.BREX | Phage_integrase+Phage_integrase-><-N6-METHYLASE\|\|N6-METHYLASE-p->N6-METHYLASE->N6-METHYLASE-p-><-N6-METHYLASE\|\|N6-METHYLASE->BrxA-DNA-binding->BrxB-BrxF-iSTAND->DUF6079*->?->PglZ-> [illegible] | 1194 | Archaea>Euryarchaeota>Methanomada_group>Methanobacteria>Methanobacteriales>Methanobacteriaceae | GCF_000016525.1 | Methanobrevibacter | MSM_RS08755 | 1809971..1813555 | 2172 | MULTISPECIES:-BREX-system-P-loop-protein-BrxC-[Methanobrevibacter] |
| WP_143744747.1 | Type1.BREX | <-SIGMA+OB-Fold+AAA-ATPase+Lon-Protease-BrxL<-SIGMA+OB-Fold+AAA-ATPase+Lon-Protease-BrxL-p<- [illegible] | 1174 | Archaea>Euryarchaeota>Methanomada_group>Methanobacteria>Methanobacteriales>Methanobacteriaceae | GCF_002208625.1 | Methanobrevibacter_sp._87.7 | BGI41_RS00855 | 14881..18405 | 387957 | BREX-system-P-loop-protein-BrxC-[Methanobrevibacter-sp.-87.7] |
| WP_100907789.1 | Type1.BREX | EVE->BrxD-AAA->SF1-SF2-Helicase+wHTH+Small-B-barrel+DUF1998->?->SbcC+SbcC->HIN-HTH-><-REase+DUF4268\|\| [illegible] | 1197 | Archaea>Euryarchaeota>Methanomada_group>Methanobacteria>Methanobacteriales>Methanobacteriaceae | GCF_002813675.1 | Methanobacterium_sp._MZ-A1 | BK008_RS11195 | 2292330..2295923 | 1911685 | BREX-system-P-loop-protein-BrxC-[Methanobacterium-sp.-MZ-A1] |
| WP_100909051.1 | Type1.BREX | BrxD-AAA->SF1-SF2-Helicase+wHTH+Small-B-barrel+DUF1998->?->SbcC+SbcC-><-TPR+TPR+TPR+TPR<-?<-REase+DUF4268\|\| [illegible] | 1197 | Archaea>Euryarchaeota>Methanomada_group>Methanobacteria>Methanobacteriales>Methanobacteriaceae | GCF_002813695.1 | Methanobacterium_subterraneum | BK009_RS03450 | 727019..730612 | 59277 | BREX-system-P-loop-protein-BrxC-[Methanobacterium-subterraneum] |
| WP_191216096.1 | Type1.BREX | <-DTXR-HTH\|\|?->?-><-?\|\|?->?-><-SIGMA+OB-Fold+AAA-ATPase+Lon-Protease-BrxL<-PglZ<-N6-METHYLASE<-DUF6079*<- [illegible] | 1183 | Archaea>Euryarchaeota>Methanomada_group>Methanobacteria>Methanobacteriales>Methanobacteriaceae | GCF_009914355.1 | Methanothermobacter_sp._THM-1 | FZP57_06355 | 1234960..1238511 | 2606911 | BREX-system-P-loop-protein-BrxC-[Methanothermobacter-sp.-THM-1] |
| WP_071906589.1 | Type1.BREX | <-McrC-NTD+REase<-McrB-NTD+McrB-AAA\|\|?-><-ParB+HNH<-ASCH<-SIGMA+OB-Fold+AAA-ATPase+Lon-Protease-BrxL<-PglZ<- [illegible] | 1198 | Archaea>Euryarchaeota>Methanomada_group>Methanobacteria>Methanobacteriales>Methanobacteriaceae | GCF_900095295.1 | Methanobacterium_congolense | MCBB_RS04215 | 888415..892011 | 118062 | BREX-system-P-loop-protein-BrxC-[Methanobacterium-congolense] |
| WP_042694044.1 | Type1.BREX | TRD-><-SIGMA+OB-Fold+AAA-ATPase+Lon-Protease-BrxL<-PglZ<-?<-DUF6079*<-BrxB-BrxF-iSTAND<-BrxA-DNA-binding<- [illegible] | 1193 | Archaea>Euryarchaeota>Methanomada_group>Methanobacteria>Methanobacteriales>Methanobacteriaceae | GCF_900289035.1 | Methanobrevibacter_oralis | C9989_RS00250 | 18115..21696 | 66851 | BREX-system-P-loop-protein-BrxC-[Methanobrevibacter-oralis] |
| WP_052331840.1 | Type1.BREX | TRD-><-SIGMA+OB-Fold+AAA-ATPase+Lon-Protease-BrxL<-PglZ<-?<-DUF6079*<-BrxB-BrxF-iSTAND<-BrxA-DNA-binding<- [illegible] | 1188 | Archaea>Euryarchaeota>Methanomada_group>Methanobacteria>Methanobacteriales>Methanobacteriaceae | GCF_902384065.1 | Methanobrevibacter_oralis | TY05_RS07165 | 191000..194566 | 66851 | BREX-system-P-loop-protein-BrxC-[Methanobrevibacter-oralis] |
| ERJ05676.1 | Type1.BREX | <-N6-METHYLASE-p\|\|viral-protein-><-?<-NTP_transf+wHTH<-HTH+ICLR-HTH<-REase<-DUF6079<-BrxB-BrxF-iSTAND<- [illegible] | 1231 | Archaea>Euryarchaeota>Stenosarchaea_group>Halobacteria>Halobacteriales>Haloarculaceae | GCA_000215915.3 | Halorhabdus_tiamatea_SARL4B | HLRTI_002298 | 10094..13789 | 1033806 | hypothetical-protein-HLRTI_002298-[Halorhabdus-tiamatea-SARL4B] |
| EMA17933.1 | Type1.BREX | <-Exonuc_V_gamma+REase<-TUDOR+SF1-SF2-Helicase+iREase+CTD<-PglZ<-N6-METHYLASE<-REase<-wHTH<-wHTH<- [illegible] | 1280 | Archaea>Euryarchaeota>Stenosarchaea_group>Halobacteria>Halobacteriales>Haloarculaceae | GCA_000336895.1 | Haloarcula_argentinensis_DSM_12282 | C443_20652 | 152002..155844 | 1230451 | hypothetical-protein-C443_20652-[Haloarcula-argentinensis-DSM-12282] |
| EMA17935.1 | Type1.BREX | <-PglZ<-N6-METHYLASE<-REase<-wHTH<-wHTH<-NTP_transf+wHTH<-HTH+ICLR-HTH<-DUF6079<-BrxB-BrxF-iSTAND<-DUF6079*<- [illegible] | 1227 | Archaea>Euryarchaeota>Stenosarchaea_group>Halobacteria>Halobacteriales>Haloarculaceae | GCA_000336895.1 | Haloarcula_argentinensis_DSM_12282 | C443_20662 | 156419..160102 | 1230451 | hypothetical-protein-C443_20662-[Haloarcula-argentinensis-DSM-12282] |
| ELZ19699.1 | Type1.BREX | <-N6-METHYLASE-p\|\|?->?->?-><-PIN\|\|?-><-REase<-?\|\|?-><-DUF6079*<-BrxB-BrxF-iSTAND<-DUF6079<-BrxA-DNA-binding<- [illegible] | 1198 | Archaea>Euryarchaeota>Stenosarchaea_group>Halobacteria>Halobacteriales>Haloarculaceae | GCA_000337455.1 | Halosimplex_carlsbadense_2-9-1 | C475_22179 | 59263..62859 | 797114 | hypothetical-protein-C475_22179-[Halosimplex-carlsbadense-2-9-1] |
| ELZ19701.1 | Type1.BREX | <-PIN\|\|?-><-REase<-?\|\|?-><-DUF6079<-BrxB-BrxF-iSTAND<-DUF6079*<-BrxA-DNA-binding<-CR-ATPase9+REase<-?<-?\|\| [illegible] | 1248 | Archaea>Euryarchaeota>Stenosarchaea_group>Halobacteria>Halobacteriales>Haloarculaceae | GCA_000337455.1 | Halosimplex_carlsbadense_2-9-1 | C475_22189 | 63665..67411 | 797114 | hypothetical-protein-C475_22189-[Halosimplex-carlsbadense-2-9-1] |
| WP_015789649.1 | Type1.BREX | <-REase<-?<-?\|\|DOC+zhth-><-BirA-HTH<-?\|\|?->BrxA-DNA-binding->DUF6079*->BrxB-BrxF-iSTAND->DUF6079-><-?<-?\|\|?-> [illegible] | 1236 | Archaea>Euryarchaeota>Stenosarchaea_group>Halobacteria>Halobacteriales>Haloarculaceae | GCF_000023945.1 | Halorhabdus_utahensis | HUTA_RS09330 | 1919731..1923441 | 146826 | hypothetical-protein-[Halorhabdus-utahensis] |
| WP_015789651.1 | Type1.BREX | DOC+zhth-><-BirA-HTH<-?\|\|?->BrxA-DNA-binding->DUF6079->BrxB-BrxF-iSTAND->DUF6079*-><-?<-?\|\|?-><-OLD-ABC+TOPRIM [illegible] | 1280 | Archaea>Euryarchaeota>Stenosarchaea_group>Halobacteria>Halobacteriales>Haloarculaceae | GCF_000023945.1 | Halorhabdus_utahensis | HUTA_RS09340 | 1924016..1927858 | 146826 | hypothetical-protein-[Halorhabdus-utahensis] |
| WP_008308279.1 | Type1.BREX | ParA-Soj-PloopNTPase->BrxA-DNA-binding->DUF6079*->BrxB-BrxF-iSTAND->DUF6079-p-> | 1227 | Archaea>Euryarchaeota>Stenosarchaea_group>Halobacteria>Halobacteriales>Haloarculaceae | GCF_000336615.1 | Haloarcula_amylolytica | C442_RS04815 | 261082..264765 | 396317 | hypothetical-protein-[Haloarcula-amylolytica] |
| WP_008308281.1 | Type1.BREX | ParA-Soj-PloopNTPase->BrxA-DNA-binding->DUF6079->BrxB-BrxF-iSTAND->DUF6079-p*-> | 959 | Archaea>Euryarchaeota>Stenosarchaea_group>Halobacteria>Halobacteriales>Haloarculaceae | GCF_000336615.1 | Haloarcula_amylolytica | C442_RS04825 | 265340..268215 | 396317 | hypothetical-protein,-partial-[Haloarcula-amylolytica] |
| WP_049931199.1 | Type1.BREX | <-PglZ<-N6-METHYLASE<-PIN<-?\|\|?-><-Peptidase_M1<-REase<-?\|\|?-><-DUF6079*<-BrxB-BrxF-iSTAND<-DUF6079<- [illegible] | 1272 | Archaea>Euryarchaeota>Stenosarchaea_group>Halobacteria>Halobacteriales>Haloarculaceae | GCF_000337455.1 | Halosimplex_carlsbadense | C475_RS21845 | 59263..63081 | 171164 | hypothetical-protein-[Halosimplex-carlsbadense] |
| WP_049931237.1 | Type1.BREX | <-PIN<-?\|\|?-><-Peptidase_M1<-REase<-?\|\|?-><-DUF6079<-BrxB-BrxF-iSTAND<-DUF6079*<-BrxA-DNA-binding<- [illegible] | 1239 | Archaea>Euryarchaeota>Stenosarchaea_group>Halobacteria>Halobacteriales>Haloarculaceae | GCF_000337455.1 | Halosimplex_carlsbadense | C475_RS21855 | 63665..67384 | 171164 | hypothetical-protein-[Halosimplex-carlsbadense] |
| WP_245545399.1 | Type1.BREX | <-SLATT<-SMODS<-SMODS\|\|CopG/RHH-><-?\|\|MarR-HTH-><-?<-?<-DUF6079-p*\|\|?-><-BrxB-BrxF-iSTAND<-DUF6079<- [illegible] | 991 | Archaea>Euryarchaeota>Stenosarchaea_group>Halobacteria>Halobacteriales>Haloarculaceae | GCF_000379085.1 | Halomicrobium_katesii | HALKA_RS0114555 | 2756089..2759064 | 437163 | hypothetical-protein-[Halomicrobium-katesii] |
| WP_018258688.1 | Type1.BREX | <-SMODS\|\|CopG/RHH-><-?\|\|MarR-HTH-><-?<-?<-DUF6079-p\|\|?-><-BrxB-BrxF-iSTAND<-DUF6079*<-BrxA-DNA-binding<- [illegible] | 1248 | Archaea>Euryarchaeota>Stenosarchaea_group>Halobacteria>Halobacteriales>Haloarculaceae | GCF_000379085.1 | Halomicrobium_katesii | HALKA_RS0114565 | 2760536..2764282 | 437163 | hypothetical-protein-[Halomicrobium-katesii] |
| WP_008528537.1 | Type1.BREX | <-SbcC<-?<-N6-METHYLASE<-zf-IS66+DDE_Tnp_IS66\|\|viral-protein-><-?<-NTP_transf+wHTH<-HTH+ICLR-HTH<-REase<- [illegible] | 1279 | Archaea>Euryarchaeota>Stenosarchaea_group>Halobacteria>Halobacteriales>Haloarculaceae | GCF_000470655.1 | Halorhabdus_tiamatea | HTIA_RS15355 | 321396..325235 | 430914 | hypothetical-protein-[Halorhabdus-tiamatea] |
| WP_008528535.1 | Type1.BREX | <-N6-METHYLASE<-zf-IS66+DDE_Tnp_IS66\|\|viral-protein-><-?<-NTP_transf+wHTH<-HTH+ICLR-HTH<-REase<-DUF6079<- [illegible] | 1226 | Archaea>Euryarchaeota>Stenosarchaea_group>Halobacteria>Halobacteriales>Haloarculaceae | GCF_000470655.1 | Halorhabdus_tiamatea | HTIA_RS15365 | 325810..329490 | 430914 | hypothetical-protein-[Halorhabdus-tiamatea] |
| WP_058991492.1 | Type1.BREX | <-TUDOR+SF1-SF2-Helicase+iREase+CTD<-N6-METHYLASE<-PglZ<-?<-N6-METHYLASE<-REase<-wHTH<-NTP_transf+wHTH<- [illegible] | 1280 | Archaea>Euryarchaeota>Stenosarchaea_group>Halobacteria>Halobacteriales>Haloarculaceae | GCF_001485575.1 | Haloarcula_sp._CBA1127 | AV059_RS01020 | 247527..251369 | 1765055 | hypothetical-protein-[Haloarcula-sp.-CBA1127] |
| WP_058991494.1 | Type1.BREX | <-PglZ<-?<-N6-METHYLASE<-REase<-wHTH<-NTP_transf+wHTH<-HTH+ICLR-HTH<-DUF6079<-BrxB-BrxF-iSTAND<-DUF6079*<- [illegible] | 1227 | Archaea>Euryarchaeota>Stenosarchaea_group>Halobacteria>Halobacteriales>Haloarculaceae | GCF_001485575.1 | Haloarcula_sp._CBA1127 | AV059_RS01030 | 251944..255627 | 1765055 | hypothetical-protein-[Haloarcula-sp.-CBA1127] |
| WP_108383896.1 | Type1.BREX | <-SF1-SF2-Helicase-p\|\|?-><-?<-?<-?<-?\|\|BrxA-DNA-binding->DUF6079*->BrxB-BrxF-iSTAND->DUF6079->REase-><-?<-?\|\| [illegible] | 1236 | Archaea>Euryarchaeota>Stenosarchaea_group>Halobacteria>Halobacteriales>Haloarculaceae | GCF_003058365.1 | Halococcoides_cellulosivorans | HARCEL1_RS12430 | 2549904..2553614 | 1679096 | hypothetical-protein-[Halococcoides-cellulosivorans] |
| WP_108383898.1 | Type1.BREX | <-SF1-SF2-Helicase-p\|\|?-><-?<-?<-?<-?\|\|BrxA-DNA-binding->DUF6079->BrxB-BrxF-iSTAND->DUF6079*->REase-><-?<-?\|\| [illegible] | 1280 | Archaea>Euryarchaeota>Stenosarchaea_group>Halobacteria>Halobacteriales>Haloarculaceae | GCF_003058365.1 | Halococcoides_cellulosivorans | HARCEL1_RS12440 | 2554189..2558031 | 1679096 | hypothetical-protein-[Halococcoides-cellulosivorans] |
| WP_129754743.1 | Type1.BREX | ParA-Soj-PloopNTPase->BrxA-DNA-binding->DUF6079*->BrxB-BrxF-iSTAND->DUF6079-><-viral-protein\|\|N6-METHYLASE-> [illegible] | 1227 | Archaea>Euryarchaeota>Stenosarchaea_group>Halobacteria>Halobacteriales>Haloarculaceae | GCF_004150385.1 | Haloarcula_hispanica | ELS20_RS00155 | 36123..39806 | 51589 | hypothetical-protein-[Haloarcula-hispanica] |
| WP_136689168.1 | Type1.BREX | <-35exo<-35exo<-MNS+REase<-?<-ARG\|\|?->?->SF1-SF2-Helicase+REase->BrxA-DNA-binding->DUF6079*->BrxB-BrxF-iSTAND-> [illegible] | 1249 | Archaea>Euryarchaeota>Stenosarchaea_group>Halobacteria>Halobacteriales>Haloarculaceae | GCF_004944975.1 | Halorhabdus_amylolytica | E3374_RS11205 | 84563..88312 | 2559573 | hypothetical-protein-[Halorhabdus-amylolytica] |
| WP_136689170.1 | Type1.BREX | <-MNS+REase<-?<-ARG\|\|?->?->SF1-SF2-Helicase+REase->BrxA-DNA-binding->DUF6079->BrxB-BrxF-iSTAND->DUF6079*-> [illegible] | 1279 | Archaea>Euryarchaeota>Stenosarchaea_group>Halobacteria>Halobacteriales>Haloarculaceae | GCF_004944975.1 | Halorhabdus_amylolytica | E3374_RS11215 | 88926..92765 | 2559573 | hypothetical-protein-[Halorhabdus-amylolytica] |
| WP_152120782.1 | Type1.BREX | BrxB-BrxF-iSTAND->DUF6079*->N6-METHYLASE->REase->PglZ->SIGMA+OB-Fold+AAA-ATPase+Lon-Protease-BrxL-> [illegible] | 1215 | Archaea>Euryarchaeota>Stenosarchaea_group>Halobacteria>Halobacteriales>Haloarculaceae | GCF_009184545.1 | Halosegnis_rubeus | DMP03_RS11345 | 583..4230 | 2212850 | BREX-system-P-loop-protein-BrxC-[Halosegnis-rubeus] |
| WP_188853474.1 | Type1.BREX | <-ABC-ATPase+SbcC+VirB11+ABC-ATPase+SbcC<-?<-Calcineurin<-?<-?\|\|?->?->?->BrxA-DNA-binding->DUF6079*-> [illegible] | 1227 | Archaea>Euryarchaeota>Stenosarchaea_group>Halobacteria>Halobacteriales>Haloarculaceae | GCF_014647095.1 | Haloarcula_salaria | IEY10_RS16005 | 45657..49340 | 575195 | hypothetical-protein-[Haloarcula-salaria] |
| WP_188853476.1 | Type1.BREX | <-Calcineurin<-?<-?\|\|?->?->?->BrxA-DNA-binding->DUF6079->BrxB-BrxF-iSTAND->DUF6079*->HTH+ICLR-HTH-> [illegible] | 1280 | Archaea>Euryarchaeota>Stenosarchaea_group>Halobacteria>Halobacteriales>Haloarculaceae | GCF_014647095.1 | Haloarcula_salaria | IEY10_RS16015 | 49915..53757 | 575195 | hypothetical-protein-[Haloarcula-salaria] |
| WP_188997387.1 | Type1.BREX | MarR-HTH-><-SF1-SF2-Helicase+REase<-Exonuc_V_gamma+REase\|\|BrxA-DNA-binding->DUF6079*->BrxB-BrxF-iSTAND-> [illegible] | 1251 | Archaea>Euryarchaeota>Stenosarchaea_group>Halobacteria>Halobacteriales>Haloarculaceae | GCF_014647235.1 | Halomicroarcula_pellucida | IEY28_RS11100 | 280416..284171 | 1427151 | hypothetical-protein-[Halomicroarcula-pellucida] |
| WP_188997391.1 | Type1.BREX | MarR-HTH-><-SF1-SF2-Helicase+REase<-Exonuc_V_gamma+REase\|\|BrxA-DNA-binding->DUF6079->BrxB-BrxF-iSTAND-> [illegible] | 1288 | Archaea>Euryarchaeota>Stenosarchaea_group>Halobacteria>Halobacteriales>Haloarculaceae | GCF_014647235.1 | Halomicroarcula_pellucida | IEY28_RS11110 | 284755..288621 | 1427151 | hypothetical-protein-[Halomicroarcula-pellucida] |
| WP_229112049.1 | Type1.BREX | <-TUDOR+SF1-SF2-Helicase+iREase+CTD<-PglZ<-N6-METHYLASE-p<-?<-DDE_Tnp_1<-N6-METHYLASE\|\|HD-><-?<-HTH+ICLR-HTH<- [illegible] | 1278 | Archaea>Euryarchaeota>Stenosarchaea_group>Halobacteria>Halobacteriales>Haloarculaceae | GCF_017094525.1 | Halapricum_desulfuricans | HSR122_RS06875 | 1337225..1341061 | 2841257 | hypothetical-protein-[Halapricum-desulfuricans] |
| WP_229112051.1 | Type1.BREX | <-N6-METHYLASE-p<-?<-DDE_Tnp_1<-N6-METHYLASE\|\|HD-><-?<-HTH+ICLR-HTH<-DUF6079<-BrxB-BrxF-iSTAND<-DUF6079*<- [illegible] | 1248 | Archaea>Euryarchaeota>Stenosarchaea_group>Halobacteria>Halobacteriales>Haloarculaceae | GCF_017094525.1 | Halapricum_desulfuricans | HSR122_RS06885 | 1341681..1345427 | 2841257 | hypothetical-protein-[Halapricum-desulfuricans] |
| WP_129754745.1 | Type1.BREX | ParA-Soj-PloopNTPase->BrxA-DNA-binding->BrxB-BrxF-iSTAND->DUF6079*-><-viral-protein\|\|N6-METHYLASE->PglZ-><-?<- [illegible] | 1280 | Archaea>Euryarchaeota>Stenosarchaea_group>Halobacteria>Halobacteriales>Haloarculaceae | GCF_022807515.1 | Haloarcula_hispanica | ELS20_RS00165 | 40381..44223 | 51589 | hypothetical-protein-[Haloarcula-hispanica] |
| WP_274325353.1 | Type1.BREX | <-PIN<-?\|\|?-><-Peptidase_M1<-REase<-?\|\|?-><-DUF6079<-BrxB-BrxF-iSTAND<-DUF6079*<-Zn_Tnp_IS1595+DUF2089+ [illegible] | 1126 | Archaea>Euryarchaeota>Stenosarchaea_group>Halobacteria>Halobacteriales>Haloarculaceae | GCF_028747785.1 | Halosimplex_sp._XZYJT29 | PSC72_RS07975 | 1604962..1608342 | 3026162 | hypothetical-protein-[Halosimplex-sp.-XZYJT29] |
| WP_135666499.1 | Type1.BREX | <-PglZ<-RelE<-?<-N6-METHYLASE<-TBP+TBP\|\|TBP+TBP->?-><-?<-?<-DUF6079*<-BrxB-BrxF-iSTAND<-DUF6079<- [illegible] | 1277 | Archaea>Euryarchaeota>Stenosarchaea_group>Halobacteria>Halobacteriales>Haloarculaceae | GCF_900880625.1 | Halorhabdus_rudnickae | BN2694_RS13550 | 37861..41694 | 1775544 | hypothetical-protein-[Halorhabdus-rudnickae] |
| WP_135666504.1 | Type1.BREX | <-N6-METHYLASE<-TBP+TBP\|\|TBP+TBP->?-><-?<-?<-DUF6079<-BrxB-BrxF-iSTAND<-DUF6079*<-BrxA-DNA-binding<- [illegible] | 1248 | Archaea>Euryarchaeota>Stenosarchaea_group>Halobacteria>Halobacteriales>Haloarculaceae | GCF_900880625.1 | Halorhabdus_rudnickae | BN2694_RS13560 | 42308..46054 | 1775544 | hypothetical-protein-[Halorhabdus-rudnickae] |
| WP_049892750.1 | Type1.BREX | <-N6-METHYLASE\|\|OLD-ABC+TOPRIM-><-DDE_Tnp_1\|\|?-><-?<-NTP_transf+wHTH<-NTP_transf+wHTH<-CopG/RHH<-HTH+ICLR-HTH<- [illegible] | 1280 | Archaea>Euryarchaeota>Stenosarchaea_group>Halobacteria>Halobacteriales>Halobacteriaceae | GCF_000069025.1 | Halobacterium_salinarum | OE_RS13050 | 230465..234307 | 2242 | hypothetical-protein-[Halobacterium-salinarum] |
| WP_049892751.1 | Type1.BREX | <-DDE_Tnp_1\|\|?-><-?<-NTP_transf+wHTH<-NTP_transf+wHTH<-CopG/RHH<-HTH+ICLR-HTH<-DUF6079<-BrxB-BrxF-iSTAND<- [illegible] | 1227 | Archaea>Euryarchaeota>Stenosarchaea_group>Halobacteria>Halobacteriales>Halobacteriaceae | GCF_000069025.1 | Halobacterium_salinarum | OE_RS13060 | 234882..238565 | 2242 | hypothetical-protein-[Halobacterium-salinarum] |
| WP_123076504.1 | Type1.BREX | <-TUDOR+SF1-SF2-Helicase+iREase+CTD<-PglZ<-N6-METHYLASE\|\|?->?-><-?\|\|?->SF1-SF2-Helicase-><-DUF6079*<- [illegible] | 1277 | Archaea>Euryarchaeota>Stenosarchaea_group>Halobacteria>Halobacteriales>Halobacteriaceae | GCF_003721175.2 | Halarchaeum_sp._CBA1220 | EFA46_RS09075 | 1680640..1684473 | 1853682 | hypothetical-protein-[Halarchaeum-sp.-CBA1220] |
| WP_123076500.1 | Type1.BREX | <-PglZ<-N6-METHYLASE\|\|?->?-><-?\|\|?->SF1-SF2-Helicase-><-DUF6079<-BrxB-BrxF-iSTAND<-DUF6079*<-BrxA-DNA-binding<- [illegible] | 1248 | Archaea>Euryarchaeota>Stenosarchaea_group>Halobacteria>Halobacteriales>Halobacteriaceae | GCF_003721175.2 | Halarchaeum_sp._CBA1220 | EFA46_RS09085 | 1685087..1688833 | 1853682 | hypothetical-protein-[Halarchaeum-sp.-CBA1220] |
| WP_135823790.1 | Type1.BREX | N6-METHYLASE->?-><-DUF6079*<-BrxB-BrxF-iSTAND<-DUF6079<-?\|\|Peptidase_M50-> | 1264 | Archaea>Euryarchaeota>Stenosarchaea_group>Halobacteria>Halobacteriales>Halobacteriaceae | GCF_004765785.1 | Halorussus_ruber | EPL01_RS03100 | 202199..205993 | 1126238 | hypothetical-protein-[Halorussus-ruber] |
| WP_135823792.1 | Type1.BREX | N6-METHYLASE->?-><-DUF6079<-BrxB-BrxF-iSTAND<-DUF6079*<-?\|\|Peptidase_M50->?->?->?->?-><-?\|\|?->CBS+CBS-> | 1304 | Archaea>Euryarchaeota>Stenosarchaea_group>Halobacteria>Halobacteriales>Halobacteriaceae | GCF_004765785.1 | Halorussus_ruber | EPL01_RS03110 | 206652..210566 | 1126238 | hypothetical-protein-[Halorussus-ruber] |
| WP_136361239.1 | Type1.BREX | <-HTH+BirA-HTH<-?<-DDE_Tnp_1\|\|Classical-AAA->?-><-DDE_Tnp_1<-N6-METHYLASE-p\|\|BrxA-DNA-binding->DUF6079*-> [illegible] | 1233 | Archaea>Euryarchaeota>Stenosarchaea_group>Halobacteria>Halobacteriales>Halobacteriaceae | GCF_004799605.1 | Halobacterium_salinarum | HBSAL_RS04975 | 932815..936516 | 2242 | hypothetical-protein-[Halobacterium-salinarum] |
| WP_136361241.1 | Type1.BREX | <-DDE_Tnp_1\|\|Classical-AAA->?-><-DDE_Tnp_1<-N6-METHYLASE-p\|\|BrxA-DNA-binding->DUF6079->BrxB-BrxF-iSTAND-> [illegible] | 1231 | Archaea>Euryarchaeota>Stenosarchaea_group>Halobacteria>Halobacteriales>Halobacteriaceae | GCF_004799605.1 | Halobacterium_salinarum | HBSAL_RS04985 | 937132..940827 | 2242 | hypothetical-protein-[Halobacterium-salinarum] |
| WP_233265034.1 | Type1.BREX | <-NTP_transf\|\|?->?-><-PIWI<-SF1-SF2-Helicase+REase<-SF1-SF2-Helicase+REase<-SF1-SF2-Helicase+wHTH+ [illegible] | 962 | Archaea>Euryarchaeota>Stenosarchaea_group>Halobacteria>Halobacteriales>Halobacteriaceae | GCF_009791395.1 | Halomarina_oriensis | GQS65_RS04705 | 84531..87418 | 671145 | hypothetical-protein,-partial-[Halomarina-oriensis] |
| WP_205252856.1 | Type1.BREX | <-Exonuc_V_gamma+REase<-TUDOR+SF1-SF2-Helicase+iREase+CTD<-HEPN<-PglZ<-?<-N6-METHYLASE\|\|?->?-><-REase<- [illegible] | 1280 | Archaea>Euryarchaeota>Stenosarchaea_group>Halobacteria>Halobacteriales>Halobacteriaceae | GCF_016917835.2 | Halobacterium_sp._BOL4-2 | JRZ79_RS24035 | 52137..55979 | 2810537 | hypothetical-protein-[Halobacterium-sp.-BOL4-2] |
| WP_205252782.1 | Type1.BREX | <-HEPN<-PglZ<-?<-N6-METHYLASE\|\|?->?-><-REase<-DUF6079<-BrxB-BrxF-iSTAND<-DUF6079*<-BrxA-DNA-binding\|\| [illegible] | 1227 | Archaea>Euryarchaeota>Stenosarchaea_group>Halobacteria>Halobacteriales>Halobacteriaceae | GCF_016917835.2 | Halobacterium_sp._BOL4-2 | JRZ79_RS24045 | 56554..60237 | 2810537 | hypothetical-protein-[Halobacterium-sp.-BOL4-2] |
| WP_227231659.1 | Type1.BREX | SF1-SF2-Helicase+REase->SF1-SF2-Helicase+REase-><-DOC+HTH<-PglZ<-DUF6079*<-BrxB-BrxF-iSTAND<-DUF6079 | 1231 | Archaea>Euryarchaeota>Stenosarchaea_group>Halobacteria>Halobacteriales>Halobacteriaceae | GCF_020567425.1 | Salinirubrum_litoreum | LI337_RS19795 | 90015..93710 | 1126234 | hypothetical-protein-[Salinirubrum-litoreum] |
| WP_227231661.1 | Type1.BREX | SF1-SF2-Helicase+REase->SF1-SF2-Helicase+REase-><-DOC+HTH<-PglZ<-DUF6079<-BrxB-BrxF-iSTAND<-DUF6079* | 1054 | Archaea>Euryarchaeota>Stenosarchaea_group>Halobacteria>Halobacteriales>Halobacteriaceae | GCF_020567425.1 | Salinirubrum_litoreum | LI337_RS19805 | 94326..97490 | 1126234 | hypothetical-protein,-partial-[Salinirubrum-litoreum] |
| WP_231187339.1 | Type1.BREX | BrxA-DNA-binding->DUF6079*->BrxB-BrxF-iSTAND->DUF6079->N6-METHYLASE-p->PglZ->?->BirA-HTH->MarR-HTH-> [illegible] | 1228 | Archaea>Euryarchaeota>Stenosarchaea_group>Halobacteria>Halobacteriales>Halobacteriaceae | GCF_021083305.1 | Haladaptatus_sp._DYF46 | LJF34_RS20420 | 52158..55844 | 2886041 | hypothetical-protein-[Haladaptatus-sp.-DYF46] |
| WP_231187343.1 | Type1.BREX | BrxA-DNA-binding->DUF6079->BrxB-BrxF-iSTAND->DUF6079*->N6-METHYLASE-p->PglZ->?->BirA-HTH->MarR-HTH-> [illegible] | 1283 | Archaea>Euryarchaeota>Stenosarchaea_group>Halobacteria>Halobacteriales>Halobacteriaceae | GCF_021083305.1 | Haladaptatus_sp._DYF46 | LJF34_RS20430 | 56419..60270 | 2886041 | hypothetical-protein-[Haladaptatus-sp.-DYF46] |
| WP_235167905.1 | Type1.BREX | <-Phage_int_SAM_5<-?<-ParA-Soj-PloopNTPase<-SF1-SF2-Helicase+REase<-Exonuc_V_gamma+REase\|\|BrxA-DNA-binding-> [illegible] | 1248 | Archaea>Euryarchaeota>Stenosarchaea_group>Halobacteria>Halobacteriales>Halobacteriaceae | GCF_021503975.1 | Halobacterium_salinarum | KI372_RS02110 | 10921..14667 | 2242 | hypothetical-protein-[Halobacterium-salinarum] |
| WP_235262714.1 | Type1.BREX | <-TUDOR+SF1-SF2-Helicase+iREase+CTD<-?<-?<-?<-PglZ<-N6-METHYLASE<-?\|\|?->?-><-DUF6079*<-BrxB-BrxF-iSTAND<- [illegible] | 1279 | Archaea>Euryarchaeota>Stenosarchaea_group>Halobacteria>Halobacteriales>Halobacteriaceae | GCF_021503975.1 | Halobacterium_salinarum | KVP02_RS02090 | 31387..35226 | 2242 | hypothetical-protein-[Halobacterium-salinarum] |
| WP_267638768.1 | Type1.BREX | <-HEPN<-Peptidase_M1\|\|?->?->Calcineurin-SbcD-><-DUF6079<-BrxB-BrxF-iSTAND<-DUF6079*<-BrxA-DNA-binding\|\|REase-> [illegible] | 1251 | Archaea>Euryarchaeota>Stenosarchaea_group>Halobacteria>Halobacteriales>Halobacteriaceae | GCF_026586665.1 | Halovenus_rubra | N5098_RS15400 | 10573..14328 | 869890 | hypothetical-protein-[Halovenus-rubra] |
| WP_121821034.1 | Type1.BREX | <-Calcineurin<-?\|\|BrxA-DNA-binding->DUF6079*->BrxB-BrxF-iSTAND->DUF6079->?-><-?<-?\|\|N6-METHYLASE->?->PglZ-> [illegible] | 1237 | Archaea>Euryarchaeota>Stenosarchaea_group>Halobacteria>Halobacteriales>Natronoarchaeaceae | GCF_003675855.1 | Halostella_salina | D8896_RS05250 | 383144..386857 | 1547897 | hypothetical-protein-[Halostella-salina] |
| WP_121821036.1 | Type1.BREX | <-Calcineurin<-?\|\|BrxA-DNA-binding->DUF6079->BrxB-BrxF-iSTAND->DUF6079*->?-><-?<-?\|\|N6-METHYLASE->?->PglZ-> [illegible] | 1280 | Archaea>Euryarchaeota>Stenosarchaea_group>Halobacteria>Halobacteriales>Natronoarchaeaceae | GCF_003675855.1 | Halostella_salina | D8896_RS05260 | 387432..391274 | 1547897 | hypothetical-protein-[Halostella-salina] |
| WP_008387699.1 | Type1.BREX | <-N6-METHYLASE<-REase<-?<-REase+METHYLASE<-DUF6079*<-BrxB-BrxF-iSTAND<-DUF6079<-BrxA-DNA-binding\|\| [illegible] | 1270 | Archaea>Euryarchaeota>Stenosarchaea_group>Halobacteria>Haloferacales>Haloferacaceae | GCF_000337095.1 | Halogeometricum_pallidum | C474_RS13405 | 6094..9906 | 411361 | hypothetical-protein-[Halogeometricum-pallidum] |
| WP_008387702.1 | Type1.BREX | <-N6-METHYLASE<-REase<-?<-REase+METHYLASE<-DUF6079<-BrxB-BrxF-iSTAND<-DUF6079*<-BrxA-DNA-binding\|\| [illegible] | 1243 | Archaea>Euryarchaeota>Stenosarchaea_group>Halobacteria>Haloferacales>Haloferacaceae | GCF_000337095.1 | Halogeometricum_pallidum | C474_RS13415 | 10487..14218 | 411361 | hypothetical-protein-[Halogeometricum-pallidum] |
| WP_049935523.1 | Type1.BREX | ABC-ATPase+SbcC->?->?->?->DTXR-HTH->tRNA-><-?<-PglZ<-N6-METHYLASE-p<-DUF6079*<-BrxB-BrxF-iSTAND<-DUF6079<- [illegible] | 1271 | Archaea>Euryarchaeota>Stenosarchaea_group>Halobacteria>Haloferacales>Haloferacaceae | GCF_000427685.1 | Haloplanus_natans | HALNA_RS06110 | 731546..735361 | 376171 | hypothetical-protein-[Haloplanus-natans] |
| WP_049935526.1 | Type1.BREX | DTXR-HTH->tRNA-><-?<-PglZ<-N6-METHYLASE-p<-DUF6079<-BrxB-BrxF-iSTAND<-DUF6079*<-BrxA-DNA-binding\|\|?-><-?<-PIN [illegible] | 1253 | Archaea>Euryarchaeota>Stenosarchaea_group>Halobacteria>Haloferacales>Haloferacaceae | GCF_000427685.1 | Haloplanus_natans | HALNA_RS06120 | 735945..739706 | 376171 | hypothetical-protein-[Haloplanus-natans] |
| WP_122106903.1 | Type1.BREX | <-DDE_Tnp_1<-?<-?\|\|MarR-HTH-><-?<-ParA-Soj-PloopNTPase<-SF1-SF2-Helicase+REase<-Exonuc_V_gamma+REase\|\| [illegible] | 1248 | Archaea>Euryarchaeota>Stenosarchaea_group>Halobacteria>Haloferacales>Haloferacaceae | GCF_003697845.1 | Halobellus_captivus | EA078_RS16655 | 16343..20089 | 2592614 | hypothetical-protein-[Halobellus-captivus] |
| WP_122106905.1 | Type1.BREX | MarR-HTH-><-?<-ParA-Soj-PloopNTPase<-SF1-SF2-Helicase+REase<-Exonuc_V_gamma+REase\|\|BrxA-DNA-binding->DUF6079-> [illegible] | 1279 | Archaea>Euryarchaeota>Stenosarchaea_group>Halobacteria>Haloferacales>Haloferacaceae | GCF_003697845.1 | Halobellus_captivus | EA078_RS16665 | 20703..24542 | 2592614 | hypothetical-protein-[Halobellus-captivus] |
| WP_248897121.1 | Type1.BREX | BirA-HTH->?-><-?<-?\|\|?->BrxA-DNA-binding->DUF6079*->BrxB-BrxF-iSTAND->DUF6079->HTH+ICLR-HTH->PglZ-> | 1247 | Archaea>Euryarchaeota>Stenosarchaea_group>Halobacteria>Haloferacales>Haloferacaceae | GCF_023276355.1 | Haloplanus_sp._XH21 | MXB53_RS09475 | 1803325..1807068 | 2934938 | hypothetical-protein-[Haloplanus-sp.-XH21] |
| WP_248897123.1 | Type1.BREX | BirA-HTH->?-><-?<-?\|\|?->BrxA-DNA-binding->DUF6079->BrxB-BrxF-iSTAND->DUF6079*->HTH+ICLR-HTH->PglZ-><-?<-?\|\|?-> [illegible] | 1277 | Archaea>Euryarchaeota>Stenosarchaea_group>Halobacteria>Haloferacales>Haloferacaceae | GCF_023276355.1 | Haloplanus_sp._XH21 | MXB53_RS09485 | 1807682..1811515 | 2934938 | hypothetical-protein-[Haloplanus-sp.-XH21] |
| WP_256422112.1 | Type1.BREX | <-N6-METHYLASE\|\|?-><-?<-?<-NTP_transf+wHTH<-HTH+ICLR-HTH<-DUF6079<-BrxB-BrxF-iSTAND<-DUF6079*<- [illegible] | 1248 | Archaea>Euryarchaeota>Stenosarchaea_group>Halobacteria>Haloferacales>Haloferacaceae | GCF_024494595.1 | Halobellus_rarus | N0876_RS13095 | 193250..196996 | 1126237 | hypothetical-protein-[Halobellus-rarus] |
| ELZ39021.1 | Type1.BREX | OrfB_IS605+OrfB_Zn_ribbon->DDE_Tnp_1->?-><-ORC-CDC6\|\|BrxA-DNA-binding->DUF6079->BrxB-BrxF-iSTAND->DUF6079*-> [illegible] | 1280 | Archaea>Euryarchaeota>Stenosarchaea_group>Halobacteria>Haloferacales>Halorubraceae | GCA_000336875.1 | Halorubrum_californiense_DSM_19288 | C463_17695 | 23296..27138 | 1227465 | hypothetical-protein-C463_17695-[Halorubrum-californiense-DSM-19288] |
| PHQ38249.1 | Type1.BREX | SF1-SF2-Helicase->SF1-SF2-Helicase+REase->HTH_28->NTP_transf_5-><-?\|\|?-><-?\|\|<-?\|\|BrxA-DNA-binding->DUF6079*-> [illegible] | 1227 | Archaea>Euryarchaeota>Stenosarchaea_group>Halobacteria>Haloferacales>Halorubraceae | GCA_002727125.1 | Halorubrum_persicum | DJ69_12930 | 85034..88717 | 1383844 | hypothetical-protein-DJ69_12930-[Halorubrum-persicum] |
| SFP79187.1 | Type1.BREX | <-PglZ\|\|ZnR+REase->OLD-ABC+TOPRIM-><-N6-METHYLASE\|\|viral-protein-><-REase\|\|?-><-DUF6079<-BrxB-BrxF-iSTAND<- [illegible] | 1232 | Archaea>Euryarchaeota>Stenosarchaea_group>Halobacteria>Haloferacales>Halorubraceae | GCA_900115675.1 | Halolamina_pelagica | SAMN05216277_10870 | 87547..91245 | 699431 | hypothetical-protein-SAMN05216277_10870-[Halolamina-pelagica] |
| WP_015911586.1 | Type1.BREX | <-N6-METHYLASE-p\|\|MazE_antitoxin->PIN-><-DUF6079*<-BrxB-BrxF-iSTAND<-DUF6079<-BrxA-DNA-binding<- [illegible] | 1270 | Archaea>Euryarchaeota>Stenosarchaea_group>Halobacteria>Haloferacales>Halorubraceae | GCF_000022205.1 | Halorubrum_lacusprofundi | CF318_RS14700 | 1993..5805 | 2247 | hypothetical-protein-[Halorubrum-lacusprofundi] |
| WP_015911588.1 | Type1.BREX | <-TUDOR+SF1-SF2-Helicase+iREase+CTD<-PglZ<-N6-METHYLASE\|\|DDE_Tnp_1-><-N6-METHYLASE-p\|\|MazE_antitoxin->PIN-> [illegible] | 1239 | Archaea>Euryarchaeota>Stenosarchaea_group>Halobacteria>Haloferacales>Halorubraceae | GCF_000022205.1 | Halorubrum_lacusprofundi | CF318_RS14710 | 6389..10108 | 2247 | hypothetical-protein-[Halorubrum-lacusprofundi] |
| WP_008446367.1 | Type1.BREX | OrfB_IS605+OrfB_Zn_ribbon->\|\|DDE_Tnp_1->?-><-ORC-CDC6\|\|BrxA-DNA-binding->DUF6079*->BrxB-BrxF-iSTAND->DUF6079-> [illegible] | 1227 | Archaea>Euryarchaeota>Stenosarchaea_group>Halobacteria>Haloferacales>Halorubraceae | GCF_000336875.1 | Halorubrum_californiense | C463_RS17190 | 19038..22721 | 416585 | hypothetical-protein-[Halorubrum-californiense] |
| WP_239638934.1 | Type1.BREX | OrfB_IS605+OrfB_Zn_ribbon->\|\|DDE_Tnp_1->?-><-ORC-CDC6\|\|BrxA-DNA-binding->DUF6079->BrxB-BrxF-iSTAND->DUF6079*-> [illegible] | 1235 | Archaea>Euryarchaeota>Stenosarchaea_group>Halobacteria>Haloferacales>Halorubraceae | GCF_000336875.1 | Halorubrum_californiense | C463_RS17200 | 23431..27138 | 416585 | hypothetical-protein-[Halorubrum-californiense] |
| WP_235271349.1 | Type1.BREX | <-Phage_integrase<-TUDOR+SF1-SF2-Helicase+iREase+CTD<-REase<-PglZ<-PglZ<-N6-METHYLASE\|\|?->?-><-?<-DUF6079*<- [illegible] | 1111 | Archaea>Euryarchaeota>Stenosarchaea_group>Halobacteria>Haloferacales>Halorubraceae | GCF_000700025.1 | Halorubrum_saccharovorum | FK85_RS03110 | 45328..48663 | 2248 | hypothetical-protein-[Halorubrum-saccharovorum] |
| WP_235271352.1 | Type1.BREX | <-PglZ<-PglZ<-N6-METHYLASE\|\|?->?-><-?<-DUF6079<-?<-BrxB-BrxF-iSTAND<-DUF6079-p*<-DUF6079-p<-BrxA-DNA-binding<- [illegible] | 692 | Archaea>Euryarchaeota>Stenosarchaea_group>Halobacteria>Haloferacales>Halorubraceae | GCF_000700025.1 | Halorubrum_saccharovorum | FK85_RS18475 | 49771..51849 | 2248 | hypothetical-protein-[Halorubrum-saccharovorum] |
| WP_094582515.1 | Type1.BREX | <-N6-METHYLASE<-DUF6079*<-BrxB-BrxF-iSTAND<-DUF6079<-BrxA-DNA-binding\|\|?->ABC-ATPase+SbcC-> | 1277 | Archaea>Euryarchaeota>Stenosarchaea_group>Halobacteria>Haloferacales>Halorubraceae | GCF_002252835.1 | Halorubrum_ezzemoulense | DJ78_RS02240 | 3110..6943 | 337243 | hypothetical-protein-[Halorubrum-ezzemoulense] |
| WP_094582519.1 | Type1.BREX | <-N6-METHYLASE<-DUF6079<-BrxB-BrxF-iSTAND<-DUF6079*<-BrxA-DNA-binding\|\|?->ABC-ATPase+SbcC->?-><-?\|\|?->?-> [illegible] | 1226 | Archaea>Euryarchaeota>Stenosarchaea_group>Halobacteria>Haloferacales>Halorubraceae | GCF_002252835.1 | Halorubrum_ezzemoulense | DJ78_RS02250 | 7518..11198 | 337243 | hypothetical-protein-[Halorubrum-ezzemoulense] |
| WP_096396418.1 | Type1.BREX | <-ParA-Soj-PloopNTPase<-PADR-HTH<-?\|\|?-><-?<-?<-?<-?\|\|BrxA-DNA-binding->DUF6079*-> | 1227 | Archaea>Euryarchaeota>Stenosarchaea_group>Halobacteria>Haloferacales>Halorubraceae | GCF_002355655.1 | Halorubrum_trapanicum | CPZ01_RS14385 | 137266..140949 | 29284 | hypothetical-protein-[Halorubrum-trapanicum] |
| WP_099288831.1 | Type1.BREX | <-GDResolvase\|\|?->?->?->?->BrxA-DNA-binding->DUF6079*->BrxB-BrxF-iSTAND->DUF6079-> | 1248 | Archaea>Euryarchaeota>Stenosarchaea_group>Halobacteria>Haloferacales>Halorubraceae | GCF_002727095.1 | Halorubrum_sp._C191 | Z052_RS03555 | 9147..12893 | 1383842 | hypothetical-protein-[Halorubrum-sp.-C191] |
| WP_099288832.1 | Type1.BREX | <-GDResolvase\|\|?->?->?->?->BrxA-DNA-binding->DUF6079->BrxB-BrxF-iSTAND->DUF6079*-> | 1280 | Archaea>Euryarchaeota>Stenosarchaea_group>Halobacteria>Haloferacales>Halorubraceae | GCF_002727095.1 | Halorubrum_sp._C191 | Z052_RS03565 | 13507..17349 | 1383842 | hypothetical-protein-[Halorubrum-sp.-C191] |
| WP_245851676.1 | Type1.BREX | SF1-SF2-Helicase->SF1-SF2-Helicase+REase->HTH_28->NTP_transf_5-><-?\|\|?-><-?\|\|<-?\|\|BrxA-DNA-binding->DUF6079*-> [illegible] | 1139 | Archaea>Euryarchaeota>Stenosarchaea_group>Halobacteria>Haloferacales>Halorubraceae | GCF_002727125.1 | Halorubrum_persicum | DJ69_RS13035 | 85298..88717 | 1383844 | hypothetical-protein-[Halorubrum-persicum] |
| WP_099255996.1 | Type1.BREX | HTH_28->NTP_transf_5-><-?\|\|?-><-?\|\|<-?\|\|BrxA-DNA-binding->DUF6079->BrxB-BrxF-iSTAND->DUF6079*->wHTH->REase-> [illegible] | 1278 | Archaea>Euryarchaeota>Stenosarchaea_group>Halobacteria>Haloferacales>Halorubraceae | GCF_002727125.1 | Halorubrum_persicum | DJ69_RS13045 | 89292..93128 | 1383844 | hypothetical-protein-[Halorubrum-persicum] |
| WP_112078641.1 | Type1.BREX | ABC-ATPase+SbcC->?->BPD_transp_2->?-><-?\|\|BrxA-DNA-binding->DUF6079*->BrxB-BrxF-iSTAND->DUF6079->N6-METHYLASE- [illegible] | 1251 | Archaea>Euryarchaeota>Stenosarchaea_group>Halobacteria>Haloferacales>Halorubraceae | GCF_003287355.1 | Halorubrum_sp._48-1-W | DQW50_RS07315 | 54126..57881 | 2249761 | hypothetical-protein-[Halorubrum-sp.-48-1-W] |
| WP_112078643.1 | Type1.BREX | ABC-ATPase+SbcC->?->BPD_transp_2->?-><-?\|\|BrxA-DNA-binding->DUF6079->BrxB-BrxF-iSTAND->DUF6079*->N6-METHYLASE- [illegible] | 1277 | Archaea>Euryarchaeota>Stenosarchaea_group>Halobacteria>Haloferacales>Halorubraceae | GCF_003287355.1 | Halorubrum_sp._48-1-W | DQW50_RS07325 | 58459..62292 | 2249761 | hypothetical-protein-[Halorubrum-sp.-48-1-W] |
| WP_137716102.1 | Type1.BREX | Phage_integrase-><-Phage_integrase\|\|?->HD->?->?-><-?\|\|BrxA-DNA-binding->DUF6079*->BrxB-BrxF-iSTAND->DUF6079-> [illegible] | 1248 | Archaea>Euryarchaeota>Stenosarchaea_group>Halobacteria>Haloferacales>Halorubraceae | GCF_005435055.1 | Halorubrum_sp._SS7 | EXE44_RS03240 | 153853..157599 | 2518119 | hypothetical-protein-[Halorubrum-sp.-SS7] |
| WP_137715939.1 | Type1.BREX | <-Phage_integrase\|\|?->HD->?->?-><-?\|\|BrxA-DNA-binding->DUF6079->BrxB-BrxF-iSTAND->DUF6079*->?-><-MarR-HTH\|\|?-> [illegible] | 1277 | Archaea>Euryarchaeota>Stenosarchaea_group>Halobacteria>Haloferacales>Halorubraceae | GCF_005435055.1 | Halorubrum_sp._SS7 | EXE44_RS03250 | 158213..162046 | 2518119 | hypothetical-protein-[Halorubrum-sp.-SS7] |
| WP_137705565.1 | Type1.BREX | <-ParA-Soj-PloopNTPase\|\|?-><-PADR-HTH<-?<-?<-?<-?\|\|BrxA-DNA-binding->DUF6079*->BrxB-BrxF-iSTAND->DUF6079-> [illegible] | 1227 | Archaea>Euryarchaeota>Stenosarchaea_group>Halobacteria>Haloferacales>Halorubraceae | GCF_005435115.1 | Halorubrum_sp._ARQ200 | EXE50_RS16210 | 7684..11367 | 1855872 | hypothetical-protein-[Halorubrum-sp.-ARQ200] |
| WP_137705567.1 | Type1.BREX | <-PADR-HTH<-?<-?<-?<-?\|\|BrxA-DNA-binding->DUF6079->BrxB-BrxF-iSTAND->DUF6079*->REase-><-?<-?\|\|N6-METHYLASE-> [illegible] | 1280 | Archaea>Euryarchaeota>Stenosarchaea_group>Halobacteria>Haloferacales>Halorubraceae | GCF_005435115.1 | Halorubrum_sp._ARQ200 | EXE50_RS16220 | 11942..15784 | 1855872 | hypothetical-protein-[Halorubrum-sp.-ARQ200] |
| WP_210114021.1 | Type1.BREX | <-N6-METHYLASE-p\|\|BrxA-DNA-binding->DUF6079*->BrxB-BrxF-iSTAND->DUF6079->PglZ->DOC+HTH-><-SF1-SF2-Helicase+ [illegible] | 1233 | Archaea>Euryarchaeota>Stenosarchaea_group>Halobacteria>Haloferacales>Halorubraceae | GCF_017873555.1 | Halorubrum_trapanicum | J2744_RS15020 | 2668..6369 | 29284 | hypothetical-protein-[Halorubrum-trapanicum] |
| WP_210114022.1 | Type1.BREX | <-N6-METHYLASE-p\|\|BrxA-DNA-binding->DUF6079->BrxB-BrxF-iSTAND->DUF6079*->PglZ->DOC+HTH-><-SF1-SF2-Helicase+ [illegible] | 1230 | Archaea>Euryarchaeota>Stenosarchaea_group>Halobacteria>Haloferacales>Halorubraceae | GCF_017873555.1 | Halorubrum_trapanicum | J2744_RS15030 | 6985..10677 | 29284 | hypothetical-protein-[Halorubrum-trapanicum] |
| WP_271958997.1 | Type1.BREX | TOPRIM->?-><-SF1-SF2-Helicase+REase<-SF1-SF2-Helicase<-?<-?\|\|HTH->?->BrxA-DNA-binding->DUF6079*-> [illegible] | 1226 | Archaea>Euryarchaeota>Stenosarchaea_group>Halobacteria>Haloferacales>Halorubraceae | GCF_028211185.1 | Halorubrum_ezzemoulense | PM020_RS15300 | 39362..43042 | 337243 | hypothetical-protein-[Halorubrum-ezzemoulense] |
| WP_271952852.1 | Type1.BREX | <-DUF6079<-BrxB-BrxF-iSTAND<-DUF6079*<-BrxA-DNA-binding\|\|?-><-?\|\|GDResolvase-> | 1248 | Archaea>Euryarchaeota>Stenosarchaea_group>Halobacteria>Haloferacales>Halorubraceae | GCF_028211255.1 | Halorubrum_ezzemoulense | PM030_RS17190 | 6805..10551 | 337243 | hypothetical-protein-[Halorubrum-ezzemoulense] |
| WP_271959521.1 | Type1.BREX | <-N6-METHYLASE\|\|BrxA-DNA-binding->DUF6079*->BrxB-BrxF-iSTAND->DUF6079->PglZ-><-?<-?<-?<-?<-?\|\|ABC-ATPase+SbcC [illegible] | 1233 | Archaea>Euryarchaeota>Stenosarchaea_group>Halobacteria>Haloferacales>Halorubraceae | GCF_028211295.1 | Halorubrum_ezzemoulense | PM026_RS14195 | 3267..6968 | 337243 | hypothetical-protein-[Halorubrum-ezzemoulense] |
| WP_271959526.1 | Type1.BREX | <-N6-METHYLASE\|\|BrxA-DNA-binding->DUF6079->BrxB-BrxF-iSTAND->DUF6079*->PglZ-><-?<-?<-?<-?<-?\|\|ABC-ATPase+SbcC [illegible] | 1230 | Archaea>Euryarchaeota>Stenosarchaea_group>Halobacteria>Haloferacales>Halorubraceae | GCF_028211295.1 | Halorubrum_ezzemoulense | PM026_RS14205 | 7584..11276 | 337243 | hypothetical-protein-[Halorubrum-ezzemoulense] |
| WP_074878712.1 | Type1.BREX | <-TUDOR+SF1-SF2-Helicase+iREase+CTD<-REase+METHYLASE<-PglZ\|\|ZnR+REase->OLD-ABC+TOPRIM-><-N6-METHYLASE\|\| [illegible] | 1279 | Archaea>Euryarchaeota>Stenosarchaea_group>Halobacteria>Haloferacales>Halorubraceae | GCF_900115675.1 | Halolamina | BMZ83_RS11295 | 83133..86972 | 1075397 | MULTISPECIES:-hypothetical-protein-[Halolamina] |
| WP_074878715.1 | Type1.BREX | <-PglZ\|\|ZnR+REase->OLD-ABC+TOPRIM-><-N6-METHYLASE\|\|viral-protein-><-REase\|\|?-><-DUF6079<-BrxB-BrxF-iSTAND<- [illegible] | 1227 | Archaea>Euryarchaeota>Stenosarchaea_group>Halobacteria>Haloferacales>Halorubraceae | GCF_900115675.1 | Halolamina | BMZ83_RS11305 | 87547..91230 | 1075397 | MULTISPECIES:-hypothetical-protein-[Halolamina] |
| WP_013878506.1 | Type1.BREX | <-ParB+N6_N4_METHYLASE\|\|BrxA-DNA-binding->DUF6079->BrxB-BrxF-iSTAND->DUF6079*->REase->HTH+ICLR-HTH-> [illegible] | 1280 | Archaea>Euryarchaeota>Stenosarchaea_group>Halobacteria>Natrialbales>Natrialbaceae | GCF_000217715.1 | Halopiger_xanaduensis | HALXA_RS01375 | 280540..284382 | 387343 | hypothetical-protein-[Halopiger-xanaduensis] |
| WP_006184162.1 | Type1.BREX | <-BrxB-BrxF-iSTAND<-DUF6079*<-BrxA-DNA-binding\|\|TBP+TBP->REase->MORC+DUF3684+REase->ORC-CDC6+HTH-><-?<- [illegible] | 1243 | Archaea>Euryarchaeota>Stenosarchaea_group>Halobacteria>Natrialbales>Natrialbaceae | GCF_000337615.1 | Natrinema_pallidum | C487_RS02820 | 457..4188 | 69527 | hypothetical-protein-[Natrinema-pallidum] |
| WP_138651678.1 | Type1.BREX | <-PglZ<-N6-METHYLASE\|\|?->?-><-REase\|\|?-><-NTP_transf+wHTH<-HTH+ICLR-HTH<-REase<-DUF6079*<-BrxB-BrxF-iSTAND<- [illegible] | 1280 | Archaea>Euryarchaeota>Stenosarchaea_group>Halobacteria>Natrialbales>Natrialbaceae | GCF_005890195.1 | Natrinema_pallidum | FGF80_RS00910 | 194921..198763 | 69527 | hypothetical-protein-[Natrinema-pallidum] |
| WP_226042478.1 | Type1.BREX | RelE-><-TUDOR+SF1-SF2-Helicase+iREase+CTD<-REase+METHYLASE\|\|?-><-PglZ<-N6-METHYLASE\|\|?->?-><-REase<-DUF6079*<- [illegible] | 1280 | Archaea>Euryarchaeota>Stenosarchaea_group>Halobacteria>Natrialbales>Natrialbaceae | GCF_020405225.1 | Natrinema_sp._DC36 | LDH74_RS10650 | 2032265..2036107 | 2878680 | hypothetical-protein-[Natrinema-sp.-DC36] |
| WP_226042480.1 | Type1.BREX | <-REase+METHYLASE\|\|?-><-PglZ<-N6-METHYLASE\|\|?->?-><-REase<-DUF6079<-BrxB-BrxF-iSTAND<-DUF6079*<- [illegible] | 1227 | Archaea>Euryarchaeota>Stenosarchaea_group>Halobacteria>Natrialbales>Natrialbaceae | GCF_020405225.1 | Natrinema_sp._DC36 | LDH74_RS10660 | 2036682..2040365 | 2878680 | hypothetical-protein-[Natrinema-sp.-DC36] |
| WP_252697945.1 | Type1.BREX | <-REase<-HTH+REase<-?<-SbcC\|\|?->?->?->?->BrxA-DNA-binding->DUF6079*->BrxB-BrxF-iSTAND->DUF6079-><-?\|\| [illegible] | 1226 | Archaea>Euryarchaeota>Stenosarchaea_group>Halobacteria>Natrialbales>Natrialbaceae | GCF_023973145.1 | Natribaculum_sp._CGA3 | NGM68_RS09840 | 2053676..2057356 | 2953881 | hypothetical-protein-[Natribaculum-sp.-CGA3] |
| WP_252697948.1 | Type1.BREX | <-SbcC\|\|?->?->?->?->BrxA-DNA-binding->DUF6079->BrxB-BrxF-iSTAND->DUF6079*-><-?\|\|N6-METHYLASE->PglZ->?->?-><- [illegible] | 1284 | Archaea>Euryarchaeota>Stenosarchaea_group>Halobacteria>Natrialbales>Natrialbaceae | GCF_023973145.1 | Natribaculum_sp._CGA3 | NGM68_RS09850 | 2057949..2061803 | 2953881 | hypothetical-protein-[Natribaculum-sp.-CGA3] |
| WP_254767378.1 | Type1.BREX | <-TUDOR+SF1-SF2-Helicase+iREase+CTD<-REase<-PglZ<-N6-METHYLASE<-SLATT<-SMODS<-?\|\|MarR-HTH-><-DUF6079*<- [illegible] | 1278 | Archaea>Euryarchaeota>Stenosarchaea_group>Halobacteria>Natrialbales>Natrialbaceae | GCF_024296665.1 | Natribaculum_sp._AD-4 | NKG98_RS17310 | 3290404..3294240 | 2961596 | hypothetical-protein-[Natribaculum-sp.-AD-4] |
| WP_254767380.1 | Type1.BREX | <-REase<-PglZ<-N6-METHYLASE<-SLATT<-SMODS<-?\|\|MarR-HTH-><-DUF6079<-BrxB-BrxF-iSTAND<-DUF6079*<- [illegible] | 1265 | Archaea>Euryarchaeota>Stenosarchaea_group>Halobacteria>Natrialbales>Natrialbaceae | GCF_024296665.1 | Natribaculum_sp._AD-4 | NKG98_RS17320 | 3294854..3298651 | 2961596 | hypothetical-protein-[Natribaculum-sp.-AD-4] |
| WP_255191907.1 | Type1.BREX | <-REC+CheB-Methylesterase<-REC+OmpR-HTH<-?\|\|?-><-?\|\|HTH_23+HTH_32+HTH_33-><-McrC-NTD+REase<-?\|\| [illegible] | 1261 | Archaea>Euryarchaeota>Stenosarchaea_group>Halobacteria>Natrialbales>Natrialbaceae | GCF_024362485.1 | Halovivax_sp._WLHSJ27 | NMQ09_RS17725 | 3493167..3496952 | 2963433 | hypothetical-protein-[Halovivax-sp.-WLHSJ27] |
| WP_255191909.1 | Type1.BREX | HTH_23+HTH_32+HTH_33-><-McrC-NTD+REase<-?\|\|BrxA-DNA-binding->DUF6079->BrxB-BrxF-iSTAND->DUF6079*-> [illegible] | 1273 | Archaea>Euryarchaeota>Stenosarchaea_group>Halobacteria>Natrialbales>Natrialbaceae | GCF_024362485.1 | Halovivax_sp._WLHSJ27 | NMQ09_RS17735 | 3497545..3501366 | 2963433 | hypothetical-protein-[Halovivax-sp.-WLHSJ27] |
| WP_090620839.1 | Type1.BREX | GDResolvase->?->?->?->?->?-><-?<-ParB+N6_N4_METHYLASE\|\|BrxA-DNA-binding->DUF6079*->BrxB-BrxF-iSTAND->DUF6079-> [illegible] | 1227 | Archaea>Euryarchaeota>Stenosarchaea_group>Halobacteria>Natrialbales>Natrialbaceae | GCF_900110865.1 | Natrinema_salaciae | BMX07_RS18655 | 46177..49860 | 1186196 | hypothetical-protein-[Natrinema-salaciae] |
| WP_090620841.1 | Type1.BREX | <-ParB+N6_N4_METHYLASE\|\|BrxA-DNA-binding->DUF6079->BrxB-BrxF-iSTAND->DUF6079*-><-?\|\|REase-><-viral-protein\|\| [illegible] | 1280 | Archaea>Euryarchaeota>Stenosarchaea_group>Halobacteria>Natrialbales>Natrialbaceae | GCF_900110865.1 | Natrinema_salaciae | BMX07_RS18665 | 50435..54277 | 1186196 | hypothetical-protein-[Natrinema-salaciae] |
| KUK70180.1 | Type1.BREX | <-SIGMA+OB-Fold+AAA-ATPase+Lon-Protease-BrxL<-PglZ<-N6-METHYLASE<-DUF6079*<-BrxB-BrxF-iSTAND<-BrxA-DNA-binding [illegible] | 1157 | Archaea>Euryarchaeota>Stenosarchaea_group>Methanomicrobia>Methanomicrobiales>Methanocalculaceae | GCA_001508455.1 | Methanocalculus_sp._52_23 | XD88_0819 | 8030..11503 | 1641382 | MAG:-Uncharacterized-protein-XD88_0819-[Methanocalculus-sp.-52_23] |
| MBR1368475.1 | Type1.BREX | <-HTH_Tnp_1\|\|OLD-ABC+TOPRIM->SF1-SF2-Helicase->BrxA-DNA-binding->BrxB-BrxF-iSTAND->DUF6079*->N6-METHYLASE->?-> [illegible] | 1155 | Archaea>Euryarchaeota>Stenosarchaea_group>Methanomicrobia>Methanomicrobiales>Methanocalculaceae | GCA_018132105.1 | Methanocalculus_chunghsingensis | RJ53_02730 | 5881..9348 | 156457 | hypothetical-protein-[Methanocalculus-chunghsingensis] |
| WP_246495968.1 | Type1.BREX | OLD-ABC+TOPRIM->SF1-SF2-Helicase->BrxA-DNA-binding->BrxB-BrxF-iSTAND->DUF6079*->N6-METHYLASE->?-> [illegible] | 1148 | Archaea>Euryarchaeota>Stenosarchaea_group>Methanomicrobia>Methanomicrobiales>Methanocalculaceae | GCF_018132105.1 | Methanocalculus_chunghsingensis | RJ53_RS02795 | 5902..9348 | 156457 | BREX-system-P-loop-protein-BrxC-[Methanocalculus-chunghsingensis] |
| WP_253460960.1 | Type1.BREX | <-TPR<-TBP+TBP<-ASCH\|\|BrxB-BrxF-iSTAND->DUF6079*->N6-METHYLASE->Schlafen+HATPase+wHTH-p->N6-METHYLASE-p-> [illegible] | 1180 | Archaea>Euryarchaeota>Stenosarchaea_group>Methanomicrobia>Methanomicrobiales>Methanocalculaceae | GCF_024170265.1 | Methanocalculus_sp._AMF5 | J2T61_RS09750 | 4070..7612 | 1198257 | BREX-system-P-loop-protein-BrxC-[Methanocalculus-sp.-AMF5] |
| WP_253488285.1 | Type1.BREX | <-SIGMA+OB-Fold+AAA-ATPase+Lon-Protease-BrxL<-PglZ<-TOPRIM<-OLD-ABC<-N6-METHYLASE<-DUF6079*<-BrxB-BrxF-iSTAND [illegible] | 1155 | Archaea>Euryarchaeota>Stenosarchaea_group>Methanomicrobia>Methanomicrobiales>Methanocalculaceae | GCF_024170505.1 | Methanocalculus_alkaliphilus | J2T58_RS06410 | 9528..12995 | 768730 | BREX-system-P-loop-protein-BrxC-[Methanocalculus-alkaliphilus] |
| WP_245241072.1 | Type1.BREX | HTH+REase->?->?-><-SIGMA+OB-Fold+AAA-ATPase+Lon-Protease-BrxL<-PglZ<-N6-METHYLASE-p<-N6-METHYLASE-p<- [illegible] | 1180 | Archaea>Euryarchaeota>Stenosarchaea_group>Methanomicrobia>Methanomicrobiales>Methanocorpusculaceae | GCF_012062385.1 | Methanocorpusculum_sp._GPch4 | EYA77_RS00170 | 16814..20356 | 2527877 | BREX-system-P-loop-protein-BrxC-[Methanocorpusculum-sp.-GPch4] |
| HIH85712.1 | Type1.BREX | <-SIGMA+OB-Fold+AAA-ATPase+Lon-Protease-BrxL<-PglZ<-N6-METHYLASE<-DUF6079*<-BrxB-BrxF-iSTAND<-BrxA-DNA-binding [illegible] | 1155 | Archaea>Euryarchaeota>Stenosarchaea_group>Methanomicrobia>Methanomicrobiales>Methanomicrobiaceae | GCA_013329555.1 | Methanoculleus_sp. | HA342_02595 | 8043..11510 | 90427 | BREX-system-P-loop-protein-BrxC-[Methanoculleus-sp.] |
| QYZ80392.1 | Type1.BREX | ArdCN+MPTase->?-><-?<-Glyco_hydro\|\|?->ORC-CDC6+HTH->?->BrxA-DNA-binding->BrxB-BrxF-iSTAND->DUF6079*-> [illegible] | 1159 | Archaea>Euryarchaeota>Stenosarchaea_group>Methanomicrobia>Methanomicrobiales>Methanomicrobiaceae | GCA_019633745.1 | Methanofollis_formosanus | E2N92_03980 | 901654..905133 | 299308 | BREX-system-P-loop-protein-BrxC-[Methanofollis-formosanus] |
| MCK4269451.1 | Type1.BREX | BrxA-DNA-binding->BrxB-BrxF-iSTAND->DUF6079*->N6-METHYLASE->N6-METHYLASE-p-> | 1207 | Archaea>Euryarchaeota>Stenosarchaea_group>Methanomicrobia>Methanomicrobiales>Methanomicrobiaceae | GCA_023128355.1 | Methanogenium_sp. | KAW93_03130 | 3124..6747 | 2196 | BREX-system-P-loop-protein-BrxC-[Methanogenium-sp.] |
| WP_004077401.1 | Type1.BREX | wHTH+WYL-><-SIGMA+OB-Fold+AAA-ATPase+Lon-Protease-BrxL<-PglZ<-N6-METHYLASE-p<-HTH+REase<-?<-?<-N6-METHYLASE<- [illegible] | 1180 | Archaea>Euryarchaeota>Stenosarchaea_group>Methanomicrobia>Methanomicrobiales>Methanomicrobiaceae | GCF_000243255.1 | Methanoplanus_limicola | METLIM_RS07750 | 1605385..1608927 | 2315 | BREX-system-P-loop-protein-BrxC-[Methanoplanus-limicola] |
| WP_004078487.1 | Type1.BREX | <-CR-ATPase5+REase\|\|MORC+REase-p-><-SIGMA+OB-Fold+AAA-ATPase+Lon-Protease-BrxL<-PglZ<-?<-HTH+REase<- [illegible] | 1155 | Archaea>Euryarchaeota>Stenosarchaea_group>Methanomicrobia>Methanomicrobiales>Methanomicrobiaceae | GCF_000243255.1 | Methanoplanus_limicola | METLIM_RS10810 | 2304713..2308180 | 2315 | BREX-system-P-loop-protein-BrxC-[Methanoplanus-limicola] |
| WP_004038977.1 | Type1.BREX | <-ASCH<-REase+DUF5655<-EVE<-SIGMA+OB-Fold+AAA-ATPase+Lon-Protease-BrxL<-PglZ<-HD<-SbcC<-?<-N6-METHYLASE<- [illegible] | 1222 | Archaea>Euryarchaeota>Stenosarchaea_group>Methanomicrobia>Methanomicrobiales>Methanomicrobiaceae | GCF_000275865.1 | Methanofollis_liminatans | METLI_RS06530 | 1309221..1312889 | 2201 | BREX-system-P-loop-protein-BrxC-[Methanofollis-liminatans] |
| WP_014866844.1 | Type1.BREX | <-HEPN<-EVE<-REase+DUF5655<-ASCH<-EVE<-SIGMA+OB-Fold+AAA-ATPase+Lon-Protease-BrxL<-PglZ<-N6-METHYLASE<- [illegible] | 1155 | Archaea>Euryarchaeota>Stenosarchaea_group>Methanomicrobia>Methanomicrobiales>Methanomicrobiaceae | GCF_000304355.2 | Methanoculleus_bourgensis | BN140_RS04760 | 992975..996442 | 83986 | BREX-system-P-loop-protein-BrxC-[Methanoculleus-bourgensis] |
| WP_048152685.1 | Type1.BREX | Schlafen+HATPase+wHTH-><-?\|\|?-><-REase+DUF5655<-EVE\|\|BrxB-BrxF-iSTAND->DUF6079*->N6-METHYLASE-> [illegible] | 1179 | Archaea>Euryarchaeota>Stenosarchaea_group>Methanomicrobia>Methanomicrobiales>Methanomicrobiaceae | GCF_000784355.1 | Methanolacinia_paynteri | METPAY_RS11475 | 99003..102542 | 230356 | BREX-system-P-loop-protein-BrxC-[Methanolacinia-paynteri] |
| WP_250317376.1 | Type1.BREX | <-PolB\|\|?-><-SbcC<-Calcineurin\|\|?->REase->BrxA-DNA-binding->BrxB-BrxF-iSTAND->DUF6079-p->DUF6079*-> [illegible] | 1034 | Archaea>Euryarchaeota>Stenosarchaea_group>Methanomicrobia>Methanomicrobiales>Methanomicrobiaceae | GCF_018495055.1 | Methanoculleus_bourgensis | KH990_RS12090 | 14004..17108 | 83986 | BREX-system-P-loop-protein-BrxC-[Methanoculleus-bourgensis] |

| Protein-ID | System | Gene-Neighbourhoods | Length | Taxonomy | Assembly-ID | Strain | Locus-Tag | Genomic-coordinates | TaxID | Fasta-define |
|---|---|---|---|---|---|---|---|---|---|---|
| WP_250317375.1 | Type1.BREX | <-PolB<-PolB\|\|?-><-SbcC<-Calcineurin\|\|?->REase->BrxA-DNA-binding->BrxB-BrxF-iSTAND->DUF6079-p*->DUF6079->N6-METHYLASE->PglZ->SIGMA+OB-Fold+AAA-ATPase+Lon-Protease-BrxL->EVE->ZDNA-HTH-[illegible] | 149 | Archaea>Euryarchaeota>Stenosarchaea_group>Methanomicrobia>Methanomicrobiales>Methanomicrobiaceae | GCF_018495055.1 | Methanoculleus_bourgensis | KH990_RS13175 | 13640..14089 | 83986 | hypothetical-protein-[Methanoculleus-bourgensis] |
| WP_246589305.1 | Type1.BREX | ArdCN+MPTase->?-><-?<-Glyco_hydro\|\|?->ORC-CDC6+HTH->?->BrxA-DNA-binding->BrxB-BrxF-iSTAND->DUF6079*->N6-METHYLASE->?->PglZ->SIGMA+OB-Fold+AAA-ATPase+Lon-Protease-BrxL-> | 1160 | Archaea>Euryarchaeota>Stenosarchaea_group>Methanomicrobia>Methanomicrobiales>Methanomicrobiaceae | GCF_019633745.1 | Methanofollis_formosanus | E2N92_RS04005 | 901651..905133 | 299308 | BREX-system-P-loop-protein-BrxC-[Methanofollis-formosanus] |
| WP_257742634.1 | Type1.BREX | <-REase\|\|PIN-><-?\|\|Peptidase_M42->BrxA-DNA-binding->BrxB-BrxF-iSTAND->DUF6079*->N6-METHYLASE->HTH+REase->DDE_Tnp_1->?->PglZ->SIGMA+OB-Fold+AAA-ATPase+Lon-Protease-BrxL->?-><-DDE_Tnp | 1155 | Archaea>Euryarchaeota>Stenosarchaea_group>Methanomicrobia>Methanomicrobiales>Methanomicrobiaceae | GCF_024662215.1 | Methanoplanus_endosymbiosus | L6E24_RS14385 | 3075748..3079215 | 33865 | BREX-system-P-loop-protein-BrxC-[Methanoplanus-endosymbiosus] |
| WP_074369579.1 | Type1.BREX | 35exo->?->HTH+REase-><-TBP+TBP<-ASCH<-EVE\|\|BrxA-DNA-binding->BrxB-BrxF-iSTAND->DUF6079*->N6-METHYLASE->?->PglZ->SIGMA+OB-Fold+AAA-ATPase+Lon-Protease-BrxL->REase-><-DOC+HTH | 1160 | Archaea>Euryarchaeota>Stenosarchaea_group>Methanomicrobia>Methanomicrobiales>Methanomicrobiaceae | GCF_900095385.1 | Methanoculleus_chikugoensis | L21_RS05950 | 37665..41147 | 118126 | BREX-system-P-loop-protein-BrxC-[Methanoculleus-chikugoensis] |
| ABD40832.1 | Type1.BREX | <-N6-METHYLASE<-?<-SIGMA+OB-Fold+AAA-ATPase+Lon-Protease-BrxL<-PglZ<-TOPRIM<-OLD-ABC<-N6-METHYLASE<-RHH<-DUF6079*<-BrxB-BrxF-iSTAND<-BrxA-DNA-binding<-?<-?<-Glyco_hydro | 1166 | Archaea>Euryarchaeota>Stenosarchaea_group>Methanomicrobia>Methanomicrobiales>Methanospirillaceae | GCA_000013445.1 | Methanospirillum_hungatei_JF-1 | Mhun_1083 | 1223800..1227300 | 323259 | hypothetical-protein-Mhun_1083-[Methanospirillum-hungatei-JF-1] |
| MBN1165665.1 | Type1.BREX | TPR+TPR+TPR->BrxA-DNA-binding->BrxB-BrxF-iSTAND->DUF6079*->N6-METHYLASE-p->N6-METHYLASE->?->?->ParA-Soj-PloopNTPase->?->HTH+REase->PglZ->SIGMA+OB-Fold+AAA-ATPase+Lon-Protease-BrxL-> | 1218 | Archaea>Euryarchaeota>Stenosarchaea_group>Methanomicrobia>Methanomicrobiales>Methanospirillaceae | GCA_016926505.1 | Methanospirillaceae_archaeon | JXA44_00885 | 48354..52010 | 2813030 | BREX-system-P-loop-protein-BrxC-[Methanospirillaceae-archaeon] |
| MCA1916608.1 | Type1.BREX | <-N6-METHYLASE-p<-Schlafen+HATPase+wHTH<-N6-METHYLASE<-DUF6079*<-BrxB-BrxF-iSTAND\|\|OLD-ABC->TPR-> | 1178 | Archaea>Euryarchaeota>Stenosarchaea_group>Methanomicrobia>Methanomicrobiales>Methanospirillaceae | GCA_020216685.1 | Methanospirillum_hungatei | LDL35_09440 | 4587..8123 | 2203 | BREX-system-P-loop-protein-BrxC-[Methanospirillum-hungatei] |
| WP_052288823.1 | Type1.BREX | <-SIGMA+OB-Fold+AAA-ATPase+Lon-Protease-BrxL<-PglZ<-TOPRIM<-OLD-ABC<-N6-METHYLASE<-?<-RHH<-DUF6079*<-BrxB-BrxF-iSTAND<-BrxA-DNA-binding<-?<-?<-?<-Glyco_hydro | 1159 | Archaea>Euryarchaeota>Stenosarchaea_group>Methanomicrobia>Methanomicrobiales>Methanospirillaceae | GCF_000013445.1 | Methanospirillum_hungatei | MHUN_RS05665 | 1223800..1227279 | 2203 | BREX-system-P-loop-protein-BrxC-[Methanospirillum-hungatei] |
| WP_011448873.1 | Type1.BREX | CR-ATPase8+REase-><-TPR+TPR<-TPR<-TOPRIM<-OLD-ABC<-?\|\|BrxB-BrxF-iSTAND->DUF6079*->N6-METHYLASE->?->?->?->HATPase+wHTH->HTH+REase->N6-METHYLASE-p-><-DDE_Tnp\|\|N6-METHYLASE-p-> | 1178 | Archaea>Euryarchaeota>Stenosarchaea_group>Methanomicrobia>Methanomicrobiales>Methanospirillaceae | GCF_000013445.1 | Methanospirillum_hungatei | MHUN_RS09850 | 2137740..2141276 | 2203 | BREX-system-P-loop-protein-BrxC-[Methanospirillum-hungatei] |
| WP_214418671.1 | Type1.BREX | <-SIGMA+OB-Fold+AAA-ATPase+Lon-Protease-BrxL<-PglZ<-N6-METHYLASE-p<-HTH+REase<-Schlafen+HATPase+wHTH<-N6-METHYLASE<-DUF6079*<-BrxB-BrxF-iSTAND<-zf-IS66<-N6-METHYLASE<-?<-RNase<-?<-REase\|\|EVE-> | 1171 | Archaea>Euryarchaeota>Stenosarchaea_group>Methanomicrobia>Methanomicrobiales>Methanospirillaceae | GCF_018502485.1 | Methanospirillum_sp._J.3.6.1-F.2.7.3 | KHC33_RS10925 | 2269426..2272941 | 2834276 | BREX-system-P-loop-protein-BrxC-[Methanospirillum-sp.-J.3.6.1-F.2.7.3] |
| WP_218608642.1 | Type1.BREX | <-TBP+TBP<-?<-HD<-Peptidase_S8<-Classical-AAA<-N6-METHYLASE\|\|?->?-><-?<-DUF6079*<-BrxB-BrxF-iSTAND<-BrxA-DNA-binding<-?\|\|?-><-HicB_lk_antitox<-HicA_toxin\|\|?->?->Schlafen+HATPase+wHTH-> | 1214 | Archaea>Euryarchaeota>Stenosarchaea_group>Methanomicrobia>Methanomicrobiales>Methanospirillaceae | GCF_019263745.1 | Methanospirillum_hungatei | KSK55_RS06665 | 1421419..1425063 | 2203 | BREX-system-P-loop-protein-BrxC-[Methanospirillum-hungatei] |
| MCE8425368.1 | Type1.BREX | <-PglZ<-Schlafen+HATPase+wHTH<-N6-METHYLASE<-DUF6079*<-BrxB-BrxF-iSTAND<-BrxA-DNA-binding\|\|SF1-SF2-Helicase+REase-><-ADH_N+ADH_zinc_N_2 | 1196 | Archaea>Euryarchaeota>Stenosarchaea_group>Methanomicrobia>Methanosarcinales>Candidatus_Methanoperedenaceae | GCA_021409295.1 | Candidatus_Methanoperedens_sp. | J5U17_06270 | 6093..9683 | 2032688 | BREX-system-P-loop-protein-BrxC-[Candidatus-Methanoperedens-sp.] |
| AAM05760.1 | Type1.BREX | <-Calcineurin<-SIGMA+OB-Fold+AAA-ATPase+Lon-Protease-BrxL<-PglZ<-MNS+REase\|\|PHD_like+HicB_lk_antitox-><-Schlafen+HATPase+wHTH<-Schlafen+HATPase+wHTH<-HDRnase<-N6-METHYLASE<-DUF6079*<-BrxB-BrxF-iSTAND\|\|?->PolB->?-><-?\|\|?-><-GT4+DSBH<-Glyco_hydro_130 | 1201 | Archaea>Euryarchaeota>Stenosarchaea_group>Methanomicrobia>Methanosarcinales>Methanosarcinaceae | GCA_000007345.1 | Methanosarcina_acetivorans_C2A | MA_2373 | 2935402..2939007 | 188937 | predicted-protein-[Methanosarcina-acetivorans-C2A] |
| AFV22360.1 | Type1.BREX | MNS->?->McrB-AAA->McrC-NTD+REase-><-DNA-glycosylase\|\|?->CarD+HTH->CarD+HTH->BrxB-BrxF-iSTAND->DUF6079*->N6-METHYLASE->Schlafen+HATPase+wHTH->N6-METHYLASE-p->ASCH->HD->PglZ->SIGMA+OB-Fold+AAA-ATPase+Lon-Protease-BrxL-><-REase<-SF1-SF2-Helicase | 1185 | Archaea>Euryarchaeota>Stenosarchaea_group>Methanomicrobia>Methanosarcinales>Methanosarcinaceae | GCA_000306725.1 | Methanolobus_psychrophilus_R15 | Mpsy_0147 | 147504..151061 | 1094980 | hypothetical-protein-Mpsy_0147-[Methanolobus-psychrophilus-R15] |
| AKB55571.1 | Type1.BREX | <-SIGMA+OB-Fold+AAA-ATPase+Lon-Protease-BrxL<-PglZ<-Schlafen-p<-TIR<-N6-METHYLASE<-DUF6079-p*<-DUF6079-p<-BrxB-BrxF-iSTAND\|\|?-><-tRNA<-tRNA<-?<-?<-?<-?<-?<-ABC-ATPase+SbcC | 797 | Archaea>Euryarchaeota>Stenosarchaea_group>Methanomicrobia>Methanosarcinales>Methanosarcinaceae | GCA_000970025.1 | Methanosarcina_barkeri_MS | MSBRM_2573 | 3399926..3402319 | 1434108 | hypothetical-protein-MSBRM_2573-[Methanosarcina-barkeri-MS] |
| AKB55572.1 | Type1.BREX | <-SIGMA+OB-Fold+AAA-ATPase+Lon-Protease-BrxL<-PglZ<-Schlafen-p<-TIR<-N6-METHYLASE<-DUF6079-p<-DUF6079-p*<-BrxB-BrxF-iSTAND\|\|?-><-tRNA<-tRNA<-?<-?<-?<-?<-?<-ABC-ATPase+SbcC | 353 | Archaea>Euryarchaeota>Stenosarchaea_group>Methanomicrobia>Methanosarcinales>Methanosarcinaceae | GCA_000970025.1 | Methanosarcina_barkeri_MS | MSBRM_2574 | 3402467..3403528 | 1434108 | ATPase-like-protein-[Methanosarcina-barkeri-MS] |
| KXS44097.1 | Type1.BREX | <-SIGMA+OB-Fold+AAA-ATPase+Lon-Protease-BrxL<-PglZ<-TOPRIM<-OLD-ABC<-?<-N6-METHYLASE-p<-Schlafen+HATPase+wHTH<-N6-METHYLASE<-DUF6079*<-BrxB-BrxF-iSTAND<-McrC-NTD+REase<-EVE+McrB-AAA<-?\|\|McrC-NTD+REase->MNS+REase->RADICAL-SAM-> | 1185 | Archaea>Euryarchaeota>Stenosarchaea_group>Methanomicrobia>Methanosarcinales>Methanosarcinaceae | GCA_001577565.1 | Methanolobus_sp._T82-4 | AWU59_730 | 16022..19579 | 1794908 | MAG:-hypothetical-protein-AWU59_730-[Methanolobus-sp.-T82-4] |
| OBZ35401.1 | Type1.BREX | BrxB-BrxF-iSTAND->DUF6079*->Schlafen+HATPase+wHTH->N6-METHYLASE->?->PglZ->SIGMA+OB-Fold+AAA-ATPase+Lon-Protease-BrxL->RNaseH-><-?<-REase<-McrB-AAA | 1187 | Archaea>Euryarchaeota>Stenosarchaea_group>Methanomicrobia>Methanosarcinales>Methanosarcinaceae | GCA_001683555.1 | Methanohalophilus_sp._DAL1 | A9957_07285 | 3150..6713 | 1864608 | MAG:-hypothetical-protein-A9957_07285-[Methanohalophilus-sp.-DAL1] |
| NPE28183.1 | Type1.BREX | <-SIGMA+OB-Fold+AAA-ATPase+Lon-Protease-BrxL<-PglZ<-TOPRIM<-OLD-ABC<-TIR<-N6-METHYLASE-p<-Schlafen+HATPase+wHTH<-N6-METHYLASE<-DUF6079*<-BrxB-BrxF-iSTAND<-URI-DNase<-?<-SF1-SF2-Helicase+REase\|\|<-?\|\|?->HNH-> | 1185 | Archaea>Euryarchaeota>Stenosarchaea_group>Methanomicrobia>Methanosarcinales>Methanosarcinaceae | GCA_013166765.1 | Methanococcoides_sp._SA1 | HNV12_09470 | 12693..16250 | 2735871 | BREX-system-P-loop-protein-BrxC-[Methanococcoides-sp.-SA1] |
| HII01702.1 | Type1.BREX | <-SF1-SF2-Helicase<-?<-OLD-ABC+TOPRIM<-?<-N6-METHYLASE<-DUF6079*<-BrxB-BrxF-iSTAND\|\|?->SAM-METHYLASE->?->MNS-><-?\|\|HAD+HAD-><-TPR+TPR+TPR+TPR<-REC+OmpR-HTH | 1196 | Archaea>Euryarchaeota>Stenosarchaea_group>Methanomicrobia>Methanosarcinales>Methanosarcinaceae | GCA_013329415.1 | Methanosarcinaceae_archaeon | HA351_08670 | 9336..12926 | 1945595 | BREX-system-P-loop-protein-BrxC-[Methanosarcinaceae-archaeon] |
| HIH93569.1 | Type1.BREX | <-PglZ<-?<-MNS+REase\|\|PHD_like+HicB_lk_antitox->HicA_toxin-><-Schlafen+HATPase+wHTH<-Schlafen-p<-HDRnase<-?<-DUF6079*<-BrxB-BrxF-iSTAND | 1124 | Archaea>Euryarchaeota>Stenosarchaea_group>Methanomicrobia>Methanosarcinales>Methanosarcinaceae | GCA_013329655.1 | Methanosarcina_acetivorans | HA338_05875 | 28858..32230 | 2214 | BREX-system-P-loop-protein-BrxC-[Methanosarcina-acetivorans] |
| MBN2489135.1 | Type1.BREX | <-ArdCN+MPTase\|\|?->URI-DNase->DNA-glycosylase-><-SF1-SF2-Helicase+REase\|\|BrxB-BrxF-iSTAND->DUF6079-p*-> | 551 | Archaea>Euryarchaeota>Stenosarchaea_group>Methanomicrobia>Methanosarcinales>Methanosarcinaceae | GCA_016935475.1 | Methanosarcinaceae_archaeon | JXA98_08910 | 15184..16838 | 1945595 | BREX-system-P-loop-protein-BrxC-[Methanosarcinaceae-archaeon] |
| MCD4820926.1 | Type1.BREX | <-ASCH\|\|Phage_int_SAM_4+Phage_integrase->?-><-SIGMA+OB-Fold+AAA-ATPase+Lon-Protease-BrxL<-PglZ<-?<-N6-METHYLASE-p<-Schlafen+HATPase+wHTH<-N6-METHYLASE<-DUF6079*<-BrxB-BrxF-iSTAND<-URI-DNase<-SF1-SF2-Helicase+REase\|\|HEPN+PAS+DUF4011+SF1-SF2-Helicase+REase->?->EVE+McrB-AAA->McrC-NTD+REase-> | 1185 | Archaea>Euryarchaeota>Stenosarchaea_group>Methanomicrobia>Methanosarcinales>Methanosarcinaceae | GCA_021108275.1 | Methanococcoides_sp. | K8R11_02360 | 106094..109651 | 1966350 | BREX-system-P-loop-protein-BrxC-[Methanococcoides-sp.] |
| WP_011032108.1 | Type1.BREX | <-D5N+viral-AAA+CDC6-HTH<-?<-?<-Phage_int_SAM_4+Phage_integrase\|\|BrxB-BrxF-iSTAND->DUF6079*-><-TPR+TPR+TPR+TPR+TPR+TPR+TPR\|\|N6-METHYLASE->Schlafen-p->Schlafen+HATPase+wHTH-p->HATPase+wHTH->PadR-HTH-><-?\|\|Mrr_N+ACET-><-HicA_toxin | 1212 | Archaea>Euryarchaeota>Stenosarchaea_group>Methanomicrobia>Methanosarcinales>Methanosarcinaceae | GCF_000007065.1 | Methanosarcina_mazei | MM_RS00835 | 194588..198226 | 2209 | BREX-system-P-loop-protein-BrxC-[Methanosarcina-mazei] |
| WP_048065383.1 | Type1.BREX | PHD_like+HicB_lk_antitox->HicA_toxin-><-?<-HATPase_c<-Schlafen+HATPase+wHTH<-Schlafen-p<-Schlafen+HATPase+wHTH<-HDRnase<-N6-METHYLASE<-DUF6079*<-BrxB-BrxF-iSTAND<-tRNA\|\|?->PolB-><-?\|\|HEPN->?-><-GT4+DSBH<-Glyco_hydro_130 | 1198 | Archaea>Euryarchaeota>Stenosarchaea_group>Methanomicrobia>Methanosarcinales>Methanosarcinaceae | GCF_000007345.1 | Methanosarcina_acetivorans | MA_RS12305 | 2935402..2938998 | 2214 | BREX-system-P-loop-protein-BrxC-[Methanosarcina-acetivorans] |
| WP_013898237.1 | Type1.BREX | <-SIGMA+OB-Fold+AAA-ATPase+Lon-Protease-BrxL<-PglZ<-HD<-N6-METHYLASE<-Schlafen+HATPase+wHTH<-DUF6079*<-BrxB-BrxF-iSTAND-p<-BrxB-BrxF-iSTAND<-HKD-DNase+MED12\|\|?->?->?->?->HAD+HAD-> | 1188 | Archaea>Euryarchaeota>Stenosarchaea_group>Methanomicrobia>Methanosarcinales>Methanosarcinaceae | GCF_000217995.1 | Methanosalsum_zhilinae | MZHIL_RS04755 | 1026278..1029844 | 39669 | BREX-system-P-loop-protein-BrxC-[Methanosalsum-zhilinae] |
| WP_015324772.1 | Type1.BREX | <-TOPRIM<-OLD-ABC<-?<-?\|\|?-><-?<-N6-METHYLASE-p<-?<-?<-N6-METHYLASE<-DUF6079*<-BrxB-BrxF-iSTAND\|\|?->?->TPR+TPR-><-?\|\|DSBH->?-><-?\|\|35exo-> | 1187 | Archaea>Euryarchaeota>Stenosarchaea_group>Methanomicrobia>Methanosarcinales>Methanosarcinaceae | GCF_000328665.1 | Methanomethylovorans_hollandica | METHO_RS06665 | 1392190..1395753 | 101192 | BREX-system-P-loop-protein-BrxC-[Methanomethylovorans-hollandica] |
| WP_015410898.1 | Type1.BREX | PRTase-><-?\|\|?->?->tRNA->BrxB-BrxF-iSTAND->DUF6079*-><-TPR+TPR+TPR+TPR+TPR+TPR+TPR\|\|N6-METHYLASE->N6-METHYLASE-p->Schlafen-p->Schlafen+HATPase+wHTH-p->HATPase+wHTH->PadR-HTH-><-?\|\|Mrr_N+ACET-> | 1212 | Archaea>Euryarchaeota>Stenosarchaea_group>Methanomicrobia>Methanosarcinales>Methanosarcinaceae | GCF_000341715.1 | Methanosarcina_mazei | MMTUC01_RS00750 | 161332..164970 | 2209 | BREX-system-P-loop-protein-BrxC-[Methanosarcina-mazei] |
| WP_048118458.1 | Type1.BREX | BrxA-DNA-binding->BrxB-BrxF-iSTAND->DUF6079*->N6-METHYLASE->PglZ->SIGMA+OB-Fold+AAA-ATPase+Lon-Protease-BrxL-><-DUF488<-Glycos_transf_1+Glyco_trans_1_2<-?<-DSBH | 1143 | Archaea>Euryarchaeota>Stenosarchaea_group>Methanomicrobia>Methanosarcinales>Methanosarcinaceae | GCF_000969905.1 | Methanosarcina_vacuolata | MSVAZ_RS04205 | 977468..980899 | 2215 | BREX-system-P-loop-protein-BrxC-[Methanosarcina-vacuolata] |
| WP_048158954.1 | Type1.BREX | DDE_Tnp_1->DDE_Tnp_1-><-Schlafen+HATPase+wHTH<-Schlafen-p<-Schlafen+HATPase+wHTH<-?<-?<-TIR<-N6-METHYLASE<-DUF6079*<-BrxB-BrxF-iSTAND\|\|?->RRXRR+HNH->PglZ->SIGMA+OB-Fold+AAA-ATPase+Lon-Protease-BrxL->Calcineurin->SbcC-><-SF1-SF2-Helicase+REase | 1223 | Archaea>Euryarchaeota>Stenosarchaea_group>Methanomicrobia>Methanosarcinales>Methanosarcinaceae | GCF_000969965.1 | Methanosarcina_sp._WWM596 | MSWHS_RS09215 | 2091991..2095662 | 1434103 | BREX-system-P-loop-protein-BrxC-[Methanosarcina-sp.-WWM596] |
| WP_048171288.1 | Type1.BREX | PHD_like+HicB_lk_antitox->HicA_toxin-><-HATPase+wHTH<-Schlafen+HATPase+wHTH<-Schlafen-p<-Schlafen+HATPase+wHTH<-SMODS<-SLATT<-N6-METHYLASE<-DUF6079*<-BrxB-BrxF-iSTAND<-tRNA<-?\|\|?->?->?->DnaB-><-REC+OmpR-HTH<-DnaB\|\|tRNA->ACET-> | 1220 | Archaea>Euryarchaeota>Stenosarchaea_group>Methanomicrobia>Methanosarcinales>Methanosarcinaceae | GCF_000970085.1 | Methanosarcina_siciliae | MSSIT_RS07430 | 1736961..1740623 | 38027 | BREX-system-P-loop-protein-BrxC-[Methanosarcina-siciliae] |
| WP_148705147.1 | Type1.BREX | <-HATPase+wHTH<-Schlafen+HATPase+wHTH<-Schlafen-p<-Schlafen+HATPase+wHTH<-?<-?<-TIR<-N6-METHYLASE\|\|TPR+TPR+TPR+TPR+TPR+TPR+TPR-><-DUF6079*<-BrxB-BrxF-iSTAND<-tRNA<-?\|\|?-><-?\|\|?->?->DnaB-><-REC+OmpR-HTH<-DnaB | 1222 | Archaea>Euryarchaeota>Stenosarchaea_group>Methanomicrobia>Methanosarcinales>Methanosarcinaceae | GCF_000970125.1 | Methanosarcina_siciliae | MSSIH_RS07150 | 1687969..1691637 | 38027 | BREX-system-P-loop-protein-BrxC-[Methanosarcina-siciliae] |
| WP_048181889.1 | Type1.BREX | SF1-SF2-Helicase-><-SbcC<-Calcineurin<-SIGMA+OB-Fold+AAA-ATPase+Lon-Protease-BrxL<-PglZ<-SF1-SF2-Helicase<-SbcC<-N6-METHYLASE<-DUF6079*<-BrxB-BrxF-iSTAND<-tRNA<-?\|\|?-><-?\|\|RVT->RVT->ABC-ATPase+SbcC->Peptidase-> | 1212 | Archaea>Euryarchaeota>Stenosarchaea_group>Methanomicrobia>Methanosarcinales>Methanosarcinaceae | GCF_000970145.1 | Methanosarcina_siciliae | MSSAC_RS09150 | 2102967..2106605 | 38027 | BREX-system-P-loop-protein-BrxC-[Methanosarcina-siciliae] |
| WP_048052927.1 | Type1.BREX | <-Mrr_N+ACET\|\|?-><-PadR-HTH<-?<-HATPase+wHTH<-Schlafen+HATPase+wHTH-p<-Schlafen-p<-N6-METHYLASE\|\|TPR+TPR+TPR+TPR+TPR+TPR+TPR-><-DUF6079*<-BrxB-BrxF-iSTAND<-tRNA<-?<-?\|\|?-><-PRTase | 1212 | Archaea>Euryarchaeota>Stenosarchaea_group>Methanomicrobia>Methanosarcinales>Methanosarcinaceae | GCF_000970165.1 | Methanosarcina_mazei | MSMAW_RS06205 | 1427636..1431274 | 2209 | BREX-system-P-loop-protein-BrxC-[Methanosarcina-mazei] |
| WP_048046369.1 | Type1.BREX | HicA_toxin-><-Mrr_N+ACET\|\|?-><-PadR-HTH<-HATPase+wHTH<-Schlafen+HATPase+wHTH-p<-Schlafen-p<-N6-METHYLASE\|\|TPR+TPR+TPR+TPR+TPR+TPR+TPR-><-DUF6079*<-BrxB-BrxF-iSTAND<-DDE_Tnp_1<-?<-OLD-ABC<-Cas_Cas1<-?\|\|?->?->HNH-> | 1212 | Archaea>Euryarchaeota>Stenosarchaea_group>Methanomicrobia>Methanosarcinales>Methanosarcinaceae | GCF_000970205.1 | Methanosarcina_mazei | MSMAS_RS06220 | 1426586..1430224 | 2209 | BREX-system-P-loop-protein-BrxC-[Methanosarcina-mazei] |
| WP_048040167.1 | Type1.BREX | <-MNS+REase<-?<-ACET<-?<-HATPase+wHTH<-Schlafen-p<-TIR<-N6-METHYLASE\|\|TPR+TPR+TPR+TPR+TPR+TPR+TPR-><-DUF6079*<-BrxB-BrxF-iSTAND<-tRNA<-?<-?\|\|?-><-PRTase | 1212 | Archaea>Euryarchaeota>Stenosarchaea_group>Methanomicrobia>Methanosarcinales>Methanosarcinaceae | GCF_000970225.1 | Methanosarcina_mazei | MSMAL_RS06195 | 1422644..1426282 | 2209 | BREX-system-P-loop-protein-BrxC-[Methanosarcina-mazei] |
| WP_048038520.1 | Type1.BREX | HicA_toxin-><-Mrr_N+ACET\|\|?-><-PadR-HTH<-HATPase+wHTH<-Schlafen+HATPase+wHTH-p<-Schlafen-p<-N6-METHYLASE\|\|TPR+TPR+TPR+TPR+TPR+TPR+TPR-><-DUF6079*<-BrxB-BrxF-iSTAND<-tRNA<-?<-?\|\|?-><-PRTase | 1212 | Archaea>Euryarchaeota>Stenosarchaea_group>Methanomicrobia>Methanosarcinales>Methanosarcinaceae | GCF_000970245.1 | Methanosarcina_mazei | MSMAC_RS13360 | 3106154..3109792 | 2209 | BREX-system-P-loop-protein-BrxC-[Methanosarcina-mazei] |
| WP_048125616.1 | Type1.BREX | RRXRR+HNH->RNase->?->PolB->HEPN-><-DDE_Tnp_1\|\|?-><-?\|\|BrxB-BrxF-iSTAND->DUF6079*->N6-METHYLASE->PglZ->SIGMA+OB-Fold+AAA-ATPase+Lon-Protease-BrxL->SF1-SF2-Helicase->?->ParB+HNH-><-BirA-HTH | 1252 | Archaea>Euryarchaeota>Stenosarchaea_group>Methanomicrobia>Methanosarcinales>Methanosarcinaceae | GCF_000970265.1 | Methanosarcina_lacustris | MSLAZ_RS07215 | 1686822..1690580 | 170861 | BREX-system-P-loop-protein-BrxC-[Methanosarcina-lacustris] |
| WP_048126494.1 | Type1.BREX | <-DDE_Tnp_1<-DDE_Tnp_1\|\|<-SIGMA+OB-Fold+AAA-ATPase+Lon-Protease-BrxL<-PglZ<-SF1-SF2-Helicase<-OLD-ABC+TOPRIM<-?<-N6-METHYLASE\|\|TPR+TPR+TPR+TPR+TPR+TPR+TPR-><-DUF6079*<-BrxB-BrxF-iSTAND<-?<-?\|\|?-><-TPR+TPR+TPR+TPR | 1211 | Archaea>Euryarchaeota>Stenosarchaea_group>Methanomicrobia>Methanosarcinales>Methanosarcinaceae | GCF_000970265.1 | Methanosarcina_lacustris | MSLAZ_RS10160 | 2470398..2474033 | 170861 | BREX-system-P-loop-protein-BrxC-[Methanosarcina-lacustris] |
| WP_048048522.1 | Type1.BREX | HicA_toxin-><-Mrr_N+ACET\|\|?-><-PadR-HTH<-HATPase+wHTH<-Schlafen+HATPase+wHTH-p<-Schlafen-p<-N6-METHYLASE\|\|TPR+TPR+TPR+TPR+TPR+TPR+TPR-><-DUF6079*<-BrxB-BrxF-iSTAND<-tRNA<-?<-?\|\|?-><-PRTase | 1212 | Archaea>Euryarchaeota>Stenosarchaea_group>Methanomicrobia>Methanosarcinales>Methanosarcinaceae | GCF_000979015.1 | Methanosarcina_mazei | DU63_RS00595 | 17697..21335 | 2209 | BREX-system-P-loop-protein-BrxC-[Methanosarcina-mazei] |
| WP_048048860.1 | Type1.BREX | <-ACET\|\|?->?->?->?->?->Glyco_hydro_130->GT4+DSBH->\|\|BrxB-BrxF-iSTAND->DUF6079*->N6-METHYLASE->Schlafen-p->Schlafen+HATPase+wHTH-p->HATPase+wHTH->?->PadR-HTH-><-?\|\|Mrr_N+ACET-><-HicA_toxin | 1212 | Archaea>Euryarchaeota>Stenosarchaea_group>Methanomicrobia>Methanosarcinales>Methanosarcinaceae | GCF_000979185.1 | Methanosarcina_mazei | DU71_RS16355 | 152756..156394 | 2209 | BREX-system-P-loop-protein-BrxC-[Methanosarcina-mazei] |
| WP_048133224.1 | Type1.BREX | <-SIGMA+OB-Fold+AAA-ATPase+Lon-Protease-BrxL<-PglZ<-HTH_Tnp_1<-?<-?<-?<-?<-TIR<-N6-METHYLASE<-DUF6079*<-BrxB-BrxF-iSTAND | 1243 | Archaea>Euryarchaeota>Stenosarchaea_group>Methanomicrobia>Methanosarcinales>Methanosarcinaceae | GCF_000979295.1 | Methanosarcina_sp._1.H.T.1A.1 | E095_RS06145 | 15439..19170 | 1483602 | BREX-system-P-loop-protein-BrxC-[Methanosarcina-sp.-1.H.T.1A.1] |
| WP_048041771.1 | Type1.BREX | PIN-><-?<-?\|\|?-><-Phage_int_SAM_4+Phage_integrase\|\|BrxB-BrxF-iSTAND->DUF6079*-><-TPR+TPR+TPR+TPR+TPR+TPR+TPR\|\|N6-METHYLASE->Schlafen-p->Schlafen+HATPase+wHTH-p->HATPase+wHTH->PadR-HTH-><-?\|\|Mrr_N+ACET-><-HicA_toxin | 1212 | Archaea>Euryarchaeota>Stenosarchaea_group>Methanomicrobia>Methanosarcinales>Methanosarcinaceae | GCF_000979335.1 | Methanosarcina_mazei | DU82_RS15310 | 140764..144402 | 2209 | BREX-system-P-loop-protein-BrxC-[Methanosarcina-mazei] |
| WP_048037387.1 | Type1.BREX | HicA_toxin-><-Mrr_N+ACET\|\|?-><-PadR-HTH<-HATPase+wHTH<-Schlafen+HATPase+wHTH-p<-Schlafen-p<-N6-METHYLASE\|\|TPR+TPR+TPR+TPR+TPR+TPR+TPR-><-DUF6079*<-BrxB-BrxF-iSTAND\|\|<-GT4+DSBH<-Glyco_hydro_130<-?<-?<-?\|\|Y1_Tnp->HTH_OrfB_IS605+OrfB_IS605+OrfB_Zn_ribbon-> | 1212 | Archaea>Euryarchaeota>Stenosarchaea_group>Methanomicrobia>Methanosarcinales>Methanosarcinaceae | GCF_000979345.1 | Methanosarcina | DU57_RS09710 | 64540..68178 | 2207 | MULTISPECIES:-BREX-system-P-loop-protein-BrxC-[Methanosarcina] |
| WP_048045678.1 | Type1.BREX | PRTase-><-?\|\|?->?->tRNA->BrxB-BrxF-iSTAND->DUF6079*-><-TPR+TPR+TPR+TPR+TPR+TPR+TPR\|\|N6-METHYLASE->Schlafen-p->HATPase+wHTH->PadR-HTH->Mrr_N+ACET-><-PHD_like+HicB_lk_antitox\|\|MNS+REase-> | 1212 | Archaea>Euryarchaeota>Stenosarchaea_group>Methanomicrobia>Methanosarcinales>Methanosarcinaceae | GCF_000979375.1 | Methanosarcina_mazei | DU40_09105 | 234397..238035 | 2209 | BREX-system-P-loop-protein-BrxC-[Methanosarcina-mazei] |
| WP_048049967.1 | Type1.BREX | PRTase-><-?\|\|?->?->tRNA->BrxB-BrxF-iSTAND->DUF6079*-><-TPR+TPR+TPR+TPR+TPR+TPR+TPR\|\|N6-METHYLASE->TIR->Schlafen-p->Schlafen+HATPase+wHTH-p->HATPase+wHTH->PadR-HTH-><-?\|\|Mrr_N+ACET-> | 1212 | Archaea>Euryarchaeota>Stenosarchaea_group>Methanomicrobia>Methanosarcinales>Methanosarcinaceae | GCF_000979685.1 | Methanosarcina_mazei | DU55_RS15175 | 41781..45419 | 2209 | BREX-system-P-loop-protein-BrxC-[Methanosarcina-mazei] |
| WP_048042209.1 | Type1.BREX | HicA_toxin-><-Mrr_N+ACET\|\|?-><-PadR-HTH<-Schlafen+HATPase+wHTH<-Schlafen+HATPase+wHTH-p<-Schlafen-p<-N6-METHYLASE\|\|TPR+TPR+TPR+TPR+TPR+TPR+TPR-><-DUF6079*<-BrxB-BrxF-iSTAND<-tRNA<-?<-NACHT<-?\|\|<-HKD-DNase<-tRNA<-?<-?\|\|?-><-PRTase | 1212 | Archaea>Euryarchaeota>Stenosarchaea_group>Methanomicrobia>Methanosarcinales>Methanosarcinaceae | GCF_000979755.1 | Methanosarcina_mazei | DU69_RS16895 | 67838..71476 | 2209 | BREX-system-P-loop-protein-BrxC-[Methanosarcina-mazei] |
| WP_048162601.1 | Type1.BREX | Glyco_hydro_130->GT4+DSBH-><-?<-?\|\|tRNA-><-NUDIX+DUF429\|\|?->BrxB-BrxF-iSTAND->DUF6079*->N6-METHYLASE->HTH+REase->N6-METHYLASE-p->?->Schlafen+HATPase+wHTH->?->ACET->Mrr_N+ACET-><-TPR+TPR+TPR | 1223 | Archaea>Euryarchaeota>Stenosarchaea_group>Methanomicrobia>Methanosarcinales>Methanosarcinaceae | GCF_000979975.1 | Methanosarcina_sp._1.H.A.2.2 | E093_RS11285 | 63696..67367 | 1483601 | BREX-system-P-loop-protein-BrxC-[Methanosarcina-sp.-1.H.A.2.2] |
| WP_096712430.1 | Type1.BREX | RADICAL-SAM->tRNA->tRNA-><-?<-?<-?\|\|?-><-SF1-SF2-Helicase\|\|BrxB-BrxF-iSTAND->DUF6079*->N6-METHYLASE->?->SbcC-><-PglZ->SIGMA+OB-Fold+AAA-ATPase+Lon-Protease-BrxL->RNaseH-><-REase<-McrB-AAA<-McrC-NTD+REase | 1188 | Archaea>Euryarchaeota>Stenosarchaea_group>Methanomicrobia>Methanosarcinales>Methanosarcinaceae | GCF_001726015.1 | Methanohalophilus_euhalobius | CP947_RS06580 | 29477..33043 | 51203 | BREX-system-P-loop-protein-BrxC-[Methanohalophilus-euhalobius] |
| WP_094227763.1 | Type1.BREX | TPR+TPR+TPR+TPR->?->DDE_Tnp<-?\|\|?->?->SF1-SF2-Helicase+REase->BrxB-BrxF-iSTAND->DUF6079*->N6-METHYLASE-p->SbcC->PglZ->SIGMA+OB-Fold+AAA-ATPase+Lon-Protease-BrxL->SF1-SF2-Helicase-><-DDE_Tnp_1\|\|HD->OLD-ABC-> | 1185 | Archaea>Euryarchaeota>Stenosarchaea_group>Methanomicrobia>Methanosarcinales>Methanosarcinaceae | GCF_002243045.1 | Methanolobus_psychrotolerans | BB781_RS06865 | 19266..22823 | 1874706 | BREX-system-P-loop-protein-BrxC-[Methanolobus-psychrotolerans] |
| WP_123137046.1 | Type1.BREX | VirB11->?->?->RADICAL-SAM->tRNA->?->?-><-?<-SF1-SF2-Helicase\|\|BrxB-BrxF-iSTAND->DUF6079*->Schlafen+HATPase+wHTH->N6-METHYLASE->?->PglZ->SIGMA+OB-Fold+AAA-ATPase+Lon-Protease-BrxL-><-?<-REase<-SF1-SF2-Helicase+REase<-RNase+DSRBD | 1187 | Archaea>Euryarchaeota>Stenosarchaea_group>Methanomicrobia>Methanosarcinales>Methanosarcinaceae | GCF_003722075.1 | Methanohalophilus_sp._RSK | EFE42_RS09705 | 37539..41102 | 2485783 | BREX-system-P-loop-protein-BrxC-[Methanohalophilus-sp.-RSK] |
| WP_128754626.1 | Type1.BREX | RADICAL-SAM->tRNA->tRNA->?-><-?<-?<-?\|\|?-><-SF1-SF2-Helicase\|\|BrxB-BrxF-iSTAND->DUF6079*->N6-METHYLASE->HD->PglZ->SIGMA+OB-Fold+AAA-ATPase+Lon-Protease-BrxL->RNaseH-><-SF1-SF2-Helicase+REase\|\|CRISPR_Cas6->?->RAMPs-> | 1188 | Archaea>Euryarchaeota>Stenosarchaea_group>Methanomicrobia>Methanosarcinales>Methanosarcinaceae | GCF_004104135.1 | Methanohalophilus_sp._WG1-DM | CI957_RS08065 | 29040..32606 | 2491675 | BREX-system-P-loop-protein-BrxC-[Methanohalophilus-sp.-WG1-DM] |
| WP_163645423.1 | Type1.BREX | HicA_toxin-><-Mrr_N+ACET\|\|?-><-?<-HATPase+wHTH<-Schlafen+HATPase+wHTH-p<-Schlafen-p<-N6-METHYLASE<-DUF6079*<-BrxB-BrxF-iSTAND\|\|<-GT4+DSBH<-Glyco_hydro_130<-?<-?<-?\|\|?->Phage_int_SAM_4+Phage_integrase-> | 1212 | Archaea>Euryarchaeota>Stenosarchaea_group>Methanomicrobia>Methanosarcinales>Methanosarcinaceae | GCF_010706455.1 | Methanosarcina_mazei | FQU78_RS06260 | 1447297..1450935 | 2209 | BREX-system-P-loop-protein-BrxC-[Methanosarcina-mazei] |
| WP_229393447.1 | Type1.BREX | DDE_Tnp_IS66->?->?->BrxB-BrxF-iSTAND->DUF6079*->N6-METHYLASE-><-Schlafen+HATPase+wHTH\|\|Mrr_N+ACET->PglZ->SIGMA+OB-Fold+AAA-ATPase+Lon-Protease-BrxL->MNS+REase-><-?\|\|Calcineurin->SbcC-> | 1202 | Archaea>Euryarchaeota>Stenosarchaea_group>Methanomicrobia>Methanosarcinales>Methanosarcinaceae | GCF_020804205.1 | Methanosarcina_sp._DH1 | FXW07_RS01525 | 42111..45719 | 2605695 | BREX-system-P-loop-protein-BrxC-[Methanosarcina-sp.-DH1] |
| WP_229395640.1 | Type1.BREX | BrxA-DNA-binding->BrxB-BrxF-iSTAND->DUF6079*->N6-METHYLASE->ParB+HNH->PglZ->SIGMA+OB-Fold+AAA-ATPase+Lon-Protease-BrxL-><-DUF488<-Glycos_transf_1+Glyco_trans_1_2<-?<-DSBH | 1143 | Archaea>Euryarchaeota>Stenosarchaea_group>Methanomicrobia>Methanosarcinales>Methanosarcinaceae | GCF_020804205.1 | Methanosarcina_sp._DH1 | FXW07_RS12025 | 174817..178248 | 2605695 | BREX-system-P-loop-protein-BrxC-[Methanosarcina-sp.-DH1] |
| WP_229391422.1 | Type1.BREX | BrxD-AAA->SF1-SF2-Helicase+wHTH+Small-B-barrel+DUF1998-><-SIGMA+OB-Fold+AAA-ATPase+Lon-Protease-BrxL<-PglZ\|\|TPR+TPR+TPR+TPR-><-PadR-HTH<-Schlafen+HATPase+wHTH<-TIR<-N6-METHYLASE<-DUF6079*<-BrxB-BrxF-iSTAND<-?\|\|\|\|?->?->?->?->?-><-?<-N6-METHYLASE-p | 1201 | Archaea>Euryarchaeota>Stenosarchaea_group>Methanomicrobia>Methanosarcinales>Methanosarcinaceae | GCF_020804225.1 | Methanosarcina_sp._DH2 | FXV91_RS14640 | 168630..172235 | 2605639 | BREX-system-P-loop-protein-BrxC-[Methanosarcina-sp.-DH2] |
| WP_250868420.1 | Type1.BREX | SF1-SF2-Helicase->?-><-URI-DNase<-?<-?<-SF1-SF2-Helicase+REase\|\|SF1-SF2-Helicase->BrxB-BrxF-iSTAND->DUF6079-p*->DUF6079-p->?->?->?->PglZ->SIGMA+OB-Fold+AAA-ATPase+Lon-Protease-BrxL-><-tRNA<-tRNA<-RADICAL-SAM | 749 | Archaea>Euryarchaeota>Stenosarchaea_group>Methanomicrobia>Methanosarcinales>Methanosarcinaceae | GCF_023684405.1 | Methanococcoides_seepicolus | KDK67_RS08780 | 13773..16022 | 2828780 | BREX-system-P-loop-protein-BrxC-[Methanococcoides-seepicolus] |
| WP_256623626.1 | Type1.BREX | EVE+McrB-AAA->McrC-NTD+REase->?->URI-DNase->BrxB-BrxF-iSTAND->DUF6079*->N6-METHYLASE->Schlafen+HATPase+wHTH->N6-METHYLASE-p->SbcC-><-?\|\|?->?->OLD-ABC->TOPRIM-> | 1185 | Archaea>Euryarchaeota>Stenosarchaea_group>Methanomicrobia>Methanosarcinales>Methanosarcinaceae | GCF_024500045.1 | Methanolobus_chelungpuianus | PV02_RS11875 | 317319..320876 | 502115 | BREX-system-P-loop-protein-BrxC-[Methanolobus-chelungpuianus] |
| WP_270096529.1 | Type1.BREX | SF1-SF2-Helicase+REase->HNH-><-SIGMA+OB-Fold+AAA-ATPase+Lon-Protease-BrxL<-PglZ<-TOPRIM<-OLD-ABC<-HD<-N6-METHYLASE-p<-N6-METHYLASE-p<-DUF6079*<-BrxB-BrxF-iSTAND<-REase<-?<-?<-RNase\|\|Classical-AAA->SF1-SF2-Helicase-> | 1190 | Archaea>Euryarchaeota>Stenosarchaea_group>Methanomicrobia>Methanosarcinales>Methanosarcinaceae | GCF_027594145.1 | Methanococcoides_alaskense | PBV53_RS07490 | 737341..740913 | 325778 | BREX-system-P-loop-protein-BrxC-[Methanococcoides-alaskense] |
| WP_091932407.1 | Type1.BREX | SF1-SF2-Helicase+REase->?-><-SIGMA+OB-Fold+AAA-ATPase+Lon-Protease-BrxL<-PglZ<-TOPRIM<-OLD-ABC<-N6-METHYLASE-p<-Schlafen+HATPase+wHTH<-N6-METHYLASE<-DUF6079*<-BrxB-BrxF-iSTAND\|\|McrC-NTD+REase->McrB-AAA->SF1-SF2-Helicase-><-DDE_Tnp_1+DDE_Tnp_1_2 | 1184 | Archaea>Euryarchaeota>Stenosarchaea_group>Methanomicrobia>Methanosarcinales>Methanosarcinaceae | GCF_900114835.1 | Methanolobus_profundi | BM008_RS01885 | 368008..371562 | 487685 | BREX-system-P-loop-protein-BrxC-[Methanolobus-profundi] |
| WP_244603415.1 | Type1.BREX | <-SIR2<-SIGMA+OB-Fold+AAA-ATPase+Lon-Protease-BrxL<-PglZ<-SF1-SF2-Helicase<-OLD-ABC+TOPRIM<-N6-METHYLASE-p<-N6-METHYLASE-p<-DUF6079*<-BrxB-BrxF-iSTAND<-URI-DNase<-?<-?\|\|?->TPR->TPR-><-ZnR+HTH_Tnp_ISL3+DDE_Tnp_ISL3 | 1181 | Archaea>Euryarchaeota>Stenosarchaea_group>Methanomicrobia>Methanosarcinales>Methanosarcinaceae | GCF_900774285.1 | Methanococcoides_sp._NM1 | E7X56_RS05470 | 155089..158634 | 1201013 | BREX-system-P-loop-protein-BrxC-[Methanococcoides-sp.-NM1] |
| WP_172363658.1 | Type1.BREX | <-SIGMA+OB-Fold+AAA-ATPase+Lon-Protease-BrxL<-PglZ<-TOPRIM<-OLD-ABC<-TIR<-N6-METHYLASE-p<-Schlafen+HATPase+wHTH<-N6-METHYLASE<-DUF6079*<-BrxB-BrxF-iSTAND<-URI-DNase<-?<-SF1-SF2-Helicase+REase\|\|<-?\|\|?->HNH-> | 1185 | Archaea>Euryarchaeota>Stenosarchaea_group>Methanomicrobia>Methanosarcinales>Methanosarcinaceae | - | Methanococcoides_sp._SA1 | HNV12_09470 | 12693..16250 | 2735871 | BREX-system-P-loop-protein-BrxC-[Methanococcoides-sp.-SA1] |
| WP_048171216.1 | Type1.BREX | Glyco_hydro_130->GT4+DSBH-><-?<-?\|\|tRNA-><-NUDIX+DUF429\|\|?->BrxB-BrxF-iSTAND->DUF6079*->N6-METHYLASE->?->OLD-ABC+TOPRIM->SF1-SF2-Helicase->Schlafen-p-> | 1204 | Archaea>Euryarchaeota>Stenosarchaea_group>Methanomicrobia>Methanosarcinales>Methanosarcinaceae | - | Methanosarcina_sp._2.H.A.1B.4 | E092_RS09675 | 12629..16243 | 1483600 | BREX-system-P-loop-protein-BrxC-[Methanosarcina-sp.-2.H.A.1B.4] |
| OFV66969.1 | Type1.BREX | <-PglZ<-N6-METHYLASE-p<-?<-?<-SbcC<-?<-N6-METHYLASE<-DUF6079*<-BrxB-BrxF-iSTAND<-BrxA-DNA-binding<-SIGMA+OB-Fold+AAA-ATPase+Lon-Protease-BrxL\|\|?-><-?<-?<-nSTAND | 1184 | Archaea>Euryarchaeota>Stenosarchaea_group>Methanomicrobia>Methanosarcinales>Methanosarcinales_incertae_sedis | GCA_001766825.1 | Candidatus_Syntrophoarchaeum_butanivorans | SBU_000262 | 234688..238242 | 1839936 | ATPase-[Candidatus-Syntrophoarchaeum-butanivorans] |
| MCD4767073.1 | Type1.BREX | <-N6-METHYLASE-p<-BirA-HTH<-DrHyd<-N6-METHYLASE<-DUF6079*<-BrxB-BrxF-iSTAND | 1179 | Archaea>Euryarchaeota>Stenosarchaea_group>Methanomicrobia>Methanosarcinales>unclassified_Methanosarcinales | GCA_021107935.1 | Methanosarcinales_archaeon | K8R34_11245 | 4319..7858 | 2250255 | BREX-system-P-loop-protein-BrxC-[Methanosarcinales-archaeon] |
| MCD4843299.1 | Type1.BREX | <-AIPR<-HD<-HTH+REase<-Schlafen-p<-TIR<-N6-METHYLASE-p<-?<-DrHyd+HTH<-N6-METHYLASE<-DUF6079*<-BrxB-BrxF-iSTAND<-?<-HicA_toxin<-HicB_lk_antitox<-?<-SAM-METHYLASE | 1180 | Archaea>Euryarchaeota>Stenosarchaea_group>Methanomicrobia>Methanosarcinales>unclassified_Methanosarcinales | GCA_021108405.1 | Methanosarcinales_archaeon | K8R25_02310 | 18778..22320 | 2250255 | BREX-system-P-loop-protein-BrxC-[Methanosarcinales-archaeon] |
| QNO42144.1 | Type1.BREX | <-SIGMA+OB-Fold+AAA-ATPase+Lon-Protease-BrxL<-PglZ<-Schlafen+HATPase+wHTH<-SF1-SF2-Helicase<-OLD-ABC+TOPRIM<-N6-METHYLASE<-DUF6079-p<-DUF6079-p*<-BrxB-BrxF-iSTAND\|\|DDE_Tnp_1-> | 488 | Archaea>Euryarchaeota>Stenosarchaea_group>Methanomicrobia>Methanosarcinales>unclassified_Methanosarcinales | - | Methanosarcinales_archaeon_ANME-2c_ERB4 | CAOPPJJI_00010 | 16805..18271 | 2759911 | hypothetical-protein-CAOPPJJI_00010-[Methanosarcinales-archaeon-ANME-2c-ERB4] |
| OPY50193.1 | Type1.BREX | DUF772+DDE_Tnp_1+DDE_Tnp_1_6->REase->?-><-?<-?<-SIGMA+OB-Fold+AAA-ATPase+Lon-Protease-BrxL<-PglZ<-N6-METHYLASE<-DUF6079*<-BrxB-BrxF-iSTAND<-BrxA-DNA-binding<-?<-?<-Phage_int_SAM_4+Phage_integrase | 1204 | Archaea>Euryarchaeota>Stenosarchaea_group>Methanomicrobia>Methanotrichales>Methanotrichaceae | GCA_002067365.1 | Methanosaeta_sp._PtaU1.Bin060 | A4E48_01974 | 19816..23430 | 1811688 | MAG:-hypothetical-protein-A4E48_01974-[Methanosaeta-sp.-PtaU1.Bin060] |
| MBN1236634.1 | Type1.BREX | <-SIGMA+OB-Fold+AAA-ATPase+Lon-Protease-BrxL<-PglZ<-?<-N6-METHYLASE\|\|SbcC->?-><-DUF6079*<-BrxB-BrxF-iSTAND<-BrxA-DNA-binding | 1217 | Archaea>Euryarchaeota>Stenosarchaea_group>Methanomicrobia>Methanotrichales>Methanotrichaceae | GCA_016927055.1 | Methanotrichaceae_archaeon | JW999_11415 | 12015..15668 | 2595067 | BREX-system-P-loop-protein-BrxC-[Methanotrichaceae-archaeon] |
| MBP7068907.1 | Type1.BREX | <-SIGMA+OB-Fold+AAA-ATPase+Lon-Protease-BrxL<-PglZ<-?<-N6-METHYLASE<-DUF6079*<-BrxB-BrxF-iSTAND<-BrxA-DNA-binding<-REase | 1215 | Archaea>Euryarchaeota>Stenosarchaea_group>Methanomicrobia>Methanotrichales>Methanotrichaceae | GCA_017994095.1 | Methanothrix_sp. | KBB04_11615 | 10819..14466 | 90426 | BREX-system-P-loop-protein-BrxC-[Methanothrix-sp.] |
| MBU4220824.1 | Type1.BREX | PAIREDC-HTH->?->?->?->?-><-?\|\|BrxB-BrxF-iSTAND->DUF6079*->N6-METHYLASE->TIR->Schlafen-p->HEPN->Schlafen+HATPase+wHTH->?->?->PglZ-p-> | 1180 | Archaea>Euryarchaeota>unclassified_Euryarchaeota | GCA_018898145.1 | Euryarchaeota_archaeon | KKA10_04250 | 6252..9794 | 2026739 | BREX-system-P-loop-protein-BrxC-[Euryarchaeota-archaeon] |
| EMR73247.1 | Type1.BREX | BrxA-DNA-binding->BrxB-BrxF-iSTAND->DUF6079*->N6-METHYLASE->PglZ->?->SIGMA+OB-Fold+AAA-ATPase+Lon-Protease-BrxL-p-> | 1207 | Archaea>TACK_group>Nitrososphaerota>unclassified_Nitrososphaerota | GCA_000349625.1 | Thaumarchaeota_archaeon_SCGC_AB-539-E09 | MCGE09_00285 | 3053..6676 | 1198115 | hypothetical-protein-MCGE09_00285-[Thaumarchaeota-archaeon-SCGC-AB-539-E09] |
| MBA3712418.1 | Type1.BREX | <-GDResolvase+Recombinase<-GDResolvase\|\|<-?<-?\|\|GspA-ATPase->?->TOPRIM+Topoisom_bac->BrxA-DNA-binding->BrxB-BrxF-iSTAND->DUF6079-p*-> | 967 | Bacteria>Acidobacteriota>Blastocatellia>Blastocatellales>Pyrinomonadaceae>unclassified_Pyrinomonadaceae | GCA_013816505.1 | Pyrinomonadaceae_bacterium | HOW76_08205 | 12993..15893 | 2283092 | BREX-system-P-loop-protein-BrxC-[Pyrinomonadaceae-bacterium] |
| MBD0371455.1 | Type1.BREX | VirB11->PSBP->?-><-SIGMA+OB-Fold+AAA-ATPase+Lon-Protease-BrxL<-PglZ<-ParB+HNH<-NERD<-N6-METHYLASE<-HEPN<-DUF6079*<-BrxB-BrxF-iSTAND<-BrxA-DNA-binding | 1218 | Bacteria>Acidobacteriota>Blastocatellia>Blastocatellales>Pyrinomonadaceae>unclassified_Pyrinomonadaceae | GCA_014534425.1 | Pyrinomonadaceae_bacterium | ICV60_11505 | 29495..33151 | 2283092 | BREX-system-P-loop-protein-BrxC-[Pyrinomonadaceae-bacterium] |
| MBK9154354.1 | Type1.BREX | Phage_integrase->DUF488-><-?<-?<-?\|\|HTH_3->Peptidase_M78->BrxA-DNA-binding->BrxB-BrxF-iSTAND->DUF6079-p*->?->DUF6079-p->AIPR->?->PglZ->PglZ->Exonuc_V_gamma+REase->SF1-SF2-Helicase->SF1-SF2-Helicase-> | 481 | Bacteria>Acidobacteriota>Blastocatellia>Chloracidobacterium>unclassified_Chloracidobacterium | GCA_016716365.1 | Chloracidobacterium_sp. | IPM25_09080 | 169835..171280 | 2099375 | BREX-system-P-loop-protein-BrxC-[Chloracidobacterium-sp.] |
| MCI0490100.1 | Type1.BREX | BrxA-DNA-binding->BrxB-BrxF-iSTAND->DUF6079*->HEPN->Four-helical-protein-><-?<-?\|\|N6-METHYLASE->Schlafen+HATPase+wHTH-p-> | 1218 | Bacteria>Acidobacteriota>Blastocatellia>unclassified_Blastocatellia | GCA_022615215.1 | Blastocatellia_bacterium | L0229_26200 | 2085..5741 | 2052146 | BREX-system-P-loop-protein-BrxC-[Blastocatellia-bacterium] |
| WP_005031444.1 | Type1.BREX | HEPN+PAS+DUF4011+SF1-SF2-Helicase+REase-><-SIGMA+OB-Fold+AAA-ATPase+Lon-Protease-BrxL<-PglZ<-Schlafen+HATPase+wHTH<-N6-METHYLASE<-Trypsin<-DUF6079*<-BrxB-BrxF-iSTAND<-BrxA-DNA-binding\|\|?->ZetaToxin->wHTH+SAD+HNH->?-><-DOC+HTH | 1205 | Bacteria>Acidobacteriota>Holophagae>Holophagales>Holophagaceae>Holophaga | GCF_000242615.2 | Holophaga_foetida | HOLFODRAFT_RS0203665 | 863160..866777 | 35839 | BREX-system-P-loop-protein-BrxC-[Holophaga-foetida] |
| MXX75209.1 | Type1.BREX | Cysteine-Desulfurase-><-?\|\|BrxA-DNA-binding->BrxB-BrxF-iSTAND->DUF6079*->HTH+MPTase->?->?->?->N6-METHYLASE->PglZ->SIGMA+OB-Fold+AAA-ATPase+Lon-Protease-BrxL->?-><-ABhydrolase | 1173 | Bacteria>Acidobacteriota>Holophagae>Holophagales>unclassified_Holophagales | GCA_009837885.1 | Holophagales_bacterium | F4Y77_07375 | 64446..67967 | 2478486 | BREX-system-P-loop-protein-BrxC-[Holophagales-bacterium] |
| MBK9374878.1 | Type1.BREX | <-NtrC-AAA+FIS-HTH\|\|?->?-><-?\|\|?->?->BrxA-DNA-binding->DUF6079*->N6-METHYLASE-p->?->PglZ->Tox-RelE->HTH_37-> | 1147 | Bacteria>Acidobacteriota>Holophagae>Holophagales>unclassified_Holophagales | GCA_016719485.1 | Holophagales_bacterium | IPN03_14410 | 526057..529500 | 2478486 | BREX-system-P-loop-protein-BrxC-[Holophagales-bacterium] |
| MBK9968089.1 | Type1.BREX | <-HAD+HAD<-?<-?<-?<-?<-?\|\|XIS-HTH->BrxA-DNA-binding->BrxB-BrxF-iSTAND->DUF6079*->N6-METHYLASE->DOC+HTH->PglZ->SIGMA+OB-Fold+AAA-ATPase+Lon-Protease-BrxL-> | 1245 | Bacteria>Acidobacteriota>Holophagae>Holophagales>unclassified_Holophagales | GCA_016722225.1 | Holophagales_bacterium | IPP07_25730 | 57868..61605 | 2478486 | BREX-system-P-loop-protein-BrxC-[Holophagales-bacterium] |
| MBZ2183842.1 | Type1.BREX | <-UbiA<-?\|\|?->ABhydrolase-><-?\|\|?-><-?\|\|BrxA-DNA-binding->BrxB-BrxF-iSTAND->DUF6079*->N6-METHYLASE->ArdCN+MPTase->Schlafen+HATPase+wHTH->ParA-Soj-PloopNTPase->PglZ->SIGMA+OB-Fold+AAA-ATPase+Lon-Protease-BrxL->?->?->ROK-HTH-> | 1166 | Bacteria>Acidobacteriota>Terriglobia>Bryobacterales>Bryobacteraceae>Bryobacter | GCA_019931185.1 | Bryobacter_sp. | K7J46_03960 | 13925..17425 | 1962911 | BREX-system-P-loop-protein-BrxC-[Bryobacter-sp.] |
| WP_238326045.1 | Type1.BREX | <-SIGMA+OB-Fold+AAA-ATPase+Lon-Protease-BrxL<-PglZ<-PHP+SbcC<-N6-METHYLASE<-DUF6079*<-BrxB-BrxF-iSTAND<-BrxA-DNA-binding\|\|?->AbiEii-> | 1159 | Bacteria>Acidobacteriota>Terriglobia>Bryobacterales>Bryobacteraceae>Bryobacter | GCF_000702445.1 | Bryobacter_aggregatus | M017_RS0125570 | 1351195..1354674 | 360054 | BREX-system-P-loop-protein-BrxC-[Bryobacter-aggregatus] |
| GIU77527.1 | Type1.BREX | Phage_int_SAM_4+Phage_integrase->HTH_21+rve-><-?<-?\|\|?->HTH_21+rve-><-PHP+SbcC<-SIGMA+OB-Fold+AAA-ATPase+Lon-Protease-BrxL<-PglZ<-N6-METHYLASE<-DUF6079*<-BrxB-BrxF-iSTAND<-BrxA-DNA-binding<-?\|\|DUF772+DDE_Tnp_1+DDE_Tnp_1_6-><-XIS-HTH\|\|MULE-TRANSPOSASE->HIN-HTH+rve+rve_3->?->DUF772+DDE_Tnp_1+DDE_Tnp_1_6-> | 1157 | Bacteria>Acidobacteriota>Terriglobia>Bryobacterales>Bryobacteraceae>unclassified_Bryobacteraceae | GCA_026002875.1 | Bryobacteraceae_bacterium | KatS3mg005_0765 | 905101..908574 | 2212468 | MAG:-hypothetical-protein-KatS3mg005_0765-[Bryobacteraceae-bacterium] |
| MBK9165938.1 | Type1.BREX | XIS-HTH-><-?<-SIGMA+OB-Fold+AAA-ATPase+Lon-Protease-BrxL<-PglZ<-PglZ<-PglZ<-?<-N6-METHYLASE<-HEPN<-DUF6079*<-BrxB-BrxF-iSTAND<-BrxA-DNA-binding<-CopG/RHH\|\|?-><-ParD_antitoxin<-HTH_37\|\|?->?->Tox-RelE-> | 1234 | Bacteria>Acidobacteriota>Terriglobia>Bryobacterales>unclassified_Bryobacterales | GCA_016716385.1 | Bryobacterales_bacterium | IPM24_00570 | 108049..111753 | 2026791 | BREX-system-P-loop-protein-BrxC-[Bryobacterales-bacterium] |
| MCX6390067.1 | Type1.BREX | ParB->ParB->TPR-><-MACRODOMAIN<-SIGMA+OB-Fold+AAA-ATPase+Lon-Protease-BrxL<-PglZ<-N6-METHYLASE-p<-DOC+HTH<-N6-METHYLASE<-DUF6079*<-BrxB-BrxF-iSTAND<-BrxA-DNA-binding<-?<-ART-DarT<-SF1-SF2-Helicase<-HTH+MPTase | 1241 | Bacteria>Acidobacteriota>Terriglobia>Bryobacterales>unclassified_Bryobacterales | GCA_020847895.1 | Bryobacterales_bacterium | IT167_05665 | 14981..18706 | 2026791 | BREX-system-P-loop-protein-BrxC-[Bryobacterales-bacterium] |
| WP_089839623.1 | Type1.BREX | ParB->TIR->Peptidase_C14-><-?<-REase<-SIGMA+OB-Fold+AAA-ATPase+Lon-Protease-BrxL<-PglZ<-N6-METHYLASE<-DUF6079*<-BrxB-BrxF-iSTAND<-BrxA-DNA-binding<-?\|\|?-><-?<-MODE-HTH+PBPII\|\|XIS-HTH->Phage_int_SAM_3+Phage_integrase-> | 1149 | Bacteria>Acidobacteriota>Terriglobia>Terriglobales>Acidobacteriaceae>Granulicella | GCF_900114625.1 | Granulicella_pectinivorans | BM400_RS13360 | 3371671..3375120 | 474950 | BREX-system-P-loop-protein-BrxC-[Granulicella-pectinivorans] |
| MBV9502366.1 | Type1.BREX | CopG/RHH->BrxA-DNA-binding->BrxB-BrxF-iSTAND->DUF6079-p*->SIGMA+OB-Fold+AAA-ATPase+Lon-Protease-BrxL->SIGMA+OB-Fold+AAA-ATPase+Lon-Protease-BrxL->ArdCN+MPTase-> | 537 | Bacteria>Acidobacteriota>Terriglobia>Terriglobales>Acidobacteriaceae>unclassified_Acidobacteriaceae | GCA_019244055.1 | Acidobacteriaceae_bacterium | J0138_23605 | 3237..4850 | 2052142 | hypothetical-protein-[Acidobacteriaceae-bacterium] |
| MBZ5499154.1 | Type1.BREX | Phage_int_SAM_1+Phage_integrase-><-?<-SIGMA+OB-Fold+AAA-ATPase+Lon-Protease-BrxL<-PglZ<-Schlafen+HATPase+wHTH<-SbcC<-N6-METHYLASE<-?<-TIR<-DUF6079*<-BrxB-BrxF-iSTAND<-BrxA-DNA-binding\|\|XIS-HTH-> | 1147 | Bacteria>Acidobacteriota>Terriglobia>unclassified_Terriglobia | GCA_020072565.1 | Acidobacteriia_bacterium | LAP85_22370 | 28879..32322 | 2081523 | BREX-system-P-loop-protein-BrxC-[Acidobacteriia-bacterium] |
| MBP7799662.1 | Type1.BREX | <-CR-ATPase5+REase<-REase+SF1-SF2-Helicase<-?\|\|DDE_Tnp-><-SIGMA+OB-Fold+AAA-ATPase+Lon-Protease-BrxL<-PglZ<-Schlafen+HATPase+wHTH<-N6-METHYLASE<-?<-DUF6079*<-BrxB-BrxF-iSTAND<-BrxA-DNA-binding | 1146 | Bacteria>Acidobacteriota>Thermoanaerobaculia>Thermoanaerobaculales>Thermoanaerobaculaceae>unclassified_Thermoanaerobaculaceae | GCA_018000955.1 | Thermoanaerobaculaceae_bacterium | KA072_13665 | 19624..23064 | 2699758 | BREX-system-P-loop-protein-BrxC-[Thermoanaerobaculaceae-bacterium] |
| OFW00070.1 | Type1.BREX | HTH_21+rve+rve_3-><-?<-?<-?\|\|DDE_Tnp_1_4-><-SIGMA+OB-Fold+AAA-ATPase+Lon-Protease-BrxL<-PglZ<-N6-METHYLASE<-DUF6079*<-BrxB-BrxF-iSTAND<-BrxA-DNA-binding<-REase<-?<-?\|\|HNH->ParB+N6_N4_METHYLASE-> | 1145 | Bacteria>Acidobacteriota>unclassified_Acidobacteriota | GCA_001767075.1 | Acidobacteria_bacterium_RIFCSPLOWO2_12_FULL_54_10 | A3F68_05125 | 19518..22955 | 1797186 | hypothetical-protein-A3F68_05125-[Acidobacteria-bacterium-RIFCSPLOWO2_12_FULL_54_10] |
| OFW17305.1 | Type1.BREX | <-SF1-SF2-Helicase<-SIGMA+OB-Fold+AAA-ATPase+Lon-Protease-BrxL<-PglZ<-HATPase+wHTH<-HATPase_c_4<-Schlafen-p<-N6-METHYLASE<-HEPN<-PolB<-DUF6079*<-BrxB-BrxF-iSTAND<-BrxA-DNA-binding\|\|wHTH+AbiEi_2->AbiEii-><-?<-?<-PIN | 1156 | Bacteria>Acidobacteriota>unclassified_Acidobacteriota | GCA_001767195.1 | Acidobacteria_bacterium_RIFCSPLOWO2_02_FULL_67_21 | A3H29_10520 | 11544..15014 | 1797183 | hypothetical-protein-A3H29_10520-[Acidobacteria-bacterium-RIFCSPLOWO2_02_FULL_67_21] |
| OFW39743.1 | Type1.BREX | PIN->?->?-><-AbiEii<-wHTH+AbiEi_2\|\|BrxA-DNA-binding->BrxB-BrxF-iSTAND->DUF6079*->PolB->HEPN->N6-METHYLASE->Schlafen+HATPase+wHTH->HATPase+wHTH->PglZ->SIGMA+OB-Fold+AAA-ATPase+Lon-Protease-BrxL->SF1-SF2-Helicase->DDE_Tnp_1_4-> | 1156 | Bacteria>Acidobacteriota>unclassified_Acidobacteriota | GCA_001767515.1 | Acidobacteria_bacterium_RIFCSPLOWO2_12_FULL_67_14 | A3F70_01760 | 7438..10908 | 1797192 | hypothetical-protein-A3F70_01760-[Acidobacteria-bacterium-RIFCSPLOWO2_12_FULL_67_14] |
| PYQ26570.1 | Type1.BREX | BrxA-DNA-binding->BrxB-BrxF-iSTAND->DUF6079*->N6-METHYLASE->DOC+HTH->PglZ->SIGMA+OB-Fold+AAA-ATPase+Lon-Protease-BrxL->?-><-?\|\|ACET+ACET->SIGMA-HTH-> | 1155 | Bacteria>Acidobacteriota>unclassified_Acidobacteriota | GCA_003223645.1 | Acidobacteriota_bacterium | DMF56_23440 | 3756..7223 | 1978231 | BREX-system-P-loop-protein-BrxC-[Acidobacteriota-bacterium] |
| MYN66269.1 | Type1.BREX | BrxA-DNA-binding->BrxB-BrxF-iSTAND->DUF6079*->OLD-ABC->TOPRIM->Schlafen+HATPase+wHTH->N6-METHYLASE->?->?->SbcC->PHP+SbcC->Schlafen+HATPase+wHTH-> | 1283 | Bacteria>Acidobacteriota>unclassified_Acidobacteriota | GCA_009861545.1 | Acidobacteriota_bacterium | F4X11_14760 | 2943..6794 | 1978231 | BREX-system-P-loop-protein-BrxC-[Acidobacteriota-bacterium] |
| NLV30311.1 | Type1.BREX | DNA-glycosylase-><-?<-SH3\|\|wHTH+WYL+WCX->SF1-SF2-Helicase->BrxA-DNA-binding->BrxB-BrxF-iSTAND->DUF6079*->Schlafen+HATPase+wHTH+HTH->N6-METHYLASE->PglZ->SIGMA+OB-Fold+AAA-ATPase+HKD-DNase-BrxL->HigB_toxin->HTH+MPTase->SF1-SF2-Helicase->?-><-DSBA+Thioredoxin | 1169 | Bacteria>Acidobacteriota>unclassified_Acidobacteriota | GCA_012729555.1 | Acidobacteriota_bacterium | GXY47_04070 | 27594..31103 | 1978231 | BREX-system-P-loop-protein-BrxC-[Acidobacteriota-bacterium] |
| MBI2837087.1 | Type1.BREX | <-SIGMA+OB-Fold+AAA-ATPase+Lon-Protease-BrxL<-PglZ<-Schlafen+HATPase+wHTH<-PHP+SbcC<-N6-METHYLASE<-Schlafen+HATPase+wHTH<-PHP+SbcC<-N6-METHYLASE<-DUF6079*<-BrxB-BrxF-iSTAND<-BrxA-DNA-binding<-XIS-HTH | 1224 | Bacteria>Acidobacteriota>unclassified_Acidobacteriota | GCA_016191015.1 | Acidobacteriota_bacterium | HYX75_02140 | 17742..21416 | 1978231 | BREX-system-P-loop-protein-BrxC-[Acidobacteriota-bacterium] |
| MBI4904787.1 | Type1.BREX | Sulfatase+DUF4994-><-?<-?<-?<-?\|\|XIS-HTH->BrxA-DNA-binding->BrxB-BrxF-iSTAND->DUF6079*->N6-METHYLASE->REase+AIPR->PglZ->SIGMA+OB-Fold+AAA-ATPase+Lon-Protease-BrxL->?->?->?-><-?<-HTH_3 | 1144 | Bacteria>Acidobacteriota>unclassified_Acidobacteriota | GCA_016213775.1 | Acidobacteriota_bacterium | HY820_14195 | 24664..28098 | 1978231 | BREX-system-P-loop-protein-BrxC-[Acidobacteriota-bacterium] |
| MBS1828808.1 | Type1.BREX | D5N+D5ATPase-><-?\|\|ParB-><-?<-Phage_integrase+Phage_integrase<-?\|\|CopG/RHH->BrxA-DNA-binding->BrxB-BrxF-iSTAND->DUF6079*->HEPN->N6-METHYLASE->PglZ->SIGMA+OB-Fold+AAA-ATPase+Lon-Protease-BrxL->REase->HEPN->?->Phage_integrase-> | 1226 | Bacteria>Acidobacteriota>unclassified_Acidobacteriota | GCA_018268935.1 | Acidobacteriota_bacterium | JST93_26130 | 24024..27704 | 1978231 | BREX-system-P-loop-protein-BrxC-[Acidobacteriota-bacterium] |
| MBV8891344.1 | Type1.BREX | HTH_Tnp_1+HTH_21+rve+rve_3->?->CopG/RHH->BrxA-DNA-binding->BrxB-BrxF-iSTAND->DUF6079-p*-> | 334 | Bacteria>Acidobacteriota>unclassified_Acidobacteriota | GCA_019243065.1 | Acidobacteriota_bacterium | JO266_05105 | 5695..6697 | 1978231 | BREX-system-P-loop-protein-BrxC-[Acidobacteriota-bacterium] |
| MCI0392588.1 | Type1.BREX | NtrC-AAA+FIS-HTH->BrxA-DNA-binding->BrxB-BrxF-iSTAND->DUF6079*->HKD-DNase+DUF3150->N6-METHYLASE-> | 1160 | Bacteria>Acidobacteriota>unclassified_Acidobacteriota | GCA_022614935.1 | Acidobacteriota_bacterium | MOB07_27970 | 3246..6728 | 1978231 | BREX-system-P-loop-protein-BrxC-[Acidobacteriota-bacterium] |
| MCI0393103.1 | Type1.BREX | BrxA-DNA-binding->BrxB-BrxF-iSTAND->DUF6079*-><-?\|\|HEPN->N6-METHYLASE->Schlafen+HATPase+wHTH+HTH->PglZ->SIGMA+OB-Fold+AAA-ATPase+Lon-Protease-BrxL-><-MPTase<-HTH_3 | 1220 | Bacteria>Acidobacteriota>unclassified_Acidobacteriota | GCA_022614935.1 | Acidobacteriota_bacterium | MOB07_30595 | 1910..5572 | 1978231 | BREX-system-P-loop-protein-BrxC-[Acidobacteriota-bacterium] |
| MCI0407271.1 | Type1.BREX | HNH->URI-DNase->?->REase->TPR+TPR->?->CHTH->BrxA-DNA-binding->BrxB-BrxF-iSTAND->DUF6079*->N6-METHYLASE->PglZ->SIGMA+OB-Fold+AAA-ATPase+Lon-Protease-BrxL->METHYLASE->Radical-SAM->?->?->XIS-HTH->Thermonuclease-> | 1222 | Bacteria>Acidobacteriota>unclassified_Acidobacteriota | GCA_022615035.1 | Acidobacteriota_bacterium | L0191_01695 | 12711..16379 | 1978231 | BREX-system-P-loop-protein-BrxC-[Acidobacteriota-bacterium] |
| MCI0624094.1 | Type1.BREX | <-HHH_5+RecR+TOPRIM<-?<-RFC-AAA+RuvB-AAA<-ncRNA<-PglZ<-PIN<-PhdYeFM_antitox<-REase<-DUF6079*<-BrxB-BrxF-iSTAND<-BrxA-DNA-binding\|\|?->?->?-><-AA_kinase+ACT+ACT_7<-NAD_binding_3+Homoserine_dh<-REC+NtrC-AAA+FIS-HTH<-GAF+PAS+HISKIN | 1197 | Bacteria>Acidobacteriota>unclassified_Acidobacteriota | GCA_022615935.1 | Acidobacteriota_bacterium | L0387_20995 | 11986..15579 | 1978231 | BREX-system-P-loop-protein-BrxC-[Acidobacteriota-bacterium] |
| MCI0628306.1 | Type1.BREX | TraJ-RHH->VirD4-TraG+VirD4-FtsK-><-Schlafen+PIWI<-SIGMA+OB-Fold+AAA-ATPase+Lon-Protease-BrxL<-PglZ<-?<-N6-METHYLASE-p<-DOC+HTH<-N6-METHYLASE<-DUF6079* | 1127 | Bacteria>Acidobacteriota>unclassified_Acidobacteriota | GCA_022615935.1 | Acidobacteriota_bacterium | L0387_42755 | 64096..67479 | 1978231 | BREX-system-P-loop-protein-BrxC-[Acidobacteriota-bacterium] |
| MCI0724954.1 | Type1.BREX | DUF6079-p*->N6-METHYLASE->DOC+HTH->N6-METHYLASE-p->PglZ->SIGMA+OB-Fold+AAA-ATPase+Lon-Protease-BrxL-p-> | 736 | Bacteria>Acidobacteriota>unclassified_Acidobacteriota | GCA_022616615.1 | Acidobacteriota_bacterium | L0338_39220 | 1..2211 | 1978231 | hypothetical-protein-[Acidobacteriota-bacterium] |
| PKP62156.1 | Type1.BREX | SbcC+SbcC->BrxA-DNA-binding->BrxB-BrxF-iSTAND->DUF6079*->N6-METHYLASE->PglZ->SIGMA+OB-Fold+AAA-ATPase+Lon-Protease-BrxL->HNH->STYKIN->TPR+TPR->TPR-> | 1193 | Bacteria>Atribacterota>unclassified_Atribacterota | GCA_002841055.1 | Candidatus_Atribacteria_bacterium_HGW-Atribacteria-1 | CVT91_00360 | 3425..7006 | 2013676 | BREX-system-P-loop-protein-BrxC-[Candidatus-Atribacteria-bacterium-HGW-Atribacteria-1] |
| PKP59611.1 | Type1.BREX | BrxB-BrxF-iSTAND-p->DUF6079-p->DUF6079*->N6-METHYLASE->PglZ-> | 1026 | Bacteria>Atribacterota>unclassified_Atribacterota | GCA_002841055.1 | Candidatus_Atribacteria_bacterium_HGW-Atribacteria-1 | CVT91_06440 | 727..3807 | 2013676 | BREX-system-P-loop-protein-BrxC-[Candidatus-Atribacteria-bacterium-HGW-Atribacteria-1] |
| PKP59550.1 | Type1.BREX | <-PglZ<-N6-METHYLASE<-DUF6079*<-BrxB-BrxF-iSTAND-p | 1170 | Bacteria>Atribacterota>unclassified_Atribacterota | GCA_002841055.1 | Candidatus_Atribacteria_bacterium_HGW-Atribacteria-1 | CVT91_06535 | 4911..8423 | 2013676 | BREX-system-P-loop-protein-BrxC-[Candidatus-Atribacteria-bacterium-HGW-Atribacteria-1] |
| PKP57580.1 | Type1.BREX | <-DUF6079*<-BrxB-BrxF-iSTAND<-DUF4007<-Mrr_N<-N6-METHYLASE<-?<-?\|\|rve-> | 1366 | Bacteria>Atribacterota>unclassified_Atribacterota | GCA_002841055.1 | Candidatus_Atribacteria_bacterium_HGW-Atribacteria-1 | CVT91_10260 | 255..4355 | 2013676 | hypothetical-protein-CVT91_10260-[Candidatus-Atribacteria-bacterium-HGW-Atribacteria-1] |
| MBN2413375.1 | Type1.BREX | BrxA-DNA-binding->BrxB-BrxF-iSTAND->DUF6079*->PglZ->N6-METHYLASE->SIR2->AbiEi_4->AbiEii->SIGMA+OB-Fold+AAA-ATPase+Lon-Protease-BrxL->?->HTH_3->HIN-HTH-> | 1212 | Bacteria>Bacteria_incertae_sedis>Bacteria_candidate_phyla>candidate_division_KSB1>unclassified_candidate_division_KSB1 | GCA_016934825.1 | candidate_division_KSB1_bacterium | JXQ31_16970 | 1525..5163 | 2172550 | BREX-system-P-loop-protein-BrxC-[candidate-division-KSB1-bacterium] |
| MCI0698477.1 | Type1.BREX | <-Schlafen+HATPase+wHTH<-HEPN<-DUF6079*<-BrxB-BrxF-iSTAND<-BrxA-DNA-binding<-AbiEii+ZnR<-AbiEi_4+AbiEi_1 | 1215 | Bacteria>Bacteria_incertae_sedis>Bacteria_candidate_phyla>candidate_division_KSB1>unclassified_candidate_division_KSB1 | GCA_022616415.1 | candidate_division_KSB1_bacterium | L0337_41575 | 2047..5694 | 2172550 | BREX-system-P-loop-protein-BrxC-[candidate-division-KSB1-bacterium] |
| KUK86067.1 | Type1.BREX | <-SIGMA+OB-Fold+AAA-ATPase+Lon-Protease-BrxL<-PglZ<-N6-METHYLASE<-DUF6079*<-BrxB-BrxF-iSTAND<-BrxA-DNA-binding | 1197 | Bacteria>Bacteria_incertae_sedis>Bacteria_candidate_phyla>candidate_division_TA06 | GCA_001508575.1 | candidate_division_TA06_bacterium_34_109 | XE03_1737 | 9245..12838 | 1635277 | hypothetical-protein-XE03_1737-[candidate-division-TA06-bacterium-34_109] |
| NIM12892.1 | Type1.BREX | BrxA-DNA-binding->BrxB-BrxF-iSTAND->DUF6079*->N6-METHYLASE->PglZ->SIGMA+OB-Fold+AAA-ATPase+Lon-Protease-BrxL->?-><-?<-?<-TPR+TPR<-?<-TPR | 1215 | Bacteria>Bacteria_incertae_sedis>Bacteria_candidate_phyla>Candidatus_Aminicenantes>unclassified_Aminicenantes | GCA_011771225.1 | Candidatus_Aminicenantes_bacterium | GT081_12995 | 3544..7191 | 2052149 | BREX-system-P-loop-protein-BrxC-[Candidatus-Aminicenantes-bacterium] |
| NIM13883.1 | Type1.BREX | <-SIGMA+OB-Fold+AAA-ATPase+Lon-Protease-BrxL<-PglZ<-N6-METHYLASE<-DUF6079*<-BrxB-BrxF-iSTAND<-BrxA-DNA-binding\|\|?->?-><-?<-?\|\|HTH_3->?->HTH_3-> | 1218 | Bacteria>Bacteria_incertae_sedis>Bacteria_candidate_phyla>Candidatus_Aminicenantes>unclassified_Aminicenantes | GCA_011771225.1 | Candidatus_Aminicenantes_bacterium | GT081_18040 | 9506..13162 | 2052149 | BREX-system-P-loop-protein-BrxC-[Candidatus-Aminicenantes-bacterium] |
| HDN19844.1 | Type1.BREX | BrxA-DNA-binding->BrxB-BrxF-iSTAND->DUF6079*->N6-METHYLASE->?->?->N6-METHYLASE->PglZ->SIGMA+OB-Fold+AAA-ATPase+Lon-Protease-BrxL->?->HTH_36-> | 1231 | Bacteria>Bacteria_incertae_sedis>Bacteria_candidate_phyla>Candidatus_Bipolaricaulota>unclassified_Candidatus_Bipolaricaulota | GCA_011042375.1 | Candidatus_Acetothermia_bacterium | ENF46_01950 | 2856..6551 | 2053493 | BREX-system-P-loop-protein-BrxC-[Candidatus-Acetothermia-bacterium] |
| MCD5408800.1 | Type1.BREX | BrxB-BrxF-iSTAND->DUF6079-p*->DUF6079-p->N6-METHYLASE->PglZ->SIGMA+OB-Fold+AAA-ATPase+Lon-Protease-BrxL->?->?-><-?\|\|FURR-HTH-> | 174 | Bacteria>Bacteria_incertae_sedis>Bacteria_candidate_phyla>Candidatus_Bipolaricaulota>unclassified_Candidatus_Bipolaricaulota | GCA_021129895.1 | Candidatus_Bipolaricaulota_bacterium | LR090_06240 | 1440..1964 | 2306012 | hypothetical-protein-[Candidatus-Bipolaricaulota-bacterium] |
| MCD5408801.1 | Type1.BREX | BrxB-BrxF-iSTAND->DUF6079-p->DUF6079-p*->N6-METHYLASE->PglZ->SIGMA+OB-Fold+AAA-ATPase+Lon-Protease-BrxL->?->?-><-?\|\|FURR-HTH->?->ABC-ATPase+SbcC-> | 995 | Bacteria>Bacteria_incertae_sedis>Bacteria_candidate_phyla>Candidatus_Bipolaricaulota>unclassified_Candidatus_Bipolaricaulota | GCA_021129895.1 | Candidatus_Bipolaricaulota_bacterium | LR090_06245 | 1904..4891 | 2306012 | BREX-system-P-loop-protein-BrxC-[Candidatus-Bipolaricaulota-bacterium] |
| MCC6649984.1 | Type1.BREX | <-BrxD-AAA<-PglZ<-Schlafen+HATPase+wHTH<-RNase+PHP\|\|?->?-><-N6-METHYLASE<-?<-DUF6079-p* | 705 | Bacteria>Bacteria_incertae_sedis>Bacteria_candidate_phyla>Candidatus_Eisenbacteria | GCA_020849095.1 | Candidatus_Eisenbacteria_bacterium | IT348_02410 | 16201..18318 | 2212470 | hypothetical-protein-[Candidatus-Eisenbacteria-bacterium] |
| MCK4303758.1 | Type1.BREX | BrxA-DNA-binding->BrxB-BrxF-iSTAND->DUF6079*->N6-METHYLASE->Schlafen+HATPase+wHTH-><-FGS<-?\|\|?-><-DDE_Tnp | 1190 | Bacteria>Bacteria_incertae_sedis>Bacteria_candidate_phyla>Candidatus_Eisenbacteria>unclassified_Candidatus_Eisenbacteria | GCA_023128935.1 | Candidatus_Eisenbacteria_sp. | KAY24_05935 | 1288..4860 | 2814261 | BREX-system-P-loop-protein-BrxC-[Candidatus-Eisenbacteria-sp.] |
| MCK4412365.1 | Type1.BREX | <-ACET\|\|?->Radical_SAM->?-><-SIGMA+OB-Fold+AAA-ATPase+Lon-Protease-BrxL<-PglZ<-N6-METHYLASE<-HATPase+wHTH<-?<-DUF6079*<-BrxB-BrxF-iSTAND<-BrxA-DNA-binding<-?\|\|?->?-><-?<-DSBH | 1148 | Bacteria>Bacteria_incertae_sedis>Bacteria_candidate_phyla>Candidatus_Eisenbacteria>unclassified_Candidatus_Eisenbacteria | GCA_023129715.1 | Candidatus_Eisenbacteria_sp. | KAY32_02355 | 86405..89851 | 2814261 | BREX-system-P-loop-protein-BrxC-[Candidatus-Eisenbacteria-sp.] |
| OGG45667.1 | Type1.BREX | <-DOC+HTH<-PIN<-wHTH+HTH_3<-N6-METHYLASE<-DUF6079*<-BrxB-BrxF-iSTAND<-BrxA-DNA-binding | 1145 | Bacteria>Bacteria_incertae_sedis>Bacteria_candidate_phyla>Candidatus_Handelsmanbacteria | GCA_001781055.1 | Candidatus_Handelsmanbacteria_bacterium_RIFCSPLOWO2_12_FULL_64_10 | A3F84_14490 | 8296..11733 | 1817868 | hypothetical-protein-A3F84_14490-[Candidatus-Handelsmanbacteria-bacterium-RIFCSPLOWO2_12_FULL_64_10] |
| MCK9474444.1 | Type1.BREX | BrxA-DNA-binding->BrxB-BrxF-iSTAND->DUF6079*->OLD-ABC->NERD->N6-METHYLASE->PglZ->SIGMA+OB-Fold+AAA-ATPase+HKD-DNase-BrxL->?->ZnR->?->PadR-HTH-> | 1170 | Bacteria>Bacteria_incertae_sedis>Bacteria_candidate_phyla>Candidatus_Muirbacteria>Candidatus_Muirbacterium | GCA_023228995.1 | Candidatus_Muirbacterium_halophilum | MOR46_00860 | 28955..32467 | 2053465 | BREX-system-P-loop-protein-BrxC-[Candidatus-Muirbacterium-halophilum] |
| MBD3180773.1 | Type1.BREX | BrxA-DNA-binding->BrxB-BrxF-iSTAND->DUF6079-p*->DUF6079-p->N6-METHYLASE-> | 557 | Bacteria>Bacteria_incertae_sedis>Bacteria_candidate_phyla>Candidatus_Poribacteria>unclassified_Candidatus_Poribacteria | GCA_014730235.1 | Candidatus_Poribacteria_bacterium | GF312_00675 | 1397..3070 | 2026781 | BREX-system-P-loop-protein-BrxC-[Candidatus-Poribacteria-bacterium] |
| MCD6506891.1 | Type1.BREX | <-N6-METHYLASE<-?<-Schlafen+HATPase+wHTH<-N6-METHYLASE<-DUF6079*<-BrxB-BrxF-iSTAND<-BrxA-DNA-binding<-SIGMA+OB-Fold+AAA-ATPase+Lon-Protease-BrxL | 1172 | Bacteria>Bacteria_incertae_sedis>Bacteria_candidate_phyla>Candidatus_Poribacteria>unclassified_Candidatus_Poribacteria | GCA_021162805.1 | Candidatus_Poribacteria_bacterium | J7M22_09745 | 5721..9239 | 2026781 | BREX-system-P-loop-protein-BrxC-[Candidatus-Poribacteria-bacterium] |
| PKL50472.1 | Type1.BREX | <-MPTase<-HTH_19<-?\|\|BrxA-DNA-binding->BrxB-BrxF-iSTAND->DUF6079*->Schlafen+HATPase+wHTH+HTH->OLD-ABC->REase->N6-METHYLASE->PglZ->SIGMA+OB-Fold+AAA-ATPase+HKD-DNase-BrxL-> | 1185 | Bacteria>Bacteria_incertae_sedis>Bacteria_candidate_phyla>Candidatus_Riflebacteria>unclassified_Candidatus_Riflebacteria | GCA_002839435.1 | Candidatus_Riflebacteria_bacterium_HGW-Riflebacteria-2 | CVV42_02960 | 62937..66494 | 2013830 | MAG:-BREX-system-P-loop-protein-BrxC-[Candidatus-Riflebacteria-bacterium-HGW-Riflebacteria-2] |
| MBF0500206.1 | Type1.BREX | IG+IG-><-ANK<-?<-ANK+ANK+ANK<-?\|\|EVE->BrxA-DNA-binding->BrxB-BrxF-iSTAND->DUF6079*->N6-METHYLASE->OLD-ABC->TOPRIM->N6-METHYLASE->PglZ->SIGMA+OB-Fold+AAA-ATPase+Lon-Protease-BrxL->DDE_Tnp_IS66-><-SbcC | 1142 | Bacteria>Bacteria_incertae_sedis>Bacteria_candidate_phyla>Candidatus_Riflebacteria>unclassified_Candidatus_Riflebacteria | GCA_015233155.1 | Candidatus_Riflebacteria_bacterium | HQM09_08735 | 64284..67712 | 2053586 | BREX-system-P-loop-protein-BrxC-[Candidatus-Riflebacteria-bacterium] |
| OGL14724.1 | Type1.BREX | REase-><-?<-ArdCN+MPTase<-SIGMA+OB-Fold+AAA-ATPase+Lon-Protease-BrxL<-PglZ<-Schlafen+HATPase+wHTH<-RNase+PHP<-N6-METHYLASE<-DUF6079*<-BrxB-BrxF-iSTAND<-BrxA-DNA-binding | 1145 | Bacteria>Bacteria_incertae_sedis>Bacteria_candidate_phyla>Candidatus_Rokubacteria>unclassified_Candidatus_Rokubacteria | GCA_001790335.1 | Candidatus_Rokubacteria_bacterium_RIFCSPLOWO2_12_FULL_71_22 | A3F92_03770 | 25373..28810 | 1802110 | MAG:-hypothetical-protein-A3F92_03770-[Candidatus-Rokubacteria-bacterium-RIFCSPLOWO2_12_FULL_71_22] |
| OIP86292.1 | Type1.BREX | SLATT-><-HerA/FtsK\|\|ZnR+HARE-HTH->ABhydrolase-><-?\|\|BrxA-DNA-binding->BrxB-BrxF-iSTAND->DUF6079*->N6-METHYLASE->PglZ->SIGMA+OB-Fold+AAA-ATPase+Lon-Protease-BrxL->?->?-><-HerA/FtsK | 1102 | Bacteria>Bacteria_incertae_sedis>Bacteria_candidate_phyla>Candidatus_Saccharibacteria>unclassified_Candidatus_Saccharibacteria | GCA_001873625.1 | Candidatus_Saccharibacteria_bacterium_CG2_30_41_52 | AUK57_00775 | 137723..141031 | 1805375 | hypothetical-protein-AUK57_00775-[Candidatus-Saccharibacteria-bacterium-CG2_30_41_52] |
| NCC19592.1 | Type1.BREX | <-SIGMA+OB-Fold+AAA-ATPase+Lon-Protease-BrxL<-PglZ<-N6-METHYLASE<-DUF6079*<-BrxB-BrxF-iSTAND<-BrxA-DNA-binding<-?<-ZnR+HARE-HTH<-HerA/FtsK | 1102 | Bacteria>Bacteria_incertae_sedis>Bacteria_candidate_phyla>Candidatus_Saccharibacteria>unclassified_Candidatus_Saccharibacteria | GCA_009929655.1 | Candidatus_Saccharibacteria_bacterium | EOM33_00865 | 8194..11502 | 2026720 | BREX-system-P-loop-protein-BrxC-[Candidatus-Saccharibacteria-bacterium] |
| MBQ3440720.1 | Type1.BREX | <-Glycos_transf_1+Glyco_trans_1_2<-?<-?\|\|?-><-?<-?<-?\|\|BrxA-DNA-binding->BrxB-BrxF-iSTAND->DUF6079*->N6-METHYLASE->PglZ->SIGMA+OB-Fold+AAA-ATPase+Lon-Protease-BrxL->?->ZnR-> | 1108 | Bacteria>Bacteria_incertae_sedis>Bacteria_candidate_phyla>Candidatus_Saccharibacteria>unclassified_Candidatus_Saccharibacteria | GCA_017406285.1 | Candidatus_Saccharibacteria_bacterium | IJG27_00165 | 27802..31128 | 2026720 | BREX-system-P-loop-protein-BrxC-[Candidatus-Saccharibacteria-bacterium] |
| MBO7131877.1 | Type1.BREX | ZnR->?->?-><-?\|\|DOC+HTH-><-SIGMA+OB-Fold+AAA-ATPase+Lon-Protease-BrxL<-PglZ<-N6-METHYLASE<-DUF6079*<-BrxB-BrxF-iSTAND<-BrxA-DNA-binding<-ParB+HNH<-?\|\|HD-> | 1109 | Bacteria>Bacteria_incertae_sedis>Bacteria_candidate_phyla>Candidatus_Saccharibacteria>unclassified_Candidatus_Saccharibacteria | GCA_017645955.1 | Candidatus_Saccharibacteria_bacterium | J6V85_01240 | 41463..44792 | 2026720 | BREX-system-P-loop-protein-BrxC-[Candidatus-Saccharibacteria-bacterium] |
| MCA9347639.1 | Type1.BREX | <-ACET<-DUF6079*<-BrxB-BrxF-iSTAND<-BrxA-DNA-binding<-?<-ZnR+HARE-HTH<-ABhydrolase\|\|ABC-ATPase+SbcC+ABC-ATPase+SbcC->AAA-ATPase-> | 1102 | Bacteria>Bacteria_incertae_sedis>Bacteria_candidate_phyla>Candidatus_Saccharibacteria>unclassified_Candidatus_Saccharibacteria | GCA_020428535.1 | Candidatus_Saccharibacteria_bacterium | KC930_03630 | 351..3659 | 2026720 | BREX-system-P-loop-protein-BrxC-[Candidatus-Saccharibacteria-bacterium] |
| MBI2566519.1 | Type1.BREX | NtrC-AAA+FIS-HTH->?->?-><-?<-?\|\|BrxA-DNA-binding->BrxB-BrxF-iSTAND->DUF6079*->N6-METHYLASE->HEPN->ParA-Soj-PloopNTPase->?->PglZ->HNH-><-?<-NtrC-AAA+FIS-HTH | 1109 | Bacteria>Bacteria_incertae_sedis>Bacteria_candidate_phyla>Candidatus_Schekmanbacteria | GCA_016188575.1 | Candidatus_Schekmanbacteria_bacterium | HYV63_05760 | 102186..105515 | 2419984 | BREX-system-P-loop-protein-BrxC-[Candidatus-Schekmanbacteria-bacterium] |
| MBI2568435.1 | Type1.BREX | RADICAL-SAM->METHYLASE->RADICAL-SAM->RADICAL-SAM->Glycos_transf_1+Glyco_trans_1_2-><-REase\|\|?->BrxA-DNA-binding->BrxB-BrxF-iSTAND->DUF6079*->REase->N6-METHYLASE->DOC+HTH->N6-METHYLASE-p->?->?->PglZ->SIGMA+OB-Fold+AAA-ATPase+Lon-Protease-BrxL-> | 1243 | Bacteria>Bacteria_incertae_sedis>Bacteria_candidate_phyla>Candidatus_Schekmanbacteria | GCA_016188575.1 | Candidatus_Schekmanbacteria_bacterium | HYV63_15550 | 53896..57627 | 2419984 | BREX-system-P-loop-protein-BrxC-[Candidatus-Schekmanbacteria-bacterium] |
| PKK90022.1 | Type1.BREX | <-Amidohydro_1<-?<-?\|\|BrxA-DNA-binding->BrxB-BrxF-iSTAND->DUF6079*->HEPN->N6-METHYLASE-p->?->HEPN-> | 1176 | Bacteria>Bacteria_incertae_sedis>Bacteria_candidate_phyla>Candidatus_Wallbacteria | GCA_002838975.1 | Candidatus_Wallbacteria_bacterium_HGW-Wallbacteria-1 | CVV64_11925 | 128140..131670 | 2013854 | BREX-system-P-loop-protein-BrxC-[Candidatus-Wallbacteria-bacterium-HGW-Wallbacteria-1] |
| MCB9789984.1 | Type1.BREX | MarR-HTH->?-><-?\|\|ArdCN+MPTase->?->?->HTH_3->BrxA-DNA-binding->BrxB-BrxF-iSTAND->DUF6079-p*->?->?->N6-METHYLASE-p->N6-METHYLASE->N6-METHYLASE-p->?->PglZ->SIGMA+OB-Fold+AAA-ATPase+Lon-Protease-BrxL->REase-> | 654 | Bacteria>Bacteria_incertae_sedis>Bacteria_candidate_phyla>Patescibacteria_group>Parcubacteria_group>Candidatus_Nomurabacteria | GCA_020632435.1 | Candidatus_Nomurabacteria_bacterium | H6762_03295 | 728705..730669 | 2052152 | BREX-system-P-loop-protein-BrxC-[Candidatus-Nomurabacteria-bacterium] |
| USN96045.1 | Type1.BREX | HNH-><-?\|\|?-><-?<-?\|\|BrxA-DNA-binding->BrxB-BrxF-iSTAND->DUF6079*->N6-METHYLASE->PglZ->SIGMA+OB-Fold+AAA-ATPase+Lon-Protease-BrxL->SbcC->YTCD-HTH-><-?\|\|?->VirB6-> | 1102 | Bacteria>Bacteria_incertae_sedis>Bacteria_candidate_phyla>Patescibacteria_group>Parcubacteria_group>Candidatus_Nomurabacteria | GCA_023898645.1 | Candidatus_Nomurabacteria_bacterium | H6797_03115 | 608928..612236 | 2052152 | MAG:-BREX-system-P-loop-protein-BrxC-[Candidatus-Nomurabacteria-bacterium] |
| MCB9093205.1 | Type1.BREX | <-NtrC-AAA+FIS-HTH\|\|?->S1COLD->N6-METHYLASE->DUF4007->LON-AAA->DUF6079*->PglZ->SIGMA+OB-Fold+AAA-ATPase+Lon-Protease-BrxL-><-REase<-ParB+HNH<-ParB+HNH | 1375 | Bacteria>Bdellovibrionota>Bacteriovoracia>Bacteriovoracales>Halobacteriovoraceae>unclassified_Halobacteriovoraceae | GCA_020635155.1 | Halobacteriovoraceae_bacterium | H6620_11660 | 12013..16140 | 2026745 | chitobiase/beta-hexosaminidase-C-terminal-domain-containing-protein-[Halobacteriovoraceae-bacterium] |
| NUM59087.1 | Type1.BREX | Phage_integrase->Phage_integrase->Phage_int_SAM_1+Phage_integrase->?-><-?<-SbcC+SbcC<-SIGMA+OB-Fold+AAA-ATPase+HKD-DNase-BrxL<-PglZ<-N6-METHYLASE<-DUF6079*<-BrxB-BrxF-iSTAND<-BrxA-DNA-binding<-XIS-HTH<-?<-?<-ABhydrolase<-REC+NtrC-AAA+FIS-HTH<-HISKIN+MORC | 1157 | Bacteria>Bdellovibrionota>Bdellovibrionia>Bdellovibrionales>Pseudobdellovibrionaceae>unclassified_Pseudobdellovibrionaceae | GCA_013359135.1 | Pseudobdellovibrionaceae_bacterium | HUU56_10670 | 15517..18990 | 2026715 | BREX-system-P-loop-protein-BrxC-[Pseudobdellovibrionaceae-bacterium] |
| PIT99446.1 | Type1.BREX | <-ABhydrolase<-REC+NtrC-AAA+FIS-HTH<-HISKIN+MORC<-?<-?\|\|XIS-HTH->BrxA-DNA-binding->BrxB-BrxF-iSTAND->DUF6079*->N6-METHYLASE->PglZ->SIGMA+OB-Fold+AAA-ATPase+HKD-DNase-BrxL->SIR2-><-MuB-ATPase<-HTH_28+HTH_32+rve+rve_3 | 1164 | Bacteria>Bdellovibrionota>Bdellovibrionia>Bdellovibrionales>unclassified_Bdellovibrionales | GCA_002778785.1 | Bdellovibrionales_bacterium_CG10_big_fil_rev_8_21_14_0_10_45_34 | COT74_10625 | 394529..398023 | 1973927 | BREX-system-P-loop-protein-BrxC-[Bdellovibrionales-bacterium-CG10_big_fil_rev_8_21_14_0_10_45_34] |
| NLI42049.1 | Type1.BREX | <-YTCD-HTH<-?<-?<-SIGMA+OB-Fold+AAA-ATPase+Lon-Protease-BrxL<-PglZ<-N6-METHYLASE<-DUF6079*<-BrxB-BrxF-iSTAND<-BrxA-DNA-binding | 1102 | Bacteria>Caldisericota/Cryosericota_group>Caldisericota>Caldisericia>Caldisericales>unclassified_Caldisericales | GCA_012517765.1 | Caldisericales_bacterium | GX421_12905 | 11000..14308 | 2614251 | BREX-system-P-loop-protein-BrxC-[Caldisericales-bacterium] |
| MCK6622791.1 | Type1.BREX | <-ABhydrolase<-?<-?<-SIGMA+OB-Fold+AAA-ATPase+Lon-Protease-BrxL<-PglZ<-DOC+HTH<-N6-METHYLASE<-Y1_Tnp<-DUF6079*<-BrxB-BrxF-iSTAND<-BrxA-DNA-binding | 1161 | Bacteria>Calditrichota>Calditrichia>unclassified_Calditrichia | GCA_023151135.1 | Calditrichia_bacterium | L6Q94_20070 | 39359..42844 | 2268178 | BREX-system-P-loop-protein-BrxC-[Calditrichia-bacterium] |
| KAA3613611.1 | Type1.BREX | <-PIN\|\|?->?->BrxA-DNA-binding->BrxB-BrxF-iSTAND->DUF6079*->N6-METHYLASE->?->PglZ->SIGMA+OB-Fold+AAA-ATPase+HKD-DNase-BrxL->REase->CARF+NtrC-AAA+FIS-HTH->REase-> | 1159 | Bacteria>Calditrichota>unclassified_Calditrichota | GCA_008501795.1 | Calditrichota_bacterium | DWQ03_12295 | 106728..110207 | 2212469 | BREX-system-P-loop-protein-BrxC-[Calditrichota-bacterium] |

<table>
<tr><th>Protein-ID</th><th>System</th><th>Gene-Neighbourhoods</th><th>Length</th><th>Taxonomy</th><th>Assembly-ID</th><th>Strain</th><th>Locus-Tag</th><th>Genomic-coordinates</th><th>TaxID</th><th>Fasta-define</th></tr>
<tr><td>WP_046997142.1</td><td>Type1.BREX</td><td>wHTH+WYL+WCX->BrxA-DNA-binding->BrxB-BrxF-iSTAND->DUF6079*->N6-METHYLASE->?->HEPN->Schlafen+HATPase+wHTH->PglZ->SIGMA+OB-Fold+AAA-ATPase+Lon-Protease-BrxL-></td><td>1202</td><td>Bacteria>Campylobacterota>Epsilonproteobacteria>Campylobacterales>Arcobacteraceae>Aliarcobacter</td><td>GCF_001010585.1</td><td>Aliarcobacter_butzleri</td><td>AA20_RS09495</td><td>10373..13981</td><td>28197</td><td>BREX-system-P-loop-protein-BrxC-[Aliarcobacter-butzleri]</td></tr>
<tr><td>WP_141050577.1</td><td>Type1.BREX</td><td><-DnaB+TnsC-ATPase<-ParB<-ParA-Soj-PloopNTPase+ORC-CDC6-like<-DDE_5+DDE_Tnp_1||BirA-HTH->?->wHTH+WYL+WCX->BrxA-DNA-binding->BrxB-BrxF-iSTAND->DUF6079*->HTH+REase->N6-METHYLASE-p->?->N6-METHYLASE-p->AbiEi_4->AbiEii->Schlafen+HATPase+wHTH->REase+SF1-SF2-Helicase-></td><td>1162</td><td>Bacteria>Campylobacterota>Epsilonproteobacteria>Campylobacterales>Arcobacteraceae>Aliarcobacter</td><td>GCF_006503615.1</td><td>Aliarcobacter_cryaerophilus</td><td>AV936_RS08795</td><td>11744..15232</td><td>28198</td><td>BREX-system-P-loop-protein-BrxC-[Aliarcobacter-cryaerophilus]</td></tr>
<tr><td>WP_187475233.1</td><td>Type1.BREX</td><td>METHYLASE->Peptidase_M3->?->?->?-><-DDE_Tnp_1_4||?->XIS-HTH->BrxA-DNA-binding->BrxB-BrxF-iSTAND->DUF6079*->OLD-ABC->REase->N6-METHYLASE->DrHyd->Schlafen+HATPase+wHTH->Trypsin->?->PglZ->SIGMA+OB-Fold+AAA-ATPase+HKD-DNase-BrxL-></td><td>1166</td><td>Bacteria>Campylobacterota>Epsilonproteobacteria>Campylobacterales>Arcobacteraceae>Aliarcobacter</td><td>GCF_014352915.1</td><td>Aliarcobacter_cryaerophilus</td><td>HOO34_RS05640</td><td>1060903..1064403</td><td>28198</td><td>BREX-system-P-loop-protein-BrxC-[Aliarcobacter-cryaerophilus]</td></tr>
<tr><td>WP_260949077.1</td><td>Type1.BREX</td><td><-Couple_hipA+HipA_C<-Calcineurin<-SIGMA+OB-Fold+AAA-ATPase+Lon-Protease-BrxL<-PglZ<-Schlafen+HATPase+wHTH<-Schlafen+HATPase+wHTH<-N6-METHYLASE-p<-?<-N6-METHYLASE-p<-DUF6079*<-BrxB-BrxF-iSTAND<-BrxA-DNA-binding<-wHTH+WYL+WCX<-?<-?<-SF1-SF2-Helicase<-REase<-McrB-AAA<-HTH+HEPN</td><td>1162</td><td>Bacteria>Campylobacterota>Epsilonproteobacteria>Campylobacterales>Arcobacteraceae>Aliarcobacter</td><td>GCF_025334565.1</td><td>Aliarcobacter_cryaerophilus</td><td>N5T87_RS09120</td><td>17819..21307</td><td>28198</td><td>BREX-system-P-loop-protein-BrxC-[Aliarcobacter-cryaerophilus]</td></tr>
<tr><td>WP_260942902.1</td><td>Type1.BREX</td><td><-GDResolvase+HIN-HTH<-Calcineurin<-SIGMA+OB-Fold+AAA-ATPase+Lon-Protease-BrxL<-PglZ<-Tox-RES<-?<-?<-N6-METHYLASE<-DUF6079*<-BrxB-BrxF-iSTAND<-BrxA-DNA-binding<-wHTH+WYL+WCX<-?||ParA-Soj-PloopNTPase+ORC-CDC6-like->ParB->SF1-SF2-Helicase-></td><td>1162</td><td>Bacteria>Campylobacterota>Epsilonproteobacteria>Campylobacterales>Arcobacteraceae>Aliarcobacter</td><td>GCF_025334745.1</td><td>Aliarcobacter_cryaerophilus</td><td>N5T78_RS08220</td><td>26893..30381</td><td>28198</td><td>BREX-system-P-loop-protein-BrxC-[Aliarcobacter-cryaerophilus]</td></tr>
<tr><td>WP_260896239.1</td><td>Type1.BREX</td><td>Peptidase->Phage_integrase->?->XIS-HTH->BrxA-DNA-binding->BrxB-BrxF-iSTAND->DUF6079*->?->N6-METHYLASE->DrHyd->Schlafen+HATPase+wHTH->REase->ACET->PglZ->SIGMA+OB-Fold+AAA-ATPase+HKD-DNase-BrxL-></td><td>1166</td><td>Bacteria>Campylobacterota>Epsilonproteobacteria>Campylobacterales>Arcobacteraceae>Aliarcobacter</td><td>GCF_025335025.1</td><td>Aliarcobacter_cryaerophilus</td><td>N5T57_RS09580</td><td>13633..17133</td><td>28198</td><td>BREX-system-P-loop-protein-BrxC-[Aliarcobacter-cryaerophilus]</td></tr>
<tr><td>WP_260916976.1</td><td>Type1.BREX</td><td>SF1-SF2-Helicase-><-Calcineurin<-SIGMA+OB-Fold+AAA-ATPase+Lon-Protease-BrxL<-PglZ<-Schlafen+HATPase+wHTH<-?<-AbiEi_4<-?<-N6-METHYLASE<-DUF6079*<-BrxB-BrxF-iSTAND<-BrxA-DNA-binding<-wHTH+WYL+WCX<-?||?->?-><-?||PRTase-></td><td>1162</td><td>Bacteria>Campylobacterota>Epsilonproteobacteria>Campylobacterales>Arcobacteraceae>Aliarcobacter</td><td>GCF_025335525.1</td><td>Aliarcobacter_butzleri</td><td>N5U17_RS09665</td><td>19417..22905</td><td>28197</td><td>BREX-system-P-loop-protein-BrxC-[Aliarcobacter-butzleri]</td></tr>
<tr><td>WP_260968564.1</td><td>Type1.BREX</td><td>ABC-ATPase+SbcC->?->LuxR-HTH-><-?||Phage_integrase->?->wHTH+WYL+WCX->BrxA-DNA-binding->BrxB-BrxF-iSTAND->DUF6079*->N6-METHYLASE-p->HEPN+RES->N6-METHYLASE-p->Schlafen+HATPase+wHTH->Schlafen+HATPase+wHTH->PglZ->SIGMA+OB-Fold+AAA-ATPase+Lon-Protease-BrxL->Calcineurin-><-SIGMA-HTH</td><td>1162</td><td>Bacteria>Campylobacterota>Epsilonproteobacteria>Campylobacterales>Arcobacteraceae>Aliarcobacter</td><td>GCF_025335945.1</td><td>Aliarcobacter_cryaerophilus</td><td>N5S77_RS10385</td><td>37284..40772</td><td>28198</td><td>BREX-system-P-loop-protein-BrxC-[Aliarcobacter-cryaerophilus]</td></tr>
<tr><td>WP_260934490.1</td><td>Type1.BREX</td><td><-ZetaToxin<-AbiEi_4<-?<-N6-METHYLASE-p<-?<-N6-METHYLASE-p<-DUF6079*<-BrxB-BrxF-iSTAND<-BrxA-DNA-binding<-wHTH+WYL+WCX||RHH->RelE-ParE->HTH+rve-><-DnaB<-?<-ParB</td><td>1162</td><td>Bacteria>Campylobacterota>Epsilonproteobacteria>Campylobacterales>Arcobacteraceae>Aliarcobacter</td><td>GCF_025336285.1</td><td>Aliarcobacter_cryaerophilus</td><td>N5T58_RS09420</td><td>6714..10202</td><td>28198</td><td>BREX-system-P-loop-protein-BrxC-[Aliarcobacter-cryaerophilus]</td></tr>
<tr><td>WP_152054592.1</td><td>Type1.BREX</td><td><-ParA-Soj-PloopNTPase<-SIGMA+OB-Fold+AAA-ATPase+Lon-Protease-BrxL<-PglZ<-Schlafen+HATPase+wHTH<-N6-METHYLASE-p<-REase<-?<-?<-N6-METHYLASE-p<-DUF6079*<-BrxB-BrxF-iSTAND<-BrxA-DNA-binding<-Phage_integrase||?-><-KAP-NTPase<-DOC+HTH<-?<-HD+HD_assoc</td><td>1162</td><td>Bacteria>Campylobacterota>Epsilonproteobacteria>Campylobacterales>Arcobacteraceae>Aliarcobacter</td><td>GCF_902500815.1</td><td>Aliarcobacter_butzleri</td><td>GDI74_RS00980</td><td>25690..29178</td><td>28197</td><td>BREX-system-P-loop-protein-BrxC-[Aliarcobacter-butzleri]</td></tr>
<tr><td>RXJ95785.1</td><td>Type1.BREX</td><td>wHTH+WYL+WCX->?->?->RADICAL-SAM->?->HIN-HTH->BrxA-DNA-binding->BrxB-BrxF-iSTAND->DUF6079*->N6-METHYLASE-p->?->?->HTH+REase->N6-METHYLASE-p->AbiEi_4->AbiEii->PglZ->SIGMA+OB-Fold+AAA-ATPase+Lon-Protease-BrxL-></td><td>1162</td><td>Bacteria>Campylobacterota>Epsilonproteobacteria>Campylobacterales>Arcobacteraceae>Arcobacter</td><td>GCA_004116575.1</td><td>Arcobacter_sp._CECT_9299</td><td>CRU94_03995</td><td>49858..53346</td><td>2044506</td><td>hypothetical-protein-CRU94_03995-[Arcobacter-sp.-CECT-9299]</td></tr>
<tr><td>WP_129105531.1</td><td>Type1.BREX</td><td>wHTH+WYL+WCX->?->?->RADICAL-SAM->?->BrxA-DNA-binding->BrxB-BrxF-iSTAND->DUF6079*->N6-METHYLASE-p->?->?->HTH+REase->N6-METHYLASE-p->AbiEi_4->AbiEii->PglZ->SIGMA+OB-Fold+AAA-ATPase+Lon-Protease-BrxL-></td><td>1162</td><td>Bacteria>Campylobacterota>Epsilonproteobacteria>Campylobacterales>Arcobacteraceae>Arcobacter</td><td>GCF_004116575.1</td><td>Arcobacter_sp._CECT_9299</td><td>CRU94_RS03995</td><td>49858..53346</td><td>2044506</td><td>BREX-system-P-loop-protein-BrxC-[Arcobacter-sp.-CECT-9299]</td></tr>
<tr><td>WP_164968523.1</td><td>Type1.BREX</td><td>BrxA-DNA-binding->BrxB-BrxF-iSTAND->DUF6079*->N6-METHYLASE-p->?->?->HTH+REase->N6-METHYLASE-p->AbiEi_4->AbiEii->PglZ->SIGMA+OB-Fold+AAA-ATPase+Lon-Protease-BrxL-></td><td>1160</td><td>Bacteria>Campylobacterota>Epsilonproteobacteria>Campylobacterales>Arcobacteraceae>Arcobacter</td><td>GCF_013201725.1</td><td>Arcobacter_defluvii</td><td>ADFLV_RS07175</td><td>1422397..1425879</td><td>873191</td><td>BREX-system-P-loop-protein-BrxC-[Arcobacter-defluvii]</td></tr>
<tr><td>MCD8478249.1</td><td>Type1.BREX</td><td>PIN->PhdYeFM_antitox->?->PhdYeFM_antitox->?->BrxA-DNA-binding->BrxB-BrxF-iSTAND->DUF6079-p*->?->?->?->?->?->N6-METHYLASE-p->N6-METHYLASE-p->N6-METHYLASE-p-></td><td>544</td><td>Bacteria>Campylobacterota>Epsilonproteobacteria>Campylobacterales>Sulfurospirillaceae>Sulfurospirillum</td><td>GCA_021206415.1</td><td>Sulfurospirillum_sp.</td><td>LRY68_10520</td><td>4304..5938</td><td>2053622</td><td>BREX-system-P-loop-protein-BrxC-[Sulfurospirillum-sp.]</td></tr>
<tr><td>OZA03558.1</td><td>Type1.BREX</td><td><-SAM-METHYLASE||?-><-TPR||?->BrxB-BrxF-iSTAND->ACET->XIS-HTH->BrxA-DNA-binding->BrxB-BrxF-iSTAND->DUF6079*->HEPN->N6-METHYLASE->?->ZnR+HEPN->?->PglZ->SIGMA+OB-Fold+AAA-ATPase+Lon-Protease-BrxL-></td><td>1158</td><td>Bacteria>Campylobacterota>Epsilonproteobacteria>Campylobacterales>Thiovulaceae>Sulfuricurvum</td><td>GCA_002282015.1</td><td>Sulfuricurvum_sp._17-40-25</td><td>B7X89_02520</td><td>35980..39456</td><td>1970439</td><td>MAG:-hypothetical-protein-B7X89_02520-[Sulfuricurvum-sp.-17-40-25]</td></tr>
<tr><td>WP_013460546.1</td><td>Type1.BREX</td><td>Phage_integrase-><-XIS-HTH<-HTH_36<-?||HTH_3-><-XIS-HTH||BrxA-DNA-binding->BrxB-BrxF-iSTAND->DUF6079*-><-?||N6-METHYLASE-p->?->?->HEPN->BrxD->N6-METHYLASE-p->CR-ATPase5+REase->PglZ-></td><td>1163</td><td>Bacteria>Campylobacterota>Epsilonproteobacteria>Campylobacterales>Thiovulaceae>Sulfuricurvum</td><td>GCF_000183725.1</td><td>Sulfuricurvum_kujiense</td><td>SULKU_RS08475</td><td>1692937..1696428</td><td>148813</td><td>BREX-system-P-loop-protein-BrxC-[Sulfuricurvum-kujiense]</td></tr>
<tr><td>PHS59105.1</td><td>Type1.BREX</td><td><-Methyltransf_30<-?<-?<-?<-?<-?||?->BrxA-DNA-binding->BrxB-BrxF-iSTAND->DUF6079*->PemK_toxin->N6-METHYLASE-p->?->?->N6-METHYLASE-p->?->HTH+REase->PglZ->SIGMA+OB-Fold+AAA-ATPase+Lon-Protease-BrxL-></td><td>1158</td><td>Bacteria>Campylobacterota>Epsilonproteobacteria>Campylobacterales>Thiovulaceae>Sulfurimonas</td><td>GCA_002733155.1</td><td>Sulfurimonas_sp.</td><td>COB17_00805</td><td>100943..104419</td><td>2022749</td><td>MAG:-hypothetical-protein-COB17_00805-[Sulfurimonas-sp.]</td></tr>
<tr><td>QFR43750.1</td><td>Type1.BREX</td><td>Glycos_transf_4->?->?->tRNA->tRNA->XIS-HTH->BrxA-DNA-binding->BrxB-BrxF-iSTAND->DUF6079*->HEPN->N6-METHYLASE-p->?->DUF3644->?->N6-METHYLASE-p->?->PglZ->SIGMA+OB-Fold+AAA-ATPase+Lon-Protease-BrxL-></td><td>1163</td><td>Bacteria>Campylobacterota>Epsilonproteobacteria>Campylobacterales>Thiovulaceae>Sulfurimonas</td><td>GCA_009258045.1</td><td>Sulfurimonas_xiamenensis</td><td>FJR47_07440</td><td>1472966..1476457</td><td>2590021</td><td>BREX-system-P-loop-protein-BrxC-[Sulfurimonas-xiamenensis]</td></tr>
<tr><td>MBT5934549.1</td><td>Type1.BREX</td><td>Arm-DNA-bind_3+Phage_integrase->?->XIS-HTH->?->?->ZnR+TOPRIM+AAA-helicase+wHTH->BrxA-DNA-binding->BrxB-BrxF-iSTAND->DUF6079-p*-></td><td>988</td><td>Bacteria>Campylobacterota>Epsilonproteobacteria>Campylobacterales>Thiovulaceae>Sulfurimonas</td><td>GCA_018698455.1</td><td>Sulfurimonas_sp.</td><td>HOG88_04280</td><td>23331..26296</td><td>2022749</td><td>BREX-system-P-loop-protein-BrxC-[Sulfurimonas-sp.]</td></tr>
<tr><td>WP_188093712.1</td><td>Type1.BREX</td><td>Glycos_transf_4->?->?->tRNA->tRNA->XIS-HTH->BrxA-DNA-binding->BrxB-BrxF-iSTAND->DUF6079*->HEPN->N6-METHYLASE-p->?->DUF3644->?->N6-METHYLASE-p->?->PglZ->SIGMA+OB-Fold+AAA-ATPase+Lon-Protease-BrxL-></td><td>1160</td><td>Bacteria>Campylobacterota>Epsilonproteobacteria>Campylobacterales>Thiovulaceae>Sulfurimonas</td><td>GCF_009258045.1</td><td>Sulfurimonas_xiamenensis</td><td>FJR47_RS07440</td><td>1472975..1476457</td><td>2590021</td><td>BREX-system-P-loop-protein-BrxC-[Sulfurimonas-xiamenensis]</td></tr>
<tr><td>WP_193150356.1</td><td>Type1.BREX</td><td><-Phage_integrase<-SIGMA+OB-Fold+AAA-ATPase+Lon-Protease-BrxL<-PglZ<-?<-SKI<-N6-METHYLASE-p<-RVT<-N6-METHYLASE-p<-HEPN<-DUF6079*<-BrxB-BrxF-iSTAND<-BrxA-DNA-binding<-DDE_Tnp<-?<-DDE_Tnp</td><td>1160</td><td>Bacteria>Campylobacterota>Epsilonproteobacteria>Campylobacterales>Thiovulaceae>Sulfurimonas</td><td>GCF_014905115.1</td><td>Sulfurimonas_sediminis</td><td>FJR45_RS09735</td><td>1812721..1816203</td><td>2590020</td><td>BREX-system-P-loop-protein-BrxC-[Sulfurimonas-sediminis]</td></tr>
<tr><td>WP_226069221.1</td><td>Type1.BREX</td><td>Phage_integrase->HTH+rve->AAA-ATPase->?->?->?->?->BrxA-DNA-binding->BrxB-BrxF-iSTAND->DUF6079*->N6-METHYLASE->OLD-ABC+TOPRIM->SF1-SF2-Helicase->?->Schlafen+HATPase+wHTH->PglZ->SIGMA+OB-Fold+AAA-ATPase+Lon-Protease-BrxL->?->Phage_integrase-></td><td>1201</td><td>Bacteria>Campylobacterota>Epsilonproteobacteria>Campylobacterales>Thiovulaceae>Sulfurimonas</td><td>GCF_020410885.1</td><td>Sulfurimonas_sp._SWIR-19</td><td>LCX93_RS05490</td><td>1055837..1059442</td><td>2878390</td><td>BREX-system-P-loop-protein-BrxC-[Sulfurimonas-sp.-SWIR-19]</td></tr>
<tr><td>OYZ34277.1</td><td>Type1.BREX</td><td><-SIGMA+OB-Fold+AAA-ATPase+Lon-Protease-BrxL<-PglZ<-?<-ZnR+HEPN<-?<-N6-METHYLASE<-HEPN<-DUF6079*<-DUF6079-p<-BrxB-BrxF-iSTAND<-BrxA-DNA-binding<-XIS-HTH<-ACET<-BrxB-BrxF-iSTAND<-?||TPR-></td><td>1013</td><td>Bacteria>Campylobacterota>Epsilonproteobacteria>Campylobacterales>unclassified_Campylobacterales</td><td>GCA_002281685.1</td><td>Campylobacterales_bacterium_16-40-21</td><td>B7Y30_07055</td><td>21520..24561</td><td>1970501</td><td>hypothetical-protein-B7Y30_07055-[Campylobacterales-bacterium-16-40-21]</td></tr>
<tr><td>MBD3789167.1</td><td>Type1.BREX</td><td>ABC-ATPase+SbcC->?-><-GGDEF+EAL||?-><-?<-?||XIS-HTH->BrxA-DNA-binding->BrxB-BrxF-iSTAND->DUF6079-p*-></td><td>671</td><td>Bacteria>Campylobacterota>Epsilonproteobacteria>Campylobacterales>unclassified_Campylobacterales</td><td>GCA_014763895.1</td><td>Campylobacterales_bacterium</td><td>IE885_02155</td><td>92297..94311</td><td>2268180</td><td>BREX-system-P-loop-protein-BrxC-[Campylobacterales-bacterium]</td></tr>
<tr><td>OQX77395.1</td><td>Type1.BREX</td><td><-Phage_integrase<-?||BrxA-DNA-binding->BrxB-BrxF-iSTAND->DUF6079*->AbiEi_4-></td><td>1159</td><td>Bacteria>Campylobacterota>unclassified_Campylobacterota</td><td>GCA_002085305.1</td><td>Epsilonproteobacteria_bacterium_4484_65</td><td>B6D54_02040</td><td>4837..8316</td><td>1970777</td><td>hypothetical-protein-B6D54_02040-[Epsilonproteobacteria-bacterium-4484_65]</td></tr>
<tr><td>WP_013452128.1</td><td>Type1.BREX</td><td>CR-ATPase8+REase-><-?<-SIGMA+OB-Fold+AAA-ATPase+Lon-Protease-BrxL<-PglZ<-N6-METHYLASE<-HEPN<-N6-METHYLASE-p<-DUF6079*<-BrxB-BrxF-iSTAND<-BrxA-DNA-binding||CR-ATPase8+REase-></td><td>1161</td><td>Bacteria>Deferribacterota>Deferribacteres>Deferribacterales>Calditerrivibrionaceae>Calditerrivibrio</td><td>GCF_000183405.1</td><td>Calditerrivibrio_nitroreducens</td><td>CALNI_RS10260</td><td>2104217..2107702</td><td>477976</td><td>BREX-system-P-loop-protein-BrxC-[Calditerrivibrio-nitroreducens]</td></tr>
<tr><td>MBZ4644075.1</td><td>Type1.BREX</td><td><-TPR<-ARG<-?||McrB-AAA->McrC-NTD+REase->BrxA-DNA-binding->BrxB-BrxF-iSTAND->DUF6079*->N6-METHYLASE-p->HEPN->N6-METHYLASE->PglZ->SIGMA+OB-Fold+AAA-ATPase+Lon-Protease-BrxL->?->SF1-SF2-Helicase-></td><td>1168</td><td>Bacteria>Deferribacterota>Deferribacteres>Deferribacterales>Deferribacteraceae>unclassified_Deferribacteraceae</td><td>GCA_020054865.1</td><td>Deferribacteraceae_bacterium</td><td>JG767_1684</td><td>7812..11318</td><td>2053543</td><td>hypothetical-protein-[Deferribacteraceae-bacterium]</td></tr>
<tr><td>WP_242820841.1</td><td>Type1.BREX</td><td>NUDIX-><-TPR+S1<-ARG<-?<-?||McrB-AAA->McrC-NTD+REase->BrxA-DNA-binding->BrxB-BrxF-iSTAND->DUF6079*->N6-METHYLASE-p->HEPN->N6-METHYLASE->PglZ->SIGMA+OB-Fold+AAA-ATPase+Lon-Protease-BrxL->?->SF1-SF2-Helicase-></td><td>1168</td><td>Bacteria>Deferribacterota>Deferribacteres>Deferribacterales>Deferrivibrionaceae>Deferrivibrio</td><td>GCA_020480685.1</td><td>Deferrivibrio_essentukiensis</td><td>LF845_RS09815</td><td>29861..33367</td><td>2880922</td><td>BREX-system-P-loop-protein-BrxC-[Deferrivibrio-essentukiensis]</td></tr>
<tr><td>MBF0239771.1</td><td>Type1.BREX</td><td><-SIGMA+OB-Fold+AAA-ATPase+Lon-Protease-BrxL-p<-REase<-PglZ<-?<-?||?-><-N6-METHYLASE-p<-REase<-DUF6079*<-BrxB-BrxF-iSTAND<-BrxA-DNA-binding</td><td>1178</td><td>Bacteria>delta/epsilon_subdivisions>Deltaproteobacteria>Deltaproteobacteria_incertae_sedis>SAR324_cluster</td><td>GCA_015231535.1</td><td>SAR324_cluster_bacterium</td><td>HQM12_18890</td><td>8068..11604</td><td>2024889</td><td>BREX-system-P-loop-protein-BrxC-[SAR324-cluster-bacterium]</td></tr>
<tr><td>EFK06105.1</td><td>Type1.BREX</td><td><-Acyltransferase<-?<-?<-SIGMA+OB-Fold+AAA-ATPase+Lon-Protease-BrxL<-PglZ<-TOPRIM<-OLD-ABC<-N6-METHYLASE<-HEPN<-DUF6079*<-BrxB-BrxF-iSTAND-p||?-><-BrxA-DNA-binding||?->?->?-><-DnaJ</td><td>1215</td><td>Bacteria>delta/epsilon_subdivisions>Deltaproteobacteria>unclassified_Deltaproteobacteria</td><td>GCA_000179315.1</td><td>delta_proteobacterium_NaphS2</td><td>NPH_2628</td><td>46790..50437</td><td>88274</td><td>conserved-hypothetical-protein-[delta-proteobacterium-NaphS2]</td></tr>
<tr><td>OGR04765.1</td><td>Type1.BREX</td><td><-LexA-protease||BrxA-DNA-binding->BrxB-BrxF-iSTAND->DUF6079-p*->PglZ->SIGMA+OB-Fold+AAA-ATPase+Lon-Protease-BrxL-></td><td>983</td><td>Bacteria>delta/epsilon_subdivisions>Deltaproteobacteria>unclassified_Deltaproteobacteria</td><td>GCA_001799225.1</td><td>Deltaproteobacteria_bacterium_RIFOXYD12_FULL_50_9</td><td>A2511_13070</td><td>2145..5095</td><td>1797902</td><td>ATPase,-partial-[Deltaproteobacteria-bacterium-RIFOXYD12_FULL_50_9]</td></tr>
<tr><td>MAE16011.1</td><td>Type1.BREX</td><td><-N6-METHYLASE-p<-DUF6079*<-BrxB-BrxF-iSTAND<-BrxA-DNA-binding<-?<-?<-RNasePH</td><td>1175</td><td>Bacteria>delta/epsilon_subdivisions>Deltaproteobacteria>unclassified_Deltaproteobacteria</td><td>GCA_002685875.1</td><td>Deltaproteobacteria_bacterium</td><td>CL911_01040</td><td>1115..4642</td><td>2026735</td><td>hypothetical-protein-[Deltaproteobacteria-bacterium]</td></tr>
<tr><td>PIV21173.1</td><td>Type1.BREX</td><td><-SIGMA+OB-Fold+AAA-ATPase+Lon-Protease-BrxL<-PglZ<-Schlafen+HATPase+wHTH+HTH<-N6-METHYLASE<-DUF6079*<-BrxB-BrxF-iSTAND<-BrxA-DNA-binding<-SIR2<-?<-DOC+HTH</td><td>1215</td><td>Bacteria>delta/epsilon_subdivisions>Deltaproteobacteria>unclassified_Deltaproteobacteria</td><td>GCA_002780715.1</td><td>Deltaproteobacteria_bacterium_CG03_land_8_20_14_0_80_45_14</td><td>COS40_08745</td><td>11135..14782</td><td>1973969</td><td>BREX-system-P-loop-protein-BrxC-[Deltaproteobacteria-bacterium-CG03_land_8_20_14_0_80_45_14]</td></tr>
<tr><td>PKN56508.1</td><td>Type1.BREX</td><td>BrxA-DNA-binding->BrxB-BrxF-iSTAND->DUF6079*->N6-METHYLASE-><-?||?-><-?<-HTH_3<-?<-?<-SopB+HNH</td><td>1206</td><td>Bacteria>delta/epsilon_subdivisions>Deltaproteobacteria>unclassified_Deltaproteobacteria</td><td>GCA_002841945.1</td><td>Deltaproteobacteria_bacterium_HGW-Deltaproteobacteria-14</td><td>CVU56_15695</td><td>1564..5184</td><td>2013741</td><td>BREX-system-P-loop-protein-BrxC-[Deltaproteobacteria-bacterium-HGW-Deltaproteobacteria-14]</td></tr>
<tr><td>PXF57197.1</td><td>Type1.BREX</td><td><-PemK_toxin<-MetJarc-HTH<-PglZ<-TOPRIM<-OLD-ABC<-N6-METHYLASE<-PIN<-Phd-YefM<-DUF6079*<-BrxB-BrxF-iSTAND<-BrxA-DNA-binding<-PIN<-?||?->?->?->?->His_biosynth-></td><td>1215</td><td>Bacteria>delta/epsilon_subdivisions>Deltaproteobacteria>unclassified_Deltaproteobacteria</td><td>GCA_003194495.1</td><td>Deltaproteobacteria_bacterium</td><td>C4B58_10725</td><td>19842..23489</td><td>2026735</td><td>BREX-system-P-loop-protein-BrxC-[Deltaproteobacteria-bacterium]</td></tr>
<tr><td>HBL23376.1</td><td>Type1.BREX</td><td><-DUF6079-p*<-BrxB-BrxF-iSTAND<-BrxA-DNA-binding<-XIS-HTH||Four-helical-protein-></td><td>888</td><td>Bacteria>delta/epsilon_subdivisions>Deltaproteobacteria>unclassified_Deltaproteobacteria</td><td>GCA_003506795.1</td><td>Deltaproteobacteria_bacterium</td><td>DDZ40_04590</td><td>1..2666</td><td>2026735</td><td>BREX-system-P-loop-protein-BrxC-[Deltaproteobacteria-bacterium]</td></tr>
<tr><td>RLC18759.1</td><td>Type1.BREX</td><td>BrxA-DNA-binding->BrxB-BrxF-iSTAND->DUF6079*->?->PemK_toxin->N6-METHYLASE-p-></td><td>1189</td><td>Bacteria>delta/epsilon_subdivisions>Deltaproteobacteria>unclassified_Deltaproteobacteria</td><td>GCA_003647425.1</td><td>Deltaproteobacteria_bacterium</td><td>DRI57_08000</td><td>870..4439</td><td>2026735</td><td>BREX-system-P-loop-protein-BrxC-[Deltaproteobacteria-bacterium]</td></tr>
<tr><td>TDB40153.1</td><td>Type1.BREX</td><td>Phage_integrase->SIR2-><-Thermonuclease<-?<-LexA-protease||?->BrxA-DNA-binding->BrxB-BrxF-iSTAND->DUF6079*->Schlafen+HATPase+wHTH+HTH->N6-METHYLASE->REase->PglZ->SIGMA+OB-Fold+AAA-ATPase+HKD-DNase-BrxL-><-?<-?<-35exo<-Calcineurin</td><td>1180</td><td>Bacteria>delta/epsilon_subdivisions>Deltaproteobacteria>unclassified_Deltaproteobacteria</td><td>GCA_004348915.1</td><td>Deltaproteobacteria_bacterium</td><td>D9V46_00535</td><td>112112..115654</td><td>2026735</td><td>BREX-system-P-loop-protein-BrxC-[Deltaproteobacteria-bacterium]</td></tr>
<tr><td>MBC8466717.1</td><td>Type1.BREX</td><td><-RAMA<-SIGMA+OB-Fold+AAA-ATPase+HKD-DNase-BrxL<-PglZ<-N6-METHYLASE<-REase<-OLD-ABC<-DUF6079*<-BrxB-BrxF-iSTAND<-BrxA-DNA-binding</td><td>1172</td><td>Bacteria>delta/epsilon_subdivisions>Deltaproteobacteria>unclassified_Deltaproteobacteria</td><td>GCA_014384065.1</td><td>Deltaproteobacteria_bacterium</td><td>H8D55_02650</td><td>10326..13844</td><td>2026735</td><td>BREX-system-P-loop-protein-BrxC-[Deltaproteobacteria-bacterium]</td></tr>
<tr><td>MBF0105032.1</td><td>Type1.BREX</td><td><-SIGMA+OB-Fold+AAA-ATPase+Lon-Protease-BrxL<-PglZ<-HTH+REase<-N6-METHYLASE<-DUF6079*<-BrxB-BrxF-iSTAND<-BrxA-DNA-binding</td><td>1166</td><td>Bacteria>delta/epsilon_subdivisions>Deltaproteobacteria>unclassified_Deltaproteobacteria</td><td>GCA_015228875.1</td><td>Deltaproteobacteria_bacterium</td><td>HQM16_06860</td><td>10380..13880</td><td>2026735</td><td>BREX-system-P-loop-protein-BrxC-[Deltaproteobacteria-bacterium]</td></tr>
<tr><td>MBF0105437.1</td><td>Type1.BREX</td><td><-CR-ATPase5+REase||?-><-?<-?||?->BrxA-DNA-binding->BrxB-BrxF-iSTAND->DUF6079*->N6-METHYLASE->HTH+REase->?->PglZ->SIGMA+OB-Fold+AAA-ATPase+Lon-Protease-BrxL->RADICAL-SAM-></td><td>1155</td><td>Bacteria>delta/epsilon_subdivisions>Deltaproteobacteria>unclassified_Deltaproteobacteria</td><td>GCA_015228875.1</td><td>Deltaproteobacteria_bacterium</td><td>HQM16_08940</td><td>21511..24978</td><td>2026735</td><td>BREX-system-P-loop-protein-BrxC-[Deltaproteobacteria-bacterium]</td></tr>
<tr><td>MBI2354427.1</td><td>Type1.BREX</td><td><-DUF6079*<-BrxB-BrxF-iSTAND<-BrxA-DNA-binding<-XIS-HTH||AAA-ToxA->?->?->REase->?->MoxR-AAA-></td><td>1182</td><td>Bacteria>delta/epsilon_subdivisions>Deltaproteobacteria>unclassified_Deltaproteobacteria</td><td>GCA_016182975.1</td><td>Deltaproteobacteria_bacterium</td><td>HYV06_05270</td><td>743..4291</td><td>2026735</td><td>BREX-system-P-loop-protein-BrxC-[Deltaproteobacteria-bacterium]</td></tr>
<tr><td>MBI4517718.1</td><td>Type1.BREX</td><td>BrxA-DNA-binding->BrxB-BrxF-iSTAND->DUF6079*->N6-METHYLASE->SbcC+ABC-ATPase->PglZ->SIGMA+OB-Fold+AAA-ATPase+Lon-Protease-BrxL-></td><td>1149</td><td>Bacteria>delta/epsilon_subdivisions>Deltaproteobacteria>unclassified_Deltaproteobacteria</td><td>GCA_016210005.1</td><td>Deltaproteobacteria_bacterium</td><td>HY699_18080</td><td>1797..5246</td><td>2026735</td><td>BREX-system-P-loop-protein-BrxC-[Deltaproteobacteria-bacterium]</td></tr>
<tr><td>MBI5514929.1</td><td>Type1.BREX</td><td>HAD+HAD-><-?<-?||MerR-HTH-><-LexA-protease||BrxA-DNA-binding->BrxB-BrxF-iSTAND->DUF6079*->N6-METHYLASE->PglZ->SIGMA+OB-Fold+AAA-ATPase+Lon-Protease-BrxL->HTH+MPTase->?->?->MarR-HTH->AbiEii-></td><td>1224</td><td>Bacteria>delta/epsilon_subdivisions>Deltaproteobacteria>unclassified_Deltaproteobacteria</td><td>GCA_016218725.1</td><td>Deltaproteobacteria_bacterium</td><td>HY909_14230</td><td>59420..63094</td><td>2026735</td><td>BREX-system-P-loop-protein-BrxC-[Deltaproteobacteria-bacterium]</td></tr>
<tr><td>MBI5606167.1</td><td>Type1.BREX</td><td><-Schlafen+HATPase+wHTH<-DUF6079*<-BrxB-BrxF-iSTAND<-BrxA-DNA-binding||?-><-PIN<-?||HicA_toxin->HicB_lk_antitox->ACET-></td><td>1147</td><td>Bacteria>delta/epsilon_subdivisions>Deltaproteobacteria>unclassified_Deltaproteobacteria</td><td>GCA_016219225.1</td><td>Deltaproteobacteria_bacterium</td><td>HY879_22775</td><td>11994..15437</td><td>2026735</td><td>BREX-system-P-loop-protein-BrxC-[Deltaproteobacteria-bacterium]</td></tr>
<tr><td>MBK7759463.1</td><td>Type1.BREX</td><td><-N6-METHYLASE-p<-DUF6079-p<-DUF6079-p<-DUF6079-p*<-BrxB-BrxF-iSTAND<-?<-DEDD_Tnp_IS110+Transposase_20</td><td>766</td><td>Bacteria>delta/epsilon_subdivisions>Deltaproteobacteria>unclassified_Deltaproteobacteria</td><td>GCA_016706815.1</td><td>Deltaproteobacteria_bacterium</td><td>IPI35_24325</td><td>2903..5203</td><td>2026735</td><td>BREX-system-P-loop-protein-BrxC-[Deltaproteobacteria-bacterium]</td></tr>
<tr><td>MBN1905588.1</td><td>Type1.BREX</td><td>PAS+NtrC-AAA+FIS-HTH-><-SIGMA+OB-Fold+AAA-ATPase+Lon-Protease-BrxL<-PglZ<-N6-METHYLASE<-N6-METHYLASE-p<-TOPRIM<-OLD-ABC<-DUF6079*<-BrxA-DNA-binding</td><td>1190</td><td>Bacteria>delta/epsilon_subdivisions>Deltaproteobacteria>unclassified_Deltaproteobacteria</td><td>GCA_016931115.1</td><td>Deltaproteobacteria_bacterium</td><td>JW927_10865</td><td>38502..42074</td><td>2026735</td><td>BREX-system-P-loop-protein-BrxC-[Deltaproteobacteria-bacterium]</td></tr>
<tr><td>MBP8650772.1</td><td>Type1.BREX</td><td><-DUF4096+DDE_Tnp_1+DDE_Tnp_1_2||?->CASPASE-><-SIGMA+OB-Fold+AAA-ATPase+Lon-Protease-BrxL<-PglZ<-Polbeta<-MarR-HTH<-DUF932<-N6-METHYLASE<-DUF6079*<-BrxB-BrxF-iSTAND<-BrxA-DNA-binding</td><td>1145</td><td>Bacteria>delta/epsilon_subdivisions>Deltaproteobacteria>unclassified_Deltaproteobacteria</td><td>GCA_018053065.1</td><td>Deltaproteobacteria_bacterium</td><td>KBI48_08700</td><td>29985..33422</td><td>2026735</td><td>BREX-system-P-loop-protein-BrxC-[Deltaproteobacteria-bacterium]</td></tr>
<tr><td>MBT3255199.1</td><td>Type1.BREX</td><td><-wHTH+WYL+WCX<-?||?->DUF932->BrxA-DNA-binding->BrxB-BrxF-iSTAND->DUF6079*->N6-METHYLASE->OLD-ABC->TOPRIM->PglZ->SIGMA+OB-Fold+AAA-ATPase+Lon-Protease-BrxL->?->?-><-SAVED<-JAB</td><td>1216</td><td>Bacteria>delta/epsilon_subdivisions>Deltaproteobacteria>unclassified_Deltaproteobacteria</td><td>GCA_018644785.1</td><td>Deltaproteobacteria_bacterium</td><td>HN366_01900</td><td>5574..9224</td><td>2026735</td><td>BREX-system-P-loop-protein-BrxC-[Deltaproteobacteria-bacterium]</td></tr>
<tr><td>MBT4264397.1</td><td>Type1.BREX</td><td><-PAS+PAS+SHELIX+LuxR-HTH<-Peptidase||?->?->?-><-REC+PAS+PAS+HISKIN||?-><-Phage_integrase<-DUF6079-p*<-BrxB-BrxF-iSTAND<-BrxA-DNA-binding||<-tRNA<-?<-?||?->CHASE2+ACYC-><-TPR+TPR+TPR<-PCMT+METHYLASE</td><td>409</td><td>Bacteria>delta/epsilon_subdivisions>Deltaproteobacteria>unclassified_Deltaproteobacteria</td><td>GCA_018651965.1</td><td>Deltaproteobacteria_bacterium</td><td>HOD85_09330</td><td>22894..24123</td><td>2026735</td><td>BREX-system-P-loop-protein-BrxC-[Deltaproteobacteria-bacterium]</td></tr>
<tr><td>MBW1741678.1</td><td>Type1.BREX</td><td><-N6-METHYLASE-p<-DUF6079*<-BrxB-BrxF-iSTAND<-BrxA-DNA-binding<-AbiEii<-AbiEi_4<-SKI<-Shikimate_dh_N+SDH_C</td><td>1106</td><td>Bacteria>delta/epsilon_subdivisions>Deltaproteobacteria>unclassified_Deltaproteobacteria</td><td>GCA_019306505.1</td><td>Deltaproteobacteria_bacterium</td><td>JRJ42_11205</td><td>451..3771</td><td>2026735</td><td>BREX-system-P-loop-protein-BrxC-[Deltaproteobacteria-bacterium]</td></tr>
<tr><td>MBW1799340.1</td><td>Type1.BREX</td><td><-DUF6079-p*<-BrxB-BrxF-iSTAND<-BrxA-DNA-binding<-?<-?<-?||ADH_N+ADH_zinc_N+METHYLASE-></td><td>188</td><td>Bacteria>delta/epsilon_subdivisions>Deltaproteobacteria>unclassified_Deltaproteobacteria</td><td>GCA_019306865.1</td><td>Deltaproteobacteria_bacterium</td><td>JRJ85_01290</td><td>1..565</td><td>2026735</td><td>BREX-system-P-loop-protein-BrxC-[Deltaproteobacteria-bacterium]</td></tr>
<tr><td>MBW1938726.1</td><td>Type1.BREX</td><td><-Schlafen+HATPase+wHTH<-N6-METHYLASE<-DUF6079*<-BrxB-BrxF-iSTAND<-BrxA-DNA-binding</td><td>1178</td><td>Bacteria>delta/epsilon_subdivisions>Deltaproteobacteria>unclassified_Deltaproteobacteria</td><td>GCA_019307905.1</td><td>Deltaproteobacteria_bacterium</td><td>JRI67_08190</td><td>4087..7623</td><td>2026735</td><td>BREX-system-P-loop-protein-BrxC-[Deltaproteobacteria-bacterium]</td></tr>
<tr><td>MBW1967272.1</td><td>Type1.BREX</td><td><-N6-METHYLASE-p<-REase<-OLD-ABC<-DUF6079*<-BrxB-BrxF-iSTAND<-BrxA-DNA-binding<-XIS-HTH</td><td>1154</td><td>Bacteria>delta/epsilon_subdivisions>Deltaproteobacteria>unclassified_Deltaproteobacteria</td><td>GCA_019308155.1</td><td>Deltaproteobacteria_bacterium</td><td>JRI48_07780</td><td>2430..5894</td><td>2026735</td><td>BREX-system-P-loop-protein-BrxC-[Deltaproteobacteria-bacterium]</td></tr>
<tr><td>MBW2031933.1</td><td>Type1.BREX</td><td>XIS-HTH->BrxA-DNA-binding-p->BrxB-BrxF-iSTAND->DUF6079-p*-></td><td>162</td><td>Bacteria>delta/epsilon_subdivisions>Deltaproteobacteria>unclassified_Deltaproteobacteria</td><td>GCA_019308745.1</td><td>Deltaproteobacteria_bacterium</td><td>JRJ31_23030</td><td>2222..2706</td><td>2026735</td><td>hypothetical-protein-[Deltaproteobacteria-bacterium]</td></tr>
<tr><td>MBW2308571.1</td><td>Type1.BREX</td><td>SF1-SF2-Helicase->||XIS-HTH->BrxA-DNA-binding->BrxB-BrxF-iSTAND->N6-METHYLASE->N6-METHYLASE-p->DUF6079*->PglZ->SIGMA+OB-Fold+AAA-ATPase+Lon-Protease-BrxL-></td><td>1220</td><td>Bacteria>delta/epsilon_subdivisions>Deltaproteobacteria>unclassified_Deltaproteobacteria</td><td>GCA_019310535.1</td><td>Deltaproteobacteria_bacterium</td><td>JRG73_16720</td><td>7753..11415</td><td>2026735</td><td>BREX-system-P-loop-protein-BrxC-[Deltaproteobacteria-bacterium]</td></tr>
<tr><td>MCC6503799.1</td><td>Type1.BREX</td><td><-SbcC<-Calcineurin+Calcineurin-SbcD<-SbcC+APATPase+SbcC<-SIGMA+OB-Fold+AAA-ATPase+HKD-DNase-BrxL<-PglZ<-N6-METHYLASE<-REase<-OLD-ABC<-DUF6079*<-BrxB-BrxF-iSTAND<-BrxA-DNA-binding<-XIS-HTH</td><td>1181</td><td>Bacteria>delta/epsilon_subdivisions>Deltaproteobacteria>unclassified_Deltaproteobacteria</td><td>GCA_020846435.1</td><td>Deltaproteobacteria_bacterium</td><td>IT362_11790</td><td>16699..20244</td><td>2026735</td><td>BREX-system-P-loop-protein-BrxC-[Deltaproteobacteria-bacterium]</td></tr>
<tr><td>MCD6185271.1</td><td>Type1.BREX</td><td><-DUF6079-p*<-BrxB-BrxF-iSTAND<-BrxA-DNA-binding<-DOC+HTH<-?<-Calcineurin<-RADICAL-SAM<-?||URI-DNase+UVR+UvrC_HhH_N+HHH-></td><td>852</td><td>Bacteria>delta/epsilon_subdivisions>Deltaproteobacteria>unclassified_Deltaproteobacteria</td><td>GCA_021157555.1</td><td>Deltaproteobacteria_bacterium</td><td>J7K84_05705</td><td>1..2556</td><td>2026735</td><td>BREX-system-P-loop-protein-BrxC-[Deltaproteobacteria-bacterium]</td></tr>
<tr><td>MCD6306396.1</td><td>Type1.BREX</td><td><-N6-METHYLASE-p||PhdYeFM_antitox->PIN-><-DUF6079*<-BrxB-BrxF-iSTAND<-BrxA-DNA-binding||wHTH+WYL+WCX-p-></td><td>1148</td><td>Bacteria>delta/epsilon_subdivisions>Deltaproteobacteria>unclassified_Deltaproteobacteria</td><td>GCA_021159525.1</td><td>Deltaproteobacteria_bacterium</td><td>J7M32_08925</td><td>1514..4960</td><td>2026735</td><td>BREX-system-P-loop-protein-BrxC-[Deltaproteobacteria-bacterium]</td></tr>
<tr><td>MCD6532985.1</td><td>Type1.BREX</td><td><-PglZ<-Schlafen+HATPase+wHTH<-PHP+SbcC<-PIN<-?<-Schlafen+HATPase+wHTH<-N6-METHYLASE<-REase<-OLD-ABC<-DUF6079-p*</td><td>691</td><td>Bacteria>delta/epsilon_subdivisions>Deltaproteobacteria>unclassified_Deltaproteobacteria</td><td>GCA_021163445.1</td><td>Deltaproteobacteria_bacterium</td><td>J7L25_02755</td><td>20972..23047</td><td>2026735</td><td>BREX-system-P-loop-protein-BrxC-[Deltaproteobacteria-bacterium]</td></tr>
<tr><td>MCK9263307.1</td><td>Type1.BREX</td><td>SIGMA-HTH+SIGMA-HTH->SIGMA-HTH->?->BrxA-DNA-binding->BrxB-BrxF-iSTAND->DUF6079*->Schlafen+HATPase+wHTH-></td><td>1179</td><td>Bacteria>delta/epsilon_subdivisions>Deltaproteobacteria>unclassified_Deltaproteobacteria</td><td>GCA_023230665.1</td><td>Deltaproteobacteria_bacterium</td><td>MOR18_05785</td><td>10308..13847</td><td>2026735</td><td>BREX-system-P-loop-protein-BrxC-[Deltaproteobacteria-bacterium]</td></tr>
<tr><td>MCP4158880.1</td><td>Type1.BREX</td><td><-CARF<-HKD-DNase+HKD-DNase<-?<-S1+SF1-SF2-Helicase+S1<-SIGMA+OB-Fold+AAA-ATPase+HKD-DNase-BrxL<-PglZ<-Couple_hipA+HipA_C<-HTH_3<-N6-METHYLASE<-DUF6079*</td><td>1006</td><td>Bacteria>delta/epsilon_subdivisions>Deltaproteobacteria>unclassified_Deltaproteobacteria</td><td>GCA_024224625.1</td><td>Deltaproteobacteria_bacterium</td><td>GY760_02295</td><td>25697..28718</td><td>2026735</td><td>BREX-system-P-loop-protein-BrxC-[Deltaproteobacteria-bacterium]</td></tr>
<tr><td>MCP4577326.1</td><td>Type1.BREX</td><td><-Thioredoxin||?->?->?-><-?||BrxA-DNA-binding->BrxB-BrxF-iSTAND->DUF6079*->RNase->?->HTH+REase->N6-METHYLASE->Schlafen+HATPase+wHTH->PglZ->SIGMA+OB-Fold+AAA-ATPase+Lon-Protease-BrxL->?->ABC-ATPase+SbcC-></td><td>1196</td><td>Bacteria>delta/epsilon_subdivisions>Deltaproteobacteria>unclassified_Deltaproteobacteria</td><td>GCA_024231575.1</td><td>Deltaproteobacteria_bacterium</td><td>GY846_13700</td><td>13452..17042</td><td>2026735</td><td>BREX-system-P-loop-protein-BrxC-[Deltaproteobacteria-bacterium]</td></tr>
<tr><td>OGR40953.1</td><td>Type1.BREX</td><td><-HTH+REase<-SMODS<-?<-SIGMA+OB-Fold+AAA-ATPase+Lon-Protease-BrxL<-PglZ<-AbiEii<-AbiEi_4<-N6-METHYLASE<-DUF6079*<-BrxB-BrxF-iSTAND<-BrxA-DNA-binding||?->?->?->?->Phage_int_SAM_1+Phage_integrase-><-?||MerR-HTH-></td><td>1147</td><td>Bacteria>Elusimicrobiota>unclassified_Elusimicrobiota</td><td>GCA_001799575.1</td><td>Elusimicrobia_bacterium_GWA2_56_46</td><td>A2X28_06390</td><td>23644..27087</td><td>1797922</td><td>nucleoside-triphosphate-hydrolase-[Elusimicrobia-bacterium-GWA2_56_46]</td></tr>
<tr><td>MBK7688407.1</td><td>Type1.BREX</td><td>Thermonuclease->BrxA-DNA-binding->BrxB-BrxF-iSTAND->DUF6079-p*->DUF6079-p->DUF6079-p->N6-METHYLASE->PglZ->SIGMA+OB-Fold+AAA-ATPase+Lon-Protease-BrxL-></td><td>415</td><td>Bacteria>Elusimicrobiota>unclassified_Elusimicrobiota</td><td>GCA_016709555.1</td><td>Elusimicrobiota_bacterium</td><td>IPJ35_05275</td><td>9900..11147</td><td>2030800</td><td>hypothetical-protein-[Elusimicrobiota-bacterium]</td></tr>
<tr><td>MBK7688408.1</td><td>Type1.BREX</td><td>Thermonuclease->BrxA-DNA-binding->BrxB-BrxF-iSTAND->DUF6079-p->DUF6079-p*->DUF6079-p->N6-METHYLASE->PglZ->SIGMA+OB-Fold+AAA-ATPase+Lon-Protease-BrxL-></td><td>466</td><td>Bacteria>Elusimicrobiota>unclassified_Elusimicrobiota</td><td>GCA_016709555.1</td><td>Elusimicrobiota_bacterium</td><td>IPJ35_05280</td><td>11117..12517</td><td>2030800</td><td>hypothetical-protein-[Elusimicrobiota-bacterium]</td></tr>
<tr><td>MXY08911.1</td><td>Type1.BREX</td><td><-MPTase||BrxA-DNA-binding->BrxB-BrxF-iSTAND->DUF6079*->N6-METHYLASE->PglZ->SIGMA+OB-Fold+AAA-ATPase+Lon-Protease-BrxL->?-><-Phage_int_SAM_4+Phage_integrase</td><td>1183</td><td>Bacteria>FCB_group>Bacteroidetes/Chlorobi_group>Bacteroidota>Bacteroidetes_incertae_sedis>Rhodothermaceae</td><td>GCA_009838245.1</td><td>Rhodothermaceae_bacterium</td><td>F4Y61_09720</td><td>3347..6898</td><td>2026787</td><td>BREX-system-P-loop-protein-BrxC-[Rhodothermaceae-bacterium]</td></tr>
<tr><td>MCS4056789.1</td><td>Type1.BREX</td><td><-Calcineurin-SbcD<-SIGMA+OB-Fold+AAA-ATPase+HKD-DNase-BrxL<-PglZ<-HEPN<-PolB||?-><-?<-N6-METHYLASE<-DOC+HTH<-DUF6079*<-BrxB-BrxF-iSTAND<-BrxA-DNA-binding<-?<-viral-protein<-?<-?||?-><-RHH</td><td>1165</td><td>Bacteria>FCB_group>Bacteroidetes/Chlorobi_group>Bacteroidota>Bacteroidetes_incertae_sedis>Rhodothermaceae</td><td>GCA_024809315.1</td><td>Salinibacter_ruber</td><td>GGP75_002131</td><td>31231..34728</td><td>146919</td><td>hypothetical-protein-[Salinibacter-ruber]</td></tr>
<tr><td>WP_119843039.1</td><td>Type1.BREX</td><td><-HD<-SbcC+SbcC<-Calcineurin-SbcD<-HORMA<-SIGMA+OB-Fold+AAA-ATPase+HKD-DNase-BrxL<-PglZ<-N6-METHYLASE<-DUF6079*<-BrxB-BrxF-iSTAND<-BrxA-DNA-binding<-REase+RNase<-viral-protein||?-><-Phage_holin_2_4<-?<-?<-PAS+PAS+PAS+GAF+HISKIN</td><td>1201</td><td>Bacteria>FCB_group>Bacteroidetes/Chlorobi_group>Bacteroidota>Bacteroidetes_incertae_sedis>Rhodothermaceae</td><td>GCF_002290345.1</td><td>Salinibacter_ruber</td><td>B7W84_RS15115</td><td>76596..80201</td><td>146919</td><td>BREX-system-P-loop-protein-BrxC-[Salinibacter-ruber]</td></tr>
<tr><td>WP_103028550.1</td><td>Type1.BREX</td><td><-N6-METHYLASE<-DUF6079*<-BrxB-BrxF-iSTAND<-BrxA-DNA-binding||Calcineurin-SbcD->SbcC+SbcC->SbcC->Phage_integrase-></td><td>1168</td><td>Bacteria>FCB_group>Bacteroidetes/Chlorobi_group>Bacteroidota>Bacteroidetes_incertae_sedis>Rhodothermaceae</td><td>GCF_002894625.1</td><td>Salinibacter_altiplanensis</td><td>B7R94_RS01140</td><td>2369..5875</td><td>1803181</td><td>BREX-system-P-loop-protein-BrxC-[Salinibacter-altiplanensis]</td></tr>
<tr><td>WP_259084851.1</td><td>Type1.BREX</td><td><-REase+DUF4268<-SbcC+SbcC<-Calcineurin-SbcD<-SIGMA+OB-Fold+AAA-ATPase+Lon-Protease-BrxL<-PglZ<-SF1-SF2-Helicase<-OLD-ABC+TOPRIM<-N6-METHYLASE<-DOC+HTH<-DUF6079*<-BrxB-BrxF-iSTAND<-BrxA-DNA-binding||REase-><-Phage_int_SAM_1+Phage_integrase<-?||HTH_3-><-TetR-HTH<-Phage_integrase<-Phage_integrase</td><td>1143</td><td>Bacteria>FCB_group>Bacteroidetes/Chlorobi_group>Bacteroidota>Bacteroidetes_incertae_sedis>Rhodothermaceae</td><td>GCF_024808015.1</td><td>Salinibacter_ruber</td><td>GGP69_RS05925</td><td>149309..152740</td><td>146919</td><td>BREX-system-P-loop-protein-BrxC-[Salinibacter-ruber]</td></tr>
<tr><td>WP_259111349.1</td><td>Type1.BREX</td><td>RHH->Phage_holin_2_4->?->viral-protein-><-REase||BrxA-DNA-binding->BrxB-BrxF-iSTAND->DUF6079*->DOC+HTH->N6-METHYLASE->?->PglZ->SIGMA+OB-Fold+AAA-ATPase+Lon-Protease-BrxL->Calcineurin-SbcD->SbcC+SbcC->HD-></td><td>1145</td><td>Bacteria>FCB_group>Bacteroidetes/Chlorobi_group>Bacteroidota>Bacteroidetes_incertae_sedis>Rhodothermaceae</td><td>GCF_024808065.1</td><td>Salinibacter_ruber</td><td>GGP79_RS14420</td><td>5761..9198</td><td>146919</td><td>BREX-system-P-loop-protein-BrxC-[Salinibacter-ruber]</td></tr>
<tr><td>WP_259049434.1</td><td>Type1.BREX</td><td><-HISKIN<-SIGMA+OB-Fold+AAA-ATPase+Lon-Protease-BrxL<-PglZ<-DrHyd<-?<-N6-METHYLASE<-DOC+HTH<-DUF6079*<-BrxB-BrxF-iSTAND<-BrxA-DNA-binding||REase-><-Arm-DNA-bind_5+Phage_int_SAM_5+Phage_integrase||?->Trypsin+PDZ-></td><td>1145</td><td>Bacteria>FCB_group>Bacteroidetes/Chlorobi_group>Bacteroidota>Bacteroidetes_incertae_sedis>Rhodothermaceae</td><td>GCF_024808355.1</td><td>Salinibacter_ruber</td><td>GGP97_RS08705</td><td>20679..24116</td><td>146919</td><td>BREX-system-P-loop-protein-BrxC-[Salinibacter-ruber]</td></tr>
<tr><td>WP_259099618.1</td><td>Type1.BREX</td><td><-HD<-SbcC+SbcC<-Calcineurin-SbcD<-SIGMA+OB-Fold+AAA-ATPase+Lon-Protease-BrxL<-PglZ<-OLD-ABC+TOPRIM<-N6-METHYLASE<-DOC+HTH<-DUF6079*<-BrxB-BrxF-iSTAND<-BrxA-DNA-binding||REase-><-?<-viral-protein</td><td>1143</td><td>Bacteria>FCB_group>Bacteroidetes/Chlorobi_group>Bacteroidota>Bacteroidetes_incertae_sedis>Rhodothermaceae</td><td>GCF_024808405.1</td><td>Salinibacter_ruber</td><td>GGP77_002965</td><td>30959..34390</td><td>146919</td><td>BREX-system-P-loop-protein-BrxC-[Salinibacter-ruber]</td></tr>
<tr><td>WP_259094322.1</td><td>Type1.BREX</td><td><-HD<-SbcC+SbcC<-Calcineurin-SbcD<-SIGMA+OB-Fold+AAA-ATPase+Lon-Protease-BrxL<-PglZ<-SF1-SF2-Helicase<-OLD-ABC+TOPRIM<-N6-METHYLASE<-DOC+HTH<-DUF6079*<-BrxB-BrxF-iSTAND<-BrxA-DNA-binding<-REase<-HKD-DNase+SF1-SF2-Helicase<-?<-?||?-><-RHH</td><td>1143</td><td>Bacteria>FCB_group>Bacteroidetes/Chlorobi_group>Bacteroidota>Bacteroidetes_incertae_sedis>Rhodothermaceae</td><td>GCF_024809025.1</td><td>Salinibacter_ruber</td><td>GGP87_RS12315</td><td>32840..36271</td><td>146919</td><td>BREX-system-P-loop-protein-BrxC-[Salinibacter-ruber]</td></tr>
<tr><td>WP_259115052.1</td><td>Type1.BREX</td><td><-SbcC+SbcC<-Calcineurin-SbcD<-SIGMA+OB-Fold+AAA-ATPase+HKD-DNase-BrxL<-PglZ<-HEPN<-PolB<-?<-N6-METHYLASE<-DOC+HTH<-DUF6079*<-BrxB-BrxF-iSTAND<-BrxA-DNA-binding<-?<-viral-protein<-?<-?||?-><-RHH||HTH_3+Peptidase_S24-></td><td>1167</td><td>Bacteria>FCB_group>Bacteroidetes/Chlorobi_group>Bacteroidota>Bacteroidetes_incertae_sedis>Rhodothermaceae</td><td>GCF_024809315.1</td><td>Salinibacter_ruber</td><td>GGP75_RS10390</td><td>31231..34734</td><td>146919</td><td>BREX-system-P-loop-protein-BrxC-[Salinibacter-ruber]</td></tr>
<tr><td>WP_259076695.1</td><td>Type1.BREX</td><td>TraJ-RHH->RHH->Phage_holin_2_4->||viral-protein->?->BrxA-DNA-binding->BrxB-BrxF-iSTAND->DUF6079*->DOC+HTH->N6-METHYLASE->?->?->PglZ->SIGMA+OB-Fold+AAA-ATPase+HKD-DNase-BrxL->Calcineurin-SbcD->SbcC+SbcC-></td><td>1167</td><td>Bacteria>FCB_group>Bacteroidetes/Chlorobi_group>Bacteroidota>Bacteroidetes_incertae_sedis>Rhodothermaceae</td><td>GCF_024809595.1</td><td>Salinibacter_ruber</td><td>GGP73_RS15155</td><td>19341..22844</td><td>146919</td><td>BREX-system-P-loop-protein-BrxC-[Salinibacter-ruber]</td></tr>
<tr><td>WP_259210815.1</td><td>Type1.BREX</td><td><-MarR-HTH<-Phage_integrase<-SbcC+SbcC<-Calcineurin-SbcD<-SIGMA+OB-Fold+AAA-ATPase+Lon-Protease-BrxL<-PglZ<-?<-N6-METHYLASE<-DOC+HTH<-DUF6079*<-BrxB-BrxF-iSTAND<-BrxA-DNA-binding||REase-><-viral-protein<-?<-RHH<-TraJ-RHH||HTH_3+Peptidase_S24-></td><td>1145</td><td>Bacteria>FCB_group>Bacteroidetes/Chlorobi_group>Bacteroidota>Bacteroidetes_incertae_sedis>Rhodothermaceae</td><td>GCF_024809615.1</td><td>Salinibacter_ruber</td><td>GGP43_RS13275</td><td>33115..36552</td><td>146919</td><td>BREX-system-P-loop-protein-BrxC-[Salinibacter-ruber]</td></tr>
<tr><td>WP_259195582.1</td><td>Type1.BREX</td><td>Phage_integrase->Phage_integrase->TetR-HTH-><-HTH_3||?->Phage_int_SAM_1+Phage_integrase-><-REase||BrxA-DNA-binding->BrxB-BrxF-iSTAND->DUF6079*->DOC+HTH->N6-METHYLASE->OLD-ABC+TOPRIM->PglZ->SIGMA+OB-Fold+AAA-ATPase+Lon-Protease-BrxL->Calcineurin-SbcD->SbcC+SbcC->REase+DUF4268-></td><td>1144</td><td>Bacteria>FCB_group>Bacteroidetes/Chlorobi_group>Bacteroidota>Bacteroidetes_incertae_sedis>Rhodothermaceae</td><td>GCF_024809635.1</td><td>Salinibacter_ruber</td><td>GGP50_RS04130</td><td>10424..13858</td><td>146919</td><td>BREX-system-P-loop-protein-BrxC-[Salinibacter-ruber]</td></tr>
<tr><td>WP_259087839.1</td><td>Type1.BREX</td><td>URI-DNase-><-SIGMA+OB-Fold+AAA-ATPase+HKD-DNase-BrxL<-PglZ<-PIN<-XIS-HTH<-?<-?<-N6-METHYLASE<-DUF6079*<-BrxB-BrxF-iSTAND<-BrxA-DNA-binding<-?||Phage_integrase-><-TOPRIM<-?||XIS-HTH->Phage_int_SAM_4+Phage_integrase-><-tRNA||BetaPropeller+ABhydrolase-></td><td>1168</td><td>Bacteria>FCB_group>Bacteroidetes/Chlorobi_group>Bacteroidota>Bacteroidetes_incertae_sedis>Rhodothermaceae</td><td>GCF_024809725.1</td><td>Salinibacter_ruber</td><td>GGQ07_RS07865</td><td>19343..22849</td><td>146919</td><td>BREX-system-P-loop-protein-BrxC-[Salinibacter-ruber]</td></tr>
<tr><td>WP_251951689.1</td><td>Type1.BREX</td><td><-SbcC+SbcC<-Calcineurin-SbcD<-SIGMA+OB-Fold+AAA-ATPase+Lon-Protease-BrxL<-PglZ<-?<-?<-N6-METHYLASE<-?<-?<-DUF6079*<-BrxB-BrxF-iSTAND<-BrxA-DNA-binding<-REase<-HKD-DNase+SF1-SF2-Helicase<-SF1-SF2-Helicase+REase<-viral-protein<-?<-?<-RHH</td><td>1143</td><td>Bacteria>FCB_group>Bacteroidetes/Chlorobi_group>Bacteroidota>Bacteroidetes_incertae_sedis>Rhodothermaceae</td><td>GCF_943908255.1</td><td>Salinibacter_ruber</td><td>NGL47_RS12295</td><td>24332..27763</td><td>146919</td><td>BREX-system-P-loop-protein-BrxC-[Salinibacter-ruber]</td></tr>
<tr><td>WP_263820582.1</td><td>Type1.BREX</td><td><-SbcC+SbcC<-Calcineurin-SbcD<-SIGMA+OB-Fold+AAA-ATPase+HKD-DNase-BrxL<-PglZ<-?<-?<-N6-METHYLASE<-DOC+HTH<-DUF6079*<-BrxB-BrxF-iSTAND<-BrxA-DNA-binding<-DUF4258<-REase+RNase<-viral-protein<-?||CHTH-><-Phage_holin_2_4<-RHH</td><td>1167</td><td>Bacteria>FCB_group>Bacteroidetes/Chlorobi_group>Bacteroidota>Bacteroidetes_incertae_sedis>Rhodothermaceae</td><td>GCF_947077735.1</td><td>Salinibacter_sp.</td><td>OJA43_RS08805</td><td>45420..48923</td><td>2065818</td><td>BREX-system-P-loop-protein-BrxC-[Salinibacter-sp.]</td></tr>
<tr><td>WP_259248797.1</td><td>Type1.BREX</td><td><-SbcC+SbcC<-Calcineurin-SbcD<-SIGMA+OB-Fold+AAA-ATPase+HKD-DNase-BrxL<-PglZ<-CHTH<-RVT<-N6-METHYLASE<-DOC+HTH<-DUF6079*<-BrxB-BrxF-iSTAND<-BrxA-DNA-binding<-DUF4258<-REase+RNase<-HKD-DNase+SF1-SF2-Helicase<-RHH||HTH_23+HTH_32+HTH_33->DDE_3-><-TraJ-RHH</td><td>1167</td><td>Bacteria>FCB_group>Bacteroidetes/Chlorobi_group>Bacteroidota>Bacteroidetes_incertae_sedis>Rhodothermaceae</td><td>-</td><td>Salinibacter_ruber</td><td>GGP55_RS14300</td><td>49306..52809</td><td>146919</td><td>BREX-system-P-loop-protein-BrxC-[Salinibacter-ruber]</td></tr>
<tr><td>HHU27149.1</td><td>Type1.BREX</td><td><-DUF6079-p*<-BrxB-BrxF-iSTAND<-BrxA-DNA-binding||DEDD_Tnp_IS110+Transposase_20-></td><td>343</td><td>Bacteria>FCB_group>Bacteroidetes/Chlorobi_group>Bacteroidota>Bacteroidia>Bacteroidales</td><td>-</td><td>Bacteroidales_bacterium</td><td>GXZ56_11115</td><td>1..1030</td><td>2030927</td><td>BREX-system-P-loop-protein-BrxC-[Bacteroidales-bacterium]</td></tr>
<tr><td>WP_204433169.1</td><td>Type1.BREX</td><td>BrxA-DNA-binding->BrxB-BrxF-iSTAND->DUF6079*->OLD-ABC->TOPRIM->N6-METHYLASE-></td><td>1195</td><td>Bacteria>FCB_group>Bacteroidetes/Chlorobi_group>Bacteroidota>Bacteroidia>Bacteroidales</td><td>-</td><td>Bacteroides_mediterraneensis</td><td>H6B28_RS04775</td><td>126749..130336</td><td>1841856</td><td>BREX-system-P-loop-protein-BrxC-[Bacteroides-mediterraneensis]</td></tr>
<tr><td>MCS3257591.1</td><td>Type1.BREX</td><td>RNase-><-?<-HipA_C<-Couple_hipA<-HTH_3||HTH_3->BrxA-DNA-binding->BrxB-BrxF-iSTAND->DUF6079-p*-><-METHYLASE<-?<-HTH_58<-?<-?<-?<-?<-?<-SIGMA-HTH</td><td>733</td><td>Bacteria>FCB_group>Bacteroidetes/Chlorobi_group>Bacteroidota>Bacteroidia>Bacteroidales</td><td>-</td><td>Bacteroides_thetaiotaomicron</td><td>NXV10_10775</td><td>1863438..1865639</td><td>818</td><td>BREX-system-P-loop-protein-BrxC-[Bacteroides-thetaiotaomicron]</td></tr>
<tr><td>WP_118696414.1</td><td>Type1.BREX</td><td><-Arm-DNA-bind_5+Phage_int_SAM_5+Phage_integrase||DUF4007->BrxB-BrxF-iSTAND->DUF6079-p->DUF6079-p*->N6-METHYLASE->PglZ->SIGMA+OB-Fold+AAA-ATPase+HKD-DNase-BrxL-><-?<-RHH+MobC||ARG->?->UDG-></td><td>938</td><td>Bacteria>FCB_group>Bacteroidetes/Chlorobi_group>Bacteroidota>Bacteroidia>Bacteroidales</td><td>-</td><td>Bacteroides_uniformis</td><td>KSX92_08935</td><td>3061..5877</td><td>820</td><td>BREX-system-P-loop-protein-BrxC-[Bacteroides-uniformis]</td></tr>
<tr><td>EEZ19147.1</td><td>Type1.BREX</td><td>AbrB-HTH-><-?||?-><-KAP-NTPase<-SIGMA+OB-Fold+AAA-ATPase+Lon-Protease-BrxL<-PglZ<-Schlafen+HATPase+wHTH<-N6-METHYLASE<-DUF6079*<-BrxB-BrxF-iSTAND<-BrxA-DNA-binding<-HTH_3||?->XIS-HTH->XIS-HTH-></td><td>1213</td><td>Bacteria>FCB_group>Bacteroidetes/Chlorobi_group>Bacteroidota>Bacteroidia>Bacteroidales</td><td>GCA_000162195.1</td><td>Bacteroides_sp._3_1_33FAA</td><td>HMPREF0105_4547</td><td>25602..29243</td><td>457391</td><td>hypothetical-protein-HMPREF0105_4547-[Bacteroides-sp.-3_1_33FAA]</td></tr>
<tr><td>CDD81792.1</td><td>Type1.BREX</td><td><-HIRAN<-RADICAL-SAM<-NADAR<-HTH_3<-SIGMA+OB-Fold+AAA-ATPase+HKD-DNase-BrxL<-PglZ<-Schlafen+HATPase+wHTH<-N6-METHYLASE<-Four-helical-protein<-DUF6079*<-BrxB-BrxF-iSTAND<-BrxA-DNA-binding<-HTH_3||?->?->?->?->?-><-XIS-HTH</td><td>1217</td><td>Bacteria>FCB_group>Bacteroidetes/Chlorobi_group>Bacteroidota>Bacteroidia>Bacteroidales</td><td>GCA_000435095.1</td><td>Bacteroides_sp._CAG:462</td><td>BN666_01223</td><td>46564..50217</td><td>1262740</td><td>putative-uncharacterized-protein-[Bacteroides-sp.-CAG:462]</td></tr>
<tr><td>CDD50103.1</td><td>Type1.BREX</td><td>OLD-ABC->?->?-><-?<-?<-?<-?||BrxA-DNA-binding->BrxB-BrxF-iSTAND->DUF6079*->SbcC->OLD-ABC->TOPRIM->N6-METHYLASE->N6-METHYLASE-p-></td><td>1195</td><td>Bacteria>FCB_group>Bacteroidetes/Chlorobi_group>Bacteroidota>Bacteroidia>Bacteroidales</td><td>GCA_000435415.1</td><td>Bacteroides_sp._CAG:875</td><td>BN800_01988</td><td>17775..21362</td><td>1262752</td><td>putative-uncharacterized-protein-[Bacteroides-sp.-CAG:875]</td></tr>
<tr><td>CDD33290.1</td><td>Type1.BREX</td><td><-TOPRIM<-OLD-ABC<-DUF6079*<-BrxB-BrxF-iSTAND<-BrxA-DNA-binding<-HTH_3<-Phage_holin<-HD</td><td>1195</td><td>Bacteria>FCB_group>Bacteroidetes/Chlorobi_group>Bacteroidota>Bacteroidia>Bacteroidales</td><td>GCA_000436055.1</td><td>Bacteroides_sp._CAG:714</td><td>BN762_00981</td><td>3639..7226</td><td>1262749</td><td>putative-uncharacterized-protein-[Bacteroides-sp.-CAG:714]</td></tr>
<tr><td>CDE09669.1</td><td>Type1.BREX</td><td>ArdA->HTH_3->BrxA-DNA-binding->BrxB-BrxF-iSTAND->DUF6079*->Pkinase->N6-METHYLASE->HEPN->?->?->PglZ->SIGMA+OB-Fold+AAA-ATPase+Lon-Protease-BrxL-></td><td>1221</td><td>Bacteria>FCB_group>Bacteroidetes/Chlorobi_group>Bacteroidota>Bacteroidia>Bacteroidales</td><td>GCA_000437495.1</td><td>Prevotella_sp._CAG:485</td><td>BN677_00784</td><td>6933..10598</td><td>1262927</td><td>putative-uncharacterized-protein-[Prevotella-sp.-CAG:485]</td></tr>
<tr><td>OCW94638.1</td><td>Type1.BREX</td><td>MarR-HTH->Polbeta->Calcineurin-><-DOC+HTH||?->HTH_3->BrxA-DNA-binding->BrxB-BrxF-iSTAND->DUF6079*->N6-METHYLASE->PglZ->SIGMA+OB-Fold+AAA-ATPase+HKD-DNase-BrxL->?->?->XIS-HTH->TOPRIM->XIS-HTH-><-AbiEii</td><td>1202</td><td>Bacteria>FCB_group>Bacteroidetes/Chlorobi_group>Bacteroidota>Bacteroidia>Bacteroidales</td><td>GCA_001704385.2</td><td>Macellibacteroides_sp._HH-ZS</td><td>A9168_06455</td><td>108865..112473</td><td>1857568</td><td>hypothetical-protein-A9168_06455-[Macellibacteroides-sp.-HH-ZS]</td></tr>
<tr><td>OKZ04598.1</td><td>Type1.BREX</td><td><-SIGMA-HTH<-?||35exo+SF1-SF2-Helicase-><-SIGMA+OB-Fold+AAA-ATPase+HKD-DNase-BrxL<-PglZ<-Schlafen+HATPase+wHTH<-SIR2+SIR2<-N6-METHYLASE<-DUF6079*<-BrxB-BrxF-iSTAND<-BrxA-DNA-binding<-?||<-REase||HTH_3->DNA_METHYLASE->McrC-NTD+REase->SbcC->REase-></td><td>1202</td><td>Bacteria>FCB_group>Bacteroidetes/Chlorobi_group>Bacteroidota>Bacteroidia>Bacteroidales</td><td>GCA_001915435.1</td><td>Bacteroides_sp._41_26</td><td>BHV71_10470</td><td>85707..89315</td><td>1896973</td><td>MAG:-hypothetical-protein-BHV71_10470-[Bacteroides-sp.-41_26]</td></tr>
<tr><td>OKZ20119.1</td><td>Type1.BREX</td><td><-REase<-?<-?<-?<-SIGMA+OB-Fold+AAA-ATPase+HKD-DNase-BrxL<-PglZ<-Schlafen+HATPase+wHTH<-N6-METHYLASE<-DUF6079*<-BrxB-BrxF-iSTAND<-BrxA-DNA-binding<-HTH_3<-?<-?<-?||?->?->SIGMA-HTH-></td><td>1185</td><td>Bacteria>FCB_group>Bacteroidetes/Chlorobi_group>Bacteroidota>Bacteroidia>Bacteroidales</td><td>GCA_001915545.1</td><td>Bacteroides_sp._43_108</td><td>BHV77_10980</td><td>38384..41941</td><td>1896974</td><td>MAG:-hypothetical-protein-BHV77_10980-[Bacteroides-sp.-43_108]</td></tr>
<tr><td>OKZ19958.1</td><td>Type1.BREX</td><td>MarR-HTH->Peptidase_M56+TonBC+TonBC-><-?<-?||BrxA-DNA-binding->BrxB-BrxF-iSTAND->DUF6079*->?->N6-METHYLASE->Schlafen+HATPase+wHTH->PglZ->SIGMA+OB-Fold+AAA-ATPase+Lon-Protease-BrxL-></td><td>1213</td><td>Bacteria>FCB_group>Bacteroidetes/Chlorobi_group>Bacteroidota>Bacteroidia>Bacteroidales</td><td>GCA_001915545.1</td><td>Bacteroides_sp._43_108</td><td>BHV77_11420</td><td>12434..16075</td><td>1896974</td><td>MAG:-hypothetical-protein-BHV77_11420-[Bacteroides-sp.-43_108]</td></tr>
<tr><td>HAH90337.1</td><td>Type1.BREX</td><td><-D5ATPase+DUF3874<-?||BrxA-DNA-binding->BrxB-BrxF-iSTAND->DUF6079*->HTH+MPTase->PIN->?->?->?->N6-METHYLASE->PglZ->SIGMA+OB-Fold+AAA-ATPase+Lon-Protease-BrxL->HAD+HAD-></td><td>1222</td><td>Bacteria>FCB_group>Bacteroidetes/Chlorobi_group>Bacteroidota>Bacteroidia>Bacteroidales</td><td>GCA_003450425.1</td><td>Prevotella_sp.</td><td>DCL96_01170</td><td>7620..11288</td><td>59823</td><td>BREX-system-P-loop-protein-BrxC-[Prevotella-sp.]</td></tr>
<tr><td>HAH91106.1</td><td>Type1.BREX</td><td><-HIRAN<-RADICAL-SAM<-ARG<-?<-?||HTH_3->wHTH+WYL+WCX->BrxA-DNA-binding->BrxB-BrxF-iSTAND->DUF6079*->OLD-ABC->TOPRIM->?->SF1-SF2-Helicase->PglZ->SIGMA+OB-Fold+AAA-ATPase+HKD-DNase-BrxL->McrB-AAA->McrC-NTD+REase-></td><td>1202</td><td>Bacteria>FCB_group>Bacteroidetes/Chlorobi_group>Bacteroidota>Bacteroidia>Bacteroidales</td><td>GCA_003450425.1</td><td>Prevotella_sp.</td><td>DCL96_05230</td><td>7787..11395</td><td>59823</td><td>BREX-system-P-loop-protein-BrxC-[Prevotella-sp.]</td></tr>
<tr><td>RHC75801.1</td><td>Type1.BREX</td><td>sGTP->Phage_int_SAM_5+Phage_integrase->?->XIS-HTH->SMS-KaiC->HTH_3->BrxA-DNA-binding->BrxB-BrxF-iSTAND->DUF6079-p*->DUF6079-p->OLD-ABC->REase->N6-METHYLASE->PglZ->SIGMA+OB-Fold+AAA-ATPase+HKD-DNase-BrxL-><-REC+AraC-HTH+AraC-HTH</td><td>691</td><td>Bacteria>FCB_group>Bacteroidetes/Chlorobi_group>Bacteroidota>Bacteroidia>Bacteroidales</td><td>GCA_003468465.1</td><td>Bacteroides_uniformis</td><td>DW831_02855</td><td>19890..21965</td><td>820</td><td>BREX-system-P-loop-protein-BrxC-[Bacteroides-uniformis]</td></tr>
<tr><td>RHK76870.1</td><td>Type1.BREX</td><td>HTH_3->wHTH+WYL+WCX->BrxA-DNA-binding->BrxB-BrxF-iSTAND->DUF6079*->N6-METHYLASE->ACET+TPR-></td><td>1208</td><td>Bacteria>FCB_group>Bacteroidetes/Chlorobi_group>Bacteroidota>Bacteroidia>Bacteroidales</td><td>GCA_003474155.1</td><td>Phocaeicola_vulgatus</td><td>DW047_18670</td><td>2653..6279</td><td>821</td><td>BREX-system-P-loop-protein-BrxC-[Phocaeicola-vulgatus]</td></tr>
<tr><td>RHR54760.1</td><td>Type1.BREX</td><td>ACET-><-?<-Cysteine-Desulfurase||tRNA-><-?<-?<-?<-?||BrxA-DNA-binding->BrxB-BrxF-iSTAND->DUF6079*->N6-METHYLASE->Schlafen+HATPase+wHTH->PglZ->SIGMA+OB-Fold+AAA-ATPase+HKD-DNase-BrxL-><-?||?->PolB->HEPN-></td><td>1216</td><td>Bacteria>FCB_group>Bacteroidetes/Chlorobi_group>Bacteroidota>Bacteroidia>Bacteroidales</td><td>GCA_003479205.1</td><td>Parabacteroides_sp._AF17-28</td><td>DWW90_15085</td><td>22918..26568</td><td>2292241</td><td>BREX-system-P-loop-protein-BrxC-[Parabacteroides-sp.-AF17-28]</td></tr>
<tr><td>HBT86696.1</td><td>Type1.BREX</td><td><-DUF6079*<-BrxB-BrxF-iSTAND<-BrxA-DNA-binding||?-><-tRNA||NTP_transferase+MannoseP_isomer-><-?<-Glyco_hydro_32N+Glyco_hydro_43<-?<-?<-Glyco_hydro_43</td><td>1062</td><td>Bacteria>FCB_group>Bacteroidetes/Chlorobi_group>Bacteroidota>Bacteroidia>Bacteroidales</td><td>GCA_003510605.1</td><td>Porphyromonadaceae_bacterium</td><td>DEB12_12440</td><td>1..3185</td><td>2049046</td><td>BREX-system-P-loop-protein-BrxC-[Porphyromonadaceae-bacterium]</td></tr>
<tr><td>HBQ57053.1</td><td>Type1.BREX</td><td><-SF1-SF2-Helicase||tRNA-><-?<-?<-SIGMA+OB-Fold+AAA-ATPase+HKD-DNase-BrxL<-PglZ<-N6-METHYLASE<-DUF6079*<-BrxB-BrxF-iSTAND<-BrxA-DNA-binding<-?<-REase</td><td>1176</td><td>Bacteria>FCB_group>Bacteroidetes/Chlorobi_group>Bacteroidota>Bacteroidia>Bacteroidales</td><td>GCA_003511405.1</td><td>Porphyromonadaceae_bacterium</td><td>DD786_07890</td><td>28635..32165</td><td>2049046</td><td>BREX-system-P-loop-protein-BrxC-[Porphyromonadaceae-bacterium]</td></tr>
<tr><td>HCI54649.1</td><td>Type1.BREX</td><td><-DUF6079-p*<-BrxB-BrxF-iSTAND<-BrxA-DNA-binding<-HTH_26||<-?<-?<-?<-HTH_3</td><td>227</td><td>Bacteria>FCB_group>Bacteroidetes/Chlorobi_group>Bacteroidota>Bacteroidia>Bacteroidales</td><td>GCA_003521565.1</td><td>Bacteroidales_bacterium</td><td>DFI01_01845</td><td>1..682</td><td>2030927</td><td>BREX-system-P-loop-protein-BrxC-[Bacteroidales-bacterium]</td></tr>
<tr><td>HCY74134.1</td><td>Type1.BREX</td><td><-SAM-METHYLASE<-N6-METHYLASE-p<-DUF6079*<-BrxB-BrxF-iSTAND<-BrxA-DNA-binding</td><td>1205</td><td>Bacteria>FCB_group>Bacteroidetes/Chlorobi_group>Bacteroidota>Bacteroidia>Bacteroidales</td><td>GCA_003537735.1</td><td>Rikenellaceae_bacterium</td><td>DHU75_08775</td><td>2145..5762</td><td>2049048</td><td>BREX-system-P-loop-protein-BrxC-[Rikenellaceae-bacterium]</td></tr>
<tr><td>HCV16199.1</td><td>Type1.BREX</td><td><-DUF4099+DUF3945+TOPRIM+TOPRIM||HTH_3->BrxA-DNA-binding->BrxB-BrxF-iSTAND->DUF6079-p*->DUF6079-p->N6-METHYLASE->Schlafen+HATPase+wHTH->PglZ-></td><td>581</td><td>Bacteria>FCB_group>Bacteroidetes/Chlorobi_group>Bacteroidota>Bacteroidia>Bacteroidales</td><td>GCA_003542255.1</td><td>Rikenellaceae_bacterium</td><td>DF637_07690</td><td>3098..4843</td><td>2049048</td><td>hypothetical-protein-[Rikenellaceae-bacterium]</td></tr>
<tr><td>ROS89081.1</td><td>Type1.BREX</td><td><-SF1-SF2-Helicase<-SIGMA+OB-Fold+AAA-ATPase+HKD-DNase-BrxL<-PglZ||?-><-Schlafen+HATPase+wHTH<-N6-METHYLASE<-DUF6079*<-BrxB-BrxF-iSTAND<-BrxA-DNA-binding<-HTH_3</td><td>1215</td><td>Bacteria>FCB_group>Bacteroidetes/Chlorobi_group>Bacteroidota>Bacteroidia>Bacteroidales</td><td>GCA_003762635.1</td><td>Muribaculaceae_bacterium_Isolate-043_(Harlan)</td><td>EEL36_13385</td><td>16553..20200</td><td>2486463</td><td>BREX-system-P-loop-protein-BrxC-[Muribaculaceae-bacterium-Isolate-043-(Harlan)]</td></tr>
<tr><td>ROT16603.1</td><td>Type1.BREX</td><td><-SIGMA+OB-Fold+AAA-ATPase+HKD-DNase-BrxL<-PglZ<-MORC+DUF3684+SF1-SF2-Helicase<-ACET+TPR<-AT-HTH-HYD2<-Tox-ART-HYD2<-AT-HTH-HYD2<-N6-METHYLASE<-DUF6079*<-BrxB-BrxF-iSTAND<-BrxA-DNA-binding<-HTH_3||SAVED->SMODS->?->?-><-Phage_lysozyme</td><td>1195</td><td>Bacteria>FCB_group>Bacteroidetes/Chlorobi_group>Bacteroidota>Bacteroidia>Bacteroidales</td><td>GCA_003762745.1</td><td>Muribaculaceae_bacterium_Isolate-105_(HZI)</td><td>EEL50_01930</td><td>18926..22513</td><td>2486472</td><td>BREX-system-P-loop-protein-BrxC-[Muribaculaceae-bacterium-Isolate-105-(HZI)]</td></tr>
<tr><td>ROT18783.1</td><td>Type1.BREX</td><td><-SIGMA+OB-Fold+AAA-ATPase+HKD-DNase-BrxL<-PglZ<-Schlafen+HATPase+wHTH<-REase<-N6-METHYLASE<-?<-Tox-ART-HYD2<-?<-DUF6079*<-Four-helical-protein<-BrxB-BrxF-iSTAND<-BrxA-DNA-binding<-HTH_3<-?<-AbiEii<-AbiEii</td><td>1223</td><td>Bacteria>FCB_group>Bacteroidetes/Chlorobi_group>Bacteroidota>Bacteroidia>Bacteroidales</td><td>GCA_003833095.1</td><td>Muribaculaceae_bacterium_Isolate-114_(HZI)</td><td>EEL53_11930</td><td>25441..29112</td><td>2486475</td><td>BREX-system-P-loop-protein-BrxC-[Muribaculaceae-bacterium-Isolate-114-(HZI)]</td></tr>
<tr><td>KAA3824840.1</td><td>Type1.BREX</td><td>DUF6079*->N6-METHYLASE->SIR2+SIR2->Schlafen+HATPase+wHTH->PglZ->SIGMA+OB-Fold+AAA-ATPase+HKD-DNase-BrxL->||BACON+Peptidase_M64-></td><td>1065</td><td>Bacteria>FCB_group>Bacteroidetes/Chlorobi_group>Bacteroidota>Bacteroidia>Bacteroidales</td><td>GCA_008565655.1</td><td>Bacteroides_ovatus</td><td>F3F58_10380</td><td>1..3200</td><td>28116</td><td>BREX-system-P-loop-protein-BrxC,-partial-[Bacteroides-ovatus]</td></tr>
<tr><td>KAA3823632.1</td><td>Type1.BREX</td><td><-DOC+HTH<-?<-N6-METHYLASE<-REase+SF1-SF2-Helicase<-HTH_3<-?<-?||BrxA-DNA-binding->BrxB-BrxF-iSTAND->DUF6079-p*-></td><td>131</td><td>Bacteria>FCB_group>Bacteroidetes/Chlorobi_group>Bacteroidota>Bacteroidia>Bacteroidales</td><td>GCA_008565655.1</td><td>Bacteroides_ovatus</td><td>F3F58_17080</td><td>102919..103310</td><td>28116</td><td>BREX-system-P-loop-protein-BrxC,-partial-[Bacteroides-ovatus]</td></tr>
<tr><td>KAA3934629.1</td><td>Type1.BREX</td><td><-DOC+HTH<-?<-N6-METHYLASE<-REase+SF1-SF2-Helicase<-HTH_3<-?<-?||BrxA-DNA-binding->BrxB-BrxF-iSTAND->DUF6079-p*-></td><td>197</td><td>Bacteria>FCB_group>Bacteroidetes/Chlorobi_group>Bacteroidota>Bacteroidia>Bacteroidales</td><td>GCA_008565965.1</td><td>Bacteroides_ovatus</td><td>F3F50_10370</td><td>48351..48941</td><td>28116</td><td>BREX-system-P-loop-protein-BrxC,-partial-[Bacteroides-ovatus]</td></tr>
<tr><td>KAA3965513.1</td><td>Type1.BREX</td><td><-REase+SF1-SF2-Helicase<-HTH_3<-?<-?||BrxA-DNA-binding->BrxB-BrxF-iSTAND->DUF6079-p*-></td><td>437</td><td>Bacteria>FCB_group>Bacteroidetes/Chlorobi_group>Bacteroidota>Bacteroidia>Bacteroidales</td><td>GCA_008566245.1</td><td>Bacteroides_ovatus</td><td>F3F43_12105</td><td>5385..6696</td><td>28116</td><td>BREX-system-P-loop-protein-BrxC,-partial-[Bacteroides-ovatus]</td></tr>
<tr><td>KAA4088264.1</td><td>Type1.BREX</td><td><-SIGMA+OB-Fold+AAA-ATPase+HKD-DNase-BrxL<-PglZ<-SWIM+DUF3670+SF1-SF2-Helicase<-?<-?<-?<-Peptidase_C39_2<-ACET+TPR<-N6-METHYLASE<-DUF6079*</td><td>1069</td><td>Bacteria>FCB_group>Bacteroidetes/Chlorobi_group>Bacteroidota>Bacteroidia>Bacteroidales</td><td>GCA_008566385.1</td><td>Bacteroides_ovatus</td><td>F3F49_00290</td><td>70210..73420</td><td>28116</td><td>BREX-system-P-loop-protein-BrxC,-partial-[Bacteroides-ovatus]</td></tr>
<tr><td>KAA4085421.1</td><td>Type1.BREX</td><td><-DUF6079-p*<-BrxB-BrxF-iSTAND<-BrxA-DNA-binding<-wHTH+WYL+WCX<-HTH_3||?->?->?->NUDIX+NUDIX-></td><td>134</td><td>Bacteria>FCB_group>Bacteroidetes/Chlorobi_group>Bacteroidota>Bacteroidia>Bacteroidales</td><td>GCA_008566385.1</td><td>Bacteroides_ovatus</td><td>F3F49_09110</td><td>1..402</td><td>28116</td><td>BREX-system-P-loop-protein-BrxC,-partial-[Bacteroides-ovatus]</td></tr>
<tr><td>KAA4007861.1</td><td>Type1.BREX</td><td><-NUDIX+NUDIX<-?<-?||HTH_3->wHTH+WYL+WCX->BrxA-DNA-binding->BrxB-BrxF-iSTAND->DUF6079*->N6-METHYLASE->ACET+TPR->Peptidase_C39_2->?->?->?->SWIM+DUF3670+SF1-SF2-Helicase->PglZ->SIGMA+OB-Fold+AAA-ATPase+HKD-DNase-BrxL-></td><td>1208</td><td>Bacteria>FCB_group>Bacteroidetes/Chlorobi_group>Bacteroidota>Bacteroidia>Bacteroidales</td><td>GCA_008566415.1</td><td>Bacteroides_ovatus</td><td>F3F37_14290</td><td>84100..87726</td><td>28116</td><td>BREX-system-P-loop-protein-BrxC-[Bacteroides-ovatus]</td></tr>
<tr><td>KAA4743085.1</td><td>Type1.BREX</td><td>AbiEi_4-><-Phage_int_SAM_5+Phage_integrase||STYKinase-like->?->?->||HTH_3->BrxA-DNA-binding->BrxB-BrxF-iSTAND->DUF6079*->DUF6079-p->N6-METHYLASE->Schlafen+HATPase+wHTH->PglZ->SIGMA+OB-Fold+AAA-ATPase+HKD-DNase-BrxL->?->URI-DNase+RAMA->DOC+HTH-></td><td>1056</td><td>Bacteria>FCB_group>Bacteroidetes/Chlorobi_group>Bacteroidota>Bacteroidia>Bacteroidales</td><td>GCA_008569225.1</td><td>Bacteroides_fragilis</td><td>F3B36_09960</td><td>278156..281326</td><td>817</td><td>BREX-system-P-loop-protein-BrxC-[Bacteroides-fragilis]</td></tr>
<tr><td>KAB5317657.1</td><td>Type1.BREX</td><td><-Phage_int_SAM_5<-Phage_integrase<-Phage_int_SAM_5+Phage_integrase<-SIGMA+OB-Fold+AAA-ATPase+HKD-DNase-BrxL<-PglZ<-N6-METHYLASE<-REase<-OLD-ABC<-DUF6079*<-BrxB-BrxF-iSTAND<-BrxA-DNA-binding<-HTH_3<-SMS-KaiC<-XIS-HTH</td><td>1196</td><td>Bacteria>FCB_group>Bacteroidetes/Chlorobi_group>Bacteroidota>Bacteroidia>Bacteroidales</td><td>GCA_009025145.1</td><td>Bacteroides_stercoris</td><td>F9949_11590</td><td>61674..65264</td><td>46506</td><td>BREX-system-P-loop-protein-BrxC-[Bacteroides-stercoris]</td></tr>
<tr><td>NBJ07681.1</td><td>Type1.BREX</td><td>LRR_5->HTH_3->PSBP->?->?->?->HTH_3->BrxA-DNA-binding->BrxB-BrxF-iSTAND->DUF6079*->N6-METHYLASE->REase->Schlafen+HATPase+wHTH->PglZ->SIGMA+OB-Fold+AAA-ATPase+HKD-DNase-BrxL->SF1-SF2-Helicase->?->?->MPTase-></td><td>1223</td><td>Bacteria>FCB_group>Bacteroidetes/Chlorobi_group>Bacteroidota>Bacteroidia>Bacteroidales</td><td>GCA_009911575.1</td><td>Alistipes_sp._Z76</td><td>D1647_16085</td><td>15947..19618</td><td>2304565</td><td>BREX-system-P-loop-protein-BrxC-[Alistipes-sp.-Z76]</td></tr>
<tr><td>NLA16135.1</td><td>Type1.BREX</td><td><-HTH_3||HTH+REase-><-SIGMA+OB-Fold+AAA-ATPase+HKD-DNase-BrxL<-PglZ<-N6-METHYLASE<-DUF6079*<-BrxB-BrxF-iSTAND<-BrxA-DNA-binding<-HTH_3<-?||?->tRNA-><-?<-?||?->LuxR-HTH-></td><td>1174</td><td>Bacteria>FCB_group>Bacteroidetes/Chlorobi_group>Bacteroidota>Bacteroidia>Bacteroidales</td><td>GCA_012510445.1</td><td>Bacteroidales_bacterium</td><td>GX877_06335</td><td>9723..13247</td><td>2030927</td><td>BREX-system-P-loop-protein-BrxC-[Bacteroidales-bacterium]</td></tr>
<tr><td>NLN95595.1</td><td>Type1.BREX</td><td>PolB->HEPN->?-><-?<-SIGMA+OB-Fold+AAA-ATPase+HKD-DNase-BrxL<-PglZ<-?<-N6-METHYLASE<-?<-DUF6079*<-BrxB-BrxF-iSTAND<-BrxA-DNA-binding<-HTH_3||?->TRD+TRD-></td><td>1176</td><td>Bacteria>FCB_group>Bacteroidetes/Chlorobi_group>Bacteroidota>Bacteroidia>Bacteroidales</td><td>GCA_012522795.1</td><td>Bacteroidales_bacterium</td><td>GX128_05430</td><td>111716..115246</td><td>2030927</td><td>BREX-system-P-loop-protein-BrxC-[Bacteroidales-bacterium]</td></tr>
<tr><td>HHU96164.1</td><td>Type1.BREX</td><td><-SIGMA+OB-Fold+AAA-ATPase+Lon-Protease-BrxL<-PglZ<-AbiEii<-AbiEi_4<-N6-METHYLASE<-DUF6079*<-BrxB-BrxF-iSTAND<-BrxA-DNA-binding<-?<-?||?-><-tRNA||?->?->?->sGTP-></td><td>1202</td><td>Bacteria>FCB_group>Bacteroidetes/Chlorobi_group>Bacteroidota>Bacteroidia>Bacteroidales</td><td>GCA_012839475.1</td><td>Petrimonas_sp.</td><td>GXX67_02725</td><td>36773..40381</td><td>2023866</td><td>BREX-system-P-loop-protein-BrxC-[Petrimonas-sp.]</td></tr>
<tr><td>QKG81082.1</td><td>Type1.BREX</td><td>ABhydrolase-><-PDZ+Peptidase<-?<-?<-?||BrxA-DNA-binding->BrxB-BrxF-iSTAND->DUF6079*->N6-METHYLASE->PglZ->SIGMA+OB-Fold+AAA-ATPase+Lon-Protease-BrxL-><-tRNA<-?||?->sGTP-><-TPR+TPR+TPR+TPR+TPR+TPR+HISKIN+REC</td><td>1206</td><td>Bacteria>FCB_group>Bacteroidetes/Chlorobi_group>Bacteroidota>Bacteroidia>Bacteroidales</td><td>GCA_013265555.1</td><td>Tenuifilum_thalassicum</td><td>FHG85_12670</td><td>3070622..3074242</td><td>2590900</td><td>BREX-system-P-loop-protein-BrxC-[Tenuifilum-thalassicum]</td></tr>
<tr><td>MBD5181931.1</td><td>Type1.BREX</td><td><-Acyltransferase<-?<-?||HISKIN+MORC->REC+OmpR-HTH->BrxA-DNA-binding->BrxB-BrxF-iSTAND->DUF6079*->N6-METHYLASE->OLD-ABC->TOPRIM->PglZ->SIGMA+OB-Fold+AAA-ATPase+HKD-DNase-BrxL-></td><td>1193</td><td>Bacteria>FCB_group>Bacteroidetes/Chlorobi_group>Bacteroidota>Bacteroidia>Bacteroidales</td><td>GCA_014800235.1</td><td>Bacteroidales_bacterium</td><td>HDS98_05210</td><td>20716..24297</td><td>2030927</td><td>BREX-system-P-loop-protein-BrxC-[Bacteroidales-bacterium]</td></tr>
<tr><td>MBD5263514.1</td><td>Type1.BREX</td><td><-Calcineurin<-?<-?<-?<-Aminotran_1_2||?-><-?||BrxA-DNA-binding->BrxB-BrxF-iSTAND->DUF6079*->N6-METHYLASE->Schlafen+HATPase+wHTH->SF1-SF2-Helicase->BirA-HTH->PglZ->SIGMA+OB-Fold+AAA-ATPase+HKD-DNase-BrxL-></td><td>1221</td><td>Bacteria>FCB_group>Bacteroidetes/Chlorobi_group>Bacteroidota>Bacteroidia>Bacteroidales</td><td>GCA_014801365.1</td><td>Bacteroides_sp.</td><td>HDS39_05520</td><td>20086..23751</td><td>29523</td><td>BREX-system-P-loop-protein-BrxC-[Bacteroides-sp.]</td></tr>
<tr><td>MBD5304899.1</td><td>Type1.BREX</td><td>Tox-RelE->HTH_3-><-Schlafen+HATPase+wHTH<-?||?-><-?||HTH_3->BrxA-DNA-binding->BrxB-BrxF-iSTAND->DUF6079*->N6-METHYLASE->Schlafen+HATPase+wHTH->PglZ->SIGMA+OB-Fold+AAA-ATPase+Lon-Protease-BrxL->LRR+LRR-><-Arm-DNA-bind_5+Phage_int_SAM_5+Phage_integrase</td><td>1182</td><td>Bacteria>FCB_group>Bacteroidetes/Chlorobi_group>Bacteroidota>Bacteroidia>Bacteroidales</td><td>GCA_014801805.1</td><td>Bacteroides_sp.</td><td>HDS12_01205</td><td>15900..19448</td><td>29523</td><td>BREX-system-P-loop-protein-BrxC-[Bacteroides-sp.]</td></tr>
<tr><td>MBD5349669.1</td><td>Type1.BREX</td><td><-ABhydrolase<-JAB<-?<-?<-SIGMA+OB-Fold+AAA-ATPase+HKD-DNase-BrxL<-Schlafen+HATPase+wHTH<-?<-N6-METHYLASE<-Four-helical-protein<-DUF6079*<-BrxB-BrxF-iSTAND<-BrxA-DNA-binding<-HTH_3<-TraJ-RHH||Zn_Tnp_IS1595+DDE_Tnp_IS1595->?-><-?<-HTH_58<-METHYLASE</td><td>1219</td><td>Bacteria>FCB_group>Bacteroidetes/Chlorobi_group>Bacteroidota>Bacteroidia>Bacteroidales</td><td>GCA_014802205.1</td><td>Bacteroides_sp.</td><td>HDR89_02130</td><td>36441..40100</td><td>29523</td><td>BREX-system-P-loop-protein-BrxC-[Bacteroides-sp.]</td></tr>
<tr><td>MBD5357714.1</td><td>Type1.BREX</td><td>AAA+->?->XIS-HTH-><-?<-?<-?<-?||BrxA-DNA-binding->BrxB-BrxF-iSTAND->DUF6079*->N6-METHYLASE->HU-HIG+REase->PglZ->SIGMA+OB-Fold+AAA-ATPase+HKD-DNase-BrxL->REase->?->CASPASE-></td><td>1203</td><td>Bacteria>FCB_group>Bacteroidetes/Chlorobi_group>Bacteroidota>Bacteroidia>Bacteroidales</td><td>GCA_014802265.1</td><td>Bacteroides_sp.</td><td>HDR88_12035</td><td>22916..26527</td><td>29523</td><td>BREX-system-P-loop-protein-BrxC-[Bacteroides-sp.]</td></tr>
<tr><td>MBD5419378.1</td><td>Type1.BREX</td><td>BrxB-BrxF-iSTAND->DUF6079*->N6-METHYLASE->Schlafen+HATPase+wHTH->PglZ->SIGMA+OB-Fold+AAA-ATPase+HKD-DNase-BrxL->?->Calcineurin->Glyco_hydro_3-></td><td>1201</td><td>Bacteria>FCB_group>Bacteroidetes/Chlorobi_group>Bacteroidota>Bacteroidia>Bacteroidales</td><td>GCA_014803765.1</td><td>Bacteroides_sp.</td><td>HDR48_05020</td><td>1391..4996</td><td>29523</td><td>BREX-system-P-loop-protein-BrxC-[Bacteroides-sp.]</td></tr>
<tr><td>MBE6254618.1</td><td>Type1.BREX</td><td><-SF1-SF2-Helicase<-?<-SIGMA+OB-Fold+AAA-ATPase+HKD-DNase-BrxL<-PglZ<-SIR2<-N6-METHYLASE<-DUF6079*<-BrxB-BrxF-iSTAND<-BrxA-DNA-binding<-wHTH+WYL+WCX</td><td>1211</td><td>Bacteria>FCB_group>Bacteroidetes/Chlorobi_group>Bacteroidota>Bacteroidia>Bacteroidales</td><td>GCA_015060555.1</td><td>Prevotella_sp.</td><td>E7105_03785</td><td>25107..28742</td><td>59823</td><td>BREX-system-P-loop-protein-BrxC-[Prevotella-sp.]</td></tr>
<tr><td>MBN2915288.1</td><td>Type1.BREX</td><td><-McrC-NTD+REase<-McrB-AAA<-SIGMA+OB-Fold+AAA-ATPase+HKD-DNase-BrxL<-PglZ<-SIR2+SIR2<-N6-METHYLASE<-TOPRIM<-OLD-ABC<-DUF6079-p*</td><td>971</td><td>Bacteria>FCB_group>Bacteroidetes/Chlorobi_group>Bacteroidota>Bacteroidia>Bacteroidales</td><td>GCA_016939035.1</td><td>Prevotella_sp.</td><td>JTJ26_04215</td><td>25012..27927</td><td>59823</td><td>BREX-system-P-loop-protein-BrxC-[Prevotella-sp.]</td></tr>
<tr><td>MBQ2195595.1</td><td>Type1.BREX</td><td>ABC-ATPase+SbcC->HTH_26->BrxA-DNA-binding->BrxB-BrxF-iSTAND->DUF6079*-><-DUF4372+DDE_Tnp_1||HEPN-><-?<-Relaxase</td><td>1009</td><td>Bacteria>FCB_group>Bacteroidetes/Chlorobi_group>Bacteroidota>Bacteroidia>Bacteroidales</td><td>GCA_017389665.1</td><td>Prevotella_sp.</td><td>II407_01515</td><td>3565..6594</td><td>59823</td><td>BREX-system-P-loop-protein-BrxC-[Prevotella-sp.]</td></tr>
<tr><td>MBQ2852763.1</td><td>Type1.BREX</td><td>AbiEi_4->AbiEii-><-Response_reg||HTH_3->BrxA-DNA-binding->BrxB-BrxF-iSTAND->DUF6079*->SbcC->N6-METHYLASE->OLD-ABC->HNH->PglZ->SIGMA+OB-Fold+AAA-ATPase+Lon-Protease-BrxL-><-?<-?<-XIS-HTH</td><td>1215</td><td>Bacteria>FCB_group>Bacteroidetes/Chlorobi_group>Bacteroidota>Bacteroidia>Bacteroidales</td><td>GCA_017409275.1</td><td>Bacteroidales_bacterium</td><td>IJE76_05160</td><td>110084..113731</td><td>2030927</td><td>hypothetical-protein-[Bacteroidales-bacterium]</td></tr>
<tr><td>MBQ6178640.1</td><td>Type1.BREX</td><td>ArdCN+PRTase->?->?->?-><-wHTH+WYL+WCX<-?||HD->BrxA-DNA-binding->BrxB-BrxF-iSTAND->DUF6079*->N6-METHYLASE->REase->PglZ->SIGMA+OB-Fold+AAA-ATPase+HKD-DNase-BrxL-></td><td>1202</td><td>Bacteria>FCB_group>Bacteroidetes/Chlorobi_group>Bacteroidota>Bacteroidia>Bacteroidales</td><td>GCA_017429185.1</td><td>Bacteroidales_bacterium</td><td>IJK32_03880</td><td>18867..22475</td><td>2030927</td><td>BREX-system-P-loop-protein-BrxC-[Bacteroidales-bacterium]</td></tr>
<tr><td>MBR0492955.1</td><td>Type1.BREX</td><td>Relaxase->?-><-Response_reg<-SIGMA+OB-Fold+AAA-ATPase+Lon-Protease-BrxL<-PglZ<-DOC+HTH<-N6-METHYLASE<-SbcC<-SbcC<-DUF6079*<-BrxB-BrxF-iSTAND<-BrxA-DNA-binding<-HTH_3||?->TPR+TPR+TPR+TPR+TPR+TPR->?-><-IG+IG</td><td>1215</td><td>Bacteria>FCB_group>Bacteroidetes/Chlorobi_group>Bacteroidota>Bacteroidia>Bacteroidales</td><td>GCA_017430125.1</td><td>Muribaculaceae_bacterium</td><td>IJJ83_04260</td><td>18614..22261</td><td>2498093</td><td>BREX-system-P-loop-protein-BrxC-[Muribaculaceae-bacterium]</td></tr>
<tr><td>MBQ6872144.1</td><td>Type1.BREX</td><td>BrxA-DNA-binding->BrxB-BrxF-iSTAND->DUF6079*->N6-METHYLASE->PglZ->SIGMA+OB-Fold+AAA-ATPase+HKD-DNase-BrxL-></td><td>1190</td><td>Bacteria>FCB_group>Bacteroidetes/Chlorobi_group>Bacteroidota>Bacteroidia>Bacteroidales</td><td>GCA_017435685.1</td><td>Bacteroidales_bacterium</td><td>IJO23_07295</td><td>3964..7536</td><td>2030927</td><td>BREX-system-P-loop-protein-BrxC-[Bacteroidales-bacterium]</td></tr>
<tr><td>MBQ6577354.1</td><td>Type1.BREX</td><td><-SIGMA+OB-Fold+AAA-ATPase+Lon-Protease-BrxL<-PglZ<-REase<-N6-METHYLASE<-DUF6079-p*</td><td>744</td><td>Bacteria>FCB_group>Bacteroidetes/Chlorobi_group>Bacteroidota>Bacteroidia>Bacteroidales</td><td>GCA_017438445.1</td><td>Bacteroidales_bacterium</td><td>IJL91_06365</td><td>8459..10695</td><td>2030927</td><td>hypothetical-protein-[Bacteroidales-bacterium]</td></tr>
<tr><td>MBR0262391.1</td><td>Type1.BREX</td><td><-DUF6079-p*<-BrxB-BrxF-iSTAND<-BrxA-DNA-binding<-wHTH+WYL+WCX<-HTH_3||HTH_3->?-><-Phage_lysozyme<-VirD4-TraG+VirD4-FtsK</td><td>992</td><td>Bacteria>FCB_group>Bacteroidetes/Chlorobi_group>Bacteroidota>Bacteroidia>Bacteroidales</td><td>GCA_017443345.1</td><td>Prevotella_sp.</td><td>IJQ60_00735</td><td>1..2978</td><td>59823</td><td>BREX-system-P-loop-protein-BrxC-[Prevotella-sp.]</td></tr>
<tr><td>MBQ9588226.1</td><td>Type1.BREX</td><td>Peptidase->?-><-HD||BrxA-DNA-binding->BrxB-BrxF-iSTAND->DUF6079*->N6-METHYLASE->HTH+REase->N6-METHYLASE-p->PglZ->SIGMA+OB-Fold+AAA-ATPase+HKD-DNase-BrxL-></td><td>1183</td><td>Bacteria>FCB_group>Bacteroidetes/Chlorobi_group>Bacteroidota>Bacteroidia>Bacteroidales</td><td>GCA_017446745.1</td><td>Bacteroidales_bacterium</td><td>IJR26_10275</td><td>19779..23330</td><td>2030927</td><td>BREX-system-P-loop-protein-BrxC-[Bacteroidales-bacterium]</td></tr>
<tr><td>MBR0254975.1</td><td>Type1.BREX</td><td><-Glyco_hydro_43+GH43_C2<-HTH_3<-?<-?||BrxA-DNA-binding->BrxB-BrxF-iSTAND->DUF6079*->N6-METHYLASE->PglZ->SIGMA+OB-Fold+AAA-ATPase+Lon-Protease-BrxL->TIR->?->STAND->HTH_3->REase->HEPN-></td><td>1217</td><td>Bacteria>FCB_group>Bacteroidetes/Chlorobi_group>Bacteroidota>Bacteroidia>Bacteroidales</td><td>GCA_017447265.1</td><td>Bacteroidales_bacterium</td><td>IJQ69_03550</td><td>66425..70078</td><td>2030927</td><td>BREX-system-P-loop-protein-BrxC-[Bacteroidales-bacterium]</td></tr>
<tr><td>MBQ7672328.1</td><td>Type1.BREX</td><td>REase+35exo->?->?->LRR_5->LRR_5->BrxA-DNA-binding->BrxB-BrxF-iSTAND->DUF6079*->N6-METHYLASE->PglZ->SIGMA+OB-Fold+AAA-ATPase+HKD-DNase-BrxL->?->DOC+HTH-></td><td>1193</td><td>Bacteria>FCB_group>Bacteroidetes/Chlorobi_group>Bacteroidota>Bacteroidia>Bacteroidales</td><td>GCA_017450385.1</td><td>Paludibacteraceae_bacterium</td><td>IJS49_04535</td><td>76772..80353</td><td>2748031</td><td>BREX-system-P-loop-protein-BrxC-[Paludibacteraceae-bacterium]</td></tr>
<tr><td>MBQ8673311.1</td><td>Type1.BREX</td><td>Peptidase_M20->?-><-?<-?<-?<-?||BrxA-DNA-binding->BrxB-BrxF-iSTAND->DUF6079*->N6-METHYLASE-p-></td><td>1180</td><td>Bacteria>FCB_group>Bacteroidetes/Chlorobi_group>Bacteroidota>Bacteroidia>Bacteroidales</td><td>GCA_017471925.1</td><td>Bacteroides_sp.</td><td>IJ511_04565</td><td>26677..30219</td><td>29523</td><td>BREX-system-P-loop-protein-BrxC-[Bacteroides-sp.]</td></tr>
<tr><td>MBR2237166.1</td><td>Type1.BREX</td><td><-Glyco_hydro_32N+Glyco_hydro_43<-?<-?<-HD<-SIGMA+OB-Fold+AAA-ATPase+Lon-Protease-BrxL<-PglZ<-Schlafen+HATPase+wHTH<-N6-METHYLASE<-DUF6079*<-BrxB-BrxF-iSTAND<-BrxA-DNA-binding<-HTH_3||?->?->DsbC+DsbD+Thioredoxin_7->PDZ+Peptidase-></td><td>1217</td><td>Bacteria>FCB_group>Bacteroidetes/Chlorobi_group>Bacteroidota>Bacteroidia>Bacteroidales</td><td>GCA_017475265.1</td><td>Prevotella_sp.</td><td>IJ887_04725</td><td>43951..47604</td><td>59823</td><td>BREX-system-P-loop-protein-BrxC-[Prevotella-sp.]</td></tr>
</table>

| Protein-ID | System | Gene-Neighbourhoods | Length | Taxonomy | Assembly-ID | Strain | Locus-Tag | Genomic-coordinates | TaxID | Fasta-define |
|---|---|---|---|---|---|---|---|---|---|---|
| MBR3491242.1 | Type1.BREX | <-CopG/RHH<-SIGMA-HTH+MA<-KAP-NTPase\|\|BrxA-DNA-binding->BrxB-BrxF-iSTAND->DUF6079*->N6-METHYLASE-p-> | 1195 | [illegible] | [illegible] | [illegible] | [illegible] | [illegible] | [illegible] | [illegible] |
| MBR6936560.1 | Type1.BREX | <-SIGMA+OB-Fold+AAA-ATPase+Lon-Protease-BrxL<-PglZ<-Schlafen+HATPase+wHTH<-N6-METHYLASE<-?<-OLD-ABC<-ParB<-DUF6079-p*<-DUF6079-p | 827 | [illegible] | [illegible] | [illegible] | [illegible] | [illegible] | [illegible] | [illegible] |
| MBR6937288.1 | Type1.BREX | HTH_3->Couple_hipA->HipA_C->Phage_integrase->?->HTH_3->BrxA-DNA-binding->BrxB-BrxF-iSTAND->DUF6079-p*-> | 288 | [illegible] | [illegible] | [illegible] | [illegible] | [illegible] | [illegible] | [illegible] |
| MBR4647494.1 | Type1.BREX | MNS+CR-ATPase5-><-?\|\|?->?->Calcineurin->SbcC+CCDC158->BrxA-DNA-binding->BrxB-BrxF-iSTAND->DUF6079*->OLD-ABC->PIN->N6-METHYLASE->PglZ->SIGMA+OB-Fold+AAA-ATPase+Lon-Protease-BrxL->?->?->?->Cysteine-Desulfurase-> | 1174 | [illegible] | [illegible] | [illegible] | [illegible] | [illegible] | [illegible] | [illegible] |
| MBR3727503.1 | Type1.BREX | <-Response_reg<-SIGMA+OB-Fold+AAA-ATPase+Lon-Protease-BrxL<-PglZ<-DOC+HTH<-N6-METHYLASE<-SbcC<-SbcC<-DUF6079*<-BrxB-BrxF-iSTAND<-BrxA-DNA-binding<-HTH_3\|\|LRR_5+LRR_5+LRR_5+LRR_5->IG+IG->wHTH+WYL->?->SSB-><-NTP_transferase+MannoseP_isomer | 1215 | [illegible] | [illegible] | [illegible] | [illegible] | [illegible] | [illegible] | [illegible] |
| MBR3711479.1 | Type1.BREX | <-CR-ATPase5+REase\|\|BrxA-DNA-binding->BrxB-BrxF-iSTAND->DUF6079*->N6-METHYLASE-p-> | 1195 | [illegible] | [illegible] | [illegible] | [illegible] | [illegible] | [illegible] | [illegible] |
| MBR6863890.1 | Type1.BREX | <-ParB+HNH<-wHTH+WYL+WCX<-?\|\|?->Couple_hipA+HIRAN->EVE->HD->BrxA-DNA-binding->BrxB-BrxF-iSTAND->DUF6079*->N6-METHYLASE->PglZ->SIGMA+OB-Fold+AAA-ATPase+HKD-DNase-BrxL->?->SF1-SF2-Helicase-> | 1202 | [illegible] | [illegible] | [illegible] | [illegible] | [illegible] | [illegible] | [illegible] |
| MBR4380326.1 | Type1.BREX | <-HTH+REase<-Phage_int_SAM_5+Phage_integrase<-NACHT+nSTAND4<-?<-SIGMA+OB-Fold+AAA-ATPase+Lon-Protease-BrxL<-PglZ<-Schlafen+HATPase+wHTH<-N6-METHYLASE<-DUF6079*<-BrxB-BrxF-iSTAND<-BrxA-DNA-binding<-HTH_3\|\|XIS-HTH->?->?-><-Phage_int_SAM_5 | 1216 | [illegible] | [illegible] | [illegible] | [illegible] | [illegible] | [illegible] | [illegible] |
| MBR5893775.1 | Type1.BREX | <-SIGMA+OB-Fold+AAA-ATPase+HKD-DNase-BrxL<-PglZ<-DOC+HTH<-N6-METHYLASE<-?<-DUF6079*<-BrxB-BrxF-iSTAND<-BrxA-DNA-binding<-?<-HAD+HAD | 1185 | [illegible] | [illegible] | [illegible] | [illegible] | [illegible] | [illegible] | [illegible] |
| MBR5630362.1 | Type1.BREX | <-CopG/RHH\|\|BrxA-DNA-binding->BrxB-BrxF-iSTAND->DUF6079*->N6-METHYLASE->PglZ->SIGMA+OB-Fold+AAA-ATPase+HKD-DNase-BrxL-> | 1171 | [illegible] | [illegible] | [illegible] | [illegible] | [illegible] | [illegible] | [illegible] |
| MBR5576788.1 | Type1.BREX | <-LRR_5+LRR_5+LRR_5+LRR_5<-?<-?<-?<-?<-?<-?\|\|BrxA-DNA-binding->BrxB-BrxF-iSTAND->DUF6079*->N6-METHYLASE->SbcC+SbcC->?->ACET+TPR->?->SF1-SF2-Helicase->PglZ->SIGMA+OB-Fold+AAA-ATPase+HKD-DNase-BrxL-> | 1209 | [illegible] | [illegible] | [illegible] | [illegible] | [illegible] | [illegible] | [illegible] |
| MBO7470104.1 | Type1.BREX | BrxA-DNA-binding->BrxB-BrxF-iSTAND->DUF6079*->N6-METHYLASE->?->?->PglZ->SIGMA+OB-Fold+AAA-ATPase+HKD-DNase-BrxL-> | 1169 | [illegible] | [illegible] | [illegible] | [illegible] | [illegible] | [illegible] | [illegible] |
| MBO7636706.1 | Type1.BREX | <-Peptidase_C39_2<-N6-METHYLASE<-SIR2<-N6-METHYLASE<-DUF6079*<-BrxB-BrxF-iSTAND<-BrxA-DNA-binding | 1214 | [illegible] | [illegible] | [illegible] | [illegible] | [illegible] | [illegible] | [illegible] |
| MBP6870975.1 | Type1.BREX | <-SF1-SF2-Helicase<-STYKinase-like<-TRPR-HTH\|\|HTH_3->Couple_hipA->HipA_C->?->BrxA-DNA-binding->BrxB-BrxF-iSTAND->DUF6079*->?->?->?->CR-Kinase+Classical-AAA->?->?->?->N6-METHYLASE-> | 1166 | [illegible] | [illegible] | [illegible] | [illegible] | [illegible] | [illegible] | [illegible] |
| MBP9983647.1 | Type1.BREX | <-N6-METHYLASE<-DUF6079*<-BrxB-BrxF-iSTAND<-BrxA-DNA-binding<-HTH_3<-SMS-KaiC<-XIS-HTH<-?<-Phage_int_SAM_5+Phage_integrase<-XIS-HTH+XIS-HTH+XIS-HTH+XIS-HTH<-sGTP | 1184 | [illegible] | [illegible] | [illegible] | [illegible] | [illegible] | [illegible] | [illegible] |
| MBS6238336.1 | Type1.BREX | <-PAS+HISKIN\|\|?-><-AraC-HTH+AraC-HTH<-SIGMA+OB-Fold+AAA-ATPase+HKD-DNase-BrxL<-PglZ<-Schlafen+HATPase+wHTH<-ParB+HNH<-ParB+HNH<-N6-METHYLASE<-DUF6079*<-BrxB-BrxF-iSTAND<-BrxA-DNA-binding<-HTH_3<-?<-?<-?<-?\|\|SIGMA-HTH-> | 1205 | [illegible] | [illegible] | [illegible] | [illegible] | [illegible] | [illegible] | [illegible] |
| MBV1818013.1 | Type1.BREX | HNH->?-><-D5N+D5ATPase<-?<-SF1-SF2-Helicase+RdgC+REase<-SIGMA+OB-Fold+AAA-ATPase+Lon-Protease-BrxL-p<-PglZ<-N6-METHYLASE<-DUF6079*<-BrxB-BrxF-iSTAND<-BrxA-DNA-binding<-?<-REase<-SF1-SF2-Helicase<-OLD-ABC+TOPRIM | 1195 | [illegible] | [illegible] | [illegible] | [illegible] | [illegible] | [illegible] | [illegible] |
| MBV1819082.1 | Type1.BREX | <-RNase<-PglZ<-N6-METHYLASE<-DUF6079*<-BrxB-BrxF-iSTAND<-BrxA-DNA-binding | 1185 | [illegible] | [illegible] | [illegible] | [illegible] | [illegible] | [illegible] | [illegible] |
| MBW6491485.1 | Type1.BREX | <-HTH_3\|\|XIS-HTH->?->?->?->BrxA-DNA-binding->BrxB-BrxF-iSTAND->DUF6079*->HTH+REase->N6-METHYLASE->AbiEii->PglZ->SIGMA+OB-Fold+AAA-ATPase+Lon-Protease-BrxL->REase->HEPN->ClpABN-AAA+ClpABC-AAA-> | 1144 | [illegible] | [illegible] | [illegible] | [illegible] | [illegible] | [illegible] | [illegible] |
| MCB9017748.1 | Type1.BREX | <-ACET\|\|?-><-SIGMA+OB-Fold+AAA-ATPase+HKD-DNase-BrxL<-PglZ<-N6-METHYLASE-p<-HTH+REase<-?<-Schlafen+HATPase+wHTH<-DUF6079*<-BrxB-BrxF-iSTAND<-BrxA-DNA-binding<-?<-PALP | 1207 | [illegible] | [illegible] | [illegible] | [illegible] | [illegible] | [illegible] | [illegible] |
| HJF82219.1 | Type1.BREX | <-SF1-SF2-Helicase-p<-HTH_3<-?<-?\|\|BrxA-DNA-binding->BrxB-BrxF-iSTAND->DUF6079*->OLD-ABC->TOPRIM->N6-METHYLASE->SIR2+SIR2->PglZ->SIGMA+OB-Fold+AAA-ATPase+HKD-DNase-BrxL-> | 1195 | [illegible] | [illegible] | [illegible] | [illegible] | [illegible] | [illegible] | [illegible] |
| HJF93068.1 | Type1.BREX | <-SIR2+SIR2<-N6-METHYLASE<-DUF6079*<-BrxB-BrxF-iSTAND<-BrxA-DNA-binding<-HTH_3 | 1217 | [illegible] | [illegible] | [illegible] | [illegible] | [illegible] | [illegible] | [illegible] |
| MCC8037087.1 | Type1.BREX | <-HAD+PRTase+PRTase+REase+SF1-SF2-Helicase<-?<-?<-?<-STYKIN<-?<-SIGMA+OB-Fold+AAA-ATPase+HKD-DNase-BrxL<-PglZ<-DUF6079*<-BrxB-BrxF-iSTAND<-BrxA-DNA-binding<-?<-?<-?<-?<-Acetate_kinase | 1219 | [illegible] | [illegible] | [illegible] | [illegible] | [illegible] | [illegible] | [illegible] |
| MCC8072421.1 | Type1.BREX | <-TOPRIM+Topoisom_bac<-?<-?<-?\|\|?->?->tRNA->BrxA-DNA-binding->BrxB-BrxF-iSTAND->DUF6079*-> | 1191 | [illegible] | [illegible] | [illegible] | [illegible] | [illegible] | [illegible] | [illegible] |
| MCC8115076.1 | Type1.BREX | PglZ->SIGMA+OB-Fold+AAA-ATPase+HKD-DNase-BrxL->BrxA-DNA-binding->BrxB-BrxF-iSTAND->DUF6079-p*->DUF6079->N6-METHYLASE-> | 159 | [illegible] | [illegible] | [illegible] | [illegible] | [illegible] | [illegible] | [illegible] |
| MCC8115077.1 | Type1.BREX | PglZ->SIGMA+OB-Fold+AAA-ATPase+HKD-DNase-BrxL->BrxA-DNA-binding->BrxB-BrxF-iSTAND->DUF6079-p->DUF6079*->N6-METHYLASE-> | 1040 | [illegible] | [illegible] | [illegible] | [illegible] | [illegible] | [illegible] | [illegible] |
| MCC8119638.1 | Type1.BREX | <-PglZ<-Schlafen+HATPase+wHTH<-?<-ParB+HNH<-AAA-ATPase-p<-Schlafen+HATPase+wHTH<-?<-N6-METHYLASE<-HTH+REase<-DUF6079*<-BrxB-BrxF-iSTAND<-BrxA-DNA-binding<-tRNA<-35exo+T7DNApol | 1189 | [illegible] | [illegible] | [illegible] | [illegible] | [illegible] | [illegible] | [illegible] |
| MCD8313154.1 | Type1.BREX | <-N6-METHYLASE-p<-DUF6079-p*<-BrxB-BrxF-iSTAND<-BrxA-DNA-binding | 762 | [illegible] | [illegible] | [illegible] | [illegible] | [illegible] | [illegible] | [illegible] |
| MCD8387916.1 | Type1.BREX | <-AbiEii<-?<-?<-?<-SIGMA+OB-Fold+AAA-ATPase+Lon-Protease-BrxL<-PglZ<-N6-METHYLASE<-DUF6079*<-BrxB-BrxF-iSTAND<-BrxA-DNA-binding<-sGTP-MglA+sGTP<-CNMP+CRP-HTH | 1201 | [illegible] | [illegible] | [illegible] | [illegible] | [illegible] | [illegible] | [illegible] |
| MCE5179211.1 | Type1.BREX | <-SIGMA+OB-Fold+AAA-ATPase+Lon-Protease-BrxL<-PglZ<-AbiEii<-AbiEi_4<-N6-METHYLASE<-?<-DUF6079*<-BrxB-BrxF-iSTAND<-BrxA-DNA-binding<-?<-?<-?<-?<-SIGMA-HTH | 1197 | [illegible] | [illegible] | [illegible] | [illegible] | [illegible] | [illegible] | [illegible] |
| MCF0219894.1 | Type1.BREX | CR-Kinase+Classical-AAA->?-><-TPR<-SIGMA+OB-Fold+AAA-ATPase+Lon-Protease-BrxL<-PglZ<-?<-Schlafen+HATPase+wHTH<-N6-METHYLASE<-DUF6079*<-BrxB-BrxF-iSTAND<-BrxA-DNA-binding<-?<-XIS-HTH<-?<-Phage_int_SAM_5+Phage_integrase\|\|LRR_5+LRR_5+LRR_5+LRR_5-> | 1215 | [illegible] | [illegible] | [illegible] | [illegible] | [illegible] | [illegible] | [illegible] |
| MCF2581772.1 | Type1.BREX | <-SIGMA+OB-Fold+AAA-ATPase+Lon-Protease-BrxL<-PglZ<-Schlafen+HATPase+wHTH<-N6-METHYLASE<-SLOG+BirA-HTH<-SF1-SF2-Helicase+PRTase<-HNH<-OLD-ABC<-DUF6079*<-BrxB-BrxF-iSTAND<-BrxA-DNA-binding | 1182 | [illegible] | [illegible] | [illegible] | [illegible] | [illegible] | [illegible] | [illegible] |
| MCF8307437.1 | Type1.BREX | <-HTH_3<-?<-?<-N6-METHYLASE-p<-DUF6079*<-BrxB-BrxF-iSTAND<-BrxA-DNA-binding<-HTH_3<-?<-tRNA<-RADICAL_SAM | 1165 | [illegible] | [illegible] | [illegible] | [illegible] | [illegible] | [illegible] | [illegible] |
| MCH4018978.1 | Type1.BREX | XIS-HTH->?->AAA+-><-?\|\|wHTH+WYL+WCX->HTH_3->BrxA-DNA-binding->BrxB-BrxF-iSTAND->DUF6079*->N6-METHYLASE->Schlafen+HATPase+wHTH->SIR2->PglZ->SIGMA+OB-Fold+AAA-ATPase+HKD-DNase-BrxL->SIR2+SIR2->ZnR->DDE_Tnp_ISL3->Tox-RelE-> | 1198 | [illegible] | [illegible] | [illegible] | [illegible] | [illegible] | [illegible] | [illegible] |
| MCI1742488.1 | Type1.BREX | <-HTH_31\|\|tRNA-><-?<-SIR2+SIR2<-SIGMA+OB-Fold+AAA-ATPase+HKD-DNase-BrxL<-PglZ<-HTH+REase<-Schlafen+HATPase+wHTH<-N6-METHYLASE<-DUF6079*<-BrxB-BrxF-iSTAND<-BrxA-DNA-binding<-HTH_3<-wHTH+WYL+WCX\|\|?-><-AAA+<-?<-XIS-HTH | 1206 | [illegible] | [illegible] | [illegible] | [illegible] | [illegible] | [illegible] | [illegible] |
| MCI2087046.1 | Type1.BREX | <-SIGMA+OB-Fold+AAA-ATPase+HKD-DNase-BrxL<-PglZ<-HTH+REase<-Schlafen+HATPase+wHTH<-N6-METHYLASE<-DUF6079*<-BrxB-BrxF-iSTAND<-BrxA-DNA-binding<-HTH_3<-wHTH+WYL+WCX\|\|?-><-AAA+<-?<-XIS-HTH | 1198 | [illegible] | [illegible] | [illegible] | [illegible] | [illegible] | [illegible] | [illegible] |
| MCI6111298.1 | Type1.BREX | ABC-ATPase+SbcC-><-?<-Phosphodiest\|\|?-><-Phos_pyr_kin\|\|?->?->BrxA-DNA-binding->BrxB-BrxF-iSTAND->DUF6079*->N6-METHYLASE-> | 1219 | [illegible] | [illegible] | [illegible] | [illegible] | [illegible] | [illegible] | [illegible] |
| MCI6081496.1 | Type1.BREX | <-METHYLASE<-?<-?<-?<-?<-?\|\|?->BrxA-DNA-binding->BrxB-BrxF-iSTAND->DUF6079*->ParB->OLD-ABC->?->N6-METHYLASE->DOC+HTH->PglZ->SIGMA+OB-Fold+AAA-ATPase+Lon-Protease-BrxL->Aminotran_1_2-> | 1215 | [illegible] | [illegible] | [illegible] | [illegible] | [illegible] | [illegible] | [illegible] |
| MCI6195664.1 | Type1.BREX | <-PglZ<-SIR2+SIR2<-Peptidase_C39_2<-?<-SIR2<-?<-N6-METHYLASE<-TOPRIM<-OLD-ABC<-DUF6079*<-BrxB-BrxF-iSTAND<-BrxA-DNA-binding<-SF1-SF2-Helicase<-?\|\|CR-ATPase5+REase-><-ARG<-?<-?\|\|SbcC-> | 1190 | [illegible] | [illegible] | [illegible] | [illegible] | [illegible] | [illegible] | [illegible] |
| MCI7017346.1 | Type1.BREX | <-SIGMA+OB-Fold+AAA-ATPase+HKD-DNase-BrxL<-PglZ<-?<-Schlafen+HATPase+wHTH<-N6-METHYLASE<-AT-HTH-HYD2<-Tox-ART-HYD2<-AT-HTH-HYD2<-DUF6079*<-BrxB-BrxF-iSTAND<-BrxA-DNA-binding<-wHTH+WYL+WCX | 1189 | [illegible] | [illegible] | [illegible] | [illegible] | [illegible] | [illegible] | [illegible] |
| MCI6403340.1 | Type1.BREX | AAA-ATPase-p-><-?\|\|BrxA-DNA-binding->BrxB-BrxF-iSTAND->DUF6079*->Schlafen+HATPase+wHTH->N6-METHYLASE-> | 1217 | [illegible] | [illegible] | [illegible] | [illegible] | [illegible] | [illegible] | [illegible] |
| MCI6372654.1 | Type1.BREX | <-N6-METHYLASE-p<-DUF6079*<-BrxB-BrxF-iSTAND<-BrxA-DNA-binding<-wHTH+WYL+WCX<-HTH_3\|\|?->?->HTH_3->?->TrwF-> | 1205 | [illegible] | [illegible] | [illegible] | [illegible] | [illegible] | [illegible] | [illegible] |
| MCI6336437.1 | Type1.BREX | <-DUF6079-p*<-BrxB-BrxF-iSTAND<-BrxA-DNA-binding<-wHTH+WYL+WCX<-HTH_3\|\|?-><-?<-PRTase<-Phage_lysozyme | 337 | [illegible] | [illegible] | [illegible] | [illegible] | [illegible] | [illegible] | [illegible] |
| MCI6336457.1 | Type1.BREX | PIWI->?->BrxA-DNA-binding->BrxB-BrxF-iSTAND->DUF6079*->ParB->OLD-ABC->?->N6-METHYLASE->OLD-ABC->HNH->PglZ->SIGMA+OB-Fold+AAA-ATPase+Lon-Protease-BrxL-> | 1215 | [illegible] | [illegible] | [illegible] | [illegible] | [illegible] | [illegible] | [illegible] |
| MCI6336758.1 | Type1.BREX | DUF6079-p*->OLD-ABC->TOPRIM->N6-METHYLASE->?->SIR2->ACET+TPR->?->Peptidase_C39_2-> | 873 | [illegible] | [illegible] | [illegible] | [illegible] | [illegible] | [illegible] | [illegible] |
| MCI7256454.1 | Type1.BREX | <-SIGMA+OB-Fold+AAA-ATPase+Lon-Protease-BrxL-p<-PglZ<-N6-METHYLASE-p<-DOC+HTH<-N6-METHYLASE<-AT-HTH-HYD2<-AT-HTH-HYD2<-DUF6079*<-BrxB-BrxF-iSTAND<-BrxA-DNA-binding<-HTH_3\|\|Thioredoxin-> | 1195 | [illegible] | [illegible] | [illegible] | [illegible] | [illegible] | [illegible] | [illegible] |
| MCI7250520.1 | Type1.BREX | ABC-ATPase+SbcC-><-?<-Phosphodiest\|\|?-><-Phos_pyr_kin\|\|?->?->BrxA-DNA-binding->BrxB-BrxF-iSTAND->DUF6079-p*-> | 275 | [illegible] | [illegible] | [illegible] | [illegible] | [illegible] | [illegible] | [illegible] |
| MCI7548156.1 | Type1.BREX | BrxA-DNA-binding->BrxB-BrxF-iSTAND->DUF6079*->N6-METHYLASE->?->PglZ->SIGMA+OB-Fold+AAA-ATPase+Lon-Protease-BrxL-> | 1222 | [illegible] | [illegible] | [illegible] | [illegible] | [illegible] | [illegible] | [illegible] |
| MCI7614186.1 | Type1.BREX | <-SF1-SF2-Helicase<-?\|\|HTH_3->BrxA-DNA-binding->BrxB-BrxF-iSTAND->DUF6079*->ParB->OLD-ABC->?->N6-METHYLASE->DOC+HTH->PglZ-> | 1215 | [illegible] | [illegible] | [illegible] | [illegible] | [illegible] | [illegible] | [illegible] |
| MCK4345284.1 | Type1.BREX | <-Phage_integrase\|\|?->HTH->BrxA-DNA-binding->BrxB-BrxF-iSTAND->DUF6079*-> | 1141 | [illegible] | [illegible] | [illegible] | [illegible] | [illegible] | [illegible] | [illegible] |
| MCK9398992.1 | Type1.BREX | PIN-><-?\|\|DDE_Tnp_1-><-?<-?\|\|HTH_3->BrxA-DNA-binding->BrxB-BrxF-iSTAND->DUF6079*->N6-METHYLASE->PglZ->SIGMA+OB-Fold+AAA-ATPase+HKD-DNase-BrxL->?->?->?-><-tRNA\|\|RADICAL-SAM-> | 1163 | [illegible] | [illegible] | [illegible] | [illegible] | [illegible] | [illegible] | [illegible] |
| MCK9310968.1 | Type1.BREX | ABC-ATPase+SbcC->?->?->?->HD->BrxA-DNA-binding->BrxB-BrxF-iSTAND->DUF6079*->N6-METHYLASE-p-> | 1143 | [illegible] | [illegible] | [illegible] | [illegible] | [illegible] | [illegible] | [illegible] |
| MCM1267126.1 | Type1.BREX | <-SbcC<-?<-HNH<-SIGMA+OB-Fold+AAA-ATPase+Lon-Protease-BrxL<-PglZ<-DOC+HTH<-?<-N6-METHYLASE<-DUF6079*<-BrxB-BrxF-iSTAND<-BrxA-DNA-binding<-?<-?<-?<-CHTH | 1173 | [illegible] | [illegible] | [illegible] | [illegible] | [illegible] | [illegible] | [illegible] |
| MCQ2071598.1 | Type1.BREX | <-TOPRIM<-?<-?\|\|Peptidase_M23->BrxA-DNA-binding->BrxB-BrxF-iSTAND->DUF6079*->N6-METHYLASE->Schlafen+HATPase+wHTH->PglZ->SIGMA+OB-Fold+AAA-ATPase+HKD-DNase-BrxL->XIS-HTH->wHTH+WYL+WCX->?->HHH_5+SF1-SF2-Helicase-> | 1190 | [illegible] | [illegible] | [illegible] | [illegible] | [illegible] | [illegible] | [illegible] |
| MCS2813929.1 | Type1.BREX | <-Peptidase_C39_2<-ACET+TPR<-N6-METHYLASE-p<-N6-METHYLASE<-N6-METHYLASE-p<-DUF6079-p<-?<-DUF6079-p<-DUF499<-DUF6079-p*<-DUF6079-p<-BrxB-BrxF-iSTAND<-BrxA-DNA-binding<-wHTH+WYL+WCX<-HTH_3 | 246 | [illegible] | [illegible] | [illegible] | [illegible] | [illegible] | [illegible] | [illegible] |
| MCS2813930.1 | Type1.BREX | <-ACET+TPR<-N6-METHYLASE-p<-N6-METHYLASE<-N6-METHYLASE-p<-DUF6079-p<-?<-DUF6079-p<-DUF499<-DUF6079-p<-DUF6079-p*<-BrxB-BrxF-iSTAND<-BrxA-DNA-binding<-wHTH+WYL+WCX<-HTH_3\|\|?->?->?->?->NUDIX+NUDIX-> | 131 | [illegible] | [illegible] | [illegible] | [illegible] | [illegible] | [illegible] | [illegible] |
| WP_156315612.1 | Type1.BREX | XIS-HTH-><-?<-?\|\|?->?->?->HTH_3->BrxA-DNA-binding->BrxB-BrxF-iSTAND->DUF6079-p*->DUF6079-p->N6-METHYLASE->PglZ-p->PglZ->SIGMA+OB-Fold+AAA-ATPase+HKD-DNase-BrxL->KAP-NTPase-> | 925 | [illegible] | [illegible] | [illegible] | [illegible] | [illegible] | [illegible] | [illegible] |
| WP_118354616.1 | Type1.BREX | <-N6-METHYLASE-p<-DOC+HTH<-?<-N6-METHYLASE-p<-N6-METHYLASE-p<-Four-helical-protein<-DUF6079*<-BrxB-BrxF-iSTAND<-BrxA-DNA-binding<-wHTH+WYL+WCX<-HTH_3\|\|?-><-?<-?<-?<-AA_kinase | 1194 | [illegible] | [illegible] | [illegible] | [illegible] | [illegible] | [illegible] | [illegible] |
| SCH71957.1 | Type1.BREX | <-SIGMA+OB-Fold+AAA-ATPase+Lon-Protease-BrxL<-PglZ<-Schlafen+HATPase+wHTH<-N6-METHYLASE<-SLOG+BirA-HTH<-SF1-SF2-Helicase+PRTase<-HNH<-OLD-ABC<-DUF6079*<-BrxB-BrxF-iSTAND<-BrxA-DNA-binding | 1179 | [illegible] | [illegible] | [illegible] | [illegible] | [illegible] | [illegible] | [illegible] |
| SCI35540.1 | Type1.BREX | <-N6-METHYLASE<-TOPRIM<-OLD-ABC<-DUF6079*<-BrxB-BrxF-iSTAND<-BrxA-DNA-binding | 1195 | [illegible] | [illegible] | [illegible] | [illegible] | [illegible] | [illegible] | [illegible] |
| SHF69941.1 | Type1.BREX | <-RADICAL-SAM<-?\|\|tRNA-><-SIGMA+OB-Fold+AAA-ATPase+Lon-Protease-BrxL<-PglZ<-Schlafen+HATPase+wHTH<-ParB+HNH<-N6-METHYLASE<-?<-DUF6079*<-BrxB-BrxF-iSTAND<-BrxA-DNA-binding | 1182 | [illegible] | [illegible] | [illegible] | [illegible] | [illegible] | [illegible] | [illegible] |
| SHK68061.1 | Type1.BREX | <-Classical-AAA<-?\|\|tRNA-><-?<-?<-?<-HTH+DOC\|\|?->HTH_3->BrxA-DNA-binding->BrxB-BrxF-iSTAND->DUF6079*->N6-METHYLASE->SbcC+SbcC->?->SIR2+SIR2->PglZ->SIGMA+OB-Fold+AAA-ATPase+HKD-DNase-BrxL->?->ParB+HNH->ParB+HNH-> | 1206 | [illegible] | [illegible] | [illegible] | [illegible] | [illegible] | [illegible] | [illegible] |
| WP_008760280.1 | Type1.BREX | D5ATPase+DUF3874-><-?\|\|ADH_N+ADH_zinc_N+METHYLASE->SPOUT-><-SIGMA+OB-Fold+AAA-ATPase+HKD-DNase-BrxL<-PglZ<-HTH+REase<-N6-METHYLASE-p<-N6-METHYLASE<-DUF6079*<-BrxB-BrxF-iSTAND<-BrxA-DNA-binding<-HTH_3<-?<-?<-?<-?<-TIR<-?<-TIR | 1193 | [illegible] | [illegible] | [illegible] | [illegible] | [illegible] | [illegible] | [illegible] |
| WP_004312187.1 | Type1.BREX | <-SIGMA+OB-Fold+AAA-ATPase+HKD-DNase-BrxL<-PglZ<-SWIM+DUF3670+SF1-SF2-Helicase<-?<-?<-?<-Peptidase_C39_2<-ACET+TPR<-N6-METHYLASE<-DUF6079*<-BrxB-BrxF-iSTAND<-BrxA-DNA-binding<-wHTH+WYL+WCX<-HTH_3\|\|?->?->?->NUDIX+NUDIX-> | 1208 | [illegible] | [illegible] | [illegible] | [illegible] | [illegible] | [illegible] | [illegible] |
| WP_006745116.1 | Type1.BREX | <-MODE-HTH+PBPII\|\|?-><-?<-NUDIX\|\|ABhydrolase->DNA-glycosylase-><-SIGMA+OB-Fold+AAA-ATPase+HKD-DNase-BrxL<-PglZ<-N6-METHYLASE<-DUF6079*<-BrxB-BrxF-iSTAND<-BrxA-DNA-binding<-?\|\|?->REase-><-?<-?<-?<-SIGMA-HTH | 1192 | [illegible] | [illegible] | [illegible] | [illegible] | [illegible] | [illegible] | [illegible] |
| WP_018109295.1 | Type1.BREX | <-FURR-HTH<-?<-Peptidase_M49<-AraC-HTH\|\|SF1-SF2-Helicase+PRTase->tRNA->HTH_3->BrxA-DNA-binding->BrxB-BrxF-iSTAND->DUF6079*->N6-METHYLASE->PglZ->SIGMA+OB-Fold+AAA-ATPase+HKD-DNase-BrxL->?->?->XIS-HTH->HTH_3->MerR-HTH->TOPRIM-> | 1202 | [illegible] | [illegible] | [illegible] | [illegible] | [illegible] | [illegible] | [illegible] |
| WP_002561973.1 | Type1.BREX | 35exo+SF1-SF2-Helicase-><-DDE_Tnp\|\|<-SIGMA+OB-Fold+AAA-ATPase+HKD-DNase-BrxL<-PglZ<-Schlafen+HATPase+wHTH<-SF1-SF2-Helicase<-OLD-ABC+TOPRIM<-N6-METHYLASE<-DUF6079*<-BrxB-BrxF-iSTAND<-BrxA-DNA-binding | 1202 | [illegible] | [illegible] | [illegible] | [illegible] | [illegible] | [illegible] | [illegible] |
| WP_024993540.1 | Type1.BREX | <-wHTH+WYL+WCX<-wHTH+WYL+WCX<-SIGMA+OB-Fold+AAA-ATPase+HKD-DNase-BrxL<-PglZ<-Schlafen+HATPase+wHTH<-N6-METHYLASE<-DUF6079*<-BrxB-BrxF-iSTAND<-BrxA-DNA-binding<-HTH_3 | 1210 | [illegible] | [illegible] | [illegible] | [illegible] | [illegible] | [illegible] | [illegible] |
| WP_042520323.1 | Type1.BREX | DDE_Tnp_ISL3-><-PARG+MACRODOMAIN<-REase<-SIGMA+OB-Fold+AAA-ATPase+HKD-DNase-BrxL<-PglZ<-N6-METHYLASE<-DUF6079*<-BrxB-BrxF-iSTAND<-BrxA-DNA-binding<-HTH_3 | 1204 | [illegible] | [illegible] | [illegible] | [illegible] | [illegible] | [illegible] | [illegible] |
| WP_044073670.1 | Type1.BREX | HTH_3->BrxA-DNA-binding->BrxB-BrxF-iSTAND->DUF6079*->N6-METHYLASE->PglZ->SIGMA+OB-Fold+AAA-ATPase+HKD-DNase-BrxL->REase->PARG+MACRODOMAIN-><-DDE_Tnp_ISL3 | 1206 | [illegible] | [illegible] | [illegible] | [illegible] | [illegible] | [illegible] | [illegible] |
| WP_094389547.1 | Type1.BREX | LRR_5->?->?->?-><-?\|\|?->HTH_3->BrxA-DNA-binding->BrxB-BrxF-iSTAND->DUF6079*->N6-METHYLASE->PglZ->SIGMA+OB-Fold+AAA-ATPase+HKD-DNase-BrxL->REase-> | 1228 | [illegible] | [illegible] | [illegible] | [illegible] | [illegible] | [illegible] | [illegible] |
| WP_094382632.1 | Type1.BREX | <-SIGMA+OB-Fold+AAA-ATPase+Lon-Protease-BrxL<-PglZ<-DOC+HTH<-N6-METHYLASE<-SbcC<-SbcC<-DUF6079*<-BrxB-BrxF-iSTAND<-BrxA-DNA-binding<-?<-PIWI<-?<-?<-?\|\|?->Glyco_hydro_32N+Glyco_hydro_43-> | 1215 | [illegible] | [illegible] | [illegible] | [illegible] | [illegible] | [illegible] | [illegible] |
| WP_094881847.1 | Type1.BREX | <-TOPRIM<-OLD-ABC<-DUF6079*<-BrxB-BrxF-iSTAND<-BrxA-DNA-binding<-wHTH+WYL+WCX<-HTH_3 | 1206 | [illegible] | [illegible] | [illegible] | [illegible] | [illegible] | [illegible] | [illegible] |
| WP_233544533.1 | Type1.BREX | ABC-ATPase+SbcC-><-?\|\|?->?->BrxA-DNA-binding->BrxB-BrxF-iSTAND->DUF6079-p->DUF6079*-> | 1083 | [illegible] | [illegible] | [illegible] | [illegible] | [illegible] | [illegible] | [illegible] |
| WP_233544532.1 | Type1.BREX | ABC-ATPase+SbcC->?-><-?\|\|?->?->BrxA-DNA-binding->BrxB-BrxF-iSTAND->DUF6079-p*->DUF6079-> | 126 | [illegible] | [illegible] | [illegible] | [illegible] | [illegible] | [illegible] | [illegible] |
| WP_118255350.1 | Type1.BREX | PRTase-><-?<-?<-?<-?\|\|HTH_3->wHTH+WYL+WCX->BrxA-DNA-binding->BrxB-BrxF-iSTAND->DUF6079*->DOC+HTH->N6-METHYLASE-><-?<-tRNA<-tRNA<-tRNA<-?<-HTH_3<-?<-?<-?<-ParA-Soj-PloopNTPase | 1206 | [illegible] | [illegible] | [illegible] | [illegible] | [illegible] | [illegible] | [illegible] |
| WP_118441918.1 | Type1.BREX | HTH_Tnp_1+HTH_21+rve+rve_3-><-?<-?\|\|BrxA-DNA-binding->BrxB-BrxF-iSTAND->DUF6079*->OLD-ABC->TOPRIM->N6-METHYLASE->?->PglZ->SIGMA+OB-Fold+AAA-ATPase+HKD-DNase-BrxL->?->?->Peptidase_M28-> | 1195 | [illegible] | [illegible] | [illegible] | [illegible] | [illegible] | [illegible] | [illegible] |
| WP_122342696.1 | Type1.BREX | Arm-DNA-bind_5+Phage_int_SAM_5+Phage_integrase->?->AbrB-HTH-><-HIN-HTH+rve+rve_3<-KAP-NTPase<-SIGMA+OB-Fold+AAA-ATPase+Lon-Protease-BrxL<-PglZ<-Schlafen+HATPase+wHTH<-N6-METHYLASE<-DUF6079*<-BrxB-BrxF-iSTAND<-BrxA-DNA-binding<-HTH_3\|\|?->XIS-HTH->XIS-HTH->Arm-DNA-bind_5+Phage_int_SAM_5+Phage_integrase->XIS-HTH->XIS-HTH-> | 1211 | [illegible] | [illegible] | [illegible] | [illegible] | [illegible] | [illegible] | [illegible] |
| WP_135993587.1 | Type1.BREX | <-SF1-SF2-Helicase<-SIGMA+OB-Fold+AAA-ATPase+HKD-DNase-BrxL<-PglZ<-Schlafen+HATPase+wHTH<-SIR2<-N6-METHYLASE<-DUF6079*<-BrxB-BrxF-iSTAND<-BrxA-DNA-binding<-HTH_3<-?\|\|Phage_int_SAM_1+Phage_integrase->Phage_integrase->Phage_int_SAM_1+Phage_integrase-><-Peptidase_M23 | 1221 | [illegible] | [illegible] | [illegible] | [illegible] | [illegible] | [illegible] | [illegible] |
| WP_172774764.1 | Type1.BREX | OLD-ABC->TOPRIM->XIS-HTH-><-?<-?<-?\|\|HTH_3->BrxA-DNA-binding->BrxB-BrxF-iSTAND->DUF6079*->N6-METHYLASE->OLD-ABC+TOPRIM->SF1-SF2-Helicase->Schlafen+HATPase+wHTH->PglZ->SIGMA+OB-Fold+AAA-ATPase+HKD-DNase-BrxL->NADAR->RADICAL-SAM->HIRAN-> | 1217 | [illegible] | [illegible] | [illegible] | [illegible] | [illegible] | [illegible] | [illegible] |
| WP_173076490.1 | Type1.BREX | ABhydrolase-><-PDZ+Peptidase<-?<-?<-?\|\|BrxA-DNA-binding->BrxB-BrxF-iSTAND->DUF6079*->N6-METHYLASE->PglZ->SIGMA+OB-Fold+AAA-ATPase+Lon-Protease-BrxL-><-tRNA<-?\|\|?->sGTP-><-TPR+TPR+TPR+TPR+TPR+TPR+HISKIN+REC | 1195 | [illegible] | [illegible] | [illegible] | [illegible] | [illegible] | [illegible] | [illegible] |
| WP_175324259.1 | Type1.BREX | <-SIGMA+OB-Fold+AAA-ATPase+HKD-DNase-BrxL<-PglZ<-SWIM+DUF3670+SF1-SF2-Helicase<-?<-?<-Peptidase_C39_2<-ACET+TPR<-N6-METHYLASE<-DUF6079*<-BrxB-BrxF-iSTAND<-BrxA-DNA-binding<-wHTH+WYL+WCX<-HTH_3\|\|?->?->NUDIX+NUDIX-> | 1208 | [illegible] | [illegible] | [illegible] | [illegible] | [illegible] | [illegible] | [illegible] |
| WP_179400143.1 | Type1.BREX | <-Phage_holin\|\|?->tRNA-><-?\|\|?->?->ABC-ATPase+SbcC-><-?\|\|BrxA-DNA-binding->BrxB-BrxF-iSTAND->DUF6079*->N6-METHYLASE->ParB->OLD-ABC->?->PglZ->SIGMA+OB-Fold+AAA-ATPase+Lon-Protease-BrxL-><-?<-HTH_Tnp_1+HTH_21+rve+rve_3<-HTH_Tnp_1 | 1192 | [illegible] | [illegible] | [illegible] | [illegible] | [illegible] | [illegible] | [illegible] |
| WP_181213825.1 | Type1.BREX | TOPRIM->?->TOPRIM->?-><-KAP-NTPase<-SIGMA+OB-Fold+AAA-ATPase+HKD-DNase-BrxL<-PglZ<-Schlafen+HATPase+wHTH<-N6-METHYLASE<-DUF6079*<-BrxB-BrxF-iSTAND<-BrxA-DNA-binding<-HTH_3\|\|?->?-><-XIS-HTH\|\|?->?->wHTH+WYL+WCX-> | 1212 | [illegible] | [illegible] | [illegible] | [illegible] | [illegible] | [illegible] | [illegible] |
| WP_115498725.1 | Type1.BREX | <-SIGMA+OB-Fold+AAA-ATPase+HKD-DNase-BrxL<-PglZ<-Schlafen+HATPase+wHTH<-N6-METHYLASE<-DUF6079*<-BrxB-BrxF-iSTAND<-BrxA-DNA-binding\|\|?->?->?->?-><-tRNA\|\|Cysteine-Desulfurase->?-><-ACET | 1211 | [illegible] | [illegible] | [illegible] | [illegible] | [illegible] | [illegible] | [illegible] |
| WP_004319896.1 | Type1.BREX | BrxA-DNA-binding->BrxB-BrxF-iSTAND->DUF6079*->N6-METHYLASE->SIR2+SIR2->Schlafen+HATPase+wHTH->PglZ->SIGMA+OB-Fold+AAA-ATPase+HKD-DNase-BrxL-><-35exo+SF1-SF2-Helicase\|\|BACON+Peptidase_M64-> | 1202 | [illegible] | [illegible] | [illegible] | [illegible] | [illegible] | [illegible] | [illegible] |
| WP_195290998.1 | Type1.BREX | Calcineurin-><-?<-SIGMA+OB-Fold+AAA-ATPase+HKD-DNase-BrxL<-PglZ<-Cas4+SF1-SF2-Helicase<-ACET+TPR<-N6-METHYLASE<-DUF6079*<-BrxB-BrxF-iSTAND<-BrxA-DNA-binding<-HTH_3<-Arm-DNA-bind_3+Phage_integrase | 1188 | [illegible] | [illegible] | [illegible] | [illegible] | [illegible] | [illegible] | [illegible] |
| WP_118403962.1 | Type1.BREX | <-Phage_int_SAM_5+Phage_integrase\|\|STYKinase-like->?->?-><-HEPN\|\|HTH_3->BrxA-DNA-binding->BrxB-BrxF-iSTAND->DUF6079*->N6-METHYLASE->Schlafen+HATPase+wHTH->PglZ->SIGMA+OB-Fold+AAA-ATPase+HKD-DNase-BrxL->?->URI-DNase+RAMA-> | 1211 | [illegible] | [illegible] | [illegible] | [illegible] | [illegible] | [illegible] | [illegible] |
| WP_195571580.1 | Type1.BREX | BrxA-DNA-binding->BrxB-BrxF-iSTAND->DUF6079*->N6-METHYLASE->SIR2+SIR2->Schlafen+HATPase+wHTH->PglZ->SIGMA+OB-Fold+AAA-ATPase+HKD-DNase-BrxL-><-35exo+SF1-SF2-Helicase\|\|?->SIGMA-HTH-> | 1202 | [illegible] | [illegible] | [illegible] | [illegible] | [illegible] | [illegible] | [illegible] |
| WP_195628684.1 | Type1.BREX | <-MORC+REase<-?<-REase+nSTAND3\|\|?->?-><-SIGMA+OB-Fold+AAA-ATPase+HKD-DNase-BrxL<-PglZ<-N6-METHYLASE<-DUF6079-p*\|\|?->?->?->?->?->?-><-?<-ACET | 769 | [illegible] | [illegible] | [illegible] | [illegible] | [illegible] | [illegible] | [illegible] |
| WP_195663259.1 | Type1.BREX | <-SIGMA+OB-Fold+AAA-ATPase+HKD-DNase-BrxL<-PglZ<-SWIM+DUF3670+SF1-SF2-Helicase<-?<-?<-?<-?<-DOC+HTH<-N6-METHYLASE<-DUF6079*<-BrxB-BrxF-iSTAND<-BrxA-DNA-binding<-HTH_3\|\|?->?-><-?<-?<-RADICAL-SAM | 1206 | [illegible] | [illegible] | [illegible] | [illegible] | [illegible] | [illegible] | [illegible] |
| WP_203041477.1 | Type1.BREX | Phage_lysozyme->PRTase-><-?<-?<-?\|\|HTH_3->wHTH+WYL+WCX->BrxA-DNA-binding->BrxB-BrxF-iSTAND->DUF6079*->N6-METHYLASE->SIR2+SIR2->PglZ->SIGMA+OB-Fold+AAA-ATPase+HKD-DNase-BrxL-> | 1207 | [illegible] | [illegible] | [illegible] | [illegible] | [illegible] | [illegible] | [illegible] |
| WP_204430319.1 | Type1.BREX | <-NADAR<-SIGMA+OB-Fold+AAA-ATPase+HKD-DNase-BrxL<-PglZ<-Schlafen+HATPase+wHTH<-SIR2<-N6-METHYLASE<-DUF6079*<-BrxB-BrxF-iSTAND<-BrxA-DNA-binding<-HTH_3\|\|?->?->?->?-><-XIS-HTH<-TOPRIM | 1217 | [illegible] | [illegible] | [illegible] | [illegible] | [illegible] | [illegible] | [illegible] |
| WP_205111283.1 | Type1.BREX | OLD-ABC->XIS-HTH-><-?<-?<-?\|\|HTH_3->BrxA-DNA-binding->BrxB-BrxF-iSTAND->DUF6079*->N6-METHYLASE->AT-HTH-HYD2->Tox-ART-HYD2->AT-HTH-HYD2->N6-METHYLASE-p->SIR2+SIR2->Schlafen+HATPase+wHTH->PglZ-> | 1217 | [illegible] | [illegible] | [illegible] | [illegible] | [illegible] | [illegible] | [illegible] |
| WP_204392825.1 | Type1.BREX | <-SbcC<-HKD-DNase<-SIGMA+OB-Fold+AAA-ATPase+HKD-DNase-BrxL<-PglZ<-N6-METHYLASE<-DUF6079*<-BrxB-BrxF-iSTAND<-BrxA-DNA-binding\|\|?-><-?<-Sulfatase+Nab6_mRNP_bdg+Phosphodiest<-Glyco_hydro_88\|\|?->?-><-LexA-protease | 1185 | [illegible] | [illegible] | [illegible] | [illegible] | [illegible] | [illegible] | [illegible] |
| WP_204506195.1 | Type1.BREX | HTH_3->BrxA-DNA-binding->BrxB-BrxF-iSTAND->DUF6079*->N6-METHYLASE->Schlafen+HATPase+wHTH->PglZ->SIGMA+OB-Fold+AAA-ATPase+HKD-DNase-BrxL->?->?->?->REase-> | 1187 | [illegible] | [illegible] | [illegible] | [illegible] | [illegible] | [illegible] | [illegible] |
| WP_204401091.1 | Type1.BREX | Calcineurin+Calcineurin-conf->?->?->?->?->BrxA-DNA-binding->BrxB-BrxF-iSTAND->DUF6079*->OLD-ABC->HNH->SF1-SF2-Helicase+PRTase->SLOG+BirA-HTH->N6-METHYLASE->Schlafen+HATPase+wHTH->PglZ->SIGMA+OB-Fold+AAA-ATPase+Lon-Protease-BrxL->SbcC-> | 1179 | [illegible] | [illegible] | [illegible] | [illegible] | [illegible] | [illegible] | [illegible] |
| WP_211451926.1 | Type1.BREX | Peptidase_M23-><-?<-?<-?<-Calcineurin<-SIGMA+OB-Fold+AAA-ATPase+HKD-DNase-BrxL<-PglZ<-SIR2<-N6-METHYLASE<-DUF6079*<-BrxB-BrxF-iSTAND<-BrxA-DNA-binding<-wHTH+WYL+WCX<-HTH_3\|\|?->?->?->NUDIX+NUDIX-> | 1208 | [illegible] | [illegible] | [illegible] | [illegible] | [illegible] | [illegible] | [illegible] |
| WP_215652320.1 | Type1.BREX | Phage_lysozyme->PRTase-><-?<-?<-?\|\|HTH_3->wHTH+WYL+WCX->BrxA-DNA-binding->BrxB-BrxF-iSTAND->DUF6079*->N6-METHYLASE->SIR2->Peptidase_C39_2->PglZ->SIGMA+OB-Fold+AAA-ATPase+HKD-DNase-BrxL->McrB-AAA->McrC-NTD+REase-> | 1206 | [illegible] | [illegible] | [illegible] | [illegible] | [illegible] | [illegible] | [illegible] |
| WP_215708904.1 | Type1.BREX | Arm-DNA-bind_5-><-UDG<-?<-ARG\|\|<-SIGMA+OB-Fold+AAA-ATPase+HKD-DNase-BrxL<-PglZ<-N6-METHYLASE<-DUF6079*<-BrxB-BrxF-iSTAND<-DUF4007\|\|Arm-DNA-bind_5+Phage_int_SAM_5+Phage_integrase->Virulence_RhuM-><-?\|\|Mob_Pre-><-TPR | 1215 | [illegible] | [illegible] | [illegible] | [illegible] | [illegible] | [illegible] | [illegible] |
| WP_216951216.1 | Type1.BREX | <-HTH_Tnp<-?<-?\|\|HTH_3-><-?<-?\|\|wHTH+WYL+WCX->BrxA-DNA-binding->BrxB-BrxF-iSTAND->DUF6079*->N6-METHYLASE->ACET+TPR->Peptidase_C39_2->?->?->SWIM+DUF3670+SF1-SF2-Helicase->PglZ->SIGMA+OB-Fold+AAA-ATPase+HKD-DNase-BrxL-> | 1208 | [illegible] | [illegible] | [illegible] | [illegible] | [illegible] | [illegible] | [illegible] |
| WP_217745553.1 | Type1.BREX | <-HIRAN<-RADICAL-SAM<-ARG<-?<-?\|\|HTH_3->wHTH+WYL+WCX->BrxA-DNA-binding->BrxB-BrxF-iSTAND->DUF6079*->OLD-ABC->TOPRIM->N6-METHYLASE->SIR2+SIR2->PglZ->SIGMA+OB-Fold+AAA-ATPase+HKD-DNase-BrxL->McrB-AAA->McrC-NTD+REase-> | 1206 | [illegible] | [illegible] | [illegible] | [illegible] | [illegible] | [illegible] | [illegible] |
| WP_217718390.1 | Type1.BREX | TOPRIM->XIS-HTH-><-?<-?<-?\|\|HTH_3->BrxA-DNA-binding->BrxB-BrxF-iSTAND->DUF6079*->N6-METHYLASE->OLD-ABC+TOPRIM->SF1-SF2-Helicase->Schlafen+HATPase+wHTH->PglZ->SIGMA+OB-Fold+AAA-ATPase+HKD-DNase-BrxL->NADAR->RADICAL-SAM->HIRAN-> | 1217 | [illegible] | [illegible] | [illegible] | [illegible] | [illegible] | [illegible] | [illegible] |
| WP_217725669.1 | Type1.BREX | <-BACON+Peptidase_M64\|\|35exo+SF1-SF2-Helicase-><-SIGMA+OB-Fold+AAA-ATPase+HKD-DNase-BrxL<-PglZ<-Schlafen+HATPase+wHTH<-SIR2+SIR2<-N6-METHYLASE<-DUF6079*<-BrxB-BrxF-iSTAND<-BrxA-DNA-binding | 1202 | [illegible] | [illegible] | [illegible] | [illegible] | [illegible] | [illegible] | [illegible] |
| WP_217719995.1 | Type1.BREX | DUF4007->BrxB-BrxF-iSTAND->DUF6079-p*->N6-METHYLASE->PglZ->SIGMA+OB-Fold+AAA-ATPase+HKD-DNase-BrxL-><-RHH+MobC\|\|ARG->?->UDG-><-Arm-DNA-bind_5 | 265 | [illegible] | [illegible] | [illegible] | [illegible] | [illegible] | [illegible] | [illegible] |
| WP_115473019.1 | Type1.BREX | <-OLD-ABC\|\|?-><-Phage_int_SAM_5+Phage_integrase\|\|?->?-><-HEPN\|\|HTH_3->BrxA-DNA-binding->BrxB-BrxF-iSTAND->DUF6079*->N6-METHYLASE->Schlafen+HATPase+wHTH->PglZ->SIGMA+OB-Fold+AAA-ATPase+HKD-DNase-BrxL->?-><-?<-MULE-TRANSPOSASE\|\|DOC+HTH-> | 1211 | [illegible] | [illegible] | [illegible] | [illegible] | [illegible] | [illegible] | [illegible] |
| WP_120489887.1 | Type1.BREX | BrxA-DNA-binding->BrxB-BrxF-iSTAND->DUF6079*->N6-METHYLASE->SIR2->Schlafen+HATPase+wHTH->PglZ->SIGMA+OB-Fold+AAA-ATPase+HKD-DNase-BrxL-><-35exo+SF1-SF2-Helicase\|\|BACON+Peptidase_M64-> | 1202 | [illegible] | [illegible] | [illegible] | [illegible] | [illegible] | [illegible] | [illegible] |
| WP_130041244.1 | Type1.BREX | Polbeta+HEPN->D5ATPase+DUF3874-><-?\|\|ADH_N+ADH_zinc_N+METHYLASE->SPOUT-><-SIGMA+OB-Fold+AAA-ATPase+HKD-DNase-BrxL<-PglZ<-HTH+REase<-N6-METHYLASE<-DUF6079*<-BrxB-BrxF-iSTAND<-BrxA-DNA-binding<-HTH_3<-sGTP\|\|?->Response_reg->NAD_binding_10-> | 1175 | [illegible] | [illegible] | [illegible] | [illegible] | [illegible] | [illegible] | [illegible] |
| WP_221729077.1 | Type1.BREX | <-DUF772+DDE_Tnp_1+DDE_Tnp_1_6<-?\|\|HTH_3->BrxA-DNA-binding->BrxB-BrxF-iSTAND->DUF6079*->N6-METHYLASE->DOC+HTH->N6-METHYLASE-p->SIR2->PglZ->SIGMA+OB-Fold+AAA-ATPase+HKD-DNase-BrxL-><-TnpB_IS66 | 1206 | [illegible] | [illegible] | [illegible] | [illegible] | [illegible] | [illegible] | [illegible] |
| WP_060386337.1 | Type1.BREX | <-SF1-SF2-Helicase<-?\|\|?->AbrB-HTH-><-KAP-NTPase<-SIGMA+OB-Fold+AAA-ATPase+Lon-Protease-BrxL<-PglZ<-Schlafen+HATPase+wHTH<-N6-METHYLASE<-DUF6079*<-BrxB-BrxF-iSTAND<-BrxA-DNA-binding<-HTH_3\|\|?->XIS-HTH->XIS-HTH-><-DEDD_Tnp_IS110+Transposase_20\|\|?->Trypsin-> | 1213 | [illegible] | [illegible] | [illegible] | [illegible] | [illegible] | [illegible] | [illegible] |
| WP_225659484.1 | Type1.BREX | Polbeta+HEPN->D5ATPase+DUF3874-><-?\|\|ADH_N+ADH_zinc_N+METHYLASE->SPOUT-><-SIGMA+OB-Fold+AAA-ATPase+HKD-DNase-BrxL<-PglZ<-HTH+REase<-N6-METHYLASE<-DUF6079*<-BrxB-BrxF-iSTAND<-BrxA-DNA-binding<-HTH_3<-sGTP\|\|?->Response_reg->NAD_binding_10-> | 1175 | [illegible] | [illegible] | [illegible] | [illegible] | [illegible] | [illegible] | [illegible] |
| WP_227088012.1 | Type1.BREX | ArdCN+MPTase->TOPRIM->?->TOPRIM->?->?->HTH_3->BrxA-DNA-binding->BrxB-BrxF-iSTAND->DUF6079*->N6-METHYLASE->Schlafen+HATPase+wHTH->PglZ->SIGMA+OB-Fold+AAA-ATPase+Lon-Protease-BrxL->RNaseH-><-?\|\|?->?-><-XIS-HTH | 1176 | [illegible] | [illegible] | [illegible] | [illegible] | [illegible] | [illegible] | [illegible] |
| WP_217720628.1 | Type1.BREX | <-NUDIX+NUDIX<-?<-?<-?\|\|HTH_3->wHTH+WYL+WCX->BrxA-DNA-binding->BrxB-BrxF-iSTAND->DUF6079*->N6-METHYLASE->ACET+TPR->Peptidase_C39_2->?->?->?->SWIM+DUF3670+SF1-SF2-Helicase->PglZ->SIGMA+OB-Fold+AAA-ATPase+HKD-DNase-BrxL-> | 1208 | [illegible] | [illegible] | [illegible] | [illegible] | [illegible] | [illegible] | [illegible] |
| WP_234117060.1 | Type1.BREX | <-BACON+Peptidase_M64\|\|35exo+SF1-SF2-Helicase-><-SIGMA+OB-Fold+AAA-ATPase+HKD-DNase-BrxL<-PglZ<-Schlafen+HATPase+wHTH<-SIR2+SIR2<-N6-METHYLASE<-DUF6079*<-BrxB-BrxF-iSTAND<-BrxA-DNA-binding | 1202 | [illegible] | [illegible] | [illegible] | [illegible] | [illegible] | [illegible] | [illegible] |
| WP_235327427.1 | Type1.BREX | <-SIGMA+OB-Fold+AAA-ATPase+Lon-Protease-BrxL<-PglZ<-Schlafen+HATPase+wHTH<-N6-METHYLASE<-SLOG+BirA-HTH<-SF1-SF2-Helicase+PRTase<-HNH<-OLD-ABC<-DUF6079*<-BrxB-BrxF-iSTAND<-BrxA-DNA-binding | 1179 | [illegible] | [illegible] | [illegible] | [illegible] | [illegible] | [illegible] | [illegible] |
| WP_235417293.1 | Type1.BREX | BrxA-DNA-binding->BrxB-BrxF-iSTAND->DUF6079*->N6-METHYLASE->Schlafen+HATPase+wHTH->PglZ->SIGMA+OB-Fold+AAA-ATPase+Lon-Protease-BrxL->?->REase+Bacteriocin_IIc-> | 1176 | [illegible] | [illegible] | [illegible] | [illegible] | [illegible] | [illegible] | [illegible] |
| WP_243323291.1 | Type1.BREX | XIS-HTH-><-?\|\|TraJ-RHH->?->?-><-?<-SIGMA+OB-Fold+AAA-ATPase+Lon-Protease-BrxL<-PglZ<-N6-METHYLASE<-DUF6079* | 1142 | [illegible] | [illegible] | [illegible] | [illegible] | [illegible] | [illegible] | [illegible] |
| WP_243326763.1 | Type1.BREX | <-DUF6079-p*<-BrxB-BrxF-iSTAND<-BrxA-DNA-binding<-HTH_3\|\|Phage_lysozyme->Arm-DNA-bind_5+Phage_int_SAM_5+Phage_integrase-> | 90 | [illegible] | [illegible] | [illegible] | [illegible] | [illegible] | [illegible] | [illegible] |
| WP_249093422.1 | Type1.BREX | TOPRIM->XIS-HTH-><-?<-?<-?\|\|HTH_3->BrxA-DNA-binding->BrxB-BrxF-iSTAND->DUF6079*->N6-METHYLASE->SIR2->Schlafen+HATPase+wHTH->PglZ->SIGMA+OB-Fold+AAA-ATPase+HKD-DNase-BrxL->NADAR->RADICAL-SAM->HIRAN->HHH_5+SF1-SF2-Helicase-> | 1217 | [illegible] | [illegible] | [illegible] | [illegible] | [illegible] | [illegible] | [illegible] |
| WP_250803006.1 | Type1.BREX | <-SIGMA+OB-Fold+AAA-ATPase+HKD-DNase-BrxL<-PglZ<-SWIM+DUF3670+SF1-SF2-Helicase<-?<-?<-?<-Peptidase_C39_2<-ACET+TPR<-N6-METHYLASE<-DUF6079*<-BrxB-BrxF-iSTAND<-BrxA-DNA-binding<-wHTH+WYL+WCX<-HTH_3\|\|?->?->?->NUDIX+NUDIX->DUF4365+REase-> | 1208 | [illegible] | [illegible] | [illegible] | [illegible] | [illegible] | [illegible] | [illegible] |
| WP_250702187.1 | Type1.BREX | <-GDResolvase+HIN-HTH<-HTH+REase<-SF1-SF2-Helicase\|\|ADH_N+ADH_zinc_N+METHYLASE-><-?<-SPOUT\|\|CR-ATPase5+REase->BrxA-DNA-binding->BrxB-BrxF-iSTAND->DUF6079*->N6-METHYLASE->PglZ->SIGMA+OB-Fold+AAA-ATPase+Lon-Protease-BrxL-><-?<-?<-TraJ-RHH<-ZnR+TOPRIM<-CR-ATPase4<-XIS-HTH | 1215 | [illegible] | [illegible] | [illegible] | [illegible] | [illegible] | [illegible] | [illegible] |
| WP_252760249.1 | Type1.BREX | <-TIR<-?<-SIGMA+OB-Fold+AAA-ATPase+HKD-DNase-BrxL<-PglZ<-N6-METHYLASE-p<-?<-N6-METHYLASE<-DUF6079*<-BrxB-BrxF-iSTAND<-BrxA-DNA-binding<-HTH_3<-wHTH+WYL+WCX\|\|?->?->?-><-?\|\|ACET-> | 1205 | [illegible] | [illegible] | [illegible] | [illegible] | [illegible] | [illegible] | [illegible] |
| WP_252760927.1 | Type1.BREX | DDE_Tnp_ISL3->XIS-HTH->?->AAA+-><-?\|\|wHTH+WYL+WCX->HTH_3->BrxA-DNA-binding->BrxB-BrxF-iSTAND->DUF6079*->N6-METHYLASE->Schlafen+HATPase+wHTH->PglZ->SIGMA+OB-Fold+AAA-ATPase+HKD-DNase-BrxL->SIR2+SIR2->?->?-><-tRNA\|\|HTH_31-> | 1206 | [illegible] | [illegible] | [illegible] | [illegible] | [illegible] | [illegible] | [illegible] |
| WP_258327305.1 | Type1.BREX | <-HHH_5+SF1-SF2-Helicase<-HIRAN<-RADICAL-SAM<-NADAR<-SIGMA+OB-Fold+AAA-ATPase+HKD-DNase-BrxL<-PglZ<-Schlafen+HATPase+wHTH<-SIR2<-N6-METHYLASE<-DUF6079*<-BrxB-BrxF-iSTAND<-BrxA-DNA-binding<-HTH_3\|\|?->?->?->?-><-XIS-HTH<-TOPRIM | 1217 | [illegible] | [illegible] | [illegible] | [illegible] | [illegible] | [illegible] | [illegible] |
| WP_258898999.1 | Type1.BREX | <-SIGMA+OB-Fold+AAA-ATPase+HKD-DNase-BrxL<-PglZ<-SWIM+DUF3670+SF1-SF2-Helicase<-?<-?<-?<-Peptidase_C39_2<-ACET+TPR<-N6-METHYLASE<-DUF6079*<-BrxB-BrxF-iSTAND<-BrxA-DNA-binding<-wHTH+WYL+WCX<-HTH_3\|\|?->?->?->NUDIX+NUDIX->DUF4365+REase-> | 1208 | [illegible] | [illegible] | [illegible] | [illegible] | [illegible] | [illegible] | [illegible] |
| WP_258900416.1 | Type1.BREX | <-PglZ<-SF1-SF2-Helicase-p<-SWIM+DUF3670+SF1-SF2-Helicase<-?<-?<-?<-Peptidase_C39_2<-ACET+TPR<-N6-METHYLASE<-DUF6079*<-BrxB-BrxF-iSTAND<-BrxA-DNA-binding<-wHTH+WYL+WCX\|\|?->?->?->NUDIX+NUDIX->?-><-Phage_lysozyme | 1208 | [illegible] | [illegible] | [illegible] | [illegible] | [illegible] | [illegible] | [illegible] |
| WP_234146092.1 | Type1.BREX | Arm-DNA-bind_5+Phage_int_SAM_5+Phage_integrase-><-Schlafen+HATPase+wHTH<-SIGMA+OB-Fold+AAA-ATPase+Lon-Protease-BrxL<-PglZ<-N6-METHYLASE<-PIN<-OLD-ABC<-DUF6079*<-BrxB-BrxF-iSTAND<-BrxA-DNA-binding<-HTH_3\|\|?->?->N6-METHYLASE-><-AAA+<-HTH_36<-XIS-HTH | 1180 | [illegible] | [illegible] | [illegible] | [illegible] | [illegible] | [illegible] | [illegible] |
| WP_258991265.1 | Type1.BREX | <-SIGMA+OB-Fold+AAA-ATPase+HKD-DNase-BrxL<-PglZ<-SWIM+DUF3670+SF1-SF2-Helicase<-?<-?<-?<-Peptidase_C39_2<-ACET+TPR<-N6-METHYLASE<-DUF6079*<-BrxB-BrxF-iSTAND<-BrxA-DNA-binding<-wHTH+WYL+WCX<-HTH_3\|\|?->?->?->NUDIX+NUDIX-> | 1208 | [illegible] | [illegible] | [illegible] | [illegible] | [illegible] | [illegible] | [illegible] |
| WP_259011403.1 | Type1.BREX | BrxA-DNA-binding->BrxB-BrxF-iSTAND->DUF6079*->N6-METHYLASE->ParB->OLD-ABC->?->Schlafen+HATPase+wHTH->PglZ->SIGMA+OB-Fold+AAA-ATPase+HKD-DNase-BrxL-><-35exo+SF1-SF2-Helicase | 1202 | [illegible] | [illegible] | [illegible] | [illegible] | [illegible] | [illegible] | [illegible] |
| WP_258973412.1 | Type1.BREX | Polbeta+HEPN->D5ATPase+DUF3874-><-?\|\|ADH_N+ADH_zinc_N+METHYLASE->SPOUT-><-SIGMA+OB-Fold+AAA-ATPase+HKD-DNase-BrxL<-PglZ<-HTH+REase<-N6-METHYLASE<-DUF6079*<-?<-BrxB-BrxF-iSTAND<-BrxA-DNA-binding<-HTH_3\|\|CR-ATPase5+REase->?->?-><-RVT | 1133 | [illegible] | [illegible] | [illegible] | [illegible] | [illegible] | [illegible] | [illegible] |
| WP_258979859.1 | Type1.BREX | BrxA-DNA-binding->BrxB-BrxF-iSTAND->DUF6079-p*->DUF6079-p->N6-METHYLASE->ParB->OLD-ABC->?->Schlafen+HATPase+wHTH->PglZ->SIGMA+OB-Fold+AAA-ATPase+HKD-DNase-BrxL-> | 780 | [illegible] | [illegible] | [illegible] | [illegible] | [illegible] | [illegible] | [illegible] |
| WP_009000708.1 | Type1.BREX | BrxA-DNA-binding->BrxB-BrxF-iSTAND->DUF6079*->N6-METHYLASE->?->?->Schlafen+HATPase+wHTH->PglZ->SIGMA+OB-Fold+AAA-ATPase+HKD-DNase-BrxL-><-35exo+SF1-SF2-Helicase\|\|BACON+Peptidase_M64-> | 1202 | [illegible] | [illegible] | [illegible] | [illegible] | [illegible] | [illegible] | [illegible] |
| WP_262301578.1 | Type1.BREX | <-SIGMA+OB-Fold+AAA-ATPase+Lon-Protease-BrxL<-PglZ<-Schlafen+HATPase+wHTH<-N6-METHYLASE<-DUF6079*<-BrxB-BrxF-iSTAND<-BrxA-DNA-binding<-REase | 1218 | [illegible] | [illegible] | [illegible] | [illegible] | [illegible] | [illegible] | [illegible] |
| WP_148330685.1 | Type1.BREX | <-N6-METHYLASE<-TOPRIM<-OLD-ABC<-DUF6079*<-BrxB-BrxF-iSTAND<-BrxA-DNA-binding | 1195 | [illegible] | [illegible] | [illegible] | [illegible] | [illegible] | [illegible] | [illegible] |
| WP_195661285.1 | Type1.BREX | BrxA-DNA-binding->BrxB-BrxF-iSTAND->DUF6079*->N6-METHYLASE->SIR2+SIR2->Schlafen+HATPase+wHTH->PglZ->SIGMA+OB-Fold+AAA-ATPase+HKD-DNase-BrxL-> | 1202 | [illegible] | [illegible] | [illegible] | [illegible] | [illegible] | [illegible] | [illegible] |
| WP_264912090.1 | Type1.BREX | <-HIRAN<-RADICAL-SAM<-ARG<-?<-?\|\|HTH_3->wHTH+WYL+WCX->BrxA-DNA-binding->BrxB-BrxF-iSTAND->DUF6079*->DOC+HTH->N6-METHYLASE->SIR2+SIR2->PglZ->SIGMA+OB-Fold+AAA-ATPase+HKD-DNase-BrxL->McrB-AAA->McrC-NTD+REase-> | 1206 | [illegible] | [illegible] | [illegible] | [illegible] | [illegible] | [illegible] | [illegible] |
| WP_264965401.1 | Type1.BREX | ABC-ATPase+ABC-ATPase+ABC-ATPase+SbcC->Peptidase_M13_N+Peptidase_M13-><-SIGMA+OB-Fold+AAA-ATPase+HKD-DNase-BrxL<-PglZ<-N6-METHYLASE-p<-Schlafen+HATPase+wHTH<-N6-METHYLASE<-DUF6079*<-BrxB-BrxF-iSTAND<-BrxA-DNA-binding<-?<-?\|\|?-><-?<-?<-ABC-ATPase+SbcC | 1194 | [illegible] | [illegible] | [illegible] | [illegible] | [illegible] | [illegible] | [illegible] |
| WP_270558549.1 | Type1.BREX | <-HISKIN<-?<-Polbeta<-MarR-HTH<-?\|\|HTH_3->wHTH+WYL+WCX->BrxA-DNA-binding->BrxB-BrxF-iSTAND->DUF6079*->N6-METHYLASE->DOC+HTH->SIR2+SIR2->N6-METHYLASE->PglZ->SIGMA+OB-Fold+AAA-ATPase+HKD-DNase-BrxL->CASPASE->SLATT_4-> | 1216 | [illegible] | [illegible] | [illegible] | [illegible] | [illegible] | [illegible] | [illegible] |
| WP_271687043.1 | Type1.BREX | <-SIGMA+OB-Fold+AAA-ATPase+HKD-DNase-BrxL<-PglZ<-SWIM+DUF3670+SF1-SF2-Helicase<-?<-?<-?<-Peptidase_C39_2<-ACET+TPR<-N6-METHYLASE<-DUF6079*<-BrxB-BrxF-iSTAND<-BrxA-DNA-binding<-wHTH+WYL+WCX<-HTH_3\|\|?->?->?->NUDIX+NUDIX-> | 1208 | [illegible] | [illegible] | [illegible] | [illegible] | [illegible] | [illegible] | [illegible] |
| WP_271704578.1 | Type1.BREX | HTH_3->MORC+DUF3684+REase->SF1-SF2-Helicase+PIN->HTH_3->BrxA-DNA-binding->BrxB-BrxF-iSTAND->DUF6079*->N6-METHYLASE->SIR2+SIR2->Schlafen+HATPase+wHTH->PglZ->SIGMA+OB-Fold+AAA-ATPase+HKD-DNase-BrxL-> | 1209 | [illegible] | [illegible] | [illegible] | [illegible] | [illegible] | [illegible] | [illegible] |
| WP_272709339.1 | Type1.BREX | <-DUF4365+REase<-NUDIX+NUDIX<-?<-?<-?\|\|HTH_3->wHTH+WYL+WCX->BrxA-DNA-binding->BrxB-BrxF-iSTAND->DUF6079*->N6-METHYLASE->Peptidase_C39_2->SIR2->PglZ->SIGMA+OB-Fold+AAA-ATPase+HKD-DNase-BrxL-> | 1208 | [illegible] | [illegible] | [illegible] | [illegible] | [illegible] | [illegible] | [illegible] |
| WP_274224871.1 | Type1.BREX | Arm-DNA-bind_5+Phage_int_SAM_5+Phage_integrase->?->AbrB-HTH-><-KAP-NTPase<-SIGMA+OB-Fold+AAA-ATPase+Lon-Protease-BrxL<-PglZ<-Schlafen+HATPase+wHTH<-N6-METHYLASE<-DUF6079*<-BrxB-BrxF-iSTAND<-BrxA-DNA-binding<-HTH_3\|\|?->XIS-HTH->XIS-HTH-> | 1213 | [illegible] | [illegible] | [illegible] | [illegible] | [illegible] | [illegible] | [illegible] |
| WP_032560932.1 | Type1.BREX | SF1-SF2-Helicase->?-><-DOC+HTH<-URI-DNase+RAMA<-?<-SIGMA+OB-Fold+AAA-ATPase+HKD-DNase-BrxL<-PglZ<-Schlafen+HATPase+wHTH<-N6-METHYLASE<-DUF6079*<-BrxB-BrxF-iSTAND<-BrxA-DNA-binding<-HTH_3\|\|HEPN-><-?<-?<-STYKinase-like\|\|Phage_int_SAM_5+Phage_integrase-> | 1211 | [illegible] | [illegible] | [illegible] | [illegible] | [illegible] | [illegible] | [illegible] |

| Protein-ID | System | Gene-Neighbourhoods | Length | Taxonomy | Assembly-ID | Strain | Locus-Tag | Genomic-coordinates | TaxID | Fasta-define |
|---|---|---|---|---|---|---|---|---|---|---|
| WP_154043608.1 | Type1.BREX | <-SbcC<-SIGMA+OB-Fold+AAA-ATPase+Lon-Protease-BrxL<-PglZ<-Schlafen+HATPase+wHTH<-N6-METHYLASE<-SLOG+BirA-HTH<-SF1-SF2-Helicase+PRTase<-HNH<-OLD-ABC<-DUF6079*<-BrxB-BrxF-iSTAND<- [illegible] | 1179 | Bacteria>FCB_group>Bacteroidetes/Chlorobi_group>Bacteroidota>Bacteroidia>Bacteroidales | GCF_900066445.1 | Bacteroides_cellulolyticus | OCV82_RS05980 | 56504..60043 | 2981780 | BREX-system-P-loop-protein-BrxC-[Bacteroides-cellulolyticus] |
| WP_074557116.1 | Type1.BREX | <-HAD+PRTase+PRTase+REase+SF1-SF2-Helicase<-?<-TPR+TPR<-?<-Schlafen+HATPase+wHTH<-TrwF\|\|BrxA-DNA-binding->BrxB-BrxF-iSTAND->DUF6079*->N6-METHYLASE->SIR2->Schlafen+HATPase+wHTH->PglZ-> [illegible] | 1202 | Bacteria>FCB_group>Bacteroidetes/Chlorobi_group>Bacteroidota>Bacteroidia>Bacteroidales | GCF_900102645.1 | Bacteroides_ovatus | BLR14_RS05840 | 106864..110472 | 28116 | BREX-system-P-loop-protein-BrxC-[Bacteroides-ovatus] |
| WP_072542576.1 | Type1.BREX | <-AAA-ATPase<-HTH+rve<-?<-SIGMA+OB-Fold+AAA-ATPase+HKD-DNase-BrxL<-PglZ<-SIR2+SIR2<-N6-METHYLASE<-TOPRIM<-OLD-ABC<-DUF6079*<-BrxB-BrxF-iSTAND<-BrxA-DNA-binding\|\|?->?->?->?-><-?<-?<- [illegible] | 1195 | Bacteria>FCB_group>Bacteroidetes/Chlorobi_group>Bacteroidota>Bacteroidia>Bacteroidales | GCF_900128455.1 | Bacteroides_mediterraneensis | BQ7480_RS08760 | 2129051..2132638 | 1841856 | BREX-system-P-loop-protein-BrxC-[Bacteroides-mediterraneensis] |
| WP_073402639.1 | Type1.BREX | <-RADICAL-SAM<-?\|\|tRNA-><-SIGMA+OB-Fold+AAA-ATPase+Lon-Protease-BrxL<-PglZ<-Schlafen+HATPase+wHTH<-ParB+HNH<-N6-METHYLASE<-?<-DUF6079*<-BrxB-BrxF-iSTAND<-BrxA-DNA-binding | 1188 | Bacteria>FCB_group>Bacteroidetes/Chlorobi_group>Bacteroidota>Bacteroidia>Bacteroidales | GCF_900128905.1 | Bacteroides_luti | BUC21_RS11790 | 81108..84674 | 1297750 | BREX-system-P-loop-protein-BrxC-[Bacteroides-luti] |
| WP_254795118.1 | Type1.BREX | <-Classical-AAA<-?\|\|tRNA-><-?<-?<-HTH+DOC\|\|?->HTH_3->BrxA-DNA-binding->BrxB-BrxF-iSTAND->DUF6079*->N6-METHYLASE->SbcC+SbcC->?->SIR2+SIR2->PglZ->SIGMA+OB-Fold+AAA-ATPase+HKD-DNase- [illegible] | 1209 | Bacteria>FCB_group>Bacteroidetes/Chlorobi_group>Bacteroidota>Bacteroidia>Bacteroidales | GCF_900142525.1 | Prevotella_ruminicola | BUB86_RS07050 | 46256..49885 | 839 | BREX-system-P-loop-protein-BrxC-[Prevotella-ruminicola] |
| WP_077197246.1 | Type1.BREX | DDE_Tnp_IS66+DDE_Tnp_IS66_C-><-?<-?<-?<-?<-?\|\|HTH_3->BrxA-DNA-binding->BrxB-BrxF-iSTAND->DUF6079*->N6-METHYLASE->Schlafen+HATPase+wHTH->PglZ->SIGMA+OB-Fold+AAA-ATPase+Lon-Protease- [illegible] | 1176 | Bacteria>FCB_group>Bacteroidetes/Chlorobi_group>Bacteroidota>Bacteroidia>Bacteroidales | GCF_900155655.1 | Prevotella_ihumii | CZ367_RS13300 | 3061778..3065308 | 1917878 | BREX-system-P-loop-protein-BrxC-[Prevotella-ihumii] |
| WP_176134075.1 | Type1.BREX | <-Phage_holin\|\|?->tRNA-><-?\|\|?->?->ABC-ATPase+SbcC-><-?\|\|BrxA-DNA-binding->BrxB-BrxF-iSTAND->DUF6079*->N6-METHYLASE->OLD-ABC+TOPRIM->SF1-SF2-Helicase->?->N6-METHYLASE-p->ParB+HNH-> [illegible] | 1192 | Bacteria>FCB_group>Bacteroidetes/Chlorobi_group>Bacteroidota>Bacteroidia>Bacteroidales | GCF_900168155.1 | Parabacteroides_chartae | B5X57_RS12575 | 48804..52382 | 1037355 | BREX-system-P-loop-protein-BrxC-[Parabacteroides-chartae] |
| WP_130829424.1 | Type1.BREX | <-SIGMA+OB-Fold+AAA-ATPase+Lon-Protease-BrxL<-PglZ<-Schlafen+HATPase+wHTH<-Schlafen<-N6-METHYLASE<-DUF6079*<-BrxB-BrxF-iSTAND<-BrxA-DNA-binding<-HTH_3<-?\|\|?-><-PTSEIIA1\|\|?-> [illegible] | 1153 | Bacteria>FCB_group>Bacteroidetes/Chlorobi_group>Bacteroidota>Bacteroidia>Bacteroidales | GCF_900607215.1 | Prevotella | ETF27_RS09605 | 9675..13136 | 838 | MULTISPECIES:-BREX-system-P-loop-protein-BrxC-[Prevotella] |
| WP_125105113.1 | Type1.BREX | <-HHH_5+SF1-SF2-Helicase<-HIRAN<-RADICAL-SAM<-NADAR<-SIGMA+OB-Fold+AAA-ATPase+HKD-DNase-BrxL<-PglZ<-Schlafen+HATPase+wHTH<-SIR2<-N6-METHYLASE<-DUF6079*<-BrxB-BrxF-iSTAND<- [illegible] | 1217 | Bacteria>FCB_group>Bacteroidetes/Chlorobi_group>Bacteroidota>Bacteroidia>Bacteroidales | GCF_900625065.1 | Prevotella_marseillensis | EJL04_RS12230 | 3058927..3062580 | 2479840 | BREX-system-P-loop-protein-BrxC-[Prevotella-marseillensis] |
| WP_144153273.1 | Type1.BREX | <-HIRAN<-RADICAL-SAM<-ARG<-?<-?\|\|HTH_3->wHTH+WYL+WCX->BrxA-DNA-binding->BrxB-BrxF-iSTAND->DUF6079*->N6-METHYLASE->SIR2+SIR2->PglZ->SIGMA+OB-Fold+AAA-ATPase+HKD-DNase-BrxL->McrB-AAA-> [illegible] | 1194 | Bacteria>FCB_group>Bacteroidetes/Chlorobi_group>Bacteroidota>Bacteroidia>Bacteroidales | GCF_902167755.1 | Prevotella_copri | BIL0145876EF_RS09275 | 107988..111572 | 165179 | BREX-system-P-loop-protein-BrxC-[Prevotella-copri] |
| WP_032543214.1 | Type1.BREX | <-METHYLASE<-ACET<-DOC+HTH<-URI-DNase+RAMA<-?<-SIGMA+OB-Fold+AAA-ATPase+HKD-DNase-BrxL<-PglZ<-Schlafen+HATPase+wHTH<-N6-METHYLASE<-DUF6079*<-BrxB-BrxF-iSTAND<-BrxA-DNA-binding<- [illegible] | 1211 | Bacteria>FCB_group>Bacteroidetes/Chlorobi_group>Bacteroidota>Bacteroidia>Bacteroidales | GCF_902364655.1 | Bacteroides_fragilis | PDN68_RS18445 | 100181..103816 | 817 | BREX-system-P-loop-protein-BrxC-[Bacteroides-fragilis] |
| WP_060407611.1 | Type1.BREX | <-SIGMA+OB-Fold+AAA-ATPase+HKD-DNase-BrxL<-PglZ<-Schlafen+HATPase+wHTH<-SIR2<-N6-METHYLASE-p<-N6-METHYLASE<-DUF6079*<-BrxB-BrxF-iSTAND<-BrxA-DNA-binding<-HTH_3<-?<-?<-?\|\|SIGMA-HTH-> | 1205 | Bacteria>FCB_group>Bacteroidetes/Chlorobi_group>Bacteroidota>Bacteroidia>Bacteroidales | GCF_902387305.1 | Bacteroides_cellulosilyticus | FYC30_RS03960 | 75039..78656 | 246787 | BREX-system-P-loop-protein-BrxC-[Bacteroides-cellulosilyticus] |
| WP_118419043.1 | Type1.BREX | BrxA-DNA-binding->BrxB-BrxF-iSTAND->DUF6079*->N6-METHYLASE->SIR2+SIR2->Schlafen+HATPase+wHTH->PglZ->SIGMA+OB-Fold+AAA-ATPase+HKD-DNase-BrxL-><-35exo+SF1-SF2-Helicase\|\|BACON+Peptidase_ [illegible] | 1202 | Bacteria>FCB_group>Bacteroidetes/Chlorobi_group>Bacteroidota>Bacteroidia>Bacteroidales | GCF_918279245.1 | Bacteroides_ovatus | PO203_RS09380 | 101158..104766 | 28116 | BREX-system-P-loop-protein-BrxC-[Bacteroides-ovatus] |
| WP_117985926.1 | Type1.BREX | XIS-HTH->SMS-KaiC-><-SF1-SF2-Helicase<-SIGMA+OB-Fold+AAA-ATPase+HKD-DNase-BrxL<-PglZ<-Schlafen+HATPase+wHTH<-SF1-SF2-Helicase<-OLD-ABC+TOPRIM<-N6-METHYLASE<-DUF6079*<-BrxB-BrxF-iSTAND [illegible] | 1210 | Bacteria>FCB_group>Bacteroidetes/Chlorobi_group>Bacteroidota>Bacteroidia>Bacteroidales | GCF_934726555.1 | Bacteroides | O1446_RS12920 | 68574..72206 | 816 | MULTISPECIES:-BREX-system-P-loop-protein-BrxC-[Bacteroides] |
| WP_158582848.1 | Type1.BREX | ACET-><-?<-Cysteine-Desulfurase\|\|tRNA-><-?<-?<-?<-?\|\|BrxA-DNA-binding->BrxB-BrxF-iSTAND->DUF6079*->N6-METHYLASE->Schlafen+HATPase+wHTH->PglZ->SIGMA+OB-Fold+AAA-ATPase+HKD-DNase-BrxL-> [illegible] | 1210 | Bacteria>FCB_group>Bacteroidetes/Chlorobi_group>Bacteroidota>Bacteroidia>Bacteroidales | GCF_938033175.1 | Parabacteroides_sp._AF17-28 | DWW90_RS15090 | 22936..26568 | 2292241 | BREX-system-P-loop-protein-BrxC-[Parabacteroides-sp.-AF17-28] |
| WP_118139864.1 | Type1.BREX | <-HIRAN<-RADICAL-SAM<-ARG<-?<-?\|\|HTH_3->wHTH+WYL+WCX->BrxA-DNA-binding->BrxB-BrxF-iSTAND->DUF6079*->OLD-ABC->TOPRIM->N6-METHYLASE->N6-METHYLASE-p->?->SF1-SF2-Helicase->PglZ-> [illegible] | 1202 | Bacteria>FCB_group>Bacteroidetes/Chlorobi_group>Bacteroidota>Bacteroidia>Bacteroidales | - | Prevotella_copri | DWV60_RS07145 | 31878..35486 | 165179 | BREX-system-P-loop-protein-BrxC-[Prevotella-copri] |
| WP_153112500.1 | Type1.BREX | HAD+HAD->?->?->?->BrxA-DNA-binding->BrxB-BrxF-iSTAND->DUF6079*->N6-METHYLASE-> [illegible] | 1188 | Bacteria>FCB_group>Bacteroidetes/Chlorobi_group>Bacteroidota>Bacteroidia>Bacteroidales | - | Prevotella_copri | F7D34_RS01710 | 38543..42109 | 165179 | BREX-system-P-loop-protein-BrxC-[Prevotella-copri] |
| WP_264909062.1 | Type1.BREX | <-HAD+HAD<-SIGMA+OB-Fold+AAA-ATPase+Lon-Protease-BrxL<-PglZ<-N6-METHYLASE<-?<-?<- [illegible] | 1222 | Bacteria>FCB_group>Bacteroidetes/Chlorobi_group>Bacteroidota>Bacteroidia>Bacteroidales | - | Prevotella_copri | ONT11_RS09740 | 2183354..2187022 | 165179 | BREX-system-P-loop-protein-BrxC-[Prevotella-copri] |
| WP_264908465.1 | Type1.BREX | <-McrC-NTD+REase<-McrB-AAA<-SIGMA+OB-Fold+AAA-ATPase+HKD-DNase-BrxL<-PglZ<- [illegible] | 1206 | Bacteria>FCB_group>Bacteroidetes/Chlorobi_group>Bacteroidota>Bacteroidia>Bacteroidales | - | Prevotella_copri | ONT12_RS10110 | 2444120..2447740 | 165179 | BREX-system-P-loop-protein-BrxC-[Prevotella-copri] |
| WP_264951499.1 | Type1.BREX | SLOG+BirA-HTH-><-?\|\|AT-HTH-HYD2->AT-HTH-HYD2-><-HNH<-OLD-ABC<-?\|\|BrxA-DNA-binding [illegible] | 1222 | Bacteria>FCB_group>Bacteroidetes/Chlorobi_group>Bacteroidota>Bacteroidia>Bacteroidales | - | Prevotella_copri | ONT17_RS13510 | 3302536..3306204 | 165179 | BREX-system-P-loop-protein-BrxC-[Prevotella-copri] |
| WP_264904325.1 | Type1.BREX | <-HIRAN<-RADICAL-SAM<-ARG<-?<-?\|\|HTH_3->wHTH+WYL+WCX->BrxA-DNA-binding-> [illegible] | 1206 | Bacteria>FCB_group>Bacteroidetes/Chlorobi_group>Bacteroidota>Bacteroidia>Bacteroidales | - | Prevotella_copri | ONT18_RS04485 | 1079147..1082767 | 165179 | BREX-system-P-loop-protein-BrxC-[Prevotella-copri] |
| WP_073207711.1 | Type1.BREX | <-Classical-AAA<-?\|\|tRNA-><-?<-?<-HTH+DOC\|\|?->HTH_3->BrxA-DNA-binding-> [illegible] | 1206 | Bacteria>FCB_group>Bacteroidetes/Chlorobi_group>Bacteroidota>Bacteroidia>Bacteroidales | - | Prevotella_ruminicola | SAMN05216463_10942 | 46265..49885 | 839 | BREX-system-P-loop-protein-BrxC-[Prevotella-ruminicola] |
| WP_181980669.1 | Type1.BREX | TOPRIM->OLD-ABC+TOPRIM->SF1-SF2-Helicase->MORC+DUF3684+REase-><- [illegible] | 1192 | Bacteria>FCB_group>Bacteroidetes/Chlorobi_group>Bacteroidota>Bacteroidia | GCF_015558535.1 | Bacteroidales | QH473_RS20245 | 26845..30423 | 171549 | MULTISPECIES:-BREX-system-P-loop-protein-BrxC-[Bacteroidales] |
| WP_057098610.1 | Type1.BREX | <-KAP-NTPase<-SIGMA+OB-Fold+AAA-ATPase+Lon-Protease-BrxL<-PglZ<-Schlafen+HATPase+ [illegible] | 1212 | Bacteria>FCB_group>Bacteroidetes/Chlorobi_group>Bacteroidota>Bacteroidia | GCF_017309675.2 | Bacteroidales | DWX60_RS07020 | 13179..16817 | 171549 | MULTISPECIES:-BREX-system-P-loop-protein-BrxC-[Bacteroidales] |
| WP_046151576.1 | Type1.BREX | <-wHTH+WYL+WCX<-ParB+HNH<-SIR2<-SIGMA+OB-Fold+AAA-ATPase+Lon-Protease-BrxL<- [illegible] | 1184 | Bacteria>FCB_group>Bacteroidetes/Chlorobi_group>Bacteroidota>Bacteroidia | GCF_900102645.1 | Bacteroidales | BLR14_RS08960 | 69320..72874 | 171549 | MULTISPECIES:-BREX-system-P-loop-protein-BrxC-[Bacteroidales] |
| WP_009275691.1 | Type1.BREX | <-NUDIX+NUDIX<-?<-?<-?\|\|HTH_3->wHTH+WYL+WCX->BrxA-DNA-binding->BrxB-BrxF-iSTAND-> [illegible] | 1208 | Bacteria>FCB_group>Bacteroidetes/Chlorobi_group>Bacteroidota>Bacteroidia | GCF_949084405.1 | Bacteroidales | KSZ59_RS23265 | 10034..13660 | 171549 | MULTISPECIES:-BREX-system-P-loop-protein-BrxC-[Bacteroidales] |
| BBE18573.1 | Type1.BREX | TPR+TPR->BrxA-DNA-binding->BrxB-BrxF-iSTAND->DUF6079*->N6-METHYLASE->?->DOC+HTH-> [illegible] | 1211 | Bacteria>FCB_group>Bacteroidetes/Chlorobi_group>Bacteroidota>Bacteroidia>Marinilabiliales | - | Aquipluma_nitroreducens | AQPE_2736 | 3049074..3052709 | 2010828 | hypothetical-protein-AQPE_2736-[Aquipluma-nitroreducens] |
| BBE20443.1 | Type1.BREX | TraJ-RHH->TraI-VirD2-RCRE-><-wHTH+WYL+WCX\|\|?->SMODS->?->HTH_3->BrxA-DNA-binding-> [illegible] | 1173 | Bacteria>FCB_group>Bacteroidetes/Chlorobi_group>Bacteroidota>Bacteroidia>Marinilabiliales | - | Aquipluma_nitroreducens | AQPE_4635 | 5186015..5189536 | 2010828 | hypothetical-protein-AQPE_4635-[Aquipluma-nitroreducens] |
| MCF8362389.1 | Type1.BREX | <-SIGMA+OB-Fold+AAA-ATPase+HKD-DNase-BrxL<-PglZ<-PglZ<-Schlafen+HATPase+wHTH<- [illegible] | 1180 | Bacteria>FCB_group>Bacteroidetes/Chlorobi_group>Bacteroidota>Bacteroidia>Marinilabiliales | GCA_021740345.1 | Prolixibacteraceae_bacterium | K9G70_07170 | 11489..15031 | 2053594 | BREX-system-P-loop-protein-BrxC-[Prolixibacteraceae-bacterium] |
| MCK9411439.1 | Type1.BREX | Peptidase_A8->Peptidase_A8-><-wHTH+WYL+WCX\|\|?->SMODS->?->HTH_3-> [illegible] | 1173 | Bacteria>FCB_group>Bacteroidetes/Chlorobi_group>Bacteroidota>Bacteroidia>Marinilabiliales | GCA_023229575.1 | Prolixibacteraceae_bacterium | M0Q53_04000 | 25065..28586 | 2053594 | BREX-system-P-loop-protein-BrxC-[Prolixibacteraceae-bacterium] |
| MCU4173933.1 | Type1.BREX | <-REase+RNase\|\|?->?->?->?->?->HTH_3->BrxA-DNA-binding->BrxB-BrxF-iSTAND-> [illegible] | 1167 | Bacteria>FCB_group>Bacteroidetes/Chlorobi_group>Bacteroidota>Bacteroidia>Marinilabiliales | GCA_025499625.1 | Marinilabiliaceae_bacterium_N1Y90 | J1N11_02510 | 10462..13965 | 2816964 | BREX-system-P-loop-protein-BrxC-[Marinilabiliaceae-bacterium-N1Y90] |
| WP_027470973.1 | Type1.BREX | wHTH+WYL+WCX->HTH_3->BrxA-DNA-binding->BrxB-BrxF-iSTAND->DUF6079*->?->SbcC->?-> [illegible] | 1172 | Bacteria>FCB_group>Bacteroidetes/Chlorobi_group>Bacteroidota>Bacteroidia>Marinilabiliales | GCF_000583675.1 | Saccharicrinis_fermentans | JCM21142_RS14985 | 5014..8532 | 982 | BREX-system-P-loop-protein-BrxC-[Saccharicrinis-fermentans] |
| WP_062122677.1 | Type1.BREX | ABhydrolase->tRNA-><-SIGMA+OB-Fold+AAA-ATPase+HKD-DNase-BrxL<-PglZ<- [illegible] | 329 | Bacteria>FCB_group>Bacteroidetes/Chlorobi_group>Bacteroidota>Bacteroidia>Marinilabiliales | GCF_000974365.1 | Geofilum_rubicundum | JCM15548_RS04470 | 156242..157231 | 472113 | hypothetical-protein-[Geofilum-rubicundum] |
| WP_075603947.1 | Type1.BREX | TPR+TPR+TPR+TPR->?->XIS-HTH->?->CHC2+TOPRIM-DNAG->?->BrxA-DNA-binding-> [illegible] | 1204 | Bacteria>FCB_group>Bacteroidetes/Chlorobi_group>Bacteroidota>Bacteroidia>Marinilabiliales | GCF_001660705.2 | Saccharicrinis_aurantiacus | A8B35_RS21005 | 65274..68888 | 1849719 | BREX-system-P-loop-protein-BrxC-[Saccharicrinis-aurantiacus] |
| WP_107823954.1 | Type1.BREX | <-STYKinase-like<-Couple_hipA<-HTH_3<-SIGMA+OB-Fold+AAA-ATPase+HKD-DNase-BrxL<- [illegible] | 1168 | Bacteria>FCB_group>Bacteroidetes/Chlorobi_group>Bacteroidota>Bacteroidia>Marinilabiliales | GCF_003046625.1 | Mangrovibacterium_marinum | C8N47_RS17935 | 11546..15052 | 1639118 | BREX-system-P-loop-protein-BrxC-[Mangrovibacterium-marinum] |
| WP_163710191.1 | Type1.BREX | HTH_3->BrxA-DNA-binding->BrxB-BrxF-iSTAND->DUF6079*->Schlafen+HATPase+wHTH-> [illegible] | 1168 | Bacteria>FCB_group>Bacteroidetes/Chlorobi_group>Bacteroidota>Bacteroidia>Marinilabiliales | GCF_010882175.1 | Mangrovibacterium_lignilyticum | GJV22_RS06145 | 1519093..1522599 | 2668052 | BREX-system-P-loop-protein-BrxC-[Mangrovibacterium-lignilyticum] |
| WP_250725046.1 | Type1.BREX | DDE_Tnp_ISL3->DUF4372+DDE_Tnp_1-><-SIGMA+OB-Fold+AAA-ATPase+HKD-DNase-BrxL<- [illegible] | 1173 | Bacteria>FCB_group>Bacteroidetes/Chlorobi_group>Bacteroidota>Bacteroidia>Marinilabiliales | GCF_023674465.1 | Alkaliflexus_sp._Ai-910 | M9189_RS04830 | 1155477..1158998 | 2949635 | BREX-system-P-loop-protein-BrxC-[Alkaliflexus-sp.-Ai-910] |
| OJV92258.1 | Type1.BREX | RNase->?->TatD_DNase->?->tRNA-><-SIGMA+OB-Fold+AAA-ATPase+HKD-DNase-BrxL<-PglZ<- [illegible] | 1200 | Bacteria>FCB_group>Bacteroidetes/Chlorobi_group>Bacteroidota>Bacteroidia>unclassified_Bacteroidia | GCA_001897595.1 | Bacteroidia_bacterium_44-10 | BGO34_14870 | 43076..46678 | 1895721 | hypothetical-protein-BGO34_14870-[Bacteroidia-bacterium-44-10] |
| OSZ81159.1 | Type1.BREX | BrxA-DNA-binding->BrxB-BrxF-iSTAND->DUF6079*->N6-METHYLASE->McrB-AAA-> [illegible] | 1144 | Bacteria>FCB_group>Bacteroidetes/Chlorobi_group>Bacteroidota>Chitinophagia>Chitinophagales | GCA_002127295.1 | Chitinophagaceae_bacterium_IBVUCB2 | CAP36_07965 | 1792100..1795534 | 1985174 | hypothetical-protein-CAP36_07965-[Chitinophagaceae-bacterium-IBVUCB2] |
| PJE46487.1 | Type1.BREX | <-DOC+HTH<-?<-?<-HerA/FtsK\|\|wHTH+WYL+WCX->?->REase+RNase->BrxA-DNA-binding-> [illegible] | 1169 | Bacteria>FCB_group>Bacteroidetes/Chlorobi_group>Bacteroidota>Chitinophagia>Chitinophagales | GCA_002786355.1 | Sediminibacterium_sp._FEMGT703S | CUR34_07570 | 66784..70293 | 2080766 | MAG:-hypothetical-protein-CUR34_07570-[Sediminibacterium-sp.] |
| QEC55281.1 | Type1.BREX | <-AbiEi_4<-MORC+SF1-SF2-Helicase<-TatD_DNase<-?<-KAP-glycosylase<-KAP-NTPase<- [illegible] | 1177 | Bacteria>FCB_group>Bacteroidetes/Chlorobi_group>Bacteroidota>Chitinophagia>Chitinophagales | GCA_007970805.1 | Flavisolibacter_ginsenosidimutans | FSB75_04955 | 1196460..1199993 | 661481 | BREX-system-P-loop-protein-BrxC-[Flavisolibacter-ginsenosidimutans] |
| NTS43012.1 | Type1.BREX | <-SIGMA+OB-Fold+AAA-ATPase+HKD-DNase-BrxL<-PglZ<-N6-METHYLASE<-DUF6079*<- [illegible] | 1173 | Bacteria>FCB_group>Bacteroidetes/Chlorobi_group>Bacteroidota>Chitinophagia>Chitinophagales | GCA_013327885.1 | Flavisolibacter_longurius | HRG84_19150 | 28734..32255 | 2740583 | BREX-system-P-loop-protein-BrxC-[Flavisolibacter-longurius] |
| MBL7764603.1 | Type1.BREX | HTH->BrxA-DNA-binding->BrxB-BrxF-iSTAND->DUF6079*->N6-METHYLASE->McrB-AAA-> [illegible] | 1146 | Bacteria>FCB_group>Bacteroidetes/Chlorobi_group>Bacteroidota>Chitinophagia>Chitinophagales | GCA_016786845.1 | Chitinophagaceae_bacterium | JNJ58_00790 | 47471..50911 | 1869212 | BREX-system-P-loop-protein-BrxC-[Chitinophagaceae-bacterium] |
| MBP8114974.1 | Type1.BREX | <-SIGMA+OB-Fold+AAA-ATPase+Lon-Protease-BrxL-p<-PglZ<-?<-N6-METHYLASE<-DUF6079* | 1136 | Bacteria>FCB_group>Bacteroidetes/Chlorobi_group>Bacteroidota>Chitinophagia>Chitinophagales | GCA_018005575.1 | Chitinophagaceae_bacterium | KAY50_06430 | 6842..10254 | 1869212 | BREX-system-P-loop-protein-BrxC-[Chitinophagaceae-bacterium] |
| MCB9043396.1 | Type1.BREX | sGTP->?->HTH->BrxA-DNA-binding->BrxB-BrxF-iSTAND->DUF6079*-> [illegible] | 1197 | Bacteria>FCB_group>Bacteroidetes/Chlorobi_group>Bacteroidota>Chitinophagia>Chitinophagales | GCA_020636475.1 | Chitinophagales_bacterium | H6554_06480 | 28579..32172 | 2448779 | BREX-system-P-loop-protein-BrxC-[Chitinophagales-bacterium] |
| MCK9404665.1 | Type1.BREX | <-HARE-HTH+REase<-SIGMA+OB-Fold+AAA-ATPase+Lon-Protease-BrxL<-PglZ<-?<- [illegible] | 1144 | Bacteria>FCB_group>Bacteroidetes/Chlorobi_group>Bacteroidota>Chitinophagia>Chitinophagales | GCA_023229465.1 | Chitinophagaceae_bacterium | M0Q26_14830 | 11931..15365 | 1869212 | BREX-system-P-loop-protein-BrxC-[Chitinophagaceae-bacterium] |
| WP_172623061.1 | Type1.BREX | <-AbiEi_4<-MORC+SF1-SF2-Helicase<-TatD_DNase<-?<-KAP-glycosylase<-KAP-NTPase<- [illegible] | 1174 | Bacteria>FCB_group>Bacteroidetes/Chlorobi_group>Bacteroidota>Chitinophagia>Chitinophagales | GCF_007970805.1 | Flavisolibacter_ginsenosidimutans | FSB75_RS04955 | 1196460..1199984 | 661481 | BREX-system-P-loop-protein-BrxC-[Flavisolibacter-ginsenosidimutans] |
| WP_072366579.1 | Type1.BREX | <-HTH_Tnp_1\|\|HTH_3->BrxA-DNA-binding->BrxB-BrxF-iSTAND->DUF6079*->?-> [illegible] | 1142 | Bacteria>FCB_group>Bacteroidetes/Chlorobi_group>Bacteroidota>Chitinophagia>Chitinophagales | GCF_900119105.1 | Chitinophaga_sancti | BT118_RS32550 | 2428..5856 | 1004 | BREX-system-P-loop-protein-BrxC-[Chitinophaga-sancti] |
| ADB40822.1 | Type1.BREX | GDResolvase+HIN-HTH-><-?\|\|?-><-?\|\|?-><-?\|\|URI-DNase->BrxA-DNA-binding-> [illegible] | 1186 | Bacteria>FCB_group>Bacteroidetes/Chlorobi_group>Bacteroidota>Cytophagia>Cytophagales | GCA_000024525.1 | Spirosoma_linguale_DSM_74 | Slin_4844 | 5894993..5898553 | 504472 | hypothetical-protein-Slin_4844-[Spirosoma-linguale-DSM-74] |
| UOQ83254.1 | Type1.BREX | <-pPIWI+pPIWI\|\|HEPN-><-REase<-?\|\|Phage_integrase+Phage_integrase->NUMOD4+HNH-> [illegible] | 1148 | Bacteria>FCB_group>Bacteroidetes/Chlorobi_group>Bacteroidota>Cytophagia>Cytophagales | GCA_022919455.1 | Hymenobacter_sp._5414T-23 | MUN83_21225 | 87571..91017 | 2932252 | BREX-system-P-loop-protein-BrxC-[Hymenobacter-sp.-5414T-23] |
| SKB69504.1 | Type1.BREX | D5ATPase->?->?->?->?->CopG/RHH->HTH->BrxA-DNA-binding->BrxB-BrxF-iSTAND-> [illegible] | 1201 | Bacteria>FCB_group>Bacteroidetes/Chlorobi_group>Bacteroidota>Cytophagia>Cytophagales | GCA_900167945.1 | Dyadobacter_psychrophilus | SAMN05660293_01532 | 106911..110516 | 651661 | hypothetical-protein-SAMN05660293_01532-[Dyadobacter-psychrophilus] |
| WP_013929909.1 | Type1.BREX | Peptidase_S51->?-><-SIGMA+OB-Fold+AAA-ATPase+Lon-Protease-BrxL<-PglZ<- [illegible] | 1173 | Bacteria>FCB_group>Bacteroidetes/Chlorobi_group>Bacteroidota>Cytophagia>Cytophagales | GCF_000218895.1 | Runella_slithyformis | RUNSL_RS20935 | 4933430..4936951 | 106 | BREX-system-P-loop-protein-BrxC-[Runella-slithyformis] |
| WP_008625904.1 | Type1.BREX | Calcineurin->?->SbcC->?-><-SF1-SF2-Helicase\|\|?->?->BrxA-DNA-binding-> [illegible] | 1146 | Bacteria>FCB_group>Bacteroidetes/Chlorobi_group>Bacteroidota>Cytophagia>Cytophagales | GCF_000330725.2 | Mariniradius_saccharolyticus | C943_RS07085 | 84904..88344 | 1245591 | BREX-system-P-loop-protein-BrxC-[Mariniradius-saccharolyticus] |
| WP_018619849.1 | Type1.BREX | <-SAM-METHYLASE\|\|?->HTH_Tnp_1+HTH_21+rve+rve_3-><-SIGMA+OB-Fold+AAA-ATPase+ [illegible] | 1142 | Bacteria>FCB_group>Bacteroidetes/Chlorobi_group>Bacteroidota>Cytophagia>Cytophagales | GCF_000374065.1 | Spirosoma_luteum | B156_RS0111470 | 131208..134636 | 431553 | BREX-system-P-loop-protein-BrxC-[Spirosoma-luteum] |
| WP_028665035.1 | Type1.BREX | DSBH+AraC-HTH->?-><-?<-CNMP+CRP-HTH<-?\|\|BrxA-DNA-binding->BrxB-BrxF-iSTAND-> [illegible] | 1171 | Bacteria>FCB_group>Bacteroidetes/Chlorobi_group>Bacteroidota>Cytophagia>Cytophagales | GCF_000423565.1 | Runella_zeae | G563_RS0108985 | 25803..29318 | 94255 | BREX-system-P-loop-protein-BrxC-[Runella-zeae] |
| WP_053405044.1 | Type1.BREX | <-SIGMA+OB-Fold+AAA-ATPase+HKD-DNase-BrxL<-PglZ\|\|ParB+HNH-><-N6-METHYLASE<- [illegible] | 1179 | Bacteria>FCB_group>Bacteroidetes/Chlorobi_group>Bacteroidota>Cytophagia>Cytophagales | GCF_001274635.1 | Persicobacter_sp._CCB-QB2 | NG22_RS06825 | 1502189..1505728 | 1561025 | BREX-system-P-loop-protein-BrxC-[Persicobacter-sp.-CCB-QB2] |
| WP_073850520.1 | Type1.BREX | <-HNH+SAVED<-?<-?<-?<-?<-?<-SIGMA+OB-Fold+AAA-ATPase+HKD-DNase-BrxL<-PglZ<- [illegible] | 1179 | Bacteria>FCB_group>Bacteroidetes/Chlorobi_group>Bacteroidota>Cytophagia>Cytophagales | GCF_001907195.1 | Pontibacter_flavimaris | A3841_RS01550 | 388287..391826 | 1797110 | BREX-system-P-loop-protein-BrxC-[Pontibacter-flavimaris] |
| WP_094816172.1 | Type1.BREX | GDResolvase->REase->?-><-MIT_C<-SIGMA+OB-Fold+AAA-ATPase+Lon-Protease-BrxL<- [illegible] | 1165 | Bacteria>FCB_group>Bacteroidetes/Chlorobi_group>Bacteroidota>Cytophagia>Cytophagales | GCF_002261125.1 | Siphonobacter_sp._BAB-5385 | BWI93_RS24020 | 10112..13609 | 1864822 | BREX-system-P-loop-protein-BrxC-[Siphonobacter-sp.-BAB-5385] |
| WP_273212250.1 | Type1.BREX | <-AbiEii<-wHTH+AbiEi_2<-SIGMA+OB-Fold+AAA-ATPase+Lon-Protease-BrxL<-PglZ<- [illegible] | 1173 | Bacteria>FCB_group>Bacteroidetes/Chlorobi_group>Bacteroidota>Cytophagia>Cytophagales | GCF_002483185.1 | Runella_zeae | B4E63_RS12460 | 307458..310979 | 94255 | BREX-system-P-loop-protein-BrxC-[Runella-zeae] |
| WP_100629834.1 | Type1.BREX | <-SbcC\|\|BrxA-DNA-binding->BrxB-BrxF-iSTAND->DUF6079-p*->DUF6079-p->N6-METHYLASE-> [illegible] | 553 | Bacteria>FCB_group>Bacteroidetes/Chlorobi_group>Bacteroidota>Cytophagia>Cytophagales | GCF_002807035.1 | Algoriphagus_formosus | CDH51_RS17530 | 460328..461989 | 2007308 | BREX-system-P-loop-protein-BrxC-[Algoriphagus-formosus] |
| WP_119666207.1 | Type1.BREX | <-IG+IG<-?<-NTP_transf_5<-MarR-HTH<-?\|\|METHYLASE->ChlI+Lon_C+AAA-> [illegible] | 1143 | Bacteria>FCB_group>Bacteroidetes/Chlorobi_group>Bacteroidota>Cytophagia>Cytophagales | GCF_003583365.1 | Fibrisoma_montanum | DYU11_RS03320 | 792110..795541 | 2305895 | BREX-system-P-loop-protein-BrxC-[Fibrisoma-montanum] |
| WP_122934661.1 | Type1.BREX | <-ParB<-ParA-Soj-PloopNTPase\|\|AbiEi_4->AbiEii->BrxA-DNA-binding->BrxB-BrxF-iSTAND [illegible] | 1142 | Bacteria>FCB_group>Bacteroidetes/Chlorobi_group>Bacteroidota>Cytophagia>Cytophagales | GCF_003711225.1 | Runella_sp._SP2 | DTQ70_RS30140 | 43566..46994 | 2268026 | BREX-system-P-loop-protein-BrxC-[Runella-sp.-SP2] |
| WP_133557368.1 | Type1.BREX | <-SF1-SF2-Helicase<-HKD-DNase<-?\|\|?->HAD+PRTase+PRTase+REase+SF1-SF2-Helicase->?- [illegible] | 1167 | Bacteria>FCB_group>Bacteroidetes/Chlorobi_group>Bacteroidota>Cytophagia>Cytophagales | GCF_004362805.1 | Algoriphagus_boseongensis | DFQ04_RS15130 | 300982..304485 | 1442587 | BREX-system-P-loop-protein-BrxC-[Algoriphagus-boseongensis] |
| WP_138994213.1 | Type1.BREX | TraN->?->?->?->?->GDResolvase+HIN-HTH->BrxA-DNA-binding->BrxB-BrxF-iSTAND-> [illegible] | 1214 | Bacteria>FCB_group>Bacteroidetes/Chlorobi_group>Bacteroidota>Cytophagia>Cytophagales | GCF_006149005.1 | Larkinella_sp._C7 | FH871_RS25315 | 15156..18800 | 2576607 | BREX-system-P-loop-protein-BrxC-[Larkinella-sp.-C7] |
| WP_146895591.1 | Type1.BREX | <-wHTH+WYL+WCX\|\|SF1-SF2-Helicase->RVT->?->?-><-SIGMA+OB-Fold+AAA-ATPase+ [illegible] | 1143 | Bacteria>FCB_group>Bacteroidetes/Chlorobi_group>Bacteroidota>Cytophagia>Cytophagales | GCF_007991635.1 | Adhaeribacter_aerolatus | AAE02nite_RS05160 | 90394..93825 | 670289 | BREX-system-P-loop-protein-BrxC-[Adhaeribacter-aerolatus] |
| WP_146899771.1 | Type1.BREX | <-SIGMA+OB-Fold+AAA-ATPase+HKD-DNase-BrxL<-PglZ<-N6-METHYLASE<-DUF6079*<- [illegible] | 1182 | Bacteria>FCB_group>Bacteroidetes/Chlorobi_group>Bacteroidota>Cytophagia>Cytophagales | GCF_007991635.1 | Adhaeribacter_aerolatus | AAE02nite_RS16235 | 111241..114789 | 670289 | BREX-system-P-loop-protein-BrxC-[Adhaeribacter-aerolatus] |
| WP_159471214.1 | Type1.BREX | <-XIS-HTH<-AbiEii<-AbiEi_4+AbiEi_1<-REase<-SIGMA+OB-Fold+AAA-ATPase+Lon-Protease- [illegible] | 1179 | Bacteria>FCB_group>Bacteroidetes/Chlorobi_group>Bacteroidota>Cytophagia>Cytophagales | GCF_009801115.1 | Dyadobacter_sp._3J3 | FZF08_RS17845 | 89009..92548 | 2606600 | BREX-system-P-loop-protein-BrxC-[Dyadobacter-sp.-3J3] |
| WP_168675169.1 | Type1.BREX | <-CASPASE<-DDE_Tnp_4+DDE_Tnp_1_2\|\|Phage_integrase+Phage_integrase-><-McrC-NTD+ [illegible] | 1172 | Bacteria>FCB_group>Bacteroidetes/Chlorobi_group>Bacteroidota>Cytophagia>Cytophagales | GCF_012275535.1 | Hymenobacter_artigasi | HBN54_RS20840 | 55199..58717 | 2719616 | BREX-system-P-loop-protein-BrxC-[Hymenobacter-artigasi] |
| WP_181308258.1 | Type1.BREX | HEPN->RNaseH-><-?<-HU-HIG+REase<-REase+N-glycosylase\|\|BrxA-DNA-binding-> [illegible] | 1144 | Bacteria>FCB_group>Bacteroidetes/Chlorobi_group>Bacteroidota>Cytophagia>Cytophagales | GCF_013623775.1 | Rufibacter_sp._XAAS-G3-1 | HH050_RS18475 | 14255..17689 | 2729134 | BREX-system-P-loop-protein-BrxC-[Rufibacter-sp.-XAAS-G3-1] |
| WP_183972373.1 | Type1.BREX | D5ATPase->?->?->HKD-DNase-><-?<-?<-?\|\|BrxA-DNA-binding->BrxB-BrxF-iSTAND-> [illegible] | 1202 | Bacteria>FCB_group>Bacteroidetes/Chlorobi_group>Bacteroidota>Cytophagia>Cytophagales | GCF_014195535.1 | Runella_defluvii | FHS57_RS08095 | 111516..115124 | 370973 | BREX-system-P-loop-protein-BrxC-[Runella-defluvii] |
| WP_185195977.1 | Type1.BREX | <-TIR<-?<-HTH+MPTase<-?\|\|AbiEi_4->AbiEii-><-SIGMA+OB-Fold+AAA-ATPase+HKD-DNase- [illegible] | 1179 | Bacteria>FCB_group>Bacteroidetes/Chlorobi_group>Bacteroidota>Cytophagia>Cytophagales | GCF_014212495.1 | Pontibacter_sp._Tf4 | H7F15_RS05560 | 22760..26299 | 2761620 | BREX-system-P-loop-protein-BrxC-[Pontibacter-sp.-Tf4] |
| WP_187455529.1 | Type1.BREX | <-N6-METHYLASE<-?\|\|XIS-HTH-><-SIGMA+OB-Fold+AAA-ATPase+HKD-DNase-BrxL<-PglZ<- [illegible] | 1178 | Bacteria>FCB_group>Bacteroidetes/Chlorobi_group>Bacteroidota>Cytophagia>Cytophagales | GCF_014349165.1 | Hymenobacter_sp._BT491 | H9I52_RS21275 | 49708..53244 | 2766779 | BREX-system-P-loop-protein-BrxC-[Hymenobacter-sp.-BT491] |
| WP_188169955.1 | Type1.BREX | <-OLD-ABC<-Arm-DNA-bind_5+Phage_int_SAM_5+Phage_integrase<-MACRODOMAIN<-ART-DarT< [illegible] | 1216 | Bacteria>FCB_group>Bacteroidetes/Chlorobi_group>Bacteroidota>Cytophagia>Cytophagales | GCF_014534745.1 | Flammeovirga_sp._EKP202 | IC221_RS16125 | 83755..87405 | 2770592 | BREX-system-P-loop-protein-BrxC-[Flammeovirga-sp.-EKP202] |
| WP_188933335.1 | Type1.BREX | <-wHTH+WYL+WCX\|\|?->TIR-><-HTH_28+HTH_32+rve+rve_3<-SF1-SF2-Helicase<- [illegible] | 1165 | Bacteria>FCB_group>Bacteroidetes/Chlorobi_group>Bacteroidota>Cytophagia>Cytophagales | GCF_014641595.1 | Dyadobacter_endophyticus | IEY24_RS14735 | 618400..621897 | 1749036 | BREX-system-P-loop-protein-BrxC-[Dyadobacter-endophyticus] |
| WP_189584875.1 | Type1.BREX | AbiGi+seqA+RAMA->BrxA-DNA-binding->BrxB-BrxF-iSTAND->DUF6079*->N6-METHYLASE-> [illegible] | 1146 | Bacteria>FCB_group>Bacteroidetes/Chlorobi_group>Bacteroidota>Cytophagia>Cytophagales | GCF_014652135.1 | Mongoliitalea_lutea | IE080_RS15850 | 2275..5715 | 849756 | BREX-system-P-loop-protein-BrxC-[Mongoliitalea-lutea] |
| WP_191042738.1 | Type1.BREX | TraJ-RHH->TraI-VirD2-RCRE-><-NtrC-AAA+FIS-HTH\|\|35exo+wHTH+SF1-SF2-Helicase+HTH-> [illegible] | 1206 | Bacteria>FCB_group>Bacteroidetes/Chlorobi_group>Bacteroidota>Cytophagia>Cytophagales | GCF_014699035.1 | Spirosoma_validum | IC230_RS29845 | 51060..54680 | 2771355 | BREX-system-P-loop-protein-BrxC-[Spirosoma-validum] |
| WP_201982535.1 | Type1.BREX | HEPN->CHTH->TOPRIM->Sp-><-ParB<-OLD-ABC+TOPRIM<-SIGMA+OB-Fold+AAA-ATPase+Lon- [illegible] | 1143 | Bacteria>FCB_group>Bacteroidetes/Chlorobi_group>Bacteroidota>Cytophagia>Cytophagales | GCF_016734815.1 | Hymenobacter_rubidus | JKL48_RS11490 | 20898..24329 | 1441626 | BREX-system-P-loop-protein-BrxC-[Hymenobacter-rubidus] |
| WP_205503392.1 | Type1.BREX | <-wHTH+WYL+WCX\|\|SF1-SF2-Helicase->RVT->?-><-SIGMA+OB-Fold+AAA-ATPase+HKD-DNase- [illegible] | 1180 | Bacteria>FCB_group>Bacteroidetes/Chlorobi_group>Bacteroidota>Cytophagia>Cytophagales | GCF_016924645.1 | Rufibacter_sp._SYSU_D00308 | JT718_RS14320 | 179098..182640 | 2812556 | BREX-system-P-loop-protein-BrxC-[Rufibacter-sp.-SYSU-D00308] |
| WP_234861365.1 | Type1.BREX | STYKIN-><-?\|\|?->GDResolvase-><-?\|\|TPR+TPR->BrxA-DNA-binding->BrxB-BrxF-iSTAND-> [illegible] | 1174 | Bacteria>FCB_group>Bacteroidetes/Chlorobi_group>Bacteroidota>Cytophagia>Cytophagales | GCF_021491425.1 | Mariniradius_sp._RY-2 | LOU89_RS09865 | 251856..255380 | 2909237 | BREX-system-P-loop-protein-BrxC-[Mariniradius-sp.-RY-2] |
| WP_241348547.1 | Type1.BREX | <-ParB+N6_N4_METHYLASE<-?<-?<-SIGMA+OB-Fold+AAA-ATPase+HKD-DNase-BrxL<-PglZ<- [illegible] | 1174 | Bacteria>FCB_group>Bacteroidetes/Chlorobi_group>Bacteroidota>Cytophagia>Cytophagales | GCF_022549775.1 | Belliella_sp._DSM_111904 | MM239_RS12315 | 24692..28216 | 2923435 | BREX-system-P-loop-protein-BrxC-[Belliella-sp.-DSM-111904] |
| WP_244713422.1 | Type1.BREX | <-XIS-HTH\|\|HAD+PRTase+PRTase+REase+SF1-SF2- [illegible] | 1145 | Bacteria>FCB_group>Bacteroidetes/Chlorobi_group>Bacteroidota>Cytophagia>Cytophagales | GCF_022919135.1 | Hymenobacter_sp._5116S-27 | MUN80_RS13030 | 3065695..3069132 | 2932249 | BREX-system-P-loop-protein-BrxC-[Hymenobacter-sp.-5116S-27] |
| WP_262922299.1 | Type1.BREX | <-pPIWI+pPIWI\|\|HEPN-><-REase<-?\|\|Phage_integrase [illegible] | 1164 | Bacteria>FCB_group>Bacteroidetes/Chlorobi_group>Bacteroidota>Cytophagia>Cytophagales | GCF_022919455.1 | Hymenobacter_sp._5414T-23 | MUN83_RS21225 | 87523..91017 | 2932252 | BREX-system-P-loop-protein-BrxC-[Hymenobacter-sp.-5414T-23] |
| WP_255036052.1 | Type1.BREX | XIS-HTH->D5ATPase->?->?->?->?->BrxA-DNA-binding- [illegible] | 1199 | Bacteria>FCB_group>Bacteroidetes/Chlorobi_group>Bacteroidota>Cytophagia>Cytophagales | GCF_024343775.1 | Lacihabitans_soyangensis | EGI31_RS04905 | 50707..54306 | 869394 | BREX-system-P-loop-protein-BrxC-[Lacihabitans-soyangensis] |
| WP_269011923.1 | Type1.BREX | HTH_3->Couple_hipA+HipA_C-><-?<-HNH+Polbeta<- [illegible] | 1166 | Bacteria>FCB_group>Bacteroidetes/Chlorobi_group>Bacteroidota>Cytophagia>Cytophagales | GCF_026938185.1 | Aquirufa_antheringensis | G9H55_RS04755 | 131184..134684 | 2516559 | BREX-system-P-loop-protein-BrxC-[Aquirufa-antheringensis] |
| WP_082214111.1 | Type1.BREX | XIS-HTH->D5ATPase->?->?->CopG/RHH->HTH->BrxA- [illegible] | 1201 | Bacteria>FCB_group>Bacteroidetes/Chlorobi_group>Bacteroidota>Cytophagia>Cytophagales | GCF_900167945.1 | Dyadobacter_psychrophilus | B5X68_RS07585 | 106911..110516 | 651661 | BREX-system-P-loop-protein-BrxC-[Dyadobacter-psychrophilus] |
| WP_084121639.1 | Type1.BREX | <-TraJ-RHH\|\|?->TPR+TPR->MazG-><-?<-SIGMA+OB-Fold [illegible] | 1146 | Bacteria>FCB_group>Bacteroidetes/Chlorobi_group>Bacteroidota>Cytophagia>Cytophagales | GCF_900176595.1 | Aquiflexum_balticum | B9A52_RS17175 | 4064780..4068220 | 280473 | BREX-system-P-loop-protein-BrxC-[Aquiflexum-balticum] |
| WP_244679416.1 | Type1.BREX | <-pPIWI+pPIWI\|\|HEPN-><-REase<-?\|\|Phage_integrase [illegible] | 1148 | Bacteria>FCB_group>Bacteroidetes/Chlorobi_group>Bacteroidota>Cytophagia>Cytophagales | - | Hymenobacter_sp._5414T-23 | MUN83_21225 | 87571..91017 | 2932252 | BREX-system-P-loop-protein-BrxC-[Hymenobacter-sp.-5414T-23] |
| MBT5528618.1 | Type1.BREX | HEPN-><-Phage_integrase\|\|BrxA-DNA-binding->BrxB- [illegible] | 473 | Bacteria>FCB_group>Bacteroidetes/Chlorobi_group>Bacteroidota>Cytophagia>unclassified_Cytophagia | GCA_018665725.1 | Cytophagia_bacterium | HOK35_05630 | 3390..4810 | 2026730 | BREX-system-P-loop-protein-BrxC-[Cytophagia-bacterium] |
| OIQ21186.1 | Type1.BREX | CHTH+LexA-protease-><-GDResolvase+Recombinase+ [illegible] | 1165 | Bacteria>FCB_group>Bacteroidetes/Chlorobi_group>Bacteroidota>Flavobacteriia>Flavobacteriales | GCA_001874145.1 | Lacinutrix_sp._MedPE-SW | BM549_09430 | 19615..23112 | 1860087 | MAG:-hypothetical-protein-BM549_09430-[Lacinutrix-sp.-MedPE-SW] |
| GAW88409.1 | Type1.BREX | XIS-HTH->HTH_36->Classical-AAA->HTH_3->BrxA-DNA- [illegible] | 385 | Bacteria>FCB_group>Bacteroidetes/Chlorobi_group>Bacteroidota>Flavobacteriia>Flavobacteriales | GCA_002207815.1 | Flavobacterium_psychrophilum | FPS14_contig00004-0074 | 61867..63024 | 96345 | hypothetical-protein-FPS14_contig00004-0074-[Flavobacterium-psychrophilum] |
| GAW88410.1 | Type1.BREX | XIS-HTH->HTH_36->Classical-AAA->HTH_3->BrxA-DNA- [illegible] | 768 | Bacteria>FCB_group>Bacteroidetes/Chlorobi_group>Bacteroidota>Flavobacteriia>Flavobacteriales | GCA_002207815.1 | Flavobacterium_psychrophilum | FPS14_contig00004-0075 | 63069..65375 | 96345 | hypothetical-protein-FPS14_contig00004-0075-[Flavobacterium-psychrophilum] |
| OYX86687.1 | Type1.BREX | <-RADICAL-SAM+TRAM\|\|?->Arm-DNA-bind_5+Phage_int_ [illegible] | 1199 | Bacteria>FCB_group>Bacteroidetes/Chlorobi_group>Bacteroidota>Flavobacteriia>Flavobacteriales | GCA_002280815.1 | Flavobacteriales_bacterium_32-34-25 | B7Y83_00570 | 11855..15454 | 1970508 | hypothetical-protein-B7Y83_00570-[Flavobacteriales-bacterium-32-34-25] |
| MAM30277.1 | Type1.BREX | <-ParB+N6_N4_METHYLASE\|\|XIS-HTH-><-SIGMA+OB-Fold [illegible] | 1165 | Bacteria>FCB_group>Bacteroidetes/Chlorobi_group>Bacteroidota>Flavobacteriia>Flavobacteriales | GCA_002693005.1 | Flavobacteriaceae_bacterium | CMC13_14765 | 19860..23357 | 1871037 | hypothetical-protein-[Flavobacteriaceae-bacterium] |
| MAL23063.1 | Type1.BREX | <-METHYLASE<-?<-SF1-SF2-Helicase<-?<-?<-REase+ [illegible] | 1157 | Bacteria>FCB_group>Bacteroidetes/Chlorobi_group>Bacteroidota>Flavobacteriia>Flavobacteriales | GCA_002694765.1 | Xanthomarina_sp. | CMP04_07455 | 107327..110800 | 1931211 | hypothetical-protein-[Xanthomarina-sp.] |
| HBR11728.1 | Type1.BREX | <-GDResolvase+Recombinase+ZnR<-?\|\|REase+HTH+35exo [illegible] | 1210 | Bacteria>FCB_group>Bacteroidetes/Chlorobi_group>Bacteroidota>Flavobacteriia>Flavobacteriales | GCA_003511565.1 | Chryseobacterium_sp. | DD740_05880 | 144786..148418 | 1871047 | BREX-system-P-loop-protein-BrxC-[Chryseobacterium-sp.] |
| HCC95656.1 | Type1.BREX | Arm-DNA-bind_5+Phage_int_SAM_5+Phage_integrase-> [illegible] | 980 | Bacteria>FCB_group>Bacteroidetes/Chlorobi_group>Bacteroidota>Flavobacteriia>Flavobacteriales | GCA_003517615.1 | Flavobacteriaceae_bacterium | DEQ26_15305 | 12785..15728 | 1871037 | BREX-system-P-loop-protein-BrxC-[Flavobacteriaceae-bacterium] |
| ROI04325.1 | Type1.BREX | <-HTH_3<-?\|\|?->?->?<-?<-SIGMA+OB-Fold+AAA-ATPase [illegible] | 1174 | Bacteria>FCB_group>Bacteroidetes/Chlorobi_group>Bacteroidota>Flavobacteriia>Flavobacteriales | GCA_003729975.1 | Kaistella_haifensis | EGH90_10140 | 55406..58930 | 421525 | BREX-system-P-loop-protein-BrxC-[Kaistella-haifensis] |
| AZB21330.1 | Type1.BREX | <-SLOG<-?\|\|?-><-ChlI+Lon_C+AAA\|\|BrxA-DNA-binding [illegible] | 808 | Bacteria>FCB_group>Bacteroidetes/Chlorobi_group>Bacteroidota>Flavobacteriia>Flavobacteriales | GCA_003816015.1 | Kaistella_haifensis | EG338_03940 | 799177..801603 | 421525 | BREX-system-P-loop-protein-BrxC-[Kaistella-haifensis] |
| RZJ34503.1 | Type1.BREX | <-GDResolvase\|\|HTH_3->BrxA-DNA-binding->BrxB- [illegible] | 1165 | Bacteria>FCB_group>Bacteroidetes/Chlorobi_group>Bacteroidota>Flavobacteriia>Flavobacteriales | GCA_004210845.1 | Flavobacterium_sp. | EOO51_09215 | 4934..8431 | 239 | MAG:-BREX-system-P-loop-protein-BrxC-[Flavobacterium-sp.] |
| GEM55955.1 | Type1.BREX | <-SIGMA+OB-Fold+AAA-ATPase+Lon-Protease-BrxL<- [illegible] | 1216 | Bacteria>FCB_group>Bacteroidetes/Chlorobi_group>Bacteroidota>Flavobacteriia>Flavobacteriales | GCA_007990815.1 | Flavobacterium_branchiophilum_NBRC_15030_=_ATCC_35035 | FB1_21760 | 14247..17897 | 1218110 | hypothetical-protein-FB1_21760-[Flavobacterium-branchiophilum-NBRC-15030-=-ATCC-35035] |
| NJY61426.1 | Type1.BREX | Arm-DNA-bind_5+Phage_int_SAM_5+Phage_integrase-> [illegible] | 1202 | Bacteria>FCB_group>Bacteroidetes/Chlorobi_group>Bacteroidota>Flavobacteriia>Flavobacteriales | GCA_012038135.1 | Salinimicrobium_nanhaiense | HC174_01475 | 311288..314896 | 2722701 | BREX-system-P-loop-protein-BrxC-[Salinimicrobium-nanhaiense] |
| NOQ74050.1 | Type1.BREX | <-REC+LuxR-HTH<-?\|\|sGTP-><-35exo+SF1-SF2-Helicase [illegible] | 1203 | Bacteria>FCB_group>Bacteroidetes/Chlorobi_group>Bacteroidota>Flavobacteriia>Flavobacteriales | GCA_013138735.1 | Crocinitomix_sp. | GQ574_18735 | 89954..93565 | 2039232 | BREX-system-P-loop-protein-BrxC-[Crocinitomix-sp.] |
| MBQ0768324.1 | Type1.BREX | <-N6-METHYLASE<-Schlafen+HATPase+wHTH<-DUF6079*<- [illegible] | 1160 | Bacteria>FCB_group>Bacteroidetes/Chlorobi_group>Bacteroidota>Flavobacteriia>Flavobacteriales | GCA_018067115.1 | Bizionia_sp. | KBT58_03470 | 3917..7399 | 1954480 | BREX-system-P-loop-protein-BrxC-[Bizionia-sp.] |
| MBW7869163.1 | Type1.BREX | HTH_3->BrxA-DNA-binding->BrxB-BrxF-iSTAND-> [illegible] | 1166 | Bacteria>FCB_group>Bacteroidetes/Chlorobi_group>Bacteroidota>Flavobacteriia>Flavobacteriales | GCA_019455205.1 | Brumimicrobium_sp. | H3C31_12675 | 1666..5166 | 2029867 | BREX-system-P-loop-protein-BrxC-[Brumimicrobium-sp.] |
| MBZ0328278.1 | Type1.BREX | <-SIGMA+OB-Fold+AAA-ATPase+HKD-DNase-BrxL<-PglZ<- [illegible] | 1164 | Bacteria>FCB_group>Bacteroidetes/Chlorobi_group>Bacteroidota>Flavobacteriia>Flavobacteriales | GCA_019913785.1 | Altibacter_sp. | K8F54_11775 | 128036..131530 | 2024823 | BREX-system-P-loop-protein-BrxC-[Altibacter-sp.] |
| WCC43686.1 | Type1.BREX | <-DUF6079-p<-DUF6079-p*<-BrxB-BrxF-iSTAND<- [illegible] | 780 | Bacteria>FCB_group>Bacteroidetes/Chlorobi_group>Bacteroidota>Flavobacteriia>Flavobacteriales | GCA_028010105.1 | Tenacibaculum_finnmarkense | PJW08_00020 | 1487..3829 | 2781243 | BREX-system-P-loop-protein-BrxC-[Tenacibaculum-finnmarkense] |
| WP_014083579.1 | Type1.BREX | <-SIGMA+OB-Fold+AAA-ATPase+Lon-Protease-BrxL<- [illegible] | 1199 | Bacteria>FCB_group>Bacteroidetes/Chlorobi_group>Bacteroidota>Flavobacteriia>Flavobacteriales | GCF_000253275.1 | Flavobacterium_branchiophilum | FBFL15_RS05125 | 1197053..1200652 | 55197 | BREX-system-P-loop-protein-BrxC-[Flavobacterium-branchiophilum] |
| WP_051224566.1 | Type1.BREX | <-TPR+TPR<-?<-SIGMA+OB-Fold+AAA-ATPase+HKD-DNase- [illegible] | 1168 | Bacteria>FCB_group>Bacteroidetes/Chlorobi_group>Bacteroidota>Flavobacteriia>Flavobacteriales | GCF_000425485.1 | Flavobacterium_tegetincola | H602_RS18955 | 62387..65893 | 150172 | BREX-system-P-loop-protein-BrxC-[Flavobacterium-tegetincola] |
| WP_028123425.1 | Type1.BREX | <-XIS-HTH\|\|?-><-?<-?\|\|HTH_3->BrxA-DNA-binding-> [illegible] | 1188 | Bacteria>FCB_group>Bacteroidetes/Chlorobi_group>Bacteroidota>Flavobacteriia>Flavobacteriales | GCF_000428485.1 | Epilithonimonas_tenax | F564_RS0115480 | 6701..10267 | 191577 | BREX-system-P-loop-protein-BrxC-[Epilithonimonas-tenax] |
| WP_026451333.1 | Type1.BREX | VirB5->TrwI->TraK->?->TraN->?->HTH_3->BrxA-DNA- [illegible] | 1160 | Bacteria>FCB_group>Bacteroidetes/Chlorobi_group>Bacteroidota>Flavobacteriia>Flavobacteriales | GCF_000429125.1 | Aequorivita_capsosiphonis | G440_RS0110860 | 108295..111777 | 487317 | BREX-system-P-loop-protein-BrxC-[Aequorivita-capsosiphonis] |
| WP_034657463.1 | Type1.BREX | REase->?->HEPN->REase->?-><-?\|\|HTH_3->BrxA-DNA- [illegible] | 1160 | Bacteria>FCB_group>Bacteroidetes/Chlorobi_group>Bacteroidota>Flavobacteriia>Flavobacteriales | GCF_000468595.1 | Cellulophaga_baltica | M665_RS0118760 | 90424..93906 | 76594 | BREX-system-P-loop-protein-BrxC-[Cellulophaga-baltica] |
| WP_025613698.1 | Type1.BREX | Relaxase->VirD4-TraG+VirD4-FtsK->MORC+DUF3684+ [illegible] | 1199 | Bacteria>FCB_group>Bacteroidetes/Chlorobi_group>Bacteroidota>Flavobacteriia>Flavobacteriales | GCF_000477035.2 | Cellulophaga_baltica | M667_RS02035 | 472716..476315 | 76594 | BREX-system-P-loop-protein-BrxC-[Cellulophaga-baltica] |
| WP_034059321.1 | Type1.BREX | <-SAM-METHYLASE<-?<-PglZ<-METHYLASE<-?<-SF1-SF2- [illegible] | 1160 | Bacteria>FCB_group>Bacteroidetes/Chlorobi_group>Bacteroidota>Flavobacteriia>Flavobacteriales | GCF_000797445.1 | Lacinutrix_jangbogonensis | RH82_RS07100 | 67712..71194 | 1469557 | BREX-system-P-loop-protein-BrxC-[Lacinutrix-jangbogonensis] |
| WP_059014619.1 | Type1.BREX | <-HEPN<-ParB+HNH<-N6-METHYLASE<-DUF6079*<- [illegible] | 1169 | Bacteria>FCB_group>Bacteroidetes/Chlorobi_group>Bacteroidota>Flavobacteriia>Flavobacteriales | GCF_001485515.1 | Flavobacterium_psychrophilum | FPK15_RS07995 | 6803..10312 | 96345 | BREX-system-P-loop-protein-BrxC-[Flavobacterium-psychrophilum] |
| WP_076662677.1 | Type1.BREX | DOC+HTH-><-?<-?<-SIGMA+OB-Fold+AAA-ATPase+Lon- [illegible] | 1201 | Bacteria>FCB_group>Bacteroidetes/Chlorobi_group>Bacteroidota>Flavobacteriia>Flavobacteriales | GCF_001975575.1 | Mangrovimonas_sp._DI_80 | BKM32_RS04295 | 828004..831609 | 1779330 | BREX-system-P-loop-protein-BrxC-[Mangrovimonas-sp.-DI-80] |
| WP_088323652.1 | Type1.BREX | <-XIS-HTH<-?<-ParB+N6_N4_Mtase<-?<-GDResolvase<- [illegible] | 1158 | Bacteria>FCB_group>Bacteroidetes/Chlorobi_group>Bacteroidota>Flavobacteriia>Flavobacteriales | GCF_002201235.1 | Polaribacter_tangerinus | CCS00_RS05195 | 350003..353479 | 1920034 | BREX-system-P-loop-protein-BrxC-[Polaribacter-tangerinus] |
| WP_094415998.1 | Type1.BREX | <-PIN<-?<-SIGMA+OB-Fold+AAA-ATPase+Lon-Protease- [illegible] | 1197 | Bacteria>FCB_group>Bacteroidetes/Chlorobi_group>Bacteroidota>Flavobacteriia>Flavobacteriales | GCF_002251835.1 | Flavobacterium_cyanobacteriorum | CHU92_RS12285 | 18327..21920 | 2022802 | BREX-system-P-loop-protein-BrxC-[Flavobacterium-cyanobacteriorum] |
| WP_107011557.1 | Type1.BREX | <-PglZ<-?<-HAD+PRTase+PRTase+REase+SF1-SF2- [illegible] | 1166 | Bacteria>FCB_group>Bacteroidetes/Chlorobi_group>Bacteroidota>Flavobacteriia>Flavobacteriales | GCF_003024155.1 | Gramella_fulva | C7S20_RS05585 | 1214513..1218013 | 2126553 | BREX-system-P-loop-protein-BrxC-[Christiangramia-fulva] |
| WP_108879107.1 | Type1.BREX | MoxR-AAA->?->wHTH+WYL+WCX->ZnR+CC+REase->BrxA- [illegible] | 1166 | Bacteria>FCB_group>Bacteroidetes/Chlorobi_group>Bacteroidota>Flavobacteriia>Flavobacteriales | GCF_003095655.1 | Dokdonia_sp._Dokd-P16 | DCS32_RS15465 | 3477371..3480871 | 2173169 | BREX-system-P-loop-protein-BrxC-[Dokdonia-sp.-Dokd-P16] |
| WP_129016554.1 | Type1.BREX | <-SIGMA+OB-Fold+AAA-ATPase+Lon-Protease-BrxL<- [illegible] | 1202 | Bacteria>FCB_group>Bacteroidetes/Chlorobi_group>Bacteroidota>Flavobacteriia>Flavobacteriales | GCF_004115975.1 | Gelidibacter_gilvus | ESZ48_RS06605 | 20961..24569 | 59602 | BREX-system-P-loop-protein-BrxC-[Gelidibacter-gilvus] |
| WP_129366655.1 | Type1.BREX | <-HNH<-SbcC<-SIGMA+OB-Fold+AAA-ATPase+HKD-DNase- [illegible] | 1168 | Bacteria>FCB_group>Bacteroidetes/Chlorobi_group>Bacteroidota>Flavobacteriia>Flavobacteriales | GCF_004121075.1 | Lutibacter_sp._HS1-25 | EC396_RS10485 | 12877..16383 | 2485000 | BREX-system-P-loop-protein-BrxC-[Lutibacter-sp.-HS1-25] |
| WP_132215847.1 | Type1.BREX | <-CARF+REase<-CARF<-HKD-DNase+HKD-DNase<-?<- [illegible] | 1169 | Bacteria>FCB_group>Bacteroidetes/Chlorobi_group>Bacteroidota>Flavobacteriia>Flavobacteriales | GCF_004341235.1 | Mariniflexile_fucanivorans | EV196_RS05490 | 443037..446546 | 264023 | BREX-system-P-loop-protein-BrxC-[Mariniflexile-fucanivorans] |
| WP_131908313.1 | Type1.BREX | <-XIS-HTH<-?\|\|?-><-SIGMA+OB-Fold+AAA-ATPase+Lon- [illegible] | 1203 | Bacteria>FCB_group>Bacteroidetes/Chlorobi_group>Bacteroidota>Flavobacteriia>Flavobacteriales | GCF_004349085.1 | Flavobacterium_caseinilyticum | E0F89_RS02690 | 598586..602197 | 2541732 | BREX-system-P-loop-protein-BrxC-[Flavobacterium-caseinilyticum] |
| WP_133546801.1 | Type1.BREX | <-XIS-HTH<-?<-?<-AraC-HTH<-Arm-DNA-bind_5+ [illegible] | 1167 | Bacteria>FCB_group>Bacteroidetes/Chlorobi_group>Bacteroidota>Flavobacteriia>Flavobacteriales | GCF_004362395.1 | Salegentibacter_sp._24 | DET49_RS03950 | 90042..93545 | 2183986 | BREX-system-P-loop-protein-BrxC-[Salegentibacter-sp.-24] |
| WP_136564624.1 | Type1.BREX | XIS-HTH->?->AbiEi_4+AbiEi_1->AbiEii+ZnR->wHTH+ [illegible] | 1164 | Bacteria>FCB_group>Bacteroidetes/Chlorobi_group>Bacteroidota>Flavobacteriia>Flavobacteriales | GCF_004916895.1 | Muricauda_alvinocaridis | EZV76_RS00375 | 77026..80520 | 2530200 | BREX-system-P-loop-protein-BrxC-[Muricauda-alvinocaridis] |
| WP_141421442.1 | Type1.BREX | <-HAD+PRTase+PRTase+REase+SF1-SF2-Helicase<-?<- [illegible] | 1165 | Bacteria>FCB_group>Bacteroidetes/Chlorobi_group>Bacteroidota>Flavobacteriia>Flavobacteriales | GCF_006546625.1 | Haloflavibacter_putidus | FKR84_RS06225 | 21354..24851 | 2576776 | BREX-system-P-loop-protein-BrxC-[Haloflavibacter-putidus] |
| WP_019974718.1 | Type1.BREX | <-SIGMA+OB-Fold+AAA-ATPase+HKD-DNase-BrxL<-PglZ<- [illegible] | 1191 | Bacteria>FCB_group>Bacteroidetes/Chlorobi_group>Bacteroidota>Flavobacteriia>Flavobacteriales | GCF_007990795.1 | Empedobacter_brevis | EB1_RS03060 | 16539..20114 | 247 | BREX-system-P-loop-protein-BrxC-[Empedobacter-brevis] |
| WP_089081509.1 | Type1.BREX | <-SIGMA+OB-Fold+AAA-ATPase+Lon-Protease-BrxL<- [illegible] | 1199 | Bacteria>FCB_group>Bacteroidetes/Chlorobi_group>Bacteroidota>Flavobacteriia>Flavobacteriales | GCF_007990815.1 | Flavobacterium_branchiophilum | FB1_RS11100 | 14247..17846 | 55197 | BREX-system-P-loop-protein-BrxC-[Flavobacterium-branchiophilum] |
| WP_153266434.1 | Type1.BREX | HTH_36->Classical-AAA->TPRR-HTH->STYKinase-like- [illegible] | 1157 | Bacteria>FCB_group>Bacteroidetes/Chlorobi_group>Bacteroidota>Flavobacteriia>Flavobacteriales | GCF_009498295.1 | Oceanihabitans_sp._IOP_32 | FEZ18_RS00165 | 31374..34847 | 2529032 | BREX-system-P-loop-protein-BrxC-[Oceanihabitans-sp.-IOP_32] |
| WP_158973850.1 | Type1.BREX | <-ANK+ANK\|\|wHTH+WYL+WCX->BrxA-DNA-binding->BrxB- [illegible] | 1170 | Bacteria>FCB_group>Bacteroidetes/Chlorobi_group>Bacteroidota>Flavobacteriia>Flavobacteriales | GCF_009797025.1 | Cellulophaga_sp._L1A9 | GQR94_RS02145 | 525105..528617 | 2686362 | BREX-system-P-loop-protein-BrxC-[Cellulophaga-sp.-L1A9] |
| WP_165150638.1 | Type1.BREX | <-ABhydrolase<-REase+RNase<-?<-wHTH+WYL+WCX\|\|?-> [illegible] | 1198 | Bacteria>FCB_group>Bacteroidetes/Chlorobi_group>Bacteroidota>Flavobacteriia>Flavobacteriales | GCF_011065475.1 | Aequorivita_sp._KMM_9714 | G3A46_RS13570 | 21223..24819 | 2707173 | BREX-system-P-loop-protein-BrxC-[Aequorivita-sp.-KMM-9714] |
| WP_177760874.1 | Type1.BREX | HTH_3-><-McrC-NTD+REase\|\|HTH+rve->AAA-ATPase-><- [illegible] | 1217 | Bacteria>FCB_group>Bacteroidetes/Chlorobi_group>Bacteroidota>Flavobacteriia>Flavobacteriales | GCF_013389595.1 | Flavobacterium_sp._I3-2 | HW119_RS00890 | 194599..198252 | 2748319 | BREX-system-P-loop-protein-BrxC-[Flavobacterium-sp.-I3-2] |
| WP_163432971.1 | Type1.BREX | <-LexA-protease<-?\|\|?->XIS-HTH->PIN->GDResolvase- [illegible] | 1189 | Bacteria>FCB_group>Bacteroidetes/Chlorobi_group>Bacteroidota>Flavobacteriia>Flavobacteriales | GCF_014050215.1 | unclassified_Flavobacterium | G7B00_RS14255 | 32052..35621 | 196869 | MULTISPECIES:-BREX-system-P-loop-protein-BrxC-[unclassified-Flavobacterium] |
| WP_073117772.1 | Type1.BREX | SLATT_1-><-METHYLASE-p<-?<-SMODS<-?<-?<-SIGMA+OB- [illegible] | 1170 | Bacteria>FCB_group>Bacteroidetes/Chlorobi_group>Bacteroidota>Flavobacteriia>Flavobacteriales | GCF_014636515.1 | Wenyingzhuangia_marina | IEV03_RS11465 | 186450..189962 | 1195760 | BREX-system-P-loop-protein-BrxC-[Wenyingzhuangia-marina] |
| WP_093139951.1 | Type1.BREX | <-SF1-SF2-Helicase<-SIGMA+OB-Fold+AAA-ATPase+Lon- [illegible] | 1202 | Bacteria>FCB_group>Bacteroidetes/Chlorobi_group>Bacteroidota>Flavobacteriia>Flavobacteriales | GCF_014651275.1 | Ulvibacter_litoralis | IE050_RS03690 | 820645..824253 | 227084 | BREX-system-P-loop-protein-BrxC-[Ulvibacter-litoralis] |
| WP_194850146.1 | Type1.BREX | <-SF1-SF2-Helicase<-OLD-ABC+TOPRIM<-SIGMA+OB-Fold [illegible] | 1160 | Bacteria>FCB_group>Bacteroidetes/Chlorobi_group>Bacteroidota>Flavobacteriia>Flavobacteriales | GCF_015356755.1 | Nonlabens_antarcticus | IQ282_RS01355 | 76947..80429 | 392714 | BREX-system-P-loop-protein-BrxC-[Nonlabens-antarcticus] |
| WP_198639224.1 | Type1.BREX | REase->SF1-SF2-Helicase+REase->NERD+SF1-SF2- [illegible] | 1201 | Bacteria>FCB_group>Bacteroidetes/Chlorobi_group>Bacteroidota>Flavobacteriia>Flavobacteriales | GCF_016236975.1 | Salegentibacter_maritimus | I6U50_RS13470 | 35086..38691 | 2794347 | BREX-system-P-loop-protein-BrxC-[Salegentibacter-maritimus] |
| WP_199600307.1 | Type1.BREX | Relaxase->VirD4-TraG+VirD4-FtsK->Classical-AAA-> [illegible] | 1160 | Bacteria>FCB_group>Bacteroidetes/Chlorobi_group>Bacteroidota>Flavobacteriia>Flavobacteriales | GCF_016469035.1 | Gelidibacter_salicanalis | JEM65_RS13235 | 170248..173730 | 291193 | BREX-system-P-loop-protein-BrxC-[Gelidibacter-salicanalis] |
| WP_203057453.1 | Type1.BREX | <-DUF4011+SF1-SF2-Helicase+REase<-SIGMA+OB-Fold+ [illegible] | 1178 | Bacteria>FCB_group>Bacteroidetes/Chlorobi_group>Bacteroidota>Flavobacteriia>Flavobacteriales | GCF_016786255.1 | Robertkochia_sediminum | IS446_RS01110 | 192082..195618 | 2785326 | BREX-system-P-loop-protein-BrxC-[Robertkochia-sediminum] |
| WP_203251454.1 | Type1.BREX | XIS-HTH-><-SIGMA+OB-Fold+AAA-ATPase+HKD-DNase- [illegible] | 1160 | Bacteria>FCB_group>Bacteroidetes/Chlorobi_group>Bacteroidota>Flavobacteriia>Flavobacteriales | GCF_016811105.1 | Aurantibacter_crassamenti | JQC67_RS03370 | 810968..814450 | 1837375 | BREX-system-P-loop-protein-BrxC-[Aurantibacter-crassamenti] |
| WP_179314673.1 | Type1.BREX | <-ParB+HNH<-PglZ<-METHYLASE<-?<-SF1-SF2- [illegible] | 1160 | Bacteria>FCB_group>Bacteroidetes/Chlorobi_group>Bacteroidota>Flavobacteriia>Flavobacteriales | GCF_019278425.1 | Winogradskyella | HOI32_RS04080 | 939044..942526 | 286104 | MULTISPECIES:-BREX-system-P-loop-protein-BrxC-[Winogradskyella] |
| WP_224137560.1 | Type1.BREX | <-REC<-N6-METHYLASE<-ASCH<-HIRAN<-SIGMA+OB-Fold+ [illegible] | 1174 | Bacteria>FCB_group>Bacteroidetes/Chlorobi_group>Bacteroidota>Flavobacteriia>Flavobacteriales | GCF_020084865.1 | Candidatus_Kaistella_beijingensis | J4771_RS06115 | 1314125..1317649 | 2820270 | BREX-system-P-loop-protein-BrxC-[Candidatus-Kaistella-beijingensis] |
| WP_224474755.1 | Type1.BREX | <-SIGMA+OB-Fold+AAA-ATPase+HKD-DNase-BrxL<-PglZ<- [illegible] | 1167 | Bacteria>FCB_group>Bacteroidetes/Chlorobi_group>Bacteroidota>Flavobacteriia>Flavobacteriales | GCF_020164485.1 | Salegentibacter_tibetensis | LB467_RS14420 | 15605..19108 | 2873600 | BREX-system-P-loop-protein-BrxC-[Salegentibacter-tibetensis] |

| Protein-ID | System | Gene-Neighbourhoods | Length | Taxonomy | Assembly-ID | Strain | Locus-Tag | Genomic-coordinates | TaxID | Fasta-define |
|---|---|---|---|---|---|---|---|---|---|---|
| WP_226063322.1 | Type1.BREX | [illegible] | 1175 | [illegible] | [illegible] | [illegible] | [illegible] | 1609850..1613377 | 2878534 | [illegible] |
| WP_232121385.1 | Type1.BREX | [illegible] | 1176 | [illegible] | [illegible] | [illegible] | [illegible] | 20484..24014 | 669041 | [illegible] |
| WP_232125551.1 | Type1.BREX | [illegible] | 1176 | [illegible] | [illegible] | [illegible] | [illegible] | 18832..22362 | 2781243 | [illegible] |
| WP_233899778.1 | Type1.BREX | [illegible] | 1176 | [illegible] | [illegible] | [illegible] | [illegible] | 464400..467930 | 1458515 | [illegible] |
| WP_232123632.1 | Type1.BREX | [illegible] | 1176 | [illegible] | [illegible] | [illegible] | [illegible] | 3814..7344 | 2781243 | [illegible] |
| WP_193709240.1 | Type1.BREX | [illegible] | 1176 | [illegible] | [illegible] | [illegible] | [illegible] | 69854..73384 | 2781243 | [illegible] |
| WP_244340081.1 | Type1.BREX | [illegible] | 1169 | [illegible] | [illegible] | [illegible] | [illegible] | 3214137..3217646 | 237 | [illegible] |
| WP_249986967.1 | Type1.BREX | [illegible] | 1176 | [illegible] | [illegible] | [illegible] | [illegible] | 308529..312059 | 2927126 | [illegible] |
| WP_264521759.1 | Type1.BREX | [illegible] | 1166 | [illegible] | [illegible] | [illegible] | [illegible] | 842577..846077 | 2986827 | [illegible] |
| WP_267609146.1 | Type1.BREX | [illegible] | 1173 | [illegible] | [illegible] | [illegible] | [illegible] | 10693..14214 | 2997334 | [illegible] |
| WP_225581981.1 | Type1.BREX | [illegible] | 1189 | [illegible] | [illegible] | [illegible] | [illegible] | 4194..7763 | 76832 | [illegible] |
| WP_089756260.1 | Type1.BREX | [illegible] | 1176 | [illegible] | [illegible] | [illegible] | [illegible] | 831929..835459 | 311333 | [illegible] |
| WP_089663492.1 | Type1.BREX | [illegible] | 1167 | [illegible] | [illegible] | [illegible] | [illegible] | 3168257..3171760 | 279359 | [illegible] |
| WP_103999283.1 | Type1.BREX | [illegible] | 1170 | [illegible] | [illegible] | [illegible] | [illegible] | 195084..198596 | 935224 | [illegible] |
| WP_074938106.1 | Type1.BREX | [illegible] | 1160 | [illegible] | [illegible] | [illegible] | [illegible] | 97299..100781 | 221126 | [illegible] |
| WP_091208763.1 | Type1.BREX | [illegible] | 1172 | [illegible] | [illegible] | [illegible] | [illegible] | 46469..49987 | 935223 | [illegible] |
| WP_090838887.1 | Type1.BREX | [illegible] | 1160 | [illegible] | [illegible] | [illegible] | [illegible] | 330310..333792 | 1144750 | [illegible] |
| WP_089819303.1 | Type1.BREX | [illegible] | 1174 | [illegible] | [illegible] | [illegible] | [illegible] | 1141726..1145250 | 631455 | [illegible] |
| WP_159447525.1 | Type1.BREX | [illegible] | 719 | [illegible] | [illegible] | [illegible] | [illegible] | 472839..474998 | 561365 | [illegible] |
| WP_094162522.1 | Type1.BREX | [illegible] | 1198 | [illegible] | [illegible] | [illegible] | [illegible] | 11651..15247 | 96345 | [illegible] |
| WP_095070899.1 | Type1.BREX | [illegible] | 1178 | [illegible] | [illegible] | [illegible] | [illegible] | 977692..981228 | 536441 | [illegible] |
| WP_089379946.1 | Type1.BREX | [illegible] | 1168 | [illegible] | [illegible] | [illegible] | [illegible] | 359460..362966 | 1109740 | [illegible] |
| WP_089376855.1 | Type1.BREX | [illegible] | 1165 | [illegible] | [illegible] | [illegible] | [illegible] | 583517..587014 | 691689 | [illegible] |
| WP_213195478.1 | Type1.BREX | [illegible] | 1176 | [illegible] | [illegible] | [illegible] | [illegible] | 263131..266661 | 2004710 | [illegible] |
| WP_271393637.1 | Type1.BREX | [illegible] | 1168 | [illegible] | [illegible] | [illegible] | [illegible] | 126765..130271 | 1382458 | [illegible] |
| WP_271393758.1 | Type1.BREX | [illegible] | 1165 | [illegible] | [illegible] | [illegible] | [illegible] | 23754..27251 | 1382458 | [illegible] |
| WP_272023608.1 | Type1.BREX | [illegible] | 1160 | [illegible] | [illegible] | [illegible] | [illegible] | 26845..30327 | 1144750 | [illegible] |
| MBK6861507.1 | Type1.BREX | [illegible] | 1144 | [illegible] | [illegible] | [illegible] | [illegible] | 1781325..1784759 | 2202734 | [illegible] |
| MBK8736252.1 | Type1.BREX | [illegible] | 919 | [illegible] | [illegible] | [illegible] | [illegible] | 1583..4342 | 2202734 | [illegible] |
| MBK8109385.1 | Type1.BREX | [illegible] | 128 | [illegible] | [illegible] | [illegible] | [illegible] | 78067..78453 | 2202734 | [illegible] |
| MBK8623789.1 | Type1.BREX | [illegible] | 1197 | [illegible] | [illegible] | [illegible] | [illegible] | 313073..316666 | 2202734 | [illegible] |
| MBK8854955.1 | Type1.BREX | [illegible] | 155 | [illegible] | [illegible] | [illegible] | [illegible] | 55975..56442 | 2202734 | [illegible] |
| MBK8854956.1 | Type1.BREX | [illegible] | 772 | [illegible] | [illegible] | [illegible] | [illegible] | 56456..58774 | 2202734 | [illegible] |
| MBK9103294.1 | Type1.BREX | [illegible] | 591 | [illegible] | [illegible] | [illegible] | [illegible] | 365771..367546 | 2202734 | [illegible] |
| MBK9689678.1 | Type1.BREX | [illegible] | 236 | [illegible] | [illegible] | [illegible] | [illegible] | 1629..2339 | 2202734 | [illegible] |
| MBL0081467.1 | Type1.BREX | [illegible] | 143 | [illegible] | [illegible] | [illegible] | [illegible] | 15607..16038 | 2202734 | [illegible] |
| MBK9994339.1 | Type1.BREX | [illegible] | 1197 | [illegible] | [illegible] | [illegible] | [illegible] | 1480..5073 | 2202734 | [illegible] |
| MBP6234856.1 | Type1.BREX | [illegible] | 1152 | [illegible] | [illegible] | [illegible] | [illegible] | 136824..140282 | 2202734 | [illegible] |
| MCB0517804.1 | Type1.BREX | [illegible] | 1174 | [illegible] | [illegible] | [illegible] | [illegible] | 9612..13136 | 2202734 | [illegible] |
| MCB0520339.1 | Type1.BREX | [illegible] | 1145 | [illegible] | [illegible] | [illegible] | [illegible] | 21267..24704 | 2202734 | [illegible] |
| MCB0524505.1 | Type1.BREX | [illegible] | 1094 | [illegible] | [illegible] | [illegible] | [illegible] | 14975..18259 | 2202734 | [illegible] |
| MCB0625399.1 | Type1.BREX | [illegible] | 1151 | [illegible] | [illegible] | [illegible] | [illegible] | 4651..8106 | 2202734 | [illegible] |
| MCB9352089.1 | Type1.BREX | [illegible] | 1176 | [illegible] | [illegible] | [illegible] | [illegible] | 43547..47077 | 2760807 | [illegible] |
| MCB9352382.1 | Type1.BREX | [illegible] | 1214 | [illegible] | [illegible] | [illegible] | [illegible] | 34149..37793 | 2760807 | [illegible] |
| MCB9053313.1 | Type1.BREX | [illegible] | 1180 | [illegible] | [illegible] | [illegible] | [illegible] | 325530..329072 | 2760807 | [illegible] |
| WP_013769021.1 | Type1.BREX | [illegible] | 1144 | [illegible] | [illegible] | [illegible] | [illegible] | 27488..30922 | 2350 | [illegible] |
| WP_020534907.1 | Type1.BREX | [illegible] | 1159 | [illegible] | [illegible] | [illegible] | [illegible] | 116498..119977 | 70995 | [illegible] |
| WP_044220103.1 | Type1.BREX | [illegible] | 1186 | [illegible] | [illegible] | [illegible] | [illegible] | 11358..14918 | 1524460 | [illegible] |
| WP_147165921.1 | Type1.BREX | [illegible] | 1186 | [illegible] | [illegible] | [illegible] | [illegible] | 87702..91262 | 1564516 | [illegible] |
| WP_147932457.1 | Type1.BREX | [illegible] | 1198 | [illegible] | [illegible] | [illegible] | [illegible] | 18155..21751 | 2602767 | [illegible] |
| WP_168040578.1 | Type1.BREX | [illegible] | 1211 | [illegible] | [illegible] | [illegible] | [illegible] | 12164..15799 | 442734 | [illegible] |
| WP_236977640.1 | Type1.BREX | [illegible] | 1159 | [illegible] | [illegible] | [illegible] | [illegible] | 21527..25006 | 2904244 | [illegible] |
| OWY22675.1 | Type1.BREX | [illegible] | 1168 | [illegible] | [illegible] | [illegible] | [illegible] | 6474..9980 | 1932005 | [illegible] |
| OYD42847.1 | Type1.BREX | [illegible] | 1175 | [illegible] | [illegible] | [illegible] | [illegible] | 557163..560690 | 1768011 | [illegible] |
| QQS30389.1 | Type1.BREX | [illegible] | 1202 | [illegible] | [illegible] | [illegible] | [illegible] | 1549850..1553458 | 2044944 | [illegible] |
| WP_157278947.1 | Type1.BREX | [illegible] | 1169 | [illegible] | [illegible] | [illegible] | [illegible] | 7227..10736 | 376470 | [illegible] |
| WP_094282244.1 | Type1.BREX | [illegible] | 1175 | [illegible] | [illegible] | [illegible] | [illegible] | 18381..21908 | 1768011 | [illegible] |
| WP_070561019.1 | Type1.BREX | [illegible] | 1187 | [illegible] | [illegible] | [illegible] | [illegible] | 95716..99279 | 1581095 | [illegible] |
| WP_165305702.1 | Type1.BREX | [illegible] | 1187 | [illegible] | [illegible] | [illegible] | [illegible] | 1244063..1247626 | 2713573 | [illegible] |
| WP_200066585.1 | Type1.BREX | [illegible] | 1199 | [illegible] | [illegible] | [illegible] | [illegible] | 547285..550884 | 2793071 | [illegible] |
| WP_090698254.1 | Type1.BREX | [illegible] | 1169 | [illegible] | [illegible] | [illegible] | [illegible] | 316504..320013 | 990371 | [illegible] |
| MCE1168906.1 | Type1.BREX | [illegible] | 1202 | [illegible] | [illegible] | [illegible] | [illegible] | 2695..6303 | 2035813 | [illegible] |
| KPP94735.1 | Type1.BREX | [illegible] | 1203 | [illegible] | [illegible] | [illegible] | [illegible] | 188157..191768 | 1666909 | [illegible] |
| OPZ99879.1 | Type1.BREX | [illegible] | 1147 | [illegible] | [illegible] | [illegible] | [illegible] | 505..3948 | 1852814 | [illegible] |
| PKP36801.1 | Type1.BREX | [illegible] | 1199 | [illegible] | [illegible] | [illegible] | [illegible] | 1837..5436 | 2013682 | [illegible] |
| PSR13219.1 | Type1.BREX | [illegible] | 1045 | [illegible] | [illegible] | [illegible] | [illegible] | 1..3135 | 1898104 | [illegible] |
| PTM08477.1 | Type1.BREX | [illegible] | 1159 | [illegible] | [illegible] | [illegible] | [illegible] | 6731..10210 | 1898104 | [illegible] |
| KAF0128492.1 | Type1.BREX | [illegible] | 1144 | [illegible] | [illegible] | [illegible] | [illegible] | 38823..42257 | 1898104 | [illegible] |
| NQT60364.1 | Type1.BREX | [illegible] | 1348 | [illegible] | [illegible] | [illegible] | [illegible] | 13554..17600 | 1898104 | [illegible] |
| NUQ82570.1 | Type1.BREX | [illegible] | 1197 | [illegible] | [illegible] | [illegible] | [illegible] | 47380..50973 | 1898104 | [illegible] |
| MBK7107567.1 | Type1.BREX | [illegible] | 1197 | [illegible] | [illegible] | [illegible] | [illegible] | 12776..16369 | 1898104 | [illegible] |
| MBK7967322.1 | Type1.BREX | [illegible] | 197 | [illegible] | [illegible] | [illegible] | [illegible] | 46949..47542 | 1898104 | [illegible] |
| MBL0279470.1 | Type1.BREX | [illegible] | 1197 | [illegible] | [illegible] | [illegible] | [illegible] | 937205..940798 | 1898104 | [illegible] |
| MBM3454071.1 | Type1.BREX | [illegible] | 1144 | [illegible] | [illegible] | [illegible] | [illegible] | 4277..7711 | 1898104 | [illegible] |
| MBN8706499.1 | Type1.BREX | [illegible] | 1173 | [illegible] | [illegible] | [illegible] | [illegible] | 8364..11885 | 1898104 | [illegible] |
| MBS1911241.1 | Type1.BREX | [illegible] | 1145 | [illegible] | [illegible] | [illegible] | [illegible] | 55920..59357 | 1898104 | [illegible] |
| MBT4727235.1 | Type1.BREX | [illegible] | 481 | [illegible] | [illegible] | [illegible] | [illegible] | 7916..9358 | 1898104 | [illegible] |
| MBT7826368.1 | Type1.BREX | [illegible] | 1165 | [illegible] | [illegible] | [illegible] | [illegible] | 2065..5562 | 1898104 | [illegible] |
| MBU2649683.1 | Type1.BREX | [illegible] | 1189 | [illegible] | [illegible] | [illegible] | [illegible] | 1..3566 | 1898104 | [illegible] |
| MCB0536651.1 | Type1.BREX | [illegible] | 1197 | [illegible] | [illegible] | [illegible] | [illegible] | 1012..4605 | 1898104 | [illegible] |
| MCH7965547.1 | Type1.BREX | [illegible] | 1161 | [illegible] | [illegible] | [illegible] | [illegible] | 28479..31964 | 1898104 | [illegible] |
| MCR9155224.1 | Type1.BREX | [illegible] | 1141 | [illegible] | [illegible] | [illegible] | [illegible] | 83336..86761 | 1898104 | [illegible] |
| MAO64836.1 | Type1.BREX | [illegible] | 1168 | [illegible] | [illegible] | [illegible] | [illegible] | 664531..668037 | 2024824 | [illegible] |
| TVQ65167.1 | Type1.BREX | [illegible] | 1164 | [illegible] | [illegible] | [illegible] | [illegible] | 1062..4556 | 2053516 | [illegible] |
| MCC5907043.1 | Type1.BREX | [illegible] | 1167 | [illegible] | [illegible] | [illegible] | [illegible] | 29362..32865 | 2053516 | [illegible] |
| MCH2449130.1 | Type1.BREX | [illegible] | 1151 | [illegible] | [illegible] | [illegible] | [illegible] | 295077..298532 | 1974203 | [illegible] |
| WP_095606906.1 | Type1.BREX | [illegible] | 1213 | [illegible] | [illegible] | [illegible] | [illegible] | 230173..233814 | 2032627 | [illegible] |
| WP_210512160.1 | Type1.BREX | [illegible] | 1141 | [illegible] | [illegible] | [illegible] | [illegible] | 86175..89600 | 2812953 | [illegible] |
| WP_237852616.1 | Type1.BREX | [illegible] | 1164 | [illegible] | [illegible] | [illegible] | [illegible] | 217944..221438 | 2911366 | [illegible] |
| WP_265763895.1 | Type1.BREX | [illegible] | 1165 | [illegible] | [illegible] | [illegible] | [illegible] | 34405..37902 | 2820877 | [illegible] |
| ANT65175.1 | Type1.BREX | [illegible] | 1144 | [illegible] | [illegible] | [illegible] | [illegible] | 1430885..1434319 | 1868325 | [illegible] |
| TNJ34090.1 | Type1.BREX | [illegible] | 1144 | [illegible] | [illegible] | [illegible] | [illegible] | 14740..18174 | 1098 | [illegible] |
| KAA6232806.1 | Type1.BREX | [illegible] | 1204 | [illegible] | [illegible] | [illegible] | [illegible] | 1409593..1413207 | 1094 | [illegible] |
| NMW20007.1 | Type1.BREX | [illegible] | 1160 | [illegible] | [illegible] | [illegible] | [illegible] | 1..3483 | 2067991 | [illegible] |
| MBV5326244.1 | Type1.BREX | [illegible] | 1179 | [illegible] | [illegible] | [illegible] | [illegible] | 2432..5971 | 1095 | [illegible] |
| MBV5326252.1 | Type1.BREX | [illegible] | 1179 | [illegible] | [illegible] | [illegible] | [illegible] | 9464..13003 | 1095 | [illegible] |
| MBV5328349.1 | Type1.BREX | [illegible] | 1208 | [illegible] | [illegible] | [illegible] | [illegible] | 10298..13924 | 1095 | [illegible] |
| MBV5329669.1 | Type1.BREX | [illegible] | 1150 | [illegible] | [illegible] | [illegible] | [illegible] | 2439..5891 | 1095 | [illegible] |
| MBV5304386.1 | Type1.BREX | [illegible] | 1138 | [illegible] | [illegible] | [illegible] | [illegible] | 1849..5265 | 1095 | [illegible] |
| MCF8215288.1 | Type1.BREX | [illegible] | 1145 | [illegible] | [illegible] | [illegible] | [illegible] | 58948..62385 | 1095 | [illegible] |
| WP_012508724.1 | Type1.BREX | [illegible] | 1211 | [illegible] | [illegible] | [illegible] | [illegible] | 2123667..2127302 | 34090 | [illegible] |
| WP_204208384.1 | Type1.BREX | [illegible] | 997 | [illegible] | [illegible] | [illegible] | [illegible] | 1057250..1060243 | 1868325 | [illegible] |
| WP_204208337.1 | Type1.BREX | [illegible] | 1145 | [illegible] | [illegible] | [illegible] | [illegible] | 1430885..1434322 | 1868325 | [illegible] |
| WP_076791024.1 | Type1.BREX | [illegible] | 1179 | [illegible] | [illegible] | [illegible] | [illegible] | 12313..15852 | 1917528 | [illegible] |
| WP_085659853.1 | Type1.BREX | [illegible] | 1209 | [illegible] | [illegible] | [illegible] | [illegible] | 1603101..1606730 | 1974213 | [illegible] |
| WP_139626761.1 | Type1.BREX | [illegible] | 1204 | [illegible] | [illegible] | [illegible] | [illegible] | 9816..13430 | 1098 | [illegible] |
| WP_211340437.1 | Type1.BREX | [illegible] | 1145 | [illegible] | [illegible] | [illegible] | [illegible] | 14740..18177 | 1098 | [illegible] |
| WP_192496514.1 | Type1.BREX | [illegible] | 1203 | [illegible] | [illegible] | [illegible] | [illegible] | 1409593..1413204 | 1094 | [illegible] |
| WP_151419733.1 | Type1.BREX | [illegible] | 1183 | [illegible] | [illegible] | [illegible] | [illegible] | 13780..17331 | 1094 | [illegible] |
| MBM3423463.1 | Type1.BREX | [illegible] | 1182 | [illegible] | [illegible] | [illegible] | [illegible] | 5435..8983 | 2268192 | [illegible] |
| QOJ30044.1 | Type1.BREX | [illegible] | 1198 | [illegible] | [illegible] | [illegible] | [illegible] | 3719207..3722803 | 2049428 | [illegible] |
| MBV6512081.1 | Type1.BREX | [illegible] | 1198 | [illegible] | [illegible] | [illegible] | [illegible] | 145578..149174 | 2651163 | [illegible] |
| MBV6422081.1 | Type1.BREX | [illegible] | 1214 | [illegible] | [illegible] | [illegible] | [illegible] | 33171..36815 | 2651163 | [illegible] |
| MCB9211065.1 | Type1.BREX | [illegible] | 1377 | [illegible] | [illegible] | [illegible] | [illegible] | 10432..14565 | 2049428 | [illegible] |
| MCK6605711.1 | Type1.BREX | [illegible] | 1179 | [illegible] | [illegible] | [illegible] | [illegible] | 25884..29423 | 2651163 | [illegible] |
| MBL0321410.1 | Type1.BREX | [illegible] | 737 | [illegible] | [illegible] | [illegible] | [illegible] | 309754..311967 | 2053306 | [illegible] |
| MBK7413718.1 | Type1.BREX | [illegible] | 1183 | [illegible] | [illegible] | [illegible] | [illegible] | 1380288..1383839 | 2053306 | [illegible] |
| WP_157621530.1 | Type1.BREX | [illegible] | 1117 | [illegible] | [illegible] | [illegible] | [illegible] | 2150..5503 | 503170 | [illegible] |
| WP_040606398.1 | Type1.BREX | [illegible] | 1247 | [illegible] | [illegible] | [illegible] | [illegible] | 6164..9907 | 503170 | [illegible] |
| WP_098060820.1 | Type1.BREX | [illegible] | 1206 | [illegible] | [illegible] | [illegible] | [illegible] | 364524..368144 | 1469170 | [illegible] |
| MBP9037128.1 | Type1.BREX | [illegible] | 1152 | [illegible] | [illegible] | [illegible] | [illegible] | 2613..6071 | 2053534 | [illegible] |
| OQC05381.1 | Type1.BREX | [illegible] | 1162 | [illegible] | [illegible] | [illegible] | [illegible] | 1812..5300 | 1852836 | [illegible] |
| OQC10188.1 | Type1.BREX | [illegible] | 1149 | [illegible] | [illegible] | [illegible] | [illegible] | 19726..23175 | 1852835 | [illegible] |
| MCB5286841.1 | Type1.BREX | [illegible] | 1145 | [illegible] | [illegible] | [illegible] | [illegible] | 9843..13280 | 2030808 | [illegible] |
| MCB5271576.1 | Type1.BREX | [illegible] | 1171 | [illegible] | [illegible] | [illegible] | [illegible] | 18816..22331 | 2030808 | [illegible] |
| KZD20046.1 | Type1.BREX | [illegible] | 1146 | [illegible] | [illegible] | [illegible] | [illegible] | 11808..15248 | 1729713 | [illegible] |
| MCD6589067.1 | Type1.BREX | [illegible] | 1169 | [illegible] | [illegible] | [illegible] | [illegible] | 26..3535 | 2774599 | [illegible] |
| NLV46851.1 | Type1.BREX | [illegible] | 1339 | [illegible] | [illegible] | [illegible] | [illegible] | 9749..13768 | 2030809 | [illegible] |
| NUM53512.1 | Type1.BREX | [illegible] | 1149 | [illegible] | [illegible] | [illegible] | [illegible] | 154693..158142 | 2030809 | [illegible] |
| MCC6145266.1 | Type1.BREX | [illegible] | 1156 | [illegible] | [illegible] | [illegible] | [illegible] | 16372..19842 | 2030809 | [illegible] |
| MCC6156024.1 | Type1.BREX | [illegible] | 1142 | [illegible] | [illegible] | [illegible] | [illegible] | 10018..13447 | 2030809 | [illegible] |
| RKY59330.1 | Type1.BREX | [illegible] | 1192 | [illegible] | [illegible] | [illegible] | [illegible] | 4031..7609 | 2053570 | [illegible] |
| MCC7260964.1 | Type1.BREX | [illegible] | 1146 | [illegible] | [illegible] | [illegible] | [illegible] | 7937..11377 | 2053570 | [illegible] |
| MBT4282223.1 | Type1.BREX | [illegible] | 1156 | [illegible] | [illegible] | [illegible] | [illegible] | 1407..4877 | 2026760 | [illegible] |
| MBT4361892.1 | Type1.BREX | [illegible] | 1148 | [illegible] | [illegible] | [illegible] | [illegible] | 1..3448 | 2026760 | [illegible] |
| MBT5758478.1 | Type1.BREX | [illegible] | 1156 | [illegible] | [illegible] | [illegible] | [illegible] | 1576..5046 | 2026760 | [illegible] |
| OWV23454.1 | Type1.BREX | [illegible] | 1174 | [illegible] | [illegible] | [illegible] | [illegible] | 365376..368900 | 1964358 | [illegible] |
| NLW30564.1 | Type1.BREX | [illegible] | 1185 | [illegible] | [illegible] | [illegible] | [illegible] | 4326..7883 | 35828 | [illegible] |
| NLW30641.1 | Type1.BREX | [illegible] | 1213 | [illegible] | [illegible] | [illegible] | [illegible] | 2375..6016 | 35828 | [illegible] |
| SHK42502.1 | Type1.BREX | [illegible] | 1186 | [illegible] | [illegible] | [illegible] | [illegible] | 444346..447906 | 1896211 | [illegible] |
| WP_212668349.1 | Type1.BREX | [illegible] | 1186 | [illegible] | [illegible] | [illegible] | [illegible] | 153736..157296 | 1964353 | [illegible] |
| WP_198953245.1 | Type1.BREX | [illegible] | 1176 | [illegible] | [illegible] | [illegible] | [illegible] | 365376..368906 | 1964358 | [illegible] |
| WP_073283173.1 | Type1.BREX | [illegible] | 1184 | [illegible] | [illegible] | [illegible] | [illegible] | 351031..354585 | 1896212 | [illegible] |

| Protein-ID | System | Gene-Neighbourhoods | Length | Taxonomy | Assembly-ID | Strain | Locus-Tag | Genomic-coordinates | TaxID | Fasta-defline |
|---|---|---|---|---|---|---|---|---|---|---|
| WP_217650997.1 | Type1.BREX | <-AbiEii<-AbiEi_4<-?\|\|?-><-MORC+REase\|\|<-PIN<-Phd-YefM\|\|BrxA-DNA-binding->BrxB-BrxF-iSTAND->DUF6079*->SbcC->SbcC->N6-METHYLASE->Schlafen+HATPase+wHTH->PglZ->SIGMA+OB-Fold+AAA-ATPase+L[illegible] | 1176 | Bacteria>FCB_group>Fibrobacterota>Fibrobacteria>Fibrobacterales>Fibrobacteraceae | GCF_900142475.1 | Fibrobacter_sp._UWH4 | BUA93_RS10220 | 85922..89452 | 1896210 | BREX-system-P-loop-protein-BrxC-[Fibrobacter-sp.-UWH4] |
| WP_073156756.1 | Type1.BREX | <-VirE<-?<-XIS-HTH\|\|?->?->?->BrxA-DNA-binding->BrxB-BrxF-iSTAND->DUF6079*->ParB+HNH->N6-METHYLASE->Schlafen+HATPase+wHTH->PglZ->SIGMA+OB-Fold+AAA-ATPase+Lon-Protease-BrxL->TIR->?->REa[illegible] | 1184 | Bacteria>FCB_group>Fibrobacterota>Fibrobacteria>Fibrobacterales>Fibrobacteraceae | GCF_900142505.1 | Fibrobacter_sp._UWH5 | BUB38_RS01950 | 444352..447906 | 1896211 | BREX-system-P-loop-protein-BrxC-[Fibrobacter-sp.-UWH5] |
| WP_073424902.1 | Type1.BREX | ABhydrolase->?-><-SIGMA+OB-Fold+AAA-ATPase+HKD-DNase-BrxL<-PglZ<-N6-METHYLASE<-DUF6079*<-BrxB-BrxF-iSTAND<-BrxA-DNA-binding<-LRR_5<-PIN<-RelB | 1212 | Bacteria>FCB_group>Fibrobacterota>Fibrobacteria>Fibrobacterales>Fibrobacteraceae | GCF_900142945.1 | Fibrobacter_sp._UWB7 | BUC30_RS10995 | 309469..313107 | 1896206 | BREX-system-P-loop-protein-BrxC-[Fibrobacter-sp.-UWB7] |
| MBK8058263.1 | Type1.BREX | PIN->DOC+zhth->DOC+HTH-><-SIGMA+OB-Fold+AAA-ATPase+Lon-Protease-BrxL<-PglZ<-N6-METHYLASE-p<-DOC+HTH<-N6-METHYLASE<-?<-DUF6079*<-BrxB-BrxF-iSTAND<-BrxA-DNA-binding\|\|?-><-?<-REase | 1035 | Bacteria>FCB_group>Gemmatimonadota>unclassified_Gemmatimonadota | GCA_016713015.1 | Gemmatimonadota_bacterium | IPK33_10390 | 16210..19317 | 2026742 | BREX-system-P-loop-protein-BrxC-[Gemmatimonadota-bacterium] |
| MBK8060292.1 | Type1.BREX | TPR+TPR+TPR+TPR+HDGYP-><-?<-?<-?\|\|?->BrxA-DNA-binding->BrxB-BrxF-iSTAND->DUF6079*->N6-METHYLASE->PglZ->SIGMA+OB-Fold+AAA-ATPase+Lon-Protease-BrxL->?-><-?\|\|?->?->O6MGT-HTH->SAM-METHYLA[illegible] | 1154 | Bacteria>FCB_group>Gemmatimonadota>unclassified_Gemmatimonadota | GCA_016713015.1 | Gemmatimonadota_bacterium | IPK33_21145 | 255603..259067 | 2026742 | BREX-system-P-loop-protein-BrxC-[Gemmatimonadota-bacterium] |
| MBK9977127.1 | Type1.BREX | TPR+TPR+TPR+TPR+HDGYP-><-?<-?\|\|?->?->?->DUF6079-p*->DUF6079-p->?->DUF6079-p->N6-METHYLASE-p->N6-METHYLASE-p->N6-METHYLASE->?->N6-METHYLASE-p->PglZ-p-> | 79 | Bacteria>FCB_group>Gemmatimonadota>unclassified_Gemmatimonadota | GCA_016721935.1 | Gemmatimonadota_bacterium | IPP20_04220 | 1050046..1050285 | 2026742 | hypothetical-protein-[Gemmatimonadota-bacterium] |
| WP_185874362.1 | Type1.BREX | MORC->?->?->SF1-SF2-Helicase->?->?->SWIM+DUF3670+SF1-SF2-Helicase->zf-ISL3->DUF6079*->SbcC->?->N6-METHYLASE->McrB-AAA->REase-><-McrB-AAA<-?<-N6-METHYLASE-p | 1243 | Bacteria>Fusobacteriota>Fusobacteriia>Fusobacteriales>Fusobacteriaceae>Cetobacterium | GCF_014250675.1 | Cetobacterium_sp._8H | H5J22_RS00820 | 148071..151802 | 2759681 | BREX-system-P-loop-protein-BrxC-[Cetobacterium-sp.-8H] |
| CDA09060.1 | Type1.BREX | <-SIGMA+OB-Fold+AAA-ATPase+Lon-Protease-BrxL<-PglZ<-CR-ATPase8+REase<-OLD-ABC+TOPRIM<-?<-?<-N6-METHYLASE<-TOPRIM<-OLD-ABC<-DUF6079*<-?<-SWIM+DUF3670+SF1-SF2-Helicase<-?<-?<-?<-SF1-SF2[illegible] | 1250 | Bacteria>Fusobacteriota>Fusobacteriia>Fusobacteriales>Fusobacteriaceae>Fusobacterium | GCA_000433695.1 | Fusobacterium_sp._CAG:649 | BN748_00884 | 23068..26820 | 1262900 | putative-uncharacterized-protein-[Fusobacterium-sp.-CAG:649] |
| ALF25292.1 | Type1.BREX | SKI->JAB+sGTP->wHTH+WYL+WCX->?->?->ABC-ATPase+SbcC->?-><-?<-?\|\|DUF6079*->OLD-ABC->TOPRIM->Schlafen+HATPase+wHTH->N6-METHYLASE->KAP-NTPase->HNH->SbcC->SIR2->HerA/FtsK-> | 1250 | Bacteria>Fusobacteriota>Fusobacteriia>Fusobacteriales>Fusobacteriaceae>Fusobacterium | GCA_001296185.1 | Fusobacterium_nucleatum_subsp._nucleatum | RN95_02150 | 455192..458944 | 76856 | cobalamin-adenosyltransferase-[Fusobacterium-nucleatum-subsp.-nucleatum] |
| KAB0576672.1 | Type1.BREX | <-RADICAL-SAM<-?\|\|\|\|<-SbcC<-SIGMA+OB-Fold+AAA-ATPase+Lon-Protease-BrxL<-PglZ<-AT-HTH-HYD2-2<-?<-Tox-ART-HYD2<-N6-METHYLASE<-DUF6079*<-BrxB-BrxF-iSTAND<-BrxA-DNA-binding<-XIS-HTH<-?<-M[illegible] | 1217 | Bacteria>Fusobacteriota>Fusobacteriia>Fusobacteriales>Fusobacteriaceae>Fusobacterium | GCA_008801825.1 | Fusobacterium_naviforme | F7P78_08540 | 57971..61624 | 77917 | BREX-system-P-loop-protein-BrxC-[Fusobacterium-naviforme] |
| MCF0162780.1 | Type1.BREX | PrmA+METHYLASE->?->?->METHYLASE->?->?->?->DUF6079*->N6-METHYLASE->?-><-ABC-ATPase\|\|AlcRN+AraC-HTH+AraC-HTH-> | 1240 | Bacteria>Fusobacteriota>Fusobacteriia>Fusobacteriales>Fusobacteriaceae>Fusobacterium | GCA_021443685.1 | Fusobacterium_necrophorum | HUJ88_RS09480 | 48208..51930 | 859 | BREX-system-P-loop-protein-BrxC-[Fusobacterium-necrophorum] |
| MCF2639759.1 | Type1.BREX | DUF6079*->N6-METHYLASE->PglZ->BrxA-DNA-binding->BrxB-BrxF-iSTAND->CR-ATPase5+REase->DOC->Calcineurin->HIN-HTH-> | 1244 | Bacteria>Fusobacteriota>Fusobacteriia>Fusobacteriales>Fusobacteriaceae>Fusobacterium | GCA_021531615.1 | Fusobacterium_varium | I6E31_07220 | 5895..9629 | 856 | BREX-system-P-loop-protein-BrxC-[Fusobacterium-varium] |
| WP_008797395.1 | Type1.BREX | <-HerA/FtsK<-SIR2<-SbcC<-HNH<-KAP-NTPase<-KAP-NTPase<-N6-METHYLASE<-TOPRIM<-OLD-ABC<-DUF6079*\|\|?->?-><-?<-ABC-ATPase+SbcC<-?<-?<-wHTH+WYL+WCX<-JAB+sGTP<-SKI | 1249 | Bacteria>Fusobacteriota>Fusobacteriia>Fusobacteriales>Fusobacteriaceae>Fusobacterium | GCF_000163915.2 | Fusobacterium_nucleatum | HMPREF0405_RS05640 | 1192926..1196675 | 851 | BREX-system-P-loop-protein-BrxC-[Fusobacterium-vincentii] |
| WP_005903411.1 | Type1.BREX | <-PglZ<-ParB+HNH+RAMA<-OLD-ABC+TOPRIM<-N6-METHYLASE<-Schlafen+HATPase+wHTH<-?<-RNase+PHP<-?<-OLD-ABC<-DUF6079*\|\|?->?-><-?<-ABC-ATPase+SbcC<-?<-?<-wHTH+WYL+WCX<-JAB+sGTP<-SKI | 1250 | Bacteria>Fusobacteriota>Fusobacteriia>Fusobacteriales>Fusobacteriaceae>Fusobacterium | GCF_000178895.1 | Fusobacterium_nucleatum | C4N14_RS08075 | 1667383..1671135 | 851 | BREX-system-P-loop-protein-BrxC-[Fusobacterium-nucleatum] |
| WP_008692035.1 | Type1.BREX | DUF3670+SF1-SF2-Helicase->?-><-?<-?<-?<-?\|\|DUF6079*->OLD-ABC->TOPRIM->N6-METHYLASE->?->OLD-ABC+TOPRIM->CR-ATPase8+REase->PglZ->SIGMA+OB-Fold+AAA-ATPase+Lon-Protease-BrxL->BrxA-DNA-bin[illegible] | 1250 | Bacteria>Fusobacteriota>Fusobacteriia>Fusobacteriales>Fusobacteriaceae>Fusobacterium | GCF_000218655.1 | Fusobacterium_nucleatum | HMPREF0401_RS10120 | 557323..561075 | 851 | BREX-system-P-loop-protein-BrxC-[Fusobacterium-animalis] |
| WP_008700610.1 | Type1.BREX | SWIM+DUF3670+SF1-SF2-Helicase->?-><-?<-?<-?<-?\|\|DUF6079*->OLD-ABC->TOPRIM->KAP-NTPase->HNH->SbcC->SIR2->HerA/FtsK->CR-ATPase8+REase->Classical-AAA-> | 1250 | Bacteria>Fusobacteriota>Fusobacteriia>Fusobacteriales>Fusobacteriaceae>Fusobacterium | GCF_000234075.2 | Fusobacterium_nucleatum | FSDG_RS04805 | 1011636..1015388 | 851 | MULTISPECIES:-BREX-system-P-loop-protein-BrxC-[Fusobacterium] |
| [illegible] | [illegible] | [illegible] | [illegible] | [illegible] | [illegible] | [illegible] | [illegible] | [illegible] | [illegible] | [illegible] |

| Protein-ID | System | Gene-Neighbourhoods | Length | Taxonomy | Assembly-ID | Strain | Locus-Tag | Genomic-coordinates | TaxID | Fasta-define |
|---|---|---|---|---|---|---|---|---|---|---|
| WP_266332120.1 | Type1.BREX | <-SF1-SF2-Helicase<-HKD-DNase+HKD-DNase<-wHTH<-SF1-SF2-Helicase+REase<-SF1-SF2-Helicase+REase<-SIGMA+OB-Fold+AAA-ATPase+HKD-DNase-BrxL<-PglZ<-N6-METHYLASE<-DUF6079*<-BrxB-BrxF-iSTAND<-BrxA-DNA-binding<-wHTH+WYL+WCX<-MODE-HTH+PBPII\|\|TetR-HTH-><-RHH<-VirB10<-VirB9<-VirB8 | 1179 | Bacteria>Pseudomonadota>Alphaproteobacteria>Hyphomicrobiales>Kaistiaceae>Kaistia | GCF_026343955.1 | Kaistia_defluvii | OSH10_RS12655 | 370045..373584 | 410841 | BREX-system-P-loop-protein-BrxC-[Kaistia-defluvii] |
| WP_056266971.1 | Type1.BREX | <-SIGMA+OB-Fold+AAA-ATPase+Lon-Protease-BrxL<-PglZ<-N6-METHYLASE<-DUF6079*<-BrxB-BrxF-iSTAND<-BrxA-DNA-binding | 1150 | Bacteria>Pseudomonadota>Alphaproteobacteria>Hyphomicrobiales>Methylobacteriaceae>Methylobacterium | GCF_001422775.1 | Methylobacterium_sp._Leaf87 | ASF22_RS18370 | 367037..370489 | 1736243 | BREX-system-P-loop-protein-BrxC-[Methylobacterium-sp.-Leaf87] |
| WP_147828804.1 | Type1.BREX | Arm-DNA-bind_3+Phage_integrase->?->Phage_integrase-><-SIGMA+OB-Fold+AAA-ATPase+Lon-Protease-BrxL<-PglZ<-N6-METHYLASE<-DUF6079*<-BrxB-BrxF-iSTAND<-BrxA-DNA-binding<-?<-SF1-SF2-Helicase<-AAA-ATPase<-REase\|\|rve+rve_3->ParB+HNH->ParB+HNH-> | 1149 | Bacteria>Pseudomonadota>Alphaproteobacteria>Hyphomicrobiales>Methylobacteriaceae>Methylobacterium | GCF_008040045.1 | Methylobacterium_sp._WL8 | FV234_RS09050 | 16496..19945 | 2603899 | BREX-system-P-loop-protein-BrxC-[Methylobacterium-sp.-WL8] |
| WP_009764060.1 | Type1.BREX | <-Phage_integrase\|\|PRTase-><-?\|\|?->?->\|\|BrxA-DNA-binding->BrxB-BrxF-iSTAND->DUF6079*->N6-METHYLASE->PglZ->SIGMA+OB-Fold+AAA-ATPase+Lon-Protease-BrxL->REase-> | 1149 | Bacteria>Pseudomonadota>Alphaproteobacteria>Hyphomicrobiales>Methylobacteriaceae>Microvirga | GCF_000262405.1 | Microvirga_lotononidis | MICLODRAFT_RS22145 | 1808535..1811984 | 864069 | BREX-system-P-loop-protein-BrxC-[Microvirga-lotononidis] |
| WP_202057660.1 | Type1.BREX | Phage_integrase+Phage_integrase->REase->DnaK->?->?->?->BrxA-DNA-binding->BrxB-BrxF-iSTAND->DUF6079*->N6-METHYLASE->PglZ->SIGMA+OB-Fold+AAA-ATPase+HKD-DNase-BrxL->?->?->?-><-?<-tRNA<-?<-CDP-OH_P_transf | 1176 | Bacteria>Pseudomonadota>Alphaproteobacteria>Hyphomicrobiales>Methylobacteriaceae>Microvirga | GCF_016743775.1 | Microvirga_aerilata | JKG68_RS07685 | 69600..73130 | 670292 | BREX-system-P-loop-protein-BrxC-[Microvirga-aerilata] |
| WP_036280608.1 | Type1.BREX | HKD-DNase->SF1-SF2-Helicase-><-Small-B-barrel+SF1-SF2-Helicase+RdgC+REase<-SF1-SF2-Helicase+REase<-35Exo+REase\|\|wHTH+WYL+WCX->BrxA-DNA-binding->BrxB-BrxF-iSTAND->DUF6079*->OLD-ABC->REase->N6-METHYLASE->PglZ->SIGMA+OB-Fold+AAA-ATPase+HKD-DNase-BrxL->HIN-HTH->?->LZ_Tnp_IS66+zf-IS66+DDE_Tnp_IS66+DDE_Tnp_IS66_C-> | 1178 | Bacteria>Pseudomonadota>Alphaproteobacteria>Hyphomicrobiales>Methylocystaceae>Methylocystis | GCF_000188155.2 | Methylocystis_sp._ATCC_49242 | MET49242_RS03150 | 256459..259995 | 622637 | BREX-system-P-loop-protein-BrxC-[Methylocystis-sp.-ATCC-49242] |
| WP_024514760.1 | Type1.BREX | HerA/FtsK->?-><-?<-?\|\|?-><-SIGMA+OB-Fold+AAA-ATPase+Lon-Protease-BrxL<-PglZ<-N6-METHYLASE<-DUF6079*<-BrxB-BrxF-iSTAND<-BrxA-DNA-binding\|\|McrB-AAA->OLD-ABC+TOPRIM->SF1-SF2-Helicase-> | 1155 | Bacteria>Pseudomonadota>Alphaproteobacteria>Hyphomicrobiales>Nitrobacteraceae>Bradyrhizobium | GCF_000472425.1 | Bradyrhizobium_sp._Tv2a-2 | A3AI_RS0106300 | 29834..33301 | 113395 | BREX-system-P-loop-protein-BrxC-[Bradyrhizobium-sp.-Tv2a-2] |
| WP_027571504.1 | Type1.BREX | <-Thermonuclease<-?\|\|ACET-><-AAA-ATPase<-REase+HTH+35exo+Small-B-barrel+SF1-SF2-Helicase\|\|BrxA-DNA-binding->BrxB-BrxF-iSTAND->DUF6079*->N6-METHYLASE->PglZ->SIGMA+OB-Fold+AAA-ATPase+Lon-Protease-BrxL-><-?<-HTH+HKD-DNase<-Pkinase+SF1-SF2-Helicase<-SF1-SF2-Helicase | 1149 | Bacteria>Pseudomonadota>Alphaproteobacteria>Hyphomicrobiales>Nitrobacteraceae>Bradyrhizobium | GCF_000473065.1 | Bradyrhizobium_sp._WSM1743 | YU9_RS0103440 | 276500..279949 | 318996 | BREX-system-P-loop-protein-BrxC-[Bradyrhizobium-sp.-WSM1743] |
| WP_027565284.1 | Type1.BREX | <-MarR-HTH<-?<-CR-ATPase<-GDResolvase+HIN-HTH<-?<-SIGMA+OB-Fold+AAA-ATPase+Lon-Protease-BrxL<-PglZ<-N6-METHYLASE<-DUF6079*<-BrxB-BrxF-iSTAND<-BrxA-DNA-binding<-?<-Phage_integrase<-tRNA<-?\|\|S4+PSYN->SIGMA-HTH+SIGMA-HTH-> | 1144 | Bacteria>Pseudomonadota>Alphaproteobacteria>Hyphomicrobiales>Nitrobacteraceae>Bradyrhizobium | GCF_000518345.1 | Bradyrhizobium_sp._URHA0013 | N551_RS0100500 | 109615..113049 | 1380352 | BREX-system-P-loop-protein-BrxC-[Bradyrhizobium-sp.-URHA0013] |
| WP_044590041.1 | Type1.BREX | <-Peptidase+SF1-SF2-Helicase+REase<-?<-SIGMA-HTH<-SbcC+CCDC158<-Calcineurin-SbcD\|\|BrxA-DNA-binding->BrxB-BrxF-iSTAND->DUF6079*->N6-METHYLASE->PglZ->SIGMA+OB-Fold+AAA-ATPase+Lon-Protease-BrxL-><-?<-?<-?<-Thermonuclease<-SF1-SF2-Helicase | 1149 | Bacteria>Pseudomonadota>Alphaproteobacteria>Hyphomicrobiales>Nitrobacteraceae>Bradyrhizobium | GCF_000938235.1 | Bradyrhizobium_sp._LTSPM299 | UP10_RS26195 | 631176..634625 | 1619233 | BREX-system-P-loop-protein-BrxC-[Bradyrhizobium-sp.-LTSPM299] |
| WP_085384424.1 | Type1.BREX | DUF772+DDE_Tnp_1+DDE_Tnp_1_6->?->?->?->?->BrxA-DNA-binding->BrxB-BrxF-iSTAND->DUF6079*->N6-METHYLASE->PglZ->SIGMA+OB-Fold+AAA-ATPase+Lon-Protease-BrxL-> | 1199 | Bacteria>Pseudomonadota>Alphaproteobacteria>Hyphomicrobiales>Nitrobacteraceae>Bradyrhizobium | GCF_002108905.1 | Bradyrhizobium_canariense | BSR47_RS02850 | 9232..12831 | 255045 | BREX-system-P-loop-protein-BrxC-[Bradyrhizobium-canariense] |
| WP_130224749.1 | Type1.BREX | <-ArdCN+MPTase<-HEPN<-Small-B-barrel+SF1-SF2-Helicase+RdgC+REase<-35Exo+REase<-?\|\|wHTH+WYL+WCX->BrxA-DNA-binding->BrxB-BrxF-iSTAND->DUF6079*->N6-METHYLASE->PglZ->SIGMA+OB-Fold+AAA-ATPase+HKD-DNase-BrxL->AbiEi_4->AbiEii->OLD-ABC+TOPRIM->SF1-SF2-Helicase->REase-><-ABhydrolase | 1181 | Bacteria>Pseudomonadota>Alphaproteobacteria>Hyphomicrobiales>Nitrobacteraceae>Bradyrhizobium | GCF_004212675.1 | Bradyrhizobium_sp._Leo121 | CWO90_RS07090 | 48469..52014 | 1571195 | BREX-system-P-loop-protein-BrxC-[Bradyrhizobium-sp.-Leo121] |
| WP_152104118.1 | Type1.BREX | BrxA-DNA-binding->BrxB-BrxF-iSTAND->DUF6079*->N6-METHYLASE->PglZ->SIGMA+OB-Fold+AAA-ATPase+Lon-Protease-BrxL->?->?->dCache_1+HisKA+HATPase_c->REC+NtrC-AAA+FIS-HTH-> | 1199 | Bacteria>Pseudomonadota>Alphaproteobacteria>Hyphomicrobiales>Nitrobacteraceae>Bradyrhizobium | GCF_009176665.1 | Bradyrhizobium_ottawaense | GDN72_RS19495 | 4098964..4102563 | 931866 | BREX-system-P-loop-protein-BrxC-[Bradyrhizobium-ottawaense] |
| WP_157343672.1 | Type1.BREX | ABC-ATPase+SbcC->?->?-><-tRNA<-Small-B-barrel+SF1-SF2-Helicase+RdgC+REase<-SF1-SF2-Helicase+REase<-35Exo+REase\|\|wHTH+WYL+WCX->BrxA-DNA-binding->BrxB-BrxF-iSTAND->DUF6079*->N6-METHYLASE->PglZ->SIGMA+OB-Fold+AAA-ATPase+HKD-DNase-BrxL-><-?<-HTH+REase<-Arm-DNA-bind_3+Phage_integrase<-?\|\|?-><-Peptidase_M24 | 1181 | Bacteria>Pseudomonadota>Alphaproteobacteria>Hyphomicrobiales>Nitrobacteraceae>Bradyrhizobium | GCF_009759865.1 | Bradyrhizobium_pachyrhizi | GPL21_13485 | 267907..271452 | 280333 | BREX-system-P-loop-protein-BrxC-[Bradyrhizobium-pachyrhizi] |
| WP_212056797.1 | Type1.BREX | <-SF1-SF2-Helicase<-MORC+DUF3684+REase<-HHH_4+SF1-SF2-Helicase+REase<-AbiEi_4<-SIGMA+OB-Fold+AAA-ATPase+HKD-DNase-BrxL<-PglZ<-N6-METHYLASE<-DUF6079*<-BrxB-BrxF-iSTAND<-BrxA-DNA-binding<-wHTH+WYL+WCX\|\|35Exo+REase->SF1-SF2-Helicase+REase->Small-B-barrel+SF1-SF2-Helicase+RdgC+REase-><-?\|\|GDResolvase+Recombinase+ZnR-> | 1180 | Bacteria>Pseudomonadota>Alphaproteobacteria>Hyphomicrobiales>Nitrobacteraceae>Bradyrhizobium | GCF_018131045.1 | Bradyrhizobium_elkanii | JQ569_RS15725 | 68040..71582 | 29448 | BREX-system-P-loop-protein-BrxC-[Bradyrhizobium-elkanii] |
| WP_225161486.1 | Type1.BREX | <-ParB+HNH+Schlafen<-?<-SIGMA+OB-Fold+AAA-ATPase+HKD-DNase-BrxL<-PglZ<-N6-METHYLASE<-DUF6079*<-BrxB-BrxF-iSTAND<-BrxA-DNA-binding<-wHTH+WYL+WCX\|\|?->35Exo+REase->SF1-SF2-Helicase+REase->35exo+SF1-SF2-Helicase-><-Phage_integrase | 1183 | Bacteria>Pseudomonadota>Alphaproteobacteria>Hyphomicrobiales>Nitrobacteraceae>Bradyrhizobium | GCF_020194035.1 | Bradyrhizobium_sp._BRP56 | I6F38_19500 | 73459..77010 | 2793819 | BREX-system-P-loop-protein-BrxC-[Bradyrhizobium-sp.-BRP56] |
| WP_247808194.1 | Type1.BREX | <-TOPRIM<-SAM-METHYLASE+SF1-SF2-Helicase\|\|?-><-ParB<-DUF932\|\|XIS-HTH-><-SIGMA+OB-Fold+AAA-ATPase+Lon-Protease-BrxL<-PglZ<-N6-METHYLASE<-DUF6079*<-BrxB-BrxF-iSTAND<-BrxA-DNA-binding\|\|Calcineurin-SbcD->SbcC+CCDC158->SIGMA-HTH->?->Peptidase+SF1-SF2-Helicase+REase-> | 1149 | Bacteria>Pseudomonadota>Alphaproteobacteria>Hyphomicrobiales>Nitrobacteraceae>Bradyrhizobium | GCF_023101025.1 | unclassified_Bradyrhizobium | IVB17_RS35670 | 7402145..7405594 | 2631580 | MULTISPECIES:-BREX-system-P-loop-protein-BrxC-[unclassified-Bradyrhizobium] |
| WP_260373497.1 | Type1.BREX | <-ParB+HNH+Schlafen<-?<-SIGMA+OB-Fold+AAA-ATPase+HKD-DNase-BrxL<-PglZ<-N6-METHYLASE<-DUF6079*<-BrxB-BrxF-iSTAND<-BrxA-DNA-binding<-wHTH+WYL+WCX\|\|?->35Exo+REase->SF1-SF2-Helicase+REase->35exo+SF1-SF2-Helicase-><-Phage_integrase | 1183 | Bacteria>Pseudomonadota>Alphaproteobacteria>Hyphomicrobiales>Nitrobacteraceae>Bradyrhizobium | GCF_025200925.1 | Bradyrhizobium_sp._CB1015 | N2604_RS01610 | 347161..350712 | 2976822 | BREX-system-P-loop-protein-BrxC-[Bradyrhizobium-sp.-CB1015] |
| WP_275169502.1 | Type1.BREX | <-Trypsin+REase\|\|?-><-?\|\|?-><-SIGMA+OB-Fold+AAA-ATPase+Lon-Protease-BrxL<-PglZ<-N6-METHYLASE<-DUF6079*<-BrxB-BrxF-iSTAND<-BrxA-DNA-binding\|\|<-?<-tRNA<-?<-METHYLASE<-?\|\|?->?->HTH_Tnp_2-> | 1145 | Bacteria>Pseudomonadota>Alphaproteobacteria>Hyphomicrobiales>Nitrobacteraceae>Bradyrhizobium | GCF_029051725.1 | Bradyrhizobium_sp._CSS354 | GWG67_RS11755 | 378554..381991 | 2699172 | BREX-system-P-loop-protein-BrxC-[Bradyrhizobium-sp.-CSS354] |
| WP_110780661.1 | Type1.BREX | <-DOC+HTH<-SF1-SF2-Helicase<-HKD-DNase+HKD-DNase<-wHTH<-SF1-SF2-Helicase+REase<-SF1-SF2-Helicase+REase<-SIGMA+OB-Fold+AAA-ATPase+HKD-DNase-BrxL<-PglZ<-N6-METHYLASE<-DUF6079*<-BrxB-BrxF-iSTAND<-BrxA-DNA-binding<-wHTH+WYL+WCX<-MODE-HTH+PBPII\|\|TetR-HTH-><-RHH<-VirB10<-VirB9<-VirB8 | 1179 | Bacteria>Pseudomonadota>Alphaproteobacteria>Hyphomicrobiales>Nitrobacteraceae>Rhodopseudomonas | GCF_003217325.1 | Rhodopseudomonas_faecalis | BJ122_RS08545 | 91514..95053 | 99655 | BREX-system-P-loop-protein-BrxC-[Rhodopseudomonas-faecalis] |
| WP_184518066.1 | Type1.BREX | HTH+MPTase-><-SIGMA+OB-Fold+AAA-ATPase+Lon-Protease-BrxL<-PglZ<-N6-METHYLASE-p<-N6-METHYLASE<-DUF6079*<-BrxB-BrxF-iSTAND<-BrxA-DNA-binding<-CopG/RHH<-TraI-VirD2-RCRE<-?<-?<-?<-ABhydrolase<-HISKIN | 1196 | Bacteria>Pseudomonadota>Alphaproteobacteria>Hyphomicrobiales>Nitrobacteraceae>Tardiphaga | GCF_014189415.1 | Tardiphaga_robiniae | HB776_RS12405 | 2595278..2598868 | 943830 | BREX-system-P-loop-protein-BrxC-[Tardiphaga-robiniae] |
| MAM94519.1 | Type1.BREX | VirB10->RHH-><-TetR-HTH\|\|MODE-HTH+PBPII-><-SbcC+SbcC<-Calcineurin+Calcineurin-SbcD\|\|wHTH+WYL+WCX->BrxA-DNA-binding->BrxB-BrxF-iSTAND->DUF6079*->N6-METHYLASE->PglZ->SIGMA+OB-Fold+AAA-ATPase+HKD-DNase-BrxL->OLD-ABC+TOPRIM->SF1-SF2-Helicase-><-Arm-DNA-bind_3+Phage_int_SAM_3+Phage_integrase<-?<-?<-MerR-HTH | 1179 | Bacteria>Pseudomonadota>Alphaproteobacteria>Hyphomicrobiales>Parvibaculaceae>Parvibaculum | GCA_002694985.1 | Parvibaculum_sp. | CMI61_07805 | 145242..148781 | 2024848 | hypothetical-protein-[Parvibaculum-sp.] |
| WP_110030326.1 | Type1.BREX | RAMPs->RAMPs->?->RVT->CRISPR_Cas6->?-><-?\|\|BrxA-DNA-binding->BrxB-BrxF-iSTAND->DUF6079*->N6-METHYLASE->PglZ->SIGMA+OB-Fold+AAA-ATPase+HKD-DNase-BrxL->?->HKD-DNase+HKD-DNase->SF1-SF2-Helicase->MORC+DUF3684+REase->REase+SF1-SF2-Helicase-> | 1173 | Bacteria>Pseudomonadota>Alphaproteobacteria>Hyphomicrobiales>Phyllobacteriaceae>Hoeflea | GCF_003182275.1 | Hoeflea_marina | DFR52_RS02490 | 542434..545955 | 274592 | BREX-system-P-loop-protein-BrxC-[Hoeflea-marina] |
| ESX24129.1 | Type1.BREX | <-SIGMA+OB-Fold+AAA-ATPase+HKD-DNase-BrxL<-PglZ<-N6-METHYLASE<-DUF6079*<-BrxB-BrxF-iSTAND<-BrxA-DNA-binding<-?\|\|REase->HIN-HTH->?->LZ_Tnp_IS66+zf-IS66+DDE_Tnp_IS66+DDE_Tnp_IS66_C-> | 1186 | Bacteria>Pseudomonadota>Alphaproteobacteria>Hyphomicrobiales>Phyllobacteriaceae>Mesorhizobium | GCA_000502755.1 | Mesorhizobium_sp._LSJC264A00 | X767_13205 | 150936..154496 | 1287321 | hypothetical-protein-X767_13205-[Mesorhizobium-sp.-LSJC264A00] |
| ESX84801.1 | Type1.BREX | PIWI->HIN-HTH->HTH_21+rve+rve_3->Y1_Tnp-><-SIGMA+OB-Fold+AAA-ATPase+HKD-DNase-BrxL<-PglZ<-N6-METHYLASE<-NERD<-OLD-ABC<-DUF6079*<-BrxB-BrxF-iSTAND<-BrxA-DNA-binding\|\|?->?-><-HEPN<-HTH_37 | 1177 | Bacteria>Pseudomonadota>Alphaproteobacteria>Hyphomicrobiales>Phyllobacteriaceae>Mesorhizobium | GCA_000503035.1 | Mesorhizobium_sp._LSHC412B00 | X756_23895 | 63877..67410 | 1287285 | hypothetical-protein-X756_23895-[Mesorhizobium-sp.-LSHC412B00] |
| RWC29997.1 | Type1.BREX | [illegible] | 1181 | Bacteria>Pseudomonadota>Alphaproteobacteria>Hyphomicrobiales>Phyllobacteriaceae>Mesorhizobium | GCA_004017395.1 | Mesorhizobium_sp. | EOS27_14215 | 67406..70951 | 1871066 | BREX-system-P-loop-protein-BrxC-[Mesorhizobium-sp.] |
| RWE30575.1 | Type1.BREX | [illegible] | 1034 | Bacteria>Pseudomonadota>Alphaproteobacteria>Hyphomicrobiales>Phyllobacteriaceae>Mesorhizobium | GCA_004017865.1 | Mesorhizobium_sp. | EOS77_19540 | 17552..20653 | 1871066 | BREX-system-P-loop-protein-BrxC,-partial-[Mesorhizobium-sp.] |
| RWI14537.1 | Type1.BREX | [illegible] | 1202 | Bacteria>Pseudomonadota>Alphaproteobacteria>Hyphomicrobiales>Phyllobacteriaceae>Mesorhizobium | GCA_004018995.1 | Mesorhizobium_sp. | EOQ92_28755 | 60098..63706 | 1871066 | BREX-system-P-loop-protein-BrxC-[Mesorhizobium-sp.] |
| RWK37907.1 | Type1.BREX | [illegible] | 1137 | Bacteria>Pseudomonadota>Alphaproteobacteria>Hyphomicrobiales>Phyllobacteriaceae>Mesorhizobium | GCA_004019695.1 | Mesorhizobium_sp. | EOR46_25140 | 24164..27577 | 1871066 | BREX-system-P-loop-protein-BrxC-[Mesorhizobium-sp.] |
| RWM07956.1 | Type1.BREX | [illegible] | 1184 | Bacteria>Pseudomonadota>Alphaproteobacteria>Hyphomicrobiales>Phyllobacteriaceae>Mesorhizobium | GCA_004020545.1 | Mesorhizobium_sp. | EOR71_14460 | 11856..15410 | 1871066 | BREX-system-P-loop-protein-BrxC-[Mesorhizobium-sp.] |
| RWO20271.1 | Type1.BREX | [illegible] | 1178 | Bacteria>Pseudomonadota>Alphaproteobacteria>Hyphomicrobiales>Phyllobacteriaceae>Mesorhizobium | GCA_004021465.1 | Mesorhizobium_sp. | EOS09_27860 | 18398..21934 | 1871066 | BREX-system-P-loop-protein-BrxC-[Mesorhizobium-sp.] |
| TIL41012.1 | Type1.BREX | [illegible] | 845 | Bacteria>Pseudomonadota>Alphaproteobacteria>Hyphomicrobiales>Phyllobacteriaceae>Mesorhizobium | GCA_004961205.1 | Mesorhizobium_sp. | E5Y82_03575 | 245762..248299 | 1871066 | BREX-system-P-loop-protein-BrxC-[Mesorhizobium-sp.] |
| TIV88924.1 | Type1.BREX | [illegible] | 1179 | Bacteria>Pseudomonadota>Alphaproteobacteria>Hyphomicrobiales>Phyllobacteriaceae>Mesorhizobium | GCA_004963965.1 | Mesorhizobium_sp. | E5V78_03705 | 4955..8494 | 1871066 | BREX-system-P-loop-protein-BrxC-[Mesorhizobium-sp.] |
| WP_156951459.1 | Type1.BREX | [illegible] | 1188 | Bacteria>Pseudomonadota>Alphaproteobacteria>Hyphomicrobiales>Phyllobacteriaceae>Mesorhizobium | GCF_000502755.1 | Mesorhizobium_sp._LSJC264A00 | X768_RS154485 | 150936..154502 | 1287321 | BREX-system-P-loop-protein-BrxC-[Mesorhizobium-sp.-LSJC264A00] |
| WP_023809232.1 | Type1.BREX | [illegible] | 1150 | Bacteria>Pseudomonadota>Alphaproteobacteria>Hyphomicrobiales>Phyllobacteriaceae>Mesorhizobium | GCF_000502875.1 | Mesorhizobium_sp._L2C084A000 | X735_RS161770 | 861519..864971 | 1287116 | BREX-system-P-loop-protein-BrxC-[Mesorhizobium-sp.-L2C084A000] |
| WP_156933469.1 | Type1.BREX | [illegible] | 1179 | Bacteria>Pseudomonadota>Alphaproteobacteria>Hyphomicrobiales>Phyllobacteriaceae>Mesorhizobium | GCF_000503035.1 | Mesorhizobium_sp._LSHC412B00 | X757_RS142950 | 63877..67416 | 1287285 | BREX-system-P-loop-protein-BrxC-[Mesorhizobium-sp.-LSHC412B00] |
| WP_071036304.1 | Type1.BREX | [illegible] | 1137 | Bacteria>Pseudomonadota>Alphaproteobacteria>Hyphomicrobiales>Phyllobacteriaceae>Mesorhizobium | GCF_001854815.1 | Mesorhizobium_sp._LCM_4576 | LCM4576_RS31865 | 30762..34175 | 1848289 | BREX-system-P-loop-protein-BrxC-[Mesorhizobium-sp.-LCM-4576] |
| WP_095085237.1 | Type1.BREX | [illegible] | 1138 | Bacteria>Pseudomonadota>Alphaproteobacteria>Hyphomicrobiales>Phyllobacteriaceae>Mesorhizobium | GCF_002270415.1 | Mesorhizobium_sophorae | CGZ66_RS16485 | 77748..81164 | 1300294 | BREX-system-P-loop-protein-BrxC-[Mesorhizobium-sophorae] |
| WP_128286158.1 | Type1.BREX | [illegible] | 1181 | Bacteria>Pseudomonadota>Alphaproteobacteria>Hyphomicrobiales>Phyllobacteriaceae>Mesorhizobium | GCF_004023115.1 | Mesorhizobium_sp._M6A.T.Cr.TU.017.01.1.1 | EOB36_RS28010 | 14606..18151 | 2496774 | BREX-system-P-loop-protein-BrxC-[Mesorhizobium-sp.-M6A.T.Cr.TU.017.01.1.1] |
| WP_136091391.1 | Type1.BREX | [illegible] | 1181 | Bacteria>Pseudomonadota>Alphaproteobacteria>Hyphomicrobiales>Phyllobacteriaceae>Mesorhizobium | GCF_004794715.1 | Mesorhizobium_sp._M8A.F.Ca.ET.207.01.1.1 | EN850_RS12645 | 56072..59617 | 2563968 | BREX-system-P-loop-protein-BrxC-[Mesorhizobium-sp.-M8A.F.Ca.ET.207.01.1.1] |
| WP_199202187.1 | Type1.BREX | [illegible] | 1185 | Bacteria>Pseudomonadota>Alphaproteobacteria>Hyphomicrobiales>Phyllobacteriaceae>Mesorhizobium | GCF_016756495.1 | Mesorhizobium_sp._113-3-3 | MesoLj113b_71250 | 318356..321913 | 2744516 | BREX-system-P-loop-protein-BrxC-[Mesorhizobium-sp.-113-3-3] |
| WP_092958042.1 | Type1.BREX | [illegible] | 1179 | Bacteria>Pseudomonadota>Alphaproteobacteria>Hyphomicrobiales>Phyllobacteriaceae>Mesorhizobium | GCF_020164785.1 | Mesorhizobium_sp._ES1-1 | LB513_RS22695 | 51971..55510 | 2876629 | BREX-system-P-loop-protein-BrxC-[Mesorhizobium-sp.-ES1-1] |
| WP_256745318.1 | Type1.BREX | [illegible] | 1211 | Bacteria>Pseudomonadota>Alphaproteobacteria>Hyphomicrobiales>Phyllobacteriaceae>Mesorhizobium | GCF_024518695.1 | Mesorhizobium_sp._LMG17149 | NP945_RS00755 | 156415..160050 | 2968497 | BREX-system-P-loop-protein-BrxC-[Mesorhizobium-sp.-LMG17149] |
| WP_009450523.1 | Type1.BREX | [illegible] | 1179 | Bacteria>Pseudomonadota>Alphaproteobacteria>Hyphomicrobiales>Phyllobacteriaceae>Nitratireductor | GCF_000300515.1 | Nitratireductor_indicus | NA8A_RS11845 | 137447..140986 | 721133 | BREX-system-P-loop-protein-BrxC-[Nitratireductor-indicus] |
| WP_080921652.1 | Type1.BREX | [illegible] | 1179 | Bacteria>Pseudomonadota>Alphaproteobacteria>Hyphomicrobiales>Phyllobacteriaceae>Pseudaminobacter | GCF_002075885.1 | Pseudaminobacter_manganicus | BFN67_RS22220 | 17195..20734 | 1873176 | BREX-system-P-loop-protein-BrxC-[Pseudaminobacter-manganicus] |
| WP_150288342.1 | Type1.BREX | [illegible] | 1173 | Bacteria>Pseudomonadota>Alphaproteobacteria>Hyphomicrobiales>Rhabdaerophilaceae>Rhabdaerophilum | GCF_008641065.1 | Rhabdaerophilum_calidifontis | FWJ24_RS13435 | 109797..113318 | 2604328 | BREX-system-P-loop-protein-BrxC-[Rhabdaerophilum-calidifontis] |
| WP_154722158.1 | Type1.BREX | [illegible] | 1181 | Bacteria>Pseudomonadota>Alphaproteobacteria>Hyphomicrobiales>Rhizobiaceae>Ciceribacter | GCF_900492205.1 | Ciceribacter_naphthalenivorans | DYZ14_RS19075 | 12704..16249 | 1118451 | BREX-system-P-loop-protein-BrxC-[Ciceribacter-naphthalenivorans] |
| WP_225246472.1 | Type1.BREX | [illegible] | 1175 | Bacteria>Pseudomonadota>Alphaproteobacteria>Hyphomicrobiales>Rhizobiaceae>Rhizobium/Agrobacterium_group | - | Agrobacterium_tumefaciens_complex | HW573_RS24905 | 44719..48246 | 1183400 | MULTISPECIES:-BREX-system-P-loop-protein-BrxC-[Agrobacterium-tumefaciens-complex] |
| WP_174060722.1 | Type1.BREX | [illegible] | 1175 | Bacteria>Pseudomonadota>Alphaproteobacteria>Hyphomicrobiales>Rhizobiaceae>Rhizobium/Agrobacterium_group | - | Agrobacterium_tumefaciens | G6M83_RS06120 | 12437..15964 | 358 | BREX-system-P-loop-protein-BrxC-[Agrobacterium-tumefaciens] |
| WP_156535776.1 | Type1.BREX | [illegible] | 1181 | Bacteria>Pseudomonadota>Alphaproteobacteria>Hyphomicrobiales>Rhizobiaceae>Rhizobium/Agrobacterium_group | - | Agrobacterium_vitis | GOZ98_21320 | 20620..24165 | 373 | BREX-system-P-loop-protein-BrxC-[Agrobacterium-vitis] |
| OJF91479.1 | Type1.BREX | [illegible] | 1163 | Bacteria>Pseudomonadota>Alphaproteobacteria>Hyphomicrobiales>Rhizobiaceae>Rhizobium/Agrobacterium_group | GCA_001885575.1 | Rhizobium_sp._58 | AX761_22345 | 8315..11806 | 1798804 | hypothetical-protein-AX761_22345-[Rhizobium-sp.-58] |
| PZU76796.1 | Type1.BREX | [illegible] | 1188 | Bacteria>Pseudomonadota>Alphaproteobacteria>Hyphomicrobiales>Rhizobiaceae>Rhizobium/Agrobacterium_group | GCA_003248675.1 | Rhizobium_sp. | DI546_06985 | 39524..43090 | 391 | BREX-system-P-loop-protein-BrxC-[Rhizobium-sp.] |
| TQN59072.1 | Type1.BREX | [illegible] | 1179 | Bacteria>Pseudomonadota>Alphaproteobacteria>Hyphomicrobiales>Rhizobiaceae>Rhizobium/Agrobacterium_group | GCA_006757705.1 | Agrobacterium_tumefaciens | FLX27_23795 | 22501..26040 | 358 | BREX-system-P-loop-protein-BrxC-[Agrobacterium-tumefaciens] |
| NEH61390.1 | Type1.BREX | [illegible] | 1174 | Bacteria>Pseudomonadota>Alphaproteobacteria>Hyphomicrobiales>Rhizobiaceae>Rhizobium/Agrobacterium_group | GCA_010668215.1 | Rhizobium_leguminosarum | GR207_02220 | 29199..32723 | 384 | BREX-system-P-loop-protein-BrxC-[Rhizobium-leguminosarum] |
| MBY3093307.1 | Type1.BREX | [illegible] | 1174 | Bacteria>Pseudomonadota>Alphaproteobacteria>Hyphomicrobiales>Rhizobiaceae>Rhizobium/Agrobacterium_group | GCA_019778505.1 | Rhizobium_laguerreae | HFO72_21270 | 353189..356713 | 1076926 | BREX-system-P-loop-protein-BrxC-[Rhizobium-laguerreae] |
| MBY5873817.1 | Type1.BREX | [illegible] | 1174 | Bacteria>Pseudomonadota>Alphaproteobacteria>Hyphomicrobiales>Rhizobiaceae>Rhizobium/Agrobacterium_group | GCA_019799745.1 | Rhizobium_leguminosarum | HFN53_17500 | 401377..404901 | 384 | BREX-system-P-loop-protein-BrxC-[Rhizobium-leguminosarum] |
| UIJ87239.1 | Type1.BREX | [illegible] | 1174 | Bacteria>Pseudomonadota>Alphaproteobacteria>Hyphomicrobiales>Rhizobiaceae>Rhizobium/Agrobacterium_group | GCA_021391615.1 | Rhizobium_leguminosarum | LZK77_04465 | 905701..909225 | 384 | BREX-system-P-loop-protein-BrxC-[Rhizobium-leguminosarum] |
| UIK00356.1 | Type1.BREX | [illegible] | 1137 | Bacteria>Pseudomonadota>Alphaproteobacteria>Hyphomicrobiales>Rhizobiaceae>Rhizobium/Agrobacterium_group | GCA_021391655.1 | Rhizobium_leguminosarum | LZK82_09445 | 1889801..1893214 | 384 | BREX-system-P-loop-protein-BrxC-[Rhizobium-leguminosarum] |
| UIK12452.1 | Type1.BREX | [illegible] | 292 | Bacteria>Pseudomonadota>Alphaproteobacteria>Hyphomicrobiales>Rhizobiaceae>Rhizobium/Agrobacterium_group | GCA_021391695.1 | Rhizobium_leguminosarum | LZK80_09475 | 1889966..1890844 | 384 | hypothetical-protein-LZK80_09475-[Rhizobium-leguminosarum] |
| UIK12453.1 | Type1.BREX | [illegible] | 533 | Bacteria>Pseudomonadota>Alphaproteobacteria>Hyphomicrobiales>Rhizobiaceae>Rhizobium/Agrobacterium_group | GCA_021391695.1 | Rhizobium_leguminosarum | LZK80_09480 | 1891035..1892636 | 384 | hypothetical-protein-LZK80_09480-[Rhizobium-leguminosarum] |
| UIL29451.1 | Type1.BREX | [illegible] | 410 | Bacteria>Pseudomonadota>Alphaproteobacteria>Hyphomicrobiales>Rhizobiaceae>Rhizobium/Agrobacterium_group | GCA_021397605.1 | Rhizobium_leguminosarum | LZK75_09525 | 1890231..1891463 | 384 | hypothetical-protein-LZK75_09525-[Rhizobium-leguminosarum] |
| UIL29452.1 | Type1.BREX | [illegible] | 662 | Bacteria>Pseudomonadota>Alphaproteobacteria>Hyphomicrobiales>Rhizobiaceae>Rhizobium/Agrobacterium_group | GCA_021397605.1 | Rhizobium_leguminosarum | LZK75_09530 | 1891654..1893642 | 384 | hypothetical-protein-LZK75_09530-[Rhizobium-leguminosarum] |
| WP_131614693.1 | Type1.BREX | [illegible] | 1174 | Bacteria>Pseudomonadota>Alphaproteobacteria>Hyphomicrobiales>Rhizobiaceae>Rhizobium/Agrobacterium_group | GCA_030820635.1 | Rhizobium_leguminosarum | HFN52_RS21030 | 83963..87487 | 384 | BREX-system-P-loop-protein-BrxC-[Rhizobium-leguminosarum] |
| CAD6617282.1 | Type1.BREX | [illegible] | 1172 | Bacteria>Pseudomonadota>Alphaproteobacteria>Hyphomicrobiales>Rhizobiaceae>Rhizobium/Agrobacterium_group | GCA_902502895.2 | Rhizobium_sp._Khangiran2 | RKHAN_03345 | 3387102..3390620 | 1435607 | BREX-system-P-loop-protein-BrxC-[Rhizobium-sp.-Khangiran2] |
| WP_018481320.1 | Type1.BREX | [illegible] | 1207 | Bacteria>Pseudomonadota>Alphaproteobacteria>Hyphomicrobiales>Rhizobiaceae>Rhizobium/Agrobacterium_group | GCF_000373285.1 | Rhizobium_leguminosarum | B052_RS0106775 | 169147..172770 | 384 | BREX-system-P-loop-protein-BrxC-[Rhizobium-leguminosarum] |
| WP_027666511.1 | Type1.BREX | [illegible] | 1174 | Bacteria>Pseudomonadota>Alphaproteobacteria>Hyphomicrobiales>Rhizobiaceae>Rhizobium/Agrobacterium_group | GCF_000427945.1 | Rhizobium_leguminosarum | A19Q_RS0117655 | 62384..65908 | 384 | BREX-system-P-loop-protein-BrxC-[Rhizobium-leguminosarum] |
| WP_046669699.1 | Type1.BREX | [illegible] | 1182 | Bacteria>Pseudomonadota>Alphaproteobacteria>Hyphomicrobiales>Rhizobiaceae>Rhizobium/Agrobacterium_group | GCF_000985915.1 | Neorhizobium_galegae | AAW20_RS29810 | 5236..8784 | 399 | BREX-system-P-loop-protein-BrxC-[Neorhizobium-galegae] |
| WP_088677288.1 | Type1.BREX | [illegible] | 1194 | Bacteria>Pseudomonadota>Alphaproteobacteria>Hyphomicrobiales>Rhizobiaceae>Rhizobium/Agrobacterium_group | GCF_002211395.1 | Rhizobium_sp._R339 | ATY76_RS24585 | 353019..356603 | 1764273 | BREX-system-P-loop-protein-BrxC-[Rhizobium-sp.-R339] |
| WP_126920114.1 | Type1.BREX | [illegible] | 1175 | Bacteria>Pseudomonadota>Alphaproteobacteria>Hyphomicrobiales>Rhizobiaceae>Rhizobium/Agrobacterium_group | GCF_003985155.1 | Rhizobium_vallis | EFQ99_RS06935 | 372119..375646 | 634290 | BREX-system-P-loop-protein-BrxC-[Rhizobium-vallis] |
| WP_128407956.1 | Type1.BREX | [illegible] | 1137 | Bacteria>Pseudomonadota>Alphaproteobacteria>Hyphomicrobiales>Rhizobiaceae>Rhizobium/Agrobacterium_group | GCF_004028055.1 | Rhizobium_leguminosarum | EHI43_RS33385 | 4123..7536 | 384 | BREX-system-P-loop-protein-BrxC-[Rhizobium-leguminosarum] |
| WP_130741058.1 | Type1.BREX | [illegible] | 1207 | Bacteria>Pseudomonadota>Alphaproteobacteria>Hyphomicrobiales>Rhizobiaceae>Rhizobium/Agrobacterium_group | GCF_004303625.1 | Rhizobium_leguminosarum | ELI01_RS08965 | 1760640..1764263 | 384 | BREX-system-P-loop-protein-BrxC-[Rhizobium-leguminosarum] |
| WP_130755622.1 | Type1.BREX | [illegible] | 1174 | Bacteria>Pseudomonadota>Alphaproteobacteria>Hyphomicrobiales>Rhizobiaceae>Rhizobium/Agrobacterium_group | GCF_004304705.1 | Rhizobium_leguminosarum | ELH79_RS02355 | 502725..506249 | 384 | BREX-system-P-loop-protein-BrxC-[Rhizobium-leguminosarum] |
| WP_130805164.1 | Type1.BREX | [illegible] | 1207 | Bacteria>Pseudomonadota>Alphaproteobacteria>Hyphomicrobiales>Rhizobiaceae>Rhizobium/Agrobacterium_group | GCF_004305935.1 | Rhizobium_leguminosarum | ELG76_RS08485 | 1639412..1643035 | 384 | BREX-system-P-loop-protein-BrxC-[Rhizobium-leguminosarum] |
| WP_130816714.1 | Type1.BREX | [illegible] | 1137 | Bacteria>Pseudomonadota>Alphaproteobacteria>Hyphomicrobiales>Rhizobiaceae>Rhizobium/Agrobacterium_group | GCF_004306815.1 | Rhizobium_leguminosarum | ELG72_RS09600 | 1900026..1903439 | 384 | BREX-system-P-loop-protein-BrxC-[Rhizobium-leguminosarum] |
| WP_131718742.1 | Type1.BREX | [illegible] | 1178 | Bacteria>Pseudomonadota>Alphaproteobacteria>Hyphomicrobiales>Rhizobiaceae>Rhizobium/Agrobacterium_group | GCF_004330655.1 | Rhizobium_leguminosarum | E0H57_RS05710 | 350884..354420 | 384 | BREX-system-P-loop-protein-BrxC-[Rhizobium-leguminosarum] |
| WP_164009387.1 | Type1.BREX | [illegible] | 1137 | Bacteria>Pseudomonadota>Alphaproteobacteria>Hyphomicrobiales>Rhizobiaceae>Rhizobium/Agrobacterium_group | GCF_010668805.1 | Rhizobium_phaseoli | GR197_RS11280 | 144008..147421 | 396 | BREX-system-P-loop-protein-BrxC-[Rhizobium-phaseoli] |
| WP_168337993.1 | Type1.BREX | [illegible] | 1174 | Bacteria>Pseudomonadota>Alphaproteobacteria>Hyphomicrobiales>Rhizobiaceae>Rhizobium/Agrobacterium_group | GCF_012275885.1 | Rhizobium_laguerreae | GFL43_RS28845 | 8439..11963 | 1076926 | BREX-system-P-loop-protein-BrxC-[Rhizobium-laguerreae] |
| WP_168349960.1 | Type1.BREX | [illegible] | 190 | Bacteria>Pseudomonadota>Alphaproteobacteria>Hyphomicrobiales>Rhizobiaceae>Rhizobium/Agrobacterium_group | GCF_012276085.1 | Rhizobium_leguminosarum | GFM28_RS29310 | 8641..9213 | 384 | hypothetical-protein-[Rhizobium-leguminosarum] |
| WP_182828337.1 | Type1.BREX | [illegible] | 1150 | Bacteria>Pseudomonadota>Alphaproteobacteria>Hyphomicrobiales>Rhizobiaceae>Rhizobium/Agrobacterium_group | GCF_014158035.1 | Rhizobium_sp._G21 | H2014_RS15370 | 3060977..3064429 | 2758439 | BREX-system-P-loop-protein-BrxC-[Rhizobium-sp.-G21] |
| WP_136542766.1 | Type1.BREX | [illegible] | 1179 | Bacteria>Pseudomonadota>Alphaproteobacteria>Hyphomicrobiales>Rhizobiaceae>Rhizobium/Agrobacterium_group | GCF_014202175.1 | Rhizobium_rosettiformans | FAA86_RS19555 | 64899..68438 | 1368430 | BREX-system-P-loop-protein-BrxC-[Rhizobium-rosettiformans] |
| WP_192117159.1 | Type1.BREX | [illegible] | 1174 | Bacteria>Pseudomonadota>Alphaproteobacteria>Hyphomicrobiales>Rhizobiaceae>Rhizobium/Agrobacterium_group | GCF_014841565.1 | Rhizobium_sp._CFBP_8762 | IFT84_RS03115 | 659482..663006 | 2775279 | BREX-system-P-loop-protein-BrxC-[Rhizobium-sp.-CFBP-8762] |
| WP_215570403.1 | Type1.BREX | [illegible] | 1175 | Bacteria>Pseudomonadota>Alphaproteobacteria>Hyphomicrobiales>Rhizobiaceae>Rhizobium/Agrobacterium_group | GCF_018760785.1 | Rhizobium_sp._CSW-27 | KLP42_RS24160 | 30886..34413 | 2839985 | BREX-system-P-loop-protein-BrxC-[Rhizobium-sp.-CSW-27] |
| WP_246638929.1 | Type1.BREX | [illegible] | 1170 | Bacteria>Pseudomonadota>Alphaproteobacteria>Hyphomicrobiales>Rhizobiaceae>Rhizobium/Agrobacterium_group | GCF_019778505.1 | Rhizobium_laguerreae | HFO72_RS21050 | 353201..356713 | 1076926 | BREX-system-P-loop-protein-BrxC-[Rhizobium-laguerreae] |
| WP_222012036.1 | Type1.BREX | [illegible] | 1141 | Bacteria>Pseudomonadota>Alphaproteobacteria>Hyphomicrobiales>Rhizobiaceae>Rhizobium/Agrobacterium_group | GCF_019778925.1 | Rhizobium_laguerreae | HFO18_RS33045 | 24769..28194 | 1076926 | BREX-system-P-loop-protein-BrxC-[Rhizobium-laguerreae] |
| WP_222013551.1 | Type1.BREX | [illegible] | 1137 | Bacteria>Pseudomonadota>Alphaproteobacteria>Hyphomicrobiales>Rhizobiaceae>Rhizobium/Agrobacterium_group | GCF_019779115.1 | Rhizobium_laguerreae | HFO05_RS34495 | 7944..11357 | 1076926 | BREX-system-P-loop-protein-BrxC-[Rhizobium-laguerreae] |
| WP_222362949.1 | Type1.BREX | [illegible] | 1174 | Bacteria>Pseudomonadota>Alphaproteobacteria>Hyphomicrobiales>Rhizobiaceae>Rhizobium/Agrobacterium_group | GCF_019798305.1 | Rhizobium_leguminosarum | HFO99_RS19360 | 191490..195014 | 384 | BREX-system-P-loop-protein-BrxC-[Rhizobium-leguminosarum] |
| WP_131591195.1 | Type1.BREX | [illegible] | 1137 | Bacteria>Pseudomonadota>Alphaproteobacteria>Hyphomicrobiales>Rhizobiaceae>Rhizobium/Agrobacterium_group | GCF_019798685.1 | Rhizobium_leguminosarum | E0H55_RS21895 | 87979..91392 | 384 | BREX-system-P-loop-protein-BrxC-[Rhizobium-leguminosarum] |
| WP_222263630.1 | Type1.BREX | [illegible] | 1141 | Bacteria>Pseudomonadota>Alphaproteobacteria>Hyphomicrobiales>Rhizobiaceae>Rhizobium/Agrobacterium_group | GCF_019798865.1 | Rhizobium_leguminosarum | HFO42_RS34260 | 4774..8199 | 384 | BREX-system-P-loop-protein-BrxC-[Rhizobium-leguminosarum] |
| WP_222338071.1 | Type1.BREX | [illegible] | 1141 | Bacteria>Pseudomonadota>Alphaproteobacteria>Hyphomicrobiales>Rhizobiaceae>Rhizobium/Agrobacterium_group | GCF_019799025.1 | Rhizobium_leguminosarum | HFO50_RS18885 | 16648..20073 | 384 | BREX-system-P-loop-protein-BrxC-[Rhizobium-leguminosarum] |
| WP_222342569.1 | Type1.BREX | [illegible] | 1174 | Bacteria>Pseudomonadota>Alphaproteobacteria>Hyphomicrobiales>Rhizobiaceae>Rhizobium/Agrobacterium_group | GCF_019799465.1 | Rhizobium_leguminosarum | HFO12_RS18165 | 9414..12938 | 384 | BREX-system-P-loop-protein-BrxC-[Rhizobium-leguminosarum] |
| WP_229574330.1 | Type1.BREX | [illegible] | 1137 | Bacteria>Pseudomonadota>Alphaproteobacteria>Hyphomicrobiales>Rhizobiaceae>Rhizobium/Agrobacterium_group | GCF_020889605.1 | Rhizobium_sp._'Codium_1' | LJE98_RS06515 | 266612..270025 | 2940484 | BREX-system-P-loop-protein-BrxC-[Rhizobium-sp.-'Codium-1'] |
| WP_231283598.1 | Type1.BREX | [illegible] | 1207 | Bacteria>Pseudomonadota>Alphaproteobacteria>Hyphomicrobiales>Rhizobiaceae>Rhizobium/Agrobacterium_group | GCF_021052345.1 | Rhizobium_leguminosarum | RlegSU303_RS06785 | 1350799..1354422 | 384 | BREX-system-P-loop-protein-BrxC-[Rhizobium-leguminosarum] |
| WP_221976415.1 | Type1.BREX | [illegible] | 1174 | Bacteria>Pseudomonadota>Alphaproteobacteria>Hyphomicrobiales>Rhizobiaceae>Rhizobium/Agrobacterium_group | GCF_021391715.1 | Rhizobium | LZK79_RS04540 | 933670..937194 | 379 | MULTISPECIES:-BREX-system-P-loop-protein-BrxC-[Rhizobium] |
| WP_239619846.1 | Type1.BREX | [illegible] | 1174 | Bacteria>Pseudomonadota>Alphaproteobacteria>Hyphomicrobiales>Rhizobiaceae>Rhizobium/Agrobacterium_group | GCF_022354505.1 | Rhizobium_sp._C104 | MF410_RS14985 | 2774142..2777666 | 2917727 | BREX-system-P-loop-protein-BrxC-[Rhizobium-sp.-C104] |
| WP_245134299.1 | Type1.BREX | [illegible] | 1177 | Bacteria>Pseudomonadota>Alphaproteobacteria>Hyphomicrobiales>Rhizobiaceae>Rhizobium/Agrobacterium_group | GCF_030369675.1 | Rhizobium_sp._SSM4.3 | MKJ03_RS01550 | 316527..320060 | 2919917 | BREX-system-P-loop-protein-BrxC-[Rhizobium-sp.-SSM4.3] |
| WP_077987199.1 | Type1.BREX | [illegible] | 1175 | Bacteria>Pseudomonadota>Alphaproteobacteria>Hyphomicrobiales>Rhizobiaceae>Rhizobium/Agrobacterium_group | GCF_030440405.1 | Rhizobium_pusense | BTE56_RS17195 | 18928..22455 | 648995 | BREX-system-P-loop-protein-BrxC-[Rhizobium-pusense] |
| WP_149748055.1 | Type1.BREX | [illegible] | 1175 | Bacteria>Pseudomonadota>Alphaproteobacteria>Hyphomicrobiales>Rhizobiaceae>Rhizobium/Agrobacterium_group | GCF_900156005.1 | Rhizobium_sp._RU35A | F3H95_RS25745 | 3768..7295 | 1907414 | BREX-system-P-loop-protein-BrxC-[Rhizobium-sp.-RU35A] |
| WP_245278427.1 | Type1.BREX | [illegible] | 1170 | Bacteria>Pseudomonadota>Alphaproteobacteria>Hyphomicrobiales>Rhizobiaceae>Rhizobium/Agrobacterium_group | - | Rhizobium_leguminosarum | A3A3_RS0116705 | 192694..196206 | 384 | BREX-system-P-loop-protein-BrxC-[Rhizobium-leguminosarum] |
| WP_131707092.1 | Type1.BREX | [illegible] | 1174 | Bacteria>Pseudomonadota>Alphaproteobacteria>Hyphomicrobiales>Rhizobiaceae>Rhizobium/Agrobacterium_group | - | Rhizobium_leguminosarum | E0H59_RS18255 | 300646..304170 | 384 | BREX-system-P-loop-protein-BrxC-[Rhizobium-leguminosarum] |
| WP_024321352.1 | Type1.BREX | [illegible] | 1174 | Bacteria>Pseudomonadota>Alphaproteobacteria>Hyphomicrobiales>Rhizobiaceae>Rhizobium/Agrobacterium_group | - | Rhizobium_leguminosarum | E0J20_RS04835 | 93871..97395 | 384 | BREX-system-P-loop-protein-BrxC-[Rhizobium-leguminosarum] |
| WP_130784075.1 | Type1.BREX | [illegible] | 1178 | Bacteria>Pseudomonadota>Alphaproteobacteria>Hyphomicrobiales>Rhizobiaceae>Rhizobium/Agrobacterium_group | - | Rhizobium_leguminosarum | ELG96_RS28915 | 11184..14720 | 384 | BREX-system-P-loop-protein-BrxC-[Rhizobium-leguminosarum] |
| WP_018070454.1 | Type1.BREX | [illegible] | 1207 | Bacteria>Pseudomonadota>Alphaproteobacteria>Hyphomicrobiales>Rhizobiaceae>Rhizobium/Agrobacterium_group | - | Rhizobium_leguminosarum | ELH14_RS07055 | 1387374..1390997 | 384 | BREX-system-P-loop-protein-BrxC-[Rhizobium-leguminosarum] |
| WP_130823534.1 | Type1.BREX | [illegible] | 1137 | Bacteria>Pseudomonadota>Alphaproteobacteria>Hyphomicrobiales>Rhizobiaceae>Rhizobium/Agrobacterium_group | - | Rhizobium_leguminosarum | ELH15_RS08895 | 1753674..1757087 | 384 | BREX-system-P-loop-protein-BrxC-[Rhizobium-leguminosarum] |
| WP_130764093.1 | Type1.BREX | [illegible] | 1174 | Bacteria>Pseudomonadota>Alphaproteobacteria>Hyphomicrobiales>Rhizobiaceae>Rhizobium/Agrobacterium_group | - | Rhizobium_leguminosarum | ELI00_RS29085 | 22850..26374 | 384 | BREX-system-P-loop-protein-BrxC-[Rhizobium-leguminosarum] |
| WP_245488667.1 | Type1.BREX | [illegible] | 1170 | Bacteria>Pseudomonadota>Alphaproteobacteria>Hyphomicrobiales>Rhizobiaceae>Rhizobium/Agrobacterium_group | - | Rhizobium_leguminosarum | ELI09_RS02630 | 538702..542214 | 384 | BREX-system-P-loop-protein-BrxC-[Rhizobium-leguminosarum] |
| WP_130672003.1 | Type1.BREX | [illegible] | 1174 | Bacteria>Pseudomonadota>Alphaproteobacteria>Hyphomicrobiales>Rhizobiaceae>Rhizobium/Agrobacterium_group | - | Rhizobium_leguminosarum | ELI10_RS02625 | 537164..540688 | 384 | BREX-system-P-loop-protein-BrxC-[Rhizobium-leguminosarum] |
| WP_165586283.1 | Type1.BREX | [illegible] | 1207 | Bacteria>Pseudomonadota>Alphaproteobacteria>Hyphomicrobiales>Rhizobiaceae>Rhizobium/Agrobacterium_group | - | Rhizobium_leguminosarum | G7039_RS08625 | 1692346..1695969 | 384 | BREX-system-P-loop-protein-BrxC-[Rhizobium-leguminosarum] |
| WP_168316672.1 | Type1.BREX | [illegible] | 1181 | Bacteria>Pseudomonadota>Alphaproteobacteria>Hyphomicrobiales>Rhizobiaceae>Rhizobium/Agrobacterium_group | - | Rhizobium_leguminosarum | GFL49_RS25835 | 227535..231080 | 384 | BREX-system-P-loop-protein-BrxC-[Rhizobium-leguminosarum] |
| WP_168308740.1 | Type1.BREX | [illegible] | 1181 | Bacteria>Pseudomonadota>Alphaproteobacteria>Hyphomicrobiales>Rhizobiaceae>Rhizobium/Agrobacterium_group | - | Rhizobium_leguminosarum | GFM13_RS06520 | 103411..106956 | 384 | BREX-system-P-loop-protein-BrxC-[Rhizobium-leguminosarum] |
| WP_246708550.1 | Type1.BREX | [illegible] | 1170 | Bacteria>Pseudomonadota>Alphaproteobacteria>Hyphomicrobiales>Rhizobiaceae>Rhizobium/Agrobacterium_group | - | Rhizobium_leguminosarum | GR207_RS02230 | 29211..32723 | 384 | BREX-system-P-loop-protein-BrxC-[Rhizobium-leguminosarum] |
| PZU19146.1 | Type1.BREX | [illegible] | 1175 | Bacteria>Pseudomonadota>Alphaproteobacteria>Hyphomicrobiales>Rhizobiaceae>Shinella | GCA_003248545.1 | Shinella_sp. | DI589_22170 | 12827..16354 | 1870904 | MAG:-BREX-system-P-loop-protein-BrxC-[Shinella-sp.] |
| WP_004434288.1 | Type1.BREX | [illegible] | 1177 | Bacteria>Pseudomonadota>Alphaproteobacteria>Hyphomicrobiales>Rhizobiaceae>Sinorhizobium/Ensifer_group | GCF_000236945.1 | Sinorhizobium_meliloti | SM0020_RS29775 | 166866..170399 | 382 | BREX-system-P-loop-protein-BrxC-[Sinorhizobium-meliloti] |
| WP_037424884.1 | Type1.BREX | [illegible] | 1181 | Bacteria>Pseudomonadota>Alphaproteobacteria>Hyphomicrobiales>Rhizobiaceae>Sinorhizobium/Ensifer_group | GCF_000261005.1 | Sinorhizobium_sp._CCBAU_05631 | SS05631_RS30325 | 18322..21867 | 794846 | BREX-system-P-loop-protein-BrxC-[Sinorhizobium-sp.-CCBAU-05631] |
| WP_244425598.1 | Type1.BREX | [illegible] | 705 | Bacteria>Pseudomonadota>Alphaproteobacteria>Hyphomicrobiales>Rhizobiaceae>Sinorhizobium/Ensifer_group | GCF_000261865.1 | Sinorhizobium_fredii | IC47_RS36510 | 5889..8006 | 380 | BREX-system-P-loop-protein-BrxC-[Sinorhizobium-fredii] |
| WP_028054052.1 | Type1.BREX | [illegible] | 1150 | Bacteria>Pseudomonadota>Alphaproteobacteria>Hyphomicrobiales>Rhizobiaceae>Sinorhizobium/Ensifer_group | GCF_000427485.1 | Sinorhizobium_medicae | SINMEL_RS0108765 | 254501..257953 | 110321 | BREX-system-P-loop-protein-BrxC-[Sinorhizobium-medicae] |
| WP_164817971.1 | Type1.BREX | [illegible] | 1141 | Bacteria>Pseudomonadota>Alphaproteobacteria>Hyphomicrobiales>Rhizobiaceae>Sinorhizobium/Ensifer_group | GCF_004001385.1 | Sinorhizobium_meliloti | CN238_RS30780 | 8247..11672 | 382 | BREX-system-P-loop-protein-BrxC-[Sinorhizobium-meliloti] |
| WP_127638472.1 | Type1.BREX | [illegible] | 1188 | Bacteria>Pseudomonadota>Alphaproteobacteria>Hyphomicrobiales>Rhizobiaceae>Sinorhizobium/Ensifer_group | GCF_004003915.1 | Sinorhizobium_meliloti | CN087_RS12760 | 106891..110457 | 382 | BREX-system-P-loop-protein-BrxC-[Sinorhizobium-meliloti] |
| CAG0989916.1 | Type1.BREX | [illegible] | 1140 | Bacteria>Pseudomonadota>Alphaproteobacteria>Hyphomicrobiales>Rhizobiaceae>unclassified_Rhizobiaceae | GCA_905339215.1 | Rhizobiaceae_bacterium | RHIZO_02194 | 11543..14965 | 1913961 | hypothetical-protein-RHIZO_02194-[Rhizobiaceae-bacterium] |
| WP_027288600.1 | Type1.BREX | [illegible] | 1150 | Bacteria>Pseudomonadota>Alphaproteobacteria>Hyphomicrobiales>Rhodovibrionaceae>Rhodovibrio | GCF_000515255.1 | Rhodovibrio_salinarum | RHOSA_RS0110320 | 2222437..2225889 | 1087 | BREX-system-P-loop-protein-BrxC-[Rhodovibrio-salinarum] |
| WP_074444907.1 | Type1.BREX | [illegible] | 1148 | Bacteria>Pseudomonadota>Alphaproteobacteria>Hyphomicrobiales>Salinarimonadaceae>Saliniramus | GCF_900094735.1 | Saliniramus_fredricksonii | GA0071312_RS10335 | 795252..798698 | 1653334 | BREX-system-P-loop-protein-BrxC-[Saliniramus-fredricksonii] |
| WP_035540926.1 | Type1.BREX | [illegible] | 1157 | Bacteria>Pseudomonadota>Alphaproteobacteria>Hyphomonadales>Hyphomonadaceae>Hyphomonas | GCF_000685295.1 | Hyphomonas_oceanitis | HOC_RS17370 | 29720..33193 | 81033 | BREX-system-P-loop-protein-BrxC-[Hyphomonas-oceanitis] |
| MCE2892634.1 | Type1.BREX | [illegible] | 1195 | Bacteria>Pseudomonadota>Alphaproteobacteria>Hyphomonadales>Hyphomonadaceae>unclassified_Hyphomonadaceae | GCA_021299265.1 | Hyphomonadaceae_bacterium | LW848_16090 | 9073..12660 | 2026748 | BREX-system-P-loop-protein-BrxC-[Hyphomonadaceae-bacterium] |
| WP_130470524.1 | Type1.BREX | [illegible] | 1193 | Bacteria>Pseudomonadota>Alphaproteobacteria>Magnetococcales>Candidatus_Magnetaquicoccaceae>Candidatus_Magnetaquicoccus | GCF_004217665.1 | Candidatus_Magnetaquicoccus_inordinatus | EJP87_RS03730 | 21320..24901 | 2496818 | BREX-system-P-loop-protein-BrxC-[Candidatus-Magnetaquicoccus-inordinatus] |
| MBF0098005.1 | Type1.BREX | [illegible] | 1144 | Bacteria>Pseudomonadota>Alphaproteobacteria>Magnetococcales>unclassified_Magnetococcales | GCA_015228825.1 | Magnetococcales_bacterium | HQM05_11780 | 14372..17806 | 2026759 | BREX-system-P-loop-protein-BrxC-[Magnetococcales-bacterium] |
| MBF0399382.1 | Type1.BREX | [illegible] | 1144 | Bacteria>Pseudomonadota>Alphaproteobacteria>Magnetococcales>unclassified_Magnetococcales | GCA_015232635.1 | Magnetococcales_bacterium | HQL90_01290 | 18375..21809 | 2026759 | BREX-system-P-loop-protein-BrxC-[Magnetococcales-bacterium] |
| ADM09203.1 | Type1.BREX | [illegible] | 1171 | Bacteria>Pseudomonadota>Alphaproteobacteria>Parvularculales>Parvularculaceae>Parvularcula | GCA_000152825.2 | Parvularcula_bermudensis_HTCC2503 | PB2503_05652 | 1076655..1080170 | 314260 | hypothetical-protein-PB2503_05652-[Parvularcula-bermudensis-HTCC2503] |
| WP_238525821.1 | Type1.BREX | [illegible] | 1168 | Bacteria>Pseudomonadota>Alphaproteobacteria>Parvularculales>Parvularculaceae>Parvularcula | GCF_000152825.2 | Parvularcula_bermudensis | PB2503_RS05170 | 1076655..1080161 | 208216 | BREX-system-P-loop-protein-BrxC-[Parvularcula-bermudensis] |
| WP_217361198.1 | Type1.BREX | [illegible] | 1168 | Bacteria>Pseudomonadota>Alphaproteobacteria>Rhodobacterales>Paracoccaceae>Aliiroseovarius | GCF_013283115.1 | unclassified_Aliiroseovarius | XMV209_RS00060 | 11614..15120 | 2623558 | MULTISPECIES:-BREX-system-P-loop-protein-BrxC-[unclassified-Aliiroseovarius] |
| WP_173486905.1 | Type1.BREX | [illegible] | 1173 | Bacteria>Pseudomonadota>Alphaproteobacteria>Rhodobacterales>Paracoccaceae>Aliiroseovarius | GCF_013283525.1 | unclassified_Aliiroseovarius | XMM309_RS11965 | 259901..263422 | 2623558 | MULTISPECIES:-BREX-system-P-loop-protein-BrxC-[unclassified-Aliiroseovarius] |
| WP_035840066.1 | Type1.BREX | [illegible] | 1208 | Bacteria>Pseudomonadota>Alphaproteobacteria>Rhodobacterales>Paracoccaceae>Allgaiera | GCF_014653075.1 | Allgaiera_indica | IE192_RS04280 | 311977..315603 | 765699 | BREX-system-P-loop-protein-BrxC-[Allgaiera-indica] |
| WP_245513828.1 | Type1.BREX | [illegible] | 659 | Bacteria>Pseudomonadota>Alphaproteobacteria>Rhodobacterales>Paracoccaceae>Antarcticimicrobium | GCF_004358185.1 | Antarcticimicrobium_luteum | E1832_RS04855 | 61315..63294 | 2547397 | BREX-system-P-loop-protein-BrxC-[Antarcticimicrobium-luteum] |
| WP_146284416.1 | Type1.BREX | [illegible] | 1173 | Bacteria>Pseudomonadota>Alphaproteobacteria>Rhodobacterales>Paracoccaceae>Gemmobacter | GCF_007859075.1 | Gemmobacter_aquaticus | FQU74_RS00760 | 54180..57701 | 490185 | BREX-system-P-loop-protein-BrxC-[Gemmobacter-aquaticus] |
| WP_076528450.1 | Type1.BREX | [illegible] | 1179 | Bacteria>Pseudomonadota>Alphaproteobacteria>Rhodobacterales>Paracoccaceae>Gemmobacter | GCF_014636395.1 | Gemmobacter_megaterium | IEV79_RS02805 | 604470..608009 | 1086013 | BREX-system-P-loop-protein-BrxC-[Gemmobacter-megaterium] |
| WP_229704463.1 | Type1.BREX | [illegible] | 1169 | Bacteria>Pseudomonadota>Alphaproteobacteria>Rhodobacterales>Paracoccaceae>Gemmobacter | GCF_014645295.1 | Gemmobacter_aquaticus | IEX45_RS17475 | 6610..10119 | 490185 | BREX-system-P-loop-protein-BrxC-[Gemmobacter-aquaticus] |
| WP_131389735.1 | Type1.BREX | [illegible] | 1179 | Bacteria>Pseudomonadota>Alphaproteobacteria>Rhodobacterales>Paracoccaceae>Haematobacter | GCF_004328555.1 | Haematobacter_massiliensis | HmaOT1_RS15910 | 65338..68877 | 195105 | BREX-system-P-loop-protein-BrxC-[Haematobacter-massiliensis] |
| WP_039019257.1 | Type1.BREX | [illegible] | 1172 | Bacteria>Pseudomonadota>Alphaproteobacteria>Rhodobacterales>Paracoccaceae>Halocynthiibacter | GCF_000812685.1 | Halocynthiibacter_namhaensis | RJ38_RS12670 | 13538..17056 | 1290553 | BREX-system-P-loop-protein-BrxC-[Halocynthiibacter-namhaensis] |
| WP_263953721.1 | Type1.BREX | [illegible] | 1172 | Bacteria>Pseudomonadota>Alphaproteobacteria>Rhodobacterales>Paracoccaceae>Halocynthiibacter | GCF_025802015.1 | unclassified_Halocynthiibacter | OH136_RS09920 | 459600..463118 | 2640401 | MULTISPECIES:-BREX-system-P-loop-protein-BrxC-[unclassified-Halocynthiibacter] |
| WP_274653192.1 | Type1.BREX | [illegible] | 1172 | Bacteria>Pseudomonadota>Alphaproteobacteria>Rhodobacterales>Paracoccaceae>Halocynthiibacter | GCF_028823805.1 | Halocynthiibacter_sp._C4 | OU789_RS03095 | 642215..645733 | 2992758 | BREX-system-P-loop-protein-BrxC-[Halocynthiibacter-sp.-C4] |
| WP_269398497.1 | Type1.BREX | [illegible] | 1171 | Bacteria>Pseudomonadota>Alphaproteobacteria>Rhodobacterales>Paracoccaceae>Limimaricola | GCF_027213785.1 | Limimaricola_sp._G21655-S1 | O4G76_RS17300 | 20007..23522 | 3014768 | BREX-system-P-loop-protein-BrxC-[Limimaricola-sp.-G21655-S1] |
| WP_094023246.1 | Type1.BREX | [illegible] | 1175 | Bacteria>Pseudomonadota>Alphaproteobacteria>Rhodobacterales>Paracoccaceae>Maliponia | GCF_900184945.1 | Maliponia_aquimaris | MAA8898_RS22355 | 47898..51425 | 1673631 | BREX-system-P-loop-protein-BrxC-[Maliponia-aquimaris] |
| MBO6774806.1 | Type1.BREX | [illegible] | 1172 | Bacteria>Pseudomonadota>Alphaproteobacteria>Rhodobacterales>Paracoccaceae>Marinibacterium | GCA_017643305.1 | Marinibacterium_sp. | JJ897_04735 | 11284..14802 | 1931201 | BREX-system-P-loop-protein-BrxC-[Marinibacterium-sp.] |
| APO88218.1 | Type1.BREX | [illegible] | 1166 | Bacteria>Pseudomonadota>Alphaproteobacteria>Rhodobacterales>Paracoccaceae>Marivivens_group | GCA_001908835.1 | Marivivens_sp._JLT3646 | BSK21_01620 | 313072..316572 | 1920883 | hypothetical-protein-BSK21_01620-[Marivivens-sp.-JLT3646] |
| WP_017998950.1 | Type1.BREX | [illegible] | 1177 | Bacteria>Pseudomonadota>Alphaproteobacteria>Rhodobacterales>Paracoccaceae>Paracoccus | GCF_000371965.1 | Paracoccus_sp._N5 | PARN5_RS0106405 | 1284773..1288306 | 1101189 | BREX-system-P-loop-protein-BrxC-[Paracoccus-sp.-N5] |

| Protein-ID | System | Gene-Neighbourhoods | Length | Taxonomy | Assembly-ID | Strain | Locus-Tag | Genomic-coordinates | TaxID | Fasta-define |
|---|---|---|---|---|---|---|---|---|---|---|
| WP_042245816.1 | Type1.BREX | DDE_Tnp->DDE_Tnp-><-HKD-DNase+Preader14<-SIGMA+OB-Fold+AAA-ATPase+HKD-DNase-BrxL<-PglZ<-PglZ-p<-N6-METHYLASE-p<-N6-METHYLASE<-DUF6079*<-BrxB-BrxF-iSTAND<-BrxA-DN [illegible] | 1174 | Bacteria>Pseudomonadota>Alphaproteobacteria>Rhodobacterales>Paracoccaceae>Paracoccus | GCF_000787695.1 | Paracoccus_sp._PAMC_22219 | TY51_RS03260 | 14723..18247 | 1569209 | BREX-system-P-loop-protein-BrxC-[Paracoccus-sp.-PAMC-22219] |
| WP_046001801.1 | Type1.BREX | TarH+HAMP+MA-><-Glyoxalase<-?<-ARG\|\|WXG-><-PIN<-SIGMA+OB-Fold+AAA-ATPase+HKD-DNase-BrxL<-PglZ<-N6-METHYLASE<-DUF6079*<-BrxB-BrxF-iSTAND<-BrxA-DNA-binding<-35exo [illegible] | 1174 | Bacteria>Pseudomonadota>Alphaproteobacteria>Rhodobacterales>Paracoccaceae>Paracoccus | GCF_000967825.1 | Paracoccus_sp._228 | SY26_RS18685 | 30060..33584 | 1192054 | BREX-system-P-loop-protein-BrxC-[Paracoccus-sp.-228] |
| WP_199257821.1 | Type1.BREX | GDResolvase+Recombinase+ZnR-><-SF1-SF2-Helicase<-HKD-DNase+HKD-DNase<-wHTH<-SF1-SF2-Helicase+REase<-SF1-SF2-Helicase+REase<-SIGMA+OB-Fold+AAA-ATPase+HKD-DNase-Br [illegible] | 1179 | Bacteria>Pseudomonadota>Alphaproteobacteria>Rhodobacterales>Paracoccaceae>Paracoccus | GCF_016446475.1 | Paracoccus_binzhouensis | I9P22_RS02450 | 23714..27253 | 2796149 | BREX-system-P-loop-protein-BrxC-[Paracoccus-binzhouensis] |
| WP_199258165.1 | Type1.BREX | <-N6-METHYLASE-p<-REase<-OLD-ABC<-DUF6079*<-BrxB-BrxF-iSTAND<-BrxA-DNA-binding<-wHTH+WYL+WCX\|\|?->Thermonuclease->TrbM-><-?<-HTH_3 | 1173 | Bacteria>Pseudomonadota>Alphaproteobacteria>Rhodobacterales>Paracoccaceae>Paracoccus | GCF_016446475.1 | Paracoccus_binzhouensis | I9P22_RS04365 | 3344..6865 | 2796149 | BREX-system-P-loop-protein-BrxC-[Paracoccus-binzhouensis] |
| WP_260276663.1 | Type1.BREX | URI-DNase+RAMA->XIS-HTH->wHTH+WYL+WCX->BrxA-DNA-binding->BrxB-BrxF-iSTAND->DUF6079*->N6-METHYLASE->PglZ->SIGMA+OB-Fold+AAA-ATPase+HKD-DNase-BrxL->URI-DNase+RAMA- [illegible] | 1173 | Bacteria>Pseudomonadota>Alphaproteobacteria>Rhodobacterales>Paracoccaceae>Paracoccus | GCF_025195095.1 | Paracoccus_sp._YLB-12 | MU516_RS07900 | 40235..43756 | 2933292 | BREX-system-P-loop-protein-BrxC-[Paracoccus-sp.-YLB-12] |
| WP_271109779.1 | Type1.BREX | MerR-HTH-><-HTH_Tnp_1+HTH_28+HTH_32+rve+rve_3\|\|BrxB-BrxF-iSTAND->DUF6079*->DDE_Tnp_ISL3-> | 1173 | Bacteria>Pseudomonadota>Alphaproteobacteria>Rhodobacterales>Paracoccaceae>Paracoccus | GCF_027912675.1 | Paracoccus_sp._SCSIO_75233 | PAF12_17375 | 77006..80527 | 3017782 | BREX-system-P-loop-protein-BrxC-[Paracoccus-sp.-SCSIO-75233] |
| WP_271118256.1 | Type1.BREX | HTH_3->?-><-TrbM<-Thermonuclease<-?\|\|wHTH+WYL+WCX->BrxA-DNA-binding->BrxB-BrxF-iSTAND->DUF6079*->N6-METHYLASE->PglZ->SIGMA+OB-Fold+AAA-ATPase+HKD-DNase-BrxL->HIN [illegible] | 1173 | Bacteria>Pseudomonadota>Alphaproteobacteria>Rhodobacterales>Paracoccaceae>Paracoccus | GCF_027912835.1 | Paracoccus_sp._SCSIO_76264 | PAF18_RS16795 | 74012..77533 | 3017783 | BREX-system-P-loop-protein-BrxC-[Paracoccus-sp.-SCSIO-76264] |
| WP_149766716.1 | Type1.BREX | <-HTH<-Classical-AAA<-?<-?\|\|?->AAA+HTH->wHTH+WYL+WCX->BrxA-DNA-binding->BrxB-BrxF-iSTAND->DUF6079*->PglZ->SIGMA+OB-Fold+AAA-ATPase+HKD-DNase-BrxL->?->ParB-> | 1177 | Bacteria>Pseudomonadota>Alphaproteobacteria>Rhodobacterales>Paracoccaceae>Paracoccus | GCF_900156255.1 | Paracoccus_thiocyanatus | F3H32_RS16120 | 13661..17194 | 34006 | BREX-system-P-loop-protein-BrxC-[Paracoccus-thiocyanatus] |
| WP_085502591.1 | Type1.BREX | <-ABhydrolase+ABhydrolase<-?\|\|MODE-HTH+PBPII-><-SbcC+SbcC<-Calcineurin+Calcineurin-SbcD\|\|wHTH+WYL+WCX->BrxA-DNA-binding->BrxB-BrxF-iSTAND->DUF6079*->N6-METHYLASE [illegible] | 1179 | Bacteria>Pseudomonadota>Alphaproteobacteria>Rhodobacterales>Paracoccaceae>Paracoccus | GCF_900177635.1 | Paracoccus_sp._J56 | B9Y79_RS14640 | 24785..28324 | 935850 | BREX-system-P-loop-protein-BrxC-[Paracoccus-sp.-J56] |
| WP_113913307.1 | Type1.BREX | SIR2-><-?<-REase<-Immank+ANK+ANK<-GreA_GreB+SF1-SF2-Helicase<-SIGMA+OB-Fold+AAA-ATPase+HKD-DNase-BrxL<-PglZ<-N6-METHYLASE<-DUF6079*<-BrxB-BrxF-iSTAND<-BrxA-DNA-b [illegible] | 1179 | Bacteria>Pseudomonadota>Alphaproteobacteria>Rhodobacterales>Paracoccaceae>Pelagivirga | GCF_003316635.1 | Pelagivirga_dicentrarchi | DQ230_RS14395 | 19757..23296 | 2250573 | BREX-system-P-loop-protein-BrxC-[Pelagivirga-dicentrarchi] |
| NYS26401.1 | Type1.BREX | <-N6-METHYLASE-p<-DUF6079*<-DUF6079-p<-BrxB-BrxF-iSTAND<-BrxA-DNA-binding<-MetJarc-HTH<-?<-GDResolvase+ZnR<-RAMA | 1025 | Bacteria>Pseudomonadota>Alphaproteobacteria>Rhodobacterales>Paracoccaceae>Rhabdonatronobacter | GCA_013415485.1 | Rhabdonatronobacter_sediminivivens | HUK65_15550 | 1855..4932 | 2743469 | BREX-system-P-loop-protein-BrxC-[Rhabdonatronobacter-sediminivivens] |
| WP_268839433.1 | Type1.BREX | <-N6-METHYLASE-p<-DUF6079*<-DUF6079-p<-BrxB-BrxF-iSTAND<-BrxA-DNA-binding<-?<-GDResolvase+ZnR<-RAMA | 1018 | Bacteria>Pseudomonadota>Alphaproteobacteria>Rhodobacterales>Paracoccaceae>Rhabdonatronobacter | GCF_013415485.1 | Rhabdonatronobacter_sediminivivens | HUK65_RS15575 | 1855..4911 | 2743469 | BREX-system-P-loop-protein-BrxC-[Rhabdonatronobacter-sediminivivens] |
| WP_179907198.1 | Type1.BREX | <-N6-METHYLASE-p<-DUF6079<-DUF6079-p*<-BrxB-BrxF-iSTAND<-BrxA-DNA-binding<-?<-GDResolvase+ZnR<-RAMA | 105 | Bacteria>Pseudomonadota>Alphaproteobacteria>Rhodobacterales>Paracoccaceae>Rhabdonatronobacter | GCF_013415485.1 | Rhabdonatronobacter_sediminivivens | HUK65_RS15580 | 4960..5277 | 2743469 | hypothetical-protein-[Rhabdonatronobacter-sediminivivens] |
| RBP80165.1 | Type1.BREX | XIS-HTH->wHTH+WYL+WCX->BrxA-DNA-binding->BrxB-BrxF-iSTAND->DUF6079*->OLD-ABC->N6-METHYLASE->PglZ->SIGMA+OB-Fold+AAA-ATPase+HKD-DNase-BrxL-><-?<-Glyco_hydro_53+Ph [illegible] | 1172 | Bacteria>Pseudomonadota>Alphaproteobacteria>Rhodobacterales>Paracoccaceae>Rhodobacter | GCA_003315475.1 | Rhodobacter_sp._140A | DFO80_1418 | 4623..8141 | 2183910 | hypothetical-protein-DFO80_1418-[Rhodobacter-sp.-140A] |
| WP_223637900.1 | Type1.BREX | MMR_HSR1+sGTP+sGTP->?-><-?\|\|?->?->SF1-SF2-Helicase+DUF1998->BrxA-DNA-binding->BrxB-BrxF-iSTAND->DUF6079*->N6-METHYLASE->PglZ->SIGMA+OB-Fold+AAA-ATPase+HKD-DNase- [illegible] | 1175 | Bacteria>Pseudomonadota>Alphaproteobacteria>Rhodobacterales>Paracoccaceae>Rhodobacter | GCF_020023995.1 | Rhodobacter_sp._TJ_12 | CKO11_RS10405 | 86480..90007 | 2029399 | BREX-system-P-loop-protein-BrxC-[Rhodobacter-sp.-TJ_12] |
| WP_243695875.1 | Type1.BREX | HTH_3->?-><-TrbM<-Thermonuclease<-?\|\|wHTH+WYL+WCX->BrxA-DNA-binding->BrxB-BrxF-iSTAND->DUF6079-p*->DDE_Tnp_1_5+DDE_Tnp_1->GDResolvase+HIN-HTH-><-TetR-HTH\|\|?->ABC [illegible] | 633 | Bacteria>Pseudomonadota>Alphaproteobacteria>Rhodobacterales>Paracoccaceae>Rhodovulum | GCF_004343075.1 | Rhodovulum_marinum | EV662_RS15600 | 23147..25048 | 320662 | BREX-system-P-loop-protein-BrxC-[Rhodovulum-marinum] |
| WP_227427259.1 | Type1.BREX | [illegible] | 1140 | Bacteria>Pseudomonadota>Alphaproteobacteria>Rhodobacterales>Paracoccaceae>Roseibaca | GCF_020667835.1 | Roseibaca_sp._Y0-43 | J1C52_RS07230 | 292113..295535 | 2816854 | BREX-system-P-loop-protein-BrxC-[Roseibaca-sp.-Y0-43] |
| WP_121096034.1 | Type1.BREX | [illegible] | 1173 | Bacteria>Pseudomonadota>Alphaproteobacteria>Rhodobacterales>Paracoccaceae>Roseibaca | GCF_900499075.1 | Roseibaca_ekhonensis | ROE7235_RS13515 | 24046..27567 | 254356 | BREX-system-P-loop-protein-BrxC-[Roseibaca-ekhonensis] |
| WP_245964025.1 | Type1.BREX | [illegible] | 637 | Bacteria>Pseudomonadota>Alphaproteobacteria>Rhodobacterales>Paracoccaceae>Roseibaca | GCF_900499075.1 | Roseibaca_ekhonensis | ROE7235_RS18160 | 88..2001 | 254356 | BREX-system-P-loop-protein-BrxC-[Roseibaca-ekhonensis] |
| WP_218631289.1 | Type1.BREX | [illegible] | 1173 | Bacteria>Pseudomonadota>Alphaproteobacteria>Rhodobacterales>Paracoccaceae>Sedimentimonas | GCF_019264375.1 | Sedimentimonas_flavescens | KTQ37_RS14800 | 8221..11742 | 2851012 | BREX-system-P-loop-protein-BrxC-[Sedimentimonas-flavescens] |
| WP_108641830.1 | Type1.BREX | [illegible] | 1173 | Bacteria>Pseudomonadota>Alphaproteobacteria>Rhodobacterales>Paracoccaceae>Thalassorhabdomicrobium | GCF_003072065.1 | Thalassorhabdomicrobium_marinisediminis | DC363_RS14250 | 78584..82105 | 2170577 | BREX-system-P-loop-protein-BrxC-[Thalassorhabdomicrobium-marinisediminis] |
| WP_058289795.1 | Type1.BREX | [illegible] | 1172 | Bacteria>Pseudomonadota>Alphaproteobacteria>Rhodobacterales>Paracoccaceae>Tritonibacter | GCF_001458415.1 | Tritonibacter_multivorans | PGB32_RS12460 | 15411..18929 | 928856 | BREX-system-P-loop-protein-BrxC-[Tritonibacter-multivorans] |
| WP_065322029.1 | Type1.BREX | [illegible] | 1172 | Bacteria>Pseudomonadota>Alphaproteobacteria>Rhodobacterales>Paracoccaceae>Tritonibacter | GCF_001681255.1 | Tritonibacter_mobilis | UB66_RS09365 | 554629..558147 | 379347 | BREX-system-P-loop-protein-BrxC-[Tritonibacter-mobilis] |
| WP_225262956.1 | Type1.BREX | [illegible] | 1172 | Bacteria>Pseudomonadota>Alphaproteobacteria>Rhodobacterales>Paracoccaceae>Tritonibacter | GCF_020216085.1 | Tritonibacter_mobilis | LC038_RS07430 | 133716..137234 | 379347 | BREX-system-P-loop-protein-BrxC-[Tritonibacter-mobilis] |
| WP_065331859.1 | Type1.BREX | [illegible] | 1204 | Bacteria>Pseudomonadota>Alphaproteobacteria>Rhodobacterales>Paracoccaceae>Tritonibacter | GCF_030159715.1 | Tritonibacter_mobilis | BLU61_RS13075 | 124334..127948 | 379347 | BREX-system-P-loop-protein-BrxC-[Tritonibacter-mobilis] |
| PHO04850.1 | Type1.BREX | [illegible] | 1166 | Bacteria>Pseudomonadota>Alphaproteobacteria>Rhodobacterales>Paracoccaceae>unclassified_Paracoccaceae | GCA_002700045.1 | Rhodobacteraceae_bacterium_4F10 | CSC82_04805 | 8196..11696 | 2045445 | hypothetical-protein-CSC82_04805-[Rhodobacteraceae-bacterium-4F10] |
| NKX29277.1 | Type1.BREX | [illegible] | 1168 | Bacteria>Pseudomonadota>Alphaproteobacteria>Rhodobacterales>Paracoccaceae>unclassified_Paracoccaceae | GCA_012395685.1 | Rhodobacteraceae_bacterium_R_SAG6 | HGE68_05220 | 14041..17547 | 2719008 | BREX-system-P-loop-protein-BrxC-[Rhodobacteraceae-bacterium-R_SAG6] |
| MBD1206697.1 | Type1.BREX | [illegible] | 1181 | Bacteria>Pseudomonadota>Alphaproteobacteria>Rhodobacterales>Paracoccaceae>unclassified_Paracoccaceae | GCA_014654775.1 | Paracoccaceae_bacterium | H9533_21535 | 9124..12669 | 1904441 | BREX-system-P-loop-protein-BrxC-[Paracoccaceae-bacterium] |
| MBL4542464.1 | Type1.BREX | [illegible] | 1173 | Bacteria>Pseudomonadota>Alphaproteobacteria>Rhodobacterales>Paracoccaceae>unclassified_Paracoccaceae | GCA_016762795.1 | Paracoccaceae_bacterium | JKP97_11775 | 15111..18632 | 1904441 | BREX-system-P-loop-protein-BrxC-[Paracoccaceae-bacterium] |
| MBL4767704.1 | Type1.BREX | [illegible] | 663 | Bacteria>Pseudomonadota>Alphaproteobacteria>Rhodobacterales>Paracoccaceae>unclassified_Paracoccaceae | GCA_016764515.1 | Paracoccaceae_bacterium | JKY94_08320 | 11908..13895 | 1904441 | BREX-system-P-loop-protein-BrxC-[Paracoccaceae-bacterium] |
| MCR9111160.1 | Type1.BREX | [illegible] | 1176 | Bacteria>Pseudomonadota>Alphaproteobacteria>Rhodobacterales>Paracoccaceae>unclassified_Paracoccaceae | GCA_024742895.1 | Paracoccaceae_bacterium | NXH94_20060 | 7296..10826 | 1904441 | BREX-system-P-loop-protein-BrxC-[Paracoccaceae-bacterium] |
| MBD3677805.1 | Type1.BREX | [illegible] | 1173 | Bacteria>Pseudomonadota>Alphaproteobacteria>Rhodobacterales>Paracoccaceae>unclassified_Paracoccaceae | - | Paracoccaceae_bacterium | HUJ27_05300 | 15046..18567 | 1904441 | BREX-system-P-loop-protein-BrxC-[Paracoccaceae-bacterium] |
| WP_089279695.1 | Type1.BREX | [illegible] | 1173 | Bacteria>Pseudomonadota>Alphaproteobacteria>Rhodobacterales>Roseobacteraceae>Antarctobacter | GCF_900188425.1 | Antarctobacter_heliothermus | CHB90_RS20470 | 19687..23208 | 74033 | BREX-system-P-loop-protein-BrxC-[Antarctobacter-heliothermus] |
| WP_089279817.1 | Type1.BREX | [illegible] | 1147 | Bacteria>Pseudomonadota>Alphaproteobacteria>Rhodobacterales>Roseobacteraceae>Antarctobacter | GCF_900188425.1 | Antarctobacter_heliothermus | CHB90_RS21205 | 7452..10895 | 74033 | BREX-system-P-loop-protein-BrxC-[Antarctobacter-heliothermus] |
| MAN15901.1 | Type1.BREX | [illegible] | 265 | Bacteria>Pseudomonadota>Alphaproteobacteria>Rhodobacterales>Roseobacteraceae>Dinoroseobacter | GCA_002693265.1 | Dinoroseobacter_sp. | CL945_14875 | 7316..8113 | 2003366 | ATPase-[Dinoroseobacter-sp.] |
| WP_106262543.1 | Type1.BREX | [illegible] | 1172 | Bacteria>Pseudomonadota>Alphaproteobacteria>Rhodobacterales>Roseobacteraceae>Donghicola | GCF_003003355.1 | Donghicola_tyrosinivorans | CLV74_RS02110 | 436162..439680 | 1652492 | BREX-system-P-loop-protein-BrxC-[Donghicola-tyrosinivorans] |
| WP_027259561.1 | Type1.BREX | [illegible] | 1172 | Bacteria>Pseudomonadota>Alphaproteobacteria>Rhodobacterales>Roseobacteraceae>Leisingera | GCF_000473165.1 | Leisingera_aquimarina | AQUI_RS0122640 | 3713027..3716545 | 476529 | BREX-system-P-loop-protein-BrxC-[Leisingera-aquimarina] |
| WP_263764345.1 | Type1.BREX | [illegible] | 1175 | Bacteria>Pseudomonadota>Alphaproteobacteria>Rhodobacterales>Roseobacteraceae>Lentibacter | GCF_025751535.1 | Lentibacter_sp._XHP0401 | OE750_RS11445 | 67112..70639 | 2984334 | BREX-system-P-loop-protein-BrxC-[Lentibacter-sp.-XHP0401] |
| WP_263765218.1 | Type1.BREX | [illegible] | 1167 | Bacteria>Pseudomonadota>Alphaproteobacteria>Rhodobacterales>Roseobacteraceae>Lentibacter | GCF_025751535.1 | Lentibacter_sp._XHP0401 | OE750_RS15480 | 13619..17122 | 2984334 | BREX-system-P-loop-protein-BrxC-[Lentibacter-sp.-XHP0401] |
| WP_088668915.1 | Type1.BREX | [illegible] | 1179 | Bacteria>Pseudomonadota>Alphaproteobacteria>Rhodobacterales>Roseobacteraceae>Mameliella | GCF_002211215.1 | Mameliella_alba | CDZ95_RS03685 | 62213..65752 | 561184 | BREX-system-P-loop-protein-BrxC-[Mameliella-alba] |
| WP_167646419.1 | Type1.BREX | [illegible] | 1175 | Bacteria>Pseudomonadota>Alphaproteobacteria>Rhodobacterales>Roseobacteraceae>Mameliella | GCF_011858205.1 | Mameliella_alba | DAA47_RS08650 | 638377..641904 | 561184 | BREX-system-P-loop-protein-BrxC-[Mameliella-alba] |
| WP_091845786.1 | Type1.BREX | [illegible] | 1172 | Bacteria>Pseudomonadota>Alphaproteobacteria>Rhodobacterales>Roseobacteraceae>Maribius | GCF_900110115.1 | Maribius_pelagius | BMX96_RS12425 | 183654..187172 | 387096 | BREX-system-P-loop-protein-BrxC-[Maribius-pelagius] |
| WP_274591702.1 | Type1.BREX | [illegible] | 1173 | Bacteria>Pseudomonadota>Alphaproteobacteria>Rhodobacterales>Roseobacteraceae>Marinovum | GCF_028807375.1 | Marinovum_sp._PR37 | PVW49_RS22970 | 3287..6808 | 3028382 | BREX-system-P-loop-protein-BrxC-[Marinovum-sp.-PR37] |
| WP_123792783.1 | Type1.BREX | [illegible] | 1172 | Bacteria>Pseudomonadota>Alphaproteobacteria>Rhodobacterales>Roseobacteraceae>Pacificibacter | GCF_003814915.1 | Pacificibacter_maritimus | EDD53_RS08420 | 394668..398186 | 762213 | BREX-system-P-loop-protein-BrxC-[Pacificibacter-maritimus] |
| MAE92616.1 | Type1.BREX | [illegible] | 1210 | Bacteria>Pseudomonadota>Alphaproteobacteria>Rhodobacterales>Roseobacteraceae>Pelagibaca | GCA_002687285.1 | Pelagibaca_sp. | CMI67_24080 | 2344..5976 | 1979400 | ATPase-[Pelagibaca-sp.] |
| MAE92765.1 | Type1.BREX | [illegible] | 1219 | Bacteria>Pseudomonadota>Alphaproteobacteria>Rhodobacterales>Roseobacteraceae>Pelagibaca | GCA_002687285.1 | Pelagibaca_sp. | CMI67_24850 | 4770..8429 | 1979400 | ATPase-[Pelagibaca-sp.] |
| WP_244686757.1 | Type1.BREX | [illegible] | 1173 | Bacteria>Pseudomonadota>Alphaproteobacteria>Rhodobacterales>Roseobacteraceae>Phaeobacter | GCF_022867865.1 | Phaeobacter_sp._J2-8 | MUB49_RS06760 | 203466..206987 | 2931394 | BREX-system-P-loop-protein-BrxC-[Phaeobacter-sp.-J2-8] |
| WP_260112390.1 | Type1.BREX | [illegible] | 1172 | Bacteria>Pseudomonadota>Alphaproteobacteria>Rhodobacterales>Roseobacteraceae>Phaeobacter | GCF_025142595.1 | Phaeobacter_inhibens | K4F86_RS11215 | 2363506..2367024 | 221822 | BREX-system-P-loop-protein-BrxC-[Phaeobacter-inhibens] |
| WP_009810044.1 | Type1.BREX | [illegible] | 1172 | Bacteria>Pseudomonadota>Alphaproteobacteria>Rhodobacterales>Roseobacteraceae>Roseobacter | GCF_000152965.1 | Roseobacter_sp._MED193 | MED193_RS12240 | 2047846..2051364 | 314262 | BREX-system-P-loop-protein-BrxC-[Roseobacter-sp.-MED193] |
| KJS41393.1 | Type1.BREX | [illegible] | 1178 | Bacteria>Pseudomonadota>Alphaproteobacteria>Rhodobacterales>Roseobacteraceae>Roseovarius | GCA_000961715.1 | Roseovarius_sp._BRH_c41 | VR71_18660 | 26779..30315 | 1629709 | hypothetical-protein-VR71_18660-[Roseovarius-sp.-BRH_c41] |
| WP_160325893.1 | Type1.BREX | [illegible] | 1110 | Bacteria>Pseudomonadota>Alphaproteobacteria>Rhodobacterales>Roseobacteraceae>Roseovarius | GCF_001441635.1 | Roseovarius_indicus | XM52_RS23920 | 112621..115950 | 540747 | BREX-system-P-loop-protein-BrxC,-partial-[Roseovarius-indicus] |
| WP_074940632.1 | Type1.BREX | [illegible] | 1119 | Bacteria>Pseudomonadota>Alphaproteobacteria>Rhodobacterales>Roseobacteraceae>Roseovarius | GCF_008728195.1 | Roseovarius_indicus | RIdsm_RS22200 | 4630317..4633676 | 540747 | BREX-system-P-loop-protein-BrxC-[Roseovarius-indicus] |
| WP_139076871.1 | Type1.BREX | [illegible] | 1175 | Bacteria>Pseudomonadota>Alphaproteobacteria>Rhodobacterales>Roseobacteraceae>Rubellimicrobium | GCF_006152115.1 | Rubellimicrobium_rubrum | FHG66_11370 | 28349..31876 | 2585369 | BREX-system-P-loop-protein-BrxC-[Rubellimicrobium-rubrum] |
| WP_068340009.1 | Type1.BREX | [illegible] | 1203 | Bacteria>Pseudomonadota>Alphaproteobacteria>Rhodobacterales>Roseobacteraceae>Ruegeria | GCF_001507545.1 | Ruegeria_profundi | AVO44_RS17790 | 11048..14659 | 1685378 | BREX-system-P-loop-protein-BrxC-[Ruegeria-profundi] |
| WP_254429641.1 | Type1.BREX | [illegible] | 1170 | Bacteria>Pseudomonadota>Alphaproteobacteria>Rhodobacterales>Roseobacteraceae>Ruegeria | GCF_013031165.1 | Ruegeria_sp._HKCCD7251 | GOX75_RS12475 | 203228..206740 | 2683011 | BREX-system-P-loop-protein-BrxC-[Ruegeria-sp.-HKCCD7251] |
| WP_171123550.1 | Type1.BREX | [illegible] | 1174 | Bacteria>Pseudomonadota>Alphaproteobacteria>Rhodobacterales>Roseobacteraceae>Ruegeria | GCF_013031185.1 | Ruegeria_sp._HKCCD7239 | GOX74_RS13410 | 208876..212400 | 2683010 | BREX-system-P-loop-protein-BrxC-[Ruegeria-sp.-HKCCD7239] |
| WP_170339699.1 | Type1.BREX | [illegible] | 1173 | Bacteria>Pseudomonadota>Alphaproteobacteria>Rhodobacterales>Roseobacteraceae>Ruegeria | GCF_013031325.1 | Ruegeria_arenilitoris | GOX56_RS07985 | 128142..131663 | 1173585 | BREX-system-P-loop-protein-BrxC-[Ruegeria-arenilitoris] |
| WP_170340900.1 | Type1.BREX | [illegible] | 1140 | Bacteria>Pseudomonadota>Alphaproteobacteria>Rhodobacterales>Roseobacteraceae>Ruegeria | GCF_013031325.1 | Ruegeria_arenilitoris | GOX56_RS11935 | 236782..240204 | 1173585 | BREX-system-P-loop-protein-BrxC-[Ruegeria-arenilitoris] |
| WP_254433969.1 | Type1.BREX | [illegible] | 1170 | Bacteria>Pseudomonadota>Alphaproteobacteria>Rhodobacterales>Roseobacteraceae>Ruegeria | GCF_013032735.1 | Ruegeria_arenilitoris | GOW80_RS09865 | 349565..353077 | 1173585 | BREX-system-P-loop-protein-BrxC-[Ruegeria-arenilitoris] |
| WP_170509651.1 | Type1.BREX | [illegible] | 1174 | Bacteria>Pseudomonadota>Alphaproteobacteria>Rhodobacterales>Roseobacteraceae>Ruegeria | GCF_013032765.1 | Ruegeria_arenilitoris | GOW79_RS10365 | 805076..808600 | 1173585 | BREX-system-P-loop-protein-BrxC-[Ruegeria-arenilitoris] |
| WP_170364516.1 | Type1.BREX | [illegible] | 1140 | Bacteria>Pseudomonadota>Alphaproteobacteria>Rhodobacterales>Roseobacteraceae>Ruegeria | GCF_013032855.1 | Ruegeria_arenilitoris | GOW72_RS01075 | 235735..239157 | 1173585 | BREX-system-P-loop-protein-BrxC-[Ruegeria-arenilitoris] |
| WP_209509473.1 | Type1.BREX | [illegible] | 1147 | Bacteria>Pseudomonadota>Alphaproteobacteria>Rhodobacterales>Roseobacteraceae>Ruegeria | GCF_017872615.1 | Ruegeria_sp._HKCCE4150 | I9183_RS11345 | 408329..411772 | 2794828 | BREX-system-P-loop-protein-BrxC-[Ruegeria-sp.-HKCCE4150] |
| WP_222457132.1 | Type1.BREX | [illegible] | 1174 | Bacteria>Pseudomonadota>Alphaproteobacteria>Rhodobacterales>Roseobacteraceae>Ruegeria | GCF_019801065.1 | Ruegeria_arenilitoris | KUW05_RS05645 | 1133713..1137237 | 1173585 | BREX-system-P-loop-protein-BrxC-[Ruegeria-arenilitoris] |
| WP_234170466.1 | Type1.BREX | [illegible] | 899 | Bacteria>Pseudomonadota>Alphaproteobacteria>Rhodobacterales>Roseobacteraceae>Ruegeria | GCF_021405425.1 | Ruegeria_pomeroyi | KBY28_RS08200 | 126596..129295 | 89184 | BREX-system-P-loop-protein-BrxC-[Ruegeria-pomeroyi] |
| WP_150282504.1 | Type1.BREX | [illegible] | 1173 | Bacteria>Pseudomonadota>Alphaproteobacteria>Rhodobacterales>Roseobacteraceae>Salipiger | GCF_008692285.1 | Salipiger_aestuarii | AL036_RS18780 | 13068..16589 | 568098 | BREX-system-P-loop-protein-BrxC-[Salipiger-aestuarii] |
| WP_089852186.1 | Type1.BREX | [illegible] | 1173 | Bacteria>Pseudomonadota>Alphaproteobacteria>Rhodobacterales>Roseobacteraceae>Salipiger | GCF_900100085.1 | Salipiger_marinus | BLR51_RS25320 | 13015..16536 | 555512 | BREX-system-P-loop-protein-BrxC-[Salipiger-marinus] |
| WP_025045421.1 | Type1.BREX | [illegible] | 1147 | Bacteria>Pseudomonadota>Alphaproteobacteria>Rhodobacterales>Roseobacteraceae>Sulfitobacter | GCF_013407805.1 | Sulfitobacter_geojensis | GGR94_RS09535 | 1982054..1985497 | 1342299 | BREX-system-P-loop-protein-BrxC-[Sulfitobacter-geojensis] |
| WP_203210578.1 | Type1.BREX | [illegible] | 1172 | Bacteria>Pseudomonadota>Alphaproteobacteria>Rhodobacterales>Roseobacteraceae>Sulfitobacter | GCF_016801775.1 | Sulfitobacter_mediterraneus | JNX03_RS00635 | 149782..153300 | 83219 | BREX-system-P-loop-protein-BrxC-[Sulfitobacter-mediterraneus] |
| WP_259953659.1 | Type1.BREX | [illegible] | 1174 | Bacteria>Pseudomonadota>Alphaproteobacteria>Rhodobacterales>Roseobacteraceae>Sulfitobacter | GCF_025140475.1 | Sulfitobacter_mediterraneus | K3753_RS16990 | 3485906..3489430 | 83219 | BREX-system-P-loop-protein-BrxC-[Sulfitobacter-mediterraneus] |
| WP_267167638.1 | Type1.BREX | [illegible] | 1173 | Bacteria>Pseudomonadota>Alphaproteobacteria>Rhodobacterales>Roseobacteraceae>Sulfitobacter | GCF_026410285.1 | Sulfitobacter_sp._F26169L | OS189_RS12380 | 41979..45500 | 2996015 | BREX-system-P-loop-protein-BrxC-[Sulfitobacter-sp.-F26169L] |
| WP_271692167.1 | Type1.BREX | [illegible] | 1174 | Bacteria>Pseudomonadota>Alphaproteobacteria>Rhodobacterales>Roseobacteraceae>Sulfitobacter | GCF_028201715.1 | Sulfitobacter_faviae | PL335_RS12830 | 2584668..2588192 | 1775881 | BREX-system-P-loop-protein-BrxC-[Sulfitobacter-faviae] |
| WP_223427796.1 | Type1.BREX | [illegible] | 1179 | Bacteria>Pseudomonadota>Alphaproteobacteria>Rhodobacterales>Roseobacteraceae>Tateyamaria | GCF_019966495.1 | Tateyamaria_pelophila | LC354_RS21665 | 9757..13296 | 328415 | BREX-system-P-loop-protein-BrxC-[Tateyamaria-pelophila] |
| WP_058242644.1 | Type1.BREX | [illegible] | 1173 | Bacteria>Pseudomonadota>Alphaproteobacteria>Rhodobacterales>Roseobacteraceae>Thalassobius | GCF_001458255.1 | Thalassobius_autumnalis | TL5120_RS05495 | 4804..8325 | 2072972 | BREX-system-P-loop-protein-BrxC-[Thalassobius-autumnalis] |
| WP_058243041.1 | Type1.BREX | [illegible] | 1172 | Bacteria>Pseudomonadota>Alphaproteobacteria>Rhodobacterales>Roseobacteraceae>Thalassobius | GCF_001458275.1 | Thalassobius_autumnalis | TL5120_RS07660 | 24794..28312 | 2072972 | BREX-system-P-loop-protein-BrxC-[Thalassobius-autumnalis] |
| OHC57392.1 | Type1.BREX | [illegible] | 1173 | Bacteria>Pseudomonadota>Alphaproteobacteria>Rhodobacterales>unclassified_Rhodobacterales | GCA_001828855.1 | Rhodobacterales_bacterium_RIFCSPHIGHO2_02_FULL_62_130 | A3D16_23615 | 14989..18510 | 1802014 | hypothetical-protein-A3D16_23615-[Rhodobacterales-bacterium-RIFCSPHIGHO2_02_FULL_62_130] |
| OUS21258.1 | Type1.BREX | [illegible] | 1173 | Bacteria>Pseudomonadota>Alphaproteobacteria>Rhodobacterales>unclassified_Rhodobacterales | GCA_002162885.1 | Rhodobacterales_bacterium_59_46_T64 | A9Q95_07015 | 5961..9482 | 1856300 | hypothetical-protein-A9Q95_07015-[Rhodobacterales-bacterium-59_46_T64] |
| WP_061508432.1 | Type1.BREX | [illegible] | 1177 | Bacteria>Pseudomonadota>Alphaproteobacteria>Rhodospirillales>Acetobacteraceae>Acetobacter | GCF_001580695.1 | Acetobacter_malorum | AD933_RS09485 | 34256..37789 | 178901 | BREX-system-P-loop-protein-BrxC-[Acetobacter-malorum] |
| WP_212335128.1 | Type1.BREX | [illegible] | 1177 | Bacteria>Pseudomonadota>Alphaproteobacteria>Rhodospirillales>Acetobacteraceae>Acetobacter | GCF_018256855.1 | Acetobacter_thailandicus | JK172_RS08930 | 62967..66500 | 1502842 | BREX-system-P-loop-protein-BrxC-[Acetobacter-thailandicus] |
| OYV89048.1 | Type1.BREX | [illegible] | 1173 | Bacteria>Pseudomonadota>Alphaproteobacteria>Rhodospirillales>Acetobacteraceae>Acidiphilium | GCA_002279355.1 | Acidiphilium_sp._37-60-79 | B7Z57_13830 | 139..3660 | 1970298 | MAG:-hypothetical-protein-B7Z57_13830-[Acidiphilium-sp.-37-60-79] |
| CAG4901729.1 | Type1.BREX | [illegible] | 459 | Bacteria>Pseudomonadota>Alphaproteobacteria>Rhodospirillales>Acetobacteraceae>Acidocella | GCA_907164655.1 | Acidocella_sp._C78 | ACIDOCELLA78_258 | 233780..235159 | 1671486 | unnamed-protein-product,-partial-[Acidocella-sp.-C78] |
| WP_164698243.1 | Type1.BREX | [illegible] | 1143 | Bacteria>Pseudomonadota>Alphaproteobacteria>Rhodospirillales>Acetobacteraceae>Falsiroseomonas | GCF_011040385.1 | Falsiroseomonas_algicola | G3576_RS30405 | 8811..12242 | 2716930 | BREX-system-P-loop-protein-BrxC-[Falsiroseomonas-algicola] |
| WP_211586184.1 | Type1.BREX | [illegible] | 1211 | Bacteria>Pseudomonadota>Alphaproteobacteria>Rhodospirillales>Acetobacteraceae>Gluconobacter | GCF_018256215.1 | Gluconobacter_sp._Dm-73 | JK176_RS02065 | 37036..40671 | 2799802 | BREX-system-P-loop-protein-BrxC-[Gluconobacter-sp.-Dm-73] |
| WP_226610978.1 | Type1.BREX | [illegible] | 1144 | Bacteria>Pseudomonadota>Alphaproteobacteria>Rhodospirillales>Acetobacteraceae>Roseicella | GCF_020523605.1 | Roseicella_aerolata | LHA35_RS18965 | 9480..12914 | 2883479 | BREX-system-P-loop-protein-BrxC-[Roseicella-aerolata] |
| MCA3353385.1 | Type1.BREX | [illegible] | 1178 | Bacteria>Pseudomonadota>Alphaproteobacteria>Rhodospirillales>Acetobacteraceae>Roseomonas | GCA_020249745.1 | Roseomonas_sp. | INF80_03760 | 34583..38119 | 1869190 | BREX-system-P-loop-protein-BrxC-[Roseomonas-sp.] |
| WP_209351251.1 | Type1.BREX | [illegible] | 1177 | Bacteria>Pseudomonadota>Alphaproteobacteria>Rhodospirillales>Acetobacteraceae>Roseomonas | GCF_017813355.1 | Roseomonas_sp._PWR1 | J5Y09_RS08145 | 78258..81791 | 2820810 | BREX-system-P-loop-protein-BrxC-[Roseomonas-sp.-PWR1] |
| WP_239419500.1 | Type1.BREX | [illegible] | 1170 | Bacteria>Pseudomonadota>Alphaproteobacteria>Rhodospirillales>Acetobacteraceae>Roseomonas | - | Roseomonas_sp._ACRSG | MHZ93_RS20975 | 7121..10633 | 2918212 | BREX-system-P-loop-protein-BrxC-[Roseomonas-sp.-ACRSG] |
| PWC40000.1 | Type1.BREX | [illegible] | 1148 | Bacteria>Pseudomonadota>Alphaproteobacteria>Rhodospirillales>Azospirillaceae>Azospirillum | GCA_003116095.1 | Azospirillum_sp._TSO35-2 | TSO352_06450 | 1114896..1118342 | 716796 | hypothetical-protein-TSO352_06450-[Azospirillum-sp.-TSO35-2] |
| WP_014187927.1 | Type1.BREX | [illegible] | 1148 | Bacteria>Pseudomonadota>Alphaproteobacteria>Rhodospirillales>Azospirillaceae>Azospirillum | GCF_000283655.1 | Azospirillum_lipoferum | AZOLI_RS14530 | 158802..162248 | 193 | BREX-system-P-loop-protein-BrxC-[Azospirillum-lipoferum] |
| WP_094302078.1 | Type1.BREX | [illegible] | 1215 | Bacteria>Pseudomonadota>Alphaproteobacteria>Rhodospirillales>Azospirillaceae>Azospirillum | GCF_002245955.1 | Azospirillum_brasilense | CHT98_04470 | 49941..53588 | 192 | BREX-system-P-loop-protein-BrxC-[Azospirillum-brasilense] |
| WP_109096920.1 | Type1.BREX | [illegible] | 1215 | Bacteria>Pseudomonadota>Alphaproteobacteria>Rhodospirillales>Azospirillaceae>Azospirillum | GCF_003116035.1 | Azospirillum_sp._TSH64 | TSH64_RS11970 | 2295743..2299390 | 652740 | BREX-system-P-loop-protein-BrxC-[Azospirillum-sp.-TSH64] |
| WP_247871928.1 | Type1.BREX | [illegible] | 1149 | Bacteria>Pseudomonadota>Alphaproteobacteria>Rhodospirillales>Azospirillaceae>Azospirillum | GCF_003116095.1 | Azospirillum_sp._TSO35-2 | TSO352_RS06555 | 1114896..1118345 | 716796 | BREX-system-P-loop-protein-BrxC-[Azospirillum-sp.-TSO35-2] |
| WP_137104425.1 | Type1.BREX | [illegible] | 1148 | Bacteria>Pseudomonadota>Alphaproteobacteria>Rhodospirillales>Azospirillaceae>Azospirillum | GCF_005222185.1 | Azospirillum_argentinense | D3867_15345 | 176394..179840 | 2970906 | BREX-system-P-loop-protein-BrxC-[Azospirillum-argentinense] |
| WP_089290578.1 | Type1.BREX | [illegible] | 1202 | Bacteria>Pseudomonadota>Alphaproteobacteria>Rhodospirillales>Azospirillaceae>Azospirillum | GCF_900188385.1 | unclassified_Azospirillum | CHB97_RS24120 | 34901..38509 | 2630922 | MULTISPECIES:-BREX-system-P-loop-protein-BrxC-[unclassified-Azospirillum] |
| MAD36284.1 | Type1.BREX | [illegible] | 971 | Bacteria>Pseudomonadota>Alphaproteobacteria>Rhodospirillales>Geminicoccaceae>Tistrella | GCA_002684955.1 | Tistrella_sp. | CMO28_06235 | 72..2987 | 2024861 | hypothetical-protein-[Tistrella-sp.] |
| MBA78722.1 | Type1.BREX | [illegible] | 971 | Bacteria>Pseudomonadota>Alphaproteobacteria>Rhodospirillales>Geminicoccaceae>Tistrella | GCA_002709065.1 | Tistrella_sp. | CMO30_25940 | 72..2987 | 2024861 | hypothetical-protein-[Tistrella-sp.] |
| BAE50844.1 | Type1.BREX | [illegible] | 794 | Bacteria>Pseudomonadota>Alphaproteobacteria>Rhodospirillales>Rhodospirillaceae>Magnetospirillum | GCA_000009985.1 | Magnetospirillum_magneticum_AMB-1 | amb2040 | 2216694..2219078 | 342108 | hypothetical-protein-amb2040-[Magnetospirillum-magneticum-AMB-1] |
| WP_043746510.1 | Type1.BREX | [illegible] | 790 | Bacteria>Pseudomonadota>Alphaproteobacteria>Rhodospirillales>Rhodospirillaceae>Magnetospirillum | GCF_000009985.1 | Magnetospirillum_magneticum | AMB_RS10300 | 2216706..2219078 | 84159 | BREX-system-P-loop-protein-BrxC,-partial-[Magnetospirillum-magneticum] |
| WP_008620174.1 | Type1.BREX | [illegible] | 1205 | Bacteria>Pseudomonadota>Alphaproteobacteria>Rhodospirillales>Rhodospirillaceae>Magnetospirillum | GCF_000342045.1 | Magnetospirillum_caucaseum | H261_RS17615 | 20424..24041 | 1244869 | BREX-system-P-loop-protein-BrxC-[Magnetospirillum-caucaseum] |
| WP_008621107.1 | Type1.BREX | [illegible] | 1208 | Bacteria>Pseudomonadota>Alphaproteobacteria>Rhodospirillales>Rhodospirillaceae>Magnetospirillum | GCF_000342045.1 | Magnetospirillum_caucaseum | H261_RS19775 | 10510..14136 | 1244869 | BREX-system-P-loop-protein-BrxC-[Magnetospirillum-caucaseum] |
| WP_068504545.1 | Type1.BREX | [illegible] | 1205 | Bacteria>Pseudomonadota>Alphaproteobacteria>Rhodospirillales>Rhodospirillaceae>Magnetospirillum | GCF_001650635.1 | Magnetospirillum_moscoviense | A6A05_RS18805 | 69930..73547 | 1437059 | BREX-system-P-loop-protein-BrxC-[Magnetospirillum-moscoviense] |
| MBA4095641.1 | Type1.BREX | [illegible] | 1149 | Bacteria>Pseudomonadota>Alphaproteobacteria>Rhodospirillales>Rhodospirillaceae>Rhodospirillum | GCA_013822765.1 | Rhodospirillum_sp. | CO484_02575 | 552126..555575 | 37792 | BREX-system-P-loop-protein-BrxC-[Rhodospirillum-sp.] |
| WP_086466635.1 | Type1.BREX | [illegible] | 1149 | Bacteria>Pseudomonadota>Alphaproteobacteria>Rhodospirillales>Thalassobaculaceae>Oceanibaculum | GCF_002148795.1 | Oceanibaculum_nanhaiense | BKM74_RS15615 | 60819..64268 | 1909734 | BREX-system-P-loop-protein-BrxC-[Oceanibaculum-nanhaiense] |
| WP_069959207.1 | Type1.BREX | [illegible] | 1190 | Bacteria>Pseudomonadota>Alphaproteobacteria>Rhodospirillales>Thalassospiraceae>Magnetovibrio | GCF_001746755.1 | Magnetovibrio_blakemorei | BEN30_RS15365 | 12021..15593 | 28181 | BREX-system-P-loop-protein-BrxC-[Magnetovibrio-blakemorei] |
| MBL4843424.1 | Type1.BREX | [illegible] | 1179 | Bacteria>Pseudomonadota>Alphaproteobacteria>Rhodospirillales>Thalassospiraceae>Thalassospira | GCA_016765055.1 | Thalassospira_sp. | JKY47_21635 | 20483..24022 | 1912094 | BREX-system-P-loop-protein-BrxC-[Thalassospira-sp.] |
| WP_085595207.1 | Type1.BREX | [illegible] | 1182 | Bacteria>Pseudomonadota>Alphaproteobacteria>Rhodospirillales>Thalassospiraceae>Thalassospira | GCF_002115825.1 | Thalassospira_sp._MCCC_1A01428 | THS27_RS23975 | 1729..5277 | 1470575 | BREX-system-P-loop-protein-BrxC-[Thalassospira-sp.-MCCC-1A01428] |
| SEP16519.1 | Type1.BREX | [illegible] | 1142 | Bacteria>Pseudomonadota>Alphaproteobacteria>Rhodospirillales>unclassified_Rhodospirillales | GCA_900110395.1 | Rhodospirillales_bacterium_URHD0017 | SAMN02990966_04223 | 161142..164570 | 1380357 | hypothetical-protein-SAMN02990966_04223-[Rhodospirillales-bacterium-URHD0017] |
| MBY0534082.1 | Type1.BREX | [illegible] | 1197 | Bacteria>Pseudomonadota>Alphaproteobacteria>Rickettsiales>Rickettsiaceae>unclassified_Rickettsiaceae | GCA_019752735.1 | Rickettsiaceae_bacterium | K2P31_03860 | 4925..8518 | 2023360 | BREX-system-P-loop-protein-BrxC-[Rickettsiaceae-bacterium] |
| WP_143251171.1 | Type1.BREX | [illegible] | 1141 | Bacteria>Pseudomonadota>Alphaproteobacteria>Sphingomonadales>Erythrobacteraceae>Altererythrobacter | GCF_007065885.1 | Altererythrobacter_sp._TH136 | C0V74_RS07000 | 1448696..1452121 | 2067415 | BREX-system-P-loop-protein-BrxC-[Altererythrobacter-sp.-TH136] |
| WP_160484443.1 | Type1.BREX | [illegible] | 1172 | Bacteria>Pseudomonadota>Alphaproteobacteria>Sphingomonadales>Erythrobacteraceae>Aurantiacibacter | GCF_009807005.1 | Aurantiacibacter_rhizosphaerae | GRF63_RS02615 | 556566..560084 | 2691582 | BREX-system-P-loop-protein-BrxC-[Aurantiacibacter-rhizosphaerae] |
| RNJ61798.1 | Type1.BREX | [illegible] | 1173 | Bacteria>Pseudomonadota>Alphaproteobacteria>Sphingomonadales>Erythrobacteraceae>Erythrobacter/Porphyrobacter_group | GCA_003724185.1 | Porphyrobacter_sp._IPPAS_B-1204 | EDM03_09690 | 47855..51376 | 2483793 | BREX-system-P-loop-protein-BrxC-[Porphyrobacter-sp.-IPPAS-B-1204] |
| WP_011415860.1 | Type1.BREX | [illegible] | 1172 | Bacteria>Pseudomonadota>Alphaproteobacteria>Sphingomonadales>Erythrobacteraceae>Erythrobacter/Porphyrobacter_group | GCF_000013005.1 | Erythrobacter_litoralis | EL2594_RS14475 | 2964416..2967934 | 39960 | BREX-system-P-loop-protein-BrxC-[Erythrobacter-litoralis] |
| WP_196847550.1 | Type1.BREX | [illegible] | 1172 | Bacteria>Pseudomonadota>Alphaproteobacteria>Sphingomonadales>Erythrobacteraceae>Erythrobacter/Porphyrobacter_group | GCF_015767595.1 | Erythrobacter_sp._A30-3 | IT881_RS05450 | 1101811..1105329 | 2785907 | BREX-system-P-loop-protein-BrxC-[Erythrobacter-sp.-A30-3] |
| WP_226699433.1 | Type1.BREX | [illegible] | 1170 | Bacteria>Pseudomonadota>Alphaproteobacteria>Sphingomonadales>Erythrobacteraceae>Qipengyuania | GCF_020524765.1 | Qipengyuania_gaetbuli | LC083_RS04940 | 959412..962924 | 266952 | BREX-system-P-loop-protein-BrxC-[Qipengyuania-gaetbuli] |
| MBK9012577.1 | Type1.BREX | [illegible] | 1212 | Bacteria>Pseudomonadota>Alphaproteobacteria>Sphingomonadales>Sphingomonadaceae>Novosphingobium | GCA_016719755.1 | Novosphingobium_sp. | IPM87_18255 | 676827..680465 | 1874826 | BREX-system-P-loop-protein-BrxC-[Novosphingobium-sp.] |
| WP_054107369.1 | Type1.BREX | [illegible] | 1214 | Bacteria>Pseudomonadota>Alphaproteobacteria>Sphingomonadales>Sphingomonadaceae>Novosphingobium | GCF_001295795.1 | Novosphingobium_sp._AAP83 | IP81_RS07510 | 164945..168589 | 1523425 | BREX-system-P-loop-protein-BrxC-[Novosphingobium-sp.-AAP83] |
| WP_240530317.1 | Type1.BREX | [illegible] | 735 | Bacteria>Pseudomonadota>Alphaproteobacteria>Sphingomonadales>Sphingomonadaceae>Novosphingobium | GCF_002556635.1 | Novosphingobium_sp._PC22D | B2G71_RS22885 | 26272..28479 | 1962403 | BREX-system-P-loop-protein-BrxC-[Novosphingobium-sp.-PC22D] |
| WP_160987331.1 | Type1.BREX | [illegible] | 675 | Bacteria>Pseudomonadota>Alphaproteobacteria>Sphingomonadales>Sphingomonadaceae>Novosphingobium | GCF_009856825.1 | Novosphingobium_silvae | GR702_RS19995 | 24258..26285 | 2692619 | BREX-system-P-loop-protein-BrxC-[Novosphingobium-silvae] |
| WP_243992543.1 | Type1.BREX | [illegible] | 1189 | Bacteria>Pseudomonadota>Alphaproteobacteria>Sphingomonadales>Sphingomonadaceae>Novosphingobium | GCF_022832385.1 | Novosphingobium_sp._B2580 | MTR64_RS07820 | 57412..60981 | 2930093 | BREX-system-P-loop-protein-BrxC-[Novosphingobium-sp.-B2580] |
| WP_243995499.1 | Type1.BREX | [illegible] | 1179 | Bacteria>Pseudomonadota>Alphaproteobacteria>Sphingomonadales>Sphingomonadaceae>Novosphingobium | GCF_022832385.1 | Novosphingobium_sp._B2580 | MTR64_RS16290 | 56132..59671 | 2930093 | BREX-system-P-loop-protein-BrxC-[Novosphingobium-sp.-B2580] |
| WP_159983052.1 | Type1.BREX | [illegible] | 1212 | Bacteria>Pseudomonadota>Alphaproteobacteria>Sphingomonadales>Sphingomonadaceae>Novosphingobium | GCF_902706195.1 | unclassified_Novosphingobium | GVL61_RS22205 | 8858..12496 | 2644732 | MULTISPECIES:-BREX-system-P-loop-protein-BrxC-[unclassified-Novosphingobium] |
| WP_022673601.1 | Type1.BREX | [illegible] | 1189 | Bacteria>Pseudomonadota>Alphaproteobacteria>Sphingomonadales>Sphingomonadaceae>Novosphingopyxis | GCF_000420305.1 | Novosphingopyxis_baekryungensis | G410_RS0110420 | 254465..258034 | 279369 | BREX-system-P-loop-protein-BrxC-[Novosphingopyxis-baekryungensis] |
| ARR56525.1 | Type1.BREX | [illegible] | 1189 | Bacteria>Pseudomonadota>Alphaproteobacteria>Sphingomonadales>Sphingomonadaceae>Rhizorhabdus | GCA_000715175.2 | Rhizorhabdus_wittichii_DC-6 | HY78_25365 | 5371615..5375184 | 1283312 | hypothetical-protein-HY78_25365-[Rhizorhabdus-wittichii-DC-6] |
| WP_209047668.1 | Type1.BREX | [illegible] | 1179 | Bacteria>Pseudomonadota>Alphaproteobacteria>Sphingomonadales>Sphingomonadaceae>Rhizorhabdus | GCF_029543145.1 | Rhizorhabdus_histidinilytica | P7B04_RS17610 | 19171..22710 | 439228 | MULTISPECIES:-BREX-system-P-loop-protein-BrxC-[Sphingomonadaceae] |
| WP_010336176.1 | Type1.BREX | [illegible] | 1179 | Bacteria>Pseudomonadota>Alphaproteobacteria>Sphingomonadales>Sphingomonadaceae>Sphingobium | GCF_000224695.1 | Sphingobium_yanoikuyae | SYA_RS0104815 | 38393..41932 | 13690 | BREX-system-P-loop-protein-BrxC-[Sphingobium-yanoikuyae] |
| WP_007688323.1 | Type1.BREX | [illegible] | 1189 | Bacteria>Pseudomonadota>Alphaproteobacteria>Sphingomonadales>Sphingomonadaceae>Sphingobium | GCF_000264945.2 | Sphingobium_indicum | SIDU_RS03565 | 700467..704036 | 332055 | BREX-system-P-loop-protein-BrxC-[Sphingobium-indicum] |
| WP_021240096.1 | Type1.BREX | [illegible] | 1177 | Bacteria>Pseudomonadota>Alphaproteobacteria>Sphingomonadales>Sphingomonadaceae>Sphingobium | GCF_000445065.1 | Sphingobium_quisquiliarum | L288_RS41575 | 32905..36438 | 538379 | BREX-system-P-loop-protein-BrxC-[Sphingobium-quisquiliarum] |
| WP_130029930.1 | Type1.BREX | [illegible] | 1140 | Bacteria>Pseudomonadota>Alphaproteobacteria>Sphingomonadales>Sphingomonadaceae>Sphingobium | GCF_014636045.1 | Sphingobium_fuliginis | EWH10_RS07560 | 584587..588009 | 336203 | BREX-system-P-loop-protein-BrxC-[Sphingobium-fuliginis] |
| WP_264583946.1 | Type1.BREX | [illegible] | 1179 | Bacteria>Pseudomonadota>Alphaproteobacteria>Sphingomonadales>Sphingomonadaceae>Sphingobium | GCF_025961895.1 | Sphingobium_sp._B7D2B | M2341_RS01365 | 293056..296595 | 2940583 | BREX-system-P-loop-protein-BrxC-[Sphingobium-sp.-B7D2B] |
| EJU13065.1 | Type1.BREX | [illegible] | 1179 | Bacteria>Pseudomonadota>Alphaproteobacteria>Sphingomonadales>Sphingomonadaceae>Sphingomonas | GCA_000293195.2 | Sphingomonas_sp._LH128 | LH128_10616 | 1306..4845 | 473781 | hypothetical-protein-LH128_10616-[Sphingomonas-sp.-LH128] |
| MBY0305717.1 | Type1.BREX | [illegible] | 1212 | Bacteria>Pseudomonadota>Alphaproteobacteria>Sphingomonadales>Sphingomonadaceae>Sphingomonas | GCA_019750015.1 | Sphingomonas_sp. | K2W86_11290 | 121509..125147 | 28214 | BREX-system-P-loop-protein-BrxC-[Sphingomonas-sp.] |
| WP_056486438.1 | Type1.BREX | [illegible] | 1214 | Bacteria>Pseudomonadota>Alphaproteobacteria>Sphingomonadales>Sphingomonadaceae>Sphingomonas | GCF_001421415.1 | Sphingomonas_sp._Leaf208 | ASE69_07675 | 294901..298545 | 1735679 | BREX-system-P-loop-protein-BrxC-[Sphingomonas-sp.-Leaf208] |
| WP_066478880.1 | Type1.BREX | [illegible] | 1179 | Bacteria>Pseudomonadota>Alphaproteobacteria>Sphingomonadales>Sphingomonadaceae>Sphingomonas | GCF_001555785.1 | unclassified_Sphingomonas | AX667_RS15610 | 42451..45990 | 196159 | MULTISPECIES:-BREX-system-P-loop-protein-BrxC-[unclassified-Sphingomonas] |
| WP_107968397.1 | Type1.BREX | [illegible] | 1186 | Bacteria>Pseudomonadota>Alphaproteobacteria>Sphingomonadales>Sphingomonadaceae>Sphingomonas | GCF_003050615.1 | Sphingomonas_oleivorans | CLG96_RS13035 | 363272..366832 | 1735121 | BREX-system-P-loop-protein-BrxC-[Sphingomonas-oleivorans] |
| WP_192976950.1 | Type1.BREX | [illegible] | 1177 | Bacteria>Pseudomonadota>Alphaproteobacteria>Sphingomonadales>Sphingomonadaceae>Sphingomonas | GCF_014892665.1 | Sphingomonas_sp._CFBP_13603 | IFR23_RS17335 | 87025..90558 | 2774040 | BREX-system-P-loop-protein-BrxC-[Sphingomonas-sp.-CFBP-13603] |
| WP_089221081.1 | Type1.BREX | [illegible] | 1179 | Bacteria>Pseudomonadota>Alphaproteobacteria>Sphingomonadales>Sphingomonadaceae>Sphingomonas | GCF_900188165.1 | Sphingomonas_laterariae | CHB74_RS18985 | 16083..19622 | 861865 | BREX-system-P-loop-protein-BrxC-[Sphingomonas-laterariae] |
| WP_079639881.1 | Type1.BREX | [illegible] | 1179 | Bacteria>Pseudomonadota>Alphaproteobacteria>Sphingomonadales>Sphingomonadaceae>Sphingopyxis | GCF_900168005.1 | Sphingopyxis_flava | B5X69_RS17550 | 26842..30381 | 1507287 | BREX-system-P-loop-protein-BrxC-[Sphingopyxis-flava] |
| WP_133494816.1 | Type1.BREX | [illegible] | 1189 | Bacteria>Pseudomonadota>Alphaproteobacteria>Sphingomonadales>Sphingomonadaceae>Stakelama | GCF_014645395.1 | Stakelama_pacifica | IEX20_RS07270 | 305060..308629 | 517720 | BREX-system-P-loop-protein-BrxC-[Stakelama-pacifica] |
| WP_121052748.1 | Type1.BREX | [illegible] | 1179 | Bacteria>Pseudomonadota>Alphaproteobacteria>Sphingomonadales>Sphingosinicellaceae>Sphingosinicella | GCF_003967095.1 | Sphingosinicella_microcystinivorans | SmB9_RS16135 | 3426933..3430472 | 335406 | BREX-system-P-loop-protein-BrxC-[Sphingosinicella-microcystinivorans] |
| OYY96513.1 | Type1.BREX | [illegible] | 1189 | Bacteria>Pseudomonadota>Alphaproteobacteria>Sphingomonadales>unclassified_Sphingomonadales | GCA_002281585.1 | Sphingomonadales_bacterium_28-56-43 | B7Y38_10875 | 3957..7526 | 1970590 | MAG:-hypothetical-protein-B7Y38_10875-[Sphingomonadales-bacterium-28-56-43] |
| TNE62528.1 | Type1.BREX | [illegible] | 1175 | Bacteria>Pseudomonadota>Alphaproteobacteria>Sphingomonadales>unclassified_Sphingomonadales | GCA_006226385.1 | Sphingomonadales_bacterium | EP341_00225 | 33441..36968 | 1978525 | MAG:-BREX-system-P-loop-protein-BrxC-[Sphingomonadales-bacterium] |
| WP_015740180.1 | Type1.BREX | [illegible] | 1179 | Bacteria>Pseudomonadota>Alphaproteobacteria>Sphingomonadales>Zymomonadaceae>Zymomonas | GCF_000024245.1 | Zymomonas_mobilis | Za10_1374 | 1591366..1594905 | 542 | BREX-system-P-loop-protein-BrxC-[Zymomonas-mobilis] |
| OQC17755.1 | Type1.BREX | [illegible] | 1145 | Bacteria>Pseudomonadota>Alphaproteobacteria>unclassified_Alphaproteobacteria | GCA_002070795.1 | Alphaproteobacteria_bacterium_ADurb.Bin100 | BWX79_00187 | 2015..5452 | 1852791 | hypothetical-protein-BWX79_00187-[Alphaproteobacteria-bacterium-ADurb.Bin100] |
| OYU71768.1 | Type1.BREX | [illegible] | 1219 | Bacteria>Pseudomonadota>Alphaproteobacteria>unclassified_Alphaproteobacteria | GCA_002256425.1 | Alphaproteobacteria_bacterium_PA2 | CFE28_16215 | 3389137..3392796 | 2015570 | hypothetical-protein-CFE28_16215-[Alphaproteobacteria-bacterium-PA2] |

| Protein-ID | System | Gene-Neighbourhoods | Length | Taxonomy | Assembly-ID | Strain | Locus-Tag | Genomic-coordinates | TaxID | Fasta-defline |
|---|---|---|---|---|---|---|---|---|---|---|
| RYH00367.1 | Type1.BREX | <-HTH_3<-XIS-HTH<-?<-ParB+HNH\|\|?->Preader17-><-SIGMA+OB-Fold+AAA-ATPase+Lon-Protease-BrxL<-PglZ<-N6-METHYLASE<-DUF6079*<-BrxB-BrxF-iSTAND<-BrxA-DNA-binding\|\|?->?->?->?->35exo+SF1-SF2- [illegible] | 1212 | Bacteria>Pseudomonadota>Alphaproteobacteria>unclassified_Alphaproteobacteria | GCA_004145575.1 | Alphaproteobacteria_bacterium | EON58_00375 | 13524..17162 | 1913988 | BREX-system-P-loop-protein-BrxC-[Alphaproteobacteria-bacterium] |
| MBN9530327.1 | Type1.BREX | <-ICLR-HTH<-?\|\|?->HTH_21+rve+rve_3->HTH_21+rve+rve_3-><-SIGMA+OB-Fold+AAA-ATPase+HKD-DNase-BrxL<-PglZ<-N6-METHYLASE<-DUF6079*<-BrxB-BrxF-iSTAND<-BrxA-DNA-binding<-wHTH+WYL+WCX\|\|35Exo+ [illegible] | 1180 | Bacteria>Pseudomonadota>Alphaproteobacteria>unclassified_Alphaproteobacteria | GCA_017307375.1 | Alphaproteobacteria_bacterium | J0H82_29215 | 188733..192275 | 1913988 | BREX-system-P-loop-protein-BrxC-[Alphaproteobacteria-bacterium] |
| MBN9589014.1 | Type1.BREX | <-S4+PSYN\|\|?->?-><-?\|\|tRNA-><-Peptidase_M29+DUF2971\|\|?-><-SIGMA+OB-Fold+AAA-ATPase+Lon-Protease-BrxL<-PglZ<-N6-METHYLASE<-DUF6079*<-BrxB-BrxF-iSTAND<-BrxA-DNA-binding<-?\|\|REase+DUF565 [illegible] | 1155 | Bacteria>Pseudomonadota>Alphaproteobacteria>unclassified_Alphaproteobacteria | GCA_017308475.1 | Alphaproteobacteria_bacterium | J0G99_08395 | 190029..193496 | 1913988 | BREX-system-P-loop-protein-BrxC-[Alphaproteobacteria-bacterium] |
| MBQ2845004.1 | Type1.BREX | Peptidase-M15->?-><-HKD-DNase<-?\|\|BrxA-DNA-binding->BrxB-BrxF-iSTAND->DUF6079*->N6-METHYLASE->PglZ->SIGMA+OB-Fold+AAA-ATPase+Lon-Protease-BrxL->ANK+ANK+ANK->?->Small-B-barrel+SF1-SF2- [illegible] | 1173 | Bacteria>Pseudomonadota>Alphaproteobacteria>unclassified_Alphaproteobacteria | GCA_017409215.1 | Alphaproteobacteria_bacterium | IJE79_03245 | 53916..57437 | 1913988 | BREX-system-P-loop-protein-BrxC-[Alphaproteobacteria-bacterium] |
| MBQ2886030.1 | Type1.BREX | AbiEi_4->AbiEii-><-Response_reg\|\|HTH_3->BrxA-DNA-binding->BrxB-BrxF-iSTAND->DUF6079*-> | 1131 | Bacteria>Pseudomonadota>Alphaproteobacteria>unclassified_Alphaproteobacteria | GCA_017409945.1 | Alphaproteobacteria_bacterium | IJE43_20100 | 7355..10747 | 1913988 | hypothetical-protein-[Alphaproteobacteria-bacterium] |
| MBQ2886288.1 | Type1.BREX | <-wHTH+WYL<-REase<-SIGMA+OB-Fold+AAA-ATPase+Lon-Protease-BrxL<-PglZ<-Schlafen+HATPase+wHTH<-N6-METHYLASE<-PIN<-OLD-ABC<-DUF6079*<-BrxB-BrxF-iSTAND<-BrxA-DNA-binding<-HTH_3 | 1180 | Bacteria>Pseudomonadota>Alphaproteobacteria>unclassified_Alphaproteobacteria | GCA_017409945.1 | Alphaproteobacteria_bacterium | IJE43_21410 | 13139..16681 | 1913988 | BREX-system-P-loop-protein-BrxC-[Alphaproteobacteria-bacterium] |
| MBQ8347028.1 | Type1.BREX | VirD4-TraG+VirD4-FtsK->?-><-TraI-VirD2-RCRE<-?<-?<-?<-SIGMA+OB-Fold+AAA-ATPase+HKD-DNase-BrxL<-PglZ<-N6-METHYLASE<-DUF6079*<-BrxB-BrxF-iSTAND<-BrxA-DNA-binding\|\|?-><-?<-ArdCN+MPTase\|\| [illegible] | 1150 | Bacteria>Pseudomonadota>Alphaproteobacteria>unclassified_Alphaproteobacteria | GCA_017462585.1 | Alphaproteobacteria_bacterium | IJY17_05445 | 46698..50150 | 1913988 | BREX-system-P-loop-protein-BrxC-[Alphaproteobacteria-bacterium] |
| MBT3788616.1 | Type1.BREX | ParA-Soj-PloopNTPase->?-><-?<-?\|\|DEDD_Tnp_IS110+Transposase_20->?->HTH_Tnp_1+HTH_21+rve+rve_3-><-DUF6079* | 1185 | Bacteria>Pseudomonadota>Alphaproteobacteria>unclassified_Alphaproteobacteria | GCA_018654745.1 | Alphaproteobacteria_bacterium | HOF94_09595 | 7711..11268 | 1913988 | BREX-system-P-loop-protein-BrxC-[Alphaproteobacteria-bacterium] |
| EXI84705.1 | Type1.BREX | DDE_Tnp_1-><-?<-MqsA_antitoxin<-HTH<-?<-Arm-DNA-bind_3+Phage_integrase\|\|BrxA-DNA-binding->BrxB-BrxF-iSTAND->DUF6079*->N6-METHYLASE->HTH+REase->PglZ->SIGMA+OB-Fold+AAA-ATPase+Lon-Prote [illegible] | 1204 | Bacteria>Pseudomonadota>Betaproteobacteria>Betaproteobacteria_incertae_sedis>Candidatus_Accumulibacter | GCA_000585075.1 | Candidatus_Accumulibacter_regalis | AW11_03760 | 6873..10487 | 522306 | MAG:-hypothetical-protein-AW11_03760-[Candidatus-Accumulibacter-regalis] |
| RDE50960.1 | Type1.BREX | <-Phage_int_SAM_4+Phage_integrase\|\|?->TPR+TPR+TPR+TPR+TPR->BrxA-DNA-binding->BrxB-BrxF-iSTAND->DUF6079*->HTH_3->N6-METHYLASE->SF1-SF2-Helicase+PRTase->SLOG+BirA-HTH->PglZ->SIGMA+OB-Fo [illegible] | 1154 | Bacteria>Pseudomonadota>Betaproteobacteria>Betaproteobacteria_incertae_sedis>Candidatus_Accumulibacter | GCA_003332265.1 | Candidatus_Accumulibacter_meliphilus | DVS81_08175 | 4944..8408 | 2211374 | MAG:-BREX-system-P-loop-protein-BrxC-[Candidatus-Accumulibacter-meliphilus] |
| RDE49751.1 | Type1.BREX | <-PIN+HTH<-SIGMA+OB-Fold+AAA-ATPase+Lon-Protease-BrxL<-PglZ<-TOPRIM<-OLD-ABC<-DUF6079*<-BrxB-BrxF-iSTAND<-BrxA-DNA-binding\|\|?->?-><-?<-?\|\|?-><-?<-Zn_Tnp_IS91+Y2_Tnp | 1071 | Bacteria>Pseudomonadota>Betaproteobacteria>Betaproteobacteria_incertae_sedis>Candidatus_Accumulibacter | GCA_003332265.1 | Candidatus_Accumulibacter_meliphilus | DVS81_14920 | 17642..20857 | 2211374 | MAG:-BREX-system-P-loop-protein-BrxC-[Candidatus-Accumulibacter-meliphilus] |
| MQM35874.1 | Type1.BREX | <-MqsA_antitoxin<-HTH<-?<-Arm-DNA-bind_3+Phage_integrase\|\|BrxA-DNA-binding->BrxB-BrxF-iSTAND->DUF6079-p*-> | 732 | Bacteria>Pseudomonadota>Betaproteobacteria>Betaproteobacteria_incertae_sedis>Candidatus_Accumulibacter | GCA_009467855.1 | Candidatus_Accumulibacter_phosphatis | CRU72_16025 | 5196..7392 | 327160 | ATPase-[Candidatus-Accumulibacter-phosphatis] |
| MQM30738.1 | Type1.BREX | PolB-><-SIGMA+OB-Fold+AAA-ATPase+Lon-Protease-BrxL<-PglZ<-?<-SbcC<-N6-METHYLASE<-HTH_3<-DUF6079*<-BrxB-BrxF-iSTAND<-BrxA-DNA-binding<-TPR+TPR+TPR+TPR+TPR<-?\|\|?->?->?->NUDIX-> | 1154 | Bacteria>Pseudomonadota>Betaproteobacteria>Betaproteobacteria_incertae_sedis>Candidatus_Accumulibacter | GCA_009467885.1 | Candidatus_Accumulibacter_phosphatis | CRU78_09485 | 11967..15431 | 327160 | hypothetical-protein-[Candidatus-Accumulibacter-phosphatis] |
| WP_169070206.1 | Type1.BREX | [illegible] | 1204 | Bacteria>Pseudomonadota>Betaproteobacteria>Betaproteobacteria_incertae_sedis>Candidatus_Accumulibacter | GCF_012940005.1 | Candidatus_Accumulibacter_contiguus | E4Q08_RS09460 | 34303..37917 | 2954381 | BREX-system-P-loop-protein-BrxC-[Candidatus-Accumulibacter-contiguus] |
| EXI72915.1 | Type1.BREX | [illegible] | 1155 | Bacteria>Pseudomonadota>Betaproteobacteria>Betaproteobacteria_incertae_sedis>Candidatus_Accumulibacter>unclassified_Candidatus_Accumulibacter | GCA_000584995.1 | Candidatus_Accumulibacter_sp._SK-11 | AW07_03051 | 55405..58872 | 1454000 | MAG:-hypothetical-protein-AW07_03051-[Candidatus-Accumulibacter-sp.-SK-11] |
| HAY27737.1 | Type1.BREX | [illegible] | 1286 | Bacteria>Pseudomonadota>Betaproteobacteria>Betaproteobacteria_incertae_sedis>Candidatus_Accumulibacter>unclassified_Candidatus_Accumulibacter | GCA_003487685.1 | Candidatus_Accumulibacter_sp. | DCY47_09605 | 6303..10163 | 2053492 | BREX-system-P-loop-protein-BrxC-[Candidatus-Accumulibacter-sp.] |
| HCN68765.1 | Type1.BREX | [illegible] | 1154 | Bacteria>Pseudomonadota>Betaproteobacteria>Betaproteobacteria_incertae_sedis>Candidatus_Accumulibacter>unclassified_Candidatus_Accumulibacter | GCA_003535635.1 | Candidatus_Accumulibacter_sp. | DIT03_10970 | 31798..35262 | 2053492 | BREX-system-P-loop-protein-BrxC-[Candidatus-Accumulibacter-sp.] |
| MBK8385355.1 | Type1.BREX | [illegible] | 1154 | Bacteria>Pseudomonadota>Betaproteobacteria>Betaproteobacteria_incertae_sedis>Candidatus_Accumulibacter>unclassified_Candidatus_Accumulibacter | GCA_016711725.1 | Candidatus_Accumulibacter_sp. | IPL11_06655 | 113668..117132 | 2053492 | BREX-system-P-loop-protein-BrxC-[Candidatus-Accumulibacter-sp.] |
| MBK8115934.1 | Type1.BREX | [illegible] | 1154 | Bacteria>Pseudomonadota>Betaproteobacteria>Betaproteobacteria_incertae_sedis>Candidatus_Accumulibacter>unclassified_Candidatus_Accumulibacter | GCA_016712935.1 | Candidatus_Accumulibacter_sp. | IPK44_16290 | 126114..129578 | 2053492 | BREX-system-P-loop-protein-BrxC-[Candidatus-Accumulibacter-sp.] |
| MBK7952886.1 | Type1.BREX | [illegible] | 1207 | Bacteria>Pseudomonadota>Betaproteobacteria>Betaproteobacteria_incertae_sedis>Candidatus_Accumulibacter>unclassified_Candidatus_Accumulibacter | GCA_016713625.1 | Candidatus_Accumulibacter_sp. | IPK02_02340 | 220445..224068 | 2053492 | BREX-system-P-loop-protein-BrxC-[Candidatus-Accumulibacter-sp.] |
| MBK8578271.1 | Type1.BREX | [illegible] | 1154 | Bacteria>Pseudomonadota>Betaproteobacteria>Betaproteobacteria_incertae_sedis>Candidatus_Accumulibacter>unclassified_Candidatus_Accumulibacter | GCA_016714935.1 | Candidatus_Accumulibacter_sp. | IPN78_08405 | 116670..120134 | 2053492 | BREX-system-P-loop-protein-BrxC-[Candidatus-Accumulibacter-sp.] |
| MBL8396239.1 | Type1.BREX | [illegible] | 1204 | Bacteria>Pseudomonadota>Betaproteobacteria>Betaproteobacteria_incertae_sedis>Candidatus_Accumulibacter>unclassified_Candidatus_Accumulibacter | GCA_016789985.1 | Candidatus_Accumulibacter_sp. | JNK99_16080 | 8411..12025 | 2053492 | BREX-system-P-loop-protein-BrxC-[Candidatus-Accumulibacter-sp.] |
| MBN8454657.1 | Type1.BREX | [illegible] | 1157 | Bacteria>Pseudomonadota>Betaproteobacteria>Betaproteobacteria_incertae_sedis>Candidatus_Accumulibacter>unclassified_Candidatus_Accumulibacter | GCA_017302415.1 | Candidatus_Accumulibacter_sp. | J0M31_16455 | 17156..20629 | 2053492 | BREX-system-P-loop-protein-BrxC-[Candidatus-Accumulibacter-sp.] |
| MBN8436515.1 | Type1.BREX | [illegible] | 1154 | Bacteria>Pseudomonadota>Betaproteobacteria>Betaproteobacteria_incertae_sedis>Candidatus_Accumulibacter>unclassified_Candidatus_Accumulibacter | GCA_017302555.1 | Candidatus_Accumulibacter_sp. | J0L95_00470 | 105300..108764 | 2053492 | BREX-system-P-loop-protein-BrxC-[Candidatus-Accumulibacter-sp.] |
| MBN8436950.1 | Type1.BREX | [illegible] | 1146 | Bacteria>Pseudomonadota>Betaproteobacteria>Betaproteobacteria_incertae_sedis>Candidatus_Accumulibacter>unclassified_Candidatus_Accumulibacter | GCA_017302555.1 | Candidatus_Accumulibacter_sp. | J0L95_02725 | 187324..190764 | 2053492 | BREX-system-P-loop-protein-BrxC-[Candidatus-Accumulibacter-sp.] |
| MCM8614222.1 | Type1.BREX | [illegible] | 1155 | Bacteria>Pseudomonadota>Betaproteobacteria>Betaproteobacteria_incertae_sedis>Candidatus_Accumulibacter>unclassified_Candidatus_Accumulibacter | GCA_023806505.1 | Candidatus_Accumulibacter_sp. | NFW17_19430 | 33411..36878 | 2053492 | BREX-system-P-loop-protein-BrxC-[Candidatus-Accumulibacter-sp.] |
| WP_050447538.1 | Type1.BREX | [illegible] | 1153 | Bacteria>Pseudomonadota>Betaproteobacteria>Burkholderiales>Alcaligenaceae>Achromobacter | GCF_001189595.1 | Achromobacter_spanius | AFM18_RS14295 | 68240..71701 | 217203 | BREX-system-P-loop-protein-BrxC-[Achromobacter-spanius] |
| WP_063584995.1 | Type1.BREX | [illegible] | 1146 | Bacteria>Pseudomonadota>Betaproteobacteria>Burkholderiales>Alcaligenaceae>Achromobacter | GCF_001637105.1 | Achromobacter_ruhlandii | AOT39_RS22745 | 36149..39589 | 72557 | BREX-system-P-loop-protein-BrxC-[Achromobacter-ruhlandii] |
| WP_076518288.1 | Type1.BREX | [illegible] | 1181 | Bacteria>Pseudomonadota>Betaproteobacteria>Burkholderiales>Alcaligenaceae>Achromobacter | GCF_900156745.1 | Achromobacter_sp._MFA1_R4 | BXA00_RS09440 | 2091384..2094929 | 1881016 | BREX-system-P-loop-protein-BrxC-[Achromobacter-sp.-MFA1-R4] |
| WP_175212942.1 | Type1.BREX | [illegible] | 1181 | Bacteria>Pseudomonadota>Betaproteobacteria>Burkholderiales>Alcaligenaceae>Achromobacter | GCF_902859965.1 | Achromobacter_aegrifaciens | HV793_RS07505 | 1613364..1616909 | 1287736 | BREX-system-P-loop-protein-BrxC-[Achromobacter-aegrifaciens] |
| WP_200235209.1 | Type1.BREX | [illegible] | 1190 | Bacteria>Pseudomonadota>Betaproteobacteria>Burkholderiales>Alcaligenaceae>Advenella | GCF_016595155.1 | Advenella_mandrilli | JHL22_RS06635 | 201531..205103 | 2800330 | BREX-system-P-loop-protein-BrxC-[Advenella-mandrilli] |
| WP_109088944.1 | Type1.BREX | [illegible] | 1146 | Bacteria>Pseudomonadota>Betaproteobacteria>Burkholderiales>Alcaligenaceae>Alcaligenes | GCF_003122065.1 | Alcaligenes_faecalis | DF183_RS09485 | 797094..800534 | 511 | BREX-system-P-loop-protein-BrxC-[Alcaligenes-faecalis] |
| WP_121740016.1 | Type1.BREX | [illegible] | 1190 | Bacteria>Pseudomonadota>Betaproteobacteria>Burkholderiales>Alcaligenaceae>Alcaligenes | GCF_003671915.1 | Alcaligenes_aquatilis | D3M96_RS02075 | 458513..462085 | 323284 | BREX-system-P-loop-protein-BrxC-[Alcaligenes-aquatilis] |
| CCJ47860.1 | Type1.BREX | [illegible] | 1194 | Bacteria>Pseudomonadota>Betaproteobacteria>Burkholderiales>Alcaligenaceae>Bordetella | GCA_000317935.1 | Bordetella_parapertussis_Bpp5 | BN117_0527 | 555038..558622 | 1208660 | conserved-hypothetical-protein-[Bordetella-parapertussis-Bpp5] |
| MBO9356876.1 | Type1.BREX | [illegible] | 1153 | Bacteria>Pseudomonadota>Betaproteobacteria>Burkholderiales>Alcaligenaceae>Bordetella | GCA_017745595.1 | Bordetella_petrii | GG851_23045 | 14450..17911 | 94624 | BREX-system-P-loop-protein-BrxC-[Bordetella-petrii] |
| WP_164926424.1 | Type1.BREX | [illegible] | 1188 | Bacteria>Pseudomonadota>Betaproteobacteria>Burkholderiales>Alcaligenaceae>Bordetella | GCF_000317935.1 | Bordetella_parapertussis | BN117_RS02695 | 555056..558622 | 519 | BREX-system-P-loop-protein-BrxC-[Bordetella-parapertussis] |
| WP_197022064.1 | Type1.BREX | [illegible] | 1150 | Bacteria>Pseudomonadota>Betaproteobacteria>Burkholderiales>Alcaligenaceae>Bordetella | GCF_000518845.1 | Bordetella_petrii | L636_RS0102150 | 157001..160453 | 94624 | BREX-system-P-loop-protein-BrxC-[Bordetella-petrii] |
| WP_043223478.1 | Type1.BREX | [illegible] | 1146 | Bacteria>Pseudomonadota>Betaproteobacteria>Burkholderiales>Alcaligenaceae>Bordetella | GCF_000657755.1 | Bordetella_hinzii | L539_RS12625 | 94736..98176 | 103855 | BREX-system-P-loop-protein-BrxC-[Bordetella-hinzii] |
| WP_165720894.1 | Type1.BREX | [illegible] | 1181 | Bacteria>Pseudomonadota>Betaproteobacteria>Burkholderiales>Alcaligenaceae>Bordetella | GCF_011386965.1 | Bordetella_trematum | G6Z49_RS19625 | 4277130..4280675 | 123899 | BREX-system-P-loop-protein-BrxC-[Bordetella-trematum] |
| WP_043683814.1 | Type1.BREX | [illegible] | 1153 | Bacteria>Pseudomonadota>Betaproteobacteria>Burkholderiales>Alcaligenaceae>Castellaniella | GCF_000612685.1 | Castellaniella_defragrans | BN940_RS13240 | 2967883..2971344 | 75697 | BREX-system-P-loop-protein-BrxC-[Castellaniella-defragrans] |
| WP_066458434.1 | Type1.BREX | [illegible] | 1196 | Bacteria>Pseudomonadota>Betaproteobacteria>Burkholderiales>Alcaligenaceae>Castellaniella | GCF_001592225.1 | Castellaniella_caeni | CUB94_RS15500 | 2114..5704 | 266123 | BREX-system-P-loop-protein-BrxC-[Castellaniella-caeni] |
| WP_085318435.1 | Type1.BREX | [illegible] | 1153 | Bacteria>Pseudomonadota>Betaproteobacteria>Burkholderiales>Alcaligenaceae>Derxia | GCF_002105195.1 | Derxia_lacustris | B9P64_RS23165 | 16077..19538 | 764842 | BREX-system-P-loop-protein-BrxC-[Derxia-lacustris] |
| WP_068373657.1 | Type1.BREX | [illegible] | 1176 | Bacteria>Pseudomonadota>Betaproteobacteria>Burkholderiales>Alcaligenaceae>Kerstersia | GCF_000988095.1 | Kerstersia_gyiorum | AAV32_RS14310 | 103983..107513 | 206506 | BREX-system-P-loop-protein-BrxC-[Kerstersia-gyiorum] |
| WP_115075260.1 | Type1.BREX | [illegible] | 1187 | Bacteria>Pseudomonadota>Betaproteobacteria>Burkholderiales>Alcaligenaceae>Oligella | GCF_900454255.1 | Oligella_urethralis | DX091_RS10510 | 2175019..2178582 | 90245 | BREX-system-P-loop-protein-BrxC-[Oligella-urethralis] |
| WP_115063824.1 | Type1.BREX | [illegible] | 1155 | Bacteria>Pseudomonadota>Betaproteobacteria>Burkholderiales>Alcaligenaceae>Oligella | GCF_900454315.1 | Oligella_ureolytica | NCTC11998_02420 | 2456151..2459618 | 90244 | BREX-system-P-loop-protein-BrxC-[Oligella-ureolytica] |
| WP_166406662.1 | Type1.BREX | [illegible] | 1190 | Bacteria>Pseudomonadota>Betaproteobacteria>Burkholderiales>Alcaligenaceae>Paenalcaligenes | GCF_011290505.1 | Paenalcaligenes_suwonensis | GY634_RS00685 | 163356..166928 | 1202713 | BREX-system-P-loop-protein-BrxC-[Paenalcaligenes-suwonensis] |
| MCK9506949.1 | Type1.BREX | [illegible] | 1069 | Bacteria>Pseudomonadota>Betaproteobacteria>Burkholderiales>Alcaligenaceae>Pigmentiphaga | GCA_023228545.1 | Pigmentiphaga_sp. | M0Q54_00780 | 41..3250 | 1977564 | BREX-system-P-loop-protein-BrxC-[Pigmentiphaga-sp.] |
| WP_132980217.1 | Type1.BREX | [illegible] | 1181 | Bacteria>Pseudomonadota>Betaproteobacteria>Burkholderiales>Alcaligenaceae>Pigmentiphaga | GCF_004352985.1 | Pigmentiphaga_sp._D-2 | EZY13_RS19720 | 185799..189344 | 1002116 | BREX-system-P-loop-protein-BrxC-[Pigmentiphaga-sp.-D-2] |
| WP_148812759.1 | Type1.BREX | [illegible] | 1153 | Bacteria>Pseudomonadota>Betaproteobacteria>Burkholderiales>Alcaligenaceae>Pigmentiphaga | GCF_008119665.1 | Pigmentiphaga_aceris | FXN63_RS03140 | 736142..739603 | 1940612 | BREX-system-P-loop-protein-BrxC-[Pigmentiphaga-aceris] |
| WP_102070953.1 | Type1.BREX | [illegible] | 1181 | Bacteria>Pseudomonadota>Betaproteobacteria>Burkholderiales>Alcaligenaceae>Pollutimonas | GCF_002849615.1 | Pollutimonas_nitritireducens | CR155_RS15505 | 30043..33588 | 2045209 | BREX-system-P-loop-protein-BrxC-[Pollutimonas-nitritireducens] |
| RIY39966.1 | Type1.BREX | [illegible] | 1145 | Bacteria>Pseudomonadota>Betaproteobacteria>Burkholderiales>Alcaligenaceae>Pusillimonas | GCA_003585905.1 | Pusillimonas_maritima | CJP73_12365 | 2891..6328 | 2026239 | hypothetical-protein-CJP73_12365-[Pusillimonas-maritima] |
| WP_218972830.1 | Type1.BREX | [illegible] | 1146 | Bacteria>Pseudomonadota>Betaproteobacteria>Burkholderiales>Alcaligenaceae>Pusillimonas | GCF_003585905.1 | Pusillimonas_maritima | CJP73_RS12395 | 2888..6328 | 2026239 | BREX-system-P-loop-protein-BrxC-[Pusillimonas-maritima] |
| NYT60502.1 | Type1.BREX | [illegible] | 1188 | Bacteria>Pseudomonadota>Betaproteobacteria>Burkholderiales>Alcaligenaceae>unclassified_Alcaligenaceae | GCA_013416405.1 | Alcaligenaceae_bacterium | H0A65_16390 | 10015..13581 | 2021368 | BREX-system-P-loop-protein-BrxC-[Alcaligenaceae-bacterium] |
| NYT60768.1 | Type1.BREX | [illegible] | 1146 | Bacteria>Pseudomonadota>Betaproteobacteria>Burkholderiales>Alcaligenaceae>unclassified_Alcaligenaceae | GCA_013416445.1 | Alcaligenaceae_bacterium | H0A66_00285 | 70051..73491 | 2021368 | BREX-system-P-loop-protein-BrxC-[Alcaligenaceae-bacterium] |
| NYT63370.1 | Type1.BREX | [illegible] | 1191 | Bacteria>Pseudomonadota>Betaproteobacteria>Burkholderiales>Alcaligenaceae>unclassified_Alcaligenaceae | GCA_013416445.1 | Alcaligenaceae_bacterium | H0A66_13705 | 63657..67232 | 2021368 | BREX-system-P-loop-protein-BrxC-[Alcaligenaceae-bacterium] |
| NYT66107.1 | Type1.BREX | [illegible] | 1193 | Bacteria>Pseudomonadota>Betaproteobacteria>Burkholderiales>Alcaligenaceae>unclassified_Alcaligenaceae | GCA_013416485.1 | Alcaligenaceae_bacterium | H0A58_09005 | 172517..176098 | 2021368 | BREX-system-P-loop-protein-BrxC-[Alcaligenaceae-bacterium] |
| WP_217193526.1 | Type1.BREX | [illegible] | 1146 | Bacteria>Pseudomonadota>Betaproteobacteria>Burkholderiales>Burkholderiaceae>Burkholderia | - | Burkholderia_multivorans | OCJ09_RS26065 | 2174..5614 | 87883 | BREX-system-P-loop-protein-BrxC-[Burkholderia-multivorans] |
| ABO55019.1 | Type1.BREX | [illegible] | 1210 | Bacteria>Pseudomonadota>Betaproteobacteria>Burkholderiales>Burkholderiaceae>Burkholderia | GCA_000016205.1 | Burkholderia_vietnamiensis_G4 | Bcep1808_2017 | 2211669..2215301 | 269482 | ATPase-like-protein-[Burkholderia-vietnamiensis-G4] |
| POM13919.1 | Type1.BREX | [illegible] | 837 | Bacteria>Pseudomonadota>Betaproteobacteria>Burkholderiales>Burkholderiaceae>Burkholderia | GCA_002906895.1 | Burkholderia_cepacia | CSX04_08370 | 7336..9849 | 292 | hypothetical-protein-CSX04_08370-[Burkholderia-cepacia] |
| POM13920.1 | Type1.BREX | [illegible] | 315 | Bacteria>Pseudomonadota>Betaproteobacteria>Burkholderiales>Burkholderiaceae>Burkholderia | GCA_002906895.1 | Burkholderia_cepacia | CSX04_08371 | 9849..10796 | 292 | hypothetical-protein-CSX04_08371-[Burkholderia-cepacia] |
| MCA8485082.1 | Type1.BREX | [illegible] | 879 | Bacteria>Pseudomonadota>Betaproteobacteria>Burkholderiales>Burkholderiaceae>Burkholderia | GCA_020421385.1 | Burkholderia_multivorans | LGM85_14180 | 202982..205621 | 87883 | BREX-system-P-loop-protein-BrxC-[Burkholderia-multivorans] |
| MCA8485083.1 | Type1.BREX | [illegible] | 266 | Bacteria>Pseudomonadota>Betaproteobacteria>Burkholderiales>Burkholderiaceae>Burkholderia | GCA_020421385.1 | Burkholderia_multivorans | LGM85_14185 | 205622..206422 | 87883 | hypothetical-protein-[Burkholderia-multivorans] |
| SAK30849.1 | Type1.BREX | [illegible] | 843 | Bacteria>Pseudomonadota>Betaproteobacteria>Burkholderiales>Burkholderiaceae>Burkholderia | GCA_900079585.1 | Burkholderia_multivorans | UA11_00037 | 36503..39034 | 87883 | ATPase-like-protein-[Burkholderia-multivorans] |
| WP_013697442.1 | Type1.BREX | [illegible] | 1223 | Bacteria>Pseudomonadota>Betaproteobacteria>Burkholderiales>Burkholderiaceae>Burkholderia | GCF_000194745.1 | Burkholderia_gladioli | BGLA_RS07070 | 1576041..1579712 | 28095 | BREX-system-P-loop-protein-BrxC-[Burkholderia-gladioli] |
| WP_025097273.1 | Type1.BREX | [illegible] | 1153 | Bacteria>Pseudomonadota>Betaproteobacteria>Burkholderiales>Burkholderiaceae>Burkholderia | GCF_000633145.1 | Burkholderia_sp._A1 | DP01_RS0105180 | 92035..95496 | 148446 | BREX-system-P-loop-protein-BrxC-[Burkholderia-sp.-A1] |
| WP_038780921.1 | Type1.BREX | [illegible] | 1152 | Bacteria>Pseudomonadota>Betaproteobacteria>Burkholderiales>Burkholderiaceae>Burkholderia | GCF_000774265.1 | Burkholderia_pseudomallei | X888_RS15495 | 3559753..3563211 | 28450 | BREX-system-P-loop-protein-BrxC-[Burkholderia-pseudomallei] |
| WP_059567309.1 | Type1.BREX | [illegible] | 1153 | Bacteria>Pseudomonadota>Betaproteobacteria>Burkholderiales>Burkholderiaceae>Burkholderia | GCF_001523965.1 | Burkholderia_cepacia_complex | C6T69_RS26905 | 12049..15510 | 87882 | MULTISPECIES:-BREX-system-P-loop-protein-BrxC-[Burkholderia-cepacia-complex] |
| WP_060225079.1 | Type1.BREX | [illegible] | 1146 | Bacteria>Pseudomonadota>Betaproteobacteria>Burkholderiales>Burkholderiaceae>Burkholderia | GCF_001531525.1 | Burkholderia_cepacia | WL34_RS17275 | 238513..241953 | 292 | BREX-system-P-loop-protein-BrxC-[Burkholderia-cepacia] |
| WP_060189440.1 | Type1.BREX | [illegible] | 1186 | Bacteria>Pseudomonadota>Betaproteobacteria>Burkholderiales>Burkholderiaceae>Burkholderia | GCF_001533155.1 | Burkholderia_diffusa | WL85_RS21680 | 47133..50693 | 488732 | BREX-system-P-loop-protein-BrxC-[Burkholderia-diffusa] |
| WP_231941297.1 | Type1.BREX | [illegible] | 1183 | Bacteria>Pseudomonadota>Betaproteobacteria>Burkholderiales>Burkholderiaceae>Burkholderia | GCF_001577465.1 | Burkholderia_sp._PAMC_28687 | AX768_RS02030 | 441347..444898 | 1795874 | BREX-system-P-loop-protein-BrxC-[Burkholderia-sp.-PAMC-28687] |
| WP_105762828.1 | Type1.BREX | [illegible] | 1146 | Bacteria>Pseudomonadota>Betaproteobacteria>Burkholderiales>Burkholderiaceae>Burkholderia | GCF_002980785.1 | Burkholderia_multivorans | C6P92_RS30780 | 36936..40376 | 87883 | BREX-system-P-loop-protein-BrxC-[Burkholderia-multivorans] |
| WP_260428004.1 | Type1.BREX | [illegible] | 929 | Bacteria>Pseudomonadota>Betaproteobacteria>Burkholderiales>Burkholderiaceae>Burkholderia | GCF_003853135.1 | Burkholderia_stagnalis | DF110_RS10485 | 163..2952 | 1503054 | BREX-system-P-loop-protein-BrxC-[Burkholderia-stagnalis] |
| WP_260431103.1 | Type1.BREX | [illegible] | 623 | Bacteria>Pseudomonadota>Betaproteobacteria>Burkholderiales>Burkholderiaceae>Burkholderia | GCF_003853195.1 | Burkholderia_stagnalis | DF031_RS07940 | 269197..271068 | 1503054 | BREX-system-P-loop-protein-BrxC-[Burkholderia-stagnalis] |
| WP_124831269.1 | Type1.BREX | [illegible] | 1200 | Bacteria>Pseudomonadota>Betaproteobacteria>Burkholderiales>Burkholderiaceae>Burkholderia | GCF_003857875.1 | Burkholderia_stagnalis | DF113_RS32940 | 12280..15882 | 1503054 | BREX-system-P-loop-protein-BrxC-[Burkholderia-stagnalis] |
| WP_205797411.1 | Type1.BREX | [illegible] | 1223 | Bacteria>Pseudomonadota>Betaproteobacteria>Burkholderiales>Burkholderiaceae>Burkholderia | GCF_017104785.1 | Burkholderia_sp._Ac-20353 | G3N94_RS31910 | 10724..14395 | 2703894 | BREX-system-P-loop-protein-BrxC-[Burkholderia-sp.-Ac-20353] |
| WP_212170318.1 | Type1.BREX | [illegible] | 1153 | Bacteria>Pseudomonadota>Betaproteobacteria>Burkholderiales>Burkholderiaceae>Burkholderia | GCF_018147655.1 | Burkholderia_vietnamiensis | KDX28_RS26080 | 57628..61089 | 60552 | BREX-system-P-loop-protein-BrxC-[Burkholderia-vietnamiensis] |
| WP_226243435.1 | Type1.BREX | [illegible] | 1200 | Bacteria>Pseudomonadota>Betaproteobacteria>Burkholderiales>Burkholderiaceae>Burkholderia | GCF_020419265.1 | Burkholderia_metallica | LGM59_RS24125 | 116704..120306 | 488729 | BREX-system-P-loop-protein-BrxC-[Burkholderia-metallica] |
| WP_269509108.1 | Type1.BREX | [illegible] | 1153 | Bacteria>Pseudomonadota>Betaproteobacteria>Burkholderiales>Burkholderiaceae>Burkholderia | GCF_027257875.1 | Burkholderia_sp._IMCC1007 | O1Q03_RS33565 | 11993..15454 | 3004104 | BREX-system-P-loop-protein-BrxC-[Burkholderia-sp.-IMCC1007] |
| WP_122833867.1 | Type1.BREX | [illegible] | 1223 | Bacteria>Pseudomonadota>Betaproteobacteria>Burkholderiales>Burkholderiaceae>Burkholderia | GCF_900597175.1 | Burkholderia_pseudomallei | EFT75_RS19485 | 195449..199120 | 28450 | BREX-system-P-loop-protein-BrxC-[Burkholderia-pseudomallei] |
| WP_175792486.1 | Type1.BREX | [illegible] | 1207 | Bacteria>Pseudomonadota>Betaproteobacteria>Burkholderiales>Burkholderiaceae>Burkholderia | GCF_902830065.1 | Burkholderia_ambifaria | HWW16_RS27390 | 202712..206335 | 152480 | BREX-system-P-loop-protein-BrxC-[Burkholderia-ambifaria] |
| WP_111998987.1 | Type1.BREX | [illegible] | 1153 | Bacteria>Pseudomonadota>Betaproteobacteria>Burkholderiales>Burkholderiaceae>Burkholderia | GCF_902830635.1 | Burkholderia_cepacia | HWX29_RS29485 | 220004..223465 | 292 | BREX-system-P-loop-protein-BrxC-[Burkholderia-cepacia] |
| WP_186109347.1 | Type1.BREX | [illegible] | 1153 | Bacteria>Pseudomonadota>Betaproteobacteria>Burkholderiales>Burkholderiaceae>Burkholderia | GCF_902831275.1 | Burkholderia_gladioli | H9D39_RS35055 | 68649..72110 | 28095 | BREX-system-P-loop-protein-BrxC-[Burkholderia-gladioli] |
| WP_186089005.1 | Type1.BREX | [illegible] | 1153 | Bacteria>Pseudomonadota>Betaproteobacteria>Burkholderiales>Burkholderiaceae>Burkholderia | GCF_902831595.1 | Burkholderia_gladioli | H9D22_RS09670 | 586813..590274 | 28095 | BREX-system-P-loop-protein-BrxC-[Burkholderia-gladioli] |
| WP_186146941.1 | Type1.BREX | [illegible] | 1153 | Bacteria>Pseudomonadota>Betaproteobacteria>Burkholderiales>Burkholderiaceae>Burkholderia | GCF_902831935.1 | Burkholderia_gladioli | H9E05_RS08890 | 201279..204740 | 28095 | BREX-system-P-loop-protein-BrxC-[Burkholderia-gladioli] |
| WP_186177578.1 | Type1.BREX | [illegible] | 1153 | Bacteria>Pseudomonadota>Betaproteobacteria>Burkholderiales>Burkholderiaceae>Burkholderia | GCF_902832085.1 | Burkholderia_gladioli | H9E56_RS05565 | 447000..450461 | 28095 | BREX-system-P-loop-protein-BrxC-[Burkholderia-gladioli] |
| WP_186266772.1 | Type1.BREX | [illegible] | 1153 | Bacteria>Pseudomonadota>Betaproteobacteria>Burkholderiales>Burkholderiaceae>Burkholderia | GCF_902832635.1 | Burkholderia_gladioli | H9H41_RS07040 | 658576..662037 | 28095 | BREX-system-P-loop-protein-BrxC-[Burkholderia-gladioli] |
| WP_176025807.1 | Type1.BREX | [illegible] | 1153 | Bacteria>Pseudomonadota>Betaproteobacteria>Burkholderiales>Burkholderiaceae>Burkholderia | GCF_902833615.1 | Burkholderia_vietnamiensis | HWX82_RS31015 | 13799..17260 | 60552 | BREX-system-P-loop-protein-BrxC-[Burkholderia-vietnamiensis] |
| WP_087670041.1 | Type1.BREX | [illegible] | 1204 | Bacteria>Pseudomonadota>Betaproteobacteria>Burkholderiales>Burkholderiaceae>Caballeronia | GCF_001544475.1 | Caballeronia_humi | AWB65_RS26815 | 12713..16327 | 326474 | BREX-system-P-loop-protein-BrxC-[Caballeronia-humi] |
| WP_061146343.1 | Type1.BREX | [illegible] | 1223 | Bacteria>Pseudomonadota>Betaproteobacteria>Burkholderiales>Burkholderiaceae>Caballeronia | GCF_001544695.2 | Caballeronia_arvi | AWB74_RS08600 | 195274..198945 | 1777135 | BREX-system-P-loop-protein-BrxC-[Caballeronia-arvi] |
| WP_087048072.1 | Type1.BREX | [illegible] | 1153 | Bacteria>Pseudomonadota>Betaproteobacteria>Burkholderiales>Burkholderiaceae>Caballeronia | GCF_001545075.1 | Caballeronia_ptereochthonis | AWB83_04706 | 123490..126951 | 1777144 | BREX-system-P-loop-protein-BrxC-[Caballeronia-ptereochthonis] |
| WP_165087512.1 | Type1.BREX | [illegible] | 1204 | Bacteria>Pseudomonadota>Betaproteobacteria>Burkholderiales>Burkholderiaceae>Caballeronia | GCF_011039955.1 | Caballeronia_sp._SBC2 | SBC2_RS04820 | 1105795..1109409 | 2705547 | BREX-system-P-loop-protein-BrxC-[Caballeronia-sp.-SBC2] |
| WP_250522809.1 | Type1.BREX | [illegible] | 1153 | Bacteria>Pseudomonadota>Betaproteobacteria>Burkholderiales>Burkholderiaceae>Caballeronia | GCF_023631205.1 | unclassified_Caballeronia | NCS65_RS00835 | 183401..186862 | 2646786 | MULTISPECIES:-BREX-system-P-loop-protein-BrxC-[unclassified-Caballeronia] |
| WP_250514397.1 | Type1.BREX | [illegible] | 1153 | Bacteria>Pseudomonadota>Betaproteobacteria>Burkholderiales>Burkholderiaceae>Caballeronia | GCF_023631325.1 | Caballeronia_sp._INDeC2 | NCS28_RS14240 | 310434..313895 | 2921747 | BREX-system-P-loop-protein-BrxC-[Caballeronia-sp.-INDeC2] |
| WP_018314059.1 | Type1.BREX | [illegible] | 1221 | Bacteria>Pseudomonadota>Betaproteobacteria>Burkholderiales>Burkholderiaceae>Cupriavidus | GCF_000379565.1 | Cupriavidus_sp._UYPR2.512 | A3A5_RS0129585 | 33129..36794 | 1080187 | BREX-system-P-loop-protein-BrxC-[Cupriavidus-sp.-UYPR2.512] |
| WP_224078061.1 | Type1.BREX | [illegible] | 1153 | Bacteria>Pseudomonadota>Betaproteobacteria>Burkholderiales>Burkholderiaceae>Cupriavidus | GCF_914271435.1 | Cupriavidus_laharis | LDO29_RS01695 | 364592..368053 | 151654 | BREX-system-P-loop-protein-BrxC-[Cupriavidus-laharis] |
| WP_125094459.1 | Type1.BREX | [illegible] | 1146 | Bacteria>Pseudomonadota>Betaproteobacteria>Burkholderiales>Burkholderiaceae>Lautropia | GCF_003892345.1 | Lautropia_dentalis | EHV23_RS01850 | 480599..484039 | 2490857 | BREX-system-P-loop-protein-BrxC-[Lautropia-dentalis] |
| WP_275272077.1 | Type1.BREX | [illegible] | 1193 | Bacteria>Pseudomonadota>Betaproteobacteria>Burkholderiales>Burkholderiaceae>Limnobacter | GCF_029087225.1 | Limnobacter_sp._P1 | PY990_RS13680 | 12633..16214 | 3031298 | BREX-system-P-loop-protein-BrxC-[Limnobacter-sp.-P1] |
| WP_087752374.1 | Type1.BREX | [illegible] | 1153 | Bacteria>Pseudomonadota>Betaproteobacteria>Burkholderiales>Burkholderiaceae>Paraburkholderia | GCF_002179925.1 | Paraburkholderia_caledonica | CDV47_RS15050 | 148364..151825 | 134536 | BREX-system-P-loop-protein-BrxC-[Paraburkholderia-caledonica] |
| WP_121308231.1 | Type1.BREX | [illegible] | 1204 | Bacteria>Pseudomonadota>Betaproteobacteria>Burkholderiales>Burkholderiaceae>Paraburkholderia | GCF_003634175.1 | Paraburkholderia_sp._BL17N1 | B0G82_RS17815 | 3937370..3940984 | 1938798 | BREX-system-P-loop-protein-BrxC-[Paraburkholderia-sp.-BL17N1] |
| WP_153135349.1 | Type1.BREX | [illegible] | 1153 | Bacteria>Pseudomonadota>Betaproteobacteria>Burkholderiales>Burkholderiaceae>Paraburkholderia | GCF_009455635.1 | Paraburkholderia_agricolaris | GH665_RS07660 | 1680328..1683789 | 2152888 | BREX-system-P-loop-protein-BrxC-[Paraburkholderia-agricolaris] |
| WP_176383829.1 | Type1.BREX | [illegible] | 1210 | Bacteria>Pseudomonadota>Betaproteobacteria>Burkholderiales>Burkholderiaceae>Paraburkholderia | GCF_013371975.1 | Paraburkholderia_youngii | FSB08_RS32050 | 68379..72011 | 2782701 | BREX-system-P-loop-protein-BrxC-[Paraburkholderia-youngii] |
| WP_254334233.1 | Type1.BREX | [illegible] | 1210 | Bacteria>Pseudomonadota>Betaproteobacteria>Burkholderiales>Burkholderiaceae>Paraburkholderia | GCF_024206475.1 | Paraburkholderia_sp._CNPSo_3274 | M3I54_RS07285 | 43504..47136 | 2940932 | BREX-system-P-loop-protein-BrxC-[Paraburkholderia-sp.-CNPSo-3274] |
| WP_167306478.1 | Type1.BREX | [illegible] | 1182 | Bacteria>Pseudomonadota>Betaproteobacteria>Burkholderiales>Burkholderiaceae>Paraburkholderia | GCF_900100565.1 | Paraburkholderia_steynii | BLT65_RS12240 | 289685..293233 | 1245441 | BREX-system-P-loop-protein-BrxC-[Paraburkholderia-steynii] |
| WP_165186500.1 | Type1.BREX | [illegible] | 1210 | Bacteria>Pseudomonadota>Betaproteobacteria>Burkholderiales>Burkholderiaceae>Paraburkholderia | GCF_902459535.1 | Paraburkholderia_dioscoreae | PDMSB3_RS12750 | 2839967..2843599 | 2604047 | BREX-system-P-loop-protein-BrxC-[Paraburkholderia-dioscoreae] |
| WP_201659285.1 | Type1.BREX | [illegible] | 1218 | Bacteria>Pseudomonadota>Betaproteobacteria>Burkholderiales>Burkholderiaceae>Paraburkholderia | GCF_904848645.1 | Paraburkholderia_sabiae | JMY25_RS35995 | 30018..33674 | 273251 | BREX-system-P-loop-protein-BrxC-[Paraburkholderia-sabiae] |
| WP_201107905.1 | Type1.BREX | [illegible] | 1179 | Bacteria>Pseudomonadota>Betaproteobacteria>Burkholderiales>Burkholderiaceae>Paraburkholderia | GCF_905220855.1 | Paraburkholderia_domus | KHO36_RS10755 | 51203..54742 | 2793075 | BREX-system-P-loop-protein-BrxC-[Paraburkholderia-domus] |
| AGW91529.1 | Type1.BREX | [illegible] | 1153 | Bacteria>Pseudomonadota>Betaproteobacteria>Burkholderiales>Burkholderiaceae>Ralstonia | GCA_000471925.1 | Ralstonia_pickettii_DTP0602 | N234_15980 | 3439152..3442613 | 1366050 | hypothetical-protein-N234_15980-[Ralstonia-pickettii-DTP0602] |
| RAA12362.1 | Type1.BREX | [illegible] | 1153 | Bacteria>Pseudomonadota>Betaproteobacteria>Burkholderiales>Burkholderiaceae>Ralstonia | GCA_003256405.1 | Ralstonia_pseudosolanacearum | DOT79_19220 | 30474..33935 | 1310165 | BREX-system-P-loop-protein-BrxC-[Ralstonia-pseudosolanacearum] |
| MBN6206497.1 | Type1.BREX | [illegible] | 1366 | Bacteria>Pseudomonadota>Betaproteobacteria>Burkholderiales>Burkholderiaceae>Ralstonia | GCA_017162415.1 | Ralstonia_pickettii | JYK21_08530 | 22216..26316 | 329 | hypothetical-protein-[Ralstonia-pickettii] |
| UGS92597.1 | Type1.BREX | [illegible] | 1153 | Bacteria>Pseudomonadota>Betaproteobacteria>Burkholderiales>Burkholderiaceae>Ralstonia | GCA_021173085.1 | Ralstonia_wenshanensis | KOL96_10360 | 529423..532884 | 2842456 | BREX-system-P-loop-protein-BrxC-[Ralstonia-wenshanensis] |
| WP_223259776.1 | Type1.BREX | [illegible] | 1156 | Bacteria>Pseudomonadota>Betaproteobacteria>Burkholderiales>Burkholderiaceae>Ralstonia | GCF_003256405.1 | Ralstonia_pseudosolanacearum | DOT79_RS19235 | 30465..33935 | 1310165 | BREX-system-P-loop-protein-BrxC-[Ralstonia-pseudosolanacearum] |
| WP_232042032.1 | Type1.BREX | [illegible] | 1156 | Bacteria>Pseudomonadota>Betaproteobacteria>Burkholderiales>Burkholderiaceae>Ralstonia | GCF_021173085.1 | Ralstonia_wenshanensis | KOL96_RS10420 | 529414..532884 | 2842456 | BREX-system-P-loop-protein-BrxC-[Ralstonia-wenshanensis] |
| MBC7381112.1 | Type1.BREX | [illegible] | 1146 | Bacteria>Pseudomonadota>Betaproteobacteria>Burkholderiales>Burkholderiaceae>unclassified_Burkholderiaceae | GCA_014376335.1 | Burkholderiaceae_bacterium | H7346_27255 | 2496..5936 | 2030806 | BREX-system-P-loop-protein-BrxC-[Burkholderiaceae-bacterium] |
| MBK8767853.1 | Type1.BREX | [illegible] | 1148 | Bacteria>Pseudomonadota>Betaproteobacteria>Burkholderiales>Burkholderiaceae>unclassified_Burkholderiaceae | GCA_016710525.1 | Burkholderiaceae_bacterium | IPM01_25890 | 21329..24775 | 2030806 | BREX-system-P-loop-protein-BrxC-[Burkholderiaceae-bacterium] |
| MBX9901152.1 | Type1.BREX | [illegible] | 1220 | Bacteria>Pseudomonadota>Betaproteobacteria>Burkholderiales>Burkholderiaceae>unclassified_Burkholderiaceae | GCA_019745005.1 | Burkholderiaceae_bacterium | K2Y28_10275 | 3751..7413 | 2030806 | BREX-system-P-loop-protein-BrxC-[Burkholderiaceae-bacterium] |
| MCD6671945.1 | Type1.BREX | [illegible] | 1148 | Bacteria>Pseudomonadota>Betaproteobacteria>Burkholderiales>Burkholderiaceae>unclassified_Burkholderiaceae | GCA_021166835.1 | Burkholderiaceae_bacterium | LT106_03745 | 32812..36258 | 2030806 | BREX-system-P-loop-protein-BrxC-[Burkholderiaceae-bacterium] |
| MCK6425125.1 | Type1.BREX | [illegible] | 1081 | Bacteria>Pseudomonadota>Betaproteobacteria>Burkholderiales>Burkholderiaceae>unclassified_Burkholderiaceae | GCA_023150375.1 | Burkholderiaceae_bacterium | L6Q75_08530 | 1698..4943 | 2030806 | BREX-system-P-loop-protein-BrxC-[Burkholderiaceae-bacterium] |
| MCK6421971.1 | Type1.BREX | [illegible] | 1153 | Bacteria>Pseudomonadota>Betaproteobacteria>Burkholderiales>Burkholderiales_genera_incertae_sedis>Aquabacterium | GCA_023150345.1 | Aquabacterium_sp. | L6Q73_13845 | 24265..27726 | 1872578 | BREX-system-P-loop-protein-BrxC-[Aquabacterium-sp.] |
| WP_243738604.1 | Type1.BREX | [illegible] | 1224 | Bacteria>Pseudomonadota>Betaproteobacteria>Burkholderiales>Burkholderiales_genera_incertae_sedis>Aquabacterium | GCF_004362855.1 | Aquabacterium_commune | EV672_RS09250 | 184197..187871 | 70586 | BREX-system-P-loop-protein-BrxC-[Aquabacterium-commune] |
| WP_008906812.1 | Type1.BREX | [illegible] | 1213 | Bacteria>Pseudomonadota>Betaproteobacteria>Burkholderiales>Comamonadaceae>Acidovorax | GCF_000238595.1 | Acidovorax_sp._NO-1 | KYG_RS20325 | 5938..9579 | 512030 | BREX-system-P-loop-protein-BrxC-[Acidovorax-sp.-NO-1] |
| WP_086927320.1 | Type1.BREX | [illegible] | 1153 | Bacteria>Pseudomonadota>Betaproteobacteria>Burkholderiales>Comamonadaceae>Acidovorax | GCF_002157125.1 | Acidovorax_carolinensis | CBP36_RS10055 | 2162910..2166371 | 553814 | BREX-system-P-loop-protein-BrxC-[Acidovorax-carolinensis] |
| WP_094097616.1 | Type1.BREX | [illegible] | 1214 | Bacteria>Pseudomonadota>Betaproteobacteria>Burkholderiales>Comamonadaceae>Acidovorax | GCF_002157145.1 | Acidovorax_carolinensis | CBP34_RS06480 | 1379414..1383058 | 553814 | BREX-system-P-loop-protein-BrxC-[Acidovorax-carolinensis] |
| WP_086912102.1 | Type1.BREX | [illegible] | 1213 | Bacteria>Pseudomonadota>Betaproteobacteria>Burkholderiales>Comamonadaceae>Acidovorax | GCF_002157165.1 | Acidovorax_carolinensis | CBP33_RS08115 | 1704915..1708556 | 553814 | BREX-system-P-loop-protein-BrxC-[Acidovorax-carolinensis] |
| WP_204247869.1 | Type1.BREX | [illegible] | 590 | Bacteria>Pseudomonadota>Betaproteobacteria>Burkholderiales>Comamonadaceae>Acidovorax | GCF_002157185.1 | Acidovorax_carolinensis | CBP35_RS21920 | 1914645..1916417 | 553814 | BREX-system-P-loop-protein-BrxC-[Acidovorax-carolinensis] |
| GAO25248.1 | Type1.BREX | [illegible] | 1205 | Bacteria>Pseudomonadota>Betaproteobacteria>Burkholderiales>Comamonadaceae>Alicycliphilus | GCA_000974645.1 | Alicycliphilus_sp._B1 | ALISP_5068 | 1538304..1541921 | 1603291 | ATPase-like-protein-[Alicycliphilus-sp.-B1] |
| WP_103018746.1 | Type1.BREX | [illegible] | 1153 | Bacteria>Pseudomonadota>Betaproteobacteria>Burkholderiales>Comamonadaceae>Alicycliphilus | GCF_002894355.1 | Alicycliphilus_denitrificans | CUB93_RS20845 | 10909..14370 | 179636 | BREX-system-P-loop-protein-BrxC-[Alicycliphilus-denitrificans] |
| WP_169107995.1 | Type1.BREX | [illegible] | 1153 | Bacteria>Pseudomonadota>Betaproteobacteria>Burkholderiales>Comamonadaceae>Comamonas | GCF_012844455.2 | Comamonas_suwonensis | HF327_RS09545 | 49446..52907 | 2606214 | BREX-system-P-loop-protein-BrxC-[Comamonas-suwonensis] |
| KEH09926.1 | Type1.BREX | [illegible] | 1185 | Bacteria>Pseudomonadota>Betaproteobacteria>Burkholderiales>Comamonadaceae>Delftia | GCA_000710165.1 | Delftia_tsuruhatensis | GY14_08945 | 17729..21286 | 180282 | hypothetical-protein-GY14_08945-[Delftia-tsuruhatensis] |
| OJX16153.1 | Type1.BREX | [illegible] | 1181 | Bacteria>Pseudomonadota>Betaproteobacteria>Burkholderiales>Comamonadaceae>Delftia | GCA_001898745.1 | Delftia_sp._67-8 | BGO79_00315 | 75959..79504 | 1895749 | hypothetical-protein-BGO79_00315-[Delftia-sp.-67-8] |
| MBL8354914.1 | Type1.BREX | [illegible] | 1153 | Bacteria>Pseudomonadota>Betaproteobacteria>Burkholderiales>Comamonadaceae>Delftia | GCA_016790545.1 | Delftia_acidovorans | JNM01_08765 | 1915..5376 | 80866 | BREX-system-P-loop-protein-BrxC-[Delftia-acidovorans] |
| TFI47144.1 | Type1.BREX | [illegible] | 1188 | Bacteria>Pseudomonadota>Betaproteobacteria>Burkholderiales>Comamonadaceae>Diaphorobacter | GCA_004563745.1 | Diaphorobacter_sp._DS2 | E4O93_14250 | 10625..14191 | 1302548 | BREX-system-P-loop-protein-BrxC-[Diaphorobacter-sp.-DS2] |
| WP_172207178.1 | Type1.BREX | [illegible] | 1213 | Bacteria>Pseudomonadota>Betaproteobacteria>Burkholderiales>Comamonadaceae>Diaphorobacter | GCF_013154915.1 | Diaphorobacter_sp._JS3050 | HND92_RS11875 | 2478331..2481972 | 2735554 | BREX-system-P-loop-protein-BrxC-[Diaphorobacter-sp.-JS3050] |
| WP_054075976.1 | Type1.BREX | [illegible] | 1209 | Bacteria>Pseudomonadota>Betaproteobacteria>Burkholderiales>Comamonadaceae | GCF_001294075.1 | Comamonas | KF712C_RS25790 | 15422..19051 | 283 | MULTISPECIES:-BREX-system-P-loop-protein-BrxC-[Comamonas] |
| MBN9372145.1 | Type1.BREX | [illegible] | 1153 | Bacteria>Pseudomonadota>Betaproteobacteria>Burkholderiales>Comamonadaceae>Hydrogenophaga | GCA_017307075.1 | Hydrogenophaga_sp. | JOI38_13780 | 21785..25246 | 1904254 | BREX-system-P-loop-protein-BrxC-[Hydrogenophaga-sp.] |
| WP_070397447.1 | Type1.BREX | [illegible] | 1153 | Bacteria>Pseudomonadota>Betaproteobacteria>Burkholderiales>Comamonadaceae>Hydrogenophaga | GCF_001777155.1 | Hydrogenophaga_sp._PML113 | BHU06_RS01940 | 415865..419326 | 1899350 | BREX-system-P-loop-protein-BrxC-[Hydrogenophaga-sp.-PML113] |
| WP_226462338.1 | Type1.BREX | [illegible] | 1204 | Bacteria>Pseudomonadota>Betaproteobacteria>Burkholderiales>Comamonadaceae>Hydrogenophaga | GCF_020510545.1 | Hydrogenophaga_taeniospiralis | KI616_RS15915 | 3311609..3315223 | 65656 | BREX-system-P-loop-protein-BrxC-[Hydrogenophaga-taeniospiralis] |
| WP_104799767.1 | Type1.BREX | [illegible] | 1202 | Bacteria>Pseudomonadota>Betaproteobacteria>Burkholderiales>Comamonadaceae>Limnohabitans | GCF_002943475.1 | Limnohabitans_sp._TS-CS-82 | C5F52_RS18160 | 199496..203104 | 2094193 | BREX-system-P-loop-protein-BrxC-[Limnohabitans-sp.-TS-CS-82] |
| WP_108373352.1 | Type1.BREX | [illegible] | 1209 | Bacteria>Pseudomonadota>Betaproteobacteria>Burkholderiales>Comamonadaceae>Limnohabitans | GCF_003063315.1 | Limnohabitans_sp._T6-5 | B9Z51_RS07570 | 412563..416192 | 1100724 | BREX-system-P-loop-protein-BrxC-[Limnohabitans-sp.-T6-5] |
| WP_108330269.1 | Type1.BREX | [illegible] | 1154 | Bacteria>Pseudomonadota>Betaproteobacteria>Burkholderiales>Comamonadaceae>Limnohabitans | GCF_003063435.1 | Limnohabitans_sp._2KL-1 | B9Z47_RS05555 | 224189..227653 | 1100699 | BREX-system-P-loop-protein-BrxC-[Limnohabitans-sp.-2KL-1] |
| WP_108311897.1 | Type1.BREX | [illegible] | 1153 | Bacteria>Pseudomonadota>Betaproteobacteria>Burkholderiales>Comamonadaceae>Limnohabitans | GCF_003063455.1 | Limnohabitans_parvus | B9Z37_RS03875 | 837390..840851 | 540061 | BREX-system-P-loop-protein-BrxC-[Limnohabitans-parvus] |
| WP_101103324.1 | Type1.BREX | [illegible] | 1153 | Bacteria>Pseudomonadota>Betaproteobacteria>Burkholderiales>Comamonadaceae>Macromonas | GCF_002837135.1 | Macromonas_bipunctata | CON76_RS10585 | 3906..7367 | 183670 | BREX-system-P-loop-protein-BrxC-[Macromonas-bipunctata] |
| WP_161125024.1 | Type1.BREX | [illegible] | 1153 | Bacteria>Pseudomonadota>Betaproteobacteria>Burkholderiales>Comamonadaceae>Malikia | GCF_009867145.1 | Malikia_spinosa | F5985_RS08150 | 68993..72454 | 86180 | BREX-system-P-loop-protein-BrxC-[Malikia-spinosa] |
| WP_161126216.1 | Type1.BREX | [illegible] | 1153 | Bacteria>Pseudomonadota>Betaproteobacteria>Burkholderiales>Comamonadaceae>Malikia | GCF_009867145.1 | Malikia_spinosa | F5985_RS16415 | 9368..12829 | 86180 | BREX-system-P-loop-protein-BrxC-[Malikia-spinosa] |

| Protein-ID | System | Gene-Neighbourhoods | Length | Taxonomy | Assembly-ID | Strain | Locus-Tag | Genomic-coordinates | TaxID | Fasta-define |
|---|---|---|---|---|---|---|---|---|---|---|
| MBK6612622.1 | Type1.BREX | TraJ-RHH-><-?<-?<-SIGMA+OB-Fold+AAA-ATPase+Lon-Protease-BrxL<-PglZ<-?<-viral-capsid<-ZnR<-N6-METHYLASE<-DUF6079*<-BrxB-BrxF-iSTAND<-BrxA-DNA-binding<-RHH_5<-MULE-TRANSPOSASE<-?<-Phage_integrase<-DUF499\|\|HEPN->HTH+REase-> | 1153 | Bacteria>Pseudomonadota>Betaproteobacteria>Burkholderiales>Comamonadaceae>Ottowia | GCA_016703855.1 | Ottowia_sp. | IPG27_00505 | 113652..117113 | 1898956 | BREX-system-P-loop-protein-BrxC-[Ottowia-sp.] |
| OOG58024.1 | Type1.BREX | <-HTH_3<-?<-?\|\|?-><-SIGMA+OB-Fold+AAA-ATPase+Lon-Protease-BrxL<-PglZ<-HEPN<-N6-METHYLASE<-DUF6079*<-BrxB-BrxF-iSTAND<-BrxA-DNA-binding\|\|?->HD+PspB-> | 1207 | Bacteria>Pseudomonadota>Betaproteobacteria>Burkholderiales>Comamonadaceae>Polaromonas | GCA_002002705.1 | Polaromonas_sp._C04 | BOE49_04110 | 64714..68337 | 1945857 | ATPase-[Polaromonas-sp.-C04] |
| TAG33475.1 | Type1.BREX | <-DUF6079*<-BrxB-BrxF-iSTAND<-BrxA-DNA-binding<-HTH_26<-?<-Peptidase_M20\|\|?->?->?->ABC-ATPase+SbcC-> | 1139 | Bacteria>Pseudomonadota>Betaproteobacteria>Burkholderiales>Comamonadaceae>Polaromonas | GCA_004293655.1 | Polaromonas_sp. | EAZ34_07410 | 1..3418 | 1869339 | MAG:-BREX-system-P-loop-protein-BrxC,-partial-[Polaromonas-sp.] |
| MBG6075367.1 | Type1.BREX | <-SWIM<-SIGMA+OB-Fold+AAA-ATPase+Lon-Protease-BrxL<-PglZ<-HTH+REase<-DOC+HTH<-?<-N6-METHYLASE<-Couple_hipA+HipA_C<-HTH_3<-DUF6079-p*\|\|?-><-BrxB-BrxF-iSTAND<-BrxA-DNA-binding<-?<-?<-Phage_integrase\|\|?->?->ABhydrolase+ABhydrolase-> | 921 | Bacteria>Pseudomonadota>Betaproteobacteria>Burkholderiales>Comamonadaceae>Polaromonas | GCA_015752225.1 | Polaromonas_sp._CG_9.11 | IWX85_001192 | 30062..32827 | 2787730 | hypothetical-protein-[Polaromonas-sp.-CG_9.11] |
| WP_011485879.1 | Type1.BREX | N6-METHYLASE-><-?<-?<-?<-REC<-HISKIN\|\|HTH_3->BrxA-DNA-binding->BrxB-BrxF-iSTAND->DUF6079*->CHTH->Tox-RES->N6-METHYLASE->HEPN->NERD+Pkinase+Pkinase+SF1-SF2-Helicase->?->PglZ->SIGMA+OB-Fold+AAA-ATPase+Lon-Protease-BrxL-><-PHP+SbcC | 1202 | Bacteria>Pseudomonadota>Betaproteobacteria>Burkholderiales>Comamonadaceae>Polaromonas | GCF_000013865.1 | Polaromonas_sp._JS666 | BPRO_RS25145 | 121747..125355 | 296591 | BREX-system-P-loop-protein-BrxC-[Polaromonas-sp.-JS666] |
| WP_231379566.1 | Type1.BREX | <-SWIM<-SIGMA+OB-Fold+AAA-ATPase+Lon-Protease-BrxL<-PglZ<-HTH+REase<-DOC+HTH<-?<-N6-METHYLASE<-Couple_hipA+HipA_C<-HTH_3<-DUF6079*<-BrxB-BrxF-iSTAND<-BrxA-DNA-binding<-?<-?<-Phage_integrase | 1220 | Bacteria>Pseudomonadota>Betaproteobacteria>Burkholderiales>Comamonadaceae>Polaromonas | GCF_015752225.1 | Polaromonas_sp._CG_9.11 | IWX85_RS05685 | 30062..33724 | 2787730 | BREX-system-P-loop-protein-BrxC-[Polaromonas-sp.-CG_9.11] |
| WP_169420224.1 | Type1.BREX | <-SIGMA+OB-Fold+AAA-ATPase+Lon-Protease-BrxL<-PglZ<-ParB+HNH<-?<-TPR+REase<-?<-?<-N6-METHYLASE<-DUF6079*<-BrxB-BrxF-iSTAND<-BrxA-DNA-binding<-RHH_5<-HAD+CHTH<-?<-ParA-Soj-PloopNTPase\|\|?->D5ATPase-> | 1153 | Bacteria>Pseudomonadota>Betaproteobacteria>Burkholderiales>Comamonadaceae>Ramlibacter | GCF_012927085.1 | Ramlibacter_agri | HHL11_RS19385 | 189581..193042 | 2728837 | BREX-system-P-loop-protein-BrxC-[Ramlibacter-agri] |
| APW46516.1 | Type1.BREX | Glyco_transf_9-><-?\|\|?->?->TPR+TPR+TPR->?->tRNA-><-?\|\|DUF6079-p*->MarR-HTH+Polbeta->HEPN->HEPN->N6-METHYLASE-p->N6-METHYLASE->?->?->?-><-DDE_Tnp_1+DDE_Tnp_1 | 269 | Bacteria>Pseudomonadota>Betaproteobacteria>Burkholderiales>Comamonadaceae>Rhodoferax | GCA_001955735.1 | Rhodoferax_antarcticus | RA876_09190 | 1894413..1895222 | 81479 | hypothetical-protein-RA876_09190-[Rhodoferax-antarcticus] |
| MBP9058688.1 | Type1.BREX | <-PglZ<-SLOG+BirA-HTH<-SF1-SF2-Helicase+PRTase<-?<-TPR+REase<-?<-N6-METHYLASE<-HEPN<-MarR-HTH+Polbeta<-DUF6079*<-BrxB-BrxF-iSTAND<-BrxA-DNA-binding<-RHH_5\|\|N6-METHYLASE->ORC-CDC6+ZnR-> | 1153 | Bacteria>Pseudomonadota>Betaproteobacteria>Burkholderiales>Comamonadaceae>Rhodoferax | GCA_018057375.1 | Rhodoferax_sp. | KBF98_00065 | 15882..19343 | 50421 | BREX-system-P-loop-protein-BrxC-[Rhodoferax-sp.] |
| MCF8208673.1 | Type1.BREX | <-DUF6079-p*<-BrxB-BrxF-iSTAND<-BrxA-DNA-binding<-Peptidase_M3_N+Peptidase_M3\|\|NTP_transf->dCache_1+HisKA+HATPase_c->TPR+TPR+TPR+TPR+TPR+TPR+HISKIN+REC-><-REC+GGDEF+EAL<-REC+OmpR-HTH | 609 | Bacteria>Pseudomonadota>Betaproteobacteria>Burkholderiales>Comamonadaceae>Rhodoferax | GCA_021738345.1 | Rhodoferax_sp. | K9K38_04605 | 1..1828 | 50421 | BREX-system-P-loop-protein-BrxC-[Rhodoferax-sp.] |
| MCF8211220.1 | Type1.BREX | BrxA-DNA-binding->BrxB-BrxF-iSTAND->DUF6079*->MarR-HTH+Polbeta->OLD-ABC->REase->HEPN->N6-METHYLASE-p-> | 1153 | Bacteria>Pseudomonadota>Betaproteobacteria>Burkholderiales>Comamonadaceae>Rhodoferax | GCA_021738345.1 | Rhodoferax_sp. | K9K38_17735 | 1925..5386 | 50421 | BREX-system-P-loop-protein-BrxC-[Rhodoferax-sp.] |
| WP_139313405.1 | Type1.BREX | ABC-ATPase+SbcC->?->?->Glyco_transf_9-><-?\|\|?->?->TPR+TPR+TPR->?->tRNA->\|\|DUF6079-p*->MarR-HTH+Polbeta->HEPN->?->HEPN->N6-METHYLASE-p->N6-METHYLASE->?->?->RAMA-> | 323 | Bacteria>Pseudomonadota>Betaproteobacteria>Burkholderiales>Comamonadaceae>Rhodoferax | GCF_001955735.1 | Rhodoferax_antarcticus | RA876_RS09190 | 1894251..1895222 | 81479 | hypothetical-protein-[Rhodoferax-antarcticus] |
| WP_264569449.1 | Type1.BREX | <-Glyco_transf_9\|\|?->?->?->TPR+TPR+TPR->?->tRNA->\|\|DUF6079-p*->MarR-HTH+Polbeta->HEPN->?->HEPN->N6-METHYLASE-p->N6-METHYLASE-p-> | 323 | Bacteria>Pseudomonadota>Betaproteobacteria>Burkholderiales>Comamonadaceae>Rhodoferax | GCF_025961535.1 | Rhodoferax_antarcticus | M2244_RS19480 | 9793..10764 | 81479 | hypothetical-protein-[Rhodoferax-antarcticus] |
| PIQ51624.1 | Type1.BREX | <-OLD-ABC<-HEPN<-?<-N6-METHYLASE<-?<-?<-N6-METHYLASE-p<-AbiEii+ZnR<-AbiEi_4+AbiEi_1<-DUF6079*<-BrxB-BrxF-iSTAND<-BrxA-DNA-binding<-wHTH+WYL+WCX<-?<-RADICAL-SAM | 1204 | Bacteria>Pseudomonadota>Betaproteobacteria>Burkholderiales>Comamonadaceae>unclassified_Comamonadaceae | GCA_002786915.1 | Comamonadaceae_bacterium_CG12_big_fil_rev_8_21_14_0_65_59_15 | COW02_13880 | 31766..35380 | 1973952 | ATPase-[Comamonadaceae-bacterium-CG12_big_fil_rev_8_21_14_0_65_59_15] |
| PIQ50876.1 | Type1.BREX | dCache_1+HisKA+HATPase_c-><-SIGMA+OB-Fold+AAA-ATPase+Lon-Protease-BrxL<-PglZ<-Schlafen+HATPase+wHTH<-SbcC<-N6-METHYLASE<-PIN<-MazE_antitoxin<-DUF6079*<-BrxB-BrxF-iSTAND<-BrxA-DNA-binding<-RHH_5\|\|?->SF1-SF2-Helicase+REase->?->RelE-ParE-> | 1153 | Bacteria>Pseudomonadota>Betaproteobacteria>Burkholderiales>Comamonadaceae>unclassified_Comamonadaceae | GCA_002786915.1 | Comamonadaceae_bacterium_CG12_big_fil_rev_8_21_14_0_65_59_15 | COW02_17785 | 48153..51614 | 1973952 | hypothetical-protein-COW02_17785-[Comamonadaceae-bacterium-CG12_big_fil_rev_8_21_14_0_65_59_15] |
| MBH2043138.1 | Type1.BREX | RHH_5->BrxA-DNA-binding->BrxB-BrxF-iSTAND->DUF6079*->MarR-HTH+Polbeta->HEPN->OLD-ABC->HNH->PglZ->SIGMA+OB-Fold+AAA-ATPase+Lon-Protease-BrxL-> | 1153 | Bacteria>Pseudomonadota>Betaproteobacteria>Burkholderiales>Comamonadaceae>unclassified_Comamonadaceae | GCA_016000285.1 | Comamonadaceae_bacterium | I8H87_06315 | 1878..5339 | 1871071 | BREX-system-P-loop-protein-BrxC-[Comamonadaceae-bacterium] |
| WP_097197978.1 | Type1.BREX | <-DOC+HTH\|\|Phage_integrase->HIN-HTH->?->LZ_Tnp_IS66+zf-IS66+DDE_Tnp_IS66+DDE_Tnp_IS66_C->?->\|\|BrxA-DNA-binding->BrxB-BrxF-iSTAND->DUF6079*->?->N6-METHYLASE->PHP+SbcC->PglZ->SIGMA+OB-Fold+AAA-ATPase+Lon-Protease-BrxL-><-RNase<-HTH+MPTase | 1153 | Bacteria>Pseudomonadota>Betaproteobacteria>Burkholderiales>Comamonadaceae>Variovorax | GCF_900215425.1 | Variovorax_sp._YR752 | CRP32_RS05955 | 1262049..1265510 | 1884383 | BREX-system-P-loop-protein-BrxC-[Variovorax-sp.-YR752] |
| WP_186915942.1 | Type1.BREX | wHTH+WYL+WCX->?->?->wHTH+WYL+WCX->ANK->XIS-HTH->BrxA-DNA-binding->BrxB-BrxF-iSTAND->DUF6079*->OLD-ABC->REase->N6-METHYLASE->ParB+HNH->PglZ->SIGMA+OB-Fold+AAA-ATPase+HKD-DNase-BrxL-> | 1183 | Bacteria>Pseudomonadota>Betaproteobacteria>Burkholderiales>Oxalobacteraceae | GCF_014284115.1 | Undibacterium | H8K35_RS08325 | 157325..160876 | 401469 | MULTISPECIES:-BREX-system-P-loop-protein-BrxC-[Undibacterium] |
| WP_253548067.1 | Type1.BREX | <-SF1-SF2-Helicase<-OLD-ABC+TOPRIM\|\|GDResolvase+HIN-HTH-><-SIGMA+OB-Fold+AAA-ATPase+Lon-Protease-BrxL<-PglZ<-N6-METHYLASE<-N6-METHYLASE-p<-DUF6079*<-BrxB-BrxF-iSTAND<-BrxA-DNA-binding<-RHH_5\|\|GDResolvase+HIN-HTH->?-><-?<-XIS-HTH\|\|XIS-HTH->REase+SF1-SF2-Helicase-> | 1153 | Bacteria>Pseudomonadota>Betaproteobacteria>Burkholderiales>Oxalobacteraceae>Herbaspirillum | GCF_024170025.1 | Herbaspirillum_rubrisubalbicans | J2S30_RS12610 | 2752801..2756262 | 80842 | BREX-system-P-loop-protein-BrxC-[Herbaspirillum-rubrisubalbicans] |
| PHV04095.1 | Type1.BREX | BrxA-DNA-binding->BrxB-BrxF-iSTAND->DUF6079*->PglZ->SIGMA+OB-Fold+AAA-ATPase+Lon-Protease-BrxL->SMODS->?->JAB->SAVED->AAA-ATPase-p->Zn_Tnp_IS1595+DDE_Tnp_IS1595-> | 1205 | Bacteria>Pseudomonadota>Betaproteobacteria>Burkholderiales>Oxalobacteraceae>Janthinobacterium | GCA_002735605.1 | Janthinobacterium_sp._BJB412 | CSQ96_27685 | 29262..32879 | 1871178 | ATPase-[Janthinobacterium-sp.-BJB412] |
| WP_071078524.1 | Type1.BREX | Phage_integrase-><-?<-?<-SIGMA+OB-Fold+AAA-ATPase+Lon-Protease-BrxL<-PglZ<-HNH<-OLD-ABC<-N6-METHYLASE<-DUF6079*<-BrxB-BrxF-iSTAND<-BrxA-DNA-binding<-RHH_5<-?<-HEPN+OST-HTH | 1153 | Bacteria>Pseudomonadota>Betaproteobacteria>Burkholderiales>Oxalobacteraceae>Janthinobacterium | GCF_001854915.1 | Janthinobacterium_lividum | AKG95_RS19180 | 51113..54574 | 29581 | BREX-system-P-loop-protein-BrxC-[Janthinobacterium-lividum] |
| WP_096234088.1 | Type1.BREX | HTH_3->?->?->?->BrxA-DNA-binding->BrxB-BrxF-iSTAND->DUF6079*->N6-METHYLASE->OLD-ABC->TOPRIM->PglZ->SIGMA+OB-Fold+AAA-ATPase+Lon-Protease-BrxL->HEPN-><-TPR | 1204 | Bacteria>Pseudomonadota>Betaproteobacteria>Burkholderiales>Oxalobacteraceae>Janthinobacterium | GCF_002327145.1 | Janthinobacterium_svalbardensis | CNX70_RS06880 | 1564323..1567937 | 368607 | BREX-system-P-loop-protein-BrxC-[Janthinobacterium-svalbardensis] |
| WP_099389566.1 | Type1.BREX | HTH+MPTase->RNase-><-SIGMA+OB-Fold+AAA-ATPase+Lon-Protease-BrxL<-PglZ<-SbcC<-N6-METHYLASE<-DUF6079*<-BrxB-BrxF-iSTAND<-BrxA-DNA-binding<-?\|\|?->?-><-ABhydrolase | 1153 | Bacteria>Pseudomonadota>Betaproteobacteria>Burkholderiales>Oxalobacteraceae>Janthinobacterium | GCF_002735685.1 | Janthinobacterium_sp._BJB301 | CSQ91_RS21655 | 521004..524465 | 1560195 | BREX-system-P-loop-protein-BrxC-[Janthinobacterium-sp.-BJB301] |
| WP_219136384.1 | Type1.BREX | REase->?-><-SIGMA+OB-Fold+AAA-ATPase+Lon-Protease-BrxL<-PglZ<-?<-?<-?<-N6-METHYLASE<-DUF6079*<-BrxB-BrxF-iSTAND<-BrxA-DNA-binding\|\|TPR-><-HEPN+OST-HTH\|\|?-><-HEPN<-?<-HTH_3 | 1153 | Bacteria>Pseudomonadota>Betaproteobacteria>Burkholderiales>Oxalobacteraceae>Janthinobacterium | GCF_900095235.1 | Janthinobacterium_sp._UMAB-60 | KOF86_RS22385 | 15275..18736 | 1365365 | BREX-system-P-loop-protein-BrxC-[Janthinobacterium-sp.-UMAB-60] |
| WP_106758218.1 | Type1.BREX | AA_kinase+ACT+ACT_7->tRNA->?->?->?->BrxA-DNA-binding->BrxB-BrxF-iSTAND->DUF6079*->sGTP+GTPase-AIG->N6-METHYLASE->ParB+HNH->ZnR->viral-capsid->?->PglZ->SIGMA+OB-Fold+AAA-ATPase+Lon-Protease-BrxL-> | 1154 | Bacteria>Pseudomonadota>Betaproteobacteria>Burkholderiales>Oxalobacteraceae>Massilia_group | GCF_003011895.2 | Massilia_glaciei | C7C56_RS15200 | 27208..30672 | 1524097 | BREX-system-P-loop-protein-BrxC-[Massilia-glaciei] |
| WP_183556379.1 | Type1.BREX | <-Peptidase_M22<-?\|\|?->?->?->?->SIGMA-HTH+SIGMA-HTH+SIGMA-HTH->tRNA->BrxA-DNA-binding->BrxB-BrxF-iSTAND->DUF6079*->N6-METHYLASE->PglZ->SIGMA+OB-Fold+AAA-ATPase+Lon-Protease-BrxL->?-><-HISKIN\|\|?-><-?<-Rrf2-HTH | 1153 | Bacteria>Pseudomonadota>Betaproteobacteria>Burkholderiales>Oxalobacteraceae>Massilia_group | GCF_014200505.1 | Massilia_aurea | HD842_RS19385 | 505200..508661 | 373040 | BREX-system-P-loop-protein-BrxC-[Massilia-aurea] |
| WP_223466304.1 | Type1.BREX | BrxA-DNA-binding->BrxB-BrxF-iSTAND->DUF6079*->N6-METHYLASE->PglZ->SIGMA+OB-Fold+AAA-ATPase+Lon-Protease-BrxL->?->SMODS->SMODS->?->JAB->SAVED-> | 1205 | Bacteria>Pseudomonadota>Betaproteobacteria>Burkholderiales>Oxalobacteraceae>Massilia_group | GCF_016809835.1 | Massilia_soli | I4X03_RS04425 | 2147..5764 | 2792854 | BREX-system-P-loop-protein-BrxC-[Massilia-soli] |
| WP_258819858.1 | Type1.BREX | Phage_integrase-><-?<-SbcC<-SIGMA+OB-Fold+AAA-ATPase+HKD-DNase-BrxL<-PglZ<-N6-METHYLASE-p<-REase<-OLD-ABC<-N6-METHYLASE<-DUF6079*<-BrxB-BrxF-iSTAND<-BrxA-DNA-binding<-wHTH+WYL+WCX | 1186 | Bacteria>Pseudomonadota>Betaproteobacteria>Burkholderiales>Oxalobacteraceae>Massilia_group | GCF_024753285.1 | Massilia_pinisoli | NX784_RS27510 | 42185..45745 | 1772194 | BREX-system-P-loop-protein-BrxC-[Massilia-pinisoli] |
| WP_090328302.1 | Type1.BREX | <-D5N+D5ATPase<-?<-?\|\|TIR->?-><-?\|\|BrxA-DNA-binding->BrxB-BrxF-iSTAND->DUF6079*->N6-METHYLASE->PglZ->SIGMA+OB-Fold+AAA-ATPase+Lon-Protease-BrxL->MORC+REase-><-HTH_21+rve+rve_3 | 1204 | Bacteria>Pseudomonadota>Betaproteobacteria>Burkholderiales>Oxalobacteraceae>Massilia_group | GCF_900110275.1 | Duganella_sp._CF517 | BMX85_RS29475 | 9688..13302 | 1881038 | BREX-system-P-loop-protein-BrxC-[Duganella-sp.-CF517] |
| WP_146328427.1 | Type1.BREX | <-STYKIN<-?<-?<-?\|\|?->HTH_3-><-?\|\|BrxA-DNA-binding->BrxB-BrxF-iSTAND->DUF6079*->N6-METHYLASE->ZnR->viral-capsid->?->PglZ->SIGMA+OB-Fold+AAA-ATPase+Lon-Protease-BrxL->?->CR-ATPase4-> | 1153 | Bacteria>Pseudomonadota>Betaproteobacteria>Burkholderiales>Oxalobacteraceae>Noviherbaspirillum | GCF_007874125.1 | Noviherbaspirillum_sp._UKPF54 | FAY22_RS00565 | 114292..117753 | 2601898 | BREX-system-P-loop-protein-BrxC-[Noviherbaspirillum-sp.-UKPF54] |
| WP_182219587.1 | Type1.BREX | <-RNaseP<-?\|\|RFC-AAA+DNAA-HTH->?->MORC+DNA_gyrase+TOPRIM->wHTH+WYL+WCX->BrxA-DNA-binding->BrxB-BrxF-iSTAND->DUF6079*->N6-METHYLASE->OLD-ABC->NERD->N6-METHYLASE-p->PglZ->SIGMA+OB-Fold+AAA-ATPase+HKD-DNase-BrxL->PHP->?->REase-> | 1186 | Bacteria>Pseudomonadota>Betaproteobacteria>Burkholderiales>Oxalobacteraceae>Rugamonas | GCF_014042365.1 | Rugamonas_fusca | H3H36_RS18580 | 34208..37768 | 2758568 | BREX-system-P-loop-protein-BrxC-[Rugamonas-fusca] |
| WP_151122046.1 | Type1.BREX | Phage_integrase+Phage_integrase->wHTH+WYL+WCX->BrxA-DNA-binding->BrxB-BrxF-iSTAND->DUF6079*->sGTP+GTPase-AIG->N6-METHYLASE->ZnR->viral-capsid->?->PglZ->SIGMA+OB-Fold+AAA-ATPase+Lon-Protease-BrxL->MukF->JetB-> | 1153 | Bacteria>Pseudomonadota>Betaproteobacteria>Burkholderiales>Sphaerotilaceae>Ideonella | GCF_021049305.1 | Ideonella_dechloratans | LRM40_RS14105 | 3039018..3042479 | 36863 | BREX-system-P-loop-protein-BrxC-[Ideonella-dechloratans] |
| EGJ09025.1 | Type1.BREX | <-DUF6079-p*<-BrxB-BrxF-iSTAND<-BrxA-DNA-binding<-RHH_5<-HTH_26\|\|Phage_AlpA->Arm-DNA-bind_2+Phage_int_SAM_3+Phage_integrase->?-><-URI-DNase-> | 280 | Bacteria>Pseudomonadota>Betaproteobacteria>Burkholderiales>Sphaerotilaceae>Rubrivivax | GCA_000190375.2 | Rubrivivax_benzoatilyticus_JA2_=_ATCC_BAA-35 | RBXJA2T_01790 | 1..841 | 987059 | hypothetical-protein-RBXJA2T_01790,-partial-[Rubrivivax-benzoatilyticus-JA2-=-ATCC-BAA-35] |
| WP_009855692.1 | Type1.BREX | <-SIGMA+OB-Fold+AAA-ATPase+Lon-Protease-BrxL<-PglZ<-SLOG+BirA-HTH<-SF1-SF2-Helicase+PRTase<-N6-METHYLASE<-DUF6079-p* | 874 | Bacteria>Pseudomonadota>Betaproteobacteria>Burkholderiales>Sphaerotilaceae>Rubrivivax | GCF_000190375.1 | Rubrivivax_benzoatilyticus | RBXJA2T_RS02255 | 28686..31310 | 316997 | BREX-system-P-loop-protein-BrxC,-partial-[Rubrivivax-benzoatilyticus] |
| WP_029719002.1 | Type1.BREX | <-URI-DNase<-?<-?<-Arm-DNA-bind_2+Phage_int_SAM_3+Phage_integrase<-Phage_AlpA\|\|HTH_26->RHH_5->BrxA-DNA-binding->BrxB-BrxF-iSTAND->DUF6079*->N6-METHYLASE->SF1-SF2-Helicase+PRTase->SLOG+BirA-HTH->PglZ->SIGMA+OB-Fold+AAA-ATPase+Lon-Protease-BrxL-> | 1153 | Bacteria>Pseudomonadota>Betaproteobacteria>Burkholderiales>Sphaerotilaceae>Rubrivivax | GCF_000420125.1 | Rubrivivax_benzoatilyticus | F799_RS0117210 | 9638..13099 | 316997 | BREX-system-P-loop-protein-BrxC-[Rubrivivax-benzoatilyticus] |
| WP_230248250.1 | Type1.BREX | GNTR-HTH+LRP-HTH->?->?->wHTH+WYL+WCX->BrxA-DNA-binding->BrxB-BrxF-iSTAND->DUF6079*->N6-METHYLASE->PglZ->SIGMA+OB-Fold+AAA-ATPase+Lon-Protease-BrxL->?->?->DUF932->?->ParA-Soj-PloopNTPase->ParB+RepB-> | 1207 | Bacteria>Pseudomonadota>Betaproteobacteria>Burkholderiales>Sphaerotilaceae>Rubrivivax | GCF_020966545.1 | unclassified_Rubrivivax | LOC64_RS00220 | 46220..49843 | 2649762 | MULTISPECIES:-BREX-system-P-loop-protein-BrxC-[unclassified-Rubrivivax] |
| MAG79942.1 | Type1.BREX | Phage_integrase->AbiEi_4->?-><-?\|\|ZnR+TOPRIM+AAA-helicase+wHTH->Phage_integrase->RHH_5->BrxA-DNA-binding->BrxB-BrxF-iSTAND->DUF6079*->N6-METHYLASE->PglZ->SIGMA+OB-Fold+AAA-ATPase+Lon-Protease-BrxL->?->XIS-HTH-><-?<-REase | 1155 | Bacteria>Pseudomonadota>Betaproteobacteria>Burkholderiales>Sutterellaceae>unclassified_Sutterellaceae | GCA_002690025.1 | Sutterellaceae_bacterium | CMN88_03370 | 89998..93465 | 2026794 | hypothetical-protein-[Sutterellaceae-bacterium] |
| RIK89376.1 | Type1.BREX | AAA-ATPase->HIN-HTH-><-?<-?<-?<-?\|\|GNTR-HTH+Aminotran_1_2->BrxA-DNA-binding->BrxB-BrxF-iSTAND->DUF6079*->Schlafen+HATPase+wHTH->ParA-Soj-PloopNTPase->PglZ->SIGMA+OB-Fold+AAA-ATPase+Lon-Protease-BrxL-><-?<-ABC-ATPase+SbcC<-ABC-ATPase+SbcC | 1146 | Bacteria>Pseudomonadota>Betaproteobacteria>Burkholderiales>unclassified_Burkholderiales | GCA_003577305.1 | Burkholderiales_bacterium | DCC70_08090 | 13043..16483 | 1891238 | BREX-system-P-loop-protein-BrxC-[Burkholderiales-bacterium] |
| MBL8529773.1 | Type1.BREX | <-Phage_integrase<-SIGMA+OB-Fold+AAA-ATPase+Lon-Protease-BrxL<-PglZ<-N6-METHYLASE<-DOC+HTH<-N6-METHYLASE<-DUF6079*<-BrxB-BrxF-iSTAND<-BrxA-DNA-binding<-TraG_N<-TraH<-?<-?<-TraH | 1244 | Bacteria>Pseudomonadota>Betaproteobacteria>Burkholderiales>unclassified_Burkholderiales | GCA_016791285.1 | Burkholderiales_bacterium | JNL68_18995 | 11051..14785 | 1891238 | BREX-system-P-loop-protein-BrxC-[Burkholderiales-bacterium] |
| MBN8486519.1 | Type1.BREX | <-SIR2\|\|?->wHTH+WYL+WCX->BrxA-DNA-binding->BrxB-BrxF-iSTAND->DUF6079*->N6-METHYLASE->OLD-ABC->NERD->RVT->N6-METHYLASE-p->HTH+REase->PglZ->SIGMA+OB-Fold+AAA-ATPase+HKD-DNase-BrxL-><-HISKIN | 1186 | Bacteria>Pseudomonadota>Betaproteobacteria>Burkholderiales>unclassified_Burkholderiales | GCA_017302485.1 | Burkholderiales_bacterium | JOM20_02125 | 4400..7960 | 1891238 | BREX-system-P-loop-protein-BrxC-[Burkholderiales-bacterium] |
| MBP7846551.1 | Type1.BREX | <-TrwC+SF1-SF2-Helicase<-SF1-SF2-Helicase+REase<-SIGMA+OB-Fold+AAA-ATPase+Lon-Protease-BrxL<-PglZ<-TOPRIM<-OLD-ABC<-N6-METHYLASE-p<-N6-METHYLASE<-DUF6079*<-BrxB-BrxF-iSTAND<-BrxA-DNA-binding<-SWIM+DUF3670+SF1-SF2-Helicase<-?<-SWIM+DUF3670+SF1-SF2-Helicase | 1193 | Bacteria>Pseudomonadota>Betaproteobacteria>Burkholderiales>unclassified_Burkholderiales | GCA_018001735.1 | Burkholderiales_bacterium | KAZ94_00285 | 68005..71586 | 1891238 | BREX-system-P-loop-protein-BrxC-[Burkholderiales-bacterium] |
| MBX9598966.1 | Type1.BREX | BrxA-DNA-binding->BrxB-BrxF-iSTAND->DUF6079*->N6-METHYLASE->TOPRIM->OLD-ABC->TOPRIM->PglZ->SIGMA+OB-Fold+AAA-ATPase+Lon-Protease-BrxL->?->Radical_SAM->PIN-> | 1193 | Bacteria>Pseudomonadota>Betaproteobacteria>Burkholderiales>unclassified_Burkholderiales | GCA_019748155.1 | Burkholderiales_bacterium | K2X04_10405 | 1415..4996 | 1891238 | BREX-system-P-loop-protein-BrxC-[Burkholderiales-bacterium] |
| MCP5252671.1 | Type1.BREX | <-HNH<-TraI-VirD2-RCRE<-TraJ-RHH<-XIS-HTH<-?\|\|DEDD_Tnp_IS110+Transposase_20->?->ParA\|\|AAA-ATPase->BrxB-BrxF-iSTAND->DUF6079*->Schlafen+HATPase+wHTH-><-?<-HTH_3<-Tox-RelE<-Arm-DNA-bind_3<-?\|\|HKD-DNase+HKD-DNase-> | 1144 | Bacteria>Pseudomonadota>Betaproteobacteria>Burkholderiales>unclassified_Burkholderiales | GCA_024235735.1 | Burkholderiales_bacterium | H6939_13335 | 16624..20058 | 1891238 | BREX-system-P-loop-protein-BrxC-[Burkholderiales-bacterium] |
| MCP5292583.1 | Type1.BREX | <-HKD-DNase+HKD-DNase\|\|?->Arm-DNA-bind_3->Tox-RelE->HTH_3->?-><-Schlafen+HATPase+wHTH<-DUF6079*<-BrxB-BrxF-iSTAND<-AAA-ATPase-p | 1144 | Bacteria>Pseudomonadota>Betaproteobacteria>Burkholderiales>unclassified_Burkholderiales | GCA_024235775.1 | Burkholderiales_bacterium | H6938_08650 | 46881..50315 | 1891238 | BREX-system-P-loop-protein-BrxC-[Burkholderiales-bacterium] |
| MCP5275500.1 | Type1.BREX | PhdYeFM_antitox->RelE-ParE-><-HTH_32+DDE_3<-?<-BrnT_toxin<-REase+TOPRIM<-HEPN+OST-HTH\|\|BrxA-DNA-binding->BrxB-BrxF-iSTAND->DUF6079*->ABC-ATPase->REase->N6-METHYLASE->ParB+HNH->OLD-ABC->?->PglZ->SIGMA+OB-Fold+AAA-ATPase+HKD-DNase-BrxL-> | 1176 | Bacteria>Pseudomonadota>Betaproteobacteria>Burkholderiales>unclassified_Burkholderiales | GCA_024235795.1 | Burkholderiales_bacterium | H6936_11775 | 653459..656989 | 1891238 | BREX-system-P-loop-protein-BrxC-[Burkholderiales-bacterium] |
| MCP5275800.1 | Type1.BREX | GDResolvase+Recombinase+ZnR->Arm-DNA-bind_3+Phage_integrase-><-REase+TOPRIM<-HEPN+OST-HTH\|\|BrxA-DNA-binding->BrxB-BrxF-iSTAND->DUF6079*->ABC-ATPase->REase->N6-METHYLASE-> | 1177 | Bacteria>Pseudomonadota>Betaproteobacteria>Burkholderiales>unclassified_Burkholderiales | GCA_024235795.1 | Burkholderiales_bacterium | H6936_13310 | 1016712..1020245 | 1891238 | BREX-system-P-loop-protein-BrxC-[Burkholderiales-bacterium] |
| WP_131354865.1 | Type1.BREX | RNA_METHYLASE->Arm-DNA-bind_3+Phage_integrase-><-?\|\|wHTH+WYL+WCX->BrxA-DNA-binding->BrxB-BrxF-iSTAND->DUF6079*->JAB->N6-METHYLASE->?->?->PglZ->DOC+HTH->SIGMA+OB-Fold+AAA-ATPase+Lon-Protease-BrxL-> | 1224 | Bacteria>Pseudomonadota>Betaproteobacteria>Neisseriales>Chromobacteriaceae>Aquitalea | GCF_004328945.1 | Aquitalea_sp._USM4 | DKK66_RS01410 | 305665..309339 | 1590041 | BREX-system-P-loop-protein-BrxC-[Aquitalea-sp.-USM4] |
| WP_205677506.1 | Type1.BREX | MerR-HTH->tRNA-><-STYKinase-like<-TRPR-HTH\|\|?->RHH_5->BrxA-DNA-binding->BrxB-BrxF-iSTAND->DUF6079*->N6-METHYLASE->PHP+SbcC->PglZ->SIGMA+OB-Fold+AAA-ATPase+Lon-Protease-BrxL-><-?<-Arm-DNA-bind_3+Phage_integrase\|\|tRNA->HKD-DNase-> | 1165 | Bacteria>Pseudomonadota>Betaproteobacteria>Neisseriales>Chromobacteriaceae>Aquitalea | GCF_009856625.1 | Aquitalea_denitrificans | GSR16_RS06460 | 1403398..1406895 | 519081 | BREX-system-P-loop-protein-BrxC-[Aquitalea-denitrificans] |
| WP_046167751.1 | Type1.BREX | BrxA-DNA-binding->BrxB-BrxF-iSTAND->DUF6079*->N6-METHYLASE->OLD-ABC->HNH->PglZ->SIGMA+OB-Fold+AAA-ATPase+Lon-Protease-BrxL->?->?-><-AraC-HTH+AraC-HTH | 1153 | Bacteria>Pseudomonadota>Betaproteobacteria>Neisseriales>Chromobacteriaceae>Chromobacterium_group | GCF_000971355.1 | Chromobacterium_vaccinii | VL68_RS13365 | 2032..5493 | 1108595 | BREX-system-P-loop-protein-BrxC-[Chromobacterium-vaccinii] |
| WP_047248551.1 | Type1.BREX | SbcC-><-SIGMA+OB-Fold+AAA-ATPase+Lon-Protease-BrxL<-PglZ<-PglZ-p<-TOPRIM<-OLD-ABC<-N6-METHYLASE<-DUF6079*<-BrxB-BrxF-iSTAND<-BrxA-DNA-binding<-wHTH+WYL+WCX\|\|DUF932->?->?->?-><-HTH_19 | 1228 | Bacteria>Pseudomonadota>Betaproteobacteria>Neisseriales>Chromobacteriaceae>Chromobacterium_group | GCF_001020585.1 | Chromobacterium_subtsugae | AA094_RS04310 | 55231..58917 | 251747 | BREX-system-P-loop-protein-BrxC-[Chromobacterium-subtsugae] |
| WP_199155307.1 | Type1.BREX | <-N6-METHYLASE-p<-DUF6079*<-BrxB-BrxF-iSTAND<-BrxA-DNA-binding<-wHTH+WYL+WCX\|\|<-?<-?<-?\|\|?->?-><-GGDEF | 1225 | Bacteria>Pseudomonadota>Betaproteobacteria>Neisseriales>Chromobacteriaceae>Chromobacterium_group | GCF_016428945.1 | Chromobacterium_sp._ASV23 | JHD59_RS18460 | 1242..4919 | 2795110 | BREX-system-P-loop-protein-BrxC-[Chromobacterium-sp.-ASV23] |
| WP_203569835.1 | Type1.BREX | <-HAD+HAD\|\|?->HTH_Tnp_1+HTH_21+rve+rve_3->wHTH+WYL+WCX->BrxA-DNA-binding->BrxB-BrxF-iSTAND->DUF6079*->HEPN->N6-METHYLASE->?->?->?->PglZ->SIGMA+OB-Fold+AAA-ATPase+Lon-Protease-BrxL-> | 1156 | Bacteria>Pseudomonadota>Betaproteobacteria>Neisseriales>Chromobacteriaceae | GCF_014042245.2 | Deefgea | H1229_RS02985 | 646302..649772 | 400947 | MULTISPECIES:-BREX-system-P-loop-protein-BrxC-[Deefgea] |
| MCG9052529.1 | Type1.BREX | HIN-HTH-><-SIGMA+OB-Fold+AAA-ATPase+Lon-Protease-BrxL<-PglZ<-TOPRIM<-OLD-ABC<-N6-METHYLASE<-DOC+HTH<-Couple_hipA+HipA_C<-TRPR-HTH<-DUF6079*<-BrxB-BrxF-iSTAND<-BrxA-DNA-binding<-wHTH+WYL+WCX-p<-?<-XIS-HTH\|\|?-><-Arm-DNA-bind_3+Phage_integrase | 1224 | Bacteria>Pseudomonadota>Betaproteobacteria>Neisseriales>Chromobacteriaceae>Laribacter | GCA_022394235.1 | Laribacter_hongkongensis | LH447_05335 | 18538..22212 | 168471 | BREX-system-P-loop-protein-BrxC-[Laribacter-hongkongensis] |
| MCG9059785.1 | Type1.BREX | DDE_Tnp_1-><-?\|\|ARG-><-SIGMA+OB-Fold+AAA-ATPase+Lon-Protease-BrxL<-PglZ<-TOPRIM<-OLD-ABC<-N6-METHYLASE<-HKD-DNase<-DUF6079*<-BrxB-BrxF-iSTAND<-BrxA-DNA-binding<-wHTH+WYL+WCX\|\|\|\|?->DOC-><-DUF772+DDE_Tnp_1\|\|?->\|\|AAA-ATPase-><-CRISPR_Cas6 | 1217 | Bacteria>Pseudomonadota>Betaproteobacteria>Neisseriales>Chromobacteriaceae>Laribacter | GCA_022394335.1 | Laribacter_hongkongensis | LH452_12755 | 30373..34026 | 168471 | BREX-system-P-loop-protein-BrxC-[Laribacter-hongkongensis] |
| MCG9118882.1 | Type1.BREX | <-SIGMA+OB-Fold+AAA-ATPase+Lon-Protease-BrxL<-PglZ<-HNH<-?<-?<-N6-METHYLASE<-?<-DUF6079*<-BrxB-BrxF-iSTAND<-BrxA-DNA-binding<-wHTH+WYL+WCX<-?<-?<-?\|\|?->HTH_Tnp_1+HTH_21+rve+rve_3->Phage_int_SAM_1+Phage_integrase-> | 1221 | Bacteria>Pseudomonadota>Betaproteobacteria>Neisseriales>Chromobacteriaceae>Laribacter | GCA_022394685.1 | Laribacter_hongkongensis | LH456_09760 | 59894..63559 | 168471 | BREX-system-P-loop-protein-BrxC-[Laribacter-hongkongensis] |
| WP_275663441.1 | Type1.BREX | wHTH+WYL+WCX-p->BrxA-DNA-binding->BrxB-BrxF-iSTAND->DUF6079*->TRPR-HTH->Couple_hipA+HipA_C->DOC+HTH->N6-METHYLASE->?->TOPRIM->PglZ->SIGMA+OB-Fold+AAA-ATPase+Lon-Protease-BrxL-><-HIN-HTH | 1229 | Bacteria>Pseudomonadota>Betaproteobacteria>Neisseriales>Chromobacteriaceae>Laribacter | GCF_022394235.1 | Laribacter_hongkongensis | LH442_RS13325 | 2333..6022 | 168471 | BREX-system-P-loop-protein-BrxC-[Laribacter-hongkongensis] |
| WP_239907373.1 | Type1.BREX | <-SIGMA+OB-Fold+AAA-ATPase+HKD-DNase-BrxL<-PglZ<-HTH+REase<-N6-METHYLASE-p<-HerA/FtsK<-SIR2<-REase<-OLD-ABC<-N6-METHYLASE<-DUF6079*<-BrxB-BrxF-iSTAND<-BrxA-DNA-binding<-wHTH+WYL+WCX<-Phage_integrase+Phage_integrase<-HTH_36<-XIS-HTH<-?<-Arm-DNA-bind_3+Phage_integrase<-sGTP | 1185 | Bacteria>Pseudomonadota>Betaproteobacteria>Neisseriales>Chromobacteriaceae>Laribacter | GCF_022394255.1 | Laribacter_hongkongensis | LH425_05855 | 38844..42401 | 168471 | BREX-system-P-loop-protein-BrxC-[Laribacter-hongkongensis] |
| WP_239898705.1 | Type1.BREX | DDE_Tnp_1-><-?\|\|ARG-><-SIGMA+OB-Fold+AAA-ATPase+Lon-Protease-BrxL<-PglZ<-TOPRIM<-OLD-ABC<-N6-METHYLASE<-HKD-DNase<-DUF6079*<-BrxB-BrxF-iSTAND<-BrxA-DNA-binding<-wHTH+WYL+WCX\|\|\|\|?->DOC-><-DUF772+DDE_Tnp_1\|\|?->\|\|AAA-ATPase-><-Cas_Cas1 | 1222 | Bacteria>Pseudomonadota>Betaproteobacteria>Neisseriales>Chromobacteriaceae>Laribacter | GCF_022394335.1 | Laribacter_hongkongensis | LH452_RS12745 | 30373..34041 | 168471 | BREX-system-P-loop-protein-BrxC-[Laribacter-hongkongensis] |
| WP_239902638.1 | Type1.BREX | ABC-ATPase+ABC-ATPase+ABC-ATPase+SbcC->ABC-ATPase+SbcC-><-?\|\|RHH_5->BrxA-DNA-binding->BrxB-BrxF-iSTAND->DUF6079*->N6-METHYLASE->OLD-ABC+TOPRIM->SF1-SF2-Helicase->PglZ->SIGMA+OB-Fold+AAA-ATPase+Lon-Protease-BrxL->HTH_19-><-?<-Phage_integrase+Phage_integrase | 1153 | Bacteria>Pseudomonadota>Betaproteobacteria>Neisseriales>Chromobacteriaceae>Laribacter | GCF_022394475.1 | Laribacter_hongkongensis | LH449_RS00385 | 78525..81986 | 168471 | BREX-system-P-loop-protein-BrxC-[Laribacter-hongkongensis] |
| WP_239909460.1 | Type1.BREX | <-CopG/RHH<-?<-?\|\|HTH_3->?->TraI-VirD2-RCRE->wHTH+WYL+WCX->BrxA-DNA-binding->BrxB-BrxF-iSTAND->DUF6079*->N6-METHYLASE->OLD-ABC->REase->SIR2->HerA/FtsK->N6-METHYLASE-p->OLD-ABC->PglZ->SIGMA+OB-Fold+AAA-ATPase+HKD-DNase-BrxL-> | 1186 | Bacteria>Pseudomonadota>Betaproteobacteria>Neisseriales>Chromobacteriaceae>Laribacter | GCF_022394655.1 | Laribacter_hongkongensis | LH423_RS01205 | 99182..102742 | 168471 | BREX-system-P-loop-protein-BrxC-[Laribacter-hongkongensis] |
| WP_275667170.1 | Type1.BREX | <-SIGMA+OB-Fold+AAA-ATPase+Lon-Protease-BrxL<-PglZ<-HNH<-?<-?<-N6-METHYLASE<-?<-DUF6079*<-BrxB-BrxF-iSTAND<-BrxA-DNA-binding<-wHTH+WYL+WCX<-?<-?<-?\|\|?->HTH_Tnp_1+HTH_21+rve+rve_3->Phage_int_SAM_1+Phage_integrase-> | 1216 | Bacteria>Pseudomonadota>Betaproteobacteria>Neisseriales>Chromobacteriaceae>Laribacter | GCF_022394685.1 | Laribacter_hongkongensis | LH456_RS09740 | 59894..63544 | 168471 | BREX-system-P-loop-protein-BrxC-[Laribacter-hongkongensis] |
| WP_239900332.1 | Type1.BREX | HIN-HTH-><-SIGMA+OB-Fold+AAA-ATPase+Lon-Protease-BrxL<-PglZ<-TOPRIM<-OLD-ABC<-N6-METHYLASE<-DOC+HTH<-Couple_hipA+HipA_C<-TRPR-HTH<-DUF6079*<-BrxB-BrxF-iSTAND<-BrxA-DNA-binding<-wHTH+WYL+WCX-p<-?<-XIS-HTH\|\|?-><-Arm-DNA-bind_3+Phage_integrase | 1224 | Bacteria>Pseudomonadota>Betaproteobacteria>Neisseriales>Chromobacteriaceae>Laribacter | - | Laribacter_hongkongensis | LH447_05335 | 18538..22212 | 168471 | BREX-system-P-loop-protein-BrxC-[Laribacter-hongkongensis] |
| WP_239891840.1 | Type1.BREX | DDE_Tnp_1-><-?\|\|ARG-><-SIGMA+OB-Fold+AAA-ATPase+Lon-Protease-BrxL<-PglZ<-TOPRIM<-OLD-ABC<-N6-METHYLASE<-HKD-DNase<-DUF6079*<-BrxB-BrxF-iSTAND<-BrxA-DNA-binding<-wHTH+WYL+WCX\|\|\|\|?->DOC-><-DUF772+DDE_Tnp_1\|\|?->\|\|AAA-ATPase-><-CRISPR_Cas6 | 1217 | Bacteria>Pseudomonadota>Betaproteobacteria>Neisseriales>Chromobacteriaceae>Laribacter | - | Laribacter_hongkongensis | LH452_12755 | 30373..34026 | 168471 | BREX-system-P-loop-protein-BrxC-[Laribacter-hongkongensis] |
| WP_239900962.1 | Type1.BREX | TOPRIM+Primase-><-SIGMA+OB-Fold+AAA-ATPase+Lon-Protease-BrxL<-PglZ<-TOPRIM<-OLD-ABC<-N6-METHYLASE<-DOC+HTH<-Couple_hipA+HipA_C<-TRPR-HTH<-DUF6079*<-BrxB-BrxF-iSTAND<-BrxA-DNA-binding<-wHTH+WYL+WCX<-?\|\|?->35exo->?->Calcineurin-><-Arm-DNA-bind_3+Phage_integrase | 1224 | Bacteria>Pseudomonadota>Betaproteobacteria>Neisseriales>Chromobacteriaceae>Laribacter | - | Laribacter_hongkongensis | LH454_02040 | 72475..76149 | 168471 | BREX-system-P-loop-protein-BrxC-[Laribacter-hongkongensis] |
| WP_239901116.1 | Type1.BREX | <-SIGMA+OB-Fold+AAA-ATPase+Lon-Protease-BrxL<-PglZ<-HNH<-?<-?<-N6-METHYLASE<-?<-DUF6079*<-BrxB-BrxF-iSTAND<-BrxA-DNA-binding<-wHTH+WYL+WCX<-?<-?<-?\|\|?->HTH_Tnp_1+HTH_21+rve+rve_3->Phage_int_SAM_1+Phage_integrase-> | 1221 | Bacteria>Pseudomonadota>Betaproteobacteria>Neisseriales>Chromobacteriaceae>Laribacter | - | Laribacter_hongkongensis | LH456_09760 | 59894..63559 | 168471 | BREX-system-P-loop-protein-BrxC-[Laribacter-hongkongensis] |
| WP_089417825.1 | Type1.BREX | <-SIGMA+OB-Fold+AAA-ATPase+Lon-Protease-BrxL<-PglZ<-REase+AIPR<-N6-METHYLASE<-REase<-OLD-ABC<-DUF6079*<-BrxB-BrxF-iSTAND<-BrxA-DNA-binding<-wHTH+WYL+WCX<-sGTP\|\|?->RHS+RHS+RHS+RHS->?->?-><-HTH_Tnp_1 | 1153 | Bacteria>Pseudomonadota>Betaproteobacteria>Neisseriales>Neisseriaceae>Vitreoscilla | GCF_002222655.1 | Vitreoscilla_filiformis | VITFI_RS15935 | 3397543..3401004 | 63 | BREX-system-P-loop-protein-BrxC-[Vitreoscilla-filiformis] |
| NCN89398.1 | Type1.BREX | BrxA-DNA-binding->BrxB-BrxF-iSTAND->DUF6079*->AbiEi_4->AbiEii->N6-METHYLASE->PglZ->SIGMA+OB-Fold+AAA-ATPase+Lon-Protease-BrxL->MarR-HTH->Polbeta-> | 1207 | Bacteria>Pseudomonadota>Betaproteobacteria>Nitrosomonadales>Gallionellaceae>Gallionella | GCA_009992105.1 | Gallionella_sp. | GW936_08180 | 1399..5022 | 2052161 | BREX-system-P-loop-protein-BrxC-[Gallionella-sp.] |
| WP_013292857.1 | Type1.BREX | VirD4-TraG+VirB4-FtsK->?->?->Peptidase_M50->?->?-><-?\|\|BrxA-DNA-binding->BrxB-BrxF-iSTAND->DUF6079*->AbiEi_4->AbiEii->N6-METHYLASE->HTH+REase->REase->?->OLD-ABC->TOPRIM->Schlafen+HATPase+wHTH-> | 1207 | Bacteria>Pseudomonadota>Betaproteobacteria>Nitrosomonadales>Gallionellaceae>Gallionella | GCF_000145255.1 | Gallionella_capsiferriformans | GALF_RS04415 | 928777..932400 | 370405 | BREX-system-P-loop-protein-BrxC-[Gallionella-capsiferriformans] |
| WP_013293480.1 | Type1.BREX | <-Aminotran_3\|\|Phage_integrase->Phage_integrase-><-ArdA\|\|BrxA-DNA-binding->BrxB-BrxF-iSTAND->DUF6079*->HTH_3->Couple_hipA+HipA_C->N6-METHYLASE->PglZ->SIGMA+OB-Fold+AAA-ATPase+Lon-Protease-BrxL-><-PadR-HTH<-HerA/FtsK | 1161 | Bacteria>Pseudomonadota>Betaproteobacteria>Nitrosomonadales>Gallionellaceae>Gallionella | GCF_000145255.1 | Gallionella_capsiferriformans | GALF_RS07565 | 1653086..1656571 | 370405 | BREX-system-P-loop-protein-BrxC-[Gallionella-capsiferriformans] |
| TXI44178.1 | Type1.BREX | <-EVE+HNH<-wHTH+WYL+WCX<-REase+HTH+35exo+Small-B-barrel+SF1-SF2-Helicase\|\|?->BrxA-DNA-binding->BrxB-BrxF-iSTAND->DUF6079*->N6-METHYLASE->SH3->OLD-ABC+TOPRIM->ZnR+RmuC->DOC+HTH->PglZ->SIGMA+OB-Fold+AAA-ATPase+HKD-DNase-BrxL-> | 1198 | Bacteria>Pseudomonadota>Betaproteobacteria>Nitrosomonadales>Methylophilaceae>Methylophilus | GCA_008015755.1 | Methylophilus_sp. | E6Q52_09965 | 12844..16440 | 29541 | MAG:-BREX-system-P-loop-protein-BrxC-[Methylophilus-sp.] |
| ROH85757.1 | Type1.BREX | Phage_holin->Phage_lysozyme->?-><-HARE-HTH+REase<-SIGMA+OB-Fold+AAA-ATPase+Lon-Protease-BrxL<-PglZ<-ParB+HNH<-N6-METHYLASE<-DUF6079*<-BrxB-BrxF-iSTAND<-BrxA-DNA-binding<-wHTH+WYL+WCX\|\|JetA->JetB->SbcC->?-><-PIN<-HTH+MPTase | 1146 | Bacteria>Pseudomonadota>Betaproteobacteria>Nitrosomonadales>Methylophilaceae>Pseudomethylobacillus | GCA_003730115.1 | Pseudomethylobacillus_aquaticus | ED236_08405 | 68439..71879 | 2676064 | BREX-system-P-loop-protein-BrxC-[Pseudomethylobacillus-aquaticus] |
| WP_205125664.1 | Type1.BREX | Phage_holin->Phage_lysozyme->?-><-HARE-HTH+REase<-SIGMA+OB-Fold+AAA-ATPase+Lon-Protease-BrxL<-PglZ<-ParB+HNH<-N6-METHYLASE<-DUF6079*<-BrxB-BrxF-iSTAND<-BrxA-DNA-binding<-wHTH+WYL+WCX\|\|JetA->JetB->SbcC->?-><-PIN<-HTH+MPTase | 1150 | Bacteria>Pseudomonadota>Betaproteobacteria>Nitrosomonadales>Methylophilaceae>Pseudomethylobacillus | GCF_003730115.1 | Pseudomethylobacillus_aquaticus | ED236_RS08405 | 68439..71891 | 2676064 | BREX-system-P-loop-protein-BrxC-[Pseudomethylobacillus-aquaticus] |
| PTQ77404.1 | Type1.BREX | rve->Glyco_tran_28_C-><-?<-Phage_integrase+Phage_integrase<-?<-?\|\|?->BrxA-DNA-binding->BrxB-BrxF-iSTAND->DUF6079*->N6-METHYLASE->DarT+Peptidase->MACRODOMAIN->PglZ->SIGMA+OB-Fold+AAA-ATPase+Lon-Protease-BrxL-><-?<-Arm-DNA-bind_3+Phage_integrase | 1208 | Bacteria>Pseudomonadota>Betaproteobacteria>Nitrosomonadales>Nitrosomonadaceae>Nitrosomonas | GCA_003050805.1 | Nitrosomonas_oligotropha | C8R26_10850 | 43938..47564 | 42354 | hypothetical-protein-C8R26_10850-[Nitrosomonas-oligotropha] |
| MBK7364915.1 | Type1.BREX | <-HIN-HTH+rve+rve_3\|\|tRNA-><-SIGMA+OB-Fold+AAA-ATPase+HKD-DNase-BrxL<-PglZ<-N6-METHYLASE<-REase<-OLD-ABC<-DUF6079*<-BrxB-BrxF-iSTAND<-BrxA-DNA-binding<-XIS-HTH\|\|ArdA->TRPR-HTH->?-><-DOC+HTH<-HNH<-TraI-VirD2-RCRE | 1187 | Bacteria>Pseudomonadota>Betaproteobacteria>Nitrosomonadales>Nitrosomonadaceae>Nitrosomonas | GCA_016705625.1 | Nitrosomonas_sp. | IPI97_07925 | 1671776..1675339 | 42353 | BREX-system-P-loop-protein-BrxC-[Nitrosomonas-sp.] |
| MBK7491769.1 | Type1.BREX | HIN-HTH->?-><-SF1-SF2-Helicase<-?\|\|?->DUF772+DDE_Tnp_1->PIN+OST-HTH->BrxA-DNA-binding->BrxB-BrxF-iSTAND->DUF6079*->N6-METHYLASE->HEPN->HTH+REase->PglZ->SIGMA+OB-Fold+AAA-ATPase+Lon-Protease-BrxL->HIN-HTH-> | 1203 | Bacteria>Pseudomonadota>Betaproteobacteria>Nitrosomonadales>Nitrosomonadaceae>Nitrosomonas | GCA_016707165.1 | Nitrosomonas_sp. | IPI17_07005 | 52164..55775 | 42353 | BREX-system-P-loop-protein-BrxC-[Nitrosomonas-sp.] |
| MBK7352718.1 | Type1.BREX | TraI-VirD2-RCRE->HNH-><-?<-TRPR-HTH<-?<-ArdA\|\|XIS-HTH->BrxA-DNA-binding->BrxB-BrxF-iSTAND->DUF6079-p*->DUF6079-p->?->?->OLD-ABC->N6-METHYLASE->PglZ->SIGMA+OB-Fold+AAA-ATPase+HKD-DNase-BrxL->HIN-HTH->AAA-ATPase-> | 705 | Bacteria>Pseudomonadota>Betaproteobacteria>Nitrosomonadales>Nitrosomonadaceae>Nitrosomonas | GCA_016710135.1 | Nitrosomonas_sp. | IPJ05_03020 | 595038..597155 | 42353 | BREX-system-P-loop-protein-BrxC-[Nitrosomonas-sp.] |
| MBV6446594.1 | Type1.BREX | TraI-VirD2-RCRE->HNH->?-><-Couple_hipA+HipA_C<-TRPR-HTH<-ArdA\|\|XIS-HTH->BrxA-DNA-binding->BrxB-BrxF-iSTAND->DUF6079*->OLD-ABC->REase->N6-METHYLASE->PglZ->SIGMA+OB-Fold+AAA-ATPase+HKD-DNase-BrxL->SbcC+SbcC-><-Peptidase_S8<-BrxB-BrxF-iSTAND\|\|DOC+HTH-> | 1182 | Bacteria>Pseudomonadota>Betaproteobacteria>Nitrosomonadales>Nitrosomonadaceae>Nitrosomonas | GCA_019187415.1 | Nitrosomonas_sp. | FCKEDINB_00187 | 17095..20643 | 42353 | hypothetical-protein-[Nitrosomonas-sp.] |
| MBX3639968.1 | Type1.BREX | <-SIGMA+OB-Fold+AAA-ATPase+HKD-DNase-BrxL<-PglZ<-REase<-N6-METHYLASE<-?<-DUF6079*<-BrxB-BrxF-iSTAND<-BrxA-DNA-binding<-XIS-HTH\|\|HIN-HTH->SIR2-> | 1182 | Bacteria>Pseudomonadota>Betaproteobacteria>Nitrosomonadales>Nitrosomonadaceae>Nitrosomonas | GCA_019635175.1 | Nitrosomonas_sp. | KF888_05565 | 120382..123930 | 42353 | BREX-system-P-loop-protein-BrxC-[Nitrosomonas-sp.] |
| MBX9637025.1 | Type1.BREX | GDResolvase+HIN-HTH->?->BrxA-DNA-binding->BrxB-BrxF-iSTAND->DUF6079*->N6-METHYLASE->HTH+REase->PglZ->SIGMA+OB-Fold+AAA-ATPase+Lon-Protease-BrxL->SIR2->HerA/FtsK->Calcineurin-SbcD->SbcC+CCDC158-> | 1212 | Bacteria>Pseudomonadota>Betaproteobacteria>Nitrosomonadales>Nitrosomonadaceae>Nitrosomonas | GCA_019748355.1 | Nitrosomonas_sp. | K2Q45_05685 | 2429..6067 | 42353 | BREX-system-P-loop-protein-BrxC-[Nitrosomonas-sp.] |
| MCB1936159.1 | Type1.BREX | <-HTH+REase<-N6-METHYLASE<-REase<-ABC-ATPase<-DUF6079*<-BrxB-BrxF-iSTAND<-BrxA-DNA-binding\|\|HEPN+OST-HTH->REase+TOPRIM-> | 1174 | Bacteria>Pseudomonadota>Betaproteobacteria>Nitrosomonadales>Nitrosomonadaceae>Nitrosomonas | GCA_020446045.1 | Nitrosomonas_sp. | KDF59_09470 | 5848..9372 | 42353 | BREX-system-P-loop-protein-BrxC-[Nitrosomonas-sp.] |
| SDW37048.1 | Type1.BREX | <-SIGMA+OB-Fold+AAA-ATPase+Lon-Protease-BrxL<-PglZ<-ParB+HNH<-PHP+SbcC<-N6-METHYLASE<-Schlafen+HATPase+wHTH<-Schlafen+HATPase+wHTH<-DUF6079*<-BrxB-BrxF-iSTAND<-BrxA-DNA-binding\|\|JetA->JetB->SbcC->REase-> | 1145 | Bacteria>Pseudomonadota>Betaproteobacteria>Nitrosomonadales>Nitrosomonadaceae>Nitrosomonas | GCA_900106545.1 | Nitrosomonas_communis | SAMN05421882_100921 | 53532..56969 | 44574 | hypothetical-protein-SAMN05421882_100921-[Nitrosomonas-communis] |
| WP_181258202.1 | Type1.BREX | <-REase+TOPRIM<-HEPN+OST-HTH\|\|BrxA-DNA-binding->BrxB-BrxF-iSTAND->DUF6079*->N6-METHYLASE->ParB+HNH->OLD-ABC->?->PglZ->SIGMA+OB-Fold+AAA-ATPase+HKD-DNase-BrxL->SbcC->Calcineurin+Calcineurin-SbcD-><-MULE-TRANSPOSASE | 1179 | Bacteria>Pseudomonadota>Betaproteobacteria>Nitrosomonadales>Nitrosomonadaceae>Nitrosomonas | GCF_003046585.1 | Nitrosomonas_aestuarii | C8R11_107137 | 171439..174978 | 52441 | BREX-system-P-loop-protein-BrxC-[Nitrosomonas-aestuarii] |
| WP_107802981.1 | Type1.BREX | Glyco_tran_28_C-><-?<-Phage_integrase+Phage_integrase<-?<-?\|\|?->BrxA-DNA-binding->BrxB-BrxF-iSTAND->DUF6079*->N6-METHYLASE->DarT+Peptidase->MACRODOMAIN->PglZ->SIGMA+OB-Fold+AAA-ATPase+Lon-Protease-BrxL-><-?<-Arm-DNA-bind_3+Phage_integrase | 1208 | Bacteria>Pseudomonadota>Betaproteobacteria>Nitrosomonadales>Nitrosomonadaceae>Nitrosomonas | GCF_003050805.1 | Nitrosomonas_oligotropha | C8R26_RS06410 | 43938..47564 | 42354 | BREX-system-P-loop-protein-BrxC-[Nitrosomonas-oligotropha] |
| WP_107790392.1 | Type1.BREX | ParB+RepB->?-><-SbcC<-Calcineurin+Calcineurin-SbcD<-SbcC<-SIGMA+OB-Fold+AAA-ATPase+HKD-DNase-BrxL<-PglZ<-N6-METHYLASE<-Schlafen+HATPase+wHTH+HTH<-DUF6079*<-BrxB-BrxF-iSTAND<-BrxA-DNA-binding\|\|?->?->PIN-> | 1180 | Bacteria>Pseudomonadota>Betaproteobacteria>Nitrosomonadales>Nitrosomonadaceae>Nitrosomonas | GCF_003051105.1 | Nitrosomonas_nitrosa | C8R30_RS13785 | 18563..22105 | 52442 | BREX-system-P-loop-protein-BrxC-[Nitrosomonas-nitrosa] |
| WP_211752264.1 | Type1.BREX | <-SIGMA+OB-Fold+AAA-ATPase+Lon-Protease-BrxL<-PglZ<-ParB+HNH<-PHP+SbcC<-N6-METHYLASE<-Schlafen+HATPase+wHTH<-Schlafen+HATPase+wHTH<-DUF6079*<-BrxB-BrxF-iSTAND<-BrxA-DNA-binding\|\|JetA->JetB->SbcC->REase->?-><-?<-ABC-ATPase+SbcC | 1146 | Bacteria>Pseudomonadota>Betaproteobacteria>Nitrosomonadales>Nitrosomonadaceae>Nitrosomonas | GCF_900106545.1 | Nitrosomonas_communis | BLV21_RS05435 | 53532..56972 | 44574 | BREX-system-P-loop-protein-BrxC-[Nitrosomonas-communis] |
| WP_090415704.1 | Type1.BREX | <-REase<-ABC-ATPase<-DUF6079*<-BrxB-BrxF-iSTAND<-BrxA-DNA-binding\|\|LexA-protease-> | 1182 | Bacteria>Pseudomonadota>Betaproteobacteria>Nitrosomonadales>Nitrosomonadaceae>Nitrosomonas | GCF_900107165.1 | Nitrosomonas_halophila | BLV85_RS13200 | 2958..6506 | 44576 | BREX-system-P-loop-protein-BrxC-[Nitrosomonas-halophila] |
| WP_204800510.1 | Type1.BREX | <-SbcC<-Calcineurin+Calcineurin-SbcD<-SbcC<-SIGMA+OB-Fold+AAA-ATPase+HKD-DNase-BrxL<-PglZ<-HTH+REase<-?<-Schlafen+HATPase+wHTH+HTH<-DUF6079*<-BrxB-BrxF-iSTAND<-BrxA-DNA-binding<-XIS-HTH<-?<-Arm-DNA-bind_3+Phage_integrase | 1180 | Bacteria>Pseudomonadota>Betaproteobacteria>Nitrosomonadales>Nitrosomonadaceae>Nitrosomonas | GCF_905220645.1 | Nitrosomonas_nitrosa | NUZ5A_RS15530 | 137140..140682 | 52442 | BREX-system-P-loop-protein-BrxC-[Nitrosomonas-nitrosa] |
| NOT17080.1 | Type1.BREX | <-PglZ<-SF1-SF2-Helicase<-REase<-HTH+REase<-TOPRIM<-OLD-ABC<-N6-METHYLASE<-AbiEii<-AbiEi_4<-DUF6079*<-BrxB-BrxF-iSTAND<-BrxA-DNA-binding<-wHTH+WYL+WCX<-SbcC+CCDC158<-SbcC+CCDC158<-Calcineurin+Calcineurin-SbcD<-LZ_Tnp_IS66+zf-IS66+DDE_Tnp_IS66+DDE_Tnp_IS66_C<-TnpB_IS66<-HIN-HTH | 1202 | Bacteria>Pseudomonadota>Betaproteobacteria>Nitrosomonadales>Sulfuricellaceae>Sulfuriferula | GCA_013140765.1 | Sulfuriferula_sp. | HOP20_03285 | 21073..24681 | 2025307 | BREX-system-P-loop-protein-BrxC-[Sulfuriferula-sp.] |
| ODV10849.1 | Type1.BREX | <-RelE-ParE<-?<-?<-DOC<-MazE_antitoxin\|\|PhdYeFM_antitox->PIN->BrxA-DNA-binding->DUF6079-p*-> | 451 | Bacteria>Pseudomonadota>Betaproteobacteria>Nitrosomonadales>Thiobacillaceae>Thiobacillus | GCA_001725535.1 | Thiobacillus_sp._SCN_64-317 | ABT22_10740 | 12066..13421 | 1660143 | MAG:-hypothetical-protein-ABT22_10740-[Thiobacillus-sp.-SCN-64-317] |
| MBN8763784.1 | Type1.BREX | <-PRTase\|\|?->?->N6-METHYLASE-p<-DUF6079-p*<-PIN<-PhdYeFM_antitox\|\|MazE_antitoxin->DOC->?->RelE-ParE-> | 865 | Bacteria>Pseudomonadota>Betaproteobacteria>Nitrosomonadales>Thiobacillaceae>Thiobacillus | GCA_017309235.1 | Thiobacillus_sp. | J0J02_08935 | 6734..9331 | 924 | hypothetical-protein-[Thiobacillus-sp.] |
| OGS68620.1 | Type1.BREX | VirD4-TraG+VirB4-FtsK->?->?->Peptidase_M50->?->?-><-?\|\|BrxA-DNA-binding->BrxB-BrxF-iSTAND->DUF6079*->AbiEi_4->AbiEii->N6-METHYLASE->HTH+REase->?->PglZ->SIGMA+OB-Fold+AAA-ATPase+Lon-Protease-BrxL->Classical-AAA->Peptidase_S8-> | 1207 | Bacteria>Pseudomonadota>Betaproteobacteria>Nitrosomonadales>unclassified_Nitrosomonadales | GCA_001801025.1 | Gallionellales_bacterium_GWA2_54_124 | A2Z87_08865 | 78784..82407 | 1798243 | ATPase-[Gallionellales-bacterium-GWA2_54_124] |
| OGT18472.1 | Type1.BREX | wHTH+WYL+WCX-><-wHTH+AbiEi_2\|\|Ntox49->HigB_toxin->HTH_3->BrxA-DNA-binding->BrxB-BrxF-iSTAND->DUF6079*->HTH_3->DOC+HTH->N6-METHYLASE->SbcC->?->?->PglZ->SIGMA+OB-Fold+AAA-ATPase+Lon-Protease-BrxL-><-DOC+HTH | 1154 | Bacteria>Pseudomonadota>Betaproteobacteria>Nitrosomonadales>unclassified_Nitrosomonadales | GCA_001801155.1 | Gallionellales_bacterium_RIFCSPHIGHO2_02_FULL_57_16 | A3J49_13100 | 5649..9113 | 1798251 | hypothetical-protein-A3J49_13100-[Gallionellales-bacterium-RIFCSPHIGHO2_02_FULL_57_16] |
| OGS98638.1 | Type1.BREX | BrxA-DNA-binding->BrxB-BrxF-iSTAND->DUF6079*->AbiEi_4->AbiEii->N6-METHYLASE->PglZ->SIGMA+OB-Fold+AAA-ATPase+Lon-Protease-BrxL-> | 1207 | Bacteria>Pseudomonadota>Betaproteobacteria>Nitrosomonadales>unclassified_Nitrosomonadales | GCA_001801215.1 | Gallionellales_bacterium_RIFCSPLOWO2_02_FULL_59_110 | A3H99_12810 | 1693..5316 | 1798255 | ATPase-[Gallionellales-bacterium-RIFCSPLOWO2_02_FULL_59_110] |
| PIX05110.1 | Type1.BREX | <-DUF6079-p*<-BrxB-BrxF-iSTAND<-BrxA-DNA-binding<-XIS-HTH<-HTH_36<-?<-?<-?<-Phage_integrase | 340 | Bacteria>Pseudomonadota>Betaproteobacteria>Nitrosomonadales>unclassified_Nitrosomonadales | GCA_002782465.1 | Gallionellales_bacterium_CG_4_8_14_3_um_filter_54_18 | COZ77_02910 | 1..1019 | 1974014 | BREX-system-P-loop-protein-BrxC,-partial-[Gallionellales-bacterium-CG_4_8_14_3_um_filter_54_18] |
| MBK9161130.1 | Type1.BREX | Phage_holin->Phage_lysozyme->?-><-HARE-HTH+REase<-SIGMA+OB-Fold+AAA-ATPase+Lon-Protease-BrxL<-PglZ<-ParB+HNH<-N6-METHYLASE<-DUF6079*<-BrxB-BrxF-iSTAND<-BrxA-DNA-binding<-GreA_GreB+SF1-SF2-Helicase<-SLOG+BirA-HTH<-SF1-SF2-Helicase+PRTase<-XIS-HTH<-?<-ARG | 1146 | Bacteria>Pseudomonadota>Betaproteobacteria>Nitrosomonadales>unclassified_Nitrosomonadales | GCA_016716325.1 | Nitrosomonadales_bacterium | IPM27_06155 | 1292283..1295723 | 2026761 | BREX-system-P-loop-protein-BrxC-[Nitrosomonadales-bacterium] |
| MBL8410223.1 | Type1.BREX | TOPRIM->?-><-ZnR+TPR<-SIGMA+OB-Fold+AAA-ATPase+Lon-Protease-BrxL<-PglZ<-HKD-DNase<-N6-METHYLASE<-Couple_hipA+HipA_C<-HTH_3<-DUF6079*<-BrxB-BrxF-iSTAND<-BrxA-DNA-binding<-?\|\|NACHT->?->HD+Tox-RelA-> | 1154 | Bacteria>Pseudomonadota>Betaproteobacteria>Rhodocyclales>Azonexaceae>Dechloromonas | GCA_016791065.1 | Dechloromonas_sp. | JNM15_02850 | 32710..36174 | 1917218 | BREX-system-P-loop-protein-BrxC-[Dechloromonas-sp.] |
| MBT9520412.1 | Type1.BREX | <-SF1-SF2-Helicase<-OLD-ABC+TOPRIM<-MORC<-SIGMA+OB-Fold+AAA-ATPase+Lon-Protease-BrxL<-PglZ<-MarR-HTH<-N6-METHYLASE<-AbiEii<-AbiEi_4<-DUF6079*<-BrxB-BrxF-iSTAND<-BrxA-DNA-binding<-XIS-HTH<-Phage_integrase<-XIS-HTH<-HTH_3 | 1208 | Bacteria>Pseudomonadota>Betaproteobacteria>Rhodocyclales>Azonexaceae>Dechloromonas | GCA_018780195.1 | Dechloromonas_sp. | IV101_05910 | 54359..57985 | 1917218 | BREX-system-P-loop-protein-BrxC-[Dechloromonas-sp.] |
| MCK0506222.1 | Type1.BREX | <-METHYLASE<-SIGMA+OB-Fold+AAA-ATPase+Lon-Protease-BrxL<-PglZ<-PIWI<-OLD-ABC<-TOPRIM<-OLD-ABC<-N6-METHYLASE\|\|RHH_6-><-DUF6079*<-BrxB-BrxF-iSTAND<-BrxA-DNA-binding<-wHTH+WYL+WCX\|\|?->?->?-><-35exo\|\|Peptidase_M48_N+Peptidase_M48->S1+GTPase-AIG-> | 1186 | Bacteria>Pseudomonadota>Betaproteobacteria>Rhodocyclales>Rhodocyclaceae>Aromatoleum | GCA_012910705.2 | Aromatoleum_anaerobium | GO606_004945 | 104237..107797 | 182180 | BREX-system-P-loop-protein-BrxC-[Aromatoleum-anaerobium] |
| NMG55067.1 | Type1.BREX | <-TraK<-TraE<-TraM<-DUF3262<-?\|\|XIS-HTH->BrxA-DNA-binding->BrxB-BrxF-iSTAND->DUF6079*->N6-METHYLASE->OLD-ABC->REase->SIR2->HerA/FtsK->N6-METHYLASE-p->PglZ->SIGMA+OB-Fold+AAA-ATPase+HKD-DNase-BrxL->Phage-HTH-> | 1189 | Bacteria>Pseudomonadota>Betaproteobacteria>Rhodocyclales>Rhodocyclaceae>Aromatoleum | GCA_012910915.1 | Aromatoleum_aromaticum | GPA23_10435 | 14208..17777 | 551760 | BREX-system-P-loop-protein-BrxC-[Aromatoleum-aromaticum] |
| WP_011238768.1 | Type1.BREX | <-Peptidase_M48_N+Peptidase_M48\|\|35exo-><-?<-?<-?\|\|?->wHTH+WYL+WCX->BrxA-DNA-binding->BrxB-BrxF-iSTAND->DUF6079*-><-PIN<-RHH_6\|\|?->N6-METHYLASE->OLD-ABC->HNH->OLD-ABC->TOPRIM->OLD-ABC-> | 1203 | Bacteria>Pseudomonadota>Betaproteobacteria>Rhodocyclales>Rhodocyclaceae>Aromatoleum | GCF_000025965.1 | Aromatoleum_aromaticum | EBN1_RS14750 | 3092506..3096117 | 551760 | BREX-system-P-loop-protein-BrxC-[Aromatoleum-aromaticum] |
| WP_169117250.1 | Type1.BREX | <-METHYLASE<-SIGMA+OB-Fold+AAA-ATPase+Lon-Protease-BrxL<-PglZ<-PIWI<-OLD-ABC<-TOPRIM<-OLD-ABC<-N6-METHYLASE\|\|RHH_6-><-DUF6079*<-BrxB-BrxF-iSTAND<-BrxA-DNA-binding<-wHTH+WYL+WCX\|\|?->?->?-><-35exo\|\|Peptidase_M48_N+Peptidase_M48->S1+GTPase-AIG-> | 1203 | Bacteria>Pseudomonadota>Betaproteobacteria>Rhodocyclales>Rhodocyclaceae>Aromatoleum | GCF_012910705.2 | Aromatoleum_anaerobium | GO606_RS04945 | 104237..107848 | 182180 | BREX-system-P-loop-protein-BrxC-[Aromatoleum-anaerobium] |
| WP_168953997.1 | Type1.BREX | <-TraK<-TraE<-TraM<-DUF3262<-?\|\|XIS-HTH->BrxA-DNA-binding->BrxB-BrxF-iSTAND->DUF6079*->N6-METHYLASE->OLD-ABC->REase->SIR2->HerA/FtsK->N6-METHYLASE-p->PglZ->SIGMA+OB-Fold+AAA-ATPase+HKD-DNase-BrxL->Phage-HTH-> | 1188 | Bacteria>Pseudomonadota>Betaproteobacteria>Rhodocyclales>Rhodocyclaceae>Aromatoleum | GCF_023704195.1 | Aromatoleum_aromaticum | L8M16_RS08665 | 1808924..1812490 | 551760 | BREX-system-P-loop-protein-BrxC-[Aromatoleum-aromaticum] |
| WP_205250204.1 | Type1.BREX | SbcC->?-><-SIGMA+OB-Fold+AAA-ATPase+Lon-Protease-BrxL<-PglZ<-SF1-SF2-Helicase<-REase<-OLD-ABC+TOPRIM<-N6-METHYLASE<-REase<-OLD-ABC<-DUF6079*<-BrxB-BrxF-iSTAND<-BrxA-DNA-binding<-RHH_5<-HTH_37\|\|Tn7_Tnp_TnsA_N+Tn7_Tnp_TnsA_C+rve+Mu-transpos_C->TnsC-ATPase-> | 1153 | Bacteria>Pseudomonadota>Betaproteobacteria>Rhodocyclales>Rhodocyclaceae>Azospira | GCF_905120365.1 | Azospira_oryzae | JYJ48_RS05820 | 17001..20462 | 146939 | BREX-system-P-loop-protein-BrxC-[Azospira-oryzae] |
| MBP6421402.1 | Type1.BREX | SF1-SF2-Helicase-><-SIGMA+OB-Fold+AAA-ATPase+HKD-DNase-BrxL<-PglZ<-ParB+HNH<-DUF488-s<-DUF488-l<-DUF4326<-N6-METHYLASE<-N6-METHYLASE<-DUF6079-p* | 923 | Bacteria>Pseudomonadota>Betaproteobacteria>Rhodocyclales>Rhodocyclaceae>Propionivibrio | GCA_017988215.1 | Propionivibrio_sp. | KA271_00765 | 12654..15427 | 2212460 | BREX-system-P-loop-protein-BrxC-[Propionivibrio-sp.] |
| MCQ3924618.1 | Type1.BREX | ARG->Ntox49->?->?->TPR->?->REase+RNase\|\|?->BrxA-DNA-binding->BrxB-BrxF-iSTAND->DUF6079*->HTH_3->Couple_hipA+HipA_C->N6-METHYLASE->SbcC->?->PglZ->SIGMA+OB-Fold+AAA-ATPase+Lon-Protease-BrxL->GrpE->GrpE-> | 1164 | Bacteria>Pseudomonadota>Betaproteobacteria>Rhodocyclales>Rhodocyclaceae>unclassified_Rhodocyclaceae | GCA_024434155.1 | Rhodocyclaceae_bacterium | DPW12_10550 | 16269..19763 | 1898103 | BREX-system-P-loop-protein-BrxC-[Rhodocyclaceae-bacterium] |
| OHC72318.1 | Type1.BREX | <-Integrase_1\|\|BrxA-DNA-binding->BrxB-BrxF-iSTAND->DUF6079*->N6-METHYLASE->GrpE->?->HTH+REase->PglZ->SIGMA+OB-Fold+AAA-ATPase+Lon-Protease-BrxL-><-TPR+TPR+TPR+TPR+TPR+TPR+TPR<-CR-Kinase<-REase | 1152 | Bacteria>Pseudomonadota>Betaproteobacteria>Rhodocyclales>unclassified_Rhodocyclales | GCA_001828935.1 | Rhodocyclales_bacterium_RIFCSPLOWO2_02_FULL_63_24 | A3H93_14585 | 39152..42610 | 1802018 | MAG:-hypothetical-protein-A3H93_14585-[Rhodocyclales-bacterium-RIFCSPLOWO2_02_FULL_63_24] |
| WP_191660805.1 | Type1.BREX | TPR-><-RADICAL-SAM\|\|ANK+ANK+ANK-><-?\|\|Phage_integrase+Phage_integrase-><-SIGMA+OB-Fold+AAA-ATPase+Lon-Protease-BrxL<-PglZ<-MORC+DUF3684+DUF3883<-N6-METHYLASE<-DUF6079*<-BrxB-BrxF-iSTAND<-BrxA-DNA-binding<-wHTH+WYL+WCX<-HTH_26\|\|?->LON-AAA-> | 1204 | Bacteria>Pseudomonadota>Betaproteobacteria>Rhodocyclales>Zoogloeaceae>Azoarcus | - | Azoarcus_sp._Aa7 | AZOA_39690 | 55571..59185 | 2303328 | BREX-system-P-loop-protein-BrxC-[Azoarcus-sp.-Aa7] |
| NMG49986.1 | Type1.BREX | <-TraK<-TraE<-TraM<-DUF3262<-?\|\|HipA_C->XIS-HTH->BrxA-DNA-binding->BrxB-BrxF-iSTAND->DUF6079*->N6-METHYLASE->OLD-ABC->REase->SIR2->HerA/FtsK->N6-METHYLASE-p->PglZ->SIGMA+OB-Fold+AAA-ATPase+HKD-DNase-BrxL->SbcC+SbcC-> | 1189 | Bacteria>Pseudomonadota>Betaproteobacteria>Rhodocyclales>Zoogloeaceae>Parazoarcus | GCA_012910825.1 | Parazoarcus_communis | GO613_17970 | 21745..25314 | 41977 | BREX-system-P-loop-protein-BrxC-[Parazoarcus-communis] |
| WP_108973925.1 | Type1.BREX | RNase->TPR->NERD+SF1-SF2-Helicase-><-SF1-SF2-Helicase<-MORC+DUF3684+REase\|\|DUF772+DDE_Tnp_1+DDE_Tnp_1_6->wHTH+WYL+WCX->BrxA-DNA-binding->BrxB-BrxF-iSTAND->DUF6079*->N6-METHYLASE->DUF3644->HTH+REase->?->PglZ->VapB_antitoxin->PIN->SIGMA+OB-Fold+AAA-ATPase+Lon-Protease-BrxL-><-Phage_integrase | 1215 | Bacteria>Pseudomonadota>Betaproteobacteria>Rhodocyclales>Zoogloeaceae>Parazoarcus | GCF_003111665.1 | Parazoarcus_communis | CEW87_RS14060 | 3075229..3078876 | 41977 | BREX-system-P-loop-protein-BrxC-[Parazoarcus-communis] |
| WP_168942691.1 | Type1.BREX | <-TraK<-TraE<-TraM<-DUF3262<-?\|\|XIS-HTH->BrxA-DNA-binding->BrxB-BrxF-iSTAND->DUF6079*->N6-METHYLASE->OLD-ABC->REase->SIR2->HerA/FtsK->N6-METHYLASE-p->PglZ->SIGMA+OB-Fold+AAA-ATPase+HKD-DNase-BrxL->SbcC+SbcC-> | 1188 | Bacteria>Pseudomonadota>Betaproteobacteria>Rhodocyclales>Zoogloeaceae>Parazoarcus | GCF_012910825.1 | Parazoarcus_communis | GO613_RS17995 | 21748..25314 | 41977 | BREX-system-P-loop-protein-BrxC-[Parazoarcus-communis] |
| TXH82461.1 | Type1.BREX | PhdYeFM_antitox->PIN->BrxA-DNA-binding->BrxB-BrxF-iSTAND->DUF6079*->OLD-ABC->REase->N6-METHYLASE-p-> | 1153 | Bacteria>Pseudomonadota>Betaproteobacteria>Rhodocyclales>Zoogloeaceae>Thauera | GCA_008015255.1 | Thauera_aminoaromatica | E6Q80_15435 | 2474..5935 | 164330 | BREX-system-P-loop-protein-BrxC-[Thauera-aminoaromatica] |
| MBP6132433.1 | Type1.BREX | ZnR->OLD-ABC+TOPRIM->SF1-SF2-Helicase->?->?->?->RHH_5->BrxA-DNA-binding->BrxB-BrxF-iSTAND->DUF6079*->?->N6-METHYLASE-p-> | 1153 | Bacteria>Pseudomonadota>Betaproteobacteria>Rhodocyclales>Zoogloeaceae>Thauera | GCA_017985115.1 | Thauera_sp. | KA496_12935 | 35100..38561 | 1905334 | BREX-system-P-loop-protein-BrxC-[Thauera-sp.] |
| CAH1748211.1 | Type1.BREX | <-DUF3584+DUF5401<-JetB<-MukF<-?\|\|Terminase_5+HTH_33+DDE_3-><-?\|\|wHTH+WYL+WCX->BrxA-DNA-binding->BrxB-BrxF-iSTAND->DUF6079*->HKD-DNase->N6-METHYLASE->TIR->SbcC+NACHT->OLD-ABC->TOPRIM->PIWI->PglZ->SIGMA+OB-Fold+AAA-ATPase+Lon-Protease-BrxL-> | 1203 | Bacteria>Pseudomonadota>Betaproteobacteria>Rhodocyclales>Zoogloeaceae>Thauera | GCA_932126575.1 | Thauera_sp._PIV-1 | C4PIVTH_3071 | 3140142..3143753 | 204461 | ATPase-like-protein-[Thauera-sp.-PIV-1] |
| WP_037984790.1 | Type1.BREX | wHTH+WYL+WCX->BrxA-DNA-binding->BrxB-BrxF-iSTAND->DUF6079*-><-PIN<-RHH_6\|\|N6-METHYLASE->UDG->REase->?->PglZ-> | 1203 | Bacteria>Pseudomonadota>Betaproteobacteria>Rhodocyclales>Zoogloeaceae>Thauera | GCF_000310145.2 | Thauera_sp._28 | C662_RS14985 | 3144..6755 | 303682 | BREX-system-P-loop-protein-BrxC-[Thauera-sp.-28] |
| WP_043743640.1 | Type1.BREX | <-SIGMA+OB-Fold+AAA-ATPase+Lon-Protease-BrxL<-PglZ<-REase+AIPR<-N6-METHYLASE<-?<-viral-capsid<-ZnR<-N6-METHYLASE<-?<-DUF6079*<-BrxB-BrxF-iSTAND<-BrxA-DNA-binding<-RHH_5<-?<-?<-TPR+TPR<-REC+OmpR-HTH | 1153 | Bacteria>Pseudomonadota>Betaproteobacteria>Rhodocyclales>Zoogloeaceae>Thauera | GCF_000831325.1 | Thauera_sp._SWB20 | PO78_RS10565 | 31733..35194 | 1572758 | BREX-system-P-loop-protein-BrxC-[Thauera-sp.-SWB20] |
| WP_068639549.1 | Type1.BREX | <-HEPN<-HTH+REase<-N6-METHYLASE<-DUF6079*<-BrxB-BrxF-iSTAND<-BrxA-DNA-binding<-XIS-HTH | 1188 | Bacteria>Pseudomonadota>Betaproteobacteria>Rhodocyclales>Zoogloeaceae>Thauera | GCF_001591165.1 | Thauera_butanivorans | TBU01S_RS21545 | 4844..8410 | 86174 | BREX-system-P-loop-protein-BrxC-[Thauera-butanivorans] |
| WP_068803137.1 | Type1.BREX | <-HTH_3\|\|?-><-?<-SWIM<-SIGMA+OB-Fold+AAA-ATPase+Lon-Protease-BrxL<-PglZ<-REase+AIPR<-N6-METHYLASE<-?<-DUF6079*<-BrxB-BrxF-iSTAND<-BrxA-DNA-binding<-RHH_5<-?<-?<-TPR+TPR<-SF1-SF2-Helicase | 1153 | Bacteria>Pseudomonadota>Betaproteobacteria>Rhodocyclales>Zoogloeaceae>Thauera | GCF_001696715.1 | Thauera_phenolivorans | BB334_RS00200 | 55015..58476 | 1792543 | BREX-system-P-loop-protein-BrxC-[Thauera-phenolivorans] |
| WP_094269325.1 | Type1.BREX | <-wHTH+WYL+WCX<-ARG<-Ntox49\|\|35exo-><-?\|\|RHH_5->BrxA-DNA-binding->BrxB-BrxF-iSTAND->DUF6079*->sGTP+GTPase-AIG->N6-METHYLASE-p-> | 1153 | Bacteria>Pseudomonadota>Betaproteobacteria>Rhodocyclales>Zoogloeaceae>Thauera | GCF_002245655.1 | Thauera_propionica | CGK74_RS15430 | 17351..20812 | 2019431 | BREX-system-P-loop-protein-BrxC-[Thauera-propionica] |
| WP_096448538.1 | Type1.BREX | <-ParB+HNH<-PHP+SbcC<-DUF4096+DDE_Tnp_1+DDE_Tnp_1_2<-N6-METHYLASE<-?<-viral-capsid<-ZnR<-N6-METHYLASE<-?<-DUF6079*<-BrxB-BrxF-iSTAND<-BrxA-DNA-binding<-RHH_5<-?\|\|?-><-?\|\|DDE_Tnp_1-> | 1153 | Bacteria>Pseudomonadota>Betaproteobacteria>Rhodocyclales>Zoogloeaceae>Thauera | GCF_002354895.1 | Thauera_sp._K11 | CCZ27_12255 | 2842153..2845614 | 2005884 | BREX-system-P-loop-protein-BrxC-[Thauera-sp.-K11] |
| WP_107221110.1 | Type1.BREX | METHYLASE->ParA-Soj-PloopNTPase-><-SIGMA+OB-Fold+AAA-ATPase+Lon-Protease-BrxL\|\|DUF772+DDE_Tnp_1+DDE_Tnp_1_6-><-N6-METHYLASE-p<-REase<-OLD-ABC<-PIN<-MazE_antitoxin<-DUF6079*<-BrxB-BrxF-iSTAND<-BrxA-DNA-binding<-SPOUT<-RnaseHII | 1153 | Bacteria>Pseudomonadota>Betaproteobacteria>Rhodocyclales>Zoogloeaceae>Thauera | GCF_003030465.1 | Thauera_aromatica | Tharo_RS10350 | 2186270..2189731 | 59405 | BREX-system-P-loop-protein-BrxC-[Thauera-aromatica] |
| WP_247724881.1 | Type1.BREX | <-PglZ-p<-REase+AIPR<-N6-METHYLASE<-PIN<-MazE_antitoxin<-DUF6079*<-BrxB-BrxF-iSTAND<-BrxA-DNA-binding<-SPOUT<-RnaseHII | 1153 | Bacteria>Pseudomonadota>Betaproteobacteria>Rhodocyclales>Zoogloeaceae>Thauera | GCF_023093595.1 | Thauera_aromatica | MXC99_RS07615 | 6533..9994 | 59405 | BREX-system-P-loop-protein-BrxC-[Thauera-aromatica] |
| WP_247725480.1 | Type1.BREX | <-SIGMA+OB-Fold+AAA-ATPase+Lon-Protease-BrxL<-PglZ<-PIWI<-TOPRIM<-OLD-ABC<-N6-METHYLASE\|\|RHH_6->PIN-><-DUF6079*<-BrxB-BrxF-iSTAND<-BrxA-DNA-binding<-wHTH+WYL+WCX | 1203 | Bacteria>Pseudomonadota>Betaproteobacteria>Rhodocyclales>Zoogloeaceae>Thauera | GCF_023093595.1 | Thauera_aromatica | MXC99_RS10910 | 53320..56931 | 59405 | BREX-system-P-loop-protein-BrxC-[Thauera-aromatica] |
| WP_247737217.1 | Type1.BREX | BrxA-DNA-binding->BrxB-BrxF-iSTAND->DUF6079*->MazE_antitoxin->PIN->OLD-ABC->REase->N6-METHYLASE-p-> | 1153 | Bacteria>Pseudomonadota>Betaproteobacteria>Rhodocyclales>Zoogloeaceae>Thauera | - | Thauera_aromatica | MX653_RS17175 | 1332..4793 | 59405 | BREX-system-P-loop-protein-BrxC-[Thauera-aromatica] |
| MBK6655412.1 | Type1.BREX | HDRnase->?-><-HTH_19\|\|?->?->PhdYeFM_antitox->PIN->BrxA-DNA-binding->BrxB-BrxF-iSTAND->DUF6079*->OLD-ABC->REase->N6-METHYLASE->ZnR->viral-capsid->?->?->TPR+REase-> | 1153 | Bacteria>Pseudomonadota>Betaproteobacteria>Rhodocyclales>Zoogloeaceae>Zoogloea | GCA_016703645.1 | Zoogloea_sp. | IPG40_16580 | 748210..751671 | 49181 | BREX-system-P-loop-protein-BrxC-[Zoogloea-sp.] |
| MBK7848388.1 | Type1.BREX | Arm-DNA-bind_2+Phage_int_SAM_3+Phage_integrase->?-><-HTH_19\|\|?->?->PhdYeFM_antitox->PIN->BrxA-DNA-binding->BrxB-BrxF-iSTAND->DUF6079*->REase->ZnR->viral-capsid->?->?->TPR+REase->?->PglZ->SIGMA+OB-Fold+AAA-ATPase+Lon-Protease-BrxL-> | 1153 | Bacteria>Pseudomonadota>Betaproteobacteria>Rhodocyclales>Zoogloeaceae>Zoogloea | GCA_016714095.1 | Zoogloea_sp. | IPJ73_15250 | 34710..38171 | 49181 | BREX-system-P-loop-protein-BrxC-[Zoogloea-sp.] |
| WP_141354347.1 | Type1.BREX | wHTH+WYL+WCX->BrxA-DNA-binding->BrxB-BrxF-iSTAND->DUF6079*-><-PIN<-RHH_6\|\|N6-METHYLASE-p->HKD-DNase->N6-METHYLASE->HTH+REase->ZnR+TOPRIM->PglZ->SIGMA+OB-Fold+AAA-ATPase+Lon-Protease-BrxL-> | 1203 | Bacteria>Pseudomonadota>Betaproteobacteria>Rhodocyclales>Zoogloeaceae>Zoogloea | GCF_006539865.1 | Zoogloea_ramigera | ZRA01_RS16820 | 2482..6093 | 350 | BREX-system-P-loop-protein-BrxC-[Zoogloea-ramigera] |
| WP_148578091.1 | Type1.BREX | Arm-DNA-bind_2+Phage_int_SAM_3+Phage_integrase->HDRnase->?-><-HTH_19\|\|?->?->wHTH+WYL+WCX->BrxA-DNA-binding->BrxB-BrxF-iSTAND->DUF6079*->N6-METHYLASE->OLD-ABC->REase->SIR2->HerA/FtsK->N6-METHYLASE-p->HTH+REase->PglZ->SIGMA+OB-Fold+AAA-ATPase+HKD-DNase-BrxL-> | 1186 | Bacteria>Pseudomonadota>Betaproteobacteria>Rhodocyclales>Zoogloeaceae>Zoogloea | GCF_008107625.1 | Zoogloea_oleivorans | ETQ85_RS05700 | 193093..196653 | 1552750 | BREX-system-P-loop-protein-BrxC-[Zoogloea-oleivorans] |
| WP_148578351.1 | Type1.BREX | <-SbcC+SbcC+ABC-ATPase+SbcC\|\|?-><-HTH_3\|\|?->\|\|wHTH+WYL+WCX->BirA-HTH->BrxA-DNA-binding->BrxB-BrxF-iSTAND->DUF6079*->N6-METHYLASE->DUF932->ParB+HNH->OLD-ABC->HNH->PglZ->SIGMA+OB-Fold+AAA-ATPase+Lon-Protease-BrxL->DUF772+DDE_Tnp_1-> | 1146 | Bacteria>Pseudomonadota>Betaproteobacteria>Rhodocyclales>Zoogloeaceae>Zoogloea | GCF_008107625.1 | Zoogloea_oleivorans | ETQ85_RS07180 | 242245..245685 | 1552750 | BREX-system-P-loop-protein-BrxC-[Zoogloea-oleivorans] |
| OQA32553.1 | Type1.BREX | Phage_integrase->?->?->ParB->?->HTH_3->BrxA-DNA-binding->BrxB-BrxF-iSTAND->DUF6079*-> | 1140 | Bacteria>Pseudomonadota>Betaproteobacteria>unclassified_Betaproteobacteria | GCA_002068855.1 | Betaproteobacteria_bacterium_ADurb.Bin341 | BWY57_01861 | 6243..9665 | 1852821 | hypothetical-protein-BWY57_01861-[Betaproteobacteria-bacterium-ADurb.Bin341] |
| PKO80619.1 | Type1.BREX | <-ZnR<-N6-METHYLASE<-URI-DNase<-DUF6079*<-BrxB-BrxF-iSTAND<-BrxA-DNA-binding<-RHH_5\|\|?->HTH-> | 1153 | Bacteria>Pseudomonadota>Betaproteobacteria>unclassified_Betaproteobacteria | GCA_002842325.1 | Betaproteobacteria_bacterium_HGW-Betaproteobacteria-13 | CVU19_11375 | 4671..8132 | 2013704 | BREX-system-P-loop-protein-BrxC-[Betaproteobacteria-bacterium-HGW-Betaproteobacteria-13] |
| PKO87722.1 | Type1.BREX | <-SIGMA+OB-Fold+AAA-ATPase+Lon-Protease-BrxL-p<-PglZ<-N6-METHYLASE<-DUF6079*<-BrxB-BrxF-iSTAND | 1149 | Bacteria>Pseudomonadota>Betaproteobacteria>unclassified_Betaproteobacteria | GCA_002842415.1 | Betaproteobacteria_bacterium_HGW-Betaproteobacteria-10 | CVU16_15240 | 6333..9782 | 2013701 | BREX-system-P-loop-protein-BrxC-[Betaproteobacteria-bacterium-HGW-Betaproteobacteria-10] |
| MBI3372694.1 | Type1.BREX | <-AAA-ATPase\|\|?->?-><-?\|\|BrxA-DNA-binding->BrxB-BrxF-iSTAND->DUF6079*->OLD-ABC->TOPRIM->?->?->SbcC->N6-METHYLASE->ArdCN+MPTase->PglZ->SIGMA+OB-Fold+AAA-ATPase+Lon-Protease-BrxL-> | 1148 | Bacteria>Pseudomonadota>Betaproteobacteria>unclassified_Betaproteobacteria | GCA_016195915.1 | Betaproteobacteria_bacterium | HYO17_13195 | 52819..56265 | 1891241 | BREX-system-P-loop-protein-BrxC-[Betaproteobacteria-bacterium] |
| WP_219254923.1 | Type1.BREX | <-PglZ<-TOPRIM<-OLD-ABC<-N6-METHYLASE-p<-HerA/FtsK<-SIR2<-N6-METHYLASE<-TOPRIM<-OLD-ABC<-DUF6079*<-BrxB-BrxF-iSTAND<-BrxA-DNA-binding<-RHS<-?\|\|wHTH+WYL+WCX-><-?\|\|LexA-protease->?-><-DOC+HTH | 1226 | Bacteria>Pseudomonadota>Gammaproteobacteria>Aeromonadales>Aeromonadaceae>Aeromonas | - | Aeromonas_hydrophila | GPV00_14065 | 286470..290150 | 644 | BREX-system-P-loop-protein-BrxC-[Aeromonas-hydrophila] |
| AUZ77883.1 | Type1.BREX | <-HIN-HTH<-SIGMA+OB-Fold+AAA-ATPase+HKD-DNase-BrxL<-PglZ<-ParB+HNH<-N6-METHYLASE<-REase<-OLD-ABC<-DUF6079*<-BrxB-BrxF-iSTAND<-BrxA-DNA-binding<-XIS-HTH\|\|<-DOC+HTH<-?<-XIS-HTH<-Arm-DNA-bind_3+Phage_integrase\|\|?-><-Sulfate_transp+STAS | 1183 | Bacteria>Pseudomonadota>Gammaproteobacteria>Aeromonadales>Aeromonadaceae>Aeromonas | GCA_002934425.1 | Aeromonas_sp._ASNIH4 | C2U40_16585 | 3601638..3605189 | 1636609 | BREX-system-P-loop-protein-BrxC-[Aeromonas-sp.-ASNIH4] |
| PZQ93954.1 | Type1.BREX | <-HNH<-DUF6079*<-BrxB-BrxF-iSTAND\|\|HIN-HTH->IstB_IS21-><-HTH_Tnp_1+HTH_21+rve+rve_3 | 1218 | Bacteria>Pseudomonadota>Gammaproteobacteria>Aeromonadales>Aeromonadaceae>Aeromonas | GCA_003243215.1 | Aeromonas_media | DI541_18955 | 3274..6930 | 651 | BREX-system-P-loop-protein-BrxC-[Aeromonas-media] |
| BBR09393.1 | Type1.BREX | Arm-DNA-bind_3+Phage_integrase-><-?\|\|?->?-><-wHTH+WYL+WCX\|\|?->RHS->BrxA-DNA-binding->BrxB-BrxF-iSTAND->DUF6079*->OLD-ABC->TOPRIM->ParB+HNH->N6-METHYLASE->OLD-ABC->TOPRIM->PglZ->SIGMA+OB-Fold+AAA-ATPase+Lon-Protease-BrxL-> | 1226 | Bacteria>Pseudomonadota>Gammaproteobacteria>Aeromonadales>Aeromonadaceae>Aeromonas | GCA_014158455.1 | Aeromonas_caviae | WP3S18E02_10540 | 1118346..1122026 | 648 | hypothetical-protein-WP3S18E02_10540-[Aeromonas-caviae] |
| HAU4888102.1 | Type1.BREX | <-N6-METHYLASE-p<-TOPRIM<-OLD-ABC<-DUF6079*<-BrxB-BrxF-iSTAND<-BrxA-DNA-binding<-RHS<-ZnR+ZnR\|\|wHTH+WYL+WCX-><-?\|\|LexA-protease->?-><-DOC+HTH | 1231 | Bacteria>Pseudomonadota>Gammaproteobacteria>Aeromonadales>Aeromonadaceae>Aeromonas | GCA_016067875.1 | Aeromonas_hydrophila | GD804_RS12725 | 3923..7618 | 644 | BREX-system-P-loop-protein-BrxC-[Aeromonas-hydrophila] |
| EGX6956813.1 | Type1.BREX | <-SIGMA+OB-Fold+AAA-ATPase+Lon-Protease-BrxL<-PglZ<-?<-N6-METHYLASE-p<-HerA/FtsK<-SIR2<-N6-METHYLASE<-ParB+HNH<-DUF6079*<-BrxB-BrxF-iSTAND<-BrxA-DNA-binding<-?<-XIS-HTH\|\|?-><-Arm-DNA-bind_3+Phage_int_SAM_3+Phage_integrase | 1218 | Bacteria>Pseudomonadota>Gammaproteobacteria>Aeromonadales>Aeromonadaceae>Aeromonas | GCA_016414985.1 | Aeromonas_hydrophila | JFQ85_000577 | 650804..654460 | 644 | BREX-system-P-loop-protein-BrxC-[Aeromonas-hydrophila] |
| MBS4697267.1 | Type1.BREX | <-SIGMA+OB-Fold+AAA-ATPase+Lon-Protease-BrxL<-PglZ<-TOPRIM<-OLD-ABC<-N6-METHYLASE<-ParB+HNH<-TOPRIM<-OLD-ABC<-DUF6079*<-BrxB-BrxF-iSTAND<-BrxA-DNA-binding<-RHS | 1226 | Bacteria>Pseudomonadota>Gammaproteobacteria>Aeromonadales>Aeromonadaceae>Aeromonas | GCA_018359965.1 | Aeromonas_allosaccharophila | J4G65_17610 | 87511..91191 | 656 | BREX-system-P-loop-protein-BrxC-[Aeromonas-allosaccharophila] |
| MBW3845327.1 | Type1.BREX | <-PglZ<-TOPRIM<-OLD-ABC<-N6-METHYLASE-p<-HerA/FtsK<-SIR2<-N6-METHYLASE<-TOPRIM<-OLD-ABC<-DUF6079*<-BrxB-BrxF-iSTAND<-BrxA-DNA-binding<-RHS<-?\|\|wHTH+WYL+WCX-><-?\|\|LexA-protease->?-><-DOC+HTH | 1226 | Bacteria>Pseudomonadota>Gammaproteobacteria>Aeromonadales>Aeromonadaceae>Aeromonas | GCA_019348915.1 | Aeromonas_hydrophila | GPV00_14065 | 286470..290150 | 644 | BREX-system-P-loop-protein-BrxC-[Aeromonas-hydrophila] |
| MBX9561629.1 | Type1.BREX | <-HIN-HTH+rve+rve_3\|\|SF1-SF2-Helicase+REase-><-HORMA<-HNH<-OLD-ABC<-N6-METHYLASE<-DUF6079*<-BrxB-BrxF-iSTAND<-BrxA-DNA-binding<-URI-DNase<-?\|\|LexA-protease->?->PAIREDC-HTH-><-MODE-HTH+PBPII | 1157 | Bacteria>Pseudomonadota>Gammaproteobacteria>Aeromonadales>Aeromonadaceae>Aeromonas | GCA_019742455.1 | Aeromonas_hydrophila | FKF73_02860 | 531067..534540 | 644 | BREX-system-P-loop-protein-BrxC-[Aeromonas-hydrophila] |
| GJA32139.1 | Type1.BREX | <-Peptidase_C39_2<-?<-N6-METHYLASE-p<-HEPN<-N6-METHYLASE-p\|\|DDE_Tnp_1_assoc+DDE_Tnp_1-><-N6-METHYLASE<-?<-HNH<-DUF6079*<-BrxB-BrxF-iSTAND<-BrxA-DNA-binding<-RHS<-?<-Ntox49\|\|wHTH+WYL+WCX-><-XIS-HTH\|\|?-><-HTH_26 | 1227 | Bacteria>Pseudomonadota>Gammaproteobacteria>Aeromonadales>Aeromonadaceae>Aeromonas | GCA_019971155.1 | Aeromonas_caviae | KAM347_17720 | 32751..36434 | 648 | hypothetical-protein-KAM341_18170-[Aeromonas-caviae] |
| GJA65768.1 | Type1.BREX | DDE_Tnp_1_assoc+DDE_Tnp_1->wHTH+WYL+WCX->BrxA-DNA-binding->BrxB-BrxF-iSTAND->DUF6079*-> | 927 | Bacteria>Pseudomonadota>Gammaproteobacteria>Aeromonadales>Aeromonadaceae>Aeromonas | GCA_019971515.1 | Aeromonas_caviae | KAM351_43790 | 4448..7231 | 648 | hypothetical-protein-KAM351_43790-[Aeromonas-caviae] |

| Protein-ID | System | Gene-Neighbourhoods | Length | Taxonomy | Assembly-ID | Strain | Locus-Tag | Genomic-coordinates | TaxID | Fasta-defline |
|---|---|---|---|---|---|---|---|---|---|---|
| BDO08311.1 | Type1.BREX | ARG->RHS->HTH_23+HTH_32+HTH_33->DDE_3->RHS->BrxA-DNA-binding->BrxB-BrxF-iSTAND->DUF6079-p-><-DDE_Tnp_1\|\|DUF6079*->ParB+HNH->N6-METHYLASE->HEPN->PAIREDC-HTH+HIN-HTH->N6-METHYLASE-p-> [illegible] | 1089 | Bacteria>Pseudomonadota>Gammaproteobacteria>Aeromonadales>Aeromonadaceae>Aeromonas | GCA_024347775.1 | Aeromonas_caviae | KAM643c_18840 | 2028863..2032132 | 648 | hypothetical-protein-KAM643c_18840-[Aeromonas-caviae] |
| EJN6954191.1 | Type1.BREX | <-LexA-protease\|\|?-><-wHTH+WYL+WCX\|\|ARG->?->ZnR->RHS->BrxA-DNA-binding->BrxB-BrxF-iSTAND->DUF6079*->N6-METHYLASE->OLD-ABC->HNH->PglZ->HORMA-><-SF1-SF2-Helicase+REase<-SF1-SF2-Helicase [illegible] | 1157 | Bacteria>Pseudomonadota>Gammaproteobacteria>Aeromonadales>Aeromonadaceae>Aeromonas | GCA_024506355.1 | Aeromonas_hydrophila | NRD12_000732 | 206979..210452 | 644 | BREX-system-P-loop-protein-BrxC-[Aeromonas-hydrophila] |
| WP_010634561.1 | Type1.BREX | DUF2787->?->?->?->SF1-SF2-Helicase+REase->SF1-SF2-Helicase+REase-><-PglZ<-ParB+HNH<-N6-METHYLASE<-DUF6079*<-BrxB-BrxF-iSTAND<-BrxA-DNA-binding<-RHS<-ZnR<-?<-ARG\|\|wHTH+WYL+WCX-><-?\|\| [illegible] | 1157 | Bacteria>Pseudomonadota>Gammaproteobacteria>Aeromonadales>Aeromonadaceae>Aeromonas | GCF_000315195.1 | Aeromonas_dhakensis | AQU_RS10070 | 136801..140274 | 196024 | BREX-system-P-loop-protein-BrxC-[Aeromonas-dhakensis] |
| WP_043124789.1 | Type1.BREX | SF1-SF2-Helicase+REase->SF1-SF2-Helicase+REase-><-HORMA<-PglZ<-HNH<-OLD-ABC<-N6-METHYLASE<-DUF6079*<-BrxB-BrxF-iSTAND<-BrxA-DNA-binding<-URI-DNase<-?\|\|LexA-protease-> | 1157 | Bacteria>Pseudomonadota>Gammaproteobacteria>Aeromonadales>Aeromonadaceae>Aeromonas | GCF_000724965.1 | Aeromonas_hydrophila | HR52_RS13935 | 3102820..3106293 | 644 | BREX-system-P-loop-protein-BrxC-[Aeromonas-hydrophila] |
| WP_043557612.1 | Type1.BREX | REase-><-?\|\|?-><-SIGMA+OB-Fold+AAA-ATPase+HKD-DNase-BrxL<-PglZ<-ParB+HNH<-N6-METHYLASE<-REase<-OLD-ABC<-DUF6079*<-BrxB-BrxF-iSTAND<-BrxA-DNA-binding<-XIS-HTH<-?<-AAA+HTH<-?<-XIS-HTH | 1182 | Bacteria>Pseudomonadota>Gammaproteobacteria>Aeromonadales>Aeromonadaceae>Aeromonas | GCF_000819745.1 | Aeromonas_bestiarum | BN1118_RS19650 | 69736..73284 | 105751 | BREX-system-P-loop-protein-BrxC-[Aeromonas-bestiarum] |
| WP_042011559.1 | Type1.BREX | <-SIGMA+OB-Fold+AAA-ATPase+Lon-Protease-BrxL<-PglZ<-TOPRIM<-OLD-ABC<-?<-DUF3944<-N6-METHYLASE<-TOPRIM<-OLD-ABC<-DUF6079*<-BrxB-BrxF-iSTAND<-BrxA-DNA-binding<-RHS<-HKD-DNase [illegible] | 1239 | Bacteria>Pseudomonadota>Gammaproteobacteria>Aeromonadales>Aeromonadaceae>Aeromonas | GCF_000819885.1 | Aeromonas_fluvialis | BN1125_RS07645 | 79425..83144 | 591962 | BREX-system-P-loop-protein-BrxC-[Aeromonas-fluvialis] |
| WP_059169685.1 | Type1.BREX | <-PglZ<-N6-METHYLASE-p<-HEPN<-RVT<-N6-METHYLASE<-?<-HNH<-TOPRIM<-OLD-ABC<-DUF6079*<-BrxB-BrxF-iSTAND<-BrxA-DNA-binding<-RHS<-ARG<-RHS\|\|LexA-protease->?-><-HDRnase [illegible] | 1240 | Bacteria>Pseudomonadota>Gammaproteobacteria>Aeromonadales>Aeromonadaceae>Aeromonas | GCF_001481535.2 | Aeromonas_salmonicida | VO70_RS19205 | 18975..22697 | 645 | BREX-system-P-loop-protein-BrxC-[Aeromonas-salmonicida] |
| WP_087756130.1 | Type1.BREX | <-SIGMA+OB-Fold+AAA-ATPase+Lon-Protease-BrxL<-PglZ<-?<-DUF3944<-N6-METHYLASE-p<-N6-METHYLASE<-ParB+HNH<-TOPRIM<-OLD-ABC<-DUF6079*<-BrxB-BrxF-iSTAND<-BrxA-DNA-binding<-RHS\|\|ZnR->HEPN-> [illegible] | 1231 | Bacteria>Pseudomonadota>Gammaproteobacteria>Aeromonadales>Aeromonadaceae>Aeromonas | GCF_002180335.1 | Aeromonas_salmonicida | O23A_RS10590 | 2337193..2340888 | 645 | BREX-system-P-loop-protein-BrxC-[Aeromonas-salmonicida] |
| WP_095593101.1 | Type1.BREX | Arm-DNA-bind_3-><-SIGMA+OB-Fold+AAA-ATPase+Lon-Protease-BrxL<-PglZ<-?<-DUF3944<-N6-METHYLASE-p<-N6-METHYLASE<-ParB+HNH<-DUF6079*<-BrxB-BrxF-iSTAND<-BrxA-DNA-binding<-RHS\|\|HEPN-><-? [illegible] | 1231 | Bacteria>Pseudomonadota>Gammaproteobacteria>Aeromonadales>Aeromonadaceae>Aeromonas | GCF_002285935.1 | Aeromonas_dhakensis | CK627_RS18865 | 4072109..4075804 | 196024 | BREX-system-P-loop-protein-BrxC-[Aeromonas-dhakensis] |
| WP_101617899.1 | Type1.BREX | <-TetR-HTH<-Phage_integrase+Phage_integrase<-Phage_integrase+Phage_integrase<-Phage_integrase<-LexA-protease\|\|?->RHS->BrxA-DNA-binding->BrxB-BrxF-iSTAND->DUF6079*->OLD-ABC->TOPRIM-> [illegible] | 1231 | Bacteria>Pseudomonadota>Gammaproteobacteria>Aeromonadales>Aeromonadaceae>Aeromonas | GCF_002850695.3 | Aeromonas_hydrophila | C1A23_RS07645 | 1215070..1218765 | 644 | BREX-system-P-loop-protein-BrxC-[Aeromonas-hydrophila] |
| WP_113721905.1 | Type1.BREX | SF1-SF2-Helicase+REase->SF1-SF2-Helicase+REase-><-HEPN+Sel1+SEC-C<-PglZ<-HNH<-OLD-ABC<-N6-METHYLASE<-DUF6079*<-BrxB-BrxF-iSTAND<-BrxA-DNA-binding<-RHS<-?<-ARG\|\|wHTH+WYL+WCX-><-?\|\| [illegible] | 1157 | Bacteria>Pseudomonadota>Gammaproteobacteria>Aeromonadales>Aeromonadaceae>Aeromonas | GCF_003265515.1 | Aeromonas_salmonicida | CPA05_RS19945 | 20643..24116 | 645 | BREX-system-P-loop-protein-BrxC-[Aeromonas-salmonicida] |
| WP_113995752.1 | Type1.BREX | <-N6-METHYLASE<-DUF6079*<-BrxB-BrxF-iSTAND<-BrxA-DNA-binding<-RHS<-?<-ARG<-ARG\|\|wHTH+WYL+WCX-><-?\|\|LexA-protease-> | 1157 | Bacteria>Pseudomonadota>Gammaproteobacteria>Aeromonadales>Aeromonadaceae>Aeromonas | GCF_003323285.1 | Aeromonas_hydrophila | C6C11_RS19255 | 2250..5723 | 644 | BREX-system-P-loop-protein-BrxC-[Aeromonas-hydrophila] |
| WP_114523382.1 | Type1.BREX | <-LexA-protease\|\|?->URI-DNase->BrxA-DNA-binding->BrxB-BrxF-iSTAND->DUF6079*->N6-METHYLASE->OLD-ABC->HNH->PglZ-> | 1157 | Bacteria>Pseudomonadota>Gammaproteobacteria>Aeromonadales>Aeromonadaceae>Aeromonas | GCF_003344485.1 | Aeromonas_sp._ARM81 | ASJ36_RS21100 | 4447..7920 | 1747384 | BREX-system-P-loop-protein-BrxC-[Aeromonas-sp.-ARM81] |
| WP_120415687.1 | Type1.BREX | <-TetR-HTH\|\|HHH+DNA_pol_B+RNase->tRNA-><-?<-LexA-protease\|\|?->RHS->BrxA-DNA-binding->BrxB-BrxF-iSTAND->DUF6079*->ParB+HNH->N6-METHYLASE->N6-METHYLASE-p->DUF3944->?->PglZ-> [illegible] | 1231 | Bacteria>Pseudomonadota>Gammaproteobacteria>Aeromonadales>Aeromonadaceae>Aeromonas | GCF_003611985.1 | Aeromonas_veronii | D6R50_RS16910 | 827539..831234 | 654 | BREX-system-P-loop-protein-BrxC-[Aeromonas-veronii] |
| WP_123173885.1 | Type1.BREX | DOC+HTH-><-?<-LexA-protease\|\|?-><-wHTH+WYL+WCX\|\|ZnR+ZnR->RHS->BrxA-DNA-binding->BrxB-BrxF-iSTAND->DUF6079*->OLD-ABC->TOPRIM->N6-METHYLASE->N6-METHYLASE->DUF3944->?-><-HTH_Tnp_1 [illegible] | 1231 | Bacteria>Pseudomonadota>Gammaproteobacteria>Aeromonadales>Aeromonadaceae>Aeromonas | GCF_003722175.1 | Aeromonas_veronii | EFI48_RS19250 | 4187979..4191674 | 654 | BREX-system-P-loop-protein-BrxC-[Aeromonas-veronii] |
| WP_128297388.1 | Type1.BREX | RHS->BrxA-DNA-binding->BrxB-BrxF-iSTAND->DUF6079*->OLD-ABC->TOPRIM->N6-METHYLASE-> | 1225 | Bacteria>Pseudomonadota>Gammaproteobacteria>Aeromonadales>Aeromonadaceae>Aeromonas | GCF_004024195.1 | Aeromonas_caviae | DN612_RS19420 | 2423..6100 | 648 | BREX-system-P-loop-protein-BrxC-[Aeromonas-caviae] |
| WP_240039238.1 | Type1.BREX | <-PglZ<-TOPRIM<-OLD-ABC<-N6-METHYLASE-p<-HerA/FtsK<-SIR2<-N6-METHYLASE<-TOPRIM<-OLD-ABC<-DUF6079*<-BrxB-BrxF-iSTAND<-BrxA-DNA-binding<-RHS<-?\|\|wHTH+WYL+WCX-><-?\|\|LexA-protease->?-> [illegible] | 1280 | Bacteria>Pseudomonadota>Gammaproteobacteria>Aeromonadales>Aeromonadaceae>Aeromonas | GCF_006243275.1 | Aeromonas_hydrophila | CF138_RS14425 | 159803..163645 | 644 | BREX-system-P-loop-protein-BrxC-[Aeromonas-hydrophila] |
| WP_160837717.1 | Type1.BREX | <-N6-METHYLASE-p<-TIR<-HEPN<-N6-METHYLASE<-TOPRIM<-OLD-ABC<-DUF6079*<-BrxB-BrxF-iSTAND<-BrxA-DNA-binding<-RHS<-ZnR<-ARG<-RHS\|\|LexA-protease->?-><-HDRnase | 1231 | Bacteria>Pseudomonadota>Gammaproteobacteria>Aeromonadales>Aeromonadaceae>Aeromonas | GCF_009831085.1 | Aeromonas_caviae | GR302_RS08560 | 7464..11159 | 648 | BREX-system-P-loop-protein-BrxC-[Aeromonas-caviae] |
| WP_159144201.1 | Type1.BREX | <-SIGMA+OB-Fold+AAA-ATPase+Lon-Protease-BrxL<-PglZ<-TOPRIM<-OLD-ABC<-?<-DUF3944<-N6-METHYLASE<-N6-METHYLASE<-ParB+HNH<-DUF6079*<-BrxB-BrxF-iSTAND<-BrxA-DNA-binding<-RHS<-?\|\| [illegible] | 1231 | Bacteria>Pseudomonadota>Gammaproteobacteria>Aeromonadales>Aeromonadaceae>Aeromonas | GCF_009834065.1 | Aeromonas_veronii | GRF56_RS18105 | 3933894..3937589 | 654 | BREX-system-P-loop-protein-BrxC-[Aeromonas-veronii] |
| WP_171280306.1 | Type1.BREX | AAA-ToxA+Classical-AAA->Classical-AAA->ARG->?->ZnR->RHS->BrxA-DNA-binding->BrxB-BrxF-iSTAND->DUF6079*->N6-METHYLASE->ParB+HNH->PglZ->HEPN+Sel1+SEC-C-><-SF1-SF2-Helicase+REase [illegible] | 1157 | Bacteria>Pseudomonadota>Gammaproteobacteria>Aeromonadales>Aeromonadaceae>Aeromonas | GCF_013085505.1 | Aeromonas_sp._2692-1 | E5E97_RS12155 | 2531425..2534898 | 2560029 | BREX-system-P-loop-protein-BrxC-[Aeromonas-sp.-2692-1] |
| WP_232090117.1 | Type1.BREX | XIS-HTH->?->?-><-wHTH+WYL+WCX\|\|?->RHS->BrxA-DNA-binding->BrxB-BrxF-iSTAND->DUF6079*->OLD-ABC->TOPRIM->ParB+HNH->N6-METHYLASE->OLD-ABC->TOPRIM->PglZ-> [illegible] | 1281 | Bacteria>Pseudomonadota>Gammaproteobacteria>Aeromonadales>Aeromonadaceae>Aeromonas | GCF_014158455.1 | Aeromonas_caviae | H7X73_RS05535 | 1118181..1122026 | 648 | BREX-system-P-loop-protein-BrxC-[Aeromonas-caviae] |
| WP_182951579.1 | Type1.BREX | <-LexA-protease\|\|?->REase+N6_METHYLASE->TRD+TRD->REase+SF1-SF2-Helicase->zf-ISL3+HTH_Tnp_ISL3+DDE_Tnp_ISL3->URI-DNase->BrxA-DNA-binding->BrxB-BrxF-iSTAND->DUF6079*->N6-METHYLASE-> [illegible] | 1157 | Bacteria>Pseudomonadota>Gammaproteobacteria>Aeromonadales>Aeromonadaceae>Aeromonas | GCF_014168615.1 | Aeromonas_sp._WP2-W18-CRE-05 | WP2W18C05_13360 | 1426228..1429701 | 2675707 | BREX-system-P-loop-protein-BrxC-[Aeromonas-sp.-WP2-W18-CRE-05] |
| WP_190283730.1 | Type1.BREX | <-PglZ<-?<-DUF3944 [illegible] | 1226 | Bacteria>Pseudomonadota>Gammaproteobacteria>Aeromonadales>Aeromonadaceae>Aeromonas | GCF_014679585.1 | Aeromonas_hydrophila | IFA22_RS17960 | 3928101..3931781 | 644 | BREX-system-P-loop-protein-BrxC-[Aeromonas-hydrophila] |
| WP_193000848.1 | Type1.BREX | <-DUF3944<-N6-METHYLASE [illegible] | 1226 | Bacteria>Pseudomonadota>Gammaproteobacteria>Aeromonadales>Aeromonadaceae>Aeromonas | GCF_014892695.1 | Aeromonas_simiae | FE240_RS10745 | 2272968..2276648 | 218936 | BREX-system-P-loop-protein-BrxC-[Aeromonas-simiae] |
| WP_201964470.1 | Type1.BREX | [illegible] | 432 | Bacteria>Pseudomonadota>Gammaproteobacteria>Aeromonadales>Aeromonadaceae>Aeromonas | GCF_016729105.1 | Aeromonas_caviae | JD512_RS20845 | 12456..13754 | 648 | BREX-system-P-loop-protein-BrxC-[Aeromonas-caviae] |
| WP_201944276.1 | Type1.BREX | [illegible] | 1231 | Bacteria>Pseudomonadota>Gammaproteobacteria>Aeromonadales>Aeromonadaceae>Aeromonas | GCF_016729595.1 | Aeromonas_veronii | JD494_RS16545 | 22676..26371 | 654 | BREX-system-P-loop-protein-BrxC-[Aeromonas-veronii] |
| WP_201932781.1 | Type1.BREX | [illegible] | 1226 | Bacteria>Pseudomonadota>Gammaproteobacteria>Aeromonadales>Aeromonadaceae>Aeromonas | GCF_016729935.1 | Aeromonas_caviae | JD546_RS01995 | 64762..68442 | 648 | BREX-system-P-loop-protein-BrxC-[Aeromonas-caviae] |
| WP_202047008.1 | Type1.BREX | [illegible] | 1226 | Bacteria>Pseudomonadota>Gammaproteobacteria>Aeromonadales>Aeromonadaceae>Aeromonas | GCF_016730065.1 | Aeromonas_media | JD497_RS12375 | 531279..534959 | 651 | BREX-system-P-loop-protein-BrxC-[Aeromonas-media] |
| WP_203130882.1 | Type1.BREX | <-wHTH+WYL+WCX\|\|ZnR+ZnR [illegible] | 935 | Bacteria>Pseudomonadota>Gammaproteobacteria>Aeromonadales>Aeromonadaceae>Aeromonas | GCF_016805325.1 | Aeromonas_jandaei | GNZ06_RS06060 | 150251..153058 | 650 | BREX-system-P-loop-protein-BrxC-[Aeromonas-jandaei] |
| WP_203130883.1 | Type1.BREX | <-wHTH+WYL+WCX\|\|ZnR+ZnR [illegible] | 226 | Bacteria>Pseudomonadota>Gammaproteobacteria>Aeromonadales>Aeromonadaceae>Aeromonas | GCF_016805325.1 | Aeromonas_jandaei | GNZ06_RS06065 | 153058..153738 | 650 | hypothetical-protein-[Aeromonas-jandaei] |
| WP_203133261.1 | Type1.BREX | [illegible] | 1157 | Bacteria>Pseudomonadota>Gammaproteobacteria>Aeromonadales>Aeromonadaceae>Aeromonas | GCF_016805365.1 | Aeromonas_jandaei | G4224_RS08640 | 376570..380043 | 650 | BREX-system-P-loop-protein-BrxC-[Aeromonas-jandaei] |
| WP_210557479.1 | Type1.BREX | [illegible] | 1218 | Bacteria>Pseudomonadota>Gammaproteobacteria>Aeromonadales>Aeromonadaceae>Aeromonas | GCF_017939245.1 | Aeromonas_sp._PrichA-15 | J9889_RS17255 | 4362..8018 | 2823360 | BREX-system-P-loop-protein-BrxC-[Aeromonas-sp.-PrichA-15] |
| WP_213387487.1 | Type1.BREX | [illegible] | 1157 | Bacteria>Pseudomonadota>Gammaproteobacteria>Aeromonadales>Aeromonadaceae>Aeromonas | GCF_018359765.1 | Aeromonas_veronii | J4G64_RS11995 | 290568..294041 | 654 | BREX-system-P-loop-protein-BrxC-[Aeromonas-veronii] |
| WP_244842130.1 | Type1.BREX | [illegible] | 1281 | Bacteria>Pseudomonadota>Gammaproteobacteria>Aeromonadales>Aeromonadaceae>Aeromonas | GCF_018359965.1 | Aeromonas_allosaccharophila | J4G65_RS17605 | 87511..91356 | 656 | BREX-system-P-loop-protein-BrxC-[Aeromonas-allosaccharophila] |
| WP_215805224.1 | Type1.BREX | [illegible] | 1157 | Bacteria>Pseudomonadota>Gammaproteobacteria>Aeromonadales>Aeromonadaceae>Aeromonas | GCF_018802365.1 | Aeromonas_jandaei | HQ398_RS10760 | 2353585..2357058 | 650 | BREX-system-P-loop-protein-BrxC-[Aeromonas-jandaei] |
| WP_216986928.1 | Type1.BREX | [illegible] | 1157 | Bacteria>Pseudomonadota>Gammaproteobacteria>Aeromonadales>Aeromonadaceae>Aeromonas | GCF_019048565.1 | Aeromonas_sp._FDAARGOS_1406 | I6L36_RS00705 | 137070..140543 | 2778055 | BREX-system-P-loop-protein-BrxC-[Aeromonas-sp.-FDAARGOS-1406] |
| WP_216990395.1 | Type1.BREX | [illegible] | 1157 | Bacteria>Pseudomonadota>Gammaproteobacteria>Aeromonadales>Aeromonadaceae>Aeromonas | GCF_019048705.1 | Aeromonas_sp._FDAARGOS_1409 | I6L39_RS17660 | 3801324..3804797 | 2778058 | BREX-system-P-loop-protein-BrxC-[Aeromonas-sp.-FDAARGOS-1409] |
| WP_216992842.1 | Type1.BREX | [illegible] | 1157 | Bacteria>Pseudomonadota>Gammaproteobacteria>Aeromonadales>Aeromonadaceae>Aeromonas | GCF_019048725.1 | Aeromonas_sp._FDAARGOS_1407 | I6L37_20730 | 4426496..4429969 | 2778056 | BREX-system-P-loop-protein-BrxC-[Aeromonas-sp.-FDAARGOS-1407] |
| WP_203133976.1 | Type1.BREX | [illegible] | 1156 | Bacteria>Pseudomonadota>Gammaproteobacteria>Aeromonadales>Aeromonadaceae>Aeromonas | GCF_019348745.1 | Aeromonas_hydrophila | PVL97_RS11265 | 2527654..2531124 | 644 | BREX-system-P-loop-protein-BrxC-[Aeromonas-hydrophila] |
| WP_257621860.1 | Type1.BREX | [illegible] | 1281 | Bacteria>Pseudomonadota>Gammaproteobacteria>Aeromonadales>Aeromonadaceae>Aeromonas | GCF_019348915.1 | Aeromonas_hydrophila | GPV00_RS14000 | 286470..290315 | 644 | BREX-system-P-loop-protein-BrxC-[Aeromonas-hydrophila] |
| WP_223938541.1 | Type1.BREX | [illegible] | 1157 | Bacteria>Pseudomonadota>Gammaproteobacteria>Aeromonadales>Aeromonadaceae>Aeromonas | GCF_019971155.1 | Aeromonas_caviae | LDN24_RS08690 | 33954..37427 | 648 | BREX-system-P-loop-protein-BrxC-[Aeromonas-caviae] |
| WP_223931722.1 | Type1.BREX | [illegible] | 1231 | Bacteria>Pseudomonadota>Gammaproteobacteria>Aeromonadales>Aeromonadaceae>Aeromonas | GCF_019972495.1 | Aeromonas_caviae | LDN23_RS02850 | 18313..22008 | 648 | BREX-system-P-loop-protein-BrxC-[Aeromonas-caviae] |
| WP_234578020.1 | Type1.BREX | [illegible] | 1231 | Bacteria>Pseudomonadota>Gammaproteobacteria>Aeromonadales>Aeromonadaceae>Aeromonas | GCF_021440865.1 | Aeromonas_salmonicida | LZ632_RS15270 | 23400..27095 | 645 | BREX-system-P-loop-protein-BrxC-[Aeromonas-salmonicida] |
| WP_235462874.1 | Type1.BREX | [illegible] | 1157 | Bacteria>Pseudomonadota>Gammaproteobacteria>Aeromonadales>Aeromonadaceae>Aeromonas | GCF_021538905.1 | Aeromonas_australiensis | EHZ86_RS12350 | 8704..12177 | 1114880 | BREX-system-P-loop-protein-BrxC-[Aeromonas-australiensis] |
| WP_236274804.1 | Type1.BREX | [illegible] | 1231 | Bacteria>Pseudomonadota>Gammaproteobacteria>Aeromonadales>Aeromonadaceae>Aeromonas | GCF_021609985.1 | Aeromonas_caviae | L1871_RS17025 | 3638140..3641835 | 648 | BREX-system-P-loop-protein-BrxC-[Aeromonas-caviae] |
| WP_139392567.1 | Type1.BREX | [illegible] | 1225 | Bacteria>Pseudomonadota>Gammaproteobacteria>Aeromonadales>Aeromonadaceae>Aeromonas | GCF_021654355.1 | Aeromonas_hydrophila | L6424_RS03515 | 743150..746827 | 644 | BREX-system-P-loop-protein-BrxC-[Aeromonas-hydrophila] |
| WP_241325388.1 | Type1.BREX | [illegible] | 1157 | Bacteria>Pseudomonadota>Gammaproteobacteria>Aeromonadales>Aeromonadaceae>Aeromonas | GCF_022549295.1 | Aeromonas_sp._MR7 | MM192_RS09020 | 75622..79095 | 2923419 | BREX-system-P-loop-protein-BrxC-[Aeromonas-sp.-MR7] |
| WP_241871100.1 | Type1.BREX | [illegible] | 1157 | Bacteria>Pseudomonadota>Gammaproteobacteria>Aeromonadales>Aeromonadaceae>Aeromonas | GCF_022631175.1 | Aeromonas_hydrophila | MOO40_RS17050 | 3752340..3755813 | 644 | BREX-system-P-loop-protein-BrxC-[Aeromonas-hydrophila] |
| WP_247824022.1 | Type1.BREX | [illegible] | 1157 | Bacteria>Pseudomonadota>Gammaproteobacteria>Aeromonadales>Aeromonadaceae>Aeromonas | GCF_023100745.1 | Aeromonas_veronii | MYF86_09055 | 1896227..1899700 | 654 | BREX-system-P-loop-protein-BrxC-[Aeromonas-veronii] |
| WP_252657151.1 | Type1.BREX | [illegible] | 1231 | Bacteria>Pseudomonadota>Gammaproteobacteria>Aeromonadales>Aeromonadaceae>Aeromonas | GCF_023958325.1 | Aeromonas_veronii | NG896_RS17505 | 21137..24832 | 654 | BREX-system-P-loop-protein-BrxC-[Aeromonas-veronii] |
| WP_261925329.1 | Type1.BREX | [illegible] | 1089 | Bacteria>Pseudomonadota>Gammaproteobacteria>Aeromonadales>Aeromonadaceae>Aeromonas | GCF_024347775.1 | Aeromonas_caviae | OCX50_RS09760 | 2028863..2032132 | 648 | BREX-system-P-loop-protein-BrxC-[Aeromonas-caviae] |
| WP_256683847.1 | Type1.BREX | [illegible] | 991 | Bacteria>Pseudomonadota>Gammaproteobacteria>Aeromonadales>Aeromonadaceae>Aeromonas | GCF_024508255.1 | Aeromonas_salmonicida | NP805_RS12070 | 2625523..2628498 | 645 | BREX-system-P-loop-protein-BrxC-[Aeromonas-salmonicida] |
| WP_257706959.1 | Type1.BREX | [illegible] | 1182 | Bacteria>Pseudomonadota>Gammaproteobacteria>Aeromonadales>Aeromonadaceae>Aeromonas | GCF_024649325.1 | Aeromonas_caviae | NUK33_RS01495 | 150155..153703 | 648 | BREX-system-P-loop-protein-BrxC-[Aeromonas-caviae] |
| WP_257688091.1 | Type1.BREX | [illegible] | 1231 | Bacteria>Pseudomonadota>Gammaproteobacteria>Aeromonadales>Aeromonadaceae>Aeromonas | GCF_024649335.1 | Aeromonas_veronii | NUK45_RS19510 | 3671..7366 | 654 | BREX-system-P-loop-protein-BrxC-[Aeromonas-veronii] |
| WP_257712004.1 | Type1.BREX | [illegible] | 1281 | Bacteria>Pseudomonadota>Gammaproteobacteria>Aeromonadales>Aeromonadaceae>Aeromonas | GCF_024649525.1 | Aeromonas_caviae | NUK31_RS14230 | 17564..21409 | 648 | BREX-system-P-loop-protein-BrxC-[Aeromonas-caviae] |
| WP_264787731.1 | Type1.BREX | [illegible] | 1157 | Bacteria>Pseudomonadota>Gammaproteobacteria>Aeromonadales>Aeromonadaceae>Aeromonas | GCF_026000055.1 | Aeromonas_caviae | OPI30_RS10860 | 2213781..2217254 | 648 | BREX-system-P-loop-protein-BrxC-[Aeromonas-caviae] |
| WP_265455065.1 | Type1.BREX | [illegible] | 1231 | Bacteria>Pseudomonadota>Gammaproteobacteria>Aeromonadales>Aeromonadaceae>Aeromonas | GCF_026174125.1 | Aeromonas_salmonicida | ORM31_RS04470 | 132358..136053 | 645 | BREX-system-P-loop-protein-BrxC-[Aeromonas-salmonicida] |
| WP_265776110.1 | Type1.BREX | [illegible] | 1231 | Bacteria>Pseudomonadota>Gammaproteobacteria>Aeromonadales>Aeromonadaceae>Aeromonas | GCF_026230455.1 | Aeromonas_veronii | OD804_RS12725 | 3923..7618 | 654 | BREX-system-P-loop-protein-BrxC-[Aeromonas-veronii] |
| WP_265783524.1 | Type1.BREX | [illegible] | 1235 | Bacteria>Pseudomonadota>Gammaproteobacteria>Aeromonadales>Aeromonadaceae>Aeromonas | GCF_026230585.1 | Aeromonas_veronii | OD810_RS03320 | 107717..111424 | 654 | BREX-system-P-loop-protein-BrxC-[Aeromonas-veronii] |
| WP_270670318.1 | Type1.BREX | [illegible] | 1157 | Bacteria>Pseudomonadota>Gammaproteobacteria>Aeromonadales>Aeromonadaceae>Aeromonas | GCF_027705165.1 | Aeromonas_sp._QDB04 | OKC37_RS13865 | 9577..13050 | 2990477 | BREX-system-P-loop-protein-BrxC-[Aeromonas-sp.-QDB04] |
| WP_270698108.1 | Type1.BREX | [illegible] | 1280 | Bacteria>Pseudomonadota>Gammaproteobacteria>Aeromonadales>Aeromonadaceae>Aeromonas | GCF_027705185.1 | Aeromonas_sp._QDB05 | OKC32_RS09690 | 4766..8608 | 2990478 | BREX-system-P-loop-protein-BrxC-[Aeromonas-sp.-QDB05] |
| WP_270661186.1 | Type1.BREX | [illegible] | 1182 | Bacteria>Pseudomonadota>Gammaproteobacteria>Aeromonadales>Aeromonadaceae>Aeromonas | GCF_027705205.1 | unclassified_Aeromonas | OKC30_RS15245 | 6003..9551 | 257493 | MULTISPECIES:-BREX-system-P-loop-protein-BrxC-[unclassified-Aeromonas] |
| WP_270793601.1 | Type1.BREX | [illegible] | 1226 | Bacteria>Pseudomonadota>Gammaproteobacteria>Aeromonadales>Aeromonadaceae>Aeromonas | GCF_027705385.1 | unclassified_Aeromonas | OKB76_RS13245 | 1893..5573 | 257493 | MULTISPECIES:-BREX-system-P-loop-protein-BrxC-[unclassified-Aeromonas] |
| WP_270805301.1 | Type1.BREX | [illegible] | 1240 | Bacteria>Pseudomonadota>Gammaproteobacteria>Aeromonadales>Aeromonadaceae>Aeromonas | GCF_027705505.1 | Aeromonas_sp._QDB56 | OKB81_RS14905 | 8131..11853 | 2990495 | BREX-system-P-loop-protein-BrxC-[Aeromonas-sp.-QDB56] |
| WP_270809144.1 | Type1.BREX | [illegible] | 1157 | Bacteria>Pseudomonadota>Gammaproteobacteria>Aeromonadales>Aeromonadaceae>Aeromonas | GCF_027705685.1 | Aeromonas_sp._QDB08 | OKB79_RS12620 | 11744..15217 | 2990480 | BREX-system-P-loop-protein-BrxC-[Aeromonas-sp.-QDB08] |
| WP_270659881.1 | Type1.BREX | [illegible] | 1157 | Bacteria>Pseudomonadota>Gammaproteobacteria>Aeromonadales>Aeromonadaceae>Aeromonas | GCF_027705965.1 | Aeromonas_sp._Y301-2 | OKB83_RS15925 | 143279..146752 | 2990505 | BREX-system-P-loop-protein-BrxC-[Aeromonas-sp.-Y301-2] |
| WP_275056847.1 | Type1.BREX | [illegible] | 1231 | Bacteria>Pseudomonadota>Gammaproteobacteria>Aeromonadales>Aeromonadaceae>Aeromonas | GCF_029024985.1 | Aeromonas_allosaccharophila | PYU98_RS18280 | 3929296..3932991 | 656 | BREX-system-P-loop-protein-BrxC-[Aeromonas-allosaccharophila] |
| WP_076362312.1 | Type1.BREX | [illegible] | 1231 | Bacteria>Pseudomonadota>Gammaproteobacteria>Aeromonadales>Aeromonadaceae>Aeromonas | GCF_900155975.1 | Aeromonas_hydrophila | BW963_RS17620 | 135140..138835 | 644 | BREX-system-P-loop-protein-BrxC-[Aeromonas-hydrophila] |
| WP_050664996.1 | Type1.BREX | [illegible] | 1231 | Bacteria>Pseudomonadota>Gammaproteobacteria>Aeromonadales>Aeromonadaceae | GCF_000820105.1 | Aeromonas | BN1133_RS02615 | 38531..42226 | 642 | MULTISPECIES:-BREX-system-P-loop-protein-BrxC-[Aeromonas] |
| WP_181014924.1 | Type1.BREX | [illegible] | 1182 | Bacteria>Pseudomonadota>Gammaproteobacteria>Aeromonadales>Aeromonadaceae | GCF_002934425.1 | Aeromonas | C2U40_RS16580 | 3601638..3605186 | 642 | MULTISPECIES:-BREX-system-P-loop-protein-BrxC-[Aeromonas] |
| WP_182788683.1 | Type1.BREX | [illegible] | 1236 | Bacteria>Pseudomonadota>Gammaproteobacteria>Aeromonadales>Aeromonadaceae | GCF_014161425.1 | Aeromonas | H7K58_RS09750 | 2083810..2087520 | 642 | MULTISPECIES:-BREX-system-P-loop-protein-BrxC-[Aeromonas] |
| WP_223915718.1 | Type1.BREX | [illegible] | 1218 | Bacteria>Pseudomonadota>Gammaproteobacteria>Aeromonadales>Aeromonadaceae | GCF_019972515.1 | Aeromonas | LDN24_RS09390 | 22833..26489 | 642 | MULTISPECIES:-BREX-system-P-loop-protein-BrxC-[Aeromonas] |
| WP_201934227.1 | Type1.BREX | [illegible] | 1231 | Bacteria>Pseudomonadota>Gammaproteobacteria>Aeromonadales>Aeromonadaceae | GCF_019972675.1 | Aeromonas | MWG22_RS17445 | 1818..5513 | 642 | MULTISPECIES:-BREX-system-P-loop-protein-BrxC-[Aeromonas] |
| WP_171273737.1 | Type1.BREX | [illegible] | 1231 | Bacteria>Pseudomonadota>Gammaproteobacteria>Aeromonadales>Aeromonadaceae | GCF_021654375.1 | Aeromonas | E4185_RS15845 | 3355159..3358854 | 642 | MULTISPECIES:-BREX-system-P-loop-protein-BrxC-[Aeromonas] |
| WP_234605103.1 | Type1.BREX | [illegible] | 1231 | Bacteria>Pseudomonadota>Gammaproteobacteria>Aeromonadales>Aeromonadaceae | GCF_026173065.1 | Aeromonas | ORM39_RS00835 | 21547..25242 | 642 | BREX-system-P-loop-protein-BrxC-[Aeromonas-salmonicida] |
| WP_270796656.1 | Type1.BREX | [illegible] | 1157 | Bacteria>Pseudomonadota>Gammaproteobacteria>Aeromonadales>Aeromonadaceae | GCF_029024985.1 | Aeromonas | OKC40_RS19420 | 7580..11053 | 642 | MULTISPECIES:-BREX-system-P-loop-protein-BrxC-[Aeromonas] |
| WP_069785631.1 | Type1.BREX | [illegible] | 1157 | Bacteria>Pseudomonadota>Gammaproteobacteria>Aeromonadales>Aeromonadaceae | GCF_029773025.1 | Aeromonas | QBS29_RS17430 | 3038195..3041668 | 642 | MULTISPECIES:-BREX-system-P-loop-protein-BrxC-[Aeromonas] |
| WP_019935798.1 | Type1.BREX | [illegible] | 1146 | Bacteria>Pseudomonadota>Gammaproteobacteria>Aeromonadales>Aeromonadaceae>Oceanimonas | GCF_000381965.1 | Oceanimonas_smirnovii | G364_RS0114130 | 4273..7713 | 264574 | BREX-system-P-loop-protein-BrxC-[Oceanimonas-smirnovii] |
| WP_228212351.1 | Type1.BREX | [illegible] | 1221 | Bacteria>Pseudomonadota>Gammaproteobacteria>Aeromonadales>Aeromonadaceae>Oceanimonas | GCF_020781865.1 | Oceanimonas_baumannii | LL240_RS06775 | 52784..56449 | 129578 | BREX-system-P-loop-protein-BrxC-[Oceanimonas-baumannii] |
| MBP8039255.1 | Type1.BREX | [illegible] | 1043 | Bacteria>Pseudomonadota>Gammaproteobacteria>Aeromonadales>Aeromonadaceae>unclassified_Aeromonadaceae | GCA_018004635.1 | Aeromonadaceae_bacterium | KAY67_01800 | 1..3131 | 1948889 | BREX-system-P-loop-protein-BrxC-[Aeromonadaceae-bacterium] |
| WP_019002750.1 | Type1.BREX | [illegible] | 1210 | Bacteria>Pseudomonadota>Gammaproteobacteria>Aeromonadales>Succinivibrionaceae>Succinimonas | GCF_000378405.1 | Succinimonas_amylolytica | F557_RS20035 | 169312..172944 | 83769 | BREX-system-P-loop-protein-BrxC-[Succinimonas-amylolytica] |
| WP_219938189.1 | Type1.BREX | [illegible] | 1210 | Bacteria>Pseudomonadota>Gammaproteobacteria>Aeromonadales>Succinivibrionaceae>Succinivibrio | GCF_019448115.1 | Succinivibrio_faecicola | J5V48_RS08685 | 2494..6126 | 2820300 | BREX-system-P-loop-protein-BrxC-[Succinivibrio-faecicola] |
| MCR4553833.1 | Type1.BREX | [illegible] | 1184 | Bacteria>Pseudomonadota>Gammaproteobacteria>Aeromonadales>Succinivibrionaceae>unclassified_Succinivibrionaceae | GCA_024683965.1 | Succinivibrionaceae_bacterium | K5752_02815 | 10310..13864 | 2053620 | BREX-system-P-loop-protein-BrxC-[Succinivibrionaceae-bacterium] |
| WP_163133170.1 | Type1.BREX | [illegible] | 1160 | Bacteria>Pseudomonadota>Gammaproteobacteria>Alteromonadales>Alteromonadaceae>Agarivorans | GCF_900537085.1 | Agarivorans_sp._Alg241-V36 | G6R11_RS11270 | 302012..305494 | 2305992 | BREX-system-P-loop-protein-BrxC-[Agarivorans-sp.-Alg241-V36] |
| WP_008844769.1 | Type1.BREX | [illegible] | 1229 | Bacteria>Pseudomonadota>Gammaproteobacteria>Alteromonadales>Alteromonadaceae>Aliiglaciecola | GCF_000314975.1 | Aliiglaciecola_lipolytica | GLIP_RS11310 | 7346..11035 | 477689 | BREX-system-P-loop-protein-BrxC-[Aliiglaciecola-lipolytica] |
| OZB42971.1 | Type1.BREX | [illegible] | 1224 | Bacteria>Pseudomonadota>Gammaproteobacteria>Alteromonadales>Alteromonadaceae>Alishewanella | GCA_002278995.1 | Alishewanella_sp._34-51-39 | B7X50_02430 | 10903..14577 | 1970316 | MAG:-hypothetical-protein-B7X50_02430-[Alishewanella-sp.-34-51-39] |
| GGW57088.1 | Type1.BREX | [illegible] | 1114 | Bacteria>Pseudomonadota>Gammaproteobacteria>Alteromonadales>Alteromonadaceae>Alishewanella | GCA_014652375.1 | Alishewanella_tabrizica | GCM10008111_11430 | 80462..83806 | 671278 | hypothetical-protein-GCM10008111_11430-[Alishewanella-tabrizica] |
| WP_057786312.1 | Type1.BREX | [illegible] | 1157 | Bacteria>Pseudomonadota>Gammaproteobacteria>Alteromonadales>Alteromonadaceae>Alishewanella | GCF_001441695.1 | Alishewanella_sp._WH16-1 | AAY72_RS06860 | 5881..9354 | 1651088 | BREX-system-P-loop-protein-BrxC-[Alishewanella-sp.-WH16-1] |
| WP_229796961.1 | Type1.BREX | [illegible] | 1125 | Bacteria>Pseudomonadota>Gammaproteobacteria>Alteromonadales>Alteromonadaceae>Alishewanella | GCF_014652375.1 | Alishewanella_tabrizica | IE070_RS05825 | 80429..83806 | 671278 | BREX-system-P-loop-protein-BrxC-[Alishewanella-tabrizica] |
| WP_265045927.1 | Type1.BREX | [illegible] | 1194 | Bacteria>Pseudomonadota>Gammaproteobacteria>Alteromonadales>Alteromonadaceae>Alkalimarinus | GCF_026016465.1 | Alkalimarinus_sp._A2M4 | NKI27_RS10060 | 2208861..2212445 | 2961619 | BREX-system-P-loop-protein-BrxC-[Alkalimarinus-sp.-A2M4] |
| WP_014998461.1 | Type1.BREX | [illegible] | 1157 | Bacteria>Pseudomonadota>Gammaproteobacteria>Alteromonadales>Alteromonadaceae>Alteromonas/Salinimonas_group | - | Alteromonas_macleodii | AMBLS11_RS09820 | 2297331..2300804 | 28108 | BREX-system-P-loop-protein-BrxC-[Alteromonas-macleodii] |
| WP_100972015.1 | Type1.BREX | [illegible] | 1157 | Bacteria>Pseudomonadota>Gammaproteobacteria>Alteromonadales>Alteromonadaceae>Alteromonas/Salinimonas_group | GCF_002831605.1 | Alteromonas_sp._MB-3u-76 | CW735_RS10480 | 2338685..2342158 | 2058133 | BREX-system-P-loop-protein-BrxC-[Alteromonas-sp.-MB-3u-76] |
| WP_189403478.1 | Type1.BREX | [illegible] | 1215 | Bacteria>Pseudomonadota>Gammaproteobacteria>Alteromonadales>Alteromonadaceae>Alteromonas/Salinimonas_group | GCF_014651815.1 | Alteromonas_halophila | IE054_RS02435 | 519507..523154 | 516698 | BREX-system-P-loop-protein-BrxC-[Alteromonas-halophila] |
| WP_207611650.1 | Type1.BREX | [illegible] | 1157 | Bacteria>Pseudomonadota>Gammaproteobacteria>Alteromonadales>Alteromonadaceae>Alteromonas/Salinimonas_group | GCF_017368475.1 | Alteromonas_sp._5E99-2 | J3L16_RS04210 | 79358..82831 | 2817683 | BREX-system-P-loop-protein-BrxC-[Alteromonas-sp.-5E99-2] |
| WP_273638258.1 | Type1.BREX | [illegible] | 1215 | Bacteria>Pseudomonadota>Gammaproteobacteria>Alteromonadales>Alteromonadaceae>Alteromonas/Salinimonas_group | GCF_028595265.1 | Alteromonas_sp._chi3 | OIK42_RS02990 | 673231..676878 | 2987522 | BREX-system-P-loop-protein-BrxC-[Alteromonas-sp.-chi3] |
| WP_006006589.1 | Type1.BREX | [illegible] | 1189 | Bacteria>Pseudomonadota>Gammaproteobacteria>Alteromonadales>Alteromonadaceae>Glaciecola | GCF_000252165.1 | Glaciecola_punicea | GPUN_RS10800 | 142812..146381 | 56804 | BREX-system-P-loop-protein-BrxC-[Glaciecola-punicea] |
| WP_136550441.1 | Type1.BREX | [illegible] | 1193 | Bacteria>Pseudomonadota>Gammaproteobacteria>Alteromonadales>Alteromonadaceae>Hydrocarboniclastica | GCF_004851605.1 | Hydrocarboniclastica_marina | soil367_RS18210 | 4131404..4134985 | 2259620 | BREX-system-P-loop-protein-BrxC-[Hydrocarboniclastica-marina] |
| PKG98979.1 | Type1.BREX | [illegible] | 1231 | Bacteria>Pseudomonadota>Gammaproteobacteria>Alteromonadales>Alteromonadaceae>Paraglaciecola | GCA_002835935.1 | Paraglaciecola_sp._MB-3u-78 | CXF95_12105 | 233468..237163 | 2058332 | BREX-system-P-loop-protein-BrxC-[Paraglaciecola-sp.-MB-3u-78] |
| WP_007105022.1 | Type1.BREX | [illegible] | 1229 | Bacteria>Pseudomonadota>Gammaproteobacteria>Alteromonadales>Alteromonadaceae>Paraglaciecola | GCF_000315055.1 | Paraglaciecola_polaris | GPLA_RS11155 | 47408..51097 | 222814 | BREX-system-P-loop-protein-BrxC-[Paraglaciecola-polaris] |
| WP_232787809.1 | Type1.BREX | [illegible] | 1241 | Bacteria>Pseudomonadota>Gammaproteobacteria>Alteromonadales>Alteromonadaceae>Paraglaciecola | GCF_002835935.1 | Paraglaciecola_sp._MB-3u-78 | CXF95_RS12105 | 233438..237163 | 2058332 | BREX-system-P-loop-protein-BrxC-[Paraglaciecola-sp.-MB-3u-78] |
| WP_158768857.1 | Type1.BREX | [illegible] | 1222 | Bacteria>Pseudomonadota>Gammaproteobacteria>Alteromonadales>Alteromonadaceae>Paraglaciecola | GCF_009796745.1 | Paraglaciecola_sp._L1A13 | GQR89_RS03920 | 955654..959322 | 2686359 | BREX-system-P-loop-protein-BrxC-[Paraglaciecola-sp.-L1A13] |
| WP_240919947.1 | Type1.BREX | [illegible] | 1180 | Bacteria>Pseudomonadota>Gammaproteobacteria>Alteromonadales>Alteromonadaceae>Paraglaciecola | GCF_011378905.1 | Paraglaciecola_sp._20A4 | GQS54_RS13935 | 91436..94978 | 2687288 | BREX-system-P-loop-protein-BrxC-[Paraglaciecola-sp.-20A4] |
| NQY33786.1 | Type1.BREX | [illegible] | 1200 | Bacteria>Pseudomonadota>Gammaproteobacteria>Alteromonadales>Alteromonadaceae>unclassified_Alteromonadaceae | GCA_013216015.1 | Alteromonadaceae_bacterium | HRT37_02225 | 108462..112064 | 1916082 | BREX-system-P-loop-protein-BrxC-[Alteromonadaceae-bacterium] |
| ASP49913.1 | Type1.BREX | [illegible] | 1229 | Bacteria>Pseudomonadota>Gammaproteobacteria>Alteromonadales>Colwelliaceae>Cognaticolwellia | GCA_002076895.1 | Cognaticolwellia_beringensis | B5D82_06290 | 1500021..1503710 | 1967665 | DNA-repair-protein-[Cognaticolwellia-beringensis] |
| WP_245807568.1 | Type1.BREX | [illegible] | 1232 | Bacteria>Pseudomonadota>Gammaproteobacteria>Alteromonadales>Colwelliaceae>Cognaticolwellia | GCF_002076895.1 | Cognaticolwellia_beringensis | B5D82_RS06290 | 1500012..1503710 | 1967665 | BREX-system-P-loop-protein-BrxC-[Cognaticolwellia-beringensis] |
| AWB59509.1 | Type1.BREX | [illegible] | 1232 | Bacteria>Pseudomonadota>Gammaproteobacteria>Alteromonadales>Colwelliaceae>Colwellia | GCA_003061515.1 | Colwellia_sp._Arc7-D | DBO93_09830 | 2239405..2243103 | 2161872 | BREX-system-P-loop-protein-BrxC-[Colwellia-sp.-Arc7-D] |
| WP_239058953.1 | Type1.BREX | [illegible] | 1235 | Bacteria>Pseudomonadota>Gammaproteobacteria>Alteromonadales>Colwelliaceae>Colwellia | GCF_003061515.1 | Colwellia_sp._Arc7-D | DBO93_RS09830 | 2239405..2243112 | 2161872 | BREX-system-P-loop-protein-BrxC-[Colwellia-sp.-Arc7-D] |
| WP_146799747.1 | Type1.BREX | [illegible] | 1165 | Bacteria>Pseudomonadota>Gammaproteobacteria>Alteromonadales>Colwelliaceae>Colwellia | GCF_007954305.1 | Colwellia_hornerae | ESZ26_RS11985 | 115858..119355 | 89402 | BREX-system-P-loop-protein-BrxC-[Colwellia-hornerae] |
| WP_085284213.1 | Type1.BREX | [illegible] | 1243 | Bacteria>Pseudomonadota>Gammaproteobacteria>Alteromonadales>Colwelliaceae>Colwellia | GCF_900109795.1 | Colwellia_chukchiensis | B6K83_RS06980 | 108494..112225 | 641665 | BREX-system-P-loop-protein-BrxC-[Colwellia-chukchiensis] |
| WP_130603913.1 | Type1.BREX | [illegible] | 1188 | Bacteria>Pseudomonadota>Gammaproteobacteria>Alteromonadales>Colwelliaceae>Litorilituus | GCF_004295665.1 | Litorilituus_sediminis | EMK97_RS16655 | 3755554..3759120 | 718192 | BREX-system-P-loop-protein-BrxC-[Litorilituus-sediminis] |
| WP_189376285.1 | Type1.BREX | [illegible] | 1235 | Bacteria>Pseudomonadota>Gammaproteobacteria>Alteromonadales>Colwelliaceae>Thalassotalea | GCF_014653195.1 | Thalassotalea_profundi | IE155_RS01345 | 297657..301364 | 2036687 | BREX-system-P-loop-protein-BrxC-[Thalassotalea-profundi] |
| KXS36208.1 | Type1.BREX | [illegible] | 1150 | Bacteria>Pseudomonadota>Gammaproteobacteria>Alteromonadales>Idiomarinaceae>Idiomarina | GCA_001577655.1 | Idiomarina_sp._T82-3 | AWU56_323 | 88359..91811 | 1794809 | hypothetical-protein-AWU56_323-[Idiomarina-sp.-T82-3] |
| WP_258240870.1 | Type1.BREX | [illegible] | 1150 | Bacteria>Pseudomonadota>Gammaproteobacteria>Alteromonadales>Idiomarinaceae>Pseudidiomarina | GCF_024723895.1 | Pseudidiomarina_homiensis | NV376_RS09715 | 149869..153321 | 364198 | BREX-system-P-loop-protein-BrxC-[Pseudidiomarina-homiensis] |
| WP_092590479.1 | Type1.BREX | [illegible] | 1224 | Bacteria>Pseudomonadota>Gammaproteobacteria>Alteromonadales>Idiomarinaceae>Pseudidiomarina | GCF_900104245.1 | Pseudidiomarina_indica | BLP64_RS00065 | 10133..13807 | 1159017 | BREX-system-P-loop-protein-BrxC-[Pseudidiomarina-indica] |
| WP_112714387.1 | Type1.BREX | [illegible] | 1199 | Bacteria>Pseudomonadota>Gammaproteobacteria>Alteromonadales>Moritellaceae>Moritella | GCF_900465055.1 | Moritella_yayanosii | MORIYA_RS08605 | 1893116..1896715 | 69539 | BREX-system-P-loop-protein-BrxC-[Moritella-yayanosii] |
| WP_010607320.1 | Type1.BREX | [illegible] | 1188 | Bacteria>Pseudomonadota>Gammaproteobacteria>Alteromonadales>Pseudoalteromonadaceae | GCF_000276645.1 | Pseudoalteromonas | UY7_RS0121200 | 90697..94263 | 53246 | MULTISPECIES:-BREX-system-P-loop-protein-BrxC-[Pseudoalteromonas] |
| CCQ10850.1 | Type1.BREX | [illegible] | 1237 | Bacteria>Pseudomonadota>Gammaproteobacteria>Alteromonadales>Pseudoalteromonadaceae>Pseudoalteromonas | GCA_000333235.1 | Pseudoalteromonas_luteoviolacea_B_=_ATCC_29581 | PALB_17240 | 21761..25474 | 1268239 | ATPase-like-protein-[Pseudoalteromonas-luteoviolacea-B-=-ATCC-29581] |
| HDY93539.1 | Type1.BREX | [illegible] | 1159 | Bacteria>Pseudomonadota>Gammaproteobacteria>Alteromonadales>Pseudoalteromonadaceae>Pseudoalteromonas | GCA_011050215.1 | Pseudoalteromonas_sp. | ENH79_15755 | 24321..27800 | 53249 | BREX-system-P-loop-protein-BrxC-[Pseudoalteromonas-sp.] |
| WP_024611453.1 | Type1.BREX | [illegible] | 1193 | Bacteria>Pseudomonadota>Gammaproteobacteria>Alteromonadales>Pseudoalteromonadaceae>Pseudoalteromonas | GCF_000498095.1 | Pseudoalteromonas_sp._TB64 | L647_RS0112855 | 131987..135568 | 1938600 | BREX-system-P-loop-protein-BrxC-[Pseudoalteromonas-sp.-TB64] |
| WP_054983666.1 | Type1.BREX | [illegible] | 1159 | Bacteria>Pseudomonadota>Gammaproteobacteria>Alteromonadales>Pseudoalteromonadaceae>Pseudoalteromonas | GCF_001400005.1 | Pseudoalteromonas_sp._P1-30 | AN395_RS16640 | 61256..64735 | 1723760 | BREX-system-P-loop-protein-BrxC-[Pseudoalteromonas-sp.-P1-30] |
| WP_058373560.1 | Type1.BREX | [illegible] | 1229 | Bacteria>Pseudomonadota>Gammaproteobacteria>Alteromonadales>Pseudoalteromonadaceae>Pseudoalteromonas | GCF_001465295.1 | Pseudoalteromonas_translucida | PTRA_a2157 | 1987847..1991536 | 166935 | BREX-system-P-loop-protein-BrxC-[Pseudoalteromonas-translucida] |
| WP_064663367.1 | Type1.BREX | [illegible] | 1242 | Bacteria>Pseudomonadota>Gammaproteobacteria>Alteromonadales>Pseudoalteromonadaceae>Pseudoalteromonas | GCF_001661605.1 | Pseudoalteromonas_sp._MQS005 | A9479_RS10285 | 16730..20458 | 1854052 | BREX-system-P-loop-protein-BrxC-[Pseudoalteromonas-sp.-MQS005] |
| WP_076923084.1 | Type1.BREX | [illegible] | 1229 | Bacteria>Pseudomonadota>Gammaproteobacteria>Alteromonadales>Pseudoalteromonadaceae>Pseudoalteromonas | GCF_001974845.1 | Pseudoalteromonas_sp._SK20 | BW140_RS17330 | 3309..6998 | 1938367 | BREX-system-P-loop-protein-BrxC-[Pseudoalteromonas-sp.-SK20] |
| WP_121955020.1 | Type1.BREX | [illegible] | 1027 | Bacteria>Pseudomonadota>Gammaproteobacteria>Alteromonadales>Pseudoalteromonadaceae>Pseudoalteromonas | GCF_002863625.1 | Pseudoalteromonas_sp._MelDa3 | CXF89_RS14715 | 1..3085 | 888435 | BREX-system-P-loop-protein-BrxC,-partial-[Pseudoalteromonas-sp.-MelDa3] |
| WP_234000034.1 | Type1.BREX | [illegible] | 1252 | Bacteria>Pseudomonadota>Gammaproteobacteria>Alteromonadales>Pseudoalteromonadaceae>Pseudoalteromonas | GCF_002964725.1 | Pseudoalteromonas_sp._T1lg76 | C2M35_RS06890 | 48486..52244 | 2077103 | BREX-system-P-loop-protein-BrxC-[Pseudoalteromonas-sp.-T1lg76] |
| WP_233998422.1 | Type1.BREX | [illegible] | 1252 | Bacteria>Pseudomonadota>Gammaproteobacteria>Alteromonadales>Pseudoalteromonadaceae>Pseudoalteromonas | GCF_002964785.1 | Pseudoalteromonas_sp._T1lg48 | C2M29_RS12220 | 655771..659529 | 2077100 | BREX-system-P-loop-protein-BrxC-[Pseudoalteromonas-sp.-T1lg48] |
| WP_130052623.1 | Type1.BREX | [illegible] | 1244 | Bacteria>Pseudomonadota>Gammaproteobacteria>Alteromonadales>Pseudoalteromonadaceae>Pseudoalteromonas | GCF_004195145.1 | Pseudoalteromonas_sp._MEBiC_03485 | EVU92_RS18430 | 351830..355564 | 2571103 | BREX-system-P-loop-protein-BrxC-[Pseudoalteromonas-sp.-MEBiC-03485] |
| WP_130146693.1 | Type1.BREX | [illegible] | 1225 | Bacteria>Pseudomonadota>Gammaproteobacteria>Alteromonadales>Pseudoalteromonadaceae>Pseudoalteromonas | GCF_004208955.1 | Pseudoalteromonas_sp._CO325X | EXT46_RS08650 | 70759..74436 | 1777262 | BREX-system-P-loop-protein-BrxC-[Pseudoalteromonas-sp.-CO325X] |
| WP_138480410.1 | Type1.BREX | [illegible] | 1233 | Bacteria>Pseudomonadota>Gammaproteobacteria>Alteromonadales>Pseudoalteromonadaceae>Pseudoalteromonas | GCF_005876835.1 | Pseudoalteromonas_phenolica | C1E24_RS08165 | 13273..16974 | 161398 | BREX-system-P-loop-protein-BrxC-[Pseudoalteromonas-phenolica] |
| WP_096038338.1 | Type1.BREX | [illegible] | 1159 | Bacteria>Pseudomonadota>Gammaproteobacteria>Alteromonadales>Pseudoalteromonadaceae>Pseudoalteromonas | GCF_007991235.1 | Pseudoalteromonas_tetraodonis | PTE01_RS15665 | 26152..29631 | 43659 | BREX-system-P-loop-protein-BrxC-[Pseudoalteromonas-tetraodonis] |
| WP_089368387.1 | Type1.BREX | [illegible] | 1229 | Bacteria>Pseudomonadota>Gammaproteobacteria>Alteromonadales>Pseudoalteromonadaceae>Pseudoalteromonas | GCF_007991255.1 | Pseudoalteromonas_nigrifaciens | PNIG_RS09950 | 2060327..2064016 | 28109 | BREX-system-P-loop-protein-BrxC-[Pseudoalteromonas-nigrifaciens] |
| WP_089368479.1 | Type1.BREX | [illegible] | 1169 | Bacteria>Pseudomonadota>Gammaproteobacteria>Alteromonadales>Pseudoalteromonadaceae>Pseudoalteromonas | GCF_007991255.1 | Pseudoalteromonas_nigrifaciens | PNIG_RS10730 | 2243615..2247124 | 28109 | BREX-system-P-loop-protein-BrxC-[Pseudoalteromonas-nigrifaciens] |
| WP_167658112.1 | Type1.BREX | [illegible] | 1188 | Bacteria>Pseudomonadota>Gammaproteobacteria>Alteromonadales>Pseudoalteromonadaceae>Pseudoalteromonas | GCF_011806445.1 | Pseudoalteromonas_sp._HF66 | HCZ11_RS07660 | 212811..216377 | 2721559 | BREX-system-P-loop-protein-BrxC-[Pseudoalteromonas-sp.-HF66] |
| WP_182739649.1 | Type1.BREX | [illegible] | 1235 | Bacteria>Pseudomonadota>Gammaproteobacteria>Alteromonadales>Pseudoalteromonadaceae>Pseudoalteromonas | GCF_014163545.1 | Pseudoalteromonas_sp._SR45-6 | H5158_RS05620 | 135410..139117 | 2760927 | BREX-system-P-loop-protein-BrxC-[Pseudoalteromonas-sp.-SR45-6] |
| WP_182752860.1 | Type1.BREX | [illegible] | 1159 | Bacteria>Pseudomonadota>Gammaproteobacteria>Alteromonadales>Pseudoalteromonadaceae>Pseudoalteromonas | GCF_014163875.1 | Pseudoalteromonas_sp._SR43-3 | H5088_RS10925 | 129286..132765 | 2760943 | BREX-system-P-loop-protein-BrxC-[Pseudoalteromonas-sp.-SR43-3] |
| WP_193038371.1 | Type1.BREX | [illegible] | 1159 | Bacteria>Pseudomonadota>Gammaproteobacteria>Alteromonadales>Pseudoalteromonadaceae>Pseudoalteromonas | GCF_014897925.1 | Pseudoalteromonas_nigrifaciens | INT11_RS00030 | 7982..11461 | 28109 | BREX-system-P-loop-protein-BrxC-[Pseudoalteromonas-nigrifaciens] |
| WP_074988793.1 | Type1.BREX | [illegible] | 1242 | Bacteria>Pseudomonadota>Gammaproteobacteria>Alteromonadales>Pseudoalteromonadaceae>Pseudoalteromonas | GCF_014925285.1 | Pseudoalteromonas_lipolytica | PLIP_RS19215 | 618195..621923 | 570156 | BREX-system-P-loop-protein-BrxC-[Pseudoalteromonas-lipolytica] |
| WP_197117904.1 | Type1.BREX | [illegible] | 1229 | Bacteria>Pseudomonadota>Gammaproteobacteria>Alteromonadales>Pseudoalteromonadaceae>Pseudoalteromonas | GCF_015864765.1 | Pseudoalteromonas_sp._NZS11 | I6E82_RS11050 | 319401..323090 | 2792049 | BREX-system-P-loop-protein-BrxC-[Pseudoalteromonas-sp.-NZS11] |
| WP_200220630.1 | Type1.BREX | [illegible] | 1193 | Bacteria>Pseudomonadota>Gammaproteobacteria>Alteromonadales>Pseudoalteromonadaceae>Pseudoalteromonas | GCF_016598755.1 | Pseudoalteromonas_sp._LC2018020214 | JG479_RS17805 | 301190..304771 | 2799564 | BREX-system-P-loop-protein-BrxC-[Pseudoalteromonas-sp.-LC2018020214] |
| WP_248533819.1 | Type1.BREX | [illegible] | 1229 | Bacteria>Pseudomonadota>Gammaproteobacteria>Alteromonadales>Pseudoalteromonadaceae>Pseudoalteromonas | GCF_023161935.1 | Pseudoalteromonas_sp._2CM28B | MTN96_RS29910 | 10677..14366 | 2929851 | BREX-system-P-loop-protein-BrxC-[Pseudoalteromonas-sp.-2CM28B] |
| WP_165723736.1 | Type1.BREX | [illegible] | 1225 | Bacteria>Pseudomonadota>Gammaproteobacteria>Alteromonadales>Pseudoalteromonadaceae>Pseudoalteromonas | GCF_943789085.1 | Pseudoalteromonas_sp._Z1A2 | QDP19_RS17640 | 3065370..3069047 | 2686350 | MULTISPECIES:-BREX-system-P-loop-protein-BrxC-[Pseudoalteromonas] |
| WP_197236686.1 | Type1.BREX | [illegible] | 1234 | Bacteria>Pseudomonadota>Gammaproteobacteria>Alteromonadales>Pseudoalteromonadaceae>Pseudoalteromonas | - | Pseudoalteromonas_sp._NZS71 | I6E85_05230 | 196559..200263 | 2792052 | BREX-system-P-loop-protein-BrxC-[Pseudoalteromonas-sp.-NZS71] |
| MCK5817827.1 | Type1.BREX | [illegible] | 582 | Bacteria>Pseudomonadota>Gammaproteobacteria>Alteromonadales>Psychromonadaceae>Psychromonas | GCA_023145575.1 | Psychromonas_sp. | KAH18_00955 | 1..1748 | 1884585 | BREX-system-P-loop-protein-BrxC-[Psychromonas-sp.] |

| Protein-ID | System | Gene-Neighbourhoods | Length | Taxonomy | Assembly-ID | Strain | Locus-Tag | Genomic-coordinates | TaxID | Fasta-define |
|---|---|---|---|---|---|---|---|---|---|---|
| WP_015465415.1 | Type1.BREX | wHTH+WYL+WCX->FGase-><-?\|\|SF1-SF2-Helicase+REase->SF1-SF2-Helicase+REase-><-TPR<-HEPN+SF1-SF2-Helicase+RNase+RNase<-PglZ<-N6-METHYLASE<-DUF6079*<-BrxB-BrxF-iSTAND<-BrxA-DNA-binding<-wHTH+WYL+WCX<-XIS-HTH\|\|?-><-tRNA\|\|?-><-?\|\|?->35exo+HRDC-> | 1159 | Bacteria>Pseudomonadota>Gammaproteobacteria>Alteromonadales>Psychromonadaceae>Psychromonas | GCF_000153405.2 | Psychromonas_sp._CNPT3 | PCNPT3_RS08205 | 1892696..1896175 | 314282 | BREX-system-P-loop-protein-BrxC-[Psychromonas-sp.-CNPT3] |
| WP_037429580.1 | Type1.BREX | <-PglZ<-ParB+HNH<-N6-METHYLASE<-DUF6079*<-BrxB-BrxF-iSTAND<-BrxA-DNA-binding<-?<-HEPN<-?<-MoxR-AAA<-NtrC-AAA+FIS-HTH<-XIS-HTH | 1157 | Bacteria>Pseudomonadota>Gammaproteobacteria>Alteromonadales>Shewanellaceae | GCF_000282755.1 | Shewanella | SHEWPOL2_RS15310 | 7136..10609 | 22 | MULTISPECIES:-BREX-system-P-loop-protein-BrxC-[Shewanella] |
| PKG74712.1 | Type1.BREX | <-SIGMA+OB-Fold+AAA-ATPase+HKD-DNase-BrxL<-PglZ<-ParB+HNH<-N6-METHYLASE<-REase<-OLD-ABC<-DUF6079*<-BrxB-BrxF-iSTAND<-BrxA-DNA-binding<-XIS-HTH<-?<-TnsC-ATPase<-rve+Mu-transpos_C<-REase+wHTH | 1191 | Bacteria>Pseudomonadota>Gammaproteobacteria>Alteromonadales>Shewanellaceae>Shewanella | GCA_002836135.1 | Shewanella_sp._GutCb | CXF86_11690 | 122253..125828 | 2058315 | BREX-system-P-loop-protein-BrxC-[Shewanella-sp.-GutCb] |
| HAY94623.1 | Type1.BREX | <-DUF2787<-CbeA_antitoxin<-?<-?\|\|XIS-HTH->wHTH+WYL+WCX-p->wHTH+WYL+WCX->BrxA-DNA-binding->BrxB-BrxF-iSTAND->DUF6079*->HNH->?->?->ParB+HNH->N6-METHYLASE->HEPN->N6-METHYLASE-p-> | 1224 | Bacteria>Pseudomonadota>Gammaproteobacteria>Alteromonadales>Shewanellaceae>Shewanella | GCA_003486485.1 | Shewanella_sp. | DCY70_12125 | 4933..8607 | 50422 | BREX-system-P-loop-protein-BrxC-[Shewanella-sp.] |
| MBB1438394.1 | Type1.BREX | <-CHTH\|\|XIS-HTH->BrxA-DNA-binding->BrxB-BrxF-iSTAND->DUF6079*->N6-METHYLASE->PglZ->SIGMA+OB-Fold+AAA-ATPase+HKD-DNase-BrxL-><-?<-MerR-HTH<-?<-HTH_26\|\|Phage_AlpA->Arm-DNA-bind_2+Phage_int_SAM_3+Phage_integrase-> | 1182 | Bacteria>Pseudomonadota>Gammaproteobacteria>Alteromonadales>Shewanellaceae>Shewanella | GCA_014164505.1 | Shewanella_sp._SG41-4 | H5202_06785 | 3542..7090 | 2760976 | BREX-system-P-loop-protein-BrxC-[Shewanella-sp.-SG41-4] |
| MBO2546554.1 | Type1.BREX | <-SIGMA+OB-Fold+AAA-ATPase+Lon-Protease-BrxL<-PglZ<-?<-N6-METHYLASE-p<-RVT<-N6-METHYLASE<-?<-HNH<-DUF6079*<-BrxB-BrxF-iSTAND<-BrxA-DNA-binding<-wHTH+WYL+WCX<-?<-?<-XIS-HTH<-XIS-HTH<-?<-Arm-DNA-bind_3+Phage_integrase | 1225 | Bacteria>Pseudomonadota>Gammaproteobacteria>Alteromonadales>Shewanellaceae>Shewanella | GCA_017580215.1 | Shewanella_algae | I6M36_02045 | 117246..120923 | 38313 | BREX-system-P-loop-protein-BrxC-[Shewanella-algae] |
| MCD8561045.1 | Type1.BREX | <-N6-METHYLASE-p<-HEPN<-N6-METHYLASE-p<-N6-METHYLASE-p<-N6-METHYLASE<-?<-HNH<-DUF6079*<-BrxB-BrxF-iSTAND<-BrxA-DNA-binding<-wHTH+WYL+WCX<-XIS-HTH | 1231 | Bacteria>Pseudomonadota>Gammaproteobacteria>Alteromonadales>Shewanellaceae>Shewanella | GCA_021209305.1 | Shewanella_xiamenensis | LRY75_19995 | 6155..9850 | 332186 | BREX-system-P-loop-protein-BrxC-[Shewanella-xiamenensis] |
| VEE64276.1 | Type1.BREX | <-Cas10\|\|Cas_NE0113-><-?<-HDRnase<-Arm-DNA-bind_2+Phage_int_SAM_3+Phage_integrase<-Phage_AlpA\|\|wHTH+WYL+WCX->BrxA-DNA-binding->BrxB-BrxF-iSTAND->DUF6079*->N6-METHYLASE->N6-METHYLASE-p->RVT->HEPN->PglZ->PglZ->SIGMA+OB-Fold+AAA-ATPase+Lon-Protease-BrxL->REase->CHTH-> | 1268 | Bacteria>Pseudomonadota>Gammaproteobacteria>Alteromonadales>Shewanellaceae>Shewanella | GCA_900636665.1 | Shewanella_putrefaciens | NCTC12093_04119 | 4646571..4650377 | 24 | Uncharacterised-protein-[Shewanella-putrefaciens] |
| WP_011622483.1 | Type1.BREX | XIS-HTH->NtrC-AAA+FIS-HTH->MoxR-AAA->?->HEPN->?->BrxA-DNA-binding->BrxB-BrxF-iSTAND->DUF6079*->N6-METHYLASE->ParB+HNH->PglZ->ZetaToxin->?-><-SF1-SF2-Helicase+REase<-SF1-SF2-Helicase+REase\|\|HerA/FtsK-> | 1157 | Bacteria>Pseudomonadota>Gammaproteobacteria>Alteromonadales>Shewanellaceae>Shewanella | GCF_000014685.1 | Shewanella_sp._MR-4 | SHEWMR4_RS09030 | 2019364..2022837 | 60480 | BREX-system-P-loop-protein-BrxC-[Shewanella-sp.-MR-4] |
| WP_011716828.1 | Type1.BREX | XIS-HTH->NtrC-AAA+FIS-HTH->MoxR-AAA->?->HEPN->?->BrxA-DNA-binding->BrxB-BrxF-iSTAND->DUF6079*->N6-METHYLASE->ParB+HNH->PglZ->ZetaToxin->?-><-SF1-SF2-Helicase+REase<-SF1-SF2-Helicase+REase\|\|HerA/FtsK-> | 1157 | Bacteria>Pseudomonadota>Gammaproteobacteria>Alteromonadales>Shewanellaceae>Shewanella | GCF_000203935.1 | Shewanella_sp._ANA-3 | SHEWANA3_RS09590 | 2144277..2147750 | 94122 | BREX-system-P-loop-protein-BrxC-[Shewanella-sp.-ANA-3] |
| WP_006086465.1 | Type1.BREX | <-TraI-VirD2-RCRE\|\|?-><-?\|\|?->wHTH+WYL+WCX->Peptidase_C47->wHTH+WYL+WCX->BrxA-DNA-binding->BrxB-BrxF-iSTAND->DUF6079*->ParB+HNH->OLD-ABC->TOPRIM->N6-METHYLASE->PglZ->SIGMA+OB-Fold+AAA-ATPase+Lon-Protease-BrxL->SF1-SF2-Helicase->?-><-DDE_Tnp_1_assoc+DDE_Tnp_1 | 1229 | Bacteria>Pseudomonadota>Gammaproteobacteria>Alteromonadales>Shewanellaceae>Shewanella | GCF_000231345.1 | Shewanella_baltica | SBAL625DRAFT_RS15525 | 212954..216643 | 62322 | BREX-system-P-loop-protein-BrxC-[Shewanella-baltica] |
| WP_028764576.1 | Type1.BREX | BrxA-DNA-binding->BrxB-BrxF-iSTAND->DUF6079*->N6-METHYLASE->ParB+HNH->PglZ->HEPN+Sel1+SEC-C-><-SF1-SF2-Helicase+REase<-SF1-SF2-Helicase+REase<-?<-MerR-HTH | 1157 | Bacteria>Pseudomonadota>Gammaproteobacteria>Alteromonadales>Shewanellaceae>Shewanella | GCF_000518705.1 | Shewanella_colwelliana | L876_RS0115155 | 4861..8334 | 23 | BREX-system-P-loop-protein-BrxC-[Shewanella-colwelliana] |
| WP_231570521.1 | Type1.BREX | TetR-HTH-><-?<-?<-?\|\|?->?-><-PglZ<-ParB+HNH<-N6-METHYLASE<-DUF6079-p*\|\|?->?-><-?<-PDZ+Peptidase<-PTSEIIA1 | 941 | Bacteria>Pseudomonadota>Gammaproteobacteria>Alteromonadales>Shewanellaceae>Shewanella | GCF_000798835.1 | Shewanella_sp._ZOR0012 | L976_RS20155 | 4209796..4212621 | 1339231 | BREX-system-P-loop-protein-BrxC-[Shewanella-sp.-ZOR0012] |
| WP_055648628.1 | Type1.BREX | <-PVC-AAA<-?\|\|SF1-SF2-Helicase+REase->SF1-SF2-Helicase+REase-><-?<-ZetaToxin<-PglZ<-ParB+HNH<-N6-METHYLASE<-DUF6079*<-BrxB-BrxF-iSTAND<-BrxA-DNA-binding<-?<-HEPN | 1157 | Bacteria>Pseudomonadota>Gammaproteobacteria>Alteromonadales>Shewanellaceae>Shewanella | GCF_001308045.1 | Shewanella_sp._Sh95 | AEA42_RS19845 | 36695..40168 | 1689868 | BREX-system-P-loop-protein-BrxC-[Shewanella-sp.-Sh95] |
| WP_244159705.1 | Type1.BREX | <-HDRnase<-DDE_Tnp_1<-Arm-DNA-bind_2+Phage_int_SAM_3+Phage_integrase<-Phage_AlpA\|\|HTH_26->wHTH+WYL+WCX->BrxA-DNA-binding->BrxB-BrxF-iSTAND->DUF6079*->HNH->?->N6-METHYLASE->N6-METHYLASE-p->HEPN->N6-METHYLASE-p->MULE-TRANSPOSASE->N6-METHYLASE-p-> | 1254 | Bacteria>Pseudomonadota>Gammaproteobacteria>Alteromonadales>Shewanellaceae>Shewanella | GCF_001723195.1 | Shewanella_xiamenensis | ABT47_RS18205 | 4085889..4089653 | 332186 | BREX-system-P-loop-protein-BrxC-[Shewanella-xiamenensis] |
| WP_069454895.1 | Type1.BREX | <-HDRnase<-Arm-DNA-bind_2+Phage_int_SAM_3+Phage_integrase<-Phage_AlpA\|\|HTH_26->wHTH+WYL+WCX->BrxA-DNA-binding->BrxB-BrxF-iSTAND->DUF6079*->HNH->?->N6-METHYLASE->N6-METHYLASE-p->HEPN->N6-METHYLASE-p->MULE-TRANSPOSASE->N6-METHYLASE-p-> | 1240 | Bacteria>Pseudomonadota>Gammaproteobacteria>Alteromonadales>Shewanellaceae>Shewanella | GCF_002074855.1 | Shewanella_xiamenensis | VO00_RS12940 | 4702..8424 | 332186 | BREX-system-P-loop-protein-BrxC-[Shewanella-xiamenensis] |
| WP_080916542.1 | Type1.BREX | <-Pkinase+STYKIN\|\|?->?->wHTH+WYL+WCX->BrxA-DNA-binding->BrxB-BrxF-iSTAND->DUF6079-p*->ParB+HNH->N6-METHYLASE->PglZ->SIGMA+OB-Fold+AAA-ATPase+Lon-Protease-BrxL-><-MerR-HTH\|\|?->?->Phage_int_SAM_4-> | 1234 | Bacteria>Pseudomonadota>Gammaproteobacteria>Alteromonadales>Shewanellaceae>Shewanella | GCF_002075795.1 | Shewanella_japonica | SJ2017_RS16905 | 3936240..3939944 | 93973 | BREX-system-P-loop-protein-BrxC-[Shewanella-japonica] |
| WP_088585687.1 | Type1.BREX | ZnR-><-HEPN-p<-HEPN-p<-SIGMA+OB-Fold+AAA-ATPase+Lon-Protease-BrxL<-PglZ<-N6-METHYLASE-p\|\|DDE_Tnp_1-><-N6-METHYLASE<-DUF6079*<-BrxB-BrxF-iSTAND<-BrxA-DNA-binding<-DDE_Tnp_1<-Prok-JAB<-wHTH+WYL+WCX<-?<-XIS-HTH | 1229 | Bacteria>Pseudomonadota>Gammaproteobacteria>Alteromonadales>Shewanellaceae>Shewanella | GCF_002209245.2 | Shewanella_sp._FDAARGOS_354 | CEQ32_RS05910 | 1298859..1302548 | 1930557 | BREX-system-P-loop-protein-BrxC-[Shewanella-sp.-FDAARGOS_354] |
| WP_257985331.1 | Type1.BREX | <-SIGMA+OB-Fold+AAA-ATPase+HKD-DNase-BrxL<-PglZ<-ParB+HNH<-N6-METHYLASE<-REase<-OLD-ABC<-DUF6079*<-BrxB-BrxF-iSTAND<-BrxA-DNA-binding<-XIS-HTH<-?<-TnsC-ATPase<-rve+Mu-transpos_C<-REase+wHTH | 1193 | Bacteria>Pseudomonadota>Gammaproteobacteria>Alteromonadales>Shewanellaceae>Shewanella | GCF_002836135.1 | Shewanella_sp._GutCb | CXF86_RS11730 | 122253..125834 | 2058315 | BREX-system-P-loop-protein-BrxC-[Shewanella-sp.-GutCb] |
| WP_107947297.1 | Type1.BREX | <-Phage_integrase<-DUF2787<-DUF2787<-CbeA_antitoxin<-?<-?\|\|wHTH+WYL+WCX->BrxA-DNA-binding->BrxB-BrxF-iSTAND->DUF6079*->HNH->?->N6-METHYLASE->SIR2->HerA/FtsK->N6-METHYLASE-p->PglZ->SIGMA+OB-Fold+AAA-ATPase+Lon-Protease-BrxL-> | 1246 | Bacteria>Pseudomonadota>Gammaproteobacteria>Alteromonadales>Shewanellaceae>Shewanella | GCF_003052765.1 | Shewanella_baltica | C7T95_RS09475 | 2180761..2184501 | 62322 | BREX-system-P-loop-protein-BrxC-[Shewanella-baltica] |
| WP_108947326.1 | Type1.BREX | <-Phage_integrase<-Phage_integrase<-?\|\|35exo->?->wHTH+WYL+WCX->BrxA-DNA-binding->BrxB-BrxF-iSTAND->DUF6079*->N6-METHYLASE->PglZ->SIGMA+OB-Fold+AAA-ATPase+Lon-Protease-BrxL->SbcC->HDRnase->VirD4-TraG+VirB4-FtsK-> | 1225 | Bacteria>Pseudomonadota>Gammaproteobacteria>Alteromonadales>Shewanellaceae>Shewanella | GCF_003112715.1 | Shewanella_halifaxensis | C6V78_RS24360 | 248833..252510 | 271098 | BREX-system-P-loop-protein-BrxC-[Shewanella-halifaxensis] |
| WP_173294709.1 | Type1.BREX | wHTH(SBD)+HNH->BrxA-DNA-binding->BrxB-BrxF-iSTAND->DUF6079*->N6-METHYLASE->PglZ-><-SF1-SF2-Helicase+REase<-SF1-SF2-Helicase+REase\|\|?-><-PAIREDC-HTH | 1157 | Bacteria>Pseudomonadota>Gammaproteobacteria>Alteromonadales>Shewanellaceae>Shewanella | GCF_013248905.1 | Shewanella_sp._VB17 | HQQ94_RS12350 | 2876249..2879722 | 2739432 | BREX-system-P-loop-protein-BrxC-[Shewanella-sp.-VB17] |
| WP_252977707.1 | Type1.BREX | XIS-HTH->NtrC-AAA+FIS-HTH->MoxR-AAA->?->?->?->BrxA-DNA-binding->BrxB-BrxF-iSTAND->DUF6079-p*-><-DEDD_Tnp_IS110+Transposase_20\|\|DUF6079-p->DUF6079-p->?->N6-METHYLASE-p->N6-METHYLASE-p->?->N6-METHYLASE-p->ParB+HNH-> | 261 | Bacteria>Pseudomonadota>Gammaproteobacteria>Alteromonadales>Shewanellaceae>Shewanella | GCF_013307415.1 | Shewanella_sp._ZOR0012 | HT094_RS07805 | 1783246..1784031 | 1339231 | hypothetical-protein-[Shewanella-sp.-ZOR0012] |
| WP_252977708.1 | Type1.BREX | NtrC-AAA+FIS-HTH->MoxR-AAA->?->?->?->BrxA-DNA-binding->BrxB-BrxF-iSTAND->DUF6079-p*-><-DEDD_Tnp_IS110+Transposase_20\|\|DUF6079-p*->DUF6079-p->?->N6-METHYLASE-p->N6-METHYLASE-p->?->N6-METHYLASE-p->ParB+HNH->PglZ->ZetaToxin-> | 746 | Bacteria>Pseudomonadota>Gammaproteobacteria>Alteromonadales>Shewanellaceae>Shewanella | GCF_013307415.1 | Shewanella_sp._ZOR0012 | HT094_RS07815 | 1785170..1787410 | 1339231 | BREX-system-P-loop-protein-BrxC-[Shewanella-sp.-ZOR0012] |
| WP_259397220.1 | Type1.BREX | <-CHTH\|\|XIS-HTH->BrxA-DNA-binding->BrxB-BrxF-iSTAND->DUF6079*->N6-METHYLASE->PglZ->SIGMA+OB-Fold+AAA-ATPase+HKD-DNase-BrxL-><-?<-MerR-HTH<-?<-HTH_26\|\|Phage_AlpA->Arm-DNA-bind_2+Phage_int_SAM_3+Phage_integrase-> | 1180 | Bacteria>Pseudomonadota>Gammaproteobacteria>Alteromonadales>Shewanellaceae>Shewanella | GCF_014164505.1 | Shewanella_sp._SG41-4 | H5202_RS06785 | 3548..7090 | 2760976 | BREX-system-P-loop-protein-BrxC-[Shewanella-sp.-SG41-4] |
| WP_203431812.1 | Type1.BREX | <-LexA-protease<-SIGMA+OB-Fold+AAA-ATPase+Lon-Protease-BrxL<-PglZ<-N6-METHYLASE<-DUF6079*<-BrxB-BrxF-iSTAND<-BrxA-DNA-binding<-wHTH+WYL+WCX<-Prok-JAB<-wHTH+WYL+WCX<-?<-XIS-HTH | 1224 | Bacteria>Pseudomonadota>Gammaproteobacteria>Alteromonadales>Shewanellaceae>Shewanella | GCF_016859145.1 | Shewanella_sp._LZH-2 | JM642_RS11340 | 2554148..2557822 | 2806008 | BREX-system-P-loop-protein-BrxC-[Shewanella-sp.-LZH-2] |
| WP_208124435.1 | Type1.BREX | XIS-HTH->NtrC-AAA+FIS-HTH->MoxR-AAA->?->HEPN->BrxA-DNA-binding->BrxB-BrxF-iSTAND->DUF6079*->N6-METHYLASE->ParB+HNH->PglZ->ZetaToxin->?-><-SF1-SF2-Helicase+REase<-SF1-SF2-Helicase+REase\|\|REase->HIN-HTH-><-GDResolvase+HIN-HTH | 1157 | Bacteria>Pseudomonadota>Gammaproteobacteria>Alteromonadales>Shewanellaceae>Shewanella | GCF_017570045.1 | Shewanella_algae | JKK45_RS07515 | 1709365..1712838 | 38313 | BREX-system-P-loop-protein-BrxC-[Shewanella-algae] |
| WP_208251007.1 | Type1.BREX | <-DUF2787<-?<-?\|\|XIS-HTH->?\|\|HTH_3->wHTH+WYL+WCX->BrxA-DNA-binding->BrxB-BrxF-iSTAND->DUF6079*->OLD-ABC->TOPRIM->N6-METHYLASE->PglZ->SIGMA+OB-Fold+AAA-ATPase+Lon-Protease-BrxL->?->?->?->ANK+ANK+ANK-> | 1225 | Bacteria>Pseudomonadota>Gammaproteobacteria>Alteromonadales>Shewanellaceae>Shewanella | GCF_017580085.1 | Shewanella_algae | I6M29_RS02105 | 9413..13090 | 38313 | BREX-system-P-loop-protein-BrxC-[Shewanella-algae] |
| WP_243891830.1 | Type1.BREX | <-SIGMA+OB-Fold+AAA-ATPase+Lon-Protease-BrxL<-PglZ<-?<-N6-METHYLASE-p<-RVT<-N6-METHYLASE<-?<-HNH<-DUF6079*<-BrxB-BrxF-iSTAND<-BrxA-DNA-binding<-wHTH+WYL+WCX<-?<-?<-XIS-HTH<-?<-Arm-DNA-bind_3+Phage_integrase | 1226 | Bacteria>Pseudomonadota>Gammaproteobacteria>Alteromonadales>Shewanellaceae>Shewanella | GCF_017580215.1 | Shewanella_algae | I6M36_RS02045 | 117246..120926 | 38313 | BREX-system-P-loop-protein-BrxC-[Shewanella-algae] |
| WP_144225094.1 | Type1.BREX | <-Phage_integrase<-DUF2787<-?<-?\|\|XIS-HTH->XIS-HTH->BrxA-DNA-binding->BrxB-BrxF-iSTAND-p->DUF6079*->OLD-ABC->PIN->N6-METHYLASE->PglZ->SIGMA+OB-Fold+AAA-ATPase+Lon-Protease-BrxL->?-><-Arm-DNA-bind_3+Phage_integrase<-?\|\|RNasePH-> | 1145 | Bacteria>Pseudomonadota>Gammaproteobacteria>Alteromonadales>Shewanellaceae>Shewanella | GCF_017580345.1 | Shewanella_algae | I6M43_RS01845 | 49386..52823 | 38313 | BREX-system-P-loop-protein-BrxC-[Shewanella-algae] |
| WP_101059463.1 | Type1.BREX | HTH_21+rve+rve_3->BrxA-DNA-binding->BrxB-BrxF-iSTAND->DUF6079*->N6-METHYLASE->ParB+HNH->PglZ->ZetaToxin->?-><-SF1-SF2-Helicase+REase<-SF1-SF2-Helicase+REase | 1157 | Bacteria>Pseudomonadota>Gammaproteobacteria>Alteromonadales>Shewanellaceae>Shewanella | GCF_017580385.1 | Shewanella_algae | CX652_RS12220 | 2312..5785 | 38313 | BREX-system-P-loop-protein-BrxC-[Shewanella-algae] |
| WP_144191359.1 | Type1.BREX | XIS-HTH->NtrC-AAA+FIS-HTH->MoxR-AAA->?->HEPN->BrxA-DNA-binding->BrxB-BrxF-iSTAND->DUF6079*->N6-METHYLASE->ParB+HNH->PglZ->ZetaToxin->?-><-SF1-SF2-Helicase+REase<-SF1-SF2-Helicase+REase | 1157 | Bacteria>Pseudomonadota>Gammaproteobacteria>Alteromonadales>Shewanellaceae>Shewanella | GCF_017580415.1 | Shewanella_algae | I6M45_RS07395 | 495996..499469 | 38313 | BREX-system-P-loop-protein-BrxC-[Shewanella-algae] |
| WP_208223521.1 | Type1.BREX | rve->BrxA-DNA-binding->BrxB-BrxF-iSTAND->DUF6079*->N6-METHYLASE->ParB+HNH->PglZ->ZetaToxin->?-><-SF1-SF2-Helicase+REase<-SF1-SF2-Helicase+REase\|\|REase-><-GDResolvase+HIN-HTH | 1157 | Bacteria>Pseudomonadota>Gammaproteobacteria>Alteromonadales>Shewanellaceae>Shewanella | GCF_017580425.1 | Shewanella_algae | I6M46_RS07445 | 1788..5261 | 38313 | BREX-system-P-loop-protein-BrxC-[Shewanella-algae] |
| WP_208162985.1 | Type1.BREX | HTH_3->wHTH+WYL+WCX->BrxA-DNA-binding->BrxB-BrxF-iSTAND->DUF6079*->OLD-ABC->TOPRIM->N6-METHYLASE->PglZ->SIGMA+OB-Fold+AAA-ATPase+Lon-Protease-BrxL-> | 1225 | Bacteria>Pseudomonadota>Gammaproteobacteria>Alteromonadales>Shewanellaceae>Shewanella | GCF_017580535.1 | Shewanella_algae | I6M49_RS22460 | 3510..7187 | 38313 | BREX-system-P-loop-protein-BrxC-[Shewanella-algae] |
| WP_220062217.1 | Type1.BREX | <-wHTH\|\|MerR-HTH->?->SF1-SF2-Helicase+REase->SF1-SF2-Helicase+REase-><-HEPN+Sel1+SEC-C<-PglZ<-ParB+HNH<-N6-METHYLASE<-DUF6079*<-BrxB-BrxF-iSTAND<-BrxA-DNA-binding\|\|sGTP->sGTP->?->sGTP->HEPN-><-XIS-HTH | 1157 | Bacteria>Pseudomonadota>Gammaproteobacteria>Alteromonadales>Shewanellaceae>Shewanella | GCF_019457515.1 | Shewanella_sp._FJAT-53870 | KO173_RS17055 | 3880275..3883748 | 2864208 | BREX-system-P-loop-protein-BrxC-[Shewanella-sp.-FJAT-53870] |
| WP_220768073.1 | Type1.BREX | MobI-><-?<-LexA-protease\|\|35exo->?->wHTH+WYL+WCX->BrxA-DNA-binding->BrxB-BrxF-iSTAND->DUF6079*->ParB+HNH->N6-METHYLASE->PglZ->SIGMA+OB-Fold+AAA-ATPase+Lon-Protease-BrxL->NACHT+HEAT+REase->HDRnase->VirD4-TraG+VirB4-FtsK-> | 1225 | Bacteria>Pseudomonadota>Gammaproteobacteria>Alteromonadales>Shewanellaceae>Shewanella | GCF_019654915.1 | Shewanella_sp._MBTL60-007 | K5Q92_RS05655 | 54742..58419 | 2815911 | BREX-system-P-loop-protein-BrxC-[Shewanella-sp.-MBTL60-007] |
| WP_221058246.1 | Type1.BREX | <-ANK+ANK+ANK<-?<-?<-?<-SIGMA+OB-Fold+AAA-ATPase+Lon-Protease-BrxL<-PglZ<-N6-METHYLASE<-ParB+HNH<-DUF6079*<-BrxB-BrxF-iSTAND<-BrxA-DNA-binding<-wHTH+WYL+WCX<-HTH_3\|\|?-><-XIS-HTH\|\|?->?->DUF2787-> | 1199 | Bacteria>Pseudomonadota>Gammaproteobacteria>Alteromonadales>Shewanellaceae>Shewanella | GCF_019670685.1 | Shewanella_algae | K5631_RS12960 | 2926333..2929932 | 38313 | BREX-system-P-loop-protein-BrxC-[Shewanella-algae] |
| WP_224020606.1 | Type1.BREX | <-HerA/FtsK\|\|SF1-SF2-Helicase+REase->SF1-SF2-Helicase+REase-><-?<-ZetaToxin<-PglZ<-ParB+HNH<-N6-METHYLASE<-DUF6079*<-BrxB-BrxF-iSTAND<-BrxA-DNA-binding<-?<-HEPN<-?<-MoxR-AAA<-NtrC-AAA+FIS-HTH<-XIS-HTH | 1157 | Bacteria>Pseudomonadota>Gammaproteobacteria>Alteromonadales>Shewanellaceae>Shewanella | GCF_019973655.1 | Shewanella_xiamenensis | LDO18_RS12250 | 2746465..2749938 | 332186 | BREX-system-P-loop-protein-BrxC-[Shewanella-xiamenensis] |
| WP_248084385.1 | Type1.BREX | SF1-SF2-Helicase-><-DUF2787<-CbeA_antitoxin<-?<-?\|\|XIS-HTH->wHTH+WYL+WCX->BrxA-DNA-binding->BrxB-BrxF-iSTAND->DUF6079*->HNH->?->ParB+HNH->N6-METHYLASE->HEPN->N6-METHYLASE-p-> | 1231 | Bacteria>Pseudomonadota>Gammaproteobacteria>Alteromonadales>Shewanellaceae>Shewanella | GCF_023159095.1 | unclassified_Shewanella | MOD77_RS02670 | 216752..220447 | 196818 | MULTISPECIES:-BREX-system-P-loop-protein-BrxC-[unclassified-Shewanella] |
| WP_249554920.1 | Type1.BREX | TetR-HTH->ABhydrolase->?-><-PRTase<-RNasePH\|\|?->Arm-DNA-bind_3+Phage_integrase->BrxA-DNA-binding->BrxB-BrxF-iSTAND->DUF6079*->N6-METHYLASE->PglZ->SIGMA+OB-Fold+AAA-ATPase+HKD-DNase-BrxL->MORC+DUF3684+REase-><-XIS-HTH\|\|?->?->DUF2787->Phage_integrase-> | 1188 | Bacteria>Pseudomonadota>Gammaproteobacteria>Alteromonadales>Shewanellaceae>Shewanella | GCF_023475945.1 | Shewanella_sp._8A | M0413_RS16385 | 25717..29283 | 2943323 | BREX-system-P-loop-protein-BrxC-[Shewanella-sp.-8A] |
| WP_249553736.1 | Type1.BREX | <-SIGMA+OB-Fold+AAA-ATPase+Lon-Protease-BrxL<-PglZ<-?<-N6-METHYLASE<-HEPN<-RVT<-N6-METHYLASE-p<-N6-METHYLASE<-DUF6079*<-BrxB-BrxF-iSTAND<-BrxA-DNA-binding<-wHTH+WYL+WCX\|\|?->?->CbeA_antitoxin->DUF2787->DUF2787->Phage_integrase-> | 1268 | Bacteria>Pseudomonadota>Gammaproteobacteria>Alteromonadales>Shewanellaceae>Shewanella | GCF_023475945.1 | Shewanella_sp._8A | M2897_RS12020 | 2756867..2760673 | 2943323 | BREX-system-P-loop-protein-BrxC-[Shewanella-sp.-8A] |
| WP_259476695.1 | Type1.BREX | XIS-HTH->?->wHTH+WYL+WCX->Prok-JAB->wHTH+WYL+WCX->BrxA-DNA-binding->BrxB-BrxF-iSTAND->DUF6079*->ParB+HNH->N6-METHYLASE->PglZ->SIGMA+OB-Fold+AAA-ATPase+Lon-Protease-BrxL->?->HerA/FtsK-> | 1225 | Bacteria>Pseudomonadota>Gammaproteobacteria>Alteromonadales>Shewanellaceae>Shewanella | GCF_024972795.1 | Shewanella_xiamenensis | K5M76_RS09815 | 2212502..2216179 | 332186 | BREX-system-P-loop-protein-BrxC-[Shewanella-xiamenensis] |
| WP_259573703.1 | Type1.BREX | <-CHTH<-REase<-SIGMA+OB-Fold+AAA-ATPase+Lon-Protease-BrxL<-PglZ<-N6-METHYLASE-p<-HEPN<-RVT<-N6-METHYLASE-p<-N6-METHYLASE<-DUF6079*<-BrxB-BrxF-iSTAND<-BrxA-DNA-binding<-wHTH+WYL+WCX<-HTH_26\|\|Phage_AlpA->Arm-DNA-bind_2+Phage_int_SAM_3+Phage_integrase->HDRnase->?-><-SF1-SF2-Helicase | 1240 | Bacteria>Pseudomonadota>Gammaproteobacteria>Alteromonadales>Shewanellaceae>Shewanella | GCF_024999165.1 | Shewanella_baltica | G7A68_RS06675 | 25637..29359 | 62322 | BREX-system-P-loop-protein-BrxC-[Shewanella-baltica] |
| WP_259506670.1 | Type1.BREX | <-Herpes_LMP1+REase<-?<-?\|\|SF1-SF2-Helicase+REase->SF1-SF2-Helicase+REase-><-HEPN+Sel1+SEC-C<-PglZ<-ParB+HNH<-N6-METHYLASE<-DUF6079*<-BrxB-BrxF-iSTAND<-BrxA-DNA-binding\|\|?->?->CbeA_antitoxin->DUF2787->Phage_integrase->?->RHH_1-> | 1157 | Bacteria>Pseudomonadota>Gammaproteobacteria>Alteromonadales>Shewanellaceae>Shewanella | GCF_024999425.1 | Shewanella_baltica | G3479_RS14595 | 93624..97097 | 62322 | BREX-system-P-loop-protein-BrxC-[Shewanella-baltica] |
| WP_261753251.1 | Type1.BREX | Arm-DNA-bind_3+Phage_integrase->SF1-SF2-Helicase+REase->SF1-SF2-Helicase+REase->?-><-?<-ZetaToxin<-PglZ<-ParB+HNH<-N6-METHYLASE<-DUF6079*<-BrxB-BrxF-iSTAND<-BrxA-DNA-binding | 1157 | Bacteria>Pseudomonadota>Gammaproteobacteria>Alteromonadales>Shewanellaceae>Shewanella | GCF_025398225.1 | Shewanella_algae | N4366_RS19550 | 70548..74021 | 38313 | BREX-system-P-loop-protein-BrxC-[Shewanella-algae] |
| WP_261732933.1 | Type1.BREX | Peptidase_M2-><-?\|\|<-HDRnase<-Arm-DNA-bind_2+Phage_int_SAM_3+Phage_integrase<-Phage_AlpA\|\|HTH_26->BrxA-DNA-binding->BrxB-BrxF-iSTAND->DUF6079*->N6-METHYLASE->N6-METHYLASE-p->OLD-ABC->?->TOPRIM->PglZ-p->PglZ->HEPN+Sel1+SEC-C-> | 1157 | Bacteria>Pseudomonadota>Gammaproteobacteria>Alteromonadales>Shewanellaceae>Shewanella | GCF_025398315.1 | Shewanella_sp._KJ10-1 | N4T56_RS08695 | 1854721..1858194 | 2978345 | BREX-system-P-loop-protein-BrxC-[Shewanella-sp.-KJ10-1] |
| WP_263145911.1 | Type1.BREX | <-N6-METHYLASE<-HerA/FtsK<-SIR2<-N6-METHYLASE<-ParB+HNH<-DUF6079*<-BrxB-BrxF-iSTAND<-BrxA-DNA-binding<-wHTH+WYL+WCX<-XIS-HTH\|\|?-><-?<-CbeA_antitoxin<-DUF2787->DUF2787-> | 1224 | Bacteria>Pseudomonadota>Gammaproteobacteria>Alteromonadales>Shewanellaceae>Shewanella | GCF_025653395.1 | Shewanella_sp._SW24 | L5M17_RS05810 | 187685..191359 | 2912805 | BREX-system-P-loop-protein-BrxC-[Shewanella-sp.-SW24] |
| WP_263212792.1 | Type1.BREX | <-McrC-NTD+REase<-Preader17<-SIGMA+OB-Fold+AAA-ATPase+Lon-Protease-BrxL<-PglZ<-N6-METHYLASE<-DUF6079*<-BrxB-BrxF-iSTAND<-BrxA-DNA-binding<-wHTH+WYL+WCX<-Peptidase_C47<-wHTH+WYL+WCX<-?<-XIS-HTH | 1224 | Bacteria>Pseudomonadota>Gammaproteobacteria>Alteromonadales>Shewanellaceae>Shewanella | GCF_025653775.1 | Shewanella_sp._SM72 | L5M36_RS22725 | 11851..15525 | 2912805 | BREX-system-P-loop-protein-BrxC-[Shewanella-sp.-SM72] |
| WP_263171306.1 | Type1.BREX | <-Phage_integrase<-DUF2787<-DUF2787<-CbeA_antitoxin<-?<-?\|\|wHTH+WYL+WCX->BrxA-DNA-binding->BrxB-BrxF-iSTAND->DUF6079*->HNH->?->N6-METHYLASE->SIR2->HerA/FtsK->N6-METHYLASE->PglZ->SIGMA+OB-Fold+AAA-ATPase+Lon-Protease-BrxL-> | 1235 | Bacteria>Pseudomonadota>Gammaproteobacteria>Alteromonadales>Shewanellaceae>Shewanella | GCF_025653795.1 | Shewanella_sp._SM73 | L5M51_RS01900 | 416485..420192 | 2912806 | BREX-system-P-loop-protein-BrxC-[Shewanella-sp.-SM73] |
| WP_263177453.1 | Type1.BREX | <-SIGMA+OB-Fold+AAA-ATPase+Lon-Protease-BrxL<-PglZ<-N6-METHYLASE<-ParB+HNH<-DUF6079*<-BrxB-BrxF-iSTAND<-BrxA-DNA-binding<-wHTH+WYL+WCX<-?<-wHTH+WYL+WCX<-?<-XIS-HTH\|\|<-?\|\|TraI-VirD2-RCRE-> | 1225 | Bacteria>Pseudomonadota>Gammaproteobacteria>Alteromonadales>Shewanellaceae>Shewanella | GCF_025654415.1 | unclassified_Shewanella | L5M16_RS21425 | 11352..15029 | 196818 | MULTISPECIES:-BREX-system-P-loop-protein-BrxC-[unclassified-Shewanella] |
| WP_263197229.1 | Type1.BREX | <-SIGMA+OB-Fold+AAA-ATPase+Lon-Protease-BrxL<-PglZ<-N6-METHYLASE-p<-HEPN<-N6-METHYLASE-p<-N6-METHYLASE<-?<-HNH<-DUF6079*<-BrxB-BrxF-iSTAND<-BrxA-DNA-binding<-wHTH+WYL+WCX<-HTH_26\|\|Phage_AlpA->Arm-DNA-bind_2+Phage_int_SAM_3+Phage_integrase->HDRnase-> | 1246 | Bacteria>Pseudomonadota>Gammaproteobacteria>Alteromonadales>Shewanellaceae>Shewanella | GCF_025654545.1 | Shewanella_sp._SM20 | L5M18_RS13095 | 42119..45859 | 2912792 | BREX-system-P-loop-protein-BrxC-[Shewanella-sp.-SM20] |
| WP_264921054.1 | Type1.BREX | XIS-HTH->NtrC-AAA+FIS-HTH->MoxR-AAA->?->HEPN->HIN-HTH->BrxA-DNA-binding->BrxB-BrxF-iSTAND->DUF6079*->N6-METHYLASE->ParB+HNH->PglZ->ZetaToxin->?-><-SF1-SF2-Helicase+REase<-SF1-SF2-Helicase+REase<-ZnR+HTH_Tnp_ISL3+DDE_Tnp_ISL3\|\|N6-N4-METHYLASE-> | 1157 | Bacteria>Pseudomonadota>Gammaproteobacteria>Alteromonadales>Shewanellaceae>Shewanella | GCF_026013945.1 | Shewanella_algae | OLL83_RS07695 | 1703466..1706939 | 38313 | BREX-system-P-loop-protein-BrxC-[Shewanella-algae] |
| WP_275482470.1 | Type1.BREX | MobI-><-?<-LexA-protease\|\|35exo->?->wHTH+WYL+WCX->BrxA-DNA-binding->BrxB-BrxF-iSTAND->DUF6079*->ParB+HNH->N6-METHYLASE->PglZ->SIGMA+OB-Fold+AAA-ATPase+Lon-Protease-BrxL->CARF+NtrC-AAA+FIS-HTH->OBFOLD-NUCLEASE-> | 1225 | Bacteria>Pseudomonadota>Gammaproteobacteria>Alteromonadales>Shewanellaceae>Shewanella | GCF_029167175.1 | Shewanella_sp._A32 | PY479_RS00080 | 16338..20015 | 3031327 | BREX-system-P-loop-protein-BrxC-[Shewanella-sp.-A32] |
| WP_076500079.1 | Type1.BREX | SF1-SF2-Helicase+REase->SF1-SF2-Helicase+REase-><-REase+SF1-SF2-Helicase+REase<-REase<-PglZ<-TOPRIM<-OLD-ABC<-N6-METHYLASE<-DUF6079*<-BrxB-BrxF-iSTAND<-BrxA-DNA-binding\|\|?->?->CbeA_antitoxin->DUF2787->DUF2787->Phage_integrase-> | 1157 | Bacteria>Pseudomonadota>Gammaproteobacteria>Alteromonadales>Shewanellaceae>Shewanella | GCF_900156405.1 | Shewanella_morhuae | BW947_RS12490 | 20495..23968 | 365591 | BREX-system-P-loop-protein-BrxC-[Shewanella-morhuae] |
| WP_112352819.1 | Type1.BREX | MerR-HTH->?-><-?<-SIGMA+OB-Fold+AAA-ATPase+HKD-DNase-BrxL<-PglZ<-N6-METHYLASE<-REase<-OLD-ABC<-DUF6079*<-BrxB-BrxF-iSTAND<-BrxA-DNA-binding<-XIS-HTH<-HNH+NERD+ZnR<-SAM-METHYLASE+wHTH+HNH<-?\|\|URI-DNase-><-MerR-HTH | 1197 | Bacteria>Pseudomonadota>Gammaproteobacteria>Alteromonadales>Shewanellaceae>Shewanella | GCF_900476435.1 | Shewanella_benthica | SHEWBE_RS12055 | 2634220..2637813 | 43661 | BREX-system-P-loop-protein-BrxC-[Shewanella-benthica] |
| WP_126513302.1 | Type1.BREX | Cas_NE0113-><-?<-HDRnase<-Arm-DNA-bind_2+Phage_int_SAM_3+Phage_integrase<-Phage_AlpA\|\|HTH_26->wHTH+WYL+WCX->BrxA-DNA-binding->BrxB-BrxF-iSTAND->DUF6079*->N6-METHYLASE->N6-METHYLASE-p->RVT->HEPN->N6-METHYLASE-p->PglZ->SIGMA+OB-Fold+AAA-ATPase+Lon-Protease-BrxL->REase->CHTH-> | 1254 | Bacteria>Pseudomonadota>Gammaproteobacteria>Alteromonadales>Shewanellaceae>Shewanella | - | Shewanella_putrefaciens | EL161_RS20440 | 4646613..4650377 | 24 | BREX-system-P-loop-protein-BrxC-[Shewanella-putrefaciens] |
| WP_102039083.1 | Type1.BREX | <-SIGMA+OB-Fold+AAA-ATPase+HKD-DNase-BrxL<-PglZ<-ParB+HNH<-N6-METHYLASE<-REase<-OLD-ABC<-DUF6079*<-BrxB-BrxF-iSTAND<-BrxA-DNA-binding<-XIS-HTH<-?<-TnsC-ATPase<-rve+Mu-transpos_C<-REase+wHTH | 1191 | Bacteria>Pseudomonadota>Gammaproteobacteria>Alteromonadales>Shewanellaceae>Shewanella | - | Shewanella_sp._GutCb | CXF86_11690 | 122253..125828 | 2058315 | BREX-system-P-loop-protein-BrxC-[Shewanella-sp.-GutCb] |
| WP_182771402.1 | Type1.BREX | <-CHTH\|\|XIS-HTH->BrxA-DNA-binding->BrxB-BrxF-iSTAND->DUF6079*->N6-METHYLASE->PglZ->SIGMA+OB-Fold+AAA-ATPase+HKD-DNase-BrxL-><-?<-MerR-HTH<-?<-HTH_26\|\|Phage_AlpA->Arm-DNA-bind_2+Phage_int_SAM_3+Phage_integrase-> | 1182 | Bacteria>Pseudomonadota>Gammaproteobacteria>Alteromonadales>Shewanellaceae>Shewanella | - | Shewanella_sp._SG41-4 | H5202_06785 | 3542..7090 | 2760976 | BREX-system-P-loop-protein-BrxC-[Shewanella-sp.-SG41-4] |
| MBV7434872.1 | Type1.BREX | <-SIGMA+OB-Fold+AAA-ATPase+HKD-DNase-BrxL<-PglZ<-ParB+HNH<-N6-METHYLASE<-DUF6079*<-BrxB-BrxF-iSTAND<-BrxA-DNA-binding<-?<-?<-?<-?<-?<-Peptidase_M20 | 1175 | Bacteria>Pseudomonadota>Gammaproteobacteria>Cardiobacteriales>Cardiobacteriaceae>unclassified_Cardiobacteriaceae | GCA_019218315.1 | Cardiobacteriaceae_bacterium_TAE3-ERU3 | KRX19_07510 | 264796..268323 | 2856340 | BREX-system-P-loop-protein-BrxC-[Cardiobacteriaceae-bacterium-TAE3-ERU3] |
| WP_096087095.1 | Type1.BREX | <-SbcC<-SIGMA+OB-Fold+AAA-ATPase+Lon-Protease-BrxL<-PglZ<-N6-METHYLASE<-TOPRIM<-OLD-ABC<-DUF6079*<-BrxB-BrxF-iSTAND<-BrxA-DNA-binding<-wHTH+WYL+WCX<-?<-35exo\|\|LexA-protease->?-><-MobI | 1225 | Bacteria>Pseudomonadota>Gammaproteobacteria>Cellvibrionales>Cellvibrionaceae>Agaribacterium | GCF_002312815.1 | Agaribacterium_haliotis | CFK45_RS14065 | 11991..15668 | 2013869 | BREX-system-P-loop-protein-BrxC-[Agaribacterium-haliotis] |
| WP_007642692.1 | Type1.BREX | <-SIGMA+OB-Fold+AAA-ATPase+Lon-Protease-BrxL<-PglZ<-TOPRIM<-OLD-ABC<-N6-METHYLASE-p<-RVT<-N6-METHYLASE<-HATPase+wHTH<-DUF6079*<-BrxB-BrxF-iSTAND<-BrxA-DNA-binding<-wHTH+WYL+WCX<-REase\|\|?-><-?\|\|?-><-MPTase | 1220 | Bacteria>Pseudomonadota>Gammaproteobacteria>Cellvibrionales>Cellvibrionaceae>Cellvibrio | GCF_000263355.1 | Cellvibrio_sp._BR | O59_RS11305 | 237548..241210 | 1134474 | BREX-system-P-loop-protein-BrxC-[Cellvibrio-sp.-BR] |
| WP_054114051.1 | Type1.BREX | DDE_Tnp_IS66+DDE_Tnp_IS66_C-><-SIGMA+OB-Fold+AAA-ATPase+Lon-Protease-BrxL<-PglZ<-N6-METHYLASE<-TOPRIM<-OLD-ABC<-DUF6079*<-BrxB-BrxF-iSTAND<-BrxA-DNA-binding<-wHTH+WYL+WCX<-?\|\|DDE_Tnp_IS66+DDE_Tnp_IS66_C-> | 1229 | Bacteria>Pseudomonadota>Gammaproteobacteria>Cellvibrionales>Cellvibrionaceae>Marinagarivorans | GCF_001296225.1 | Marinagarivorans_algicola | AC526_RS14090 | 30886..34575 | 1513270 | BREX-system-P-loop-protein-BrxC-[Marinagarivorans-algicola] |
| WP_265366287.1 | Type1.BREX | Arm-DNA-bind_3+Phage_integrase-><-XIS-HTH<-AAA+HTH<-XIS-HTH\|\|MobI->XIS-HTH->BrxA-DNA-binding->BrxB-BrxF-iSTAND->DUF6079*->Schlafen+HATPase+wHTH+HTH->OLD-ABC->REase->N6-METHYLASE->ParB+HNH->PglZ->SIGMA+OB-Fold+AAA-ATPase+HKD-DNase-BrxL->SbcC+APATPase+SbcC-> | 1193 | Bacteria>Pseudomonadota>Gammaproteobacteria>Cellvibrionales>Cellvibrionaceae>Marinimicrobium | GCF_026153455.1 | Marinimicrobium_koreense | OOT55_RS13025 | 3024461..3028042 | 306545 | BREX-system-P-loop-protein-BrxC-[Marinimicrobium-sp.-C6131] |
| NHN36564.1 | Type1.BREX | <-SIGMA+OB-Fold+AAA-ATPase+Lon-Protease-BrxL<-PglZ<-Polbeta<-MarR-HTH<-N6-METHYLASE<-?<-DUF6079*<-BrxB-BrxF-iSTAND<-BrxA-DNA-binding<-tRNA\|\|?-><-?-><-MerR-HTH | 1151 | Bacteria>Pseudomonadota>Gammaproteobacteria>Cellvibrionales>Cellvibrionaceae>Pseudomaricurvus | GCA_011440395.1 | Pseudomaricurvus_alcaniphilus | G8764_04575 | 1080165..1083620 | 1166482 | BREX-system-P-loop-protein-BrxC-[Pseudomaricurvus-alcaniphilus] |
| WP_011469234.1 | Type1.BREX | HNH-><-TraJ-RHH<-RelE-ParE<-PhdYeFM_antitox<-SIGMA+OB-Fold+AAA-ATPase+Lon-Protease-BrxL<-PglZ<-TOPRIM<-OLD-ABC<-N6-METHYLASE<-DUF6079*<-BrxB-BrxF-iSTAND<-BrxA-DNA-binding<-XIS-HTH\|\|?->GDResolvase+ZnR->?-><-ACET | 1145 | Bacteria>Pseudomonadota>Gammaproteobacteria>Cellvibrionales>Cellvibrionaceae>Saccharophagus | GCF_000013665.1 | Saccharophagus_degradans | SDE_RS14425 | 3492242..3495679 | 86304 | BREX-system-P-loop-protein-BrxC-[Saccharophagus-degradans] |
| WP_187410606.1 | Type1.BREX | Phage_AlpA->DnaB->?->TraJ-RHH->?->PIN->\|\|wHTH+WYL+WCX->BrxA-DNA-binding->BrxB-BrxF-iSTAND->DUF6079*->N6-METHYLASE->PHP+SbcC->ParB+HNH->PglZ->SIGMA+OB-Fold+AAA-ATPase+Lon-Protease-BrxL->?->ABC-ATPase+SbcC-> | 1146 | Bacteria>Pseudomonadota>Gammaproteobacteria>Cellvibrionales>Cellvibrionaceae>Saccharophagus | GCF_014336955.1 | Saccharophagus_sp._K07 | DWB84_RS14900 | 71821..75261 | 2283636 | BREX-system-P-loop-protein-BrxC-[Saccharophagus-sp.-K07] |
| WP_009576096.1 | Type1.BREX | Calcineurin-><-?<-?<-?<-HD<-SLATT\|\|nSTAND1+HNH->BrxA-DNA-binding->BrxB-BrxF-iSTAND->DUF6079*->N6-METHYLASE->PglZ->SIGMA+OB-Fold+AAA-ATPase+Lon-Protease-BrxL->URI-DNase+RAMA->?->?-><-?<-?<-RADICAL-SAM | 1178 | Bacteria>Pseudomonadota>Gammaproteobacteria>Cellvibrionales>Halieaceae>Aequoribacter | GCF_000204315.1 | Aequoribacter_fuscus | EYZ66_RS04240 | 942285..945821 | 2518989 | BREX-system-P-loop-protein-BrxC-[Aequoribacter-fuscus] |
| WP_084199902.1 | Type1.BREX | CHTH->Calcineurin-><-?<-TnsC-ATPase<-?<-?\|\|XIS-HTH->BrxA-DNA-binding->BrxB-BrxF-iSTAND->DUF6079*->OLD-ABC->REase->N6-METHYLASE->PglZ->SIGMA+OB-Fold+AAA-ATPase+HKD-DNase-BrxL-> | 1184 | Bacteria>Pseudomonadota>Gammaproteobacteria>Cellvibrionales>Halieaceae>Halioglobus | GCF_014652255.1 | Halioglobus_japonicus | COO29_RS05530 | 1195751..1199305 | 930805 | BREX-system-P-loop-protein-BrxC-[Halioglobus-japonicus] |
| WP_101520955.1 | Type1.BREX | HEPN->HNH+Polbeta->?->?->Peptidase_M29+DUF2971->XIS-HTH->BrxA-DNA-binding->BrxB-BrxF-iSTAND->DUF6079*->N6-METHYLASE->MarR-HTH->Polbeta->PglZ->SIGMA+OB-Fold+AAA-ATPase+Lon-Protease-BrxL->HEPN+SF1-SF2-Helicase+RNase+RNase->TPR-><-?<-Arm-DNA-bind_3+Phage_integrase | 1151 | Bacteria>Pseudomonadota>Gammaproteobacteria>Cellvibrionales>Halieaceae>Kineobactrum | GCF_002868715.1 | Kineobactrum_sediminis | CWI75_RS08140 | 627029..630484 | 1905677 | BREX-system-P-loop-protein-BrxC-[Kineobactrum-sediminis] |
| WP_148062597.1 | Type1.BREX | XIS-HTH->TOPRIM->AAA+wHTH->?->?->XIS-HTH->BrxA-DNA-binding->BrxB-BrxF-iSTAND->DUF6079*->SF1-SF2-Helicase+PRTase->SLOG+BirA-HTH->N6-METHYLASE->RVT->PglZ->Immank+ANK+ANK->REase->SIGMA+OB-Fold+AAA-ATPase+HKD-DNase-BrxL-> | 1180 | Bacteria>Pseudomonadota>Gammaproteobacteria>Cellvibrionales>Halieaceae>Parahaliea | GCF_008064635.1 | Parahaliea_aestuarii | FVW59_RS02260 | 491803..495345 | 1852021 | BREX-system-P-loop-protein-BrxC-[Parahaliea-aestuarii] |
| WP_043319021.1 | Type1.BREX | <-TetR-HTH<-?\|\|?-><-?<-SIGMA+OB-Fold+AAA-ATPase+Lon-Protease-BrxL<-PglZ<-ParB+HNH<-N6-METHYLASE<-Schlafen+HATPase+wHTH<-DUF6079*<-BrxB-BrxF-iSTAND<-BrxA-DNA-binding<-tRNA<-?<-?<-?<-?<-Classical-AAA<-SAM-METHYLASE | 1145 | Bacteria>Pseudomonadota>Gammaproteobacteria>Cellvibrionales>Microbulbiferaceae>Microbulbifer | GCF_000708675.1 | Microbulbifer_sp._HZ11 | AU11_RS09640 | 451911..455348 | 1453501 | BREX-system-P-loop-protein-BrxC-[Microbulbifer-sp.-HZ11] |
| WP_265965539.1 | Type1.BREX | ABhydrolase->MORC+DNA_gyrase+TOPRIM->HTH_3->?->BrxA-DNA-binding->BrxB-BrxF-iSTAND->DUF6079*->N6-METHYLASE->SIR2->HerA/FtsK->N6-METHYLASE-p->?->PglZ->SIGMA+OB-Fold+AAA-ATPase+Lon-Protease-BrxL-><-?<-ACET+ACET | 1223 | Bacteria>Pseudomonadota>Gammaproteobacteria>Cellvibrionales>Microbulbiferaceae>Microbulbifer | GCF_026241645.1 | Microbulbifer_thermotolerans | OQJ62_RS10425 | 90626..94297 | 252514 | BREX-system-P-loop-protein-BrxC-[Microbulbifer-thermotolerans] |
| WP_036859475.1 | Type1.BREX | Classical-AAA->?->?->?->?->tRNA->Arm-DNA-bind_3+Phage_integrase->XIS-HTH->BrxA-DNA-binding->BrxB-BrxF-iSTAND->DUF6079*->SF1-SF2-Helicase+PRTase->SLOG+BirA-HTH->N6-METHYLASE->ParB+HNH->PglZ->SIGMA+OB-Fold+AAA-ATPase+HKD-DNase-BrxL->?->XIS-HTH-> | 1187 | Bacteria>Pseudomonadota>Gammaproteobacteria>Cellvibrionales>Porticoccaceae>Porticoccus | GCF_000744735.1 | Porticoccus_hydrocarbonoclasticus | U740_RS05135 | 1073957..1077520 | 1073414 | BREX-system-P-loop-protein-BrxC-[Porticoccus-hydrocarbonoclasticus] |
| MAT50826.1 | Type1.BREX | <-N6-METHYLASE<-SLOG+BirA-HTH<-SF1-SF2-Helicase+PRTase<-Schlafen+HATPase+wHTH+HTH<-DUF6079*<-BrxB-BrxF-iSTAND<-BrxA-DNA-binding<-XIS-HTH\|\|?->?-><-GDResolvase | 1182 | Bacteria>Pseudomonadota>Gammaproteobacteria>Cellvibrionales>Porticoccaceae>unclassified_Porticoccaceae | GCA_002702725.1 | Porticoccaceae_bacterium | CMK32_06560 | 7929..11477 | 2026782 | hypothetical-protein-[Porticoccaceae-bacterium] |
| WP_019531408.1 | Type1.BREX | <-DUF6079*<-BrxB-BrxF-iSTAND<-BrxA-DNA-binding<-SF1-SF2-Helicase+REase<-Exonuc_V_gamma+REase<-MULE-TRANSPOSASE<-?<-TraJ-RHH\|\|?->Peptidase_M50+PDZ_2+Lon_C-> | 1156 | Bacteria>Pseudomonadota>Gammaproteobacteria>Cellvibrionales>Spongiibacteraceae>Dasania | GCF_000373485.1 | Dasania_marina | B067_RS0117580 | 1458..4928 | 471499 | BREX-system-P-loop-protein-BrxC-[Dasania-marina] |
| WP_268905143.1 | Type1.BREX | <-PHP+SbcC<-REase+HTH+35exo+Small-B-barrel+SF1-SF2-Helicase<-REase+iREase<-PIWI<-REase+PNTD1+PNTD2<-SIGMA+OB-Fold+AAA-ATPase+Lon-Protease-BrxL<-PglZ<-N6-METHYLASE<-DUF6079*<-BrxB-BrxF-iSTAND<-BrxA-DNA-binding<-wHTH+WYL+WCX\|\|Thioredoxin_4-> | 1219 | Bacteria>Pseudomonadota>Gammaproteobacteria>Cellvibrionales>Spongiibacteraceae>Dasania | GCF_026891035.1 | Dasania_sp._GY-19 | OOV09_RS14670 | 22695..26354 | 2965066 | BREX-system-P-loop-protein-BrxC-[Dasania-sp.-GY-19] |
| WP_268905395.1 | Type1.BREX | Phage_integrase-><-?<-SMODS<-SLATT<-SIGMA+OB-Fold+AAA-ATPase+Lon-Protease-BrxL<-PglZ<-ParB+HNH<-N6-METHYLASE<-Schlafen+HATPase+wHTH<-DUF6079*<-BrxB-BrxF-iSTAND<-BrxA-DNA-binding<-wHTH+WYL+WCX\|\|GGDEF+EAL-><-HTH_3 | 1220 | Bacteria>Pseudomonadota>Gammaproteobacteria>Cellvibrionales>Spongiibacteraceae>Dasania | GCF_026891035.1 | Dasania_sp._GY-19 | OOV09_RS18305 | 21121..24783 | 2965066 | BREX-system-P-loop-protein-BrxC-[Dasania-sp.-GY-19] |
| WP_269621356.1 | Type1.BREX | wHTH+WYL+WCX->BrxA-DNA-binding->BrxB-BrxF-iSTAND->DUF6079*->MarR-HTH->AbiEii->?->N6-METHYLASE->DUF4158+DDE_Tnp_Tn3-> | 1219 | Bacteria>Pseudomonadota>Gammaproteobacteria>Cellvibrionales>Spongiibacteraceae>Marortus | GCF_027257955.1 | Marortus_sp._BJYM1 | O9X78_RS19220 | 4639..8298 | 2975599 | BREX-system-P-loop-protein-BrxC-[Marortus-sp.-BJYM1] |
| MBU73272.1 | Type1.BREX | <-N6-METHYLASE-p<-DUF6079*<-BrxB-BrxF-iSTAND<-BrxA-DNA-binding<-?<-?<-TraI-VirD2-RCRE<-TraJ-RHH | 1221 | Bacteria>Pseudomonadota>Gammaproteobacteria>Cellvibrionales>Spongiibacteraceae>Spongiibacter | GCA_002730815.1 | Spongiibacter_sp. | CMN81_14305 | 814..4479 | 2024860 | ATPase-[Spongiibacter-sp.] |
| WP_190766547.1 | Type1.BREX | ARG-><-SIGMA+OB-Fold+AAA-ATPase+Lon-Protease-BrxL<-DOC+HTH<-?<-ZNR+RmuC<-?<-?<-N6-METHYLASE<-ParB+HNH<-DUF6079*<-BrxB-BrxF-iSTAND<-BrxA-DNA-binding\|\|wHTH+WYL+WCX-><-?<-SF1-SF2-Helicase | 1192 | Bacteria>Pseudomonadota>Gammaproteobacteria>Cellvibrionales>Spongiibacteraceae>Spongiibacter | GCF_014705705.1 | Spongiibacter_pelagi | IB286_RS13890 | 27960..31538 | 2760804 | BREX-system-P-loop-protein-BrxC-[Spongiibacter-pelagi] |
| WP_273033648.1 | Type1.BREX | <-SIGMA+OB-Fold+AAA-ATPase+HKD-DNase-BrxL<-PglZ<-ParB+HNH<-N6-METHYLASE-p<-HerA/FtsK<-?<-CR-ATPase3<-N6-METHYLASE<-Schlafen+HATPase+wHTH+HTH<-DUF6079*<-BrxB-BrxF-iSTAND<-BrxA-DNA-binding<-XIS-HTH<-MobI\|\|XIS-HTH->AAA+HTH->XIS-HTH-><-Arm-DNA-bind_3+Phage_integrase<-SIGMA-HTH | 1193 | Bacteria>Pseudomonadota>Gammaproteobacteria>Cellvibrionales>Spongiibacteraceae>Spongiibacter | GCF_950061765.1 | Spongiibacter_tropicus | B4C78_RS03795 | 102796..106377 | 454602 | BREX-system-P-loop-protein-BrxC-[Spongiibacter-tropicus] |
| EIF42662.1 | Type1.BREX | wHTH+WYL+WCX->BrxA-DNA-binding->BrxB-BrxF-iSTAND->DUF6079*->MarR-HTH->AbiEii->?->N6-METHYLASE->PglZ->SIGMA+OB-Fold+AAA-ATPase+Lon-Protease-BrxL->SbcC->REase+iREase->HEPN-> | 1220 | Bacteria>Pseudomonadota>Gammaproteobacteria>Cellvibrionales>Spongiibacteraceae>unclassified_Spongiibacteraceae | GCA_000259575.1 | gamma_proteobacterium_BDW918 | DOK_12571 | 2372..6034 | 1168065 | putative-ATPase-like-protein-[gamma-proteobacterium-BDW918] |
| MBN49711.1 | Type1.BREX | <-SIGMA+OB-Fold+AAA-ATPase+Lon-Protease-BrxL<-PglZ<-DOC+HTH<-?<-ZNR+RmuC<-HEPN<-?<-N6-METHYLASE<-?<-DUF6079*<-BrxB-BrxF-iSTAND<-BrxA-DNA-binding\|\|wHTH+WYL+WCX->?-><-?<-SF1-SF2-Helicase\|\|?->?->Phage_integrase-> | 1193 | Bacteria>Pseudomonadota>Gammaproteobacteria>Cellvibrionales>Spongiibacteraceae>unclassified_Spongiibacteraceae | GCA_002723235.1 | Spongiibacteraceae_bacterium | CMN85_09225 | 177592..181173 | 2026793 | hypothetical-protein-[Spongiibacteraceae-bacterium] |
| MTW21824.1 | Type1.BREX | <-SAM-METHYLASE<-SIGMA+OB-Fold+AAA-ATPase+HKD-DNase-BrxL<-PglZ<-ParB+HNH<-Schlafen+HATPase+wHTH<-PhdYeFM_antitox<-N6-METHYLASE<-REase<-OLD-ABC<-DUF6079*<-BrxB-BrxF-iSTAND<-BrxA-DNA-binding<-DnaB+HINT+DnaB+HINT | 1173 | Bacteria>Pseudomonadota>Gammaproteobacteria>Chromatiales>Chromatiaceae>Allochromatium | GCA_009720725.1 | Allochromatium_palmeri | GJ668_12065 | 37225..40746 | 231048 | BREX-system-P-loop-protein-BrxC-[Allochromatium-palmeri] |
| WP_186343071.1 | Type1.BREX | <-SAM-METHYLASE<-SIGMA+OB-Fold+AAA-ATPase+HKD-DNase-BrxL<-PglZ<-ParB+HNH<-Schlafen+HATPase+wHTH<-PhdYeFM_antitox<-N6-METHYLASE<-REase<-OLD-ABC<-DUF6079*<-BrxB-BrxF-iSTAND<-BrxA-DNA-binding<-DnaB+HINT+DnaB+HINT | 1172 | Bacteria>Pseudomonadota>Gammaproteobacteria>Chromatiales>Chromatiaceae>Allochromatium | GCF_009720725.1 | Allochromatium_palmeri | GJ668_RS12065 | 37225..40743 | 231048 | BREX-system-P-loop-protein-BrxC-[Allochromatium-palmeri] |
| WP_176976536.1 | Type1.BREX | <-SF1-SF2-Helicase<-?<-HD+HD_assoc<-SIGMA+OB-Fold+AAA-ATPase+Lon-Protease-BrxL<-PglZ<-TOPRIM<-OLD-ABC<-N6-METHYLASE<-DUF6079*<-BrxB-BrxF-iSTAND<-BrxA-DNA-binding<-?<-Arm-DNA-bind_3+Phage_integrase<-CHTH<-RelE | 1198 | Bacteria>Pseudomonadota>Gammaproteobacteria>Chromatiales>Chromatiaceae>Allochromatium | GCF_013385175.1 | Allochromatium_humboldtianum | HW932_RS10995 | 221965..225561 | 504901 | BREX-system-P-loop-protein-BrxC-[Allochromatium-humboldtianum] |
| WP_200158841.1 | Type1.BREX | <-SIGMA+OB-Fold+AAA-ATPase+Lon-Protease-BrxL<-PglZ<-HNH<-OLD-ABC<-SF1-SF2-Helicase<-OLD-ABC+TOPRIM<-N6-METHYLASE<-PIN<-OLD-ABC<-DUF6079*<-BrxB-BrxF-iSTAND<-BrxA-DNA-binding<-GDResolvase+HIN-HTH<-ZnR | 1154 | Bacteria>Pseudomonadota>Gammaproteobacteria>Chromatiales>Chromatiaceae>Allochromatium | GCF_016584135.1 | Allochromatium_vinosum | CKO29_RS16480 | 18892..22356 | 1049 | BREX-system-P-loop-protein-BrxC-[Allochromatium-vinosum] |
| NEX15944.1 | Type1.BREX | <-NTP_transf<-?<-SIGMA+OB-Fold+AAA-ATPase+HKD-DNase-BrxL<-PglZ-p<-PglZ<-ParB+HNH<-N6-METHYLASE<-REase<-OLD-ABC<-DUF6079*<-BrxB-BrxF-iSTAND<-BrxA-DNA-binding<-?<-?<-ABC-ATPase+SbcC<-?<-?\|\|PALP-> | 1173 | Bacteria>Pseudomonadota>Gammaproteobacteria>Chromatiales>Chromatiaceae>Halochromatium | GCA_010912705.1 | Halochromatium_sp. | C1943_04720 | 67001..70522 | 2049430 | BREX-system-P-loop-protein-BrxC-[Halochromatium-sp.] |
| WP_200346460.1 | Type1.BREX | CR-ATPase5+REase->MazE_antitoxin->PemK_toxin->VapB_antitoxin->BrxA-DNA-binding->BrxB-BrxF-iSTAND->DUF6079*->RHH_6->N6-METHYLASE->OLD-ABC+TOPRIM->SF1-SF2-Helicase->Schlafen+HATPase+wHTH->MerR-HTH+GDResolvase->HTH_OrfB_IS605+OrfB_IS605+OrfB_Zn_ribbon->HTH_3->PhdYeFM_antitox-> | 1152 | Bacteria>Pseudomonadota>Gammaproteobacteria>Chromatiales>Chromatiaceae>Halochromatium | GCF_016584085.1 | Halochromatium_glycolicum | CKO40_RS12030 | 27931..31389 | 85075 | BREX-system-P-loop-protein-BrxC-[Halochromatium-glycolicum] |
| WP_201220083.1 | Type1.BREX | <-OLD-ABC<-SF1-SF2-Helicase<-OLD-ABC+TOPRIM<-N6-METHYLASE<-VapB_antitoxin<-PemK_toxin<-MazE_antitoxin<-PemK_toxin<-RHH<-DUF6079*<-BrxB-BrxF-iSTAND<-BrxA-DNA-binding<-?<-?<-?\|\|ABhydrolase->MlaE->ABC-ATPase+SbcC-> | 1154 | Bacteria>Pseudomonadota>Gammaproteobacteria>Chromatiales>Chromatiaceae>Halochromatium | GCF_016653315.1 | Halochromatium_roseum | CCR96_RS10495 | 53560..57024 | 391920 | BREX-system-P-loop-protein-BrxC-[Halochromatium-roseum] |
| WP_201222419.1 | Type1.BREX | <-PCMT+METHYLASE<-?<-?<-?\|\|?->tRNA->?->BrxA-DNA-binding->BrxB-BrxF-iSTAND->DUF6079*->OLD-ABC->N6-METHYLASE->Schlafen+HATPase+wHTH->ParB+HNH->PglZ->REase->MIT_C-> | 1174 | Bacteria>Pseudomonadota>Gammaproteobacteria>Chromatiales>Chromatiaceae>Halochromatium | GCF_016653315.1 | Halochromatium_roseum | CCR96_RS19410 | 137258..140782 | 391920 | BREX-system-P-loop-protein-BrxC-[Halochromatium-roseum] |
| WP_200250838.1 | Type1.BREX | <-ParB+HNH<-HNH<-OLD-ABC<-N6-METHYLASE<-REase<-OLD-ABC<-DUF6079*<-BrxB-BrxF-iSTAND<-BrxA-DNA-binding | 1173 | Bacteria>Pseudomonadota>Gammaproteobacteria>Chromatiales>Chromatiaceae>Lamprobacter | GCF_016583905.1 | Lamprobacter_modestohalophilus | CKO42_RS24315 | 8724..12245 | 1064514 | BREX-system-P-loop-protein-BrxC-[Lamprobacter-modestohalophilus] |
| MBV5272983.1 | Type1.BREX | <-PglZ<-?<-REase+AIPR<-N6-METHYLASE-p<-N6-METHYLASE-p<-PIN<-MazE_antitoxin<-DUF6079*<-BrxB-BrxF-iSTAND<-BrxA-DNA-binding<-Aminotran_3<-FURR-HTH\|\|?-><-?<-?<-?<-Classical-AAA | 1143 | Bacteria>Pseudomonadota>Gammaproteobacteria>Chromatiales>Chromatiaceae>Lamprocystis | GCA_019163055.1 | Lamprocystis_purpurea | JZU52_04840 | 9550..12981 | 61598 | BREX-system-P-loop-protein-BrxC-[Lamprocystis-purpurea] |
| WP_020505452.1 | Type1.BREX | RNase-><-?<-?\|\|BrxA-DNA-binding->BrxB-BrxF-iSTAND->DUF6079*-><-?\|\|N6-METHYLASE->Schlafen+HATPase+wHTH->PglZ->SIGMA+OB-Fold+AAA-ATPase+Lon-Protease-BrxL-> | 1146 | Bacteria>Pseudomonadota>Gammaproteobacteria>Chromatiales>Chromatiaceae>Lamprocystis | GCF_000379525.1 | Lamprocystis_purpurea | A39O_RS0114850 | 122427..125867 | 61598 | BREX-system-P-loop-protein-BrxC-[Lamprocystis-purpurea] |
| WP_020507234.1 | Type1.BREX | <-PIN<-MazE_antitoxin<-DUF6079*<-BrxB-BrxF-iSTAND<-BrxA-DNA-binding<-Aminotran_3<-FURR-HTH | 1153 | Bacteria>Pseudomonadota>Gammaproteobacteria>Chromatiales>Chromatiaceae>Lamprocystis | GCF_000379525.1 | Lamprocystis_purpurea | A39O_RS0124120 | 1671..5132 | 61598 | BREX-system-P-loop-protein-BrxC-[Lamprocystis-purpurea] |
| WP_062271501.1 | Type1.BREX | <-OLD-ABC<-RHH<-SF1-SF2-Helicase<-OLD-ABC+TOPRIM<-N6-METHYLASE<-PIN<-OLD-ABC<-PIN<-?<-DUF6079*<-BrxB-BrxF-iSTAND<-BrxA-DNA-binding<-tRNA<-Aminotran_1_2\|\|SF1-SF2-Helicase-> | 1153 | Bacteria>Pseudomonadota>Gammaproteobacteria>Chromatiales>Chromatiaceae>Marichromatium | GCF_001583505.1 | Marichromatium_gracile | AY586_RS00820 | 23071..26532 | 1048 | BREX-system-P-loop-protein-BrxC-[Marichromatium-gracile] |
| WP_134356376.1 | Type1.BREX | <-DUF488-l<-DUF4326\|\|?-><-?\|\|?->DDE_Tnp_1_assoc+DDE_Tnp_1-><-?\|\|BrxA-DNA-binding->BrxB-BrxF-iSTAND->DUF6079*->HicB_lk_antitox->HicA_toxin->N6-METHYLASE->PglZ->SIGMA+OB-Fold+AAA-ATPase+Lon-Protease-BrxL-><-?<-DDE_3<-HTH_23+HTH_32+HTH_33 | 1147 | Bacteria>Pseudomonadota>Gammaproteobacteria>Chromatiales>Chromatiaceae>Nitrosococcus | GCF_004421105.1 | Nitrosococcus_wardiae | E3U44_RS01695 | 339932..343375 | 1814290 | BREX-system-P-loop-protein-BrxC-[Nitrosococcus-wardiae] |
| TXH98297.1 | Type1.BREX | <-DUF6079*<-BrxB-BrxF-iSTAND<-BrxA-DNA-binding<-tRNA<-?\|\|?->?-><-Glyco_hydro_3+Glyco_hydro_3_C+Fn3-like | 1197 | Bacteria>Pseudomonadota>Gammaproteobacteria>Chromatiales>Chromatiaceae>Rheinheimera | GCA_008015325.1 | Rheinheimera_sp. | E6Q75_01850 | 1..3593 | 1869214 | BREX-system-P-loop-protein-BrxC,-partial-[Rheinheimera-sp.] |
| WP_264346732.1 | Type1.BREX | HEPN->?->PVC-AAA->?->GreA_GreB+SF1-SF2-Helicase->BrxA-DNA-binding->BrxB-BrxF-iSTAND->DUF6079*->N6-METHYLASE->OLD-ABC->HNH->ParB+HNH->PglZ->HEPN+Sel1+SEC-C-><-?<-NERD+SF1-SF2-Helicase<-?<-SF1-SF2-Helicase+REase | 1157 | Bacteria>Pseudomonadota>Gammaproteobacteria>Chromatiales>Chromatiaceae>Rheinheimera | GCF_947090785.1 | Rheinheimera_sp. | OM978_RS09960 | 2096767..2100240 | 1869214 | BREX-system-P-loop-protein-BrxC-[Rheinheimera-sp.-MM224] |
| QVL48954.1 | Type1.BREX | <-PglZ<-PIN<-ParD_antitoxin<-wHTH<-TOPRIM<-OLD-ABC<-N6-METHYLASE<-ACET+ACET<-TraJ-RHH<-DUF6079*<-BrxB-BrxF-iSTAND<-BrxA-DNA-binding<-tRNA\|\|RADICAL-SAM->Glycos_transf_2->?<-?\|\|?-><-tRNA\|\|NTP_transf+HDRnase-> | 1136 | Bacteria>Pseudomonadota>Gammaproteobacteria>Chromatiales>Chromatiaceae>Thiocapsa | GCA_018399035.1 | Thiocapsa_sp. | KFB96_26095 | 5699625..5703035 | 2024551 | MAG:-BREX-system-P-loop-protein-BrxC-[Thiocapsa-sp.] |
| WP_120797099.1 | Type1.BREX | <-HTH_3<-DDE_Tnp\|\|?->?<-?\|\|?->?->BrxA-DNA-binding->BrxB-BrxF-iSTAND->DUF6079*->N6-METHYLASE-p->?<-DDE_Tnp\|\|ParB+HNH->PglZ->?->SIGMA+OB-Fold+AAA-ATPase+HKD-DNase-BrxL-> | 1173 | Bacteria>Pseudomonadota>Gammaproteobacteria>Chromatiales>Chromatiaceae>Thiocapsa | GCF_003634315.1 | Thiocapsa_rosea | BDD21_RS10290 | 2320412..2323933 | 69360 | BREX-system-P-loop-protein-BrxC-[Thiocapsa-rosea] |
| WP_200385867.1 | Type1.BREX | DUF6079*->?->PIN-><-?\|\|N6-METHYLASE->OLD-ABC->HNH->PglZ->AbrB-HTH->RHH-> | 1179 | Bacteria>Pseudomonadota>Gammaproteobacteria>Chromatiales>Chromatiaceae>Thiocapsa | GCF_016583985.1 | Thiocapsa_imhoffii | CKO25_RS00005 | 26..3565 | 382777 | BREX-system-P-loop-protein-BrxC-[Thiocapsa-imhoffii] |
| WP_200387145.1 | Type1.BREX | <-SIGMA+OB-Fold+AAA-ATPase+Lon-Protease-BrxL<-PglZ<-wHTH<-TOPRIM<-OLD-ABC<-N6-METHYLASE<-ACET+ACET<-TraJ-RHH<-DUF6079*<-BrxB-BrxF-iSTAND<-BrxA-DNA-binding<-tRNA\|\|RADICAL-SAM->Glycos_transf_2-><-?\|\|?-><-tRNA\|\|NTP_transf+HDRnase-> | 1136 | Bacteria>Pseudomonadota>Gammaproteobacteria>Chromatiales>Chromatiaceae>Thiocapsa | GCF_016583985.1 | Thiocapsa_imhoffii | CKO25_RS06645 | 47680..51090 | 382777 | BREX-system-P-loop-protein-BrxC-[Thiocapsa-imhoffii] |
| WP_200389392.1 | Type1.BREX | <-PAS+PAS+PAS+GGDEF<-?<-RuvB-AAA+RuvB-HTH<-?\|\|tRNA->BrxA-DNA-binding->BrxB-BrxF-iSTAND->DUF6079*->MazE_antitoxin->AbrB-HTH->PIN->?->?->?->OLD-ABC->TOPRIM-> | 1193 | Bacteria>Pseudomonadota>Gammaproteobacteria>Chromatiales>Chromatiaceae>Thiocapsa | GCF_016583985.1 | Thiocapsa_imhoffii | CKO25_RS18260 | 8675..12256 | 382777 | BREX-system-P-loop-protein-BrxC-[Thiocapsa-imhoffii] |
| WP_232409231.1 | Type1.BREX | <-Tox-RelE<-TOPRIM<-OLD-ABC<-?<-PIN<-VapB_antitoxin<-VapB_antitoxin<-PemK_toxin<-MazE_antitoxin<-DUF6079-p*\|\|DDE_Tnp_1_4-><-DUF6079-p<-BrxB-BrxF-iSTAND<-BrxA-DNA-binding<-?<-HEPN<-Ntox45<-HTH_Tnp_1+HTH_21+rve+rve_3 | 909 | Bacteria>Pseudomonadota>Gammaproteobacteria>Chromatiales>Chromatiaceae>Thiocapsa | GCF_021228795.1 | Thiocapsa_bogorovii | LT988_RS05545 | 1229820..1232549 | 521689 | BREX-system-P-loop-protein-BrxC-[Thiocapsa-bogorovii] |
| WP_232409232.1 | Type1.BREX | <-OLD-ABC<-?<-PIN<-VapB_antitoxin<-VapB_antitoxin<-PemK_toxin<-MazE_antitoxin<-DUF6079-p\|\|DDE_Tnp_1_4-><-DUF6079-p*<-BrxB-BrxF-iSTAND<-BrxA-DNA-binding<-?<-HEPN<-Ntox45<-HTH_Tnp_1+HTH_21+rve+rve_3<-?\|\|ICLR-HTH->MoxR-AAA-> | 245 | Bacteria>Pseudomonadota>Gammaproteobacteria>Chromatiales>Chromatiaceae>Thiocapsa | GCF_021228795.1 | Thiocapsa_bogorovii | LT988_RS05555 | 1234280..1235017 | 521689 | hypothetical-protein-[Thiocapsa-bogorovii] |
| WP_100919677.1 | Type1.BREX | <-PspA\|\|REC+OmpR-HTH->HISKIN->?-><-?<-CASPASE\|\|BrxA-DNA-binding->BrxB-BrxF-iSTAND->DUF6079*->?->TraJ-RHH->N6-METHYLASE->OLD-ABC->TOPRIM->wHTH->PglZ->SIGMA+OB-Fold+AAA-ATPase+Lon-Protease-BrxL-><-ROK-HTH | 1156 | Bacteria>Pseudomonadota>Gammaproteobacteria>Chromatiales>Chromatiaceae>Thiodictyon | GCF_002813775.1 | Candidatus_Thiodictyon_syntrophicum | THSYN_RS13895 | 3302661..3306131 | 1166950 | BREX-system-P-loop-protein-BrxC-[Candidatus-Thiodictyon-syntrophicum] |
| WP_100920747.1 | Type1.BREX | BrxA-DNA-binding->BrxB-BrxF-iSTAND->?->DUF6079*->PhdYeFM_antitox->PIN->MqsA_antitoxin->N6-METHYLASE-p->N6-METHYLASE->Transposase_31->OLD-ABC->Polbeta->PglZ-> | 1194 | Bacteria>Pseudomonadota>Gammaproteobacteria>Chromatiales>Chromatiaceae>Thiodictyon | GCF_002813775.1 | Candidatus_Thiodictyon_syntrophicum | THSYN_RS20415 | 4833326..4836910 | 1166950 | BREX-system-P-loop-protein-BrxC-[Candidatus-Thiodictyon-syntrophicum] |
| MBK1629582.1 | Type1.BREX | <-SbcC<-?<-RFC-AAA+DNAA-HTH\|\|?->RNaseP->?->sGTP->BrxA-DNA-binding->BrxB-BrxF-iSTAND->DUF6079*->MazE_antitoxin->PemK_toxin->DOC+HTH->N6-METHYLASE-p->PHP+SbcC->Schlafen+HATPase+wHTH->PglZ->SIGMA+OB-Fold+AAA-ATPase+Lon-Protease-BrxL-> | 1152 | Bacteria>Pseudomonadota>Gammaproteobacteria>Chromatiales>Chromatiaceae>Thiohalocapsa | GCA_016583825.1 | Thiohalocapsa_halophila | CKO31_02265 | 100295..103753 | 69359 | hypothetical-protein-[Thiohalocapsa-halophila] |
| MCG6940983.1 | Type1.BREX | <-N6-METHYLASE-p<-PIN<-VapB_antitoxin<-DUF6079-p*<-tRNA<-?<-AA_kinase+ACT+ACT_7 | 251 | Bacteria>Pseudomonadota>Gammaproteobacteria>Chromatiales>Chromatiaceae>Thiohalocapsa | GCA_022340605.1 | Thiohalocapsa_sp. | LJE69_07005 | 1995..2750 | 2497641 | hypothetical-protein-[Thiohalocapsa-sp.] |
| WP_058554877.1 | Type1.BREX | DDE_Tnp_1->?->?->?->?->AA_kinase+ACT+ACT_7->?->tRNA->BrxA-DNA-binding->BrxB-BrxF-iSTAND->DUF6079*->VapB_antitoxin->VapB_antitoxin->PIN-><-?\|\|N6-METHYLASE-> | 1152 | Bacteria>Pseudomonadota>Gammaproteobacteria>Chromatiales>Chromatiaceae>Thiohalocapsa | GCF_001469165.1 | Thiohalocapsa_sp._ML1 | ML01_RS08025 | 156057..159515 | 1431688 | BREX-system-P-loop-protein-BrxC-[Thiohalocapsa-sp.-ML1] |
| WP_242475077.1 | Type1.BREX | HAD+HAD-><-MORC+DNA_gyrase+TOPRIM<-SbcC<-RFC-AAA+DNAA-HTH\|\|?->?->sGTP->BrxA-DNA-binding->BrxB-BrxF-iSTAND->DUF6079*->MazE_antitoxin->PemK_toxin->RHH_6-><-?\|\|N6-METHYLASE->DOC+HTH->N6-METHYLASE-p->PHP+SbcC->Schlafen+HATPase+wHTH-> | 1154 | Bacteria>Pseudomonadota>Gammaproteobacteria>Chromatiales>Chromatiaceae>Thiohalocapsa | GCF_016583825.1 | Thiohalocapsa_halophila | CKO31_RS02315 | 100289..103753 | 69359 | BREX-system-P-loop-protein-BrxC-[Thiohalocapsa-halophila] |
| WP_200239121.1 | Type1.BREX | <-PglZ<-PIN<-PhdYeFM_antitox<-HicA_toxin<-HicB_lk_antitox<-MazE_antitoxin<-PIN<-MazE_antitoxin<-DUF6079*<-BrxB-BrxF-iSTAND<-BrxA-DNA-binding<-?<-ABhydrolase<-?<-?\|\|?->?-><-REase | 1197 | Bacteria>Pseudomonadota>Gammaproteobacteria>Chromatiales>Chromatiaceae>Thiohalocapsa | GCF_016583825.1 | Thiohalocapsa_halophila | CKO31_RS15145 | 13443..17036 | 69359 | BREX-system-P-loop-protein-BrxC-[Thiohalocapsa-halophila] |
| EIC19850.1 | Type1.BREX | <-SIGMA+OB-Fold+AAA-ATPase+Lon-Protease-BrxL<-PglZ<-RelE-ParE<-PhdYeFM_antitox<-Schlafen+HATPase+wHTH<-N6-METHYLASE<-PIN<-VapB_antitoxin<-DUF6079*<-BrxB-BrxF-iSTAND<-BrxA-DNA-binding<-?<-HISKIN<-?<-?<-Thioredoxin | 1152 | Bacteria>Pseudomonadota>Gammaproteobacteria>Chromatiales>Chromatiaceae>Thiorhodovibrio | GCA_000228725.3 | Thiorhodovibrio_sp._970 | Thi970DRAFT_03455 | 369329..372787 | 631362 | hypothetical-protein-Thi970DRAFT_03455-[Thiorhodovibrio-sp.-970] |
| WP_223296802.1 | Type1.BREX | <-SIGMA+OB-Fold+AAA-ATPase+Lon-Protease-BrxL<-PglZ<-RelE-ParE<-PhdYeFM_antitox<-Schlafen+HATPase+wHTH<-N6-METHYLASE<-PIN<-VapB_antitoxin<-DUF6079*<-BrxB-BrxF-iSTAND<-BrxA-DNA-binding<-?<-HISKIN<-?<-?<-Thioredoxin\|\|?->REC+GGDEF+EAL-> | 1154 | Bacteria>Pseudomonadota>Gammaproteobacteria>Chromatiales>Chromatiaceae>Thiorhodovibrio | GCF_000228725.2 | Thiorhodovibrio_sp._970 | THI970DRAFT_RS16355 | 369329..372793 | 631362 | BREX-system-P-loop-protein-BrxC-[Thiorhodovibrio-sp.-970] |
| WP_201066887.1 | Type1.BREX | <-wHTH<-?<-PIN<-?<-N6-METHYLASE<-HicB_lk_antitox<-MazE_antitoxin<-?<-PIN<-DUF6079*<-BrxB-BrxF-iSTAND<-BrxA-DNA-binding<-tRNA<-Sulfate_transp+STAS | 1203 | Bacteria>Pseudomonadota>Gammaproteobacteria>Chromatiales>Chromatiaceae>Thiorhodovibrio | GCF_016653415.1 | Thiorhodovibrio_winogradskyi | CCR91_RS10560 | 25140..28751 | 77007 | BREX-system-P-loop-protein-BrxC-[Thiorhodovibrio-winogradskyi] |
| MBK8638067.1 | Type1.BREX | AAA-ATPase->?->ZnR->?->?->?->DUF6079*->PhdYeFM_antitox->AbrB-HTH->PIN->N6-METHYLASE->PIN->?->PglZ->SIGMA+OB-Fold+AAA-ATPase+Lon-Protease-BrxL-><-SIR2 | 1067 | Bacteria>Pseudomonadota>Gammaproteobacteria>Chromatiales>Chromatiaceae>unclassified_Chromatiaceae | GCA_016714645.1 | Chromatiaceae_bacterium | IPN92_07155 | 42706..45909 | 2049432 | BREX-system-P-loop-protein-BrxC-[Chromatiaceae-bacterium] |
| MBP6581931.1 | Type1.BREX | <-SIGMA+OB-Fold+AAA-ATPase+Lon-Protease-BrxL<-PglZ<-?<-PglZ-p<-SbcC+CCDC158<-Calcineurin-SbcD<-N6-METHYLASE<-DUF6079*<-BrxB-BrxF-iSTAND<-BrxA-DNA-binding<-tRNA\|\|?-><-?<-?<-?<-?<-ABC-ATPase+SbcC | 1144 | Bacteria>Pseudomonadota>Gammaproteobacteria>Chromatiales>Chromatiaceae>unclassified_Chromatiaceae | GCA_017989555.1 | Chromatiaceae_bacterium | KA204_00415 | 17018..20452 | 2049432 | BREX-system-P-loop-protein-BrxC-[Chromatiaceae-bacterium] |
| MBP6733459.1 | Type1.BREX | BrxA-DNA-binding->BrxB-BrxF-iSTAND->DUF6079*-><-BrxB-BrxF-iSTAND\|\|?->PhdYeFM_antitox->PIN->N6-METHYLASE->HTH+REase->REase->N6-METHYLASE-p-> | 1171 | Bacteria>Pseudomonadota>Gammaproteobacteria>Chromatiales>Chromatiaceae>unclassified_Chromatiaceae | GCA_017990755.1 | Chromatiaceae_bacterium | KA142_01990 | 1791..5306 | 2049432 | BREX-system-P-loop-protein-BrxC-[Chromatiaceae-bacterium] |
| MBP6806725.1 | Type1.BREX | <-SIGMA+OB-Fold+AAA-ATPase+Lon-Protease-BrxL<-PglZ<-?<-?<-PglZ-p<-SbcC+CCDC158<-Calcineurin-SbcD<-N6-METHYLASE<-DUF6079-p* | 782 | Bacteria>Pseudomonadota>Gammaproteobacteria>Chromatiales>Chromatiaceae>unclassified_Chromatiaceae | GCA_017991215.1 | Chromatiaceae_bacterium | KA125_00340 | 14653..17002 | 2049432 | BREX-system-P-loop-protein-BrxC-[Chromatiaceae-bacterium] |
| MBP6807296.1 | Type1.BREX | BrxA-DNA-binding->BrxB-BrxF-iSTAND->DUF6079*->MazE_antitoxin->PIN->N6-METHYLASE-p->AbrB-HTH-> | 1194 | Bacteria>Pseudomonadota>Gammaproteobacteria>Chromatiales>Chromatiaceae>unclassified_Chromatiaceae | GCA_017991215.1 | Chromatiaceae_bacterium | KA125_03275 | 984..4568 | 2049432 | BREX-system-P-loop-protein-BrxC-[Chromatiaceae-bacterium] |
| MCF7978332.1 | Type1.BREX | <-AbrB-HTH<-PIN<-AbrB-HTH<-N6-METHYLASE<-DOC<-MazE_antitoxin<-MqsA_antitoxin<-?<-PIN<-DUF6079* | 1223 | Bacteria>Pseudomonadota>Gammaproteobacteria>Chromatiales>Chromatiaceae>unclassified_Chromatiaceae | GCA_021736685.1 | Chromatiaceae_bacterium | K9L82_09985 | 9865..13536 | 2049432 | BREX-system-P-loop-protein-BrxC-[Chromatiaceae-bacterium] |
| MCF8004221.1 | Type1.BREX | <-DUF6079-p*<-BrxB-BrxF-iSTAND<-BrxA-DNA-binding<-ABC-ATPase+SbcC<-?<-?<-?\|\|PALP-> | 370 | Bacteria>Pseudomonadota>Gammaproteobacteria>Chromatiales>Chromatiaceae>unclassified_Chromatiaceae | GCA_021736845.1 | Chromatiaceae_bacterium | K9L32_08450 | 1..1111 | 2049432 | BREX-system-P-loop-protein-BrxC-[Chromatiaceae-bacterium] |
| MCF8016509.1 | Type1.BREX | <-AbrB-HTH<-PIN<-AbrB-HTH<-N6-METHYLASE<-DOC<-MazE_antitoxin<-MqsA_antitoxin<-?<-PIN<-DUF6079-p* | 835 | Bacteria>Pseudomonadota>Gammaproteobacteria>Chromatiales>Chromatiaceae>unclassified_Chromatiaceae | GCA_021736865.1 | Chromatiaceae_bacterium | K9L65_12595 | 6416..8923 | 2049432 | BREX-system-P-loop-protein-BrxC-[Chromatiaceae-bacterium] |
| MCP5407916.1 | Type1.BREX | ABhydrolase->?->TraU->?->TraG_N-><-?\|\|BrxA-DNA-binding->BrxB-BrxF-iSTAND->DUF6079*->MazE_antitoxin->PIN->N6-METHYLASE->SIR2->?->PglZ->SIGMA+OB-Fold+AAA-ATPase+Lon-Protease-BrxL->XIS-HTH-> | 1194 | Bacteria>Pseudomonadota>Gammaproteobacteria>Chromatiales>Chromatiaceae>unclassified_Chromatiaceae | GCA_024235335.1 | Chromatiaceae_bacterium | H6963_01295 | 248686..252270 | 2049432 | BREX-system-P-loop-protein-BrxC-[Chromatiaceae-bacterium] |
| WP_018716649.1 | Type1.BREX | HerA/FtsK->Relaxase+SF1-SF2-Helicase->?->XIS-HTH->BrxA-DNA-binding->BrxB-BrxF-iSTAND->DUF6079*->Schlafen+HATPase+wHTH->N6-METHYLASE->ParB+HNH->PglZ->SIGMA+OB-Fold+AAA-ATPase+Lon-Protease-BrxL-> | 1144 | Bacteria>Pseudomonadota>Gammaproteobacteria>Chromatiales>Ectothiorhodospiraceae>Arhodomonas | GCF_000374645.1 | Arhodomonas_aquaeolei | C516_RS0101825 | 422257..425691 | 2369 | BREX-system-P-loop-protein-BrxC-[Arhodomonas-aquaeolei] |
| WP_063464066.1 | Type1.BREX | ZnR->RADICAL-SAM->?->?->?->GGDEF->XIS-HTH->BrxA-DNA-binding->BrxB-BrxF-iSTAND->DUF6079*->Schlafen+HATPase+wHTH+HTH->N6-METHYLASE-p->OLD-ABC->REase->SIR2->HerA/FtsK->N6-METHYLASE-p->OLD-ABC->HNH-> | 1189 | Bacteria>Pseudomonadota>Gammaproteobacteria>Chromatiales>Ectothiorhodospiraceae>Ectothiorhodospira | GCF_001632845.1 | Ectothiorhodospira_sp._BSL-9 | ECTOBSL9_RS04500 | 914827..918396 | 1442136 | BREX-system-P-loop-protein-BrxC-[Ectothiorhodospira-sp.-BSL-9] |
| WP_238627878.1 | Type1.BREX | <-DUF6079-p*<-BrxB-BrxF-iSTAND<-BrxA-DNA-binding<-XIS-HTH<-?<-?<-?<-RADICAL-SAM<-ZnR\|\|SWACOS-> | 688 | Bacteria>Pseudomonadota>Gammaproteobacteria>Chromatiales>Ectothiorhodospiraceae>Ectothiorhodospira | GCF_022227565.1 | Ectothiorhodospira_sp._9100 | LPW29_RS09995 | 1..2064 | 2897388 | BREX-system-P-loop-protein-BrxC,-partial-[Ectothiorhodospira-sp.-9100] |
| WP_238631208.1 | Type1.BREX | <-PglZ<-TOPRIM<-OLD-ABC<-N6-METHYLASE-p<-HerA/FtsK<-?<-CR-ATPase3<-N6-METHYLASE<-Schlafen+HATPase+wHTH+HTH<-DUF6079*<-BrxB-BrxF-iSTAND<-BrxA-DNA-binding<-XIS-HTH<-?<-?<-?<-RADICAL-SAM<-ZnR\|\|SWACOS-> | 1189 | Bacteria>Pseudomonadota>Gammaproteobacteria>Chromatiales>Ectothiorhodospiraceae>Ectothiorhodospira | GCF_022227695.1 | Ectothiorhodospira_sp._9905 | LPW28_RS03070 | 43824..47393 | 2897387 | BREX-system-P-loop-protein-BrxC-[Ectothiorhodospira-sp.-9905] |
| WP_238620557.1 | Type1.BREX | sGTP-><-35exo+T7DNApol\|\|?->?->BrxA-DNA-binding->BrxB-BrxF-iSTAND->DUF6079*->N6-METHYLASE->OLD-ABC->HNH->OLD-ABC->HNH->PglZ->SIGMA+OB-Fold+AAA-ATPase+Lon-Protease-BrxL-><-Schlafen+HATPase+wHTH<-Schlafen+HATPase+wHTH | 1154 | Bacteria>Pseudomonadota>Gammaproteobacteria>Chromatiales>Ectothiorhodospiraceae>Ectothiorhodospira | GCF_022227785.1 | Ectothiorhodospira_sp._B14B | LQ952_00335 | 78448..81912 | 2899127 | BREX-system-P-loop-protein-BrxC-[Ectothiorhodospira-sp.-B14B] |
| WP_090207826.1 | Type1.BREX | SbcC->REase->OrfB_IS605+OrfB_Zn_ribbon-><-SIGMA+OB-Fold+AAA-ATPase+Lon-Protease-BrxL-p<-Phage_AlpA<-PglZ<-TOPRIM<-OLD-ABC<-N6-METHYLASE<-DUF6079*<-BrxB-BrxF-iSTAND<-BrxA-DNA-binding\|\|tRNA-><-?<-REC+REC<-tRNA<-tRNA<-?<-?<-?<-?<-GAF+PAS+GGDEF+EAL | 1231 | Bacteria>Pseudomonadota>Gammaproteobacteria>Chromatiales>Ectothiorhodospiraceae>Ectothiorhodospira | GCF_900110965.1 | Ectothiorhodospira_magna | BMX62_RS09270 | 49857..53552 | 867345 | BREX-system-P-loop-protein-BrxC-[Ectothiorhodospira-magna] |
| MBK5942672.1 | Type1.BREX | <-ParB+HNH<-?<-N6-METHYLASE<-DUF6079*<-BrxB-BrxF-iSTAND<-BrxA-DNA-binding<-SbcC+CCDC158<-Calcineurin<-?<-STAS<-HDRnase<-?<-Arm-DNA-bind_2+Phage_int_SAM_3+Phage_integrase | 1146 | Bacteria>Pseudomonadota>Gammaproteobacteria>Chromatiales>Ectothiorhodospiraceae>Halorhodospira | GCA_016653405.1 | Halorhodospira_halophila | CCR79_02095 | 7976..11416 | 1053 | hypothetical-protein-[Halorhodospira-halophila] |
| WP_242510795.1 | Type1.BREX | <-ParB+HNH<-?<-N6-METHYLASE<-DUF6079*<-BrxB-BrxF-iSTAND<-BrxA-DNA-binding<-Calcineurin<-?<-STAS<-HDRnase<-?<-Arm-DNA-bind_2+Phage_int_SAM_3+Phage_integrase<-Phage_AlpA | 1148 | Bacteria>Pseudomonadota>Gammaproteobacteria>Chromatiales>Ectothiorhodospiraceae>Halorhodospira | GCF_016653405.1 | Halorhodospira_halophila | CCR79_RS02135 | 7976..11422 | 1053 | BREX-system-P-loop-protein-BrxC-[Halorhodospira-halophila] |
| AGM41053.1 | Type1.BREX | PhdYeFM_antitox->PIN-><-?<-nSTAND1+HNH\|\|?-><-ParB+HNH<-SIGMA+OB-Fold+AAA-ATPase+Lon-Protease-BrxL<-PglZ<-N6-METHYLASE<-DUF6079*<-BrxB-BrxF-iSTAND<-BrxA-DNA-binding<-CopG/RHH<-?\|\|?->?->SAM-METHYLASE-><-MerR-HTH | 1145 | Bacteria>Pseudomonadota>Gammaproteobacteria>Chromatiales>Ectothiorhodospiraceae>Spiribacter | GCA_000319575.2 | Spiribacter_salinus_M19-40 | SPISAL_04795 | 979011..982448 | 1260251 | hypothetical-protein-SPISAL_04795-[Spiribacter-salinus-M19-40] |
| TQE98798.1 | Type1.BREX | BrxA-DNA-binding->BrxB-BrxF-iSTAND->DUF6079*->Schlafen+HATPase+wHTH->N6-METHYLASE-p-> | 1193 | Bacteria>Pseudomonadota>Gammaproteobacteria>Chromatiales>Ectothiorhodospiraceae>Spiribacter | GCA_006569195.1 | Spiribacter_salinus | FKY71_11930 | 1820..5401 | 1335746 | BREX-system-P-loop-protein-BrxC-[Spiribacter-salinus] |
| KAF0285942.1 | Type1.BREX | <-Phage_AlpA\|\|?->?->?->TPR->?->CopG/RHH->BrxA-DNA-binding->BrxB-BrxF-iSTAND->DUF6079*->N6-METHYLASE->PglZ->SIGMA+OB-Fold+AAA-ATPase+Lon-Protease-BrxL->ParB+HNH-> | 1145 | Bacteria>Pseudomonadota>Gammaproteobacteria>Chromatiales>Ectothiorhodospiraceae>Spiribacter | GCA_009805925.1 | Spiribacter_sp._SSL99 | BA899_01280 | 102096..105533 | 1866884 | hypothetical-protein-BA899_01280-[Spiribacter-sp.-SSL99] |
| WP_245539889.1 | Type1.BREX | PhdYeFM_antitox->PIN-><-?<-nSTAND1+HNH<-?<-ParB+HNH<-SIGMA+OB-Fold+AAA-ATPase+Lon-Protease-BrxL<-PglZ<-N6-METHYLASE<-DUF6079*<-BrxB-BrxF-iSTAND<-BrxA-DNA-binding<-CopG/RHH<-?\|\|?->?->SAM-METHYLASE->?-><-MerR-HTH | 1151 | Bacteria>Pseudomonadota>Gammaproteobacteria>Chromatiales>Ectothiorhodospiraceae>Spiribacter | GCF_000319575.2 | Spiribacter_salinus | SPISAL_RS04875 | 979011..982466 | 1335746 | BREX-system-P-loop-protein-BrxC-[Spiribacter-salinus] |
| WP_154297058.1 | Type1.BREX | <-ParB+HNH<-SIGMA+OB-Fold+AAA-ATPase+Lon-Protease-BrxL<-PglZ<-Polbeta<-MarR-HTH<-N6-METHYLASE<-DUF6079*<-BrxB-BrxF-iSTAND<-BrxA-DNA-binding\|\|SIGMA-HTH+HIN-HTH-><-?<-MerR-HTH<-HTH_Tnp_1+HTH_21+rve+rve_3<-?<-DNA_METHYLASE | 1145 | Bacteria>Pseudomonadota>Gammaproteobacteria>Chromatiales>Ectothiorhodospiraceae>Spiribacter | GCF_009676705.1 | Spiribacter_sp._2438 | GJ672_RS05635 | 1128117..1131554 | 2666185 | BREX-system-P-loop-protein-BrxC-[Spiribacter-sp.-2438] |
| WP_235013293.1 | Type1.BREX | TPR->?->CopG/RHH->BrxA-DNA-binding->BrxB-BrxF-iSTAND->DUF6079*->N6-METHYLASE->PglZ->SIGMA+OB-Fold+AAA-ATPase+Lon-Protease-BrxL->ParB+HNH-> | 1151 | Bacteria>Pseudomonadota>Gammaproteobacteria>Chromatiales>Ectothiorhodospiraceae>Spiribacter | GCF_009805925.1 | Spiribacter_sp._SSL99 | BA899_RS04165 | 102078..105533 | 1866884 | BREX-system-P-loop-protein-BrxC-[Spiribacter-sp.-SSL99] |
| WP_244406389.1 | Type1.BREX | <-PVC-AAA+Classical-AAA<-?\|\|wHTH+WYL+WCX->?->?->BrxA-DNA-binding->BrxB-BrxF-iSTAND->DUF6079*->MULE-TRANSPOSASE->DUF6079-p->REase->N6-METHYLASE->MarR-HTH->Polbeta->PglZ->SIGMA+OB-Fold+AAA-ATPase+Lon-Protease-BrxL->HIN-HTH-> | 1041 | Bacteria>Pseudomonadota>Gammaproteobacteria>Chromatiales>Ectothiorhodospiraceae>Thioalkalivibrio | GCF_000025545.1 | Thioalkalivibrio_sp._K90mix | TK90_RS05670 | 1196495..1199620 | 396595 | BREX-system-P-loop-protein-BrxC-[Thioalkalivibrio-sp.-K90mix] |
| WP_019570731.1 | Type1.BREX | BrxA-DNA-binding->BrxB-BrxF-iSTAND->DUF6079*->?->N6-METHYLASE->MarR-HTH->Polbeta->PglZ->SIGMA+OB-Fold+AAA-ATPase+Lon-Protease-BrxL-><-AAA-ToxA+Classical-AAA | 1145 | Bacteria>Pseudomonadota>Gammaproteobacteria>Chromatiales>Ectothiorhodospiraceae>Thioalkalivibrio | GCF_000381205.1 | Thioalkalivibrio_sp._ALE11 | F620_RS0111965 | 2244..5681 | 1265494 | BREX-system-P-loop-protein-BrxC-[Thioalkalivibrio-sp.-ALE11] |
| WP_019570748.1 | Type1.BREX | PglZ->SIGMA+OB-Fold+AAA-ATPase+Lon-Protease-BrxL-><-AAA-ToxA+Classical-AAA\|\|?->?->?->wHTH+WYL+WCX->BrxA-DNA-binding->BrxB-BrxF-iSTAND->DUF6079*->SF1-SF2-Helicase+PRTase->SLOG+BirA-HTH->N6-METHYLASE->ParB+HNH->PglZ->SIGMA+OB-Fold+AAA-ATPase+Lon-Protease-BrxL->?->Phage_integrase-> | 1225 | Bacteria>Pseudomonadota>Gammaproteobacteria>Chromatiales>Ectothiorhodospiraceae>Thioalkalivibrio | GCF_000381205.1 | Thioalkalivibrio_sp._ALE11 | F620_RS0112040 | 26049..29726 | 1265494 | BREX-system-P-loop-protein-BrxC-[Thioalkalivibrio-sp.-ALE11] |
| QEP43020.1 | Type1.BREX | <-DDE_Tnp_1<-HTH_Tnp_1+HTH_21+rve+rve_3<-HTH_Tnp_1<-TOPRIM<-OLD-ABC<-RNase<-N6-METHYLASE<-HNH<-OLD-ABC<-DUF6079*<-BrxB-BrxF-iSTAND<-BrxA-DNA-binding<-Phage_holin<-?<-?\|\|GDResolvase+ZnR->?->DDE_Tnp_IS66->HD+SF1-SF2-Helicase-> | 1145 | Bacteria>Pseudomonadota>Gammaproteobacteria>Chromatiales>Ectothiorhodospiraceae>unclassified_Ectothiorhodospiraceae | GCA_008375315.1 | Ectothiorhodospiraceae_bacterium_BW-2 | D5085_07760 | 1613185..1616622 | 947515 | BREX-system-P-loop-protein-BrxC-[Ectothiorhodospiraceae-bacterium-BW-2] |

| Protein-ID | System | Gene-Neighbourhoods | Length | Taxonomy | Assembly-ID | Strain | Locus-Tag | Genomic-coordinates | TaxID | Fasta-define |
|---|---|---|---|---|---|---|---|---|---|---|
| MCB1864346.1 | Type1.BREX | <-DNA_METHYLASE\|\|?->TraU->TraH->?->TraG_N-><-?\|\|BrxA-DNA-binding->BrxB-BrxF-iSTAND->DUF6079*->N6-METHYLASE->Pg [illegible] | 1147 | Bacteria>Pseudomonadota>Gammaproteobacteria>Chromatiales>unclassified_Chromatiales | GCA_020445605.1 | Chromatiales_bacterium | KDG50_02880 | 27912..31355 | 2026725 | BREX-system-P-loop-protein-BrxC-[Chromatiales-bacterium] |
| MCK7578720.1 | Type1.BREX | BrxA-DNA-binding->BrxB-BrxF-iSTAND->DUF6079-p->DUF6079-p*->DUF6079-p->PhdYeFM_antitox->RelE-ParE->N6-METHYLASE [illegible] | 647 | Bacteria>Pseudomonadota>Gammaproteobacteria>Chromatiales>unclassified_Chromatiales | GCA_023156155.1 | Chromatiales_bacterium | MZV65_25270 | 40289..42232 | 2026725 | BREX-system-P-loop-protein-BrxC-[Chromatiales-bacterium] |
| MCK7579547.1 | Type1.BREX | <-OLD-ABC+TOPRIM<-?<-?<-N6-METHYLASE<-PIN<-OLD-ABC\|\|?-><-DUF6079-p<-DUF6079-p*<-?<-DUF6079-p<-BrxB-BrxF-iSTAND [illegible] | 299 | Bacteria>Pseudomonadota>Gammaproteobacteria>Chromatiales>unclassified_Chromatiales | GCA_023156155.1 | Chromatiales_bacterium | MZV65_30115 | 14491..15390 | 2026725 | hypothetical-protein-[Chromatiales-bacterium] |
| MCK7579549.1 | Type1.BREX | <-N6-METHYLASE<-PIN<-OLD-ABC\|\|?-><-DUF6079-p<-DUF6079-p<-?<-DUF6079-p*<-BrxB-BrxF-iSTAND<-BrxA-DNA-binding-p<- [illegible] | 648 | Bacteria>Pseudomonadota>Gammaproteobacteria>Chromatiales>unclassified_Chromatiales | GCA_023156155.1 | Chromatiales_bacterium | MZV65_30125 | 15595..17541 | 2026725 | BREX-system-P-loop-protein-BrxC-[Chromatiales-bacterium] |
| RFF27207.1 | Type1.BREX | <-SbcC+SbcC<-?<-HEPN<-?\|\|Integrase_1-><-?<-SIGMA+OB-Fold+AAA-ATPase+Lon-Protease-BrxL<-PglZ<-N6-METHYLASE<-DUF [illegible] | 1166 | Bacteria>Pseudomonadota>Gammaproteobacteria>Chromatiales>Wenzhouxiangellaceae>Wenzhouxiangella | GCA_003410035.1 | Wenzhouxiangella_sp._15181 | DZK25_09525 | 145045..148545 | 2301224 | BREX-system-P-loop-protein-BrxC-[Wenzhouxiangella-sp.-15181] |
| WP_181918413.1 | Type1.BREX | <-SbcC+SbcC<-?<-HEPN<-?\|\|Integrase_1-><-?<-SIGMA+OB-Fold+AAA-ATPase+Lon-Protease-BrxL<-PglZ<-N6-METHYLASE<-DUF [illegible] | 1149 | Bacteria>Pseudomonadota>Gammaproteobacteria>Chromatiales>Wenzhouxiangellaceae>Wenzhouxiangella | GCF_003410035.1 | unclassified_Wenzhouxiangella | DZK25_RS09530 | 145045..148494 | 2613841 | MULTISPECIES:-BREX-system-P-loop-protein-BrxC-[unclassified-Wenzhouxiangella] |
| RJT19525.1 | Type1.BREX | <-PglZ<-RVT<-HEPN<-N6-METHYLASE<-ParB+HNH<-DUF6079*<-BrxB-BrxF-iSTAND<-BrxA-DNA- [illegible] | 1233 | Bacteria>Pseudomonadota>Gammaproteobacteria>Enterobacterales>Enterobacteriaceae>Buttiauxella | GCA_003601925.1 | Buttiauxella_izardii | D6029_18525 | 11062..14763 | 82991 | BREX-system-P-loop-protein-BrxC-[Buttiauxella-izardii] |
| WP_244212427.1 | Type1.BREX | <-PglZ<-N6-METHYLASE-p<-N6-METHYLASE-p<-RVT<-HEPN<-N6-METHYLASE<-ParB+HNH<-DUF607 [illegible] | 1237 | Bacteria>Pseudomonadota>Gammaproteobacteria>Enterobacterales>Enterobacteriaceae>Buttiauxella | GCF_003601925.1 | Buttiauxella_izardii | D6029_RS18475 | 11062..14775 | 82991 | BREX-system-P-loop-protein-BrxC-[Buttiauxella-izardii] |
| WP_124023337.1 | Type1.BREX | <-sGTP<-wHTH+WYL<-NACHT+SF1-SF2-Helicase+HKD-DNase\|\|SF1-SF2-Helicase+REase->SF1- [illegible] | 1157 | Bacteria>Pseudomonadota>Gammaproteobacteria>Enterobacterales>Enterobacteriaceae>Buttiauxella | GCF_003818135.1 | Buttiauxella_warmboldiae | EHN07_RS06350 | 61248..64721 | 82993 | BREX-system-P-loop-protein-BrxC-[Buttiauxella-warmboldiae] |
| WP_167614622.1 | Type1.BREX | <-HerA/FtsK+A32\|\|?-><-?<-SAM-METHYLASE\|\|tRNA->Arm-DNA-bind_3+Phage_integrase-><-? [illegible] | 1213 | Bacteria>Pseudomonadota>Gammaproteobacteria>Enterobacterales>Enterobacteriaceae>Cedecea | GCF_011808225.1 | Cedecea_colo | E2L00_RS19595 | 113572..117213 | 2552946 | BREX-system-P-loop-protein-BrxC-[Cedecea-colo] |
| WP_094760992.1 | Type1.BREX | XIS-HTH->?-><-Sulfatase<-?<-?<-?<-PHP+SbcC\|\|BrxA-DNA-binding->BrxB-BrxF-iSTAND-> [illegible] | 1233 | Bacteria>Pseudomonadota>Gammaproteobacteria>Enterobacterales>Enterobacteriaceae>Citrobacter | - | Citrobacter_braakii | B9P84_05295 | 62196..65897 | 57706 | MULTISPECIES:-BREX-system-P-loop-protein-BrxC-[Citrobacter] |
| WP_181218785.1 | Type1.BREX | CbeA_antitoxin->?->METHYLASE->HTH_3->?->REase-><-SIGMA+OB-Fold+AAA-ATPase+Lon-Pro [illegible] | 1213 | Bacteria>Pseudomonadota>Gammaproteobacteria>Enterobacterales>Enterobacteriaceae>Citrobacter | - | Citrobacter_freundii | HV183_02915 | 608608..612249 | 546 | BREX-system-P-loop-protein-BrxC-[Citrobacter-freundii] |
| EJM8694624.1 | Type1.BREX | DOC+HTH-><-?\|\|Phage_Cox->XIS-HTH->AAA+wHTH->?-><-SIGMA+OB-Fold+AAA-ATPase+Lon-Pro [illegible] | 1213 | Bacteria>Pseudomonadota>Gammaproteobacteria>Enterobacterales>Enterobacteriaceae>Citrobacter | - | Citrobacter_freundii | NO746_003414 | 23548..27189 | 546 | BREX-system-P-loop-protein-BrxC-[Citrobacter-freundii] |
| WP_182347709.1 | Type1.BREX | <-Schlafen+HATPase+wHTH<-PHP+SbcC<-?\|\|BrxA-DNA-binding->BrxB-BrxF-iSTAND->DUF6079 [illegible] | 1233 | Bacteria>Pseudomonadota>Gammaproteobacteria>Enterobacterales>Enterobacteriaceae>Citrobacter | - | Citrobacter_sp._RHBSTW-00013 | HV034_17810 | 3734345..3738046 | 2742628 | BREX-system-P-loop-protein-BrxC-[Citrobacter-sp.-RHBSTW-00013] |
| WP_181630678.1 | Type1.BREX | <-Sulfatase<-?<-?<-?<-Schlafen+HATPase+wHTH<-PHP+SbcC<-?\|\|BrxA-DNA-binding->BrxB- [illegible] | 1233 | Bacteria>Pseudomonadota>Gammaproteobacteria>Enterobacterales>Enterobacteriaceae>Citrobacter | - | Citrobacter_sp._RHBSTW-01013 | HV350_21470 | 4470372..4474073 | 2742677 | BREX-system-P-loop-protein-BrxC-[Citrobacter-sp.-RHBSTW-01013] |
| OXU12831.1 | Type1.BREX | XIS-HTH->?-><-Sulfatase<-?<-?<-?<-PHP+SbcC\|\|BrxA-DNA-binding->BrxB-BrxF-iSTAND-> [illegible] | 1233 | Bacteria>Pseudomonadota>Gammaproteobacteria>Enterobacterales>Enterobacteriaceae>Citrobacter | GCA_002239605.1 | Citrobacter_braakii | B9P84_05295 | 62196..65897 | 57706 | DNA-repair-protein-[Citrobacter-braakii] |
| QLR26611.1 | Type1.BREX | <-Sulfatase<-?<-?<-?<-Schlafen+HATPase+wHTH<-PHP+SbcC<-?\|\|BrxA-DNA-binding->BrxB- [illegible] | 1233 | Bacteria>Pseudomonadota>Gammaproteobacteria>Enterobacterales>Enterobacteriaceae>Citrobacter | GCA_013781985.1 | Citrobacter_sp._RHBSTW-01013 | HV350_21470 | 4470372..4474073 | 2742677 | BREX-system-P-loop-protein-BrxC-[Citrobacter-sp.-RHBSTW-01013] |
| MBA8131541.1 | Type1.BREX | <-Schlafen+HATPase+wHTH<-PHP+SbcC<-?\|\|BrxA-DNA-binding->BrxB-BrxF-iSTAND->DUF6079 [illegible] | 1233 | Bacteria>Pseudomonadota>Gammaproteobacteria>Enterobacterales>Enterobacteriaceae>Citrobacter | GCA_014104375.1 | Citrobacter_sp._RHBSTW-00013 | HV034_17810 | 3734345..3738046 | 2742628 | BREX-system-P-loop-protein-BrxC-[Citrobacter-sp.-RHBSTW-00013] |
| MBM9568541.1 | Type1.BREX | <-wHTH+WYL<-DUF932<-nSTAND1+GTPase-AIG+sGTP<-CopG/RHH\|\|XIS-HTH->?->?->BrxA-DNA-bi [illegible] | 1233 | Bacteria>Pseudomonadota>Gammaproteobacteria>Enterobacterales>Enterobacteriaceae>Citrobacter | GCA_014931625.2 | Citrobacter_sedlakii | INP82_14060 | 158441..162142 | 67826 | BREX-system-P-loop-protein-BrxC-[Citrobacter-sedlakii] |
| EGT0661408.1 | Type1.BREX | REase+SPF-cons->?->?-><-?<-Schlafen+HATPase+wHTH<-PHP+SbcC<-?\|\|BrxA-DNA-binding-> [illegible] | 1233 | Bacteria>Pseudomonadota>Gammaproteobacteria>Enterobacterales>Enterobacteriaceae>Citrobacter | GCA_015942525.1 | Citrobacter_werkmanii | JAF83_001575 | 366383..370084 | 67827 | BREX-system-P-loop-protein-BrxC-[Citrobacter-werkmanii] |
| HAT7499716.1 | Type1.BREX | <-Peptidase_M17_N+Peptidase_M17\|\|?->?-><-HerA/FtsK+A32<-?\|\|tRNA->Arm-DNA-bind_3+P [illegible] | 1157 | Bacteria>Pseudomonadota>Gammaproteobacteria>Enterobacterales>Enterobacteriaceae>Citrobacter | GCA_015956985.1 | Citrobacter_freundii | JAV71_002653 | 147091..150564 | 546 | BREX-system-P-loop-protein-BrxC-[Citrobacter-freundii] |
| HAU4366733.1 | Type1.BREX | SKI-><-?\|\|tRNA->Arm-DNA-bind_3+Phage_integrase-><-?<-SIGMA+OB-Fold+AAA-ATPase+Lon [illegible] | 1213 | Bacteria>Pseudomonadota>Gammaproteobacteria>Enterobacterales>Enterobacteriaceae>Citrobacter | GCA_016081455.1 | Citrobacter_amalonaticus | F6W26_02175 | 132901..136542 | 35703 | BREX-system-P-loop-protein-BrxC-[Citrobacter-amalonaticus] |
| HBC2890663.1 | Type1.BREX | BrxA-DNA-binding->BrxB-BrxF-iSTAND->DUF6079*->N6-METHYLASE->OLD-ABC->TOPRIM->PglZ [illegible] | 1204 | Bacteria>Pseudomonadota>Gammaproteobacteria>Enterobacterales>Enterobacteriaceae>Citrobacter | GCA_018089985.1 | Citrobacter_freundii | JD321_004755 | 1401..5015 | 546 | BREX-system-P-loop-protein-BrxC-[Citrobacter-freundii] |
| HCL6754282.1 | Type1.BREX | <-HerA/FtsK+A32<-?\|\|tRNA->Arm-DNA-bind_3+Phage_integrase-><-HKD-DNase+OGG-like<- [illegible] | 1213 | Bacteria>Pseudomonadota>Gammaproteobacteria>Enterobacterales>Enterobacteriaceae>Citrobacter | GCA_025039155.1 | Citrobacter_freundii | N2R09_003052 | 112932..116573 | 546 | BREX-system-P-loop-protein-BrxC-[Citrobacter-freundii] |
| SYX30363.1 | Type1.BREX | GDResolvase+HIN-HTH->MPTase-><-SIGMA+OB-Fold+AAA-ATPase+Lon-Protease-BrxL<-PglZ<- [illegible] | 1233 | Bacteria>Pseudomonadota>Gammaproteobacteria>Enterobacterales>Enterobacteriaceae>Citrobacter | GCA_900520375.1 | Citrobacter_freundii | CFU2785_04435 | 4465123..4468824 | 546 | Uncharacterised-protein-[Citrobacter-freundii] |
| WP_044264596.1 | Type1.BREX | DDE_Tnp_IS240->BrxA-DNA-binding->BrxB-BrxF-iSTAND->DUF6079*->N6-METHYLASE->OLD-AB [illegible] | 1213 | Bacteria>Pseudomonadota>Gammaproteobacteria>Enterobacterales>Enterobacteriaceae>Citrobacter | GCF_000731055.1 | Citrobacter_amalonaticus | DQ02_RS05315 | 398614..402255 | 35703 | BREX-system-P-loop-protein-BrxC-[Citrobacter-amalonaticus] |
| WP_046487315.1 | Type1.BREX | <-HIN-HTH\|\|<-?\|\|?->?->BrxA-DNA-binding->BrxB-BrxF-iSTAND->DUF6079*->PglZ->SIGMA+O [illegible] | 1213 | Bacteria>Pseudomonadota>Gammaproteobacteria>Enterobacterales>Enterobacteriaceae>Citrobacter | GCF_000981805.1 | Citrobacter_amalonaticus | F384_RS16675 | 3635289..3638930 | 35703 | BREX-system-P-loop-protein-BrxC-[Citrobacter-amalonaticus] |
| WP_231587318.1 | Type1.BREX | <-wHTH+WYL<-nSTAND1+sGTP<-?\|\|XIS-HTH->?->?->BrxA-DNA-binding->BrxB-BrxF-iSTAND-> [illegible] | 1240 | Bacteria>Pseudomonadota>Gammaproteobacteria>Enterobacterales>Enterobacteriaceae>Citrobacter | GCF_001037515.1 | Citrobacter_sp._MGH110 | SK38_RS14780 | 3143856..3147578 | 1686383 | BREX-system-P-loop-protein-BrxC-[Citrobacter-sp.-MGH110] |
| WP_121581750.1 | Type1.BREX | SF1-SF2-Helicase-><-?<-?<-?<-wHTH+WYL<-DUF932<-nSTAND1+GTPase-AIG+sGTP\|\|BrxA-DNA- [illegible] | 1204 | Bacteria>Pseudomonadota>Gammaproteobacteria>Enterobacterales>Enterobacteriaceae>Citrobacter | GCF_003665675.1 | Citrobacter_freundii | CUC48_RS18615 | 3736317..3739931 | 546 | BREX-system-P-loop-protein-BrxC-[Citrobacter-freundii] |
| WP_135912646.1 | Type1.BREX | <-PAS+AraC-HTH+AraC-HTH\|\|?->?->?->?->SymE_toxin-><-Glyoxalase_8\|\|BrxA-DNA-binding [illegible] | 1213 | Bacteria>Pseudomonadota>Gammaproteobacteria>Enterobacterales>Enterobacteriaceae>Citrobacter | GCF_004792475.1 | Citrobacter_freundii | E5284_RS20140 | 4206675..4210316 | 546 | BREX-system-P-loop-protein-BrxC-[Citrobacter-freundii] |
| WP_149691146.1 | Type1.BREX | <-HTH_36<-HTH+HTH<-ParB+BAM<-XIS-HTH<-wHTH<-ZnR\|\|BrxA-DNA-binding->BrxB-BrxF-iSTA [illegible] | 1204 | Bacteria>Pseudomonadota>Gammaproteobacteria>Enterobacterales>Enterobacteriaceae>Citrobacter | GCF_008373795.1 | Citrobacter_pasteurii | DXF85_RS00930 | 181362..184976 | 1563222 | BREX-system-P-loop-protein-BrxC-[Citrobacter-pasteurii] |
| WP_129695037.1 | Type1.BREX | Glyoxalase_8->URI-DNase-><-RHH_antitoxin\|\|?->XIS-HTH->?->ZnR->BrxA-DNA-binding-> [illegible] | 1213 | Bacteria>Pseudomonadota>Gammaproteobacteria>Enterobacterales>Enterobacteriaceae>Citrobacter | GCF_013618825.1 | Citrobacter_werkmanii | EVY00_RS24285 | 8302..11943 | 67827 | BREX-system-P-loop-protein-BrxC-[Citrobacter-werkmanii] |
| WP_249418019.1 | Type1.BREX | <-Sulfatase<-?<-?<-?<-Schlafen+HATPase+wHTH<-PHP+SbcC<-?\|\|BrxA-DNA-binding->BrxB- [illegible] | 1240 | Bacteria>Pseudomonadota>Gammaproteobacteria>Enterobacterales>Enterobacteriaceae>Citrobacter | GCF_013781985.1 | Citrobacter_sp._RHBSTW-01013 | HV350_RS21465 | 4470351..4474073 | 2742677 | BREX-system-P-loop-protein-BrxC-[Citrobacter-sp.-RHBSTW-01013] |
| WP_181621180.1 | Type1.BREX | SF1-SF2-Helicase-><-?<-?<-?<-wHTH+WYL<-DUF932<-nSTAND1+GTPase-AIG+sGTP\|\|BrxA-DNA- [illegible] | 1204 | Bacteria>Pseudomonadota>Gammaproteobacteria>Enterobacterales>Enterobacteriaceae>Citrobacter | GCF_013791615.1 | Citrobacter_freundii | HV083_RS19895 | 4141726..4145340 | 546 | BREX-system-P-loop-protein-BrxC-[Citrobacter-freundii] |
| WP_249424276.1 | Type1.BREX | <-Schlafen+HATPase+wHTH<-PHP+SbcC<-?\|\|BrxA-DNA-binding->BrxB-BrxF-iSTAND->DUF6079 [illegible] | 1240 | Bacteria>Pseudomonadota>Gammaproteobacteria>Enterobacterales>Enterobacteriaceae>Citrobacter | GCF_014104375.1 | Citrobacter_sp._RHBSTW-00013 | HV034_RS17810 | 3734324..3738046 | 2742628 | BREX-system-P-loop-protein-BrxC-[Citrobacter-sp.-RHBSTW-00013] |
| WP_192466357.1 | Type1.BREX | <-HerA/FtsK+A32<-?\|\|tRNA->Arm-DNA-bind_3+Phage_integrase-><-HKD-DNase+OGG-like<- [illegible] | 1213 | Bacteria>Pseudomonadota>Gammaproteobacteria>Enterobacterales>Enterobacteriaceae>Citrobacter | GCF_014855915.1 | Citrobacter_koseri | FOT44_RS18785 | 338007..341648 | 545 | BREX-system-P-loop-protein-BrxC-[Citrobacter-koseri] |
| WP_200025358.1 | Type1.BREX | Glyoxalase_8->URI-DNase-><-RHH_antitoxin\|\|?->XIS-HTH->?->ZnR->BrxA-DNA-binding-> [illegible] | 1213 | Bacteria>Pseudomonadota>Gammaproteobacteria>Enterobacterales>Enterobacteriaceae>Citrobacter | GCF_016500865.1 | Citrobacter_freundii | I5388_RS15310 | 359245..362886 | 546 | BREX-system-P-loop-protein-BrxC-[Citrobacter-freundii] |
| WP_200031143.1 | Type1.BREX | <-HerA/FtsK+A32<-?\|\|tRNA->Arm-DNA-bind_3+Phage_integrase-><-?<-SIGMA+OB-Fold+AAA- [illegible] | 1213 | Bacteria>Pseudomonadota>Gammaproteobacteria>Enterobacterales>Enterobacteriaceae>Citrobacter | GCF_016501985.1 | Citrobacter_koseri | I5424_RS14480 | 22464..26105 | 545 | BREX-system-P-loop-protein-BrxC-[Citrobacter-koseri] |
| WP_200054816.1 | Type1.BREX | <-Peptidase_M17_N+Peptidase_M17\|\|?->?-><-HerA/FtsK+A32<-?\|\|tRNA->Arm-DNA-bind_3+P [illegible] | 1157 | Bacteria>Pseudomonadota>Gammaproteobacteria>Enterobacterales>Enterobacteriaceae>Citrobacter | GCF_016504925.1 | Citrobacter_freundii | GHT45_RS09240 | 237227..240700 | 546 | BREX-system-P-loop-protein-BrxC-[Citrobacter-freundii] |
| WP_203457741.1 | Type1.BREX | <-ArdB<-DUF932<-CopG/RHH\|\|?->XIS-HTH->ZnR->BrxA-DNA-binding->BrxB-BrxF-iSTAND-> [illegible] | 1204 | Bacteria>Pseudomonadota>Gammaproteobacteria>Enterobacterales>Enterobacteriaceae>Citrobacter | GCF_016859695.1 | Citrobacter_braakii | GF710_RS09095 | 85023..88637 | 57706 | BREX-system-P-loop-protein-BrxC-[Citrobacter-braakii] |
| WP_205688206.1 | Type1.BREX | <-MAD-NTD+HNH<-HTH_Tnp_1+HTH_21+rve+rve_3\|\|?->?->SF1-SF2-Helicase+REase->SF1-SF2- [illegible] | 1157 | Bacteria>Pseudomonadota>Gammaproteobacteria>Enterobacterales>Enterobacteriaceae>Citrobacter | GCF_017086505.1 | Citrobacter_freundii | GHT30_RS20265 | 4262904..4266377 | 546 | BREX-system-P-loop-protein-BrxC-[Citrobacter-freundii] |
| WP_241511105.1 | Type1.BREX | <-HerA/FtsK+A32<-?\|\|tRNA->Arm-DNA-bind_3+Phage_integrase-><-HKD-DNase+OGG-like<- [illegible] | 1197 | Bacteria>Pseudomonadota>Gammaproteobacteria>Enterobacterales>Enterobacteriaceae>Citrobacter | GCF_022585175.1 | Citrobacter_freundii | MMQ01_RS10405 | 144077..147670 | 546 | BREX-system-P-loop-protein-BrxC-[Citrobacter-freundii] |
| WP_252170698.1 | Type1.BREX | <-DUF905<-?<-?<-?<-wHTH+WYL<-DUF932<-nSTAND1+GTPase-AIG+sGTP\|\|BrxA-DNA-binding-> [illegible] | 1204 | Bacteria>Pseudomonadota>Gammaproteobacteria>Enterobacterales>Enterobacteriaceae>Citrobacter | GCF_023821945.1 | Citrobacter_cronae | M4W13_RS12100 | 107491..111105 | 1748967 | BREX-system-P-loop-protein-BrxC-[Citrobacter-cronae] |
| WP_275379687.1 | Type1.BREX | Arm-DNA-bind_3+Phage_integrase-><-SIGMA+OB-Fold+AAA-ATPase+Lon-Protease-BrxL<-Pgl [illegible] | 1240 | Bacteria>Pseudomonadota>Gammaproteobacteria>Enterobacterales>Enterobacteriaceae>Citrobacter | GCF_029104295.1 | Citrobacter_freundii | L2W90_RS09965 | 166217..169939 | 546 | BREX-system-P-loop-protein-BrxC-[Citrobacter-freundii] |
| WP_275399266.1 | Type1.BREX | <-DUF932<-nSTAND1+GTPase-AIG+sGTP<-?\|\|?->XIS-HTH->?->?->BrxA-DNA-binding->BrxB- [illegible] | 1095 | Bacteria>Pseudomonadota>Gammaproteobacteria>Enterobacterales>Enterobacteriaceae>Citrobacter | GCF_029104415.1 | Citrobacter_freundii | L2W96_RS12535 | 133812..137099 | 546 | BREX-system-P-loop-protein-BrxC-[Citrobacter-freundii] |
| WP_196010428.1 | Type1.BREX | Glyoxalase_8->URI-DNase-><-RHH_antitoxin\|\|?->XIS-HTH->?->ZnR->BrxA-DNA-binding-> [illegible] | 1213 | Bacteria>Pseudomonadota>Gammaproteobacteria>Enterobacterales>Enterobacteriaceae>Citrobacter | GCF_029717585.1 | Citrobacter_amalonaticus | NFJ54_RS22965 | 4694172..4697813 | 35703 | MULTISPECIES:-BREX-system-P-loop-protein-BrxC-[Enterobacteriaceae] |
| WP_148977713.1 | Type1.BREX | ZnR-><-NACHT+SF1-SF2-Helicase+HKD-DNase\|\|BrxA-DNA-binding->BrxB-BrxF-iSTAND->DUF6 [illegible] | 1204 | Bacteria>Pseudomonadota>Gammaproteobacteria>Enterobacterales>Enterobacteriaceae>Citrobacter | GCF_902807175.1 | Citrobacter_freundii | FYK57_RS19440 | 6782..10396 | 546 | BREX-system-P-loop-protein-BrxC-[Citrobacter-freundii] |
| PUV47226.1 | Type1.BREX | <-wHTH+WYL<-sGTP<-?\|\|XIS-HTH->?->?->BrxA-DNA-binding->BrxB-BrxF-iSTAND->DUF6079*- [illegible] | 1233 | Bacteria>Pseudomonadota>Gammaproteobacteria>Enterobacterales>Enterobacteriaceae>Cronobacter | GCA_002532375.1 | Cronobacter_sakazakii | CDU02_01545 | 336773..340474 | 28141 | BREX-system-P-loop-protein-BrxC-[Cronobacter-sakazakii] |
| KAB0815149.1 | Type1.BREX | <-wHTH(SBD)+HNH\|\|tRNA->Arm-DNA-bind_3+Phage_integrase-><-SF1-SF2-Helicase\|\|OLD-AB [illegible] | 1233 | Bacteria>Pseudomonadota>Gammaproteobacteria>Enterobacterales>Enterobacteriaceae>Cronobacter | GCA_008805175.1 | Cronobacter_sakazakii | FZI24_17905 | 37972..41673 | 28141 | BREX-system-P-loop-protein-BrxC-[Cronobacter-sakazakii] |
| EGT4327973.1 | Type1.BREX | BrxA-DNA-binding->BrxB-BrxF-iSTAND->DUF6079*->N6-METHYLASE->PglZ->SIGMA+OB-Fold+ [illegible] | 1213 | Bacteria>Pseudomonadota>Gammaproteobacteria>Enterobacterales>Enterobacteriaceae>Cronobacter | GCA_016076655.1 | Cronobacter_sakazakii | AUM89_20300 | 1368..5009 | 28141 | BREX-system-P-loop-protein-BrxC-[Cronobacter-sakazakii] |
| EGT4509729.1 | Type1.BREX | URI-DNase-><-RHH_antitoxin\|\|?->XIS-HTH->?-><-?\|\|ZnR->BrxA-DNA-binding->BrxB-BrxF- [illegible] | 1213 | Bacteria>Pseudomonadota>Gammaproteobacteria>Enterobacterales>Enterobacteriaceae>Cronobacter | GCA_016077415.1 | Cronobacter_sakazakii | AUN08_13950 | 41596..45237 | 28141 | BREX-system-P-loop-protein-BrxC-[Cronobacter-sakazakii] |
| WP_007729758.1 | Type1.BREX | <-REC+LuxR-HTH<-SAM-METHYLASE<-wHTH+HNH\|\|tRNA->Arm-DNA-bind_3+Phage_integrase-><- [illegible] | 1213 | Bacteria>Pseudomonadota>Gammaproteobacteria>Enterobacterales>Enterobacteriaceae>Cronobacter | GCF_000409225.1 | Cronobacter_dublinensis | AFK67_RS02785 | 592165..595806 | 413497 | BREX-system-P-loop-protein-BrxC-[Cronobacter-dublinensis] |
| WP_075184066.1 | Type1.BREX | <-TarH+HAMP+MA\|\|ACET->DOC+HTH-><-?\|\|BrxA-DNA-binding->BrxB-BrxF-iSTAND->DUF6079*- [illegible] | 1213 | Bacteria>Pseudomonadota>Gammaproteobacteria>Enterobacterales>Enterobacteriaceae>Cronobacter | GCF_001922925.1 | Cronobacter_sakazakii | BS410_RS19680 | 61425..65066 | 28141 | BREX-system-P-loop-protein-BrxC-[Cronobacter-sakazakii] |
| WP_239984508.1 | Type1.BREX | <-wHTH+WYL<-GTPase-AIG<-?\|\|XIS-HTH->?-><-TIR\|\|BrxA-DNA-binding->BrxB-BrxF-iSTAND- [illegible] | 1240 | Bacteria>Pseudomonadota>Gammaproteobacteria>Enterobacterales>Enterobacteriaceae>Cronobacter | GCF_002942265.1 | Cronobacter_sakazakii | C3D67_RS10955 | 271432..275154 | 28141 | BREX-system-P-loop-protein-BrxC-[Cronobacter-sakazakii] |
| WP_105547575.1 | Type1.BREX | Phage_Cox->XIS-HTH->AAA+wHTH->?-><-Zn_Tnp_IS1+HTH_Tnp_IS1+DDE_Tnp_IS1<-HKD-DNase+ [illegible] | 1213 | Bacteria>Pseudomonadota>Gammaproteobacteria>Enterobacterales>Enterobacteriaceae>Cronobacter | GCF_002974775.1 | Cronobacter_sakazakii | C5959_RS02070 | 327806..331447 | 28141 | BREX-system-P-loop-protein-BrxC-[Cronobacter-sakazakii] |
| WP_105579035.1 | Type1.BREX | Glyoxalase_8->URI-DNase-><-RHH_antitoxin\|\|?->XIS-HTH->?->ZnR->BrxA-DNA-binding-> [illegible] | 1213 | Bacteria>Pseudomonadota>Gammaproteobacteria>Enterobacterales>Enterobacteriaceae>Cronobacter | GCF_002977115.1 | Cronobacter_sakazakii | CI818_RS20655 | 12068..15709 | 28141 | BREX-system-P-loop-protein-BrxC-[Cronobacter-sakazakii] |
| WP_110812233.1 | Type1.BREX | XIS-HTH->?->AAA+HTH-><-SIGMA+OB-Fold+AAA-ATPase+Lon-Protease-BrxL<-PglZ<-N6-METHY [illegible] | 1213 | Bacteria>Pseudomonadota>Gammaproteobacteria>Enterobacterales>Enterobacteriaceae>Cronobacter | GCF_003206925.1 | Cronobacter_sakazakii | DOM37_RS13070 | 492836..496477 | 28141 | BREX-system-P-loop-protein-BrxC-[Cronobacter-sakazakii] |
| WP_161575571.1 | Type1.BREX | <-Peptidase_M17_N+Peptidase_M17\|\|?->?->?->?-><-?\|\|tRNA->Arm-DNA-bind_3+Phage_inte [illegible] | 1157 | Bacteria>Pseudomonadota>Gammaproteobacteria>Enterobacterales>Enterobacteriaceae>Cronobacter | GCF_009938355.1 | Cronobacter_sakazakii | EHJ08_RS11755 | 205127..208600 | 28141 | BREX-system-P-loop-protein-BrxC-[Cronobacter-sakazakii] |
| WP_166500158.1 | Type1.BREX | BrxA-DNA-binding->DUF6079*->N6-METHYLASE->PglZ->SIGMA+OB-Fold+AAA-ATPase+Lon-Prot [illegible] | 1213 | Bacteria>Pseudomonadota>Gammaproteobacteria>Enterobacterales>Enterobacteriaceae>Cronobacter | GCF_011605595.1 | Cronobacter_sakazakii | G9X52_RS02005 | 2082..5723 | 28141 | BREX-system-P-loop-protein-BrxC-[Cronobacter-sakazakii] |
| WP_166501056.1 | Type1.BREX | MarR-HTH->?-><-?<-?<-?<-HTH_19<-REase\|\|BrxA-DNA-binding->BrxB-BrxF-iSTAND->DUF607 [illegible] | 1213 | Bacteria>Pseudomonadota>Gammaproteobacteria>Enterobacterales>Enterobacteriaceae>Cronobacter | GCF_011605635.1 | Cronobacter_dublinensis | G9X53_RS16040 | 318383..322024 | 413497 | BREX-system-P-loop-protein-BrxC-[Cronobacter-dublinensis] |
| WP_239978191.1 | Type1.BREX | <-SAM-METHYLASE<-wHTH(SBD)+HNH\|\|tRNA->Arm-DNA-bind_3+Phage_integrase-><-SIGMA+OB- [illegible] | 1240 | Bacteria>Pseudomonadota>Gammaproteobacteria>Enterobacterales>Enterobacteriaceae>Cronobacter | GCF_017845055.1 | Cronobacter_sakazakii | KAR72_RS09245 | 217726..221448 | 28141 | BREX-system-P-loop-protein-BrxC-[Cronobacter-sakazakii] |
| WP_269506681.1 | Type1.BREX | <-wHTH+WYL<-DUF932<-nSTAND1+GTPase-AIG+sGTP\|\|BrxA-DNA-binding->BrxB-BrxF-iSTAND-> [illegible] | 1204 | Bacteria>Pseudomonadota>Gammaproteobacteria>Enterobacterales>Enterobacteriaceae>Cronobacter | GCF_027277125.1 | Cronobacter_sakazakii | O6D31_RS12675 | 364123..367737 | 28141 | BREX-system-P-loop-protein-BrxC-[Cronobacter-sakazakii] |
| CAF2998376.1 | Type1.BREX | Arm-DNA-bind_3+Phage_integrase-><-SIGMA+OB-Fold+AAA-ATPase+Lon-Protease-BrxL<-Pgl [illegible] | 1240 | Bacteria>Pseudomonadota>Gammaproteobacteria>Enterobacterales>Enterobacteriaceae>Enterobacter | - | Enterobacter_cloacae | AI2972V1_0508 | 557695..561417 | 550 | hypothetical-protein-AI2972V1_0508-[Enterobacter-cloacae] |
| WP_129258922.1 | Type1.BREX | <-CopG/RHH\|\|XIS-HTH->?->?->SF1-SF2-Helicase+REase->SF1-SF2-Helicase+REase-><-PglZ [illegible] | 1157 | Bacteria>Pseudomonadota>Gammaproteobacteria>Enterobacterales>Enterobacteriaceae>Enterobacter | - | Enterobacter_cloacae_complex_sp._GF14B | DD606_17375 | 24450..27923 | 2511982 | BREX-system-P-loop-protein-BrxC-[Enterobacter-cloacae-complex-sp.-GF14B] |
| WP_163363286.1 | Type1.BREX | Arm-DNA-bind_3+Phage_integrase-><-HKD-DNase+OGG-like<-SIGMA+OB-Fold+AAA-ATPase+Lo [illegible] | 1197 | Bacteria>Pseudomonadota>Gammaproteobacteria>Enterobacterales>Enterobacteriaceae>Enterobacter | - | Enterobacter_cloacae | G4556_RS12910 | 16910..20503 | 550 | BREX-system-P-loop-protein-BrxC-[Enterobacter-cloacae] |
| WP_203465577.1 | Type1.BREX | nSTAND1+HNH->RNaseH->NACHT+SF1-SF2-Helicase+HKD-DNase->REase-><-HKD-DNase+OGG-lik [illegible] | 1213 | Bacteria>Pseudomonadota>Gammaproteobacteria>Enterobacterales>Enterobacteriaceae>Enterobacter | - | Enterobacter_cloacae | GEA59_RS02615 | 551852..555493 | 550 | BREX-system-P-loop-protein-BrxC-[Enterobacter-cloacae] |
| WP_039268698.1 | Type1.BREX | Arm-DNA-bind_3+Phage_integrase-><-SIGMA+OB-Fold+AAA-ATPase+Lon-Protease-BrxL<-Pgl [illegible] | 1233 | Bacteria>Pseudomonadota>Gammaproteobacteria>Enterobacterales>Enterobacteriaceae>Enterobacter | - | Enterobacter_hormaechei | LI66_02640 | 557969..561670 | 158836 | MULTISPECIES:-BREX-system-P-loop-protein-BrxC-[Enterobacter] |
| WP_181507052.1 | Type1.BREX | REase+SPF-cons->?->?-><-?<-Schlafen+HATPase+wHTH<-PHP+SbcC<-?\|\|BrxA-DNA-binding-> [illegible] | 1233 | Bacteria>Pseudomonadota>Gammaproteobacteria>Enterobacterales>Enterobacteriaceae>Enterobacter | - | Enterobacter_roggenkampii | JAF83_001575 | 366383..370084 | 1812935 | MULTISPECIES:-BREX-system-P-loop-protein-BrxC-[Enterobacteriaceae] |
| WP_212478308.1 | Type1.BREX | SymE_toxin->wHTH+HNH->SAM-METHYLASE->?->DUF2754->?->DOC+HTH->BrxA-DNA-binding->Br [illegible] | 1213 | Bacteria>Pseudomonadota>Gammaproteobacteria>Enterobacterales>Enterobacteriaceae>Enterobacter | - | Enterobacter_sp._JGM127 | JK158_RS17050 | 116414..120055 | 2799790 | BREX-system-P-loop-protein-BrxC-[Enterobacter-sp.-JGM127] |
| AFP68425.1 | Type1.BREX | Arm-DNA-bind_3+Phage_integrase-><-SIGMA+OB-Fold+AAA-ATPase+Lon-Protease-BrxL<-Pgl [illegible] | 1233 | Bacteria>Pseudomonadota>Gammaproteobacteria>Enterobacterales>Enterobacteriaceae>Enterobacter | GCA_000286275.1 | Enterobacter_kobei | ECENHK_02660 | 581253..584954 | 208224 | hypothetical-protein-ECENHK_02660-[Enterobacter-kobei] |
| ESM83248.1 | Type1.BREX | CbeA_antitoxin->?-><-?<-SIGMA+OB-Fold+AAA-ATPase+Lon-Protease-BrxL<-PglZ<-N6-METH [illegible] | 611 | Bacteria>Pseudomonadota>Gammaproteobacteria>Enterobacterales>Enterobacteriaceae>Enterobacter | GCA_000492895.1 | Enterobacter_roggenkampii_MGH_34 | L380_02580 | 2783361..2785196 | 1329832 | hypothetical-protein-L380_02580-[Enterobacter-roggenkampii-MGH-34] |
| AJB80307.1 | Type1.BREX | Arm-DNA-bind_3+Phage_integrase-><-SIGMA+OB-Fold+AAA-ATPase+Lon-Protease-BrxL<-Pgl [illegible] | 1233 | Bacteria>Pseudomonadota>Gammaproteobacteria>Enterobacterales>Enterobacteriaceae>Enterobacter | GCA_000814225.1 | Enterobacter_hormaechei_subsp._xiangfangensis | LI66_02640 | 557969..561670 | 1296536 | DNA-repair-protein-[Enterobacter-hormaechei-subsp.-xiangfangensis] |
| KZR28106.1 | Type1.BREX | <-wHTH+WYL<-sGTP<-?\|\|XIS-HTH->?->?-><-?\|\|BrxA-DNA-binding->BrxB-BrxF-iSTAND->DUF6 [illegible] | 1233 | Bacteria>Pseudomonadota>Gammaproteobacteria>Enterobacterales>Enterobacteriaceae>Enterobacter | GCA_001631925.1 | Enterobacter_hormaechei_subsp._steigerwaltii | A3N67_04030 | 129053..132754 | 299766 | DNA-repair-protein-[Enterobacter-hormaechei-subsp.-steigerwaltii] |
| AWF32098.1 | Type1.BREX | Arm-DNA-bind_3+Phage_integrase-><-SIGMA+OB-Fold+AAA-ATPase+Lon-Protease-BrxL<-Pgl [illegible] | 1233 | Bacteria>Pseudomonadota>Gammaproteobacteria>Enterobacterales>Enterobacteriaceae>Enterobacter | GCA_003073995.1 | Enterobacter_hormaechei | CSC19_3711 | 3849441..3853142 | 158836 | hypothetical-protein-CSC19_3711-[Enterobacter-hormaechei] |
| PWG69505.1 | Type1.BREX | NACHT+SF1-SF2-Helicase+HKD-DNase->REase-><-HKD-DNase+OGG-like<-SIGMA+OB-Fold+AAA- [illegible] | 1213 | Bacteria>Pseudomonadota>Gammaproteobacteria>Enterobacterales>Enterobacteriaceae>Enterobacter | GCA_003129945.1 | Enterobacter_mori | DEM28_19810 | 168758..172399 | 539813 | BREX-system-P-loop-protein-BrxC-[Enterobacter-mori] |
| RVR09283.1 | Type1.BREX | <-DUF6079-p*<-BrxB-BrxF-iSTAND<-BrxA-DNA-binding<-Glyoxalase<-ZnR<-METHYLASE | 859 | Bacteria>Pseudomonadota>Gammaproteobacteria>Enterobacterales>Enterobacteriaceae>Enterobacter | GCA_004004525.1 | Enterobacter_hormaechei | EOL34_23625 | 1..2579 | 158836 | BREX-system-P-loop-protein-BrxC,-partial-[Enterobacter-hormaechei] |
| MBG0523800.1 | Type1.BREX | Arm-DNA-bind_3+Phage_integrase-><-SIGMA+OB-Fold+AAA-ATPase+Lon-Protease-BrxL<-Pgl [illegible] | 1233 | Bacteria>Pseudomonadota>Gammaproteobacteria>Enterobacterales>Enterobacteriaceae>Enterobacter | GCA_015704905.1 | Enterobacter_cloacae | IOR24_17915 | 3774454..3778155 | 550 | BREX-system-P-loop-protein-BrxC-[Enterobacter-cloacae] |
| HAV2283855.1 | Type1.BREX | <-DUF6079-p*<-BrxB-BrxF-iSTAND<-BrxA-DNA-binding<-SAM-METHYLASE<-wHTH+HNH\|\|?->?-> [illegible] | 932 | Bacteria>Pseudomonadota>Gammaproteobacteria>Enterobacterales>Enterobacteriaceae>Enterobacter | GCA_016445985.1 | Enterobacter_cloacae | JHK37_004627 | 1..2796 | 550 | BREX-system-P-loop-protein-BrxC-[Enterobacter-cloacae] |
| MBK3145473.1 | Type1.BREX | Glyoxalase->ACET->YjhX_toxin->SAM-METHYLASE->ZnR->Glyoxalase->BrxA-DNA-binding->B [illegible] | 1233 | Bacteria>Pseudomonadota>Gammaproteobacteria>Enterobacterales>Enterobacteriaceae>Enterobacter | GCA_016616795.1 | Enterobacter_hormaechei | FKK11_12155 | 273388..277089 | 158836 | BREX-system-P-loop-protein-BrxC-[Enterobacter-hormaechei] |
| MBT2005130.1 | Type1.BREX | wHTH+SAD+HNH-><-NACHT+SF1-SF2-Helicase+HKD-DNase\|\|McrB-AAA->McrC-NTD+REase-><-SIG [illegible] | 1204 | Bacteria>Pseudomonadota>Gammaproteobacteria>Enterobacterales>Enterobacteriaceae>Enterobacter | GCA_018597745.1 | Enterobacter_hormaechei_subsp._xiangfangensis | KKZ39_07415 | 340677..344291 | 1296536 | BREX-system-P-loop-protein-BrxC-[Enterobacter-hormaechei-subsp.-xiangfangensis] |
| MBX8934563.1 | Type1.BREX | DEDD_Tnp_IS110+Transposase_20-><-SIGMA+OB-Fold+AAA-ATPase+Lon-Protease-BrxL<-PglZ [illegible] | 1225 | Bacteria>Pseudomonadota>Gammaproteobacteria>Enterobacterales>Enterobacteriaceae>Enterobacter | GCA_019731165.1 | Enterobacter_sp._K62_1 | HCH30_08625 | 10503..14180 | 2718933 | BREX-system-P-loop-protein-BrxC-[Enterobacter-sp.-K62_1] |
| HBL8846881.1 | Type1.BREX | DUF6079*->ParB+HNH->N6-METHYLASE->PglZ->SIGMA+OB-Fold+AAA-ATPase+Lon-Protease-Brx [illegible] | 1089 | Bacteria>Pseudomonadota>Gammaproteobacteria>Enterobacterales>Enterobacteriaceae>Enterobacter | GCA_021095225.1 | Enterobacter_hormaechei | LTW48_004335 | 1..3271 | 158836 | BREX-system-P-loop-protein-BrxC-[Enterobacter-hormaechei] |
| HBM2523095.1 | Type1.BREX | <-DUF6079-p*<-BrxB-BrxF-iSTAND<-BrxA-DNA-binding<-URI-DNase\|\|<-EVE+HNH\|\|SymE_toxi [illegible] | 599 | Bacteria>Pseudomonadota>Gammaproteobacteria>Enterobacterales>Enterobacteriaceae>Enterobacter | GCA_021175225.1 | Enterobacter_hormaechei | LUX61_004833 | 1..1799 | 158836 | BREX-system-P-loop-protein-BrxC-[Enterobacter-hormaechei] |
| HBM2647612.1 | Type1.BREX | <-DUF6079-p*<-BrxB-BrxF-iSTAND<-BrxA-DNA-binding<-URI-DNase\|\|<-EVE+HNH\|\|SymE_toxi [illegible] | 586 | Bacteria>Pseudomonadota>Gammaproteobacteria>Enterobacterales>Enterobacteriaceae>Enterobacter | GCA_021176145.1 | Enterobacter_hormaechei | LUX77_003527 | 1..1758 | 158836 | BREX-system-P-loop-protein-BrxC-[Enterobacter-hormaechei] |
| MCE1563408.1 | Type1.BREX | wHTH+SAD+HNH-><-NACHT+SF1-SF2-Helicase+HKD-DNase\|\|McrB-AAA->McrC-NTD+REase-><-SIG [illegible] | 1204 | Bacteria>Pseudomonadota>Gammaproteobacteria>Enterobacterales>Enterobacteriaceae>Enterobacter | GCA_021285765.1 | Enterobacter_hormaechei | LWT81_04630 | 337849..341463 | 158836 | BREX-system-P-loop-protein-BrxC-[Enterobacter-hormaechei] |
| MCI9498943.1 | Type1.BREX | wHTH+SAD+HNH-><-NACHT+SF1-SF2-Helicase+HKD-DNase\|\|McrB-AAA->McrC-NTD+REase-><-SIG [illegible] | 1204 | Bacteria>Pseudomonadota>Gammaproteobacteria>Enterobacterales>Enterobacteriaceae>Enterobacter | GCA_022800955.1 | Enterobacter_hormaechei_subsp._steigerwaltii | HFE69_15020 | 342834..346448 | 299766 | BREX-system-P-loop-protein-BrxC-[Enterobacter-hormaechei-subsp.-steigerwaltii] |
| HCM9270259.1 | Type1.BREX | Glyco_hydro_43+GH43_C2-><-ICLR-HTH<-?\|\|?->?->?->tRNA->Arm-DNA-bind_3+Phage_integr [illegible] | 1157 | Bacteria>Pseudomonadota>Gammaproteobacteria>Enterobacterales>Enterobacteriaceae>Enterobacter | GCA_025212085.1 | Enterobacter_cloacae_subsp._cloacae | N5L33_003913 | 155329..158802 | 336306 | BREX-system-P-loop-protein-BrxC-[Enterobacter-cloacae-subsp.-cloacae] |
| HCM9559898.1 | Type1.BREX | Glyco_hydro_43+GH43_C2-><-ICLR-HTH<-?\|\|?->?->?->tRNA->BrxA-DNA-binding->BrxB-BrxF [illegible] | 1157 | Bacteria>Pseudomonadota>Gammaproteobacteria>Enterobacterales>Enterobacteriaceae>Enterobacter | GCA_025214055.1 | Enterobacter_cloacae_subsp._cloacae | N5O13_002966 | 176473..179946 | 336306 | BREX-system-P-loop-protein-BrxC-[Enterobacter-cloacae-subsp.-cloacae] |
| WP_228371847.1 | Type1.BREX | Arm-DNA-bind_3+Phage_integrase-><-SIGMA+OB-Fold+AAA-ATPase+Lon-Protease-BrxL<-Pgl [illegible] | 1240 | Bacteria>Pseudomonadota>Gammaproteobacteria>Enterobacterales>Enterobacteriaceae>Enterobacter | GCF_000286275.1 | Enterobacter_kobei | ECENHK_RS02760 | 581253..584975 | 208224 | BREX-system-P-loop-protein-BrxC-[Enterobacter-kobei] |
| WP_021240656.1 | Type1.BREX | <-SymE_toxin<-?<-?<-?\|\|wHTH+HNH->SAM-METHYLASE->BrxA-DNA-binding->BrxB-BrxF-iSTAN [illegible] | 1213 | Bacteria>Pseudomonadota>Gammaproteobacteria>Enterobacterales>Enterobacteriaceae>Enterobacter | GCF_000467655.2 | Enterobacter_roggenkampii | L799_RS145635 | 328329..331970 | 1812935 | BREX-system-P-loop-protein-BrxC-[Enterobacter-roggenkampii] |
| WP_234704204.1 | Type1.BREX | Arm-DNA-bind_3+Phage_integrase-><-SIGMA+OB-Fold+AAA-ATPase+Lon-Protease-BrxL<-Pgl [illegible] | 1240 | Bacteria>Pseudomonadota>Gammaproteobacteria>Enterobacterales>Enterobacteriaceae>Enterobacter | GCF_000814225.1 | Enterobacter_cloacae_complex | LI66_RS02640 | 557969..561691 | 354276 | MULTISPECIES:-BREX-system-P-loop-protein-BrxC-[Enterobacter-cloacae-complex] |
| WP_058675376.1 | Type1.BREX | wHTH+SAD+HNH-><-NACHT+SF1-SF2-Helicase+HKD-DNase\|\|McrB-AAA->McrC-NTD+REase-><-SIG [illegible] | 1204 | Bacteria>Pseudomonadota>Gammaproteobacteria>Enterobacterales>Enterobacteriaceae>Enterobacter | GCF_001473175.1 | Enterobacter_hormaechei | ASV30_RS20360 | 113168..116782 | 158836 | BREX-system-P-loop-protein-BrxC-[Enterobacter-hormaechei] |
| WP_059514247.1 | Type1.BREX | BrxA-DNA-binding->BrxB-BrxF-iSTAND->DUF6079*->N6-METHYLASE->OLD-ABC->TOPRIM->PglZ [illegible] | 1204 | Bacteria>Pseudomonadota>Gammaproteobacteria>Enterobacterales>Enterobacteriaceae>Enterobacter | GCF_001525475.1 | Enterobacter_hormaechei | CI630_RS14005 | 1978..5592 | 158836 | BREX-system-P-loop-protein-BrxC-[Enterobacter-hormaechei] |
| WP_230373248.1 | Type1.BREX | <-wHTH+WYL<-sGTP<-?\|\|XIS-HTH->?->?->BrxA-DNA-binding->BrxB-BrxF-iSTAND->DUF6079*- [illegible] | 1240 | Bacteria>Pseudomonadota>Gammaproteobacteria>Enterobacterales>Enterobacteriaceae>Enterobacter | GCF_001631925.1 | Enterobacter_hormaechei | A3N67_RS02925 | 129032..132754 | 158836 | BREX-system-P-loop-protein-BrxC-[Enterobacter-hormaechei] |
| WP_069597812.1 | Type1.BREX | <-ICLR-HTH<-?<-?\|\|?->?->?->tRNA-><-ZnR\|\|BrxA-DNA-binding->BrxB-BrxF-iSTAND->DUF60 [illegible] | 1204 | Bacteria>Pseudomonadota>Gammaproteobacteria>Enterobacterales>Enterobacteriaceae>Enterobacter | GCF_001729805.1 | Enterobacter_roggenkampii | BFV67_RS02540 | 543370..546984 | 1812935 | BREX-system-P-loop-protein-BrxC-[Enterobacter-roggenkampii] |
| WP_095453563.1 | Type1.BREX | <-SymE_toxin<-?<-?<-?\|\|wHTH+HNH->SAM-METHYLASE->BrxA-DNA-binding->BrxB-BrxF-iSTAN [illegible] | 1213 | Bacteria>Pseudomonadota>Gammaproteobacteria>Enterobacterales>Enterobacteriaceae>Enterobacter | GCF_002278355.1 | Enterobacter_cloacae | CIW57_RS16410 | 135960..139601 | 550 | BREX-system-P-loop-protein-BrxC-[Enterobacter-cloacae] |
| WP_100165592.1 | Type1.BREX | <-SIGMA+OB-Fold+AAA-ATPase+Lon-Protease-BrxL<-PglZ<-TOPRIM<-OLD-ABC<-N6-METHYLASE [illegible] | 1204 | Bacteria>Pseudomonadota>Gammaproteobacteria>Enterobacterales>Enterobacteriaceae>Enterobacter | GCF_002785605.1 | Enterobacter_mori | B9Q20_RS04890 | 379505..383119 | 539813 | BREX-system-P-loop-protein-BrxC-[Enterobacter-mori] |
| WP_237764241.1 | Type1.BREX | Arm-DNA-bind_3+Phage_integrase-><-SIGMA+OB-Fold+AAA-ATPase+Lon-Protease-BrxL<-Pgl [illegible] | 1240 | Bacteria>Pseudomonadota>Gammaproteobacteria>Enterobacterales>Enterobacteriaceae>Enterobacter | GCF_003073995.1 | Enterobacter_hormaechei | CSC19_RS19005 | 3849441..3853163 | 158836 | BREX-system-P-loop-protein-BrxC-[Enterobacter-hormaechei] |
| WP_110108109.1 | Type1.BREX | XIS-HTH->PIWI-><-?<-Phage_integrase<-ZnR\|\|BrxA-DNA-binding->BrxB-BrxF-iSTAND->DUF [illegible] | 1197 | Bacteria>Pseudomonadota>Gammaproteobacteria>Enterobacterales>Enterobacteriaceae>Enterobacter | GCF_003186415.1 | Enterobacter_hormaechei | BET69_RS02790 | 559852..563445 | 158836 | BREX-system-P-loop-protein-BrxC-[Enterobacter-hormaechei] |
| WP_110875386.1 | Type1.BREX | <-CopG/RHH\|\|XIS-HTH->?->?->SF1-SF2-Helicase+REase->SF1-SF2-Helicase+REase-><-PglZ [illegible] | 1157 | Bacteria>Pseudomonadota>Gammaproteobacteria>Enterobacterales>Enterobacteriaceae>Enterobacter | GCF_003207215.1 | Enterobacter_hormaechei | DOM70_RS06250 | 341163..344636 | 158836 | BREX-system-P-loop-protein-BrxC-[Enterobacter-hormaechei] |
| WP_123923007.1 | Type1.BREX | TOPRIM-DNAG+D5N+D5ATPase+P4PRIMASE-HTH->BrxB-BrxF-iSTAND->ACET->?->BrxA-DNA-bindi [illegible] | 1204 | Bacteria>Pseudomonadota>Gammaproteobacteria>Enterobacterales>Enterobacteriaceae>Enterobacter | GCF_003812145.1 | Enterobacter_roggenkampii | EGY04_RS13705 | 2428639..2432253 | 1812935 | BREX-system-P-loop-protein-BrxC-[Enterobacter-roggenkampii] |
| WP_125357895.1 | Type1.BREX | <-SIGMA+OB-Fold+AAA-ATPase+Lon-Protease-BrxL<-PglZ<-N6-METHYLASE<-ParB+HNH<-DUF60 [illegible] | 1225 | Bacteria>Pseudomonadota>Gammaproteobacteria>Enterobacterales>Enterobacteriaceae>Enterobacter | GCF_003937295.2 | Enterobacter_hormaechei | EGK38_RS19825 | 4108658..4112335 | 158836 | BREX-system-P-loop-protein-BrxC-[Enterobacter-hormaechei] |
| WP_126349337.1 | Type1.BREX | Glyco_hydro_43+GH43_C2-><-ICLR-HTH<-?<-?\|\|?->?->?->tRNA->BrxA-DNA-binding->BrxB-B [illegible] | 1157 | Bacteria>Pseudomonadota>Gammaproteobacteria>Enterobacterales>Enterobacteriaceae>Enterobacter | GCF_003965695.1 | Enterobacter_roggenkampii | EKN52_RS20255 | 23415..26888 | 1812935 | BREX-system-P-loop-protein-BrxC-[Enterobacter-roggenkampii] |
| WP_126800556.1 | Type1.BREX | SF1-SF2-Helicase+REase->SF1-SF2-Helicase+REase-><-PglZ<-ParB+HNH<-N6-METHYLASE<-D [illegible] | 1157 | Bacteria>Pseudomonadota>Gammaproteobacteria>Enterobacterales>Enterobacteriaceae>Enterobacter | GCF_003986875.1 | Enterobacter_sp._WCHEn090032 | EKN83_RS06700 | 53011..56484 | 2497435 | BREX-system-P-loop-protein-BrxC-[Enterobacter-sp.-WCHEn090032] |
| WP_241243342.1 | Type1.BREX | <-DUF6079-p*<-BrxB-BrxF-iSTAND<-BrxA-DNA-binding<-Glyoxalase<-ZnR<-METHYLASE | 866 | Bacteria>Pseudomonadota>Gammaproteobacteria>Enterobacterales>Enterobacteriaceae>Enterobacter | GCF_004004525.1 | Enterobacter_hormaechei | EOL34_RS23625 | 1..2600 | 158836 | BREX-system-P-loop-protein-BrxC,-partial-[Enterobacter-hormaechei] |
| WP_129328058.1 | Type1.BREX | Phage_Cox->XIS-HTH->AAA+wHTH->?-><-SIGMA+OB-Fold+AAA-ATPase+Lon-Protease-BrxL<-Pg [illegible] | 1204 | Bacteria>Pseudomonadota>Gammaproteobacteria>Enterobacterales>Enterobacteriaceae>Enterobacter | GCF_004127365.1 | Enterobacter_cloacae | DD604_RS20965 | 29802..33416 | 550 | BREX-system-P-loop-protein-BrxC-[Enterobacter-cloacae] |
| WP_129694025.1 | Type1.BREX | <-REase<-?<-HTH_Tnp_1+HTH_21+rve+rve_3\|\|SF1-SF2-Helicase+REase->SF1-SF2-Helicase+ [illegible] | 1157 | Bacteria>Pseudomonadota>Gammaproteobacteria>Enterobacterales>Enterobacteriaceae>Enterobacter | GCF_004138605.1 | Enterobacter_roggenkampii | D8675_RS06390 | 1085829..1089302 | 1812935 | BREX-system-P-loop-protein-BrxC-[Enterobacter-roggenkampii] |
| WP_148769092.1 | Type1.BREX | PAAR_motif+Pyocin_S+ColE7+HNH->?->?-><-SymE_toxin\|\|wHTH+HNH->SAM-METHYLASE->BrxA- [illegible] | 1213 | Bacteria>Pseudomonadota>Gammaproteobacteria>Enterobacterales>Enterobacteriaceae>Enterobacter | GCF_008122455.1 | Enterobacter_sp._C6 | DOU50_RS06770 | 7671..11312 | 1299469 | BREX-system-P-loop-protein-BrxC-[Enterobacter-sp.-C6] |
| WP_164474334.1 | Type1.BREX | SF1-SF2-Helicase+REase->SF1-SF2-Helicase+REase-><-PglZ<-ParB+HNH<-N6-METHYLASE<-N [illegible] | 1157 | Bacteria>Pseudomonadota>Gammaproteobacteria>Enterobacterales>Enterobacteriaceae>Enterobacter | GCF_011045335.1 | Enterobacter_hormaechei | G5C52_RS10105 | 2135134..2138607 | 158836 | BREX-system-P-loop-protein-BrxC-[Enterobacter-hormaechei] |
| WP_168980132.1 | Type1.BREX | LACI-HTH+PBPI-><-ASCH+Glyoxalase\|\|?->?-><-SymE_toxin\|\|wHTH+HNH->SAM-METHYLASE->Br [illegible] | 1213 | Bacteria>Pseudomonadota>Gammaproteobacteria>Enterobacterales>Enterobacteriaceae>Enterobacter | GCF_012843805.1 | Enterobacter_asburiae | HF856_RS13430 | 88040..91681 | 61645 | BREX-system-P-loop-protein-BrxC-[Enterobacter-asburiae] |
| WP_181236548.1 | Type1.BREX | <-DUF905<-?<-?<-?<-wHTH+WYL<-DUF932<-nSTAND1+GTPase-AIG+sGTP\|\|BrxA-DNA-binding->B [illegible] | 1204 | Bacteria>Pseudomonadota>Gammaproteobacteria>Enterobacterales>Enterobacteriaceae>Enterobacter | GCF_013624605.1 | Enterobacter_hormaechei | HV094_RS19965 | 4127702..4131316 | 158836 | BREX-system-P-loop-protein-BrxC-[Enterobacter-hormaechei] |
| WP_182062523.1 | Type1.BREX | OLD-ABC+TOPRIM->SF1-SF2-Helicase-><-?<-?<-?<-wHTH+WYL<-nSTAND1+GTPase-AIG+sGTP\|\|B [illegible] | 1202 | Bacteria>Pseudomonadota>Gammaproteobacteria>Enterobacterales>Enterobacteriaceae>Enterobacter | GCF_013927005.1 | Enterobacter_sp._RHBSTW-00175 | HV107_RS06435 | 1368672..1372280 | 2742639 | BREX-system-P-loop-protein-BrxC-[Enterobacter-sp.-RHBSTW-00175] |
| WP_182014101.1 | Type1.BREX | Glyoxalase-><-HTH_Tnp_1+HTH_21+rve+rve_3\|\|McrC-NTD+REase-><-SIGMA+OB-Fold+AAA-ATP [illegible] | 1204 | Bacteria>Pseudomonadota>Gammaproteobacteria>Enterobacterales>Enterobacteriaceae>Enterobacter | GCF_013927185.1 | Enterobacter_roggenkampii | HV097_RS13200 | 2641536..2645150 | 1812935 | BREX-system-P-loop-protein-BrxC-[Enterobacter-roggenkampii] |
| WP_182278818.1 | Type1.BREX | <-SymE_toxin\|\|HIN-HTH+PAIREDC-HTH-><-?<-?<-?\|\|wHTH+HNH->SAM-METHYLASE->BrxA-DNA-b [illegible] | 1213 | Bacteria>Pseudomonadota>Gammaproteobacteria>Enterobacterales>Enterobacteriaceae>Enterobacter | GCF_014101255.1 | Enterobacter_roggenkampii | HV177_RS13220 | 2740666..2744307 | 1812935 | BREX-system-P-loop-protein-BrxC-[Enterobacter-roggenkampii] |
| WP_100126890.1 | Type1.BREX | <-DUF905<-?<-?<-?<-wHTH+WYL<-DUF932<-nSTAND1+GTPase-AIG+sGTP\|\|BrxA-DNA-binding->B [illegible] | 1204 | Bacteria>Pseudomonadota>Gammaproteobacteria>Enterobacterales>Enterobacteriaceae>Enterobacter | GCF_014103055.1 | Enterobacter_cloacae_complex | HV354_RS20740 | 74806..78420 | 354276 | MULTISPECIES:-BREX-system-P-loop-protein-BrxC-[Enterobacter-cloacae-complex] |
| WP_057055966.1 | Type1.BREX | Arm-DNA-bind_3+Phage_integrase-><-HKD-DNase+OGG-like<-SIGMA+OB-Fold+AAA-ATPase+Lo [illegible] | 1197 | Bacteria>Pseudomonadota>Gammaproteobacteria>Enterobacterales>Enterobacteriaceae>Enterobacter | GCF_014169295.1 | Enterobacter_cloacae_complex | H7R61_RS03370 | 684757..688350 | 354276 | MULTISPECIES:-BREX-system-P-loop-protein-BrxC-[Enterobacter-cloacae-complex] |
| WP_230863208.1 | Type1.BREX | Arm-DNA-bind_3+Phage_integrase-><-SIGMA+OB-Fold+AAA-ATPase+Lon-Protease-BrxL<-Pgl [illegible] | 1240 | Bacteria>Pseudomonadota>Gammaproteobacteria>Enterobacterales>Enterobacteriaceae>Enterobacter | GCF_015704905.1 | Enterobacter_cloacae | IOR24_RS17915 | 3774454..3778176 | 550 | BREX-system-P-loop-protein-BrxC-[Enterobacter-cloacae] |
| WP_234450174.1 | Type1.BREX | Glycoside_Hydrolase->?->Glyoxalase->YjhX_toxin->SAM-METHYLASE->ZnR->Glyoxalase->B [illegible] | 1240 | Bacteria>Pseudomonadota>Gammaproteobacteria>Enterobacterales>Enterobacteriaceae>Enterobacter | GCF_016616795.1 | Enterobacter_hormaechei | FKK11_RS12010 | 273367..277089 | 158836 | BREX-system-P-loop-protein-BrxC-[Enterobacter-hormaechei] |
| WP_211449384.1 | Type1.BREX | <-SymE_toxin\|\|wHTH(SBD)+HNH-><-NACHT+SF1-SF2-Helicase+HKD-DNase\|\|SF1-SF2-Helicase [illegible] | 1157 | Bacteria>Pseudomonadota>Gammaproteobacteria>Enterobacterales>Enterobacteriaceae>Enterobacter | GCF_018299605.1 | Enterobacter_mori | GUY16_RS21845 | 58984..62457 | 539813 | BREX-system-P-loop-protein-BrxC-[Enterobacter-mori] |
| WP_006812019.1 | Type1.BREX | Phage_Cox->XIS-HTH->AAA+wHTH->?-><-SIGMA+OB-Fold+AAA-ATPase+Lon-Protease-BrxL<-Pg [illegible] | 1204 | Bacteria>Pseudomonadota>Gammaproteobacteria>Enterobacterales>Enterobacteriaceae>Enterobacter | GCF_019048245.1 | Enterobacter_hormaechei | I6L63_RS02015 | 410151..413765 | 158836 | BREX-system-P-loop-protein-BrxC-[Enterobacter-hormaechei] |
| WP_219280521.1 | Type1.BREX | <-HNH<-OLD-ABC<-N6-N4-METHYLASE<-HTH_3<-?<-ZnR\|\|BrxA-DNA-binding->BrxB-BrxF-iSTAN [illegible] | 1197 | Bacteria>Pseudomonadota>Gammaproteobacteria>Enterobacterales>Enterobacteriaceae>Enterobacter | GCF_019351595.1 | Enterobacter_cloacae | JW319_RS20560 | 173870..177463 | 550 | BREX-system-P-loop-protein-BrxC-[Enterobacter-cloacae] |
| WP_222665203.1 | Type1.BREX | Phage_Cox->XIS-HTH->AAA+wHTH->?-><-SIGMA+OB-Fold+AAA-ATPase+Lon-Protease-BrxL<-Pg [illegible] | 1204 | Bacteria>Pseudomonadota>Gammaproteobacteria>Enterobacterales>Enterobacteriaceae>Enterobacter | GCF_019839635.1 | Enterobacter_roggenkampii | K4X45_RS16780 | 84245..87859 | 1812935 | BREX-system-P-loop-protein-BrxC-[Enterobacter-roggenkampii] |
| WP_032634492.1 | Type1.BREX | BrxA-DNA-binding->BrxB-BrxF-iSTAND->DUF6079*->N6-METHYLASE->OLD-ABC->TOPRIM->PglZ [illegible] | 1204 | Bacteria>Pseudomonadota>Gammaproteobacteria>Enterobacterales>Enterobacteriaceae>Enterobacter | GCF_019974415.1 | Enterobacter_cloacae_complex | DDA68_RS23710 | 1545..5159 | 354276 | MULTISPECIES:-BREX-system-P-loop-protein-BrxC-[Enterobacter-cloacae-complex] |
| WP_230238963.1 | Type1.BREX | Glyco_hydro_43+GH43_C2-><-ICLR-HTH<-?<-?\|\|?->?->?->tRNA->BrxA-DNA-binding->BrxB-B [illegible] | 1157 | Bacteria>Pseudomonadota>Gammaproteobacteria>Enterobacterales>Enterobacteriaceae>Enterobacter | GCF_020971765.1 | Enterobacter_sp._AN-K1 | LOC22_RS13480 | 2816173..2819646 | 2894203 | BREX-system-P-loop-protein-BrxC-[Enterobacter-sp.-AN-K1] |
| WP_241512123.1 | Type1.BREX | Arm-DNA-bind_3+Phage_integrase-><-SIGMA+OB-Fold+AAA-ATPase+Lon-Protease-BrxL<-Pgl [illegible] | 1240 | Bacteria>Pseudomonadota>Gammaproteobacteria>Enterobacterales>Enterobacteriaceae>Enterobacter | GCF_022585395.1 | Enterobacter_hormaechei | MMO50_RS10500 | 125167..128889 | 158836 | BREX-system-P-loop-protein-BrxC-[Enterobacter-hormaechei] |
| WP_248201800.1 | Type1.BREX | Glyoxalase_8->URI-DNase-><-RHH_antitoxin\|\|?->XIS-HTH->?->ZnR->BrxA-DNA-binding-> [illegible] | 1213 | Bacteria>Pseudomonadota>Gammaproteobacteria>Enterobacterales>Enterobacteriaceae>Enterobacter | GCF_023151575.1 | Enterobacter_asburiae | L8T09_RS18420 | 27571..31212 | 61645 | BREX-system-P-loop-protein-BrxC-[Enterobacter-asburiae] |
| WP_248198186.1 | Type1.BREX | XIS-HTH->?->?->?->TOPRIM-DNAG+D5N+D5ATPase+P4PRIMASE-HTH-><-?<-SIGMA+OB-Fold+AAA- [illegible] | 1213 | Bacteria>Pseudomonadota>Gammaproteobacteria>Enterobacterales>Enterobacteriaceae>Enterobacter | GCF_023151985.1 | Enterobacter_bugandensis | L8S35_RS17040 | 181002..184643 | 881260 | BREX-system-P-loop-protein-BrxC-[Enterobacter-bugandensis] |
| WP_248241159.1 | Type1.BREX | <-NACHT+SF1-SF2-Helicase+HKD-DNase\|\|McrB-AAA->McrC-NTD+REase-><-SIGMA+OB-Fold+AAA [illegible] | 1204 | Bacteria>Pseudomonadota>Gammaproteobacteria>Enterobacterales>Enterobacteriaceae>Enterobacter | GCF_023153155.1 | Enterobacter_roggenkampii | L8Q49_RS17945 | 130096..133710 | 1812935 | BREX-system-P-loop-protein-BrxC-[Enterobacter-roggenkampii] |
| WP_248076370.1 | Type1.BREX | YjhX_toxin->SAM-METHYLASE->ZnR->Glyoxalase->BrxA-DNA-binding->BrxB-BrxF-iSTAND->D [illegible] | 1240 | Bacteria>Pseudomonadota>Gammaproteobacteria>Enterobacterales>Enterobacteriaceae>Enterobacter | GCF_023153855.1 | Enterobacter_cloacae | L8P05_RS09265 | 348723..352445 | 550 | BREX-system-P-loop-protein-BrxC-[Enterobacter-cloacae] |
| WP_248053563.1 | Type1.BREX | <-ZnR\|\|BrxA-DNA-binding->BrxB-BrxF-iSTAND->DUF6079*->N6-METHYLASE->OLD-ABC->TOPRI [illegible] | 1197 | Bacteria>Pseudomonadota>Gammaproteobacteria>Enterobacterales>Enterobacteriaceae>Enterobacter | GCF_023154935.1 | Enterobacter_chengduensis | L8P32_RS20960 | 6466..10059 | 2494701 | BREX-system-P-loop-protein-BrxC-[Enterobacter-chengduensis] |
| WP_032621388.1 | Type1.BREX | <-SymE_toxin\|\|wHTH(SBD)+HNH-><-SIGMA+OB-Fold+AAA-ATPase+Lon-Protease-BrxL<-PglZ<- [illegible] | 1204 | Bacteria>Pseudomonadota>Gammaproteobacteria>Enterobacterales>Enterobacteriaceae>Enterobacter | GCF_023734955.1 | Enterobacter_hormaechei | NVR78_RS04785 | 326171..329785 | 158836 | BREX-system-P-loop-protein-BrxC-[Enterobacter-hormaechei] |
| WP_192468115.1 | Type1.BREX | URI-DNase-><-HIN-HTH<-RHH_antitoxin\|\|?->XIS-HTH->?->ZnR->BrxA-DNA-binding->BrxB-B [illegible] | 1213 | Bacteria>Pseudomonadota>Gammaproteobacteria>Enterobacterales>Enterobacteriaceae>Enterobacter | GCF_023749245.1 | Enterobacter_asburiae | M8N38_RS13285 | 26000..29641 | 61645 | BREX-system-P-loop-protein-BrxC-[Enterobacter-asburiae] |
| WP_252070049.1 | Type1.BREX | Arm-DNA-bind_3+Phage_integrase-><-?<-SIGMA+OB-Fold+AAA-ATPase+Lon-Protease-BrxL<- [illegible] | 1213 | Bacteria>Pseudomonadota>Gammaproteobacteria>Enterobacterales>Enterobacteriaceae>Enterobacter | GCF_023750345.1 | Enterobacter_ludwigii | M8O29_RS15875 | 136440..140081 | 299767 | BREX-system-P-loop-protein-BrxC-[Enterobacter-ludwigii] |
| WP_253179320.1 | Type1.BREX | wHTH+SAD+HNH-><-NACHT+SF1-SF2-Helicase+HKD-DNase\|\|McrB-AAA->McrC-NTD+REase-><-SIG [illegible] | 1204 | Bacteria>Pseudomonadota>Gammaproteobacteria>Enterobacterales>Enterobacteriaceae>Enterobacter | GCF_024128585.1 | Enterobacter_mori | NJA11_RS17930 | 132016..135630 | 539813 | BREX-system-P-loop-protein-BrxC-[Enterobacter-mori] |
| WP_257273059.1 | Type1.BREX | LACI-HTH+PBPI-><-ASCH+Glyoxalase\|\|?-><-SymE_toxin\|\|wHTH+HNH->SAM-METHYLASE->SF1-S [illegible] | 398 | Bacteria>Pseudomonadota>Gammaproteobacteria>Enterobacterales>Enterobacteriaceae>Enterobacter | GCF_024607735.1 | Enterobacter_soli | NR352_RS15925 | 54991..56187 | 885040 | BREX-system-P-loop-protein-BrxC-[Enterobacter-soli] |
| WP_257246766.1 | Type1.BREX | <-ASCH+Glyoxalase\|\|?->DUF2754->?-><-SymE_toxin\|\|wHTH+SAD+HNH-><-NACHT+SF1-SF2-Hel [illegible] | 1197 | Bacteria>Pseudomonadota>Gammaproteobacteria>Enterobacterales>Enterobacteriaceae>Enterobacter | GCF_024607805.1 | unclassified_Enterobacter | NR293_RS15365 | 69785..73378 | 2608935 | MULTISPECIES:-BREX-system-P-loop-protein-BrxC-[unclassified-Enterobacter] |
| WP_058688926.1 | Type1.BREX | nSTAND1+HNH->RNaseH->NACHT+SF1-SF2-Helicase+HKD-DNase->REase-><-HKD-DNase+OGG-lik [illegible] | 1213 | Bacteria>Pseudomonadota>Gammaproteobacteria>Enterobacterales>Enterobacteriaceae>Enterobacter | GCF_024655335.1 | Enterobacter_hormaechei | ASU86_RS07715 | 183720..187361 | 158836 | BREX-system-P-loop-protein-BrxC-[Enterobacter-hormaechei] |
| WP_258247898.1 | Type1.BREX | <-ZnR\|\|BrxA-DNA-binding->BrxB-BrxF-iSTAND->DUF6079*->N6-METHYLASE->OLD-ABC->TOPRI [illegible] | 1204 | Bacteria>Pseudomonadota>Gammaproteobacteria>Enterobacterales>Enterobacteriaceae>Enterobacter | GCF_024733415.1 | Enterobacter_hormaechei | NUJ48_RS00150 | 29183..32797 | 158836 | BREX-system-P-loop-protein-BrxC-[Enterobacter-hormaechei] |
| WP_265188117.1 | Type1.BREX | wHTH+SAD+HNH-><-NACHT+SF1-SF2-Helicase+HKD-DNase\|\|McrB-AAA->McrC-NTD+REase-><-SIG [illegible] | 1204 | Bacteria>Pseudomonadota>Gammaproteobacteria>Enterobacterales>Enterobacteriaceae>Enterobacter | GCF_026115025.1 | Enterobacter_hormaechei | M7778_RS15260 | 340339..343953 | 158836 | BREX-system-P-loop-protein-BrxC-[Enterobacter-hormaechei] |
| WP_271139057.1 | Type1.BREX | <-ICLR-HTH<-?<-?\|\|?->?->?->tRNA-><-ZnR\|\|BrxA-DNA-binding->BrxB-BrxF-iSTAND->DUF60 [illegible] | 1204 | Bacteria>Pseudomonadota>Gammaproteobacteria>Enterobacterales>Enterobacteriaceae>Enterobacter | GCF_027920365.1 | Enterobacter_hormaechei | PBS85_RS02750 | 569121..572735 | 158836 | BREX-system-P-loop-protein-BrxC-[Enterobacter-hormaechei] |
| WP_274416986.1 | Type1.BREX | XIS-HTH->ZnR->ZnR->?-><-NACHT+SF1-SF2-Helicase+HKD-DNase\|\|BrxA-DNA-binding->BrxB- [illegible] | 1204 | Bacteria>Pseudomonadota>Gammaproteobacteria>Enterobacterales>Enterobacteriaceae>Enterobacter | GCF_028765355.1 | Enterobacter_kobei | PVM09_RS00190 | 32375..35989 | 208224 | BREX-system-P-loop-protein-BrxC-[Enterobacter-kobei] |
| WP_111946923.1 | Type1.BREX | wHTH+SAD+HNH-><-NACHT+SF1-SF2-Helicase+HKD-DNase\|\|McrB-AAA->McrC-NTD+REase-><-SIG [illegible] | 1204 | Bacteria>Pseudomonadota>Gammaproteobacteria>Enterobacterales>Enterobacteriaceae>Enterobacter | GCF_030324195.1 | Enterobacter_hormaechei | J9785_RS10685 | 292266..295880 | 158836 | BREX-system-P-loop-protein-BrxC-[Enterobacter-hormaechei] |
| WP_180264800.1 | Type1.BREX | <-HTH_36<-HTH+HTH<-ParB+BAM<-XIS-HTH<-wHTH<-ZnR\|\|BrxA-DNA-binding->BrxB-BrxF-iSTA [illegible] | 1204 | Bacteria>Pseudomonadota>Gammaproteobacteria>Enterobacterales>Enterobacteriaceae>Enterobacter | GCF_900558725.1 | Enterobacter_hormaechei | H0848_RS18555 | 27730..31344 | 158836 | BREX-system-P-loop-protein-BrxC-[Enterobacter-hormaechei] |

| Protein-ID | System | Gene-Neighbourhoods | Length | Taxonomy | Assembly-ID | Strain | Locus-Tag | Genomic-coordinates | TaxID | Fasta-define |
|---|---|---|---|---|---|---|---|---|---|---|
| WP_063851018.1 | Type1.BREX | <-HTH_36<-HTH+HTH<-ParB+BAM<-XIS-HTH<-wHTH<-ZnR\|\|BrxA-DNA-binding->BrxB-BrxF-iSTAND->DUF6079*->N6-METHYLASE->OLD-ABC->TOPRIM->PglZ->SIGMA+OB-Fold+AAA-ATPase+Lon-Protease-BrxL-><-wHTH(SBD)+HNH\|\|SymE_toxin-> | 1204 | Bacteria>Pseudomonadota>Gammaproteobacteria>Enterobacterales>Enterobacteriaceae>Enterobacter | GCF_902160625.1 | Enterobacter_hormaechei | FN214_RS22220 | 26549..30163 | 158836 | BREX-system-P-loop-protein-BrxC-[Enterobacter-hormaechei] |
| WP_112781824.1 | Type1.BREX | <-SIGMA+OB-Fold+AAA-ATPase+Lon-Protease-BrxL<-PglZ<-TOPRIM<-OLD-ABC<-N6-METHYLASE<-DUF6079*<-BrxB-BrxF-iSTAND<-BrxA-DNA-binding<-XIS-HTH<-?\|\|CopG/RHH->DUF932->ArdB-> | 1204 | Bacteria>Pseudomonadota>Gammaproteobacteria>Enterobacterales>Enterobacteriaceae>Enterobacter | GCF_902387755.1 | Enterobacter_cloacae | FYC20_RS23065 | 9489..13103 | 550 | MULTISPECIES:-BREX-system-P-loop-protein-BrxC-[Enterobacteriaceae] |
| EHX5558535.1 | Type1.BREX | <-HerA/FtsK+A32<-?\|\|tRNA->Arm-DNA-bind_3+Phage_integrase->EVE+HNH-><-REase+iREase\|\|BrxA-DNA-binding->BrxB-BrxF-iSTAND->DUF6079*->N6-METHYLASE->PglZ->SIGMA+OB-Fold+AAA-ATPase+Lon-Protease-BrxL-><-ZnR<-ZnR<-?<-?\|\|RHH_antitoxin->XIS-HTH-> | 1213 | Bacteria>Pseudomonadota>Gammaproteobacteria>Enterobacterales>Enterobacteriaceae>Escherichia | - | Escherichia_albertii | K2D51_002305 | 108873..112514 | 208962 | BREX-system-P-loop-protein-BrxC-[Escherichia-albertii] |
| WP_095526333.1 | Type1.BREX | <-35exo<-SIGMA+OB-Fold+AAA-ATPase+Lon-Protease-BrxL<-PglZ<-N6-METHYLASE<-ParB+HNH<-DUF6079*<-BrxB-BrxF-iSTAND<-BrxA-DNA-binding<-wHTH+WYL+WCX<-XIS-HTH<-ZnR\|\|?-><-?\|\|?->ArdB-> | 1225 | Bacteria>Pseudomonadota>Gammaproteobacteria>Enterobacterales>Enterobacteriaceae>Escherichia | - | Escherichia_coli | CDN88_RS21130 | 10739..14416 | 562 | BREX-system-P-loop-protein-BrxC-[Escherichia-coli] |
| WP_121868993.1 | Type1.BREX | <-HerA/FtsK+A32<-?\|\|tRNA->Arm-DNA-bind_3+Phage_integrase-><-SIGMA+OB-Fold+AAA-ATPase+Lon-Protease-BrxL<-PglZ<-N6-METHYLASE-p<-N6-METHYLASE<-ParB+HNH<-DUF6079*<-BrxB-BrxF-iSTAND<-BrxA-DNA-binding<-?<-?<-XIS-HTH\|\|?->nSTAND1+sGTP->wHTH+WYL-> | 1233 | Bacteria>Pseudomonadota>Gammaproteobacteria>Enterobacterales>Enterobacteriaceae>Escherichia | - | Escherichia_coli | EAI39_15065 | 84115..87816 | 562 | BREX-system-P-loop-protein-BrxC-[Escherichia-coli] |
| EFH4523723.1 | Type1.BREX | <-METHYLASE\|\|SKI-><-?\|\|tRNA->Arm-DNA-bind_3+Phage_integrase-><-REase+iREase\|\|BrxA-DNA-binding->BrxB-BrxF-iSTAND->DUF6079*->N6-METHYLASE->PglZ->SIGMA+OB-Fold+AAA-ATPase+Lon-Protease-BrxL-><-ZnR<-ZnR | 1213 | Bacteria>Pseudomonadota>Gammaproteobacteria>Enterobacterales>Enterobacteriaceae>Escherichia | - | Escherichia_coli | F9608_003902 | 25000..28641 | 562 | BREX-system-P-loop-protein-BrxC-[Escherichia-coli] |
| WP_181716349.1 | Type1.BREX | XIS-HTH->PIWI-><-DDE_Tnp_1_5+DDE_Tnp_1\|\|PIWI-><-?<-Phage_integrase\|\|Arm-DNA-bind_3+Phage_integrase->BrxA-DNA-binding->BrxB-BrxF-iSTAND->DUF6079*->N6-METHYLASE->OLD-ABC->TOPRIM->SIGMA+OB-Fold+AAA-ATPase+Lon-Protease-BrxL-><-DDE_Tnp_1_5+DDE_Tnp_1<-?\|\|DDE_Tnp_1->RNase-><-HKD-DNase+MED12 | 1204 | Bacteria>Pseudomonadota>Gammaproteobacteria>Enterobacterales>Enterobacteriaceae>Escherichia | - | Escherichia_coli | H5E40_RS01425 | 278751..282365 | 562 | BREX-system-P-loop-protein-BrxC-[Escherichia-coli] |
| WP_174580485.1 | Type1.BREX | DDE_Tnp_IS1->?->ZnR->BrxA-DNA-binding->BrxB-BrxF-iSTAND->DUF6079*->ParB+HNH->N6-METHYLASE->HEPN->RVT->N6-METHYLASE-p->PglZ->SIGMA+OB-Fold+AAA-ATPase+Lon-Protease-BrxL-><-Arm-DNA-bind_3+Phage_integrase\|\|Phage_integrase-> | 1233 | Bacteria>Pseudomonadota>Gammaproteobacteria>Enterobacterales>Enterobacteriaceae>Escherichia | - | Escherichia_coli | HUC40_15465 | 2945..6646 | 562 | BREX-system-P-loop-protein-BrxC-[Escherichia-coli] |
| WP_227303300.1 | Type1.BREX | <-Phage_integrase<-Phage_integrase\|\|?->?->?->\|\|wHTH+SAD+HNH-><-REase+iREase\|\|BrxA-DNA-binding->BrxB-BrxF-iSTAND->DUF6079*->N6-METHYLASE->PglZ->SIGMA+OB-Fold+AAA-ATPase+Lon-Protease-BrxL->HKD-DNase+OGG-like->\|\|AbiEi_3_N+AbiEi_3->AbiEii->?-><-DUF6216 | 1213 | Bacteria>Pseudomonadota>Gammaproteobacteria>Enterobacterales>Enterobacteriaceae>Escherichia | - | Escherichia_coli | LJD42_RS04270 | 217680..221321 | 562 | BREX-system-P-loop-protein-BrxC-[Escherichia-coli] |
| MCN1699418.1 | Type1.BREX | <-LACI-HTH+PBPI<-?<-?<-METHYLASE\|\|SKI-><-?\|\|tRNA->Arm-DNA-bind_3+Phage_integrase->BrxA-DNA-binding->BrxB-BrxF-iSTAND->DUF6079*->N6-METHYLASE->OLD-ABC->TOPRIM->PglZ->SIGMA+OB-Fold+AAA-ATPase+Lon-Protease-BrxL-><-URI-DNase+RAMA | 1204 | Bacteria>Pseudomonadota>Gammaproteobacteria>Enterobacterales>Enterobacteriaceae>Escherichia | - | Escherichia_coli | MLI78_16455 | 116212..119826 | 562 | BREX-system-P-loop-protein-BrxC-[Escherichia-coli] |
| MCN1334053.1 | Type1.BREX | <-LACI-HTH+PBPI\|\|?->?-><-Zn_Tnp_IS1+HTH_Tnp_IS1+DDE_Tnp_IS1<-REase<-REase\|\|BrxA-DNA-binding->BrxB-BrxF-iSTAND->DUF6079*->N6-METHYLASE->PglZ->SIGMA+OB-Fold+AAA-ATPase+Lon-Protease-BrxL-><-ZnR<-HTH_Tnp_1+HTH_21+rve+rve_3 | 1213 | Bacteria>Pseudomonadota>Gammaproteobacteria>Enterobacterales>Enterobacteriaceae>Escherichia | - | Escherichia_coli | MLR45_18135 | 8747..12388 | 562 | BREX-system-P-loop-protein-BrxC-[Escherichia-coli] |
| MCN3919819.1 | Type1.BREX | <-METHYLASE\|\|SKI-><-?\|\|tRNA->Arm-DNA-bind_3+Phage_integrase-><-SIGMA+OB-Fold+AAA-ATPase+Lon-Protease-BrxL<-PglZ<-N6-METHYLASE<-ParB+HNH<-DUF6079*<-BrxB-BrxF-iSTAND<-BrxA-DNA-binding<-?<-?<-XIS-HTH\|\|?->wHTH+WYL-> | 1240 | Bacteria>Pseudomonadota>Gammaproteobacteria>Enterobacterales>Enterobacteriaceae>Escherichia | - | Escherichia_coli | MLZ57_18615 | 66742..70464 | 562 | BREX-system-P-loop-protein-BrxC-[Escherichia-coli] |
| EIH98049.1 | Type1.BREX | REase+N6_METHYLASE->?->ParB->?-><-?<-SIGMA+OB-Fold+AAA-ATPase+Lon-Protease-BrxL<-PglZ<-N6-METHYLASE<-DUF6079-p*<-DUF6079-p<-BrxB-BrxF-iSTAND<-BrxA-DNA-binding\|\|REase+iREase-><-wHTH+SAD+HNH\|\|HNH-> | 804 | Bacteria>Pseudomonadota>Gammaproteobacteria>Enterobacterales>Enterobacteriaceae>Escherichia | GCA_000215245.2 | Escherichia_coli_96.154 | EC96154_4669 | 254732..257146 | 869683 | hypothetical-protein-EC96154_4669-[Escherichia-coli-96.154] |
| EHW21311.1 | Type1.BREX | <-AbiEii<-AbiEi_3_N+AbiEi_3<-HKD-DNase+OGG-like\|\|?-><-SIGMA+OB-Fold+AAA-ATPase+Lon-Protease-BrxL<-Zn_Tnp_IS91+Y2_Tnp<-?\|\|?-><-N6-METHYLASE<-DUF6079*\|\|HIN-HTH->HTH_Tnp_1+HTH_21+rve+rve_3-><-BrxA-DNA-binding\|\|REase+iREase-><-wHTH+SAD+HNH\|\|HNH-> | 1001 | Bacteria>Pseudomonadota>Gammaproteobacteria>Enterobacterales>Enterobacteriaceae>Escherichia | GCA_000249875.2 | Escherichia_coli_DEC8E | ECDEC8E_5320 | 144024..147029 | 868174 | putative-DNA-repair-ATPase-[Escherichia-coli-DEC8E] |
| EHX52156.1 | Type1.BREX | DUF6216-><-?<-AbiEii<-AbiEi_3_N+AbiEi_3<-HKD-DNase+OGG-like<-SIGMA+OB-Fold+AAA-ATPase+Lon-Protease-BrxL<-PglZ<-N6-METHYLASE-p<-N6-METHYLASE<-DUF6079-p*<-DUF6079-p<-BrxB-BrxF-iSTAND<-BrxA-DNA-binding\|\|?->REase+iREase-><-wHTH+SAD+HNH\|\|HNH-> | 788 | Bacteria>Pseudomonadota>Gammaproteobacteria>Enterobacterales>Enterobacteriaceae>Escherichia | GCA_000250315.2 | Escherichia_coli_DEC13A | ECDEC13A_0072 | 76997..79363 | 868196 | putative-DNA-repair-ATPase-[Escherichia-coli-DEC13A] |
| EHX55090.1 | Type1.BREX | <-AbiEii<-AbiEi_3_N+AbiEi_3<-HKD-DNase+OGG-like<-SIGMA+OB-Fold+AAA-ATPase+Lon-Protease-BrxL<-PglZ<-N6-METHYLASE-p<-N6-METHYLASE-p<-N6-METHYLASE<-DUF6079-p*<-DUF6079-p<-BrxB-BrxF-iSTAND<-BrxA-DNA-binding\|\|REase+iREase-><-wHTH+SAD+HNH\|\|HNH-> | 804 | Bacteria>Pseudomonadota>Gammaproteobacteria>Enterobacterales>Enterobacteriaceae>Escherichia | GCA_000250335.2 | Escherichia_coli_DEC13B | ECDEC13B_4632 | 100761..103175 | 868197 | putative-DNA-repair-ATPase-[Escherichia-coli-DEC13B] |
| ESD29615.1 | Type1.BREX | ZnR-><-SIGMA+OB-Fold+AAA-ATPase+Lon-Protease-BrxL<-PglZ<-N6-METHYLASE<-DUF6079*<-BrxB-BrxF-iSTAND<-BrxA-DNA-binding<-?\|\|REase->REase-> | 1218 | Bacteria>Pseudomonadota>Gammaproteobacteria>Enterobacterales>Enterobacteriaceae>Escherichia | GCA_000488435.1 | Escherichia_coli_907715 | HMPREF1600_01130 | 9186..12842 | 1268987 | hypothetical-protein-HMPREF1600_01130-[Escherichia-coli-907715] |
| ESD66367.1 | Type1.BREX | wHTH+SAD+HNH-><-REase+iREase\|\|?->BrxA-DNA-binding->BrxB-BrxF-iSTAND->DUF6079*->N6-METHYLASE-> | 1218 | Bacteria>Pseudomonadota>Gammaproteobacteria>Enterobacterales>Enterobacteriaceae>Escherichia | GCA_000488595.1 | Escherichia_coli_908541 | HMPREF1609_05038 | 9764..13420 | 1268996 | hypothetical-protein-HMPREF1609_05038-[Escherichia-coli-908541] |
| CDL44502.1 | Type1.BREX | <-DDE_Tnp_IS1<-?<-REase<-REase<-?\|\|BrxA-DNA-binding->BrxB-BrxF-iSTAND->DUF6079*->DUF6079-p->N6-METHYLASE->N6-METHYLASE-p->PglZ->SIGMA+OB-Fold+AAA-ATPase+Lon-Protease-BrxL-><-ZnR<-HTH_Tnp_1+HTH_21+rve+rve_3<-RVT | 1116 | Bacteria>Pseudomonadota>Gammaproteobacteria>Enterobacterales>Enterobacteriaceae>Escherichia | GCA_000529555.1 | Escherichia_coli_ISC41 | . | 3336..6686 | 1432559 | putative-DNA-repair-ATPase-[Escherichia-coli-ISC41] |
| KEM88219.1 | Type1.BREX | <-HTH_Tnp_1+HTH_21+rve+rve_3<-HTH_Tnp_1\|\|wHTH+WYL+WCX->BrxA-DNA-binding->BrxB-BrxF-iSTAND->DUF6079-p->DUF6079-p*->ParB+HNH->N6-METHYLASE->PglZ->SIGMA+OB-Fold+AAA-ATPase+Lon-Protease-BrxL->35exo->?-><-HTH_21+rve+rve_3<-HIN-HTH\|\|DDE_Tnp_IS1-> | 947 | Bacteria>Pseudomonadota>Gammaproteobacteria>Enterobacterales>Enterobacteriaceae>Escherichia | GCA_000713255.1 | Escherichia_coli_6-537-08_S4_C1 | AC92_4028 | 7171..10014 | 1444227 | putative-aTPase-like-protein-[Escherichia-coli-6-537-08_S4_C1] |
| KGM58490.1 | Type1.BREX | <-DDE_Tnp_IS1<-TetR-HTH<-?<-?<-?<-REase+iREase<-?\|\|BrxA-DNA-binding->BrxB-BrxF-iSTAND->DUF6079*->N6-METHYLASE->PglZ->SIGMA+OB-Fold+AAA-ATPase+Lon-Protease-BrxL-><-LZ_Tnp_IS66+zf-IS66+DDE_Tnp_IS66+DDE_Tnp_IS66_C<-?<-HIN-HTH<-DUF772+DDE_Tnp_1<-DDE_Tnp_IS1 | 1218 | Bacteria>Pseudomonadota>Gammaproteobacteria>Enterobacterales>Enterobacteriaceae>Escherichia | GCA_000768425.1 | Escherichia_coli_G3/10 | EL76_4293 | 581374..585030 | 1455601 | hypothetical-protein-EL76_4293-[Escherichia-coli-G3/10] |
| CTT26978.1 | Type1.BREX | DUF6079*->N6-METHYLASE->N6-METHYLASE-p->PglZ->SIGMA+OB-Fold+AAA-ATPase+Lon-Protease-BrxL->HKD-DNase+OGG-like->AbiEi_3_N+AbiEi_3->AbiEii-><-DUF6216<-Arm-DNA-bind_3 | 1088 | Bacteria>Pseudomonadota>Gammaproteobacteria>Enterobacterales>Enterobacteriaceae>Escherichia | GCA_001283645.1 | Escherichia_coli | ERS085343_03077 | 214..3480 | 562 | ATPase-like-protein-[Escherichia-coli] |
| CTW38723.1 | Type1.BREX | DUF6079-p*->N6-METHYLASE->PglZ->SIGMA+OB-Fold+AAA-ATPase+Lon-Protease-BrxL->HKD-DNase+OGG-like->AbiEi_3_N+AbiEi_3->AbiEii-><-DUF6216<-Arm-DNA-bind_3 | 788 | Bacteria>Pseudomonadota>Gammaproteobacteria>Enterobacterales>Enterobacteriaceae>Escherichia | GCA_001283765.1 | Escherichia_coli | ERS085413_04497 | 243..2609 | 562 | ATPase-like-protein-[Escherichia-coli] |
| CUA46950.1 | Type1.BREX | DUF6079*->N6-METHYLASE->N6-METHYLASE-p->PglZ->SIGMA+OB-Fold+AAA-ATPase+Lon-Protease-BrxL->HKD-DNase+OGG-like->AbiEi_3_N+AbiEi_3->AbiEii-><-DUF6216<-Arm-DNA-bind_3 | 1088 | Bacteria>Pseudomonadota>Gammaproteobacteria>Enterobacterales>Enterobacteriaceae>Escherichia | GCA_001286505.1 | Escherichia_coli | ERS139261_03500 | 135..3401 | 562 | ATPase-like-protein-[Escherichia-coli] |
| OWF25643.1 | Type1.BREX | DUF6079-p*->ParB+HNH->N6-METHYLASE-p->PglZ->SIGMA+OB-Fold+AAA-ATPase+Lon-Protease-BrxL-><-HTH_Tnp_1+HTH_21+rve+rve_3 | 902 | Bacteria>Pseudomonadota>Gammaproteobacteria>Enterobacterales>Enterobacteriaceae>Escherichia | GCA_002190095.1 | Escherichia_coli | A8M76_10400 | 1..2709 | 562 | DNA-repair-protein,-partial-[Escherichia-coli] |
| POU04217.1 | Type1.BREX | HNH-><-?\|\|?->XIS-HTH->SF1-SF2-Helicase+REase->SF1-SF2-Helicase+REase-><-PglZ<-ParB+HNH<-N6-METHYLASE<-DUF6079*<-BrxB-BrxF-iSTAND<-BrxA-DNA-binding<-Arm-DNA-bind_3+Phage_integrase<-tRNA\|\|wHTH(SBD)+HNH->SAM-METHYLASE-> | 1157 | Bacteria>Pseudomonadota>Gammaproteobacteria>Enterobacterales>Enterobacteriaceae>Escherichia | GCA_002918705.1 | Escherichia_sp._ESNIH1 | C3369_02380 | 500495..503968 | 1985876 | BREX-system-P-loop-protein-BrxC-[Escherichia-sp.-ESNIH1] |
| AUZ10893.1 | Type1.BREX | <-35exo\|\|?->?->?-><-HNH\|\|wHTH+SAD+HNH-><-REase+iREase\|\|BrxA-DNA-binding->BrxB-BrxF-iSTAND->\|\|DUF6079-p*->N6-METHYLASE->PglZ->SIGMA+OB-Fold+AAA-ATPase+Lon-Protease-BrxL->?-><-?<-ParB<-?\|\|<-REase+N6_METHYLASE<-TRD | 788 | Bacteria>Pseudomonadota>Gammaproteobacteria>Enterobacterales>Enterobacteriaceae>Escherichia | GCA_002968935.1 | Escherichia_coli | BWI88_05955 | 1182148..1184514 | 562 | DNA-repair-ATPase-[Escherichia-coli] |
| RLY30110.1 | Type1.BREX | <-HerA/FtsK+A32<-?\|\|tRNA->Arm-DNA-bind_3+Phage_integrase-><-SIGMA+OB-Fold+AAA-ATPase+Lon-Protease-BrxL<-PglZ<-N6-METHYLASE-p<-N6-METHYLASE<-ParB+HNH<-DUF6079*<-BrxB-BrxF-iSTAND<-BrxA-DNA-binding<-?<-?<-XIS-HTH\|\|?->nSTAND1+sGTP->wHTH+WYL-> | 1233 | Bacteria>Pseudomonadota>Gammaproteobacteria>Enterobacterales>Enterobacteriaceae>Escherichia | GCA_003687125.1 | Escherichia_coli | EAI39_15065 | 84115..87816 | 562 | BREX-system-P-loop-protein-BrxC-[Escherichia-coli] |
| EEG9579271.1 | Type1.BREX | DUF6079*->N6-METHYLASE->PglZ->SIGMA+OB-Fold+AAA-ATPase+Lon-Protease-BrxL->HKD-DNase+OGG-like->\|\|AbiEi_3_N+AbiEi_3->AbiEii->?-><-DUF6216<-Arm-DNA-bind_3 | 1111 | Bacteria>Pseudomonadota>Gammaproteobacteria>Enterobacterales>Enterobacteriaceae>Escherichia | GCA_003736085.2 | Escherichia_coli | EB576_RS15540 | 1..3336 | 562 | BREX-system-P-loop-protein-BrxC-[Escherichia-coli] |
| EED0771052.1 | Type1.BREX | BrxA-DNA-binding->BrxB-BrxF-iSTAND->DUF6079*->N6-METHYLASE->PglZ->SIGMA+OB-Fold+AAA-ATPase+Lon-Protease-BrxL->HKD-DNase+OGG-like->\|\|AbiEi_3_N+AbiEi_3->AbiEii-><-DUF6216<-Arm-DNA-bind_3 | 1213 | Bacteria>Pseudomonadota>Gammaproteobacteria>Enterobacterales>Enterobacteriaceae>Escherichia | GCA_003747865.2 | Escherichia_coli | ED219_RS15230 | 935..4576 | 562 | BREX-system-P-loop-protein-BrxC-[Escherichia-coli] |
| EED1762068.1 | Type1.BREX | DUF6079*->PglZ->SIGMA+OB-Fold+AAA-ATPase+Lon-Protease-BrxL->HKD-DNase+OGG-like->\|\|AbiEi_3_N+AbiEi_3->AbiEii-><-DUF6216<-Arm-DNA-bind_3<-tRNA\|\|?->HerA/FtsK+A32-> | 1073 | Bacteria>Pseudomonadota>Gammaproteobacteria>Enterobacterales>Enterobacteriaceae>Escherichia | GCA_003786715.2 | Escherichia_coli | EEP34_RS13370 | 1..3223 | 562 | BREX-system-P-loop-protein-BrxC-[Escherichia-coli] |
| EEC9685943.1 | Type1.BREX | DUF6079-p*->N6-METHYLASE->PglZ->SIGMA+OB-Fold+AAA-ATPase+Lon-Protease-BrxL->HKD-DNase+OGG-like->\|\|AbiEi_3_N+AbiEi_3->AbiEii-><-DUF6216<-Arm-DNA-bind_3 | 989 | Bacteria>Pseudomonadota>Gammaproteobacteria>Enterobacterales>Enterobacteriaceae>Escherichia | GCA_004231765.2 | Escherichia_coli | D8K54_RS17535 | 1..2971 | 562 | BREX-system-P-loop-protein-BrxC-[Escherichia-coli] |
| EEC7405250.1 | Type1.BREX | <-Phage_integrase\|\|?->?->?-><-HNH\|\|wHTH+SAD+HNH-><-REase+iREase\|\|BrxA-DNA-binding->BrxB-BrxF-iSTAND->DUF6079*->N6-METHYLASE->PglZ->SIGMA+OB-Fold+AAA-ATPase+Lon-Protease-BrxL->HKD-DNase+OGG-like->\|\|AbiEi_3_N+AbiEi_3->AbiEii-><-DUF6216<-Arm-DNA-bind_3 | 1213 | Bacteria>Pseudomonadota>Gammaproteobacteria>Enterobacterales>Enterobacteriaceae>Escherichia | GCA_004272605.2 | Escherichia_coli | D4L89_RS08005 | 71644..75285 | 562 | BREX-system-P-loop-protein-BrxC-[Escherichia-coli] |
| GCK20492.1 | Type1.BREX | <-Fimbrial<-Phage_integrase<-Phage_integrase\|\|?->?->?->wHTH(SBD)+HNH->BrxA-DNA-binding->BrxB-BrxF-iSTAND->DUF6079-p*-> | 621 | Bacteria>Pseudomonadota>Gammaproteobacteria>Enterobacterales>Enterobacteriaceae>Escherichia | GCA_005401165.1 | Escherichia_coli | BvCmsA58A_02120 | 111143..113008 | 562 | ATPase-like-protein-[Escherichia-coli] |
| TZF09961.1 | Type1.BREX | RVT->DEDD_Tnp_IS110+Transposase_20-><-SIGMA+OB-Fold+AAA-ATPase+Lon-Protease-BrxL<-PglZ<-N6-METHYLASE<-ParB+HNH<-DUF6079* | 1149 | Bacteria>Pseudomonadota>Gammaproteobacteria>Enterobacterales>Enterobacteriaceae>Escherichia | GCA_008273745.1 | Escherichia_coli | EOI42_22665 | 16149..19600 | 562 | BREX-system-P-loop-protein-BrxC,-partial-[Escherichia-coli] |
| TZE19045.1 | Type1.BREX | RVT->DEDD_Tnp_IS110+Transposase_20-><-SIGMA+OB-Fold+AAA-ATPase+Lon-Protease-BrxL<-PglZ<-N6-METHYLASE<-DUF6079*<-BrxB-BrxF-iSTAND<-BrxA-DNA-binding<-wHTH+WYL+WCX<-?\|\|RHH->\|\|CopG/RHH->?-><-DDE_Tnp_IS1 | 1225 | Bacteria>Pseudomonadota>Gammaproteobacteria>Enterobacterales>Enterobacteriaceae>Escherichia | GCA_008273875.1 | Escherichia_coli | EOI13_22465 | 16132..19809 | 562 | BREX-system-P-loop-protein-BrxC-[Escherichia-coli] |
| TZE05560.1 | Type1.BREX | <-CopG/RHH\|\|<-RHH\|\|?->wHTH+WYL+WCX->BrxA-DNA-binding->BrxB-BrxF-iSTAND->DUF6079*->ParB+HNH->N6-METHYLASE->PglZ->SIGMA+OB-Fold+AAA-ATPase+Lon-Protease-BrxL-><-DEDD_Tnp_IS110+Transposase_20<-RVT | 1225 | Bacteria>Pseudomonadota>Gammaproteobacteria>Enterobacterales>Enterobacteriaceae>Escherichia | GCA_008273905.1 | Escherichia_coli | EOI14_22605 | 9953..13630 | 562 | BREX-system-P-loop-protein-BrxC-[Escherichia-coli] |
| KAA9812768.1 | Type1.BREX | DUF6079-p*->N6-METHYLASE->PglZ->SIGMA+OB-Fold+AAA-ATPase+Lon-Protease-BrxL->HKD-DNase+OGG-like->\|\|AbiEi_3_N+AbiEi_3->AbiEii-><-DUF6216<-Arm-DNA-bind_3 | 891 | Bacteria>Pseudomonadota>Gammaproteobacteria>Enterobacterales>Enterobacteriaceae>Escherichia | GCA_008755545.1 | Escherichia_coli | F7F73_23785 | 1..2676 | 562 | BREX-system-P-loop-protein-BrxC,-partial-[Escherichia-coli] |
| KAA9812739.1 | Type1.BREX | DUF6079-p*->N6-METHYLASE->PglZ->SIGMA+OB-Fold+AAA-ATPase+Lon-Protease-BrxL-><-ZnR<-ZnR<-?\|\|HIN-HTH+HIN-HTH-> | 891 | Bacteria>Pseudomonadota>Gammaproteobacteria>Enterobacterales>Enterobacteriaceae>Escherichia | GCA_008755545.1 | Escherichia_coli | F7F73_23930 | 1..2676 | 562 | BREX-system-P-loop-protein-BrxC,-partial-[Escherichia-coli] |
| KAB2983059.1 | Type1.BREX | <-Phage_integrase\|\|?->?->?-><-HNH\|\|wHTH+SAD+HNH-><-REase+iREase\|\|BrxA-DNA-binding->BrxB-BrxF-iSTAND->DUF6079*->N6-METHYLASE->PglZ->SIGMA+OB-Fold+AAA-ATPase+Lon-Protease-BrxL->?-><-?<-ParB<-?\|\|<-REase+N6_METHYLASE<-TRD | 1213 | Bacteria>Pseudomonadota>Gammaproteobacteria>Enterobacterales>Enterobacteriaceae>Escherichia | GCA_009014785.1 | Escherichia_coli | F9179_13170 | 9843..13484 | 562 | BREX-system-P-loop-protein-BrxC-[Escherichia-coli] |
| EEQ1797639.1 | Type1.BREX | DUF6079-p*->N6-METHYLASE->PglZ->SIGMA+OB-Fold+AAA-ATPase+Lon-Protease-BrxL->HKD-DNase+OGG-like->\|\|AbiEi_3_N+AbiEi_3->AbiEii-><-DUF6216<-Arm-DNA-bind_3 | 861 | Bacteria>Pseudomonadota>Gammaproteobacteria>Enterobacterales>Enterobacteriaceae>Escherichia | GCA_011802225.1 | Escherichia_coli | GUY43_002901 | 1..2587 | 562 | BREX-system-P-loop-protein-BrxC-[Escherichia-coli] |
| EEQ2509736.1 | Type1.BREX | BrxA-DNA-binding->BrxB-BrxF-iSTAND->DUF6079*->N6-METHYLASE->PglZ->SIGMA+OB-Fold+AAA-ATPase+Lon-Protease-BrxL->HKD-DNase+OGG-like->\|\|AbiEi_3_N+AbiEi_3->AbiEii-><-DUF6216<-Arm-DNA-bind_3 | 1213 | Bacteria>Pseudomonadota>Gammaproteobacteria>Enterobacterales>Enterobacteriaceae>Escherichia | GCA_011806325.1 | Escherichia_coli | GSE75_002859 | 1085..4726 | 562 | BREX-system-P-loop-protein-BrxC-[Escherichia-coli] |
| EEQ6465042.1 | Type1.BREX | <-XIS-HTH<-RHH_antitoxin\|\|?->?->ZnR->ZnR-><-SIGMA+OB-Fold+AAA-ATPase+Lon-Protease-BrxL<-PglZ<-N6-METHYLASE<-DUF6079*<-BrxB-BrxF-iSTAND<-BrxA-DNA-binding\|\|<-HTH_Tnp_1+HTH_21+rve+rve_3 | 1213 | Bacteria>Pseudomonadota>Gammaproteobacteria>Enterobacterales>Enterobacteriaceae>Escherichia | GCA_011825175.1 | Escherichia_coli | GMM94_000612 | 221240..224881 | 562 | BREX-system-P-loop-protein-BrxC-[Escherichia-coli] |
| EEQ7780471.1 | Type1.BREX | <-HNH\|\|wHTH+SAD+HNH-><-REase+iREase\|\|BrxA-DNA-binding->BrxB-BrxF-iSTAND->DUF6079*->N6-METHYLASE->PglZ->SIGMA+OB-Fold+AAA-ATPase+Lon-Protease-BrxL->HKD-DNase+OGG-like->\|\|AbiEi_3_N+AbiEi_3->AbiEii-><-DUF6216<-Arm-DNA-bind_3 | 1213 | Bacteria>Pseudomonadota>Gammaproteobacteria>Enterobacterales>Enterobacteriaceae>Escherichia | GCA_011830195.1 | Escherichia_coli | GLS89_002414 | 7714..11355 | 562 | BREX-system-P-loop-protein-BrxC-[Escherichia-coli] |
| EER7097196.1 | Type1.BREX | DUF6079*->N6-METHYLASE->PglZ->SIGMA+OB-Fold+AAA-ATPase+Lon-Protease-BrxL->HKD-DNase+OGG-like->\|\|AbiEi_3_N+AbiEi_3->AbiEii-><-DUF6216<-Arm-DNA-bind_3 | 1064 | Bacteria>Pseudomonadota>Gammaproteobacteria>Enterobacterales>Enterobacteriaceae>Escherichia | GCA_011867305.1 | Escherichia_coli | F2Z67_003270 | 1..3197 | 562 | BREX-system-P-loop-protein-BrxC-[Escherichia-coli] |
| EER7518149.1 | Type1.BREX | DUF6079-p*->N6-METHYLASE->PglZ->SIGMA+OB-Fold+AAA-ATPase+Lon-Protease-BrxL->HKD-DNase+OGG-like->\|\|AbiEi_3_N+AbiEi_3->AbiEii-><-DUF6216<-Arm-DNA-bind_3 | 990 | Bacteria>Pseudomonadota>Gammaproteobacteria>Enterobacterales>Enterobacteriaceae>Escherichia | GCA_011869725.1 | Escherichia_coli | F1Z98_003251 | 1..2974 | 562 | BREX-system-P-loop-protein-BrxC-[Escherichia-coli] |
| EER8632679.1 | Type1.BREX | DUF6079*->N6-METHYLASE->PglZ->SIGMA+OB-Fold+AAA-ATPase+Lon-Protease-BrxL->HKD-DNase+OGG-like->\|\|AbiEi_3_N+AbiEi_3->AbiEii-><-DUF6216<-Arm-DNA-bind_3 | 1080 | Bacteria>Pseudomonadota>Gammaproteobacteria>Enterobacterales>Enterobacteriaceae>Escherichia | GCA_011873745.1 | Escherichia_coli | F0E18_004248 | 1..3245 | 562 | BREX-system-P-loop-protein-BrxC-[Escherichia-coli] |
| MBC1090136.1 | Type1.BREX | <-HerA/FtsK+A32<-?\|\|tRNA->Arm-DNA-bind_3+Phage_integrase->EVE+HNH-><-REase+iREase\|\|BrxA-DNA-binding->BrxB-BrxF-iSTAND->DUF6079*->N6-METHYLASE->PglZ->SIGMA+OB-Fold+AAA-ATPase+Lon-Protease-BrxL-><-ZnR<-ZnR<-?<-?\|\|RHH_antitoxin->XIS-HTH-> | 1213 | Bacteria>Pseudomonadota>Gammaproteobacteria>Enterobacterales>Enterobacteriaceae>Escherichia | GCA_011890865.2 | Escherichia_coli | FSG28_003850 | 139502..143143 | 562 | BREX-system-P-loop-protein-BrxC-[Escherichia-coli] |
| EES3817132.1 | Type1.BREX | DUF6079*->N6-METHYLASE->PglZ->SIGMA+OB-Fold+AAA-ATPase+Lon-Protease-BrxL->HKD-DNase+OGG-like->\|\|AbiEi_3_N+AbiEi_3->AbiEii-><-DUF6216<-Arm-DNA-bind_3 | 952 | Bacteria>Pseudomonadota>Gammaproteobacteria>Enterobacterales>Enterobacteriaceae>Escherichia | GCA_011896505.1 | Escherichia_coli | FS433_003751 | 1..2860 | 562 | BREX-system-P-loop-protein-BrxC-[Escherichia-coli] |
| EES8430772.1 | Type1.BREX | DUF6079-p*->N6-METHYLASE->PglZ->SIGMA+OB-Fold+AAA-ATPase+Lon-Protease-BrxL->HKD-DNase+OGG-like->\|\|AbiEi_3_N+AbiEi_3->AbiEii-><-DUF6216<-Arm-DNA-bind_3 | 988 | Bacteria>Pseudomonadota>Gammaproteobacteria>Enterobacterales>Enterobacteriaceae>Escherichia | GCA_011915225.1 | Escherichia_coli | FU162_003513 | 1..2968 | 562 | BREX-system-P-loop-protein-BrxC-[Escherichia-coli] |
| EET5616501.1 | Type1.BREX | <-RHH_antitoxin\|\|?->?->ZnR->ZnR-><-SIGMA+OB-Fold+AAA-ATPase+Lon-Protease-BrxL<-PglZ<-N6-METHYLASE<-DUF6079*<-BrxB-BrxF-iSTAND<-BrxA-DNA-binding\|\|<-EVE+HNH<-Arm-DNA-bind_3+Phage_integrase<-tRNA\|\|?->HerA/FtsK+A32-><-?<-?\|\|Peptidase_M17_N+Peptidase_M17-> | 1213 | Bacteria>Pseudomonadota>Gammaproteobacteria>Enterobacterales>Enterobacteriaceae>Escherichia | GCA_011940945.1 | Escherichia_coli | FFE61_003335 | 12923..16564 | 562 | BREX-system-P-loop-protein-BrxC-[Escherichia-coli] |
| EET5967869.1 | Type1.BREX | DUF6079*->N6-METHYLASE->PglZ->SIGMA+OB-Fold+AAA-ATPase+Lon-Protease-BrxL->HKD-DNase+OGG-like->\|\|AbiEi_3_N+AbiEi_3->AbiEii-><-DUF6216<-Arm-DNA-bind_3 | 1065 | Bacteria>Pseudomonadota>Gammaproteobacteria>Enterobacterales>Enterobacteriaceae>Escherichia | GCA_011942865.1 | Escherichia_coli | FE570_001648 | 1..3198 | 562 | BREX-system-P-loop-protein-BrxC-[Escherichia-coli] |
| EEU4094847.1 | Type1.BREX | DUF6079-p*->N6-METHYLASE->PglZ->SIGMA+OB-Fold+AAA-ATPase+Lon-Protease-BrxL->HKD-DNase+OGG-like->\|\|AbiEi_3_N+AbiEi_3->AbiEii-><-DUF6216<-Arm-DNA-bind_3 | 779 | Bacteria>Pseudomonadota>Gammaproteobacteria>Enterobacterales>Enterobacteriaceae>Escherichia | GCA_011974935.1 | Escherichia_coli | E8H53_003141 | 1..2340 | 562 | BREX-system-P-loop-protein-BrxC-[Escherichia-coli] |
| EEU4096206.1 | Type1.BREX | <-DUF6079-p*<-BrxB-BrxF-iSTAND<-BrxA-DNA-binding\|\|<-wHTH(SBD)+HNH\|\|HNH-><-?<-?<-?\|\|Phage_integrase->Phage_integrase-> | 433 | Bacteria>Pseudomonadota>Gammaproteobacteria>Enterobacterales>Enterobacteriaceae>Escherichia | GCA_011974935.1 | Escherichia_coli | E8H53_004589 | 1..1299 | 562 | BREX-system-P-loop-protein-BrxC-[Escherichia-coli] |
| EEV0352856.1 | Type1.BREX | DUF6079*->N6-METHYLASE->PglZ->SIGMA+OB-Fold+AAA-ATPase+Lon-Protease-BrxL->HKD-DNase+OGG-like->\|\|AbiEi_3_N+AbiEi_3->AbiEii-><-DUF6216<-Arm-DNA-bind_3 | 1194 | Bacteria>Pseudomonadota>Gammaproteobacteria>Enterobacterales>Enterobacteriaceae>Escherichia | GCA_012011485.1 | Escherichia_coli | EYE11_003099 | 1..3587 | 562 | BREX-system-P-loop-protein-BrxC-[Escherichia-coli] |
| EEV1146578.1 | Type1.BREX | <-Phage_integrase\|\|?->?->?-><-HNH\|\|wHTH(SBD)+HNH->\|\|BrxA-DNA-binding->BrxB-BrxF-iSTAND->DUF6079*->N6-METHYLASE-> | 1213 | Bacteria>Pseudomonadota>Gammaproteobacteria>Enterobacterales>Enterobacteriaceae>Escherichia | GCA_012013845.1 | Escherichia_coli | EUB49_004354 | 8713..12354 | 562 | BREX-system-P-loop-protein-BrxC-[Escherichia-coli] |
| EEV8721794.1 | Type1.BREX | <-DUF6079-p*<-BrxB-BrxF-iSTAND<-BrxA-DNA-binding\|\|REase+iREase-><-wHTH+SAD+HNH\|\|HNH-><-?<-?<-?\|\|Phage_integrase-> | 634 | Bacteria>Pseudomonadota>Gammaproteobacteria>Enterobacterales>Enterobacteriaceae>Escherichia | GCA_012057805.1 | Escherichia_coli | EHU66_22975 | 1..1904 | 562 | BREX-system-P-loop-protein-BrxC-[Escherichia-coli] |
| EEV8657615.1 | Type1.BREX | DUF6079*->N6-METHYLASE->PglZ->SIGMA+OB-Fold+AAA-ATPase+Lon-Protease-BrxL->HKD-DNase+OGG-like->\|\|AbiEi_3_N+AbiEi_3->AbiEii-><-DUF6216<-Arm-DNA-bind_3 | 1180 | Bacteria>Pseudomonadota>Gammaproteobacteria>Enterobacterales>Enterobacteriaceae>Escherichia | GCA_012058765.1 | Escherichia_coli | EHJ50_001183 | 1..3543 | 562 | BREX-system-P-loop-protein-BrxC-[Escherichia-coli] |
| EEV9926442.1 | Type1.BREX | DUF772+DDE_Tnp_1-><-REase+iREase\|\|BrxA-DNA-binding->BrxB-BrxF-iSTAND->DUF6079*->N6-METHYLASE->PglZ->SIGMA+OB-Fold+AAA-ATPase+Lon-Protease-BrxL-><-ZnR<-ZnR<-?<-?<-?\|\|RHH_antitoxin-> | 1213 | Bacteria>Pseudomonadota>Gammaproteobacteria>Enterobacterales>Enterobacteriaceae>Escherichia | GCA_012061085.1 | Escherichia_coli | EG211_22240 | 4333..7974 | 562 | BREX-system-P-loop-protein-BrxC-[Escherichia-coli] |
| EEW0980710.1 | Type1.BREX | <-Phage_integrase\|\|?->?->?-><-HNH\|\|wHTH+SAD+HNH-><-REase+iREase\|\|BrxA-DNA-binding->BrxB-BrxF-iSTAND->DUF6079*->N6-METHYLASE->PglZ->SIGMA+OB-Fold+AAA-ATPase+Lon-Protease-BrxL->HKD-DNase+OGG-like->\|\|AbiEi_3_N+AbiEi_3->AbiEii-><-DUF6216<-Arm-DNA-bind_3 | 1213 | Bacteria>Pseudomonadota>Gammaproteobacteria>Enterobacterales>Enterobacteriaceae>Escherichia | GCA_012068555.1 | Escherichia_coli | ED289_06180 | 111258..114899 | 562 | BREX-system-P-loop-protein-BrxC-[Escherichia-coli] |
| EEW0945476.1 | Type1.BREX | <-Phage_integrase\|\|?->?->?-><-HNH\|\|wHTH+SAD+HNH-><-REase+iREase\|\|BrxA-DNA-binding->BrxB-BrxF-iSTAND->DUF6079*->PglZ->SIGMA+OB-Fold+AAA-ATPase+Lon-Protease-BrxL->?-><-?<-ParB<-?\|\|<-REase+N6_METHYLASE<-TRD<-REase | 1213 | Bacteria>Pseudomonadota>Gammaproteobacteria>Enterobacterales>Enterobacteriaceae>Escherichia | GCA_012068805.1 | Escherichia_coli | BCB75_000479 | 153764..157405 | 562 | BREX-system-P-loop-protein-BrxC-[Escherichia-coli] |
| EEW0874914.1 | Type1.BREX | <-Phage_integrase\|\|?->?->?-><-HNH\|\|wHTH+SAD+HNH-><-REase+iREase\|\|BrxA-DNA-binding->BrxB-BrxF-iSTAND->DUF6079*->N6-METHYLASE->PglZ->SIGMA+OB-Fold+AAA-ATPase+Lon-Protease-BrxL->?-><-?<-ParB | 1213 | Bacteria>Pseudomonadota>Gammaproteobacteria>Enterobacterales>Enterobacteriaceae>Escherichia | GCA_012068885.1 | Escherichia_coli | ED728_000829 | 111230..114871 | 562 | BREX-system-P-loop-protein-BrxC-[Escherichia-coli] |
| EEW2565584.1 | Type1.BREX | Arm-DNA-bind_3+Phage_integrase->N6-METHYLASE->ORC-CDC6+ZnR->HEPN-><-REase<-REase\|\|BrxA-DNA-binding->BrxB-BrxF-iSTAND->DUF6079*->N6-METHYLASE->PglZ->SIGMA+OB-Fold+AAA-ATPase+Lon-Protease-BrxL-><-ZnR<-ZnR<-?<-?\|\|RHH_antitoxin->XIS-HTH-> | 1213 | Bacteria>Pseudomonadota>Gammaproteobacteria>Enterobacterales>Enterobacteriaceae>Escherichia | GCA_012076045.1 | Escherichia_coli | D9E67_18610 | 72989..76630 | 562 | BREX-system-P-loop-protein-BrxC-[Escherichia-coli] |
| EEW3089662.1 | Type1.BREX | <-Phage_integrase<-Phage_integrase\|\|?->?->?-><-HNH\|\|wHTH(SBD)+HNH->\|\|BrxA-DNA-binding->BrxB-BrxF-iSTAND->DUF6079*->N6-METHYLASE->PglZ->SIGMA+OB-Fold+AAA-ATPase+Lon-Protease-BrxL->HKD-DNase+OGG-like->\|\|AbiEi_3_N+AbiEi_3->AbiEii-><-DUF6216<-Arm-DNA-bind_3 | 1213 | Bacteria>Pseudomonadota>Gammaproteobacteria>Enterobacterales>Enterobacteriaceae>Escherichia | GCA_012078325.1 | Escherichia_coli | EBA98_002639 | 14697..18338 | 562 | BREX-system-P-loop-protein-BrxC-[Escherichia-coli] |
| EEW3885152.1 | Type1.BREX | <-HNH\|\|wHTH(SBD)+HNH->\|\|BrxA-DNA-binding->BrxB-BrxF-iSTAND->DUF6079*->N6-METHYLASE->PglZ->SIGMA+OB-Fold+AAA-ATPase+Lon-Protease-BrxL->HKD-DNase+OGG-like->\|\|AbiEi_3_N+AbiEi_3->AbiEii-><-DUF6216<-Arm-DNA-bind_3 | 1213 | Bacteria>Pseudomonadota>Gammaproteobacteria>Enterobacterales>Enterobacteriaceae>Escherichia | GCA_012080795.1 | Escherichia_coli | D9Y07_001392 | 7166..10807 | 562 | BREX-system-P-loop-protein-BrxC-[Escherichia-coli] |
| EEW4281070.1 | Type1.BREX | DUF6079*->N6-METHYLASE->PglZ->SIGMA+OB-Fold+AAA-ATPase+Lon-Protease-BrxL->HKD-DNase+OGG-like->\|\|AbiEi_3_N+AbiEi_3->AbiEii-><-DUF6216<-Arm-DNA-bind_3 | 1085 | Bacteria>Pseudomonadota>Gammaproteobacteria>Enterobacterales>Enterobacteriaceae>Escherichia | GCA_012082755.1 | Escherichia_coli | D9T01_23345 | 1..3259 | 562 | BREX-system-P-loop-protein-BrxC-[Escherichia-coli] |
| EEW6030677.1 | Type1.BREX | DUF6079*->PglZ->SIGMA+OB-Fold+AAA-ATPase+Lon-Protease-BrxL->HKD-DNase+OGG-like->\|\|AbiEi_3_N+AbiEi_3->AbiEii-><-DUF6216<-Arm-DNA-bind_3<-tRNA\|\|?->HerA/FtsK+A32-> | 1178 | Bacteria>Pseudomonadota>Gammaproteobacteria>Enterobacterales>Enterobacteriaceae>Escherichia | GCA_012088895.1 | Escherichia_coli | D7V62_16715 | 1..3539 | 562 | BREX-system-P-loop-protein-BrxC-[Escherichia-coli] |
| EEW5973241.1 | Type1.BREX | DUF6079*->N6-METHYLASE->PglZ->SIGMA+OB-Fold+AAA-ATPase+Lon-Protease-BrxL-><-ZnR<-ZnR<-?\|\|RHH_antitoxin->XIS-HTH-> | 1050 | Bacteria>Pseudomonadota>Gammaproteobacteria>Enterobacterales>Enterobacteriaceae>Escherichia | GCA_012088995.1 | Escherichia_coli | D7W18_24725 | 1..3153 | 562 | BREX-system-P-loop-protein-BrxC-[Escherichia-coli] |
| EEW8735903.1 | Type1.BREX | <-HIN-HTH\|\|<-35exo<-SIGMA+OB-Fold+AAA-ATPase+Lon-Protease-BrxL<-PglZ<-N6-METHYLASE<-ParB+HNH<-DUF6079*<-BrxB-BrxF-iSTAND<-BrxA-DNA-binding<-wHTH+WYL+WCX<-ZnR | 1221 | Bacteria>Pseudomonadota>Gammaproteobacteria>Enterobacterales>Enterobacteriaceae>Escherichia | GCA_012099745.1 | Escherichia_coli | D5Q61_02350 | 121702..125367 | 562 | BREX-system-P-loop-protein-BrxC-[Escherichia-coli] |
| EEX1719682.1 | Type1.BREX | <-HNH\|\|wHTH(SBD)+HNH->\|\|BrxA-DNA-binding->BrxB-BrxF-iSTAND->DUF6079*->N6-METHYLASE->PglZ->SIGMA+OB-Fold+AAA-ATPase+Lon-Protease-BrxL->HKD-DNase+OGG-like->\|\|AbiEi_3_N+AbiEi_3->AbiEii-><-DUF6216<-Arm-DNA-bind_3 | 1213 | Bacteria>Pseudomonadota>Gammaproteobacteria>Enterobacterales>Enterobacteriaceae>Escherichia | GCA_012115095.1 | Escherichia_coli | D3S79_003407 | 6685..10326 | 562 | BREX-system-P-loop-protein-BrxC-[Escherichia-coli] |
| EEX4954053.1 | Type1.BREX | BrxB-BrxF-iSTAND->DUF6079*->N6-METHYLASE->PglZ->SIGMA+OB-Fold+AAA-ATPase+Lon-Protease-BrxL->HKD-DNase+OGG-like->\|\|AbiEi_3_N+AbiEi_3->AbiEii-> | 1213 | Bacteria>Pseudomonadota>Gammaproteobacteria>Enterobacterales>Enterobacteriaceae>Escherichia | GCA_012126795.1 | Escherichia_coli | D0791_003833 | 606..4247 | 562 | BREX-system-P-loop-protein-BrxC-[Escherichia-coli] |
| EEX5072170.1 | Type1.BREX | <-N6-METHYLASE<-DUF6079*<-BrxB-BrxF-iSTAND<-BrxA-DNA-binding\|\|REase+iREase-><-wHTH+SAD+HNH\|\|HNH-><-?<-?<-?\|\|Phage_integrase-> | 1213 | Bacteria>Pseudomonadota>Gammaproteobacteria>Enterobacterales>Enterobacteriaceae>Escherichia | GCA_012127315.1 | Escherichia_coli | DYU47_004374 | 2562..6203 | 562 | BREX-system-P-loop-protein-BrxC-[Escherichia-coli] |
| EEX6773445.1 | Type1.BREX | DUF6079-p*->N6-METHYLASE->PglZ->SIGMA+OB-Fold+AAA-ATPase+Lon-Protease-BrxL->HKD-DNase+OGG-like->\|\|AbiEi_3_N+AbiEi_3->AbiEii-><-DUF6216<-Arm-DNA-bind_3 | 987 | Bacteria>Pseudomonadota>Gammaproteobacteria>Enterobacterales>Enterobacteriaceae>Escherichia | GCA_012133985.1 | Escherichia_coli | DYL55_002403 | 1..2965 | 562 | BREX-system-P-loop-protein-BrxC-[Escherichia-coli] |
| EEX6733972.1 | Type1.BREX | <-DUF6079*<-BrxB-BrxF-iSTAND<-BrxA-DNA-binding\|\|<-wHTH(SBD)+HNH\|\|HNH-><-?<-?<-?\|\|Phage_integrase->Phage_integrase-> | 1175 | Bacteria>Pseudomonadota>Gammaproteobacteria>Enterobacterales>Enterobacteriaceae>Escherichia | GCA_012135265.1 | Escherichia_coli | L0765_004693 | 1..3524 | 562 | BREX-system-P-loop-protein-BrxC-[Escherichia-coli] |
| EEX7673116.1 | Type1.BREX | DUF6079*->N6-METHYLASE->PglZ->SIGMA+OB-Fold+AAA-ATPase+Lon-Protease-BrxL->HKD-DNase+OGG-like->\|\|AbiEi_3_N+AbiEi_3->AbiEii-><-DUF6216<-Arm-DNA-bind_3 | 1156 | Bacteria>Pseudomonadota>Gammaproteobacteria>Enterobacterales>Enterobacteriaceae>Escherichia | GCA_012137685.1 | Escherichia_coli | DX034_003680 | 1..3471 | 562 | BREX-system-P-loop-protein-BrxC-[Escherichia-coli] |
| EEY1277168.1 | Type1.BREX | DUF6079*->N6-METHYLASE->PglZ->SIGMA+OB-Fold+AAA-ATPase+Lon-Protease-BrxL->HKD-DNase+OGG-like->\|\|AbiEi_3_N+AbiEi_3->AbiEii->?-><-DUF6216<-Arm-DNA-bind_3 | 1189 | Bacteria>Pseudomonadota>Gammaproteobacteria>Enterobacterales>Enterobacteriaceae>Escherichia | GCA_012153105.1 | Escherichia_coli | BFU70_002920 | 1..3572 | 562 | BREX-system-P-loop-protein-BrxC-[Escherichia-coli] |
| EEY1600790.1 | Type1.BREX | <-Phage_integrase\|\|?->?->?-><-HNH\|\|wHTH+SAD+HNH-><-REase+iREase\|\|BrxA-DNA-binding->BrxB-BrxF-iSTAND->DUF6079*->N6-METHYLASE->PglZ->SIGMA+OB-Fold+AAA-ATPase+Lon-Protease-BrxL->HKD-DNase+OGG-like->\|\|AbiEi_3_N+AbiEi_3->AbiEii-><-DUF6216<-Arm-DNA-bind_3 | 1213 | Bacteria>Pseudomonadota>Gammaproteobacteria>Enterobacterales>Enterobacteriaceae>Escherichia | GCA_012154025.1 | Escherichia_coli | DP662_003727 | 9743..13384 | 562 | BREX-system-P-loop-protein-BrxC-[Escherichia-coli] |
| EEY2596885.1 | Type1.BREX | DUF6079*->N6-METHYLASE->PglZ->SIGMA+OB-Fold+AAA-ATPase+Lon-Protease-BrxL->HKD-DNase+OGG-like->\|\|AbiEi_3_N+AbiEi_3->AbiEii-><-DUF6216<-tRNA\|\|?->HerA/FtsK+A32-> | 1032 | Bacteria>Pseudomonadota>Gammaproteobacteria>Enterobacterales>Enterobacteriaceae>Escherichia | GCA_012158085.1 | Escherichia_coli | E9W20_002908 | 1..3101 | 562 | BREX-system-P-loop-protein-BrxC-[Escherichia-coli] |
| EEY3542786.1 | Type1.BREX | <-DUF6079*<-BrxB-BrxF-iSTAND<-BrxA-DNA-binding\|\|REase+iREase-><-wHTH+SAD+HNH\|\|HNH-><-?<-?<-?\|\|Phage_integrase-> | 1157 | Bacteria>Pseudomonadota>Gammaproteobacteria>Enterobacterales>Enterobacteriaceae>Escherichia | GCA_012162105.1 | Escherichia_coli | EQJ55_17955 | 1..3471 | 562 | BREX-system-P-loop-protein-BrxC-[Escherichia-coli] |
| EEY3544735.1 | Type1.BREX | <-Phage_integrase\|\|?->?->?-><-HNH\|\|wHTH+SAD+HNH-><-REase+iREase\|\|BrxA-DNA-binding->BrxB-BrxF-iSTAND->DUF6079*->N6-METHYLASE->PglZ->SIGMA+OB-Fold+AAA-ATPase+Lon-Protease-BrxL->?-><-?<-ParB<-?\|\|<-REase+N6_METHYLASE<-TRD | 1213 | Bacteria>Pseudomonadota>Gammaproteobacteria>Enterobacterales>Enterobacteriaceae>Escherichia | GCA_012162185.1 | Escherichia_coli | EQJ57_01200 | 8758..12399 | 562 | BREX-system-P-loop-protein-BrxC-[Escherichia-coli] |
| EEY3950320.1 | Type1.BREX | <-RHH_antitoxin\|\|?->?->?->ZnR->ZnR-><-SIGMA+OB-Fold+AAA-ATPase+Lon-Protease-BrxL<-PglZ<-N6-METHYLASE<-DUF6079*<-BrxB-BrxF-iSTAND<-BrxA-DNA-binding\|\|REase+iREase-><-EVE+HNH<-Arm-DNA-bind_3+Phage_integrase<-tRNA\|\|?->HerA/FtsK+A32-> | 1213 | Bacteria>Pseudomonadota>Gammaproteobacteria>Enterobacterales>Enterobacteriaceae>Escherichia | GCA_012163455.1 | Escherichia_coli | DFX67_15375 | 24900..28541 | 562 | BREX-system-P-loop-protein-BrxC-[Escherichia-coli] |
| EEY5880446.1 | Type1.BREX | <-Phage_integrase\|\|?->?->?-><-HNH\|\|wHTH+SAD+HNH-><-REase+iREase\|\|BrxA-DNA-binding->BrxB-BrxF-iSTAND->DUF6079*->N6-METHYLASE->PglZ->SIGMA+OB-Fold+AAA-ATPase+Lon-Protease-BrxL->HKD-DNase+OGG-like->\|\|AbiEi_3_N+AbiEi_3->AbiEii-><-DUF6216<-Arm-DNA-bind_3 | 1213 | Bacteria>Pseudomonadota>Gammaproteobacteria>Enterobacterales>Enterobacteriaceae>Escherichia | GCA_012171595.1 | Escherichia_coli | EU637_10820 | 71590..75231 | 562 | BREX-system-P-loop-protein-BrxC-[Escherichia-coli] |
| EEZ0357268.1 | Type1.BREX | <-Phage_integrase\|\|?->?->?-><-HNH\|\|wHTH+SAD+HNH-><-REase+iREase\|\|BrxA-DNA-binding->BrxB-BrxF-iSTAND->DUF6079*->N6-METHYLASE->PglZ->SIGMA+OB-Fold+AAA-ATPase+Lon-Protease-BrxL->?-><-?<-ParB<-?\|\|<-REase+N6_METHYLASE<-TRD | 1213 | Bacteria>Pseudomonadota>Gammaproteobacteria>Enterobacterales>Enterobacteriaceae>Escherichia | GCA_012192205.1 | Escherichia_coli | E9E86_001508 | 71741..75382 | 562 | BREX-system-P-loop-protein-BrxC-[Escherichia-coli] |
| EEZ3165536.1 | Type1.BREX | DUF6079*->N6-METHYLASE->PglZ->SIGMA+OB-Fold+AAA-ATPase+Lon-Protease-BrxL->HKD-DNase+OGG-like->\|\|AbiEi_3_N+AbiEi_3->AbiEii-><-DUF6216<-Arm-DNA-bind_3 | 1136 | Bacteria>Pseudomonadota>Gammaproteobacteria>Enterobacterales>Enterobacteriaceae>Escherichia | GCA_012200425.1 | Escherichia_coli | DIN68_003954 | 1..3411 | 562 | BREX-system-P-loop-protein-BrxC-[Escherichia-coli] |
| EEZ2582139.1 | Type1.BREX | <-DUF6079-p*<-BrxB-BrxF-iSTAND<-BrxA-DNA-binding\|\|<-wHTH(SBD)+HNH\|\|HNH-><-?<-?<-?\|\|Phage_integrase->Phage_integrase-> | 800 | Bacteria>Pseudomonadota>Gammaproteobacteria>Enterobacterales>Enterobacteriaceae>Escherichia | GCA_012201865.1 | Escherichia_coli | DMH54_003958 | 1..2402 | 562 | BREX-system-P-loop-protein-BrxC-[Escherichia-coli] |
| EEZ3215399.1 | Type1.BREX | <-HNH\|\|wHTH(SBD)+HNH->\|\|BrxA-DNA-binding->BrxB-BrxF-iSTAND->DUF6079*->N6-METHYLASE-> | 1213 | Bacteria>Pseudomonadota>Gammaproteobacteria>Enterobacterales>Enterobacteriaceae>Escherichia | GCA_012202545.1 | Escherichia_coli | DXV48_004298 | 6695..10336 | 562 | BREX-system-P-loop-protein-BrxC-[Escherichia-coli] |
| EEZ5289151.1 | Type1.BREX | <-Phage_integrase\|\|?->?->?-><-HNH\|\|wHTH+SAD+HNH-><-REase+iREase\|\|BrxA-DNA-binding->BrxB-BrxF-iSTAND->DUF6079*->N6-METHYLASE->PglZ->SIGMA+OB-Fold+AAA-ATPase+Lon-Protease-BrxL->HKD-DNase+OGG-like->\|\|AbiEi_3_N+AbiEi_3->AbiEii-><-DUF6216<-Arm-DNA-bind_3 | 1213 | Bacteria>Pseudomonadota>Gammaproteobacteria>Enterobacterales>Enterobacteriaceae>Escherichia | GCA_012210255.1 | Escherichia_coli | DDL85_000894 | 111149..114790 | 562 | BREX-system-P-loop-protein-BrxC-[Escherichia-coli] |
| EEZ5688036.1 | Type1.BREX | <-Phage_integrase\|\|?->?->?-><-HNH\|\|wHTH+SAD+HNH-><-REase+iREase\|\|BrxA-DNA-binding->BrxB-BrxF-iSTAND->DUF6079*->N6-METHYLASE->PglZ->SIGMA+OB-Fold+AAA-ATPase+Lon-Protease-BrxL->HKD-DNase+OGG-like->\|\|AbiEi_3_N+AbiEi_3->AbiEii-><-DUF6216<-Arm-DNA-bind_3 | 1213 | Bacteria>Pseudomonadota>Gammaproteobacteria>Enterobacterales>Enterobacteriaceae>Escherichia | GCA_012215265.1 | Escherichia_coli_O25 | DBU11_001474 | 113366..117007 | 1095709 | BREX-system-P-loop-protein-BrxC-[Escherichia-coli-O25] |
| EEZ5714529.1 | Type1.BREX | <-wHTH+WYL<-nSTAND1+sGTP<-?\|\|XIS-HTH->?->?->ZnR->BrxA-DNA-binding->BrxB-BrxF-iSTAND->DUF6079*->ParB+HNH->N6-METHYLASE->HEPN->RVT->N6-METHYLASE-p->PglZ->SIGMA+OB-Fold+AAA-ATPase+Lon-Protease-BrxL-><-Arm-DNA-bind_3+Phage_integrase | 1233 | Bacteria>Pseudomonadota>Gammaproteobacteria>Enterobacterales>Enterobacteriaceae>Escherichia | GCA_012215525.1 | Escherichia_coli_O25 | DBU27_000026 | 20065..23766 | 1095709 | BREX-system-P-loop-protein-BrxC-[Escherichia-coli-O25] |
| EEZ7212406.1 | Type1.BREX | <-REase+iREase\|\|BrxA-DNA-binding->BrxB-BrxF-iSTAND->DUF6079*->N6-METHYLASE->PglZ->SIGMA+OB-Fold+AAA-ATPase+Lon-Protease-BrxL->HKD-DNase+OGG-like->\|\|AbiEi_3_N+AbiEi_3->AbiEii-><-DUF6216<-Arm-DNA-bind_3 | 1213 | Bacteria>Pseudomonadota>Gammaproteobacteria>Enterobacterales>Enterobacteriaceae>Escherichia | GCA_012218605.1 | Escherichia_coli | DAB40_000420 | 2198..5839 | 562 | BREX-system-P-loop-protein-BrxC-[Escherichia-coli] |
| EEZ7669684.1 | Type1.BREX | <-N6-METHYLASE<-DUF6079*<-BrxB-BrxF-iSTAND<-BrxA-DNA-binding\|\|REase+iREase-><-wHTH+SAD+HNH\|\|HNH-><-?<-?<-?\|\|Phage_integrase-> | 1213 | Bacteria>Pseudomonadota>Gammaproteobacteria>Enterobacterales>Enterobacteriaceae>Escherichia | GCA_012222835.1 | Escherichia_coli | C7P55_004830 | 2589..6230 | 562 | BREX-system-P-loop-protein-BrxC-[Escherichia-coli] |
| EEZ9804432.1 | Type1.BREX | <-Phage_integrase\|\|?->?->?-><-HNH\|\|wHTH+SAD+HNH-><-REase+iREase\|\|BrxA-DNA-binding->BrxB-BrxF-iSTAND->DUF6079*->N6-METHYLASE->PglZ->SIGMA+OB-Fold+AAA-ATPase+Lon-Protease-BrxL->?-><-?<-ParB<-?\|\|<-REase+N6_METHYLASE<-TRD | 1213 | Bacteria>Pseudomonadota>Gammaproteobacteria>Enterobacterales>Enterobacteriaceae>Escherichia | GCA_012226505.1 | Escherichia_coli_O25 | DF264_000712 | 59918..63559 | 1095709 | BREX-system-P-loop-protein-BrxC-[Escherichia-coli-O25] |
| EEZ9156819.1 | Type1.BREX | DUF6079*->N6-METHYLASE->PglZ->SIGMA+OB-Fold+AAA-ATPase+Lon-Protease-BrxL->HKD-DNase+OGG-like->\|\|AbiEi_3_N+AbiEi_3->AbiEii-><-DUF6216<-Arm-DNA-bind_3 | 1010 | Bacteria>Pseudomonadota>Gammaproteobacteria>Enterobacterales>Enterobacteriaceae>Escherichia | GCA_012228465.1 | Escherichia_coli | DDB94_002970 | 1..3035 | 562 | BREX-system-P-loop-protein-BrxC-[Escherichia-coli] |
| EEZ8952063.1 | Type1.BREX | <-Phage_integrase\|\|?->?->?-><-HNH\|\|wHTH+SAD+HNH-><-REase+iREase\|\|BrxA-DNA-binding->BrxB-BrxF-iSTAND->DUF6079*->PglZ->SIGMA+OB-Fold+AAA-ATPase+Lon-Protease-BrxL->HKD-DNase+OGG-like->\|\|AbiEi_3_N+AbiEi_3->AbiEii-><-DUF6216<-Arm-DNA-bind_3<-TOPRIM-DNAG+D5N+D5ATPase+P4PRIMASE-HTH | 1171 | Bacteria>Pseudomonadota>Gammaproteobacteria>Enterobacterales>Enterobacteriaceae>Escherichia | GCA_012229475.1 | Escherichia_coli_O35 | C2D51_003183 | 9758..13273 | 2072460 | BREX-system-P-loop-protein-BrxC-[Escherichia-coli-O35] |
| EFA3420990.1 | Type1.BREX | DUF6079*->N6-METHYLASE->PglZ->SIGMA+OB-Fold+AAA-ATPase+Lon-Protease-BrxL-><-ZnR | 1196 | Bacteria>Pseudomonadota>Gammaproteobacteria>Enterobacterales>Enterobacteriaceae>Escherichia | GCA_012244525.1 | Escherichia_coli | E2K00_23170 | 1..3593 | 562 | BREX-system-P-loop-protein-BrxC-[Escherichia-coli] |
| EFA4240016.1 | Type1.BREX | <-HNH\|\|wHTH+SAD+HNH-><-REase+iREase\|\|BrxA-DNA-binding->BrxB-BrxF-iSTAND->DUF6079*->N6-METHYLASE->PglZ->SIGMA+OB-Fold+AAA-ATPase+Lon-Protease-BrxL->?-><-?<-ParB<-?\|\|<-REase+N6_METHYLASE<-TRD | 1213 | Bacteria>Pseudomonadota>Gammaproteobacteria>Enterobacterales>Enterobacteriaceae>Escherichia | GCA_012246545.1 | Escherichia_coli_O36:H5 | C5879_000979 | 4688..8329 | 2100480 | BREX-system-P-loop-protein-BrxC-[Escherichia-coli-O36:H5] |
| EFA5158822.1 | Type1.BREX | <-Phage_integrase\|\|?->?->?-><-HNH\|\|wHTH+SAD+HNH-><-REase+iREase\|\|BrxA-DNA-binding->BrxB-BrxF-iSTAND->DUF6079*->N6-METHYLASE->PglZ->SIGMA+OB-Fold+AAA-ATPase+Lon-Protease-BrxL->HKD-DNase+OGG-like->\|\|AbiEi_3_N+AbiEi_3->AbiEii-><-DUF6216<-Arm-DNA-bind_3 | 1213 | Bacteria>Pseudomonadota>Gammaproteobacteria>Enterobacterales>Enterobacteriaceae>Escherichia | GCA_012250845.1 | Escherichia_coli | E2930_09975 | 121051..124692 | 562 | BREX-system-P-loop-protein-BrxC-[Escherichia-coli] |
| EFA5480296.1 | Type1.BREX | <-XIS-HTH<-RHH_antitoxin\|\|?->ZnR->ZnR-><-SIGMA+OB-Fold+AAA-ATPase+Lon-Protease-BrxL<-PglZ<-N6-METHYLASE<-DUF6079*<-BrxB-BrxF-iSTAND<-BrxA-DNA-binding\|\|REase->REase-><-HEPN<-ORC-CDC6+ZnR<-N6-METHYLASE<-Arm-DNA-bind_3+Phage_integrase | 1213 | Bacteria>Pseudomonadota>Gammaproteobacteria>Enterobacterales>Enterobacteriaceae>Escherichia | GCA_012252925.1 | Escherichia_coli_O8 | EIG07_17100 | 60763..64404 | 1010796 | BREX-system-P-loop-protein-BrxC-[Escherichia-coli-O8] |
| EFA6389818.1 | Type1.BREX | <-nSTAND1+GTPase-AIG+sGTP<-wHTH+WYL\|\|ZnR->ZnR-><-SIGMA+OB-Fold+AAA-ATPase+Lon-Protease-BrxL<-PglZ<-TOPRIM<-OLD-ABC<-N6-METHYLASE<-DUF6079*<-BrxB-BrxF-iSTAND<-BrxA-DNA-binding<-Arm-DNA-bind_3+Phage_integrase<-tRNA\|\|?->HerA/FtsK+A32-><-?<-?\|\|Peptidase_M17_N+Peptidase_M17-> | 1204 | Bacteria>Pseudomonadota>Gammaproteobacteria>Enterobacterales>Enterobacteriaceae>Escherichia | GCA_012256985.1 | Escherichia_coli | CIW09_001013 | 351923..355537 | 562 | BREX-system-P-loop-protein-BrxC-[Escherichia-coli] |
| EFA8105886.1 | Type1.BREX | <-REase+iREase\|\|BrxA-DNA-binding->BrxB-BrxF-iSTAND->DUF6079*->N6-METHYLASE->PglZ->SIGMA+OB-Fold+AAA-ATPase+Lon-Protease-BrxL-><-ZnR<-AAA-ATPase<-HIN-HTH<-?<-?\|\|XIS-HTH-> | 1213 | Bacteria>Pseudomonadota>Gammaproteobacteria>Enterobacterales>Enterobacteriaceae>Escherichia | GCA_012266535.1 | Escherichia_coli | C3Z38_001327 | 2540..6181 | 562 | BREX-system-P-loop-protein-BrxC-[Escherichia-coli] |
| EFA9203627.1 | Type1.BREX | DUF6079*->N6-METHYLASE->SIGMA+OB-Fold+AAA-ATPase+Lon-Protease-BrxL->?-><-?<-ParB<-?\|\|<-REase+N6_METHYLASE<-TRD<-REase | 1112 | Bacteria>Pseudomonadota>Gammaproteobacteria>Enterobacterales>Enterobacteriaceae>Escherichia | GCA_012268365.1 | Escherichia_coli | C2702_002033 | 1..3339 | 562 | BREX-system-P-loop-protein-BrxC-[Escherichia-coli] |
| EFB1479888.1 | Type1.BREX | <-DUF6079-p*<-BrxB-BrxF-iSTAND<-BrxA-DNA-binding\|\|REase+iREase-><-wHTH+SAD+HNH\|\|HNH-><-?<-?<-?\|\|Phage_integrase-> | 428 | Bacteria>Pseudomonadota>Gammaproteobacteria>Enterobacterales>Enterobacteriaceae>Escherichia | GCA_012288905.1 | Escherichia_coli | FJ807_23450 | 1..1284 | 562 | BREX-system-P-loop-protein-BrxC-[Escherichia-coli] |
| EFB2626049.1 | Type1.BREX | <-HKD-DNase+OGG-like<-SIGMA+OB-Fold+AAA-ATPase+Lon-Protease-BrxL<-PglZ<-N6-METHYLASE<-DUF6079*<-BrxB-BrxF-iSTAND<-BrxA-DNA-binding\|\|REase+iREase-><-wHTH+SAD+HNH\|\|HNH-><-?<-?<-?\|\|Phage_integrase-> | 1213 | Bacteria>Pseudomonadota>Gammaproteobacteria>Enterobacterales>Enterobacteriaceae>Escherichia | GCA_012293805.1 | Escherichia_coli | FHL63_20525 | 10185..13826 | 562 | BREX-system-P-loop-protein-BrxC-[Escherichia-coli] |
| HAG7757106.1 | Type1.BREX | <-wHTH+WYL\|\|ZnR->ZnR-><-SIGMA+OB-Fold+AAA-ATPase+Lon-Protease-BrxL<-PglZ<-TOPRIM<-OLD-ABC<-N6-METHYLASE<-DUF6079*<-BrxB-BrxF-iSTAND<-BrxA-DNA-binding<-Arm-DNA-bind_3+Phage_integrase<-tRNA\|\|?->HerA/FtsK+A32-><-?<-?\|\|Peptidase_M17_N+Peptidase_M17-> | 1204 | Bacteria>Pseudomonadota>Gammaproteobacteria>Enterobacterales>Enterobacteriaceae>Escherichia | GCA_012297605.1 | Escherichia_coli | HGR18_09675 | 351902..355516 | 562 | BREX-system-P-loop-protein-BrxC-[Escherichia-coli] |
| EFB3218130.1 | Type1.BREX | <-Zn_Tnp_IS1+HTH_Tnp_IS1\|\|wHTH+WYL+WCX->BrxA-DNA-binding->BrxB-BrxF-iSTAND->DUF6079*->ParB+HNH->N6-METHYLASE->PglZ->SIGMA+OB-Fold+AAA-ATPase+Lon-Protease-BrxL-><-HTH_Tnp_1+HTH_21+rve+rve_3<-HIN-HTH<-PIN<-MazE_antitoxin | 1225 | Bacteria>Pseudomonadota>Gammaproteobacteria>Enterobacterales>Enterobacteriaceae>Escherichia | GCA_012299345.1 | Escherichia_coli | FLO28_20235 | 2574..6251 | 562 | BREX-system-P-loop-protein-BrxC-[Escherichia-coli] |
| EFB4448575.1 | Type1.BREX | <-SIGMA+OB-Fold+AAA-ATPase+Lon-Protease-BrxL<-PglZ<-N6-METHYLASE<-DUF6079*<-BrxB-BrxF-iSTAND<-BrxA-DNA-binding\|\|?->?->TetR-HTH-><-Arm-DNA-bind_3+Phage_integrase | 1213 | Bacteria>Pseudomonadota>Gammaproteobacteria>Enterobacterales>Enterobacteriaceae>Escherichia | GCA_012307385.1 | Escherichia_coli | FQB94_07765 | 8545..12186 | 562 | BREX-system-P-loop-protein-BrxC-[Escherichia-coli] |
| EFB4439448.1 | Type1.BREX | Arm-DNA-bind_3+Phage_integrase-><-?<-?<-REase+iREase\|\|BrxA-DNA-binding->BrxB-BrxF-iSTAND->DUF6079*->N6-METHYLASE->PglZ->SIGMA+OB-Fold+AAA-ATPase+Lon-Protease-BrxL-><-ZnR<-ZnR<-?\|\|RHH_antitoxin-> | 1213 | Bacteria>Pseudomonadota>Gammaproteobacteria>Enterobacterales>Enterobacteriaceae>Escherichia | GCA_012309125.1 | Escherichia_coli | COW91_001589 | 30727..34368 | 562 | BREX-system-P-loop-protein-BrxC-[Escherichia-coli] |
| EFB6629151.1 | Type1.BREX | <-SIGMA-HTH<-?<-ArsR-HTH\|\|?-><-DDE_Tnp_1_5+DDE_Tnp_1\|\|\|\|wHTH+WYL+WCX->BrxA-DNA-binding->BrxB-BrxF-iSTAND->DUF6079*->ParB+HNH->N6-METHYLASE->PglZ->SIGMA+OB-Fold+AAA-ATPase+Lon-Protease-BrxL-><-HTH_Tnp_1+HTH_21+rve+rve_3<-HIN-HTH<-PIN<-MazE_antitoxin | 1225 | Bacteria>Pseudomonadota>Gammaproteobacteria>Enterobacterales>Enterobacteriaceae>Escherichia | GCA_012316585.1 | Escherichia_coli | FQF26_18830 | 72071..75748 | 562 | BREX-system-P-loop-protein-BrxC-[Escherichia-coli] |
| EFB7539253.1 | Type1.BREX | <-Phage_integrase\|\|?->?->?-><-HNH\|\|wHTH+SAD+HNH-><-REase+iREase\|\|BrxA-DNA-binding->BrxB-BrxF-iSTAND->DUF6079*->N6-METHYLASE->PglZ->SIGMA+OB-Fold+AAA-ATPase+Lon-Protease-BrxL->?-><-?<-ParB<-?\|\|<-REase+N6_METHYLASE<-TRD | 1213 | Bacteria>Pseudomonadota>Gammaproteobacteria>Enterobacterales>Enterobacteriaceae>Escherichia | GCA_012318925.1 | Escherichia_coli | E4J12_05355 | 111245..114886 | 562 | BREX-system-P-loop-protein-BrxC-[Escherichia-coli] |
| EFB9770829.1 | Type1.BREX | <-REase<-REase\|\|BrxA-DNA-binding->BrxB-BrxF-iSTAND->DUF6079*->N6-METHYLASE->PglZ->SIGMA+OB-Fold+AAA-ATPase+Lon-Protease-BrxL-><-ZnR<-ZnR<-?\|\|RHH_antitoxin->XIS-HTH-> | 1213 | Bacteria>Pseudomonadota>Gammaproteobacteria>Enterobacterales>Enterobacteriaceae>Escherichia | GCA_012329995.1 | Escherichia_coli | FAD49_21945 | 2845..6486 | 562 | BREX-system-P-loop-protein-BrxC-[Escherichia-coli] |
| EFB9701597.1 | Type1.BREX | ParB->?-><-?<-SIGMA+OB-Fold+AAA-ATPase+Lon-Protease-BrxL<-DUF6079*<-BrxB-BrxF-iSTAND<-BrxA-DNA-binding\|\|REase+iREase-> | 1385 | Bacteria>Pseudomonadota>Gammaproteobacteria>Enterobacterales>Enterobacteriaceae>Escherichia | GCA_012330505.1 | Escherichia_coli | E5H11_23300 | 6377..10534 | 562 | BREX-system-P-loop-protein-BrxC-[Escherichia-coli] |
| EFC0631243.1 | Type1.BREX | <-Peptidase_M17_N+Peptidase_M17\|\|?->?-><-HerA/FtsK+A32<-?\|\|tRNA->Arm-DNA-bind_3+Phage_integrase->BrxA-DNA-binding->BrxB-BrxF-iSTAND->DUF6079*->N6-METHYLASE->OLD-ABC->TOPRIM->PglZ->SIGMA+OB-Fold+AAA-ATPase+Lon-Protease-BrxL-><-?<-ZnR<-ZnR<-wHTH | 1204 | Bacteria>Pseudomonadota>Gammaproteobacteria>Enterobacterales>Enterobacteriaceae>Escherichia | GCA_012334465.1 | Escherichia_coli | FA817_13710 | 286581..290195 | 562 | BREX-system-P-loop-protein-BrxC-[Escherichia-coli] |
| EFC1641769.1 | Type1.BREX | <-HerA/FtsK+A32<-?\|\|tRNA->DUF6216-><-AbiEii<-AbiEi_3_N+AbiEi_3\|\|<-HKD-DNase+OGG-like<-SIGMA+OB-Fold+AAA-ATPase+Lon-Protease-BrxL<-N6-METHYLASE<-DUF6079* | 1140 | Bacteria>Pseudomonadota>Gammaproteobacteria>Enterobacterales>Enterobacteriaceae>Escherichia | GCA_012339895.1 | Escherichia_coli | F2I36_20020 | 26288..29710 | 562 | BREX-system-P-loop-protein-BrxC-[Escherichia-coli] |
| EFC2245710.1 | Type1.BREX | <-Phage_integrase\|\|?->?->?-><-HNH\|\|wHTH+SAD+HNH-><-REase+iREase\|\|BrxA-DNA-binding->BrxB-BrxF-iSTAND->DUF6079*->SIGMA+OB-Fold+AAA-ATPase+Lon-Protease-BrxL->?-><-?<-ParB<-?\|\|<-REase+N6_METHYLASE<-TRD<-REase | 1385 | Bacteria>Pseudomonadota>Gammaproteobacteria>Enterobacterales>Enterobacteriaceae>Escherichia | GCA_012340135.1 | Escherichia_coli | E5H86_07830 | 9758..13915 | 562 | BREX-system-P-loop-protein-BrxC-[Escherichia-coli] |
| EFC3446201.1 | Type1.BREX | <-DUF6079-p*<-BrxB-BrxF-iSTAND<-BrxA-DNA-binding\|\|<-wHTH(SBD)+HNH\|\|HNH-><-?<-?<-?\|\|Phage_integrase->Phage_integrase-> | 844 | Bacteria>Pseudomonadota>Gammaproteobacteria>Enterobacterales>Enterobacteriaceae>Escherichia | GCA_012346015.1 | Escherichia_coli | CO751_004598 | 1..2534 | 562 | BREX-system-P-loop-protein-BrxC-[Escherichia-coli] |
| MBB9587758.1 | Type1.BREX | TRD->REase+N6_METHYLASE->\|\|?->ParB->?-><-?<-SIGMA+OB-Fold+AAA-ATPase+Lon-Protease-BrxL<-PglZ<-N6-METHYLASE<-DUF6079*<-BrxB-BrxF-iSTAND<-BrxA-DNA-binding\|\|REase+iREase-><-wHTH+SAD+HNH\|\|HNH-><-?<-?<-?\|\|Phage_integrase-> | 1213 | Bacteria>Pseudomonadota>Gammaproteobacteria>Enterobacterales>Enterobacteriaceae>Escherichia | GCA_012370635.2 | Escherichia_coli | FSJ86_010580 | 16354..19995 | 562 | BREX-system-P-loop-protein-BrxC-[Escherichia-coli] |
| EFD0914260.1 | Type1.BREX | <-DUF6079*<-BrxB-BrxF-iSTAND<-BrxA-DNA-binding\|\|REase+iREase-><-wHTH+SAD+HNH\|\|HNH-> | 1198 | Bacteria>Pseudomonadota>Gammaproteobacteria>Enterobacterales>Enterobacteriaceae>Escherichia | GCA_012379395.1 | Escherichia_coli | KO979_004413 | 1..3594 | 562 | BREX-system-P-loop-protein-BrxC-[Escherichia-coli] |
| EFD0312067.1 | Type1.BREX | GDResolvase+HIN-HTH-><-SIGMA+OB-Fold+AAA-ATPase+Lon-Protease-BrxL<-PglZ<-N6-METHYLASE-p<-RVT<-HEPN<-N6-METHYLASE<-ParB+HNH<-DUF6079*<-BrxB-BrxF-iSTAND<-BrxA-DNA-binding<-?<-?<-XIS-HTH<-?\|\|?->nSTAND1+GTPase-AIG+sGTP->DUF932-> | 1233 | Bacteria>Pseudomonadota>Gammaproteobacteria>Enterobacterales>Enterobacteriaceae>Escherichia | GCA_012380475.1 | Escherichia_coli | E6D44_20535 | 19474..23175 | 562 | BREX-system-P-loop-protein-BrxC-[Escherichia-coli] |
| MBC0786087.1 | Type1.BREX | <-Phage_integrase\|\|?->?->?-><-HNH\|\|wHTH+SAD+HNH-><-REase+iREase\|\|BrxA-DNA-binding->BrxB-BrxF-iSTAND->DUF6079*->N6-METHYLASE->PglZ->SIGMA+OB-Fold+AAA-ATPase+Lon-Protease-BrxL->HKD-DNase+OGG-like->\|\|AbiEi_3_N+AbiEi_3->AbiEii->?-><-DUF6216<-Arm-DNA-bind_3 | 1213 | Bacteria>Pseudomonadota>Gammaproteobacteria>Enterobacterales>Enterobacteriaceae>Escherichia | GCA_012382335.2 | Escherichia_coli | FSG57_002365 | 111024..114665 | 562 | BREX-system-P-loop-protein-BrxC-[Escherichia-coli] |
| EFD1659139.1 | Type1.BREX | <-Phage_integrase\|\|?->?->?-><-HNH\|\|wHTH+SAD+HNH-><-REase+iREase\|\|BrxA-DNA-binding->BrxB-BrxF-iSTAND->DUF6079*->N6-METHYLASE->PglZ->SIGMA+OB-Fold+AAA-ATPase+Lon-Protease-BrxL->HKD-DNase+OGG-like->\|\|AbiEi_3_N+AbiEi_3->AbiEii-><-DUF6216<-Arm-DNA-bind_3 | 1213 | Bacteria>Pseudomonadota>Gammaproteobacteria>Enterobacterales>Enterobacteriaceae>Escherichia | GCA_012382975.1 | Escherichia_coli | FGU32_03035 | 69619..73260 | 562 | BREX-system-P-loop-protein-BrxC-[Escherichia-coli] |
| EFD3459874.1 | Type1.BREX | <-HNH\|\|wHTH(SBD)+HNH->\|\|BrxA-DNA-binding->BrxB-BrxF-iSTAND->DUF6079*->N6-METHYLASE->PglZ->SIGMA+OB-Fold+AAA-ATPase+Lon-Protease-BrxL-> | 1213 | Bacteria>Pseudomonadota>Gammaproteobacteria>Enterobacterales>Enterobacteriaceae>Escherichia | GCA_012389335.1 | Escherichia_coli | CL211_004302 | 6759..10400 | 562 | BREX-system-P-loop-protein-BrxC-[Escherichia-coli] |
| EFE1965719.1 | Type1.BREX | <-Phage_integrase<-Phage_integrase\|\|?->?->?-><-HNH\|\|wHTH(SBD)+HNH->\|\|BrxA-DNA-binding->BrxB-BrxF-iSTAND->DUF6079*->N6-METHYLASE->PglZ->SIGMA+OB-Fold+AAA-ATPase+Lon-Protease-BrxL->HKD-DNase+OGG-like->\|\|AbiEi_3_N+AbiEi_3->AbiEii-><-DUF6216<-Arm-DNA-bind_3 | 1213 | Bacteria>Pseudomonadota>Gammaproteobacteria>Enterobacterales>Enterobacteriaceae>Escherichia | GCA_012428445.1 | Escherichia_coli | CJV81_002667 | 14696..18337 | 562 | BREX-system-P-loop-protein-BrxC-[Escherichia-coli] |
| EFE8676226.1 | Type1.BREX | BrxA-DNA-binding->BrxB-BrxF-iSTAND->DUF6079*->N6-METHYLASE->PglZ->SIGMA+OB-Fold+AAA-ATPase+Lon-Protease-BrxL-><-ZnR<-ZnR<-?\|\|RHH_antitoxin-> | 1213 | Bacteria>Pseudomonadota>Gammaproteobacteria>Enterobacterales>Enterobacteriaceae>Escherichia | GCA_012457165.1 | Escherichia_coli | CAA49_004703 | 1709..5350 | 562 | BREX-system-P-loop-protein-BrxC-[Escherichia-coli] |
| EFF2616657.1 | Type1.BREX | <-Phage_integrase\|\|?->?->?-><-HNH\|\|wHTH+SAD+HNH-><-REase+iREase\|\|BrxA-DNA-binding->BrxB-BrxF-iSTAND->DUF6079*->N6-METHYLASE->PglZ->SIGMA+OB-Fold+AAA-ATPase+Lon-Protease-BrxL->?-><-?<-ParB<-?\|\|<-REase+N6_METHYLASE<-TRD | 1213 | Bacteria>Pseudomonadota>Gammaproteobacteria>Enterobacterales>Enterobacteriaceae>Escherichia | GCA_012476875.1 | Escherichia_coli | B9L79_001313 | 111240..114881 | 562 | BREX-system-P-loop-protein-BrxC-[Escherichia-coli] |
| EFF3077603.1 | Type1.BREX | <-Phage_integrase\|\|?->?->?-><-HNH\|\|wHTH+SAD+HNH-><-REase+iREase\|\|BrxA-DNA-binding->BrxB-BrxF-iSTAND->DUF6079*->N6-METHYLASE->PglZ->SIGMA+OB-Fold+AAA-ATPase+Lon-Protease-BrxL->?-><-?<-ParB<-?\|\|<-REase+N6_METHYLASE<-TRD | 1213 | Bacteria>Pseudomonadota>Gammaproteobacteria>Enterobacterales>Enterobacteriaceae>Escherichia | GCA_012480115.1 | Escherichia_coli | BOR91_001555 | 57086..60727 | 562 | BREX-system-P-loop-protein-BrxC-[Escherichia-coli] |
| EFF3363917.1 | Type1.BREX | <-Phage_integrase\|\|?->?->?-><-HNH\|\|wHTH+SAD+HNH-><-REase+iREase\|\|BrxA-DNA-binding->BrxB-BrxF-iSTAND->DUF6079*->N6-METHYLASE->PglZ->SIGMA+OB-Fold+AAA-ATPase+Lon-Protease-BrxL->?-><-?<-ParB<-?\|\|<-REase+N6_METHYLASE<-TRD | 1213 | Bacteria>Pseudomonadota>Gammaproteobacteria>Enterobacterales>Enterobacteriaceae>Escherichia | GCA_012481335.1 | Escherichia_coli | B7M15_000791 | 9758..13399 | 562 | BREX-system-P-loop-protein-BrxC-[Escherichia-coli] |
| EFF3839116.1 | Type1.BREX | <-Phage_integrase\|\|?->?->?-><-HNH\|\|wHTH+SAD+HNH-><-REase+iREase\|\|BrxA-DNA-binding->BrxB-BrxF-iSTAND->DUF6079-p*-> | 473 | Bacteria>Pseudomonadota>Gammaproteobacteria>Enterobacterales>Enterobacteriaceae>Escherichia | GCA_012482765.1 | Escherichia_coli | B6310_003036 | 111336..112756 | 562 | BREX-system-P-loop-protein-BrxC-[Escherichia-coli] |
| EFF3839660.1 | Type1.BREX | DUF6079-p*->N6-METHYLASE->PglZ->SIGMA+OB-Fold+AAA-ATPase+Lon-Protease-BrxL->HKD-DNase+OGG-like->\|\|AbiEi_3_N+AbiEi_3->AbiEii-><-DUF6216<-Arm-DNA-bind_3 | 739 | Bacteria>Pseudomonadota>Gammaproteobacteria>Enterobacterales>Enterobacteriaceae>Escherichia | GCA_012482765.1 | Escherichia_coli | B6310_003615 | 1..2221 | 562 | BREX-system-P-loop-protein-BrxC-[Escherichia-coli] |
| EFF7960190.1 | Type1.BREX | <-Phage_integrase\|\|?->?->?-><-HNH\|\|wHTH+SAD+HNH-><-REase+iREase\|\|BrxA-DNA-binding->BrxB-BrxF-iSTAND->DUF6079*->N6-METHYLASE->PglZ->SIGMA+OB-Fold+AAA-ATPase+Lon-Protease-BrxL->?-><-?<-ParB<-?\|\|<-REase+N6_METHYLASE<-TRD | 1213 | Bacteria>Pseudomonadota>Gammaproteobacteria>Enterobacterales>Enterobacteriaceae>Escherichia | GCA_012505155.1 | Escherichia_coli | BUX12_001709 | 111293..114934 | 562 | BREX-system-P-loop-protein-BrxC-[Escherichia-coli] |
| HAH2520952.1 | Type1.BREX | <-Phage_integrase\|\|?->?->?-><-HNH\|\|wHTH+SAD+HNH-><-REase+iREase\|\|BrxA-DNA-binding->BrxB-BrxF-iSTAND->DUF6079*->N6-METHYLASE->PglZ->SIGMA+OB-Fold+AAA-ATPase+Lon-Protease-BrxL->HKD-DNase+OGG-like->\|\|AbiEi_3_N+AbiEi_3->AbiEii-> | 1162 | Bacteria>Pseudomonadota>Gammaproteobacteria>Enterobacterales>Enterobacteriaceae>Escherichia | GCA_012530295.1 | Escherichia_coli | GF659_15740 | 71446..74934 | 562 | BREX-system-P-loop-protein-BrxC-[Escherichia-coli] |
| EFG1482424.1 | Type1.BREX | <-Zn_Tnp_IS1+HTH_Tnp_IS1<-REase<-REase\|\|BrxA-DNA-binding->BrxB-BrxF-iSTAND->DUF6079*->N6-METHYLASE->PglZ->SIGMA+OB-Fold+AAA-ATPase+Lon-Protease-BrxL-><-ZnR<-ZnR | 1213 | Bacteria>Pseudomonadota>Gammaproteobacteria>Enterobacterales>Enterobacteriaceae>Escherichia | GCA_012544495.1 | Escherichia_coli | BRV97_004207 | 3158..6799 | 562 | BREX-system-P-loop-protein-BrxC-[Escherichia-coli] |
| EFG3005928.1 | Type1.BREX | DUF6079-p*->N6-METHYLASE->PglZ->SIGMA+OB-Fold+AAA-ATPase+Lon-Protease-BrxL->HKD-DNase+OGG-like->\|\|AbiEi_3_N+AbiEi_3->AbiEii-><-DUF6216<-Arm-DNA-bind_3 | 897 | Bacteria>Pseudomonadota>Gammaproteobacteria>Enterobacterales>Enterobacteriaceae>Escherichia | GCA_012550755.1 | Escherichia_coli | BOZ89_003087 | 1..2694 | 562 | BREX-system-P-loop-protein-BrxC-[Escherichia-coli] |
| EFG4404338.1 | Type1.BREX | <-Phage_integrase<-Phage_integrase\|\|?->?->?-><-HNH\|\|wHTH(SBD)+HNH->\|\|BrxA-DNA-binding->BrxB-BrxF-iSTAND->DUF6079*->N6-METHYLASE->PglZ->SIGMA+OB-Fold+AAA-ATPase+Lon-Protease-BrxL->HKD-DNase+OGG-like->\|\|AbiEi_3_N+AbiEi_3->AbiEii-><-DUF6216<-Arm-DNA-bind_3 | 1213 | Bacteria>Pseudomonadota>Gammaproteobacteria>Enterobacterales>Enterobacteriaceae>Escherichia | GCA_012556825.1 | Escherichia_coli | BM772_002472 | 14697..18338 | 562 | BREX-system-P-loop-protein-BrxC-[Escherichia-coli] |
| EFH0548056.1 | Type1.BREX | DUF6079-p*->N6-METHYLASE->PglZ->SIGMA+OB-Fold+AAA-ATPase+Lon-Protease-BrxL->HKD-DNase+OGG-like->\|\|AbiEi_3_N+AbiEi_3->AbiEii-><-DUF6216<-Arm-DNA-bind_3 | 789 | Bacteria>Pseudomonadota>Gammaproteobacteria>Enterobacterales>Enterobacteriaceae>Escherichia | GCA_012582395.1 | Escherichia_coli | BJE23_002820 | 1..2370 | 562 | BREX-system-P-loop-protein-BrxC-[Escherichia-coli] |
| EFH0549024.1 | Type1.BREX | <-Phage_integrase<-Phage_integrase\|\|?->?->?-><-HNH\|\|wHTH(SBD)+HNH->\|\|BrxA-DNA-binding->BrxB-BrxF-iSTAND->DUF6079-p*-> | 422 | Bacteria>Pseudomonadota>Gammaproteobacteria>Enterobacterales>Enterobacteriaceae>Escherichia | GCA_012582395.1 | Escherichia_coli | BJE23_003837 | 39992..41258 | 562 | BREX-system-P-loop-protein-BrxC-[Escherichia-coli] |
| EFH3554397.1 | Type1.BREX | <-Phage_integrase\|\|?->?->?-><-HNH\|\|wHTH+SAD+HNH-><-REase+iREase\|\|BrxA-DNA-binding->BrxB-BrxF-iSTAND->DUF6079*->N6-METHYLASE-p-> | 1213 | Bacteria>Pseudomonadota>Gammaproteobacteria>Enterobacterales>Enterobacteriaceae>Escherichia | GCA_012596955.1 | Escherichia_coli | GQW80_16270 | 69288..72929 | 562 | BREX-system-P-loop-protein-BrxC-[Escherichia-coli] |
| EFH3789843.1 | Type1.BREX | <-TetR-HTH<-?<-?\|\|BrxA-DNA-binding->BrxB-BrxF-iSTAND->DUF6079*->N6-METHYLASE->PglZ->SIGMA+OB-Fold+AAA-ATPase+Lon-Protease-BrxL-> | 1213 | Bacteria>Pseudomonadota>Gammaproteobacteria>Enterobacterales>Enterobacteriaceae>Escherichia | GCA_012598785.1 | Escherichia_coli | F9418_21630 | 6934..10575 | 562 | BREX-system-P-loop-protein-BrxC-[Escherichia-coli] |
| EFH4793016.1 | Type1.BREX | BrxA-DNA-binding->BrxB-BrxF-iSTAND->DUF6079*->N6-METHYLASE->PglZ->SIGMA+OB-Fold+AAA-ATPase+Lon-Protease-BrxL-> | 1213 | Bacteria>Pseudomonadota>Gammaproteobacteria>Enterobacterales>Enterobacteriaceae>Escherichia | GCA_012602795.1 | Escherichia_coli | F9381_22830 | 5106..8747 | 562 | BREX-system-P-loop-protein-BrxC-[Escherichia-coli] |
| EFH4936891.1 | Type1.BREX | <-Phage_integrase\|\|?->?->?-><-HNH\|\|wHTH+SAD+HNH-><-REase+iREase\|\|BrxA-DNA-binding->BrxB-BrxF-iSTAND->DUF6079*->N6-METHYLASE->PglZ->SIGMA+OB-Fold+AAA-ATPase+Lon-Protease-BrxL->HKD-DNase+OGG-like->\|\|AbiEi_3_N+AbiEi_3->AbiEii-><-DUF6216<-Arm-DNA-bind_3 | 1213 | Bacteria>Pseudomonadota>Gammaproteobacteria>Enterobacterales>Enterobacteriaceae>Escherichia | GCA_012603315.1 | Escherichia_coli | F9408_05585 | 110160..113801 | 562 | BREX-system-P-loop-protein-BrxC-[Escherichia-coli] |
| EFH7369082.1 | Type1.BREX | <-RelE-ParE<-CopG/RHH\|\|<-RHH\|\|?->wHTH+WYL+WCX->BrxA-DNA-binding->BrxB-BrxF-iSTAND->DUF6079*->ParB+HNH->N6-METHYLASE-p-> | 1225 | Bacteria>Pseudomonadota>Gammaproteobacteria>Enterobacterales>Enterobacteriaceae>Escherichia | GCA_012614485.1 | Escherichia_coli | GKQ19_21210 | 5665..9342 | 562 | BREX-system-P-loop-protein-BrxC-[Escherichia-coli] |
| EFH8432482.1 | Type1.BREX | ParB->?-><-?<-SIGMA+OB-Fold+AAA-ATPase+Lon-Protease-BrxL<-PglZ<-N6-METHYLASE<-DUF6079*<-BrxB-BrxF-iSTAND<-BrxA-DNA-binding\|\|REase+iREase-> | 1213 | Bacteria>Pseudomonadota>Gammaproteobacteria>Enterobacterales>Enterobacteriaceae>Escherichia | GCA_012619365.1 | Escherichia_coli | GSM49_21410 | 10956..14597 | 562 | BREX-system-P-loop-protein-BrxC-[Escherichia-coli] |
| EFH9688867.1 | Type1.BREX | ZnR->ZnR-><-SIGMA+OB-Fold+AAA-ATPase+Lon-Protease-BrxL<-PglZ<-N6-METHYLASE<-DUF6079*<-BrxB-BrxF-iSTAND<-BrxA-DNA-binding\|\|REase->REase-><-HEPN<-ORC-CDC6+ZnR<-N6-METHYLASE<-Arm-DNA-bind_3+Phage_integrase | 1213 | Bacteria>Pseudomonadota>Gammaproteobacteria>Enterobacterales>Enterobacteriaceae>Escherichia | GCA_012624895.1 | Escherichia_coli | F8Z35_13525 | 11254..14895 | 562 | BREX-system-P-loop-protein-BrxC-[Escherichia-coli] |
| EFH9748630.1 | Type1.BREX | <-Phage_integrase\|\|?->?->?-><-HNH\|\|wHTH+SAD+HNH-><-REase+iREase\|\|BrxA-DNA-binding->BrxB-BrxF-iSTAND->DUF6079*->N6-METHYLASE->PglZ->SIGMA+OB-Fold+AAA-ATPase+Lon-Protease-BrxL->?-><-?<-ParB<-?\|\|<-REase+N6_METHYLASE<-TRD | 1213 | Bacteria>Pseudomonadota>Gammaproteobacteria>Enterobacterales>Enterobacteriaceae>Escherichia | GCA_012625865.1 | Escherichia_coli | F7N48_00500 | 90362..94003 | 562 | BREX-system-P-loop-protein-BrxC-[Escherichia-coli] |

<table>
<tr><th>Protein-ID</th><th>System</th><th>Gene-Neighbourhoods</th><th>Length</th><th>Taxonomy</th><th>Assembly-ID</th><th>Strain</th><th>Locus-Tag</th><th>Genomic-coordinates</th><th>TaxID</th><th>Fasta-define</th></tr>
<tr><td>EFI1580322.1</td><td>Type1.BREX</td><td>DUF6079*->N6-METHYLASE->PglZ->SIGMA+OB-Fold+AAA-ATPase+Lon-Protease-BrxL->HKD-DNase+OGG-like->||AbiEi_3_N+AbiEi_3->AbiEii-><-DUF6216<-Arm-DNA-bind_3</td><td>1071</td><td>Bacteria>Pseudomonadota>Gammaproteobacteria>Enterobacterales>Enterobacteriaceae>Escherichia</td><td>GCA_012634445.1</td><td>Escherichia_coli</td><td>BFN79_002999</td><td>1..3216</td><td>562</td><td>BREX-system-P-loop-protein-BrxC-[Escherichia-coli]</td></tr>
<tr><td>EFI3320066.1</td><td>Type1.BREX</td><td>DDE_3-><-HKD-DNase+OGG-like<-SIGMA+OB-Fold+AAA-ATPase+Lon-Protease-BrxL<-PglZ<-N6-METHYLASE<-DUF6079*<-BrxB-BrxF-iSTAND<-BrxA-DNA-binding||REase+iREase-><-wHTH+SAD+HNH||HNH-><-?<-?<-?||Phage_integrase-></td><td>1213</td><td>Bacteria>Pseudomonadota>Gammaproteobacteria>Enterobacterales>Enterobacteriaceae>Escherichia</td><td>GCA_012643725.1</td><td>Escherichia_coli</td><td>BCC72_001616</td><td>10313..13954</td><td>562</td><td>BREX-system-P-loop-protein-BrxC-[Escherichia-coli]</td></tr>
<tr><td>EFI8237360.1</td><td>Type1.BREX</td><td>DUF6079*->N6-METHYLASE->PglZ->SIGMA+OB-Fold+AAA-ATPase+Lon-Protease-BrxL-><-DDE_Tnp_IS66+DDE_Tnp_IS66_C</td><td>1179</td><td>Bacteria>Pseudomonadota>Gammaproteobacteria>Enterobacterales>Enterobacteriaceae>Escherichia</td><td>GCA_012674225.1</td><td>Escherichia_coli</td><td>HIB78_004536</td><td>1..3541</td><td>562</td><td>BREX-system-P-loop-protein-BrxC-[Escherichia-coli]</td></tr>
<tr><td>EFI9515006.1</td><td>Type1.BREX</td><td><-XIS-HTH<-RHH_antitoxin||?->?->ZnR->ZnR-><-SIGMA+OB-Fold+AAA-ATPase+Lon-Protease-BrxL<-PglZ<-N6-METHYLASE<-DUF6079*<-BrxB-BrxF-iSTAND<-BrxA-DNA-binding||REase+iREase-></td><td>1213</td><td>Bacteria>Pseudomonadota>Gammaproteobacteria>Enterobacterales>Enterobacteriaceae>Escherichia</td><td>GCA_012683285.1</td><td>Escherichia_coli</td><td>G4U18_004344</td><td>13351..16992</td><td>562</td><td>BREX-system-P-loop-protein-BrxC-[Escherichia-coli]</td></tr>
<tr><td>EFJ2123507.1</td><td>Type1.BREX</td><td>DDE_Tnp_1_5+DDE_Tnp_1-><-?||DDE_Tnp_1_5+DDE_Tnp_1->||?-><-RHH||?->wHTH+WYL+WCX->BrxA-DNA-binding->BrxB-BrxF-iSTAND->DUF6079-p*-></td><td>422</td><td>Bacteria>Pseudomonadota>Gammaproteobacteria>Enterobacterales>Enterobacteriaceae>Escherichia</td><td>GCA_012696185.1</td><td>Escherichia_coli</td><td>G6K59_004265</td><td>24578..25844</td><td>562</td><td>BREX-system-P-loop-protein-BrxC-[Escherichia-coli]</td></tr>
<tr><td>EFJ5167524.1</td><td>Type1.BREX</td><td>DUF6079*->N6-METHYLASE->PglZ->SIGMA+OB-Fold+AAA-ATPase+Lon-Protease-BrxL->HKD-DNase+OGG-like->||AbiEi_3_N+AbiEi_3->AbiEii-><-DUF6216<-Arm-DNA-bind_3</td><td>1181</td><td>Bacteria>Pseudomonadota>Gammaproteobacteria>Enterobacterales>Enterobacteriaceae>Escherichia</td><td>GCA_012707625.1</td><td>Escherichia_coli</td><td>A5T96_003401</td><td>1..3546</td><td>562</td><td>BREX-system-P-loop-protein-BrxC-[Escherichia-coli]</td></tr>
<tr><td>EFJ6798268.1</td><td>Type1.BREX</td><td><-XIS-HTH<-RHH_antitoxin||?->?->ZnR->ZnR-><-SIGMA+OB-Fold+AAA-ATPase+Lon-Protease-BrxL<-PglZ<-N6-METHYLASE<-DUF6079*<-BrxB-BrxF-iSTAND<-BrxA-DNA-binding||REase+iREase-><-EVE+HNH<-Arm-DNA-bind_3+Phage_integrase<-tRNA||?->HerA/FtsK+A32-></td><td>1213</td><td>Bacteria>Pseudomonadota>Gammaproteobacteria>Enterobacterales>Enterobacteriaceae>Escherichia</td><td>GCA_012714605.1</td><td>Escherichia_coli</td><td>A2G00_000895</td><td>22019..25660</td><td>562</td><td>BREX-system-P-loop-protein-BrxC-[Escherichia-coli]</td></tr>
<tr><td>HAH8430180.1</td><td>Type1.BREX</td><td>DUF6079-p*->N6-METHYLASE->PglZ->SIGMA+OB-Fold+AAA-ATPase+Lon-Protease-BrxL->HKD-DNase+OGG-like->||AbiEi_3_N+AbiEi_3->AbiEii-><-DUF6216<-Arm-DNA-bind_3</td><td>846</td><td>Bacteria>Pseudomonadota>Gammaproteobacteria>Enterobacterales>Enterobacteriaceae>Escherichia</td><td>GCA_012716735.1</td><td>Escherichia_coli</td><td>HIR43_003168</td><td>1..2542</td><td>562</td><td>BREX-system-P-loop-protein-BrxC-[Escherichia-coli]</td></tr>
<tr><td>EFK2991875.1</td><td>Type1.BREX</td><td><-REase<-REase||BrxA-DNA-binding->BrxB-BrxF-iSTAND->DUF6079*->N6-METHYLASE->PglZ->SIGMA+OB-Fold+AAA-ATPase+Lon-Protease-BrxL-><-ZnR<-ZnR<-?||RHH_antitoxin->XIS-HTH-></td><td>1213</td><td>Bacteria>Pseudomonadota>Gammaproteobacteria>Enterobacterales>Enterobacteriaceae>Escherichia</td><td>GCA_012752375.1</td><td>Escherichia_coli</td><td>A8514_004288</td><td>2939..6580</td><td>562</td><td>BREX-system-P-loop-protein-BrxC-[Escherichia-coli]</td></tr>
<tr><td>HAH9940666.1</td><td>Type1.BREX</td><td><-Phage_integrase||?->?->?-><-HNH||wHTH+SAD+HNH-><-REase+iREase||BrxA-DNA-binding->BrxB-BrxF-iSTAND->DUF6079*->N6-METHYLASE->PglZ->SIGMA+OB-Fold+AAA-ATPase+Lon-Protease-BrxL->HKD-DNase+OGG-like->||AbiEi_3_N+AbiEi_3->AbiEii-><-DUF6216<-Arm-DNA-bind_3<-REase</td><td>1213</td><td>Bacteria>Pseudomonadota>Gammaproteobacteria>Enterobacterales>Enterobacteriaceae>Escherichia</td><td>GCA_012753595.1</td><td>Escherichia_coli</td><td>HIP13_004044</td><td>9744..13385</td><td>562</td><td>BREX-system-P-loop-protein-BrxC-[Escherichia-coli]</td></tr>
<tr><td>EFK7079585.1</td><td>Type1.BREX</td><td><-Phage_integrase||?->?->?-><-HNH||wHTH+SAD+HNH-><-REase+iREase||BrxA-DNA-binding->BrxB-BrxF-iSTAND->DUF6079*->PglZ->SIGMA+OB-Fold+AAA-ATPase+Lon-Protease-BrxL->?-><-?<-?||<-REase+N6_METHYLASE<-TRD<-REase</td><td>1213</td><td>Bacteria>Pseudomonadota>Gammaproteobacteria>Enterobacterales>Enterobacteriaceae>Escherichia</td><td>GCA_012771275.1</td><td>Escherichia_coli</td><td>AS076_001188</td><td>111240..114881</td><td>562</td><td>BREX-system-P-loop-protein-BrxC-[Escherichia-coli]</td></tr>
<tr><td>EFK7446827.1</td><td>Type1.BREX</td><td>DDE_Tnp_IS1-><-SIGMA+OB-Fold+AAA-ATPase+Lon-Protease-BrxL<-PglZ<-N6-METHYLASE<-DUF6079*<-BrxB-BrxF-iSTAND<-BrxA-DNA-binding||REase->?->||<-Arm-DNA-bind_3+Phage_integrase<-tRNA||?->HerA/FtsK+A32-></td><td>1213</td><td>Bacteria>Pseudomonadota>Gammaproteobacteria>Enterobacterales>Enterobacteriaceae>Escherichia</td><td>GCA_012772735.1</td><td>Escherichia_coli</td><td>BBQ60_004521</td><td>8855..12496</td><td>562</td><td>BREX-system-P-loop-protein-BrxC-[Escherichia-coli]</td></tr>
<tr><td>EFK8341906.1</td><td>Type1.BREX</td><td><-Phage_integrase<-Phage_integrase||?->?->?->wHTH+SAD+HNH-><-REase+iREase||BrxA-DNA-binding->BrxB-BrxF-iSTAND->DUF6079-p*-></td><td>800</td><td>Bacteria>Pseudomonadota>Gammaproteobacteria>Enterobacterales>Enterobacteriaceae>Escherichia</td><td>GCA_012777335.1</td><td>Escherichia_coli</td><td>A0793_004522</td><td>36430..38831</td><td>562</td><td>BREX-system-P-loop-protein-BrxC-[Escherichia-coli]</td></tr>
<tr><td>EFL0019973.1</td><td>Type1.BREX</td><td><-Phage_integrase||?->?->?-><-HNH||wHTH+SAD+HNH-><-REase+iREase||BrxA-DNA-binding->BrxB-BrxF-iSTAND->DUF6079*->N6-METHYLASE->PglZ->SIGMA+OB-Fold+AAA-ATPase+Lon-Protease-BrxL->?-><-?<-ParB<-?||<-REase+N6_METHYLASE<-TRD</td><td>1213</td><td>Bacteria>Pseudomonadota>Gammaproteobacteria>Enterobacterales>Enterobacteriaceae>Escherichia</td><td>GCA_012786995.1</td><td>Escherichia_coli</td><td>ACG50_000782</td><td>71932..75573</td><td>562</td><td>BREX-system-P-loop-protein-BrxC-[Escherichia-coli]</td></tr>
<tr><td>EFL0544862.1</td><td>Type1.BREX</td><td>DUF6079-p*->N6-METHYLASE->PglZ->SIGMA+OB-Fold+AAA-ATPase+Lon-Protease-BrxL->HKD-DNase+OGG-like->||AbiEi_3_N+AbiEi_3->AbiEii-><-DUF6216<-Arm-DNA-bind_3<-TOPRIM-DNAG+D5N+D5ATPase+P4PRIMASE-HTH</td><td>803</td><td>Bacteria>Pseudomonadota>Gammaproteobacteria>Enterobacterales>Enterobacteriaceae>Escherichia</td><td>GCA_012789875.1</td><td>Escherichia_coli</td><td>HIZ25_002081</td><td>1..2412</td><td>562</td><td>BREX-system-P-loop-protein-BrxC-[Escherichia-coli]</td></tr>
<tr><td>EFL4011089.1</td><td>Type1.BREX</td><td><-Phage_integrase||?->?->?-><-HNH||wHTH+SAD+HNH-><-REase+iREase||BrxA-DNA-binding->BrxB-BrxF-iSTAND->DUF6079*->N6-METHYLASE->PglZ->SIGMA+OB-Fold+AAA-ATPase+Lon-Protease-BrxL->HKD-DNase+OGG-like->||AbiEi_3_N+AbiEi_3->AbiEii-><-DUF6216<-Arm-DNA-bind_3</td><td>1213</td><td>Bacteria>Pseudomonadota>Gammaproteobacteria>Enterobacterales>Enterobacteriaceae>Escherichia</td><td>GCA_012809955.1</td><td>Escherichia_coli</td><td>AC312_001861</td><td>91319..94960</td><td>562</td><td>BREX-system-P-loop-protein-BrxC-[Escherichia-coli]</td></tr>
<tr><td>EFL7364898.1</td><td>Type1.BREX</td><td>DUF6079*->N6-METHYLASE->PglZ->SIGMA+OB-Fold+AAA-ATPase+Lon-Protease-BrxL->?-><-?<-ParB<-?||<-REase+N6_METHYLASE<-TRD</td><td>1211</td><td>Bacteria>Pseudomonadota>Gammaproteobacteria>Enterobacterales>Enterobacteriaceae>Escherichia</td><td>GCA_012822795.1</td><td>Escherichia_coli</td><td>VT16_000316</td><td>1..3636</td><td>562</td><td>BREX-system-P-loop-protein-BrxC-[Escherichia-coli]</td></tr>
<tr><td>EFL7041959.1</td><td>Type1.BREX</td><td><-XIS-HTH<-RHH_antitoxin||?->?->ZnR->ZnR-><-SIGMA+OB-Fold+AAA-ATPase+Lon-Protease-BrxL<-PglZ<-N6-METHYLASE<-DUF6079*<-BrxB-BrxF-iSTAND<-BrxA-DNA-binding||<-EVE+HNH<-Arm-DNA-bind_3+Phage_integrase<-tRNA||?->HerA/FtsK+A32-><-?<-?||Peptidase_M17_N+Peptidase_M17-></td><td>1213</td><td>Bacteria>Pseudomonadota>Gammaproteobacteria>Enterobacterales>Enterobacteriaceae>Escherichia</td><td>GCA_012822855.1</td><td>Escherichia_coli</td><td>X077_001368</td><td>13786..17427</td><td>562</td><td>BREX-system-P-loop-protein-BrxC-[Escherichia-coli]</td></tr>
<tr><td>EFM0326192.1</td><td>Type1.BREX</td><td><-Phage_integrase||?->?->?-><-HNH||wHTH+SAD+HNH-><-REase+iREase||BrxA-DNA-binding->BrxB-BrxF-iSTAND->DUF6079*->N6-METHYLASE->PglZ->SIGMA+OB-Fold+AAA-ATPase+Lon-Protease-BrxL->?-><-?<-ParB<-?||<-REase+N6_METHYLASE<-TRD</td><td>1213</td><td>Bacteria>Pseudomonadota>Gammaproteobacteria>Enterobacterales>Enterobacteriaceae>Escherichia</td><td>GCA_012835895.1</td><td>Escherichia_coli</td><td>CM47_002122</td><td>16867..20508</td><td>562</td><td>BREX-system-P-loop-protein-BrxC-[Escherichia-coli]</td></tr>
<tr><td>MBB7947572.1</td><td>Type1.BREX</td><td>BrxA-DNA-binding->BrxB-BrxF-iSTAND->DUF6079*->N6-METHYLASE->PglZ->SIGMA+OB-Fold+AAA-ATPase+Lon-Protease-BrxL-><-ZnR<-AAA-ATPase<-HIN-HTH</td><td>1213</td><td>Bacteria>Pseudomonadota>Gammaproteobacteria>Enterobacterales>Enterobacteriaceae>Escherichia</td><td>GCA_012847135.2</td><td>Escherichia_coli</td><td>HEL38_020820</td><td>1537..5178</td><td>562</td><td>BREX-system-P-loop-protein-BrxC-[Escherichia-coli]</td></tr>
<tr><td>HAI2941917.1</td><td>Type1.BREX</td><td><-METHYLASE||SKI-><-?||tRNA->Arm-DNA-bind_3+Phage_integrase->EVE+HNH-><-REase+iREase||BrxA-DNA-binding->BrxB-BrxF-iSTAND->DUF6079*->N6-METHYLASE->PglZ->SIGMA+OB-Fold+AAA-ATPase+Lon-Protease-BrxL-><-ZnR<-ZnR<-?<-?||XIS-HTH-></td><td>1213</td><td>Bacteria>Pseudomonadota>Gammaproteobacteria>Enterobacterales>Enterobacteriaceae>Escherichia</td><td>GCA_012856395.1</td><td>Escherichia_coli</td><td>HJL49_003091</td><td>149607..153248</td><td>562</td><td>BREX-system-P-loop-protein-BrxC-[Escherichia-coli]</td></tr>
<tr><td>HAI3030635.1</td><td>Type1.BREX</td><td><-Zn_Tnp_IS1+HTH_Tnp_IS1<-TetR-HTH-><-?||BrxA-DNA-binding->BrxB-BrxF-iSTAND->DUF6079*->N6-METHYLASE->PglZ->SIGMA+OB-Fold+AAA-ATPase+Lon-Protease-BrxL-><-?<-HIN-HTH<-ParB+HNH<-ParB+HNH||RVT->DEDD_Tnp_IS110+Transposase_20-></td><td>1213</td><td>Bacteria>Pseudomonadota>Gammaproteobacteria>Enterobacterales>Enterobacteriaceae>Escherichia</td><td>GCA_012856755.1</td><td>Escherichia_coli</td><td>HJL73_003515</td><td>6743..10384</td><td>562</td><td>BREX-system-P-loop-protein-BrxC-[Escherichia-coli]</td></tr>
<tr><td>HAI3857478.1</td><td>Type1.BREX</td><td>DEDD_Tnp_IS110+Transposase_20-><-RHH||?->wHTH+WYL+WCX->BrxA-DNA-binding->BrxB-BrxF-iSTAND->DUF6079*->ParB+HNH->N6-METHYLASE->PglZ->SIGMA+OB-Fold+AAA-ATPase+Lon-Protease-BrxL-><-DEDD_Tnp_IS110+Transposase_20<-PIN<-MazE_antitoxin</td><td>1225</td><td>Bacteria>Pseudomonadota>Gammaproteobacteria>Enterobacterales>Enterobacteriaceae>Escherichia</td><td>GCA_012860255.1</td><td>Escherichia_coli</td><td>HJN25_004326</td><td>5261..8938</td><td>562</td><td>BREX-system-P-loop-protein-BrxC-[Escherichia-coli]</td></tr>
<tr><td>HAI4157988.1</td><td>Type1.BREX</td><td><-METHYLASE||SKI-><-?||tRNA->Arm-DNA-bind_3+Phage_integrase->EVE+HNH-><-REase+iREase||BrxA-DNA-binding->BrxB-BrxF-iSTAND->DUF6079*->N6-METHYLASE->PglZ->SIGMA+OB-Fold+AAA-ATPase+Lon-Protease-BrxL-><-ZnR<-ZnR<-?<-?||XIS-HTH-></td><td>1213</td><td>Bacteria>Pseudomonadota>Gammaproteobacteria>Enterobacterales>Enterobacteriaceae>Escherichia</td><td>GCA_012863435.1</td><td>Escherichia_coli</td><td>HJD09_003308</td><td>150572..154213</td><td>562</td><td>BREX-system-P-loop-protein-BrxC-[Escherichia-coli]</td></tr>
<tr><td>EFM7055934.1</td><td>Type1.BREX</td><td>DUF6079*->N6-METHYLASE->PglZ->SIGMA+OB-Fold+AAA-ATPase+Lon-Protease-BrxL->HKD-DNase+OGG-like->||AbiEi_3_N+AbiEi_3->AbiEii-><-DUF6216<-Arm-DNA-bind_3</td><td>1050</td><td>Bacteria>Pseudomonadota>Gammaproteobacteria>Enterobacterales>Enterobacteriaceae>Escherichia</td><td>GCA_012894495.1</td><td>Escherichia_coli</td><td>BAU66_002360</td><td>1..3155</td><td>562</td><td>BREX-system-P-loop-protein-BrxC-[Escherichia-coli]</td></tr>
<tr><td>EFN0218458.1</td><td>Type1.BREX</td><td><-Zn_Tnp_IS1+HTH_Tnp_IS1<-REase+iREase||BrxA-DNA-binding->BrxB-BrxF-iSTAND->DUF6079*->N6-METHYLASE->PglZ->SIGMA+OB-Fold+AAA-ATPase+Lon-Protease-BrxL-><-ZnR<-ZnR<-?<-?||RHH_antitoxin->XIS-HTH-></td><td>1213</td><td>Bacteria>Pseudomonadota>Gammaproteobacteria>Enterobacterales>Enterobacteriaceae>Escherichia</td><td>GCA_012909925.1</td><td>Escherichia_fergusonii</td><td>HNH53_002980</td><td>3165..6806</td><td>564</td><td>BREX-system-P-loop-protein-BrxC-[Escherichia-fergusonii]</td></tr>
<tr><td>EFN0586557.1</td><td>Type1.BREX</td><td><-REase<-REase||BrxA-DNA-binding->BrxB-BrxF-iSTAND->DUF6079*->N6-METHYLASE->PglZ->SIGMA+OB-Fold+AAA-ATPase+Lon-Protease-BrxL-></td><td>1213</td><td>Bacteria>Pseudomonadota>Gammaproteobacteria>Enterobacterales>Enterobacteriaceae>Escherichia</td><td>GCA_012914285.1</td><td>Escherichia_coli</td><td>HIK38_004176</td><td>2784..6425</td><td>562</td><td>BREX-system-P-loop-protein-BrxC-[Escherichia-coli]</td></tr>
<tr><td>MBB7654234.1</td><td>Type1.BREX</td><td><-XIS-HTH<-RHH_antitoxin||?->?->ZnR->ZnR-><-SIGMA+OB-Fold+AAA-ATPase+Lon-Protease-BrxL<-PglZ<-N6-METHYLASE<-DUF6079*<-BrxB-BrxF-iSTAND<-BrxA-DNA-binding||REase+iREase-><-EVE+HNH<-Arm-DNA-bind_3+Phage_integrase<-tRNA||?->HerA/FtsK+A32-></td><td>1213</td><td>Bacteria>Pseudomonadota>Gammaproteobacteria>Enterobacterales>Enterobacteriaceae>Escherichia</td><td>GCA_012916875.2</td><td>Escherichia_coli</td><td>HEM24_008890</td><td>24657..28298</td><td>562</td><td>BREX-system-P-loop-protein-BrxC-[Escherichia-coli]</td></tr>
<tr><td>EFN2130505.1</td><td>Type1.BREX</td><td>ParB->?-><-?<-SIGMA+OB-Fold+AAA-ATPase+Lon-Protease-BrxL<-PglZ<-N6-METHYLASE<-DUF6079*<-BrxB-BrxF-iSTAND<-BrxA-DNA-binding||REase+iREase-><-wHTH+SAD+HNH||MAD-NTD+HNH-></td><td>1213</td><td>Bacteria>Pseudomonadota>Gammaproteobacteria>Enterobacterales>Enterobacteriaceae>Escherichia</td><td>GCA_012923035.1</td><td>Escherichia_coli</td><td>HG367_004308</td><td>11483..15124</td><td>562</td><td>BREX-system-P-loop-protein-BrxC-[Escherichia-coli]</td></tr>
<tr><td>HAI9090450.1</td><td>Type1.BREX</td><td>DUF6079*->N6-METHYLASE->PglZ->SIGMA+OB-Fold+AAA-ATPase+Lon-Protease-BrxL->HKD-DNase+OGG-like->||AbiEi_3_N+AbiEi_3->AbiEii-><-DUF6216<-Arm-DNA-bind_3</td><td>1180</td><td>Bacteria>Pseudomonadota>Gammaproteobacteria>Enterobacterales>Enterobacteriaceae>Escherichia</td><td>GCA_012925565.1</td><td>Escherichia_coli</td><td>HKC72_003106</td><td>1..3544</td><td>562</td><td>BREX-system-P-loop-protein-BrxC-[Escherichia-coli]</td></tr>
<tr><td>EFN3304096.1</td><td>Type1.BREX</td><td>ZnR-><-SIGMA+OB-Fold+AAA-ATPase+Lon-Protease-BrxL<-PglZ<-N6-METHYLASE<-DUF6079*<-BrxB-BrxF-iSTAND<-BrxA-DNA-binding||REase+iREase-><-EVE+HNH</td><td>1213</td><td>Bacteria>Pseudomonadota>Gammaproteobacteria>Enterobacterales>Enterobacteriaceae>Escherichia</td><td>GCA_012969905.1</td><td>Escherichia_coli</td><td>HKH56_004291</td><td>12014..15655</td><td>562</td><td>BREX-system-P-loop-protein-BrxC-[Escherichia-coli]</td></tr>
<tr><td>EFN3922650.1</td><td>Type1.BREX</td><td><-Phage_integrase||?->?->?-><-HNH||wHTH+SAD+HNH-><-REase+iREase||BrxA-DNA-binding->BrxB-BrxF-iSTAND->DUF6079*->N6-METHYLASE->PglZ->SIGMA+OB-Fold+AAA-ATPase+Lon-Protease-BrxL->?-><-?<-ParB<-?||<-REase+N6_METHYLASE<-TRD</td><td>1213</td><td>Bacteria>Pseudomonadota>Gammaproteobacteria>Enterobacterales>Enterobacteriaceae>Escherichia</td><td>GCA_012987865.1</td><td>Escherichia_coli</td><td>FQJ23_02905</td><td>87001..90642</td><td>562</td><td>BREX-system-P-loop-protein-BrxC-[Escherichia-coli]</td></tr>
<tr><td>EFN5198232.1</td><td>Type1.BREX</td><td><-TIR||BrxA-DNA-binding->BrxB-BrxF-iSTAND->DUF6079*->ParB+HNH->N6-METHYLASE->HEPN->RVT->N6-METHYLASE-p->PglZ->SIGMA+OB-Fold+AAA-ATPase+Lon-Protease-BrxL->wHTH+WYL+WCX-></td><td>1233</td><td>Bacteria>Pseudomonadota>Gammaproteobacteria>Enterobacterales>Enterobacteriaceae>Escherichia</td><td>GCA_012993025.1</td><td>Escherichia_coli</td><td>FQG10_00765</td><td>2767..6468</td><td>562</td><td>BREX-system-P-loop-protein-BrxC-[Escherichia-coli]</td></tr>
<tr><td>EFN5705755.1</td><td>Type1.BREX</td><td><-RHH_antitoxin||?->?->?->ZnR->ZnR-><-SIGMA+OB-Fold+AAA-ATPase+Lon-Protease-BrxL<-PglZ<-N6-METHYLASE<-DUF6079*<-BrxB-BrxF-iSTAND<-BrxA-DNA-binding||REase+iREase-><-EVE+HNH<-Arm-DNA-bind_3+Phage_integrase<-tRNA||?->HerA/FtsK+A32-></td><td>1213</td><td>Bacteria>Pseudomonadota>Gammaproteobacteria>Enterobacterales>Enterobacteriaceae>Escherichia</td><td>GCA_012995165.1</td><td>Escherichia_coli</td><td>FQH62_10005</td><td>24813..28454</td><td>562</td><td>BREX-system-P-loop-protein-BrxC-[Escherichia-coli]</td></tr>
<tr><td>EFN5734792.1</td><td>Type1.BREX</td><td><-Phage_integrase||?->?->?-><-HNH||wHTH+SAD+HNH-><-REase+iREase||BrxA-DNA-binding->BrxB-BrxF-iSTAND->DUF6079*->N6-METHYLASE->PglZ->SIGMA+OB-Fold+AAA-ATPase+Lon-Protease-BrxL->HKD-DNase+OGG-like->||AbiEi_3_N+AbiEi_3->AbiEii-><-DUF6216<-Arm-DNA-bind_3</td><td>1213</td><td>Bacteria>Pseudomonadota>Gammaproteobacteria>Enterobacterales>Enterobacteriaceae>Escherichia</td><td>GCA_012995705.1</td><td>Escherichia_coli</td><td>FQJ28_11185</td><td>111254..114895</td><td>562</td><td>BREX-system-P-loop-protein-BrxC-[Escherichia-coli]</td></tr>
<tr><td>HAJ1412079.1</td><td>Type1.BREX</td><td>DUF6079*->PglZ->SIGMA+OB-Fold+AAA-ATPase+Lon-Protease-BrxL->HKD-DNase+OGG-like->||AbiEi_3_N+AbiEi_3->AbiEii-></td><td>1046</td><td>Bacteria>Pseudomonadota>Gammaproteobacteria>Enterobacterales>Enterobacteriaceae>Escherichia</td><td>GCA_013000285.1</td><td>Escherichia_coli</td><td>HMG43_22065</td><td>1..3142</td><td>562</td><td>BREX-system-P-loop-protein-BrxC-[Escherichia-coli]</td></tr>
<tr><td>HAJ4265401.1</td><td>Type1.BREX</td><td>BrxA-DNA-binding->BrxB-BrxF-iSTAND->DUF6079*->N6-METHYLASE->PglZ->SIGMA+OB-Fold+AAA-ATPase+Lon-Protease-BrxL-></td><td>1213</td><td>Bacteria>Pseudomonadota>Gammaproteobacteria>Enterobacterales>Enterobacteriaceae>Escherichia</td><td>GCA_013021445.1</td><td>Escherichia_coli</td><td>HLU44_19895</td><td>1490..5131</td><td>562</td><td>BREX-system-P-loop-protein-BrxC-[Escherichia-coli]</td></tr>
<tr><td>HAJ6497178.1</td><td>Type1.BREX</td><td>DsbC+DsbD+Thioredoxin_7->TetR-HTH->tRNA->Arm-DNA-bind_3+Phage_integrase-><-Sulfatase<-?||DDE_Tnp_1_5+DDE_Tnp_1-><-REase+iREase||BrxA-DNA-binding->BrxB-BrxF-iSTAND->DUF6079*->N6-METHYLASE->PglZ->SIGMA+OB-Fold+AAA-ATPase+Lon-Protease-BrxL-><-ZnR<-ZnR</td><td>1213</td><td>Bacteria>Pseudomonadota>Gammaproteobacteria>Enterobacterales>Enterobacteriaceae>Escherichia</td><td>GCA_013038245.1</td><td>Escherichia_coli</td><td>HNC37_02435</td><td>510930..514571</td><td>562</td><td>BREX-system-P-loop-protein-BrxC-[Escherichia-coli]</td></tr>
<tr><td>HAJ6661538.1</td><td>Type1.BREX</td><td><-RHH_antitoxin||?->?->?->ZnR->ZnR-><-SIGMA+OB-Fold+AAA-ATPase+Lon-Protease-BrxL<-PglZ<-N6-METHYLASE<-DUF6079*<-BrxB-BrxF-iSTAND<-BrxA-DNA-binding||REase+iREase-></td><td>1213</td><td>Bacteria>Pseudomonadota>Gammaproteobacteria>Enterobacterales>Enterobacteriaceae>Escherichia</td><td>GCA_013038825.1</td><td>Escherichia_coli</td><td>HNC51_24215</td><td>13272..16913</td><td>562</td><td>BREX-system-P-loop-protein-BrxC-[Escherichia-coli]</td></tr>
<tr><td>HAJ6872573.1</td><td>Type1.BREX</td><td><-Phage_integrase||?->?->?-><-HNH||wHTH+SAD+HNH-><-REase+iREase||BrxA-DNA-binding->BrxB-BrxF-iSTAND->DUF6079-p*-></td><td>576</td><td>Bacteria>Pseudomonadota>Gammaproteobacteria>Enterobacterales>Enterobacteriaceae>Escherichia</td><td>GCA_013039665.1</td><td>Escherichia_coli</td><td>HND09_18460</td><td>71676..73402</td><td>562</td><td>BREX-system-P-loop-protein-BrxC-[Escherichia-coli]</td></tr>
<tr><td>EFN7556473.1</td><td>Type1.BREX</td><td><-REase+iREase||BrxA-DNA-binding->BrxB-BrxF-iSTAND->DUF6079*->N6-METHYLASE->PglZ->SIGMA+OB-Fold+AAA-ATPase+Lon-Protease-BrxL-><-ZnR<-ZnR<-?<-?<-?||RHH_antitoxin-></td><td>1213</td><td>Bacteria>Pseudomonadota>Gammaproteobacteria>Enterobacterales>Enterobacteriaceae>Escherichia</td><td>GCA_013041485.1</td><td>Escherichia_coli</td><td>EHH76_16080</td><td>2195..5836</td><td>562</td><td>BREX-system-P-loop-protein-BrxC-[Escherichia-coli]</td></tr>
<tr><td>EFO0007667.1</td><td>Type1.BREX</td><td><-N6-METHYLASE<-DUF6079*<-BrxB-BrxF-iSTAND<-BrxA-DNA-binding||REase+iREase-></td><td>1213</td><td>Bacteria>Pseudomonadota>Gammaproteobacteria>Enterobacterales>Enterobacteriaceae>Escherichia</td><td>GCA_013057425.1</td><td>Escherichia_coli</td><td>D1356_22460</td><td>2492..6133</td><td>562</td><td>BREX-system-P-loop-protein-BrxC-[Escherichia-coli]</td></tr>
<tr><td>EFO0695969.1</td><td>Type1.BREX</td><td><-Phage_integrase||?->?->?-><-HNH||wHTH+SAD+HNH-><-REase+iREase||BrxA-DNA-binding->BrxB-BrxF-iSTAND->DUF6079*->N6-METHYLASE->PglZ->SIGMA+OB-Fold+AAA-ATPase+Lon-Protease-BrxL-></td><td>1213</td><td>Bacteria>Pseudomonadota>Gammaproteobacteria>Enterobacterales>Enterobacteriaceae>Escherichia</td><td>GCA_013059045.1</td><td>Escherichia_coli</td><td>KI087_004400</td><td>8657..12298</td><td>562</td><td>BREX-system-P-loop-protein-BrxC-[Escherichia-coli]</td></tr>
<tr><td>EFO0712882.1</td><td>Type1.BREX</td><td><-METHYLASE||SKI-><-?||tRNA->Arm-DNA-bind_3+Phage_integrase->EVE+HNH-><-REase<-REase+iREase||BrxA-DNA-binding->BrxB-BrxF-iSTAND->DUF6079*->N6-METHYLASE->PglZ->SIGMA+OB-Fold+AAA-ATPase+Lon-Protease-BrxL-><-ZnR<-ZnR<-?<-?<-?||RHH_antitoxin-></td><td>1213</td><td>Bacteria>Pseudomonadota>Gammaproteobacteria>Enterobacterales>Enterobacteriaceae>Escherichia</td><td>GCA_013059225.1</td><td>Escherichia_coli</td><td>DYR48_06815</td><td>148369..152010</td><td>562</td><td>BREX-system-P-loop-protein-BrxC-[Escherichia-coli]</td></tr>
<tr><td>EFO1505998.1</td><td>Type1.BREX</td><td><-DUF6079-p*<-BrxB-BrxF-iSTAND<-BrxA-DNA-binding||REase+iREase-><-wHTH+SAD+HNH||HNH-><-?<-?<-?||35exo-></td><td>919</td><td>Bacteria>Pseudomonadota>Gammaproteobacteria>Enterobacterales>Enterobacteriaceae>Escherichia</td><td>GCA_013063585.1</td><td>Escherichia_coli</td><td>DWG87_23060</td><td>1..2758</td><td>562</td><td>BREX-system-P-loop-protein-BrxC-[Escherichia-coli]</td></tr>
<tr><td>EFO2091244.1</td><td>Type1.BREX</td><td><-35exo||?->?->?-><-HNH||wHTH+SAD+HNH-><-REase+iREase||BrxA-DNA-binding->BrxB-BrxF-iSTAND->DUF6079*->N6-METHYLASE->PglZ->SIGMA+OB-Fold+AAA-ATPase+Lon-Protease-BrxL->HKD-DNase+OGG-like->||AbiEi_3_N+AbiEi_3->AbiEii-><-DUF6216<-Arm-DNA-bind_3</td><td>1213</td><td>Bacteria>Pseudomonadota>Gammaproteobacteria>Enterobacterales>Enterobacteriaceae>Escherichia</td><td>GCA_013066245.1</td><td>Escherichia_coli_O54</td><td>DP047_19880</td><td>9767..13408</td><td>2184075</td><td>BREX-system-P-loop-protein-BrxC-[Escherichia-coli-O54]</td></tr>
<tr><td>EFO2510526.1</td><td>Type1.BREX</td><td>ZnR->ZnR-><-SIGMA+OB-Fold+AAA-ATPase+Lon-Protease-BrxL<-PglZ<-N6-METHYLASE<-DUF6079*<-BrxB-BrxF-iSTAND<-BrxA-DNA-binding||REase->REase-><-HEPN<-ORC-CDC6+ZnR<-N6-METHYLASE<-Arm-DNA-bind_3+Phage_integrase</td><td>1213</td><td>Bacteria>Pseudomonadota>Gammaproteobacteria>Enterobacterales>Enterobacteriaceae>Escherichia</td><td>GCA_013067425.1</td><td>Escherichia_coli</td><td>B6Q63_13725</td><td>11254..14895</td><td>562</td><td>BREX-system-P-loop-protein-BrxC-[Escherichia-coli]</td></tr>
<tr><td>HAJ7467856.1</td><td>Type1.BREX</td><td><-RelE-ParE<-CopG/RHH||<-RHH||?->wHTH+WYL+WCX->BrxA-DNA-binding->BrxB-BrxF-iSTAND->DUF6079*->ParB+HNH->N6-METHYLASE->PglZ->SIGMA+OB-Fold+AAA-ATPase+Lon-Protease-BrxL-><-RVT<-PIN<-MazE_antitoxin||?->Phage_holin-></td><td>1225</td><td>Bacteria>Pseudomonadota>Gammaproteobacteria>Enterobacterales>Enterobacteriaceae>Escherichia</td><td>GCA_013077315.1</td><td>Escherichia_coli</td><td>HNV92_21025</td><td>13565..17242</td><td>562</td><td>BREX-system-P-loop-protein-BrxC-[Escherichia-coli]</td></tr>
<tr><td>EFP0188320.1</td><td>Type1.BREX</td><td>DUF772+DDE_Tnp_1->BrxA-DNA-binding->BrxB-BrxF-iSTAND->DUF6079*->N6-METHYLASE->PglZ->SIGMA+OB-Fold+AAA-ATPase+Lon-Protease-BrxL-><-ZnR<-ZnR|||nSTAND1+sGTP-></td><td>1213</td><td>Bacteria>Pseudomonadota>Gammaproteobacteria>Enterobacterales>Enterobacteriaceae>Escherichia</td><td>GCA_013129375.1</td><td>Escherichia_coli</td><td>HLX13_004311</td><td>1679..5320</td><td>562</td><td>BREX-system-P-loop-protein-BrxC-[Escherichia-coli]</td></tr>
<tr><td>EFP2415668.1</td><td>Type1.BREX</td><td><-Phage_integrase||?->?->?-><-HNH||wHTH+SAD+HNH-><-REase+iREase||BrxA-DNA-binding->BrxB-BrxF-iSTAND->DUF6079*->ZnR->ZnR->?-><-wHTH+WYL||?->?->?->HIN-HTH-></td><td>1160</td><td>Bacteria>Pseudomonadota>Gammaproteobacteria>Enterobacterales>Enterobacteriaceae>Escherichia</td><td>GCA_013157045.1</td><td>Escherichia_coli</td><td>HP898_000185</td><td>8665..12147</td><td>562</td><td>BREX-system-P-loop-protein-BrxC-[Escherichia-coli]</td></tr>
<tr><td>NUC65042.1</td><td>Type1.BREX</td><td>DDE_Tnp_IS1->?->ZnR->BrxA-DNA-binding->BrxB-BrxF-iSTAND->DUF6079*->ParB+HNH->N6-METHYLASE->HEPN->RVT->N6-METHYLASE-p->PglZ->SIGMA+OB-Fold+AAA-ATPase+Lon-Protease-BrxL-><-Arm-DNA-bind_3+Phage_integrase||Phage_integrase-></td><td>1233</td><td>Bacteria>Pseudomonadota>Gammaproteobacteria>Enterobacterales>Enterobacteriaceae>Escherichia</td><td>GCA_013342565.1</td><td>Escherichia_coli</td><td>HUC40_15465</td><td>2945..6646</td><td>562</td><td>BREX-system-P-loop-protein-BrxC-[Escherichia-coli]</td></tr>
<tr><td>EFR2049109.1</td><td>Type1.BREX</td><td>DUF6079-p*->N6-METHYLASE->PglZ->SIGMA+OB-Fold+AAA-ATPase+Lon-Protease-BrxL->HKD-DNase+OGG-like->||AbiEi_3_N+AbiEi_3->AbiEii-><-DUF6216<-Arm-DNA-bind_3</td><td>884</td><td>Bacteria>Pseudomonadota>Gammaproteobacteria>Enterobacterales>Enterobacteriaceae>Escherichia</td><td>GCA_013546455.1</td><td>Escherichia_coli</td><td>HZ098_003737</td><td>1..2655</td><td>562</td><td>BREX-system-P-loop-protein-BrxC-[Escherichia-coli]</td></tr>
<tr><td>EFR9656371.1</td><td>Type1.BREX</td><td>DUF6079-p*->N6-METHYLASE->PglZ->SIGMA+OB-Fold+AAA-ATPase+Lon-Protease-BrxL->HKD-DNase+OGG-like->||AbiEi_3_N+AbiEi_3->AbiEii-><-DUF6216<-Arm-DNA-bind_3</td><td>991</td><td>Bacteria>Pseudomonadota>Gammaproteobacteria>Enterobacterales>Enterobacteriaceae>Escherichia</td><td>GCA_013588925.1</td><td>Escherichia_coli</td><td>H2N11_003246</td><td>1..2976</td><td>562</td><td>BREX-system-P-loop-protein-BrxC-[Escherichia-coli]</td></tr>
<tr><td>EFS8463113.1</td><td>Type1.BREX</td><td><-Phage_integrase||?->?->?-><-HNH||wHTH+SAD+HNH-><-REase+iREase||BrxA-DNA-binding->BrxB-BrxF-iSTAND->DUF6079*->N6-METHYLASE->PglZ->SIGMA+OB-Fold+AAA-ATPase+Lon-Protease-BrxL->HKD-DNase+OGG-like->||AbiEi_3_N+AbiEi_3->AbiEii-><-DUF6216<-Arm-DNA-bind_3</td><td>1213</td><td>Bacteria>Pseudomonadota>Gammaproteobacteria>Enterobacterales>Enterobacteriaceae>Escherichia</td><td>GCA_013669455.1</td><td>Escherichia_coli</td><td>HWT39_002319</td><td>97196..100837</td><td>562</td><td>BREX-system-P-loop-protein-BrxC-[Escherichia-coli]</td></tr>
<tr><td>EFT2808154.1</td><td>Type1.BREX</td><td>ZnR->wHTH+WYL+WCX->BrxA-DNA-binding->BrxB-BrxF-iSTAND->DUF6079*->ParB+HNH->N6-METHYLASE->PglZ->SIGMA+OB-Fold+AAA-ATPase+Lon-Protease-BrxL->35exo-></td><td>1221</td><td>Bacteria>Pseudomonadota>Gammaproteobacteria>Enterobacterales>Enterobacteriaceae>Escherichia</td><td>GCA_013688275.1</td><td>Escherichia_coli</td><td>HVL53_003849</td><td>3457..7122</td><td>562</td><td>BREX-system-P-loop-protein-BrxC-[Escherichia-coli]</td></tr>
<tr><td>EFT2742093.1</td><td>Type1.BREX</td><td>TRD->REase+N6_METHYLASE->||?->ParB->?-><-?<-SIGMA+OB-Fold+AAA-ATPase+Lon-Protease-BrxL<-PglZ<-N6-METHYLASE<-DUF6079*<-BrxB-BrxF-iSTAND<-BrxA-DNA-binding||REase+iREase-><-wHTH+SAD+HNH||HNH-><-?<-?<-?||Phage_integrase-></td><td>1213</td><td>Bacteria>Pseudomonadota>Gammaproteobacteria>Enterobacterales>Enterobacteriaceae>Escherichia</td><td>GCA_013688615.1</td><td>Escherichia_coli</td><td>HVL39_003223</td><td>27812..31453</td><td>562</td><td>BREX-system-P-loop-protein-BrxC-[Escherichia-coli]</td></tr>
<tr><td>EFU0098527.1</td><td>Type1.BREX</td><td>DUF6079*->N6-METHYLASE->PglZ->SIGMA+OB-Fold+AAA-ATPase+Lon-Protease-BrxL->HKD-DNase+OGG-like->||AbiEi_3_N+AbiEi_3->AbiEii->?-><-DUF6216<-Arm-DNA-bind_3</td><td>1009</td><td>Bacteria>Pseudomonadota>Gammaproteobacteria>Enterobacterales>Enterobacteriaceae>Escherichia</td><td>GCA_013728855.1</td><td>Escherichia_coli</td><td>HVV04_003689</td><td>1..3030</td><td>562</td><td>BREX-system-P-loop-protein-BrxC-[Escherichia-coli]</td></tr>
<tr><td>HAL0559649.1</td><td>Type1.BREX</td><td>DUF6079*->N6-METHYLASE->PglZ->SIGMA+OB-Fold+AAA-ATPase+Lon-Protease-BrxL-><-ZnR<-HTH_Tnp_1+HTH_21+rve+rve_3</td><td>1111</td><td>Bacteria>Pseudomonadota>Gammaproteobacteria>Enterobacterales>Enterobacteriaceae>Escherichia</td><td>GCA_013967645.1</td><td>Escherichia_coli</td><td>H4D61_004677</td><td>1..3338</td><td>562</td><td>BREX-system-P-loop-protein-BrxC-[Escherichia-coli]</td></tr>
<tr><td>HAL0969191.1</td><td>Type1.BREX</td><td>DUF6079*->N6-METHYLASE->PglZ->SIGMA+OB-Fold+AAA-ATPase+Lon-Protease-BrxL-><-ZnR<-HTH_Tnp_1+HTH_21+rve+rve_3</td><td>1186</td><td>Bacteria>Pseudomonadota>Gammaproteobacteria>Enterobacterales>Enterobacteriaceae>Escherichia</td><td>GCA_013984825.1</td><td>Escherichia_coli</td><td>H4H65_005028</td><td>1..3562</td><td>562</td><td>BREX-system-P-loop-protein-BrxC-[Escherichia-coli]</td></tr>
<tr><td>HAL9997354.1</td><td>Type1.BREX</td><td>DUF6079*->N6-METHYLASE->PglZ->SIGMA+OB-Fold+AAA-ATPase+Lon-Protease-BrxL->HKD-DNase+OGG-like->||AbiEi_3_N+AbiEi_3->AbiEii-><-DUF6216<-Arm-DNA-bind_3</td><td>1081</td><td>Bacteria>Pseudomonadota>Gammaproteobacteria>Enterobacterales>Enterobacteriaceae>Escherichia</td><td>GCA_014189205.1</td><td>Escherichia_coli</td><td>H7W48_003266</td><td>1..3246</td><td>562</td><td>BREX-system-P-loop-protein-BrxC-[Escherichia-coli]</td></tr>
<tr><td>EGC7303784.1</td><td>Type1.BREX</td><td>HIN-HTH->AAA-ATPase->?->ZnR->ZnR-><-SIGMA+OB-Fold+AAA-ATPase+Lon-Protease-BrxL<-PglZ<-N6-METHYLASE<-DUF6079*<-BrxB-BrxF-iSTAND<-BrxA-DNA-binding||REase+iREase-><-EVE+HNH<-Arm-DNA-bind_3+Phage_integrase<-tRNA||?->HerA/FtsK+A32-></td><td>1213</td><td>Bacteria>Pseudomonadota>Gammaproteobacteria>Enterobacterales>Enterobacteriaceae>Escherichia</td><td>GCA_014304595.1</td><td>Escherichia_coli</td><td>JTG03_003848</td><td>38080..41721</td><td>562</td><td>BREX-system-P-loop-protein-BrxC-[Escherichia-coli]</td></tr>
<tr><td>EGD5155863.1</td><td>Type1.BREX</td><td><-35exo||?->?->?-><-HNH||wHTH+SAD+HNH-><-REase+iREase||BrxA-DNA-binding->BrxB-BrxF-iSTAND->DUF6079*->N6-METHYLASE->PglZ->SIGMA+OB-Fold+AAA-ATPase+Lon-Protease-BrxL->HKD-DNase+OGG-like->||AbiEi_3_N+AbiEi_3->AbiEii-><-DUF6216<-Arm-DNA-bind_3</td><td>1213</td><td>Bacteria>Pseudomonadota>Gammaproteobacteria>Enterobacterales>Enterobacteriaceae>Escherichia</td><td>GCA_014387005.1</td><td>Escherichia_coli</td><td>DTY31_07100</td><td>71746..75387</td><td>562</td><td>BREX-system-P-loop-protein-BrxC-[Escherichia-coli]</td></tr>
<tr><td>HAM5125585.1</td><td>Type1.BREX</td><td>DUF6079-p*->N6-METHYLASE->PglZ->SIGMA+OB-Fold+AAA-ATPase+Lon-Protease-BrxL-><-ZnR<-RVT</td><td>770</td><td>Bacteria>Pseudomonadota>Gammaproteobacteria>Enterobacterales>Enterobacteriaceae>Escherichia</td><td>GCA_014468575.1</td><td>Escherichia_coli</td><td>IB005_004014</td><td>1..2313</td><td>562</td><td>BREX-system-P-loop-protein-BrxC-[Escherichia-coli]</td></tr>
<tr><td>EGF7347429.1</td><td>Type1.BREX</td><td>DUF6079*->HKD-DNase+OGG-like->||AbiEi_3_N+AbiEi_3->AbiEii-><-DUF6216<-tRNA||?->HerA/FtsK+A32-><-?<-?||Peptidase_M17_N+Peptidase_M17-></td><td>1213</td><td>Bacteria>Pseudomonadota>Gammaproteobacteria>Enterobacterales>Enterobacteriaceae>Escherichia</td><td>GCA_014521725.1</td><td>Escherichia_coli</td><td>HJV04_003560</td><td>44..3685</td><td>562</td><td>BREX-system-P-loop-protein-BrxC-[Escherichia-coli]</td></tr>
<tr><td>HAN3626817.1</td><td>Type1.BREX</td><td><-N6-METHYLASE<-DUF6079*<-BrxB-BrxF-iSTAND<-BrxA-DNA-binding||REase+iREase-><-wHTH+SAD+HNH||HNH-><-?<-?<-?||Phage_integrase-></td><td>1213</td><td>Bacteria>Pseudomonadota>Gammaproteobacteria>Enterobacterales>Enterobacteriaceae>Escherichia</td><td>GCA_014691525.1</td><td>Escherichia_coli</td><td>IE966_004457</td><td>2731..6372</td><td>562</td><td>BREX-system-P-loop-protein-BrxC-[Escherichia-coli]</td></tr>
<tr><td>HAN6349201.1</td><td>Type1.BREX</td><td><-Zn_Tnp_IS1+HTH_Tnp_IS1<-REase||BrxA-DNA-binding->BrxB-BrxF-iSTAND->DUF6079*->N6-METHYLASE->PglZ->SIGMA+OB-Fold+AAA-ATPase+Lon-Protease-BrxL-><-ZnR<-ZnR<-?||XIS-HTH-></td><td>1213</td><td>Bacteria>Pseudomonadota>Gammaproteobacteria>Enterobacterales>Enterobacteriaceae>Escherichia</td><td>GCA_014709895.1</td><td>Escherichia_coli</td><td>IF023_004329</td><td>2523..6164</td><td>562</td><td>BREX-system-P-loop-protein-BrxC-[Escherichia-coli]</td></tr>
<tr><td>EGI2318416.1</td><td>Type1.BREX</td><td><-Phage_integrase||?->?->?-><-HNH||wHTH+SAD+HNH-><-REase+iREase||BrxA-DNA-binding->BrxB-BrxF-iSTAND->DUF6079*-></td><td>1027</td><td>Bacteria>Pseudomonadota>Gammaproteobacteria>Enterobacterales>Enterobacteriaceae>Escherichia</td><td>GCA_014736875.1</td><td>Escherichia_coli</td><td>IGA36_003675</td><td>97131..100211</td><td>562</td><td>BREX-system-P-loop-protein-BrxC-[Escherichia-coli]</td></tr>
<tr><td>EGI6859711.1</td><td>Type1.BREX</td><td>DUF6079*->N6-METHYLASE->PglZ->SIGMA+OB-Fold+AAA-ATPase+Lon-Protease-BrxL-><-ZnR<-ZnR<-?||RHH_antitoxin->XIS-HTH-></td><td>1173</td><td>Bacteria>Pseudomonadota>Gammaproteobacteria>Enterobacterales>Enterobacteriaceae>Escherichia</td><td>GCA_014766135.1</td><td>Escherichia_coli</td><td>IGX90_003813</td><td>1..3522</td><td>562</td><td>BREX-system-P-loop-protein-BrxC-[Escherichia-coli]</td></tr>
<tr><td>HAO0324358.1</td><td>Type1.BREX</td><td><-HNH||wHTH+SAD+HNH-><-REase||BrxA-DNA-binding->BrxB-BrxF-iSTAND->DUF6079*->N6-METHYLASE-p-></td><td>1213</td><td>Bacteria>Pseudomonadota>Gammaproteobacteria>Enterobacterales>Enterobacteriaceae>Escherichia</td><td>GCA_014771515.1</td><td>Escherichia_coli</td><td>IGZ64_004245</td><td>6773..10414</td><td>562</td><td>BREX-system-P-loop-protein-BrxC-[Escherichia-coli]</td></tr>
<tr><td>HAO2821705.1</td><td>Type1.BREX</td><td>DUF6079*->PglZ->SIGMA+OB-Fold+AAA-ATPase+Lon-Protease-BrxL->HKD-DNase+OGG-like->||AbiEi_3_N+AbiEi_3->AbiEii-><-DUF6216<-Arm-DNA-bind_3<-tRNA||?->HerA/FtsK+A32-></td><td>1111</td><td>Bacteria>Pseudomonadota>Gammaproteobacteria>Enterobacterales>Enterobacteriaceae>Escherichia</td><td>GCA_014823275.1</td><td>Escherichia_coli</td><td>IHL91_001055</td><td>1..3336</td><td>562</td><td>BREX-system-P-loop-protein-BrxC-[Escherichia-coli]</td></tr>
<tr><td>HAO3163381.1</td><td>Type1.BREX</td><td><-HIN-HTH<-RelE-ParE<-CopG/RHH||<-RHH||?->wHTH+WYL+WCX->BrxA-DNA-binding->BrxB-BrxF-iSTAND->DUF6079*-></td><td>1136</td><td>Bacteria>Pseudomonadota>Gammaproteobacteria>Enterobacterales>Enterobacteriaceae>Escherichia</td><td>GCA_014826135.1</td><td>Escherichia_coli</td><td>IHN66_004302</td><td>5400..8808</td><td>562</td><td>BREX-system-P-loop-protein-BrxC-[Escherichia-coli]</td></tr>
<tr><td>EGK3607993.1</td><td>Type1.BREX</td><td>Arm-DNA-bind_3+Phage_integrase->XIS-HTH->?->?->?->BrxA-DNA-binding->BrxB-BrxF-iSTAND->DUF6079*->ParB+HNH->N6-METHYLASE->SIR2->HerA/FtsK->N6-METHYLASE-p-></td><td>1238</td><td>Bacteria>Pseudomonadota>Gammaproteobacteria>Enterobacterales>Enterobacteriaceae>Escherichia</td><td>GCA_014957385.1</td><td>Escherichia_coli</td><td>IOT31_003182</td><td>92049..95765</td><td>562</td><td>BREX-system-P-loop-protein-BrxC-[Escherichia-coli]</td></tr>
<tr><td>EGK3603428.1</td><td>Type1.BREX</td><td>DUF6079*->PglZ->SIGMA+OB-Fold+AAA-ATPase+Lon-Protease-BrxL->HKD-DNase+OGG-like->||AbiEi_3_N+AbiEi_3->AbiEii-><-DUF6216<-Arm-DNA-bind_3<-tRNA||?->HerA/FtsK+A32-></td><td>1179</td><td>Bacteria>Pseudomonadota>Gammaproteobacteria>Enterobacterales>Enterobacteriaceae>Escherichia</td><td>GCA_014957425.1</td><td>Escherichia_coli</td><td>IOT28_003449</td><td>1..3540</td><td>562</td><td>BREX-system-P-loop-protein-BrxC-[Escherichia-coli]</td></tr>
<tr><td>EGJ9762745.1</td><td>Type1.BREX</td><td><-Phage_integrase||?->?->?->||wHTH+SAD+HNH-><-REase+iREase||BrxA-DNA-binding->BrxB-BrxF-iSTAND->DUF6079*->N6-METHYLASE->PglZ->SIGMA+OB-Fold+AAA-ATPase+Lon-Protease-BrxL->?-><-?<-ParB<-?||<-REase+N6_METHYLASE<-TRD</td><td>1213</td><td>Bacteria>Pseudomonadota>Gammaproteobacteria>Enterobacterales>Enterobacteriaceae>Escherichia</td><td>GCA_014978025.1</td><td>Escherichia_coli</td><td>LZF81_002887</td><td>55534..59175</td><td>562</td><td>BREX-system-P-loop-protein-BrxC-[Escherichia-coli]</td></tr>
<tr><td>HAO5539965.1</td><td>Type1.BREX</td><td><-REase<-REase||BrxA-DNA-binding->BrxB-BrxF-iSTAND->DUF6079*->N6-METHYLASE->PglZ->SIGMA+OB-Fold+AAA-ATPase+Lon-Protease-BrxL-><-ZnR<-RVT</td><td>1213</td><td>Bacteria>Pseudomonadota>Gammaproteobacteria>Enterobacterales>Enterobacteriaceae>Escherichia</td><td>GCA_015013895.1</td><td>Escherichia_coli</td><td>IP613_004381</td><td>2370..6011</td><td>562</td><td>BREX-system-P-loop-protein-BrxC-[Escherichia-coli]</td></tr>
<tr><td>EGN4319649.1</td><td>Type1.BREX</td><td>DUF6079*->N6-METHYLASE->PglZ->SIGMA+OB-Fold+AAA-ATPase+Lon-Protease-BrxL->?-><-?<-ParB<-?||<-REase+N6_METHYLASE</td><td>1081</td><td>Bacteria>Pseudomonadota>Gammaproteobacteria>Enterobacterales>Enterobacteriaceae>Escherichia</td><td>GCA_015200385.1</td><td>Escherichia_coli</td><td>IH255_004211</td><td>1..3247</td><td>562</td><td>BREX-system-P-loop-protein-BrxC-[Escherichia-coli]</td></tr>
<tr><td>HAU8224518.1</td><td>Type1.BREX</td><td><-Phage_integrase||?->?->?-><-HNH||wHTH+SAD+HNH-><-REase+iREase||BrxA-DNA-binding->BrxB-BrxF-iSTAND->DUF6079*->N6-METHYLASE->PglZ->SIGMA+OB-Fold+AAA-ATPase+Lon-Protease-BrxL->?-><-?<-ParB<-?||<-REase+N6_METHYLASE<-TRD</td><td>1213</td><td>Bacteria>Pseudomonadota>Gammaproteobacteria>Enterobacterales>Enterobacteriaceae>Escherichia</td><td>GCA_016109325.1</td><td>Escherichia_coli</td><td>IG138_000798</td><td>111239..114880</td><td>562</td><td>BREX-system-P-loop-protein-BrxC-[Escherichia-coli]</td></tr>
<tr><td>HAU8802979.1</td><td>Type1.BREX</td><td>DDE_Tnp_IS1-><-SIGMA+OB-Fold+AAA-ATPase+Lon-Protease-BrxL<-PglZ<-N6-METHYLASE<-DUF6079*<-BrxB-BrxF-iSTAND<-BrxA-DNA-binding||REase->?->||<-Arm-DNA-bind_3+Phage_integrase<-tRNA||?->HerA/FtsK+A32-></td><td>1213</td><td>Bacteria>Pseudomonadota>Gammaproteobacteria>Enterobacterales>Enterobacteriaceae>Escherichia</td><td>GCA_016115265.1</td><td>Escherichia_coli</td><td>JEW39_004409</td><td>9027..12668</td><td>562</td><td>BREX-system-P-loop-protein-BrxC-[Escherichia-coli]</td></tr>
<tr><td>EGV9472870.1</td><td>Type1.BREX</td><td><-Phage_integrase||?->?->?-><-HNH||wHTH+SAD+HNH-><-REase+iREase||BrxA-DNA-binding->BrxB-BrxF-iSTAND->DUF6079*->N6-METHYLASE->PglZ->SIGMA+OB-Fold+AAA-ATPase+Lon-Protease-BrxL->?-><-?<-ParB<-?||<-REase+N6_METHYLASE<-TRD</td><td>1213</td><td>Bacteria>Pseudomonadota>Gammaproteobacteria>Enterobacterales>Enterobacteriaceae>Escherichia</td><td>GCA_016298225.1</td><td>Escherichia_coli</td><td>JF871_001476</td><td>111292..114933</td><td>562</td><td>BREX-system-P-loop-protein-BrxC-[Escherichia-coli]</td></tr>
<tr><td>EGY5636086.1</td><td>Type1.BREX</td><td>DUF6079*->N6-METHYLASE->PglZ->SIGMA+OB-Fold+AAA-ATPase+Lon-Protease-BrxL->HKD-DNase+OGG-like->||AbiEi_3_N+AbiEi_3->AbiEii-><-DUF6216<-Arm-DNA-bind_3</td><td>1171</td><td>Bacteria>Pseudomonadota>Gammaproteobacteria>Enterobacterales>Enterobacteriaceae>Escherichia</td><td>GCA_016519105.1</td><td>Escherichia_coli</td><td>JI450_003022</td><td>1..3517</td><td>562</td><td>BREX-system-P-loop-protein-BrxC-[Escherichia-coli]</td></tr>
<tr><td>EGY5620863.1</td><td>Type1.BREX</td><td><-Phage_integrase<-Phage_integrase||?->?->?-><-HNH||wHTH(SBD)+HNH->||BrxA-DNA-binding->BrxB-BrxF-iSTAND->DUF6079*->N6-METHYLASE->PglZ->SIGMA+OB-Fold+AAA-ATPase+Lon-Protease-BrxL->HKD-DNase+OGG-like->||AbiEi_3_N+AbiEi_3->AbiEii-><-DUF6216<-Arm-DNA-bind_3</td><td>1213</td><td>Bacteria>Pseudomonadota>Gammaproteobacteria>Enterobacterales>Enterobacteriaceae>Escherichia</td><td>GCA_016519195.1</td><td>Escherichia_coli</td><td>JI433_002890</td><td>14696..18337</td><td>562</td><td>BREX-system-P-loop-protein-BrxC-[Escherichia-coli]</td></tr>
<tr><td>EGY7451904.1</td><td>Type1.BREX</td><td><-DUF6079-p*<-BrxB-BrxF-iSTAND<-BrxA-DNA-binding||<-wHTH(SBD)+HNH||HNH-><-?<-?<-?||Phage_integrase->Phage_integrase-></td><td>968</td><td>Bacteria>Pseudomonadota>Gammaproteobacteria>Enterobacterales>Enterobacteriaceae>Escherichia</td><td>GCA_016533805.1</td><td>Escherichia_coli</td><td>JI519_004377</td><td>1..2904</td><td>562</td><td>BREX-system-P-loop-protein-BrxC-[Escherichia-coli]</td></tr>
<tr><td>EGY8947056.1</td><td>Type1.BREX</td><td>DUF6079-p*->N6-METHYLASE->PglZ->SIGMA+OB-Fold+AAA-ATPase+Lon-Protease-BrxL->HKD-DNase+OGG-like->||AbiEi_3_N+AbiEi_3->AbiEii-><-DUF6216<-Arm-DNA-bind_3</td><td>763</td><td>Bacteria>Pseudomonadota>Gammaproteobacteria>Enterobacterales>Enterobacteriaceae>Escherichia</td><td>GCA_016541865.1</td><td>Escherichia_coli</td><td>JI613_002965</td><td>1..2292</td><td>562</td><td>BREX-system-P-loop-protein-BrxC-[Escherichia-coli]</td></tr>
<tr><td>EGY8948619.1</td><td>Type1.BREX</td><td><-DUF6079-p*<-BrxB-BrxF-iSTAND<-BrxA-DNA-binding||<-wHTH(SBD)+HNH||HNH-><-?<-?<-?||Phage_integrase->Phage_integrase-></td><td>450</td><td>Bacteria>Pseudomonadota>Gammaproteobacteria>Enterobacterales>Enterobacteriaceae>Escherichia</td><td>GCA_016541865.1</td><td>Escherichia_coli</td><td>JI613_004635</td><td>1..1349</td><td>562</td><td>BREX-system-P-loop-protein-BrxC-[Escherichia-coli]</td></tr>
<tr><td>EGY9082468.1</td><td>Type1.BREX</td><td>DUF6079*->N6-METHYLASE->PglZ->SIGMA+OB-Fold+AAA-ATPase+Lon-Protease-BrxL->HKD-DNase+OGG-like->||AbiEi_3_N+AbiEi_3->AbiEii-><-DUF6216<-Arm-DNA-bind_3</td><td>1175</td><td>Bacteria>Pseudomonadota>Gammaproteobacteria>Enterobacterales>Enterobacteriaceae>Escherichia</td><td>GCA_016542525.1</td><td>Escherichia_coli</td><td>JI646_002303</td><td>1..3528</td><td>562</td><td>BREX-system-P-loop-protein-BrxC-[Escherichia-coli]</td></tr>
<tr><td>EGY9306082.1</td><td>Type1.BREX</td><td><-HNH||wHTH(SBD)+HNH->||BrxA-DNA-binding->BrxB-BrxF-iSTAND->DUF6079*->N6-METHYLASE->PglZ-></td><td>1213</td><td>Bacteria>Pseudomonadota>Gammaproteobacteria>Enterobacterales>Enterobacteriaceae>Escherichia</td><td>GCA_016543505.1</td><td>Escherichia_coli</td><td>HZ096_004640</td><td>4089..7730</td><td>562</td><td>BREX-system-P-loop-protein-BrxC-[Escherichia-coli]</td></tr>
<tr><td>EGY9336114.1</td><td>Type1.BREX</td><td>DUF6079*->N6-METHYLASE->PglZ->SIGMA+OB-Fold+AAA-ATPase+Lon-Protease-BrxL->HKD-DNase+OGG-like->||AbiEi_3_N+AbiEi_3->AbiEii-><-DUF6216<-Arm-DNA-bind_3</td><td>1177</td><td>Bacteria>Pseudomonadota>Gammaproteobacteria>Enterobacterales>Enterobacteriaceae>Escherichia</td><td>GCA_016543525.1</td><td>Escherichia_coli</td><td>HZ078_003642</td><td>1..3535</td><td>562</td><td>BREX-system-P-loop-protein-BrxC-[Escherichia-coli]</td></tr>
<tr><td>EGY9928133.1</td><td>Type1.BREX</td><td>DUF6079-p*->N6-METHYLASE->PglZ->SIGMA+OB-Fold+AAA-ATPase+Lon-Protease-BrxL-><-ZnR<-ZnR<-?||HIN-HTH+HIN-HTH-></td><td>865</td><td>Bacteria>Pseudomonadota>Gammaproteobacteria>Enterobacterales>Enterobacteriaceae>Escherichia</td><td>GCA_016546565.1</td><td>Escherichia_coli</td><td>HVV03_004451</td><td>1..2599</td><td>562</td><td>BREX-system-P-loop-protein-BrxC-[Escherichia-coli]</td></tr>
<tr><td>EGZ0051356.1</td><td>Type1.BREX</td><td>DUF6079-p*->N6-METHYLASE->PglZ->SIGMA+OB-Fold+AAA-ATPase+Lon-Protease-BrxL->HKD-DNase+OGG-like->||AbiEi_3_N+AbiEi_3->AbiEii-><-DUF6216<-Arm-DNA-bind_3</td><td>834</td><td>Bacteria>Pseudomonadota>Gammaproteobacteria>Enterobacterales>Enterobacteriaceae>Escherichia</td><td>GCA_016546855.1</td><td>Escherichia_coli</td><td>JJA24_002218</td><td>1..2505</td><td>562</td><td>BREX-system-P-loop-protein-BrxC-[Escherichia-coli]</td></tr>
<tr><td>EGZ1085523.1</td><td>Type1.BREX</td><td>DUF6079-p*->N6-METHYLASE->PglZ->SIGMA+OB-Fold+AAA-ATPase+Lon-Protease-BrxL-><-ZnR<-ZnR</td><td>964</td><td>Bacteria>Pseudomonadota>Gammaproteobacteria>Enterobacterales>Enterobacteriaceae>Escherichia</td><td>GCA_016551725.1</td><td>Escherichia_coli</td><td>FO146_004816</td><td>1..2896</td><td>562</td><td>BREX-system-P-loop-protein-BrxC-[Escherichia-coli]</td></tr>
<tr><td>EGZ1391640.1</td><td>Type1.BREX</td><td>DUF6079-p*->N6-METHYLASE->PglZ->SIGMA+OB-Fold+AAA-ATPase+Lon-Protease-BrxL->HKD-DNase+OGG-like->||AbiEi_3_N+AbiEi_3->AbiEii-><-DUF6216<-Arm-DNA-bind_3</td><td>789</td><td>Bacteria>Pseudomonadota>Gammaproteobacteria>Enterobacterales>Enterobacteriaceae>Escherichia</td><td>GCA_016552825.1</td><td>Escherichia_coli</td><td>FG160_002854</td><td>1..2370</td><td>562</td><td>BREX-system-P-loop-protein-BrxC-[Escherichia-coli]</td></tr>
<tr><td>EGZ1415768.1</td><td>Type1.BREX</td><td>Arm-DNA-bind_3->DUF6216-><-AbiEii<-AbiEi_3_N+AbiEi_3||<-HKD-DNase+OGG-like<-SIGMA+OB-Fold+AAA-ATPase+Lon-Protease-BrxL<-PglZ<-N6-METHYLASE<-DUF6079*<-BrxB-BrxF-iSTAND<-BrxA-DNA-binding||<-wHTH+SAD+HNH||HNH-></td><td>1213</td><td>Bacteria>Pseudomonadota>Gammaproteobacteria>Enterobacterales>Enterobacteriaceae>Escherichia</td><td>GCA_016552885.1</td><td>Escherichia_coli</td><td>FFU33_001578</td><td>154175..157816</td><td>562</td><td>BREX-system-P-loop-protein-BrxC-[Escherichia-coli]</td></tr>
<tr><td>EGZ1559467.1</td><td>Type1.BREX</td><td>DUF6079-p*->N6-METHYLASE->PglZ->SIGMA+OB-Fold+AAA-ATPase+Lon-Protease-BrxL->HKD-DNase+OGG-like->||AbiEi_3_N+AbiEi_3->AbiEii-><-DUF6216<-Arm-DNA-bind_3</td><td>768</td><td>Bacteria>Pseudomonadota>Gammaproteobacteria>Enterobacterales>Enterobacteriaceae>Escherichia</td><td>GCA_016553465.1</td><td>Escherichia_coli</td><td>FAV16_003764</td><td>1..2307</td><td>562</td><td>BREX-system-P-loop-protein-BrxC-[Escherichia-coli]</td></tr>
<tr><td>EGZ1560300.1</td><td>Type1.BREX</td><td><-DUF6079-p*<-BrxB-BrxF-iSTAND<-BrxA-DNA-binding||<-wHTH(SBD)+HNH||HNH-><-?<-?<-?||Phage_integrase->Phage_integrase-></td><td>444</td><td>Bacteria>Pseudomonadota>Gammaproteobacteria>Enterobacterales>Enterobacteriaceae>Escherichia</td><td>GCA_016553465.1</td><td>Escherichia_coli</td><td>FAV16_004661</td><td>1..1332</td><td>562</td><td>BREX-system-P-loop-protein-BrxC-[Escherichia-coli]</td></tr>
<tr><td>EGZ1869331.1</td><td>Type1.BREX</td><td>DUF6079-p*->N6-METHYLASE->PglZ->SIGMA+OB-Fold+AAA-ATPase+Lon-Protease-BrxL->HKD-DNase+OGG-like->||AbiEi_3_N+AbiEi_3->AbiEii-><-DUF6216<-Arm-DNA-bind_3</td><td>796</td><td>Bacteria>Pseudomonadota>Gammaproteobacteria>Enterobacterales>Enterobacteriaceae>Escherichia</td><td>GCA_016554585.1</td><td>Escherichia_coli</td><td>EYE54_001992</td><td>1..2391</td><td>562</td><td>BREX-system-P-loop-protein-BrxC-[Escherichia-coli]</td></tr>
<tr><td>EGZ1871551.1</td><td>Type1.BREX</td><td><-DUF6079-p*<-BrxB-BrxF-iSTAND<-BrxA-DNA-binding||<-wHTH(SBD)+HNH||HNH-><-?<-?<-?||Phage_integrase->Phage_integrase-></td><td>416</td><td>Bacteria>Pseudomonadota>Gammaproteobacteria>Enterobacterales>Enterobacteriaceae>Escherichia</td><td>GCA_016554585.1</td><td>Escherichia_coli</td><td>EYE54_004341</td><td>1..1248</td><td>562</td><td>BREX-system-P-loop-protein-BrxC-[Escherichia-coli]</td></tr>
<tr><td>EGZ2189929.1</td><td>Type1.BREX</td><td><-DUF6079-p*<-BrxB-BrxF-iSTAND<-BrxA-DNA-binding||<-wHTH(SBD)+HNH||HNH-><-?<-?<-?||Phage_integrase-></td><td>628</td><td>Bacteria>Pseudomonadota>Gammaproteobacteria>Enterobacterales>Enterobacteriaceae>Escherichia</td><td>GCA_016555885.1</td><td>Escherichia_coli</td><td>ED609_004465</td><td>1..1884</td><td>562</td><td>BREX-system-P-loop-protein-BrxC-[Escherichia-coli]</td></tr>
<tr><td>EGZ2354229.1</td><td>Type1.BREX</td><td><-Phage_integrase<-Phage_integrase||?->?->?-><-HNH||wHTH(SBD)+HNH->||BrxA-DNA-binding->BrxB-BrxF-iSTAND->DUF6079*->N6-METHYLASE->PglZ->SIGMA+OB-Fold+AAA-ATPase+Lon-Protease-BrxL->HKD-DNase+OGG-like->||AbiEi_3_N+AbiEi_3->AbiEii-><-DUF6216<-Arm-DNA-bind_3</td><td>1213</td><td>Bacteria>Pseudomonadota>Gammaproteobacteria>Enterobacterales>Enterobacteriaceae>Escherichia</td><td>GCA_016556665.1</td><td>Escherichia_coli</td><td>D7J47_002044</td><td>14697..18338</td><td>562</td><td>BREX-system-P-loop-protein-BrxC-[Escherichia-coli]</td></tr>
<tr><td>EGZ2497683.1</td><td>Type1.BREX</td><td>DUF6079-p*->N6-METHYLASE->PglZ->SIGMA+OB-Fold+AAA-ATPase+Lon-Protease-BrxL->HKD-DNase+OGG-like->||AbiEi_3_N+AbiEi_3->AbiEii-><-DUF6216<-Arm-DNA-bind_3</td><td>969</td><td>Bacteria>Pseudomonadota>Gammaproteobacteria>Enterobacterales>Enterobacteriaceae>Escherichia</td><td>GCA_016557105.1</td><td>Escherichia_coli</td><td>D6W78_003631</td><td>1..2912</td><td>562</td><td>BREX-system-P-loop-protein-BrxC-[Escherichia-coli]</td></tr>
<tr><td>EGZ2497894.1</td><td>Type1.BREX</td><td>DUF6079*->N6-METHYLASE->PglZ->SIGMA+OB-Fold+AAA-ATPase+Lon-Protease-BrxL-><-ZnR<-ZnR<-?||HIN-HTH+PAIREDC-HTH->||RHH_antitoxin->XIS-HTH-></td><td>1012</td><td>Bacteria>Pseudomonadota>Gammaproteobacteria>Enterobacterales>Enterobacteriaceae>Escherichia</td><td>GCA_016557105.1</td><td>Escherichia_coli</td><td>D6W78_003856</td><td>1..3041</td><td>562</td><td>BREX-system-P-loop-protein-BrxC-[Escherichia-coli]</td></tr>
<tr><td>EGZ2674120.1</td><td>Type1.BREX</td><td>DUF6079*->N6-METHYLASE->PglZ->SIGMA+OB-Fold+AAA-ATPase+Lon-Protease-BrxL->HKD-DNase+OGG-like->||AbiEi_3_N+AbiEi_3->AbiEii-><-DUF6216<-Arm-DNA-bind_3</td><td>1096</td><td>Bacteria>Pseudomonadota>Gammaproteobacteria>Enterobacterales>Enterobacteriaceae>Escherichia</td><td>GCA_016557745.1</td><td>Escherichia_coli</td><td>D1H08_002340</td><td>1..3291</td><td>562</td><td>BREX-system-P-loop-protein-BrxC-[Escherichia-coli]</td></tr>
<tr><td>EGZ2777080.1</td><td>Type1.BREX</td><td><-Phage_integrase<-Phage_integrase||?->?->?-><-HNH||wHTH(SBD)+HNH->||BrxA-DNA-binding->BrxB-BrxF-iSTAND->DUF6079*->N6-METHYLASE->PglZ->SIGMA+OB-Fold+AAA-ATPase+Lon-Protease-BrxL->HKD-DNase+OGG-like->||AbiEi_3_N+AbiEi_3->AbiEii-><-DUF6216<-Arm-DNA-bind_3</td><td>1213</td><td>Bacteria>Pseudomonadota>Gammaproteobacteria>Enterobacterales>Enterobacteriaceae>Escherichia</td><td>GCA_016558225.1</td><td>Escherichia_coli</td><td>DVD41_002389</td><td>14697..18338</td><td>562</td><td>BREX-system-P-loop-protein-BrxC-[Escherichia-coli]</td></tr>
<tr><td>EGZ3242782.1</td><td>Type1.BREX</td><td><-Phage_integrase<-Phage_integrase||?->?->?-><-HNH||wHTH(SBD)+HNH->||BrxA-DNA-binding->BrxB-BrxF-iSTAND->DUF6079*->N6-METHYLASE->PglZ->SIGMA+OB-Fold+AAA-ATPase+Lon-Protease-BrxL->HKD-DNase+OGG-like->||AbiEi_3_N+AbiEi_3->AbiEii-><-DUF6216<-Arm-DNA-bind_3</td><td>1213</td><td>Bacteria>Pseudomonadota>Gammaproteobacteria>Enterobacterales>Enterobacteriaceae>Escherichia</td><td>GCA_016560025.1</td><td>Escherichia_coli</td><td>C5N70_002404</td><td>39991..43632</td><td>562</td><td>BREX-system-P-loop-protein-BrxC-[Escherichia-coli]</td></tr>
<tr><td>EGZ4145352.1</td><td>Type1.BREX</td><td>DUF6079*->N6-METHYLASE->PglZ->SIGMA+OB-Fold+AAA-ATPase+Lon-Protease-BrxL->HKD-DNase+OGG-like->||AbiEi_3_N+AbiEi_3->AbiEii-><-DUF6216<-Arm-DNA-bind_3</td><td>1095</td><td>Bacteria>Pseudomonadota>Gammaproteobacteria>Enterobacterales>Enterobacteriaceae>Escherichia</td><td>GCA_016563925.1</td><td>Escherichia_coli</td><td>CKY95_002785</td><td>1..3290</td><td>562</td><td>BREX-system-P-loop-protein-BrxC-[Escherichia-coli]</td></tr>
<tr><td>EGZ5166758.1</td><td>Type1.BREX</td><td><-DUF6079-p*<-BrxB-BrxF-iSTAND<-BrxA-DNA-binding||<-wHTH(SBD)+HNH||HNH-><-?<-?<-?||Phage_integrase->Phage_integrase-></td><td>624</td><td>Bacteria>Pseudomonadota>Gammaproteobacteria>Enterobacterales>Enterobacteriaceae>Escherichia</td><td>GCA_016568175.1</td><td>Escherichia_coli</td><td>CDS35_004366</td><td>1..1874</td><td>562</td><td>BREX-system-P-loop-protein-BrxC-[Escherichia-coli]</td></tr>
<tr><td>EGZ5520004.1</td><td>Type1.BREX</td><td>DUF6079-p*->N6-METHYLASE->PglZ->SIGMA+OB-Fold+AAA-ATPase+Lon-Protease-BrxL->HKD-DNase+OGG-like->||AbiEi_3_N+AbiEi_3->AbiEii->?-><-DUF6216<-Arm-DNA-bind_3</td><td>815</td><td>Bacteria>Pseudomonadota>Gammaproteobacteria>Enterobacterales>Enterobacteriaceae>Escherichia</td><td>GCA_016569715.1</td><td>Escherichia_coli</td><td>LF100_002181</td><td>1..2450</td><td>562</td><td>BREX-system-P-loop-protein-BrxC-[Escherichia-coli]</td></tr>
<tr><td>EGZ5591347.1</td><td>Type1.BREX</td><td><-DUF6079-p*<-BrxB-BrxF-iSTAND<-BrxA-DNA-binding||<-wHTH(SBD)+HNH||HNH-><-?<-?<-?||Phage_integrase->Phage_integrase-></td><td>707</td><td>Bacteria>Pseudomonadota>Gammaproteobacteria>Enterobacterales>Enterobacteriaceae>Escherichia</td><td>GCA_016569835.1</td><td>Escherichia_coli</td><td>BZZ56_004473</td><td>1..2121</td><td>562</td><td>BREX-system-P-loop-protein-BrxC-[Escherichia-coli]</td></tr>
<tr><td>EGZ5731917.1</td><td>Type1.BREX</td><td><-DUF6079-p*<-BrxB-BrxF-iSTAND<-BrxA-DNA-binding||<-wHTH(SBD)+HNH||HNH-><-?<-?<-?||Phage_integrase->Phage_integrase-></td><td>885</td><td>Bacteria>Pseudomonadota>Gammaproteobacteria>Enterobacterales>Enterobacteriaceae>Escherichia</td><td>GCA_016570515.1</td><td>Escherichia_coli</td><td>BL334_004670</td><td>1..2655</td><td>562</td><td>BREX-system-P-loop-protein-BrxC-[Escherichia-coli]</td></tr>
<tr><td>EGZ5978150.1</td><td>Type1.BREX</td><td><-Phage_integrase<-Phage_integrase||?->?->?-><-HNH||wHTH(SBD)+HNH->||BrxA-DNA-binding->BrxB-BrxF-iSTAND->DUF6079-p*-></td><td>421</td><td>Bacteria>Pseudomonadota>Gammaproteobacteria>Enterobacterales>Enterobacteriaceae>Escherichia</td><td>GCA_016571515.1</td><td>Escherichia_coli</td><td>BIB10_004104</td><td>39992..41255</td><td>562</td><td>BREX-system-P-loop-protein-BrxC-[Escherichia-coli]</td></tr>
<tr><td>EGZ7391807.1</td><td>Type1.BREX</td><td>DUF6079-p*->N6-METHYLASE->PglZ->SIGMA+OB-Fold+AAA-ATPase+Lon-Protease-BrxL->HKD-DNase+OGG-like->||AbiEi_3_N+AbiEi_3->AbiEii-><-DUF6216<-Arm-DNA-bind_3</td><td>970</td><td>Bacteria>Pseudomonadota>Gammaproteobacteria>Enterobacterales>Enterobacteriaceae>Escherichia</td><td>GCA_016578925.1</td><td>Escherichia_coli</td><td>BAU88_004031</td><td>1..2913</td><td>562</td><td>BREX-system-P-loop-protein-BrxC-[Escherichia-coli]</td></tr>
<tr><td>EGZ7392043.1</td><td>Type1.BREX</td><td>DUF6079-p*->N6-METHYLASE->PglZ->SIGMA+OB-Fold+AAA-ATPase+Lon-Protease-BrxL-><-ZnR<-ZnR<-?||HIN-HTH+PAIREDC-HTH->||RHH_antitoxin->XIS-HTH-></td><td>928</td><td>Bacteria>Pseudomonadota>Gammaproteobacteria>Enterobacterales>Enterobacteriaceae>Escherichia</td><td>GCA_016578925.1</td><td>Escherichia_coli</td><td>BAU88_004280</td><td>1..2788</td><td>562</td><td>BREX-system-P-loop-protein-BrxC-[Escherichia-coli]</td></tr>
<tr><td>EGZ7370259.1</td><td>Type1.BREX</td><td>DUF6079-p*->N6-METHYLASE->PglZ->SIGMA+OB-Fold+AAA-ATPase+Lon-Protease-BrxL->HKD-DNase+OGG-like->||AbiEi_3_N+AbiEi_3->AbiEii-><-DUF6216<-Arm-DNA-bind_3</td><td>880</td><td>Bacteria>Pseudomonadota>Gammaproteobacteria>Enterobacterales>Enterobacteriaceae>Escherichia</td><td>GCA_016578935.1</td><td>Escherichia_coli</td><td>BAU61_002957</td><td>1..2643</td><td>562</td><td>BREX-system-P-loop-protein-BrxC-[Escherichia-coli]</td></tr>
<tr><td>EGZ7468561.1</td><td>Type1.BREX</td><td>DUF6079-p*->N6-METHYLASE->PglZ->SIGMA+OB-Fold+AAA-ATPase+Lon-Protease-BrxL->HKD-DNase+OGG-like->||AbiEi_3_N+AbiEi_3->AbiEii-><-DUF6216<-Arm-DNA-bind_3</td><td>931</td><td>Bacteria>Pseudomonadota>Gammaproteobacteria>Enterobacterales>Enterobacteriaceae>Escherichia</td><td>GCA_016579325.1</td><td>Escherichia_coli</td><td>A6F02_004042</td><td>1..2798</td><td>562</td><td>BREX-system-P-loop-protein-BrxC-[Escherichia-coli]</td></tr>
<tr><td>EGZ7468749.1</td><td>Type1.BREX</td><td>DUF6079-p*->N6-METHYLASE->PglZ->SIGMA+OB-Fold+AAA-ATPase+Lon-Protease-BrxL-><-ZnR<-ZnR<-?||HIN-HTH+PAIREDC-HTH->||RHH_antitoxin->XIS-HTH-></td><td>931</td><td>Bacteria>Pseudomonadota>Gammaproteobacteria>Enterobacterales>Enterobacteriaceae>Escherichia</td><td>GCA_016579325.1</td><td>Escherichia_coli</td><td>A6F02_004243</td><td>1..2798</td><td>562</td><td>BREX-system-P-loop-protein-BrxC-[Escherichia-coli]</td></tr>
<tr><td>EGZ7821797.1</td><td>Type1.BREX</td><td>DUF6079-p*->N6-METHYLASE->PglZ->SIGMA+OB-Fold+AAA-ATPase+Lon-Protease-BrxL->HKD-DNase+OGG-like->||AbiEi_3_N+AbiEi_3->AbiEii-><-DUF6216<-Arm-DNA-bind_3</td><td>729</td><td>Bacteria>Pseudomonadota>Gammaproteobacteria>Enterobacterales>Enterobacteriaceae>Escherichia</td><td>GCA_016581105.1</td><td>Escherichia_coli</td><td>AWC87_001490</td><td>1..2190</td><td>562</td><td>BREX-system-P-loop-protein-BrxC-[Escherichia-coli]</td></tr>
<tr><td>EGZ8838308.1</td><td>Type1.BREX</td><td><-Phage_integrase||?->?->?-><-HNH||wHTH(SBD)+HNH->||BrxA-DNA-binding->BrxB-BrxF-iSTAND->DUF6079*->N6-METHYLASE->PglZ->SIGMA+OB-Fold+AAA-ATPase+Lon-Protease-BrxL->HKD-DNase+OGG-like->||AbiEi_3_N+AbiEi_3->AbiEii-><-DUF6216<-Arm-DNA-bind_3</td><td>1213</td><td>Bacteria>Pseudomonadota>Gammaproteobacteria>Enterobacterales>Enterobacteriaceae>Escherichia</td><td>GCA_016590495.1</td><td>Escherichia_coli</td><td>AT823_000493</td><td>8719..12360</td><td>562</td><td>BREX-system-P-loop-protein-BrxC-[Escherichia-coli]</td></tr>
<tr><td>EGZ9024440.1</td><td>Type1.BREX</td><td>DUF6079-p*->N6-METHYLASE->PglZ->SIGMA+OB-Fold+AAA-ATPase+Lon-Protease-BrxL->HKD-DNase+OGG-like->||AbiEi_3_N+AbiEi_3->AbiEii-><-DUF6216<-Arm-DNA-bind_3</td><td>963</td><td>Bacteria>Pseudomonadota>Gammaproteobacteria>Enterobacterales>Enterobacteriaceae>Escherichia</td><td>GCA_016591155.1</td><td>Escherichia_coli</td><td>AFE50_004176</td><td>1..2892</td><td>562</td><td>BREX-system-P-loop-protein-BrxC-[Escherichia-coli]</td></tr>
<tr><td>EGZ9024631.1</td><td>Type1.BREX</td><td>DUF6079-p*->N6-METHYLASE->PglZ->SIGMA+OB-Fold+AAA-ATPase+Lon-Protease-BrxL-><-ZnR<-ZnR<-?||HIN-HTH+PAIREDC-HTH->||RHH_antitoxin->XIS-HTH-></td><td>980</td><td>Bacteria>Pseudomonadota>Gammaproteobacteria>Enterobacterales>Enterobacteriaceae>Escherichia</td><td>GCA_016591155.1</td><td>Escherichia_coli</td><td>AFE50_004379</td><td>1..2945</td><td>562</td><td>BREX-system-P-loop-protein-BrxC-[Escherichia-coli]</td></tr>
<tr><td>EHA0949388.1</td><td>Type1.BREX</td><td>DUF6079-p*->N6-METHYLASE->PglZ->SIGMA+OB-Fold+AAA-ATPase+Lon-Protease-BrxL->HKD-DNase+OGG-like->||AbiEi_3_N+AbiEi_3->AbiEii-><-DUF6216<-Arm-DNA-bind_3</td><td>762</td><td>Bacteria>Pseudomonadota>Gammaproteobacteria>Enterobacterales>Enterobacteriaceae>Escherichia</td><td>GCA_016610545.1</td><td>Escherichia_coli</td><td>JI650_002930</td><td>1..2291</td><td>562</td><td>BREX-system-P-loop-protein-BrxC-[Escherichia-coli]</td></tr>
<tr><td>HAW1448903.1</td><td>Type1.BREX</td><td><-Phage_integrase||?->?->?-><-HNH||wHTH+SAD+HNH-><-REase+iREase||BrxA-DNA-binding->BrxB-BrxF-iSTAND->DUF6079*->N6-METHYLASE->PglZ->SIGMA+OB-Fold+AAA-ATPase+Lon-Protease-BrxL->HKD-DNase+OGG-like->||AbiEi_3_N+AbiEi_3->AbiEii-><-DUF6216<-Arm-DNA-bind_3</td><td>1213</td><td>Bacteria>Pseudomonadota>Gammaproteobacteria>Enterobacterales>Enterobacteriaceae>Escherichia</td><td>GCA_016674355.1</td><td>Escherichia_coli</td><td>JLM49_000126</td><td>135704..139345</td><td>562</td><td>BREX-system-P-loop-protein-BrxC-[Escherichia-coli]</td></tr>
<tr><td>HAW2754029.1</td><td>Type1.BREX</td><td><-HerA/FtsK+A32<-?||tRNA->Arm-DNA-bind_3+Phage_integrase->EVE+HNH-><-REase+iREase||BrxA-DNA-binding->BrxB-BrxF-iSTAND->DUF6079*->N6-METHYLASE->PglZ->SIGMA+OB-Fold+AAA-ATPase+Lon-Protease-BrxL-><-ZnR<-ZnR</td><td>1213</td><td>Bacteria>Pseudomonadota>Gammaproteobacteria>Enterobacterales>Enterobacteriaceae>Escherichia</td><td>GCA_016680635.1</td><td>Escherichia_coli</td><td>JLQ48_003685</td><td>19788..23429</td><td>562</td><td>BREX-system-P-loop-protein-BrxC-[Escherichia-coli]</td></tr>
<tr><td>HAW6060333.1</td><td>Type1.BREX</td><td><-DUF6079-p*<-BrxB-BrxF-iSTAND<-BrxA-DNA-binding||REase+iREase-><-wHTH+SAD+HNH</td><td>613</td><td>Bacteria>Pseudomonadota>Gammaproteobacteria>Enterobacterales>Enterobacteriaceae>Escherichia</td><td>GCA_016697785.1</td><td>Escherichia_coli</td><td>JL413_004248</td><td>1..1839</td><td>562</td><td>BREX-system-P-loop-protein-BrxC-[Escherichia-coli]</td></tr>
<tr><td>HAW6308706.1</td><td>Type1.BREX</td><td>ZnR->ZnR-><-SIGMA+OB-Fold+AAA-ATPase+Lon-Protease-BrxL<-PglZ<-N6-METHYLASE<-DUF6079*<-BrxB-BrxF-iSTAND<-BrxA-DNA-binding||REase+iREase->?->?-><-DDE_Tnp_IS1</td><td>1213</td><td>Bacteria>Pseudomonadota>Gammaproteobacteria>Enterobacterales>Enterobacteriaceae>Escherichia</td><td>GCA_016734185.1</td><td>Escherichia_coli</td><td>JLP42_004227</td><td>9916..13557</td><td>562</td><td>BREX-system-P-loop-protein-BrxC-[Escherichia-coli]</td></tr>
<tr><td>EHA9087556.1</td><td>Type1.BREX</td><td><-Phage_integrase<-Phage_integrase||?->?->?-><-HNH||wHTH(SBD)+HNH->||BrxA-DNA-binding->BrxB-BrxF-iSTAND->DUF6079*->N6-METHYLASE->PglZ->SIGMA+OB-Fold+AAA-ATPase+Lon-Protease-BrxL->HKD-DNase+OGG-like->||AbiEi_3_N+AbiEi_3->AbiEii-><-DUF6216<-Arm-DNA-bind_3</td><td>1213</td><td>Bacteria>Pseudomonadota>Gammaproteobacteria>Enterobacterales>Enterobacteriaceae>Escherichia</td><td>GCA_016846225.1</td><td>Escherichia_coli</td><td>D4094_002230</td><td>14699..18340</td><td>562</td><td>BREX-system-P-loop-protein-BrxC-[Escherichia-coli]</td></tr>
<tr><td>HAW9526082.1</td><td>Type1.BREX</td><td><-N6-METHYLASE<-DUF6079*<-BrxB-BrxF-iSTAND<-BrxA-DNA-binding||REase+iREase-><-wHTH+SAD+HNH||HNH-><-?<-?<-?||Phage_integrase-></td><td>1213</td><td>Bacteria>Pseudomonadota>Gammaproteobacteria>Enterobacterales>Enterobacteriaceae>Escherichia</td><td>GCA_016946495.1</td><td>Escherichia_coli</td><td>JY346_004480</td><td>2578..6219</td><td>562</td><td>BREX-system-P-loop-protein-BrxC-[Escherichia-coli]</td></tr>
<tr><td>HAX1741345.1</td><td>Type1.BREX</td><td>MobI->?-><-?<-LexA-protease||35exo->?->wHTH+WYL+WCX->BrxA-DNA-binding->BrxB-BrxF-iSTAND->DUF6079-p*-></td><td>414</td><td>Bacteria>Pseudomonadota>Gammaproteobacteria>Enterobacterales>Enterobacteriaceae>Escherichia</td><td>GCA_016984655.1</td><td>Escherichia_coli</td><td>JXS76_004269</td><td>14153..15393</td><td>562</td><td>BREX-system-P-loop-protein-BrxC-[Escherichia-coli]</td></tr>
<tr><td>HAX1741551.1</td><td>Type1.BREX</td><td>DUF6079*->ParB+HNH->N6-METHYLASE->PglZ->SIGMA+OB-Fold+AAA-ATPase+Lon-Protease-BrxL->SbcC-></td><td>793</td><td>Bacteria>Pseudomonadota>Gammaproteobacteria>Enterobacterales>Enterobacteriaceae>Escherichia</td><td>GCA_016984655.1</td><td>Escherichia_coli</td><td>JXS76_004485</td><td>1..2383</td><td>562</td><td>BREX-system-P-loop-protein-BrxC-[Escherichia-coli]</td></tr>
<tr><td>HAX2048855.1</td><td>Type1.BREX</td><td>ZnR->ZnR-><-SIGMA+OB-Fold+AAA-ATPase+Lon-Protease-BrxL<-PglZ<-N6-METHYLASE<-DUF6079*<-BrxB-BrxF-iSTAND<-BrxA-DNA-binding||REase+iREase-></td><td>1213</td><td>Bacteria>Pseudomonadota>Gammaproteobacteria>Enterobacterales>Enterobacteriaceae>Escherichia</td><td>GCA_016988055.1</td><td>Escherichia_coli</td><td>JW528_004279</td><td>9399..13040</td><td>562</td><td>BREX-system-P-loop-protein-BrxC-[Escherichia-coli]</td></tr>
<tr><td>HAX3725917.1</td><td>Type1.BREX</td><td>Glyoxalase_8-><-?<-?<-VirD4-TraG+VirB4-FtsK<-HDRnase<-SbcC<-SIGMA+OB-Fold+AAA-ATPase+Lon-Protease-BrxL<-PglZ<-N6-METHYLASE<-DUF6079-p*</td><td>928</td><td>Bacteria>Pseudomonadota>Gammaproteobacteria>Enterobacterales>Enterobacteriaceae>Escherichia</td><td>GCA_017002415.1</td><td>Escherichia_coli</td><td>JV380_003192</td><td>66035..68823</td><td>562</td><td>BREX-system-P-loop-protein-BrxC-[Escherichia-coli]</td></tr>
<tr><td>HAX3800328.1</td><td>Type1.BREX</td><td><-REase<-REase||BrxA-DNA-binding->BrxB-BrxF-iSTAND->DUF6079*->N6-METHYLASE->PglZ->SIGMA+OB-Fold+AAA-ATPase+Lon-Protease-BrxL-><-ZnR<-ZnR<-?||RHH_antitoxin->XIS-HTH-></td><td>1213</td><td>Bacteria>Pseudomonadota>Gammaproteobacteria>Enterobacterales>Enterobacteriaceae>Escherichia</td><td>GCA_017002675.1</td><td>Escherichia_coli</td><td>JV366_004401</td><td>2784..6425</td><td>562</td><td>BREX-system-P-loop-protein-BrxC-[Escherichia-coli]</td></tr>
<tr><td>HAX4031531.1</td><td>Type1.BREX</td><td><-Phage_integrase||?->?->?-><-HNH||wHTH+SAD+HNH-><-REase+iREase||BrxA-DNA-binding->BrxB-BrxF-iSTAND->DUF6079*->N6-METHYLASE->PglZ->SIGMA+OB-Fold+AAA-ATPase+Lon-Protease-BrxL->HKD-DNase+OGG-like->||AbiEi_3_N+AbiEi_3->AbiEii-><-DUF6216<-Arm-DNA-bind_3</td><td>1213</td><td>Bacteria>Pseudomonadota>Gammaproteobacteria>Enterobacterales>Enterobacteriaceae>Escherichia</td><td>GCA_017003755.1</td><td>Escherichia_coli</td><td>JV397_001890</td><td>160694..164335</td><td>562</td><td>BREX-system-P-loop-protein-BrxC-[Escherichia-coli]</td></tr>
<tr><td>HAX4880267.1</td><td>Type1.BREX</td><td><-wHTH+WYL||wHTH->ZnR->ZnR-><-SIGMA+OB-Fold+AAA-ATPase+Lon-Protease-BrxL<-PglZ<-TOPRIM<-OLD-ABC<-N6-METHYLASE<-DUF6079*<-BrxB-BrxF-iSTAND<-BrxA-DNA-binding<-tRNA||?-><-SKI||METHYLASE->?->?->LACI-HTH+PBPI->HerA/FtsK+A32-></td><td>1204</td><td>Bacteria>Pseudomonadota>Gammaproteobacteria>Enterobacterales>Enterobacteriaceae>Escherichia</td><td>GCA_017016975.1</td><td>Escherichia_coli</td><td>JZF83_001472</td><td>109249..112863</td><td>562</td><td>BREX-system-P-loop-protein-BrxC-[Escherichia-coli]</td></tr>
<tr><td>EHC1662562.1</td><td>Type1.BREX</td><td><-N6-METHYLASE<-DUF6079*<-BrxB-BrxF-iSTAND<-BrxA-DNA-binding||REase+iREase-><-wHTH+SAD+HNH||HNH-></td><td>1213</td><td>Bacteria>Pseudomonadota>Gammaproteobacteria>Enterobacterales>Enterobacteriaceae>Escherichia</td><td>GCA_017030065.1</td><td>Escherichia_coli</td><td>JTZ37_004160</td><td>2115..5756</td><td>562</td><td>BREX-system-P-loop-protein-BrxC-[Escherichia-coli]</td></tr>
<tr><td>EHC2076257.1</td><td>Type1.BREX</td><td><-Phage_integrase||?->?->?-><-HNH||wHTH+SAD+HNH-><-REase+iREase||BrxA-DNA-binding->BrxB-BrxF-iSTAND->DUF6079*->N6-METHYLASE->PglZ->SIGMA+OB-Fold+AAA-ATPase+Lon-Protease-BrxL->HKD-DNase+OGG-like->||AbiEi_3_N+AbiEi_3->AbiEii-><-DUF6216<-Arm-DNA-bind_3</td><td>1213</td><td>Bacteria>Pseudomonadota>Gammaproteobacteria>Enterobacterales>Enterobacteriaceae>Escherichia</td><td>GCA_017031715.1</td><td>Escherichia_coli</td><td>JT651_000259</td><td>8773..12414</td><td>562</td><td>BREX-system-P-loop-protein-BrxC-[Escherichia-coli]</td></tr>
<tr><td>EHC2846675.1</td><td>Type1.BREX</td><td>Arm-DNA-bind_3+Phage_integrase-><-Sulfatase<-?||DDE_Tnp_1_5+DDE_Tnp_1-><-REase+iREase||BrxA-DNA-binding->BrxB-BrxF-iSTAND->DUF6079*->N6-METHYLASE->PglZ->SIGMA+OB-Fold+AAA-ATPase+Lon-Protease-BrxL-><-ZnR<-ZnR<-?<-?||RHH_antitoxin->XIS-HTH-></td><td>1213</td><td>Bacteria>Pseudomonadota>Gammaproteobacteria>Enterobacterales>Enterobacteriaceae>Escherichia</td><td>GCA_017035095.1</td><td>Escherichia_coli</td><td>JTY00_000601</td><td>175379..179020</td><td>562</td><td>BREX-system-P-loop-protein-BrxC-[Escherichia-coli]</td></tr>
<tr><td>HAX6962711.1</td><td>Type1.BREX</td><td><-XIS-HTH<-RHH_antitoxin||?->ZnR->ZnR-><-SIGMA+OB-Fold+AAA-ATPase+Lon-Protease-BrxL<-PglZ<-N6-METHYLASE<-DUF6079*<-BrxB-BrxF-iSTAND<-BrxA-DNA-binding||REase+iREase-><-EVE+HNH<-Arm-DNA-bind_3+Phage_integrase<-tRNA||?->HerA/FtsK+A32-></td><td>1213</td><td>Bacteria>Pseudomonadota>Gammaproteobacteria>Enterobacterales>Enterobacteriaceae>Escherichia</td><td>GCA_017045635.1</td><td>Escherichia_coli</td><td>JTN21_000929</td><td>60484..64125</td><td>562</td><td>BREX-system-P-loop-protein-BrxC-[Escherichia-coli]</td></tr>
<tr><td>HAX7126694.1</td><td>Type1.BREX</td><td><-TetR-HTH<-?<-?<-REase+iREase||BrxA-DNA-binding->BrxB-BrxF-iSTAND->DUF6079*->N6-METHYLASE->PglZ->SIGMA+OB-Fold+AAA-ATPase+Lon-Protease-BrxL-><-ZnR<-ZnR<-?||RHH_antitoxin->XIS-HTH-></td><td>1213</td><td>Bacteria>Pseudomonadota>Gammaproteobacteria>Enterobacterales>Enterobacteriaceae>Escherichia</td><td>GCA_017046015.1</td><td>Escherichia_coli</td><td>JTN94_003485</td><td>7595..11236</td><td>562</td><td>BREX-system-P-loop-protein-BrxC-[Escherichia-coli]</td></tr>
<tr><td>EHC4928901.1</td><td>Type1.BREX</td><td>BrxA-DNA-binding->BrxB-BrxF-iSTAND->DUF6079*->N6-METHYLASE->PglZ->SIGMA+OB-Fold+AAA-ATPase+Lon-Protease-BrxL-><-ZnR<-ZnR</td><td>1213</td><td>Bacteria>Pseudomonadota>Gammaproteobacteria>Enterobacterales>Enterobacteriaceae>Escherichia</td><td>GCA_017057795.1</td><td>Escherichia_coli</td><td>JS667_004217</td><td>1769..5410</td><td>562</td><td>BREX-system-P-loop-protein-BrxC-[Escherichia-coli]</td></tr>
<tr><td>HAX9732773.1</td><td>Type1.BREX</td><td><-Phage_integrase||?->?->?-><-HNH||wHTH+SAD+HNH-><-REase+iREase||BrxA-DNA-binding->BrxB-BrxF-iSTAND->DUF6079*->N6-METHYLASE->PglZ->SIGMA+OB-Fold+AAA-ATPase+Lon-Protease-BrxL->HKD-DNase+OGG-like->||AbiEi_3_N+AbiEi_3->AbiEii-><-DUF6216<-Arm-DNA-bind_3</td><td>1213</td><td>Bacteria>Pseudomonadota>Gammaproteobacteria>Enterobacterales>Enterobacteriaceae>Escherichia</td><td>GCA_017073235.1</td><td>Escherichia_coli</td><td>JS163_001163</td><td>104185..107826</td><td>562</td><td>BREX-system-P-loop-protein-BrxC-[Escherichia-coli]</td></tr>
<tr><td>HAZ7757191.1</td><td>Type1.BREX</td><td>Glyoxalase_8-><-?<-?<-VirD4-TraG+VirB4-FtsK<-HDRnase<-SbcC<-SIGMA+OB-Fold+AAA-ATPase+Lon-Protease-BrxL<-PglZ<-HTH_Tnp_1+HTH_21+rve+rve_3<-DUF6079*<-BrxB-BrxF-iSTAND<-BrxA-DNA-binding||DDE_Tnp_IS240-><-wHTH+WYL+WCX<-?<-35exo||LexA-protease->?-><-MobI</td><td>1225</td><td>Bacteria>Pseudomonadota>Gammaproteobacteria>Enterobacterales>Enterobacteriaceae>Escherichia</td><td>GCA_017680345.1</td><td>Escherichia_coli</td><td>J7089_002325</td><td>28074..31751</td><td>562</td><td>BREX-system-P-loop-protein-BrxC-[Escherichia-coli]</td></tr>
<tr><td>HBA3405375.1</td><td>Type1.BREX</td><td>ZnR->ZnR-><-SIGMA+OB-Fold+AAA-ATPase+Lon-Protease-BrxL<-PglZ<-N6-METHYLASE<-DUF6079*<-BrxB-BrxF-iSTAND<-BrxA-DNA-binding||REase+iREase->?->?->TetR-HTH-></td><td>1213</td><td>Bacteria>Pseudomonadota>Gammaproteobacteria>Enterobacterales>Enterobacteriaceae>Escherichia</td><td>GCA_017728455.1</td><td>Escherichia_coli</td><td>J5T69_004101</td><td>10657..14298</td><td>562</td><td>BREX-system-P-loop-protein-BrxC-[Escherichia-coli]</td></tr>
<tr><td>HBA3375039.1</td><td>Type1.BREX</td><td><-Phage_integrase||?->?->?-><-HNH||wHTH+SAD+HNH-><-REase+iREase||BrxA-DNA-binding->BrxB-BrxF-iSTAND->DUF6079*->N6-METHYLASE->PglZ->SIGMA+OB-Fold+AAA-ATPase+Lon-Protease-BrxL->HKD-DNase+OGG-like->||AbiEi_3_N+AbiEi_3->AbiEii-><-DUF6216<-Arm-DNA-bind_3</td><td>1213</td><td>Bacteria>Pseudomonadota>Gammaproteobacteria>Enterobacterales>Enterobacteriaceae>Escherichia</td><td>GCA_017728515.1</td><td>Escherichia_coli</td><td>J8Z50_001951</td><td>111486..115127</td><td>562</td><td>BREX-system-P-loop-protein-BrxC-[Escherichia-coli]</td></tr>
<tr><td>HBA6912638.1</td><td>Type1.BREX</td><td><-Phage_integrase||?->?->?-><-HNH||wHTH+SAD+HNH-><-REase+iREase||BrxA-DNA-binding->BrxB-BrxF-iSTAND->DUF6079*->N6-METHYLASE->PglZ->SIGMA+OB-Fold+AAA-ATPase+Lon-Protease-BrxL->HKD-DNase+OGG-like->||AbiEi_3_N+AbiEi_3->AbiEii-><-DUF6216<-Arm-DNA-bind_3</td><td>1213</td><td>Bacteria>Pseudomonadota>Gammaproteobacteria>Enterobacterales>Enterobacteriaceae>Escherichia</td><td>GCA_017762865.1</td><td>Escherichia_coli</td><td>J1V61_002563</td><td>8737..12378</td><td>562</td><td>BREX-system-P-loop-protein-BrxC-[Escherichia-coli]</td></tr>
</table>

| Protein-ID | System | Gene-Neighbourhoods | Length | Taxonomy | Assembly-ID | Strain | Locus-Tag | Genomic-coordinates | TaxID | Fasta-defline |
|---|---|---|---|---|---|---|---|---|---|---|
| HBA9095564.1 | Type1.BREX | AraC-HTH+AraC-HTH-><-DDE_Tnp_IS240\|\|?->ZnR->ZnR-><-SIGMA+OB-Fold+AAA-ATPase+Lon-Protease-BrxL<-PglZ<-N6-METHYLASE<-DUF6079*<-BrxB-BrxF-iSTAND<-BrxA-DNA-binding\|\|REase->REase-><-HEPN<-ORC-CDC6+ZnR<-N6-METHYLASE<-Arm-DNA-bind_3+Phage_integrase | 1213 | Bacteria>Pseudomonadota>Gammaproteobacteria>Enterobacterales>Enterobacteriaceae>Escherichia | GCA_017772965.1 | Escherichia_coli | J1298_002132 | 65423..69064 | 562 | BREX-system-P-loop-protein-BrxC-[Escherichia-coli] |
| HBB1458964.1 | Type1.BREX | <-XIS-HTH\|\|?->?->ZnR->ZnR-><-SIGMA+OB-Fold+AAA-ATPase+Lon-Protease-BrxL<-PglZ<-N6-METHYLASE<-DUF6079*<-BrxB-BrxF-iSTAND<-BrxA-DNA-binding\|\|REase+iREase-><-EVE+HNH<-Arm-DNA-bind_3+Phage_integrase<-tRNA\|\|?->HerA/FtsK+A32-> | 1213 | Bacteria>Pseudomonadota>Gammaproteobacteria>Enterobacterales>Enterobacteriaceae>Escherichia | GCA_017785795.1 | Escherichia_coli | J8712_001728 | 57793..61434 | 562 | BREX-system-P-loop-protein-BrxC-[Escherichia-coli] |
| HBB1702364.1 | Type1.BREX | <-Phage_integrase\|\|?->?->?-><-HNH\|\|wHTH+SAD+HNH-><-REase+iREase\|\|BrxA-DNA-binding->BrxB-BrxF-iSTAND->DUF6079*-><-?<-?<-?\|\|HTH_6+SIS->\|\|?->?->?->DndB-> | 1213 | Bacteria>Pseudomonadota>Gammaproteobacteria>Enterobacterales>Enterobacteriaceae>Escherichia | GCA_017786745.1 | Escherichia_coli | J0606_001985 | 110955..114596 | 562 | BREX-system-P-loop-protein-BrxC-[Escherichia-coli] |
| HBB3467020.1 | Type1.BREX | DUF6079-p*->N6-METHYLASE->PglZ->SIGMA+OB-Fold+AAA-ATPase+Lon-Protease-BrxL-><-ZnR<-HTH_Tnp_1+HTH_21+rve+rve_3 | 834 | Bacteria>Pseudomonadota>Gammaproteobacteria>Enterobacterales>Enterobacteriaceae>Escherichia | GCA_017841335.1 | Escherichia_coli | KAD37_004247 | 1..2507 | 562 | BREX-system-P-loop-protein-BrxC-[Escherichia-coli] |
| HBB4280095.1 | Type1.BREX | DUF6079*->N6-METHYLASE->PglZ->SIGMA+OB-Fold+AAA-ATPase+Lon-Protease-BrxL->HKD-DNase+OGG-like-> | 1060 | Bacteria>Pseudomonadota>Gammaproteobacteria>Enterobacterales>Enterobacteriaceae>Escherichia | GCA_017899135.1 | Escherichia_coli | KAW34_004647 | 1..3185 | 562 | BREX-system-P-loop-protein-BrxC-[Escherichia-coli] |
| EHJ8640897.1 | Type1.BREX | <-Phage_integrase<-Phage_integrase\|\|?->?->?-><-HNH\|\|wHTH(SBD)+HNH->\|\|BrxA-DNA-binding->BrxB-BrxF-iSTAND->DUF6079*->N6-METHYLASE->PglZ->SIGMA+OB-Fold+AAA-ATPase+Lon-Protease-BrxL->HKD-DNase+OGG-like->\|\|AbiEi_3_N+AbiEi_3->AbiEii->?-><-DUF6216<-Arm-DNA-bind_3 | 1213 | Bacteria>Pseudomonadota>Gammaproteobacteria>Enterobacterales>Enterobacteriaceae>Escherichia | GCA_017945505.1 | Escherichia_coli | KB366_000307 | 330961..334602 | 562 | BREX-system-P-loop-protein-BrxC-[Escherichia-coli] |
| EHK0633846.1 | Type1.BREX | <-Phage_integrase\|\|?->?->?-><-HNH\|\|wHTH+SAD+HNH-><-REase+iREase\|\|BrxA-DNA-binding->BrxB-BrxF-iSTAND->DUF6079*->N6-METHYLASE->PglZ->SIGMA+OB-Fold+AAA-ATPase+Lon-Protease-BrxL->HKD-DNase+OGG-like->\|\|AbiEi_3_N+AbiEi_3->AbiEii->?-><-DUF6216<-Arm-DNA-bind_3 | 1213 | Bacteria>Pseudomonadota>Gammaproteobacteria>Enterobacterales>Enterobacteriaceae>Escherichia | GCA_017971565.1 | Escherichia_coli | KCG30_002478 | 111267..114908 | 562 | BREX-system-P-loop-protein-BrxC-[Escherichia-coli] |
| EHK3416136.1 | Type1.BREX | DUF6079*->N6-METHYLASE->PglZ->SIGMA+OB-Fold+AAA-ATPase+Lon-Protease-BrxL->HKD-DNase+OGG-like->\|\|AbiEi_3_N+AbiEi_3->AbiEii->?-><-DUF6216<-Arm-DNA-bind_3 | 1126 | Bacteria>Pseudomonadota>Gammaproteobacteria>Enterobacterales>Enterobacteriaceae>Escherichia | GCA_018020775.1 | Escherichia_coli | JYE89_003262 | 1..3383 | 562 | BREX-system-P-loop-protein-BrxC-[Escherichia-coli] |
| EHK3567501.1 | Type1.BREX | AAA+wHTH->?-><-SWIM+DUF3670+SF1-SF2-Helicase<-?<-?<-?\|\|BrxA-DNA-binding->BrxB-BrxF-iSTAND->DUF6079*->N6-METHYLASE->PglZ->SIGMA+OB-Fold+AAA-ATPase+Lon-Protease-BrxL-><-DDE_Tnp_IS1 | 1213 | Bacteria>Pseudomonadota>Gammaproteobacteria>Enterobacterales>Enterobacteriaceae>Escherichia | GCA_018022115.1 | Escherichia_coli | JT624_003007 | 41653..45294 | 562 | BREX-system-P-loop-protein-BrxC-[Escherichia-coli] |
| EHK3830299.1 | Type1.BREX | HTH_Tnp_1+HTH_21+rve+rve_3->wHTH+WYL+WCX->BrxA-DNA-binding->BrxB-BrxF-iSTAND->DUF6079*->ParB+HNH->N6-METHYLASE->PglZ->SIGMA+OB-Fold+AAA-ATPase+Lon-Protease-BrxL-><-DEDD_Tnp_IS110+Transposase_20 | 1225 | Bacteria>Pseudomonadota>Gammaproteobacteria>Enterobacterales>Enterobacteriaceae>Escherichia | GCA_018024135.1 | Escherichia_coli | JR760_003915 | 2875..6552 | 562 | BREX-system-P-loop-protein-BrxC-[Escherichia-coli] |
| EHL0175571.1 | Type1.BREX | <-DUF6079*<-BrxB-BrxF-iSTAND<-BrxA-DNA-binding\|\|<-wHTH(SBD)+HNH\|\|HNH-><-?<-?<-?\|\|Phage_integrase->Phage_integrase-> | 1016 | Bacteria>Pseudomonadota>Gammaproteobacteria>Enterobacterales>Enterobacteriaceae>Escherichia | GCA_018090545.1 | Escherichia_coli | D7K57_004368 | 1..3050 | 562 | BREX-system-P-loop-protein-BrxC-[Escherichia-coli] |
| EHL0363096.1 | Type1.BREX | DUF6079-p*->N6-METHYLASE->PglZ->SIGMA+OB-Fold+AAA-ATPase+Lon-Protease-BrxL->HKD-DNase+OGG-like->\|\|AbiEi_3_N+AbiEi_3->AbiEii->?-><-DUF6216<-Arm-DNA-bind_3 | 921 | Bacteria>Pseudomonadota>Gammaproteobacteria>Enterobacterales>Enterobacteriaceae>Escherichia | GCA_018090865.1 | Escherichia_coli | D1H14_003269 | 1..2766 | 562 | BREX-system-P-loop-protein-BrxC-[Escherichia-coli] |
| EHL0676199.1 | Type1.BREX | <-Phage_integrase<-Phage_integrase\|\|?->?->?-><-HNH\|\|wHTH(SBD)+HNH->\|\|BrxA-DNA-binding->BrxB-BrxF-iSTAND->DUF6079*->N6-METHYLASE-> | 1213 | Bacteria>Pseudomonadota>Gammaproteobacteria>Enterobacterales>Enterobacteriaceae>Escherichia | GCA_018092345.1 | Escherichia_coli | CP667_004253 | 9829..13470 | 562 | BREX-system-P-loop-protein-BrxC-[Escherichia-coli] |
| EHL0920537.1 | Type1.BREX | DUF6079-p*->N6-METHYLASE->PglZ->SIGMA+OB-Fold+AAA-ATPase+Lon-Protease-BrxL->HKD-DNase+OGG-like->\|\|AbiEi_3_N+AbiEi_3->AbiEii->?-><-DUF6216<-Arm-DNA-bind_3 | 767 | Bacteria>Pseudomonadota>Gammaproteobacteria>Enterobacterales>Enterobacteriaceae>Escherichia | GCA_018093285.1 | Escherichia_coli | CFF49_003158 | 1..2306 | 562 | BREX-system-P-loop-protein-BrxC-[Escherichia-coli] |
| EHL1132628.1 | Type1.BREX | DUF6079-p*->N6-METHYLASE->PglZ->SIGMA+OB-Fold+AAA-ATPase+Lon-Protease-BrxL->HKD-DNase+OGG-like->\|\|AbiEi_3_N+AbiEi_3->AbiEii->?-><-DUF6216<-Arm-DNA-bind_3 | 752 | Bacteria>Pseudomonadota>Gammaproteobacteria>Enterobacterales>Enterobacteriaceae>Escherichia | GCA_018094045.1 | Escherichia_coli | BIB01_002904 | 1..2259 | 562 | BREX-system-P-loop-protein-BrxC-[Escherichia-coli] |
| EHL1134570.1 | Type1.BREX | <-DUF6079-p*<-BrxB-BrxF-iSTAND<-BrxA-DNA-binding\|\|<-wHTH(SBD)+HNH\|\|HNH-><-?<-?<-?\|\|Phage_integrase->Phage_integrase-> | 460 | Bacteria>Pseudomonadota>Gammaproteobacteria>Enterobacterales>Enterobacteriaceae>Escherichia | GCA_018094045.1 | Escherichia_coli | BIB01_004961 | 1..1380 | 562 | BREX-system-P-loop-protein-BrxC-[Escherichia-coli] |
| EHL1162648.1 | Type1.BREX | <-Phage_integrase<-Phage_integrase\|\|?->?->?-><-HNH\|\|wHTH(SBD)+HNH->\|\|BrxA-DNA-binding->BrxB-BrxF-iSTAND->DUF6079*->N6-METHYLASE->PglZ->SIGMA+OB-Fold+AAA-ATPase+Lon-Protease-BrxL->HKD-DNase+OGG-like->\|\|AbiEi_3_N+AbiEi_3->AbiEii->?-><-DUF6216<-Arm-DNA-bind_3 | 1213 | Bacteria>Pseudomonadota>Gammaproteobacteria>Enterobacterales>Enterobacteriaceae>Escherichia | GCA_018094265.1 | Escherichia_coli | BE892_002471 | 14698..18339 | 562 | BREX-system-P-loop-protein-BrxC-[Escherichia-coli] |
| EHL3508970.1 | Type1.BREX | DUF6079*->N6-METHYLASE->PglZ->SIGMA+OB-Fold+AAA-ATPase+Lon-Protease-BrxL->?-><-?<-ParB<-?\|\|<-REase+N6_METHYLASE<-TRD | 1212 | Bacteria>Pseudomonadota>Gammaproteobacteria>Enterobacterales>Enterobacteriaceae>Escherichia | GCA_018111065.1 | Escherichia_coli | KDM69_001438 | 1..3639 | 562 | BREX-system-P-loop-protein-BrxC-[Escherichia-coli] |
| EHL6853247.1 | Type1.BREX | <-Phage_integrase\|\|?->?->?-><-HNH\|\|wHTH+SAD+HNH-><-REase+iREase\|\|BrxA-DNA-binding->BrxB-BrxF-iSTAND->DUF6079*->N6-METHYLASE-p-> | 1213 | Bacteria>Pseudomonadota>Gammaproteobacteria>Enterobacterales>Enterobacteriaceae>Escherichia | GCA_018143355.1 | Escherichia_coli | KEN32_003203 | 111251..114892 | 562 | BREX-system-P-loop-protein-BrxC-[Escherichia-coli] |
| HBC6897266.1 | Type1.BREX | DEDD_Tnp_IS110+Transposase_20-><-RHH\|\|?->wHTH+WYL+WCX->BrxA-DNA-binding->BrxB-BrxF-iSTAND->DUF6079*->ParB+HNH->N6-METHYLASE->PglZ-p-> | 1225 | Bacteria>Pseudomonadota>Gammaproteobacteria>Enterobacterales>Enterobacteriaceae>Escherichia | GCA_018163035.1 | Escherichia_coli | KEZ94_003915 | 5261..8938 | 562 | BREX-system-P-loop-protein-BrxC-[Escherichia-coli] |
| HBC7950668.1 | Type1.BREX | <-Phage_integrase\|\|?->?->?-><-HNH\|\|wHTH+SAD+HNH-><-REase+iREase\|\|BrxA-DNA-binding->BrxB-BrxF-iSTAND->DUF6079*->N6-METHYLASE->PglZ->SIGMA+OB-Fold+AAA-ATPase+Lon-Protease-BrxL->?-><-?<-ParB<-?\|\|<-REase+N6_METHYLASE<-TRD | 1213 | Bacteria>Pseudomonadota>Gammaproteobacteria>Enterobacterales>Enterobacteriaceae>Escherichia | GCA_018168015.1 | Escherichia_coli | KEZ88_001190 | 111008..114649 | 562 | BREX-system-P-loop-protein-BrxC-[Escherichia-coli] |
| EHM5922455.1 | Type1.BREX | <-HNH\|\|wHTH+SAD+HNH-><-REase+iREase\|\|BrxA-DNA-binding->BrxB-BrxF-iSTAND->DUF6079*->N6-METHYLASE->PglZ->SIGMA+OB-Fold+AAA-ATPase+Lon-Protease-BrxL->HKD-DNase+OGG-like->\|\|AbiEi_3_N+AbiEi_3->AbiEii->?-><-DUF6216<-Arm-DNA-bind_3 | 1213 | Bacteria>Pseudomonadota>Gammaproteobacteria>Enterobacterales>Enterobacteriaceae>Escherichia | GCA_018270935.1 | Escherichia_coli | KGR67_002708 | 6748..10389 | 562 | BREX-system-P-loop-protein-BrxC-[Escherichia-coli] |
| HBD2985690.1 | Type1.BREX | <-REase+iREase\|\|BrxA-DNA-binding->BrxB-BrxF-iSTAND->DUF6079*->N6-METHYLASE->PglZ->SIGMA+OB-Fold+AAA-ATPase+Lon-Protease-BrxL-><-ZnR<-ZnR<-?<-?\|\|RHH_antitoxin->XIS-HTH-> | 1213 | Bacteria>Pseudomonadota>Gammaproteobacteria>Enterobacterales>Enterobacteriaceae>Escherichia | GCA_018351255.1 | Escherichia_coli | NP403_001237 | 2672..6313 | 562 | BREX-system-P-loop-protein-BrxC-[Escherichia-coli] |
| EHN3342070.1 | Type1.BREX | <-Phage_integrase<-Phage_integrase\|\|?->?->?-><-HNH\|\|wHTH(SBD)+HNH->\|\|BrxA-DNA-binding->BrxB-BrxF-iSTAND->DUF6079*->N6-METHYLASE->PglZ->SIGMA+OB-Fold+AAA-ATPase+Lon-Protease-BrxL->HKD-DNase+OGG-like->\|\|AbiEi_3_N+AbiEi_3->AbiEii->?-><-DUF6216<-Arm-DNA-bind_3 | 1213 | Bacteria>Pseudomonadota>Gammaproteobacteria>Enterobacterales>Enterobacteriaceae>Escherichia | GCA_018356405.1 | Escherichia_coli | KIR71_003047 | 14697..18338 | 562 | BREX-system-P-loop-protein-BrxC-[Escherichia-coli] |
| HBD3449422.1 | Type1.BREX | <-N6-METHYLASE-p<-ParB+HNH<-DUF6079*<-BrxB-BrxF-iSTAND<-BrxA-DNA-binding<-wHTH+WYL+WCX<-Arm-DNA-bind_3+Phage_integrase\|\|?-><-?\|\|?->METHYLASE-> | 1225 | Bacteria>Pseudomonadota>Gammaproteobacteria>Enterobacterales>Enterobacteriaceae>Escherichia | GCA_018390165.1 | Escherichia_coli | KMB60_001824 | 3370..7047 | 562 | BREX-system-P-loop-protein-BrxC-[Escherichia-coli] |
| GIP82493.1 | Type1.BREX | <-Peptidase_M17_N+Peptidase_M17\|\|?->?-><-HerA/FtsK+A32\|\|tRNA->Arm-DNA-bind_3+Phage_integrase->EVE+HNH-><-REase+iREase\|\|BrxA-DNA-binding->BrxB-BrxF-iSTAND->DUF6079*->N6-METHYLASE->PglZ->SIGMA+OB-Fold+AAA-ATPase+Lon-Protease-BrxL-><-ZnR<-ZnR<-?\|\|RHH_antitoxin->XIS-HTH-> | 1213 | Bacteria>Pseudomonadota>Gammaproteobacteria>Enterobacterales>Enterobacteriaceae>Escherichia | GCA_018403555.1 | Escherichia_coli | pm083_35810 | 54245..57886 | 562 | ATPase-like-protein-[Escherichia-coli] |
| HBD4982009.1 | Type1.BREX | <-XIS-HTH<-RHH_antitoxin\|\|?->?->ZnR->ZnR-><-SIGMA+OB-Fold+AAA-ATPase+Lon-Protease-BrxL<-PglZ<-N6-METHYLASE<-DUF6079*<-BrxB-BrxF-iSTAND<-BrxA-DNA-binding\|\|REase+iREase-> | 1213 | Bacteria>Pseudomonadota>Gammaproteobacteria>Enterobacterales>Enterobacteriaceae>Escherichia | GCA_018458945.1 | Escherichia_coli | KKK67_004276 | 14430..18071 | 562 | BREX-system-P-loop-protein-BrxC-[Escherichia-coli] |
| HBD5250199.1 | Type1.BREX | <-Phage_integrase\|\|Arm-DNA-bind_3->DUF6216-><-AbiEii<-AbiEi_3_N+AbiEi_3\|\|<-HKD-DNase+OGG-like<-SIGMA+OB-Fold+AAA-ATPase+Lon-Protease-BrxL<-PglZ<-N6-METHYLASE<-DUF6079* | 1060 | Bacteria>Pseudomonadota>Gammaproteobacteria>Enterobacterales>Enterobacteriaceae>Escherichia | GCA_018460085.1 | Escherichia_coli | KKL26_003784 | 28719..31903 | 562 | BREX-system-P-loop-protein-BrxC-[Escherichia-coli] |
| HBD5388086.1 | Type1.BREX | <-HIN-HTH<-RHH_antitoxin\|\|?->?->ZnR->ZnR-><-SIGMA+OB-Fold+AAA-ATPase+Lon-Protease-BrxL<-PglZ<-N6-METHYLASE<-DUF6079*<-BrxB-BrxF-iSTAND<-BrxA-DNA-binding\|\|REase->REase-> | 1213 | Bacteria>Pseudomonadota>Gammaproteobacteria>Enterobacterales>Enterobacteriaceae>Escherichia | GCA_018460465.1 | Escherichia_coli | KKL44_003112 | 108799..112440 | 562 | BREX-system-P-loop-protein-BrxC-[Escherichia-coli] |
| HBE5478594.1 | Type1.BREX | DUF6079*->N6-METHYLASE->PglZ->SIGMA+OB-Fold+AAA-ATPase+Lon-Protease-BrxL-> | 1064 | Bacteria>Pseudomonadota>Gammaproteobacteria>Enterobacterales>Enterobacteriaceae>Escherichia | GCA_018566785.1 | Escherichia_coli | KMB17_004111 | 1..3196 | 562 | BREX-system-P-loop-protein-BrxC-[Escherichia-coli] |
| HBE6046090.1 | Type1.BREX | <-Phage_integrase\|\|?->?->?-><-HNH\|\|wHTH+SAD+HNH-><-REase+iREase\|\|BrxA-DNA-binding->BrxB-BrxF-iSTAND->DUF6079*->N6-METHYLASE->PglZ->SIGMA+OB-Fold+AAA-ATPase+Lon-Protease-BrxL->?-><-?<-ParB<-?\|\|<-REase+N6_METHYLASE<-TRD | 1213 | Bacteria>Pseudomonadota>Gammaproteobacteria>Enterobacterales>Enterobacteriaceae>Escherichia | GCA_018577025.1 | Escherichia_coli | KMG52_000552 | 111009..114650 | 562 | BREX-system-P-loop-protein-BrxC-[Escherichia-coli] |
| EHQ7823790.1 | Type1.BREX | DUF6079-p*->N6-METHYLASE->PglZ->SIGMA+OB-Fold+AAA-ATPase+Lon-Protease-BrxL->HKD-DNase+OGG-like->\|\|AbiEi_3_N+AbiEi_3->AbiEii->?-><-DUF6216<-Arm-DNA-bind_3 | 927 | Bacteria>Pseudomonadota>Gammaproteobacteria>Enterobacterales>Enterobacteriaceae>Escherichia | GCA_018975885.1 | Escherichia_coli | KSS40_002990 | 1..2786 | 562 | BREX-system-P-loop-protein-BrxC-[Escherichia-coli] |
| HBH4475334.1 | Type1.BREX | <-Zn_Tnp_IS1+HTH_Tnp_IS1\|\|BrxA-DNA-binding->BrxB-BrxF-iSTAND->DUF6079*->PglZ->SIGMA+OB-Fold+AAA-ATPase+Lon-Protease-BrxL-> | 1213 | Bacteria>Pseudomonadota>Gammaproteobacteria>Enterobacterales>Enterobacteriaceae>Escherichia | GCA_019001705.1 | Escherichia_coli | KTE98_004875 | 1931..5572 | 562 | BREX-system-P-loop-protein-BrxC-[Escherichia-coli] |
| EHR8027111.1 | Type1.BREX | <-REase+iREase\|\|BrxA-DNA-binding->BrxB-BrxF-iSTAND->DUF6079*->N6-METHYLASE->PglZ->SIGMA+OB-Fold+AAA-ATPase+Lon-Protease-BrxL-> | 1213 | Bacteria>Pseudomonadota>Gammaproteobacteria>Enterobacterales>Enterobacteriaceae>Escherichia | GCA_019077505.1 | Escherichia_coli | KUT12_004099 | 2676..6317 | 562 | BREX-system-P-loop-protein-BrxC-[Escherichia-coli] |
| HBH7337791.1 | Type1.BREX | BrxA-DNA-binding->BrxB-BrxF-iSTAND->DUF6079*->N6-METHYLASE->PglZ->SIGMA+OB-Fold+AAA-ATPase+Lon-Protease-BrxL-><-ZnR<-ZnR<-HIN-HTH<-?<-?\|\|RHH_antitoxin-> | 1213 | Bacteria>Pseudomonadota>Gammaproteobacteria>Enterobacterales>Enterobacteriaceae>Escherichia | GCA_019178095.1 | Escherichia_coli | KWY89_004355 | 1906..5547 | 562 | BREX-system-P-loop-protein-BrxC-[Escherichia-coli] |
| EHW5636696.1 | Type1.BREX | <-Phage_integrase\|\|?->?->?-><-HNH\|\|wHTH+SAD+HNH-><-REase+iREase\|\|BrxA-DNA-binding->BrxB-BrxF-iSTAND->DUF6079*->N6-METHYLASE->PglZ->SIGMA+OB-Fold+AAA-ATPase+Lon-Protease-BrxL->HKD-DNase+OGG-like->\|\|AbiEi_3_N+AbiEi_3->AbiEii->?-><-DUF6216<-Arm-DNA-bind_3 | 1213 | Bacteria>Pseudomonadota>Gammaproteobacteria>Enterobacterales>Enterobacteriaceae>Escherichia | GCA_019482305.1 | Escherichia_coli | K2955_003127 | 35708..39349 | 562 | BREX-system-P-loop-protein-BrxC-[Escherichia-coli] |
| EHX2153194.1 | Type1.BREX | BrxA-DNA-binding->BrxB-BrxF-iSTAND->DUF6079*->N6-METHYLASE->PglZ->SIGMA+OB-Fold+AAA-ATPase+Lon-Protease-BrxL->HKD-DNase+OGG-like->\|\|AbiEi_3_N+AbiEi_3->AbiEii->?-><-DUF6216<-Arm-DNA-bind_3 | 1213 | Bacteria>Pseudomonadota>Gammaproteobacteria>Enterobacterales>Enterobacteriaceae>Escherichia | GCA_019516635.1 | Escherichia_coli | K3P68_001530 | 1396..5037 | 562 | BREX-system-P-loop-protein-BrxC-[Escherichia-coli] |
| EHX8392339.1 | Type1.BREX | Arm-DNA-bind_3->DUF6216-><-?<-AbiEii<-AbiEi_3_N+AbiEi_3\|\|<-HKD-DNase+OGG-like<-SIGMA+OB-Fold+AAA-ATPase+Lon-Protease-BrxL<-PglZ<-DUF6079*<-BrxB-BrxF-iSTAND<-BrxA-DNA-binding\|\|REase+iREase-><-wHTH+SAD+HNH\|\|HNH-><-?<-?<-?\|\|Phage_integrase-> | 1213 | Bacteria>Pseudomonadota>Gammaproteobacteria>Enterobacterales>Enterobacteriaceae>Escherichia | GCA_019550135.1 | Escherichia_coli | K3368_004098 | 15146..18787 | 562 | BREX-system-P-loop-protein-BrxC-[Escherichia-coli] |
| EHX8476217.1 | Type1.BREX | <-Phage_integrase\|\|?->?->?-><-HNH\|\|wHTH+SAD+HNH-><-REase+iREase\|\|BrxA-DNA-binding->BrxB-BrxF-iSTAND->DUF6079*->N6-METHYLASE->PglZ->SIGMA+OB-Fold+AAA-ATPase+Lon-Protease-BrxL->HKD-DNase+OGG-like->\|\|AbiEi_3_N+AbiEi_3->AbiEii->?-><-DUF6216<-Arm-DNA-bind_3 | 1213 | Bacteria>Pseudomonadota>Gammaproteobacteria>Enterobacterales>Enterobacteriaceae>Escherichia | GCA_019550495.1 | Escherichia_coli | K3324_002225 | 23193..26834 | 562 | BREX-system-P-loop-protein-BrxC-[Escherichia-coli] |
| EHY5030619.1 | Type1.BREX | <-wHTH+WYL<-?\|\|XIS-HTH->?->BrxA-DNA-binding->BrxB-BrxF-iSTAND->DUF6079*->ParB+HNH->N6-METHYLASE->PglZ->SIGMA+OB-Fold+AAA-ATPase+Lon-Protease-BrxL-><-Arm-DNA-bind_3+Phage_integrase\|\|Phage_integrase-> | 1233 | Bacteria>Pseudomonadota>Gammaproteobacteria>Enterobacterales>Enterobacteriaceae>Escherichia | GCA_019582855.1 | Escherichia_coli | K4I20_002754 | 15002..18703 | 562 | BREX-system-P-loop-protein-BrxC-[Escherichia-coli] |
| EHY5298658.1 | Type1.BREX | DUF6079-p*->N6-METHYLASE->PglZ->SIGMA+OB-Fold+AAA-ATPase+Lon-Protease-BrxL->HKD-DNase+OGG-like->\|\|AbiEi_3_N+AbiEi_3->AbiEii->?-><-DUF6216<-Arm-DNA-bind_3 | 861 | Bacteria>Pseudomonadota>Gammaproteobacteria>Enterobacterales>Enterobacteriaceae>Escherichia | GCA_019588505.1 | Escherichia_coli | K4I57_002534 | 1..2588 | 562 | BREX-system-P-loop-protein-BrxC-[Escherichia-coli] |
| EHY5690081.1 | Type1.BREX | <-Phage_integrase<-Phage_integrase\|\|?->?->?-><-HNH\|\|wHTH(SBD)+HNH->\|\|BrxA-DNA-binding->BrxB-BrxF-iSTAND->DUF6079*->N6-METHYLASE->PglZ->SIGMA+OB-Fold+AAA-ATPase+Lon-Protease-BrxL->HKD-DNase+OGG-like->\|\|AbiEi_3_N+AbiEi_3->AbiEii->?-><-DUF6216<-Arm-DNA-bind_3 | 1213 | Bacteria>Pseudomonadota>Gammaproteobacteria>Enterobacterales>Enterobacteriaceae>Escherichia | GCA_019590565.1 | Escherichia_coli | K4J54_002559 | 14696..18337 | 562 | BREX-system-P-loop-protein-BrxC-[Escherichia-coli] |
| EHZ2564701.1 | Type1.BREX | DUF6079*->N6-METHYLASE->PglZ->SIGMA+OB-Fold+AAA-ATPase+Lon-Protease-BrxL->HKD-DNase+OGG-like->\|\|AbiEi_3_N+AbiEi_3->AbiEii->?-><-DUF6216<-Arm-DNA-bind_3 | 1130 | Bacteria>Pseudomonadota>Gammaproteobacteria>Enterobacterales>Enterobacteriaceae>Escherichia | GCA_019681855.1 | Escherichia_coli | K5N16_003075 | 1..3394 | 562 | BREX-system-P-loop-protein-BrxC-[Escherichia-coli] |
| EHZ4708255.1 | Type1.BREX | <-Phage_integrase<-Phage_integrase\|\|?->?->?->\|\|wHTH+SAD+HNH-><-REase+iREase\|\|BrxA-DNA-binding->BrxB-BrxF-iSTAND->DUF6079*->N6-METHYLASE->PglZ->SIGMA+OB-Fold+AAA-ATPase+Lon-Protease-BrxL-> | 1213 | Bacteria>Pseudomonadota>Gammaproteobacteria>Enterobacterales>Enterobacteriaceae>Escherichia | GCA_019701545.1 | Escherichia_coli | K5X72_003965 | 35776..39417 | 562 | BREX-system-P-loop-protein-BrxC-[Escherichia-coli] |
| EIB0293606.1 | Type1.BREX | <-DUF6079-p*<-BrxB-BrxF-iSTAND<-BrxA-DNA-binding\|\|<-wHTH(SBD)+HNH\|\|HNH-><-?<-?<-?\|\|Phage_integrase->Phage_integrase-> | 529 | Bacteria>Pseudomonadota>Gammaproteobacteria>Enterobacterales>Enterobacteriaceae>Escherichia | GCA_019848435.1 | Escherichia_coli | K7644_004610 | 1..1589 | 562 | BREX-system-P-loop-protein-BrxC-[Escherichia-coli] |
| HBI8276592.1 | Type1.BREX | <-METHYLASE\|\|SKI-><-?\|\|tRNA->Arm-DNA-bind_3+Phage_integrase->EVE+HNH-><-REase<-REase+iREase\|\|BrxA-DNA-binding->BrxB-BrxF-iSTAND->DUF6079*->N6-METHYLASE->PglZ->SIGMA+OB-Fold+AAA-ATPase+Lon-Protease-BrxL-><-ZnR<-ZnR<-?<-?\|\|RHH_antitoxin->XIS-HTH-> | 1213 | Bacteria>Pseudomonadota>Gammaproteobacteria>Enterobacterales>Enterobacteriaceae>Escherichia | GCA_019886465.1 | Escherichia_coli | K8321_003292 | 148369..152010 | 562 | BREX-system-P-loop-protein-BrxC-[Escherichia-coli] |
| HBJ0235562.1 | Type1.BREX | <-HKD-DNase+OGG-like<-SIGMA+OB-Fold+AAA-ATPase+Lon-Protease-BrxL<-PglZ<-N6-METHYLASE<-DUF6079*<-BrxB-BrxF-iSTAND<-BrxA-DNA-binding\|\|REase+iREase-><-wHTH+SAD+HNH\|\|HNH-> | 1213 | Bacteria>Pseudomonadota>Gammaproteobacteria>Enterobacterales>Enterobacteriaceae>Escherichia | GCA_019980105.1 | Escherichia_coli | LAM28_003686 | 10586..14227 | 562 | BREX-system-P-loop-protein-BrxC-[Escherichia-coli] |
| HBJ1150970.1 | Type1.BREX | BrxA-DNA-binding->BrxB-BrxF-iSTAND->DUF6079*->N6-METHYLASE->PglZ->SIGMA+OB-Fold+AAA-ATPase+Lon-Protease-BrxL-><-ZnR<-ZnR<-?<-?\|\|RHH_antitoxin->XIS-HTH-> | 1213 | Bacteria>Pseudomonadota>Gammaproteobacteria>Enterobacterales>Enterobacteriaceae>Escherichia | GCA_019988955.1 | Escherichia_coli | LAQ35_004277 | 1344..4985 | 562 | BREX-system-P-loop-protein-BrxC-[Escherichia-coli] |
| HBJ1306075.1 | Type1.BREX | <-METHYLASE\|\|SKI-><-?\|\|tRNA->Arm-DNA-bind_3+Phage_integrase->EVE+HNH-><-REase+iREase\|\|BrxA-DNA-binding->BrxB-BrxF-iSTAND->DUF6079*->N6-METHYLASE->PglZ->SIGMA+OB-Fold+AAA-ATPase+Lon-Protease-BrxL-><-ZnR | 1213 | Bacteria>Pseudomonadota>Gammaproteobacteria>Enterobacterales>Enterobacteriaceae>Escherichia | GCA_019996045.1 | Escherichia_coli | LAS09_001693 | 95554..99195 | 562 | BREX-system-P-loop-protein-BrxC-[Escherichia-coli] |
| EID3638522.1 | Type1.BREX | <-Phage_integrase\|\|?->?->?-><-HNH\|\|wHTH+SAD+HNH-><-REase+iREase\|\|BrxA-DNA-binding->BrxB-BrxF-iSTAND->DUF6079*->N6-METHYLASE->PglZ->SIGMA+OB-Fold+AAA-ATPase+Lon-Protease-BrxL->HKD-DNase+OGG-like->\|\|AbiEi_3_N+AbiEi_3->AbiEii->?-><-DUF6216<-Arm-DNA-bind_3 | 1213 | Bacteria>Pseudomonadota>Gammaproteobacteria>Enterobacterales>Enterobacteriaceae>Escherichia | GCA_020041725.1 | Escherichia_coli | LB206_001712 | 80689..84330 | 562 | BREX-system-P-loop-protein-BrxC-[Escherichia-coli] |
| EID8685857.1 | Type1.BREX | Arm-DNA-bind_3->DUF6216-><-?<-AbiEii<-AbiEi_3_N+AbiEi_3\|\|<-HKD-DNase+OGG-like<-SIGMA+OB-Fold+AAA-ATPase+Lon-Protease-BrxL<-PglZ<-N6-METHYLASE<-DUF6079-p* | 949 | Bacteria>Pseudomonadota>Gammaproteobacteria>Enterobacterales>Enterobacteriaceae>Escherichia | GCA_020094635.1 | Escherichia_coli | LCI88_002092 | 151835..154685 | 562 | BREX-system-P-loop-protein-BrxC-[Escherichia-coli] |
| EID8792051.1 | Type1.BREX | wHTH+WYL+WCX->BrxA-DNA-binding->BrxB-BrxF-iSTAND->DUF6079*->ParB+HNH->N6-METHYLASE->PglZ->SIGMA+OB-Fold+AAA-ATPase+Lon-Protease-BrxL->HTH_Tnp_1+HTH_21+rve+rve_3<-HIN-HTH<-PIN<-MazE_antitoxin | 1225 | Bacteria>Pseudomonadota>Gammaproteobacteria>Enterobacterales>Enterobacteriaceae>Escherichia | GCA_020095335.1 | Escherichia_coli | LCJ17_004431 | 3670..7347 | 562 | BREX-system-P-loop-protein-BrxC-[Escherichia-coli] |
| MBZ8679488.1 | Type1.BREX | <-Phage_integrase\|\|?->?->?-><-HNH\|\|wHTH+SAD+HNH-><-REase+iREase\|\|BrxA-DNA-binding->BrxB-BrxF-iSTAND->DUF6079*->N6-METHYLASE->PglZ->SIGMA+OB-Fold+AAA-ATPase+Lon-Protease-BrxL->?-><-?<-ParB<-?\|\|<-REase+N6_METHYLASE<-TRD | 1213 | Bacteria>Pseudomonadota>Gammaproteobacteria>Enterobacterales>Enterobacteriaceae>Escherichia | GCA_020145545.1 | Escherichia_coli | H5S60_20510 | 9779..13420 | 562 | BREX-system-P-loop-protein-BrxC-[Escherichia-coli] |
| EIF2649115.1 | Type1.BREX | DUF6079*->N6-METHYLASE->PglZ->SIGMA+OB-Fold+AAA-ATPase+Lon-Protease-BrxL-><-ZnR<-ZnR<-?\|\|HIN-HTH+PAIREDC-HTH->\|\|RHH_antitoxin->XIS-HTH-> | 1005 | Bacteria>Pseudomonadota>Gammaproteobacteria>Enterobacterales>Enterobacteriaceae>Escherichia | GCA_020238205.1 | Escherichia_coli | LFR91_004210 | 1..3019 | 562 | BREX-system-P-loop-protein-BrxC-[Escherichia-coli] |
| EIG2625391.1 | Type1.BREX | DUF6079-p*->N6-METHYLASE->PglZ->SIGMA+OB-Fold+AAA-ATPase+Lon-Protease-BrxL->HKD-DNase+OGG-like->\|\|AbiEi_3_N+AbiEi_3->AbiEii->?-><-DUF6216<-Arm-DNA-bind_3 | 771 | Bacteria>Pseudomonadota>Gammaproteobacteria>Enterobacterales>Enterobacteriaceae>Escherichia | GCA_020303335.1 | Escherichia_coli | GLS16_002890 | 1..2318 | 562 | BREX-system-P-loop-protein-BrxC-[Escherichia-coli] |
| EIG4139450.1 | Type1.BREX | <-REase+iREase\|\|BrxA-DNA-binding->BrxB-BrxF-iSTAND->DUF6079*->N6-METHYLASE->PglZ->SIGMA+OB-Fold+AAA-ATPase+Lon-Protease-BrxL-><-ZnR<-ZnR<-?<-?\|\|XIS-HTH-> | 1213 | Bacteria>Pseudomonadota>Gammaproteobacteria>Enterobacterales>Enterobacteriaceae>Escherichia | GCA_020309325.1 | Escherichia_coli | FPR33_004782 | 2597..6238 | 562 | BREX-system-P-loop-protein-BrxC-[Escherichia-coli] |
| EIG6470759.1 | Type1.BREX | <-DUF6079-p*<-BrxB-BrxF-iSTAND<-BrxA-DNA-binding\|\|<-wHTH(SBD)+HNH\|\|HNH-><-?<-?<-?\|\|Phage_integrase->Phage_integrase-> | 634 | Bacteria>Pseudomonadota>Gammaproteobacteria>Enterobacterales>Enterobacteriaceae>Escherichia | GCA_020319175.1 | Escherichia_coli | EG665_005071 | 1..1903 | 562 | BREX-system-P-loop-protein-BrxC-[Escherichia-coli] |
| EIG7860374.1 | Type1.BREX | DUF6079*->N6-METHYLASE->PglZ->SIGMA+OB-Fold+AAA-ATPase+Lon-Protease-BrxL->HKD-DNase+OGG-like->\|\|AbiEi_3_N+AbiEi_3->AbiEii->?-><-DUF6216<-Arm-DNA-bind_3 | 1077 | Bacteria>Pseudomonadota>Gammaproteobacteria>Enterobacterales>Enterobacteriaceae>Escherichia | GCA_020324895.1 | Escherichia_coli | D0584_004201 | 1..3234 | 562 | BREX-system-P-loop-protein-BrxC-[Escherichia-coli] |
| HBK2401145.1 | Type1.BREX | <-Phage_integrase\|\|?->?->?-><-HNH\|\|wHTH+SAD+HNH-><-REase+iREase\|\|BrxA-DNA-binding->BrxB-BrxF-iSTAND->DUF6079*-> | 1149 | Bacteria>Pseudomonadota>Gammaproteobacteria>Enterobacterales>Enterobacteriaceae>Escherichia | GCA_020359375.1 | Escherichia_coli | LGJ14_002758 | 111240..114688 | 562 | BREX-system-P-loop-protein-BrxC-[Escherichia-coli] |
| HBK2662992.1 | Type1.BREX | <-Phage_integrase\|\|?->?->?-><-HNH\|\|wHTH+SAD+HNH-><-REase+iREase\|\|BrxA-DNA-binding->BrxB-BrxF-iSTAND->DUF6079-p*-> | 626 | Bacteria>Pseudomonadota>Gammaproteobacteria>Enterobacterales>Enterobacteriaceae>Escherichia | GCA_020362085.1 | Escherichia_coli | LGK41_002196 | 111240..113117 | 562 | BREX-system-P-loop-protein-BrxC-[Escherichia-coli] |
| EIH7117202.1 | Type1.BREX | DUF6079*->N6-METHYLASE->PglZ->SIGMA+OB-Fold+AAA-ATPase+Lon-Protease-BrxL->HKD-DNase+OGG-like->\|\|AbiEi_3_N+AbiEi_3->AbiEii->?-><-DUF6216<-Arm-DNA-bind_3 | 1174 | Bacteria>Pseudomonadota>Gammaproteobacteria>Enterobacterales>Enterobacteriaceae>Escherichia | GCA_020376505.1 | Escherichia_coli | BB077_003959 | 1..3527 | 562 | BREX-system-P-loop-protein-BrxC-[Escherichia-coli] |
| MCA7636345.1 | Type1.BREX | <-XIS-HTH<-RHH_antitoxin\|\|?->?->ZnR->ZnR-><-SIGMA+OB-Fold+AAA-ATPase+Lon-Protease-BrxL<-PglZ<-N6-METHYLASE<-DUF6079* | 1132 | Bacteria>Pseudomonadota>Gammaproteobacteria>Enterobacterales>Enterobacteriaceae>Escherichia | GCA_020418475.1 | Escherichia_coli | LEZ98_21685 | 17948..21346 | 562 | BREX-system-P-loop-protein-BrxC-[Escherichia-coli] |
| MCA7852868.1 | Type1.BREX | <-Phage_integrase<-Phage_integrase\|\|?->?->?->\|\|wHTH+SAD+HNH-><-REase+iREase\|\|BrxA-DNA-binding->BrxB-BrxF-iSTAND->DUF6079-p*-> | 920 | Bacteria>Pseudomonadota>Gammaproteobacteria>Enterobacterales>Enterobacteriaceae>Escherichia | GCA_020420865.1 | Escherichia_coli | LE030_10780 | 111187..113946 | 562 | BREX-system-P-loop-protein-BrxC-[Escherichia-coli] |
| EIL3113271.1 | Type1.BREX | <-HerA/FtsK+A32<-?\|\|tRNA->Arm-DNA-bind_3+Phage_integrase->EVE+HNH-><-REase+iREase\|\|BrxA-DNA-binding->BrxB-BrxF-iSTAND->DUF6079*->N6-METHYLASE->PglZ->SIGMA+OB-Fold+AAA-ATPase+Lon-Protease-BrxL-><-ZnR<-ZnR<-?<-?\|\|RHH_antitoxin->XIS-HTH-> | 1213 | Bacteria>Pseudomonadota>Gammaproteobacteria>Enterobacterales>Enterobacteriaceae>Escherichia | GCA_020689665.1 | Escherichia_coli | LLT94_003844 | 18999..22640 | 562 | BREX-system-P-loop-protein-BrxC-[Escherichia-coli] |
| EIN0528288.1 | Type1.BREX | <-GDResolvase+HIN-HTH<-?<-RelE-ParE<-CopG/RHH\|\|<-RHH\|\|?->wHTH+WYL+WCX->BrxA-DNA-binding->BrxB-BrxF-iSTAND->DUF6079-p*-> | 671 | Bacteria>Pseudomonadota>Gammaproteobacteria>Enterobacterales>Enterobacteriaceae>Escherichia | GCA_020842825.1 | Escherichia_coli | LMH12_004589 | 11532..13543 | 562 | BREX-system-P-loop-protein-BrxC-[Escherichia-coli] |
| EIN2019349.1 | Type1.BREX | <-Phage_integrase\|\|?->?->?-><-HNH\|\|wHTH+SAD+HNH-><-REase+iREase\|\|BrxA-DNA-binding->BrxB-BrxF-iSTAND->DUF6079*->N6-METHYLASE->PglZ->SIGMA+OB-Fold+AAA-ATPase+Lon-Protease-BrxL->?-><-?<-ParB<-?\|\|<-REase+N6_METHYLASE<-TRD | 1213 | Bacteria>Pseudomonadota>Gammaproteobacteria>Enterobacterales>Enterobacteriaceae>Escherichia | GCA_020867105.1 | Escherichia_coli | LOJ72_000111 | 111240..114881 | 562 | BREX-system-P-loop-protein-BrxC-[Escherichia-coli] |
| EIO0019318.1 | Type1.BREX | DUF6079-p*->N6-METHYLASE->PglZ->SIGMA+OB-Fold+AAA-ATPase+Lon-Protease-BrxL->HKD-DNase+OGG-like->\|\|AbiEi_3_N+AbiEi_3->AbiEii->?-><-DUF6216<-Arm-DNA-bind_3 | 816 | Bacteria>Pseudomonadota>Gammaproteobacteria>Enterobacterales>Enterobacteriaceae>Escherichia | GCA_020946425.1 | Escherichia_coli | LO099_004443 | 1..2453 | 562 | BREX-system-P-loop-protein-BrxC-[Escherichia-coli] |
| EIO1138690.1 | Type1.BREX | <-Phage_integrase\|\|?->?->?-><-HNH\|\|wHTH+SAD+HNH-><-REase+iREase\|\|BrxA-DNA-binding->BrxB-BrxF-iSTAND->DUF6079*->N6-METHYLASE->PglZ->SIGMA+OB-Fold+AAA-ATPase+Lon-Protease-BrxL->?-><-?<-ParB<-?\|\|<-REase+N6_METHYLASE<-TRD | 1213 | Bacteria>Pseudomonadota>Gammaproteobacteria>Enterobacterales>Enterobacteriaceae>Escherichia | GCA_020952215.1 | Escherichia_coli | LQD99_001531 | 111293..114934 | 562 | BREX-system-P-loop-protein-BrxC-[Escherichia-coli] |
| HBL7859045.1 | Type1.BREX | DUF6079*->N6-METHYLASE->PglZ->SIGMA+OB-Fold+AAA-ATPase+Lon-Protease-BrxL-><-ZnR<-HTH_Tnp_1+HTH_21+rve+rve_3 | 1097 | Bacteria>Pseudomonadota>Gammaproteobacteria>Enterobacterales>Enterobacteriaceae>Escherichia | GCA_021087345.1 | Escherichia_coli | LTB56_004870 | 1..3294 | 562 | BREX-system-P-loop-protein-BrxC-[Escherichia-coli] |
| EIS3583963.1 | Type1.BREX | <-N6-METHYLASE<-DUF6079*<-BrxB-BrxF-iSTAND<-BrxA-DNA-binding\|\|REase+iREase-><-wHTH+SAD+HNH\|\|HNH-><-?<-?<-?\|\|Phage_integrase-> | 1213 | Bacteria>Pseudomonadota>Gammaproteobacteria>Enterobacterales>Enterobacteriaceae>Escherichia | GCA_021351055.1 | Escherichia_coli | LZO81_004279 | 2281..5922 | 562 | BREX-system-P-loop-protein-BrxC-[Escherichia-coli] |
| EIS4879897.1 | Type1.BREX | Arm-DNA-bind_3+Phage_integrase->N6-METHYLASE->ORC-CDC6+ZnR->HEPN-><-REase<-REase\|\|BrxA-DNA-binding->BrxB-BrxF-iSTAND->DUF6079*->N6-METHYLASE->PglZ->SIGMA+OB-Fold+AAA-ATPase+Lon-Protease-BrxL-><-ZnR<-ZnR<-?\|\|RHH_antitoxin->XIS-HTH-> | 1213 | Bacteria>Pseudomonadota>Gammaproteobacteria>Enterobacterales>Enterobacteriaceae>Escherichia | GCA_021362895.1 | Escherichia_coli | LZX86_002736 | 36303..39944 | 562 | BREX-system-P-loop-protein-BrxC-[Escherichia-coli] |
| HBM7577416.1 | Type1.BREX | <-REase+iREase\|\|BrxA-DNA-binding->BrxB-BrxF-iSTAND->DUF6079*->N6-METHYLASE->PglZ->SIGMA+OB-Fold+AAA-ATPase+Lon-Protease-BrxL-> | 1213 | Bacteria>Pseudomonadota>Gammaproteobacteria>Enterobacterales>Enterobacteriaceae>Escherichia | GCA_021378865.1 | Escherichia_coli | LZR97_004339 | 2197..5838 | 562 | BREX-system-P-loop-protein-BrxC-[Escherichia-coli] |
| HBM8747261.1 | Type1.BREX | <-Phage_integrase\|\|?->?->?-><-HNH\|\|wHTH+SAD+HNH-><-REase+iREase\|\|BrxA-DNA-binding->BrxB-BrxF-iSTAND->DUF6079*->N6-METHYLASE->PglZ->SIGMA+OB-Fold+AAA-ATPase+Lon-Protease-BrxL->HKD-DNase+OGG-like->\|\|AbiEi_3_N+AbiEi_3->AbiEii->?-><-DUF6216<-Arm-DNA-bind_3 | 1213 | Bacteria>Pseudomonadota>Gammaproteobacteria>Enterobacterales>Enterobacteriaceae>Escherichia | GCA_021417505.1 | Escherichia_coli | LOV31_003260 | 39874..43515 | 562 | BREX-system-P-loop-protein-BrxC-[Escherichia-coli] |
| HBM8228633.1 | Type1.BREX | TRD->REase+N6_METHYLASE->\|\|?->ParB->?-><-?<-SIGMA+OB-Fold+AAA-ATPase+Lon-Protease-BrxL<-PglZ<-N6-METHYLASE<-DUF6079*<-BrxB-BrxF-iSTAND<-BrxA-DNA-binding\|\|REase+iREase-><-wHTH+SAD+HNH\|\|HNH-><-?<-?<-?\|\|Phage_integrase-> | 1213 | Bacteria>Pseudomonadota>Gammaproteobacteria>Enterobacterales>Enterobacteriaceae>Escherichia | GCA_021429145.1 | Escherichia_coli | LOW06_003442 | 20140..23781 | 562 | BREX-system-P-loop-protein-BrxC-[Escherichia-coli] |
| HBN0250039.1 | Type1.BREX | <-DUF932<-ArdB<-?\|\|?->XIS-HTH->ZnR->wHTH+WYL+WCX->BrxA-DNA-binding->BrxB-BrxF-iSTAND->DUF6079*->ParB+HNH->N6-METHYLASE->PglZ->SIGMA+OB-Fold+AAA-ATPase+Lon-Protease-BrxL->35exo->DUF772+DDE_Tnp_1->OBFOLD-NUCLEASE-> | 1225 | Bacteria>Pseudomonadota>Gammaproteobacteria>Enterobacterales>Enterobacteriaceae>Escherichia | GCA_021454405.1 | Escherichia_coli | L1L95_003624 | 57124..60801 | 562 | BREX-system-P-loop-protein-BrxC-[Escherichia-coli] |
| HBN0573063.1 | Type1.BREX | Arm-DNA-bind_3->DUF6216-><-?<-AbiEii<-AbiEi_3_N+AbiEi_3\|\|<-HKD-DNase+OGG-like<-SIGMA+OB-Fold+AAA-ATPase+Lon-Protease-BrxL<-PglZ<-N6-METHYLASE<-DUF6079*<-BrxB-BrxF-iSTAND<-BrxA-DNA-binding\|\|REase+iREase-><-wHTH+SAD+HNH\|\|HNH-><-?<-?<-?\|\|Phage_integrase-> | 1213 | Bacteria>Pseudomonadota>Gammaproteobacteria>Enterobacterales>Enterobacteriaceae>Escherichia | GCA_021454505.1 | Escherichia_coli | L1N12_002093 | 35071..38712 | 562 | BREX-system-P-loop-protein-BrxC-[Escherichia-coli] |
| HBN0877260.1 | Type1.BREX | <-XIS-HTH\|\|?->?->ZnR->ZnR-><-SIGMA+OB-Fold+AAA-ATPase+Lon-Protease-BrxL<-PglZ<-N6-METHYLASE<-DUF6079*<-BrxB-BrxF-iSTAND<-BrxA-DNA-binding\|\|REase+iREase-> | 1213 | Bacteria>Pseudomonadota>Gammaproteobacteria>Enterobacterales>Enterobacteriaceae>Escherichia | GCA_021468605.1 | Escherichia_coli | L1V71_000763 | 365927..369568 | 562 | BREX-system-P-loop-protein-BrxC-[Escherichia-coli] |
| HBN1947859.1 | Type1.BREX | BrxA-DNA-binding->BrxB-BrxF-iSTAND->DUF6079*->N6-METHYLASE->PglZ->SIGMA+OB-Fold+AAA-ATPase+Lon-Protease-BrxL-> | 1213 | Bacteria>Pseudomonadota>Gammaproteobacteria>Enterobacterales>Enterobacteriaceae>Escherichia | GCA_021489855.1 | Escherichia_coli | L1401_003657 | 2289..5930 | 562 | BREX-system-P-loop-protein-BrxC-[Escherichia-coli] |
| HBP9919082.1 | Type1.BREX | <-Phage_integrase<-Phage_integrase\|\|?->?->?-><-HNH\|\|wHTH(SBD)+HNH->\|\|BrxA-DNA-binding->BrxB-BrxF-iSTAND->DUF6079*->N6-METHYLASE->PglZ->SIGMA+OB-Fold+AAA-ATPase+Lon-Protease-BrxL->HKD-DNase+OGG-like->\|\|AbiEi_3_N+AbiEi_3->AbiEii->?-><-DUF6216<-Arm-DNA-bind_3 | 1213 | Bacteria>Pseudomonadota>Gammaproteobacteria>Enterobacterales>Enterobacteriaceae>Escherichia | GCA_021766545.1 | Escherichia_coli | L7G70_002290 | 14696..18337 | 562 | BREX-system-P-loop-protein-BrxC-[Escherichia-coli] |
| HBQ4241594.1 | Type1.BREX | <-N6-METHYLASE<-DUF6079*<-BrxB-BrxF-iSTAND<-BrxA-DNA-binding\|\|REase+iREase-><-EVE+HNH<-Arm-DNA-bind_3+Phage_integrase<-tRNA\|\|?-><-SKI\|\|METHYLASE-> | 1213 | Bacteria>Pseudomonadota>Gammaproteobacteria>Enterobacterales>Enterobacteriaceae>Escherichia | GCA_021857785.1 | Escherichia_coli | L7222_003680 | 2636..6277 | 562 | BREX-system-P-loop-protein-BrxC-[Escherichia-coli] |
| HBV0129404.1 | Type1.BREX | <-DUF6079-p*<-BrxB-BrxF-iSTAND<-BrxA-DNA-binding\|\|REase+iREase->?->?->TetR-HTH-><-AAA-ATPase<-HIN-HTH<-AraC-HTH | 804 | Bacteria>Pseudomonadota>Gammaproteobacteria>Enterobacterales>Enterobacteriaceae>Escherichia | GCA_022118015.1 | Escherichia_coli | L4R27_004066 | 1..2413 | 562 | BREX-system-P-loop-protein-BrxC-[Escherichia-coli] |
| EIW5817588.1 | Type1.BREX | <-Phage_integrase\|\|?->?->?-><-HNH\|\|wHTH+SAD+HNH-><-REase+iREase\|\|BrxA-DNA-binding->BrxB-BrxF-iSTAND->DUF6079*->N6-METHYLASE->PglZ->SIGMA+OB-Fold+AAA-ATPase+Lon-Protease-BrxL->?-><-?<-ParB<-?\|\|<-REase+N6_METHYLASE<-TRD | 1213 | Bacteria>Pseudomonadota>Gammaproteobacteria>Enterobacterales>Enterobacteriaceae>Escherichia | GCA_022185325.1 | Escherichia_coli | ME061_001720 | 111299..114940 | 562 | BREX-system-P-loop-protein-BrxC-[Escherichia-coli] |
| EIX9838476.1 | Type1.BREX | DUF6079*->N6-METHYLASE->PglZ->SIGMA+OB-Fold+AAA-ATPase+Lon-Protease-BrxL->HKD-DNase+OGG-like->\|\|AbiEi_3_N+AbiEi_3->AbiEii->?-><-DUF6216<-Arm-DNA-bind_3 | 1208 | Bacteria>Pseudomonadota>Gammaproteobacteria>Enterobacterales>Enterobacteriaceae>Escherichia | GCA_022433775.1 | Escherichia_coli | ML439_002959 | 1..3628 | 562 | BREX-system-P-loop-protein-BrxC-[Escherichia-coli] |
| EIZ4505608.1 | Type1.BREX | <-XIS-HTH<-RHH_antitoxin\|\|?->?->ZnR->ZnR-><-SIGMA+OB-Fold+AAA-ATPase+Lon-Protease-BrxL<-PglZ<-N6-METHYLASE<-DUF6079*<-BrxB-BrxF-iSTAND<-BrxA-DNA-binding\|\|REase+iREase->?->?->TetR-HTH-> | 1213 | Bacteria>Pseudomonadota>Gammaproteobacteria>Enterobacterales>Enterobacteriaceae>Escherichia | GCA_022636095.1 | Escherichia_coli | MPU38_003973 | 70705..74346 | 562 | BREX-system-P-loop-protein-BrxC-[Escherichia-coli] |
| EJA4563483.1 | Type1.BREX | <-Phage_integrase\|\|?->?->?-><-HNH\|\|wHTH+SAD+HNH-><-REase+iREase\|\|BrxA-DNA-binding->BrxB-BrxF-iSTAND->DUF6079*->N6-METHYLASE->PglZ->SIGMA+OB-Fold+AAA-ATPase+Lon-Protease-BrxL->HKD-DNase+OGG-like->\|\|AbiEi_3_N+AbiEi_3->AbiEii->?-><-DUF6216<-Arm-DNA-bind_3 | 1213 | Bacteria>Pseudomonadota>Gammaproteobacteria>Enterobacterales>Enterobacteriaceae>Escherichia | GCA_022888875.1 | Escherichia_coli | MVR17_003830 | 10074..13715 | 562 | BREX-system-P-loop-protein-BrxC-[Escherichia-coli] |
| EJB1753906.1 | Type1.BREX | ZnR->ZnR-><-SIGMA+OB-Fold+AAA-ATPase+Lon-Protease-BrxL<-PglZ<-N6-METHYLASE<-DUF6079*<-BrxB-BrxF-iSTAND<-BrxA-DNA-binding\|\|REase+iREase-><-EVE+HNH<-Arm-DNA-bind_3+Phage_integrase<-tRNA\|\|?->HerA/FtsK+A32-> | 1213 | Bacteria>Pseudomonadota>Gammaproteobacteria>Enterobacterales>Enterobacteriaceae>Escherichia | GCA_022927665.1 | Escherichia_coli | MWZ83_002631 | 9550..13191 | 562 | BREX-system-P-loop-protein-BrxC-[Escherichia-coli] |
| EJB4937501.1 | Type1.BREX | <-XIS-HTH<-RHH_antitoxin\|\|?->?->ZnR->ZnR-><-SIGMA+OB-Fold+AAA-ATPase+Lon-Protease-BrxL<-PglZ<-N6-METHYLASE<-DUF6079*<-BrxB-BrxF-iSTAND<-BrxA-DNA-binding\|\|REase+iREase->?->?->TetR-HTH-><-AAA-ATPase | 1213 | Bacteria>Pseudomonadota>Gammaproteobacteria>Enterobacterales>Enterobacteriaceae>Escherichia | GCA_022947155.1 | Escherichia_coli | MTZ30_000457 | 137240..140881 | 562 | BREX-system-P-loop-protein-BrxC-[Escherichia-coli] |
| HCB2140406.1 | Type1.BREX | <-Zn_Tnp_IS91+Y2_Tnp\|\|<-?<-Phage_integrase\|\|Arm-DNA-bind_3+Phage_integrase->BrxA-DNA-binding->BrxB-BrxF-iSTAND->DUF6079*->N6-METHYLASE-> | 1204 | Bacteria>Pseudomonadota>Gammaproteobacteria>Enterobacterales>Enterobacteriaceae>Escherichia | GCA_023036775.1 | Escherichia_coli | MYQ63_004000 | 17767..21381 | 562 | BREX-system-P-loop-protein-BrxC-[Escherichia-coli] |
| EJC4972778.1 | Type1.BREX | <-XIS-HTH<-RHH_antitoxin\|\|?->?->ZnR->ZnR-><-SIGMA+OB-Fold+AAA-ATPase+Lon-Protease-BrxL<-PglZ<-N6-METHYLASE<-DUF6079*<-BrxB-BrxF-iSTAND<-BrxA-DNA-binding\|\|Zn_Tnp_IS1+HTH_Tnp_IS1-> | 1213 | Bacteria>Pseudomonadota>Gammaproteobacteria>Enterobacterales>Enterobacteriaceae>Escherichia | GCA_023055825.1 | Escherichia_coli | MZE40_000805 | 401924..405565 | 562 | BREX-system-P-loop-protein-BrxC-[Escherichia-coli] |
| MCK3244657.1 | Type1.BREX | <-HIN-HTH<-RelE-ParE<-CopG/RHH\|\|<-RHH\|\|?->wHTH+WYL+WCX->BrxA-DNA-binding->BrxB-BrxF-iSTAND->DUF6079*->ParB+HNH-> | 1225 | Bacteria>Pseudomonadota>Gammaproteobacteria>Enterobacterales>Enterobacteriaceae>Escherichia | GCA_023098955.1 | Escherichia_coli | MZH07_24520 | 5688..9365 | 562 | BREX-system-P-loop-protein-BrxC-[Escherichia-coli] |
| EJD7925982.1 | Type1.BREX | <-REC+CITB-HTH<-HIN-HTH\|\|DDE_Tnp_1_5+DDE_Tnp_1->?-><-REase+iREase\|\|BrxA-DNA-binding->BrxB-BrxF-iSTAND->DUF6079*->N6-METHYLASE->PglZ->SIGMA+OB-Fold+AAA-ATPase+Lon-Protease-BrxL-><-ZnR<-ZnR<-?<-?\|\|RHH_antitoxin->XIS-HTH-> | 1213 | Bacteria>Pseudomonadota>Gammaproteobacteria>Enterobacterales>Enterobacteriaceae>Escherichia | GCA_023216415.1 | Escherichia_coli | M2Y98_004402 | 13269..16910 | 562 | BREX-system-P-loop-protein-BrxC-[Escherichia-coli] |
| EJE3980761.1 | Type1.BREX | <-RelE-ParE<-CopG/RHH\|\|<-RHH\|\|?->wHTH+WYL+WCX->BrxA-DNA-binding->BrxB-BrxF-iSTAND->DUF6079*->ParB+HNH->N6-METHYLASE->PglZ->SIGMA+OB-Fold+AAA-ATPase+Lon-Protease-BrxL-><-DEDD_Tnp_IS110+Transposase_20<-RVT | 1225 | Bacteria>Pseudomonadota>Gammaproteobacteria>Enterobacterales>Enterobacteriaceae>Escherichia | GCA_023310495.1 | Escherichia_coli | M4B16_004238 | 5667..9344 | 562 | BREX-system-P-loop-protein-BrxC-[Escherichia-coli] |
| EJE7557557.1 | Type1.BREX | <-DUF6079-p*<-BrxB-BrxF-iSTAND<-BrxA-DNA-binding\|\|REase+iREase->?->?->TetR-HTH-><-AAA-ATPase<-HIN-HTH | 625 | Bacteria>Pseudomonadota>Gammaproteobacteria>Enterobacterales>Enterobacteriaceae>Escherichia | GCA_023344655.1 | Escherichia_coli | M5O73_003716 | 1..1876 | 562 | BREX-system-P-loop-protein-BrxC-[Escherichia-coli] |
| EJE7788338.1 | Type1.BREX | <-Phage_integrase\|\|?->?->?-><-HNH\|\|wHTH+SAD+HNH-><-REase+iREase\|\|BrxA-DNA-binding->BrxB-BrxF-iSTAND->DUF6079-p*-> | 859 | Bacteria>Pseudomonadota>Gammaproteobacteria>Enterobacterales>Enterobacteriaceae>Escherichia | GCA_023345825.1 | Escherichia_coli | M5R54_003441 | 111186..113763 | 562 | BREX-system-P-loop-protein-BrxC-[Escherichia-coli] |
| EJH0789763.1 | Type1.BREX | DUF6079*->N6-METHYLASE->PglZ->SIGMA+OB-Fold+AAA-ATPase+Lon-Protease-BrxL->HKD-DNase+OGG-like->\|\|AbiEi_3_N+AbiEi_3->AbiEii->?-><-DUF6216<-Arm-DNA-bind_3 | 1107 | Bacteria>Pseudomonadota>Gammaproteobacteria>Enterobacterales>Enterobacteriaceae>Escherichia | GCA_023637565.1 | Escherichia_coli | NCI57_002418 | 1..3324 | 562 | BREX-system-P-loop-protein-BrxC-[Escherichia-coli] |
| MCM4369403.1 | Type1.BREX | <-HerA/FtsK+A32<-?\|\|tRNA->Arm-DNA-bind_3+Phage_integrase->EVE+HNH-><-REase+iREase\|\|BrxA-DNA-binding->BrxB-BrxF-iSTAND->DUF6079*->N6-METHYLASE->PglZ->SIGMA+OB-Fold+AAA-ATPase+Lon-Protease-BrxL-><-ZnR<-ZnR<-?\|\|?-><-?\|\|XIS-HTH-> | 1213 | Bacteria>Pseudomonadota>Gammaproteobacteria>Enterobacterales>Enterobacteriaceae>Escherichia | GCA_023723445.1 | Escherichia_coli | MMS14_20555 | 25622..29263 | 562 | BREX-system-P-loop-protein-BrxC-[Escherichia-coli] |
| EJH6809378.1 | Type1.BREX | <-DUF6079-p*<-BrxB-BrxF-iSTAND<-BrxA-DNA-binding\|\|REase+iREase-><-wHTH+SAD+HNH\|\|HNH-> | 851 | Bacteria>Pseudomonadota>Gammaproteobacteria>Enterobacterales>Enterobacteriaceae>Escherichia | GCA_023728085.1 | Escherichia_coli | NFZ65_004801 | 1..2554 | 562 | BREX-system-P-loop-protein-BrxC-[Escherichia-coli] |
| EJH8598536.1 | Type1.BREX | <-XIS-HTH<-RHH_antitoxin\|\|?->?->ZnR->ZnR-><-SIGMA+OB-Fold+AAA-ATPase+Lon-Protease-BrxL<-PglZ<-N6-METHYLASE<-DUF6079*<-BrxB-BrxF-iSTAND<-BrxA-DNA-binding\|\|REase+iREase-><-EVE+HNH<-Arm-DNA-bind_3+Phage_integrase<-tRNA\|\|?->HerA/FtsK+A32-> | 1213 | Bacteria>Pseudomonadota>Gammaproteobacteria>Enterobacterales>Enterobacteriaceae>Escherichia | GCA_023738335.1 | Escherichia_coli | NF498_001054 | 22625..26266 | 562 | BREX-system-P-loop-protein-BrxC-[Escherichia-coli] |
| HCE0292887.1 | Type1.BREX | <-REase+iREase\|\|BrxA-DNA-binding->BrxB-BrxF-iSTAND->DUF6079*->N6-METHYLASE->PglZ->SIGMA+OB-Fold+AAA-ATPase+Lon-Protease-BrxL-><-ZnR<-XIS-HTH<-?\|\|?-><-?\|\|CopG/RHH-> | 1213 | Bacteria>Pseudomonadota>Gammaproteobacteria>Enterobacterales>Enterobacteriaceae>Escherichia | GCA_023779205.1 | Escherichia_coli | NEO28_002917 | 2558..6199 | 562 | BREX-system-P-loop-protein-BrxC-[Escherichia-coli] |
| EJI6887231.1 | Type1.BREX | <-REase+iREase\|\|BrxA-DNA-binding->BrxB-BrxF-iSTAND->DUF6079*->N6-METHYLASE->PglZ->SIGMA+OB-Fold+AAA-ATPase+Lon-Protease-BrxL-><-ZnR<-XIS-HTH | 1213 | Bacteria>Pseudomonadota>Gammaproteobacteria>Enterobacterales>Enterobacteriaceae>Escherichia | GCA_023803175.1 | Escherichia_coli | NFV42_004498 | 5105..8746 | 562 | BREX-system-P-loop-protein-BrxC-[Escherichia-coli] |
| HCE4109215.1 | Type1.BREX | <-XIS-HTH\|\|?->?->ZnR->ZnR-><-SIGMA+OB-Fold+AAA-ATPase+Lon-Protease-BrxL<-PglZ<-N6-METHYLASE<-DUF6079* | 1120 | Bacteria>Pseudomonadota>Gammaproteobacteria>Enterobacterales>Enterobacteriaceae>Escherichia | GCA_023826315.1 | Escherichia_coli | NES30_001638 | 158765..162129 | 562 | BREX-system-P-loop-protein-BrxC-[Escherichia-coli] |
| MCN5345816.1 | Type1.BREX | <-DUF932<-nSTAND1+GTPase-AIG+sGTP<-?\|\|?->XIS-HTH-><-?<-ZnR\|\|BrxA-DNA-binding->BrxB-BrxF-iSTAND->DUF6079*->N6-METHYLASE->OLD-ABC->TOPRIM->PglZ->SIGMA+OB-Fold+AAA-ATPase+Lon-Protease-BrxL->Zn_Tnp_IS1+HTH_Tnp_IS1-> | 1204 | Bacteria>Pseudomonadota>Gammaproteobacteria>Enterobacterales>Enterobacteriaceae>Escherichia | GCA_023869305.1 | Escherichia_coli | MLU97_13810 | 69828..73442 | 562 | BREX-system-P-loop-protein-BrxC-[Escherichia-coli] |
| EJM6441909.1 | Type1.BREX | DUF6079*->N6-METHYLASE->PglZ->SIGMA+OB-Fold+AAA-ATPase+Lon-Protease-BrxL->HKD-DNase+OGG-like->\|\|AbiEi_3_N+AbiEi_3->AbiEii->?-><-DUF6216<-Arm-DNA-bind_3 | 1006 | Bacteria>Pseudomonadota>Gammaproteobacteria>Enterobacterales>Enterobacteriaceae>Escherichia | GCA_024340725.1 | Escherichia_coli | NOR68_003993 | 1..3023 | 562 | BREX-system-P-loop-protein-BrxC-[Escherichia-coli] |
| EJN1844348.1 | Type1.BREX | <-RHH\|\|?->wHTH+WYL+WCX->BrxA-DNA-binding->BrxB-BrxF-iSTAND->DUF6079*->ParB+HNH->N6-METHYLASE->PglZ->SIGMA+OB-Fold+AAA-ATPase+Lon-Protease-BrxL-><-DEDD_Tnp_IS110+Transposase_20 | 1225 | Bacteria>Pseudomonadota>Gammaproteobacteria>Enterobacterales>Enterobacteriaceae>Escherichia | GCA_024422675.1 | Escherichia_coli | NPJ45_004376 | 3256..6933 | 562 | BREX-system-P-loop-protein-BrxC-[Escherichia-coli] |
| MCQ4517261.1 | Type1.BREX | DUF6216-><-?<-AbiEii<-AbiEi_3_N+AbiEi_3\|\|<-HKD-DNase+OGG-like<-SIGMA+OB-Fold+AAA-ATPase+Lon-Protease-BrxL<-PglZ<-N6-METHYLASE<-DUF6079*<-BrxB-BrxF-iSTAND<-BrxA-DNA-binding\|\|REase+iREase-> | 1213 | Bacteria>Pseudomonadota>Gammaproteobacteria>Enterobacterales>Enterobacteriaceae>Escherichia | GCA_024450325.1 | Escherichia_coli | NO030_20220 | 28424..32065 | 562 | BREX-system-P-loop-protein-BrxC-[Escherichia-coli] |
| HCJ2602063.1 | Type1.BREX | <-TetR-HTH<-?<-?<-REase+iREase\|\|BrxA-DNA-binding->BrxB-BrxF-iSTAND->DUF6079*->N6-METHYLASE->PglZ->SIGMA+OB-Fold+AAA-ATPase+Lon-Protease-BrxL-> | 1213 | Bacteria>Pseudomonadota>Gammaproteobacteria>Enterobacterales>Enterobacteriaceae>Escherichia | GCA_024493375.1 | Escherichia_coli | NQS58_004405 | 6692..10333 | 562 | BREX-system-P-loop-protein-BrxC-[Escherichia-coli] |
| MCQ6644900.1 | Type1.BREX | <-CbeA_antitoxin<-?<-?<-DUF932<-nSTAND1+sGTP<-?<-XIS-HTH\|\|\|\|Zn_Tnp_IS1+HTH_Tnp_IS1+DDE_Tnp_IS1-><-N6-METHYLASE-p<-DUF6079*<-BrxB-BrxF-iSTAND<-BrxA-DNA-binding\|\|REase+iREase-><-Zn_Tnp_IS1+HTH_Tnp_IS1+DDE_Tnp_IS1\|\|HIN-HTH->?->?->?->PhdYeFM_antitox-> | 1213 | Bacteria>Pseudomonadota>Gammaproteobacteria>Enterobacterales>Enterobacteriaceae>Escherichia | GCA_024496475.1 | Escherichia_coli | NO371_23560 | 27231..30872 | 562 | BREX-system-P-loop-protein-BrxC-[Escherichia-coli] |
| EJN8021038.1 | Type1.BREX | <-REase+iREase\|\|BrxA-DNA-binding->BrxB-BrxF-iSTAND->DUF6079*->N6-METHYLASE->PglZ->SIGMA+OB-Fold+AAA-ATPase+Lon-Protease-BrxL->HKD-DNase+OGG-like->\|\|AbiEi_3_N+AbiEi_3->AbiEii->?-><-DUF6216<-Arm-DNA-bind_3 | 1213 | Bacteria>Pseudomonadota>Gammaproteobacteria>Enterobacterales>Enterobacteriaceae>Escherichia | GCA_024524615.1 | Escherichia_coli | NRW59_000779 | 2207..5848 | 562 | BREX-system-P-loop-protein-BrxC-[Escherichia-coli] |
| EJO9498073.1 | Type1.BREX | <-HNH\|\|wHTH+SAD+HNH-><-REase+iREase\|\|BrxA-DNA-binding->BrxB-BrxF-iSTAND->DUF6079*->N6-METHYLASE->PglZ->SIGMA+OB-Fold+AAA-ATPase+Lon-Protease-BrxL->HKD-DNase+OGG-like->\|\|AbiEi_3_N+AbiEi_3->AbiEii->?-><-DUF6216<-Arm-DNA-bind_3 | 1213 | Bacteria>Pseudomonadota>Gammaproteobacteria>Enterobacterales>Enterobacteriaceae>Escherichia | GCA_024612475.1 | Escherichia_coli | NUZ09_002875 | 7894..11535 | 562 | BREX-system-P-loop-protein-BrxC-[Escherichia-coli] |
| HCJ9453756.1 | Type1.BREX | <-DDE_Tnp_IS240<-?\|\|XIS-HTH->PIWI-><-?<-Phage_integrase\|\|Arm-DNA-bind_3+Phage_integrase->BrxA-DNA-binding->BrxB-BrxF-iSTAND->DUF6079*->N6-METHYLASE->OLD-ABC->TOPRIM->PglZ->SIGMA+OB-Fold+AAA-ATPase+Lon-Protease-BrxL-> | 1204 | Bacteria>Pseudomonadota>Gammaproteobacteria>Enterobacterales>Enterobacteriaceae>Escherichia | GCA_024682335.1 | Escherichia_coli | NV934_004004 | 9250..12864 | 562 | BREX-system-P-loop-protein-BrxC-[Escherichia-coli] |
| HCJ9689412.1 | Type1.BREX | <-SWIM+DUF3670+SF1-SF2-Helicase<-?<-?<-?\|\|BrxA-DNA-binding->BrxB-BrxF-iSTAND->DUF6079*->N6-METHYLASE->PglZ->SIGMA+OB-Fold+AAA-ATPase+Lon-Protease-BrxL-> | 1213 | Bacteria>Pseudomonadota>Gammaproteobacteria>Enterobacterales>Enterobacteriaceae>Escherichia | GCA_024685835.1 | Escherichia_coli | NV971_003963 | 13318..16959 | 562 | BREX-system-P-loop-protein-BrxC-[Escherichia-coli] |
| MCS2047788.1 | Type1.BREX | <-GntR-HTH+TrmB+ZDNA-HTH<-?<-?\|\|?->?-><-?<-Fimbrial<-Fimbrial<-Zn_Tnp_IS1+HTH_Tnp_IS1+DDE_Tnp_IS1\|\|DUF6079*->N6-METHYLASE->PglZ->SIGMA+OB-Fold+AAA-ATPase+Lon-Protease-BrxL->?-><-?<-ParB<-?\|\|<-REase+N6_METHYLASE | 1213 | Bacteria>Pseudomonadota>Gammaproteobacteria>Enterobacterales>Enterobacteriaceae>Escherichia | GCA_024768515.1 | Escherichia_coli | MWK73_15380 | 3165783..3169424 | 562 | BREX-system-P-loop-protein-BrxC-[Escherichia-coli] |
| EJQ5102253.1 | Type1.BREX | DUF6079-p*->N6-METHYLASE->PglZ->SIGMA+OB-Fold+AAA-ATPase+Lon-Protease-BrxL-><-ZnR<-ZnR<-?\|\|HIN-HTH+HIN-HTH-> | 973 | Bacteria>Pseudomonadota>Gammaproteobacteria>Enterobacterales>Enterobacteriaceae>Escherichia | GCA_024841725.1 | Escherichia_coli | N0K27_004784 | 1..2924 | 562 | BREX-system-P-loop-protein-BrxC-[Escherichia-coli] |
| EJQ6147017.1 | Type1.BREX | <-XIS-HTH<-RHH_antitoxin\|\|?->?->ZnR->ZnR-><-SIGMA+OB-Fold+AAA-ATPase+Lon-Protease-BrxL<-PglZ<-N6-METHYLASE<-DUF6079*<-BrxB-BrxF-iSTAND<-BrxA-DNA-binding\|\|REase+iREase-><-EVE+HNH<-Arm-DNA-bind_3+Phage_integrase<-tRNA\|\|?->HerA/FtsK+A32-> | 1213 | Bacteria>Pseudomonadota>Gammaproteobacteria>Enterobacterales>Enterobacteriaceae>Escherichia | GCA_024846805.1 | Escherichia_albertii | NZZ21_002663 | 93437..97078 | 208962 | BREX-system-P-loop-protein-BrxC-[Escherichia-albertii] |
| HCL8518718.1 | Type1.BREX | <-Phage_integrase\|\|?->?->?-><-HNH\|\|wHTH+SAD+HNH-><-REase+iREase\|\|BrxA-DNA-binding->BrxB-BrxF-iSTAND->DUF6079-p*-> | 605 | Bacteria>Pseudomonadota>Gammaproteobacteria>Enterobacterales>Enterobacteriaceae>Escherichia | GCA_025048635.1 | Escherichia_coli | N2025_002882 | 80925..82740 | 562 | BREX-system-P-loop-protein-BrxC-[Escherichia-coli] |
| EJT7594528.1 | Type1.BREX | <-N6-METHYLASE<-DUF6079*<-BrxB-BrxF-iSTAND<-BrxA-DNA-binding\|\|REase+iREase-><-wHTH+SAD+HNH\|\|HNH-><-?<-?<-?\|\|Phage_integrase-> | 1213 | Bacteria>Pseudomonadota>Gammaproteobacteria>Enterobacterales>Enterobacteriaceae>Escherichia | GCA_025102805.1 | Escherichia_coli | N3T17_004670 | 1985..5626 | 562 | BREX-system-P-loop-protein-BrxC-[Escherichia-coli] |
| EJU9647232.1 | Type1.BREX | <-DDE_Tnp_1_5+DDE_Tnp_1\|\|\|\|wHTH+WYL+WCX->BrxA-DNA-binding->BrxB-BrxF-iSTAND->DUF6079*->ParB+HNH->N6-METHYLASE->PglZ->SIGMA+OB-Fold+AAA-ATPase+Lon-Protease-BrxL-><-HTH_Tnp_1+HTH_21+rve+rve_3<-HIN-HTH<-PIN<-MazE_antitoxin | 1225 | Bacteria>Pseudomonadota>Gammaproteobacteria>Enterobacterales>Enterobacteriaceae>Escherichia | GCA_025224895.1 | Escherichia_coli | N5N25_003986 | 5462..9139 | 562 | BREX-system-P-loop-protein-BrxC-[Escherichia-coli] |
| HCN4580231.1 | Type1.BREX | <-Phage_integrase\|\|?->?->?-><-HNH\|\|wHTH+SAD+HNH-><-REase+iREase\|\|BrxA-DNA-binding->BrxB-BrxF-iSTAND->DUF6079*->wHTH+WYL | 1213 | Bacteria>Pseudomonadota>Gammaproteobacteria>Enterobacterales>Enterobacteriaceae>Escherichia | GCA_025275895.1 | Escherichia_coli | N6V60_002026 | 111191..114832 | 562 | BREX-system-P-loop-protein-BrxC-[Escherichia-coli] |
| HCN4662824.1 | Type1.BREX | <-REase+iREase\|\|BrxA-DNA-binding->BrxB-BrxF-iSTAND->DUF6079*->N6-METHYLASE->PglZ->SIGMA+OB-Fold+AAA-ATPase+Lon-Protease-BrxL->HKD-DNase+OGG-like->\|\|AbiEi_3_N+AbiEi_3->AbiEii->?-><-DUF6216<-Arm-DNA-bind_3 | 1213 | Bacteria>Pseudomonadota>Gammaproteobacteria>Enterobacterales>Enterobacteriaceae>Escherichia | GCA_025276155.1 | Escherichia_coli | N6W02_002919 | 2176..5817 | 562 | BREX-system-P-loop-protein-BrxC-[Escherichia-coli] |
| HCN5442922.1 | Type1.BREX | DUF6079*->N6-METHYLASE->PglZ->SIGMA+OB-Fold+AAA-ATPase+Lon-Protease-BrxL->?-><-?<-ParB<-?\|\|<-REase+N6_METHYLASE<-TRD | 1088 | Bacteria>Pseudomonadota>Gammaproteobacteria>Enterobacterales>Enterobacteriaceae>Escherichia | GCA_025279415.1 | Escherichia_coli | N6X60_000278 | 1..3269 | 562 | BREX-system-P-loop-protein-BrxC-[Escherichia-coli] |
| HCN5870003.1 | Type1.BREX | DUF6079*->N6-METHYLASE->PglZ->SIGMA+OB-Fold+AAA-ATPase+Lon-Protease-BrxL-><-ZnR<-ZnR<-?\|\|RHH_antitoxin->XIS-HTH-> | 1016 | Bacteria>Pseudomonadota>Gammaproteobacteria>Enterobacterales>Enterobacteriaceae>Escherichia | GCA_025281175.1 | Escherichia_coli | N6Y80_004176 | 1..3051 | 562 | BREX-system-P-loop-protein-BrxC-[Escherichia-coli] |
| HCN7834282.1 | Type1.BREX | <-N6-METHYLASE<-DUF6079*<-BrxB-BrxF-iSTAND<-BrxA-DNA-binding<-tRNA\|\|?-><-SKI\|\|METHYLASE->?->?->LACI-HTH+PBPI->HerA/FtsK+A32-> | 1204 | Bacteria>Pseudomonadota>Gammaproteobacteria>Enterobacterales>Enterobacteriaceae>Escherichia | GCA_025289555.1 | Escherichia_coli | N6366_003464 | 2753..6367 | 562 | BREX-system-P-loop-protein-BrxC-[Escherichia-coli] |
| HCO1420187.1 | Type1.BREX | <-MAD-NTD+HNH\|\|wHTH+SAD+HNH-><-REase+iREase\|\|BrxA-DNA-binding->BrxB-BrxF-iSTAND->DUF6079*->N6-METHYLASE-> | 1213 | Bacteria>Pseudomonadota>Gammaproteobacteria>Enterobacterales>Enterobacteriaceae>Escherichia | GCA_025325225.1 | Escherichia_coli | N7L77_004122 | 7756..11397 | 562 | BREX-system-P-loop-protein-BrxC-[Escherichia-coli] |
| EJV7137460.1 | Type1.BREX | <-Phage_integrase\|\|?->?->?-><-HNH\|\|wHTH+SAD+HNH-><-REase+iREase\|\|BrxA-DNA-binding->BrxB-BrxF-iSTAND->DUF6079*->N6-METHYLASE->PglZ->SIGMA+OB-Fold+AAA-ATPase+Lon-Protease-BrxL->?-><-?<-ParB<-?\|\|<-REase+N6_METHYLASE<-TRD | 1213 | Bacteria>Pseudomonadota>Gammaproteobacteria>Enterobacterales>Enterobacteriaceae>Escherichia | GCA_025337425.1 | Escherichia_coli | FHP62_000281 | 290632..294273 | 562 | BREX-system-P-loop-protein-BrxC-[Escherichia-coli] |
| HCO5071633.1 | Type1.BREX | <-RelE-ParE<-CopG/RHH\|\|<-RHH\|\|?->wHTH+WYL+WCX->BrxA-DNA-binding->BrxB-BrxF-iSTAND->DUF6079*->ParB+HNH-> | 1225 | Bacteria>Pseudomonadota>Gammaproteobacteria>Enterobacterales>Enterobacteriaceae>Escherichia | GCA_025390635.1 | Escherichia_coli | N8039_004383 | 5665..9342 | 562 | BREX-system-P-loop-protein-BrxC-[Escherichia-coli] |
| HCO6583160.1 | Type1.BREX | <-Phage_integrase\|\|?->?->?-><-HNH\|\|wHTH+SAD+HNH-><-REase+iREase\|\|BrxA-DNA-binding->BrxB-BrxF-iSTAND->DUF6079*->N6-METHYLASE->PglZ->SIGMA+OB-Fold+AAA-ATPase+Lon-Protease-BrxL->HKD-DNase+OGG-like->\|\|AbiEi_3_N+AbiEi_3->AbiEii-><-DUF6216<-Arm-DNA-bind_3 | 1213 | Bacteria>Pseudomonadota>Gammaproteobacteria>Enterobacterales>Enterobacteriaceae>Escherichia | GCA_025412985.1 | Escherichia_coli | OBR12_001368 | 111262..114903 | 562 | BREX-system-P-loop-protein-BrxC-[Escherichia-coli] |
| HCO7023289.1 | Type1.BREX | <-Phage_integrase\|\|?->?->?-><-HNH\|\|wHTH+SAD+HNH-><-REase+iREase\|\|BrxA-DNA-binding->BrxB-BrxF-iSTAND->DUF6079-p*-> | 886 | Bacteria>Pseudomonadota>Gammaproteobacteria>Enterobacterales>Enterobacteriaceae>Escherichia | GCA_025415015.1 | Escherichia_coli | OBS61_002730 | 83196..85855 | 562 | BREX-system-P-loop-protein-BrxC-[Escherichia-coli] |
| EJW3212697.1 | Type1.BREX | <-DUF6079-p*<-BrxB-BrxF-iSTAND<-BrxA-DNA-binding\|\|<-wHTH(SBD)+HNH\|\|HNH-><-?<-?<-?\|\|Phage_integrase->Phage_integrase-> | 418 | Bacteria>Pseudomonadota>Gammaproteobacteria>Enterobacterales>Enterobacteriaceae>Escherichia | GCA_025418235.1 | Escherichia_coli | OBZ04_004372 | 1..1254 | 562 | BREX-system-P-loop-protein-BrxC-[Escherichia-coli] |
| HCP0312404.1 | Type1.BREX | <-METHYLASE\|\|SKI-><-?\|\|tRNA->Arm-DNA-bind_3+Phage_integrase->EVE+HNH-><-REase+iREase\|\|BrxA-DNA-binding->BrxB-BrxF-iSTAND->DUF6079*->N6-METHYLASE->PglZ->SIGMA+OB-Fold+AAA-ATPase+Lon-Protease-BrxL-><-ZnR<-ZnR<-?<-?\|\|XIS-HTH-> | 1213 | Bacteria>Pseudomonadota>Gammaproteobacteria>Enterobacterales>Enterobacteriaceae>Escherichia | GCA_025467455.1 | Escherichia_coli | OCM84_003824 | 150572..154213 | 562 | BREX-system-P-loop-protein-BrxC-[Escherichia-coli] |
| HCP1392420.1 | Type1.BREX | Arm-DNA-bind_3->DUF6216-><-AbiEii<-AbiEi_3_N+AbiEi_3\|\|<-HKD-DNase+OGG-like<-SIGMA+OB-Fold+AAA-ATPase+Lon-Protease-BrxL<-PglZ<-N6-METHYLASE<-DUF6079*<-BrxB-BrxF-iSTAND<-BrxA-DNA-binding\|\|REase+iREase-><-wHTH+SAD+HNH\|\|HNH-><-?<-?<-?\|\|Phage_integrase-> | 1213 | Bacteria>Pseudomonadota>Gammaproteobacteria>Enterobacterales>Enterobacteriaceae>Escherichia | GCA_025491965.1 | Escherichia_coli | OCX38_000210 | 218312..221953 | 562 | BREX-system-P-loop-protein-BrxC-[Escherichia-coli] |
| WP_250907975.1 | Type1.BREX | DDE_Tnp_IS1->?->ZnR->BrxA-DNA-binding->BrxB-BrxF-iSTAND->DUF6079*->ParB+HNH->N6-METHYLASE->HEPN->RVT->N6-METHYLASE-p->PglZ->SIGMA+OB-Fold+AAA-ATPase+Lon-Protease-BrxL-><-Arm-DNA-bind_3+Phage_integrase\|\|Phage_integrase-> | 1240 | Bacteria>Pseudomonadota>Gammaproteobacteria>Enterobacterales>Enterobacteriaceae>Escherichia | GCA_030820635.1 | Escherichia_coli | HUC40_RS15465 | 2924..6646 | 562 | BREX-system-P-loop-protein-BrxC-[Escherichia-coli] |
| CEK08497.1 | Type1.BREX | DUF6216-><-?<-AbiEii<-AbiEi_3_N+AbiEi_3<-HKD-DNase+OGG-like<-SIGMA+OB-Fold+AAA-ATPase+Lon-Protease-BrxL<-PglZ<-N6-METHYLASE<-N6-METHYLASE-p<-DUF6079*<-BrxB-BrxF-iSTAND<-BrxA-DNA-binding\|\|REase+iREase-><-wHTH+SAD+HNH<-?<-?<-?\|\|Phage_integrase-> | 1218 | Bacteria>Pseudomonadota>Gammaproteobacteria>Enterobacterales>Enterobacteriaceae>Escherichia | GCA_900000205.1 | Escherichia_coli_O26:H11 | ECO26H__800033 | 35711..39367 | 244319 | conserved-hypothetical-protein-[Escherichia-coli-O26:H11] |
| STH24241.1 | Type1.BREX | <-DUF772+DDE_Tnp_1\|\|?->DiS_P_DiS+Peptidase_A24->?-><-DDE_Tnp_IS1<-DUF6079-p*<-BrxB-BrxF-iSTAND<-BrxA-DNA-binding\|\|REase+iREase->?->?->?-><-DUF772+DDE_Tnp_1\|\|?->TetR-HTH-> | 778 | Bacteria>Pseudomonadota>Gammaproteobacteria>Enterobacterales>Enterobacteriaceae>Escherichia | GCA_900448945.1 | Escherichia_coli | NCTC11122_00780 | 798799..801135 | 562 | ATPase-like-protein-[Escherichia-coli] |
| STI20016.1 | Type1.BREX | <-Phage_integrase\|\|?->?->?->wHTH+SAD+HNH-><-REase+iREase\|\|BrxA-DNA-binding->BrxB-BrxF-iSTAND->DUF6079-p*->DUF6079-p->DUF6079-p->DUF6079-p->N6-METHYLASE-p->N6-METHYLASE-p->N6-METHYLASE-p->PglZ-> | 488 | Bacteria>Pseudomonadota>Gammaproteobacteria>Enterobacterales>Enterobacteriaceae>Escherichia | GCA_900448955.1 | Escherichia_coli | NCTC10865_05411 | 4206168..4207634 | 562 | ATPase-like-protein-[Escherichia-coli] |
| STN20139.1 | Type1.BREX | DDE_Tnp_1_5+DDE_Tnp_1-><-?\|\|HTH_21+rve+rve_3-><-RHH\|\|?->wHTH+WYL+WCX->BrxA-DNA-binding->BrxB-BrxF-iSTAND->DUF6079-p*->DUF6079-p*->DUF6079-p->?->DUF6079-p->ParB+HNH->ParB+HNH->N6-METHYLASE->?->PglZ-p-> | 353 | Bacteria>Pseudomonadota>Gammaproteobacteria>Enterobacterales>Enterobacteriaceae>Escherichia | GCA_900450565.1 | Escherichia_fergusonii | NCTC12128_00054 | 38373..39434 | 564 | ATPase-(modular-protein)-[Escherichia-fergusonii] |
| WP_001019654.1 | Type1.BREX | LexA-protease->\|\|Arm-DNA-bind_3+Phage_integrase->N6-METHYLASE->ORC-CDC6+ZnR->HEPN-><-Zn_Tnp_IS1+HTH_Tnp_IS1+DDE_Tnp_IS1<-REase\|\|BrxA-DNA-binding->BrxB-BrxF-iSTAND->DUF6079*->N6-METHYLASE->PglZ->SIGMA+OB-Fold+AAA-ATPase+Lon-Protease-BrxL-><-ZnR<-ZnR<-?\|\|HTH_Tnp_1+HTH_21+rve+rve_3-><-LuxR-HTH | 1213 | Bacteria>Pseudomonadota>Gammaproteobacteria>Enterobacterales>Enterobacteriaceae>Escherichia | GCF_000017765.1 | Escherichia_coli | ECHS_RS01710 | 342111..345752 | 562 | BREX-system-P-loop-protein-BrxC-[Escherichia-coli] |
| WP_001019642.1 | Type1.BREX | DUF6216-><-?<-AbiEii<-AbiEi_3_N+AbiEi_3\|\|<-HKD-DNase+OGG-like<-SIGMA+OB-Fold+AAA-ATPase+Lon-Protease-BrxL<-PglZ<-N6-METHYLASE\|\|HTH_Tnp_1+HTH_21+rve+rve_3-><-DUF6079*<-BrxB-BrxF-iSTAND<-BrxA-DNA-binding\|\|<-wHTH+SAD+HNH\|\|<-?<-?<-?\|\|Phage_integrase->Phage_integrase-> | 1188 | Bacteria>Pseudomonadota>Gammaproteobacteria>Enterobacterales>Enterobacteriaceae>Escherichia | GCF_000194355.1 | Escherichia_coli | ECJB195_RS15905 | 80241..83807 | 562 | BREX-system-P-loop-protein-BrxC-[Escherichia-coli] |
| WP_001019646.1 | Type1.BREX | REase+N6_METHYLASE-><-?\|\|?->ParB->?-><-?<-SIGMA+OB-Fold+AAA-ATPase+Lon-Protease-BrxL<-PglZ<-N6-METHYLASE<-DUF6079*<-BrxB-BrxF-iSTAND<-BrxA-DNA-binding\|\|REase+iREase-><-wHTH+SAD+HNH\|\|<-?<-?<-?\|\|Phage_integrase->Phage_integrase-> | 1213 | Bacteria>Pseudomonadota>Gammaproteobacteria>Enterobacterales>Enterobacteriaceae>Escherichia | GCF_000234235.1 | Escherichia_coli | KEM_RS0123875 | 120239..123880 | 562 | BREX-system-P-loop-protein-BrxC-[Escherichia-coli] |
| WP_001019647.1 | Type1.BREX | Arm-DNA-bind_3+Phage_integrase->N6-METHYLASE->ORC-CDC6+ZnR->HEPN-><-Zn_Tnp_IS1+HTH_Tnp_IS1+DDE_Tnp_IS1<-REase<-REase\|\|BrxA-DNA-binding->BrxB-BrxF-iSTAND->DUF6079*->N6-METHYLASE->PglZ->SIGMA+OB-Fold+AAA-ATPase+Lon-Protease-BrxL-><-ZnR<-ZnR<-?\|\|RHH_antitoxin->XIS-HTH-><-HTH_Tnp_1+HTH_21+rve+rve_3 | 1213 | Bacteria>Pseudomonadota>Gammaproteobacteria>Enterobacterales>Enterobacteriaceae>Escherichia | GCF_000249715.1 | Escherichia_coli | ECDEC7B_RS22125 | 12544..16185 | 562 | BREX-system-P-loop-protein-BrxC-[Escherichia-coli] |
| WP_001019640.1 | Type1.BREX | TRD->REase+N6_METHYLASE-><-?\|\|?->ParB->?-><-?<-SIGMA+OB-Fold+AAA-ATPase+Lon-Protease-BrxL<-PglZ<-DUF6079*<-BrxB-BrxF-iSTAND<-BrxA-DNA-binding\|\|REase+iREase-><-wHTH+SAD+HNH\|\|<-?<-?<-?\|\|Phage_integrase->Fimbrial-> | 1213 | Bacteria>Pseudomonadota>Gammaproteobacteria>Enterobacterales>Enterobacteriaceae>Escherichia | GCF_000250075.1 | Escherichia_coli | BE83_RS00870 | 326860..330501 | 562 | BREX-system-P-loop-protein-BrxC-[Escherichia-coli] |
| WP_001019639.1 | Type1.BREX | REase+N6_METHYLASE-><-?\|\|?->ParB->?-><-?<-SIGMA+OB-Fold+AAA-ATPase+Lon-Protease-BrxL<-PglZ<-DUF6079*<-BrxB-BrxF-iSTAND<-BrxA-DNA-binding\|\|REase+iREase-><-wHTH+SAD+HNH\|\|<-?<-?<-?\|\|Phage_integrase->Phage_integrase-> | 1213 | Bacteria>Pseudomonadota>Gammaproteobacteria>Enterobacterales>Enterobacteriaceae>Escherichia | GCF_000250475.1 | Escherichia_coli | BE27_RS00915 | 17908..21549 | 562 | BREX-system-P-loop-protein-BrxC-[Escherichia-coli] |
| WP_001019650.1 | Type1.BREX | wHTH->ZnR->ZnR->URI-DNase+RAMA-><-SIGMA+OB-Fold+AAA-ATPase+Lon-Protease-BrxL<-PglZ<-TOPRIM<-OLD-ABC<-N6-METHYLASE<-DUF6079*<-BrxB-BrxF-iSTAND<-BrxA-DNA-binding | 1204 | Bacteria>Pseudomonadota>Gammaproteobacteria>Enterobacterales>Enterobacteriaceae>Escherichia | GCF_000264195.1 | Escherichia_coli | ECKD2_RS09380 | 24128..27742 | 562 | BREX-system-P-loop-protein-BrxC-[Escherichia-coli] |
| WP_001019643.1 | Type1.BREX | ZnR-><-SIGMA+OB-Fold+AAA-ATPase+Lon-Protease-BrxL<-PglZ<-N6-METHYLASE<-DUF6079*<-BrxB-BrxF-iSTAND<-BrxA-DNA-binding\|\|REase->REase-> | 1213 | Bacteria>Pseudomonadota>Gammaproteobacteria>Enterobacterales>Enterobacteriaceae>Escherichia | GCF_000326345.1 | Escherichia_coli | WK3_RS01055 | 9311..12952 | 562 | BREX-system-P-loop-protein-BrxC-[Escherichia-coli] |

| Protein-ID | System | Gene-Neighbourhoods | Length | Taxonomy | Assembly-ID | Strain | Locus-Tag | Genomic-coordinates | TaxID | Fasta-define |
|---|---|---|---|---|---|---|---|---|---|---|
| WP_001698869.1 | Type1.BREX | RVT->DEDD_Tnp_IS110+Transposase_20-><-SIGMA+OB-Fold+AAA-ATPase+Lon-Protease-BrxL<-PglZ<-N6-METHYLASE<-ParB+HNH<-DUF6079*<-BrxB-BrxF-iSTAND<-BrxA-DNA-binding<-wHTH+WYL+WCX<-?\|\|RHH-><-DEDD_Tnp_IS110+Transposase_20<-?\|\|MerR-HTH+ParA-Soj-PloopNTPase->ParB-> | 1225 | Bacteria>Pseudomonadota>Gammaproteobacteria>Enterobacterales>Enterobacteriaceae>Escherichia | GCF_000340255.1 | Escherichia_coli | C201_RS01205 | 14823..18500 | 562 | BREX-system-P-loop-protein-BrxC-[Escherichia-coli] |
| WP_001704019.1 | Type1.BREX | <-HerA/FtsK+A32<-?\|\|tRNA->DUF6216-><-AbiEii<-AbiEi_3_N+AbiEi_3\|\|<-HKD-DNase+OGG-like<-SIGMA+OB-Fold+AAA-ATPase+Lon-Protease-BrxL<-PglZ<-N6-METHYLASE<-DUF6079*<-BrxB-BrxF-iSTAND<-BrxA-DNA-binding\|\|REase+iREase-><-wHTH+SAD+HNH\|\|<-?<-?<-?\|\|Phage_integrase-> | 1213 | Bacteria>Pseudomonadota>Gammaproteobacteria>Enterobacterales>Enterobacteriaceae>Escherichia | GCF_000354275.2 | Escherichia_coli | EC2875000_RS00745 | 314498..318139 | 562 | BREX-system-P-loop-protein-BrxC-[Escherichia-coli] |
| WP_001704755.1 | Type1.BREX | DUF6216-><-?<-AbiEii<-AbiEi_3_N+AbiEi_3\|\|<-HKD-DNase+OGG-like<-SIGMA+OB-Fold+AAA-ATPase+Lon-Protease-BrxL<-PglZ<-N6-METHYLASE<-DUF6079*<-BrxB-BrxF-iSTAND<-BrxA-DNA-binding\|\|REase+iREase-><-wHTH+SAD+HNH\|\|<-?<-?<-?\|\|Phage_integrase->Phage_integrase-> | 1213 | Bacteria>Pseudomonadota>Gammaproteobacteria>Enterobacterales>Enterobacteriaceae>Escherichia | GCF_000354455.1 | Escherichia_coli | EC2872800_RS00650 | 314338..317979 | 562 | BREX-system-P-loop-protein-BrxC-[Escherichia-coli] |
| WP_001707406.1 | Type1.BREX | DUF6216-><-?<-AbiEii<-AbiEi_3_N+AbiEi_3\|\|<-HKD-DNase+OGG-like<-SIGMA+OB-Fold+AAA-ATPase+Lon-Protease-BrxL<-PglZ<-N6-METHYLASE<-DUF6079*<-BrxB-BrxF-iSTAND<-BrxA-DNA-binding\|\|REase+iREase-><-wHTH+SAD+HNH\|\|<-?<-?<-?\|\|Phage_integrase->Phage_integrase-> | 1213 | Bacteria>Pseudomonadota>Gammaproteobacteria>Enterobacterales>Enterobacteriaceae>Escherichia | GCF_000354815.2 | Escherichia_coli | EC2867750_RS24725 | 314336..317977 | 562 | BREX-system-P-loop-protein-BrxC-[Escherichia-coli] |
| WP_004019909.1 | Type1.BREX | <-HerA/FtsK+A32<-?\|\|tRNA->\|\|<-REase+iREase\|\|BrxA-DNA-binding->BrxB-BrxF-iSTAND->DUF6079*->N6-METHYLASE->PglZ->SIGMA+OB-Fold+AAA-ATPase+Lon-Protease-BrxL-><-ZnR\|\|<-DEDD_Tnp_IS110+Transposase_20\|\|<-LACI-HTH+PBPI | 1213 | Bacteria>Pseudomonadota>Gammaproteobacteria>Enterobacterales>Enterobacteriaceae>Escherichia | GCF_000354915.1 | Escherichia_coli | EC174750_RS00780 | 7732..11373 | 562 | BREX-system-P-loop-protein-BrxC-[Escherichia-coli] |
| WP_003969750.1 | Type1.BREX | DUF6216-><-?<-AbiEii<-AbiEi_3_N+AbiEi_3\|\|<-HKD-DNase+OGG-like<-SIGMA+OB-Fold+AAA-ATPase+Lon-Protease-BrxL<-PglZ<-N6-METHYLASE<-DUF6079*<-BrxB-BrxF-iSTAND<-BrxA-DNA-binding\|\|REase+iREase-><-wHTH+SAD+HNH\|\|<-?<-?<-?\|\|Phage_integrase->Phage_integrase-> | 1213 | Bacteria>Pseudomonadota>Gammaproteobacteria>Enterobacterales>Enterobacteriaceae>Escherichia | GCF_000354935.2 | Escherichia_coli | EC2866450_RS25180 | 218299..221940 | 562 | BREX-system-P-loop-protein-BrxC-[Escherichia-coli] |
| WP_001709521.1 | Type1.BREX | DUF6216-><-?<-AbiEii<-AbiEi_3_N+AbiEi_3\|\|<-HKD-DNase+OGG-like<-SIGMA+OB-Fold+AAA-ATPase+Lon-Protease-BrxL<-PglZ<-N6-METHYLASE<-DUF6079*<-BrxB-BrxF-iSTAND<-BrxA-DNA-binding\|\|REase+iREase-><-wHTH+SAD+HNH\|\|<-?<-?<-?\|\|Phage_integrase->Phage_integrase-> | 1213 | Bacteria>Pseudomonadota>Gammaproteobacteria>Enterobacterales>Enterobacteriaceae>Escherichia | GCF_000354955.1 | Escherichia_coli | EC2866550_RS00550 | 55545..59186 | 562 | BREX-system-P-loop-protein-BrxC-[Escherichia-coli] |
| WP_001713779.1 | Type1.BREX | DUF6216-><-?<-AbiEii<-AbiEi_3_N+AbiEi_3\|\|<-HKD-DNase+OGG-like<-SIGMA+OB-Fold+AAA-ATPase+Lon-Protease-BrxL<-PglZ<-N6-METHYLASE<-DUF6079*<-BrxB-BrxF-iSTAND<-BrxA-DNA-binding\|\|REase+iREase-><-wHTH+SAD+HNH\|\|<-?<-?<-?\|\|Phage_integrase-> | 1213 | Bacteria>Pseudomonadota>Gammaproteobacteria>Enterobacterales>Enterobacteriaceae>Escherichia | GCF_000355015.2 | Escherichia_coli | EC2851500_RS25090 | 314221..317862 | 562 | BREX-system-P-loop-protein-BrxC-[Escherichia-coli] |
| WP_001715918.1 | Type1.BREX | DUF6216-><-?<-AbiEii<-AbiEi_3_N+AbiEi_3\|\|<-HKD-DNase+OGG-like<-SIGMA+OB-Fold+AAA-ATPase+Lon-Protease-BrxL<-PglZ<-N6-METHYLASE<-DUF6079*<-BrxB-BrxF-iSTAND<-BrxA-DNA-binding\|\|REase+iREase-><-wHTH+SAD+HNH\|\|<-?<-?<-?\|\|Phage_integrase->Phage_integrase-> | 1213 | Bacteria>Pseudomonadota>Gammaproteobacteria>Enterobacterales>Enterobacteriaceae>Escherichia | GCF_000355055.1 | Escherichia_coli | EC2850400_RS01425 | 314375..318016 | 562 | BREX-system-P-loop-protein-BrxC-[Escherichia-coli] |
| WP_001714681.1 | Type1.BREX | DUF6216-><-?<-AbiEii<-AbiEi_3_N+AbiEi_3\|\|<-HKD-DNase+OGG-like<-SIGMA+OB-Fold+AAA-ATPase+Lon-Protease-BrxL<-PglZ<-N6-METHYLASE<-DUF6079*<-BrxB-BrxF-iSTAND<-BrxA-DNA-binding\|\|REase+iREase-><-wHTH+SAD+HNH\|\|<-?<-?<-?\|\|Phage_integrase-> | 1213 | Bacteria>Pseudomonadota>Gammaproteobacteria>Enterobacterales>Enterobacteriaceae>Escherichia | GCF_000355335.2 | Escherichia_coli | EC2850750_RS25235 | 55686..59327 | 562 | BREX-system-P-loop-protein-BrxC-[Escherichia-coli] |
| WP_001462984.1 | Type1.BREX | DUF6216-><-?<-AbiEii<-AbiEi_3_N+AbiEi_3\|\|<-HKD-DNase+OGG-like<-SIGMA+OB-Fold+AAA-ATPase+Lon-Protease-BrxL<-PglZ<-N6-METHYLASE<-DUF6079*<-BrxB-BrxF-iSTAND<-BrxA-DNA-binding\|\|REase+iREase-><-wHTH+SAD+HNH\|\|<-?<-?<-?\|\|Phage_integrase->Phage_integrase-> | 1213 | Bacteria>Pseudomonadota>Gammaproteobacteria>Enterobacterales>Enterobacteriaceae>Escherichia | GCF_000357005.2 | Escherichia_coli | ECP02994383_RS24020 | 76495..80136 | 562 | BREX-system-P-loop-protein-BrxC-[Escherichia-coli] |
| WP_001464881.1 | Type1.BREX | DUF6216-><-?<-AbiEii<-AbiEi_3_N+AbiEi_3\|\|<-HKD-DNase+OGG-like<-SIGMA+OB-Fold+AAA-ATPase+Lon-Protease-BrxL<-PglZ<-N6-METHYLASE<-DUF6079*<-BrxB-BrxF-iSTAND<-BrxA-DNA-binding\|\|REase+iREase-><-wHTH+SAD+HNH\|\|<-?<-?<-?\|\|Phage_integrase->Phage_integrase-> | 1213 | Bacteria>Pseudomonadota>Gammaproteobacteria>Enterobacterales>Enterobacteriaceae>Escherichia | GCF_000357045.2 | Escherichia_coli | ECP02994386_RS23895 | 144466..148107 | 562 | BREX-system-P-loop-protein-BrxC-[Escherichia-coli] |
| WP_001466204.1 | Type1.BREX | <-Phage_integrase<-Phage_integrase\|\|?->?->?->\|\|wHTH+SAD+HNH-><-REase+iREase\|\|BrxA-DNA-binding->BrxB-BrxF-iSTAND->DUF6079*->N6-METHYLASE->PglZ->SIGMA+OB-Fold+AAA-ATPase+Lon-Protease-BrxL->HKD-DNase+OGG-like->\|\|AbiEi_3_N+AbiEi_3->AbiEii->?-><-DUF6216 | 1213 | Bacteria>Pseudomonadota>Gammaproteobacteria>Enterobacterales>Enterobacteriaceae>Escherichia | GCF_000357245.1 | Escherichia_coli | ECP02994387_RS14340 | 10081..13722 | 562 | BREX-system-P-loop-protein-BrxC-[Escherichia-coli] |
| WP_001505004.1 | Type1.BREX | DUF6216-><-?<-AbiEii<-AbiEi_3_N+AbiEi_3\|\|<-HKD-DNase+OGG-like<-SIGMA+OB-Fold+AAA-ATPase+Lon-Protease-BrxL<-PglZ<-N6-METHYLASE<-DUF6079*<-BrxB-BrxF-iSTAND<-BrxA-DNA-binding\|\|REase+iREase-><-wHTH+SAD+HNH\|\|<-?<-?<-?\|\|Phage_integrase->Phage_integrase-> | 1213 | Bacteria>Pseudomonadota>Gammaproteobacteria>Enterobacterales>Enterobacteriaceae>Escherichia | GCF_000446625.2 | Escherichia_coli | ECC52794_RS10870 | 64245..67886 | 562 | BREX-system-P-loop-protein-BrxC-[Escherichia-coli] |
| WP_024222704.1 | Type1.BREX | <-Fimbrial<-Phage_integrase\|\|?->?->?->\|\|wHTH+SAD+HNH-><-REase+iREase\|\|BrxA-DNA-binding->BrxB-BrxF-iSTAND->DUF6079*->PglZ->SIGMA+OB-Fold+AAA-ATPase+Lon-Protease-BrxL->HKD-DNase+OGG-like->\|\|AbiEi_3_N+AbiEi_3->AbiEii->?-><-DUF6216<-?<-TOPRIM-DNAG+D5N+D5ATPase+P4PRIMASE-HTH | 1213 | Bacteria>Pseudomonadota>Gammaproteobacteria>Enterobacterales>Enterobacteriaceae>Escherichia | GCF_000446725.2 | Escherichia_coli | CTH77_RS23215 | 13991..17632 | 562 | BREX-system-P-loop-protein-BrxC-[Escherichia-coli] |
| WP_024236748.1 | Type1.BREX | DUF6079*->PglZ->SIGMA+OB-Fold+AAA-ATPase+Lon-Protease-BrxL->HKD-DNase+OGG-like->\|\|AbiEi_3_N+AbiEi_3->AbiEii->?-><-DUF6216<-tRNA\|\|?->HerA/FtsK+A32-> | 1129 | Bacteria>Pseudomonadota>Gammaproteobacteria>Enterobacterales>Enterobacteriaceae>Escherichia | GCF_000446745.2 | Escherichia_coli | ECC71710_RS03330 | 1..3392 | 562 | BREX-system-P-loop-protein-BrxC,-partial-[Escherichia-coli] |
| WP_024243520.1 | Type1.BREX | DUF6216-><-AbiEii<-AbiEi_3_N+AbiEi_3\|\|<-HKD-DNase+OGG-like<-SIGMA+OB-Fold+AAA-ATPase+Lon-Protease-BrxL<-PglZ<-N6-METHYLASE<-DUF6079*<-BrxB-BrxF-iSTAND<-BrxA-DNA-binding\|\|REase+iREase-><-wHTH+SAD+HNH | 1213 | Bacteria>Pseudomonadota>Gammaproteobacteria>Enterobacterales>Enterobacteriaceae>Escherichia | GCF_000446965.2 | Escherichia_coli | C3I80_RS17935 | 35165..38806 | 562 | BREX-system-P-loop-protein-BrxC-[Escherichia-coli] |
| WP_001019649.1 | Type1.BREX | ZnR->ZnR->URI-DNase+RAMA-><-SIGMA+OB-Fold+AAA-ATPase+Lon-Protease-BrxL<-PglZ<-TOPRIM<-OLD-ABC<-N6-METHYLASE<-DUF6079*<-BrxB-BrxF-iSTAND<-BrxA-DNA-binding<-tRNA\|\|?-><-SKI\|\|METHYLASE->?->?->LACI-HTH+PBPI-> | 1204 | Bacteria>Pseudomonadota>Gammaproteobacteria>Enterobacterales>Enterobacteriaceae>Escherichia | GCF_000503455.1 | Escherichia_coli | U109_RS0110040 | 13723..17337 | 562 | BREX-system-P-loop-protein-BrxC-[Escherichia-coli] |
| WP_025670435.1 | Type1.BREX | <-XIS-HTH\|\|?-><-?\|\|?->ZnR->ZnR-><-SIGMA+OB-Fold+AAA-ATPase+Lon-Protease-BrxL<-PglZ<-N6-METHYLASE<-DUF6079*<-BrxB-BrxF-iSTAND<-BrxA-DNA-binding\|\|REase+iREase-><-EVE+HNH<-Arm-DNA-bind_3+Phage_integrase<-tRNA\|\|?-><-SKI\|\|METHYLASE-> | 1213 | Bacteria>Pseudomonadota>Gammaproteobacteria>Enterobacterales>Enterobacteriaceae>Escherichia | GCF_000511525.1 | Escherichia_coli | A7216_RS07695 | 16662..20303 | 562 | BREX-system-P-loop-protein-BrxC-[Escherichia-coli] |
| WP_029488486.1 | Type1.BREX | TOPRIM-DNAG+D5N+D5ATPase+P4PRIMASE-HTH->DUF6216-><-?<-AbiEii<-AbiEi_3_N+AbiEi_3\|\|<-HKD-DNase+OGG-like<-SIGMA+OB-Fold+AAA-ATPase+Lon-Protease-BrxL<-PglZ<-N6-METHYLASE<-DUF6079*<-BrxB-BrxF-iSTAND<-BrxA-DNA-binding\|\|REase+iREase-><-wHTH+SAD+HNH\|\|<-?<-?<-?\|\|Phage_integrase->Phage_integrase-> | 1213 | Bacteria>Pseudomonadota>Gammaproteobacteria>Enterobacterales>Enterobacteriaceae>Escherichia | GCF_000511545.1 | Escherichia_coli | U124_RS0103885 | 150968..154609 | 562 | BREX-system-P-loop-protein-BrxC-[Escherichia-coli] |
| WP_032271764.1 | Type1.BREX | <-HerA/FtsK+A32<-?\|\|tRNA->DUF6216-><-AbiEii<-AbiEi_3_N+AbiEi_3\|\|<-HKD-DNase+OGG-like<-SIGMA+OB-Fold+AAA-ATPase+Lon-Protease-BrxL<-PglZ<-N6-METHYLASE<-DUF6079-p* | 996 | Bacteria>Pseudomonadota>Gammaproteobacteria>Enterobacterales>Enterobacteriaceae>Escherichia | GCF_000614035.2 | Escherichia_coli | BX93_RS13740 | 78498..81489 | 562 | BREX-system-P-loop-protein-BrxC,-partial-[Escherichia-coli] |
| WP_032272074.1 | Type1.BREX | DUF6079*->N6-METHYLASE->PglZ->SIGMA+OB-Fold+AAA-ATPase+Lon-Protease-BrxL->HKD-DNase+OGG-like->\|\|AbiEi_3_N+AbiEi_3->AbiEii-><-DUF6216<-tRNA\|\|?->HerA/FtsK+A32-> | 1188 | Bacteria>Pseudomonadota>Gammaproteobacteria>Enterobacterales>Enterobacteriaceae>Escherichia | GCF_000615605.2 | Escherichia_coli | BX59_RS22390 | 1..3568 | 562 | BREX-system-P-loop-protein-BrxC,-partial-[Escherichia-coli] |
| WP_032353829.1 | Type1.BREX | DUF6216-><-?<-AbiEii<-AbiEi_3_N+AbiEi_3\|\|<-HKD-DNase+OGG-like<-SIGMA+OB-Fold+AAA-ATPase+Lon-Protease-BrxL<-PglZ<-N6-METHYLASE<-DUF6079*<-BrxB-BrxF-iSTAND<-BrxA-DNA-binding\|\|<-wHTH+SAD+HNH\|\|<-?<-?<-?\|\|Phage_integrase->Phage_integrase->Fimbrial-> | 1213 | Bacteria>Pseudomonadota>Gammaproteobacteria>Enterobacterales>Enterobacteriaceae>Escherichia | GCF_000632675.2 | Escherichia_coli | BX99_RS21625 | 78780..82421 | 562 | BREX-system-P-loop-protein-BrxC-[Escherichia-coli] |
| WP_032216088.1 | Type1.BREX | XIS-HTH-><-HTH_Tnp_1+HTH_21+rve+rve_3\|\|wHTH+WYL+WCX->BrxA-DNA-binding->BrxB-BrxF-iSTAND->DUF6079*->ParB+HNH->N6-METHYLASE->PglZ->SIGMA+OB-Fold+AAA-ATPase+Lon-Protease-BrxL->35exo->\|\|HIN-HTH-> | 1225 | Bacteria>Pseudomonadota>Gammaproteobacteria>Enterobacterales>Enterobacteriaceae>Escherichia | GCF_000713175.1 | Escherichia_coli | AD46_RS16460 | 5265..8942 | 562 | BREX-system-P-loop-protein-BrxC-[Escherichia-coli] |
| WP_032305083.1 | Type1.BREX | XIS-HTH-><-HTH_Tnp_1+HTH_21+rve+rve_3\|\|wHTH+WYL+WCX->BrxA-DNA-binding->BrxB-BrxF-iSTAND->DUF6079*->ParB+HNH->N6-METHYLASE->PglZ->SIGMA+OB-Fold+AAA-ATPase+Lon-Protease-BrxL->35exo-> | 1225 | Bacteria>Pseudomonadota>Gammaproteobacteria>Enterobacterales>Enterobacteriaceae>Escherichia | GCF_000714055.1 | Escherichia_coli | AC36_RS01690 | 6338..10015 | 562 | BREX-system-P-loop-protein-BrxC-[Escherichia-coli] |
| WP_032222683.1 | Type1.BREX | XIS-HTH-><-HTH_Tnp_1+HTH_21+rve+rve_3\|\|wHTH+WYL+WCX->BrxA-DNA-binding->BrxB-BrxF-iSTAND->DUF6079*->ParB+HNH->N6-METHYLASE->PglZ->SIGMA+OB-Fold+AAA-ATPase+Lon-Protease-BrxL->35exo-> | 1225 | Bacteria>Pseudomonadota>Gammaproteobacteria>Enterobacterales>Enterobacteriaceae>Escherichia | GCF_000714385.1 | Escherichia_coli | AC91_RS16385 | 5217..8894 | 562 | BREX-system-P-loop-protein-BrxC-[Escherichia-coli] |
| WP_033804924.1 | Type1.BREX | <-DSBH+AraC-HTH+AraC-HTH\|\|?->DUF488-><-?\|\|GAF+GGDEF->?->?-><-?<-OrfB_IS605+OrfB_Zn_ribbon\|\|DUF6079-p*->N6-METHYLASE->PglZ->SIGMA+OB-Fold+AAA-ATPase+Lon-Protease-BrxL->HKD-DNase+OGG-like->\|\|AbiEi_3_N+AbiEi_3->AbiEii->?-><-DUF6216 | 862 | Bacteria>Pseudomonadota>Gammaproteobacteria>Enterobacterales>Enterobacteriaceae>Escherichia | GCF_000752535.1 | Escherichia_coli | MY85_RS14095 | 76836..79424 | 562 | BREX-system-P-loop-protein-BrxC,-partial-[Escherichia-coli] |
| WP_042106733.1 | Type1.BREX | wHTH->ZnR->ZnR->URI-DNase+RAMA-><-SIGMA+OB-Fold+AAA-ATPase+Lon-Protease-BrxL<-PglZ<-TOPRIM<-OLD-ABC<-N6-METHYLASE<-DUF6079-p* | 895 | Bacteria>Pseudomonadota>Gammaproteobacteria>Enterobacterales>Enterobacteriaceae>Escherichia | GCF_000777265.1 | Escherichia_coli | PP93_RS24150 | 26949..29632 | 562 | BREX-system-P-loop-protein-BrxC,-partial-[Escherichia-coli] |
| WP_033566562.1 | Type1.BREX | <-HerA/FtsK+A32<-?\|\|tRNA->Arm-DNA-bind_3+Phage_integrase->\|\|<-?\|\|BrxA-DNA-binding->BrxB-BrxF-iSTAND->DUF6079-p*-> | 781 | Bacteria>Pseudomonadota>Gammaproteobacteria>Enterobacterales>Enterobacteriaceae>Escherichia | GCF_000783335.1 | Escherichia_coli | LW92_RS03930 | 20620..22963 | 562 | BREX-system-P-loop-protein-BrxC,-partial-[Escherichia-coli] |
| WP_034167068.1 | Type1.BREX | <-DnaB+HerA/FtsK<-HDRnase<-SbcC<-SIGMA+OB-Fold+AAA-ATPase+Lon-Protease-BrxL<-PglZ<-N6-METHYLASE<-DUF6079*<-BrxB-BrxF-iSTAND<-BrxA-DNA-binding<-wHTH+WYL+WCX<-?\|\|<-Couple_hipA+HipA_C<-HTH_3 | 1225 | Bacteria>Pseudomonadota>Gammaproteobacteria>Enterobacterales>Enterobacteriaceae>Escherichia | GCF_000804325.1 | Escherichia_coli | RK25_RS05450 | 34844..38521 | 562 | BREX-system-P-loop-protein-BrxC-[Escherichia-coli] |
| WP_038989008.1 | Type1.BREX | wHTH->ZnR->ZnR->URI-DNase+RAMA-><-SIGMA+OB-Fold+AAA-ATPase+Lon-Protease-BrxL<-PglZ<-TOPRIM<-OLD-ABC<-N6-METHYLASE<-DUF6079*<-BrxB-BrxF-iSTAND<-BrxA-DNA-binding<-Arm-DNA-bind_3+Phage_integrase<-tRNA\|\|?->HerA/FtsK+A32-><-?<-?<-?\|\|Peptidase_M17_N+Peptidase_M17-> | 1204 | Bacteria>Pseudomonadota>Gammaproteobacteria>Enterobacterales>Enterobacteriaceae>Escherichia | GCF_000812725.1 | Escherichia_coli | RK29_RS16985 | 15030..18644 | 562 | BREX-system-P-loop-protein-BrxC-[Escherichia-coli] |
| WP_040091953.1 | Type1.BREX | DUF6079-p*->N6-METHYLASE->PglZ->SIGMA+OB-Fold+AAA-ATPase+Lon-Protease-BrxL-> | 837 | Bacteria>Pseudomonadota>Gammaproteobacteria>Enterobacterales>Enterobacteriaceae>Escherichia | GCF_000819205.1 | Escherichia_coli | PU45_RS22665 | 1..2514 | 562 | BREX-system-P-loop-protein-BrxC,-partial-[Escherichia-coli] |
| WP_050867845.1 | Type1.BREX | <-Phage_integrase<-Phage_integrase\|\|?->?->?->\|\|wHTH+SAD+HNH-><-REase+iREase\|\|BrxA-DNA-binding->BrxB-BrxF-iSTAND->DUF6079*->PglZ->SIGMA+OB-Fold+AAA-ATPase+Lon-Protease-BrxL->?-><-?<-ParB<-?\|\|<-REase+N6_METHYLASE<-TRD<-REase | 1213 | Bacteria>Pseudomonadota>Gammaproteobacteria>Enterobacterales>Enterobacteriaceae>Escherichia | GCF_001191345.1 | Escherichia_coli | WQ87_RS09225 | 9779..13420 | 562 | BREX-system-P-loop-protein-BrxC-[Escherichia-coli] |
| WP_052909519.1 | Type1.BREX | <-Phage_integrase<-Phage_integrase\|\|?->?->?->\|\|wHTH+SAD+HNH-><-REase+iREase\|\|BrxA-DNA-binding->BrxB-BrxF-iSTAND->DUF6079*->N6-METHYLASE->PglZ->SIGMA+OB-Fold+AAA-ATPase+Lon-Protease-BrxL->?-><-?<-ParB<-?\|\|<-REase+N6_METHYLASE<-TRD | 1213 | Bacteria>Pseudomonadota>Gammaproteobacteria>Enterobacterales>Enterobacteriaceae>Escherichia | GCF_001262895.1 | Escherichia_coli | AGA27_RS05565 | 9779..13420 | 562 | BREX-system-P-loop-protein-BrxC-[Escherichia-coli] |
| WP_053916754.1 | Type1.BREX | DUF6216-><-?<-AbiEii<-AbiEi_3_N+AbiEi_3\|\|<-HKD-DNase+OGG-like<-SIGMA+OB-Fold+AAA-ATPase+Lon-Protease-BrxL<-PglZ<-N6-METHYLASE<-DUF6079*<-BrxB-BrxF-iSTAND<-BrxA-DNA-binding\|\|REase+iREase-><-wHTH+SAD+HNH\|\|<-?<-?<-?\|\|Phage_integrase-> | 1213 | Bacteria>Pseudomonadota>Gammaproteobacteria>Enterobacterales>Enterobacteriaceae>Escherichia | GCF_001283285.1 | Escherichia_coli | AOH25_RS20055 | 35212..38853 | 562 | BREX-system-P-loop-protein-BrxC-[Escherichia-coli] |
| WP_053892086.1 | Type1.BREX | <-Fimbrial<-Phage_integrase\|\|?->?->?->\|\|wHTH+SAD+HNH-><-REase+iREase\|\|BrxA-DNA-binding->BrxB-BrxF-iSTAND->DUF6079*->N6-METHYLASE->PglZ->SIGMA+OB-Fold+AAA-ATPase+Lon-Protease-BrxL->HKD-DNase+OGG-like->\|\|AbiEi_3_N+AbiEi_3->AbiEii->?-><-DUF6216 | 1213 | Bacteria>Pseudomonadota>Gammaproteobacteria>Enterobacterales>Enterobacteriaceae>Escherichia | GCF_001285285.1 | Escherichia_coli | AOH78_RS17135 | 37162..40803 | 562 | BREX-system-P-loop-protein-BrxC-[Escherichia-coli] |
| WP_021568966.1 | Type1.BREX | DUF6216-><-?<-AbiEii<-AbiEi_3_N+AbiEi_3\|\|<-HKD-DNase+OGG-like<-SIGMA+OB-Fold+AAA-ATPase+Lon-Protease-BrxL<-PglZ<-N6-METHYLASE<-DUF6079*<-BrxB-BrxF-iSTAND<-BrxA-DNA-binding\|\|REase+iREase-><-wHTH+SAD+HNH\|\|<-?<-?<-?\|\|Phage_integrase->Phage_integrase-> | 1213 | Bacteria>Pseudomonadota>Gammaproteobacteria>Enterobacterales>Enterobacteriaceae>Escherichia | GCF_001285965.1 | Escherichia_coli | AOJ37_RS12845 | 35181..38822 | 562 | BREX-system-P-loop-protein-BrxC-[Escherichia-coli] |
| WP_032257286.1 | Type1.BREX | <-Phage_integrase<-Phage_integrase\|\|?->?->?->\|\|wHTH+SAD+HNH-><-REase+iREase\|\|BrxA-DNA-binding->BrxB-BrxF-iSTAND->DUF6079*->N6-METHYLASE->PglZ->SIGMA+OB-Fold+AAA-ATPase+Lon-Protease-BrxL->HKD-DNase+OGG-like->\|\|AbiEi_3_N+AbiEi_3->AbiEii->?-><-DUF6216 | 1213 | Bacteria>Pseudomonadota>Gammaproteobacteria>Enterobacterales>Enterobacteriaceae>Escherichia | GCF_001286385.1 | Escherichia_coli | AOJ18_RS01745 | 37207..40848 | 562 | BREX-system-P-loop-protein-BrxC-[Escherichia-coli] |
| WP_053897162.1 | Type1.BREX | DUF6079*->PglZ->SIGMA+OB-Fold+AAA-ATPase+Lon-Protease-BrxL->HKD-DNase+OGG-like->\|\|AbiEi_3_N+AbiEi_3->AbiEii->?-><-DUF6216<-tRNA\|\|?->HerA/FtsK+A32-> | 1132 | Bacteria>Pseudomonadota>Gammaproteobacteria>Enterobacterales>Enterobacteriaceae>Escherichia | GCF_001286505.1 | Escherichia_coli | AOJ58_RS17615 | 1..3401 | 562 | BREX-system-P-loop-protein-BrxC,-partial-[Escherichia-coli] |
| WP_060616116.1 | Type1.BREX | <-SIGMA+OB-Fold+AAA-ATPase+Lon-Protease-BrxL<-PglZ<-TOPRIM<-OLD-ABC<-N6-METHYLASE<-DUF6079*<-BrxB-BrxF-iSTAND<-BrxA-DNA-binding\|\|ZnR->?-><-XIS-HTH | 1204 | Bacteria>Pseudomonadota>Gammaproteobacteria>Enterobacterales>Enterobacteriaceae>Escherichia | GCF_001484475.1 | Escherichia_coli | DT570_RS09710 | 10713..14327 | 562 | BREX-system-P-loop-protein-BrxC-[Escherichia-coli] |
| WP_059338855.1 | Type1.BREX | ZnR-><-SIGMA+OB-Fold+AAA-ATPase+Lon-Protease-BrxL<-PglZ<-N6-METHYLASE<-DUF6079-p* | 991 | Bacteria>Pseudomonadota>Gammaproteobacteria>Enterobacterales>Enterobacteriaceae>Escherichia | GCF_001519915.1 | Escherichia_coli | AWF46_RS03555 | 9408..12384 | 562 | BREX-system-P-loop-protein-BrxC,-partial-[Escherichia-coli] |
| WP_059322911.1 | Type1.BREX | DUF6079-p*->N6-METHYLASE->PglZ->SIGMA+OB-Fold+AAA-ATPase+Lon-Protease-BrxL-><-ZnR<-RVT | 992 | Bacteria>Pseudomonadota>Gammaproteobacteria>Enterobacterales>Enterobacteriaceae>Escherichia | GCF_001520675.1 | Escherichia_coli | AWF39_RS02795 | 1..2981 | 562 | BREX-system-P-loop-protein-BrxC,-partial-[Escherichia-coli] |
| WP_061359766.1 | Type1.BREX | <-REase+iREase\|\|BrxA-DNA-binding->BrxB-BrxF-iSTAND->DUF6079*->N6-METHYLASE->PglZ->SIGMA+OB-Fold+AAA-ATPase+Lon-Protease-BrxL-><-ZnR<-ZnR<-?<-?\|\|RHH_antitoxin-> | 1213 | Bacteria>Pseudomonadota>Gammaproteobacteria>Enterobacterales>Enterobacteriaceae>Escherichia | GCF_001575355.1 | Escherichia_coli | AUP84_RS21650 | 2336..5977 | 562 | BREX-system-P-loop-protein-BrxC-[Escherichia-coli] |
| WP_062863778.1 | Type1.BREX | DUF6079-p*->N6-METHYLASE->PglZ->SIGMA+OB-Fold+AAA-ATPase+Lon-Protease-BrxL->HKD-DNase+OGG-like->\|\|AbiEi_3_N+AbiEi_3->AbiEii->?-><-DUF6216 | 998 | Bacteria>Pseudomonadota>Gammaproteobacteria>Enterobacterales>Enterobacteriaceae>Escherichia | GCF_001607105.1 | Escherichia_coli | AML45_RS08440 | 1..2999 | 562 | BREX-system-P-loop-protein-BrxC,-partial-[Escherichia-coli] |
| WP_063100938.1 | Type1.BREX | <-XIS-HTH<-RHH_antitoxin\|\|?->?->ZnR->ZnR-><-SIGMA+OB-Fold+AAA-ATPase+Lon-Protease-BrxL<-PglZ<-N6-METHYLASE<-DUF6079*<-BrxB-BrxF-iSTAND<-BrxA-DNA-binding\|\|REase+iREase->?->?->TetR-HTH-><-?<-AAA-ATPase<-HIN-HTH | 1213 | Bacteria>Pseudomonadota>Gammaproteobacteria>Enterobacterales>Enterobacteriaceae>Escherichia | GCF_001615735.1 | Escherichia_coli | AVN29_RS04305 | 18241..21882 | 562 | BREX-system-P-loop-protein-BrxC-[Escherichia-coli] |
| WP_069924879.1 | Type1.BREX | <-Phage_integrase<-Phage_integrase\|\|?->?->?->\|\|wHTH+SAD+HNH-><-REase+iREase\|\|BrxA-DNA-binding->BrxB-BrxF-iSTAND->DUF6079*->N6-METHYLASE->PglZ->SIGMA+OB-Fold+AAA-ATPase+Lon-Protease-BrxL->?-><-?<-ParB<-?\|\|<-REase+N6_METHYLASE<-TRD | 1213 | Bacteria>Pseudomonadota>Gammaproteobacteria>Enterobacterales>Enterobacteriaceae>Escherichia | GCF_001748825.1 | Escherichia_coli | BHF18_RS13015 | 262302..265943 | 562 | BREX-system-P-loop-protein-BrxC-[Escherichia-coli] |
| WP_072679154.1 | Type1.BREX | <-XIS-HTH<-RHH_antitoxin\|\|?->?->ZnR->ZnR-><-SIGMA+OB-Fold+AAA-ATPase+Lon-Protease-BrxL<-PglZ<-N6-METHYLASE<-DUF6079*<-BrxB-BrxF-iSTAND<-BrxA-DNA-binding\|\|REase+iREase->?->\|\|<-Arm-DNA-bind_3+Phage_integrase<-tRNA\|\|?->HerA/FtsK+A32-> | 1213 | Bacteria>Pseudomonadota>Gammaproteobacteria>Enterobacterales>Enterobacteriaceae>Escherichia | GCF_001894085.1 | Escherichia_coli | BK377_RS08410 | 37601..41242 | 562 | BREX-system-P-loop-protein-BrxC-[Escherichia-coli] |
| WP_072752451.1 | Type1.BREX | DDE_Tnp_1_5+DDE_Tnp_1-><-35exo<-SIGMA+OB-Fold+AAA-ATPase+Lon-Protease-BrxL<-PglZ<-N6-METHYLASE<-ParB+HNH<-DUF6079*<-BrxB-BrxF-iSTAND<-BrxA-DNA-binding<-wHTH+WYL+WCX | 1225 | Bacteria>Pseudomonadota>Gammaproteobacteria>Enterobacterales>Enterobacteriaceae>Escherichia | GCF_001894415.1 | Escherichia_coli | BK380_RS19185 | 23163..26840 | 562 | BREX-system-P-loop-protein-BrxC-[Escherichia-coli] |
| WP_072748589.1 | Type1.BREX | <-HerA/FtsK+A32\|\|tRNA->Arm-DNA-bind_3+Phage_integrase->EVE+HNH-><-REase+iREase\|\|BrxA-DNA-binding->BrxB-BrxF-iSTAND->DUF6079*->N6-METHYLASE->PglZ->SIGMA+OB-Fold+AAA-ATPase+Lon-Protease-BrxL-><-ZnR<-DEOR-HTH+ISOCOT | 1213 | Bacteria>Pseudomonadota>Gammaproteobacteria>Enterobacterales>Enterobacteriaceae>Escherichia | GCF_001894585.1 | Escherichia_coli | BK398_RS07575 | 128185..131826 | 562 | BREX-system-P-loop-protein-BrxC-[Escherichia-coli] |
| WP_074525756.1 | Type1.BREX | DUF6216-><-?<-AbiEii<-AbiEi_3_N+AbiEi_3\|\|<-HKD-DNase+OGG-like<-SIGMA+OB-Fold+AAA-ATPase+Lon-Protease-BrxL<-PglZ<-N6-METHYLASE<-DUF6079*<-BrxB-BrxF-iSTAND<-BrxA-DNA-binding\|\|REase+iREase-><-wHTH+SAD+HNH\|\|<-?<-?<-?\|\|Phage_integrase->Phage_integrase-> | 1213 | Bacteria>Pseudomonadota>Gammaproteobacteria>Enterobacterales>Enterobacteriaceae>Escherichia | GCF_001911555.1 | Escherichia_coli | ACN68_RS20915 | 50623..54264 | 562 | BREX-system-P-loop-protein-BrxC-[Escherichia-coli] |
| WP_085314183.1 | Type1.BREX | <-Phage_integrase<-Phage_integrase\|\|?->?->?->\|\|wHTH+SAD+HNH-><-REase+iREase\|\|BrxA-DNA-binding->BrxB-BrxF-iSTAND->DUF6079*->DUF6079-p->N6-METHYLASE-><-wHTH+WYL\|\|?->?->?->HIN-HTH-> | 1078 | Bacteria>Pseudomonadota>Gammaproteobacteria>Enterobacterales>Enterobacteriaceae>Escherichia | GCF_002104645.1 | Escherichia_coli | B8A27_RS24555 | 59111..62345 | 562 | BREX-system-P-loop-protein-BrxC,-partial-[Escherichia-coli] |
| WP_244571630.1 | Type1.BREX | <-LexA-protease\|\|35exo->?->wHTH+WYL+WCX->BrxA-DNA-binding->BrxB-BrxF-iSTAND->DUF6079-p*-> | 673 | Bacteria>Pseudomonadota>Gammaproteobacteria>Enterobacterales>Enterobacteriaceae>Escherichia | GCF_002231255.1 | Escherichia_coli | ATB30_RS25725 | 4607..6624 | 562 | BREX-system-P-loop-protein-BrxC,-partial-[Escherichia-coli] |
| WP_089613509.1 | Type1.BREX | <-Phage_integrase<-Phage_integrase\|\|?->?->?->\|\|wHTH+SAD+HNH-><-REase+iREase\|\|BrxA-DNA-binding->BrxB-BrxF-iSTAND->DUF6079*->N6-METHYLASE->PglZ->SIGMA+OB-Fold+AAA-ATPase+Lon-Protease-BrxL->HKD-DNase+OGG-like->\|\|AbiEi_3_N+AbiEi_3->AbiEii->?-><-DUF6216 | 1213 | Bacteria>Pseudomonadota>Gammaproteobacteria>Enterobacterales>Enterobacteriaceae>Escherichia | GCF_002231555.1 | Escherichia_coli | ATB14_RS09425 | 78080..81721 | 562 | BREX-system-P-loop-protein-BrxC-[Escherichia-coli] |
| WP_089592460.1 | Type1.BREX | <-XIS-HTH<-RHH_antitoxin\|\|?->ZnR->ZnR-><-SIGMA+OB-Fold+AAA-ATPase+Lon-Protease-BrxL<-PglZ<-N6-METHYLASE<-DUF6079*<-BrxB-BrxF-iSTAND<-BrxA-DNA-binding\|\|REase+iREase-> | 1213 | Bacteria>Pseudomonadota>Gammaproteobacteria>Enterobacterales>Enterobacteriaceae>Escherichia | GCF_002232915.1 | Escherichia_coli | AS998_RS25490 | 17146..20787 | 562 | BREX-system-P-loop-protein-BrxC-[Escherichia-coli] |
| WP_096933204.1 | Type1.BREX | <-REase+iREase\|\|BrxA-DNA-binding->BrxB-BrxF-iSTAND->DUF6079*->N6-METHYLASE->PglZ->SIGMA+OB-Fold+AAA-ATPase+Lon-Protease-BrxL-><-ZnR<-ZnR<-?<-?\|\|RHH_antitoxin->XIS-HTH-> | 1213 | Bacteria>Pseudomonadota>Gammaproteobacteria>Enterobacterales>Enterobacteriaceae>Escherichia | GCF_002442365.1 | Escherichia_coli | BMR57_RS14870 | 5774..9415 | 562 | BREX-system-P-loop-protein-BrxC-[Escherichia-coli] |
| WP_096939573.1 | Type1.BREX | <-HerA/FtsK+A32<-?\|\|tRNA->Arm-DNA-bind_3+Phage_integrase->EVE+HNH-><-REase+iREase\|\|BrxA-DNA-binding->BrxB-BrxF-iSTAND->DUF6079*->N6-METHYLASE->PglZ->SIGMA+OB-Fold+AAA-ATPase+Lon-Protease-BrxL-><-ZnR<-ZnR | 1213 | Bacteria>Pseudomonadota>Gammaproteobacteria>Enterobacterales>Enterobacteriaceae>Escherichia | GCF_002464005.1 | Escherichia_coli | A7578_RS18845 | 20590..24231 | 562 | BREX-system-P-loop-protein-BrxC-[Escherichia-coli] |
| WP_096969144.1 | Type1.BREX | <-METHYLASE<-?\|\|?-><-?\|\|Arm-DNA-bind_3+Phage_integrase->wHTH+WYL+WCX->BrxA-DNA-binding->BrxB-BrxF-iSTAND->DUF6079*->ParB+HNH->N6-METHYLASE->PglZ->SIGMA+OB-Fold+AAA-ATPase+Lon-Protease-BrxL->35exo-><-DDE_Tnp_1_5+DDE_Tnp_1 | 1225 | Bacteria>Pseudomonadota>Gammaproteobacteria>Enterobacterales>Enterobacteriaceae>Escherichia | GCF_002466035.1 | Escherichia_coli | BSN44_RS10850 | 73237..76914 | 562 | BREX-system-P-loop-protein-BrxC-[Escherichia-coli] |
| WP_097339933.1 | Type1.BREX | <-SIGMA+OB-Fold+AAA-ATPase+Lon-Protease-BrxL<-PglZ<-N6-METHYLASE<-DUF6079*<-BrxB-BrxF-iSTAND<-BrxA-DNA-binding\|\|REase+iREase-><-wHTH+SAD+HNH | 1213 | Bacteria>Pseudomonadota>Gammaproteobacteria>Enterobacterales>Enterobacteriaceae>Escherichia | GCF_002484685.1 | Escherichia_coli | A7197_RS22885 | 8895..12536 | 562 | BREX-system-P-loop-protein-BrxC-[Escherichia-coli] |
| WP_097426405.1 | Type1.BREX | <-AbiEii<-AbiEi_3_N+AbiEi_3\|\|<-HKD-DNase+OGG-like<-SIGMA+OB-Fold+AAA-ATPase+Lon-Protease-BrxL<-PglZ<-N6-METHYLASE<-DUF6079*<-BrxB-BrxF-iSTAND<-BrxA-DNA-binding\|\|REase+iREase-><-wHTH+SAD+HNH\|\|<-?<-?<-?\|\|Phage_integrase->Phage_integrase-> | 1213 | Bacteria>Pseudomonadota>Gammaproteobacteria>Enterobacterales>Enterobacteriaceae>Escherichia | GCF_002510325.1 | Escherichia_coli | A7660_RS13795 | 13148..16789 | 562 | BREX-system-P-loop-protein-BrxC-[Escherichia-coli] |
| WP_097429067.1 | Type1.BREX | <-LACI-HTH+PBPI\|\|<-Zn_Tnp_IS1+HTH_Tnp_IS1+DDE_Tnp_IS1\|\|BrxA-DNA-binding->BrxB-BrxF-iSTAND->DUF6079*->N6-METHYLASE->PglZ->SIGMA+OB-Fold+AAA-ATPase+Lon-Protease-BrxL-><-ZnR<-ZnR<-?\|\|RHH_antitoxin->HIN-HTH->XIS-HTH-> | 1213 | Bacteria>Pseudomonadota>Gammaproteobacteria>Enterobacterales>Enterobacteriaceae>Escherichia | GCF_002510645.1 | Escherichia_coli | A7614_RS02560 | 133569..137210 | 562 | BREX-system-P-loop-protein-BrxC-[Escherichia-coli] |
| WP_244587252.1 | Type1.BREX | DUF6079*->PglZ->SIGMA+OB-Fold+AAA-ATPase+Lon-Protease-BrxL->HKD-DNase+OGG-like->\|\|AbiEi_3_N+AbiEi_3->AbiEii->?-><-DUF6216<-tRNA\|\|?->HerA/FtsK+A32-> | 1137 | Bacteria>Pseudomonadota>Gammaproteobacteria>Enterobacterales>Enterobacteriaceae>Escherichia | GCF_002519325.1 | Escherichia_coli | A7034_RS17685 | 1..3416 | 562 | BREX-system-P-loop-protein-BrxC,-partial-[Escherichia-coli] |
| WP_097760627.1 | Type1.BREX | ZnR->ZnR-><-SIGMA+OB-Fold+AAA-ATPase+Lon-Protease-BrxL<-PglZ<-N6-METHYLASE<-DUF6079*<-BrxB-BrxF-iSTAND<-BrxA-DNA-binding\|\|REase+iREase-><-EVE+HNH<-Arm-DNA-bind_3+Phage_integrase<-tRNA\|\|?->HerA/FtsK+A32-> | 1213 | Bacteria>Pseudomonadota>Gammaproteobacteria>Enterobacterales>Enterobacteriaceae>Escherichia | GCF_002534675.1 | Escherichia_coli | BEA63_RS10320 | 9803..13444 | 562 | BREX-system-P-loop-protein-BrxC-[Escherichia-coli] |
| WP_099139404.1 | Type1.BREX | <-Phage_integrase<-Phage_integrase\|\|?->?->?->\|\|wHTH+SAD+HNH-><-REase+iREase\|\|BrxA-DNA-binding->BrxB-BrxF-iSTAND->DUF6079*->N6-METHYLASE->PglZ->SIGMA+OB-Fold+AAA-ATPase+Lon-Protease-BrxL->HKD-DNase+OGG-like->\|\|AbiEi_3_N+AbiEi_3->AbiEii->?-><-DUF6216 | 1213 | Bacteria>Pseudomonadota>Gammaproteobacteria>Enterobacterales>Enterobacteriaceae>Escherichia | GCF_002631765.2 | Escherichia_coli | CQR85_RS13645 | 71476..75117 | 562 | BREX-system-P-loop-protein-BrxC-[Escherichia-coli] |
| WP_096974508.1 | Type1.BREX | DUF6216-><-?<-AbiEii<-AbiEi_3_N+AbiEi_3\|\|<-HKD-DNase+OGG-like<-SIGMA+OB-Fold+AAA-ATPase+Lon-Protease-BrxL<-PglZ<-N6-METHYLASE<-DUF6079*<-BrxB-BrxF-iSTAND<-BrxA-DNA-binding\|\|REase+iREase-><-wHTH+SAD+HNH\|\|<-?<-?<-?\|\|Phage_integrase->Phage_integrase-> | 1213 | Bacteria>Pseudomonadota>Gammaproteobacteria>Enterobacterales>Enterobacteriaceae>Escherichia | GCF_002741355.1 | Escherichia_coli | BFZ27_RS06875 | 15053..18694 | 562 | BREX-system-P-loop-protein-BrxC-[Escherichia-coli] |
| WP_102812546.1 | Type1.BREX | ZnR->ZnR-><-SIGMA+OB-Fold+AAA-ATPase+Lon-Protease-BrxL<-PglZ<-N6-METHYLASE<-DUF6079*<-BrxB-BrxF-iSTAND<-BrxA-DNA-binding\|\|REase->REase-> | 1213 | Bacteria>Pseudomonadota>Gammaproteobacteria>Enterobacterales>Enterobacteriaceae>Escherichia | GCF_002888475.1 | Escherichia_coli | C1T15_RS24495 | 11055..14696 | 562 | BREX-system-P-loop-protein-BrxC-[Escherichia-coli] |
| WP_148116075.1 | Type1.BREX | <-HIN-HTH<-?<-?<-?\|\|wHTH+WYL-><-N6-METHYLASE<-DUF6079*<-BrxB-BrxF-iSTAND<-BrxA-DNA-binding\|\|REase+iREase-><-wHTH+SAD+HNH\|\|<-?<-?<-?\|\|Phage_integrase->Phage_integrase-> | 1213 | Bacteria>Pseudomonadota>Gammaproteobacteria>Enterobacterales>Enterobacteriaceae>Escherichia | GCF_002951795.1 | Escherichia_coli | BWI87_RS06400 | 1249242..1252883 | 562 | BREX-system-P-loop-protein-BrxC-[Escherichia-coli] |
| WP_106879201.1 | Type1.BREX | DUF6216-><-?<-AbiEii<-AbiEi_3_N+AbiEi_3\|\|<-HKD-DNase+OGG-like<-SIGMA+OB-Fold+AAA-ATPase+Lon-Protease-BrxL<-PglZ<-N6-METHYLASE<-DUF6079*<-BrxB-BrxF-iSTAND<-BrxA-DNA-binding\|\|REase+iREase-><-wHTH+SAD+HNH\|\|<-?<-?<-?\|\|Phage_integrase->Phage_integrase-> | 1213 | Bacteria>Pseudomonadota>Gammaproteobacteria>Enterobacterales>Enterobacteriaceae>Escherichia | GCF_003018215.1 | Escherichia_coli | C6P67_RS24165 | 4665409..4669050 | 562 | BREX-system-P-loop-protein-BrxC-[Escherichia-coli] |
| WP_044805810.1 | Type1.BREX | DUF6216-><-?<-AbiEii<-AbiEi_3_N+AbiEi_3\|\|<-HKD-DNase+OGG-like<-SIGMA+OB-Fold+AAA-ATPase+Lon-Protease-BrxL<-PglZ<-DUF6079*<-BrxB-BrxF-iSTAND<-BrxA-DNA-binding\|\|REase+iREase-><-wHTH+SAD+HNH\|\|<-?<-?<-?\|\|Phage_integrase->Phage_integrase-> | 1213 | Bacteria>Pseudomonadota>Gammaproteobacteria>Enterobacterales>Enterobacteriaceae>Escherichia | GCF_003018495.1 | Escherichia_coli | C6953_RS06925 | 1324145..1327786 | 562 | BREX-system-P-loop-protein-BrxC-[Escherichia-coli] |
| WP_109548991.1 | Type1.BREX | <-Fimbrial<-Phage_integrase<-Phage_integrase\|\|?->?->?->\|\|wHTH(SBD)+HNH->\|\|BrxA-DNA-binding->BrxB-BrxF-iSTAND->DUF6079*->N6-METHYLASE->PglZ->SIGMA+OB-Fold+AAA-ATPase+Lon-Protease-BrxL->HKD-DNase+OGG-like->\|\|AbiEi_3_N+AbiEi_3->AbiEii-><-DUF6216 | 1213 | Bacteria>Pseudomonadota>Gammaproteobacteria>Enterobacterales>Enterobacteriaceae>Escherichia | GCF_003145675.1 | Escherichia_coli | B6S34_RS22850 | 50490..54131 | 562 | BREX-system-P-loop-protein-BrxC-[Escherichia-coli] |
| WP_097502528.1 | Type1.BREX | DUF772+DDE_Tnp_1->?-><-TetR-HTH<-?<-?<-REase+iREase\|\|BrxA-DNA-binding->BrxB-BrxF-iSTAND->DUF6079*->N6-METHYLASE->PglZ->SIGMA+OB-Fold+AAA-ATPase+Lon-Protease-BrxL-><-ZnR\|\|?->METHYLASE-><-tRNA\|\|?->HHH+DNA_pol_B+RNase-> | 1213 | Bacteria>Pseudomonadota>Gammaproteobacteria>Enterobacterales>Enterobacteriaceae>Escherichia | GCF_003176235.1 | Escherichia_coli | DDA49_RS07225 | 8727..12368 | 562 | BREX-system-P-loop-protein-BrxC-[Escherichia-coli] |
| WP_113316937.1 | Type1.BREX | <-REase<-REase\|\|BrxA-DNA-binding->BrxB-BrxF-iSTAND->DUF6079*->N6-METHYLASE->PglZ->SIGMA+OB-Fold+AAA-ATPase+Lon-Protease-BrxL-> | 1213 | Bacteria>Pseudomonadota>Gammaproteobacteria>Enterobacterales>Enterobacteriaceae>Escherichia | GCF_003299815.1 | Escherichia_coli | DUK12_RS25275 | 2498..6139 | 562 | BREX-system-P-loop-protein-BrxC-[Escherichia-coli] |
| WP_121351411.1 | Type1.BREX | <-Fimbrial<-Phage_integrase<-Phage_integrase\|\|?->?->\|\|wHTH+SAD+HNH-><-REase+iREase\|\|BrxA-DNA-binding->BrxB-BrxF-iSTAND->DUF6079*->N6-METHYLASE->PglZ->SIGMA+OB-Fold+AAA-ATPase+Lon-Protease-BrxL->HKD-DNase+OGG-like->\|\|AbiEi_3_N+AbiEi_3->AbiEii->?-><-DUF6216 | 1213 | Bacteria>Pseudomonadota>Gammaproteobacteria>Enterobacterales>Enterobacteriaceae>Escherichia | GCF_003651285.1 | Escherichia_coli | EAO63_RS07885 | 97180..100821 | 562 | BREX-system-P-loop-protein-BrxC-[Escherichia-coli] |
| WP_252363318.1 | Type1.BREX | <-HerA/FtsK+A32\|\|tRNA->Arm-DNA-bind_3+Phage_integrase-><-SIGMA+OB-Fold+AAA-ATPase+Lon-Protease-BrxL<-PglZ<-N6-METHYLASE-p<-N6-METHYLASE<-ParB+HNH<-DUF6079*<-BrxB-BrxF-iSTAND<-BrxA-DNA-binding<-?<-?<-XIS-HTH\|\|?->nSTAND1+sGTP->wHTH+WYL-> | 1240 | Bacteria>Pseudomonadota>Gammaproteobacteria>Enterobacterales>Enterobacteriaceae>Escherichia | GCF_003687125.1 | Escherichia_coli | EAI39_RS15070 | 84115..87837 | 562 | BREX-system-P-loop-protein-BrxC-[Escherichia-coli] |
| WP_122927386.1 | Type1.BREX | Zn_Tnp_IS1+HTH_Tnp_IS1+DDE_Tnp_IS1-><-?<-?<-HNH<-?<-wHTH+WYL+WCX\|\|DUF772+DDE_Tnp_1->BrxA-DNA-binding->BrxB-BrxF-iSTAND->DUF6079*->N6-METHYLASE->PglZ->SIGMA+OB-Fold+AAA-ATPase+Lon-Protease-BrxL->HTH_Tnp_1+HTH_21+rve+rve_3->?->?-><-Zn_Tnp_IS1+HTH_Tnp_IS1+DDE_Tnp_IS1 | 1213 | Bacteria>Pseudomonadota>Gammaproteobacteria>Enterobacterales>Enterobacteriaceae>Escherichia | GCF_003711005.1 | Escherichia_coli | EB670_RS21355 | 4330120..4333761 | 562 | BREX-system-P-loop-protein-BrxC-[Escherichia-coli] |
| WP_128448083.1 | Type1.BREX | <-HerA/FtsK+A32<-?\|\|tRNA->Arm-DNA-bind_3+Phage_integrase->EVE+HNH-><-REase+iREase\|\|BrxA-DNA-binding->BrxB-BrxF-iSTAND->DUF6079*->N6-METHYLASE->PglZ->SIGMA+OB-Fold+AAA-ATPase+Lon-Protease-BrxL-><-ZnR<-ZnR | 1213 | Bacteria>Pseudomonadota>Gammaproteobacteria>Enterobacterales>Enterobacteriaceae>Escherichia | GCF_004054205.1 | Escherichia_coli | EQH54_15620 | 68233..71874 | 562 | BREX-system-P-loop-protein-BrxC-[Escherichia-coli] |
| WP_135129151.1 | Type1.BREX | <-LACI-HTH+PBPI<-?<-?<-METHYLASE\|\|SKI-><-?\|\|tRNA->Arm-DNA-bind_3+Phage_integrase->BrxA-DNA-binding->BrxB-BrxF-iSTAND->DUF6079*->N6-METHYLASE->OLD-ABC-> | 1204 | Bacteria>Pseudomonadota>Gammaproteobacteria>Enterobacterales>Enterobacteriaceae>Escherichia | GCF_004566475.1 | Escherichia_coli | ELY39_RS16045 | 71744..75358 | 562 | BREX-system-P-loop-protein-BrxC-[Escherichia-coli] |
| WP_135497797.1 | Type1.BREX | Arm-DNA-bind_3+Phage_integrase->N6-METHYLASE->ORC-CDC6+ZnR->HEPN-><-Zn_Tnp_IS1+HTH_Tnp_IS1+DDE_Tnp_IS1<-REase<-REase\|\|BrxA-DNA-binding->BrxB-BrxF-iSTAND->DUF6079*->N6-METHYLASE->PglZ->SIGMA+OB-Fold+AAA-ATPase+Lon-Protease-BrxL-><-ZnR<-ZnR<-?\|\|RHH_antitoxin->XIS-HTH-> | 1213 | Bacteria>Pseudomonadota>Gammaproteobacteria>Enterobacterales>Enterobacteriaceae>Escherichia | GCF_004767105.1 | Escherichia_coli | DAH29_RS19080 | 36143..39784 | 562 | BREX-system-P-loop-protein-BrxC-[Escherichia-coli] |
| WP_016246797.1 | Type1.BREX | TetR-HTH->tRNA->Arm-DNA-bind_3+Phage_integrase-><-Zn_Tnp_IS1+HTH_Tnp_IS1+DDE_Tnp_IS1<-TetR-HTH<-?<-?<-REase+iREase\|\|BrxA-DNA-binding->BrxB-BrxF-iSTAND->DUF6079*->N6-METHYLASE->PglZ->SIGMA+OB-Fold+AAA-ATPase+Lon-Protease-BrxL-><-DDE_Tnp | 1213 | Bacteria>Pseudomonadota>Gammaproteobacteria>Enterobacterales>Enterobacteriaceae>Escherichia | GCF_005384525.1 | Escherichia_coli | FFB03_RS10735 | 79495..83136 | 562 | BREX-system-P-loop-protein-BrxC-[Escherichia-coli] |
| WP_044782573.1 | Type1.BREX | <-Fimbrial<-Phage_integrase\|\|?->?->?->\|\|wHTH+SAD+HNH-><-REase+iREase\|\|BrxA-DNA-binding->BrxB-BrxF-iSTAND->DUF6079*->N6-METHYLASE->PglZ->SIGMA+OB-Fold+AAA-ATPase+Lon-Protease-BrxL->HKD-DNase+OGG-like->\|\|AbiEi_3_N+AbiEi_3->AbiEii->?-><-DUF6216 | 1213 | Bacteria>Pseudomonadota>Gammaproteobacteria>Enterobacterales>Enterobacteriaceae>Escherichia | GCF_005390485.1 | Escherichia_coli | FFG15_RS13655 | 95922..99563 | 562 | BREX-system-P-loop-protein-BrxC-[Escherichia-coli] |
| WP_136724038.1 | Type1.BREX | <-Phage_integrase<-Phage_integrase\|\|?->?->?->\|\|wHTH+SAD+HNH-><-REase+iREase\|\|BrxA-DNA-binding->BrxB-BrxF-iSTAND->DUF6079*->N6-METHYLASE->PglZ->SIGMA+OB-Fold+AAA-ATPase+Lon-Protease-BrxL->?-><-?<-ParB<-?\|\|<-REase+N6_METHYLASE<-TRD | 1213 | Bacteria>Pseudomonadota>Gammaproteobacteria>Enterobacterales>Enterobacteriaceae>Escherichia | GCF_005390625.1 | Escherichia_coli | FFG35_RS20705 | 9778..13419 | 562 | BREX-system-P-loop-protein-BrxC-[Escherichia-coli] |
| WP_097512272.1 | Type1.BREX | DUF6216-><-?<-AbiEii<-AbiEi_3_N+AbiEi_3\|\|<-HKD-DNase+OGG-like<-SIGMA+OB-Fold+AAA-ATPase+Lon-Protease-BrxL<-PglZ<-N6-METHYLASE<-DUF6079*<-BrxB-BrxF-iSTAND<-BrxA-DNA-binding\|\|REase+iREase-><-wHTH(SBD)+HNH\|\|<-?<-?<-?\|\|Phage_integrase->Phage_integrase-> | 1213 | Bacteria>Pseudomonadota>Gammaproteobacteria>Enterobacterales>Enterobacteriaceae>Escherichia | GCF_005391145.1 | Escherichia_coli | A7301_RS09385 | 138963..142604 | 562 | BREX-system-P-loop-protein-BrxC-[Escherichia-coli] |
| WP_137536909.1 | Type1.BREX | <-Fimbrial<-Phage_integrase\|\|?->?->?->\|\|wHTH+SAD+HNH-><-REase+iREase\|\|BrxA-DNA-binding->BrxB-BrxF-iSTAND->DUF6079*->N6-METHYLASE->PglZ->SIGMA+OB-Fold+AAA-ATPase+Lon-Protease-BrxL->HKD-DNase+OGG-like->\|\|AbiEi_3_N+AbiEi_3->AbiEii->?-><-DUF6216 | 1213 | Bacteria>Pseudomonadota>Gammaproteobacteria>Enterobacterales>Enterobacteriaceae>Escherichia | GCF_005392025.1 | Escherichia_coli | FFI00_RS00495 | 95917..99558 | 562 | BREX-system-P-loop-protein-BrxC-[Escherichia-coli] |
| WP_063115883.1 | Type1.BREX | DUF6216-><-?<-AbiEii<-AbiEi_3_N+AbiEi_3\|\|<-HKD-DNase+OGG-like<-SIGMA+OB-Fold+AAA-ATPase+Lon-Protease-BrxL<-PglZ<-N6-METHYLASE<-DUF6079*<-BrxB-BrxF-iSTAND<-BrxA-DNA-binding\|\|REase+iREase-><-wHTH+SAD+HNH\|\|<-?<-?<-?\|\|Phage_integrase->Phage_integrase-> | 1213 | Bacteria>Pseudomonadota>Gammaproteobacteria>Enterobacterales>Enterobacteriaceae>Escherichia | GCF_005393265.1 | Escherichia_coli | MMU60_RS13225 | 76316..79957 | 562 | BREX-system-P-loop-protein-BrxC-[Escherichia-coli] |
| WP_137551577.1 | Type1.BREX | <-Fimbrial<-Phage_integrase\|\|?->?->?->\|\|wHTH+SAD+HNH-><-REase+iREase\|\|BrxA-DNA-binding->BrxB-BrxF-iSTAND->DUF6079*->N6-METHYLASE->PglZ->SIGMA+OB-Fold+AAA-ATPase+Lon-Protease-BrxL->HKD-DNase+OGG-like->\|\|AbiEi_3_N+AbiEi_3->AbiEii->?-><-DUF6216 | 1213 | Bacteria>Pseudomonadota>Gammaproteobacteria>Enterobacterales>Enterobacteriaceae>Escherichia | GCF_005394845.1 | Escherichia_coli | FFJ65_RS02650 | 22335..25976 | 562 | BREX-system-P-loop-protein-BrxC-[Escherichia-coli] |
| WP_137573648.1 | Type1.BREX | <-XIS-HTH\|\|?-><-?\|\|?->ZnR->ZnR-><-SIGMA+OB-Fold+AAA-ATPase+Lon-Protease-BrxL<-PglZ<-N6-METHYLASE<-DUF6079*<-BrxB-BrxF-iSTAND<-BrxA-DNA-binding\|\|REase+iREase-> | 1213 | Bacteria>Pseudomonadota>Gammaproteobacteria>Enterobacterales>Enterobacteriaceae>Escherichia | GCF_005397085.1 | Escherichia_coli | FFK68_RS25060 | 21797..25438 | 562 | BREX-system-P-loop-protein-BrxC-[Escherichia-coli] |
| WP_137585959.1 | Type1.BREX | <-Phage_integrase<-Phage_integrase\|\|?->?->?->\|\|wHTH+SAD+HNH-><-REase+iREase\|\|BrxA-DNA-binding->BrxB-BrxF-iSTAND->DUF6079*->N6-METHYLASE->PglZ->SIGMA+OB-Fold+AAA-ATPase+Lon-Protease-BrxL->HKD-DNase+OGG-like->\|\|AbiEi_3_N+AbiEi_3->AbiEii->?-><-DUF6216<-TOPRIM-DNAG+D5N+D5ATPase+P4PRIMASE-HTH | 1213 | Bacteria>Pseudomonadota>Gammaproteobacteria>Enterobacterales>Enterobacteriaceae>Escherichia | GCF_005399505.1 | Escherichia_coli | FFL73_RS10880 | 97640..101281 | 562 | BREX-system-P-loop-protein-BrxC-[Escherichia-coli] |
| WP_137590371.1 | Type1.BREX | DUF6216-><-?<-AbiEii<-AbiEi_3_N+AbiEi_3\|\|<-HKD-DNase+OGG-like<-SIGMA+OB-Fold+AAA-ATPase+Lon-Protease-BrxL<-PglZ<-N6-METHYLASE<-DUF6079*<-BrxB-BrxF-iSTAND<-BrxA-DNA-binding\|\|REase+iREase-><-wHTH+SAD+HNH\|\|<-?<-?\|\|Phage_integrase->Phage_integrase->Fimbrial-> | 1213 | Bacteria>Pseudomonadota>Gammaproteobacteria>Enterobacterales>Enterobacteriaceae>Escherichia | GCF_005399685.1 | Escherichia_coli | FFM00_RS14520 | 19836..23477 | 562 | BREX-system-P-loop-protein-BrxC-[Escherichia-coli] |
| WP_139060210.1 | Type1.BREX | <-DUF6079-p*<-BrxB-BrxF-iSTAND<-BrxA-DNA-binding\|\|REase+iREase-><-wHTH+SAD+HNH\|\|<-?<-?<-?\|\|Phage_integrase->Phage_integrase-> | 635 | Bacteria>Pseudomonadota>Gammaproteobacteria>Enterobacterales>Enterobacteriaceae>Escherichia | GCF_006151715.1 | Escherichia_coli | FD592_RS22960 | 1..1905 | 562 | BREX-system-P-loop-protein-BrxC,-partial-[Escherichia-coli] |
| WP_147697439.1 | Type1.BREX | DEDD_Tnp_IS110+Transposase_20-><-SIGMA+OB-Fold+AAA-ATPase+Lon-Protease-BrxL<-PglZ<-N6-METHYLASE<-ParB+HNH<-DUF6079*<-BrxB-BrxF-iSTAND<-BrxA-DNA-binding<-wHTH+WYL+WCX<-?\|\|RHH->\|\|CopG/RHH->RelE-ParE->HIN-HTH-> | 1225 | Bacteria>Pseudomonadota>Gammaproteobacteria>Enterobacterales>Enterobacteriaceae>Escherichia | GCF_008041475.1 | Escherichia_coli | FV285_RS21415 | 12602..16279 | 562 | BREX-system-P-loop-protein-BrxC-[Escherichia-coli] |
| WP_032192228.1 | Type1.BREX | <-Phage_integrase\|\|?->?->?->\|\|wHTH+SAD+HNH-><-REase+iREase\|\|BrxA-DNA-binding->BrxB-BrxF-iSTAND->DUF6079*-> | 1213 | Bacteria>Pseudomonadota>Gammaproteobacteria>Enterobacterales>Enterobacteriaceae>Escherichia | GCF_008326465.1 | Escherichia_coli | F1555_RS25960 | 8938..12579 | 562 | BREX-system-P-loop-protein-BrxC-[Escherichia-coli] |
| WP_149488190.1 | Type1.BREX | DDE_Tnp_1_5+DDE_Tnp_1-><-35exo<-SIGMA+OB-Fold+AAA-ATPase+Lon-Protease-BrxL<-PglZ<-N6-METHYLASE<-ParB+HNH<-DUF6079*<-BrxB-BrxF-iSTAND<-BrxA-DNA-binding<-wHTH+WYL+WCX<-Arm-DNA-bind_3+Phage_integrase\|\|?-><-?\|\|?->METHYLASE-> | 1225 | Bacteria>Pseudomonadota>Gammaproteobacteria>Enterobacterales>Enterobacteriaceae>Escherichia | GCF_008364435.1 | Escherichia_coli | F0318_RS09425 | 23572..27249 | 562 | BREX-system-P-loop-protein-BrxC-[Escherichia-coli] |
| WP_023063795.1 | Type1.BREX | <-35exo<-SIGMA+OB-Fold+AAA-ATPase+Lon-Protease-BrxL<-PglZ<-N6-METHYLASE<-ParB+HNH<-DUF6079*<-BrxB-BrxF-iSTAND<-BrxA-DNA-binding<-wHTH+WYL+WCX | 1225 | Bacteria>Pseudomonadota>Gammaproteobacteria>Enterobacterales>Enterobacteriaceae>Escherichia | GCF_008370755.1 | Escherichia_coli | L913_RS00645 | 11831..15508 | 562 | BREX-system-P-loop-protein-BrxC-[Escherichia-coli] |
| WP_050939406.1 | Type1.BREX | DUF6216-><-?<-AbiEii<-AbiEi_3_N+AbiEi_3\|\|<-HKD-DNase+OGG-like<-SIGMA+OB-Fold+AAA-ATPase+Lon-Protease-BrxL<-PglZ<-N6-METHYLASE<-DUF6079*<-BrxB-BrxF-iSTAND<-BrxA-DNA-binding\|\|REase+iREase-><-wHTH+SAD+HNH\|\|<-?<-?<-?\|\|Phage_integrase->Phage_integrase-> | 1213 | Bacteria>Pseudomonadota>Gammaproteobacteria>Enterobacterales>Enterobacteriaceae>Escherichia | GCF_009618015.1 | Escherichia_coli | DTB71_RS01510 | 297715..301356 | 562 | BREX-system-P-loop-protein-BrxC-[Escherichia-coli] |
| WP_154139479.1 | Type1.BREX | <-SIGMA+OB-Fold+AAA-ATPase+Lon-Protease-BrxL<-PglZ<-N6-METHYLASE<-DUF6079*<-BrxB-BrxF-iSTAND<-BrxA-DNA-binding | 1213 | Bacteria>Pseudomonadota>Gammaproteobacteria>Enterobacterales>Enterobacteriaceae>Escherichia | GCF_009671045.1 | Escherichia_coli | ML316_RS21570 | 8749..12390 | 562 | BREX-system-P-loop-protein-BrxC-[Escherichia-coli] |
| WP_160518859.1 | Type1.BREX | <-Phage_integrase<-Phage_integrase\|\|?->?->?->\|\|wHTH(SBD)+HNH-><-REase+iREase\|\|BrxA-DNA-binding->BrxB-BrxF-iSTAND->DUF6079*->N6-METHYLASE->PglZ->SIGMA+OB-Fold+AAA-ATPase+Lon-Protease-BrxL->HKD-DNase+OGG-like->\|\|AbiEi_3_N+AbiEi_3->AbiEii->?-><-DUF6216\|\|XIS-HTH-> | 1213 | Bacteria>Pseudomonadota>Gammaproteobacteria>Enterobacterales>Enterobacteriaceae>Escherichia | GCF_009820915.1 | Escherichia_coli | FQ032_RS06170 | 111352..114993 | 562 | BREX-system-P-loop-protein-BrxC-[Escherichia-coli] |
| WP_163451335.1 | Type1.BREX | <-DUF932<-nSTAND1+GTPase-AIG+sGTP\|\|BrxA-DNA-binding->BrxB-BrxF-iSTAND->DUF6079*->N6-METHYLASE->OLD-ABC->TOPRIM->PglZ->SIGMA+OB-Fold+AAA-ATPase+Lon-Protease-BrxL-><-?<-AAA+wHTH<-XIS-HTH\|\|<-Phage_Cox | 1204 | Bacteria>Pseudomonadota>Gammaproteobacteria>Enterobacterales>Enterobacteriaceae>Escherichia | GCF_011008705.1 | Escherichia_coli | G4V04_RS14425 | 2939..6553 | 562 | BREX-system-P-loop-protein-BrxC-[Escherichia-coli] |
| WP_164476769.1 | Type1.BREX | <-Fimbrial<-Phage_integrase<-Phage_integrase\|\|?->?->?->\|\|<-REase+iREase\|\|BrxA-DNA-binding->BrxB-BrxF-iSTAND->DUF6079*->N6-METHYLASE->PglZ->SIGMA+OB-Fold+AAA-ATPase+Lon-Protease-BrxL->HKD-DNase+OGG-like->\|\|AbiEi_3_N+AbiEi_3->AbiEii-><-DUF6216<-REase | 1213 | Bacteria>Pseudomonadota>Gammaproteobacteria>Enterobacterales>Enterobacteriaceae>Escherichia | GCF_011045235.1 | Escherichia_coli | G6Z93_RS19880 | 4188029..4191670 | 562 | BREX-system-P-loop-protein-BrxC-[Escherichia-coli] |
| WP_136516625.1 | Type1.BREX | <-DUF772+DDE_Tnp_1<-35exo<-SIGMA+OB-Fold+AAA-ATPase+Lon-Protease-BrxL<-PglZ<-N6-METHYLASE<-ParB+HNH<-DUF6079*<-BrxB-BrxF-iSTAND<-BrxA-DNA-binding<-wHTH+WYL+WCX<-XIS-HTH<-ZnR\|\|?-><-XIS-HTH | 1225 | Bacteria>Pseudomonadota>Gammaproteobacteria>Enterobacterales>Enterobacteriaceae>Escherichia | GCF_011045875.1 | Escherichia_coli | EIZ78_RS19775 | 10945..14622 | 562 | BREX-system-P-loop-protein-BrxC-[Escherichia-coli] |
| WP_072657276.1 | Type1.BREX | wHTH+WYL+WCX->BrxA-DNA-binding->BrxB-BrxF-iSTAND->DUF6079*->ParB+HNH->N6-METHYLASE->PglZ->SIGMA+OB-Fold+AAA-ATPase+Lon-Protease-BrxL->35exo->\|\|<-ParB+HNH | 1225 | Bacteria>Pseudomonadota>Gammaproteobacteria>Enterobacterales>Enterobacteriaceae>Escherichia | GCF_011077045.1 | Escherichia_coli | G6Q90_RS23590 | 2674..6351 | 562 | BREX-system-P-loop-protein-BrxC-[Escherichia-coli] |
| WP_152922120.1 | Type1.BREX | <-XIS-HTH<-?\|\|?->ZnR->ZnR-><-SIGMA+OB-Fold+AAA-ATPase+Lon-Protease-BrxL<-PglZ<-N6-METHYLASE<-DUF6079*<-BrxB-BrxF-iSTAND<-BrxA-DNA-binding\|\|REase+iREase-><-EVE+HNH<-Arm-DNA-bind_3+Phage_integrase<-tRNA\|\|?-><-SKI\|\|METHYLASE-> | 1213 | Bacteria>Pseudomonadota>Gammaproteobacteria>Enterobacterales>Enterobacteriaceae>Escherichia | GCF_011404475.1 | Escherichia_coli | HRR20_RS19820 | 4109090..4112731 | 562 | BREX-system-P-loop-protein-BrxC-[Escherichia-coli] |
| WP_167824001.1 | Type1.BREX | <-HIN-HTH<-?<-?<-?\|\|wHTH+WYL-><-N6-METHYLASE<-DUF6079*<-BrxB-BrxF-iSTAND<-BrxA-DNA-binding\|\|REase+iREase-><-wHTH+SAD+HNH\|\|<-?<-?<-?\|\|Phage_integrase->Phage_integrase-> | 1213 | Bacteria>Pseudomonadota>Gammaproteobacteria>Enterobacterales>Enterobacteriaceae>Escherichia | GCF_012063235.1 | Escherichia_coli | GTJ82_RS11140 | 125491..129132 | 562 | BREX-system-P-loop-protein-BrxC-[Escherichia-coli] |
| WP_250854521.1 | Type1.BREX | DUF6079-p*->N6-METHYLASE->PglZ->SIGMA+OB-Fold+AAA-ATPase+Lon-Protease-BrxL->HKD-DNase+OGG-like->\|\|AbiEi_3_N+AbiEi_3->AbiEii->?-><-DUF6216 | 887 | Bacteria>Pseudomonadota>Gammaproteobacteria>Enterobacterales>Enterobacteriaceae>Escherichia | GCF_012556305.1 | Escherichia_coli | MLIZ_RS15475 | 1..2666 | 562 | BREX-system-P-loop-protein-BrxC,-partial-[Escherichia-coli] |
| WP_170969386.1 | Type1.BREX | <-Phage_integrase\|\|?->?->?-><-HNH\|\|wHTH+SAD+HNH-><-REase+iREase\|\|BrxA-DNA-binding->BrxB-BrxF-iSTAND->DUF6079*->N6-METHYLASE->PglZ->SIGMA+OB-Fold+AAA-ATPase+Lon-Protease-BrxL->?-><-?<-ParB<-?\|\|<-REase+N6_METHYLASE<-TRD | 1213 | Bacteria>Pseudomonadota>Gammaproteobacteria>Enterobacterales>Enterobacteriaceae>Escherichia | GCF_013085185.1 | Escherichia_coli | HHJ24_09500 | 1928934..1932575 | 562 | BREX-system-P-loop-protein-BrxC-[Escherichia-coli] |
| WP_174563320.1 | Type1.BREX | <-REase+iREase\|\|BrxA-DNA-binding->BrxB-BrxF-iSTAND->DUF6079*->N6-METHYLASE->PglZ->SIGMA+OB-Fold+AAA-ATPase+Lon-Protease-BrxL-><-ZnR | 1213 | Bacteria>Pseudomonadota>Gammaproteobacteria>Enterobacterales>Enterobacteriaceae>Escherichia | GCF_013341485.1 | Escherichia_coli | HUC04_RS15605 | 6608..10249 | 562 | BREX-system-P-loop-protein-BrxC-[Escherichia-coli] |
| WP_097472690.1 | Type1.BREX | <-HerA/FtsK+A32<-?\|\|tRNA->Arm-DNA-bind_3+Phage_integrase->EVE+HNH-><-REase+iREase\|\|BrxA-DNA-binding->BrxB-BrxF-iSTAND->DUF6079*-> | 1213 | Bacteria>Pseudomonadota>Gammaproteobacteria>Enterobacterales>Enterobacteriaceae>Escherichia | GCF_013341925.1 | Escherichia_coli | MMN47_RS21975 | 16744..20385 | 562 | BREX-system-P-loop-protein-BrxC-[Escherichia-coli] |
| WP_032218952.1 | Type1.BREX | <-Peptidase_M17_N+Peptidase_M17\|\|?->?->?-><-HerA/FtsK+A32<-?\|\|tRNA->Arm-DNA-bind_3+Phage_integrase->BrxA-DNA-binding->BrxB-BrxF-iSTAND->DUF6079*->N6-METHYLASE->OLD-ABC->TOPRIM->PglZ->SIGMA+OB-Fold+AAA-ATPase+Lon-Protease-BrxL-><-URI-DNase+RAMA<-ZnR<-ZnR<-wHTH | 1204 | Bacteria>Pseudomonadota>Gammaproteobacteria>Enterobacterales>Enterobacteriaceae>Escherichia | GCF_013370905.1 | Escherichia_coli | DNN97_RS07945 | 182755..186369 | 562 | BREX-system-P-loop-protein-BrxC-[Escherichia-coli] |
| WP_176670934.1 | Type1.BREX | <-XIS-HTH<-?<-?\|\|HTH_Tnp_1+HTH_21+rve+rve_3-><-?<-SIGMA+OB-Fold+AAA-ATPase+Lon-Protease-BrxL<-PglZ<-N6-METHYLASE<-DUF6079*<-BrxB-BrxF-iSTAND<-BrxA-DNA-binding\|\|REase+iREase-><-wHTH+SAD+HNH\|\|<-?<-?<-?\|\|Phage_integrase->Phage_integrase-> | 1213 | Bacteria>Pseudomonadota>Gammaproteobacteria>Enterobacterales>Enterobacteriaceae>Escherichia | GCF_013376895.1 | Escherichia_coli | A7A13_RS22790 | 4759790..4763431 | 562 | BREX-system-P-loop-protein-BrxC-[Escherichia-coli] |
| WP_179778150.1 | Type1.BREX | <-Phage_integrase<-Phage_integrase\|\|?->?->?->\|\|wHTH+SAD+HNH-><-REase+iREase\|\|BrxA-DNA-binding->BrxB-BrxF-iSTAND->DUF6079*->N6-METHYLASE->PglZ->SIGMA+OB-Fold+AAA-ATPase+Lon-Protease-BrxL->?-><-?<-ParB<-?\|\|<-REase+N6_METHYLASE<-TRD | 1213 | Bacteria>Pseudomonadota>Gammaproteobacteria>Enterobacterales>Enterobacteriaceae>Escherichia | GCF_013413745.1 | Escherichia_coli | HZT14_RS03845 | 97447..101088 | 562 | BREX-system-P-loop-protein-BrxC-[Escherichia-coli] |
| WP_032228177.1 | Type1.BREX | wHTH+WYL+WCX->BrxA-DNA-binding->BrxB-BrxF-iSTAND->DUF6079*->ParB+HNH->N6-METHYLASE->PglZ->SIGMA+OB-Fold+AAA-ATPase+Lon-Protease-BrxL->35exo->\|\|HIN-HTH->\|\|HIN-HTH+PAIREDC-HTH-><-Arm-DNA-bind_3+Phage_integrase<-tRNA<-TetR-HTH | 1225 | Bacteria>Pseudomonadota>Gammaproteobacteria>Enterobacterales>Enterobacteriaceae>Escherichia | GCF_013424245.1 | Escherichia_coli | HOP07_RS08595 | 2755..6432 | 562 | BREX-system-P-loop-protein-BrxC-[Escherichia-coli] |
| WP_180489728.1 | Type1.BREX | <-REase<-REase\|\|BrxA-DNA-binding->BrxB-BrxF-iSTAND->DUF6079*->N6-METHYLASE->PglZ->SIGMA+OB-Fold+AAA-ATPase+Lon-Protease-BrxL-><-ZnR<-ZnR<-?\|\|RHH_antitoxin->XIS-HTH-> | 1213 | Bacteria>Pseudomonadota>Gammaproteobacteria>Enterobacterales>Enterobacteriaceae>Escherichia | GCF_013424415.1 | Escherichia_coli | HOO79_RS20980 | 3102..6743 | 562 | BREX-system-P-loop-protein-BrxC-[Escherichia-coli] |
| WP_181678095.1 | Type1.BREX | <-REase<-REase\|\|BrxA-DNA-binding->BrxB-BrxF-iSTAND->DUF6079*->N6-METHYLASE->PglZ->SIGMA+OB-Fold+AAA-ATPase+Lon-Protease-BrxL-><-ZnR<-ZnR | 1213 | Bacteria>Pseudomonadota>Gammaproteobacteria>Enterobacterales>Enterobacteriaceae>Escherichia | GCF_013851105.1 | Escherichia_coli | H4962_RS13940 | 2269..5910 | 562 | BREX-system-P-loop-protein-BrxC-[Escherichia-coli] |
| WP_182913085.1 | Type1.BREX | <-HIN-HTH<-RHH_antitoxin\|\|?->?->ZnR->ZnR-><-SIGMA+OB-Fold+AAA-ATPase+Lon-Protease-BrxL<-PglZ<-N6-METHYLASE<-DUF6079*<-BrxB-BrxF-iSTAND<-BrxA-DNA-binding\|\|REase+iREase->?->?->?-><-DDE_Tnp_IS1<-Zn_Tnp_IS1+HTH_Tnp_IS1<-Arm-DNA-bind_3+Phage_integrase | 1213 | Bacteria>Pseudomonadota>Gammaproteobacteria>Enterobacterales>Enterobacteriaceae>Escherichia | GCF_014168755.1 | Escherichia_coli | WP2S18E08_08260 | 876105..879746 | 562 | BREX-system-P-loop-protein-BrxC-[Escherichia-coli] |
| WP_025653408.1 | Type1.BREX | <-Phage_integrase<-Phage_integrase\|\|?->?->?->\|\|wHTH(SBD)+HNH-><-REase+iREase\|\|BrxA-DNA-binding->BrxB-BrxF-iSTAND->DUF6079*->N6-METHYLASE->PglZ->SIGMA+OB-Fold+AAA-ATPase+Lon-Protease-BrxL->HKD-DNase+OGG-like->\|\|AbiEi_3_N+AbiEi_3->AbiEii->?-><-DUF6216 | 1213 | Bacteria>Pseudomonadota>Gammaproteobacteria>Enterobacterales>Enterobacteriaceae>Escherichia | GCF_014169595.1 | Escherichia_coli | H7R13_RS19900 | 4165197..4168838 | 562 | BREX-system-P-loop-protein-BrxC-[Escherichia-coli] |
| WP_192481452.1 | Type1.BREX | <-Zn_Tnp_IS1+HTH_Tnp_IS1+DDE_Tnp_IS1\|\|?->?->ZnR->ZnR-><-SIGMA+OB-Fold+AAA-ATPase+Lon-Protease-BrxL<-PglZ<-N6-METHYLASE<-DUF6079*<-BrxB-BrxF-iSTAND<-BrxA-DNA-binding\|\|REase+iREase-><-EVE+HNH<-Arm-DNA-bind_3+Phage_integrase<-tRNA\|\|?->HerA/FtsK+A32-> | 1213 | Bacteria>Pseudomonadota>Gammaproteobacteria>Enterobacterales>Enterobacteriaceae>Escherichia | GCF_014858085.1 | Escherichia_coli | IHV13_RS01935 | 12078..15719 | 562 | BREX-system-P-loop-protein-BrxC-[Escherichia-coli] |
| WP_192960469.1 | Type1.BREX | <-REase+iREase\|\|BrxA-DNA-binding->BrxB-BrxF-iSTAND->DUF6079*->N6-METHYLASE-p-> | 1213 | Bacteria>Pseudomonadota>Gammaproteobacteria>Enterobacterales>Enterobacteriaceae>Escherichia | GCF_014892435.1 | Escherichia_coli | INS74_RS22970 | 2229..5870 | 562 | BREX-system-P-loop-protein-BrxC-[Escherichia-coli] |
| WP_193098973.1 | Type1.BREX | <-DUF6079-p*<-BrxB-BrxF-iSTAND<-BrxA-DNA-binding\|\|REase+iREase-><-EVE+HNH<-Arm-DNA-bind_3+Phage_integrase<-tRNA\|\|?->HerA/FtsK+A32-> | 684 | Bacteria>Pseudomonadota>Gammaproteobacteria>Enterobacterales>Enterobacteriaceae>Escherichia | GCF_014900935.1 | Escherichia_coli | ICK29_RS00005 | 1..2052 | 562 | BREX-system-P-loop-protein-BrxC,-partial-[Escherichia-coli] |
| WP_193097043.1 | Type1.BREX | <-N6-METHYLASE-p<-DUF6079*<-BrxB-BrxF-iSTAND<-BrxA-DNA-binding\|\|REase+iREase-><-EVE+HNH<-Arm-DNA-bind_3+Phage_integrase<-tRNA\|\|?->HerA/FtsK+A32-> | 1213 | Bacteria>Pseudomonadota>Gammaproteobacteria>Enterobacterales>Enterobacteriaceae>Escherichia | GCF_014900955.1 | Escherichia_coli | ICK28_RS00010 | 345..3986 | 562 | BREX-system-P-loop-protein-BrxC-[Escherichia-coli] |
| WP_087888065.1 | Type1.BREX | ABhydrolase-><-LZ_Tnp_IS66+zf-IS66+DDE_Tnp_IS66+DDE_Tnp_IS66_C\|\|?-><-Zn_Tnp_IS1+HTH_Tnp_IS1+DDE_Tnp_IS1\|\|wHTH+WYL+WCX->BrxA-DNA-binding->BrxB-BrxF-iSTAND->DUF6079*->ParB+HNH->N6-METHYLASE->PglZ->SIGMA+OB-Fold+AAA-ATPase+Lon-Protease-BrxL-><-DEDD_Tnp_IS110+Transposase_20<-RVT | 1225 | Bacteria>Pseudomonadota>Gammaproteobacteria>Enterobacterales>Enterobacteriaceae>Escherichia | GCF_015229345.1 | Escherichia_coli | GVI22_RS02065 | 20896..24573 | 562 | BREX-system-P-loop-protein-BrxC-[Escherichia-coli] |
| WP_097450867.1 | Type1.BREX | <-XIS-HTH<-RHH_antitoxin\|\|?->?->ZnR->ZnR-><-SIGMA+OB-Fold+AAA-ATPase+Lon-Protease-BrxL<-PglZ<-N6-METHYLASE<-DUF6079*<-BrxB-BrxF-iSTAND<-BrxA-DNA-binding\|\|<-EVE+HNH<-Arm-DNA-bind_3+Phage_integrase<-tRNA\|\|?-><-SKI\|\|METHYLASE-> | 1213 | Bacteria>Pseudomonadota>Gammaproteobacteria>Enterobacterales>Enterobacteriaceae>Escherichia | GCF_015476745.1 | Escherichia_coli | IOB98_RS17950 | 606603..610244 | 562 | BREX-system-P-loop-protein-BrxC-[Escherichia-coli] |
| WP_195390166.1 | Type1.BREX | <-XIS-HTH\|\|?-><-?\|\|?->ZnR->ZnR-><-SIGMA+OB-Fold+AAA-ATPase+Lon-Protease-BrxL<-PglZ<-N6-METHYLASE<-DUF6079*<-BrxB-BrxF-iSTAND<-BrxA-DNA-binding\|\|REase+iREase-><-EVE+HNH<-Arm-DNA-bind_3+Phage_integrase<-tRNA\|\|?-><-SKI\|\|METHYLASE-> | 1213 | Bacteria>Pseudomonadota>Gammaproteobacteria>Enterobacterales>Enterobacteriaceae>Escherichia | GCF_015553825.1 | Escherichia_coli | I2D81_RS15750 | 15887..19528 | 562 | BREX-system-P-loop-protein-BrxC-[Escherichia-coli] |
| WP_195745416.1 | Type1.BREX | <-REase+iREase\|\|BrxA-DNA-binding->BrxB-BrxF-iSTAND->DUF6079*->N6-METHYLASE->PglZ->SIGMA+OB-Fold+AAA-ATPase+Lon-Protease-BrxL-><-ZnR<-ZnR | 1213 | Bacteria>Pseudomonadota>Gammaproteobacteria>Enterobacterales>Enterobacteriaceae>Escherichia | GCF_015620345.1 | Escherichia_coli | I3C17_RS21905 | 2642..6283 | 562 | BREX-system-P-loop-protein-BrxC-[Escherichia-coli] |
| WP_197364841.1 | Type1.BREX | BrxB-BrxF-iSTAND->DUF6079*->N6-METHYLASE->PglZ->SIGMA+OB-Fold+AAA-ATPase+Lon-Protease-BrxL->HKD-DNase+OGG-like->\|\|AbiEi_3_N+AbiEi_3->AbiEii-><-DUF6216<-tRNA\|\|?->HerA/FtsK+A32-> | 1213 | Bacteria>Pseudomonadota>Gammaproteobacteria>Enterobacterales>Enterobacteriaceae>Escherichia | GCF_015912555.1 | Escherichia_coli | F2X22_RS13815 | 740..4381 | 562 | BREX-system-P-loop-protein-BrxC-[Escherichia-coli] |
| WP_201358321.1 | Type1.BREX | XIS-HTH->?-><-?<-PemK_toxin\|\|?->?-><-Zn_Tnp_IS1+HTH_Tnp_IS1+DDE_Tnp_IS1\|\|BrxA-DNA-binding->BrxB-BrxF-iSTAND->DUF6079*->PglZ->SIGMA+OB-Fold+AAA-ATPase+Lon-Protease-BrxL-><-ZnR<-ZnR<-?<-?\|\|RHH_antitoxin-><-HTH_Tnp_1+HTH_21+rve+rve_3 | 1213 | Bacteria>Pseudomonadota>Gammaproteobacteria>Enterobacterales>Enterobacteriaceae>Escherichia | GCF_016587155.1 | Escherichia_coli | JMH48_RS18845 | 518753..522394 | 562 | BREX-system-P-loop-protein-BrxC-[Escherichia-coli] |
| WP_202845940.1 | Type1.BREX | <-PemK_toxin<-HIN-HTH\|\|?->?-><-Zn_Tnp_IS1+HTH_Tnp_IS1+DDE_Tnp_IS1\|\|BrxA-DNA-binding->BrxB-BrxF-iSTAND->DUF6079*->N6-METHYLASE->PglZ->SIGMA+OB-Fold+AAA-ATPase+Lon-Protease-BrxL-><-ZnR<-ZnR<-?<-?\|\|RHH_antitoxin-><-HTH_Tnp_1+HTH_21+rve+rve_3 | 1213 | Bacteria>Pseudomonadota>Gammaproteobacteria>Enterobacterales>Enterobacteriaceae>Escherichia | GCF_016776325.1 | Escherichia_coli | JNN51_RS16875 | 3525018..3528659 | 562 | BREX-system-P-loop-protein-BrxC-[Escherichia-coli] |
| WP_032343442.1 | Type1.BREX | REase+N6_METHYLASE-><-?\|\|?->ParB-><-?<-SIGMA+OB-Fold+AAA-ATPase+Lon-Protease-BrxL<-PglZ<-N6-METHYLASE<-DUF6079*<-BrxB-BrxF-iSTAND<-BrxA-DNA-binding\|\|REase+iREase-><-wHTH+SAD+HNH\|\|<-?<-?<-?\|\|Phage_integrase->Phage_integrase-> | 1213 | Bacteria>Pseudomonadota>Gammaproteobacteria>Enterobacterales>Enterobacteriaceae>Escherichia | GCF_016858785.1 | Escherichia_coli | BEA16_RS14875 | 18248..21889 | 562 | BREX-system-P-loop-protein-BrxC-[Escherichia-coli] |
| WP_208634821.1 | Type1.BREX | <-HIN-HTH<-?<-XIS-HTH\|\|HIN-HTH->ZnR->ZnR-><-SIGMA+OB-Fold+AAA-ATPase+Lon-Protease-BrxL<-PglZ<-N6-METHYLASE<-DUF6079*<-BrxB-BrxF-iSTAND<-BrxA-DNA-binding\|\|REase+iREase->?->?-><-Zn_Tnp_IS1+HTH_Tnp_IS1+DDE_Tnp_IS1<-Arm-DNA-bind_3+Phage_integrase<-RelE-ParE<-MetJarc-HTH | 1213 | Bacteria>Pseudomonadota>Gammaproteobacteria>Enterobacterales>Enterobacteriaceae>Escherichia | GCF_016904795.1 | Escherichia_albertii | JRC43_RS04070 | 837246..840887 | 208962 | BREX-system-P-loop-protein-BrxC-[Escherichia-albertii] |
| WP_072666953.1 | Type1.BREX | Arm-DNA-bind_3+Phage_integrase-><-Sulfatase<-?\|\|DDE_Tnp_1_5+DDE_Tnp_1-><-Zn_Tnp_IS1+HTH_Tnp_IS1+DDE_Tnp_IS1<-?<-REase+iREase\|\|BrxA-DNA-binding->BrxB-BrxF-iSTAND->DUF6079*->N6-METHYLASE->PglZ->SIGMA+OB-Fold+AAA-ATPase+Lon-Protease-BrxL-><-ZnR<-ZnR<-?<-?\|\|RHH_antitoxin->XIS-HTH-> | 1213 | Bacteria>Pseudomonadota>Gammaproteobacteria>Enterobacterales>Enterobacteriaceae>Escherichia | GCF_017088485.1 | Escherichia_coli | JR883_RS20005 | 4162845..4166486 | 562 | BREX-system-P-loop-protein-BrxC-[Escherichia-coli] |
| WP_205937296.1 | Type1.BREX | DUF6216-><-?<-AbiEii<-AbiEi_3_N+AbiEi_3\|\|<-HKD-DNase+OGG-like<-SIGMA+OB-Fold+AAA-ATPase+Lon-Protease-BrxL<-PglZ<-N6-METHYLASE<-DUF6079*<-BrxB-BrxF-iSTAND<-BrxA-DNA-binding\|\|REase+iREase-><-wHTH+SAD+HNH\|\|<-?<-?<-?\|\|Phage_integrase->Fimbrial-> | 1213 | Bacteria>Pseudomonadota>Gammaproteobacteria>Enterobacterales>Enterobacteriaceae>Escherichia | GCF_017117165.1 | Escherichia_coli | C3E82_RS10505 | 78550..82191 | 562 | BREX-system-P-loop-protein-BrxC-[Escherichia-coli] |
| WP_207516242.1 | Type1.BREX | <-Phage_integrase<-Phage_integrase\|\|?->?->?->\|\|wHTH+SAD+HNH-><-REase+iREase\|\|BrxA-DNA-binding->BrxB-BrxF-iSTAND->DUF6079*->N6-METHYLASE->PglZ->SIGMA+OB-Fold+AAA-ATPase+Lon-Protease-BrxL->HKD-DNase+OGG-like->\|\|AbiEi_3_N+AbiEi_3->AbiEii->?-><-DUF6216 | 1213 | Bacteria>Pseudomonadota>Gammaproteobacteria>Enterobacterales>Enterobacteriaceae>Escherichia | GCF_017356105.1 | Escherichia_coli | J4Q20_RS14625 | 71539..75180 | 562 | BREX-system-P-loop-protein-BrxC-[Escherichia-coli] |
| WP_208482710.1 | Type1.BREX | <-RHH_antitoxin\|\|HIN-HTH->?->?->ZnR->ZnR-><-SIGMA+OB-Fold+AAA-ATPase+Lon-Protease-BrxL<-PglZ<-N6-METHYLASE<-DUF6079*<-BrxB-BrxF-iSTAND<-BrxA-DNA-binding\|\|REase+iREase-> | 1213 | Bacteria>Pseudomonadota>Gammaproteobacteria>Enterobacterales>Enterobacteriaceae>Escherichia | GCF_017598785.1 | Escherichia_coli | J7R96_RS22835 | 18064..21705 | 562 | BREX-system-P-loop-protein-BrxC-[Escherichia-coli] |
| WP_210173219.1 | Type1.BREX | <-Phage_integrase<-Phage_integrase\|\|?->?->?->\|\|wHTH+SAD+HNH-><-REase+iREase\|\|BrxA-DNA-binding->BrxB-BrxF-iSTAND->DUF6079*->N6-METHYLASE->PglZ->SIGMA+OB-Fold+AAA-ATPase+Lon-Protease-BrxL->HKD-DNase+OGG-like->\|\|AbiEi_3_N+AbiEi_3->AbiEii->?-><-DUF6216 | 1213 | Bacteria>Pseudomonadota>Gammaproteobacteria>Enterobacterales>Enterobacteriaceae>Escherichia | GCF_017897155.1 | Escherichia_coli | J8352_RS05700 | 22726..26367 | 562 | BREX-system-P-loop-protein-BrxC-[Escherichia-coli] |
| WP_186240487.1 | Type1.BREX | <-Phage_integrase<-Phage_integrase\|\|?->?->?->\|\|wHTH+SAD+HNH-><-REase+iREase\|\|BrxA-DNA-binding->BrxB-BrxF-iSTAND->DUF6079*->N6-METHYLASE->PglZ->SIGMA+OB-Fold+AAA-ATPase+Lon-Protease-BrxL->HKD-DNase+OGG-like->\|\|AbiEi_3_N+AbiEi_3->AbiEii->?-><-DUF6216\|\|XIS-HTH-> | 1213 | Bacteria>Pseudomonadota>Gammaproteobacteria>Enterobacterales>Enterobacteriaceae>Escherichia | GCF_018141145.1 | Escherichia_coli | HUS69_RS19530 | 4048381..4052022 | 562 | BREX-system-P-loop-protein-BrxC-[Escherichia-coli] |
| WP_097470372.1 | Type1.BREX | <-Peptidase_M17_N+Peptidase_M17\|\|?->?->?-><-HerA/FtsK+A32<-?\|\|tRNA->Arm-DNA-bind_3+Phage_integrase->BrxA-DNA-binding->BrxB-BrxF-iSTAND->DUF6079*->N6-METHYLASE->OLD-ABC->TOPRIM->PglZ->SIGMA+OB-Fold+AAA-ATPase+Lon-Protease-BrxL-><-URI-DNase+RAMA<-ZnR<-ZnR<-wHTH | 1204 | Bacteria>Pseudomonadota>Gammaproteobacteria>Enterobacterales>Enterobacteriaceae>Escherichia | GCF_018682695.1 | Escherichia_coli | BEG75_RS10330 | 67748..71362 | 562 | BREX-system-P-loop-protein-BrxC-[Escherichia-coli] |
| WP_216317187.1 | Type1.BREX | <-Phage_integrase<-Phage_integrase\|\|?->?->?->\|\|wHTH+SAD+HNH-><-REase+iREase\|\|BrxA-DNA-binding->BrxB-BrxF-iSTAND->DUF6079*->N6-METHYLASE->PglZ->SIGMA+OB-Fold+AAA-ATPase+Lon-Protease-BrxL->HKD-DNase+OGG-like->\|\|AbiEi_3_N+AbiEi_3->AbiEii->?-><-DUF6216<-TOPRIM-DNAG+D5N+D5ATPase+P4PRIMASE-HTH | 1213 | Bacteria>Pseudomonadota>Gammaproteobacteria>Enterobacterales>Enterobacteriaceae>Escherichia | GCF_018884345.1 | Escherichia_coli | EXA04_RS21875 | 4479425..4483066 | 562 | BREX-system-P-loop-protein-BrxC-[Escherichia-coli] |
| WP_217800889.1 | Type1.BREX | <-Phage_integrase<-Phage_integrase\|\|?->?->?->\|\|wHTH+SAD+HNH-><-REase+iREase\|\|BrxA-DNA-binding->BrxB-BrxF-iSTAND->DUF6079*->N6-METHYLASE->PglZ->SIGMA+OB-Fold+AAA-ATPase+Lon-Protease-BrxL->?-><-?<-ParB<-?\|\|?-><-REase+N6_METHYLASE | 1213 | Bacteria>Pseudomonadota>Gammaproteobacteria>Enterobacterales>Enterobacteriaceae>Escherichia | GCF_019120875.1 | Escherichia_coli | KR452_RS02525 | 132614..136255 | 562 | BREX-system-P-loop-protein-BrxC-[Escherichia-coli] |
| WP_220298575.1 | Type1.BREX | <-SIGMA-HTH\|\|?-><-REase+iREase\|\|BrxA-DNA-binding->BrxB-BrxF-iSTAND->DUF6079*->N6-METHYLASE->PglZ->SIGMA+OB-Fold+AAA-ATPase+Lon-Protease-BrxL-><-ZnR<-ZnR<-?<-?\|\|RHH_antitoxin->XIS-HTH-> | 1213 | Bacteria>Pseudomonadota>Gammaproteobacteria>Enterobacterales>Enterobacteriaceae>Escherichia | GCF_019509425.1 | Escherichia_coli | KZA94_RS20415 | 22821..26462 | 562 | BREX-system-P-loop-protein-BrxC-[Escherichia-coli] |
| WP_222694585.1 | Type1.BREX | <-METHYLASE\|\|SKI-><-?\|\|tRNA->Arm-DNA-bind_3+Phage_integrase->EVE+HNH-><-REase+iREase\|\|BrxA-DNA-binding->BrxB-BrxF-iSTAND->DUF6079-p*-> | 849 | Bacteria>Pseudomonadota>Gammaproteobacteria>Enterobacterales>Enterobacteriaceae>Escherichia | GCF_019839475.1 | Escherichia_coli | K6681_RS00785 | 145766..148312 | 562 | BREX-system-P-loop-protein-BrxC,-partial-[Escherichia-coli] |
| WP_222687482.1 | Type1.BREX | BrxA-DNA-binding->BrxB-BrxF-iSTAND->DUF6079*->N6-METHYLASE->PglZ->SIGMA+OB-Fold+AAA-ATPase+Lon-Protease-BrxL-><-ZnR<-ZnR | 1213 | Bacteria>Pseudomonadota>Gammaproteobacteria>Enterobacterales>Enterobacteriaceae>Escherichia | GCF_019840005.1 | Escherichia_coli | K6684_RS15390 | 1591..5232 | 562 | BREX-system-P-loop-protein-BrxC-[Escherichia-coli] |
| WP_222698798.1 | Type1.BREX | <-REase+iREase\|\|BrxA-DNA-binding->BrxB-BrxF-iSTAND->DUF6079*->N6-METHYLASE-p-> | 1213 | Bacteria>Pseudomonadota>Gammaproteobacteria>Enterobacterales>Enterobacteriaceae>Escherichia | GCF_019840195.1 | Escherichia_coli | K6666_RS17905 | 2229..5870 | 562 | BREX-system-P-loop-protein-BrxC-[Escherichia-coli] |
| WP_220680248.1 | Type1.BREX | <-RHH_antitoxin\|\|?->?->ZnR->ZnR-><-SIGMA+OB-Fold+AAA-ATPase+Lon-Protease-BrxL<-PglZ<-N6-METHYLASE<-DUF6079*<-BrxB-BrxF-iSTAND<-BrxA-DNA-binding\|\|REase->REase-> | 1213 | Bacteria>Pseudomonadota>Gammaproteobacteria>Enterobacterales>Enterobacteriaceae>Escherichia | GCF_019926805.1 | Escherichia_coli | JOW43_RS22215 | 13140..16781 | 562 | BREX-system-P-loop-protein-BrxC-[Escherichia-coli] |
| WP_219370674.1 | Type1.BREX | <-XIS-HTH<-RHH_antitoxin\|\|?->?->ZnR->ZnR-><-SIGMA+OB-Fold+AAA-ATPase+Lon-Protease-BrxL<-PglZ<-DDE_Tnp_1<-DUF6079*<-BrxB-BrxF-iSTAND<-BrxA-DNA-binding\|\|REase+iREase-><-EVE+HNH<-Arm-DNA-bind_3+Phage_integrase<-tRNA\|\|?->HerA/FtsK+A32-><-?<-NAD_binding_10 | 1213 | Bacteria>Pseudomonadota>Gammaproteobacteria>Enterobacterales>Enterobacteriaceae>Escherichia | GCF_020002285.1 | Escherichia_fergusonii | KYC43_RS19615 | 4024013..4027654 | 564 | BREX-system-P-loop-protein-BrxC-[Escherichia-fergusonii] |
| WP_224452732.1 | Type1.BREX | <-HTH_OrfB_IS605+OrfB_IS605+OrfB_Zn_ribbon<-ParB<-MerR-HTH+ParA-Soj-PloopNTPase\|\|?->DEDD_Tnp_IS110+Transposase_20->\|\|wHTH+WYL+WCX->BrxA-DNA-binding->BrxB-BrxF-iSTAND->DUF6079*->ParB+HNH->N6-METHYLASE->PglZ->SIGMA+OB-Fold+AAA-ATPase+Lon-Protease-BrxL-> | 1225 | Bacteria>Pseudomonadota>Gammaproteobacteria>Enterobacterales>Enterobacteriaceae>Escherichia | GCF_020162075.2 | Escherichia_coli | KD159_RS16650 | 41518..45195 | 562 | BREX-system-P-loop-protein-BrxC-[Escherichia-coli] |
| WP_089559427.1 | Type1.BREX | <-XIS-HTH<-RHH_antitoxin\|\|?->?->ZnR->ZnR-><-SIGMA+OB-Fold+AAA-ATPase+Lon-Protease-BrxL<-PglZ<-N6-METHYLASE<-DUF6079*<-BrxB-BrxF-iSTAND<-BrxA-DNA-binding<-DDE_Tnp_1_5+DDE_Tnp_1\|\|?->Sulfatase-><-Arm-DNA-bind_3+Phage_integrase<-tRNA<-DsbC+DsbD+Thioredoxin_7 | 1213 | Bacteria>Pseudomonadota>Gammaproteobacteria>Enterobacterales>Enterobacteriaceae>Escherichia | GCF_020230375.1 | Escherichia_coli | LF808_RS18060 | 3738120..3741761 | 562 | BREX-system-P-loop-protein-BrxC-[Escherichia-coli] |

| Protein-ID | System | Gene-Neighbourhoods | Length | Taxonomy | Assembly-ID | Strain | Locus-Tag | Genomic-coordinates | TaxID | Fasta-define |
|---|---|---|---|---|---|---|---|---|---|---|
| WP_063122485.1 | Type1.BREX | <-RHH_antitoxin\|\|?->?->ZnR->ZnR-><-SIGMA+OB-Fold+AAA-ATPase+Lon-Protease-BrxL<-PglZ<-N6-METHYLASE<-DUF6079*<-BrxB-BrxF-iSTAND<-BrxA-DNA-binding\|\|REase+iREase->?->?-><-Arm-DNA-bind_3+Phage_integrase<-RelE-ParE<-MetJarc-HTH<-METHYLASE | 1213 | Bacteria>Pseudomonadota>Gammaproteobacteria>Enterobacterales>Enterobacteriaceae>Escherichia | GCF_020230435.1 | Escherichia_coli | LF812_RS09505 | 1992870..1996511 | 562 | BREX-system-P-loop-protein-BrxC-[Escherichia-coli] |
| WP_130560426.1 | Type1.BREX | <-XIS-HTH\|\|?->?->ZnR->ZnR-><-SIGMA+OB-Fold+AAA-ATPase+Lon-Protease-BrxL<-PglZ<-N6-METHYLASE<-DUF6079*<-BrxB-BrxF-iSTAND<-BrxA-DNA-binding<-DDE_Tnp_IS1 | 1213 | Bacteria>Pseudomonadota>Gammaproteobacteria>Enterobacterales>Enterobacteriaceae>Escherichia | GCF_020282105.1 | Escherichia_coli | IHV37_RS24260 | 16243..19884 | 562 | BREX-system-P-loop-protein-BrxC-[Escherichia-coli] |
| WP_227262353.1 | Type1.BREX | <-Peptidase_M17_N+Peptidase_M17\|\|?->?-><-HerA/FtsK+A32\|\|tRNA->Arm-DNA-bind_3+Phage_integrase->EVE+HNH-><-REase+iREase\|\|BrxA-DNA-binding->BrxB-BrxF-iSTAND->DUF6079*->N6-METHYLASE->PglZ->SIGMA+OB-Fold+AAA-ATPase+Lon-Protease-BrxL-><-ZnR<-ZnR<-?<-?\|\|RHH_antitoxin->XIS-HTH-> | 1213 | Bacteria>Pseudomonadota>Gammaproteobacteria>Enterobacterales>Enterobacteriaceae>Escherichia | GCF_020594775.1 | Escherichia_coli | LJD39_RS10205 | 66062..69703 | 562 | BREX-system-P-loop-protein-BrxC-[Escherichia-coli] |
| WP_229322431.1 | Type1.BREX | <-HerA/FtsK+A32<-?\|\|tRNA->Arm-DNA-bind_3+Phage_integrase->EVE+HNH-><-REase+iREase\|\|BrxA-DNA-binding->BrxB-BrxF-iSTAND->DUF6079*->N6-METHYLASE->PglZ->SIGMA+OB-Fold+AAA-ATPase+Lon-Protease-BrxL-><-ZnR<-ZnR | 1213 | Bacteria>Pseudomonadota>Gammaproteobacteria>Enterobacterales>Enterobacteriaceae>Escherichia | GCF_020883515.1 | Escherichia_coli | L2W85_RS01370 | 290770..294411 | 562 | BREX-system-P-loop-protein-BrxC-[Escherichia-coli] |
| WP_231833501.1 | Type1.BREX | <-ParB+HNH<-DUF6079*<-BrxB-BrxF-iSTAND<-BrxA-DNA-binding<-wHTH+WYL+WCX\|\|DUF4158+DDE_Tnp_Tn3-> | 1225 | Bacteria>Pseudomonadota>Gammaproteobacteria>Enterobacterales>Enterobacteriaceae>Escherichia | GCF_021168085.1 | Escherichia_coli | LTT78_RS21790 | 2950..6627 | 562 | BREX-system-P-loop-protein-BrxC-[Escherichia-coli] |
| WP_097495602.1 | Type1.BREX | <-TetR-HTH<-?<-?<-REase<-REase\|\|BrxA-DNA-binding->BrxB-BrxF-iSTAND->DUF6079*->N6-METHYLASE->PglZ->SIGMA+OB-Fold+AAA-ATPase+Lon-Protease-BrxL-> | 1213 | Bacteria>Pseudomonadota>Gammaproteobacteria>Enterobacterales>Enterobacteriaceae>Escherichia | GCF_021547795.1 | Escherichia_coli | BEC04_RS22745 | 6957..10598 | 562 | BREX-system-P-loop-protein-BrxC-[Escherichia-coli] |
| WP_236265313.1 | Type1.BREX | <-XIS-HTH<-RHH_antitoxin\|\|?->?->ZnR->ZnR-><-SIGMA+OB-Fold+AAA-ATPase+Lon-Protease-BrxL<-PglZ<-N6-METHYLASE<-DUF6079*<-BrxB-BrxF-iSTAND<-BrxA-DNA-binding\|\|REase+iREase-> | 1213 | Bacteria>Pseudomonadota>Gammaproteobacteria>Enterobacterales>Enterobacteriaceae>Escherichia | GCF_021647095.1 | Escherichia_coli | L6H61_RS22580 | 19345..22986 | 562 | BREX-system-P-loop-protein-BrxC-[Escherichia-coli] |
| WP_236486706.1 | Type1.BREX | BrxA-DNA-binding->BrxB-BrxF-iSTAND->DUF6079*->N6-METHYLASE->PglZ->SIGMA+OB-Fold+AAA-ATPase+Lon-Protease-BrxL-><-ZnR<-ZnR | 1213 | Bacteria>Pseudomonadota>Gammaproteobacteria>Enterobacterales>Enterobacteriaceae>Escherichia | GCF_021648945.1 | Escherichia_coli | L6E82_RS22710 | 1925..5566 | 562 | BREX-system-P-loop-protein-BrxC-[Escherichia-coli] |
| WP_112885923.1 | Type1.BREX | <-HerA/FtsK+A32<-?\|\|tRNA->Arm-DNA-bind_3+Phage_integrase->EVE+HNH-><-REase+iREase\|\|BrxA-DNA-binding->BrxB-BrxF-iSTAND->DUF6079*->N6-METHYLASE->PglZ->SIGMA+OB-Fold+AAA-ATPase+Lon-Protease-BrxL-><-ZnR<-ZnR<-?<-?\|\|DDE_Tnp_1_5+DDE_Tnp_1-> | 1213 | Bacteria>Pseudomonadota>Gammaproteobacteria>Enterobacterales>Enterobacteriaceae>Escherichia | GCF_021697115.1 | Escherichia_coli | IHV65_RS22435 | 19394..23035 | 562 | BREX-system-P-loop-protein-BrxC-[Escherichia-coli] |
| WP_236936469.1 | Type1.BREX | <-Phage_integrase<-Phage_integrase\|\|?->?->?->\|\|wHTH+SAD+HNH-><-REase+iREase\|\|BrxA-DNA-binding->BrxB-BrxF-iSTAND->DUF6079*->N6-METHYLASE->PglZ->SIGMA+OB-Fold+AAA-ATPase+Lon-Protease-BrxL->HKD-DNase+OGG-like->\|\|AbiEi_3_N+AbiEi_3->AbiEii->?-><-DUF6216 | 1213 | Bacteria>Pseudomonadota>Gammaproteobacteria>Enterobacterales>Enterobacteriaceae>Escherichia | GCF_021698335.1 | Escherichia_coli | IHV39_RS14485 | 12915..16556 | 562 | BREX-system-P-loop-protein-BrxC-[Escherichia-coli] |
| WP_089663046.1 | Type1.BREX | <-Peptidase_M17_N+Peptidase_M17\|\|?->?->?-><-HerA/FtsK+A32<-?\|\|tRNA->Arm-DNA-bind_3+Phage_integrase->BrxA-DNA-binding->BrxB-BrxF-iSTAND->DUF6079*->N6-METHYLASE->OLD-ABC->TOPRIM->PglZ->SIGMA+OB-Fold+AAA-ATPase+Lon-Protease-BrxL-><-URI-DNase+RAMA<-ZnR<-ZnR<-wHTH | 1204 | Bacteria>Pseudomonadota>Gammaproteobacteria>Enterobacterales>Enterobacteriaceae>Escherichia | GCF_022489725.1 | Escherichia_coli | ML322_RS02965 | 174957..178571 | 562 | BREX-system-P-loop-protein-BrxC-[Escherichia-coli] |
| WP_213838985.1 | Type1.BREX | <-RHH_antitoxin\|\|?->?->?->ZnR->ZnR-><-SIGMA+OB-Fold+AAA-ATPase+Lon-Protease-BrxL<-PglZ<-N6-METHYLASE<-DUF6079*<-BrxB-BrxF-iSTAND<-BrxA-DNA-binding\|\|REase+iREase->?->?-><-Zn_Tnp_IS1+HTH_Tnp_IS1+DDE_Tnp_IS1<-Arm-DNA-bind_3+Phage_integrase | 1213 | Bacteria>Pseudomonadota>Gammaproteobacteria>Enterobacterales>Enterobacteriaceae>Escherichia | GCF_022493455.1 | Escherichia_coli | AOY76_RS04040 | 833620..837261 | 562 | BREX-system-P-loop-protein-BrxC-[Escherichia-coli] |
| WP_241227056.1 | Type1.BREX | XIS-HTH->AAA+wHTH->?-><-SWIM+DUF3670+SF1-SF2-Helicase<-?<-?\|\|BrxA-DNA-binding->DUF6079*->N6-METHYLASE->PglZ->SIGMA+OB-Fold+AAA-ATPase+Lon-Protease-BrxL-><-DDE_Tnp_IS1 | 1213 | Bacteria>Pseudomonadota>Gammaproteobacteria>Enterobacterales>Enterobacteriaceae>Escherichia | GCF_022547975.1 | Escherichia_coli | EFK08_RS14245 | 31309..34950 | 562 | BREX-system-P-loop-protein-BrxC-[Escherichia-coli] |
| WP_109114033.1 | Type1.BREX | ZnR->ZnR-><-SIGMA+OB-Fold+AAA-ATPase+Lon-Protease-BrxL<-PglZ<-N6-METHYLASE<-DUF6079*<-BrxB-BrxF-iSTAND<-BrxA-DNA-binding | 1213 | Bacteria>Pseudomonadota>Gammaproteobacteria>Enterobacterales>Enterobacteriaceae>Escherichia | GCF_022602835.1 | Escherichia_coli | JEU19_RS23720 | 11740..15381 | 562 | BREX-system-P-loop-protein-BrxC-[Escherichia-coli] |
| WP_247153716.1 | Type1.BREX | <-Phage_integrase<-Phage_integrase\|\|?->?->?->\|\|wHTH+SAD+HNH-><-REase+iREase\|\|BrxA-DNA-binding->BrxB-BrxF-iSTAND->DUF6079-p*-> | 427 | Bacteria>Pseudomonadota>Gammaproteobacteria>Enterobacterales>Enterobacteriaceae>Escherichia | GCF_022968215.1 | Escherichia_coli | MWG73_RS16615 | 71483..72763 | 562 | BREX-system-P-loop-protein-BrxC,-partial-[Escherichia-coli] |
| WP_247178533.1 | Type1.BREX | <-Phage_integrase<-Phage_integrase\|\|?->?->?->\|\|wHTH+SAD+HNH-><-REase+iREase\|\|BrxA-DNA-binding->BrxB-BrxF-iSTAND->DUF6079-p*-> | 625 | Bacteria>Pseudomonadota>Gammaproteobacteria>Enterobacterales>Enterobacteriaceae>Escherichia | GCF_022970855.1 | Escherichia_coli | MZQ30_RS16040 | 111241..113117 | 562 | BREX-system-P-loop-protein-BrxC,-partial-[Escherichia-coli] |
| WP_069898759.1 | Type1.BREX | <-Phage_integrase<-Phage_integrase\|\|?->?->?->\|\|wHTH+SAD+HNH-><-REase+iREase\|\|BrxA-DNA-binding->BrxB-BrxF-iSTAND->DUF6079*->N6-METHYLASE->PglZ->SIGMA+OB-Fold+AAA-ATPase+Lon-Protease-BrxL-> | 1213 | Bacteria>Pseudomonadota>Gammaproteobacteria>Enterobacterales>Enterobacteriaceae>Escherichia | GCF_022971155.1 | Escherichia_coli | EAI54_RS17715 | 71532..75173 | 562 | BREX-system-P-loop-protein-BrxC-[Escherichia-coli] |
| WP_247149149.1 | Type1.BREX | <-METHYLASE<-?\|\|?-><-?\|\|<-AAA-ATPase<-HIN-HTH\|\|wHTH+WYL+WCX->BrxA-DNA-binding->BrxB-BrxF-iSTAND->DUF6079*->ParB+HNH->N6-METHYLASE->PglZ->SIGMA+OB-Fold+AAA-ATPase+Lon-Protease-BrxL->35exo-><-DDE_Tnp_1_5+DDE_Tnp_1 | 1225 | Bacteria>Pseudomonadota>Gammaproteobacteria>Enterobacterales>Enterobacteriaceae>Escherichia | GCF_022975115.1 | Escherichia_coli | MZS66_RS10030 | 63457..67134 | 562 | BREX-system-P-loop-protein-BrxC-[Escherichia-coli] |
| WP_247168067.1 | Type1.BREX | <-XIS-HTH<-RHH_antitoxin\|\|?->?->ZnR->ZnR-><-SIGMA+OB-Fold+AAA-ATPase+Lon-Protease-BrxL<-PglZ<-N6-METHYLASE<-DUF6079*<-BrxB-BrxF-iSTAND<-BrxA-DNA-binding | 1213 | Bacteria>Pseudomonadota>Gammaproteobacteria>Enterobacterales>Enterobacteriaceae>Escherichia | GCF_022975475.1 | Escherichia_coli | MZT24_RS24480 | 15782..19423 | 562 | BREX-system-P-loop-protein-BrxC-[Escherichia-coli] |
| WP_247151822.1 | Type1.BREX | <-SIGMA+OB-Fold+AAA-ATPase+Lon-Protease-BrxL<-PglZ<-N6-METHYLASE<-DUF6079*<-BrxB-BrxF-iSTAND<-BrxA-DNA-binding\|\|REase+iREase-><-wHTH+SAD+HNH\|\|<-?<-?\|\|Phage_integrase->Phage_integrase->Fimbrial-> | 1213 | Bacteria>Pseudomonadota>Gammaproteobacteria>Enterobacterales>Enterobacteriaceae>Escherichia | GCF_022976225.1 | Escherichia_coli | MZU12_RS18965 | 11408..15049 | 562 | BREX-system-P-loop-protein-BrxC-[Escherichia-coli] |
| WP_247169646.1 | Type1.BREX | DUF6079-p*->N6-METHYLASE->PglZ->SIGMA+OB-Fold+AAA-ATPase+Lon-Protease-BrxL-><-ZnR<-ZnR<-?<-?\|\|RHH_antitoxin->XIS-HTH-> | 956 | Bacteria>Pseudomonadota>Gammaproteobacteria>Enterobacterales>Enterobacteriaceae>Escherichia | GCF_022977435.1 | Escherichia_coli | MZU72_RS23215 | 1..2871 | 562 | BREX-system-P-loop-protein-BrxC,-partial-[Escherichia-coli] |
| WP_152933333.1 | Type1.BREX | BrxA-DNA-binding->BrxB-BrxF-iSTAND->DUF6079*->N6-METHYLASE->PglZ->SIGMA+OB-Fold+AAA-ATPase+Lon-Protease-BrxL-><-ZnR<-ZnR | 1213 | Bacteria>Pseudomonadota>Gammaproteobacteria>Enterobacterales>Enterobacteriaceae>Escherichia | GCF_023482315.1 | Escherichia_coli | M5X22_RS20110 | 1744..5385 | 562 | BREX-system-P-loop-protein-BrxC-[Escherichia-coli] |
| WP_249573915.1 | Type1.BREX | wHTH+WYL+WCX->BrxA-DNA-binding->BrxB-BrxF-iSTAND->DUF6079*->ParB+HNH->N6-METHYLASE->PglZ->SIGMA+OB-Fold+AAA-ATPase+Lon-Protease-BrxL->35exo-> | 1225 | Bacteria>Pseudomonadota>Gammaproteobacteria>Enterobacterales>Enterobacteriaceae>Escherichia | GCF_023483525.1 | Escherichia_coli | M5X65_RS22540 | 2590..6267 | 562 | BREX-system-P-loop-protein-BrxC-[Escherichia-coli] |
| WP_250239414.1 | Type1.BREX | MobI-><-?<-LexA-protease\|\|35exo->?->wHTH+WYL+WCX->BrxA-DNA-binding->BrxB-BrxF-iSTAND->DUF6079*->ParB+HNH->N6-METHYLASE->PglZ->SIGMA+OB-Fold+AAA-ATPase+Lon-Protease-BrxL->SbcC->HDRnase->VirD4-TraG+VirB4-FtsK-> | 1225 | Bacteria>Pseudomonadota>Gammaproteobacteria>Enterobacterales>Enterobacteriaceae>Escherichia | GCF_023599595.1 | Escherichia_coli | NFF73_RS04910 | 234938..238615 | 562 | MULTISPECIES:-BREX-system-P-loop-protein-BrxC-[Enterobacterales] |
| WP_097733507.1 | Type1.BREX | <-HerA/FtsK+A32<-?\|\|tRNA->Arm-DNA-bind_3+Phage_integrase->EVE+HNH-><-REase+iREase\|\|BrxA-DNA-binding->BrxB-BrxF-iSTAND->DUF6079*->N6-METHYLASE->PglZ->SIGMA+OB-Fold+AAA-ATPase+Lon-Protease-BrxL-><-ZnR<-ZnR<-?<-?\|\|RHH_antitoxin->XIS-HTH-> | 1213 | Bacteria>Pseudomonadota>Gammaproteobacteria>Enterobacterales>Enterobacteriaceae>Escherichia | GCF_023600485.1 | Escherichia_coli | LL309_RS17435 | 16120..19761 | 562 | BREX-system-P-loop-protein-BrxC-[Escherichia-coli] |
| WP_089551103.1 | Type1.BREX | <-XIS-HTH<-RHH_antitoxin\|\|?->?->ZnR->ZnR-><-SIGMA+OB-Fold+AAA-ATPase+Lon-Protease-BrxL<-PglZ<-N6-METHYLASE<-DUF6079*<-BrxB-BrxF-iSTAND<-BrxA-DNA-binding\|\|REase-> | 1213 | Bacteria>Pseudomonadota>Gammaproteobacteria>Enterobacterales>Enterobacteriaceae>Escherichia | GCF_023953245.1 | Escherichia_coli | M1V37_RS24515 | 16803..20444 | 562 | BREX-system-P-loop-protein-BrxC-[Escherichia-coli] |
| WP_254825569.1 | Type1.BREX | XIS-HTH->PIWI-><-?<-Phage_integrase\|\|Arm-DNA-bind_3+Phage_integrase->BrxA-DNA-binding->BrxB-BrxF-iSTAND->DUF6079*->N6-METHYLASE->OLD-ABC->TOPRIM->PglZ->SIGMA+OB-Fold+AAA-ATPase+Lon-Protease-BrxL-><-DDE_Tnp_1_5+DDE_Tnp_1<-?\|\|?->HEPN-> | 1204 | Bacteria>Pseudomonadota>Gammaproteobacteria>Enterobacterales>Enterobacteriaceae>Escherichia | GCF_024298945.1 | Escherichia_coli | LCR13_RS23605 | 136646..140260 | 562 | BREX-system-P-loop-protein-BrxC-[Escherichia-coli] |
| WP_255397240.1 | Type1.BREX | <-Phage_integrase<-Phage_integrase\|\|?->?->?->\|\|wHTH+SAD+HNH-><-REase+iREase\|\|BrxA-DNA-binding->BrxB-BrxF-iSTAND->DUF6079-p*-> | 919 | Bacteria>Pseudomonadota>Gammaproteobacteria>Enterobacterales>Enterobacteriaceae>Escherichia | GCF_024380595.1 | Escherichia_coli | NOE18_RS13095 | 111191..113948 | 562 | BREX-system-P-loop-protein-BrxC,-partial-[Escherichia-coli] |
| WP_255944649.1 | Type1.BREX | DUF6216-><-?<-AbiEii<-AbiEi_3_N+AbiEi_3\|\|<-HKD-DNase+OGG-like<-SIGMA+OB-Fold+AAA-ATPase+Lon-Protease-BrxL<-PglZ<-N6-METHYLASE<-DUF6079*<-BrxB-BrxF-iSTAND<-BrxA-DNA-binding\|\|REase+iREase-> | 1213 | Bacteria>Pseudomonadota>Gammaproteobacteria>Enterobacterales>Enterobacteriaceae>Escherichia | GCF_024450325.1 | Escherichia_coli | NO030_RS20220 | 28424..32065 | 562 | BREX-system-P-loop-protein-BrxC-[Escherichia-coli] |
| WP_256168548.1 | Type1.BREX | DsbC+DsbD+Thioredoxin_7->TetR-HTH->tRNA->Arm-DNA-bind_3+Phage_integrase-><-DOC<-PhdYeFM_antitox<-Zn_Tnp_IS1+HTH_Tnp_IS1+DDE_Tnp_IS1<-REase+iREase\|\|BrxA-DNA-binding->BrxB-BrxF-iSTAND->DUF6079*->N6-METHYLASE->PglZ->SIGMA+OB-Fold+AAA-ATPase+Lon-Protease-BrxL-><-ZnR<-?<-XIS-HTH | 1213 | Bacteria>Pseudomonadota>Gammaproteobacteria>Enterobacterales>Enterobacteriaceae>Escherichia | GCF_024467035.1 | Escherichia_coli | NO473_RS12000 | 2442717..2446358 | 562 | BREX-system-P-loop-protein-BrxC-[Escherichia-coli] |
| WP_256876221.1 | Type1.BREX | AraC-HTH->?-><-TetR-HTH<-?<-?<-REase<-REase\|\|BrxA-DNA-binding->BrxB-BrxF-iSTAND->DUF6079*->N6-METHYLASE->PglZ->SIGMA+OB-Fold+AAA-ATPase+Lon-Protease-BrxL-><-ZnR | 1213 | Bacteria>Pseudomonadota>Gammaproteobacteria>Enterobacterales>Enterobacteriaceae>Escherichia | GCF_024532055.1 | Escherichia_albertii | NHH32_RS22595 | 12425..16066 | 208962 | BREX-system-P-loop-protein-BrxC-[Escherichia-albertii] |
| WP_262043070.1 | Type1.BREX | RVT->ZnR-><-SIGMA+OB-Fold+AAA-ATPase+Lon-Protease-BrxL<-PglZ<-N6-METHYLASE<-DUF6079*<-BrxB-BrxF-iSTAND<-BrxA-DNA-binding\|\|REase->REase-> | 1213 | Bacteria>Pseudomonadota>Gammaproteobacteria>Enterobacterales>Enterobacteriaceae>Escherichia | GCF_024546885.1 | Escherichia_coli | NO383_RS24480 | 18816..22457 | 562 | BREX-system-P-loop-protein-BrxC-[Escherichia-coli] |
| WP_257249285.1 | Type1.BREX | HNH->?->?->\|\|wHTH+SAD+HNH-><-REase+iREase\|\|BrxA-DNA-binding->BrxB-BrxF-iSTAND->DUF6079*->N6-METHYLASE->PglZ->SIGMA+OB-Fold+AAA-ATPase+Lon-Protease-BrxL->HKD-DNase+OGG-like->\|\|AbiEi_3_N+AbiEi_3->AbiEii->?-><-DUF6216 | 1213 | Bacteria>Pseudomonadota>Gammaproteobacteria>Enterobacterales>Enterobacteriaceae>Escherichia | GCF_024608215.1 | Escherichia_coli | NO078_RS04965 | 41836..45477 | 562 | BREX-system-P-loop-protein-BrxC-[Escherichia-coli] |
| WP_257766119.1 | Type1.BREX | Arm-DNA-bind_3+Phage_integrase-><-Zn_Tnp_IS1+HTH_Tnp_IS1+DDE_Tnp_IS1<-TetR-HTH<-?<-?<-REase+iREase\|\|BrxA-DNA-binding->BrxB-BrxF-iSTAND->DUF6079*->N6-METHYLASE-> | 1213 | Bacteria>Pseudomonadota>Gammaproteobacteria>Enterobacterales>Enterobacteriaceae>Escherichia | GCF_024655425.1 | Escherichia_coli | NUU44_RS01945 | 122932..126573 | 562 | BREX-system-P-loop-protein-BrxC-[Escherichia-coli] |
| WP_257797852.1 | Type1.BREX | <-XIS-HTH<-RHH_antitoxin\|\|?->ZnR->ZnR-><-SIGMA+OB-Fold+AAA-ATPase+Lon-Protease-BrxL<-PglZ<-N6-METHYLASE<-DUF6079*<-BrxB-BrxF-iSTAND<-BrxA-DNA-binding\|\|REase+iREase-> | 1213 | Bacteria>Pseudomonadota>Gammaproteobacteria>Enterobacterales>Enterobacteriaceae>Escherichia | GCF_024700395.1 | Escherichia_coli | NU286_RS11325 | 47264..50905 | 562 | BREX-system-P-loop-protein-BrxC-[Escherichia-coli] |
| WP_260253294.1 | Type1.BREX | Zn_Tnp_IS1+HTH_Tnp_IS1+DDE_Tnp_IS1-><-?<-?\|\|PemK_toxin->PemK_toxin-><-Zn_Tnp_IS1+HTH_Tnp_IS1+DDE_Tnp_IS1<-SIGMA+OB-Fold+AAA-ATPase+Lon-Protease-BrxL<-PglZ<-N6-METHYLASE<-DUF6079*<-BrxB-BrxF-iSTAND<-BrxA-DNA-binding<-Zn_Tnp_IS1+HTH_Tnp_IS1+DDE_Tnp_IS1<-Zn_Tnp_IS1+HTH_Tnp_IS1+DDE_Tnp_IS1 | 1213 | Bacteria>Pseudomonadota>Gammaproteobacteria>Enterobacterales>Enterobacteriaceae>Escherichia | GCF_025200615.1 | Escherichia_coli | N1707_RS18325 | 3745633..3749274 | 562 | BREX-system-P-loop-protein-BrxC-[Escherichia-coli] |
| WP_261629031.1 | Type1.BREX | <-HIN-HTH<-RelE-ParE<-CopG/RHH\|\|<-RHH\|\|?->wHTH+WYL+WCX->BrxA-DNA-binding->BrxB-BrxF-iSTAND->DUF6079*->ParB+HNH->N6-METHYLASE->PglZ->SIGMA+OB-Fold+AAA-ATPase+Lon-Protease-BrxL-><-DEDD_Tnp_IS110+Transposase_20\|\|Zn_Tnp_IS1+HTH_Tnp_IS1+DDE_Tnp_IS1-> | 1225 | Bacteria>Pseudomonadota>Gammaproteobacteria>Enterobacterales>Enterobacteriaceae>Escherichia | GCF_025404235.1 | Escherichia_coli | J0W24_RS20355 | 5362..9039 | 562 | BREX-system-P-loop-protein-BrxC-[Escherichia-coli] |
| WP_262931584.1 | Type1.BREX | ZnR->ZnR-><-SIGMA+OB-Fold+AAA-ATPase+Lon-Protease-BrxL<-PglZ<-N6-METHYLASE<-DUF6079*<-BrxB-BrxF-iSTAND<-BrxA-DNA-binding\|\|REase+iREase-><-EVE+HNH<-Arm-DNA-bind_3+Phage_integrase<-tRNA\|\|?->HerA/FtsK+A32-> | 1213 | Bacteria>Pseudomonadota>Gammaproteobacteria>Enterobacterales>Enterobacteriaceae>Escherichia | GCF_025599895.1 | Escherichia_albertii | N6N71_RS13920 | 11398..15039 | 208962 | BREX-system-P-loop-protein-BrxC-[Escherichia-albertii] |
| WP_263767786.1 | Type1.BREX | DUF6216-><-?<-AbiEii<-AbiEi_3_N+AbiEi_3\|\|<-HKD-DNase+OGG-like<-SIGMA+OB-Fold+AAA-ATPase+Lon-Protease-BrxL<-PglZ<-N6-METHYLASE<-DUF6079* | 1072 | Bacteria>Pseudomonadota>Gammaproteobacteria>Enterobacterales>Enterobacteriaceae>Escherichia | GCF_025750965.1 | Escherichia_coli | OHF01_RS16785 | 78632..81850 | 562 | BREX-system-P-loop-protein-BrxC,-partial-[Escherichia-coli] |
| WP_171854687.1 | Type1.BREX | <-XIS-HTH<-RHH_antitoxin\|\|?->?->ZnR->ZnR-><-SIGMA+OB-Fold+AAA-ATPase+Lon-Protease-BrxL<-PglZ<-N6-METHYLASE<-DUF6079*<-BrxB-BrxF-iSTAND<-BrxA-DNA-binding\|\|<-EVE+HNH<-Arm-DNA-bind_3+Phage_integrase<-tRNA\|\|?-><-SKI\|\|METHYLASE-> | 1213 | Bacteria>Pseudomonadota>Gammaproteobacteria>Enterobacterales>Enterobacteriaceae>Escherichia | GCF_025946525.1 | Escherichia_coli | N7L43_RS20535 | 4233126..4236767 | 562 | BREX-system-P-loop-protein-BrxC-[Escherichia-coli] |
| WP_265188955.1 | Type1.BREX | <-XIS-HTH<-RHH_antitoxin\|\|?->ZnR->ZnR-><-SIGMA+OB-Fold+AAA-ATPase+Lon-Protease-BrxL<-PglZ<-N6-METHYLASE<-DUF6079*<-BrxB-BrxF-iSTAND<-BrxA-DNA-binding\|\|REase+iREase->Zn_Tnp_IS1+HTH_Tnp_IS1+DDE_Tnp_IS1-><-HEPN<-ORC-CDC6+ZnR<-N6-METHYLASE<-Arm-DNA-bind_3+Phage_integrase | 1213 | Bacteria>Pseudomonadota>Gammaproteobacteria>Enterobacterales>Enterobacteriaceae>Escherichia | GCF_026134335.1 | Escherichia_coli | OHD11_RS10990 | 2114745..2118386 | 562 | BREX-system-P-loop-protein-BrxC-[Escherichia-coli] |
| WP_265199179.1 | Type1.BREX | Arm-DNA-bind_3+Phage_integrase->N6-METHYLASE->ORC-CDC6+ZnR->HEPN-><-Zn_Tnp_IS1+HTH_Tnp_IS1+DDE_Tnp_IS1<-REase<-REase\|\|BrxA-DNA-binding->BrxB-BrxF-iSTAND->DUF6079*->N6-METHYLASE->PglZ->SIGMA+OB-Fold+AAA-ATPase+Lon-Protease-BrxL-><-HTH_Tnp_1+HTH_21+rve+rve_3\|\|RHH_antitoxin->XIS-HTH-> | 1213 | Bacteria>Pseudomonadota>Gammaproteobacteria>Enterobacterales>Enterobacteriaceae>Escherichia | GCF_026135205.1 | Escherichia_coli | OHD45_RS08935 | 1859993..1863634 | 562 | BREX-system-P-loop-protein-BrxC-[Escherichia-coli] |
| WP_265209809.1 | Type1.BREX | TPR-><-LACI-HTH+PBPI\|\|?->?-><-Zn_Tnp_IS1+HTH_Tnp_IS1+DDE_Tnp_IS1<-REase<-REase\|\|BrxA-DNA-binding->BrxB-BrxF-iSTAND->DUF6079*->N6-METHYLASE->PglZ->SIGMA+OB-Fold+AAA-ATPase+Lon-Protease-BrxL-><-ZnR<-RVT | 1213 | Bacteria>Pseudomonadota>Gammaproteobacteria>Enterobacterales>Enterobacteriaceae>Escherichia | GCF_026135825.1 | Escherichia_coli | NYV54_RS21460 | 4297326..4300967 | 562 | BREX-system-P-loop-protein-BrxC-[Escherichia-coli] |
| WP_265931224.1 | Type1.BREX | <-Peptidase_M17_N+Peptidase_M17\|\|?->?->?-><-HerA/FtsK+A32<-?\|\|tRNA->Arm-DNA-bind_3+Phage_integrase->BrxA-DNA-binding->BrxB-BrxF-iSTAND->DUF6079*->N6-METHYLASE->OLD-ABC->TOPRIM->PglZ->SIGMA+OB-Fold+AAA-ATPase+Lon-Protease-BrxL-><-URI-DNase+RAMA<-ZnR<-ZnR<-wHTH | 1204 | Bacteria>Pseudomonadota>Gammaproteobacteria>Enterobacterales>Enterobacteriaceae>Escherichia | GCF_026228105.1 | Escherichia_coli | NMW20_RS04715 | 287437..291051 | 562 | BREX-system-P-loop-protein-BrxC-[Escherichia-coli] |
| WP_267833950.1 | Type1.BREX | BrxA-DNA-binding->BrxB-BrxF-iSTAND->DUF6079*->N6-METHYLASE->PglZ->SIGMA+OB-Fold+AAA-ATPase+Lon-Protease-BrxL-><-ZnR | 1213 | Bacteria>Pseudomonadota>Gammaproteobacteria>Enterobacterales>Enterobacteriaceae>Escherichia | GCF_026612595.1 | Escherichia_coli | IGP49_RS06410 | 1748..5389 | 562 | BREX-system-P-loop-protein-BrxC-[Escherichia-coli] |
| WP_268646541.1 | Type1.BREX | <-VirD4-TraG+VirB4-FtsK<-HDRnase<-SbcC<-SIGMA+OB-Fold+AAA-ATPase+Lon-Protease-BrxL<-PglZ<-N6-METHYLASE<-ParB+HNH<-DUF6079*<-BrxB-BrxF-iSTAND<-BrxA-DNA-binding<-wHTH+WYL+WCX<-?<-35exo\|\|LexA-protease->?-><-?<-MobI | 1225 | Bacteria>Pseudomonadota>Gammaproteobacteria>Enterobacterales>Enterobacteriaceae>Escherichia | GCF_026801415.1 | Escherichia_coli | OZI01_RS14450 | 44846..48523 | 562 | BREX-system-P-loop-protein-BrxC-[Escherichia-coli] |
| WP_270658579.1 | Type1.BREX | <-Fimbrial<-Phage_integrase<-Phage_integrase\|\|?->?->\|\|wHTH+SAD+HNH-><-REase+iREase\|\|BrxA-DNA-binding->BrxB-BrxF-iSTAND->DUF6079*->N6-METHYLASE->PglZ->SIGMA+OB-Fold+AAA-ATPase+Lon-Protease-BrxL->HKD-DNase+OGG-like->\|\|AbiEi_3_N+AbiEi_3->AbiEii->?-><-DUF6216\|\|XIS-HTH-> | 1213 | Bacteria>Pseudomonadota>Gammaproteobacteria>Enterobacterales>Enterobacteriaceae>Escherichia | GCF_027696005.1 | Escherichia_coli | PG805_RS04695 | 85075..88716 | 562 | BREX-system-P-loop-protein-BrxC-[Escherichia-coli] |
| WP_272434363.1 | Type1.BREX | <-DUF772+DDE_Tnp_1<-HIN-HTH<-RelE-ParE<-CopG/RHH\|\|<-RHH\|\|?->wHTH+WYL+WCX->BrxA-DNA-binding->BrxB-BrxF-iSTAND->DUF6079*->ParB+HNH->N6-METHYLASE->PglZ->SIGMA+OB-Fold+AAA-ATPase+Lon-Protease-BrxL-><-DEDD_Tnp_IS110+Transposase_20\|\|Zn_Tnp_IS1+HTH_Tnp_IS1+DDE_Tnp_IS1->?->?->HEPN+SF1-SF2-Helicase+RNase+RNase-> | 1225 | Bacteria>Pseudomonadota>Gammaproteobacteria>Enterobacterales>Enterobacteriaceae>Escherichia | GCF_028436275.1 | Escherichia_coli | IXW67_RS19930 | 12892..16569 | 562 | BREX-system-P-loop-protein-BrxC-[Escherichia-coli] |
| WP_071702268.1 | Type1.BREX | <-TetR-HTH<-?<-?<-REase+iREase\|\|BrxA-DNA-binding->BrxB-BrxF-iSTAND->DUF6079*->N6-METHYLASE->PglZ->SIGMA+OB-Fold+AAA-ATPase+Lon-Protease-BrxL-><-ZnR<-ZnR<-?<-?\|\|RHH_antitoxin->XIS-HTH-> | 1213 | Bacteria>Pseudomonadota>Gammaproteobacteria>Enterobacterales>Enterobacteriaceae>Escherichia | GCF_029351265.1 | Escherichia_coli | P3474_RS20935 | 8656..12297 | 562 | BREX-system-P-loop-protein-BrxC-[Escherichia-coli] |
| WP_115723901.1 | Type1.BREX | ZnR->ZnR->URI-DNase+RAMA-><-SIGMA+OB-Fold+AAA-ATPase+Lon-Protease-BrxL<-PglZ<-TOPRIM<-OLD-ABC<-N6-METHYLASE<-DUF6079*<-BrxB-BrxF-iSTAND<-BrxA-DNA-binding<-Arm-DNA-bind_3+Phage_integrase<-tRNA\|\|?->HerA/FtsK+A32-><-?<-?<-?\|\|Peptidase_M17_N+Peptidase_M17-> | 1204 | Bacteria>Pseudomonadota>Gammaproteobacteria>Enterobacterales>Enterobacteriaceae>Escherichia | GCF_029542125.1 | Escherichia_coli | OM087_RS11620 | 2419674..2423288 | 562 | BREX-system-P-loop-protein-BrxC-[Escherichia-coli] |
| WP_023157617.1 | Type1.BREX | Phage_Cox->\|\|XIS-HTH->AAA+wHTH->?-><-SIGMA+OB-Fold+AAA-ATPase+Lon-Protease-BrxL<-PglZ<-TOPRIM<-OLD-ABC<-N6-METHYLASE<-DUF6079*<-BrxB-BrxF-iSTAND<-BrxA-DNA-binding\|\|nSTAND1+GTPase-AIG+sGTP->DUF932-> | 1204 | Bacteria>Pseudomonadota>Gammaproteobacteria>Enterobacterales>Enterobacteriaceae>Escherichia | GCF_030163335.1 | Escherichia_coli | HMPREF1620_RS03645 | 19303..22917 | 562 | BREX-system-P-loop-protein-BrxC-[Escherichia-coli] |
| WP_112915232.1 | Type1.BREX | <-SIGMA+OB-Fold+AAA-ATPase+Lon-Protease-BrxL<-PglZ<-N6-METHYLASE<-DUF6079*<-BrxB-BrxF-iSTAND<-BrxA-DNA-binding\|\|REase+iREase-><-wHTH+SAD+HNH\|\|<-?<-?<-?\|\|Phage_integrase->Phage_integrase-> | 1213 | Bacteria>Pseudomonadota>Gammaproteobacteria>Enterobacterales>Enterobacteriaceae>Escherichia | GCF_030324795.1 | Escherichia_coli | EFJ90_RS23235 | 8334..11975 | 562 | BREX-system-P-loop-protein-BrxC-[Escherichia-coli] |
| WP_087894339.1 | Type1.BREX | <-35exo<-SIGMA+OB-Fold+AAA-ATPase+Lon-Protease-BrxL<-PglZ<-N6-METHYLASE<-ParB+HNH<-DUF6079*<-BrxB-BrxF-iSTAND<-BrxA-DNA-binding<-wHTH+WYL+WCX | 1225 | Bacteria>Pseudomonadota>Gammaproteobacteria>Enterobacterales>Enterobacteriaceae>Escherichia | GCF_030716655.1 | Escherichia_coli | A8C62_RS24175 | 11745..15422 | 562 | BREX-system-P-loop-protein-BrxC-[Escherichia-coli] |
| WP_096309279.1 | Type1.BREX | <-DUF6079-p*<-BrxB-BrxF-iSTAND<-BrxA-DNA-binding\|\|REase+iREase-><-wHTH+SAD+HNH | 918 | Bacteria>Pseudomonadota>Gammaproteobacteria>Enterobacterales>Enterobacteriaceae>Escherichia | GCF_900195535.1 | Escherichia_coli | CPH42_RS29515 | 1..2754 | 562 | BREX-system-P-loop-protein-BrxC,-partial-[Escherichia-coli] |
| WP_096262508.1 | Type1.BREX | <-XIS-HTH<-RHH_antitoxin\|\|?->?->ZnR->ZnR-><-SIGMA+OB-Fold+AAA-ATPase+Lon-Protease-BrxL<-PglZ<-N6-METHYLASE<-DUF6079*<-BrxB-BrxF-iSTAND<-BrxA-DNA-binding\|\|REase-> | 1213 | Bacteria>Pseudomonadota>Gammaproteobacteria>Enterobacterales>Enterobacteriaceae>Escherichia | GCF_900196515.1 | Escherichia_coli | CPH64_RS18250 | 76258..79899 | 562 | BREX-system-P-loop-protein-BrxC-[Escherichia-coli] |
| WP_096317297.1 | Type1.BREX | HTH_21+rve+rve_3\|\|?->wHTH+WYL+WCX->BrxA-DNA-binding->BrxB-BrxF-iSTAND->DUF6079*->ParB+HNH->N6-METHYLASE->PglZ->SIGMA+OB-Fold+AAA-ATPase+Lon-Protease-BrxL-><-RVT | 1225 | Bacteria>Pseudomonadota>Gammaproteobacteria>Enterobacterales>Enterobacteriaceae>Escherichia | GCF_900196545.1 | Escherichia_coli | LLQ47_RS24465 | 3245..6922 | 562 | BREX-system-P-loop-protein-BrxC-[Escherichia-coli] |
| WP_110093138.1 | Type1.BREX | BrxA-DNA-binding->BrxB-BrxF-iSTAND->DUF6079*->N6-METHYLASE->PglZ->SIGMA+OB-Fold+AAA-ATPase+Lon-Protease-BrxL-><-ZnR\|\|RVT-><-?<-?<-DDE_Tnp_IS1 | 1213 | Bacteria>Pseudomonadota>Gammaproteobacteria>Enterobacterales>Enterobacteriaceae>Escherichia | GCF_900196585.1 | Escherichia_coli | CPH25_RS23340 | 1849..5490 | 562 | BREX-system-P-loop-protein-BrxC-[Escherichia-coli] |
| WP_115738447.1 | Type1.BREX | REase+N6_METHYLASE+REase<-?<-?\|\|?->?->?-><-SIGMA+OB-Fold+AAA-ATPase+Lon-Protease-BrxL<-PglZ<-N6-METHYLASE<-DUF6079*<-BrxB-BrxF-iSTAND<-BrxA-DNA-binding\|\|REase+iREase-><-wHTH+SAD+HNH\|\|<-?<-?<-?\|\|Phage_integrase->Phage_integrase-> | 1213 | Bacteria>Pseudomonadota>Gammaproteobacteria>Enterobacterales>Enterobacteriaceae>Escherichia | GCF_900499355.1 | Escherichia_coli | DY383_RS04210 | 180625..184266 | 562 | BREX-system-P-loop-protein-BrxC-[Escherichia-coli] |
| WP_126509145.1 | Type1.BREX | <-XIS-HTH<-RHH_antitoxin\|\|?->?->ZnR->ZnR-><-SIGMA+OB-Fold+AAA-ATPase+Lon-Protease-BrxL<-PglZ<-N6-METHYLASE<-DUF6079*<-BrxB-BrxF-iSTAND<-BrxA-DNA-binding\|\|REase+iREase->REase->Zn_Tnp_IS1+HTH_Tnp_IS1+DDE_Tnp_IS1->?->Sulfatase-><-Arm-DNA-bind_3+Phage_integrase | 1213 | Bacteria>Pseudomonadota>Gammaproteobacteria>Enterobacterales>Enterobacteriaceae>Escherichia | GCF_900636125.1 | Escherichia_coli | EL118_RS04180 | 856584..860225 | 562 | BREX-system-P-loop-protein-BrxC-[Escherichia-coli] |
| WP_001548950.1 | Type1.BREX | TetR-HTH->tRNA->Arm-DNA-bind_3+Phage_integrase-><-Zn_Tnp_IS1+HTH_Tnp_IS1+DDE_Tnp_IS1<-TetR-HTH<-?<-?<-REase+iREase\|\|BrxA-DNA-binding->BrxB-BrxF-iSTAND->DUF6079*->N6-METHYLASE->PglZ->SIGMA+OB-Fold+AAA-ATPase+Lon-Protease-BrxL-> | 1213 | Bacteria>Pseudomonadota>Gammaproteobacteria>Enterobacterales>Enterobacteriaceae>Escherichia | GCF_900706885.1 | Escherichia_coli | LF679_RS14520 | 38542..42183 | 562 | BREX-system-P-loop-protein-BrxC-[Escherichia-coli] |
| WP_152072595.1 | Type1.BREX | <-Phage_integrase<-Phage_integrase\|\|?->?->?->\|\|wHTH+SAD+HNH-><-REase+iREase\|\|BrxA-DNA-binding->BrxB-BrxF-iSTAND->DUF6079*->N6-METHYLASE->PglZ->SIGMA+OB-Fold+AAA-ATPase+Lon-Protease-BrxL->HKD-DNase+OGG-like->\|\|AbiEi_3_N+AbiEi_3->AbiEii->?-><-DUF6216 | 1213 | Bacteria>Pseudomonadota>Gammaproteobacteria>Enterobacterales>Enterobacteriaceae>Escherichia | GCF_902501575.1 | Escherichia_coli | GDK01_RS16655 | 24181..27822 | 562 | BREX-system-P-loop-protein-BrxC-[Escherichia-coli] |
| WP_001552307.1 | Type1.BREX | wHTH->ZnR->ZnR->URI-DNase+RAMA-><-SIGMA+OB-Fold+AAA-ATPase+Lon-Protease-BrxL<-PglZ<-TOPRIM<-OLD-ABC<-N6-METHYLASE<-DUF6079*<-BrxB-BrxF-iSTAND<-BrxA-DNA-binding<-Arm-DNA-bind_3+Phage_integrase<-tRNA\|\|?-><-SKI\|\|METHYLASE->?->?->LACI-HTH+PBPI-> | 1204 | Bacteria>Pseudomonadota>Gammaproteobacteria>Enterobacterales>Enterobacteriaceae>Escherichia | GCF_902505225.1 | Escherichia_coli | GDK13_RS03595 | 266015..269629 | 562 | BREX-system-P-loop-protein-BrxC-[Escherichia-coli] |
| WP_096955310.1 | Type1.BREX | DUF6216-><-?<-AbiEii<-AbiEi_3_N+AbiEi_3\|\|<-HKD-DNase+OGG-like<-SIGMA+OB-Fold+AAA-ATPase+Lon-Protease-BrxL<-PglZ<-N6-METHYLASE<-DUF6079*<-BrxB-BrxF-iSTAND<-BrxA-DNA-binding\|\|REase+iREase-><-wHTH+SAD+HNH\|\|<-?<-?<-?\|\|Phage_integrase->Phage_integrase-> | 1213 | Bacteria>Pseudomonadota>Gammaproteobacteria>Enterobacterales>Enterobacteriaceae>Escherichia | GCF_904831825.1 | Escherichia_coli | A7198_RS13140 | 15059..18700 | 562 | BREX-system-P-loop-protein-BrxC-[Escherichia-coli] |
| WP_001702945.1 | Type1.BREX | Arm-DNA-bind_3+Phage_integrase->N6-METHYLASE->ORC-CDC6+ZnR->HEPN-><-Zn_Tnp_IS1+HTH_Tnp_IS1+DDE_Tnp_IS1<-REase<-REase\|\|BrxA-DNA-binding->BrxB-BrxF-iSTAND->DUF6079*->N6-METHYLASE->PglZ->SIGMA+OB-Fold+AAA-ATPase+Lon-Protease-BrxL-><-ZnR | 1213 | Bacteria>Pseudomonadota>Gammaproteobacteria>Enterobacterales>Enterobacteriaceae>Escherichia | GCF_907176105.1 | Escherichia_coli | A7133_RS15540 | 52191..55832 | 562 | BREX-system-P-loop-protein-BrxC-[Escherichia-coli] |
| WP_187798258.1 | Type1.BREX | <-Sulfatase<-?\|\|DDE_Tnp_1_5+DDE_Tnp_1->?-><-Zn_Tnp_IS1+HTH_Tnp_IS1+DDE_Tnp_IS1<-?<-REase+iREase\|\|BrxA-DNA-binding->BrxB-BrxF-iSTAND->DUF6079*->N6-METHYLASE->PglZ->SIGMA+OB-Fold+AAA-ATPase+Lon-Protease-BrxL-><-ZnR<-ZnR\|\|HIN-HTH-><-RVT\|\|<-ZnR\|\|<-ZnR | 1213 | Bacteria>Pseudomonadota>Gammaproteobacteria>Enterobacterales>Enterobacteriaceae>Escherichia | GCF_929617055.1 | Escherichia_coli | MJ085_RS02910 | 12212..15853 | 562 | BREX-system-P-loop-protein-BrxC-[Escherichia-coli] |
| WP_088209948.1 | Type1.BREX | <-RNase<-DEOR-HTH\|\|?->DDE_Tnp_1_5+DDE_Tnp_1-><-SIGMA+OB-Fold+AAA-ATPase+Lon-Protease-BrxL<-PglZ<-TOPRIM<-OLD-ABC<-N6-METHYLASE<-DUF6079*<-BrxB-BrxF-iSTAND<-BrxA-DNA-binding<-Arm-DNA-bind_3+Phage_integrase\|\|Phage_integrase->?-><-PIWI<-XIS-HTH\|\|?->DDE_Tnp_IS240-> | 1204 | Bacteria>Pseudomonadota>Gammaproteobacteria>Enterobacterales>Enterobacteriaceae>Escherichia | GCF_943733315.1 | Escherichia_coli | OVQ79_RS01305 | 107612..111226 | 562 | BREX-system-P-loop-protein-BrxC-[Escherichia-coli] |
| WP_096964704.1 | Type1.BREX | DUF6216-><-?<-AbiEii<-AbiEi_3_N+AbiEi_3\|\|<-HKD-DNase+OGG-like<-SIGMA+OB-Fold+AAA-ATPase+Lon-Protease-BrxL<-PglZ<-N6-METHYLASE<-DUF6079*<-BrxB-BrxF-iSTAND<-BrxA-DNA-binding\|\|REase+iREase-><-wHTH+SAD+HNH | 1213 | Bacteria>Pseudomonadota>Gammaproteobacteria>Enterobacterales>Enterobacteriaceae>Escherichia | GCF_948144475.1 | Escherichia_coli | A7257_RS18210 | 50280..53921 | 562 | BREX-system-P-loop-protein-BrxC-[Escherichia-coli] |
| WP_052920517.1 | Type1.BREX | ZnR->ZnR->URI-DNase+RAMA-><-SIGMA+OB-Fold+AAA-ATPase+Lon-Protease-BrxL<-PglZ<-TOPRIM<-OLD-ABC<-N6-METHYLASE<-DUF6079*<-BrxB-BrxF-iSTAND<-BrxA-DNA-binding<-Arm-DNA-bind_3+Phage_integrase<-tRNA\|\|?->HerA/FtsK+A32-><-?<-?<-?\|\|Peptidase_M17_N+Peptidase_M17-> | 1204 | Bacteria>Pseudomonadota>Gammaproteobacteria>Enterobacterales>Enterobacteriaceae>Escherichia | GCF_948144575.1 | Escherichia_coli | AGA26_RS17910 | 13907..17521 | 562 | BREX-system-P-loop-protein-BrxC-[Escherichia-coli] |
| WP_244586021.1 | Type1.BREX | <-wHTH+WYL<-?\|\|XIS-HTH->?->BrxA-DNA-binding->BrxB-BrxF-iSTAND->DUF6079*->ParB+HNH->N6-METHYLASE->PglZ-> | 1240 | Bacteria>Pseudomonadota>Gammaproteobacteria>Enterobacterales>Enterobacteriaceae>Escherichia | GCF_948144585.1 | Escherichia_coli | A7Y91_RS22675 | 15059..18781 | 562 | BREX-system-P-loop-protein-BrxC-[Escherichia-coli] |
| WP_160450150.1 | Type1.BREX | <-ParB+HNH<-?\|\|GNTR-HTH+Aminotran_1_2-><-REase+HTH\|\|?->?-><-REase+iREase\|\|BrxA-DNA-binding->BrxB-BrxF-iSTAND->DUF6079*->N6-METHYLASE-><-?\|\|PglZ->SIGMA+OB-Fold+AAA-ATPase+Lon-Protease-BrxL->HKD-DNase+OGG-like->\|\|AbiEi_3_N+AbiEi_3->AbiEii->?-><-DUF6216 | 1213 | Bacteria>Pseudomonadota>Gammaproteobacteria>Enterobacterales>Enterobacteriaceae | - | Escherichia | H1X95_RS06210 | 181594..185235 | 561 | MULTISPECIES:-BREX-system-P-loop-protein-BrxC-[Escherichia] |
| WP_024559724.1 | Type1.BREX | <-SF1-SF2-Helicase<-SF1-SF2-Helicase<-?<-?<-?<-?\|\|<-SIGMA+OB-Fold+AAA-ATPase+Lon-Protease-BrxL<-PglZ<-N6-METHYLASE<-DUF6079*<-BrxB-BrxF-iSTAND<-BrxA-DNA-binding\|\|TIR-><-HTH_35\|\|Peptidase_M20->?->ParB-> | 1213 | Bacteria>Pseudomonadota>Gammaproteobacteria>Enterobacterales>Enterobacteriaceae>Franconibacter | GCF_000485985.1 | Franconibacter_pulveris | U709_RS0101725 | 38623..42264 | 435910 | MULTISPECIES:-BREX-system-P-loop-protein-BrxC-[Franconibacter-pulveris] |
| WP_001019652.1 | Type1.BREX | Arm-DNA-bind_3->NACHT+SF1-SF2-Helicase+HKD-DNase->REase-><-SIGMA+OB-Fold+AAA-ATPase+Lon-Protease-BrxL<-PglZ<-TOPRIM<-OLD-ABC<-N6-METHYLASE<-DUF6079*<-BrxB-BrxF-iSTAND<-BrxA-DNA-binding<-?<-SAM-METHYLASE<-YjhX_toxin\|\|?->REase+iREase->HEPN-> | 1213 | Bacteria>Pseudomonadota>Gammaproteobacteria>Enterobacterales>Enterobacteriaceae | GCF_000006945.2 | Salmonella | SL1344_RS23060 | 4767568..4771209 | 590 | MULTISPECIES:-BREX-system-P-loop-protein-BrxC-[Salmonella] |
| WP_008501367.1 | Type1.BREX | <-SymE_toxin<-?<-?<-?\|\|wHTH+HNH->SAM-METHYLASE->BrxA-DNA-binding->BrxB-BrxF-iSTAND->DUF6079*->N6-METHYLASE->PglZ->SIGMA+OB-Fold+AAA-ATPase+Lon-Protease-BrxL-> | 1213 | Bacteria>Pseudomonadota>Gammaproteobacteria>Enterobacterales>Enterobacteriaceae | GCF_000286655.1 | Enterobacter | B498_RS11105 | 63158..66799 | 547 | MULTISPECIES:-BREX-system-P-loop-protein-BrxC-[Enterobacter] |
| WP_052939500.1 | Type1.BREX | <-HEPN<-?\|\|YjhX_toxin->SAM-METHYLASE->?->\|\|?-><-ZnR\|\|BrxA-DNA-binding->BrxB-BrxF-iSTAND->DUF6079*->N6-METHYLASE->OLD-ABC->TOPRIM->PglZ->SIGMA+OB-Fold+AAA-ATPase+Lon-Protease-BrxL->HKD-DNase+OGG-like-><-REase<-NACHT+SF1-SF2-Helicase+HKD-DNase | 1204 | Bacteria>Pseudomonadota>Gammaproteobacteria>Enterobacterales>Enterobacteriaceae | GCF_001569545.1 | Salmonella | MA624_RS20100 | 4127586..4131200 | 590 | MULTISPECIES:-BREX-system-P-loop-protein-BrxC-[Salmonella] |
| WP_063308429.1 | Type1.BREX | Glyoxalase_8->URI-DNase-><-RHH_antitoxin\|\|?->XIS-HTH->?->ZnR->BrxA-DNA-binding->BrxB-BrxF-iSTAND->DUF6079*->N6-METHYLASE->PglZ->SIGMA+OB-Fold+AAA-ATPase+Lon-Protease-BrxL->HKD-DNase+OGG-like->\|\|<-REase<-Arm-DNA-bind_3+Phage_integrase | 1213 | Bacteria>Pseudomonadota>Gammaproteobacteria>Enterobacterales>Enterobacteriaceae | GCF_001623605.1 | Enterobacter | A4308_RS00205 | 35188..38829 | 547 | MULTISPECIES:-BREX-system-P-loop-protein-BrxC-[Enterobacter] |
| WP_241755053.1 | Type1.BREX | <-METHYLASE\|\|SKI-><-?\|\|tRNA->Arm-DNA-bind_3+Phage_integrase-><-SIGMA+OB-Fold+AAA-ATPase+Lon-Protease-BrxL<-PglZ<-N6-METHYLASE<-ParB+HNH<-DUF6079*<-BrxB-BrxF-iSTAND<-BrxA-DNA-binding\|\|?->PHP+SbcC->Schlafen+HATPase+wHTH->?->?->?->Sulfatase-> | 1240 | Bacteria>Pseudomonadota>Gammaproteobacteria>Enterobacterales>Enterobacteriaceae | GCF_013791465.1 | Citrobacter | HV084_RS03275 | 673516..677238 | 544 | MULTISPECIES:-BREX-system-P-loop-protein-BrxC-[Citrobacter] |
| WP_233470897.1 | Type1.BREX | <-DUF932<-nSTAND1+GTPase-AIG+sGTP<-CopG/RHH\|\|?->XIS-HTH->?->?->BrxA-DNA-binding->BrxB-BrxF-iSTAND->DUF6079*->ParB+HNH->N6-METHYLASE->HEPN->RVT->N6-METHYLASE->PglZ->SIGMA+OB-Fold+AAA-ATPase+Lon-Protease-BrxL-><-Arm-DNA-bind_3+Phage_integrase | 1240 | Bacteria>Pseudomonadota>Gammaproteobacteria>Enterobacterales>Enterobacteriaceae | GCF_014931625.2 | Citrobacter | INP82_RS14360 | 158420..162142 | 544 | MULTISPECIES:-BREX-system-P-loop-protein-BrxC-[Citrobacter] |
| WP_129336016.1 | Type1.BREX | <-HEPN<-REase+iREase<-?\|\|YjhX_toxin->SAM-METHYLASE->?->BrxA-DNA-binding->BrxB-BrxF-iSTAND->DUF6079*->N6-METHYLASE->OLD-ABC->TOPRIM->PglZ->SIGMA+OB-Fold+AAA-ATPase+Lon-Protease-BrxL-><-REase<-NACHT+SF1-SF2-Helicase+HKD-DNase<-Arm-DNA-bind_3 | 1213 | Bacteria>Pseudomonadota>Gammaproteobacteria>Enterobacterales>Enterobacteriaceae | GCF_019670325.1 | Salmonella | K5676_RS20565 | 4193364..4197005 | 590 | MULTISPECIES:-BREX-system-P-loop-protein-BrxC-[Salmonella] |
| WP_080247850.1 | Type1.BREX | <-HEPN<-REase+iREase<-?\|\|YjhX_toxin->SAM-METHYLASE->?->BrxA-DNA-binding->BrxB-BrxF-iSTAND->DUF6079*->N6-METHYLASE->OLD-ABC->TOPRIM->PglZ->SIGMA+OB-Fold+AAA-ATPase+Lon-Protease-BrxL-><-REase\|\|<-?<-?<-wHTH+WYL | 1213 | Bacteria>Pseudomonadota>Gammaproteobacteria>Enterobacterales>Enterobacteriaceae | GCF_020221795.1 | Salmonella | LDD68_RS20280 | 4140919..4144560 | 590 | MULTISPECIES:-BREX-system-P-loop-protein-BrxC-[Salmonella] |
| WP_070534016.1 | Type1.BREX | LACI-HTH+PBPI<-?\|\|<-ASCH+Glyoxalase\|\|?->SymE_toxin\|\|<-CASPASE<-?\|\|BrxA-DNA-binding->BrxB-BrxF-iSTAND->DUF6079*->N6-METHYLASE->PglZ->SIGMA+OB-Fold+AAA-ATPase+Lon-Protease-BrxL->HKD-DNase+OGG-like-><-REase\|\|SF1-SF2-Helicase-> | 1213 | Bacteria>Pseudomonadota>Gammaproteobacteria>Enterobacterales>Enterobacteriaceae | GCF_024104775.1 | Enterobacter | KGY84_RS21865 | 326949..330590 | 547 | MULTISPECIES:-BREX-system-P-loop-protein-BrxC-[Enterobacter] |
| WP_047722721.1 | Type1.BREX | <-LuxR-HTH<-?\|\|\|\|BrxA-DNA-binding->BrxB-BrxF-iSTAND->DUF6079*->N6-METHYLASE->PglZ->SIGMA+OB-Fold+AAA-ATPase+Lon-Protease-BrxL-><-DnaJ<-?<-Arm-DNA-bind_3+Phage_integrase | 1213 | Bacteria>Pseudomonadota>Gammaproteobacteria>Enterobacterales>Enterobacteriaceae | GCF_026008675.1 | Klebsiella/Raoultella_group | OPC88_RS09775 | 105880..109521 | 2890311 | MULTISPECIES:-BREX-system-P-loop-protein-BrxC-[Klebsiella/Raoultella-group] |
| WP_192856165.1 | Type1.BREX | TetR-HTH->tRNA->Arm-DNA-bind_3+Phage_integrase->Sulfatase-><-?\|\|DDE_Tnp_1_5+DDE_Tnp_1-><-?<-REase+iREase\|\|BrxA-DNA-binding->BrxB-BrxF-iSTAND->DUF6079*->N6-METHYLASE->PglZ->SIGMA+OB-Fold+AAA-ATPase+Lon-Protease-BrxL-><-ZnR<-ZnR<-?<-?\|\|RHH_antitoxin->XIS-HTH-> | 1213 | Bacteria>Pseudomonadota>Gammaproteobacteria>Enterobacterales>Enterobacteriaceae | GCF_029962465.1 | Escherichia | H5I61_RS19040 | 73585..77226 | 561 | MULTISPECIES:-BREX-system-P-loop-protein-BrxC-[Escherichia] |
| WP_050858669.1 | Type1.BREX | <-Peptidase_M17_N+Peptidase_M17\|\|?->?-><-HerA/FtsK+A32<-?\|\|tRNA->Arm-DNA-bind_3+Phage_integrase->BrxA-DNA-binding->BrxB-BrxF-iSTAND->DUF6079*->N6-METHYLASE->OLD-ABC->TOPRIM->PglZ->SIGMA+OB-Fold+AAA-ATPase+Lon-Protease-BrxL-><-URI-DNase+RAMA<-ZnR<-ZnR<-wHTH | 1204 | Bacteria>Pseudomonadota>Gammaproteobacteria>Enterobacterales>Enterobacteriaceae | GCF_030295055.1 | Escherichia | AST78_RS16735 | 106796..110410 | 561 | MULTISPECIES:-BREX-system-P-loop-protein-BrxC-[Escherichia] |
| WP_048999825.1 | Type1.BREX | <-NACHT+SF1-SF2-Helicase+HKD-DNase\|\|BrxA-DNA-binding->BrxB-BrxF-iSTAND->DUF6079*->N6-METHYLASE->OLD-ABC->TOPRIM->PglZ->SIGMA+OB-Fold+AAA-ATPase+Lon-Protease-BrxL->HKD-DNase+OGG-like->HKD-DNase+OGG-like-><-Arm-DNA-bind_3+Phage_integrase | 1204 | Bacteria>Pseudomonadota>Gammaproteobacteria>Enterobacterales>Enterobacteriaceae | GCF_900775105.1 | Klebsiella/Raoultella_group | E7V87_RS13635 | 63633..67247 | 2890311 | MULTISPECIES:-BREX-system-P-loop-protein-BrxC-[Klebsiella/Raoultella-group] |
| WP_225990544.1 | Type1.BREX | Glyoxalase->ACET->YjhX_toxin->SAM-METHYLASE->ZnR->Glyoxalase->BrxA-DNA-binding->BrxB-BrxF-iSTAND->DUF6079*->ParB+HNH->N6-METHYLASE->HEPN->RVT->N6-METHYLASE-p->PglZ->SIGMA+OB-Fold+AAA-ATPase+Lon-Protease-BrxL-><-Arm-DNA-bind_3+Phage_integrase | 1240 | Bacteria>Pseudomonadota>Gammaproteobacteria>Enterobacterales>Enterobacteriaceae | GCF_954736865.1 | Enterobacter | CVO81_RS00925 | 173954..177676 | 547 | MULTISPECIES:-BREX-system-P-loop-protein-BrxC-[Enterobacter] |
| CDL15658.1 | Type1.BREX | Arm-DNA-bind_3+Phage_integrase->EVE+HNH-><-REase+iREase\|\|BrxA-DNA-binding->BrxB-BrxF-iSTAND->DUF6079-p->DUF6079-p*->N6-METHYLASE-p->N6-METHYLASE-> | 950 | Bacteria>Pseudomonadota>Gammaproteobacteria>Enterobacterales>Enterobacteriaceae>Klebsiella/Raoultella_group | GCA_000529945.1 | Klebsiella_pneumoniae_IS46 | - | 6252..9104 | 1432553 | putative-DNA-repair-ATPase-[Klebsiella-pneumoniae-IS46] |
| PLL44193.1 | Type1.BREX | <-N6-METHYLASE-p<-N6-METHYLASE<-ParB+HNH<-DUF6079*<-BrxB-BrxF-iSTAND<-BrxA-DNA-binding\|\|<-XIS-HTH | 1233 | Bacteria>Pseudomonadota>Gammaproteobacteria>Enterobacterales>Enterobacteriaceae>Klebsiella/Raoultella_group | GCA_002856085.1 | Klebsiella_michiganensis | CWN50_01600 | 4530..8231 | 1134687 | BREX-system-P-loop-protein-BrxC-[Klebsiella-michiganensis] |
| AUN75366.1 | Type1.BREX | Arm-DNA-bind_3+Phage_integrase-><-SIGMA+OB-Fold+AAA-ATPase+Lon-Protease-BrxL<-PglZ<-N6-METHYLASE-p<-RVT<-HEPN<-N6-METHYLASE<-ParB+HNH<-DUF6079*<-BrxB-BrxF-iSTAND<-BrxA-DNA-binding<-?<-?<-XIS-HTH\|\|?->nSTAND1+sGTP->wHTH+WYL-> | 1233 | Bacteria>Pseudomonadota>Gammaproteobacteria>Enterobacterales>Enterobacteriaceae>Klebsiella/Raoultella_group | GCA_002870985.1 | Klebsiella_pneumoniae | C0077_01580 | 326913..330614 | 573 | BREX-system-P-loop-protein-BrxC-[Klebsiella-pneumoniae] |
| QHX00137.1 | Type1.BREX | YjhX_toxin->ACET->SAM-METHYLASE->ZnR->Glyoxalase->BrxA-DNA-binding->BrxB-BrxF-iSTAND->DUF6079*->ParB+HNH->N6-METHYLASE->HEPN->RVT->N6-METHYLASE-p->PglZ->SIGMA+OB-Fold+AAA-ATPase+Lon-Protease-BrxL-><-Arm-DNA-bind_3+Phage_integrase | 1233 | Bacteria>Pseudomonadota>Gammaproteobacteria>Enterobacterales>Enterobacteriaceae>Klebsiella/Raoultella_group | GCA_010365565.1 | Klebsiella_variicola | GZS05_24295 | 4949707..4953408 | 244366 | BREX-system-P-loop-protein-BrxC-[Klebsiella-variicola] |
| QJK24913.1 | Type1.BREX | DUF2754-><-?<-NUDIX\|\|ColE7+HNH->?->BrxA-DNA-binding->BrxB-BrxF-iSTAND->DUF6079*->N6-METHYLASE->PglZ->SIGMA+OB-Fold+AAA-ATPase+Lon-Protease-BrxL->HKD-DNase+OGG-like-><-REase<-NACHT+SF1-SF2-Helicase+HKD-DNase<-RNaseH<-nSTAND1+HNH | 1213 | Bacteria>Pseudomonadota>Gammaproteobacteria>Enterobacterales>Enterobacteriaceae>Klebsiella/Raoultella_group | GCA_012967125.1 | Klebsiella_pneumoniae | HJX28_24000 | 4748106..4751747 | 573 | BREX-system-P-loop-protein-BrxC-[Klebsiella-pneumoniae] |
| QJM45197.1 | Type1.BREX | <-wHTH+WYL<-sGTP<-?\|\|XIS-HTH->?->?->BrxA-DNA-binding->BrxB-BrxF-iSTAND->DUF6079*->ParB+HNH->N6-METHYLASE->HEPN->RVT->N6-METHYLASE-p->PglZ->SIGMA+OB-Fold+AAA-ATPase+Lon-Protease-BrxL-><-Arm-DNA-bind_3+Phage_integrase | 1233 | Bacteria>Pseudomonadota>Gammaproteobacteria>Enterobacterales>Enterobacteriaceae>Klebsiella/Raoultella_group | GCA_012970665.1 | Klebsiella_pneumoniae | HJW71_23035 | 4620464..4624165 | 573 | BREX-system-P-loop-protein-BrxC-[Klebsiella-pneumoniae] |
| HAT1683007.1 | Type1.BREX | Peptidase_M20->?->?->DDE_Tnp-><-SIGMA+OB-Fold+AAA-ATPase+Lon-Protease-BrxL<-PglZ<-N6-METHYLASE<-DUF6079*<-BrxB-BrxF-iSTAND<-BrxA-DNA-binding | 1213 | Bacteria>Pseudomonadota>Gammaproteobacteria>Enterobacterales>Enterobacteriaceae>Klebsiella/Raoultella_group | GCA_015893305.1 | Klebsiella_oxytoca | I8Y21_003722 | 15957..19598 | 571 | BREX-system-P-loop-protein-BrxC-[Klebsiella-oxytoca] |
| EGT0044780.1 | Type1.BREX | <-DUF905<-DUF932<-CopG/RHH\|\|?->XIS-HTH->?-><-TIR<-TIR\|\|BrxA-DNA-binding->BrxB-BrxF-iSTAND->DUF6079*->ParB+HNH->N6-METHYLASE->N6-METHYLASE-p->PglZ->SIGMA+OB-Fold+AAA-ATPase+Lon-Protease-BrxL->wHTH+WYL+WCX->?->HNH-> | 1233 | Bacteria>Pseudomonadota>Gammaproteobacteria>Enterobacterales>Enterobacteriaceae>Klebsiella/Raoultella_group | GCA_015929385.1 | Klebsiella_oxytoca | I9Y61_001542 | 37432..41133 | 571 | BREX-system-P-loop-protein-BrxC-[Klebsiella-oxytoca] |
| EGT0068692.1 | Type1.BREX | HIN-HTH->?->LZ_Tnp_IS66+zf-IS66+DDE_Tnp_IS66+DDE_Tnp_IS66_C-><-HIN-HTH\|\|BrxA-DNA-binding->BrxB-BrxF-iSTAND->DUF6079*->N6-METHYLASE->PglZ->SIGMA+OB-Fold+AAA-ATPase+Lon-Protease-BrxL-><-?<-DnaJ | 1213 | Bacteria>Pseudomonadota>Gammaproteobacteria>Enterobacterales>Enterobacteriaceae>Klebsiella/Raoultella_group | GCA_015929475.1 | Klebsiella_michiganensis | I9Y62_005522 | 4489..8130 | 1134687 | BREX-system-P-loop-protein-BrxC-[Klebsiella-michiganensis] |
| EJG2380286.1 | Type1.BREX | <-DUF905<-DUF932<-CopG/RHH\|\|?->XIS-HTH->?-><-TIR<-TIR\|\|BrxA-DNA-binding->BrxB-BrxF-iSTAND->DUF6079*->ParB+HNH->N6-METHYLASE->HEPN->RVT->N6-METHYLASE-p->PglZ->SIGMA+OB-Fold+AAA-ATPase+Lon-Protease-BrxL->wHTH+WYL+WCX-> | 1233 | Bacteria>Pseudomonadota>Gammaproteobacteria>Enterobacterales>Enterobacteriaceae>Klebsiella/Raoultella_group | GCA_016088515.2 | Raoultella_ornithinolytica | AM381_RS08835 | 19475..23176 | 54291 | BREX-system-P-loop-protein-BrxC-[Raoultella-ornithinolytica] |
| HAV2047149.1 | Type1.BREX | <-Calcineurin<-?<-?<-?\|\|tRNA-><-SIGMA+OB-Fold+AAA-ATPase+Lon-Protease-BrxL<-PglZ<-N6-METHYLASE-p<-N6-METHYLASE<-ParB+HNH<-DUF6079*<-BrxB-BrxF-iSTAND<-BrxA-DNA-binding\|\|?-><-Glyco_hydro_43+GH43_C2<-?\|\|AP_endonuc_2-> | 1233 | Bacteria>Pseudomonadota>Gammaproteobacteria>Enterobacterales>Enterobacteriaceae>Klebsiella/Raoultella_group | GCA_016445365.1 | Raoultella_ornithinolytica | JHJ87_003574 | 41911..45612 | 54291 | BREX-system-P-loop-protein-BrxC-[Raoultella-ornithinolytica] |
| MBL3233196.1 | Type1.BREX | <-HKD-DNase+OGG-like<-SIGMA+OB-Fold+AAA-ATPase+Lon-Protease-BrxL<-PglZ<-N6-METHYLASE<-DUF6079*<-BrxB-BrxF-iSTAND<-BrxA-DNA-binding<-?<-ColE7+HNH\|\|?-><-DUF2754 | 1213 | Bacteria>Pseudomonadota>Gammaproteobacteria>Enterobacterales>Enterobacteriaceae>Klebsiella/Raoultella_group | GCA_016755095.1 | Klebsiella_pneumoniae | ELP31_24695 | 10226..13867 | 573 | BREX-system-P-loop-protein-BrxC-[Klebsiella-pneumoniae] |
| MBT7733855.1 | Type1.BREX | <-wHTH+WYL<-sGTP<-?\|\|XIS-HTH->?->?->BrxA-DNA-binding->BrxB-BrxF-iSTAND->DUF6079*->ParB+HNH->N6-METHYLASE->HEPN->RVT->N6-METHYLASE-p->PglZ->SIGMA+OB-Fold+AAA-ATPase+Lon-Protease-BrxL-><-Arm-DNA-bind_3+Phage_integrase | 1233 | Bacteria>Pseudomonadota>Gammaproteobacteria>Enterobacterales>Enterobacteriaceae>Klebsiella/Raoultella_group | GCA_017255495.1 | Klebsiella_pneumoniae | JX616_18220 | 31914..35615 | 573 | BREX-system-P-loop-protein-BrxC-[Klebsiella-pneumoniae] |
| MBZ6545107.1 | Type1.BREX | <-wHTH+WYL<-nSTAND1+sGTP<-?\|\|XIS-HTH->?->?->BrxA-DNA-binding->BrxB-BrxF-iSTAND->DUF6079*->ParB+HNH-> | 1233 | Bacteria>Pseudomonadota>Gammaproteobacteria>Enterobacterales>Enterobacteriaceae>Klebsiella/Raoultella_group | GCA_020115175.1 | Klebsiella_pneumoniae | FMJ66_23200 | 19841..23542 | 573 | BREX-system-P-loop-protein-BrxC-[Klebsiella-pneumoniae] |
| MBZ7196032.1 | Type1.BREX | ACET->?->YjhX_toxin->ACET->SAM-METHYLASE->ZnR->Glyoxalase->BrxA-DNA-binding->BrxB-BrxF-iSTAND->DUF6079*->ParB+HNH->N6-METHYLASE->N6-METHYLASE-p->PglZ->SIGMA+OB-Fold+AAA-ATPase+Lon-Protease-BrxL-><-Arm-DNA-bind_3+Phage_integrase | 1233 | Bacteria>Pseudomonadota>Gammaproteobacteria>Enterobacterales>Enterobacteriaceae>Klebsiella/Raoultella_group | GCA_020118595.1 | Klebsiella_oxytoca | FMX61_04485 | 39995..43696 | 571 | BREX-system-P-loop-protein-BrxC-[Klebsiella-oxytoca] |
| MCB3335667.1 | Type1.BREX | <-DDE_Tnp_1_5+DDE_Tnp_1\|\|?->?->ZnR->ZnR-><-SIGMA+OB-Fold+AAA-ATPase+Lon-Protease-BrxL<-PglZ<-N6-METHYLASE<-DUF6079*<-BrxB-BrxF-iSTAND<-BrxA-DNA-binding\|\|REase+iREase-><-EVE+HNH<-Arm-DNA-bind_3+Phage_integrase | 1213 | Bacteria>Pseudomonadota>Gammaproteobacteria>Enterobacterales>Enterobacteriaceae>Klebsiella/Raoultella_group | GCA_020478865.1 | Klebsiella_pneumoniae | H5849_24975 | 11785..15426 | 573 | BREX-system-P-loop-protein-BrxC-[Klebsiella-pneumoniae] |
| HBQ2245107.1 | Type1.BREX | <-DUF6079*<-BrxB-BrxF-iSTAND<-BrxA-DNA-binding<-URI-DNase\|\|Phage_integrase-> | 1164 | Bacteria>Pseudomonadota>Gammaproteobacteria>Enterobacterales>Enterobacteriaceae>Klebsiella/Raoultella_group | GCA_021823445.1 | Klebsiella_pneumoniae | L7082_005482 | 1..3492 | 573 | BREX-system-P-loop-protein-BrxC-[Klebsiella-pneumoniae] |
| HBQ2805341.1 | Type1.BREX | <-DUF6079-p*<-BrxB-BrxF-iSTAND<-BrxA-DNA-binding<-URI-DNase\|\|Phage_integrase-> | 606 | Bacteria>Pseudomonadota>Gammaproteobacteria>Enterobacterales>Enterobacteriaceae>Klebsiella/Raoultella_group | GCA_021848835.1 | Klebsiella_pneumoniae | L7V14_005501 | 1..1820 | 573 | BREX-system-P-loop-protein-BrxC-[Klebsiella-pneumoniae] |
| EIV3098057.1 | Type1.BREX | <-wHTH+WYL<-nSTAND1+sGTP<-?\|\|XIS-HTH->?->?->BrxA-DNA-binding->BrxB-BrxF-iSTAND->DUF6079*->ParB+HNH->N6-METHYLASE->HEPN->RVT->N6-METHYLASE-p->PglZ->SIGMA+OB-Fold+AAA-ATPase+Lon-Protease-BrxL-><-Arm-DNA-bind_3+Phage_integrase | 1233 | Bacteria>Pseudomonadota>Gammaproteobacteria>Enterobacterales>Enterobacteriaceae>Klebsiella/Raoultella_group | GCA_021854435.1 | Klebsiella_pneumoniae | L7W77_004838 | 45573..49274 | 573 | BREX-system-P-loop-protein-BrxC-[Klebsiella-pneumoniae] |
| HBQ4025835.1 | Type1.BREX | <-ABC-ATPase+SbcC<-?<-?<-?\|\|MODE-HTH+PBPII->?->DDE_Tnp_1_5+DDE_Tnp_1->BrxA-DNA-binding->BrxB-BrxF-iSTAND->DUF6079*->N6-METHYLASE->PglZ->SIGMA+OB-Fold+AAA-ATPase+Lon-Protease-BrxL-><-RVT<-HIN-HTH\|\|DEDD_Tnp_IS110+Transposase_20-><-TPR<-?<-DnaJ | 1213 | Bacteria>Pseudomonadota>Gammaproteobacteria>Enterobacterales>Enterobacteriaceae>Klebsiella/Raoultella_group | GCA_021857085.1 | Klebsiella_pneumoniae | L7127_004405 | 18391..22032 | 573 | BREX-system-P-loop-protein-BrxC-[Klebsiella-pneumoniae] |
| HBQ4939450.1 | Type1.BREX | XIS-HTH->SF1-SF2-Helicase+REase->SF1-SF2-Helicase+REase-><-PglZ<-ParB+HNH<-N6-METHYLASE<-DUF6079*<-BrxB-BrxF-iSTAND<-BrxA-DNA-binding<-Arm-DNA-bind_3+Phage_integrase | 1157 | Bacteria>Pseudomonadota>Gammaproteobacteria>Enterobacterales>Enterobacteriaceae>Klebsiella/Raoultella_group | GCA_021861305.1 | Klebsiella_pneumoniae | L7366_003904 | 52372..55845 | 573 | BREX-system-P-loop-protein-BrxC-[Klebsiella-pneumoniae] |
| EIV3902456.1 | Type1.BREX | Arm-DNA-bind_3+Phage_integrase-><-SIGMA+OB-Fold+AAA-ATPase+Lon-Protease-BrxL<-PglZ<-TOPRIM<-OLD-ABC<-N6-METHYLASE<-DUF6079*<-BrxB-BrxF-iSTAND<-BrxA-DNA-binding\|\|NACHT+SF1-SF2-Helicase+HKD-DNase-> | 1204 | Bacteria>Pseudomonadota>Gammaproteobacteria>Enterobacterales>Enterobacteriaceae>Klebsiella/Raoultella_group | GCA_021871195.1 | Klebsiella_pneumoniae | L8E75_001739 | 326019..329633 | 573 | BREX-system-P-loop-protein-BrxC-[Klebsiella-pneumoniae] |
| HBQ6953848.1 | Type1.BREX | XIS-HTH->XIS-HTH->?->REase+SPF-cons-><-SF1-SF2-Helicase<-HKD-DNase+HKD-DNase<-?\|\|BrxA-DNA-binding->BrxB-BrxF-iSTAND->DUF6079*-> | 1043 | Bacteria>Pseudomonadota>Gammaproteobacteria>Enterobacterales>Enterobacteriaceae>Klebsiella/Raoultella_group | GCA_021875055.1 | Klebsiella_pneumoniae | L8079_001073 | 40057..43187 | 573 | BREX-system-P-loop-protein-BrxC-[Klebsiella-pneumoniae] |
| HBQ7765849.1 | Type1.BREX | <-DUF6079-p*<-BrxB-BrxF-iSTAND<-BrxA-DNA-binding<-URI-DNase\|\|Phage_integrase-> | 741 | Bacteria>Pseudomonadota>Gammaproteobacteria>Enterobacterales>Enterobacteriaceae>Klebsiella/Raoultella_group | GCA_021878555.1 | Klebsiella_pneumoniae | L8R48_006010 | 1..2224 | 573 | BREX-system-P-loop-protein-BrxC-[Klebsiella-pneumoniae] |
| HBQ7782691.1 | Type1.BREX | <-DUF6079*<-BrxB-BrxF-iSTAND<-BrxA-DNA-binding<-URI-DNase\|\|Phage_integrase-> | 1042 | Bacteria>Pseudomonadota>Gammaproteobacteria>Enterobacterales>Enterobacteriaceae>Klebsiella/Raoultella_group | GCA_021878575.1 | Klebsiella_pneumoniae | L8R50_005941 | 1..3128 | 573 | BREX-system-P-loop-protein-BrxC-[Klebsiella-pneumoniae] |
| HBR1207023.1 | Type1.BREX | <-DUF905<-?<-?<-?<-wHTH+WYL<-DUF932<-nSTAND1+GTPase-AIG+sGTP\|\|BrxA-DNA-binding->BrxB-BrxF-iSTAND->DUF6079*->N6-METHYLASE->OLD-ABC->TOPRIM->PglZ->SIGMA+OB-Fold+AAA-ATPase+Lon-Protease-BrxL-><-?<-AAA+wHTH<-XIS-HTH<-Phage_Cox | 1204 | Bacteria>Pseudomonadota>Gammaproteobacteria>Enterobacterales>Enterobacteriaceae>Klebsiella/Raoultella_group | GCA_021909055.1 | Klebsiella_pneumoniae | L9K92_004097 | 80517..84131 | 573 | BREX-system-P-loop-protein-BrxC-[Klebsiella-pneumoniae] |
| HBR1207682.1 | Type1.BREX | Glyco_hydro_43+GH43_C2->wHTH+SAD+HNH->wHTH+HNH-><-HIN-HTH<-SIGMA+OB-Fold+AAA-ATPase+Lon-Protease-BrxL<-PglZ<-TOPRIM<-OLD-ABC<-N6-METHYLASE<-DUF6079*<-BrxB-BrxF-iSTAND<-BrxA-DNA-binding\|\|ZnR-><-tRNA\|\|?->?->?->?->?->Calcineurin-> | 1204 | Bacteria>Pseudomonadota>Gammaproteobacteria>Enterobacterales>Enterobacteriaceae>Klebsiella/Raoultella_group | GCA_021909055.1 | Klebsiella_pneumoniae | L9K92_004777 | 40693..44307 | 573 | BREX-system-P-loop-protein-BrxC-[Klebsiella-pneumoniae] |
| HBR1265003.1 | Type1.BREX | <-AP_endonuc_2\|\|?->Glyco_hydro_43+GH43_C2-><-MAD-NTD+HNH\|\|wHTH+SAD+HNH-><-AbiEii<-AbiEi_4\|\|BrxA-DNA-binding->BrxB-BrxF-iSTAND->DUF6079*->N6-METHYLASE->PglZ->SIGMA+OB-Fold+AAA-ATPase+Lon-Protease-BrxL->HKD-DNase+OGG-like-><-REase<-NACHT+SF1-SF2-Helicase+HKD-DNase<-RNaseH<-nSTAND1+HNH | 1213 | Bacteria>Pseudomonadota>Gammaproteobacteria>Enterobacterales>Enterobacteriaceae>Klebsiella/Raoultella_group | GCA_021909255.1 | Klebsiella_quasipneumoniae_subsp._quasipneumoniae | L9L24_003166 | 12201..15842 | 1667327 | BREX-system-P-loop-protein-BrxC-[Klebsiella-quasipneumoniae-subsp.-quasipneumoniae] |
| HBR1743014.1 | Type1.BREX | wHTH+SAD+HNH-><-?\|\|?->DUF2754-><-?\|\|ColE7+HNH->?->BrxA-DNA-binding->BrxB-BrxF-iSTAND->DUF6079*->N6-METHYLASE->PglZ->SIGMA+OB-Fold+AAA-ATPase+Lon-Protease-BrxL->HKD-DNase+OGG-like-><-REase<-NACHT+SF1-SF2-Helicase+HKD-DNase<-RNaseH<-nSTAND1+HNH | 1213 | Bacteria>Pseudomonadota>Gammaproteobacteria>Enterobacterales>Enterobacteriaceae>Klebsiella/Raoultella_group | GCA_021911065.1 | Klebsiella_pneumoniae | L9M23_004127 | 37623..41264 | 573 | BREX-system-P-loop-protein-BrxC-[Klebsiella-pneumoniae] |
| HBR1749211.1 | Type1.BREX | REase+iREase-><-RHH\|\|?->wHTH+WYL+WCX->BrxA-DNA-binding->BrxB-BrxF-iSTAND->DUF6079*->ParB+HNH->N6-METHYLASE->PglZ->SIGMA+OB-Fold+AAA-ATPase+Lon-Protease-BrxL-><-DEDD_Tnp_IS110+Transposase_20<-PIN<-MazE_antitoxin\|\|?-><-EAL | 1225 | Bacteria>Pseudomonadota>Gammaproteobacteria>Enterobacterales>Enterobacteriaceae>Klebsiella/Raoultella_group | GCA_021911085.1 | Klebsiella_pneumoniae | L9M14_005022 | 6747..10424 | 573 | BREX-system-P-loop-protein-BrxC-[Klebsiella-pneumoniae] |
| HBR1784209.1 | Type1.BREX | <-wHTH+WYL<-sGTP<-?\|\|XIS-HTH->?->?->BrxA-DNA-binding->BrxB-BrxF-iSTAND->DUF6079*->ParB+HNH->N6-METHYLASE->HEPN->RVT->N6-METHYLASE-p->PglZ->SIGMA+OB-Fold+AAA-ATPase+Lon-Protease-BrxL-><-Arm-DNA-bind_3+Phage_integrase | 1233 | Bacteria>Pseudomonadota>Gammaproteobacteria>Enterobacterales>Enterobacteriaceae>Klebsiella/Raoultella_group | GCA_021911385.1 | Klebsiella_quasipneumoniae_subsp._quasipneumoniae | L9M28_003408 | 20306..24007 | 1667327 | BREX-system-P-loop-protein-BrxC-[Klebsiella-quasipneumoniae-subsp.-quasipneumoniae] |
| HBR3944992.1 | Type1.BREX | <-HTH_36<-HTH+HTH<-ParB+BAM<-XIS-HTH<-wHTH\|\|BrxA-DNA-binding->BrxB-BrxF-iSTAND->DUF6079*->N6-METHYLASE-p-> | 1157 | Bacteria>Pseudomonadota>Gammaproteobacteria>Enterobacterales>Enterobacteriaceae>Klebsiella/Raoultella_group | GCA_021923985.1 | Klebsiella_pneumoniae | L9T85_005193 | 16003..19476 | 573 | BREX-system-P-loop-protein-BrxC-[Klebsiella-pneumoniae] |
| HBR4416044.1 | Type1.BREX | wHTH+SAD+HNH-><-?\|\|?->DUF2754-><-?\|\|ColE7+HNH->?->BrxA-DNA-binding->BrxB-BrxF-iSTAND->DUF6079*->N6-METHYLASE->PglZ->SIGMA+OB-Fold+AAA-ATPase+Lon-Protease-BrxL->HKD-DNase+OGG-like-><-REase<-NACHT+SF1-SF2-Helicase+HKD-DNase<-RNaseH<-nSTAND1+HNH | 1213 | Bacteria>Pseudomonadota>Gammaproteobacteria>Enterobacterales>Enterobacteriaceae>Klebsiella/Raoultella_group | GCA_021925925.1 | Klebsiella_pneumoniae | L9V39_001889 | 16089..19730 | 573 | BREX-system-P-loop-protein-BrxC-[Klebsiella-pneumoniae] |
| HBR4497442.1 | Type1.BREX | REase-><-SIGMA+OB-Fold+AAA-ATPase+Lon-Protease-BrxL<-PglZ<-N6-METHYLASE<-DUF6079*<-BrxB-BrxF-iSTAND<-BrxA-DNA-binding<-URI-DNase\|\|<-?<-?\|\|\|\|URI-DNase-> | 1213 | Bacteria>Pseudomonadota>Gammaproteobacteria>Enterobacterales>Enterobacteriaceae>Klebsiella/Raoultella_group | GCA_021926505.1 | Klebsiella_pneumoniae | L9V75_004938 | 11616..15257 | 573 | BREX-system-P-loop-protein-BrxC-[Klebsiella-pneumoniae] |
| HBR5241810.1 | Type1.BREX | DUF2754-><-?<-NUDIX\|\|ColE7+HNH->?->BrxA-DNA-binding->BrxB-BrxF-iSTAND->DUF6079*->N6-METHYLASE->PglZ->SIGMA+OB-Fold+AAA-ATPase+Lon-Protease-BrxL-><-REase<-NACHT+SF1-SF2-Helicase+HKD-DNase<-RNaseH<-nSTAND1+HNH | 1213 | Bacteria>Pseudomonadota>Gammaproteobacteria>Enterobacterales>Enterobacteriaceae>Klebsiella/Raoultella_group | GCA_021929495.1 | Klebsiella_pneumoniae | L9Y21_004664 | 37685..41326 | 573 | BREX-system-P-loop-protein-BrxC-[Klebsiella-pneumoniae] |
| HBR6062602.1 | Type1.BREX | ColE7+HNH->?->BrxA-DNA-binding->BrxB-BrxF-iSTAND->DUF6079*->N6-METHYLASE->PglZ->SIGMA+OB-Fold+AAA-ATPase+Lon-Protease-BrxL->HKD-DNase+OGG-like-><-REase<-NACHT+SF1-SF2-Helicase+HKD-DNase<-RNaseH | 1213 | Bacteria>Pseudomonadota>Gammaproteobacteria>Enterobacterales>Enterobacteriaceae>Klebsiella/Raoultella_group | GCA_021932975.1 | Klebsiella_pneumoniae | L9O24_005225 | 4087..7728 | 573 | BREX-system-P-loop-protein-BrxC-[Klebsiella-pneumoniae] |
| HBR8923219.1 | Type1.BREX | DUF6079*->N6-METHYLASE->PglZ->SIGMA+OB-Fold+AAA-ATPase+Lon-Protease-BrxL-><-ZnR<-ZnR | 1005 | Bacteria>Pseudomonadota>Gammaproteobacteria>Enterobacterales>Enterobacteriaceae>Klebsiella/Raoultella_group | GCA_021945235.1 | Klebsiella_pneumoniae | L9729_005179 | 1..3020 | 573 | BREX-system-P-loop-protein-BrxC-[Klebsiella-pneumoniae] |
| HBS2746978.1 | Type1.BREX | <-wHTH+WYL<-nSTAND1+sGTP<-?\|\|XIS-HTH->?->?->BrxA-DNA-binding->BrxB-BrxF-iSTAND->DUF6079*->ParB+HNH-> | 1233 | Bacteria>Pseudomonadota>Gammaproteobacteria>Enterobacterales>Enterobacteriaceae>Klebsiella/Raoultella_group | GCA_021960165.1 | Klebsiella_quasipneumoniae_subsp._quasipneumoniae | MAE70_005431 | 15539..19240 | 1667327 | BREX-system-P-loop-protein-BrxC-[Klebsiella-quasipneumoniae-subsp.-quasipneumoniae] |
| HBS5818882.1 | Type1.BREX | Arm-DNA-bind_3+Phage_integrase->BrxA-DNA-binding->BrxB-BrxF-iSTAND->DUF6079*->ParB+HNH->PglZ-><-SF1-SF2-Helicase+REase<-SF1-SF2-Helicase+REase<-XIS-HTH | 1157 | Bacteria>Pseudomonadota>Gammaproteobacteria>Enterobacterales>Enterobacteriaceae>Klebsiella/Raoultella_group | GCA_021971955.1 | Klebsiella_pneumoniae | MAK71_003676 | 3132..6605 | 573 | BREX-system-P-loop-protein-BrxC-[Klebsiella-pneumoniae] |
| HBS6040887.1 | Type1.BREX | Glyoxalase_8->URI-DNase-><-RHH_antitoxin\|\|?->XIS-HTH->?->ZnR->BrxA-DNA-binding->BrxB-BrxF-iSTAND->DUF6079*->N6-METHYLASE->PglZ->SIGMA+OB-Fold+AAA-ATPase+Lon-Protease-BrxL->?-><-Arm-DNA-bind_3+Phage_integrase | 1213 | Bacteria>Pseudomonadota>Gammaproteobacteria>Enterobacterales>Enterobacteriaceae>Klebsiella/Raoultella_group | GCA_021973155.1 | Klebsiella_aerogenes | MAL38_002165 | 35308..38949 | 548 | BREX-system-P-loop-protein-BrxC-[Klebsiella-aerogenes] |
| HBS6654372.1 | Type1.BREX | <-HNH-p\|\|wHTH+SAD+HNH->wHTH+HNH-><-HIN-HTH<-SIGMA+OB-Fold+AAA-ATPase+Lon-Protease-BrxL<-PglZ<-TOPRIM<-OLD-ABC<-N6-METHYLASE<-DUF6079*<-BrxB-BrxF-iSTAND<-BrxA-DNA-binding\|\|ZnR-><-tRNA\|\|?->?->?->?->Calcineurin-> | 1204 | Bacteria>Pseudomonadota>Gammaproteobacteria>Enterobacterales>Enterobacteriaceae>Klebsiella/Raoultella_group | GCA_021975495.1 | Klebsiella_pneumoniae | MAN10_004114 | 53582..57196 | 573 | BREX-system-P-loop-protein-BrxC-[Klebsiella-pneumoniae] |
| HBS6827429.1 | Type1.BREX | <-DUF932<-nSTAND1+GTPase-AIG+sGTP<-CopG/RHH\|\|?->XIS-HTH->?->?->BrxA-DNA-binding->BrxB-BrxF-iSTAND->DUF6079*-> | 1021 | Bacteria>Pseudomonadota>Gammaproteobacteria>Enterobacterales>Enterobacteriaceae>Klebsiella/Raoultella_group | GCA_021976135.1 | Klebsiella_pneumoniae | MAN94_004909 | 32044..35105 | 573 | BREX-system-P-loop-protein-BrxC-[Klebsiella-pneumoniae] |
| HBS7652074.1 | Type1.BREX | <-Calcineurin<-?<-?<-?<-?<-?\|\|tRNA-><-ZnR\|\|BrxA-DNA-binding->BrxB-BrxF-iSTAND->DUF6079*->N6-METHYLASE->OLD-ABC->TOPRIM->PglZ->SIGMA+OB-Fold+AAA-ATPase+Lon-Protease-BrxL->HIN-HTH-><-wHTH+HNH<-wHTH+SAD+HNH\|\|MAD-NTD+HNH-> | 1204 | Bacteria>Pseudomonadota>Gammaproteobacteria>Enterobacterales>Enterobacteriaceae>Klebsiella/Raoultella_group | GCA_021979555.1 | Klebsiella_pneumoniae | MAR21_002848 | 317524..321138 | 573 | BREX-system-P-loop-protein-BrxC-[Klebsiella-pneumoniae] |
| HBS9599072.1 | Type1.BREX | <-HNH-p\|\|wHTH+SAD+HNH->wHTH+HNH-><-HIN-HTH<-SIGMA+OB-Fold+AAA-ATPase+Lon-Protease-BrxL<-PglZ<-TOPRIM<-OLD-ABC<-N6-METHYLASE<-DUF6079*<-BrxB-BrxF-iSTAND<-BrxA-DNA-binding\|\|ZnR-><-tRNA\|\|?->?->?->?->Calcineurin-> | 1204 | Bacteria>Pseudomonadota>Gammaproteobacteria>Enterobacterales>Enterobacteriaceae>Klebsiella/Raoultella_group | GCA_021986995.1 | Klebsiella_pneumoniae | MAW98_001666 | 53581..57195 | 573 | BREX-system-P-loop-protein-BrxC-[Klebsiella-pneumoniae] |
| EIV7918660.1 | Type1.BREX | DUF2754-><-HTH_Tnp_1+HTH_21+rve+rve_3\|\|ColE7+HNH->?->BrxA-DNA-binding->BrxB-BrxF-iSTAND->DUF6079*->N6-METHYLASE->PglZ->SIGMA+OB-Fold+AAA-ATPase+Lon-Protease-BrxL->HKD-DNase+OGG-like-><-REase<-NACHT+SF1-SF2-Helicase+HKD-DNase<-RNaseH<-nSTAND1+HNH | 1213 | Bacteria>Pseudomonadota>Gammaproteobacteria>Enterobacterales>Enterobacteriaceae>Klebsiella/Raoultella_group | GCA_022009395.1 | Klebsiella_pneumoniae | MA315_005123 | 14682..18323 | 573 | BREX-system-P-loop-protein-BrxC-[Klebsiella-pneumoniae] |
| EJD7072427.1 | Type1.BREX | Arm-DNA-bind_3+Phage_integrase->EVE+HNH-><-REase+iREase\|\|BrxA-DNA-binding->BrxB-BrxF-iSTAND->DUF6079-p*-> | 584 | Bacteria>Pseudomonadota>Gammaproteobacteria>Enterobacterales>Enterobacteriaceae>Klebsiella/Raoultella_group | GCA_022024375.2 | Klebsiella_pneumoniae | MOJ36_RS17010 | 155077..156830 | 573 | BREX-system-P-loop-protein-BrxC-[Klebsiella-pneumoniae] |
| HBT6124508.1 | Type1.BREX | <-DUF6079-p*<-BrxB-BrxF-iSTAND<-BrxA-DNA-binding<-URI-DNase\|\|Phage_integrase-> | 662 | Bacteria>Pseudomonadota>Gammaproteobacteria>Enterobacterales>Enterobacteriaceae>Klebsiella/Raoultella_group | GCA_022055945.1 | Klebsiella_pneumoniae | MCD08_005761 | 1..1988 | 573 | BREX-system-P-loop-protein-BrxC-[Klebsiella-pneumoniae] |
| HBU3475135.1 | Type1.BREX | DUF2754->\|\|ColE7+HNH->?->BrxA-DNA-binding->BrxB-BrxF-iSTAND->DUF6079*->N6-METHYLASE->PglZ->SIGMA+OB-Fold+AAA-ATPase+Lon-Protease-BrxL->HKD-DNase+OGG-like-><-REase<-NACHT+SF1-SF2-Helicase+HKD-DNase<-RNaseH<-nSTAND1+HNH | 1213 | Bacteria>Pseudomonadota>Gammaproteobacteria>Enterobacterales>Enterobacteriaceae>Klebsiella/Raoultella_group | GCA_022085545.1 | Klebsiella_pneumoniae | MCW21_004534 | 37275..40916 | 573 | BREX-system-P-loop-protein-BrxC-[Klebsiella-pneumoniae] |
| HBV1956328.1 | Type1.BREX | URI-DNase->BrxA-DNA-binding->BrxB-BrxF-iSTAND->DUF6079*->N6-METHYLASE->PglZ->SIGMA+OB-Fold+AAA-ATPase+Lon-Protease-BrxL->HKD-DNase+OGG-like-><-REase | 1213 | Bacteria>Pseudomonadota>Gammaproteobacteria>Enterobacterales>Enterobacteriaceae>Klebsiella/Raoultella_group | GCA_022126105.1 | Klebsiella_pneumoniae | MDQ85_004847 | 2358..5999 | 573 | BREX-system-P-loop-protein-BrxC-[Klebsiella-pneumoniae] |
| HBV3572283.1 | Type1.BREX | DUF6079*->N6-METHYLASE->PglZ->SIGMA+OB-Fold+AAA-ATPase+Lon-Protease-BrxL-><-ZnR<-ZnR | 1061 | Bacteria>Pseudomonadota>Gammaproteobacteria>Enterobacterales>Enterobacteriaceae>Klebsiella/Raoultella_group | GCA_022140925.1 | Klebsiella_pneumoniae | MDV19_005297 | 1..3186 | 573 | BREX-system-P-loop-protein-BrxC-[Klebsiella-pneumoniae] |
| HBV3618888.1 | Type1.BREX | DUF6079*->N6-METHYLASE->PglZ->SIGMA+OB-Fold+AAA-ATPase+Lon-Protease-BrxL-><-ZnR<-ZnR | 1060 | Bacteria>Pseudomonadota>Gammaproteobacteria>Enterobacterales>Enterobacteriaceae>Klebsiella/Raoultella_group | GCA_022141165.1 | Klebsiella_pneumoniae | MDV14_005210 | 1..3185 | 573 | BREX-system-P-loop-protein-BrxC-[Klebsiella-pneumoniae] |
| HBV8971411.1 | Type1.BREX | <-LACI-HTH+PBPI<-?\|\|HNH->SymE_toxin->Glyoxalase->?->\|\|URI-DNase->BrxA-DNA-binding->BrxB-BrxF-iSTAND->DUF6079*->N6-METHYLASE->PglZ->SIGMA+OB-Fold+AAA-ATPase+Lon-Protease-BrxL->HKD-DNase+OGG-like-><-REase<-NACHT+SF1-SF2-Helicase+HKD-DNase<-RNaseH<-nSTAND1+HNH\|\|XIS-HTH-> | 1213 | Bacteria>Pseudomonadota>Gammaproteobacteria>Enterobacterales>Enterobacteriaceae>Klebsiella/Raoultella_group | GCA_022162485.1 | Klebsiella_oxytoca | MD887_005073 | 22560..26201 | 571 | BREX-system-P-loop-protein-BrxC-[Klebsiella-oxytoca] |
| HBW3476175.1 | Type1.BREX | nSTAND1+HNH->RNaseH->NACHT+SF1-SF2-Helicase+HKD-DNase->REase-><-HKD-DNase+OGG-like<-SIGMA+OB-Fold+AAA-ATPase+Lon-Protease-BrxL<-PglZ<-N6-METHYLASE<-DUF6079* | 1162 | Bacteria>Pseudomonadota>Gammaproteobacteria>Enterobacterales>Enterobacteriaceae>Klebsiella/Raoultella_group | GCA_022184305.1 | Klebsiella_pneumoniae | MEZ45_004510 | 80809..84297 | 573 | BREX-system-P-loop-protein-BrxC-[Klebsiella-pneumoniae] |
| HBW3608486.1 | Type1.BREX | DUF2754-><-?\|\|ColE7+HNH->?->BrxA-DNA-binding->BrxB-BrxF-iSTAND->DUF6079*->N6-METHYLASE->PglZ->SIGMA+OB-Fold+AAA-ATPase+Lon-Protease-BrxL->HKD-DNase+OGG-like-><-REase<-NACHT+SF1-SF2-Helicase+HKD-DNase<-RNaseH<-nSTAND1+HNH | 1213 | Bacteria>Pseudomonadota>Gammaproteobacteria>Enterobacterales>Enterobacteriaceae>Klebsiella/Raoultella_group | GCA_022184865.1 | Klebsiella_pneumoniae | MEZ81_004956 | 16088..19729 | 573 | BREX-system-P-loop-protein-BrxC-[Klebsiella-pneumoniae] |
| HBW4624689.1 | Type1.BREX | ZnR->ZnR-><-SIGMA+OB-Fold+AAA-ATPase+Lon-Protease-BrxL<-PglZ<-N6-METHYLASE<-DUF6079*<-BrxB-BrxF-iSTAND<-BrxA-DNA-binding\|\|REase+iREase-><-EVE+HNH<-Arm-DNA-bind_3+Phage_integrase | 1213 | Bacteria>Pseudomonadota>Gammaproteobacteria>Enterobacterales>Enterobacteriaceae>Klebsiella/Raoultella_group | GCA_022189205.1 | Klebsiella_pneumoniae | MES66_005166 | 11368..15009 | 573 | BREX-system-P-loop-protein-BrxC-[Klebsiella-pneumoniae] |
| HBW6426293.1 | Type1.BREX | BrxA-DNA-binding->BrxB-BrxF-iSTAND->DUF6079*->N6-METHYLASE->PglZ->SIGMA+OB-Fold+AAA-ATPase+Lon-Protease-BrxL-> | 1213 | Bacteria>Pseudomonadota>Gammaproteobacteria>Enterobacterales>Enterobacteriaceae>Klebsiella/Raoultella_group | GCA_022195805.1 | Klebsiella_pneumoniae | MFA43_005255 | 1622..5263 | 573 | BREX-system-P-loop-protein-BrxC-[Klebsiella-pneumoniae] |
| HBW7740799.1 | Type1.BREX | DUF2754-><-?\|\|ColE7+HNH->?->BrxA-DNA-binding->BrxB-BrxF-iSTAND->DUF6079*->N6-METHYLASE->PglZ->SIGMA+OB-Fold+AAA-ATPase+Lon-Protease-BrxL->HKD-DNase+OGG-like-><-REase<-NACHT+SF1-SF2-Helicase+HKD-DNase<-RNaseH<-nSTAND1+HNH | 1213 | Bacteria>Pseudomonadota>Gammaproteobacteria>Enterobacterales>Enterobacteriaceae>Klebsiella/Raoultella_group | GCA_022200625.1 | Klebsiella_pneumoniae | MFE66_003732 | 16104..19745 | 573 | BREX-system-P-loop-protein-BrxC-[Klebsiella-pneumoniae] |
| HBW8426142.1 | Type1.BREX | wHTH+SAD+HNH-><-?\|\|?->DUF2754-><-?\|\|ColE7+HNH->?->BrxA-DNA-binding->BrxB-BrxF-iSTAND->DUF6079*->N6-METHYLASE->PglZ->SIGMA+OB-Fold+AAA-ATPase+Lon-Protease-BrxL-><-REase<-NACHT+SF1-SF2-Helicase+HKD-DNase<-RNaseH<-nSTAND1+HNH | 1213 | Bacteria>Pseudomonadota>Gammaproteobacteria>Enterobacterales>Enterobacteriaceae>Klebsiella/Raoultella_group | GCA_022203225.1 | Klebsiella_pneumoniae | MFL89_004730 | 37523..41164 | 573 | BREX-system-P-loop-protein-BrxC-[Klebsiella-pneumoniae] |

| Protein-ID | System | Gene-Neighbourhoods | Length | Taxonomy | Assembly-ID | Strain | Locus-Tag | Genomic-coordinates | TaxID | Fasta-defline |
|---|---|---|---|---|---|---|---|---|---|---|
| HBW9808854.1 | Type1.BREX | <-DUF6079*<-BrxB-BrxF-iSTAND<-BrxA-DNA-binding\|\|REase+iREase-><-EVE+HNH<-Arm-DNA-bind_3+Phage_integrase | 1056 | Bacteria>Pseudomonadota>Gammaproteobacteria>Enterobacterales>Enterobacteriaceae>Klebsiella/Raoultella_group | GCA_022208425.1 | Klebsiella_pneumoniae | MFX42_22985 | 1..3169 | 573 | BREX-system-P-loop-protein-BrxC-[Klebsiella-pneumoniae] |
| HBX0790077.1 | Type1.BREX | wHTH+SAD+HNH-><-?<-?\|\|?->DUF2754->\|\|ColE7+HNH->?->BrxA-DNA-binding->BrxB-BrxF-iSTAND->DUF6079*->N6-METHYLASE->PglZ->SIGMA+OB-Fold+AAA-ATPase+Lon-Protease-BrxL->HKD-DNase+OGG-like-><-REase<-NACHT+SF1-SF2-Helicase+HKD-DNase<-RNaseH<-nSTAND1+HNH | 1213 | Bacteria>Pseudomonadota>Gammaproteobacteria>Enterobacterales>Enterobacteriaceae>Klebsiella/Raoultella_group | GCA_022212165.1 | Klebsiella_pneumoniae | MF115_18880 | 50016..53657 | 573 | BREX-system-P-loop-protein-BrxC-[Klebsiella-pneumoniae] |
| HBX5767913.1 | Type1.BREX | MULE-TRANSPOSASE-><-AAA-ATPase<-rve\|\|<-SF1-SF2-Helicase<-HKD-DNase+HKD-DNase<-?<-?\|\|BrxA-DNA-binding->BrxB-BrxF-iSTAND->DUF6079*->N6-METHYLASE->PglZ->SIGMA+OB-Fold+AAA-ATPase+Lon-Protease-BrxL-><-RVT\|\|?-><-TPR\|\|<-?<-DnaJ | 1213 | Bacteria>Pseudomonadota>Gammaproteobacteria>Enterobacterales>Enterobacteriaceae>Klebsiella/Raoultella_group | GCA_022248915.1 | Klebsiella_pneumoniae | MH553_25875 | 12227..15868 | 573 | BREX-system-P-loop-protein-BrxC-[Klebsiella-pneumoniae] |
| HBX5876367.1 | Type1.BREX | <-SF1-SF2-Helicase\|\|BrxA-DNA-binding->BrxB-BrxF-iSTAND->DUF6079*->N6-METHYLASE->OLD-ABC->TOPRIM->PglZ->SIGMA+OB-Fold+AAA-ATPase+Lon-Protease-BrxL-><-Arm-DNA-bind_3+Phage_integrase | 1204 | Bacteria>Pseudomonadota>Gammaproteobacteria>Enterobacterales>Enterobacteriaceae>Klebsiella/Raoultella_group | GCA_022249175.1 | Klebsiella_pneumoniae | MH570_26215 | 1979..5593 | 573 | BREX-system-P-loop-protein-BrxC-[Klebsiella-pneumoniae] |
| HBX7751797.1 | Type1.BREX | ColE7+HNH->?-><-SIGMA+OB-Fold+AAA-ATPase+Lon-Protease-BrxL<-PglZ<-TOPRIM<-OLD-ABC<-N6-METHYLASE<-DUF6079*<-BrxB-BrxF-iSTAND<-BrxA-DNA-binding\|\|ZnR-> | 1204 | Bacteria>Pseudomonadota>Gammaproteobacteria>Enterobacterales>Enterobacteriaceae>Klebsiella/Raoultella_group | GCA_022258155.1 | Klebsiella_pneumoniae | MIG23_15250 | 14841..18455 | 573 | BREX-system-P-loop-protein-BrxC-[Klebsiella-pneumoniae] |
| HBY0194703.1 | Type1.BREX | <-DUF6079-p*<-BrxB-BrxF-iSTAND<-BrxA-DNA-binding<-URI-DNase\|\|ASCH+Glyoxalase->Phage_integrase-> | 704 | Bacteria>Pseudomonadota>Gammaproteobacteria>Enterobacterales>Enterobacteriaceae>Klebsiella/Raoultella_group | GCA_022267575.1 | Klebsiella_pneumoniae | MIN90_27480 | 1..2112 | 573 | BREX-system-P-loop-protein-BrxC-[Klebsiella-pneumoniae] |
| HBY1687637.1 | Type1.BREX | Glyoxalase_8->URI-DNase-><-RHH_antitoxin\|\|?->XIS-HTH->?->ZnR->BrxA-DNA-binding->BrxB-BrxF-iSTAND->DUF6079*->N6-METHYLASE->PglZ->SIGMA+OB-Fold+AAA-ATPase+Lon-Protease-BrxL->HKD-DNase+OGG-like-><-Arm-DNA-bind_3+Phage_integrase | 1213 | Bacteria>Pseudomonadota>Gammaproteobacteria>Enterobacterales>Enterobacteriaceae>Klebsiella/Raoultella_group | GCA_022272455.1 | Klebsiella_pneumoniae | MIQ44_23625 | 54198..57839 | 573 | BREX-system-P-loop-protein-BrxC-[Klebsiella-pneumoniae] |
| HBY1974060.1 | Type1.BREX | <-DUF6079*<-BrxB-BrxF-iSTAND<-BrxA-DNA-binding\|\|REase+iREase-><-EVE+HNH<-Arm-DNA-bind_3+Phage_integrase | 1065 | Bacteria>Pseudomonadota>Gammaproteobacteria>Enterobacterales>Enterobacteriaceae>Klebsiella/Raoultella_group | GCA_022274115.1 | Klebsiella_pneumoniae | MIR32_28085 | 1..3194 | 573 | BREX-system-P-loop-protein-BrxC-[Klebsiella-pneumoniae] |
| HBY5777257.1 | Type1.BREX | <-DUF6079*<-BrxB-BrxF-iSTAND<-BrxA-DNA-binding<-URI-DNase\|\|ASCH+Glyoxalase->Phage_integrase-> | 1075 | Bacteria>Pseudomonadota>Gammaproteobacteria>Enterobacterales>Enterobacteriaceae>Klebsiella/Raoultella_group | GCA_022292055.1 | Klebsiella_pneumoniae | MIZ42_28895 | 1..3225 | 573 | BREX-system-P-loop-protein-BrxC-[Klebsiella-pneumoniae] |
| HBY8309079.1 | Type1.BREX | DUF6079*->ParB+HNH->N6-METHYLASE->PglZ->SIGMA+OB-Fold+AAA-ATPase+Lon-Protease-BrxL-><-HTH_Tnp_1+HTH_21+rve+rve_3 | 1167 | Bacteria>Pseudomonadota>Gammaproteobacteria>Enterobacterales>Enterobacteriaceae>Klebsiella/Raoultella_group | GCA_022301875.1 | Klebsiella_pneumoniae | MIZ37_25465 | 1..3506 | 573 | BREX-system-P-loop-protein-BrxC-[Klebsiella-pneumoniae] |
| HBZ4474772.1 | Type1.BREX | <-DUF6079-p*<-BrxB-BrxF-iSTAND<-BrxA-DNA-binding<-URI-DNase\|\|Phage_integrase-> | 737 | Bacteria>Pseudomonadota>Gammaproteobacteria>Enterobacterales>Enterobacteriaceae>Klebsiella/Raoultella_group | GCA_022336725.1 | Klebsiella_pneumoniae | L8R61_005966 | 1..2211 | 573 | BREX-system-P-loop-protein-BrxC-[Klebsiella-pneumoniae] |
| EIX9162689.1 | Type1.BREX | <-MODE-HTH+PBPII\|\|ANK+ANK+ANK->?->?->?->?->ACET->\|\|BrxA-DNA-binding->BrxB-BrxF-iSTAND->DUF6079*->N6-METHYLASE->PglZ->SIGMA+OB-Fold+AAA-ATPase+Lon-Protease-BrxL-><-ZnR<-ZnR<-?<-?\|\|RHH_antitoxin->XIS-HTH-> | 1213 | Bacteria>Pseudomonadota>Gammaproteobacteria>Enterobacterales>Enterobacteriaceae>Klebsiella/Raoultella_group | GCA_022430005.1 | Klebsiella_pneumoniae | ML399_002334 | 77875..81516 | 573 | BREX-system-P-loop-protein-BrxC-[Klebsiella-pneumoniae] |
| EIX9370752.1 | Type1.BREX | <-wHTH+WYL<-nSTAND1+sGTP<-?\|\|XIS-HTH->?->BrxA-DNA-binding->BrxB-BrxF-iSTAND->DUF6079*->ParB+HNH->N6-METHYLASE->PglZ->SIGMA+OB-Fold+AAA-ATPase+Lon-Protease-BrxL-><-Arm-DNA-bind_3+Phage_integrase | 1233 | Bacteria>Pseudomonadota>Gammaproteobacteria>Enterobacterales>Enterobacteriaceae>Klebsiella/Raoultella_group | GCA_022430905.1 | Klebsiella_pneumoniae | L7649_004474 | 53291..56992 | 573 | BREX-system-P-loop-protein-BrxC-[Klebsiella-pneumoniae] |
| EIX9551660.1 | Type1.BREX | <-MAD-NTD+HNH\|\|<-?\|\|?->\|\|?->?-><-Phage_integrase\|\|URI-DNase->BrxA-DNA-binding->BrxB-BrxF-iSTAND->DUF6079*->N6-METHYLASE->PglZ->SIGMA+OB-Fold+AAA-ATPase+Lon-Protease-BrxL->HKD-DNase+OGG-like-><-REase | 1213 | Bacteria>Pseudomonadota>Gammaproteobacteria>Enterobacterales>Enterobacteriaceae>Klebsiella/Raoultella_group | GCA_022431205.1 | Klebsiella_pneumoniae | ML061_004130 | 17308..20949 | 573 | BREX-system-P-loop-protein-BrxC-[Klebsiella-pneumoniae] |
| EIX9552648.1 | Type1.BREX | <-DUF6079-p*<-BrxB-BrxF-iSTAND<-BrxA-DNA-binding<-URI-DNase\|\|Phage_integrase-> | 891 | Bacteria>Pseudomonadota>Gammaproteobacteria>Enterobacterales>Enterobacteriaceae>Klebsiella/Raoultella_group | GCA_022431205.1 | Klebsiella_pneumoniae | ML061_005170 | 1..2675 | 573 | BREX-system-P-loop-protein-BrxC-[Klebsiella-pneumoniae] |
| HBZ8012775.1 | Type1.BREX | <-MAD-NTD+HNH\|\|wHTH+SAD+HNH->\|\|?->DUF2754-><-?\|\|ColE7+HNH->?->BrxA-DNA-binding->BrxB-BrxF-iSTAND->DUF6079*->N6-METHYLASE->PglZ->SIGMA+OB-Fold+AAA-ATPase+Lon-Protease-BrxL->HKD-DNase+OGG-like-><-REase<-NACHT+SF1-SF2-Helicase+HKD-DNase<-RNaseH<-nSTAND1+HNH | 1213 | Bacteria>Pseudomonadota>Gammaproteobacteria>Enterobacterales>Enterobacteriaceae>Klebsiella/Raoultella_group | GCA_022500545.1 | Klebsiella_pneumoniae | MM158_002941 | 16412..20053 | 573 | BREX-system-P-loop-protein-BrxC-[Klebsiella-pneumoniae] |
| EIY5117082.1 | Type1.BREX | <-Calcineurin<-?<-?<-?<-?<-?\|\|tRNA-><-ZnR\|\|BrxA-DNA-binding->BrxB-BrxF-iSTAND->DUF6079*->N6-METHYLASE->OLD-ABC->TOPRIM->PglZ->SIGMA+OB-Fold+AAA-ATPase+Lon-Protease-BrxL-><-wHTH(SBD)+HNH\|\|MAD-NTD+HNH-><-Glyco_hydro_43+GH43_C2 | 1197 | Bacteria>Pseudomonadota>Gammaproteobacteria>Enterobacterales>Enterobacteriaceae>Klebsiella/Raoultella_group | GCA_022503905.1 | Klebsiella_quasipneumoniae | ML076_004996 | 20031..23624 | 1463165 | BREX-system-P-loop-protein-BrxC-[Klebsiella-quasipneumoniae] |
| HCA0942642.1 | Type1.BREX | Glyco_hydro_43+GH43_C2-><-MAD-NTD+HNH\|\|<-?\|\|\|\|?->?-><-Phage_integrase\|\|URI-DNase->BrxA-DNA-binding->BrxB-BrxF-iSTAND->DUF6079*->N6-METHYLASE->PglZ->SIGMA+OB-Fold+AAA-ATPase+Lon-Protease-BrxL->HKD-DNase+OGG-like-><-REase | 1213 | Bacteria>Pseudomonadota>Gammaproteobacteria>Enterobacterales>Enterobacteriaceae>Klebsiella/Raoultella_group | GCA_022528725.1 | Klebsiella_pneumoniae | MNJ11_004335 | 17307..20948 | 573 | BREX-system-P-loop-protein-BrxC-[Klebsiella-pneumoniae] |
| HCA3733838.1 | Type1.BREX | DUF6079*->N6-METHYLASE->PglZ->SIGMA+OB-Fold+AAA-ATPase+Lon-Protease-BrxL-><-ZnR<-ZnR | 1102 | Bacteria>Pseudomonadota>Gammaproteobacteria>Enterobacterales>Enterobacteriaceae>Klebsiella/Raoultella_group | GCA_022599315.1 | Klebsiella_pneumoniae | MPI41_005488 | 1..3309 | 573 | BREX-system-P-loop-protein-BrxC-[Klebsiella-pneumoniae] |
| HCB0434648.1 | Type1.BREX | DUF2754-><-?\|\|ColE7+HNH->?->BrxA-DNA-binding->BrxB-BrxF-iSTAND->DUF6079*->N6-METHYLASE->PglZ->SIGMA+OB-Fold+AAA-ATPase+Lon-Protease-BrxL->HKD-DNase+OGG-like-><-REase<-NACHT+SF1-SF2-Helicase+HKD-DNase<-RNaseH<-nSTAND1+HNH | 1213 | Bacteria>Pseudomonadota>Gammaproteobacteria>Enterobacterales>Enterobacteriaceae>Klebsiella/Raoultella_group | GCA_023030065.1 | Klebsiella_variicola_subsp._variicola | MYM82_004336 | 18617..22258 | 2590157 | BREX-system-P-loop-protein-BrxC-[Klebsiella-variicola-subsp.-variicola] |
| HCB0569860.1 | Type1.BREX | YjhX_toxin->ACET->METHYLASE->ZnR->Glyoxalase->BrxA-DNA-binding->BrxB-BrxF-iSTAND->DUF6079*->OLD-ABC->TOPRIM->N6-METHYLASE->PglZ->SIGMA+OB-Fold+AAA-ATPase+Lon-Protease-BrxL-><-Arm-DNA-bind_3+Phage_integrase | 1233 | Bacteria>Pseudomonadota>Gammaproteobacteria>Enterobacterales>Enterobacteriaceae>Klebsiella/Raoultella_group | GCA_023030625.1 | Klebsiella_quasipneumoniae_subsp._similipneumoniae | MYL58_003257 | 37179..40880 | 1463164 | BREX-system-P-loop-protein-BrxC-[Klebsiella-quasipneumoniae-subsp.-similipneumoniae] |
| HCD1392381.1 | Type1.BREX | <-wHTH+WYL+WCX<-SIGMA+OB-Fold+AAA-ATPase+Lon-Protease-BrxL<-PglZ<-N6-METHYLASE-p<-RVT<-HEPN<-N6-METHYLASE<-ParB+HNH<-DUF6079*<-BrxB-BrxF-iSTAND<-BrxA-DNA-binding\|\|TIR->TIR-><-XIS-HTH<-?\|\|CopG/RHH->DUF932->DUF905-> | 1233 | Bacteria>Pseudomonadota>Gammaproteobacteria>Enterobacterales>Enterobacteriaceae>Klebsiella/Raoultella_group | GCA_023530865.1 | Klebsiella_pneumoniae_subsp._pneumoniae | M9022_004645 | 23383..27084 | 72407 | BREX-system-P-loop-protein-BrxC-[Klebsiella-pneumoniae-subsp.-pneumoniae] |
| HCD5426239.1 | Type1.BREX | <-DUF4400<-XIS-HTH<-wHTH<-ZnR\|\|BrxA-DNA-binding->BrxB-BrxF-iSTAND->DUF6079*->N6-METHYLASE-> | 1204 | Bacteria>Pseudomonadota>Gammaproteobacteria>Enterobacterales>Enterobacteriaceae>Klebsiella/Raoultella_group | GCA_023613715.1 | Klebsiella_aerogenes | NBX34_004630 | 15041..18655 | 548 | BREX-system-P-loop-protein-BrxC-[Klebsiella-aerogenes] |
| MCM6196114.1 | Type1.BREX | <-Calcineurin<-?<-?<-?<-?<-?\|\|tRNA->Arm-DNA-bind_3+Phage_integrase->BrxA-DNA-binding->BrxB-BrxF-iSTAND->DUF6079*->N6-METHYLASE->ParB+HNH->PglZ-><-SF1-SF2-Helicase+REase<-SF1-SF2-Helicase+REase<-XIS-HTH | 1157 | Bacteria>Pseudomonadota>Gammaproteobacteria>Enterobacterales>Enterobacteriaceae>Klebsiella/Raoultella_group | GCA_023741115.1 | Klebsiella_pneumoniae | M4C47_13300 | 309323..312796 | 573 | BREX-system-P-loop-protein-BrxC-[Klebsiella-pneumoniae] |
| HCF8566127.1 | Type1.BREX | <-HNH-p\|\|wHTH+SAD+HNH-><-HIN-HTH<-SIGMA+OB-Fold+AAA-ATPase+Lon-Protease-BrxL<-PglZ<-TOPRIM<-OLD-ABC<-N6-METHYLASE<-DUF6079*<-BrxB-BrxF-iSTAND<-BrxA-DNA-binding\|\|ZnR-><-tRNA\|\|?->?->?->?->?->Calcineurin-> | 1204 | Bacteria>Pseudomonadota>Gammaproteobacteria>Enterobacterales>Enterobacteriaceae>Klebsiella/Raoultella_group | GCA_023980665.1 | Klebsiella_pneumoniae | NIV99_001798 | 32314..35928 | 573 | BREX-system-P-loop-protein-BrxC-[Klebsiella-pneumoniae] |
| MCP6627038.1 | Type1.BREX | nSTAND1+HNH->RNaseH->NACHT+SF1-SF2-Helicase+HKD-DNase->REase-><-HKD-DNase+OGG-like<-SIGMA+OB-Fold+AAA-ATPase+Lon-Protease-BrxL<-PglZ<-N6-METHYLASE<-DUF6079*<-BrxB-BrxF-iSTAND<-BrxA-DNA-binding<-?<-ColE7+HNH<-EVE+HNH\|\|NUDIX->?-><-DUF2754 | 1213 | Bacteria>Pseudomonadota>Gammaproteobacteria>Enterobacterales>Enterobacteriaceae>Klebsiella/Raoultella_group | GCA_024240675.1 | Klebsiella_pneumoniae | NL507_17490 | 188287..191928 | 573 | BREX-system-P-loop-protein-BrxC-[Klebsiella-pneumoniae] |
| HCI4589554.1 | Type1.BREX | XIS-HTH->PIWI->?-><-Phage_integrase<-ZnR\|\|BrxA-DNA-binding->BrxB-BrxF-iSTAND->DUF6079*->N6-METHYLASE->OLD-ABC->TOPRIM->PglZ->SIGMA+OB-Fold+AAA-ATPase+Lon-Protease-BrxL-><-SF1-SF2-Helicase<-?\|\|HIN-HTH->URI-DNase-> | 1204 | Bacteria>Pseudomonadota>Gammaproteobacteria>Enterobacterales>Enterobacteriaceae>Klebsiella/Raoultella_group | GCA_024379455.1 | Klebsiella_quasipneumoniae_subsp._quasipneumoniae | N0899_004582 | 16188..19802 | 1667327 | BREX-system-P-loop-protein-BrxC-[Klebsiella-quasipneumoniae-subsp.-quasipneumoniae] |
| HCI5816641.1 | Type1.BREX | XIS-HTH->?->REase+SPF-cons-><-SF1-SF2-Helicase<-HKD-DNase+HKD-DNase<-?\|\|BrxA-DNA-binding->BrxB-BrxF-iSTAND->DUF6079*->N6-METHYLASE->PglZ->SIGMA+OB-Fold+AAA-ATPase+Lon-Protease-BrxL-><-RVT<-TPR<-?<-DnaJ<-Arm-DNA-bind_3+Phage_integrase | 1213 | Bacteria>Pseudomonadota>Gammaproteobacteria>Enterobacterales>Enterobacteriaceae>Klebsiella/Raoultella_group | GCA_024431185.1 | Klebsiella_pneumoniae | NPN32_003719 | 27788..31429 | 573 | BREX-system-P-loop-protein-BrxC-[Klebsiella-pneumoniae] |
| HCI5938461.1 | Type1.BREX | <-ZnR\|\|BrxA-DNA-binding->BrxB-BrxF-iSTAND->DUF6079*->N6-METHYLASE->OLD-ABC->TOPRIM->PglZ->SIGMA+OB-Fold+AAA-ATPase+Lon-Protease-BrxL-> | 1197 | Bacteria>Pseudomonadota>Gammaproteobacteria>Enterobacterales>Enterobacteriaceae>Klebsiella/Raoultella_group | GCA_024431715.1 | Klebsiella_quasipneumoniae_subsp._quasipneumoniae | NPN45_004672 | 2243..5836 | 1667327 | BREX-system-P-loop-protein-BrxC-[Klebsiella-quasipneumoniae-subsp.-quasipneumoniae] |
| MCS5967892.1 | Type1.BREX | <-nSTAND1+sGTP<-?<-?\|\|?->BrxA-DNA-binding->BrxB-BrxF-iSTAND->DUF6079-p*->DUF6079-p->?->DUF6079-p->ParB+HNH->ParB+HNH->N6-METHYLASE->HEPN->?->RVT-> | 389 | Bacteria>Pseudomonadota>Gammaproteobacteria>Enterobacterales>Enterobacteriaceae>Klebsiella/Raoultella_group | GCA_024990005.1 | Klebsiella_variicola_subsp._variicola | LNP26_07865 | 1516930..1518099 | 2590157 | BREX-system-P-loop-protein-BrxC-[Klebsiella-variicola-subsp.-variicola] |
| GKN17816.1 | Type1.BREX | Arm-DNA-bind_3+Phage_integrase-><-HKD-DNase+OGG-like<-SIGMA+OB-Fold+AAA-ATPase+Lon-Protease-BrxL<-PglZ<-N6-METHYLASE<-DUF6079-p*<-DUF6079-p<-BrxB-BrxF-iSTAND<-BrxA-DNA-binding<-?<-XIS-HTH<-?\|\|RHH_antitoxin-><-URI-DNase<-Glyoxalase_8 | 725 | Bacteria>Pseudomonadota>Gammaproteobacteria>Enterobacterales>Enterobacteriaceae>Klebsiella/Raoultella_group | GCA_026006635.1 | Klebsiella_pneumoniae | NUBL17184_45010 | 12144..14321 | 573 | hypothetical-protein-NUBL17184_45010-[Klebsiella-pneumoniae] |
| GKN17819.1 | Type1.BREX | Arm-DNA-bind_3+Phage_integrase-><-HKD-DNase+OGG-like<-SIGMA+OB-Fold+AAA-ATPase+Lon-Protease-BrxL<-PglZ<-N6-METHYLASE<-DUF6079-p<-DUF6079-p*<-BrxB-BrxF-iSTAND<-BrxA-DNA-binding<-?<-XIS-HTH<-?\|\|RHH_antitoxin-><-URI-DNase<-Glyoxalase_8<-RNase | 453 | Bacteria>Pseudomonadota>Gammaproteobacteria>Enterobacterales>Enterobacteriaceae>Klebsiella/Raoultella_group | GCA_026006635.1 | Klebsiella_pneumoniae | NUBL17184_45020 | 14424..15785 | 573 | hypothetical-protein-NUBL17184_45020-[Klebsiella-pneumoniae] |
| GKP62583.1 | Type1.BREX | <-HIN-HTH<-HIN-HTH\|\|HIN-HTH->?->LZ_Tnp_IS66+zf-IS66+DDE_Tnp_IS66+DDE_Tnp_IS66_C->BrxA-DNA-binding->BrxB-BrxF-iSTAND->DUF6079*->N6-METHYLASE->OLD-ABC->TOPRIM->PglZ->SIGMA+OB-Fold+AAA-ATPase+Lon-Protease-BrxL-><-?<-?<-Arm-DNA-bind_3+Phage_integrase | 1204 | Bacteria>Pseudomonadota>Gammaproteobacteria>Enterobacterales>Enterobacteriaceae>Klebsiella/Raoultella_group | GCA_026008315.1 | Klebsiella_quasipneumoniae | NUKP47_17630 | 173713..177327 | 1463165 | hypothetical-protein-NUKP47_17630-[Klebsiella-quasipneumoniae] |
| WEJ89011.1 | Type1.BREX | <-wHTH(SBD)+HNH\|\|tRNA->Arm-DNA-bind_3+Phage_integrase-><-SIGMA+OB-Fold+AAA-ATPase+Lon-Protease-BrxL<-PglZ<-N6-METHYLASE-p<-RVT<-HEPN<-N6-METHYLASE<-ParB+HNH<-DUF6079*<-BrxB-BrxF-iSTAND<-BrxA-DNA-binding<-ZnR<-XIS-HTH<-?\|\|HIN-HTH->?->LZ_Tnp_IS66+zf-IS66+DDE_Tnp_IS66+DDE_Tnp_IS66_C-> | 1240 | Bacteria>Pseudomonadota>Gammaproteobacteria>Enterobacterales>Enterobacteriaceae>Klebsiella/Raoultella_group | GCA_029202365.1 | Klebsiella_huaxiensis | P0Y63_27795 | 5833350..5837072 | 2153354 | MAG:-BREX-system-P-loop-protein-BrxC-[Klebsiella-huaxiensis] |
| SAY23702.1 | Type1.BREX | RVT->HTH_Tnp_1+HTH_21+rve+rve_3-><-SIGMA+OB-Fold+AAA-ATPase+Lon-Protease-BrxL<-PglZ\|\|?->?-><-DUF6079*<-BrxB-BrxF-iSTAND<-BrxA-DNA-binding | 1052 | Bacteria>Pseudomonadota>Gammaproteobacteria>Enterobacterales>Enterobacteriaceae>Klebsiella/Raoultella_group | GCA_900086305.1 | Klebsiella_quasipneumoniae | SAMEA2273696_05187 | 11957..15115 | 1463165 | Uncharacterised-protein-[Klebsiella-quasipneumoniae] |
| STR42120.1 | Type1.BREX | <-SF1-SF2-Helicase<-HKD-DNase+HKD-DNase<-?<-?<-?<-?\|\|BrxA-DNA-binding->BrxB-BrxF-iSTAND->DUF6079-p*->DUF6079-p->DUF6079-p->N6-METHYLASE-p->?->ACET->N6-METHYLASE-p->N6-METHYLASE-p->N6-METHYLASE-p->PglZ-p-> | 800 | Bacteria>Pseudomonadota>Gammaproteobacteria>Enterobacterales>Enterobacteriaceae>Klebsiella/Raoultella_group | GCA_900451375.1 | Klebsiella_michiganensis | NCTC11694_03329 | 2640700..2643102 | 1134687 | Uncharacterised-protein-[Klebsiella-michiganensis] |
| SYJ62179.1 | Type1.BREX | <-DUF6079*<-BrxB-BrxF-iSTAND<-BrxA-DNA-binding<-ZnR<-?<-XIS-HTH<-?\|\|RHH_antitoxin-><-URI-DNase<-Glyoxalase_8 | 1186 | Bacteria>Pseudomonadota>Gammaproteobacteria>Enterobacterales>Enterobacteriaceae>Klebsiella/Raoultella_group | GCA_900515475.1 | Klebsiella_pneumoniae | SAMEA3538642_04756 | 88..3648 | 573 | Uncharacterised-protein-[Klebsiella-pneumoniae] |
| VAO27126.1 | Type1.BREX | DUF2754-><-?<-?\|\|ACET->ColE7+HNH->?->BrxA-DNA-binding->BrxB-BrxF-iSTAND->DUF6079*->N6-METHYLASE->PglZ->SIGMA+OB-Fold+AAA-ATPase+Lon-Protease-BrxL-><-DDE_Tnp_1_5+DDE_Tnp_1<-?<-DnaJ<-Arm-DNA-bind_3+Phage_integrase | 1213 | Bacteria>Pseudomonadota>Gammaproteobacteria>Enterobacterales>Enterobacteriaceae>Klebsiella/Raoultella_group | GCA_900559435.1 | Klebsiella_pneumoniae | SAMEA2053954_04091 | 16530..20171 | 573 | Uncharacterised-protein-[Klebsiella-pneumoniae] |
| VTO17843.1 | Type1.BREX | RNaseH->NACHT+SF1-SF2-Helicase+HKD-DNase->REase-><-HKD-DNase+OGG-like<-HKD-DNase+OGG-like<-SIGMA+OB-Fold+AAA-ATPase+Lon-Protease-BrxL<-SIGMA+OB-Fold+AAA-ATPase+Lon-Protease-BrxL<-PglZ<-N6-METHYLASE<-DUF6079-p*<-DUF6079-p<-BrxB-BrxF-iSTAND<-BrxA-DNA-binding<-?<-ColE7+HNH<-ACET\|\|?->?-><-DUF2754 | 864 | Bacteria>Pseudomonadota>Gammaproteobacteria>Enterobacterales>Enterobacteriaceae>Klebsiella/Raoultella_group | GCA_901421915.1 | Klebsiella_variicola | NCTC9178_02481 | 2086491..2089085 | 244366 | Uncharacterised-protein-[Klebsiella-variicola] |
| WP_014837241.1 | Type1.BREX | <-wHTH+HNH\|\|tRNA->Arm-DNA-bind_3+Phage_integrase->DnaJ->?->TPR-><-DEDD_Tnp_IS110+Transposase_20\|\|RVT-><-SIGMA+OB-Fold+AAA-ATPase+Lon-Protease-BrxL<-PglZ<-DUF6079*<-BrxB-BrxF-iSTAND<-BrxA-DNA-binding<-DDE_Tnp_1_5+DDE_Tnp_1\|\|?-><-?<-MODE-HTH+PBPII | 1213 | Bacteria>Pseudomonadota>Gammaproteobacteria>Enterobacterales>Enterobacteriaceae>Klebsiella/Raoultella_group | GCF_000276705.2 | Klebsiella | A225_RS02820 | 612873..616514 | 570 | MULTISPECIES:-BREX-system-P-loop-protein-BrxC-[Klebsiella] |
| WP_228134609.1 | Type1.BREX | Glyoxalase->ACET->YjhX_toxin->METHYLASE->ZnR->Glyoxalase->BrxA-DNA-binding->BrxB-BrxF-iSTAND->DUF6079*->ParB+HNH->N6-METHYLASE->N6-METHYLASE->PglZ->SIGMA+OB-Fold+AAA-ATPase+Lon-Protease-BrxL-><-Arm-DNA-bind_3+Phage_integrase<-tRNA\|\|SAM-METHYLASE-> | 1240 | Bacteria>Pseudomonadota>Gammaproteobacteria>Enterobacterales>Enterobacteriaceae>Klebsiella/Raoultella_group | GCF_000632415.1 | Klebsiella_michiganensis | J415_RS00260 | 45760..49482 | 1134687 | BREX-system-P-loop-protein-BrxC-[Klebsiella-michiganensis] |
| WP_038432196.1 | Type1.BREX | KAP-NTPase->?->?->TatD_DNase->REase-><-HKD-DNase+OGG-like<-SIGMA+OB-Fold+AAA-ATPase+Lon-Protease-BrxL<-PglZ<-N6-METHYLASE<-DUF6079*<-BrxB-BrxF-iSTAND<-BrxA-DNA-binding<-URI-DNase\|\|<-?<-?\|\|<-?\|\|?->\|\|MAD-NTD+HNH-><-Glyco_hydro_43+GH43_C2 | 1213 | Bacteria>Pseudomonadota>Gammaproteobacteria>Enterobacterales>Enterobacteriaceae>Klebsiella/Raoultella_group | GCF_000742755.1 | Klebsiella_pneumoniae | PF543_RS25520 | 27597..31238 | 573 | BREX-system-P-loop-protein-BrxC-[Klebsiella-pneumoniae] |
| WP_047066027.1 | Type1.BREX | ColE7+HNH->?->NACHT+SF1-SF2-Helicase+HKD-DNase->REase-><-SIGMA+OB-Fold+AAA-ATPase+Lon-Protease-BrxL<-PglZ<-DUF6079*<-BrxB-BrxF-iSTAND<-BrxA-DNA-binding<-wHTH+SAD+HNH\|\|REase+iREase-><-?<-HNH | 1213 | Bacteria>Pseudomonadota>Gammaproteobacteria>Enterobacterales>Enterobacteriaceae>Klebsiella/Raoultella_group | GCF_001011795.1 | Klebsiella_aerogenes | G4873_RS08800 | 147034..150675 | 548 | BREX-system-P-loop-protein-BrxC-[Klebsiella-aerogenes] |
| WP_064406990.1 | Type1.BREX | <-HTH_23+HTH_32+HTH_33+DDE_3<-RHH_antitoxin\|\|?->XIS-HTH->?->ZnR->BrxA-DNA-binding->BrxB-BrxF-iSTAND->DUF6079*->N6-METHYLASE->PglZ->SIGMA+OB-Fold+AAA-ATPase+Lon-Protease-BrxL->HKD-DNase+OGG-like-><-REase<-Arm-DNA-bind_3+Phage_integrase | 1213 | Bacteria>Pseudomonadota>Gammaproteobacteria>Enterobacterales>Enterobacteriaceae>Klebsiella/Raoultella_group | GCF_001598695.1 | Klebsiella_oxytoca | K01_RS15010 | 40025..43666 | 571 | BREX-system-P-loop-protein-BrxC-[Klebsiella-oxytoca] |
| WP_094487773.1 | Type1.BREX | DUF2754-><-?<-NUDIX\|\|ACET->ColE7+HNH->?->BrxA-DNA-binding->BrxB-BrxF-iSTAND->DUF6079*->N6-METHYLASE->PglZ->SIGMA+OB-Fold+AAA-ATPase+Lon-Protease-BrxL->HKD-DNase+OGG-like-><-REase<-NACHT+SF1-SF2-Helicase+HKD-DNase<-RNaseH<-nSTAND1+HNH | 1213 | Bacteria>Pseudomonadota>Gammaproteobacteria>Enterobacterales>Enterobacteriaceae>Klebsiella/Raoultella_group | GCF_002251875.2 | Klebsiella_pneumoniae | CI667_RS22335 | 4584289..4587930 | 573 | BREX-system-P-loop-protein-BrxC-[Klebsiella-pneumoniae] |
| WP_256991583.1 | Type1.BREX | <-DUF905<-DUF932<-CopG/RHH\|\|?->XIS-HTH->ZnR-><-TIR\|\|BrxA-DNA-binding->BrxB-BrxF-iSTAND->DUF6079*->ParB+HNH->N6-METHYLASE->HEPN->RVT->N6-METHYLASE-p->PglZ->SIGMA+OB-Fold+AAA-ATPase+Lon-Protease-BrxL->wHTH+WYL+WCX-> | 1240 | Bacteria>Pseudomonadota>Gammaproteobacteria>Enterobacterales>Enterobacteriaceae>Klebsiella/Raoultella_group | GCF_002264145.1 | Raoultella_planticola | CIG23_RS11860 | 59081..62803 | 575 | BREX-system-P-loop-protein-BrxC-[Raoultella-planticola] |
| WP_096925291.1 | Type1.BREX | URI-DNase->BrxA-DNA-binding->BrxB-BrxF-iSTAND->DUF6079*->N6-METHYLASE->PglZ->SIGMA+OB-Fold+AAA-ATPase+Lon-Protease-BrxL->HKD-DNase+OGG-like-><-REase | 1213 | Bacteria>Pseudomonadota>Gammaproteobacteria>Enterobacterales>Enterobacteriaceae>Klebsiella/Raoultella_group | GCF_002411925.1 | Klebsiella_pneumoniae | CP905_RS22210 | 5854..9495 | 573 | BREX-system-P-loop-protein-BrxC-[Klebsiella-pneumoniae] |
| WP_096927069.1 | Type1.BREX | ZnR->ZnR-><-SIGMA+OB-Fold+AAA-ATPase+Lon-Protease-BrxL<-PglZ<-N6-METHYLASE<-DUF6079*<-BrxB-BrxF-iSTAND<-BrxA-DNA-binding\|\|REase+iREase-><-Arm-DNA-bind_3+Phage_integrase | 1213 | Bacteria>Pseudomonadota>Gammaproteobacteria>Enterobacterales>Enterobacteriaceae>Klebsiella/Raoultella_group | GCF_002417545.1 | Klebsiella_pneumoniae | CQA60_RS24305 | 11428..15069 | 573 | BREX-system-P-loop-protein-BrxC-[Klebsiella-pneumoniae] |
| WP_102012827.1 | Type1.BREX | <-DUF6079-p*<-BrxB-BrxF-iSTAND<-BrxA-DNA-binding<-?<-ColE7+HNH<-ACET\|\|NUDIX-> | 986 | Bacteria>Pseudomonadota>Gammaproteobacteria>Enterobacterales>Enterobacteriaceae>Klebsiella/Raoultella_group | GCF_002856315.1 | Klebsiella_pneumoniae | CWN62_RS18100 | 1..2959 | 573 | BREX-system-P-loop-protein-BrxC,-partial-[Klebsiella-pneumoniae] |
| WP_105178007.1 | Type1.BREX | DUF2754-><-?<-NUDIX\|\|EVE+HNH->ColE7+HNH->?->BrxA-DNA-binding->BrxB-BrxF-iSTAND->DUF6079*->N6-METHYLASE->PglZ->SIGMA+OB-Fold+AAA-ATPase+Lon-Protease-BrxL->HKD-DNase+OGG-like-><-REase<-NACHT+SF1-SF2-Helicase+HKD-DNase<-RNaseH<-nSTAND1+HNH | 1213 | Bacteria>Pseudomonadota>Gammaproteobacteria>Enterobacterales>Enterobacteriaceae>Klebsiella/Raoultella_group | GCF_002934865.1 | Klebsiella_pneumoniae | CSC91_RS11390 | 28681..32322 | 573 | BREX-system-P-loop-protein-BrxC-[Klebsiella-pneumoniae] |
| WP_104443227.1 | Type1.BREX | <-ZnR\|\|BrxA-DNA-binding->BrxB-BrxF-iSTAND->DUF6079*->N6-METHYLASE->OLD-ABC->TOPRIM->PglZ->SIGMA+OB-Fold+AAA-ATPase+Lon-Protease-BrxL-><-?<-ColE7+HNH | 1204 | Bacteria>Pseudomonadota>Gammaproteobacteria>Enterobacterales>Enterobacteriaceae>Klebsiella/Raoultella_group | GCF_002934985.1 | Klebsiella_pneumoniae | CSC89_RS24445 | 2230..5844 | 573 | BREX-system-P-loop-protein-BrxC-[Klebsiella-pneumoniae] |
| WP_108704264.1 | Type1.BREX | MPTase->?-><-REase\|\|BrxA-DNA-binding->BrxB-BrxF-iSTAND->DUF6079*->N6-METHYLASE->PglZ->SIGMA+OB-Fold+AAA-ATPase+Lon-Protease-BrxL-><-TPR<-?<-DnaJ<-?<-Arm-DNA-bind_3+Phage_integrase<-tRNA\|\|wHTH+HNH-> | 1213 | Bacteria>Pseudomonadota>Gammaproteobacteria>Enterobacterales>Enterobacteriaceae>Klebsiella/Raoultella_group | GCF_003073975.1 | Klebsiella_oxytoca | CSC17_RS12015 | 2422340..2425981 | 571 | BREX-system-P-loop-protein-BrxC-[Klebsiella-oxytoca] |
| WP_108986419.1 | Type1.BREX | <-DUF6079*<-BrxB-BrxF-iSTAND<-BrxA-DNA-binding\|\|REase+iREase-><-Arm-DNA-bind_3+Phage_integrase | 1064 | Bacteria>Pseudomonadota>Gammaproteobacteria>Enterobacterales>Enterobacteriaceae>Klebsiella/Raoultella_group | GCF_003115315.1 | Klebsiella_pneumoniae | DB703_RS28015 | 1..3192 | 573 | BREX-system-P-loop-protein-BrxC,-partial-[Klebsiella-pneumoniae] |
| WP_023313883.1 | Type1.BREX | Glyco_hydro_43+GH43_C2-><-MAD-NTD+HNH\|\|<-?\|\|?->\|\|?->?->\|\|URI-DNase->BrxA-DNA-binding->BrxB-BrxF-iSTAND->DUF6079*->N6-METHYLASE->PglZ->SIGMA+OB-Fold+AAA-ATPase+Lon-Protease-BrxL->HKD-DNase+OGG-like-><-REase | 1213 | Bacteria>Pseudomonadota>Gammaproteobacteria>Enterobacterales>Enterobacteriaceae>Klebsiella/Raoultella_group | GCF_003427195.1 | Klebsiella_pneumoniae | D2N15_RS14250 | 17171..20812 | 573 | BREX-system-P-loop-protein-BrxC-[Klebsiella-pneumoniae] |
| WP_087652989.1 | Type1.BREX | DUF2754-><-?<-DUF772+DDE_Tnp_1<-?\|\|ColE7+HNH->?->BrxA-DNA-binding->BrxB-BrxF-iSTAND->DUF6079*->N6-METHYLASE->PglZ->SIGMA+OB-Fold+AAA-ATPase+Lon-Protease-BrxL->HKD-DNase+OGG-like-><-REase<-NACHT+SF1-SF2-Helicase+HKD-DNase<-RNaseH<-nSTAND1+HNH | 1213 | Bacteria>Pseudomonadota>Gammaproteobacteria>Enterobacterales>Enterobacteriaceae>Klebsiella/Raoultella_group | GCF_003429625.1 | Klebsiella_variicola | BC497_RS18300 | 3693589..3697230 | 244366 | BREX-system-P-loop-protein-BrxC-[Klebsiella-variicola] |
| WP_264757344.1 | Type1.BREX | <-wHTH+WYL<-nSTAND1+sGTP<-?\|\|XIS-HTH->?->?->BrxA-DNA-binding->BrxB-BrxF-iSTAND-><-DDE_Tnp_1_5+DDE_Tnp_1\|\|DUF6079-p*->ParB+HNH->N6-METHYLASE->HEPN->RVT->N6-METHYLASE-p->PglZ->SIGMA+OB-Fold+AAA-ATPase+Lon-Protease-BrxL-><-Arm-DNA-bind_3+Phage_integrase | 995 | Bacteria>Pseudomonadota>Gammaproteobacteria>Enterobacterales>Enterobacteriaceae>Klebsiella/Raoultella_group | GCF_003660185.1 | Klebsiella_pneumoniae | D9K63_RS23205 | 4667642..4670629 | 573 | BREX-system-P-loop-protein-BrxC-[Klebsiella-pneumoniae] |
| WP_049088891.1 | Type1.BREX | Arm-DNA-bind_3+Phage_integrase->DnaJ->?->TPR-><-SIGMA+OB-Fold+AAA-ATPase+Lon-Protease-BrxL<-PglZ<-N6-METHYLASE<-DUF6079*<-BrxB-BrxF-iSTAND<-BrxA-DNA-binding\|\|?->HKD-DNase+HKD-DNase->SF1-SF2-Helicase-><-REase+SPF-cons<-?<-XIS-HTH | 1213 | Bacteria>Pseudomonadota>Gammaproteobacteria>Enterobacterales>Enterobacteriaceae>Klebsiella/Raoultella_group | GCF_003851825.1 | Klebsiella | KPPSC5_RS01070 | 215107..218748 | 570 | MULTISPECIES:-BREX-system-P-loop-protein-BrxC-[Klebsiella] |
| WP_206615409.1 | Type1.BREX | <-ABC-ATPase+SbcC<-Calcineurin<-?<-?<-?\|\|tRNA->Arm-DNA-bind_3+Phage_integrase->wHTH+WYL+WCX->BrxA-DNA-binding->BrxB-BrxF-iSTAND->DUF6079-p*-> | 129 | Bacteria>Pseudomonadota>Gammaproteobacteria>Enterobacterales>Enterobacteriaceae>Klebsiella/Raoultella_group | GCF_004024155.1 | Raoultella_planticola | DN603_RS10420 | 221178..221567 | 575 | DUF6079-family-protein-[Raoultella-planticola] |
| WP_087867252.1 | Type1.BREX | TOPRIM-DNAG+D5N+D5ATPase+P4PRIMASE-HTH->nSTAND1+HNH->RNaseH->NACHT+SF1-SF2-Helicase+HKD-DNase->REase-><-HKD-DNase+OGG-like<-SIGMA+OB-Fold+AAA-ATPase+Lon-Protease-BrxL<-PglZ<-N6-METHYLASE<-DUF6079*<-BrxB-BrxF-iSTAND<-BrxA-DNA-binding<-?<-ColE7+HNH<-ACET\|\|?-><-DUF2754 | 1213 | Bacteria>Pseudomonadota>Gammaproteobacteria>Enterobacterales>Enterobacteriaceae>Klebsiella/Raoultella_group | GCF_004311705.1 | Klebsiella_pneumoniae | E0M06_RS18340 | 199992..203633 | 573 | BREX-system-P-loop-protein-BrxC-[Klebsiella-pneumoniae] |
| WP_130973675.1 | Type1.BREX | <-SIGMA+OB-Fold+AAA-ATPase+Lon-Protease-BrxL<-PglZ<-N6-METHYLASE<-ParB+HNH<-DUF6079*<-BrxB-BrxF-iSTAND<-BrxA-DNA-binding<-wHTH+WYL+WCX<-?\|\|RHH-> | 1225 | Bacteria>Pseudomonadota>Gammaproteobacteria>Enterobacterales>Enterobacteriaceae>Klebsiella/Raoultella_group | GCF_004311945.1 | Klebsiella_pneumoniae | E0M14_RS25420 | 10722..14399 | 573 | BREX-system-P-loop-protein-BrxC-[Klebsiella-pneumoniae] |
| WP_131033948.1 | Type1.BREX | nSTAND1+HNH->RNaseH->NACHT+SF1-SF2-Helicase+HKD-DNase->REase-><-HKD-DNase+OGG-like<-SIGMA+OB-Fold+AAA-ATPase+Lon-Protease-BrxL<-PglZ<-N6-METHYLASE<-DUF6079*<-BrxB-BrxF-iSTAND<-BrxA-DNA-binding<-?<-ColE7+HNH\|\|NUDIX->?->?-><-DUF2754 | 1213 | Bacteria>Pseudomonadota>Gammaproteobacteria>Enterobacterales>Enterobacteriaceae>Klebsiella/Raoultella_group | GCF_004312565.1 | Klebsiella_variicola | E0N46_RS24005 | 106854..110495 | 244366 | BREX-system-P-loop-protein-BrxC-[Klebsiella-variicola] |
| WP_032410729.1 | Type1.BREX | Glyoxalase_8->URI-DNase-><-RHH_antitoxin\|\|?->XIS-HTH->?->ZnR->BrxA-DNA-binding->BrxB-BrxF-iSTAND->DUF6079*->N6-METHYLASE->PglZ->SIGMA+OB-Fold+AAA-ATPase+Lon-Protease-BrxL->HKD-DNase+OGG-like-><-Arm-DNA-bind_3+Phage_integrase | 1213 | Bacteria>Pseudomonadota>Gammaproteobacteria>Enterobacterales>Enterobacteriaceae>Klebsiella/Raoultella_group | GCF_004312665.1 | Klebsiella_pneumoniae | E0N31_RS20140 | 33140..36781 | 573 | BREX-system-P-loop-protein-BrxC-[Klebsiella-pneumoniae] |
| WP_223814682.1 | Type1.BREX | <-wHTH+WYL<-nSTAND1+sGTP<-?\|\|XIS-HTH->?->?->BrxA-DNA-binding->BrxB-BrxF-iSTAND->DUF6079*->ParB+HNH->N6-METHYLASE->HEPN->RVT->N6-METHYLASE-p->PglZ->SIGMA+OB-Fold+AAA-ATPase+Lon-Protease-BrxL-><-Arm-DNA-bind_3+Phage_integrase | 1240 | Bacteria>Pseudomonadota>Gammaproteobacteria>Enterobacterales>Enterobacteriaceae>Klebsiella/Raoultella_group | GCF_004313365.1 | Klebsiella | E0O18_RS24630 | 24494..28216 | 570 | MULTISPECIES:-BREX-system-P-loop-protein-BrxC-[Klebsiella] |
| WP_048334010.1 | Type1.BREX | DUF2754-><-?<-NUDIX\|\|EVE+HNH->ColE7+HNH->?->BrxA-DNA-binding->BrxB-BrxF-iSTAND->DUF6079*->N6-METHYLASE->PglZ->SIGMA+OB-Fold+AAA-ATPase+Lon-Protease-BrxL->HKD-DNase+OGG-like-><-REase | 1213 | Bacteria>Pseudomonadota>Gammaproteobacteria>Enterobacterales>Enterobacteriaceae>Klebsiella/Raoultella_group | GCF_004313665.1 | Klebsiella_pneumoniae | E0O19_RS25945 | 16153..19794 | 573 | BREX-system-P-loop-protein-BrxC-[Klebsiella-pneumoniae] |
| WP_131930858.1 | Type1.BREX | MPTase->?-><-REase<-HTH_23+HTH_32+HTH_33+DDE_3\|\|BrxA-DNA-binding->BrxB-BrxF-iSTAND->DUF6079*->N6-METHYLASE->PglZ->SIGMA+OB-Fold+AAA-ATPase+Lon-Protease-BrxL-><-?<-DnaJ<-Arm-DNA-bind_3+Phage_integrase<-tRNA\|\|wHTH+HNH->SAM-METHYLASE-> | 1213 | Bacteria>Pseudomonadota>Gammaproteobacteria>Enterobacterales>Enterobacteriaceae>Klebsiella/Raoultella_group | GCF_004343645.1 | Klebsiella_grimontii | E0D83_RS25820 | 59532..63173 | 2058152 | BREX-system-P-loop-protein-BrxC-[Klebsiella-grimontii] |
| WP_147075822.1 | Type1.BREX | <-HNH-p\|\|wHTH+SAD+HNH->wHTH+HNH-><-HIN-HTH<-SIGMA+OB-Fold+AAA-ATPase+Lon-Protease-BrxL<-PglZ<-TOPRIM<-OLD-ABC<-N6-METHYLASE<-DUF6079*<-BrxB-BrxF-iSTAND<-BrxA-DNA-binding\|\|ZnR-><-tRNA\|\|?->?->?->?->?->Calcineurin-> | 1204 | Bacteria>Pseudomonadota>Gammaproteobacteria>Enterobacterales>Enterobacteriaceae>Klebsiella/Raoultella_group | GCF_007955295.1 | Klebsiella_pneumoniae | FTO41_25020 | 32326..35940 | 573 | BREX-system-P-loop-protein-BrxC-[Klebsiella-pneumoniae] |
| WP_096812545.1 | Type1.BREX | Arm-DNA-bind_3+Phage_integrase->BrxA-DNA-binding->BrxB-BrxF-iSTAND->DUF6079*->N6-METHYLASE->ParB+HNH->PglZ-><-SF1-SF2-Helicase+REase<-SF1-SF2-Helicase+REase<-XIS-HTH | 1157 | Bacteria>Pseudomonadota>Gammaproteobacteria>Enterobacterales>Enterobacteriaceae>Klebsiella/Raoultella_group | GCF_009907625.1 | Klebsiella_pneumoniae | GV742_RS00800 | 3194..6667 | 573 | BREX-system-P-loop-protein-BrxC-[Klebsiella-pneumoniae] |
| WP_227665788.1 | Type1.BREX | XIS-HTH->ZnR-><-TIR<-TIR\|\|BrxA-DNA-binding->BrxB-BrxF-iSTAND->DUF6079*->ParB+HNH->N6-METHYLASE->?->N6-METHYLASE-p->PglZ->SIGMA+OB-Fold+AAA-ATPase+Lon-Protease-BrxL-><-Arm-DNA-bind_3+Phage_integrase | 1240 | Bacteria>Pseudomonadota>Gammaproteobacteria>Enterobacterales>Enterobacteriaceae>Klebsiella/Raoultella_group | GCF_010602405.1 | Klebsiella_variicola | G4552_RS17215 | 23636..27358 | 244366 | BREX-system-P-loop-protein-BrxC-[Klebsiella-variicola] |
| WP_181250259.1 | Type1.BREX | <-LuxR-HTH<-?\|\|\|\|BrxA-DNA-binding->BrxB-BrxF-iSTAND->DUF6079*->N6-METHYLASE->PglZ->SIGMA+OB-Fold+AAA-ATPase+Lon-Protease-BrxL-><-?<-DnaJ<-?<-Arm-DNA-bind_3+Phage_integrase | 1213 | Bacteria>Pseudomonadota>Gammaproteobacteria>Enterobacterales>Enterobacteriaceae>Klebsiella/Raoultella_group | GCF_013624375.1 | Klebsiella_michiganensis | HV105_RS26495 | 5508481..5512122 | 1134687 | BREX-system-P-loop-protein-BrxC-[Klebsiella-michiganensis] |
| WP_032425511.1 | Type1.BREX | DUF2754-><-?<-?<-NUDIX\|\|EVE+HNH->ColE7+HNH->?->BrxA-DNA-binding->BrxB-BrxF-iSTAND->DUF6079*->N6-METHYLASE->PglZ->SIGMA+OB-Fold+AAA-ATPase+Lon-Protease-BrxL->HKD-DNase+OGG-like-><-REase<-NACHT+SF1-SF2-Helicase+HKD-DNase<-RNaseH<-nSTAND1+HNH | 1213 | Bacteria>Pseudomonadota>Gammaproteobacteria>Enterobacterales>Enterobacteriaceae>Klebsiella/Raoultella_group | GCF_013735155.1 | Klebsiella_pneumoniae | HV247_RS22800 | 4632201..4635842 | 573 | BREX-system-P-loop-protein-BrxC-[Klebsiella-pneumoniae] |
| WP_182012819.1 | Type1.BREX | HIN-HTH-><-SIGMA+OB-Fold+AAA-ATPase+Lon-Protease-BrxL<-PglZ<-N6-METHYLASE<-ParB+HNH<-DUF6079*<-BrxB-BrxF-iSTAND<-BrxA-DNA-binding<-wHTH+WYL+WCX<-Phage_int_SAM_4+Phage_integrase<-?<-ArdCN+MPTase<-?\|\|HEPN-><-Phage_int_SAM_2+Integrase_1 | 1225 | Bacteria>Pseudomonadota>Gammaproteobacteria>Enterobacterales>Enterobacteriaceae>Klebsiella/Raoultella_group | GCF_013926025.1 | Klebsiella_michiganensis | HV264_RS11965 | 2488431..2492108 | 1134687 | BREX-system-P-loop-protein-BrxC-[Klebsiella-michiganensis] |
| WP_182409424.1 | Type1.BREX | HerA/FtsK->HNH->?-><-SIGMA+OB-Fold+AAA-ATPase+Lon-Protease-BrxL<-PglZ<-TOPRIM<-OLD-ABC<-N6-METHYLASE<-DUF6079*<-BrxB-BrxF-iSTAND<-BrxA-DNA-binding<-XIS-HTH<-?\|\|CopG/RHH->DUF932->ArdB-> | 1204 | Bacteria>Pseudomonadota>Gammaproteobacteria>Enterobacterales>Enterobacteriaceae>Klebsiella/Raoultella_group | GCF_014129405.1 | Klebsiella_michiganensis | H0753_RS11150 | 98138..101752 | 1134687 | BREX-system-P-loop-protein-BrxC-[Klebsiella-michiganensis] |
| WP_088607455.1 | Type1.BREX | <-XIS-HTH<-RHH_antitoxin\|\|?->?->ZnR->ZnR-><-SIGMA+OB-Fold+AAA-ATPase+Lon-Protease-BrxL<-PglZ<-N6-METHYLASE<-DUF6079*<-BrxB-BrxF-iSTAND<-BrxA-DNA-binding\|\|REase+iREase-><-Arm-DNA-bind_3+Phage_integrase | 1213 | Bacteria>Pseudomonadota>Gammaproteobacteria>Enterobacterales>Enterobacteriaceae>Klebsiella/Raoultella_group | GCF_014169335.1 | Klebsiella_pneumoniae | H7R18_RS22570 | 4678469..4682110 | 573 | BREX-system-P-loop-protein-BrxC-[Klebsiella-pneumoniae] |
| WP_187950301.1 | Type1.BREX | wHTH+WYL+WCX->BrxA-DNA-binding->BrxB-BrxF-iSTAND->DUF6079*->ParB+HNH->N6-METHYLASE->PglZ->SIGMA+OB-Fold+AAA-ATPase+Lon-Protease-BrxL-> | 1225 | Bacteria>Pseudomonadota>Gammaproteobacteria>Enterobacterales>Enterobacteriaceae>Klebsiella/Raoultella_group | GCF_014500865.1 | Klebsiella_pneumoniae | HKB44_RS26575 | 2539..6216 | 573 | BREX-system-P-loop-protein-BrxC-[Klebsiella-pneumoniae] |
| WP_196942032.1 | Type1.BREX | HNH->SymE_toxin->Glyoxalase->?->\|\|URI-DNase->?->BrxA-DNA-binding->BrxB-BrxF-iSTAND->DUF6079*->N6-METHYLASE->PglZ->SIGMA+OB-Fold+AAA-ATPase+Lon-Protease-BrxL->HKD-DNase+OGG-like-><-REase<-NACHT+SF1-SF2-Helicase+HKD-DNase<-nSTAND1+HNH\|\|wHTH->XIS-HTH-> | 1213 | Bacteria>Pseudomonadota>Gammaproteobacteria>Enterobacterales>Enterobacteriaceae>Klebsiella/Raoultella_group | GCF_015773165.1 | Klebsiella_michiganensis | I5M64_RS05730 | 22758..26399 | 1134687 | BREX-system-P-loop-protein-BrxC-[Klebsiella-michiganensis] |
| WP_200794272.1 | Type1.BREX | DUF2754-><-?<-NUDIX\|\|EVE+HNH->ColE7+HNH->?->BrxA-DNA-binding->BrxB-BrxF-iSTAND->DUF6079*->N6-METHYLASE->PglZ->SIGMA+OB-Fold+AAA-ATPase+Lon-Protease-BrxL->HKD-DNase+OGG-like-><-REase<-NACHT+SF1-SF2-Helicase+HKD-DNase<-nSTAND1+HNH\|\|RVT-> | 1213 | Bacteria>Pseudomonadota>Gammaproteobacteria>Enterobacterales>Enterobacteriaceae>Klebsiella/Raoultella_group | GCF_016635835.1 | Klebsiella_pneumoniae | HZ910_RS23410 | 16106..19747 | 573 | BREX-system-P-loop-protein-BrxC-[Klebsiella-pneumoniae] |
| WP_202707962.1 | Type1.BREX | wHTH+WYL+WCX->BrxA-DNA-binding->BrxB-BrxF-iSTAND->DUF6079*->ParB+HNH->N6-METHYLASE->PglZ->SIGMA+OB-Fold+AAA-ATPase+Lon-Protease-BrxL-> | 1225 | Bacteria>Pseudomonadota>Gammaproteobacteria>Enterobacterales>Enterobacteriaceae>Klebsiella/Raoultella_group | GCF_016761975.1 | Klebsiella_pneumoniae | H9R49_RS24845 | 3004..6681 | 573 | BREX-system-P-loop-protein-BrxC-[Klebsiella-pneumoniae] |
| WP_117118908.1 | Type1.BREX | HTH_Tnp_1+HTH_21+rve+rve_3-><-?\|\|?->MPTase->?-><-REase<-HTH_23+HTH_32+HTH_33+DDE_3\|\|BrxA-DNA-binding->BrxB-BrxF-iSTAND->DUF6079*->N6-METHYLASE->PglZ->SIGMA+OB-Fold+AAA-ATPase+Lon-Protease-BrxL-><-?<-DnaJ<-?<-Arm-DNA-bind_3+Phage_integrase | 1213 | Bacteria>Pseudomonadota>Gammaproteobacteria>Enterobacterales>Enterobacteriaceae>Klebsiella/Raoultella_group | GCF_017310385.1 | Klebsiella_pneumoniae | H4I72_RS22460 | 4645304..4648945 | 573 | BREX-system-P-loop-protein-BrxC-[Klebsiella-pneumoniae] |
| WP_211540916.1 | Type1.BREX | <-SIGMA+OB-Fold+AAA-ATPase+Lon-Protease-BrxL<-PglZ<-N6-METHYLASE<-DUF6079*<-BrxB-BrxF-iSTAND | 1213 | Bacteria>Pseudomonadota>Gammaproteobacteria>Enterobacterales>Enterobacteriaceae>Klebsiella/Raoultella_group | GCF_018108615.1 | Klebsiella_pneumoniae | FD796_RS26745 | 8816..12457 | 573 | BREX-system-P-loop-protein-BrxC-[Klebsiella-pneumoniae] |
| WP_099719181.1 | Type1.BREX | nSTAND1+HNH->RNaseH->NACHT+SF1-SF2-Helicase+HKD-DNase->REase-><-HKD-DNase+OGG-like<-SIGMA+OB-Fold+AAA-ATPase+Lon-Protease-BrxL<-PglZ<-N6-METHYLASE<-DUF6079*<-BrxB-BrxF-iSTAND<-BrxA-DNA-binding<-?<-HNH<-ACET\|\|?-><-DUF2754 | 1213 | Bacteria>Pseudomonadota>Gammaproteobacteria>Enterobacterales>Enterobacteriaceae>Klebsiella/Raoultella_group | GCF_018137755.1 | Klebsiella_pneumoniae | KFB18_RS20840 | 34480..38121 | 573 | BREX-system-P-loop-protein-BrxC-[Klebsiella-pneumoniae] |
| WP_211586105.1 | Type1.BREX | DUF2754-><-?\|\|ACET->ColE7+HNH->?->BrxA-DNA-binding->BrxB-BrxF-iSTAND->DUF6079*->N6-METHYLASE->PglZ->SIGMA+OB-Fold+AAA-ATPase+Lon-Protease-BrxL->HKD-DNase+OGG-like-><-REase<-NACHT+SF1-SF2-Helicase+HKD-DNase<-RNaseH<-nSTAND1+HNH | 1213 | Bacteria>Pseudomonadota>Gammaproteobacteria>Enterobacterales>Enterobacteriaceae>Klebsiella/Raoultella_group | GCF_018137855.1 | Klebsiella_pneumoniae | G3W03_RS23040 | 37519..41160 | 573 | BREX-system-P-loop-protein-BrxC-[Klebsiella-pneumoniae] |
| WP_211776395.1 | Type1.BREX | <-ZnR\|\|BrxA-DNA-binding->BrxB-BrxF-iSTAND->DUF6079*->N6-METHYLASE->OLD-ABC->TOPRIM->PglZ->SIGMA+OB-Fold+AAA-ATPase+Lon-Protease-BrxL-><-?<-ColE7+HNH<-ACET | 1204 | Bacteria>Pseudomonadota>Gammaproteobacteria>Enterobacterales>Enterobacteriaceae>Klebsiella/Raoultella_group | GCF_018156605.1 | Klebsiella_pneumoniae | J5D66_RS24885 | 2349..5963 | 573 | BREX-system-P-loop-protein-BrxC-[Klebsiella-pneumoniae] |
| WP_049123912.1 | Type1.BREX | Arm-DNA-bind_3+Phage_integrase->BrxA-DNA-binding->BrxB-BrxF-iSTAND->DUF6079*->N6-METHYLASE->ParB+HNH->PglZ-><-?<-XIS-HTH\|\|CopG/RHH->nSTAND1+sGTP->?->DUF932-> | 1157 | Bacteria>Pseudomonadota>Gammaproteobacteria>Enterobacterales>Enterobacteriaceae>Klebsiella/Raoultella_group | GCF_018223925.1 | Klebsiella | KHQ26_RS17845 | 3287..6760 | 570 | MULTISPECIES:-BREX-system-P-loop-protein-BrxC-[Klebsiella] |
| WP_217046681.1 | Type1.BREX | <-XIS-HTH<-RHH_antitoxin\|\|?->ZnR->ZnR-><-SIGMA+OB-Fold+AAA-ATPase+Lon-Protease-BrxL<-PglZ<-N6-METHYLASE<-DUF6079*<-BrxB-BrxF-iSTAND<-BrxA-DNA-binding\|\|Zn_Tnp_IS1+HTH_Tnp_IS1+DDE_Tnp_IS1-><-ACET<-?<-?<-?<-?<-ANK+ANK+ANK | 1213 | Bacteria>Pseudomonadota>Gammaproteobacteria>Enterobacterales>Enterobacteriaceae>Klebsiella/Raoultella_group | GCF_019038575.1 | Klebsiella_pneumoniae | KRP25_RS23525 | 4832898..4836539 | 573 | BREX-system-P-loop-protein-BrxC-[Klebsiella-pneumoniae] |
| WP_217074485.1 | Type1.BREX | <-REase+iREase\|\|wHTH(SBD)+HNH-><-?<-?<-SIGMA+OB-Fold+AAA-ATPase+Lon-Protease-BrxL<-PglZ<-TOPRIM<-OLD-ABC<-N6-METHYLASE<-DUF6079*<-BrxB-BrxF-iSTAND<-BrxA-DNA-binding\|\|ZnR->\|\|?->?->?->ParB+HNH-><-Arm-DNA-bind_3+Phage_integrase | 1204 | Bacteria>Pseudomonadota>Gammaproteobacteria>Enterobacterales>Enterobacteriaceae>Klebsiella/Raoultella_group | GCF_019048885.1 | Klebsiella_aerogenes | KSL85_RS22185 | 4652934..4656548 | 548 | BREX-system-P-loop-protein-BrxC-[Klebsiella-aerogenes] |
| WP_219665441.1 | Type1.BREX | <-Phage_holin<-?\|\|MazE_antitoxin->PIN->DEDD_Tnp_IS110+Transposase_20-><-SIGMA+OB-Fold+AAA-ATPase+Lon-Protease-BrxL<-PglZ<-N6-METHYLASE<-ParB+HNH<-DUF6079*<-BrxB-BrxF-iSTAND<-BrxA-DNA-binding<-wHTH+WYL+WCX | 1225 | Bacteria>Pseudomonadota>Gammaproteobacteria>Enterobacterales>Enterobacteriaceae>Klebsiella/Raoultella_group | GCF_019401235.1 | Klebsiella_pneumoniae | GQQ64_RS25415 | 25134..28811 | 573 | BREX-system-P-loop-protein-BrxC-[Klebsiella-pneumoniae] |
| WP_219810043.1 | Type1.BREX | <-Zn_Tnp_IS1+HTH_Tnp_IS1<-SIGMA+OB-Fold+AAA-ATPase+Lon-Protease-BrxL<-PglZ<-TOPRIM<-OLD-ABC<-N6-METHYLASE<-DUF6079*<-BrxB-BrxF-iSTAND<-BrxA-DNA-binding\|\|<-DEDD_Tnp_IS110+Transposase_20\|\|Phage_integrase->Arm-DNA-bind_3+Phage_integrase->XIS-HTH->DnaB->?->HTH_37-> | 1189 | Bacteria>Pseudomonadota>Gammaproteobacteria>Enterobacterales>Enterobacteriaceae>Klebsiella/Raoultella_group | GCF_019401565.1 | Klebsiella_pneumoniae | GQQ45_RS24745 | 10483..14052 | 573 | BREX-system-P-loop-protein-BrxC-[Klebsiella-pneumoniae] |
| WP_219712976.1 | Type1.BREX | DUF2754-><-?\|\|ACET->ColE7+HNH->?->BrxA-DNA-binding->BrxB-BrxF-iSTAND->DUF6079*->N6-METHYLASE-> | 1213 | Bacteria>Pseudomonadota>Gammaproteobacteria>Enterobacterales>Enterobacteriaceae>Klebsiella/Raoultella_group | GCF_019426525.1 | Klebsiella_pneumoniae | CFM93_RS25160 | 7914..11555 | 573 | BREX-system-P-loop-protein-BrxC-[Klebsiella-pneumoniae] |
| WP_221013059.1 | Type1.BREX | ColE7+HNH->?->BrxA-DNA-binding->BrxB-BrxF-iSTAND->DUF6079*->N6-METHYLASE-> | 1213 | Bacteria>Pseudomonadota>Gammaproteobacteria>Enterobacterales>Enterobacteriaceae>Klebsiella/Raoultella_group | GCF_019679075.1 | Klebsiella_pneumoniae | C6952_RS26355 | 3277..6918 | 573 | BREX-system-P-loop-protein-BrxC-[Klebsiella-pneumoniae] |
| WP_223098448.1 | Type1.BREX | DUF2754-><-?\|\|EVE+HNH->ColE7+HNH->?->BrxA-DNA-binding->BrxB-BrxF-iSTAND->DUF6079*->N6-METHYLASE->PglZ->SIGMA+OB-Fold+AAA-ATPase+Lon-Protease-BrxL->HKD-DNase+OGG-like-><-REase<-NACHT+SF1-SF2-Helicase+HKD-DNase<-RNaseH<-nSTAND1+HNH | 1213 | Bacteria>Pseudomonadota>Gammaproteobacteria>Enterobacterales>Enterobacteriaceae>Klebsiella/Raoultella_group | GCF_019900805.2 | Klebsiella_pneumoniae | KZ657_RS22900 | 4647952..4651593 | 573 | BREX-system-P-loop-protein-BrxC-[Klebsiella-pneumoniae] |
| WP_223065518.1 | Type1.BREX | <-AP_endonuc_2\|\|?->Glyco_hydro_43+GH43_C2-><-MAD-NTD+HNH\|\|wHTH+SAD+HNH-><-AbiEii<-AbiEi_4\|\|DDE_Tnp_1_5+DDE_Tnp_1->BrxB-BrxF-iSTAND->DUF6079*->N6-METHYLASE->PglZ->SIGMA+OB-Fold+AAA-ATPase+Lon-Protease-BrxL->?-><-Arm-DNA-bind_3+Phage_integrase | 1213 | Bacteria>Pseudomonadota>Gammaproteobacteria>Enterobacterales>Enterobacteriaceae>Klebsiella/Raoultella_group | GCF_019900825.1 | Klebsiella_quasipneumoniae | KZ658_RS22630 | 4681396..4685037 | 1463165 | BREX-system-P-loop-protein-BrxC-[Klebsiella-quasipneumoniae] |
| WP_053064557.1 | Type1.BREX | Arm-DNA-bind_3+Phage_integrase-><-HKD-DNase+OGG-like<-SIGMA+OB-Fold+AAA-ATPase+Lon-Protease-BrxL<-PglZ<-N6-METHYLASE<-DUF6079*<-BrxB-BrxF-iSTAND<-BrxA-DNA-binding<-ZnR<-?<-XIS-HTH<-?\|\|RHH_antitoxin-><-URI-DNase<-Glyoxalase_8 | 1213 | Bacteria>Pseudomonadota>Gammaproteobacteria>Enterobacterales>Enterobacteriaceae>Klebsiella/Raoultella_group | GCF_019930185.1 | Klebsiella | KFB82_RS19625 | 81811..85452 | 570 | MULTISPECIES:-BREX-system-P-loop-protein-BrxC-[Klebsiella] |
| WP_194863478.1 | Type1.BREX | MPTase->?-><-REase\|\|DEDD_Tnp_IS110+Transposase_20->BrxA-DNA-binding->BrxB-BrxF-iSTAND->DUF6079*->N6-METHYLASE-> | 1213 | Bacteria>Pseudomonadota>Gammaproteobacteria>Enterobacterales>Enterobacteriaceae>Klebsiella/Raoultella_group | GCF_020119055.1 | Klebsiella_michiganensis | FMK49_RS27050 | 40834..44475 | 1134687 | BREX-system-P-loop-protein-BrxC-[Klebsiella-michiganensis] |
| WP_101858977.1 | Type1.BREX | DUF2754-><-?<-NUDIX\|\|ColE7+HNH->?->BrxA-DNA-binding->BrxB-BrxF-iSTAND->DUF6079*->N6-METHYLASE->PglZ->SIGMA+OB-Fold+AAA-ATPase+Lon-Protease-BrxL->HKD-DNase+OGG-like-><-NACHT+SF1-SF2-Helicase+HKD-DNase<-RNaseH<-nSTAND1+HNH | 1213 | Bacteria>Pseudomonadota>Gammaproteobacteria>Enterobacterales>Enterobacteriaceae>Klebsiella/Raoultella_group | GCF_020479405.1 | Klebsiella_variicola | B6I40_RS13715 | 18030..21671 | 244366 | BREX-system-P-loop-protein-BrxC-[Klebsiella-variicola] |
| WP_116956119.1 | Type1.BREX | <-XIS-HTH<-RHH_antitoxin\|\|?->ZnR->ZnR-><-SIGMA+OB-Fold+AAA-ATPase+Lon-Protease-BrxL<-PglZ<-N6-METHYLASE<-DUF6079*<-BrxB-BrxF-iSTAND<-BrxA-DNA-binding\|\|REase+iREase->?->?->TetR-HTH-><-?<-AAA-ATPase<-HIN-HTH | 1213 | Bacteria>Pseudomonadota>Gammaproteobacteria>Enterobacterales>Enterobacteriaceae>Klebsiella/Raoultella_group | GCF_020576615.1 | Klebsiella_pneumoniae | KAF60_RS20580 | 68875..72516 | 573 | BREX-system-P-loop-protein-BrxC-[Klebsiella-pneumoniae] |
| WP_211639658.1 | Type1.BREX | <-wHTH+WYL<-nSTAND1+GTPase-AIG+sGTP\|\|HIN-HTH->?->XIS-HTH-><-AAA+wHTH\|\|BrxA-DNA-binding->BrxB-BrxF-iSTAND->DUF6079*->N6-METHYLASE->PglZ->SIGMA+OB-Fold+AAA-ATPase+Lon-Protease-BrxL-> | 1213 | Bacteria>Pseudomonadota>Gammaproteobacteria>Enterobacterales>Enterobacteriaceae>Klebsiella/Raoultella_group | GCF_020576935.1 | Klebsiella_pneumoniae | KAF51_RS13395 | 161239..164880 | 573 | BREX-system-P-loop-protein-BrxC-[Klebsiella-pneumoniae] |
| WP_100751493.1 | Type1.BREX | nSTAND1+HNH->RNaseH->NACHT+SF1-SF2-Helicase+HKD-DNase->REase-><-HKD-DNase+OGG-like<-SIGMA+OB-Fold+AAA-ATPase+Lon-Protease-BrxL<-PglZ<-N6-METHYLASE<-DUF6079*<-BrxB-BrxF-iSTAND<-BrxA-DNA-binding<-?<-ColE7+HNH\|\|?-><-DUF2754 | 1213 | Bacteria>Pseudomonadota>Gammaproteobacteria>Enterobacterales>Enterobacteriaceae>Klebsiella/Raoultella_group | GCF_021261545.1 | Klebsiella | CWM56_RS26125 | 34746..38387 | 570 | MULTISPECIES:-BREX-system-P-loop-protein-BrxC-[Klebsiella] |
| WP_023329155.1 | Type1.BREX | <-SAM-METHYLASE<-wHTH(SBD)+HNH\|\|tRNA->Arm-DNA-bind_3+Phage_integrase->wHTH+WYL+WCX->BrxA-DNA-binding->BrxB-BrxF-iSTAND->DUF6079*->N6-METHYLASE->ParB+HNH->PglZ-><-SF1-SF2-Helicase+REase<-SF1-SF2-Helicase+REase\|\|NACHT+SF1-SF2-Helicase+HKD-DNase-> | 1157 | Bacteria>Pseudomonadota>Gammaproteobacteria>Enterobacterales>Enterobacteriaceae>Klebsiella/Raoultella_group | GCF_021264605.1 | Klebsiella_oxytoca | LVW29_RS13105 | 76515..79988 | 571 | BREX-system-P-loop-protein-BrxC-[Klebsiella-oxytoca] |
| WP_234580820.1 | Type1.BREX | <-DUF932<-CopG/RHH\|\|?->XIS-HTH-><-TIR<-TIR\|\|BrxA-DNA-binding->BrxB-BrxF-iSTAND->DUF6079-p*-> | 744 | Bacteria>Pseudomonadota>Gammaproteobacteria>Enterobacterales>Enterobacteriaceae>Klebsiella/Raoultella_group | GCF_021440775.1 | Raoultella_ornithinolytica | LZ676_RS28690 | 5491..7723 | 54291 | BREX-system-P-loop-protein-BrxC,-partial-[Raoultella-ornithinolytica] |
| WP_234545002.1 | Type1.BREX | <-DUF6079-p*<-BrxB-BrxF-iSTAND<-BrxA-DNA-binding\|\|TIR->TIR-><-XIS-HTH<-?\|\|CopG/RHH->DUF932->DUF905-> | 741 | Bacteria>Pseudomonadota>Gammaproteobacteria>Enterobacterales>Enterobacteriaceae>Klebsiella/Raoultella_group | GCF_021441325.1 | Raoultella_ornithinolytica | LZ678_RS28780 | 1..2223 | 54291 | BREX-system-P-loop-protein-BrxC,-partial-[Raoultella-ornithinolytica] |
| WP_235410418.1 | Type1.BREX | <-wHTH+WYL<-sGTP<-?\|\|XIS-HTH->?->?->BrxA-DNA-binding->BrxB-BrxF-iSTAND->DUF6079*->ParB+HNH->N6-METHYLASE->HEPN->RVT->N6-METHYLASE-p->PglZ->SIGMA+OB-Fold+AAA-ATPase+Lon-Protease-BrxL-><-Arm-DNA-bind_3+Phage_integrase | 1240 | Bacteria>Pseudomonadota>Gammaproteobacteria>Enterobacterales>Enterobacteriaceae>Klebsiella/Raoultella_group | GCF_021532695.1 | Klebsiella_pneumoniae | LZ602_RS12600 | 31915..35637 | 573 | BREX-system-P-loop-protein-BrxC-[Klebsiella-pneumoniae] |
| WP_233435617.1 | Type1.BREX | PIWI-><-Phage_integrase\|\|tRNA-><-SIGMA+OB-Fold+AAA-ATPase+Lon-Protease-BrxL<-PglZ<-N6-METHYLASE-p<-ParB+HNH<-DUF6079*<-BrxB-BrxF-iSTAND<-BrxA-DNA-binding\|\|?-><-Glyco_hydro_43+GH43_C2<-?\|\|AP_endonuc_2->DDE_Tnp_1_5+DDE_Tnp_1-> | 1240 | Bacteria>Pseudomonadota>Gammaproteobacteria>Enterobacterales>Enterobacteriaceae>Klebsiella/Raoultella_group | GCF_021699015.1 | Raoultella_ornithinolytica | INS59_RS15575 | 205995..209717 | 54291 | BREX-system-P-loop-protein-BrxC-[Raoultella-ornithinolytica] |
| WP_238640446.1 | Type1.BREX | Arm-DNA-bind_3+Phage_integrase-><-SIGMA+OB-Fold+AAA-ATPase+Lon-Protease-BrxL<-PglZ<-N6-METHYLASE<-ParB+HNH<-DUF6079*<-BrxB-BrxF-iSTAND<-BrxA-DNA-binding<-?<-XIS-HTH\|\|?->nSTAND1+sGTP->wHTH+WYL-> | 1240 | Bacteria>Pseudomonadota>Gammaproteobacteria>Enterobacterales>Enterobacteriaceae>Klebsiella/Raoultella_group | GCF_022229165.1 | Klebsiella_pneumoniae | LLG23_RS24005 | 30146..33868 | 573 | BREX-system-P-loop-protein-BrxC-[Klebsiella-pneumoniae] |
| WP_223339928.1 | Type1.BREX | <-wHTH+WYL<-sGTP<-?\|\|XIS-HTH->?->?->BrxA-DNA-binding->BrxB-BrxF-iSTAND->DUF6079*->ParB+HNH->N6-METHYLASE->HEPN->RVT->N6-METHYLASE-p->PglZ->SIGMA+OB-Fold+AAA-ATPase+Lon-Protease-BrxL-><-Arm-DNA-bind_3+Phage_integrase | 1240 | Bacteria>Pseudomonadota>Gammaproteobacteria>Enterobacterales>Enterobacteriaceae>Klebsiella/Raoultella_group | GCF_022353805.1 | Klebsiella_pneumoniae | JX616_RS18220 | 31893..35615 | 573 | BREX-system-P-loop-protein-BrxC-[Klebsiella-pneumoniae] |
| WP_241187802.1 | Type1.BREX | <-XIS-HTH<-wHTH\|\|nSTAND1+HNH->NACHT+SF1-SF2-Helicase+HKD-DNase->REase-><-HKD-DNase+OGG-like<-SIGMA+OB-Fold+AAA-ATPase+Lon-Protease-BrxL<-PglZ<-N6-METHYLASE<-DUF6079*<-BrxB-BrxF-iSTAND<-BrxA-DNA-binding<-URI-DNase\|\|<-?<-Glyoxalase<-SymE_toxin<-HNH\|\|?->LACI-HTH+PBPI-> | 1213 | Bacteria>Pseudomonadota>Gammaproteobacteria>Enterobacterales>Enterobacteriaceae>Klebsiella/Raoultella_group | GCF_022543825.1 | Klebsiella_michiganensis | MOA98_RS25025 | 250948..254589 | 1134687 | BREX-system-P-loop-protein-BrxC-[Klebsiella-michiganensis] |
| WP_241506431.1 | Type1.BREX | nSTAND1+HNH->RNaseH->NACHT+SF1-SF2-Helicase+HKD-DNase->REase-><-HKD-DNase+OGG-like<-SIGMA+OB-Fold+AAA-ATPase+Lon-Protease-BrxL<-PglZ<-N6-METHYLASE<-DUF6079*<-BrxB-BrxF-iSTAND<-BrxA-DNA-binding<-URI-DNase\|\|<-?<-?\|\|<-?\|\|?->\|\|MAD-NTD+HNH-><-Glyco_hydro_43+GH43_C2 | 1213 | Bacteria>Pseudomonadota>Gammaproteobacteria>Enterobacterales>Enterobacteriaceae>Klebsiella/Raoultella_group | GCF_022585755.1 | Klebsiella_pneumoniae | MMP84_RS15875 | 154032..157673 | 573 | BREX-system-P-loop-protein-BrxC-[Klebsiella-pneumoniae] |
| WP_157261982.1 | Type1.BREX | Arm-DNA-bind_3+Phage_integrase->?->DnaJ->?-><-SIGMA+OB-Fold+AAA-ATPase+Lon-Protease-BrxL<-PglZ<-N6-METHYLASE<-DUF6079*<-BrxB-BrxF-iSTAND<-BrxA-DNA-binding\|\|?->HKD-DNase+HKD-DNase->SF1-SF2-Helicase-><-REase+SPF-cons<-?<-XIS-HTH | 1213 | Bacteria>Pseudomonadota>Gammaproteobacteria>Enterobacterales>Enterobacteriaceae>Klebsiella/Raoultella_group | GCF_022759445.1 | Klebsiella | GSE35_RS20710 | 46310..49951 | 570 | MULTISPECIES:-BREX-system-P-loop-protein-BrxC-[Klebsiella] |
| WP_202181851.1 | Type1.BREX | <-XIS-HTH\|\|nSTAND1+HNH->RNaseH->NACHT+SF1-SF2-Helicase+HKD-DNase->REase-><-HKD-DNase+OGG-like<-SIGMA+OB-Fold+AAA-ATPase+Lon-Protease-BrxL<-PglZ<-N6-METHYLASE<-DUF6079*<-BrxB-BrxF-iSTAND<-BrxA-DNA-binding<-?<-ColE7+HNH<-ACET\|\|?-><-DUF2754 | 1213 | Bacteria>Pseudomonadota>Gammaproteobacteria>Enterobacterales>Enterobacteriaceae>Klebsiella/Raoultella_group | GCF_023703955.1 | Klebsiella_pneumoniae | KU670_RS11590 | 2422249..2425890 | 573 | BREX-system-P-loop-protein-BrxC-[Klebsiella-pneumoniae] |
| WP_254541266.1 | Type1.BREX | <-Calcineurin<-?<-?<-?<-?<-?\|\|tRNA->Arm-DNA-bind_3+Phage_integrase->BrxA-DNA-binding->BrxB-BrxF-iSTAND->DUF6079*->N6-METHYLASE->ParB+HNH->PglZ-><-SF1-SF2-Helicase+REase<-SF1-SF2-Helicase+REase\|\|HIN-HTH-><-XIS-HTH | 1157 | Bacteria>Pseudomonadota>Gammaproteobacteria>Enterobacterales>Enterobacteriaceae>Klebsiella/Raoultella_group | GCF_024224875.1 | Klebsiella_pneumoniae | NLZ12_RS02805 | 582600..586073 | 573 | BREX-system-P-loop-protein-BrxC-[Klebsiella-pneumoniae] |
| WP_255075067.1 | Type1.BREX | nSTAND1+HNH->RNaseH->NACHT+SF1-SF2-Helicase+HKD-DNase->REase-><-HKD-DNase+OGG-like<-SIGMA+OB-Fold+AAA-ATPase+Lon-Protease-BrxL<-PglZ<-N6-METHYLASE<-DUF6079*<-BrxB-BrxF-iSTAND<-BrxA-DNA-binding<-?<-ColE7+HNH | 1213 | Bacteria>Pseudomonadota>Gammaproteobacteria>Enterobacterales>Enterobacteriaceae>Klebsiella/Raoultella_group | GCF_024357365.1 | Klebsiella_pneumoniae | L1C80_RS20005 | 166541..170182 | 573 | BREX-system-P-loop-protein-BrxC-[Klebsiella-pneumoniae] |
| WP_250099520.1 | Type1.BREX | Zn_Tnp_IS1+HTH_Tnp_IS1+DDE_Tnp_IS1->BrxA-DNA-binding->BrxB-BrxF-iSTAND->DUF6079*->N6-METHYLASE->PglZ->SIGMA+OB-Fold+AAA-ATPase+Lon-Protease-BrxL-><-ZnR | 1213 | Bacteria>Pseudomonadota>Gammaproteobacteria>Enterobacterales>Enterobacteriaceae>Klebsiella/Raoultella_group | GCF_024516295.1 | Klebsiella_pneumoniae | JEK46_RS25555 | 2056..5697 | 573 | BREX-system-P-loop-protein-BrxC-[Klebsiella-pneumoniae] |
| WP_256768592.1 | Type1.BREX | <-MAD-NTD+HNH\|\|wHTH+SAD+HNH->wHTH+HNH-><-HIN-HTH<-SIGMA+OB-Fold+AAA-ATPase+Lon-Protease-BrxL<-PglZ<-TOPRIM<-OLD-ABC<-N6-METHYLASE<-DUF6079*<-BrxB-BrxF-iSTAND<-BrxA-DNA-binding\|\|ZnR-><-tRNA\|\|?->?->?->?->?->Calcineurin-> | 1204 | Bacteria>Pseudomonadota>Gammaproteobacteria>Enterobacterales>Enterobacteriaceae>Klebsiella/Raoultella_group | GCF_024517905.1 | Klebsiella_pneumoniae | JEK00_RS18440 | 19794..23408 | 573 | BREX-system-P-loop-protein-BrxC-[Klebsiella-pneumoniae] |
| WP_101881469.1 | Type1.BREX | Arm-DNA-bind_3+Phage_integrase->?->DnaJ->?-><-SIGMA+OB-Fold+AAA-ATPase+Lon-Protease-BrxL<-PglZ<-N6-METHYLASE<-DUF6079*<-BrxB-BrxF-iSTAND<-BrxA-DNA-binding\|\|?->HKD-DNase+HKD-DNase->SF1-SF2-Helicase-><-REase+SPF-cons<-?<-XIS-HTH<-XIS-HTH | 1213 | Bacteria>Pseudomonadota>Gammaproteobacteria>Enterobacterales>Enterobacteriaceae>Klebsiella/Raoultella_group | GCF_024813835.1 | Klebsiella_quasipneumoniae | B6J71_RS18320 | 31229..34870 | 1463165 | BREX-system-P-loop-protein-BrxC-[Klebsiella-quasipneumoniae] |
| WP_196588765.1 | Type1.BREX | <-ArdB<-DUF932<-CopG/RHH\|\|?->XIS-HTH->BrxA-DNA-binding->BrxB-BrxF-iSTAND->DUF6079*->N6-METHYLASE->OLD-ABC->TOPRIM->PglZ->SIGMA+OB-Fold+AAA-ATPase+Lon-Protease-BrxL-><-?<-HNH<-HerA/FtsK | 1204 | Bacteria>Pseudomonadota>Gammaproteobacteria>Enterobacterales>Enterobacteriaceae>Klebsiella/Raoultella_group | GCF_024917295.1 | Klebsiella | M5R35_RS24840 | 5265325..5268939 | 570 | MULTISPECIES:-BREX-system-P-loop-protein-BrxC-[Klebsiella] |
| WP_259574830.1 | Type1.BREX | REase-><-HKD-DNase+OGG-like<-SIGMA+OB-Fold+AAA-ATPase+Lon-Protease-BrxL<-PglZ<-N6-METHYLASE<-DUF6079*<-BrxB-BrxF-iSTAND<-BrxA-DNA-binding<-URI-DNase | 1213 | Bacteria>Pseudomonadota>Gammaproteobacteria>Enterobacterales>Enterobacteriaceae>Klebsiella/Raoultella_group | GCF_024998145.1 | Klebsiella_pneumoniae | KQ276_RS10525 | 179571..183212 | 573 | BREX-system-P-loop-protein-BrxC-[Klebsiella-pneumoniae] |
| WP_064144681.1 | Type1.BREX | <-N6-METHYLASE<-DUF6079*<-BrxB-BrxF-iSTAND<-BrxA-DNA-binding<-?<-ColE7+HNH | 1213 | Bacteria>Pseudomonadota>Gammaproteobacteria>Enterobacterales>Enterobacteriaceae>Klebsiella/Raoultella_group | GCF_026007595.1 | Klebsiella_variicola | OPB18_RS25725 | 2398..6039 | 244366 | BREX-system-P-loop-protein-BrxC-[Klebsiella-variicola] |
| WP_264975991.1 | Type1.BREX | HIN-HTH->?->LZ_Tnp_IS66+zf-IS66+DDE_Tnp_IS66+DDE_Tnp_IS66_C->BrxA-DNA-binding->BrxB-BrxF-iSTAND->DUF6079*->N6-METHYLASE->OLD-ABC->TOPRIM->PglZ->SIGMA+OB-Fold+AAA-ATPase+Lon-Protease-BrxL-><-?<-Arm-DNA-bind_3+Phage_integrase<-tRNA<-?<-TetR-HTH | 1204 | Bacteria>Pseudomonadota>Gammaproteobacteria>Enterobacterales>Enterobacteriaceae>Klebsiella/Raoultella_group | GCF_026008315.1 | Klebsiella_quasipneumoniae | OPC12_RS09035 | 173713..177327 | 1463165 | BREX-system-P-loop-protein-BrxC-[Klebsiella-quasipneumoniae] |
| WP_265815522.1 | Type1.BREX | <-wHTH\|\|nSTAND1+HNH->RNaseH->NACHT+SF1-SF2-Helicase+HKD-DNase->REase-><-HKD-DNase+OGG-like<-SIGMA+OB-Fold+AAA-ATPase+Lon-Protease-BrxL<-PglZ<-N6-METHYLASE<-DUF6079*<-BrxB-BrxF-iSTAND<-BrxA-DNA-binding<-URI-DNase\|\|<-?<-Glyoxalase<-SymE_toxin<-HNH\|\|?->LACI-HTH+PBPI-> | 1213 | Bacteria>Pseudomonadota>Gammaproteobacteria>Enterobacterales>Enterobacteriaceae>Klebsiella/Raoultella_group | GCF_026224285.1 | Klebsiella_oxytoca | ORM16_RS17370 | 107113..110754 | 571 | BREX-system-P-loop-protein-BrxC-[Klebsiella-oxytoca] |
| WP_224260387.1 | Type1.BREX | <-RelE-ParE<-RHH<-DUF4400<-ZnR\|\|BrxA-DNA-binding->BrxB-BrxF-iSTAND->DUF6079*->N6-METHYLASE->OLD-ABC->TOPRIM->PglZ->SIGMA+OB-Fold+AAA-ATPase+Lon-Protease-BrxL-><-?<-Glyoxalase<-SymE_toxin<-HNH | 1204 | Bacteria>Pseudomonadota>Gammaproteobacteria>Enterobacterales>Enterobacteriaceae>Klebsiella/Raoultella_group | GCF_026224455.1 | Klebsiella_michiganensis | ORL69_RS11440 | 318057..321671 | 1134687 | BREX-system-P-loop-protein-BrxC-[Klebsiella-michiganensis] |
| WP_064161342.1 | Type1.BREX | DUF4158+DDE_Tnp_Tn3->DEDD_Tnp_IS110+Transposase_20->\|\|DEDD_Tnp_IS110+Transposase_20-><-REase+ZnR\|\|DUF772+DDE_Tnp_1-><-Zn_Tnp_IS1+HTH_Tnp_IS1+DDE_Tnp_IS1\|\|wHTH+WYL+WCX->BrxA-DNA-binding->BrxB-BrxF-iSTAND->DUF6079*->ParB+HNH->N6-METHYLASE->PglZ->SIGMA+OB-Fold+AAA-ATPase+Lon-Protease-BrxL->\|\|GDResolvase+HIN-HTH->?->?->DDE_Tnp_1_5+DDE_Tnp_1-> | 1225 | Bacteria>Pseudomonadota>Gammaproteobacteria>Enterobacterales>Enterobacteriaceae>Klebsiella/Raoultella_group | GCF_027577885.1 | Klebsiella | OEE45_RS29300 | 43964..47641 | 570 | MULTISPECIES:-BREX-system-P-loop-protein-BrxC-[Klebsiella] |
| WP_274901739.1 | Type1.BREX | nSTAND1+HNH->RNaseH->NACHT+SF1-SF2-Helicase+HKD-DNase->REase-><-HKD-DNase+OGG-like<-SIGMA+OB-Fold+AAA-ATPase+Lon-Protease-BrxL<-PglZ<-N6-METHYLASE<-DUF6079*<-BrxB-BrxF-iSTAND<-BrxA-DNA-binding<-?<-ColE7+HNH\|\|HTH_Tnp_1+HTH_21+rve+rve_3-><-DUF2754 | 1213 | Bacteria>Pseudomonadota>Gammaproteobacteria>Enterobacterales>Enterobacteriaceae>Klebsiella/Raoultella_group | GCF_028894275.1 | Klebsiella_pneumoniae | PXW84_RS25015 | 34732..38373 | 573 | BREX-system-P-loop-protein-BrxC-[Klebsiella-pneumoniae] |
| WP_275268565.1 | Type1.BREX | DUF6079*->N6-METHYLASE->PglZ->SIGMA+OB-Fold+AAA-ATPase+Lon-Protease-BrxL->HKD-DNase+OGG-like-><-NACHT+SF1-SF2-Helicase+HKD-DNase<-RNaseH<-nSTAND1+HNH | 1196 | Bacteria>Pseudomonadota>Gammaproteobacteria>Enterobacterales>Enterobacteriaceae>Klebsiella/Raoultella_group | GCF_029079245.1 | Klebsiella_variicola | PZY30_RS10230 | 1..3591 | 244366 | BREX-system-P-loop-protein-BrxC,-partial-[Klebsiella-variicola] |
| WP_064171377.1 | Type1.BREX | DUF2754-><-?\|\|ColE7+HNH->?->BrxA-DNA-binding->BrxB-BrxF-iSTAND->DUF6079*->N6-METHYLASE->PglZ->SIGMA+OB-Fold+AAA-ATPase+Lon-Protease-BrxL->HKD-DNase+OGG-like-><-REase<-NACHT+SF1-SF2-Helicase+HKD-DNase<-RNaseH<-nSTAND1+HNH<-ParB+HNH | 1213 | Bacteria>Pseudomonadota>Gammaproteobacteria>Enterobacterales>Enterobacteriaceae>Klebsiella/Raoultella_group | GCF_900084795.1 | Klebsiella_variicola | GJ687_RS21830 | 17638..21279 | 244366 | BREX-system-P-loop-protein-BrxC-[Klebsiella-variicola] |
| WP_064182275.1 | Type1.BREX | MazE_antitoxin->PIN->DEDD_Tnp_IS110+Transposase_20->DEDD_Tnp_IS110+Transposase_20-><-SIGMA+OB-Fold+AAA-ATPase+Lon-Protease-BrxL<-PglZ<-N6-METHYLASE<-ParB+HNH<-DUF6079*<-BrxB-BrxF-iSTAND<-BrxA-DNA-binding<-wHTH+WYL+WCX\|\|Zn_Tnp_IS1+HTH_Tnp_IS1+DDE_Tnp_IS1->DDE_Tnp-><-HEPN<-RelB\|\|HTH_3-> | 1225 | Bacteria>Pseudomonadota>Gammaproteobacteria>Enterobacterales>Enterobacteriaceae>Klebsiella/Raoultella_group | GCF_900084835.1 | Klebsiella_pneumoniae | A8U42_RS24700 | 50900..54577 | 573 | BREX-system-P-loop-protein-BrxC-[Klebsiella-pneumoniae] |
| WP_064143068.1 | Type1.BREX | <-Calcineurin<-?<-?<-?<-?<-?\|\|tRNA-><-ZnR\|\|BrxA-DNA-binding->BrxB-BrxF-iSTAND->DUF6079*->N6-METHYLASE->OLD-ABC->TOPRIM->PglZ->SIGMA+OB-Fold+AAA-ATPase+Lon-Protease-BrxL-> | 1204 | Bacteria>Pseudomonadota>Gammaproteobacteria>Enterobacterales>Enterobacteriaceae>Klebsiella/Raoultella_group | GCF_900085185.1 | Klebsiella_variicola | A8S52_RS23955 | 19840..23454 | 244366 | BREX-system-P-loop-protein-BrxC-[Klebsiella-variicola] |
| WP_064158594.1 | Type1.BREX | RVT-><-SIGMA+OB-Fold+AAA-ATPase+Lon-Protease-BrxL<-PglZ<-N6-METHYLASE<-DUF6079*<-DUF6079-p<-BrxB-BrxF-iSTAND<-BrxA-DNA-binding | 1058 | Bacteria>Pseudomonadota>Gammaproteobacteria>Enterobacterales>Enterobacteriaceae>Klebsiella/Raoultella_group | GCF_900086305.1 | Klebsiella_quasipneumoniae | A8T30_RS26420 | 11957..15133 | 1463165 | BREX-system-P-loop-protein-BrxC,-partial-[Klebsiella-quasipneumoniae] |
| WP_065808708.1 | Type1.BREX | DUF2754-><-?\|\|ACET->ColE7+HNH->?->BrxA-DNA-binding->BrxB-BrxF-iSTAND->DUF6079*->N6-METHYLASE->PglZ->SIGMA+OB-Fold+AAA-ATPase+Lon-Protease-BrxL->HKD-DNase+OGG-like-><-REase<-NACHT+SF1-SF2-Helicase+HKD-DNase<-RNaseH<-nSTAND1+HNH | 1213 | Bacteria>Pseudomonadota>Gammaproteobacteria>Enterobacterales>Enterobacteriaceae>Klebsiella/Raoultella_group | GCF_900093245.1 | Klebsiella_pneumoniae | BC433_RS15565 | 37858..41499 | 573 | BREX-system-P-loop-protein-BrxC-[Klebsiella-pneumoniae] |
| WP_080916293.1 | Type1.BREX | <-HNH+NERD+ZnR<-Peptidase_S49+Peptidase_S49<-?<-DDE_Tnp_1_5+DDE_Tnp_1\|\|?->?->\|\|URI-DNase->BrxA-DNA-binding->BrxB-BrxF-iSTAND->DUF6079*->N6-METHYLASE->PglZ->SIGMA+OB-Fold+AAA-ATPase+Lon-Protease-BrxL->HKD-DNase+OGG-like-><-REase | 1213 | Bacteria>Pseudomonadota>Gammaproteobacteria>Enterobacterales>Enterobacteriaceae>Klebsiella/Raoultella_group | GCF_900172605.1 | Klebsiella_pneumoniae | B7F35_RS00110 | 14289..17930 | 573 | BREX-system-P-loop-protein-BrxC-[Klebsiella-pneumoniae] |
| WP_227643422.1 | Type1.BREX | Arm-DNA-bind_3+Phage_integrase-><-SIGMA+OB-Fold+AAA-ATPase+Lon-Protease-BrxL<-PglZ<-N6-METHYLASE-p<-RVT<-HEPN<-N6-METHYLASE<-ParB+HNH<-DUF6079*<-BrxB-BrxF-iSTAND<-BrxA-DNA-binding<-?<-?<-XIS-HTH<-?\|\|CopG/RHH->nSTAND1+GTPase-AIG+sGTP->DUF932-> | 1240 | Bacteria>Pseudomonadota>Gammaproteobacteria>Enterobacterales>Enterobacteriaceae>Klebsiella/Raoultella_group | GCF_900173735.1 | Klebsiella_pneumoniae | B7E22_RS15515 | 109572..113294 | 573 | BREX-system-P-loop-protein-BrxC-[Klebsiella-pneumoniae] |
| WP_109220871.1 | Type1.BREX | Glyoxalase_8->URI-DNase-><-RHH_antitoxin\|\|?->XIS-HTH->?->ZnR->BrxA-DNA-binding->BrxB-BrxF-iSTAND->DUF6079*->PglZ->SIGMA+OB-Fold+AAA-ATPase+Lon-Protease-BrxL->HKD-DNase+OGG-like-><-Arm-DNA-bind_3+Phage_integrase<-tRNA\|\|wHTH(SBD)+HNH->SAM-METHYLASE-> | 1213 | Bacteria>Pseudomonadota>Gammaproteobacteria>Enterobacterales>Enterobacteriaceae>Klebsiella/Raoultella_group | GCF_900407115.1 | Klebsiella_oxytoca | DK336_RS07845 | 45020..48661 | 571 | BREX-system-P-loop-protein-BrxC-[Klebsiella-oxytoca] |
| WP_115658225.1 | Type1.BREX | nSTAND1+HNH->RNaseH->NACHT+SF1-SF2-Helicase+HKD-DNase->REase-><-HKD-DNase+OGG-like<-SIGMA+OB-Fold+AAA-ATPase+Lon-Protease-BrxL<-PglZ<-N6-METHYLASE<-DUF6079*<-BrxB-BrxF-iSTAND<-BrxA-DNA-binding<-?<-ColE7+HNH<-ACET\|\|?-><-DUF2754 | 1213 | Bacteria>Pseudomonadota>Gammaproteobacteria>Enterobacterales>Enterobacteriaceae>Klebsiella/Raoultella_group | GCF_900498955.1 | Klebsiella_pneumoniae | DY137_RS22260 | 323150..326791 | 573 | BREX-system-P-loop-protein-BrxC-[Klebsiella-pneumoniae] |
| WP_117104235.1 | Type1.BREX | nSTAND1+HNH->RNaseH->NACHT+SF1-SF2-Helicase+HKD-DNase->REase-><-HKD-DNase+OGG-like<-SIGMA+OB-Fold+AAA-ATPase+Lon-Protease-BrxL<-PglZ<-N6-METHYLASE<-DUF6079*<-BrxB-BrxF-iSTAND<-BrxA-DNA-binding<-?<-ColE7+HNH\|\|?-><-DUF2754\|\|?-><-wHTH+SAD+HNH\|\|MAD-NTD+HNH-> | 1213 | Bacteria>Pseudomonadota>Gammaproteobacteria>Enterobacterales>Enterobacteriaceae>Klebsiella/Raoultella_group | GCF_900502405.1 | Klebsiella_pneumoniae | D2Z44_RS21380 | 236554..240195 | 573 | BREX-system-P-loop-protein-BrxC-[Klebsiella-pneumoniae] |
| WP_117044181.1 | Type1.BREX | DUF2754-><-?<-?\|\|EVE+HNH->ColE7+HNH->?->BrxA-DNA-binding->BrxB-BrxF-iSTAND->DUF6079*->N6-METHYLASE->PglZ->SIGMA+OB-Fold+AAA-ATPase+Lon-Protease-BrxL->HKD-DNase+OGG-like-><-REase<-NACHT+SF1-SF2-Helicase+HKD-DNase<-RNaseH<-nSTAND1+HNH | 1213 | Bacteria>Pseudomonadota>Gammaproteobacteria>Enterobacterales>Enterobacteriaceae>Klebsiella/Raoultella_group | GCF_900503165.1 | Klebsiella_pneumoniae | D2094_RS08665 | 16725..20366 | 573 | BREX-system-P-loop-protein-BrxC-[Klebsiella-pneumoniae] |
| WP_117268636.1 | Type1.BREX | HNH->?->HIN-HTH->?->LZ_Tnp_IS66+zf-IS66+DDE_Tnp_IS66+DDE_Tnp_IS66_C-><-SIGMA+OB-Fold+AAA-ATPase+Lon-Protease-BrxL<-PglZ<-TOPRIM<-OLD-ABC<-DUF6079*<-BrxB-BrxF-iSTAND<-BrxA-DNA-binding<-XIS-HTH<-?\|\|CopG/RHH->DUF932->ArdB-> | 1204 | Bacteria>Pseudomonadota>Gammaproteobacteria>Enterobacterales>Enterobacteriaceae>Klebsiella/Raoultella_group | GCF_900509065.1 | Klebsiella_pneumoniae | D2728_RS15300 | 268407..272021 | 573 | BREX-system-P-loop-protein-BrxC-[Klebsiella-pneumoniae] |
| WP_117133599.1 | Type1.BREX | nSTAND1+HNH->RNaseH->NACHT+SF1-SF2-Helicase+HKD-DNase->REase-><-HKD-DNase+OGG-like<-SIGMA+OB-Fold+AAA-ATPase+Lon-Protease-BrxL<-PglZ<-N6-METHYLASE<-DUF6079*<-BrxB-BrxF-iSTAND<-BrxA-DNA-binding<-?<-ColE7+HNH\|\|?-><-DUF2754 | 1213 | Bacteria>Pseudomonadota>Gammaproteobacteria>Enterobacterales>Enterobacteriaceae>Klebsiella/Raoultella_group | GCF_900509525.1 | Klebsiella_variicola | D2R55_RS21945 | 231466..235107 | 244366 | BREX-system-P-loop-protein-BrxC-[Klebsiella-variicola] |
| WP_117082329.1 | Type1.BREX | REase-><-HKD-DNase+OGG-like<-SIGMA+OB-Fold+AAA-ATPase+Lon-Protease-BrxL<-PglZ<-N6-METHYLASE<-DUF6079*<-BrxB-BrxF-iSTAND<-BrxA-DNA-binding\|\|?-><-URI-DNase<-HIN-HTH | 1213 | Bacteria>Pseudomonadota>Gammaproteobacteria>Enterobacterales>Enterobacteriaceae>Klebsiella/Raoultella_group | GCF_900514375.1 | Klebsiella_pneumoniae | SAMEA3515241_04816 | 29985..33626 | 573 | BREX-system-P-loop-protein-BrxC-[Klebsiella-pneumoniae] |
| WP_117102453.1 | Type1.BREX | Arm-DNA-bind_3+Phage_integrase->?->DnaJ->?-><-SIGMA+OB-Fold+AAA-ATPase+Lon-Protease-BrxL<-PglZ<-N6-METHYLASE<-DUF6079*<-BrxB-BrxF-iSTAND<-BrxA-DNA-binding\|\|HTH_23+HTH_32+HTH_33+DDE_3->REase-><-?<-MPTase<-?\|\|\|\|?->HKD-DNase+HKD-DNase-> | 1213 | Bacteria>Pseudomonadota>Gammaproteobacteria>Enterobacterales>Enterobacteriaceae>Klebsiella/Raoultella_group | GCF_900517475.1 | Klebsiella_pneumoniae | D2Y27_RS13960 | 242329..245970 | 573 | BREX-system-P-loop-protein-BrxC-[Klebsiella-pneumoniae] |
| WP_122533088.1 | Type1.BREX | DUF2754-><-?\|\|ColE7+HNH->?->BrxA-DNA-binding->BrxB-BrxF-iSTAND->DUF6079*->N6-METHYLASE->PglZ->SIGMA+OB-Fold+AAA-ATPase+Lon-Protease-BrxL->HKD-DNase+OGG-like-><-REase<-NACHT+SF1-SF2-Helicase+HKD-DNase<-RNaseH<-nSTAND1+HNH | 1213 | Bacteria>Pseudomonadota>Gammaproteobacteria>Enterobacterales>Enterobacteriaceae>Klebsiella/Raoultella_group | GCF_900576735.1 | Klebsiella_variicola | EES49_RS00110 | 18527..22168 | 244366 | BREX-system-P-loop-protein-BrxC-[Klebsiella-variicola] |
| WP_085803383.1 | Type1.BREX | nSTAND1+HNH->RNaseH->NACHT+SF1-SF2-Helicase+HKD-DNase->REase-><-HKD-DNase+OGG-like<-SIGMA+OB-Fold+AAA-ATPase+Lon-Protease-BrxL<-PglZ<-N6-METHYLASE<-DUF6079*<-BrxB-BrxF-iSTAND<-BrxA-DNA-binding<-?<-ColE7+HNH<-EVE+HNH\|\|?->?-><-DUF2754 | 1213 | Bacteria>Pseudomonadota>Gammaproteobacteria>Enterobacterales>Enterobacteriaceae>Klebsiella/Raoultella_group | GCF_900774735.1 | Klebsiella_pneumoniae | EF470_RS14885 | 16936..20577 | 573 | BREX-system-P-loop-protein-BrxC-[Klebsiella-pneumoniae] |
| WP_058836894.1 | Type1.BREX | XIS-HTH-><-AAA+wHTH<-Zn_Tnp_IS1+HTH_Tnp_IS1+DDE_Tnp_IS1<-?<-?<-DUF772+DDE_Tnp_1\|\|BrxA-DNA-binding->BrxB-BrxF-iSTAND->DUF6079*->N6-METHYLASE->PglZ->SIGMA+OB-Fold+AAA-ATPase+Lon-Protease-BrxL-><-?<-DnaJ<-?<-Arm-DNA-bind_3+Phage_integrase | 1213 | Bacteria>Pseudomonadota>Gammaproteobacteria>Enterobacterales>Enterobacteriaceae>Klebsiella/Raoultella_group | GCF_900774805.1 | Klebsiella | E7U83_RS18910 | 22123..25764 | 570 | MULTISPECIES:-BREX-system-P-loop-protein-BrxC-[Klebsiella] |
| WP_031591770.1 | Type1.BREX | DUF2754-><-?\|\|ACET->ColE7+HNH->?->BrxA-DNA-binding->BrxB-BrxF-iSTAND->DUF6079*->N6-METHYLASE->PglZ->SIGMA+OB-Fold+AAA-ATPase+Lon-Protease-BrxL->HKD-DNase+OGG-like-><-REase<-NACHT+SF1-SF2-Helicase+HKD-DNase<-RNaseH<-nSTAND1+HNH | 1213 | Bacteria>Pseudomonadota>Gammaproteobacteria>Enterobacterales>Enterobacteriaceae>Klebsiella/Raoultella_group | GCF_900774955.1 | Klebsiella_pneumoniae | F3L21_RS14210 | 37429..41070 | 573 | BREX-system-P-loop-protein-BrxC-[Klebsiella-pneumoniae] |
| WP_135729167.1 | Type1.BREX | wHTH+SAD+HNH->wHTH+HNH-><-HIN-HTH<-SIGMA+OB-Fold+AAA-ATPase+Lon-Protease-BrxL<-PglZ<-TOPRIM<-OLD-ABC<-N6-METHYLASE<-DUF6079*<-BrxB-BrxF-iSTAND<-BrxA-DNA-binding\|\|ZnR-><-tRNA\|\|?->?->?->?->?->Calcineurin-> | 1204 | Bacteria>Pseudomonadota>Gammaproteobacteria>Enterobacterales>Enterobacteriaceae>Klebsiella/Raoultella_group | GCF_900775295.1 | Klebsiella_pneumoniae | E7W89_RS19490 | 55176..58790 | 573 | BREX-system-P-loop-protein-BrxC-[Klebsiella-pneumoniae] |
| WP_114690464.1 | Type1.BREX | nSTAND1+HNH->RNaseH->NACHT+SF1-SF2-Helicase+HKD-DNase->REase-><-HKD-DNase+OGG-like<-SIGMA+OB-Fold+AAA-ATPase+Lon-Protease-BrxL<-PglZ<-N6-METHYLASE<-DUF6079*<-BrxB-BrxF-iSTAND<-BrxA-DNA-binding<-?<-ColE7+HNH<-ACET\|\|NUDIX->?->?-><-DUF2754 | 1213 | Bacteria>Pseudomonadota>Gammaproteobacteria>Enterobacterales>Enterobacteriaceae>Klebsiella/Raoultella_group | GCF_900775385.1 | Klebsiella_pneumoniae | MC274_RS12705 | 209352..212993 | 573 | BREX-system-P-loop-protein-BrxC-[Klebsiella-pneumoniae] |

<table>
<tr><th>Protein-ID</th><th>System</th><th>Gene-Neighbourhoods</th><th>Length</th><th>Taxonomy</th><th>Assembly-ID</th><th>Strain</th><th>Locus-Tag</th><th>Genomic-coordinates</th><th>TaxID</th><th>Fasta-define</th></tr>
<tr><td>WP_023285784.1</td><td>Type1.BREX</td><td><-Calcineurin<-?<-?<-?<-?<-?||tRNA-><-ZnR||BrxA-DNA-binding->BrxB-BrxF-iSTAND->DUF6079*->N6-METHYLASE->OLD-ABC->TOPRIM->PglZ->SIGMA+OB-Fold+AAA-ATPase+Lon-Protease-BrxL->HIN-HTH-><-wH[illegible]</td><td>1204</td><td>Bacteria>Pseudomonadota>Gammaproteobacteria>Enterobacterales>Enterobacteriaceae>Klebsiella/Raoultella_group</td><td>GCF_900775395.1</td><td>Klebsiella_pneumoniae</td><td>PXW47_RS20390</td><td>324730..328344</td><td>573</td><td>BREX-system-P-loop-protein-BrxC-[Klebsiella-pneumoniae]</td></tr>
<tr><td>WP_064153723.1</td><td>Type1.BREX</td><td>nSTAND1+HNH->RNaseH->NACHT+SF1-SF2-Helicase+HKD-DNase->REase-><-HKD-DNase+OGG-like<-SIGMA+OB-Fold+AAA-ATPase+Lon-Protease-BrxL<-PglZ<-N6-METHYLASE<-DUF6079*<-BrxB-BrxF-iSTAND<-BrxA-DN[illegible]</td><td>1213</td><td>Bacteria>Pseudomonadota>Gammaproteobacteria>Enterobacterales>Enterobacteriaceae>Klebsiella/Raoultella_group</td><td>GCF_900775555.1</td><td>Klebsiella_pneumoniae</td><td>E7W23_RS23250</td><td>82036..85677</td><td>573</td><td>BREX-system-P-loop-protein-BrxC-[Klebsiella-pneumoniae]</td></tr>
<tr><td>WP_135732751.1</td><td>Type1.BREX</td><td>XIS-HTH->?->?->?->TOPRIM-DNAG+D5N+D5ATPase+P4PRIMASE-HTH-><-ZnR||BrxA-DNA-binding->BrxB-BrxF-iSTAND->DUF6079*->N6-METHYLASE->OLD-ABC->TOPRIM->PglZ->SIGMA+OB-Fold+AAA-ATPase+Lon-Protea[illegible]</td><td>1204</td><td>Bacteria>Pseudomonadota>Gammaproteobacteria>Enterobacterales>Enterobacteriaceae>Klebsiella/Raoultella_group</td><td>GCF_900775595.1</td><td>Klebsiella_quasipneumoniae</td><td>ICC30_RS25220</td><td>11695..15309</td><td>1463165</td><td>BREX-system-P-loop-protein-BrxC-[Klebsiella-quasipneumoniae]</td></tr>
<tr><td>WP_135723013.1</td><td>Type1.BREX</td><td>Arm-DNA-bind_3+Phage_integrase-><-HKD-DNase+OGG-like<-HKD-DNase+OGG-like<-SIGMA+OB-Fold+AAA-ATPase+Lon-Protease-BrxL<-PglZ<-TOPRIM<-OLD-ABC<-N6-METHYLASE<-DUF6079*<-BrxB-BrxF-iSTAND<-[illegible]</td><td>1204</td><td>Bacteria>Pseudomonadota>Gammaproteobacteria>Enterobacterales>Enterobacteriaceae>Klebsiella/Raoultella_group</td><td>GCF_900775875.1</td><td>Klebsiella_pneumoniae</td><td>E7X40_RS22910</td><td>78837..82451</td><td>573</td><td>BREX-system-P-loop-protein-BrxC-[Klebsiella-pneumoniae]</td></tr>
<tr><td>WP_136086191.1</td><td>Type1.BREX</td><td>nSTAND1+HNH->RNaseH->NACHT+SF1-SF2-Helicase+HKD-DNase->REase-><-HKD-DNase+OGG-like<-SIGMA+OB-Fold+AAA-ATPase+Lon-Protease-BrxL<-PglZ<-N6-METHYLASE<-DUF6079*<-BrxB-BrxF-iSTAND<-BrxA-DN[illegible]</td><td>1213</td><td>Bacteria>Pseudomonadota>Gammaproteobacteria>Enterobacterales>Enterobacteriaceae>Klebsiella/Raoultella_group</td><td>GCF_900978435.1</td><td>Klebsiella_variicola</td><td>E9559_RS11315</td><td>84067..87708</td><td>244366</td><td>BREX-system-P-loop-protein-BrxC-[Klebsiella-variicola]</td></tr>
<tr><td>WP_142470399.1</td><td>Type1.BREX</td><td><-SAM-METHYLASE<-wHTH+HNH||tRNA->Arm-DNA-bind_3+Phage_integrase-><-HKD-DNase+OGG-like<-SIGMA+OB-Fold+AAA-ATPase+Lon-Protease-BrxL<-PglZ<-N6-METHYLASE<-DUF6079*<-BrxB-BrxF-iSTAND<-BrxA[illegible]</td><td>1213</td><td>Bacteria>Pseudomonadota>Gammaproteobacteria>Enterobacterales>Enterobacteriaceae>Klebsiella/Raoultella_group</td><td>GCF_902158685.1</td><td>Klebsiella_pasteurii</td><td>SB6424_04698</td><td>319733..323374</td><td>2587529</td><td>BREX-system-P-loop-protein-BrxC-[Klebsiella-pasteurii]</td></tr>
<tr><td>WP_224385469.1</td><td>Type1.BREX</td><td><-HTH_Tnp_1+HTH_21+rve+rve_3<-PglZ<-N6-METHYLASE-p<-N6-METHYLASE<-ParB+HNH<-DUF6079*<-BrxB-BrxF-iSTAND<-BrxA-DNA-binding<-Glyoxalase<-ZnR<-METHYLASE<-YjhX_toxin<-ACET<-Glyoxalase</td><td>1240</td><td>Bacteria>Pseudomonadota>Gammaproteobacteria>Enterobacterales>Enterobacteriaceae>Klebsiella/Raoultella_group</td><td>GCF_902158715.1</td><td>Klebsiella</td><td>FN110_RS17010</td><td>11054..14776</td><td>570</td><td>MULTISPECIES:-BREX-system-P-loop-protein-BrxC-[Klebsiella]</td></tr>
<tr><td>WP_256682067.1</td><td>Type1.BREX</td><td><-DUF905<-DUF932<-CopG/RHH||?->XIS-HTH-><-TIR<-TIR||BrxA-DNA-binding->BrxB-BrxF-iSTAND->DUF6079*->ParB+HNH->N6-METHYLASE->?->N6-METHYLASE-p->PglZ->SIGMA+OB-Fold+AAA-ATPase+Lon-Proteas[illegible]</td><td>1240</td><td>Bacteria>Pseudomonadota>Gammaproteobacteria>Enterobacterales>Enterobacteriaceae>Klebsiella/Raoultella_group</td><td>GCF_902158845.1</td><td>Klebsiella_michiganensis</td><td>FN096_RS13265</td><td>22370..26092</td><td>1134687</td><td>BREX-system-P-loop-protein-BrxC-[Klebsiella-michiganensis]</td></tr>
<tr><td>WP_154062666.1</td><td>Type1.BREX</td><td><-HTH_36<-HTH+HTH<-ParB+BAM<-XIS-HTH<-wHTH<-ZnR||BrxA-DNA-binding->BrxB-BrxF-iSTAND->DUF6079*->N6-METHYLASE->OLD-ABC->TOPRIM->PglZ->SIGMA+OB-Fold+AAA-ATPase+Lon-Protease-BrxL-><-?<-Gl[illegible]</td><td>1204</td><td>Bacteria>Pseudomonadota>Gammaproteobacteria>Enterobacterales>Enterobacteriaceae>Klebsiella/Raoultella_group</td><td>GCF_902159535.1</td><td>Klebsiella_michiganensis</td><td>FN635_RS25500</td><td>5342283..5345897</td><td>1134687</td><td>BREX-system-P-loop-protein-BrxC-[Klebsiella-michiganensis]</td></tr>
<tr><td>WP_227635281.1</td><td>Type1.BREX</td><td>XIS-HTH-><-TIR<-TIR||BrxA-DNA-binding->BrxB-BrxF-iSTAND->DUF6079*->ParB+HNH->N6-METHYLASE->HEPN->RVT->N6-METHYLASE-p->PglZ->SIGMA+OB-Fold+AAA-ATPase+Lon-Protease-BrxL->wHTH+WYL+WCX-></td><td>1240</td><td>Bacteria>Pseudomonadota>Gammaproteobacteria>Enterobacterales>Enterobacteriaceae>Klebsiella/Raoultella_group</td><td>GCF_902159715.1</td><td>Klebsiella</td><td>PIK69_RS17535</td><td>2737..6459</td><td>570</td><td>MULTISPECIES:-BREX-system-P-loop-protein-BrxC-[Klebsiella]</td></tr>
<tr><td>WP_048266247.1</td><td>Type1.BREX</td><td>REase-><-HKD-DNase+OGG-like<-SIGMA+OB-Fold+AAA-ATPase+Lon-Protease-BrxL<-PglZ<-N6-METHYLASE<-DUF6079*<-BrxB-BrxF-iSTAND<-BrxA-DNA-binding<-URI-DNase</td><td>1213</td><td>Bacteria>Pseudomonadota>Gammaproteobacteria>Enterobacterales>Enterobacteriaceae>Klebsiella/Raoultella_group</td><td>GCF_902161325.1</td><td>Klebsiella_pneumoniae</td><td>FN306_RS22675</td><td>100097..103738</td><td>573</td><td>BREX-system-P-loop-protein-BrxC-[Klebsiella-pneumoniae]</td></tr>
<tr><td>WP_085820485.1</td><td>Type1.BREX</td><td>REase-><-HKD-DNase+OGG-like<-SIGMA+OB-Fold+AAA-ATPase+Lon-Protease-BrxL<-PglZ<-N6-METHYLASE<-DUF6079*<-BrxB-BrxF-iSTAND<-BrxA-DNA-binding<-URI-DNase||<-?<-?||<-?||?->||MAD-NTD+HNH-><-[illegible]</td><td>1213</td><td>Bacteria>Pseudomonadota>Gammaproteobacteria>Enterobacterales>Enterobacteriaceae>Klebsiella/Raoultella_group</td><td>GCF_902508285.1</td><td>Klebsiella_pneumoniae</td><td>GVM87_RS22970</td><td>29420..33061</td><td>573</td><td>BREX-system-P-loop-protein-BrxC-[Klebsiella-pneumoniae]</td></tr>
<tr><td>WP_177340625.1</td><td>Type1.BREX</td><td><-OBFOLD-NUCLEASE<-CARF+NtrC-AAA+FIS-HTH<-SIGMA+OB-Fold+AAA-ATPase+Lon-Protease-BrxL<-PglZ<-N6-METHYLASE<-DUF6079*<-BrxB-BrxF-iSTAND<-BrxA-DNA-binding<-wHTH+WYL+WCX<-?<-35exo<-RVT||Le[illegible]</td><td>1225</td><td>Bacteria>Pseudomonadota>Gammaproteobacteria>Enterobacterales>Enterobacteriaceae>Klebsiella/Raoultella_group</td><td>GCF_902754085.1</td><td>Klebsiella_michiganensis</td><td>HV187_RS24210</td><td>5087544..5091221</td><td>1134687</td><td>BREX-system-P-loop-protein-BrxC-[Klebsiella-michiganensis]</td></tr>
<tr><td>WP_224423598.1</td><td>Type1.BREX</td><td>DUF2754-><-?||ACET->ColE7+HNH->?->BrxA-DNA-binding->BrxB-BrxF-iSTAND->DUF6079*->N6-METHYLASE->PglZ->SIGMA+OB-Fold+AAA-ATPase+Lon-Protease-BrxL->?-><-Arm-DNA-bind_3+Phage_integrase||Pa[illegible]</td><td>1213</td><td>Bacteria>Pseudomonadota>Gammaproteobacteria>Enterobacterales>Enterobacteriaceae>Klebsiella/Raoultella_group</td><td>GCF_903166255.1</td><td>Klebsiella_quasivariicola</td><td>QJU95_RS24650</td><td>36202..39843</td><td>2026240</td><td>BREX-system-P-loop-protein-BrxC-[Klebsiella-quasivariicola]</td></tr>
<tr><td>WP_225377085.1</td><td>Type1.BREX</td><td><-NACHT+SF1-SF2-Helicase+HKD-DNase||BrxA-DNA-binding->BrxB-BrxF-iSTAND->DUF6079*->N6-METHYLASE->OLD-ABC->TOPRIM->PglZ->SIGMA+OB-Fold+AAA-ATPase+Lon-Protease-BrxL->HKD-DNase+OGG-like->[illegible]</td><td>1204</td><td>Bacteria>Pseudomonadota>Gammaproteobacteria>Enterobacterales>Enterobacteriaceae>Klebsiella/Raoultella_group</td><td>GCF_916172235.1</td><td>Klebsiella_pneumoniae</td><td>LF666_RS16195</td><td>10713..14327</td><td>573</td><td>BREX-system-P-loop-protein-BrxC-[Klebsiella-pneumoniae]</td></tr>
<tr><td>WP_225386868.1</td><td>Type1.BREX</td><td>DUF2754-><-?<-?||ColE7+HNH->?->BrxA-DNA-binding->BrxB-BrxF-iSTAND->DUF6079*->N6-METHYLASE->PglZ->SIGMA+OB-Fold+AAA-ATPase+Lon-Protease-BrxL->HKD-DNase+OGG-like-><-REase<-NACHT+SF1-SF2[illegible]</td><td>1213</td><td>Bacteria>Pseudomonadota>Gammaproteobacteria>Enterobacterales>Enterobacteriaceae>Klebsiella/Raoultella_group</td><td>GCF_916173145.1</td><td>Klebsiella_variicola</td><td>LF736_RS25380</td><td>17904..21545</td><td>244366</td><td>BREX-system-P-loop-protein-BrxC-[Klebsiella-variicola]</td></tr>
<tr><td>WP_048289717.1</td><td>Type1.BREX</td><td><-NUDIX||EVE+HNH->ColE7+HNH->?-><-SIGMA+OB-Fold+AAA-ATPase+Lon-Protease-BrxL<-PglZ<-TOPRIM<-OLD-ABC<-N6-METHYLASE<-DUF6079*<-BrxB-BrxF-iSTAND<-BrxA-DNA-binding||ZnR-></td><td>1204</td><td>Bacteria>Pseudomonadota>Gammaproteobacteria>Enterobacterales>Enterobacteriaceae>Klebsiella/Raoultella_group</td><td>GCF_934247105.1</td><td>Klebsiella_pneumoniae</td><td>A8T29_RS25610</td><td>25582..29196</td><td>573</td><td>BREX-system-P-loop-protein-BrxC-[Klebsiella-pneumoniae]</td></tr>
<tr><td>WP_101867996.1</td><td>Type1.BREX</td><td><-SAM-METHYLASE<-wHTH(SBD)+HNH||tRNA->Arm-DNA-bind_3+Phage_integrase->wHTH+WYL+WCX->BrxA-DNA-binding->BrxB-BrxF-iSTAND->DUF6079*->N6-METHYLASE->ParB+HNH-></td><td>1157</td><td>Bacteria>Pseudomonadota>Gammaproteobacteria>Enterobacterales>Enterobacteriaceae>Klebsiella/Raoultella_group</td><td>-</td><td>Klebsiella_michiganensis</td><td>CWN04_RS12955</td><td>12917..16390</td><td>1134687</td><td>BREX-system-P-loop-protein-BrxC-[Klebsiella-michiganensis]</td></tr>
<tr><td>WP_154904474.1</td><td>Type1.BREX</td><td><-N6-METHYLASE-p<-N6-METHYLASE<-ParB+HNH<-DUF6079*<-BrxB-BrxF-iSTAND<-BrxA-DNA-binding||<-XIS-HTH</td><td>1233</td><td>Bacteria>Pseudomonadota>Gammaproteobacteria>Enterobacterales>Enterobacteriaceae>Klebsiella/Raoultella_group</td><td>-</td><td>Klebsiella_michiganensis</td><td>CWN50_01600</td><td>4530..8231</td><td>1134687</td><td>BREX-system-P-loop-protein-BrxC-[Klebsiella-michiganensis]</td></tr>
<tr><td>WP_227627429.1</td><td>Type1.BREX</td><td><-wHTH+WYL+WCX<-?<-SIGMA+OB-Fold+AAA-ATPase+Lon-Protease-BrxL<-PglZ<-N6-METHYLASE[illegible]</td><td>1240</td><td>Bacteria>Pseudomonadota>Gammaproteobacteria>Enterobacterales>Enterobacteriaceae>Klebsiella/Raoultella_group</td><td>-</td><td>Klebsiella_michiganensis</td><td>G4403_RS10440</td><td>98242..101964</td><td>1134687</td><td>BREX-system-P-loop-protein-BrxC-[Klebsiella-michiganensis]</td></tr>
<tr><td>WP_097365540.1</td><td>Type1.BREX</td><td><-DUF905<-DUF932<-CopG/RHH||?->XIS-HTH-><-TIR<-TIR||BrxA-DNA-binding->BrxB-BrxF-i[illegible]</td><td>1233</td><td>Bacteria>Pseudomonadota>Gammaproteobacteria>Enterobacterales>Enterobacteriaceae>Klebsiella/Raoultella_group</td><td>-</td><td>Klebsiella_oxytoca</td><td>I9Y61_001542</td><td>37432..41133</td><td>571</td><td>MULTISPECIES:-BREX-system-P-loop-protein-BrxC-[Klebsiella]</td></tr>
<tr><td>ADJ51152.1</td><td>Type1.BREX</td><td><-DUF6079-p*<-BrxB-BrxF-iSTAND<-BrxA-DNA-binding<-wHTH+WYL+WCX||Zn_Tnp_IS1+HTH_Tn[illegible]</td><td>484</td><td>Bacteria>Pseudomonadota>Gammaproteobacteria>Enterobacterales>Enterobacteriaceae>Klebsiella/Raoultella_group</td><td>-</td><td>Klebsiella_pneumoniae</td><td>.</td><td>1..1453</td><td>573</td><td>ATPase-like-protein,-partial-[Klebsiella-pneumoniae]</td></tr>
<tr><td>WP_117042583.1</td><td>Type1.BREX</td><td><-DUF905<-DUF932<-CopG/RHH||?->XIS-HTH-><-TIR<-TIR||BrxA-DNA-binding->BrxB-BrxF-i[illegible]</td><td>1233</td><td>Bacteria>Pseudomonadota>Gammaproteobacteria>Enterobacterales>Enterobacteriaceae>Klebsiella/Raoultella_group</td><td>-</td><td>Klebsiella_pneumoniae</td><td>AM381_RS08835</td><td>19475..23176</td><td>573</td><td>MULTISPECIES:-BREX-system-P-loop-protein-BrxC-[Klebsiella]</td></tr>
<tr><td>WP_114263049.1</td><td>Type1.BREX</td><td><-DUF932<-GspA-ATPase+GTPase<-CopG/RHH||?->XIS-HTH->?->?->BrxA-DNA-binding->BrxB-[illegible]</td><td>1233</td><td>Bacteria>Pseudomonadota>Gammaproteobacteria>Enterobacterales>Enterobacteriaceae>Klebsiella/Raoultella_group</td><td>-</td><td>Klebsiella_pneumoniae</td><td>AN2353V1_3685</td><td>3967252..3970953</td><td>573</td><td>MULTISPECIES:-BREX-system-P-loop-protein-BrxC-[Enterobacteriaceae]</td></tr>
<tr><td>WP_062729754.1</td><td>Type1.BREX</td><td>Arm-DNA-bind_3+Phage_integrase-><-SIGMA+OB-Fold+AAA-ATPase+Lon-Protease-BrxL<-Pgl[illegible]</td><td>1233</td><td>Bacteria>Pseudomonadota>Gammaproteobacteria>Enterobacterales>Enterobacteriaceae>Klebsiella/Raoultella_group</td><td>-</td><td>Klebsiella_pneumoniae</td><td>C0077_01580</td><td>326913..330614</td><td>573</td><td>MULTISPECIES:-BREX-system-P-loop-protein-BrxC-[Enterobacteriaceae]</td></tr>
<tr><td>WP_073511310.1</td><td>Type1.BREX</td><td><-wHTH+WYL<-sGTP<-?||XIS-HTH->?->?->BrxA-DNA-binding->BrxB-BrxF-iSTAND->DUF6079*-[illegible]</td><td>1233</td><td>Bacteria>Pseudomonadota>Gammaproteobacteria>Enterobacterales>Enterobacteriaceae>Klebsiella/Raoultella_group</td><td>-</td><td>Klebsiella_pneumoniae</td><td>L7V03_004584</td><td>32186..35887</td><td>573</td><td>BREX-system-P-loop-protein-BrxC-[Klebsiella-pneumoniae]</td></tr>
<tr><td>WP_117247358.1</td><td>Type1.BREX</td><td><-wHTH+WYL<-nSTAND1+sGTP<-?||XIS-HTH->?->?->BrxA-DNA-binding->BrxB-BrxF-iSTAND->D[illegible]</td><td>1233</td><td>Bacteria>Pseudomonadota>Gammaproteobacteria>Enterobacterales>Enterobacteriaceae>Klebsiella/Raoultella_group</td><td>-</td><td>Klebsiella_pneumoniae</td><td>L7W77_004838</td><td>45573..49274</td><td>573</td><td>MULTISPECIES:-BREX-system-P-loop-protein-BrxC-[Klebsiella]</td></tr>
<tr><td>EIV5715275.1</td><td>Type1.BREX</td><td>METHYLASE->nSTAND1+HNH->RNaseH->NACHT+SF1-SF2-Helicase+HKD-DNase->REase-><-HKD-DN[illegible]</td><td>1213</td><td>Bacteria>Pseudomonadota>Gammaproteobacteria>Enterobacterales>Enterobacteriaceae>Klebsiella/Raoultella_group</td><td>-</td><td>Klebsiella_pneumoniae</td><td>L9F37_004806</td><td>56067..59708</td><td>573</td><td>BREX-system-P-loop-protein-BrxC-[Klebsiella-pneumoniae]</td></tr>
<tr><td>HCB8952633.1</td><td>Type1.BREX</td><td>DUF2754-><-?<-NUDIX||ColE7+HNH->?->BrxA-DNA-binding->BrxB-BrxF-iSTAND->DUF6079*->[illegible]</td><td>1213</td><td>Bacteria>Pseudomonadota>Gammaproteobacteria>Enterobacterales>Enterobacteriaceae>Klebsiella/Raoultella_group</td><td>-</td><td>Klebsiella_pneumoniae</td><td>M2O84_005147</td><td>37298..40939</td><td>573</td><td>BREX-system-P-loop-protein-BrxC-[Klebsiella-pneumoniae]</td></tr>
<tr><td>HCC8216583.1</td><td>Type1.BREX</td><td>GDResolvase+HIN-HTH->ZnR-><-SIGMA+OB-Fold+AAA-ATPase+Lon-Protease-BrxL<-PglZ<-N6-[illegible]</td><td>1213</td><td>Bacteria>Pseudomonadota>Gammaproteobacteria>Enterobacterales>Enterobacteriaceae>Klebsiella/Raoultella_group</td><td>-</td><td>Klebsiella_pneumoniae</td><td>M2P48_005497</td><td>10102..13743</td><td>573</td><td>BREX-system-P-loop-protein-BrxC-[Klebsiella-pneumoniae]</td></tr>
<tr><td>WP_117076897.1</td><td>Type1.BREX</td><td><-wHTH+WYL<-nSTAND1+sGTP<-?||XIS-HTH->?->BrxA-DNA-binding->BrxB-BrxF-iSTAND->DUF6[illegible]</td><td>1233</td><td>Bacteria>Pseudomonadota>Gammaproteobacteria>Enterobacterales>Enterobacteriaceae>Klebsiella/Raoultella_group</td><td>-</td><td>Klebsiella_pneumoniae</td><td>ML041_004566</td><td>31807..35508</td><td>573</td><td>BREX-system-P-loop-protein-BrxC-[Klebsiella-pneumoniae]</td></tr>
<tr><td>HCA5579377.1</td><td>Type1.BREX</td><td>XIS-HTH->?->REase+SPF-cons-><-SF1-SF2-Helicase<-HKD-DNase+HKD-DNase<-?||BrxA-DNA-[illegible]</td><td>1213</td><td>Bacteria>Pseudomonadota>Gammaproteobacteria>Enterobacterales>Enterobacteriaceae>Klebsiella/Raoultella_group</td><td>-</td><td>Klebsiella_pneumoniae</td><td>MXD08_004593</td><td>70758..74399</td><td>573</td><td>BREX-system-P-loop-protein-BrxC-[Klebsiella-pneumoniae]</td></tr>
<tr><td>WP_168234029.1</td><td>Type1.BREX</td><td>ParB+HNH->nSTAND1+HNH->RNaseH->NACHT+SF1-SF2-Helicase+HKD-DNase->REase-><-HKD-DNa[illegible]</td><td>1213</td><td>Bacteria>Pseudomonadota>Gammaproteobacteria>Enterobacterales>Enterobacteriaceae>Klebsiella/Raoultella_group</td><td>-</td><td>Klebsiella_variicola</td><td>HGK55_RS15775</td><td>337116..340757</td><td>244366</td><td>BREX-system-P-loop-protein-BrxC-[Klebsiella-variicola]</td></tr>
<tr><td>EJN1576646.1</td><td>Type1.BREX</td><td>Glyoxalase->ACET->YjhX_toxin->ACET->METHYLASE->ZnR->Glyoxalase->BrxA-DNA-binding-[illegible]</td><td>1233</td><td>Bacteria>Pseudomonadota>Gammaproteobacteria>Enterobacterales>Enterobacteriaceae>Klebsiella/Raoultella_group</td><td>-</td><td>Klebsiella_variicola</td><td>NPH96_004932</td><td>32414..36115</td><td>244366</td><td>BREX-system-P-loop-protein-BrxC-[Klebsiella-variicola]</td></tr>
<tr><td>URQ56457.1</td><td>Type1.BREX</td><td>HTH_21+rve+rve_3-><-RHH||?->wHTH+WYL+WCX->BrxA-DNA-binding->BrxB-BrxF-iSTAND->DUF[illegible]</td><td>769</td><td>Bacteria>Pseudomonadota>Gammaproteobacteria>Enterobacterales>Enterobacteriaceae>Klebsiella/Raoultella_group</td><td>-</td><td>Raoultella_ornithinolytica</td><td>.</td><td>79835..82144</td><td>54291</td><td>Hypothetical-protein-[Raoultella-ornithinolytica]</td></tr>
<tr><td>EHT9899814.1</td><td>Type1.BREX</td><td><-Calcineurin<-?<-?<-?||tRNA-><-SIGMA+OB-Fold+AAA-ATPase+Lon-Protease-BrxL<-PglZ<-N6-METHYLASE-p<-N6-METHYLASE<-ParB+HNH<-DUF6079*<-BrxB-BrxF-iSTAND<-BrxA-DNA-binding||?-><-Glyco_hydr[illegible]</td><td>1233</td><td>Bacteria>Pseudomonadota>Gammaproteobacteria>Enterobacterales>Enterobacteriaceae>Klebsiella/Raoultella_group</td><td>-</td><td>Raoultella_ornithinolytica</td><td>KY209_001651</td><td>277271..280972</td><td>54291</td><td>BREX-system-P-loop-protein-BrxC-[Raoultella-ornithinolytica]</td></tr>
<tr><td>WP_094898295.1</td><td>Type1.BREX</td><td><-TIR||BrxA-DNA-binding->BrxB-BrxF-iSTAND->DUF6079*->ParB+HNH->N6-METHYLASE->HEPN->RVT->N6-METHYLASE-p->PglZ->SIGMA+OB-Fold+AAA-ATPase+Lon-Protease-BrxL->wHTH+WYL+WCX-></td><td>1233</td><td>Bacteria>Pseudomonadota>Gammaproteobacteria>Enterobacterales>Enterobacteriaceae>Klebsiella/Raoultella_group</td><td>-</td><td>Raoultella_planticola</td><td>FQG10_00765</td><td>2767..6468</td><td>575</td><td>BREX-system-P-loop-protein-BrxC-[Raoultella-planticola]</td></tr>
<tr><td>HAT1572918.1</td><td>Type1.BREX</td><td>MarR-HTH->?->?->BZIP+SF1-SF2-Helicase->||HIN-HTH->HTH_Tnp_1+HTH_21+rve+rve_3->||wHTH+WYL+WCX->BrxA-DNA-binding->BrxB-BrxF-iSTAND->DUF6079*->ParB+HNH->N6-METHYLASE->PglZ->SIGMA+OB-Fold[illegible]</td><td>1225</td><td>Bacteria>Pseudomonadota>Gammaproteobacteria>Enterobacterales>Enterobacteriaceae>Kluyvera</td><td>GCA_015892965.1</td><td>Kluyvera_cryocrescens</td><td>I8Y10_004331</td><td>41743..45420</td><td>580</td><td>BREX-system-P-loop-protein-BrxC-[Kluyvera-cryocrescens]</td></tr>
<tr><td>WP_271458914.1</td><td>Type1.BREX</td><td><-N6-METHYLASE<-DUF6079*<-BrxB-BrxF-iSTAND<-BrxA-DNA-binding||ZnR->wHTH->XIS-HTH->ParB+BAM->HTH+HTH->HTH_36-></td><td>1204</td><td>Bacteria>Pseudomonadota>Gammaproteobacteria>Enterobacterales>Enterobacteriaceae>Kluyvera</td><td>GCF_028069715.1</td><td>Kluyvera_sp._Awk_3</td><td>NNN71_RS16325</td><td>2505..6119</td><td>2963956</td><td>BREX-system-P-loop-protein-BrxC-[Kluyvera-sp.-Awk-3]</td></tr>
<tr><td>WP_112204418.1</td><td>Type1.BREX</td><td>GDResolvase+HIN-HTH-><-REase<-?||HTH_3->Arm-DNA-bind_3+Phage_integrase-><-?<-PglZ<-N6-METHYLASE<-DUF6079*<-BrxB-BrxF-iSTAND<-BrxA-DNA-binding||ArdB->?->CbeA_antitoxin->?->REase-></td><td>1213</td><td>Bacteria>Pseudomonadota>Gammaproteobacteria>Enterobacterales>Enterobacteriaceae>Kluyvera</td><td>GCF_900452275.1</td><td>Kluyvera_cryocrescens</td><td>DS850_RS16305</td><td>27927..31568</td><td>580</td><td>BREX-system-P-loop-protein-BrxC-[Kluyvera-cryocrescens]</td></tr>
<tr><td>WP_064568621.1</td><td>Type1.BREX</td><td>ABC-ATPase+SbcC->?-><-DSBH+AraC-HTH+AraC-HTH||?->?->TarH+HAMP+MA->BrxA-DNA-binding->BrxB-BrxF-iSTAND->DUF6079*->N6-METHYLASE->PglZ->SIGMA+OB-Fold+AAA-ATPase+Lon-Protease-BrxL->?-><-tR[illegible]</td><td>1216</td><td>Bacteria>Pseudomonadota>Gammaproteobacteria>Enterobacterales>Enterobacteriaceae>Kosakonia</td><td>GCF_001658025.2</td><td>Kosakonia_oryzae</td><td>AWR26_RS22590</td><td>4759672..4763322</td><td>497725</td><td>BREX-system-P-loop-protein-BrxC-[Kosakonia-oryzae]</td></tr>
<tr><td>WP_144838631.1</td><td>Type1.BREX</td><td>Aminotran_3->GNTR-HTH->SymE_toxin-><-?||wHTH+HNH->SAM-METHYLASE->BrxA-DNA-binding->BrxB-BrxF-iSTAND->DUF6079*->N6-METHYLASE->PglZ->SIGMA+OB-Fold+AAA-ATPase+Lon-Protease-BrxL->HKD-DNas[illegible]</td><td>1213</td><td>Bacteria>Pseudomonadota>Gammaproteobacteria>Enterobacterales>Enterobacteriaceae>Leclercia</td><td>GCF_007680555.1</td><td>Leclercia_adecarboxylata</td><td>FS784_RS12375</td><td>119273..122914</td><td>83655</td><td>BREX-system-P-loop-protein-BrxC-[Leclercia-adecarboxylata]</td></tr>
<tr><td>QXB23949.1</td><td>Type1.BREX</td><td>XIS-HTH->?->ZnR->ZnR->?->||BrxA-DNA-binding->BrxB-BrxF-iSTAND->DUF6079*->ParB+HNH->N6-METHYLASE->HEPN->RVT->N6-METHYLASE-><-HIN-HTH||PglZ->SIGMA+OB-Fold+AAA-ATPase+Lon-Protease-BrxL->[illegible]</td><td>1233</td><td>Bacteria>Pseudomonadota>Gammaproteobacteria>Enterobacterales>Enterobacteriaceae>Lelliottia</td><td>GCA_019048185.1</td><td>Lelliottia_amnigena</td><td>I6L76_15755</td><td>3242307..3246008</td><td>61646</td><td>BREX-system-P-loop-protein-BrxC-[Lelliottia-amnigena]</td></tr>
<tr><td>WP_102104498.1</td><td>Type1.BREX</td><td><-HKD-DNase+OGG-like<-SIGMA+OB-Fold+AAA-ATPase+Lon-Protease-BrxL<-PglZ<-TOPRIM<-OLD-ABC<-N6-METHYLASE<-DUF6079*<-BrxB-BrxF-iSTAND<-BrxA-DNA-binding||NACHT+SF1-SF2-Helicase+HKD-DNase->[illegible]</td><td>1204</td><td>Bacteria>Pseudomonadota>Gammaproteobacteria>Enterobacterales>Enterobacteriaceae>Lelliottia</td><td>GCF_002870005.1</td><td>unclassified_Lelliottia</td><td>F159LOC_RS08780</td><td>168198..171812</td><td>2642424</td><td>MULTISPECIES:-BREX-system-P-loop-protein-BrxC-[unclassified-Lelliottia]</td></tr>
<tr><td>WP_254208058.1</td><td>Type1.BREX</td><td>XIS-HTH->?->ZnR->ZnR->?->||BrxA-DNA-binding->BrxB-BrxF-iSTAND->DUF6079*->ParB+HNH->N6-METHYLASE->HEPN->RVT->N6-METHYLASE-><-HIN-HTH||PglZ->SIGMA+OB-Fold+AAA-ATPase+Lon-Protease-BrxL->[illegible]</td><td>1237</td><td>Bacteria>Pseudomonadota>Gammaproteobacteria>Enterobacterales>Enterobacteriaceae>Lelliottia</td><td>GCF_019048185.1</td><td>Lelliottia_amnigena</td><td>I6L76_RS15755</td><td>3242295..3246008</td><td>61646</td><td>BREX-system-P-loop-protein-BrxC-[Lelliottia-amnigena]</td></tr>
<tr><td>WP_216981896.1</td><td>Type1.BREX</td><td>XIS-HTH->?->ZnR->ZnR->?->||BrxA-DNA-binding->BrxB-BrxF-iSTAND->DUF6079*->ParB+HNH->N6-METHYLASE->HEPN->RVT->N6-METHYLASE-><-HIN-HTH||PglZ->SIGMA+OB-Fold+AAA-ATPase+Lon-Protease-BrxL->[illegible]</td><td>1233</td><td>Bacteria>Pseudomonadota>Gammaproteobacteria>Enterobacterales>Enterobacteriaceae>Lelliottia</td><td>-</td><td>Lelliottia_amnigena</td><td>I6L76_15755</td><td>3242307..3246008</td><td>61646</td><td>BREX-system-P-loop-protein-BrxC-[Lelliottia-amnigena]</td></tr>
<tr><td>WP_077737804.1</td><td>Type1.BREX</td><td>ZnR->ZnR-><-SIGMA+OB-Fold+AAA-ATPase+Lon-Protease-BrxL<-PglZ<-N6-METHYLASE<-DUF6079*<-BrxB-BrxF-iSTAND<-BrxA-DNA-binding||REase+iREase->Zn_Tnp_IS1+HTH_Tnp_IS1+DDE_Tnp_IS1-><-?<-DDE_Tn[illegible]</td><td>1213</td><td>Bacteria>Pseudomonadota>Gammaproteobacteria>Enterobacterales</td><td>-</td><td>Enterobacteriaceae</td><td>NFO83_RS22575</td><td>9834..13475</td><td>543</td><td>MULTISPECIES:-BREX-system-P-loop-protein-BrxC-[Enterobacteriaceae]</td></tr>
<tr><td>WP_233672193.1</td><td>Type1.BREX</td><td><-SIGMA+OB-Fold+AAA-ATPase+Lon-Protease-BrxL<-PglZ<-TOPRIM<-OLD-ABC<-N6-METHYLASE-p<-TIR<-N6-METHYLASE<-TOPRIM<-OLD-ABC<-DUF6079*<-BrxB-BrxF-iSTAND<-BrxA-DNA-binding<-?<-?<-?<-XIS-HTH[illegible]</td><td>1238</td><td>Bacteria>Pseudomonadota>Gammaproteobacteria>Enterobacterales>Enterobacteriaceae>Plesiomonas</td><td>GCF_021368255.1</td><td>Plesiomonas_sp._PI-19</td><td>LQ901_RS09600</td><td>2145541..2149257</td><td>2898798</td><td>BREX-system-P-loop-protein-BrxC-[Plesiomonas-sp.-PI-19]</td></tr>
<tr><td>WP_265846293.1</td><td>Type1.BREX</td><td><-RAMPs<-Cas10||GTPase-AIG->GTPase-AIG->XIS-HTH->?->?->BrxA-DNA-binding->BrxB-BrxF-iSTAND->DUF6079*->ParB+HNH->TIR->N6-METHYLASE-p->?->PglZ-></td><td>1238</td><td>Bacteria>Pseudomonadota>Gammaproteobacteria>Enterobacterales>Enterobacteriaceae>Plesiomonas</td><td>GCF_026240155.1</td><td>Plesiomonas_shigelloides</td><td>OQ486_RS15170</td><td>39648..43364</td><td>703</td><td>BREX-system-P-loop-protein-BrxC-[Plesiomonas-shigelloides]</td></tr>
<tr><td>EHC84522.1</td><td>Type1.BREX</td><td><-SIGMA+OB-Fold+AAA-ATPase+Lon-Protease-BrxL<-PglZ<-N6-METHYLASE-p<-N6-METHYLASE<-ParB+HNH<-DUF6079*<-BrxB-BrxF-iSTAND<-BrxA-DNA-binding<-ZnR<-Glyoxalase<-METHYLASE</td><td>1247</td><td>Bacteria>Pseudomonadota>Gammaproteobacteria>Enterobacterales>Enterobacteriaceae>Salmonella</td><td>GCA_000231705.2</td><td>Salmonella_enterica_subsp._enterica_serovar_Uganda_str._R8-3404</td><td>LTSEUGA_5994</td><td>10868..14611</td><td>913083</td><td>hypothetical-protein-LTSEUGA_5994-[Salmonella-enterica-subsp.-enterica-serovar-Uganda-str.-R8-3404]</td></tr>
<tr><td>ESG75014.1</td><td>Type1.BREX</td><td><-HEPN<-?||YjhX_toxin->METHYLASE->?-><-wHTH+SAD+HNH||?->BrxA-DNA-binding->DUF6079*->?->N6-METHYLASE->OLD-ABC->TOPRIM->PglZ->SIGMA+OB-Fold+AAA-ATPase+Lon-Protease-BrxL->HKD-DNase+OGG-l[illegible]</td><td>1142</td><td>Bacteria>Pseudomonadota>Gammaproteobacteria>Enterobacterales>Enterobacteriaceae>Salmonella</td><td>GCA_000487055.1</td><td>Salmonella_enterica_subsp._enterica_serovar_Kentucky_str._ATCC_9263</td><td>SEEK9263_16857</td><td>19218..22646</td><td>984236</td><td>ATPase-like-protein-[Salmonella-enterica-subsp.-enterica-serovar-Kentucky-str.-ATCC-9263]</td></tr>
<tr><td>APV85542.1</td><td>Type1.BREX</td><td>Glyoxalase->ACET->YjhX_toxin->ACET->METHYLASE->ZnR->Glyoxalase->BrxA-DNA-binding->BrxB-BrxF-iSTAND->DUF6079*->ParB+HNH->N6-METHYLASE->PglZ->SIGMA+OB-Fold+AAA-ATPase+Lon-Protease-BrxL-[illegible]</td><td>1233</td><td>Bacteria>Pseudomonadota>Gammaproteobacteria>Enterobacterales>Enterobacteriaceae>Salmonella</td><td>GCA_000487155.2</td><td>Salmonella_enterica_subsp._enterica_serovar_Inverness_str._ATCC_10720</td><td>SEEI0720_021515</td><td>4182897..4186598</td><td>941187</td><td>DNA-repair-protein-[Salmonella-enterica-subsp.-enterica-serovar-Inverness-str.-ATCC-10720]</td></tr>
<tr><td>AXD95438.1</td><td>Type1.BREX</td><td><-METHYLASE||SKI-><-?||tRNA->Arm-DNA-bind_3+Phage_integrase-><-SIGMA+OB-Fold+AAA-ATPase+Lon-Protease-BrxL<-PglZ<-N6-METHYLASE<-ParB+HNH<-DUF6079*<-BrxB-BrxF-iSTAND<-BrxA-DNA-binding<-[illegible]</td><td>1233</td><td>Bacteria>Pseudomonadota>Gammaproteobacteria>Enterobacterales>Enterobacteriaceae>Salmonella</td><td>GCA_003325015.1</td><td>Salmonella_enterica</td><td>CHC75_12745</td><td>2557798..2561499</td><td>28901</td><td>BREX-system-P-loop-protein-BrxC-[Salmonella-enterica]</td></tr>
<tr><td>MDG46363.1</td><td>Type1.BREX</td><td><-HEPN<-REase+iREase<-?||YjhX_toxin->SAM-METHYLASE->?->BrxA-DNA-binding->BrxB-BrxF-iSTAND->DUF6079*->N6-METHYLASE->OLD-ABC->TOPRIM->PglZ->SIGMA+OB-Fold+AAA-ATPase+Lon-Protease-BrxL-><[illegible]</td><td>1213</td><td>Bacteria>Pseudomonadota>Gammaproteobacteria>Enterobacterales>Enterobacteriaceae>Salmonella</td><td>GCA_003733245.1</td><td>Salmonella_enterica</td><td>EB289_07920</td><td>38672..42313</td><td>28901</td><td>BREX-system-P-loop-protein-BrxC-[Salmonella-enterica]</td></tr>
<tr><td>MIX28204.1</td><td>Type1.BREX</td><td>YjhX_toxin->SAM-METHYLASE-><-?||?->?->BrxA-DNA-binding->BrxB-BrxF-iSTAND->DUF6079*->N6-METHYLASE->OLD-ABC->TOPRIM->PglZ->SIGMA+OB-Fold+AAA-ATPase+Lon-Protease-BrxL-><-REase<-NACHT+SF1[illegible]</td><td>1213</td><td>Bacteria>Pseudomonadota>Gammaproteobacteria>Enterobacterales>Enterobacteriaceae>Salmonella</td><td>GCA_003876625.1</td><td>Salmonella_enterica_subsp._enterica_serovar_Livingstone</td><td>CRQ34_16010</td><td>159179..162820</td><td>260680</td><td>BREX-system-P-loop-protein-BrxC-[Salmonella-enterica-subsp.-enterica-serovar-Livingstone]</td></tr>
<tr><td>MLQ35368.1</td><td>Type1.BREX</td><td><-MORC+HISKIN<-DNA_METHYLASE<-?||HIN-HTH-><-SIGMA+OB-Fold+AAA-ATPase+Lon-Protease-BrxL<-PglZ<-TOPRIM<-OLD-ABC<-N6-METHYLASE<-DUF6079*<-BrxB-BrxF-iSTAND<-BrxA-DNA-binding<-?<-?<-SAM-ME[illegible]</td><td>1213</td><td>Bacteria>Pseudomonadota>Gammaproteobacteria>Enterobacterales>Enterobacteriaceae>Salmonella</td><td>GCA_003910355.1</td><td>Salmonella_enterica_subsp._enterica</td><td>DRM12_04740</td><td>261903..265544</td><td>59201</td><td>BREX-system-P-loop-protein-BrxC-[Salmonella-enterica-subsp.-enterica]</td></tr>
<tr><td>MLX95744.1</td><td>Type1.BREX</td><td><-HEPN<-REase+iREase<-?||||YjhX_toxin->SAM-METHYLASE->?->?->BrxA-DNA-binding->BrxB-BrxF-iSTAND->DUF6079-p*-></td><td>669</td><td>Bacteria>Pseudomonadota>Gammaproteobacteria>Enterobacterales>Enterobacteriaceae>Salmonella</td><td>GCA_003913195.1</td><td>Salmonella_enterica_subsp._enterica</td><td>DSF42_09165</td><td>170394..172402</td><td>59201</td><td>BREX-system-P-loop-protein-BrxC-[Salmonella-enterica-subsp.-enterica]</td></tr>
<tr><td>AZT73891.1</td><td>Type1.BREX</td><td><-wHTH||wHTH+WYL+WCX-><-MarR-wHTH<-SIGMA-HTH+REase<-wHTH+SAD+HNH||?-><-ZnR||BrxA-DNA-binding->BrxB-BrxF-iSTAND->DUF6079*->N6-METHYLASE->OLD-ABC->TOPRIM->PglZ->SIGMA+OB-Fold+AAA-ATPase[illegible]</td><td>1204</td><td>Bacteria>Pseudomonadota>Gammaproteobacteria>Enterobacterales>Enterobacteriaceae>Salmonella</td><td>GCA_003994735.1</td><td>Salmonella_enterica_subsp._enterica_serovar_Waycross</td><td>ELZ91_19520</td><td>3951400..3955014</td><td>58101</td><td>BREX-system-P-loop-protein-BrxC-[Salmonella-enterica-subsp.-enterica-serovar-Waycross]</td></tr>
<tr><td>EAA2289696.1</td><td>Type1.BREX</td><td><-HEPN<-REase+iREase<-?||||YjhX_toxin->SAM-METHYLASE->?->?->BrxA-DNA-binding->BrxB-BrxF-iSTAND->DUF6079*-></td><td>1103</td><td>Bacteria>Pseudomonadota>Gammaproteobacteria>Enterobacterales>Enterobacteriaceae>Salmonella</td><td>GCA_004165775.1</td><td>Salmonella_enterica_subsp._enterica</td><td>DNI00_14480</td><td>60001..63308</td><td>59201</td><td>BREX-system-P-loop-protein-BrxC-[Salmonella-enterica-subsp.-enterica]</td></tr>
<tr><td>EAA3385003.1</td><td>Type1.BREX</td><td><-HEPN<-?||||YjhX_toxin->SAM-METHYLASE-><-?||?->?->BrxA-DNA-binding->BrxB-BrxF-iSTAND->DUF6079*->N6-METHYLASE->OLD-ABC->TOPRIM->PglZ->SIGMA+OB-Fold+AAA-ATPase+Lon-Protease-BrxL-><-REa[illegible]</td><td>1213</td><td>Bacteria>Pseudomonadota>Gammaproteobacteria>Enterobacterales>Enterobacteriaceae>Salmonella</td><td>GCA_004174495.1</td><td>Salmonella_enterica</td><td>DMV54_10710</td><td>131940..135581</td><td>28901</td><td>BREX-system-P-loop-protein-BrxC-[Salmonella-enterica]</td></tr>
<tr><td>EAA3988072.1</td><td>Type1.BREX</td><td>Arm-DNA-bind_3->NACHT+SF1-SF2-Helicase+HKD-DNase->REase-><-SIGMA+OB-Fold+AAA-ATPase+Lon-Protease-BrxL<-PglZ<-TOPRIM<-OLD-ABC<-N6-METHYLASE<-DUF6079*<-BrxB-BrxF-iSTAND<-BrxA-DNA-bindin[illegible]</td><td>1213</td><td>Bacteria>Pseudomonadota>Gammaproteobacteria>Enterobacterales>Enterobacteriaceae>Salmonella</td><td>GCA_004177075.1</td><td>Salmonella_enterica_subsp._enterica</td><td>DO557_10295</td><td>132678..136319</td><td>59201</td><td>BREX-system-P-loop-protein-BrxC-[Salmonella-enterica-subsp.-enterica]</td></tr>
<tr><td>EAA4435938.1</td><td>Type1.BREX</td><td>XIS-HTH->?->?-><-?<-ZnR||BrxA-DNA-binding->BrxB-BrxF-iSTAND->DUF6079*->ParB+HNH->N6-METHYLASE->HEPN->RVT->N6-METHYLASE-p->PglZ->SIGMA+OB-Fold+AAA-ATPase+Lon-Protease-BrxL-><-tRNA||?->[illegible]</td><td>1233</td><td>Bacteria>Pseudomonadota>Gammaproteobacteria>Enterobacterales>Enterobacteriaceae>Salmonella</td><td>GCA_004179155.1</td><td>Salmonella_enterica_subsp._salamae</td><td>DPE44_05505</td><td>71371..75072</td><td>59202</td><td>BREX-system-P-loop-protein-BrxC-[Salmonella-enterica-subsp.-salamae]</td></tr>
<tr><td>EAA4290852.1</td><td>Type1.BREX</td><td><-HEPN<-REase+iREase<-?||||YjhX_toxin->SAM-METHYLASE->?->?->BrxA-DNA-binding->BrxB-BrxF-iSTAND->DUF6079-p*-></td><td>989</td><td>Bacteria>Pseudomonadota>Gammaproteobacteria>Enterobacterales>Enterobacteriaceae>Salmonella</td><td>GCA_004180125.1</td><td>Salmonella_enterica_subsp._enterica</td><td>DO981_06670</td><td>47529..50496</td><td>59201</td><td>BREX-system-P-loop-protein-BrxC-[Salmonella-enterica-subsp.-enterica]</td></tr>
<tr><td>EAA4442641.1</td><td>Type1.BREX</td><td><-HEPN<-REase+iREase<-?||||YjhX_toxin->SAM-METHYLASE->?->?->BrxA-DNA-binding->BrxB-BrxF-iSTAND->DUF6079-p*-></td><td>833</td><td>Bacteria>Pseudomonadota>Gammaproteobacteria>Enterobacterales>Enterobacteriaceae>Salmonella</td><td>GCA_004180805.1</td><td>Salmonella_enterica_subsp._enterica</td><td>DPE76_16250</td><td>60221..62721</td><td>59201</td><td>BREX-system-P-loop-protein-BrxC-[Salmonella-enterica-subsp.-enterica]</td></tr>
<tr><td>EAA4605168.1</td><td>Type1.BREX</td><td><-HEPN<-REase+iREase<-?||||YjhX_toxin->SAM-METHYLASE-><-?||?->||BrxA-DNA-binding->BrxB-BrxF-iSTAND->DUF6079*->N6-METHYLASE->PglZ->SIGMA+OB-Fold+AAA-ATPase+Lon-Protease-BrxL-></td><td>1213</td><td>Bacteria>Pseudomonadota>Gammaproteobacteria>Enterobacterales>Enterobacteriaceae>Salmonella</td><td>GCA_004181295.1</td><td>Salmonella_enterica_subsp._enterica</td><td>DPJ21_07760</td><td>164877..168518</td><td>59201</td><td>BREX-system-P-loop-protein-BrxC-[Salmonella-enterica-subsp.-enterica]</td></tr>
<tr><td>EAA4842519.1</td><td>Type1.BREX</td><td><-HEPN<-REase+iREase<-?||||YjhX_toxin->SAM-METHYLASE->?->?->BrxA-DNA-binding->BrxB-BrxF-iSTAND->DUF6079*->OLD-ABC->TOPRIM->PglZ->SIGMA+OB-Fold+AAA-ATPase+Lon-Protease-BrxL-><-REase<-N[illegible]</td><td>1213</td><td>Bacteria>Pseudomonadota>Gammaproteobacteria>Enterobacterales>Enterobacteriaceae>Salmonella</td><td>GCA_004183005.1</td><td>Salmonella_enterica_subsp._enterica</td><td>DPP49_10705</td><td>60116..63757</td><td>59201</td><td>BREX-system-P-loop-protein-BrxC-[Salmonella-enterica-subsp.-enterica]</td></tr>
<tr><td>EAA5451231.1</td><td>Type1.BREX</td><td><-REase+iREase<-?||||YjhX_toxin->SAM-METHYLASE-><-?||?->?->BrxA-DNA-binding->BrxB-BrxF-iSTAND->DUF6079*->N6-METHYLASE->OLD-ABC->TOPRIM->PglZ->SIGMA+OB-Fold+AAA-ATPase+Lon-Protease-Brx[illegible]</td><td>1213</td><td>Bacteria>Pseudomonadota>Gammaproteobacteria>Enterobacterales>Enterobacteriaceae>Salmonella</td><td>GCA_004184965.1</td><td>Salmonella_enterica_subsp._enterica</td><td>DQC86_00800</td><td>160528..164169</td><td>59201</td><td>BREX-system-P-loop-protein-BrxC-[Salmonella-enterica-subsp.-enterica]</td></tr>
<tr><td>EAA5642506.1</td><td>Type1.BREX</td><td><-HEPN<-?||?->||YjhX_toxin->SAM-METHYLASE->?->?->BrxA-DNA-binding->BrxB-BrxF-iSTAND->DUF6079*->N6-METHYLASE->OLD-ABC->TOPRIM->PglZ->SIGMA+OB-Fold+AAA-ATPase+Lon-Protease-BrxL->?-><-RE[illegible]</td><td>1213</td><td>Bacteria>Pseudomonadota>Gammaproteobacteria>Enterobacterales>Enterobacteriaceae>Salmonella</td><td>GCA_004185395.1</td><td>Salmonella_enterica_subsp._enterica</td><td>DQJ71_10390</td><td>149103..152744</td><td>59201</td><td>BREX-system-P-loop-protein-BrxC-[Salmonella-enterica-subsp.-enterica]</td></tr>
<tr><td>EAA5580610.1</td><td>Type1.BREX</td><td><-HEPN<-?||?->||YjhX_toxin->SAM-METHYLASE->?->?->BrxA-DNA-binding->BrxB-BrxF-iSTAND->DUF6079*->N6-METHYLASE->OLD-ABC->TOPRIM->PglZ->SIGMA+OB-Fold+AAA-ATPase+Lon-Protease-BrxL->HKD-DNa[illegible]</td><td>1213</td><td>Bacteria>Pseudomonadota>Gammaproteobacteria>Enterobacterales>Enterobacteriaceae>Salmonella</td><td>GCA_004185805.1</td><td>Salmonella_enterica_subsp._enterica</td><td>DQD90_06840</td><td>58937..62578</td><td>59201</td><td>BREX-system-P-loop-protein-BrxC-[Salmonella-enterica-subsp.-enterica]</td></tr>
<tr><td>EAA6071870.1</td><td>Type1.BREX</td><td>YjhX_toxin->SAM-METHYLASE-><-?||?->?->BrxA-DNA-binding->BrxB-BrxF-iSTAND->DUF6079*->N6-METHYLASE->OLD-ABC->TOPRIM->PglZ->SIGMA+OB-Fold+AAA-ATPase+Lon-Protease-BrxL-><-REase<-Couple_hi[illegible]</td><td>1213</td><td>Bacteria>Pseudomonadota>Gammaproteobacteria>Enterobacterales>Enterobacteriaceae>Salmonella</td><td>GCA_004187475.1</td><td>Salmonella_enterica_subsp._enterica</td><td>DQZ25_06110</td><td>158264..161905</td><td>59201</td><td>BREX-system-P-loop-protein-BrxC-[Salmonella-enterica-subsp.-enterica]</td></tr>
<tr><td>EAA6168130.1</td><td>Type1.BREX</td><td>HTH_3->Couple_hipA+HipA_C->REase-><-SIGMA+OB-Fold+AAA-ATPase+Lon-Protease-BrxL<-PglZ<-TOPRIM<-OLD-ABC<-N6-METHYLASE<-DUF6079*<-BrxB-BrxF-iSTAND<-BrxA-DNA-binding<-?<-?||?-><-SAM-METHY[illegible]</td><td>1213</td><td>Bacteria>Pseudomonadota>Gammaproteobacteria>Enterobacterales>Enterobacteriaceae>Salmonella</td><td>GCA_004188635.1</td><td>Salmonella_enterica_subsp._enterica</td><td>DQZ68_14465</td><td>24635..28276</td><td>59201</td><td>BREX-system-P-loop-protein-BrxC-[Salmonella-enterica-subsp.-enterica]</td></tr>
<tr><td>EAA7005308.1</td><td>Type1.BREX</td><td><-HEPN<-?||||YjhX_toxin->SAM-METHYLASE-><-?||?->?->BrxA-DNA-binding->BrxB-BrxF-iSTAND->DUF6079*->N6-METHYLASE-></td><td>1213</td><td>Bacteria>Pseudomonadota>Gammaproteobacteria>Enterobacterales>Enterobacteriaceae>Salmonella</td><td>GCA_004191915.1</td><td>Salmonella_enterica</td><td>AVW22_12590</td><td>47439..51080</td><td>28901</td><td>BREX-system-P-loop-protein-BrxC-[Salmonella-enterica]</td></tr>
<tr><td>EAA7109615.1</td><td>Type1.BREX</td><td>YjhX_toxin->SAM-METHYLASE-><-?||?->?->BrxA-DNA-binding->BrxB-BrxF-iSTAND->DUF6079*->N6-METHYLASE->OLD-ABC->TOPRIM->PglZ->SIGMA+OB-Fold+AAA-ATPase+Lon-Protease-BrxL-><-REase</td><td>1213</td><td>Bacteria>Pseudomonadota>Gammaproteobacteria>Enterobacterales>Enterobacteriaceae>Salmonella</td><td>GCA_004192185.1</td><td>Salmonella_enterica_subsp._enterica</td><td>DSE94_16630</td><td>52127..55768</td><td>59201</td><td>BREX-system-P-loop-protein-BrxC-[Salmonella-enterica-subsp.-enterica]</td></tr>
<tr><td>EAA7310017.1</td><td>Type1.BREX</td><td><-REase+iREase<-?||||YjhX_toxin->SAM-METHYLASE-><-?||?->?->BrxA-DNA-binding->BrxB-BrxF-iSTAND->DUF6079*->N6-METHYLASE->OLD-ABC->TOPRIM->PglZ->SIGMA+OB-Fold+AAA-ATPase+Lon-Protease-Brx[illegible]</td><td>1213</td><td>Bacteria>Pseudomonadota>Gammaproteobacteria>Enterobacterales>Enterobacteriaceae>Salmonella</td><td>GCA_004192645.1</td><td>Salmonella_enterica_subsp._enterica</td><td>TL61_10425</td><td>156005..159646</td><td>59201</td><td>BREX-system-P-loop-protein-BrxC-[Salmonella-enterica-subsp.-enterica]</td></tr>
<tr><td>EAA7604354.1</td><td>Type1.BREX</td><td><-HEPN<-?||||YjhX_toxin->SAM-METHYLASE->?->?->BrxA-DNA-binding->BrxB-BrxF-iSTAND->DUF6079*->N6-METHYLASE->OLD-ABC->TOPRIM->PglZ->SIGMA+OB-Fold+AAA-ATPase+Lon-Protease-BrxL-></td><td>1213</td><td>Bacteria>Pseudomonadota>Gammaproteobacteria>Enterobacterales>Enterobacteriaceae>Salmonella</td><td>GCA_004218255.1</td><td>Salmonella_enterica</td><td>ADQ66_03555</td><td>130477..134118</td><td>28901</td><td>BREX-system-P-loop-protein-BrxC-[Salmonella-enterica]</td></tr>
<tr><td>EAA7639151.1</td><td>Type1.BREX</td><td>BrxB-BrxF-iSTAND->ACET->||?-><-Phage_integrase||BrxA-DNA-binding->BrxB-BrxF-iSTAND->DUF6079*->N6-METHYLASE->PglZ->SIGMA+OB-Fold+AAA-ATPase+Lon-Protease-BrxL-><-AAA+HTH<-?<-XIS-HTH</td><td>1213</td><td>Bacteria>Pseudomonadota>Gammaproteobacteria>Enterobacterales>Enterobacteriaceae>Salmonella</td><td>GCA_004218325.1</td><td>Salmonella_enterica_subsp._enterica</td><td>AAV76_19225</td><td>212134..215775</td><td>59201</td><td>BREX-system-P-loop-protein-BrxC-[Salmonella-enterica-subsp.-enterica]</td></tr>
<tr><td>EAA8317754.1</td><td>Type1.BREX</td><td><-HEPN<-?||?->||YjhX_toxin->SAM-METHYLASE->?->?->BrxA-DNA-binding->BrxB-BrxF-iSTAND->DUF6079*->N6-METHYLASE->OLD-ABC->TOPRIM->PglZ->SIGMA+OB-Fold+AAA-ATPase+Lon-Protease-BrxL->?-><-RE[illegible]</td><td>1213</td><td>Bacteria>Pseudomonadota>Gammaproteobacteria>Enterobacterales>Enterobacteriaceae>Salmonella</td><td>GCA_004221325.1</td><td>Salmonella_enterica</td><td>AOS09_04965</td><td>162610..166251</td><td>28901</td><td>BREX-system-P-loop-protein-BrxC-[Salmonella-enterica]</td></tr>
<tr><td>EAA8523361.1</td><td>Type1.BREX</td><td><-HEPN<-?||||YjhX_toxin->SAM-METHYLASE-><-?||?->?->BrxA-DNA-binding->BrxB-BrxF-iSTAND->DUF6079*->N6-METHYLASE->OLD-ABC->TOPRIM->PglZ->SIGMA+OB-Fold+AAA-ATPase+Lon-Protease-BrxL-><-REa[illegible]</td><td>1213</td><td>Bacteria>Pseudomonadota>Gammaproteobacteria>Enterobacterales>Enterobacteriaceae>Salmonella</td><td>GCA_004222265.1</td><td>Salmonella_enterica_subsp._enterica</td><td>ADQ53_00815</td><td>161051..164692</td><td>59201</td><td>BREX-system-P-loop-protein-BrxC-[Salmonella-enterica-subsp.-enterica]</td></tr>
<tr><td>EAA8875705.1</td><td>Type1.BREX</td><td><-HEPN<-?||YjhX_toxin->SAM-METHYLASE->?-><-wHTH+SAD+HNH||URI-DNase->BrxA-DNA-binding->BrxB-BrxF-iSTAND->DUF6079*->N6-METHYLASE->OLD-ABC->TOPRIM->PglZ->SIGMA+OB-Fold+AAA-ATPase+Lon-Pro[illegible]</td><td>1213</td><td>Bacteria>Pseudomonadota>Gammaproteobacteria>Enterobacterales>Enterobacteriaceae>Salmonella</td><td>GCA_004223005.1</td><td>Salmonella_enterica</td><td>CIU45_09625</td><td>381857..385498</td><td>28901</td><td>BREX-system-P-loop-protein-BrxC-[Salmonella-enterica]</td></tr>
<tr><td>EAA8954969.1</td><td>Type1.BREX</td><td>YjhX_toxin->SAM-METHYLASE-><-?||?->?->BrxA-DNA-binding->BrxB-BrxF-iSTAND->DUF6079*->N6-METHYLASE->PglZ->SIGMA+OB-Fold+AAA-ATPase+Lon-Protease-BrxL->||PIN-><-HTH_21+rve+rve_3</td><td>1213</td><td>Bacteria>Pseudomonadota>Gammaproteobacteria>Enterobacterales>Enterobacteriaceae>Salmonella</td><td>GCA_004224545.1</td><td>Salmonella_enterica</td><td>CBJ10_12720</td><td>157324..160965</td><td>28901</td><td>BREX-system-P-loop-protein-BrxC-[Salmonella-enterica]</td></tr>
<tr><td>EAA9319736.1</td><td>Type1.BREX</td><td>YjhX_toxin->SAM-METHYLASE-><-?||?->?->BrxA-DNA-binding->BrxB-BrxF-iSTAND->DUF6079*->N6-METHYLASE->OLD-ABC->TOPRIM->PglZ->SIGMA+OB-Fold+AAA-ATPase+Lon-Protease-BrxL-><-REase<-Couple_hi[illegible]</td><td>1213</td><td>Bacteria>Pseudomonadota>Gammaproteobacteria>Enterobacterales>Enterobacteriaceae>Salmonella</td><td>GCA_004226405.1</td><td>Salmonella_enterica</td><td>CRQ39_09900</td><td>167436..171077</td><td>28901</td><td>BREX-system-P-loop-protein-BrxC-[Salmonella-enterica]</td></tr>
<tr><td>EAA9627982.1</td><td>Type1.BREX</td><td><-HEPN<-REase+iREase<-?||||YjhX_toxin->SAM-METHYLASE->?->?->BrxA-DNA-binding->BrxB-BrxF-iSTAND->DUF6079*->N6-METHYLASE->OLD-ABC->TOPRIM->PglZ->SIGMA+OB-Fold+AAA-ATPase+Lon-Protease-Br[illegible]</td><td>1213</td><td>Bacteria>Pseudomonadota>Gammaproteobacteria>Enterobacterales>Enterobacteriaceae>Salmonella</td><td>GCA_004227565.1</td><td>Salmonella_enterica_subsp._enterica</td><td>DVO1_07255</td><td>170919..174560</td><td>59201</td><td>BREX-system-P-loop-protein-BrxC-[Salmonella-enterica-subsp.-enterica]</td></tr>
<tr><td>EAB1768963.1</td><td>Type1.BREX</td><td><-HEPN<-REase+iREase<-?||YjhX_toxin->SAM-METHYLASE->?->BrxA-DNA-binding->BrxB-BrxF-iSTAND->DUF6079*->N6-METHYLASE->OLD-ABC->TOPRIM->PglZ->SIGMA+OB-Fold+AAA-ATPase+Lon-Protease-BrxL-><[illegible]</td><td>1213</td><td>Bacteria>Pseudomonadota>Gammaproteobacteria>Enterobacterales>Enterobacteriaceae>Salmonella</td><td>GCA_004236005.1</td><td>Salmonella_enterica</td><td>EIS13_11220</td><td>18269..21910</td><td>28901</td><td>BREX-system-P-loop-protein-BrxC-[Salmonella-enterica]</td></tr>
<tr><td>EAB2294236.1</td><td>Type1.BREX</td><td><-HEPN<-?||?->YjhX_toxin->SAM-METHYLASE->?->||BrxA-DNA-binding->BrxB-BrxF-iSTAND->DUF6079*->N6-METHYLASE-p-></td><td>1213</td><td>Bacteria>Pseudomonadota>Gammaproteobacteria>Enterobacterales>Enterobacteriaceae>Salmonella</td><td>GCA_004236865.1</td><td>Salmonella_enterica</td><td>EHO06_05115</td><td>132073..135714</td><td>28901</td><td>BREX-system-P-loop-protein-BrxC-[Salmonella-enterica]</td></tr>
<tr><td>EAB2013162.1</td><td>Type1.BREX</td><td>XIS-HTH->?->?-><-ZnR||?->BrxA-DNA-binding->BrxB-BrxF-iSTAND->DUF6079*->N6-METHYLASE->OLD-ABC->TOPRIM->PglZ->SIGMA+OB-Fold+AAA-ATPase+Lon-Protease-BrxL-></td><td>1204</td><td>Bacteria>Pseudomonadota>Gammaproteobacteria>Enterobacterales>Enterobacteriaceae>Salmonella</td><td>GCA_004236885.1</td><td>Salmonella_enterica</td><td>D7J64_23090</td><td>5108..8722</td><td>28901</td><td>BREX-system-P-loop-protein-BrxC-[Salmonella-enterica]</td></tr>
<tr><td>EAB2831578.1</td><td>Type1.BREX</td><td><-REase+iREase<-?||?->||YjhX_toxin->SAM-METHYLASE->?->?->BrxA-DNA-binding->BrxB-BrxF-iSTAND->DUF6079*->N6-METHYLASE->OLD-ABC->TOPRIM->PglZ->SIGMA+OB-Fold+AAA-ATPase+Lon-Protease-BrxL-[illegible]</td><td>1213</td><td>Bacteria>Pseudomonadota>Gammaproteobacteria>Enterobacterales>Enterobacteriaceae>Salmonella</td><td>GCA_004240605.1</td><td>Salmonella_enterica</td><td>EAV48_05270</td><td>157994..161635</td><td>28901</td><td>BREX-system-P-loop-protein-BrxC-[Salmonella-enterica]</td></tr>
<tr><td>EAB2910642.1</td><td>Type1.BREX</td><td>Arm-DNA-bind_3->NACHT+SF1-SF2-Helicase+HKD-DNase->REase-><-SIGMA+OB-Fold+AAA-ATPase+Lon-Protease-BrxL<-PglZ<-TOPRIM<-OLD-ABC<-N6-METHYLASE<-DUF6079*<-BrxB-BrxF-iSTAND<-BrxA-DNA-bindin[illegible]</td><td>1213</td><td>Bacteria>Pseudomonadota>Gammaproteobacteria>Enterobacterales>Enterobacteriaceae>Salmonella</td><td>GCA_004240965.1</td><td>Salmonella_enterica</td><td>EHO38_04055</td><td>132608..136249</td><td>28901</td><td>BREX-system-P-loop-protein-BrxC-[Salmonella-enterica]</td></tr>
<tr><td>EAB6380447.1</td><td>Type1.BREX</td><td>HTH_3->Couple_hipA+HipA_C->REase-><-SIGMA+OB-Fold+AAA-ATPase+Lon-Protease-BrxL<-PglZ<-TOPRIM<-OLD-ABC<-N6-METHYLASE<-DUF6079*<-BrxB-BrxF-iSTAND<-BrxA-DNA-binding<-?<-?||?-><-SAM-METHY[illegible]</td><td>1213</td><td>Bacteria>Pseudomonadota>Gammaproteobacteria>Enterobacterales>Enterobacteriaceae>Salmonella</td><td>GCA_004255205.1</td><td>Salmonella_enterica_subsp._enterica</td><td>CWG97_20720</td><td>30069..33710</td><td>59201</td><td>BREX-system-P-loop-protein-BrxC-[Salmonella-enterica-subsp.-enterica]</td></tr>
<tr><td>EAB6416573.1</td><td>Type1.BREX</td><td>YjhX_toxin->SAM-METHYLASE-><-?||?->||?->BrxA-DNA-binding->BrxB-BrxF-iSTAND->DUF6079*->N6-METHYLASE->OLD-ABC->TOPRIM->PglZ->SIGMA+OB-Fold+AAA-ATPase+Lon-Protease-BrxL-><-REase<-NACHT+S[illegible]</td><td>1213</td><td>Bacteria>Pseudomonadota>Gammaproteobacteria>Enterobacterales>Enterobacteriaceae>Salmonella</td><td>GCA_004256685.1</td><td>Salmonella_enterica_subsp._enterica</td><td>D6S13_10910</td><td>52351..55992</td><td>59201</td><td>BREX-system-P-loop-protein-BrxC-[Salmonella-enterica-subsp.-enterica]</td></tr>
<tr><td>EAB6711531.1</td><td>Type1.BREX</td><td>HTH_3->Couple_hipA+HipA_C->REase-><-SIGMA+OB-Fold+AAA-ATPase+Lon-Protease-BrxL<-PglZ<-N6-METHYLASE<-DUF6079*<-BrxB-BrxF-iSTAND<-BrxA-DNA-binding<-?<-?||?-><-SAM-METHYLASE<-YjhX_toxin|[illegible]</td><td>1213</td><td>Bacteria>Pseudomonadota>Gammaproteobacteria>Enterobacterales>Enterobacteriaceae>Salmonella</td><td>GCA_004258665.1</td><td>Salmonella_enterica_subsp._enterica</td><td>DU754_17280</td><td>25213..28854</td><td>59201</td><td>BREX-system-P-loop-protein-BrxC-[Salmonella-enterica-subsp.-enterica]</td></tr>
<tr><td>EAB6843525.1</td><td>Type1.BREX</td><td><-GntR-HTH++TrmB+ZDNA-HTH||<-?<-HEPN<-?||YjhX_toxin->METHYLASE->?->BrxA-DNA-binding->BrxB-BrxF-iSTAND->DUF6079*->N6-METHYLASE->OLD-ABC->TOPRIM->PglZ->SIGMA+OB-Fold+AAA-ATPase+Lon-Prot[illegible]</td><td>1213</td><td>Bacteria>Pseudomonadota>Gammaproteobacteria>Enterobacterales>Enterobacteriaceae>Salmonella</td><td>GCA_004258725.1</td><td>Salmonella_enterica_subsp._salamae</td><td>XT66_14185</td><td>50997..54638</td><td>59202</td><td>BREX-system-P-loop-protein-BrxC-[Salmonella-enterica-subsp.-salamae]</td></tr>
<tr><td>EAB7363624.1</td><td>Type1.BREX</td><td><-HEPN<-REase+iREase<-?||YjhX_toxin->SAM-METHYLASE->?->BrxA-DNA-binding->BrxB-BrxF-iSTAND->DUF6079*->N6-METHYLASE->OLD-ABC->TOPRIM->PglZ->SIGMA+OB-Fold+AAA-ATPase+Lon-Protease-BrxL-><[illegible]</td><td>1213</td><td>Bacteria>Pseudomonadota>Gammaproteobacteria>Enterobacterales>Enterobacteriaceae>Salmonella</td><td>GCA_004260605.1</td><td>Salmonella_enterica_subsp._enterica</td><td>EBO89_06925</td><td>58191..61832</td><td>59201</td><td>BREX-system-P-loop-protein-BrxC-[Salmonella-enterica-subsp.-enterica]</td></tr>
<tr><td>EAB9078796.1</td><td>Type1.BREX</td><td>YjhX_toxin->SAM-METHYLASE-><-?||?->?->BrxA-DNA-binding->BrxB-BrxF-iSTAND->DUF6079*->N6-METHYLASE->OLD-ABC->TOPRIM->PglZ->SIGMA+OB-Fold+AAA-ATPase+Lon-Protease-BrxL->HKD-DNase+OGG-like[illegible]</td><td>1213</td><td>Bacteria>Pseudomonadota>Gammaproteobacteria>Enterobacterales>Enterobacteriaceae>Salmonella</td><td>GCA_004265565.1</td><td>Salmonella_enterica_subsp._enterica</td><td>AAL03_01510</td><td>52372..56013</td><td>59201</td><td>BREX-system-P-loop-protein-BrxC-[Salmonella-enterica-subsp.-enterica]</td></tr>
<tr><td>EAB9374475.1</td><td>Type1.BREX</td><td><-REase+iREase<-?||?->||YjhX_toxin->SAM-METHYLASE->?->?->BrxA-DNA-binding->BrxB-BrxF-iSTAND->DUF6079*->N6-METHYLASE->OLD-ABC->TOPRIM->PglZ->SIGMA+OB-Fold+AAA-ATPase+Lon-Protease-BrxL-[illegible]</td><td>1213</td><td>Bacteria>Pseudomonadota>Gammaproteobacteria>Enterobacterales>Enterobacteriaceae>Salmonella</td><td>GCA_004270415.1</td><td>Salmonella_enterica_subsp._enterica</td><td>D7N03_15200</td><td>60704..64345</td><td>59201</td><td>BREX-system-P-loop-protein-BrxC-[Salmonella-enterica-subsp.-enterica]</td></tr>
<tr><td>EAC1009595.1</td><td>Type1.BREX</td><td>YjhX_toxin->SAM-METHYLASE-><-?||?->?->BrxA-DNA-binding->BrxB-BrxF-iSTAND->DUF6079*->N6-METHYLASE->OLD-ABC->TOPRIM->PglZ->SIGMA+OB-Fold+AAA-ATPase+Lon-Protease-BrxL->?-><-REase<-NACHT+[illegible]</td><td>1213</td><td>Bacteria>Pseudomonadota>Gammaproteobacteria>Enterobacterales>Enterobacteriaceae>Salmonella</td><td>GCA_004275715.1</td><td>Salmonella_enterica_subsp._enterica</td><td>YR14_13800</td><td>150440..154081</td><td>59201</td><td>BREX-system-P-loop-protein-BrxC-[Salmonella-enterica-subsp.-enterica]</td></tr>
<tr><td>EAC1236045.1</td><td>Type1.BREX</td><td><-REase+iREase<-?||?->||YjhX_toxin->SAM-METHYLASE->?->?->BrxA-DNA-binding->BrxB-BrxF-iSTAND->DUF6079*->N6-METHYLASE->OLD-ABC->TOPRIM->PglZ->SIGMA+OB-Fold+AAA-ATPase+Lon-Protease-BrxL-[illegible]</td><td>1213</td><td>Bacteria>Pseudomonadota>Gammaproteobacteria>Enterobacterales>Enterobacteriaceae>Salmonella</td><td>GCA_004276135.1</td><td>Salmonella_enterica_subsp._enterica</td><td>DTG84_00810</td><td>157931..161572</td><td>59201</td><td>BREX-system-P-loop-protein-BrxC-[Salmonella-enterica-subsp.-enterica]</td></tr>
<tr><td>EAM1956840.1</td><td>Type1.BREX</td><td><-HEPN<-REase+iREase<-?||||YjhX_toxin->?->?->BrxA-DNA-binding->BrxB-BrxF-iSTAND->DUF6079*->N6-METHYLASE-></td><td>1213</td><td>Bacteria>Pseudomonadota>Gammaproteobacteria>Enterobacterales>Enterobacteriaceae>Salmonella</td><td>GCA_005406505.1</td><td>Salmonella_enterica</td><td>D7U14_11535</td><td>170827..174468</td><td>28901</td><td>BREX-system-P-loop-protein-BrxC-[Salmonella-enterica]</td></tr>
<tr><td>EAM3013482.1</td><td>Type1.BREX</td><td><-HEPN<-?||?->||YjhX_toxin->SAM-METHYLASE->?->?->BrxA-DNA-binding->BrxB-BrxF-iSTAND->DUF6079*->N6-METHYLASE->OLD-ABC->TOPRIM->PglZ->SIGMA+OB-Fold+AAA-ATPase+Lon-Protease-BrxL->HKD-DNa[illegible]</td><td>1213</td><td>Bacteria>Pseudomonadota>Gammaproteobacteria>Enterobacterales>Enterobacteriaceae>Salmonella</td><td>GCA_005412095.1</td><td>Salmonella_enterica</td><td>CBJ93_01865</td><td>394707..398348</td><td>28901</td><td>BREX-system-P-loop-protein-BrxC-[Salmonella-enterica]</td></tr>
<tr><td>EAM3141100.1</td><td>Type1.BREX</td><td>Arm-DNA-bind_3->NACHT+SF1-SF2-Helicase+HKD-DNase->REase-><-HKD-DNase+OGG-like<-SIGMA+OB-Fold+AAA-ATPase+Lon-Protease-BrxL<-PglZ<-OLD-ABC<-N6-METHYLASE<-DUF6079*<-BrxB-BrxF-iSTAND<-Brx[illegible]</td><td>1204</td><td>Bacteria>Pseudomonadota>Gammaproteobacteria>Enterobacterales>Enterobacteriaceae>Salmonella</td><td>GCA_005412435.1</td><td>Salmonella_enterica</td><td>DMT78_01470</td><td>307802..311416</td><td>28901</td><td>BREX-system-P-loop-protein-BrxC-[Salmonella-enterica]</td></tr>
<tr><td>EAM5567470.1</td><td>Type1.BREX</td><td><-SIGMA+OB-Fold+AAA-ATPase+Lon-Protease-BrxL<-PglZ<-TOPRIM<-OLD-ABC<-N6-METHYLASE<-DUF6079*<-BrxB-BrxF-iSTAND<-BrxA-DNA-binding</td><td>1213</td><td>Bacteria>Pseudomonadota>Gammaproteobacteria>Enterobacterales>Enterobacteriaceae>Salmonella</td><td>GCA_005423155.1</td><td>Salmonella_enterica</td><td>EK556_21780</td><td>11069..14710</td><td>28901</td><td>BREX-system-P-loop-protein-BrxC-[Salmonella-enterica]</td></tr>
<tr><td>EAM6081349.1</td><td>Type1.BREX</td><td>Arm-DNA-bind_3->NACHT+SF1-SF2-Helicase+HKD-DNase->REase-><-?<-SIGMA+OB-Fold+AAA-ATPase+Lon-Protease-BrxL<-PglZ<-TOPRIM<-OLD-ABC<-N6-METHYLASE<-DUF6079*<-BrxB-BrxF-iSTAND<-BrxA-DNA-bin[illegible]</td><td>1213</td><td>Bacteria>Pseudomonadota>Gammaproteobacteria>Enterobacterales>Enterobacteriaceae>Salmonella</td><td>GCA_005424535.1</td><td>Salmonella_enterica</td><td>ERR94_15350</td><td>17401..21042</td><td>28901</td><td>BREX-system-P-loop-protein-BrxC-[Salmonella-enterica]</td></tr>
<tr><td>EAM5877058.1</td><td>Type1.BREX</td><td>Arm-DNA-bind_3->NACHT+SF1-SF2-Helicase+HKD-DNase->REase-><-SIGMA+OB-Fold+AAA-ATPase+Lon-Protease-BrxL<-PglZ<-TOPRIM<-OLD-ABC<-N6-METHYLASE<-DUF6079*<-BrxB-BrxF-iSTAND<-BrxA-DNA-bindin[illegible]</td><td>1213</td><td>Bacteria>Pseudomonadota>Gammaproteobacteria>Enterobacterales>Enterobacteriaceae>Salmonella</td><td>GCA_005424775.1</td><td>Salmonella_enterica</td><td>EPF42_07745</td><td>346357..349998</td><td>28901</td><td>BREX-system-P-loop-protein-BrxC-[Salmonella-enterica]</td></tr>
<tr><td>EAM9976205.1</td><td>Type1.BREX</td><td><-HEPN<-REase+iREase<-?||YjhX_toxin->SAM-METHYLASE->?->BrxA-DNA-binding->BrxB-BrxF-iSTAND->DUF6079*->N6-METHYLASE-p-></td><td>1213</td><td>Bacteria>Pseudomonadota>Gammaproteobacteria>Enterobacterales>Enterobacteriaceae>Salmonella</td><td>GCA_005442735.1</td><td>Salmonella_enterica</td><td>EU643_14555</td><td>35351..38992</td><td>28901</td><td>BREX-system-P-loop-protein-BrxC-[Salmonella-enterica]</td></tr>
<tr><td>EAN1187097.1</td><td>Type1.BREX</td><td><-HEPN<-REase+iREase<-?||||YjhX_toxin->SAM-METHYLASE->?->?->BrxA-DNA-binding->BrxB-BrxF-iSTAND->DUF6079-p*-></td><td>797</td><td>Bacteria>Pseudomonadota>Gammaproteobacteria>Enterobacterales>Enterobacteriaceae>Salmonella</td><td>GCA_005448045.1</td><td>Salmonella_enterica</td><td>D7G02_16655</td><td>60001..62392</td><td>28901</td><td>BREX-system-P-loop-protein-BrxC-[Salmonella-enterica]</td></tr>
<tr><td>EAN9691870.1</td><td>Type1.BREX</td><td><-HEPN<-?||?->YjhX_toxin->SAM-METHYLASE->?->BrxA-DNA-binding->BrxB-BrxF-iSTAND->DUF6079*->N6-METHYLASE-></td><td>1213</td><td>Bacteria>Pseudomonadota>Gammaproteobacteria>Enterobacterales>Enterobacteriaceae>Salmonella</td><td>GCA_005483095.1</td><td>Salmonella_enterica</td><td>ETI19_18710</td><td>73016..76657</td><td>28901</td><td>BREX-system-P-loop-protein-BrxC-[Salmonella-enterica]</td></tr>
<tr><td>EAO1947720.1</td><td>Type1.BREX</td><td><-HEPN<-REase+iREase<-?||YjhX_toxin->SAM-METHYLASE->?->BrxA-DNA-binding->BrxB-BrxF-iSTAND->DUF6079*->N6-METHYLASE->OLD-ABC->TOPRIM->PglZ->SIGMA+OB-Fold+AAA-ATPase+Lon-Protease-BrxL-><[illegible]</td><td>1213</td><td>Bacteria>Pseudomonadota>Gammaproteobacteria>Enterobacterales>Enterobacteriaceae>Salmonella</td><td>GCA_005492965.1</td><td>Salmonella_enterica</td><td>E2X64_08235</td><td>25607..29248</td><td>28901</td><td>BREX-system-P-loop-protein-BrxC-[Salmonella-enterica]</td></tr>
<tr><td>EAO2684614.1</td><td>Type1.BREX</td><td><-GntR-HTH++TrmB+ZDNA-HTH||<-?<-HEPN<-?||YjhX_toxin->SAM-METHYLASE->?->||BrxA-DNA-binding->BrxB-BrxF-iSTAND->DUF6079*->N6-METHYLASE->OLD-ABC->TOPRIM->PglZ->SIGMA+OB-Fold+AAA-ATPase+Lo[illegible]</td><td>1213</td><td>Bacteria>Pseudomonadota>Gammaproteobacteria>Enterobacterales>Enterobacteriaceae>Salmonella</td><td>GCA_005496785.1</td><td>Salmonella_enterica</td><td>E2X65_05650</td><td>67584..71225</td><td>28901</td><td>BREX-system-P-loop-protein-BrxC-[Salmonella-enterica]</td></tr>
<tr><td>EAO4363249.1</td><td>Type1.BREX</td><td><-HEPN<-REase+iREase<-?||?->YjhX_toxin->SAM-METHYLASE->?->||BrxA-DNA-binding->BrxB-BrxF-iSTAND->DUF6079*->N6-METHYLASE->OLD-ABC->TOPRIM->PglZ->SIGMA+OB-Fold+AAA-ATPase+Lon-Protease-Br[illegible]</td><td>1213</td><td>Bacteria>Pseudomonadota>Gammaproteobacteria>Enterobacterales>Enterobacteriaceae>Salmonella</td><td>GCA_005505735.1</td><td>Salmonella_enterica</td><td>E5197_09130</td><td>175421..179062</td><td>28901</td><td>BREX-system-P-loop-protein-BrxC-[Salmonella-enterica]</td></tr>
<tr><td>EAO8797691.1</td><td>Type1.BREX</td><td><-HEPN<-REase+iREase<-?||YjhX_toxin->SAM-METHYLASE->?->BrxA-DNA-binding->BrxB-BrxF-iSTAND->DUF6079*->N6-METHYLASE->OLD-ABC->TOPRIM->PglZ->SIGMA+OB-Fold+AAA-ATPase+Lon-Protease-BrxL-><[illegible]</td><td>1213</td><td>Bacteria>Pseudomonadota>Gammaproteobacteria>Enterobacterales>Enterobacteriaceae>Salmonella</td><td>GCA_005530015.1</td><td>Salmonella_enterica</td><td>E0391_14535</td><td>25872..29513</td><td>28901</td><td>BREX-system-P-loop-protein-BrxC-[Salmonella-enterica]</td></tr>
<tr><td>EGG4120336.1</td><td>Type1.BREX</td><td><-HEPN<-REase+iREase<-?||YjhX_toxin->SAM-METHYLASE->?->BrxA-DNA-binding->BrxB-BrxF-iSTAND->DUF6079*->N6-METHYLASE->OLD-ABC->TOPRIM->PglZ->SIGMA+OB-Fold+AAA-ATPase+Lon-Protease-BrxL-><[illegible]</td><td>1213</td><td>Bacteria>Pseudomonadota>Gammaproteobacteria>Enterobacterales>Enterobacteriaceae>Salmonella</td><td>GCA_005532775.2</td><td>Salmonella_enterica</td><td>SI35_20740</td><td>157325..160966</td><td>28901</td><td>BREX-system-P-loop-protein-BrxC-[Salmonella-enterica]</td></tr>
<tr><td>EAP0128501.1</td><td>Type1.BREX</td><td>YjhX_toxin->SAM-METHYLASE-><-?||?->?->BrxA-DNA-binding->BrxB-BrxF-iSTAND->DUF6079*->N6-METHYLASE->OLD-ABC->TOPRIM->PglZ->SIGMA+OB-Fold+AAA-ATPase+Lon-Protease-BrxL->?-><-REase<-NACHT+[illegible]</td><td>1213</td><td>Bacteria>Pseudomonadota>Gammaproteobacteria>Enterobacterales>Enterobacteriaceae>Salmonella</td><td>GCA_005535515.1</td><td>Salmonella_enterica</td><td>DKP73_00815</td><td>161390..165031</td><td>28901</td><td>BREX-system-P-loop-protein-BrxC-[Salmonella-enterica]</td></tr>
<tr><td>EAP0644643.1</td><td>Type1.BREX</td><td>Arm-DNA-bind_3->NACHT+SF1-SF2-Helicase+HKD-DNase->REase-><-SIGMA+OB-Fold+AAA-ATPase+Lon-Protease-BrxL<-PglZ<-TOPRIM<-OLD-ABC<-N6-METHYLASE<-DUF6079*<-BrxB-BrxF-iSTAND<-BrxA-DNA-bindin[illegible]</td><td>1213</td><td>Bacteria>Pseudomonadota>Gammaproteobacteria>Enterobacterales>Enterobacteriaceae>Salmonella</td><td>GCA_005538295.1</td><td>Salmonella_enterica</td><td>C6277_11605</td><td>108063..111704</td><td>28901</td><td>BREX-system-P-loop-protein-BrxC-[Salmonella-enterica]</td></tr>
<tr><td>EAP5601609.1</td><td>Type1.BREX</td><td>wHTH+WYL+WCX-><-MarR-wHTH<-SIGMA-HTH+REase<-wHTH+SAD+HNH||?-><-?<-ZnR||BrxA-DNA-binding->BrxB-BrxF-iSTAND->DUF6079*->N6-METHYLASE-></td><td>1204</td><td>Bacteria>Pseudomonadota>Gammaproteobacteria>Enterobacterales>Enterobacteriaceae>Salmonella</td><td>GCA_005559495.1</td><td>Salmonella_enterica</td><td>EHT80_06400</td><td>135199..138813</td><td>28901</td><td>BREX-system-P-loop-protein-BrxC-[Salmonella-enterica]</td></tr>
<tr><td>EAP9950624.1</td><td>Type1.BREX</td><td>XIS-HTH->?->?-><-ZnR||?->BrxA-DNA-binding->BrxB-BrxF-iSTAND->DUF6079*->N6-METHYLASE->OLD-ABC->TOPRIM->PglZ->SIGMA+OB-Fold+AAA-ATPase+Lon-Protease-BrxL-></td><td>1204</td><td>Bacteria>Pseudomonadota>Gammaproteobacteria>Enterobacterales>Enterobacteriaceae>Salmonella</td><td>GCA_005576875.1</td><td>Salmonella_enterica</td><td>D3G27_06335</td><td>176089..179703</td><td>28901</td><td>BREX-system-P-loop-protein-BrxC-[Salmonella-enterica]</td></tr>
<tr><td>EAQ0497027.1</td><td>Type1.BREX</td><td><-HEPN<-REase+iREase<-?||YjhX_toxin->SAM-METHYLASE->?->BrxA-DNA-binding->BrxB-BrxF-iSTAND->DUF6079*->N6-METHYLASE->OLD-ABC->TOPRIM->PglZ->SIGMA+OB-Fold+AAA-ATPase+Lon-Protease-BrxL-><[illegible]</td><td>1213</td><td>Bacteria>Pseudomonadota>Gammaproteobacteria>Enterobacterales>Enterobacteriaceae>Salmonella</td><td>GCA_005582475.1</td><td>Salmonella_enterica</td><td>EW683_07755</td><td>36904..40545</td><td>28901</td><td>BREX-system-P-loop-protein-BrxC-[Salmonella-enterica]</td></tr>
<tr><td>EAQ5906792.1</td><td>Type1.BREX</td><td><-HEPN<-REase+iREase<-?||YjhX_toxin->SAM-METHYLASE->?->BrxA-DNA-binding->BrxB-BrxF-iSTAND->DUF6079*->N6-METHYLASE-></td><td>1213</td><td>Bacteria>Pseudomonadota>Gammaproteobacteria>Enterobacterales>Enterobacteriaceae>Salmonella</td><td>GCA_005605705.1</td><td>Salmonella_enterica</td><td>FEF66_01945</td><td>118005..121646</td><td>28901</td><td>BREX-system-P-loop-protein-BrxC-[Salmonella-enterica]</td></tr>
<tr><td>EAQ6130543.1</td><td>Type1.BREX</td><td><-HEPN<-?||||YjhX_toxin->SAM-METHYLASE-><-?||?->?->BrxA-DNA-binding->BrxB-BrxF-iSTAND->DUF6079*->N6-METHYLASE->OLD-ABC->TOPRIM->PglZ->SIGMA+OB-Fold+AAA-ATPase+Lon-Protease-BrxL->?-><-[illegible]</td><td>1213</td><td>Bacteria>Pseudomonadota>Gammaproteobacteria>Enterobacterales>Enterobacteriaceae>Salmonella</td><td>GCA_005607075.1</td><td>Salmonella_enterica</td><td>AXA91_18730</td><td>62340..65981</td><td>28901</td><td>BREX-system-P-loop-protein-BrxC-[Salmonella-enterica]</td></tr>
<tr><td>EAQ7293575.1</td><td>Type1.BREX</td><td>YjhX_toxin->SAM-METHYLASE-><-?||?->?->BrxA-DNA-binding->BrxB-BrxF-iSTAND->DUF6079*->N6-METHYLASE->OLD-ABC->TOPRIM->PglZ->SIGMA+OB-Fold+AAA-ATPase+Lon-Protease-BrxL-><-REase<-Couple_hi[illegible]</td><td>1213</td><td>Bacteria>Pseudomonadota>Gammaproteobacteria>Enterobacterales>Enterobacteriaceae>Salmonella</td><td>GCA_005612115.1</td><td>Salmonella_enterica</td><td>CRQ46_08445</td><td>406739..410380</td><td>28901</td><td>BREX-system-P-loop-protein-BrxC-[Salmonella-enterica]</td></tr>
<tr><td>EAS1883622.1</td><td>Type1.BREX</td><td><-REase+iREase<-?||||YjhX_toxin->SAM-METHYLASE-><-?||?->?->BrxA-DNA-binding->BrxB-BrxF-iSTAND->DUF6079*->N6-METHYLASE->PglZ->SIGMA+OB-Fold+AAA-ATPase+Lon-Protease-BrxL-><-REase<-Coupl[illegible]</td><td>1213</td><td>Bacteria>Pseudomonadota>Gammaproteobacteria>Enterobacterales>Enterobacteriaceae>Salmonella</td><td>GCA_005675995.1</td><td>Salmonella_enterica</td><td>AIH17_08775</td><td>402397..406038</td><td>28901</td><td>BREX-system-P-loop-protein-BrxC-[Salmonella-enterica]</td></tr>
<tr><td>EAS9365410.1</td><td>Type1.BREX</td><td><-HEPN<-REase+iREase<-?||YjhX_toxin->SAM-METHYLASE->?->BrxA-DNA-binding->BrxB-BrxF-iSTAND->DUF6079*->N6-METHYLASE->OLD-ABC->TOPRIM->PglZ->SIGMA+OB-Fold+AAA-ATPase+Lon-Protease-BrxL-><[illegible]</td><td>1213</td><td>Bacteria>Pseudomonadota>Gammaproteobacteria>Enterobacterales>Enterobacteriaceae>Salmonella</td><td>GCA_005712675.1</td><td>Salmonella_enterica</td><td>EPA30_12270</td><td>170842..174483</td><td>28901</td><td>BREX-system-P-loop-protein-BrxC-[Salmonella-enterica]</td></tr>
<tr><td>EAT1838141.1</td><td>Type1.BREX</td><td><-HEPN<-REase+iREase<-?||YjhX_toxin->SAM-METHYLASE->?->BrxA-DNA-binding->BrxB-BrxF-iSTAND->DUF6079*->N6-METHYLASE->OLD-ABC->TOPRIM->PglZ->SIGMA+OB-Fold+AAA-ATPase+Lon-Protease-BrxL-><[illegible]</td><td>1213</td><td>Bacteria>Pseudomonadota>Gammaproteobacteria>Enterobacterales>Enterobacteriaceae>Salmonella</td><td>GCA_005717435.1</td><td>Salmonella_enterica</td><td>EJR21_08315</td><td>102985..106626</td><td>28901</td><td>BREX-system-P-loop-protein-BrxC-[Salmonella-enterica]</td></tr>
<tr><td>EAT1639063.1</td><td>Type1.BREX</td><td><-HEPN<-?||?->YjhX_toxin->SAM-METHYLASE->?->BrxA-DNA-binding->BrxB-BrxF-iSTAND->DUF6079*->N6-METHYLASE->OLD-ABC->TOPRIM->PglZ->SIGMA+OB-Fold+AAA-ATPase+Lon-Protease-BrxL->HKD-DNase+OG[illegible]</td><td>1213</td><td>Bacteria>Pseudomonadota>Gammaproteobacteria>Enterobacterales>Enterobacteriaceae>Salmonella</td><td>GCA_005720135.1</td><td>Salmonella_enterica</td><td>EKF56_11740</td><td>25006..28647</td><td>28901</td><td>BREX-system-P-loop-protein-BrxC-[Salmonella-enterica]</td></tr>
<tr><td>EAT2430707.1</td><td>Type1.BREX</td><td><-HEPN<-REase+iREase<-?||YjhX_toxin->SAM-METHYLASE->?->BrxA-DNA-binding->BrxB-BrxF-iSTAND->DUF6079*->N6-METHYLASE->OLD-ABC->TOPRIM->PglZ->SIGMA+OB-Fold+AAA-ATPase+Lon-Protease-BrxL->H[illegible]</td><td>1213</td><td>Bacteria>Pseudomonadota>Gammaproteobacteria>Enterobacterales>Enterobacteriaceae>Salmonella</td><td>GCA_005726115.1</td><td>Salmonella_enterica</td><td>EI207_04325</td><td>177142..180783</td><td>28901</td><td>BREX-system-P-loop-protein-BrxC-[Salmonella-enterica]</td></tr>
<tr><td>EAU0116267.1</td><td>Type1.BREX</td><td>YjhX_toxin->SAM-METHYLASE-><-?||?->?->BrxA-DNA-binding->BrxB-BrxF-iSTAND->DUF6079*->N6-METHYLASE->OLD-ABC->TOPRIM->PglZ->SIGMA+OB-Fold+AAA-ATPase+Lon-Protease-BrxL->?-><-REase<-NACHT+[illegible]</td><td>1213</td><td>Bacteria>Pseudomonadota>Gammaproteobacteria>Enterobacterales>Enterobacteriaceae>Salmonella</td><td>GCA_005756415.1</td><td>Salmonella_enterica</td><td>ASR58_09430</td><td>25917..29558</td><td>28901</td><td>BREX-system-P-loop-protein-BrxC-[Salmonella-enterica]</td></tr>
<tr><td>EAU0152691.1</td><td>Type1.BREX</td><td>YjhX_toxin->SAM-METHYLASE-><-?||?->?->BrxA-DNA-binding->BrxB-BrxF-iSTAND->DUF6079*->N6-METHYLASE->OLD-ABC->TOPRIM->PglZ->SIGMA+OB-Fold+AAA-ATPase+Lon-Protease-BrxL->?-><-REase<-NACHT+[illegible]</td><td>1213</td><td>Bacteria>Pseudomonadota>Gammaproteobacteria>Enterobacterales>Enterobacteriaceae>Salmonella</td><td>GCA_005756695.1</td><td>Salmonella_enterica</td><td>AZG01_01865</td><td>390716..394357</td><td>28901</td><td>BREX-system-P-loop-protein-BrxC-[Salmonella-enterica]</td></tr>
<tr><td>EAU0181515.1</td><td>Type1.BREX</td><td>Glyoxalase->ACET->||YjhX_toxin->ACET->METHYLASE->ZnR->Glyoxalase->BrxA-DNA-binding->BrxB-BrxF-iSTAND->DUF6079*->ParB+HNH->N6-METHYLASE->PglZ->SIGMA+OB-Fold+AAA-ATPase+Lon-Protease-Brx[illegible]</td><td>1233</td><td>Bacteria>Pseudomonadota>Gammaproteobacteria>Enterobacterales>Enterobacteriaceae>Salmonella</td><td>GCA_005756755.1</td><td>Salmonella_enterica</td><td>A4I02_04465</td><td>211497..215198</td><td>28901</td><td>BREX-system-P-loop-protein-BrxC-[Salmonella-enterica]</td></tr>
<tr><td>EAV4850774.1</td><td>Type1.BREX</td><td>Response_reg->NACHT+SF1-SF2-Helicase+HKD-DNase->REase-><-?<-SIGMA+OB-Fold+AAA-ATPase+Lon-Protease-BrxL<-PglZ<-TOPRIM<-OLD-ABC<-N6-METHYLASE<-DUF6079*<-BrxB-BrxF-iSTAND<-BrxA-DNA-bindi[illegible]</td><td>1213</td><td>Bacteria>Pseudomonadota>Gammaproteobacteria>Enterobacterales>Enterobacteriaceae>Salmonella</td><td>GCA_005847515.1</td><td>Salmonella_enterica</td><td>EGE73_10390</td><td>37027..40668</td><td>28901</td><td>BREX-system-P-loop-protein-BrxC-[Salmonella-enterica]</td></tr>
<tr><td>EAW0893908.1</td><td>Type1.BREX</td><td><-HEPN<-?||?->||YjhX_toxin->SAM-METHYLASE->?->BrxA-DNA-binding->BrxB-BrxF-iSTAND->DUF6079*->N6-METHYLASE->OLD-ABC->TOPRIM->PglZ->SIGMA+OB-Fold+AAA-ATPase+Lon-Protease-BrxL->?-><-REase[illegible]</td><td>1213</td><td>Bacteria>Pseudomonadota>Gammaproteobacteria>Enterobacterales>Enterobacteriaceae>Salmonella</td><td>GCA_005897045.1</td><td>Salmonella_enterica</td><td>EA086_11660</td><td>157064..160705</td><td>28901</td><td>BREX-system-P-loop-protein-BrxC-[Salmonella-enterica]</td></tr>
<tr><td>EAW3206019.1</td><td>Type1.BREX</td><td>Arm-DNA-bind_3->NACHT+SF1-SF2-Helicase+HKD-DNase->REase-><-HKD-DNase+OGG-like<-SIGMA+OB-Fold+AAA-ATPase+Lon-Protease-BrxL<-PglZ<-TOPRIM<-OLD-ABC<-N6-METHYLASE<-DUF6079*<-BrxB-BrxF-iST[illegible]</td><td>1204</td><td>Bacteria>Pseudomonadota>Gammaproteobacteria>Enterobacterales>Enterobacteriaceae>Salmonella</td><td>GCA_005905825.1</td><td>Salmonella_enterica</td><td>FEL99_04750</td><td>232863..236477</td><td>28901</td><td>BREX-system-P-loop-protein-BrxC-[Salmonella-enterica]</td></tr>
<tr><td>EAW5830197.1</td><td>Type1.BREX</td><td><-HEPN<-REase+iREase<-?||YjhX_toxin->SAM-METHYLASE->?->BrxA-DNA-binding->BrxB-BrxF-iSTAND->DUF6079*->N6-METHYLASE->OLD-ABC->TOPRIM->PglZ->SIGMA+OB-Fold+AAA-ATPase+Lon-Protease-BrxL-><[illegible]</td><td>1213</td><td>Bacteria>Pseudomonadota>Gammaproteobacteria>Enterobacterales>Enterobacteriaceae>Salmonella</td><td>GCA_005917905.1</td><td>Salmonella_enterica</td><td>FF074_06105</td><td>170712..174353</td><td>28901</td><td>BREX-system-P-loop-protein-BrxC-[Salmonella-enterica]</td></tr>
<tr><td>EAW8145855.1</td><td>Type1.BREX</td><td><-HEPN<-?||YjhX_toxin->SAM-METHYLASE->?->BrxA-DNA-binding->BrxB-BrxF-iSTAND->DUF6079*->N6-METHYLASE->OLD-ABC->TOPRIM->PglZ->SIGMA+OB-Fold+AAA-ATPase+Lon-Protease-BrxL->?->||<-tRNA||?-[illegible]</td><td>1213</td><td>Bacteria>Pseudomonadota>Gammaproteobacteria>Enterobacterales>Enterobacteriaceae>Salmonella</td><td>GCA_005934015.1</td><td>Salmonella_enterica</td><td>HM10_03825</td><td>163517..167158</td><td>28901</td><td>BREX-system-P-loop-protein-BrxC-[Salmonella-enterica]</td></tr>
<tr><td>EAW9396387.1</td><td>Type1.BREX</td><td><-wHTH||wHTH+WYL+WCX-><-MarR-wHTH<-SIGMA-HTH+REase<-wHTH+SAD+HNH||?-><-ZnR||BrxA-DNA-binding->BrxB-BrxF-iSTAND->DUF6079*->N6-METHYLASE->OLD-ABC->TOPRIM->PglZ->SIGMA+OB-Fold+AAA-ATPase[illegible]</td><td>1204</td><td>Bacteria>Pseudomonadota>Gammaproteobacteria>Enterobacterales>Enterobacteriaceae>Salmonella</td><td>GCA_005940865.1</td><td>Salmonella_enterica</td><td>QE38_19735</td><td>48409..52023</td><td>28901</td><td>BREX-system-P-loop-protein-BrxC-[Salmonella-enterica]</td></tr>
<tr><td>EAW9404815.1</td><td>Type1.BREX</td><td>NACHT+SF1-SF2-Helicase+HKD-DNase->REase-><-?<-SIGMA+OB-Fold+AAA-ATPase+Lon-Protease-BrxL<-PglZ<-TOPRIM<-OLD-ABC<-N6-METHYLASE<-DUF6079*<-BrxB-BrxF-iSTAND<-BrxA-DNA-binding<-?<-SAM-MET[illegible]</td><td>1213</td><td>Bacteria>Pseudomonadota>Gammaproteobacteria>Enterobacterales>Enterobacteriaceae>Salmonella</td><td>GCA_005940875.1</td><td>Salmonella_enterica</td><td>QD58_16790</td><td>37524..41165</td><td>28901</td><td>BREX-system-P-loop-protein-BrxC-[Salmonella-enterica]</td></tr>
<tr><td>EAX4456710.1</td><td>Type1.BREX</td><td><-HEPN<-REase+iREase<-?||YjhX_toxin->SAM-METHYLASE->?->BrxA-DNA-binding->BrxB-BrxF-iSTAND->DUF6079*->N6-METHYLASE->OLD-ABC->TOPRIM->PglZ->SIGMA+OB-Fold+AAA-ATPase+Lon-Protease-BrxL-><[illegible]</td><td>1213</td><td>Bacteria>Pseudomonadota>Gammaproteobacteria>Enterobacterales>Enterobacteriaceae>Salmonella</td><td>GCA_005963445.1</td><td>Salmonella_enterica</td><td>AC467_14820</td><td>18038..21679</td><td>28901</td><td>BREX-system-P-loop-protein-BrxC-[Salmonella-enterica]</td></tr>
<tr><td>EAX8471827.1</td><td>Type1.BREX</td><td><-GntR-HTH++TrmB+ZDNA-HTH||<-?<-HEPN<-?||YjhX_toxin->SAM-METHYLASE->?->BrxA-DNA-binding->BrxB-BrxF-iSTAND->DUF6079*->N6-METHYLASE->OLD-ABC->TOPRIM->PglZ->SIGMA+OB-Fold+AAA-ATPase+Lon-[illegible]</td><td>1213</td><td>Bacteria>Pseudomonadota>Gammaproteobacteria>Enterobacterales>Enterobacteriaceae>Salmonella</td><td>GCA_005980785.1</td><td>Salmonella_enterica</td><td>AH421_04140</td><td>246530..250171</td><td>28901</td><td>BREX-system-P-loop-protein-BrxC-[Salmonella-enterica]</td></tr>
<tr><td>EAY3206374.1</td><td>Type1.BREX</td><td><-HEPN<-?||YjhX_toxin->SAM-METHYLASE->?->BrxA-DNA-binding->BrxB-BrxF-iSTAND->DUF6079*->N6-METHYLASE-></td><td>1213</td><td>Bacteria>Pseudomonadota>Gammaproteobacteria>Enterobacterales>Enterobacteriaceae>Salmonella</td><td>GCA_006001705.1</td><td>Salmonella_enterica</td><td>EAO91_18695</td><td>11389..15030</td><td>28901</td><td>BREX-system-P-loop-protein-BrxC-[Salmonella-enterica]</td></tr>
<tr><td>EAY5330993.1</td><td>Type1.BREX</td><td>ACET->METHYLASE->ZnR->Glyoxalase->BrxA-DNA-binding->BrxB-BrxF-iSTAND->DUF6079*->ParB+HNH->N6-METHYLASE->PglZ->SIGMA+OB-Fold+AAA-ATPase+Lon-Protease-BrxL-><-Arm-DNA-bind_3+Phage_integr[illegible]</td><td>1233</td><td>Bacteria>Pseudomonadota>Gammaproteobacteria>Enterobacterales>Enterobacteriaceae>Salmonella</td><td>GCA_006011505.1</td><td>Salmonella_enterica</td><td>LS33_08700</td><td>4073..7774</td><td>28901</td><td>BREX-system-P-loop-protein-BrxC-[Salmonella-enterica]</td></tr>
<tr><td>EAY5508618.1</td><td>Type1.BREX</td><td><-wHTH||wHTH+WYL+WCX-><-MarR-wHTH<-?<-SIGMA-HTH+REase<-wHTH(SBD)+HNH||?->BrxA-DNA-binding->BrxB-BrxF-iSTAND->DUF6079*->N6-METHYLASE->OLD-ABC->TOPRIM->PglZ->SIGMA+OB-Fold+AAA-ATPase+Lo[illegible]</td><td>1204</td><td>Bacteria>Pseudomonadota>Gammaproteobacteria>Enterobacterales>Enterobacteriaceae>Salmonella</td><td>GCA_006012725.1</td><td>Salmonella_enterica</td><td>NE47_11015</td><td>49392..53006</td><td>28901</td><td>BREX-system-P-loop-protein-BrxC-[Salmonella-enterica]</td></tr>
<tr><td>EAY8201055.1</td><td>Type1.BREX</td><td>Glyoxalase->ACET->YjhX_toxin->ACET->METHYLASE->ZnR->Glyoxalase->BrxA-DNA-binding->BrxB-BrxF-iSTAND->DUF6079*->ParB+HNH->N6-METHYLASE->PglZ->SIGMA+OB-Fold+AAA-ATPase+Lon-Protease-BrxL-[illegible]</td><td>1233</td><td>Bacteria>Pseudomonadota>Gammaproteobacteria>Enterobacterales>Enterobacteriaceae>Salmonella</td><td>GCA_006024175.1</td><td>Salmonella_enterica</td><td>ATQ84_14875</td><td>10975..14676</td><td>28901</td><td>BREX-system-P-loop-protein-BrxC-[Salmonella-enterica]</td></tr>
<tr><td>EAZ0388355.1</td><td>Type1.BREX</td><td><-HEPN<-REase+iREase<-?||YjhX_toxin->SAM-METHYLASE->?->BrxA-DNA-binding->BrxB-BrxF-iSTAND->DUF6079*-></td><td>1023</td><td>Bacteria>Pseudomonadota>Gammaproteobacteria>Enterobacterales>Enterobacteriaceae>Salmonella</td><td>GCA_006033355.1</td><td>Salmonella_enterica</td><td>A7762_09675</td><td>30532..33600</td><td>28901</td><td>BREX-system-P-loop-protein-BrxC-[Salmonella-enterica]</td></tr>
<tr><td>EAZ5773634.1</td><td>Type1.BREX</td><td>Arm-DNA-bind_3->NACHT+SF1-SF2-Helicase+HKD-DNase->REase-><-SIGMA+OB-Fold+AAA-ATPase+Lon-Protease-BrxL<-PglZ<-TOPRIM<-OLD-ABC<-N6-METHYLASE<-DUF6079*<-BrxB-BrxF-iSTAND<-BrxA-DNA-bindin[illegible]</td><td>1213</td><td>Bacteria>Pseudomonadota>Gammaproteobacteria>Enterobacterales>Enterobacteriaceae>Salmonella</td><td>GCA_006057755.1</td><td>Salmonella_enterica</td><td>CCZ51_07205</td><td>108064..111705</td><td>28901</td><td>BREX-system-P-loop-protein-BrxC-[Salmonella-enterica]</td></tr>
<tr><td>EAZ9311961.1</td><td>Type1.BREX</td><td><-HEPN<-?||YjhX_toxin->SAM-METHYLASE->?->BrxA-DNA-binding->BrxB-BrxF-iSTAND->DUF6079*->N6-METHYLASE->OLD-ABC->TOPRIM->PglZ->SIGMA+OB-Fold+AAA-ATPase+Lon-Protease-BrxL->HKD-DNase+OGG-l[illegible]</td><td>1213</td><td>Bacteria>Pseudomonadota>Gammaproteobacteria>Enterobacterales>Enterobacteriaceae>Salmonella</td><td>GCA_006073635.1</td><td>Salmonella_enterica</td><td>CBJ28_03025</td><td>158675..162316</td><td>28901</td><td>BREX-system-P-loop-protein-BrxC-[Salmonella-enterica]</td></tr>
<tr><td>EAZ9943132.1</td><td>Type1.BREX</td><td><-HEPN<-REase+iREase<-?||YjhX_toxin->SAM-METHYLASE->?->BrxA-DNA-binding->BrxB-BrxF-iSTAND->DUF6079*->N6-METHYLASE-></td><td>1213</td><td>Bacteria>Pseudomonadota>Gammaproteobacteria>Enterobacterales>Enterobacteriaceae>Salmonella</td><td>GCA_006074375.1</td><td>Salmonella_enterica_subsp._enterica_serovar_Typhimurium</td><td>EP721_03820</td><td>139949..143590</td><td>90371</td><td>BREX-system-P-loop-protein-BrxC-[Salmonella-enterica-subsp.-enterica-serovar-Typhimurium]</td></tr>
</table>

| Protein-ID | System | Gene-Neighbourhoods | Length | Taxonomy | Assembly-ID | Strain | Locus-Tag | Genomic-coordinates | TaxID | Fasta-define |
|---|---|---|---|---|---|---|---|---|---|---|
| EBA4497717.1 | Type1.BREX | <-HEPN<-?\|\|YjhX_toxin->SAM-METHYLASE->?->BrxA-DNA-binding->BrxB-BrxF-iSTAND->DUF6079*->N6-METHYLASE->OLD-ABC->TOPRIM->PglZ->SIGMA+OB-Fold+AAA-ATPase+Lon-Protease-BrxL-><-Couple_hipA+HipA_C<-HTH_3 | 1213 | Bacteria>Pseudomonadota>Gammaproteobacteria>Enterobacterales>Enterobacteriaceae>Salmonella | GCA_006097275.1 | Salmonella_enterica | A7G53_12020 | 208894..212535 | 28901 | BREX-system-P-loop-protein-BrxC-[Salmonella-enterica] |
| EBB0684615.1 | Type1.BREX | HTH_3->Couple_hipA+HipA_C->REase-><-SIGMA+OB-Fold+AAA-ATPase+Lon-Protease-BrxL<-PglZ<-TOPRIM<-OLD-ABC<-N6-METHYLASE<-DUF6079*<-BrxB-BrxF-iSTAND<-BrxA-DNA-binding<-?<-SAM-METHYLASE<-YjhX_toxin\|\|?->REase+iREase->HEPN-> | 1213 | Bacteria>Pseudomonadota>Gammaproteobacteria>Enterobacterales>Enterobacteriaceae>Salmonella | GCA_006125875.1 | Salmonella_enterica | CG859_01610 | 345657..349298 | 28901 | BREX-system-P-loop-protein-BrxC-[Salmonella-enterica] |
| EBB1941184.1 | Type1.BREX | <-HEPN<-REase+iREase<-?\|\|YjhX_toxin->SAM-METHYLASE->?->BrxA-DNA-binding->BrxB-BrxF-iSTAND->DUF6079*->N6-METHYLASE->OLD-ABC->TOPRIM->PglZ->SIGMA+OB-Fold+AAA-ATPase+Lon-Protease-BrxL-><-REase<-NACHT+SF1-SF2-Helicase+HKD-DNase<-Arm-DNA-bind_3 | 1213 | Bacteria>Pseudomonadota>Gammaproteobacteria>Enterobacterales>Enterobacteriaceae>Salmonella | GCA_006131775.1 | Salmonella_enterica | CJP21_07730 | 38679..42320 | 28901 | BREX-system-P-loop-protein-BrxC-[Salmonella-enterica] |
| EBB3832829.1 | Type1.BREX | <-HEPN<-?\|\|YjhX_toxin->SAM-METHYLASE->?->BrxA-DNA-binding->BrxB-BrxF-iSTAND->DUF6079*->N6-METHYLASE->OLD-ABC->TOPRIM->PglZ->SIGMA+OB-Fold+AAA-ATPase+Lon-Protease-BrxL->?-><-REase<-NACHT+SF1-SF2-Helicase+HKD-DNase | 1213 | Bacteria>Pseudomonadota>Gammaproteobacteria>Enterobacterales>Enterobacteriaceae>Salmonella | GCA_006140415.1 | Salmonella_enterica | CJ140_05520 | 149724..153365 | 28901 | BREX-system-P-loop-protein-BrxC-[Salmonella-enterica] |
| EBB4495255.1 | Type1.BREX | <-HEPN<-REase+iREase<-?\|\|\|\|YjhX_toxin->SAM-METHYLASE->?->?->BrxA-DNA-binding->BrxB-BrxF-iSTAND->DUF6079*->N6-METHYLASE->OLD-ABC->TOPRIM->PglZ->SIGMA+OB-Fold+AAA-ATPase+Lon-Protease-BrxL-><-REase<-NACHT+SF1-SF2-Helicase+HKD-DNase<-Arm-DNA-bind_3 | 1213 | Bacteria>Pseudomonadota>Gammaproteobacteria>Enterobacterales>Enterobacteriaceae>Salmonella | GCA_006143655.1 | Salmonella_enterica_subsp._enterica_serovar_Typhimurium | CET45_10230 | 170918..174559 | 90371 | BREX-system-P-loop-protein-BrxC-[Salmonella-enterica-subsp.-enterica-serovar-Typhimurium] |
| EBB5938894.1 | Type1.BREX | <-GntR-HTH++TrmB+ZDNA-HTH\|\|?-><-?<-HEPN<-?\|\|YjhX_toxin->?->BrxA-DNA-binding->BrxB-BrxF-iSTAND->DUF6079*->N6-METHYLASE->OLD-ABC->PglZ->SIGMA+OB-Fold+AAA-ATPase+Lon-Protease-BrxL->?-><-REase<-NACHT+SF1-SF2-Helicase+HKD-DNase | 1213 | Bacteria>Pseudomonadota>Gammaproteobacteria>Enterobacterales>Enterobacteriaceae>Salmonella | GCA_006151465.1 | Salmonella_enterica | CMR97_09925 | 156823..160464 | 28901 | BREX-system-P-loop-protein-BrxC-[Salmonella-enterica] |
| EBC0774151.1 | Type1.BREX | HTH_3->Couple_hipA+HipA_C->REase-><-SIGMA+OB-Fold+AAA-ATPase+Lon-Protease-BrxL<-PglZ<-TOPRIM<-OLD-ABC<-N6-METHYLASE<-DUF6079*<-BrxB-BrxF-iSTAND<-BrxA-DNA-binding<-?<-SAM-METHYLASE<-YjhX_toxin\|\|?->REase+iREase->HEPN-> | 1213 | Bacteria>Pseudomonadota>Gammaproteobacteria>Enterobacterales>Enterobacteriaceae>Salmonella | GCA_006172705.1 | Salmonella_enterica | CS696_13045 | 23540..27181 | 28901 | BREX-system-P-loop-protein-BrxC-[Salmonella-enterica] |
| EBC4250051.1 | Type1.BREX | <-HEPN<-REase+iREase<-?\|\|YjhX_toxin->SAM-METHYLASE->?->BrxA-DNA-binding->BrxB-BrxF-iSTAND->DUF6079*->N6-METHYLASE->OLD-ABC->TOPRIM->PglZ->SIGMA+OB-Fold+AAA-ATPase+Lon-Protease-BrxL-><-REase<-NACHT+SF1-SF2-Helicase+HKD-DNase<-Arm-DNA-bind_3 | 1213 | Bacteria>Pseudomonadota>Gammaproteobacteria>Enterobacterales>Enterobacteriaceae>Salmonella | GCA_006188785.1 | Salmonella_enterica | CZB00_07690 | 213870..217511 | 28901 | BREX-system-P-loop-protein-BrxC-[Salmonella-enterica] |
| EBD9210544.1 | Type1.BREX | Arm-DNA-bind_3->NACHT+SF1-SF2-Helicase+HKD-DNase->REase-><-SIGMA+OB-Fold+AAA-ATPase+Lon-Protease-BrxL<-PglZ<-TOPRIM<-OLD-ABC<-N6-METHYLASE<-DUF6079*<-BrxB-BrxF-iSTAND<-BrxA-DNA-binding<-?<-SAM-METHYLASE<-YjhX_toxin\|\|?->REase+iREase->HEPN-> | 1213 | Bacteria>Pseudomonadota>Gammaproteobacteria>Enterobacterales>Enterobacteriaceae>Salmonella | GCA_006272795.1 | Salmonella_enterica | DA198_05755 | 187002..190643 | 28901 | BREX-system-P-loop-protein-BrxC-[Salmonella-enterica] |
| EBE1252189.1 | Type1.BREX | <-HEPN<-REase+iREase<-?\|\|YjhX_toxin->SAM-METHYLASE->?->BrxA-DNA-binding->BrxB-BrxF-iSTAND->DUF6079*->N6-METHYLASE->OLD-ABC->TOPRIM->PglZ->SIGMA+OB-Fold+AAA-ATPase+Lon-Protease-BrxL-><-REase<-NACHT+SF1-SF2-Helicase+HKD-DNase<-Arm-DNA-bind_3 | 1213 | Bacteria>Pseudomonadota>Gammaproteobacteria>Enterobacterales>Enterobacteriaceae>Salmonella | GCA_006282055.1 | Salmonella_enterica | DDQ39_02640 | 124417..128058 | 28901 | BREX-system-P-loop-protein-BrxC-[Salmonella-enterica] |
| EBE3451720.1 | Type1.BREX | <-HEPN<-REase+iREase<-?\|\|YjhX_toxin->SAM-METHYLASE->?->BrxA-DNA-binding->BrxB-BrxF-iSTAND->DUF6079*-> | 1011 | Bacteria>Pseudomonadota>Gammaproteobacteria>Enterobacterales>Enterobacteriaceae>Salmonella | GCA_006291715.1 | Salmonella_enterica | DKG97_14175 | 60236..63269 | 28901 | BREX-system-P-loop-protein-BrxC-[Salmonella-enterica] |
| EBE5553332.1 | Type1.BREX | Response_reg->NACHT+SF1-SF2-Helicase+HKD-DNase->REase-><-?<-SIGMA+OB-Fold+AAA-ATPase+Lon-Protease-BrxL<-PglZ<-TOPRIM<-OLD-ABC<-N6-METHYLASE<-DUF6079*<-BrxB-BrxF-iSTAND<-BrxA-DNA-binding<-?<-SAM-METHYLASE<-YjhX_toxin\|\|?->REase+iREase->HEPN-> | 1213 | Bacteria>Pseudomonadota>Gammaproteobacteria>Enterobacterales>Enterobacteriaceae>Salmonella | GCA_006301715.1 | Salmonella_enterica | CP597_01400 | 309928..313569 | 28901 | BREX-system-P-loop-protein-BrxC-[Salmonella-enterica] |
| EBE9623448.1 | Type1.BREX | <-HEPN<-REase+iREase<-?\|\|YjhX_toxin->SAM-METHYLASE->?->BrxA-DNA-binding->BrxB-BrxF-iSTAND->DUF6079*->N6-METHYLASE->OLD-ABC->TOPRIM->PglZ->SIGMA+OB-Fold+AAA-ATPase+Lon-Protease-BrxL-><-REase<-NACHT+SF1-SF2-Helicase+HKD-DNase<-Arm-DNA-bind_3 | 1213 | Bacteria>Pseudomonadota>Gammaproteobacteria>Enterobacterales>Enterobacteriaceae>Salmonella | GCA_006320265.1 | Salmonella_enterica | DA967_03425 | 170963..174604 | 28901 | BREX-system-P-loop-protein-BrxC-[Salmonella-enterica] |
| EBF0840956.1 | Type1.BREX | <-HEPN<-?\|\|YjhX_toxin->SAM-METHYLASE->?->BrxA-DNA-binding->BrxB-BrxF-iSTAND->DUF6079*->N6-METHYLASE->OLD-ABC->TOPRIM->PglZ->SIGMA+OB-Fold+AAA-ATPase+Lon-Protease-BrxL->?-><-REase<-NACHT+SF1-SF2-Helicase+HKD-DNase | 1213 | Bacteria>Pseudomonadota>Gammaproteobacteria>Enterobacterales>Enterobacteriaceae>Salmonella | GCA_006325425.1 | Salmonella_enterica | DDQ86_07950 | 150304..153945 | 28901 | BREX-system-P-loop-protein-BrxC-[Salmonella-enterica] |
| EBF3214239.1 | Type1.BREX | Glyoxalase->ACET->YjhX_toxin->ACET->METHYLASE->ZnR->Glyoxalase->BrxA-DNA-binding->BrxB-BrxF-iSTAND->DUF6079*->ParB+HNH->N6-METHYLASE->PglZ->SIGMA+OB-Fold+AAA-ATPase+Lon-Protease-BrxL-><-Arm-DNA-bind_3+Phage_integrase<-tRNA\|\|?-><-SKI\|\|METHYLASE-> | 1233 | Bacteria>Pseudomonadota>Gammaproteobacteria>Enterobacterales>Enterobacteriaceae>Salmonella | GCA_006336125.1 | Salmonella_enterica | FHK34_03265 | 183037..186738 | 28901 | BREX-system-P-loop-protein-BrxC-[Salmonella-enterica] |
| EBF4455779.1 | Type1.BREX | YjhX_toxin->SAM-METHYLASE->?->\|\|?-><-?<-ZnR<-ZnR\|\|BrxA-DNA-binding->BrxB-BrxF-iSTAND->DUF6079*->N6-METHYLASE->OLD-ABC->TOPRIM->PglZ->SIGMA+OB-Fold+AAA-ATPase+Lon-Protease-BrxL->HKD-DNase+OGG-like-><-REase<-NACHT+SF1-SF2-Helicase+HKD-DNase<-Arm-DNA-bind_3 | 1204 | Bacteria>Pseudomonadota>Gammaproteobacteria>Enterobacterales>Enterobacteriaceae>Salmonella | GCA_006343165.1 | Salmonella_enterica | FH443_09905 | 412115..415729 | 28901 | BREX-system-P-loop-protein-BrxC-[Salmonella-enterica] |
| EBF8311517.1 | Type1.BREX | <-HEPN<-REase+iREase<-?\|\|YjhX_toxin->SAM-METHYLASE->?->\|\|BrxA-DNA-binding->BrxB-BrxF-iSTAND->DUF6079*->N6-METHYLASE->OLD-ABC->TOPRIM->PglZ->SIGMA+OB-Fold+AAA-ATPase+Lon-Protease-BrxL->\|\|<-HTH_Tnp_1+HTH_21+rve+rve_3 | 1213 | Bacteria>Pseudomonadota>Gammaproteobacteria>Enterobacterales>Enterobacteriaceae>Salmonella | GCA_006394955.1 | Salmonella_enterica_subsp._enterica | FGE90_18580 | 53285..56926 | 59201 | BREX-system-P-loop-protein-BrxC-[Salmonella-enterica-subsp.-enterica] |
| EBG0770112.1 | Type1.BREX | BrxA-DNA-binding->BrxB-BrxF-iSTAND->DUF6079*->N6-METHYLASE->OLD-ABC->TOPRIM->PglZ->SIGMA+OB-Fold+AAA-ATPase+Lon-Protease-BrxL-><-REase<-Couple_hipA+HipA_C<-HTH_3 | 1213 | Bacteria>Pseudomonadota>Gammaproteobacteria>Enterobacterales>Enterobacteriaceae>Salmonella | GCA_006406395.1 | Salmonella_enterica | FH890_08325 | 3073..6714 | 28901 | BREX-system-P-loop-protein-BrxC-[Salmonella-enterica] |
| EBG3411773.1 | Type1.BREX | <-HEPN<-REase+iREase<-?\|\|YjhX_toxin->SAM-METHYLASE->?->BrxA-DNA-binding->BrxB-BrxF-iSTAND->DUF6079*->N6-METHYLASE->OLD-ABC->TOPRIM->PglZ->SIGMA+OB-Fold+AAA-ATPase+Lon-Protease-BrxL-><-REase<-NACHT+SF1-SF2-Helicase+HKD-DNase<-Arm-DNA-bind_3 | 1213 | Bacteria>Pseudomonadota>Gammaproteobacteria>Enterobacterales>Enterobacteriaceae>Salmonella | GCA_006423935.1 | Salmonella_enterica | FIW39_06205 | 171028..174669 | 28901 | BREX-system-P-loop-protein-BrxC-[Salmonella-enterica] |
| EBG5655091.1 | Type1.BREX | Arm-DNA-bind_3->NACHT+SF1-SF2-Helicase+HKD-DNase->REase-><-SIGMA+OB-Fold+AAA-ATPase+Lon-Protease-BrxL<-PglZ<-TOPRIM<-OLD-ABC<-N6-METHYLASE<-DUF6079*<-BrxB-BrxF-iSTAND<-BrxA-DNA-binding | 1213 | Bacteria>Pseudomonadota>Gammaproteobacteria>Enterobacterales>Enterobacteriaceae>Salmonella | GCA_006430455.1 | Salmonella_enterica_subsp._enterica | FI236_11975 | 108064..111705 | 59201 | BREX-system-P-loop-protein-BrxC-[Salmonella-enterica-subsp.-enterica] |
| EBG7129666.1 | Type1.BREX | <-HEPN<-?\|\|YjhX_toxin->SAM-METHYLASE->?->BrxA-DNA-binding->BrxB-BrxF-iSTAND->DUF6079*->N6-METHYLASE-> | 1213 | Bacteria>Pseudomonadota>Gammaproteobacteria>Enterobacterales>Enterobacteriaceae>Salmonella | GCA_006457905.1 | Salmonella_enterica_subsp._enterica | FJA05_19325 | 25635..29276 | 59201 | BREX-system-P-loop-protein-BrxC-[Salmonella-enterica-subsp.-enterica] |
| EBH0811989.1 | Type1.BREX | <-HEPN<-REase+iREase<-?\|\|YjhX_toxin->SAM-METHYLASE->?->BrxA-DNA-binding->BrxB-BrxF-iSTAND->DUF6079*->N6-METHYLASE->OLD-ABC->TOPRIM->PglZ->SIGMA+OB-Fold+AAA-ATPase+Lon-Protease-BrxL-><-REase<-NACHT+SF1-SF2-Helicase+HKD-DNase<-Arm-DNA-bind_3 | 1213 | Bacteria>Pseudomonadota>Gammaproteobacteria>Enterobacterales>Enterobacteriaceae>Salmonella | GCA_006475165.1 | Salmonella_enterica | FJ896_02670 | 170932..174573 | 28901 | BREX-system-P-loop-protein-BrxC-[Salmonella-enterica] |
| EBH4970956.1 | Type1.BREX | <-HEPN<-REase+iREase<-?\|\|YjhX_toxin->SAM-METHYLASE->?->BrxA-DNA-binding->BrxB-BrxF-iSTAND->DUF6079*->N6-METHYLASE->OLD-ABC->TOPRIM->PglZ->SIGMA+OB-Fold+AAA-ATPase+Lon-Protease-BrxL-><-REase<-NACHT+SF1-SF2-Helicase+HKD-DNase<-Arm-DNA-bind_3 | 1213 | Bacteria>Pseudomonadota>Gammaproteobacteria>Enterobacterales>Enterobacteriaceae>Salmonella | GCA_006499395.1 | Salmonella_enterica | CKR35_08925 | 170347..173988 | 28901 | BREX-system-P-loop-protein-BrxC-[Salmonella-enterica] |
| EBH5838355.1 | Type1.BREX | Arm-DNA-bind_3->NACHT+SF1-SF2-Helicase+HKD-DNase->REase-><-SIGMA+OB-Fold+AAA-ATPase+Lon-Protease-BrxL<-PglZ<-TOPRIM<-OLD-ABC<-N6-METHYLASE<-DUF6079*<-BrxB-BrxF-iSTAND<-BrxA-DNA-binding\|\|ZnR->?-><-?\|\|wHTH+SAD+HNH->SIGMA-HTH+REase->?->MarR-wHTH-> | 1204 | Bacteria>Pseudomonadota>Gammaproteobacteria>Enterobacterales>Enterobacteriaceae>Salmonella | GCA_006503215.1 | Salmonella_enterica | CQL58_06375 | 305201..308815 | 28901 | BREX-system-P-loop-protein-BrxC-[Salmonella-enterica] |
| EBI6018519.1 | Type1.BREX | YjhX_toxin->SAM-METHYLASE-><-?\|\|?->?->BrxA-DNA-binding->BrxB-BrxF-iSTAND->DUF6079*->N6-METHYLASE->PglZ->SIGMA+OB-Fold+AAA-ATPase+Lon-Protease-BrxL->\|\|HIN-HTH-> | 1213 | Bacteria>Pseudomonadota>Gammaproteobacteria>Enterobacterales>Enterobacteriaceae>Salmonella | GCA_006562665.1 | Salmonella_enterica | DW438_24035 | 31966..35607 | 28901 | BREX-system-P-loop-protein-BrxC-[Salmonella-enterica] |
| EBI8027930.1 | Type1.BREX | <-REase+iREase<-?\|\|\|\|YjhX_toxin->SAM-METHYLASE-><-?\|\|?->?->BrxA-DNA-binding->BrxB-BrxF-iSTAND->DUF6079*->N6-METHYLASE->OLD-ABC->TOPRIM->PglZ->SIGMA+OB-Fold+AAA-ATPase+Lon-Protease-BrxL-><-REase<-Arm-DNA-bind_3<-tRNA\|\|?-><-SKI | 1213 | Bacteria>Pseudomonadota>Gammaproteobacteria>Enterobacterales>Enterobacteriaceae>Salmonella | GCA_006571625.1 | Salmonella_enterica | CHM23_07390 | 170919..174560 | 28901 | BREX-system-P-loop-protein-BrxC-[Salmonella-enterica] |
| EBK5803035.1 | Type1.BREX | <-HEPN<-?\|\|?->\|\|YjhX_toxin->SAM-METHYLASE->?->\|\|?->BrxA-DNA-binding->BrxB-BrxF-iSTAND->DUF6079*->N6-METHYLASE->OLD-ABC->TOPRIM->PglZ->SIGMA+OB-Fold+AAA-ATPase+Lon-Protease-BrxL->?-><-REase<-NACHT+SF1-SF2-Helicase+HKD-DNase<-Arm-DNA-bind_3 | 1213 | Bacteria>Pseudomonadota>Gammaproteobacteria>Enterobacterales>Enterobacteriaceae>Salmonella | GCA_006648525.1 | Salmonella_enterica | DPM88_04065 | 158403..162044 | 28901 | BREX-system-P-loop-protein-BrxC-[Salmonella-enterica] |
| EBK7108450.1 | Type1.BREX | <-HEPN<-REase+iREase<-?\|\|\|\|YjhX_toxin->SAM-METHYLASE->?->?->BrxA-DNA-binding->BrxB-BrxF-iSTAND->DUF6079*->N6-METHYLASE->OLD-ABC->TOPRIM->PglZ->SIGMA+OB-Fold+AAA-ATPase+Lon-Protease-BrxL-><-REase<-NACHT+SF1-SF2-Helicase+HKD-DNase<-Arm-DNA-bind_3 | 1213 | Bacteria>Pseudomonadota>Gammaproteobacteria>Enterobacterales>Enterobacteriaceae>Salmonella | GCA_006653645.1 | Salmonella_enterica | DSC93_06685 | 170929..174570 | 28901 | BREX-system-P-loop-protein-BrxC-[Salmonella-enterica] |
| EBK9871839.1 | Type1.BREX | YjhX_toxin->SAM-METHYLASE-><-?\|\|?->?->BrxA-DNA-binding->BrxB-BrxF-iSTAND->DUF6079*->N6-METHYLASE->OLD-ABC->TOPRIM->PglZ->SIGMA+OB-Fold+AAA-ATPase+Lon-Protease-BrxL->?-><-REase<-NACHT+SF1-SF2-Helicase+HKD-DNase | 1213 | Bacteria>Pseudomonadota>Gammaproteobacteria>Enterobacterales>Enterobacteriaceae>Salmonella | GCA_006668805.1 | Salmonella_enterica | DYD89_19150 | 25917..29558 | 28901 | BREX-system-P-loop-protein-BrxC-[Salmonella-enterica] |
| EBL1260729.1 | Type1.BREX | <-HEPN<-REase+iREase<-?\|\|\|\|YjhX_toxin->SAM-METHYLASE->?->?->BrxA-DNA-binding->BrxB-BrxF-iSTAND->DUF6079*->N6-METHYLASE->OLD-ABC->TOPRIM->PglZ->SIGMA+OB-Fold+AAA-ATPase+Lon-Protease-BrxL-><-REase<-NACHT+SF1-SF2-Helicase+HKD-DNase<-Arm-DNA-bind_3 | 1213 | Bacteria>Pseudomonadota>Gammaproteobacteria>Enterobacterales>Enterobacteriaceae>Salmonella | GCA_006672445.1 | Salmonella_enterica | DO790_07470 | 171130..174771 | 28901 | BREX-system-P-loop-protein-BrxC-[Salmonella-enterica] |
| EBL2857392.1 | Type1.BREX | Arm-DNA-bind_3->NACHT+SF1-SF2-Helicase+HKD-DNase->REase-><-SIGMA+OB-Fold+AAA-ATPase+Lon-Protease-BrxL<-PglZ<-TOPRIM<-OLD-ABC<-N6-METHYLASE<-DUF6079*<-BrxB-BrxF-iSTAND<-BrxA-DNA-binding<-?<-?<-SAM-METHYLASE<-YjhX_toxin\|\|\|\|?->REase+iREase->HEPN-> | 1213 | Bacteria>Pseudomonadota>Gammaproteobacteria>Enterobacterales>Enterobacteriaceae>Salmonella | GCA_006679365.1 | Salmonella_enterica | D3Q95_12500 | 132391..136032 | 28901 | BREX-system-P-loop-protein-BrxC-[Salmonella-enterica] |
| EBL4178053.1 | Type1.BREX | <-HEPN<-REase+iREase<-?\|\|YjhX_toxin->SAM-METHYLASE->?->BrxA-DNA-binding->BrxB-BrxF-iSTAND->DUF6079*->N6-METHYLASE->OLD-ABC->TOPRIM->PglZ->SIGMA+OB-Fold+AAA-ATPase+Lon-Protease-BrxL-><-REase<-NACHT+SF1-SF2-Helicase+HKD-DNase<-Arm-DNA-bind_3 | 1213 | Bacteria>Pseudomonadota>Gammaproteobacteria>Enterobacterales>Enterobacteriaceae>Salmonella | GCA_006685525.1 | Salmonella_enterica_subsp._enterica_serovar_Typhimurium | E5L78_02825 | 18164..21805 | 90371 | BREX-system-P-loop-protein-BrxC-[Salmonella-enterica-subsp.-enterica-serovar-Typhimurium] |
| EBL6294791.1 | Type1.BREX | Arm-DNA-bind_3->NACHT+SF1-SF2-Helicase+HKD-DNase->REase-><-SIGMA+OB-Fold+AAA-ATPase+Lon-Protease-BrxL<-PglZ<-TOPRIM<-OLD-ABC<-N6-METHYLASE<-DUF6079-p* | 846 | Bacteria>Pseudomonadota>Gammaproteobacteria>Enterobacterales>Enterobacteriaceae>Salmonella | GCA_006694585.1 | Salmonella_enterica_subsp._enterica_serovar_Typhimurium | AC864_13995 | 181813..184355 | 90371 | BREX-system-P-loop-protein-BrxC-[Salmonella-enterica-subsp.-enterica-serovar-Typhimurium] |
| EBL7150924.1 | Type1.BREX | SKI-><-?\|\|tRNA->Arm-DNA-bind_3->REase-><-SIGMA+OB-Fold+AAA-ATPase+Lon-Protease-BrxL<-PglZ<-TOPRIM<-OLD-ABC<-N6-METHYLASE<-DUF6079*<-BrxB-BrxF-iSTAND<-BrxA-DNA-binding<-?<-?<-SAM-METHYLASE<-YjhX_toxin\|\|\|\|?->REase+iREase->HEPN-> | 1213 | Bacteria>Pseudomonadota>Gammaproteobacteria>Enterobacterales>Enterobacteriaceae>Salmonella | GCA_006698245.1 | Salmonella_enterica | CFL44_11230 | 80176..83817 | 28901 | BREX-system-P-loop-protein-BrxC-[Salmonella-enterica] |
| EBM2234242.1 | Type1.BREX | <-REase+iREase<-?\|\|\|\|YjhX_toxin->SAM-METHYLASE-><-?\|\|?->?->BrxA-DNA-binding->BrxB-BrxF-iSTAND->DUF6079*->N6-METHYLASE-> | 1213 | Bacteria>Pseudomonadota>Gammaproteobacteria>Enterobacterales>Enterobacteriaceae>Salmonella | GCA_006721345.1 | Salmonella_enterica | DWD21_19365 | 25859..29500 | 28901 | BREX-system-P-loop-protein-BrxC-[Salmonella-enterica] |
| EBM6187520.1 | Type1.BREX | <-REase+iREase<-?\|\|\|\|YjhX_toxin->SAM-METHYLASE-><-?\|\|?->?->BrxA-DNA-binding->BrxB-BrxF-iSTAND->DUF6079*->N6-METHYLASE->OLD-ABC->TOPRIM->PglZ->SIGMA+OB-Fold+AAA-ATPase+Lon-Protease-BrxL->HKD-DNase+OGG-like-><-REase<-NACHT+SF1-SF2-Helicase+HKD-DNase<-Arm-DNA-bind_3 | 1213 | Bacteria>Pseudomonadota>Gammaproteobacteria>Enterobacterales>Enterobacteriaceae>Salmonella | GCA_006738745.1 | Salmonella_enterica | D1943_04230 | 100576..104217 | 28901 | BREX-system-P-loop-protein-BrxC-[Salmonella-enterica] |
| EBN5015145.1 | Type1.BREX | <-HEPN<-?\|\|\|\|YjhX_toxin->SAM-METHYLASE-><-?\|\|?->?->BrxA-DNA-binding->BrxB-BrxF-iSTAND->DUF6079*->N6-METHYLASE-> | 1213 | Bacteria>Pseudomonadota>Gammaproteobacteria>Enterobacterales>Enterobacteriaceae>Salmonella | GCA_006773125.1 | Salmonella_enterica | DT736_14590 | 144280..147921 | 28901 | BREX-system-P-loop-protein-BrxC-[Salmonella-enterica] |
| EBQ4102575.1 | Type1.BREX | <-N6-METHYLASE<-DUF6079*<-BrxB-BrxF-iSTAND<-BrxA-DNA-binding<-?<-SAM-METHYLASE<-YjhX_toxin\|\|?->HEPN-> | 1213 | Bacteria>Pseudomonadota>Gammaproteobacteria>Enterobacterales>Enterobacteriaceae>Salmonella | GCA_006909045.1 | Salmonella_enterica | BFN90_18795 | 2722..6363 | 28901 | BREX-system-P-loop-protein-BrxC-[Salmonella-enterica] |
| EBQ6490464.1 | Type1.BREX | <-HEPN<-?\|\|YjhX_toxin->SAM-METHYLASE->?->BrxA-DNA-binding->BrxB-BrxF-iSTAND->DUF6079*->N6-METHYLASE->PglZ->SIGMA+OB-Fold+AAA-ATPase+Lon-Protease-BrxL-><-Arm-DNA-bind_3+Phage_integrase<-TOPRIM-DNAG+D5N+D5ATPase+P4PRIMASE-HTH<-?<-?<-?<-XIS-HTH | 1213 | Bacteria>Pseudomonadota>Gammaproteobacteria>Enterobacterales>Enterobacteriaceae>Salmonella | GCA_006919175.1 | Salmonella_enterica | BU548_03115 | 149047..152688 | 28901 | BREX-system-P-loop-protein-BrxC-[Salmonella-enterica] |
| EBR0226079.1 | Type1.BREX | wHTH+WYL+WCX-><-MarR-wHTH<-SIGMA-HTH+REase<-wHTH+SAD+HNH\|\|?-><-?<-ZnR\|\|BrxA-DNA-binding->BrxB-BrxF-iSTAND->DUF6079*->N6-METHYLASE->OLD-ABC->TOPRIM->PglZ->SIGMA+OB-Fold+AAA-ATPase+Lon-Protease-BrxL->HKD-DNase+OGG-like-><-REase<-NACHT+SF1-SF2-Helicase+HKD-DNase | 1204 | Bacteria>Pseudomonadota>Gammaproteobacteria>Enterobacterales>Enterobacteriaceae>Salmonella | GCA_006936115.1 | Salmonella_enterica_subsp._enterica_serovar_Monschaui | GPN34_002360 | 410647..414261 | 29480 | BREX-system-P-loop-protein-BrxC-[Salmonella-enterica-subsp.-enterica-serovar-Monschaui] |
| EBR4438819.1 | Type1.BREX | Arm-DNA-bind_3->NACHT+SF1-SF2-Helicase+HKD-DNase->REase-><-SIGMA+OB-Fold+AAA-ATPase+Lon-Protease-BrxL<-PglZ<-TOPRIM<-OLD-ABC<-N6-METHYLASE<-DUF6079*<-BrxB-BrxF-iSTAND<-BrxA-DNA-binding<-?<-SAM-METHYLASE<-YjhX_toxin\|\|?->REase+iREase->HEPN-> | 1213 | Bacteria>Pseudomonadota>Gammaproteobacteria>Enterobacterales>Enterobacteriaceae>Salmonella | GCA_006955825.1 | Salmonella_enterica | BSE18_11505 | 108064..111705 | 28901 | BREX-system-P-loop-protein-BrxC-[Salmonella-enterica] |
| EBR5149893.1 | Type1.BREX | <-MarR-wHTH<-?<-SIGMA-HTH+REase<-wHTH+SAD+HNH\|\|?-><-?<-ZnR\|\|BrxA-DNA-binding->BrxB-BrxF-iSTAND->DUF6079*->N6-METHYLASE->OLD-ABC->TOPRIM->PglZ->SIGMA+OB-Fold+AAA-ATPase+Lon-Protease-BrxL->HKD-DNase+OGG-like-><-REase<-NACHT+SF1-SF2-Helicase+HKD-DNase<-Arm-DNA-bind_3 | 1204 | Bacteria>Pseudomonadota>Gammaproteobacteria>Enterobacterales>Enterobacteriaceae>Salmonella | GCA_006958505.1 | Salmonella_enterica | B5W64_09845 | 350811..354425 | 28901 | BREX-system-P-loop-protein-BrxC-[Salmonella-enterica] |
| EBR9945161.1 | Type1.BREX | <-HEPN<-REase+iREase<-?\|\|\|\|YjhX_toxin->SAM-METHYLASE-><-?\|\|?->BrxA-DNA-binding->BrxB-BrxF-iSTAND->DUF6079*->N6-METHYLASE->OLD-ABC->TOPRIM->PglZ->SIGMA+OB-Fold+AAA-ATPase+Lon-Protease-BrxL-><-Couple_hipA+HipA_C<-HTH_3 | 1213 | Bacteria>Pseudomonadota>Gammaproteobacteria>Enterobacterales>Enterobacteriaceae>Salmonella | GCA_006983125.1 | Salmonella_enterica_subsp._enterica_serovar_Krefeld | DTF03_11145 | 153391..157032 | 1242084 | BREX-system-P-loop-protein-BrxC-[Salmonella-enterica-subsp.-enterica-serovar-Krefeld] |
| EBS4429082.1 | Type1.BREX | YjhX_toxin->SAM-METHYLASE-><-?\|\|?->?->BrxA-DNA-binding->BrxB-BrxF-iSTAND->DUF6079*->N6-METHYLASE->OLD-ABC->TOPRIM->PglZ->SIGMA+OB-Fold+AAA-ATPase+Lon-Protease-BrxL->?-><-REase<-NACHT+SF1-SF2-Helicase+HKD-DNase | 1213 | Bacteria>Pseudomonadota>Gammaproteobacteria>Enterobacterales>Enterobacteriaceae>Salmonella | GCA_007008305.1 | Salmonella_enterica_subsp._enterica_serovar_Poona | DQJ97_07365 | 25927..29568 | 436295 | BREX-system-P-loop-protein-BrxC-[Salmonella-enterica-subsp.-enterica-serovar-Poona] |
| EBS7702178.1 | Type1.BREX | <-MORC+HISKIN<-DNA_METHYLASE<-?\|\|HIN-HTH-><-SIGMA+OB-Fold+AAA-ATPase+Lon-Protease-BrxL<-PglZ<-TOPRIM<-OLD-ABC<-N6-METHYLASE<-DUF6079*<-BrxB-BrxF-iSTAND<-BrxA-DNA-binding<-?<-SAM-METHYLASE<-YjhX_toxin\|\|?->REase+iREase->HEPN-> | 1213 | Bacteria>Pseudomonadota>Gammaproteobacteria>Enterobacterales>Enterobacteriaceae>Salmonella | GCA_007022165.1 | Salmonella_enterica | CD603_08310 | 64360..68001 | 28901 | BREX-system-P-loop-protein-BrxC-[Salmonella-enterica] |
| EBS7807126.1 | Type1.BREX | Glyoxalase->ACET->YjhX_toxin->ACET->METHYLASE->ZnR->Glyoxalase->BrxA-DNA-binding->BrxB-BrxF-iSTAND->DUF6079*->ParB+HNH->N6-METHYLASE->PglZ->SIGMA+OB-Fold+AAA-ATPase+Lon-Protease-BrxL-><-Arm-DNA-bind_3+Phage_integrase<-tRNA\|\|?-><-SKI\|\|METHYLASE-> | 1233 | Bacteria>Pseudomonadota>Gammaproteobacteria>Enterobacterales>Enterobacteriaceae>Salmonella | GCA_007023485.1 | Salmonella_enterica | CD664_12540 | 43837..47538 | 28901 | BREX-system-P-loop-protein-BrxC-[Salmonella-enterica] |
| EBS9243151.1 | Type1.BREX | BrxB-BrxF-iSTAND->ACET->\|\|?-><-Phage_integrase\|\|DOC+HTH->BrxA-DNA-binding->BrxB-BrxF-iSTAND->DUF6079*->N6-METHYLASE->PglZ->SIGMA+OB-Fold+AAA-ATPase+Lon-Protease-BrxL-><-AAA+HTH<-?<-XIS-HTH | 1213 | Bacteria>Pseudomonadota>Gammaproteobacteria>Enterobacterales>Enterobacteriaceae>Salmonella | GCA_007029675.1 | Salmonella_enterica | CF713_13020 | 212136..215777 | 28901 | BREX-system-P-loop-protein-BrxC-[Salmonella-enterica] |
| EBT4520742.1 | Type1.BREX | Arm-DNA-bind_3->NACHT+SF1-SF2-Helicase+HKD-DNase->REase-><-SIGMA+OB-Fold+AAA-ATPase+Lon-Protease-BrxL<-PglZ<-TOPRIM<-OLD-ABC<-N6-METHYLASE<-DUF6079-p* | 962 | Bacteria>Pseudomonadota>Gammaproteobacteria>Enterobacterales>Enterobacteriaceae>Salmonella | GCA_007056145.1 | Salmonella_enterica | CXQ08_17680 | 108064..110954 | 28901 | BREX-system-P-loop-protein-BrxC-[Salmonella-enterica] |
| EBU3113248.1 | Type1.BREX | <-DUF6079-p*<-BrxB-BrxF-iSTAND<-BrxA-DNA-binding<-Glyoxalase<-ZnR<-METHYLASE<-ACET<-YjhX_toxin | 778 | Bacteria>Pseudomonadota>Gammaproteobacteria>Enterobacterales>Enterobacteriaceae>Salmonella | GCA_007098435.1 | Salmonella_enterica_subsp._enterica | CSQ72_21675 | 1..2335 | 59201 | BREX-system-P-loop-protein-BrxC-[Salmonella-enterica-subsp.-enterica] |
| EBU3750854.1 | Type1.BREX | <-HEPN<-?\|\|YjhX_toxin->SAM-METHYLASE->?->\|\|BrxA-DNA-binding->BrxB-BrxF-iSTAND->DUF6079*->N6-METHYLASE->OLD-ABC->TOPRIM->PglZ->SIGMA+OB-Fold+AAA-ATPase+Lon-Protease-BrxL->?-><-REase<-NACHT+SF1-SF2-Helicase+HKD-DNase<-Arm-DNA-bind_3 | 1213 | Bacteria>Pseudomonadota>Gammaproteobacteria>Enterobacterales>Enterobacteriaceae>Salmonella | GCA_007101645.1 | Salmonella_enterica | CVX24_10925 | 25914..29555 | 28901 | BREX-system-P-loop-protein-BrxC-[Salmonella-enterica] |
| EBU6515329.1 | Type1.BREX | SAM-METHYLASE->?->?->BrxA-DNA-binding->BrxB-BrxF-iSTAND->DUF6079*->N6-METHYLASE->OLD-ABC->TOPRIM->PglZ->SIGMA+OB-Fold+AAA-ATPase+Lon-Protease-BrxL-><-REase<-NACHT+SF1-SF2-Helicase+HKD-DNase<-Arm-DNA-bind_3 | 1213 | Bacteria>Pseudomonadota>Gammaproteobacteria>Enterobacterales>Enterobacteriaceae>Salmonella | GCA_007137465.1 | Salmonella_enterica_subsp._enterica_serovar_Typhimurium | DKJ21_01470 | 4814..8455 | 90371 | BREX-system-P-loop-protein-BrxC-[Salmonella-enterica-subsp.-enterica-serovar-Typhimurium] |
| EBU6749119.1 | Type1.BREX | DUF6079-p*->N6-METHYLASE->OLD-ABC->TOPRIM->PglZ->SIGMA+OB-Fold+AAA-ATPase+Lon-Protease-BrxL-><-REase<-NACHT+SF1-SF2-Helicase+HKD-DNase<-Arm-DNA-bind_3 | 874 | Bacteria>Pseudomonadota>Gammaproteobacteria>Enterobacterales>Enterobacteriaceae>Salmonella | GCA_007137985.1 | Salmonella_enterica_subsp._enterica_serovar_Typhimurium | DXS55_01035 | 1..2627 | 90371 | BREX-system-P-loop-protein-BrxC-[Salmonella-enterica-subsp.-enterica-serovar-Typhimurium] |
| EBU7641088.1 | Type1.BREX | YjhX_toxin->SAM-METHYLASE-><-?\|\|?->?->BrxA-DNA-binding->BrxB-BrxF-iSTAND->DUF6079*->N6-METHYLASE->OLD-ABC->TOPRIM->PglZ->SIGMA+OB-Fold+AAA-ATPase+Lon-Protease-BrxL->HKD-DNase+OGG-like-><-REase<-NACHT+SF1-SF2-Helicase+HKD-DNase | 1213 | Bacteria>Pseudomonadota>Gammaproteobacteria>Enterobacterales>Enterobacteriaceae>Salmonella | GCA_007142065.1 | Salmonella_enterica_subsp._enterica_serovar_Brevik | DK709_13715 | 59395..63036 | 2564275 | BREX-system-P-loop-protein-BrxC-[Salmonella-enterica-subsp.-enterica-serovar-Brevik] |
| EBU8677660.1 | Type1.BREX | HTH_3->Couple_hipA+HipA_C->REase-><-SIGMA+OB-Fold+AAA-ATPase+Lon-Protease-BrxL<-PglZ<-TOPRIM<-OLD-ABC<-N6-METHYLASE<-DUF6079*<-BrxB-BrxF-iSTAND<-BrxA-DNA-binding | 1213 | Bacteria>Pseudomonadota>Gammaproteobacteria>Enterobacterales>Enterobacteriaceae>Salmonella | GCA_007146445.1 | Salmonella_enterica_subsp._enterica_serovar_Parkroyal | DLR30_20715 | 39960..43601 | 2564909 | BREX-system-P-loop-protein-BrxC-[Salmonella-enterica-subsp.-enterica-serovar-Parkroyal] |
| EBU9347433.1 | Type1.BREX | Arm-DNA-bind_3->NACHT+SF1-SF2-Helicase+HKD-DNase->REase-><-HKD-DNase+OGG-like<-SIGMA+OB-Fold+AAA-ATPase+Lon-Protease-BrxL<-PglZ<-TOPRIM<-OLD-ABC<-N6-METHYLASE<-DUF6079*<-BrxB-BrxF-iSTAND<-BrxA-DNA-binding\|\|ZnR->?-><-?\|\|wHTH+SAD+HNH-> | 1204 | Bacteria>Pseudomonadota>Gammaproteobacteria>Enterobacterales>Enterobacteriaceae>Salmonella | GCA_007149485.1 | Salmonella_enterica_subsp._enterica_serovar_Angoda | DMU95_04690 | 163241..166855 | 2564159 | BREX-system-P-loop-protein-BrxC-[Salmonella-enterica-subsp.-enterica-serovar-Angoda] |
| EBU9954738.1 | Type1.BREX | YjhX_toxin->SAM-METHYLASE-><-?\|\|?->?->BrxA-DNA-binding->BrxB-BrxF-iSTAND->DUF6079*->N6-METHYLASE->OLD-ABC->TOPRIM->PglZ->SIGMA+OB-Fold+AAA-ATPase+Lon-Protease-BrxL->\|\|<-HTH_Tnp_1+HTH_21+rve+rve_3 | 1213 | Bacteria>Pseudomonadota>Gammaproteobacteria>Enterobacterales>Enterobacteriaceae>Salmonella | GCA_007151145.1 | Salmonella_enterica_subsp._enterica_serovar_Tamberma | DSB42_14900 | 163634..167275 | 2565079 | BREX-system-P-loop-protein-BrxC-[Salmonella-enterica-subsp.-enterica-serovar-Tamberma] |
| EBU9775078.1 | Type1.BREX | BrxA-DNA-binding->BrxB-BrxF-iSTAND->DUF6079*->N6-METHYLASE->OLD-ABC->TOPRIM->PglZ->SIGMA+OB-Fold+AAA-ATPase+Lon-Protease-BrxL->HKD-DNase+OGG-like-> | 1213 | Bacteria>Pseudomonadota>Gammaproteobacteria>Enterobacterales>Enterobacteriaceae>Salmonella | GCA_007151525.1 | Salmonella_enterica_subsp._enterica_serovar_Kumasi | DM304_21645 | 3074..6715 | 2493647 | BREX-system-P-loop-protein-BrxC-[Salmonella-enterica-subsp.-enterica-serovar-Kumasi] |
| EBU9878489.1 | Type1.BREX | <-HEPN<-REase+iREase<-?\|\|\|\|YjhX_toxin->SAM-METHYLASE->?->?->BrxA-DNA-binding->BrxB-BrxF-iSTAND->DUF6079*->N6-METHYLASE->OLD-ABC->TOPRIM->PglZ->SIGMA+OB-Fold+AAA-ATPase+Lon-Protease-BrxL-><-REase<-NACHT+SF1-SF2-Helicase+HKD-DNase<-Arm-DNA-bind_3 | 1213 | Bacteria>Pseudomonadota>Gammaproteobacteria>Enterobacterales>Enterobacteriaceae>Salmonella | GCA_007152025.1 | Salmonella_enterica_subsp._enterica_serovar_Typhimurium | DM361_04530 | 60215..63856 | 90371 | BREX-system-P-loop-protein-BrxC-[Salmonella-enterica-subsp.-enterica-serovar-Typhimurium] |
| EBV0268830.1 | Type1.BREX | <-HEPN<-REase+iREase<-?\|\|\|\|YjhX_toxin->SAM-METHYLASE->?->?->BrxA-DNA-binding->BrxB-BrxF-iSTAND->DUF6079*->N6-METHYLASE->OLD-ABC->TOPRIM->PglZ->SIGMA+OB-Fold+AAA-ATPase+Lon-Protease-BrxL-><-REase<-NACHT+SF1-SF2-Helicase+HKD-DNase<-Arm-DNA-bind_3 | 1213 | Bacteria>Pseudomonadota>Gammaproteobacteria>Enterobacterales>Enterobacteriaceae>Salmonella | GCA_007152865.1 | Salmonella_enterica_subsp._enterica_serovar_Typhimurium | DNN74_05345 | 60001..63642 | 90371 | BREX-system-P-loop-protein-BrxC-[Salmonella-enterica-subsp.-enterica-serovar-Typhimurium] |
| EBV2735692.1 | Type1.BREX | <-HEPN<-?\|\|?->\|\|YjhX_toxin->SAM-METHYLASE->?->BrxA-DNA-binding->BrxB-BrxF-iSTAND->DUF6079*->N6-METHYLASE->OLD-ABC->TOPRIM->PglZ->SIGMA+OB-Fold+AAA-ATPase+Lon-Protease-BrxL->HKD-DNase+OGG-like-><-REase<-NACHT+SF1-SF2-Helicase+HKD-DNase | 1213 | Bacteria>Pseudomonadota>Gammaproteobacteria>Enterobacterales>Enterobacteriaceae>Salmonella | GCA_007164145.1 | Salmonella_enterica_subsp._enterica_serovar_Stourbridge | DOA65_09445 | 58944..62585 | 1160766 | BREX-system-P-loop-protein-BrxC-[Salmonella-enterica-subsp.-enterica-serovar-Stourbridge] |
| EBV4609756.1 | Type1.BREX | BrxA-DNA-binding->BrxB-BrxF-iSTAND->DUF6079*->N6-METHYLASE->OLD-ABC->TOPRIM->PglZ->SIGMA+OB-Fold+AAA-ATPase+Lon-Protease-BrxL->HKD-DNase+OGG-like-><-REase<-NACHT+SF1-SF2-Helicase+HKD-DNase | 1213 | Bacteria>Pseudomonadota>Gammaproteobacteria>Enterobacterales>Enterobacteriaceae>Salmonella | GCA_007170635.1 | Salmonella_enterica_subsp._enterica_serovar_Solt | DOX24_02175 | 3073..6714 | 2565040 | BREX-system-P-loop-protein-BrxC-[Salmonella-enterica-subsp.-enterica-serovar-Solt] |
| EBV4672043.1 | Type1.BREX | YjhX_toxin->SAM-METHYLASE-><-?\|\|?->?->BrxA-DNA-binding->BrxB-BrxF-iSTAND->DUF6079*->N6-METHYLASE->OLD-ABC->TOPRIM->PglZ->SIGMA+OB-Fold+AAA-ATPase+Lon-Protease-BrxL-> | 1213 | Bacteria>Pseudomonadota>Gammaproteobacteria>Enterobacterales>Enterobacteriaceae>Salmonella | GCA_007170935.1 | Salmonella_enterica_subsp._enterica_serovar_Tamberma | DOW22_12520 | 149175..152816 | 2565079 | BREX-system-P-loop-protein-BrxC-[Salmonella-enterica-subsp.-enterica-serovar-Tamberma] |
| EBV5450231.1 | Type1.BREX | <-HEPN<-REase+iREase<-?\|\|\|\|YjhX_toxin->SAM-METHYLASE->?->?->BrxA-DNA-binding->BrxB-BrxF-iSTAND->DUF6079*->N6-METHYLASE->OLD-ABC->TOPRIM->PglZ->SIGMA+OB-Fold+AAA-ATPase+Lon-Protease-BrxL-><-REase<-NACHT+SF1-SF2-Helicase+HKD-DNase<-Arm-DNA-bind_3 | 1213 | Bacteria>Pseudomonadota>Gammaproteobacteria>Enterobacterales>Enterobacteriaceae>Salmonella | GCA_007175875.1 | Salmonella_enterica_subsp._enterica_serovar_Typhimurium | DO810_06270 | 60219..63860 | 90371 | BREX-system-P-loop-protein-BrxC-[Salmonella-enterica-subsp.-enterica-serovar-Typhimurium] |
| EBV7110754.1 | Type1.BREX | <-REase+iREase<-?\|\|\|\|YjhX_toxin->SAM-METHYLASE-><-?\|\|?->?->BrxA-DNA-binding->BrxB-BrxF-iSTAND->DUF6079*->N6-METHYLASE->OLD-ABC->TOPRIM->PglZ->SIGMA+OB-Fold+AAA-ATPase+Lon-Protease-BrxL-><-REase<-Couple_hipA+HipA_C<-HTH_3 | 1213 | Bacteria>Pseudomonadota>Gammaproteobacteria>Enterobacterales>Enterobacteriaceae>Salmonella | GCA_007180535.1 | Salmonella_enterica_subsp._enterica_serovar_Ohio | YY60_02790 | 154724..158365 | 117541 | BREX-system-P-loop-protein-BrxC-[Salmonella-enterica-subsp.-enterica-serovar-Ohio] |
| EBW0065987.1 | Type1.BREX | <-HEPN<-REase+iREase<-?\|\|YjhX_toxin->SAM-METHYLASE->?->BrxA-DNA-binding->BrxB-BrxF-iSTAND->DUF6079*->N6-METHYLASE-> | 1213 | Bacteria>Pseudomonadota>Gammaproteobacteria>Enterobacterales>Enterobacteriaceae>Salmonella | GCA_007195055.1 | Salmonella_enterica_subsp._enterica_serovar_Typhimurium_var._5- | AR716_20745 | 18317..21958 | 1620419 | BREX-system-P-loop-protein-BrxC-[Salmonella-enterica-subsp.-enterica-serovar-Typhimurium-var.-5-] |
| EBW2326418.1 | Type1.BREX | <-HEPN<-?\|\|YjhX_toxin->SAM-METHYLASE->?->\|\|BrxA-DNA-binding->BrxB-BrxF-iSTAND->DUF6079*->N6-METHYLASE->OLD-ABC->TOPRIM->PglZ->SIGMA+OB-Fold+AAA-ATPase+Lon-Protease-BrxL-><-REase<-NACHT+SF1-SF2-Helicase+HKD-DNase | 1213 | Bacteria>Pseudomonadota>Gammaproteobacteria>Enterobacterales>Enterobacteriaceae>Salmonella | GCA_007201245.1 | Salmonella_enterica_subsp._enterica_serovar_Agoueve | CBJ63_000885 | 165096..168737 | 1505961 | BREX-system-P-loop-protein-BrxC-[Salmonella-enterica-subsp.-enterica-serovar-Agoueve] |
| EBW1953456.1 | Type1.BREX | <-HEPN<-?\|\|\|\|YjhX_toxin->SAM-METHYLASE-><-?\|\|?->?->BrxA-DNA-binding->BrxB-BrxF-iSTAND->DUF6079*->N6-METHYLASE->OLD-ABC->TOPRIM->PglZ->SIGMA+OB-Fold+AAA-ATPase+Lon-Protease-BrxL-><-REase<-NACHT+SF1-SF2-Helicase+HKD-DNase<-Arm-DNA-bind_3 | 1213 | Bacteria>Pseudomonadota>Gammaproteobacteria>Enterobacterales>Enterobacteriaceae>Salmonella | GCA_007201375.1 | Salmonella_enterica_subsp._enterica_serovar_Cerro | C3N96_11410 | 44119..47760 | 340188 | BREX-system-P-loop-protein-BrxC-[Salmonella-enterica-subsp.-enterica-serovar-Cerro] |
| EBW2731416.1 | Type1.BREX | <-HEPN<-REase+iREase<-?\|\|\|\|YjhX_toxin->SAM-METHYLASE->?->?->BrxA-DNA-binding->BrxB-BrxF-iSTAND->DUF6079*->N6-METHYLASE->OLD-ABC->TOPRIM->PglZ->SIGMA+OB-Fold+AAA-ATPase+Lon-Protease-BrxL-><-REase<-NACHT+SF1-SF2-Helicase+HKD-DNase<-Arm-DNA-bind_3 | 1213 | Bacteria>Pseudomonadota>Gammaproteobacteria>Enterobacterales>Enterobacteriaceae>Salmonella | GCA_007205375.1 | Salmonella_enterica_subsp._enterica_serovar_Typhimurium | DPA65_05335 | 170935..174576 | 90371 | BREX-system-P-loop-protein-BrxC-[Salmonella-enterica-subsp.-enterica-serovar-Typhimurium] |
| EBW3295108.1 | Type1.BREX | <-HEPN<-?\|\|\|\|YjhX_toxin->SAM-METHYLASE-><-?\|\|?->?->BrxA-DNA-binding->BrxB-BrxF-iSTAND->DUF6079*->N6-METHYLASE->OLD-ABC->TOPRIM->PglZ->SIGMA+OB-Fold+AAA-ATPase+Lon-Protease-BrxL-><-REase<-NACHT+SF1-SF2-Helicase+HKD-DNase<-Arm-DNA-bind_3 | 1213 | Bacteria>Pseudomonadota>Gammaproteobacteria>Enterobacterales>Enterobacteriaceae>Salmonella | GCA_007207035.1 | Salmonella_enterica_subsp._enterica_serovar_Cerro | DPE09_02360 | 44118..47759 | 340188 | BREX-system-P-loop-protein-BrxC-[Salmonella-enterica-subsp.-enterica-serovar-Cerro] |
| EBW4591360.1 | Type1.BREX | <-HEPN<-REase+iREase<-?\|\|\|\|YjhX_toxin->SAM-METHYLASE->?->?->BrxA-DNA-binding->BrxB-BrxF-iSTAND->DUF6079*-> | 1144 | Bacteria>Pseudomonadota>Gammaproteobacteria>Enterobacterales>Enterobacteriaceae>Salmonella | GCA_007212435.1 | Salmonella_enterica_subsp._enterica_serovar_Typhimurium | DPP13_05415 | 60001..63431 | 90371 | BREX-system-P-loop-protein-BrxC-[Salmonella-enterica-subsp.-enterica-serovar-Typhimurium] |
| EBW5570928.1 | Type1.BREX | HTH_3->Couple_hipA+HipA_C->REase-><-SIGMA+OB-Fold+AAA-ATPase+Lon-Protease-BrxL<-PglZ<-TOPRIM<-OLD-ABC<-N6-METHYLASE<-DUF6079*<-BrxB-BrxF-iSTAND<-BrxA-DNA-binding<-?<-?\|\|?-><-SAM-METHYLASE<-YjhX_toxin\|\|\|\|?->REase+iREase-> | 1213 | Bacteria>Pseudomonadota>Gammaproteobacteria>Enterobacterales>Enterobacteriaceae>Salmonella | GCA_007216555.1 | Salmonella_enterica_subsp._enterica_serovar_Emek | DPY54_19345 | 30061..33702 | 286784 | BREX-system-P-loop-protein-BrxC-[Salmonella-enterica-subsp.-enterica-serovar-Emek] |
| EBW6592762.1 | Type1.BREX | SAM-METHYLASE-><-?\|\|?->\|\|?-><-?<-ZnR\|\|?->BrxA-DNA-binding->BrxB-BrxF-iSTAND->DUF6079-p*-> | 907 | Bacteria>Pseudomonadota>Gammaproteobacteria>Enterobacterales>Enterobacteriaceae>Salmonella | GCA_007223235.1 | Salmonella_enterica_subsp._enterica_serovar_Kentucky | DP741_24395 | 20426..23147 | 192955 | BREX-system-P-loop-protein-BrxC-[Salmonella-enterica-subsp.-enterica-serovar-Kentucky] |
| EBW8181804.1 | Type1.BREX | Glyoxalase->ACET->YjhX_toxin->ACET->METHYLASE->ZnR->Glyoxalase->BrxA-DNA-binding->BrxB-BrxF-iSTAND->DUF6079*->ParB+HNH->N6-METHYLASE->PglZ->SIGMA+OB-Fold+AAA-ATPase+Lon-Protease-BrxL-><-Arm-DNA-bind_3+Phage_integrase<-tRNA\|\|?-><-SKI\|\|METHYLASE-> | 1233 | Bacteria>Pseudomonadota>Gammaproteobacteria>Enterobacterales>Enterobacteriaceae>Salmonella | GCA_007227235.1 | Salmonella_enterica_subsp._enterica_serovar_Uganda | AR874_12565 | 406191..409892 | 487004 | BREX-system-P-loop-protein-BrxC-[Salmonella-enterica-subsp.-enterica-serovar-Uganda] |
| EBW8394988.1 | Type1.BREX | BrxA-DNA-binding->BrxB-BrxF-iSTAND->DUF6079*->N6-METHYLASE->OLD-ABC->TOPRIM->PglZ->SIGMA+OB-Fold+AAA-ATPase+Lon-Protease-BrxL->?-><-REase<-NACHT+SF1-SF2-Helicase+HKD-DNase<-Arm-DNA-bind_3 | 1213 | Bacteria>Pseudomonadota>Gammaproteobacteria>Enterobacterales>Enterobacteriaceae>Salmonella | GCA_007228915.1 | Salmonella_enterica_subsp._enterica_serovar_Florida | A3X38_19915 | 3074..6715 | 2564436 | BREX-system-P-loop-protein-BrxC-[Salmonella-enterica-subsp.-enterica-serovar-Florida] |
| EBX0519400.1 | Type1.BREX | <-HEPN<-?\|\|\|\|YjhX_toxin->SAM-METHYLASE-><-?\|\|?->?->BrxA-DNA-binding->BrxB-BrxF-iSTAND->DUF6079*->PglZ->SIGMA+OB-Fold+AAA-ATPase+Lon-Protease-BrxL->HKD-DNase+OGG-like-><-REase<-NACHT+SF1-SF2-Helicase+HKD-DNase<-Arm-DNA-bind_3<-tRNA\|\|?-><-SKI\|\|METHYLASE-> | 1213 | Bacteria>Pseudomonadota>Gammaproteobacteria>Enterobacterales>Enterobacteriaceae>Salmonella | GCA_007236875.1 | Salmonella_enterica_subsp._enterica_serovar_Saintpaul | AHL42_03320 | 144059..147700 | 90105 | BREX-system-P-loop-protein-BrxC-[Salmonella-enterica-subsp.-enterica-serovar-Saintpaul] |
| EBX6018293.1 | Type1.BREX | <-SIGMA+OB-Fold+AAA-ATPase+Lon-Protease-BrxL<-PglZ<-TOPRIM<-OLD-ABC<-N6-METHYLASE<-DUF6079*<-BrxB-BrxF-iSTAND<-BrxA-DNA-binding<-?<-?<-SAM-METHYLASE<-YjhX_toxin\|\|\|\|?->HEPN-> | 1213 | Bacteria>Pseudomonadota>Gammaproteobacteria>Enterobacterales>Enterobacteriaceae>Salmonella | GCA_007258805.1 | Salmonella_enterica_subsp._enterica_serovar_Dortmund | DSR74_22245 | 12037..15678 | 2564368 | BREX-system-P-loop-protein-BrxC-[Salmonella-enterica-subsp.-enterica-serovar-Dortmund] |
| EBX6302551.1 | Type1.BREX | <-HEPN<-?\|\|?->\|\|YjhX_toxin->SAM-METHYLASE->?->?->BrxA-DNA-binding->BrxB-BrxF-iSTAND->DUF6079*->N6-METHYLASE->OLD-ABC->TOPRIM->PglZ->SIGMA+OB-Fold+AAA-ATPase+Lon-Protease-BrxL->?-><-REase<-NACHT+SF1-SF2-Helicase+HKD-DNase | 1213 | Bacteria>Pseudomonadota>Gammaproteobacteria>Enterobacterales>Enterobacteriaceae>Salmonella | GCA_007260765.1 | Salmonella_enterica_subsp._enterica_serovar_Enugu | DS191_04270 | 25805..29446 | 2564400 | BREX-system-P-loop-protein-BrxC-[Salmonella-enterica-subsp.-enterica-serovar-Enugu] |
| EBX8836172.1 | Type1.BREX | <-HEPN<-?\|\|\|\|YjhX_toxin->SAM-METHYLASE-><-?\|\|?->?->BrxA-DNA-binding->BrxB-BrxF-iSTAND->DUF6079*->N6-METHYLASE->OLD-ABC->TOPRIM->PglZ->SIGMA+OB-Fold+AAA-ATPase+Lon-Protease-BrxL->?-><-REase<-NACHT+SF1-SF2-Helicase+HKD-DNase<-Arm-DNA-bind_3 | 1213 | Bacteria>Pseudomonadota>Gammaproteobacteria>Enterobacterales>Enterobacteriaceae>Salmonella | GCA_007270555.1 | Salmonella_enterica_subsp._enterica_serovar_Urbana | DTU76_05370 | 52211..55852 | 260677 | BREX-system-P-loop-protein-BrxC-[Salmonella-enterica-subsp.-enterica-serovar-Urbana] |
| EBY1390424.1 | Type1.BREX | NACHT+SF1-SF2-Helicase+HKD-DNase->REase-><-?<-SIGMA+OB-Fold+AAA-ATPase+Lon-Protease-BrxL<-PglZ<-TOPRIM<-OLD-ABC<-N6-METHYLASE<-DUF6079*<-BrxB-BrxF-iSTAND<-BrxA-DNA-binding<-?<-?<-SAM-METHYLASE<-YjhX_toxin\|\|<-?\|\|?->HEPN-> | 1213 | Bacteria>Pseudomonadota>Gammaproteobacteria>Enterobacterales>Enterobacteriaceae>Salmonella | GCA_007281935.1 | Salmonella_enterica_subsp._enterica_serovar_Wagenia | DUW06_08905 | 152900..156541 | 322833 | BREX-system-P-loop-protein-BrxC-[Salmonella-enterica-subsp.-enterica-serovar-Wagenia] |
| EBY1767815.1 | Type1.BREX | <-HEPN<-?\|\|?->\|\|YjhX_toxin->SAM-METHYLASE->?->?->BrxA-DNA-binding->BrxB-BrxF-iSTAND->DUF6079*->N6-METHYLASE->OLD-ABC->TOPRIM->PglZ->SIGMA+OB-Fold+AAA-ATPase+Lon-Protease-BrxL->HKD-DNase+OGG-like-><-REase<-NACHT+SF1-SF2-Helicase+HKD-DNase<-Arm-DNA-bind_3 | 1213 | Bacteria>Pseudomonadota>Gammaproteobacteria>Enterobacterales>Enterobacteriaceae>Salmonella | GCA_007283275.1 | Salmonella_enterica_subsp._enterica_serovar_Georgia | DU102_16105 | 50651..54292 | 2564467 | BREX-system-P-loop-protein-BrxC-[Salmonella-enterica-subsp.-enterica-serovar-Georgia] |
| EBY2665620.1 | Type1.BREX | YjhX_toxin->SAM-METHYLASE-><-?\|\|?->?->BrxA-DNA-binding->BrxB-BrxF-iSTAND->DUF6079*->N6-METHYLASE->OLD-ABC->TOPRIM->PglZ->SIGMA+OB-Fold+AAA-ATPase+Lon-Protease-BrxL->?-><-REase<-NACHT+SF1-SF2-Helicase+HKD-DNase | 1213 | Bacteria>Pseudomonadota>Gammaproteobacteria>Enterobacterales>Enterobacteriaceae>Salmonella | GCA_007287115.1 | Salmonella_enterica_subsp._enterica_serovar_Teko | DVF23_02315 | 17137..20778 | 2079198 | BREX-system-P-loop-protein-BrxC-[Salmonella-enterica-subsp.-enterica-serovar-Teko] |
| EBY3147217.1 | Type1.BREX | <-HEPN<-?\|\|YjhX_toxin->SAM-METHYLASE->?->BrxA-DNA-binding->BrxB-BrxF-iSTAND->DUF6079*->N6-METHYLASE->OLD-ABC->TOPRIM->PglZ->SIGMA+OB-Fold+AAA-ATPase+Lon-Protease-BrxL-><-REase<-NACHT+SF1-SF2-Helicase+HKD-DNase<-tRNA\|\|?-><-SKI | 1213 | Bacteria>Pseudomonadota>Gammaproteobacteria>Enterobacterales>Enterobacteriaceae>Salmonella | GCA_007287795.1 | Salmonella_enterica_subsp._enterica_serovar_Teshie | KZE80_000349 | 384366..388007 | 2565098 | BREX-system-P-loop-protein-BrxC-[Salmonella-enterica-subsp.-enterica-serovar-Teshie] |
| EBY3006707.1 | Type1.BREX | YjhX_toxin->SAM-METHYLASE-><-?\|\|?->?->BrxA-DNA-binding->BrxB-BrxF-iSTAND->DUF6079*->N6-METHYLASE-> | 1213 | Bacteria>Pseudomonadota>Gammaproteobacteria>Enterobacterales>Enterobacteriaceae>Salmonella | GCA_007287955.1 | Salmonella_enterica_subsp._enterica_serovar_Ekpoui | DVG04_05655 | 135015..138656 | 2564392 | BREX-system-P-loop-protein-BrxC-[Salmonella-enterica-subsp.-enterica-serovar-Ekpoui] |
| EBZ6440064.1 | Type1.BREX | DUF6079-p*->N6-METHYLASE->OLD-ABC->TOPRIM->PglZ->SIGMA+OB-Fold+AAA-ATPase+Lon-Protease-BrxL-><-REase<-NACHT+SF1-SF2-Helicase+HKD-DNase<-Arm-DNA-bind_3 | 897 | Bacteria>Pseudomonadota>Gammaproteobacteria>Enterobacterales>Enterobacteriaceae>Salmonella | GCA_007344875.1 | Salmonella_enterica_subsp._enterica_serovar_Typhimurium | ED482_12665 | 1..2696 | 90371 | BREX-system-P-loop-protein-BrxC-[Salmonella-enterica-subsp.-enterica-serovar-Typhimurium] |
| EBZ6491792.1 | Type1.BREX | <-HEPN<-?\|\|?->YjhX_toxin->SAM-METHYLASE->?->BrxA-DNA-binding->BrxB-BrxF-iSTAND->DUF6079*->N6-METHYLASE->OLD-ABC->TOPRIM->PglZ->SIGMA+OB-Fold+AAA-ATPase+Lon-Protease-BrxL->HKD-DNase+OGG-like-><-REase<-NACHT+SF1-SF2-Helicase+HKD-DNase<-Arm-DNA-bind_3 | 1213 | Bacteria>Pseudomonadota>Gammaproteobacteria>Enterobacterales>Enterobacteriaceae>Salmonella | GCA_007345015.1 | Salmonella_enterica_subsp._enterica_serovar_Dugbe | ED480_20060 | 16934..20575 | 2564376 | BREX-system-P-loop-protein-BrxC-[Salmonella-enterica-subsp.-enterica-serovar-Dugbe] |
| EBZ8208146.1 | Type1.BREX | Glyoxalase->ACET->YjhX_toxin->ACET->METHYLASE->ZnR->Glyoxalase->BrxA-DNA-binding->BrxB-BrxF-iSTAND->DUF6079*->ParB+HNH->N6-METHYLASE->PglZ->SIGMA+OB-Fold+AAA-ATPase+Lon-Protease-BrxL-><-Arm-DNA-bind_3+Phage_integrase<-tRNA\|\|?-><-SKI\|\|METHYLASE-> | 1233 | Bacteria>Pseudomonadota>Gammaproteobacteria>Enterobacterales>Enterobacteriaceae>Salmonella | GCA_007351255.1 | Salmonella_enterica_subsp._enterica_serovar_Zanzibar | EGK94_05900 | 411162..414863 | 399588 | BREX-system-P-loop-protein-BrxC-[Salmonella-enterica-subsp.-enterica-serovar-Zanzibar] |
| EBZ9408825.1 | Type1.BREX | <-HEPN<-?\|\|YjhX_toxin->SAM-METHYLASE->?->\|\|BrxA-DNA-binding->BrxB-BrxF-iSTAND->DUF6079*->N6-METHYLASE-> | 1213 | Bacteria>Pseudomonadota>Gammaproteobacteria>Enterobacterales>Enterobacteriaceae>Salmonella | GCA_007357915.1 | Salmonella_enterica_subsp._enterica_serovar_Saintpaul | EHQ15_16500 | 82377..86018 | 90105 | BREX-system-P-loop-protein-BrxC-[Salmonella-enterica-subsp.-enterica-serovar-Saintpaul] |
| ECA8969862.1 | Type1.BREX | <-HEPN<-?\|\|YjhX_toxin->SAM-METHYLASE->?->BrxA-DNA-binding->BrxB-BrxF-iSTAND->DUF6079*->N6-METHYLASE->OLD-ABC->TOPRIM->PglZ->SIGMA+OB-Fold+AAA-ATPase+Lon-Protease-BrxL->?->\|\|HIN-HTH-><-tRNA\|\|?-><-SKI | 1213 | Bacteria>Pseudomonadota>Gammaproteobacteria>Enterobacterales>Enterobacteriaceae>Salmonella | GCA_007399965.1 | Salmonella_enterica_subsp._enterica_serovar_Omuna | ETB55_05725 | 57828..61469 | 2564887 | BREX-system-P-loop-protein-BrxC-[Salmonella-enterica-subsp.-enterica-serovar-Omuna] |
| ECB3141175.1 | Type1.BREX | <-HEPN<-REase+iREase<-?\|\|YjhX_toxin->SAM-METHYLASE->?->BrxA-DNA-binding->BrxB-BrxF-iSTAND->DUF6079-p*-> | 876 | Bacteria>Pseudomonadota>Gammaproteobacteria>Enterobacterales>Enterobacteriaceae>Salmonella | GCA_007419865.1 | Salmonella_enterica_subsp._enterica_serovar_Typhimurium | EWE34_19370 | 60221..62849 | 90371 | BREX-system-P-loop-protein-BrxC-[Salmonella-enterica-subsp.-enterica-serovar-Typhimurium] |
| ECB3805264.1 | Type1.BREX | <-HEPN<-?\|\|YjhX_toxin->SAM-METHYLASE->?->BrxA-DNA-binding->BrxB-BrxF-iSTAND->DUF6079*->N6-METHYLASE->OLD-ABC->TOPRIM->PglZ->SIGMA+OB-Fold+AAA-ATPase+Lon-Protease-BrxL->HKD-DNase+OGG-like-><-REase<-NACHT+SF1-SF2-Helicase+HKD-DNase | 1213 | Bacteria>Pseudomonadota>Gammaproteobacteria>Enterobacterales>Enterobacteriaceae>Salmonella | GCA_007422765.1 | Salmonella_enterica_subsp._enterica_serovar_Fufu | EW367_11360 | 52142..55783 | 2564449 | BREX-system-P-loop-protein-BrxC-[Salmonella-enterica-subsp.-enterica-serovar-Fufu] |
| ECB4380457.1 | Type1.BREX | <-DUF6079-p*<-BrxB-BrxF-iSTAND<-BrxA-DNA-binding<-?<-SAM-METHYLASE<-YjhX_toxin\|\|?->REase+iREase->HEPN-> | 685 | Bacteria>Pseudomonadota>Gammaproteobacteria>Enterobacterales>Enterobacteriaceae>Salmonella | GCA_007426465.1 | Salmonella_enterica_subsp._enterica_serovar_Typhimurium | EXA28_15315 | 1..2055 | 90371 | BREX-system-P-loop-protein-BrxC-[Salmonella-enterica-subsp.-enterica-serovar-Typhimurium] |
| ECC0743076.1 | Type1.BREX | Arm-DNA-bind_3->NACHT+SF1-SF2-Helicase+HKD-DNase->REase-><-SIGMA+OB-Fold+AAA-ATPase+Lon-Protease-BrxL<-PglZ<-TOPRIM<-OLD-ABC<-N6-METHYLASE<-DUF6079*<-BrxB-BrxF-iSTAND<-BrxA-DNA-binding<-?<-SAM-METHYLASE<-YjhX_toxin\|\|?->REase+iREase->HEPN-> | 1213 | Bacteria>Pseudomonadota>Gammaproteobacteria>Enterobacterales>Enterobacteriaceae>Salmonella | GCA_007457215.1 | Salmonella_enterica_subsp._enterica | FMV06_07395 | 108064..111705 | 59201 | BREX-system-P-loop-protein-BrxC-[Salmonella-enterica-subsp.-enterica] |
| ECC1667299.1 | Type1.BREX | <-HEPN<-REase+iREase<-?\|\|YjhX_toxin->METHYLASE->?->\|\|BrxA-DNA-binding->BrxB-BrxF-iSTAND->DUF6079*->N6-METHYLASE->OLD-ABC->TOPRIM->PglZ->SIGMA+OB-Fold+AAA-ATPase+Lon-Protease-BrxL->HKD-DNase+OGG-like-><-REase<-NACHT+SF1-SF2-Helicase+HKD-DNase<-nSTAND1+HNH | 1213 | Bacteria>Pseudomonadota>Gammaproteobacteria>Enterobacterales>Enterobacteriaceae>Salmonella | GCA_007463295.1 | Salmonella_enterica_subsp._salamae | FNN95_01255 | 267827..271468 | 59202 | BREX-system-P-loop-protein-BrxC-[Salmonella-enterica-subsp.-salamae] |
| ECC3171797.1 | Type1.BREX | <-HEPN<-REase+iREase<-?\|\|YjhX_toxin->SAM-METHYLASE->?->BrxA-DNA-binding->BrxB-BrxF-iSTAND->DUF6079*->N6-METHYLASE->OLD-ABC->TOPRIM->PglZ->SIGMA+OB-Fold+AAA-ATPase+Lon-Protease-BrxL-><-REase<-NACHT+SF1-SF2-Helicase+HKD-DNase<-Arm-DNA-bind_3 | 1213 | Bacteria>Pseudomonadota>Gammaproteobacteria>Enterobacterales>Enterobacteriaceae>Salmonella | GCA_007472855.1 | Salmonella_enterica_subsp._enterica | R914_02595 | 77607..81248 | 59201 | BREX-system-P-loop-protein-BrxC-[Salmonella-enterica-subsp.-enterica] |
| ECC8603900.1 | Type1.BREX | <-HEPN<-REase+iREase<-?\|\|\|\|YjhX_toxin->SAM-METHYLASE->?->?->BrxA-DNA-binding->BrxB-BrxF-iSTAND->DUF6079-p*-> | 987 | Bacteria>Pseudomonadota>Gammaproteobacteria>Enterobacterales>Enterobacteriaceae>Salmonella | GCA_007497185.1 | Salmonella_enterica_subsp._enterica | DK044_12730 | 60001..62961 | 59201 | BREX-system-P-loop-protein-BrxC-[Salmonella-enterica-subsp.-enterica] |
| ECD1705587.1 | Type1.BREX | Arm-DNA-bind_3->NACHT+SF1-SF2-Helicase+HKD-DNase->REase-><-SIGMA+OB-Fold+AAA-ATPase+Lon-Protease-BrxL<-PglZ<-TOPRIM<-OLD-ABC<-N6-METHYLASE<-DUF6079*<-BrxB-BrxF-iSTAND<-BrxA-DNA-binding<-?<-SAM-METHYLASE<-YjhX_toxin\|\|?->REase+iREase->HEPN-> | 1213 | Bacteria>Pseudomonadota>Gammaproteobacteria>Enterobacterales>Enterobacteriaceae>Salmonella | GCA_007509785.1 | Salmonella_enterica_subsp._enterica | EWD17_08950 | 108064..111705 | 59201 | BREX-system-P-loop-protein-BrxC-[Salmonella-enterica-subsp.-enterica] |
| ECD2821735.1 | Type1.BREX | <-HEPN<-?\|\|?->YjhX_toxin->SAM-METHYLASE->?->BrxA-DNA-binding->BrxB-BrxF-iSTAND->DUF6079*->N6-METHYLASE->OLD-ABC->TOPRIM->PglZ->SIGMA+OB-Fold+AAA-ATPase+Lon-Protease-BrxL->HKD-DNase+OGG-like-><-REase<-NACHT+SF1-SF2-Helicase+HKD-DNase<-Arm-DNA-bind_3 | 1213 | Bacteria>Pseudomonadota>Gammaproteobacteria>Enterobacterales>Enterobacteriaceae>Salmonella | GCA_007513645.1 | Salmonella_enterica_subsp._enterica_serovar_Fanti | EXA45_11150 | 52318..55959 | 2564418 | BREX-system-P-loop-protein-BrxC-[Salmonella-enterica-subsp.-enterica-serovar-Fanti] |
| ECD3895174.1 | Type1.BREX | <-HEPN<-?\|\|YjhX_toxin->SAM-METHYLASE->?->BrxA-DNA-binding->BrxB-BrxF-iSTAND->DUF6079*->N6-METHYLASE->OLD-ABC->TOPRIM->PglZ->SIGMA+OB-Fold+AAA-ATPase+Lon-Protease-BrxL-><-REase<-NACHT+SF1-SF2-Helicase+HKD-DNase | 1213 | Bacteria>Pseudomonadota>Gammaproteobacteria>Enterobacterales>Enterobacteriaceae>Salmonella | GCA_007520185.1 | Salmonella_enterica_subsp._enterica_serovar_Gloucester | EOU70_05950 | 164775..168416 | 2564474 | BREX-system-P-loop-protein-BrxC-[Salmonella-enterica-subsp.-enterica-serovar-Gloucester] |
| ECD4420808.1 | Type1.BREX | Glyoxalase->ACET->YjhX_toxin->ACET->METHYLASE->ZnR->Glyoxalase->BrxA-DNA-binding->BrxB-BrxF-iSTAND->DUF6079*->ParB+HNH->N6-METHYLASE->PglZ->SIGMA+OB-Fold+AAA-ATPase+Lon-Protease-BrxL-><-Arm-DNA-bind_3+Phage_integrase<-tRNA\|\|?-><-SKI\|\|METHYLASE-> | 1233 | Bacteria>Pseudomonadota>Gammaproteobacteria>Enterobacterales>Enterobacteriaceae>Salmonella | GCA_007522325.1 | Salmonella_enterica_subsp._enterica_serovar_Berkeley | EO910_10430 | 8115..11816 | 1965103 | BREX-system-P-loop-protein-BrxC-[Salmonella-enterica-subsp.-enterica-serovar-Berkeley] |
| ECD9608662.1 | Type1.BREX | <-HEPN<-REase+iREase<-?\|\|YjhX_toxin->METHYLASE->?->\|\|BrxA-DNA-binding->BrxB-BrxF-iSTAND->DUF6079*->N6-METHYLASE->OLD-ABC->TOPRIM->PglZ->SIGMA+OB-Fold+AAA-ATPase+Lon-Protease-BrxL->HKD-DNase+OGG-like-> | 1213 | Bacteria>Pseudomonadota>Gammaproteobacteria>Enterobacterales>Enterobacteriaceae>Salmonella | GCA_007542335.1 | Salmonella_enterica_subsp._salamae | FNZ18_04560 | 256517..260158 | 59202 | BREX-system-P-loop-protein-BrxC-[Salmonella-enterica-subsp.-salamae] |
| ECE0383737.1 | Type1.BREX | <-HEPN<-?\|\|?->\|\|YjhX_toxin->SAM-METHYLASE-><-?\|\|?->BrxA-DNA-binding->BrxB-BrxF-iSTAND->DUF6079*->N6-METHYLASE->OLD-ABC->TOPRIM->PglZ->SIGMA+OB-Fold+AAA-ATPase+Lon-Protease-BrxL->?-><-REase<-NACHT+SF1-SF2-Helicase+HKD-DNase<-Arm-DNA-bind_3 | 1213 | Bacteria>Pseudomonadota>Gammaproteobacteria>Enterobacterales>Enterobacteriaceae>Salmonella | GCA_007548695.1 | Salmonella_enterica_subsp._enterica | XZ45_12010 | 25704..29345 | 59201 | BREX-system-P-loop-protein-BrxC-[Salmonella-enterica-subsp.-enterica] |
| ECE6154798.1 | Type1.BREX | <-HEPN<-?\|\|?->\|\|YjhX_toxin->SAM-METHYLASE->?->?->BrxA-DNA-binding->BrxB-BrxF-iSTAND->DUF6079*->N6-METHYLASE-> | 1213 | Bacteria>Pseudomonadota>Gammaproteobacteria>Enterobacterales>Enterobacteriaceae>Salmonella | GCA_007577415.1 | Salmonella_enterica_subsp._enterica | DPK65_21905 | 11265..14906 | 59201 | BREX-system-P-loop-protein-BrxC-[Salmonella-enterica-subsp.-enterica] |
| ECE6725605.1 | Type1.BREX | <-RHH<-?<-?<-SIGMA+OB-Fold+AAA-ATPase+Lon-Protease-BrxL<-PglZ<-N6-METHYLASE<-DUF6079*<-BrxB-BrxF-iSTAND<-BrxA-DNA-binding<-wHTH+WYL+WCX\|\|DUF4236-><-CR-ATPase5+REase<-DDE_Tnp_IS1 | 1226 | Bacteria>Pseudomonadota>Gammaproteobacteria>Enterobacterales>Enterobacteriaceae>Salmonella | GCA_007580505.1 | Salmonella_enterica_subsp._enterica | DSQ87_23555 | 10655..14335 | 59201 | BREX-system-P-loop-protein-BrxC-[Salmonella-enterica-subsp.-enterica] |
| ECE7382814.1 | Type1.BREX | <-DUF6079-p*<-BrxB-BrxF-iSTAND<-BrxA-DNA-binding<-?<-?<-SAM-METHYLASE<-YjhX_toxin\|\|\|\|?->REase+iREase->HEPN-> | 693 | Bacteria>Pseudomonadota>Gammaproteobacteria>Enterobacterales>Enterobacteriaceae>Salmonella | GCA_007582365.1 | Salmonella_enterica_subsp._enterica | D7M99_18000 | 1..2079 | 59201 | BREX-system-P-loop-protein-BrxC-[Salmonella-enterica-subsp.-enterica] |
| ECF0142189.1 | Type1.BREX | <-MarR-wHTH<-?<-SIGMA-HTH+REase<-wHTH+SAD+HNH\|\|?-><-?<-ZnR\|\|BrxA-DNA-binding->BrxB-BrxF-iSTAND->DUF6079*->N6-METHYLASE->OLD-ABC->TOPRIM->PglZ->SIGMA+OB-Fold+AAA-ATPase+Lon-Protease-BrxL->HKD-DNase+OGG-like-><-REase<-NACHT+SF1-SF2-Helicase+HKD-DNase<-AAA+HTH | 1204 | Bacteria>Pseudomonadota>Gammaproteobacteria>Enterobacterales>Enterobacteriaceae>Salmonella | GCA_007595955.1 | Salmonella_enterica_subsp._enterica_serovar_Monschaui | EX931_14155 | 69682..73296 | 29480 | BREX-system-P-loop-protein-BrxC-[Salmonella-enterica-subsp.-enterica-serovar-Monschaui] |
| ECF4176041.1 | Type1.BREX | Arm-DNA-bind_3->NACHT+SF1-SF2-Helicase+HKD-DNase->REase-><-SIGMA+OB-Fold+AAA-ATPase+Lon-Protease-BrxL<-PglZ<-TOPRIM<-OLD-ABC<-N6-METHYLASE<-DUF6079-p* | 864 | Bacteria>Pseudomonadota>Gammaproteobacteria>Enterobacterales>Enterobacteriaceae>Salmonella | GCA_007613175.1 | Salmonella_enterica_subsp._enterica_serovar_Typhimurium | E5U03_12955 | 108064..110659 | 90371 | BREX-system-P-loop-protein-BrxC-[Salmonella-enterica-subsp.-enterica-serovar-Typhimurium] |
| ECF4172399.1 | Type1.BREX | <-HEPN<-REase+iREase<-?\|\|YjhX_toxin->SAM-METHYLASE->?->BrxA-DNA-binding->BrxB-BrxF-iSTAND->DUF6079*-> | 1076 | Bacteria>Pseudomonadota>Gammaproteobacteria>Enterobacterales>Enterobacteriaceae>Salmonella | GCA_007613215.1 | Salmonella_enterica_subsp._enterica | E5100_18995 | 60001..63229 | 59201 | BREX-system-P-loop-protein-BrxC-[Salmonella-enterica-subsp.-enterica] |
| ECG1483122.1 | Type1.BREX | YjhX_toxin->SAM-METHYLASE-><-?\|\|?->\|\|?->BrxA-DNA-binding->BrxB-BrxF-iSTAND->DUF6079*->N6-METHYLASE->OLD-ABC->TOPRIM->PglZ->SIGMA+OB-Fold+AAA-ATPase+Lon-Protease-BrxL-><-REase<-Couple_hipA+HipA_C<-HTH_3 | 1213 | Bacteria>Pseudomonadota>Gammaproteobacteria>Enterobacterales>Enterobacteriaceae>Salmonella | GCA_007649655.1 | Salmonella_enterica_subsp._enterica | DT144_20820 | 52449..56090 | 59201 | BREX-system-P-loop-protein-BrxC-[Salmonella-enterica-subsp.-enterica] |
| ECG1893117.1 | Type1.BREX | <-HEPN<-?\|\|?->YjhX_toxin->SAM-METHYLASE->?->BrxA-DNA-binding->BrxB-BrxF-iSTAND->DUF6079*->N6-METHYLASE->OLD-ABC->TOPRIM->PglZ->SIGMA+OB-Fold+AAA-ATPase+Lon-Protease-BrxL->?-><-REase<-NACHT+SF1-SF2-Helicase+HKD-DNase | 1213 | Bacteria>Pseudomonadota>Gammaproteobacteria>Enterobacterales>Enterobacteriaceae>Salmonella | GCA_007650535.1 | Salmonella_enterica_subsp._enterica | EI460_05105 | 155639..159280 | 59201 | BREX-system-P-loop-protein-BrxC-[Salmonella-enterica-subsp.-enterica] |
| ECG3781073.1 | Type1.BREX | <-HEPN<-?\|\|?->YjhX_toxin->SAM-METHYLASE->?->BrxA-DNA-binding->BrxB-BrxF-iSTAND->DUF6079*->N6-METHYLASE->OLD-ABC->TOPRIM->PglZ->SIGMA+OB-Fold+AAA-ATPase+Lon-Protease-BrxL-><-REase<-Couple_hipA+HipA_C<-HTH_3 | 1213 | Bacteria>Pseudomonadota>Gammaproteobacteria>Enterobacterales>Enterobacteriaceae>Salmonella | GCA_007660485.1 | Salmonella_enterica_subsp._enterica_serovar_Breukelen | EXA83_17820 | 52108..55749 | 2564274 | BREX-system-P-loop-protein-BrxC-[Salmonella-enterica-subsp.-enterica-serovar-Breukelen] |
| ECG5099189.1 | Type1.BREX | <-HEPN<-?\|\|YjhX_toxin->SAM-METHYLASE->?->BrxA-DNA-binding->BrxB-BrxF-iSTAND->DUF6079*->N6-METHYLASE->OLD-ABC->TOPRIM->PglZ->SIGMA+OB-Fold+AAA-ATPase+Lon-Protease-BrxL-><-REase | 1213 | Bacteria>Pseudomonadota>Gammaproteobacteria>Enterobacterales>Enterobacteriaceae>Salmonella | GCA_007668905.1 | Salmonella_enterica_subsp._enterica | E0547_20045 | 59933..63574 | 59201 | BREX-system-P-loop-protein-BrxC-[Salmonella-enterica-subsp.-enterica] |
| ECG6954637.1 | Type1.BREX | <-HEPN<-REase+iREase<-?\|\|YjhX_toxin->SAM-METHYLASE->?->BrxA-DNA-binding->BrxB-BrxF-iSTAND->DUF6079*->N6-METHYLASE-> | 1213 | Bacteria>Pseudomonadota>Gammaproteobacteria>Enterobacterales>Enterobacteriaceae>Salmonella | GCA_007687065.1 | Salmonella_enterica_subsp._enterica_serovar_Typhimurium | E5150_19655 | 60001..63642 | 90371 | BREX-system-P-loop-protein-BrxC-[Salmonella-enterica-subsp.-enterica-serovar-Typhimurium] |
| ECG8301814.1 | Type1.BREX | <-HEPN<-?\|\|?->YjhX_toxin->SAM-METHYLASE->?->BrxA-DNA-binding->BrxB-BrxF-iSTAND->DUF6079*->N6-METHYLASE->OLD-ABC->TOPRIM->PglZ->SIGMA+OB-Fold+AAA-ATPase+Lon-Protease-BrxL->HKD-DNase+OGG-like-><-REase<-NACHT+SF1-SF2-Helicase+HKD-DNase | 1213 | Bacteria>Pseudomonadota>Gammaproteobacteria>Enterobacterales>Enterobacteriaceae>Salmonella | GCA_007694625.1 | Salmonella_enterica_subsp._enterica_serovar_Glostrup | E1853_02135 | 58935..62576 | 1151180 | BREX-system-P-loop-protein-BrxC-[Salmonella-enterica-subsp.-enterica-serovar-Glostrup] |
| ECG9327787.1 | Type1.BREX | Arm-DNA-bind_3->NACHT+SF1-SF2-Helicase+HKD-DNase->REase-><-HKD-DNase+OGG-like<-SIGMA+OB-Fold+AAA-ATPase+Lon-Protease-BrxL<-PglZ<-TOPRIM<-OLD-ABC<-N6-METHYLASE<-DUF6079*<-BrxB-BrxF-iSTAND<-BrxA-DNA-binding<-?<-SAM-METHYLASE<-YjhX_toxin\|\|?->HEPN-> | 1213 | Bacteria>Pseudomonadota>Gammaproteobacteria>Enterobacterales>Enterobacteriaceae>Salmonella | GCA_007700105.1 | Salmonella_enterica | FD199_03415 | 131756..135397 | 28901 | BREX-system-P-loop-protein-BrxC-[Salmonella-enterica] |
| ECH3013710.1 | Type1.BREX | Arm-DNA-bind_3->NACHT+SF1-SF2-Helicase+HKD-DNase->REase-><-SIGMA+OB-Fold+AAA-ATPase+Lon-Protease-BrxL<-PglZ<-TOPRIM<-OLD-ABC<-N6-METHYLASE<-DUF6079*<-BrxB-BrxF-iSTAND<-BrxA-DNA-binding<-?<-SAM-METHYLASE<-YjhX_toxin\|\|?->REase+iREase->HEPN-> | 1213 | Bacteria>Pseudomonadota>Gammaproteobacteria>Enterobacterales>Enterobacteriaceae>Salmonella | GCA_007716945.1 | Salmonella_enterica | FOV18_07420 | 343098..346739 | 28901 | BREX-system-P-loop-protein-BrxC-[Salmonella-enterica] |
| ECI2073658.1 | Type1.BREX | Arm-DNA-bind_3->NACHT+SF1-SF2-Helicase+HKD-DNase->REase-><-SIGMA+OB-Fold+AAA-ATPase+Lon-Protease-BrxL<-PglZ<-TOPRIM<-OLD-ABC<-N6-METHYLASE<-DUF6079-p* | 851 | Bacteria>Pseudomonadota>Gammaproteobacteria>Enterobacterales>Enterobacteriaceae>Salmonella | GCA_007762095.1 | Salmonella_enterica_subsp._enterica_serovar_Kentucky | AUD11_09545 | 108064..110621 | 192955 | BREX-system-P-loop-protein-BrxC-[Salmonella-enterica-subsp.-enterica-serovar-Kentucky] |
| ECI3716680.1 | Type1.BREX | YjhX_toxin->SAM-METHYLASE-><-?\|\|?->?->BrxA-DNA-binding->BrxB-BrxF-iSTAND->DUF6079*->N6-METHYLASE->OLD-ABC->TOPRIM->PglZ->SIGMA+OB-Fold+AAA-ATPase+Lon-Protease-BrxL-><-REase<-NACHT+SF1-SF2-Helicase+HKD-DNase<-Arm-DNA-bind_3 | 1213 | Bacteria>Pseudomonadota>Gammaproteobacteria>Enterobacterales>Enterobacteriaceae>Salmonella | GCA_007768905.1 | Salmonella_enterica_subsp._enterica | DK077_11320 | 44493..48134 | 59201 | BREX-system-P-loop-protein-BrxC-[Salmonella-enterica-subsp.-enterica] |
| ECI5247229.1 | Type1.BREX | <-HEPN<-REase+iREase<-?\|\|\|\|YjhX_toxin->SAM-METHYLASE->?->?->BrxA-DNA-binding->BrxB-BrxF-iSTAND->DUF6079-p*-> | 849 | Bacteria>Pseudomonadota>Gammaproteobacteria>Enterobacterales>Enterobacteriaceae>Salmonella | GCA_007773965.1 | Salmonella_enterica_subsp._enterica | DUU01_16740 | 60001..62549 | 59201 | BREX-system-P-loop-protein-BrxC-[Salmonella-enterica-subsp.-enterica] |
| ECI6492267.1 | Type1.BREX | <-HEPN<-?\|\|?->YjhX_toxin->SAM-METHYLASE->?->BrxA-DNA-binding->BrxB-BrxF-iSTAND->DUF6079*->N6-METHYLASE->OLD-ABC->TOPRIM->PglZ->SIGMA+OB-Fold+AAA-ATPase+Lon-Protease-BrxL-><-REase<-Couple_hipA+HipA_C<-HTH_3<-HTH_21+rve+rve_3 | 1213 | Bacteria>Pseudomonadota>Gammaproteobacteria>Enterobacterales>Enterobacteriaceae>Salmonella | GCA_007781245.1 | Salmonella_enterica_subsp._enterica | EIL18_20695 | 55793..59434 | 59201 | BREX-system-P-loop-protein-BrxC-[Salmonella-enterica-subsp.-enterica] |
| ECI6612371.1 | Type1.BREX | <-HEPN<-?\|\|?->YjhX_toxin->SAM-METHYLASE->?->BrxA-DNA-binding->BrxB-BrxF-iSTAND->DUF6079*->N6-METHYLASE->OLD-ABC->TOPRIM->PglZ->SIGMA+OB-Fold+AAA-ATPase+Lon-Protease-BrxL-><-REase | 1213 | Bacteria>Pseudomonadota>Gammaproteobacteria>Enterobacterales>Enterobacteriaceae>Salmonella | GCA_007781585.1 | Salmonella_enterica_subsp._enterica | EU408_19955 | 62734..66375 | 59201 | BREX-system-P-loop-protein-BrxC-[Salmonella-enterica-subsp.-enterica] |
| ECI6683601.1 | Type1.BREX | <-HEPN<-?\|\|?->YjhX_toxin->SAM-METHYLASE->?->\|\|BrxA-DNA-binding->BrxB-BrxF-iSTAND->DUF6079*->N6-METHYLASE->OLD-ABC->TOPRIM->PglZ->SIGMA+OB-Fold+AAA-ATPase+Lon-Protease-BrxL-><-REase<-Couple_hipA+HipA_C<-HTH_3 | 1213 | Bacteria>Pseudomonadota>Gammaproteobacteria>Enterobacterales>Enterobacteriaceae>Salmonella | GCA_007781825.1 | Salmonella_enterica_subsp._enterica | EHE65_20725 | 52107..55748 | 59201 | BREX-system-P-loop-protein-BrxC-[Salmonella-enterica-subsp.-enterica] |
| ECI6882011.1 | Type1.BREX | <-HEPN<-REase+iREase<-?\|\|YjhX_toxin->SAM-METHYLASE->?->BrxA-DNA-binding->BrxB-BrxF-iSTAND->DUF6079-p*-> | 929 | Bacteria>Pseudomonadota>Gammaproteobacteria>Enterobacterales>Enterobacteriaceae>Salmonella | GCA_007782565.1 | Salmonella_enterica_subsp._enterica | EWD50_16145 | 60001..62787 | 59201 | BREX-system-P-loop-protein-BrxC-[Salmonella-enterica-subsp.-enterica] |
| ECI6944166.1 | Type1.BREX | <-HEPN<-REase+iREase<-?\|\|YjhX_toxin->SAM-METHYLASE->?->BrxA-DNA-binding->BrxB-BrxF-iSTAND->DUF6079-p*-> | 979 | Bacteria>Pseudomonadota>Gammaproteobacteria>Enterobacterales>Enterobacteriaceae>Salmonella | GCA_007783905.1 | Salmonella_enterica_subsp._enterica | EWD54_17660 | 60227..63165 | 59201 | BREX-system-P-loop-protein-BrxC-[Salmonella-enterica-subsp.-enterica] |
| ECI7333736.1 | Type1.BREX | Arm-DNA-bind_3->NACHT+SF1-SF2-Helicase+HKD-DNase->REase-><-SIGMA+OB-Fold+AAA-ATPase+Lon-Protease-BrxL<-PglZ<-TOPRIM<-OLD-ABC<-N6-METHYLASE<-DUF6079*<-BrxB-BrxF-iSTAND<-BrxA-DNA-binding<-?<-SAM-METHYLASE<-YjhX_toxin\|\|?->REase+iREase->HEPN-> | 1213 | Bacteria>Pseudomonadota>Gammaproteobacteria>Enterobacterales>Enterobacteriaceae>Salmonella | GCA_007783945.1 | Salmonella_enterica_subsp._enterica | EZ611_10915 | 108064..111705 | 59201 | BREX-system-P-loop-protein-BrxC-[Salmonella-enterica-subsp.-enterica] |
| ECJ2539588.1 | Type1.BREX | YjhX_toxin->METHYLASE->?-><-wHTH(SBD)+HNH\|\|SF1-SF2-Helicase+REase-><-PglZ<-ParB+HNH<-N6-METHYLASE<-DUF6079*<-BrxB-BrxF-iSTAND<-BrxA-DNA-binding\|\|wHTH->XIS-HTH->?->?-><-MAD-NTD+HNH<-?\|\|DOC+HTH-> | 1157 | Bacteria>Pseudomonadota>Gammaproteobacteria>Enterobacterales>Enterobacteriaceae>Salmonella | GCA_007808255.1 | Salmonella_enterica_subsp._salamae | FNI64_03990 | 384975..388448 | 59202 | BREX-system-P-loop-protein-BrxC-[Salmonella-enterica-subsp.-salamae] |
| ECJ4523530.1 | Type1.BREX | <-HEPN<-REase+iREase<-?\|\|\|\|YjhX_toxin->SAM-METHYLASE->?->?->BrxA-DNA-binding->BrxB-BrxF-iSTAND->DUF6079-p*-> | 599 | Bacteria>Pseudomonadota>Gammaproteobacteria>Enterobacterales>Enterobacteriaceae>Salmonella | GCA_007819955.1 | Salmonella_enterica_subsp._enterica | DPQ09_03955 | 170822..172620 | 59201 | BREX-system-P-loop-protein-BrxC-[Salmonella-enterica-subsp.-enterica] |
| ECJ6092446.1 | Type1.BREX | <-GntR-HTH++TrmB+ZDNA-HTH\|\|<-?<-HEPN<-?\|\|YjhX_toxin->METHYLASE->?->\|\|BrxA-DNA-binding->BrxB-BrxF-iSTAND->DUF6079*->N6-METHYLASE->OLD-ABC->TOPRIM->PglZ->SIGMA+OB-Fold+AAA-ATPase+Lon-Protease-BrxL->HKD-DNase+OGG-like-><-REase<-NACHT+SF1-SF2-Helicase+HKD-DNase\|\|wHTH-> | 1213 | Bacteria>Pseudomonadota>Gammaproteobacteria>Enterobacterales>Enterobacteriaceae>Salmonella | GCA_007834095.1 | Salmonella_enterica | GR78_06760 | 372350..375991 | 28901 | BREX-system-P-loop-protein-BrxC-[Salmonella-enterica] |
| ECJ9595725.1 | Type1.BREX | <-HEPN<-REase+iREase<-?\|\|YjhX_toxin->SAM-METHYLASE->?->BrxA-DNA-binding->BrxB-BrxF-iSTAND->DUF6079*->N6-METHYLASE-> | 1213 | Bacteria>Pseudomonadota>Gammaproteobacteria>Enterobacterales>Enterobacteriaceae>Salmonella | GCA_007852735.1 | Salmonella_enterica | FQP47_21215 | 18123..21764 | 28901 | BREX-system-P-loop-protein-BrxC-[Salmonella-enterica] |
| ECK3339581.1 | Type1.BREX | Glyoxalase->ACET->YjhX_toxin->ACET->METHYLASE->ZnR->Glyoxalase->BrxA-DNA-binding->BrxB-BrxF-iSTAND->DUF6079*->ParB+HNH->N6-METHYLASE->PglZ->SIGMA+OB-Fold+AAA-ATPase+Lon-Protease-BrxL-><-Arm-DNA-bind_3+Phage_integrase<-tRNA\|\|?-><-SKI\|\|METHYLASE-> | 1233 | Bacteria>Pseudomonadota>Gammaproteobacteria>Enterobacterales>Enterobacteriaceae>Salmonella | GCA_007873945.1 | Salmonella_enterica | FRH68_09850 | 220275..223976 | 28901 | BREX-system-P-loop-protein-BrxC-[Salmonella-enterica] |
| ECK5773200.1 | Type1.BREX | <-HEPN<-REase+iREase<-?\|\|YjhX_toxin->SAM-METHYLASE->?->BrxA-DNA-binding->BrxB-BrxF-iSTAND->DUF6079*->N6-METHYLASE->OLD-ABC->TOPRIM->PglZ->SIGMA+OB-Fold+AAA-ATPase+Lon-Protease-BrxL-><-REase<-NACHT+SF1-SF2-Helicase+HKD-DNase<-Arm-DNA-bind_3 | 1213 | Bacteria>Pseudomonadota>Gammaproteobacteria>Enterobacterales>Enterobacteriaceae>Salmonella | GCA_007886645.1 | Salmonella_enterica_subsp._enterica_serovar_Typhimurium | AXS32_00815 | 172577..176218 | 90371 | BREX-system-P-loop-protein-BrxC-[Salmonella-enterica-subsp.-enterica-serovar-Typhimurium] |
| ECM2518301.1 | Type1.BREX | <-DUF6079-p*<-BrxB-BrxF-iSTAND<-BrxA-DNA-binding<-?<-SAM-METHYLASE<-YjhX_toxin\|\|?->REase+iREase->HEPN-> | 636 | Bacteria>Pseudomonadota>Gammaproteobacteria>Enterobacterales>Enterobacteriaceae>Salmonella | GCA_008018705.1 | Salmonella_enterica_subsp._enterica_serovar_Typhimurium | PZ44_21595 | 1..1907 | 90371 | BREX-system-P-loop-protein-BrxC-[Salmonella-enterica-subsp.-enterica-serovar-Typhimurium] |
| ECM3029728.1 | Type1.BREX | <-HEPN<-REase+iREase<-?\|\|YjhX_toxin->SAM-METHYLASE->?->BrxA-DNA-binding->BrxB-BrxF-iSTAND->DUF6079*->N6-METHYLASE->OLD-ABC->TOPRIM->PglZ->SIGMA+OB-Fold+AAA-ATPase+Lon-Protease-BrxL-><-REase<-NACHT+SF1-SF2-Helicase+HKD-DNase<-Arm-DNA-bind_3 | 1213 | Bacteria>Pseudomonadota>Gammaproteobacteria>Enterobacterales>Enterobacteriaceae>Salmonella | GCA_008020915.1 | Salmonella_enterica_subsp._enterica_serovar_Typhimurium | QB31_08405 | 60227..63868 | 90371 | BREX-system-P-loop-protein-BrxC-[Salmonella-enterica-subsp.-enterica-serovar-Typhimurium] |
| ECM3363968.1 | Type1.BREX | SKI-><-?\|\|tRNA->Arm-DNA-bind_3->NACHT+SF1-SF2-Helicase+HKD-DNase->REase-><-SIGMA+OB-Fold+AAA-ATPase+Lon-Protease-BrxL<-PglZ<-TOPRIM<-OLD-ABC<-DUF6079*<-BrxB-BrxF-iSTAND<-BrxA-DNA-binding<-?<-SAM-METHYLASE<-YjhX_toxin\|\|?->REase+iREase->HEPN-> | 1213 | Bacteria>Pseudomonadota>Gammaproteobacteria>Enterobacterales>Enterobacteriaceae>Salmonella | GCA_008021815.1 | Salmonella_enterica_subsp._enterica_serovar_Typhimurium | PW85_07020 | 108063..111704 | 90371 | BREX-system-P-loop-protein-BrxC-[Salmonella-enterica-subsp.-enterica-serovar-Typhimurium] |
| ECN0902565.1 | Type1.BREX | <-HEPN<-REase+iREase<-?\|\|YjhX_toxin->SAM-METHYLASE->?->BrxA-DNA-binding->BrxB-BrxF-iSTAND->DUF6079*->OLD-ABC->TOPRIM->PglZ->SIGMA+OB-Fold+AAA-ATPase+Lon-Protease-BrxL-><-REase<-NACHT+SF1-SF2-Helicase+HKD-DNase<-Arm-DNA-bind_3<-tRNA\|\|?-><-SKI | 1213 | Bacteria>Pseudomonadota>Gammaproteobacteria>Enterobacterales>Enterobacteriaceae>Salmonella | GCA_008061715.1 | Salmonella_enterica_subsp._enterica_serovar_Typhimurium | YR82_02405 | 60002..63643 | 90371 | BREX-system-P-loop-protein-BrxC-[Salmonella-enterica-subsp.-enterica-serovar-Typhimurium] |
| ECO0332142.1 | Type1.BREX | Arm-DNA-bind_3->NACHT+SF1-SF2-Helicase+HKD-DNase->REase-><-SIGMA+OB-Fold+AAA-ATPase+Lon-Protease-BrxL<-PglZ<-TOPRIM<-OLD-ABC<-N6-METHYLASE<-DUF6079*<-BrxB-BrxF-iSTAND<-BrxA-DNA-binding<-?<-SAM-METHYLASE<-YjhX_toxin\|\|?->REase+iREase->HEPN-> | 1213 | Bacteria>Pseudomonadota>Gammaproteobacteria>Enterobacterales>Enterobacteriaceae>Salmonella | GCA_008112305.1 | Salmonella_enterica_subsp._enterica_serovar_Typhimurium | ZS66_00475 | 108064..111705 | 90371 | BREX-system-P-loop-protein-BrxC-[Salmonella-enterica-subsp.-enterica-serovar-Typhimurium] |
| ECO1322281.1 | Type1.BREX | <-HEPN<-REase+iREase<-?\|\|YjhX_toxin->SAM-METHYLASE->?->BrxA-DNA-binding->BrxB-BrxF-iSTAND->DUF6079-p*-> | 631 | Bacteria>Pseudomonadota>Gammaproteobacteria>Enterobacterales>Enterobacteriaceae>Salmonella | GCA_008116905.1 | Salmonella_enterica_subsp._enterica_serovar_Typhimurium | AC815_12045 | 38663..40557 | 90371 | BREX-system-P-loop-protein-BrxC-[Salmonella-enterica-subsp.-enterica-serovar-Typhimurium] |
| ECO5284314.1 | Type1.BREX | <-HEPN<-?\|\|YjhX_toxin->SAM-METHYLASE->?->BrxA-DNA-binding->BrxB-BrxF-iSTAND->DUF6079*->N6-METHYLASE->OLD-ABC->TOPRIM->PglZ->SIGMA+OB-Fold+AAA-ATPase+Lon-Protease-BrxL-> | 1213 | Bacteria>Pseudomonadota>Gammaproteobacteria>Enterobacterales>Enterobacteriaceae>Salmonella | GCA_008153185.1 | Salmonella_enterica | FYE58_11845 | 398014..401655 | 28901 | BREX-system-P-loop-protein-BrxC-[Salmonella-enterica] |
| ECO7189041.1 | Type1.BREX | BrxB-BrxF-iSTAND->ACET->\|\|?-><-?<-?<-Phage_integrase\|\|DOC+HTH->BrxA-DNA-binding->BrxB-BrxF-iSTAND->DUF6079*->N6-METHYLASE->PglZ->SIGMA+OB-Fold+AAA-ATPase+Lon-Protease-BrxL-><-AAA+HTH<-?<-XIS-HTH\|\|?->?-><-Phage_integrase | 1213 | Bacteria>Pseudomonadota>Gammaproteobacteria>Enterobacterales>Enterobacteriaceae>Salmonella | GCA_008162045.1 | Salmonella_enterica | F6029_01875 | 16689..20330 | 28901 | BREX-system-P-loop-protein-BrxC-[Salmonella-enterica] |
| ECQ9432768.1 | Type1.BREX | BrxA-DNA-binding->BrxB-BrxF-iSTAND->DUF6079*->N6-METHYLASE->OLD-ABC->TOPRIM->PglZ->SIGMA+OB-Fold+AAA-ATPase+Lon-Protease-BrxL->?-><-REase<-NACHT+SF1-SF2-Helicase+HKD-DNase<-Arm-DNA-bind_3 | 1213 | Bacteria>Pseudomonadota>Gammaproteobacteria>Enterobacterales>Enterobacteriaceae>Salmonella | GCA_008312495.1 | Salmonella_enterica | F0371_05165 | 2795..6436 | 28901 | BREX-system-P-loop-protein-BrxC-[Salmonella-enterica] |
| ECR2528674.1 | Type1.BREX | Arm-DNA-bind_3->NACHT+SF1-SF2-Helicase+HKD-DNase->REase-><-SIGMA+OB-Fold+AAA-ATPase+Lon-Protease-BrxL<-PglZ<-TOPRIM<-OLD-ABC<-N6-METHYLASE<-DUF6079*<-BrxB-BrxF-iSTAND<-BrxA-DNA-binding<-?<-SAM-METHYLASE<-YjhX_toxin\|\|?->REase+iREase->HEPN-> | 1213 | Bacteria>Pseudomonadota>Gammaproteobacteria>Enterobacterales>Enterobacteriaceae>Salmonella | GCA_008341925.1 | Salmonella_enterica_subsp._enterica | F1J42_15865 | 108064..111705 | 59201 | BREX-system-P-loop-protein-BrxC-[Salmonella-enterica-subsp.-enterica] |
| ECS4740325.1 | Type1.BREX | Arm-DNA-bind_3->NACHT+SF1-SF2-Helicase+HKD-DNase->REase-><-SIGMA+OB-Fold+AAA-ATPase+Lon-Protease-BrxL<-PglZ<-TOPRIM<-OLD-ABC<-N6-METHYLASE<-DUF6079-p* | 907 | Bacteria>Pseudomonadota>Gammaproteobacteria>Enterobacterales>Enterobacteriaceae>Salmonella | GCA_008423865.1 | Salmonella_enterica_subsp._enterica_serovar_Typhimurium_var._5- | A9R80_15960 | 85998..88723 | 1620419 | BREX-system-P-loop-protein-BrxC-[Salmonella-enterica-subsp.-enterica-serovar-Typhimurium-var.-5-] |
| ECT8081314.1 | Type1.BREX | YjhX_toxin->SAM-METHYLASE-><-?\|\|?->?->BrxA-DNA-binding->BrxB-BrxF-iSTAND->DUF6079*->N6-METHYLASE->OLD-ABC->TOPRIM->PglZ->SIGMA+OB-Fold+AAA-ATPase+Lon-Protease-BrxL->?-><-REase<-NACHT+SF1-SF2-Helicase+HKD-DNase | 1213 | Bacteria>Pseudomonadota>Gammaproteobacteria>Enterobacterales>Enterobacteriaceae>Salmonella | GCA_008481645.1 | Salmonella_enterica_subsp._enterica_serovar_Carrau | BL796_09860 | 150076..153717 | 1160739 | BREX-system-P-loop-protein-BrxC-[Salmonella-enterica-subsp.-enterica-serovar-Carrau] |
| ECU0288387.1 | Type1.BREX | <-HEPN<-REase+iREase<-?\|\|\|\|YjhX_toxin->SAM-METHYLASE->?->?->BrxA-DNA-binding->BrxB-BrxF-iSTAND->DUF6079*->N6-METHYLASE->OLD-ABC->TOPRIM->PglZ->SIGMA+OB-Fold+AAA-ATPase+Lon-Protease-BrxL-><-REase<-NACHT+SF1-SF2-Helicase+HKD-DNase<-Arm-DNA-bind_3 | 1213 | Bacteria>Pseudomonadota>Gammaproteobacteria>Enterobacterales>Enterobacteriaceae>Salmonella | GCA_008492025.1 | Salmonella_enterica_subsp._enterica_serovar_Typhimurium | C7B88_00710 | 140025..143666 | 90371 | BREX-system-P-loop-protein-BrxC-[Salmonella-enterica-subsp.-enterica-serovar-Typhimurium] |

| Protein-ID | System | Gene-Neighbourhoods | Length | Taxonomy | Assembly-ID | Strain | Locus-Tag | Genomic-coordinates | TaxID | Fasta-defline |
|---|---|---|---|---|---|---|---|---|---|---|
| ECU9351200.1 | Type1.BREX | [illegible] | 1213 | Bacteria>Pseudomonadota>Gammaproteobacteria>Enterobacterales>Enterobacteriaceae>Salmonella | GCA_008530195.1 | Salmonella_enterica_subsp._enterica_serovar_Typhimurium | CIH13_12040 | 171029..174670 | 90371 | BREX-system-P-loop-protein-BrxC-[Salmonella-enterica-subsp.-enterica-serovar-Typhimurium] |
| ECV7778489.1 | Type1.BREX | [illegible] | 1213 | Bacteria>Pseudomonadota>Gammaproteobacteria>Enterobacterales>Enterobacteriaceae>Salmonella | GCA_008576505.1 | Salmonella_enterica_subsp._enterica_serovar_Typhimurium | ZX52_02590 | 60002..63643 | 90371 | BREX-system-P-loop-protein-BrxC-[Salmonella-enterica-subsp.-enterica-serovar-Typhimurium] |
| ECY3255615.1 | Type1.BREX | [illegible] | 1204 | Bacteria>Pseudomonadota>Gammaproteobacteria>Enterobacterales>Enterobacteriaceae>Salmonella | GCA_008731355.1 | Salmonella_enterica_subsp._enterica_serovar_Alachua | AU577_03140 | 17059..20673 | 913240 | BREX-system-P-loop-protein-BrxC-[Salmonella-enterica-subsp.-enterica-serovar-Alachua] |
| ECY8229161.1 | Type1.BREX | [illegible] | 1213 | Bacteria>Pseudomonadota>Gammaproteobacteria>Enterobacterales>Enterobacteriaceae>Salmonella | GCA_008752395.1 | Salmonella_enterica_subsp._enterica_serovar_Stanley | AU695_08280 | 160733..164374 | 192953 | BREX-system-P-loop-protein-BrxC-[Salmonella-enterica-subsp.-enterica-serovar-Stanley] |
| ECZ9372117.1 | Type1.BREX | [illegible] | 1213 | Bacteria>Pseudomonadota>Gammaproteobacteria>Enterobacterales>Enterobacteriaceae>Salmonella | GCA_008834045.1 | Salmonella_enterica_subsp._enterica_serovar_Typhimurium | A3U95_10245 | 108064..111705 | 90371 | BREX-system-P-loop-protein-BrxC-[Salmonella-enterica-subsp.-enterica-serovar-Typhimurium] |
| EDA5898036.1 | Type1.BREX | [illegible] | 1213 | Bacteria>Pseudomonadota>Gammaproteobacteria>Enterobacterales>Enterobacteriaceae>Salmonella | GCA_008864045.1 | Salmonella_enterica_subsp._enterica | F9X73_05935 | 25917..29558 | 59201 | BREX-system-P-loop-protein-BrxC-[Salmonella-enterica-subsp.-enterica] |
| EDA7420100.1 | Type1.BREX | [illegible] | 1213 | Bacteria>Pseudomonadota>Gammaproteobacteria>Enterobacterales>Enterobacteriaceae>Salmonella | GCA_008872905.1 | Salmonella_enterica_subsp._enterica_serovar_Typhimurium | A3U84_08760 | 60219..63860 | 90371 | BREX-system-P-loop-protein-BrxC-[Salmonella-enterica-subsp.-enterica-serovar-Typhimurium] |
| EDB0845244.1 | Type1.BREX | [illegible] | 1213 | Bacteria>Pseudomonadota>Gammaproteobacteria>Enterobacterales>Enterobacteriaceae>Salmonella | GCA_008888525.1 | Salmonella_enterica | F9C59_12685 | 144464..148105 | 28901 | BREX-system-P-loop-protein-BrxC-[Salmonella-enterica] |
| EDB2044587.1 | Type1.BREX | [illegible] | 1233 | Bacteria>Pseudomonadota>Gammaproteobacteria>Enterobacterales>Enterobacteriaceae>Salmonella | GCA_008894145.1 | Salmonella_enterica_subsp._enterica | F9P84_01850 | 405984..409685 | 59201 | BREX-system-P-loop-protein-BrxC-[Salmonella-enterica-subsp.-enterica] |
| EDB9656968.1 | Type1.BREX | [illegible] | 1213 | Bacteria>Pseudomonadota>Gammaproteobacteria>Enterobacterales>Enterobacteriaceae>Salmonella | GCA_008938335.1 | Salmonella_enterica_subsp._enterica_serovar_Typhimurium | BEO39_09545 | 27455..31096 | 90371 | BREX-system-P-loop-protein-BrxC-[Salmonella-enterica-subsp.-enterica-serovar-Typhimurium] |
| EDC2707334.1 | Type1.BREX | <-HEPN<-REase+iREase<-?\|\|YjhX_toxin->SAM-METHYLASE->?->BrxA-DNA-binding->BrxB-BrxF-iSTAND->DUF6079*->N6-METHYLASE-> | 1213 | Bacteria>Pseudomonadota>Gammaproteobacteria>Enterobacterales>Enterobacteriaceae>Salmonella | GCA_008954065.1 | Salmonella_enterica | GAD24_16260 | 80632..84273 | 28901 | BREX-system-P-loop-protein-BrxC-[Salmonella-enterica] |
| EDC2377951.1 | Type1.BREX | [illegible] | 1213 | Bacteria>Pseudomonadota>Gammaproteobacteria>Enterobacterales>Enterobacteriaceae>Salmonella | GCA_008954485.1 | Salmonella_enterica | GAA61_09305 | 25917..29558 | 28901 | BREX-system-P-loop-protein-BrxC-[Salmonella-enterica] |
| EDE2293304.1 | Type1.BREX | [illegible] | 1213 | Bacteria>Pseudomonadota>Gammaproteobacteria>Enterobacterales>Enterobacteriaceae>Salmonella | GCA_009089805.1 | Salmonella_enterica | GBE85_07255 | 108064..111705 | 28901 | BREX-system-P-loop-protein-BrxC-[Salmonella-enterica] |
| EDH2237970.1 | Type1.BREX | [illegible] | 1213 | Bacteria>Pseudomonadota>Gammaproteobacteria>Enterobacterales>Enterobacteriaceae>Salmonella | GCA_009231185.1 | Salmonella_enterica | GC580_06310 | 170513..174154 | 28901 | BREX-system-P-loop-protein-BrxC-[Salmonella-enterica] |
| EDH5631745.1 | Type1.BREX | <-HEPN<-?\|\|YjhX_toxin->SAM-METHYLASE->?->\|\|BrxA-DNA-binding->BrxB-BrxF-iSTAND->DUF6079*->N6-METHYLASE->OLD-ABC->TOPRIM->PglZ->SIGMA+OB-Fold+AAA-ATPase+Lon-Protease-BrxL-> | 1213 | Bacteria>Pseudomonadota>Gammaproteobacteria>Enterobacterales>Enterobacteriaceae>Salmonella | GCA_009247085.1 | Salmonella_enterica_subsp._enterica_serovar_Claibornei | CB172_15080 | 158451..162092 | 2572709 | BREX-system-P-loop-protein-BrxC-[Salmonella-enterica-subsp.-enterica-serovar-Claibornei] |
| EDH5972513.1 | Type1.BREX | <-HEPN<-REase+iREase<-?\|\|YjhX_toxin->SAM-METHYLASE->?->BrxA-DNA-binding->BrxB-BrxF-iSTAND->DUF6079*-> | 1176 | Bacteria>Pseudomonadota>Gammaproteobacteria>Enterobacterales>Enterobacteriaceae>Salmonella | GCA_009248125.1 | Salmonella_enterica_subsp._enterica_serovar_Typhimurium | CB276_16685 | 60001..63528 | 90371 | BREX-system-P-loop-protein-BrxC-[Salmonella-enterica-subsp.-enterica-serovar-Typhimurium] |
| EDH6190230.1 | Type1.BREX | [illegible] | 1213 | Bacteria>Pseudomonadota>Gammaproteobacteria>Enterobacterales>Enterobacteriaceae>Salmonella | GCA_009249585.1 | Salmonella_enterica_subsp._enterica_serovar_Typhi | CB347_02510 | 200361..204002 | 90370 | BREX-system-P-loop-protein-BrxC-[Salmonella-enterica-subsp.-enterica-serovar-Typhi] |
| EDH5995080.1 | Type1.BREX | [illegible] | 958 | Bacteria>Pseudomonadota>Gammaproteobacteria>Enterobacterales>Enterobacteriaceae>Salmonella | GCA_009250025.1 | Salmonella_enterica_subsp._enterica_serovar_Typhimurium | CB280_15355 | 60001..62876 | 90371 | BREX-system-P-loop-protein-BrxC-[Salmonella-enterica-subsp.-enterica-serovar-Typhimurium] |
| EDH6721610.1 | Type1.BREX | [illegible] | 691 | Bacteria>Pseudomonadota>Gammaproteobacteria>Enterobacterales>Enterobacteriaceae>Salmonella | GCA_009251865.1 | Salmonella_enterica_subsp._enterica_serovar_Heidelberg | CB494_23570 | 5530..7603 | 611 | BREX-system-P-loop-protein-BrxC-[Salmonella-enterica-subsp.-enterica-serovar-Heidelberg] |
| EDH9429027.1 | Type1.BREX | [illegible] | 1069 | Bacteria>Pseudomonadota>Gammaproteobacteria>Enterobacterales>Enterobacteriaceae>Salmonella | GCA_009288565.1 | Salmonella_enterica_subsp._enterica_serovar_Typhimurium | CC452_16960 | 59990..63196 | 90371 | BREX-system-P-loop-protein-BrxC-[Salmonella-enterica-subsp.-enterica-serovar-Typhimurium] |
| EDH9929876.1 | Type1.BREX | [illegible] | 1213 | Bacteria>Pseudomonadota>Gammaproteobacteria>Enterobacterales>Enterobacteriaceae>Salmonella | GCA_009291205.1 | Salmonella_enterica_subsp._enterica_serovar_Typhimurium | CC611_10195 | 170364..174005 | 90371 | BREX-system-P-loop-protein-BrxC-[Salmonella-enterica-subsp.-enterica-serovar-Typhimurium] |
| EDH9999209.1 | Type1.BREX | [illegible] | 978 | Bacteria>Pseudomonadota>Gammaproteobacteria>Enterobacterales>Enterobacteriaceae>Salmonella | GCA_009291525.1 | Salmonella_enterica_subsp._enterica_serovar_Typhimurium | CC636_17605 | 60002..62935 | 90371 | BREX-system-P-loop-protein-BrxC-[Salmonella-enterica-subsp.-enterica-serovar-Typhimurium] |
| EDH9995453.1 | Type1.BREX | [illegible] | 837 | Bacteria>Pseudomonadota>Gammaproteobacteria>Enterobacterales>Enterobacteriaceae>Salmonella | GCA_009292885.1 | Salmonella_enterica_subsp._enterica_serovar_Java | CC627_22205 | 1..2514 | 224729 | BREX-system-P-loop-protein-BrxC-[Salmonella-enterica-subsp.-enterica-serovar-Java] |
| EDI1182989.1 | Type1.BREX | [illegible] | 1060 | Bacteria>Pseudomonadota>Gammaproteobacteria>Enterobacterales>Enterobacteriaceae>Salmonella | GCA_009300115.1 | Salmonella_enterica_subsp._enterica_serovar_Typhimurium | CC995_04970 | 170835..174015 | 90371 | BREX-system-P-loop-protein-BrxC-[Salmonella-enterica-subsp.-enterica-serovar-Typhimurium] |
| EDI5281730.1 | Type1.BREX | [illegible] | 1213 | Bacteria>Pseudomonadota>Gammaproteobacteria>Enterobacterales>Enterobacteriaceae>Salmonella | GCA_009327915.1 | Salmonella_enterica_subsp._enterica_serovar_Typhimurium | CEM90_10175 | 170818..174459 | 90371 | BREX-system-P-loop-protein-BrxC-[Salmonella-enterica-subsp.-enterica-serovar-Typhimurium] |
| EDI9669678.1 | Type1.BREX | [illegible] | 1213 | Bacteria>Pseudomonadota>Gammaproteobacteria>Enterobacterales>Enterobacteriaceae>Salmonella | GCA_009349595.1 | Salmonella_enterica | GEV75_12805 | 38669..42310 | 28901 | BREX-system-P-loop-protein-BrxC-[Salmonella-enterica] |
| EDL1929631.1 | Type1.BREX | [illegible] | 1213 | Bacteria>Pseudomonadota>Gammaproteobacteria>Enterobacterales>Enterobacteriaceae>Salmonella | GCA_009508775.1 | Salmonella_enterica_subsp._enterica_serovar_Typhimurium | CQD92_11450 | 108064..111705 | 90371 | BREX-system-P-loop-protein-BrxC-[Salmonella-enterica-subsp.-enterica-serovar-Typhimurium] |
| EDL2092523.1 | Type1.BREX | [illegible] | 1213 | Bacteria>Pseudomonadota>Gammaproteobacteria>Enterobacterales>Enterobacteriaceae>Salmonella | GCA_009509795.1 | Salmonella_enterica_subsp._enterica_serovar_Typhimurium | CQF25_02855 | 72413..76054 | 90371 | BREX-system-P-loop-protein-BrxC-[Salmonella-enterica-subsp.-enterica-serovar-Typhimurium] |
| EDL6685014.1 | Type1.BREX | [illegible] | 1213 | Bacteria>Pseudomonadota>Gammaproteobacteria>Enterobacterales>Enterobacteriaceae>Salmonella | GCA_009531865.1 | Salmonella_enterica | CXC07_02670 | 170913..174554 | 28901 | BREX-system-P-loop-protein-BrxC-[Salmonella-enterica] |
| EDL7421472.1 | Type1.BREX | [illegible] | 1213 | Bacteria>Pseudomonadota>Gammaproteobacteria>Enterobacterales>Enterobacteriaceae>Salmonella | GCA_009534895.1 | Salmonella_enterica | CU840_19785 | 23186..26827 | 28901 | BREX-system-P-loop-protein-BrxC-[Salmonella-enterica] |
| EDM4126305.1 | Type1.BREX | [illegible] | 1213 | Bacteria>Pseudomonadota>Gammaproteobacteria>Enterobacterales>Enterobacteriaceae>Salmonella | GCA_009562515.1 | Salmonella_enterica_subsp._enterica_serovar_Typhimurium | CR810_06735 | 172578..176219 | 90371 | BREX-system-P-loop-protein-BrxC-[Salmonella-enterica-subsp.-enterica-serovar-Typhimurium] |
| EDM6640280.1 | Type1.BREX | [illegible] | 1213 | Bacteria>Pseudomonadota>Gammaproteobacteria>Enterobacterales>Enterobacteriaceae>Salmonella | GCA_009573765.1 | Salmonella_enterica_subsp._enterica_serovar_Typhimurium | CUM00_00810 | 170918..174559 | 90371 | BREX-system-P-loop-protein-BrxC-[Salmonella-enterica-subsp.-enterica-serovar-Typhimurium] |
| EDN6133533.1 | Type1.BREX | [illegible] | 1204 | Bacteria>Pseudomonadota>Gammaproteobacteria>Enterobacterales>Enterobacteriaceae>Salmonella | GCA_010008765.1 | Salmonella_enterica_subsp._enterica_serovar_Kentucky | GJQ78_12610 | 62378..65992 | 192955 | BREX-system-P-loop-protein-BrxC-[Salmonella-enterica-subsp.-enterica-serovar-Kentucky] |
| HAC8921685.1 | Type1.BREX | [illegible] | 778 | Bacteria>Pseudomonadota>Gammaproteobacteria>Enterobacterales>Enterobacteriaceae>Salmonella | GCA_010188005.1 | Salmonella_enterica | G0J09_01290 | 58631..60969 | 28901 | BREX-system-P-loop-protein-BrxC-[Salmonella-enterica] |
| HAD0448194.1 | Type1.BREX | [illegible] | 1213 | Bacteria>Pseudomonadota>Gammaproteobacteria>Enterobacterales>Enterobacteriaceae>Salmonella | GCA_010193865.1 | Salmonella_enterica_subsp._enterica_serovar_Typhimurium | G0N57_00710 | 150026..153667 | 90371 | BREX-system-P-loop-protein-BrxC-[Salmonella-enterica-subsp.-enterica-serovar-Typhimurium] |
| HAD1283638.1 | Type1.BREX | [illegible] | 1107 | Bacteria>Pseudomonadota>Gammaproteobacteria>Enterobacterales>Enterobacteriaceae>Salmonella | GCA_010197825.1 | Salmonella_enterica_subsp._enterica_serovar_Typhimurium | GTH76_16775 | 108064..111389 | 90371 | BREX-system-P-loop-protein-BrxC-[Salmonella-enterica-subsp.-enterica-serovar-Typhimurium] |
| EDP8828062.1 | Type1.BREX | [illegible] | 793 | Bacteria>Pseudomonadota>Gammaproteobacteria>Enterobacterales>Enterobacteriaceae>Salmonella | GCA_010369285.1 | Salmonella_enterica_subsp._enterica | S669_001969 | 1..2380 | 59201 | BREX-system-P-loop-protein-BrxC-[Salmonella-enterica-subsp.-enterica] |
| EDQ1995876.1 | Type1.BREX | [illegible] | 509 | Bacteria>Pseudomonadota>Gammaproteobacteria>Enterobacterales>Enterobacteriaceae>Salmonella | GCA_010387725.1 | Salmonella_enterica | GT143_004731 | 5665..7190 | 28901 | BREX-system-P-loop-protein-BrxC-[Salmonella-enterica] |
| EDQ2665795.1 | Type1.BREX | [illegible] | 1213 | Bacteria>Pseudomonadota>Gammaproteobacteria>Enterobacterales>Enterobacteriaceae>Salmonella | GCA_010390975.1 | Salmonella_enterica_subsp._enterica | ZH92_003658 | 52353..55994 | 59201 | BREX-system-P-loop-protein-BrxC-[Salmonella-enterica-subsp.-enterica] |
| EDQ2405063.1 | Type1.BREX | [illegible] | 1213 | Bacteria>Pseudomonadota>Gammaproteobacteria>Enterobacterales>Enterobacteriaceae>Salmonella | GCA_010391095.1 | Salmonella_enterica_subsp._enterica_serovar_Saintpaul | DQ96_001225 | 144464..148105 | 90105 | BREX-system-P-loop-protein-BrxC-[Salmonella-enterica-subsp.-enterica-serovar-Saintpaul] |
| EDR5087378.1 | Type1.BREX | [illegible] | 1213 | Bacteria>Pseudomonadota>Gammaproteobacteria>Enterobacterales>Enterobacteriaceae>Salmonella | GCA_010449355.1 | Salmonella_enterica_subsp._enterica | GRZ33_003685 | 38268..41909 | 59201 | BREX-system-P-loop-protein-BrxC-[Salmonella-enterica-subsp.-enterica] |
| HAE3790823.1 | Type1.BREX | [illegible] | 1213 | Bacteria>Pseudomonadota>Gammaproteobacteria>Enterobacterales>Enterobacteriaceae>Salmonella | GCA_010485735.1 | Salmonella_enterica_subsp._enterica_serovar_Typhimurium | G4A23_000143 | 146702..150343 | 90371 | BREX-system-P-loop-protein-BrxC-[Salmonella-enterica-subsp.-enterica-serovar-Typhimurium] |
| EDS3575399.1 | Type1.BREX | [illegible] | 1204 | Bacteria>Pseudomonadota>Gammaproteobacteria>Enterobacterales>Enterobacteriaceae>Salmonella | GCA_010490095.1 | Salmonella_enterica_subsp._enterica | GR179_002518 | 3402..7016 | 59201 | BREX-system-P-loop-protein-BrxC-[Salmonella-enterica-subsp.-enterica] |
| EDS5421358.1 | Type1.BREX | [illegible] | 1213 | Bacteria>Pseudomonadota>Gammaproteobacteria>Enterobacterales>Enterobacteriaceae>Salmonella | GCA_010498755.1 | Salmonella_enterica_subsp._enterica | ZH17_000156 | 158281..161922 | 59201 | BREX-system-P-loop-protein-BrxC-[Salmonella-enterica-subsp.-enterica] |
| EDT1456935.1 | Type1.BREX | [illegible] | 1213 | Bacteria>Pseudomonadota>Gammaproteobacteria>Enterobacterales>Enterobacteriaceae>Salmonella | GCA_010529175.1 | Salmonella_enterica_subsp._enterica_serovar_Ramatgan | CE894_001670 | 153160..156801 | 2564942 | BREX-system-P-loop-protein-BrxC-[Salmonella-enterica-subsp.-enterica-serovar-Ramatgan] |
| EDT2838946.1 | Type1.BREX | [illegible] | 1213 | Bacteria>Pseudomonadota>Gammaproteobacteria>Enterobacterales>Enterobacteriaceae>Salmonella | GCA_010536715.1 | Salmonella_enterica_subsp._enterica | S958_003413 | 14802..18443 | 59201 | BREX-system-P-loop-protein-BrxC-[Salmonella-enterica-subsp.-enterica] |
| EDT5820088.1 | Type1.BREX | [illegible] | 1213 | Bacteria>Pseudomonadota>Gammaproteobacteria>Enterobacterales>Enterobacteriaceae>Salmonella | GCA_010551565.1 | Salmonella_enterica_subsp._enterica_serovar_4,[5],12:i:- | B7921_002250 | 108064..111705 | 440524 | BREX-system-P-loop-protein-BrxC-[Salmonella-enterica-subsp.-enterica-serovar-4,[5] |
| EDT6691760.1 | Type1.BREX | [illegible] | 1213 | Bacteria>Pseudomonadota>Gammaproteobacteria>Enterobacterales>Enterobacteriaceae>Salmonella | GCA_010555745.1 | Salmonella_enterica_subsp._enterica | H2V91_002192 | 153095..156736 | 59201 | BREX-system-P-loop-protein-BrxC-[Salmonella-enterica-subsp.-enterica] |
| EDT7239824.1 | Type1.BREX | [illegible] | 1213 | Bacteria>Pseudomonadota>Gammaproteobacteria>Enterobacterales>Enterobacteriaceae>Salmonella | GCA_010557585.1 | Salmonella_enterica_subsp._enterica | AHZ96_001967 | 93382..97023 | 59201 | BREX-system-P-loop-protein-BrxC-[Salmonella-enterica-subsp.-enterica] |
| EDT7223673.1 | Type1.BREX | [illegible] | 1204 | Bacteria>Pseudomonadota>Gammaproteobacteria>Enterobacterales>Enterobacteriaceae>Salmonella | GCA_010559045.1 | Salmonella_enterica_subsp._enterica | AH268_003334 | 169288..172902 | 59201 | BREX-system-P-loop-protein-BrxC-[Salmonella-enterica-subsp.-enterica] |
| EDT8878002.1 | Type1.BREX | [illegible] | 1213 | Bacteria>Pseudomonadota>Gammaproteobacteria>Enterobacterales>Enterobacteriaceae>Salmonella | GCA_010565975.1 | Salmonella_enterica_subsp._enterica | EK51_003288 | 149002..152643 | 59201 | BREX-system-P-loop-protein-BrxC-[Salmonella-enterica-subsp.-enterica] |
| EDT9358547.1 | Type1.BREX | [illegible] | 1213 | Bacteria>Pseudomonadota>Gammaproteobacteria>Enterobacterales>Enterobacteriaceae>Salmonella | GCA_010568345.1 | Salmonella_enterica_subsp._enterica_serovar_Miami | A9571_000728 | 158750..162391 | 286780 | BREX-system-P-loop-protein-BrxC-[Salmonella-enterica-subsp.-enterica-serovar-Miami] |
| EDU3766992.1 | Type1.BREX | [illegible] | 1213 | Bacteria>Pseudomonadota>Gammaproteobacteria>Enterobacterales>Enterobacteriaceae>Salmonella | GCA_010595665.1 | Salmonella_enterica_subsp._enterica_serovar_Minnesota | C4O09_000357 | 390744..394385 | 70803 | BREX-system-P-loop-protein-BrxC-[Salmonella-enterica-subsp.-enterica-serovar-Minnesota] |
| EDU7470137.1 | Type1.BREX | [illegible] | 1213 | Bacteria>Pseudomonadota>Gammaproteobacteria>Enterobacterales>Enterobacteriaceae>Salmonella | GCA_010625705.1 | Salmonella_enterica_subsp._enterica | CQO18_002340 | 108064..111705 | 59201 | BREX-system-P-loop-protein-BrxC-[Salmonella-enterica-subsp.-enterica] |
| EDU9161212.1 | Type1.BREX | [illegible] | 1213 | Bacteria>Pseudomonadota>Gammaproteobacteria>Enterobacterales>Enterobacteriaceae>Salmonella | GCA_010634945.1 | Salmonella_enterica_subsp._enterica | IW54_004329 | 11544..15185 | 59201 | BREX-system-P-loop-protein-BrxC-[Salmonella-enterica-subsp.-enterica] |
| EDU9624695.1 | Type1.BREX | [illegible] | 1213 | Bacteria>Pseudomonadota>Gammaproteobacteria>Enterobacterales>Enterobacteriaceae>Salmonella | GCA_010637145.1 | Salmonella_enterica_subsp._enterica | XW65_002877 | 59802..63443 | 59201 | BREX-system-P-loop-protein-BrxC-[Salmonella-enterica-subsp.-enterica] |
| EDV2155932.1 | Type1.BREX | [illegible] | 1213 | Bacteria>Pseudomonadota>Gammaproteobacteria>Enterobacterales>Enterobacteriaceae>Salmonella | GCA_010649605.1 | Salmonella_enterica | JR65_004155 | 25186..28827 | 28901 | BREX-system-P-loop-protein-BrxC-[Salmonella-enterica] |
| EDV2418012.1 | Type1.BREX | [illegible] | 1213 | Bacteria>Pseudomonadota>Gammaproteobacteria>Enterobacterales>Enterobacteriaceae>Salmonella | GCA_010650645.1 | Salmonella_enterica_subsp._enterica_serovar_Ohio | CQ222_000233 | 225731..229372 | 117541 | BREX-system-P-loop-protein-BrxC-[Salmonella-enterica-subsp.-enterica-serovar-Ohio] |
| HAE7521102.1 | Type1.BREX | [illegible] | 1213 | Bacteria>Pseudomonadota>Gammaproteobacteria>Enterobacterales>Enterobacteriaceae>Salmonella | GCA_010655205.1 | Salmonella_enterica_subsp._enterica_serovar_Carrau | G4P13_000881 | 149047..152688 | 1160739 | BREX-system-P-loop-protein-BrxC-[Salmonella-enterica-subsp.-enterica-serovar-Carrau] |
| EDV4604931.1 | Type1.BREX | [illegible] | 1213 | Bacteria>Pseudomonadota>Gammaproteobacteria>Enterobacterales>Enterobacteriaceae>Salmonella | GCA_010662535.1 | Salmonella_enterica_subsp._enterica | ZP62_000973 | 72422..76063 | 59201 | BREX-system-P-loop-protein-BrxC-[Salmonella-enterica-subsp.-enterica] |
| EDV5116990.1 | Type1.BREX | [illegible] | 1213 | Bacteria>Pseudomonadota>Gammaproteobacteria>Enterobacterales>Enterobacteriaceae>Salmonella | GCA_010665025.1 | Salmonella_enterica_subsp._enterica | ZJ36_000825 | 28924..32565 | 59201 | BREX-system-P-loop-protein-BrxC-[Salmonella-enterica-subsp.-enterica] |
| EDV6600966.1 | Type1.BREX | [illegible] | 1213 | Bacteria>Pseudomonadota>Gammaproteobacteria>Enterobacterales>Enterobacteriaceae>Salmonella | GCA_010670385.1 | Salmonella_enterica_subsp._enterica_serovar_Typhimurium_var._5- | BH535_001152 | 81343..84984 | 1620419 | BREX-system-P-loop-protein-BrxC-[Salmonella-enterica-subsp.-enterica-serovar-Typhimurium-var.-5-] |
| EDV9644204.1 | Type1.BREX | [illegible] | 1213 | Bacteria>Pseudomonadota>Gammaproteobacteria>Enterobacterales>Enterobacteriaceae>Salmonella | GCA_010680145.1 | Salmonella_enterica_subsp._enterica_serovar_Carno | HM25_003997 | 164513..168154 | 2024268 | BREX-system-P-loop-protein-BrxC-[Salmonella-enterica-subsp.-enterica-serovar-Carno] |
| EDV9109398.1 | Type1.BREX | [illegible] | 1213 | Bacteria>Pseudomonadota>Gammaproteobacteria>Enterobacterales>Enterobacteriaceae>Salmonella | GCA_010691445.1 | Salmonella_enterica_subsp._enterica | ZK68_001600 | 51998..55639 | 59201 | BREX-system-P-loop-protein-BrxC-[Salmonella-enterica-subsp.-enterica] |
| EDW0696245.1 | Type1.BREX | [illegible] | 1213 | Bacteria>Pseudomonadota>Gammaproteobacteria>Enterobacterales>Enterobacteriaceae>Salmonella | GCA_010695935.1 | Salmonella_enterica_subsp._enterica | KV01_000535 | 150350..153991 | 59201 | BREX-system-P-loop-protein-BrxC-[Salmonella-enterica-subsp.-enterica] |
| EDW2626492.1 | Type1.BREX | [illegible] | 1213 | Bacteria>Pseudomonadota>Gammaproteobacteria>Enterobacterales>Enterobacteriaceae>Salmonella | GCA_010704865.1 | Salmonella_enterica_subsp._enterica | XX60_002382 | 156709..160350 | 59201 | BREX-system-P-loop-protein-BrxC-[Salmonella-enterica-subsp.-enterica] |
| EDW4488615.1 | Type1.BREX | [illegible] | 1213 | Bacteria>Pseudomonadota>Gammaproteobacteria>Enterobacterales>Enterobacteriaceae>Salmonella | GCA_010711495.1 | Salmonella_enterica_subsp._enterica | YV39_000487 | 166105..169746 | 59201 | BREX-system-P-loop-protein-BrxC-[Salmonella-enterica-subsp.-enterica] |
| EDW7722488.1 | Type1.BREX | [illegible] | 1213 | Bacteria>Pseudomonadota>Gammaproteobacteria>Enterobacterales>Enterobacteriaceae>Salmonella | GCA_010729405.1 | Salmonella_enterica | GTR06_001871 | 397975..401616 | 28901 | BREX-system-P-loop-protein-BrxC-[Salmonella-enterica] |
| EDX0369803.1 | Type1.BREX | [illegible] | 1213 | Bacteria>Pseudomonadota>Gammaproteobacteria>Enterobacterales>Enterobacteriaceae>Salmonella | GCA_010742735.1 | Salmonella_enterica | GTL21_000361 | 386606..390247 | 28901 | BREX-system-P-loop-protein-BrxC-[Salmonella-enterica] |
| EDX7895849.1 | Type1.BREX | [illegible] | 1213 | Bacteria>Pseudomonadota>Gammaproteobacteria>Enterobacterales>Enterobacteriaceae>Salmonella | GCA_010782335.1 | Salmonella_enterica_subsp._enterica_serovar_4,[5],12:i:- | C2Y98_001768 | 38662..42303 | 440524 | BREX-system-P-loop-protein-BrxC-[Salmonella-enterica-subsp.-enterica-serovar-4,[5] |
| EDZ7084220.1 | Type1.BREX | [illegible] | 1213 | Bacteria>Pseudomonadota>Gammaproteobacteria>Enterobacterales>Enterobacteriaceae>Salmonella | GCA_010862365.1 | Salmonella_enterica | GS086_003556 | 2815..6456 | 28901 | BREX-system-P-loop-protein-BrxC-[Salmonella-enterica] |
| EEB8473036.1 | Type1.BREX | [illegible] | 1213 | Bacteria>Pseudomonadota>Gammaproteobacteria>Enterobacterales>Enterobacteriaceae>Salmonella | GCA_010964635.1 | Salmonella_enterica | GN471_06045 | 170718..174359 | 28901 | BREX-system-P-loop-protein-BrxC-[Salmonella-enterica] |
| EEC0292871.1 | Type1.BREX | [illegible] | 1213 | Bacteria>Pseudomonadota>Gammaproteobacteria>Enterobacterales>Enterobacteriaceae>Salmonella | GCA_010970835.1 | Salmonella_enterica_subsp._enterica | YV76_000179 | 186389..190030 | 59201 | BREX-system-P-loop-protein-BrxC-[Salmonella-enterica-subsp.-enterica] |
| EEC1070526.1 | Type1.BREX | [illegible] | 1213 | Bacteria>Pseudomonadota>Gammaproteobacteria>Enterobacterales>Enterobacteriaceae>Salmonella | GCA_010973255.1 | Salmonella_enterica_subsp._enterica | AH545_000138 | 139989..143630 | 59201 | BREX-system-P-loop-protein-BrxC-[Salmonella-enterica-subsp.-enterica] |
| EEC4816191.1 | Type1.BREX | [illegible] | 1213 | Bacteria>Pseudomonadota>Gammaproteobacteria>Enterobacterales>Enterobacteriaceae>Salmonella | GCA_010993355.1 | Salmonella_enterica | G4L83_001633 | 157032..160673 | 28901 | BREX-system-P-loop-protein-BrxC-[Salmonella-enterica] |
| EED3636775.1 | Type1.BREX | [illegible] | 1213 | Bacteria>Pseudomonadota>Gammaproteobacteria>Enterobacterales>Enterobacteriaceae>Salmonella | GCA_011081015.1 | Salmonella_enterica_subsp._enterica | S719_003281 | 157579..161220 | 59201 | BREX-system-P-loop-protein-BrxC-[Salmonella-enterica-subsp.-enterica] |
| EEE1152714.1 | Type1.BREX | [illegible] | 1213 | Bacteria>Pseudomonadota>Gammaproteobacteria>Enterobacterales>Enterobacteriaceae>Salmonella | GCA_011117415.1 | Salmonella_enterica_subsp._enterica_serovar_Teko | CAE49_001975 | 330756..334397 | 2079198 | BREX-system-P-loop-protein-BrxC-[Salmonella-enterica-subsp.-enterica-serovar-Teko] |
| HAF0812803.1 | Type1.BREX | [illegible] | 1213 | Bacteria>Pseudomonadota>Gammaproteobacteria>Enterobacterales>Enterobacteriaceae>Salmonella | GCA_011125695.1 | Salmonella_enterica | G9G17_001783 | 108064..111705 | 28901 | BREX-system-P-loop-protein-BrxC-[Salmonella-enterica] |
| EEE6742962.1 | Type1.BREX | [illegible] | 1213 | Bacteria>Pseudomonadota>Gammaproteobacteria>Enterobacterales>Enterobacteriaceae>Salmonella | GCA_011144815.1 | Salmonella_enterica_subsp._enterica_serovar_Westhampton | CBJ05_002020 | 408792..412433 | 356621 | BREX-system-P-loop-protein-BrxC-[Salmonella-enterica-subsp.-enterica-serovar-Westhampton] |
| EEE9192570.1 | Type1.BREX | [illegible] | 555 | Bacteria>Pseudomonadota>Gammaproteobacteria>Enterobacterales>Enterobacteriaceae>Salmonella | GCA_011156215.1 | Salmonella_enterica_subsp._enterica_serovar_Typhimurium | DA558_21335 | 48506..50170 | 90371 | BREX-system-P-loop-protein-BrxC-[Salmonella-enterica-subsp.-enterica-serovar-Typhimurium] |
| EEH0007757.1 | Type1.BREX | [illegible] | 1213 | Bacteria>Pseudomonadota>Gammaproteobacteria>Enterobacterales>Enterobacteriaceae>Salmonella | GCA_011260745.1 | Salmonella_enterica | G3P26_001654 | 368950..372591 | 28901 | BREX-system-P-loop-protein-BrxC-[Salmonella-enterica] |
| EEH2707246.1 | Type1.BREX | [illegible] | 1213 | Bacteria>Pseudomonadota>Gammaproteobacteria>Enterobacterales>Enterobacteriaceae>Salmonella | GCA_011274245.1 | Salmonella_enterica_subsp._enterica | GYE71_000437 | 54012..57653 | 59201 | BREX-system-P-loop-protein-BrxC-[Salmonella-enterica-subsp.-enterica] |
| EEH3158636.1 | Type1.BREX | [illegible] | 1213 | Bacteria>Pseudomonadota>Gammaproteobacteria>Enterobacterales>Enterobacteriaceae>Salmonella | GCA_011275545.1 | Salmonella_enterica | GZB32_002496 | 108064..111705 | 28901 | BREX-system-P-loop-protein-BrxC-[Salmonella-enterica] |
| EEH9995094.1 | Type1.BREX | [illegible] | 1213 | Bacteria>Pseudomonadota>Gammaproteobacteria>Enterobacterales>Enterobacteriaceae>Salmonella | GCA_011315605.1 | Salmonella_enterica_subsp._enterica_serovar_4,[5],12:i:- | G7H85_000159 | 170419..174060 | 440524 | BREX-system-P-loop-protein-BrxC-[Salmonella-enterica-subsp.-enterica-serovar-4,[5] |
| EEI0202361.1 | Type1.BREX | [illegible] | 908 | Bacteria>Pseudomonadota>Gammaproteobacteria>Enterobacterales>Enterobacteriaceae>Salmonella | GCA_011321375.1 | Salmonella_enterica | GNM15_17120 | 108064..110791 | 28901 | BREX-system-P-loop-protein-BrxC-[Salmonella-enterica] |
| EEI5362885.1 | Type1.BREX | [illegible] | 1204 | Bacteria>Pseudomonadota>Gammaproteobacteria>Enterobacterales>Enterobacteriaceae>Salmonella | GCA_011357245.1 | Salmonella_enterica_subsp._enterica_serovar_Kentucky | GXM95_001563 | 173373..176987 | 192955 | BREX-system-P-loop-protein-BrxC-[Salmonella-enterica-subsp.-enterica-serovar-Kentucky] |
| EEJ3565315.1 | Type1.BREX | [illegible] | 1213 | Bacteria>Pseudomonadota>Gammaproteobacteria>Enterobacterales>Enterobacteriaceae>Salmonella | GCA_011417115.1 | Salmonella_enterica_subsp._enterica | AA437_001844 | 125694..129335 | 59201 | BREX-system-P-loop-protein-BrxC-[Salmonella-enterica-subsp.-enterica] |
| EEJ7235528.1 | Type1.BREX | [illegible] | 1213 | Bacteria>Pseudomonadota>Gammaproteobacteria>Enterobacterales>Enterobacteriaceae>Salmonella | GCA_011437855.1 | Salmonella_enterica_subsp._salamae | S483_003519 | 152399..156040 | 59202 | BREX-system-P-loop-protein-BrxC-[Salmonella-enterica-subsp.-salamae] |
| EEJ7298184.1 | Type1.BREX | [illegible] | 1213 | Bacteria>Pseudomonadota>Gammaproteobacteria>Enterobacterales>Enterobacteriaceae>Salmonella | GCA_011438135.1 | Salmonella_enterica_subsp._enterica_serovar_Livingstone | DV24_002305 | 312113..315754 | 260680 | BREX-system-P-loop-protein-BrxC-[Salmonella-enterica-subsp.-enterica-serovar-Livingstone] |
| EEJ9771966.1 | Type1.BREX | [illegible] | 1213 | Bacteria>Pseudomonadota>Gammaproteobacteria>Enterobacterales>Enterobacteriaceae>Salmonella | GCA_011448615.1 | Salmonella_enterica_subsp._enterica_serovar_Typhimurium | HL79_09630 | 170931..174572 | 90371 | BREX-system-P-loop-protein-BrxC-[Salmonella-enterica-subsp.-enterica-serovar-Typhimurium] |
| EEK0538631.1 | Type1.BREX | [illegible] | 1213 | Bacteria>Pseudomonadota>Gammaproteobacteria>Enterobacterales>Enterobacteriaceae>Salmonella | GCA_011452795.1 | Salmonella_enterica_subsp._enterica_serovar_Typhimurium | IH96_05510 | 150023..153664 | 90371 | BREX-system-P-loop-protein-BrxC-[Salmonella-enterica-subsp.-enterica-serovar-Typhimurium] |
| EEK5480526.1 | Type1.BREX | [illegible] | 1213 | Bacteria>Pseudomonadota>Gammaproteobacteria>Enterobacterales>Enterobacteriaceae>Salmonella | GCA_011476295.1 | Salmonella_enterica_subsp._enterica_serovar_Typhimurium | K6Z50_000356 | 395891..399532 | 90371 | BREX-system-P-loop-protein-BrxC-[Salmonella-enterica-subsp.-enterica-serovar-Typhimurium] |
| EEL0135869.1 | Type1.BREX | [illegible] | 1213 | Bacteria>Pseudomonadota>Gammaproteobacteria>Enterobacterales>Enterobacteriaceae>Salmonella | GCA_011496945.1 | Salmonella_enterica | G4L63_001980 | 126120..129761 | 28901 | BREX-system-P-loop-protein-BrxC-[Salmonella-enterica] |
| EEL6995865.1 | Type1.BREX | [illegible] | 1213 | Bacteria>Pseudomonadota>Gammaproteobacteria>Enterobacterales>Enterobacteriaceae>Salmonella | GCA_011533325.1 | Salmonella_enterica | FPY45_19715 | 27496..31137 | 28901 | BREX-system-P-loop-protein-BrxC-[Salmonella-enterica] |
| EEL7097503.1 | Type1.BREX | [illegible] | 1213 | Bacteria>Pseudomonadota>Gammaproteobacteria>Enterobacterales>Enterobacteriaceae>Salmonella | GCA_011533985.1 | Salmonella_enterica | GKH84_08575 | 158871..162512 | 28901 | BREX-system-P-loop-protein-BrxC-[Salmonella-enterica] |
| EEM2455575.1 | Type1.BREX | [illegible] | 1204 | Bacteria>Pseudomonadota>Gammaproteobacteria>Enterobacterales>Enterobacteriaceae>Salmonella | GCA_011558015.1 | Salmonella_enterica_subsp._enterica_serovar_Kentucky | CWG70_02775 | 173706..177320 | 192955 | BREX-system-P-loop-protein-BrxC-[Salmonella-enterica-subsp.-enterica-serovar-Kentucky] |
| EEO3564960.1 | Type1.BREX | [illegible] | 1213 | Bacteria>Pseudomonadota>Gammaproteobacteria>Enterobacterales>Enterobacteriaceae>Salmonella | GCA_011665275.1 | Salmonella_enterica_subsp._enterica_serovar_Poona | DEI68_08995 | 52368..56009 | 436295 | BREX-system-P-loop-protein-BrxC-[Salmonella-enterica-subsp.-enterica-serovar-Poona] |
| EEP2127992.1 | Type1.BREX | [illegible] | 1213 | Bacteria>Pseudomonadota>Gammaproteobacteria>Enterobacterales>Enterobacteriaceae>Salmonella | GCA_011714235.1 | Salmonella_enterica | HA780_002410 | 111301..114942 | 28901 | BREX-system-P-loop-protein-BrxC-[Salmonella-enterica] |
| EEP5746981.1 | Type1.BREX | [illegible] | 1213 | Bacteria>Pseudomonadota>Gammaproteobacteria>Enterobacterales>Enterobacteriaceae>Salmonella | GCA_011732675.1 | Salmonella_enterica | HBB29_002275 | 108064..111705 | 28901 | BREX-system-P-loop-protein-BrxC-[Salmonella-enterica] |
| EEP8318663.1 | Type1.BREX | [illegible] | 1213 | Bacteria>Pseudomonadota>Gammaproteobacteria>Enterobacterales>Enterobacteriaceae>Salmonella | GCA_011746585.1 | Salmonella_enterica_subsp._enterica | HCM12_000954 | 52440..56081 | 59201 | BREX-system-P-loop-protein-BrxC-[Salmonella-enterica-subsp.-enterica] |
| EEP9788441.1 | Type1.BREX | [illegible] | 1213 | Bacteria>Pseudomonadota>Gammaproteobacteria>Enterobacterales>Enterobacteriaceae>Salmonella | GCA_011756305.1 | Salmonella_enterica_subsp._enterica_serovar_Saintpaul | HCP67_002105 | 64937..68578 | 90105 | BREX-system-P-loop-protein-BrxC-[Salmonella-enterica-subsp.-enterica-serovar-Saintpaul] |
| HAF9991656.1 | Type1.BREX | [illegible] | 1213 | Bacteria>Pseudomonadota>Gammaproteobacteria>Enterobacterales>Enterobacteriaceae>Salmonella | GCA_011779145.1 | Salmonella_enterica | G8Q07_002648 | 170744..174385 | 28901 | BREX-system-P-loop-protein-BrxC-[Salmonella-enterica] |
| HAG2842216.1 | Type1.BREX | [illegible] | 1213 | Bacteria>Pseudomonadota>Gammaproteobacteria>Enterobacterales>Enterobacteriaceae>Salmonella | GCA_011791385.1 | Salmonella_enterica | G8Z41_002592 | 114973..118614 | 28901 | BREX-system-P-loop-protein-BrxC-[Salmonella-enterica] |
| HAG5552356.1 | Type1.BREX | [illegible] | 1213 | Bacteria>Pseudomonadota>Gammaproteobacteria>Enterobacterales>Enterobacteriaceae>Salmonella | GCA_011798725.1 | Salmonella_enterica | G8Q25_001512 | 186770..190411 | 28901 | BREX-system-P-loop-protein-BrxC-[Salmonella-enterica] |
| EFP5236617.1 | Type1.BREX | [illegible] | 1213 | Bacteria>Pseudomonadota>Gammaproteobacteria>Enterobacterales>Enterobacteriaceae>Salmonella | GCA_013442935.1 | Salmonella_enterica | H1F05_002027 | 170348..173989 | 28901 | BREX-system-P-loop-protein-BrxC-[Salmonella-enterica] |
| EFP6288854.1 | Type1.BREX | [illegible] | 755 | Bacteria>Pseudomonadota>Gammaproteobacteria>Enterobacterales>Enterobacteriaceae>Salmonella | GCA_013446435.1 | Salmonella_enterica | H1E36_001915 | 171036..173302 | 28901 | BREX-system-P-loop-protein-BrxC-[Salmonella-enterica] |
| EFQ1578017.1 | Type1.BREX | [illegible] | 1213 | Bacteria>Pseudomonadota>Gammaproteobacteria>Enterobacterales>Enterobacteriaceae>Salmonella | GCA_013479125.1 | Salmonella_enterica | H1L81_001832 | 38672..42313 | 28901 | BREX-system-P-loop-protein-BrxC-[Salmonella-enterica] |
| HAK0222817.1 | Type1.BREX | [illegible] | 805 | Bacteria>Pseudomonadota>Gammaproteobacteria>Enterobacterales>Enterobacteriaceae>Salmonella | GCA_013485925.1 | Salmonella_enterica | H1O47_004541 | 10945..13359 | 28901 | BREX-system-P-loop-protein-BrxC-[Salmonella-enterica] |
| HAK0744369.1 | Type1.BREX | [illegible] | 982 | Bacteria>Pseudomonadota>Gammaproteobacteria>Enterobacterales>Enterobacteriaceae>Salmonella | GCA_013487565.1 | Salmonella_enterica | H1P65_002115 | 170411..173355 | 28901 | BREX-system-P-loop-protein-BrxC-[Salmonella-enterica] |
| EFQ3650906.1 | Type1.BREX | [illegible] | 1213 | Bacteria>Pseudomonadota>Gammaproteobacteria>Enterobacterales>Enterobacteriaceae>Salmonella | GCA_013488685.1 | Salmonella_enterica | HZ593_001598 | 149045..152686 | 28901 | BREX-system-P-loop-protein-BrxC-[Salmonella-enterica] |
| EFR0716065.1 | Type1.BREX | [illegible] | 720 | Bacteria>Pseudomonadota>Gammaproteobacteria>Enterobacterales>Enterobacteriaceae>Salmonella | GCA_013531855.1 | Salmonella_enterica | HYQ91_004636 | 1..2164 | 28901 | BREX-system-P-loop-protein-BrxC-[Salmonella-enterica] |
| EFQ9845222.1 | Type1.BREX | [illegible] | 1213 | Bacteria>Pseudomonadota>Gammaproteobacteria>Enterobacterales>Enterobacteriaceae>Salmonella | GCA_013535435.1 | Salmonella_enterica | HZL63_001579 | 150017..153658 | 28901 | BREX-system-P-loop-protein-BrxC-[Salmonella-enterica] |
| EFS3421932.1 | Type1.BREX | [illegible] | 1213 | Bacteria>Pseudomonadota>Gammaproteobacteria>Enterobacterales>Enterobacteriaceae>Salmonella | GCA_013632355.1 | Salmonella_enterica | HX201_001605 | 150173..153814 | 28901 | BREX-system-P-loop-protein-BrxC-[Salmonella-enterica] |
| HAK8810847.1 | Type1.BREX | [illegible] | 1213 | Bacteria>Pseudomonadota>Gammaproteobacteria>Enterobacterales>Enterobacteriaceae>Salmonella | GCA_013646795.1 | Salmonella_enterica | H2X42_001962 | 158401..162042 | 28901 | BREX-system-P-loop-protein-BrxC-[Salmonella-enterica] |
| EFS5965938.1 | Type1.BREX | [illegible] | 1213 | Bacteria>Pseudomonadota>Gammaproteobacteria>Enterobacterales>Enterobacteriaceae>Salmonella | GCA_013657295.1 | Salmonella_enterica | HW654_000972 | 195304..198945 | 28901 | BREX-system-P-loop-protein-BrxC-[Salmonella-enterica] |
| EFS7669868.1 | Type1.BREX | [illegible] | 1078 | Bacteria>Pseudomonadota>Gammaproteobacteria>Enterobacterales>Enterobacteriaceae>Salmonella | GCA_013666175.1 | Salmonella_enterica | HWZ11_001253 | 1..3239 | 28901 | BREX-system-P-loop-protein-BrxC-[Salmonella-enterica] |
| EFU4335446.1 | Type1.BREX | [illegible] | 1213 | Bacteria>Pseudomonadota>Gammaproteobacteria>Enterobacterales>Enterobacteriaceae>Salmonella | GCA_013751665.1 | Salmonella_enterica | HUC58_001813 | 170349..173990 | 28901 | BREX-system-P-loop-protein-BrxC-[Salmonella-enterica] |
| EGA9157240.1 | Type1.BREX | [illegible] | 1213 | Bacteria>Pseudomonadota>Gammaproteobacteria>Enterobacterales>Enterobacteriaceae>Salmonella | GCA_014125285.1 | Salmonella_enterica_subsp._enterica_serovar_Kirkee | H6M98_002631 | 52334..55975 | 2564624 | BREX-system-P-loop-protein-BrxC-[Salmonella-enterica-subsp.-enterica-serovar-Kirkee] |
| EGB5523462.1 | Type1.BREX | [illegible] | 1213 | Bacteria>Pseudomonadota>Gammaproteobacteria>Enterobacterales>Enterobacteriaceae>Salmonella | GCA_014209965.1 | Salmonella_enterica | H7074_002104 | 159039..162680 | 28901 | BREX-system-P-loop-protein-BrxC-[Salmonella-enterica] |
| EGB5697881.1 | Type1.BREX | [illegible] | 1204 | Bacteria>Pseudomonadota>Gammaproteobacteria>Enterobacterales>Enterobacteriaceae>Salmonella | GCA_014213495.1 | Salmonella_enterica_subsp._enterica | H7531_003071 | 17313..20927 | 59201 | BREX-system-P-loop-protein-BrxC-[Salmonella-enterica-subsp.-enterica] |
| EGC3767042.1 | Type1.BREX | [illegible] | 1213 | Bacteria>Pseudomonadota>Gammaproteobacteria>Enterobacterales>Enterobacteriaceae>Salmonella | GCA_014295145.1 | Salmonella_enterica | H9A68_002330 | 38672..42313 | 28901 | BREX-system-P-loop-protein-BrxC-[Salmonella-enterica] |
| EGC8584468.1 | Type1.BREX | [illegible] | 1213 | Bacteria>Pseudomonadota>Gammaproteobacteria>Enterobacterales>Enterobacteriaceae>Salmonella | GCA_014317295.1 | Salmonella_enterica | H9103_003699 | 1964..5605 | 28901 | BREX-system-P-loop-protein-BrxC-[Salmonella-enterica] |
| EGD1472167.1 | Type1.BREX | [illegible] | 1213 | Bacteria>Pseudomonadota>Gammaproteobacteria>Enterobacterales>Enterobacteriaceae>Salmonella | GCA_014351875.1 | Salmonella_enterica | IAV68_002850 | 108064..111705 | 28901 | BREX-system-P-loop-protein-BrxC-[Salmonella-enterica] |
| EGF1249453.1 | Type1.BREX | [illegible] | 1213 | Bacteria>Pseudomonadota>Gammaproteobacteria>Enterobacterales>Enterobacteriaceae>Salmonella | GCA_014468995.1 | Salmonella_enterica | IB037_001321 | 170919..174560 | 28901 | BREX-system-P-loop-protein-BrxC-[Salmonella-enterica] |
| EGH1765314.1 | Type1.BREX | [illegible] | 1213 | Bacteria>Pseudomonadota>Gammaproteobacteria>Enterobacterales>Enterobacteriaceae>Salmonella | GCA_014604855.1 | Salmonella_enterica | ID459_000689 | 170687..174328 | 28901 | BREX-system-P-loop-protein-BrxC-[Salmonella-enterica] |
| EGH2238321.1 | Type1.BREX | [illegible] | 1213 | Bacteria>Pseudomonadota>Gammaproteobacteria>Enterobacterales>Enterobacteriaceae>Salmonella | GCA_014610595.1 | Salmonella_enterica | ID764_000354 | 390871..394512 | 28901 | BREX-system-P-loop-protein-BrxC-[Salmonella-enterica] |
| EGH8508053.1 | Type1.BREX | [illegible] | 1213 | Bacteria>Pseudomonadota>Gammaproteobacteria>Enterobacterales>Enterobacteriaceae>Salmonella | GCA_014678145.1 | Salmonella_enterica | IFE76_001736 | 26096..29737 | 28901 | BREX-system-P-loop-protein-BrxC-[Salmonella-enterica] |
| EGH9456388.1 | Type1.BREX | [illegible] | 1213 | Bacteria>Pseudomonadota>Gammaproteobacteria>Enterobacterales>Enterobacteriaceae>Salmonella | GCA_014692265.1 | Salmonella_enterica_subsp._enterica_serovar_Leeuwarden | HKU92_000347 | 389340..392981 | 2564678 | BREX-system-P-loop-protein-BrxC-[Salmonella-enterica-subsp.-enterica-serovar-Leeuwarden] |
| EGI8941862.1 | Type1.BREX | [illegible] | 1213 | Bacteria>Pseudomonadota>Gammaproteobacteria>Enterobacterales>Enterobacteriaceae>Salmonella | GCA_014780755.1 | Salmonella_enterica | IG687_002353 | 272986..276627 | 28901 | BREX-system-P-loop-protein-BrxC-[Salmonella-enterica] |
| EGJ1340019.1 | Type1.BREX | [illegible] | 1213 | Bacteria>Pseudomonadota>Gammaproteobacteria>Enterobacterales>Enterobacteriaceae>Salmonella | GCA_014793405.1 | Salmonella_enterica_subsp._enterica_serovar_Typhimurium | IHB80_004023 | 18317..21958 | 90371 | BREX-system-P-loop-protein-BrxC-[Salmonella-enterica-subsp.-enterica-serovar-Typhimurium] |
| EGK3406186.1 | Type1.BREX | [illegible] | 1213 | Bacteria>Pseudomonadota>Gammaproteobacteria>Enterobacterales>Enterobacteriaceae>Salmonella | GCA_014949985.1 | Salmonella_enterica | IOS83_004393 | 25212..28853 | 28901 | BREX-system-P-loop-protein-BrxC-[Salmonella-enterica] |
| EGL7108184.1 | Type1.BREX | [illegible] | 1213 | Bacteria>Pseudomonadota>Gammaproteobacteria>Enterobacterales>Enterobacteriaceae>Salmonella | GCA_015048385.1 | Salmonella_enterica | IMT80_001261 | 170929..174570 | 28901 | BREX-system-P-loop-protein-BrxC-[Salmonella-enterica] |
| EGP1242555.1 | Type1.BREX | [illegible] | 1213 | Bacteria>Pseudomonadota>Gammaproteobacteria>Enterobacterales>Enterobacteriaceae>Salmonella | GCA_015377445.1 | Salmonella_enterica | IV995_001915 | 60140..63781 | 28901 | BREX-system-P-loop-protein-BrxC-[Salmonella-enterica] |
| HAS9574207.1 | Type1.BREX | [illegible] | 1213 | Bacteria>Pseudomonadota>Gammaproteobacteria>Enterobacterales>Enterobacteriaceae>Salmonella | GCA_015871885.1 | Salmonella_enterica_subsp._enterica_serovar_Typhimurium | I8M64_002289 | 108065..111706 | 90371 | BREX-system-P-loop-protein-BrxC-[Salmonella-enterica-subsp.-enterica-serovar-Typhimurium] |
| EGS3682914.1 | Type1.BREX | [illegible] | 912 | Bacteria>Pseudomonadota>Gammaproteobacteria>Enterobacterales>Enterobacteriaceae>Salmonella | GCA_015875385.1 | Salmonella_enterica | I5X77_001841 | 392372..395108 | 28901 | BREX-system-P-loop-protein-BrxC-[Salmonella-enterica] |
| HAT0208696.1 | Type1.BREX | [illegible] | 773 | Bacteria>Pseudomonadota>Gammaproteobacteria>Enterobacterales>Enterobacteriaceae>Salmonella | GCA_015882745.1 | Salmonella_enterica_subsp._enterica | I8Q75_000460 | 75065..77386 | 59201 | BREX-system-P-loop-protein-BrxC-[Salmonella-enterica-subsp.-enterica] |
| HAT0211425.1 | Type1.BREX | [illegible] | 408 | Bacteria>Pseudomonadota>Gammaproteobacteria>Enterobacterales>Enterobacteriaceae>Salmonella | GCA_015882745.1 | Salmonella_enterica_subsp._enterica | I8Q75_003248 | 1..1225 | 59201 | BREX-system-P-loop-protein-BrxC-[Salmonella-enterica-subsp.-enterica] |
| EGT0819117.1 | Type1.BREX | [illegible] | 1213 | Bacteria>Pseudomonadota>Gammaproteobacteria>Enterobacterales>Enterobacteriaceae>Salmonella | GCA_015953925.1 | Salmonella_enterica_subsp._enterica_serovar_Ohio | I3I53_003218 | 25100..28741 | 117541 | BREX-system-P-loop-protein-BrxC-[Salmonella-enterica-subsp.-enterica-serovar-Ohio] |
| EGT1190058.1 | Type1.BREX | [illegible] | 1213 | Bacteria>Pseudomonadota>Gammaproteobacteria>Enterobacterales>Enterobacteriaceae>Salmonella | GCA_015955585.1 | Salmonella_enterica | I3G58_002375 | 108064..111705 | 28901 | BREX-system-P-loop-protein-BrxC-[Salmonella-enterica] |
| MBH5277084.1 | Type1.BREX | [illegible] | 1213 | Bacteria>Pseudomonadota>Gammaproteobacteria>Enterobacterales>Enterobacteriaceae>Salmonella | GCA_016019715.1 | Salmonella_enterica | I3A81_01650 | 358597..362238 | 28901 | BREX-system-P-loop-protein-BrxC-[Salmonella-enterica] |
| HAU6657706.1 | Type1.BREX | [illegible] | 1157 | Bacteria>Pseudomonadota>Gammaproteobacteria>Enterobacterales>Enterobacteriaceae>Salmonella | GCA_016095655.1 | Salmonella_enterica | JEL20_001876 | 30497..33967 | 28901 | BREX-system-P-loop-protein-BrxC-[Salmonella-enterica] |
| EGU3680633.1 | Type1.BREX | [illegible] | 1213 | Bacteria>Pseudomonadota>Gammaproteobacteria>Enterobacterales>Enterobacteriaceae>Salmonella | GCA_016165525.1 | Salmonella_enterica | HOR40_002395 | 27396..31037 | 28901 | BREX-system-P-loop-protein-BrxC-[Salmonella-enterica] |
| EGU5502543.1 | Type1.BREX | [illegible] | 1213 | Bacteria>Pseudomonadota>Gammaproteobacteria>Enterobacterales>Enterobacteriaceae>Salmonella | GCA_016174335.1 | Salmonella_enterica | HGK88_001891 | 343296..346937 | 28901 | BREX-system-P-loop-protein-BrxC-[Salmonella-enterica] |

| Protein-ID | System | Gene-Neighbourhoods | Length | Taxonomy | Assembly-ID | Strain | Locus-Tag | Genomic-coordinates | TaxID | Fasta-define |
|---|---|---|---|---|---|---|---|---|---|---|
| HAV0565027.1 | Type1.BREX | Arm-DNA-bind_3->NACHT+SF1-SF2-Helicase+HKD-DNase->REase-><-SIGMA+OB-Fold+AAA-ATPase+Lon-Protease-BrxL<-PglZ<-TOPRIM<-OLD-ABC<-N6-METHYLASE<-DUF6079*<-BrxB-BrxF-iSTAND<-BrxA-DNA-bindin[illegible] | 1213 | Bacteria>Pseudomonadota>Gammaproteobacteria>Enterobacterales>Enterobacteriaceae>Salmonella | GCA_016249375.1 | Salmonella_enterica_subsp._enterica | JF300_000748 | 89373..93014 | 59201 | BREX-system-P-loop-protein-BrxC-[Salmonella-enterica-subsp.-enterica] |
| EGV8384248.1 | Type1.BREX | <-HEPN<-REase+iREase<-?\|\|YjhX_toxin->SAM-METHYLASE->?->BrxA-DNA-binding->BrxB-BrxF-iSTAND->DUF6079*->N6-METHYLASE->OLD-ABC->TOPRIM->PglZ->SIGMA+OB-Fold+AAA-ATPase+Lon-Protease-BrxL->[illegible] | 1214 | Bacteria>Pseudomonadota>Gammaproteobacteria>Enterobacterales>Enterobacteriaceae>Salmonella | GCA_016279875.1 | Salmonella_enterica | EHN43_00830 | 170919..174563 | 28901 | BREX-system-P-loop-protein-BrxC-[Salmonella-enterica] |
| EGZ8568475.1 | Type1.BREX | <-HEPN<-REase+iREase<-?\|\|YjhX_toxin->SAM-METHYLASE->?->BrxA-DNA-binding->BrxB-BrxF-iSTAND->DUF6079*->N6-METHYLASE->OLD-ABC->TOPRIM->PglZ->SIGMA+OB-Fold+AAA-ATPase+Lon-Protease-BrxL->[illegible] | 1213 | Bacteria>Pseudomonadota>Gammaproteobacteria>Enterobacterales>Enterobacteriaceae>Salmonella | GCA_016588995.1 | Salmonella_enterica_subsp._enterica_serovar_Typhimurium | JJM06_003392 | 12585..16226 | 90371 | BREX-system-P-loop-protein-BrxC-[Salmonella-enterica-subsp.-enterica-serovar-Typhimurium] |
| EHA2523073.1 | Type1.BREX | <-HEPN<-?\|\|YjhX_toxin->SAM-METHYLASE->?->BrxA-DNA-binding->BrxB-BrxF-iSTAND->DUF6079*->N6-METHYLASE->OLD-ABC->TOPRIM->PglZ->SIGMA+OB-Fold+AAA-ATPase+Lon-Protease-BrxL-><-REase<-SF1-SF[illegible] | 1213 | Bacteria>Pseudomonadota>Gammaproteobacteria>Enterobacterales>Enterobacteriaceae>Salmonella | GCA_016628375.1 | Salmonella_enterica | JKI82_003469 | 165981..169622 | 28901 | BREX-system-P-loop-protein-BrxC-[Salmonella-enterica] |
| EHA8894993.1 | Type1.BREX | <-HEPN<-?\|\|YjhX_toxin->SAM-METHYLASE->?->BrxA-DNA-binding->BrxB-BrxF-iSTAND->DUF6079*->N6-METHYLASE->OLD-ABC->TOPRIM->PglZ->SIGMA+OB-Fold+AAA-ATPase+Lon-Protease-BrxL-><-REase<-Couple[illegible] | 1213 | Bacteria>Pseudomonadota>Gammaproteobacteria>Enterobacterales>Enterobacteriaceae>Salmonella | GCA_016843845.1 | Salmonella_enterica_subsp._enterica | JHW15_001254 | 44404..48045 | 59201 | BREX-system-P-loop-protein-BrxC-[Salmonella-enterica-subsp.-enterica] |
| EHB1109136.1 | Type1.BREX | <-HEPN<-REase+iREase<-?\|\|YjhX_toxin->SAM-METHYLASE->?->BrxA-DNA-binding->BrxB-BrxF-iSTAND->DUF6079*->N6-METHYLASE-> | 1213 | Bacteria>Pseudomonadota>Gammaproteobacteria>Enterobacterales>Enterobacteriaceae>Salmonella | GCA_016950725.1 | Salmonella_enterica_subsp._enterica_serovar_Typhimurium | JYP19_004114 | 18317..21958 | 90371 | BREX-system-P-loop-protein-BrxC-[Salmonella-enterica-subsp.-enterica-serovar-Typhimurium] |
| EHC3781321.1 | Type1.BREX | <-HEPN<-REase+iREase<-?\|\|YjhX_toxin->SAM-METHYLASE->?->BrxA-DNA-binding->BrxB-BrxF-iSTAND->DUF6079*->N6-METHYLASE->OLD-ABC->TOPRIM->PglZ->SIGMA+OB-Fold+AAA-ATPase+Lon-Protease-BrxL->[illegible] | 1213 | Bacteria>Pseudomonadota>Gammaproteobacteria>Enterobacterales>Enterobacteriaceae>Salmonella | GCA_017044955.1 | Salmonella_enterica_subsp._enterica_serovar_Typhimurium | JS923_000750 | 173044..176685 | 90371 | BREX-system-P-loop-protein-BrxC-[Salmonella-enterica-subsp.-enterica-serovar-Typhimurium] |
| QSG17817.1 | Type1.BREX | <-Peptidase_M17_N+Peptidase_M17\|\|?-><-?\|\|SKI-><-?\|\|tRNA->Arm-DNA-bind_3->NACHT+SF1-SF2-Helicase+HKD-DNase-><-TOPRIM<-DUF6079*<-BrxA-DNA-binding<-?<-YjhX_toxin\|\|REase+iREase->?-><-?\|\|[illegible] | 1213 | Bacteria>Pseudomonadota>Gammaproteobacteria>Enterobacterales>Enterobacteriaceae>Salmonella | GCA_017094545.1 | Salmonella_enterica_subsp._enterica_serovar_Typhimurium | IRY60_23155 | 4744141..4747782 | 90371 | BREX-system-P-loop-protein-BrxC-[Salmonella-enterica-subsp.-enterica-serovar-Typhimurium] |
| QSG23651.1 | Type1.BREX | <-LACI-HTH+PBPI\|\|SKI-><-?\|\|tRNA->Arm-DNA-bind_3->REase-><-SIGMA+OB-Fold+AAA-ATPase+Lon-Protease-BrxL<-TOPRIM<-OLD-ABC<-DUF6079*<-?<-YjhX_toxin\|\|REase+iREase->?-><-?\|\|<-?\|\|<-MODE-HTH+P[illegible] | 1213 | Bacteria>Pseudomonadota>Gammaproteobacteria>Enterobacterales>Enterobacteriaceae>Salmonella | GCA_017094585.1 | Salmonella_enterica_subsp._enterica_serovar_Typhimurium | IRY58_23115 | 4749695..4753336 | 90371 | BREX-system-P-loop-protein-BrxC-[Salmonella-enterica-subsp.-enterica-serovar-Typhimurium] |
| EHD7670518.1 | Type1.BREX | <-HEPN<-REase+iREase<-?\|\|YjhX_toxin->SAM-METHYLASE->?->BrxA-DNA-binding->BrxB-BrxF-iSTAND->DUF6079*->N6-METHYLASE->OLD-ABC->TOPRIM->PglZ->SIGMA+OB-Fold+AAA-ATPase+Lon-Protease-BrxL->[illegible] | 1213 | Bacteria>Pseudomonadota>Gammaproteobacteria>Enterobacterales>Enterobacteriaceae>Salmonella | GCA_017183075.1 | Salmonella_enterica | JQD35_001754 | 170582..174223 | 28901 | BREX-system-P-loop-protein-BrxC-[Salmonella-enterica] |
| EHD9459645.1 | Type1.BREX | <-HEPN<-REase+iREase<-?\|\|YjhX_toxin->SAM-METHYLASE->?->BrxA-DNA-binding->BrxB-BrxF-iSTAND->DUF6079*->N6-METHYLASE->OLD-ABC->TOPRIM->PglZ->SIGMA+OB-Fold+AAA-ATPase+Lon-Protease-BrxL->[illegible] | 1213 | Bacteria>Pseudomonadota>Gammaproteobacteria>Enterobacterales>Enterobacteriaceae>Salmonella | GCA_017198155.1 | Salmonella_enterica_subsp._enterica_serovar_Typhimurium | JYL48_000531 | 170932..174573 | 90371 | BREX-system-P-loop-protein-BrxC-[Salmonella-enterica-subsp.-enterica-serovar-Typhimurium] |
| EHE5127202.1 | Type1.BREX | ACET->YjhX_toxin->ACET->METHYLASE->ZnR->Glyoxalase->ZnR->BrxA-DNA-binding->BrxB-BrxF-iSTAND->DUF6079*->ParB+HNH->N6-METHYLASE->PglZ->SIGMA+OB-Fold+AAA-ATPase+Lon-Protease-BrxL-><-Arm-[illegible] | 1233 | Bacteria>Pseudomonadota>Gammaproteobacteria>Enterobacterales>Enterobacteriaceae>Salmonella | GCA_017241415.1 | Salmonella_enterica | JMY69_002120 | 181649..185350 | 28901 | BREX-system-P-loop-protein-BrxC-[Salmonella-enterica] |
| EHH9023834.1 | Type1.BREX | <-HEPN<-REase+iREase<-?\|\|YjhX_toxin->SAM-METHYLASE->?->BrxA-DNA-binding->BrxB-BrxF-iSTAND->DUF6079*->N6-METHYLASE->OLD-ABC->TOPRIM->PglZ->SIGMA+OB-Fold+AAA-ATPase+Lon-Protease-BrxL->[illegible] | 1213 | Bacteria>Pseudomonadota>Gammaproteobacteria>Enterobacterales>Enterobacteriaceae>Salmonella | GCA_017748765.1 | Salmonella_enterica | J8439_003629 | 18317..21958 | 28901 | BREX-system-P-loop-protein-BrxC-[Salmonella-enterica] |
| EHI1197052.1 | Type1.BREX | Arm-DNA-bind_3->NACHT+SF1-SF2-Helicase+HKD-DNase->REase-><-SIGMA+OB-Fold+AAA-ATPase+Lon-Protease-BrxL<-PglZ<-TOPRIM<-OLD-ABC<-N6-METHYLASE<-DUF6079*<-BrxB-BrxF-iSTAND<-BrxA-DNA-bindin[illegible] | 1213 | Bacteria>Pseudomonadota>Gammaproteobacteria>Enterobacterales>Enterobacteriaceae>Salmonella | GCA_017785955.1 | Salmonella_enterica | J8551_000470 | 108064..111705 | 28901 | BREX-system-P-loop-protein-BrxC-[Salmonella-enterica] |
| EHI4209832.1 | Type1.BREX | <-HEPN<-REase+iREase<-?\|\|YjhX_toxin->SAM-METHYLASE->?->BrxA-DNA-binding->BrxB-BrxF-iSTAND->DUF6079*->N6-METHYLASE-> | 1213 | Bacteria>Pseudomonadota>Gammaproteobacteria>Enterobacterales>Enterobacteriaceae>Salmonella | GCA_017802295.1 | Salmonella_enterica | J9Y57_000957 | 124173..127814 | 28901 | BREX-system-P-loop-protein-BrxC-[Salmonella-enterica] |
| EHJ0877869.1 | Type1.BREX | <-HEPN<-REase+iREase<-?\|\|YjhX_toxin->SAM-METHYLASE->?->BrxA-DNA-binding->BrxB-BrxF-iSTAND->DUF6079*->N6-METHYLASE->OLD-ABC->TOPRIM->PglZ->SIGMA+OB-Fold+AAA-ATPase+Lon-Protease-BrxL->[illegible] | 1213 | Bacteria>Pseudomonadota>Gammaproteobacteria>Enterobacterales>Enterobacteriaceae>Salmonella | GCA_017846115.1 | Salmonella_enterica | KAB51_002697 | 18125..21766 | 28901 | BREX-system-P-loop-protein-BrxC-[Salmonella-enterica] |
| EHJ1242128.1 | Type1.BREX | Arm-DNA-bind_3->NACHT+S[illegible] | 1213 | Bacteria>Pseudomonadota>Gammaproteobacteria>Enterobacterales>Enterobacteriaceae>Salmonella | GCA_017851875.1 | Salmonella_enterica_subsp._enterica_serovar_Typhimurium | KAC06_001466 | 81358..84999 | 90371 | BREX-system-P-loop-protein-BrxC-[Salmonella-enterica-subsp.-enterica-serovar-Typhimurium] |
| EHJ7117733.1 | Type1.BREX | <-HEPN<-?\|\|YjhX_toxin->[illegible] | 1204 | Bacteria>Pseudomonadota>Gammaproteobacteria>Enterobacterales>Enterobacteriaceae>Salmonella | GCA_017926475.1 | Salmonella_enterica_subsp._enterica_serovar_Kentucky | KAP12_002965 | 412329..415943 | 192955 | BREX-system-P-loop-protein-BrxC-[Salmonella-enterica-subsp.-enterica-serovar-Kentucky] |
| EHJ9839635.1 | Type1.BREX | HTH_3->Couple_hipA+HipA[illegible] | 1213 | Bacteria>Pseudomonadota>Gammaproteobacteria>Enterobacterales>Enterobacteriaceae>Salmonella | GCA_017967145.1 | Salmonella_enterica | KEH66_000998 | 398114..401755 | 28901 | BREX-system-P-loop-protein-BrxC-[Salmonella-enterica] |
| EHN4465874.1 | Type1.BREX | Arm-DNA-bind_3->NACHT+S[illegible] | 1213 | Bacteria>Pseudomonadota>Gammaproteobacteria>Enterobacterales>Enterobacteriaceae>Salmonella | GCA_018387865.1 | Salmonella_enterica | KIZ21_001044 | 348400..352041 | 28901 | BREX-system-P-loop-protein-BrxC-[Salmonella-enterica] |
| EHN4525913.1 | Type1.BREX | <-HEPN<-REase+iREase<-?[illegible] | 1213 | Bacteria>Pseudomonadota>Gammaproteobacteria>Enterobacterales>Enterobacteriaceae>Salmonella | GCA_018388245.1 | Salmonella_enterica_subsp._enterica_serovar_Ohio | KI018_001081 | 92040..95681 | 117541 | BREX-system-P-loop-protein-BrxC-[Salmonella-enterica-subsp.-enterica-serovar-Ohio] |
| EHO4506760.1 | Type1.BREX | <-HEPN<-?\|\|YjhX_toxin->[illegible] | 1213 | Bacteria>Pseudomonadota>Gammaproteobacteria>Enterobacterales>Enterobacteriaceae>Salmonella | GCA_018524125.1 | Salmonella_enterica_subsp._enterica_serovar_Carrau | KD666_001066 | 149262..152903 | 1160739 | BREX-system-P-loop-protein-BrxC-[Salmonella-enterica-subsp.-enterica-serovar-Carrau] |
| MBT0981085.1 | Type1.BREX | <-HEPN<-REase+iREase<-?[illegible] | 1213 | Bacteria>Pseudomonadota>Gammaproteobacteria>Enterobacterales>Enterobacteriaceae>Salmonella | GCA_018531305.1 | Salmonella_enterica_subsp._enterica_serovar_Typhimurium | KJ032_08020 | 170949..174590 | 90371 | BREX-system-P-loop-protein-BrxC-[Salmonella-enterica-subsp.-enterica-serovar-Typhimurium] |
| EHO5101500.1 | Type1.BREX | HTH_Tnp_1+HTH_21+rve+rv[illegible] | 1213 | Bacteria>Pseudomonadota>Gammaproteobacteria>Enterobacterales>Enterobacteriaceae>Salmonella | GCA_018532585.1 | Salmonella_enterica | KLP74_002745 | 12418..16059 | 28901 | BREX-system-P-loop-protein-BrxC-[Salmonella-enterica] |
| EHO5062413.1 | Type1.BREX | <-HEPN<-?\|\|YjhX_toxin->[illegible] | 1213 | Bacteria>Pseudomonadota>Gammaproteobacteria>Enterobacterales>Enterobacteriaceae>Salmonella | GCA_018532705.1 | Salmonella_enterica | KLP81_000353 | 388533..392174 | 28901 | BREX-system-P-loop-protein-BrxC-[Salmonella-enterica] |
| EHO5648342.1 | Type1.BREX | <-HEPN<-?\|\|YjhX_toxin->[illegible] | 1213 | Bacteria>Pseudomonadota>Gammaproteobacteria>Enterobacterales>Enterobacteriaceae>Salmonella | GCA_018535125.1 | Salmonella_enterica | KLR08_000690 | 144464..148105 | 28901 | BREX-system-P-loop-protein-BrxC-[Salmonella-enterica] |
| EHO5627401.1 | Type1.BREX | <-HEPN<-?\|\|YjhX_toxin->[illegible] | 1213 | Bacteria>Pseudomonadota>Gammaproteobacteria>Enterobacterales>Enterobacteriaceae>Salmonella | GCA_018535225.1 | Salmonella_enterica | KLQ90_002307 | 30800..34441 | 28901 | BREX-system-P-loop-protein-BrxC-[Salmonella-enterica] |
| EHP4799940.1 | Type1.BREX | <-HEPN<-?\|\|YjhX_toxin->[illegible] | 1213 | Bacteria>Pseudomonadota>Gammaproteobacteria>Enterobacterales>Enterobacteriaceae>Salmonella | GCA_018776965.1 | Salmonella_enterica_subsp._enterica_serovar_Sangera | KM018_002015 | 44674..48315 | 1173574 | BREX-system-P-loop-protein-BrxC-[Salmonella-enterica-subsp.-enterica-serovar-Sangera] |
| EHQ7935671.1 | Type1.BREX | Arm-DNA-bind_3->NACHT+S[illegible] | 1213 | Bacteria>Pseudomonadota>Gammaproteobacteria>Enterobacterales>Enterobacteriaceae>Salmonella | GCA_018976465.1 | Salmonella_enterica | KSM73_002317 | 108064..111705 | 28901 | BREX-system-P-loop-protein-BrxC-[Salmonella-enterica] |
| EHS1468535.1 | Type1.BREX | <-HEPN<-REase+iREase<-?[illegible] | 1213 | Bacteria>Pseudomonadota>Gammaproteobacteria>Enterobacterales>Enterobacteriaceae>Salmonella | GCA_019106545.1 | Salmonella_enterica_subsp._enterica_serovar_Typhimurium | KVU04_001403 | 171297..174938 | 90371 | BREX-system-P-loop-protein-BrxC-[Salmonella-enterica-subsp.-enterica-serovar-Typhimurium] |
| EHS3976100.1 | Type1.BREX | BrxA-DNA-binding->BrxB-[illegible] | 1213 | Bacteria>Pseudomonadota>Gammaproteobacteria>Enterobacterales>Enterobacteriaceae>Salmonella | GCA_019140515.1 | Salmonella_enterica_subsp._enterica_serovar_Poona | KV430_003532 | 2009..5650 | 436295 | BREX-system-P-loop-protein-BrxC-[Salmonella-enterica-subsp.-enterica-serovar-Poona] |
| EHS4662101.1 | Type1.BREX | Arm-DNA-bind_3->NACHT+S[illegible] | 1213 | Bacteria>Pseudomonadota>Gammaproteobacteria>Enterobacterales>Enterobacteriaceae>Salmonella | GCA_019144395.1 | Salmonella_enterica | KV604_003674 | 73553..77194 | 28901 | BREX-system-P-loop-protein-BrxC-[Salmonella-enterica] |
| EHS5456435.1 | Type1.BREX | <-HEPN<-?\|\|YjhX_toxin->[illegible] | 1213 | Bacteria>Pseudomonadota>Gammaproteobacteria>Enterobacterales>Enterobacteriaceae>Salmonella | GCA_019151305.1 | Salmonella_enterica_subsp._enterica_serovar_Corvallis | KV802_001386 | 165237..168878 | 593905 | BREX-system-P-loop-protein-BrxC-[Salmonella-enterica-subsp.-enterica-serovar-Corvallis] |
| EHT1034281.1 | Type1.BREX | <-GntR-HTH++TrmB+ZDNA-H[illegible] | 1213 | Bacteria>Pseudomonadota>Gammaproteobacteria>Enterobacterales>Enterobacteriaceae>Salmonella | GCA_019186465.1 | Salmonella_enterica | KW750_001284 | 149102..152743 | 28901 | BREX-system-P-loop-protein-BrxC-[Salmonella-enterica] |
| EHT6941866.1 | Type1.BREX | <-HEPN<-REase+iREase<-?[illegible] | 1213 | Bacteria>Pseudomonadota>Gammaproteobacteria>Enterobacterales>Enterobacteriaceae>Salmonella | GCA_019250575.1 | Salmonella_enterica | KYH33_001187 | 170918..174559 | 28901 | BREX-system-P-loop-protein-BrxC-[Salmonella-enterica] |
| EHU5044814.1 | Type1.BREX | SKI-><-?\|\|tRNA->NACHT+S[illegible] | 1213 | Bacteria>Pseudomonadota>Gammaproteobacteria>Enterobacterales>Enterobacteriaceae>Salmonella | GCA_019291225.1 | Salmonella_enterica | KY920_001891 | 99712..103353 | 28901 | BREX-system-P-loop-protein-BrxC-[Salmonella-enterica] |
| EHU7518552.1 | Type1.BREX | <-HEPN<-?\|\|YjhX_toxin->[illegible] | 1213 | Bacteria>Pseudomonadota>Gammaproteobacteria>Enterobacterales>Enterobacteriaceae>Salmonella | GCA_019318345.1 | Salmonella_enterica | KZL07_001983 | 273071..276712 | 28901 | BREX-system-P-loop-protein-BrxC-[Salmonella-enterica] |
| EHV2030459.1 | Type1.BREX | <-HEPN<-?\|\|YjhX_toxin->[illegible] | 1213 | Bacteria>Pseudomonadota>Gammaproteobacteria>Enterobacterales>Enterobacteriaceae>Salmonella | GCA_019356775.1 | Salmonella_enterica | KZ138_001531 | 389696..393337 | 28901 | BREX-system-P-loop-protein-BrxC-[Salmonella-enterica] |
| EHW6237841.1 | Type1.BREX | <-HEPN<-?\|\|YjhX_toxin->[illegible] | 1213 | Bacteria>Pseudomonadota>Gammaproteobacteria>Enterobacterales>Enterobacteriaceae>Salmonella | GCA_019485205.1 | Salmonella_enterica | K2495_002337 | 30739..34380 | 28901 | BREX-system-P-loop-protein-BrxC-[Salmonella-enterica] |
| EHW8503808.1 | Type1.BREX | <-HEPN<-?\|\|YjhX_toxin->[illegible] | 1213 | Bacteria>Pseudomonadota>Gammaproteobacteria>Enterobacterales>Enterobacteriaceae>Salmonella | GCA_019496265.1 | Salmonella_enterica | K3U91_002209 | 149047..152688 | 28901 | BREX-system-P-loop-protein-BrxC-[Salmonella-enterica] |
| EHX4782365.1 | Type1.BREX | <-HEPN<-?\|\|YjhX_toxin->[illegible] | 1213 | Bacteria>Pseudomonadota>Gammaproteobacteria>Enterobacterales>Enterobacteriaceae>Salmonella | GCA_019533715.1 | Salmonella_enterica | K3Y02_002640 | 144286..147927 | 28901 | BREX-system-P-loop-protein-BrxC-[Salmonella-enterica] |
| EHZ3116575.1 | Type1.BREX | <-HEPN<-REase+iREase<-?[illegible] | 957 | Bacteria>Pseudomonadota>Gammaproteobacteria>Enterobacterales>Enterobacteriaceae>Salmonella | GCA_019692095.1 | Salmonella_enterica_subsp._enterica_serovar_Typhimurium | K5O64_003315 | 60042..62911 | 90371 | BREX-system-P-loop-protein-BrxC-[Salmonella-enterica-subsp.-enterica-serovar-Typhimurium] |
| EHZ3783988.1 | Type1.BREX | <-HEPN<-REase+iREase<-?[illegible] | 1213 | Bacteria>Pseudomonadota>Gammaproteobacteria>Enterobacterales>Enterobacteriaceae>Salmonella | GCA_019697575.1 | Salmonella_enterica | K5U30_001272 | 170919..174560 | 28901 | BREX-system-P-loop-protein-BrxC-[Salmonella-enterica] |
| EIA3194373.1 | Type1.BREX | Arm-DNA-bind_3->NACHT+S[illegible] | 1213 | Bacteria>Pseudomonadota>Gammaproteobacteria>Enterobacterales>Enterobacteriaceae>Salmonella | GCA_019788365.1 | Salmonella_enterica | K7D38_001961 | 187001..190642 | 28901 | BREX-system-P-loop-protein-BrxC-[Salmonella-enterica] |
| EIA3335811.1 | Type1.BREX | <-HEPN<-REase+iREase<-?[illegible] | 1213 | Bacteria>Pseudomonadota>Gammaproteobacteria>Enterobacterales>Enterobacteriaceae>Salmonella | GCA_019789135.1 | Salmonella_enterica | K7D78_001298 | 400174..403815 | 28901 | BREX-system-P-loop-protein-BrxC-[Salmonella-enterica] |
| EIB1064378.1 | Type1.BREX | Arm-DNA-bind_3->NACHT+S[illegible] | 1213 | Bacteria>Pseudomonadota>Gammaproteobacteria>Enterobacterales>Enterobacteriaceae>Salmonella | GCA_019855255.1 | Salmonella_enterica | K7907_001965 | 108064..111705 | 28901 | BREX-system-P-loop-protein-BrxC-[Salmonella-enterica] |
| EIB7346154.1 | Type1.BREX | BrxA-DNA-binding->BrxB-[illegible] | 1213 | Bacteria>Pseudomonadota>Gammaproteobacteria>Enterobacterales>Enterobacteriaceae>Salmonella | GCA_019904735.1 | Salmonella_enterica | K9H00_003649 | 2808..6449 | 28901 | BREX-system-P-loop-protein-BrxC-[Salmonella-enterica] |
| EIB7574746.1 | Type1.BREX | ACET->YjhX_toxin->ACET-[illegible] | 1233 | Bacteria>Pseudomonadota>Gammaproteobacteria>Enterobacterales>Enterobacteriaceae>Salmonella | GCA_019905755.1 | Salmonella_enterica | K9I12_000373 | 406371..410072 | 28901 | BREX-system-P-loop-protein-BrxC-[Salmonella-enterica] |
| EIC4097432.1 | Type1.BREX | <-HEPN<-?\|\|YjhX_toxin->[illegible] | 1213 | Bacteria>Pseudomonadota>Gammaproteobacteria>Enterobacterales>Enterobacteriaceae>Salmonella | GCA_019951695.1 | Salmonella_enterica | K9796_003003 | 22191..25832 | 28901 | BREX-system-P-loop-protein-BrxC-[Salmonella-enterica] |
| EIC8525255.1 | Type1.BREX | <-REase+iREase<-?\|\|YjhX[illegible] | 1204 | Bacteria>Pseudomonadota>Gammaproteobacteria>Enterobacterales>Enterobacteriaceae>Salmonella | GCA_019993965.1 | Salmonella_enterica | LAR10_002215 | 30447..34061 | 28901 | BREX-system-P-loop-protein-BrxC-[Salmonella-enterica] |
| HBJ3502651.1 | Type1.BREX | <-HEPN<-REase+iREase<-?[illegible] | 1213 | Bacteria>Pseudomonadota>Gammaproteobacteria>Enterobacterales>Enterobacteriaceae>Salmonella | GCA_020048005.1 | Salmonella_enterica_subsp._enterica_serovar_Typhimurium | LB282_001657 | 170703..174344 | 90371 | BREX-system-P-loop-protein-BrxC-[Salmonella-enterica-subsp.-enterica-serovar-Typhimurium] |
| MBZ4773899.1 | Type1.BREX | <-DUF6079*<-BrxB-BrxF-i[illegible] | 1129 | Bacteria>Pseudomonadota>Gammaproteobacteria>Enterobacterales>Enterobacteriaceae>Salmonella | GCA_020070065.1 | Salmonella_enterica_subsp._enterica_serovar_Typhimurium | HU130_03520 | 1..3389 | 90371 | BREX-system-P-loop-protein-BrxC-[Salmonella-enterica-subsp.-enterica-serovar-Typhimurium] |
| MBZ4755449.1 | Type1.BREX | <-HEPN<-REase+iREase<-?[illegible] | 1174 | Bacteria>Pseudomonadota>Gammaproteobacteria>Enterobacterales>Enterobacteriaceae>Salmonella | GCA_020070115.1 | Salmonella_enterica_subsp._enterica_serovar_Typhimurium | HU026_04360 | 60999..64520 | 90371 | BREX-system-P-loop-protein-BrxC-[Salmonella-enterica-subsp.-enterica-serovar-Typhimurium] |
| EID5777526.1 | Type1.BREX | <-HEPN<-?\|\|YjhX_toxin->[illegible] | 1213 | Bacteria>Pseudomonadota>Gammaproteobacteria>Enterobacterales>Enterobacteriaceae>Salmonella | GCA_020070325.1 | Salmonella_enterica | LB861_002368 | 82497..86138 | 28901 | BREX-system-P-loop-protein-BrxC-[Salmonella-enterica] |
| MBZ4898657.1 | Type1.BREX | Arm-DNA-bind_3->NACHT+S[illegible] | 1213 | Bacteria>Pseudomonadota>Gammaproteobacteria>Enterobacterales>Enterobacteriaceae>Salmonella | GCA_020070625.1 | Salmonella_enterica_subsp._enterica_serovar_Typhimurium | HU117_01215 | 58948..62589 | 90371 | BREX-system-P-loop-protein-BrxC-[Salmonella-enterica-subsp.-enterica-serovar-Typhimurium] |
| MBZ5058250.1 | Type1.BREX | <-HEPN<-REase+iREase<-?[illegible] | 1213 | Bacteria>Pseudomonadota>Gammaproteobacteria>Enterobacterales>Enterobacteriaceae>Salmonella | GCA_020071235.1 | Salmonella_enterica_subsp._enterica_serovar_Typhimurium | HU058_13730 | 15756..19397 | 90371 | BREX-system-P-loop-protein-BrxC-[Salmonella-enterica-subsp.-enterica-serovar-Typhimurium] |
| MBZ5141240.1 | Type1.BREX | <-HEPN<-REase+iREase<-?[illegible] | 601 | Bacteria>Pseudomonadota>Gammaproteobacteria>Enterobacterales>Enterobacteriaceae>Salmonella | GCA_020071745.1 | Salmonella_enterica_subsp._enterica_serovar_Typhimurium | HU084_13180 | 16232..18035 | 90371 | BREX-system-P-loop-protein-BrxC-[Salmonella-enterica-subsp.-enterica-serovar-Typhimurium] |
| EIE3342236.1 | Type1.BREX | <-HEPN<-REase+iREase<-?[illegible] | 1213 | Bacteria>Pseudomonadota>Gammaproteobacteria>Enterobacterales>Enterobacteriaceae>Salmonella | GCA_020155375.1 | Salmonella_enterica_subsp._enterica_serovar_Typhimurium | LDC84_002298 | 60209..63850 | 90371 | BREX-system-P-loop-protein-BrxC-[Salmonella-enterica-subsp.-enterica-serovar-Typhimurium] |
| EIF4998474.1 | Type1.BREX | <-HEPN<-REase+iREase<-?[illegible] | 848 | Bacteria>Pseudomonadota>Gammaproteobacteria>Enterobacterales>Enterobacteriaceae>Salmonella | GCA_020260065.1 | Salmonella_enterica_subsp._enterica_serovar_Typhimurium | LFO03_001522 | 60001..62543 | 90371 | BREX-system-P-loop-protein-BrxC-[Salmonella-enterica-subsp.-enterica-serovar-Typhimurium] |
| EIF5096942.1 | Type1.BREX | <-HEPN<-REase+iREase<-?[illegible] | 899 | Bacteria>Pseudomonadota>Gammaproteobacteria>Enterobacterales>Enterobacteriaceae>Salmonella | GCA_020260525.1 | Salmonella_enterica_subsp._enterica_serovar_Typhimurium | LFF62_002381 | 60001..62698 | 90371 | BREX-system-P-loop-protein-BrxC-[Salmonella-enterica-subsp.-enterica-serovar-Typhimurium] |
| EIF5593860.1 | Type1.BREX | <-HEPN<-REase+iREase<-?[illegible] | 1058 | Bacteria>Pseudomonadota>Gammaproteobacteria>Enterobacterales>Enterobacteriaceae>Salmonella | GCA_020262845.1 | Salmonella_enterica_subsp._enterica_serovar_Typhimurium | LFF21_002838 | 60001..63175 | 90371 | BREX-system-P-loop-protein-BrxC-[Salmonella-enterica-subsp.-enterica-serovar-Typhimurium] |
| EIF9427095.1 | Type1.BREX | <-HEPN<-?\|\|YjhX_toxin->[illegible] | 1213 | Bacteria>Pseudomonadota>Gammaproteobacteria>Enterobacterales>Enterobacteriaceae>Salmonella | GCA_020283115.1 | Salmonella_enterica | HW154_002103 | 25917..29558 | 28901 | BREX-system-P-loop-protein-BrxC-[Salmonella-enterica] |
| EIF9623260.1 | Type1.BREX | <-HEPN<-?\|\|YjhX_toxin->[illegible] | 1213 | Bacteria>Pseudomonadota>Gammaproteobacteria>Enterobacterales>Enterobacteriaceae>Salmonella | GCA_020284415.1 | Salmonella_enterica | HQ981_002275 | 25917..29558 | 28901 | BREX-system-P-loop-protein-BrxC-[Salmonella-enterica] |
| EII7462484.1 | Type1.BREX | DUF6079-p*->N6-METHYLAS[illegible] | 865 | Bacteria>Pseudomonadota>Gammaproteobacteria>Enterobacterales>Enterobacteriaceae>Salmonella | GCA_020462505.1 | Salmonella_enterica_subsp._enterica_serovar_Typhimurium | LHR57_000982 | 1..2599 | 90371 | BREX-system-P-loop-protein-BrxC-[Salmonella-enterica-subsp.-enterica-serovar-Typhimurium] |
| EIJ0372099.1 | Type1.BREX | <-HEPN<-REase+iREase<-?[illegible] | 1213 | Bacteria>Pseudomonadota>Gammaproteobacteria>Enterobacterales>Enterobacteriaceae>Salmonella | GCA_020490715.1 | Salmonella_enterica_subsp._enterica_serovar_Typhimurium | LH081_001559 | 170919..174560 | 90371 | BREX-system-P-loop-protein-BrxC-[Salmonella-enterica-subsp.-enterica-serovar-Typhimurium] |
| EIJ0643398.1 | Type1.BREX | Arm-DNA-bind_3->NACHT+S[illegible] | 1101 | Bacteria>Pseudomonadota>Gammaproteobacteria>Enterobacterales>Enterobacteriaceae>Salmonella | GCA_020492735.1 | Salmonella_enterica | LH177_001939 | 72298..75605 | 28901 | BREX-system-P-loop-protein-BrxC-[Salmonella-enterica] |
| EIJ2988432.1 | Type1.BREX | <-HEPN<-REase+iREase<-?[illegible] | 1213 | Bacteria>Pseudomonadota>Gammaproteobacteria>Enterobacterales>Enterobacteriaceae>Salmonella | GCA_020513955.1 | Salmonella_enterica | LIS40_000161 | 170919..174560 | 28901 | BREX-system-P-loop-protein-BrxC-[Salmonella-enterica] |
| EIJ6143992.1 | Type1.BREX | <-HEPN<-?\|\|YjhX_toxin->[illegible] | 1213 | Bacteria>Pseudomonadota>Gammaproteobacteria>Enterobacterales>Enterobacteriaceae>Salmonella | GCA_020548845.1 | Salmonella_enterica_subsp._enterica_serovar_4:a:- | LI678_001014 | 394746..398387 | 2885150 | BREX-system-P-loop-protein-BrxC-[Salmonella-enterica-subsp.-enterica-serovar-4:a:-] |
| EIM3792263.1 | Type1.BREX | <-HEPN<-REase+iREase<-?[illegible] | 1213 | Bacteria>Pseudomonadota>Gammaproteobacteria>Enterobacterales>Enterobacteriaceae>Salmonella | GCA_020780095.1 | Salmonella_enterica | LM721_003554 | 27348..30989 | 28901 | BREX-system-P-loop-protein-BrxC-[Salmonella-enterica] |
| EIN7178581.1 | Type1.BREX | <-HEPN<-REase+iREase<-?[illegible] | 1137 | Bacteria>Pseudomonadota>Gammaproteobacteria>Enterobacterales>Enterobacteriaceae>Salmonella | GCA_020912625.1 | Salmonella_enterica_subsp._enterica_serovar_Typhimurium | LPQ66_002313 | 59987..63398 | 90371 | BREX-system-P-loop-protein-BrxC-[Salmonella-enterica-subsp.-enterica-serovar-Typhimurium] |
| EIO7765743.1 | Type1.BREX | <-HEPN<-REase+iREase<-?[illegible] | 1213 | Bacteria>Pseudomonadota>Gammaproteobacteria>Enterobacterales>Enterobacteriaceae>Salmonella | GCA_020997775.1 | Salmonella_enterica | LQR80_000699 | 170919..174560 | 28901 | BREX-system-P-loop-protein-BrxC-[Salmonella-enterica] |
| HBL9882169.1 | Type1.BREX | <-HEPN<-?\|\|YjhX_toxin->[illegible] | 1213 | Bacteria>Pseudomonadota>Gammaproteobacteria>Enterobacterales>Enterobacteriaceae>Salmonella | GCA_021103155.1 | Salmonella_enterica_subsp._enterica_serovar_Jangwani | LT029_003490 | 155244..158885 | 2564577 | BREX-system-P-loop-protein-BrxC-[Salmonella-enterica-subsp.-enterica-serovar-Jangwani] |
| HBM1480881.1 | Type1.BREX | <-DUF6079-p*<-BrxB-BrxF[illegible] | 700 | Bacteria>Pseudomonadota>Gammaproteobacteria>Enterobacterales>Enterobacteriaceae>Salmonella | GCA_021143585.1 | Salmonella_enterica_subsp._enterica | LUI79_002478 | 1..2100 | 59201 | BREX-system-P-loop-protein-BrxC-[Salmonella-enterica-subsp.-enterica] |
| EIQ4281221.1 | Type1.BREX | <-HEPN<-REase+iREase<-?[illegible] | 1213 | Bacteria>Pseudomonadota>Gammaproteobacteria>Enterobacterales>Enterobacteriaceae>Salmonella | GCA_021189835.1 | Salmonella_enterica | LU454_001105 | 102830..106471 | 28901 | BREX-system-P-loop-protein-BrxC-[Salmonella-enterica] |
| EIQ7941123.1 | Type1.BREX | <-DUF6079*<-BrxB-BrxF-i[illegible] | 1180 | Bacteria>Pseudomonadota>Gammaproteobacteria>Enterobacterales>Enterobacteriaceae>Salmonella | GCA_021224035.1 | Salmonella_enterica_subsp._enterica_serovar_Typhimurium | LVZ53_003070 | 1..3541 | 90371 | BREX-system-P-loop-protein-BrxC-[Salmonella-enterica-subsp.-enterica-serovar-Typhimurium] |
| EIR7569906.1 | Type1.BREX | Arm-DNA-bind_3->NACHT+S[illegible] | 873 | Bacteria>Pseudomonadota>Gammaproteobacteria>Enterobacterales>Enterobacteriaceae>Salmonella | GCA_021281445.1 | Salmonella_enterica_subsp._enterica_serovar_Agbeni | LXI61_003348 | 30366..32989 | 1967642 | BREX-system-P-loop-protein-BrxC-[Salmonella-enterica-subsp.-enterica-serovar-Agbeni] |
| EIR8875059.1 | Type1.BREX | BrxA-DNA-binding->BrxB-[illegible] | 1213 | Bacteria>Pseudomonadota>Gammaproteobacteria>Enterobacterales>Enterobacteriaceae>Salmonella | GCA_021305935.1 | Salmonella_enterica | LXE31_003492 | 2812..6453 | 28901 | BREX-system-P-loop-protein-BrxC-[Salmonella-enterica] |
| EIS9096680.1 | Type1.BREX | <-N6-METHYLASE<-DUF6079[illegible] | 1213 | Bacteria>Pseudomonadota>Gammaproteobacteria>Enterobacterales>Enterobacteriaceae>Salmonella | GCA_021435525.1 | Salmonella_enterica | LO705_004818 | 2605..6246 | 28901 | BREX-system-P-loop-protein-BrxC-[Salmonella-enterica] |
| HBN2343839.1 | Type1.BREX | <-HEPN<-REase+iREase<-?[illegible] | 1083 | Bacteria>Pseudomonadota>Gammaproteobacteria>Enterobacterales>Enterobacteriaceae>Salmonella | GCA_021498555.1 | Salmonella_enterica_subsp._enterica_serovar_Typhimurium | L2A29_001994 | 59989..63237 | 90371 | BREX-system-P-loop-protein-BrxC-[Salmonella-enterica-subsp.-enterica-serovar-Typhimurium] |
| EIT9287885.1 | Type1.BREX | <-HEPN<-REase+iREase<-?[illegible] | 960 | Bacteria>Pseudomonadota>Gammaproteobacteria>Enterobacterales>Enterobacteriaceae>Salmonella | GCA_021558855.1 | Salmonella_enterica_subsp._enterica_serovar_Derby | L3N85_002601 | 53297..56175 | 28144 | BREX-system-P-loop-protein-BrxC-[Salmonella-enterica-subsp.-enterica-serovar-Derby] |
| EIU6597714.1 | Type1.BREX | <-HEPN<-?\|\|YjhX_toxin->[illegible] | 1213 | Bacteria>Pseudomonadota>Gammaproteobacteria>Enterobacterales>Enterobacteriaceae>Salmonella | GCA_021692515.1 | Salmonella_enterica | L5W02_001568 | 377290..380931 | 28901 | BREX-system-P-loop-protein-BrxC-[Salmonella-enterica] |
| HBZ6150869.1 | Type1.BREX | <-HEPN<-?\|\|YjhX_toxin->[illegible] | 1213 | Bacteria>Pseudomonadota>Gammaproteobacteria>Enterobacterales>Enterobacteriaceae>Salmonella | GCA_022432985.1 | Salmonella_enterica | ML330_001731 | 63563..67204 | 28901 | BREX-system-P-loop-protein-BrxC-[Salmonella-enterica] |
| EJB4514208.1 | Type1.BREX | <-HEPN<-?\|\|YjhX_toxin->[illegible] | 1213 | Bacteria>Pseudomonadota>Gammaproteobacteria>Enterobacterales>Enterobacteriaceae>Salmonella | GCA_022945175.1 | Salmonella_enterica | MT088_002306 | 380173..383814 | 28901 | BREX-system-P-loop-protein-BrxC-[Salmonella-enterica] |
| EJC3638306.1 | Type1.BREX | <-HEPN<-?\|\|YjhX_toxin->[illegible] | 1213 | Bacteria>Pseudomonadota>Gammaproteobacteria>Enterobacterales>Enterobacteriaceae>Salmonella | GCA_023047235.1 | Salmonella_enterica | MYX88_003393 | 151139..154780 | 28901 | BREX-system-P-loop-protein-BrxC-[Salmonella-enterica] |
| HCB4841064.1 | Type1.BREX | <-HEPN<-?\|\|YjhX_toxin->[illegible] | 1213 | Bacteria>Pseudomonadota>Gammaproteobacteria>Enterobacterales>Enterobacteriaceae>Salmonella | GCA_023082205.1 | Salmonella_enterica_subsp._enterica_serovar_Bredeney | MOF44_000351 | 390385..394026 | 134047 | BREX-system-P-loop-protein-BrxC-[Salmonella-enterica-subsp.-enterica-serovar-Bredeney] |
| EJF4683727.1 | Type1.BREX | Arm-DNA-bind_3->NACHT+S[illegible] | 1213 | Bacteria>Pseudomonadota>Gammaproteobacteria>Enterobacterales>Enterobacteriaceae>Salmonella | GCA_023476405.1 | Salmonella_enterica | M8N25_002793 | 108064..111705 | 28901 | BREX-system-P-loop-protein-BrxC-[Salmonella-enterica] |
| MCL8965752.1 | Type1.BREX | <-HEPN<-REase+iREase<-?[illegible] | 1213 | Bacteria>Pseudomonadota>Gammaproteobacteria>Enterobacterales>Enterobacteriaceae>Salmonella | GCA_023608175.1 | Salmonella_enterica_subsp._enterica_serovar_Enteritidis | MU637_15175 | 170549..174190 | 149539 | BREX-system-P-loop-protein-BrxC-[Salmonella-enterica-subsp.-enterica-serovar-Enteritidis] |
| MCM9949031.1 | Type1.BREX | Arm-DNA-bind_3->NACHT+S[illegible] | 1213 | Bacteria>Pseudomonadota>Gammaproteobacteria>Enterobacterales>Enterobacteriaceae>Salmonella | GCA_023843835.1 | Salmonella_enterica | M5G05_07280 | 348323..351964 | 28901 | BREX-system-P-loop-protein-BrxC-[Salmonella-enterica] |
| EJK0228892.1 | Type1.BREX | Arm-DNA-bind_3->NACHT+S[illegible] | 1213 | Bacteria>Pseudomonadota>Gammaproteobacteria>Enterobacterales>Enterobacteriaceae>Salmonella | GCA_024068175.1 | Salmonella_enterica | NKK90_001819 | 323151..326792 | 28901 | BREX-system-P-loop-protein-BrxC-[Salmonella-enterica] |
| HCH7501034.1 | Type1.BREX | REC-><-HIN-HTH\|\|<-RHH\|\|[illegible] | 633 | Bacteria>Pseudomonadota>Gammaproteobacteria>Enterobacterales>Enterobacteriaceae>Salmonella | GCA_024265105.1 | Salmonella_enterica | NM874_003883 | 7331..9230 | 28901 | BREX-system-P-loop-protein-BrxC-[Salmonella-enterica] |
| EJM7985883.1 | Type1.BREX | <-GntR-HTH++TrmB+ZDNA-H[illegible] | 1213 | Bacteria>Pseudomonadota>Gammaproteobacteria>Enterobacterales>Enterobacteriaceae>Salmonella | GCA_024373005.1 | Salmonella_enterica | NO486_001112 | 369092..372733 | 28901 | BREX-system-P-loop-protein-BrxC-[Salmonella-enterica] |
| EJN2869328.1 | Type1.BREX | <-HEPN<-REase+iREase<-?[illegible] | 1213 | Bacteria>Pseudomonadota>Gammaproteobacteria>Enterobacterales>Enterobacteriaceae>Salmonella | GCA_024442015.1 | Salmonella_enterica_subsp._enterica_serovar_Derby | NPT71_003578 | 165015..168656 | 28144 | BREX-system-P-loop-protein-BrxC-[Salmonella-enterica-subsp.-enterica-serovar-Derby] |
| UTZ95782.1 | Type1.BREX | <-HEPN<-?\|\|YjhX_toxin->[illegible] | 1204 | Bacteria>Pseudomonadota>Gammaproteobacteria>Enterobacterales>Enterobacteriaceae>Salmonella | GCA_024442275.1 | Salmonella_enterica_subsp._enterica_serovar_Kentucky | NOQ95_21560 | 4448475..4452089 | 192955 | BREX-system-P-loop-protein-BrxC-[Salmonella-enterica-subsp.-enterica-serovar-Kentucky] |
| EJN9121676.1 | Type1.BREX | <-HEPN<-REase+iREase<-?[illegible] | 1213 | Bacteria>Pseudomonadota>Gammaproteobacteria>Enterobacterales>Enterobacteriaceae>Salmonella | GCA_024535515.1 | Salmonella_enterica | NR521_001803 | 170929..174570 | 28901 | BREX-system-P-loop-protein-BrxC-[Salmonella-enterica] |
| EJO5979497.1 | Type1.BREX | <-HEPN<-?\|\|YjhX_toxin->[illegible] | 1213 | Bacteria>Pseudomonadota>Gammaproteobacteria>Enterobacterales>Enterobacteriaceae>Salmonella | GCA_024573615.1 | Salmonella_enterica | NRR85_002625 | 159039..162680 | 28901 | BREX-system-P-loop-protein-BrxC-[Salmonella-enterica] |
| EJS8182320.1 | Type1.BREX | <-HEPN<-?\|\|YjhX_toxin->[illegible] | 1213 | Bacteria>Pseudomonadota>Gammaproteobacteria>Enterobacterales>Enterobacteriaceae>Salmonella | GCA_024988615.1 | Salmonella_enterica | NWJ04_001330 | 25917..29558 | 28901 | BREX-system-P-loop-protein-BrxC-[Salmonella-enterica] |
| HCL5270538.1 | Type1.BREX | <-N6-METHYLASE<-DUF6079[illegible] | 1213 | Bacteria>Pseudomonadota>Gammaproteobacteria>Enterobacterales>Enterobacteriaceae>Salmonella | GCA_025012985.1 | Salmonella_enterica | N2G33_002753 | 3666..7307 | 28901 | BREX-system-P-loop-protein-BrxC-[Salmonella-enterica] |
| HCM2417898.1 | Type1.BREX | Arm-DNA-bind_3->NACHT+S[illegible] | 1213 | Bacteria>Pseudomonadota>Gammaproteobacteria>Enterobacterales>Enterobacteriaceae>Salmonella | GCA_025074775.1 | Salmonella_enterica_subsp._enterica_serovar_Typhimurium_var._monophasic_4,[5],12:i:- | N3D22_002716 | 108064..111705 | 1461749 | BREX-system-P-loop-protein-BrxC-[Salmonella-enterica-subsp.-enterica-serovar-Typhimurium-var.-monophasic-4,[5] |
| HCM2697231.1 | Type1.BREX | <-HEPN<-?\|\|YjhX_toxin->[illegible] | 1213 | Bacteria>Pseudomonadota>Gammaproteobacteria>Enterobacterales>Enterobacteriaceae>Salmonella | GCA_025075935.1 | Salmonella_enterica_subsp._enterica_serovar_Yaba | N3D89_002099 | 63778..67419 | 2036205 | BREX-system-P-loop-protein-BrxC-[Salmonella-enterica-subsp.-enterica-serovar-Yaba] |
| EJU7772063.1 | Type1.BREX | <-HEPN<-REase+iREase<-?[illegible] | 1213 | Bacteria>Pseudomonadota>Gammaproteobacteria>Enterobacterales>Enterobacteriaceae>Salmonella | GCA_025208445.1 | Salmonella_enterica_subsp._salamae_serovar_4,12:e,n,x:1,6 | KAB52_003008 | 279421..283062 | 2978379 | BREX-system-P-loop-protein-BrxC-[Salmonella-enterica-subsp.-salamae-serovar-4,12:e,n,x:1,6] |
| EJV9378859.1 | Type1.BREX | wHTH+WYL+WCX->BrxA-DNA-[illegible] | 1225 | Bacteria>Pseudomonadota>Gammaproteobacteria>Enterobacterales>Enterobacteriaceae>Salmonella | GCA_025367915.1 | Salmonella_enterica | N7887_004537 | 2923..6600 | 28901 | BREX-system-P-loop-protein-BrxC-[Salmonella-enterica] |
| EJW7283568.1 | Type1.BREX | Arm-DNA-bind_3->NACHT+S[illegible] | 1213 | Bacteria>Pseudomonadota>Gammaproteobacteria>Enterobacterales>Enterobacteriaceae>Salmonella | GCA_025474975.1 | Salmonella_enterica_subsp._enterica_serovar_Typhimurium | OCR56_001321 | 81267..84908 | 90371 | BREX-system-P-loop-protein-BrxC-[Salmonella-enterica-subsp.-enterica-serovar-Typhimurium] |
| WP_096117035.1 | Type1.BREX | <-DUF6079*<-BrxB-BrxF-i[illegible] | 1166 | Bacteria>Pseudomonadota>Gammaproteobacteria>Enterobacterales>Enterobacteriaceae>Salmonella | GCA_030820635.1 | Salmonella_enterica | CDC58_RS01455 | 1..3498 | 28901 | BREX-system-P-loop-protein-BrxC,-partial-[Salmonella-enterica] |
| WP_111761031.1 | Type1.BREX | nSTAND1+HNH->NACHT+SF1-[illegible] | 1213 | Bacteria>Pseudomonadota>Gammaproteobacteria>Enterobacterales>Enterobacteriaceae>Salmonella | GCA_030820635.1 | Salmonella_enterica | KXD81_RS02520 | 21979..25620 | 28901 | BREX-system-P-loop-protein-BrxC-[Salmonella-enterica] |
| WP_256863860.1 | Type1.BREX | <-DUF6079-p*<-BrxB-BrxF[illegible] | 840 | Bacteria>Pseudomonadota>Gammaproteobacteria>Enterobacterales>Enterobacteriaceae>Salmonella | GCA_030820635.1 | Salmonella_enterica | SO19_RS21855 | 1..2520 | 28901 | BREX-system-P-loop-protein-BrxC,-partial-[Salmonella-enterica] |
| SUG73766.1 | Type1.BREX | <-SIGMA+OB-Fold+AAA-ATP[illegible] | 900 | Bacteria>Pseudomonadota>Gammaproteobacteria>Enterobacterales>Enterobacteriaceae>Salmonella | GCA_900456435.1 | Salmonella_enterica_subsp._enterica | NCTC5798_05055 | 4086397..4089099 | 59201 | DNA-repair-ATPase-[Salmonella-enterica-subsp.-enterica] |
| SUI30166.1 | Type1.BREX | YjhX_toxin->SAM-METHYLA[illegible] | 864 | Bacteria>Pseudomonadota>Gammaproteobacteria>Enterobacterales>Enterobacteriaceae>Salmonella | GCA_900456825.1 | Salmonella_enterica_subsp._enterica_serovar_Derby | NCTC5722_04070 | 4063548..4066142 | 28144 | DNA-repair-ATPase-[Salmonella-enterica-subsp.-enterica-serovar-Derby] |
| SUH94769.1 | Type1.BREX | YjhX_toxin->SAM-METHYLA[illegible] | 530 | Bacteria>Pseudomonadota>Gammaproteobacteria>Enterobacterales>Enterobacteriaceae>Salmonella | GCA_900456925.1 | Salmonella_enterica_subsp._enterica_serovar_Typhimurium | NCTC13347_04212 | 4172999..4174591 | 90371 | DNA-repair-ATPase-[Salmonella-enterica-subsp.-enterica-serovar-Typhimurium] |
| VUC90716.1 | Type1.BREX | YjhX_toxin->SAM-METHYLA[illegible] | 532 | Bacteria>Pseudomonadota>Gammaproteobacteria>Enterobacterales>Enterobacteriaceae>Salmonella | GCA_902109495.1 | Salmonella_sp._NCTC_11881 | NCTC11881_05559 | 4058427..4060025 | 2583579 | DNA-repair-ATPase-[Salmonella-sp.-NCTC-11881] |
| WP_001523601.1 | Type1.BREX | <-HEPN<-?\|\|YjhX_toxin->[illegible] | 1213 | Bacteria>Pseudomonadota>Gammaproteobacteria>Enterobacterales>Enterobacteriaceae>Salmonella | GCF_000170235.1 | Salmonella_enterica | SESPB_RS10230 | 40505..44146 | 28901 | BREX-system-P-loop-protein-BrxC-[Salmonella-enterica] |
| WP_001019653.1 | Type1.BREX | HTH_3->Couple_hipA+HipA[illegible] | 1213 | Bacteria>Pseudomonadota>Gammaproteobacteria>Enterobacterales>Enterobacteriaceae>Salmonella | GCF_000230875.1 | Salmonella_enterica | SEENINOB_RS28275 | 4768987..4772628 | 28901 | BREX-system-P-loop-protein-BrxC-[Salmonella-enterica] |
| WP_001019651.1 | Type1.BREX | Arm-DNA-bind_3->NACHT+S[illegible] | 1213 | Bacteria>Pseudomonadota>Gammaproteobacteria>Enterobacterales>Enterobacteriaceae>Salmonella | GCF_000252875.1 | Salmonella_enterica | JM982_RS03985 | 854415..858056 | 28901 | BREX-system-P-loop-protein-BrxC-[Salmonella-enterica] |
| WP_024154967.1 | Type1.BREX | NACHT+SF1-SF2-Helicase+[illegible] | 1213 | Bacteria>Pseudomonadota>Gammaproteobacteria>Enterobacterales>Enterobacteriaceae>Salmonella | GCF_000412855.1 | Salmonella_enterica | AP074_RS03115 | 646291..649932 | 28901 | BREX-system-P-loop-protein-BrxC-[Salmonella-enterica] |
| WP_071055527.1 | Type1.BREX | <-HEPN<-?\|\|YjhX_toxin->[illegible] | 1213 | Bacteria>Pseudomonadota>Gammaproteobacteria>Enterobacterales>Enterobacteriaceae>Salmonella | GCF_000486165.2 | Salmonella_enterica | SES26_RS19315 | 4009469..4013110 | 28901 | BREX-system-P-loop-protein-BrxC-[Salmonella-enterica] |
| WP_023217629.1 | Type1.BREX | <-HEPN<-?\|\|YjhX_toxin->[illegible] | 1213 | Bacteria>Pseudomonadota>Gammaproteobacteria>Enterobacterales>Enterobacteriaceae>Salmonella | GCF_000486855.2 | Salmonella_enterica | SEEM9284_RS19920 | 3931825..3935466 | 28901 | BREX-system-P-loop-protein-BrxC-[Salmonella-enterica] |
| WP_077956997.1 | Type1.BREX | <-METHYLASE\|\|SKI-><-?\|\|[illegible] | 1240 | Bacteria>Pseudomonadota>Gammaproteobacteria>Enterobacterales>Enterobacteriaceae>Salmonella | GCF_000487155.2 | Salmonella_enterica | A7T10_RS09345 | 418937..422659 | 28901 | BREX-system-P-loop-protein-BrxC-[Salmonella-enterica] |
| WP_023213578.1 | Type1.BREX | <-HEPN<-?\|\|YjhX_toxin->[illegible] | 1213 | Bacteria>Pseudomonadota>Gammaproteobacteria>Enterobacterales>Enterobacteriaceae>Salmonella | GCF_000493295.2 | Salmonella_enterica | SEEP1673_RS21585 | 4142040..4145681 | 28901 | BREX-system-P-loop-protein-BrxC-[Salmonella-enterica] |
| WP_050956242.1 | Type1.BREX | Arm-DNA-bind_3->NACHT+S[illegible] | 1213 | Bacteria>Pseudomonadota>Gammaproteobacteria>Enterobacterales>Enterobacteriaceae>Salmonella | GCF_001217085.1 | Salmonella_enterica | AYW49_RS05955 | 256224..259865 | 28901 | BREX-system-P-loop-protein-BrxC-[Salmonella-enterica] |
| WP_052925311.1 | Type1.BREX | Arm-DNA-bind_3->NACHT+S[illegible] | 1213 | Bacteria>Pseudomonadota>Gammaproteobacteria>Enterobacterales>Enterobacteriaceae>Salmonella | GCF_001243555.1 | Salmonella_enterica | AEV74_RS03660 | 108085..111726 | 28901 | BREX-system-P-loop-protein-BrxC-[Salmonella-enterica] |
| WP_057523065.1 | Type1.BREX | <-HEPN<-REase+iREase<-?[illegible] | 1213 | Bacteria>Pseudomonadota>Gammaproteobacteria>Enterobacterales>Enterobacteriaceae>Salmonella | GCF_001326235.1 | Salmonella_enterica | HG82_RS07610 | 171678..175319 | 28901 | BREX-system-P-loop-protein-BrxC-[Salmonella-enterica] |
| WP_058799135.1 | Type1.BREX | <-HEPN<-REase+iREase<-?[illegible] | 1213 | Bacteria>Pseudomonadota>Gammaproteobacteria>Enterobacterales>Enterobacteriaceae>Salmonella | GCF_001478065.1 | Salmonella_enterica | DD74_RS03635 | 38743..42384 | 28901 | BREX-system-P-loop-protein-BrxC-[Salmonella-enterica] |
| WP_061383284.1 | Type1.BREX | NACHT+SF1-SF2-Helicase+[illegible] | 1204 | Bacteria>Pseudomonadota>Gammaproteobacteria>Enterobacterales>Enterobacteriaceae>Salmonella | GCF_001570045.1 | Salmonella_enterica | AW140_RS06880 | 353892..357506 | 28901 | BREX-system-P-loop-protein-BrxC-[Salmonella-enterica] |
| WP_061380447.1 | Type1.BREX | <-HEPN<-?\|\|YjhX_toxin->[illegible] | 1213 | Bacteria>Pseudomonadota>Gammaproteobacteria>Enterobacterales>Enterobacteriaceae>Salmonella | GCF_001570285.1 | Salmonella_enterica | AW153_RS18915 | 158422..162063 | 28901 | BREX-system-P-loop-protein-BrxC-[Salmonella-enterica] |
| WP_061584452.1 | Type1.BREX | METHYLASE->Arm-DNA-bind[illegible] | 1213 | Bacteria>Pseudomonadota>Gammaproteobacteria>Enterobacterales>Enterobacteriaceae>Salmonella | GCF_001588495.1 | Salmonella_enterica | AF388_RS19360 | 33009..36650 | 28901 | BREX-system-P-loop-protein-BrxC-[Salmonella-enterica] |
| WP_064013329.1 | Type1.BREX | BrxA-DNA-binding->BrxB-[illegible] | 1204 | Bacteria>Pseudomonadota>Gammaproteobacteria>Enterobacterales>Enterobacteriaceae>Salmonella | GCF_001643815.1 | Salmonella_enterica | IJ96_RS19570 | 3320..6934 | 28901 | BREX-system-P-loop-protein-BrxC-[Salmonella-enterica] |
| WP_069106895.1 | Type1.BREX | <-HEPN<-REase+iREase<-?[illegible] | 1213 | Bacteria>Pseudomonadota>Gammaproteobacteria>Enterobacterales>Enterobacteriaceae>Salmonella | GCF_001714545.1 | Salmonella_enterica | AHW33_RS11985 | 172706..176347 | 28901 | BREX-system-P-loop-protein-BrxC-[Salmonella-enterica] |
| WP_070132690.1 | Type1.BREX | SF1-SF2-Helicase+HKD-DN[illegible] | 1213 | Bacteria>Pseudomonadota>Gammaproteobacteria>Enterobacterales>Enterobacteriaceae>Salmonella | GCF_001756155.1 | Salmonella_enterica | BG484_RS05620 | 12664..16305 | 28901 | BREX-system-P-loop-protein-BrxC-[Salmonella-enterica] |
| WP_070134780.1 | Type1.BREX | DUF6079*->N6-METHYLASE-[illegible] | 1201 | Bacteria>Pseudomonadota>Gammaproteobacteria>Enterobacterales>Enterobacteriaceae>Salmonella | GCF_001756205.1 | Salmonella_enterica | BG487_15355 | 1..3606 | 28901 | BREX-system-P-loop-protein-BrxC,-partial-[Salmonella-enterica] |
| WP_070793518.1 | Type1.BREX | SKI-><-?\|\|tRNA->Arm-DNA[illegible] | 1213 | Bacteria>Pseudomonadota>Gammaproteobacteria>Enterobacterales>Enterobacteriaceae>Salmonella | GCF_001832595.1 | Salmonella_enterica | A7T54_RS11885 | 259865..263506 | 28901 | BREX-system-P-loop-protein-BrxC-[Salmonella-enterica] |
| WP_076937571.1 | Type1.BREX | <-HEPN<-REase+iREase<-?[illegible] | 1213 | Bacteria>Pseudomonadota>Gammaproteobacteria>Enterobacterales>Enterobacteriaceae>Salmonella | GCF_001975325.1 | Salmonella_enterica | LFZ3_RS17405 | 3406419..3410060 | 28901 | BREX-system-P-loop-protein-BrxC-[Salmonella-enterica] |
| WP_076736168.1 | Type1.BREX | HTH_3->Couple_hipA+HipA[illegible] | 1213 | Bacteria>Pseudomonadota>Gammaproteobacteria>Enterobacterales>Enterobacteriaceae>Salmonella | GCF_001975445.1 | Salmonella_enterica | LFZ24_RS24380 | 4843050..4846691 | 28901 | BREX-system-P-loop-protein-BrxC-[Salmonella-enterica] |
| WP_078367156.1 | Type1.BREX | <-HEPN<-REase+iREase<-?[illegible] | 1213 | Bacteria>Pseudomonadota>Gammaproteobacteria>Enterobacterales>Enterobacteriaceae>Salmonella | GCF_002018325.1 | Salmonella_enterica | B1S11_RS05200 | 170490..174131 | 28901 | BREX-system-P-loop-protein-BrxC-[Salmonella-enterica] |
| WP_078351828.1 | Type1.BREX | SKI-><-?\|\|tRNA->Arm-DNA[illegible] | 1213 | Bacteria>Pseudomonadota>Gammaproteobacteria>Enterobacterales>Enterobacteriaceae>Salmonella | GCF_002018355.1 | Salmonella_enterica | B1211_RS18830 | 150004..153645 | 28901 | BREX-system-P-loop-protein-BrxC-[Salmonella-enterica] |
| WP_001729064.1 | Type1.BREX | <-HKD-DNase+OGG-like<-S[illegible] | 1213 | Bacteria>Pseudomonadota>Gammaproteobacteria>Enterobacterales>Enterobacteriaceae>Salmonella | GCF_002034005.1 | Salmonella_enterica | AJH59_RS23280 | 9917..13558 | 28901 | BREX-system-P-loop-protein-BrxC-[Salmonella-enterica] |
| WP_079808688.1 | Type1.BREX | nSTAND1+HNH->NACHT+SF1-[illegible] | 1213 | Bacteria>Pseudomonadota>Gammaproteobacteria>Enterobacterales>Enterobacteriaceae>Salmonella | GCF_002035875.1 | Salmonella_enterica | Y985_RS08730 | 120043..123684 | 28901 | BREX-system-P-loop-protein-BrxC-[Salmonella-enterica] |
| WP_079814448.1 | Type1.BREX | XIS-HTH->?->?-><-ZnR<-Z[illegible] | 1204 | Bacteria>Pseudomonadota>Gammaproteobacteria>Enterobacterales>Enterobacteriaceae>Salmonella | GCF_002036095.1 | Salmonella_enterica | S481_13075 | 121983..125597 | 28901 | BREX-system-P-loop-protein-BrxC-[Salmonella-enterica] |
| WP_079897969.1 | Type1.BREX | NACHT+SF1-SF2-Helicase+[illegible] | 1204 | Bacteria>Pseudomonadota>Gammaproteobacteria>Enterobacterales>Enterobacteriaceae>Salmonella | GCF_002045805.1 | Salmonella_enterica | VY62_RS13065 | 181728..185342 | 28901 | BREX-system-P-loop-protein-BrxC-[Salmonella-enterica] |
| WP_079972900.1 | Type1.BREX | Arm-DNA-bind_3->NACHT+S[illegible] | 1213 | Bacteria>Pseudomonadota>Gammaproteobacteria>Enterobacterales>Enterobacteriaceae>Salmonella | GCF_002046615.1 | Salmonella_enterica | VX95_RS09560 | 163568..167209 | 28901 | BREX-system-P-loop-protein-BrxC-[Salmonella-enterica] |
| WP_079965457.1 | Type1.BREX | <-HEPN<-?\|\|YjhX_toxin->[illegible] | 1213 | Bacteria>Pseudomonadota>Gammaproteobacteria>Enterobacterales>Enterobacteriaceae>Salmonella | GCF_002046745.1 | Salmonella_enterica | VX54_RS10650 | 52174..55815 | 28901 | BREX-system-P-loop-protein-BrxC-[Salmonella-enterica] |
| WP_079902376.1 | Type1.BREX | <-GntR-HTH++TrmB+ZDNA-H[illegible] | 1213 | Bacteria>Pseudomonadota>Gammaproteobacteria>Enterobacterales>Enterobacteriaceae>Salmonella | GCF_002048195.1 | Salmonella_enterica | VX33_RS03275 | 157928..161569 | 28901 | BREX-system-P-loop-protein-BrxC-[Salmonella-enterica] |
| WP_080094962.1 | Type1.BREX | <-HEPN<-?\|\|?->\|\|YjhX_to[illegible] | 1213 | Bacteria>Pseudomonadota>Gammaproteobacteria>Enterobacterales>Enterobacteriaceae>Salmonella | GCF_002052895.1 | Salmonella_enterica | DL104_05730 | 29230..32871 | 28901 | BREX-system-P-loop-protein-BrxC-[Salmonella-enterica] |
| WP_079944402.1 | Type1.BREX | <-HEPN<-?\|\|YjhX_toxin->[illegible] | 1213 | Bacteria>Pseudomonadota>Gammaproteobacteria>Enterobacterales>Enterobacteriaceae>Salmonella | GCF_002058485.1 | Salmonella_enterica | AJM56_RS05775 | 30789..34430 | 28901 | BREX-system-P-loop-protein-BrxC-[Salmonella-enterica] |
| WP_080162863.1 | Type1.BREX | <-HEPN<-?\|\|YjhX_toxin->[illegible] | 1213 | Bacteria>Pseudomonadota>Gammaproteobacteria>Enterobacterales>Enterobacteriaceae>Salmonella | GCF_002062115.1 | Salmonella_enterica | SO66_RS04055 | 158804..162445 | 28901 | BREX-system-P-loop-protein-BrxC-[Salmonella-enterica] |

| Protein-ID | System | Gene-Neighbourhoods | Length | Taxonomy | Assembly-ID | Strain | Locus-Tag | Genomic-coordinates | TaxID | Fasta-define |
|---|---|---|---|---|---|---|---|---|---|---|
| WP_139780228.1 | Type1.BREX | <-DUF6079*<-BrxB-BrxF-iSTAND<-BrxA-DNA-binding<-?<-SAM-METHYLASE<-YjhX_toxin\|\|?->HEPN-> | 1015 | Bacteria>Pseudomonadota>Gammaproteobacteria>Enterobacterales>Enterobacteriaceae>Salmonella | GCF_002064225.1 | Salmonella_enterica | SN68_RS12600 | 1..3045 | 28901 | BREX-system-P-loop-protein-BrxC,-partial-[Salmonella-enterica] |
| WP_079905548.1 | Type1.BREX | <-MukE<-wHTH\|\|wHTH+WYL+WCX-><-HKD-DNase<-MarR-wHTH<-SIGMA-HTH+REase\|\|?->BrxA-DNA-binding->BrxB-BrxF-iSTAND->DUF6079*->N6-METHYLASE->OLD-ABC->TOPRIM->PglZ->SIGMA+OB-Fold+AAA-ATPase+Lon [illegible] | 1204 | Bacteria>Pseudomonadota>Gammaproteobacteria>Enterobacterales>Enterobacteriaceae>Salmonella | GCF_002065435.1 | Salmonella_enterica | SN48_RS13425 | 18385..21999 | 28901 | BREX-system-P-loop-protein-BrxC-[Salmonella-enterica] |
| WP_080250285.1 | Type1.BREX | <-HEPN<-?\|\|YjhX_toxin->SAM-METHYLASE->?->BrxA-DNA-binding->BrxB-BrxF-iSTAND->DUF6079*->N6-METHYLASE->OLD-ABC->TOPRIM->PglZ->SIGMA+OB-Fold+AAA-ATPase+Lon-Protease-BrxL->HKD-DNase+OGG-l [illegible] | 1213 | Bacteria>Pseudomonadota>Gammaproteobacteria>Enterobacterales>Enterobacteriaceae>Salmonella | GCF_002066965.1 | Salmonella_enterica | RZ13_RS02295 | 164882..168523 | 28901 | BREX-system-P-loop-protein-BrxC-[Salmonella-enterica] |
| WP_111769566.1 | Type1.BREX | NACHT+SF1-SF2-Helicase+HKD-DNase->REase-><-?<-SIGMA+OB-Fold+AAA-ATPase+Lon-Protease-BrxL<-PglZ<-TOPRIM<-OLD-ABC<-N6-METHYLASE<-DUF6079*<-BrxB-BrxF-iSTAND<-BrxA-DNA-binding<-?<-SAM-MET [illegible] | 1213 | Bacteria>Pseudomonadota>Gammaproteobacteria>Enterobacterales>Enterobacteriaceae>Salmonella | GCF_002106425.1 | Salmonella_enterica | R537_RS06030 | 154441..158082 | 28901 | BREX-system-P-loop-protein-BrxC-[Salmonella-enterica] |
| WP_086805524.1 | Type1.BREX | NACHT+SF1-SF2-Helicase+HKD-DNase->REase-><-HKD-DNase+OGG-like<-SIGMA+OB-Fold+AAA-ATPase+Lon-Protease-BrxL<-PglZ<-TOPRIM<-OLD-ABC<-N6-METHYLASE<-DUF6079*<-BrxB-BrxF-iSTAND<-BrxA-DNA-bi [illegible] | 1204 | Bacteria>Pseudomonadota>Gammaproteobacteria>Enterobacterales>Enterobacteriaceae>Salmonella | GCF_002156225.1 | Salmonella_enterica | CCL33_RS10730 | 52147..55761 | 28901 | BREX-system-P-loop-protein-BrxC-[Salmonella-enterica] |
| WP_065348243.1 | Type1.BREX | <-HEPN<-?\|\|YjhX_toxin->SAM-METHYLASE->?->BrxA-DNA-binding->BrxB-BrxF-iSTAND->DUF6079*->N6-METHYLASE->OLD-ABC->TOPRIM->PglZ->SIGMA+OB-Fold+AAA-ATPase+Lon-Protease-BrxL->?-><-REase<-NAC [illegible] | 1213 | Bacteria>Pseudomonadota>Gammaproteobacteria>Enterobacterales>Enterobacteriaceae>Salmonella | GCF_002196535.1 | Salmonella_enterica | GX95_RS20905 | 4055067..4058708 | 28901 | BREX-system-P-loop-protein-BrxC-[Salmonella-enterica] |
| WP_088546528.1 | Type1.BREX | <-GntR-HTH++TrmB+ZDNA-HTH\|\|?-><-?<-HEPN<-?\|\|YjhX_toxin->?->BrxA-DNA-binding->BrxB-BrxF-iSTAND->DUF6079*->N6-METHYLASE->OLD-ABC->TOPRIM->PglZ->SIGMA+OB-Fold+AAA-ATPase+Lon-Protease-Brx [illegible] | 1213 | Bacteria>Pseudomonadota>Gammaproteobacteria>Enterobacterales>Enterobacteriaceae>Salmonella | GCF_002208255.1 | Salmonella_enterica | LFZ35_RS21090 | 4061970..4065611 | 28901 | BREX-system-P-loop-protein-BrxC-[Salmonella-enterica] |
| WP_088759215.1 | Type1.BREX | <-MukB<-MukE<-wHTH\|\|wHTH+WYL+WCX-><-MarR-wHTH<-SIGMA-HTH+REase\|\|?->BrxA-DNA-binding->BrxB-BrxF-iSTAND->DUF6079*->N6-METHYLASE->OLD-ABC->TOPRIM->PglZ->SIGMA+OB-Fold+AAA-ATPase+Lon-Prot [illegible] | 1204 | Bacteria>Pseudomonadota>Gammaproteobacteria>Enterobacterales>Enterobacteriaceae>Salmonella | GCF_002211845.1 | Salmonella_enterica | CE137_RS21050 | 4095775..4099389 | 28901 | BREX-system-P-loop-protein-BrxC-[Salmonella-enterica] |
| WP_080090697.1 | Type1.BREX | <-HEPN<-REase+iREase<-?\|\|YjhX_toxin->SAM-METHYLASE->?->BrxA-DNA-binding->BrxB-BrxF-iSTAND->DUF6079*->N6-METHYLASE->OLD-ABC->TOPRIM->PglZ->SIGMA+OB-Fold+AAA-ATPase+Lon-Protease-BrxL-><-[illegible] | 1213 | Bacteria>Pseudomonadota>Gammaproteobacteria>Enterobacterales>Enterobacteriaceae>Salmonella | GCF_002234755.2 | Salmonella_enterica | CHD02_RS05505 | 1125404..1129045 | 28901 | BREX-system-P-loop-protein-BrxC-[Salmonella-enterica] |
| WP_053445538.1 | Type1.BREX | <-HEPN<-?\|\|YjhX_toxin->SAM-METHYLASE->?->BrxA-DNA-binding->BrxB-BrxF-iSTAND->DUF6079*->N6-METHYLASE->OLD-ABC->TOPRIM->PglZ->SIGMA+OB-Fold+AAA-ATPase+Lon-Protease-BrxL-><-REase | 1213 | Bacteria>Pseudomonadota>Gammaproteobacteria>Enterobacterales>Enterobacteriaceae>Salmonella | GCF_002238275.1 | Salmonella_enterica | F1E41_RS17160 | 150695..154336 | 28901 | BREX-system-P-loop-protein-BrxC-[Salmonella-enterica] |
| WP_094797764.1 | Type1.BREX | <-HEPN<-REase+iREase<-?\|\|YjhX_toxin->SAM-METHYLASE->?->BrxA-DNA-binding->BrxB-BrxF-iSTAND->DUF6079*->N6-METHYLASE->OLD-ABC->TOPRIM->PglZ->SIGMA+OB-Fold+AAA-ATPase+Lon-Protease-BrxL-><-[illegible] | 1213 | Bacteria>Pseudomonadota>Gammaproteobacteria>Enterobacterales>Enterobacteriaceae>Salmonella | GCF_002260565.1 | Salmonella_enterica | CFB03_RS03285 | 15872..19513 | 28901 | BREX-system-P-loop-protein-BrxC-[Salmonella-enterica] |
| WP_094887773.1 | Type1.BREX | REase-><-SIGMA+OB-Fold+AAA-ATPase+Lon-Protease-BrxL<-PglZ<-TOPRIM<-OLD-ABC<-N6-METHYLASE<-DUF6079*<-BrxB-BrxF-iSTAND<-BrxA-DNA-binding | 1213 | Bacteria>Pseudomonadota>Gammaproteobacteria>Enterobacterales>Enterobacteriaceae>Salmonella | GCF_002265975.1 | Salmonella_enterica | CC043_RS23255 | 12854..16495 | 28901 | BREX-system-P-loop-protein-BrxC-[Salmonella-enterica] |
| WP_094964979.1 | Type1.BREX | <-HEPN<-?\|\|YjhX_toxin->SAM-METHYLASE->?->BrxA-DNA-binding->BrxB-BrxF-iSTAND->DUF6079*->N6-METHYLASE->OLD-ABC->TOPRIM->PglZ->SIGMA+OB-Fold+AAA-ATPase+Lon-Protease-BrxL->\|\|<-HTH_Tnp_1+H [illegible] | 1213 | Bacteria>Pseudomonadota>Gammaproteobacteria>Enterobacterales>Enterobacteriaceae>Salmonella | GCF_002266055.1 | Salmonella_enterica | CC049_RS14335 | 399067..402708 | 28901 | BREX-system-P-loop-protein-BrxC-[Salmonella-enterica] |
| WP_094946975.1 | Type1.BREX | <-HEPN<-?\|\|YjhX_toxin->SAM-METHYLASE->?->BrxA-DNA-binding->BrxB-BrxF-iSTAND->DUF6079*->N6-METHYLASE->OLD-ABC->TOPRIM->PglZ->SIGMA+OB-Fold+AAA-ATPase+Lon-Protease-BrxL-><-REase<-NACHT+[illegible] | 1213 | Bacteria>Pseudomonadota>Gammaproteobacteria>Enterobacterales>Enterobacteriaceae>Salmonella | GCF_002266315.1 | Salmonella_enterica | CC036_RS01880 | 387042..390683 | 28901 | BREX-system-P-loop-protein-BrxC-[Salmonella-enterica] |
| WP_097369131.1 | Type1.BREX | <-HEPN<-REase+iREase<-?\|\|YjhX_toxin->SAM-METHYLASE->?->BrxA-DNA-binding->BrxB-BrxF-iSTAND->DUF6079*->N6-METHYLASE->OLD-ABC->TOPRIM->PglZ->SIGMA+OB-Fold+AAA-ATPase+Lon-Protease-BrxL-><[illegible] | 1213 | Bacteria>Pseudomonadota>Gammaproteobacteria>Enterobacterales>Enterobacteriaceae>Salmonella | GCF_002507575.1 | Salmonella_enterica | CBI65_RS08825 | 170892..174533 | 28901 | BREX-system-P-loop-protein-BrxC-[Salmonella-enterica] |
| WP_097368036.1 | Type1.BREX | <-HEPN<-REase+iREase<-?\|\|YjhX_toxin->SAM-METHYLASE->?->BrxA-DNA-binding->BrxB-BrxF-iSTAND->DUF6079*->N6-METHYLASE->OLD-ABC->TOPRIM->PglZ->SIGMA+OB-Fold+AAA-ATPase+Lon-Protease-BrxL-><[illegible] | 1213 | Bacteria>Pseudomonadota>Gammaproteobacteria>Enterobacterales>Enterobacteriaceae>Salmonella | GCF_002507625.1 | Salmonella_enterica | CBI66_RS11180 | 171059..174700 | 28901 | BREX-system-P-loop-protein-BrxC-[Salmonella-enterica] |
| WP_097375567.1 | Type1.BREX | <-HEPN<-REase+iREa[illegible] | 1213 | Bacteria>Pseudomonadota>Gammaproteobacteria>Enterobacterales>Enterobacteriaceae>Salmonella | GCF_002507635.1 | Salmonella_enterica | CBI67_RS08835 | 170640..174281 | 28901 | BREX-system-P-loop-protein-BrxC-[Salmonella-enterica] |
| WP_107326618.1 | Type1.BREX | Arm-DNA-bind_3->N[illegible] | 1213 | Bacteria>Pseudomonadota>Gammaproteobacteria>Enterobacterales>Enterobacteriaceae>Salmonella | GCF_003032235.1 | Salmonella_enterica | C5I04_RS23495 | 346379..350020 | 28901 | BREX-system-P-loop-protein-BrxC-[Salmonella-enterica] |
| WP_116574272.1 | Type1.BREX | TOPRIM-DNAG+D5N+D5[illegible] | 1213 | Bacteria>Pseudomonadota>Gammaproteobacteria>Enterobacterales>Enterobacteriaceae>Salmonella | GCF_003077035.1 | Salmonella_enterica | C4874_RS10725 | 43609..47250 | 28901 | BREX-system-P-loop-protein-BrxC-[Salmonella-enterica] |
| WP_058114204.1 | Type1.BREX | <-GntR-HTH++TrmB+Z[illegible] | 1213 | Bacteria>Pseudomonadota>Gammaproteobacteria>Enterobacterales>Enterobacteriaceae>Salmonella | GCF_003324795.1 | Salmonella_enterica | DOE60_RS06770 | 1364315..1367956 | 28901 | BREX-system-P-loop-protein-BrxC-[Salmonella-enterica] |
| WP_079798429.1 | Type1.BREX | Glyoxalase->ACET->[illegible] | 1240 | Bacteria>Pseudomonadota>Gammaproteobacteria>Enterobacterales>Enterobacteriaceae>Salmonella | GCF_003325015.1 | Salmonella_enterica | HFU03_RS19925 | 4101513..4105235 | 28901 | BREX-system-P-loop-protein-BrxC-[Salmonella-enterica] |
| WP_114051212.1 | Type1.BREX | HTH_3->Couple_hip[illegible] | 1213 | Bacteria>Pseudomonadota>Gammaproteobacteria>Enterobacterales>Enterobacteriaceae>Salmonella | GCF_003325275.1 | Salmonella_enterica | CHE40_RS23180 | 4543503..4547144 | 28901 | BREX-system-P-loop-protein-BrxC-[Salmonella-enterica] |
| WP_114052981.1 | Type1.BREX | <-GntR-HTH++TrmB+Z[illegible] | 1213 | Bacteria>Pseudomonadota>Gammaproteobacteria>Enterobacterales>Enterobacteriaceae>Salmonella | GCF_003325295.1 | Salmonella_enterica | CHD06_RS08895 | 1727033..1730674 | 28901 | BREX-system-P-loop-protein-BrxC-[Salmonella-enterica] |
| WP_133207218.1 | Type1.BREX | XIS-HTH->?->AAA+HT[illegible] | 1213 | Bacteria>Pseudomonadota>Gammaproteobacteria>Enterobacterales>Enterobacteriaceae>Salmonella | GCF_004354245.1 | Salmonella_enterica | EIM31_RS05595 | 1133023..1136664 | 28901 | BREX-system-P-loop-protein-BrxC-[Salmonella-enterica] |
| WP_140237021.1 | Type1.BREX | BrxA-DNA-binding-[illegible] | 1213 | Bacteria>Pseudomonadota>Gammaproteobacteria>Enterobacterales>Enterobacteriaceae>Salmonella | GCF_006379925.2 | Salmonella_enterica | FG677_RS21910 | 2844..6485 | 28901 | BREX-system-P-loop-protein-BrxC-[Salmonella-enterica] |
| WP_080166838.1 | Type1.BREX | <-HEPN<-?\|\|YjhX_to[illegible] | 1213 | Bacteria>Pseudomonadota>Gammaproteobacteria>Enterobacterales>Enterobacteriaceae>Salmonella | GCF_012273475.1 | Salmonella_enterica | FOC17_RS02975 | 512521..516162 | 28901 | BREX-system-P-loop-protein-BrxC-[Salmonella-enterica] |
| WP_181299515.1 | Type1.BREX | <-HEPN<-?\|\|\|\|YjhX[illegible] | 1213 | Bacteria>Pseudomonadota>Gammaproteobacteria>Enterobacterales>Enterobacteriaceae>Salmonella | GCF_013624525.1 | Salmonella_enterica | D6413_08715 | 52330..55971 | 28901 | BREX-system-P-loop-protein-BrxC-[Salmonella-enterica] |
| WP_080213758.1 | Type1.BREX | <-GntR-HTH++TrmB+Z[illegible] | 1213 | Bacteria>Pseudomonadota>Gammaproteobacteria>Enterobacterales>Enterobacteriaceae>Salmonella | GCF_015180625.1 | Salmonella_enterica | AS04_RS02250 | 44431..48072 | 28901 | BREX-system-P-loop-protein-BrxC-[Salmonella-enterica] |
| WP_079788274.1 | Type1.BREX | <-HEPN<-REase+iREa[illegible] | 1213 | Bacteria>Pseudomonadota>Gammaproteobacteria>Enterobacterales>Enterobacteriaceae>Salmonella | GCF_016772815.1 | Salmonella_enterica | JM978_RS19965 | 4115108..4118749 | 28901 | BREX-system-P-loop-protein-BrxC-[Salmonella-enterica] |
| WP_079805580.1 | Type1.BREX | SLOG->HAD-><-SIGM[illegible] | 1240 | Bacteria>Pseudomonadota>Gammaproteobacteria>Enterobacterales>Enterobacteriaceae>Salmonella | GCF_016903075.1 | Salmonella_enterica | AJH70_RS01330 | 291078..294800 | 28901 | BREX-system-P-loop-protein-BrxC-[Salmonella-enterica] |
| WP_210138875.1 | Type1.BREX | SKI-><-?\|\|tRNA->A[illegible] | 1213 | Bacteria>Pseudomonadota>Gammaproteobacteria>Enterobacterales>Enterobacteriaceae>Salmonella | GCF_017893505.1 | Salmonella_enterica | JJB80_RS22605 | 4685033..4688674 | 28901 | BREX-system-P-loop-protein-BrxC-[Salmonella-enterica] |
| WP_241575978.1 | Type1.BREX | Arm-DNA-bind_3->N[illegible] | 1213 | Bacteria>Pseudomonadota>Gammaproteobacteria>Enterobacterales>Enterobacteriaceae>Salmonella | GCF_022603615.1 | Salmonella_enterica | K5800_RS10145 | 108176..111817 | 28901 | BREX-system-P-loop-protein-BrxC-[Salmonella-enterica] |
| WP_247258912.1 | Type1.BREX | NACHT+SF1-SF2-Heli[illegible] | 1204 | Bacteria>Pseudomonadota>Gammaproteobacteria>Enterobacterales>Enterobacteriaceae>Salmonella | GCF_023007725.1 | Salmonella_enterica | MA804_RS14880 | 357716..361330 | 28901 | BREX-system-P-loop-protein-BrxC-[Salmonella-enterica] |
| WP_247223752.1 | Type1.BREX | <-HEPN<-REase+iREa[illegible] | 1213 | Bacteria>Pseudomonadota>Gammaproteobacteria>Enterobacterales>Enterobacteriaceae>Salmonella | GCF_023016865.1 | Salmonella_sp._L-S2638 | MVT37_RS15260 | 170948..174589 | 2933299 | BREX-system-P-loop-protein-BrxC-[Salmonella-sp.-L-S2638] |
| WP_198899489.1 | Type1.BREX | <-HEPN<-REase+iREa[illegible] | 1213 | Bacteria>Pseudomonadota>Gammaproteobacteria>Enterobacterales>Enterobacteriaceae>Salmonella | GCF_023017625.1 | Salmonella_sp._14ESS1484 | MVA85_RS11375 | 170949..174590 | 2933314 | BREX-system-P-loop-protein-BrxC-[Salmonella-sp.-14ESS1484] |
| WP_254499948.1 | Type1.BREX | Arm-DNA-bind_3->N[illegible] | 1213 | Bacteria>Pseudomonadota>Gammaproteobacteria>Enterobacterales>Enterobacteriaceae>Salmonella | GCF_024261625.1 | Salmonella_enterica | NNJ27_RS08465 | 108147..111788 | 28901 | BREX-system-P-loop-protein-BrxC-[Salmonella-enterica] |
| WP_255223204.1 | Type1.BREX | <-HEPN<-REase+iREa[illegible] | 1213 | Bacteria>Pseudomonadota>Gammaproteobacteria>Enterobacterales>Enterobacteriaceae>Salmonella | GCF_024364985.1 | Salmonella_enterica | NMU37_RS20780 | 4227543..4231184 | 28901 | BREX-system-P-loop-protein-BrxC-[Salmonella-enterica] |
| WP_255929194.1 | Type1.BREX | <-HEPN<-?\|\|YjhX_to[illegible] | 1204 | Bacteria>Pseudomonadota>Gammaproteobacteria>Enterobacterales>Enterobacteriaceae>Salmonella | GCF_024442275.1 | Salmonella_enterica | NOQ95_RS21560 | 4448475..4452089 | 28901 | BREX-system-P-loop-protein-BrxC-[Salmonella-enterica] |
| WP_259423574.1 | Type1.BREX | <-METHYLASE\|\|SKI-[illegible] | 921 | Bacteria>Pseudomonadota>Gammaproteobacteria>Enterobacterales>Enterobacteriaceae>Salmonella | GCF_024919285.1 | Salmonella_enterica | NXM87_RS15320 | 123096..125861 | 28901 | BREX-system-P-loop-protein-BrxC-[Salmonella-enterica] |
| WP_260776303.1 | Type1.BREX | <-HEPN<-?\|\|YjhX_to[illegible] | 1213 | Bacteria>Pseudomonadota>Gammaproteobacteria>Enterobacterales>Enterobacteriaceae>Salmonella | GCF_025291975.1 | Salmonella_enterica | M1709_RS12435 | 17008..20649 | 28901 | BREX-system-P-loop-protein-BrxC-[Salmonella-enterica] |
| WP_080092317.1 | Type1.BREX | <-HEPN<-?\|\|YjhX_to[illegible] | 1213 | Bacteria>Pseudomonadota>Gammaproteobacteria>Enterobacterales>Enterobacteriaceae>Salmonella | GCF_025594515.1 | Salmonella_enterica | J3T40_RS14905 | 380423..384064 | 28901 | BREX-system-P-loop-protein-BrxC-[Salmonella-enterica] |
| WP_267272757.1 | Type1.BREX | <-SIGMA+OB-Fold+A[illegible] | 1213 | Bacteria>Pseudomonadota>Gammaproteobacteria>Enterobacterales>Enterobacteriaceae>Salmonella | GCF_026460795.1 | Salmonella_enterica | OWO74_RS14195 | 11359..15000 | 28901 | BREX-system-P-loop-protein-BrxC-[Salmonella-enterica] |
| WP_274877667.1 | Type1.BREX | Glyoxalase->ACET->[illegible] | 806 | Bacteria>Pseudomonadota>Gammaproteobacteria>Enterobacterales>Enterobacteriaceae>Salmonella | GCF_028892685.1 | Salmonella_enterica | PQP98_RS20365 | 4148316..4150736 | 28901 | BREX-system-P-loop-protein-BrxC-[Salmonella-enterica] |
| WP_274894925.1 | Type1.BREX | Glyoxalase->ACET->[illegible] | 1240 | Bacteria>Pseudomonadota>Gammaproteobacteria>Enterobacterales>Enterobacteriaceae>Salmonella | GCF_028892745.1 | Salmonella_enterica | PQQ04_RS20380 | 4148528..4152250 | 28901 | BREX-system-P-loop-protein-BrxC-[Salmonella-enterica] |
| WP_097173203.1 | Type1.BREX | Arm-DNA-bind_3->N[illegible] | 1213 | Bacteria>Pseudomonadota>Gammaproteobacteria>Enterobacterales>Enterobacteriaceae>Salmonella | GCF_900003195.1 | Salmonella_enterica | CRP09_RS12230 | 81403..85044 | 28901 | BREX-system-P-loop-protein-BrxC-[Salmonella-enterica] |
| WP_261983090.1 | Type1.BREX | <-METHYLASE\|\|SKI-[illegible] | 1240 | Bacteria>Pseudomonadota>Gammaproteobacteria>Enterobacterales>Enterobacteriaceae>Salmonella | GCF_901212535.1 | Salmonella_enterica | ECOMC1_RS14180 | 234231..237953 | 28901 | BREX-system-P-loop-protein-BrxC-[Salmonella-enterica] |
| WP_094914772.1 | Type1.BREX | <-HEPN<-?\|\|YjhX_to[illegible] | 1213 | Bacteria>Pseudomonadota>Gammaproteobacteria>Enterobacterales>Enterobacteriaceae>Salmonella | GCF_903930985.1 | Salmonella_enterica | DQN83_RS20180 | 3946680..3950321 | 28901 | BREX-system-P-loop-protein-BrxC-[Salmonella-enterica] |
| WP_088524188.1 | Type1.BREX | NACHT+SF1-SF2-Heli[illegible] | 1204 | Bacteria>Pseudomonadota>Gammaproteobacteria>Enterobacterales>Enterobacteriaceae>Salmonella | GCF_903931065.1 | Salmonella_enterica | LFZ40_19410 | 3810154..3813768 | 28901 | BREX-system-P-loop-protein-BrxC-[Salmonella-enterica] |
| WP_183072251.1 | Type1.BREX | <-HEPN<-REase+iREa[illegible] | 1213 | Bacteria>Pseudomonadota>Gammaproteobacteria>Enterobacterales>Enterobacteriaceae>Salmonella | GCF_903931105.1 | Salmonella_enterica | H7T17_RS07610 | 54171..57812 | 28901 | BREX-system-P-loop-protein-BrxC-[Salmonella-enterica] |
| WP_058685106.1 | Type1.BREX | Arm-DNA-bind_3->N[illegible] | 1213 | Bacteria>Pseudomonadota>Gammaproteobacteria>Enterobacterales>Enterobacteriaceae>Salmonella | GCF_940677185.1 | Salmonella_enterica | NBW86_RS10200 | 108141..111782 | 28901 | BREX-system-P-loop-protein-BrxC-[Salmonella-enterica] |
| WP_080205562.1 | Type1.BREX | <-HEPN<-REase+iREa[illegible] | 1213 | Bacteria>Pseudomonadota>Gammaproteobacteria>Enterobacterales>Enterobacteriaceae>Salmonella | GCF_943114325.1 | Salmonella_enterica | OCU44_RS00985 | 212044..215685 | 28901 | BREX-system-P-loop-protein-BrxC-[Salmonella-enterica] |
| WP_039590210.1 | Type1.BREX | <-HEPN<-REase+iREa[illegible] | 1213 | Bacteria>Pseudomonadota>Gammaproteobacteria>Enterobacterales>Enterobacteriaceae>Salmonella | GCF_943119195.1 | Salmonella_enterica | OCU40_RS05645 | 1155326..1158967 | 28901 | BREX-system-P-loop-protein-BrxC-[Salmonella-enterica] |
| WP_154023496.1 | Type1.BREX | Glyoxalase->ACET->[illegible] | 1233 | Bacteria>Pseudomonadota>Gammaproteobacteria>Enterobacterales>Enterobacteriaceae>Salmonella | - | Salmonella_enterica | A4I02_04465 | 211497..215198 | 28901 | BREX-system-P-loop-protein-BrxC-[Salmonella-enterica] |
| WP_097363546.1 | Type1.BREX | <-METHYLASE\|\|SKI-[illegible] | 1233 | Bacteria>Pseudomonadota>Gammaproteobacteria>Enterobacterales>Enterobacteriaceae>Salmonella | - | Salmonella_enterica | CHC75_12745 | 2557798..2561499 | 28901 | BREX-system-P-loop-protein-BrxC-[Salmonella-enterica] |
| EJF5860346.1 | Type1.BREX | <-HEPN<-REase+iREa[illegible] | 1213 | Bacteria>Pseudomonadota>Gammaproteobacteria>Enterobacterales>Enterobacteriaceae>Salmonella | - | Salmonella_enterica | M8X04_001881 | 170514..174155 | 28901 | BREX-system-P-loop-protein-BrxC-[Salmonella-enterica] |
| WP_023227805.1 | Type1.BREX | Glyoxalase->ACET->[illegible] | 1233 | Bacteria>Pseudomonadota>Gammaproteobacteria>Enterobacterales>Enterobacteriaceae>Salmonella | - | Salmonella_enterica | SEEI0720_021515 | 4182897..4186598 | 28901 | BREX-system-P-loop-protein-BrxC-[Salmonella-enterica] |
| ECG0705015.1 | Type1.BREX | <-HEPN<-REase+iREa[illegible] | 1213 | Bacteria>Pseudomonadota>Gammaproteobacteria>Enterobacterales>Enterobacteriaceae>Salmonella | - | Salmonella_enterica_subsp._salamae | DL122_04700 | 35596..39237 | 59202 | BREX-system-P-loop-protein-BrxC-[Salmonella-enterica-subsp.-salamae] |
| HCC77786.1 | Type1.BREX | AAA+HTH-><-SIGMA+[illegible] | 1216 | Bacteria>Pseudomonadota>Gammaproteobacteria>Enterobacterales>Enterobacteriaceae>Shigella | GCA_003517755.1 | Shigella_sp. | DEQ42_19255 | 11300..14950 | 625 | BREX-system-P-loop-protein-BrxC-[Shigella-sp.] |
| MRS92356.1 | Type1.BREX | DUF932-><-?\|\|ArdB-[illegible] | 1157 | Bacteria>Pseudomonadota>Gammaproteobacteria>Enterobacterales>Enterobacteriaceae>unclassified_Enterobacteriaceae | GCA_009668015.1 | Enterobacteriaceae_bacterium_RIT714 | GJV04_20305 | 30577..34050 | 2666193 | BREX-system-P-loop-protein-BrxC-[Enterobacteriaceae-bacterium-RIT714] |
| NUU66316.1 | Type1.BREX | BrxA-DNA-binding-[illegible] | 1213 | Bacteria>Pseudomonadota>Gammaproteobacteria>Enterobacterales>Enterobacteriaceae>unclassified_Enterobacteriaceae | GCA_013359925.1 | Enterobacteriaceae_bacterium_BIT-123 | HQN64_09360 | 363314..366955 | 2742797 | BREX-system-P-loop-protein-BrxC-[Enterobacteriaceae-bacterium-BIT-123] |
| UCB31815.1 | Type1.BREX | <-wHTH+WYL<-?\|\|XI[illegible] | 1240 | Bacteria>Pseudomonadota>Gammaproteobacteria>Enterobacterales>Erwiniaceae>Duffyella | GCA_020342335.1 | Duffyella_gerundensis | J9874_02366 | 2569295..2573017 | 1619313 | hypothetical-protein-J9874_02366-[Duffyella-gerundensis] |
| WP_225700910.1 | Type1.BREX | <-wHTH+WYL<-?\|\|XI[illegible] | 1237 | Bacteria>Pseudomonadota>Gammaproteobacteria>Enterobacterales>Erwiniaceae>Duffyella | GCF_020342335.1 | Duffyella_gerundensis | J9874_RS11820 | 2569304..2573017 | 1619313 | BREX-system-P-loop-protein-BrxC-[Duffyella-gerundensis] |
| WP_012669046.1 | Type1.BREX | HTH_Tnp_1+HTH_21+[illegible] | 1214 | Bacteria>Pseudomonadota>Gammaproteobacteria>Enterobacterales>Erwiniaceae>Erwinia | GCF_000027265.1 | Erwinia_pyrifoliae | EPC_RS14330 | 3076453..3080097 | 79967 | BREX-system-P-loop-protein-BrxC-[Erwinia-pyrifoliae] |
| WP_014543466.1 | Type1.BREX | PCMT+METHYLASE->L[illegible] | 1214 | Bacteria>Pseudomonadota>Gammaproteobacteria>Enterobacterales>Erwiniaceae>Erwinia | GCF_000165815.1 | Erwinia_sp._Ejp617 | EJP617_RS09645 | 2138156..2141800 | 215689 | BREX-system-P-loop-protein-BrxC-[Erwinia-sp.-Ejp617] |
| WP_253458939.1 | Type1.BREX | XIS-HTH->?->AAA+H[illegible] | 1216 | Bacteria>Pseudomonadota>Gammaproteobacteria>Enterobacterales>Erwiniaceae>Erwinia | GCF_024168495.1 | Erwinia_persicina | J3D56_RS16965 | 3646326..3649976 | 55211 | BREX-system-P-loop-protein-BrxC-[Erwinia-persicina] |
| WP_259816994.1 | Type1.BREX | SIGMA-HTH+SIGMA-H[illegible] | 1214 | Bacteria>Pseudomonadota>Gammaproteobacteria>Enterobacterales>Erwiniaceae>Erwinia | GCF_025153555.1 | Erwinia_pyrifoliae | NYP82_RS06320 | 1343257..1346901 | 79967 | BREX-system-P-loop-protein-BrxC-[Erwinia-pyrifoliae] |
| WP_259826113.1 | Type1.BREX | PCMT+METHYLASE->L[illegible] | 1214 | Bacteria>Pseudomonadota>Gammaproteobacteria>Enterobacterales>Erwiniaceae>Erwinia | GCF_025159015.1 | Erwinia_pyrifoliae | NYP84_RS03425 | 759232..762876 | 79967 | BREX-system-P-loop-protein-BrxC-[Erwinia-pyrifoliae] |
| WP_261642253.1 | Type1.BREX | <-SIGMA+OB-Fold+A[illegible] | 1213 | Bacteria>Pseudomonadota>Gammaproteobacteria>Enterobacterales>Erwiniaceae>Erwinia | GCF_025388925.1 | Erwinia_mallotivora | OCH37_RS12855 | 149371..153012 | 69222 | BREX-system-P-loop-protein-BrxC-[Erwinia-mallotivora] |
| WP_159378885.1 | Type1.BREX | HTH_Tnp_1+HTH_21+[illegible] | 1213 | Bacteria>Pseudomonadota>Gammaproteobacteria>Enterobacterales>Erwiniaceae>Mixta | GCF_009866505.1 | Mixta_calida | Y888_RS19575 | 9519..13160 | 665913 | BREX-system-P-loop-protein-BrxC-[Mixta-calida] |
| HCP27301.1 | Type1.BREX | Phage_integrase->[illegible] | 1233 | Bacteria>Pseudomonadota>Gammaproteobacteria>Enterobacterales>Erwiniaceae>Pantoea | GCA_003533915.1 | Pantoea_ananatis | DIT85_12780 | 28453..32154 | 553 | BREX-system-P-loop-protein-BrxC-[Pantoea-ananatis] |
| RZK06926.1 | Type1.BREX | <-DEDD_Tnp_IS110<[illegible] | 1213 | Bacteria>Pseudomonadota>Gammaproteobacteria>Enterobacterales>Erwiniaceae>Pantoea | GCA_004295855.1 | Pantoea_sp. | EO084_12150 | 162361..166002 | 69393 | MAG:-BREX-system-P-loop-protein-BrxC-[Pantoea-sp.] |
| NEG80386.1 | Type1.BREX | <-wHTH(SBD)+HNH\|\|[illegible] | 1233 | Bacteria>Pseudomonadota>Gammaproteobacteria>Enterobacterales>Erwiniaceae>Pantoea | GCA_010667945.1 | Pantoea_agglomerans | GQQ17_05160 | 300399..304100 | 549 | BREX-system-P-loop-protein-BrxC-[Pantoea-agglomerans] |
| WP_007890763.1 | Type1.BREX | ABhydrolase+METHY[illegible] | 1202 | Bacteria>Pseudomonadota>Gammaproteobacteria>Enterobacterales>Erwiniaceae>Pantoea | GCF_000282675.1 | Pantoea_sp._GM01 | PMI17_RS16905 | 182397..186005 | 1144320 | BREX-system-P-loop-protein-BrxC-[Pantoea-sp.-GM01] |
| WP_137241123.1 | Type1.BREX | <-SymE_toxin<-CHT[illegible] | 1202 | Bacteria>Pseudomonadota>Gammaproteobacteria>Enterobacterales>Erwiniaceae>Pantoea | GCF_005233495.1 | Pantoea_agglomerans | PagCFBP13505_RS08740 | 40890..44498 | 549 | BREX-system-P-loop-protein-BrxC-[Pantoea-agglomerans] |
| WP_145889319.1 | Type1.BREX | <-SAM-METHYLASE<-[illegible] | 1213 | Bacteria>Pseudomonadota>Gammaproteobacteria>Enterobacterales>Erwiniaceae>Pantoea | GCF_007833795.1 | Pantoea_dispersa | D8B20_RS13380 | 2861795..2865436 | 59814 | BREX-system-P-loop-protein-BrxC-[Pantoea-dispersa] |
| WP_154209858.1 | Type1.BREX | <-SIGMA+OB-Fold+A[illegible] | 1202 | Bacteria>Pseudomonadota>Gammaproteobacteria>Enterobacterales>Erwiniaceae>Pantoea | GCF_009668105.1 | Pantoea_agglomerans | GJV05_RS04780 | 299589..303197 | 549 | BREX-system-P-loop-protein-BrxC-[Pantoea-agglomerans] |
| WP_158149594.1 | Type1.BREX | <-CHTH\|\|ColE7+HNH-[illegible] | 1157 | Bacteria>Pseudomonadota>Gammaproteobacteria>Enterobacterales>Erwiniaceae>Pantoea | GCF_009765475.1 | Pantoea_agglomerans | GSQ69_RS16285 | 3495115..3498588 | 549 | BREX-system-P-loop-protein-BrxC-[Pantoea-agglomerans] |
| WP_163841538.1 | Type1.BREX | <-DEDD_Tnp_IS110<[illegible] | 1213 | Bacteria>Pseudomonadota>Gammaproteobacteria>Enterobacterales>Erwiniaceae>Pantoea | GCF_010667775.1 | Pantoea_agglomerans | GQQ23_RS01345 | 302336..305977 | 549 | BREX-system-P-loop-protein-BrxC-[Pantoea-agglomerans] |
| WP_242462783.1 | Type1.BREX | <-wHTH(SBD)+HNH\|\|[illegible] | 1240 | Bacteria>Pseudomonadota>Gammaproteobacteria>Enterobacterales>Erwiniaceae>Pantoea | GCF_010667945.1 | Pantoea_agglomerans | GQQ17_RS05165 | 300399..304121 | 549 | BREX-system-P-loop-protein-BrxC-[Pantoea-agglomerans] |
| WP_167070220.1 | Type1.BREX | <-wHTH+WYL+WCX\|\|?[illegible] | 1184 | Bacteria>Pseudomonadota>Gammaproteobacteria>Enterobacterales>Erwiniaceae>Pantoea | GCF_011752565.1 | Pantoea_sp._Ap-967 | F3J45_27210 | 36650..40204 | 2608362 | BREX-system-P-loop-protein-BrxC-[Pantoea-sp.-Ap-967] |
| WP_182687314.1 | Type1.BREX | <-DEDD_Tnp_IS110<[illegible] | 1213 | Bacteria>Pseudomonadota>Gammaproteobacteria>Enterobacterales>Erwiniaceae>Pantoea | GCF_014156615.1 | Pantoea_pleuroti | EPG66_RS14530 | 313158..316799 | 1592631 | BREX-system-P-loop-protein-BrxC-[Pantoea-pleuroti] |
| WP_187492467.1 | Type1.BREX | <-SymE_toxin\|\|?->[illegible] | 1157 | Bacteria>Pseudomonadota>Gammaproteobacteria>Enterobacterales>Erwiniaceae>Pantoea | GCF_014353895.1 | Pantoea_agglomerans | IBU46_RS01740 | 23077..26550 | 549 | BREX-system-P-loop-protein-BrxC-[Pantoea-agglomerans] |
| WP_191919036.1 | Type1.BREX | <-SymE_toxin<-CHT[illegible] | 1157 | Bacteria>Pseudomonadota>Gammaproteobacteria>Enterobacterales>Erwiniaceae>Pantoea | GCF_014838975.1 | Pantoea_agglomerans | IFU26_RS12730 | 289507..292980 | 549 | BREX-system-P-loop-protein-BrxC-[Pantoea-agglomerans] |
| WP_202940066.1 | Type1.BREX | <-DEDD_Tnp_IS110<[illegible] | 1021 | Bacteria>Pseudomonadota>Gammaproteobacteria>Enterobacterales>Erwiniaceae>Pantoea | GCF_014839325.1 | Pantoea_agglomerans | IFT41_RS17295 | 170215..173280 | 549 | BREX-system-P-loop-protein-BrxC-[Pantoea-agglomerans] |
| WP_222233110.1 | Type1.BREX | <-HKD-DNase+OGG-l[illegible] | 1202 | Bacteria>Pseudomonadota>Gammaproteobacteria>Enterobacterales>Erwiniaceae>Pantoea | GCF_019795975.1 | Pantoea_sp._DY-17 | K5Y33_RS01225 | 252070..255678 | 2871490 | BREX-system-P-loop-protein-BrxC-[Pantoea-sp.-DY-17] |
| WP_245115608.1 | Type1.BREX | <-SymE_toxin<-CHT[illegible] | 1204 | Bacteria>Pseudomonadota>Gammaproteobacteria>Enterobacterales>Erwiniaceae>Pantoea | GCF_022966105.1 | Pantoea_agglomerans | LZ609_RS16055 | 2632558..2636172 | 549 | BREX-system-P-loop-protein-BrxC-[Pantoea-agglomerans] |
| WP_264214894.1 | Type1.BREX | <-DEDD_Tnp_IS110<[illegible] | 1213 | Bacteria>Pseudomonadota>Gammaproteobacteria>Enterobacterales>Erwiniaceae>Pantoea | GCF_025881995.1 | Pantoea_ananatis | NB695_RS01430 | 332354..335995 | 553 | BREX-system-P-loop-protein-BrxC-[Pantoea-ananatis] |
| WP_272844016.1 | Type1.BREX | CbeA_antitoxin->?[illegible] | 1157 | Bacteria>Pseudomonadota>Gammaproteobacteria>Enterobacterales>Erwiniaceae>Pantoea | GCF_028555165.1 | Pantoea_ananatis | SZ66_RS15300 | 342532..346005 | 553 | BREX-system-P-loop-protein-BrxC-[Pantoea-ananatis] |
| PIJ55036.1 | Type1.BREX | <-N6-METHYLASE<-D[illegible] | 1202 | Bacteria>Pseudomonadota>Gammaproteobacteria>Enterobacterales>Erwiniaceae>unclassified_Erwiniaceae | GCA_002752575.1 | Erwinia_sp._OLMDLW33 | BOM23_20680 | 2080..5688 | 1928656 | DNA-repair-protein-[Erwinia-sp.-OLMDLW33] |
| WP_001019641.1 | Type1.BREX | <-Phage_integrase[illegible] | 1213 | Bacteria>Pseudomonadota>Gammaproteobacteria>Enterobacterales | GCF_000010765.1 | Enterobacteriaceae | PYB84_RS07565 | 16979..20620 | 543 | MULTISPECIES:-BREX-system-P-loop-protein-BrxC-[Enterobacteriaceae] |
| WP_000687847.1 | Type1.BREX | <-LexA-protease\|\|[illegible] | 1225 | Bacteria>Pseudomonadota>Gammaproteobacteria>Enterobacterales | GCF_000026225.1 | Enterobacteriaceae | EFER_RS00085 | 15017..18694 | 543 | MULTISPECIES:-BREX-system-P-loop-protein-BrxC-[Enterobacteriaceae] |
| WP_001019638.1 | Type1.BREX | <-Phage_integrase[illegible] | 1213 | Bacteria>Pseudomonadota>Gammaproteobacteria>Enterobacterales | GCF_000166575.1 | Enterobacteriaceae | EC33884_RS23090 | 1218224..1221865 | 543 | MULTISPECIES:-BREX-system-P-loop-protein-BrxC-[Enterobacteriaceae] |
| WP_001019648.1 | Type1.BREX | Arm-DNA-bind_3+Ph[illegible] | 1213 | Bacteria>Pseudomonadota>Gammaproteobacteria>Enterobacterales | GCF_000194705.1 | Enterobacteriaceae | ECB41_RS05715 | 178680..182321 | 543 | MULTISPECIES:-BREX-system-P-loop-protein-BrxC-[Enterobacteriaceae] |
| WP_001019636.1 | Type1.BREX | <-METHYLASE\|\|SKI-[illegible] | 1213 | Bacteria>Pseudomonadota>Gammaproteobacteria>Enterobacterales | GCF_000233875.1 | Enterobacteriaceae | IO2_RS24015 | 4889707..4893348 | 543 | MULTISPECIES:-BREX-system-P-loop-protein-BrxC-[Enterobacteriaceae] |
| WP_001019644.1 | Type1.BREX | <-Fimbrial<-Fimbr[illegible] | 1213 | Bacteria>Pseudomonadota>Gammaproteobacteria>Enterobacterales | GCF_000282855.1 | Enterobacteriaceae | DTB66_RS23695 | 24279..27920 | 543 | MULTISPECIES:-BREX-system-P-loop-protein-BrxC-[Enterobacteriaceae] |
| WP_006788201.1 | Type1.BREX | <-SIGMA+OB-Fold+A[illegible] | 1225 | Bacteria>Pseudomonadota>Gammaproteobacteria>Enterobacterales | GCF_000388555.1 | Enterobacteriaceae | MUG00_RS24960 | 10799..14476 | 543 | MULTISPECIES:-BREX-system-P-loop-protein-BrxC-[Enterobacteriaceae] |
| WP_032187199.1 | Type1.BREX | HEPN+SF1-SF2-Heli[illegible] | 1225 | Bacteria>Pseudomonadota>Gammaproteobacteria>Enterobacterales | GCF_000703445.1 | Enterobacteriaceae | AD25_RS20800 | 23140..26817 | 543 | MULTISPECIES:-BREX-system-P-loop-protein-BrxC-[Enterobacteriaceae] |
| WP_053287615.1 | Type1.BREX | <-Zn_Tnp_IS1+HTH_[illegible] | 1213 | Bacteria>Pseudomonadota>Gammaproteobacteria>Enterobacterales | GCF_001268905.1 | Enterobacteriaceae | AMM63_RS24455 | 5812..9453 | 543 | MULTISPECIES:-BREX-system-P-loop-protein-BrxC-[Enterobacteriaceae] |
| WP_054528932.1 | Type1.BREX | DDE_Tnp_1_5+DDE_T[illegible] | 1225 | Bacteria>Pseudomonadota>Gammaproteobacteria>Enterobacterales | GCF_002788295.1 | Enterobacteriaceae | AM338_RS26100 | 11174..14851 | 543 | MULTISPECIES:-BREX-system-P-loop-protein-BrxC-[Enterobacteriaceae] |
| WP_032408831.1 | Type1.BREX | Arm-DNA-bind_3+Ph[illegible] | 1213 | Bacteria>Pseudomonadota>Gammaproteobacteria>Enterobacterales | GCF_002897415.1 | Enterobacteriaceae | HED41_RS19555 | 135079..138720 | 543 | MULTISPECIES:-BREX-system-P-loop-protein-BrxC-[Enterobacteriaceae] |
| WP_097333547.1 | Type1.BREX | <-HIN-HTH\|\|?->?->[illegible] | 1213 | Bacteria>Pseudomonadota>Gammaproteobacteria>Enterobacterales | GCF_004194615.2 | Enterobacteriaceae | DBZ52_RS03520 | 719980..723621 | 543 | MULTISPECIES:-BREX-system-P-loop-protein-BrxC-[Enterobacteriaceae] |
| WP_223177211.1 | Type1.BREX | <-wHTH+WYL<-nSTAN[illegible] | 1240 | Bacteria>Pseudomonadota>Gammaproteobacteria>Enterobacterales | GCF_004312085.1 | Enterobacteriaceae | EOM53_RS14900 | 53286..57008 | 543 | MULTISPECIES:-BREX-system-P-loop-protein-BrxC-[Enterobacteriaceae] |
| WP_004249342.1 | Type1.BREX | <-N6-METHYLASE<-D[illegible] | 1225 | Bacteria>Pseudomonadota>Gammaproteobacteria>Enterobacterales | GCF_008041895.1 | Morganellaceae | E4V60_RS00205 | 3632..7309 | 1903414 | MULTISPECIES:-BREX-system-P-loop-protein-BrxC-[Morganellaceae] |
| WP_019078269.1 | Type1.BREX | <-HerA/FtsK+A32<-[illegible] | 1213 | Bacteria>Pseudomonadota>Gammaproteobacteria>Enterobacterales | GCF_011030295.1 | Enterobacteriaceae | QFD02_RS07350 | 209926..213567 | 543 | MULTISPECIES:-BREX-system-P-loop-protein-BrxC-[Enterobacteriaceae] |
| WP_080865889.1 | Type1.BREX | GDResolvase+HIN-H[illegible] | 1213 | Bacteria>Pseudomonadota>Gammaproteobacteria>Enterobacterales | GCF_011604705.1 | Enterobacteriaceae | HA514_RS19415 | 4027238..4030879 | 543 | MULTISPECIES:-BREX-system-P-loop-protein-BrxC-[Enterobacteriaceae] |
| WP_049194743.1 | Type1.BREX | MobI-><-?<-LexA-p[illegible] | 1225 | Bacteria>Pseudomonadota>Gammaproteobacteria>Enterobacterales | GCF_013357405.1 | Morganellaceae | HO271_RS14475 | 3106185..3109862 | 1903414 | MULTISPECIES:-BREX-system-P-loop-protein-BrxC-[Morganellaceae] |
| WP_117259785.1 | Type1.BREX | <-Peptidase_M17_N[illegible] | 1157 | Bacteria>Pseudomonadota>Gammaproteobacteria>Enterobacterales | GCF_013395115.1 | Enterobacteriaceae | HYG96_RS13000 | 239421..242894 | 543 | MULTISPECIES:-BREX-system-P-loop-protein-BrxC-[Enterobacteriaceae] |
| WP_115625931.1 | Type1.BREX | HIN-HTH->\|\|?->LZ_[illegible] | 1204 | Bacteria>Pseudomonadota>Gammaproteobacteria>Enterobacterales | GCF_013635755.1 | Enterobacteriaceae | HV027_RS18510 | 3797222..3800836 | 543 | MULTISPECIES:-BREX-system-P-loop-protein-BrxC-[Enterobacteriaceae] |
| WP_242461228.1 | Type1.BREX | Zn_Tnp_IS1+HTH_Tn[illegible] | 1240 | Bacteria>Pseudomonadota>Gammaproteobacteria>Enterobacterales | GCF_013705425.1 | Enterobacteriaceae | HV198_RS20030 | 4234756..4238478 | 543 | MULTISPECIES:-BREX-system-P-loop-protein-BrxC-[Enterobacteriaceae] |
| WP_232051962.1 | Type1.BREX | <-DUF932<-nSTAND1[illegible] | 1240 | Bacteria>Pseudomonadota>Gammaproteobacteria>Enterobacterales | GCF_018421715.1 | Enterobacteriaceae | L6492_RS02235 | 461320..465042 | 543 | MULTISPECIES:-BREX-system-P-loop-protein-BrxC-[Enterobacteriaceae] |
| WP_222691531.1 | Type1.BREX | BrxA-DNA-binding-[illegible] | 1204 | Bacteria>Pseudomonadota>Gammaproteobacteria>Enterobacterales | GCF_019841255.1 | Enterobacteriaceae | K7A35_RS15405 | 1428..5042 | 543 | MULTISPECIES:-BREX-system-P-loop-protein-BrxC-[Enterobacteriaceae] |
| WP_200541075.1 | Type1.BREX | <-SIGMA+OB-Fold+A[illegible] | 1213 | Bacteria>Pseudomonadota>Gammaproteobacteria>Enterobacterales | GCF_019927965.1 | Enterobacteriaceae | KFB01_RS26200 | 8731..12372 | 543 | MULTISPECIES:-BREX-system-P-loop-protein-BrxC-[Enterobacteriaceae] |
| WP_224243997.1 | Type1.BREX | <-SymE_toxin<-Gly[illegible] | 1157 | Bacteria>Pseudomonadota>Gammaproteobacteria>Enterobacterales | GCF_023752605.1 | Enterobacteriaceae | M8S83_RS11470 | 342574..346047 | 543 | MULTISPECIES:-BREX-system-P-loop-protein-BrxC-[Enterobacteriaceae] |
| WP_088474912.1 | Type1.BREX | Phage_integrase->[illegible] | 1213 | Bacteria>Pseudomonadota>Gammaproteobacteria>Enterobacterales | GCF_024713235.1 | Enterobacteriaceae | CD912_RS19430 | 9686..13327 | 543 | MULTISPECIES:-BREX-system-P-loop-protein-BrxC-[Enterobacteriaceae] |
| WP_080816097.1 | Type1.BREX | Phage_integrase->[illegible] | 1213 | Bacteria>Pseudomonadota>Gammaproteobacteria>Enterobacterales | GCF_026460795.1 | Enterobacteriaceae | OWO74_RS12055 | 662264..665905 | 543 | MULTISPECIES:-BREX-system-P-loop-protein-BrxC-[Enterobacteriaceae] |
| WP_224293199.1 | Type1.BREX | <-DUF905<-DUF932<[illegible] | 1240 | Bacteria>Pseudomonadota>Gammaproteobacteria>Enterobacterales | GCF_029226505.1 | Enterobacteriaceae | PVK22_RS25565 | 5368784..5372506 | 543 | MULTISPECIES:-BREX-system-P-loop-protein-BrxC-[Enterobacteriaceae] |
| WP_047644001.1 | Type1.BREX | <-XIS-HTH<-RHH_an[illegible] | 1213 | Bacteria>Pseudomonadota>Gammaproteobacteria>Enterobacterales | GCF_030296855.1 | Enterobacteriaceae | CPH82_RS06655 | 82494..86135 | 543 | MULTISPECIES:-BREX-system-P-loop-protein-BrxC-[Enterobacteriaceae] |
| WP_063149682.1 | Type1.BREX | <-SIGMA+OB-Fold+A[illegible] | 1225 | Bacteria>Pseudomonadota>Gammaproteobacteria>Enterobacterales | GCF_900075035.1 | Enterobacteriaceae | K1J36_RS23220 | 10587..14264 | 543 | MULTISPECIES:-BREX-system-P-loop-protein-BrxC-[Enterobacteriaceae] |
| WP_072662972.1 | Type1.BREX | DDE_Tnp_IS1-><-SI[illegible] | 1213 | Bacteria>Pseudomonadota>Gammaproteobacteria>Enterobacterales | GCF_900775105.1 | Enterobacteriaceae | B5I96_RS21955 | 8761..12402 | 543 | MULTISPECIES:-BREX-system-P-loop-protein-BrxC-[Enterobacteriaceae] |
| WP_045345932.1 | Type1.BREX | XIS-HTH->?->?->SF[illegible] | 1157 | Bacteria>Pseudomonadota>Gammaproteobacteria>Enterobacterales | GCF_902160145.1 | Enterobacteriaceae | FN179_RS14130 | 343492..346965 | 543 | MULTISPECIES:-BREX-system-P-loop-protein-BrxC-[Enterobacteriaceae] |
| WP_044349975.1 | Type1.BREX | DEDD_Tnp_IS110+Tr[illegible] | 1213 | Bacteria>Pseudomonadota>Gammaproteobacteria>Enterobacterales | GCF_902162255.1 | Enterobacteriaceae | FN407_RS22090 | 106116..109757 | 543 | MULTISPECIES:-BREX-system-P-loop-protein-BrxC-[Enterobacteriaceae] |
| WP_110225552.1 | Type1.BREX | BrxA-DNA-binding-[illegible] | 1213 | Bacteria>Pseudomonadota>Gammaproteobacteria>Enterobacterales | GCF_902499505.1 | Enterobacteriaceae | G4V40_RS25520 | 1772..5413 | 543 | MULTISPECIES:-BREX-system-P-loop-protein-BrxC-[Enterobacteriaceae] |
| WP_117108864.1 | Type1.BREX | <-METHYLASE\|\|SKI-[illegible] | 1189 | Bacteria>Pseudomonadota>Gammaproteobacteria>Enterobacterales | GCF_903684385.1 | Enterobacteriaceae | MJ801_RS10050 | 169597..173166 | 543 | MULTISPECIES:-BREX-system-P-loop-protein-BrxC-[Enterobacteriaceae] |
| WP_039021203.1 | Type1.BREX | rve-><-RHH_antito[illegible] | 1213 | Bacteria>Pseudomonadota>Gammaproteobacteria>Enterobacterales | GCF_904387335.1 | Enterobacteriaceae | G3M52_RS00045 | 12929..16570 | 543 | MULTISPECIES:-BREX-system-P-loop-protein-BrxC-[Enterobacteriaceae] |
| WP_047929056.1 | Type1.BREX | MazE_antitoxin->P[illegible] | 1225 | Bacteria>Pseudomonadota>Gammaproteobacteria>Enterobacterales | GCF_904799765.1 | Enterobacteriaceae | PCN061_RS24650 | 104442..108119 | 543 | MULTISPECIES:-BREX-system-P-loop-protein-BrxC-[Enterobacteriaceae] |
| WP_064143490.1 | Type1.BREX | <-Phage_integrase[illegible] | 1213 | Bacteria>Pseudomonadota>Gammaproteobacteria>Enterobacterales | GCF_910592795.1 | Enterobacteriaceae | KXI98_RS20085 | 95010..98651 | 543 | MULTISPECIES:-BREX-system-P-loop-protein-BrxC-[Enterobacteriaceae] |
| WP_175320436.1 | Type1.BREX | <-DUF932<-nSTAND1[illegible] | 1233 | Bacteria>Pseudomonadota>Gammaproteobacteria>Enterobacterales>Hafniaceae>Hafnia | GCF_013360205.1 | Hafnia_paralvei | G4F99_RS14350 | 55379..59080 | 546367 | BREX-system-P-loop-protein-BrxC-[Hafnia-paralvei] |
| WP_226228481.1 | Type1.BREX | GNTR-HTH+Aminotra[illegible] | 1202 | Bacteria>Pseudomonadota>Gammaproteobacteria>Enterobacterales>Hafniaceae>Hafnia | GCF_020450285.1 | Hafnia_paralvei | LA328_RS17860 | 3833328..3836936 | 546367 | BREX-system-P-loop-protein-BrxC-[Hafnia-paralvei] |
| WP_234619586.1 | Type1.BREX | <-nSTAND1+GTPase-[illegible] | 1237 | Bacteria>Pseudomonadota>Gammaproteobacteria>Enterobacterales>Hafniaceae>Hafnia | GCF_021440285.1 | Hafnia_alvei | LZ667_RS17495 | 58169..61882 | 569 | BREX-system-P-loop-protein-BrxC-[Hafnia-alvei] |
| WP_108479632.1 | Type1.BREX | MobI-><-?<-LexA-p[illegible] | 1225 | Bacteria>Pseudomonadota>Gammaproteobacteria>Enterobacterales>Morganellaceae | GCF_003057415.1 | Providencia | C6581_RS17855 | 39720..43397 | 586 | MULTISPECIES:-BREX-system-P-loop-protein-BrxC-[Providencia] |
| WP_241543889.1 | Type1.BREX | <-HTH_37<-?\|\|GDRe[illegible] | 1225 | Bacteria>Pseudomonadota>Gammaproteobacteria>Enterobacterales>Morganellaceae>Moellerella | GCF_022591895.1 | Moellerella_wisconsensis | MNY64_RS16755 | 120814..124491 | 158849 | BREX-system-P-loop-protein-BrxC-[Moellerella-wisconsensis] |
| WP_241542954.1 | Type1.BREX | <-SIGMA+OB-Fold+A[illegible] | 1225 | Bacteria>Pseudomonadota>Gammaproteobacteria>Enterobacterales>Morganellaceae>Moellerella | GCF_022592015.1 | Moellerella_wisconsensis | MNY72_RS15945 | 14916..18593 | 158849 | BREX-system-P-loop-protein-BrxC-[Moellerella-wisconsensis] |
| WP_241501907.1 | Type1.BREX | <-HTH_37<-?\|\|GDRe[illegible] | 1225 | Bacteria>Pseudomonadota>Gammaproteobacteria>Enterobacterales>Morganellaceae>Moellerella | GCF_022592395.1 | Moellerella_wisconsensis | MNY66_RS17045 | 197034..200711 | 158849 | BREX-system-P-loop-protein-BrxC-[Moellerella-wisconsensis] |
| AUR32853.1 | Type1.BREX | HTH->Pertactin+Au[illegible] | 1233 | Bacteria>Pseudomonadota>Gammaproteobacteria>Enterobacterales>Morganellaceae>Morganella | GCA_002891475.1 | Morganella_morganii | C1O70_15730 | 3342273..3345974 | 582 | BREX-system-P-loop-protein-BrxC-[Morganella-morganii] |
| MBS5193172.1 | Type1.BREX | REase->wHTH+WYL+W[illegible] | 1157 | Bacteria>Pseudomonadota>Gammaproteobacteria>Enterobacterales>Morganellaceae>Morganella | GCA_018366325.1 | Morganella_morganii | KHY92_05180 | 18454..21927 | 582 | BREX-system-P-loop-protein-BrxC-[Morganella-morganii] |
| WP_225973345.1 | Type1.BREX | HTH->Pertactin+Au[illegible] | 1240 | Bacteria>Pseudomonadota>Gammaproteobacteria>Enterobacterales>Morganellaceae>Morganella | GCF_002891475.1 | Morganella_morganii | C1O70_RS15730 | 3342252..3345974 | 582 | BREX-system-P-loop-protein-BrxC-[Morganella-morganii] |
| WP_214179407.1 | Type1.BREX | XIS-HTH->BrxA-DNA[illegible] | 1225 | Bacteria>Pseudomonadota>Gammaproteobacteria>Enterobacterales>Morganellaceae>Morganella | GCF_018474625.1 | Morganella_morganii | IS084_RS18050 | 5738..9415 | 582 | BREX-system-P-loop-protein-BrxC-[Morganella-morganii] |
| WP_274765494.1 | Type1.BREX | <-Peptidase_M17_N[illegible] | 1157 | Bacteria>Pseudomonadota>Gammaproteobacteria>Enterobacterales>Morganellaceae>Morganella | GCF_028867915.1 | Morganella_morganii | PS169_RS13665 | 158962..162435 | 582 | BREX-system-P-loop-protein-BrxC-[Morganella-morganii] |
| WP_081119725.1 | Type1.BREX | <-wHTH+WYL<-?<-?\|\|[illegible] | 1225 | Bacteria>Pseudomonadota>Gammaproteobacteria>Enterobacterales>Morganellaceae>Morganella | - | Morganella_morganii | B2H99_RS12750 | 8810..12487 | 582 | BREX-system-P-loop-protein-BrxC-[Morganella-morganii] |
| CAQ82523.1 | Type1.BREX | Peptidase_M28->AB[illegible] | 1216 | Bacteria>Pseudomonadota>Gammaproteobacteria>Enterobacterales>Morganellaceae>Photorhabdus | GCA_000196475.1 | Photorhabdus_asymbiotica | PAU_00430 | 471573..475223 | 291112 | putative-atpase-involved-in-dna-repair-[Photorhabdus-asymbiotica] |
| WP_041381498.1 | Type1.BREX | Peptidase_M28->AB[illegible] | 1215 | Bacteria>Pseudomonadota>Gammaproteobacteria>Enterobacterales>Morganellaceae>Photorhabdus | GCF_000196475.1 | Photorhabdus_asymbiotica | PAU_RS02125 | 471576..475223 | 291112 | BREX-system-P-loop-protein-BrxC-[Photorhabdus-asymbiotica] |
| WP_036771207.1 | Type1.BREX | ABhydrolase+ABhyd[illegible] | 1213 | Bacteria>Pseudomonadota>Gammaproteobacteria>Enterobacterales>Morganellaceae>Photorhabdus | GCF_000711895.1 | Photorhabdus_australis | BO26_RS10370 | 20693..24334 | 286156 | BREX-system-P-loop-protein-BrxC-[Photorhabdus-australis] |
| WP_054478994.1 | Type1.BREX | <-PglZ<-N6-METHYL[illegible] | 1213 | Bacteria>Pseudomonadota>Gammaproteobacteria>Enterobacterales>Morganellaceae>Photorhabdus | GCF_001280945.1 | Photorhabdus_heterorhabditis | AM629_RS11280 | 27281..30922 | 880156 | BREX-system-P-loop-protein-BrxC-[Photorhabdus-heterorhabditis] |

| Protein-ID | System | Gene-Neighbourhoods | Length | Taxonomy | Assembly-ID | Strain | Locus-Tag | Genomic-coordinates | TaxID | Fasta-define |
|---|---|---|---|---|---|---|---|---|---|---|
| WP_065824091.1 | Type1.BREX | [illegible] | 1213 | Bacteria>Pseudomonadota>Gammaproteobacteria>Enterobacterales>Morganellaceae>Photorhabdus | GCF_001696805.1 | Photorhabdus_australis | Ppb6_RS16855 | 33737..37378 | 286156 | BREX-system-P-loop-protein-BrxC-[Photorhabdus-australis] |
| WP_149617320.1 | Type1.BREX | [illegible] | 1213 | Bacteria>Pseudomonadota>Gammaproteobacteria>Enterobacterales>Morganellaceae>Photorhabdus | GCF_008370335.1 | Photorhabdus_heterorhabditis | FOL16_RS17495 | 23615..27256 | 880156 | BREX-system-P-loop-protein-BrxC-[Photorhabdus-heterorhabditis] |
| WP_166309887.1 | Type1.BREX | [illegible] | 1213 | Bacteria>Pseudomonadota>Gammaproteobacteria>Enterobacterales>Morganellaceae>Photorhabdus | GCF_011189575.1 | Photorhabdus_cinerea | C5469_RS18920 | 23326..26967 | 471575 | BREX-system-P-loop-protein-BrxC-[Photorhabdus-cinerea] |
| WP_172909759.1 | Type1.BREX | [illegible] | 1213 | Bacteria>Pseudomonadota>Gammaproteobacteria>Enterobacterales>Morganellaceae>Photorhabdus | GCF_013277735.1 | Photorhabdus_heterorhabditis | HHJ84_RS11995 | 18661..22302 | 880156 | BREX-system-P-loop-protein-BrxC-[Photorhabdus-heterorhabditis] |
| WP_214086333.1 | Type1.BREX | [illegible] | 1213 | Bacteria>Pseudomonadota>Gammaproteobacteria>Enterobacterales>Morganellaceae>Photorhabdus | GCF_018448905.1 | Photorhabdus_heterorhabditis | EAE89_RS17790 | 25052..28693 | 880156 | BREX-system-P-loop-protein-BrxC-[Photorhabdus-heterorhabditis] |
| QIG07578.1 | Type1.BREX | [illegible] | 1183 | Bacteria>Pseudomonadota>Gammaproteobacteria>Enterobacterales>Morganellaceae>Proteus | GCA_011046025.1 | Proteus_sp._ZN5 | GTK47_19170 | 4181966..4185517 | 2697019 | BREX-system-P-loop-protein-BrxC-[Proteus-sp.-ZN5] |
| HAT2164885.1 | Type1.BREX | [illegible] | 1225 | Bacteria>Pseudomonadota>Gammaproteobacteria>Enterobacterales>Morganellaceae>Proteus | GCA_015896365.1 | Proteus_mirabilis | I8052_002910 | 10079..13756 | 584 | BREX-system-P-loop-protein-BrxC-[Proteus-mirabilis] |
| MBI6417160.1 | Type1.BREX | [illegible] | 1225 | Bacteria>Pseudomonadota>Gammaproteobacteria>Enterobacterales>Morganellaceae>Proteus | GCA_016238675.1 | Proteus_mirabilis | JEP37_15655 | 69055..72732 | 584 | BREX-system-P-loop-protein-BrxC-[Proteus-mirabilis] |
| HBM9710663.1 | Type1.BREX | [illegible] | 1225 | Bacteria>Pseudomonadota>Gammaproteobacteria>Enterobacterales>Morganellaceae>Proteus | GCA_021439885.1 | Proteus_mirabilis | L1C35_000984 | 2327..6004 | 584 | BREX-system-P-loop-protein-BrxC-[Proteus-mirabilis] |
| WP_087726352.1 | Type1.BREX | [illegible] | 1229 | Bacteria>Pseudomonadota>Gammaproteobacteria>Enterobacterales>Morganellaceae>Proteus | GCF_002180235.1 | Proteus_mirabilis | NFG13_RS03655 | 19477..23166 | 584 | BREX-system-P-loop-protein-BrxC-[Proteus-mirabilis] |
| WP_152118569.1 | Type1.BREX | [illegible] | 1225 | Bacteria>Pseudomonadota>Gammaproteobacteria>Enterobacterales>Morganellaceae>Proteus | GCF_009183735.1 | Proteus_mirabilis | GBN13_RS03935 | 69067..72744 | 584 | BREX-system-P-loop-protein-BrxC-[Proteus-mirabilis] |
| WP_206535871.1 | Type1.BREX | [illegible] | 1185 | Bacteria>Pseudomonadota>Gammaproteobacteria>Enterobacterales>Morganellaceae>Proteus | GCF_011046025.1 | Proteus_sp._ZN5 | GTK47_RS19230 | 4181960..4185517 | 2697019 | BREX-system-P-loop-protein-BrxC-[Proteus-sp.-ZN5] |
| WP_165544872.1 | Type1.BREX | [illegible] | 1225 | Bacteria>Pseudomonadota>Gammaproteobacteria>Enterobacterales>Morganellaceae>Proteus | GCF_011149675.1 | Proteus_mirabilis | G9C79_RS07990 | 1716006..1719683 | 584 | BREX-system-P-loop-protein-BrxC-[Proteus-mirabilis] |
| WP_209124420.1 | Type1.BREX | [illegible] | 761 | Bacteria>Pseudomonadota>Gammaproteobacteria>Enterobacterales>Morganellaceae>Proteus | GCF_017693315.1 | Proteus_mirabilis | IS513_RS13265 | 1..2284 | 584 | BREX-system-P-loop-protein-BrxC,-partial-[Proteus-mirabilis] |
| WP_220699168.1 | Type1.BREX | [illegible] | 1225 | Bacteria>Pseudomonadota>Gammaproteobacteria>Enterobacterales>Morganellaceae>Proteus | GCF_019660825.1 | Proteus_mirabilis | K2V98_RS14125 | 5608..9285 | 584 | BREX-system-P-loop-protein-BrxC-[Proteus-mirabilis] |
| WP_210849185.1 | Type1.BREX | [illegible] | 1225 | Bacteria>Pseudomonadota>Gammaproteobacteria>Enterobacterales>Morganellaceae>Proteus | GCF_022349905.1 | Proteus_mirabilis | MJ636_RS02915 | 47390..51067 | 584 | BREX-system-P-loop-protein-BrxC-[Proteus-mirabilis] |
| WP_243213747.1 | Type1.BREX | [illegible] | 1225 | Bacteria>Pseudomonadota>Gammaproteobacteria>Enterobacterales>Morganellaceae>Proteus | GCF_022802355.1 | Proteus_mirabilis | JNG69_RS16550 | 64025..67702 | 584 | BREX-system-P-loop-protein-BrxC-[Proteus-mirabilis] |
| WP_261229609.1 | Type1.BREX | [illegible] | 1225 | Bacteria>Pseudomonadota>Gammaproteobacteria>Enterobacterales>Morganellaceae>Proteus | GCF_025379495.1 | Proteus_mirabilis | NFG18_RS15790 | 23870..27547 | 584 | BREX-system-P-loop-protein-BrxC-[Proteus-mirabilis] |
| WP_261598777.1 | Type1.BREX | [illegible] | 1225 | Bacteria>Pseudomonadota>Gammaproteobacteria>Enterobacterales>Morganellaceae>Proteus | GCF_025399735.1 | Proteus_mirabilis | NMC40_RS18460 | 15597..19274 | 584 | BREX-system-P-loop-protein-BrxC-[Proteus-mirabilis] |
| WP_271972693.1 | Type1.BREX | [illegible] | 1225 | Bacteria>Pseudomonadota>Gammaproteobacteria>Enterobacterales>Morganellaceae>Proteus | GCF_028335785.1 | Proteus_terrae | ONR71_RS01795 | 360053..363730 | 1574161 | BREX-system-P-loop-protein-BrxC-[Proteus-terrae] |
| WP_209036130.1 | Type1.BREX | [illegible] | 1225 | Bacteria>Pseudomonadota>Gammaproteobacteria>Enterobacterales>Morganellaceae>Proteus | GCF_029542405.1 | Proteus_mirabilis | IGB04_RS13130 | 82611..86288 | 584 | BREX-system-P-loop-protein-BrxC-[Proteus-mirabilis] |
| UVX22921.1 | Type1.BREX | [illegible] | 789 | Bacteria>Pseudomonadota>Gammaproteobacteria>Enterobacterales>Morganellaceae>Proteus | - | Proteus_terrae_subsp._cibarius | . | 73397..75766 | 626774 | hypothetical-protein-[Proteus-terrae-subsp.-cibarius] |
| OBY34180.1 | Type1.BREX | [illegible] | 1225 | Bacteria>Pseudomonadota>Gammaproteobacteria>Enterobacterales>Morganellaceae>Providencia | GCA_001680125.1 | Providencia_rettgeri | PR729_03900 | 44936..48613 | 587 | DNA-repair-protein-[Providencia-rettgeri] |
| EJD6671300.1 | Type1.BREX | [illegible] | 1225 | Bacteria>Pseudomonadota>Gammaproteobacteria>Enterobacterales>Morganellaceae>Providencia | GCA_020680725.2 | Providencia_rettgeri | MOK80_RS10540 | 473799..477476 | 587 | BREX-system-P-loop-protein-BrxC-[Providencia-rettgeri] |
| MCL0017528.1 | Type1.BREX | [illegible] | 1225 | Bacteria>Pseudomonadota>Gammaproteobacteria>Enterobacterales>Morganellaceae>Providencia | GCA_022387615.2 | Providencia_rettgeri | MKT11_000360 | 78084..81761 | 587 | BREX-system-P-loop-protein-BrxC-[Providencia-rettgeri] |
| MCK9997621.1 | Type1.BREX | [illegible] | 1225 | Bacteria>Pseudomonadota>Gammaproteobacteria>Enterobacterales>Morganellaceae>Providencia | GCA_022387885.2 | Providencia_rettgeri | MKT39_000560 | 19282..22959 | 587 | BREX-system-P-loop-protein-BrxC-[Providencia-rettgeri] |
| MCL0006618.1 | Type1.BREX | [illegible] | 1225 | Bacteria>Pseudomonadota>Gammaproteobacteria>Enterobacterales>Morganellaceae>Providencia | GCA_022388475.2 | Providencia_rettgeri | MKT60_007090 | 64410..68087 | 587 | BREX-system-P-loop-protein-BrxC-[Providencia-rettgeri] |
| WP_137022260.1 | Type1.BREX | [illegible] | 1204 | Bacteria>Pseudomonadota>Gammaproteobacteria>Enterobacterales>Morganellaceae>Providencia | GCF_005157325.1 | Providencia_heimbachae | C9446_RS17890 | 3853703..3857317 | 333962 | BREX-system-P-loop-protein-BrxC-[Providencia-heimbachae] |
| WP_166185547.1 | Type1.BREX | [illegible] | 1225 | Bacteria>Pseudomonadota>Gammaproteobacteria>Enterobacterales>Morganellaceae>Providencia | GCF_011078165.1 | Providencia_sp._M-8 | G6Q30_RS18775 | 21019..24696 | 2713151 | BREX-system-P-loop-protein-BrxC-[Providencia-sp.-M-8] |
| WP_265497329.1 | Type1.BREX | [illegible] | 1225 | Bacteria>Pseudomonadota>Gammaproteobacteria>Enterobacterales>Morganellaceae>Providencia | GCF_026173365.1 | Providencia_rustigianii | ORM28_RS01805 | 9364..13041 | 158850 | BREX-system-P-loop-protein-BrxC-[Providencia-rustigianii] |
| WP_269723554.1 | Type1.BREX | [illegible] | 1225 | Bacteria>Pseudomonadota>Gammaproteobacteria>Enterobacterales>Morganellaceae>Providencia | GCF_027362595.1 | Providencia_sp._210H12SH02B-Prov | O7C57_RS02370 | 500730..504407 | 3015951 | BREX-system-P-loop-protein-BrxC-[Providencia-sp.-210H12SH02B-Prov] |
| WP_272572142.1 | Type1.BREX | [illegible] | 1225 | Bacteria>Pseudomonadota>Gammaproteobacteria>Enterobacterales>Morganellaceae>Providencia | GCF_028477905.1 | unclassified_Providencia | PR658_RS19440 | 8800..12477 | 2633465 | MULTISPECIES:-BREX-system-P-loop-protein-BrxC-[unclassified-Providencia] |
| WP_272670012.1 | Type1.BREX | [illegible] | 1185 | Bacteria>Pseudomonadota>Gammaproteobacteria>Enterobacterales>Morganellaceae>Providencia | GCF_028480605.1 | unclassified_Providencia | PSE59_RS21125 | 8138..11695 | 2633465 | MULTISPECIES:-BREX-system-P-loop-protein-BrxC-[unclassified-Providencia] |
| WP_272668877.1 | Type1.BREX | [illegible] | 1225 | Bacteria>Pseudomonadota>Gammaproteobacteria>Enterobacterales>Morganellaceae>Providencia | GCF_028481005.1 | Providencia_sp._PROV151 | PSE52_RS00990 | 220703..224380 | 2949861 | BREX-system-P-loop-protein-BrxC-[Providencia-sp.-PROV151] |
| WP_272687949.1 | Type1.BREX | [illegible] | 1225 | Bacteria>Pseudomonadota>Gammaproteobacteria>Enterobacterales>Morganellaceae>Providencia | GCF_028484835.1 | Providencia_sp._PROV035 | PSF71_RS08200 | 5027..8704 | 2949766 | BREX-system-P-loop-protein-BrxC-[Providencia-sp.-PROV035] |
| AAM07998.1 | Type1.BREX | [illegible] | 947 | Bacteria>Pseudomonadota>Gammaproteobacteria>Enterobacterales>Morganellaceae>Providencia | - | Providencia_rettgeri | . | 15799..18642 | 587 | putative-ATPase-[Providencia-rettgeri] |
| CAD27664.1 | Type1.BREX | [illegible] | 485 | Bacteria>Pseudomonadota>Gammaproteobacteria>Enterobacterales>Morganellaceae>Providencia | - | Providencia_rettgeri | . | 6214..7668 | 587 | hypothetical-protein,-partial-[Providencia-rettgeri] |
| EIP8110719.1 | Type1.BREX | [illegible] | 1225 | Bacteria>Pseudomonadota>Gammaproteobacteria>Enterobacterales>Morganellaceae>Providencia | - | Providencia_rettgeri | LT185_001364 | 559434..563111 | 587 | BREX-system-P-loop-protein-BrxC-[Providencia-rettgeri] |
| WP_045967689.1 | Type1.BREX | [illegible] | 1212 | Bacteria>Pseudomonadota>Gammaproteobacteria>Enterobacterales>Morganellaceae>Xenorhabdus | GCF_000968195.1 | Xenorhabdus_doucetiae | LY16_RS12985 | 24233..27871 | 351671 | BREX-system-P-loop-protein-BrxC-[Xenorhabdus-doucetiae] |
| WP_121588635.1 | Type1.BREX | [illegible] | 1213 | Bacteria>Pseudomonadota>Gammaproteobacteria>Enterobacterales>Pectobacteriaceae>Brenneria | GCF_022358645.1 | Brenneria_goodwinii | JMY82_RS09645 | 95315..98956 | 1109412 | BREX-system-P-loop-protein-BrxC-[Brenneria-goodwinii] |
| WP_102802217.1 | Type1.BREX | [illegible] | 1213 | Bacteria>Pseudomonadota>Gammaproteobacteria>Enterobacterales>Pectobacteriaceae>Dickeya | GCF_002887555.1 | Dickeya_zeae | C1O30_RS16230 | 3581616..3585257 | 204042 | BREX-system-P-loop-protein-BrxC-[Dickeya-zeae] |
| WP_107760228.1 | Type1.BREX | [illegible] | 1213 | Bacteria>Pseudomonadota>Gammaproteobacteria>Enterobacterales>Pectobacteriaceae>Dickeya | GCF_003049805.1 | Dickeya_sp._Secpp_1600 | CPA59_RS17580 | 3939970..3943611 | 2037915 | BREX-system-P-loop-protein-BrxC-[Dickeya-sp.-Secpp-1600] |
| WP_040031759.1 | Type1.BREX | [illegible] | 1202 | Bacteria>Pseudomonadota>Gammaproteobacteria>Enterobacterales>Pectobacteriaceae | GCF_026723725.1 | Pectobacterium | GWC53_RS08230 | 1782306..1785914 | 122277 | MULTISPECIES:-BREX-system-P-loop-protein-BrxC-[Pectobacterium] |
| GKV76958.1 | Type1.BREX | [illegible] | 1157 | Bacteria>Pseudomonadota>Gammaproteobacteria>Enterobacterales>Pectobacteriaceae>Pectobacterium | GCA_024342785.1 | Pectobacterium_carotovorum_subsp._carotovorum | PEC106568_21320 | 196499..199972 | 555 | hypothetical-protein-PEC106568_21320-[Pectobacterium-carotovorum-subsp.-carotovorum] |
| GKX39595.1 | Type1.BREX | [illegible] | 1202 | Bacteria>Pseudomonadota>Gammaproteobacteria>Enterobacterales>Pectobacteriaceae>Pectobacterium | GCA_024343445.1 | Pectobacterium_carotovorum_subsp._carotovorum | SOASR014_33340 | 124830..128438 | 555 | hypothetical-protein-SOASR014_33340-[Pectobacterium-carotovorum-subsp.-carotovorum] |
| WP_005974434.1 | Type1.BREX | [illegible] | 1204 | Bacteria>Pseudomonadota>Gammaproteobacteria>Enterobacterales>Pectobacteriaceae>Pectobacterium | GCF_000291725.1 | Pectobacterium_wasabiae | Y17_RS17690 | 22336..25950 | 55208 | BREX-system-P-loop-protein-BrxC-[Pectobacterium-wasabiae] |
| WP_014916385.1 | Type1.BREX | [illegible] | 1213 | Bacteria>Pseudomonadota>Gammaproteobacteria>Enterobacterales>Pectobacteriaceae>Pectobacterium | GCF_000294535.1 | Pectobacterium_carotovorum | PCC21_RS16225 | 3704007..3707648 | 554 | BREX-system-P-loop-protein-BrxC-[Pectobacterium-carotovorum] |
| WP_103160989.1 | Type1.BREX | [illegible] | 1202 | Bacteria>Pseudomonadota>Gammaproteobacteria>Enterobacterales>Pectobacteriaceae>Pectobacterium | GCF_002904045.1 | Pectobacterium_odoriferum | BVY05_RS19610 | 74039..77647 | 78398 | BREX-system-P-loop-protein-BrxC-[Pectobacterium-odoriferum] |
| WP_103199124.1 | Type1.BREX | [illegible] | 1202 | Bacteria>Pseudomonadota>Gammaproteobacteria>Enterobacterales>Pectobacteriaceae>Pectobacterium | GCF_002904085.1 | Pectobacterium_odoriferum | BV921_RS15390 | 113976..117584 | 78398 | BREX-system-P-loop-protein-BrxC-[Pectobacterium-odoriferum] |
| WP_116155987.1 | Type1.BREX | [illegible] | 1202 | Bacteria>Pseudomonadota>Gammaproteobacteria>Enterobacterales>Pectobacteriaceae>Pectobacterium | GCF_003382585.2 | Pectobacterium_aquaticum | DMB81_RS10735 | 4900..8508 | 2204145 | BREX-system-P-loop-protein-BrxC-[Pectobacterium-aquaticum] |
| WP_181847885.1 | Type1.BREX | [illegible] | 1204 | Bacteria>Pseudomonadota>Gammaproteobacteria>Enterobacterales>Pectobacteriaceae>Pectobacterium | GCF_013913645.1 | Pectobacterium_aroidearum | H2Y54_RS11285 | 236602..240216 | 1201031 | BREX-system-P-loop-protein-BrxC-[Pectobacterium-aroidearum] |
| WP_211024717.1 | Type1.BREX | [illegible] | 1157 | Bacteria>Pseudomonadota>Gammaproteobacteria>Enterobacterales>Pectobacteriaceae>Pectobacterium | GCF_018094745.1 | Pectobacterium_versatile | F9U42_RS09295 | 89992..93465 | 2488639 | BREX-system-P-loop-protein-BrxC-[Pectobacterium-versatile] |
| WP_249681848.1 | Type1.BREX | [illegible] | 1157 | Bacteria>Pseudomonadota>Gammaproteobacteria>Enterobacterales>Pectobacteriaceae>Pectobacterium | - | Pectobacterium_polaris | EXT50_03115 | 82893..86366 | 2042057 | BREX-system-P-loop-protein-BrxC-[Pectobacterium-polaris] |
| WP_060426782.1 | Type1.BREX | [illegible] | 1213 | Bacteria>Pseudomonadota>Gammaproteobacteria>Enterobacterales>Yersiniaceae | GCF_001537885.1 | Serratia | AWX06_RS15845 | 182718..186359 | 613 | MULTISPECIES:-BREX-system-P-loop-protein-BrxC-[Serratia] |
| WP_050135880.1 | Type1.BREX | [illegible] | 1225 | Bacteria>Pseudomonadota>Gammaproteobacteria>Enterobacterales>Yersiniaceae | GCF_025758435.1 | Yersinia | N4W04_RS19100 | 4124889..4128566 | 629 | MULTISPECIES:-BREX-system-P-loop-protein-BrxC-[Yersinia] |
| WP_138775748.1 | Type1.BREX | [illegible] | 1202 | Bacteria>Pseudomonadota>Gammaproteobacteria>Enterobacterales>Yersiniaceae | GCF_902168625.1 | Yersinia | FFE93_018510 | 3893626..3897234 | 629 | MULTISPECIES:-BREX-system-P-loop-protein-BrxC-[Yersinia] |
| WP_112197427.1 | Type1.BREX | [illegible] | 1204 | Bacteria>Pseudomonadota>Gammaproteobacteria>Enterobacterales>Yersiniaceae>Rahnella | GCF_003263525.1 | Rahnella_sp._NRRL_B-41462 | UC14_RS06670 | 31474..35088 | 1610579 | BREX-system-P-loop-protein-BrxC-[Rahnella-sp.-NRRL-B-41462] |
| WP_217147154.1 | Type1.BREX | [illegible] | 1157 | Bacteria>Pseudomonadota>Gammaproteobacteria>Enterobacterales>Yersiniaceae>Rahnella | GCF_019049515.1 | Rahnella_aceris | J1789_RS04120 | 11901..15374 | 2703885 | BREX-system-P-loop-protein-BrxC-[Rahnella-aceris] |
| WP_232770162.1 | Type1.BREX | [illegible] | 1157 | Bacteria>Pseudomonadota>Gammaproteobacteria>Enterobacterales>Yersiniaceae>Rahnella | GCF_021276285.1 | Rahnella_victoriana | J9880_RS04875 | 1051789..1055262 | 1510570 | BREX-system-P-loop-protein-BrxC-[Rahnella-victoriana] |
| WP_266057125.1 | Type1.BREX | [illegible] | 1157 | Bacteria>Pseudomonadota>Gammaproteobacteria>Enterobacterales>Yersiniaceae>Rahnella | GCF_026241995.1 | Rahnella_perminowiae | ORG37_RS00070 | 21089..24562 | 2816244 | BREX-system-P-loop-protein-BrxC-[Rahnella-perminowiae] |
| NYA44684.1 | Type1.BREX | [illegible] | 1158 | Bacteria>Pseudomonadota>Gammaproteobacteria>Enterobacterales>Yersiniaceae>Serratia | GCA_013401715.1 | Serratia_fonticola | HZI31_15400 | 48979..52455 | 47917 | BREX-system-P-loop-protein-BrxC-[Serratia-fonticola] |
| EGS9996488.1 | Type1.BREX | [illegible] | 1233 | Bacteria>Pseudomonadota>Gammaproteobacteria>Enterobacterales>Yersiniaceae>Serratia | GCA_015929585.1 | Serratia_marcescens | I9Y15_003567 | 178923..182624 | 615 | BREX-system-P-loop-protein-BrxC-[Serratia-marcescens] |
| HBE9089660.1 | Type1.BREX | [illegible] | 1204 | Bacteria>Pseudomonadota>Gammaproteobacteria>Enterobacterales>Yersiniaceae>Serratia | GCA_018627075.1 | Serratia_fonticola | KNG84_001684 | 34672..38286 | 47917 | BREX-system-P-loop-protein-BrxC-[Serratia-fonticola] |
| HBK4794038.1 | Type1.BREX | [illegible] | 1233 | Bacteria>Pseudomonadota>Gammaproteobacteria>Enterobacterales>Yersiniaceae>Serratia | GCA_020682225.1 | Serratia_marcescens | LLT15_004323 | 35577..39278 | 615 | BREX-system-P-loop-protein-BrxC-[Serratia-marcescens] |
| WP_089185250.1 | Type1.BREX | [illegible] | 1213 | Bacteria>Pseudomonadota>Gammaproteobacteria>Enterobacterales>Yersiniaceae>Serratia | GCF_002220535.1 | Serratia_marcescens | BVG90_RS23850 | 5002559..5006200 | 615 | BREX-system-P-loop-protein-BrxC-[Serratia-marcescens] |
| WP_218833283.1 | Type1.BREX | [illegible] | 1197 | Bacteria>Pseudomonadota>Gammaproteobacteria>Enterobacterales>Yersiniaceae>Serratia | GCF_013401715.1 | Serratia_fonticola | HZI31_RS15400 | 48979..52572 | 47917 | BREX-system-P-loop-protein-BrxC-[Serratia-fonticola] |
| WP_197785670.1 | Type1.BREX | [illegible] | 1213 | Bacteria>Pseudomonadota>Gammaproteobacteria>Enterobacterales>Yersiniaceae>Serratia | GCF_016007555.1 | Serratia_ureilytica | I5P92_RS12860 | 24369..28010 | 300181 | BREX-system-P-loop-protein-BrxC-[Serratia-ureilytica] |
| WP_206196473.1 | Type1.BREX | [illegible] | 1157 | Bacteria>Pseudomonadota>Gammaproteobacteria>Enterobacterales>Yersiniaceae>Serratia | GCF_017152235.1 | Serratia_ureilytica | JY544_RS23375 | 37448..40921 | 300181 | BREX-system-P-loop-protein-BrxC-[Serratia-ureilytica] |
| WP_212602139.1 | Type1.BREX | [illegible] | 1213 | Bacteria>Pseudomonadota>Gammaproteobacteria>Enterobacterales>Yersiniaceae>Serratia | GCF_018257315.1 | Serratia_rubidaea | JK228_RS02460 | 34162..37803 | 61652 | BREX-system-P-loop-protein-BrxC-[Serratia-rubidaea] |
| WP_222398282.1 | Type1.BREX | [illegible] | 1213 | Bacteria>Pseudomonadota>Gammaproteobacteria>Enterobacterales>Yersiniaceae>Serratia | GCF_019795995.1 | Serratia_marcescens | L8N14_16510 | 93200..96841 | 615 | BREX-system-P-loop-protein-BrxC-[Serratia-marcescens] |
| WP_223536794.1 | Type1.BREX | [illegible] | 1158 | Bacteria>Pseudomonadota>Gammaproteobacteria>Enterobacterales>Yersiniaceae>Serratia | GCF_019968705.1 | unclassified_Serratia_(in:_enterobacteria) | KGP26_RS00385 | 95452..98928 | 2647522 | MULTISPECIES:-BREX-system-P-loop-protein-BrxC-[unclassified-Serratia-(in:-enterobacteria)] |
| WP_244757920.1 | Type1.BREX | [illegible] | 1213 | Bacteria>Pseudomonadota>Gammaproteobacteria>Enterobacterales>Yersiniaceae>Serratia | GCF_022870585.1 | Serratia_marcescens | HW877_RS02460 | 535877..539518 | 615 | BREX-system-P-loop-protein-BrxC-[Serratia-marcescens] |
| WP_260965787.1 | Type1.BREX | [illegible] | 1240 | Bacteria>Pseudomonadota>Gammaproteobacteria>Enterobacterales>Yersiniaceae>Serratia | GCF_025336405.1 | Serratia_marcescens | N7751_RS05105 | 1087639..1091361 | 615 | BREX-system-P-loop-protein-BrxC-[Serratia-marcescens] |
| WP_273866014.1 | Type1.BREX | [illegible] | 1157 | Bacteria>Pseudomonadota>Gammaproteobacteria>Enterobacterales>Yersiniaceae>Serratia | GCF_028616515.1 | Serratia_ureilytica | PVU68_RS17145 | 216123..219596 | 300181 | BREX-system-P-loop-protein-BrxC-[Serratia-ureilytica] |
| WP_261139451.1 | Type1.BREX | [illegible] | 1213 | Bacteria>Pseudomonadota>Gammaproteobacteria>Enterobacterales>Yersiniaceae>Serratia | GCF_946226675.1 | Serratia_liquefaciens | N8M82_RS16050 | 322435..326076 | 614 | BREX-system-P-loop-protein-BrxC-[Serratia-liquefaciens] |
| WP_261135874.1 | Type1.BREX | [illegible] | 1213 | Bacteria>Pseudomonadota>Gammaproteobacteria>Enterobacterales>Yersiniaceae>Serratia | GCF_946226775.1 | Serratia_liquefaciens | N8M94_RS14775 | 322685..326326 | 614 | BREX-system-P-loop-protein-BrxC-[Serratia-liquefaciens] |
| WP_261427666.1 | Type1.BREX | [illegible] | 1213 | Bacteria>Pseudomonadota>Gammaproteobacteria>Enterobacterales>Yersiniaceae>Serratia | GCF_946227925.1 | Serratia_proteamaculans | OCH17_RS14295 | 357572..361213 | 28151 | BREX-system-P-loop-protein-BrxC-[Serratia-proteamaculans] |
| WP_262254721.1 | Type1.BREX | [illegible] | 1240 | Bacteria>Pseudomonadota>Gammaproteobacteria>Enterobacterales>Yersiniaceae>Serratia | GCF_946228755.1 | Serratia_proteamaculans | OC575_RS10145 | 2133622..2137344 | 28151 | BREX-system-P-loop-protein-BrxC-[Serratia-proteamaculans] |
| WP_195869502.1 | Type1.BREX | [illegible] | 1202 | Bacteria>Pseudomonadota>Gammaproteobacteria>Enterobacterales>Yersiniaceae>Serratia | - | Serratia_liquefaciens | I1A46_RS14725 | 213767..217375 | 614 | BREX-system-P-loop-protein-BrxC-[Serratia-liquefaciens] |
| WP_004722527.1 | Type1.BREX | [illegible] | 1202 | Bacteria>Pseudomonadota>Gammaproteobacteria>Enterobacterales>Yersiniaceae>Yersinia | GCF_000173755.1 | Yersinia_ruckeri | YRUCK0001_RS14080 | 15308..18916 | 29486 | BREX-system-P-loop-protein-BrxC-[Yersinia-ruckeri] |
| WP_046051668.1 | Type1.BREX | [illegible] | 1157 | Bacteria>Pseudomonadota>Gammaproteobacteria>Enterobacterales>Yersiniaceae>Yersinia | GCF_000968115.1 | Yersinia_enterocolitica | YE5303_RS19630 | 4225906..4229379 | 630 | BREX-system-P-loop-protein-BrxC-[Yersinia-enterocolitica] |
| WP_050073003.1 | Type1.BREX | [illegible] | 1157 | Bacteria>Pseudomonadota>Gammaproteobacteria>Enterobacterales>Yersiniaceae>Yersinia | GCF_001088505.1 | Yersinia_intermedia | AE858_RS05265 | 31386..34859 | 631 | BREX-system-P-loop-protein-BrxC-[Yersinia-intermedia] |
| WP_050124850.1 | Type1.BREX | [illegible] | 1202 | Bacteria>Pseudomonadota>Gammaproteobacteria>Enterobacterales>Yersiniaceae>Yersinia | GCF_001115185.1 | Yersinia_aleksiciae | AEB18_RS00865 | 216657..220265 | 263819 | BREX-system-P-loop-protein-BrxC-[Yersinia-aleksiciae] |
| WP_053011787.1 | Type1.BREX | [illegible] | 1202 | Bacteria>Pseudomonadota>Gammaproteobacteria>Enterobacterales>Yersiniaceae>Yersinia | GCF_001244675.1 | Yersinia_intermedia | ALC37_RS11575 | 459968..463576 | 631 | BREX-system-P-loop-protein-BrxC-[Yersinia-intermedia] |
| WP_128451204.1 | Type1.BREX | [illegible] | 1202 | Bacteria>Pseudomonadota>Gammaproteobacteria>Enterobacterales>Yersiniaceae>Yersinia | GCF_004100405.1 | Yersinia_sp._2105_StPb_PI | EQP49_RS07200 | 292011..295619 | 2507058 | BREX-system-P-loop-protein-BrxC-[Yersinia-sp.-2105-StPb-PI] |
| WP_145931270.1 | Type1.BREX | [illegible] | 1202 | Bacteria>Pseudomonadota>Gammaproteobacteria>Enterobacterales>Yersiniaceae>Yersinia | GCF_902168565.1 | Yersinia_bercovieri | FS908_RS01775 | 53708..57316 | 634 | BREX-system-P-loop-protein-BrxC-[Yersinia-bercovieri] |
| WP_145532653.1 | Type1.BREX | [illegible] | 1202 | Bacteria>Pseudomonadota>Gammaproteobacteria>Enterobacterales>Yersiniaceae>Yersinia | GCF_902170345.1 | Yersinia_alsatica | FTB07_RS01240 | 22698..26306 | 2890317 | BREX-system-P-loop-protein-BrxC-[Yersinia-alsatica] |
| WP_145580439.1 | Type1.BREX | [illegible] | 1202 | Bacteria>Pseudomonadota>Gammaproteobacteria>Enterobacterales>Yersiniaceae>Yersinia | GCF_902170795.1 | Yersinia_mollaretii | FTB33_RS16505 | 6738..10346 | 33060 | BREX-system-P-loop-protein-BrxC-[Yersinia-mollaretii] |
| MBZ4194014.1 | Type1.BREX | [illegible] | 1138 | Bacteria>Pseudomonadota>Gammaproteobacteria>Gammaproteobacteria_incertae_sedis>Candidatus_Competibacteraceae>Candidatus_Contendobacter | GCA_020035205.1 | Candidatus_Contendobacter_sp. | LAE24_06875 | 5150..8566 | 2651161 | BREX-system-P-loop-protein-BrxC-[Candidatus-Contendobacter-sp.] |
| WP_034430846.1 | Type1.BREX | [illegible] | 1138 | Bacteria>Pseudomonadota>Gammaproteobacteria>Gammaproteobacteria_incertae_sedis>Candidatus_Competibacteraceae>Candidatus_Contendobacter | GCF_000531125.1 | Candidatus_Contendobacter_odensis | BN874_RS03435 | 764..4180 | 1400860 | BREX-system-P-loop-protein-BrxC-[Candidatus-Contendobacter-odensis] |
| WP_132542053.1 | Type1.BREX | [illegible] | 1153 | Bacteria>Pseudomonadota>Gammaproteobacteria>Gammaproteobacteria_incertae_sedis>Candidatus_Competibacteraceae>Plasticicumulans | GCF_004341245.1 | Plasticicumulans_lactativorans | EV699_RS09620 | 184945..188406 | 1133106 | BREX-system-P-loop-protein-BrxC-[Plasticicumulans-lactativorans] |
| MBK8752263.1 | Type1.BREX | [illegible] | 1139 | Bacteria>Pseudomonadota>Gammaproteobacteria>Gammaproteobacteria_incertae_sedis>Candidatus_Competibacteraceae>unclassified_Candidatus_Competibacteraceae | GCA_016710535.1 | Candidatus_Competibacteraceae_bacterium | IPL99_11760 | 9123..12542 | 2053538 | BREX-system-P-loop-protein-BrxC-[Candidatus-Competibacteraceae-bacterium] |
| MBK8753015.1 | Type1.BREX | [illegible] | 1154 | Bacteria>Pseudomonadota>Gammaproteobacteria>Gammaproteobacteria_incertae_sedis>Candidatus_Competibacteraceae>unclassified_Candidatus_Competibacteraceae | GCA_016710535.1 | Candidatus_Competibacteraceae_bacterium | IPL99_15935 | 14692..18156 | 2053538 | BREX-system-P-loop-protein-BrxC-[Candidatus-Competibacteraceae-bacterium] |
| MBK8536094.1 | Type1.BREX | [illegible] | 491 | Bacteria>Pseudomonadota>Gammaproteobacteria>Gammaproteobacteria_incertae_sedis>Candidatus_Competibacteraceae>unclassified_Candidatus_Competibacteraceae | GCA_016710825.1 | Candidatus_Competibacteraceae_bacterium | IPL59_13760 | 11734..13209 | 2053538 | BREX-system-P-loop-protein-BrxC-[Candidatus-Competibacteraceae-bacterium] |
| MCB1769904.1 | Type1.BREX | [illegible] | 760 | Bacteria>Pseudomonadota>Gammaproteobacteria>Gammaproteobacteria_incertae_sedis>Candidatus_Competibacteraceae>unclassified_Candidatus_Competibacteraceae | GCA_020445005.1 | Candidatus_Competibacteraceae_bacterium | KDJ31_09455 | 11685..13969 | 2053538 | BREX-system-P-loop-protein-BrxC-[Candidatus-Competibacteraceae-bacterium] |
| WP_271768605.1 | Type1.BREX | [illegible] | 1154 | Bacteria>Pseudomonadota>Gammaproteobacteria>Gammaproteobacteria_incertae_sedis>Candidatus_Pseudothioglobus | GCF_028273125.1 | Candidatus_Pseudothioglobus_singularis | OAH81_RS01105 | 228598..232062 | 1427364 | BREX-system-P-loop-protein-BrxC-[Candidatus-Pseudothioglobus-singularis] |
| WP_248566606.1 | Type1.BREX | [illegible] | 1145 | Bacteria>Pseudomonadota>Gammaproteobacteria>Gammaproteobacteria_incertae_sedis>Methylonatrum | GCF_023195885.1 | Methylonatrum_kenyense | MO534_RS01350 | 290769..294206 | 455253 | BREX-system-P-loop-protein-BrxC-[Methylonatrum-kenyense] |
| HDZ09174.1 | Type1.BREX | [illegible] | 1208 | Bacteria>Pseudomonadota>Gammaproteobacteria>Gammaproteobacteria_incertae_sedis>Pseudohongiella>unclassified_Pseudohongiella | GCA_011050365.1 | Pseudohongiella_sp. | ENH71_08845 | 47068..50694 | 1979412 | BREX-system-P-loop-protein-BrxC-[Pseudohongiella-sp.] |
| WP_029131813.1 | Type1.BREX | [illegible] | 1147 | Bacteria>Pseudomonadota>Gammaproteobacteria>Gammaproteobacteria_incertae_sedis>Sedimenticola | GCF_000428045.1 | Sedimenticola_selenatireducens | A3GD_RS0101005 | 196433..199876 | 191960 | BREX-system-P-loop-protein-BrxC-[Sedimenticola-selenatireducens] |
| MBT4669056.1 | Type1.BREX | [illegible] | 1167 | Bacteria>Pseudomonadota>Gammaproteobacteria>Gammaproteobacteria_incertae_sedis>sulfur-oxidizing_symbionts>Candidatus_Ruthia | GCA_018657745.1 | Candidatus_Ruthia_sp. | HN615_06440 | 8657..12160 | 2735566 | BREX-system-P-loop-protein-BrxC-[Candidatus-Ruthturnera-sp.] |
| MCG7867606.1 | Type1.BREX | [illegible] | 1172 | Bacteria>Pseudomonadota>Gammaproteobacteria>Gammaproteobacteria_incertae_sedis>sulfur-oxidizing_symbionts>Candidatus_Thiodiazotropha | GCA_016842585.1 | Candidatus_Thiodiazotropha_taylori | JAY74_14755 | 23622..27140 | 2792791 | BREX-system-P-loop-protein-BrxC-[Candidatus-Thiodiazotropha-taylori] |
| MCG8047516.1 | Type1.BREX | [illegible] | 1147 | Bacteria>Pseudomonadota>Gammaproteobacteria>Gammaproteobacteria_incertae_sedis>sulfur-oxidizing_symbionts>Candidatus_Thiodiazotropha | GCA_016843075.1 | Candidatus_Thiodiazotropha_taylori | JAY66_17825 | 23755..27198 | 2792791 | BREX-system-P-loop-protein-BrxC-[Candidatus-Thiodiazotropha-taylori] |
| MCG7871843.1 | Type1.BREX | [illegible] | 1174 | Bacteria>Pseudomonadota>Gammaproteobacteria>Gammaproteobacteria_incertae_sedis>sulfur-oxidizing_symbionts>Candidatus_Thiodiazotropha | GCA_016848925.1 | Candidatus_Thiodiazotropha_lotti | JAZ11_07035 | 23024..26548 | 2792787 | BREX-system-P-loop-protein-BrxC-[Candidatus-Thiodiazotropha-lotti] |
| MCG7872379.1 | Type1.BREX | [illegible] | 1170 | Bacteria>Pseudomonadota>Gammaproteobacteria>Gammaproteobacteria_incertae_sedis>sulfur-oxidizing_symbionts>Candidatus_Thiodiazotropha | GCA_016848925.1 | Candidatus_Thiodiazotropha_lotti | JAZ11_09800 | 178889..182401 | 2792787 | BREX-system-P-loop-protein-BrxC-[Candidatus-Thiodiazotropha-lotti] |
| MCG7872685.1 | Type1.BREX | [illegible] | 1151 | Bacteria>Pseudomonadota>Gammaproteobacteria>Gammaproteobacteria_incertae_sedis>sulfur-oxidizing_symbionts>Candidatus_Thiodiazotropha | GCA_016848925.1 | Candidatus_Thiodiazotropha_lotti | JAZ11_11345 | 100935..104390 | 2792787 | BREX-system-P-loop-protein-BrxC-[Candidatus-Thiodiazotropha-lotti] |
| WP_177417917.1 | Type1.BREX | [illegible] | 1188 | Bacteria>Pseudomonadota>Gammaproteobacteria>Gammaproteobacteria_incertae_sedis>sulfur-oxidizing_symbionts | GCF_011947365.2 | endosymbiont_of_Lamellibrachia_barhami | G9Q73_RS00010 | 458..4024 | 205975 | BREX-system-P-loop-protein-BrxC-[endosymbiont-of-Lamellibrachia-barhami] |
| WP_044865655.1 | Type1.BREX | [illegible] | 1213 | Bacteria>Pseudomonadota>Gammaproteobacteria | GCF_001540705.1 | Enterobacterales | TJ11_RS03570 | 21952..25593 | 91347 | MULTISPECIES:-BREX-system-P-loop-protein-BrxC-[Enterobacterales] |
| WP_223374968.1 | Type1.BREX | [illegible] | 1240 | Bacteria>Pseudomonadota>Gammaproteobacteria | GCF_003051945.2 | Enterobacterales | N5D94_RS08195 | 82368..86090 | 91347 | MULTISPECIES:-BREX-system-P-loop-protein-BrxC-[Enterobacterales] |
| WP_060554888.1 | Type1.BREX | [illegible] | 1225 | Bacteria>Pseudomonadota>Gammaproteobacteria | GCF_003176495.1 | Enterobacterales | DDA63_RS11920 | 2661..6338 | 91347 | MULTISPECIES:-BREX-system-P-loop-protein-BrxC-[Enterobacterales] |
| WP_052921406.1 | Type1.BREX | [illegible] | 1225 | Bacteria>Pseudomonadota>Gammaproteobacteria | GCF_003402955.1 | Enterobacterales | DS732_RS04425 | 849664..853341 | 91347 | MULTISPECIES:-BREX-system-P-loop-protein-BrxC-[Enterobacterales] |
| WP_112020499.1 | Type1.BREX | [illegible] | 1204 | Bacteria>Pseudomonadota>Gammaproteobacteria | GCF_013927185.1 | Enterobacterales | DP180_RS12665 | 36252..39866 | 91347 | MULTISPECIES:-BREX-system-P-loop-protein-BrxC-[Enterobacterales] |
| WP_099433763.1 | Type1.BREX | [illegible] | 1233 | Bacteria>Pseudomonadota>Gammaproteobacteria | GCF_014168815.1 | Enterobacterales | H7R56_RS21645 | 4484852..4488553 | 91347 | MULTISPECIES:-BREX-system-P-loop-protein-BrxC-[Enterobacterales] |
| WP_192176748.1 | Type1.BREX | [illegible] | 1225 | Bacteria>Pseudomonadota>Gammaproteobacteria | GCF_014843115.1 | Enterobacterales | H6P66_RS01965 | 387207..390884 | 91347 | MULTISPECIES:-BREX-system-P-loop-protein-BrxC-[Enterobacterales] |
| MAZ66828.1 | Type1.BREX | [illegible] | 1234 | Bacteria>Pseudomonadota>Gammaproteobacteria>Kangiellales>Kangiellaceae>unclassified_Kangiellaceae | GCA_002707725.1 | Kangiellaceae_bacterium | CMF25_06935 | 12477..16181 | 2026751 | DNA-repair-protein-[Kangiellaceae-bacterium] |
| WP_207387236.1 | Type1.BREX | [illegible] | 1194 | Bacteria>Pseudomonadota>Gammaproteobacteria>Legionellales>Legionellaceae>Fluoribacter | GCF_900640105.1 | Fluoribacter_gormanii | E1P13_RS01655 | 70546..74130 | 464 | BREX-system-P-loop-protein-BrxC-[Fluoribacter-gormanii] |
| TIG67286.1 | Type1.BREX | [illegible] | 1185 | Bacteria>Pseudomonadota>Gammaproteobacteria>Legionellales>Legionellaceae>Legionella | GCA_004921705.1 | Legionella_pneumophila | JAN10_12325 | 31417..34974 | 446 | BREX-system-P-loop-protein-BrxC-[Legionella-pneumophila] |
| HAT1978996.1 | Type1.BREX | [illegible] | 1184 | Bacteria>Pseudomonadota>Gammaproteobacteria>Legionellales>Legionellaceae>Legionella | GCA_015895665.1 | Legionella_pneumophila | I8Z83_002252 | 23760..27314 | 446 | BREX-system-P-loop-protein-BrxC-[Legionella-pneumophila] |
| HAT6348316.1 | Type1.BREX | [illegible] | 1185 | Bacteria>Pseudomonadota>Gammaproteobacteria>Legionellales>Legionellaceae>Legionella | GCA_015945075.1 | Legionella_pneumophila | JAJ33_001368 | 159147..162704 | 446 | BREX-system-P-loop-protein-BrxC-[Legionella-pneumophila] |
| HAT6334489.1 | Type1.BREX | [illegible] | 1185 | Bacteria>Pseudomonadota>Gammaproteobacteria>Legionellales>Legionellaceae>Legionella | GCA_015946305.1 | Legionella_pneumophila | JAJ38_001743 | 27953..31510 | 446 | BREX-system-P-loop-protein-BrxC-[Legionella-pneumophila] |
| HAT9211453.1 | Type1.BREX | [illegible] | 1185 | Bacteria>Pseudomonadota>Gammaproteobacteria>Legionellales>Legionellaceae>Legionella | GCA_015973865.1 | Legionella_pneumophila_subsp._pneumophila | JBB81_06050 | 33487..37044 | 91891 | BREX-system-P-loop-protein-BrxC-[Legionella-pneumophila-subsp.-pneumophila] |
| HAU0774874.1 | Type1.BREX | [illegible] | 1185 | Bacteria>Pseudomonadota>Gammaproteobacteria>Legionellales>Legionellaceae>Legionella | GCA_015983005.1 | Legionella_pneumophila | JBH09_07400 | 2447..6004 | 446 | BREX-system-P-loop-protein-BrxC-[Legionella-pneumophila] |
| HAU2155475.1 | Type1.BREX | [illegible] | 1185 | Bacteria>Pseudomonadota>Gammaproteobacteria>Legionellales>Legionellaceae>Legionella | GCA_016020315.1 | Legionella_pneumophila | JBK36_03075 | 164784..168341 | 446 | BREX-system-P-loop-protein-BrxC-[Legionella-pneumophila] |
| HAU3861597.1 | Type1.BREX | [illegible] | 1185 | Bacteria>Pseudomonadota>Gammaproteobacteria>Legionellales>Legionellaceae>Legionella | GCA_016062615.1 | Legionella_pneumophila | F7Z80_07285 | 265287..268844 | 446 | BREX-system-P-loop-protein-BrxC-[Legionella-pneumophila] |
| HCJ1122636.1 | Type1.BREX | [illegible] | 1184 | Bacteria>Pseudomonadota>Gammaproteobacteria>Legionellales>Legionellaceae>Legionella | GCA_024486645.1 | Legionella_pneumophila | NQN52_000248 | 14141..17695 | 446 | BREX-system-P-loop-protein-BrxC-[Legionella-pneumophila] |
| HCJ1069247.1 | Type1.BREX | [illegible] | 1225 | Bacteria>Pseudomonadota>Gammaproteobacteria>Legionellales>Legionellaceae>Legionella | GCA_024486765.1 | Legionella_pneumophila | NQN29_000516 | 40432..44109 | 446 | BREX-system-P-loop-protein-BrxC-[Legionella-pneumophila] |
| WP_061693758.1 | Type1.BREX | [illegible] | 1184 | Bacteria>Pseudomonadota>Gammaproteobacteria>Legionellales>Legionellaceae>Legionella | GCF_002002625.1 | Legionella_pneumophila | LPM_RS06580 | 16051..19605 | 446 | BREX-system-P-loop-protein-BrxC-[Legionella-pneumophila] |
| WP_192875104.1 | Type1.BREX | [illegible] | 1184 | Bacteria>Pseudomonadota>Gammaproteobacteria>Legionellales>Legionellaceae>Legionella | GCF_003004175.1 | Legionella_pneumophila | CA922_RS00900 | 198017..201571 | 446 | BREX-system-P-loop-protein-BrxC-[Legionella-pneumophila] |
| WP_061721013.1 | Type1.BREX | [illegible] | 1184 | Bacteria>Pseudomonadota>Gammaproteobacteria>Legionellales>Legionellaceae>Legionella | GCF_900054435.1 | Legionella_pneumophila | A2E52_RS14865 | 12190..15744 | 446 | BREX-system-P-loop-protein-BrxC-[Legionella-pneumophila] |
| WP_061637858.1 | Type1.BREX | [illegible] | 1184 | Bacteria>Pseudomonadota>Gammaproteobacteria>Legionellales>Legionellaceae>Legionella | GCF_900061645.1 | Legionella_pneumophila | A1L82_RS06820 | 180647..184201 | 446 | BREX-system-P-loop-protein-BrxC-[Legionella-pneumophila] |
| WP_027222687.1 | Type1.BREX | [illegible] | 1184 | Bacteria>Pseudomonadota>Gammaproteobacteria>Legionellales>Legionellaceae>Legionella | GCF_900639945.1 | Legionella_pneumophila | E1076_RS05405 | 246005..249559 | 446 | BREX-system-P-loop-protein-BrxC-[Legionella-pneumophila] |
| WP_165474921.1 | Type1.BREX | [illegible] | 1184 | Bacteria>Pseudomonadota>Gammaproteobacteria>Legionellales>Legionellaceae>Legionella | GCF_900640175.1 | Legionella_sp._W10-070 | E1O65_RS06915 | 46772..50326 | 1117709 | BREX-system-P-loop-protein-BrxC-[Legionella-sp.-W10-070] |
| MCD2450192.1 | Type1.BREX | [illegible] | 1213 | Bacteria>Pseudomonadota>Gammaproteobacteria>Methylococcales>Methylococcaceae>Methylicorpusculum | GCA_009828925.2 | Methylicorpusculum_oleiharenae | GD003_007310 | 1557698..1561339 | 1338687 | BREX-system-P-loop-protein-BrxC-[Methylicorpusculum-oleiharenae] |
| WP_159653266.1 | Type1.BREX | [illegible] | 1184 | Bacteria>Pseudomonadota>Gammaproteobacteria>Methylococcales>Methylococcaceae>Methylicorpusculum | GCF_009828925.2 | Methylicorpusculum_oleiharenae | GD003_RS20515 | 4601128..4604682 | 1338687 | BREX-system-P-loop-protein-BrxC-[Methylicorpusculum-oleiharenae] |
| WP_159652892.1 | Type1.BREX | [illegible] | 1213 | Bacteria>Pseudomonadota>Gammaproteobacteria>Methylococcales>Methylococcaceae>Methylicorpusculum | - | Methylicorpusculum_oleiharenae | GD003_007310 | 1557698..1561339 | 1338687 | BREX-system-P-loop-protein-BrxC-[Methylicorpusculum-oleiharenae] |
| MCL7421787.1 | Type1.BREX | [illegible] | 1146 | Bacteria>Pseudomonadota>Gammaproteobacteria>Methylococcales>Methylococcaceae>Methylobacter | GCA_023544625.1 | Methylobacter_sp. | M8364_12865 | 42290..45730 | 2051955 | BREX-system-P-loop-protein-BrxC-[Methylobacter-sp.] |
| WP_031438510.1 | Type1.BREX | [illegible] | 1218 | Bacteria>Pseudomonadota>Gammaproteobacteria>Methylococcales>Methylococcaceae>Methylobacter | GCF_000733835.1 | Methylobacter_tundripaludum | T449_RS0121480 | 4638570..4642226 | 173365 | BREX-system-P-loop-protein-BrxC-[Methylobacter-tundripaludum] |
| WP_262965888.1 | Type1.BREX | [illegible] | 1212 | Bacteria>Pseudomonadota>Gammaproteobacteria>Methylococcales>Methylococcaceae>Methylobacter | GCF_025583945.1 | Methylobacter_psychrophilus | N7974_RS12230 | 10707..14345 | 96941 | BREX-system-P-loop-protein-BrxC-[Methylobacter-psychrophilus] |
| WP_174625954.1 | Type1.BREX | [illegible] | 1149 | Bacteria>Pseudomonadota>Gammaproteobacteria>Methylococcales>Methylococcaceae>Methylobacter | GCF_902806695.1 | Candidatus_Methylobacter_favarea | HUX64_RS09295 | 14302..17751 | 2707345 | BREX-system-P-loop-protein-BrxC-[Candidatus-Methylobacter-favarea] |
| WP_210024304.1 | Type1.BREX | [illegible] | 1199 | Bacteria>Pseudomonadota>Gammaproteobacteria>Methylococcales>Methylococcaceae>Methylocaldum | GCF_017832375.1 | Methylocaldum_sp._RMAD-M | JOD69_RS22430 | 17805..21404 | 2806557 | BREX-system-P-loop-protein-BrxC-[Methylocaldum-sp.-RMAD-M] |
| MBE0434438.1 | Type1.BREX | [illegible] | 1177 | Bacteria>Pseudomonadota>Gammaproteobacteria>Methylococcales>Methylococcaceae>Methylomicrobium | GCA_014859425.1 | Methylomicrobium_sp. | IBX56_01390 | 2720..6253 | 2303549 | BREX-system-P-loop-protein-BrxC-[Methylomicrobium-sp.] |
| MBE0436274.1 | Type1.BREX | [illegible] | 251 | Bacteria>Pseudomonadota>Gammaproteobacteria>Methylococcales>Methylococcaceae>Methylomicrobium | GCA_014859425.1 | Methylomicrobium_sp. | IBX56_10780 | 2140..2891 | 2303549 | BREX-system-P-loop-protein-BrxC-[Methylomicrobium-sp.] |
| MBS3952115.1 | Type1.BREX | [illegible] | 1206 | Bacteria>Pseudomonadota>Gammaproteobacteria>Methylococcales>Methylococcaceae>Methylomicrobium | GCA_018333785.1 | Methylomicrobium_sp. | KGZ88_04105 | 40122..43742 | 2303549 | BREX-system-P-loop-protein-BrxC-[Methylomicrobium-sp.] |
| MBS3952424.1 | Type1.BREX | [illegible] | 1178 | Bacteria>Pseudomonadota>Gammaproteobacteria>Methylococcales>Methylococcaceae>Methylomicrobium | GCA_018333785.1 | Methylomicrobium_sp. | KGZ88_05695 | 2093..5629 | 2303549 | BREX-system-P-loop-protein-BrxC-[Methylomicrobium-sp.] |

| Protein-ID | System | Gene-Neighbourhoods | Length | Taxonomy | Assembly-ID | Strain | Locus-Tag | Genomic-coordinates | TaxID | Fasta-define |
|---|---|---|---|---|---|---|---|---|---|---|
| WP_024296674.1 | Type1.BREX | [illegible] | 1202 | [illegible] | GCF_000527095.1 | [illegible] | METLA_RS0100460 | 95992..99600 | 136992 | [illegible] |
| MBS3964846.1 | Type1.BREX | [illegible] | 1209 | [illegible] | GCA_018333915.1 | [illegible] | KGZ80_10205 | 1968..5597 | 418 | [illegible] |
| WP_256360610.1 | Type1.BREX | [illegible] | 1145 | [illegible] | GCF_001312005.1 | [illegible] | JCM16701_RS06940 | 49293..52730 | 702114 | [illegible] |
| WP_071159674.1 | Type1.BREX | [illegible] | 1153 | [illegible] | GCF_001856095.1 | [illegible] | BJL95_RS17210 | 210511..213972 | 1905845 | [illegible] |
| WP_096876853.1 | Type1.BREX | [illegible] | 1180 | [illegible] | GCF_002441955.1 | [illegible] | MKLM6_RS22285 | 58076..61618 | 702114 | [illegible] |
| WP_221054675.1 | Type1.BREX | [illegible] | 1176 | [illegible] | GCF_019669905.1 | [illegible] | MKFW12EY_RS21895 | 239254..1338 | 702114 | [illegible] |
| WP_255188315.1 | Type1.BREX | [illegible] | 1206 | [illegible] | GCF_024360925.2 | [illegible] | NM686_RS13235 | 2825288..2828908 | 2963939 | [illegible] |
| WP_255189595.1 | Type1.BREX | [illegible] | 1209 | [illegible] | GCF_024360925.2 | [illegible] | NM686_RS20105 | 4241783..4245412 | 2963939 | [illegible] |
| WP_192375581.1 | Type1.BREX | [illegible] | 1207 | [illegible] | GCF_903064715.1 | [illegible] | EBA_RS15675 | 3420343..3423966 | 1854563 | [illegible] |
| MBE0471466.1 | Type1.BREX | [illegible] | 1214 | [illegible] | GCA_014859465.1 | [illegible] | IBX55_18395 | 2520..6164 | 2020875 | [illegible] |
| WP_080523593.1 | Type1.BREX | [illegible] | 1206 | [illegible] | GCF_002072955.1 | [illegible] | AU255_RS14210 | 277762..281382 | 1420851 | [illegible] |
| MBZ4201977.1 | Type1.BREX | [illegible] | 909 | [illegible] | GCA_020035215.1 | [illegible] | LAC70_02510 | 1..2727 | 1916980 | [illegible] |
| MCF8007296.1 | Type1.BREX | [illegible] | 781 | [illegible] | GCA_021736885.1 | [illegible] | K9K84_07785 | 12351..14696 | 1916980 | [illegible] |
| WP_088619247.1 | Type1.BREX | [illegible] | 1201 | [illegible] | GCF_002209385.1 | [illegible] | CEK71_RS09965 | 2189788..2193393 | 1704499 | [illegible] |
| WP_103975698.1 | Type1.BREX | [illegible] | 1201 | [illegible] | GCF_002923755.1 | [illegible] | AADEFJLK_RS21345 | 25888..29493 | 1704499 | [illegible] |
| WP_215479507.1 | Type1.BREX | [illegible] | 1201 | [illegible] | GCF_018722605.1 | [illegible] | KFZ76_RS21360 | 105012..108617 | 1704499 | [illegible] |
| TSA38613.1 | Type1.BREX | [illegible] | 1218 | [illegible] | GCA_007280895.1 | [illegible] | D4R63_10680 | 1521..5177 | 1933926 | [illegible] |
| MCF7971355.1 | Type1.BREX | [illegible] | 1151 | [illegible] | GCA_021736705.1 | [illegible] | K9L22_09360 | 3141..6596 | 1933926 | [illegible] |
| RIZ69585.1 | Type1.BREX | [illegible] | 799 | [illegible] | GCA_003584865.1 | [illegible] | D0528_03735 | 16784..19183 | 2304002 | [illegible] |
| WP_104129248.1 | Type1.BREX | [illegible] | 739 | [illegible] | - | [illegible] | AbaMCR10126_RS06085 | 1..2216 | 470 | [illegible] |
| EGY2544509.1 | Type1.BREX | [illegible] | 942 | [illegible] | - | [illegible] | JHZ71_003464 | 4150..6977 | 470 | [illegible] |
| EHN8630147.1 | Type1.BREX | [illegible] | 561 | [illegible] | - | [illegible] | KKI61_003746 | 8069..9753 | 470 | [illegible] |
| EJC5774375.1 | Type1.BREX | [illegible] | 881 | [illegible] | - | [illegible] | MZL57_003344 | 1..2643 | 470 | [illegible] |
| WP_099476278.1 | Type1.BREX | [illegible] | 1226 | [illegible] | - | [illegible] | HPG40_RS00205 | 37613..41293 | 48296 | [illegible] |
| WP_252316424.1 | Type1.BREX | [illegible] | 1212 | [illegible] | - | [illegible] | J687_RS15310 | 12712..16350 | 1310788 | [illegible] |
| WP_019836725.1 | Type1.BREX | [illegible] | 1226 | [illegible] | - | [illegible] | H048_RS0104285 | 13386..17066 | 1209355 | [illegible] |
| WP_005303500.1 | Type1.BREX | [illegible] | 1226 | [illegible] | - | [illegible] | F886_RS07830 | 464704..468384 | 1217688 | [illegible] |
| CAM84763.1 | Type1.BREX | [illegible] | 1229 | [illegible] | GCA_000069245.1 | [illegible] | p3ABAYE0070 | 63476..67165 | 509173 | [illegible] |
| EEY88530.1 | Type1.BREX | [illegible] | 793 | [illegible] | GCA_000162095.1 | [illegible] | HMPREF0017_02853 | 41683..44064 | 575588 | [illegible] |
| ENX11167.1 | Type1.BREX | [illegible] | 911 | [illegible] | GCA_000369645.1 | [illegible] | F895_03860 | 13889..16624 | 1217694 | [illegible] |
| EXS00102.1 | Type1.BREX | [illegible] | 1190 | [illegible] | GCA_000588415.1 | [illegible] | J687_2205 | 12778..16350 | 1310788 | [illegible] |
| AZB97741.1 | Type1.BREX | [illegible] | 728 | [illegible] | GCA_003359275.2 | [illegible] | DKE42_019155 | 44169..46355 | 48296 | [illegible] |
| HAV3580163.1 | Type1.BREX | [illegible] | 1226 | [illegible] | GCA_016485765.1 | [illegible] | JIE63_001443 | 180845..184525 | 470 | [illegible] |
| HAV4485227.1 | Type1.BREX | [illegible] | 759 | [illegible] | GCA_016498165.1 | [illegible] | JIR97_15920 | 19224..21501 | 470 | [illegible] |
| HAV4527427.1 | Type1.BREX | [illegible] | 759 | [illegible] | GCA_016498325.1 | [illegible] | JIR72_17730 | 19224..21501 | 470 | [illegible] |
| HAV4540658.1 | Type1.BREX | [illegible] | 1226 | [illegible] | GCA_016498465.1 | [illegible] | JIS19_16850 | 25447..29127 | 470 | [illegible] |
| HAV4891449.1 | Type1.BREX | [illegible] | 676 | [illegible] | GCA_016503425.1 | [illegible] | JIU86_16385 | 19224..21252 | 470 | [illegible] |
| HAV4979216.1 | Type1.BREX | [illegible] | 1226 | [illegible] | GCA_016507565.1 | [illegible] | JIZ05_14385 | 7251..10931 | 470 | [illegible] |
| HAV5318348.1 | Type1.BREX | [illegible] | 1226 | [illegible] | GCA_016510035.1 | [illegible] | JI043_16030 | 3866..7546 | 470 | [illegible] |
| HAV6134262.1 | Type1.BREX | [illegible] | 1226 | [illegible] | GCA_016514395.1 | [illegible] | JI295_12625 | 9155..12835 | 470 | [illegible] |
| EHU1604380.1 | Type1.BREX | [illegible] | 1226 | [illegible] | GCA_016522135.2 | [illegible] | A1Z38_RS18510 | 4859..8539 | 470 | [illegible] |
| EHU2434196.1 | Type1.BREX | [illegible] | 1226 | [illegible] | GCA_016527835.2 | [illegible] | A0165_RS18865 | 10464..14144 | 470 | [illegible] |
| GIT84604.1 | Type1.BREX | [illegible] | 1224 | [illegible] | GCA_018403785.1 | [illegible] | DSM16313_23860 | 1741..5415 | 281376 | [illegible] |
| BCZ16231.1 | Type1.BREX | [illegible] | 1142 | [illegible] | GCA_019703285.1 | [illegible] | OCUAc18_37710 | 57364..60792 | 470 | [illegible] |
| UDM39811.1 | Type1.BREX | [illegible] | 1226 | [illegible] | GCA_020616635.1 | [illegible] | LIS44_16805 | 284403..288083 | 29430 | [illegible] |
| MCE1270820.1 | Type1.BREX | [illegible] | 1226 | [illegible] | GCA_021287125.1 | [illegible] | LWW88_04505 | 5070..8750 | 472 | [illegible] |
| HCA4923662.1 | Type1.BREX | [illegible] | 1226 | [illegible] | GCA_022931145.1 | [illegible] | MW112_001902 | 190856..194536 | 470 | [illegible] |
| USI88235.1 | Type1.BREX | [illegible] | 1226 | [illegible] | GCA_023822225.1 | [illegible] | LZ086_18665 | 10226..13906 | 40214 | [illegible] |
| WP_136143645.1 | Type1.BREX | [illegible] | 1226 | [illegible] | GCA_030820635.1 | [illegible] | EQG52_RS02450 | 14374..18054 | 202956 | [illegible] |
| CAI3142244.1 | Type1.BREX | [illegible] | 416 | [illegible] | GCA_947045015.1 | [illegible] | MWMV16_MWMV16_03034 | 31325..32575 | 470 | [illegible] |
| WP_001021844.1 | Type1.BREX | [illegible] | 1226 | [illegible] | GCF_000069245.1 | [illegible] | ABAYE_RS00380 | 63476..67156 | 909768 | [illegible] |
| WP_005401825.1 | Type1.BREX | [illegible] | 1226 | [illegible] | GCF_000162055.1 | [illegible] | HMPREF0016_RS15525 | 47644..51324 | 40214 | [illegible] |
| WP_010116441.1 | Type1.BREX | [illegible] | 1226 | [illegible] | GCF_000214135.1 | [illegible] | GGG_RS0115320 | 12069..15749 | 1029823 | [illegible] |
| WP_017400109.1 | Type1.BREX | [illegible] | 1226 | [illegible] | GCF_000302375.1 | [illegible] | E8E07_RS19610 | 55428..59108 | 909768 | [illegible] |
| WP_001021843.1 | Type1.BREX | [illegible] | 1226 | [illegible] | GCF_000305235.1 | [illegible] | ACIN5098_RS09930 | 47279..50959 | 470 | [illegible] |
| WP_019383579.1 | Type1.BREX | [illegible] | 1226 | [illegible] | GCF_000308235.1 | [illegible] | A489_RS0100450 | 77471..81151 | 52133 | [illegible] |
| WP_019384245.1 | Type1.BREX | [illegible] | 1225 | [illegible] | GCF_000308235.1 | [illegible] | A489_RS0105135 | 855128..858805 | 52133 | [illegible] |
| WP_004758928.1 | Type1.BREX | [illegible] | 1226 | [illegible] | GCF_000368305.1 | [illegible] | F973_RS14130 | 48127..51807 | 1144664 | [illegible] |
| WP_004811070.1 | Type1.BREX | [illegible] | 1225 | [illegible] | GCF_000368465.1 | [illegible] | F965_RS04725 | 393977..397654 | 108981 | [illegible] |
| WP_004827026.1 | Type1.BREX | [illegible] | 1226 | [illegible] | GCF_000368505.1 | [illegible] | F963_00144 | 48426..52106 | 106648 | [illegible] |
| WP_004984466.1 | Type1.BREX | [illegible] | 1190 | [illegible] | GCF_000368805.1 | [illegible] | F946_RS15710 | 148915..152487 | 40214 | [illegible] |
| WP_005245153.1 | Type1.BREX | [illegible] | 1226 | [illegible] | GCF_000369665.1 | [illegible] | F894_RS00155 | 31081..34761 | 1144670 | [illegible] |
| WP_005247073.1 | Type1.BREX | [illegible] | 1226 | [illegible] | GCF_000369665.1 | [illegible] | F894_RS08220 | 945028..948708 | 1144670 | [illegible] |
| WP_005253593.1 | Type1.BREX | [illegible] | 1226 | [illegible] | GCF_000369685.1 | [illegible] | F893_RS16420 | 115413..119093 | 1144665 | [illegible] |
| WP_005264232.1 | Type1.BREX | [illegible] | 1226 | [illegible] | GCF_000369725.1 | [illegible] | F891_RS14635 | 23007..26687 | 1144662 | [illegible] |
| WP_016539314.1 | Type1.BREX | [illegible] | 1248 | [illegible] | GCF_000414055.1 | [illegible] | Agu01S_RS05845 | 113550..117296 | 106649 | [illegible] |
| WP_023013360.1 | Type1.BREX | [illegible] | 1226 | [illegible] | GCF_000475335.1 | [illegible] | IO68_RS25385 | 11690..15370 | 1397525 | [illegible] |
| WP_031946304.1 | Type1.BREX | [illegible] | 1226 | [illegible] | GCF_000516575.2 | [illegible] | UN97_RS12380 | 246428..250108 | 470 | [illegible] |
| WP_034686569.1 | Type1.BREX | [illegible] | 1226 | [illegible] | GCF_000581035.1 | [illegible] | J520_RS12320 | 4406..8086 | 1310621 | [illegible] |
| WP_034680334.1 | Type1.BREX | [illegible] | 1226 | [illegible] | GCF_000582135.1 | [illegible] | J559_RS13455 | 17998..21678 | 1310660 | [illegible] |
| WP_032016259.1 | Type1.BREX | [illegible] | 1226 | [illegible] | GCF_000584335.1 | [illegible] | J609_RS12655 | 131163..134843 | 470 | [illegible] |
| WP_025095052.1 | Type1.BREX | [illegible] | 1226 | [illegible] | GCF_000633005.1 | [illegible] | EG58_RS0106955 | 8205..11885 | 487316 | [illegible] |
| WP_034584422.1 | Type1.BREX | [illegible] | 1226 | [illegible] | GCF_000773685.1 | [illegible] | GW12_RS05240 | 10375..14055 | 1509403 | [illegible] |
| WP_039048213.1 | Type1.BREX | [illegible] | 1226 | [illegible] | GCF_000813495.1 | [illegible] | RP26_RS13815 | 17614..21294 | 40215 | [illegible] |
| WP_045544683.1 | Type1.BREX | [illegible] | 1226 | [illegible] | GCF_000828935.1 | [illegible] | IOMTU433_RS19345 | 66852..70532 | 470 | [illegible] |
| WP_016658783.1 | Type1.BREX | [illegible] | 1226 | [illegible] | GCF_000830155.1 | [illegible] | AIN01_RS11690 | 13692..17372 | 756892 | [illegible] |
| WP_043972752.1 | Type1.BREX | [illegible] | 1226 | [illegible] | GCF_000876115.1 | [illegible] | Aba26S_RS13795 | 7444..11124 | 1550715 | [illegible] |
| WP_228136291.1 | Type1.BREX | [illegible] | 613 | [illegible] | GCF_000949675.1 | [illegible] | Aba28S_RS17090 | 3475..5316 | 106654 | [illegible] |
| WP_032059948.1 | Type1.BREX | [illegible] | 1226 | [illegible] | GCF_001077555.2 | [illegible] | AB895_RS13070 | 2715747..2719427 | 470 | [illegible] |
| WP_058952662.1 | Type1.BREX | [illegible] | 1220 | [illegible] | GCF_001484935.1 | [illegible] | EHS42_RS19740 | 21243..24905 | 40214 | [illegible] |
| WP_058971759.1 | Type1.BREX | [illegible] | 1226 | [illegible] | GCF_001500155.2 | [illegible] | NDM229_RS00350 | 66308..69988 | 106648 | [illegible] |
| WP_065184483.1 | Type1.BREX | [illegible] | 1226 | [illegible] | GCF_001674695.1 | [illegible] | GKP36_RS06470 | 16105..19785 | 470 | [illegible] |
| WP_068540167.1 | Type1.BREX | [illegible] | 1226 | [illegible] | GCF_001706415.2 | [illegible] | BFR83_RS18935 | 77132..80812 | 48296 | [illegible] |
| WP_069122561.1 | Type1.BREX | [illegible] | 1226 | [illegible] | GCF_001716945.1 | [illegible] | AXH23_RS05625 | 247550..251230 | 48296 | [illegible] |
| WP_075040289.1 | Type1.BREX | [illegible] | 1226 | [illegible] | GCF_001917365.1 | [illegible] | SA188iCDA_RS03905 | 54372..58052 | 40216 | [illegible] |
| WP_078390063.1 | Type1.BREX | [illegible] | 1224 | [illegible] | GCF_002018895.1 | [illegible] | MF4640_RS14775 | 20084..23758 | 1960826 | [illegible] |
| WP_086231303.1 | Type1.BREX | [illegible] | 1226 | [illegible] | GCF_002137255.1 | [illegible] | B9Y01_RS02195 | 14367..18047 | 470 | [illegible] |
| WP_143224472.1 | Type1.BREX | [illegible] | 593 | [illegible] | GCF_002145235.1 | [illegible] | CAT65_RS01675 | 11536..13316 | 48296 | [illegible] |
| WP_087537459.1 | Type1.BREX | [illegible] | 1226 | [illegible] | GCF_002165255.2 | [illegible] | CDG55_RS00865 | 132887..136567 | 2004644 | [illegible] |
| WP_087528528.1 | Type1.BREX | [illegible] | 1226 | [illegible] | GCF_002165305.2 | [illegible] | CDG62_RS00815 | 35375..39055 | 2004646 | [illegible] |
| WP_087554207.1 | Type1.BREX | [illegible] | 1226 | [illegible] | GCF_002165345.2 | [illegible] | CDG68_RS02780 | 35949..39629 | 1879050 | [illegible] |
| WP_096902445.1 | Type1.BREX | [illegible] | 1222 | [illegible] | GCF_002412355.1 | [illegible] | CF596_RS10765 | 15331..18999 | 2018564 | [illegible] |
| WP_099349802.1 | Type1.BREX | [illegible] | 1226 | [illegible] | GCF_002727055.2 | [illegible] | CPI82_RS19405 | 19969..23649 | 470 | [illegible] |
| WP_099475573.1 | Type1.BREX | [illegible] | 1226 | [illegible] | GCF_002740745.1 | [illegible] | CT134_RS17990 | 28430..32110 | 48296 | [illegible] |
| WP_100834299.1 | Type1.BREX | [illegible] | 1226 | [illegible] | GCF_002813065.1 | [illegible] | BSR56_RS16115 | 29817..33497 | 29430 | [illegible] |
| WP_104123635.1 | Type1.BREX | [illegible] | 1226 | [illegible] | GCF_002927775.1 | [illegible] | AbaMCR9238_RS08400 | 60585..64265 | 470 | [illegible] |
| WP_104442411.1 | Type1.BREX | [illegible] | 1226 | [illegible] | GCF_002934965.1 | [illegible] | C3F34_RS18690 | 72475..76155 | 1758189 | [illegible] |
| WP_104500994.1 | Type1.BREX | [illegible] | 1226 | [illegible] | GCF_002938745.1 | [illegible] | C5505_RS10055 | 19022..22702 | 756892 | [illegible] |
| WP_104852217.1 | Type1.BREX | [illegible] | 1247 | [illegible] | GCF_002946595.1 | [illegible] | C3Y93_RS04635 | 951564..955307 | 2079596 | [illegible] |
| WP_109291743.1 | Type1.BREX | [illegible] | 1226 | [illegible] | GCF_003131095.1 | [illegible] | DF172_RS10505 | 4712..8392 | 470 | [illegible] |
| WP_110975939.1 | Type1.BREX | [illegible] | 1226 | [illegible] | GCF_003231195.1 | [illegible] | DPG01_RS17630 | 3221..6901 | 2213016 | [illegible] |
| WP_111281982.1 | Type1.BREX | [illegible] | 1226 | [illegible] | GCF_003258335.1 | [illegible] | NVV25_RS10895 | 13399..17079 | 40216 | [illegible] |
| WP_119025213.1 | Type1.BREX | [illegible] | 1226 | [illegible] | GCF_003569585.1 | [illegible] | D5I70_RS01315 | 8436..12116 | 487316 | [illegible] |
| WP_119047908.1 | Type1.BREX | [illegible] | 1226 | [illegible] | GCF_003569645.1 | [illegible] | D5I76_RS05685 | 52847..56527 | 470 | [illegible] |
| WP_119518033.1 | Type1.BREX | [illegible] | 817 | [illegible] | GCF_003583605.1 | [illegible] | D3X29_RS17285 | 1..2451 | 470 | [illegible] |
| WP_119879428.1 | Type1.BREX | [illegible] | 1226 | [illegible] | GCF_003594775.1 | [illegible] | D5055_RS12040 | 10913..14593 | 40216 | [illegible] |
| WP_120366206.1 | Type1.BREX | [illegible] | 1226 | [illegible] | GCF_003611535.1 | [illegible] | D7V68_RS14625 | 28683..32363 | 2136182 | [illegible] |
| WP_121523424.1 | Type1.BREX | [illegible] | 1226 | [illegible] | GCF_003664645.1 | [illegible] | D9K81_RS12845 | 89350..93030 | 2420890 | [illegible] |
| WP_121533276.1 | Type1.BREX | [illegible] | 1226 | [illegible] | GCF_003664655.1 | [illegible] | D9K79_RS15510 | 4801..8481 | 2136182 | [illegible] |
| WP_123785432.1 | Type1.BREX | [illegible] | 1226 | [illegible] | GCF_003812065.1 | [illegible] | LH406_RS10280 | 68046..71726 | 470 | [illegible] |
| WP_125269804.1 | Type1.BREX | [illegible] | 1512 | [illegible] | GCF_003938865.1 | [illegible] | EGS47_RS18730 | 3064192..3068730 | 2420307 | [illegible] |
| WP_125274900.1 | Type1.BREX | [illegible] | 1226 | [illegible] | GCF_003940285.1 | [illegible] | EGT73_RS16195 | 9004..12684 | 40214 | [illegible] |
| WP_125318492.1 | Type1.BREX | [illegible] | 1226 | [illegible] | GCF_003940305.1 | [illegible] | EGT64_RS07935 | 8553..12233 | 40215 | [illegible] |
| WP_125525774.1 | Type1.BREX | [illegible] | 1226 | [illegible] | GCF_003947765.1 | [illegible] | EA755_RS16870 | 59824..63504 | 48296 | [illegible] |
| WP_127800367.1 | Type1.BREX | [illegible] | 1226 | [illegible] | GCF_004005445.1 | [illegible] | ENC21_RS11140 | 10000..13680 | 756892 | [illegible] |
| WP_129260966.1 | Type1.BREX | [illegible] | 1226 | [illegible] | GCF_004123275.1 | [illegible] | ETZ13_RS00160 | 34893..38573 | 40215 | [illegible] |
| WP_131273640.1 | Type1.BREX | [illegible] | 1154 | [illegible] | GCF_004331195.1 | [illegible] | E0H88_RS07250 | 81102..84566 | 2529840 | [illegible] |
| WP_131214141.1 | Type1.BREX | [illegible] | 1190 | [illegible] | GCF_004331285.1 | [illegible] | E0H89_RS01980 | 72363..75935 | 2529835 | [illegible] |
| WP_138000997.1 | Type1.BREX | [illegible] | 1226 | [illegible] | GCF_005519305.1 | [illegible] | E3H47_RS14125 | 6771..10451 | 40216 | [illegible] |
| WP_140948309.1 | Type1.BREX | [illegible] | 1226 | [illegible] | GCF_006491975.1 | [illegible] | FJV17_RS05945 | 24470..28150 | 470 | [illegible] |
| WP_140967408.1 | Type1.BREX | [illegible] | 1226 | [illegible] | GCF_006492985.1 | [illegible] | FJU65_RS09860 | 11803..15483 | 470 | [illegible] |
| WP_005021608.1 | Type1.BREX | [illegible] | 1226 | [illegible] | GCF_006757745.1 | [illegible] | ACRAD_RS15795 | 43375..47055 | 40216 | [illegible] |
| WP_144583841.1 | Type1.BREX | [illegible] | 1226 | [illegible] | GCF_007713425.1 | [illegible] | FPV60_RS17750 | 3413..7093 | 280145 | [illegible] |
| WP_148728183.1 | Type1.BREX | [illegible] | 1226 | [illegible] | GCF_008121475.1 | [illegible] | FX017_RS20770 | 81542..85222 | 48296 | [illegible] |
| WP_149931258.1 | Type1.BREX | [illegible] | 1220 | [illegible] | GCF_008580825.1 | [illegible] | F6C27_RS15450 | 2605..6267 | 487316 | [illegible] |
| WP_151708311.1 | Type1.BREX | [illegible] | 1226 | [illegible] | GCF_008979805.1 | [illegible] | GBZ45_RS13505 | 5536..9216 | 40215 | [illegible] |
| WP_151725173.1 | Type1.BREX | [illegible] | 1226 | [illegible] | GCF_008981805.1 | [illegible] | GBJ57_RS12485 | 1752..5432 | 108980 | [illegible] |
| WP_151726277.1 | Type1.BREX | [illegible] | 1226 | [illegible] | GCF_008982065.1 | [illegible] | GBJ55_RS11105 | 8766..12446 | 108980 | [illegible] |
| WP_151732002.1 | Type1.BREX | [illegible] | 1226 | [illegible] | GCF_008982325.1 | [illegible] | GBJ63_RS14825 | 20101..23781 | 108980 | [illegible] |
| WP_151742861.1 | Type1.BREX | [illegible] | 1226 | [illegible] | GCF_008982465.1 | [illegible] | GBL32_RS17905 | 1894..5574 | 196816 | [illegible] |
| WP_151792238.1 | Type1.BREX | [illegible] | 1249 | [illegible] | GCF_008995635.1 | [illegible] | IC766_RS15995 | 3376176..3379925 | 1530123 | [illegible] |
| WP_151817454.1 | Type1.BREX | [illegible] | 1226 | [illegible] | GCF_009004075.1 | [illegible] | GBR91_RS15640 | 8514..12194 | 108980 | [illegible] |
| WP_151834043.1 | Type1.BREX | [illegible] | 1190 | [illegible] | GCF_009012055.1 | [illegible] | GBX89_RS12550 | 5375..8947 | 108980 | [illegible] |
| WP_151837873.1 | Type1.BREX | [illegible] | 1226 | [illegible] | GCF_009013055.1 | [illegible] | GBZ97_RS11405 | 21380..25060 | 196816 | [illegible] |
| WP_151959064.1 | Type1.BREX | [illegible] | 1226 | [illegible] | GCF_009013895.1 | [illegible] | GB010_RS14460 | 6225..9905 | 40215 | [illegible] |
| WP_153389475.1 | Type1.BREX | [illegible] | 1224 | [illegible] | GCF_009601085.1 | [illegible] | GHJ48_RS13160 | 17934..21608 | 2662362 | [illegible] |
| WP_153569296.1 | Type1.BREX | [illegible] | 1226 | [illegible] | GCF_009645555.1 | [illegible] | F4T96_RS05690 | 68625..72305 | 106654 | [illegible] |
| WP_159516812.1 | Type1.BREX | [illegible] | 1226 | [illegible] | GCF_009829025.1 | [illegible] | GVV58_RS14230 | 9426..13106 | 2692137 | [illegible] |
| WP_160126737.1 | Type1.BREX | [illegible] | 1226 | [illegible] | GCF_009892245.1 | [illegible] | AhaeAN59_RS17345 | 50610..54290 | 29430 | [illegible] |
| WP_160255370.1 | Type1.BREX | [illegible] | 1226 | [illegible] | GCF_009914395.1 | [illegible] | GA646_RS00530 | 100361..104041 | 756892 | [illegible] |
| WP_163071913.1 | Type1.BREX | [illegible] | 1226 | [illegible] | GCF_010499775.1 | [illegible] | G3N14_RS16470 | 10817..14497 | 470 | [illegible] |
| WP_163146588.1 | Type1.BREX | [illegible] | 1226 | [illegible] | GCF_010918375.1 | [illegible] | FSC09_RS16655 | 95015..98695 | 756892 | [illegible] |
| WP_164543101.1 | Type1.BREX | [illegible] | 1224 | [illegible] | GCF_011058205.1 | [illegible] | G4V72_RS07960 | 7784..11458 | 28090 | [illegible] |

| Protein-ID | System | Gene-Neighbourhoods | Length | Taxonomy | Assembly-ID | Strain | Locus-Tag | Genomic-coordinates | TaxID | Fasta-defline |
|---|---|---|---|---|---|---|---|---|---|---|
| WP_166137070.1 | Type1.BREX | <-N6-METHYLASE<-DUF6079*<-BrxB-BrxF-iSTAND<-BrxA-DNA-binding<-wHTH+WYL+WCX\|\|DUF4236-><-CR-ATPase5+REase<-ParB+HNH<-Phage_integrase<-?<-SbcC+SbcC+ABC-ATPase+SbcC | 1226 | Bacteria>Pseudomonadota>Gammaproteobacteria>Moraxellales>Moraxellaceae>Acinetobacter | GCF_011192225.1 | unclassified_Acinetobacter | G5T41_RS07850 | 4541..8221 | 196816 | MULTISPECIES:-BREX-system-P-loop-protein-BrxC-[unclassified-Acinetobacter] |
| WP_168370063.1 | Type1.BREX | DUF932->?->?->XIS-HTH->SAM-METHYLASE->wHTH(SBD)+HNH->wHTH+WYL+WCX->BrxA-DNA-binding->BrxB-BrxF-iSTAND->DUF6079*->N6-METHYLASE->ParB+HNH->PglZ->SIGMA+OB-Fold+AAA-ATPase+Lon-Protease-BrxL->?-><-DOC+HTH | 1224 | Bacteria>Pseudomonadota>Gammaproteobacteria>Moraxellales>Moraxellaceae>Acinetobacter | GCF_012278715.1 | Acinetobacter_indicus | HHC64_RS10945 | 2259038..2262712 | 756892 | BREX-system-P-loop-protein-BrxC-[Acinetobacter-indicus] |
| WP_168382103.1 | Type1.BREX | Phage_integrase+Phage_integrase->Phage_integrase+Phage_integrase->?->ParB+HNH->CR-ATPase5+REase-><-DUF4236\|\|wHTH+WYL+WCX->BrxA-DNA-binding->BrxB-BrxF-iSTAND->DUF6079*->N6-METHYLASE->PglZ->SIGMA+OB-Fold+AAA-ATPase+Lon-Protease-BrxL->\|\|?->?->?->?->?->GDResolvase+HIN-HTH-> | 1226 | Bacteria>Pseudomonadota>Gammaproteobacteria>Moraxellales>Moraxellaceae>Acinetobacter | GCF_012366635.1 | Acinetobacter_indicus | HHC47_RS11970 | 17406..21086 | 756892 | BREX-system-P-loop-protein-BrxC-[Acinetobacter-indicus] |
| WP_168396480.1 | Type1.BREX | Phage_integrase+Phage_integrase->Phage_integrase+Phage_integrase->?->ParB+HNH->CR-ATPase5+REase-><-DUF4236\|\|wHTH+WYL+WCX->BrxA-DNA-binding->BrxB-BrxF-iSTAND->DUF6079*->N6-METHYLASE->SF1-SF2-Helicase+PRTase->SLOG+BirA-HTH->PglZ->SIGMA+OB-Fold+AAA-ATPase+Lon-Protease-BrxL->SF1-SF2-Helicase+REase-> | 1224 | Bacteria>Pseudomonadota>Gammaproteobacteria>Moraxellales>Moraxellaceae>Acinetobacter | GCF_012366875.1 | Acinetobacter_indicus | HG607_RS04385 | 142310..145984 | 756892 | BREX-system-P-loop-protein-BrxC-[Acinetobacter-indicus] |
| WP_168386656.1 | Type1.BREX | HTH_Tnp_1+HTH_21+rve+rve_3-><-?<-ABhydrolase\|\|MerR-HTH->DUF4096+DDE_Tnp_1+DDE_Tnp_1_2->MerR-HTH-><-SIGMA+OB-Fold+AAA-ATPase+Lon-Protease-BrxL<-PglZ<-N6-METHYLASE<-DUF6079*<-BrxB-BrxF-iSTAND<-BrxA-DNA-binding<-wHTH+WYL+WCX\|\|DUF4236-><-?<-ParB+HNH-p<-CR-ATPase5+REase<-ParB+HNH<-HTH_Tnp_1+HTH_21+rve+rve_3 | 1226 | Bacteria>Pseudomonadota>Gammaproteobacteria>Moraxellales>Moraxellaceae>Acinetobacter | GCF_012393445.1 | Acinetobacter_lwoffii | FGB28_RS00230 | 39022..42702 | 28090 | BREX-system-P-loop-protein-BrxC-[Acinetobacter-lwoffii] |
| WP_169025139.1 | Type1.BREX | Phage_integrase+Phage_integrase->Phage_integrase->HTH_Tnp_1+HTH_21+rve+rve_3->CR-ATPase5+REase-><-DUF4236\|\|wHTH+WYL+WCX->BrxA-DNA-binding->BrxB-BrxF-iSTAND->DUF6079*->N6-METHYLASE->ParB+HNH->PglZ->SIGMA+OB-Fold+AAA-ATPase+Lon-Protease-BrxL-><-?<-ABhydrolase<-Classical-AAA<-?<-SMODS | 1226 | Bacteria>Pseudomonadota>Gammaproteobacteria>Moraxellales>Moraxellaceae>Acinetobacter | GCF_012931665.1 | Acinetobacter_baumannii | HIN88_RS18590 | 12016..15696 | 470 | BREX-system-P-loop-protein-BrxC-[Acinetobacter-baumannii] |
| WP_171530296.1 | Type1.BREX | <-ACET<-?\|\|<-SIGMA+OB-Fold+AAA-ATPase+Lon-Protease-BrxL<-PglZ<-ParB+HNH<-N6-METHYLASE<-DUF6079*<-BrxB-BrxF-iSTAND<-BrxA-DNA-binding<-wHTH+WYL+WCX\|\|DUF4236-><-CR-ATPase5+REase<-ParB+HNH<-Phage_integrase+Phage_integrase<-Phage_integrase+Phage_integrase | 1226 | Bacteria>Pseudomonadota>Gammaproteobacteria>Moraxellales>Moraxellaceae>Acinetobacter | GCF_013072655.1 | Acinetobacter_defluvii | A7P53_RS01880 | 14398..18078 | 1871111 | BREX-system-P-loop-protein-BrxC-[Acinetobacter-defluvii] |
| WP_171431587.1 | Type1.BREX | ParB+HNH->CR-ATPase5+REase-><-DUF4236\|\|wHTH+WYL+WCX->BrxA-DNA-binding->BrxB-BrxF-iSTAND->DUF6079*->N6-METHYLASE->PglZ->SIGMA+OB-Fold+AAA-ATPase+Lon-Protease-BrxL->\|\|?->?->?->?->?->AAA-ATPase-p-> | 1228 | Bacteria>Pseudomonadota>Gammaproteobacteria>Moraxellales>Moraxellaceae>Acinetobacter | GCF_013116605.1 | Acinetobacter_indicus | HMI53_RS01195 | 262958..266644 | 756892 | BREX-system-P-loop-protein-BrxC-[Acinetobacter-indicus] |
| WP_174660656.1 | Type1.BREX | <-GDResolvase+HIN-HTH<-DDE_Tnp_1_5+DDE_Tnp_1\|\|?->?->?-><-ICLR-HTH<-SIGMA+OB-Fold+AAA-ATPase+Lon-Protease-BrxL<-PglZ<-N6-METHYLASE<-DUF6079*<-BrxB-BrxF-iSTAND<-BrxA-DNA-binding<-wHTH+WYL+WCX\|\|DUF4236-><-CR-ATPase5+REase<-ParB+HNH<-Phage_integrase+Phage_integrase<-Phage_integrase+Phage_integrase | 1226 | Bacteria>Pseudomonadota>Gammaproteobacteria>Moraxellales>Moraxellaceae>Acinetobacter | GCF_013343215.1 | Acinetobacter_sp._10FS3-1 | E5Y90_RS16040 | 29443..33123 | 2563897 | BREX-system-P-loop-protein-BrxC-[Acinetobacter-sp.-10FS3-1] |
| WP_174660691.1 | Type1.BREX | <-ACET<-?\|\|TRD+TRD-><-?<-SIGMA+OB-Fold+AAA-ATPase+Lon-Protease-BrxL<-PglZ<-SbcC<-N6-METHYLASE<-DUF6079*<-BrxB-BrxF-iSTAND<-BrxA-DNA-binding<-wHTH+WYL+WCX\|\|DUF4236-><-DOC+HTH<-?<-ParB+HNH<-Phage_integrase+Phage_integrase<-Phage_integrase+Phage_integrase | 1226 | Bacteria>Pseudomonadota>Gammaproteobacteria>Moraxellales>Moraxellaceae>Acinetobacter | GCF_013343215.1 | Acinetobacter_sp._10FS3-1 | E5Y90_RS16730 | 19958..23638 | 2563897 | BREX-system-P-loop-protein-BrxC-[Acinetobacter-sp.-10FS3-1] |
| WP_174727826.1 | Type1.BREX | BrxA-DNA-binding->BrxB-BrxF-iSTAND->DUF6079*->N6-METHYLASE->ParB+HNH->PglZ->?->?->?->?->?->ZnR+ZnR-> | 1224 | Bacteria>Pseudomonadota>Gammaproteobacteria>Moraxellales>Moraxellaceae>Acinetobacter | GCF_013344765.1 | Acinetobacter_lactucae | HUN22_RS07895 | 1711..5385 | 1785128 | BREX-system-P-loop-protein-BrxC-[Acinetobacter-lactucae] |
| WP_174761284.1 | Type1.BREX | <-N6-METHYLASE<-DUF6079*<-BrxB-BrxF-iSTAND<-BrxA-DNA-binding<-wHTH+WYL+WCX\|\|DUF4236-> | 1226 | Bacteria>Pseudomonadota>Gammaproteobacteria>Moraxellales>Moraxellaceae>Acinetobacter | GCF_013346575.1 | Acinetobacter_bereziniae | HUN33_RS09130 | 1857..5537 | 106648 | BREX-system-P-loop-protein-BrxC-[Acinetobacter-bereziniae] |
| WP_176538601.1 | Type1.BREX | <-DUF4372+DDE_Tnp_1<-LZ_Tnp_IS66+zf-IS66+DDE_Tnp_IS66+DDE_Tnp_IS66_C<-?<-HIN-HTH<-SIGMA+OB-Fold+AAA-ATPase+Lon-Protease-BrxL<-PglZ<-N6-METHYLASE<-DUF6079*<-BrxB-BrxF-iSTAND<-BrxA-DNA-binding<-wHTH+WYL+WCX\|\|DUF4236-><-?\|\|<-Phage_integrase | 1226 | Bacteria>Pseudomonadota>Gammaproteobacteria>Moraxellales>Moraxellaceae>Acinetobacter | GCF_013374795.1 | Acinetobacter_sp._NEB_394 | HUK62_RS18295 | 47703..51383 | 2743575 | BREX-system-P-loop-protein-BrxC-[Acinetobacter-sp.-NEB-394] |
| WP_179994566.1 | Type1.BREX | Arm-DNA-bind_3+Phage_integrase-><-?<-?<-?<-?<-TetR-HTH\|\|wHTH+WYL+WCX->BrxA-DNA-binding->BrxB-BrxF-iSTAND->DUF6079*->N6-METHYLASE->ParB+HNH->PglZ->SIGMA+OB-Fold+AAA-ATPase+Lon-Protease-BrxL->SF1-SF2-Helicase+REase->DarT+Peptidase->MACRODOMAIN-> | 1226 | Bacteria>Pseudomonadota>Gammaproteobacteria>Moraxellales>Moraxellaceae>Acinetobacter | GCF_013416255.1 | Acinetobacter_baumannii | HLG75_RS04210 | 863150..866830 | 470 | BREX-system-P-loop-protein-BrxC-[Acinetobacter-baumannii] |
| WP_180070068.1 | Type1.BREX | Phage_integrase->\|\|Zn_Tnp_IS1+DDE_Tnp_IS1->\|\|Y1_Tnp->ParB+HNH->CR-ATPase5+REase-><-DUF4236\|\|wHTH+WYL+WCX->BrxA-DNA-binding->BrxB-BrxF-iSTAND->DUF6079*->N6-METHYLASE-p-> | 1226 | Bacteria>Pseudomonadota>Gammaproteobacteria>Moraxellales>Moraxellaceae>Acinetobacter | GCF_013417275.1 | Acinetobacter_sp._YH16038 | FTV34_RS10085 | 73238..76918 | 2601183 | BREX-system-P-loop-protein-BrxC-[Acinetobacter-sp.-YH16038] |
| WP_180001763.1 | Type1.BREX | <-SIGMA+OB-Fold+AAA-ATPase+Lon-Protease-BrxL<-PglZ<-N6-METHYLASE<-DUF6079*<-BrxB-BrxF-iSTAND<-BrxA-DNA-binding<-wHTH+WYL+WCX<-ABC-ATPase+SbcC+ABC-ATPase+SbcC | 1226 | Bacteria>Pseudomonadota>Gammaproteobacteria>Moraxellales>Moraxellaceae>Acinetobacter | GCF_013417355.1 | Acinetobacter_sp._YH12255 | FTV30_RS13425 | 9473..13153 | 2601179 | BREX-system-P-loop-protein-BrxC-[Acinetobacter-sp.-YH12255] |
| WP_180012151.1 | Type1.BREX | CR-ATPase5+REase-><-DUF4236\|\|wHTH+WYL+WCX->BrxA-DNA-binding->BrxB-BrxF-iSTAND->DUF6079*->N6-METHYLASE->PglZ->SIGMA+OB-Fold+AAA-ATPase+Lon-Protease-BrxL->\|\|?->?->?->?->?->GDResolvase+HIN-HTH-> | 1226 | Bacteria>Pseudomonadota>Gammaproteobacteria>Moraxellales>Moraxellaceae>Acinetobacter | GCF_013417705.1 | Acinetobacter_sp._YH12235 | FTV13_RS12105 | 5418..9098 | 2601162 | BREX-system-P-loop-protein-BrxC-[Acinetobacter-sp.-YH12235] |
| WP_180058343.1 | Type1.BREX | <-HTH_3<-HigB_toxin\|\|<-SF1-SF2-Helicase+REase<-SIGMA+OB-Fold+AAA-ATPase+Lon-Protease-BrxL<-PglZ<-N6-METHYLASE<-DUF6079*<-BrxB-BrxF-iSTAND<-BrxA-DNA-binding<-wHTH+WYL+WCX<-wHTH+HNH\|\|<-SF1-SF2-Helicase<-?<-SAM-METHYLASE | 1223 | Bacteria>Pseudomonadota>Gammaproteobacteria>Moraxellales>Moraxellaceae>Acinetobacter | GCF_013417785.1 | Acinetobacter_sp._YH12227 | FTV09_RS11090 | 15051..18722 | 2601158 | BREX-system-P-loop-protein-BrxC-[Acinetobacter-sp.-YH12227] |
| WP_180041457.1 | Type1.BREX | ParB+HNH-><-?\|\|SAM-METHYLASE->?->SF1-SF2-Helicase->wHTH+HNH->wHTH+WYL+WCX->BrxA-DNA-binding->BrxB-BrxF-iSTAND->DUF6079*->N6-METHYLASE->HKD-DNase+DUF3150->PglZ->SIGMA+OB-Fold+AAA-ATPase+Lon-Protease-BrxL->SF1-SF2-Helicase+REase-> | 1221 | Bacteria>Pseudomonadota>Gammaproteobacteria>Moraxellales>Moraxellaceae>Acinetobacter | GCF_013417925.1 | Acinetobacter_sp._YH12218 | FTV03_RS11390 | 13993..17658 | 2601152 | BREX-system-P-loop-protein-BrxC-[Acinetobacter-sp.-YH12218] |
| WP_180082253.1 | Type1.BREX | DEOR-HTH-><-SIGMA+OB-Fold+AAA-ATPase+Lon-Protease-BrxL<-PglZ<-?<-N6-METHYLASE<-DUF6079*<-BrxB-BrxF-iSTAND<-BrxA-DNA-binding<-wHTH+WYL+WCX\|\|DUF4236-><-CR-ATPase5+REase | 1226 | Bacteria>Pseudomonadota>Gammaproteobacteria>Moraxellales>Moraxellaceae>Acinetobacter | GCF_013418145.1 | Acinetobacter_sp._YH12201 | FTU91_RS11740 | 12103..15783 | 2601140 | BREX-system-P-loop-protein-BrxC-[Acinetobacter-sp.-YH12201] |
| WP_180106519.1 | Type1.BREX | MarR-HTH->AbiEii->?-><-SIGMA+OB-Fold+AAA-ATPase+Lon-Protease-BrxL<-PglZ<-N6-METHYLASE<-DUF6079*<-BrxB-BrxF-iSTAND<-BrxA-DNA-binding<-wHTH+WYL+WCX | 1226 | Bacteria>Pseudomonadota>Gammaproteobacteria>Moraxellales>Moraxellaceae>Acinetobacter | GCF_013418285.1 | unclassified_Acinetobacter | FTU83_RS11060 | 37089..40769 | 196816 | MULTISPECIES:-BREX-system-P-loop-protein-BrxC-[unclassified-Acinetobacter] |
| WP_180056096.1 | Type1.BREX | CR-ATPase5+REase-><-DUF4236\|\|wHTH+WYL+WCX->BrxA-DNA-binding->BrxB-BrxF-iSTAND->DUF6079*->N6-METHYLASE->PglZ->SIGMA+OB-Fold+AAA-ATPase+Lon-Protease-BrxL-><-?<-AbiEii<-MarR-HTH | 1226 | Bacteria>Pseudomonadota>Gammaproteobacteria>Moraxellales>Moraxellaceae>Acinetobacter | GCF_013418435.1 | Acinetobacter_sp._YH12142 | FTU77_RS14435 | 5429..9109 | 2601126 | BREX-system-P-loop-protein-BrxC-[Acinetobacter-sp.-YH12142] |
| WP_180058089.1 | Type1.BREX | wHTH+WYL+WCX->BrxA-DNA-binding->BrxB-BrxF-iSTAND->DUF6079*->N6-METHYLASE->PglZ-> | 1226 | Bacteria>Pseudomonadota>Gammaproteobacteria>Moraxellales>Moraxellaceae>Acinetobacter | GCF_013418465.1 | Acinetobacter_sp._YH12141 | FTU76_RS14840 | 1801..5481 | 2601125 | BREX-system-P-loop-protein-BrxC-[Acinetobacter-sp.-YH12141] |
| WP_180052746.1 | Type1.BREX | <-SIGMA+OB-Fold+AAA-ATPase+Lon-Protease-BrxL<-PglZ<-?<-N6-METHYLASE<-DUF6079*<-BrxB-BrxF-iSTAND<-BrxA-DNA-binding<-wHTH+WYL+WCX\|\|DUF4236-><-CR-ATPase5+REase<-ParB+HNH<-Y1_Tnp\|\|HTH_OrfB_IS605+OrfB_IS605+OrfB_Zn_ribbon-><-Phage_integrase | 1226 | Bacteria>Pseudomonadota>Gammaproteobacteria>Moraxellales>Moraxellaceae>Acinetobacter | GCF_013418535.1 | unclassified_Acinetobacter | FTU72_RS09265 | 17103..20783 | 196816 | MULTISPECIES:-BREX-system-P-loop-protein-BrxC-[unclassified-Acinetobacter] |
| WP_180045992.1 | Type1.BREX | LZ_Tnp_IS66+zf-IS66+DDE_Tnp_IS66+DDE_Tnp_IS66_C->ParB+HNH->SAM-METHYLASE->LexA-protease->?->wHTH(SBD)+HNH->wHTH+WYL+WCX->BrxA-DNA-binding->BrxB-BrxF-iSTAND->DUF6079*->N6-METHYLASE->ParB+HNH->?->PglZ->SIGMA+OB-Fold+AAA-ATPase+Lon-Protease-BrxL->?->?->HIN-HTH-> | 1218 | Bacteria>Pseudomonadota>Gammaproteobacteria>Moraxellales>Moraxellaceae>Acinetobacter | GCF_013418865.1 | Acinetobacter_sp._YH12117 | FTU55_RS01900 | 159420..163076 | 2601104 | BREX-system-P-loop-protein-BrxC-[Acinetobacter-sp.-YH12117] |
| WP_180047260.1 | Type1.BREX | CR-ATPase5+REase-><-DUF4236\|\|wHTH+WYL+WCX->BrxA-DNA-binding->BrxB-BrxF-iSTAND->DUF6079*->N6-METHYLASE->PglZ->SIGMA+OB-Fold+AAA-ATPase+Lon-Protease-BrxL->\|\|SbcC-><-HTH_3<-RelE-ParE\|\|?->GDResolvase+HIN-HTH-> | 1226 | Bacteria>Pseudomonadota>Gammaproteobacteria>Moraxellales>Moraxellaceae>Acinetobacter | GCF_013418865.1 | Acinetobacter_sp._YH12117 | FTU55_RS12625 | 5420..9100 | 2601104 | BREX-system-P-loop-protein-BrxC-[Acinetobacter-sp.-YH12117] |
| WP_180074827.1 | Type1.BREX | <-SIGMA+OB-Fold+AAA-ATPase+Lon-Protease-BrxL<-PglZ<-N6-METHYLASE<-DUF6079*<-BrxB-BrxF-iSTAND<-BrxA-DNA-binding<-wHTH+WYL+WCX\|\|DUF4236-><-CR-ATPase5+REase<-ParB+HNH | 1226 | Bacteria>Pseudomonadota>Gammaproteobacteria>Moraxellales>Moraxellaceae>Acinetobacter | GCF_013418895.1 | Acinetobacter_sp._YH12116 | FTU54_RS13385 | 8809..12489 | 2601103 | BREX-system-P-loop-protein-BrxC-[Acinetobacter-sp.-YH12116] |
| WP_180076555.1 | Type1.BREX | Phage_integrase+Phage_integrase->?->ParB+HNH->?->DOC+HTH-><-DUF4236\|\|wHTH+WYL+WCX->BrxA-DNA-binding->BrxB-BrxF-iSTAND->DUF6079*->N6-METHYLASE->PglZ->SIGMA+OB-Fold+AAA-ATPase+Lon-Protease-BrxL-> | 1224 | Bacteria>Pseudomonadota>Gammaproteobacteria>Moraxellales>Moraxellaceae>Acinetobacter | GCF_013418995.1 | Acinetobacter_sp._YH12106 | FTU45_RS13255 | 18925..22599 | 2601094 | BREX-system-P-loop-protein-BrxC-[Acinetobacter-sp.-YH12106] |
| WP_180086491.1 | Type1.BREX | <-DUF6079*<-BrxB-BrxF-iSTAND<-BrxA-DNA-binding<-wHTH+WYL+WCX\|\|DUF4236-><-DOC+HTH<-?<-?\|\|?-><-ParB | 1224 | Bacteria>Pseudomonadota>Gammaproteobacteria>Moraxellales>Moraxellaceae>Acinetobacter | GCF_013419115.1 | Acinetobacter_sp._YH12103 | FTU43_RS13620 | 1696..5370 | 2601092 | BREX-system-P-loop-protein-BrxC-[Acinetobacter-sp.-YH12103] |
| WP_180036170.1 | Type1.BREX | <-SIGMA+OB-Fold+AAA-ATPase+Lon-Protease-BrxL<-PglZ<-N6-METHYLASE<-DUF6079*<-BrxB-BrxF-iSTAND<-BrxA-DNA-binding<-wHTH+WYL+WCX\|\|DUF4236-> | 1226 | Bacteria>Pseudomonadota>Gammaproteobacteria>Moraxellales>Moraxellaceae>Acinetobacter | GCF_013419295.1 | unclassified_Acinetobacter | FTU34_RS16295 | 9372..13052 | 196816 | MULTISPECIES:-BREX-system-P-loop-protein-BrxC-[unclassified-Acinetobacter] |
| WP_180108075.1 | Type1.BREX | <-SIGMA+OB-Fold+AAA-ATPase+HKD-DNase-BrxL<-PglZ<-N6-METHYLASE<-REase<-OLD-ABC<-DUF6079*<-BrxB-BrxF-iSTAND<-XIS-HTH<-?<-wHTH+HNH<-SF1-SF2-Helicase<-?<-SAM-METHYLASE\|\|?-><-ParB+HNH | 1190 | Bacteria>Pseudomonadota>Gammaproteobacteria>Moraxellales>Moraxellaceae>Acinetobacter | GCF_013419415.1 | Acinetobacter_sp._YH12085 | FTU28_RS13915 | 49013..52585 | 2601077 | BREX-system-P-loop-protein-BrxC-[Acinetobacter-sp.-YH12085] |
| WP_180131186.1 | Type1.BREX | Phage_integrase-><-HTH_OrfB_IS605+OrfB_IS605+OrfB_Zn_ribbon\|\|Y1_Tnp->ParB+HNH->CR-ATPase5+REase-><-DUF4236\|\|wHTH+WYL+WCX->BrxA-DNA-binding->BrxB-BrxF-iSTAND->DUF6079*->N6-METHYLASE->HKD-DNase+DUF3150->PglZ->SIGMA+OB-Fold+AAA-ATPase+Lon-Protease-BrxL-> | 1226 | Bacteria>Pseudomonadota>Gammaproteobacteria>Moraxellales>Moraxellaceae>Acinetobacter | GCF_013419565.1 | Acinetobacter_sp._YH12072 | FTU19_RS11080 | 35259..38939 | 2601068 | BREX-system-P-loop-protein-BrxC-[Acinetobacter-sp.-YH12072] |
| WP_180114868.1 | Type1.BREX | CR-ATPase5+REase-><-DUF4236\|\|wHTH+WYL+WCX->BrxA-DNA-binding->BrxB-BrxF-iSTAND->DUF6079*->N6-METHYLASE->PglZ->SIGMA+OB-Fold+AAA-ATPase+Lon-Protease-BrxL->\|\|?->?->?->?-><-DDE_Tnp_IS240 | 1226 | Bacteria>Pseudomonadota>Gammaproteobacteria>Moraxellales>Moraxellaceae>Acinetobacter | GCF_013419715.1 | Acinetobacter_sp._YH12063 | FTU12_RS15130 | 5418..9098 | 2601061 | BREX-system-P-loop-protein-BrxC-[Acinetobacter-sp.-YH12063] |
| WP_180160550.1 | Type1.BREX | <-SIGMA+OB-Fold+AAA-ATPase+Lon-Protease-BrxL<-PglZ<-?<-N6-METHYLASE<-DUF6079*<-BrxB-BrxF-iSTAND<-BrxA-DNA-binding<-wHTH+WYL+WCX\|\|DUF4236-><-CR-ATPase5+REase | 1226 | Bacteria>Pseudomonadota>Gammaproteobacteria>Moraxellales>Moraxellaceae>Acinetobacter | GCF_013419795.1 | Acinetobacter_sp._YH12054 | FTU07_RS15240 | 20642..24322 | 2601056 | BREX-system-P-loop-protein-BrxC-[Acinetobacter-sp.-YH12054] |
| WP_180165638.1 | Type1.BREX | <-SIGMA+OB-Fold+AAA-ATPase+Lon-Protease-BrxL<-PglZ<-?<-N6-METHYLASE<-DUF6079*<-BrxB-BrxF-iSTAND<-BrxA-DNA-binding<-wHTH+WYL+WCX\|\|DUF4236-><-CR-ATPase5+REase | 1226 | Bacteria>Pseudomonadota>Gammaproteobacteria>Moraxellales>Moraxellaceae>Acinetobacter | GCF_013419875.1 | Acinetobacter_sp._YH12049 | FTU05_RS13040 | 31963..35643 | 2601054 | BREX-system-P-loop-protein-BrxC-[Acinetobacter-sp.-YH12049] |
| WP_180174851.1 | Type1.BREX | <-DUF4096+DDE_Tnp_1+DDE_Tnp_1_2<-?\|\|?-><-MACRODOMAIN<-REase\|\|<-SF1-SF2-Helicase+REase<-SIGMA+OB-Fold+AAA-ATPase+Lon-Protease-BrxL<-PglZ<-N6-METHYLASE<-DUF6079*<-BrxB-BrxF-iSTAND<-BrxA-DNA-binding<-wHTH+WYL+WCX<-wHTH+HNH<-DDE_Tnp | 1226 | Bacteria>Pseudomonadota>Gammaproteobacteria>Moraxellales>Moraxellaceae>Acinetobacter | GCF_013420235.1 | Acinetobacter_sp._YH01022 | FTT87_RS10805 | 36696..40376 | 2601036 | BREX-system-P-loop-protein-BrxC-[Acinetobacter-sp.-YH01022] |
| WP_180087504.1 | Type1.BREX | Phage_integrase+Phage_integrase->ParB+HNH->CR-ATPase5+REase->\|\|wHTH+WYL+WCX->BrxA-DNA-binding->BrxB-BrxF-iSTAND->DUF6079*->N6-METHYLASE->PglZ->SIGMA+OB-Fold+AAA-ATPase+Lon-Protease-BrxL->?-><-DDE_Tnp_IS240 | 1226 | Bacteria>Pseudomonadota>Gammaproteobacteria>Moraxellales>Moraxellaceae>Acinetobacter | GCF_013420255.1 | unclassified_Acinetobacter | FTT86_RS12000 | 35844..39524 | 196816 | MULTISPECIES:-BREX-system-P-loop-protein-BrxC-[unclassified-Acinetobacter] |
| WP_180178840.1 | Type1.BREX | <-SIGMA+OB-Fold+AAA-ATPase+Lon-Protease-BrxL<-PglZ<-N6-METHYLASE<-DUF6079*<-BrxB-BrxF-iSTAND<-BrxA-DNA-binding<-wHTH+WYL+WCX\|\|DUF4236-><-CR-ATPase5+REase | 1226 | Bacteria>Pseudomonadota>Gammaproteobacteria>Moraxellales>Moraxellaceae>Acinetobacter | GCF_013420795.1 | Acinetobacter_sp._YH01013 | FTT80_RS13290 | 20368..24048 | 2601029 | BREX-system-P-loop-protein-BrxC-[Acinetobacter-sp.-YH01013] |
| WP_180006134.1 | Type1.BREX | DDE_Tnp_IS240-><-?<-?<-?<-?\|\|<-SIGMA+OB-Fold+AAA-ATPase+Lon-Protease-BrxL<-PglZ<-N6-METHYLASE<-DUF6079*<-BrxB-BrxF-iSTAND<-BrxA-DNA-binding<-wHTH+WYL+WCX\|\|DUF4236-><-CR-ATPase5+REase | 1226 | Bacteria>Pseudomonadota>Gammaproteobacteria>Moraxellales>Moraxellaceae>Acinetobacter | GCF_013420805.1 | unclassified_Acinetobacter | FTT79_RS10555 | 27794..31474 | 196816 | MULTISPECIES:-BREX-system-P-loop-protein-BrxC-[unclassified-Acinetobacter] |
| WP_182925313.1 | Type1.BREX | <-SF1-SF2-Helicase+REase<-SIGMA+OB-Fold+AAA-ATPase+Lon-Protease-BrxL<-PglZ<-SbcC<-N6-METHYLASE<-DUF6079*<-BrxB-BrxF-iSTAND<-BrxA-DNA-binding<-wHTH+WYL+WCX<-wHTH+HNH<-SAM-METHYLASE<-XIS-HTH | 1226 | Bacteria>Pseudomonadota>Gammaproteobacteria>Moraxellales>Moraxellaceae>Acinetobacter | GCF_014169655.1 | Acinetobacter_baumannii | H7R41_RS13110 | 2778553..2782233 | 470 | BREX-system-P-loop-protein-BrxC-[Acinetobacter-baumannii] |
| WP_185218776.1 | Type1.BREX | CR-ATPase5+REase-><-DUF4236\|\|wHTH+WYL+WCX->BrxA-DNA-binding->BrxB-BrxF-iSTAND->DUF6079*->N6-METHYLASE->ParB+HNH->PglZ->SIGMA+OB-Fold+AAA-ATPase+Lon-Protease-BrxL->\|\|?->?->?->?-><-DDE_Tnp_IS240 | 1226 | Bacteria>Pseudomonadota>Gammaproteobacteria>Moraxellales>Moraxellaceae>Acinetobacter | GCF_014218985.1 | Acinetobacter_baumannii | H6W49_RS17735 | 6511..10191 | 470 | BREX-system-P-loop-protein-BrxC-[Acinetobacter-baumannii] |
| WP_188199768.1 | Type1.BREX | <-RelE-ParE<-RelB\|\|?->ParB+HNH->CR-ATPase5+REase-><-DUF4236\|\|wHTH+WYL+WCX->BrxA-DNA-binding->BrxB-BrxF-iSTAND->DUF6079*->N6-METHYLASE->REase->PglZ->SIGMA+OB-Fold+AAA-ATPase+Lon-Protease-BrxL->\|\|SIR2-><-HTH_3<-RelE-ParE\|\|HEPN->DDE_Tnp_IS240-> | 1226 | Bacteria>Pseudomonadota>Gammaproteobacteria>Moraxellales>Moraxellaceae>Acinetobacter | GCF_014560035.1 | Acinetobacter_baumannii | HPJ37_RS18500 | 29485..33165 | 470 | BREX-system-P-loop-protein-BrxC-[Acinetobacter-baumannii] |
| WP_193893631.1 | Type1.BREX | HigB_toxin->HTH_3->CR-ATPase5+REase-><-DUF4236\|\|wHTH+WYL+WCX->BrxA-DNA-binding->BrxB-BrxF-iSTAND->DUF6079*->N6-METHYLASE->ParB+HNH->PglZ->SIGMA+OB-Fold+AAA-ATPase+Lon-Protease-BrxL-> | 1226 | Bacteria>Pseudomonadota>Gammaproteobacteria>Moraxellales>Moraxellaceae>Acinetobacter | GCF_015209685.1 | Acinetobacter_albensis | IQR32_RS12760 | 7183..10863 | 1673609 | BREX-system-P-loop-protein-BrxC-[Acinetobacter-albensis] |
| WP_180025627.1 | Type1.BREX | DDE_Tnp_IS240-><-?<-?<-?<-?\|\|<-SIGMA+OB-Fold+AAA-ATPase+Lon-Protease-BrxL<-PglZ<-N6-METHYLASE<-DUF6079*<-BrxB-BrxF-iSTAND<-BrxA-DNA-binding<-wHTH+WYL+WCX\|\|DUF4236-><-CR-ATPase5+REase\|\|?->HATPase+wHTH-><-?<-Zn_Tnp_IS91+Y2_Tnp | 1226 | Bacteria>Pseudomonadota>Gammaproteobacteria>Moraxellales>Moraxellaceae>Acinetobacter | GCF_015218525.1 | unclassified_Acinetobacter | G0029_RS17570 | 86789..90469 | 196816 | MULTISPECIES:-BREX-system-P-loop-protein-BrxC-[unclassified-Acinetobacter] |
| WP_136512845.1 | Type1.BREX | <-SIGMA+OB-Fold+AAA-ATPase+Lon-Protease-BrxL<-PglZ<-ParB+HNH<-N6-METHYLASE<-DUF6079*<-BrxB-BrxF-iSTAND<-BrxA-DNA-binding<-wHTH+WYL+WCX<-DUF932\|\|PIN+OST-HTH-><-ZnR+DUF3482+ZnR | 1226 | Bacteria>Pseudomonadota>Gammaproteobacteria>Moraxellales>Moraxellaceae>Acinetobacter | GCF_015265775.1 | Acinetobacter_baumannii | O9X70_RS06810 | 133723..137403 | 470 | BREX-system-P-loop-protein-BrxC-[Acinetobacter-baumannii] |
| WP_196075355.1 | Type1.BREX | AbiEii->\|\|ParB+HNH->CR-ATPase5+REase-><-DUF4236\|\|wHTH+WYL+WCX->BrxA-DNA-binding->BrxB-BrxF-iSTAND->DUF6079*->ParB+HNH->PglZ->?->Arm-DNA-bind_3+Phage_integrase->HicB-><-?\|\|HTH_26->\|\|DEOR-HTH-> | 1204 | Bacteria>Pseudomonadota>Gammaproteobacteria>Moraxellales>Moraxellaceae>Acinetobacter | GCF_015676315.1 | unclassified_Acinetobacter | IM039_RS19270 | 9799..13413 | 196816 | MULTISPECIES:-BREX-system-P-loop-protein-BrxC-[unclassified-Acinetobacter] |
| WP_198080379.1 | Type1.BREX | <-SF1-SF2-Helicase+REase<-SIGMA+OB-Fold+AAA-ATPase+Lon-Protease-BrxL<-PglZ<-ParB+HNH<-N6-METHYLASE<-DUF6079*<-BrxB-BrxF-iSTAND<-wHTH+WYL+WCX | 1222 | Bacteria>Pseudomonadota>Gammaproteobacteria>Moraxellales>Moraxellaceae>Acinetobacter | GCF_016082245.1 | Acinetobacter_calcoaceticus | BL242_RS20750 | 15792..19460 | 471 | BREX-system-P-loop-protein-BrxC-[Acinetobacter-calcoaceticus] |
| WP_199973361.1 | Type1.BREX | AraC-HTH->?-><-?<-REase<-?\|\|tRNA-><-?\|\|HTH_19-><-PglZ<-N6-METHYLASE<-DUF6079*<-BrxB-BrxF-iSTAND<-BrxA-DNA-binding\|\|?->tRNA-><-?<-SAM-METHYLASE<-?<-DSBH\|\|?->MODE-HTH+PBPII-> | 1249 | Bacteria>Pseudomonadota>Gammaproteobacteria>Moraxellales>Moraxellaceae>Acinetobacter | GCF_016502305.1 | Acinetobacter_courvalinii | I6M62_RS06800 | 311191..314940 | 280147 | BREX-system-P-loop-protein-BrxC-[Acinetobacter-courvalinii] |
| WP_032001825.1 | Type1.BREX | PhdYeFM_antitox-><-OLD-ABC+TOPRIM<-SF1-SF2-Helicase+REase<-SIGMA+OB-Fold+AAA-ATPase+Lon-Protease-BrxL<-PglZ<-?<-N6-METHYLASE<-DUF6079*<-BrxB-BrxF-iSTAND<-BrxA-DNA-binding<-wHTH+WYL+WCX\|\|<-DOC+HTH<-?\|\|DDE_Tnp_1_5+DDE_Tnp_1-> | 1226 | Bacteria>Pseudomonadota>Gammaproteobacteria>Moraxellales>Moraxellaceae>Acinetobacter | GCF_016502765.1 | Acinetobacter_calcoaceticus/baumannii_complex | I6M92_RS14775 | 66667..70347 | 909768 | MULTISPECIES:-BREX-system-P-loop-protein-BrxC-[Acinetobacter-calcoaceticus/baumannii-complex] |
| WP_199939733.1 | Type1.BREX | <-Phage_GPD+Phage_base_V+T6SS_Vgr+DUF2345<-TetR-HTH\|\|?-><-?<-SIGMA+OB-Fold+AAA-ATPase+Lon-Protease-BrxL<-PglZ<-ParB+HNH<-N6-METHYLASE<-DUF6079*<-BrxB-BrxF-iSTAND<-BrxA-DNA-binding<-wHTH+WYL+WCX<-DUF932\|\|PIN+OST-HTH-><-ZnR+DUF3482+ZnR | 1226 | Bacteria>Pseudomonadota>Gammaproteobacteria>Moraxellales>Moraxellaceae>Acinetobacter | GCF_016506315.1 | Acinetobacter_baumannii | I5555_RS13010 | 139911..143591 | 470 | BREX-system-P-loop-protein-BrxC-[Acinetobacter-baumannii] |
| WP_200230916.1 | Type1.BREX | SbcC+SbcC+ABC-ATPase+SbcC->?->Phage_integrase->ParB+HNH->CR-ATPase5+REase-><-DUF4236\|\|wHTH+WYL+WCX->BrxA-DNA-binding->BrxB-BrxF-iSTAND->DUF6079*->N6-METHYLASE->PglZ->SIGMA+OB-Fold+AAA-ATPase+Lon-Protease-BrxL->\|<-?<-?<-?<-?<-?<-ArsR-HTH | 1226 | Bacteria>Pseudomonadota>Gammaproteobacteria>Moraxellales>Moraxellaceae>Acinetobacter | GCF_016607565.1 | Acinetobacter_variabilis | IAQ69_RS15330 | 32669..36349 | 70346 | BREX-system-P-loop-protein-BrxC-[Acinetobacter-variabilis] |
| WP_201704335.1 | Type1.BREX | SbcC+SbcC+ABC-ATPase+SbcC->DDE_Tnp_1_5+DDE_Tnp_1-><-ADH_N+ADH_zinc_N<-?\|\|<-GDResolvase+HIN-HTH<-HEPN+RES<-SIGMA+OB-Fold+AAA-ATPase+Lon-Protease-BrxL<-PglZ<-N6-METHYLASE<-DUF6079*<-BrxB-BrxF-iSTAND<-BrxA-DNA-binding<-wHTH+WYL+WCX\|\|DUF4236-><-?\|\|<-Phage_integrase\|\|\|\|<-LZ_Tnp_IS66+zf-IS66+DDE_Tnp_IS66+DDE_Tnp_IS66_C<-?<-HIN-HTH | 1223 | Bacteria>Pseudomonadota>Gammaproteobacteria>Moraxellales>Moraxellaceae>Acinetobacter | GCF_016726625.1 | Acinetobacter_johnsonii | I6I51_RS17025 | 51852..55523 | 40214 | BREX-system-P-loop-protein-BrxC-[Acinetobacter-johnsonii] |
| WP_203080037.1 | Type1.BREX | TRD->Cap4_nuclease->?->?->?-><-TetR-HTH\|\|BrxA-DNA-binding->BrxB-BrxF-iSTAND->DUF6079*->N6-METHYLASE->ParB+HNH->PglZ->SIGMA+OB-Fold+AAA-ATPase+Lon-Protease-BrxL->SF1-SF2-Helicase+REase-><-DUF772+DDE_Tnp_1+DDE_Tnp_1_6\|\|DarT+Peptidase->MACRODOMAIN-> | 1226 | Bacteria>Pseudomonadota>Gammaproteobacteria>Moraxellales>Moraxellaceae>Acinetobacter | GCF_016804045.1 | Acinetobacter_pittii | HRJ47_RS04290 | 884929..888609 | 48296 | BREX-system-P-loop-protein-BrxC-[Acinetobacter-pittii] |
| WP_204086201.1 | Type1.BREX | PhdYeFM_antitox-><-OLD-ABC+TOPRIM<-SF1-SF2-Helicase+REase<-SIGMA+OB-Fold+AAA-ATPase+Lon-Protease-BrxL<-PglZ<-?<-N6-METHYLASE<-DUF6079*<-BrxB-BrxF-iSTAND<-BrxA-DNA-binding<-wHTH+WYL+WCX\|\|<-DOC+HTH<-?\|\|DDE_Tnp_1_5+DDE_Tnp_1-> | 1226 | Bacteria>Pseudomonadota>Gammaproteobacteria>Moraxellales>Moraxellaceae>Acinetobacter | GCF_016889825.1 | Acinetobacter_pittii | I6J44_RS19355 | 45796..49476 | 48296 | BREX-system-P-loop-protein-BrxC-[Acinetobacter-pittii] |
| WP_205779680.1 | Type1.BREX | Phage_integrase+Phage_integrase->Phage_integrase+Phage_integrase->?->ParB+HNH->CR-ATPase5+REase-><-DUF4236\|\|wHTH+WYL+WCX->BrxA-DNA-binding->BrxB-BrxF-iSTAND->DUF6079*->N6-METHYLASE->SF1-SF2-Helicase+PRTase->SLOG+BirA-HTH->ParB+HNH->PglZ->SIGMA+OB-Fold+AAA-ATPase+Lon-Protease-BrxL->SF1-SF2-Helicase+REase-><-LZ_Tnp_IS66+zf-IS66+DDE_Tnp_IS66+DDE_Tnp_IS66_C | 1226 | Bacteria>Pseudomonadota>Gammaproteobacteria>Moraxellales>Moraxellaceae>Acinetobacter | GCF_017132735.1 | Acinetobacter_indicus | JYB86_RS06365 | 1326455..1330135 | 756892 | BREX-system-P-loop-protein-BrxC-[Acinetobacter-indicus] |
| WP_206863034.1 | Type1.BREX | CR-ATPase5+REase-><-DUF4236\|\|wHTH+WYL+WCX->BrxA-DNA-binding->BrxB-BrxF-iSTAND->DUF6079*->N6-METHYLASE->PglZ->SIGMA+OB-Fold+AAA-ATPase+Lon-Protease-BrxL->\|\|?->?->?->?->?->GDResolvase+HIN-HTH-> | 1226 | Bacteria>Pseudomonadota>Gammaproteobacteria>Moraxellales>Moraxellaceae>Acinetobacter | GCF_017309425.1 | Acinetobacter_indicus | JOW32_RS14080 | 4215..7895 | 756892 | BREX-system-P-loop-protein-BrxC-[Acinetobacter-indicus] |
| WP_206896797.1 | Type1.BREX | Phage_integrase+Phage_integrase->Phage_integrase+Phage_integrase->?->ParB+HNH->CR-ATPase5+REase-><-DUF4236\|\|wHTH+WYL+WCX->BrxA-DNA-binding->BrxB-BrxF-iSTAND->DUF6079*->N6-METHYLASE->PglZ->SIGMA+OB-Fold+AAA-ATPase+Lon-Protease-BrxL->SF1-SF2-Helicase+REase-><-HIN-HTH+rve+rve_3<-DOC+HTH\|\|SAM-METHYLASE->SAM-METHYLASE-> | 1226 | Bacteria>Pseudomonadota>Gammaproteobacteria>Moraxellales>Moraxellaceae>Acinetobacter | GCF_017309445.1 | Acinetobacter_indicus | JOW33_RS13795 | 2887588..2891268 | 756892 | BREX-system-P-loop-protein-BrxC-[Acinetobacter-indicus] |
| WP_208464845.1 | Type1.BREX | <-DUF6079-p*<-BrxB-BrxF-iSTAND<-BrxA-DNA-binding<-wHTH+WYL+WCX\|\|DUF4236-><-CR-ATPase5+REase<-ParB+HNH | 537 | Bacteria>Pseudomonadota>Gammaproteobacteria>Moraxellales>Moraxellaceae>Acinetobacter | GCF_017592465.1 | Acinetobacter_haemolyticus | J5N55_RS15595 | 1..1612 | 29430 | BREX-system-P-loop-protein-BrxC,-partial-[Acinetobacter-haemolyticus] |
| WP_109426631.1 | Type1.BREX | DUF932->?->?->?-><-?<-TetR-HTH\|\|wHTH+WYL+WCX->BrxA-DNA-binding->BrxB-BrxF-iSTAND->DUF6079*->N6-METHYLASE->ParB+HNH->PglZ->SIGMA+OB-Fold+AAA-ATPase+Lon-Protease-BrxL->SF1-SF2-Helicase+REase->DarT+Peptidase->MACRODOMAIN-> | 1226 | Bacteria>Pseudomonadota>Gammaproteobacteria>Moraxellales>Moraxellaceae>Acinetobacter | GCF_017726575.1 | Acinetobacter_baumannii | J6I88_RS04260 | 873693..877373 | 470 | BREX-system-P-loop-protein-BrxC-[Acinetobacter-baumannii] |
| WP_212272673.1 | Type1.BREX | <-DDE_Tnp_1_5+DDE_Tnp_1\|\|?->DOC+HTH->\|\|wHTH+WYL+WCX->BrxA-DNA-binding->BrxB-BrxF-iSTAND->DUF6079*->N6-METHYLASE->?->PglZ->SIGMA+OB-Fold+AAA-ATPase+Lon-Protease-BrxL->SF1-SF2-Helicase+REase->OLD-ABC+TOPRIM-> | 1226 | Bacteria>Pseudomonadota>Gammaproteobacteria>Moraxellales>Moraxellaceae>Acinetobacter | GCF_018139165.1 | Acinetobacter_nosocomialis | KC103_RS15085 | 69521..73201 | 106654 | BREX-system-P-loop-protein-BrxC-[Acinetobacter-nosocomialis] |
| WP_213705174.1 | Type1.BREX | Phage_integrase+Phage_integrase->Phage_integrase+Phage_integrase->ParB+HNH->?->DOC+HTH-><-DUF4236\|\|wHTH+WYL+WCX->BrxA-DNA-binding->BrxB-BrxF-iSTAND->DUF6079*->N6-METHYLASE->PglZ->SIGMA+OB-Fold+AAA-ATPase+Lon-Protease-BrxL-><-?<-?<-?\|\|?->?-><-LuxR-HTH | 1226 | Bacteria>Pseudomonadota>Gammaproteobacteria>Moraxellales>Moraxellaceae>Acinetobacter | GCF_018446675.1 | Acinetobacter_baumannii | KKV18_RS14925 | 28314..31994 | 470 | BREX-system-P-loop-protein-BrxC-[Acinetobacter-baumannii] |
| WP_215276195.1 | Type1.BREX | <-ADH_N+ADH_zinc_N_2\|\|?->Phage_integrase+Phage_integrase->Phage_integrase->wHTH(SBD)+HNH->wHTH+WYL+WCX->BrxA-DNA-binding->BrxB-BrxF-iSTAND->DUF6079*->N6-METHYLASE->Aminotran_3->?->?->?->?-><-MODE-HTH+PBPII | 1226 | Bacteria>Pseudomonadota>Gammaproteobacteria>Moraxellales>Moraxellaceae>Acinetobacter | GCF_018678125.1 | Acinetobacter_baumannii | KMP96_RS12165 | 16275..19955 | 470 | BREX-system-P-loop-protein-BrxC-[Acinetobacter-baumannii] |
| WP_216174886.1 | Type1.BREX | DUF932->?->?->?-><-?<-TetR-HTH\|\|wHTH+WYL+WCX->BrxA-DNA-binding->BrxB-BrxF-iSTAND->DUF6079*->N6-METHYLASE->ParB+HNH->PglZ->SIGMA+OB-Fold+AAA-ATPase+Lon-Protease-BrxL->SF1-SF2-Helicase+REase->DarT+Peptidase->MACRODOMAIN-> | 1226 | Bacteria>Pseudomonadota>Gammaproteobacteria>Moraxellales>Moraxellaceae>Acinetobacter | GCF_018863255.1 | Acinetobacter_baumannii | KPE97_RS15710 | 83323..87003 | 470 | BREX-system-P-loop-protein-BrxC-[Acinetobacter-baumannii] |
| WP_216116819.1 | Type1.BREX | Peptidase_M48-><-MACRODOMAIN<-DarT+Peptidase<-SF1-SF2-Helicase+REase<-SIGMA+OB-Fold+AAA-ATPase+Lon-Protease-BrxL<-PglZ<-ParB+HNH<-N6-METHYLASE<-DUF6079*<-BrxB-BrxF-iSTAND<-BrxA-DNA-binding<-wHTH+WYL+WCX\|\|TetR-HTH-><-?<-?<-Cap4_nuclease<-TRD | 1226 | Bacteria>Pseudomonadota>Gammaproteobacteria>Moraxellales>Moraxellaceae>Acinetobacter | GCF_018883565.1 | Acinetobacter_baumannii | KPF69_RS05265 | 35520..39200 | 470 | BREX-system-P-loop-protein-BrxC-[Acinetobacter-baumannii] |
| WP_216936951.1 | Type1.BREX | <-HTH_3\|\|?-><-Phage_integrase<-Phage_integrase+Phage_integrase<-HTH\|\|XIS-HTH->BrxA-DNA-binding->BrxB-BrxF-iSTAND->DUF6079*->OLD-ABC->REase->N6-METHYLASE->PglZ->SIGMA+OB-Fold+AAA-ATPase+HKD-DNase-BrxL-><-?<-REase\|\|ZnR-> | 1190 | Bacteria>Pseudomonadota>Gammaproteobacteria>Moraxellales>Moraxellaceae>Acinetobacter | GCF_019038255.1 | unclassified_Acinetobacter | KUG40_RS10660 | 10626..14198 | 196816 | MULTISPECIES:-BREX-system-P-loop-protein-BrxC-[unclassified-Acinetobacter] |
| WP_218698253.1 | Type1.BREX | <-MarR-HTH<-SF1-SF2-Helicase<-STYKIN+CR-Kinase<-?<-SIGMA+OB-Fold+AAA-ATPase+Lon-Protease-BrxL<-PglZ<-HKD-DNase+DUF3150<-ParB+HNH<-N6-METHYLASE<-DUF6079*<-BrxB-BrxF-iSTAND<-BrxA-DNA-binding<-wHTH+WYL+WCX\|\|DUF4236-><-RelE-ParE<-ParD_antitoxin\|\|<-?<-HIN-HTH | 1224 | Bacteria>Pseudomonadota>Gammaproteobacteria>Moraxellales>Moraxellaceae>Acinetobacter | GCF_019279015.1 | Acinetobacter_harbinensis | KZA13_RS08660 | 34272..37946 | 1353941 | BREX-system-P-loop-protein-BrxC-[Acinetobacter-harbinensis] |
| WP_221260378.1 | Type1.BREX | <-sGTP<-wHTH+WYL+WCX<-?<-?<-?<-SIGMA+OB-Fold+AAA-ATPase+Lon-Protease-BrxL<-PglZ<-ParB+HNH<-N6-METHYLASE<-DUF6079*<-BrxB-BrxF-iSTAND<-BrxA-DNA-binding<-wHTH+WYL+WCX<-wHTH(SBD)+HNH<-SF1-SF2-Helicase<-?<-SAM-METHYLASE<-ParB+HNH\|\|Calcineurin-> | 1226 | Bacteria>Pseudomonadota>Gammaproteobacteria>Moraxellales>Moraxellaceae>Acinetobacter | GCF_019702905.1 | Acinetobacter_sp._Tol_5 | TOL5_RS12615 | 2696813..2700493 | 710648 | BREX-system-P-loop-protein-BrxC-[Acinetobacter-sp.-Tol-5] |
| WP_223106928.1 | Type1.BREX | <-DEDD_Tnp_IS110+Transposase_20\|\|?->ADH_N+ADH_zinc_N+METHYLASE-><-HTH_Tnp_4+DDE_Tnp_4\|\|ABhydrolase->HTH_Tnp_1+HTH_21+rve+rve_3-><-SIGMA+OB-Fold+AAA-ATPase+Lon-Protease-BrxL<-PglZ<-N6-METHYLASE<-DUF6079*<-BrxB-BrxF-iSTAND<-BrxA-DNA-binding<-wHTH+WYL+WCX\|\|DUF4236-><-CR-ATPase5+REase\|\|?->HIN-HTH->AAA-ATPase->Couple_hipA+HipA_C-> | 1226 | Bacteria>Pseudomonadota>Gammaproteobacteria>Moraxellales>Moraxellaceae>Acinetobacter | GCF_019903235.1 | Acinetobacter_baumannii | H2787_RS17960 | 80252..83932 | 470 | BREX-system-P-loop-protein-BrxC-[Acinetobacter-baumannii] |
| WP_223239187.1 | Type1.BREX | <-DUF4236\|\|wHTH+WYL+WCX->BrxA-DNA-binding->BrxB-BrxF-iSTAND->DUF6079*->N6-METHYLASE->?->PglZ->SIGMA+OB-Fold+AAA-ATPase+Lon-Protease-BrxL-> | 1226 | Bacteria>Pseudomonadota>Gammaproteobacteria>Moraxellales>Moraxellaceae>Acinetobacter | GCF_019920865.1 | Acinetobacter_baumannii | HLI16_RS17115 | 2695..6375 | 470 | BREX-system-P-loop-protein-BrxC-[Acinetobacter-baumannii] |
| WP_207272916.1 | Type1.BREX | <-MACRODOMAIN<-DarT+Peptidase<-SF1-SF2-Helicase+REase<-SIGMA+OB-Fold+AAA-ATPase+Lon-Protease-BrxL<-PglZ<-ParB+HNH<-N6-METHYLASE<-DUF6079*<-BrxB-BrxF-iSTAND<-BrxA-DNA-binding<-wHTH+WYL+WCX\|\|TetR-HTH-> | 1226 | Bacteria>Pseudomonadota>Gammaproteobacteria>Moraxellales>Moraxellaceae>Acinetobacter | GCF_019921105.1 | Acinetobacter_baumannii | HLH91_RS14165 | 18027..21707 | 470 | BREX-system-P-loop-protein-BrxC-[Acinetobacter-baumannii] |
| WP_005236280.1 | Type1.BREX | <-DUF4096+DDE_Tnp_1+DDE_Tnp_1_2\|\|?->?->SbcC+ABC-ATPase-><-SF1-SF2-Helicase+REase<-SIGMA+OB-Fold+AAA-ATPase+Lon-Protease-BrxL<-PglZ<-ParB+HNH<-N6-METHYLASE<-DUF6079*<-BrxB-BrxF-iSTAND<-BrxA-DNA-binding<-wHTH+WYL+WCX<-wHTH(SBD)+HNH<-SF1-SF2-Helicase<-?<-SAM-METHYLASE<-ParB+HNH\|\|Calcineurin-> | 1226 | Bacteria>Pseudomonadota>Gammaproteobacteria>Moraxellales>Moraxellaceae>Acinetobacter | GCF_020099375.1 | Acinetobacter_variabilis | LA331_RS04655 | 963646..967326 | 70346 | BREX-system-P-loop-protein-BrxC-[Acinetobacter-variabilis] |
| WP_225554349.1 | Type1.BREX | DUF6079*->N6-METHYLASE->ParB+HNH->PglZ->SIGMA+OB-Fold+AAA-ATPase+Lon-Protease-BrxL->?-><-?<-DEOR-HTH<-?\|\|MerR-HTH-> | 1226 | Bacteria>Pseudomonadota>Gammaproteobacteria>Moraxellales>Moraxellaceae>Acinetobacter | GCF_020271765.1 | Acinetobacter_pseudolwoffii | LD052_RS14245 | 193616..1867 | 2053287 | BREX-system-P-loop-protein-BrxC-[Acinetobacter-pseudolwoffii] |
| WP_225554403.1 | Type1.BREX | <-GDResolvase+HIN-HTH<-?<-?<-?<-?<-?\|\|<-SIGMA+OB-Fold+AAA-ATPase+Lon-Protease-BrxL<-PglZ<-N6-METHYLASE<-DUF6079*<-BrxB-BrxF-iSTAND<-BrxA-DNA-binding<-wHTH+WYL+WCX\|\|DUF4236-><-DOC+HTH<-?\|\|?-><-ParB<-ParA-Soj-PloopNTPase | 1226 | Bacteria>Pseudomonadota>Gammaproteobacteria>Moraxellales>Moraxellaceae>Acinetobacter | GCF_020271765.1 | Acinetobacter_pseudolwoffii | LD052_RS15430 | 47806..51486 | 2053287 | BREX-system-P-loop-protein-BrxC-[Acinetobacter-pseudolwoffii] |
| WP_228414014.1 | Type1.BREX | ZnR+DUF3482+ZnR->\|\|DUF932->wHTH+WYL+WCX->BrxA-DNA-binding->BrxB-BrxF-iSTAND->DUF6079*->N6-METHYLASE->ParB+HNH->PglZ->SIGMA+OB-Fold+AAA-ATPase+Lon-Protease-BrxL-><-Peptidase_M48<-?\|\|?->TetR-HTH-> | 1226 | Bacteria>Pseudomonadota>Gammaproteobacteria>Moraxellales>Moraxellaceae>Acinetobacter | GCF_020810655.1 | Acinetobacter_baumannii | NLHDIDDJ_RS06545 | 1352098..1355778 | 470 | BREX-system-P-loop-protein-BrxC-[Acinetobacter-baumannii] |
| WP_230245698.1 | Type1.BREX | DDE_Tnp_1_5+DDE_Tnp_1->CR-ATPase5+REase-><-DUF4236\|\|wHTH+WYL+WCX->BrxA-DNA-binding->BrxB-BrxF-iSTAND->DUF6079*->N6-METHYLASE->PglZ->SIGMA+OB-Fold+AAA-ATPase+Lon-Protease-BrxL-><-DDE_Tnp_1<-?\|\|?->?->?->RNase-> | 1226 | Bacteria>Pseudomonadota>Gammaproteobacteria>Moraxellales>Moraxellaceae>Acinetobacter | GCF_020982655.1 | Acinetobacter_sp._PW68 | K1T76_RS14000 | 121130..124810 | 2865162 | BREX-system-P-loop-protein-BrxC-[Acinetobacter-sp.-PW68] |
| WP_233751176.1 | Type1.BREX | <-DDE_Tnp_1_5+DDE_Tnp_1\|\|?->DOC+HTH->\|\|wHTH+WYL+WCX->BrxA-DNA-binding->BrxB-BrxF-iSTAND->DUF6079*->N6-METHYLASE->?->PglZ->SIGMA+OB-Fold+AAA-ATPase+Lon-Protease-BrxL-><-ADH_N+ADH_zinc_N\|\|DDE_Tnp_1_5+DDE_Tnp_1-><-HTH_26\|\|?-><-Arm-DNA-bind_3+Phage_integrase | 1226 | Bacteria>Pseudomonadota>Gammaproteobacteria>Moraxellales>Moraxellaceae>Acinetobacter | GCF_021386775.1 | Acinetobacter_pittii | KR570_RS16050 | 9603..13283 | 48296 | BREX-system-P-loop-protein-BrxC-[Acinetobacter-pittii] |
| WP_233945509.1 | Type1.BREX | <-SF1-SF2-Helicase+REase<-SIGMA+OB-Fold+AAA-ATPase+Lon-Protease-BrxL<-PglZ<-ParB+HNH<-SLOG+BirA-HTH<-SF1-SF2-Helicase+PRTase<-N6-METHYLASE<-DUF6079*<-BrxB-BrxF-iSTAND<-BrxA-DNA-binding<-wHTH+WYL+WCX<-wHTH(SBD)+HNH<-SAM-METHYLASE<-ParB+HNH\|\|Calcineurin-> | 1226 | Bacteria>Pseudomonadota>Gammaproteobacteria>Moraxellales>Moraxellaceae>Acinetobacter | GCF_021391555.1 | Acinetobacter_sp._SH20PTE14 | LXF01_RS17045 | 120215..123895 | 2905879 | BREX-system-P-loop-protein-BrxC-[Acinetobacter-sp.-SH20PTE14] |
| WP_234115082.1 | Type1.BREX | Phage_integrase+Phage_integrase->Phage_integrase+Phage_integrase->ParB+HNH->?->DOC+HTH-><-DUF4236\|\|wHTH+WYL+WCX->BrxA-DNA-binding->BrxB-BrxF-iSTAND->DUF6079*->N6-METHYLASE->HKD-DNase+DUF3150->PglZ->SIGMA+OB-Fold+AAA-ATPase+Lon-Protease-BrxL-><-?<-ABhydrolase<-Classical-AAA<-?<-SMODS | 1226 | Bacteria>Pseudomonadota>Gammaproteobacteria>Moraxellales>Moraxellaceae>Acinetobacter | GCF_021432045.1 | Acinetobacter_towneri | LZG54_RS14455 | 23931..27611 | 202956 | BREX-system-P-loop-protein-BrxC-[Acinetobacter-towneri] |
| WP_234693498.1 | Type1.BREX | Phage_integrase+Phage_integrase->Phage_integrase+Phage_integrase->ParB+HNH->?->DOC+HTH-><-DUF4236\|\|wHTH+WYL+WCX->BrxA-DNA-binding->BrxB-BrxF-iSTAND->DUF6079*->N6-METHYLASE->HKD-DNase+DUF3150->PglZ->SIGMA+OB-Fold+AAA-ATPase+Lon-Protease-BrxL-><-?<-ABhydrolase<-Classical-AAA<-?<-SMODS | 1226 | Bacteria>Pseudomonadota>Gammaproteobacteria>Moraxellales>Moraxellaceae>Acinetobacter | GCF_021460555.1 | Acinetobacter_baumannii | LH676_RS19735 | 26787..30467 | 470 | BREX-system-P-loop-protein-BrxC-[Acinetobacter-baumannii] |
| WP_236095241.1 | Type1.BREX | <-PglZ<-ParB+HNH<-N6-METHYLASE<-DUF6079*<-BrxB-BrxF-iSTAND<-BrxA-DNA-binding<-wHTH+WYL+WCX\|\|DUF4236-><-CR-ATPase5+REase<-HTH_Tnp_1+HTH_21+rve+rve_3<-ParB+HNH<-Phage_integrase | 1226 | Bacteria>Pseudomonadota>Gammaproteobacteria>Moraxellales>Moraxellaceae>Acinetobacter | GCF_021560715.1 | Acinetobacter_sp._SCLZS86 | LZP46_RS14905 | 6362..10042 | 2908637 | BREX-system-P-loop-protein-BrxC-[Acinetobacter-sp.-SCLZS86] |
| WP_236163089.1 | Type1.BREX | Phage_integrase->ParB+HNH->HTH_Tnp_1+HTH_21+rve+rve_3->CR-ATPase5+REase-><-DUF4236\|\|wHTH+WYL+WCX->BrxA-DNA-binding->BrxB-BrxF-iSTAND->DUF6079*->N6-METHYLASE->ParB+HNH->PglZ->SIGMA+OB-Fold+AAA-ATPase+Lon-Protease-BrxL->\|\|?->?->?->SbcC+ABC-ATPase-> | 1226 | Bacteria>Pseudomonadota>Gammaproteobacteria>Moraxellales>Moraxellaceae>Acinetobacter | GCF_021602685.1 | unclassified_Acinetobacter | L6B80_RS11000 | 14595..18275 | 196816 | MULTISPECIES:-BREX-system-P-loop-protein-BrxC-[unclassified-Acinetobacter] |
| WP_239046817.1 | Type1.BREX | Phage_integrase+Phage_integrase->Phage_integrase+Phage_integrase->ParB+HNH->?->DOC+HTH-><-DUF4236\|\|wHTH+WYL+WCX->BrxA-DNA-binding->BrxB-BrxF-iSTAND->DUF6079*->N6-METHYLASE->PglZ->SIGMA+OB-Fold+AAA-ATPase+Lon-Protease-BrxL-><-?<-ABhydrolase<-Classical-AAA<-?<-SMODS\|\|ZnR+HTH_Tnp_ISL3+DDE_Tnp_ISL3-> | 1226 | Bacteria>Pseudomonadota>Gammaproteobacteria>Moraxellales>Moraxellaceae>Acinetobacter | GCF_022343575.1 | Acinetobacter_sp._AG3 | L3W46_RS22595 | 25482..29162 | 2912245 | BREX-system-P-loop-protein-BrxC-[Acinetobacter-sp.-AG3] |
| WP_241305487.1 | Type1.BREX | <-ABhydrolase<-ADH_N+ADH_zinc_N+METHYLASE<-?\|\|ADH_N+ADH_zinc_N_2->ADH_N+ADH_zinc_N_2->?-><-?<-PglZ<-N6-METHYLASE<-DUF6079*<-BrxB-BrxF-iSTAND<-BrxA-DNA-binding<-?<-?<-DUF932<-ZnR+ZnR | 1248 | Bacteria>Pseudomonadota>Gammaproteobacteria>Moraxellales>Moraxellaceae>Acinetobacter | GCF_022549325.1 | Acinetobacter_sp._NIPH_605 | MMP73_RS14775 | 133059..136805 | 2923434 | BREX-system-P-loop-protein-BrxC-[Acinetobacter-sp.-NIPH-605] |
| WP_241338465.1 | Type1.BREX | <-SIGMA+OB-Fold+AAA-ATPase+Lon-Protease-BrxL<-PglZ<-ParB+HNH<-SLOG+BirA-HTH<-SF1-SF2-Helicase+PRTase<-N6-METHYLASE<-DUF6079*<-BrxB-BrxF-iSTAND<-BrxA-DNA-binding<-wHTH+WYL+WCX<-wHTH(SBD)+HNH<-SF1-SF2-Helicase<-?<-SAM-METHYLASE<-ParB+HNH\|\|Calcineurin-> | 1226 | Bacteria>Pseudomonadota>Gammaproteobacteria>Moraxellales>Moraxellaceae>Acinetobacter | GCF_022549435.1 | Acinetobacter_sp._ANC_5659 | MMP64_RS09425 | 122256..125936 | 2923427 | BREX-system-P-loop-protein-BrxC-[Acinetobacter-sp.-ANC-5659] |
| WP_243475070.1 | Type1.BREX | DOC-><-MACRODOMAIN\|\|?->DEOR-HTH-><-SIGMA+OB-Fold+AAA-ATPase+Lon-Protease-BrxL<-PglZ<-N6-METHYLASE<-DUF6079*<-BrxB-BrxF-iSTAND<-BrxA-DNA-binding<-wHTH+WYL+WCX\|\|DUF4236-><-CR-ATPase5+REase<-ParB+HNH<-?<-AAA-ATPase<-HIN-HTH | 1226 | Bacteria>Pseudomonadota>Gammaproteobacteria>Moraxellales>Moraxellaceae>Acinetobacter | GCF_022809915.1 | Acinetobacter_lwoffii | MTX11_RS10505 | 45498..49178 | 28090 | BREX-system-P-loop-protein-BrxC-[Acinetobacter-lwoffii] |
| WP_248102847.1 | Type1.BREX | Phage_integrase->ParB+HNH-p->CR-ATPase5+REase-><-DUF4236\|\|wHTH+WYL+WCX->BrxA-DNA-binding->BrxB-BrxF-iSTAND->DUF6079*->N6-METHYLASE->PglZ->SIGMA+OB-Fold+AAA-ATPase+Lon-Protease-BrxL->ICLR-HTH-> | 1226 | Bacteria>Pseudomonadota>Gammaproteobacteria>Moraxellales>Moraxellaceae>Acinetobacter | GCF_023158935.1 | Acinetobacter_portensis | MZ021_RS15025 | 45372..49052 | 1839785 | BREX-system-P-loop-protein-BrxC-[Acinetobacter-portensis] |
| WP_249468201.1 | Type1.BREX | PAIREDC-HTH-><-?<-HTH_3<-?<-SIGMA+OB-Fold+AAA-ATPase+Lon-Protease-BrxL<-PglZ<-N6-METHYLASE<-DUF6079*<-BrxB-BrxF-iSTAND<-BrxA-DNA-binding<-wHTH+WYL+WCX\|\|DUF4236-><-CR-ATPase5+REase | 1226 | Bacteria>Pseudomonadota>Gammaproteobacteria>Moraxellales>Moraxellaceae>Acinetobacter | GCF_023371125.1 | Acinetobacter_baumannii | M4045_RS18565 | 26114..29794 | 470 | BREX-system-P-loop-protein-BrxC-[Acinetobacter-baumannii] |
| WP_249571539.1 | Type1.BREX | MarR-HTH->AbiEii->?-><-SIGMA+OB-Fold+AAA-ATPase+Lon-Protease-BrxL<-PglZ<-N6-METHYLASE<-DUF6079*<-BrxB-BrxF-iSTAND<-BrxA-DNA-binding<-wHTH+WYL+WCX\|\|DUF4236-><-CR-ATPase5+REase | 1226 | Bacteria>Pseudomonadota>Gammaproteobacteria>Moraxellales>Moraxellaceae>Acinetobacter | GCF_023488045.1 | Acinetobacter_sp._ANC5681 | M5121_RS14220 | 11775..15455 | 2929504 | BREX-system-P-loop-protein-BrxC-[Acinetobacter-sp.-ANC5681] |
| WP_250180441.1 | Type1.BREX | HTH_Tnp_1+HTH_21+rve+rve_3->CR-ATPase5+REase-><-DUF4236\|\|wHTH+WYL+WCX->BrxA-DNA-binding->BrxB-BrxF-iSTAND->DUF6079-p*-> | 386 | Bacteria>Pseudomonadota>Gammaproteobacteria>Moraxellales>Moraxellaceae>Acinetobacter | GCF_023572305.1 | Acinetobacter_baumannii | M9Y19_RS19145 | 4860..6017 | 470 | BREX-system-P-loop-protein-BrxC,-partial-[Acinetobacter-baumannii] |
| WP_128202440.1 | Type1.BREX | <-TPR+TPR<-?<-MACRODOMAIN<-DarT+Peptidase<-SF1-SF2-Helicase+REase<-SIGMA+OB-Fold+AAA-ATPase+Lon-Protease-BrxL<-PglZ<-ParB+HNH<-N6-METHYLASE<-DUF6079*<-BrxB-BrxF-iSTAND<-BrxA-DNA-binding<-wHTH+WYL+WCX\|\|TetR-HTH-><-?\|\|?-><-Arm-DNA-bind_3+Phage_integrase | 1226 | Bacteria>Pseudomonadota>Gammaproteobacteria>Moraxellales>Moraxellaceae>Acinetobacter | GCF_023896505.1 | Acinetobacter_baumannii | O9X56_RS14280 | 107190..110870 | 470 | BREX-system-P-loop-protein-BrxC-[Acinetobacter-baumannii] |
| WP_253153041.1 | Type1.BREX | ParB+HNH->CR-ATPase5+REase-><-DUF4236\|\|wHTH+WYL+WCX->BrxA-DNA-binding->BrxB-BrxF-iSTAND->DUF6079*->N6-METHYLASE->PglZ->SIGMA+OB-Fold+AAA-ATPase+Lon-Protease-BrxL->\|\|?-><-DDE_Tnp_IS240<-?\|\|MerR-HTH-><-HTH_37 | 1226 | Bacteria>Pseudomonadota>Gammaproteobacteria>Moraxellales>Moraxellaceae>Acinetobacter | GCF_024129285.1 | Acinetobacter_sp._S4397-1 | NI467_RS11480 | 6279..9959 | 2972915 | BREX-system-P-loop-protein-BrxC-[Acinetobacter-sp.-S4397-1] |
| WP_253097522.1 | Type1.BREX | <-SIGMA+OB-Fold+AAA-ATPase+Lon-Protease-BrxL<-PglZ<-N6-METHYLASE<-DUF6079*<-BrxB-BrxF-iSTAND<-BrxA-DNA-binding<-wHTH+WYL+WCX\|\|DUF4236-><-CR-ATPase5+REase<-ParB+HNH<-Phage_integrase<-?<-rve | 1226 | Bacteria>Pseudomonadota>Gammaproteobacteria>Moraxellales>Moraxellaceae>Acinetobacter | GCF_024129755.1 | Acinetobacter_indicus | NI393_RS13245 | 9324..13004 | 756892 | BREX-system-P-loop-protein-BrxC-[Acinetobacter-indicus] |
| WP_253167193.1 | Type1.BREX | ParB+HNH->SAM-METHYLASE->wHTH+HNH->wHTH+WYL+WCX->BrxA-DNA-binding->BrxB-BrxF-iSTAND->DUF6079*->N6-METHYLASE->PglZ->SIGMA+OB-Fold+AAA-ATPase+Lon-Protease-BrxL-><-?<-RADICAL-SAM | 1226 | Bacteria>Pseudomonadota>Gammaproteobacteria>Moraxellales>Moraxellaceae>Acinetobacter | GCF_024129795.1 | Acinetobacter_indicus | NI451_RS01020 | 208268..211948 | 756892 | BREX-system-P-loop-protein-BrxC-[Acinetobacter-indicus] |
| WP_253133338.1 | Type1.BREX | RADICAL-SAM->?-><-SIGMA+OB-Fold+AAA-ATPase+Lon-Protease-BrxL<-PglZ<-N6-METHYLASE<-DUF6079*<-BrxB-BrxF-iSTAND<-BrxA-DNA-binding<-wHTH+WYL+WCX<-wHTH+HNH<-SAM-METHYLASE<-ParB+HNH | 1224 | Bacteria>Pseudomonadota>Gammaproteobacteria>Moraxellales>Moraxellaceae>Acinetobacter | GCF_024129835.1 | Acinetobacter_indicus | NI394_RS03575 | 37848..41522 | 756892 | BREX-system-P-loop-protein-BrxC-[Acinetobacter-indicus] |
| WP_253145790.1 | Type1.BREX | DDE_Tnp_IS66+DDE_Tnp_IS66_C-><-HEPN<-?<-SIGMA+OB-Fold+AAA-ATPase+Lon-Protease-BrxL<-PglZ<-?<-N6-METHYLASE<-DUF6079*<-BrxB-BrxF-iSTAND<-BrxA-DNA-binding<-wHTH+WYL+WCX\|\|DUF4236-><-?<-ParB+HNH-p<-CR-ATPase5+REase\|\|HicA->HicB-> | 1222 | Bacteria>Pseudomonadota>Gammaproteobacteria>Moraxellales>Moraxellaceae>Acinetobacter | GCF_024129855.1 | Acinetobacter_lwoffii | NI465_RS14465 | 11210..14878 | 28090 | BREX-system-P-loop-protein-BrxC-[Acinetobacter-lwoffii] |
| WP_257259040.1 | Type1.BREX | <-MACRODOMAIN<-DarT+Peptidase<-SF1-SF2-Helicase+REase<-SIGMA+OB-Fold+AAA-ATPase+Lon-Protease-BrxL<-PglZ<-ParB+HNH<-N6-METHYLASE<-DUF6079*<-BrxB-BrxF-iSTAND<-BrxA-DNA-binding<-wHTH+WYL+WCX\|\|TetR-HTH->?-><-?<-?<-?<-DUF932 | 1226 | Bacteria>Pseudomonadota>Gammaproteobacteria>Moraxellales>Moraxellaceae>Acinetobacter | GCF_024589115.1 | Acinetobacter_baumannii | L2615_RS07865 | 180845..184525 | 470 | BREX-system-P-loop-protein-BrxC-[Acinetobacter-baumannii] |
| WP_180004195.1 | Type1.BREX | Zn_Tnp_IS91+Y2_Tnp->?-><-HATPase+wHTH<-?\|\|CR-ATPase5+REase-><-DUF4236\|\|wHTH+WYL+WCX->BrxA-DNA-binding->BrxB-BrxF-iSTAND->DUF6079*->N6-METHYLASE->PglZ->SIGMA+OB-Fold+AAA-ATPase+Lon-Protease-BrxL->\|\|?->?->?->?->?->GDResolvase+HIN-HTH-> | 1226 | Bacteria>Pseudomonadota>Gammaproteobacteria>Moraxellales>Moraxellaceae>Acinetobacter | GCF_024605785.1 | unclassified_Acinetobacter | MST16_RS17335 | 65383..69063 | 196816 | MULTISPECIES:-BREX-system-P-loop-protein-BrxC-[unclassified-Acinetobacter] |
| WP_257995145.1 | Type1.BREX | <-TPR+TPR<-MACRODOMAIN<-DarT+Peptidase<-SF1-SF2-Helicase+REase<-SIGMA+OB-Fold+AAA-ATPase+Lon-Protease-BrxL<-PglZ<-ParB+HNH<-N6-METHYLASE<-DUF6079*<-BrxB-BrxF-iSTAND<-BrxA-DNA-binding<-wHTH+WYL+WCX\|\|TetR-HTH->?-><-?<-?<-?<-DUF932 | 1226 | Bacteria>Pseudomonadota>Gammaproteobacteria>Moraxellales>Moraxellaceae>Acinetobacter | GCF_024705725.1 | Acinetobacter_baumannii | MSH04_RS15115 | 32588..36268 | 470 | BREX-system-P-loop-protein-BrxC-[Acinetobacter-baumannii] |
| WP_238640059.1 | Type1.BREX | <-SIGMA+OB-Fold+AAA-ATPase+Lon-Protease-BrxL<-PglZ<-ParB+HNH<-N6-METHYLASE<-DUF6079*<-BrxB-BrxF-iSTAND<-BrxA-DNA-binding<-wHTH+WYL+WCX\|\|DUF4236-><-CR-ATPase5+REase | 1226 | Bacteria>Pseudomonadota>Gammaproteobacteria>Moraxellales>Moraxellaceae>Acinetobacter | GCF_025264005.1 | Acinetobacter_baumannii | NT399_RS17670 | 10742..14422 | 470 | BREX-system-P-loop-protein-BrxC-[Acinetobacter-baumannii] |
| WP_262463188.1 | Type1.BREX | <-SIGMA+OB-Fold+AAA-ATPase+Lon-Protease-BrxL<-PglZ<-?<-N6-METHYLASE<-DUF6079*<-BrxB-BrxF-iSTAND<-BrxA-DNA-binding<-wHTH+WYL+WCX\|\|DUF4236-><-CR-ATPase5+REase | 1226 | Bacteria>Pseudomonadota>Gammaproteobacteria>Moraxellales>Moraxellaceae>Acinetobacter | GCF_025513035.1 | Acinetobacter_pittii | KTG74_RS19040 | 10083..13763 | 48296 | BREX-system-P-loop-protein-BrxC-[Acinetobacter-pittii] |
| WP_262456128.1 | Type1.BREX | CR-ATPase5+REase-><-DUF4236\|\|wHTH+WYL+WCX->BrxA-DNA-binding->BrxB-BrxF-iSTAND->DUF6079*->N6-METHYLASE->?->PglZ->SIGMA+OB-Fold+AAA-ATPase+Lon-Protease-BrxL-><-?<-AbiEii<-MarR-HTH | 1226 | Bacteria>Pseudomonadota>Gammaproteobacteria>Moraxellales>Moraxellaceae>Acinetobacter | GCF_025513465.1 | Acinetobacter_bereziniae | KTH44_RS20855 | 4679..8359 | 106648 | BREX-system-P-loop-protein-BrxC-[Acinetobacter-bereziniae] |
| WP_262468026.1 | Type1.BREX | RVT->?-><-?<-SIGMA+OB-Fold+AAA-ATPase+Lon-Protease-BrxL<-PglZ<-?<-N6-METHYLASE<-DUF6079*<-BrxB-BrxF-iSTAND<-BrxA-DNA-binding<-wHTH+WYL+WCX\|\|DUF4236-><-?<-ParB+HNH-p<-CR-ATPase5+REase<-ParB+HNH | 1226 | Bacteria>Pseudomonadota>Gammaproteobacteria>Moraxellales>Moraxellaceae>Acinetobacter | GCF_025513595.1 | Acinetobacter_schindleri | KTH34_RS14665 | 11459..15139 | 108981 | BREX-system-P-loop-protein-BrxC-[Acinetobacter-schindleri] |
| WP_262737085.1 | Type1.BREX | ParA-Soj-PloopNTPase->?-><-DDE_Tnp_IS240\|\|<-SIGMA+OB-Fold+AAA-ATPase+Lon-Protease-BrxL<-PglZ<-N6-METHYLASE<-DUF6079*<-BrxB-BrxF-iSTAND<-BrxA-DNA-binding<-wHTH+WYL+WCX\|\|DUF4236-><-CR-ATPase5+REase | 1226 | Bacteria>Pseudomonadota>Gammaproteobacteria>Moraxellales>Moraxellaceae>Acinetobacter | GCF_025514155.1 | Acinetobacter_pittii | KTI11_RS17510 | 13692..17372 | 48296 | BREX-system-P-loop-protein-BrxC-[Acinetobacter-pittii] |
| WP_086355030.1 | Type1.BREX | DEOR-HTH->ParB->ParB+HNH->CR-ATPase5+REase-><-DUF4236<-DUF772+DDE_Tnp_1+DDE_Tnp_1_6\|\|wHTH+WYL+WCX->BrxA-DNA-binding->BrxB-BrxF-iSTAND->DUF6079*->N6-METHYLASE->PglZ->SIGMA+OB-Fold+AAA-ATPase+Lon-Protease-BrxL-><-DDE_Tnp_IS240\|\|Sulfate_transp+STAS->NERD+Pkinase+Pkinase+SF1-SF2-Helicase->?->HIN-HTH->AAA-ATPase-> | 1226 | Bacteria>Pseudomonadota>Gammaproteobacteria>Moraxellales>Moraxellaceae>Acinetobacter | GCF_025514625.1 | Acinetobacter_calcoaceticus/baumannii_complex | KTI87_RS17630 | 13118..16798 | 909768 | MULTISPECIES:-BREX-system-P-loop-protein-BrxC-[Acinetobacter-calcoaceticus/baumannii-complex] |
| WP_262737771.1 | Type1.BREX | <-DDE_Tnp_1_5+DDE_Tnp_1\|\|MarR-HTH->AbiEii->?-><-SIGMA+OB-Fold+AAA-ATPase+Lon-Protease-BrxL<-PglZ<-N6-METHYLASE<-DUF6079*<-BrxB-BrxF-iSTAND<-BrxA-DNA-binding<-wHTH+WYL+WCX\|\|DUF4236-><-CR-ATPase5+REase<-ParB+HNH<-Y1_Tnp\|\|HTH_OrfB_IS605+OrfB_IS605+OrfB_Zn_ribbon-><-Phage_integrase | 1226 | Bacteria>Pseudomonadota>Gammaproteobacteria>Moraxellales>Moraxellaceae>Acinetobacter | GCF_025515095.1 | Acinetobacter_variabilis | KTJ53_RS03615 | 103263..106943 | 70346 | BREX-system-P-loop-protein-BrxC-[Acinetobacter-variabilis] |
| WP_262579272.1 | Type1.BREX | <-SIGMA+OB-Fold+AAA-ATPase+Lon-Protease-BrxL<-PglZ<-ParB+HNH<-N6-METHYLASE<-DUF6079*<-BrxB-BrxF-iSTAND<-BrxA-DNA-binding<-wHTH+WYL+WCX\|\|DUF4236-><-CR-ATPase5+REase | 1226 | Bacteria>Pseudomonadota>Gammaproteobacteria>Moraxellales>Moraxellaceae>Acinetobacter | GCF_025562355.1 | Acinetobacter_junii | KTH64_RS13010 | 10561..14241 | 40215 | BREX-system-P-loop-protein-BrxC-[Acinetobacter-junii] |
| WP_262558821.1 | Type1.BREX | BrxA-DNA-binding->BrxB-BrxF-iSTAND->DUF6079*->N6-METHYLASE->PglZ-p-> | 1224 | Bacteria>Pseudomonadota>Gammaproteobacteria>Moraxellales>Moraxellaceae>Acinetobacter | GCF_025562515.1 | Acinetobacter_bereziniae | KTH91_RS20770 | 1047..4721 | 106648 | BREX-system-P-loop-protein-BrxC-[Acinetobacter-bereziniae] |
| WP_263513455.1 | Type1.BREX | DDE_Tnp_1_5+DDE_Tnp_1->?->ADH_N+ADH_zinc_N+METHYLASE-><-DDE_Tnp_IS240<-PglZ<-N6-METHYLASE<-DUF6079*<-BrxB-BrxF-iSTAND<-BrxA-DNA-binding<-wHTH+WYL+WCX\|\|DUF4236-><-DOC+HTH<-?<-ParB+HNH-p\|\|Couple_hipA+HipA_C-><-HTH_26 | 1226 | Bacteria>Pseudomonadota>Gammaproteobacteria>Moraxellales>Moraxellaceae>Acinetobacter | GCF_025723145.1 | Acinetobacter_ursingii | LSD60_RS18250 | 102062..105742 | 108980 | BREX-system-P-loop-protein-BrxC-[Acinetobacter-ursingii] |
| WP_265466015.1 | Type1.BREX | <-Calcineurin\|\|ParB+HNH->SAM-METHYLASE->?->SF1-SF2-Helicase->wHTH(SBD)+HNH->wHTH+WYL+WCX->BrxA-DNA-binding->BrxB-BrxF-iSTAND->DUF6079*->N6-METHYLASE->ParB+HNH->PglZ->SIGMA+OB-Fold+AAA-ATPase+Lon-Protease-BrxL->SF1-SF2-Helicase+REase-><-?<-AA_permease | 1226 | Bacteria>Pseudomonadota>Gammaproteobacteria>Moraxellales>Moraxellaceae>Acinetobacter | GCF_026168575.1 | Acinetobacter_sp._BIT-DXN8 | OKC24_RS16365 | 19397..23077 | 2989714 | BREX-system-P-loop-protein-BrxC-[Acinetobacter-sp.-BIT-DXN8] |
| WP_267982262.1 | Type1.BREX | GDResolvase+HIN-HTH->CR-ATPase5+REase-><-DUF4236\|\|wHTH+WYL+WCX->BrxA-DNA-binding->BrxB-BrxF-iSTAND->DUF6079*->N6-METHYLASE->PglZ->SIGMA+OB-Fold+AAA-ATPase+Lon-Protease-BrxL-><-?<-AbiEii<-MarR-HTH<-?\|\|DEOR-HTH->HTH_37-> | 1226 | Bacteria>Pseudomonadota>Gammaproteobacteria>Moraxellales>Moraxellaceae>Acinetobacter | GCF_026736615.1 | Acinetobacter_sp._VNH17 | OYK78_RS17990 | 7578..11258 | 2998078 | BREX-system-P-loop-protein-BrxC-[Acinetobacter-sp.-VNH17] |
| WP_071225618.1 | Type1.BREX | ADH_N+ADH_zinc_N-><-MODE-HTH+PBPII\|\|MarR-HTH->AbiEii->?-><-SIGMA+OB-Fold+AAA-ATPase+Lon-Protease-BrxL<-PglZ<-ParB+HNH<-N6-METHYLASE<-DUF6079*<-BrxB-BrxF-iSTAND<-BrxA-DNA-binding<-wHTH+WYL+WCX<-wHTH(SBD)+HNH<-Phage_integrase<-Phage_integrase+Phage_integrase<-?<-DDE_Tnp_1 | 1226 | Bacteria>Pseudomonadota>Gammaproteobacteria>Moraxellales>Moraxellaceae>Acinetobacter | GCF_027854895.1 | Acinetobacter_baumannii | MJ031_RS19545 | 43789..47469 | 470 | BREX-system-P-loop-protein-BrxC-[Acinetobacter-baumannii] |
| WP_274579352.1 | Type1.BREX | <-DDE_Tnp_1_5+DDE_Tnp_1\|\|?->DOC+HTH-><-DUF4236\|\|wHTH+WYL+WCX->BrxA-DNA-binding->BrxB-BrxF-iSTAND->DUF6079*->N6-METHYLASE->ParB+HNH->PglZ->SIGMA+OB-Fold+AAA-ATPase+Lon-Protease-BrxL->SF1-SF2-Helicase+REase->OLD-ABC+TOPRIM-p-> | 1226 | Bacteria>Pseudomonadota>Gammaproteobacteria>Moraxellales>Moraxellaceae>Acinetobacter | GCF_028774645.1 | Acinetobacter_lactucae | MO054_RS17975 | 24360..28040 | 1785128 | BREX-system-P-loop-protein-BrxC-[Acinetobacter-lactucae] |
| WP_275290158.1 | Type1.BREX | <-HTH_Tnp_1+HTH_21+rve+rve_3\|\|<-HTH_3<-HigB_toxin\|\|CR-ATPase5+REase-><-DUF4236\|\|wHTH+WYL+WCX->BrxA-DNA-binding->BrxB-BrxF-iSTAND->DUF6079*->N6-METHYLASE->SF1-SF2-Helicase+PRTase->SLOG+BirA-HTH->ParB+HNH->PglZ->DDE_Tnp_IS240-><-?<-PAS+NtrC-AAA+FIS-HTH | 1226 | Bacteria>Pseudomonadota>Gammaproteobacteria>Moraxellales>Moraxellaceae>Acinetobacter | GCF_029100495.1 | Acinetobacter_nosocomialis | JIZ05_14385 | 7251..10931 | 106654 | BREX-system-P-loop-protein-BrxC-[Acinetobacter-nosocomialis] |
| WP_161409476.1 | Type1.BREX | wHTH+WYL+WCX->BrxA-DNA-binding->BrxB-BrxF-iSTAND->DUF6079*->N6-METHYLASE->SF1-SF2-Helicase+PRTase->SLOG+BirA-HTH->ParB+HNH->PglZ-> | 1226 | Bacteria>Pseudomonadota>Gammaproteobacteria>Moraxellales>Moraxellaceae>Acinetobacter | GCF_030329445.1 | Acinetobacter_haemolyticus | GPS61_RS07925 | 49241..52921 | 29430 | BREX-system-P-loop-protein-BrxC-[Acinetobacter-haemolyticus] |
| WP_005115116.1 | Type1.BREX | <-SIGMA+OB-Fold+AAA-ATPase+Lon-Protease-BrxL<-PglZ<-ParB+HNH<-N6-METHYLASE<-DUF6079*<-BrxB-BrxF-iSTAND<-BrxA-DNA-binding<-wHTH+WYL+WCX\|\|DUF4236-><-CR-ATPase5+REase | 1226 | Bacteria>Pseudomonadota>Gammaproteobacteria>Moraxellales>Moraxellaceae>Acinetobacter | GCF_030413705.1 | Acinetobacter_baumannii | AWM09_RS06130 | 10901..14581 | 470 | BREX-system-P-loop-protein-BrxC-[Acinetobacter-baumannii] |
| WP_112987434.1 | Type1.BREX | <-SIGMA+OB-Fold+AAA-ATPase+Lon-Protease-BrxL<-PglZ<-N6-METHYLASE<-DUF6079*<-BrxB-BrxF-iSTAND<-BrxA-DNA-binding<-wHTH+WYL+WCX\|\|DUF4236-><-CR-ATPase5+REase\|\|HigB_toxin->HTH_3-> | 1226 | Bacteria>Pseudomonadota>Gammaproteobacteria>Moraxellales>Moraxellaceae>Acinetobacter | GCF_030665425.1 | Acinetobacter_junii | DDF86_RS15435 | 8518..12198 | 40215 | BREX-system-P-loop-protein-BrxC-[Acinetobacter-junii] |
| WP_092692667.1 | Type1.BREX | ParB-><-?\|\|CR-ATPase5+REase->ParB+HNH-p->?-><-DUF4236\|\|wHTH+WYL+WCX->BrxA-DNA-binding->BrxB-BrxF-iSTAND->DUF6079*-> | 1224 | Bacteria>Pseudomonadota>Gammaproteobacteria>Moraxellales>Moraxellaceae>Acinetobacter | GCF_900107285.1 | Acinetobacter_kyonggiensis | BLV99_RS16485 | 12458..16132 | 595670 | BREX-system-P-loop-protein-BrxC-[Acinetobacter-kyonggiensis] |
| WP_074947383.1 | Type1.BREX | ParB+HNH-p->ParB+HNH-p->?-><-DUF4236\|\|wHTH+WYL+WCX->BrxA-DNA-binding->BrxB-BrxF-iSTAND->DUF6079*->Phage_integrase->Phage_integrase+Phage_integrase->Phage_integrase+Phage_integrase->TetR-HTH-><-SF1-SF2-Helicase\|\|HTH_Tnp_1+HTH_21+rve+rve_3-><-?<-SF1-SF2-Helicase\|\|ParB+HNH-> | 1226 | Bacteria>Pseudomonadota>Gammaproteobacteria>Moraxellales>Moraxellaceae>Acinetobacter | GCF_900116265.1 | Acinetobacter_bohemicus | F7P73_RS14505 | 5974..9654 | 1435036 | BREX-system-P-loop-protein-BrxC-[Acinetobacter-bohemicus] |
| WP_005403809.1 | Type1.BREX | Peptidase_A8->zf-ISL3+HTH_Tnp_ISL3+DDE_Tnp_ISL3->?->?->RNase-><-DUF4236\|\|wHTH+WYL+WCX->BrxA-DNA-binding->BrxB-BrxF-iSTAND->DUF6079*->N6-METHYLASE->PglZ->SIGMA+OB-Fold+AAA-ATPase+Lon-Protease-BrxL-><-?<-AbiEii<-MarR-HTH | 1226 | Bacteria>Pseudomonadota>Gammaproteobacteria>Moraxellales>Moraxellaceae>Acinetobacter | GCF_902374105.1 | Acinetobacter_junii | FXX57_RS02410 | 133086..136766 | 40215 | BREX-system-P-loop-protein-BrxC-[Acinetobacter-junii] |
| WP_002120185.1 | Type1.BREX | PhdYeFM_antitox-><-OLD-ABC+TOPRIM\|\|HIN-HTH+PAIREDC-HTH-><-OLD-ABC+TOPRIM-p<-SIGMA+OB-Fold+AAA-ATPase+Lon-Protease-BrxL<-PglZ<-?<-N6-METHYLASE<-DUF6079*<-BrxB-BrxF-iSTAND<-BrxA-DNA-binding<-wHTH+WYL+WCX\|\|<-DOC+HTH<-?\|\|DDE_Tnp_1_5+DDE_Tnp_1-> | 1226 | Bacteria>Pseudomonadota>Gammaproteobacteria>Moraxellales>Moraxellaceae | - | Acinetobacter | LG653_RS19250 | 36203..39883 | 469 | MULTISPECIES:-BREX-system-P-loop-protein-BrxC-[Acinetobacter] |
| WP_115736701.1 | Type1.BREX | <-DDE_Tnp_1_5+DDE_Tnp_1\|\|MarR-HTH->HTH_Tnp_1+HTH_21+rve+rve_3-><-SIGMA+OB-Fold+AAA-ATPase+Lon-Protease-BrxL<-PglZ<-N6-METHYLASE<-DUF6079*<-BrxB-BrxF-iSTAND<-BrxA-DNA-binding<-wHTH+WYL+WCX\|\|DUF4236-><-CR-ATPase5+REase<-DDE_Tnp_1_5+DDE_Tnp_1<-RNase | 1226 | Bacteria>Pseudomonadota>Gammaproteobacteria>Moraxellales>Moraxellaceae | - | Acinetobacter | MK935_RS00110 | 25389..29069 | 469 | MULTISPECIES:-BREX-system-P-loop-protein-BrxC-[Acinetobacter] |
| WP_180169606.1 | Type1.BREX | DDE_Tnp_IS240-><-?<-?<-?<-?\|\|<-SIGMA+OB-Fold+AAA-ATPase+Lon-Protease-BrxL<-PglZ<-N6-METHYLASE<-DUF6079*<-BrxB-BrxF-iSTAND<-BrxA-DNA-binding<-wHTH+WYL+WCX\|\|DUF4236-><-CR-ATPase5+REase<-DUF772+DDE_Tnp_1+DDE_Tnp_1_6\|\|DSBH->\|\|?-><-Zn_Tnp_IS91+Y2_Tnp | 1226 | Bacteria>Pseudomonadota>Gammaproteobacteria>Moraxellales>Moraxellaceae>Acinetobacter | - | unclassified_Acinetobacter | FSJ21_RS00665 | 124577..128257 | 196816 | MULTISPECIES:-BREX-system-P-loop-protein-BrxC-[unclassified-Acinetobacter] |
| WP_180172593.1 | Type1.BREX | <-SIGMA+OB-Fold+AAA-ATPase+Lon-Protease-BrxL<-PglZ<-?<-N6-METHYLASE<-DUF6079*<-BrxB-BrxF-iSTAND<-BrxA-DNA-binding<-wHTH+WYL+WCX\|\|DUF4236-><-CR-ATPase5+REase\|\|?->HATPase+wHTH-><-?<-Zn_Tnp_IS91+Y2_Tnp | 1226 | Bacteria>Pseudomonadota>Gammaproteobacteria>Moraxellales>Moraxellaceae>Acinetobacter | - | unclassified_Acinetobacter | LW346_RS00160 | 40718..44398 | 196816 | MULTISPECIES:-BREX-system-P-loop-protein-BrxC-[unclassified-Acinetobacter] |
| WP_107866915.1 | Type1.BREX | <-SIGMA+OB-Fold+AAA-ATPase+Lon-Protease-BrxL<-PglZ<-TOPRIM<-?<-OLD-ABC<-N6-METHYLASE<-Couple_hipA+HipA_C<-HTH_3<-DUF6079*<-BrxB-BrxF-iSTAND<-BrxA-DNA-binding<-wHTH+WYL+WCX | 1226 | Bacteria>Pseudomonadota>Gammaproteobacteria>Moraxellales>Moraxellaceae>Agitococcus | GCF_003051055.1 | Agitococcus_lubricus | C8N29_RS15175 | 21839..25489 | 1077255 | BREX-system-P-loop-protein-BrxC-[Agitococcus-lubricus] |
| WP_134244556.1 | Type1.BREX | rve->?->RelE-ParE->\|\|CR-ATPase5+REase-><-DUF4236\|\|wHTH+WYL+WCX->BrxA-DNA-binding->BrxB-BrxF-iSTAND->DUF6079*->N6-METHYLASE->?->PglZ->SIGMA+OB-Fold+AAA-ATPase+Lon-Protease-BrxL-><-AAA-ATPase-p | 1216 | Bacteria>Pseudomonadota>Gammaproteobacteria>Moraxellales>Moraxellaceae>Alkanindiges | GCF_004377485.1 | Alkanindiges_illinoisensis | E2B99_RS08410 | 8100..11780 | 197183 | BREX-system-P-loop-protein-BrxC-[Alkanindiges-illinoisensis] |
| WP_002010901.1 | Type1.BREX | <-MACRODOMAIN<-DarT+Peptidase<-SF1-SF2-Helicase+REase<-SIGMA+OB-Fold+AAA-ATPase+Lon-Protease-BrxL<-PglZ<-ParB+HNH<-N6-METHYLASE<-DUF6079*<-BrxB-BrxF-iSTAND<-BrxA-DNA-binding<-wHTH+WYL+WCX\|\|TetR-HTH->?-><-?<-?<-?<-DUF932 | 1226 | Bacteria>Pseudomonadota>Gammaproteobacteria>Moraxellales>Moraxellaceae | GCF_000214965.1 | Acinetobacter | ACINNAV18_RS19255 | 65599..69279 | 469 | MULTISPECIES:-BREX-system-P-loop-protein-BrxC-[Acinetobacter] |
| WP_001021842.1 | Type1.BREX | <-AbiGi+seqA+RAMA<-CHTH<-RelE<-DDE_Tnp_1_5+DDE_Tnp_1\|\|CR-ATPase5+REase-><-DUF4236\|\|wHTH+WYL+WCX->BrxA-DNA-binding->BrxB-BrxF-iSTAND->DUF6079*->N6-METHYLASE->PglZ->SIGMA+OB-Fold+AAA-ATPase+Lon-Protease-BrxL-><-DDE_Tnp_IS240\|\|Sulfate_transp+STAS->NERD+Pkinase+Pkinase+SF1-SF2-Helicase->?-><-GDResolvase+HIN-HTH\|\|DDE_Tnp_IS240-> | 1226 | Bacteria>Pseudomonadota>Gammaproteobacteria>Moraxellales>Moraxellaceae | GCF_000241705.1 | Acinetobacter | HO22_RS0119175 | 11899..15579 | 469 | MULTISPECIES:-BREX-system-P-loop-protein-BrxC-[Acinetobacter] |
| WP_111885889.1 | Type1.BREX | <-SIGMA+OB-Fold+AAA-ATPase+Lon-Protease-BrxL<-PglZ<-ParB+HNH<-N6-METHYLASE<-DUF6079*<-BrxB-BrxF-iSTAND<-BrxA-DNA-binding<-wHTH+WYL+WCX\|\|DUF4236-><-CR-ATPase5+REase<-ParB+HNH<-Phage_integrase+Phage_integrase<-Phage_integrase+Phage_integrase | 1226 | Bacteria>Pseudomonadota>Gammaproteobacteria>Moraxellales>Moraxellaceae | GCF_002339435.1 | Acinetobacter | NVV26_RS12530 | 20211..23891 | 469 | MULTISPECIES:-BREX-system-P-loop-protein-BrxC-[Acinetobacter] |
| WP_108675357.1 | Type1.BREX | Arm-DNA-bind_3+Phage_integrase-><-?\|\|?->ZnR+ZnR->?->?->XIS-HTH->BrxA-DNA-binding->BrxB-BrxF-iSTAND->DUF6079*->ParB+HNH->OLD-ABC->TOPRIM->N6-METHYLASE->PglZ->SIGMA+OB-Fold+AAA-ATPase+Lon-Protease-BrxL->\|\|?->?->Thioredoxin_3-> | 1225 | Bacteria>Pseudomonadota>Gammaproteobacteria>Moraxellales>Moraxellaceae | GCF_003072585.2 | Acinetobacter | CO119_RS05575 | 1118908..1122585 | 469 | MULTISPECIES:-BREX-system-P-loop-protein-BrxC-[Acinetobacter] |
| WP_071320614.1 | Type1.BREX | Phage_integrase+Phage_integrase->ParB+HNH->CR-ATPase5+REase->ParB+HNH-p->?-><-DUF4236\|\|wHTH+WYL+WCX->BrxA-DNA-binding->BrxB-BrxF-iSTAND->DUF6079*->N6-METHYLASE-> | 1224 | Bacteria>Pseudomonadota>Gammaproteobacteria>Moraxellales>Moraxellaceae | GCF_007833615.1 | Acinetobacter | DPR03_RS15465 | 213962..217636 | 469 | MULTISPECIES:-BREX-system-P-loop-protein-BrxC-[Acinetobacter] |
| WP_075316556.1 | Type1.BREX | <-N6-METHYLASE<-DUF6079*<-BrxB-BrxF-iSTAND<-BrxA-DNA-binding<-wHTH+WYL+WCX\|\|DUF4236-><-CR-ATPase5+REase | 1226 | Bacteria>Pseudomonadota>Gammaproteobacteria>Moraxellales>Moraxellaceae | GCF_008993855.1 | Acinetobacter | GBN82_RS18495 | 2355..6035 | 469 | MULTISPECIES:-BREX-system-P-loop-protein-BrxC-[Acinetobacter] |
| WP_152630429.1 | Type1.BREX | PhdYeFM_antitox->?->Phage_integrase+Phage_integrase->Phage_integrase->\|\|wHTH+WYL+WCX->BrxA-DNA-binding->BrxB-BrxF-iSTAND->DUF6079*->N6-METHYLASE->PglZ->SIGMA+OB-Fold+AAA-ATPase+Lon-Protease-BrxL->SF1-SF2-Helicase+REase-><-?<-AA_permease<-?<-?\|\|GNTR-HTH+Aminotran_1_2-> | 1226 | Bacteria>Pseudomonadota>Gammaproteobacteria>Moraxellales>Moraxellaceae | GCF_009014155.1 | Acinetobacter | QJ110_RS14020 | 22608..26288 | 469 | MULTISPECIES:-BREX-system-P-loop-protein-BrxC-[Acinetobacter] |
| WP_152342636.1 | Type1.BREX | <-GDResolvase+HIN-HTH<-?<-?<-?<-?<-?\|\|<-SIGMA+OB-Fold+AAA-ATPase+Lon-Protease-BrxL<-PglZ<-N6-METHYLASE<-DUF6079*<-BrxB-BrxF-iSTAND<-BrxA-DNA-binding<-wHTH+WYL+WCX\|\|DUF4236-><-CR-ATPase5+REase | 1226 | Bacteria>Pseudomonadota>Gammaproteobacteria>Moraxellales>Moraxellaceae | GCF_009296145.1 | Acinetobacter | MOW12_RS15415 | 73736..77416 | 469 | MULTISPECIES:-BREX-system-P-loop-protein-BrxC-[Acinetobacter] |
| WP_100355920.1 | Type1.BREX | <-NTP_transf\|\|<-BrxA-DNA-binding\|\|<-?\|\|HTH_26->CR-ATPase5+REase-><-DUF4236\|\|wHTH+WYL+WCX->BrxA-DNA-binding->BrxB-BrxF-iSTAND->DUF6079*->N6-METHYLASE-> | 1226 | Bacteria>Pseudomonadota>Gammaproteobacteria>Moraxellales>Moraxellaceae | GCF_009914395.1 | Acinetobacter | GA646_RS15620 | 96338..100018 | 469 | MULTISPECIES:-BREX-system-P-loop-protein-BrxC-[Acinetobacter] |
| WP_163167826.1 | Type1.BREX | <-GDResolvase+HIN-HTH<-?<-?<-?<-?<-?\|\|<-SIGMA+OB-Fold+AAA-ATPase+Lon-Protease-BrxL<-PglZ<-N6-METHYLASE<-DUF6079*<-BrxB-BrxF-iSTAND<-BrxA-DNA-binding<-wHTH+WYL+WCX\|\|DUF4236-><-CR-ATPase5+REase\|\|?->?->Pkinase-> | 1226 | Bacteria>Pseudomonadota>Gammaproteobacteria>Moraxellales>Moraxellaceae | GCF_010917815.1 | Acinetobacter | FSC02_RS15715 | 99085..102765 | 469 | MULTISPECIES:-BREX-system-P-loop-protein-BrxC-[Acinetobacter] |
| WP_178319334.1 | Type1.BREX | <-DUF4096+DDE_Tnp_1+DDE_Tnp_1_2\|\|DDE_5+DDE_Tnp_1-><-?<-?\|\|?-><-DUF4236\|\|wHTH+WYL+WCX->BrxA-DNA-binding->BrxB-BrxF-iSTAND->DUF6079*->N6-METHYLASE->ParB+HNH->PglZ->SIGMA+OB-Fold+AAA-ATPase+Lon-Protease-BrxL-> | 1226 | Bacteria>Pseudomonadota>Gammaproteobacteria>Moraxellales>Moraxellaceae | GCF_013390985.1 | Acinetobacter | HX743_RS11980 | 35327..39007 | 469 | MULTISPECIES:-BREX-system-P-loop-protein-BrxC-[Acinetobacter] |
| WP_159138903.1 | Type1.BREX | <-SIGMA+OB-Fold+AAA-ATPase+Lon-Protease-BrxL<-PglZ<-N6-METHYLASE<-DUF6079*<-BrxB-BrxF-iSTAND<-BrxA-DNA-binding<-wHTH+WYL+WCX\|\|DUF4236-><-DOC+HTH | 1226 | Bacteria>Pseudomonadota>Gammaproteobacteria>Moraxellales>Moraxellaceae | GCF_013417415.1 | Acinetobacter | FTV29_RS13965 | 21709..25389 | 469 | MULTISPECIES:-BREX-system-P-loop-protein-BrxC-[Acinetobacter] |
| WP_159153679.1 | Type1.BREX | DDE_Tnp_1->CR-ATPase5+REase-><-DUF4236<-?\|\|wHTH+WYL+WCX->BrxA-DNA-binding->BrxB-BrxF-iSTAND->DUF6079*->N6-METHYLASE->PglZ->SIGMA+OB-Fold+AAA-ATPase+Lon-Protease-BrxL-><-?<-AbiEii<-MarR-HTH | 1226 | Bacteria>Pseudomonadota>Gammaproteobacteria>Moraxellales>Moraxellaceae | GCF_013420015.1 | Acinetobacter | FTT98_RS13935 | 5674..9354 | 469 | MULTISPECIES:-BREX-system-P-loop-protein-BrxC-[Acinetobacter] |
| WP_180192273.1 | Type1.BREX | <-GDResolvase+HIN-HTH<-?<-?<-?<-?<-?\|\|<-SIGMA+OB-Fold+AAA-ATPase+Lon-Protease-BrxL<-PglZ<-N6-METHYLASE<-DUF6079*<-BrxB-BrxF-iSTAND<-BrxA-DNA-binding<-wHTH+WYL+WCX\|\|DUF4236-><-CR-ATPase5+REase<-ParB+HNH<-?<-Phage_integrase+Phage_integrase | 1224 | Bacteria>Pseudomonadota>Gammaproteobacteria>Moraxellales>Moraxellaceae | GCF_015217905.1 | Acinetobacter | G0027_RS16200 | 28364..32038 | 469 | MULTISPECIES:-BREX-system-P-loop-protein-BrxC-[Acinetobacter] |
| WP_180047180.1 | Type1.BREX | Phage_integrase+Phage_integrase->Zn_Tnp_IS1+DDE_Tnp_IS1->ParB+HNH->CR-ATPase5+REase-><-DUF4236<-?\|\|wHTH+WYL+WCX->BrxA-DNA-binding->BrxB-BrxF-iSTAND->DUF6079*->N6-METHYLASE->PglZ->SIGMA+OB-Fold+AAA-ATPase+Lon-Protease-BrxL-><-HTH_Tnp_1+HTH_21+rve+rve_3<-?<-OLD-ABC<-ParB\|\|GDResolvase+HIN-HTH-><-DUF1705+Sulfatase | 1226 | Bacteria>Pseudomonadota>Gammaproteobacteria>Moraxellales>Moraxellaceae | GCF_015218165.1 | Acinetobacter | G0028_RS19990 | 40652..44332 | 469 | MULTISPECIES:-BREX-system-P-loop-protein-BrxC-[Acinetobacter] |
| WP_199980844.1 | Type1.BREX | <-DDE_Tnp_1_5+DDE_Tnp_1\|\|MarR-HTH->AbiEii->?-><-DUF4372+DDE_Tnp_1<-SIGMA+OB-Fold+AAA-ATPase+Lon-Protease-BrxL<-PglZ<-N6-METHYLASE<-DUF6079*<-BrxB-BrxF-iSTAND<-BrxA-DNA-binding<-wHTH+WYL+WCX\|\|DUF4236-><-CR-ATPase5+REase<-ParB+HNH<-Phage_integrase<-Phage_integrase+Phage_integrase | 1225 | Bacteria>Pseudomonadota>Gammaproteobacteria>Moraxellales>Moraxellaceae | GCF_019702905.1 | Acinetobacter | TOL5_RS22165 | 83634..87311 | 469 | MULTISPECIES:-BREX-system-P-loop-protein-BrxC-[Acinetobacter] |
| WP_252172191.1 | Type1.BREX | ParB+HNH-><-DUF4236\|\|wHTH+WYL+WCX->BrxA-DNA-binding->BrxB-BrxF-iSTAND->DUF6079*->N6-METHYLASE->PglZ->SIGMA+OB-Fold+AAA-ATPase+Lon-Protease-BrxL-><-?<-ABhydrolase<-Classical-AAA<-?<-SMODS<-DDE_Tnp_IS240 | 1226 | Bacteria>Pseudomonadota>Gammaproteobacteria>Moraxellales>Moraxellaceae | GCF_023809625.1 | Acinetobacter | N5980_RS16655 | 54910..58590 | 469 | MULTISPECIES:-BREX-system-P-loop-protein-BrxC-[Acinetobacter] |
| WP_180000666.1 | Type1.BREX | ParB+HNH->?->DOC+HTH-><-DUF4236\|\|wHTH+WYL+WCX->BrxA-DNA-binding->BrxB-BrxF-iSTAND->DUF6079*->PglZ->SIGMA+OB-Fold+AAA-ATPase+Lon-Protease-BrxL-><-HTH_28+rve+rve_3 | 1224 | Bacteria>Pseudomonadota>Gammaproteobacteria>Moraxellales>Moraxellaceae | GCF_024159345.1 | Acinetobacter | MUB06_RS13820 | 43929..47603 | 469 | MULTISPECIES:-BREX-system-P-loop-protein-BrxC-[Acinetobacter] |

| Protein-ID | System | Gene-Neighbourhoods | Length | Taxonomy | Assembly-ID | Strain | Locus-Tag | Genomic-coordinates | TaxID | Fasta-defline |
|---|---|---|---|---|---|---|---|---|---|---|
| ABE75386.1 | Type1.BREX | ABC-ATPase+ABC-ATPase->?->?->?->tRNA->tRNA->Arm-DNA-bind_3+Phage_integrase-><-ParB+HNH<-PglZ<-N6-METHYLASE<-DUF6079*<-BrxB-BrxF-iSTAND<-BrxA-DNA-binding<-TPR+TPR<-ZnR\|\|REase->REase->[illegible] | 1201 | Bacteria>Pseudomonadota>Gammaproteobacteria>Moraxellales>Moraxellaceae>Psychrobacter | GCA_000013905.1 | Psychrobacter_cryohalolentis_K5 | Pcryo_1609 | 1979457..1983062 | 335284 | conserved-hypothetical-protein-[Psychrobacter-cryohalolentis-K5] |
| GAF52839.1 | Type1.BREX | DUF932->?->?->XIS-HTH-><-?<-?\|\|BrxB-BrxF-iSTAND->DUF6079*->N6-METHYLASE->PglZ->?-><-Arm-DNA-bind_3+Phage_integrase<-tRNA<-tRNA<-?<-?<-?<-ABC-ATPase+ABC-ATPase | 1200 | Bacteria>Pseudomonadota>Gammaproteobacteria>Moraxellales>Moraxellaceae>Psychrobacter | GCA_000586415.1 | Psychrobacter_sp._JCM_18900 | JCM18900_11381 | 174807..178411 | 1298608 | LOW-QUALITY-PROTEIN:-hypothetical-protein-JCM18900_11381-[Psychrobacter-sp.-JCM-18900] |
| OLF41388.1 | Type1.BREX | Cas_Cas1->?-><-REase\|\|BrxA-DNA-binding->BrxB-BrxF-iSTAND->DUF6079*->N6-METHYLASE->DUF3644->OLD-ABC->TOPRIM->DUF932-> | 1154 | Bacteria>Pseudomonadota>Gammaproteobacteria>Moraxellales>Moraxellaceae>Psychrobacter | GCA_001921755.1 | Psychrobacter_sp._Rd_27.2 | BTV99_04785 | 263905..267369 | 1926479 | hypothetical-protein-BTV99_04785-[Psychrobacter-sp.-Rd-27.2] |
| HCI75257.1 | Type1.BREX | <-TOPRIM<-SbcC<-?<-REase<-REase\|\|ZnR->TPR+TPR->BrxA-DNA-binding->BrxB-BrxF-iSTAND->DUF6079*->N6-METHYLASE->PglZ->?-><-Arm-DNA-bind_3+Phage_integrase<-tRNA<-tRNA<-?<-?<-HTH_OrfB_IS605+[illegible] | 1201 | Bacteria>Pseudomonadota>Gammaproteobacteria>Moraxellales>Moraxellaceae>Psychrobacter | GCA_003530515.1 | Psychrobacter_sp. | DHV27_03225 | 245755..249360 | 56811 | BREX-system-P-loop-protein-BrxC-[Psychrobacter-sp.] |
| MCG3880112.1 | Type1.BREX | <-wHTH+REase+ZnR<-McrB-AAA<-MarR-HTH<-SF1-SF2-Helicase<-STYKIN+CR-Kinase<-?<-?\|\|BrxA-DNA-binding->BrxB-BrxF-iSTAND->DUF6079*->N6-METHYLASE->PHP+SbcC->DUF932->ParB+HNH->PglZ->SIGMA+OB-[illegible] | 1154 | Bacteria>Pseudomonadota>Gammaproteobacteria>Moraxellales>Moraxellaceae>Psychrobacter | GCA_022129235.1 | Psychrobacter_sp._Ps6 | I3255_11455 | 48655..52119 | 2790960 | BREX-system-P-loop-protein-BrxC-[Psychrobacter-sp.-Ps6] |
| WP_226939352.1 | Type1.BREX | ABC-ATPase+ABC-ATPase->?->?->?->tRNA->tRNA->Arm-DNA-bind_3+Phage_integrase-><-ParB+HNH<-PglZ<-N6-METHYLASE<-DUF6079*<-BrxB-BrxF-iSTAND<-BrxA-DNA-binding\|\|<-TPR+TPR<-ZnR\|\|REase->REase-[illegible] | 1221 | Bacteria>Pseudomonadota>Gammaproteobacteria>Moraxellales>Moraxellaceae>Psychrobacter | GCF_000013905.1 | Psychrobacter_cryohalolentis | CEP87_RS02910 | 676881..680546 | 330922 | BREX-system-P-loop-protein-BrxC-[Psychrobacter-cryohalolentis] |
| WP_261863080.1 | Type1.BREX | XIS-HTH-><-?<-?<-?<-REase\|\|BrxA-DNA-binding->BrxB-BrxF-iSTAND-p->DUF6079-p*->DUF6079-p->?->N6-METHYLASE-p->N6-METHYLASE-p->N6-METHYLASE->N6-METHYLASE-p->PglZ-> | 801 | Bacteria>Pseudomonadota>Gammaproteobacteria>Moraxellales>Moraxellaceae>Psychrobacter | GCF_000586415.1 | Psychrobacter_sp._JCM_18900 | TT10_RS07330 | 174747..177152 | 1298608 | BREX-system-P-loop-protein-BrxC-[Psychrobacter-sp.-JCM-18900] |
| WP_261869530.1 | Type1.BREX | Phage_integrase+Phage_integrase->Phage_integrase+Phage_integrase->Phage_integrase+Phage_integrase->Phage_integrase->Phage_integrase->?->BrxA-DNA-binding->BrxB-BrxF-iSTAND->DUF6079-p->[illegible] | 642 | Bacteria>Pseudomonadota>Gammaproteobacteria>Moraxellales>Moraxellaceae>Psychrobacter | GCF_000586435.1 | Psychrobacter_sp._JCM_18901 | TT11_RS05300 | 11820..13748 | 1298609 | BREX-system-P-loop-protein-BrxC-[Psychrobacter-sp.-JCM-18901] |
| WP_218607358.1 | Type1.BREX | Cas_Cas1->?-><-REase\|\|BrxA-DNA-binding->BrxB-BrxF-iSTAND->DUF6079*->N6-METHYLASE->DUF3644->OLD-ABC->TOPRIM->DUF932->ParB+HNH-> | 1162 | Bacteria>Pseudomonadota>Gammaproteobacteria>Moraxellales>Moraxellaceae>Psychrobacter | GCF_001921755.1 | Psychrobacter_sp._Rd_27.2 | BTV99_RS04780 | 263881..267369 | 1926479 | BREX-system-P-loop-protein-BrxC-[Psychrobacter-sp.-Rd-27.2] |
| WP_089554755.1 | Type1.BREX | Phage_integrase->?->XIS-HTH->AAA+HTH-><-?<-PglZ<-N6-METHYLASE<-ParB+HNH<-DUF6079*<-BrxB-BrxF-iSTAND<-BrxA-DNA-binding<-Phage_integrase<-Phage_integrase<-Phage_integrase+Phage_integras[illegible] | 1223 | Bacteria>Pseudomonadota>Gammaproteobacteria>Moraxellales>Moraxellaceae>Psychrobacter | GCF_002224105.1 | Psychrobacter_sp._DAB_AL32B | CAN34_RS05290 | 13543..17214 | 1028414 | BREX-system-P-loop-protein-BrxC-[Psychrobacter-sp.-DAB_AL32B] |
| WP_226950212.1 | Type1.BREX | XIS-HTH-><-TOPRIM<-SbcC<-REase<-REase\|\|ZnR->TPR+TPR->\|\|BrxA-DNA-binding->BrxB-BrxF-iSTAND->DUF6079*->N6-METHYLASE->PglZ->?-><-Arm-DNA-bind_3+Phage_integrase<-tRNA<-tRNA<-?<-?<-HTH_Orf[illegible] | 1221 | Bacteria>Pseudomonadota>Gammaproteobacteria>Moraxellales>Moraxellaceae>Psychrobacter | GCF_003530515.1 | unclassified_Psychrobacter | INS97_RS03560 | 152962..156627 | 196806 | MULTISPECIES:-BREX-system-P-loop-protein-BrxC-[unclassified-Psychrobacter] |
| WP_191691193.1 | Type1.BREX | <-Phage_integrase<-Phage_integrase+Phage_integrase<-ParB\|\|ZnR-><-?\|\|?-><-SIGMA+OB-Fold+AAA-ATPase+Lon-Protease-BrxL<-PglZ<-N6-METHYLASE<-DUF6079*<-BrxB-BrxF-iSTAND<-BrxA-DNA-binding<-[illegible] | 1222 | Bacteria>Pseudomonadota>Gammaproteobacteria>Moraxellales>Moraxellaceae>Psychrobacter | GCF_014836505.1 | Psychrobacter_communis | H9653_RS05990 | 24340..28008 | 2762238 | BREX-system-P-loop-protein-BrxC-[Psychrobacter-communis] |
| WP_192529491.1 | Type1.BREX | PAIREDC-HTH->HTH_26->ParB->Phage_integrase+Phage_integrase->Phage_integrase+Phage_integrase->Phage_integrase->?->BrxA-DNA-binding->BrxB-BrxF-iSTAND->DUF6079*-> | 1218 | Bacteria>Pseudomonadota>Gammaproteobacteria>Moraxellales>Moraxellaceae>Psychrobacter | GCF_014861375.1 | Psychrobacter_sp._FME5 | EI161_09970 | 10554..14210 | 2487706 | BREX-system-P-loop-protein-BrxC-[Psychrobacter-sp.-FME5] |
| WP_207991848.1 | Type1.BREX | <-TOPRIM<-SbcC<-REase<-REase\|\|ZnR->BrxA-DNA-binding->BrxB-BrxF-iSTAND->DUF6079*->N6-METHYLASE->ParB+HNH->PglZ->SIGMA+OB-Fold+AAA-ATPase+Lon-Protease-BrxL-><-Arm-DNA-bind_3+Phage_integ[illegible] | 1221 | Bacteria>Pseudomonadota>Gammaproteobacteria>Moraxellales>Moraxellaceae>Psychrobacter | GCF_017498085.1 | Psychrobacter_coccoides | J3492_RS09680 | 48612..52277 | 2818440 | BREX-system-P-loop-protein-BrxC-[Psychrobacter-coccoides] |
| WP_238038933.1 | Type1.BREX | <-wHTH+REase+ZnR<-McrB-AAA<-MarR-HTH<-SF1-SF2-Helicase<-STYKIN+CR-Kinase<-?\|\|BrxA-DNA-binding->BrxB-BrxF-iSTAND->DUF6079*->N6-METHYLASE->PHP+SbcC->DUF932->ParB+HNH->PglZ->wHTH+WYL+WCX[illegible] | 1162 | Bacteria>Pseudomonadota>Gammaproteobacteria>Moraxellales>Moraxellaceae>Psychrobacter | GCF_022129235.1 | Psychrobacter_sp._Ps6 | I3255_RS11455 | 48631..52119 | 2790960 | BREX-system-P-loop-protein-BrxC-[Psychrobacter-sp.-Ps6] |
| WP_241878421.1 | Type1.BREX | <-MIT_C<-MIT_C<-SIGMA+OB-Fold+AAA-ATPase+Lon-Protease-BrxL<-PglZ<-N6-METHYLASE-p<-N6-METHYLASE-p<-N6-METHYLASE-p<-N6-METHYLASE-p<-?<-DUF6079-p*<-DUF6079-p<-?<-BrxB-BrxF-iSTAND<-BrxA-D[illegible] | 709 | Bacteria>Pseudomonadota>Gammaproteobacteria>Moraxellales>Moraxellaceae>Psychrobacter | GCF_022631235.2 | Psychrobacter_sp._PraFG1 | MN210_RS12710 | 2340484..2342613 | 2565531 | BREX-system-P-loop-protein-BrxC-[Psychrobacter-sp.-PraFG1] |
| [illegible] | [illegible] | [illegible] | [illegible] | [illegible] | [illegible] | [illegible] | [illegible] | [illegible] | [illegible] | [illegible] |

| Protein-ID | System | Gene-Neighbourhoods | Length | Taxonomy | Assembly-ID | Strain | Locus-Tag | Genomic-coordinates | TaxID | Fasta-define |
|---|---|---|---|---|---|---|---|---|---|---|
| WP_027706208.1 | Type1.BREX | <-METHYLASE<-SAM-METHYLASE\|\|tRNA-><-?\|\|?-><-SIGMA+OB-Fold+AAA-ATPase+Lon-Protease-BrxL<-PglZ<-N6-METHYLASE<-DUF6079*<-BrxB-BrxF-iSTAND<-BrxA-DNA-binding\|\|?->?->?->ZnR-> | 1213 | Bacteria>Pseudomonadota>Gammaproteobacteria>Oceanospirillales>Halomonadaceae>Zymobacter_group | GCF_003610015.1 | Zymobacter_palmae | ZBT109_RS10990 | 2474695..2478336 | 33074 | BREX-system-P-loop-protein-BrxC-[Zymobacter-palmae] |
| WP_261843643.1 | Type1.BREX | <-LexA-protease\|\|?->35exo->?->wHTH+WYL+WCX->BrxA-DNA-binding->BrxB-BrxF-iSTAND->DUF6079*->ParB+HNH->N6-METHYLASE->PglZ->SIGMA+OB-Fold+AAA-ATPase+Lon-Protease-BrxL->SbcC->HDRnase->VirD4-TraG+VirB4-FtsK-> | 1225 | Bacteria>Pseudomonadota>Gammaproteobacteria>Oceanospirillales>Oceanospirillaceae>Aliamphritea | GCF_024347215.1 | Aliamphritea_ceti | OCU49_RS03510 | 764184..767861 | 1524258 | BREX-system-P-loop-protein-BrxC-[Aliamphritea-ceti] |
| WP_027853089.1 | Type1.BREX | <-SIGMA+OB-Fold+AAA-ATPase+Lon-Protease-BrxL<-PglZ<-HNH<-OLD-ABC<-TOPRIM<-OLD-ABC<-ParB+HNH<-N6-METHYLASE<-DUF6079*<-BrxB-BrxF-iSTAND<-BrxA-DNA-binding<-wHTH+WYL+WCX<-TIR<-?<-?<-wHTH+WYL+WCX | 1154 | Bacteria>Pseudomonadota>Gammaproteobacteria>Oceanospirillales>Oceanospirillaceae>Marinobacterium | GCF_000428985.1 | Marinobacterium_litorale | G403_RS0105385 | 191863..195327 | 404770 | BREX-system-P-loop-protein-BrxC-[Marinobacterium-litorale] |
| WP_027854332.1 | Type1.BREX | GDResolvase+HIN-HTH->?->?-><-SIGMA+OB-Fold+AAA-ATPase+HKD-DNase-BrxL<-PglZ<-N6-METHYLASE<-REase<-OLD-ABC<-Schlafen+HATPase+wHTH+HTH<-DUF6079*<-BrxB-BrxF-iSTAND<-BrxA-DNA-binding<-XIS-HTH<-MobI\|\|XIS-HTH->AAA+HTH->XIS-HTH-><-Arm-DNA-bind_3+Phage_integrase | 1193 | Bacteria>Pseudomonadota>Gammaproteobacteria>Oceanospirillales>Oceanospirillaceae>Marinobacterium | GCF_000428985.1 | Marinobacterium_litorale | G403_RS0112740 | 74962..78543 | 404770 | BREX-system-P-loop-protein-BrxC-[Marinobacterium-litorale] |
| WP_181736357.1 | Type1.BREX | XIS-HTH->?->?->TIR->wHTH+WYL+WCX->BrxA-DNA-binding->BrxB-BrxF-iSTAND->DUF6079*->N6-METHYLASE->ParB+HNH->OLD-ABC->HNH->PglZ->SIGMA+OB-Fold+AAA-ATPase+Lon-Protease-BrxL->ParB+HNH-> | 1154 | Bacteria>Pseudomonadota>Gammaproteobacteria>Oceanospirillales>Oceanospirillaceae>Marinobacterium | GCF_013868415.1 | Marinobacterium_sp._3-1745 | HIS06_RS00670 | 30655..34119 | 2756129 | BREX-system-P-loop-protein-BrxC-[Marinobacterium-sp.-3-1745] |
| WP_206838922.1 | Type1.BREX | MODE-HTH+PBPII-><-ADH_N+ADH_zinc_N<-?\|\|?->?->wHTH+WYL+WCX->BrxA-DNA-binding->BrxB-BrxF-iSTAND->DUF6079*->N6-METHYLASE->DUF932->OLD-ABC->TOPRIM->OLD-ABC->HNH->PglZ->SIGMA+OB-Fold+AAA-ATPase+Lon-Protease-BrxL->Integrase_1-> | 1154 | Bacteria>Pseudomonadota>Gammaproteobacteria>Oceanospirillales>Oceanospirillaceae>Marinobacterium | GCF_017310015.1 | Marinobacterium_georgiense | CFI10_RS02405 | 480710..484174 | 48076 | BREX-system-P-loop-protein-BrxC-[Marinobacterium-iners] |
| WP_217336191.1 | Type1.BREX | DUF772+DDE_Tnp_1+DDE_Tnp_1_6->wHTH+WYL+WCX->BrxA-DNA-binding->BrxB-BrxF-iSTAND->DUF6079*->N6-METHYLASE->DUF932->OLD-ABC->HNH->OLD-ABC->HNH->PglZ->SIGMA+OB-Fold+AAA-ATPase+Lon-Protease-BrxL-> | 1154 | Bacteria>Pseudomonadota>Gammaproteobacteria>Oceanospirillales>Oceanospirillaceae>Marinobacterium | GCF_019061305.1 | Marinobacterium_sp._A346 | KTN04_RS15750 | 3266..6730 | 2851016 | BREX-system-P-loop-protein-BrxC-[Marinobacterium-sp.-A346] |
| WP_239040425.1 | Type1.BREX | <-SF1-SF2-Helicase<-ParB+HNH<-SF1-SF2-Helicase<-N6-METHYLASE-p<-RVT<-N6-METHYLASE<-?<-AbiEii<-MarR-HTH<-DUF6079*<-BrxB-BrxF-iSTAND<-BrxA-DNA-binding<-?<-?<-wHTH+WYL+WCX<-?<-?<-?<-HTH_26 | 1223 | Bacteria>Pseudomonadota>Gammaproteobacteria>Oceanospirillales>Oceanospirillaceae>Marinobacterium | GCF_022343685.1 | Marinobacterium_sediminicola | LN244_RS14165 | 3103586..3107257 | 518898 | BREX-system-P-loop-protein-BrxC-[Marinobacterium-sediminicola] |
| AWY02455.1 | Type1.BREX | <-HTH_26\|\|Phage_AlpA->Arm-DNA-bind_2+Phage_int_SAM_3+Phage_integrase->?-><-SIGMA+OB-Fold+AAA-ATPase+HKD-DNase-BrxL<-PglZ<-ParB+HNH<-N6-METHYLASE<-REase<-DUF6079*<-BrxB-BrxF-iSTAND<-BrxA-DNA-binding<-HTH_26\|\|DUF772+DDE_Tnp_1->DUF772+DDE_Tnp_1-><-HTH_Tnp_1+HTH_21+rve+rve_3<-HIN-HTH\|\|?->SF1-SF2-Helicase-> | 1197 | Bacteria>Pseudomonadota>Gammaproteobacteria>Oceanospirillales>Oceanospirillaceae>Marinomonas | GCA_003285205.1 | Marinomonas_primoryensis | A8139_14855 | 3171018..3174611 | 178399 | hypothetical-protein-A8139_14855-[Marinomonas-primoryensis] |
| RUM53671.1 | Type1.BREX | wHTH+WYL+WCX->AAA-ToxA+Classical-AAA-><-?<-?<-?<-?<-SIGMA+OB-Fold+AAA-ATPase+Lon-Protease-BrxL<-PglZ<-N6-METHYLASE<-DUF6079*<-BrxB-BrxF-iSTAND<-BrxA-DNA-binding<-HTH_26\|\|Phage_AlpA->Arm-DNA-bind_2+Phage_int_SAM_3+Phage_integrase-> | 1229 | Bacteria>Pseudomonadota>Gammaproteobacteria>Oceanospirillales>Oceanospirillaceae>Marinomonas | GCA_003978655.1 | Marinomonas_sp. | DSY85_08610 | 29236..32925 | 1904862 | BREX-system-P-loop-protein-BrxC-[Marinomonas-sp.] |
| WP_036163554.1 | Type1.BREX | <-SIGMA+OB-Fold+AAA-ATPase+Lon-Protease-BrxL<-PglZ<-GP+REase<-McrC-NTD+REase<-McrB-AAA<-N6-METHYLASE<-SLOG+BirA-HTH<-SF1-SF2-Helicase+PRTase<-DUF6079*<-BrxB-BrxF-iSTAND<-BrxA-DNA-binding<-ABhydrolase<-HTH_26\|\|Phage_AlpA->Arm-DNA-bind_2+Phage_int_SAM_3+Phage_integrase->?-><-?<-LexA-protease | 1246 | Bacteria>Pseudomonadota>Gammaproteobacteria>Oceanospirillales>Oceanospirillaceae>Marinomonas | GCF_000521805.1 | Marinomonas_ushuaiensis | MUS1_RS14035 | 47481..51221 | 263818 | BREX-system-P-loop-protein-BrxC-[Marinomonas-ushuaiensis] |
| WP_100635909.1 | Type1.BREX | XIS-HTH->?->BrxA-DNA-binding->BrxB-BrxF-iSTAND->DUF6079*->N6-METHYLASE->ParB+HNH->PglZ->ZetaToxin->?-><-SF1-SF2-Helicase+REase<-SF1-SF2-Helicase+REase<-URI-DNase+RAMA | 1157 | Bacteria>Pseudomonadota>Gammaproteobacteria>Oceanospirillales>Oceanospirillaceae>Marinomonas | GCF_002806845.1 | Marinomonas_sp._ef1 | CCS02_RS04315 | 398110..401583 | 2005043 | BREX-system-P-loop-protein-BrxC-[Marinomonas-sp.-ef1] |
| WP_239495510.1 | Type1.BREX | Phage_AlpA->Arm-DNA-bind_2+Phage_int_SAM_3+Phage_integrase->?-><-SIGMA+OB-Fold+AAA-ATPase+HKD-DNase-BrxL<-PglZ<-ParB+HNH<-N6-METHYLASE<-REase<-OLD-ABC<-DUF6079*<-BrxB-BrxF-iSTAND<-BrxA-DNA-binding<-HTH_26\|\|<-HTH_Tnp_1+HTH_21+rve+rve_3<-HTH_19\|\|?->SF1-SF2-Helicase->TPR+HEPN-><-SF1-SF2-Helicase+wHTH+Small-B-barrel+DUF1998 | 1199 | Bacteria>Pseudomonadota>Gammaproteobacteria>Oceanospirillales>Oceanospirillaceae>Marinomonas | GCF_003285205.1 | Marinomonas_primoryensis | A8139_RS14980 | 3171018..3174617 | 178399 | BREX-system-P-loop-protein-BrxC-[Marinomonas-primoryensis] |
| WP_205113926.1 | Type1.BREX | TPR+TPR+TPR+TPR->?->?-><-Sulfate_transp+STAS\|\|SF1-SF2-Helicase->?->tRNA->BrxA-DNA-binding->BrxB-BrxF-iSTAND->DUF6079*->ABC-ATPase->REase->N6-METHYLASE->PglZ->SIGMA+OB-Fold+AAA-ATPase+HKD-DNase-BrxL-><-HTH_26\|\|Phage_AlpA->Arm-DNA-bind_2+Phage_int_SAM_3+Phage_integrase-> | 1191 | Bacteria>Pseudomonadota>Gammaproteobacteria>Oceanospirillales>Oceanospirillaceae>Marinomonas | GCF_016906065.1 | Marinomonas_foliarum | JSY38_RS15135 | 3246777..3250352 | 491950 | BREX-system-P-loop-protein-BrxC-[Marinomonas-foliarum] |
| WP_226752736.1 | Type1.BREX | <-SIGMA+OB-Fold+AAA-ATPase+Lon-Protease-BrxL<-PglZ<-Peptidase_S8<-Classical-AAA<-SbcC<-N6-METHYLASE<-TOPRIM<-OLD-ABC<-DUF6079*<-BrxB-BrxF-iSTAND<-BrxA-DNA-binding<-HTH_3\|\|wHTH+WYL->SF1-SF2-Helicase->Phage_integrase-><-CACHE+GGDEF | 1237 | Bacteria>Pseudomonadota>Gammaproteobacteria>Oceanospirillales>Oceanospirillaceae>Marinomonas | GCF_020532605.1 | Marinomonas_algarum | LG368_RS00225 | 46423..50136 | 2883105 | BREX-system-P-loop-protein-BrxC-[Marinomonas-algarum] |
| WP_067206359.1 | Type1.BREX | <-VirD4-TraG+VirB4-FtsK<-HDRnase<-SbcC<-SIGMA+OB-Fold+AAA-ATPase+Lon-Protease-BrxL<-PglZ<-N6-METHYLASE<-TOPRIM<-OLD-ABC<-DUF6079*<-BrxB-BrxF-iSTAND<-BrxA-DNA-binding<-wHTH+WYL+WCX<-?<-35exo\|\|LexA-protease->?-><-MobI | 1225 | Bacteria>Pseudomonadota>Gammaproteobacteria>Oceanospirillales>Oceanospirillaceae>Marinomonas | GCF_900089755.1 | Marinomonas_aquimarina | MAQ5080_RS04935 | 83930..87607 | 295068 | BREX-system-P-loop-protein-BrxC-[Marinomonas-aquimarina] |
| MBE0507005.1 | Type1.BREX | BrxA-DNA-binding->BrxB-BrxF-iSTAND->DUF6079*->OLD-ABC->REase->PglZ->SIGMA+OB-Fold+AAA-ATPase+Lon-Protease-BrxL-> | 1183 | Bacteria>Pseudomonadota>Gammaproteobacteria>Oceanospirillales>Oceanospirillaceae>Marinospirillum | GCA_014859565.1 | Marinospirillum_sp. | IBX50_09850 | 1716..5267 | 2183934 | BREX-system-P-loop-protein-BrxC-[Marinospirillum-sp.] |
| MBE0507394.1 | Type1.BREX | <-N6-METHYLASE<-DUF6079*<-BrxB-BrxF-iSTAND<-BrxA-DNA-binding<-wHTH+WYL+WCX\|\|?-><-Mrr-N\|\|LexA-protease-> | 1159 | Bacteria>Pseudomonadota>Gammaproteobacteria>Oceanospirillales>Oceanospirillaceae>Marinospirillum | GCA_014859565.1 | Marinospirillum_sp. | IBX50_11860 | 2713..6192 | 2183934 | BREX-system-P-loop-protein-BrxC-[Marinospirillum-sp.] |
| WP_138986811.1 | Type1.BREX | <-SIGMA+OB-Fold+AAA-ATPase+HKD-DNase-BrxL<-PglZ<-HTH+REase<-N6-METHYLASE<-DUF6079*<-BrxB-BrxF-iSTAND<-BrxA-DNA-binding<-XIS-HTH | 1190 | Bacteria>Pseudomonadota>Gammaproteobacteria>Oceanospirillales>Oceanospirillaceae>Neptunomonas | GCF_008630635.1 | Neptunomonas_concharum | FOU83_RS12940 | 2741645..2745217 | 1031538 | BREX-system-P-loop-protein-BrxC-[Neptunomonas-concharum] |
| WP_202806908.1 | Type1.BREX | <-PglZ<-TOPRIM<-OLD-ABC<-SbcC<-N6-METHYLASE-p<-N6-METHYLASE-p<-PIN<-OLD-ABC<-DUF6079-p*<-DUF6079-p<-BrxB-BrxF-iSTAND<-BrxA-DNA-binding<-XIS-HTH<-?<-HDRnase<-Phage_int_SAM<-Integrase_1\|\|XIS-HTH-> | 460 | Bacteria>Pseudomonadota>Gammaproteobacteria>Oceanospirillales>Oceanospirillaceae>Nitrincola | GCF_000585235.1 | Nitrincola_nitratireducens | D791_RS20495 | 12347..13729 | 1229521 | hypothetical-protein-[Nitrincola-nitratireducens] |
| WP_202806909.1 | Type1.BREX | <-TOPRIM<-OLD-ABC<-SbcC<-N6-METHYLASE-p<-N6-METHYLASE-p<-PIN<-OLD-ABC<-DUF6079-p<-DUF6079-p*<-BrxB-BrxF-iSTAND<-BrxA-DNA-binding<-XIS-HTH<-?<-HDRnase<-Phage_int_SAM<-Integrase_1\|\|XIS-HTH->Phage_integrase-> | 697 | Bacteria>Pseudomonadota>Gammaproteobacteria>Oceanospirillales>Oceanospirillaceae>Nitrincola | GCF_000585235.1 | Nitrincola_nitratireducens | D791_RS20500 | 13689..15782 | 1229521 | BREX-system-P-loop-protein-BrxC-[Nitrincola-nitratireducens] |
| MBL4799472.1 | Type1.BREX | <-N6-METHYLASE<-DUF6079*<-BrxB-BrxF-iSTAND<-BrxA-DNA-binding<-wHTH+WYL+WCX<-SF1-SF2-Helicase<-SbcC<-?<-SIR2 | 1239 | Bacteria>Pseudomonadota>Gammaproteobacteria>Oceanospirillales>Oceanospirillaceae>Oleispira | GCA_016764675.1 | Oleispira_sp. | JXX82_09395 | 2259..5978 | 2024846 | BREX-system-P-loop-protein-BrxC-[Oleispira-sp.] |
| WP_225677254.1 | Type1.BREX | URI-DNase-><-AAA-ATPase\|\|HEPN-><-UDG-conflict<-REase+iREase<-SIGMA+OB-Fold+AAA-ATPase+Lon-Protease-BrxL<-PglZ<-N6-METHYLASE<-DUF6079*<-BrxB-BrxF-iSTAND<-BrxA-DNA-binding<-wHTH+WYL+WCX | 1219 | Bacteria>Pseudomonadota>Gammaproteobacteria>Oceanospirillales>Oceanospirillaceae>Thalassolituus | GCF_020320395.1 | Thalassolituus_marinus | I9W95_RS17495 | 45926..49585 | 671053 | BREX-system-P-loop-protein-BrxC-[Thalassolituus-marinus] |
| WP_260997651.1 | Type1.BREX | <-Arm-DNA-bind_2+Phage_int_SAM_3+Phage_integrase<-Phage_AlpA\|\|HTH_26-><-SIGMA+OB-Fold+AAA-ATPase+Lon-Protease-BrxL<-PglZ<-TOPRIM<-OLD-ABC<-ParB+HNH<-N6-METHYLASE<-DUF6079*<-BrxB-BrxF-iSTAND<-BrxA-DNA-binding<-wHTH+WYL+WCX | 1156 | Bacteria>Pseudomonadota>Gammaproteobacteria>Oceanospirillales>Oceanospirillaceae>Thalassolituus | GCF_025345565.1 | Thalassolituus_hydrocarbonoclasticus | HUF19_RS16715 | 3723205..3726675 | 2742796 | BREX-system-P-loop-protein-BrxC-[Thalassolituus-hydrocarbonoclasticus] |
| EPJ48519.1 | Type1.BREX | <-DUF772+DDE_Tnp_1<-DUF772+DDE_Tnp_1\|\|?->MULE-TRANSPOSASE->wHTH+WYL+WCX->BrxA-DNA-binding-><-?\|\|BrxB-BrxF-iSTAND->DUF6079*->MarR-HTH->AbiEii->?->N6-METHYLASE->TIR->?->DOC+HTH->PglZ->SIGMA+OB-Fold+AAA-ATPase+Lon-Protease-BrxL-> | 1226 | Bacteria>Pseudomonadota>Gammaproteobacteria>Oceanospirillales>Oceanospirillaceae>unclassified_Oceanospirillaceae | GCA_000416275.1 | Osedax_symbiont_Rs1 | OFPII_05770 | 35450..39130 | 1330036 | MAG:-putative-ATPase-like-protein-[Osedax-symbiont-Rs1] |
| MAD46040.1 | Type1.BREX | <-SIGMA+OB-Fold+AAA-ATPase+HKD-DNase-BrxL<-PglZ<-HTH+REase<-N6-METHYLASE<-DUF6079*<-BrxB-BrxF-iSTAND<-BrxA-DNA-binding | 1189 | Bacteria>Pseudomonadota>Gammaproteobacteria>Oceanospirillales>Oceanospirillaceae>unclassified_Oceanospirillaceae | GCA_002685055.1 | Oceanospirillaceae_bacterium | CMH98_13650 | 9512..13081 | 1899355 | hypothetical-protein-[Oceanospirillaceae-bacterium] |
| MAS25385.1 | Type1.BREX | <-SIGMA+OB-Fold+AAA-ATPase+Lon-Protease-BrxL<-PglZ<-ParB+HNH<-PHP+SbcC<-N6-METHYLASE<-Schlafen+HATPase+wHTH<-DUF6079*<-BrxB-BrxF-iSTAND<-BrxA-DNA-binding<-XIS-HTH | 1145 | Bacteria>Pseudomonadota>Gammaproteobacteria>Oceanospirillales>Oceanospirillaceae>unclassified_Oceanospirillaceae | GCA_002698675.1 | Oceanospirillaceae_bacterium | CMI08_10750 | 15208..18645 | 1899355 | hypothetical-protein-[Oceanospirillaceae-bacterium] |
| MAR01289.1 | Type1.BREX | BrxB-BrxF-iSTAND-><-AbiEii<-wHTH+AbiEi_2\|\|?-><-SIGMA+OB-Fold+AAA-ATPase+HKD-DNase-BrxL<-REase<-Immank+ANK+ANK<-PglZ<-N6-METHYLASE<-DUF6079*<-BrxB-BrxF-iSTAND<-BrxA-DNA-binding\|\|HTH+REase-><-?<-?<-?<-PBPI | 1185 | Bacteria>Pseudomonadota>Gammaproteobacteria>Oceanospirillales>Oceanospirillaceae>unclassified_Oceanospirillaceae | GCA_002701445.1 | Oceanospirillaceae_bacterium | CMI00_12230 | 13782..17339 | 1899355 | hypothetical-protein-[Oceanospirillaceae-bacterium] |
| MBM94539.1 | Type1.BREX | CRISPR_Cas6->?-><-SIGMA+OB-Fold+AAA-ATPase+Lon-Protease-BrxL<-PglZ<-TOPRIM<-OLD-ABC<-N6-METHYLASE<-TOPRIM<-OLD-ABC<-DUF6079*<-BrxB-BrxF-iSTAND<-BrxA-DNA-binding\|\|?-><-?<-?<-Arm-DNA-bind_2+Phage_int_SAM_3+Phage_integrase<-Phage_AlpA\|\|HTH_26-> | 1225 | Bacteria>Pseudomonadota>Gammaproteobacteria>Oceanospirillales>Oceanospirillaceae>unclassified_Oceanospirillaceae | GCA_002723385.1 | Oceanospirillaceae_bacterium | CMI09_01635 | 47843..51520 | 1899355 | DNA-repair-protein-[Oceanospirillaceae-bacterium] |
| MBN56612.1 | Type1.BREX | <-PglZ<-?<-N6-METHYLASE<-?<-AbiEii<-MarR-HTH<-RNase<-DUF3584+DUF5401<-MukF<-DUF6079*<-BrxB-BrxF-iSTAND<-BrxA-DNA-binding<-wHTH+WYL+WCX\|\|Thioredoxin_4-> | 1216 | Bacteria>Pseudomonadota>Gammaproteobacteria>Oceanospirillales>Oceanospirillaceae>unclassified_Oceanospirillaceae | GCA_002724925.1 | Oceanospirillaceae_bacterium | CMI04_02830 | 19450..23100 | 1899355 | ATPase-[Oceanospirillaceae-bacterium] |
| NVK73702.1 | Type1.BREX | DUF4158+DDE_Tnp_Tn3->wHTH+WYL+WCX->BrxA-DNA-binding->BrxB-BrxF-iSTAND->DUF6079*->N6-METHYLASE->PglZ->SIGMA+OB-Fold+AAA-ATPase+Lon-Protease-BrxL->SbcC->REase+iREase->REase+HTH+35exo+Small-B-barrel+SF1-SF2-Helicase-><-Phage_integrase\|\|?-><-GGDEF | 1219 | Bacteria>Pseudomonadota>Gammaproteobacteria>Oceanospirillales>Oceanospirillaceae>unclassified_Oceanospirillaceae | GCA_013373235.1 | Oceanospirillaceae_bacterium | HWE24_09495 | 6489..10148 | 1899355 | BREX-system-P-loop-protein-BrxC-[Oceanospirillaceae-bacterium] |
| WP_116688140.1 | Type1.BREX | DDE_Tnp-><-HTH_3\|\|?-><-?\|\|?->DUF4096+DDE_Tnp_1+DDE_Tnp_1_2->HTH_26->BrxA-DNA-binding->BrxB-BrxF-iSTAND->DUF6079*->ParB+HNH->N6-METHYLASE->OLD-ABC->TOPRIM->OLD-ABC->TOPRIM->PglZ->SIGMA+OB-Fold+AAA-ATPase+Lon-Protease-BrxL-><-ANK+ANK | 1236 | Bacteria>Pseudomonadota>Gammaproteobacteria>Oceanospirillales>Oceanospirillales_incertae_sedis>Pelagibaculum | GCF_003097315.1 | Pelagibaculum_spongiae | DC094_RS16040 | 158438..162148 | 2080658 | BREX-system-P-loop-protein-BrxC-[Pelagibaculum-spongiae] |
| WP_100257386.1 | Type1.BREX | <-NERD+SF1-SF2-Helicase<-SIGMA+OB-Fold+AAA-ATPase+Lon-Protease-BrxL<-PglZ<-ParB+HNH<-N6-METHYLASE<-N6-METHYLASE<-DUF6079*<-BrxB-BrxF-iSTAND<-BrxA-DNA-binding<-tRNA<-MerR-HTH | 1215 | Bacteria>Pseudomonadota>Gammaproteobacteria>Oceanospirillales>Saccharospirillaceae>Reinekea | GCF_002795845.1 | Reinekea_forsetii | REIFOR_RS09825 | 2117568..2121215 | 1336806 | BREX-system-P-loop-protein-BrxC-[Reinekea-forsetii] |
| WP_119395059.1 | Type1.BREX | SF1-SF2-Helicase->HTH+HEPN->McrB-AAA->wHTH+REase+ZnR->HKD-DNase->OLD-ABC->HNH->BrxA-DNA-binding->BrxB-BrxF-iSTAND->DUF6079*->N6-METHYLASE->sGTP->sGTP->?->HerA/FtsK->HNH->?->HNH->PglZ-> | 1155 | Bacteria>Pseudomonadota>Gammaproteobacteria>Oceanospirillales>Saccharospirillaceae>Salinibius | GCF_003577475.1 | Salinibius_halmophilus | DUW71_RS09715 | 145555..149022 | 1853216 | BREX-system-P-loop-protein-BrxC-[Salinibius-halmophilus] |
| NRB78692.1 | Type1.BREX | <-sGTP<-sGTP<-?<-SIGMA+OB-Fold+AAA-ATPase+Lon-Protease-BrxL<-PglZ<-N6-METHYLASE<-DUF6079*<-BrxB-BrxF-iSTAND<-BrxA-DNA-binding<-HNH<-?\|\|?->?-><-SF1-SF2-Helicase<-HKD-DNase+HKD-Rease | 1233 | Bacteria>Pseudomonadota>Gammaproteobacteria>Oceanospirillales>Saccharospirillaceae>unclassified_Saccharospirillaceae | GCA_013214075.1 | Saccharospirillaceae_bacterium | HRU38_08495 | 18482..22183 | 2497634 | BREX-system-P-loop-protein-BrxC-[Saccharospirillaceae-bacterium] |
| NRB80667.1 | Type1.BREX | <-N6-METHYLASE<-DUF6079*<-BrxB-BrxF-iSTAND<-BrxA-DNA-binding<-wHTH+WYL+WCX<-HTH_19<-TnsC-ATPase<-Tn7_Tnp_TnsA_N+rve+Mu-transpos_C<-ncRNA<-GGDEF+EAL | 1225 | Bacteria>Pseudomonadota>Gammaproteobacteria>Oceanospirillales>Saccharospirillaceae>unclassified_Saccharospirillaceae | GCA_013214075.1 | Saccharospirillaceae_bacterium | HRU38_18705 | 4651..8328 | 2497634 | BREX-system-P-loop-protein-BrxC-[Saccharospirillaceae-bacterium] |
| MBR9867112.1 | Type1.BREX | <-SbcC<-SIGMA+OB-Fold+AAA-ATPase+Lon-Protease-BrxL<-PglZ<-TOPRIM<-OLD-ABC<-N6-METHYLASE<-ParB+HNH<-DUF6079*<-BrxB-BrxF-iSTAND<-BrxA-DNA-binding<-wHTH+WYL+WCX<-?<-?<-SLOG+BirA-HTH<-SF1-SF2-Helicase+PRTase\|\|DnaJ-> | 1225 | Bacteria>Pseudomonadota>Gammaproteobacteria>Oceanospirillales>unclassified_Oceanospirillales | GCA_018223265.1 | Oceanospirillales_bacterium | GYB18_08415 | 21169..24846 | 1914300 | BREX-system-P-loop-protein-BrxC-[Oceanospirillales-bacterium] |
| WP_241468927.1 | Type1.BREX | <-VirD4-TraG+VirB4-FtsK<-HDRnase<-SbcC<-SIGMA+OB-Fold+AAA-ATPase+Lon-Protease-BrxL<-PglZ<-N6-METHYLASE<-ParB+HNH<-DUF6079*<-BrxB-BrxF-iSTAND<-BrxA-DNA-binding<-wHTH+WYL+WCX<-?<-35exo<-?\|\|LexA-protease->?->Phage_integrase-> | 1225 | Bacteria>Pseudomonadota>Gammaproteobacteria>Pasteurellales>Pasteurellaceae>Pasteurella | GCF_022559605.1 | Pasteurella_multocida | MMZ64_RS09105 | 1895707..1899384 | 747 | BREX-system-P-loop-protein-BrxC-[Pasteurella-multocida] |
| WP_119518392.1 | Type1.BREX | Arm-DNA-bind_3+Phage_integrase->?-><-SIGMA+OB-Fold+AAA-ATPase+Lon-Protease-BrxL<-PglZ<-ParB+HNH<-?<-HNH+GCN1<-?<-N6-METHYLASE<-DUF6079*<-BrxB-BrxF-iSTAND<-BrxA-DNA-binding<-?<-Peptidase_M15<-?\|\|wHTH+WYL+WCX->?-><-PhdYeFM_antitox<-NERD | 1154 | Bacteria>Pseudomonadota>Gammaproteobacteria>Pseudomonadales | GCF_021610005.1 | Pseudomonadaceae | L1776_RS13375 | 2929669..2933133 | 135621 | MULTISPECIES:-BREX-system-P-loop-protein-BrxC-[Pseudomonadaceae] |
| KXS54417.1 | Type1.BREX | BrxA-DNA-binding->BrxB-BrxF-iSTAND->DUF6079*->DOC+HTH->N6-METHYLASE->OLD-ABC+TOPRIM->OLD-ABC-> | 1221 | Bacteria>Pseudomonadota>Gammaproteobacteria>Pseudomonadales>Marinobacteraceae>Marinobacter | GCA_001577575.1 | Marinobacter_sp._T13-3 | AWU57_1199 | 1913..5578 | 1794906 | hypothetical-protein-AWU57_1199-[Marinobacter-sp.-T13-3] |
| KXS53697.1 | Type1.BREX | <-HNH<-?<-OLD-ABC<-N6-METHYLASE<-HTH+HTH<-Schlafen+HATPase+wHTH<-DUF6079-p\|\|?-><-DUF6079-p*<-BrxB-BrxF-iSTAND<-BrxA-DNA-binding<-XIS-HTH<-MobI\|\|XIS-HTH->AAA+HTH->XIS-HTH-><-Arm-DNA-bind_3+Phage_integrase | 539 | Bacteria>Pseudomonadota>Gammaproteobacteria>Pseudomonadales>Marinobacteraceae>Marinobacter | GCA_001577575.1 | Marinobacter_sp._T13-3 | AWU57_1918 | 9032..10651 | 1794906 | hypothetical-protein-AWU57_1918-[Marinobacter-sp.-T13-3] |
| MAH31343.1 | Type1.BREX | <-Schlafen+HATPase+wHTH<-DUF6079*<-BrxB-BrxF-iSTAND<-BrxA-DNA-binding<-XIS-HTH<-MobI\|\|XIS-HTH->AAA+HTH->XIS-HTH-><-Arm-DNA-bind_3+Phage_integrase | 1193 | Bacteria>Pseudomonadota>Gammaproteobacteria>Pseudomonadales>Marinobacteraceae>Marinobacter | GCA_002690365.1 | Marinobacter_sp. | CMG78_04665 | 1378..4959 | 50741 | hypothetical-protein-[Marinobacter-sp.] |
| MBE95557.1 | Type1.BREX | BrxB-BrxF-iSTAND->DUF6079*->Schlafen+HATPase+wHTH+HTH->SF1-SF2-Helicase+PRTase->SLOG+BirA-HTH->N6-METHYLASE->ParB+HNH->PglZ->SIGMA+OB-Fold+AAA-ATPase+HKD-DNase-BrxL-><-?<-IstB_IS21 | 1193 | Bacteria>Pseudomonadota>Gammaproteobacteria>Pseudomonadales>Marinobacteraceae>Marinobacter | GCA_002714765.1 | Marinobacter_sp. | CMG89_11180 | 696..4277 | 50741 | hypothetical-protein-[Marinobacter-sp.] |
| PTB94103.1 | Type1.BREX | <-Phage_integrase\|\|BrxA-DNA-binding->BrxB-BrxF-iSTAND->DUF6079*->DOC+HTH->N6-METHYLASE-> | 1221 | Bacteria>Pseudomonadota>Gammaproteobacteria>Pseudomonadales>Marinobacteraceae>Marinobacter | GCA_003030785.1 | Marinobacter_sp._B9-2 | C9974_06180 | 29766..33431 | 2137198 | BREX-system-P-loop-protein-BrxC-[Marinobacter-sp.-B9-2] |
| HBC35650.1 | Type1.BREX | BrxB-BrxF-iSTAND->BrxB-BrxF-iSTAND-p->DUF6079-p->DUF6079-p*->DUF6079-p->DOC+HTH->DOC+HTH->N6-METHYLASE-> | 726 | Bacteria>Pseudomonadota>Gammaproteobacteria>Pseudomonadales>Marinobacteraceae>Marinobacter | GCA_003487885.1 | Marinobacter_adhaerens | DC045_15360 | 1248..3427 | 1033846 | BREX-system-P-loop-protein-BrxC-[Marinobacter-adhaerens] |
| TVT30739.1 | Type1.BREX | GDResolvase+HIN-HTH->?-><-REase<-SIGMA+OB-Fold+AAA-ATPase+Lon-Protease-BrxL<-PglZ<-Schlafen+HATPase+wHTH<-PHP+SbcC<-N6-METHYLASE<-Schlafen+HATPase+wHTH<-DUF6079*<-BrxB-BrxF-iSTAND<-BrxA-DNA-binding<-XIS-HTH\|\|XIS-HTH->APATPase->?-><-Phage_int_SAM_5+Phage_integrase | 1145 | Bacteria>Pseudomonadota>Gammaproteobacteria>Pseudomonadales>Marinobacteraceae>Marinobacter | GCA_007713625.1 | Marinobacter_vinifirmus | FHK81_16390 | 18158..21595 | 355591 | BREX-system-P-loop-protein-BrxC-[Marinobacter-vinifirmus] |
| MBC7192650.1 | Type1.BREX | <-SIGMA+OB-Fold+AAA-ATPase+Lon-Protease-BrxL<-PglZ<-HNH<-OLD-ABC<-TOPRIM<-OLD-ABC<-HTH+REase<-N6-METHYLASE<-Schlafen+HATPase+wHTH<-DUF6079*<-BrxB-BrxF-iSTAND<-BrxA-DNA-binding<-XIS-HTH\|\|HTH_Tnp_1+HTH_21+rve+rve_3->XIS-HTH->DnaB->?-><-Phage_int_SAM_5+Phage_integrase<-sGTP | 1145 | Bacteria>Pseudomonadota>Gammaproteobacteria>Pseudomonadales>Marinobacteraceae>Marinobacter | GCA_014359475.1 | Marinobacter_sp. | HSU31_09890 | 22634..26071 | 50741 | BREX-system-P-loop-protein-BrxC-[Marinobacter-sp.] |
| WBU42642.1 | Type1.BREX | <-Mrr-N\|\|?->?->?->SWIM+DUF3670+SF1-SF2-Helicase->HTH_3->BrxA-DNA-binding->BrxB-BrxF-iSTAND->DUF6079*->DOC+HTH->N6-METHYLASE->?->N6-METHYLASE->HEPN->?->TIR->?->TIR-> | 1221 | Bacteria>Pseudomonadota>Gammaproteobacteria>Pseudomonadales>Marinobacteraceae>Marinobacter | GCA_027912435.1 | Marinobacter_alkaliphilus | PBN92_07055 | 1547680..1551345 | 254719 | BREX-system-P-loop-protein-BrxC-[Marinobacter-alkaliphilus] |
| WP_011784101.1 | Type1.BREX | Arm-DNA-bind_3+Phage_integrase-><-XIS-HTH<-AAA+HTH<-XIS-HTH\|\|MobI->XIS-HTH->BrxA-DNA-binding->BrxB-BrxF-iSTAND->DUF6079*->Schlafen+HATPase+wHTH+HTH->SF1-SF2-Helicase+PRTase->SLOG+BirA-HTH->N6-METHYLASE->OLD-ABC->TOPRIM->PglZ->SIGMA+OB-Fold+AAA-ATPase+HKD-DNase-BrxL->SbcC-> | 1193 | Bacteria>Pseudomonadota>Gammaproteobacteria>Pseudomonadales>Marinobacteraceae>Marinobacter | GCF_000015365.1 | Marinobacter_nauticus | MAQU_RS02805 | 628948..632529 | 2743 | BREX-system-P-loop-protein-BrxC-[Marinobacter-nauticus] |
| WP_007349110.1 | Type1.BREX | <-Phage_int_SAM_1+Phage_integrase<-REase+N6_Mtase<-HTH_26<-TetR-HTH<-Phage_integrase+Phage_integrase<-Phage_integrase+Phage_integrase<-Phage_integrase<-N6-METHYLASE-p<-Schlafen+HATPase+wHTH<-DUF6079*<-BrxB-BrxF-iSTAND<-BrxA-DNA-binding<-XIS-HTH\|\|Arm-DNA-bind_3->?->DDE_Tnp_1_4-><-?<-?<-DDE_Tnp_IS66 | 1145 | Bacteria>Pseudomonadota>Gammaproteobacteria>Pseudomonadales>Marinobacteraceae>Marinobacter | GCF_000169375.1 | Marinobacter_sp._ELB17 | MELB17_RS05280 | 142493..145930 | 270374 | BREX-system-P-loop-protein-BrxC-[Marinobacter-sp.-ELB17] |
| WP_008174228.1 | Type1.BREX | Arm-DNA-bind_3+Phage_integrase-><-XIS-HTH<-AAA+HTH<-XIS-HTH\|\|MobI->XIS-HTH->BrxA-DNA-binding->BrxB-BrxF-iSTAND->DUF6079*->Schlafen+HATPase+wHTH+HTH->N6-METHYLASE->OLD-ABC->TOPRIM-> | 1193 | Bacteria>Pseudomonadota>Gammaproteobacteria>Pseudomonadales>Marinobacteraceae>Marinobacter | GCF_000235625.1 | Marinobacter_manganoxydans | KYE_13690 | 42273..45854 | 1150997 | BREX-system-P-loop-protein-BrxC-[Marinobacter-manganoxydans] |
| WP_018403235.1 | Type1.BREX | CHTH->GDResolvase->ZnR-><-GDResolvase+HIN-HTH<-SIGMA+OB-Fold+AAA-ATPase+Lon-Protease-BrxL<-PglZ<-REase+AIPR<-N6-METHYLASE<-Schlafen+HATPase+wHTH<-DUF6079*<-BrxB-BrxF-iSTAND<-BrxA-DNA-binding<-XIS-HTH<-tRNA<-HAD+HAD<-HDRnase<-?<-?<-DNA_METHYLASE | 1145 | Bacteria>Pseudomonadota>Gammaproteobacteria>Pseudomonadales>Marinobacteraceae>Marinobacter | GCF_000372805.1 | Marinobacter_gelidimuriae | A3OI_RS0105075 | 19872..23309 | 2739064 | BREX-system-P-loop-protein-BrxC-[Marinobacter-gelidimuriae] |
| WP_018404435.1 | Type1.BREX | OrfB_Zn_ribbon->BrxA-DNA-binding->BrxB-BrxF-iSTAND->DUF6079*->Schlafen+HATPase+wHTH+HTH->N6-METHYLASE->OLD-ABC->HNH->PglZ->SIGMA+OB-Fold+AAA-ATPase+HKD-DNase-BrxL->N6-METHYLASE-> | 1194 | Bacteria>Pseudomonadota>Gammaproteobacteria>Pseudomonadales>Marinobacteraceae>Marinobacter | GCF_000372805.1 | Marinobacter_gelidimuriae | A3OI_RS0111615 | 2203..5787 | 2739064 | BREX-system-P-loop-protein-BrxC-[Marinobacter-gelidimuriae] |
| WP_022988850.1 | Type1.BREX | <-PglZ<-TOPRIM<-OLD-ABC<-HNH<-OLD-ABC<-N6-METHYLASE<-REase<-OLD-ABC<-Schlafen+HATPase+wHTH+HTH<-DUF6079*<-BrxB-BrxF-iSTAND<-BrxA-DNA-binding<-XIS-HTH<-MobI | 1193 | Bacteria>Pseudomonadota>Gammaproteobacteria>Pseudomonadales>Marinobacteraceae>Marinobacter | GCF_000475255.1 | Marinobacter_sp._ES-1 | Q668_RS06800 | 88554..92135 | 1396858 | BREX-system-P-loop-protein-BrxC-[Marinobacter-sp.-ES-1] |
| WP_022990203.1 | Type1.BREX | <-DnaB<-XIS-HTH\|\|XIS-HTH->BrxA-DNA-binding->BrxB-BrxF-iSTAND->DUF6079*->Schlafen+HATPase+wHTH+HTH->N6-METHYLASE->REase->SIR2->HerA/FtsK->N6-METHYLASE-p->PglZ->Immank+ANK+ANK->REase-> | 1189 | Bacteria>Pseudomonadota>Gammaproteobacteria>Pseudomonadales>Marinobacteraceae>Marinobacter | GCF_000475255.1 | Marinobacter_sp._ES-1 | Q668_RS13780 | 2583..6152 | 1396858 | BREX-system-P-loop-protein-BrxC-[Marinobacter-sp.-ES-1] |
| WP_023011723.1 | Type1.BREX | <-OLD-ABC<-N6-METHYLASE-p<-HerA/FtsK<-SIR2<-REase<-N6-METHYLASE<-SLOG+BirA-HTH<-SF1-SF2-Helicase+PRTase<-Schlafen+HATPase+wHTH+HTH<-DUF6079*<-BrxB-BrxF-iSTAND<-BrxA-DNA-binding<-XIS-HTH<-MobI\|\|XIS-HTH->AAA+HTH->XIS-HTH-><-Arm-DNA-bind_3+Phage_integrase | 1193 | Bacteria>Pseudomonadota>Gammaproteobacteria>Pseudomonadales>Marinobacteraceae>Marinobacter | GCF_000475315.1 | Marinobacter_sp._EN3 | Q675_RS32885 | 62164..65745 | 1397533 | BREX-system-P-loop-protein-BrxC-[Marinobacter-sp.-EN3] |
| WP_022992169.1 | Type1.BREX | Arm-DNA-bind_3+Phage_integrase-><-XIS-HTH<-AAA+HTH<-XIS-HTH\|\|MobI->XIS-HTH->BrxA-DNA-binding->BrxB-BrxF-iSTAND->DUF6079*->Schlafen+HATPase+wHTH+HTH->N6-METHYLASE->OLD-ABC->HNH->OLD-ABC->TOPRIM->PglZ->SIGMA+OB-Fold+AAA-ATPase+HKD-DNase-BrxL-> | 1193 | Bacteria>Pseudomonadota>Gammaproteobacteria>Pseudomonadales>Marinobacteraceae>Marinobacter | GCF_000475355.1 | Marinobacter_sp._C1S70 | Q669_RS10410 | 149187..152768 | 1396859 | BREX-system-P-loop-protein-BrxC-[Marinobacter-sp.-C1S70] |
| WP_023007675.1 | Type1.BREX | sGTP->Phage_int_SAM_5+Phage_integrase-><-?<-DnaB<-XIS-HTH\|\|XIS-HTH->BrxA-DNA-binding->BrxB-BrxF-iSTAND->DUF6079*->Schlafen+HATPase+wHTH->N6-METHYLASE-p-> | 1145 | Bacteria>Pseudomonadota>Gammaproteobacteria>Pseudomonadales>Marinobacteraceae>Marinobacter | GCF_000475375.1 | Marinobacter_sp._EVN1 | Q674_RS21825 | 56461..59898 | 1397532 | BREX-system-P-loop-protein-BrxC-[Marinobacter-sp.-EVN1] |
| WP_023008352.1 | Type1.BREX | <-SbcC<-SIGMA+OB-Fold+AAA-ATPase+HKD-DNase-BrxL<-PglZ<-TOPRIM<-OLD-ABC<-HNH<-OLD-ABC<-N6-METHYLASE<-Schlafen+HATPase+wHTH+HTH<-DUF6079*<-BrxB-BrxF-iSTAND<-BrxA-DNA-binding<-XIS-HTH<-MobI\|\|XIS-HTH->AAA+HTH->XIS-HTH-><-Arm-DNA-bind_3+Phage_integrase | 1193 | Bacteria>Pseudomonadota>Gammaproteobacteria>Pseudomonadales>Marinobacteraceae>Marinobacter | GCF_000475375.1 | Marinobacter_sp._EVN1 | Q674_RS26680 | 52773..56354 | 1397532 | BREX-system-P-loop-protein-BrxC-[Marinobacter-sp.-EVN1] |
| WP_048384700.1 | Type1.BREX | Arm-DNA-bind_3+Phage_integrase-><-AbiGi\|\|?->CHTH->XIS-HTH->?->XIS-HTH->BrxA-DNA-binding->BrxB-BrxF-iSTAND->DUF6079*->Schlafen+HATPase+wHTH->N6-METHYLASE->ParB+HNH->?->DUF3644->PglZ->SIGMA+OB-Fold+AAA-ATPase+Lon-Protease-BrxL->Integrase_1-><-CHTH | 1145 | Bacteria>Pseudomonadota>Gammaproteobacteria>Pseudomonadales>Marinobacteraceae>Marinobacter | GCF_001043175.1 | Marinobacter_psychrophilus | ABA45_RS05800 | 1298938..1302375 | 330734 | BREX-system-P-loop-protein-BrxC-[Marinobacter-psychrophilus] |
| WP_058091394.1 | Type1.BREX | wHTH+WYL+WCX->?-><-Mrr-N\|\|HTH_3->BrxA-DNA-binding->BrxB-BrxF-iSTAND->DUF6079*->DOC+HTH->SF1-SF2-Helicase->N6-METHYLASE->HEPN->?->TIR->OLD-ABC->TOPRIM-> | 1221 | Bacteria>Pseudomonadota>Gammaproteobacteria>Pseudomonadales>Marinobacteraceae>Marinobacter | GCF_001447155.2 | Marinobacter_sp._LQ44 | ASQ50_RS18570 | 4052781..4056446 | 1749259 | BREX-system-P-loop-protein-BrxC-[Marinobacter-sp.-LQ44] |
| WP_071262841.1 | Type1.BREX | <-DrsE<-?\|\|tRNA->BrxA-DNA-binding->BrxB-BrxF-iSTAND->DUF6079*->MarR-HTH->AbiEii->AbiEii->?->N6-METHYLASE-p->ParB+HNH->HKD-DNase+DUF3150-> | 1217 | Bacteria>Pseudomonadota>Gammaproteobacteria>Pseudomonadales>Marinobacteraceae>Marinobacter | GCF_001858325.1 | Marinobacter_sp._AC-23 | BCA33_RS00460 | 104149..107802 | 1879031 | BREX-system-P-loop-protein-BrxC-[Marinobacter-sp.-AC-23] |
| WP_104416052.1 | Type1.BREX | Arm-DNA-bind_3+Phage_integrase-><-XIS-HTH<-AAA+HTH<-XIS-HTH\|\|MobI->XIS-HTH->BrxA-DNA-binding->BrxB-BrxF-iSTAND->DUF6079*->Schlafen+HATPase+wHTH+HTH->N6-METHYLASE->OLD-ABC->HNH->OLD-ABC->TOPRIM->PglZ->SIGMA+OB-Fold+AAA-ATPase+HKD-DNase-BrxL-> | 1193 | Bacteria>Pseudomonadota>Gammaproteobacteria>Pseudomonadales>Marinobacteraceae>Marinobacter | GCF_002934485.1 | Marinobacter_persicus | BY455_RS08980 | 65339..68920 | 930118 | BREX-system-P-loop-protein-BrxC-[Marinobacter-persicus] |
| WP_113863603.1 | Type1.BREX | Arm-DNA-bind_3+Phage_integrase-><-XIS-HTH<-AAA+HTH<-XIS-HTH\|\|MobI->XIS-HTH->BrxA-DNA-binding->BrxB-BrxF-iSTAND->DUF6079*->Schlafen+HATPase+wHTH+HTH->N6-METHYLASE->OLD-ABC->HNH->OLD-ABC->TOPRIM->PglZ->SIGMA+OB-Fold+AAA-ATPase+HKD-DNase-BrxL-> | 1193 | Bacteria>Pseudomonadota>Gammaproteobacteria>Pseudomonadales>Marinobacteraceae>Marinobacter | GCF_003315345.1 | Marinobacter_pelagius | DET50_RS14950 | 40642..44223 | 379482 | BREX-system-P-loop-protein-BrxC-[Marinobacter-pelagius] |
| WP_113879482.1 | Type1.BREX | <-N6-METHYLASE<-Schlafen+HATPase+wHTH+HTH<-DUF6079*<-BrxB-BrxF-iSTAND<-BrxA-DNA-binding<-XIS-HTH<-MobI\|\|XIS-HTH->AAA+HTH->XIS-HTH-><-Arm-DNA-bind_3+Phage_integrase | 1193 | Bacteria>Pseudomonadota>Gammaproteobacteria>Pseudomonadales>Marinobacteraceae>Marinobacter | GCF_003315555.1 | Marinobacter_nauticus | DET64_RS06865 | 4494..8075 | 2743 | BREX-system-P-loop-protein-BrxC-[Marinobacter-nauticus] |
| WP_114434424.1 | Type1.BREX | <-ParB<-HerA/FtsK<-?<-LZ_Tnp_IS66+DDE_Tnp_IS66+DDE_Tnp_IS66_C<-?<-HIN-HTH<-SIR2<-N6-METHYLASE<-SF1-SF2-Helicase<-DUF6079*<-BrxB-BrxF-iSTAND<-BrxA-DNA-binding\|\|ANK+ANK->SF1-SF2-Helicase+REase->PIWI-><-?\|\|?-><-TraI-VirD2-RCRE<-TraJ-RHH | 1217 | Bacteria>Pseudomonadota>Gammaproteobacteria>Pseudomonadales>Marinobacteraceae>Marinobacter | GCF_003337515.1 | Marinobacter_nauticus | DET61_RS09095 | 118513..122166 | 2743 | BREX-system-P-loop-protein-BrxC-[Marinobacter-nauticus] |
| WP_141068072.1 | Type1.BREX | Arm-DNA-bind_3+Phage_integrase-><-XIS-HTH<-AAA+HTH<-XIS-HTH\|\|MobI->XIS-HTH->BrxA-DNA-binding->BrxB-BrxF-iSTAND->DUF6079*->Schlafen+HATPase+wHTH+HTH->OLD-ABC->REase->N6-METHYLASE->ParB+HNH->PglZ->SIGMA+OB-Fold+AAA-ATPase+HKD-DNase-BrxL->SbcC+APATPase+SbcC-> | 1193 | Bacteria>Pseudomonadota>Gammaproteobacteria>Pseudomonadales>Marinobacteraceae>Marinobacter | GCF_006516615.1 | Marinobacter_nauticus | FH712_RS17655 | 33566..37147 | 2743 | BREX-system-P-loop-protein-BrxC-[Marinobacter-nauticus] |
| WP_273135111.1 | Type1.BREX | GDResolvase+HIN-HTH->?-><-?<-REase<-SIGMA+OB-Fold+AAA-ATPase+Lon-Protease-BrxL<-PglZ<-Schlafen+HATPase+wHTH<-PHP+SbcC<-N6-METHYLASE<-Schlafen+HATPase+wHTH<-DUF6079*<-BrxB-BrxF-iSTAND<-BrxA-DNA-binding<-XIS-HTH\|\|XIS-HTH->APATPase->?-><-Phage_int_SAM_5+Phage_integrase | 1145 | Bacteria>Pseudomonadota>Gammaproteobacteria>Pseudomonadales>Marinobacteraceae>Marinobacter | GCF_007713625.1 | Marinobacter_vinifirmus | FHK81_RS16410 | 18158..21595 | 355591 | BREX-system-P-loop-protein-BrxC-[Marinobacter-vinifirmus] |
| WP_213480159.1 | Type1.BREX | <-Couple_hipA<-SIGMA+OB-Fold+AAA-ATPase+Lon-Protease-BrxL<-PglZ<-ParB+HNH<-PHP+SbcC<-N6-METHYLASE<-DUF6079*<-BrxB-BrxF-iSTAND<-BrxA-DNA-binding<-XIS-HTH\|\|ANK+ANK->TPR-><-wHTH+WYL+WCX\|\|?->?->SbcC-> | 1145 | Bacteria>Pseudomonadota>Gammaproteobacteria>Pseudomonadales>Marinobacteraceae>Marinobacter | GCF_018424725.1 | Marinobacter_salarius | DYI41_RS19390 | 15297..18734 | 1420917 | BREX-system-P-loop-protein-BrxC-[Marinobacter-salarius] |
| WP_216008134.1 | Type1.BREX | <-PglZ<-TIR<-?<-ParB+HNH<-?<-N6-METHYLASE-p<-RVT<-N6-METHYLASE<-SF1-SF2-Helicase<-DUF6079*<-BrxB-BrxF-iSTAND<-BrxA-DNA-binding\|\|ANK+ANK->?->?->?->?->JetB-> | 1217 | Bacteria>Pseudomonadota>Gammaproteobacteria>Pseudomonadales>Marinobacteraceae>Marinobacter | GCF_018860765.1 | Marinobacter_salexigens | K0508_RS09770 | 175903..179556 | 1925763 | BREX-system-P-loop-protein-BrxC-[Marinobacter-salexigens] |
| WP_228224785.1 | Type1.BREX | TraJ-RHH->TraI-VirD2-RCRE-><-?\|\|?-><-PIWI<-SF1-SF2-Helicase+REase<-ANK+ANK\|\|BrxA-DNA-binding->BrxB-BrxF-iSTAND->DUF6079*->MarR-HTH->AbiEii->?->N6-METHYLASE->RVT->N6-METHYLASE-p->?->GP+REase-> | 1217 | Bacteria>Pseudomonadota>Gammaproteobacteria>Pseudomonadales>Marinobacteraceae>Marinobacter | GCF_020781935.1 | Marinobacter_nauticus | LL254_RS15890 | 125400..129053 | 2743 | BREX-system-P-loop-protein-BrxC-[Marinobacter-nauticus] |
| WP_247116152.1 | Type1.BREX | <-SIGMA+OB-Fold+AAA-ATPase+HKD-DNase-BrxL<-REase<-Immank+ANK+ANK<-PglZ<-?<-OLD-ABC<-ParB+HNH<-N6-METHYLASE<-DUF6079*<-BrxB-BrxF-iSTAND<-BrxA-DNA-binding\|\|?-><-?\|\|?-><-?\|\|?->\|\|?->Arm-DNA-bind_2+Phage_integrase-> | 1189 | Bacteria>Pseudomonadota>Gammaproteobacteria>Pseudomonadales>Marinobacteraceae>Marinobacter | GCF_023016045.1 | Marinobacter_sp._S6332 | MWU62_RS17920 | 19915..23484 | 2926403 | BREX-system-P-loop-protein-BrxC-[Marinobacter-sp.-S6332] |
| WP_248185657.1 | Type1.BREX | <-OLD-ABC<-Schlafen+HATPase+wHTH+HTH<-DUF6079*<-BrxB-BrxF-iSTAND<-BrxA-DNA-binding<-XIS-HTH<-MobI<-?\|\|XIS-HTH->AAA+HTH->XIS-HTH-><-Phage_integrase | 1193 | Bacteria>Pseudomonadota>Gammaproteobacteria>Pseudomonadales>Marinobacteraceae>Marinobacter | GCF_023156235.1 | Marinobacter_goseongensis | MLC70_RS17480 | 2864..6445 | 453838 | BREX-system-P-loop-protein-BrxC-[Marinobacter-goseongensis] |
| WP_248236652.1 | Type1.BREX | <-Integrase_1<-SIGMA+OB-Fold+AAA-ATPase+Lon-Protease-BrxL<-PglZ<-DUF3644<-?<-ParB+HNH<-N6-METHYLASE<-Schlafen+HATPase+wHTH<-DUF6079*<-BrxB-BrxF-iSTAND<-BrxA-DNA-binding<-XIS-HTH<-Phage_AlpA<-Arm-DNA-bind_3+Phage_integrase | 1145 | Bacteria>Pseudomonadota>Gammaproteobacteria>Pseudomonadales>Marinobacteraceae>Marinobacter | GCF_023156245.1 | Marinobacter_bryozoorum | MLC59_RS17300 | 17294..20731 | 256324 | BREX-system-P-loop-protein-BrxC-[Marinobacter-bryozoorum] |
| WP_249013861.1 | Type1.BREX | <-TIR<-?<-ParB+HNH<-?<-N6-METHYLASE-p<-RVT<-N6-METHYLASE<-DOC+HTH<-DUF6079*<-BrxB-BrxF-iSTAND<-BrxA-DNA-binding<-?\|\|?->?->DUF4096+DDE_Tnp_1_2->?->TOPRIM->JetC-> | 1217 | Bacteria>Pseudomonadota>Gammaproteobacteria>Pseudomonadales>Marinobacteraceae>Marinobacter | GCF_023311895.1 | Marinobacter_sp._M3C | MIH18_RS02775 | 634198..637851 | 2917715 | BREX-system-P-loop-protein-BrxC-[Marinobacter-sp.-M3C] |
| WP_092033189.1 | Type1.BREX | Arm-DNA-bind_3+Phage_integrase-><-XIS-HTH<-AAA+HTH<-XIS-HTH\|\|MobI->XIS-HTH->BrxA-DNA-binding->BrxB-BrxF-iSTAND->DUF6079*->Schlafen+HATPase+wHTH+HTH->OLD-ABC->REase->N6-METHYLASE->PglZ->SIGMA+OB-Fold+AAA-ATPase+HKD-DNase-BrxL->SbcC->Calcineurin+Calcineurin-SbcD->SbcC-> | 1193 | Bacteria>Pseudomonadota>Gammaproteobacteria>Pseudomonadales>Marinobacteraceae>Marinobacter | GCF_900112835.1 | Marinobacter_sp._DSM_26671 | BM035_RS16335 | 75345..78926 | 1761793 | BREX-system-P-loop-protein-BrxC-[Marinobacter-sp.-DSM-26671] |
| WP_092021229.1 | Type1.BREX | DUF932->?->?-><-HTH_3\|\|?->?->wHTH+WYL+WCX->BrxA-DNA-binding->BrxB-BrxF-iSTAND->DUF6079*->N6-METHYLASE->ParB+HNH->OLD-ABC->TOPRIM->PglZ->SIGMA+OB-Fold+AAA-ATPase+Lon-Protease-BrxL->REase+PNTD1+PNTD2->PIWI->TOPRIM-> | 1219 | Bacteria>Pseudomonadota>Gammaproteobacteria>Pseudomonadales>Marinobacteraceae>Marinobacter | GCF_900114775.1 | Marinobacter_zhejiangensis | BMZ67_RS06955 | 334318..337977 | 488535 | BREX-system-P-loop-protein-BrxC-[Marinobacter-zhejiangensis] |
| BBJ05051.1 | Type1.BREX | Arm-DNA-bind_3+Phage_integrase-><-XIS-HTH<-AAA+HTH<-XIS-HTH\|\|MobI->XIS-HTH->BrxA-DNA-binding->BrxB-BrxF-iSTAND->DUF6079*->Schlafen+HATPase+wHTH+HTH->N6-METHYLASE->RVT->HerA/FtsK->N6-METHYLASE-p->OLD-ABC->TOPRIM->PglZ->SIGMA+OB-Fold+AAA-ATPase+HKD-DNase-BrxL-> | 1193 | Bacteria>Pseudomonadota>Gammaproteobacteria>Pseudomonadales>Marinobacteraceae>Marinobacter | - | Marinobacter_nauticus | YBY_29000 | 3185607..3189188 | 2743 | hypothetical-protein-YBY_29000-[Marinobacter-nauticus] |
| WP_116918937.1 | Type1.BREX | <-XIS-HTH<-?<-?<-Arm-DNA-bind_2+Phage_int_SAM_3+Phage_integrase<-Phage_AlpA\|\|HTH_26->BrxA-DNA-binding->BrxB-BrxF-iSTAND->DUF6079*->Schlafen+HATPase+wHTH+HTH->OLD-ABC->REase->N6-METHYLASE->PglZ->Immank+ANK+ANK->REase->SIGMA+OB-Fold+AAA-ATPase+HKD-DNase-BrxL-> | 1193 | Bacteria>Pseudomonadota>Gammaproteobacteria>Pseudomonadales>Marinobacteraceae>Tamilnaduibacter | GCF_003096655.1 | Tamilnaduibacter_salinus | C8D92_104115 | 123558..127139 | 1484056 | BREX-system-P-loop-protein-BrxC-[Tamilnaduibacter-salinus] |
| WP_152390158.1 | Type1.BREX | VirB10->?-><-SIGMA+OB-Fold+AAA-ATPase+Lon-Protease-BrxL<-PglZ<-TOPRIM<-OLD-ABC<-N6-METHYLASE<-Schlafen+HATPase+wHTH<-DUF6079*<-BrxB-BrxF-iSTAND<-BrxA-DNA-binding<-wHTH+WYL+WCX<-Tox-RES<-CHTH\|\|?->?->DnaB+HerA/FtsK-> | 1224 | Bacteria>Pseudomonadota>Gammaproteobacteria>Pseudomonadales>Pseudomonadaceae>Azotobacter_group | GCF_009363155.1 | Azotobacter_salinestris | GCU53_RS24425 | 150496..154170 | 69964 | BREX-system-P-loop-protein-BrxC-[Azotobacter-salinestris] |
| WP_022966741.1 | Type1.BREX | DDE_Tnp_IS66+DDE_Tnp_IS66_C->PVC-AAA-><-?<-?<-NACHT<-SIGMA+OB-Fold+AAA-ATPase+HKD-DNase-BrxL<-PglZ<-N6-METHYLASE<-DUF6079*<-BrxB-BrxF-iSTAND<-BrxA-DNA-binding<-?<-AbiEii<-AbiEi_3_N+AbiEi_3<-35exo<-?<-?<-LexA-protease | 1184 | Bacteria>Pseudomonadota>Gammaproteobacteria>Pseudomonadales>Pseudomonadaceae>Denitrificimonas | GCF_000421765.1 | Denitrificimonas_caeni | H591_RS0109630 | 17243..20797 | 521720 | BREX-system-P-loop-protein-BrxC-[Denitrificimonas-caeni] |
| WP_003439533.1 | Type1.BREX | <-PglZ\|\|ZnR-><-TPR<-NERD+GspA-ATPase<-?<-?<-?<-N6-METHYLASE<-DOC+HTH<-DUF6079*<-BrxB-BrxF-iSTAND<-BrxA-DNA-binding<-wHTH+WYL+WCX<-?<-Phage_AlpA<-?<-Arm-DNA-bind_3+Phage_integrase | 1225 | Bacteria>Pseudomonadota>Gammaproteobacteria>Pseudomonadales>Pseudomonadaceae | GCF_000346225.2 | Pseudomonas | HBN87_RS22825 | 103169..106846 | 286 | MULTISPECIES:-BREX-system-P-loop-protein-BrxC-[Pseudomonas] |
| WP_033898350.1 | Type1.BREX | <-wHTH+WYL+WCX\|\|?->ARG->XIS-HTH->BrxA-DNA-binding->BrxB-BrxF-iSTAND->DUF6079*->N6-METHYLASE-p->?-><-?\|\|N6-METHYLASE-p->OLD-ABC->HNH->SIGMA+OB-Fold+AAA-ATPase+HKD-DNase-BrxL-> | 1185 | Bacteria>Pseudomonadota>Gammaproteobacteria>Pseudomonadales>Pseudomonadaceae | GCF_000510895.2 | Pseudomonas | GIW75_RS00865 | 8119..11676 | 286 | MULTISPECIES:-BREX-system-P-loop-protein-BrxC-[Pseudomonas] |
| WP_057711677.1 | Type1.BREX | <-Ntox49<-SF1-SF2-Helicase<-HKD-DNase+HKD-DNase<-?\|\|wHTH+WYL+WCX->?->\|\|wHTH+WYL+WCX->BrxA-DNA-binding->BrxB-BrxF-iSTAND->DUF6079*->N6-METHYLASE->NERD+GspA-ATPase->TPR->PglZ->SIGMA+OB-Fold+AAA-ATPase+Lon-Protease-BrxL->?->?-><-tRNA<-MerR-HTH | 1222 | Bacteria>Pseudomonadota>Gammaproteobacteria>Pseudomonadales>Pseudomonadaceae | GCF_001439845.1 | Pseudomonas | TX24_RS25940 | 84684..88352 | 286 | MULTISPECIES:-BREX-system-P-loop-protein-BrxC-[Pseudomonas] |
| WP_057024539.1 | Type1.BREX | ABhydrolase->?->?-><-?\|\|XIS-HTH->?->wHTH+WYL+WCX->BrxA-DNA-binding->BrxB-BrxF-iSTAND->DUF6079*->N6-METHYLASE->ParB+HNH->PglZ->SIGMA+OB-Fold+AAA-ATPase+Lon-Protease-BrxL-><-DUF4969 | 1154 | Bacteria>Pseudomonadota>Gammaproteobacteria>Pseudomonadales>Pseudomonadaceae | GCF_002091795.1 | Pseudomonas | TU77_RS22280 | 179418..182882 | 286 | MULTISPECIES:-BREX-system-P-loop-protein-BrxC-[Pseudomonas] |
| WP_096426782.1 | Type1.BREX | ZnR-><-MA+PspA+PspA+sGTP<-sGTP<-sGTP\|\|?->\|\|XIS-HTH->BrxA-DNA-binding->BrxB-BrxF-iSTAND->DUF6079*->N6-METHYLASE->ParB+HNH->PglZ->SIGMA+OB-Fold+AAA-ATPase+HKD-DNase-BrxL-><-?<-?<-DUF932 | 1185 | Bacteria>Pseudomonadota>Gammaproteobacteria>Pseudomonadales>Pseudomonadaceae | GCF_002356095.1 | Pseudomonas | J3A71_RS03870 | 842777..846334 | 286 | MULTISPECIES:-BREX-system-P-loop-protein-BrxC-[Pseudomonas] |
| WP_105523318.1 | Type1.BREX | <-SIGMA+OB-Fold+AAA-ATPase+HKD-DNase-BrxL<-PglZ<-N6-METHYLASE-p<-?<-REase<-OLD-ABC<-N6-METHYLASE<-DUF6079*<-BrxB-BrxF-iSTAND<-BrxA-DNA-binding<-XIS-HTH<-ARG<-?\|\|wHTH+WYL+WCX-><-sGTP<-sGTP | 1185 | Bacteria>Pseudomonadota>Gammaproteobacteria>Pseudomonadales>Pseudomonadaceae | GCF_002975295.1 | Pseudomonas | CQ048_RS15780 | 97047..100604 | 286 | MULTISPECIES:-BREX-system-P-loop-protein-BrxC-[Pseudomonas] |
| WP_106120190.1 | Type1.BREX | LZ_Tnp_IS66+zf-IS66+DDE_Tnp_IS66+DDE_Tnp_IS66_C-><-?<-GAF+PAS+GGDEF+EAL\|\|<-SIGMA+OB-Fold+AAA-ATPase+Lon-Protease-BrxL<-PglZ<-TPR<-NERD+GspA-ATPase<-N6-METHYLASE<-DUF6079*<-BrxB-BrxF-iSTAND<-BrxA-DNA-binding<-wHTH+WYL+WCX\|\|SF1-SF2-Helicase->?->SF1-SF2-Helicase+REase->?->MORC+REase-> | 1223 | Bacteria>Pseudomonadota>Gammaproteobacteria>Pseudomonadales>Pseudomonadaceae | GCF_003001535.1 | Pseudomonas | FHL09_RS12740 | 68451..72122 | 286 | MULTISPECIES:-BREX-system-P-loop-protein-BrxC-[Pseudomonas] |
| WP_170037970.1 | Type1.BREX | GAF+PAS+GGDEF+EAL->?-><-LZ_Tnp_IS66+zf-IS66+DDE_Tnp_IS66+DDE_Tnp_IS66_C<-?\|\|?-><-SIGMA+OB-Fold+AAA-ATPase+HKD-DNase-BrxL<-PglZ<-?<-N6-METHYLASE<-DUF6079*<-BrxB-BrxF-iSTAND<-BrxA-DNA-binding<-XIS-HTH<-ARG<-?\|\|wHTH+WYL+WCX-><-sGTP<-sGTP | 1185 | Bacteria>Pseudomonadota>Gammaproteobacteria>Pseudomonadales>Pseudomonadaceae | GCF_012985445.1 | Pseudomonas | HBO05_RS12455 | 196790..200347 | 286 | MULTISPECIES:-BREX-system-P-loop-protein-BrxC-[Pseudomonas] |
| WP_176516884.1 | Type1.BREX | <-SIGMA+OB-Fold+AAA-ATPase+Lon-Protease-BrxL<-PglZ<-REase<-N6-METHYLASE-p<-?<-HerA/FtsK<-SIR2<-N6-METHYLASE<-DUF6079*<-BrxB-BrxF-iSTAND<-BrxA-DNA-binding<-wHTH+WYL+WCX\|\|?->RVT-> | 1223 | Bacteria>Pseudomonadota>Gammaproteobacteria>Pseudomonadales>Pseudomonadaceae | GCF_013373895.1 | Pseudomonas | HXS67_RS28935 | 22447..26118 | 286 | MULTISPECIES:-BREX-system-P-loop-protein-BrxC-[Pseudomonas] |
| WP_196184087.1 | Type1.BREX | DUF932->?->?->?->DUF4969-><-SIGMA+OB-Fold+AAA-ATPase+Lon-Protease-BrxL<-PglZ<-ParB+HNH<-N6-METHYLASE<-DUF6079*<-BrxB-BrxF-iSTAND<-BrxA-DNA-binding<-wHTH+WYL+WCX<-?<-XIS-HTH\|\|RHH-><-tRNA<-MerR-HTH | 1154 | Bacteria>Pseudomonadota>Gammaproteobacteria>Pseudomonadales>Pseudomonadaceae | GCF_015680805.1 | Pseudomonas | IR013_RS15060 | 51443..54907 | 286 | MULTISPECIES:-BREX-system-P-loop-protein-BrxC-[Pseudomonas] |
| WP_198830848.1 | Type1.BREX | HNH->?->JetC-><-SIGMA+OB-Fold+AAA-ATPase+Lon-Protease-BrxL<-PglZ<-?<-?<-?<-N6-METHYLASE<-DUF6079*<-BrxB-BrxF-iSTAND<-BrxA-DNA-binding<-wHTH+WYL+WCX\|\|?-><-wHTH+WYL+WCX\|\|?->ARG-> | 1223 | Bacteria>Pseudomonadota>Gammaproteobacteria>Pseudomonadales>Pseudomonadaceae | GCF_016887925.1 | Pseudomonas | JQF37_RS21260 | 352403..356074 | 286 | MULTISPECIES:-BREX-system-P-loop-protein-BrxC-[Pseudomonas] |
| WP_218430053.1 | Type1.BREX | Phage_integrase->?->?->TraH->?-><-?<-SIGMA+OB-Fold+AAA-ATPase+HKD-DNase-BrxL<-PglZ<-N6-METHYLASE<-DUF6079*<-BrxB-BrxF-iSTAND<-BrxA-DNA-binding<-XIS-HTH\|\|DUF772+DDE_Tnp_1+DDE_Tnp_1_6-><-?<-?\|\|?->?-><-ParB | 1189 | Bacteria>Pseudomonadota>Gammaproteobacteria>Pseudomonadales>Pseudomonadaceae | GCF_019219185.1 | Pseudomonas | KW849_RS14820 | 325397..328966 | 286 | MULTISPECIES:-BREX-system-P-loop-protein-BrxC-[Pseudomonas] |
| WP_064314265.1 | Type1.BREX | <-RHH\|\|XIS-HTH->?->wHTH+WYL+WCX->BrxA-DNA-binding->BrxB-BrxF-iSTAND->DUF6079*->N6-METHYLASE->ParB+HNH->OLD-ABC->HNH->PglZ->SIGMA+OB-Fold+AAA-ATPase+Lon-Protease-BrxL-><-DUF4969 | 1154 | Bacteria>Pseudomonadota>Gammaproteobacteria>Pseudomonadales>Pseudomonadaceae | GCF_021601465.1 | Pseudomonas | L6D68_RS06035 | 31411..34875 | 286 | MULTISPECIES:-BREX-system-P-loop-protein-BrxC-[Pseudomonas] |
| WP_021219096.1 | Type1.BREX | <-SIGMA+OB-Fold+AAA-ATPase+Lon-Protease-BrxL<-PglZ<-AbiEii<-AbiEi_4<-?<-?<-N6-METHYLASE<-DNA-glycosylase<-ParB+HNH<-DUF6079*<-BrxB-BrxF-iSTAND<-BrxA-DNA-binding<-?\|\|wHTH+WYL+WCX-> | 1220 | Bacteria>Pseudomonadota>Gammaproteobacteria>Pseudomonadales>Pseudomonadaceae | GCF_023093875.1 | Pseudomonas | MY055_RS15275 | 3320590..3324252 | 286 | MULTISPECIES:-BREX-system-P-loop-protein-BrxC-[Pseudomonas] |
| WP_060765715.1 | Type1.BREX | <-HKD-DNase+HKD-DNase<-?\|\|<-SIGMA+OB-Fold+AAA-ATPase+HKD-DNase-BrxL<-PglZ<-ParB+HNH<-N6-METHYLASE<-REase<-?<-DUF6079*<-BrxB-BrxF-iSTAND<-BrxA-DNA-binding<-XIS-HTH<-?<-?<-XIS-HTH\|\|RHH-><-tRNA<-MerR-HTH | 1186 | Bacteria>Pseudomonadota>Gammaproteobacteria>Pseudomonadales>Pseudomonadaceae | GCF_029959765.1 | Pseudomonas | PFL603g_RS03160 | 383988..387548 | 286 | MULTISPECIES:-BREX-system-P-loop-protein-BrxC-[Pseudomonas] |
| WP_034126881.1 | Type1.BREX | RHH_5->BrxA-DNA-binding->BrxB-BrxF-iSTAND->DUF6079*->N6-METHYLASE->SbcC->Schlafen+HATPase+wHTH->PglZ->SIGMA+OB-Fold+AAA-ATPase+Lon-Protease-BrxL-> | 1153 | Bacteria>Pseudomonadota>Gammaproteobacteria>Pseudomonadales>Pseudomonadaceae | GCF_030505435.1 | Pseudomonas | FHL14_RS24020 | 3439..6900 | 286 | MULTISPECIES:-BREX-system-P-loop-protein-BrxC-[Pseudomonas] |
| WP_023115268.1 | Type1.BREX | <-wHTH+WYL+WCX\|\|?->Peptidase_M15->?->BrxA-DNA-binding->BrxB-BrxF-iSTAND->DUF6079*->N6-METHYLASE->?->HNH+GCN1->?->ParB+HNH->PglZ->SIGMA+OB-Fold+AAA-ATPase+Lon-Protease-BrxL->REase-> | 1154 | Bacteria>Pseudomonadota>Gammaproteobacteria>Pseudomonadales>Pseudomonadaceae | GCF_900707185.1 | Pseudomonas | E4Y87_RS04340 | 6868..10332 | 286 | MULTISPECIES:-BREX-system-P-loop-protein-BrxC-[Pseudomonas] |
| WP_025297447.1 | Type1.BREX | <-DUF6079*<-BrxB-BrxF-iSTAND<-BrxA-DNA-binding<-XIS-HTH<-?<-Peptidase_M15 | 1184 | Bacteria>Pseudomonadota>Gammaproteobacteria>Pseudomonadales>Pseudomonadaceae | GCF_900707275.1 | Pseudomonas | E4Z27_RS28920 | 187..3741 | 286 | MULTISPECIES:-BREX-system-P-loop-protein-BrxC-[Pseudomonas] |
| WP_198417420.1 | Type1.BREX | <-NERD<-?<-Integrase_1<-?\|\|XIS-HTH->?->wHTH+WYL+WCX->BrxA-DNA-binding->BrxB-BrxF-iSTAND->DUF6079*->N6-METHYLASE->ParB+HNH->PglZ->SIGMA+OB-Fold+AAA-ATPase+Lon-Protease-BrxL-><-DUF4969<-?<-?<-HIN-HTH+rve+rve_3 | 1159 | Bacteria>Pseudomonadota>Gammaproteobacteria>Pseudomonadales>Pseudomonadaceae | GCF_902329585.1 | Pseudomonas | PPTH39_RS22285 | 114113..117592 | 286 | MULTISPECIES:-BREX-system-P-loop-protein-BrxC-[Pseudomonas] |
| WP_104739193.1 | Type1.BREX | <-wHTH+WYL+WCX<-DUF772+DDE_Tnp_1+DDE_Tnp_1_6\|\|?->DUF6361->HKD-DNase->SF1-SF2-Helicase->\|\|XIS-HTH->BrxA-DNA-binding->BrxB-BrxF-iSTAND->DUF6079*->N6-METHYLASE->SbcC-p->REase->RVT->N6-METHYLASE-p->ParB+HNH->PglZ->SIGMA+OB-Fold+AAA-ATPase+HKD-DNase-BrxL-><-NtrC-AAA | 1180 | Bacteria>Pseudomonadota>Gammaproteobacteria>Pseudomonadales>Pseudomonadaceae>Halopseudomonas | GCF_014641295.1 | Halopseudomonas_oceani | IEX06_RS08390 | 366595..370137 | 1708783 | BREX-system-P-loop-protein-BrxC-[Halopseudomonas-oceani] |
| WP_269686934.1 | Type1.BREX | SF1-SF2-Helicase->SF1-SF2-Helicase->?-><-?<-RADICAL-SAM<-?<-MazE_antitoxin\|\|wHTH+WYL+WCX->BrxA-DNA-binding->BrxB-BrxF-iSTAND->DUF6079*->N6-METHYLASE->?->?->OLD-ABC->HNH->PglZ->SIGMA+OB-Fold+AAA-ATPase+Lon-Protease-BrxL-><-HTH_3\|\|PadR-HTH-> | 1223 | Bacteria>Pseudomonadota>Gammaproteobacteria>Pseudomonadales>Pseudomonadaceae>Halopseudomonas | GCF_030295685.1 | Halopseudomonas_sp._MFKK-1 | OSO48_RS10320 | 2203460..2207131 | 3015994 | BREX-system-P-loop-protein-BrxC-[Halopseudomonas-sp.-MFKK-1] |
| AUB77630.1 | Type1.BREX | Arm-DNA-bind_3+Phage_integrase->?->Phage_AlpA->?->wHTH+WYL+WCX->BrxA-DNA-binding->BrxB-BrxF-iSTAND->DUF6079*->DOC+HTH->N6-METHYLASE->?->?->NERD+GspA-ATPase->TPR->PglZ->SIGMA+OB-Fold+AAA-ATPase+Lon-Protease-BrxL-> | 1223 | Bacteria>Pseudomonadota>Gammaproteobacteria>Pseudomonadales>Pseudomonadaceae>Pseudomonas | GCA_000346225.2 | Pseudomonas_sp._Lz4W | B195_019235 | 4255709..4259380 | 1206777 | ATPase-[Pseudomonas-sp.-Lz4W] |
| ERV97546.1 | Type1.BREX | LZ_Tnp_IS66+zf-IS66+DDE_Tnp_IS66+DDE_Tnp_IS66_C->?->DDE_Tnp_1-><-?<-?<-DUF6079*<-BrxB-BrxF-iSTAND<-BrxA-DNA-binding<-wHTH+WYL+WCX<-?\|\|OLD-ABC+TOPRIM->SF1-SF2-Helicase->SF1-SF2-Helicase->?-><-Phage_integrase | 1223 | Bacteria>Pseudomonadota>Gammaproteobacteria>Pseudomonadales>Pseudomonadaceae>Pseudomonas | GCA_000481225.1 | Pseudomonas_aeruginosa_BWHPSA024 | Q037_05156 | 13246..16917 | 1402524 | hypothetical-protein-Q037_05156-[Pseudomonas-aeruginosa-BWHPSA024] |
| KPX73888.1 | Type1.BREX | PIN+OST-HTH->HNH->?->?->DUF6079-p*->PglZ->SIGMA+OB-Fold+AAA-ATPase+Lon-Protease-BrxL->HIN-HTH->HTH_Tnp_1+HTH_21+rve+rve_3-> | 889 | Bacteria>Pseudomonadota>Gammaproteobacteria>Pseudomonadales>Pseudomonadaceae>Pseudomonas | GCA_001400855.1 | Pseudomonas_syringae_pv._maculicola | ALO84_01403 | 56181..58850 | 59511 | Uncharacterized-protein-ALO84_01403-[Pseudomonas-syringae-pv.-maculicola] |
| KSG52894.1 | Type1.BREX | <-ABC-ATPase+SbcC<-?<-TIR<-?<-N6-METHYLASE-p<-RVT<-N6-METHYLASE<-DOC+HTH<-DUF6079*<-BrxB-BrxF-iSTAND<-BrxA-DNA-binding<-wHTH+WYL+WCX<-?<-Phage_AlpA<-?\|\|PAIREDC-HTH-> | 1223 | Bacteria>Pseudomonadota>Gammaproteobacteria>Pseudomonadales>Pseudomonadaceae>Pseudomonas | GCA_001450115.1 | Pseudomonas_aeruginosa | A0953_18465 | 48360..52031 | 287 | ATPase-[Pseudomonas-aeruginosa] |
| ARQ76275.1 | Type1.BREX | DUF4969-><-SIGMA+OB-Fold+AAA-ATPase+Lon-Protease-BrxL<-PglZ<-TOPRIM<-OLD-ABC<-ParB+HNH<-N6-METHYLASE<-DUF6079*<-BrxB-BrxF-iSTAND<-BrxA-DNA-binding<-?<-XIS-HTH\|\|?-><-?<-?<-?\|\|Aminotran_1_2-> | 1156 | Bacteria>Pseudomonadota>Gammaproteobacteria>Pseudomonadales>Pseudomonadaceae>Pseudomonas | GCA_002128325.1 | Pseudomonas_fragi | B6D87_19590 | 4227178..4230648 | 296 | hypothetical-protein-B6D87_19590-[Pseudomonas-fragi] |
| PAA41060.1 | Type1.BREX | HIN-HTH->AAA-ATPase-><-?\|\|XIS-HTH->?->BrxA-DNA-binding->BrxB-BrxF-iSTAND->DUF6079*->N6-METHYLASE->ParB+HNH->PglZ->SIGMA+OB-Fold+AAA-ATPase+Lon-Protease-BrxL-><-DUF4969<-?<-?<-HIN-HTH+rve+rve_3 | 1156 | Bacteria>Pseudomonadota>Gammaproteobacteria>Pseudomonadales>Pseudomonadaceae>Pseudomonas | GCA_002269445.1 | Pseudomonas_fragi | CJU80_12345 | 107560..111030 | 296 | hypothetical-protein-CJU80_12345-[Pseudomonas-fragi] |
| ATN11139.1 | Type1.BREX | <-TraI-VirD2-RCRE<-ParD_antitoxin\|\|HNH->?->\|\|BrxA-DNA-binding->BrxB-BrxF-iSTAND->DUF6079*->DNA-glycosylase->N6-METHYLASE->ParB+HNH->AbiEi_3_N+AbiEi_3->AbiEii->PglZ->SIGMA+OB-Fold+AAA-ATPase+Lon-Protease-BrxL-> | 1222 | Bacteria>Pseudomonadota>Gammaproteobacteria>Pseudomonadales>Pseudomonadaceae>Pseudomonas | GCA_002591235.1 | Pseudomonas_sp._FDAARGOS_380 | CRN80_16460 | 3507093..3510761 | 2018067 | hypothetical-protein-CRN80_16460-[Pseudomonas-sp.-FDAARGOS_380] |
| MBB59705.1 | Type1.BREX | <-DUF3584+DUF5401<-MukF<-SIGMA+OB-Fold+AAA-ATPase+Lon-Protease-BrxL<-PglZ<-TOPRIM<-OLD-ABC<-ParB+HNH<-DUF932<-N6-METHYLASE<-DUF6079*<-BrxB-BrxF-iSTAND<-BrxA-DNA-binding<-wHTH+WYL+WCX<-?<-ARG\|\|wHTH+WYL+WCX->LexA-protease-> | 1154 | Bacteria>Pseudomonadota>Gammaproteobacteria>Pseudomonadales>Pseudomonadaceae>Pseudomonas | GCA_002709345.1 | Pseudomonas_sp. | CML00_00850 | 39770..43234 | 306 | hypothetical-protein-[Pseudomonas-sp.] |
| PPB00131.1 | Type1.BREX | <-TraE<-TraM<-TraM<-DUF3262<-?\|\|HipA_C->XIS-HTH->BrxA-DNA-binding->BrxB-BrxF-iSTAND->DUF6079*->N6-METHYLASE->OLD-ABC->REase->SIR2->HerA/FtsK->N6-METHYLASE->PglZ->SIGMA+OB-Fold+AAA-ATPase+HKD-DNase-BrxL->SbcC+SbcC-> | 1194 | Bacteria>Pseudomonadota>Gammaproteobacteria>Pseudomonadales>Pseudomonadaceae>Pseudomonas | GCA_002927135.1 | Pseudomonas_aeruginosa | HV84_25295 | 2273516..2277100 | 287 | BREX-system-P-loop-protein-BrxC-[Pseudomonas-aeruginosa] |
| PPV34665.1 | Type1.BREX | <-RVT<-N6-METHYLASE<-DOC+HTH<-DUF6079*<-BrxB-BrxF-iSTAND<-BrxA-DNA-binding<-wHTH+WYL+WCX | 1223 | Bacteria>Pseudomonadota>Gammaproteobacteria>Pseudomonadales>Pseudomonadaceae>Pseudomonas | GCA_002941105.1 | Pseudomonas_oleovorans | C5L43_18395 | 7326..10997 | 301 | BREX-system-P-loop-protein-BrxC-[Pseudomonas-oleovorans] |
| HBC00740.1 | Type1.BREX | TOPRIM+Primase->?-><-?<-SIGMA+OB-Fold+AAA-ATPase+Lon-Protease-BrxL<-PglZ<-Schlafen+HATPase+wHTH<-SbcC<-N6-METHYLASE<-?<-DUF6079*<-BrxB-BrxF-iSTAND<-BrxA-DNA-binding<-RHH_5<-?<-?\|\|DEDD_Tnp_IS110+Transposase_20-><-Arm-DNA-bind_3+Phage_integrase<-tRNA<-N6-METHYLASE | 1153 | Bacteria>Pseudomonadota>Gammaproteobacteria>Pseudomonadales>Pseudomonadaceae>Pseudomonas | GCA_003488145.1 | Pseudomonas_sp. | DC032_09195 | 21916..25377 | 306 | BREX-system-P-loop-protein-BrxC-[Pseudomonas-sp.] |
| HBM65158.1 | Type1.BREX | <-ANK+ANK+ANK\|\|RADICAL-SAM->HicA_toxin->BrxA-DNA-binding->BrxB-BrxF-iSTAND->DUF6079*->N6-METHYLASE-> | 1154 | Bacteria>Pseudomonadota>Gammaproteobacteria>Pseudomonadales>Pseudomonadaceae>Pseudomonas | GCA_003504465.1 | Pseudomonas_sp. | DD418_16300 | 5931..9395 | 306 | BREX-system-P-loop-protein-BrxC-[Pseudomonas-sp.] |
| RPD95236.1 | Type1.BREX | ZnR-><-wHTH+WYL+WCX\|\|XIS-HTH->BrxA-DNA-binding->BrxB-BrxF-iSTAND->DUF6079*->N6-METHYLASE-p->PglZ->SIGMA+OB-Fold+AAA-ATPase+HKD-DNase-BrxL-><-?<-Arm-DNA-bind_3+Phage_integrase<-?\|\|GNTR-HTH+LRP-HTH-> | 1191 | Bacteria>Pseudomonadota>Gammaproteobacteria>Pseudomonadales>Pseudomonadaceae>Pseudomonas | GCA_003813925.1 | Pseudomonas_monteilii | EGN69_04400 | 958521..962096 | 76759 | BREX-system-P-loop-protein-BrxC-[Pseudomonas-monteilii] |
| RPM42463.1 | Type1.BREX | <-ABhydrolase<-TetR-HTH\|\|?->Arm-DNA-bind_3+Phage_integrase->Phage_AlpA->?->wHTH+WYL+WCX->BrxA-DNA-binding->BrxB-BrxF-iSTAND->DUF6079*->DOC+HTH->N6-METHYLASE->RVT->N6-METHYLASE-p->?-><-Y1_Tnp\|\|TIR->?->ABC-ATPase+SbcC-> | 1223 | Bacteria>Pseudomonadota>Gammaproteobacteria>Pseudomonadales>Pseudomonadaceae>Pseudomonas | GCA_003833645.1 | Pseudomonas_aeruginosa | IPC1294_03635 | 745407..749078 | 287 | ATPase-[Pseudomonas-aeruginosa] |
| TKK14312.1 | Type1.BREX | <-TraI-VirD2-RCRE<-ParD_antitoxin\|\|HNH->?->\|\|BrxA-DNA-binding->BrxB-BrxF-iSTAND->DUF6079*->DNA-glycosylase->N6-METHYLASE->ParB+HNH->AbiEi_3_N+AbiEi_3->AbiEii->PglZ->SIGMA+OB-Fold+AAA-ATPase+Lon-Protease-BrxL-> | 1222 | Bacteria>Pseudomonadota>Gammaproteobacteria>Pseudomonadales>Pseudomonadaceae>Pseudomonas | GCA_005233695.1 | Pseudomonas_fluorescens | PflCFBP13510_04710 | 495926..499594 | 294 | BREX-system-P-loop-protein-BrxC-[Pseudomonas-fluorescens] |
| TXH92079.1 | Type1.BREX | <-RelE-ParE<-RHH_1<-PglZ<-HNH<-OLD-ABC<-N6-METHYLASE-p<-NERD<-OLD-ABC<-N6-METHYLASE<-DUF6079*<-BrxB-BrxF-iSTAND<-BrxA-DNA-binding<-XIS-HTH<-?\|\|wHTH+WYL+WCX-> | 1186 | Bacteria>Pseudomonadota>Gammaproteobacteria>Pseudomonadales>Pseudomonadaceae>Pseudomonas | GCA_008015405.1 | Pseudomonas_sp. | E6Q71_01680 | 13269..16829 | 306 | BREX-system-P-loop-protein-BrxC-[Pseudomonas-sp.] |
| MRJ36970.1 | Type1.BREX | <-XIS-HTH\|\|?->DUF932->?->?->?->wHTH+WYL+WCX->BrxA-DNA-binding->BrxB-BrxF-iSTAND->DUF6079*->N6-METHYLASE->NERD+GspA-ATPase->TPR->PglZ->SIGMA+OB-Fold+AAA-ATPase+Lon-Protease-BrxL-> | 1223 | Bacteria>Pseudomonadota>Gammaproteobacteria>Pseudomonadales>Pseudomonadaceae>Pseudomonas | GCA_009659625.1 | Pseudomonas_haemolytica | FRT59_08305 | 447566..451237 | 2600065 | BREX-system-P-loop-protein-BrxC-[Pseudomonas-haemolytica] |
| MVW77192.1 | Type1.BREX | <-PglZ<-AbiEii<-AbiEi_3_N+AbiEi_3<-?<-?<-ParB+HNH<-N6-METHYLASE<-DNA-glycosylase<-ParB+HNH<-DUF6079*<-BrxB-BrxF-iSTAND<-BrxA-DNA-binding<-?\|\|wHTH+WYL+WCX->?-><-?\|\|Arm-DNA-bind_2+Phage_int_SAM_3+Phage_integrase-><-REase\|\|ZnR-> | 1221 | Bacteria>Pseudomonadota>Gammaproteobacteria>Pseudomonadales>Pseudomonadaceae>Pseudomonas | GCA_009763245.1 | Pseudomonas_xionganensis | GJV18_17880 | 24794..28459 | 2654845 | BREX-system-P-loop-protein-BrxC-[Pseudomonas-xionganensis] |
| NLU01326.1 | Type1.BREX | <-XIS-HTH\|\|?->DUF932->?->?->?->wHTH+WYL+WCX->BrxA-DNA-binding->BrxB-BrxF-iSTAND->DUF6079*->N6-METHYLASE->ParB+HNH->PglZ->SIGMA+OB-Fold+AAA-ATPase+Lon-Protease-BrxL->?-><-tRNA<-MerR-HTH | 1222 | Bacteria>Pseudomonadota>Gammaproteobacteria>Pseudomonadales>Pseudomonadaceae>Pseudomonas | GCA_012720005.1 | Pseudomonas_lundensis | GXW88_11975 | 7910..11578 | 86185 | BREX-system-P-loop-protein-BrxC-[Pseudomonas-lundensis] |
| NMY25069.1 | Type1.BREX | PIN+OST-HTH->HNH->?->?-><-wHTH+WYL+WCX\|\|?->ARG->BrxA-DNA-binding->BrxB-BrxF-iSTAND->DUF6079*->DNA-glycosylase->N6-METHYLASE->ParB+HNH->AbiEi_3_N+AbiEi_3->AbiEii->PglZ->SIGMA+OB-Fold+AAA-ATPase+Lon-Protease-BrxL->JAB-><-MerR-HTH | 1222 | Bacteria>Pseudomonadota>Gammaproteobacteria>Pseudomonadales>Pseudomonadaceae>Pseudomonas | GCA_012985535.1 | Pseudomonas_sp._WS_5021 | HBO19_03685 | 794090..797758 | 2717490 | BREX-system-P-loop-protein-BrxC-[Pseudomonas-sp.-WS-5021] |
| NMY20845.1 | Type1.BREX | <-SIGMA+OB-Fold+AAA-ATPase+Lon-Protease-BrxL<-PglZ<-TPR<-NERD+GspA-ATPase<-N6-METHYLASE<-DUF6079*<-BrxB-BrxF-iSTAND<-BrxA-DNA-binding<-wHTH+WYL+WCX\|\|?->HKD-DNase+HKD-DNase->SF1-SF2-Helicase->?->SF1-SF2-Helicase+REase-> | 1223 | Bacteria>Pseudomonadota>Gammaproteobacteria>Pseudomonadales>Pseudomonadaceae>Pseudomonas | GCA_012985645.1 | Pseudomonas_sp._WS_5410 | HBO24_11430 | 134586..138257 | 2717485 | BREX-system-P-loop-protein-BrxC-[Pseudomonas-sp.-WS-5410] |
| NMY53287.1 | Type1.BREX | <-N6-METHYLASE<-?<-DUF6079*<-BrxB-BrxF-iSTAND<-BrxA-DNA-binding<-?\|\|wHTH+WYL+WCX-><-SF1-SF2-Helicase\|\|?->?-><-Phage_integrase | 1222 | Bacteria>Pseudomonadota>Gammaproteobacteria>Pseudomonadales>Pseudomonadaceae>Pseudomonas | GCA_012985775.1 | Pseudomonas_sp._WS_5011 | HBN63_20215 | 4647..8315 | 2717477 | BREX-system-P-loop-protein-BrxC-[Pseudomonas-sp.-WS-5011] |
| NWF08290.1 | Type1.BREX | <-SIGMA+OB-Fold+AAA-ATPase+Lon-Protease-BrxL<-PglZ<-AbiEii<-AbiEi_3_N+AbiEi_3<-ParB+HNH<-N6-METHYLASE<-DNA-glycosylase<-DUF6079*<-BrxB-BrxF-iSTAND<-BrxA-DNA-binding<-?\|\|wHTH+WYL+WCX-><-tRNA<-MerR-HTH | 1222 | Bacteria>Pseudomonadota>Gammaproteobacteria>Pseudomonadales>Pseudomonadaceae>Pseudomonas | GCA_013386885.1 | Pseudomonas_salomonii | HX810_11520 | 509017..512685 | 191391 | BREX-system-P-loop-protein-BrxC-[Pseudomonas-salomonii] |
| MBD9399119.1 | Type1.BREX | <-wHTH+WYL+WCX\|\|?->JetB->?->RNase->?->BrxA-DNA-binding->BrxB-BrxF-iSTAND->DUF6079*->?->N6-METHYLASE->?->AbiEi_4->AbiEii->PglZ->SIGMA+OB-Fold+AAA-ATPase+Lon-Protease-BrxL-> | 1222 | Bacteria>Pseudomonadota>Gammaproteobacteria>Pseudomonadales>Pseudomonadaceae>Pseudomonas | GCA_014851505.1 | Pseudomonas_sp._PDM11 | IB224_16075 | 119844..123512 | 2769309 | BREX-system-P-loop-protein-BrxC-[Pseudomonas-sp.-PDM11] |
| MBF9242733.1 | Type1.BREX | SSB->Phage_integrase->Phage_integrase->?-><-HEPN\|\|BrxA-DNA-binding->BrxB-BrxF-iSTAND->DUF6079*->ParB+HNH->DNA-glycosylase->N6-METHYLASE->PglZ-> | 1222 | Bacteria>Pseudomonadota>Gammaproteobacteria>Pseudomonadales>Pseudomonadaceae>Pseudomonas | GCA_015694365.1 | Pseudomonas_syringae_pv._tomato | I4I80_00295 | 46308..49976 | 323 | BREX-system-P-loop-protein-BrxC-[Pseudomonas-syringae-pv.-tomato] |
| MBG4373384.1 | Type1.BREX | <-HIN-HTH<-REase<-wHTH+WYL+WCX\|\|?->Peptidase_M15->?->Ntox49->BrxA-DNA-binding->BrxB-BrxF-iSTAND->DUF6079*->N6-METHYLASE->?->HNH+GCN1->?->ParB+HNH->PglZ->SIGMA+OB-Fold+AAA-ATPase+Lon-Protease-BrxL-><-GreA_GreB+SF1-SF2-Helicase<-Arm-DNA-bind_3+Phage_integrase | 1154 | Bacteria>Pseudomonadota>Gammaproteobacteria>Pseudomonadales>Pseudomonadaceae>Pseudomonas | GCA_015734445.1 | Pseudomonas_aeruginosa | ISF50_23085 | 8082..11546 | 287 | BREX-system-P-loop-protein-BrxC-[Pseudomonas-aeruginosa] |
| MBG4395801.1 | Type1.BREX | <-wHTH+WYL+WCX\|\|?->Peptidase_M15->?->Ntox49->BrxA-DNA-binding->BrxB-BrxF-iSTAND->DUF6079*->N6-METHYLASE->?->HNH+GCN1->?->ParB+HNH->PglZ->SIGMA+OB-Fold+AAA-ATPase+Lon-Protease-BrxL-><-GreA_GreB+SF1-SF2-Helicase<-Arm-DNA-bind_3+Phage_integrase | 1154 | Bacteria>Pseudomonadota>Gammaproteobacteria>Pseudomonadales>Pseudomonadaceae>Pseudomonas | GCA_015734545.1 | Pseudomonas_aeruginosa | GHT73_12640 | 7113..10577 | 287 | BREX-system-P-loop-protein-BrxC-[Pseudomonas-aeruginosa] |
| MBG4463093.1 | Type1.BREX | <-HIN-HTH<-REase<-wHTH+WYL+WCX\|\|?->Peptidase_M15->?->Ntox49->BrxA-DNA-binding->BrxB-BrxF-iSTAND->DUF6079*->N6-METHYLASE->?->HNH+GCN1->?->ParB+HNH->PglZ->SIGMA+OB-Fold+AAA-ATPase+Lon-Protease-BrxL-><-GreA_GreB+SF1-SF2-Helicase<-Arm-DNA-bind_3+Phage_integrase | 1154 | Bacteria>Pseudomonadota>Gammaproteobacteria>Pseudomonadales>Pseudomonadaceae>Pseudomonas | GCA_015734755.1 | Pseudomonas_aeruginosa | GHU50_17045 | 10342..13806 | 287 | BREX-system-P-loop-protein-BrxC-[Pseudomonas-aeruginosa] |
| MBG4530231.1 | Type1.BREX | MerR-HTH->tRNA->XIS-HTH->BrxA-DNA-binding->BrxB-BrxF-iSTAND->DUF6079*->N6-METHYLASE->OLD-ABC->REase->SIR2->HerA/FtsK->N6-METHYLASE->PglZ->SIGMA+OB-Fold+AAA-ATPase+HKD-DNase-BrxL->XIS-HTH-> | 1181 | Bacteria>Pseudomonadota>Gammaproteobacteria>Pseudomonadales>Pseudomonadaceae>Pseudomonas | GCA_015734885.1 | Pseudomonas_aeruginosa | I4952_20115 | 27108..30653 | 287 | BREX-system-P-loop-protein-BrxC-[Pseudomonas-aeruginosa] |
| MBG4863325.1 | Type1.BREX | <-HIN-HTH\|\|XIS-HTH->BrxA-DNA-binding->BrxB-BrxF-iSTAND->DUF6079*->N6-METHYLASE->OLD-ABC->NERD->N6-METHYLASE-p->PglZ->SIGMA+OB-Fold+AAA-ATPase+HKD-DNase-BrxL->HIN-HTH-> | 1184 | Bacteria>Pseudomonadota>Gammaproteobacteria>Pseudomonadales>Pseudomonadaceae>Pseudomonas | GCA_015735985.1 | Pseudomonas_aeruginosa | GHU23_28970 | 1726..5280 | 287 | BREX-system-P-loop-protein-BrxC-[Pseudomonas-aeruginosa] |
| MBG5216128.1 | Type1.BREX | <-wHTH+WYL+WCX\|\|?->Peptidase_M15->?->BrxA-DNA-binding->BrxB-BrxF-iSTAND->DUF6079*->N6-METHYLASE->?->HNH+GCN1->?->ParB+HNH->PglZ->SIGMA+OB-Fold+AAA-ATPase+Lon-Protease-BrxL->REase-> | 1154 | Bacteria>Pseudomonadota>Gammaproteobacteria>Pseudomonadales>Pseudomonadaceae>Pseudomonas | GCA_015737185.1 | Pseudomonas_aeruginosa | GHT75_27125 | 7080..10544 | 287 | BREX-system-P-loop-protein-BrxC-[Pseudomonas-aeruginosa] |
| MBG6730505.1 | Type1.BREX | <-LexA-protease<-wHTH+WYL+WCX\|\|?->XIS-HTH->BrxA-DNA-binding->BrxB-BrxF-iSTAND->DUF6079*->N6-METHYLASE->SbcC-p->REase->N6-METHYLASE-p->OLD-ABC->TOPRIM->PglZ->SIGMA+OB-Fold+AAA-ATPase+HKD-DNase-BrxL-><-DUF4969 | 1184 | Bacteria>Pseudomonadota>Gammaproteobacteria>Pseudomonadales>Pseudomonadaceae>Pseudomonas | GCA_015764285.1 | Pseudomonas_aeruginosa | ISI92_11320 | 4371..7925 | 287 | BREX-system-P-loop-protein-BrxC-[Pseudomonas-aeruginosa] |
| MBH9015057.1 | Type1.BREX | <-TetR-HTH\|\|?->Arm-DNA-bind_3+Phage_integrase->?->\|\|Phage_AlpA->?->wHTH+WYL+WCX->BrxA-DNA-binding->BrxB-BrxF-iSTAND->DUF6079*->?->?->DOC+HTH->N6-METHYLASE->?->\|\|MODE-HTH+PBPII-> | 1223 | Bacteria>Pseudomonadota>Gammaproteobacteria>Pseudomonadales>Pseudomonadaceae>Pseudomonas | GCA_016090735.1 | Pseudomonas_aeruginosa | ISK34_26080 | 19802..23473 | 287 | BREX-system-P-loop-protein-BrxC-[Pseudomonas-aeruginosa] |
| MBI7168904.1 | Type1.BREX | DDE_Tnp_1-><-?<-?<-N6-METHYLASE<-DOC+HTH<-?<-DUF6079*<-BrxB-BrxF-iSTAND<-BrxA-DNA-binding<-wHTH+WYL+WCX | 1223 | Bacteria>Pseudomonadota>Gammaproteobacteria>Pseudomonadales>Pseudomonadaceae>Pseudomonas | GCA_016338125.1 | Pseudomonas_aeruginosa | JEU09_31365 | 9152..12823 | 287 | BREX-system-P-loop-protein-BrxC-[Pseudomonas-aeruginosa] |
| MBI8737743.1 | Type1.BREX | RelE-ParE-><-?<-wHTH+WYL+WCX\|\|?->Peptidase_M15->?->Ntox49->BrxA-DNA-binding->BrxB-BrxF-iSTAND->DUF6079*->N6-METHYLASE->?->HNH+GCN1->?->ParB+HNH->PglZ->SIGMA+OB-Fold+AAA-ATPase+Lon-Protease-BrxL-> | 1154 | Bacteria>Pseudomonadota>Gammaproteobacteria>Pseudomonadales>Pseudomonadaceae>Pseudomonas | GCA_016342945.1 | Pseudomonas_aeruginosa | JEZ70_02820 | 55146..58610 | 287 | BREX-system-P-loop-protein-BrxC-[Pseudomonas-aeruginosa] |
| MBM1201834.1 | Type1.BREX | XIS-HTH->?->wHTH+WYL+WCX->BrxA-DNA-binding->BrxB-BrxF-iSTAND->DUF6079*->N6-METHYLASE->ParB+HNH->PglZ->SIGMA+OB-Fold+AAA-ATPase+Lon-Protease-BrxL-><-DUF4969<-?<-?<-DUF932 | 1154 | Bacteria>Pseudomonadota>Gammaproteobacteria>Pseudomonadales>Pseudomonadaceae>Pseudomonas | GCA_016812215.1 | Pseudomonas_fragi | GYM96_18820 | 91266..94730 | 296 | BREX-system-P-loop-protein-BrxC-[Pseudomonas-fragi] |
| MBM2633269.1 | Type1.BREX | <-ABC-ATPase+SbcC<-?<-MODE-HTH+PBPII\|\|<-?<-N6-METHYLASE<-DOC+HTH<-?<-?<-DUF6079*<-BrxB-BrxF-iSTAND<-BrxA-DNA-binding<-wHTH+WYL+WCX\|\|OLD-ABC+TOPRIM->SF1-SF2-Helicase->?->SF1-SF2-Helicase-><-Phage_integrase | 1223 | Bacteria>Pseudomonadota>Gammaproteobacteria>Pseudomonadales>Pseudomonadaceae>Pseudomonas | GCA_016836445.1 | Pseudomonas_aeruginosa | JQ480_07705 | 1618759..1622430 | 287 | BREX-system-P-loop-protein-BrxC-[Pseudomonas-aeruginosa] |
| BCR26098.1 | Type1.BREX | Phage_integrase-><-?<-SF1-SF2-Helicase<-?<-SbcC\|\|?->wHTH+WYL+WCX->BrxA-DNA-binding->BrxB-BrxF-iSTAND->DUF6079*->?->?->DOC+HTH->N6-METHYLASE->RVT->N6-METHYLASE-p-><-DUF772+DDE_Tnp_1\|\|N6-METHYLASE-p-> | 1223 | Bacteria>Pseudomonadota>Gammaproteobacteria>Pseudomonadales>Pseudomonadaceae>Pseudomonas | GCA_016861485.1 | Pseudomonas_alcaligenes | KAM426_36250 | 3797295..3800966 | 43263 | hypothetical-protein-KAM426_36250-[Pseudomonas-alcaligenes] |
| MBN0979708.1 | Type1.BREX | <-DNA-glycosylase<-DUF6079*<-BrxB-BrxF-iSTAND<-BrxA-DNA-binding\|\|<-?<-GTPase-AIG<-GTPase-AIG | 1222 | Bacteria>Pseudomonadota>Gammaproteobacteria>Pseudomonadales>Pseudomonadaceae>Pseudomonas | GCA_016924785.1 | Pseudomonas_sp._SDM007_2 | JV174_20995 | 2384..6052 | 2812898 | BREX-system-P-loop-protein-BrxC-[Pseudomonas-sp.-SDM007_2] |
| MBV5724753.1 | Type1.BREX | DUF4969-><-SIGMA+OB-Fold+AAA-ATPase+HKD-DNase-BrxL<-PglZ<-ParB+HNH<-N6-METHYLASE-p<-RVT<-NERD<-OLD-ABC<-N6-METHYLASE<-DUF6079*<-BrxB-BrxF-iSTAND<-BrxA-DNA-binding<-XIS-HTH<-?\|\|wHTH+WYL+WCX-><-DUF4158+DDE_Tnp_Tn3<-GDResolvase+HIN-HTH<-GDResolvase+HIN-HTH<-Tox-RelE | 1184 | Bacteria>Pseudomonadota>Gammaproteobacteria>Pseudomonadales>Pseudomonadaceae>Pseudomonas | GCA_019166825.1 | Pseudomonas_aeruginosa | KUT37_26485 | 44544..48098 | 287 | BREX-system-P-loop-protein-BrxC-[Pseudomonas-aeruginosa] |
| GIZ69010.1 | Type1.BREX | <-DUF6079*<-BrxB-BrxF-iSTAND<-BrxA-DNA-binding<-wHTH+WYL+WCX<-?\|\|SbcC->?->SF1-SF2-Helicase->?-><-Phage_integrase | 1226 | Bacteria>Pseudomonadota>Gammaproteobacteria>Pseudomonadales>Pseudomonadaceae>Pseudomonas | GCA_019656515.1 | Pseudomonas_alcaligenes | KAM428_40950 | 2224..5904 | 43263 | hypothetical-protein-KAM428_40950-[Pseudomonas-alcaligenes] |

| Protein-ID | System | Gene-Neighbourhoods | Length | Taxonomy | Assembly-ID | Strain | Locus-Tag | Genomic-coordinates | TaxID | Fasta-defline |
|---|---|---|---|---|---|---|---|---|---|---|
| MBX6150345.1 | Type1.BREX | <-TetR-HTH\|\|?-><-SIGMA+OB-Fold+AAA-ATPase+Lon-Protease-BrxL<-PglZ<-ParB+HNH<-?<-HNH+GCN1<-?<-N6-METHYLASE<-DUF6079*<-BrxB-BrxF-iSTAND<-BrxA-DNA-binding<-?<- [illegible] | 1154 | Bacteria>Pseudomonadota>Gammaproteobacteria>Pseudomonadales>Pseudomonadaceae>Pseudomonas | GCA_019686265.1 | Pseudomonas_aeruginosa | ISD36_12550 | 248185..251649 | 287 | BREX-system-P-loop-protein-BrxC-[Pseudomonas-aeruginosa] |
| MBX5763959.1 | Type1.BREX | Phage_integrase-><-?<-SIGMA+OB-Fold+AAA-ATPase+Lon-Protease-BrxL<-PglZ<-HNH<-OLD-ABC<-?<-?<-N6-METHYLASE<-DUF6079*<-BrxB-BrxF-iSTAND<-BrxA-DNA-binding<- [illegible] | 1223 | Bacteria>Pseudomonadota>Gammaproteobacteria>Pseudomonadales>Pseudomonadaceae>Pseudomonas | GCA_019687095.1 | Pseudomonas_aeruginosa | ISD89_03975 | 862812..866483 | 287 | BREX-system-P-loop-protein-BrxC-[Pseudomonas-aeruginosa] |
| MBX5915342.1 | Type1.BREX | <-wHTH+WYL+WCX\|\|?->Peptidase_M15->?->BrxA-DNA-binding->BrxB-BrxF-iSTAND->DUF6079*->N6-METHYLASE->?->HNH+GCN1->?->ParB+HNH->PglZ->SIGMA+OB-Fold+AAA-ATPase+ [illegible] | 1154 | Bacteria>Pseudomonadota>Gammaproteobacteria>Pseudomonadales>Pseudomonadaceae>Pseudomonas | GCA_019688355.1 | Pseudomonas_aeruginosa | ISE50_29115 | 6954..10418 | 287 | BREX-system-P-loop-protein-BrxC-[Pseudomonas-aeruginosa] |
| MBY8973665.1 | Type1.BREX | <-SIGMA+OB-Fold+AAA-ATPase+Lon-Protease-BrxL<-PglZ<-AbiEii<-AbiEi_3_N+AbiEi_3<-ParB+HNH<-N6-METHYLASE<-DNA-glycosylase<-DUF6079*<-BrxB-BrxF-iSTAND<- [illegible] | 1222 | Bacteria>Pseudomonadota>Gammaproteobacteria>Pseudomonadales>Pseudomonadaceae>Pseudomonas | GCA_019891455.1 | Pseudomonas_sp._P867 | J1G33_25110 | 13708..17376 | 2816050 | BREX-system-P-loop-protein-BrxC-[Pseudomonas-sp.-P867] |
| HBN8454461.1 | Type1.BREX | XIS-HTH-><-?<-tRNA\|\|HTH_36->Phage_int_SAM_2+Integrase_1-><-SIGMA+OB-Fold+AAA-ATPase+Lon-Protease-BrxL<-PglZ<-HNH<-OLD-ABC<-N6-METHYLASE<-DUF6079*<- [illegible] | 1146 | Bacteria>Pseudomonadota>Gammaproteobacteria>Pseudomonadales>Pseudomonadaceae>Pseudomonas | GCA_021587585.1 | Pseudomonas_aeruginosa | L3711_005718 | 39513..42953 | 287 | BREX-system-P-loop-protein-BrxC-[Pseudomonas-aeruginosa] |
| HBN9708921.1 | Type1.BREX | PhdYeFM_antitox->RelE-ParE-><-?<-wHTH+WYL+WCX\|\|?->Peptidase_M15->?->BrxA-DNA-binding->BrxB-BrxF-iSTAND->DUF6079*->N6-METHYLASE->MNS+ORC-CDC6->MarR-HTH->?-> [illegible] | 1154 | Bacteria>Pseudomonadota>Gammaproteobacteria>Pseudomonadales>Pseudomonadaceae>Pseudomonas | GCA_021591785.1 | Pseudomonas_aeruginosa | L3920_001787 | 275283..278747 | 287 | BREX-system-P-loop-protein-BrxC-[Pseudomonas-aeruginosa] |
| HBO1619179.1 | Type1.BREX | <-DUF3584+DUF5401<-JetB<-MukF<-SIGMA+OB-Fold+AAA-ATPase+Lon-Protease-BrxL<-PglZ<-TOPRIM<-OLD-ABC<-ParB+HNH<-N6-METHYLASE<-DUF6079*<-BrxB-BrxF-iSTAND<- [illegible] | 1154 | Bacteria>Pseudomonadota>Gammaproteobacteria>Pseudomonadales>Pseudomonadaceae>Pseudomonas | GCA_021598085.1 | Pseudomonas_aeruginosa | L4F62_005712 | 16995..20459 | 287 | BREX-system-P-loop-protein-BrxC-[Pseudomonas-aeruginosa] |
| EIU3464059.1 | Type1.BREX | <-DUF3584+DUF5401<-JetB<-MukF<-SIGMA+OB-Fold+AAA-ATPase+Lon-Protease-BrxL<-PglZ<-HNH<-OLD-ABC<-ParB+HNH<-N6-METHYLASE<-DUF6079*<-BrxB-BrxF-iSTAND<- [illegible] | 1154 | Bacteria>Pseudomonadota>Gammaproteobacteria>Pseudomonadales>Pseudomonadaceae>Pseudomonas | GCA_021644165.1 | Pseudomonas_aeruginosa | L4P80_001287 | 38375..41839 | 287 | BREX-system-P-loop-protein-BrxC-[Pseudomonas-aeruginosa] |
| HBO3135817.1 | Type1.BREX | <-OLD-ABC<-N6-METHYLASE-p<-RVT<-NERD<-OLD-ABC<-SbcC-p<-N6-METHYLASE<-REase<-ABC-ATPase<-DUF6079*<-BrxB-BrxF-iSTAND<-BrxA-DNA-binding<-XIS-HTH | 1184 | Bacteria>Pseudomonadota>Gammaproteobacteria>Pseudomonadales>Pseudomonadaceae>Pseudomonas | GCA_021651475.1 | Pseudomonas_aeruginosa | L4Q96_001363 | 40663..44217 | 287 | BREX-system-P-loop-protein-BrxC-[Pseudomonas-aeruginosa] |
| HBO3110869.1 | Type1.BREX | Tox-RelE->HTH_3-><-tRNA\|\|HTH_36->Phage_int_SAM_2+Integrase_1-><-SIGMA+OB-Fold+AAA-ATPase+Lon-Protease-BrxL<-PglZ<-HNH<-OLD-ABC<-N6-METHYLASE<-DUF6079*<- [illegible] | 1146 | Bacteria>Pseudomonadota>Gammaproteobacteria>Pseudomonadales>Pseudomonadaceae>Pseudomonas | GCA_021651555.1 | Pseudomonas_aeruginosa | L4R01_001445 | 31462..34902 | 287 | BREX-system-P-loop-protein-BrxC-[Pseudomonas-aeruginosa] |
| HBO2945375.1 | Type1.BREX | REase+SF1-SF2-Helicase->Arm-DNA-bind_3+Phage_integrase-><-SIGMA+OB-Fold+AAA-ATPase+HKD-DNase-BrxL<-PglZ<-N6-METHYLASE-p<-RVT<-REase<-SbcC-p<-N6-METHYLASE<- [illegible] | 1184 | Bacteria>Pseudomonadota>Gammaproteobacteria>Pseudomonadales>Pseudomonadaceae>Pseudomonas | GCA_021651875.1 | Pseudomonas_aeruginosa | L4R51_003866 | 17556..21110 | 287 | BREX-system-P-loop-protein-BrxC-[Pseudomonas-aeruginosa] |
| HBO6336105.1 | Type1.BREX | <-wHTH+WYL+WCX\|\|?->Peptidase_M15->?->BrxA-DNA-binding->BrxB-BrxF-iSTAND->DUF6079*->N6-METHYLASE->?->HNH+GCN1->?->ParB+HNH->PglZ->SIGMA+OB-Fold+AAA-ATPase+ [illegible] | 1154 | Bacteria>Pseudomonadota>Gammaproteobacteria>Pseudomonadales>Pseudomonadaceae>Pseudomonas | GCA_021670735.1 | Pseudomonas_aeruginosa | L4747_005548 | 6948..10412 | 287 | BREX-system-P-loop-protein-BrxC-[Pseudomonas-aeruginosa] |
| HBO7186446.1 | Type1.BREX | DUF6079-p*->N6-METHYLASE->SbcC-p->REase->N6-METHYLASE-p->PglZ->SIGMA+OB-Fold+AAA-ATPase+HKD-DNase-BrxL-><-Arm-DNA-bind_3+Phage_integrase<-REase+SF1-SF2-Helicase<- [illegible] | 978 | Bacteria>Pseudomonadota>Gammaproteobacteria>Pseudomonadales>Pseudomonadaceae>Pseudomonas | GCA_021674675.1 | Pseudomonas_aeruginosa | L4968_002122 | 1..2938 | 287 | BREX-system-P-loop-protein-BrxC-[Pseudomonas-aeruginosa] |
| HBO6991083.1 | Type1.BREX | [illegible] | 567 | Bacteria>Pseudomonadota>Gammaproteobacteria>Pseudomonadales>Pseudomonadaceae>Pseudomonas | GCA_021674875.1 | Pseudomonas_aeruginosa | L4955_003120 | 106733..108434 | 287 | BREX-system-P-loop-protein-BrxC-[Pseudomonas-aeruginosa] |
| HBO7921132.1 | Type1.BREX | [illegible] | 1184 | Bacteria>Pseudomonadota>Gammaproteobacteria>Pseudomonadales>Pseudomonadaceae>Pseudomonas | GCA_021678155.1 | Pseudomonas_aeruginosa | L5C37_005426 | 13663..17217 | 287 | BREX-system-P-loop-protein-BrxC-[Pseudomonas-aeruginosa] |
| HBP6330513.1 | Type1.BREX | [illegible] | 1154 | Bacteria>Pseudomonadota>Gammaproteobacteria>Pseudomonadales>Pseudomonadaceae>Pseudomonas | GCA_021710255.1 | Pseudomonas_aeruginosa | L6A37_21805 | 72354..75818 | 287 | BREX-system-P-loop-protein-BrxC-[Pseudomonas-aeruginosa] |
| HBP6726062.1 | Type1.BREX | [illegible] | 1223 | Bacteria>Pseudomonadota>Gammaproteobacteria>Pseudomonadales>Pseudomonadaceae>Pseudomonas | GCA_021711795.1 | Pseudomonas_aeruginosa | L6B36_03955 | 73272..76943 | 287 | BREX-system-P-loop-protein-BrxC-[Pseudomonas-aeruginosa] |
| MCH2341807.1 | Type1.BREX | [illegible] | 1186 | Bacteria>Pseudomonadota>Gammaproteobacteria>Pseudomonadales>Pseudomonadaceae>Pseudomonas | GCA_022448005.1 | Pseudomonas_sp. | MK363_18230 | 17987..21547 | 306 | BREX-system-P-loop-protein-BrxC-[Pseudomonas-sp.] |
| HCA6672119.1 | Type1.BREX | [illegible] | 1088 | Bacteria>Pseudomonadota>Gammaproteobacteria>Pseudomonadales>Pseudomonadaceae>Pseudomonas | GCA_022988605.1 | Pseudomonas_aeruginosa | MXV24_006573 | 2961..6226 | 287 | BREX-system-P-loop-protein-BrxC-[Pseudomonas-aeruginosa] |
| MCK9534930.1 | Type1.BREX | [illegible] | 1154 | Bacteria>Pseudomonadota>Gammaproteobacteria>Pseudomonadales>Pseudomonadaceae>Pseudomonas | GCA_023228505.1 | Pseudomonas_sp. | MOQ98_09595 | 5981..9445 | 306 | BREX-system-P-loop-protein-BrxC-[Pseudomonas-sp.] |
| MCO1735128.1 | Type1.BREX | [illegible] | 1189 | Bacteria>Pseudomonadota>Gammaproteobacteria>Pseudomonadales>Pseudomonadaceae>Pseudomonas | GCA_023905245.1 | Pseudomonas_aeruginosa | FA456_28435 | 3288..6857 | 287 | BREX-system-P-loop-protein-BrxC-[Pseudomonas-aeruginosa] |
| MCO2467177.1 | Type1.BREX | [illegible] | 1146 | Bacteria>Pseudomonadota>Gammaproteobacteria>Pseudomonadales>Pseudomonadaceae>Pseudomonas | GCA_023907445.1 | Pseudomonas_aeruginosa | FA343_15170 | 67465..70905 | 287 | BREX-system-P-loop-protein-BrxC-[Pseudomonas-aeruginosa] |
| MCO2679293.1 | Type1.BREX | [illegible] | 638 | Bacteria>Pseudomonadota>Gammaproteobacteria>Pseudomonadales>Pseudomonadaceae>Pseudomonas | GCA_023908095.1 | Pseudomonas_aeruginosa | FA310_30470 | 1..1913 | 287 | BREX-system-P-loop-protein-BrxC-[Pseudomonas-aeruginosa] |
| MCO2922446.1 | Type1.BREX | [illegible] | 1189 | Bacteria>Pseudomonadota>Gammaproteobacteria>Pseudomonadales>Pseudomonadaceae>Pseudomonas | GCA_023909145.1 | Pseudomonas_aeruginosa | FA260_04940 | 145579..149148 | 287 | BREX-system-P-loop-protein-BrxC-[Pseudomonas-aeruginosa] |
| HCE7036149.1 | Type1.BREX | [illegible] | 945 | Bacteria>Pseudomonadota>Gammaproteobacteria>Pseudomonadales>Pseudomonadaceae>Pseudomonas | GCA_023918585.1 | Pseudomonas_aeruginosa | NHP64_005580 | 1..2836 | 287 | BREX-system-P-loop-protein-BrxC-[Pseudomonas-aeruginosa] |
| HCE7261574.1 | Type1.BREX | [illegible] | 1154 | Bacteria>Pseudomonadota>Gammaproteobacteria>Pseudomonadales>Pseudomonadaceae>Pseudomonas | GCA_023919205.1 | Pseudomonas_aeruginosa | NHQ13_005985 | 6842..10306 | 287 | BREX-system-P-loop-protein-BrxC-[Pseudomonas-aeruginosa] |
| HCF1762884.1 | Type1.BREX | [illegible] | 1168 | Bacteria>Pseudomonadota>Gammaproteobacteria>Pseudomonadales>Pseudomonadaceae>Pseudomonas | GCA_023941915.1 | Pseudomonas_aeruginosa | NH889_002155 | 1..3508 | 287 | BREX-system-P-loop-protein-BrxC-[Pseudomonas-aeruginosa] |
| HCF5867680.1 | Type1.BREX | [illegible] | 1181 | Bacteria>Pseudomonadota>Gammaproteobacteria>Pseudomonadales>Pseudomonadaceae>Pseudomonas | GCA_023959235.1 | Pseudomonas_aeruginosa | NIH49_002287 | 171515..175060 | 287 | BREX-system-P-loop-protein-BrxC-[Pseudomonas-aeruginosa] |
| HCF9860952.1 | Type1.BREX | [illegible] | 1223 | Bacteria>Pseudomonadota>Gammaproteobacteria>Pseudomonadales>Pseudomonadaceae>Pseudomonas | GCA_023993215.1 | Pseudomonas_aeruginosa | NI798_001205 | 427888..431559 | 287 | BREX-system-P-loop-protein-BrxC-[Pseudomonas-aeruginosa] |
| HCH0453160.1 | Type1.BREX | [illegible] | 1154 | Bacteria>Pseudomonadota>Gammaproteobacteria>Pseudomonadales>Pseudomonadaceae>Pseudomonas | GCA_024072695.1 | Pseudomonas_aeruginosa | NKO07_001951 | 228071..231535 | 287 | BREX-system-P-loop-protein-BrxC-[Pseudomonas-aeruginosa] |
| BDM24922.1 | Type1.BREX | [illegible] | 1154 | Bacteria>Pseudomonadota>Gammaproteobacteria>Pseudomonadales>Pseudomonadaceae>Pseudomonas | GCA_024349685.1 | Pseudomonas_sp._LRP2-20 | KMS_R46790 | 5328463..5331927 | 2944234 | BREX-system-P-loop-protein-BrxC-[Pseudomonas-sp.-LRP2-20] |
| WP_132538218.1 | Type1.BREX | [illegible] | 1154 | Bacteria>Pseudomonadota>Gammaproteobacteria>Pseudomonadales>Pseudomonadaceae>Pseudomonas | GCA_030820635.1 | Pseudomonas_aeruginosa | EQ548_RS01360 | 6947..10411 | 287 | BREX-system-P-loop-protein-BrxC-[Pseudomonas-aeruginosa] |
| WP_257648015.1 | Type1.BREX | [illegible] | 1073 | Bacteria>Pseudomonadota>Gammaproteobacteria>Pseudomonadales>Pseudomonadaceae>Pseudomonas | GCA_030820635.1 | Pseudomonas_aeruginosa | K3Z95_RS14360 | 10117..13340 | 287 | BREX-system-P-loop-protein-BrxC,-partial-[Pseudomonas-aeruginosa] |
| WP_230691811.1 | Type1.BREX | [illegible] | 1225 | Bacteria>Pseudomonadota>Gammaproteobacteria>Pseudomonadales>Pseudomonadaceae>Pseudomonas | GCA_030820635.1 | Pseudomonas_syringae_group_genomosp._3 | I4I80_RS00295 | 46299..49976 | 251701 | BREX-system-P-loop-protein-BrxC-[Pseudomonas-syringae-group-genomosp.-3] |
| WP_241090907.1 | Type1.BREX | [illegible] | 1225 | Bacteria>Pseudomonadota>Gammaproteobacteria>Pseudomonadales>Pseudomonadaceae>Pseudomonas | GCA_030820635.1 | Pseudomonas_viridiflava | EE220_RS21960 | 64975..68652 | 33069 | BREX-system-P-loop-protein-BrxC-[Pseudomonas-viridiflava] |
| SBW85146.1 | Type1.BREX | [illegible] | 1154 | Bacteria>Pseudomonadota>Gammaproteobacteria>Pseudomonadales>Pseudomonadaceae>Pseudomonas | GCA_900092355.1 | Pseudomonas_veronii_1YdBTEX2 | PVE_P0103 | 88310..91774 | 1295141 | hypothetical-protein-PVE_P0103-[Pseudomonas-veronii-1YdBTEX2] |
| CAB5551672.1 | Type1.BREX | [illegible] | 1154 | Bacteria>Pseudomonadota>Gammaproteobacteria>Pseudomonadales>Pseudomonadaceae>Pseudomonas | GCA_903684265.1 | Pseudomonas_putida | GHA_03446 | 127342..130806 | 303 | Uncharacterised-protein-[Pseudomonas-putida] |
| CAB5576233.1 | Type1.BREX | [illegible] | 1223 | Bacteria>Pseudomonadota>Gammaproteobacteria>Pseudomonadales>Pseudomonadaceae>Pseudomonas | GCA_903684465.1 | Pseudomonas_putida | GHA_00144 | 159878..163549 | 303 | ATPase-involved-in-DNA-repair-[Pseudomonas-putida] |
| WP_010463595.1 | Type1.BREX | [illegible] | 1185 | Bacteria>Pseudomonadota>Gammaproteobacteria>Pseudomonadales>Pseudomonadaceae>Pseudomonas | GCF_000257545.3 | Pseudomonas_mandelii | OU5_RS13450 | 3003724..3007281 | 75612 | BREX-system-P-loop-protein-BrxC-[Pseudomonas-mandelii] |
| WP_019485333.1 | Type1.BREX | [illegible] | 1193 | Bacteria>Pseudomonadota>Gammaproteobacteria>Pseudomonadales>Pseudomonadaceae>Pseudomonas | GCF_000290555.2 | Pseudomonas_aeruginosa | E4028_RS01350 | 3998..7579 | 287 | BREX-system-P-loop-protein-BrxC-[Pseudomonas-aeruginosa] |
| WP_228382933.1 | Type1.BREX | [illegible] | 1225 | Bacteria>Pseudomonadota>Gammaproteobacteria>Pseudomonadales>Pseudomonadaceae>Pseudomonas | GCF_000481225.1 | Pseudomonas_aeruginosa | Q037_RS32345 | 325963..329640 | 287 | BREX-system-P-loop-protein-BrxC-[Pseudomonas-aeruginosa] |
| WP_023113968.1 | Type1.BREX | [illegible] | 1146 | Bacteria>Pseudomonadota>Gammaproteobacteria>Pseudomonadales>Pseudomonadaceae>Pseudomonas | GCF_000481245.1 | Pseudomonas_aeruginosa | Q036_RS16830 | 2151526..2154966 | 287 | BREX-system-P-loop-protein-BrxC-[Pseudomonas-aeruginosa] |
| WP_042135808.1 | Type1.BREX | [illegible] | 1184 | Bacteria>Pseudomonadota>Gammaproteobacteria>Pseudomonadales>Pseudomonadaceae>Pseudomonas | GCF_000730625.1 | Pseudomonas_oryzihabitans | POR01S_RS10925 | 121811..125365 | 47885 | BREX-system-P-loop-protein-BrxC-[Pseudomonas-oryzihabitans] |
| WP_238579794.1 | Type1.BREX | [illegible] | 1150 | Bacteria>Pseudomonadota>Gammaproteobacteria>Pseudomonadales>Pseudomonadaceae>Pseudomonas | GCF_000789745.1 | Pseudomonas_aeruginosa | NR64_RS21585 | 177..3629 | 287 | BREX-system-P-loop-protein-BrxC-[Pseudomonas-aeruginosa] |
| WP_034081877.1 | Type1.BREX | [illegible] | 1184 | Bacteria>Pseudomonadota>Gammaproteobacteria>Pseudomonadales>Pseudomonadaceae>Pseudomonas | GCF_000793185.1 | Pseudomonas_aeruginosa | NT71_RS08285 | 193065..196619 | 287 | BREX-system-P-loop-protein-BrxC-[Pseudomonas-aeruginosa] |
| WP_033954069.1 | Type1.BREX | [illegible] | 1154 | Bacteria>Pseudomonadota>Gammaproteobacteria>Pseudomonadales>Pseudomonadaceae>Pseudomonas | GCF_000793385.1 | Pseudomonas_aeruginosa | NT88_RS29575 | 15006..18470 | 287 | BREX-system-P-loop-protein-BrxC-[Pseudomonas-aeruginosa] |
| WP_043106421.1 | Type1.BREX | [illegible] | 1223 | Bacteria>Pseudomonadota>Gammaproteobacteria>Pseudomonadales>Pseudomonadaceae>Pseudomonas | GCF_000794515.1 | Pseudomonas_aeruginosa | NQ84_RS27950 | 382..4053 | 287 | BREX-system-P-loop-protein-BrxC-[Pseudomonas-aeruginosa] |
| WP_202808417.1 | Type1.BREX | [illegible] | 937 | Bacteria>Pseudomonadota>Gammaproteobacteria>Pseudomonadales>Pseudomonadaceae>Pseudomonas | GCF_000794515.1 | Pseudomonas_aeruginosa | NQ84_RS28370 | 16579..19392 | 287 | BREX-system-P-loop-protein-BrxC-[Pseudomonas-aeruginosa] |
| WP_228480203.1 | Type1.BREX | [illegible] | 619 | Bacteria>Pseudomonadota>Gammaproteobacteria>Pseudomonadales>Pseudomonadaceae>Pseudomonas | GCF_000829885.1 | Pseudomonas_aeruginosa | EG09_RS07505 | 1715440..1717299 | 287 | BREX-system-P-loop-protein-BrxC-[Pseudomonas-aeruginosa] |
| WP_041506104.1 | Type1.BREX | [illegible] | 1184 | Bacteria>Pseudomonadota>Gammaproteobacteria>Pseudomonadales>Pseudomonadaceae>Pseudomonas | GCF_000831585.1 | Pseudomonas_plecoglossicida | RK21_RS13525 | 2976546..2980100 | 70775 | BREX-system-P-loop-protein-BrxC-[Pseudomonas-plecoglossicida] |
| WP_052507173.1 | Type1.BREX | [illegible] | 886 | Bacteria>Pseudomonadota>Gammaproteobacteria>Pseudomonadales>Pseudomonadaceae>Pseudomonas | GCF_000935725.1 | Pseudomonas_syringae_group_genomosp._3 | AC506_RS17885 | 8350..11010 | 251701 | BREX-system-P-loop-protein-BrxC-[Pseudomonas-syringae-group-genomosp.-3] |
| WP_044727109.1 | Type1.BREX | [illegible] | 1184 | Bacteria>Pseudomonadota>Gammaproteobacteria>Pseudomonadales>Pseudomonadaceae>Pseudomonas | GCF_000937465.2 | Pseudomonas_aeruginosa | TN45_RS09710 | 568754..572308 | 287 | BREX-system-P-loop-protein-BrxC-[Pseudomonas-aeruginosa] |
| WP_047303167.1 | Type1.BREX | [illegible] | 1223 | Bacteria>Pseudomonadota>Gammaproteobacteria>Pseudomonadales>Pseudomonadaceae>Pseudomonas | GCF_001020715.1 | Pseudomonas_fluorescens | AA061_RS10335 | 96304..99975 | 294 | BREX-system-P-loop-protein-BrxC-[Pseudomonas-fluorescens] |
| WP_047306816.1 | Type1.BREX | [illegible] | 1185 | Bacteria>Pseudomonadota>Gammaproteobacteria>Pseudomonadales>Pseudomonadaceae>Pseudomonas | GCF_001020715.1 | Pseudomonas_fluorescens | AA061_RS32150 | 16461..20018 | 294 | BREX-system-P-loop-protein-BrxC-[Pseudomonas-fluorescens] |
| WP_047283135.1 | Type1.BREX | [illegible] | 1154 | Bacteria>Pseudomonadota>Gammaproteobacteria>Pseudomonadales>Pseudomonadaceae>Pseudomonas | GCF_001020725.2 | Pseudomonas_lundensis | AA042_RS07115 | 1582722..1586186 | 86185 | BREX-system-P-loop-protein-BrxC-[Pseudomonas-lundensis] |
| WP_048385719.1 | Type1.BREX | [illegible] | 1185 | Bacteria>Pseudomonadota>Gammaproteobacteria>Pseudomonadales>Pseudomonadaceae>Pseudomonas | GCF_001043065.1 | Pseudomonas_helleri | TU85_RS02975 | 88982..92539 | 1608996 | BREX-system-P-loop-protein-BrxC-[Pseudomonas-helleri] |
| WP_053154299.1 | Type1.BREX | [illegible] | 1154 | Bacteria>Pseudomonadota>Gammaproteobacteria>Pseudomonadales>Pseudomonadaceae>Pseudomonas | GCF_001269805.1 | Pseudomonas_sp._P1.8 | AD668_RS02725 | 49573..53037 | 1699310 | BREX-system-P-loop-protein-BrxC-[Pseudomonas-sp.-P1.8] |
| WP_057391032.1 | Type1.BREX | [illegible] | 1154 | Bacteria>Pseudomonadota>Gammaproteobacteria>Pseudomonadales>Pseudomonadaceae>Pseudomonas | GCF_001374115.1 | Pseudomonas_aeruginosa | AYU10_RS17350 | 11442..14906 | 287 | BREX-system-P-loop-protein-BrxC-[Pseudomonas-aeruginosa] |
| WP_057429193.1 | Type1.BREX | [illegible] | 1156 | Bacteria>Pseudomonadota>Gammaproteobacteria>Pseudomonadales>Pseudomonadaceae>Pseudomonas | GCF_001401385.1 | Pseudomonas_syringae_group_genomosp._3 | AL087_RS04320 | 127987..131457 | 251701 | BREX-system-P-loop-protein-BrxC-[Pseudomonas-syringae-group-genomosp.-3] |
| WP_057978570.1 | Type1.BREX | [illegible] | 1223 | Bacteria>Pseudomonadota>Gammaproteobacteria>Pseudomonadales>Pseudomonadaceae>Pseudomonas | GCF_001439745.1 | Pseudomonas_lactis | TX25_RS28475 | 27732..31403 | 1615674 | BREX-system-P-loop-protein-BrxC-[Pseudomonas-lactis] |
| WP_235565439.1 | Type1.BREX | [illegible] | 1225 | Bacteria>Pseudomonadota>Gammaproteobacteria>Pseudomonadales>Pseudomonadaceae>Pseudomonas | GCF_001450115.1 | Pseudomonas_aeruginosa | AO953_RS18460 | 48360..52037 | 287 | BREX-system-P-loop-protein-BrxC-[Pseudomonas-aeruginosa] |
| WP_058174899.1 | Type1.BREX | [illegible] | 1184 | Bacteria>Pseudomonadota>Gammaproteobacteria>Pseudomonadales>Pseudomonadaceae>Pseudomonas | GCF_001452485.1 | Pseudomonas_aeruginosa | APA89_RS03075 | 652482..656036 | 287 | BREX-system-P-loop-protein-BrxC-[Pseudomonas-aeruginosa] |
| WP_059319043.1 | Type1.BREX | [illegible] | 1154 | Bacteria>Pseudomonadota>Gammaproteobacteria>Pseudomonadales>Pseudomonadaceae>Pseudomonas | GCF_001516265.1 | Pseudomonas_aeruginosa | HV84_RS13920 | 329043..332507 | 287 | BREX-system-P-loop-protein-BrxC-[Pseudomonas-aeruginosa] |
| WP_031639867.1 | Type1.BREX | [illegible] | 1184 | Bacteria>Pseudomonadota>Gammaproteobacteria>Pseudomonadales>Pseudomonadaceae>Pseudomonas | GCF_001548335.1 | Pseudomonas_aeruginosa | IOMTU133_RS23320 | 5069423..5072977 | 287 | BREX-system-P-loop-protein-BrxC-[Pseudomonas-aeruginosa] |
| WP_065615729.1 | Type1.BREX | [illegible] | 1223 | Bacteria>Pseudomonadota>Gammaproteobacteria>Pseudomonadales>Pseudomonadaceae>Pseudomonas | GCF_001691585.1 | Pseudomonas_moraviensis | BBA88_RS08910 | 1046415..1050086 | 321662 | BREX-system-P-loop-protein-BrxC-[Pseudomonas-moraviensis] |
| WP_069953193.1 | Type1.BREX | [illegible] | 828 | Bacteria>Pseudomonadota>Gammaproteobacteria>Pseudomonadales>Pseudomonadaceae>Pseudomonas | GCF_001750225.1 | Pseudomonas_aeruginosa | A7R77_RS00815 | 21562..24049 | 287 | BREX-system-P-loop-protein-BrxC,-partial-[Pseudomonas-aeruginosa] |
| WP_070166840.1 | Type1.BREX | [illegible] | 1182 | Bacteria>Pseudomonadota>Gammaproteobacteria>Pseudomonadales>Pseudomonadaceae>Pseudomonas | GCF_001756585.1 | Pseudomonas_aeruginosa | AN464_RS32885 | 78098..81646 | 287 | BREX-system-P-loop-protein-BrxC-[Pseudomonas-aeruginosa] |
| WP_070624003.1 | Type1.BREX | [illegible] | 1154 | Bacteria>Pseudomonadota>Gammaproteobacteria>Pseudomonadales>Pseudomonadaceae>Pseudomonas | GCF_001810465.1 | Pseudomonas_aeruginosa | HMPREF2746_RS21475 | 11855..15319 | 287 | BREX-system-P-loop-protein-BrxC-[Pseudomonas-aeruginosa] |
| WP_073844625.1 | Type1.BREX | [illegible] | 1217 | Bacteria>Pseudomonadota>Gammaproteobacteria>Pseudomonadales>Pseudomonadaceae>Pseudomonas | GCF_001907715.1 | Pseudomonas_sp._BTN1 | BMH52_RS18610 | 18591..22244 | 1750647 | BREX-system-P-loop-protein-BrxC-[Pseudomonas-sp.-BTN1] |
| WP_079783554.1 | Type1.BREX | [illegible] | 1154 | Bacteria>Pseudomonadota>Gammaproteobacteria>Pseudomonadales>Pseudomonadaceae>Pseudomonas | GCF_002037565.1 | Pseudomonas_sp._LPH1 | BHQ29_RS09350 | 2029984..2033448 | 1898684 | BREX-system-P-loop-protein-BrxC-[Pseudomonas-sp.-LPH1] |
| WP_084308437.1 | Type1.BREX | [illegible] | 1224 | Bacteria>Pseudomonadota>Gammaproteobacteria>Pseudomonadales>Pseudomonadaceae>Pseudomonas | GCF_002091575.1 | Pseudomonas_flavescens | PFL01S_RS23715 | 11165..14839 | 29435 | BREX-system-P-loop-protein-BrxC-[Pseudomonas-flavescens] |
| WP_074859114.1 | Type1.BREX | [illegible] | 1184 | Bacteria>Pseudomonadota>Gammaproteobacteria>Pseudomonadales>Pseudomonadaceae>Pseudomonas | GCF_002091815.1 | Pseudomonas_oleovorans | POL01S_RS18595 | 25960..29514 | 301 | BREX-system-P-loop-protein-BrxC-[Pseudomonas-oleovorans] |
| WP_084929168.1 | Type1.BREX | [illegible] | 1184 | Bacteria>Pseudomonadota>Gammaproteobacteria>Pseudomonadales>Pseudomonadaceae>Pseudomonas | GCF_002094805.1 | Pseudomonas_aeruginosa | B7H20_RS24080 | 14270..17824 | 287 | BREX-system-P-loop-protein-BrxC-[Pseudomonas-aeruginosa] |
| WP_205881761.1 | Type1.BREX | [illegible] | 1159 | Bacteria>Pseudomonadota>Gammaproteobacteria>Pseudomonadales>Pseudomonadaceae>Pseudomonas | GCF_002128325.1 | Pseudomonas_fragi | B6D87_RS19610 | 4227178..4230657 | 296 | BREX-system-P-loop-protein-BrxC-[Pseudomonas-fragi] |
| WP_086230732.1 | Type1.BREX | [illegible] | 1184 | Bacteria>Pseudomonadota>Gammaproteobacteria>Pseudomonadales>Pseudomonadaceae>Pseudomonas | GCF_002135805.1 | Pseudomonas_aeruginosa | CAZ03_RS13145 | 128061..131615 | 287 | BREX-system-P-loop-protein-BrxC-[Pseudomonas-aeruginosa] |
| WP_088135363.1 | Type1.BREX | [illegible] | 1154 | Bacteria>Pseudomonadota>Gammaproteobacteria>Pseudomonadales>Pseudomonadaceae>Pseudomonas | GCF_002193745.1 | Pseudomonas_aeruginosa | GNQ21_RS29160 | 3852..7316 | 287 | BREX-system-P-loop-protein-BrxC-[Pseudomonas-aeruginosa] |
| WP_095025594.1 | Type1.BREX | [illegible] | 1154 | Bacteria>Pseudomonadota>Gammaproteobacteria>Pseudomonadales>Pseudomonadaceae>Pseudomonas | GCF_002269515.1 | Pseudomonas_fragi | CJU79_RS17655 | 17310..20774 | 296 | BREX-system-P-loop-protein-BrxC-[Pseudomonas-fragi] |
| WP_095626344.1 | Type1.BREX | [illegible] | 1154 | Bacteria>Pseudomonadota>Gammaproteobacteria>Pseudomonadales>Pseudomonadaceae>Pseudomonas | GCF_002287035.1 | Pseudomonas_sp._WN033 | CK507_RS15715 | 130328..133792 | 2032629 | BREX-system-P-loop-protein-BrxC-[Pseudomonas-sp.-WN033] |
| WP_256588486.1 | Type1.BREX | [illegible] | 1225 | Bacteria>Pseudomonadota>Gammaproteobacteria>Pseudomonadales>Pseudomonadaceae>Pseudomonas | GCF_002591235.1 | Pseudomonas_sp._FDAARGOS_380 | CRN80_RS16460 | 3507084..3510761 | 2018067 | BREX-system-P-loop-protein-BrxC-[Pseudomonas-sp.-FDAARGOS_380] |
| WP_101196767.1 | Type1.BREX | [illegible] | 1180 | Bacteria>Pseudomonadota>Gammaproteobacteria>Pseudomonadales>Pseudomonadaceae>Pseudomonas | GCF_002835905.1 | Pseudomonas_monteilii | CXB65_RS25690 | 173341..176883 | 76759 | BREX-system-P-loop-protein-BrxC-[Pseudomonas-monteilii] |
| WP_102082564.1 | Type1.BREX | [illegible] | 1184 | Bacteria>Pseudomonadota>Gammaproteobacteria>Pseudomonadales>Pseudomonadaceae>Pseudomonas | GCF_002864915.1 | Pseudomonas_guariconensis | CXG53_RS23735 | 60064..63618 | 1288410 | BREX-system-P-loop-protein-BrxC-[Pseudomonas-guariconensis] |
| WP_102368059.1 | Type1.BREX | [illegible] | 1188 | Bacteria>Pseudomonadota>Gammaproteobacteria>Pseudomonadales>Pseudomonadaceae>Pseudomonas | GCF_002874965.1 | Pseudomonas_sp._NC02 | C0058_RS03795 | 845324..848890 | 2067572 | BREX-system-P-loop-protein-BrxC-[Pseudomonas-sp.-NC02] |
| WP_102684491.1 | Type1.BREX | [illegible] | 1222 | Bacteria>Pseudomonadota>Gammaproteobacteria>Pseudomonadales>Pseudomonadaceae>Pseudomonas | GCF_002883975.1 | Pseudomonas_sp._FW305-42 | C1X79_RS19065 | 47194..50862 | 2070677 | BREX-system-P-loop-protein-BrxC-[Pseudomonas-sp.-FW305-42] |
| WP_102886089.1 | Type1.BREX | [illegible] | 1184 | Bacteria>Pseudomonadota>Gammaproteobacteria>Pseudomonadales>Pseudomonadaceae>Pseudomonas | GCF_002891885.1 | Pseudomonas_putida | CBL13_04161 | 1040459..1044013 | 303 | BREX-system-P-loop-protein-BrxC-[Pseudomonas-putida] |
| WP_104727392.1 | Type1.BREX | [illegible] | 1222 | Bacteria>Pseudomonadota>Gammaproteobacteria>Pseudomonadales>Pseudomonadaceae>Pseudomonas | GCF_002941105.1 | Pseudomonas_oleovorans | C5L43_RS04465 | 3008..6676 | 301 | BREX-system-P-loop-protein-BrxC-[Pseudomonas-oleovorans] |
| WP_242445006.1 | Type1.BREX | [illegible] | 1225 | Bacteria>Pseudomonadota>Gammaproteobacteria>Pseudomonadales>Pseudomonadaceae>Pseudomonas | GCF_002941105.1 | Pseudomonas_oleovorans | C5L43_RS18405 | 7326..11003 | 301 | BREX-system-P-loop-protein-BrxC-[Pseudomonas-oleovorans] |
| WP_106118416.1 | Type1.BREX | [illegible] | 1223 | Bacteria>Pseudomonadota>Gammaproteobacteria>Pseudomonadales>Pseudomonadaceae>Pseudomonas | GCF_003001555.1 | Pseudomonas_fluorescens | C7A10_RS23710 | 130630..134301 | 294 | BREX-system-P-loop-protein-BrxC-[Pseudomonas-fluorescens] |
| WP_109933363.1 | Type1.BREX | [illegible] | 1154 | Bacteria>Pseudomonadota>Gammaproteobacteria>Pseudomonadales>Pseudomonadaceae>Pseudomonas | GCF_003176655.1 | Pseudomonas_sp._RW407 | DK254_RS08115 | 332506..335970 | 2202894 | BREX-system-P-loop-protein-BrxC-[Pseudomonas-sp.-RW407] |
| WP_116801381.1 | Type1.BREX | [illegible] | 1184 | Bacteria>Pseudomonadota>Gammaproteobacteria>Pseudomonadales>Pseudomonadaceae>Pseudomonas | GCF_003410555.2 | Pseudomonas_aeruginosa | DZ982_RS28380 | 35413..38967 | 287 | BREX-system-P-loop-protein-BrxC-[Pseudomonas-aeruginosa] |
| WP_116843887.1 | Type1.BREX | [illegible] | 549 | Bacteria>Pseudomonadota>Gammaproteobacteria>Pseudomonadales>Pseudomonadaceae>Pseudomonas | GCF_003411125.2 | Pseudomonas_aeruginosa | DZ918_RS18285 | 1..1646 | 287 | BREX-system-P-loop-protein-BrxC,-partial-[Pseudomonas-aeruginosa] |
| WP_116805555.1 | Type1.BREX | [illegible] | 1154 | Bacteria>Pseudomonadota>Gammaproteobacteria>Pseudomonadales>Pseudomonadaceae>Pseudomonas | GCF_003412045.2 | Pseudomonas_aeruginosa | L4E78_005908 | 8239..11703 | 287 | BREX-system-P-loop-protein-BrxC-[Pseudomonas-aeruginosa] |
| WP_119519035.1 | Type1.BREX | [illegible] | 1220 | Bacteria>Pseudomonadota>Gammaproteobacteria>Pseudomonadales>Pseudomonadaceae>Pseudomonas | GCF_003585225.1 | Pseudomonas_aeruginosa | AXX04_RS18775 | 17983..21645 | 287 | BREX-system-P-loop-protein-BrxC-[Pseudomonas-aeruginosa] |
| WP_121398877.1 | Type1.BREX | [illegible] | 1180 | Bacteria>Pseudomonadota>Gammaproteobacteria>Pseudomonadales>Pseudomonadaceae>Pseudomonas | GCF_003631245.1 | Pseudomonas_aeruginosa | DQH61_RS02890 | 22908..26450 | 287 | BREX-system-P-loop-protein-BrxC-[Pseudomonas-aeruginosa] |
| WP_128551186.1 | Type1.BREX | [illegible] | 1184 | Bacteria>Pseudomonadota>Gammaproteobacteria>Pseudomonadales>Pseudomonadaceae>Pseudomonas | GCF_003698635.1 | Pseudomonas_aeruginosa | IPC82_RS03385 | 45130..48684 | 287 | BREX-system-P-loop-protein-BrxC-[Pseudomonas-aeruginosa] |
| WP_236079235.1 | Type1.BREX | [illegible] | 1225 | Bacteria>Pseudomonadota>Gammaproteobacteria>Pseudomonadales>Pseudomonadaceae>Pseudomonas | GCF_003833645.1 | Pseudomonas_aeruginosa | IPC1294_RS03655 | 745401..749078 | 287 | BREX-system-P-loop-protein-BrxC-[Pseudomonas-aeruginosa] |
| WP_124138810.1 | Type1.BREX | [illegible] | 1223 | Bacteria>Pseudomonadota>Gammaproteobacteria>Pseudomonadales>Pseudomonadaceae>Pseudomonas | GCF_003833965.1 | Pseudomonas_aeruginosa | IPC1352_RS13795 | 491665..495336 | 287 | BREX-system-P-loop-protein-BrxC-[Pseudomonas-aeruginosa] |
| WP_124127279.1 | Type1.BREX | [illegible] | 1223 | Bacteria>Pseudomonadota>Gammaproteobacteria>Pseudomonadales>Pseudomonadaceae>Pseudomonas | GCF_003836525.1 | Pseudomonas_aeruginosa | IPC776_RS25325 | 73240..76911 | 287 | BREX-system-P-loop-protein-BrxC-[Pseudomonas-aeruginosa] |
| WP_201271146.1 | Type1.BREX | [illegible] | 599 | Bacteria>Pseudomonadota>Gammaproteobacteria>Pseudomonadales>Pseudomonadaceae>Pseudomonas | GCF_003838365.1 | Pseudomonas_aeruginosa | IPC642_RS23000 | 207947..209746 | 287 | BREX-system-P-loop-protein-BrxC-[Pseudomonas-aeruginosa] |
| WP_124187492.1 | Type1.BREX | [illegible] | 1154 | Bacteria>Pseudomonadota>Gammaproteobacteria>Pseudomonadales>Pseudomonadaceae>Pseudomonas | GCF_003840445.1 | Pseudomonas_aeruginosa | IPC460_RS14135 | 71105..74569 | 287 | BREX-system-P-loop-protein-BrxC-[Pseudomonas-aeruginosa] |
| WP_125881186.1 | Type1.BREX | [illegible] | 1154 | Bacteria>Pseudomonadota>Gammaproteobacteria>Pseudomonadales>Pseudomonadaceae>Pseudomonas | GCF_003935455.1 | Pseudomonas_sp._o96-267 | EGJ23_RS18845 | 351..3815 | 2479853 | BREX-system-P-loop-protein-BrxC-[Pseudomonas-sp.-o96-267] |
| WP_126190092.1 | Type1.BREX | [illegible] | 1185 | Bacteria>Pseudomonadota>Gammaproteobacteria>Pseudomonadales>Pseudomonadaceae>Pseudomonas | GCF_003957495.3 | Pseudomonas_songnenensis | EJA06_RS16615 | 27316..30873 | 1176259 | BREX-system-P-loop-protein-BrxC-[Pseudomonas-songnenensis] |
| WP_126525533.1 | Type1.BREX | [illegible] | 1154 | Bacteria>Pseudomonadota>Gammaproteobacteria>Pseudomonadales>Pseudomonadaceae>Pseudomonas | GCF_003969495.1 | Pseudomonas_aeruginosa | DZA06_RS08835 | 36272..39736 | 287 | BREX-system-P-loop-protein-BrxC-[Pseudomonas-aeruginosa] |
| WP_126532550.1 | Type1.BREX | [illegible] | 1223 | Bacteria>Pseudomonadota>Gammaproteobacteria>Pseudomonadales>Pseudomonadaceae>Pseudomonas | GCF_003969645.1 | Pseudomonas_aeruginosa | DY969_RS10905 | 72707..76378 | 287 | BREX-system-P-loop-protein-BrxC-[Pseudomonas-aeruginosa] |
| WP_126588230.1 | Type1.BREX | [illegible] | 1223 | Bacteria>Pseudomonadota>Gammaproteobacteria>Pseudomonadales>Pseudomonadaceae>Pseudomonas | GCF_003970115.1 | Pseudomonas_veronii | EKA83_RS07390 | 206636..210307 | 76761 | BREX-system-P-loop-protein-BrxC-[Pseudomonas-veronii] |
| WP_126634432.1 | Type1.BREX | [illegible] | 904 | Bacteria>Pseudomonadota>Gammaproteobacteria>Pseudomonadales>Pseudomonadaceae>Pseudomonas | GCF_003970625.1 | Pseudomonas_aeruginosa | DZA20_RS16095 | 55154..57868 | 287 | BREX-system-P-loop-protein-BrxC-[Pseudomonas-aeruginosa] |
| WP_260478239.1 | Type1.BREX | [illegible] | 1225 | Bacteria>Pseudomonadota>Gammaproteobacteria>Pseudomonadales>Pseudomonadaceae>Pseudomonas | GCF_003973785.1 | Pseudomonas_aeruginosa | IPC1623_RS31675 | 17836..21513 | 287 | BREX-system-P-loop-protein-BrxC-[Pseudomonas-aeruginosa] |
| WP_128704208.1 | Type1.BREX | [illegible] | 1184 | Bacteria>Pseudomonadota>Gammaproteobacteria>Pseudomonadales>Pseudomonadaceae>Pseudomonas | GCF_003974605.1 | Pseudomonas_aeruginosa | IPC1390_RS17360 | 217632..221186 | 287 | BREX-system-P-loop-protein-BrxC-[Pseudomonas-aeruginosa] |
| WP_129388875.1 | Type1.BREX | [illegible] | 1156 | Bacteria>Pseudomonadota>Gammaproteobacteria>Pseudomonadales>Pseudomonadaceae>Pseudomonas | GCF_004124275.1 | Pseudomonas_sp._DTU12.3 | C2E19_RS05070 | 1143497..1146967 | 2073078 | BREX-system-P-loop-protein-BrxC-[Pseudomonas-sp.-DTU12.3] |
| WP_130899395.1 | Type1.BREX | [illegible] | 1154 | Bacteria>Pseudomonadota>Gammaproteobacteria>Pseudomonadales>Pseudomonadaceae>Pseudomonas | GCF_004307505.1 | Pseudomonas_sp._Sample_11 | D8U60_RS02310 | 26340..29804 | 2448261 | BREX-system-P-loop-protein-BrxC-[Pseudomonas-sp.-Sample_11] |
| WP_134242485.1 | Type1.BREX | [illegible] | 1154 | Bacteria>Pseudomonadota>Gammaproteobacteria>Pseudomonadales>Pseudomonadaceae>Pseudomonas | GCF_004370105.1 | Pseudomonas_aeruginosa | IPC1592_RS30135 | 15141..18605 | 287 | BREX-system-P-loop-protein-BrxC-[Pseudomonas-aeruginosa] |
| WP_202876546.1 | Type1.BREX | [illegible] | 531 | Bacteria>Pseudomonadota>Gammaproteobacteria>Pseudomonadales>Pseudomonadaceae>Pseudomonas | GCF_004373095.1 | Pseudomonas_aeruginosa | IPC174_RS30040 | 16693..18288 | 287 | BREX-system-P-loop-protein-BrxC-[Pseudomonas-aeruginosa] |
| WP_202876713.1 | Type1.BREX | [illegible] | 539 | Bacteria>Pseudomonadota>Gammaproteobacteria>Pseudomonadales>Pseudomonadaceae>Pseudomonas | GCF_004373095.1 | Pseudomonas_aeruginosa | IPC175_RS30340 | 15284..16903 | 287 | hypothetical-protein-[Pseudomonas-aeruginosa] |
| WP_134282343.1 | Type1.BREX | [illegible] | 1154 | Bacteria>Pseudomonadota>Gammaproteobacteria>Pseudomonadales>Pseudomonadaceae>Pseudomonas | GCF_004373575.1 | Pseudomonas_aeruginosa | IPC135_RS30160 | 15285..18749 | 287 | BREX-system-P-loop-protein-BrxC-[Pseudomonas-aeruginosa] |
| WP_202876545.1 | Type1.BREX | [illegible] | 624 | Bacteria>Pseudomonadota>Gammaproteobacteria>Pseudomonadales>Pseudomonadaceae>Pseudomonas | GCF_004373665.1 | Pseudomonas_aeruginosa | IPC174_RS30035 | 14826..16700 | 287 | hypothetical-protein-[Pseudomonas-aeruginosa] |
| WP_136489742.1 | Type1.BREX | [illegible] | 1154 | Bacteria>Pseudomonadota>Gammaproteobacteria>Pseudomonadales>Pseudomonadaceae>Pseudomonas | GCF_004801855.1 | Pseudomonas_sp._A-1 | E5198_RS09425 | 20467..23931 | 1821274 | BREX-system-P-loop-protein-BrxC-[Pseudomonas-sp.-A-1] |
| WP_155677773.1 | Type1.BREX | [illegible] | 1155 | Bacteria>Pseudomonadota>Gammaproteobacteria>Pseudomonadales>Pseudomonadaceae>Pseudomonas | GCF_004919535.4 | Pseudomonas_veronii | E4167_12595 | 2651456..2654923 | 76761 | BREX-system-P-loop-protein-BrxC-[Pseudomonas-veronii] |
| WP_155678129.1 | Type1.BREX | [illegible] | 1221 | Bacteria>Pseudomonadota>Gammaproteobacteria>Pseudomonadales>Pseudomonadaceae>Pseudomonas | GCF_004919535.4 | Pseudomonas_veronii | E4167_RS17855 | 3896046..3899711 | 76761 | BREX-system-P-loop-protein-BrxC-[Pseudomonas-veronii] |
| WP_249351844.1 | Type1.BREX | [illegible] | 1225 | Bacteria>Pseudomonadota>Gammaproteobacteria>Pseudomonadales>Pseudomonadaceae>Pseudomonas | GCF_005223285.1 | Pseudomonas_asiatica | FD498_RS21455 | 9358..13035 | 2219225 | BREX-system-P-loop-protein-BrxC-[Pseudomonas-asiatica] |
| WP_256658147.1 | Type1.BREX | [illegible] | 1220 | Bacteria>Pseudomonadota>Gammaproteobacteria>Pseudomonadales>Pseudomonadaceae>Pseudomonas | GCF_005502555.1 | Pseudomonas_sp._D(2018) | FFS60_RS25250 | 8931..12593 | 2502238 | BREX-system-P-loop-protein-BrxC-[Pseudomonas-sp.-D(2018)] |
| WP_240721840.1 | Type1.BREX | [illegible] | 947 | Bacteria>Pseudomonadota>Gammaproteobacteria>Pseudomonadales>Pseudomonadaceae>Pseudomonas | GCF_005863515.1 | Pseudomonas_aeruginosa | FEM26_RS29610 | 9..2852 | 287 | BREX-system-P-loop-protein-BrxC-[Pseudomonas-aeruginosa] |
| WP_144958058.1 | Type1.BREX | [illegible] | 1223 | Bacteria>Pseudomonadota>Gammaproteobacteria>Pseudomonadales>Pseudomonadaceae>Pseudomonas | GCF_007680465.1 | Pseudomonas_oryzihabitans | FS730_RS21810 | 185497..189168 | 47885 | BREX-system-P-loop-protein-BrxC-[Pseudomonas-oryzihabitans] |
| WP_115288849.1 | Type1.BREX | [illegible] | 1184 | Bacteria>Pseudomonadota>Gammaproteobacteria>Pseudomonadales>Pseudomonadaceae>Pseudomonas | GCF_007833895.1 | Pseudomonas_aeruginosa | FF962_RS31550 | 6589881..6593435 | 287 | BREX-system-P-loop-protein-BrxC-[Pseudomonas-aeruginosa] |
| WP_150073350.1 | Type1.BREX | [illegible] | 1186 | Bacteria>Pseudomonadota>Gammaproteobacteria>Pseudomonadales>Pseudomonadaceae>Pseudomonas | GCF_008632325.1 | Pseudomonas_lactis | F3K52_RS19340 | 4156..7716 | 1615674 | BREX-system-P-loop-protein-BrxC-[Pseudomonas-lactis] |
| WP_151183841.1 | Type1.BREX | [illegible] | 1181 | Bacteria>Pseudomonadota>Gammaproteobacteria>Pseudomonadales>Pseudomonadaceae>Pseudomonas | GCF_008807395.1 | Pseudomonas_sp._C27(2019) | FXF61_RS02795 | 588989..592534 | 2604941 | BREX-system-P-loop-protein-BrxC-[Pseudomonas-sp.-C27(2019)] |
| WP_152745526.1 | Type1.BREX | [illegible] | 1183 | Bacteria>Pseudomonadota>Gammaproteobacteria>Pseudomonadales>Pseudomonadaceae>Pseudomonas | GCF_009296165.1 | Pseudomonas_kitaguniensis | FO169_RS03300 | 21652..25203 | 2607908 | BREX-system-P-loop-protein-BrxC-[Pseudomonas-kitaguniensis] |
| WP_153329486.1 | Type1.BREX | [illegible] | 1184 | Bacteria>Pseudomonadota>Gammaproteobacteria>Pseudomonadales>Pseudomonadaceae>Pseudomonas | GCF_009600225.1 | Pseudomonas_helleri | GHO39_RS16765 | 34385..37939 | 1608996 | BREX-system-P-loop-protein-BrxC-[Pseudomonas-helleri] |
| WP_153386820.1 | Type1.BREX | [illegible] | 1183 | Bacteria>Pseudomonadota>Gammaproteobacteria>Pseudomonadales>Pseudomonadaceae>Pseudomonas | GCF_009600315.1 | Pseudomonas_helleri | GHN86_RS14975 | 48582..52133 | 1608996 | BREX-system-P-loop-protein-BrxC-[Pseudomonas-helleri] |
| WP_153350777.1 | Type1.BREX | [illegible] | 1146 | Bacteria>Pseudomonadota>Gammaproteobacteria>Pseudomonadales>Pseudomonadaceae>Pseudomonas | GCF_009600425.1 | Pseudomonas_helleri | GHO30_RS02550 | 6514..9954 | 1608996 | BREX-system-P-loop-protein-BrxC-[Pseudomonas-helleri] |
| WP_153384138.1 | Type1.BREX | [illegible] | 1186 | Bacteria>Pseudomonadota>Gammaproteobacteria>Pseudomonadales>Pseudomonadaceae>Pseudomonas | GCF_009600565.1 | unclassified_Pseudomonas | GHO24_RS06625 | 35218..38778 | 196821 | MULTISPECIES:-BREX-system-P-loop-protein-BrxC-[unclassified-Pseudomonas] |
| WP_233280221.1 | Type1.BREX | [illegible] | 1216 | Bacteria>Pseudomonadota>Gammaproteobacteria>Pseudomonadales>Pseudomonadaceae>Pseudomonas | GCF_009763245.1 | Pseudomonas_xionganensis | GJV18_RS17875 | 24794..28444 | 2654845 | BREX-system-P-loop-protein-BrxC-[Pseudomonas-xionganensis] |
| WP_166590589.1 | Type1.BREX | [illegible] | 1223 | Bacteria>Pseudomonadota>Gammaproteobacteria>Pseudomonadales>Pseudomonadaceae>Pseudomonas | GCF_009905775.1 | Pseudomonas_sp._BC115LW | GVN25_RS11920 | 39380..43051 | 2683267 | BREX-system-P-loop-protein-BrxC-[Pseudomonas-sp.-BC115LW] |
| WP_161892243.1 | Type1.BREX | [illegible] | 1154 | Bacteria>Pseudomonadota>Gammaproteobacteria>Pseudomonadales>Pseudomonadaceae>Pseudomonas | GCF_009932375.1 | Pseudomonas_juntendi | GYA94_RS10075 | 73827..77291 | 2666183 | BREX-system-P-loop-protein-BrxC-[Pseudomonas-juntendi] |
| WP_262377456.1 | Type1.BREX | [illegible] | 787 | Bacteria>Pseudomonadota>Gammaproteobacteria>Pseudomonadales>Pseudomonadaceae>Pseudomonas | GCF_012985365.1 | Pseudomonas_veronii | HBO38_RS22345 | 78028..80391 | 76761 | BREX-system-P-loop-protein-BrxC-[Pseudomonas-veronii] |
| WP_262386226.1 | Type1.BREX | [illegible] | 1225 | Bacteria>Pseudomonadota>Gammaproteobacteria>Pseudomonadales>Pseudomonadaceae>Pseudomonas | GCF_012985535.1 | Pseudomonas_sp._WS_5021 | HBO19_RS03675 | 794081..797758 | 2717490 | BREX-system-P-loop-protein-BrxC-[Pseudomonas-sp.-WS-5021] |
| WP_169989748.1 | Type1.BREX | [illegible] | 1186 | Bacteria>Pseudomonadota>Gammaproteobacteria>Pseudomonadales>Pseudomonadaceae>Pseudomonas | GCF_012985575.1 | Pseudomonas_sp._WS_5413 | HBO21_RS11490 | 97465..101025 | 2717488 | BREX-system-P-loop-protein-BrxC-[Pseudomonas-sp.-WS-5413] |
| WP_170050473.1 | Type1.BREX | [illegible] | 1154 | Bacteria>Pseudomonadota>Gammaproteobacteria>Pseudomonadales>Pseudomonadaceae>Pseudomonas | GCF_012985775.1 | Pseudomonas_sp._WS_5011 | HBN63_RS18875 | 35814..39278 | 2717477 | BREX-system-P-loop-protein-BrxC-[Pseudomonas-sp.-WS-5011] |
| WP_262499066.1 | Type1.BREX | [illegible] | 1217 | Bacteria>Pseudomonadota>Gammaproteobacteria>Pseudomonadales>Pseudomonadaceae>Pseudomonas | GCF_012985775.1 | Pseudomonas_sp._WS_5011 | HBN63_RS20190 | 4647..8300 | 2717477 | BREX-system-P-loop-protein-BrxC-[Pseudomonas-sp.-WS-5011] |
| WP_094987826.1 | Type1.BREX | [illegible] | 1154 | Bacteria>Pseudomonadota>Gammaproteobacteria>Pseudomonadales>Pseudomonadaceae>Pseudomonas | GCF_012986325.1 | Pseudomonas_lundensis | HBN80_RS14475 | 101799..105263 | 86185 | BREX-system-P-loop-protein-BrxC-[Pseudomonas-lundensis] |
| WP_169839752.1 | Type1.BREX | [illegible] | 1154 | Bacteria>Pseudomonadota>Gammaproteobacteria>Pseudomonadales>Pseudomonadaceae>Pseudomonas | GCF_012986505.1 | Pseudomonas_lundensis | HBN72_RS14625 | 129337..132801 | 86185 | BREX-system-P-loop-protein-BrxC-[Pseudomonas-lundensis] |
| WP_223184021.1 | Type1.BREX | [illegible] | 122 | Bacteria>Pseudomonadota>Gammaproteobacteria>Pseudomonadales>Pseudomonadaceae>Pseudomonas | GCF_012986825.1 | Pseudomonas_fragi | HBN89_RS04420 | 39782..40150 | 296 | hypothetical-protein-[Pseudomonas-fragi] |
| WP_172792831.1 | Type1.BREX | [illegible] | 1154 | Bacteria>Pseudomonadota>Gammaproteobacteria>Pseudomonadales>Pseudomonadaceae>Pseudomonas | GCF_013260445.1 | Pseudomonas_aeruginosa | HQN50_RS11915 | 2493948..2497412 | 287 | BREX-system-P-loop-protein-BrxC-[Pseudomonas-aeruginosa] |
| WP_173406509.1 | Type1.BREX | [illegible] | 1154 | Bacteria>Pseudomonadota>Gammaproteobacteria>Pseudomonadales>Pseudomonadaceae>Pseudomonas | GCF_013283895.1 | Pseudomonas_sp._MPDS | HQ912_RS04605 | 1048127..1051591 | 2762896 | BREX-system-P-loop-protein-BrxC-[Pseudomonas-sp.-MPDS] |
| WP_177031785.1 | Type1.BREX | [illegible] | 1185 | Bacteria>Pseudomonadota>Gammaproteobacteria>Pseudomonadales>Pseudomonadaceae>Pseudomonas | GCF_013386345.1 | Pseudomonas_veronii | HX878_RS08640 | 6154..9711 | 76761 | BREX-system-P-loop-protein-BrxC-[Pseudomonas-veronii] |

| Protein-ID | System | Gene-Neighbourhoods | Length | Taxonomy | Assembly-ID | Strain | Locus-Tag | Genomic-coordinates | TaxID | Fasta-define |
|---|---|---|---|---|---|---|---|---|---|---|
| WP_178910878.1 | Type1.BREX | <-SIGMA+OB-Fold+AAA-ATPase+HKD-DNase-BrxL<-PglZ<-N6-METHYLASE-p<-RVT<-NERD<-OLD-ABC<-N6-METHYLASE<-DUF6079*<-BrxB-BrxF-iSTAND<-BrxA-DNA-binding<-XIS-HTH<-?\|\|?->?->\|\|Aminotran_1_2-> | 1186 | Bacteria>Pseudomonadota>Gammaproteobacteria>Pseudomonadales>Pseudomonadaceae>Pseudomonas | GCF_013386495.1 | Pseudomonas_tolaasii | HX799_RS03165 | 122791..126351 | 29442 | BREX-system-P-loop-protein-BrxC-[Pseudomonas-tolaasii] |
| WP_244972677.1 | Type1.BREX | <-SIGMA+OB-Fold+AAA-ATPase+Lon-Protease-BrxL<-PglZ<-AbiEii<-AbiEi_3_N+AbiEi_3<-HNH<-N6-METHYLASE<-DNA-glycosylase<-DUF6079*<-BrxB-BrxF-iSTAND<-BrxA-DNA-binding<-?\|\|wHTH+WYL+WCX-><-tRNA<-MerR-HTH | 1225 | Bacteria>Pseudomonadota>Gammaproteobacteria>Pseudomonadales>Pseudomonadaceae>Pseudomonas | GCF_013386885.1 | Pseudomonas_salomonii | HX810_RS11525 | 509017..512694 | 191391 | BREX-system-P-loop-protein-BrxC-[Pseudomonas-salomonii] |
| WP_177095197.1 | Type1.BREX | <-N6-METHYLASE-p<-REase<-OLD-ABC<-N6-METHYLASE<-DUF6079*<-BrxB-BrxF-iSTAND<-BrxA-DNA-binding<-XIS-HTH<-ARG<-?\|\|wHTH+WYL+WCX-><-sGTP<-sGTP | 1185 | Bacteria>Pseudomonadota>Gammaproteobacteria>Pseudomonadales>Pseudomonadaceae>Pseudomonas | GCF_013387065.1 | Pseudomonas_reactans | HX796_RS11550 | 4461..8018 | 117680 | BREX-system-P-loop-protein-BrxC-[Pseudomonas-reactans] |
| WP_180689220.1 | Type1.BREX | ABC-ATPase+SbcC->?->?->?->?->DUF772+DDE_Tnp_1->XIS-HTH->BrxA-DNA-binding->BrxB-BrxF-iSTAND->DUF6079*->N6-METHYLASE->ParB+HNH->PglZ->SIGMA+OB-Fold+AAA-ATPase+HKD-DNase-BrxL-><-Phage_integrase<-Integrase_1<-wHTH<-AAA+wHTH<-Arm-DNA-bind_3+Phage_integrase | 1185 | Bacteria>Pseudomonadota>Gammaproteobacteria>Pseudomonadales>Pseudomonadaceae>Pseudomonas | GCF_013425825.1 | Pseudomonas_putida | HOH12_RS03375 | 759269..762826 | 303 | BREX-system-P-loop-protein-BrxC-[Pseudomonas-putida] |
| WP_182367101.1 | Type1.BREX | Arm-DNA-bind_3+Phage_integrase->GreA_GreB+SF1-SF2-Helicase-><-SIGMA+OB-Fold+AAA-ATPase+Lon-Protease-BrxL<-PglZ<-ParB+HNH<-?<-MarR-HTH<-MNS+ORC-CDC6<-N6-METHYLASE<-DUF6079*<-BrxB-BrxF-iSTAND<-BrxA-DNA-binding<-Ntox49<-?<-Peptidase_M15<-?\|\|wHTH+WYL+WCX-><-?<-RelE-ParE | 1154 | Bacteria>Pseudomonadota>Gammaproteobacteria>Pseudomonadales>Pseudomonadaceae>Pseudomonas | GCF_014109765.1 | Pseudomonas_sp._MSPm1 | HS968_RS15945 | 3439076..3442540 | 2726956 | BREX-system-P-loop-protein-BrxC-[Pseudomonas-sp.-MSPm1] |
| WP_192985630.1 | Type1.BREX | HTH_19-><-SIGMA+OB-Fold+AAA-ATPase+Lon-Protease-BrxL<-PglZ<-AbiEii<-AbiEi_4<-?<-N6-METHYLASE<-DNA-glycosylase<-ParB+HNH<-DUF6079*<-BrxB-BrxF-iSTAND<-BrxA-DNA-binding | 1220 | Bacteria>Pseudomonadota>Gammaproteobacteria>Pseudomonadales>Pseudomonadaceae>Pseudomonas | GCF_014892995.1 | Pseudomonas_sp._EggHat1 | INS93_RS22765 | 65820..69482 | 2761624 | BREX-system-P-loop-protein-BrxC-[Pseudomonas-sp.-EggHat1] |
| WP_230091331.1 | Type1.BREX | <-SIGMA+OB-Fold+AAA-ATPase+Lon-Protease-BrxL<-PglZ<-AbiEii<-AbiEi_4<-?<-?<-N6-METHYLASE<-DNA-glycosylase<-ParB+HNH<-DUF6079*<-BrxB-BrxF-iSTAND<-BrxA-DNA-binding | 1215 | Bacteria>Pseudomonadota>Gammaproteobacteria>Pseudomonadales>Pseudomonadaceae>Pseudomonas | GCF_015666575.1 | Pseudomonas_aeruginosa | I1957_RS22560 | 16962..20609 | 287 | BREX-system-P-loop-protein-BrxC-[Pseudomonas-aeruginosa] |
| WP_196169752.1 | Type1.BREX | DUF4969-><-SIGMA+OB-Fold+AAA-ATPase+Lon-Protease-BrxL<-PglZ<-HNH<-OLD-ABC<-DUF932<-N6-METHYLASE<-DUF6079*<-BrxB-BrxF-iSTAND<-BrxA-DNA-binding<-wHTH+WYL+WCX<-?<-XIS-HTH\|\|CopG/RHH-><-MODE-HTH+PBPII | 1154 | Bacteria>Pseudomonadota>Gammaproteobacteria>Pseudomonadales>Pseudomonadaceae>Pseudomonas | GCF_015680295.1 | Pseudomonas_guariconensis | IRZ59_RS14965 | 185250..188714 | 1288410 | BREX-system-P-loop-protein-BrxC-[Pseudomonas-guariconensis] |
| WP_196476665.1 | Type1.BREX | PhdYeFM_antitox->RelE-ParE-><-?<-wHTH+WYL+WCX\|\|?->Peptidase_M15->?->BrxA-DNA-binding->BrxB-BrxF-iSTAND->DUF6079*->N6-METHYLASE->?->HNH+GCN1->?->ParB+HNH->PglZ->SIGMA+OB-Fold+AAA-ATPase+Lon-Protease-BrxL->REase->SF1-SF2-Helicase-> | 1154 | Bacteria>Pseudomonadota>Gammaproteobacteria>Pseudomonadales>Pseudomonadaceae>Pseudomonas | GCF_015712105.2 | Pseudomonas_sp._LMG_31766 | H3221_RS24535 | 30166..33630 | 2758695 | BREX-system-P-loop-protein-BrxC-[Pseudomonas-chaetocerotis] |
| WP_198726217.1 | Type1.BREX | <-TetR-HTH<-wHTH+WYL+WCX\|\|?->ARG->BrxA-DNA-binding->BrxB-BrxF-iSTAND->DUF6079*->N6-METHYLASE->MNS+ORC-CDC6->?->ParB+HNH->PglZ-><-Phage_integrase<-Integrase_1<-wHTH<-AAA+wHTH | 1154 | Bacteria>Pseudomonadota>Gammaproteobacteria>Pseudomonadales>Pseudomonadaceae>Pseudomonas | GCF_016307475.1 | Pseudomonas_corrugata | YA0788_RS19970 | 31466..34930 | 47879 | BREX-system-P-loop-protein-BrxC-[Pseudomonas-corrugata] |
| WP_198790924.1 | Type1.BREX | <-CopG/RHH\|\|XIS-HTH->?->?->XIS-HTH->BrxA-DNA-binding->BrxB-BrxF-iSTAND->DUF6079*->N6-METHYLASE-p->OLD-ABC->REase->?->N6-METHYLASE-p->ParB+HNH->PglZ->SIGMA+OB-Fold+AAA-ATPase+HKD-DNase-BrxL-> | 1185 | Bacteria>Pseudomonadota>Gammaproteobacteria>Pseudomonadales>Pseudomonadaceae>Pseudomonas | GCF_016405175.1 | Pseudomonas_sp._MF7451 | JFT65_RS10900 | 194530..198087 | 2797538 | BREX-system-P-loop-protein-BrxC-[Pseudomonas-sp.-MF7451] |
| WP_198832884.1 | Type1.BREX | <-SIGMA+OB-Fold+AAA-ATPase+HKD-DNase-BrxL<-PglZ<-?<-N6-METHYLASE<-DUF6079*<-BrxB-BrxF-iSTAND<-BrxA-DNA-binding<-XIS-HTH<-ARG<-?\|\|wHTH+WYL+WCX-><-sGTP | 1185 | Bacteria>Pseudomonadota>Gammaproteobacteria>Pseudomonadales>Pseudomonadaceae>Pseudomonas | GCF_016405445.1 | Pseudomonas_haemolytica | JFT42_RS28230 | 10084..13641 | 2600065 | BREX-system-P-loop-protein-BrxC-[Pseudomonas-haemolytica] |
| WP_198849248.1 | Type1.BREX | wHTH+WYL+WCX-><-SIGMA+OB-Fold+AAA-ATPase+HKD-DNase-BrxL<-HNH<-OLD-ABC<-N6-METHYLASE-p<-HerA/FtsK<-SIR2<-N6-METHYLASE-p<-DUF6079*<-BrxB-BrxF-iSTAND<-BrxA-DNA-binding<-XIS-HTH<-?<-tRNA<-MerR-HTH | 1186 | Bacteria>Pseudomonadota>Gammaproteobacteria>Pseudomonadales>Pseudomonadaceae>Pseudomonas | GCF_016405505.1 | Pseudomonas_sp._MF6772 | JFT60_RS22275 | 24346..27906 | 2797533 | BREX-system-P-loop-protein-BrxC-[Pseudomonas-sp.-MF6772] |
| WP_203327703.1 | Type1.BREX | XIS-HTH->?->wHTH+WYL+WCX->BrxA-DNA-binding->BrxB-BrxF-iSTAND->DUF6079*->N6-METHYLASE->ParB+HNH->PglZ->SIGMA+OB-Fold+AAA-ATPase+Lon-Protease-BrxL-><-DUF4969<-?<-?<-DUF932 | 1157 | Bacteria>Pseudomonadota>Gammaproteobacteria>Pseudomonadales>Pseudomonadaceae>Pseudomonas | GCF_016812215.1 | Pseudomonas_fragi | GYM96_RS18895 | 91257..94730 | 296 | BREX-system-P-loop-protein-BrxC-[Pseudomonas-fragi] |
| WP_108110611.1 | Type1.BREX | <-HIN-HTH<-REase<-?<-wHTH+WYL+WCX<-?\|\|Peptidase_M15->?->BrxA-DNA-binding->BrxB-BrxF-iSTAND->DUF6079*->N6-METHYLASE->MNS+ORC-CDC6->MarR-HTH->?->ParB+HNH->PglZ->SIGMA+OB-Fold+AAA-ATPase+Lon-Protease-BrxL-><-GreA_GreB+SF1-SF2-Helicase<-HTH_Tnp_1+HTH_21+rve+rve_3 | 1154 | Bacteria>Pseudomonadota>Gammaproteobacteria>Pseudomonadales>Pseudomonadaceae>Pseudomonas | GCF_016820125.1 | Pseudomonas_aeruginosa | JQS52_RS11310 | 2388468..2391932 | 287 | BREX-system-P-loop-protein-BrxC-[Pseudomonas-aeruginosa] |
| WP_239000709.1 | Type1.BREX | <-ABC-ATPase+SbcC<-?<-?<-MODE-HTH+PBPII\|\|<-?<-N6-METHYLASE<-DOC+HTH<-?<-?<-DUF6079*<-BrxB-BrxF-iSTAND<-BrxA-DNA-binding<-wHTH+WYL+WCX\|\|OLD-ABC+TOPRIM->SF1-SF2-Helicase->?->SF1-SF2-Helicase-><-Phage_integrase | 1225 | Bacteria>Pseudomonadota>Gammaproteobacteria>Pseudomonadales>Pseudomonadaceae>Pseudomonas | GCF_016836415.1 | Pseudomonas_aeruginosa | JQ482_RS07395 | 1563823..1567500 | 287 | BREX-system-P-loop-protein-BrxC-[Pseudomonas-aeruginosa] |
| WP_239068292.1 | Type1.BREX | Phage_integrase->Phage_integrase->?->Phage_integrase-><-SF1-SF2-Helicase<-?\|\|wHTH+WYL+WCX->BrxA-DNA-binding->BrxB-BrxF-iSTAND->DUF6079*->?->?-><-?<-AAA-ATPase<-HTH+rve\|\|DOC+HTH->N6-METHYLASE->RVT->N6-METHYLASE-p-> | 1225 | Bacteria>Pseudomonadota>Gammaproteobacteria>Pseudomonadales>Pseudomonadaceae>Pseudomonas | GCF_016861485.1 | Pseudomonas_alcaligenes | JUH08_RS19055 | 3949819..3953496 | 43263 | BREX-system-P-loop-protein-BrxC-[Pseudomonas-alcaligenes] |
| WP_203542487.1 | Type1.BREX | RHH_5->SbcC->Schlafen+HATPase+wHTH-><-?\|\|NERD->wHTH+WYL+WCX->BrxA-DNA-binding->BrxB-BrxF-iSTAND->DUF6079*->N6-METHYLASE->ParB+HNH->PglZ->SIGMA+OB-Fold+AAA-ATPase+Lon-Protease-BrxL-> | 1146 | Bacteria>Pseudomonadota>Gammaproteobacteria>Pseudomonadales>Pseudomonadaceae>Pseudomonas | GCF_016865555.1 | Pseudomonas_arcuscaelestis | IIE18_RS12195 | 346577..350017 | 2710591 | BREX-system-P-loop-protein-BrxC-[Pseudomonas-arcuscaelestis] |
| WP_240344493.1 | Type1.BREX | <-N6-METHYLASE-p<-DNA-glycosylase<-DUF6079*<-BrxB-BrxF-iSTAND<-BrxA-DNA-binding\|\|<-?<-GTPase-AIG<-GTPase-AIG | 1225 | Bacteria>Pseudomonadota>Gammaproteobacteria>Pseudomonadales>Pseudomonadaceae>Pseudomonas | GCF_016924785.1 | Pseudomonas_sp._SDM007_2 | JV174_RS20995 | 2384..6061 | 2812898 | BREX-system-P-loop-protein-BrxC-[Pseudomonas-sp.-SDM007_2] |
| WP_205518899.1 | Type1.BREX | <-PglZ<-AbiEii<-AbiEi_3_N+AbiEi_3<-HNH<-ParB+HNH<-N6-METHYLASE\|\|HIN-HTH+PAIREDC-HTH->?-><-N6-METHYLASE-p<-DNA-glycosylase<-DUF6079*<-DUF6079-p<-BrxB-BrxF-iSTAND<-BrxA-DNA-binding\|\|<-?<-GTPase-AIG<-GTPase-AIG | 1107 | Bacteria>Pseudomonadota>Gammaproteobacteria>Pseudomonadales>Pseudomonadaceae>Pseudomonas | GCF_016925675.1 | Pseudomonas_sp._SDM007 | JTY93_RS19480 | 4366691..4370014 | 2812000 | BREX-system-P-loop-protein-BrxC-[Pseudomonas-sp.-SDM007] |
| WP_206408616.1 | Type1.BREX | ARG->?->wHTH+WYL+WCX->BrxA-DNA-binding->BrxB-BrxF-iSTAND->DUF6079*->N6-METHYLASE->OLD-ABC->HNH->ParB+HNH->OLD-ABC->TOPRIM->PglZ->SIGMA+OB-Fold+AAA-ATPase+Lon-Protease-BrxL->MukF-> | 1154 | Bacteria>Pseudomonadota>Gammaproteobacteria>Pseudomonadales>Pseudomonadaceae>Pseudomonas | GCF_017168035.1 | Pseudomonas_oleovorans | ACP89_RS12320 | 3896..7360 | 301 | BREX-system-P-loop-protein-BrxC-[Pseudomonas-oleovorans] |
| WP_219730400.1 | Type1.BREX | METHYLASE->BrxA-DNA-binding->BrxB-BrxF-iSTAND->DUF6079*->ParB+HNH->DNA-glycosylase->N6-METHYLASE->\|\|PglZ->SIGMA+OB-Fold+AAA-ATPase+Lon-Protease-BrxL-><-Phage_integrase\|\|DUF4096+DDE_Tnp_1+DDE_Tnp_1_2-><-wHTH<-AAA+wHTH | 1225 | Bacteria>Pseudomonadota>Gammaproteobacteria>Pseudomonadales>Pseudomonadaceae>Pseudomonas | GCF_017350895.1 | unclassified_Pseudomonas | J4E12_RS18510 | 3053..6730 | 196821 | MULTISPECIES:-BREX-system-P-loop-protein-BrxC-[unclassified-Pseudomonas] |
| WP_210224970.1 | Type1.BREX | SF1-SF2-Helicase-><-HTH_3\|\|?->CHTH-><-?<-REase+NACHT\|\|wHTH+WYL+WCX->BrxA-DNA-binding->BrxB-BrxF-iSTAND->DUF6079*->N6-METHYLASE->SIR2->HerA/FtsK->?->N6-METHYLASE-p->sGTP->?->PglZ->SIGMA+OB-Fold+AAA-ATPase+Lon-Protease-BrxL-> | 1225 | Bacteria>Pseudomonadota>Gammaproteobacteria>Pseudomonadales>Pseudomonadaceae>Pseudomonas | GCF_017900915.1 | Pseudomonas_aeruginosa | J9247_RS05115 | 1084735..1088412 | 287 | BREX-system-P-loop-protein-BrxC-[Pseudomonas-aeruginosa] |
| WP_125970731.1 | Type1.BREX | DUF4969-><-SLOG+BirA-HTH<-SF1-SF2-Helicase+PRTase<-SIGMA+OB-Fold+AAA-ATPase+Lon-Protease-BrxL<-PglZ<-ParB+HNH<-DUF932<-N6-METHYLASE<-DUF6079*<-BrxB-BrxF-iSTAND<-BrxA-DNA-binding<-wHTH+WYL+WCX<-?<-XIS-HTH<-BirA-HTH\|\|CopG/RHH-> | 1154 | Bacteria>Pseudomonadota>Gammaproteobacteria>Pseudomonadales>Pseudomonadaceae>Pseudomonas | GCF_018336155.1 | Pseudomonas_fluorescens_group | EJJ21_RS24495 | 5255523..5258987 | 136843 | MULTISPECIES:-BREX-system-P-loop-protein-BrxC-[Pseudomonas-fluorescens-group] |
| WP_213336039.1 | Type1.BREX | DUF4969-><-SIGMA+OB-Fold+AAA-ATPase+Lon-Protease-BrxL<-PglZ<-TOPRIM<-OLD-ABC<-ParB+HNH<-N6-METHYLASE<-DUF6079*<-BrxB-BrxF-iSTAND<-BrxA-DNA-binding<-wHTH+WYL+WCX<-?<-XIS-HTH<-BirA-HTH\|\|CopG/RHH-><-Pkinase+CR-Kinase | 1157 | Bacteria>Pseudomonadota>Gammaproteobacteria>Pseudomonadales>Pseudomonadaceae>Pseudomonas | GCF_018449005.1 | Pseudomonas_lundensis | KIY13_RS15530 | 3400509..3403982 | 86185 | BREX-system-P-loop-protein-BrxC-[Pseudomonas-lundensis] |
| WP_198712541.1 | Type1.BREX | <-MORC+REase<-?<-SF1-SF2-Helicase+REase<-?\|\|?->wHTH+WYL+WCX->BrxA-DNA-binding->BrxB-BrxF-iSTAND->DUF6079*->N6-METHYLASE->NERD+GspA-ATPase->TPR->PglZ->SIGMA+OB-Fold+AAA-ATPase+Lon-Protease-BrxL-><-JetC<-?<-HNH<-tRNA<-MerR-HTH | 1217 | Bacteria>Pseudomonadota>Gammaproteobacteria>Pseudomonadales>Pseudomonadaceae>Pseudomonas | GCF_019464415.1 | Pseudomonas_sp._So3.2b | K1X80_RS19120 | 4293365..4297018 | 2864101 | BREX-system-P-loop-protein-BrxC-[Pseudomonas-sp.-So3.2b] |
| WP_222018162.1 | Type1.BREX | N6-METHYLASE->?-><-SIGMA+OB-Fold+AAA-ATPase+Lon-Protease-BrxL<-PglZ\|\|?-><-N6-METHYLASE<-TOPRIM<-OLD-ABC<-DUF6079*<-BrxB-BrxF-iSTAND<-BrxA-DNA-binding<-wHTH+WYL+WCX<-?<-?<-?<-DUF932 | 1223 | Bacteria>Pseudomonadota>Gammaproteobacteria>Pseudomonadales>Pseudomonadaceae>Pseudomonas | GCF_019775695.1 | Pseudomonas_sp._AO-1 | KVQ82_RS04465 | 959176..962847 | 2855434 | BREX-system-P-loop-protein-BrxC-[Pseudomonas-sp.-AO-1] |
| WP_223023201.1 | Type1.BREX | <-REase\|\|DNA_METHYLASE-><-CopG/RHH\|\|XIS-HTH->?->XIS-HTH->BrxA-DNA-binding->BrxB-BrxF-iSTAND->DUF6079*->OLD-ABC->REase->N6-METHYLASE->PglZ->SIGMA+OB-Fold+AAA-ATPase+HKD-DNase-BrxL-><-?<-?<-?\|\|HTH_3-> | 1183 | Bacteria>Pseudomonadota>Gammaproteobacteria>Pseudomonadales>Pseudomonadaceae>Pseudomonas | GCF_019891425.1 | Pseudomonas_carnis | J1G36_06600 | 89435..92986 | 2487355 | BREX-system-P-loop-protein-BrxC-[Pseudomonas-carnis] |
| WP_262416944.1 | Type1.BREX | <-SIGMA+OB-Fold+AAA-ATPase+Lon-Protease-BrxL<-PglZ<-AbiEii<-AbiEi_3_N+AbiEi_3<-ParB+HNH<-N6-METHYLASE<-DNA-glycosylase<-DUF6079*<-BrxB-BrxF-iSTAND<-BrxA-DNA-binding<-?\|\|wHTH+WYL+WCX-><-tRNA<-MerR-HTH | 1225 | Bacteria>Pseudomonadota>Gammaproteobacteria>Pseudomonadales>Pseudomonadaceae>Pseudomonas | GCF_019891455.1 | Pseudomonas_sp._P867 | J1G33_RS25105 | 13708..17385 | 2816050 | BREX-system-P-loop-protein-BrxC-[Pseudomonas-sp.-P867] |
| WP_231661141.1 | Type1.BREX | <-RHH\|\|XIS-HTH->?->wHTH+WYL+WCX->BrxA-DNA-binding->BrxB-BrxF-iSTAND->DUF6079*->N6-METHYLASE->ParB+HNH->PglZ->SIGMA+OB-Fold+AAA-ATPase+Lon-Protease-BrxL-><-DUF4969<-?<-?\|\|ABhydrolase-><-RNase | 1156 | Bacteria>Pseudomonadota>Gammaproteobacteria>Pseudomonadales>Pseudomonadaceae>Pseudomonas | GCF_021147865.1 | Pseudomonas_sp._CDFA_610 | KDX39_RS16865 | 240745..244215 | 2829825 | BREX-system-P-loop-protein-BrxC-[Pseudomonas-sp.-CDFA-610] |
| WP_196706085.1 | Type1.BREX | <-SIGMA+OB-Fold+AAA-ATPase+HKD-DNase-BrxL<-PglZ<-N6-METHYLASE-p<-HerA/FtsK<-SIR2<-REase<-OLD-ABC<-N6-METHYLASE<-DUF6079*<-BrxB-BrxF-iSTAND<-BrxA-DNA-binding<-XIS-HTH<-?\|\|<-?\|\|sGTP->sGTP->MA+PspA+PspA+sGTP-><-ARG | 1184 | Bacteria>Pseudomonadota>Gammaproteobacteria>Pseudomonadales>Pseudomonadaceae>Pseudomonas | GCF_021460775.1 | Pseudomonas_putida | LH705_RS23300 | 5093270..5096824 | 303 | BREX-system-P-loop-protein-BrxC-[Pseudomonas-putida] |
| WP_236166438.1 | Type1.BREX | DUF4969-><-SIGMA+OB-Fold+AAA-ATPase+Lon-Protease-BrxL<-PglZ<-HNH<-OLD-ABC<-ParB+HNH<-N6-METHYLASE<-DUF6079*<-BrxB-BrxF-iSTAND<-BrxA-DNA-binding<-wHTH+WYL+WCX<-?<-XIS-HTH | 1154 | Bacteria>Pseudomonadota>Gammaproteobacteria>Pseudomonadales>Pseudomonadaceae>Pseudomonas | GCF_021601865.1 | Pseudomonas_juntendi | L6D32_RS09730 | 148894..152358 | 2666183 | BREX-system-P-loop-protein-BrxC-[Pseudomonas-juntendi] |
| WP_236238652.1 | Type1.BREX | <-HIN-HTH<-SIGMA+OB-Fold+AAA-ATPase+HKD-DNase-BrxL<-PglZ<-ParB+HNH<-N6-METHYLASE-p<-REase<-OLD-ABC<-N6-METHYLASE-p<-DUF6079*<-BrxB-BrxF-iSTAND<-BrxA-DNA-binding<-XIS-HTH | 1185 | Bacteria>Pseudomonadota>Gammaproteobacteria>Pseudomonadales>Pseudomonadaceae>Pseudomonas | GCF_021603345.1 | Pseudomonas_rhodesiae | L6E14_RS13730 | 266364..269921 | 76760 | BREX-system-P-loop-protein-BrxC-[Pseudomonas-rhodesiae] |
| WP_236310112.1 | Type1.BREX | <-HTH+rve<-?\|\|XIS-HTH->?->wHTH+WYL+WCX->BrxA-DNA-binding->BrxB-BrxF-iSTAND->DUF6079*->N6-METHYLASE->DUF932->ParB+HNH->OLD-ABC->TOPRIM->PglZ->SIGMA+OB-Fold+AAA-ATPase+Lon-Protease-BrxL-><-DUF4969 | 1154 | Bacteria>Pseudomonadota>Gammaproteobacteria>Pseudomonadales>Pseudomonadaceae>Pseudomonas | GCF_021605645.1 | Pseudomonas_gessardii | GIW56_RS19725 | 10406..13870 | 78544 | BREX-system-P-loop-protein-BrxC-[Pseudomonas-gessardii] |
| WP_236471270.1 | Type1.BREX | XIS-HTH->?-><-REase+ZnR\|\|NERD->NERD->wHTH+WYL+WCX->BrxA-DNA-binding->BrxB-BrxF-iSTAND->DUF6079*->N6-METHYLASE->ParB+HNH->PglZ->SIGMA+OB-Fold+AAA-ATPase+Lon-Protease-BrxL-><-DUF4969<-?<-?<-?<-DUF932 | 1156 | Bacteria>Pseudomonadota>Gammaproteobacteria>Pseudomonadales>Pseudomonadaceae>Pseudomonas | GCF_021606115.1 | Pseudomonas_syringae | GIW32_RS04305 | 7736..11206 | 317 | BREX-system-P-loop-protein-BrxC-[Pseudomonas-syringae] |
| WP_243242998.1 | Type1.BREX | Amidohydro_2->ABhydrolase->AraC-HTH->?->ABC-ATPase+SbcC->?->?->BrxA-DNA-binding->BrxB-BrxF-iSTAND->DUF6079*->N6-METHYLASE->MNS+ORC-CDC6->MarR-HTH->?->ParB+HNH->PglZ->SIGMA+OB-Fold+AAA-ATPase+Lon-Protease-BrxL-><-Integrase_1<-Arm-DNA-bind_3+Phage_integrase | 1154 | Bacteria>Pseudomonadota>Gammaproteobacteria>Pseudomonadales>Pseudomonadaceae>Pseudomonas | GCF_022802935.1 | Pseudomonas_atacamensis | PSD_RS21520 | 46330..49794 | 2565368 | BREX-system-P-loop-protein-BrxC-[Pseudomonas-atacamensis] |
| WP_253132516.1 | Type1.BREX | XIS-HTH->?-><-?<-LexA-protease<-wHTH+WYL+WCX\|\|?->XIS-HTH->BrxA-DNA-binding->BrxB-BrxF-iSTAND->DUF6079*->N6-METHYLASE->OLD-ABC->REase->SIR2->HerA/FtsK->?->N6-METHYLASE-p->ParB+HNH->PglZ-> | 1184 | Bacteria>Pseudomonadota>Gammaproteobacteria>Pseudomonadales>Pseudomonadaceae>Pseudomonas | GCF_024128115.1 | Pseudomonas_guariconensis | NJC29_RS17685 | 128726..132280 | 1288410 | BREX-system-P-loop-protein-BrxC-[Pseudomonas-guariconensis] |
| WP_043087434.1 | Type1.BREX | <-HIN-HTH<-wHTH+WYL+WCX<-?\|\|?->Peptidase_M15->?->BrxA-DNA-binding->BrxB-BrxF-iSTAND->DUF6079*->N6-METHYLASE->MNS+ORC-CDC6->MarR-HTH->?->ParB+HNH->PglZ->SIGMA+OB-Fold+AAA-ATPase+Lon-Protease-BrxL-><-GreA_GreB+SF1-SF2-Helicase<-HTH_Tnp_1+HTH_21+rve+rve_3 | 1154 | Bacteria>Pseudomonadota>Gammaproteobacteria>Pseudomonadales>Pseudomonadaceae>Pseudomonas | GCF_024300845.1 | Pseudomonas_aeruginosa | MUG61_RS11500 | 2420957..2424421 | 287 | BREX-system-P-loop-protein-BrxC-[Pseudomonas-aeruginosa] |
| WP_261941654.1 | Type1.BREX | DUF932->?->?->?->DUF4969-><-SIGMA+OB-Fold+AAA-ATPase+Lon-Protease-BrxL<-PglZ<-ParB+HNH<-N6-METHYLASE<-DUF6079*<-BrxB-BrxF-iSTAND<-BrxA-DNA-binding<-wHTH+WYL+WCX<-?<-XIS-HTH\|\|RHH->?-><-?<-AraC-HTH+AraC-HTH | 1154 | Bacteria>Pseudomonadota>Gammaproteobacteria>Pseudomonadales>Pseudomonadaceae>Pseudomonas | GCF_024349685.1 | Pseudomonas_sp._LRP2-20 | OCX61_RS23895 | 5328463..5331927 | 2944234 | BREX-system-P-loop-protein-BrxC-[Pseudomonas-sp.-LRP2-20] |
| WP_010445526.1 | Type1.BREX | <-MULE-TRANSPOSASE<-?<-?\|\|HEPN-><-SIGMA+OB-Fold+AAA-ATPase+Lon-Protease-BrxL<-PglZ<-?<-sGTP<-N6-METHYLASE<-DUF6079*<-BrxB-BrxF-iSTAND<-BrxA-DNA-binding<-wHTH+WYL+WCX<-HTH+DOC<-PadR-HTH\|\|?-><-ABhydrolase | 1223 | Bacteria>Pseudomonadota>Gammaproteobacteria>Pseudomonadales>Pseudomonadaceae>Pseudomonas | GCF_024585345.2 | Pseudomonas_fuscovaginae | NLK61_RS17300 | 3830482..3834153 | 50340 | BREX-system-P-loop-protein-BrxC-[Pseudomonas-fuscovaginae] |
| WP_257780136.1 | Type1.BREX | <-LZ_Tnp_IS66+zf-IS66+DDE_Tnp_IS66+DDE_Tnp_IS66_C<-?<-HIN-HTH<-SIGMA+OB-Fold+AAA-ATPase+Lon-Protease-BrxL<-PglZ<-N6-METHYLASE<-DNA-glycosylase<-ParB+HNH<-DUF6079*<-BrxB-BrxF-iSTAND<-BrxA-DNA-binding<-?<-?<-?<-SSB<-?\|\|nSTAND1+SbcC+ABC-ATPase+SbcC-> | 1225 | Bacteria>Pseudomonadota>Gammaproteobacteria>Pseudomonadales>Pseudomonadaceae>Pseudomonas | GCF_024655625.1 | Pseudomonas_sp._18.1.10 | NUV89_RS02215 | 18654..22331 | 2969302 | BREX-system-P-loop-protein-BrxC-[Pseudomonas-sp.-18.1.10] |
| WP_258640408.1 | Type1.BREX | ABC-ATPase+SbcC->?->?->?<-RHH\|\|XIS-HTH->?->wHTH+WYL+WCX->BrxA-DNA-binding->BrxB-BrxF-iSTAND->DUF6079*->N6-METHYLASE-p->N6-METHYLASE->ParB+HNH->PglZ->SIGMA+OB-Fold+AAA-ATPase+Lon-Protease-BrxL-><-DUF4969<-?<-?\|\|ABhydrolase-> | 1154 | Bacteria>Pseudomonadota>Gammaproteobacteria>Pseudomonadales>Pseudomonadaceae>Pseudomonas | GCF_024748715.1 | Pseudomonas_moraviensis | LOY57_RS03675 | 870131..873595 | 321662 | BREX-system-P-loop-protein-BrxC-[Pseudomonas-moraviensis] |
| WP_258626742.1 | Type1.BREX | <-SIGMA+OB-Fold+AAA-ATPase+Lon-Protease-BrxL<-PglZ<-AbiEii<-AbiEi_3_N+AbiEi_3<-ParB+HNH<-N6-METHYLASE<-DNA-glycosylase<-DUF6079*<-BrxB-BrxF-iSTAND<-BrxA-DNA-binding<-ARG<-?\|\|wHTH+WYL+WCX-><-?<-GTPase-AIG<-GTPase-AIG | 1225 | Bacteria>Pseudomonadota>Gammaproteobacteria>Pseudomonadales>Pseudomonadaceae>Pseudomonas | GCF_024749045.1 | Pseudomonas_sp._B21-028 | LOY35_RS21325 | 4957302..4960979 | 2895480 | BREX-system-P-loop-protein-BrxC-[Pseudomonas-sp.-B21-028] |
| WP_259155464.1 | Type1.BREX | DUF932->?->?->?->DUF4969-><-SIGMA+OB-Fold+AAA-ATPase+Lon-Protease-BrxL<-PglZ<-ParB+HNH<-N6-METHYLASE<-DUF6079*<-BrxB-BrxF-iSTAND<-BrxA-DNA-binding<-wHTH+WYL+WCX<-?<-XIS-HTH\|\|?-><-Cysteine-Desulfurase | 1154 | Bacteria>Pseudomonadota>Gammaproteobacteria>Pseudomonadales>Pseudomonadaceae>Pseudomonas | GCF_024809425.1 | Pseudomonas_sp._JAI111 | HNR03_RS20420 | 285382..288846 | 2735913 | BREX-system-P-loop-protein-BrxC-[Pseudomonas-sp.-JAI111] |
| WP_260092499.1 | Type1.BREX | <-SF1-SF2-Helicase<-REase<-SIGMA+OB-Fold+AAA-ATPase+Lon-Protease-BrxL<-PglZ<-ParB+HNH<-?<-HNH+GCN1<-?<-N6-METHYLASE<-DUF6079*<-BrxB-BrxF-iSTAND<-BrxA-DNA-binding<-Ntox49<-?<-Peptidase_M15<-?\|\|wHTH+WYL+WCX->?-><-RelE-ParE | 1154 | Bacteria>Pseudomonadota>Gammaproteobacteria>Pseudomonadales>Pseudomonadaceae>Pseudomonas | GCF_025152895.1 | Pseudomonas_aeruginosa | MNU23_RS16440 | 110369..113833 | 287 | BREX-system-P-loop-protein-BrxC-[Pseudomonas-aeruginosa] |
| WP_261753794.1 | Type1.BREX | <-SIGMA+OB-Fold+AAA-ATPase+Lon-Protease-BrxL<-PglZ\|\|ZnR-><-TOPRIM<-OLD-ABC<-N6-METHYLASE<-DOC+HTH<-DUF6079*<-BrxB-BrxF-iSTAND<-BrxA-DNA-binding<-wHTH+WYL+WCX<-?<-Phage_AlpA<-?<-Arm-DNA-bind_3+Phage_integrase | 1225 | Bacteria>Pseudomonadota>Gammaproteobacteria>Pseudomonadales>Pseudomonadaceae>Pseudomonas | GCF_025397875.1 | Pseudomonas_veronii | MN113_RS13530 | 508981..512658 | 76761 | BREX-system-P-loop-protein-BrxC-[Pseudomonas-veronii] |
| WP_261744726.1 | Type1.BREX | HISKIN->REC+PAS+HISKIN+REC-><-?<-?<-SIGMA+OB-Fold+AAA-ATPase+Lon-Protease-BrxL<-PglZ<-?<-sGTP<-N6-METHYLASE<-DUF6079*<-BrxB-BrxF-iSTAND<-BrxA-DNA-binding<-XIS-HTH<-?\|\|<-?\|\|wHTH+WYL+WCX<-Phage_integrase<-Integrase_1<-wHTH<-AAA+wHTH | 1223 | Bacteria>Pseudomonadota>Gammaproteobacteria>Pseudomonadales>Pseudomonadaceae>Pseudomonas | GCF_025398015.1 | Pseudomonas_promysalinigenes | N5C08_RS03555 | 755414..759085 | 485898 | BREX-system-P-loop-protein-BrxC-[Pseudomonas-promysalinigenes] |
| WP_170059731.1 | Type1.BREX | <-SF1-SF2-Helicase+REase<-?<-SF1-SF2-Helicase<-HKD-DNase+HKD-DNase<-?\|\|wHTH+WYL+WCX->BrxA-DNA-binding->BrxB-BrxF-iSTAND->DUF6079*->?->N6-METHYLASE->NERD+GspA-ATPase->TPR->PglZ->SIGMA+OB-Fold+AAA-ATPase+Lon-Protease-BrxL->?-><-tRNA<-MerR-HTH | 1223 | Bacteria>Pseudomonadota>Gammaproteobacteria>Pseudomonadales>Pseudomonadaceae>Pseudomonas | GCF_025447475.1 | Pseudomonas_shahriarae | N8A88_RS04290 | 981426..985097 | 2745512 | BREX-system-P-loop-protein-BrxC-[Pseudomonas-shahriarae] |
| WP_023127678.1 | Type1.BREX | NUDIX-><-?<-wHTH+WYL+WCX\|\|ARG->?->wHTH+WYL+WCX->BrxA-DNA-binding->BrxB-BrxF-iSTAND->DUF6079*->N6-METHYLASE->DUF932->ParB+HNH->OLD-ABC->TOPRIM->PglZ->SIGMA+OB-Fold+AAA-ATPase+Lon-Protease-BrxL->MukF->JetB-> | 1154 | Bacteria>Pseudomonadota>Gammaproteobacteria>Pseudomonadales>Pseudomonadaceae>Pseudomonas | GCF_025696785.1 | Pseudomonas_aeruginosa | KUC66_RS22970 | 76351..79815 | 287 | BREX-system-P-loop-protein-BrxC-[Pseudomonas-aeruginosa] |
| WP_124171099.1 | Type1.BREX | RelE-ParE-><-?<-wHTH+WYL+WCX\|\|?->Peptidase_M15->?->BrxA-DNA-binding->BrxB-BrxF-iSTAND->DUF6079*->N6-METHYLASE->SbcC-p->REase->RVT->N6-METHYLASE-p->PglZ->SIGMA+OB-Fold+AAA-ATPase+HKD-DNase-BrxL-><-Arm-DNA-bind_3+Phage_integrase<-REase+SF1-SF2-Helicase | 1184 | Bacteria>Pseudomonadota>Gammaproteobacteria>Pseudomonadales>Pseudomonadaceae>Pseudomonas | GCF_025697325.1 | Pseudomonas_aeruginosa | IPC368_RS11155 | 506506..510060 | 287 | BREX-system-P-loop-protein-BrxC-[Pseudomonas-aeruginosa] |
| WP_265072431.1 | Type1.BREX | <-SIGMA+OB-Fold+AAA-ATPase+HKD-DNase-BrxL<-PglZ<-ParB+HNH<-N6-METHYLASE-p<-REase<-OLD-ABC<-N6-METHYLASE-p<-DUF6079*<-BrxB-BrxF-iSTAND<-BrxA-DNA-binding<-XIS-HTH\|\|HTH+REase-><-?<-?<-?<-Glyco_hydro_68 | 1186 | Bacteria>Pseudomonadota>Gammaproteobacteria>Pseudomonadales>Pseudomonadaceae>Pseudomonas | GCF_026016265.1 | Pseudomonas_viciae | LOY66_RS25465 | 5798638..5802198 | 2505979 | BREX-system-P-loop-protein-BrxC-[Pseudomonas-viciae] |
| WP_265535081.1 | Type1.BREX | <-SIGMA+OB-Fold+AAA-ATPase+HKD-DNase-BrxL<-PglZ<-N6-METHYLASE-p<-HerA/FtsK<-SIR2<-REase<-OLD-ABC<-N6-METHYLASE<-DUF6079*<-BrxB-BrxF-iSTAND<-BrxA-DNA-binding<-XIS-HTH<-?\|\|<-?\|\|sGTP->sGTP->MA+PspA+PspA+sGTP-><-ZnR | 1184 | Bacteria>Pseudomonadota>Gammaproteobacteria>Pseudomonadales>Pseudomonadaceae>Pseudomonas | GCF_026173695.1 | Pseudomonas_saponiphila | ORM81_RS30140 | 14715..18269 | 556534 | BREX-system-P-loop-protein-BrxC-[Pseudomonas-saponiphila] |
| WP_265980027.1 | Type1.BREX | Phage_integrase-><-SIGMA+OB-Fold+AAA-ATPase+HKD-DNase-BrxL<-PglZ<-ParB+HNH<-N6-METHYLASE-p<-?<-NERD<-OLD-ABC<-N6-METHYLASE<-DUF6079*<-BrxB-BrxF-iSTAND<-BrxA-DNA-binding<-XIS-HTH\|\|<-?<-?\|\|wHTH+WYL+WCX-><-JAB | 1184 | Bacteria>Pseudomonadota>Gammaproteobacteria>Pseudomonadales>Pseudomonadaceae>Pseudomonas | GCF_026241005.1 | Pseudomonas_sp._DCB_BG | OD254_RS23990 | 64105..67659 | 2993595 | BREX-system-P-loop-protein-BrxC-[Pseudomonas-sp.-DCB_BG] |
| WP_270945767.1 | Type1.BREX | MerR-HTH->\|\|?->?->DUF772+DDE_Tnp_1-><-?\|\|wHTH+WYL+WCX->BrxA-DNA-binding->BrxB-BrxF-iSTAND->DUF6079*->N6-METHYLASE->?->?->OLD-ABC->HNH->PglZ->SIGMA+OB-Fold+AAA-ATPase+Lon-Protease-BrxL->?->HTH_3-> | 1223 | Bacteria>Pseudomonadota>Gammaproteobacteria>Pseudomonadales>Pseudomonadaceae>Pseudomonas | GCF_027856115.1 | Pseudomonas_aeruginosa | KKQ36_RS16450 | 3486481..3490152 | 287 | BREX-system-P-loop-protein-BrxC-[Pseudomonas-aeruginosa] |
| WP_196620444.1 | Type1.BREX | <-XIS-HTH<-SIGMA+OB-Fold+AAA-ATPase+HKD-DNase-BrxL<-PglZ<-N6-METHYLASE-p<-HerA/FtsK<-SIR2<-REase<-OLD-ABC<-N6-METHYLASE<-DUF6079*<-BrxB-BrxF-iSTAND<-BrxA-DNA-binding<-XIS-HTH<-tRNA<-MerR-HTH | 1181 | Bacteria>Pseudomonadota>Gammaproteobacteria>Pseudomonadales>Pseudomonadaceae>Pseudomonas | GCF_027857235.1 | Pseudomonas_aeruginosa | OFL78_RS02500 | 39831..43376 | 287 | BREX-system-P-loop-protein-BrxC-[Pseudomonas-aeruginosa] |
| WP_271674697.1 | Type1.BREX | GTPase-AIG->GTPase-AIG->?-><-?\|\|wHTH+WYL+WCX\|\|?->ARG->BrxA-DNA-binding->BrxB-BrxF-iSTAND->DUF6079*->DNA-glycosylase->N6-METHYLASE-p-> | 1225 | Bacteria>Pseudomonadota>Gammaproteobacteria>Pseudomonadales>Pseudomonadaceae>Pseudomonas | GCF_028200675.1 | Pseudomonas_extremaustralis | PL026_RS31670 | 20252..23929 | 359110 | BREX-system-P-loop-protein-BrxC-[Pseudomonas-extremaustralis] |
| WP_272218090.1 | Type1.BREX | RelE-ParE-><-?<-wHTH+WYL+WCX\|\|?->Peptidase_M15->Peptidase_M15-><-?<-AAA-ATPase<-HIN-HTH\|\|DUF6079-p*->HIN-HTH->?->LZ_Tnp_IS66+zf-IS66+DDE_Tnp_IS66+DDE_Tnp_IS66_C->N6-METHYLASE-p->OLD-ABC->NERD->RVT->N6-METHYLASE-p->ParB+HNH-> | 932 | Bacteria>Pseudomonadota>Gammaproteobacteria>Pseudomonadales>Pseudomonadaceae>Pseudomonas | GCF_028404085.1 | Pseudomonas_aeruginosa | PMJ90_RS12960 | 2696481..2699279 | 287 | BREX-system-P-loop-protein-BrxC-[Pseudomonas-aeruginosa] |
| WP_273917578.1 | Type1.BREX | <-SIGMA+OB-Fold+AAA-ATPase+Lon-Protease-BrxL<-PglZ<-HEPN<-TPR<-NERD+GspA-ATPase<-N6-METHYLASE<-N6-METHYLASE<-TOPRIM<-OLD-ABC<-DUF6079*<-BrxB-BrxF-iSTAND<-BrxA-DNA-binding<-wHTH+WYL+WCX<-wHTH+WYL+WCX\|\|?->ARG-> | 1223 | Bacteria>Pseudomonadota>Gammaproteobacteria>Pseudomonadales>Pseudomonadaceae>Pseudomonas | GCF_028656895.1 | Pseudomonas_sp._TNT2022_ID1044 | M5G20_RS01570 | 347595..351266 | 2942636 | BREX-system-P-loop-protein-BrxC-[Pseudomonas-sp.-TNT2022-ID1044] |
| WP_273916456.1 | Type1.BREX | XIS-HTH->BrxA-DNA-binding->BrxB-BrxF-iSTAND->DUF6079*->N6-METHYLASE->OLD-ABC->NERD->?->?->N6-METHYLASE-p->PglZ->SIGMA+OB-Fold+AAA-ATPase+HKD-DNase-BrxL-> | 1186 | Bacteria>Pseudomonadota>Gammaproteobacteria>Pseudomonadales>Pseudomonadaceae>Pseudomonas | GCF_028656995.1 | Pseudomonas_sp._TNT2022_ID642 | M5G26_RS21745 | 58505..62065 | 2942632 | BREX-system-P-loop-protein-BrxC-[Pseudomonas-sp.-TNT2022-ID642] |
| WP_274102063.1 | Type1.BREX | Arm-DNA-bind_3+Phage_integrase->AAA+wHTH->wHTH->Integrase_1->Phage_integrase->wHTH+WYL+WCX->BrxA-DNA-binding->BrxB-BrxF-iSTAND->DUF6079*->N6-METHYLASE->PglZ->SIGMA+OB-Fold+AAA-ATPase+Lon-Protease-BrxL-><-HTH+REase<-ANK\|\|sGTP->?->sGTP->sGTP-> | 1222 | Bacteria>Pseudomonadota>Gammaproteobacteria>Pseudomonadales>Pseudomonadaceae>Pseudomonas | GCF_028682345.1 | Pseudomonas_putida | NPS33_RS09465 | 85396..89064 | 303 | BREX-system-P-loop-protein-BrxC-[Pseudomonas-putida] |
| WP_274151131.1 | Type1.BREX | Glyoxalase+Sel1+Sel1-><-NERD+ZnR<-REase+SF1-SF2-Helicase<-SIGMA+OB-Fold+AAA-ATPase+Lon-Protease-BrxL<-PglZ<-N6-METHYLASE<-DUF6079*<-BrxB-BrxF-iSTAND<-BrxA-DNA-binding<-wHTH+WYL+WCX<-Phage_integrase<-Integrase_1<-wHTH<-AAA+wHTH<-Arm-DNA-bind_3+Phage_integrase | 1223 | Bacteria>Pseudomonadota>Gammaproteobacteria>Pseudomonadales>Pseudomonadaceae>Pseudomonas | GCF_028682495.1 | Pseudomonas_sp._25571 | NPS27_RS02645 | 592231..595902 | 2967216 | BREX-system-P-loop-protein-BrxC-[Pseudomonas-sp.-25571] |
| WP_023131618.1 | Type1.BREX | <-wHTH+WYL+WCX\|\|?->BrxA-DNA-binding->BrxB-BrxF-iSTAND->DUF6079*->ParB+HNH->DNA-glycosylase->N6-METHYLASE->?->?->AbiEi_4->AbiEii->PglZ->SIGMA+OB-Fold+AAA-ATPase+Lon-Protease-BrxL-> | 1223 | Bacteria>Pseudomonadota>Gammaproteobacteria>Pseudomonadales>Pseudomonadaceae>Pseudomonas | GCF_029842935.1 | Pseudomonas_aeruginosa | N5I19_RS16015 | 16585..20256 | 287 | MULTISPECIES:-BREX-system-P-loop-protein-BrxC-[Pseudomonas] |
| WP_121324126.1 | Type1.BREX | <-REase<-wHTH+WYL+WCX\|\|?->Peptidase_M15->?->Ntox49->BrxA-DNA-binding->BrxB-BrxF-iSTAND->DUF6079*->N6-METHYLASE->?->HNH+GCN1->?->ParB+HNH->PglZ->SIGMA+OB-Fold+AAA-ATPase+Lon-Protease-BrxL-><-GreA_GreB+SF1-SF2-Helicase<-Arm-DNA-bind_3+Phage_integrase | 1154 | Bacteria>Pseudomonadota>Gammaproteobacteria>Pseudomonadales>Pseudomonadaceae>Pseudomonas | GCF_030121855.1 | Pseudomonas_aeruginosa | DQH07_RS08025 | 7041..10505 | 287 | BREX-system-P-loop-protein-BrxC-[Pseudomonas-aeruginosa] |
| WP_153334718.1 | Type1.BREX | <-SF1-SF2-Helicase+REase<-?<-?<-HKD-DNase+HKD-DNase<-?\|\|wHTH+WYL+WCX->BrxA-DNA-binding->BrxB-BrxF-iSTAND->DUF6079*->N6-METHYLASE->NERD+GspA-ATPase->TPR->PglZ->SIGMA+OB-Fold+AAA-ATPase+Lon-Protease-BrxL-> | 1223 | Bacteria>Pseudomonadota>Gammaproteobacteria>Pseudomonadales>Pseudomonadaceae>Pseudomonas | GCF_030144705.1 | Pseudomonas_sp._FSL_R10-2398 | GHO36_RS09020 | 68086..71757 | 2662201 | BREX-system-P-loop-protein-BrxC-[Pseudomonas-sp.-FSL-R10-2398] |
| WP_065945478.1 | Type1.BREX | <-AAA+wHTH<-?<-SIGMA+OB-Fold+AAA-ATPase+Lon-Protease-BrxL<-PglZ<-ParB+HNH<-?<-MarR-HTH<-MNS+ORC-CDC6<-N6-METHYLASE<-DUF6079*<-BrxB-BrxF-iSTAND<-BrxA-DNA-binding<-ARG<-?\|\|wHTH+WYL+WCX-> | 1154 | Bacteria>Pseudomonadota>Gammaproteobacteria>Pseudomonadales>Pseudomonadaceae>Pseudomonas | GCF_900004795.1 | Pseudomonas_sp._24_R_17 | BFP98_RS06365 | 99601..103065 | 1844096 | BREX-system-P-loop-protein-BrxC-[Pseudomonas-sp.-24-R-17] |
| WP_176947249.1 | Type1.BREX | <-TraK<-TraE<-TraM<-TraM<-DUF3262<-?\|\|XIS-HTH->BrxA-DNA-binding->BrxB-BrxF-iSTAND->DUF6079*->N6-METHYLASE->OLD-ABC->REase->SIR2->HerA/FtsK->N6-METHYLASE-p->PglZ->SIGMA+OB-Fold+AAA-ATPase+HKD-DNase-BrxL->Phage-HTH-> | 1188 | Bacteria>Pseudomonadota>Gammaproteobacteria>Pseudomonadales>Pseudomonadaceae>Pseudomonas | GCF_900100495.1 | Pseudomonas_benzenivorans | BLS63_RS10310 | 86902..90468 | 556533 | BREX-system-P-loop-protein-BrxC-[Pseudomonas-benzenivorans] |
| WP_092146146.1 | Type1.BREX | <-AAA+wHTH<-?<-SIGMA+OB-Fold+AAA-ATPase+Lon-Protease-BrxL<-PglZ<-ParB+HNH<-?<-MarR-HTH<-MNS+ORC-CDC6<-N6-METHYLASE<-DUF6079*<-BrxB-BrxF-iSTAND<-BrxA-DNA-binding<-ARG<-?\|\|wHTH+WYL+WCX->?->sGTP->sGTP->sGTP-> | 1154 | Bacteria>Pseudomonadota>Gammaproteobacteria>Pseudomonadales>Pseudomonadaceae>Pseudomonas | GCF_900102655.1 | unclassified_Pseudomonas | BLP71_RS14905 | 103193..106657 | 196821 | MULTISPECIES:-BREX-system-P-loop-protein-BrxC-[unclassified-Pseudomonas] |
| WP_090347579.1 | Type1.BREX | <-wHTH+WYL+WCX<-?\|\|ARG-><-ANK+ANK+ANK\|\|RADICAL-SAM->HicA_toxin->BrxA-DNA-binding->BrxB-BrxF-iSTAND->DUF6079*->N6-METHYLASE->?->HNH+GCN1->?->?->?->?->ParB+HNH->OLD-ABC-> | 1154 | Bacteria>Pseudomonadota>Gammaproteobacteria>Pseudomonadales>Pseudomonadaceae>Pseudomonas | GCF_900104805.1 | Pseudomonas_oryzae | BLT78_RS03155 | 660554..664018 | 1392877 | BREX-system-P-loop-protein-BrxC-[Pseudomonas-oryzae] |
| WP_090214592.1 | Type1.BREX | REC+PP2C->STAS-><-?<-?<-HTH_Tnp_1+HTH_21+rve+rve_3\|\|?->BrxA-DNA-binding->BrxB-BrxF-iSTAND->DUF6079*->ParB+HNH->?->?->N6-METHYLASE->?->?->?->?<-ZnR\|\|REase-> | 1218 | Bacteria>Pseudomonadota>Gammaproteobacteria>Pseudomonadales>Pseudomonadaceae>Pseudomonas | GCF_900105885.1 | Pseudomonas_guangdongensis | BLU22_RS11505 | 2422793..2426449 | 1245526 | BREX-system-P-loop-protein-BrxC-[Pseudomonas-guangdongensis] |
| WP_090192550.1 | Type1.BREX | <-35exo+SF1-SF2-Helicase\|\|XIS-HTH->BrxA-DNA-binding->BrxB-BrxF-iSTAND->DUF6079*->N6-METHYLASE->?->TPR+REase->?->SF1-SF2-Helicase+PRTase->SLOG+BirA-HTH->PglZ->SIGMA+OB-Fold+AAA-ATPase+Lon-Protease-BrxL-> | 1153 | Bacteria>Pseudomonadota>Gammaproteobacteria>Pseudomonadales>Pseudomonadaceae>Pseudomonas | GCF_900105995.1 | Pseudomonas_pohangensis | BLT89_RS00315 | 68625..72086 | 364197 | BREX-system-P-loop-protein-BrxC-[Pseudomonas-pohangensis] |
| WP_076583447.1 | Type1.BREX | DUF4969-><-SIGMA+OB-Fold+AAA-ATPase+HKD-DNase-BrxL<-PglZ<-ParB+HNH<-N6-METHYLASE-p<-RVT<-REase<-OLD-ABC<-N6-METHYLASE<-DUF6079*<-BrxB-BrxF-iSTAND<-BrxA-DNA-binding<-XIS-HTH<-?\|\|wHTH+WYL+WCX-><-?<-?<-KAP-NTPase\|\|LexA-protease-> | 1184 | Bacteria>Pseudomonadota>Gammaproteobacteria>Pseudomonadales>Pseudomonadaceae>Pseudomonas | GCF_900156545.1 | Pseudomonas_alcaligenes | BW984_RS17050 | 273058..276612 | 43263 | BREX-system-P-loop-protein-BrxC-[Pseudomonas-alcaligenes] |
| WP_095181954.1 | Type1.BREX | <-SIGMA+OB-Fold+AAA-ATPase+Lon-Protease-BrxL<-PglZ<-N6-METHYLASE<-DNA-glycosylase<-ParB+HNH<-DUF6079*<-BrxB-BrxF-iSTAND<-BrxA-DNA-binding\|\|?->Phage_GPD+Phage_base_V+DUF2345-> | 1222 | Bacteria>Pseudomonadota>Gammaproteobacteria>Pseudomonadales>Pseudomonadaceae>Pseudomonas | GCF_900187595.1 | Pseudomonas_sp._Irchel_3F6 | CKW30_RS22375 | 287368..291036 | 2009003 | BREX-system-P-loop-protein-BrxC-[Pseudomonas-sp.-Irchel-3F6] |
| WP_115284295.1 | Type1.BREX | AAA-ATPase->?->?->DUF4969-><-SIGMA+OB-Fold+AAA-ATPase+Lon-Protease-BrxL<-PglZ<-ParB+HNH<-DUF932<-N6-METHYLASE<-DUF6079*<-BrxB-BrxF-iSTAND<-BrxA-DNA-binding<-wHTH+WYL+WCX\|\|<-?<-HKD-DNase<-?\|\|?-><-?<-NADAR | 1154 | Bacteria>Pseudomonadota>Gammaproteobacteria>Pseudomonadales>Pseudomonadaceae>Pseudomonas | GCF_900455465.1 | Pseudomonas_fluorescens | DYC60_RS18930 | 4157779..4161243 | 294 | BREX-system-P-loop-protein-BrxC-[Pseudomonas-fluorescens] |
| WP_122521120.1 | Type1.BREX | <-RHH\|\|XIS-HTH->?->wHTH+WYL+WCX->BrxA-DNA-binding->BrxB-BrxF-iSTAND->DUF6079*->N6-METHYLASE->ParB+HNH->PglZ->SIGMA+OB-Fold+AAA-ATPase+Lon-Protease-BrxL-><-DUF4969<-?<-?\|\|ABhydrolase->SAM-METHYLASE-> | 1154 | Bacteria>Pseudomonadota>Gammaproteobacteria>Pseudomonadales>Pseudomonadaceae>Pseudomonas | GCF_900576635.1 | Pseudomonas_viridiflava | EET02_RS10950 | 375629..379093 | 33069 | BREX-system-P-loop-protein-BrxC-[Pseudomonas-viridiflava] |
| WP_127881958.1 | Type1.BREX | DUF932->?->?->?-><-SIGMA+OB-Fold+AAA-ATPase+Lon-Protease-BrxL<-PglZ<-ParB+HNH<-N6-METHYLASE<-DUF6079*<-BrxB-BrxF-iSTAND<-BrxA-DNA-binding<-wHTH+WYL+WCX | 1146 | Bacteria>Pseudomonadota>Gammaproteobacteria>Pseudomonadales>Pseudomonadaceae>Pseudomonas | GCF_900618835.1 | Pseudomonas_carnis | PCSC_RS23730 | 94995..98435 | 2487355 | BREX-system-P-loop-protein-BrxC-[Pseudomonas-carnis] |
| WP_150549236.1 | Type1.BREX | <-DUF3584+DUF5401<-JetB<-MukF<-SIGMA+OB-Fold+AAA-ATPase+Lon-Protease-BrxL<-PglZ<-TOPRIM<-OLD-ABC<-ParB+HNH<-N6-METHYLASE<-DUF6079*<-BrxB-BrxF-iSTAND<-BrxA-DNA-binding<-wHTH+WYL+WCX<-?<-?<-?<-XIS-HTH\|\|?-><-tRNA<-MerR-HTH | 1154 | Bacteria>Pseudomonadota>Gammaproteobacteria>Pseudomonadales>Pseudomonadaceae>Pseudomonas | GCF_900682675.2 | Pseudomonas_marincola | PMYSY11_RS11215 | 2416219..2419683 | 437900 | BREX-system-P-loop-protein-BrxC-[Pseudomonas-marincola] |
| WP_043107115.1 | Type1.BREX | GreA_GreB+SF1-SF2-Helicase-><-SIGMA+OB-Fold+AAA-ATPase+Lon-Protease-BrxL<-PglZ<-ParB+HNH<-?<-?<-HNH+GCN1<-?<-N6-METHYLASE<-DUF6079*<-BrxB-BrxF-iSTAND<-BrxA-DNA-binding<-Ntox49<-?<-Peptidase_M15<-?\|\|wHTH+WYL+WCX->REase->HIN-HTH-> | 1154 | Bacteria>Pseudomonadota>Gammaproteobacteria>Pseudomonadales>Pseudomonadaceae>Pseudomonas | GCF_900706985.1 | Pseudomonas_aeruginosa_group | E4Z08_RS29645 | 1460204..1463668 | 136841 | MULTISPECIES:-BREX-system-P-loop-protein-BrxC-[Pseudomonas-aeruginosa-group] |
| WP_134576235.1 | Type1.BREX | <-AbiEi_4<-?<-?<-N6-METHYLASE<-DNA-glycosylase<-ParB+HNH<-DUF6079-p* | 769 | Bacteria>Pseudomonadota>Gammaproteobacteria>Pseudomonadales>Pseudomonadaceae>Pseudomonas | GCF_900707355.1 | Pseudomonas_aeruginosa | E4Z10_RS26780 | 10282..12591 | 287 | BREX-system-P-loop-protein-BrxC,-partial-[Pseudomonas-aeruginosa] |
| WP_134454974.1 | Type1.BREX | <-DUF6079-p*<-BrxB-BrxF-iSTAND<-BrxA-DNA-binding<-?<-Peptidase_M15<-?\|\|wHTH+WYL+WCX-> | 704 | Bacteria>Pseudomonadota>Gammaproteobacteria>Pseudomonadales>Pseudomonadaceae>Pseudomonas | GCF_900707495.1 | Pseudomonas_aeruginosa | E4Y88_RS09690 | 1..2111 | 287 | BREX-system-P-loop-protein-BrxC,-partial-[Pseudomonas-aeruginosa] |
| WP_176605727.1 | Type1.BREX | <-MODE-HTH+PBPII\|\|<-?<-N6-METHYLASE<-DOC+HTH<-?<-?<-DUF6079*<-BrxB-BrxF-iSTAND<-BrxA-DNA-binding<-wHTH+WYL+WCX\|\|OLD-ABC+TOPRIM->SF1-SF2-Helicase->?->SF1-SF2-Helicase-><-Phage_integrase | 1225 | Bacteria>Pseudomonadota>Gammaproteobacteria>Pseudomonadales>Pseudomonadaceae>Pseudomonas | GCF_900708415.1 | Pseudomonas_aeruginosa | BUG32_RS23730 | 10964..14641 | 287 | BREX-system-P-loop-protein-BrxC-[Pseudomonas-aeruginosa] |
| WP_031684509.1 | Type1.BREX | PhdYeFM_antitox->?->?-><-wHTH+WYL+WCX\|\|?->Peptidase_M15->?->XIS-HTH->BrxA-DNA-binding->BrxB-BrxF-iSTAND->DUF6079*->N6-METHYLASE->SbcC-p->REase->?->N6-METHYLASE-p->PglZ->SIGMA+OB-Fold+AAA-ATPase+HKD-DNase-BrxL-><-Arm-DNA-bind_3+Phage_integrase\|\|SF1-SF2-Helicase-> | 1184 | Bacteria>Pseudomonadota>Gammaproteobacteria>Pseudomonadales>Pseudomonadaceae>Pseudomonas | GCF_900708485.1 | Pseudomonas_aeruginosa | IDO19_RS13300 | 2848972..2852526 | 287 | BREX-system-P-loop-protein-BrxC-[Pseudomonas-aeruginosa] |
| WP_150597441.1 | Type1.BREX | DUF4969-><-SIGMA+OB-Fold+AAA-ATPase+Lon-Protease-BrxL<-PglZ<-ParB+HNH<-N6-METHYLASE<-DUF6079*<-BrxB-BrxF-iSTAND<-BrxA-DNA-binding<-wHTH+WYL+WCX<-?<-TPR<-SF1-SF2-Helicase+RNase+RNase<-XIS-HTH\|\|RHH-> | 1154 | Bacteria>Pseudomonadota>Gammaproteobacteria>Pseudomonadales>Pseudomonadaceae>Pseudomonas | GCF_902497985.1 | Pseudomonas_fluorescens | F8N16_RS17980 | 132187..135651 | 294 | BREX-system-P-loop-protein-BrxC-[Pseudomonas-fluorescens] |
| WP_236582582.1 | Type1.BREX | <-TetR-HTH\|\|?->Arm-DNA-bind_3+Phage_integrase->?->Phage_AlpA->?->wHTH+WYL+WCX->BrxA-DNA-binding->BrxB-BrxF-iSTAND->DUF6079*->?->?->DOC+HTH->N6-METHYLASE->NERD+GspA-ATPase->TPR-><-ZnR\|\|PglZ->SIGMA+OB-Fold+AAA-ATPase+Lon-Protease-BrxL-> | 1225 | Bacteria>Pseudomonadota>Gammaproteobacteria>Pseudomonadales>Pseudomonadaceae>Pseudomonas | GCF_905071885.1 | Pseudomonas_aeruginosa | JMX72_RS23520 | 4985087..4988764 | 287 | BREX-system-P-loop-protein-BrxC-[Pseudomonas-aeruginosa] |
| WP_229999483.1 | Type1.BREX | XIS-HTH->?->URI-DNase+RAMA->Schlafen+HATPase+wHTH->DnaB+McrB-AAA->McrC-NTD+REase+ZnR->wHTH+WYL+WCX->BrxA-DNA-binding->BrxB-BrxF-iSTAND->DUF6079*->N6-METHYLASE->ParB+HNH->OLD-ABC->TOPRIM->PglZ->SIGMA+OB-Fold+AAA-ATPase+Lon-Protease-BrxL-><-DUF4969 | 1156 | Bacteria>Pseudomonadota>Gammaproteobacteria>Pseudomonadales>Pseudomonadaceae>Pseudomonas | GCF_918697895.1 | Pseudomonas_carnis | LQ224_RS08485 | 110382..113852 | 2487355 | BREX-system-P-loop-protein-BrxC-[Pseudomonas-carnis] |
| WP_033991202.1 | Type1.BREX | <-HNH<-SIGMA+OB-Fold+AAA-ATPase+Lon-Protease-BrxL<-PglZ<-MNS+REase<-?<-HNH+GCN1<-?<-N6-METHYLASE<-DUF6079*<-BrxB-BrxF-iSTAND<-BrxA-DNA-binding<-RNase<-?<-JetB<-?<-Peptidase_M15<-?\|\|wHTH+WYL+WCX-> | 1154 | Bacteria>Pseudomonadota>Gammaproteobacteria>Pseudomonadales>Pseudomonadaceae>Pseudomonas | GCF_929614685.1 | Pseudomonas_aeruginosa | MJP43_RS27275 | 28461..31925 | 287 | BREX-system-P-loop-protein-BrxC-[Pseudomonas-aeruginosa] |
| WP_033944490.1 | Type1.BREX | Arm-DNA-bind_3+Phage_integrase-><-SIGMA+OB-Fold+AAA-ATPase+HKD-DNase-BrxL<-PglZ<-N6-METHYLASE-p<-RVT<-NERD<-OLD-ABC<-N6-METHYLASE<-DUF6079*<-BrxB-BrxF-iSTAND<-BrxA-DNA-binding<-XIS-HTH<-?<-?\|\|?->?->?->HIN-HTH-> | 1184 | Bacteria>Pseudomonadota>Gammaproteobacteria>Pseudomonadales>Pseudomonadaceae>Pseudomonas | GCF_947387505.1 | Pseudomonas_aeruginosa | GPV34_RS03115 | 17071..20625 | 287 | BREX-system-P-loop-protein-BrxC-[Pseudomonas-aeruginosa] |
| WP_128722024.1 | Type1.BREX | <-TetR-HTH\|\|?->Arm-DNA-bind_3+Phage_integrase->?->?\|\|Phage_AlpA->?->BrxA-DNA-binding->BrxB-BrxF-iSTAND->DUF6079*->?->?->DOC+HTH->N6-METHYLASE->?->\|\|MODE-HTH+PBPII-> | 1223 | Bacteria>Pseudomonadota>Gammaproteobacteria>Pseudomonadales>Pseudomonadaceae>Pseudomonas | - | Pseudomonas_aeruginosa | I5K34_26080 | 19802..23473 | 287 | BREX-system-P-loop-protein-BrxC-[Pseudomonas-aeruginosa] |
| EIU5588796.1 | Type1.BREX | PhdYeFM_antitox->RelE-ParE-><-?<-wHTH+WYL+WCX\|\|?->Peptidase_M15->?->BrxA-DNA-binding->BrxB-BrxF-iSTAND->DUF6079*->N6-METHYLASE->?->HNH+GCN1->?->ParB+HNH->PglZ->SIGMA+OB-Fold+AAA-ATPase+Lon-Protease-BrxL->REase->SF1-SF2-Helicase-> | 1154 | Bacteria>Pseudomonadota>Gammaproteobacteria>Pseudomonadales>Pseudomonadaceae>Pseudomonas | - | Pseudomonas_aeruginosa | L4689_001050 | 82930..86394 | 287 | BREX-system-P-loop-protein-BrxC-[Pseudomonas-aeruginosa] |
| EIU3152871.1 | Type1.BREX | Phage_integrase->DDE_Tnp_1-><-SF1-SF2-Helicase<-?<-SF1-SF2-Helicase<-OLD-ABC+TOPRIM\|\|wHTH+WYL+WCX->BrxA-DNA-binding->BrxB-BrxF-iSTAND->DUF6079*->?->?->DOC+HTH->N6-METHYLASE-p-> | 1223 | Bacteria>Pseudomonadota>Gammaproteobacteria>Pseudomonadales>Pseudomonadaceae>Pseudomonas | - | Pseudomonas_aeruginosa | L4P27_006182 | 20932..24603 | 287 | BREX-system-P-loop-protein-BrxC-[Pseudomonas-aeruginosa] |
| WP_023124092.1 | Type1.BREX | Phage_integrase->DDE_Tnp_1-><-SF1-SF2-Helicase<-?<-SF1-SF2-Helicase<-OLD-ABC+TOPRIM\|\|wHTH+WYL+WCX->BrxA-DNA-binding->BrxB-BrxF-iSTAND->DUF6079*->?->?->DOC+HTH->N6-METHYLASE-p-> | 1223 | Bacteria>Pseudomonadota>Gammaproteobacteria>Pseudomonadales>Pseudomonadaceae>Pseudomonas | - | Pseudomonas_aeruginosa | L4P27_006182 | 20932..24603 | 287 | BREX-system-P-loop-protein-BrxC-[Pseudomonas-aeruginosa] |
| EIU3153299.1 | Type1.BREX | GreA_GreB+SF1-SF2-Helicase-p-><-SIGMA+OB-Fold+AAA-ATPase+Lon-Protease-BrxL<-PglZ<-?<-HNH+GCN1<-?<-N6-METHYLASE<-DUF6079-p* | 512 | Bacteria>Pseudomonadota>Gammaproteobacteria>Pseudomonadales>Pseudomonadaceae>Pseudomonas | - | Pseudomonas_aeruginosa | L4P27_006629 | 14238..15778 | 287 | hypothetical-protein-[Pseudomonas-aeruginosa] |
| EIU3153568.1 | Type1.BREX | <-wHTH+WYL+WCX\|\|?->Peptidase_M15->?->Ntox49->BrxA-DNA-binding->BrxB-BrxF-iSTAND->DUF6079-p*-> | 641 | Bacteria>Pseudomonadota>Gammaproteobacteria>Pseudomonadales>Pseudomonadaceae>Pseudomonas | - | Pseudomonas_aeruginosa | L4P27_006902 | 6274..8196 | 287 | BREX-system-P-loop-protein-BrxC-[Pseudomonas-aeruginosa] |
| EIU4198817.1 | Type1.BREX | DUF932->?->?->?-><-SIGMA+OB-Fold+AAA-ATPase+HKD-DNase-BrxL<-PglZ<-ParB+HNH<-N6-METHYLASE<-DUF6079*<-BrxB-BrxF-iSTAND<-BrxA-DNA-binding<-XIS-HTH\|\|<-?\|\|sGTP->sGTP->MA+PspA+PspA+sGTP-><-ZnR<-ARG | 1180 | Bacteria>Pseudomonadota>Gammaproteobacteria>Pseudomonadales>Pseudomonadaceae>Pseudomonas | - | Pseudomonas_aeruginosa | L4V49_002912 | 27945..31487 | 287 | BREX-system-P-loop-protein-BrxC-[Pseudomonas-aeruginosa] |
| WP_203788563.1 | Type1.BREX | Phage_integrase-><-?<-SF1-SF2-Helicase<-?<-SbcC\|\|?->wHTH+WYL+WCX->BrxA-DNA-binding->BrxB-BrxF-iSTAND->DUF6079*->?->?->DOC+HTH->N6-METHYLASE->RVT->N6-METHYLASE-p-><-DUF772+DDE_Tnp_1\|\|N6-METHYLASE-p-> | 1223 | Bacteria>Pseudomonadota>Gammaproteobacteria>Pseudomonadales>Pseudomonadaceae>Pseudomonas | - | Pseudomonas_alcaligenes | KAM426_36250 | 3797295..3800966 | 43263 | BREX-system-P-loop-protein-BrxC-[Pseudomonas-alcaligenes] |
| WP_154949571.1 | Type1.BREX | DUF932->?->?->DUF4969-><-SIGMA+OB-Fold+AAA-ATPase+Lon-Protease-BrxL<-PglZ<-ParB+HNH<-N6-METHYLASE<-DUF6079*<-BrxB-BrxF-iSTAND<-BrxA-DNA-binding<-wHTH+WYL+WCX<-?<-XIS-HTH\|\|RHH-><-?<-tRNA<-MerR-HTH | 1154 | Bacteria>Pseudomonadota>Gammaproteobacteria>Pseudomonadales>Pseudomonadaceae>Pseudomonas | - | Pseudomonas_fluorescens | F8N22_RS18775 | 172273..175737 | 294 | BREX-system-P-loop-protein-BrxC-[Pseudomonas-fluorescens] |
| WP_154851174.1 | Type1.BREX | <-SF1-SF2-Helicase+REase<-?<-SF1-SF2-Helicase<-HKD-DNase+HKD-DNase<-?\|\|wHTH+WYL+WCX->BrxA-DNA-binding->BrxB-BrxF-iSTAND->DUF6079*->?->N6-METHYLASE->NERD+GspA-ATPase->TPR->PglZ->SIGMA+OB-Fold+AAA-ATPase+Lon-Protease-BrxL-><-XIS-HTH | 1224 | Bacteria>Pseudomonadota>Gammaproteobacteria>Pseudomonadales>Pseudomonadaceae>Pseudomonas | - | Pseudomonas_fluorescens | F8N48_RS14725 | 264016..267690 | 294 | BREX-system-P-loop-protein-BrxC-[Pseudomonas-fluorescens] |
| WP_032902483.1 | Type1.BREX | <-AAA+wHTH<-?<-SIGMA+OB-Fold+AAA-ATPase+Lon-Protease-BrxL<-PglZ<-ParB+HNH<-?<-MarR-HTH<-MNS+ORC-CDC6<-N6-METHYLASE<-DUF6079*<-BrxB-BrxF-iSTAND<-BrxA-DNA-binding<-ARG<-?\|\|wHTH+WYL+WCX->Arm-DNA-bind_3+Phage_integrase->SF1-SF2-Helicase+REase->XIS-HTH->TOPRIM-> | 1154 | Bacteria>Pseudomonadota>Gammaproteobacteria>Pseudomonadales>Pseudomonadaceae>Pseudomonas | - | Pseudomonas_fluorescens | HO98_RS08890 | 14970..18434 | 294 | BREX-system-P-loop-protein-BrxC-[Pseudomonas-fluorescens] |
| WP_241666011.1 | Type1.BREX | <-TraI-VirD2-RCRE<-ParD_antitoxin\|\|HNH->?-><-wHTH+WYL\|\|ARG->BrxA-DNA-binding->BrxB-BrxF-iSTAND->DUF6079*->DNA-glycosylase->N6-METHYLASE->ParB+HNH->AbiEi_3_N+AbiEi_3->AbiEii->PglZ->SIGMA+OB-Fold+AAA-ATPase+Lon-Protease-BrxL-> | 1225 | Bacteria>Pseudomonadota>Gammaproteobacteria>Pseudomonadales>Pseudomonadaceae>Pseudomonas | - | Pseudomonas_fluorescens | Pf1CFBP13510_RS04715 | 495917..499594 | 294 | BREX-system-P-loop-protein-BrxC-[Pseudomonas-fluorescens] |
| WP_010656375.1 | Type1.BREX | <-SF1-SF2-Helicase+REase<-?<-Ntox49<-?<-SF1-SF2-Helicase<-HKD-DNase+HKD-DNase<-?\|\|wHTH+WYL+WCX->BrxA-DNA-binding->BrxB-BrxF-iSTAND->DUF6079*->N6-METHYLASE->NERD+GspA-ATPase->TPR->PglZ->SIGMA+OB-Fold+AAA-ATPase+Lon-Protease-BrxL->?-><-tRNA<-MerR-HTH | 1223 | Bacteria>Pseudomonadota>Gammaproteobacteria>Pseudomonadales>Pseudomonadaceae>Pseudomonas | - | Pseudomonas_fragi | AV641_RS14580 | 3283222..3286893 | 296 | BREX-system-P-loop-protein-BrxC-[Pseudomonas-fragi] |
| WP_112249959.1 | Type1.BREX | wHTH+WYL+WCX->BrxA-DNA-binding->BrxB-BrxF-iSTAND->DUF6079*->N6-METHYLASE->ParB+HNH->PglZ->SIGMA+OB-Fold+AAA-ATPase+Lon-Protease-BrxL-><-DUF4969<-?<-?<-?<-DUF932 | 1154 | Bacteria>Pseudomonadota>Gammaproteobacteria>Pseudomonadales>Pseudomonadaceae>Pseudomonas | - | Pseudomonas_putida | GT37_RS03100 | 3180..6644 | 303 | BREX-system-P-loop-protein-BrxC-[Pseudomonas-putida] |
| WP_046164318.1 | Type1.BREX | <-SIGMA+OB-Fold+AAA-ATPase+HKD-DNase-BrxL<-PglZ<-ParB+HNH<-N6-METHYLASE-p<-HerA/FtsK<-SIR2<-REase<-OLD-ABC<-N6-METHYLASE<-DUF6079*<-BrxB-BrxF-iSTAND<-BrxA-DNA-binding<-XIS-HTH<-?\|\|wHTH+WYL+WCX->sGTP->sGTP-> | 1186 | Bacteria>Pseudomonadota>Gammaproteobacteria>Pseudomonadales>Pseudomonadaceae>Pseudomonas | - | Pseudomonas_sp._10B238 | RT21_RS21355 | 15645..19205 | 1586417 | BREX-system-P-loop-protein-BrxC-[Pseudomonas-sp.-10B238] |
| WP_099493679.1 | Type1.BREX | <-SIGMA+OB-Fold+AAA-ATPase+Lon-Protease-BrxL<-PglZ<-AbiEii<-AbiEi_3_N+AbiEi_3<-ParB+HNH<-N6-METHYLASE<-DNA-glycosylase<-DUF6079*<-BrxB-BrxF-iSTAND<-BrxA-DNA-binding<-ARG<-?\|\|wHTH+WYL+WCX-><-?<-?\|\|HIN-HTH-> | 1222 | Bacteria>Pseudomonadota>Gammaproteobacteria>Pseudomonadales>Pseudomonadaceae>Pseudomonas | - | Pseudomonas_sp._2588-5 | AOA57_22445 | 341678..345346 | 1712676 | BREX-system-P-loop-protein-BrxC-[Pseudomonas-sp.-2588-5] |
| WP_098950596.1 | Type1.BREX | <-TraI-VirD2-RCRE<-ParD_antitoxin\|\|HNH->?-><-?\|\|BrxA-DNA-binding->BrxB-BrxF-iSTAND->DUF6079*->DNA-glycosylase->N6-METHYLASE->ParB+HNH->AbiEi_3_N+AbiEi_3->AbiEii->PglZ->SIGMA+OB-Fold+AAA-ATPase+Lon-Protease-BrxL-> | 1222 | Bacteria>Pseudomonadota>Gammaproteobacteria>Pseudomonadales>Pseudomonadaceae>Pseudomonas | - | Pseudomonas_sp._FDAARGOS_380 | CRN80_16460 | 3507093..3510761 | 2018067 | BREX-system-P-loop-protein-BrxC-[Pseudomonas-sp.-FDAARGOS_380] |
| WP_223019012.1 | Type1.BREX | <-SIGMA+OB-Fold+AAA-ATPase+Lon-Protease-BrxL<-PglZ<-AbiEii<-AbiEi_3_N+AbiEi_3<-ParB+HNH<-N6-METHYLASE<-DNA-glycosylase<-DUF6079*<-BrxB-BrxF-iSTAND<-BrxA-DNA-binding<-ARG<-?\|\|wHTH+WYL+WCX-><-tRNA<-MerR-HTH | 1222 | Bacteria>Pseudomonadota>Gammaproteobacteria>Pseudomonadales>Pseudomonadaceae>Pseudomonas | - | Pseudomonas_sp._P867 | J1G33_25110 | 13708..17376 | 2816050 | BREX-system-P-loop-protein-BrxC-[Pseudomonas-sp.-P867] |
| WP_225609641.1 | Type1.BREX | <-wHTH+WYL+WCX\|\|?->JetB->?->RNase->?->BrxA-DNA-binding->BrxB-BrxF-iSTAND->DUF6079*->?->N6-METHYLASE->?->AbiEi_4->AbiEii->PglZ->SIGMA+OB-Fold+AAA-ATPase+Lon-Protease-BrxL-> | 1225 | Bacteria>Pseudomonadota>Gammaproteobacteria>Pseudomonadales>Pseudomonadaceae>Pseudomonas | - | Pseudomonas_sp._PDM11 | IB224_RS16070 | 119835..123512 | 2769309 | BREX-system-P-loop-protein-BrxC-[Pseudomonas-sp.-PDM11] |
| WP_170056913.1 | Type1.BREX | PIN+OST-HTH->HNH->?->?-><-wHTH+WYL+WCX\|\|?->ARG->BrxA-DNA-binding->BrxB-BrxF-iSTAND->DUF6079*->DNA-glycosylase->N6-METHYLASE->ParB+HNH->AbiEi_3_N+AbiEi_3->AbiEii->PglZ->SIGMA+OB-Fold+AAA-ATPase+Lon-Protease-BrxL->JAB-><-MerR-HTH | 1222 | Bacteria>Pseudomonadota>Gammaproteobacteria>Pseudomonadales>Pseudomonadaceae>Pseudomonas | - | Pseudomonas_sp._WS_5021 | HBO19_03685 | 794090..797758 | 2717490 | BREX-system-P-loop-protein-BrxC-[Pseudomonas-sp.-WS-5021] |
| WP_240033455.1 | Type1.BREX | <-wHTH+WYL+WCX\|\|?->ARG->BrxA-DNA-binding->BrxB-BrxF-iSTAND->DUF6079*->DNA-glycosylase->N6-METHYLASE->ParB+HNH->AbiEi_3_N+AbiEi_3->AbiEii->PglZ->SIGMA+OB-Fold+AAA-ATPase+Lon-Protease-BrxL->?->SMODS-> | 1225 | Bacteria>Pseudomonadota>Gammaproteobacteria>Pseudomonadales>Pseudomonadaceae>Pseudomonas | - | Pseudomonas_viridiflava | EE173_RS11530 | 5062..8739 | 33069 | BREX-system-P-loop-protein-BrxC-[Pseudomonas-viridiflava] |
| WP_240043383.1 | Type1.BREX | <-Thymidylate_kin<-?\|\|ZnR-><-?<-?<-TOPRIM<-OLD-ABC<-N6-METHYLASE<-DOC+HTH<-DUF6079*<-BrxB-BrxF-iSTAND<-BrxA-DNA-binding<-wHTH+WYL+WCX | 1225 | Bacteria>Pseudomonadota>Gammaproteobacteria>Pseudomonadales>Pseudomonadaceae>Pseudomonas | - | Pseudomonas_viridiflava | EFA20_RS09810 | 85838..89515 | 33069 | BREX-system-P-loop-protein-BrxC-[Pseudomonas-viridiflava] |
| AEJ06724.1 | Type1.BREX | HTH+DOC-><-?\|\|Arm-DNA-bind_3+Phage_integrase->wHTH+WYL+WCX->BrxA-DNA-binding->BrxB-BrxF-iSTAND->DUF6079*->?-><-AAA-ATPase<-HIN-HTH\|\|REC-><-ABC-ATPase+SbcC | 1223 | Bacteria>Pseudomonadota>Gammaproteobacteria>Pseudomonadales>Pseudomonadaceae>Stutzerimonas | GCA_000219605.1 | Stutzerimonas_stutzeri | PSTAB_3443 | 3706589..3710260 | 316 | conserved-hypothetical-protein-[Stutzerimonas-stutzeri] |
| EPL61008.1 | Type1.BREX | <-Phage-HTH<-SIGMA+OB-Fold+AAA-ATPase+HKD-DNase-BrxL<-PglZ<-N6-METHYLASE-p<-HerA/FtsK<-SIR2<-REase<-OLD-ABC<-N6-METHYLASE<-DUF6079*<-BrxB-BrxF-iSTAND<-BrxA-DNA-binding<-XIS-HTH\|\|?->DUF3262->TraM->TraM->TraE->TraK-> | 1188 | Bacteria>Pseudomonadota>Gammaproteobacteria>Pseudomonadales>Pseudomonadaceae>Stutzerimonas | GCA_000416345.1 | Stutzerimonas_stutzeri_B1SMN1 | B382_18428 | 65079..68645 | 1212549 | hypothetical-protein-B382_18428-[Stutzerimonas-stutzeri-B1SMN1] |
| OWG36865.1 | Type1.BREX | <-RNase<-?<-JetB<-?<-wHTH+WYL+WCX\|\|XIS-HTH->BrxA-DNA-binding->BrxB-BrxF-iSTAND->DUF6079*->N6-METHYLASE->ParB+HNH->PglZ->SIGMA+OB-Fold+AAA-ATPase+HKD-DNase-BrxL-><-?<-?<-?<-DUF932 | 1185 | Bacteria>Pseudomonadota>Gammaproteobacteria>Pseudomonadales>Pseudomonadaceae>Stutzerimonas | GCA_002193135.1 | Stutzerimonas_stutzeri | CAQ69_18770 | 20186..23743 | 316 | hypothetical-protein-CAQ69_18770-[Stutzerimonas-stutzeri] |
| PZR87477.1 | Type1.BREX | RHH-><-TetR-HTH\|\|MODE-HTH+PBPII->wHTH+WYL+WCX->BrxA-DNA-binding->BrxB-BrxF-iSTAND->DUF6079*->PglZ->SIGMA+OB-Fold+AAA-ATPase+HKD-DNase-BrxL-><-?<-HAD+HAD | 1179 | Bacteria>Pseudomonadota>Gammaproteobacteria>Pseudomonadales>Pseudomonadaceae>Stutzerimonas | GCA_003243385.1 | Stutzerimonas_stutzeri | DI537_26065 | 4408..7947 | 316 | MAG:-BREX-system-P-loop-protein-BrxC-[Stutzerimonas-stutzeri] |
| QOZ94737.1 | Type1.BREX | <-HIRAN\|\|HIN-HTH+PAIREDC-HTH-><-?<-N6-METHYLASE-p<-N6-METHYLASE<-DOC+HTH<-?<-?<-DUF6079*<-BrxB-BrxF-iSTAND<-BrxA-DNA-binding<-wHTH+WYL+WCX<-?<-Phage_AlpA<-?<-Arm-DNA-bind_3+Phage_integrase | 1223 | Bacteria>Pseudomonadota>Gammaproteobacteria>Pseudomonadales>Pseudomonadaceae>Stutzerimonas | GCA_015291885.1 | Stutzerimonas_stutzeri | Pstu14405_04940 | 1078194..1081865 | 316 | BREX-system-P-loop-protein-BrxC-[Stutzerimonas-stutzeri] |
| MCQ2029875.1 | Type1.BREX | <-sGTP<-sGTP<-sGTP<-wHTH+WYL+WCX\|\|?-><-?\|\|XIS-HTH->BrxA-DNA-binding->BrxB-BrxF-iSTAND->DUF6079*->N6-METHYLASE->OLD-ABC->NERD->RVT->N6-METHYLASE-p->ParB+HNH->PglZ->SIGMA+OB-Fold+AAA-ATPase+HKD-DNase-BrxL->RHH_1-> | 1186 | Bacteria>Pseudomonadota>Gammaproteobacteria>Pseudomonadales>Pseudomonadaceae>Stutzerimonas | GCA_024397535.1 | [Pseudomonas]_zhaodongensis | NA613_09780 | 17231..20791 | 1176257 | BREX-system-P-loop-protein-BrxC-[[Pseudomonas] |
| MCQ4256509.1 | Type1.BREX | <-SF1-SF2-Helicase<-REase<-SIGMA+OB-Fold+AAA-ATPase+Lon-Protease-BrxL<-PglZ<-ParB+HNH<-?<-MarR-HTH<-MNS+ORC-CDC6<-N6-METHYLASE<-DUF6079*<-BrxB-BrxF-iSTAND<-BrxA-DNA-binding<-REase<-NERD\|\|?->DUF932-> | 1154 | Bacteria>Pseudomonadota>Gammaproteobacteria>Pseudomonadales>Pseudomonadaceae>Stutzerimonas | GCA_024448895.1 | Stutzerimonas_stutzeri | NA619_00645 | 147157..150621 | 316 | BREX-system-P-loop-protein-BrxC-[Stutzerimonas-stutzeri] |
| MCQ4319996.1 | Type1.BREX | Phage_integrase->?->?->?<-?<-?\|\|wHTH+WYL+WCX->BrxA-DNA-binding->BrxB-BrxF-iSTAND->DUF6079*->N6-METHYLASE->OLD-ABC->TOPRIM->N6-METHYLASE->sGTP->?->PglZ->SIGMA+OB-Fold+AAA-ATPase+Lon-Protease-BrxL-> | 1223 | Bacteria>Pseudomonadota>Gammaproteobacteria>Pseudomonadales>Pseudomonadaceae>Stutzerimonas | GCA_024448935.1 | Stutzerimonas_stutzeri | NA637_07215 | 17580..21251 | 316 | BREX-system-P-loop-protein-BrxC-[Stutzerimonas-stutzeri] |
| WP_232244016.1 | Type1.BREX | MazE_antitoxin->Arm-DNA-bind_3+Phage_integrase->?->Phage_AlpA->?->wHTH+WYL+WCX->BrxA-DNA-binding->BrxB-BrxF-iSTAND->DUF6079*->?-><-AAA-ATPase<-HIN-HTH\|\|REC-><-ABC-ATPase+SbcC | 1225 | Bacteria>Pseudomonadota>Gammaproteobacteria>Pseudomonadales>Pseudomonadaceae>Stutzerimonas | GCF_000219605.1 | Stutzerimonas_stutzeri | I6G32_RS07530 | 1635442..1639119 | 316 | BREX-system-P-loop-protein-BrxC-[Stutzerimonas-stutzeri] |
| WP_003283611.1 | Type1.BREX | <-N6-METHYLASE-p<-RVT<-N6-METHYLASE<-DOC+HTH<-?<-?<-DUF6079*<-BrxB-BrxF-iSTAND<-BrxA-DNA-binding<-wHTH+WYL+WCX<-?<-Phage_AlpA | 1225 | Bacteria>Pseudomonadota>Gammaproteobacteria>Pseudomonadales>Pseudomonadaceae>Stutzerimonas | GCF_000237885.1 | Stutzerimonas_stutzeri | PSTZOBELL_RS12590 | 11831..15508 | 316 | BREX-system-P-loop-protein-BrxC-[Stutzerimonas-stutzeri] |
| WP_014819601.1 | Type1.BREX | <-Phage-HTH<-SIGMA+OB-Fold+AAA-ATPase+HKD-DNase-BrxL<-PglZ<-N6-METHYLASE-p<-HerA/FtsK<-SIR2<-REase<-OLD-ABC<-N6-METHYLASE<-DUF6079*<-BrxB-BrxF-iSTAND<-BrxA-DNA-binding<-XIS-HTH\|\|?->DUF3262->TraM->TraM->TraE->TraK-> | 1188 | Bacteria>Pseudomonadota>Gammaproteobacteria>Pseudomonadales>Pseudomonadaceae>Stutzerimonas | GCF_000267545.1 | Stutzerimonas_stutzeri_subgroup | A458_RS28085 | 1395712..1399278 | 578833 | MULTISPECIES:-BREX-system-P-loop-protein-BrxC-[Stutzerimonas-stutzeri-subgroup] |
| WP_017245161.1 | Type1.BREX | <-PglZ<-AbiEii<-AbiEi_4<-?<-?<-ParB+HNH<-N6-METHYLASE<-DNA-glycosylase<-ParB+HNH<-DUF6079*<-BrxB-BrxF-iSTAND<-BrxA-DNA-binding\|\|?->Arm-DNA-bind_2+Phage_int_SAM_3+Phage_integrase-><-?<-?\|\|HTH_3->XIS-HTH-> | 1220 | Bacteria>Pseudomonadota>Gammaproteobacteria>Pseudomonadales>Pseudomonadaceae>Stutzerimonas | GCF_000280555.1 | Stutzerimonas_stutzeri | A598_RS0110895 | 41008..44670 | 316 | BREX-system-P-loop-protein-BrxC-[Stutzerimonas-stutzeri] |
| WP_053526945.1 | Type1.BREX | <-HTH_3<-?<-SIGMA+OB-Fold+AAA-ATPase+Lon-Protease-BrxL<-PglZ<-HNH<-OLD-ABC<-?<-?<-N6-METHYLASE<-DUF6079*<-BrxB-BrxF-iSTAND<-BrxA-DNA-binding<-wHTH+WYL+WCX\|\|?->RVT->?-><-SF1-SF2-Helicase | 1223 | Bacteria>Pseudomonadota>Gammaproteobacteria>Pseudomonadales>Pseudomonadaceae>Stutzerimonas | GCF_001276475.1 | Stutzerimonas_stutzeri | ABW53_RS00980 | 14361..18032 | 316 | BREX-system-P-loop-protein-BrxC-[Stutzerimonas-stutzeri] |
| WP_245890967.1 | Type1.BREX | JetB->?->RNase->?->BrxA-DNA-binding->BrxB-BrxF-iSTAND->DUF6079*->AbiGi+seqA+RAMA->?->REase+Bacteriocin_IIc->?->AbiEi_3_N+AbiEi_3->AbiEii->PglZ->SIGMA+OB-Fold+AAA-ATPase+Lon-Protease-BrxL-> | 1225 | Bacteria>Pseudomonadota>Gammaproteobacteria>Pseudomonadales>Pseudomonadaceae>Stutzerimonas | GCF_002929225.1 | Stutzerimonas_kunmingensis | CW305_RS04090 | 634964..638641 | 1211807 | BREX-system-P-loop-protein-BrxC-[Stutzerimonas-kunmingensis] |
| WP_125460588.1 | Type1.BREX | <-PglZ<-AbiEii<-AbiEi_3_N+AbiEi_3<-SF1-SF2-Helicase<-HKD-DNase+HKD-REase<-?<-?<-?<-N6-METHYLASE<-AbiGi+seqA+RAMA<-DUF6079*<-BrxB-BrxF-iSTAND<-BrxA-DNA-binding<-DUF772+DDE_Tnp_1\|\|?->Arm-DNA-bind_2+Phage_int_SAM_3+Phage_integrase-> | 1220 | Bacteria>Pseudomonadota>Gammaproteobacteria>Pseudomonadales>Pseudomonadaceae>Stutzerimonas | GCF_003936745.1 | Stutzerimonas_stutzeri | EGJ43_RS19065 | 24827..28489 | 316 | BREX-system-P-loop-protein-BrxC-[Stutzerimonas-stutzeri] |
| WP_125856771.1 | Type1.BREX | <-LexA-protease<-wHTH+WYL+WCX\|\|ARG->?->wHTH+WYL+WCX->BrxA-DNA-binding->BrxB-BrxF-iSTAND->DUF6079*->N6-METHYLASE->ParB+HNH->OLD-ABC->HNH->PglZ->SIGMA+OB-Fold+AAA-ATPase+Lon-Protease-BrxL->MukF->JetB->DUF3584+DUF5401-> | 1154 | Bacteria>Pseudomonadota>Gammaproteobacteria>Pseudomonadales>Pseudomonadaceae>Stutzerimonas | GCF_003940515.1 | Stutzerimonas_stutzeri | EGJ36_RS18105 | 6035..9499 | 316 | BREX-system-P-loop-protein-BrxC-[Stutzerimonas-stutzeri] |
| WP_200291399.1 | Type1.BREX | Peptidase_A8-><-?<-?\|\|LexA-protease->?-><-Phage_int_SAM<-Integrase_1\|\|BrxA-DNA-binding->BrxB-BrxF-iSTAND->DUF6079*->CR-ATPase5+REase->N6-METHYLASE->GP+REase->N6-METHYLASE-p->SIR2->HerA/FtsK->?->N6-METHYLASE-p->S1COLD-> | 1223 | Bacteria>Pseudomonadota>Gammaproteobacteria>Pseudomonadales>Pseudomonadaceae>Stutzerimonas | GCF_016599775.1 | Stutzerimonas_balearica | I6H70_RS19230 | 4083458..4087129 | 74829 | BREX-system-P-loop-protein-BrxC-[Stutzerimonas-balearica] |
| WP_218422415.1 | Type1.BREX | <-HTH_Tnp_1+HTH_21+rve+rve_3<-?\|\|?-><-ANK+ANK+ANK\|\|RADICAL-SAM->HicA_toxin->BrxA-DNA-binding->BrxB-BrxF-iSTAND->DUF6079*->N6-METHYLASE->?->HNH+GCN1->?->?->?->?->ParB+HNH->MNS+REase-> | 1154 | Bacteria>Pseudomonadota>Gammaproteobacteria>Pseudomonadales>Pseudomonadaceae>Stutzerimonas | GCF_019203165.1 | Stutzerimonas_stutzeri | KUD99_RS07220 | 1491661..1495125 | 316 | BREX-system-P-loop-protein-BrxC-[Stutzerimonas-stutzeri] |
| WP_218421912.1 | Type1.BREX | <-SIGMA+OB-Fold+AAA-ATPase+Lon-Protease-BrxL<-PglZ<-REase<-OLD-ABC<-N6-METHYLASE-p<-?<-HerA/FtsK<-SIR2<-REase<-N6-METHYLASE<-DUF6079*<-BrxB-BrxF-iSTAND<-BrxA-DNA-binding<-wHTH+WYL+WCX\|\|?->MORC->RVT-> | 1223 | Bacteria>Pseudomonadota>Gammaproteobacteria>Pseudomonadales>Pseudomonadaceae>Stutzerimonas | GCF_019203165.1 | Stutzerimonas_stutzeri | KUD99_RS18220 | 3895806..3899477 | 316 | BREX-system-P-loop-protein-BrxC-[Stutzerimonas-stutzeri] |
| WP_125870426.1 | Type1.BREX | <-DUF932<-?\|\|NERD->REase->BrxA-DNA-binding->BrxB-BrxF-iSTAND->DUF6079*->N6-METHYLASE->?->HNH+GCN1->?->PglZ->SIGMA+OB-Fold+AAA-ATPase+Lon-Protease-BrxL->REase->SF1-SF2-Helicase-><-Glycos_transf | 1154 | Bacteria>Pseudomonadota>Gammaproteobacteria>Pseudomonadales>Pseudomonadaceae>Stutzerimonas | GCF_024199075.1 | Stutzerimonas_stutzeri | NLG33_RS12680 | 207505..210969 | 316 | BREX-system-P-loop-protein-BrxC-[Stutzerimonas-stutzeri] |
| WP_255830664.1 | Type1.BREX | <-sGTP<-sGTP<-sGTP<-wHTH+WYL+WCX\|\|?-><-?\|\|XIS-HTH->BrxA-DNA-binding->BrxB-BrxF-iSTAND->DUF6079*->N6-METHYLASE->OLD-ABC->NERD->RVT->N6-METHYLASE-p->ParB+HNH->PglZ->SIGMA+OB-Fold+AAA-ATPase+HKD-DNase-BrxL->RHH_1-> | 1186 | Bacteria>Pseudomonadota>Gammaproteobacteria>Pseudomonadales>Pseudomonadaceae>Stutzerimonas | GCF_024397535.1 | [Pseudomonas]_zhaodongensis | NA613_RS09805 | 17231..20791 | 1176257 | BREX-system-P-loop-protein-BrxC-[[Pseudomonas] |
| WP_255958979.1 | Type1.BREX | <-SF1-SF2-Helicase<-REase<-SIGMA+OB-Fold+AAA-ATPase+Lon-Protease-BrxL<-PglZ<-ParB+HNH<-?<-MarR-HTH<-MNS+ORC-CDC6<-N6-METHYLASE<-DUF6079*<-BrxB-BrxF-iSTAND<-BrxA-DNA-binding<-REase<-NERD\|\|?->DUF932-> | 1154 | Bacteria>Pseudomonadota>Gammaproteobacteria>Pseudomonadales>Pseudomonadaceae>Stutzerimonas | GCF_024448895.1 | Stutzerimonas_stutzeri | NA619_RS00645 | 147157..150621 | 316 | BREX-system-P-loop-protein-BrxC-[Stutzerimonas-stutzeri] |
| WP_269263428.1 | Type1.BREX | <-HerA/FtsK<-SIR2<-N6-METHYLASE-p<-GP+REase<-N6-METHYLASE<-DOC+HTH<-?<-?<-DUF6079*<-BrxB-BrxF-iSTAND<-BrxA-DNA-binding\|\|Integrase_1->Phage_int_SAM-><-?\|\|DSBH->?-><-Phage_integrase\|\|DUF772+DDE_Tnp_1-> | 1223 | Bacteria>Pseudomonadota>Gammaproteobacteria>Pseudomonadales>Pseudomonadaceae>Stutzerimonas | GCF_027158265.1 | Stutzerimonas_balearica | O3V18_RS18735 | 22059..25730 | 74829 | BREX-system-P-loop-protein-BrxC-[Stutzerimonas-balearica] |
| WP_238586968.1 | Type1.BREX | <-TraK<-TraE<-TraM<-TraM<-DUF3262<-?\|\|XIS-HTH->BrxA-DNA-binding->BrxB-BrxF-iSTAND->DUF6079-p*->PglZ->SIGMA+OB-Fold+AAA-ATPase+HKD-DNase-BrxL->Phage-HTH-><-DUF4400<-VirD4-TraG | 718 | Bacteria>Pseudomonadota>Gammaproteobacteria>Pseudomonadales>Pseudomonadaceae>Stutzerimonas | - | Stutzerimonas_degradans | . | 57411..59567 | 2968968 | BREX-system-P-loop-protein-BrxC-[Stutzerimonas-degradans] |
| BBJ02099.1 | Type1.BREX | <-TraK<-TraE<-TraM<-TraM<-DUF3262<-?\|\|XIS-HTH->BrxA-DNA-binding->BrxB-BrxF-iSTAND->DUF6079-p*->PglZ->SIGMA+OB-Fold+AAA-ATPase+HKD-DNase-BrxL->Phage-HTH-><-DUF4400<-VirD4-TraG | 718 | Bacteria>Pseudomonadota>Gammaproteobacteria>Pseudomonadales>Pseudomonadaceae>Stutzerimonas | - | Stutzerimonas_degradans | . | 57411..59567 | 2968968 | uncharacterized-protein-sporadically-distributed-in-bacteria-and-archaea-[Stutzerimonas-degradans] |
| WP_020308058.1 | Type1.BREX | <-Phage-HTH<-SIGMA+OB-Fold+AAA-ATPase+HKD-DNase-BrxL<-PglZ<-N6-METHYLASE-p<-HerA/FtsK<-SIR2<-REase<-OLD-ABC<-N6-METHYLASE<-DUF6079*<-BrxB-BrxF-iSTAND<-BrxA-DNA-binding<-XIS-HTH\|\|?->DUF3262->TraM->TraM->TraE->TraK-> | 1188 | Bacteria>Pseudomonadota>Gammaproteobacteria>Pseudomonadales>Pseudomonadaceae>Stutzerimonas | - | Stutzerimonas_stutzeri | B382_18428 | 65079..68645 | 316 | BREX-system-P-loop-protein-BrxC-[Stutzerimonas-stutzeri] |
| WP_095461351.1 | Type1.BREX | <-wHTH+WYL+WCX-p\|\|?->wHTH+WYL+WCX->BrxA-DNA-binding->BrxB-BrxF-iSTAND->DUF6079*->N6-METHYLASE->DUF932->ParB+HNH->OLD-ABC->TOPRIM->PglZ->SIGMA+OB-Fold+AAA-ATPase+Lon-Protease-BrxL->MukF->JetB-> | 1154 | Bacteria>Pseudomonadota>Gammaproteobacteria>Pseudomonadales>Pseudomonadaceae>Stutzerimonas | - | Stutzerimonas_stutzeri | BV581_21185 | 4303..7767 | 316 | BREX-system-P-loop-protein-BrxC-[Stutzerimonas-stutzeri] |
| WP_045633496.1 | Type1.BREX | <-RNase<-?<-JetB<-?<-wHTH+WYL+WCX\|\|XIS-HTH->BrxA-DNA-binding->BrxB-BrxF-iSTAND->DUF6079*->N6-METHYLASE->ParB+HNH->PglZ->SIGMA+OB-Fold+AAA-ATPase+HKD-DNase-BrxL-><-?<-?<-?<-DUF932 | 1185 | Bacteria>Pseudomonadota>Gammaproteobacteria>Pseudomonadales>Pseudomonadaceae>Stutzerimonas | - | Stutzerimonas_stutzeri | CAQ69_18770 | 20186..23743 | 316 | MULTISPECIES:-BREX-system-P-loop-protein-BrxC-[Gammaproteobacteria] |
| WP_053105412.1 | Type1.BREX | METHYLASE->?->?->?<-?<-?<-?\|\|wHTH+WYL+WCX->BrxA-DNA-binding->BrxB-BrxF-iSTAND->DUF6079*->N6-METHYLASE->ParB+HNH->OLD-ABC->TOPRIM->HEPN->PglZ->SIGMA+OB-Fold+AAA-ATPase+Lon-Protease-BrxL->Arm-DNA-bind_3+Phage_integrase-><-SIGMA+OB-Fold+AAA-ATPase+Lon-Protease-BrxL | 1155 | Bacteria>Pseudomonadota>Gammaproteobacteria>Pseudomonadales>Pseudomonadaceae>Thiopseudomonas | GCF_001267275.1 | Thiopseudomonas_alkaliphila | AKN90_RS00615 | 123321..126788 | 1697053 | BREX-system-P-loop-protein-BrxC-[Thiopseudomonas-alkaliphila] |
| MBX9914998.1 | Type1.BREX | <-N6-METHYLASE<-Schlafen+HATPase+wHTH<-DUF6079*<-BrxB-BrxF-iSTAND<-BrxA-DNA-binding<-wHTH+WYL+WCX | 1220 | Bacteria>Pseudomonadota>Gammaproteobacteria>Pseudomonadales>Pseudomonadaceae>unclassified_Pseudomonadaceae | GCA_019745275.1 | Pseudomonadaceae_bacterium | K2Y25_13980 | 5013..8675 | 1914538 | BREX-system-P-loop-protein-BrxC-[Pseudomonadaceae-bacterium] |
| MBX9762031.1 | Type1.BREX | <-N6-METHYLASE<-DUF6079*<-BrxB-BrxF-iSTAND<-BrxA-DNA-binding<-wHTH+WYL+WCX\|\|?->MORC-> | 1223 | Bacteria>Pseudomonadota>Gammaproteobacteria>Pseudomonadales>Pseudomonadaceae>unclassified_Pseudomonadaceae | GCA_019745805.1 | Pseudomonadaceae_bacterium | K2Y24_03255 | 4991..8662 | 1914538 | BREX-system-P-loop-protein-BrxC-[Pseudomonadaceae-bacterium] |
| MBX9763927.1 | Type1.BREX | <-LexA-protease<-SF1-SF2-Helicase<-?<-wHTH+WYL+WCX\|\|ARG->?->wHTH+WYL+WCX->BrxA-DNA-binding->BrxB-BrxF-iSTAND->DUF6079*->N6-METHYLASE-> | 1154 | Bacteria>Pseudomonadota>Gammaproteobacteria>Pseudomonadales>Pseudomonadaceae>unclassified_Pseudomonadaceae | GCA_019745805.1 | Pseudomonadaceae_bacterium | K2Y24_13005 | 12458..15922 | 1914538 | BREX-system-P-loop-protein-BrxC-[Pseudomonadaceae-bacterium] |
| MAB42111.1 | Type1.BREX | <-SIGMA+OB-Fold+AAA-ATPase+Lon-Protease-BrxL<-PglZ<-TOPRIM<-OLD-ABC<-ParB+HNH<-N6-METHYLASE<-DUF6079*<-BrxB-BrxF-iSTAND<-BrxA-DNA-binding\|\|wHTH+WYL+WCX-> | 1154 | Bacteria>Pseudomonadota>Gammaproteobacteria>Pseudomonadales>unclassified_Pseudomonadales | GCA_002684155.1 | Pseudomonadales_bacterium | CMK77_07490 | 13688..17152 | 1891229 | hypothetical-protein-[Pseudomonadales-bacterium] |
| MAQ23221.1 | Type1.BREX | <-Calcineurin+Calcineurin-SbcD<-PglZ<-ParB+HNH<-N6-METHYLASE<-REase<-OLD-ABC<-Schlafen+HATPase+wHTH+HTH<-DUF6079*<-BrxB-BrxF-iSTAND<-BrxA-DNA-binding<-XIS-HTH<-MobI\|\|XIS-HTH->AAA+HTH->XIS-HTH-><-Arm-DNA-bind_3+Phage_integrase | 1193 | Bacteria>Pseudomonadota>Gammaproteobacteria>Pseudomonadales>unclassified_Pseudomonadales | GCA_002701315.1 | Pseudomonadales_bacterium | CMK83_03295 | 18015..21596 | 1891229 | hypothetical-protein-[Pseudomonadales-bacterium] |
| MBI27695.1 | Type1.BREX | Arm-DNA-bind_3+Phage_integrase-><-XIS-HTH<-AAA+HTH<-XIS-HTH\|\|MobI->XIS-HTH->BrxA-DNA-binding->BrxB-BrxF-iSTAND->DUF6079*-> | 1009 | Bacteria>Pseudomonadota>Gammaproteobacteria>Pseudomonadales>unclassified_Pseudomonadales | GCA_002718075.1 | Pseudomonadales_bacterium | CMK90_12665 | 612690..615718 | 1891229 | hypothetical-protein-[Pseudomonadales-bacterium] |

| Protein-ID | System | Gene-Neighbourhoods | Length | Taxonomy | Assembly-ID | Strain | Locus-Tag | Genomic-coordinates | TaxID | Fasta-define |
|---|---|---|---|---|---|---|---|---|---|---|
| MBV1884607.1 | Type1.BREX | <-N6-METHYLASE<-?<-AbiEii<-MarR-HTH<-DUF6079*<-BrxB-BrxF-iSTAND<-BrxA-DNA-binding<-wHTH+WYL+WCX | 1218 | Bacteria>Pseudomonadota>Gammaproteobacteria>Pseudomonadales>unclassified_Pseudomonadales | GCA_019090455.1 | Pseudomonadales_bacterium | KUG82_23410 | 5916..9572 | 1891229 | BREX-system-P-loop-protein-BrxC-[Pseudomonadales-bacterium] |
| MCB1656877.1 | Type1.BREX | <-N6-METHYLASE-p<-Couple_hipA+HipA_C<-HTH_3<-DUF6079*<-BrxB-BrxF-iSTAND<-BrxA-DNA-binding<-wHTH+WYL+WCX<-?<-HTH_Tnp_1+HTH_21+rve+rve_3 | 1216 | Bacteria>Pseudomonadota>Gammaproteobacteria>Pseudomonadales>unclassified_Pseudomonadales | GCA_020443205.1 | Pseudomonadales_bacterium | KDI39_01465 | 3417..7067 | 1891229 | BREX-system-P-loop-protein-BrxC-[Pseudomonadales-bacterium] |
| MCG8312832.1 | Type1.BREX | GDResolvase+HIN-HTH-><-?\|\|?->?->XIS-HTH->BrxA-DNA-binding->BrxB-BrxF-iSTAND->DUF6079-p*-> | 468 | Bacteria>Pseudomonadota>Gammaproteobacteria>Pseudomonadales>unclassified_Pseudomonadales | GCA_022360035.1 | Pseudomonadales_bacterium | MI976_06400 | 7277..8681 | 1891229 | BREX-system-P-loop-protein-BrxC-[Pseudomonadales-bacterium] |
| MCG8610925.1 | Type1.BREX | <-N6-METHYLASE<-HTH+REase<-DUF6079*<-BrxB-BrxF-iSTAND<-BrxA-DNA-binding<-wHTH+WYL+WCX\|\|ParA-Soj-PloopNTPase-> | 1229 | Bacteria>Pseudomonadota>Gammaproteobacteria>Pseudomonadales>unclassified_Pseudomonadales | GCA_022361325.1 | Pseudomonadales_bacterium | MI864_10365 | 4826..8515 | 1891229 | BREX-system-P-loop-protein-BrxC-[Pseudomonadales-bacterium] |
| MCJ8311755.1 | Type1.BREX | <-sGTP<-sGTP<-?<-SIGMA+OB-Fold+AAA-ATPase+Lon-Protease-BrxL<-PglZ<-N6-METHYLASE<-DUF6079*<-BrxB-BrxF-iSTAND<-BrxA-DNA-binding<-HNH<-?\|\|?->?-><-SF1-SF2-Helicase<-HKD-DNase+HKD-REase | 1233 | Bacteria>Pseudomonadota>Gammaproteobacteria>Pseudomonadales>unclassified_Pseudomonadales | GCA_022968985.1 | Pseudomonadales_bacterium | HRU38_08495 | 18482..22183 | 1891229 | BREX-system-P-loop-protein-BrxC-[Pseudomonadales-bacterium] |
| MCP5320940.1 | Type1.BREX | <-SbcC+CCDC158<-Calcineurin-SbcD<-PglZ<-ParA-Soj-PloopNTPase<-Schlafen+HATPase+wHTH<-Schlafen+HATPase+wHTH<-N6-METHYLASE<-TOPRIM<-OLD-ABC<-DUF6079*<-BrxB-BrxF-iSTAND<-BrxA-DNA-binding\|\|MerR-HTH->DUF772+DDE_Tnp_1+DDE_Tnp_1_6->?->?-><-SF1-SF2-Helicase<-SF1-SF2-Helicase<-AbiEii+ZnR | 1148 | Bacteria>Pseudomonadota>Gammaproteobacteria>Pseudomonadales>unclassified_Pseudomonadales | GCA_024234165.1 | Pseudomonadales_bacterium | H7A12_08980 | 738054..741500 | 1891229 | BREX-system-P-loop-protein-BrxC-[Pseudomonadales-bacterium] |
| MCP5344826.1 | Type1.BREX | <-PALP<-SIGMA+OB-Fold+AAA-ATPase+Lon-Protease-BrxL<-PglZ<-PHP+SbcC<-N6-METHYLASE<-REase<-OLD-ABC<-DUF6079-p* | 834 | Bacteria>Pseudomonadota>Gammaproteobacteria>Pseudomonadales>unclassified_Pseudomonadales | GCA_024234345.1 | Pseudomonadales_bacterium | H7A05_09410 | 50993..53498 | 1891229 | BREX-system-P-loop-protein-BrxC-[Pseudomonadales-bacterium] |
| MCP5331432.1 | Type1.BREX | UDG->?-><-?\|\|CHASE2+ACYC->tRNA->tRNA-><-Phage_integrase\|\|XIS-HTH->BrxA-DNA-binding->BrxB-BrxF-iSTAND->DUF6079*->OLD-ABC->REase->PHP+SbcC->PglZ->SIGMA+OB-Fold+AAA-ATPase+Lon-Protease-BrxL->PALP-> | 1145 | Bacteria>Pseudomonadota>Gammaproteobacteria>Pseudomonadales>unclassified_Pseudomonadales | GCA_024234365.1 | Pseudomonadales_bacterium | H7A07_10830 | 172054..175491 | 1891229 | BREX-system-P-loop-protein-BrxC-[Pseudomonadales-bacterium] |
| MCP5171278.1 | Type1.BREX | TraG_N-><-?<-?<-?<-?<-?\|\|wHTH+WYL+WCX->BrxA-DNA-binding->BrxB-BrxF-iSTAND->DUF6079*->MukF->JetB->DUF3584+DUF5401->RNase->MarR-HTH->AbiEii->?->N6-METHYLASE->ParB+HNH-> | 1216 | Bacteria>Pseudomonadota>Gammaproteobacteria>Pseudomonadales>unclassified_Pseudomonadales | GCA_024234435.1 | Pseudomonadales_bacterium | H7A02_03290 | 695235..698885 | 1891229 | BREX-system-P-loop-protein-BrxC-[Pseudomonadales-bacterium] |
| MCP5179014.1 | Type1.BREX | <-DUF772+DDE_Tnp_1+DDE_Tnp_1_6<-PglZ<-ParB+HNH<-?<-viral-capsid<-ZnR<-N6-METHYLASE<-HKD-DNase<-sGTP+GTPase-AIG<-DUF6079*<-BrxB-BrxF-iSTAND<-BrxA-DNA-binding<-?\|\|?-><-rve<-rve<-Couple_hipA+HipA_C<-?<-HTH_Tnp_1+HTH_21+rve+rve_3 | 1153 | Bacteria>Pseudomonadota>Gammaproteobacteria>Pseudomonadales>unclassified_Pseudomonadales | - | Pseudomonadales_bacterium | H6994_03785 | 340860..344321 | 1891229 | BREX-system-P-loop-protein-BrxC-[Pseudomonadales-bacterium] |
| WP_101840169.1 | Type1.BREX | ZnR+TOPRIM+TOPRIM->SIGMA-HTH+SIGMA-HTH+SIGMA-HTH-><-?<-wHTH+WYL+WCX\|\|?->HEPN->BrxA-DNA-binding->BrxB-BrxF-iSTAND->DUF6079*->N6-METHYLASE-> | 1210 | Bacteria>Pseudomonadota>Gammaproteobacteria>Thiotrichales>Piscirickettsiaceae>Hydrogenovibrio | GCF_002848135.1 | Hydrogenovibrio_sp._SC-1 | CYQ88_RS04565 | 151062..154694 | 2065820 | BREX-system-P-loop-protein-BrxC-[Hydrogenovibrio-sp.-SC-1] |
| WP_007146327.1 | Type1.BREX | UbiA-><-HNH<-tRNA<-?<-?<-SIGMA+OB-Fold+AAA-ATPase+HKD-DNase-BrxL<-PglZ<-REase<-ParB+HNH<-N6-METHYLASE<-DUF6079*<-BrxB-BrxF-iSTAND<-BrxA-DNA-binding<-NADAR<-ART-DarT<-?<-?<-Calcineurin | 1179 | Bacteria>Pseudomonadota>Gammaproteobacteria>Thiotrichales>Piscirickettsiaceae>Methylophaga | GCF_000214595.1 | Methylophaga_aminisulfidivorans | MAMP_RS12010 | 87136..90675 | 230105 | BREX-system-P-loop-protein-BrxC-[Methylophaga-aminisulfidivorans] |
| CAA6806547.1 | Type1.BREX | <-ParB+HNH<-N6-METHYLASE<-N6-METHYLASE-p<-?<-REase<-ABC-ATPase<-DUF6079-p* | 849 | Bacteria>Pseudomonadota>Gammaproteobacteria>Thiotrichales>Thiotrichaceae>environmental_samples | - | uncultured_Thiotrichaceae_bacterium | HELGO_WM31114 | 6431..8980 | 298394 | FIG00607287:-hypothetical-protein,-partial-[uncultured-Thiotrichaceae-bacterium] |
| WP_022954802.1 | Type1.BREX | CR-ATPase5+REase-><-?\|\|wHTH+WYL+WCX->BrxA-DNA-binding->BrxB-BrxF-iSTAND->DUF6079*->N6-METHYLASE->OLD-ABC->TOPRIM->PglZ->SIGMA+OB-Fold+AAA-ATPase+Lon-Protease-BrxL-> | 1222 | Bacteria>Pseudomonadota>Gammaproteobacteria>Thiotrichales>Thiotrichaceae>Leucothrix | GCF_000419525.1 | Leucothrix_mucor | LEUMU_RS0123780 | 65328..68996 | 45248 | BREX-system-P-loop-protein-BrxC-[Leucothrix-mucor] |
| OQX14377.1 | Type1.BREX | <-TRPR-HTH<-?<-DUF3670+SF1-SF2-Helicase\|\|?-><-SIGMA+OB-Fold+AAA-ATPase+Lon-Protease-BrxL<-PglZ<-TOPRIM<-OLD-ABC<-N6-METHYLASE<-DUF6079*<-BrxB-BrxF-iSTAND<-BrxA-DNA-binding<-TIR<-?<-?\|\|XIS-HTH-> | 1145 | Bacteria>Pseudomonadota>Gammaproteobacteria>Thiotrichales>Thiotrichaceae>Thiothrix | GCA_002083875.1 | Thiothrix_lacustris | BWK73_09690 | 20114..23551 | 525917 | hypothetical-protein-BWK73_09690-[Thiothrix-lacustris] |
| QLQ31990.1 | Type1.BREX | DrsE-><-?\|\|REase->?->Arm-DNA-bind_3+Phage_integrase-><-HTH+DOC<-?<-LexA-protease\|\|BrxB-BrxF-iSTAND->DUF6079*->AbiEi_4+AbiEi_1->AbiEii->OLD-ABC->OLD-ABC->TOPRIM->?\|\|N6-METHYLASE->N6-METHYLASE-p->HIN-HTH-> | 1209 | Bacteria>Pseudomonadota>Gammaproteobacteria>Thiotrichales>Thiotrichaceae>Thiothrix | GCA_013693955.1 | Candidatus_Thiothrix_singaporensis | HZT40_10770 | 2133663..2137292 | 2799669 | BREX-system-P-loop-protein-BrxC-[Candidatus-Thiothrix-singaporensis] |
| WP_002710590.1 | Type1.BREX | HAMP+PAS+HISKIN->REC+NtrC-AAA+FIS-HTH->BrxA-DNA-binding->BrxB-BrxF-iSTAND->DUF6079*->AbiEi_4+AbiEi_1->AbiEii->OLD-ABC->TOPRIM->N6-METHYLASE->CR-ATPase8+REase->PglZ->SIGMA+OB-Fold+AAA-ATPase+Lon-Protease-BrxL-> | 1209 | Bacteria>Pseudomonadota>Gammaproteobacteria>Thiotrichales>Thiotrichaceae>Thiothrix | GCF_000260135.1 | Thiothrix_nivea | THINI_RS21415 | 4300442..4304071 | 1031 | BREX-system-P-loop-protein-BrxC-[Thiothrix-nivea] |
| WP_210225661.1 | Type1.BREX | BrxA-DNA-binding->BrxB-BrxF-iSTAND->DUF6079*->ABC-ATPase->?->N6-METHYLASE->N6-METHYLASE-p-><-REase\|\|HATPase+wHTH->PglZ->SIGMA+OB-Fold+AAA-ATPase+HKD-DNase-BrxL-> | 1173 | Bacteria>Pseudomonadota>Gammaproteobacteria>Thiotrichales>Thiotrichaceae>Thiothrix | GCF_017901155.1 | Candidatus_Thiothrix_anitrata | JB380_RS10820 | 2094889..2098410 | 2823902 | BREX-system-P-loop-protein-BrxC-[Candidatus-Thiothrix-anitrata] |
| MBL1259972.1 | Type1.BREX | <-SIGMA+OB-Fold+AAA-ATPase+Lon-Protease-BrxL<-PglZ<-?<-RVT<-Schlafen+HATPase+wHTH<-N6-METHYLASE<-DUF6079*<-BrxB-BrxF-iSTAND<-BrxA-DNA-binding<-wHTH+WYL+WCX | 1219 | Bacteria>Pseudomonadota>Gammaproteobacteria>Thiotrichales>Thiotrichaceae>unclassified_Thiotrichaceae | GCA_002401765.2 | Thiotrichaceae_bacterium | COB33_005510 | 47175..50834 | 2030882 | BREX-system-P-loop-protein-BrxC-[Thiotrichaceae-bacterium] |
| PCJ88241.1 | Type1.BREX | <-RelE-ParE<-MetJarc-HTH\|\|?-><-HKD-DNase+DUF3150<-SIGMA+OB-Fold+AAA-ATPase+HKD-DNase-BrxL<-PglZ<-N6-METHYLASE<-REase<-OLD-ABC<-DUF6079*<-BrxB-BrxF-iSTAND<-BrxA-DNA-binding<-XIS-HTH<-?<-?\|\|RHH_5->?-><-HISKIN | 1177 | Bacteria>Pseudomonadota>Gammaproteobacteria>Thiotrichales>Thiotrichaceae>unclassified_Thiotrichaceae | GCA_002746365.1 | Thiotrichaceae_bacterium | COA54_02955 | 15853..19386 | 2030882 | hypothetical-protein-COA54_02955-[Thiotrichaceae-bacterium] |
| MBD3820859.1 | Type1.BREX | wHTH+WYL+WCX-p->BrxA-DNA-binding->BrxB-BrxF-iSTAND->DUF6079*->N6-METHYLASE->HTH+REase->REase->PglZ->SIGMA+OB-Fold+AAA-ATPase+HKD-DNase-BrxL-> | 1191 | Bacteria>Pseudomonadota>Gammaproteobacteria>Thiotrichales>unclassified_Thiotrichales | GCA_014764155.1 | Thiotrichales_bacterium | IE914_01145 | 1632..5207 | 2026796 | BREX-system-P-loop-protein-BrxC-[Thiotrichales-bacterium] |
| MBT3348501.1 | Type1.BREX | <-SIGMA+OB-Fold+AAA-ATPase+HKD-DNase-BrxL<-PglZ<-DOC+HTH<-N6-METHYLASE-p<-HerA/FtsK<-SIR2<-Schlafen+HATPase+wHTH<-N6-METHYLASE<-DUF6079*<-BrxB-BrxF-iSTAND<-BrxA-DNA-binding | 1170 | Bacteria>Pseudomonadota>Gammaproteobacteria>Thiotrichales>unclassified_Thiotrichales | GCA_018648495.1 | Thiotrichales_bacterium | HN398_08700 | 13875..17387 | 2026796 | BREX-system-P-loop-protein-BrxC-[Thiotrichales-bacterium] |
| MBK85426.1 | Type1.BREX | HHH_5+SF1-SF2-Helicase->BrxA-DNA-binding->BrxB-BrxF-iSTAND->DUF6079*->N6-METHYLASE->HEPN->Schlafen+HATPase+wHTH->TIR->PglZ->SIGMA+OB-Fold+AAA-ATPase+Lon-Protease-BrxL->?->RNase-><-SWIM+DUF3670+SF1-SF2-Helicase | 1217 | Bacteria>Pseudomonadota>Gammaproteobacteria>unclassified_Gammaproteobacteria | GCA_002721025.1 | Gammaproteobacteria_bacterium | CMD81_16515 | 7659..11312 | 1913989 | ATPase-[Gammaproteobacteria-bacterium] |
| HAG71474.1 | Type1.BREX | <-DUF6079-p*<-DUF6079-p<-BrxB-BrxF-iSTAND-p<-BrxB-BrxF-iSTAND-p<-BrxA-DNA-binding | 416 | Bacteria>Pseudomonadota>Gammaproteobacteria>unclassified_Gammaproteobacteria | GCA_003450175.1 | Gammaproteobacteria_bacterium | DCL66_04645 | 1..1249 | 1913989 | BREX-system-P-loop-protein-BrxC-[Gammaproteobacteria-bacterium] |
| RLA52302.1 | Type1.BREX | XIS-HTH->BrxA-DNA-binding->BrxB-BrxF-iSTAND->DUF6079*->N6-METHYLASE->SF1-SF2-Helicase+PRTase->SLOG+BirA-HTH-> | 1210 | Bacteria>Pseudomonadota>Gammaproteobacteria>unclassified_Gammaproteobacteria | GCA_003646365.1 | Gammaproteobacteria_bacterium | DRR42_07940 | 3455..7087 | 1913989 | BREX-system-P-loop-protein-BrxC-[Gammaproteobacteria-bacterium] |
| TXH19452.1 | Type1.BREX | <-DUF6079*<-BrxB-BrxF-iSTAND<-BrxA-DNA-binding<-XIS-HTH<-HipA_C\|\|?->DUF3262-> | 1027 | Bacteria>Pseudomonadota>Gammaproteobacteria>unclassified_Gammaproteobacteria | GCA_008012265.1 | Gammaproteobacteria_bacterium | E6R00_01845 | 1..3080 | 1913989 | BREX-system-P-loop-protein-BrxC,-partial-[Gammaproteobacteria-bacterium] |
| QFY90721.2 | Type1.BREX | TRD+TRD->DOC+HTH->?->RNase->SF1-SF2-Helicase-><-rve_3\|\|XIS-HTH->BrxA-DNA-binding->BrxB-BrxF-iSTAND->DUF6079*->?->REase->N6-METHYLASE->HTH+REase->PglZ->REase->SIGMA+OB-Fold+AAA-ATPase+HKD-DNase-BrxL->SbcC->Calcineurin+Calcineurin-SbcD-> | 1179 | Bacteria>Pseudomonadota>Gammaproteobacteria>unclassified_Gammaproteobacteria | GCA_009497875.2 | gamma_proteobacterium_SS-5 | D5125_15330 | 2185771..2189310 | 947516 | BREX-system-P-loop-protein-BrxC-[gamma-proteobacterium-SS-5] |
| MXY51046.1 | Type1.BREX | <-HEPN\|\|?-><-Phage_integrase\|\|BrxA-DNA-binding->BrxB-BrxF-iSTAND->DUF6079*->N6-METHYLASE->DOC+HTH->N6-METHYLASE-p->PglZ->SIGMA+OB-Fold+AAA-ATPase+Lon-Protease-BrxL-><-HEPN<-XIS-HTH | 1251 | Bacteria>Pseudomonadota>Gammaproteobacteria>unclassified_Gammaproteobacteria | GCA_009837725.1 | Gammaproteobacteria_bacterium | F4Y86_00740 | 6119..9874 | 1913989 | BREX-system-P-loop-protein-BrxC-[Gammaproteobacteria-bacterium] |
| MYB36894.1 | Type1.BREX | <-Glyoxalase<-SIGMA+OB-Fold+AAA-ATPase+Lon-Protease-BrxL<-PglZ<-PHP+SbcC<-N6-METHYLASE<-DUF6079*<-BrxB-BrxF-iSTAND<-BrxA-DNA-binding\|\|?-><-SAM-METHYLASE | 1146 | Bacteria>Pseudomonadota>Gammaproteobacteria>unclassified_Gammaproteobacteria | GCA_009838905.1 | Gammaproteobacteria_bacterium | F4Y26_05795 | 15436..18876 | 1913989 | BREX-system-P-loop-protein-BrxC-[Gammaproteobacteria-bacterium] |
| NCA89504.1 | Type1.BREX | <-PIN<-?<-MqsA_antitoxin\|\|Schlafen+HATPase+wHTH->BrxA-DNA-binding->BrxB-BrxF-iSTAND->DUF6079*->VapB_antitoxin->N6-METHYLASE-> | 1152 | Bacteria>Pseudomonadota>Gammaproteobacteria>unclassified_Gammaproteobacteria | GCA_009928645.1 | Gammaproteobacteria_bacterium | EOM92_11510 | 6384..9842 | 1913989 | BREX-system-P-loop-protein-BrxC-[Gammaproteobacteria-bacterium] |
| HFD80332.1 | Type1.BREX | <-SIGMA+OB-Fold+AAA-ATPase+Lon-Protease-BrxL<-PglZ<-Schlafen+HATPase+wHTH<-PHP+SbcC<-N6-METHYLASE<-PIN<-DUF6079*<-BrxB-BrxF-iSTAND<-BrxA-DNA-binding | 1156 | Bacteria>Pseudomonadota>Gammaproteobacteria>unclassified_Gammaproteobacteria | GCA_011321775.1 | Gammaproteobacteria_bacterium | ENK05_08090 | 12367..15837 | 1913989 | BREX-system-P-loop-protein-BrxC-[Gammaproteobacteria-bacterium] |
| NLY14765.1 | Type1.BREX | <-SIGMA+OB-Fold+AAA-ATPase+Lon-Protease-BrxL<-PglZ<-HEPN<-TOPRIM<-OLD-ABC<-ParB+HNH<-N6-METHYLASE<-DUF6079-p* | 438 | Bacteria>Pseudomonadota>Gammaproteobacteria>unclassified_Gammaproteobacteria | GCA_012728415.1 | Gammaproteobacteria_bacterium | GXZ05_00075 | 12795..14111 | 1913989 | hypothetical-protein-[Gammaproteobacteria-bacterium] |
| HIJ47516.1 | Type1.BREX | REC->?->PAS+PAS+HISKIN->?->BrxA-DNA-binding->BrxB-BrxF-iSTAND->DUF6079-p*-> | 470 | Bacteria>Pseudomonadota>Gammaproteobacteria>unclassified_Gammaproteobacteria | GCA_013349775.1 | Gammaproteobacteria_bacterium | HPP72_05790 | 9159..10569 | 1913989 | BREX-system-P-loop-protein-BrxC-[Gammaproteobacteria-bacterium] |
| HIJ32929.1 | Type1.BREX | BrxA-DNA-binding->BrxB-BrxF-iSTAND->DUF6079*->OLD-ABC->REase->N6-METHYLASE-p-> | 1200 | Bacteria>Pseudomonadota>Gammaproteobacteria>unclassified_Gammaproteobacteria | GCA_013349805.1 | Gammaproteobacteria_bacterium | HPP65_00765 | 772..4374 | 1913989 | BREX-system-P-loop-protein-BrxC-[Gammaproteobacteria-bacterium] |
| HIJ28180.1 | Type1.BREX | REC->?->PAS+PAS+HISKIN->?->BrxA-DNA-binding->BrxB-BrxF-iSTAND->DUF6079-p*-> | 466 | Bacteria>Pseudomonadota>Gammaproteobacteria>unclassified_Gammaproteobacteria | GCA_013349855.1 | Gammaproteobacteria_bacterium | HPP91_06560 | 8328..9724 | 1913989 | BREX-system-P-loop-protein-BrxC-[Gammaproteobacteria-bacterium] |
| NVK29385.1 | Type1.BREX | MetJarc-HTH-><-SIGMA+OB-Fold+AAA-ATPase+HKD-DNase-BrxL<-PglZ<-N6-METHYLASE<-DUF6079*<-BrxB-BrxF-iSTAND<-BrxA-DNA-binding<-wHTH+WYL+WCX\|\|?->SF1-SF2-Helicase+DUF1998-><-PIN<-MazE_antitoxin | 1180 | Bacteria>Pseudomonadota>Gammaproteobacteria>unclassified_Gammaproteobacteria | GCA_013373515.1 | Gammaproteobacteria_bacterium | HWE20_00180 | 17762..21304 | 1913989 | BREX-system-P-loop-protein-BrxC-[Gammaproteobacteria-bacterium] |
| MBK7728002.1 | Type1.BREX | Peptidase-><-DDE_Tnp<-SIGMA+OB-Fold+AAA-ATPase+Lon-Protease-BrxL<-PglZ<-HEPN<-?\|\|?-><-DUF6079*<-BrxB-BrxF-iSTAND<-BrxA-DNA-binding\|\|URI-DNase->?-><-tRNA<-MerR-HTH | 1048 | Bacteria>Pseudomonadota>Gammaproteobacteria>unclassified_Gammaproteobacteria | GCA_016709565.1 | Gammaproteobacteria_bacterium | IPJ33_05745 | 303360..306506 | 1913989 | BREX-system-P-loop-protein-BrxC-[Gammaproteobacteria-bacterium] |
| MBR9924470.1 | Type1.BREX | <-TetR-HTH\|\|?->tRNA->ncRNA->REase+wHTH->rve->TnsC-ATPase->HTH_19->BrxA-DNA-binding->BrxB-BrxF-iSTAND->DUF6079*->HTH+REase->Schlafen+HATPase+wHTH-> | 1227 | Bacteria>Pseudomonadota>Gammaproteobacteria>unclassified_Gammaproteobacteria | GCA_018223005.1 | Gammaproteobacteria_bacterium | GYB28_05690 | 375823..379506 | 1913989 | BREX-system-P-loop-protein-BrxC-[Gammaproteobacteria-bacterium] |
| MBT3489237.1 | Type1.BREX | <-DUF6079-p*<-BrxB-BrxF-iSTAND<-BrxA-DNA-binding<-?<-PAS+PAS+HISKIN<-?<-REC | 499 | Bacteria>Pseudomonadota>Gammaproteobacteria>unclassified_Gammaproteobacteria | GCA_018647365.1 | Gammaproteobacteria_bacterium | HN455_06145 | 1..1498 | 1913989 | BREX-system-P-loop-protein-BrxC-[Gammaproteobacteria-bacterium] |
| MBT3403047.1 | Type1.BREX | REC->?->PAS+PAS+HISKIN->?->BrxA-DNA-binding->BrxB-BrxF-iSTAND->DUF6079-p*-> | 701 | Bacteria>Pseudomonadota>Gammaproteobacteria>unclassified_Gammaproteobacteria | GCA_018648065.1 | Gammaproteobacteria_bacterium | HN421_04975 | 34194..36297 | 1913989 | BREX-system-P-loop-protein-BrxC-[Gammaproteobacteria-bacterium] |
| MBT3892721.1 | Type1.BREX | <-DUF6079-p*<-BrxB-BrxF-iSTAND<-BrxA-DNA-binding<-?<-PAS+PAS+HISKIN<-?<-REC | 742 | Bacteria>Pseudomonadota>Gammaproteobacteria>unclassified_Gammaproteobacteria | GCA_018649125.1 | Gammaproteobacteria_bacterium | HDC39_05580 | 1..2226 | 1913989 | BREX-system-P-loop-protein-BrxC-[Gammaproteobacteria-bacterium] |
| MBT4196271.1 | Type1.BREX | <-SIGMA+OB-Fold+AAA-ATPase+HKD-DNase-BrxL<-REase<-Immank+ANK+ANK<-PglZ<-N6-METHYLASE<-SLOG+BirA-HTH<-SF1-SF2-Helicase+PRTase<-DUF6079*<-BrxB-BrxF-iSTAND<-BrxA-DNA-binding<-HTH_26\|\|?->?->REase->rve+Mu-transpos_C->TnsC-ATPase->ZnR-> | 1178 | Bacteria>Pseudomonadota>Gammaproteobacteria>unclassified_Gammaproteobacteria | GCA_018651435.1 | Gammaproteobacteria_bacterium | HOE12_18240 | 31044..34580 | 1913989 | BREX-system-P-loop-protein-BrxC-[Gammaproteobacteria-bacterium] |
| MBT3718524.1 | Type1.BREX | <-DUF6079-p*<-BrxB-BrxF-iSTAND<-BrxA-DNA-binding<-?<-PAS+PAS+HISKIN<-?<-REC | 570 | Bacteria>Pseudomonadota>Gammaproteobacteria>unclassified_Gammaproteobacteria | GCA_018653885.1 | Gammaproteobacteria_bacterium | HOG37_06305 | 1..1711 | 1913989 | BREX-system-P-loop-protein-BrxC-[Gammaproteobacteria-bacterium] |
| MBT3846012.1 | Type1.BREX | <-DUF6079-p*<-BrxB-BrxF-iSTAND<-BrxA-DNA-binding<-?<-PAS+PAS+HISKIN<-?<-REC | 730 | Bacteria>Pseudomonadota>Gammaproteobacteria>unclassified_Gammaproteobacteria | GCA_018655435.1 | Gammaproteobacteria_bacterium | HOF59_12730 | 1..2190 | 1913989 | BREX-system-P-loop-protein-BrxC-[Gammaproteobacteria-bacterium] |
| MBT5223355.1 | Type1.BREX | BrxA-DNA-binding->BrxB-BrxF-iSTAND->DUF6079*->MazE_antitoxin->PIN->N6-METHYLASE->N6-METHYLASE->HEPN->GrpE->?->?->RelE-ParE-> | 1192 | Bacteria>Pseudomonadota>Gammaproteobacteria>unclassified_Gammaproteobacteria | GCA_018661125.1 | Gammaproteobacteria_bacterium | HOM15_11165 | 1377..4955 | 1913989 | BREX-system-P-loop-protein-BrxC-[Gammaproteobacteria-bacterium] |
| MBT5688340.1 | Type1.BREX | <-DUF6079-p*<-BrxB-BrxF-iSTAND<-BrxA-DNA-binding<-?<-PAS+PAS+HISKIN<-?<-REC | 960 | Bacteria>Pseudomonadota>Gammaproteobacteria>unclassified_Gammaproteobacteria | GCA_018667295.1 | Gammaproteobacteria_bacterium | HOI61_07455 | 1..2880 | 1913989 | BREX-system-P-loop-protein-BrxC-[Gammaproteobacteria-bacterium] |
| MBT7480622.1 | Type1.BREX | <-DUF6079-p*<-BrxB-BrxF-iSTAND<-BrxA-DNA-binding<-?<-PAS+PAS+HISKIN<-?<-REC | 580 | Bacteria>Pseudomonadota>Gammaproteobacteria>unclassified_Gammaproteobacteria | GCA_018691505.1 | Gammaproteobacteria_bacterium | HN661_14095 | 1..1740 | 1913989 | BREX-system-P-loop-protein-BrxC-[Gammaproteobacteria-bacterium] |
| MBT6879379.1 | Type1.BREX | <-DUF6079-p*<-BrxB-BrxF-iSTAND<-BrxA-DNA-binding<-?<-PAS+PAS+HISKIN<-?<-REC | 757 | Bacteria>Pseudomonadota>Gammaproteobacteria>unclassified_Gammaproteobacteria | GCA_018695175.1 | Gammaproteobacteria_bacterium | HOA22_07610 | 1..2271 | 1913989 | BREX-system-P-loop-protein-BrxC-[Gammaproteobacteria-bacterium] |
| MBU0810851.1 | Type1.BREX | DUF4969-><-SIGMA+OB-Fold+AAA-ATPase+HKD-DNase-BrxL<-PglZ<-N6-METHYLASE-p<-RVT<-NERD<-OLD-ABC<-N6-METHYLASE<-DUF6079*<-BrxB-BrxF-iSTAND<-BrxA-DNA-binding<-XIS-HTH<-?\|\|wHTH+WYL+WCX->?-><-?<-XIS-HTH<-tRNA<-MerR-HTH | 1184 | Bacteria>Pseudomonadota>Gammaproteobacteria>unclassified_Gammaproteobacteria | GCA_018815345.1 | Gammaproteobacteria_bacterium | KKH69_04595 | 28957..32511 | 1913989 | BREX-system-P-loop-protein-BrxC-[Gammaproteobacteria-bacterium] |
| MBU1286063.1 | Type1.BREX | BrxA-DNA-binding->BrxB-BrxF-iSTAND->DUF6079*->?->N6-METHYLASE->ParB+HNH->?->AbiEi_3_N+AbiEi_3->AbiEii->PglZ-> | 1222 | Bacteria>Pseudomonadota>Gammaproteobacteria>unclassified_Gammaproteobacteria | GCA_018819505.1 | Gammaproteobacteria_bacterium | KJ794_21910 | 1033..4701 | 1913989 | BREX-system-P-loop-protein-BrxC-[Gammaproteobacteria-bacterium] |
| MBZ0070158.1 | Type1.BREX | <-Peptidase+SF1-SF2-Helicase+REase<-HTH_3<-SIGMA-HTH<-SIGMA+OB-Fold+AAA-ATPase+Lon-Protease-BrxL<-PglZ<-N6-METHYLASE<-DUF6079*<-BrxB-BrxF-iSTAND<-BrxA-DNA-binding\|\|?-><-TraG_N<-?<-TraH<-TraU<-?\|\|DUF772+DDE_Tnp_1-> | 1147 | Bacteria>Pseudomonadota>Gammaproteobacteria>unclassified_Gammaproteobacteria | GCA_019911805.1 | Gammaproteobacteria_bacterium | KBI04_00285 | 50684..54127 | 1913989 | BREX-system-P-loop-protein-BrxC-[Gammaproteobacteria-bacterium] |
| MCG3145056.1 | Type1.BREX | HTH_23+HTH_32+HTH_33->DDE_3-><-SIGMA+OB-Fold+AAA-ATPase+Lon-Protease-BrxL<-PglZ<-Schlafen+HATPase+wHTH+HTH<-N6-METHYLASE<-HEPN<-DUF6079*<-BrxB-BrxF-iSTAND<-BrxA-DNA-binding<-?\|\|?-><-ParB<-ParA-Soj-PloopNTPase\|\|Glyoxalase-><-?\|\|AbiEi_4+AbiEi_1-> | 1220 | Bacteria>Pseudomonadota>Gammaproteobacteria>unclassified_Gammaproteobacteria | GCA_022072255.1 | Gammaproteobacteria_bacterium | HONDAALG_02597 | 52539..56201 | 1913989 | hypothetical-protein-[Gammaproteobacteria-bacterium] |
| MCP4410534.1 | Type1.BREX | HTH_26->BrxA-DNA-binding->BrxB-BrxF-iSTAND->DUF6079-p*->OLD-ABC->TOPRIM->Schlafen+HATPase+wHTH->PglZ->SIGMA+OB-Fold+AAA-ATPase+Lon-Protease-BrxL-> | 791 | Bacteria>Pseudomonadota>Gammaproteobacteria>unclassified_Gammaproteobacteria | GCA_024222475.1 | Gammaproteobacteria_bacterium | GY807_22880 | 5382..7757 | 1913989 | BREX-system-P-loop-protein-BrxC-[Gammaproteobacteria-bacterium] |
| MCP5451255.1 | Type1.BREX | <-ParB+HTH<-ParA-Soj-PloopNTPase\|\|?->BrxA-DNA-binding->BrxB-BrxF-iSTAND->DUF6079*->ParD_antitoxin->RelE-ParE->N6-METHYLASE->OLD-ABC->TOPRIM->REase+AIPR->PglZ->SIGMA+OB-Fold+AAA-ATPase+Lon-Protease-BrxL-><-Thermonuclease | 1154 | Bacteria>Pseudomonadota>Gammaproteobacteria>unclassified_Gammaproteobacteria | GCA_024235115.1 | Gammaproteobacteria_bacterium | H6972_12070 | 41829..45293 | 1913989 | BREX-system-P-loop-protein-BrxC-[Gammaproteobacteria-bacterium] |
| WP_245395138.1 | Type1.BREX | LON-AAA-><-Calcineurin-SbcD<-Calcineurin-><-?\|\|?->\|\|BrxA-DNA-binding->DUF6079-p-><-REase<-OLD-ABC<-DUF6079*<-HDRnase\|\|<-Arm-DNA-bind_2+Phage_int_SAM_3+Phage_integrase<-Phage_AlpA\|\|HTH_26-><-?<-tRNA\|\|RADICAL-SAM->?->DDE_Tnp_1-> | 1115 | Bacteria>Pseudomonadota>Gammaproteobacteria>unclassified_Gammaproteobacteria | GCF_016097405.1 | methane-oxidizing_endosymbiont_of_Gigantopelta_aegis | JEU20_RS06805 | 1356880..1360227 | 2794938 | BREX-system-P-loop-protein-BrxC-[methane-oxidizing-endosymbiont-of-Gigantopelta-aegis] |
| WP_005419548.1 | Type1.BREX | Peptidase_M32->SF1-SF2-Helicase+REase->SF1-SF2-Helicase+REase-><-TPR<-HEPN+SF1-SF2-Helicase+RNase+RNase<-PglZ<-ParB+HNH<-N6-METHYLASE<-DUF6079*<-BrxB-BrxF-iSTAND<-BrxA-DNA-binding<-wHTH+WYL+WCX\|\|PALP->RNase-><-?\|\|HTH_3->Couple_hipA+HipA_C-><-HNH | 1157 | Bacteria>Pseudomonadota>Gammaproteobacteria>Vibrionales>Vibrionaceae>Aliivibrio | GCF_000241785.1 | Aliivibrio_fischeri | VFSR5_RS07520 | 1604629..1608102 | 668 | BREX-system-P-loop-protein-BrxC-[Aliivibrio-fischeri] |
| WP_065641602.1 | Type1.BREX | <-DSBH+AraC-HTH+AraC-HTH\|\|?-><-?<-Glyoxalase<-?\|\|MODE-HTH+PBPII->wHTH+WYL+WCX->BrxA-DNA-binding->BrxB-BrxF-iSTAND->DUF6079*->Schlafen+HATPase+wHTH->N6-METHYLASE-><-SICOLD\|\|ParB+HNH->PglZ->SIGMA+OB-Fold+AAA-ATPase+Lon-Protease-BrxL-><-Acetate_kinase\|\|NtrC-AAA+FIS-HTH->GGDEF-> | 1227 | Bacteria>Pseudomonadota>Gammaproteobacteria>Vibrionales>Vibrionaceae>Aliivibrio | GCF_001690915.1 | Aliivibrio_fischeri | A6E13_RS16565 | 253649..257332 | 668 | BREX-system-P-loop-protein-BrxC-[Aliivibrio-fischeri] |
| WP_130049303.1 | Type1.BREX | HEPN-><-N6-METHYLASE-p<-ParB+HNH<-?<-N6-METHYLASE<-Schlafen+HATPase+wHTH<-DUF6079* | 1142 | Bacteria>Pseudomonadota>Gammaproteobacteria>Vibrionales>Vibrionaceae>Aliivibrio | GCF_004168445.1 | Aliivibrio_finisterrensis | ERW57_RS17830 | 13668..17098 | 511998 | BREX-system-P-loop-protein-BrxC,-partial-[Aliivibrio-finisterrensis] |
| WP_133416081.1 | Type1.BREX | <-DSBH+AraC-HTH+AraC-HTH\|\|?-><-?<-Glyoxalase<-?\|\|MODE-HTH+PBPII->wHTH+WYL+WCX->BrxA-DNA-binding->BrxB-BrxF-iSTAND->DUF6079*->Schlafen+HATPase+wHTH->N6-METHYLASE->ParB+HNH->ZnR+3C-ABC+TPR-> | 1227 | Bacteria>Pseudomonadota>Gammaproteobacteria>Vibrionales>Vibrionaceae>Aliivibrio | GCF_004359415.1 | Aliivibrio_fischeri | VFFQA001_RS16045 | 453579..457262 | 668 | BREX-system-P-loop-protein-BrxC-[Aliivibrio-fischeri] |
| WP_155683402.1 | Type1.BREX | SF1-SF2-Helicase-><-ZnR<-?\|\|viral-protein-><-SF1-SF2-Helicase+HTH\|\|Trypsin+PDZ->wHTH+WYL+WCX->BrxA-DNA-binding->BrxB-BrxF-iSTAND->DUF6079*->ParB+HNH->N6-METHYLASE->?->?->TOPRIM->PglZ->SIGMA+OB-Fold+AAA-ATPase+Lon-Protease-BrxL-> | 1233 | Bacteria>Pseudomonadota>Gammaproteobacteria>Vibrionales>Vibrionaceae>Aliivibrio | GCF_009727645.1 | Aliivibrio_fischeri | GNP73_RS13855 | 802390..806091 | 668 | BREX-system-P-loop-protein-BrxC-[Aliivibrio-fischeri] |
| WP_188863624.1 | Type1.BREX | MODE-HTH+PBPII->wHTH+WYL+WCX->BrxA-DNA-binding->BrxB-BrxF-iSTAND->DUF6079*->?->N6-METHYLASE->HEPN+Sel1+SEC-C->GrpE-> | 1227 | Bacteria>Pseudomonadota>Gammaproteobacteria>Vibrionales>Vibrionaceae>Aliivibrio | GCF_014647395.1 | Aliivibrio_fischeri | IEY04_RS10160 | 4332..8015 | 668 | BREX-system-P-loop-protein-BrxC-[Aliivibrio-fischeri] |
| WP_190395653.1 | Type1.BREX | <-Couple_hipA+HipA_C<-HTH_3\|\|?-><-RNase<-PALP\|\|wHTH+WYL+WCX->BrxA-DNA-binding->BrxB-BrxF-iSTAND->DUF6079*->N6-METHYLASE->OLD-ABC->TOPRIM->PglZ->HEPN+SF1-SF2-Helicase+RNase+RNase->TPR-><-SF1-SF2-Helicase+REase<-SF1-SF2-Helicase+REase<-Peptidase_M32 | 1157 | Bacteria>Pseudomonadota>Gammaproteobacteria>Vibrionales>Vibrionaceae>Aliivibrio | GCF_014694495.1 | Aliivibrio_sp._S10_S31 | HC728_RS13680 | 57939..61412 | 2720224 | BREX-system-P-loop-protein-BrxC-[Aliivibrio-sp.-S10_S31] |
| WP_199414880.1 | Type1.BREX | Peptidase_M32->SF1-SF2-Helicase+REase->SF1-SF2-Helicase+REase-><-TPR<-HEPN+SF1-SF2-Helicase+RNase+RNase<-PglZ<-TOPRIM<-OLD-ABC<-N6-METHYLASE<-DUF6079*<-BrxB-BrxF-iSTAND<-BrxA-DNA-binding<-wHTH+WYL+WCX\|\|PALP->RNase-><-?\|\|HTH_3->Couple_hipA+HipA_C-><-HNH | 1157 | Bacteria>Pseudomonadota>Gammaproteobacteria>Vibrionales>Vibrionaceae>Aliivibrio | GCF_023983475.1 | Aliivibrio_fischeri | AVFI_RS07680 | 1646065..1649538 | 668 | BREX-system-P-loop-protein-BrxC-[Aliivibrio-fischeri] |
| WP_102260244.1 | Type1.BREX | Arm-DNA-bind_3+Phage_integrase-><-XIS-HTH\|\|SF1-SF2-Helicase+REase->SF1-SF2-Helicase+REase-><-PglZ<-ParB+HNH<-N6-METHYLASE<-DUF6079*<-BrxB-BrxF-iSTAND<-BrxA-DNA-binding<-wHTH+WYL+WCX\|\|?->NERD+SF1-SF2-Helicase-><-ANK+ANK<-HicA_toxin\|\|HipA_C->HTH_19-> | 1157 | Bacteria>Pseudomonadota>Gammaproteobacteria>Vibrionales>Vibrionaceae>Enterovibrio | GCF_002874375.1 | Enterovibrio_norvegicus | BCU62_16195 | 25204..28677 | 188144 | BREX-system-P-loop-protein-BrxC-[Enterovibrio-norvegicus] |
| WP_150138110.1 | Type1.BREX | wHTH+WYL+WCX->BrxA-DNA-binding->BrxB-BrxF-iSTAND->DUF6079*->N6-METHYLASE->OLD-ABC->HNH->PglZ-> | 1159 | Bacteria>Pseudomonadota>Gammaproteobacteria>Vibrionales>Vibrionaceae>Enterovibrio | GCF_008692425.1 | Candidatus_Enterovibrio_escacola | EDM10_RS06800 | 2982..6461 | 1927127 | BREX-system-P-loop-protein-BrxC-[Candidatus-Enterovibrio-escacola] |
| WP_006071674.1 | Type1.BREX | <-DUF4113\|\|wHTH->?-><-SIGMA+OB-Fold+AAA-ATPase+Lon-Protease-BrxL<-PglZ<-Schlafen+HATPase+wHTH<-SbcC<-N6-METHYLASE<-DUF6079*<-BrxB-BrxF-iSTAND<-BrxA-DNA-binding<-wHTH+WYL+WCX<-viral-protein<-AbiEi<-SF1-SF2-Helicase<-?\|\|?->ZnR-> | 1225 | Bacteria>Pseudomonadota>Gammaproteobacteria>Vibrionales>Vibrionaceae | GCF_000181535.1 | Vibrio | VSAK1_RS11985 | 54730..58407 | 662 | MULTISPECIES:-BREX-system-P-loop-protein-BrxC-[Vibrio] |
| WP_025612391.1 | Type1.BREX | SF1-SF2-Helicase+REase->SF1-SF2-Helicase+REase-><-?<-?<-sGTP<-PglZ<-ParB+HNH<-N6-METHYLASE<-DUF6079*<-BrxB-BrxF-iSTAND<-BrxA-DNA-binding<-wHTH+WYL+WCX\|\|<-?<-CR-ATPase4 | 1157 | Bacteria>Pseudomonadota>Gammaproteobacteria>Vibrionales>Vibrionaceae | GCF_000489615.1 | Vibrio | DX35_RS0111130 | 15561..19034 | 662 | MULTISPECIES:-BREX-system-P-loop-protein-BrxC-[Vibrio] |
| WP_103201396.1 | Type1.BREX | <-VirD4-TraG+VirB4-FtsK<-HDRnase<-SbcC<-SIGMA+OB-Fold+AAA-ATPase+Lon-Protease-BrxL<-PglZ<-N6-METHYLASE<-TOPRIM<-OLD-ABC<-DUF6079*<-BrxB-BrxF-iSTAND<-BrxA-DNA-binding<-wHTH+WYL+WCX<-?<-35exo<-?\|\|LexA-protease->?-><-MobI | 1225 | Bacteria>Pseudomonadota>Gammaproteobacteria>Vibrionales>Vibrionaceae | GCF_003047085.1 | Vibrio | DA442_RS13700 | 2691122..2694799 | 662 | MULTISPECIES:-BREX-system-P-loop-protein-BrxC-[Vibrio] |
| WP_159406353.1 | Type1.BREX | <-GspA-ATPase<-?\|\|CR-ATPase4->?-><-?<-LexA-protease\|\|wHTH+WYL+WCX->BrxA-DNA-binding->BrxB-BrxF-iSTAND->DUF6079*->N6-METHYLASE->ParB+HNH->PglZ->HEPN+SF1-SF2-Helicase+RNase+RNase->TPR-><-SF1-SF2-Helicase+REase<-SF1-SF2-Helicase+REase<-viral-protein<-AbiEi | 1157 | Bacteria>Pseudomonadota>Gammaproteobacteria>Vibrionales>Vibrionaceae | GCF_009883895.1 | Vibrio | O8X55_RS08175 | 512098..515571 | 662 | MULTISPECIES:-BREX-system-P-loop-protein-BrxC-[Vibrio] |
| WP_206969396.1 | Type1.BREX | <-HEPN\|\|CR-ATPase4->?->wHTH-><-SIGMA+OB-Fold+AAA-ATPase+Lon-Protease-BrxL<-PglZ<-?<-?<-N6-METHYLASE<-DUF6079*<-BrxB-BrxF-iSTAND<-BrxA-DNA-binding<-wHTH+WYL+WCX<-viral-protein<-AbiEi<-SF1-SF2-Helicase\|\|?->ZnR-> | 1225 | Bacteria>Pseudomonadota>Gammaproteobacteria>Vibrionales>Vibrionaceae | GCF_017313425.1 | Vibrio | JO673_RS01390 | 328257..331934 | 662 | MULTISPECIES:-BREX-system-P-loop-protein-BrxC-[Vibrio] |
| WP_228075537.1 | Type1.BREX | <-PIN+OST-HTH<-CARF+NtrC-AAA<-HKD-DNase+HKD-DNase<-MarR-HTH<-SF1-SF2-Helicase\|\|wHTH+WYL+WCX->BrxA-DNA-binding->BrxB-BrxF-iSTAND->DUF6079*->N6-METHYLASE->ParB+HNH->ZnR+3C-ABC+TPR->?->?->PglZ->SIGMA+OB-Fold+AAA-ATPase+Lon-Protease-BrxL-><-?<-CR-ATPase4 | 1248 | Bacteria>Pseudomonadota>Gammaproteobacteria>Vibrionales>Vibrionaceae | GCF_020740925.1 | Vibrio | IB290_RS12355 | 161430..165176 | 662 | MULTISPECIES:-BREX-system-P-loop-protein-BrxC-[Vibrio] |
| WP_180208765.1 | Type1.BREX | AbiEi->viral-protein->SF1-SF2-Helicase+REase->SF1-SF2-Helicase+REase-><-?<-ZetaToxin<-PglZ<-ParB+HNH<-N6-METHYLASE<-DUF6079*<-BrxB-BrxF-iSTAND<-BrxA-DNA-binding<-wHTH+WYL+WCX\|\|<-?<-CR-ATPase4\|\|?->HEPN-><-sGTP | 1157 | Bacteria>Pseudomonadota>Gammaproteobacteria>Vibrionales>Vibrionaceae | GCF_021725735.1 | Vibrio | L3V27_RS03750 | 866566..870039 | 662 | MULTISPECIES:-BREX-system-P-loop-protein-BrxC-[Vibrio] |
| WP_086713494.1 | Type1.BREX | <-GrpE<-ANK+ANK+ANK\|\|?->SF1-SF2-Helicase->?->?->wHTH+WYL+WCX->BrxA-DNA-binding->BrxB-BrxF-iSTAND->DUF6079*->N6-METHYLASE->ParB+HNH->PglZ->HEPN+SF1-SF2-Helicase+RNase+RNase->TPR-><-SF1-SF2-Helicase+REase<-SF1-SF2-Helicase+REase<-?\|\|AbiEi_4-> | 1157 | Bacteria>Pseudomonadota>Gammaproteobacteria>Vibrionales>Vibrionaceae | GCF_024347515.1 | Vibrio | EDB41_RS15380 | 67742..71215 | 662 | MULTISPECIES:-BREX-system-P-loop-protein-BrxC-[Vibrio] |
| WP_198107480.1 | Type1.BREX | AbiEi->viral-protein->SF1-SF2-Helicase+REase->SF1-SF2-Helicase+REase-><-?<-ZetaToxin<-PglZ<-ParB+HNH<-N6-METHYLASE<-DUF6079*<-BrxB-BrxF-iSTAND<-BrxA-DNA-binding<-wHTH+WYL+WCX\|\|DUF4113-><-?<-CR-ATPase4 | 1157 | Bacteria>Pseudomonadota>Gammaproteobacteria>Vibrionales>Vibrionaceae | GCF_028615395.1 | Vibrio | O8Z82_RS06535 | 628630..632103 | 662 | MULTISPECIES:-BREX-system-P-loop-protein-BrxC-[Vibrio] |
| WP_130066623.1 | Type1.BREX | HEPN-><-N6-METHYLASE-p<-ParB+HNH<-?<-N6-METHYLASE<-Schlafen+HATPase+wHTH<-DUF6079*<-BrxB-BrxF-iSTAND<-BrxA-DNA-binding<-wHTH+WYL+WCX<-?<-viral-protein\|\|?->ZnR-><-?\|\|CHTH-> | 1227 | Bacteria>Pseudomonadota>Gammaproteobacteria>Vibrionales>Vibrionaceae | GCF_028765545.1 | Aliivibrio | ERW52_RS16080 | 26875..30558 | 511678 | MULTISPECIES:-BREX-system-P-loop-protein-BrxC-[Aliivibrio] |
| PSU45803.1 | Type1.BREX | HTH_8->XIS-HTH->BrxA-DNA-binding->BrxB-BrxF-iSTAND->DUF6079*->N6-METHYLASE->PglZ->SIGMA+OB-Fold+AAA-ATPase+HKD-DNase-BrxL-><-DDE_Tnp<-?\|\|?->?-><-?<-LuxR-HTH | 1185 | Bacteria>Pseudomonadota>Gammaproteobacteria>Vibrionales>Vibrionaceae>Photobacterium | GCA_003025615.1 | Photobacterium_frigidiphilum | C9J12_20970 | 5140..8697 | 264736 | BREX-system-P-loop-protein-BrxC-[Photobacterium-frigidiphilum] |
| NAW85774.1 | Type1.BREX | wHTH+WYL+WCX-p->BrxA-DNA-binding->BrxB-BrxF-iSTAND->DUF6079*->N6-METHYLASE-p-> | 1154 | Bacteria>Pseudomonadota>Gammaproteobacteria>Vibrionales>Vibrionaceae>Photobacterium | GCA_009904625.1 | Photobacterium_halotolerans | CAG71_04765 | 1726..5190 | 265726 | BREX-system-P-loop-protein-BrxC-[Photobacterium-halotolerans] |
| QDQ70156.1 | Type1.BREX | SF1-SF2-Helicase+REase->SF1-SF2-Helicase+REase-><-PglZ<-TOPRIM<-OLD-ABC<-N6-METHYLASE<-DUF6079*<-BrxB-BrxF-iSTAND<-BrxA-DNA-binding<-wHTH+WYL+WCX<-XIS-HTH<-?<-REase+SF1-SF2-Helicase\|\|SAM-METHYLASE->HKD-DNase+SF1-SF2-Helicase+HTH+HTH+SRA-> | 1154 | Bacteria>Pseudomonadota>Gammaproteobacteria>Vibrionales>Vibrionaceae>Photobacterium | GCA_014930975.1 | Photobacterium_damselae_subsp._damselae | IL982_10195 | 2142038..2145502 | 85581 | BREX-system-P-loop-protein-BrxC-[Photobacterium-damselae-subsp.-damselae] |
| WP_006647488.1 | Type1.BREX | HTH_Tnp_1+HTH_21+rve+rve_3-><-SIGMA+OB-Fold+AAA-ATPase+HKD-DNase-BrxL<-PglZ<-ParB+HNH<-N6-METHYLASE<-REase<-OLD-ABC<-HTH_Tnp_1+HTH_21+rve+rve_3<-OLD-ABC<-DUF6079*<-BrxB-BrxF-iSTAND<-BrxA-DNA-binding<-XIS-HTH<-?<-TnsC-ATPase\|\|HTH_Tnp_1+HTH_21+rve+rve_3-><-rve+Mu-transpos_C<-REase+wHTH | 1193 | Bacteria>Pseudomonadota>Gammaproteobacteria>Vibrionales>Vibrionaceae>Photobacterium | GCF_000153325.1 | Photobacterium_sp._SKA34 | SKA34_RS21260 | 87534..91115 | 121723 | BREX-system-P-loop-protein-BrxC-[Photobacterium-sp.-SKA34] |
| WP_227876424.1 | Type1.BREX | HTH_8->XIS-HTH->BrxA-DNA-binding->BrxB-BrxF-iSTAND->DUF6079*->SbcC->N6-METHYLASE->PglZ->SIGMA+OB-Fold+AAA-ATPase+HKD-DNase-BrxL-><-?\|\|?-><-?<-LuxR-HTH | 1187 | Bacteria>Pseudomonadota>Gammaproteobacteria>Vibrionales>Vibrionaceae>Photobacterium | GCF_003025615.1 | Photobacterium_frigidiphilum | C9J12_RS20995 | 5134..8697 | 264736 | BREX-system-P-loop-protein-BrxC-[Photobacterium-frigidiphilum] |
| WP_115426815.1 | Type1.BREX | <-SIGMA+OB-Fold+AAA-ATPase+Lon-Protease-BrxL<-PglZ\|\|RADICAL-SAM->Radical_SAM-><-?<-?<-ParB+HNH<-N6-METHYLASE<-?<-DUF6079*<-BrxB-BrxF-iSTAND<-BrxA-DNA-binding<-wHTH+WYL+WCX<-SF1-SF2-Helicase\|\|?-><-ACET | 1232 | Bacteria>Pseudomonadota>Gammaproteobacteria>Vibrionales>Vibrionaceae>Photobacterium | GCF_003355625.1 | Photobacterium_damselae | BC461_RS14900 | 129759..133457 | 38293 | BREX-system-P-loop-protein-BrxC-[Photobacterium-damselae] |
| WP_138222247.1 | Type1.BREX | <-wHTH+WYL+WCX<-?<-?<-?<-SIGMA+OB-Fold+AAA-ATPase+Lon-Protease-BrxL<-PglZ<-ParB+HNH<-N6-METHYLASE<-DUF6079*<-BrxB-BrxF-iSTAND<-BrxA-DNA-binding<-wHTH+WYL+WCX\|\|HTH_Tnp_1+HTH_21+rve+rve_3-><-ABC-ATPase+SbcC | 1232 | Bacteria>Pseudomonadota>Gammaproteobacteria>Vibrionales>Vibrionaceae>Photobacterium | GCF_005819835.1 | Photobacterium_damselae | FD717_RS16650 | 68908..72606 | 38293 | BREX-system-P-loop-protein-BrxC-[Photobacterium-damselae] |
| WP_138976661.1 | Type1.BREX | ACET-><-?\|\|SF1-SF2-Helicase->wHTH+WYL+WCX->BrxA-DNA-binding->BrxB-BrxF-iSTAND->DUF6079*->?->N6-METHYLASE->ParB+HNH->?->?-><-Radical_SAM<-RADICAL-SAM\|\|PglZ->SIGMA+OB-Fold+AAA-ATPase+Lon-Protease-BrxL-> | 1232 | Bacteria>Pseudomonadota>Gammaproteobacteria>Vibrionales>Vibrionaceae>Photobacterium | GCF_006124915.1 | Photobacterium_damselae | DA092_RS00785 | 169698..173396 | 38293 | BREX-system-P-loop-protein-BrxC-[Photobacterium-damselae] |
| WP_176285150.1 | Type1.BREX | <-SF1-SF2-Helicase\|\|?->sGTP+sGTP\|\|XIS-HTH-><-?->ANK+ANK->BrxA-DNA-binding->BrxB-BrxF-iSTAND->DUF6079*->N6-METHYLASE->OLD-ABC->TOPRIM->PglZ->ZetaToxin->?-><-SF1-SF2-Helicase+REase<-SF1-SF2-Helicase+REase | 1157 | Bacteria>Pseudomonadota>Gammaproteobacteria>Vibrionales>Vibrionaceae>Photobacterium | GCF_013371045.1 | Photobacterium_damselae | HU831_RS17605 | 57170..60643 | 38293 | BREX-system-P-loop-protein-BrxC-[Photobacterium-damselae] |
| WP_233092144.1 | Type1.BREX | SF1-SF2-Helicase+REase->SF1-SF2-Helicase+REase-><-PglZ<-TOPRIM<-OLD-ABC<-N6-METHYLASE<-DUF6079*<-BrxB-BrxF-iSTAND<-BrxA-DNA-binding<-wHTH+WYL+WCX<-XIS-HTH<-?<-REase+SF1-SF2-Helicase\|\|?-><-?\|\|HTH_26-> | 1153 | Bacteria>Pseudomonadota>Gammaproteobacteria>Vibrionales>Vibrionaceae>Photobacterium | GCF_014930975.1 | Photobacterium_damselae | IL982_RS10195 | 2142038..2145499 | 38293 | BREX-system-P-loop-protein-BrxC-[Photobacterium-damselae] |
| WP_207103965.1 | Type1.BREX | CR-ATPase4->?-><-SIGMA+OB-Fold+AAA-ATPase+Lon-Protease-BrxL<-PglZ<-?<-GrpE<-ParB+HNH<-N6-METHYLASE<-DUF6079*<-BrxB-BrxF-iSTAND<-BrxA-DNA-binding<-wHTH+WYL+WCX<-viral-protein<-AbiEi<-SF1-SF2-Helicase<-?\|\|?->ZnR-> | 1225 | Bacteria>Pseudomonadota>Gammaproteobacteria>Vibrionales>Vibrionaceae>Photobacterium | GCF_017329545.1 | Photobacterium_ganghwense | FH974_RS17035 | 241897..245574 | 320778 | BREX-system-P-loop-protein-BrxC-[Photobacterium-ganghwense] |
| WP_232606822.1 | Type1.BREX | ROK-HTH-><-?<-sGTP+sGTP\|\|?->ANK+ANK->wHTH+WYL+WCX-p->BrxA-DNA-binding->BrxB-BrxF-iSTAND->DUF6079*->N6-METHYLASE->N6-METHYLASE->PglZ->HEPN+SF1-SF2-Helicase+RNase+RNase->TPR-><-SF1-SF2-Helicase+REase<-SF1-SF2-Helicase+REase\|\|McrB-AAA->McrC-NTD+REase-> | 1157 | Bacteria>Pseudomonadota>Gammaproteobacteria>Vibrionales>Vibrionaceae>Photobacterium | GCF_021237045.1 | Photobacterium_phosphoreum | CJF26_RS18915 | 30954..34427 | 659 | BREX-system-P-loop-protein-BrxC-[Photobacterium-phosphoreum] |
| WP_232580803.1 | Type1.BREX | GDResolvase+HIN-HTH->Phage_integrase->Phage_integrase+Phage_integrase->Phage_integrase->?->SF1-SF2-Helicase->wHTH+WYL+WCX->BrxA-DNA-binding->BrxB-BrxF-iSTAND->DUF6079*->N6-METHYLASE->?->ParB+HNH->HTH_Tnp_1+HTH_21+rve+rve_3->ParB+HNH->?->?->PglZ->SIGMA+OB-Fold+AAA-ATPase+Lon-Protease-BrxL-> | 1238 | Bacteria>Pseudomonadota>Gammaproteobacteria>Vibrionales>Vibrionaceae>Photobacterium | GCF_021504545.1 | Photobacterium_phosphoreum | GLP29_RS03805 | 135359..139075 | 659 | BREX-system-P-loop-protein-BrxC-[Photobacterium-phosphoreum] |
| WP_239972118.1 | Type1.BREX | <-DUF2787<-?\|\|?->?\|\|?->XIS-HTH->wHTH+WYL+WCX->BrxA-DNA-binding->BrxB-BrxF-iSTAND->DUF6079*->SF1-SF2-Helicase+PRTase->SLOG+BirA-HTH->N6-METHYLASE->?->?->ParB+HNH->GrpE->?->PglZ-> | 1232 | Bacteria>Pseudomonadota>Gammaproteobacteria>Vibrionales>Vibrionaceae>Photobacterium | GCF_022398535.1 | Photobacterium_damselae | L1D45_RS07310 | 258079..261777 | 38293 | BREX-system-P-loop-protein-BrxC-[Photobacterium-damselae] |
| WP_101770358.1 | Type1.BREX | <-SIGMA+OB-Fold+AAA-ATPase+Lon-Protease-BrxL<-PglZ<-N6-METHYLASE<-DUF6079*<-TOPRIM<-OLD-ABC<-DUF6079*<-BrxB-BrxF-iSTAND<-BrxA-DNA-binding<-Recombinase<-?<-Arm-DNA-bind_2+Phage_int_SAM_3+Phage_integrase<-Phage_AlpA\|\|HTH_26-> | 1223 | Bacteria>Pseudomonadota>Gammaproteobacteria>Vibrionales>Vibrionaceae>Photobacterium | GCF_027923345.1 | Photobacterium_carnosum | CIK00_20185 | 17344..21015 | 2023717 | BREX-system-P-loop-protein-BrxC-[Photobacterium-carnosum] |
| WP_101767576.1 | Type1.BREX | <-SF1-SF2-Helicase+RNase<-SF1-SF2-Helicase+RNase<-PadR-HTH\|\|HTH_19-><-SIGMA+OB-Fold+AAA-ATPase+Lon-Protease-BrxL<-PglZ<-ParB+HNH<-?<-N6-METHYLASE<-DUF6079*<-BrxB-BrxF-iSTAND<-BrxA-DNA-binding<-wHTH+WYL+WCX<-SF1-SF2-Helicase<-?<-Phage_integrase<-Phage_integrase+Phage_integrase<-Phage_integrase<-GDResolvase+HIN-HTH | 1238 | Bacteria>Pseudomonadota>Gammaproteobacteria>Vibrionales>Vibrionaceae>Photobacterium | GCF_027923345.1 | Photobacterium_carnosum | CIK00_RS03640 | 195656..199372 | 2023717 | BREX-system-P-loop-protein-BrxC-[Photobacterium-carnosum] |
| WP_087820331.1 | Type1.BREX | SAM-METHYLASE->?-><-?<-?<-?\|\|wHTH+WYL+WCX->BrxA-DNA-binding->BrxB-BrxF-iSTAND->DUF6079*->?->N6-METHYLASE->ParB+HNH->?->?->PglZ->SIGMA+OB-Fold+AAA-ATPase+Lon-Protease-BrxL-> | 1238 | Bacteria>Pseudomonadota>Gammaproteobacteria>Vibrionales>Vibrionaceae>Photobacterium | GCF_900185565.1 | Photobacterium_aquimaris | PAQU9191_RS07280 | 444867..448583 | 512643 | BREX-system-P-loop-protein-BrxC-[Photobacterium-aquimaris] |
| CAQ34953.1 | Type1.BREX | HIN-HTH->HTH_Tnp_1+HTH_21+rve+rve_3-><-Phage_integrase<-LexA-protease\|\|35exo->?->wHTH+WYL+WCX->BrxA-DNA-binding->BrxB-BrxF-iSTAND->DUF6079*->OLD-ABC->TOPRIM->N6-METHYLASE-p-><-DDE_Tnp\|\|PglZ->SIGMA+OB-Fold+AAA-ATPase+Lon-Protease-BrxL->SbcC->HDRnase->VirD4-TraG+VirB4-FtsK-> | 1225 | Bacteria>Pseudomonadota>Gammaproteobacteria>Vibrionales>Vibrionaceae>Photobacterium | - | Photobacterium_damselae_subsp._piscicida | - | 21165..24842 | 38294 | hypothetical-protein-[Photobacterium-damselae-subsp.-piscicida] |
| WP_077485085.1 | Type1.BREX | Glyco_hydro_4+Glyco_hydro_4C-><-wHTH+WYL+WCX\|\|SF1-SF2-Helicase+REase->?->wHTH+WYL+WCX->BrxA-DNA-binding->BrxB-BrxF-iSTAND->DUF6079*->N6-METHYLASE->OLD-ABC->TOPRIM->PglZ-><-SF1-SF2-Helicase+REase<-SF1-SF2-Helicase+REase\|\|ZnR-> | 1157 | Bacteria>Pseudomonadota>Gammaproteobacteria>Vibrionales>Vibrionaceae>Salinivibrio | GCF_001995745.1 | Salinivibrio_sp._ML323 | BZG13_RS03555 | 125341..128814 | 1909474 | BREX-system-P-loop-protein-BrxC-[Salinivibrio-sp.-ML323] |
| WP_077678370.1 | Type1.BREX | <-wHTH+WYL+WCX\|\|SF1-SF2-Helicase+REase->?->wHTH+WYL+WCX->BrxA-DNA-binding->BrxB-BrxF-iSTAND->DUF6079*->N6-METHYLASE-> | 1229 | Bacteria>Pseudomonadota>Gammaproteobacteria>Vibrionales>Vibrionaceae>Salinivibrio | GCF_001996125.1 | Salinivibrio_sp._PR932 | BZG84_RS14215 | 25023..28712 | 1909492 | BREX-system-P-loop-protein-BrxC-[Salinivibrio-sp.-PR932] |
| WP_077677746.1 | Type1.BREX | Glyco_hydro_4+Glyco_hydro_4C-><-wHTH+WYL+WCX\|\|SF1-SF2-Helicase+REase->?->wHTH+WYL+WCX->BrxA-DNA-binding->BrxB-BrxF-iSTAND->DUF6079*->N6-METHYLASE-> | 1232 | Bacteria>Pseudomonadota>Gammaproteobacteria>Vibrionales>Vibrionaceae>Salinivibrio | GCF_001996245.1 | Salinivibrio_sp._MA607 | BZG81_RS14680 | 10387..14085 | 1909457 | BREX-system-P-loop-protein-BrxC-[Salinivibrio-sp.-MA607] |
| TNF18455.1 | Type1.BREX | Phage_holin->Phage_lysozyme->XIS-HTH->BrxA-DNA-binding->BrxB-BrxF-iSTAND->DUF6079*->N6-METHYLASE->OLD-ABC->TOPRIM->PglZ->SIGMA+OB-Fold+AAA-ATPase+Lon-Protease-BrxL-> | 1145 | Bacteria>Pseudomonadota>Gammaproteobacteria>Vibrionales>Vibrionaceae>unclassified_Vibrionaceae | GCA_006226665.1 | Vibrionaceae_bacterium | EP325_05025 | 51838..55275 | 1889773 | BREX-system-P-loop-protein-BrxC-[Vibrionaceae-bacterium] |
| EKK94843.1 | Type1.BREX | <-Arm-DNA-bind_3+Phage_integrase<-?\|\|MobI-><-?<-LexA-protease\|\|35exo->?->wHTH+WYL+WCX->BrxA-DNA-binding->BrxB-BrxF-iSTAND->DUF6079*->N6-METHYLASE->PglZ->SIGMA+OB-Fold+AAA-ATPase+Lon-Protease-BrxL->MarR-HTH->HEPN->HDRnase->VirD4-TraG+VirB4-FtsK-> | 1151 | Bacteria>Pseudomonadota>Gammaproteobacteria>Vibrionales>Vibrionaceae>Vibrio | GCA_000304775.1 | Vibrio_cholerae_HC-1A2 | VCHC1A2_1591 | 58635..62090 | 991937 | hypothetical-protein-VCHC1A2_1591-[Vibrio-cholerae-HC-1A2] |
| GAJ78762.1 | Type1.BREX | <-GrpE<-ParB+HNH<-ParB+HNH<-N6-METHYLASE-p<-N6-METHYLASE-p<-N6-METHYLASE-p<-DUF6079-p*<-DUF6079-p*<-BrxB-BrxF-iSTAND<-BrxA-DNA-binding<-wHTH+WYL+WCX<-viral-protein<-AbiEi<-SF1-SF2-Helicase<-SF1-SF2-Helicase | 800 | Bacteria>Pseudomonadota>Gammaproteobacteria>Vibrionales>Vibrionaceae>Vibrio | GCA_000695055.1 | Vibrio_sp._JCM_18905 | JCM18905_4752 | 16842..19244 | 1298600 | hypothetical-protein-JCM18905_4752-[Vibrio-sp.-JCM-18905] |
| GAJ78763.1 | Type1.BREX | <-GrpE<-ParB+HNH<-ParB+HNH<-N6-METHYLASE-p<-N6-METHYLASE-p<-N6-METHYLASE-p<-DUF6079-p<-DUF6079-p*<-BrxB-BrxF-iSTAND<-BrxA-DNA-binding<-wHTH+WYL+WCX<-viral-protein<-AbiEi<-SF1-SF2-Helicase<-SF1-SF2-Helicase | 425 | Bacteria>Pseudomonadota>Gammaproteobacteria>Vibrionales>Vibrionaceae>Vibrio | GCA_000695055.1 | Vibrio_sp._JCM_18905 | JCM18905_4753 | 19241..20518 | 1298600 | hypothetical-protein-JCM18905_4753-[Vibrio-sp.-JCM-18905] |
| AJZ99161.1 | Type1.BREX | <-MULE-TRANSPOSASE<-Phage_integrase<-?<-LexA-protease\|\|35exo->?->wHTH+WYL+WCX->BrxA-DNA-binding->BrxB-BrxF-iSTAND->DUF6079*->N6-METHYLASE->PglZ->SIGMA+OB-Fold+AAA-ATPase+Lon-Protease-BrxL->MarR-HTH->HEPN->HDRnase->VirD4-TraG+VirB4-FtsK-> | 1151 | Bacteria>Pseudomonadota>Gammaproteobacteria>Vibrionales>Vibrionaceae>Vibrio | GCA_000963555.1 | Vibrio_cholerae_O1_biovar_El_Tor | IR04_08395 | 718003..721458 | 686 | hypothetical-protein-IR04_08395-[Vibrio-cholerae-O1-biovar-El-Tor] |
| CSC57771.1 | Type1.BREX | <-VirD4-TraG+VirB4-FtsK<-HDRnase<-HEPN<-MarR-HTH<-SIGMA+OB-Fold+AAA-ATPase+Lon-Protease-BrxL<-PglZ<-N6-METHYLASE<-DUF6079*<-BrxB-BrxF-iSTAND<-BrxA-DNA-binding<-wHTH+WYL+WCX<-?<-35exo\|\|LexA-protease-> | 1237 | Bacteria>Pseudomonadota>Gammaproteobacteria>Vibrionales>Vibrionaceae>Vibrio | GCA_001247885.1 | Vibrio_cholerae | ERS013147_02837 | 124661..128374 | 666 | Uncharacterised-protein-[Vibrio-cholerae] |
| CSC91133.1 | Type1.BREX | <-Phage_integrase<-?<-LexA-protease\|\|?->35exo->?->BrxA-DNA-binding-p->BrxB-BrxF-iSTAND->DUF6079-p*->DUF6079-p->N6-METHYLASE-> | 268 | Bacteria>Pseudomonadota>Gammaproteobacteria>Vibrionales>Vibrionaceae>Vibrio | GCA_001254955.1 | Vibrio_cholerae | ERS013216_03026 | 6147..6953 | 666 | Uncharacterised-protein-[Vibrio-cholerae] |
| CSC91159.1 | Type1.BREX | <-Phage_integrase<-?<-LexA-protease\|\|?->35exo->?->BrxA-DNA-binding-p->BrxB-BrxF-iSTAND->DUF6079-p*->DUF6079-p*->N6-METHYLASE-> | 933 | Bacteria>Pseudomonadota>Gammaproteobacteria>Vibrionales>Vibrionaceae>Vibrio | GCA_001254955.1 | Vibrio_cholerae | ERS013216_03027 | 7037..9838 | 666 | Uncharacterised-protein-[Vibrio-cholerae] |
| CSA58950.1 | Type1.BREX | <-Phage_integrase<-?<-LexA-protease\|\|35exo->35exo->?->wHTH+WYL+WCX->BrxA-DNA-binding->BrxB-BrxF-iSTAND->DUF6079-p*->DUF6079-p->N6-METHYLASE->PglZ->SIGMA+OB-Fold+AAA-ATPase+Lon-Protease-BrxL->MarR-HTH->HEPN->HDRnase-><-?\|\|VirD4-TraG+VirB4-FtsK-> | 324 | Bacteria>Pseudomonadota>Gammaproteobacteria>Vibrionales>Vibrionaceae>Vibrio | GCA_001257255.1 | Vibrio_cholerae | ERS013131_02346 | 7278..8252 | 666 | Uncharacterised-protein-[Vibrio-cholerae] |
| CSA58983.1 | Type1.BREX | <-LexA-protease\|\|35exo->35exo->?->wHTH+WYL+WCX->BrxA-DNA-binding->BrxB-BrxF-iSTAND->DUF6079-p->DUF6079-p*->N6-METHYLASE->PglZ->SIGMA+OB-Fold+AAA-ATPase+Lon-Protease-BrxL->MarR-HTH->HEPN->HDRnase-><-?\|\|VirD4-TraG+VirB4-FtsK-> | 820 | Bacteria>Pseudomonadota>Gammaproteobacteria>Vibrionales>Vibrionaceae>Vibrio | GCA_001257255.1 | Vibrio_cholerae | ERS013131_02347 | 8501..10963 | 666 | Uncharacterised-protein-[Vibrio-cholerae] |
| CSD13814.1 | Type1.BREX | <-VirD4-TraG+VirB4-FtsK<-HDRnase<-HEPN<-MarR-HTH<-SIGMA+OB-Fold+AAA-ATPase+Lon-Protease-BrxL<-PglZ<-N6-METHYLASE<-DUF6079-p<-DUF6079-p*<-BrxB-BrxF-iSTAND<-BrxA-DNA-binding<-wHTH+WYL+WCX<-?<-35exo\|\|LexA-protease->?->Phage_integrase-> | 327 | Bacteria>Pseudomonadota>Gammaproteobacteria>Vibrionales>Vibrionaceae>Vibrio | GCA_001258535.1 | Vibrio_cholerae | ERS013225_02589 | 127512..128495 | 666 | Uncharacterised-protein-[Vibrio-cholerae] |
| KOY19349.1 | Type1.BREX | <-GDResolvase+HIN-HTH\|\|?->Arm-DNA-bind_3+Phage_integrase->SF1-SF2-Helicase+REase->SF1-SF2-Helicase+REase-><-PglZ<-TOPRIM<-OLD-ABC<-N6-METHYLASE<-DUF6079*<-BrxB-BrxF-iSTAND<-BrxA-DNA-binding<-wHTH+WYL+WCX<-XIS-HTH\|\|HTH_Tnp_1-> | 1154 | Bacteria>Pseudomonadota>Gammaproteobacteria>Vibrionales>Vibrionaceae>Vibrio | GCA_001280635.1 | Vibrio_parahaemolyticus | ACX06_21960 | 167502..170966 | 670 | hypothetical-protein-ACX06_21960-[Vibrio-parahaemolyticus] |
| ODY54465.1 | Type1.BREX | <-HTH_Tnp_1\|\|XIS-HTH->wHTH+WYL+WCX->BrxA-DNA-binding->BrxB-BrxF-iSTAND->DUF6079*->N6-METHYLASE->OLD-ABC->TOPRIM->PglZ-><-SF1-SF2-Helicase+REase<-SF1-SF2-Helicase+REase<-Arm-DNA-bind_3+Phage_integrase | 1154 | Bacteria>Pseudomonadota>Gammaproteobacteria>Vibrionales>Vibrionaceae>Vibrio | GCA_001727265.1 | Vibrio_parahaemolyticus | BBM25_06240 | 3424..6888 | 670 | hypothetical-protein-BBM25_06240-[Vibrio-parahaemolyticus] |
| ASG04983.1 | Type1.BREX | <-DeoR-HTH+ISOCOT<-DeoR-HTH+ISOCOT<-DUF2787<-?<-?\|\|XIS-HTH->XIS-HTH->BrxA-DNA-binding->BrxB-BrxF-iSTAND->DUF6079*->OLD-ABC->REase->N6-METHYLASE->PglZ->SIGMA+OB-Fold+AAA-ATPase+HKD-DNase-BrxL-><-Arm-DNA-bind_3+Phage_integrase | 1185 | Bacteria>Pseudomonadota>Gammaproteobacteria>Vibrionales>Vibrionaceae>Vibrio | GCA_002211985.1 | Vibrio_anguillarum | CEJ46_04105 | 862780..866337 | 55601 | hypothetical-protein-CEJ46_04105-[Vibrio-anguillarum] |
| ASI98027.1 | Type1.BREX | <-SIGMA+OB-Fold+AAA-ATPase+HKD-DNase-BrxL<-PglZ<-N6-METHYLASE<-NERD<-OLD-ABC<-DUF6079*<-BrxB-BrxF-iSTAND<-BrxA-DNA-binding<-XIS-HTH<-RNase<-DUF3584+DUF5401<-JetB<-MukF\|\|wHTH+WYL+WCX-> | 1183 | Bacteria>Pseudomonadota>Gammaproteobacteria>Vibrionales>Vibrionaceae>Vibrio | GCA_002214395.1 | Vibrio_rotiferianus | BSZ04_17745 | 1918666..1922217 | 190895 | hypothetical-protein-BSZ04_17745-[Vibrio-rotiferianus] |
| MVB73516.1 | Type1.BREX | GAF+PAS+GGDEF+EAL->?->Peptidase_M48->Trypsin+PDZ->35exo->?->wHTH+WYL+WCX->BrxA-DNA-binding->BrxB-BrxF-iSTAND->DUF6079*->ParB+HNH->N6-METHYLASE->PglZ->SIGMA+OB-Fold+AAA-ATPase+Lon-Protease-BrxL->SbcC->HDRnase->VirD4-TraG+VirB4-FtsK-> | 1225 | Bacteria>Pseudomonadota>Gammaproteobacteria>Vibrionales>Vibrionaceae>Vibrio | GCA_009747185.1 | Vibrio_cholerae | D6U90_04200 | 143741..147418 | 666 | BREX-system-P-loop-protein-BrxC-[Vibrio-cholerae] |
| MVC74394.1 | Type1.BREX | MobI-><-?<-LexA-protease\|\|35exo->?->wHTH+WYL+WCX->BrxA-DNA-binding->BrxB-BrxF-iSTAND->DUF6079*->ParB+HNH->N6-METHYLASE->PglZ->SIGMA+OB-Fold+AAA-ATPase+Lon-Protease-BrxL->NACHT+HEAT+REase->HDRnase->VirD4-TraG+VirB4-FtsK-> | 1225 | Bacteria>Pseudomonadota>Gammaproteobacteria>Vibrionales>Vibrionaceae>Vibrio | GCA_009747245.1 | Vibrio_cholerae | D6U22_06265 | 184143..187820 | 666 | BREX-system-P-loop-protein-BrxC-[Vibrio-cholerae] |
| NAZ69809.1 | Type1.BREX | <-SIGMA+OB-Fold+AAA-ATPase+HKD-DNase-BrxL<-PglZ<-N6-METHYLASE<-REase<-OLD-ABC<-DUF6079*<-BrxB-BrxF-iSTAND<-BrxA-DNA-binding<-XIS-HTH<-HNH<-?<-XIS-HTH\|\|?->DDE_Tnp_1_5+DDE_Tnp_1-> | 1185 | Bacteria>Pseudomonadota>Gammaproteobacteria>Vibrionales>Vibrionaceae>Vibrio | GCA_009906175.1 | Vibrio_toranzoniae | GL180_08045 | 10464..14021 | 1194427 | BREX-system-P-loop-protein-BrxC-[Vibrio-toranzoniae] |
| MBE3824873.1 | Type1.BREX | <-VirD4-TraG+VirB4-FtsK<-HDRnase<-SbcC<-SIGMA+OB-Fold+AAA-ATPase+Lon-Protease-BrxL<-PglZ<-TOPRIM<-OLD-ABC<-N6-METHYLASE<-ParB+HNH<-DUF6079*<-BrxB-BrxF-iSTAND<-BrxA-DNA-binding<-wHTH+WYL+WCX<-?<-35exo\|\|LexA-protease->?-><-MobI | 1225 | Bacteria>Pseudomonadota>Gammaproteobacteria>Vibrionales>Vibrionaceae>Vibrio | GCA_014918955.1 | Vibrio_parahaemolyticus | HJ179_16470 | 255963..259640 | 670 | BREX-system-P-loop-protein-BrxC-[Vibrio-parahaemolyticus] |
| MBE3881970.1 | Type1.BREX | MobI-><-?<-LexA-protease\|\|35exo->?->wHTH+WYL+WCX->BrxA-DNA-binding->BrxB-BrxF-iSTAND->DUF6079*->ParB+HNH->N6-METHYLASE->PglZ->SIGMA+OB-Fold+AAA-ATPase+Lon-Protease-BrxL->NACHT+HEAT+REase->HDRnase->VirD4-TraG+VirB4-FtsK-> | 1225 | Bacteria>Pseudomonadota>Gammaproteobacteria>Vibrionales>Vibrionaceae>Vibrio | GCA_014919165.1 | Vibrio_parahaemolyticus | HJ168_09865 | 226253..229930 | 670 | BREX-system-P-loop-protein-BrxC-[Vibrio-parahaemolyticus] |
| MBE4310404.1 | Type1.BREX | <-PIN+OST-HTH\|\|CR-ATPase4->?-><-?<-DUF4113\|\|wHTH-><-HTH_Tnp_1+HTH_21+rve+rve_3\|\|wHTH+WYL+WCX->BrxA-DNA-binding->BrxB-BrxF-iSTAND->DUF6079*->ParB+HNH->PglZ->HEPN+SF1-SF2-Helicase+RNase+RNase->TPR-><-SF1-SF2-Helicase+REase<-SF1-SF2-Helicase+REase<-viral-protein<-AbiEi | 1157 | Bacteria>Pseudomonadota>Gammaproteobacteria>Vibrionales>Vibrionaceae>Vibrio | GCA_014921165.1 | Vibrio_parahaemolyticus | HJ064_04290 | 963533..967006 | 670 | BREX-system-P-loop-protein-BrxC-[Vibrio-parahaemolyticus] |
| MBE4432200.1 | Type1.BREX | <-ZnR<-?\|\|?->SF1-SF2-Helicase->AbiEi->viral-protein->wHTH+WYL+WCX->BrxA-DNA-binding->BrxB-BrxF-iSTAND->DUF6079*->ParB+HNH->GrpE->?->PglZ->SIGMA+OB-Fold+AAA-ATPase+Lon-Protease-BrxL-><-?\|\|DUF4113-><-?<-CR-ATPase4 | 1225 | Bacteria>Pseudomonadota>Gammaproteobacteria>Vibrionales>Vibrionaceae>Vibrio | GCA_014921655.1 | Vibrio_parahaemolyticus | HJ022_02625 | 614361..618038 | 670 | BREX-system-P-loop-protein-BrxC-[Vibrio-parahaemolyticus] |
| MBE5192784.1 | Type1.BREX | <-ZnR<-?\|\|AbiEi<-viral-protein<-SF1-SF2-Helicase+REase-><-?<-ZetaToxin<-PglZ<-ParB+HNH<-DUF6079*<-BrxB-BrxF-iSTAND<-BrxA-DNA-binding<-wHTH+WYL+WCX\|\|<-?<-CR-ATPase4\|\|HEPN->AAA-ATPase-p-> | 1157 | Bacteria>Pseudomonadota>Gammaproteobacteria>Vibrionales>Vibrionaceae>Vibrio | GCA_014963905.1 | Vibrio_parahaemolyticus | HJ051_04870 | 1092166..1095639 | 670 | BREX-system-P-loop-protein-BrxC-[Vibrio-parahaemolyticus] |
| EGQ7646156.1 | Type1.BREX | Calcineurin-><-ZnR<-?\|\|AbiEi->viral-protein->wHTH+WYL+WCX->BrxA-DNA-binding->BrxB-BrxF-iSTAND->DUF6079*->N6-METHYLASE->ParB+HNH->GrpE->?->PglZ->SIGMA+OB-Fold+AAA-ATPase+Lon-Protease-BrxL-><-TPR<-HEPN+SF1-SF2-Helicase+RNase+RNase\|\|LexA-protease-> | 1225 | Bacteria>Pseudomonadota>Gammaproteobacteria>Vibrionales>Vibrionaceae>Vibrio | GCA_015741395.1 | Vibrio_alginolyticus | I2703_000228 | 284534..288211 | 663 | BREX-system-P-loop-protein-BrxC-[Vibrio-alginolyticus] |
| EGQ7901736.1 | Type1.BREX | <-ZnR<-?\|\|?->SF1-SF2-Helicase->AbiEi->viral-protein->wHTH+WYL+WCX->BrxA-DNA-binding->BrxB-BrxF-iSTAND->DUF6079*->N6-METHYLASE->?->?->PglZ->SIGMA+OB-Fold+AAA-ATPase+Lon-Protease-BrxL-><-wHTH\|\|<-?<-CR-ATPase4 | 1225 | Bacteria>Pseudomonadota>Gammaproteobacteria>Vibrionales>Vibrionaceae>Vibrio | GCA_015746405.1 | Vibrio_alginolyticus | H5D28_000763 | 854695..858372 | 663 | BREX-system-P-loop-protein-BrxC-[Vibrio-alginolyticus] |
| HAS3953742.1 | Type1.BREX | <-VirD4-TraG+VirB4-FtsK<-HDRnase<-HEPN<-MarR-HTH<-SIGMA+OB-Fold+AAA-ATPase+Lon-Protease-BrxL<-PglZ<-N6-METHYLASE<-DUF6079* | 1210 | Bacteria>Pseudomonadota>Gammaproteobacteria>Vibrionales>Vibrionaceae>Vibrio | GCA_015750645.1 | Vibrio_cholerae | I6306_000460 | 124735..128369 | 666 | BREX-system-P-loop-protein-BrxC-[Vibrio-cholerae] |
| QPL56152.1 | Type1.BREX | <-MULE-TRANSPOSASE<-?<-ZnR<-?\|\|AbiEi->viral-protein->wHTH+WYL+WCX->BrxA-DNA-binding->BrxB-BrxF-iSTAND->DUF6079*->N6-METHYLASE->SbcC->Schlafen+HATPase+wHTH->PglZ->SIGMA+OB-Fold+AAA-ATPase+Lon-Protease-BrxL-> | 1225 | Bacteria>Pseudomonadota>Gammaproteobacteria>Vibrionales>Vibrionaceae>Vibrio | GCA_015767675.1 | Vibrio_navarrensis | I3X05_18825 | 378934..382611 | 29495 | BREX-system-P-loop-protein-BrxC-[Vibrio-navarrensis] |
| EGQ8448095.1 | Type1.BREX | <-SIGMA+OB-Fold+AAA-ATPase+HKD-DNase-BrxL<-PglZ<-N6-METHYLASE<-REase<-OLD-ABC<-DUF6079*<-BrxB-BrxF-iSTAND<-BrxA-DNA-binding<-XIS-HTH<-HNH<-XIS-HTH\|\|?->?->DUF2787->wHTH+WYL+WCX-> | 1185 | Bacteria>Pseudomonadota>Gammaproteobacteria>Vibrionales>Vibrionaceae>Vibrio | GCA_015775815.1 | Vibrio_alginolyticus | GTP59_14930 | 64166..67723 | 663 | BREX-system-P-loop-protein-BrxC-[Vibrio-alginolyticus] |
| HAS6096890.1 | Type1.BREX | Phage_integrase->Phage_int_SAM_5->Phage_integrase->XIS-HTH-><-RVT\|\|wHTH+WYL+WCX->BrxA-DNA-binding->BrxB-BrxF-iSTAND->DUF6079*->N6-METHYLASE->SbcC->Schlafen+HATPase+wHTH->PglZ->SIGMA+OB-Fold+AAA-ATPase+Lon-Protease-BrxL-><-ZDNA-HTH<-ParB+RepB<-ParA-Soj-PloopNTPase\|\|REase+URI-DNase-> | 1229 | Bacteria>Pseudomonadota>Gammaproteobacteria>Vibrionales>Vibrionaceae>Vibrio | GCA_015776315.1 | Vibrio_vulnificus | I7145_19865 | 48069..51758 | 672 | BREX-system-P-loop-protein-BrxC-[Vibrio-vulnificus] |
| HAS6061420.1 | Type1.BREX | <-DUF6079*<-BrxB-BrxF-iSTAND<-BrxA-DNA-binding<-wHTH+WYL+WCX<-viral-protein\|\|?->ZnR-><-SF1-SF2-Helicase<-Couple_hipA+HipA_C<-HTH_3 | 1232 | Bacteria>Pseudomonadota>Gammaproteobacteria>Vibrionales>Vibrionaceae>Vibrio | GCA_015776335.1 | Vibrio_vulnificus | I7135_04535 | 448..4146 | 672 | BREX-system-P-loop-protein-BrxC-[Vibrio-vulnificus] |
| HAS6187654.1 | Type1.BREX | <-OBFOLD-NUCLEASE<-CARF+NtrC-AAA+FIS-HTH<-SIGMA+OB-Fold+AAA-ATPase+Lon-Protease-BrxL<-PglZ<-N6-METHYLASE<-ParB+HNH<-DUF6079*<-BrxB-BrxF-iSTAND<-BrxA-DNA-binding<-wHTH+WYL+WCX<-?<-35exo\|\|LexA-protease->?-><-MobI | 1225 | Bacteria>Pseudomonadota>Gammaproteobacteria>Vibrionales>Vibrionaceae>Vibrio | GCA_015777055.1 | Vibrio_vulnificus | I7170_07515 | 257971..261648 | 672 | BREX-system-P-loop-protein-BrxC-[Vibrio-vulnificus] |
| HAS6259595.1 | Type1.BREX | <-VirD4-TraG+VirB4-FtsK<-HDRnase<-NACHT+HEAT+REase<-SIGMA+OB-Fold+AAA-ATPase+Lon-Protease-BrxL<-PglZ<-TOPRIM<-OLD-ABC<-N6-METHYLASE<-ParB+HNH<-DUF6079*<-BrxB-BrxF-iSTAND<-BrxA-DNA-binding<-wHTH+WYL+WCX<-?<-35exo\|\|LexA-protease->?-><-MobI | 1225 | Bacteria>Pseudomonadota>Gammaproteobacteria>Vibrionales>Vibrionaceae>Vibrio | GCA_015777345.1 | Vibrio_vulnificus | I7226_03370 | 263913..267590 | 672 | BREX-system-P-loop-protein-BrxC-[Vibrio-vulnificus] |
| HAS6362912.1 | Type1.BREX | CR-ATPase4->?-><-SIGMA+OB-Fold+AAA-ATPase+Lon-Protease-BrxL<-PglZ<-?<-?<-ParB+HNH<-N6-METHYLASE<-DUF6079*<-BrxB-BrxF-iSTAND<-BrxA-DNA-binding<-wHTH+WYL+WCX\|\|SF1-SF2-Helicase->MarR-HTH->HKD-DNase->CARF+NtrC-AAA->PIN+OST-HTH-> | 1248 | Bacteria>Pseudomonadota>Gammaproteobacteria>Vibrionales>Vibrionaceae>Vibrio | GCA_015777565.1 | Vibrio_vulnificus | I7230_15365 | 52785..56531 | 672 | BREX-system-P-loop-protein-BrxC-[Vibrio-vulnificus] |
| HAS6390651.1 | Type1.BREX | <-HTH\|\|RNase_H-><-DUF2787<-?<-?\|\|XIS-HTH->wHTH+WYL+WCX->BrxA-DNA-binding->BrxB-BrxF-iSTAND->DUF6079*->N6-METHYLASE->OLD-ABC->HNH->PglZ->ZetaToxin-><-SF1-SF2-Helicase+REase<-SF1-SF2-Helicase+REase | 1157 | Bacteria>Pseudomonadota>Gammaproteobacteria>Vibrionales>Vibrionaceae>Vibrio | GCA_015777605.1 | Vibrio_vulnificus | I7251_16610 | 90517..93990 | 672 | BREX-system-P-loop-protein-BrxC-[Vibrio-vulnificus] |
| HAS6345170.1 | Type1.BREX | SF1-SF2-Helicase-><-ZnR<-?\|\|viral-protein-><-?<-?\|\|wHTH+WYL+WCX->BrxA-DNA-binding->BrxB-BrxF-iSTAND->DUF6079*->?->N6-METHYLASE->?->?->ParB+HNH->ZnR+3C-ABC+TPR->?->?->PglZ-> | 1232 | Bacteria>Pseudomonadota>Gammaproteobacteria>Vibrionales>Vibrionaceae>Vibrio | GCA_015778705.1 | Vibrio_vulnificus | I7243_17845 | 83102..86800 | 672 | BREX-system-P-loop-protein-BrxC-[Vibrio-vulnificus] |
| HAS6333998.1 | Type1.BREX | <-MORC<-PIN+OST-HTH<-CARF+NtrC-AAA<-HKD-DNase+HKD-DNase<-MarR-HTH<-SF1-SF2-Helicase\|\|wHTH+WYL+WCX->BrxA-DNA-binding->BrxB-BrxF-iSTAND->DUF6079*->N6-METHYLASE->?->ParB+HNH->ZnR+3C-ABC+TPR->?->?->PglZ->SIGMA+OB-Fold+AAA-ATPase+Lon-Protease-BrxL-> | 1248 | Bacteria>Pseudomonadota>Gammaproteobacteria>Vibrionales>Vibrionaceae>Vibrio | GCA_015778755.1 | Vibrio_vulnificus | I7245_08190 | 199206..202952 | 672 | BREX-system-P-loop-protein-BrxC-[Vibrio-vulnificus] |
| HAS6310370.1 | Type1.BREX | AbiEi->viral-protein->SF1-SF2-Helicase+REase->SF1-SF2-Helicase+REase-><-ZetaToxin<-PglZ<-ParB+HNH<-N6-METHYLASE<-DUF6079*<-BrxB-BrxF-iSTAND<-BrxA-DNA-binding<-wHTH+WYL+WCX\|\|LexA-protease->?-><-?<-CR-ATPase4 | 1157 | Bacteria>Pseudomonadota>Gammaproteobacteria>Vibrionales>Vibrionaceae>Vibrio | GCA_015778905.1 | Vibrio_vulnificus | I7207_04135 | 43135..46608 | 672 | BREX-system-P-loop-protein-BrxC-[Vibrio-vulnificus] |
| EGQ8649255.1 | Type1.BREX | <-VirD4-TraG+VirB4-FtsK<-HDRnase<-SbcC<-SIGMA+OB-Fold+AAA-ATPase+Lon-Protease-BrxL<-PglZ<-N6-METHYLASE<-TOPRIM<-OLD-ABC<-DUF6079*<-BrxB-BrxF-iSTAND<-BrxA-DNA-binding<-wHTH+WYL+WCX<-35exo<-?\|\|LexA-protease-> | 1225 | Bacteria>Pseudomonadota>Gammaproteobacteria>Vibrionales>Vibrionaceae>Vibrio | GCA_015781355.1 | Vibrio_cholerae | GNS53_08430 | 57189..60866 | 666 | BREX-system-P-loop-protein-BrxC-[Vibrio-cholerae] |
| EGQ8730608.1 | Type1.BREX | <-ZnR<-?\|\|?->SF1-SF2-Helicase->AbiEi->viral-protein->wHTH+WYL+WCX->BrxA-DNA-binding->BrxB-BrxF-iSTAND->DUF6079*->N6-METHYLASE->?->?->PglZ->SIGMA+OB-Fold+AAA-ATPase+Lon-Protease-BrxL-><-TPR<-HEPN+SF1-SF2-Helicase+RNase+RNase\|\|LexA-protease-> | 1225 | Bacteria>Pseudomonadota>Gammaproteobacteria>Vibrionales>Vibrionaceae>Vibrio | GCA_015781475.1 | Vibrio_parahaemolyticus | GKA80_01950 | 455980..459657 | 670 | BREX-system-P-loop-protein-BrxC-[Vibrio-parahaemolyticus] |
| EGQ8898350.1 | Type1.BREX | viral-protein->wHTH+WYL+WCX->BrxA-DNA-binding->BrxB-BrxF-iSTAND->DUF6079*->N6-METHYLASE->ParB+HNH->GrpE->?->PglZ->SIGMA+OB-Fold+AAA-ATPase+Lon-Protease-BrxL-><-TPR<-HEPN+SF1-SF2-Helicase+RNase+RNase\|\|LexA-protease-> | 1225 | Bacteria>Pseudomonadota>Gammaproteobacteria>Vibrionales>Vibrionaceae>Vibrio | GCA_015781875.1 | Vibrio_parahaemolyticus | DMW33_24570 | 2854..6531 | 670 | BREX-system-P-loop-protein-BrxC-[Vibrio-parahaemolyticus] |
| EGQ9599053.1 | Type1.BREX | sGTP-><-HEPN<-?\|\|CR-ATPase4->?->\|\|wHTH+WYL+WCX->BrxA-DNA-binding->BrxB-BrxF-iSTAND->DUF6079*->N6-METHYLASE->ParB+HNH->PglZ->ZetaToxin-><-SF1-SF2-Helicase+REase<-SF1-SF2-Helicase+REase<-viral-protein | 1157 | Bacteria>Pseudomonadota>Gammaproteobacteria>Vibrionales>Vibrionaceae>Vibrio | GCA_015785155.1 | Vibrio_parahaemolyticus | F3R80_21565 | 8289..11762 | 670 | BREX-system-P-loop-protein-BrxC-[Vibrio-parahaemolyticus] |
| EGR0072241.1 | Type1.BREX | wHTH+WYL+WCX->BrxA-DNA-binding->BrxB-BrxF-iSTAND->DUF6079*->N6-METHYLASE->ParB+HNH->?->?->PglZ->SIGMA+OB-Fold+AAA-ATPase+Lon-Protease-BrxL-><-?<-CR-ATPase4 | 1248 | Bacteria>Pseudomonadota>Gammaproteobacteria>Vibrionales>Vibrionaceae>Vibrio | GCA_015787395.1 | Vibrio_vulnificus | FR269_18510 | 2637..6383 | 672 | BREX-system-P-loop-protein-BrxC-[Vibrio-vulnificus] |
| EGR0489752.1 | Type1.BREX | AbiEi->viral-protein->SF1-SF2-Helicase+REase->SF1-SF2-Helicase+REase-><-ZetaToxin<-PglZ<-ParB+HNH<-N6-METHYLASE<-DUF6079*<-BrxB-BrxF-iSTAND<-BrxA-DNA-binding<-wHTH+WYL+WCX\|\|LexA-protease->?-><-?<-CR-ATPase4\|\|<-RVT\|\|HigA-> | 1157 | Bacteria>Pseudomonadota>Gammaproteobacteria>Vibrionales>Vibrionaceae>Vibrio | GCA_015788475.1 | Vibrio_cholerae | FG089_09200 | 79189..82662 | 666 | BREX-system-P-loop-protein-BrxC-[Vibrio-cholerae] |
| EGR0719595.1 | Type1.BREX | HTH_20-><-HEPN<-SbcC\|\|CR-ATPase4->?-><-DUF4113\|\|wHTH+WYL+WCX->BrxA-DNA-binding->BrxB-BrxF-iSTAND->DUF6079*->N6-METHYLASE->ParB+HNH->PglZ->HEPN+SF1-SF2-Helicase+RNase+RNase->TPR-><-SF1-SF2-Helicase+REase<-SF1-SF2-Helicase+REase<-viral-protein<-AbiEi | 1157 | Bacteria>Pseudomonadota>Gammaproteobacteria>Vibrionales>Vibrionaceae>Vibrio | GCA_015789455.1 | Vibrio_alginolyticus | EX464_05795 | 8586..12059 | 663 | BREX-system-P-loop-protein-BrxC-[Vibrio-alginolyticus] |
| EGR0286978.1 | Type1.BREX | Arm-DNA-bind_3+Phage_integrase->SF1-SF2-Helicase+REase->SF1-SF2-Helicase+REase-><-PglZ<-TOPRIM<-OLD-ABC<-N6-METHYLASE<-DUF6079*<-BrxB-BrxF-iSTAND<-BrxA-DNA-binding<-wHTH+WYL+WCX<-XIS-HTH\|\|?->?->DUF2787->HTH->wHTH+WYL+WCX-> | 1154 | Bacteria>Pseudomonadota>Gammaproteobacteria>Vibrionales>Vibrionaceae>Vibrio | GCA_015789595.1 | Vibrio_cholerae | FK206_11700 | 85283..88747 | 666 | BREX-system-P-loop-protein-BrxC-[Vibrio-cholerae] |
| EGR0983685.1 | Type1.BREX | MobI-><-?<-LexA-protease\|\|?->35exo->?->wHTH+WYL+WCX->BrxA-DNA-binding->BrxB-BrxF-iSTAND->DUF6079*->ParB+HNH->N6-METHYLASE->PglZ->SIGMA+OB-Fold+AAA-ATPase+Lon-Protease-BrxL->NACHT+HEAT+REase->HDRnase->VirD4-TraG+VirB4-FtsK-> | 1225 | Bacteria>Pseudomonadota>Gammaproteobacteria>Vibrionales>Vibrionaceae>Vibrio | GCA_015797355.1 | Vibrio_parahaemolyticus | EH301_01885 | 134210..137887 | 670 | BREX-system-P-loop-protein-BrxC-[Vibrio-parahaemolyticus] |
| EGR1144248.1 | Type1.BREX | SF1-SF2-Helicase+REase->SF1-SF2-Helicase+REase-><-PglZ<-TOPRIM<-OLD-ABC<-N6-METHYLASE<-DUF6079*<-BrxB-BrxF-iSTAND<-BrxA-DNA-binding<-wHTH+WYL+WCX<-XIS-HTH\|\|?->?->DUF2787->\|\|HTH->wHTH+WYL+WCX-> | 1154 | Bacteria>Pseudomonadota>Gammaproteobacteria>Vibrionales>Vibrionaceae>Vibrio | GCA_015798095.1 | Vibrio_parahaemolyticus | EC562_15505 | 41063..44527 | 670 | BREX-system-P-loop-protein-BrxC-[Vibrio-parahaemolyticus] |

| Protein-ID | System | Gene-Neighbourhoods | Length | Taxonomy | Assembly-ID | Strain | Locus-Tag | Genomic-coordinates | TaxID | Fasta-define |
|---|---|---|---|---|---|---|---|---|---|---|
| EGR1548466.1 | Type1.BREX | Arm-DNA-bind_3+Phage_integrase->SF1-SF2-Helicase+REase->SF1-SF2-Helicase+REase-><-PglZ<-TOPRIM<-OLD-ABC<-N6-METHYLASE<-DUF6079*<-BrxB-BrxF-iSTAND<-BrxA-DNA-binding<-wHTH+WYL+WCX<- [illegible] | 1154 | Bacteria>Pseudomonadota>Gammaproteobacteria>Vibrionales>Vibrionaceae>Vibrio | GCA_015798755.1 | Vibrio_parahaemolyticus | D6Y11_16950 | 161246..164710 | 670 | BREX-system-P-loop-protein-BrxC-[Vibrio-parahaemolyticus] |
| EGR2430327.1 | Type1.BREX | <-wHTH+WYL+WCX<-HTH<-DUF2787<-?<-?\|\|XIS-HTH->wHTH+WYL+WCX->BrxA-DNA-binding->BrxB-BrxF-iSTAND->DUF6079*->N6-METHYLASE->OLD-ABC->TOPRIM->PglZ-><-SF1-SF2-Helicase+REase<- [illegible] | 1154 | Bacteria>Pseudomonadota>Gammaproteobacteria>Vibrionales>Vibrionaceae>Vibrio | GCA_015802095.1 | Vibrio_cholerae | DYB83_10140 | 10866..14330 | 666 | BREX-system-P-loop-protein-BrxC-[Vibrio-cholerae] |
| EGR3302280.1 | Type1.BREX | XIS-HTH->wHTH+WYL+WCX->BrxA-DNA-binding->BrxB-BrxF-iSTAND->DUF6079*->N6-METHYLASE->OLD-ABC->TOPRIM->PglZ-><-SF1-SF2-Helicase+REase<-SF1-SF2-Helicase+REase<- [illegible] | 1154 | Bacteria>Pseudomonadota>Gammaproteobacteria>Vibrionales>Vibrionaceae>Vibrio | GCA_015805535.1 | Vibrio_parahaemolyticus | DMJ21_13130 | 8098..11562 | 670 | BREX-system-P-loop-protein-BrxC-[Vibrio-parahaemolyticus] |
| EGR3003176.1 | Type1.BREX | viral-protein->SF1-SF2-Helicase+REase->SF1-SF2-Helicase+REase-><-ZetaToxin<-PglZ<-ParB+HNH<-N6-METHYLASE<-DUF6079*<-BrxB-BrxF-iSTAND<-BrxA-DNA-binding<-wHTH+WYL+WCX\|\|<-?<-CR-ATPase4 | 1157 | Bacteria>Pseudomonadota>Gammaproteobacteria>Vibrionales>Vibrionaceae>Vibrio | GCA_015807135.1 | Vibrio_parahaemolyticus | DMW06_23030 | 12898..16371 | 670 | BREX-system-P-loop-protein-BrxC-[Vibrio-parahaemolyticus] |
| EGR4188113.1 | Type1.BREX | AbiEi->viral-protein->SF1-SF2-Helicase+REase->SF1-SF2-Helicase+REase-><-TPR<-HEPN+SF1-SF2-Helicase+RNase+RNase<-PglZ<-ParB+HNH<-N6-METHYLASE<-DUF6079*<-BrxB-BrxF-iSTAND<- [illegible] | 1157 | Bacteria>Pseudomonadota>Gammaproteobacteria>Vibrionales>Vibrionaceae>Vibrio | GCA_015809855.1 | Vibrio_cholerae | DDM99_15775 | 19819..23292 | 666 | BREX-system-P-loop-protein-BrxC-[Vibrio-cholerae] |
| EJG0966117.1 | Type1.BREX | <-ZnR<-?\|\|?->SF1-SF2-Helicase->AbiEi->viral-protein->wHTH+WYL+WCX->BrxA-DNA-binding->BrxB-BrxF-iSTAND->DUF6079*->N6-METHYLASE->ParB+HNH->GrpE->?->PglZ->SIGMA+OB-Fold+AAA-ATPase+ [illegible] | 1225 | Bacteria>Pseudomonadota>Gammaproteobacteria>Vibrionales>Vibrionaceae>Vibrio | GCA_015813265.2 | Vibrio_parahaemolyticus | C4G65_RS20915 | 24307..27984 | 670 | BREX-system-P-loop-protein-BrxC-[Vibrio-parahaemolyticus] |
| EJG0997619.1 | Type1.BREX | AbiEi->viral-protein->SF1-SF2-Helicase+REase->SF1-SF2-Helicase+REase-><-?<-ZetaToxin<-PglZ<-ParB+HNH<-N6-METHYLASE<-DUF6079*<-BrxB-BrxF-iSTAND<-BrxA-DNA-binding<-wHTH+WYL+WCX\|\| [illegible] | 1157 | Bacteria>Pseudomonadota>Gammaproteobacteria>Vibrionales>Vibrionaceae>Vibrio | GCA_015813305.2 | Vibrio_parahaemolyticus | C4G52_RS11760 | 21329..24802 | 670 | BREX-system-P-loop-protein-BrxC-[Vibrio-parahaemolyticus] |
| EJG0998764.1 | Type1.BREX | <-ZnR<-?\|\|?->SF1-SF2-Helicase->AbiEi->viral-protein->wHTH+WYL+WCX->BrxA-DNA-binding->BrxB-BrxF-iSTAND->DUF6079*->N6-METHYLASE->?->?->PglZ->SIGMA+OB-Fold+AAA-ATPase+Lon-Protease-BrxL-> [illegible] | 1225 | Bacteria>Pseudomonadota>Gammaproteobacteria>Vibrionales>Vibrionaceae>Vibrio | GCA_015813305.2 | Vibrio_parahaemolyticus | C4G52_RS17670 | 119644..123321 | 670 | BREX-system-P-loop-protein-BrxC-[Vibrio-parahaemolyticus] |
| EJG0981293.1 | Type1.BREX | Arm-DNA-bind_3+Phage_integrase->SF1-SF2-Helicase+REase->SF1-SF2-Helicase+REase-><-PglZ<-TOPRIM<-OLD-ABC<-N6-METHYLASE<-DUF6079*<-BrxB-BrxF-iSTAND<-BrxA-DNA-binding<-wHTH+WYL+WCX<- [illegible] | 1154 | Bacteria>Pseudomonadota>Gammaproteobacteria>Vibrionales>Vibrionaceae>Vibrio | GCA_015813375.2 | Vibrio_parahaemolyticus | C4G85_RS03500 | 776707..780171 | 670 | BREX-system-P-loop-protein-BrxC-[Vibrio-parahaemolyticus] |
| EJG0638704.1 | Type1.BREX | <-ZnR<-?\|\|?->SF1-SF2-Helicase->AbiEi->viral-protein->wHTH+WYL+WCX->BrxA-DNA-binding->BrxB-BrxF-iSTAND->DUF6079*->N6-METHYLASE->ParB+HNH->GrpE->?->PglZ->SIGMA+OB-Fold+AAA-ATPase+ [illegible] | 1225 | Bacteria>Pseudomonadota>Gammaproteobacteria>Vibrionales>Vibrionaceae>Vibrio | GCA_015813785.2 | Vibrio_parahaemolyticus | C4G37_RS13070 | 147789..151466 | 670 | BREX-system-P-loop-protein-BrxC-[Vibrio-parahaemolyticus] |
| EJG0936611.1 | Type1.BREX | <-GDResolvase+HIN-HTH<-?\|\|Arm-DNA-bind_3+Phage_integrase->SF1-SF2-Helicase+REase->SF1-SF2-Helicase+REase-><-PglZ<-TOPRIM<-OLD-ABC<-N6-METHYLASE<-DUF6079*<-BrxB-BrxF-iSTAND<- [illegible] | 1154 | Bacteria>Pseudomonadota>Gammaproteobacteria>Vibrionales>Vibrionaceae>Vibrio | GCA_015814185.2 | Vibrio_parahaemolyticus | C4G97_RS14675 | 118301..121765 | 670 | BREX-system-P-loop-protein-BrxC-[Vibrio-parahaemolyticus] |
| EGR4671255.1 | Type1.BREX | AbiEi->viral-protein->SF1-SF2-Helicase+REase->SF1-SF2-Helicase+REase-><-TPR<-HEPN+SF1-SF2-Helicase+RNase+RNase<-PglZ<-ParB+HNH<-N6-METHYLASE<-DUF6079*<-BrxB-BrxF-iSTAND<- [illegible] | 1157 | Bacteria>Pseudomonadota>Gammaproteobacteria>Vibrionales>Vibrionaceae>Vibrio | GCA_015814545.1 | Vibrio_parahaemolyticus | C4G45_02530 | 555237..558710 | 670 | BREX-system-P-loop-protein-BrxC-[Vibrio-parahaemolyticus] |
| EJG1873897.1 | Type1.BREX | AbiEi->viral-protein->SF1-SF2-Helicase+REase->SF1-SF2-Helicase+REase-><-?<-ZetaToxin<-PglZ<-ParB+HNH<-N6-METHYLASE<-DUF6079*<-BrxB-BrxF-iSTAND<-BrxA-DNA-binding<-wHTH+WYL+WCX\|\| [illegible] | 1157 | Bacteria>Pseudomonadota>Gammaproteobacteria>Vibrionales>Vibrionaceae>Vibrio | GCA_015817955.2 | Vibrio_parahaemolyticus | BS053_RS09090 | 170796..174269 | 670 | BREX-system-P-loop-protein-BrxC-[Vibrio-parahaemolyticus] |
| HAS8442986.1 | Type1.BREX | MobI-><-LexA-protease\|\|?->35exo->?->wHTH+WYL+WCX->BrxA-DNA-binding->BrxB-BrxF-iSTAND->DUF6079*->ParB+HNH->N6-METHYLASE->PglZ->SIGMA+OB-Fold+AAA-ATPase+Lon-Protease-BrxL->NACHT+HEAT+ [illegible] | 1225 | Bacteria>Pseudomonadota>Gammaproteobacteria>Vibrionales>Vibrionaceae>Vibrio | GCA_015820485.1 | Vibrio_vulnificus | I7740_20130 | 17039..20716 | 672 | BREX-system-P-loop-protein-BrxC-[Vibrio-vulnificus] |
| HAS8502529.1 | Type1.BREX | <-ZnR<-?\|\|?->SF1-SF2-Helicase->AbiEi->viral-protein->wHTH+WYL+WCX->BrxA-DNA-binding->BrxB-BrxF-iSTAND->DUF6079*->N6-METHYLASE->ParB+HNH->GrpE->?->PglZ->SIGMA+OB-Fold+AAA-ATPase+ [illegible] | 1225 | Bacteria>Pseudomonadota>Gammaproteobacteria>Vibrionales>Vibrionaceae>Vibrio | GCA_015820675.1 | Vibrio_vulnificus | I7764_01770 | 430591..434268 | 672 | BREX-system-P-loop-protein-BrxC-[Vibrio-vulnificus] |
| HAT7627508.1 | Type1.BREX | <-VirD4-TraG+VirB4-FtsK<-HDRnase<-HEPN<-MarR-HTH<-SIGMA+OB-Fold+AAA-ATPase+Lon-Protease-BrxL<-PglZ<-N6-METHYLASE<-DUF6079* | 1216 | Bacteria>Pseudomonadota>Gammaproteobacteria>Vibrionales>Vibrionaceae>Vibrio | GCA_015958965.1 | Vibrio_cholerae_O1 | JAW37_002070 | 69756..73407 | 127906 | BREX-system-P-loop-protein-BrxC-[Vibrio-cholerae-O1] |
| MBH9740666.1 | Type1.BREX | SF1-SF2-Helicase+HTH->MukF->JetB->XIS-HTH->BrxA-DNA-binding->BrxB-BrxF-iSTAND->DUF6079*->OLD-ABC->REase->N6-METHYLASE->PglZ->SIGMA+OB-Fold+AAA-ATPase+HKD-DNase-BrxL-> | 1185 | Bacteria>Pseudomonadota>Gammaproteobacteria>Vibrionales>Vibrionaceae>Vibrio | GCA_016094345.1 | Vibrio_navarrensis | CJ419_13180 | 152917..156474 | 29495 | hypothetical-protein-[Vibrio-navarrensis] |
| MBM4838763.1 | Type1.BREX | AbiEi->viral-protein->SF1-SF2-Helicase+REase->SF1-SF2-Helicase+REase-><-TPR<-HEPN+SF1-SF2-Helicase+RNase+RNase<-PglZ<-ParB+HNH<-N6-METHYLASE<-DUF6079*<-BrxB-BrxF-iSTAND<- [illegible] | 1157 | Bacteria>Pseudomonadota>Gammaproteobacteria>Vibrionales>Vibrionaceae>Vibrio | GCA_016876235.1 | Vibrio_parahaemolyticus | HYO55_06530 | 138658..142131 | 670 | BREX-system-P-loop-protein-BrxC-[Vibrio-parahaemolyticus] |
| MBM4875012.1 | Type1.BREX | <-AAA-ATPase-p<-HEPN\|\|CR-ATPase4->?->\|\|wHTH+WYL+WCX->BrxA-DNA-binding->BrxB-BrxF-iSTAND->DUF6079*->N6-METHYLASE->ParB+HNH->PglZ->HEPN+SF1-SF2-Helicase+RNase+RNase->TPR-><- [illegible] | 1157 | Bacteria>Pseudomonadota>Gammaproteobacteria>Vibrionales>Vibrionaceae>Vibrio | GCA_016876415.1 | Vibrio_parahaemolyticus | HYO45_05440 | 517620..521093 | 670 | BREX-system-P-loop-protein-BrxC-[Vibrio-parahaemolyticus] |
| EHD2253057.1 | Type1.BREX | viral-protein->wHTH+WYL+WCX->BrxA-DNA-binding->BrxB-BrxF-iSTAND->DUF6079*->N6-METHYLASE->ParB+HNH->GrpE->?->PglZ->SIGMA+OB-Fold+AAA-ATPase+Lon-Protease-BrxL-><-wHTH\|\|DUF4113-> | 1225 | Bacteria>Pseudomonadota>Gammaproteobacteria>Vibrionales>Vibrionaceae>Vibrio | GCA_017126235.1 | Vibrio_vulnificus | JRT24_004016 | 2833..6510 | 672 | BREX-system-P-loop-protein-BrxC-[Vibrio-vulnificus] |
| EHH1264392.1 | Type1.BREX | <-MORC<-PIN+OST-HTH<-CARF+NtrC-AAA<-HKD-DNase+HKD-DNase<-MarR-HTH<-SF1-SF2-Helicase\|\|wHTH+WYL+WCX->BrxA-DNA-binding->BrxB-BrxF-iSTAND->DUF6079*->N6-METHYLASE->ParB+HNH->?->?->PglZ-> [illegible] | 1248 | Bacteria>Pseudomonadota>Gammaproteobacteria>Vibrionales>Vibrionaceae>Vibrio | GCA_017604425.1 | Vibrio_parahaemolyticus | J7I02_003660 | 14110..17856 | 670 | BREX-system-P-loop-protein-BrxC-[Vibrio-parahaemolyticus] |
| EHH2507631.1 | Type1.BREX | Arm-DNA-bind_3+Phage_integrase->SF1-SF2-Helicase+REase->SF1-SF2-Helicase+REase-><-PglZ<-TOPRIM<-OLD-ABC<-N6-METHYLASE<-DUF6079*<-BrxB-BrxF-iSTAND<-BrxA-DNA-binding<-wHTH+WYL+WCX<- [illegible] | 1154 | Bacteria>Pseudomonadota>Gammaproteobacteria>Vibrionales>Vibrionaceae>Vibrio | GCA_017637525.1 | Vibrio_parahaemolyticus | J7X10_003794 | 109543..113007 | 670 | BREX-system-P-loop-protein-BrxC-[Vibrio-parahaemolyticus] |
| EHK0048698.1 | Type1.BREX | <-DUF4113\|\|wHTH-><-SIGMA+OB-Fold [illegible] | 1225 | Bacteria>Pseudomonadota>Gammaproteobacteria>Vibrionales>Vibrionaceae>Vibrio | GCA_017968185.1 | Vibrio_parahaemolyticus | KCA08_003292 | 85751..89428 | 670 | BREX-system-P-loop-protein-BrxC-[Vibrio-parahaemolyticus] |
| HBC2045456.1 | Type1.BREX | <-VirD4-TraG+VirB4-FtsK<-HDRnase [illegible] | 1184 | Bacteria>Pseudomonadota>Gammaproteobacteria>Vibrionales>Vibrionaceae>Vibrio | GCA_018073665.1 | Vibrio_cholerae | I6Z89_000440 | 124693..128249 | 666 | BREX-system-P-loop-protein-BrxC-[Vibrio-cholerae] |
| HBC2117287.1 | Type1.BREX | DUF6079*->N6-METHYLASE->PglZ-> [illegible] | 1191 | Bacteria>Pseudomonadota>Gammaproteobacteria>Vibrionales>Vibrionaceae>Vibrio | GCA_018074265.1 | Vibrio_cholerae | I6207_001264 | 1..3578 | 666 | BREX-system-P-loop-protein-BrxC-[Vibrio-cholerae] |
| EHK9043270.1 | Type1.BREX | <-MORC<-PIN+OST-HTH<-CARF+NtrC- [illegible] | 1248 | Bacteria>Pseudomonadota>Gammaproteobacteria>Vibrionales>Vibrionaceae>Vibrio | GCA_018080905.1 | Vibrio_vulnificus | KCU37_001412 | 195245..198991 | 672 | BREX-system-P-loop-protein-BrxC-[Vibrio-vulnificus] |
| HBC3376470.1 | Type1.BREX | <-ZnR<-?\|\|?->SF1-SF2-Helicase-> [illegible] | 1225 | Bacteria>Pseudomonadota>Gammaproteobacteria>Vibrionales>Vibrionaceae>Vibrio | GCA_018118685.1 | Vibrio_cholerae | KDX67_000916 | 90395..94072 | 666 | BREX-system-P-loop-protein-BrxC-[Vibrio-cholerae] |
| HBC3399177.1 | Type1.BREX | <-LexA-protease\|\|HEPN+SF1-SF2- [illegible] | 1225 | Bacteria>Pseudomonadota>Gammaproteobacteria>Vibrionales>Vibrionaceae>Vibrio | GCA_018118805.1 | Vibrio_parahaemolyticus | KDX79_000987 | 173788..177465 | 670 | BREX-system-P-loop-protein-BrxC-[Vibrio-parahaemolyticus] |
| HBC3428133.1 | Type1.BREX | <-PglZ<-?<-?<-?<-?<-ZnR+3C-ABC+ [illegible] | 1248 | Bacteria>Pseudomonadota>Gammaproteobacteria>Vibrionales>Vibrionaceae>Vibrio | GCA_018118885.1 | Vibrio_parahaemolyticus | KDY17_000940 | 500367..504113 | 670 | BREX-system-P-loop-protein-BrxC-[Vibrio-parahaemolyticus] |
| HBC3445072.1 | Type1.BREX | <-ZnR<-?\|\|AbiEi->viral-protein-> [illegible] | 1225 | Bacteria>Pseudomonadota>Gammaproteobacteria>Vibrionales>Vibrionaceae>Vibrio | GCA_018119025.1 | Vibrio_parahaemolyticus | KDY48_003788 | 6147..9824 | 670 | BREX-system-P-loop-protein-BrxC-[Vibrio-parahaemolyticus] |
| HBC3481459.1 | Type1.BREX | REase+SPF-cons->?-><-PIN+OST-HTH [illegible] | 1157 | Bacteria>Pseudomonadota>Gammaproteobacteria>Vibrionales>Vibrionaceae>Vibrio | GCA_018119205.1 | Vibrio_parahaemolyticus | KDY88_002916 | 65183..68656 | 670 | BREX-system-P-loop-protein-BrxC-[Vibrio-parahaemolyticus] |
| HBC3540120.1 | Type1.BREX | <-ZnR+3C-ABC+TPR<-ParB+HNH<-?<- [illegible] | 1248 | Bacteria>Pseudomonadota>Gammaproteobacteria>Vibrionales>Vibrionaceae>Vibrio | GCA_018119555.1 | Vibrio_parahaemolyticus | KDZ97_004679 | 8277..12023 | 670 | BREX-system-P-loop-protein-BrxC-[Vibrio-parahaemolyticus] |
| HBC3921009.1 | Type1.BREX | <-HigA\|\|CR-ATPase4->?-><-?<-LexA [illegible] | 1157 | Bacteria>Pseudomonadota>Gammaproteobacteria>Vibrionales>Vibrionaceae>Vibrio | GCA_018126325.1 | Vibrio_parahaemolyticus | KD834_003320 | 38197..41670 | 670 | BREX-system-P-loop-protein-BrxC-[Vibrio-parahaemolyticus] |
| HBC3861549.1 | Type1.BREX | <-VirD4-TraG+VirB4-FtsK<-HDRnase [illegible] | 1225 | Bacteria>Pseudomonadota>Gammaproteobacteria>Vibrionales>Vibrionaceae>Vibrio | GCA_018126725.1 | Vibrio_parahaemolyticus | KD871_001930 | 256177..259854 | 670 | BREX-system-P-loop-protein-BrxC-[Vibrio-parahaemolyticus] |
| HBC3986622.1 | Type1.BREX | <-DUF4113\|\|wHTH-><-SIGMA+OB-Fold [illegible] | 1225 | Bacteria>Pseudomonadota>Gammaproteobacteria>Vibrionales>Vibrionaceae>Vibrio | GCA_018126825.1 | Vibrio_parahaemolyticus | KD867_003188 | 79006..82683 | 670 | BREX-system-P-loop-protein-BrxC-[Vibrio-parahaemolyticus] |
| HBC3940910.1 | Type1.BREX | <-HigA\|\|CR-ATPase4->?-><-?<-LexA [illegible] | 1157 | Bacteria>Pseudomonadota>Gammaproteobacteria>Vibrionales>Vibrionaceae>Vibrio | GCA_018127045.1 | Vibrio_parahaemolyticus | KD880_004471 | 11650..15123 | 670 | BREX-system-P-loop-protein-BrxC-[Vibrio-parahaemolyticus] |
| MBT0011870.1 | Type1.BREX | <-SIGMA+OB-Fold+AAA-ATPase+HKD- [illegible] | 1187 | Bacteria>Pseudomonadota>Gammaproteobacteria>Vibrionales>Vibrionaceae>Vibrio | GCA_018457755.1 | Vibrio_alginolyticus | N7I24_002738 | 64007..67570 | 663 | BREX-system-P-loop-protein-BrxC-[Vibrio-alginolyticus] |
| EHW0635156.1 | Type1.BREX | wHTH-><-SIGMA+OB-Fold+AAA-ATPase [illegible] | 1225 | Bacteria>Pseudomonadota>Gammaproteobacteria>Vibrionales>Vibrionaceae>Vibrio | GCA_019442565.1 | Vibrio_vulnificus | K1B29_001386 | 71922..75599 | 672 | BREX-system-P-loop-protein-BrxC-[Vibrio-vulnificus] |
| EHW0654650.1 | Type1.BREX | HTH_20-><-HEPN<-SbcC\|\|CR-ATPase4 [illegible] | 1156 | Bacteria>Pseudomonadota>Gammaproteobacteria>Vibrionales>Vibrionaceae>Vibrio | GCA_019442625.1 | Vibrio_parahaemolyticus | K1B36_003414 | 61374..64844 | 670 | BREX-system-P-loop-protein-BrxC-[Vibrio-parahaemolyticus] |
| EHZ2755060.1 | Type1.BREX | <-ZnR<-?\|\|?->SF1-SF2-Helicase-> [illegible] | 1225 | Bacteria>Pseudomonadota>Gammaproteobacteria>Vibrionales>Vibrionaceae>Vibrio | GCA_019682715.1 | Vibrio_vulnificus | K5N53_003177 | 34547..38224 | 672 | BREX-system-P-loop-protein-BrxC-[Vibrio-vulnificus] |
| MBY7947240.1 | Type1.BREX | CR-ATPase4->?->\|\|?->wHTH-><- [illegible] | 1225 | Bacteria>Pseudomonadota>Gammaproteobacteria>Vibrionales>Vibrionaceae>Vibrio | GCA_019853335.1 | Vibrio_fluvialis | KW461_01980 | 454560..458237 | 676 | BREX-system-P-loop-protein-BrxC-[Vibrio-fluvialis] |
| MBY8044957.1 | Type1.BREX | <-HTH_Tnp_1+HTH_21+rve+rve_3\|\|<- [illegible] | 1157 | Bacteria>Pseudomonadota>Gammaproteobacteria>Vibrionales>Vibrionaceae>Vibrio | GCA_019853775.1 | Vibrio_fluvialis | KW486_21175 | 14278..17751 | 676 | BREX-system-P-loop-protein-BrxC-[Vibrio-fluvialis] |
| MBY8148861.1 | Type1.BREX | AbiEi->viral-protein->SF1-SF2- [illegible] | 1157 | Bacteria>Pseudomonadota>Gammaproteobacteria>Vibrionales>Vibrionaceae>Vibrio | GCA_019854195.1 | Vibrio_fluvialis | KW504_05155 | 39947..43420 | 676 | BREX-system-P-loop-protein-BrxC-[Vibrio-fluvialis] |
| MBY8207859.1 | Type1.BREX | <-DUF4113\|\|wHTH-><-SIGMA+OB-Fold [illegible] | 1225 | Bacteria>Pseudomonadota>Gammaproteobacteria>Vibrionales>Vibrionaceae>Vibrio | GCA_019854355.1 | Vibrio_fluvialis | KW531_20835 | 16453..20130 | 676 | BREX-system-P-loop-protein-BrxC-[Vibrio-fluvialis] |
| EIE1271992.1 | Type1.BREX | <-ZnR<-?\|\|?->SF1-SF2-Helicase-> [illegible] | 1225 | Bacteria>Pseudomonadota>Gammaproteobacteria>Vibrionales>Vibrionaceae>Vibrio | GCA_020124895.1 | Vibrio_parahaemolyticus | LCX75_001329 | 151524..155201 | 670 | BREX-system-P-loop-protein-BrxC-[Vibrio-parahaemolyticus] |
| EII2403787.1 | Type1.BREX | <-GDResolvase+HIN-HTH<-?\|\| [illegible] | 1154 | Bacteria>Pseudomonadota>Gammaproteobacteria>Vibrionales>Vibrionaceae>Vibrio | GCA_020407245.1 | Vibrio_parahaemolyticus | LG159_002760 | 116849..120313 | 670 | BREX-system-P-loop-protein-BrxC-[Vibrio-parahaemolyticus] |
| EII3725635.1 | Type1.BREX | <-VirD4-TraG+VirB4-FtsK<-HDRnase [illegible] | 1225 | Bacteria>Pseudomonadota>Gammaproteobacteria>Vibrionales>Vibrionaceae>Vibrio | GCA_020414365.1 | Vibrio_cholerae | LG468_000348 | 129198..132875 | 666 | BREX-system-P-loop-protein-BrxC-[Vibrio-cholerae] |
| HBK3330408.1 | Type1.BREX | <-TPR+HEPN\|\|<-PIN+OST-HTH<- [illegible] | 1248 | Bacteria>Pseudomonadota>Gammaproteobacteria>Vibrionales>Vibrionaceae>Vibrio | GCA_020473965.1 | Vibrio_parahaemolyticus | KD833_003429 | 17712..21458 | 670 | BREX-system-P-loop-protein-BrxC-[Vibrio-parahaemolyticus] |
| EIT6975184.1 | Type1.BREX | Couple_hipA+HipA_C->SF1-SF2- [illegible] | 1232 | Bacteria>Pseudomonadota>Gammaproteobacteria>Vibrionales>Vibrionaceae>Vibrio | GCA_021537035.1 | Vibrio_vulnificus | L2596_001303 | 248208..251906 | 672 | BREX-system-P-loop-protein-BrxC-[Vibrio-vulnificus] |
| EIU6866063.1 | Type1.BREX | CR-ATPase4->?-><-DUF4113\|\|wHTH-> [illegible] | 1225 | Bacteria>Pseudomonadota>Gammaproteobacteria>Vibrionales>Vibrionaceae>Vibrio | GCA_021704015.1 | Vibrio_parahaemolyticus | L5169_000073 | 92916..96593 | 670 | BREX-system-P-loop-protein-BrxC-[Vibrio-parahaemolyticus] |
| EIU7849849.1 | Type1.BREX | AbiEi->viral-protein->SF1-SF2- [illegible] | 1157 | Bacteria>Pseudomonadota>Gammaproteobacteria>Vibrionales>Vibrionaceae>Vibrio | GCA_021716735.1 | Vibrio_parahaemolyticus | L3I74_001636 | 175648..179121 | 670 | BREX-system-P-loop-protein-BrxC-[Vibrio-parahaemolyticus] |
| EIV8505581.1 | Type1.BREX | <-DUF4113\|\|wHTH-><-SIGMA+OB-Fold [illegible] | 1225 | Bacteria>Pseudomonadota>Gammaproteobacteria>Vibrionales>Vibrionaceae>Vibrio | GCA_022019715.1 | Vibrio_parahaemolyticus | M5229_003413 | 80224..83901 | 670 | BREX-system-P-loop-protein-BrxC-[Vibrio-parahaemolyticus] |
| MCG9593371.1 | Type1.BREX | CR-ATPase4->?-><-DUF4113\|\|wHTH-> [illegible] | 1171 | Bacteria>Pseudomonadota>Gammaproteobacteria>Vibrionales>Vibrionaceae>Vibrio | GCA_022398055.1 | Vibrio_diabolicus | L1D58_17270 | 13232..16749 | 50719 | BREX-system-P-loop-protein-BrxC-[Vibrio-diabolicus] |
| EIY9800890.1 | Type1.BREX | <-ZnR<-?\|\|?->SF1-SF2-Helicase-> [illegible] | 1225 | Bacteria>Pseudomonadota>Gammaproteobacteria>Vibrionales>Vibrionaceae>Vibrio | GCA_022594155.1 | Vibrio_parahaemolyticus | MNO17_002467 | 349786..353463 | 670 | BREX-system-P-loop-protein-BrxC-[Vibrio-parahaemolyticus] |
| EIZ1010574.1 | Type1.BREX | HTH_3->Couple_hipA+HipA_C-> [illegible] | 1187 | Bacteria>Pseudomonadota>Gammaproteobacteria>Vibrionales>Vibrionaceae>Vibrio | GCA_022609845.1 | Vibrio_vulnificus | MOS09_002154 | 288384..291947 | 672 | BREX-system-P-loop-protein-BrxC-[Vibrio-vulnificus] |
| EJB1796336.1 | Type1.BREX | SbcC+SbcC->?-><-?<-?\|\|?-> [illegible] | 1221 | Bacteria>Pseudomonadota>Gammaproteobacteria>Vibrionales>Vibrionaceae>Vibrio | GCA_022927965.1 | Vibrio_parahaemolyticus | MUE05_003019 | 106094..109759 | 670 | BREX-system-P-loop-protein-BrxC-[Vibrio-parahaemolyticus] |
| EJB8347940.1 | Type1.BREX | <-VirD4-TraG+VirB4-FtsK<-HDRnase [illegible] | 1225 | Bacteria>Pseudomonadota>Gammaproteobacteria>Vibrionales>Vibrionaceae>Vibrio | GCA_022966375.1 | Vibrio_cholerae | MW328_000282 | 40595..44272 | 666 | BREX-system-P-loop-protein-BrxC-[Vibrio-cholerae] |
| EJB8581804.1 | Type1.BREX | <-VirD4-TraG+VirB4-FtsK<-HDRnase [illegible] | 1229 | Bacteria>Pseudomonadota>Gammaproteobacteria>Vibrionales>Vibrionaceae>Vibrio | GCA_022967455.1 | Vibrio_cholerae | MW361_003123 | 34970..38659 | 666 | BREX-system-P-loop-protein-BrxC-[Vibrio-cholerae] |
| EJC1210117.1 | Type1.BREX | <-DUF2787<-?<-?\|\|XIS-HTH-> [illegible] | 1154 | Bacteria>Pseudomonadota>Gammaproteobacteria>Vibrionales>Vibrionaceae>Vibrio | GCA_022997325.1 | Vibrio_parahaemolyticus | MX350_002594 | 5689..9153 | 670 | BREX-system-P-loop-protein-BrxC-[Vibrio-parahaemolyticus] |
| EJC6937227.1 | Type1.BREX | SF1-SF2-Helicase->AbiEi-> [illegible] | 1225 | Bacteria>Pseudomonadota>Gammaproteobacteria>Vibrionales>Vibrionaceae>Vibrio | GCA_023080515.1 | Vibrio_parahaemolyticus | MX856_004599 | 4649..8326 | 670 | BREX-system-P-loop-protein-BrxC-[Vibrio-parahaemolyticus] |
| GHW31175.1 | Type1.BREX | <-HTH_Tnp_1\|\|wHTH-><-SIGMA+ [illegible] | 1225 | Bacteria>Pseudomonadota>Gammaproteobacteria>Vibrionales>Vibrionaceae>Vibrio | GCA_023163105.1 | Vibrio_cholerae | VCSRO150_3322 | 15322..18999 | 666 | DNA-repair-protein-[Vibrio-cholerae] |
| EJE4207362.1 | Type1.BREX | <-VirD4-TraG+VirB4-FtsK<-HDRnase [illegible] | 1225 | Bacteria>Pseudomonadota>Gammaproteobacteria>Vibrionales>Vibrionaceae>Vibrio | GCA_023313315.1 | Vibrio_parahaemolyticus | M3899_001773 | 267109..270786 | 670 | BREX-system-P-loop-protein-BrxC-[Vibrio-parahaemolyticus] |
| EJE8514353.1 | Type1.BREX | CR-ATPase4->?-><-DUF4113\|\|wHTH-> [illegible] | 1225 | Bacteria>Pseudomonadota>Gammaproteobacteria>Vibrionales>Vibrionaceae>Vibrio | GCA_023352155.1 | Vibrio_parahaemolyticus | M5244_002608 | 72363..76040 | 670 | BREX-system-P-loop-protein-BrxC-[Vibrio-parahaemolyticus] |
| EJE8545413.1 | Type1.BREX | <-SIGMA+OB-Fold+AAA-ATPase+ [illegible] | 1232 | Bacteria>Pseudomonadota>Gammaproteobacteria>Vibrionales>Vibrionaceae>Vibrio | GCA_023352275.1 | Vibrio_vulnificus | M5238_000046 | 65248..68946 | 672 | BREX-system-P-loop-protein-BrxC-[Vibrio-vulnificus] |
| EJE8564157.1 | Type1.BREX | AbiEi->viral-protein->SF1-SF2- [illegible] | 1157 | Bacteria>Pseudomonadota>Gammaproteobacteria>Vibrionales>Vibrionaceae>Vibrio | GCA_023352375.1 | Vibrio_parahaemolyticus | M5224_000723 | 57760..61233 | 670 | BREX-system-P-loop-protein-BrxC-[Vibrio-parahaemolyticus] |
| EJF4458718.1 | Type1.BREX | <-DUF4113\|\|wHTH-><-SIGMA+OB-Fold [illegible] | 1225 | Bacteria>Pseudomonadota>Gammaproteobacteria>Vibrionales>Vibrionaceae>Vibrio | GCA_023466975.1 | Vibrio_parahaemolyticus | MT303_002826 | 138363..142040 | 670 | BREX-system-P-loop-protein-BrxC-[Vibrio-parahaemolyticus] |
| HCD5120999.1 | Type1.BREX | AbiEi->viral-protein-> [illegible] | 1225 | Bacteria>Pseudomonadota>Gammaproteobacteria>Vibrionales>Vibrionaceae>Vibrio | GCA_023611065.1 | Vibrio_parahaemolyticus | NB556_01245 | 3462..7139 | 670 | BREX-system-P-loop-protein-BrxC-[Vibrio-parahaemolyticus] |
| HCE1978782.1 | Type1.BREX | <-ZnR<-?\|\|?->SF1-SF2-Helicase-> [illegible] | 1225 | Bacteria>Pseudomonadota>Gammaproteobacteria>Vibrionales>Vibrionaceae>Vibrio | GCA_023810845.1 | Vibrio_parahaemolyticus | NGS68_002079 | 171773..175450 | 670 | BREX-system-P-loop-protein-BrxC-[Vibrio-parahaemolyticus] |
| HCE2271447.1 | Type1.BREX | <-HEPN\|\|CR-ATPase4->?->\|\|wHTH-> [illegible] | 1225 | Bacteria>Pseudomonadota>Gammaproteobacteria>Vibrionales>Vibrionaceae>Vibrio | GCA_023811985.1 | Vibrio_parahaemolyticus | NGT56_000627 | 721824..725501 | 670 | BREX-system-P-loop-protein-BrxC-[Vibrio-parahaemolyticus] |
| HCE2679243.1 | Type1.BREX | <-SIGMA+OB-Fold+AAA-ATPase+ [illegible] | 1248 | Bacteria>Pseudomonadota>Gammaproteobacteria>Vibrionales>Vibrionaceae>Vibrio | GCA_023813525.1 | Vibrio_parahaemolyticus | NGU92_000683 | 148376..152122 | 670 | BREX-system-P-loop-protein-BrxC-[Vibrio-parahaemolyticus] |
| HCE3362112.1 | Type1.BREX | <-VirD4-TraG+VirB4-FtsK<-HDRnase [illegible] | 1225 | Bacteria>Pseudomonadota>Gammaproteobacteria>Vibrionales>Vibrionaceae>Vibrio | GCA_023821665.1 | Vibrio_parahaemolyticus | NG589_000244 | 272459..276136 | 670 | BREX-system-P-loop-protein-BrxC-[Vibrio-parahaemolyticus] |
| HCG5108676.1 | Type1.BREX | AbiEi->viral-protein->SF1-SF2- [illegible] | 1157 | Bacteria>Pseudomonadota>Gammaproteobacteria>Vibrionales>Vibrionaceae>Vibrio | GCA_024034975.1 | Vibrio_parahaemolyticus | NJY97_002344 | 346649..350122 | 670 | BREX-system-P-loop-protein-BrxC-[Vibrio-parahaemolyticus] |
| HCG6132052.1 | Type1.BREX | viral-protein->SF1-SF2- [illegible] | 1157 | Bacteria>Pseudomonadota>Gammaproteobacteria>Vibrionales>Vibrionaceae>Vibrio | GCA_024038995.1 | Vibrio_parahaemolyticus | NJ068_002907 | 67806..71279 | 670 | BREX-system-P-loop-protein-BrxC-[Vibrio-parahaemolyticus] |
| HCG7048824.1 | Type1.BREX | <-VirD4-TraG+VirB4-FtsK<-HDRnase [illegible] | 1225 | Bacteria>Pseudomonadota>Gammaproteobacteria>Vibrionales>Vibrionaceae>Vibrio | GCA_024042595.1 | Vibrio_parahaemolyticus | NJ298_000555 | 545999..549676 | 670 | BREX-system-P-loop-protein-BrxC-[Vibrio-parahaemolyticus] |
| HCG7393723.1 | Type1.BREX | McrB-AAA->McrC-NTD+REase->?-> [illegible] | 1187 | Bacteria>Pseudomonadota>Gammaproteobacteria>Vibrionales>Vibrionaceae>Vibrio | GCA_024044075.1 | Vibrio_parahaemolyticus | NJ383_004129 | 13694..17257 | 670 | BREX-system-P-loop-protein-BrxC-[Vibrio-parahaemolyticus] |
| HCG7769728.1 | Type1.BREX | <-SIGMA+OB-Fold+AAA-ATPase+ [illegible] | 1248 | Bacteria>Pseudomonadota>Gammaproteobacteria>Vibrionales>Vibrionaceae>Vibrio | GCA_024045435.1 | Vibrio_parahaemolyticus | NJ450_003037 | 12821..16567 | 670 | BREX-system-P-loop-protein-BrxC-[Vibrio-parahaemolyticus] |
| HCG8025121.1 | Type1.BREX | <-ZnR<-?\|\|?->SF1-SF2-Helicase-> [illegible] | 1225 | Bacteria>Pseudomonadota>Gammaproteobacteria>Vibrionales>Vibrionaceae>Vibrio | GCA_024046515.1 | Vibrio_parahaemolyticus | NJ909_002125 | 181233..184910 | 670 | BREX-system-P-loop-protein-BrxC-[Vibrio-parahaemolyticus] |
| HCG8164047.1 | Type1.BREX | <-PIN+OST-HTH\|\|CR-ATPase4->?->\|\| [illegible] | 1225 | Bacteria>Pseudomonadota>Gammaproteobacteria>Vibrionales>Vibrionaceae>Vibrio | GCA_024047235.1 | Vibrio_parahaemolyticus | NJ496_004031 | 74363..78040 | 670 | BREX-system-P-loop-protein-BrxC-[Vibrio-parahaemolyticus] |
| HCG8269540.1 | Type1.BREX | <-ZnR<-?\|\|?->SF1-SF2-Helicase-> [illegible] | 1225 | Bacteria>Pseudomonadota>Gammaproteobacteria>Vibrionales>Vibrionaceae>Vibrio | GCA_024047655.1 | Vibrio_parahaemolyticus | NJ590_004433 | 15397..19074 | 670 | BREX-system-P-loop-protein-BrxC-[Vibrio-parahaemolyticus] |
| HCH5919840.1 | Type1.BREX | <-ZnR<-?\|\|?->SF1-SF2-Helicase-> [illegible] | 1225 | Bacteria>Pseudomonadota>Gammaproteobacteria>Vibrionales>Vibrionaceae>Vibrio | GCA_024097015.1 | Vibrio_parahaemolyticus | NK048_004352 | 19128..22805 | 670 | BREX-system-P-loop-protein-BrxC-[Vibrio-parahaemolyticus] |
| EJK2098984.1 | Type1.BREX | <-VirD4-TraG+VirB4-FtsK<-HDRnase [illegible] | 1225 | Bacteria>Pseudomonadota>Gammaproteobacteria>Vibrionales>Vibrionaceae>Vibrio | GCA_024103575.1 | Vibrio_cholerae | NK421_001793 | 214069..217746 | 666 | BREX-system-P-loop-protein-BrxC-[Vibrio-cholerae] |
| EJL6432478.1 | Type1.BREX | <-VirD4-TraG+VirB4-FtsK<-HDRnase [illegible] | 1225 | Bacteria>Pseudomonadota>Gammaproteobacteria>Vibrionales>Vibrionaceae>Vibrio | GCA_024216615.1 | Vibrio_cholerae | NMR72_000938 | 126901..130578 | 666 | BREX-system-P-loop-protein-BrxC-[Vibrio-cholerae] |
| EJL6578344.1 | Type1.BREX | Arm-DNA-bind_3+Phage_integrase-> [illegible] | 1154 | Bacteria>Pseudomonadota>Gammaproteobacteria>Vibrionales>Vibrionaceae>Vibrio | GCA_024217525.1 | Vibrio_cholerae | NMS50_002630 | 71340..74804 | 666 | BREX-system-P-loop-protein-BrxC-[Vibrio-cholerae] |
| EJL6824798.1 | Type1.BREX | <-VirD4-TraG+VirB4-FtsK<-HDRnase [illegible] | 1202 | Bacteria>Pseudomonadota>Gammaproteobacteria>Vibrionales>Vibrionaceae>Vibrio | GCA_024218935.1 | Vibrio_cholerae | NMT01_000642 | 124775..128383 | 666 | BREX-system-P-loop-protein-BrxC-[Vibrio-cholerae] |
| UTZ28051.1 | Type1.BREX | ABhydrolase-><-HTH_18+HTH_AraC<- [illegible] | 1183 | Bacteria>Pseudomonadota>Gammaproteobacteria>Vibrionales>Vibrionaceae>Vibrio | GCA_024442175.1 | Vibrio_campbellii | HB761_02515 | 550346..553897 | 680 | BREX-system-P-loop-protein-BrxC-[Vibrio-campbellii] |
| EJO3860683.1 | Type1.BREX | AbiEi->viral-protein->SF1-SF2- [illegible] | 1157 | Bacteria>Pseudomonadota>Gammaproteobacteria>Vibrionales>Vibrionaceae>Vibrio | GCA_024563785.1 | Vibrio_parahaemolyticus | NRI58_000919 | 1072634..1076107 | 670 | BREX-system-P-loop-protein-BrxC-[Vibrio-parahaemolyticus] |
| HCJ6872077.1 | Type1.BREX | <-OBFOLD-NUCLEASE<-CARF+NtrC- [illegible] | 1225 | Bacteria>Pseudomonadota>Gammaproteobacteria>Vibrionales>Vibrionaceae>Vibrio | GCA_024621265.1 | Vibrio_cholerae | NU929_001221 | 142702..146379 | 666 | BREX-system-P-loop-protein-BrxC-[Vibrio-cholerae] |
| HCJ7299597.1 | Type1.BREX | <-OBFOLD-NUCLEASE<-CARF+NtrC- [illegible] | 1225 | Bacteria>Pseudomonadota>Gammaproteobacteria>Vibrionales>Vibrionaceae>Vibrio | GCA_024632775.1 | Vibrio_cholerae | NVH34_000348 | 35814..39491 | 666 | BREX-system-P-loop-protein-BrxC-[Vibrio-cholerae] |
| MCR9580722.1 | Type1.BREX | viral-protein->wHTH+WYL+WCX-> [illegible] | 1225 | Bacteria>Pseudomonadota>Gammaproteobacteria>Vibrionales>Vibrionaceae>Vibrio | GCA_024744095.1 | Vibrio_antiquarius | NB519_11830 | 2837..6514 | 150340 | BREX-system-P-loop-protein-BrxC-[Vibrio-antiquarius] |
| HCM0920431.1 | Type1.BREX | <-wHTH+WYL+WCX<-HTH\|\|<-DUF2787<- [illegible] | 1154 | Bacteria>Pseudomonadota>Gammaproteobacteria>Vibrionales>Vibrionaceae>Vibrio | GCA_025068375.1 | Vibrio_parahaemolyticus | N2828_003369 | 93414..96878 | 670 | BREX-system-P-loop-protein-BrxC-[Vibrio-parahaemolyticus] |
| EJU9535832.1 | Type1.BREX | AbiEi->viral-protein->SF1-SF2- [illegible] | 1157 | Bacteria>Pseudomonadota>Gammaproteobacteria>Vibrionales>Vibrionaceae>Vibrio | GCA_025224335.1 | Vibrio_alginolyticus | N5M46_001148 | 14652..18125 | 663 | BREX-system-P-loop-protein-BrxC-[Vibrio-alginolyticus] |
| EJV5950193.1 | Type1.BREX | <-SIGMA+OB-Fold+AAA-ATPase+HKD- [illegible] | 1187 | Bacteria>Pseudomonadota>Gammaproteobacteria>Vibrionales>Vibrionaceae>Vibrio | GCA_025320405.1 | Vibrio_alginolyticus | N7I24_002738 | 64007..67570 | 663 | BREX-system-P-loop-protein-BrxC-[Vibrio-alginolyticus] |
| GLT15886.1 | Type1.BREX | <-wHTH+WYL+WCX\|\|MukF->JetB-> [illegible] | 1186 | Bacteria>Pseudomonadota>Gammaproteobacteria>Vibrionales>Vibrionaceae>Vibrio | GCA_030161215.1 | Vibrio_algivorus | GCM10007931_28610 | 496776..500336 | 1667024 | hypothetical-protein-GCM10007931_28610-[Vibrio-algivorus] |
| WP_053056240.1 | Type1.BREX | <-VirD4-TraG+VirB4-FtsK<-HDRnase [illegible] | 721 | Bacteria>Pseudomonadota>Gammaproteobacteria>Vibrionales>Vibrionaceae>Vibrio | GCA_030820635.1 | Vibrio_cholerae | ALE43_RS03710 | 109450..111613 | 666 | BREX-system-P-loop-protein-BrxC,-partial-[Vibrio-cholerae] |
| WP_242567369.1 | Type1.BREX | <-Phage_integrase<-?<- [illegible] | 994 | Bacteria>Pseudomonadota>Gammaproteobacteria>Vibrionales>Vibrionaceae>Vibrio | GCA_030820635.1 | Vibrio_cholerae | ALG09_RS16130 | 6852..9838 | 666 | BREX-system-P-loop-protein-BrxC,-partial-[Vibrio-cholerae] |
| WP_114867813.1 | Type1.BREX | <-SbcC<-SIGMA+OB-Fold+AAA-ATPase [illegible] | 1225 | Bacteria>Pseudomonadota>Gammaproteobacteria>Vibrionales>Vibrionaceae>Vibrio | GCA_030820635.1 | Vibrio_parahaemolyticus | CFH89_RS14230 | 2819482..2823159 | 670 | BREX-system-P-loop-protein-BrxC-[Vibrio-parahaemolyticus] |
| WP_178326691.1 | Type1.BREX | CR-ATPase4->?->wHTH-><-SIGMA+ [illegible] | 952 | Bacteria>Pseudomonadota>Gammaproteobacteria>Vibrionales>Vibrionaceae>Vibrio | GCA_030820635.1 | Vibrio_parahaemolyticus | HX770_RS05795 | 37869..40727 | 670 | BREX-system-P-loop-protein-BrxC,-partial-[Vibrio-parahaemolyticus] |
| WP_005376166.1 | Type1.BREX | sGTP-><-HEPN\|\|CR-ATPase4->?->\|\| [illegible] | 1157 | Bacteria>Pseudomonadota>Gammaproteobacteria>Vibrionales>Vibrionaceae>Vibrio | GCF_000153505.1 | Vibrio_alginolyticus | V12G01_RS31815 | 1598723..1602196 | 663 | BREX-system-P-loop-protein-BrxC-[Vibrio-alginolyticus] |
| WP_000152592.1 | Type1.BREX | <-MULE-TRANSPOSASE<- [illegible] | 1229 | Bacteria>Pseudomonadota>Gammaproteobacteria>Vibrionales>Vibrionaceae>Vibrio | GCF_000166455.1 | Vibrio_cholerae | HJ37_RS00485 | 111831..115520 | 666 | BREX-system-P-loop-protein-BrxC-[Vibrio-cholerae] |
| WP_000934396.1 | Type1.BREX | TraI-VirD2-RCRE->?->?->CHTH->\|\| [illegible] | 1157 | Bacteria>Pseudomonadota>Gammaproteobacteria>Vibrionales>Vibrionaceae>Vibrio | GCF_000220785.1 | Vibrio_cholerae | VCHE48_RS12985 | 863732..867205 | 666 | BREX-system-P-loop-protein-BrxC-[Vibrio-cholerae] |
| WP_004748886.1 | Type1.BREX | <-Thioredoxin_4\|\|wHTH+WYL+WCX-> [illegible] | 1219 | Bacteria>Pseudomonadota>Gammaproteobacteria>Vibrionales>Vibrionaceae>Vibrio | GCF_000222665.1 | Vibrio_tubiashii | RH87_RS24955 | 79516..83175 | 29498 | BREX-system-P-loop-protein-BrxC-[Vibrio-tubiashii] |
| WP_016790631.1 | Type1.BREX | XIS-HTH->?->SF1-SF2-Helicase+HTH [illegible] | 1157 | Bacteria>Pseudomonadota>Gammaproteobacteria>Vibrionales>Vibrionaceae>Vibrio | GCF_000256605.2 | Vibrio_cyclitrophicus | OAI_RS19775 | 259622..263095 | 47951 | BREX-system-P-loop-protein-BrxC-[Vibrio-cyclitrophicus] |
| WP_005444576.1 | Type1.BREX | AbiEi->viral-protein-> [illegible] | 1225 | Bacteria>Pseudomonadota>Gammaproteobacteria>Vibrionales>Vibrionaceae>Vibrio | GCF_000259935.1 | Vibrio_harveyi | MUQ_RS25205 | 3434..7111 | 669 | BREX-system-P-loop-protein-BrxC-[Vibrio-harveyi] |
| WP_017044128.1 | Type1.BREX | HTH_19-><-ParE<-MobI<-MobI<- [illegible] | 1225 | Bacteria>Pseudomonadota>Gammaproteobacteria>Vibrionales>Vibrionaceae>Vibrio | GCF_000287075.2 | Vibrio_ordalii | A1QQ_08605 | 37533..41210 | 28174 | BREX-system-P-loop-protein-BrxC-[Vibrio-ordalii] |
| WP_000152596.1 | Type1.BREX | MobI-><-?<-LexA-protease\|\|35exo- [illegible] | 1229 | Bacteria>Pseudomonadota>Gammaproteobacteria>Vibrionales>Vibrionaceae>Vibrio | GCF_000304775.1 | Vibrio_cholerae | VCHC1A2_RS09815 | 58401..62090 | 666 | BREX-system-P-loop-protein-BrxC-[Vibrio-cholerae] |
| WP_001963844.1 | Type1.BREX | MobI-><-?<-LexA-protease\|\|35exo- [illegible] | 1225 | Bacteria>Pseudomonadota>Gammaproteobacteria>Vibrionales>Vibrionaceae>Vibrio | GCF_000304915.2 | Vibrio_cholerae | VCCP1035_RS13195 | 133177..136854 | 666 | BREX-system-P-loop-protein-BrxC-[Vibrio-cholerae] |
| WP_023585137.1 | Type1.BREX | CR-ATPase4->?-><-SIGMA+ [illegible] | 1248 | Bacteria>Pseudomonadota>Gammaproteobacteria>Vibrionales>Vibrionaceae>Vibrio | GCF_000347495.1 | Vibrio_parahaemolyticus | VPPCV08_RS0113765 | 243193..246939 | 670 | BREX-system-P-loop-protein-BrxC-[Vibrio-parahaemolyticus] |
| WP_021709035.1 | Type1.BREX | <-Thioredoxin<-?<- [illegible] | 1229 | Bacteria>Pseudomonadota>Gammaproteobacteria>Vibrionales>Vibrionaceae>Vibrio | GCF_000400265.1 | Vibrio_azureus | VAN_RS15510 | 14465..18154 | 512649 | BREX-system-P-loop-protein-BrxC-[Vibrio-azureus] |
| WP_025605138.1 | Type1.BREX | <-ZnR<-?\|\|?->SF1-SF2-Helicase-> [illegible] | 1225 | Bacteria>Pseudomonadota>Gammaproteobacteria>Vibrionales>Vibrionaceae>Vibrio | GCF_000489075.1 | Vibrio_parahaemolyticus | DX10_RS0106510 | 11492..15169 | 670 | BREX-system-P-loop-protein-BrxC-[Vibrio-parahaemolyticus] |
| WP_025525716.1 | Type1.BREX | Calcineurin->?-><-ZnR<-?\|\|AbiEi- [illegible] | 1225 | Bacteria>Pseudomonadota>Gammaproteobacteria>Vibrionales>Vibrionaceae>Vibrio | GCF_000489535.1 | Vibrio_parahaemolyticus | DX89_RS0109235 | 6826..10503 | 670 | BREX-system-P-loop-protein-BrxC-[Vibrio-parahaemolyticus] |
| WP_029810016.1 | Type1.BREX | Calcineurin->?-><-ZnR<-?\|\|AbiEi- [illegible] | 1225 | Bacteria>Pseudomonadota>Gammaproteobacteria>Vibrionales>Vibrionaceae>Vibrio | GCF_000490615.1 | Vibrio_parahaemolyticus | DX93_RS0104660 | 6873..10550 | 670 | BREX-system-P-loop-protein-BrxC-[Vibrio-parahaemolyticus] |
| WP_029847728.1 | Type1.BREX | ANK+ANK->XIS-HTH-><-?<-?<-?\|\| [illegible] | 621 | Bacteria>Pseudomonadota>Gammaproteobacteria>Vibrionales>Vibrionaceae>Vibrio | GCF_000491375.1 | Vibrio_parahaemolyticus | DY29_RS0118160 | 8133..9996 | 670 | BREX-system-P-loop-protein-BrxC,-partial-[Vibrio-parahaemolyticus] |
| WP_029830481.1 | Type1.BREX | wHTH-><-SIGMA+OB-Fold+AAA-ATPase [illegible] | 1225 | Bacteria>Pseudomonadota>Gammaproteobacteria>Vibrionales>Vibrionaceae>Vibrio | GCF_000491395.1 | Vibrio_parahaemolyticus | DY37_RS0104880 | 15763..19440 | 670 | BREX-system-P-loop-protein-BrxC-[Vibrio-parahaemolyticus] |
| WP_029805255.1 | Type1.BREX | <-ZnR<-?\|\|SF1-SF2-Helicase-> [illegible] | 1225 | Bacteria>Pseudomonadota>Gammaproteobacteria>Vibrionales>Vibrionaceae>Vibrio | GCF_000492075.1 | Vibrio_parahaemolyticus | UF29_RS13050 | 359665..363342 | 670 | BREX-system-P-loop-protein-BrxC-[Vibrio-parahaemolyticus] |
| WP_024701189.1 | Type1.BREX | CR-ATPase4->?->\|\|wHTH+WYL+WCX-> [illegible] | 1157 | Bacteria>Pseudomonadota>Gammaproteobacteria>Vibrionales>Vibrionaceae>Vibrio | GCF_000500485.1 | Vibrio_parahaemolyticus | T247_RS0121000 | 24076..27549 | 670 | BREX-system-P-loop-protein-BrxC-[Vibrio-parahaemolyticus] |
| WP_025796292.1 | Type1.BREX | AbiEi->viral-protein->SF1-SF2- [illegible] | 1157 | Bacteria>Pseudomonadota>Gammaproteobacteria>Vibrionales>Vibrionaceae>Vibrio | GCF_000591555.1 | Vibrio_parahaemolyticus | T6V_RS20315 | 13981..17454 | 670 | BREX-system-P-loop-protein-BrxC-[Vibrio-parahaemolyticus] |
| WP_032482144.1 | Type1.BREX | <-LexA-protease\|\|35exo->?-> [illegible] | 1225 | Bacteria>Pseudomonadota>Gammaproteobacteria>Vibrionales>Vibrionaceae>Vibrio | GCF_000736875.1 | Vibrio_cholerae | DN38_RS05545 | 8784..12461 | 666 | BREX-system-P-loop-protein-BrxC-[Vibrio-cholerae] |
| WP_033931157.1 | Type1.BREX | <-VirD4-TraG+VirB4-FtsK<-HDRnase [illegible] | 1225 | Bacteria>Pseudomonadota>Gammaproteobacteria>Vibrionales>Vibrionaceae>Vibrio | GCF_000788515.1 | Vibrio_cholerae | NH59_RS10860 | 62787..66464 | 666 | BREX-system-P-loop-protein-BrxC-[Vibrio-cholerae] |
| WP_045378182.1 | Type1.BREX | <-SIGMA+OB-Fold+AAA-ATPase+ [illegible] | 1248 | Bacteria>Pseudomonadota>Gammaproteobacteria>Vibrionales>Vibrionaceae>Vibrio | GCF_000818335.1 | Vibrio_campbellii | TY56_RS07080 | 532127..535873 | 680 | BREX-system-P-loop-protein-BrxC-[Vibrio-campbellii] |
| WP_041154134.1 | Type1.BREX | MobI-><-?<-LexA-protease\|\|35exo- [illegible] | 1227 | Bacteria>Pseudomonadota>Gammaproteobacteria>Vibrionales>Vibrionaceae>Vibrio | GCF_000830505.1 | Vibrio_mytili | SU60_RS02235 | 11952..15635 | 50718 | BREX-system-P-loop-protein-BrxC-[Vibrio-mytili] |
| WP_253277773.1 | Type1.BREX | <-wHTH+WYL+WCX<-DUF2787<-?<-?\|\| [illegible] | 1153 | Bacteria>Pseudomonadota>Gammaproteobacteria>Vibrionales>Vibrionaceae>Vibrio | GCF_000959745.1 | Vibrio_vulnificus | WG98_RS00450 | 91740..95201 | 672 | BREX-system-P-loop-protein-BrxC-[Vibrio-vulnificus] |
| WP_047110486.1 | Type1.BREX | sGTP-><-HEPN\|\|?->?-><-SIGMA+ [illegible] | 1232 | Bacteria>Pseudomonadota>Gammaproteobacteria>Vibrionales>Vibrionaceae>Vibrio | GCF_001013325.1 | Vibrio_vulnificus | VVYB158_12225 | 27343..31041 | 672 | BREX-system-P-loop-protein-BrxC-[Vibrio-vulnificus] |
| WP_050646850.1 | Type1.BREX | SF1-SF2-Helicase+REase->SF1-SF2- [illegible] | 1157 | Bacteria>Pseudomonadota>Gammaproteobacteria>Vibrionales>Vibrionaceae>Vibrio | GCF_001243675.1 | Vibrio_sp._J2-4 | VCR4J2_RS00160 | 35844..39317 | 1507977 | BREX-system-P-loop-protein-BrxC-[Vibrio-sp.-J2-4] |
| WP_240182084.1 | Type1.BREX | <-VirD4-TraG+VirB4-FtsK<-HDRnase [illegible] | 1072 | Bacteria>Pseudomonadota>Gammaproteobacteria>Vibrionales>Vibrionaceae>Vibrio | GCF_001247885.1 | Vibrio_cholerae | ALC59_RS14100 | 125159..128374 | 666 | BREX-system-P-loop-protein-BrxC,-partial-[Vibrio-cholerae] |
| WP_053044233.1 | Type1.BREX | BrxB-BrxF-iSTAND->DUF6079-p*-> [illegible] | 822 | Bacteria>Pseudomonadota>Gammaproteobacteria>Vibrionales>Vibrionaceae>Vibrio | GCF_001248645.1 | Vibrio_cholerae | ALC75_RS05685 | 604..3070 | 666 | BREX-system-P-loop-protein-BrxC,-partial-[Vibrio-cholerae] |
| WP_114025876.1 | Type1.BREX | DUF6079*->N6-METHYLASE->PglZ-> [illegible] | 1166 | Bacteria>Pseudomonadota>Gammaproteobacteria>Vibrionales>Vibrionaceae>Vibrio | GCF_001254815.1 | Vibrio_cholerae | ALE22_RS06100 | 1004..4506 | 666 | BREX-system-P-loop-protein-BrxC,-partial-[Vibrio-cholerae] |
| WP_053027640.1 | Type1.BREX | DUF6079-p*->N6-METHYLASE->PglZ-> [illegible] | 848 | Bacteria>Pseudomonadota>Gammaproteobacteria>Vibrionales>Vibrionaceae>Vibrio | GCF_001259875.1 | Vibrio_cholerae | ALV71_RS07765 | 696..3240 | 666 | BREX-system-P-loop-protein-BrxC,-partial-[Vibrio-cholerae] |
| WP_241772680.1 | Type1.BREX | <-GDResolvase+HIN-HTH\|\|?-> [illegible] | 1153 | Bacteria>Pseudomonadota>Gammaproteobacteria>Vibrionales>Vibrionaceae>Vibrio | GCF_001280635.1 | Vibrio_parahaemolyticus | ACX06_RS21960 | 167502..170963 | 670 | BREX-system-P-loop-protein-BrxC-[Vibrio-parahaemolyticus] |
| WP_054541287.1 | Type1.BREX | BPD_transp_2->?->?->?-> [illegible] | 1235 | Bacteria>Pseudomonadota>Gammaproteobacteria>Vibrionales>Vibrionaceae>Vibrio | GCF_001306215.1 | Vibrio_splendidus | AN167_RS02930 | 103766..107473 | 29497 | BREX-system-P-loop-protein-BrxC-[Vibrio-splendidus] |
| WP_061039701.1 | Type1.BREX | <-SLOG+BirA-HTH<-SF1-SF2- [illegible] | 1247 | Bacteria>Pseudomonadota>Gammaproteobacteria>Vibrionales>Vibrionaceae>Vibrio | GCF_001557855.1 | Vibrio_coralliirubri | ATB73_RS08125 | 549280..553023 | 1516159 | BREX-system-P-loop-protein-BrxC-[Vibrio-coralliirubri] |
| WP_061065627.1 | Type1.BREX | <-wHTH+WYL+WCX<-DUF2787<-?<-?\|\| [illegible] | 1157 | Bacteria>Pseudomonadota>Gammaproteobacteria>Vibrionales>Vibrionaceae>Vibrio | GCF_001558435.2 | Vibrio_harveyi | AL538_RS13470 | 2675377..2678850 | 669 | BREX-system-P-loop-protein-BrxC-[Vibrio-harveyi] |
| WP_241782177.1 | Type1.BREX | <-HTH\|\|RNase-><-DUF2787<-?<-?\|\| [illegible] | 1153 | Bacteria>Pseudomonadota>Gammaproteobacteria>Vibrionales>Vibrionaceae>Vibrio | GCF_001608855.1 | Vibrio_parahaemolyticus | AWQ17_RS02375 | 93824..97285 | 670 | BREX-system-P-loop-protein-BrxC-[Vibrio-parahaemolyticus] |
| WP_065090484.1 | Type1.BREX | AbiEi->viral-protein->SF1-SF2- [illegible] | 1157 | Bacteria>Pseudomonadota>Gammaproteobacteria>Vibrionales>Vibrionaceae>Vibrio | GCF_001675245.1 | Vibrio_vulnificus | FORC17_RS08380 | 1790096..1793569 | 672 | BREX-system-P-loop-protein-BrxC-[Vibrio-vulnificus] |
| WP_065091082.1 | Type1.BREX | CR-ATPase4->?-><-DUF4113\|\|wHTH-> [illegible] | 1225 | Bacteria>Pseudomonadota>Gammaproteobacteria>Vibrionales>Vibrionaceae>Vibrio | GCF_001675245.1 | Vibrio_vulnificus | FORC17_RS18340 | 777696..781373 | 672 | BREX-system-P-loop-protein-BrxC-[Vibrio-vulnificus] |
| WP_256093930.1 | Type1.BREX | XIS-HTH->wHTH+WYL+WCX-> [illegible] | 1153 | Bacteria>Pseudomonadota>Gammaproteobacteria>Vibrionales>Vibrionaceae>Vibrio | GCF_001727265.1 | Vibrio_parahaemolyticus | BBM25_RS06240 | 3427..6888 | 670 | BREX-system-P-loop-protein-BrxC-[Vibrio-parahaemolyticus] |
| WP_069530140.1 | Type1.BREX | wHTH-><-SIGMA+OB-Fold+AAA-ATPase [illegible] | 1225 | Bacteria>Pseudomonadota>Gammaproteobacteria>Vibrionales>Vibrionaceae>Vibrio | GCF_001728565.1 | Vibrio_parahaemolyticus | BBN06_RS08190 | 27008..30685 | 670 | BREX-system-P-loop-protein-BrxC-[Vibrio-parahaemolyticus] |
| WP_074189993.1 | Type1.BREX | <-DUF4113\|\|wHTH-><-SIGMA+OB-Fold [illegible] | 1225 | Bacteria>Pseudomonadota>Gammaproteobacteria>Vibrionales>Vibrionaceae>Vibrio | GCF_001909305.1 | Vibrio_harveyi_group | H058_RS04230 | 543659..547336 | 717610 | MULTISPECIES:-BREX-system-P-loop-protein-BrxC-[Vibrio-harveyi-group] |
| WP_215796802.1 | Type1.BREX | wHTH+WYL+WCX->BrxA-DNA-binding-> [illegible] | 1229 | Bacteria>Pseudomonadota>Gammaproteobacteria>Vibrionales>Vibrionaceae>Vibrio | GCF_001989775.2 | Vibrio_anguillarum | PN39_RS13080 | 3578..7267 | 55601 | BREX-system-P-loop-protein-BrxC-[Vibrio-anguillarum] |
| WP_080540649.1 | Type1.BREX | HTH_3->?->?-><-SIGMA+OB-Fold+ [illegible] | 1232 | Bacteria>Pseudomonadota>Gammaproteobacteria>Vibrionales>Vibrionaceae>Vibrio | GCF_002073135.1 | Vibrio_vulnificus | XM77_RS07295 | 72638..76336 | 672 | BREX-system-P-loop-protein-BrxC-[Vibrio-vulnificus] |
| WP_139834421.1 | Type1.BREX | HTH_19-><-ParE<-MobI<-MobI<- [illegible] | 1085 | Bacteria>Pseudomonadota>Gammaproteobacteria>Vibrionales>Vibrionaceae>Vibrio | GCF_002114605.1 | Vibrio_parahaemolyticus | B5C31_RS21495 | 50871..54130 | 670 | BREX-system-P-loop-protein-BrxC,-partial-[Vibrio-parahaemolyticus] |
| WP_198300681.1 | Type1.BREX | <-DeoR-HTH+ISOCOT<-DeoR-HTH+ [illegible] | 1187 | Bacteria>Pseudomonadota>Gammaproteobacteria>Vibrionales>Vibrionaceae>Vibrio | GCF_002211985.1 | Vibrio_anguillarum | CEJ46_RS04105 | 862774..866337 | 55601 | BREX-system-P-loop-protein-BrxC-[Vibrio-anguillarum] |
| WP_088731321.1 | Type1.BREX | HTH_19-><-ParE<-MobI<-MobI<- [illegible] | 1225 | Bacteria>Pseudomonadota>Gammaproteobacteria>Vibrionales>Vibrionaceae>Vibrio | GCF_002212025.1 | Vibrio_anguillarum | CEQ50_RS00060 | 15615..19292 | 55601 | BREX-system-P-loop-protein-BrxC-[Vibrio-anguillarum] |
| WP_197697128.1 | Type1.BREX | <-SIGMA+OB-Fold+AAA-ATPase+ [illegible] | 1185 | Bacteria>Pseudomonadota>Gammaproteobacteria>Vibrionales>Vibrionaceae>Vibrio | GCF_002214395.1 | Vibrio_rotiferianus | BSZ04_RS17965 | 1918666..1922223 | 190895 | BREX-system-P-loop-protein-BrxC-[Vibrio-rotiferianus] |

| Protein-ID | System | Gene-Neighbourhoods | Length | Taxonomy | Assembly-ID | Strain | Locus-Tag | Genomic-coordinates | TaxID | Fasta-defline |
|---|---|---|---|---|---|---|---|---|---|---|
| WP_089123599.1 | Type1.BREX | <-Couple_hipA+HipA_C<-HTH_3<-?<-SIGMA+OB-Fold+AAA-ATPase+HKD-DNase-BrxL<-PglZ<-N6-METHYLASE<-REase<-OLD-ABC<-[illegible] | 1184 | Bacteria>Pseudomonadota>Gammaproteobacteria>Vibrionales>Vibrionaceae>Vibrio | GCF_002218005.2 | Vibrio_algivorus | AVR_RS07075 | 1502603..1506157 | 1667024 | BREX-system-P-loop-protein-BrxC-[Vibrio-algivorus] |
| WP_095475146.1 | Type1.BREX | Arm-DNA-bind_3+Phage_integrase->SF1-SF2-Helicase+REase->SF1-SF2-Helicase+REase-><-?<-ZetaToxin<-PglZ<-ParB+[illegible] | 1157 | Bacteria>Pseudomonadota>Gammaproteobacteria>Vibrionales>Vibrionaceae>Vibrio | GCF_002283835.1 | Vibrio_metoecus | CGT99_RS14700 | 71972..75445 | 1481663 | BREX-system-P-loop-protein-BrxC-[Vibrio-metoecus] |
| WP_095470311.1 | Type1.BREX | Arm-DNA-bind_3+Phage_integrase->SF1-SF2-Helicase+REase->SF1-SF2-Helicase+REase-><-?<-ZetaToxin<-PglZ<-ParB+[illegible] | 1157 | Bacteria>Pseudomonadota>Gammaproteobacteria>Vibrionales>Vibrionaceae>Vibrio | GCF_002283955.1 | Vibrio_metoecus | CGT93_RS13890 | 71728..75201 | 1481663 | BREX-system-P-loop-protein-BrxC-[Vibrio-metoecus] |
| WP_096070868.1 | Type1.BREX | HTH_20->CR-ATPase4->?-><-?<-LexA-protease\|\|wHTH+WYL+WCX->BrxA-DNA-binding->BrxB-BrxF-iSTAND->DUF6079*->[illegible] | 1157 | Bacteria>Pseudomonadota>Gammaproteobacteria>Vibrionales>Vibrionaceae>Vibrio | GCF_002313025.1 | Vibrio_cholerae | FORC55_RS15555 | 276972..280445 | 666 | BREX-system-P-loop-protein-BrxC-[Vibrio-cholerae] |
| WP_100267429.1 | Type1.BREX | MobI-><-?<-LexA-protease\|\|35exo->?->wHTH+WYL+WCX->BrxA-DNA-binding->BrxB-BrxF-iSTAND->DUF6079*->ParB+HNH->[illegible] | 1225 | Bacteria>Pseudomonadota>Gammaproteobacteria>Vibrionales>Vibrionaceae>Vibrio | GCF_002796765.1 | Vibrio_fluvialis | CV024_RS02745 | 224853..228530 | 676 | BREX-system-P-loop-protein-BrxC-[Vibrio-fluvialis] |
| WP_100754281.1 | Type1.BREX | <-ParB+HNH<-N6-METHYLASE<-DUF6079*<-BrxB-BrxF-iSTAND<-BrxA-DNA-binding<-wHTH+WYL+WCX\|\|?-><-ParB\|\|RNase->?-><-[illegible] | 1248 | Bacteria>Pseudomonadota>Gammaproteobacteria>Vibrionales>Vibrionaceae>Vibrio | GCF_002811245.1 | Vibrio_salilacus | CDZ87_RS15130 | 5916..9662 | 1323749 | BREX-system-P-loop-protein-BrxC-[Vibrio-salilacus] |
| WP_102465509.1 | Type1.BREX | <-BrxD<-SIGMA+OB-Fold+AAA-ATPase+Lon-Protease-BrxL<-PglZ<-?<-?<-ZnR+3C-ABC+TPR<-ANK+ANK+ANK<-ParB+HNH<-[illegible] | 1242 | Bacteria>Pseudomonadota>Gammaproteobacteria>Vibrionales>Vibrionaceae>Vibrio | GCF_002873515.1 | Vibrio_cyclitrophicus | BCU82_RS15555 | 51743..55471 | 47951 | BREX-system-P-loop-protein-BrxC-[Vibrio-cyclitrophicus] |
| WP_102402203.1 | Type1.BREX | <-SF1-SF2-Helicase+HTH\|\|wHTH+WYL+WCX->BrxA-DNA-binding->BrxB-BrxF-iSTAND->DUF6079*->N6-METHYLASE->ParB+HNH->[illegible] | 1243 | Bacteria>Pseudomonadota>Gammaproteobacteria>Vibrionales>Vibrionaceae>Vibrio | GCF_002873825.1 | Vibrio_cyclitrophicus | BCV09_RS12225 | 6080..9811 | 47951 | BREX-system-P-loop-protein-BrxC-[Vibrio-cyclitrophicus] |
| WP_102420196.1 | Type1.BREX | TraI-VirD2-RCRE-><-?<-SF1-SF2-Helicase+HTH\|\|wHTH+WYL+WCX->BrxA-DNA-binding->BrxB-BrxF-iSTAND->DUF6079*->[illegible] | 1247 | Bacteria>Pseudomonadota>Gammaproteobacteria>Vibrionales>Vibrionaceae>Vibrio | GCF_002875525.1 | Vibrio_sp._10N.222.55.C12 | BCT20_08060 | 8620..12363 | 1884470 | BREX-system-P-loop-protein-BrxC-[Vibrio-sp.-10N.222.55.C12] |
| WP_102376359.1 | Type1.BREX | ParB-><-SIGMA+OB-Fold+AAA-ATPase+Lon-Protease-BrxL<-PglZ<-TOPRIM<-OLD-ABC<-?<-?<-N6-METHYLASE<-ParB+HNH<-[illegible] | 1233 | Bacteria>Pseudomonadota>Gammaproteobacteria>Vibrionales>Vibrionaceae>Vibrio | GCF_002876115.1 | Vibrio_breoganii | BCT61_RS03015 | 196035..199736 | 553239 | BREX-system-P-loop-protein-BrxC-[Vibrio-breoganii] |
| WP_102582639.1 | Type1.BREX | REase->Calcineurin-SbcD-><-?\|\|?->?-><-SF1-SF2-Helicase+HTH\|\|wHTH+WYL+WCX->BrxA-DNA-binding->BrxB-BrxF-iSTAND->[illegible] | 1242 | Bacteria>Pseudomonadota>Gammaproteobacteria>Vibrionales>Vibrionaceae>Vibrio | GCF_002876245.1 | Vibrio_tasmaniensis | BCT76_RS12975 | 27484..31212 | 212663 | BREX-system-P-loop-protein-BrxC-[Vibrio-tasmaniensis] |
| WP_102250444.1 | Type1.BREX | <-SF1-SF2-Helicase+PRTase-p<-?<-GrpE<-ANK+ANK+ANK<-ParB+HNH<-HEPN+Sel1+SEC-C<-N6-METHYLASE<-?<-DUF6079*<-[illegible] | 1247 | Bacteria>Pseudomonadota>Gammaproteobacteria>Vibrionales>Vibrionaceae>Vibrio | GCF_002876335.1 | Vibrio_tasmaniensis | BCT83_RS20570 | 15143..18886 | 212663 | BREX-system-P-loop-protein-BrxC-[Vibrio-tasmaniensis] |
| WP_102573392.1 | Type1.BREX | REase->Calcineurin-SbcD-><-?\|\|?->?-><-SF1-SF2-Helicase+HTH\|\|wHTH+WYL+WCX->BrxA-DNA-binding->BrxB-BrxF-iSTAND->[illegible] | 1247 | Bacteria>Pseudomonadota>Gammaproteobacteria>Vibrionales>Vibrionaceae>Vibrio | GCF_002876905.1 | Vibrio_splendidus | BCT62_RS21755 | 9396..13139 | 29497 | BREX-system-P-loop-protein-BrxC-[Vibrio-splendidus] |
| WP_102456769.1 | Type1.BREX | [illegible] | 1247 | Bacteria>Pseudomonadota>Gammaproteobacteria>Vibrionales>Vibrionaceae>Vibrio | GCF_002877315.1 | Vibrio_splendidus | BCU08_RS20225 | 19418..23161 | 29497 | BREX-system-P-loop-protein-BrxC-[Vibrio-splendidus] |
| WP_102457574.1 | Type1.BREX | [illegible] | 468 | Bacteria>Pseudomonadota>Gammaproteobacteria>Vibrionales>Vibrionaceae>Vibrio | GCF_002877525.1 | Vibrio_splendidus | BCU23_RS04325 | 6659..8062 | 29497 | BREX-system-P-loop-protein-BrxC,-partial-[Vibrio-splendidus] |
| WP_102887270.1 | Type1.BREX | [illegible] | 1225 | Bacteria>Pseudomonadota>Gammaproteobacteria>Vibrionales>Vibrionaceae>Vibrio | GCF_002891755.1 | Vibrio_vulnificus | TI06_RS01685 | 47134..50811 | 672 | BREX-system-P-loop-protein-BrxC-[Vibrio-vulnificus] |
| WP_104040811.1 | Type1.BREX | [illegible] | 1232 | Bacteria>Pseudomonadota>Gammaproteobacteria>Vibrionales>Vibrionaceae>Vibrio | GCF_002929885.1 | Vibrio_hyugaensis | COQ55_RS11480 | 5964..9662 | 1534743 | BREX-system-P-loop-protein-BrxC-[Vibrio-hyugaensis] |
| WP_104969987.1 | Type1.BREX | [illegible] | 1225 | Bacteria>Pseudomonadota>Gammaproteobacteria>Vibrionales>Vibrionaceae>Vibrio | GCF_002953335.1 | Vibrio_diabolicus | AL552_RS16285 | 3102231..3105908 | 50719 | BREX-system-P-loop-protein-BrxC-[Vibrio-diabolicus] |
| WP_040122842.1 | Type1.BREX | [illegible] | 1157 | Bacteria>Pseudomonadota>Gammaproteobacteria>Vibrionales>Vibrionaceae>Vibrio | GCF_003031205.1 | Vibrio_anguillarum | CEA93_RS18125 | 702795..706268 | 55601 | BREX-system-P-loop-protein-BrxC-[Vibrio-anguillarum] |
| WP_108187618.1 | Type1.BREX | [illegible] | 1233 | Bacteria>Pseudomonadota>Gammaproteobacteria>Vibrionales>Vibrionaceae>Vibrio | GCF_003049965.1 | Vibrio_splendidus | CWO07_RS09240 | 50503..54204 | 29497 | BREX-system-P-loop-protein-BrxC-[Vibrio-splendidus] |
| WP_108693859.1 | Type1.BREX | [illegible] | 1225 | Bacteria>Pseudomonadota>Gammaproteobacteria>Vibrionales>Vibrionaceae>Vibrio | GCF_003072005.1 | Vibrio_vulnificus | DC364_RS00725 | 56102..59779 | 672 | BREX-system-P-loop-protein-BrxC-[Vibrio-vulnificus] |
| WP_108745199.1 | Type1.BREX | [illegible] | 1225 | Bacteria>Pseudomonadota>Gammaproteobacteria>Vibrionales>Vibrionaceae>Vibrio | GCF_003076895.1 | Vibrio_parahaemolyticus | C9I78_RS18225 | 191992..195669 | 670 | BREX-system-P-loop-protein-BrxC-[Vibrio-parahaemolyticus] |
| WP_113603075.1 | Type1.BREX | [illegible] | 1225 | Bacteria>Pseudomonadota>Gammaproteobacteria>Vibrionales>Vibrionaceae>Vibrio | GCF_003311885.1 | Vibrio_tarriae | DLR65_RS02205 | 60103..63780 | 2014742 | BREX-system-P-loop-protein-BrxC-[Vibrio-tarriae] |
| WP_235868160.1 | Type1.BREX | [illegible] | 1153 | Bacteria>Pseudomonadota>Gammaproteobacteria>Vibrionales>Vibrionaceae>Vibrio | GCF_003311905.1 | Vibrio_paracholerae | DLR66_RS03590 | 70731..74192 | 650003 | BREX-system-P-loop-protein-BrxC-[Vibrio-paracholerae] |
| WP_116869874.1 | Type1.BREX | [illegible] | 1225 | Bacteria>Pseudomonadota>Gammaproteobacteria>Vibrionales>Vibrionaceae>Vibrio | GCF_003408655.1 | Vibrio_splendidus | D0583_RS01655 | 344855..348532 | 29497 | BREX-system-P-loop-protein-BrxC-[Vibrio-splendidus] |
| WP_119471206.1 | Type1.BREX | [illegible] | 1229 | Bacteria>Pseudomonadota>Gammaproteobacteria>Vibrionales>Vibrionaceae>Vibrio | GCF_003585465.1 | Vibrio_cholerae | CG489_RS12450 | 5990..9679 | 666 | BREX-system-P-loop-protein-BrxC-[Vibrio-cholerae] |
| WP_119561011.1 | Type1.BREX | [illegible] | 1071 | Bacteria>Pseudomonadota>Gammaproteobacteria>Vibrionales>Vibrionaceae>Vibrio | GCF_003585485.1 | Vibrio_cholerae | CG490_RS09140 | 1..3217 | 666 | BREX-system-P-loop-protein-BrxC,-partial-[Vibrio-cholerae] |
| WP_122015998.1 | Type1.BREX | [illegible] | 1248 | Bacteria>Pseudomonadota>Gammaproteobacteria>Vibrionales>Vibrionaceae>Vibrio | GCF_003691525.1 | Vibrio_parahaemolyticus | D0871_RS08265 | 1554661..1558407 | 670 | BREX-system-P-loop-protein-BrxC-[Vibrio-parahaemolyticus] |
| WP_130200308.1 | Type1.BREX | [illegible] | 1157 | Bacteria>Pseudomonadota>Gammaproteobacteria>Vibrionales>Vibrionaceae>Vibrio | GCF_004214625.1 | Vibrio_vulnificus | D8T51_RS18945 | 13637..17110 | 672 | BREX-system-P-loop-protein-BrxC-[Vibrio-vulnificus] |
| WP_130241041.1 | Type1.BREX | [illegible] | 803 | Bacteria>Pseudomonadota>Gammaproteobacteria>Vibrionales>Vibrionaceae>Vibrio | GCF_004214665.1 | Vibrio_vulnificus | D8T54_RS22380 | 4485..6896 | 672 | BREX-system-P-loop-protein-BrxC-[Vibrio-vulnificus] |
| WP_130241697.1 | Type1.BREX | [illegible] | 1231 | Bacteria>Pseudomonadota>Gammaproteobacteria>Vibrionales>Vibrionaceae>Vibrio | GCF_004214765.1 | Vibrio_vulnificus | D8T65_RS00475 | 112647..116342 | 672 | BREX-system-P-loop-protein-BrxC-[Vibrio-vulnificus] |
| WP_130361357.1 | Type1.BREX | [illegible] | 1231 | Bacteria>Pseudomonadota>Gammaproteobacteria>Vibrionales>Vibrionaceae>Vibrio | GCF_004215315.1 | Vibrio_vulnificus | D8T58_RS22585 | 4068..7763 | 672 | BREX-system-P-loop-protein-BrxC-[Vibrio-vulnificus] |
| WP_137027181.1 | Type1.BREX | [illegible] | 1225 | Bacteria>Pseudomonadota>Gammaproteobacteria>Vibrionales>Vibrionaceae>Vibrio | GCF_005145785.1 | Vibrio_kanaloae | FCV44_20755 | 6017..9694 | 170673 | BREX-system-P-loop-protein-BrxC-[Vibrio-kanaloae] |
| WP_136997941.1 | Type1.BREX | [illegible] | 1225 | Bacteria>Pseudomonadota>Gammaproteobacteria>Vibrionales>Vibrionaceae>Vibrio | GCF_005146415.1 | Vibrio_kanaloae | FCV52_RS09450 | 5982..9659 | 170673 | BREX-system-P-loop-protein-BrxC-[Vibrio-kanaloae] |
| WP_140128967.1 | Type1.BREX | [illegible] | 1225 | Bacteria>Pseudomonadota>Gammaproteobacteria>Vibrionales>Vibrionaceae>Vibrio | GCF_006368845.1 | Vibrio_parahaemolyticus | CGJ25_RS23760 | 3581..7258 | 670 | BREX-system-P-loop-protein-BrxC-[Vibrio-parahaemolyticus] |
| WP_140244484.1 | Type1.BREX | [illegible] | 1221 | Bacteria>Pseudomonadota>Gammaproteobacteria>Vibrionales>Vibrionaceae>Vibrio | GCF_006371105.1 | Vibrio_parahaemolyticus | CGJ91_RS11715 | 157645..161310 | 670 | BREX-system-P-loop-protein-BrxC-[Vibrio-parahaemolyticus] |
| WP_140244622.1 | Type1.BREX | [illegible] | 1225 | Bacteria>Pseudomonadota>Gammaproteobacteria>Vibrionales>Vibrionaceae>Vibrio | GCF_006371105.1 | Vibrio_parahaemolyticus | CGJ91_RS14635 | 88177..91854 | 670 | BREX-system-P-loop-protein-BrxC-[Vibrio-parahaemolyticus] |
| WP_140247448.1 | Type1.BREX | [illegible] | 1157 | Bacteria>Pseudomonadota>Gammaproteobacteria>Vibrionales>Vibrionaceae>Vibrio | GCF_006371495.1 | Vibrio_parahaemolyticus | CGJ41_RS23680 | 13693..17166 | 670 | BREX-system-P-loop-protein-BrxC-[Vibrio-parahaemolyticus] |
| WP_140344322.1 | Type1.BREX | [illegible] | 1157 | Bacteria>Pseudomonadota>Gammaproteobacteria>Vibrionales>Vibrionaceae>Vibrio | GCF_006372045.1 | Vibrio_parahaemolyticus | CGJ19_RS21455 | 3824..7297 | 670 | BREX-system-P-loop-protein-BrxC-[Vibrio-parahaemolyticus] |
| WP_140256559.1 | Type1.BREX | [illegible] | 1225 | Bacteria>Pseudomonadota>Gammaproteobacteria>Vibrionales>Vibrionaceae>Vibrio | GCF_006372155.1 | Vibrio_parahaemolyticus | D9A28_03775 | 149461..153138 | 670 | BREX-system-P-loop-protein-BrxC-[Vibrio-parahaemolyticus] |
| WP_140272583.1 | Type1.BREX | [illegible] | 1248 | Bacteria>Pseudomonadota>Gammaproteobacteria>Vibrionales>Vibrionaceae>Vibrio | GCF_006372715.1 | Vibrio_parahaemolyticus | CGI87_RS22525 | 16069..19815 | 670 | BREX-system-P-loop-protein-BrxC-[Vibrio-parahaemolyticus] |
| WP_148491305.1 | Type1.BREX | [illegible] | 1229 | Bacteria>Pseudomonadota>Gammaproteobacteria>Vibrionales>Vibrionaceae>Vibrio | GCF_008083215.1 | Vibrio_cholerae | FXF11_RS16935 | 299548..303237 | 666 | BREX-system-P-loop-protein-BrxC-[Vibrio-cholerae] |
| WP_148513618.1 | Type1.BREX | [illegible] | 1157 | Bacteria>Pseudomonadota>Gammaproteobacteria>Vibrionales>Vibrionaceae>Vibrio | GCF_008083315.1 | Vibrio_cholerae | FXF10_RS00340 | 29414..32887 | 666 | BREX-system-P-loop-protein-BrxC-[Vibrio-cholerae] |
| WP_148532770.1 | Type1.BREX | [illegible] | 1225 | Bacteria>Pseudomonadota>Gammaproteobacteria>Vibrionales>Vibrionaceae>Vibrio | GCF_008084025.1 | Vibrio_cholerae | FXE74_RS08220 | 132887..136564 | 666 | BREX-system-P-loop-protein-BrxC-[Vibrio-cholerae] |
| WP_148490856.1 | Type1.BREX | [illegible] | 1157 | Bacteria>Pseudomonadota>Gammaproteobacteria>Vibrionales>Vibrionaceae>Vibrio | GCF_008084645.1 | Vibrio_cholerae | FXE54_RS16715 | 26974..30447 | 666 | BREX-system-P-loop-protein-BrxC-[Vibrio-cholerae] |
| WP_148536810.1 | Type1.BREX | [illegible] | 1225 | Bacteria>Pseudomonadota>Gammaproteobacteria>Vibrionales>Vibrionaceae>Vibrio | GCF_008085615.1 | Vibrio_cholerae | FXE25_RS13680 | 35779..39456 | 666 | BREX-system-P-loop-protein-BrxC-[Vibrio-cholerae] |
| WP_152820636.1 | Type1.BREX | [illegible] | 1247 | Bacteria>Pseudomonadota>Gammaproteobacteria>Vibrionales>Vibrionaceae>Vibrio | GCF_009372095.1 | Vibrio_sp._B1Z05 | GCU72_RS07750 | 793260..797003 | 2654980 | BREX-system-P-loop-protein-BrxC-[Vibrio-sp.-B1Z05] |
| WP_158108820.1 | Type1.BREX | [illegible] | 1225 | Bacteria>Pseudomonadota>Gammaproteobacteria>Vibrionales>Vibrionaceae>Vibrio | GCF_009763225.1 | Vibrio_furnissii | GPY43_RS22260 | 1428276..1431953 | 29494 | BREX-system-P-loop-protein-BrxC-[Vibrio-furnissii] |
| WP_123161578.1 | Type1.BREX | [illegible] | 1225 | Bacteria>Pseudomonadota>Gammaproteobacteria>Vibrionales>Vibrionaceae>Vibrio | GCF_009763665.1 | Vibrio_cholerae | GPY01_RS14950 | 2265324..2269001 | 666 | BREX-system-P-loop-protein-BrxC-[Vibrio-cholerae] |
| WP_158155131.1 | Type1.BREX | [illegible] | 1225 | Bacteria>Pseudomonadota>Gammaproteobacteria>Vibrionales>Vibrionaceae>Vibrio | GCF_009763745.1 | Vibrio_alginolyticus | GPY47_RS00180 | 30700..34377 | 663 | BREX-system-P-loop-protein-BrxC-[Vibrio-alginolyticus] |
| WP_159408537.1 | Type1.BREX | [illegible] | 1229 | Bacteria>Pseudomonadota>Gammaproteobacteria>Vibrionales>Vibrionaceae>Vibrio | GCF_009883875.1 | Vibrio_parahaemolyticus | EHC69_RS27010 | 189487..53 | 670 | BREX-system-P-loop-protein-BrxC-[Vibrio-parahaemolyticus] |
| WP_202601719.1 | Type1.BREX | [illegible] | 1187 | Bacteria>Pseudomonadota>Gammaproteobacteria>Vibrionales>Vibrionaceae>Vibrio | GCF_009906175.1 | Vibrio_toranzoniae | GL180_RS08040 | 10464..14027 | 1194427 | BREX-system-P-loop-protein-BrxC-[Vibrio-toranzoniae] |
| WP_171981218.1 | Type1.BREX | [illegible] | 1157 | Bacteria>Pseudomonadota>Gammaproteobacteria>Vibrionales>Vibrionaceae>Vibrio | GCF_011090175.1 | Vibrio_aestuarianus | G6Z92_RS15410 | 17195..20668 | 28171 | BREX-system-P-loop-protein-BrxC-[Vibrio-aestuarianus] |
| WP_171981504.1 | Type1.BREX | [illegible] | 1225 | Bacteria>Pseudomonadota>Gammaproteobacteria>Vibrionales>Vibrionaceae>Vibrio | GCF_011090175.1 | Vibrio_aestuarianus | G6Z92_RS17720 | 1620..5297 | 28171 | BREX-system-P-loop-protein-BrxC-[Vibrio-aestuarianus] |
| WP_176314025.1 | Type1.BREX | [illegible] | 1157 | Bacteria>Pseudomonadota>Gammaproteobacteria>Vibrionales>Vibrionaceae>Vibrio | GCF_011090275.1 | Vibrio_aestuarianus | G6Z96_RS18110 | 255..3728 | 28171 | BREX-system-P-loop-protein-BrxC-[Vibrio-aestuarianus] |
| WP_172566988.1 | Type1.BREX | [illegible] | 1157 | Bacteria>Pseudomonadota>Gammaproteobacteria>Vibrionales>Vibrionaceae>Vibrio | GCF_012274945.1 | Vibrio_fluvialis | HF303_RS09150 | 434217..437690 | 676 | BREX-system-P-loop-protein-BrxC-[Vibrio-fluvialis] |
| WP_169592914.1 | Type1.BREX | [illegible] | 1225 | Bacteria>Pseudomonadota>Gammaproteobacteria>Vibrionales>Vibrionaceae>Vibrio | GCF_012956215.1 | Vibrio_parahaemolyticus | HKB29_RS26535 | 221026..224703 | 670 | BREX-system-P-loop-protein-BrxC-[Vibrio-parahaemolyticus] |
| WP_170894125.1 | Type1.BREX | [illegible] | 1225 | Bacteria>Pseudomonadota>Gammaproteobacteria>Vibrionales>Vibrionaceae>Vibrio | GCF_013048535.1 | unclassified_Vibrio | FKN13_RS12350 | 70010..73687 | 2614977 | MULTISPECIES:-BREX-system-P-loop-protein-BrxC-[unclassified-Vibrio] |
| WP_171375390.1 | Type1.BREX | [illegible] | 1225 | Bacteria>Pseudomonadota>Gammaproteobacteria>Vibrionales>Vibrionaceae>Vibrio | GCF_013114495.1 | Vibrio_aestuarianus | F0230_RS00325 | 78960..82637 | 28171 | BREX-system-P-loop-protein-BrxC-[Vibrio-aestuarianus] |
| WP_179877426.1 | Type1.BREX | [illegible] | 1169 | Bacteria>Pseudomonadota>Gammaproteobacteria>Vibrionales>Vibrionaceae>Vibrio | GCF_013414485.1 | Vibrio_cholerae | EYV61_RS05685 | 69787..73298 | 666 | BREX-system-P-loop-protein-BrxC,-partial-[Vibrio-cholerae] |
| WP_182291541.1 | Type1.BREX | [illegible] | 1157 | Bacteria>Pseudomonadota>Gammaproteobacteria>Vibrionales>Vibrionaceae>Vibrio | GCF_014107515.1 | Vibrio_vulnificus | F6X00_RS18205 | 373031..376504 | 672 | BREX-system-P-loop-protein-BrxC-[Vibrio-vulnificus] |
| WP_238930029.1 | Type1.BREX | [illegible] | 779 | Bacteria>Pseudomonadota>Gammaproteobacteria>Vibrionales>Vibrionaceae>Vibrio | GCF_014525155.1 | Vibrio_vulnificus | GO186_RS04075 | 575963..578302 | 672 | BREX-system-P-loop-protein-BrxC-[Vibrio-vulnificus] |
| WP_238930031.1 | Type1.BREX | [illegible] | 366 | Bacteria>Pseudomonadota>Gammaproteobacteria>Vibrionales>Vibrionaceae>Vibrio | GCF_014525155.1 | Vibrio_vulnificus | GO186_RS25670 | 578332..579432 | 672 | hypothetical-protein-[Vibrio-vulnificus] |
| WP_191116552.1 | Type1.BREX | [illegible] | 1248 | Bacteria>Pseudomonadota>Gammaproteobacteria>Vibrionales>Vibrionaceae>Vibrio | GCF_014769095.1 | Vibrio_parahaemolyticus | EUV06_RS14935 | 117914..121660 | 670 | BREX-system-P-loop-protein-BrxC-[Vibrio-parahaemolyticus] |
| WP_054579585.1 | Type1.BREX | [illegible] | 1157 | Bacteria>Pseudomonadota>Gammaproteobacteria>Vibrionales>Vibrionaceae>Vibrio | GCF_014769135.1 | Vibrio_alginolyticus | AOR09_RS18625 | 85180..88653 | 663 | BREX-system-P-loop-protein-BrxC-[Vibrio-alginolyticus] |
| WP_191786376.1 | Type1.BREX | [illegible] | 1225 | Bacteria>Pseudomonadota>Gammaproteobacteria>Vibrionales>Vibrionaceae>Vibrio | GCF_014836305.1 | Vibrio_parahaemolyticus | G3220_RS19475 | 66422..70099 | 670 | BREX-system-P-loop-protein-BrxC-[Vibrio-parahaemolyticus] |
| WP_194436055.1 | Type1.BREX | [illegible] | 1219 | Bacteria>Pseudomonadota>Gammaproteobacteria>Vibrionales>Vibrionaceae>Vibrio | GCF_015278125.1 | Vibrio_sp._A621 | IRQ95_RS05790 | 62806..66465 | 2783614 | BREX-system-P-loop-protein-BrxC-[Vibrio-sp.-A621] |
| WP_198106729.1 | Type1.BREX | [illegible] | 1187 | Bacteria>Pseudomonadota>Gammaproteobacteria>Vibrionales>Vibrionaceae>Vibrio | GCF_016094345.1 | Vibrio_navarrensis | CJ419_RS12980 | 152911..156474 | 29495 | BREX-system-P-loop-protein-BrxC-[Vibrio-navarrensis] |
| WP_206630456.1 | Type1.BREX | [illegible] | 1232 | Bacteria>Pseudomonadota>Gammaproteobacteria>Vibrionales>Vibrionaceae>Vibrio | GCF_017291155.1 | Vibrio_vulnificus | JOJ25_RS14690 | 90356..94054 | 672 | BREX-system-P-loop-protein-BrxC-[Vibrio-vulnificus] |
| WP_206946493.1 | Type1.BREX | [illegible] | 1157 | Bacteria>Pseudomonadota>Gammaproteobacteria>Vibrionales>Vibrionaceae>Vibrio | GCF_017313505.1 | Vibrio_sp._Vb2424 | JO679_RS01025 | 227908..231381 | 2816074 | BREX-system-P-loop-protein-BrxC-[Vibrio-sp.-Vb2424] |
| WP_210499744.1 | Type1.BREX | [illegible] | 1185 | Bacteria>Pseudomonadota>Gammaproteobacteria>Vibrionales>Vibrionaceae>Vibrio | GCF_017916775.1 | Vibrio_crassostreae | KB594_RS01450 | 322447..326004 | 246167 | BREX-system-P-loop-protein-BrxC-[Vibrio-crassostreae] |
| WP_210447442.1 | Type1.BREX | [illegible] | 1157 | Bacteria>Pseudomonadota>Gammaproteobacteria>Vibrionales>Vibrionaceae>Vibrio | GCF_017917465.1 | Vibrio_crassostreae | KB624_RS13660 | 42369..45842 | 246167 | BREX-system-P-loop-protein-BrxC-[Vibrio-crassostreae] |
| WP_210942219.1 | Type1.BREX | [illegible] | 1157 | Bacteria>Pseudomonadota>Gammaproteobacteria>Vibrionales>Vibrionaceae>Vibrio | GCF_018035165.1 | Vibrio_paracholerae | GSG51_RS14590 | 9402..12875 | 650003 | BREX-system-P-loop-protein-BrxC-[Vibrio-paracholerae] |
| WP_250156418.1 | Type1.BREX | [illegible] | 1185 | Bacteria>Pseudomonadota>Gammaproteobacteria>Vibrionales>Vibrionaceae>Vibrio | GCF_018457845.1 | Vibrio_alginolyticus | J4H60_RS12980 | 321648..325205 | 663 | BREX-system-P-loop-protein-BrxC-[Vibrio-alginolyticus] |
| WP_213895854.1 | Type1.BREX | [illegible] | 1225 | Bacteria>Pseudomonadota>Gammaproteobacteria>Vibrionales>Vibrionaceae>Vibrio | GCF_018458125.1 | Vibrio_alginolyticus | J4H64_RS03750 | 118729..122406 | 663 | BREX-system-P-loop-protein-BrxC-[Vibrio-alginolyticus] |
| WP_213968024.1 | Type1.BREX | [illegible] | 1248 | Bacteria>Pseudomonadota>Gammaproteobacteria>Vibrionales>Vibrionaceae>Vibrio | GCF_018458215.1 | Vibrio_alginolyticus | J4H41_RS21340 | 12665..16411 | 663 | BREX-system-P-loop-protein-BrxC-[Vibrio-alginolyticus] |
| WP_140279956.1 | Type1.BREX | [illegible] | 1225 | Bacteria>Pseudomonadota>Gammaproteobacteria>Vibrionales>Vibrionaceae>Vibrio | GCF_018458225.1 | Vibrio_harveyi_group | O8042_RS07565 | 631131..634808 | 717610 | MULTISPECIES:-BREX-system-P-loop-protein-BrxC-[Vibrio-harveyi-group] |
| WP_213970810.1 | Type1.BREX | [illegible] | 1157 | Bacteria>Pseudomonadota>Gammaproteobacteria>Vibrionales>Vibrionaceae>Vibrio | GCF_018458405.1 | Vibrio_alginolyticus | J4H58_RS14535 | 108443..111916 | 663 | BREX-system-P-loop-protein-BrxC-[Vibrio-alginolyticus] |
| WP_213899843.1 | Type1.BREX | [illegible] | 1157 | Bacteria>Pseudomonadota>Gammaproteobacteria>Vibrionales>Vibrionaceae>Vibrio | GCF_018458525.1 | Vibrio_alginolyticus | J4H25_RS19085 | 22843..26316 | 663 | BREX-system-P-loop-protein-BrxC-[Vibrio-alginolyticus] |
| WP_213874127.1 | Type1.BREX | [illegible] | 1157 | Bacteria>Pseudomonadota>Gammaproteobacteria>Vibrionales>Vibrionaceae>Vibrio | GCF_018458545.1 | Vibrio_alginolyticus | J4H28_RS04435 | 1018248..1021721 | 663 | BREX-system-P-loop-protein-BrxC-[Vibrio-alginolyticus] |
| WP_216506285.1 | Type1.BREX | [illegible] | 1229 | Bacteria>Pseudomonadota>Gammaproteobacteria>Vibrionales>Vibrionaceae>Vibrio | GCF_018920395.1 | Vibrio_cholerae | IN947_RS12990 | 4223..7912 | 666 | BREX-system-P-loop-protein-BrxC-[Vibrio-cholerae] |
| WP_217555808.1 | Type1.BREX | [illegible] | 1157 | Bacteria>Pseudomonadota>Gammaproteobacteria>Vibrionales>Vibrionaceae>Vibrio | GCF_019091765.1 | Vibrio_metschnikovii | KVT01_RS15810 | 26769..30242 | 28172 | BREX-system-P-loop-protein-BrxC-[Vibrio-metschnikovii] |
| WP_222133201.1 | Type1.BREX | [illegible] | 1248 | Bacteria>Pseudomonadota>Gammaproteobacteria>Vibrionales>Vibrionaceae>Vibrio | GCF_019793245.1 | Vibrio_alginolyticus | K6L11_RS14305 | 27139..30885 | 663 | BREX-system-P-loop-protein-BrxC-[Vibrio-alginolyticus] |
| WP_222134563.1 | Type1.BREX | [illegible] | 1225 | Bacteria>Pseudomonadota>Gammaproteobacteria>Vibrionales>Vibrionaceae>Vibrio | GCF_019793275.1 | Vibrio_cholerae | NB536_RS17220 | 131235..134912 | 666 | BREX-system-P-loop-protein-BrxC-[Vibrio-cholerae] |
| WP_224214086.1 | Type1.BREX | [illegible] | 1225 | Bacteria>Pseudomonadota>Gammaproteobacteria>Vibrionales>Vibrionaceae>Vibrio | GCF_020103255.1 | Vibrio_metschnikovii | LDQ06_RS03825 | 838008..841685 | 28172 | BREX-system-P-loop-protein-BrxC-[Vibrio-metschnikovii] |
| WP_224640811.1 | Type1.BREX | [illegible] | 1225 | Bacteria>Pseudomonadota>Gammaproteobacteria>Vibrionales>Vibrionaceae>Vibrio | GCF_020169625.1 | Vibrio_vulnificus | KUO11_RS09480 | 205814..209491 | 672 | BREX-system-P-loop-protein-BrxC-[Vibrio-vulnificus] |
| WP_225491794.1 | Type1.BREX | [illegible] | 1225 | Bacteria>Pseudomonadota>Gammaproteobacteria>Vibrionales>Vibrionaceae>Vibrio | GCF_020223715.1 | Vibrio_alginolyticus | LAZ44_RS12495 | 20548..24225 | 663 | BREX-system-P-loop-protein-BrxC-[Vibrio-alginolyticus] |
| WP_225482458.1 | Type1.BREX | [illegible] | 1231 | Bacteria>Pseudomonadota>Gammaproteobacteria>Vibrionales>Vibrionaceae>Vibrio | GCF_020251885.1 | Vibrio_vulnificus | JKP23_RS02685 | 250869..254564 | 672 | BREX-system-P-loop-protein-BrxC-[Vibrio-vulnificus] |
| WP_225454267.1 | Type1.BREX | [illegible] | 1248 | Bacteria>Pseudomonadota>Gammaproteobacteria>Vibrionales>Vibrionaceae>Vibrio | GCF_020252125.1 | Vibrio_vulnificus | JKP08_RS08925 | 222135..225881 | 672 | BREX-system-P-loop-protein-BrxC-[Vibrio-vulnificus] |
| WP_228096077.1 | Type1.BREX | [illegible] | 1157 | Bacteria>Pseudomonadota>Gammaproteobacteria>Vibrionales>Vibrionaceae>Vibrio | GCF_020741035.1 | Vibrio_parahaemolyticus | IB294_RS01075 | 246074..249547 | 670 | BREX-system-P-loop-protein-BrxC-[Vibrio-parahaemolyticus] |
| WP_228070153.1 | Type1.BREX | [illegible] | 1225 | Bacteria>Pseudomonadota>Gammaproteobacteria>Vibrionales>Vibrionaceae>Vibrio | GCF_020741165.1 | Vibrio_parahaemolyticus | IB301_RS18305 | 37667..41344 | 670 | BREX-system-P-loop-protein-BrxC-[Vibrio-parahaemolyticus] |
| WP_236110294.1 | Type1.BREX | [illegible] | 1157 | Bacteria>Pseudomonadota>Gammaproteobacteria>Vibrionales>Vibrionaceae>Vibrio | GCF_021568915.1 | Vibrio_sp._McD22-P3 | L2W36_RS13785 | 194928..198401 | 2724880 | BREX-system-P-loop-protein-BrxC-[Vibrio-sp.-McD22-P3] |
| WP_238795743.1 | Type1.BREX | [illegible] | 1225 | Bacteria>Pseudomonadota>Gammaproteobacteria>Vibrionales>Vibrionaceae>Vibrio | GCF_022289115.1 | Vibrio_furnissii | K6U56_RS11800 | 143910..147587 | 29494 | BREX-system-P-loop-protein-BrxC-[Vibrio-furnissii] |
| WP_238812493.1 | Type1.BREX | [illegible] | 1153 | Bacteria>Pseudomonadota>Gammaproteobacteria>Vibrionales>Vibrionaceae>Vibrio | GCF_022290755.1 | Vibrio_parahaemolyticus | K6U44_RS14230 | 7820..11281 | 670 | BREX-system-P-loop-protein-BrxC-[Vibrio-parahaemolyticus] |
| WP_140273790.1 | Type1.BREX | [illegible] | 1157 | Bacteria>Pseudomonadota>Gammaproteobacteria>Vibrionales>Vibrionaceae>Vibrio | GCF_022291015.1 | Vibrio_parahaemolyticus | K6U15_RS27495 | 32395..35868 | 670 | BREX-system-P-loop-protein-BrxC-[Vibrio-parahaemolyticus] |
| WP_238967171.1 | Type1.BREX | [illegible] | 1157 | Bacteria>Pseudomonadota>Gammaproteobacteria>Vibrionales>Vibrionaceae>Vibrio | GCF_022343145.1 | Vibrio_alginolyticus | K6750_RS21880 | 1365022..1368495 | 663 | BREX-system-P-loop-protein-BrxC-[Vibrio-alginolyticus] |
| WP_256963697.1 | Type1.BREX | [illegible] | 1171 | Bacteria>Pseudomonadota>Gammaproteobacteria>Vibrionales>Vibrionaceae>Vibrio | GCF_022398055.1 | Vibrio_diabolicus | L1D58_RS17270 | 13232..16749 | 50719 | BREX-system-P-loop-protein-BrxC,-partial-[Vibrio-diabolicus] |
| WP_243579240.1 | Type1.BREX | [illegible] | 1247 | Bacteria>Pseudomonadota>Gammaproteobacteria>Vibrionales>Vibrionaceae>Vibrio | GCF_022812045.1 | Vibrio_splendidus | LTQ02_RS02805 | 621543..625286 | 29497 | BREX-system-P-loop-protein-BrxC-[Vibrio-splendidus] |
| WP_253644856.1 | Type1.BREX | [illegible] | 1248 | Bacteria>Pseudomonadota>Gammaproteobacteria>Vibrionales>Vibrionaceae>Vibrio | GCF_024181785.1 | Vibrio_sp._Y42_MX_L11 | NL643_RS16680 | 61256..65002 | 2957765 | BREX-system-P-loop-protein-BrxC-[Vibrio-sp.-Y42_MX_L11] |
| WP_253658043.1 | Type1.BREX | [illegible] | 1248 | Bacteria>Pseudomonadota>Gammaproteobacteria>Vibrionales>Vibrionaceae>Vibrio | GCF_024181795.1 | unclassified_Vibrio | NL644_RS04175 | 947816..951562 | 2614977 | MULTISPECIES:-BREX-system-P-loop-protein-BrxC-[unclassified-Vibrio] |
| WP_261872478.1 | Type1.BREX | [illegible] | 1239 | Bacteria>Pseudomonadota>Gammaproteobacteria>Vibrionales>Vibrionaceae>Vibrio | GCF_024347075.1 | Vibrio_rarus | OCU56_RS06785 | 1441704..1445423 | 413403 | BREX-system-P-loop-protein-BrxC-[Vibrio-rarus] |
| WP_261872676.1 | Type1.BREX | [illegible] | 1151 | Bacteria>Pseudomonadota>Gammaproteobacteria>Vibrionales>Vibrionaceae>Vibrio | GCF_024347075.1 | Vibrio_rarus | OCU56_RS07815 | 1705601..1709056 | 413403 | BREX-system-P-loop-protein-BrxC-[Vibrio-rarus] |
| WP_086712895.1 | Type1.BREX | [illegible] | 1157 | Bacteria>Pseudomonadota>Gammaproteobacteria>Vibrionales>Vibrionaceae>Vibrio | GCF_024347515.1 | Vibrio_gigantis | B1026_RS08100 | 182974..186447 | 296199 | BREX-system-P-loop-protein-BrxC-[Vibrio-gigantis] |
| WP_255937424.1 | Type1.BREX | [illegible] | 1185 | Bacteria>Pseudomonadota>Gammaproteobacteria>Vibrionales>Vibrionaceae>Vibrio | GCF_024442175.1 | Vibrio_campbellii | HB761_RS02525 | 550346..553903 | 680 | BREX-system-P-loop-protein-BrxC-[Vibrio-campbellii] |
| WP_256897643.1 | Type1.BREX | [illegible] | 1248 | Bacteria>Pseudomonadota>Gammaproteobacteria>Vibrionales>Vibrionaceae>Vibrio | GCF_024530955.1 | Vibrio_parahaemolyticus | H2C82_RS04830 | 209368..213114 | 670 | BREX-system-P-loop-protein-BrxC-[Vibrio-parahaemolyticus] |
| WP_258495358.1 | Type1.BREX | [illegible] | 1157 | Bacteria>Pseudomonadota>Gammaproteobacteria>Vibrionales>Vibrionaceae>Vibrio | GCF_024744595.1 | Vibrio_alginolyticus | NB477_RS23070 | 13774..17247 | 663 | BREX-system-P-loop-protein-BrxC-[Vibrio-alginolyticus] |
| WP_248354663.1 | Type1.BREX | [illegible] | 1157 | Bacteria>Pseudomonadota>Gammaproteobacteria>Vibrionales>Vibrionaceae>Vibrio | GCF_024745175.1 | Vibrio_parahaemolyticus | DDM99_15775 | 19819..23292 | 670 | BREX-system-P-loop-protein-BrxC-[Vibrio-parahaemolyticus] |
| WP_258484886.1 | Type1.BREX | [illegible] | 1225 | Bacteria>Pseudomonadota>Gammaproteobacteria>Vibrionales>Vibrionaceae>Vibrio | GCF_024745305.1 | Vibrio_parahaemolyticus | NB589_RS16770 | 223237..226914 | 670 | BREX-system-P-loop-protein-BrxC-[Vibrio-parahaemolyticus] |
| WP_258517485.1 | Type1.BREX | [illegible] | 1225 | Bacteria>Pseudomonadota>Gammaproteobacteria>Vibrionales>Vibrionaceae>Vibrio | GCF_024745395.1 | Vibrio_parahaemolyticus | NB556_RS01245 | 3462..7139 | 670 | BREX-system-P-loop-protein-BrxC-[Vibrio-parahaemolyticus] |
| WP_140132901.1 | Type1.BREX | [illegible] | 1225 | Bacteria>Pseudomonadota>Gammaproteobacteria>Vibrionales>Vibrionaceae>Vibrio | GCF_024748295.1 | Vibrio_harveyi_group | ND930_RS02780 | 146702..150379 | 717610 | MULTISPECIES:-BREX-system-P-loop-protein-BrxC-[Vibrio-harveyi-group] |
| WP_264912148.1 | Type1.BREX | [illegible] | 1248 | Bacteria>Pseudomonadota>Gammaproteobacteria>Vibrionales>Vibrionaceae>Vibrio | GCF_026000495.1 | Vibrio_sp._STUT-A16 | OO778_RS24845 | 1838056..1841802 | 2976237 | BREX-system-P-loop-protein-BrxC-[Vibrio-sp.-STUT-A16] |
| WP_267365291.1 | Type1.BREX | [illegible] | 1248 | Bacteria>Pseudomonadota>Gammaproteobacteria>Vibrionales>Vibrionaceae>Vibrio | GCF_026537625.1 | Vibrio_parahaemolyticus | NOL37_RS23780 | 19146..22892 | 670 | BREX-system-P-loop-protein-BrxC-[Vibrio-parahaemolyticus] |
| WP_267363047.1 | Type1.BREX | [illegible] | 1187 | Bacteria>Pseudomonadota>Gammaproteobacteria>Vibrionales>Vibrionaceae>Vibrio | GCF_026537625.1 | Vibrio_parahaemolyticus | NOL37_RS24175 | 9284..12847 | 670 | BREX-system-P-loop-protein-BrxC-[Vibrio-parahaemolyticus] |
| WP_140298951.1 | Type1.BREX | [illegible] | 1225 | Bacteria>Pseudomonadota>Gammaproteobacteria>Vibrionales>Vibrionaceae>Vibrio | GCF_026538205.1 | Vibrio_parahaemolyticus | CGI26_02815 | 152399..156076 | 670 | BREX-system-P-loop-protein-BrxC-[Vibrio-parahaemolyticus] |
| WP_268670710.1 | Type1.BREX | [illegible] | 1185 | Bacteria>Pseudomonadota>Gammaproteobacteria>Vibrionales>Vibrionaceae>Vibrio | GCF_026798035.1 | Vibrio_coralliirubri | OTK49_RS06670 | 229636..233193 | 1516159 | BREX-system-P-loop-protein-BrxC-[Vibrio-coralliirubri] |
| WP_269646266.1 | Type1.BREX | [illegible] | 1225 | Bacteria>Pseudomonadota>Gammaproteobacteria>Vibrionales>Vibrionaceae>Vibrio | GCF_027277845.1 | Vibrio_parahaemolyticus | O4N70_RS12810 | 220363..224040 | 670 | BREX-system-P-loop-protein-BrxC-[Vibrio-parahaemolyticus] |
| WP_269665659.1 | Type1.BREX | [illegible] | 1225 | Bacteria>Pseudomonadota>Gammaproteobacteria>Vibrionales>Vibrionaceae>Vibrio | GCF_027277905.1 | Vibrio_parahaemolyticus | O4M94_RS09585 | 134179..137856 | 670 | BREX-system-P-loop-protein-BrxC-[Vibrio-parahaemolyticus] |
| WP_272236739.1 | Type1.BREX | [illegible] | 1229 | Bacteria>Pseudomonadota>Gammaproteobacteria>Vibrionales>Vibrionaceae>Vibrio | GCF_028446685.1 | Vibrio_europaeus | OPW18_RS21520 | 351695..355384 | 300876 | BREX-system-P-loop-protein-BrxC-[Vibrio-europaeus] |
| WP_272245950.1 | Type1.BREX | [illegible] | 1229 | Bacteria>Pseudomonadota>Gammaproteobacteria>Vibrionales>Vibrionaceae>Vibrio | GCF_028446755.1 | Vibrio_europaeus | OPW38_RS04390 | 2959..6648 | 300876 | BREX-system-P-loop-protein-BrxC-[Vibrio-europaeus] |
| WP_274056407.1 | Type1.BREX | [illegible] | 1225 | Bacteria>Pseudomonadota>Gammaproteobacteria>Vibrionales>Vibrionaceae>Vibrio | GCF_028616275.1 | Vibrio_parahaemolyticus | O8051_RS20180 | 64443..68120 | 670 | BREX-system-P-loop-protein-BrxC-[Vibrio-parahaemolyticus] |
| WP_274020835.1 | Type1.BREX | [illegible] | 1248 | Bacteria>Pseudomonadota>Gammaproteobacteria>Vibrionales>Vibrionaceae>Vibrio | GCF_028617275.1 | Vibrio_parahaemolyticus | O8254_RS13605 | 106668..110414 | 670 | BREX-system-P-loop-protein-BrxC-[Vibrio-parahaemolyticus] |
| WP_274674022.1 | Type1.BREX | [illegible] | 1187 | Bacteria>Pseudomonadota>Gammaproteobacteria>Vibrionales>Vibrionaceae>Vibrio | GCF_028857845.1 | Vibrio_aestuarianus | L9W73_RS08980 | 42250..45813 | 28171 | BREX-system-P-loop-protein-BrxC-[Vibrio-aestuarianus] |
| WP_274695775.1 | Type1.BREX | [illegible] | 1225 | Bacteria>Pseudomonadota>Gammaproteobacteria>Vibrionales>Vibrionaceae>Vibrio | GCF_028858205.1 | Vibrio_aestuarianus | MEG05_RS14945 | 3088024..3091701 | 28171 | BREX-system-P-loop-protein-BrxC-[Vibrio-aestuarianus] |
| WP_274702484.1 | Type1.BREX | [illegible] | 1228 | Bacteria>Pseudomonadota>Gammaproteobacteria>Vibrionales>Vibrionaceae>Vibrio | GCF_028858475.1 | Vibrio_aestuarianus | MCT07_RS16955 | 5869..9555 | 28171 | BREX-system-P-loop-protein-BrxC-[Vibrio-aestuarianus] |
| WP_253254707.1 | Type1.BREX | [illegible] | 1153 | Bacteria>Pseudomonadota>Gammaproteobacteria>Vibrionales>Vibrionaceae>Vibrio | GCF_030238805.1 | Vibrio_parahaemolyticus | CSR03_RS22350 | 121618..125079 | 670 | BREX-system-P-loop-protein-BrxC-[Vibrio-parahaemolyticus] |
| WP_182025746.1 | Type1.BREX | [illegible] | 1233 | Bacteria>Pseudomonadota>Gammaproteobacteria>Vibrionales>Vibrionaceae>Vibrio | GCF_903986985.1 | Vibrio_rotiferianus | H7A65_RS05210 | 396251..399952 | 190895 | BREX-system-P-loop-protein-BrxC-[Vibrio-rotiferianus] |
| WP_069535824.1 | Type1.BREX | [illegible] | 1157 | Bacteria>Pseudomonadota>Gammaproteobacteria>Vibrionales>Vibrionaceae>Vibrio | GCF_905331795.1 | Vibrio_parahaemolyticus | LPR62_RS04710 | 409258..412731 | 670 | BREX-system-P-loop-protein-BrxC-[Vibrio-parahaemolyticus] |
| WP_229611425.1 | Type1.BREX | [illegible] | 1157 | Bacteria>Pseudomonadota>Gammaproteobacteria>Vibrionales>Vibrionaceae>Vibrio | GCF_905331955.1 | Vibrio_parahaemolyticus | LPR99_RS16740 | 18511..21984 | 670 | BREX-system-P-loop-protein-BrxC-[Vibrio-parahaemolyticus] |
| WP_237468615.1 | Type1.BREX | [illegible] | 1231 | Bacteria>Pseudomonadota>Gammaproteobacteria>Vibrionales>Vibrionaceae>Vibrio | GCF_921293875.1 | Vibrio_stylophorae | L9P36_RS15955 | 492812..496507 | 659351 | BREX-system-P-loop-protein-BrxC-[Vibrio-stylophorae] |
| WP_168780466.1 | Type1.BREX | [illegible] | 1248 | Bacteria>Pseudomonadota>Gammaproteobacteria>Vibrionales>Vibrionaceae>Vibrio | GCF_944738875.1 | Vibrio_aestuarianus | OCX59_RS00150 | 25401..29147 | 28171 | BREX-system-P-loop-protein-BrxC-[Vibrio-aestuarianus] |
| WP_272166069.1 | Type1.BREX | [illegible] | 1248 | Bacteria>Pseudomonadota>Gammaproteobacteria>Vibrionales>Vibrionaceae>Vibrio | GCF_947503745.1 | Vibrio_diabolicus | PQF93_RS21015 | 28188..31934 | 50719 | BREX-system-P-loop-protein-BrxC-[Vibrio-diabolicus] |
| WP_241766700.1 | Type1.BREX | [illegible] | 1153 | Bacteria>Pseudomonadota>Gammaproteobacteria>Vibrionales>Vibrionaceae>Vibrio | GCF_948193335.1 | Vibrio_parahaemolyticus | PO77_RS19450 | 167850..171311 | 670 | BREX-system-P-loop-protein-BrxC-[Vibrio-parahaemolyticus] |
| WP_213868350.1 | Type1.BREX | [illegible] | 1187 | Bacteria>Pseudomonadota>Gammaproteobacteria>Vibrionales>Vibrionaceae>Vibrio | - | Vibrio_alginolyticus | N7I24_002738 | 64007..67570 | 663 | BREX-system-P-loop-protein-BrxC-[Vibrio-alginolyticus] |
| WP_222717221.1 | Type1.BREX | [illegible] | 1233 | Bacteria>Pseudomonadota>Gammaproteobacteria>Vibrionales>Vibrionaceae>Vibrio | - | Vibrio_atlanticus | JFJ08_RS20295 | 521..4222 | 693153 | BREX-system-P-loop-protein-BrxC-[Vibrio-atlanticus] |
| AAL59733.1 | Type1.BREX | [illegible] | 631 | Bacteria>Pseudomonadota>Gammaproteobacteria>Vibrionales>Vibrionaceae>Vibrio | - | Vibrio_cholerae | . | 28378..30273 | 666 | unknown-[Vibrio-cholerae] |
| WP_142738823.1 | Type1.BREX | [illegible] | 1224 | Bacteria>Pseudomonadota>Gammaproteobacteria>Vibrionales>Vibrionaceae>Vibrio | - | Vibrio_cholerae | FLL79_RS01410 | 36024..39698 | 666 | BREX-system-P-loop-protein-BrxC-[Vibrio-cholerae] |
| WP_069490026.1 | Type1.BREX | [illegible] | 1154 | Bacteria>Pseudomonadota>Gammaproteobacteria>Vibrionales>Vibrionaceae>Vibrio | - | Vibrio_parahaemolyticus | BBM25_06240 | 3424..6888 | 670 | BREX-system-P-loop-protein-BrxC-[Vibrio-parahaemolyticus] |
| WP_254664663.1 | Type1.BREX | [illegible] | 1153 | Bacteria>Pseudomonadota>Gammaproteobacteria>Vibrionales>Vibrionaceae>Vibrio | - | Vibrio_parahaemolyticus | BED52_RS16125 | 3457..6918 | 670 | BREX-system-P-loop-protein-BrxC-[Vibrio-parahaemolyticus] |
| WP_047508417.1 | Type1.BREX | [illegible] | 1154 | Bacteria>Pseudomonadota>Gammaproteobacteria>Vibrionales>Vibrionaceae>Vibrio | - | Vibrio_parahaemolyticus | DMJ21_13130 | 8098..11562 | 670 | BREX-system-P-loop-protein-BrxC-[Vibrio-parahaemolyticus] |
| WP_062854373.1 | Type1.BREX | [illegible] | 1154 | Bacteria>Pseudomonadota>Gammaproteobacteria>Vibrionales>Vibrionaceae>Vibrio | - | Vibrio_parahaemolyticus | LZX80_001782 | 93413..96877 | 670 | BREX-system-P-loop-protein-BrxC-[Vibrio-parahaemolyticus] |
| WP_226420703.1 | Type1.BREX | [illegible] | 1248 | Bacteria>Pseudomonadota>Gammaproteobacteria>Vibrionales>Vibrionaceae>Vibrio | - | Vibrio_sp._E14 | LIX84_RS04085 | 940499..944245 | 2849869 | BREX-system-P-loop-protein-BrxC-[Vibrio-sp.-E14] |
| WP_211003808.1 | Type1.BREX | [illegible] | 1215 | Bacteria>Pseudomonadota>Gammaproteobacteria>Xanthomonadales>Rhodanobacteraceae>Rhodanobacter | GCF_018101315.1 | Rhodanobacter_sp._B2A1Ga4 | IMW82_RS02800 | 624703..628350 | 2778647 | BREX-system-P-loop-protein-BrxC-[Rhodanobacter-sp.-B2A1Ga4] |
| MBK7210551.1 | Type1.BREX | [illegible] | 1144 | Bacteria>Pseudomonadota>Gammaproteobacteria>Xanthomonadales>unclassified_Xanthomonadales | GCA_016707485.1 | Xanthomonadales_bacterium | IPH99_09390 | 1022709..1026143 | 2006849 | BREX-system-P-loop-protein-BrxC-[Xanthomonadales-bacterium] |

| Protein-ID | System | Gene-Neighbourhoods | Length | Taxonomy | Assembly-ID | Strain | Locus-Tag | Genomic-coordinates | TaxID | Fasta-define |
|---|---|---|---|---|---|---|---|---|---|---|
| MBK9494691.1 | Type1.BREX | <-RVT<-HTH_3<-SIGMA+OB-Fold+AAA-ATPase+Lon-Protease-BrxL<-PglZ<-DUF932<-DUF6079*<-BrxB-BrxF-iSTAND<-BrxA-DNA-binding<-?<-?<-?<-?<-?||XIS-HTH-> | 1145 | Bacteria>Pseudomonadota>Gammaproteobacteria>Xanthomonadales>unclassified_Xanthomonadales | GCA_016718605.1 | Xanthomonadales_bacterium | IPO08_09120 | 1864889..1868326 | 2006849 | BREX-system-P-loop-protein-BrxC-[Xanthomonadales-bacterium] |
| WP_194339884.1 | Type1.BREX | 35exo->REase-><-SIGMA+OB-Fold+AAA-ATPase+HKD-DNase-BrxL<-PglZ<-N6-METHYLASE-p<-NERD<-OLD-ABC<-N6-METHYLASE-p<-?<-DUF6079*<-BrxB-BrxF-iSTAND<-BrxA-DNA-binding<-XIS-HTH<-?<-SbcC+SbcC<-Calcineurin+Calcineurin-SbcD | 1190 | Bacteria>Pseudomonadota>Gammaproteobacteria>Xanthomonadales>Xanthomonadaceae>Lysobacter | GCF_015277535.1 | Lysobacter_sp._H21R4 | INQ40_RS07385 | 1620041..1623613 | 2781021 | BREX-system-P-loop-protein-BrxC-[Lysobacter-sp.-H21R4] |
| WP_162376315.1 | Type1.BREX | <-NTP_transf_3+Hexapep+Hexapep||?-><-?<-SIGMA+OB-Fold+AAA-ATPase+Lon-Protease-BrxL<-PglZ<-Polbeta<-MarR-HTH<-ParB+HNH<-N6-METHYLASE<-DUF6079*<-BrxB-BrxF-iSTAND<-BrxA-DNA-binding||REase-><-?<-?<-TOPRIM<-XIS-HTH | 1148 | Bacteria>Pseudomonadota>Gammaproteobacteria>Xanthomonadales>Xanthomonadaceae>Pseudoxanthomonas | GCF_010211765.1 | Pseudoxanthomonas_kaohsiungensis | CSC66_RS05120 | 191406..194852 | 283923 | BREX-system-P-loop-protein-BrxC-[Pseudoxanthomonas-kaohsiungensis] |
| WP_271250641.1 | Type1.BREX | <-SIGMA+OB-Fold+AAA-ATPase+Lon-Protease-BrxL<-PglZ<-Polbeta<-MarR-HTH<-ParB+HNH<-N6-METHYLASE<-DUF6079*<-BrxB-BrxF-iSTAND<-BrxA-DNA-binding||REase-><-?<-?<-TOPRIM<-?<-XIS-HTH | 1148 | Bacteria>Pseudomonadota>Gammaproteobacteria>Xanthomonadales>Xanthomonadaceae>Pseudoxanthomonas | GCF_027941875.1 | Pseudoxanthomonas_mexicana | PE064_RS03375 | 689805..693251 | 128785 | BREX-system-P-loop-protein-BrxC-[Pseudoxanthomonas-mexicana] |
| MBH1702634.1 | Type1.BREX | <-RFC-AAA+RuvB-AAA||tRNA->?->AAA+wHTH-><-SIGMA+OB-Fold+AAA-ATPase+Lon-Protease-BrxL<-PglZ<-HNH<-OLD-ABC<-ParB+HNH<-N6-METHYLASE<-DUF6079*<-BrxB-BrxF-iSTAND<-BrxA-DNA-binding<-wHTH+WYL+WCX<-?<-?<-?<-DUF932||XIS-HTH-> | 1146 | Bacteria>Pseudomonadota>Gammaproteobacteria>Xanthomonadales>Xanthomonadaceae>Stenotrophomonas | GCA_015998095.1 | Stenotrophomonas_maltophilia | I5U93_05090 | 24102..27542 | 40324 | BREX-system-P-loop-protein-BrxC-[Stenotrophomonas-maltophilia] |
| WP_125428654.1 | Type1.BREX | <-RFC-AAA+RuvB-AAA||tRNA->?->AAA+wHTH-><-SIGMA+OB-Fold+AAA-ATPase+Lon-Protease-BrxL<-PglZ<-HNH<-OLD-ABC<-ParB+HNH<-N6-METHYLASE<-DUF6079*<-BrxB-BrxF-iSTAND<-BrxA-DNA-binding<-wHTH+WYL+WCX<-?<-?<-?<-DUF932||XIS-HTH-> | 1146 | Bacteria>Pseudomonadota>Gammaproteobacteria>Xanthomonadales>Xanthomonadaceae>Stenotrophomonas | GCF_003935615.1 | Stenotrophomonas_maltophilia | EGJ89_RS09210 | 127004..130444 | 40324 | BREX-system-P-loop-protein-BrxC-[Stenotrophomonas-maltophilia] |
| WP_192288052.1 | Type1.BREX | SF1-SF2-Helicase->tRNA->tRNA->BrxA-DNA-binding->BrxB-BrxF-iSTAND->DUF6079*->N6-METHYLASE->ParB+HNH->MarR-HTH->Polbeta->PglZ->SIGMA+OB-Fold+AAA-ATPase+Lon-Protease-BrxL->?-><-HTH_3||TPR-> | 1148 | Bacteria>Pseudomonadota>Gammaproteobacteria>Xanthomonadales>Xanthomonadaceae>Stenotrophomonas | GCF_014852135.1 | Stenotrophomonas_sp._STM01 | IB227_RS14890 | 150260..153706 | 2769278 | BREX-system-P-loop-protein-BrxC-[Stenotrophomonas-sp.-STM01] |
| WP_256643585.1 | Type1.BREX | <-STYKIN+Pkinase+CR-Kinase||Glyco_transf_9-><-?<-?<-SIGMA+OB-Fold+AAA-ATPase+Lon-Protease-BrxL<-PglZ<-N6-METHYLASE<-DUF6079*<-BrxB-BrxF-iSTAND<-BrxA-DNA-binding<-?<-tRNA||?-><-35exo<-RNaseH<-METHYLASE | 1219 | Bacteria>Pseudomonadota>Gammaproteobacteria>Xanthomonadales>Xanthomonadaceae>Stenotrophomonas | GCF_024499815.1 | Stenotrophomonas_sp._9(2022) | NQ204_RS08610 | 96146..99805 | 2950153 | BREX-system-P-loop-protein-BrxC-[Stenotrophomonas-sp.-9(2022)] |
| WP_225763940.1 | Type1.BREX | <-TRPR-HTH||?->PRTase->HTH_Tnp_1+HTH_21+rve+rve_3->XIS-HTH->BrxA-DNA-binding->BrxB-BrxF-iSTAND->DUF6079*->?->N6-METHYLASE->OLD-ABC->REase->?->N6-METHYLASE-p->PglZ->SIGMA+OB-Fold+AAA-ATPase+HKD-DNase-BrxL-> | 1181 | Bacteria>Pseudomonadota>Gammaproteobacteria>Xanthomonadales>Xanthomonadaceae>Stenotrophomonas | GCF_916618915.1 | Stenotrophomonas_sp._Marseille-Q4652 | LG380_RS05370 | 1171221..1174766 | 2866595 | BREX-system-P-loop-protein-BrxC-[Stenotrophomonas-sp.-Marseille-Q4652] |
| WP_139716037.1 | Type1.BREX | ARG-><-REase||HIN-HTH->?->ZnR+TOPRIM+AAA-helicase+wHTH->XIS-HTH->BrxA-DNA-binding->BrxB-BrxF-iSTAND->DUF6079*->N6-METHYLASE->PglZ->SIGMA+OB-Fold+AAA-ATPase+Lon-Protease-BrxL->?-><-?<-?<-?<-?<-DUF932 | 1205 | Bacteria>Pseudomonadota>Gammaproteobacteria>Xanthomonadales>Xanthomonadaceae>Thermomonas | GCF_006337105.1 | Thermomonas_aquatica | FHQ07_RS06475 | 1333468..1337085 | 2202149 | BREX-system-P-loop-protein-BrxC-[Thermomonas-aquatica] |
| ODU42543.1 | Type1.BREX | <-SIGMA+OB-Fold+AAA-ATPase+Lon-Protease-BrxL<-PglZ<-Polbeta<-MarR-HTH<-HNH<-OLD-ABC<-N6-METHYLASE<-DUF6079*<-BrxB-BrxF-iSTAND<-BrxA-DNA-binding||?-><-?||HKD-DNase->?->HNH->N6-METHYLASE-> | 1148 | Bacteria>Pseudomonadota>Gammaproteobacteria>Xanthomonadales>Xanthomonadaceae>unclassified_Xanthomonadaceae | GCA_001725645.1 | Xanthomonadaceae_bacterium_SCN_69-123 | ABS96_27585 | 191713..195159 | 1660170 | hypothetical-protein-ABS96_27585-[Xanthomonadaceae-bacterium-SCN-69-123] |
| RYD14825.1 | Type1.BREX | <-35exo<-SF1-SF2-Helicase+wHTH+Small-B-barrel+DUF1998||HTH_3->Couple_hipA+HipA_C->BrxA-DNA-binding->BrxB-BrxF-iSTAND->DUF6079*->N6-METHYLASE->PglZ->SIGMA+OB-Fold+AAA-ATPase+Lon-Protease-BrxL->REase-><-tRNA||NUDIX->?->?-><-REC+GGDEF+EAL | 1207 | Bacteria>Pseudomonadota>Gammaproteobacteria>Xanthomonadales>Xanthomonadaceae>unclassified_Xanthomonadaceae | GCA_004143945.1 | Xanthomonadaceae_bacterium | EOP90_11180 | 76143..79766 | 1926873 | BREX-system-P-loop-protein-BrxC-[Xanthomonadaceae-bacterium] |
| EGD07603.1 | Type1.BREX | <-RFC-AAA+RuvB-AAA||tRNA->?->AAA+wHTH-><-SIGMA+OB-Fold+AAA-ATPase+Lon-Protease-BrxL<-PglZ<-ParB+HNH<-N6-METHYLASE<-DUF6079*<-BrxB-BrxF-iSTAND<-BrxA-DNA-binding<-wHTH+WYL+WCX<-DUF772+DDE_Tnp_1||XIS-HTH-> | 1152 | Bacteria>Pseudomonadota>Gammaproteobacteria>Xanthomonadales>Xanthomonadaceae>Xanthomonas | GCA_000192025.2 | Xanthomonas_vesicatoria_ATCC_35937 | XVE_4193 | 13272..16730 | 925775 | hypothetical-protein-XVE_4193-[Xanthomonas-vesicatoria-ATCC-35937] |
| WP_042828568.1 | Type1.BREX | <-RFC-AAA+RuvB-AAA||tRNA->?->AAA+wHTH-><-SIGMA+OB-Fold+AAA-ATPase+Lon-Protease-BrxL<-PglZ<-ParB+HNH<-N6-METHYLASE<-DUF6079*<-BrxB-BrxF-iSTAND<-BrxA-DNA-binding<-wHTH+WYL+WCX||<-DUF772+DDE_Tnp_1||XIS-HTH-> | 1146 | Bacteria>Pseudomonadota>Gammaproteobacteria>Xanthomonadales>Xanthomonadaceae>Xanthomonas | GCF_000192025.1 | Xanthomonas_vesicatoria | XVE_RS20000 | 13272..16712 | 56460 | BREX-system-P-loop-protein-BrxC-[Xanthomonas-vesicatoria] |
| WP_068573669.1 | Type1.BREX | <-STYKIN+Pkinase+CR-Kinase||Glyco_transf_9-><-?||Arm-DNA-bind_3+Phage_integrase->?-><-SIGMA+OB-Fold+AAA-ATPase+Lon-Protease-BrxL<-PglZ<-N6-METHYLASE<-DUF6079*<-BrxB-BrxF-iSTAND<-BrxA-DNA-binding<-tRNA||?-><-35exo<-RNaseH<-METHYLASE | 1215 | Bacteria>Pseudomonadota>Gammaproteobacteria>Xanthomonadales>Xanthomonadaceae>Xanthomonas | GCF_000589895.1 | Xanthomonas_campestris | XCCCFBP1712_RS16305 | 3805862..3809509 | 339 | BREX-system-P-loop-protein-BrxC-[Xanthomonas-campestris] |
| WP_212590115.1 | Type1.BREX | <-STYKIN+Pkinase+CR-Kinase||Glyco_transf_9-><-?||Arm-DNA-bind_3+Phage_integrase-><-SIGMA+OB-Fold+AAA-ATPase+Lon-Protease-BrxL<-PglZ<-N6-METHYLASE<-N6-METHYLASE-p<-DUF6079*<-BrxB-BrxF-iSTAND<-BrxA-DNA-binding<-tRNA||?-><-35exo<-RNaseH<-METHYLASE | 1219 | Bacteria>Pseudomonadota>Gammaproteobacteria>Xanthomonadales>Xanthomonadaceae>Xanthomonas | GCF_000589955.1 | Xanthomonas_campestris | XCCCFBP4955_RS16075 | 3750413..3754072 | 339 | BREX-system-P-loop-protein-BrxC-[Xanthomonas-campestris] |
| WP_218196661.1 | Type1.BREX | Arm-DNA-bind_3+Phage_integrase-><-SIGMA+OB-Fold+AAA-ATPase+HKD-DNase-BrxL<-PglZ<-N6-METHYLASE-p<-?<-REase<-OLD-ABC<-N6-METHYLASE<-?<-DUF6079*<-BrxB-BrxF-iSTAND<-BrxA-DNA-binding<-XIS-HTH<-Phage_int_SAM_2+Integrase_1<-Phage_rep | 1182 | Bacteria>Pseudomonadota>Gammaproteobacteria>Xanthomonadales>Xanthomonadaceae>Xanthomonas | GCF_019193005.1 | Xanthomonas_euvesicatoria | KV691_RS15985 | 182541..186089 | 456327 | BREX-system-P-loop-protein-BrxC-[Xanthomonas-euvesicatoria] |
| WP_227971952.1 | Type1.BREX | <-N6-METHYLASE<-DUF6079*<-BrxB-BrxF-iSTAND<-BrxA-DNA-binding<-tRNA||?-><-35exo<-RNaseH<-METHYLASE | 1219 | Bacteria>Pseudomonadota>Gammaproteobacteria>Xanthomonadales>Xanthomonadaceae>Xanthomonas | GCF_020731405.1 | Xanthomonas_campestris | LLE63_RS17440 | 2847..6506 | 339 | BREX-system-P-loop-protein-BrxC-[Xanthomonas-campestris] |
| WP_228944544.1 | Type1.BREX | HTH_Tnp_1+HTH_21+rve+rve_3-><-SIGMA+OB-Fold+AAA-ATPase+Lon-Protease-BrxL<-PglZ<-N6-METHYLASE<-DUF6079*<-BrxB-BrxF-iSTAND<-BrxA-DNA-binding<-Arm-DNA-bind_3+Phage_integrase<-tRNA<-SIGMA-HTH+SIGMA-HTH+SIGMA-HTH<-?||?-><-ACET | 1217 | Bacteria>Pseudomonadota>Gammaproteobacteria>Xanthomonadales>Xanthomonadaceae>Xanthomonas | GCF_020879405.1 | Xanthomonas_campestris | LN454_RS12865 | 9123..12776 | 339 | BREX-system-P-loop-protein-BrxC-[Xanthomonas-campestris] |
| WP_260816965.1 | Type1.BREX | DnaJ->?->?->?-><-?<-XIS-HTH||wHTH+WYL+WCX->BrxA-DNA-binding->BrxB-BrxF-iSTAND->DUF6079*->N6-METHYLASE->ParB+HNH->PglZ->SIGMA+OB-Fold+AAA-ATPase+Lon-Protease-BrxL-><-AAA+wHTH<-?<-tRNA||RFC-AAA+RuvB-AAA->?->HHH_5+RecR+TOPRIM-> | 1146 | Bacteria>Pseudomonadota>Gammaproteobacteria>Xanthomonadales>Xanthomonadaceae>Xanthomonas | GCF_025263745.1 | Xanthomonas_prunicola | MOD45_RS16865 | 3940549..3943989 | 2053930 | BREX-system-P-loop-protein-BrxC-[Xanthomonas-prunicola] |
| WP_274376279.1 | Type1.BREX | METHYLASE->RNaseH->35exo-><-?||tRNA->BrxA-DNA-binding->BrxB-BrxF-iSTAND->DUF6079*->N6-METHYLASE-><-DUF772+DDE_Tnp_1+DDE_Tnp_1_6||N6-METHYLASE-p->PglZ->SIGMA+OB-Fold+AAA-ATPase+Lon-Protease-BrxL-><-?||?-><-HIN-HTH | 1219 | Bacteria>Pseudomonadota>Gammaproteobacteria>Xanthomonadales>Xanthomonadaceae>Xanthomonas | GCF_028749825.1 | Xanthomonas_campestris | JH286_RS05255 | 1188606..1192265 | 339 | BREX-system-P-loop-protein-BrxC-[Xanthomonas-campestris] |
| WP_274375850.1 | Type1.BREX | Arm-DNA-bind_3+Phage_integrase-><-HIN-HTH||<-SF1-SF2-Helicase-p<-?||?-><-SIGMA+OB-Fold+AAA-ATPase+Lon-Protease-BrxL<-PglZ<-N6-METHYLASE<-DUF6079*<-BrxB-BrxF-iSTAND<-BrxA-DNA-binding<-tRNA||?-><-35exo<-RNaseH<-METHYLASE | 1218 | Bacteria>Pseudomonadota>Gammaproteobacteria>Xanthomonadales>Xanthomonadaceae>Xanthomonas | GCF_028749885.1 | Xanthomonas_campestris | JH293_RS16145 | 3775277..3778933 | 339 | BREX-system-P-loop-protein-BrxC-[Xanthomonas-campestris] |
| WP_274507195.1 | Type1.BREX | <-RADICAL-SAM<-?<-STYKIN+Pkinase+CR-Kinase||Glyco_transf_9-><-?||Arm-DNA-bind_3+Phage_integrase-><-HIN-HTH||<-DUF6079-p*<-BrxB-BrxF-iSTAND<-BrxA-DNA-binding||?-><-?<-?<-?||HTH_19->?->REase-> | 482 | Bacteria>Pseudomonadota>Gammaproteobacteria>Xanthomonadales>Xanthomonadaceae>Xanthomonas | GCF_028750015.1 | Xanthomonas_campestris | JH282_RS16380 | 3813238..3814686 | 339 | BREX-system-P-loop-protein-BrxC-[Xanthomonas-campestris] |
| WP_000687848.1 | Type1.BREX | <-VirD4-TraG+VirB4-FtsK<-HDRnase<-SbcC<-SIGMA+OB-Fold+AAA-ATPase+Lon-Protease-BrxL<-PglZ<-N6-METHYLASE<-ParB+HNH<-DUF6079*<-BrxB-BrxF-iSTAND<-BrxA-DNA-binding<-wHTH+WYL+WCX<-?<-35exo<-?||LexA-protease->?->Phage_integrase-> | 1225 | Bacteria>Pseudomonadota | GCF_000022585.1 | Gammaproteobacteria | ALD23_RS14425 | 130597..134274 | 1236 | MULTISPECIES:-BREX-system-P-loop-protein-BrxC-[Gammaproteobacteria] |
| WP_000687849.1 | Type1.BREX | <-LexA-protease||?->35exo->?->wHTH+WYL+WCX->BrxA-DNA-binding->BrxB-BrxF-iSTAND->DUF6079*->OLD-ABC->TOPRIM->N6-METHYLASE->PglZ->SIGMA+OB-Fold+AAA-ATPase+Lon-Protease-BrxL->SbcC->HDRnase->VirD4-TraG+VirB4-FtsK-> | 1225 | Bacteria>Pseudomonadota | GCF_000299535.2 | Gammaproteobacteria | B934_RS0100870 | 5534..9211 | 1236 | MULTISPECIES:-BREX-system-P-loop-protein-BrxC-[Gammaproteobacteria] |
| WP_069709528.1 | Type1.BREX | VirB8->VirB9->VirB10->RHH-><-TetR-HTH||MODE-HTH+PBPII->wHTH+WYL+WCX->BrxA-DNA-binding->BrxB-BrxF-iSTAND->DUF6079*->N6-METHYLASE->PglZ->SIGMA+OB-Fold+AAA-ATPase+HKD-DNase-BrxL->SF1-SF2-Helicase+REase->SF1-SF2-Helicase+REase->wHTH->?-><-AAA-ATPase<-HIN-HTH | 1179 | Bacteria>Pseudomonadota | GCF_001742225.1 | Alphaproteobacteria | BES08_RS24075 | 1524283..1527822 | 28211 | MULTISPECIES:-BREX-system-P-loop-protein-BrxC-[Alphaproteobacteria] |
| WP_107394298.1 | Type1.BREX | <-HisKin<-SF1-SF2-Helicase+REase<-35Exo+REase||wHTH+WYL+WCX->BrxA-DNA-binding->BrxB-BrxF-iSTAND->DUF6079*->N6-METHYLASE->HEPN->PglZ->SIGMA+OB-Fold+AAA-ATPase+Lon-Protease-BrxL-><-AbiEii | 1140 | Bacteria>Pseudomonadota | GCF_003034225.1 | Alphaproteobacteria | CV103_RS05945 | 27177..30599 | 28211 | MULTISPECIES:-BREX-system-P-loop-protein-BrxC-[Alphaproteobacteria] |
| WP_066700609.1 | Type1.BREX | VirB8->VirB9->VirB10->RHH-><-TetR-HTH||MODE-HTH+PBPII->wHTH+WYL+WCX->BrxA-DNA-binding->BrxB-BrxF-iSTAND->DUF6079*->N6-METHYLASE->PglZ->SIGMA+OB-Fold+AAA-ATPase+HKD-DNase-BrxL->SF1-SF2-Helicase+REase->SF1-SF2-Helicase+REase->wHTH->HKD-DNase+HKD-DNase->SF1-SF2-Helicase->ABC-ATPase+SbcC-> | 1179 | Bacteria>Pseudomonadota | GCF_003967075.1 | Alphaproteobacteria | SAMIE_RS17570 | 3625371..3628910 | 28211 | MULTISPECIES:-BREX-system-P-loop-protein-BrxC-[Alphaproteobacteria] |
| WP_156734107.1 | Type1.BREX | <-LexA-protease||?->35exo->?->wHTH+WYL+WCX->BrxA-DNA-binding->BrxB-BrxF-iSTAND->DUF6079*->ParB+HNH->N6-METHYLASE->PglZ->SIGMA+OB-Fold+AAA-ATPase+Lon-Protease-BrxL->NACHT+HEAT+REase->HDRnase->VirD4-TraG+VirB4-FtsK-> | 1225 | Bacteria>Pseudomonadota | GCF_009740265.1 | Gammaproteobacteria | GTH23_RS00050 | 5219..8896 | 1236 | MULTISPECIES:-BREX-system-P-loop-protein-BrxC-[Gammaproteobacteria] |
| WP_119567031.1 | Type1.BREX | <-OBFOLD-NUCLEASE<-CARF+NtrC-AAA+FIS-HTH<-SIGMA+OB-Fold+AAA-ATPase+Lon-Protease-BrxL<-PglZ<-N6-METHYLASE<-ParB+HNH<-DUF6079*<-BrxB-BrxF-iSTAND<-BrxA-DNA-binding<-wHTH+WYL+WCX<-?<-35exo||LexA-protease->?-><-MobI | 1225 | Bacteria>Pseudomonadota | GCF_013602615.1 | Gammaproteobacteria | G9G97_RS25355 | 26642..30319 | 1236 | MULTISPECIES:-BREX-system-P-loop-protein-BrxC-[Gammaproteobacteria] |
| WP_125941663.1 | Type1.BREX | GDResolvase+HIN-HTH-><-?<-?<-SIGMA+OB-Fold+AAA-ATPase+Lon-Protease-BrxL<-PglZ<-ParB+HNH<-N6-METHYLASE-p<-HTH_23+HTH_32+HTH_33+DDE_3<-N6-METHYLASE<-DUF6079*<-BrxB-BrxF-iSTAND<-BrxA-DNA-binding<-wHTH+WYL+WCX||XIS-HTH->?->?-><-?<-Phage_int_SAM_5+Phage_integrase<-sGTP | 1154 | Bacteria>Pseudomonadota | GCF_019046925.1 | Gammaproteobacteria | I6L32_RS07295 | 1524921..1528385 | 1236 | MULTISPECIES:-BREX-system-P-loop-protein-BrxC-[Gammaproteobacteria] |
| WP_110076276.1 | Type1.BREX | <-VirD4-TraG+VirB4-FtsK<-HDRnase<-SbcC<-SIGMA+OB-Fold+AAA-ATPase+Lon-Protease-BrxL<-PglZ<-N6-METHYLASE<-ParB+HNH<-DUF6079*<-BrxB-BrxF-iSTAND<-BrxA-DNA-binding<-wHTH+WYL+WCX<-?<-35exo||LexA-protease->||?->?->Pkinase-> | 1225 | Bacteria>Pseudomonadota | GCF_021431945.1 | Gammaproteobacteria | LY388_RS16340 | 2424979..2428656 | 1236 | MULTISPECIES:-BREX-system-P-loop-protein-BrxC-[Gammaproteobacteria] |
| WP_111805008.1 | Type1.BREX | <-VirD4-TraG+VirB4-FtsK<-HDRnase<-SbcC<-SIGMA+OB-Fold+AAA-ATPase+Lon-Protease-BrxL<-PglZ<-N6-METHYLASE<-ParB+HNH<-DUF6079*<-BrxB-BrxF-iSTAND<-BrxA-DNA-binding<-wHTH+WYL+WCX<-?<-35exo||LexA-protease->?-><-MobI | 1225 | Bacteria>Pseudomonadota | GCF_028482435.1 | Gammaproteobacteria | DR130_RS11140 | 144958..148635 | 1236 | MULTISPECIES:-BREX-system-P-loop-protein-BrxC-[Gammaproteobacteria] |
| OYW40486.1 | Type1.BREX | <-SF1-SF2-Helicase<-SF1-SF2-Helicase<-BrxB-BrxF-iSTAND<-BrxA-DNA-binding<-wHTH+WYL+WCX<-MORC+DNA_gyrase+TOPRIM<-?<-RFC-AAA+DNAA-HTH||?->RNaseP-> | 1200 | Bacteria>Pseudomonadota>Hydrogenophilia>Hydrogenophilales>unclassified_Hydrogenophilales | GCA_002279795.1 | Hydrogenophilales_bacterium_12-61-10 | B7Z35_01010 | 8163..11765 | 1970515 | ATPase-[Hydrogenophilales-bacterium-12-61-10] |
| OYX29919.1 | Type1.BREX | <-ABC-ATPase+ABC-ATPase<-sGTP<-?<-RNaseP<-?||?->wHTH+WYL+WCX->BrxA-DNA-binding->BrxB-BrxF-iSTAND->DUF6079*->SF1-SF2-Helicase->SF1-SF2-Helicase->PglZ->SIGMA+OB-Fold+AAA-ATPase+Lon-Protease-BrxL->REase-> | 1200 | Bacteria>Pseudomonadota>Hydrogenophilia>Hydrogenophilales>unclassified_Hydrogenophilales | GCA_002280425.1 | Hydrogenophilales_bacterium_32-62-9 | B7Z03_07680 | 35609..39211 | 1970531 | ATPase-[Hydrogenophilales-bacterium-32-62-9] |
| PIQ12823.1 | Type1.BREX | <-HARE-HTH+REase<-SIGMA+OB-Fold+AAA-ATPase+Lon-Protease-BrxL<-PglZ<-HNH<-OLD-ABC<-N6-METHYLASE<-DUF6079*<-BrxB-BrxF-iSTAND<-BrxA-DNA-binding<-SLOG+BirA-HTH<-SF1-SF2-Helicase+PRTase<-wHTH+WYL+WCX||<-ParB<-ParB<-GDResolvase+PPP1-HTH | 1146 | Bacteria>Pseudomonadota>Hydrogenophilia>Hydrogenophilales>unclassified_Hydrogenophilales | GCA_002770925.1 | Hydrogenophilales_bacterium_CG18_big_fil_WC_8_21_14_2_50_58_12 | COW70_07940 | 10882..14322 | 1974036 | hypothetical-protein-COW70_07940-[Hydrogenophilales-bacterium-CG18_big_fil_WC_8_21_14_2_50_58_12] |
| PIY00903.1 | Type1.BREX | Peptidase_M50->?->?-><-ZDNA-HTH<-?||BrxA-DNA-binding->BrxB-BrxF-iSTAND->DUF6079*->AbiEi_4->AbiEii->N6-METHYLASE->HTH+REase-> | 1207 | Bacteria>Pseudomonadota>Hydrogenophilia>Hydrogenophilales>unclassified_Hydrogenophilales | GCA_002784065.1 | Hydrogenophilales_bacterium_CG_4_10_14_3_um_filter_58_23 | COZ23_06050 | 25268..28891 | 1974027 | BREX-system-P-loop-protein-BrxC-[Hydrogenophilales-bacterium-CG_4_10_14_3_um_filter_58_23] |
| KAA0218648.1 | Type1.BREX | HIN-HTH-><-?<-?<-?<-?||GNTR-HTH+Aminotran_1_2->BrxA-DNA-binding->BrxB-BrxF-iSTAND->DUF6079*->N6-METHYLASE->Schlafen+HATPase+wHTH->ParA-Soj-PloopNTPase->PglZ->SIGMA+OB-Fold+AAA-ATPase+Lon-Protease-BrxL-><-?<-ABC-ATPase+SbcC<-ABC-ATPase+SbcC | 1148 | Bacteria>Pseudomonadota>unclassified_Pseudomonadota | GCA_008363105.1 | Pseudomonadota_bacterium | EDM78_03970 | 14965..18411 | 1977087 | BREX-system-P-loop-protein-BrxC-[Pseudomonadota-bacterium] |
| MYJ95828.1 | Type1.BREX | PIWI->REase->ArdCN+MPTase-><-?||BrxA-DNA-binding->BrxB-BrxF-iSTAND->DUF6079-p*-> | 751 | Bacteria>Pseudomonadota>unclassified_Pseudomonadota | GCA_009845555.1 | Pseudomonadota_bacterium | F4053_09660 | 8125..10376 | 1977087 | BREX-system-P-loop-protein-BrxC-[Pseudomonadota-bacterium] |
| MBK8482432.1 | Type1.BREX | <-SIGMA+OB-Fold+AAA-ATPase+Lon-Protease-BrxL<-PglZ<-?<-N6-METHYLASE-p<-DOC+HTH<-N6-METHYLASE-p||?-><-SF1-SF2-Helicase+REase-p<-OLD-ABC+TOPRIM<-DUF6079-p*<-DUF6079-p<-BrxB-BrxF-iSTAND<-BrxA-DNA-binding<-XIS-HTH<-?<-GDResolvase+ZnR<-GDResolvase<-RAMA | 896 | Bacteria>Pseudomonadota>unclassified_Pseudomonadota | GCA_016711215.1 | Pseudomonadota_bacterium | IPL40_14915 | 141091..143781 | 1977087 | hypothetical-protein-[Pseudomonadota-bacterium] |
| MBQ1925249.1 | Type1.BREX | <-TPR<-tRNA||?->?->?->?->BrxA-DNA-binding->BrxB-BrxF-iSTAND->DUF6079-p*-> | 984 | Bacteria>Pseudomonadota>unclassified_Pseudomonadota | GCA_017392365.1 | Pseudomonadota_bacterium | II180_03915 | 15942..18894 | 1977087 | BREX-system-P-loop-protein-BrxC-[Pseudomonadota-bacterium] |
| MBQ9395285.1 | Type1.BREX | <-N6-METHYLASE-p<-nSTAND3<-?<-DUF6079*<-?<-SPOUT<-?||?->?->?->?-><-HD | 1183 | Bacteria>Pseudomonadota>unclassified_Pseudomonadota | GCA_017455165.1 | Pseudomonadota_bacterium | IJU23_07180 | 2296..5847 | 1977087 | BREX-system-P-loop-protein-BrxC-[Pseudomonadota-bacterium] |
| MBS0598081.1 | Type1.BREX | <-DUF6079*<-BrxB-BrxF-iSTAND<-BrxA-DNA-binding<-SLOG+BirA-HTH<-SF1-SF2-Helicase+PRTase<-wHTH+WYL+WCX | 1146 | Bacteria>Pseudomonadota>unclassified_Pseudomonadota | GCA_018240335.1 | Pseudomonadota_bacterium | JSR92_12305 | 1723..5163 | 1977087 | BREX-system-P-loop-protein-BrxC-[Pseudomonadota-bacterium] |
| MBS0349076.1 | Type1.BREX | RHH_5->BrxA-DNA-binding->BrxB-BrxF-iSTAND->DUF6079*->?->N6-METHYLASE->PHP+SbcC->SF1-SF2-Helicase+PRTase->SLOG+BirA-HTH->PglZ->SIGMA+OB-Fold+AAA-ATPase+Lon-Protease-BrxL->REase+TIR->PIWI-> | 1153 | Bacteria>Pseudomonadota>unclassified_Pseudomonadota | GCA_018241665.1 | Pseudomonadota_bacterium | JSR69_21735 | 2072..5533 | 1977087 | BREX-system-P-loop-protein-BrxC-[Pseudomonadota-bacterium] |
| MBS1228526.1 | Type1.BREX | Arm-DNA-bind_3+Phage_integrase-><-Phage_int_SAM_2+Integrase_1||?->?->?->?->wHTH+WYL+WCX->BrxA-DNA-binding->BrxB-BrxF-iSTAND->DUF6079*->N6-METHYLASE->HNH->?->HTH+REase->DUF3644->PglZ->SIGMA+OB-Fold+AAA-ATPase+Lon-Protease-BrxL->DOC+HTH-> | 1213 | Bacteria>Pseudomonadota>unclassified_Pseudomonadota | GCA_018262095.1 | Pseudomonadota_bacterium | H6R17_1803 | 75999..79640 | 1977087 | ATPase-like-protein-[Pseudomonadota-bacterium] |
| MBU0673651.1 | Type1.BREX | RADICAL-SAM-><-?||?->BrxA-DNA-binding->BrxB-BrxF-iSTAND->DUF6079*->Schlafen+HATPase+wHTH->?->N6-METHYLASE->REase->ParB+HNH->OLD-ABC->?->PglZ->SIGMA+OB-Fold+AAA-ATPase+HKD-DNase-BrxL-> | 1167 | Bacteria>Pseudomonadota>unclassified_Pseudomonadota | GCA_018814305.1 | Pseudomonadota_bacterium | KJ950_03300 | 48865..52368 | 1977087 | BREX-system-P-loop-protein-BrxC-[Pseudomonadota-bacterium] |
| MBU1208417.1 | Type1.BREX | SbcC->HNH->?->?-><-SIGMA+OB-Fold+AAA-ATPase+Lon-Protease-BrxL<-PglZ<-DUF6079*<-BrxB-BrxF-iSTAND<-BrxA-DNA-binding||?->PSBP-><-?<-?<-?<-?<-HTH+DSBH | 1147 | Bacteria>Pseudomonadota>unclassified_Pseudomonadota | GCA_018819135.1 | Pseudomonadota_bacterium | KKH04_16070 | 11316..14759 | 1977087 | BREX-system-P-loop-protein-BrxC-[Pseudomonadota-bacterium] |
| MBU1744600.1 | Type1.BREX | <-BrxD-AAA<-BrxD-AAA<-?<-SIGMA+OB-Fold+AAA-ATPase+Lon-Protease-BrxL<-PglZ<-N6-METHYLASE<-Schlafen+HATPase+wHTH<-DUF6079-p* | 992 | Bacteria>Pseudomonadota>unclassified_Pseudomonadota | GCA_018822765.1 | Pseudomonadota_bacterium | KJ649_06870 | 12750..15730 | 1977087 | BREX-system-P-loop-protein-BrxC-[Pseudomonadota-bacterium] |
| MBU2055247.1 | Type1.BREX | RADICAL-SAM->?->?->?->sGTP->?->SbcC->BrxA-DNA-binding->BrxB-BrxF-iSTAND->DUF6079*->N6-METHYLASE-p-> | 1139 | Bacteria>Pseudomonadota>unclassified_Pseudomonadota | GCA_018825885.1 | Pseudomonadota_bacterium | KKC25_09760 | 17486..20905 | 1977087 | BREX-system-P-loop-protein-BrxC-[Pseudomonadota-bacterium] |
| MBU2228449.1 | Type1.BREX | <-DUF6079*<-BrxB-BrxF-iSTAND<-BrxA-DNA-binding<-SbcC<-?<-sGTP<-?<-?<-?<-RADICAL-SAM | 1139 | Bacteria>Pseudomonadota>unclassified_Pseudomonadota | GCA_018827585.1 | Pseudomonadota_bacterium | KJ936_12455 | 147..3566 | 1977087 | BREX-system-P-loop-protein-BrxC-[Pseudomonadota-bacterium] |
| MBU2261623.1 | Type1.BREX | <-DUF6079-p*<-BrxB-BrxF-iSTAND<-BrxA-DNA-binding<-SbcC<-?<-sGTP<-?<-?<-?<-RADICAL-SAM | 120 | Bacteria>Pseudomonadota>unclassified_Pseudomonadota | GCA_018827665.1 | Pseudomonadota_bacterium | KKF02_06820 | 1..360 | 1977087 | BREX-system-P-loop-protein-BrxC-[Pseudomonadota-bacterium] |
| MBU4371661.1 | Type1.BREX | SIR2->BrxA-DNA-binding->BrxB-BrxF-iSTAND->DUF6079*->N6-METHYLASE->PglZ->SIGMA+OB-Fold+AAA-ATPase+Lon-Protease-BrxL-> | 1146 | Bacteria>Pseudomonadota>unclassified_Pseudomonadota | GCA_018896355.1 | Pseudomonadota_bacterium | KKD68_07180 | 16992..20432 | 1977087 | BREX-system-P-loop-protein-BrxC-[Pseudomonadota-bacterium] |
| MBU4294441.1 | Type1.BREX | <-HAMP+MA+MA<-?<-HPT+HISKIN+CheW+REC<-?<-?<-?||XIS-HTH->BrxA-DNA-binding->BrxB-BrxF-iSTAND->DUF6079*->N6-METHYLASE->Schlafen+HATPase+wHTH->PglZ->SIGMA+OB-Fold+AAA-ATPase+HKD-DNase-BrxL-> | 1176 | Bacteria>Pseudomonadota>unclassified_Pseudomonadota | GCA_018897105.1 | Pseudomonadota_bacterium | KJ862_00965 | 66976..70506 | 1977087 | BREX-system-P-loop-protein-BrxC-[Pseudomonadota-bacterium] |
| MBU4235493.1 | Type1.BREX | <-N6-METHYLASE<-REase<-OLD-ABC<-Schlafen+HATPase+wHTH<-DUF6079*<-BrxB-BrxF-iSTAND<-BrxA-DNA-binding<-XIS-HTH | 1147 | Bacteria>Pseudomonadota>unclassified_Pseudomonadota | GCA_018897885.1 | Pseudomonadota_bacterium | KKI01_04080 | 7059..10502 | 1977087 | BREX-system-P-loop-protein-BrxC-[Pseudomonadota-bacterium] |
| MBU3982443.1 | Type1.BREX | <-DUF6079-p*<-BrxB-BrxF-iSTAND<-BrxA-DNA-binding<-XIS-HTH<-Phage_integrase | 851 | Bacteria>Pseudomonadota>unclassified_Pseudomonadota | GCA_018900775.1 | Pseudomonadota_bacterium | KJ985_02955 | 1..2554 | 1977087 | BREX-system-P-loop-protein-BrxC-[Pseudomonadota-bacterium] |
| MCG8416875.1 | Type1.BREX | ZnR->?-><-?<-ParB+HTH<-ParA-Soj-PloopNTPase||?->?->DUF6079*->N6-METHYLASE-p-><-?||?-><-?<-?<-RVT | 1253 | Bacteria>Pseudomonadota>unclassified_Pseudomonadota | GCA_022360315.1 | Pseudomonadota_bacterium | MJE77_02885 | 60070..63831 | 1977087 | BREX-system-P-loop-protein-BrxC-[Pseudomonadota-bacterium] |
| MCG8417945.1 | Type1.BREX | <-TPR+TPR+TPR+TPR<-?<-Pkinase<-TPR+TPR<-Aminotran_4<-?<-REC+OmpR-HTH||DUF6079*-> | 1203 | Bacteria>Pseudomonadota>unclassified_Pseudomonadota | GCA_022360315.1 | Pseudomonadota_bacterium | MJE77_08390 | 23306..26917 | 1977087 | BREX-system-P-loop-protein-BrxC-[Pseudomonadota-bacterium] |
| WP_053322564.1 | Type1.BREX | HNH->?->?->LON-AAA->Calcineurin->?->BrxA-DNA-binding->BrxB-BrxF-iSTAND->DUF6079*->N6-METHYLASE-p->Schlafen+HATPase+wHTH->HEPN->REase->N6-METHYLASE-p->HerA/FtsK->PglZ->SIGMA+OB-Fold+AAA-ATPase+HKD-DNase-BrxL-> | 1181 | Bacteria>Pseudomonadota>Zetaproteobacteria>Mariprofundales>Mariprofundaceae>Mariprofundus | GCF_001273615.1 | Mariprofundus_ferrooxydans | AL013_RS02410 | 88551..92096 | 314344 | BREX-system-P-loop-protein-BrxC-[Mariprofundus-ferrooxydans] |
| MBN2327413.1 | Type1.BREX | <-N6-METHYLASE-p<-HicA_toxin<-?<-DUF6079*<-BrxB-BrxF-iSTAND<-BrxA-DNA-binding | 1173 | Bacteria>PVC_group>Candidatus_Omnitrophica>unclassified_Candidatus_Omnitrophica | GCA_016934335.1 | Candidatus_Omnitrophica_bacterium | JXR73_09680 | 2054..5575 | 2035772 | BREX-system-P-loop-protein-BrxC-[Candidatus-Omnitrophota-bacterium] |
| MBU1810879.1 | Type1.BREX | <-N6-METHYLASE<-Schlafen+HATPase+wHTH<-DUF6079*<-BrxB-BrxF-iSTAND<-BrxA-DNA-binding<-XIS-HTH | 1179 | Bacteria>PVC_group>Candidatus_Omnitrophica>unclassified_Candidatus_Omnitrophica | GCA_018823915.1 | Candidatus_Omnitrophica_bacterium | KJ629_07045 | 3409..6948 | 2035772 | BREX-system-P-loop-protein-BrxC-[Candidatus-Omnitrophota-bacterium] |
| MBU4477606.1 | Type1.BREX | <-SIGMA+OB-Fold+AAA-ATPase+Lon-Protease-BrxL<-PglZ<-Schlafen+HATPase+wHTH<-?<-DUF6079*<-BrxB-BrxF-iSTAND<-BrxA-DNA-binding<-XIS-HTH<-?<-?<-?<-RADICAL-SAM<-?<-ABC-ATPase+SbcC | 1181 | Bacteria>PVC_group>Candidatus_Omnitrophica>unclassified_Candidatus_Omnitrophica | GCA_018895275.1 | Candidatus_Omnitrophica_bacterium | KKH34_00715 | 10161..13706 | 2035772 | BREX-system-P-loop-protein-BrxC-[Candidatus-Omnitrophota-bacterium] |
| MCG2712035.1 | Type1.BREX | <-Schlafen+HATPase+wHTH-p<-N6-METHYLASE<-DUF6079*<-BrxB-BrxF-iSTAND<-BrxA-DNA-binding | 1181 | Bacteria>PVC_group>Candidatus_Omnitrophica>unclassified_Candidatus_Omnitrophica | GCA_022013205.1 | Candidatus_Omnitrophica_bacterium | L6416_06895 | 3795..7340 | 2035772 | BREX-system-P-loop-protein-BrxC-[Candidatus-Omnitrophota-bacterium] |
| MBQ6007751.1 | Type1.BREX | <-SIGMA+OB-Fold+AAA-ATPase+HKD-DNase-BrxL<-PglZ<-?<-N6-METHYLASE<-DUF6079*<-BrxB-BrxF-iSTAND<-BrxA-DNA-binding | 1146 | Bacteria>PVC_group>Kiritimatiellota>Kiritimatiellia>unclassified_Kiritimatiellia | GCA_017427485.1 | Kiritimatiellia_bacterium | IJL17_04385 | 19209..22649 | 2212473 | BREX-system-P-loop-protein-BrxC-[Kiritimatiellia-bacterium] |
| MBQ6248038.1 | Type1.BREX | Ferredoxin-RRM+Radical_SAM->?->Ferredoxin-RRM+Radical_SAM->?->?->BrxA-DNA-binding->BrxB-BrxF-iSTAND->DUF6079*-> | 1069 | Bacteria>PVC_group>Kiritimatiellota>Kiritimatiellia>unclassified_Kiritimatiellia | GCA_017429725.1 | Kiritimatiellia_bacterium | IJK04_14290 | 11611..14817 | 2212473 | BREX-system-P-loop-protein-BrxC-[Kiritimatiellia-bacterium] |
| MBQ6141563.1 | Type1.BREX | <-METHYLASE<-N6-METHYLASE<-?<-HAD<-?<-ABhydrolase<-?<-Schlafen+HATPase+wHTH<-N6-METHYLASE<-DUF6079*<-BrxB-BrxF-iSTAND<-BrxA-DNA-binding||NtrC-AAA-><-tRNA<-tRNA<-tRNA<-?<-NtrC-AAA+FIS-HTH<-?<-?<-HISKIN<-PHP | 1188 | Bacteria>PVC_group>Kiritimatiellota>Kiritimatiellia>unclassified_Kiritimatiellia | GCA_017432735.1 | Kiritimatiellia_bacterium | IJI54_09780 | 25512..29078 | 2212473 | BREX-system-P-loop-protein-BrxC-[Kiritimatiellia-bacterium] |
| MBQ7190494.1 | Type1.BREX | BrxA-DNA-binding->BrxB-BrxF-iSTAND->DUF6079*->N6-METHYLASE->Y1_Tnp->PglZ->SIGMA+OB-Fold+AAA-ATPase+HKD-DNase-BrxL->?->?->?->?-><-ArsR-HTH | 1172 | Bacteria>PVC_group>Kiritimatiellota>Kiritimatiellia>unclassified_Kiritimatiellia | GCA_017450695.1 | Kiritimatiellia_bacterium | IJR99_13895 | 2360..5878 | 2212473 | BREX-system-P-loop-protein-BrxC-[Kiritimatiellia-bacterium] |
| MBQ9739436.1 | Type1.BREX | <-N6-METHYLASE<-Schlafen+HATPase+wHTH<-DUF6079*<-BrxB-BrxF-iSTAND<-BrxA-DNA-binding | 1148 | Bacteria>PVC_group>Kiritimatiellota>Kiritimatiellia>unclassified_Kiritimatiellia | GCA_017457135.1 | Kiritimatiellia_bacterium | IJV91_00715 | 10471..13917 | 2212473 | BREX-system-P-loop-protein-BrxC-[Kiritimatiellia-bacterium] |
| MBR4615907.1 | Type1.BREX | <-REC+OmpR-HTH||?->BrxA-DNA-binding->BrxB-BrxF-iSTAND->DUF6079*->N6-METHYLASE-p->?->N6-METHYLASE-p-> | 1178 | Bacteria>PVC_group>Kiritimatiellota>Kiritimatiellia>unclassified_Kiritimatiellia | GCA_017525665.1 | Kiritimatiellia_bacterium | IKD55_09905 | 7558..11094 | 2212473 | BREX-system-P-loop-protein-BrxC-[Kiritimatiellia-bacterium] |
| MBQ9367250.1 | Type1.BREX | HATPase+wHTH->BrxA-DNA-binding->BrxB-BrxF-iSTAND->DUF6079*->N6-METHYLASE-p->SIR2-> | 1177 | Bacteria>PVC_group>Lentisphaerota>Lentisphaeria>Victivallales>unclassified_Victivallales | GCA_017458805.1 | Victivallales_bacterium | IJT83_05670 | 3304..6837 | 2212475 | BREX-system-P-loop-protein-BrxC-[Victivallales-bacterium] |
| MBR6059872.1 | Type1.BREX | <-SIGMA+OB-Fold+AAA-ATPase+Lon-Protease-BrxL<-PglZ<-HTH+REase<-N6-METHYLASE-p<-?<-N6-METHYLASE<-nSTAND3<-?<-DUF6079*<-BrxB-BrxF-iSTAND<-BrxA-DNA-binding | 1178 | Bacteria>PVC_group>Lentisphaerota>Lentisphaeria>Victivallales>unclassified_Victivallales | GCA_017542865.1 | Victivallales_bacterium | IKP58_17025 | 13303..16839 | 2212475 | BREX-system-P-loop-protein-BrxC-[Victivallales-bacterium] |
| MCF7790541.1 | Type1.BREX | <-HEPN<-SIGMA+OB-Fold+AAA-ATPase+HKD-DNase-BrxL<-PglZ<-TOPRIM<-OLD-ABC<-?<-?<-?<-DUF6079*<-BrxB-BrxF-iSTAND<-BrxA-DNA-binding<-wHTH+WYL+WCX<-?||?-><-?<-?||S2P+PDZ-> | 1176 | Bacteria>PVC_group>Lentisphaerota>Lentisphaeria>Victivallales>unclassified_Victivallales | GCA_021733985.1 | Victivallales_bacterium | K9M56_00960 | 14578..18108 | 2212475 | BREX-system-P-loop-protein-BrxC-[Victivallales-bacterium] |
| OGV31172.1 | Type1.BREX | XIS-HTH->BrxA-DNA-binding->BrxB-BrxF-iSTAND->DUF6079*->HTH+REase->N6-METHYLASE-p-> | 1201 | Bacteria>PVC_group>Lentisphaerota>unclassified_Lentisphaerota | GCA_001804785.1 | Lentisphaerae_bacterium_GWF2_45_14 | A2020_01685 | 2431..6036 | 1798572 | MAG:-hypothetical-protein-A2020_01685-[Lentisphaerae-bacterium-GWF2_45_14] |
| CAJ71903.1 | Type1.BREX | Unstab_antitox->RelE-ParE->RelE-ParE->HEPN->ABhydrolase-><-HEPN<-Polbeta<-SIGMA+OB-Fold+AAA-ATPase+Lon-Protease-BrxL<-PglZ<-DUF6079*<-?<-?<-?<-N6-METHYLASE-p||?-><-DDE_Tnp_1_4<-N6-METHYLASE<-BrxB-BrxF-iSTAND<-BrxA-DNA-binding | 1220 | Bacteria>PVC_group>Planctomycetota>Candidatus_Brocadiia>Candidatus_Brocadiales>Candidatus_Brocadiaceae | - | Candidatus_Kuenenia_stuttgartiensis | kustc1158 | 789101..792763 | 174633 | hypothetical-protein-kustc1158-[Candidatus-Kuenenia-stuttgartiensis] |
| TVL98972.1 | Type1.BREX | TPR+TPR->BrxA-DNA-binding->BrxB-BrxF-iSTAND->DUF6079*->HTH+REase->HTH+REase->PglZ->SIGMA+OB-Fold+AAA-ATPase+Lon-Protease-BrxL->REase+DUF5655->ParB->SIGMA-HTH-> | 1144 | Bacteria>PVC_group>Planctomycetota>Candidatus_Brocadiia>Candidatus_Brocadiales>Candidatus_Brocadiaceae | GCA_007618135.1 | Candidatus_Brocadia_sp._WS118 | CV087_19775 | 5481..8915 | 2040625 | MAG:-BREX-system-P-loop-protein-BrxC-[Candidatus-Brocadia-sp.-WS118] |
| WP_164995431.1 | Type1.BREX | BrxA-DNA-binding->BrxB-BrxF-iSTAND->N6-METHYLASE->?->?->DUF6079*->PglZ->SIGMA+OB-Fold+AAA-ATPase+Lon-Protease-BrxL->Polbeta->HEPN->?->?-><-?<-RelE-ParE<-Unstab_antitox | 1217 | Bacteria>PVC_group>Planctomycetota>Candidatus_Brocadiia>Candidatus_Brocadiales>Candidatus_Brocadiaceae | GCF_011066545.1 | Candidatus_Kuenenia_stuttgartiensis | KsCSTR_RS16670 | 3735232..3738885 | 174633 | BREX-system-P-loop-protein-BrxC-[Candidatus-Kuenenia-stuttgartiensis] |
| WP_099325977.1 | Type1.BREX | Unstab_antitox->RelE-ParE->HEPN->ABhydrolase-><-HEPN||GNAT+Tnp_DNA_bind+DDE_Tnp_1+Dimer_Tnp_Tn5-><-Polbeta<-SIGMA+OB-Fold+AAA-ATPase+Lon-Protease-BrxL<-PglZ<-DUF6079*<-?<-?<-?<-N6-METHYLASE<-BrxB-BrxF-iSTAND<-BrxA-DNA-binding | 1217 | Bacteria>PVC_group>Planctomycetota>Candidatus_Brocadiia>Candidatus_Brocadiales>Candidatus_Brocadiaceae | GCF_027493735.1 | Candidatus_Kuenenia_stuttgartiensis | KSMBR1_RS14595 | 3124365..3128018 | 174633 | BREX-system-P-loop-protein-BrxC-[Candidatus-Kuenenia-stuttgartiensis] |
| WP_169703663.1 | Type1.BREX | BrxA-DNA-binding->BrxB-BrxF-iSTAND->N6-METHYLASE->DDE_Tnp_1_4->N6-METHYLASE-p->?->?->DUF6079*->PglZ->SIGMA+OB-Fold+AAA-ATPase+Lon-Protease-BrxL->Polbeta->HEPN-><-ABhydrolase<-HEPN<-RelE-ParE<-RelE-ParE<-Unstab_antitox | 1217 | Bacteria>PVC_group>Planctomycetota>Candidatus_Brocadiia>Candidatus_Brocadiales>Candidatus_Brocadiaceae | GCF_900232175.1 | Candidatus_Kuenenia_stuttgartiensis | HLO46_RS19030 | 82646..86299 | 174633 | BREX-system-P-loop-protein-BrxC-[Candidatus-Kuenenia-stuttgartiensis] |
| ODS31202.1 | Type1.BREX | N6-METHYLASE->DUF6079-p*->?->?->PglZ->SIGMA+OB-Fold+AAA-ATPase+Lon-Protease-BrxL-> | 986 | Bacteria>PVC_group>Planctomycetota>Candidatus_Brocadiia>Candidatus_Brocadiales>Candidatus_Scalinduaceae | GCA_001723765.1 | Candidatus_Scalindua_rubra | SCARUB_03673 | 2450..5410 | 1872076 | hypothetical-protein-SCARUB_03673-[Candidatus-Scalindua-rubra] |
| MBC8551961.1 | Type1.BREX | BrxA-DNA-binding->BrxB-BrxF-iSTAND->DUF6079*->Schlafen+HATPase+wHTH->?->N6-METHYLASE-p-> | 1206 | Bacteria>PVC_group>Planctomycetota>Candidatus_Brocadiia>Candidatus_Brocadiales>unclassified_Candidatus_Brocadiales | GCA_014384865.1 | Candidatus_Brocadiales_bacterium | H8D23_20125 | 1912..5532 | 2760054 | BREX-system-P-loop-protein-BrxC-[Candidatus-Brocadiales-bacterium] |
| MBX3384511.1 | Type1.BREX | <-ORC-CDC6<-?<-?<-SIGMA+OB-Fold+AAA-ATPase+Lon-Protease-BrxL<-PglZ<-?<-?<-N6-METHYLASE<-DUF6079*<-BrxB-BrxF-iSTAND<-BrxA-DNA-binding<-?||?-><-?<-SAM-METHYLASE | 1151 | Bacteria>PVC_group>Planctomycetota>Phycisphaerae>Phycisphaerales>Phycisphaeraceae | GCA_019636615.1 | Phycisphaeraceae_bacterium | KF864_13495 | 1423896..1427351 | 2026777 | BREX-system-P-loop-protein-BrxC-[Phycisphaeraceae-bacterium] |
| MCC6426045.1 | Type1.BREX | <-HTH_3||ACET->?-><-?<-ParA-Soj-PloopNTPase<-?||BrxA-DNA-binding->BrxB-BrxF-iSTAND->DUF6079*->N6-METHYLASE->?->ArdCN+MPTase->PglZ->SIGMA+OB-Fold+AAA-ATPase+Lon-Protease-BrxL->TIR+PNTD2->PIWI-><-?<-HEPN | 1159 | Bacteria>PVC_group>Planctomycetota>Phycisphaerae>Phycisphaerales>unclassified_Phycisphaerales | GCA_020847535.1 | Phycisphaerales_bacterium | IT435_04420 | 30643..34122 | 2052180 | BREX-system-P-loop-protein-BrxC-[Phycisphaerales-bacterium] |
| MCL4743298.1 | Type1.BREX | <-Phage_integrase<-XIS-HTH<-Primpol+D5ATPase<-XIS-HTH<-?<-?||BrxA-DNA-binding->BrxB-BrxF-iSTAND->DUF6079*->N6-METHYLASE->PglZ->SIGMA+OB-Fold+AAA-ATPase+Lon-Protease-BrxL-><-Phage_integrase<-REase<-Phage_integrase | 1150 | Bacteria>PVC_group>Planctomycetota>Phycisphaerae>Phycisphaerales>unclassified_Phycisphaerales | GCA_023384195.1 | Phycisphaerales_bacterium | KJZ54_13970 | 8587..12039 | 2052180 | BREX-system-P-loop-protein-BrxC-[Phycisphaerales-bacterium] |
| WP_206292407.1 | Type1.BREX | <-HTH_3<-?<-ZnR+TOPRIM+AAA-helicase+wHTH<-XIS-HTH<-XIS-HTH<-?||BrxA-DNA-binding->BrxB-BrxF-iSTAND->DUF6079*->N6-METHYLASE->REase+AIPR->PglZ->SIGMA+OB-Fold+AAA-ATPase+Lon-Protease-BrxL-> | 1147 | Bacteria>PVC_group>Planctomycetota>Phycisphaerae>Tepidisphaerales>Tepidisphaeraceae | GCF_015169395.1 | Humisphaera_borealis | IPV69_RS24760 | 6565905..6569348 | 2807512 | BREX-system-P-loop-protein-BrxC-[Humisphaera-borealis] |
| QDV91803.1 | Type1.BREX | AAA-ATPase->?->?->?->?->BrxA-DNA-binding->BrxB-BrxF-iSTAND->DUF6079*->?->N6-METHYLASE->DNA-glycosylase->N6-METHYLASE-p->PHP+SbcC->Schlafen+HATPase+wHTH->PglZ->SIGMA+OB-Fold+AAA-ATPase+Lon-Protease-BrxL-> | 1149 | Bacteria>PVC_group>Planctomycetota>Phycisphaerae>unclassified_Phycisphaerae | GCA_007753915.1 | Phycisphaerae_bacterium_RAS2 | RAS2_29090 | 3426719..3430168 | 2528035 | hypothetical-protein-RAS2_29090-[Phycisphaerae-bacterium-RAS2] |
| MBI9018588.1 | Type1.BREX | BrxA-DNA-binding->BrxB-BrxF-iSTAND->DUF6079*->?->N6-METHYLASE->Schlafen+HATPase+wHTH->PglZ->SIGMA+OB-Fold+AAA-ATPase+Lon-Protease-BrxL->?->XIS-HTH-> | 1191 | Bacteria>PVC_group>Planctomycetota>Phycisphaerae>unclassified_Phycisphaerae | GCA_016342495.1 | Phycisphaerae_bacterium | JEZ07_15150 | 1153..4728 | 2026778 | BREX-system-P-loop-protein-BrxC-[Phycisphaerae-bacterium] |
| MBP7937491.1 | Type1.BREX | Peptidase->Phage_int_SAM_1+Phage_integrase-><-SIGMA+OB-Fold+AAA-ATPase+Lon-Protease-BrxL<-PglZ<-TOPRIM<-OLD-ABC<-N6-METHYLASE<-DUF6079*<-BrxB-BrxF-iSTAND<-BrxA-DNA-binding||Polbeta-> | 1149 | Bacteria>PVC_group>Planctomycetota>Phycisphaerae>unclassified_Phycisphaerae | GCA_018003015.1 | Phycisphaerae_bacterium | KA354_22855 | 13009..16458 | 2026778 | BREX-system-P-loop-protein-BrxC-[Phycisphaerae-bacterium] |
| MBP8911432.1 | Type1.BREX | <-Four-helical-protein||?->XIS-HTH->BrxA-DNA-binding->BrxB-BrxF-iSTAND->DUF6079*->?->N6-METHYLASE->PglZ->SIGMA+OB-Fold+AAA-ATPase+Lon-Protease-BrxL-p-> | 1214 | Bacteria>PVC_group>Planctomycetota>Phycisphaerae>unclassified_Phycisphaerae | GCA_018055905.1 | Phycisphaerae_bacterium | KBI32_08005 | 5154..8798 | 2026778 | BREX-system-P-loop-protein-BrxC-[Phycisphaerae-bacterium] |
| MCS7166358.1 | Type1.BREX | <-SIGMA+OB-Fold+AAA-ATPase+Lon-Protease-BrxL<-PglZ<-PglZ<-N6-METHYLASE<-N6-METHYLASE-p<-DUF6079-p<-DUF6079-p*<-N6-METHYLASE-p<-N6-METHYLASE<-BrxB-BrxF-iSTAND<-BrxA-DNA-binding | 483 | Bacteria>PVC_group>Planctomycetota>Planctomycetia>Gemmatales>unclassified_Gemmatales | GCA_025060195.1 | Gemmatales_bacterium | NZ914_01855 | 14978..16429 | 2838789 | hypothetical-protein-[Gemmatales-bacterium] |
| WP_145275550.1 | Type1.BREX | <-N6-N4-METHYLASE<-ParB<-Peptidase<-?<-SAM-METHYLASE||BrxA-DNA-binding->BrxB-BrxF-iSTAND->DUF6079*->N6-METHYLASE->?->PglZ->SIGMA+OB-Fold+AAA-ATPase+Lon-Protease-BrxL->?->SIGMA+OB-Fold+AAA-ATPase+Lon-Protease-BrxL->?->SAM-METHYLASE-> | 1197 | Bacteria>PVC_group>Planctomycetota>Planctomycetia>Isosphaerales>Isosphaeraceae | GCF_007752535.1 | Tautonia_plasticadhaerens | ElP_RS26955 | 6757809..6761402 | 2527974 | BREX-system-P-loop-protein-BrxC-[Tautonia-plasticadhaerens] |
| WP_152053060.1 | Type1.BREX | <-TrwC+SF1-SF2-Helicase||?->?->?->HTH_3->BrxA-DNA-binding->BrxB-BrxF-iSTAND->DUF6079*->N6-METHYLASE->PglZ->SIGMA+OB-Fold+AAA-ATPase+Lon-Protease-BrxL-><-RelE-ParE<-PhdYeFM_antitox||GDResolvase+HIN-HTH->?->ParA-Soj-PloopNTPase->CopG/RHH-> | 1200 | Bacteria>PVC_group>Planctomycetota>Planctomycetia>Isosphaerales>Isosphaeraceae | GCF_009177065.1 | Tautonia_marina | GA615_RS19855 | 162087..165689 | 2653855 | BREX-system-P-loop-protein-BrxC-[Tautonia-marina] |
| WP_146571387.1 | Type1.BREX | Pkinase-><-?||PspA->TOPRIM->?-><-SIGMA+OB-Fold+AAA-ATPase+Lon-Protease-BrxL<-PglZ<-N6-METHYLASE<-DUF6079*<-BrxB-BrxF-iSTAND<-BrxA-DNA-binding<-ArdCN+MPTase||?->?->?->?->McrB-AAA-><-HTH_32 | 1144 | Bacteria>PVC_group>Planctomycetota>Planctomycetia>Pirellulales>Lacipirellulaceae | GCF_007859815.1 | Botrimarina_hoheduenensis | Pla111_RS03510 | 890428..893862 | 2528000 | BREX-system-P-loop-protein-BrxC-[Botrimarina-hoheduenensis] |
| NLF73010.1 | Type1.BREX | XIS-HTH->BrxA-DNA-binding->BrxB-BrxF-iSTAND->DUF6079*->N6-METHYLASE->OLD-ABC->HNH->PglZ->SIGMA+OB-Fold+AAA-ATPase+Lon-Protease-BrxL-> | 1145 | Bacteria>PVC_group>Planctomycetota>Planctomycetia>Pirellulales>Pirellulaceae | GCA_012515235.1 | Candidatus_Anammoximicrobium_sp. | GX575_28570 | 2105..5542 | 2699742 | BREX-system-P-loop-protein-BrxC-[Candidatus-Anammoximicrobium-sp.] |
| NLY01510.1 | Type1.BREX | <-N6-METHYLASE<-DUF6079*<-BrxB-BrxF-iSTAND<-BrxA-DNA-binding<-XIS-HTH<-?<-?<-?<-?<-?||HEPN-> | 1215 | Bacteria>PVC_group>Planctomycetota>Planctomycetia>Pirellulales>Pirellulaceae | GCA_012728855.1 | Rhodopirellula_sp. | GXY83_35915 | 2404..6051 | 2024855 | BREX-system-P-loop-protein-BrxC-[Rhodopirellula-sp.] |
| MCC7337722.1 | Type1.BREX | SF1-SF2-Helicase+wHTH+Small-B-barrel+DUF1998->?->?->PIN->?->BrxA-DNA-binding->BrxB-BrxF-iSTAND->DUF6079*->?->N6-METHYLASE->REase+TPR+GreAB-C+PIN->MORC+DUF3684+REase->SF1-SF2-Helicase->?->PIN->PglZ->SIGMA+OB-Fold+AAA-ATPase+HKD-DNase-BrxL-> | 1171 | Bacteria>PVC_group>Planctomycetota>Planctomycetia>Pirellulales>Pirellulaceae | GCA_020852965.1 | Pirellulaceae_bacterium | IT422_21750 | 7905..11420 | 2699754 | BREX-system-P-loop-protein-BrxC-[Pirellulaceae-bacterium] |
| WP_145350724.1 | Type1.BREX | <-SIR2<-?<-SIGMA+OB-Fold+AAA-ATPase+HKD-DNase-BrxL<-PglZ<-REase+TPR+GreAB-C+PIN<-N6-METHYLASE<-Schlafen+HATPase+wHTH<-REase<-OLD-ABC<-DUF6079*<-BrxB-BrxF-iSTAND<-BrxA-DNA-binding<-?||TrwC+SF1-SF2-Helicase->?-><-?<-?<-ArdCN+MPTase | 1169 | Bacteria>PVC_group>Planctomycetota>Planctomycetia>Pirellulales>Pirellulaceae | GCF_007741495.1 | Roseimaritima_multifibrata | FF011L_RS05780 | 1542279..1545788 | 1930274 | BREX-system-P-loop-protein-BrxC-[Roseimaritima-multifibrata] |
| WP_145095869.1 | Type1.BREX | <-TPR+TPR<-Peptidase<-?<-?<-SIGMA+OB-Fold+AAA-ATPase+Lon-Protease-BrxL<-PglZ<-RAMA<-N6-METHYLASE<-ParB+HNH<-DUF6079*<-BrxB-BrxF-iSTAND<-BrxA-DNA-binding<-XIS-HTH||Thermonuclease->OLD-ABC->?->ABhydrolase-> | 1192 | Bacteria>PVC_group>Planctomycetota>Planctomycetia>Pirellulales>Pirellulaceae | GCF_007753095.1 | Rosistilla_carotiformis | Poly24_RS13225 | 3653187..3656765 | 2528017 | BREX-system-P-loop-protein-BrxC-[Rosistilla-carotiformis] |
| WP_145388328.1 | Type1.BREX | HerA/FtsK->?-><-?<-?<-?||BrxA-DNA-binding->BrxB-BrxF-iSTAND->DUF6079*->?->N6-METHYLASE->?->PglZ->SIGMA+OB-Fold+AAA-ATPase+Lon-Protease-BrxL-><-?<-?||?->Phage_int_SAM_1+Phage_integrase-> | 1202 | Bacteria>PVC_group>Planctomycetota>Planctomycetia>Pirellulales>Pirellulaceae | GCF_007754155.1 | Stieleria_neptunia | Enr13x_RS17960 | 5195275..5198883 | 2527979 | BREX-system-P-loop-protein-BrxC-[Stieleria-neptunia] |
| WP_197454902.1 | Type1.BREX | <-JAB<-?<-SMODS<-Cap4_nuclease+SAVED||BrxA-DNA-binding->BrxB-BrxF-iSTAND->DUF6079*->N6-METHYLASE->PglZ->SIGMA+OB-Fold+AAA-ATPase+Lon-Protease-BrxL->ANK+ANK+ANK-><-HTH_Tnp_1+HTH_21+rve+rve_3<-HTH_Tnp_1+HTH_21+rve+rve_3<-HIN-HTH | 1149 | Bacteria>PVC_group>Planctomycetota>Planctomycetia>Pirellulales>Pirellulaceae | GCF_007860045.1 | Stieleria_varia | Pla52n_RS23260 | 441169..444618 | 2528005 | BREX-system-P-loop-protein-BrxC-[Stieleria-varia] |
| WP_197171628.1 | Type1.BREX | <-ParB||?-><-?<-?<-?<-?<-?||BrxA-DNA-binding->BrxB-BrxF-iSTAND->DUF6079*->N6-METHYLASE->PHP+SbcC->Schlafen+HATPase+wHTH->PglZ->SIGMA+OB-Fold+AAA-ATPase+Lon-Protease-BrxL->?->RAMA+McrB-AAA->McrC-NTD+REase-> | 1149 | Bacteria>PVC_group>Planctomycetota>Planctomycetia>Pirellulales>Pirellulaceae | GCF_007860185.1 | Novipirellula_aureliae | Q31b_RS15730 | 214901..218350 | 2527966 | BREX-system-P-loop-protein-BrxC-[Novipirellula-aureliae] |
| WP_068129828.1 | Type1.BREX | <-SIGMA+OB-Fold+AAA-ATPase+HKD-DNase-BrxL<-Four-helical-protein<-PglZ<-PIN<-HTH+MPTase<-N6-METHYLASE<-DUF6079*<-BrxB-BrxF-iSTAND<-BrxA-DNA-binding<-XIS-HTH<-URI-DNase||?-><-Vrr-REase | 1169 | Bacteria>PVC_group>Planctomycetota>Planctomycetia>Pirellulales>Pirellulaceae | GCF_008065135.1 | Roseimaritima_ulvae | UC8_RS27055 | 7584147..7587656 | 980254 | BREX-system-P-loop-protein-BrxC-[Roseimaritima-ulvae] |
| WP_153558877.1 | Type1.BREX | <-SAM-METHYLASE<-?||?->DUF772+DDE_Tnp_1_6-><-?||XIS-HTH->BrxA-DNA-binding->BrxB-BrxF-iSTAND->DUF6079*->Schlafen+HATPase+wHTH->N6-METHYLASE-> | 1174 | Bacteria>PVC_group>Planctomycetota>Planctomycetia>Pirellulales>Pirellulaceae | GCF_009618275.1 | Roseimaritima_sediminicola | GE027_RS19390 | 33307..36831 | 2662066 | BREX-system-P-loop-protein-BrxC-[Roseimaritima-sediminicola] |
| WP_207395558.1 | Type1.BREX | BrxB-BrxF-iSTAND->N6-N4-METHYLASE->Peptidase->?->?->?-><-SIGMA+OB-Fold+AAA-ATPase+Lon-Protease-BrxL<-PglZ<-N6-METHYLASE<-DUF6079*<-BrxB-BrxF-iSTAND<-BrxA-DNA-binding||HTH->OLD-ABC+TOPRIM->SF1-SF2-Helicase->ArdCN+MPTase->?->?->RFC-AAA-> | 1152 | Bacteria>PVC_group>Planctomycetota>Planctomycetia>Pirellulales>Pirellulaceae | GCF_013618625.1 | Bremerella_alba | HOV93_RS05900 | 45922..49380 | 980252 | BREX-system-P-loop-protein-BrxC-[Bremerella-alba] |
| WP_207398110.1 | Type1.BREX | <-Phage_int_SAM_1+Phage_integrase<-MerR-HTH<-RFC-AAA<-?<-PIWI<-REase+PNTD1+PNTD2||XIS-HTH->BrxA-DNA-binding->BrxB-BrxF-iSTAND->DUF6079*->ParB+HNH->N6-METHYLASE->REase->?->REase->?->PglZ->SIGMA+OB-Fold+AAA-ATPase+Lon-Protease-BrxL->KAP-NTPase-> | 1192 | Bacteria>PVC_group>Planctomycetota>Planctomycetia>Pirellulales>Pirellulaceae | GCF_013618625.1 | Bremerella_alba | HOV93_RS18975 | 110238..113816 | 980252 | BREX-system-P-loop-protein-BrxC-[Bremerella-alba] |
| WP_184301662.1 | Type1.BREX | <-nSTAND1+HNH<-URI-DNase+RAMA<-SIGMA+OB-Fold+AAA-ATPase+HKD-DNase-BrxL<-PglZ<-REase+TPR+GreAB-C+PIN<-N6-METHYLASE<-?<-ABC-ATPase<-DUF6079*<-BrxB-BrxF-iSTAND<-BrxA-DNA-binding<-XIS-HTH||OLD-ABC+TOPRIM->SF1-SF2-Helicase-> | 1169 | Bacteria>PVC_group>Planctomycetota>Planctomycetia>Pirellulales>Pirellulaceae | GCF_014192335.1 | Rhodopirellula_rubra | FHS27_RS03210 | 253910..257419 | 980271 | BREX-system-P-loop-protein-BrxC-[Rhodopirellula-rubra] |
| WP_164102234.1 | Type1.BREX | BrxA-DNA-binding->BrxB-BrxF-iSTAND->DUF6079*->RNase->N6-METHYLASE->TOPRIM->PglZ->SIGMA+OB-Fold+AAA-ATPase+Lon-Protease-BrxL->SMODS-> | 1196 | Bacteria>PVC_group>Planctomycetota>Planctomycetia>Pirellulales>Pirellulaceae | GCF_016837145.1 | Candidatus_Laterigemmans_baculatus | ICJ56_RS09300 | 149679..153269 | 2770505 | BREX-system-P-loop-protein-BrxC-[Candidatus-Laterigemmans-baculatus] |
| WP_259631594.1 | Type1.BREX | <-SF1-SF2-Helicase<-?<-?||XIS-HTH->BrxA-DNA-binding->BrxB-BrxF-iSTAND->DUF6079*->N6-METHYLASE->PIN->ParB+HNH->TIR->MarR-HTH->Polbeta->PglZ->SIGMA+OB-Fold+AAA-ATPase+Lon-Protease-BrxL-> | 1147 | Bacteria>PVC_group>Planctomycetota>Planctomycetia>Pirellulales>Pirellulaceae | GCF_025060895.1 | Stieleria_sedimenti | NZK35_RS05520 | 42893..46336 | 2976331 | BREX-system-P-loop-protein-BrxC-[Stieleria-sedimenti] |
| WP_274403450.1 | Type1.BREX | <-McrB-AAA<-SIGMA+OB-Fold+AAA-ATPase+Lon-Protease-BrxL<-PglZ<-Schlafen+HATPase+wHTH<-PHP+SbcC<-N6-METHYLASE||?-><-ParA-Soj-PloopNTPase<-Schlafen+HATPase+wHTH<-DUF6079*<-BrxB-BrxF-iSTAND<-BrxA-DNA-binding||REC->HATPase_c->?-><-?||?-><-tRNA<-Calcineurin | 1149 | Bacteria>PVC_group>Planctomycetota>Planctomycetia>Pirellulales>Pirellulaceae | GCF_028768465.1 | Rhodopirellula_sp._P2 | PSR62_RS13060 | 3699964..3703413 | 2127060 | BREX-system-P-loop-protein-BrxC-[Rhodopirellula-sp.-P2] |
| MBQ9812620.1 | Type1.BREX | OLD-ABC->HEPN->BrxA-DNA-binding->BrxB-BrxF-iSTAND->DUF6079-p*->DUF6079-p->N6-METHYLASE->?->PglZ->SIGMA+OB-Fold+AAA-ATPase+Lon-Protease-BrxL-><-HNH<-OLD-ABC | 596 | Bacteria>PVC_group>Planctomycetota>Planctomycetia>Pirellulales>Thermoguttaceae | GCA_017441335.1 | Thermoguttaceae_bacterium | IJM54_04830 | 4203..5993 | 2691551 | BREX-system-P-loop-protein-BrxC-[Thermoguttaceae-bacterium] |
| MBP3694027.1 | Type1.BREX | <-DUF6079*<-BrxB-BrxF-iSTAND<-BrxA-DNA-binding<-HTH_26<-?<-Peptidase_M16+Peptidase_M16_C | 1201 | Bacteria>PVC_group>Planctomycetota>Planctomycetia>Pirellulales>Thermoguttaceae | GCA_017939795.1 | Thermoguttaceae_bacterium | J6J31_06290 | 52..3657 | 2691551 | BREX-system-P-loop-protein-BrxC-[Thermoguttaceae-bacterium] |
| MBV11173.1 | Type1.BREX | GDResolvase+PPP1-HTH->ParB->ParB-><-?<-DDE_Tnp_1_assoc+DDE_Tnp_1<-DDE_Tnp_IS66||XIS-HTH->BrxA-DNA-binding->BrxB-BrxF-iSTAND->DUF6079-p*-> | 800 | Bacteria>PVC_group>Planctomycetota>Planctomycetia>Planctomycetales>Planctomycetaceae | GCA_002731545.1 | Rubinisphaera_sp. | CMN21_18375 | 8990..11389 | 2024857 | hypothetical-protein-[Rubinisphaera-sp.] |
| HBJ35663.1 | Type1.BREX | <-SIGMA+OB-Fold+AAA-ATPase+HKD-DNase-BrxL<-PglZ<-PIN<-HTH+MPTase||ZDNA-HTH-><-?<-Pkinase+CR-Kinase<-N6-METHYLASE<-Schlafen+HATPase+wHTH<-DUF6079*<-Four-helical-protein<-BrxB-BrxF-iSTAND<-BrxA-DNA-binding<-XIS-HTH | 1188 | Bacteria>PVC_group>Planctomycetota>Planctomycetia>Planctomycetales>Planctomycetaceae | GCA_003506095.1 | Planctomycetaceae_bacterium | DDZ51_13145 | 16532..20098 | 2026779 | BREX-system-P-loop-protein-BrxC-[Planctomycetaceae-bacterium] |
| HCS52258.1 | Type1.BREX | <-SIGMA+OB-Fold+AAA-ATPase+Lon-Protease-BrxL<-PglZ<-TOPRIM<-SbcC<-N6-METHYLASE<-N6-METHYLASE-p<-DUF6079-p*<-DUF6079-p<-DUF6079-p<-BrxB-BrxF-iSTAND<-BrxA-DNA-binding<-XIS-HTH||DDE_Tnp_IS66->DDE_Tnp_1_assoc+DDE_Tnp_1->?-><-ParB | 340 | Bacteria>PVC_group>Planctomycetota>Planctomycetia>Planctomycetales>Planctomycetaceae | GCA_003531135.1 | Planctomycetaceae_bacterium | DIW81_11810 | 12228..13250 | 2026779 | BREX-system-P-loop-protein-BrxC-[Planctomycetaceae-bacterium] |
| HCS52259.1 | Type1.BREX | <-SIGMA+OB-Fold+AAA-ATPase+Lon-Protease-BrxL<-PglZ<-TOPRIM<-SbcC<-N6-METHYLASE<-N6-METHYLASE-p<-DUF6079-p<-DUF6079-p*<-DUF6079-p<-BrxB-BrxF-iSTAND<-BrxA-DNA-binding<-XIS-HTH||DDE_Tnp_IS66->DDE_Tnp_1_assoc+DDE_Tnp_1->?-><-ParB<-ParB | 393 | Bacteria>PVC_group>Planctomycetota>Planctomycetia>Planctomycetales>Planctomycetaceae | GCA_003531135.1 | Planctomycetaceae_bacterium | DIW81_11815 | 13400..14582 | 2026779 | BREX-system-P-loop-protein-BrxC-[Planctomycetaceae-bacterium] |
| HCS52260.1 | Type1.BREX | <-SIGMA+OB-Fold+AAA-ATPase+Lon-Protease-BrxL<-PglZ<-TOPRIM<-SbcC<-N6-METHYLASE<-N6-METHYLASE-p<-DUF6079-p<-DUF6079-p<-DUF6079-p*<-BrxB-BrxF-iSTAND<-BrxA-DNA-binding<-XIS-HTH||DDE_Tnp_IS66->DDE_Tnp_1_assoc+DDE_Tnp_1->?-><-ParB<-ParB<-GDResolvase+PPP1-HTH | 395 | Bacteria>PVC_group>Planctomycetota>Planctomycetia>Planctomycetales>Planctomycetaceae | GCA_003531135.1 | Planctomycetaceae_bacterium | DIW81_11820 | 14698..15884 | 2026779 | BREX-system-P-loop-protein-BrxC-[Planctomycetaceae-bacterium] |
| MBA4188926.1 | Type1.BREX | <-Sulfatase+DUF4994||GDResolvase+HIN-HTH-><-HTH_3||?->?-><-?||BrxA-DNA-binding->BrxB-BrxF-iSTAND->DUF6079*->TOPRIM->PglZ->SIGMA+OB-Fold+AAA-ATPase+Lon-Protease-BrxL-><-?<-?<-GDResolvase<-Sulfatase||PAS+NtrC-AAA+FIS-HTH-> | 1142 | Bacteria>PVC_group>Planctomycetota>Planctomycetia>Planctomycetales>Planctomycetaceae | GCA_013823135.1 | Planctomycetaceae_bacterium | C0467_13090 | 97209..100637 | 2026779 | BREX-system-P-loop-protein-BrxC-[Planctomycetaceae-bacterium] |
| MCA9037354.1 | Type1.BREX | MULE-TRANSPOSASE->?->BrxA-DNA-binding->BrxB-BrxF-iSTAND->DUF6079*->N6-METHYLASE-p-> | 1154 | Bacteria>PVC_group>Planctomycetota>Planctomycetia>Planctomycetales>Planctomycetaceae | GCA_020426345.1 | Planctomycetaceae_bacterium | KDA91_19595 | 2587..6051 | 2026779 | BREX-system-P-loop-protein-BrxC-[Planctomycetaceae-bacterium] |
| MCR9202528.1 | Type1.BREX | <-SF1-SF2-Helicase+wHTH+Small-B-barrel+DUF1998<-BrxD-AAA<-PglZ<-N6-METHYLASE<-DUF6079* | 1218 | Bacteria>PVC_group>Planctomycetota>Planctomycetia>Planctomycetales>Planctomycetaceae | GCA_024742875.1 | Planctomycetaceae_bacterium | NXI04_28130 | 14549..18205 | 2026779 | BREX-system-P-loop-protein-BrxC-[Planctomycetaceae-bacterium] |
| GIX04293.1 | Type1.BREX | <-SIGMA+OB-Fold+AAA-ATPase+Lon-Protease-BrxL<-PglZ<-N6-METHYLASE<-REase<-DUF6079*<-BrxB-BrxF-iSTAND<-BrxA-DNA-binding<-XIS-HTH<-SF1-SF2-Helicase | 1221 | Bacteria>PVC_group>Planctomycetota>Planctomycetia>Planctomycetales>Planctomycetaceae | GCA_026004855.1 | Planctomycetaceae_bacterium | KatS3mg114_0162 | 186824..190489 | 2026779 | MAG:-hypothetical-protein-KatS3mg114_0162-[Planctomycetaceae-bacterium] |
| WP_145192991.1 | Type1.BREX | <-SIGMA+OB-Fold+AAA-ATPase+Lon-Protease-BrxL<-PglZ<-?<-NTP_transf<-N6-METHYLASE<-HEPN<-DUF6079*<-BrxB-BrxF-iSTAND<-BrxA-DNA-binding<-XIS-HTH||ABhydrolase-><-?<-?<-SIGMA-HTH||GDResolvase+HIN-HTH-><-SIR2 | 1220 | Bacteria>PVC_group>Planctomycetota>Planctomycetia>Planctomycetales>Planctomycetaceae | GCF_007744075.1 | Gimesia_chilikensis | HG66A1_RS30060 | 7903021..7906683 | 2605989 | BREX-system-P-loop-protein-BrxC-[Gimesia-chilikensis] |
| HBI47071.1 | Type1.BREX | <-HTH_3<-?<-TOPRIM+Primase<-?<-TOPRIM<-XIS-HTH<-?||BrxA-DNA-binding->BrxB-BrxF-iSTAND->DUF6079*->?->N6-METHYLASE->OLD-ABC->TOPRIM->PglZ->SIGMA+OB-Fold+AAA-ATPase+Lon-Protease-BrxL-><-?<-HTH_Tnp | 1147 | Bacteria>PVC_group>Planctomycetota>Planctomycetia>Planctomycetales>unclassified_Planctomycetales | GCA_003508275.1 | Planctomycetales_bacterium | DDY78_30090 | 20346..23789 | 2053591 | BREX-system-P-loop-protein-BrxC-[Planctomycetales-bacterium] |
| MCA9126638.1 | Type1.BREX | SKI->?-><-?<-?<-?<-SMODS||BrxA-DNA-binding->BrxB-BrxF-iSTAND->DUF6079*->Schlafen+HATPase+wHTH->N6-METHYLASE->?->DOC+HTH->OLD-ABC->TOPRIM->PglZ->SIGMA+OB-Fold+AAA-ATPase+Lon-Protease-BrxL->ANK+ANK+ANK-> | 1149 | Bacteria>PVC_group>Planctomycetota>Planctomycetia>Planctomycetales>unclassified_Planctomycetales | GCA_020426625.1 | Planctomycetales_bacterium | KDB22_06125 | 7854..11303 | 2053591 | BREX-system-P-loop-protein-BrxC-[Planctomycetales-bacterium] |
| GDX95758.1 | Type1.BREX | <-35exo+T7DNApol<-?<-?<-?<-?<-?||BrxA-DNA-binding->BrxB-BrxF-iSTAND->DUF6079*->N6-METHYLASE->PglZ->SIGMA+OB-Fold+AAA-ATPase+Lon-Protease-BrxL->SIR2+SIR2-> | 1161 | Bacteria>PVC_group>Planctomycetota>Planctomycetia>unclassified_Planctomycetia | GCA_014192425.1 | Planctomycetia_bacterium | LBMAG47_14220 | 155405..158890 | 2052181 | hypothetical-protein-LBMAG47_14220-[Planctomycetia-bacterium] |
| MBE7444731.1 | Type1.BREX | REC+OmpR-HTH-><-?<-?<-?<-HEPN<-?<-?||BrxA-DNA-binding->BrxB-BrxF-iSTAND->DUF6079*-><-?<-?<-DDE_Tnp<-DDE_Tnp<-?||PglZ->PglZ-p->SIGMA+OB-Fold+AAA-ATPase+Lon-Protease-BrxL-> | 1144 | Bacteria>PVC_group>Planctomycetota>Planctomycetia>unclassified_Planctomycetia | GCA_015075005.1 | Planctomycetia_bacterium | HS132_05580 | 1084624..1088058 | 2052181 | BREX-system-P-loop-protein-BrxC-[Planctomycetia-bacterium] |
| MBL8798552.1 | Type1.BREX | <-Classical-AAA<-?<-?<-?<-?<-?||BrxA-DNA-binding->BrxB-BrxF-iSTAND->DUF6079*-> | 1139 | Bacteria>PVC_group>Planctomycetota>Planctomycetia>unclassified_Planctomycetia | GCA_016795185.1 | Planctomycetia_bacterium | JNM56_31945 | 4217..7633 | 2052181 | BREX-system-P-loop-protein-BrxC-[Planctomycetia-bacterium] |
| OQA03646.1 | Type1.BREX | HTH_7+rve+rve_3-><-?||?->BrxA-DNA-binding->BrxB-BrxF-iSTAND->DUF6079*->N6-METHYLASE->OLD-ABC->TOPRIM->PglZ->SIGMA+OB-Fold+AAA-ATPase+Lon-Protease-BrxL-> | 1145 | Bacteria>PVC_group>Planctomycetota>unclassified_Planctomycetota | GCA_002068635.1 | Planctomycetes_bacterium_ADurb.Bin401 | BWY69_00331 | 3976..7413 | 1852904 | hypothetical-protein-BWY69_00331-[Planctomycetes-bacterium-ADurb.Bin401] |
| RIK67118.1 | Type1.BREX | <-XIS-HTH<-MPTase<-?<-?<-HTH_3||ACET-><-?||BrxA-DNA-binding->BrxB-BrxF-iSTAND->DUF6079*->N6-METHYLASE->OLD-ABC->TOPRIM->PglZ->SIGMA+OB-Fold+AAA-ATPase+Lon-Protease-BrxL-><-Phage_int_SAM_1+Phage_integrase||?-><-SIGMA-HTH | 1149 | Bacteria>PVC_group>Planctomycetota>unclassified_Planctomycetota | GCA_003576905.1 | Planctomycetota_bacterium | DCC65_07820 | 141257..144706 | 2026780 | MAG:-BREX-system-P-loop-protein-BrxC-[Planctomycetota-bacterium] |
| TET34868.1 | Type1.BREX | <-HNH||?-><-?||XIS-HTH->DUF4007->LON-AAA->N6-METHYLASE->ParB+HNH->HEPN->DUF6079*->PglZ->SIGMA+OB-Fold+AAA-ATPase-BrxL->SbcC->HISKIN->wHTH+WYL->RADICAL-SAM->DEOR-HTH-><-GDResolvase+Recombinase+Zn_ribbon_recom | 1347 | Bacteria>PVC_group>Planctomycetota>unclassified_Planctomycetota | GCA_004376375.1 | Planctomycetota_bacterium | E3J72_12085 | 41890..45933 | 2026780 | MAG:-hypothetical-protein-E3J72_12085-[Planctomycetota-bacterium] |
| QDV07299.1 | Type1.BREX | <-Phage_int_SAM_1+Phage_integrase<-CHTH<-ZnR+TOPRIM+AAA-helicase+wHTH||?-><-?||Phage_integrase-><-SIGMA+OB-Fold+AAA-ATPase+Lon-Protease-BrxL<-PglZ<-N6-METHYLASE<-DUF6079*<-BrxB-BrxF-iSTAND<-BrxA-DNA-binding||REase+HTH+35exo+Small-B-barrel+SF1-SF2-Helicase->SbcC-><-HEPN||?->SIGMA-HTH->?->CR-Kinase+pi3KINASE-> | 1155 | Bacteria>PVC_group>Planctomycetota>unclassified_Planctomycetota | GCA_007751475.1 | Planctomycetes_bacterium_Poly30 | Poly30_28220 | 3862242..3865709 | 2528018 | hypothetical-protein-Poly30_28220-[Planctomycetes-bacterium-Poly30] |
| TWU10529.1 | Type1.BREX | <-SF1-SF2-Helicase+DUF1998<-SF1-SF2-Helicase<-?<-?||?-><-SF1-SF2-Helicase+REase<-Exonuc_V_gamma+REase<-SF1-SF2-Helicase+wHTH+Small-B-barrel+DUF1998<-PglZ<-DUF6079-p*<-SIR2||ABhydrolase-> | 765 | Bacteria>PVC_group>Planctomycetota>unclassified_Planctomycetota | GCA_007860105.1 | Planctomycetes_bacterium_Poly21 | Poly21_44340 | 182735..185032 | 2528014 | hypothetical-protein-Poly21_44340-[Planctomycetes-bacterium-Poly21] |
| TWU09954.1 | Type1.BREX | <-ABhydrolase<-?||?->Calcineurin->SbcC+SbcC->?->XIS-HTH->BrxA-DNA-binding->BrxB-BrxF-iSTAND->DUF6079*->ParB+HNH->N6-METHYLASE->REase->PglZ->SKI->?->SIGMA+OB-Fold+AAA-ATPase+Lon-Protease-BrxL-> | 1205 | Bacteria>PVC_group>Planctomycetota>unclassified_Planctomycetota | GCA_007860105.1 | Planctomycetes_bacterium_Poly21 | Poly21_52830 | 31284..34901 | 2528014 | hypothetical-protein-Poly21_52830-[Planctomycetes-bacterium-Poly21] |
| MBI3097943.1 | Type1.BREX | <-SIGMA+OB-Fold+AAA-ATPase+Lon-Protease-BrxL<-PglZ<-DOC+HTH<-N6-METHYLASE<-PIN<-DUF6079*<-BrxB-BrxF-iSTAND<-BrxA-DNA-binding||?->?-><-tRNA||ParA-Soj-PloopNTPase-> | 1104 | Bacteria>PVC_group>Planctomycetota>unclassified_Planctomycetota | GCA_016192525.1 | Planctomycetota_bacterium | HYY93_06795 | 72529..75843 | 2026780 | BREX-system-P-loop-protein-BrxC-[Planctomycetota-bacterium] |
| MBI3468946.1 | Type1.BREX | <-N6-METHYLASE-p<-HEPN<-DUF6079*<-BrxB-BrxF-iSTAND<-BrxA-DNA-binding||?->Amidohydro_2->?-><-Phage_integrase<-wHTH+WYL+WCX||SIGMA-HTH->Pkinase+CR-Kinase-> | 1222 | Bacteria>PVC_group>Planctomycetota>unclassified_Planctomycetota | GCA_016196255.1 | Planctomycetota_bacterium | HY000_38570 | 1768..5436 | 2026780 | BREX-system-P-loop-protein-BrxC-[Planctomycetota-bacterium] |
| MBI4584108.1 | Type1.BREX | ENDO3+NUDIX-><-?<-?<-?<-SbcC+SbcC<-Calcineurin<-SIGMA+OB-Fold+AAA-ATPase+Lon-Protease-BrxL<-PglZ<-Schlafen+HATPase+wHTH<-DUF6079*<-BrxB-BrxF-iSTAND<-BrxA-DNA-binding | 1148 | Bacteria>PVC_group>Planctomycetota>unclassified_Planctomycetota | GCA_016209605.1 | Planctomycetota_bacterium | HY717_08805 | 43440..46886 | 2026780 | BREX-system-P-loop-protein-BrxC-[Planctomycetota-bacterium] |
| MBI4586093.1 | Type1.BREX | Peptidase_M28->PALP-><-Pkinase+CR-Kinase||?-><-SIGMA+OB-Fold+AAA-ATPase+Lon-Protease-BrxL<-PglZ<-Schlafen+HATPase+wHTH+HTH<-N6-METHYLASE<-DUF6079*<-BrxB-BrxF-iSTAND<-BrxA-DNA-binding||OLD-ABC->TOPRIM->TetR-HTH->?->ABC-ATPase+SbcC-> | 1226 | Bacteria>PVC_group>Planctomycetota>unclassified_Planctomycetota | GCA_016209605.1 | Planctomycetota_bacterium | HY717_18950 | 18652..22332 | 2026780 | BREX-system-P-loop-protein-BrxC-[Planctomycetota-bacterium] |
| MBK9385761.1 | Type1.BREX | <-CASPASE||Pkinase+CR-Kinase->Sigma70_r2+Sigma70_r4_2-><-TPR+TPR+TPR<-?||BrxA-DNA-binding->BrxB-BrxF-iSTAND->DUF6079*->N6-METHYLASE->SIR2->HerA/FtsK->Schlafen+HATPase+wHTH->PglZ->SIGMA+OB-Fold+AAA-ATPase+Lon-Protease-BrxL->?->?->LZ_Tnp_IS66+DDE_Tnp_IS66+DDE_Tnp_IS66_C-> | 1149 | Bacteria>PVC_group>Planctomycetota>unclassified_Planctomycetota | GCA_016719295.1 | Planctomycetota_bacterium | IPN34_13205 | 3513017..3516466 | 2026780 | BREX-system-P-loop-protein-BrxC-[Planctomycetota-bacterium] |
| MBL1217602.1 | Type1.BREX | BrxA-DNA-binding->BrxB-BrxF-iSTAND->DUF6079*->HEPN->N6-METHYLASE->RNase->PglZ->SIGMA+OB-Fold+AAA-ATPase+Lon-Protease-BrxL-><-?<-?<-Phage_integrase | 1165 | Bacteria>PVC_group>Planctomycetota>unclassified_Planctomycetota | GCA_016743455.1 | Planctomycetota_bacterium | D8M59_08910 | 1591..5088 | 2026780 | BREX-system-P-loop-protein-BrxC-[Planctomycetota-bacterium] |
| MBL4771871.1 | Type1.BREX | <-Schlafen+HATPase+wHTH<-N6-METHYLASE<-DUF6079*<-BrxB-BrxF-iSTAND<-BrxA-DNA-binding<-XIS-HTH||?-><-?||HTH_3-><-Phage_integrase<-tRNA||FGS-> | 1192 | Bacteria>PVC_group>Planctomycetota>unclassified_Planctomycetota | GCA_016764635.1 | Planctomycetota_bacterium | JKY61_12205 | 4696..8274 | 2026780 | BREX-system-P-loop-protein-BrxC-[Planctomycetota-bacterium] |
| MBL8695658.1 | Type1.BREX | BrxB-BrxF-iSTAND->?->RADICAL-SAM->BrxA-DNA-binding->BrxB-BrxF-iSTAND->DUF6079*->N6-METHYLASE->PglZ->SIGMA+OB-Fold+AAA-ATPase+Lon-Protease-BrxL->Pkinase-> | 1150 | Bacteria>PVC_group>Planctomycetota>unclassified_Planctomycetota | GCA_016792805.1 | Planctomycetota_bacterium | JNJ88_16320 | 208992..212444 | 2026780 | BREX-system-P-loop-protein-BrxC-[Planctomycetota-bacterium] |
| MBM4067775.1 | Type1.BREX | <-HKD-DNase<-ABhydrolase<-REase<-?<-HEPN<-?<-ABhydrolase||BrxA-DNA-binding->BrxB-BrxF-iSTAND->DUF6079*->RNase->N6-METHYLASE-p-> | 1193 | Bacteria>PVC_group>Planctomycetota>unclassified_Planctomycetota | GCA_016873095.1 | Planctomycetota_bacterium | FJ271_02350 | 79784..83365 | 2026780 | BREX-system-P-loop-protein-BrxC-[Planctomycetota-bacterium] |
| MBW3538721.1 | Type1.BREX | <-SIGMA+OB-Fold+AAA-ATPase+Lon-Protease-BrxL<-PglZ<-TOPRIM<-OLD-ABC<-?<-N6-METHYLASE<-N6-METHYLASE-p||?-><-DUF6079* | 1230 | Bacteria>PVC_group>Planctomycetota>unclassified_Planctomycetota | GCA_019347905.1 | Planctomycetota_bacterium | KY476_00475 | 36012..39705 | 2026780 | BREX-system-P-loop-protein-BrxC-[Planctomycetota-bacterium] |
| MBW8002871.1 | Type1.BREX | <-Phage_int_SAM_1+Phage_integrase<-SIGMA+OB-Fold+AAA-ATPase+Lon-Protease-BrxL<-PglZ<-Schlafen+HATPase+wHTH<-NACHT<-?<-N6-METHYLASE<-DUF6079*<-BrxB-BrxF-iSTAND<-BrxA-DNA-binding<-XIS-HTH | 1214 | Bacteria>PVC_group>Planctomycetota>unclassified_Planctomycetota | GCA_019456255.1 | Planctomycetota_bacterium | FVQ80_12760 | 16380..20024 | 2026780 | BREX-system-P-loop-protein-BrxC-[Planctomycetota-bacterium] |
| MBW8002946.1 | Type1.BREX | BrxA-DNA-binding->BrxB-BrxF-iSTAND->DUF6079*->N6-METHYLASE->Ntox45->HEPN->PglZ->SIGMA+OB-Fold+AAA-ATPase+HKD-DNase-BrxL-><-HEPN<-DDE_Tnp_ISAZ013<-DDE_Tnp_ISAZ013 | 1164 | Bacteria>PVC_group>Planctomycetota>unclassified_Planctomycetota | GCA_019456255.1 | Planctomycetota_bacterium | FVQ80_13155 | 4626..8120 | 2026780 | BREX-system-P-loop-protein-BrxC-[Planctomycetota-bacterium] |
| MCC6785237.1 | Type1.BREX | <-RAMPs<-?<-?<-RAMPs<-HEPN<-Schlafen+HATPase+wHTH<-N6-METHYLASE<-SbcC<-DUF6079*<-BrxB-BrxF-iSTAND<-BrxA-DNA-binding | 1150 | Bacteria>PVC_group>Planctomycetota>unclassified_Planctomycetota | GCA_020848735.1 | Planctomycetota_bacterium | IT457_20475 | 22318..25770 | 2026780 | BREX-system-P-loop-protein-BrxC-[Planctomycetota-bacterium] |

| Protein-ID | System | Gene-Neighbourhoods | Length | Taxonomy | Assembly-ID | Strain | Locus-Tag | Genomic-coordinates | TaxID | Fasta-defline |
|---|---|---|---|---|---|---|---|---|---|---|
| MCC7315424.1 | Type1.BREX | <-SIGMA+OB-Fold+AAA-ATPase+Lon-Protease-BrxL<-PglZ<-ArdCN+MPTase<-N6-METHYLASE<-?<-TIR<-DUF6079*<-BrxB-BrxF-iSTAND<-BrxA-DNA-binding\|\|HTH_23+HTH_32-><-Peptidase_S8<-Classical-AAA<-BirA-HTH<-?\|\|MarR-HTH->Peptidase_S74-> | 1147 | Bacteria>PVC_group>Planctomycetota>unclassified_Planctomycetota | GCA_020853195.1 | Planctomycetota_bacterium | IT419_11445 | 15860..19303 | 2026780 | BREX-system-P-loop-protein-BrxC-[Planctomycetota-bacterium] |
| MCD6405191.1 | Type1.BREX | <-Schlafen+HATPase+wHTH-p<-PHP+SbcC<-SIGMA+OB-Fold+AAA-ATPase+Lon-Protease-BrxL<-PglZ<-ParB+HNH<-N6-METHYLASE<-DUF6079*<-BrxB-BrxF-iSTAND<-BrxA-DNA-binding<-XIS-HTH | 1143 | Bacteria>PVC_group>Planctomycetota>unclassified_Planctomycetota | GCA_021160625.1 | Planctomycetota_bacterium | J7M19_05155 | 12563..15994 | 2026780 | BREX-system-P-loop-protein-BrxC-[Planctomycetota-bacterium] |
| MCE9589095.1 | Type1.BREX | <-XIS-HTH<-XIS-HTH<-?<-URI-DNase+RAMA<-SIGMA+OB-Fold+AAA-ATPase+Lon-Protease-BrxL<-PglZ<-REase+AIPR<-N6-METHYLASE<-DUF6079*<-BrxB-BrxF-iSTAND<-BrxA-DNA-binding<-DDE_Tnp | 1146 | Bacteria>PVC_group>Planctomycetota>unclassified_Planctomycetota | GCA_021413865.1 | Planctomycetota_bacterium | K8S99_01060 | 75553..78993 | 2026780 | BREX-system-P-loop-protein-BrxC-[Planctomycetota-bacterium] |
| MCL5279432.1 | Type1.BREX | DUF6079-p*->N6-METHYLASE->AbiEi_4->AbiEii->?->PglZ->SIGMA+OB-Fold+AAA-ATPase+Lon-Protease-BrxL->TIR-> | 661 | Bacteria>PVC_group>Planctomycetota>unclassified_Planctomycetota | GCA_023481165.1 | Planctomycetota_bacterium | M1376_05960 | 1..1987 | 2026780 | BREX-system-P-loop-protein-BrxC-[Planctomycetota-bacterium] |
| AHF94674.1 | Type1.BREX | <-PglZ<-TOPRIM<-OLD-ABC<-PIWI<-REase+PNTD1+PNTD2<-N6-METHYLASE-p<-DOC+HTH<-N6-METHYLASE<-?<-DUF6079*<-BrxB-BrxF-iSTAND<-BrxA-DNA-binding<-?\|\|TrwC+SF1-SF2-Helicase->?->?->ClpABC-AAA-> | 1248 | Bacteria>PVC_group>Verrucomicrobiota>Opitutae>Opitutales>Opitutaceae | GCA_000242935.3 | Opitutaceae_bacterium_TAV5 | OPIT5_23275 | 5485357..5489103 | 794903 | hypothetical-protein-OPIT5_23275-[Opitutaceae-bacterium-TAV5] |
| MBC7388090.1 | Type1.BREX | DUF6079-p*->REase->N6-METHYLASE->?->PglZ->SIGMA+OB-Fold+AAA-ATPase+Lon-Protease-BrxL-> | 828 | Bacteria>PVC_group>Verrucomicrobiota>Opitutae>Opitutales>Opitutaceae | GCA_014376535.1 | Opitutaceae_bacterium | H7329_02660 | 1..2487 | 2006848 | hypothetical-protein-[Opitutaceae-bacterium] |
| MBX3736102.1 | Type1.BREX | GDResolvase+Recombinase+ZnR-><-HTH_3<-Couple_hipA+HipA_C<-HTH_3\|\|BrxA-DNA-binding->BrxB-BrxF-iSTAND->DUF6079*->TIR->N6-METHYLASE->?->TIR->TIR->PglZ->SIGMA+OB-Fold+AAA-ATPase+Lon-Protease-BrxL-> | 1149 | Bacteria>PVC_group>Verrucomicrobiota>Opitutae>Opitutales>Opitutaceae | GCA_019634575.1 | Candidatus_Didemnitutus_sp. | KF715_05395 | 1290269..1293718 | 2814259 | BREX-system-P-loop-protein-BrxC-[Candidatus-Didemnitutus-sp.] |
| MCC5808698.1 | Type1.BREX | Phage_integrase-><-SIGMA+OB-Fold+AAA-ATPase+HKD-DNase-BrxL<-PglZ<-HNH<-OLD-ABC<-N6-METHYLASE<-REase<-OLD-ABC<-DUF6079*<-BrxB-BrxF-iSTAND<-BrxA-DNA-binding<-XIS-HTH | 1183 | Bacteria>PVC_group>Verrucomicrobiota>Opitutae>Opitutales>unclassified_Opitutales | GCA_020831525.1 | Opitutales_bacterium | JJU00_20390 | 12178..15729 | 2026772 | BREX-system-P-loop-protein-BrxC-[Opitutales-bacterium] |
| MCH8474945.1 | Type1.BREX | <-DUF4236+TPR\|\|tRNA->BrxA-DNA-binding->BrxB-BrxF-iSTAND->DUF6079*->SF1-SF2-Helicase->?->?->N6-METHYLASE->N6-METHYLASE-p->SbcC->Schlafen+HATPase+wHTH->MarR-HTH->Polbeta-> | 1147 | Bacteria>PVC_group>Verrucomicrobiota>Opitutae>Opitutales>unclassified_Opitutales | GCA_022565335.1 | Opitutales_bacterium | LAT55_06930 | 5092..8535 | 2026772 | BREX-system-P-loop-protein-BrxC-[Opitutales-bacterium] |
| MBT62430.1 | Type1.BREX | <-KAP-NTPase<-N6-METHYLASE-p<-DDE_Tnp<-N6-METHYLASE-p<-RVT<-N6-METHYLASE-p<-?\|\|?-><-Schlafen+HATPase+wHTH<-DUF6079*<-BrxB-BrxF-iSTAND<-BrxA-DNA-binding<-HTH_3 | 1165 | Bacteria>PVC_group>Verrucomicrobiota>Opitutae>Puniceicoccales>Puniceicoccaceae | GCA_002729725.1 | Puniceicoccaceae_bacterium | CML13_04370 | 15109..18606 | 2026784 | hypothetical-protein-[Puniceicoccaceae-bacterium] |
| MBU3665708.1 | Type1.BREX | ABhydrolase-><-?<-?<-SIGMA+OB-Fold+AAA-ATPase+HKD-DNase-BrxL<-PglZ<-KAP-NTPase<-N6-METHYLASE<-Polbeta<-MarR-HTH<-DUF6079*<-BrxB-BrxF-iSTAND<-BrxA-DNA-binding<-ZnR<-?<-?<-?<-?<-VIT+VWA<-MerR-HTH | 1169 | Bacteria>PVC_group>Verrucomicrobiota>Spartobacteria>Chthoniobacterales>unclassified_Chthoniobacterales | GCA_018883245.1 | Chthoniobacterales_bacterium | FGM15_07520 | 29744..33253 | 2201154 | BREX-system-P-loop-protein-BrxC-[Chthoniobacterales-bacterium] |
| PTY02769.1 | Type1.BREX | RAMA->GDResolvase+ZnR->PglZ->SIGMA+OB-Fold+AAA-ATPase+HKD-DNase-BrxL-><-N6-METHYLASE<-Polbeta<-MarR-HTH<-DUF6079*<-BrxB-BrxF-iSTAND<-BrxA-DNA-binding\|\|DUF4096+DDE_Tnp_1+DDE_Tnp_1_2-> | 1172 | Bacteria>PVC_group>Verrucomicrobiota>unclassified_Verrucomicrobiota | GCA_003054695.1 | Verrucomicrobia_bacterium_LW23 | DB346_08320 | 46382..49900 | 2161867 | BREX-system-P-loop-protein-BrxC-[Verrucomicrobia-bacterium-LW23] |
| HCF96075.1 | Type1.BREX | <-N6-METHYLASE-p<-N6-METHYLASE<-REase<-OLD-ABC<-DUF6079-p<-DUF6079-p*<-DUF6079-p<-DUF6079-p<-DUF6079-p<-BrxB-BrxF-iSTAND<-BrxA-DNA-binding | 268 | Bacteria>PVC_group>Verrucomicrobiota>unclassified_Verrucomicrobiota | GCA_003534995.1 | Verrucomicrobiota_bacterium | DEW46_13535 | 6770..7576 | 2026799 | BREX-system-P-loop-protein-BrxC-[Verrucomicrobiota-bacterium] |
| HCF96078.1 | Type1.BREX | <-N6-METHYLASE-p<-N6-METHYLASE<-REase<-OLD-ABC<-DUF6079-p<-DUF6079-p<-DUF6079-p<-DUF6079-p<-DUF6079-p*<-BrxB-BrxF-iSTAND<-BrxA-DNA-binding | 531 | Bacteria>PVC_group>Verrucomicrobiota>unclassified_Verrucomicrobiota | GCA_003534995.1 | Verrucomicrobiota_bacterium | DEW46_13550 | 8732..10325 | 2026799 | BREX-system-P-loop-protein-BrxC-[Verrucomicrobiota-bacterium] |
| KAB2639241.1 | Type1.BREX | <-Y1_Tnp<-N6-METHYLASE-p<-Four-helical-protein<-Schlafen+HATPase+wHTH<-DUF6079*<-BrxB-BrxF-iSTAND<-BrxA-DNA-binding | 1168 | Bacteria>PVC_group>Verrucomicrobiota>unclassified_Verrucomicrobiota | GCA_008932935.1 | Verrucomicrobiota_bacterium | DVB25_06025 | 8272..11778 | 2026799 | BREX-system-P-loop-protein-BrxC-[Verrucomicrobiota-bacterium] |
| NBB80453.1 | Type1.BREX | HTH_3->BrxA-DNA-binding->BrxB-BrxF-iSTAND->DUF6079*->OLD-ABC->PIN->N6-METHYLASE->OLD-ABC->HNH->ParB+HNH->Schlafen+HATPase+wHTH->PglZ->SIGMA+OB-Fold+AAA-ATPase+Lon-Protease-BrxL-> | 1147 | Bacteria>PVC_group>Verrucomicrobiota>unclassified_Verrucomicrobiota | GCA_009908325.1 | Verrucomicrobiota_bacterium | GVY36_13595 | 2227..5670 | 2026799 | BREX-system-P-loop-protein-BrxC-[Verrucomicrobiota-bacterium] |
| MBI4023965.1 | Type1.BREX | CBS->BrxA-DNA-binding->BrxB-BrxF-iSTAND->DUF6079*->HEPN->N6-METHYLASE->REase+AIPR->PglZ->SIGMA+OB-Fold+AAA-ATPase+Lon-Protease-BrxL-> | 1148 | Bacteria>PVC_group>Verrucomicrobiota>unclassified_Verrucomicrobiota | GCA_016198745.1 | Verrucomicrobiota_bacterium | HY360_03230 | 10391..13837 | 2026799 | BREX-system-P-loop-protein-BrxC-[Verrucomicrobiota-bacterium] |
| MBI4624648.1 | Type1.BREX | <-PadR-HTH\|\|?->?->?->?->HTH_Tnp->BrxA-DNA-binding->BrxB-BrxF-iSTAND->DUF6079*->OLD-ABC->TOPRIM-> | 1155 | Bacteria>PVC_group>Verrucomicrobiota>unclassified_Verrucomicrobiota | GCA_016209255.1 | Verrucomicrobiota_bacterium | HY736_15695 | 9824..13291 | 2026799 | BREX-system-P-loop-protein-BrxC-[Verrucomicrobiota-bacterium] |
| MBM3880197.1 | Type1.BREX | XIS-HTH->BrxA-DNA-binding->BrxB-BrxF-iSTAND->DUF6079*->N6-METHYLASE->SbcC->PIN->?->DOC+HTH->N6-METHYLASE-p->OLD-ABC->TOPRIM->PglZ-> | 1246 | Bacteria>PVC_group>Verrucomicrobiota>unclassified_Verrucomicrobiota | GCA_016871715.1 | Verrucomicrobiota_bacterium | FJ387_10840 | 14385..18125 | 2026799 | BREX-system-P-loop-protein-BrxC-[Verrucomicrobiota-bacterium] |
| MBN9688605.1 | Type1.BREX | <-SIGMA+OB-Fold+AAA-ATPase+Lon-Protease-BrxL<-PglZ<-Schlafen+HATPase+wHTH<-PHP+SbcC<-N6-METHYLASE<-TIR<-?<-SMODS<-DUF6079*<-BrxB-BrxF-iSTAND<-BrxA-DNA-binding<-TPR<-Peptidase\|\|?-><-wHTH+HNH<-DOC+HTH<-?<-HTH+REase | 1142 | Bacteria>PVC_group>Verrucomicrobiota>unclassified_Verrucomicrobiota | GCA_017309115.1 | Verrucomicrobiota_bacterium | JOM24_00070 | 18647..22075 | 2026799 | BREX-system-P-loop-protein-BrxC-[Verrucomicrobiota-bacterium] |
| MBP7946078.1 | Type1.BREX | <-N6-METHYLASE<-Schlafen<-DUF6079*<-BrxB-BrxF-iSTAND<-BrxA-DNA-binding<-?<-?<-?<-?\|\|?->?-><-TPR+TPR+TPR+TPR+TPR | 1195 | Bacteria>PVC_group>Verrucomicrobiota>unclassified_Verrucomicrobiota | GCA_018003215.1 | Verrucomicrobiota_bacterium | KA191_08410 | 4464..8051 | 2026799 | BREX-system-P-loop-protein-BrxC-[Verrucomicrobiota-bacterium] |
| MBP7946757.1 | Type1.BREX | AbiEi_4+AbiEi_1->AbiEii+ZnR->?-><-DDE_Tnp\|\|?->?->?->BrxA-DNA-binding->BrxB-BrxF-iSTAND->DUF6079*->N6-METHYLASE-> | 1214 | Bacteria>PVC_group>Verrucomicrobiota>unclassified_Verrucomicrobiota | GCA_018003215.1 | Verrucomicrobiota_bacterium | KA191_11880 | 29346..32990 | 2026799 | BREX-system-P-loop-protein-BrxC-[Verrucomicrobiota-bacterium] |
| MCC5847925.1 | Type1.BREX | XIS-HTH->AAA-ATPase->?-><-SIGMA+OB-Fold+AAA-ATPase+Lon-Protease-BrxL<-PglZ<-nSTAND<-N6-METHYLASE<-TIR<-Schlafen+HATPase+wHTH<-DUF6079*<-BrxB-BrxF-iSTAND<-BrxA-DNA-binding\|\|?-><-?<-Phage_integrase\|\|Peptidase-> | 1154 | Bacteria>PVC_group>Verrucomicrobiota>unclassified_Verrucomicrobiota | GCA_020831635.1 | Verrucomicrobiota_bacterium | JJU29_07500 | 21995..25459 | 2026799 | BREX-system-P-loop-protein-BrxC-[Verrucomicrobiota-bacterium] |
| MCF7764832.1 | Type1.BREX | <-REC+NtrC-AAA+FIS-HTH<-HISKIN\|\|?->?-><-?<-?\|\|tRNA->?->BrxA-DNA-binding->BrxB-BrxF-iSTAND->DUF6079*->MarR-HTH->Polbeta->N6-METHYLASE-p-> | 1167 | Bacteria>PVC_group>Verrucomicrobiota>unclassified_Verrucomicrobiota | GCA_021733945.1 | Verrucomicrobiota_bacterium | K9N62_14270 | 13767..17270 | 2026799 | BREX-system-P-loop-protein-BrxC-[Verrucomicrobiota-bacterium] |
| MCU0772111.1 | Type1.BREX | <-SIGMA+OB-Fold+AAA-ATPase+Lon-Protease-BrxL<-PglZ<-?<-N6-METHYLASE<-RVT<-DUF6079*<-BrxB-BrxF-iSTAND<-BrxA-DNA-binding\|\|?->?-><-NACHT<-?<-?\|\|?->S1+GTPase-AIG-> | 1144 | Bacteria>PVC_group>Verrucomicrobiota>unclassified_Verrucomicrobiota | GCA_025457615.1 | Verrucomicrobiota_bacterium | MUE94_10150 | 10422..13856 | 2026799 | BREX-system-P-loop-protein-BrxC-[Verrucomicrobiota-bacterium] |
| SPE51781.1 | Type1.BREX | <-SIGMA+OB-Fold+AAA-ATPase+Lon-Protease-BrxL<-PglZ-p<-PglZ<-HTH+REase<-N6-METHYLASE<-?<-SMODS<-DUF6079*<-BrxB-BrxF-iSTAND<-BrxB-BrxF-iSTAND<-BrxA-DNA-binding | 1149 | Bacteria>PVC_group>Verrucomicrobiota>unclassified_Verrucomicrobiota | GCA_900290355.1 | Verrucomicrobiota_bacterium | SBV1_1420061 | 72835..76284 | 2026799 | putative-P-loop-containing-nucleoside-triphosphate-hydrolase-[Verrucomicrobiota-bacterium] |
| PAW64461.1 | Type1.BREX | BrxB-BrxF-iSTAND->DUF6079*->N6-METHYLASE->HTH+REase->Calcineurin-SbcD->SbcC+CCDC158-> | 1145 | Bacteria>PVC_group>Verrucomicrobiota>Verrucomicrobiae>unclassified_Verrucomicrobiae | GCA_002304315.1 | Verrucomicrobiae_bacterium_Tous-C4TDCM | B9S38_16345 | 386..3823 | 1982332 | MAG:-hypothetical-protein-B9S38_16345-[Verrucomicrobiae-bacterium-Tous-C4TDCM] |
| MCB1062255.1 | Type1.BREX | <-N6-METHYLASE-p<-DUF6079*<-BrxB-BrxF-iSTAND<-BrxA-DNA-binding<-BrxA-DNA-binding<-?\|\|MODE-HTH+PBPII-> | 1172 | Bacteria>PVC_group>Verrucomicrobiota>Verrucomicrobiae>unclassified_Verrucomicrobiae | GCA_020440025.1 | Verrucomicrobiae_bacterium | KDN20_04945 | 1016..4534 | 2499141 | BREX-system-P-loop-protein-BrxC-[Verrucomicrobiae-bacterium] |
| MBE6418907.1 | Type1.BREX | <-PglZ<-N6-METHYLASE<-DUF6079*<-BrxB-BrxF-iSTAND<-BrxA-DNA-binding\|\|?-><-ACET\|\|BirA-HTH->?->Glycos_transf_1+Glyco_trans_1_2-> | 1189 | Bacteria>PVC_group>Verrucomicrobiota>Verrucomicrobiae>Verrucomicrobiales>Akkermansiaceae | GCA_015062055.1 | Akkermansiaceae_bacterium | E7031_02125 | 17805..21374 | 2562705 | BREX-system-P-loop-protein-BrxC-[Akkermansiaceae-bacterium] |
| MBQ8375884.1 | Type1.BREX | <-SF1-SF2-Helicase+DUF3418\|\|?->Calcineurin->?->HAD+HAD->tRNA->tRNA->TRPR-HTH->STYKinase-like->BrxA-DNA-binding->BrxB-BrxF-iSTAND->DUF6079*->N6-METHYLASE->PglZ->SIGMA+OB-Fold+AAA-ATPase+Lon-Protease-BrxL->Tox-ART-HYD3-><-?<-?<-BirA-HTH<-Phage_integrase | 1185 | Bacteria>PVC_group>Verrucomicrobiota>Verrucomicrobiae>Verrucomicrobiales>Akkermansiaceae | GCA_017451815.1 | Akkermansia_sp. | IJX33_02385 | 35942..39499 | 1872421 | BREX-system-P-loop-protein-BrxC-[Akkermansia-sp.] |
| WP_215427518.1 | Type1.BREX | ANK+ANK+ANK->HNH->HTH_3->Couple_hipA->HipA_C->?->XIS-HTH->BrxA-DNA-binding->BrxB-BrxF-iSTAND->DUF6079*->N6-METHYLASE->PglZ->SIGMA+OB-Fold+AAA-ATPase+Lon-Protease-BrxL->HNH-><-ParB+HNH\|\|DUF932->PadR-HTH-> | 1141 | Bacteria>PVC_group>Verrucomicrobiota>Verrucomicrobiae>Verrucomicrobiales>Akkermansiaceae | GCF_018709685.1 | Akkermansia_muciniphila | HHJ06_RS03145 | 88477..91902 | 239935 | BREX-system-P-loop-protein-BrxC-[Akkermansia-muciniphila] |
| MBR01191.1 | Type1.BREX | BrxA-DNA-binding->BrxB-BrxF-iSTAND->DUF6079*->N6-METHYLASE->PglZ->REase+iREase-><-SF1-SF2-Helicase | 1162 | Bacteria>PVC_group>Verrucomicrobiota>Verrucomicrobiae>Verrucomicrobiales>unclassified_Verrucomicrobiales | GCA_002729295.1 | Verrucomicrobiales_bacterium | CMD60_10085 | 1798..5286 | 2026801 | hypothetical-protein-[Verrucomicrobiales-bacterium] |
| MBL9158389.1 | Type1.BREX | <-SIGMA+OB-Fold+AAA-ATPase+Lon-Protease-BrxL<-PglZ<-HTH+REase<-N6-METHYLASE-p<-SbcC+ABC-ATPase<-N6-METHYLASE<-TIR<-DUF6079*<-BrxB-BrxF-iSTAND<-BrxA-DNA-binding\|\|?->?->?->?->XIS-HTH->RVT-> | 1152 | Bacteria>PVC_group>Verrucomicrobiota>Verrucomicrobiae>Verrucomicrobiales>unclassified_Verrucomicrobiales | GCA_016795385.1 | Verrucomicrobiales_bacterium | JNJ70_13005 | 50985..54443 | 2026801 | BREX-system-P-loop-protein-BrxC-[Verrucomicrobiales-bacterium] |
| MCI0536692.1 | Type1.BREX | TIR-><-?\|\|BrxA-DNA-binding->BrxB-BrxF-iSTAND->DUF6079*->OLD-ABC->TOPRIM->N6-METHYLASE-p-> | 1149 | Bacteria>PVC_group>Verrucomicrobiota>Verrucomicrobiae>Verrucomicrobiales>unclassified_Verrucomicrobiales | GCA_022615925.1 | Verrucomicrobiales_bacterium | LOZ50_15855 | 3424..6873 | 2026801 | BREX-system-P-loop-protein-BrxC-[Verrucomicrobiales-bacterium] |
| MBL9184319.1 | Type1.BREX | DUF932->BrxA-DNA-binding->BrxB-BrxF-iSTAND->DUF6079*->PIN->TIR->N6-METHYLASE->?->PglZ->SIGMA+OB-Fold+AAA-ATPase+Lon-Protease-BrxL->REase+PNTD1+PNTD2->PIWI-> | 1153 | Bacteria>PVC_group>Verrucomicrobiota>Verrucomicrobiae>Verrucomicrobiales>Verrucomicrobiaceae | GCA_016795465.1 | Verrucomicrobiaceae_bacterium | JNN17_19395 | 2113..5574 | 2026800 | BREX-system-P-loop-protein-BrxC-[Verrucomicrobiaceae-bacterium] |
| MCF7788313.1 | Type1.BREX | BrxA-DNA-binding->BrxB-BrxF-iSTAND->DUF6079*->HKD-DNase+DUF3150->MarR-HTH->Polbeta->N6-METHYLASE->?->SbcC->PglZ-p->REase-> | 1167 | Bacteria>PVC_group>Verrucomicrobiota>Verrucomicrobiae>Verrucomicrobiales>Verrucomicrobiaceae | GCA_021733925.1 | Prosthecobacter_sp. | K9N47_19475 | 2530..6033 | 1965333 | BREX-system-P-loop-protein-BrxC-[Prosthecobacter-sp.] |
| MCH7225050.1 | Type1.BREX | <-SIGMA+OB-Fold+AAA-ATPase+HKD-DNase-BrxL<-PglZ<-URI-DNase<-ZnR<-?<-N6-METHYLASE<-?<-DUF6079*<-BrxB-BrxF-iSTAND<-BrxA-DNA-binding<-TPR<-Peptidase\|\|?->?->TPR+TPR+TPR+TPR+TPR-> | 1166 | Bacteria>PVC_group>Verrucomicrobiota>Verrucomicrobiae>Verrucomicrobiales>Verrucomicrobiaceae | GCA_022548915.1 | Verrucomicrobiaceae_bacterium_E54 | MLD59_02825 | 188898..192398 | 2922395 | BREX-system-P-loop-protein-BrxC-[Verrucomicrobiaceae-bacterium-E54] |
| WP_035610953.1 | Type1.BREX | <-SIGMA+OB-Fold+AAA-ATPase+Lon-Protease-BrxL<-PglZ<-N6-METHYLASE<-DUF6079*<-BrxB-BrxF-iSTAND<-BrxA-DNA-binding<-tRNA\|\|N6-METHYLASE->HTH_3-> | 1145 | Bacteria>PVC_group>Verrucomicrobiota>Verrucomicrobiae>Verrucomicrobiales>Verrucomicrobiaceae | GCF_000739615.1 | Haloferula_sp._BvORR071 | BV071_RS19610 | 47391..50828 | 1396141 | BREX-system-P-loop-protein-BrxC-[Haloferula-sp.-BvORR071] |
| TGL58697.1 | Type1.BREX | <-N6-METHYLASE<-DUF6079*<-BrxB-BrxF-iSTAND<-BrxA-DNA-binding<-?<-Classical-AAA<-?<-?<-?\|\|?->ParA-Soj-PloopNTPase-> | 1155 | Bacteria>Spirochaetota>Spirochaetia>Leptospirales>Leptospiraceae>Leptospira | GCA_004770745.1 | Leptospira_ognonensis | EHQ58_10020 | 3017..6484 | 2484945 | BREX-system-P-loop-protein-BrxC-[Leptospira-ognonensis] |
| WP_210412803.1 | Type1.BREX | <-N6-METHYLASE<-DUF6079*<-BrxB-BrxF-iSTAND<-BrxA-DNA-binding<-?<-Classical-AAA<-?<-?<-?\|\|?->ParA-Soj-PloopNTPase-> | 1157 | Bacteria>Spirochaetota>Spirochaetia>Leptospirales>Leptospiraceae>Leptospira | GCF_004770745.1 | Leptospira_ognonensis | EHQ58_RS10025 | 3017..6490 | 2484945 | BREX-system-P-loop-protein-BrxC-[Leptospira-ognonensis] |
| WP_135627473.1 | Type1.BREX | DUF6079*->N6-METHYLASE->ParB+HNH->PglZ->SIGMA+OB-Fold+AAA-ATPase+Lon-Protease-BrxL-> | 1122 | Bacteria>Spirochaetota>Spirochaetia>Leptospirales>Leptospiraceae>Leptospira | GCF_004770835.1 | Leptospira_selangorensis | EHQ82_RS10885 | 1..3369 | 2484982 | BREX-system-P-loop-protein-BrxC,-partial-[Leptospira-selangorensis] |
| MCB1192727.1 | Type1.BREX | <-SecA+SF1-SF2-Helicase<-ArdCN\|\|BrxA-DNA-binding->BrxB-BrxF-iSTAND->DUF6079*->REase+HTH->N6-METHYLASE->PglZ-> | 1148 | Bacteria>Spirochaetota>Spirochaetia>Leptospirales>Leptospiraceae>unclassified_Leptospiraceae | GCA_020440835.1 | Leptospiraceae_bacterium | KDX90_19935 | 18936..22382 | 2026757 | BREX-system-P-loop-protein-BrxC-[Leptospiraceae-bacterium] |
| MCP5494239.1 | Type1.BREX | CR-ATPase8+REase-><-SIGMA+OB-Fold+AAA-ATPase+Lon-Protease-BrxL<-?<-?<-?<-PglZ<-N6-METHYLASE<-REase+HTH<-DUF6079*<-BrxB-BrxF-iSTAND<-BrxA-DNA-binding\|\|ArdCN->SecA+SF1-SF2-Helicase-> | 1148 | Bacteria>Spirochaetota>Spirochaetia>Leptospirales>Leptospiraceae>unclassified_Leptospiraceae | GCA_024233865.1 | Leptospiraceae_bacterium | H7A23_06750 | 342984..346430 | 2026757 | BREX-system-P-loop-protein-BrxC-[Leptospiraceae-bacterium] |
| MCH3907669.1 | Type1.BREX | GspA-ATPase+REase-><-Peptidase_S58<-RHH<-?<-?<-?\|\|XIS-HTH->BrxA-DNA-binding->BrxB-BrxF-iSTAND->DUF6079*->N6-METHYLASE->Schlafen+HATPase+wHTH->PglZ->SIGMA+OB-Fold+AAA-ATPase+HKD-DNase-BrxL-><-SF1-SF2-Helicase<-SbcC<-DDE_Tnp_1 | 1157 | Bacteria>Spirochaetota>Spirochaetia>Spirochaetales>Sphaerochaetaceae>Sphaerochaeta | GCA_022482495.1 | Sphaerochaeta_sp. | LKE28_05345 | 253773..257246 | 1972642 | BREX-system-P-loop-protein-BrxC-[Sphaerochaeta-sp.] |
| WP_114628929.1 | Type1.BREX | <-Thymidylate_kin\|\|ParB+HNH->ParB+HNH-><-?<-D5N+D5ATPase\|\|XIS-HTH->BrxA-DNA-binding->BrxB-BrxF-iSTAND->DUF6079*->OLD-ABC->REase->N6-METHYLASE->SIGMA+OB-Fold+AAA-ATPase+HKD-DNase-BrxL-><-AbiEii<-AbiEi_3_N+AbiEi_3 | 1173 | Bacteria>Spirochaetota>Spirochaetia>Spirochaetales>Spirochaetaceae>Oceanispirochaeta | GCF_013166645.2 | unclassified_Oceanispirochaeta | HNV09_RS06050 | 178318..181839 | 2635722 | MULTISPECIES:-BREX-system-P-loop-protein-BrxC-[unclassified-Oceanispirochaeta] |
| OQY41116.1 | Type1.BREX | REase->BrxA-DNA-binding->BrxB-BrxF-iSTAND->DUF6079*->OLD-ABC->TOPRIM->N6-METHYLASE->ParB+HNH->ParB+HNH->SIGMA+OB-Fold+AAA-ATPase+Lon-Protease-BrxL-> | 1207 | Bacteria>Spirochaetota>Spirochaetia>Spirochaetales>Spirochaetaceae>unclassified_Spirochaetaceae | GCA_002084785.1 | Spirochaetaceae_bacterium_4572_7 | B6229_00020 | 2034..5657 | 1972456 | hypothetical-protein-B6229_00020-[Spirochaetaceae-bacterium-4572_7] |
| OQY33741.1 | Type1.BREX | <-REase<-OLD-ABC<-DUF6079*<-BrxB-BrxF-iSTAND<-BrxA-DNA-binding<-?<-?\|\|NtrC-AAA+FIS-HTH->HAMP+HisKin+HATPase_c->REC+AraC-HTH+AraC-HTH->?->ABC-ATPase+SbcC+ABC-ATPase+SbcC-> | 1173 | Bacteria>Spirochaetota>Spirochaetia>Spirochaetales>Spirochaetaceae>unclassified_Spirochaetaceae | GCA_002084805.1 | Spirochaetaceae_bacterium_4572_59 | B6241_06720 | 4557..8078 | 1971727 | hypothetical-protein-B6241_06720-[Spirochaetaceae-bacterium-4572_59] |
| TVQ28583.1 | Type1.BREX | DSBH+AraC-HTH+AraC-HTH-><-YTCD-HTH\|\|?-><-?<-Tox-RelE<-?\|\|BrxA-DNA-binding->BrxB-BrxF-iSTAND->DUF6079*->N6-METHYLASE-p-> | 1185 | Bacteria>Spirochaetota>Spirochaetia>Spirochaetales>Spirochaetaceae>unclassified_Spirochaetaceae | GCA_007694005.1 | Spirochaetaceae_bacterium | EA383_00220 | 6436..9993 | 1898206 | BREX-system-P-loop-protein-BrxC-[Spirochaetaceae-bacterium] |
| MCP5452900.1 | Type1.BREX | <-DUF6079*<-BrxB-BrxF-iSTAND<-BrxA-DNA-binding<-XIS-HTH<-?<-?<-?\|\|?->MoxR-AAA-> | 1147 | Bacteria>Spirochaetota>Spirochaetia>Spirochaetales>Spirochaetaceae>unclassified_Spirochaetaceae | - | Spirochaetaceae_bacterium | H7A27_02910 | 204..3647 | 1898206 | BREX-system-P-loop-protein-BrxC-[Spirochaetaceae-bacterium] |
| MBQ5908482.1 | Type1.BREX | <-TPR+TPR<-?\|\|BrxA-DNA-binding->BrxB-BrxF-iSTAND->DUF6079*->Schlafen+HATPase+wHTH->N6-METHYLASE->?->PglZ->SIGMA+OB-Fold+AAA-ATPase+Lon-Protease-BrxL->HTH_19-> | 1172 | Bacteria>Spirochaetota>Spirochaetia>Spirochaetales>Treponemataceae>Treponema | GCA_017381365.1 | Treponema_sp. | IIW80_10580 | 5234..8752 | 166 | BREX-system-P-loop-protein-BrxC-[Treponema-sp.] |
| MCR5123681.1 | Type1.BREX | <-N6-METHYLASE<-Schlafen+HATPase+wHTH<-DUF6079*<-BrxB-BrxF-iSTAND<-BrxA-DNA-binding<-HTH_26\|\|?->?-><-ZetaToxin<-ABC-3<-ABC-ATPase+SbcC | 1172 | Bacteria>Spirochaetota>Spirochaetia>Spirochaetales>Treponemataceae>Treponema | GCA_024697755.1 | Treponema_sp. | K6B43_00615 | 5680..9198 | 166 | BREX-system-P-loop-protein-BrxC-[Treponema-sp.] |
| NLK13330.1 | Type1.BREX | <-PglZ-p<-N6-METHYLASE<-DUF6079*<-BrxB-BrxF-iSTAND<-BrxA-DNA-binding<-?<-?\|\|SF1-SF2-Helicase->?->sGTP-> | 1160 | Bacteria>Spirochaetota>Spirochaetia>Spirochaetales>unclassified_Spirochaetales | GCA_012519115.1 | Spirochaetales_bacterium | GX313_01060 | 4407..7889 | 2026792 | BREX-system-P-loop-protein-BrxC-[Spirochaetales-bacterium] |
| NLV85368.1 | Type1.BREX | DUF6079*->N6-METHYLASE->HEPN->N6-METHYLASE-p->PglZ->SIGMA+OB-Fold+AAA-ATPase+Lon-Protease-BrxL-> | 1160 | Bacteria>Spirochaetota>Spirochaetia>Spirochaetales>unclassified_Spirochaetales | GCA_012729295.1 | Spirochaetales_bacterium | GXY60_12460 | 48..3530 | 2026792 | BREX-system-P-loop-protein-BrxC-[Spirochaetales-bacterium] |
| MBR5915376.1 | Type1.BREX | <-SH3\|\|Peptidase_M20-><-?\|\|?->REase-><-SIGMA+OB-Fold+AAA-ATPase+Lon-Protease-BrxL<-PglZ<-Schlafen+HATPase+wHTH<-N6-METHYLASE<-DUF6079*<-BrxB-BrxF-iSTAND<-BrxA-DNA-binding<-CR-ATPase5+REase<-ACET<-?<-?<-?<-HD | 1170 | Bacteria>Spirochaetota>Spirochaetia>unclassified_Spirochaetia | GCA_017554185.1 | Spirochaetia_bacterium | IXZ57_02880 | 16752..20264 | 2053615 | BREX-system-P-loop-protein-BrxC-[Spirochaetia-bacterium] |
| OQB04958.1 | Type1.BREX | <-DUF6079*<-BrxB-BrxF-iSTAND<-BrxA-DNA-binding<-XIS-HTH<-?<-wHTH+WYL+WCX\|\|?->Mrr_N+ACET-> | 1147 | Bacteria>Spirochaetota>unclassified_Spirochaetota | GCA_002070025.1 | Spirochaetes_bacterium_ADurb.Bin215 | BWY20_00149 | 30..3473 | 1852910 | hypothetical-protein-BWY20_00149-[Spirochaetes-bacterium-ADurb.Bin215] |
| RKX87882.1 | Type1.BREX | <-MarR-HTH\|\|?->tRNA->BrxA-DNA-binding->BrxB-BrxF-iSTAND->DUF6079*->PolB->HEPN->Polbeta-><-?\|\|N6-METHYLASE-p-> | 1168 | Bacteria>Spirochaetota>unclassified_Spirochaetota | GCA_003644105.1 | Spirochaetota_bacterium | DRP70_07570 | 3107..6613 | 2202144 | BREX-system-P-loop-protein-BrxC-[Spirochaetota-bacterium] |
| TXT43486.1 | Type1.BREX | <-SIGMA+OB-Fold+AAA-ATPase+HKD-DNase-BrxL<-?\|\|?-><-HTH+REase<-N6-METHYLASE-p<-N6-METHYLASE-p<-N6-METHYLASE-p<-Schlafen+HATPase+wHTH-p<-DUF6079*<-BrxB-BrxF-iSTAND<-BrxA-DNA-binding<-XIS-HTH | 1162 | Bacteria>Spirochaetota>unclassified_Spirochaetota | GCA_008080685.1 | Spirochaetota_bacterium | FD137_1824 | 16028..19516 | 2202144 | hypothetical-protein-FD137_1824-[Spirochaetota-bacterium] |
| MCI5863824.1 | Type1.BREX | <-REC+OmpR-HTH<-?<-?\|\|?-><-MACRODOMAIN\|\|?->ANK+ANK+ANK->BrxA-DNA-binding->BrxB-BrxF-iSTAND->DUF6079-p*->DUF6079-p->N6-METHYLASE->?->PglZ->SIGMA+OB-Fold+AAA-ATPase+Lon-Protease-BrxL-> | 875 | Bacteria>Synergistota>Synergistia>Synergistales>Synergistaceae>Cloacibacillus | GCA_022771785.1 | Cloacibacillus_porcorum | MR000_01150 | 159552..162179 | 1197717 | BREX-system-P-loop-protein-BrxC-[Cloacibacillus-porcorum] |
| WP_051485152.1 | Type1.BREX | MODE-HTH+PBPII->?->?-><-?<-MULE-TRANSPOSASE\|\|BrxA-DNA-binding->BrxB-BrxF-iSTAND->DUF6079*->N6-METHYLASE->SbcC->Schlafen+HATPase+wHTH->PglZ->SIGMA+OB-Fold+AAA-ATPase+Lon-Protease-BrxL->?->?->transposase-><-GNTR-HTH | 1216 | Bacteria>Synergistota>Synergistia>Synergistales>Synergistaceae>Cloacibacillus | GCF_000585335.1 | Cloacibacillus_evryensis | CLOEV_RS14965 | 3354177..3357827 | 508460 | BREX-system-P-loop-protein-BrxC-[Cloacibacillus-evryensis] |
| WP_273308638.1 | Type1.BREX | <-REC+OmpR-HTH<-?<-?\|\|?-><-MACRODOMAIN\|\|?->ANK+ANK+ANK->BrxA-DNA-binding->BrxB-BrxF-iSTAND->DUF6079-p*->DUF6079-p->N6-METHYLASE->?->PglZ->SIGMA+OB-Fold+AAA-ATPase+Lon-Protease-BrxL-> | 875 | Bacteria>Synergistota>Synergistia>Synergistales>Synergistaceae>Cloacibacillus | GCF_022771785.1 | Cloacibacillus_porcorum | MR000_RS01170 | 159552..162179 | 1197717 | BREX-system-P-loop-protein-BrxC-[Cloacibacillus-porcorum] |
| WP_006582937.1 | Type1.BREX | Amidohydro_2->?->ABhydrolase->?->?->tRNA->tRNA->tRNA->tRNA->tRNA->tRNA->XIS-HTH->BrxB-BrxF-iSTAND->DUF6079-p*->REase+N6_METHYLASE->SIR2->HerA/FtsK->TRD+TRD->DOC+HTH->REase+SF1-SF2-Helicase->PglZ-p->BrxA-DNA-binding-> | 532 | Bacteria>Synergistota>Synergistia>Synergistales>Synergistaceae>Thermanaerovibrio | GCF_000237825.1 | Thermanaerovibrio_velox | THEVEDRAFT_RS01385 | 298726..300324 | 108007 | DUF6079-family-protein-[Thermanaerovibrio-velox] |
| MBL3593011.1 | Type1.BREX | <-CHTH\|\|DUF488-1->DUF488-s->?->SF1-SF2-Helicase->?->XIS-HTH->BrxA-DNA-binding->BrxB-BrxF-iSTAND->DUF6079*->Schlafen+HATPase+wHTH->N6-METHYLASE->PglZ->SIGMA+OB-Fold+AAA-ATPase+Lon-Protease-BrxL->HARE-HTH+REase->?->GGDEF->Aminotran_4->DUF488-> | 1149 | Bacteria>Synergistota>Synergistia>Synergistales>Synergistaceae>unclassified_Synergistaceae | GCA_016756725.1 | Synergistaceae_bacterium | JMJ93_05810 | 374580..378029 | 2053624 | BREX-system-P-loop-protein-BrxC-[Synergistaceae-bacterium] |
| MCK9436263.1 | Type1.BREX | <-SIGMA+OB-Fold+AAA-ATPase+Lon-Protease-BrxL<-PglZ<-N6-METHYLASE<-DUF6079-p* | 474 | Bacteria>Synergistota>Synergistia>Synergistales>Synergistaceae>unclassified_Synergistaceae | GCA_023229345.1 | Synergistaceae_bacterium | MOQ12_03525 | 7446..8870 | 2053624 | hypothetical-protein-[Synergistaceae-bacterium] |
| GHV28822.1 | Type1.BREX | <-HicA<-?<-REase<-?<-?<-N6-METHYLASE<-N6-METHYLASE-p<-NERD<-OLD-ABC<-DUF6079*<-BrxB-BrxF-iSTAND<-BrxA-DNA-binding | 1195 | Bacteria>Synergistota>Synergistia>Synergistales>unclassified_Synergistales | GCA_026002695.1 | Synergistales_bacterium | FACS1894167_06880 | 20211..23798 | 2268201 | hypothetical-protein-FACS1894167_06880-[Synergistales-bacterium] |
| BCM94553.1 | Type1.BREX | <-HTH_28+rve<-HTH_3\|\|HTH+REase-><-GDResolvase+Recombinase+Zn_ribbon_recom<-?<-ArdCN<-SIGMA+OB-Fold+AAA-ATPase-BrxL<-PglZ<-DUF6079*<-BrxB-BrxF-iSTAND<-DUF4007<-N6-METHYLASE<-SIGMA-HTH | 1371 | Bacteria>Terrabacteria_group>Abditibacteriota>unclassified_Abditibacteriota | GCA_023169805.1 | Abditibacteriota_bacterium | IAD21_06460 | 92337..96452 | 2783868 | hypothetical-protein-IAD21_06460-[Abditibacteriota-bacterium] |
| MCB0987052.1 | Type1.BREX | <-N6-METHYLASE-p<-DUF6079*<-BrxB-BrxF-iSTAND<-BrxA-DNA-binding<-?\|\|?-><-Phage_integrase<-PIN<-HTH+MPTase<-?<-PolB | 1154 | Bacteria>Terrabacteria_group>Actinomycetota>Acidimicrobiia>Acidimicrobiales>unclassified_Acidimicrobiales | GCA_020439505.1 | Acidimicrobiales_bacterium | KDB09_04685 | 1462..4926 | 2201156 | BREX-system-P-loop-protein-BrxC-[Acidimicrobiales-bacterium] |
| WP_075374702.1 | Type1.BREX | TetR-HTH->ABC-ATPase+SbcC->?->?-><-SIGMA+OB-Fold+AAA-ATPase+HKD-DNase-BrxL<-PglZ<-Schlafen+HATPase+wHTH<-SbcC<-N6-METHYLASE<-DUF6079*<-BrxB-BrxF-iSTAND<-BrxA-DNA-binding\|\|PhoH->?->?->sGTP->?->MetJarc-HTH->PemK_toxin-> | 1172 | Bacteria>Terrabacteria_group>Actinomycetota>Actinomycetes>Actinomycetales>Actinomycetaceae | - | Actinomyces_oris | BKH18_06090 | 96462..99980 | 544580 | BREX-system-P-loop-protein-BrxC-[Actinomyces-oris] |
| WP_075392896.1 | Type1.BREX | TetR-HTH->ABC-ATPase+SbcC->?->?-><-SIGMA+OB-Fold+AAA-ATPase+HKD-DNase-BrxL<-PglZ<-Schlafen+HATPase+wHTH<-N6-METHYLASE<-DUF6079*<-BrxB-BrxF-iSTAND<-BrxA-DNA-binding\|\|PhoH->?->?->sGTP->?->MetJarc-HTH->PemK_toxin-> | 1172 | Bacteria>Terrabacteria_group>Actinomycetota>Actinomycetes>Actinomycetales>Actinomycetaceae | - | Actinomyces_oris | BKH22_09300 | 69704..73222 | 544580 | BREX-system-P-loop-protein-BrxC-[Actinomyces-oris] |
| WP_075371911.1 | Type1.BREX | TetR-HTH->ABC-ATPase+SbcC->?->?-><-SIGMA+OB-Fold+AAA-ATPase+HKD-DNase-BrxL<-PglZ<-Schlafen+HATPase+wHTH<-SbcC<-N6-METHYLASE<-DUF6079*<-BrxB-BrxF-iSTAND<-BrxA-DNA-binding\|\|PhoH->?->?->sGTP->?->MetJarc-HTH->PemK_toxin-> | 1172 | Bacteria>Terrabacteria_group>Actinomycetota>Actinomycetes>Actinomycetales>Actinomycetaceae | - | Actinomyces_oris | BKH23_02505 | 283433..286951 | 544580 | BREX-system-P-loop-protein-BrxC-[Actinomyces-oris] |
| WP_075400926.1 | Type1.BREX | <-PemK_toxin<-MetJarc-HTH<-?<-sGTP<-?<-?<-PhoH\|\|BrxA-DNA-binding->BrxB-BrxF-iSTAND->DUF6079*->N6-METHYLASE->SbcC->Schlafen+HATPase+wHTH->PglZ->SIGMA+OB-Fold+AAA-ATPase+HKD-DNase-BrxL-><-?<-?<-ABC-ATPase+SbcC<-TetR-HTH | 1171 | Bacteria>Terrabacteria_group>Actinomycetota>Actinomycetes>Actinomycetales>Actinomycetaceae | - | Actinomyces_oris | BKH24_00995 | 75611..79126 | 544580 | BREX-system-P-loop-protein-BrxC-[Actinomyces-oris] |
| WP_075550941.1 | Type1.BREX | TetR-HTH->ABC-ATPase+SbcC->?->?-><-SIGMA+OB-Fold+AAA-ATPase+HKD-DNase-BrxL<-PglZ<-Schlafen+HATPase+wHTH<-SbcC<-N6-METHYLASE<-DUF6079*<-BrxB-BrxF-iSTAND<-BrxA-DNA-binding\|\|PhoH->?->?->sGTP->?->MetJarc-HTH->PemK_toxin-> | 1172 | Bacteria>Terrabacteria_group>Actinomycetota>Actinomycetes>Actinomycetales>Actinomycetaceae | - | Actinomyces_oris | BKH26_09165 | 68268..71786 | 544580 | BREX-system-P-loop-protein-BrxC-[Actinomyces-oris] |
| WP_075411517.1 | Type1.BREX | TetR-HTH->ABC-ATPase+SbcC->?->?-><-SIGMA+OB-Fold+AAA-ATPase+HKD-DNase-BrxL<-PglZ<-Schlafen+HATPase+wHTH<-SbcC<-N6-METHYLASE<-DUF6079*<-BrxB-BrxF-iSTAND<-BrxA-DNA-binding\|\|PhoH->?->?->sGTP->?->MetJarc-HTH->PemK_toxin-> | 1172 | Bacteria>Terrabacteria_group>Actinomycetota>Actinomycetes>Actinomycetales>Actinomycetaceae | - | Actinomyces_oris | BKH31_05960 | 55043..58561 | 544580 | BREX-system-P-loop-protein-BrxC-[Actinomyces-oris] |
| WP_141407194.1 | Type1.BREX | HTH_3-><-PemK_toxin<-?<-sGTP<-?<-?<-PhoH\|\|BrxA-DNA-binding->BrxB-BrxF-iSTAND->DUF6079*->N6-METHYLASE->SbcC->Schlafen+HATPase+wHTH->PglZ->SIGMA+OB-Fold+AAA-ATPase+HKD-DNase-BrxL-><-?<-?<-ABC-ATPase+SbcC<-TetR-HTH | 1172 | Bacteria>Terrabacteria_group>Actinomycetota>Actinomycetes>Actinomycetales>Actinomycetaceae | - | Actinomyces_oris | I6I08_06235 | 1496012..1499530 | 544580 | BREX-system-P-loop-protein-BrxC-[Actinomyces-oris] |
| WP_119835580.1 | Type1.BREX | HMA+HAD+HAD-><-MarR-HTH+PADR-HTH<-?\|\|?->BrxA-DNA-binding->BrxB-BrxF-iSTAND->DUF6079*->N6-METHYLASE->SbcC->Schlafen+HATPase+wHTH->PglZ->SIGMA+OB-Fold+AAA-ATPase+HKD-DNase-BrxL->ABC-ATPase+SbcC-> | 1171 | Bacteria>Terrabacteria_group>Actinomycetota>Actinomycetes>Actinomycetales>Actinomycetaceae | - | Actinomyces_sp._2119 | D4740_02085 | 49121..52636 | 2321393 | BREX-system-P-loop-protein-BrxC-[Actinomyces-sp.-2119] |
| AOS47539.1 | Type1.BREX | <-LACI-HTH+PBPI<-?<-?\|\|?->DOC+HTH->BrxA-DNA-binding->BrxB-BrxF-iSTAND->DUF6079*->N6-METHYLASE->OLD-ABC->TOPRIM->PglZ->SIGMA+OB-Fold+AAA-ATPase+HKD-DNase-BrxL-><-ABC-3<-ABC-ATPase+SbcC | 1176 | Bacteria>Terrabacteria_group>Actinomycetota>Actinomycetes>Actinomycetales>Actinomycetaceae | GCA_001746855.1 | Pauljensenia_hongkongensis | BH719_06520 | 1584234..1587764 | 178339 | hypothetical-protein-BH719_06520-[Pauljensenia-hongkongensis] |
| OFQ22233.1 | Type1.BREX | <-ABC-ATPase+SbcC+ABC-ATPase+SbcC<-?<-SIGMA+OB-Fold+AAA-ATPase+HKD-DNase-BrxL<-PglZ<-TOPRIM<-OLD-ABC<-N6-METHYLASE<-DUF6079*<-BrxB-BrxF-iSTAND<-BrxA-DNA-binding | 1173 | Bacteria>Terrabacteria_group>Actinomycetota>Actinomycetes>Actinomycetales>Actinomycetaceae | GCA_001813985.1 | Actinomyces_sp._HMSC062G12 | HMPREF2946_04765 | 260177..263698 | 1739450 | hypothetical-protein-HMPREF2946_04765-[Actinomyces-sp.-HMSC062G12] |
| OLO46937.1 | Type1.BREX | TetR-HTH->ABC-ATPase+SbcC->?->?-><-SIGMA+OB-Fold+AAA-ATPase+HKD-DNase-BrxL<-PglZ<-Schlafen+HATPase+wHTH<-SbcC<-N6-METHYLASE<-DUF6079*<-BrxB-BrxF-iSTAND<-BrxA-DNA-binding\|\|PhoH->?->?->sGTP->?->MetJarc-HTH->PemK_toxin-> | 1172 | Bacteria>Terrabacteria_group>Actinomycetota>Actinomycetes>Actinomycetales>Actinomycetaceae | GCA_001937365.1 | Actinomyces_oris | BKH31_05960 | 55043..58561 | 544580 | hypothetical-protein-BKH31_05960-[Actinomyces-oris] |
| OLO54967.1 | Type1.BREX | TetR-HTH->ABC-ATPase+SbcC->?->?-><-SIGMA+OB-Fold+AAA-ATPase+HKD-DNase-BrxL<-PglZ<-Schlafen+HATPase+wHTH<-SbcC<-N6-METHYLASE<-DUF6079*<-BrxB-BrxF-iSTAND<-BrxA-DNA-binding\|\|PhoH->?->?->sGTP->?->MetJarc-HTH->PemK_toxin-> | 1172 | Bacteria>Terrabacteria_group>Actinomycetota>Actinomycetes>Actinomycetales>Actinomycetaceae | GCA_001937415.1 | Actinomyces_oris | BKH26_09165 | 68268..71786 | 544580 | hypothetical-protein-BKH26_09165-[Actinomyces-oris] |
| OLO62906.1 | Type1.BREX | <-PemK_toxin<-MetJarc-HTH<-?<-sGTP<-?<-?<-PhoH\|\|BrxA-DNA-binding->BrxB-BrxF-iSTAND->DUF6079*->N6-METHYLASE->SbcC->Schlafen+HATPase+wHTH->PglZ->SIGMA+OB-Fold+AAA-ATPase+HKD-DNase-BrxL-><-?<-?<-ABC-ATPase+SbcC<-TetR-HTH | 1171 | Bacteria>Terrabacteria_group>Actinomycetota>Actinomycetes>Actinomycetales>Actinomycetaceae | GCA_001937425.1 | Actinomyces_oris | BKH24_00995 | 75611..79126 | 544580 | hypothetical-protein-BKH24_00995-[Actinomyces-oris] |
| OLO63895.1 | Type1.BREX | TetR-HTH->ABC-ATPase+SbcC->?->?-><-SIGMA+OB-Fold+AAA-ATPase+HKD-DNase-BrxL<-PglZ<-Schlafen+HATPase+wHTH<-SbcC<-N6-METHYLASE<-DUF6079*<-BrxB-BrxF-iSTAND<-BrxA-DNA-binding\|\|PhoH->?->?->sGTP->?->MetJarc-HTH->PemK_toxin-> | 1172 | Bacteria>Terrabacteria_group>Actinomycetota>Actinomycetes>Actinomycetales>Actinomycetaceae | GCA_001937485.1 | Actinomyces_oris | BKH23_02505 | 283433..286951 | 544580 | hypothetical-protein-BKH23_02505-[Actinomyces-oris] |
| OLO77585.1 | Type1.BREX | TetR-HTH->ABC-ATPase+SbcC->?->?-><-SIGMA+OB-Fold+AAA-ATPase+HKD-DNase-BrxL<-PglZ<-Schlafen+HATPase+wHTH<-SbcC<-N6-METHYLASE<-DUF6079*<-BrxB-BrxF-iSTAND<-BrxA-DNA-binding\|\|PhoH->?->?->sGTP->?->MetJarc-HTH->PemK_toxin-> | 1172 | Bacteria>Terrabacteria_group>Actinomycetota>Actinomycetes>Actinomycetales>Actinomycetaceae | GCA_001937535.1 | Actinomyces_oris | BKH18_06090 | 96462..99980 | 544580 | hypothetical-protein-BKH18_06090-[Actinomyces-oris] |
| OLO63014.1 | Type1.BREX | TetR-HTH->ABC-ATPase+SbcC->?->?-><-SIGMA+OB-Fold+AAA-ATPase+HKD-DNase-BrxL<-PglZ<-Schlafen+HATPase+wHTH<-N6-METHYLASE<-DUF6079*<-BrxB-BrxF-iSTAND<-BrxA-DNA-binding\|\|PhoH->?->?->sGTP->?->MetJarc-HTH->PemK_toxin-> | 1172 | Bacteria>Terrabacteria_group>Actinomycetota>Actinomycetes>Actinomycetales>Actinomycetaceae | GCA_001937725.1 | Actinomyces_oris | BKH22_09300 | 69704..73222 | 544580 | hypothetical-protein-BKH22_09300-[Actinomyces-oris] |
| RJF44065.1 | Type1.BREX | HMA+HAD+HAD-><-MarR-HTH+PADR-HTH<-?\|\|?->BrxA-DNA-binding->BrxB-BrxF-iSTAND->DUF6079*->N6-METHYLASE->SbcC->Schlafen+HATPase+wHTH->PglZ->SIGMA+OB-Fold+AAA-ATPase+HKD-DNase-BrxL->ABC-ATPase+SbcC-> | 1171 | Bacteria>Terrabacteria_group>Actinomycetota>Actinomycetes>Actinomycetales>Actinomycetaceae | GCA_003591215.1 | Actinomyces_sp._2119 | D4740_02085 | 49121..52636 | 2321393 | BREX-system-P-loop-protein-BrxC-[Actinomyces-sp.-2119] |
| MBF0967807.1 | Type1.BREX | BrxB-BrxF-iSTAND->DUF6079*->N6-METHYLASE->OLD-ABC->TOPRIM->PglZ->SIGMA+OB-Fold+AAA-ATPase+HKD-DNase-BrxL-><-?\|\|?->ABC-ATPase+SbcC+ABC-ATPase+SbcC-> | 1169 | Bacteria>Terrabacteria_group>Actinomycetota>Actinomycetes>Actinomycetales>Actinomycetaceae | GCA_015259445.1 | Actinomyces_sp. | HXK08_00790 | 718..4227 | 29317 | BREX-system-P-loop-protein-BrxC-[Actinomyces-sp.] |
| QQC41011.1 | Type1.BREX | HTH_3-><-PemK_toxin<-?<-sGTP<-?<-?<-PhoH\|\|BrxA-DNA-binding->BrxB-BrxF-iSTAND->DUF6079*->N6-METHYLASE->SbcC->Schlafen+HATPase+wHTH->PglZ->SIGMA+OB-Fold+AAA-ATPase+HKD-DNase-BrxL-><-?<-?<-ABC-ATPase+SbcC<-TetR-HTH | 1172 | Bacteria>Terrabacteria_group>Actinomycetota>Actinomycetes>Actinomycetales>Actinomycetaceae | GCA_016127955.1 | Actinomyces_oris | I6I08_06235 | 1496012..1499530 | 544580 | BREX-system-P-loop-protein-BrxC-[Actinomyces-oris] |
| MBS6968676.1 | Type1.BREX | DEOR-HTH+ISOCOT->?-><-SIGMA+OB-Fold+AAA-ATPase+HKD-DNase-BrxL<-PglZ<-TOPRIM<-OLD-ABC<-N6-METHYLASE<-DUF6079*<-BrxB-BrxF-iSTAND<-BrxA-DNA-binding | 1169 | Bacteria>Terrabacteria_group>Actinomycetota>Actinomycetes>Actinomycetales>Actinomycetaceae | GCA_018381115.1 | Actinomyces_sp. | KH140_06985 | 13628..17137 | 29317 | BREX-system-P-loop-protein-BrxC-[Actinomyces-sp.] |
| MCI1675766.1 | Type1.BREX | ABhydrolase-><-?<-?<-?\|\|?->Calcineurin->tRNA-><-MerR-HTH+TipAS\|\|BrxA-DNA-binding->BrxB-BrxF-iSTAND->DUF6079*->N6-METHYLASE-> | 1164 | Bacteria>Terrabacteria_group>Actinomycetota>Actinomycetes>Actinomycetales>Actinomycetaceae | GCA_022648165.1 | Ancrocorticia_sp. | LKI28_05730 | 185696..189190 | 2593684 | BREX-system-P-loop-protein-BrxC-[Ancrocorticia-sp.] |
| MCI1641321.1 | Type1.BREX | sGTP->?->?->?->His_Phos_1->BrxA-DNA-binding->BrxB-BrxF-iSTAND->DUF6079*->?->N6-METHYLASE-p-> | 1167 | Bacteria>Terrabacteria_group>Actinomycetota>Actinomycetes>Actinomycetales>Actinomycetaceae | GCA_022648365.1 | Actinomyces_sp. | LKI38_02150 | 177491..180994 | 29317 | BREX-system-P-loop-protein-BrxC-[Actinomyces-sp.] |
| MCI6557834.1 | Type1.BREX | ABhydrolase-><-?<-SIGMA+OB-Fold+AAA-ATPase+HKD-DNase-BrxL<-PglZ<-?<-N6-METHYLASE-p<-Phage_int_SAM_3+Phage_integrase\|\|N6-METHYLASE-p-><-N6-METHYLASE<-DUF6079*<-BrxB-BrxF-iSTAND<-BrxA-DNA-binding\|\|SF1-SF2-Helicase+DUF3418-> | 1160 | Bacteria>Terrabacteria_group>Actinomycetota>Actinomycetes>Actinomycetales>Actinomycetaceae | GCA_022780845.1 | Schaalia_hyovaginalis | MR422_09445 | 69432..72914 | 29316 | BREX-system-P-loop-protein-BrxC-[Schaalia-hyovaginalis] |
| MCI7671412.1 | Type1.BREX | <-SIGMA+OB-Fold+AAA-ATPase+HKD-DNase-BrxL<-PglZ<-?<-N6-METHYLASE-p<-Phage_int_SAM_3+Phage_integrase\|\|N6-METHYLASE-p->DDE_Tnp_IS240->ParA-Soj-PloopNTPase-><-N6-METHYLASE-p<-DUF6079*<-BrxB-BrxF-iSTAND<-BrxA-DNA-binding\|\|SF1-SF2-Helicase+DUF3418-><-Glyco_hydro_36N+Melibiase<-?\|\|?->?->?->LACI-HTH+PBPI-> | 1161 | Bacteria>Terrabacteria_group>Actinomycetota>Actinomycetes>Actinomycetales>Actinomycetaceae | GCA_022786785.1 | Schaalia_hyovaginalis | MSS20_03495 | 24103..27588 | 29316 | BREX-system-P-loop-protein-BrxC-[Schaalia-hyovaginalis] |
| WP_009648402.1 | Type1.BREX | REase-><-?<-DDE_Tnp<-DDE_Tnp<-?\|\|?->BrxA-DNA-binding->BrxB-BrxF-iSTAND->DUF6079*->N6-METHYLASE->RVT->PglZ->SIGMA+OB-Fold+AAA-ATPase+HKD-DNase-BrxL->HEPN-> | 1169 | Bacteria>Terrabacteria_group>Actinomycetota>Actinomycetes>Actinomycetales>Actinomycetaceae | GCF_000278725.1 | Actinomyces_sp._ICM47 | HMPREF1136_RS05845 | 15295..18804 | 936548 | BREX-system-P-loop-protein-BrxC-[Actinomyces-sp.-ICM47] |
| WP_024332155.1 | Type1.BREX | HTH+REase->?->35exo->BrxA-DNA-binding->BrxB-BrxF-iSTAND->DUF6079*->N6-METHYLASE->PglZ->SIGMA+OB-Fold+AAA-ATPase+HKD-DNase-BrxL->?->?->?-><-?\|\|?-><-ABhydrolase | 1184 | Bacteria>Terrabacteria_group>Actinomycetota>Actinomycetes>Actinomycetales>Actinomycetaceae | GCF_000420405.1 | Winkia_neuii | G437_RS0108590 | 20470..24024 | 33007 | BREX-system-P-loop-protein-BrxC-[Winkia-neuii] |
| WP_026406950.1 | Type1.BREX | TetR-HTH->ABC-ATPase+SbcC->?->?-><-SIGMA+OB-Fold+AAA-ATPase+HKD-DNase-BrxL<-PglZ<-Schlafen+HATPase+wHTH<-SbcC<-N6-METHYLASE<-DUF6079*<-BrxB-BrxF-iSTAND<-BrxA-DNA-binding\|\|PhoH->?->?->sGTP->Peptidase_S9-> | 1174 | Bacteria>Terrabacteria_group>Actinomycetota>Actinomycetes>Actinomycetales>Actinomycetaceae | GCF_000429265.1 | Actinomyces_gerencseriae | G448_RS0104040 | 223321..226845 | 52769 | BREX-system-P-loop-protein-BrxC-[Actinomyces-gerencseriae] |
| WP_173647185.1 | Type1.BREX | <-LACI-HTH+PBPI<-?<-?\|\|?->DOC+HTH->BrxA-DNA-binding->BrxB-BrxF-iSTAND->DUF6079*->N6-METHYLASE->OLD-ABC->TOPRIM->PglZ->SIGMA+OB-Fold+AAA-ATPase+HKD-DNase-BrxL->?-><-ABC-3<-ABC-ATPase+SbcC | 1174 | Bacteria>Terrabacteria_group>Actinomycetota>Actinomycetes>Actinomycetales>Actinomycetaceae | GCF_001746855.1 | Pauljensenia_hongkongensis | BH719_RS06520 | 1584240..1587764 | 178339 | BREX-system-P-loop-protein-BrxC-[Pauljensenia-hongkongensis] |
| WP_256693306.1 | Type1.BREX | TetR-HTH->ABC-ATPase+SbcC->?->?-><-SIGMA+OB-Fold+AAA-ATPase+HKD-DNase-BrxL<-PglZ<-Schlafen+HATPase+wHTH<-SbcC<-N6-METHYLASE<-DUF6079*<-BrxB-BrxF-iSTAND<-BrxA-DNA-binding\|\|PhoH->?->?->sGTP->?->PemK_toxin-><-HTH_3 | 1175 | Bacteria>Terrabacteria_group>Actinomycetota>Actinomycetes>Actinomycetales>Actinomycetaceae | GCF_001937365.1 | Actinomyces_oris | BKH31_RS05960 | 55043..58570 | 544580 | BREX-system-P-loop-protein-BrxC-[Actinomyces-oris] |
| WP_256693847.1 | Type1.BREX | TetR-HTH->ABC-ATPase+SbcC->?->?-><-SIGMA+OB-Fold+AAA-ATPase+HKD-DNase-BrxL<-PglZ<-Schlafen+HATPase+wHTH<-SbcC<-N6-METHYLASE<-DUF6079*<-BrxB-BrxF-iSTAND<-BrxA-DNA-binding\|\|PhoH->?->?->sGTP->?->MetJarc-HTH->PemK_toxin-> | 1175 | Bacteria>Terrabacteria_group>Actinomycetota>Actinomycetes>Actinomycetales>Actinomycetaceae | GCF_001937415.1 | Actinomyces_oris | BKH26_RS09145 | 68268..71795 | 544580 | BREX-system-P-loop-protein-BrxC-[Actinomyces-oris] |
| WP_256699678.1 | Type1.BREX | HTH_3-><-PemK_toxin<-?<-sGTP<-?<-?<-PhoH\|\|BrxA-DNA-binding->BrxB-BrxF-iSTAND->DUF6079*->N6-METHYLASE->SbcC->Schlafen+HATPase+wHTH->PglZ->SIGMA+OB-Fold+AAA-ATPase+HKD-DNase-BrxL-><-?<-?<-ABC-ATPase+SbcC<-TetR-HTH | 1174 | Bacteria>Terrabacteria_group>Actinomycetota>Actinomycetes>Actinomycetales>Actinomycetaceae | GCF_001937425.1 | Actinomyces_oris | BKH24_RS00995 | 75602..79126 | 544580 | BREX-system-P-loop-protein-BrxC-[Actinomyces-oris] |
| WP_256699054.1 | Type1.BREX | TetR-HTH->ABC-ATPase+SbcC->?->?-><-SIGMA+OB-Fold+AAA-ATPase+HKD-DNase-BrxL<-PglZ<-Schlafen+HATPase+wHTH<-SbcC<-N6-METHYLASE<-DUF6079*<-BrxB-BrxF-iSTAND<-BrxA-DNA-binding\|\|PhoH->?->?->sGTP->?->PemK_toxin-><-HTH_3 | 1175 | Bacteria>Terrabacteria_group>Actinomycetota>Actinomycetes>Actinomycetales>Actinomycetaceae | GCF_001937485.1 | Actinomyces_oris | BKH23_RS02505 | 283433..286960 | 544580 | BREX-system-P-loop-protein-BrxC-[Actinomyces-oris] |
| WP_256694375.1 | Type1.BREX | TetR-HTH->ABC-ATPase+SbcC->?->?-><-SIGMA+OB-Fold+AAA-ATPase+HKD-DNase-BrxL<-PglZ<-Schlafen+HATPase+wHTH<-SbcC<-N6-METHYLASE<-DUF6079*<-BrxB-BrxF-iSTAND<-BrxA-DNA-binding\|\|PhoH->?->?->sGTP->?->MetJarc-HTH->PemK_toxin-> | 1175 | Bacteria>Terrabacteria_group>Actinomycetota>Actinomycetes>Actinomycetales>Actinomycetaceae | GCF_001937535.1 | Actinomyces_oris | BKH18_RS06085 | 96462..99989 | 544580 | BREX-system-P-loop-protein-BrxC-[Actinomyces-oris] |
| WP_256697766.1 | Type1.BREX | <-MetJarc-HTH\|\|TetR-HTH->ABC-ATPase+SbcC->?->?-><-SIGMA+OB-Fold+AAA-ATPase+HKD-DNase-BrxL<-PglZ<-Schlafen+HATPase+wHTH<-N6-METHYLASE<-DUF6079*<-BrxB-BrxF-iSTAND<-BrxA-DNA-binding\|\|PhoH->?->?->sGTP->?->PemK_toxin-><-HTH_3 | 1175 | Bacteria>Terrabacteria_group>Actinomycetota>Actinomycetes>Actinomycetales>Actinomycetaceae | GCF_001937725.1 | Actinomyces_oris | BKH22_RS09310 | 69704..73231 | 544580 | BREX-system-P-loop-protein-BrxC-[Actinomyces-oris] |
| WP_076066523.1 | Type1.BREX | <-AraC-HTH+AraC-HTH\|\|?->NAD_binding_10-><-SIGMA+OB-Fold+AAA-ATPase+HKD-DNase-BrxL<-PglZ<-Schlafen+HATPase+wHTH<-SbcC<-N6-METHYLASE<-DUF6079*<-BrxB-BrxF-iSTAND<-BrxA-DNA-binding\|\|PhoH->?->?->sGTP->?-><-?\|\|TetR-HTH-> | 1174 | Bacteria>Terrabacteria_group>Actinomycetota>Actinomycetes>Actinomycetales>Actinomycetaceae | GCF_001956485.1 | Actinomyces_naeslundii | BKH36_RS03225 | 74556..78080 | 1655 | BREX-system-P-loop-protein-BrxC-[Actinomyces-naeslundii] |
| WP_108716712.1 | Type1.BREX | <-SIGMA+OB-Fold+AAA-ATPase+HKD-DNase-BrxL<-PglZ<-?<-Schlafen-p<-SbcC<-N6-METHYLASE-p<-N6-METHYLASE<-DUF6079*<-BrxB-BrxF-iSTAND<-BrxA-DNA-binding\|\|Schlafen+HATPase+wHTH-p->Schlafen+HATPase+wHTH-p-><-tRNA<-?<-?<-ABC-ATPase+SbcC\|\|Sensor+HisKA_3+HATPase_c->REC+LuxR-HTH-> | 1174 | Bacteria>Terrabacteria_group>Actinomycetota>Actinomycetes>Actinomycetales>Actinomycetaceae | GCF_003073475.1 | Actinobaculum_sp._313 | DDD63_RS08615 | 1982694..1986218 | 2495645 | BREX-system-P-loop-protein-BrxC-[Actinobaculum-sp.-313] |
| WP_109093701.1 | Type1.BREX | <-PIN<-PhdYeFM_antitox\|\|?->REase-><-ParB+HNH<-SIGMA+OB-Fold+AAA-ATPase+HKD-DNase-BrxL<-PglZ<-Schlafen+HATPase+wHTH<-N6-METHYLASE<-DUF6079*<-BrxB-BrxF-iSTAND<-BrxA-DNA-binding\|\|?-><-?<-?<-LACI-HTH+PBPI<-Glyco_hydro_1 | 1170 | Bacteria>Terrabacteria_group>Actinomycetota>Actinomycetes>Actinomycetales>Actinomycetaceae | GCF_003123745.1 | Ancrocorticia_populi | DD236_RS06730 | 275849..279361 | 2175228 | BREX-system-P-loop-protein-BrxC-[Ancrocorticia-populi] |
| WP_259390601.1 | Type1.BREX | HMA+HAD+HAD-><-MarR-HTH+PADR-HTH<-?\|\|?->BrxA-DNA-binding->BrxB-BrxF-iSTAND->DUF6079*->N6-METHYLASE->SbcC->Schlafen+HATPase+wHTH->PglZ->SIGMA+OB-Fold+AAA-ATPase+HKD-DNase-BrxL->ABC-ATPase+SbcC-> | 1174 | Bacteria>Terrabacteria_group>Actinomycetota>Actinomycetes>Actinomycetales>Actinomycetaceae | GCF_003591215.1 | Actinomyces_sp._2119 | D4740_RS02085 | 49112..52636 | 2321393 | BREX-system-P-loop-protein-BrxC-[Actinomyces-sp.-2119] |
| WP_124851247.1 | Type1.BREX | <-SIGMA+OB-Fold+AAA-ATPase+HKD-DNase-BrxL<-PglZ<-HTH+REase<-N6-METHYLASE-p<-Phage_int_SAM_3+Phage_integrase\|\|N6-METHYLASE-p-><-N6-METHYLASE<-DUF6079*<-BrxB-BrxF-iSTAND<-BrxA-DNA-binding<-?\|\|?->?->?->SbcC+SbcC-> | 1170 | Bacteria>Terrabacteria_group>Actinomycetota>Actinomycetes>Actinomycetales>Actinomycetaceae | GCF_003858495.1 | Buchananella_hordeovulneris | EII12_RS02040 | 65131..68643 | 52770 | BREX-system-P-loop-protein-BrxC-[Buchananella-hordeovulneris] |
| WP_126145915.1 | Type1.BREX | <-SIGMA+OB-Fold+AAA-ATPase+HKD-DNase-BrxL<-PglZ<-?<-SbcC<-N6-METHYLASE<-DUF6079*<-BrxB-BrxF-iSTAND<-BrxA-DNA-binding\|\|Schlafen+HATPase+wHTH-><-MPTase<-HTH_3\|\|HTH_3-> | 1174 | Bacteria>Terrabacteria_group>Actinomycetota>Actinomycetes>Actinomycetales>Actinomycetaceae | GCF_003957265.1 | Actinobaculum_sp._352 | EXN07_RS06065 | 220063..223587 | 2490946 | BREX-system-P-loop-protein-BrxC-[Actinobaculum-sp.-352] |
| WP_243118374.1 | Type1.BREX | <-ABC-ATPase+SbcC<-SIGMA+OB-Fold+AAA-ATPase+HKD-DNase-BrxL<-PglZ<-Schlafen+HATPase+wHTH<-?<-N6-METHYLASE<-DUF6079*<-BrxB-BrxF-iSTAND<-BrxA-DNA-binding<-?\|\|?->MarR-HTH+PADR-HTH-><-HMA+HAD+HAD | 1174 | Bacteria>Terrabacteria_group>Actinomycetota>Actinomycetes>Actinomycetales>Actinomycetaceae | GCF_004010955.1 | Actinomyces_wuliandei | CWS50_RS00410 | 93588..97112 | 2057743 | BREX-system-P-loop-protein-BrxC-[Actinomyces-wuliandei] |
| WP_247595282.1 | Type1.BREX | ADH_N+ADH_zinc_N->TetR-HTH->ABC-ATPase+SbcC-><-SIGMA+OB-Fold+AAA-ATPase+HKD-DNase-BrxL<-PglZ<-Schlafen+HATPase+wHTH<-REase<-N6-METHYLASE<-DUF6079*<-BrxB-BrxF-iSTAND<-BrxA-DNA-binding\|\|?->PhoH->?->?->sGTP->Peptidase_S9-> | 1174 | Bacteria>Terrabacteria_group>Actinomycetota>Actinomycetes>Actinomycetales>Actinomycetaceae | GCF_004796555.1 | Actinomyces_procaprae | FAQ94_RS07530 | 102106..105630 | 2560010 | BREX-system-P-loop-protein-BrxC-[Actinomyces-procaprae] |
| WP_167202365.1 | Type1.BREX | <-GNTR-HTH+UTRA\|\|?->?->?->?->SIR2->BrxA-DNA-binding->BrxB-BrxF-iSTAND->DUF6079*->N6-METHYLASE->SbcC->Schlafen+HATPase+wHTH->PglZ->SIGMA+OB-Fold+AAA-ATPase+HKD-DNase-BrxL-> | 1171 | Bacteria>Terrabacteria_group>Actinomycetota>Actinomycetes>Actinomycetales>Actinomycetaceae | GCF_011751985.1 | Actinomyces_respiraculi | HCR20_RS03210 | 129007..132522 | 2744574 | BREX-system-P-loop-protein-BrxC-[Actinomyces-respiraculi] |
| WP_182050348.1 | Type1.BREX | HTH->Arm-DNA-bind_4+Phage_int_SAM_3+Phage_integrase->TraJ-RHH-><-?<-ZnR+ZnR<-SIGMA+OB-Fold+AAA-ATPase+Lon-Protease-BrxL<-PglZ<-DOC+HTH<-N6-METHYLASE<-DUF6079*<-BrxB-BrxF-iSTAND<-BrxA-DNA-binding<-TraJ-RHH\|\|?->?->XIS-HTH-><-REase+Bacteriocin_IIc<-REase | 1193 | Bacteria>Terrabacteria_group>Actinomycetota>Actinomycetes>Actinomycetales>Actinomycetaceae | GCF_014041995.1 | Changpingibacter_yushuensis | H2017_RS03430 | 798134..801715 | 2758440 | BREX-system-P-loop-protein-BrxC-[Changpingibacter-yushuensis] |
| WP_184451418.1 | Type1.BREX | Glyco_hydro_36N+Melibiase-><-SF1-SF2-Helicase+DUF3418\|\|?->BrxA-DNA-binding->BrxB-BrxF-iSTAND->DUF6079*->N6-METHYLASE-><-N6-METHYLASE-p\|\|Phage_int_SAM_3+Phage_integrase->N6-METHYLASE-p->?->PglZ->SIGMA+OB-Fold+AAA-ATPase+HKD-DNase-BrxL-> | 1160 | Bacteria>Terrabacteria_group>Actinomycetota>Actinomycetes>Actinomycetales>Actinomycetaceae | GCF_014208035.1 | Schaalia_hyovaginalis | HD592_RS01490 | 338165..341647 | 29316 | BREX-system-P-loop-protein-BrxC-[Schaalia-hyovaginalis] |
| WP_187994004.1 | Type1.BREX | <-NAD_binding_10\|\|?->ADH_N+ADH_zinc_N->?\|\|TetR-HTH-><-HipA_C<-?\|\|BrxA-DNA-binding->BrxB-BrxF-iSTAND->DUF6079*->N6-METHYLASE-><-N6-METHYLASE-p\|\|Phage_int_SAM_3+Phage_integrase->N6-METHYLASE-p->Schlafen+HATPase+wHTH->HATPase+wHTH->PglZ->SIGMA+OB-Fold+AAA-ATPase+HKD-DNase-BrxL-> | 1173 | Bacteria>Terrabacteria_group>Actinomycetota>Actinomycetes>Actinomycetales>Actinomycetaceae | GCF_014525425.1 | Schaalia_sp._JY-X159 | IDL90_RS00760 | 170975..174496 | 2758575 | BREX-system-P-loop-protein-BrxC-[Schaalia-sp.-JY-X159] |
| WP_166855918.1 | Type1.BREX | <-SIGMA+OB-Fold+AAA-ATPase+HKD-DNase-BrxL<-PglZ<-Schlafen+HATPase+wHTH<-SbcC<-N6-METHYLASE<-DUF6079*<-BrxB-BrxF-iSTAND<-BrxA-DNA-binding<-SIR2<-?<-?<-?<-?\|\|GNTR-HTH+UTRA-> | 1171 | Bacteria>Terrabacteria_group>Actinomycetota>Actinomycetes>Actinomycetales>Actinomycetaceae | GCF_014595995.2 | Actinomyces | ID810_RS09115 | 2190565..2194080 | 1654 | MULTISPECIES:-BREX-system-P-loop-protein-BrxC-[Actinomyces] |
| WP_232622595.1 | Type1.BREX | HTH_3-><-PemK_toxin<-?<-sGTP<-?<-?<-PhoH\|\|BrxA-DNA-binding->BrxB-BrxF-iSTAND->DUF6079*->N6-METHYLASE->SbcC->Schlafen+HATPase+wHTH->PglZ->SIGMA+OB-Fold+AAA-ATPase+HKD-DNase-BrxL-><-?<-?<-ABC-ATPase+SbcC<-TetR-HTH | 1175 | Bacteria>Terrabacteria_group>Actinomycetota>Actinomycetes>Actinomycetales>Actinomycetaceae | GCF_016127955.1 | Actinomyces_oris | I6I08_RS06235 | 1496003..1499530 | 544580 | BREX-system-P-loop-protein-BrxC-[Actinomyces-oris] |
| WP_271289309.1 | Type1.BREX | TetR-HTH->ABC-ATPase+SbcC->?->?-><-SIGMA+OB-Fold+AAA-ATPase+HKD-DNase-BrxL<-PglZ<-Schlafen+HATPase+wHTH<-?<-N6-METHYLASE<-DUF6079*<-BrxB-BrxF-iSTAND<-BrxA-DNA-binding\|\|PhoH->?->?->sGTP->?->MetJarc-HTH->PemK_toxin-> | 1175 | Bacteria>Terrabacteria_group>Actinomycetota>Actinomycetes>Actinomycetales>Actinomycetaceae | GCF_027945475.1 | Actinomyces_oris | PGE45_RS05470 | 1261783..1265310 | 544580 | BREX-system-P-loop-protein-BrxC-[Actinomyces-oris] |
| WP_272432709.1 | Type1.BREX | ABhydrolase-><-?<-SIGMA+OB-Fold+AAA-ATPase+HKD-DNase-BrxL<-PglZ<-?<-N6-METHYLASE-p<-Phage_int_SAM_3+Phage_integrase\|\|N6-METHYLASE-p-><-N6-METHYLASE<-DUF6079*<-BrxB-BrxF-iSTAND<-BrxA-DNA-binding\|\|SF1-SF2-Helicase+DUF3418-><-Glyco_hydro_36N+Melibiase<-?\|\|?->?->?->LACI-HTH+PBPI-> | 1160 | Bacteria>Terrabacteria_group>Actinomycetota>Actinomycetes>Actinomycetales>Actinomycetaceae | GCF_028437565.1 | Actinomyces_sp._B33 | M3T53_RS01725 | 406019..409501 | 2942131 | BREX-system-P-loop-protein-BrxC-[Actinomyces-sp.-B33] |
| WP_126382192.1 | Type1.BREX | <-SIGMA+OB-Fold+AAA-ATPase+HKD-DNase-BrxL<-PglZ<-Schlafen+HATPase+wHTH<-REase<-N6-METHYLASE<-DUF6079*<-BrxB-BrxF-iSTAND<-BrxA-DNA-binding<-SIR2<-?<-?<-?\|\|GNTR-HTH+UTRA-> | 1170 | Bacteria>Terrabacteria_group>Actinomycetota>Actinomycetes>Actinomycetales>Actinomycetaceae | GCF_900637165.1 | Actinomyces_howellii | GYB74_RS04555 | 58123..61635 | 52771 | BREX-system-P-loop-protein-BrxC-[Actinomyces-howellii] |
| WP_004808447.1 | Type1.BREX | <-RNase<-?\|\|?->?->?->?->HTH+REase->BrxA-DNA-binding->BrxB-BrxF-iSTAND->DUF6079*->N6-METHYLASE->PglZ->SIGMA+OB-Fold+AAA-ATPase+HKD-DNase-BrxL-><-?<-?<-ABhydrolase<-DOC+HTH<-?\|\|ZnR-> | 1184 | Bacteria>Terrabacteria_group>Actinomycetota>Actinomycetes>Actinomycetales | GCF_028201595.1 | Actinomycetaceae | PIG85_RS08980 | 1926978..1930532 | 2049 | MULTISPECIES:-BREX-system-P-loop-protein-BrxC-[Actinomycetaceae] |
| MBK6301657.1 | Type1.BREX | <-VirB11+MuB-ATPase\|\|?-><-Phage_integrase<-Phage_integrase<-HTH_26<-Phage_integrase<-PHP+SbcC<-SIGMA+OB-Fold+AAA-ATPase+Lon-Protease-BrxL<-PglZ<-N6-METHYLASE<-DUF6079*<-BrxB-BrxF-iSTAND<-BrxA-DNA-binding<-XIS-HTH<-ABhydrolase<-?<-MPTase<-HTH\|\|ArdCN+MPTase->Primpol+PriCT1-> | 1152 | Bacteria>Terrabacteria_group>Actinomycetota>Actinomycetes>Actinomycetales>unclassified_Actinomycetales | GCA_016705565.1 | Actinomycetales_bacterium | IPF40_11645 | 85880..89338 | 1911520 | BREX-system-P-loop-protein-BrxC-[Actinomycetales-bacterium] |
| MBK6956672.1 | Type1.BREX | <-PglZ<-N6-METHYLASE<-DUF6079-p*<-DUF6079-p<-BrxB-BrxF-iSTAND<-BrxA-DNA-binding<-tRNA\|\|?->REC+OmpR-HTH->HAMP+HISKIN+ST-Hiskin-> | 557 | Bacteria>Terrabacteria_group>Actinomycetota>Actinomycetes>Actinomycetales>unclassified_Actinomycetales | GCA_016708855.1 | Actinomycetales_bacterium | IPH27_14930 | 7874..9547 | 1911520 | hypothetical-protein-[Actinomycetales-bacterium] |
| MBK6956673.1 | Type1.BREX | <-PglZ<-N6-METHYLASE<-DUF6079-p<-DUF6079-p*<-BrxB-BrxF-iSTAND<-BrxA-DNA-binding<-tRNA\|\|?->REC+OmpR-HTH->HAMP+HISKIN+ST-Hiskin-> | 595 | Bacteria>Terrabacteria_group>Actinomycetota>Actinomycetes>Actinomycetales>unclassified_Actinomycetales | GCA_016708855.1 | Actinomycetales_bacterium | IPH27_14935 | 9553..11340 | 1911520 | BREX-system-P-loop-protein-BrxC-[Actinomycetales-bacterium] |
| WP_129869215.1 | Type1.BREX | MODE-HTH+PBPII->YTCD-HTH-><-HTH_3<-RelE-ParE<-ACET+ACET<-ZnR<-ZnR\|\|BrxA-DNA-binding->BrxB-BrxF-iSTAND->DUF6079*->N6-METHYLASE->PglZ->SIGMA+OB-Fold+AAA-ATPase+HKD-DNase-BrxL-><-PRTase | 1214 | Bacteria>Terrabacteria_group>Actinomycetota>Actinomycetes>Bifidobacteriales>Bifidobacteriaceae | - | Bifidobacterium_pseudolongum | PG1791B_RS04610 | 183618..187262 | 1694 | BREX-system-P-loop-protein-BrxC-[Bifidobacterium-pseudolongum] |
| KFI62072.1 | Type1.BREX | <-XIS-HTH+GDResolvase\|\|REase->?->BrxA-DNA-binding->Schlafen+HATPase+wHTH+HTH-><-XIS-HTH\|\|BrxA-DNA-binding->BrxB-BrxF-iSTAND->DUF6079*->HEPN->N6-METHYLASE->PglZ->SIGMA+OB-Fold+AAA-ATPase+HKD-DNase-BrxL-><-?<-ZnR+GP<-?<-MPTase | 1219 | Bacteria>Terrabacteria_group>Actinomycetota>Actinomycetes>Bifidobacteriales>Bifidobacteriaceae | GCA_000741575.1 | Bifidobacterium_cuniculi | BCUN_1388 | 51093..54752 | 1688 | putative-ATP-binding-protein-[Bifidobacterium-cuniculi] |
| OQM55032.1 | Type1.BREX | <-HIN-HTH<-N6-METHYLASE-p<-DUF6079*<-DUF6079-p<-BrxB-BrxF-iSTAND<-BrxA-DNA-binding<-?\|\|ACET-><-?\|\|HTH-><-35exo+BRCT | 1059 | Bacteria>Terrabacteria_group>Actinomycetota>Actinomycetes>Bifidobacteriales>Bifidobacteriaceae | GCA_002075955.1 | Bifidobacterium_dentium | B5790_1954 | 1122..4301 | 1689 | hypothetical-protein-B5790_1954-[Bifidobacterium-dentium] |
| PJM72971.1 | Type1.BREX | MACRODOMAIN-><-?<-SIGMA+OB-Fold+AAA-ATPase+HKD-DNase-BrxL<-PglZ<-?\|\|OLD-ABC->TOPRIM-><-N6-METHYLASE<-DUF6079*<-BrxB-BrxF-iSTAND<-BrxA-DNA-binding\|\|?->?-><-?<-?<-JetB | 1243 | Bacteria>Terrabacteria_group>Actinomycetota>Actinomycetes>Bifidobacteriales>Bifidobacteriaceae | GCA_002802875.1 | Bifidobacterium_primatium | CS006_06875 | 170776..174507 | 2045438 | hypothetical-protein-CS006_06875-[Bifidobacterium-primatium] |
| PKC83363.1 | Type1.BREX | <-HTH_Tnp_1\|\|RelB->N6-METHYLASE->CR-ATPase5+REase-p-><-CR-Kinase<-?\|\|BrxA-DNA-binding->BrxB-BrxF-iSTAND->DUF6079-p->DUF6079-p*->N6-METHYLASE->N6-METHYLASE->SLATT->SMODS->Schlafen+wHTH->PglZ->Lon_2->SIGMA+OB-Fold+AAA-ATPase+HKD-DNase-BrxL->TIR-> | 977 | Bacteria>Terrabacteria_group>Actinomycetota>Actinomycetes>Bifidobacteriales>Bifidobacteriaceae | GCA_002832945.1 | Bifidobacterium_longum | DPC6317_0876 | 237584..240517 | 216816 | hypothetical-protein-DPC6317_0876-[Bifidobacterium-longum] |
| PKV03451.1 | Type1.BREX | <-DUF6079-p\|\|DUF6079-p*->N6-METHYLASE->PglZ->SIGMA+OB-Fold+AAA-ATPase+HKD-DNase-BrxL-><-HEPN<-PRTase | 839 | Bacteria>Terrabacteria_group>Actinomycetota>Actinomycetes>Bifidobacteriales>Bifidobacteriaceae | GCA_002846775.1 | Bifidobacterium_pseudolongum_subsp._globosum | CQR50_1159 | 592..3111 | 1690 | hypothetical-protein-CQR50_1159-[Bifidobacterium-pseudolongum-subsp.-globosum] |
| PLS30252.1 | Type1.BREX | <-ABC-ATPase+SbcC<-?<-?<-?<-SIGMA+OB-Fold+AAA-ATPase+HKD-DNase-BrxL<-PglZ<-Schlafen+HATPase+wHTH<-N6-METHYLASE<-?<-DUF6079*<-BrxB-BrxF-iSTAND<-BrxA-DNA-binding<-?<-?<-?\|\|Peptidase_A24-> | 1244 | Bacteria>Terrabacteria_group>Actinomycetota>Actinomycetes>Bifidobacteriales>Bifidobacteriaceae | GCA_002860355.1 | Bifidobacterium_margollesii | Uis1B_1922 | 37676..41410 | 2020964 | hypothetical-protein-Uis1B_1922-[Bifidobacterium-margollesii] |
| RDX32761.1 | Type1.BREX | RelB->N6-METHYLASE->REase-><-CR-Kinase<-?\|\|BrxA-DNA-binding->BrxB-BrxF-iSTAND->DUF6079-p->DUF6079-p*->N6-METHYLASE->PglZ->PglZ-p->SIGMA+OB-Fold+AAA-ATPase+HKD-DNase-BrxL->TIR-> | 727 | Bacteria>Terrabacteria_group>Actinomycetota>Actinomycetes>Bifidobacteriales>Bifidobacteriaceae | GCA_003370195.1 | Bifidobacterium_breve | CE167_03785 | 16097..18282 | 1685 | hypothetical-protein-CE167_03785,-partial-[Bifidobacterium-breve] |
| RGY77464.1 | Type1.BREX | ABC-ATPase+SbcC->Pkinase+Pkinase->?->REC+LuxR-HTH->HISKIN-><-SbcD\|\|BrxA-DNA-binding->DUF6079*->N6-METHYLASE->SIR2->HerA/FtsK->N6-METHYLASE-p->SIR2->PglZ->SIGMA+OB-Fold+AAA-ATPase+HKD-DNase-BrxL->ZnR+GP->ZnR-> | 1219 | Bacteria>Terrabacteria_group>Actinomycetota>Actinomycetes>Bifidobacteriales>Bifidobacteriaceae | GCA_003463615.1 | Bifidobacterium_pseudocatenulatum | DXA22_03235 | 61960..65619 | 28026 | BREX-system-P-loop-protein-BrxC-[Bifidobacterium-pseudocatenulatum] |
| KAB5743305.1 | Type1.BREX | <-Abhydrolase<-ParB+HNH+RAMA<-ZnR<-ZnR+GP<-SIGMA+OB-Fold+AAA-ATPase+HKD-DNase-BrxL<-PglZ<-Schlafen+HATPase+wHTH<-SIR2<-N6-METHYLASE<-DUF6079*<-BrxB-BrxF-iSTAND<-BrxA-DNA-binding<-Pentapeptide+Pentapeptide<-RelE-ParE<-HAD\|\|SbcD-><-?<-?<-ACET+ACET | 1218 | Bacteria>Terrabacteria_group>Actinomycetota>Actinomycetes>Bifidobacteriales>Bifidobacteriaceae | GCA_009075895.1 | Bifidobacterium_adolescentis | GA749_07780 | 122966..126622 | 1680 | BREX-system-P-loop-protein-BrxC-[Bifidobacterium-adolescentis] |
| KAB5930818.1 | Type1.BREX | PemK_toxin->PolB->HEPN-><-?<-?\|\|?->\|\|Pentapeptide+Pentapeptide+Pentapeptide->BrxA-DNA-binding->BrxB-BrxF-iSTAND->DUF6079*->N6-METHYLASE->SIR2->Schlafen+HATPase+wHTH->PglZ->SIGMA+OB-Fold+AAA-ATPase+HKD-DNase-BrxL->?->?-><-REase | 1219 | Bacteria>Terrabacteria_group>Actinomycetota>Actinomycetes>Bifidobacteriales>Bifidobacteriaceae | GCA_009094485.1 | Bifidobacterium_adolescentis | GA595_06005 | 91928..95587 | 1680 | BREX-system-P-loop-protein-BrxC-[Bifidobacterium-adolescentis] |
| KAB6780790.1 | Type1.BREX | HTH+MPTase-><-?<-HEPN<-?\|\|?->BrxA-DNA-binding->BrxB-BrxF-iSTAND->DUF6079*->EVE+McrB-AAA->McrC-NTD+REase->METHYLASE-p->N6-METHYLASE-p-> | 1229 | Bacteria>Terrabacteria_group>Actinomycetota>Actinomycetes>Bifidobacteriales>Bifidobacteriaceae | GCA_009153345.1 | Bifidobacterium_longum | GBL14_01920 | 66538..70227 | 216816 | BREX-system-P-loop-protein-BrxC-[Bifidobacterium-longum] |
| KAB6790527.1 | Type1.BREX | <-N6-METHYLASE-p<-METHYLASE-p<-McrC-NTD+REase<-EVE+McrB-AAA<-DUF6079*<-BrxB-BrxF-iSTAND<-BrxA-DNA-binding | 1229 | Bacteria>Terrabacteria_group>Actinomycetota>Actinomycetes>Bifidobacteriales>Bifidobacteriaceae | GCA_009153545.1 | Bifidobacterium_longum | GBK34_07935 | 4820..8509 | 216816 | BREX-system-P-loop-protein-BrxC-[Bifidobacterium-longum] |
| KAB6796648.1 | Type1.BREX | <-N6-METHYLASE-p<-METHYLASE-p<-McrC-NTD+REase<-EVE+McrB-AAA<-DUF6079*<-BrxB-BrxF-iSTAND<-BrxA-DNA-binding<-?\|\|?->HEPN->?-><-HTH+MPTase | 1229 | Bacteria>Terrabacteria_group>Actinomycetota>Actinomycetes>Bifidobacteriales>Bifidobacteriaceae | GCA_009154515.1 | Bifidobacterium_longum | GBK79_05090 | 4820..8509 | 216816 | BREX-system-P-loop-protein-BrxC-[Bifidobacterium-longum] |
| KAB7145752.1 | Type1.BREX | HTH+MPTase-><-?<-HEPN<-?\|\|?->BrxA-DNA-binding->BrxB-BrxF-iSTAND->DUF6079*->EVE+McrB-AAA->McrC-NTD+REase->METHYLASE-p->N6-METHYLASE-p-> | 1229 | Bacteria>Terrabacteria_group>Actinomycetota>Actinomycetes>Bifidobacteriales>Bifidobacteriaceae | GCA_009156155.1 | Bifidobacterium_longum | GBC99_05955 | 66538..70227 | 216816 | BREX-system-P-loop-protein-BrxC-[Bifidobacterium-longum] |
| MBE5064977.1 | Type1.BREX | ACET->ACET+ACET->?->Thioredoxin_3->?-><-Glycoside_Hydrolase\|\|OLD-ABC->BrxA-DNA-binding->BrxB-BrxF-iSTAND->DUF6079*->N6-METHYLASE->REase->?->Schlafen+HATPase+wHTH->PglZ->SIGMA+OB-Fold+AAA-ATPase+HKD-DNase-BrxL-> | 1221 | Bacteria>Terrabacteria_group>Actinomycetota>Actinomycetes>Bifidobacteriales>Bifidobacteriaceae | GCA_014983005.1 | Bifidobacterium_pullorum_subsp._saeculare | INF27_02890 | 184023..187688 | 78257 | BREX-system-P-loop-protein-BrxC-[Bifidobacterium-pullorum-subsp.-saeculare] |
| GDY90547.1 | Type1.BREX | <-Sigma-HTH<-ParB+HNH<-ZnR<-ZnR+GP<-SIGMA+OB-Fold+AAA-ATPase+HKD-DNase-BrxL<-PglZ<-Schlafen+HATPase+wHTH<-SIR2<-N6-METHYLASE<-DUF6079*<-BrxB-BrxF-iSTAND<-BrxA-DNA-binding<-HTH_28 | 1219 | Bacteria>Terrabacteria_group>Actinomycetota>Actinomycetes>Bifidobacteriales>Bifidobacteriaceae | GCA_018326505.1 | Bifidobacteriaceae_bacterium_MCC01971 | MCC01971_21650 | 16049..19708 | 2569485 | hypothetical-protein-MCC01971_21650-[Bifidobacteriaceae-bacterium-MCC01971] |
| GDY95791.1 | Type1.BREX | <-HTH_3<-tRNA<-ACET\|\|Acyl_transf_3->Acyl_transf_3->?-><-?\|\|?->BrxA-DNA-binding->BrxB-BrxF-iSTAND->DUF6079*->N6-METHYLASE->HEPN->Schlafen+HATPase+wHTH->PglZ->SIGMA+OB-Fold+AAA-ATPase+HKD-DNase-BrxL->ZnR+GP->ZnR->ParB+HNH+RAMA-> | 1223 | Bacteria>Terrabacteria_group>Actinomycetota>Actinomycetes>Bifidobacteriales>Bifidobacteriaceae | GCA_018326565.1 | Bifidobacteriaceae_bacterium_MCC01943 | MCC01943_10780 | 5409..9080 | 2569488 | hypothetical-protein-MCC01943_10780-[Bifidobacteriaceae-bacterium-MCC01943] |
| GDZ58121.1 | Type1.BREX | <-SIR2<-?\|\|BrxA-DNA-binding->BrxB-BrxF-iSTAND->DUF6079*->N6-METHYLASE->?->Schlafen+HATPase+wHTH->PglZ->SIGMA+OB-Fold+AAA-ATPase+HKD-DNase-BrxL->?-><-?\|\|GDResolvase+HIN-HTH-> | 1219 | Bacteria>Terrabacteria_group>Actinomycetota>Actinomycetes>Bifidobacteriales>Bifidobacteriaceae | GCA_018327225.1 | Bifidobacteriaceae_bacterium_MCC01967 | MCC01967_09770 | 127200..130859 | 2569520 | hypothetical-protein-MCC01967_09770-[Bifidobacteriaceae-bacterium-MCC01967] |
| MBS6134848.1 | Type1.BREX | <-N6-METHYLASE-p<-HerA/FtsK<-SIR2<-N6-METHYLASE<-DUF6079*<-BrxB-BrxF-iSTAND<-BrxA-DNA-binding<-CR-ATPase5+REase-p<-?<-ARG | 1229 | Bacteria>Terrabacteria_group>Actinomycetota>Actinomycetes>Bifidobacteriales>Bifidobacteriaceae | GCA_018374015.1 | Bifidobacterium_longum | KH705_09795 | 10639..14328 | 216816 | BREX-system-P-loop-protein-BrxC-[Bifidobacterium-longum] |
| MBS6717590.1 | Type1.BREX | HTH_28->BrxA-DNA-binding->BrxB-BrxF-iSTAND->DUF6079*->REase->Schlafen+HATPase+wHTH->PglZ->SIGMA+OB-Fold+AAA-ATPase+HKD-DNase-BrxL->SLATT->SMODS-> | 1218 | Bacteria>Terrabacteria_group>Actinomycetota>Actinomycetes>Bifidobacteriales>Bifidobacteriaceae | GCA_018378875.1 | Bifidobacterium_longum | KH250_10240 | 6375..10031 | 216816 | BREX-system-P-loop-protein-BrxC-[Bifidobacterium-longum] |
| MBS7047476.1 | Type1.BREX | RelB->PemK_toxin->PolB->HEPN-><-?<-?<-REase\|\|BrxA-DNA-binding->BrxB-BrxF-iSTAND->DUF6079*->N6-METHYLASE->SIR2->Schlafen+HATPase+wHTH->PglZ->SIGMA+OB-Fold+AAA-ATPase+HKD-DNase-BrxL->ZnR+GP->ZnR->ParB+HNH+RAMA->Abhydrolase-> | 1218 | Bacteria>Terrabacteria_group>Actinomycetota>Actinomycetes>Bifidobacteriales>Bifidobacteriaceae | GCA_018381815.1 | Bifidobacterium_adolescentis | KH096_03850 | 341650..345306 | 1680 | BREX-system-P-loop-protein-BrxC-[Bifidobacterium-adolescentis] |

<table>
<tr><th>Protein-ID</th><th>System</th><th>Gene-Neighbourhoods</th><th>Length</th><th>Taxonomy</th><th>Assembly-ID</th><th>Strain</th><th>Locus-Tag</th><th>Genomic-coordinates</th><th>TaxID</th><th>Fasta-define</th></tr>
<tr><td>MCH4189981.1</td><td>Type1.BREX</td><td><-SIGMA+OB-Fold+AAA-ATPase+HKD-DNase-BrxL<-PglZ<-N6-METHYLASE-p<-Phage_int_SAM_3+Phage_integrase||N6-METHYLASE-p-><-N6-METHYLASE<-?<-DUF6079*<-BrxB-BrxF-iSTAND<-BrxA-DNA-binding<-?<- [illegible]</td><td>1173</td><td>Bacteria>Terrabacteria_group>Actinomycetota>Actinomycetes>Bifidobacteriales>Bifidobacteriaceae</td><td>GCA_022486485.1</td><td>Bifidobacterium_tibiigranuli</td><td>LKF45_06440</td><td>46152..49673</td><td>2172043</td><td>[illegible]</td></tr>
<tr><td>MCI1218827.1</td><td>Type1.BREX</td><td><-N6-METHYLASE-p<-?<-DUF6079*<-BrxB-BrxF-iSTAND<-BrxA-DNA-binding<-?<-?<-?<-Aminotran_1_2||?-><-?<-MoxR-AAA</td><td>1170</td><td>Bacteria>Terrabacteria_group>Actinomycetota>Actinomycetes>Bifidobacteriales>Bifidobacteriaceae</td><td>GCA_022640875.1</td><td>Bifidobacterium_crudilactis</td><td>LKG00_08945</td><td>2590..6102</td><td>327277</td><td>[illegible]</td></tr>
<tr><td>MCI2117722.1</td><td>Type1.BREX</td><td><-RNase<-BirA-HTH<-Schlafen+HATPase+wHTH<-SMODS<-SLATT<-N6-METHYLASE-p<-HerA/FtsK<-SIR2<-N6-METHYLASE<-DUF6079*<-BrxB-BrxF-iSTAND<-BrxA-DNA-binding<-CR-ATPase5+REase-p<-?||?-><-?<-?<- [illegible]</td><td>1229</td><td>Bacteria>Terrabacteria_group>Actinomycetota>Actinomycetes>Bifidobacteriales>Bifidobacteriaceae</td><td>GCA_022653935.1</td><td>Bifidobacterium_breve</td><td>LKK17_00135</td><td>29818..33507</td><td>1685</td><td>[illegible]</td></tr>
<tr><td>MCK6131816.1</td><td>Type1.BREX</td><td>35exo+HRDC->?->35exo+BRCT-><-HTH||?-><-ACET||?->BrxA-DNA-binding->BrxB-BrxF-iSTAND->DUF6079*->N6-METHYLASE->?->Schlafen+HATPase+wHTH->PglZ->SIGMA+OB-Fold+AAA-ATPase+HKD-DNase-BrxL-> [illegible]</td><td>1217</td><td>Bacteria>Terrabacteria_group>Actinomycetota>Actinomycetes>Bifidobacteriales>Bifidobacteriaceae</td><td>GCA_023147995.1</td><td>Bifidobacterium_dentium</td><td>KZX63_03525</td><td>164396..168049</td><td>1689</td><td>[illegible]</td></tr>
<tr><td>WP_014697718.1</td><td>Type1.BREX</td><td><-MODE-HTH+PBPII<-HAD+HAD||REase->ARG->YTCD-HTH->NUDIX-><-ACET+ACET||BrxA-DNA-binding->BrxB-BrxF-iSTAND->DUF6079*->N6-METHYLASE->PglZ->SIGMA+OB-Fold+AAA-ATPase+HKD-DNase-BrxL->SIR2->< [illegible]</td><td>1212</td><td>Bacteria>Terrabacteria_group>Actinomycetota>Actinomycetes>Bifidobacteriales>Bifidobacteriaceae</td><td>GCF_000260715.1</td><td>Bifidobacterium_animalis</td><td>CNCMI4602_RS04575</td><td>1015719..1019357</td><td>28025</td><td>[illegible]</td></tr>
<tr><td>WP_003812955.1</td><td>Type1.BREX</td><td>ZnR->?->ABC-ATPase+SbcC->?->?->?->BrxA-DNA-binding->BrxB-BrxF-iSTAND->DUF6079*->N6-METHYLASE->SIR2->Schlafen+HATPase+wHTH->PglZ->SIGMA+OB-Fold+AAA-ATPase+HKD-DNase-BrxL->?-><-REase|| [illegible]</td><td>1218</td><td>Bacteria>Terrabacteria_group>Actinomycetota>Actinomycetes>Bifidobacteriales>Bifidobacteriaceae</td><td>GCF_000273525.1</td><td>Bifidobacterium_bifidum</td><td>BBNG_RS0103505</td><td>833890..837546</td><td>1681</td><td>[illegible]</td></tr>
<tr><td>WP_003819016.1</td><td>Type1.BREX</td><td>ZnR->?->ABC-ATPase+SbcC->?->?->?->BrxA-DNA-binding->BrxB-BrxF-iSTAND->DUF6079*->N6-METHYLASE->REase->Schlafen+HATPase+wHTH->PglZ->SIGMA+OB-Fold+AAA-ATPase+HKD-DNase-BrxL->?-><-REase|| [illegible]</td><td>1217</td><td>Bacteria>Terrabacteria_group>Actinomycetota>Actinomycetes>Bifidobacteriales>Bifidobacteriaceae</td><td>GCF_000300215.1</td><td>Bifidobacterium_bifidum</td><td>IF1010_RS06520</td><td>82392..86045</td><td>1681</td><td>[illegible]</td></tr>
<tr><td>WP_022857559.1</td><td>Type1.BREX</td><td>HAD+HAD->MODE-HTH+PBPII->YTCD-HTH-><-ACET+ACET<-ZnR||BrxA-DNA-binding->BrxB-BrxF-iSTAND->DUF6079*->N6-METHYLASE->PglZ->SIGMA+OB-Fold+AAA-ATPase+HKD-DNase-BrxL->SIR2-><-PRTase</td><td>1214</td><td>Bacteria>Terrabacteria_group>Actinomycetota>Actinomycetes>Bifidobacteriales>Bifidobacteriaceae</td><td>GCF_000421365.1</td><td>Bifidobacterium_pseudolongum</td><td>G627_RS0100670</td><td>128741..132385</td><td>1694</td><td>[illegible]</td></tr>
<tr><td>WP_029679903.1</td><td>Type1.BREX</td><td>HTH_28->BrxA-DNA-binding->BrxB-BrxF-iSTAND->DUF6079*->N6-METHYLASE->SIR2->Schlafen+HATPase+wHTH->PglZ->SIGMA+OB-Fold+AAA-ATPase+HKD-DNase-BrxL->?-><-rve</td><td>1218</td><td>Bacteria>Terrabacteria_group>Actinomycetota>Actinomycetes>Bifidobacteriales>Bifidobacteriaceae</td><td>GCF_000421385.1</td><td>Bifidobacterium_longum</td><td>G629_RS0105465</td><td>6869..10525</td><td>216816</td><td>[illegible]</td></tr>
<tr><td>WP_026645358.1</td><td>Type1.BREX</td><td>YTCD-HTH->?-><-?||ARG-><-ACET+ACET<-ZnR||BrxA-DNA-binding->BrxB-BrxF-iSTAND->DUF6079*->N6-METHYLASE->PglZ->SIGMA+OB-Fold+AAA-ATPase+HKD-DNase-BrxL->SIR2-><-HEPN<-PRTase</td><td>1221</td><td>Bacteria>Terrabacteria_group>Actinomycetota>Actinomycetes>Bifidobacteriales>Bifidobacteriaceae</td><td>GCF_000424225.1</td><td>Bifidobacterium_sp._AGR2158</td><td>G606_RS0106975</td><td>126292..129957</td><td>1280675</td><td>[illegible]</td></tr>
<tr><td>WP_032745990.1</td><td>Type1.BREX</td><td><-Sigma-HTH<-ParB+HNH<-ZnR<-ZnR+GP<-SIGMA+OB-Fold+AAA-ATPase+HKD-DNase-BrxL<-PglZ<-Schlafen+HATPase+wHTH<-SIR2<-N6-METHYLASE<-DUF6079*<-BrxB-BrxF-iSTAND<-BrxA-DNA-binding<-HTH_28</td><td>1219</td><td>Bacteria>Terrabacteria_group>Actinomycetota>Actinomycetes>Bifidobacteriales>Bifidobacteriaceae</td><td>GCF_000730035.1</td><td>Bifidobacterium_longum</td><td>BL171B_RS06775</td><td>278302..281961</td><td>216816</td><td>[illegible]</td></tr>
<tr><td>WP_032738770.1</td><td>Type1.BREX</td><td>HTH_28->BrxA-DNA-binding->BrxB-BrxF-iSTAND->DUF6079*->N6-METHYLASE->HEPN->?->Schlafen+HATPase+wHTH->PglZ->SIGMA+OB-Fold+AAA-ATPase+HKD-DNase-BrxL-></td><td>1218</td><td>Bacteria>Terrabacteria_group>Actinomycetota>Actinomycetes>Bifidobacteriales>Bifidobacteriaceae</td><td>GCF_000730055.1</td><td>Bifidobacterium_longum</td><td>BL71B_RS01495</td><td>365977..369633</td><td>216816</td><td>[illegible]</td></tr>
<tr><td>WP_032735726.1</td><td>Type1.BREX</td><td>HTH_28->BrxA-DNA-binding->BrxB-BrxF-iSTAND->DUF6079*->N6-METHYLASE->SIR2->Schlafen+HATPase+wHTH->PglZ->SIGMA+OB-Fold+AAA-ATPase+HKD-DNase-BrxL-></td><td>1218</td><td>Bacteria>Terrabacteria_group>Actinomycetota>Actinomycetes>Bifidobacteriales>Bifidobacteriaceae</td><td>GCF_000730105.1</td><td>Bifidobacterium_longum</td><td>BL15B_RS01745</td><td>289359..293015</td><td>216816</td><td>[illegible]</td></tr>
<tr><td>WP_051922710.1</td><td>Type1.BREX</td><td><-His_Phos_1<-?||ZnR->?->ABC-ATPase+SbcC-><-?||BrxA-DNA-binding->BrxB-BrxF-iSTAND->DUF6079*->N6-METHYLASE->?->HEPN+RES->Schlafen+HATPase+wHTH->PglZ->SIGMA+OB-Fold+AAA-ATPase+HKD-DNase [illegible]</td><td>1249</td><td>Bacteria>Terrabacteria_group>Actinomycetota>Actinomycetes>Bifidobacteriales>Bifidobacteriaceae</td><td>GCF_000741785.1</td><td>Bifidobacterium_stellenboschense</td><td>BSTEL_1465</td><td>61247..64996</td><td>762211</td><td>[illegible]</td></tr>
<tr><td>WP_026646062.1</td><td>Type1.BREX</td><td><-RelB<-ParB+HNH+RAMA<-ZnR<-ZnR+GP<-SIGMA+OB-Fold+AAA-ATPase+HKD-DNase-BrxL<-PglZ<-Schlafen+HATPase+wHTH<-?<-N6-METHYLASE<-DUF6079*<-BrxB-BrxF-iSTAND<-BrxA-DNA-binding||REase->?->?-> [illegible]</td><td>1219</td><td>Bacteria>Terrabacteria_group>Actinomycetota>Actinomycetes>Bifidobacteriales>Bifidobacteriaceae</td><td>GCF_000770925.1</td><td>Bifidobacterium_ruminantium</td><td>T534_RS0102115</td><td>18145..21804</td><td>78346</td><td>[illegible]</td></tr>
<tr><td>WP_033509374.1</td><td>Type1.BREX</td><td><-SIGMA+OB-Fold+AAA-ATPase+HKD-DNase-BrxL<-PglZ<-Schlafen+HATPase+wHTH<-N6-METHYLASE-p<-HerA/FtsK<-SIR2<-N6-METHYLASE<-DUF6079*<-BrxB-BrxF-iSTAND<-BrxA-DNA-binding<-TOPRIM<-OLD-ABC|| [illegible]</td><td>1221</td><td>Bacteria>Terrabacteria_group>Actinomycetota>Actinomycetes>Bifidobacteriales>Bifidobacteriaceae</td><td>GCF_000770965.1</td><td>Bifidobacterium_pullorum</td><td>OU28_RS04650</td><td>22611..26276</td><td>78448</td><td>[illegible]</td></tr>
<tr><td>WP_160257632.1</td><td>Type1.BREX</td><td><-XIS-HTH+GDResolvase||REase->?->BrxA-DNA-binding->Schlafen+HATPase+wHTH+HTH-><-XIS-HTH||BrxA-DNA-binding->BrxB-BrxF-iSTAND-> [illegible]</td><td>1215</td><td>Bacteria>Terrabacteria_group>Actinomycetota>Actinomycetes>Bifidobacteriales>Bifidobacteriaceae</td><td>GCF_000771465.1</td><td>Bifidobacterium_cuniculi</td><td>BCUN_RS07325</td><td>51105..54752</td><td>1688</td><td>[illegible]</td></tr>
<tr><td>WP_051923722.1</td><td>Type1.BREX</td><td><-METHYLASE<-SIGMA+OB-Fold+AAA-ATPase+HKD-DNase-BrxL<-PglZ<-Schlafen+HATPase+wHTH<-SMODS<-ABhydrolase<-N6-METHYLASE<-CR-ATPase [illegible]</td><td>1247</td><td>Bacteria>Terrabacteria_group>Actinomycetota>Actinomycetes>Bifidobacteriales>Bifidobacteriaceae</td><td>GCF_000771645.1</td><td>Bifidobacterium_biavatii</td><td>OU23_RS06270</td><td>17800..21543</td><td>762212</td><td>[illegible]</td></tr>
<tr><td>WP_052382006.1</td><td>Type1.BREX</td><td><-N6-METHYLASE<-?<-?<-?<-TPR+TPR<-CR-ATPase5+REase<-DUF6079*<-BrxB-BrxF-iSTAND<-BrxA-DNA-binding||ABhydrolase->?-><-?<-HAD+S6P [illegible]</td><td>1264</td><td>Bacteria>Terrabacteria_group>Actinomycetota>Actinomycetes>Bifidobacteriales>Bifidobacteriaceae</td><td>GCF_000771685.1</td><td>Bifidobacterium_reuteri</td><td>OU25_RS00505</td><td>118725..122519</td><td>983706</td><td>[illegible]</td></tr>
<tr><td>WP_038426289.1</td><td>Type1.BREX</td><td>HTH_28->BrxA-DNA-binding->BrxB-BrxF-iSTAND->DUF6079*->N6-METHYLASE->?->?->Schlafen+HATPase+wHTH->PglZ->SIGMA+OB-Fold+AAA-ATPase [illegible]</td><td>1218</td><td>Bacteria>Terrabacteria_group>Actinomycetota>Actinomycetes>Bifidobacteriales>Bifidobacteriaceae</td><td>GCF_000772485.1</td><td>Bifidobacterium_longum</td><td>BLGT_RS02155</td><td>511827..515483</td><td>216816</td><td>[illegible]</td></tr>
<tr><td>WP_041080188.1</td><td>Type1.BREX</td><td>HTH_28->BrxA-DNA-binding->BrxB-BrxF-iSTAND->DUF6079*->N6-METHYLASE->?->Schlafen+HATPase+wHTH->PglZ->SIGMA+OB-Fold+AAA-ATPase+ [illegible]</td><td>1219</td><td>Bacteria>Terrabacteria_group>Actinomycetota>Actinomycetes>Bifidobacteriales>Bifidobacteriaceae</td><td>GCF_000829295.1</td><td>Bifidobacterium_longum</td><td>BL105A_RS01810</td><td>418620..422279</td><td>216816</td><td>[illegible]</td></tr>
<tr><td>WP_046999386.1</td><td>Type1.BREX</td><td>REase-><-ZnR<-ZnR+GP<-SIGMA+OB-Fold+AAA-ATPase+HKD-DNase-BrxL<-PglZ<-Schlafen+HATPase+wHTH<-REase<-N6-METHYLASE<-DUF6079*<- [illegible]</td><td>1217</td><td>Bacteria>Terrabacteria_group>Actinomycetota>Actinomycetes>Bifidobacteriales>Bifidobacteriaceae</td><td>GCF_001010915.1</td><td>Bifidobacterium_adolescentis</td><td>AAX71_RS05490</td><td>58570..62223</td><td>1680</td><td>[illegible]</td></tr>
<tr><td>WP_047298571.1</td><td>Type1.BREX</td><td><-Sigma-HTH<-ParB+HNH<-?<-ZnR+GP<-SIGMA+OB-Fold+AAA-ATPase+HKD-DNase-BrxL<-PglZ<-Schlafen+HATPase+wHTH<-?<-N6-METHYLASE<- [illegible]</td><td>1217</td><td>Bacteria>Terrabacteria_group>Actinomycetota>Actinomycetes>Bifidobacteriales>Bifidobacteriaceae</td><td>GCF_001020335.1</td><td>Bifidobacterium_bifidum</td><td>LMG11583_RS03370</td><td>14804..18457</td><td>1681</td><td>[illegible]</td></tr>
<tr><td>WP_003826153.1</td><td>Type1.BREX</td><td><-SbcD||HAD->PolB->HEPN-><-?<-?<-?<-REase||BrxA-DNA-binding->BrxB-BrxF-iSTAND->DUF6079*->N6-METHYLASE->SIR2->HerA/FtsK-> [illegible]</td><td>1218</td><td>Bacteria>Terrabacteria_group>Actinomycetota>Actinomycetes>Bifidobacteriales>Bifidobacteriaceae</td><td>GCF_001025155.1</td><td>Bifidobacterium_angulatum</td><td>BBAG_RS03465</td><td>863700..867356</td><td>1683</td><td>[illegible]</td></tr>
<tr><td>WP_050824906.1</td><td>Type1.BREX</td><td><-SIGMA+OB-Fold+AAA-ATPase+HKD-DNase-BrxL<-PglZ<-Schlafen+wHTH<-Schlafen+wHTH<-SMODS<-SLATT<-N6-METHYLASE<-DUF6079*<-BrxB-BrxF [illegible]</td><td>1229</td><td>Bacteria>Terrabacteria_group>Actinomycetota>Actinomycetes>Bifidobacteriales>Bifidobacteriaceae</td><td>GCF_001189355.1</td><td>Bifidobacterium_breve</td><td>BBRI4_RS05550</td><td>137028..140717</td><td>1685</td><td>[illegible]</td></tr>
<tr><td>WP_052828891.1</td><td>Type1.BREX</td><td><-TIR<-SIGMA+OB-Fold+AAA-ATPase+HKD-DNase-BrxL<-PglZ<-Schlafen+wHTH<-SMODS<-SLATT<-N6-METHYLASE<-DUF6079*<-BrxB-BrxF-iSTAND<- [illegible]</td><td>1229</td><td>Bacteria>Terrabacteria_group>Actinomycetota>Actinomycetes>Bifidobacteriales>Bifidobacteriaceae</td><td>GCF_001264045.1</td><td>Bifidobacterium_breve</td><td>BAD03_RS05925</td><td>144131..147820</td><td>1685</td><td>[illegible]</td></tr>
<tr><td>WP_052792432.1</td><td>Type1.BREX</td><td><-SIR2||BrxA-DNA-binding->BrxB-BrxF-iSTAND->DUF6079*->N6-METHYLASE->SIR2->HerA/FtsK->N6-METHYLASE-p->SIR2->Schlafen+HATPase+ [illegible]</td><td>1218</td><td>Bacteria>Terrabacteria_group>Actinomycetota>Actinomycetes>Bifidobacteriales>Bifidobacteriaceae</td><td>GCF_001264105.1</td><td>Bifidobacterium</td><td>BBM1340_RS06775</td><td>7664..11320</td><td>1678</td><td>[illegible]</td></tr>
<tr><td>WP_053495781.1</td><td>Type1.BREX</td><td><-CbiA<-?<-?<-ZDNA-HTH<-?||ACET-><-?<-HTH_3||HTH_3-><-DUF6079-p*<-BrxB-BrxF-iSTAND<-BrxA-DNA-binding||?->CR-Kinase-><-CR-ATPase [illegible]</td><td>531</td><td>Bacteria>Terrabacteria_group>Actinomycetota>Actinomycetes>Bifidobacteriales>Bifidobacteriaceae</td><td>GCF_001275745.1</td><td>Bifidobacterium_longum</td><td>BLOI2_RS03170</td><td>185000..186595</td><td>216816</td><td>[illegible]</td></tr>
<tr><td>WP_053824992.1</td><td>Type1.BREX</td><td>ZnR->?->ABC-ATPase+SbcC->?->?->?->BrxA-DNA-binding->BrxB-BrxF-iSTAND->DUF6079*->N6-METHYLASE->SIR2->Schlafen+HATPase+wHTH-> [illegible]</td><td>1217</td><td>Bacteria>Terrabacteria_group>Actinomycetota>Actinomycetes>Bifidobacteriales>Bifidobacteriaceae</td><td>GCF_001281345.1</td><td>Bifidobacterium_bifidum</td><td>RY70_RS04415</td><td>1112532..1116185</td><td>1681</td><td>[illegible]</td></tr>
<tr><td>WP_253276104.1</td><td>Type1.BREX</td><td><-SbcD||RelB->PemK_toxin->PolB->HEPN-><-?<-AIPR||BrxA-DNA-binding->BrxB-BrxF-iSTAND->DUF6079-p*->DUF6079-p->N6-METHYLASE->SIR2 [illegible]</td><td>561</td><td>Bacteria>Terrabacteria_group>Actinomycetota>Actinomycetes>Bifidobacteriales>Bifidobacteriaceae</td><td>GCF_001406455.1</td><td>Bifidobacterium_adolescentis</td><td>ARB73_RS10570</td><td>418734..420419</td><td>1680</td><td>[illegible]</td></tr>
<tr><td>WP_065465686.1</td><td>Type1.BREX</td><td><-TIR<-SIGMA+OB-Fold+AAA-ATPase+HKD-DNase-BrxL<-PglZ<-Schlafen+wHTH<-?<-N6-METHYLASE<-DUF6079*<-BrxB-BrxF-iSTAND<- [illegible]</td><td>1230</td><td>Bacteria>Terrabacteria_group>Actinomycetota>Actinomycetes>Bifidobacteriales>Bifidobacteriaceae</td><td>GCF_001685805.1</td><td>Bifidobacterium_breve</td><td>BAD00_RS06410</td><td>112826..116518</td><td>1685</td><td>[illegible]</td></tr>
<tr><td>WP_065435300.1</td><td>Type1.BREX</td><td>Pkinase+Pkinase->REC+LuxR-HTH->HISKIN-><-?<-SbcD<-?<-REase||BrxA-DNA-binding->BrxB-BrxF-iSTAND->DUF6079*->N6-METHYLASE->SIR2-> [illegible]</td><td>1219</td><td>Bacteria>Terrabacteria_group>Actinomycetota>Actinomycetes>Bifidobacteriales>Bifidobacteriaceae</td><td>GCF_001686025.1</td><td>Bifidobacterium_pseudocatenulatum</td><td>BAD11_RS06840</td><td>26725..30384</td><td>28026</td><td>[illegible]</td></tr>
<tr><td>WP_065445635.1</td><td>Type1.BREX</td><td>HTH_28->BrxA-DNA-binding->BrxB-BrxF-iSTAND->DUF6079*->N6-METHYLASE->REase->Schlafen+HATPase+wHTH->PglZ->SIGMA+OB-Fold+AAA- [illegible]</td><td>1219</td><td>Bacteria>Terrabacteria_group>Actinomycetota>Actinomycetes>Bifidobacteriales>Bifidobacteriaceae</td><td>GCF_001686165.1</td><td>Bifidobacterium_longum</td><td>MCC10077_RS10215</td><td>2512..6171</td><td>216816</td><td>[illegible]</td></tr>
<tr><td>WP_065453246.1</td><td>Type1.BREX</td><td>HTH_28->BrxA-DNA-binding->BrxB-BrxF-iSTAND->DUF6079*->N6-METHYLASE->SIR2->Schlafen+HATPase+wHTH->PglZ->SIGMA+OB-Fold+AAA-ATPase [illegible]</td><td>1219</td><td>Bacteria>Terrabacteria_group>Actinomycetota>Actinomycetes>Bifidobacteriales>Bifidobacteriaceae</td><td>GCF_001686185.1</td><td>Bifidobacterium_longum</td><td>BAD19_RS10570</td><td>392770..396429</td><td>216816</td><td>[illegible]</td></tr>
<tr><td>WP_071474421.1</td><td>Type1.BREX</td><td><-Sigma-HTH<-ParB+HNH<-ZnR<-ZnR+GP<-SIGMA+OB-Fold+AAA-ATPase+HKD-DNase-BrxL<-PglZ<-Schlafen+HATPase+wHTH<-SIR2<-N6-METHYLASE<- [illegible]</td><td>1218</td><td>Bacteria>Terrabacteria_group>Actinomycetota>Actinomycetes>Bifidobacteriales>Bifidobacteriaceae</td><td>GCF_001870705.1</td><td>Bifidobacterium_longum</td><td>BFS26_RS01875</td><td>17972..21628</td><td>216816</td><td>[illegible]</td></tr>
<tr><td>WP_071477881.1</td><td>Type1.BREX</td><td><-SIGMA+OB-Fold+AAA-ATPase+HKD-DNase-BrxL<-PglZ<-RNase<-Schlafen+HATPase+wHTH<-SMODS<-SLATT<-N6-METHYLASE<-DUF6079*<-BrxB-BrxF- [illegible]</td><td>1229</td><td>Bacteria>Terrabacteria_group>Actinomycetota>Actinomycetes>Bifidobacteriales>Bifidobacteriaceae</td><td>GCF_001870755.1</td><td>Bifidobacterium_longum</td><td>BFS25_RS03445</td><td>115725..119414</td><td>216816</td><td>[illegible]</td></tr>
<tr><td>WP_072727063.1</td><td>Type1.BREX</td><td><-OLD-ABC+TOPRIM<-N6-METHYLASE<-N6-METHYLASE-p<-DUF6079*<-BrxB-BrxF-iSTAND<-BrxA-DNA-binding||?->NTP_transf->?-><-CR-ATPase5+ [illegible]</td><td>1208</td><td>Bacteria>Terrabacteria_group>Actinomycetota>Actinomycetes>Bifidobacteriales>Bifidobacteriaceae</td><td>GCF_001895165.1</td><td>Bifidobacterium_lemurum</td><td>BRTL_RS11125</td><td>5981..9607</td><td>1603886</td><td>[illegible]</td></tr>
<tr><td>WP_094668334.1</td><td>Type1.BREX</td><td>DDE_Tnp_ISL3-><-ABhydrolase||BrxA-DNA-binding->BrxB-BrxF-iSTAND->DUF6079*->CR-ATPase5+REase->TPR+TPR->N6-METHYLASE->Schlafen+ [illegible]</td><td>1270</td><td>Bacteria>Terrabacteria_group>Actinomycetota>Actinomycetes>Bifidobacteriales>Bifidobacteriaceae</td><td>GCF_002259745.1</td><td>Bifidobacterium_myosotis</td><td>BMYO_RS10065</td><td>3739..7551</td><td>1630166</td><td>[illegible]</td></tr>
<tr><td>WP_095613737.1</td><td>Type1.BREX</td><td>MODE-HTH+PBPII->YTCD-HTH->?-><-?||ARG-><-ACET+ACET||?->BrxA-DNA-binding->BrxB-BrxF-iSTAND->DUF6079*->N6-METHYLASE-p-> [illegible]</td><td>1215</td><td>Bacteria>Terrabacteria_group>Actinomycetota>Actinomycetes>Bifidobacteriales>Bifidobacteriaceae</td><td>GCF_002286915.1</td><td>Bifidobacterium_italicum</td><td>B1400_RS07185</td><td>20228..23875</td><td>1960968</td><td>[illegible]</td></tr>
<tr><td>WP_100495666.1</td><td>Type1.BREX</td><td><-ABhydrolase||TPR+TPR+TPR+TPR->BrxA-DNA-binding->BrxB-BrxF-iSTAND->DUF6079*-><-?<-?<-?||HNH->?->SbcC->N6-METHYLASE->HEPN-> [illegible]</td><td>1240</td><td>Bacteria>Terrabacteria_group>Actinomycetota>Actinomycetes>Bifidobacteriales>Bifidobacteriaceae</td><td>GCF_002802865.1</td><td>Bifidobacterium_scaligerum</td><td>CUU80_RS01790</td><td>434556..438278</td><td>2052656</td><td>[illegible]</td></tr>
<tr><td>WP_100511074.1</td><td>Type1.BREX</td><td>MACRODOMAIN-><-?<-SIGMA+OB-Fold+AAA-ATPase+HKD-DNase-BrxL<-PglZ<-?||OLD-ABC->TOPRIM-><-N6-METHYLASE<-DUF6079*<-BrxB-BrxF- [illegible]</td><td>1251</td><td>Bacteria>Terrabacteria_group>Actinomycetota>Actinomycetes>Bifidobacteriales>Bifidobacteriaceae</td><td>GCF_002802875.1</td><td>Bifidobacterium_primatium</td><td>CS006_RS06855</td><td>170776..174531</td><td>2045438</td><td>[illegible]</td></tr>
<tr><td>WP_101025546.1</td><td>Type1.BREX</td><td>HTH_28->BrxA-DNA-binding->BrxB-BrxF-iSTAND->DUF6079*->N6-METHYLASE->?->Schlafen+HATPase+wHTH->PglZ->SIGMA+OB-Fold+AAA-ATPase+ [illegible]</td><td>1217</td><td>Bacteria>Terrabacteria_group>Actinomycetota>Actinomycetes>Bifidobacteriales>Bifidobacteriaceae</td><td>GCF_002832945.1</td><td>Bifidobacterium_longum</td><td>DPC6317_RS01855</td><td>212185..215838</td><td>216816</td><td>[illegible]</td></tr>
<tr><td>WP_101025682.1</td><td>Type1.BREX</td><td>RelB->N6-METHYLASE->CR-ATPase5+REase-p-><-CR-Kinase<-?||BrxA-DNA-binding->BrxB-BrxF-iSTAND->DUF6079-p->DUF6079*->N6-METHYLASE- [illegible]</td><td>1011</td><td>Bacteria>Terrabacteria_group>Actinomycetota>Actinomycetes>Bifidobacteriales>Bifidobacteriaceae</td><td>GCF_002832945.1</td><td>Bifidobacterium_longum</td><td>DPC6317_RS04670</td><td>237482..240517</td><td>216816</td><td>[illegible]</td></tr>
<tr><td>WP_106628753.1</td><td>Type1.BREX</td><td>rve->AAA-ATPase-><-?<-?<-?<-?<-SIR2||BrxA-DNA-binding->BrxB-BrxF-iSTAND->DUF6079*->N6-METHYLASE->SIR2->HerA/FtsK->N6-METHYLASE [illegible]</td><td>1218</td><td>Bacteria>Terrabacteria_group>Actinomycetota>Actinomycetes>Bifidobacteriales>Bifidobacteriaceae</td><td>GCF_002838305.1</td><td>Bifidobacterium_breve</td><td>NRBB11_RS04990</td><td>1136892..1140548</td><td>1685</td><td>[illegible]</td></tr>
<tr><td>WP_106620813.1</td><td>Type1.BREX</td><td><-rve||RelB->REase->N6-METHYLASE->CR-ATPase5+REase-p-><-CR-Kinase<-?||BrxA-DNA-binding->BrxB-BrxF-iSTAND->DUF6079*-> [illegible]</td><td>1229</td><td>Bacteria>Terrabacteria_group>Actinomycetota>Actinomycetes>Bifidobacteriales>Bifidobacteriaceae</td><td>GCF_002838345.1</td><td>Bifidobacterium_breve</td><td>NRBB57_RS05005</td><td>1130879..1134568</td><td>1685</td><td>[illegible]</td></tr>
<tr><td>WP_101454290.1</td><td>Type1.BREX</td><td>HAD+HAD->MODE-HTH+PBPII->YTCD-HTH-><-HTH_3<-RelE-ParE<-ACET+ACET<-ZnR||BrxA-DNA-binding->BrxB-BrxF-iSTAND->DUF6079*-> [illegible]</td><td>1214</td><td>Bacteria>Terrabacteria_group>Actinomycetota>Actinomycetes>Bifidobacteriales>Bifidobacteriaceae</td><td>GCF_002846685.1</td><td>Bifidobacterium_pseudolongum</td><td>CQR55_RS04465</td><td>180810..184454</td><td>1694</td><td>[illegible]</td></tr>
<tr><td>WP_101428597.1</td><td>Type1.BREX</td><td>HAD+HAD->MODE-HTH+PBPII->YTCD-HTH-><-ACET+ACET<-ZnR||BrxA-DNA-binding->BrxB-BrxF-iSTAND->DUF6079*->N6-METHYLASE->PglZ->SIGMA+ [illegible]</td><td>1212</td><td>Bacteria>Terrabacteria_group>Actinomycetota>Actinomycetes>Bifidobacteriales>Bifidobacteriaceae</td><td>GCF_002846755.1</td><td>Bifidobacterium_pseudolongum</td><td>CQR49_RS04725</td><td>132480..136118</td><td>1694</td><td>[illegible]</td></tr>
<tr><td>WP_180326342.1</td><td>Type1.BREX</td><td>DUF6079-p*->N6-METHYLASE->PglZ->SIGMA+OB-Fold+AAA-ATPase+HKD-DNase-BrxL-><-HEPN<-PRTase</td><td>865</td><td>Bacteria>Terrabacteria_group>Actinomycetota>Actinomycetes>Bifidobacteriales>Bifidobacteriaceae</td><td>GCF_002846775.1</td><td>Bifidobacterium_pseudolongum</td><td>CQR50_RS05955</td><td>514..3111</td><td>1694</td><td>[illegible]</td></tr>
<tr><td>WP_165782845.1</td><td>Type1.BREX</td><td><-SIGMA+OB-Fold+AAA-ATPase+HKD-DNase-BrxL<-PglZ<-Schlafen+HATPase+wHTH<-N6-METHYLASE<-?<-TPR+TPR<-DUF6079*<-BrxB-BrxF-iSTAND<- [illegible]</td><td>1259</td><td>Bacteria>Terrabacteria_group>Actinomycetota>Actinomycetes>Bifidobacteriales>Bifidobacteriaceae</td><td>GCF_002860355.1</td><td>Bifidobacterium_margollesii</td><td>Uis1B_RS09895</td><td>37676..41455</td><td>2020964</td><td>[illegible]</td></tr>
<tr><td>WP_101629209.1</td><td>Type1.BREX</td><td>ABC-ATPase+SbcC->?->?-><-?<-SIGMA+OB-Fold+AAA-ATPase+HKD-DNase-BrxL<-PglZ<-Schlafen+HATPase+wHTH<-?<-N6-METHYLASE<-DUF6079*<- [illegible]</td><td>1218</td><td>Bacteria>Terrabacteria_group>Actinomycetota>Actinomycetes>Bifidobacteriales>Bifidobacteriaceae</td><td>GCF_002861445.1</td><td>Bifidobacterium_longum</td><td>CYJ39_RS09195</td><td>68952..72608</td><td>216816</td><td>[illegible]</td></tr>
<tr><td>WP_109056521.1</td><td>Type1.BREX</td><td><-ABhydrolase||BrxA-DNA-binding->BrxB-BrxF-iSTAND->DUF6079*->N6-METHYLASE->HEPN->Schlafen+wHTH->CR-ATPase5+REase->PglZ->SIGMA+ [illegible]</td><td>1246</td><td>Bacteria>Terrabacteria_group>Actinomycetota>Actinomycetes>Bifidobacteriales>Bifidobacteriaceae</td><td>GCF_003129905.1</td><td>Bifidobacterium_callitrichidarum</td><td>DF196_RS03600</td><td>108404..112144</td><td>2052941</td><td>[illegible]</td></tr>
<tr><td>WP_115779240.1</td><td>Type1.BREX</td><td><-DUF6079-p*<-BrxB-BrxF-iSTAND<-BrxA-DNA-binding||?->CR-Kinase-><-CR-ATPase5+REase-p<-REase<-HTH_3</td><td>544</td><td>Bacteria>Terrabacteria_group>Actinomycetota>Actinomycetes>Bifidobacteriales>Bifidobacteriaceae</td><td>GCF_003370005.1</td><td>Bifidobacterium_longum</td><td>CE169_RS11735</td><td>1..1633</td><td>216816</td><td>[illegible]</td></tr>
<tr><td>WP_134859198.1</td><td>Type1.BREX</td><td>RelB->N6-METHYLASE->REase->CR-ATPase5+REase-p-><-CR-Kinase<-?||BrxA-DNA-binding->BrxB-BrxF-iSTAND->DUF6079-p*->N6-METHYLASE-> [illegible]</td><td>480</td><td>Bacteria>Terrabacteria_group>Actinomycetota>Actinomycetes>Bifidobacteriales>Bifidobacteriaceae</td><td>GCF_003370195.1</td><td>Bifidobacterium_breve</td><td>CE167_RS03750</td><td>14573..16012</td><td>1685</td><td>[illegible]</td></tr>
<tr><td>WP_115955260.1</td><td>Type1.BREX</td><td><-OLD-ABC+TOPRIM<-ZnR<-ZnR+GP<-SIGMA+OB-Fold+AAA-ATPase+HKD-DNase-BrxL<-PglZ<-Schlafen+HATPase+wHTH<-?<-HEPN<-N6-METHYLASE<- [illegible]</td><td>1217</td><td>Bacteria>Terrabacteria_group>Actinomycetota>Actinomycetes>Bifidobacteriales>Bifidobacteriaceae</td><td>GCF_003390735.1</td><td>Bifidobacterium_bifidum</td><td>CJD48_RS06210</td><td>1417204..1420857</td><td>1681</td><td>[illegible]</td></tr>
<tr><td>WP_117658188.1</td><td>Type1.BREX</td><td>Glyoxalase-><-?||REase-><-?<-SIGMA+OB-Fold+AAA-ATPase+HKD-DNase-BrxL<-PglZ<-Schlafen+HATPase+wHTH<-SIR2<-N6-METHYLASE<- [illegible]</td><td>1218</td><td>Bacteria>Terrabacteria_group>Actinomycetota>Actinomycetes>Bifidobacteriales>Bifidobacteriaceae</td><td>GCF_003436575.1</td><td>Bifidobacterium_bifidum</td><td>DXD34_RS01735</td><td>367842..371498</td><td>1681</td><td>[illegible]</td></tr>
<tr><td>WP_117656287.1</td><td>Type1.BREX</td><td><-SIGMA+OB-Fold+AAA-ATPase+HKD-DNase-BrxL<-PglZ<-Schlafen+HATPase+wHTH<-SIR2<-N6-METHYLASE-p<-HerA/FtsK<-SIR2<-N6-METHYLASE<- [illegible]</td><td>1218</td><td>Bacteria>Terrabacteria_group>Actinomycetota>Actinomycetes>Bifidobacteriales>Bifidobacteriaceae</td><td>GCF_003437075.1</td><td>Bifidobacterium_pseudocatenulatum</td><td>DXC75_RS03300</td><td>120423..124079</td><td>28026</td><td>[illegible]</td></tr>
<tr><td>WP_117664462.1</td><td>Type1.BREX</td><td><-ZnR+GP<-SIGMA+OB-Fold+AAA-ATPase+HKD-DNase-BrxL<-PglZ<-Schlafen+HATPase+wHTH<-SIR2<-SIR2<-N6-METHYLASE-p<-N6-METHYLASE-p<- [illegible]</td><td>1218</td><td>Bacteria>Terrabacteria_group>Actinomycetota>Actinomycetes>Bifidobacteriales>Bifidobacteriaceae</td><td>GCF_003437835.1</td><td>Bifidobacterium_pseudocatenulatum</td><td>DXC45_RS01980</td><td>419525..423181</td><td>28026</td><td>[illegible]</td></tr>
<tr><td>WP_117759621.1</td><td>Type1.BREX</td><td><-ParB+HNH+RAMA<-ZnR<-ZnR+GP<-SIGMA+OB-Fold+AAA-ATPase+HKD-DNase-BrxL<-PglZ<-Schlafen+HATPase+wHTH<-HEPN<-N6-METHYLASE<- [illegible]</td><td>1223</td><td>Bacteria>Terrabacteria_group>Actinomycetota>Actinomycetes>Bifidobacteriales>Bifidobacteriaceae</td><td>GCF_003458805.1</td><td>Bifidobacterium_adolescentis</td><td>DWX79_RS04615</td><td>1022537..1026208</td><td>1680</td><td>[illegible]</td></tr>
<tr><td>WP_118279738.1</td><td>Type1.BREX</td><td>REase-><-?<-SIGMA+OB-Fold+AAA-ATPase+HKD-DNase-BrxL<-PglZ<-Schlafen+HATPase+wHTH<-REase<-N6-METHYLASE<-DUF6079*<-BrxB-BrxF- [illegible]</td><td>1219</td><td>Bacteria>Terrabacteria_group>Actinomycetota>Actinomycetes>Bifidobacteriales>Bifidobacteriaceae</td><td>GCF_003472685.1</td><td>Bifidobacterium_pseudocatenulatum</td><td>DW102_RS06345</td><td>114145..117804</td><td>28026</td><td>[illegible]</td></tr>
<tr><td>WP_118269378.1</td><td>Type1.BREX</td><td>Glyoxalase-><-?||REase-><-?<-SIGMA+OB-Fold+AAA-ATPase+HKD-DNase-BrxL<-PglZ<-Schlafen+HATPase+wHTH<-SIR2<-N6-METHYLASE<- [illegible]</td><td>1218</td><td>Bacteria>Terrabacteria_group>Actinomycetota>Actinomycetes>Bifidobacteriales>Bifidobacteriaceae</td><td>GCF_003473145.1</td><td>Bifidobacterium_bifidum</td><td>DW145_RS03305</td><td>289418..293074</td><td>1681</td><td>[illegible]</td></tr>
<tr><td>WP_117641243.1</td><td>Type1.BREX</td><td>HTH_28->BrxA-DNA-binding->BrxB-BrxF-iSTAND->DUF6079*->N6-METHYLASE->?->Schlafen+HATPase+wHTH->PglZ->SIGMA+OB-Fold+AAA-ATPase+ [illegible]</td><td>1219</td><td>Bacteria>Terrabacteria_group>Actinomycetota>Actinomycetes>Bifidobacteriales>Bifidobacteriaceae</td><td>GCF_003475835.1</td><td>Bifidobacterium_longum</td><td>DWZ18_RS09025</td><td>34331..37990</td><td>216816</td><td>[illegible]</td></tr>
<tr><td>WP_124879770.1</td><td>Type1.BREX</td><td>RNasePH->?-><-?<-tRNA<-tRNA||?->HTH_28->BrxA-DNA-binding->BrxB-BrxF-iSTAND->DUF6079*->N6-METHYLASE->SIR2->Schlafen+HATPase+ [illegible]</td><td>1219</td><td>Bacteria>Terrabacteria_group>Actinomycetota>Actinomycetes>Bifidobacteriales>Bifidobacteriaceae</td><td>GCF_003860285.1</td><td>Bifidobacterium_breve</td><td>EH245_RS01900</td><td>428223..431882</td><td>1685</td><td>[illegible]</td></tr>
<tr><td>WP_126029122.1</td><td>Type1.BREX</td><td><-SIGMA+OB-Fold+AAA-ATPase+HKD-DNase-BrxL<-PglZ<-HEPN<-N6-METHYLASE<-DUF6079*<-BrxB-BrxF-iSTAND<-BrxA-DNA-binding<-?||SIGMA- [illegible]</td><td>1235</td><td>Bacteria>Terrabacteria_group>Actinomycetota>Actinomycetes>Bifidobacteriales>Bifidobacteriaceae</td><td>GCF_003952005.1</td><td>Bifidobacterium_callimiconis</td><td>D2E23_RS00850</td><td>182499..186206</td><td>2306973</td><td>[illegible]</td></tr>
<tr><td>WP_126016404.1</td><td>Type1.BREX</td><td><-ParB+HNH+RAMA<-ParB+HNH<-ZnR<-ZnR+GP<-SIGMA+OB-Fold+AAA-ATPase+HKD-DNase-BrxL<-PglZ<-Schlafen+HATPase+wHTH<-REase<- [illegible]</td><td>1217</td><td>Bacteria>Terrabacteria_group>Actinomycetota>Actinomycetes>Bifidobacteriales>Bifidobacteriaceae</td><td>GCF_003952825.1</td><td>Bifidobacterium_pseudocatenulatum</td><td>CWS99_RS05770</td><td>1339669..1343322</td><td>28026</td><td>[illegible]</td></tr>
<tr><td>WP_129237752.1</td><td>Type1.BREX</td><td><-SIGMA+OB-Fold+AAA-ATPase+HKD-DNase-BrxL<-PglZ<-Schlafen+HATPase+wHTH<-?<-N6-METHYLASE<-DUF6079*<-BrxB-BrxF-iSTAND<- [illegible]</td><td>1221</td><td>Bacteria>Terrabacteria_group>Actinomycetota>Actinomycetes>Bifidobacteriales>Bifidobacteriaceae</td><td>GCF_004135085.1</td><td>Bifidobacterium_pullorum</td><td>ESN35_RS07340</td><td>1680432..1684097</td><td>78448</td><td>[illegible]</td></tr>
<tr><td>WP_130013100.1</td><td>Type1.BREX</td><td>MODE-HTH+PBPII->YTCD-HTH-><-HTH_3<-RelE-ParE<-ACET+ACET<-ZnR<-ZnR||BrxA-DNA-binding->BrxB-BrxF-iSTAND->DUF6079*->PglZ->SIGMA+ [illegible]</td><td>1214</td><td>Bacteria>Terrabacteria_group>Actinomycetota>Actinomycetes>Bifidobacteriales>Bifidobacteriaceae</td><td>GCF_004155015.1</td><td>Bifidobacterium_pseudolongum</td><td>PG2054B_RS04700</td><td>182910..186554</td><td>1694</td><td>[illegible]</td></tr>
<tr><td>WP_129902658.1</td><td>Type1.BREX</td><td>HAD+HAD->MODE-HTH+PBPII->YTCD-HTH-><-HTH_3<-RelE-ParE<-ACET+ACET<-ZnR||BrxA-DNA-binding->BrxB-BrxF-iSTAND->DUF6079*-> [illegible]</td><td>1214</td><td>Bacteria>Terrabacteria_group>Actinomycetota>Actinomycetes>Bifidobacteriales>Bifidobacteriaceae</td><td>GCF_004155045.1</td><td>Bifidobacterium_pseudolongum</td><td>PG2083B_RS04940</td><td>90141..93785</td><td>1694</td><td>[illegible]</td></tr>
<tr><td>WP_129866281.1</td><td>Type1.BREX</td><td>MODE-HTH+PBPII->YTCD-HTH-><-HTH_3<-RelE-ParE<-ACET+ACET<-ZnR<-ZnR||BrxA-DNA-binding->BrxB-BrxF-iSTAND->DUF6079*->N6-METHYLASE- [illegible]</td><td>1214</td><td>Bacteria>Terrabacteria_group>Actinomycetota>Actinomycetes>Bifidobacteriales>Bifidobacteriaceae</td><td>GCF_004155285.1</td><td>Bifidobacterium_pseudolongum</td><td>PG2000B_RS04795</td><td>96491..100135</td><td>1694</td><td>[illegible]</td></tr>
<tr><td>WP_129867688.1</td><td>Type1.BREX</td><td>HAD+HAD->MODE-HTH+PBPII->YTCD-HTH-><-HTH_3<-RelE-ParE<-ACET+ACET<-ZnR||BrxA-DNA-binding->BrxB-BrxF-iSTAND->DUF6079*-> [illegible]</td><td>1212</td><td>Bacteria>Terrabacteria_group>Actinomycetota>Actinomycetes>Bifidobacteriales>Bifidobacteriaceae</td><td>GCF_004155405.1</td><td>Bifidobacterium_pseudolongum</td><td>PG102017_RS04060</td><td>111812..115450</td><td>1694</td><td>[illegible]</td></tr>
<tr><td>WP_129914280.1</td><td>Type1.BREX</td><td>MODE-HTH+PBPII->YTCD-HTH-><-HTH_3<-RelE-ParE<-ACET+ACET<-ZnR<-ZnR||BrxA-DNA-binding->BrxB-BrxF-iSTAND->DUF6079*->N6-METHYLASE- [illegible]</td><td>1212</td><td>Bacteria>Terrabacteria_group>Actinomycetota>Actinomycetes>Bifidobacteriales>Bifidobacteriaceae</td><td>GCF_004155645.1</td><td>Bifidobacterium_pseudolongum</td><td>PG2001B_RS04910</td><td>52519..56157</td><td>1694</td><td>[illegible]</td></tr>
<tr><td>WP_129869778.1</td><td>Type1.BREX</td><td>HAD+HAD->MODE-HTH+PBPII->YTCD-HTH-><-ACET+ACET<-ZnR||BrxA-DNA-binding->BrxB-BrxF-iSTAND->DUF6079*->N6-METHYLASE->PglZ->SIGMA+ [illegible]</td><td>1214</td><td>Bacteria>Terrabacteria_group>Actinomycetota>Actinomycetes>Bifidobacteriales>Bifidobacteriaceae</td><td>GCF_004155805.1</td><td>Bifidobacterium_pseudolongum</td><td>PG1550B_RS04805</td><td>1102192..1105836</td><td>1694</td><td>[illegible]</td></tr>
<tr><td>WP_129900679.1</td><td>Type1.BREX</td><td><-HTH_3<-RelE-ParE<-ACET+ACET<-ZnR||BrxA-DNA-binding->BrxB-BrxF-iSTAND->DUF6079*->N6-METHYLASE->PglZ->SIGMA+OB-Fold+AAA-ATPase [illegible]</td><td>1214</td><td>Bacteria>Terrabacteria_group>Actinomycetota>Actinomycetes>Bifidobacteriales>Bifidobacteriaceae</td><td>GCF_004155855.1</td><td>Bifidobacterium_pseudolongum</td><td>PG112206_RS04240</td><td>4644..8288</td><td>1694</td><td>[illegible]</td></tr>
<tr><td>WP_129917395.1</td><td>Type1.BREX</td><td>BrxA-DNA-binding->BrxB-BrxF-iSTAND->DUF6079*->N6-METHYLASE->PglZ->SIGMA+OB-Fold+AAA-ATPase+HKD-DNase-BrxL->SIR2-><-PRTase</td><td>1214</td><td>Bacteria>Terrabacteria_group>Actinomycetota>Actinomycetes>Bifidobacteriales>Bifidobacteriaceae</td><td>GCF_004156175.1</td><td>Bifidobacterium_pseudolongum</td><td>PG2103B_RS04555</td><td>1564..5208</td><td>1694</td><td>[illegible]</td></tr>
<tr><td>WP_131226808.1</td><td>Type1.BREX</td><td><-METHYLASE-p<-McrC-NTD+REase<-EVE+McrB-AAA<-?<-DUF6079*<-BrxB-BrxF-iSTAND<-BrxA-DNA-binding<-CR-ATPase5+REase-p<-?||HTH_38->? [illegible]</td><td>1229</td><td>Bacteria>Terrabacteria_group>Actinomycetota>Actinomycetes>Bifidobacteriales>Bifidobacteriaceae</td><td>GCF_004333385.1</td><td>Bifidobacterium_longum</td><td>MCC10066_RS04685</td><td>5338..9027</td><td>216816</td><td>[illegible]</td></tr>
<tr><td>WP_131210356.1</td><td>Type1.BREX</td><td><-DUF6079-p*<-BrxB-BrxF-iSTAND<-BrxA-DNA-binding<-CR-ATPase5+REase-p<-?||HTH_38->?->HEPN->?-><-HTH+MPTase</td><td>442</td><td>Bacteria>Terrabacteria_group>Actinomycetota>Actinomycetes>Bifidobacteriales>Bifidobacteriaceae</td><td>GCF_004333565.1</td><td>Bifidobacterium_longum</td><td>MCC10048_RS04670</td><td>1..1326</td><td>216816</td><td>[illegible]</td></tr>
<tr><td>WP_131292550.1</td><td>Type1.BREX</td><td><-Schlafen+HATPase+wHTH<-?<-N6-METHYLASE<-DUF6079*<-BrxB-BrxF-iSTAND<-BrxA-DNA-binding</td><td>1218</td><td>Bacteria>Terrabacteria_group>Actinomycetota>Actinomycetes>Bifidobacteriales>Bifidobacteriaceae</td><td>GCF_004333845.1</td><td>Bifidobacterium_longum</td><td>MCC10126_RS01985</td><td>6613..10269</td><td>216816</td><td>[illegible]</td></tr>
<tr><td>WP_131220402.1</td><td>Type1.BREX</td><td>HTH_28->BrxA-DNA-binding->BrxB-BrxF-iSTAND->DUF6079*->N6-METHYLASE->?->Schlafen+HATPase+wHTH->PglZ->SIGMA+OB-Fold+AAA-ATPase+ [illegible]</td><td>1219</td><td>Bacteria>Terrabacteria_group>Actinomycetota>Actinomycetes>Bifidobacteriales>Bifidobacteriaceae</td><td>GCF_004334345.1</td><td>Bifidobacterium_longum</td><td>MCC10040_RS02080</td><td>100674..104333</td><td>216816</td><td>[illegible]</td></tr>
<tr><td>WP_131308639.1</td><td>Type1.BREX</td><td><-Schlafen+wHTH<-SMODS<-SLATT<-N6-METHYLASE<-N6-METHYLASE-p<-METHYLASE-p<-McrC-NTD+REase<-EVE+McrB-AAA<-?<-DUF6079*<-BrxB-BrxF- [illegible]</td><td>1229</td><td>Bacteria>Terrabacteria_group>Actinomycetota>Actinomycetes>Bifidobacteriales>Bifidobacteriaceae</td><td>GCF_004334545.1</td><td>Bifidobacterium_longum</td><td>MCC10002_RS05185</td><td>64814..68503</td><td>216816</td><td>[illegible]</td></tr>
<tr><td>WP_136510830.1</td><td>Type1.BREX</td><td>BrxA-DNA-binding->BrxB-BrxF-iSTAND->DUF6079*->N6-METHYLASE->PglZ->SIGMA+OB-Fold+AAA-ATPase+HKD-DNase-BrxL->SIR2-><-PRTase</td><td>1214</td><td>Bacteria>Terrabacteria_group>Actinomycetota>Actinomycetes>Bifidobacteriales>Bifidobacteriaceae</td><td>GCF_004801625.1</td><td>Bifidobacterium_pseudolongum</td><td>E5991_RS00930</td><td>3050..6694</td><td>1694</td><td>[illegible]</td></tr>
<tr><td>WP_140556589.1</td><td>Type1.BREX</td><td><-Cysteine-Desulfurase<-ABC-ATPase+SbcC<-?<-?<-?<-SIGMA+OB-Fold+AAA-ATPase+HKD-DNase-BrxL<-PglZ<-Schlafen+HATPase+wHTH<- [illegible]</td><td>1232</td><td>Bacteria>Terrabacteria_group>Actinomycetota>Actinomycetes>Bifidobacteriales>Bifidobacteriaceae</td><td>GCF_006408715.1</td><td>Bifidobacterium_sp._UTCIF-39</td><td>EP30_RS04700</td><td>102584..106282</td><td>1465359</td><td>[illegible]</td></tr>
<tr><td>WP_140489867.1</td><td>Type1.BREX</td><td>ABC-ATPase+SbcC->HAD+S6PP->?-><-?<-ABhydrolase||BrxA-DNA-binding->BrxB-BrxF-iSTAND->DUF6079*->CR-ATPase5+REase->TPR+TPR-></td><td>1264</td><td>Bacteria>Terrabacteria_group>Actinomycetota>Actinomycetes>Bifidobacteriales>Bifidobacteriaceae</td><td>GCF_006408855.1</td><td>Bifidobacterium_sp._UTBIF-68</td><td>BW14_RS07205</td><td>86138..89932</td><td>1465262</td><td>[illegible]</td></tr>
<tr><td>WP_140558979.1</td><td>Type1.BREX</td><td>ABC-ATPase+SbcC->HAD+S6PP->?-><-?<-ABhydrolase||BrxA-DNA-binding->BrxB-BrxF-iSTAND->DUF6079*->CR-ATPase5+REase->TPR+TPR->?->?-> [illegible]</td><td>1264</td><td>Bacteria>Terrabacteria_group>Actinomycetota>Actinomycetes>Bifidobacteriales>Bifidobacteriaceae</td><td>GCF_006408865.1</td><td>Bifidobacterium_sp._UTBIF-56</td><td>BW10_RS07695</td><td>47972..51766</td><td>1465261</td><td>[illegible]</td></tr>
<tr><td>WP_140463094.1</td><td>Type1.BREX</td><td>ABC-ATPase+SbcC->HAD+S6PP->?-><-?<-ABhydrolase||BrxA-DNA-binding->BrxB-BrxF-iSTAND->DUF6079*->CR-ATPase5+REase->TPR+TPR-><-?|| [illegible]</td><td>1264</td><td>Bacteria>Terrabacteria_group>Actinomycetota>Actinomycetes>Bifidobacteriales>Bifidobacteriaceae</td><td>GCF_006408915.1</td><td>unclassified_Bifidobacterium</td><td>BW09_RS08245</td><td>76569..80363</td><td>2608897</td><td>[illegible]</td></tr>
<tr><td>WP_142421439.1</td><td>Type1.BREX</td><td>RNaseH-><-?<-SIGMA+OB-Fold+AAA-ATPase+HKD-DNase-BrxL<-PglZ<-Schlafen+HATPase+wHTH<-REase<-N6-METHYLASE-p<-N6-METHYLASE<- [illegible]</td><td>1218</td><td>Bacteria>Terrabacteria_group>Actinomycetota>Actinomycetes>Bifidobacteriales>Bifidobacteriaceae</td><td>GCF_006828885.1</td><td>Bifidobacterium_pseudocatenulatum</td><td>GH753_RS08045</td><td>1981980..1985636</td><td>28026</td><td>[illegible]</td></tr>
<tr><td>WP_150335795.1</td><td>Type1.BREX</td><td><-TPR+TPR<-CR-ATPase5+REase<-DUF6079*<-BrxB-BrxF-iSTAND<-BrxA-DNA-binding||ABhydrolase->?-><-?<-HAD+S6PP<-ABC-ATPase+SbcC</td><td>1264</td><td>Bacteria>Terrabacteria_group>Actinomycetota>Actinomycetes>Bifidobacteriales>Bifidobacteriaceae</td><td>GCF_008698125.1</td><td>Bifidobacterium_reuteri</td><td>EM092_RS08290</td><td>2843..6637</td><td>983706</td><td>[illegible]</td></tr>
<tr><td>WP_150379196.1</td><td>Type1.BREX</td><td><-SbcD||?-><-ABhydrolase||BrxA-DNA-binding->BrxB-BrxF-iSTAND->DUF6079*->CR-ATPase5+REase->TPR+TPR->N6-METHYLASE->HEPN-> [illegible]</td><td>1267</td><td>Bacteria>Terrabacteria_group>Actinomycetota>Actinomycetes>Bifidobacteriales>Bifidobacteriaceae</td><td>GCF_008698155.1</td><td>Bifidobacterium_myosotis</td><td>EM091_RS06000</td><td>83624..87427</td><td>1630166</td><td>[illegible]</td></tr>
<tr><td>WP_150382415.1</td><td>Type1.BREX</td><td>ABC-ATPase+SbcC-><-CR-ATPase5+REase||?-><-ZnR||MazE_antitoxin->PemK_toxin->BrxA-DNA-binding->BrxB-BrxF-iSTAND->DUF6079*-> [illegible]</td><td>1244</td><td>Bacteria>Terrabacteria_group>Actinomycetota>Actinomycetes>Bifidobacteriales>Bifidobacteriaceae</td><td>GCF_008698235.1</td><td>Bifidobacterium_rousetti</td><td>CSQ85_RS00640</td><td>119339..123073</td><td>2045439</td><td>[illegible]</td></tr>
<tr><td>WP_151916615.1</td><td>Type1.BREX</td><td>HNH->||DOC+HTH-><-sGTP||?->?->?-><-HAD+HAD||BrxA-DNA-binding->BrxB-BrxF-iSTAND->DUF6079*->CR-ATPase5+REase->TPR+TPR-> [illegible]</td><td>1269</td><td>Bacteria>Terrabacteria_group>Actinomycetota>Actinomycetes>Bifidobacteriales>Bifidobacteriaceae</td><td>GCF_009078285.1</td><td>Bifidobacterium_jacchi</td><td>EHS19_RS04610</td><td>26586..30395</td><td>2490545</td><td>[illegible]</td></tr>
<tr><td>WP_152233646.1</td><td>Type1.BREX</td><td>REase+Bacteriocin_IIc->?->?->?->?->?-><-?||BrxA-DNA-binding->BrxB-BrxF-iSTAND->DUF6079*-><-?||CR-ATPase5+REase->N6-METHYLASE-> [illegible]</td><td>1235</td><td>Bacteria>Terrabacteria_group>Actinomycetota>Actinomycetes>Bifidobacteriales>Bifidobacteriaceae</td><td>GCF_009193355.1</td><td>Bifidobacterium_leontopitheci</td><td>F7D09_RS01345</td><td>123122..126829</td><td>2650774</td><td>[illegible]</td></tr>
<tr><td>WP_117762529.1</td><td>Type1.BREX</td><td><-ParB+HNH+RAMA<-ZnR<-ZnR+GP<-SIGMA+OB-Fold+AAA-ATPase+HKD-DNase-BrxL<-PglZ<-Schlafen+HATPase+wHTH<-HEPN<-N6-METHYLASE<- [illegible]</td><td>1223</td><td>Bacteria>Terrabacteria_group>Actinomycetota>Actinomycetes>Bifidobacteriales>Bifidobacteriaceae</td><td>GCF_009832825.1</td><td>Bifidobacterium_adolescentis</td><td>DWZ00_RS05085</td><td>1174565..1178236</td><td>1680</td><td>[illegible]</td></tr>
<tr><td>WP_163196475.1</td><td>Type1.BREX</td><td><-SIGMA+OB-Fold+AAA-ATPase+HKD-DNase-BrxL<-PglZ<-Schlafen+HATPase+wHTH<-?<-N6-METHYLASE<-DUF6079*<-BrxB-BrxF-iSTAND<- [illegible]</td><td>1248</td><td>Bacteria>Terrabacteria_group>Actinomycetota>Actinomycetes>Bifidobacteriales>Bifidobacteriaceae</td><td>GCF_010667645.1</td><td>Bifidobacterium_platyrrhinorum</td><td>GFD21_RS03310</td><td>757948..761694</td><td>2661628</td><td>[illegible]</td></tr>
<tr><td>WP_163113202.1</td><td>Type1.BREX</td><td>MULE-TRANSPOSASE-><-ParB+HNH<-ZnR<-ZnR+GP<-SIGMA+OB-Fold+AAA-ATPase+HKD-DNase-BrxL<-PglZ<-Schlafen+HATPase+wHTH<-SIR2<- [illegible]</td><td>1218</td><td>Bacteria>Terrabacteria_group>Actinomycetota>Actinomycetes>Bifidobacteriales>Bifidobacteriaceae</td><td>GCF_011029205.1</td><td>Bifidobacterium_bifidum</td><td>G5T23_RS07350</td><td>43765..47421</td><td>1681</td><td>[illegible]</td></tr>
<tr><td>WP_168445148.1</td><td>Type1.BREX</td><td><-Glyoxalase||DSBH+AraC-HTH+AraC-HTH-><-ZnR<-SIGMA+OB-Fold+AAA-ATPase+HKD-DNase-BrxL<-PglZ<-PglZ<-Schlafen+HATPase+wHTH<-?<- [illegible]</td><td>1217</td><td>Bacteria>Terrabacteria_group>Actinomycetota>Actinomycetes>Bifidobacteriales>Bifidobacteriaceae</td><td>GCF_012310535.1</td><td>Bifidobacterium_dentium</td><td>PG133_RS04075</td><td>130679..134332</td><td>1689</td><td>[illegible]</td></tr>
<tr><td>WP_169274836.1</td><td>Type1.BREX</td><td><-PIN<-RelB<-?||HTH_3->?->PIN->BrxA-DNA-binding->BrxB-BrxF-iSTAND->DUF6079*->CR-ATPase5+REase->N6-METHYLASE->Schlafen+HATPase+ [illegible]</td><td>1226</td><td>Bacteria>Terrabacteria_group>Actinomycetota>Actinomycetes>Bifidobacteriales>Bifidobacteriaceae</td><td>GCF_012932365.1</td><td>Bifidobacterium_sp._DSM_109958</td><td>G1C96_RS01295</td><td>288582..292262</td><td>2705430</td><td>[illegible]</td></tr>
<tr><td>WP_193020253.1</td><td>Type1.BREX</td><td>HTH+MPTase-><-?<-HEPN<-?<-?||?->BrxA-DNA-binding->BrxB-BrxF-iSTAND->DUF6079*->N6-METHYLASE->REase->Schlafen+wHTH->PglZ->SIGMA+ [illegible]</td><td>1229</td><td>Bacteria>Terrabacteria_group>Actinomycetota>Actinomycetes>Bifidobacteriales>Bifidobacteriaceae</td><td>GCF_014898135.1</td><td>Bifidobacterium_longum</td><td>BL5934_RS04380</td><td>976256..979945</td><td>216816</td><td>[illegible]</td></tr>
<tr><td>WP_200406515.1</td><td>Type1.BREX</td><td>HTH_28+HTH_28->HTH_Tnp_1+HTH_21+rve+rve_3->HTH_28->DEDD_Tnp_IS110+Transposase_20->BrxA-DNA-binding->BrxB-BrxF-iSTAND->DUF6079* [illegible]</td><td>1219</td><td>Bacteria>Terrabacteria_group>Actinomycetota>Actinomycetes>Bifidobacteriales>Bifidobacteriaceae</td><td>GCF_014898255.1</td><td>Bifidobacterium_longum</td><td>BL7055_RS03240</td><td>790763..794422</td><td>216816</td><td>[illegible]</td></tr>
<tr><td>WP_227411048.1</td><td>Type1.BREX</td><td>ACET->ACET+ACET->?->Thioredoxin_3->ArsR-HTH-><-Glycoside_Hydrolase||OLD-ABC->BrxA-DNA-binding->BrxB-BrxF-iSTAND->DUF6079*-> [illegible]</td><td>1242</td><td>Bacteria>Terrabacteria_group>Actinomycetota>Actinomycetes>Bifidobacteriales>Bifidobacteriaceae</td><td>GCF_014983005.1</td><td>Bifidobacterium_pullorum</td><td>INF27_RS02890</td><td>183960..187688</td><td>78448</td><td>[illegible]</td></tr>
<tr><td>WP_195393981.1</td><td>Type1.BREX</td><td>PemK_toxin->?->PolB->HEPN-><-?<-?||?->BrxA-DNA-binding->BrxB-BrxF-iSTAND->DUF6079*->N6-METHYLASE->SIR2->Schlafen+HATPase+wHTH- [illegible]</td><td>1218</td><td>Bacteria>Terrabacteria_group>Actinomycetota>Actinomycetes>Bifidobacteriales>Bifidobacteriaceae</td><td>GCF_015553845.1</td><td>Bifidobacterium_adolescentis</td><td>I2D91_RS01775</td><td>423737..427393</td><td>1680</td><td>[illegible]</td></tr>
<tr><td>WP_195451643.1</td><td>Type1.BREX</td><td><-ABhydrolase<-?<-Glyoxalase||DSBH+AraC-HTH+AraC-HTH-><-ZnR<-SIGMA+OB-Fold+AAA-ATPase+HKD-DNase-BrxL<-PglZ<-Schlafen+HATPase+ [illegible]</td><td>1217</td><td>Bacteria>Terrabacteria_group>Actinomycetota>Actinomycetes>Bifidobacteriales>Bifidobacteriaceae</td><td>GCF_015555105.1</td><td>Bifidobacterium_dentium</td><td>I2E42_RS03170</td><td>226129..229782</td><td>1689</td><td>[illegible]</td></tr>
<tr><td>WP_195482034.1</td><td>Type1.BREX</td><td><-ParB+HNH+RAMA<-ZnR<-ZnR+GP<-SIGMA+OB-Fold+AAA-ATPase+HKD-DNase-BrxL<-PglZ<-Schlafen+HATPase+wHTH<-SIR2<-N6-METHYLASE<- [illegible]</td><td>1218</td><td>Bacteria>Terrabacteria_group>Actinomycetota>Actinomycetes>Bifidobacteriales>Bifidobacteriaceae</td><td>GCF_015555595.1</td><td>Bifidobacterium_adolescentis</td><td>I2E65_RS01565</td><td>362593..366249</td><td>1680</td><td>[illegible]</td></tr>
<tr><td>WP_195506526.1</td><td>Type1.BREX</td><td><-Schlafen+HATPase+wHTH<-REase<-N6-METHYLASE<-DUF6079*<-BrxB-BrxF-iSTAND<-BrxA-DNA-binding<-Pentapeptide+Pentapeptide+ [illegible]</td><td>1218</td><td>Bacteria>Terrabacteria_group>Actinomycetota>Actinomycetes>Bifidobacteriales>Bifidobacteriaceae</td><td>GCF_015556885.1</td><td>Bifidobacterium_pseudocatenulatum</td><td>I2F64_RS01000</td><td>6102..9758</td><td>28026</td><td>[illegible]</td></tr>
<tr><td>WP_195537417.1</td><td>Type1.BREX</td><td><-SbcD||RelB->PemK_toxin->HEPN-><-?<-?||?->BrxA-DNA-binding->BrxB-BrxF-iSTAND->DUF6079*->N6-METHYLASE-><-Integrase_DNA+Phage_ [illegible]</td><td>1217</td><td>Bacteria>Terrabacteria_group>Actinomycetota>Actinomycetes>Bifidobacteriales>Bifidobacteriaceae</td><td>GCF_015558565.1</td><td>Bifidobacterium_adolescentis</td><td>I2G53_RS06810</td><td>88696..92349</td><td>1680</td><td>[illegible]</td></tr>
<tr><td>WP_229088463.1</td><td>Type1.BREX</td><td>35exo+HRDC->35exo+BRCT-><-HTH||?-><-ACET||?->BrxA-DNA-binding->BrxB-BrxF-iSTAND->DUF6079-p->DUF6079-p*->DUF6079-p-></td><td>863</td><td>Bacteria>Terrabacteria_group>Actinomycetota>Actinomycetes>Bifidobacteriales>Bifidobacteriaceae</td><td>GCF_015559405.1</td><td>Bifidobacterium_dentium</td><td>I2H02_RS10335</td><td>55205..57796</td><td>1689</td><td>[illegible]</td></tr>
<tr><td>WP_195936831.1</td><td>Type1.BREX</td><td>REase-><-?<-SIGMA+OB-Fold+AAA-ATPase+HKD-DNase-BrxL<-PglZ<-Schlafen+HATPase+wHTH<-?<-N6-METHYLASE<-DUF6079*<-BrxB-BrxF-iSTAND<- [illegible]</td><td>1218</td><td>Bacteria>Terrabacteria_group>Actinomycetota>Actinomycetes>Bifidobacteriales>Bifidobacteriaceae</td><td>GCF_015667555.1</td><td>Bifidobacterium_bifidum</td><td>I4U71_RS07235</td><td>13396..17052</td><td>1681</td><td>[illegible]</td></tr>
<tr><td>WP_195938980.1</td><td>Type1.BREX</td><td>35exo+HRDC-><-HTH||ACET+ACET-><-?<-SbcD<-?<-REase||BrxA-DNA-binding->BrxB-BrxF-iSTAND->DUF6079*->N6-METHYLASE->SIR2->Schlafen+ [illegible]</td><td>1217</td><td>Bacteria>Terrabacteria_group>Actinomycetota>Actinomycetes>Bifidobacteriales>Bifidobacteriaceae</td><td>GCF_015667675.1</td><td>Bifidobacterium_pseudocatenulatum</td><td>I4U64_RS06630</td><td>371599..375252</td><td>28026</td><td>[illegible]</td></tr>
<tr><td>WP_241734738.1</td><td>Type1.BREX</td><td><-DUF6079*<-DUF6079-p<-BrxB-BrxF-iSTAND<-BrxA-DNA-binding<-?||ACET-><-?||HTH-><-35exo+BRCT<-35exo+HRDC</td><td>1076</td><td>Bacteria>Terrabacteria_group>Actinomycetota>Actinomycetes>Bifidobacteriales>Bifidobacteriaceae</td><td>GCF_015703785.1</td><td>Bifidobacterium_dentium</td><td>IHQ87_RS06495</td><td>579..3809</td><td>1689</td><td>[illegible]</td></tr>
<tr><td>WP_196336889.1</td><td>Type1.BREX</td><td>35exo+HRDC->35exo+BRCT-><-HTH||?-><-ACET||?->BrxA-DNA-binding->BrxB-BrxF-iSTAND->DUF6079*->N6-METHYLASE->HEPN->?->Schlafen+ [illegible]</td><td>1217</td><td>Bacteria>Terrabacteria_group>Actinomycetota>Actinomycetes>Bifidobacteriales>Bifidobacteriaceae</td><td>GCF_015703835.1</td><td>Bifidobacterium_dentium</td><td>IHQ82_RS06050</td><td>179848..183501</td><td>1689</td><td>[illegible]</td></tr>
<tr><td>WP_202574831.1</td><td>Type1.BREX</td><td><-Peptidase_S26<-?||ABC-ATPase+SbcC->?-><-?<-SIGMA+OB-Fold+AAA-ATPase+HKD-DNase-BrxL<-PglZ<-Schlafen+HATPase+wHTH<- [illegible]</td><td>1219</td><td>Bacteria>Terrabacteria_group>Actinomycetota>Actinomycetes>Bifidobacteriales>Bifidobacteriaceae</td><td>GCF_016759645.1</td><td>Bifidobacterium_longum</td><td>JL737_RS05465</td><td>24879..28538</td><td>216816</td><td>[illegible]</td></tr>
<tr><td>WP_204464696.1</td><td>Type1.BREX</td><td>ACET-><-O6MGT-HTH||Phage_integrase-><-?<-SIGMA+OB-Fold+AAA-ATPase+HKD-DNase-BrxL<-PglZ<-Schlafen+HATPase+wHTH<-N6-METHYLASE<- [illegible]</td><td>1221</td><td>Bacteria>Terrabacteria_group>Actinomycetota>Actinomycetes>Bifidobacteriales>Bifidobacteriaceae</td><td>GCF_016899675.1</td><td>Bifidobacterium_pullorum</td><td>H6A68_RS03865</td><td>164290..167955</td><td>78448</td><td>[illegible]</td></tr>
<tr><td>WP_204389272.1</td><td>Type1.BREX</td><td><-IstB_IS21||AAA-ATPase-><-CMP-hydrolase<-SIGMA+OB-Fold+AAA-ATPase+HKD-DNase-BrxL<-PglZ<-Schlafen+HATPase+wHTH<-N6-METHYLASE<- [illegible]</td><td>1221</td><td>Bacteria>Terrabacteria_group>Actinomycetota>Actinomycetes>Bifidobacteriales>Bifidobacteriaceae</td><td>GCF_016900615.1</td><td>Bifidobacterium_pullorum</td><td>H6A14_RS02700</td><td>186110..189775</td><td>78448</td><td>[illegible]</td></tr>
<tr><td>WP_077384433.1</td><td>Type1.BREX</td><td>HTH+MPTase-><-?<-HEPN<-?<-HTH_38||?->CR-ATPase5+REase-p->BrxA-DNA-binding->BrxB-BrxF-iSTAND->DUF6079*->?->EVE+McrB-AAA-> [illegible]</td><td>1229</td><td>Bacteria>Terrabacteria_group>Actinomycetota>Actinomycetes>Bifidobacteriales>Bifidobacteriaceae</td><td>GCF_017312625.1</td><td>Bifidobacterium_longum</td><td>MCC10017_RS04835</td><td>46545..50234</td><td>216816</td><td>[illegible]</td></tr>
<tr><td>WP_206874047.1</td><td>Type1.BREX</td><td>BrxA-DNA-binding->BrxB-BrxF-iSTAND->DUF6079*->?->McrB-AAA->McrC-NTD+REase->METHYLASE-p->N6-METHYLASE->SLATT->SMODS-> [illegible]</td><td>1229</td><td>Bacteria>Terrabacteria_group>Actinomycetota>Actinomycetes>Bifidobacteriales>Bifidobacteriaceae</td><td>GCF_017312685.1</td><td>Bifidobacterium_longum</td><td>J3464_RS07555</td><td>29014..32703</td><td>216816</td><td>[illegible]</td></tr>
<tr><td>WP_214374600.1</td><td>Type1.BREX</td><td>ZnR->?->ABC-ATPase+SbcC-><-?||?-><-ABhydrolase||BrxA-DNA-binding->BrxB-BrxF-iSTAND->DUF6079*->CR-ATPase5+REase->N6-METHYLASE-> [illegible]</td><td>1249</td><td>Bacteria>Terrabacteria_group>Actinomycetota>Actinomycetes>Bifidobacteriales>Bifidobacteriaceae</td><td>GCF_018555335.1</td><td>Bifidobacterium</td><td>JS541_RS11260</td><td>52363..56112</td><td>1678</td><td>[illegible]</td></tr>
<tr><td>WP_217054570.1</td><td>Type1.BREX</td><td><-SIGMA+OB-Fold+AAA-ATPase+HKD-DNase-BrxL<-PglZ<-HEPN+RES<-?<-?<-REase<-N6-METHYLASE<-DUF6079*<-BrxB-BrxF-iSTAND<- [illegible]</td><td>1229</td><td>Bacteria>Terrabacteria_group>Actinomycetota>Actinomycetes>Bifidobacteriales>Bifidobacteriaceae</td><td>GCF_019041665.1</td><td>Bifidobacterium_longum</td><td>KTF47_RS04030</td><td>15252..18941</td><td>216816</td><td>[illegible]</td></tr>
<tr><td>WP_219100165.1</td><td>Type1.BREX</td><td><-ABhydrolase||?->MACRODOMAIN-><-?<-SIGMA+OB-Fold+AAA-ATPase+HKD-DNase-BrxL<-PglZ<-?<-?<-N6-METHYLASE<-DUF6079*<-BrxB-BrxF- [illegible]</td><td>1253</td><td>Bacteria>Terrabacteria_group>Actinomycetota>Actinomycetes>Bifidobacteriales>Bifidobacteriaceae</td><td>GCF_019331695.1</td><td>Bifidobacterium_pluvialisilvae</td><td>KIH77_RS05905</td><td>203323..207084</td><td>2834436</td><td>[illegible]</td></tr>
<tr><td>WP_035010822.1</td><td>Type1.BREX</td><td><-N6-METHYLASE<-DUF6079*<-BrxB-BrxF-iSTAND<-BrxA-DNA-binding<-?<-?||?-><-Peptidase_M78<-HigA</td><td>1218</td><td>Bacteria>Terrabacteria_group>Actinomycetota>Actinomycetes>Bifidobacteriales>Bifidobacteriaceae</td><td>GCF_019972965.1</td><td>Bifidobacterium_adolescentis</td><td>BIFAD42_RS07975</td><td>2916..6572</td><td>1680</td><td>[illegible]</td></tr>
<tr><td>WP_226586661.1</td><td>Type1.BREX</td><td>REC+LuxR-HTH->HISKIN->?->?-><-SbcD<-?<-REase||BrxA-DNA-binding->BrxB-BrxF-iSTAND->DUF6079*->N6-METHYLASE->?->Schlafen+HATPase+ [illegible]</td><td>1219</td><td>Bacteria>Terrabacteria_group>Actinomycetota>Actinomycetes>Bifidobacteriales>Bifidobacteriaceae</td><td>GCF_020525765.1</td><td>Bifidobacterium_pseudocatenulatum</td><td>PG776_RS01085</td><td>258368..262027</td><td>28026</td><td>[illegible]</td></tr>
<tr><td>WP_226650300.1</td><td>Type1.BREX</td><td><-SIGMA+OB-Fold+AAA-ATPase+HKD-DNase-BrxL<-PglZ<-Schlafen+HATPase+wHTH<-REase<-N6-METHYLASE<-DUF6079*<-BrxB-BrxF-iSTAND<- [illegible]</td><td>1219</td><td>Bacteria>Terrabacteria_group>Actinomycetota>Actinomycetes>Bifidobacteriales>Bifidobacteriaceae</td><td>GCF_020526585.1</td><td>Bifidobacterium_pseudocatenulatum</td><td>KZP10_RS05695</td><td>1065058..1068717</td><td>28026</td><td>[illegible]</td></tr>
<tr><td>WP_235341612.1</td><td>Type1.BREX</td><td>METHYLASE->?->Glycos_transf_4-><-?||?->?->BrxA-DNA-binding->BrxB-BrxF-iSTAND->DUF6079*->PglZ->SIGMA+OB-Fold+AAA-ATPase+HKD- [illegible]</td><td>1238</td><td>Bacteria>Terrabacteria_group>Actinomycetota>Actinomycetes>Bifidobacteriales>Bifidobacteriaceae</td><td>GCF_021531155.1</td><td>Bifidobacterium_boum</td><td>I6E06_RS03585</td><td>150899..154615</td><td>78343</td><td>[illegible]</td></tr>
<tr><td>WP_237919300.1</td><td>Type1.BREX</td><td>ACET+ACET->?-><-?||?-><-SbcD||?-><-?||BrxA-DNA-binding->BrxB-BrxF-iSTAND->DUF6079*->N6-METHYLASE->SIR2->Schlafen+HATPase+wHTH- [illegible]</td><td>1219</td><td>Bacteria>Terrabacteria_group>Actinomycetota>Actinomycetes>Bifidobacteriales>Bifidobacteriaceae</td><td>GCF_022135125.1</td><td>Bifidobacterium_pseudocatenulatum</td><td>L0M88_RS00205</td><td>52697..56356</td><td>28026</td><td>[illegible]</td></tr>
<tr><td>WP_237970585.1</td><td>Type1.BREX</td><td>HTH_28->BrxA-DNA-binding->BrxB-BrxF-iSTAND->DUF6079*->N6-METHYLASE->?->Schlafen+HATPase+wHTH->PglZ->SIGMA+OB-Fold+AAA-ATPase+ [illegible]</td><td>1218</td><td>Bacteria>Terrabacteria_group>Actinomycetota>Actinomycetes>Bifidobacteriales>Bifidobacteriaceae</td><td>GCF_022136875.1</td><td>Bifidobacterium_longum</td><td>L0O84_RS04690</td><td>81950..85606</td><td>216816</td><td>[illegible]</td></tr>
<tr><td>WP_273192946.1</td><td>Type1.BREX</td><td><-SF1-SF2-Helicase||?->?-><-SPOUT||ENDO3+NUDIX->?->BrxA-DNA-binding->BrxB-BrxF-iSTAND->DUF6079*->?->N6-METHYLASE-><- [illegible]</td><td>1173</td><td>Bacteria>Terrabacteria_group>Actinomycetota>Actinomycetes>Bifidobacteriales>Bifidobacteriaceae</td><td>GCF_022487345.1</td><td>Bifidobacterium_tibiigranuli</td><td>LKF49_RS06765</td><td>121598..125119</td><td>2172043</td><td>[illegible]</td></tr>
<tr><td>WP_240833140.1</td><td>Type1.BREX</td><td><-Abhydrolase<-ParB+HNH+RAMA<-ZnR<-ZnR+GP<-SIGMA+OB-Fold+AAA-ATPase+HKD-DNase-BrxL<-PglZ<-Schlafen+HATPase+wHTH<-?<- [illegible]</td><td>1217</td><td>Bacteria>Terrabacteria_group>Actinomycetota>Actinomycetes>Bifidobacteriales>Bifidobacteriaceae</td><td>GCF_022496395.1</td><td>Bifidobacterium_pseudocatenulatum</td><td>LF914_RS07635</td><td>54595..58248</td><td>28026</td><td>[illegible]</td></tr>
<tr><td>WP_273228543.1</td><td>Type1.BREX</td><td><-N6-METHYLASE-p<-?<-DUF6079*<-BrxB-BrxF-iSTAND<-BrxA-DNA-binding<-?<-?<-?<-Aminotran_1_2||?-><-?<-MoxR-AAA</td><td>1170</td><td>Bacteria>Terrabacteria_group>Actinomycetota>Actinomycetes>Bifidobacteriales>Bifidobacteriaceae</td><td>GCF_022640875.1</td><td>Bifidobacterium_crudilactis</td><td>LKG00_RS08965</td><td>2590..6102</td><td>327277</td><td>[illegible]</td></tr>
<tr><td>WP_242075534.1</td><td>Type1.BREX</td><td>HTH_28->BrxA-DNA-binding->BrxB-BrxF-iSTAND->DUF6079*->N6-METHYLASE->Schlafen+HATPase+wHTH->PglZ->SIGMA+OB-Fold+AAA-ATPase+HKD- [illegible]</td><td>1218</td><td>Bacteria>Terrabacteria_group>Actinomycetota>Actinomycetes>Bifidobacteriales>Bifidobacteriaceae</td><td>GCF_022643745.1</td><td>Bifidobacterium_longum</td><td>G8B15_RS07005</td><td>1527026..1530682</td><td>216816</td><td>[illegible]</td></tr>
<tr><td>WP_032684176.1</td><td>Type1.BREX</td><td><-SIGMA+OB-Fold+AAA-ATPase+HKD-DNase-BrxL<-PglZ<-Schlafen+HATPase+wHTH<-N6-METHYLASE-p<-HerA/FtsK<-SIR2<-N6-METHYLASE<- [illegible]</td><td>1219</td><td>Bacteria>Terrabacteria_group>Actinomycetota>Actinomycetes>Bifidobacteriales>Bifidobacteriaceae</td><td>GCF_022701235.1</td><td>Bifidobacterium_longum</td><td>LMY38_RS03520</td><td>757375..761034</td><td>216816</td><td>[illegible]</td></tr>
<tr><td>WP_248110663.1</td><td>Type1.BREX</td><td>35exo+HRDC->?->35exo+BRCT-><-HTH||?-><-ACET||?->BrxA-DNA-binding->BrxB-BrxF-iSTAND->DUF6079*->N6-METHYLASE->?->Schlafen+ [illegible]</td><td>1219</td><td>Bacteria>Terrabacteria_group>Actinomycetota>Actinomycetes>Bifidobacteriales>Bifidobacteriaceae</td><td>GCF_023147995.1</td><td>Bifidobacterium_dentium</td><td>KZX63_RS03535</td><td>164390..168049</td><td>1689</td><td>[illegible]</td></tr>
<tr><td>WP_269374243.1</td><td>Type1.BREX</td><td><-Sigma-HTH<-ParB+HNH<-ZnR<-ZnR+GP<-SIGMA+OB-Fold+AAA-ATPase+HKD-DNase-BrxL<-PglZ<-Schlafen+HATPase+wHTH<-SIR2<-N6-METHYLASE<- [illegible]</td><td>1219</td><td>Bacteria>Terrabacteria_group>Actinomycetota>Actinomycetes>Bifidobacteriales>Bifidobacteriaceae</td><td>GCF_027214485.1</td><td>Bifidobacterium_breve</td><td>O4C76_RS09020</td><td>75244..78903</td><td>1685</td><td>[illegible]</td></tr>
<tr><td>WP_269360645.1</td><td>Type1.BREX</td><td><-RNase<-BirA-HTH<-Schlafen+HATPase+wHTH<-SMODS<-SLATT<-N6-METHYLASE-p<-HerA/FtsK<-SIR2<-N6-METHYLASE<-DUF6079*<-BrxB-BrxF- [illegible]</td><td>1229</td><td>Bacteria>Terrabacteria_group>Actinomycetota>Actinomycetes>Bifidobacteriales>Bifidobacteriaceae</td><td>GCF_027215095.1</td><td>Bifidobacterium_breve</td><td>O4D08_RS00920</td><td>188382..192071</td><td>1685</td><td>[illegible]</td></tr>
<tr><td>WP_269355165.1</td><td>Type1.BREX</td><td><-Sigma-HTH<-ParB+HNH<-ZnR<-ZnR+GP<-SIGMA+OB-Fold+AAA-ATPase+HKD-DNase-BrxL<-PglZ<-Schlafen+HATPase+wHTH<-SIR2<-N6-METHYLASE<- [illegible]</td><td>1218</td><td>Bacteria>Terrabacteria_group>Actinomycetota>Actinomycetes>Bifidobacteriales>Bifidobacteriaceae</td><td>GCF_027215185.1</td><td>Bifidobacterium_breve</td><td>O4C27_RS08995</td><td>67566..71222</td><td>1685</td><td>[illegible]</td></tr>
<tr><td>WP_269338054.1</td><td>Type1.BREX</td><td><-MULE-TRANSPOSASE||?-><-?||?->CR-ATPase5+REase-p->BrxA-DNA-binding->BrxB-BrxF-iSTAND->DUF6079-p*->DUF6079-p->N6-METHYLASE-> [illegible]</td><td>924</td><td>Bacteria>Terrabacteria_group>Actinomycetota>Actinomycetes>Bifidobacteriales>Bifidobacteriaceae</td><td>GCF_027215525.1</td><td>Bifidobacterium_breve</td><td>O4C30_RS07905</td><td>20359..23133</td><td>1685</td><td>[illegible]</td></tr>
<tr><td>WP_270294263.1</td><td>Type1.BREX</td><td><-SIGMA+OB-Fold+AAA-ATPase+HKD-DNase-BrxL<-PglZ<-Schlafen+HATPase+wHTH<-SIR2<-N6-METHYLASE-p<-HerA/FtsK<-SIR2<-N6-METHYLASE<- [illegible]</td><td>1219</td><td>Bacteria>Terrabacteria_group>Actinomycetota>Actinomycetes>Bifidobacteriales>Bifidobacteriaceae</td><td>GCF_027659585.1</td><td>Bifidobacterium_adolescentis</td><td>PGD90_RS01995</td><td>459377..463036</td><td>1680</td><td>[illegible]</td></tr>
<tr><td>WP_270267401.1</td><td>Type1.BREX</td><td><-SIGMA+OB-Fold+AAA-ATPase+HKD-DNase-BrxL<-PglZ<-RNase<-BirA-HTH<-Schlafen+HATPase+wHTH<-SMODS<-SLATT<-N6-METHYLASE<-DUF6079*<- [illegible]</td><td>1229</td><td>Bacteria>Terrabacteria_group>Actinomycetota>Actinomycetes>Bifidobacteriales>Bifidobacteriaceae</td><td>GCF_027663845.1</td><td>Bifidobacterium_breve</td><td>PGQ90_RS07415</td><td>209217..212906</td><td>1685</td><td>[illegible]</td></tr>
<tr><td>WP_117779802.1</td><td>Type1.BREX</td><td><-SIGMA+OB-Fold+AAA-ATPase+HKD-DNase-BrxL<-PglZ<-Schlafen+HATPase+wHTH<-SIR2<-N6-METHYLASE-p<-HerA/FtsK<-SIR2<-N6-METHYLASE<- [illegible]</td><td>1219</td><td>Bacteria>Terrabacteria_group>Actinomycetota>Actinomycetes>Bifidobacteriales>Bifidobacteriaceae</td><td>GCF_027664445.1</td><td>Bifidobacterium_adolescentis</td><td>PGQ86_RS05915</td><td>1356414..1360073</td><td>1680</td><td>[illegible]</td></tr>
<tr><td>WP_270271788.1</td><td>Type1.BREX</td><td>REase-><-?||?->?->HTH+HisKin-><-REase||BrxA-DNA-binding->BrxB-BrxF-iSTAND->DUF6079*->N6-METHYLASE->?->Schlafen+HATPase+wHTH-> [illegible]</td><td>1218</td><td>Bacteria>Terrabacteria_group>Actinomycetota>Actinomycetes>Bifidobacteriales>Bifidobacteriaceae</td><td>GCF_027667785.1</td><td>Bifidobacterium_bifidum</td><td>PGS22_RS05345</td><td>127181..130837</td><td>1681</td><td>[illegible]</td></tr>
<tr><td>WP_117715835.1</td><td>Type1.BREX</td><td>ZnR->?->ABC-ATPase+SbcC->?->?->?->BrxA-DNA-binding->BrxB-BrxF-iSTAND->DUF6079*->N6-METHYLASE->SIR2->Schlafen+HATPase+wHTH->PglZ [illegible]</td><td>1217</td><td>Bacteria>Terrabacteria_group>Actinomycetota>Actinomycetes>Bifidobacteriales>Bifidobacteriaceae</td><td>GCF_027669765.1</td><td>Bifidobacterium_bifidum</td><td>PHA16_RS02375</td><td>586926..590579</td><td>1681</td><td>[illegible]</td></tr>
<tr><td>WP_270200767.1</td><td>Type1.BREX</td><td>REC+LuxR-HTH->HISKIN-><-?<-SbcD<-?<-?||Pentapeptide+Pentapeptide+Pentapeptide+Pentapeptide->BrxA-DNA-binding->BrxB-BrxF-iSTAND- [illegible]</td><td>467</td><td>Bacteria>Terrabacteria_group>Actinomycetota>Actinomycetes>Bifidobacteriales>Bifidobacteriaceae</td><td>GCF_027671795.1</td><td>Bifidobacterium_pseudocatenulatum</td><td>PGT11_RS00265</td><td>67391..68794</td><td>28026</td><td>[illegible]</td></tr>
<tr><td>WP_270199955.1</td><td>Type1.BREX</td><td><-SbcD<-?<-?||?->||Pentapeptide+Pentapeptide+Pentapeptide+Pentapeptide->BrxA-DNA-binding->BrxB-BrxF-iSTAND->DUF6079*->DUF6079- [illegible]</td><td>1044</td><td>Bacteria>Terrabacteria_group>Actinomycetota>Actinomycetes>Bifidobacteriales>Bifidobacteriaceae</td><td>GCF_027675485.1</td><td>Bifidobacterium_pseudocatenulatum</td><td>PGV18_RS06405</td><td>41954..45088</td><td>28026</td><td>[illegible]</td></tr>
<tr><td>WP_270271488.1</td><td>Type1.BREX</td><td><-ZnR<-ZnR+GP<-SIGMA+OB-Fold+AAA-ATPase+HKD-DNase-BrxL<-PglZ<-Schlafen+HATPase+wHTH<-SIR2<-N6-METHYLASE<-DUF6079*<-BrxB-BrxF- [illegible]</td><td>1219</td><td>Bacteria>Terrabacteria_group>Actinomycetota>Actinomycetes>Bifidobacteriales>Bifidobacteriaceae</td><td>GCF_027679525.1</td><td>Bifidobacterium_adolescentis</td><td>PGW19_RS08965</td><td>15037..18696</td><td>1680</td><td>[illegible]</td></tr>
<tr><td>WP_270447025.1</td><td>Type1.BREX</td><td>HISKIN-><-?||<-SbcD<-?<-?||?->||Pentapeptide+Pentapeptide+Pentapeptide+Pentapeptide->BrxA-DNA-binding->BrxB-BrxF-iSTAND-> [illegible]</td><td>958</td><td>Bacteria>Terrabacteria_group>Actinomycetota>Actinomycetes>Bifidobacteriales>Bifidobacteriaceae</td><td>GCF_027682665.1</td><td>Bifidobacterium_pseudocatenulatum</td><td>PGX68_RS01375</td><td>343601..346477</td><td>28026</td><td>[illegible]</td></tr>
<tr><td>WP_270479593.1</td><td>Type1.BREX</td><td><-ZnR+GP<-SIGMA+OB-Fold+AAA-ATPase+HKD-DNase-BrxL<-PglZ<-Schlafen+HATPase+wHTH<-SIR2<-N6-METHYLASE-p<-HerA/FtsK<-SIR2<- [illegible]</td><td>1219</td><td>Bacteria>Terrabacteria_group>Actinomycetota>Actinomycetes>Bifidobacteriales>Bifidobacteriaceae</td><td>GCF_027683145.1</td><td>Bifidobacterium_adolescentis</td><td>PGX62_RS02115</td><td>472627..476286</td><td>1680</td><td>[illegible]</td></tr>
<tr><td>WP_117726345.1</td><td>Type1.BREX</td><td>ZnR->?->ABC-ATPase+SbcC->?->?->?->BrxA-DNA-binding->BrxB-BrxF-iSTAND->DUF6079*->N6-METHYLASE->?->Schlafen+HATPase+wHTH->PglZ-> [illegible]</td><td>1217</td><td>Bacteria>Terrabacteria_group>Actinomycetota>Actinomycetes>Bifidobacteriales>Bifidobacteriaceae</td><td>GCF_027685785.1</td><td>Bifidobacterium_bifidum</td><td>PG427_RS02260</td><td>555349..559002</td><td>1681</td><td>[illegible]</td></tr>
<tr><td>WP_270266433.1</td><td>Type1.BREX</td><td><-SIGMA+OB-Fold+AAA-ATPase+HKD-DNase-BrxL<-PglZ<-Schlafen+HATPase+wHTH<-?<-N6-METHYLASE<-DUF6079*<-BrxB-BrxF-iSTAND<- [illegible]</td><td>1219</td><td>Bacteria>Terrabacteria_group>Actinomycetota>Actinomycetes>Bifidobacteriales>Bifidobacteriaceae</td><td>GCF_027687815.1</td><td>Bifidobacterium_bifidum</td><td>PG972_RS08475</td><td>12443..16102</td><td>1681</td><td>[illegible]</td></tr>
</table>

| Protein-ID | System | Gene-Neighbourhoods | Length | Taxonomy | Assembly-ID | Strain | Locus-Tag | Genomic-coordinates | TaxID | Fasta-define |
|---|---|---|---|---|---|---|---|---|---|---|
| WP_270539149.1 | Type1.BREX | [illegible] | 1217 | Bacteria>Terrabacteria_group>Actinomycetota>Actinomycetes>Bifidobacteriales>Bifidobacteriaceae | GCF_027687915.1 | Bifidobacterium_bifidum | PGZ56_RS09175 | 4870..8523 | 1681 | BREX-system-P-loop-protein-BrxC-[Bifidobacterium-bifidum] |
| WP_117501722.1 | Type1.BREX | [illegible] | 1219 | Bacteria>Terrabacteria_group>Actinomycetota>Actinomycetes>Bifidobacteriales>Bifidobacteriaceae | GCF_027689935.1 | Bifidobacterium_pseudocatenulatum | PG028_RS01370 | 345834..349493 | 28026 | BREX-system-P-loop-protein-BrxC-[Bifidobacterium-pseudocatenulatum] |
| WP_270558503.1 | Type1.BREX | [illegible] | 1219 | Bacteria>Terrabacteria_group>Actinomycetota>Actinomycetes>Bifidobacteriales>Bifidobacteriaceae | GCF_027690155.1 | Bifidobacterium_adolescentis | PG057_RS05020 | 457971..461630 | 1680 | BREX-system-P-loop-protein-BrxC-[Bifidobacterium-adolescentis] |
| WP_270554212.1 | Type1.BREX | [illegible] | 1219 | Bacteria>Terrabacteria_group>Actinomycetota>Actinomycetes>Bifidobacteriales>Bifidobacteriaceae | GCF_027695195.1 | Bifidobacterium_adolescentis | PG742_RS01120 | 252838..256497 | 1680 | BREX-system-P-loop-protein-BrxC-[Bifidobacterium-adolescentis] |
| WP_270561184.1 | Type1.BREX | [illegible] | 1217 | Bacteria>Terrabacteria_group>Actinomycetota>Actinomycetes>Bifidobacteriales>Bifidobacteriaceae | GCF_027698005.1 | Bifidobacterium_dentium | PG823_RS09640 | 38589..42242 | 1689 | BREX-system-P-loop-protein-BrxC-[Bifidobacterium-dentium] |
| WP_118376243.1 | Type1.BREX | [illegible] | 1219 | Bacteria>Terrabacteria_group>Actinomycetota>Actinomycetes>Bifidobacteriales>Bifidobacteriaceae | GCF_027698785.1 | Bifidobacterium_pseudocatenulatum | PG820_RS05645 | 117035..120694 | 28026 | BREX-system-P-loop-protein-BrxC-[Bifidobacterium-pseudocatenulatum] |
| WP_271768971.1 | Type1.BREX | [illegible] | 1219 | Bacteria>Terrabacteria_group>Actinomycetota>Actinomycetes>Bifidobacteriales>Bifidobacteriaceae | GCF_028314485.1 | Bifidobacterium_pseudocatenulatum | PMN34_RS05085 | 218989..222648 | 28026 | BREX-system-P-loop-protein-BrxC-[Bifidobacterium-pseudocatenulatum] |
| WP_116463714.1 | Type1.BREX | [illegible] | 1219 | Bacteria>Terrabacteria_group>Actinomycetota>Actinomycetes>Bifidobacteriales>Bifidobacteriaceae | GCF_028898865.1 | Bifidobacterium_longum | PWA56_RS02155 | 487318..490977 | 216816 | BREX-system-P-loop-protein-BrxC-[Bifidobacterium-longum] |
| WP_077383013.1 | Type1.BREX | [illegible] | 1219 | Bacteria>Terrabacteria_group>Actinomycetota>Actinomycetes>Bifidobacteriales>Bifidobacteriaceae | GCF_900157155.1 | Bifidobacterium_longum | BQ8631_RS08415 | 45904..49563 | 216816 | BREX-system-P-loop-protein-BrxC-[Bifidobacterium-longum] |
| WP_154050436.1 | Type1.BREX | [illegible] | 1217 | Bacteria>Terrabacteria_group>Actinomycetota>Actinomycetes>Bifidobacteriales>Bifidobacteriaceae | GCF_901212475.1 | Bifidobacterium_longum | FGK88_RS06805 | 85798..89451 | 216816 | BREX-system-P-loop-protein-BrxC-[Bifidobacterium-longum] |
| WP_154035315.1 | Type1.BREX | [illegible] | 1218 | Bacteria>Terrabacteria_group>Actinomycetota>Actinomycetes>Bifidobacteriales>Bifidobacteriaceae | GCF_901212515.1 | Bifidobacterium_catenulatum | FGK79_RS07205 | 243456..247112 | 1686 | BREX-system-P-loop-protein-BrxC-[Bifidobacterium-catenulatum] |
| WP_143724686.1 | Type1.BREX | [illegible] | 1229 | Bacteria>Terrabacteria_group>Actinomycetota>Actinomycetes>Bifidobacteriales>Bifidobacteriaceae | GCF_902167555.1 | Bifidobacterium_longum | FP566_RS07260 | 1570713..1574402 | 216816 | BREX-system-P-loop-protein-BrxC-[Bifidobacterium-longum] |
| WP_143246218.1 | Type1.BREX | [illegible] | 1218 | Bacteria>Terrabacteria_group>Actinomycetota>Actinomycetes>Bifidobacteriales>Bifidobacteriaceae | GCF_902167575.1 | Bifidobacterium_breve | FPG44_RS02615 | 449270..452926 | 1685 | BREX-system-P-loop-protein-BrxC-[Bifidobacterium-breve] |
| WP_200406352.1 | Type1.BREX | [illegible] | 1229 | Bacteria>Terrabacteria_group>Actinomycetota>Actinomycetes>Bifidobacteriales>Bifidobacteriaceae | GCF_934675445.1 | Bifidobacterium_longum | BL7055_RS05750 | 1406658..1410347 | 216816 | BREX-system-P-loop-protein-BrxC-[Bifidobacterium-longum] |
| WP_245206784.1 | Type1.BREX | [illegible] | 1073 | Bacteria>Terrabacteria_group>Actinomycetota>Actinomycetes>Bifidobacteriales>Bifidobacteriaceae | GCF_937997665.1 | Bifidobacterium_dentium | J7M36_RS06065 | 1427241..1430462 | 1689 | BREX-system-P-loop-protein-BrxC-[Bifidobacterium-dentium] |
| WP_250246699.1 | Type1.BREX | [illegible] | 1229 | Bacteria>Terrabacteria_group>Actinomycetota>Actinomycetes>Bifidobacteriales>Bifidobacteriaceae | GCF_939148185.1 | Bifidobacterium_longum | NBW58_RS09955 | 18627..22316 | 216816 | BREX-system-P-loop-protein-BrxC-[Bifidobacterium-longum] |
| WP_196330425.1 | Type1.BREX | [illegible] | 1217 | Bacteria>Terrabacteria_group>Actinomycetota>Actinomycetes>Bifidobacteriales>Bifidobacteriaceae | GCF_958433885.1 | Bifidobacterium_dentium | J7M35_RS04855 | 1219062..1222715 | 1689 | BREX-system-P-loop-protein-BrxC-[Bifidobacterium-dentium] |
| WP_085380386.1 | Type1.BREX | [illegible] | 1219 | Bacteria>Terrabacteria_group>Actinomycetota>Actinomycetes>Bifidobacteriales>Bifidobacteriaceae | GCF_959020145.1 | Bifidobacterium_adolescentis | LMG10734_RS05600 | 489007..492666 | 1680 | BREX-system-P-loop-protein-BrxC-[Bifidobacterium-adolescentis] |
| WP_196335743.1 | Type1.BREX | [illegible] | 1217 | Bacteria>Terrabacteria_group>Actinomycetota>Actinomycetes>Bifidobacteriales>Bifidobacteriaceae | GCF_959600175.1 | Bifidobacterium_dentium | IHQ76_RS06180 | 228522..232175 | 1689 | BREX-system-P-loop-protein-BrxC-[Bifidobacterium-dentium] |
| WP_165367554.1 | Type1.BREX | [illegible] | 1157 | Bacteria>Terrabacteria_group>Actinomycetota>Actinomycetes>Jiangellales>Jiangellaceae | GCF_004138495.1 | Phytoactinopolyspora_endophytica | ELR45_RS07100 | 275319..278792 | 1642495 | BREX-system-P-loop-protein-BrxC-[Phytoactinopolyspora-endophytica] |
| EST38900.1 | Type1.BREX | [illegible] | 571 | Bacteria>Terrabacteria_group>Actinomycetota>Actinomycetes>Kitasatosporales>Streptomycetaceae | GCA_000497405.1 | Streptomycetaceae_bacterium_MP113-05 | N566_04930 | 13788..15503 | 1380770 | hypothetical-protein-N566_04930-[Streptomycetaceae-bacterium-MP113-05] |
| MCG7205148.1 | Type1.BREX | [illegible] | 1161 | Bacteria>Terrabacteria_group>Actinomycetota>Actinomycetes>Kitasatosporales>Streptomycetaceae | GCA_022343545.1 | Streptomyces_arenae | MB828_15195 | 11459..14944 | 29301 | BREX-system-P-loop-protein-BrxC-[Streptomyces-arenae] |
| WP_030399158.1 | Type1.BREX | [illegible] | 1161 | Bacteria>Terrabacteria_group>Actinomycetota>Actinomycetes>Kitasatosporales>Streptomycetaceae | GCF_000719665.1 | unclassified_Streptomyces | ADK87_RS19375 | 27273..30758 | 2593676 | MULTISPECIES:-BREX-system-P-loop-protein-BrxC-[unclassified-Streptomyces] |
| WP_042419815.1 | Type1.BREX | [illegible] | 1161 | Bacteria>Terrabacteria_group>Actinomycetota>Actinomycetes>Kitasatosporales>Streptomycetaceae | GCF_000787855.1 | Streptacidiphilus_anmyonensis | TR50_RS13090 | 101724..105209 | 405782 | BREX-system-P-loop-protein-BrxC-[Streptacidiphilus-anmyonensis] |
| WP_062022136.1 | Type1.BREX | [illegible] | 1161 | Bacteria>Terrabacteria_group>Actinomycetota>Actinomycetes>Kitasatosporales>Streptomycetaceae | GCF_001513985.1 | Streptomyces_griseorubiginosus | AQI84_07455 | 596850..600335 | 67304 | BREX-system-P-loop-protein-BrxC-[Streptomyces-griseorubiginosus] |
| WP_129286571.1 | Type1.BREX | [illegible] | 1163 | Bacteria>Terrabacteria_group>Actinomycetota>Actinomycetes>Kitasatosporales>Streptomycetaceae | GCF_004123705.1 | Streptomyces_sp._GZWMJZ-114 | EJA11_RS17595 | 541597..545088 | 2494734 | BREX-system-P-loop-protein-BrxC-[Streptomyces-sp.-GZWMJZ-114] |
| WP_138352701.1 | Type1.BREX | [illegible] | 1186 | Bacteria>Terrabacteria_group>Actinomycetota>Actinomycetes>Kitasatosporales>Streptomycetaceae | GCF_005860785.2 | Streptomyces_sp._So13.3 | C8250_RS11580 | 2549516..2553076 | 2136173 | BREX-system-P-loop-protein-BrxC-[Streptomyces-sp.-So13.3] |
| WP_161225978.1 | Type1.BREX | [illegible] | 1165 | Bacteria>Terrabacteria_group>Actinomycetota>Actinomycetes>Kitasatosporales>Streptomycetaceae | GCF_009862265.1 | Streptomyces_sp._SID8352 | GTY84_RS10560 | 297618..301115 | 2690338 | BREX-system-P-loop-protein-BrxC-[Streptomyces-sp.-SID8352] |
| WP_164340211.1 | Type1.BREX | [illegible] | 1163 | Bacteria>Terrabacteria_group>Actinomycetota>Actinomycetes>Kitasatosporales>Streptomycetaceae | GCF_010548045.1 | Streptomyces_anulatus | G3I47_RS13230 | 74640..78131 | 1892 | BREX-system-P-loop-protein-BrxC-[Streptomyces-anulatus] |
| WP_182441002.1 | Type1.BREX | [illegible] | 1159 | Bacteria>Terrabacteria_group>Actinomycetota>Actinomycetes>Kitasatosporales>Streptomycetaceae | GCF_014084125.1 | Streptacidiphilus_sp._PB12-B1b | GXW83_RS01100 | 242781..246260 | 2705012 | BREX-system-P-loop-protein-BrxC-[Streptacidiphilus-sp.-PB12-B1b] |
| WP_073800314.1 | Type1.BREX | [illegible] | 1163 | Bacteria>Terrabacteria_group>Actinomycetota>Actinomycetes>Kitasatosporales>Streptomycetaceae | GCF_014495705.1 | unclassified_Streptomyces | AMK20_RS03875 | 898838..902329 | 2593676 | MULTISPECIES:-BREX-system-P-loop-protein-BrxC-[unclassified-Streptomyces] |
| WP_265293811.1 | Type1.BREX | [illegible] | 1168 | Bacteria>Terrabacteria_group>Actinomycetota>Actinomycetes>Kitasatosporales>Streptomycetaceae | GCF_026126445.1 | Streptomyces_sp._SHP_1-2 | IBX28_RS14860 | 97274..100780 | 2769489 | BREX-system-P-loop-protein-BrxC-[Streptomyces-sp.-SHP-1-2] |
| WP_266948097.1 | Type1.BREX | [illegible] | 1186 | Bacteria>Terrabacteria_group>Actinomycetota>Actinomycetes>Kitasatosporales>Streptomycetaceae | GCF_026342255.1 | Streptomyces_sp._NBC_00291 | OG914_RS06430 | 1462666..1466226 | 2975704 | BREX-system-P-loop-protein-BrxC-[Streptomyces-sp.-NBC_00291] |
| WP_093695822.1 | Type1.BREX | [illegible] | 1162 | Bacteria>Terrabacteria_group>Actinomycetota>Actinomycetes>Kitasatosporales>Streptomycetaceae | GCF_900105605.1 | Streptomyces_sp._2231.1 | BLV73_RS10010 | 2127135..2130623 | 1855347 | BREX-system-P-loop-protein-BrxC-[Streptomyces-sp.-2231.1] |
| WP_242897357.1 | Type1.BREX | [illegible] | 590 | Bacteria>Terrabacteria_group>Actinomycetota>Actinomycetes>Micrococcales>Bogoriellaceae | GCF_022759505.1 | Georgenia_sp._TF02-10 | MF406_RS07595 | 1686930..1688702 | 2917725 | hypothetical-protein-[Georgenia-sp.-TF02-10] |
| WP_242897362.1 | Type1.BREX | [illegible] | 561 | Bacteria>Terrabacteria_group>Actinomycetota>Actinomycetes>Micrococcales>Bogoriellaceae | GCF_022759505.1 | Georgenia_sp._TF02-10 | MF406_RS07600 | 1688699..1690384 | 2917725 | hypothetical-protein-[Georgenia-sp.-TF02-10] |
| QZE24962.1 | Type1.BREX | [illegible] | 1152 | Bacteria>Terrabacteria_group>Actinomycetota>Actinomycetes>Micrococcales>Brevibacteriaceae | GCA_019720815.1 | Brevibacterium_casei | K4X33_12630 | 2739675..2743133 | 33889 | BREX-system-P-loop-protein-BrxC-[Brevibacterium-casei] |
| WP_005885100.1 | Type1.BREX | [illegible] | 1178 | Bacteria>Terrabacteria_group>Actinomycetota>Actinomycetes>Micrococcales>Brevibacteriaceae | GCF_000178455.1 | Brevibacterium_mcbrellneri | HMPREF0183_RS08780 | 50551..54087 | 53363 | BREX-system-P-loop-protein-BrxC-[Brevibacterium-mcbrellneri] |
| WP_096157188.1 | Type1.BREX | [illegible] | 1151 | Bacteria>Terrabacteria_group>Actinomycetota>Actinomycetes>Micrococcales>Brevibacteriaceae | GCF_002332225.1 | Brevibacterium_aurantiacum | CIK79_RS00860 | 211132..214587 | 273384 | BREX-system-P-loop-protein-BrxC-[Brevibacterium-aurantiacum] |
| WP_186372589.1 | Type1.BREX | [illegible] | 1152 | Bacteria>Terrabacteria_group>Actinomycetota>Actinomycetes>Micrococcales>Brevibacteriaceae | GCF_002339175.1 | Brevibacterium_casei | B3D11_RS04595 | 80759..84217 | 33889 | BREX-system-P-loop-protein-BrxC-[Brevibacterium-casei] |
| WP_125240568.1 | Type1.BREX | [illegible] | 1170 | Bacteria>Terrabacteria_group>Actinomycetota>Actinomycetes>Micrococcales>Brevibacteriaceae | GCF_003932995.1 | Brevibacterium_aurantiacum | CXR25_RS11695 | 2538315..2541827 | 273384 | BREX-system-P-loop-protein-BrxC-[Brevibacterium-aurantiacum] |
| WP_201670680.1 | Type1.BREX | [illegible] | 1160 | Bacteria>Terrabacteria_group>Actinomycetota>Actinomycetes>Micrococcales>Brevibacteriaceae | GCF_016726245.1 | Brevibacterium_casei | I6I57_RS09610 | 2195903..2199385 | 33889 | BREX-system-P-loop-protein-BrxC-[Brevibacterium-casei] |
| WP_259801020.1 | Type1.BREX | [illegible] | 1173 | Bacteria>Terrabacteria_group>Actinomycetota>Actinomycetes>Micrococcales>Brevibacteriaceae | GCF_025144065.1 | Brevibacterium_casei | M3G43_RS08140 | 29205..32726 | 33889 | BREX-system-P-loop-protein-BrxC-[Brevibacterium-casei] |
| WP_259820848.1 | Type1.BREX | [illegible] | 1152 | Bacteria>Terrabacteria_group>Actinomycetota>Actinomycetes>Micrococcales>Brevibacteriaceae | GCF_025152225.1 | Brevibacterium_casei | M3G00_10225 | 31944..35402 | 33889 | BREX-system-P-loop-protein-BrxC-[Brevibacterium-casei] |
| WP_264799000.1 | Type1.BREX | [illegible] | 1152 | Bacteria>Terrabacteria_group>Actinomycetota>Actinomycetes>Micrococcales>Brevibacteriaceae | GCF_026013965.1 | Brevibacterium_sp._JSBI002 | LJ362_RS10440 | 2291200..2294658 | 2886045 | BREX-system-P-loop-protein-BrxC-[Brevibacterium-sp.-JSBI002] |
| WP_101598836.1 | Type1.BREX | [illegible] | 1160 | Bacteria>Terrabacteria_group>Actinomycetota>Actinomycetes>Micrococcales>Brevibacteriaceae | GCF_900169315.1 | Brevibacterium_aurantiacum | BAURA63_03577 | 9324..12806 | 273384 | BREX-system-P-loop-protein-BrxC-[Brevibacterium-aurantiacum] |
| TGJ96612.1 | Type1.BREX | [illegible] | 969 | Bacteria>Terrabacteria_group>Actinomycetota>Actinomycetes>Micrococcales>Cellulomonadaceae | GCA_004769145.1 | Actinotalea_fermentans_ATCC_43279_=_JCM_9966_=_DSM_3133 | DLJ96_00440 | 117898..120803 | 862422 | BREX-system-P-loop-protein-BrxC,-partial-[Actinotalea-fermentans-ATCC-43279-=-JCM-9966-=-DSM-3133] |
| NMM30393.1 | Type1.BREX | [illegible] | 1157 | Bacteria>Terrabacteria_group>Actinomycetota>Actinomycetes>Micrococcales>Cellulomonadaceae | GCA_012927595.1 | Cellulomonas_sp. | HHJ10_04945 | 1662..5135 | 40001 | BREX-system-P-loop-protein-BrxC-[Cellulomonas-sp.] |
| WP_029253807.1 | Type1.BREX | [illegible] | 1160 | Bacteria>Terrabacteria_group>Actinomycetota>Actinomycetes>Micrococcales>Cellulomonadaceae | GCF_000701465.1 | Paraoerskovia_marina | Q343_RS0111945 | 114651..118133 | 545619 | BREX-system-P-loop-protein-BrxC-[Paraoerskovia-marina] |
| WP_030150650.1 | Type1.BREX | [illegible] | 1156 | Bacteria>Terrabacteria_group>Actinomycetota>Actinomycetes>Micrococcales>Cellulomonadaceae | GCF_000718325.1 | Oerskovia_turbata | IE95_RS0105585 | 234727..238197 | 1713 | BREX-system-P-loop-protein-BrxC-[Oerskovia-turbata] |
| WP_094180988.1 | Type1.BREX | [illegible] | 947 | Bacteria>Terrabacteria_group>Actinomycetota>Actinomycetes>Micrococcales>Cellulomonadaceae | GCF_002240455.1 | Cellulomonas_sp._PSBB021 | CBP52_RS11045 | 2415707..2418550 | 2003551 | BREX-system-P-loop-protein-BrxC-[Cellulomonas-sp.-PSBB021] |
| WP_130780297.1 | Type1.BREX | [illegible] | 1159 | Bacteria>Terrabacteria_group>Actinomycetota>Actinomycetes>Micrococcales>Cellulomonadaceae | GCF_004306155.1 | Cellulomonas_biazotea | CBZ_RS03705 | 43649..47128 | 1709 | BREX-system-P-loop-protein-BrxC-[Cellulomonas-biazotea] |
| WP_259803835.1 | Type1.BREX | [illegible] | 1152 | Bacteria>Terrabacteria_group>Actinomycetota>Actinomycetes>Micrococcales>Dermabacteraceae | - | Brachybacterium_muris | M3F59_04025 | 42519..45977 | 219301 | BREX-system-P-loop-protein-BrxC-[Brachybacterium-muris] |
| WP_010552220.1 | Type1.BREX | [illegible] | 1160 | Bacteria>Terrabacteria_group>Actinomycetota>Actinomycetes>Micrococcales>Dermabacteraceae | GCF_000233655.1 | Brachybacterium_paraconglomeratum | I7Y_RS0116025 | 54770..58252 | 173362 | BREX-system-P-loop-protein-BrxC-[Brachybacterium-paraconglomeratum] |
| WP_114305671.1 | Type1.BREX | [illegible] | 1160 | Bacteria>Terrabacteria_group>Actinomycetota>Actinomycetes>Micrococcales>Dermabacteraceae | GCF_003335355.1 | Brachybacterium_alimentarium | CIK67_RS09225 | 107631..111113 | 47845 | BREX-system-P-loop-protein-BrxC-[Brachybacterium-alimentarium] |
| WP_249738141.1 | Type1.BREX | [illegible] | 1173 | Bacteria>Terrabacteria_group>Actinomycetota>Actinomycetes>Micrococcales>Dermabacteraceae | GCF_023509525.1 | Brachybacterium_sp._JHP9 | Bequi_RS11780 | 103104..106625 | 2910770 | BREX-system-P-loop-protein-BrxC-[Brachybacterium-sp.-JHP9] |
| WP_179478704.1 | Type1.BREX | [illegible] | 1156 | Bacteria>Terrabacteria_group>Actinomycetota>Actinomycetes>Micrococcales>Dermacoccaceae | GCF_013410455.1 | Allobranchiibius_huperziae | HNR15_RS01985 | 409066..412536 | 1874116 | BREX-system-P-loop-protein-BrxC-[Allobranchiibius-huperziae] |
| WP_233763981.1 | Type1.BREX | [illegible] | 1156 | Bacteria>Terrabacteria_group>Actinomycetota>Actinomycetes>Micrococcales>Dermacoccaceae | GCF_021391335.1 | Allobranchiibius_sp._GilTou73 | LVQ62_RS01295 | 263639..267109 | 2904523 | BREX-system-P-loop-protein-BrxC-[Allobranchiibius-sp.-GilTou73] |
| WP_259879962.1 | Type1.BREX | [illegible] | 1173 | Bacteria>Terrabacteria_group>Actinomycetota>Actinomycetes>Micrococcales>Dermacoccaceae | GCF_025149825.1 | Dermacoccus_nishinomiyaensis | M3E15_RS12270 | 1013..4534 | 1274 | BREX-system-P-loop-protein-BrxC-[Dermacoccus-nishinomiyaensis] |
| WP_040162373.1 | Type1.BREX | [illegible] | 1178 | Bacteria>Terrabacteria_group>Actinomycetota>Actinomycetes>Micrococcales>Dermatophilaceae | GCF_902375455.1 | Mobilicoccus_massiliensis | FX004_RS16815 | 289663..293199 | 1522310 | BREX-system-P-loop-protein-BrxC-[Mobilicoccus-massiliensis] |
| WP_056922704.1 | Type1.BREX | [illegible] | 1156 | Bacteria>Terrabacteria_group>Actinomycetota>Actinomycetes>Micrococcales>Intrasporangiaceae | GCF_001429605.1 | Phycicoccus_sp._Soil802 | ASG91_RS11220 | 440237..443707 | 1736414 | BREX-system-P-loop-protein-BrxC-[Phycicoccus-sp.-Soil802] |
| WP_215519875.1 | Type1.BREX | [illegible] | 1156 | Bacteria>Terrabacteria_group>Actinomycetota>Actinomycetes>Micrococcales>Intrasporangiaceae | GCF_018729305.1 | Phycicoccus_mangrovi | KMZ30_RS04765 | 132164..135634 | 2840470 | BREX-system-P-loop-protein-BrxC-[Phycicoccus-mangrovi] |
| WP_265231741.1 | Type1.BREX | [illegible] | 1154 | Bacteria>Terrabacteria_group>Actinomycetota>Actinomycetes>Micrococcales>Intrasporangiaceae | GCF_026114115.1 | Janibacter_hoylei | ON003_RS16090 | 3001704..3005168 | 364298 | BREX-system-P-loop-protein-BrxC-[Janibacter-hoylei] |
| WP_270886872.1 | Type1.BREX | [illegible] | 1154 | Bacteria>Terrabacteria_group>Actinomycetota>Actinomycetes>Micrococcales>Intrasporangiaceae | GCF_027812955.1 | Pedococcus_sp._5OH_020 | PHA84_RS04350 | 172901..176365 | 2989814 | BREX-system-P-loop-protein-BrxC-[Pedococcus-sp.-5OH_020] |
| WP_272461397.1 | Type1.BREX | [illegible] | 1154 | Bacteria>Terrabacteria_group>Actinomycetota>Actinomycetes>Micrococcales>Intrasporangiaceae | GCF_028446585.1 | Intrasporangium_calvum | OO014_RS06100 | 11826..15290 | 53358 | BREX-system-P-loop-protein-BrxC-[Intrasporangium-calvum] |
| PPG29161.1 | Type1.BREX | [illegible] | 1183 | Bacteria>Terrabacteria_group>Actinomycetota>Actinomycetes>Micrococcales>Microbacteriaceae | GCA_002931555.1 | Pseudoclavibacter_sp._RFBB5 | C5B97_09025 | 222441..225992 | 2080574 | BREX-system-P-loop-protein-BrxC-[Pseudoclavibacter-sp.-RFBB5] |
| PPH14625.1 | Type1.BREX | [illegible] | 1175 | Bacteria>Terrabacteria_group>Actinomycetota>Actinomycetes>Micrococcales>Microbacteriaceae | GCA_002932165.1 | Rathayibacter_sp._AY1F8 | C5C35_14425 | 14771..18298 | 2080562 | BREX-system-P-loop-protein-BrxC-[Rathayibacter-sp.-AY1F8] |
| QAY61399.1 | Type1.BREX | [illegible] | 1176 | Bacteria>Terrabacteria_group>Actinomycetota>Actinomycetes>Micrococcales>Microbacteriaceae | GCA_004135285.1 | Microbacterium_protaetiae | ET475_16440 | 3529806..3533336 | 2509458 | BREX-system-P-loop-protein-BrxC-[Microbacterium-protaetiae] |
| TFC74553.1 | Type1.BREX | [illegible] | 1175 | Bacteria>Terrabacteria_group>Actinomycetota>Actinomycetes>Micrococcales>Microbacteriaceae | GCA_004402355.1 | Cryobacterium_sp._TMS1-20-1 | E3045_10185 | 26834..30361 | 1259223 | BREX-system-P-loop-protein-BrxC-[Cryobacterium-sp.-TMS1-20-1] |
| TFB71537.1 | Type1.BREX | [illegible] | 1170 | Bacteria>Terrabacteria_group>Actinomycetota>Actinomycetes>Micrococcales>Microbacteriaceae | GCA_004402535.1 | Cryobacterium_sp._HLT2-23 | E3006_11880 | 40394..43906 | 1259145 | BREX-system-P-loop-protein-BrxC-[Cryobacterium-glaciale] |
| QDZ14672.1 | Type1.BREX | [illegible] | 1176 | Bacteria>Terrabacteria_group>Actinomycetota>Actinomycetes>Micrococcales>Microbacteriaceae | GCA_007859675.1 | Humibacter_ginsenosidimutans | FPZ11_07810 | 1665101..1668631 | 2599293 | BREX-system-P-loop-protein-BrxC-[Humibacter-ginsenosidimutans] |
| QIM17270.1 | Type1.BREX | [illegible] | 1193 | Bacteria>Terrabacteria_group>Actinomycetota>Actinomycetes>Micrococcales>Microbacteriaceae | GCA_011382965.1 | Leucobacter_insecticola | G7067_02015 | 442468..446049 | 2714934 | BREX-system-P-loop-protein-BrxC-[Leucobacter-insecticola] |
| NYD70997.1 | Type1.BREX | [illegible] | 1151 | Bacteria>Terrabacteria_group>Actinomycetota>Actinomycetes>Micrococcales>Microbacteriaceae | GCA_013409865.1 | Herbiconiux_flava | BJ984_002155 | 2240959..2244414 | 881268 | hypothetical-protein-[Herbiconiux-flava] |
| MBM7754090.1 | Type1.BREX | [illegible] | 1152 | Bacteria>Terrabacteria_group>Actinomycetota>Actinomycetes>Micrococcales>Microbacteriaceae | GCA_016907555.1 | Microbacterium_laevaniformans | JOE53_002810 | 2910166..2913624 | 36807 | hypothetical-protein-[Microbacterium-laevaniformans] |
| MBO1903164.1 | Type1.BREX | [illegible] | 1152 | Bacteria>Terrabacteria_group>Actinomycetota>Actinomycetes>Micrococcales>Microbacteriaceae | GCA_017565745.1 | Leucobacter_weissii | J4H92_14565 | 1515..4973 | 1983706 | BREX-system-P-loop-protein-BrxC-[Leucobacter-weissii] |
| MBX3194872.1 | Type1.BREX | [illegible] | 1181 | Bacteria>Terrabacteria_group>Actinomycetota>Actinomycetes>Micrococcales>Microbacteriaceae | GCA_019638805.1 | Microbacteriaceae_bacterium | KF727_07200 | 15940..19485 | 1873462 | BREX-system-P-loop-protein-BrxC-[Microbacteriaceae-bacterium] |
| MBX3116430.1 | Type1.BREX | [illegible] | 1182 | Bacteria>Terrabacteria_group>Actinomycetota>Actinomycetes>Micrococcales>Microbacteriaceae | GCA_019638875.1 | Cryobacterium_sp. | KF808_03680 | 24849..28397 | 1926290 | BREX-system-P-loop-protein-BrxC-[Cryobacterium-sp.] |
| MBX3086848.1 | Type1.BREX | [illegible] | 1160 | Bacteria>Terrabacteria_group>Actinomycetota>Actinomycetes>Micrococcales>Microbacteriaceae | GCA_019638955.1 | Cryobacterium_sp. | KF742_00030 | 12107..15589 | 1926290 | BREX-system-P-loop-protein-BrxC-[Cryobacterium-sp.] |
| MBY6061973.1 | Type1.BREX | [illegible] | 1152 | Bacteria>Terrabacteria_group>Actinomycetota>Actinomycetes>Micrococcales>Microbacteriaceae | GCA_019801485.1 | Microbacterium_esteraromaticum | KUV70_11015 | 269249..272707 | 57043 | BREX-system-P-loop-protein-BrxC-[Microbacterium-esteraromaticum] |
| WP_029153784.1 | Type1.BREX | [illegible] | 1160 | Bacteria>Terrabacteria_group>Actinomycetota>Actinomycetes>Micrococcales>Microbacteriaceae | GCF_000422745.1 | Microbacterium_gubbeenense | G510_RS0111780 | 9571..13053 | 159896 | BREX-system-P-loop-protein-BrxC-[Microbacterium-gubbeenense] |
| WP_028245009.1 | Type1.BREX | [illegible] | 1170 | Bacteria>Terrabacteria_group>Actinomycetota>Actinomycetes>Micrococcales>Microbacteriaceae | GCF_000423505.1 | Pseudoclavibacter_soli | G557_RS0103810 | 190122..193634 | 452623 | BREX-system-P-loop-protein-BrxC-[Pseudoclavibacter-soli] |
| WP_053098414.1 | Type1.BREX | [illegible] | 1186 | Bacteria>Terrabacteria_group>Actinomycetota>Actinomycetes>Micrococcales>Microbacteriaceae | GCF_001266755.1 | Microbacterium_sp._CGR1 | AKG07_RS13635 | 2816924..2820484 | 1696072 | BREX-system-P-loop-protein-BrxC-[Microbacterium-sp.-CGR1] |
| WP_053352376.1 | Type1.BREX | [illegible] | 1182 | Bacteria>Terrabacteria_group>Actinomycetota>Actinomycetes>Micrococcales>Microbacteriaceae | GCF_001273845.1 | Leucobacter_musarum | CH42_RS06350 | 60565..64113 | 1930747 | BREX-system-P-loop-protein-BrxC-[Leucobacter-musarum] |
| WP_085605911.1 | Type1.BREX | [illegible] | 1160 | Bacteria>Terrabacteria_group>Actinomycetota>Actinomycetes>Micrococcales>Microbacteriaceae | GCF_002114135.1 | Microbacterium_sp._LEMMJ01 | B7W94_RS08480 | 394..3876 | 1978350 | BREX-system-P-loop-protein-BrxC-[Microbacterium-sp.-LEMMJ01] |
| WP_104291918.1 | Type1.BREX | [illegible] | 1170 | Bacteria>Terrabacteria_group>Actinomycetota>Actinomycetes>Micrococcales>Microbacteriaceae | GCF_002930935.1 | Rathayibacter_sp._AY1A2 | C5B93_RS04570 | 113948..117460 | 2080520 | BREX-system-P-loop-protein-BrxC-[Rathayibacter-sp.-AY1A2] |
| WP_104323594.1 | Type1.BREX | [illegible] | 1160 | Bacteria>Terrabacteria_group>Actinomycetota>Actinomycetes>Micrococcales>Microbacteriaceae | GCF_002931415.1 | unclassified_Rathayibacter | C5C74_RS01635 | 26455..29937 | 2609250 | MULTISPECIES:-BREX-system-P-loop-protein-BrxC-[unclassified-Rathayibacter] |
| WP_181026963.1 | Type1.BREX | [illegible] | 1151 | Bacteria>Terrabacteria_group>Actinomycetota>Actinomycetes>Micrococcales>Microbacteriaceae | GCF_002931555.1 | Pseudoclavibacter_sp._RFBB5 | C5B97_RS09025 | 222537..225992 | 2080574 | BREX-system-P-loop-protein-BrxC-[Pseudoclavibacter-sp.-RFBB5] |
| WP_104309935.1 | Type1.BREX | [illegible] | 1162 | Bacteria>Terrabacteria_group>Actinomycetota>Actinomycetes>Micrococcales>Microbacteriaceae | GCF_002932095.1 | Rathayibacter_sp._AY1G1 | C5C36_RS14015 | 21701..25189 | 2080564 | BREX-system-P-loop-protein-BrxC-[Rathayibacter-sp.-AY1G1] |
| WP_181134950.1 | Type1.BREX | [illegible] | 1151 | Bacteria>Terrabacteria_group>Actinomycetota>Actinomycetes>Micrococcales>Microbacteriaceae | GCF_002932165.1 | Rathayibacter_sp._AY1F8 | C5C35_RS14415 | 14843..18298 | 2080562 | BREX-system-P-loop-protein-BrxC-[Rathayibacter-sp.-AY1F8] |
| WP_104347518.1 | Type1.BREX | [illegible] | 1182 | Bacteria>Terrabacteria_group>Actinomycetota>Actinomycetes>Micrococcales>Microbacteriaceae | GCF_002932695.1 | Rathayibacter_sp._AY1D2 | C5D04_RS07575 | 19648..23196 | 2080543 | BREX-system-P-loop-protein-BrxC-[Rathayibacter-sp.-AY1D2] |
| WP_104281612.1 | Type1.BREX | [illegible] | 1151 | Bacteria>Terrabacteria_group>Actinomycetota>Actinomycetes>Micrococcales>Microbacteriaceae | GCF_002932795.1 | Rathayibacter_sp._AY1B5 | C5D44_RS10395 | 68293..71748 | 2080530 | BREX-system-P-loop-protein-BrxC-[Rathayibacter-sp.-AY1B5] |
| WP_104354073.1 | Type1.BREX | [illegible] | 876 | Bacteria>Terrabacteria_group>Actinomycetota>Actinomycetes>Micrococcales>Microbacteriaceae | GCF_002933035.1 | Rathayibacter_iranicus | C5E08_RS01490 | 139521..142147 | 59737 | BREX-system-P-loop-protein-BrxC,-partial-[Rathayibacter-iranicus] |
| WP_108963418.1 | Type1.BREX | [illegible] | 1151 | Bacteria>Terrabacteria_group>Actinomycetota>Actinomycetes>Micrococcales>Microbacteriaceae | GCF_003113395.1 | Mycetocola_zhujimingii | DF223_RS12635 | 389879..393334 | 2079792 | BREX-system-P-loop-protein-BrxC-[Mycetocola-zhujimingii] |
| WP_124973569.1 | Type1.BREX | [illegible] | 1170 | Bacteria>Terrabacteria_group>Actinomycetota>Actinomycetes>Micrococcales>Microbacteriaceae | GCF_003865375.1 | Gulosibacter_macacae | EG850_RS11335 | 21958..25470 | 2488791 | BREX-system-P-loop-protein-BrxC-[Gulosibacter-macacae] |
| WP_237399358.1 | Type1.BREX | [illegible] | 878 | Bacteria>Terrabacteria_group>Actinomycetota>Actinomycetes>Micrococcales>Microbacteriaceae | GCF_004011115.1 | Rathayibacter_iranicus | C7V51_RS01465 | 309166..311802 | 59737 | BREX-system-P-loop-protein-BrxC-[Rathayibacter-iranicus] |
| WP_129231959.1 | Type1.BREX | [illegible] | 1145 | Bacteria>Terrabacteria_group>Actinomycetota>Actinomycetes>Micrococcales>Microbacteriaceae | GCF_004134865.1 | Agromyces_fucosus | ESP57_RS13570 | 442497..445934 | 41985 | BREX-system-P-loop-protein-BrxC-[Agromyces-fucosus] |
| WP_129387833.1 | Type1.BREX | [illegible] | 1162 | Bacteria>Terrabacteria_group>Actinomycetota>Actinomycetes>Micrococcales>Microbacteriaceae | GCF_004135285.1 | Microbacterium_protaetiae | ET475_07150 | 1526355..1529843 | 2509458 | BREX-system-P-loop-protein-BrxC-[Microbacterium-protaetiae] |
| WP_165310971.1 | Type1.BREX | [illegible] | 1152 | Bacteria>Terrabacteria_group>Actinomycetota>Actinomycetes>Micrococcales>Microbacteriaceae | GCF_004135285.1 | Microbacterium_protaetiae | ET475_RS16440 | 3529806..3533264 | 2509458 | BREX-system-P-loop-protein-BrxC-[Microbacterium-protaetiae] |
| WP_130108838.1 | Type1.BREX | [illegible] | 1160 | Bacteria>Terrabacteria_group>Actinomycetota>Actinomycetes>Micrococcales>Microbacteriaceae | GCF_004208635.1 | Leucobacter_triazinivorans | EVS81_RS01585 | 346722..350204 | 1784719 | BREX-system-P-loop-protein-BrxC-[Leucobacter-triazinivorans] |
| WP_130279999.1 | Type1.BREX | [illegible] | 364 | Bacteria>Terrabacteria_group>Actinomycetota>Actinomycetes>Micrococcales>Microbacteriaceae | GCF_004216855.1 | Microcella_alkaliphila | EV140_RS00455 | 61..1155 | 279828 | BREX-system-P-loop-protein-BrxC-[Microcella-alkaliphila] |
| WP_166789459.1 | Type1.BREX | [illegible] | 1151 | Bacteria>Terrabacteria_group>Actinomycetota>Actinomycetes>Micrococcales>Microbacteriaceae | GCF_004402355.1 | Cryobacterium_sp._TMS1-20-1 | E3045_RS10185 | 26834..30289 | 1259223 | BREX-system-P-loop-protein-BrxC-[Cryobacterium-sp.-TMS1-20-1] |
| WP_134415956.1 | Type1.BREX | [illegible] | 1170 | Bacteria>Terrabacteria_group>Actinomycetota>Actinomycetes>Micrococcales>Microbacteriaceae | GCF_004402465.1 | Cryobacterium_sp._TMT2-14 | E3028_RS10860 | 27948..31460 | 1259245 | BREX-system-P-loop-protein-BrxC-[Cryobacterium-sp.-TMT2-14] |
| WP_241980122.1 | Type1.BREX | [illegible] | 1172 | Bacteria>Terrabacteria_group>Actinomycetota>Actinomycetes>Micrococcales>Microbacteriaceae | GCF_004402535.1 | Cryobacterium_sp._HLT2-23 | E3006_RS11880 | 40394..43912 | 1259145 | BREX-system-P-loop-protein-BrxC-[Cryobacterium-glaciale] |
| WP_134434730.1 | Type1.BREX | [illegible] | 1164 | Bacteria>Terrabacteria_group>Actinomycetota>Actinomycetes>Micrococcales>Microbacteriaceae | GCF_004403545.1 | Cryobacterium_sp._TmT2-59 | E3T24_RS13895 | 41136..44630 | 1259264 | BREX-system-P-loop-protein-BrxC-[Cryobacterium-sp.-TmT2-59] |
| WP_168203764.1 | Type1.BREX | [illegible] | 1152 | Bacteria>Terrabacteria_group>Actinomycetota>Actinomycetes>Micrococcales>Microbacteriaceae | GCF_007859675.1 | Humibacter_ginsenosidimutans | FPZ11_RS07810 | 1665101..1668559 | 2599293 | BREX-system-P-loop-protein-BrxC-[Humibacter-ginsenosidimutans] |
| WP_146322974.1 | Type1.BREX | [illegible] | 1162 | Bacteria>Terrabacteria_group>Actinomycetota>Actinomycetes>Micrococcales>Microbacteriaceae | GCF_007859675.1 | Humibacter_ginsenosidimutans | FPZ11_RS17385 | 3786036..3789524 | 2599293 | BREX-system-P-loop-protein-BrxC-[Humibacter-ginsenosidimutans] |
| WP_200838962.1 | Type1.BREX | [illegible] | 1182 | Bacteria>Terrabacteria_group>Actinomycetota>Actinomycetes>Micrococcales>Microbacteriaceae | GCF_008271365.1 | Microbacterium | FY549_RS13635 | 2819312..2822860 | 33882 | MULTISPECIES:-BREX-system-P-loop-protein-BrxC-[Microbacterium] |
| WP_198165662.1 | Type1.BREX | [illegible] | 1152 | Bacteria>Terrabacteria_group>Actinomycetota>Actinomycetes>Micrococcales>Microbacteriaceae | GCF_009749485.1 | Agromyces_laixinhei | GQQ88_RS02955 | 115313..118771 | 2585717 | BREX-system-P-loop-protein-BrxC-[Agromyces-laixinhei] |
| WP_163473715.1 | Type1.BREX | [illegible] | 1162 | Bacteria>Terrabacteria_group>Actinomycetota>Actinomycetes>Micrococcales>Microbacteriaceae | GCF_010686725.1 | Galbitalea_soli | HNR17_RS04680 | 978511..981999 | 1268042 | BREX-system-P-loop-protein-BrxC-[Galbitalea-soli] |
| WP_244301194.1 | Type1.BREX | [illegible] | 1194 | Bacteria>Terrabacteria_group>Actinomycetota>Actinomycetes>Micrococcales>Microbacteriaceae | GCF_011382965.1 | Leucobacter_insecticola | G7067_RS02010 | 442468..446052 | 2714934 | BREX-system-P-loop-protein-BrxC-[Leucobacter-insecticola] |
| WP_166866044.1 | Type1.BREX | [illegible] | 1160 | Bacteria>Terrabacteria_group>Actinomycetota>Actinomycetes>Micrococcales>Microbacteriaceae | GCF_011751865.2 | Salinibacterium_sp._ZJ70 | HCR12_RS10200 | 2144642..2148124 | 2708084 | BREX-system-P-loop-protein-BrxC-[Salinibacterium-sp.-ZJ70] |
| WP_169077984.1 | Type1.BREX | [illegible] | 1162 | Bacteria>Terrabacteria_group>Actinomycetota>Actinomycetes>Micrococcales>Microbacteriaceae | GCF_012845865.1 | Yonghaparkia | HFV16_RS05520 | 206976..210464 | 355929 | MULTISPECIES:-BREX-system-P-loop-protein-BrxC-[Yonghaparkia] |
| WP_121673638.1 | Type1.BREX | [illegible] | 1161 | Bacteria>Terrabacteria_group>Actinomycetota>Actinomycetes>Micrococcales>Microbacteriaceae | GCF_014651755.1 | Mycetocola_manganoxydans | IE057_RS13450 | 51714..55199 | 699879 | BREX-system-P-loop-protein-BrxC-[Mycetocola-manganoxydans] |
| WP_197132648.1 | Type1.BREX | [illegible] | 1171 | Bacteria>Terrabacteria_group>Actinomycetota>Actinomycetes>Micrococcales>Microbacteriaceae | GCF_015864675.1 | Salinibacterium_sp._SWN1162 | I6E58_RS09540 | 2021923..2025438 | 2792053 | BREX-system-P-loop-protein-BrxC-[Salinibacterium-sp.-SWN1162] |
| WP_206395457.1 | Type1.BREX | [illegible] | 1158 | Bacteria>Terrabacteria_group>Actinomycetota>Actinomycetes>Micrococcales>Microbacteriaceae | GCF_017167395.1 | Pseudoclavibacter_alba | JRG19_RS09065 | 120280..123756 | 272241 | BREX-system-P-loop-protein-BrxC-[Pseudoclavibacter-alba] |
| WP_207484738.1 | Type1.BREX | [illegible] | 1176 | Bacteria>Terrabacteria_group>Actinomycetota>Actinomycetes>Micrococcales>Microbacteriaceae | GCF_017355365.1 | Microbacterium_sp._SD291 | IPV10_RS13675 | 26562..30092 | 2782007 | BREX-system-P-loop-protein-BrxC-[Microbacterium-sp.-SD291] |
| WP_261393916.1 | Type1.BREX | [illegible] | 1146 | Bacteria>Terrabacteria_group>Actinomycetota>Actinomycetes>Micrococcales>Microbacteriaceae | GCF_019801485.1 | Microbacterium_esteraromaticum | KUV70_RS11015 | 269267..272707 | 57043 | BREX-system-P-loop-protein-BrxC-[Microbacterium-esteraromaticum] |
| WP_240240880.1 | Type1.BREX | [illegible] | 1178 | Bacteria>Terrabacteria_group>Actinomycetota>Actinomycetes>Micrococcales>Microbacteriaceae | GCF_022436485.1 | Agrococcus_sp._ARC_14 | MKD51_RS07980 | 1621402..1624938 | 2919927 | BREX-system-P-loop-protein-BrxC-[Agrococcus-sp.-ARC_14] |
| WP_242086581.1 | Type1.BREX | [illegible] | 1160 | Bacteria>Terrabacteria_group>Actinomycetota>Actinomycetes>Micrococcales>Microbacteriaceae | GCF_022662255.1 | Microbacterium_lacticum | L8F16_RS04085 | 1822..5304 | 33885 | BREX-system-P-loop-protein-BrxC-[Microbacterium-lacticum] |
| WP_248598773.1 | Type1.BREX | [illegible] | 1144 | Bacteria>Terrabacteria_group>Actinomycetota>Actinomycetes>Micrococcales>Microbacteriaceae | GCF_023217995.1 | Agromyces_sp._C10 | M1L56_RS07685 | 214029..217463 | 2935077 | BREX-system-P-loop-protein-BrxC-[Agromyces-sp.-C10] |
| WP_156999272.1 | Type1.BREX | [illegible] | 1173 | Bacteria>Terrabacteria_group>Actinomycetota>Actinomycetes>Micrococcales>Microbacteriaceae | GCF_024172065.1 | Agromyces_terreus | BJ978_RS07845 | 1788329..1791850 | 424795 | BREX-system-P-loop-protein-BrxC-[Agromyces-terreus] |
| WP_257523879.1 | Type1.BREX | [illegible] | 1160 | Bacteria>Terrabacteria_group>Actinomycetota>Actinomycetes>Micrococcales>Microbacteriaceae | GCF_024623505.1 | Microbacterium_sp._zg.B48 | NQ152_RS10060 | 193646..197128 | 2969408 | BREX-system-P-loop-protein-BrxC-[Microbacterium-sp.-zg.B48] |
| WP_259980691.1 | Type1.BREX | [illegible] | 1162 | Bacteria>Terrabacteria_group>Actinomycetota>Actinomycetes>Micrococcales>Microbacteriaceae | GCF_025151145.1 | Microbacterium_paraoxydans | M3D59_RS02760 | 52380..55868 | 199592 | BREX-system-P-loop-protein-BrxC-[Microbacterium-paraoxydans] |
| WP_261607502.1 | Type1.BREX | [illegible] | 1186 | Bacteria>Terrabacteria_group>Actinomycetota>Actinomycetes>Micrococcales>Microbacteriaceae | GCF_025398275.1 | Microbacterium_sp._PMIC_1C1B | N4R40_RS11380 | 57181..60741 | 2978350 | BREX-system-P-loop-protein-BrxC-[Microbacterium-sp.-PMIC_1C1B] |
| WP_204948079.1 | Type1.BREX | [illegible] | 1152 | Bacteria>Terrabacteria_group>Actinomycetota>Actinomycetes>Micrococcales>Microbacteriaceae | GCF_027921785.1 | Microbacterium_laevaniformans | PKE70_RS13835 | 19020..22478 | 36807 | BREX-system-P-loop-protein-BrxC-[Microbacterium-laevaniformans] |
| WP_179548032.1 | Type1.BREX | [illegible] | 1151 | Bacteria>Terrabacteria_group>Actinomycetota>Actinomycetes>Micrococcales>Microbacteriaceae | GCF_027922025.1 | Herbiconiux_flava | PKE81_RS17675 | 1519620..1523075 | 881268 | BREX-system-P-loop-protein-BrxC-[Herbiconiux-flava] |
| WP_263798144.1 | Type1.BREX | [illegible] | 1186 | Bacteria>Terrabacteria_group>Actinomycetota>Actinomycetes>Micrococcales>Microbacteriaceae | GCF_027943945.1 | Microbacterium_terricola | OAU46_RS04855 | 990438..993998 | 344163 | BREX-system-P-loop-protein-BrxC-[Microbacterium-terricola] |
| WP_275092901.1 | Type1.BREX | [illegible] | 1152 | Bacteria>Terrabacteria_group>Actinomycetota>Actinomycetes>Micrococcales>Microbacteriaceae | GCF_029027945.1 | Microbacterium_liquefaciens | PWF71_RS12370 | 2612718..2616176 | 33918 | BREX-system-P-loop-protein-BrxC-[Microbacterium-liquefaciens] |
| WP_257508277.1 | Type1.BREX | [illegible] | 1186 | Bacteria>Terrabacteria_group>Actinomycetota>Actinomycetes>Micrococcales>Microbacteriaceae | GCF_030246865.1 | Microbacterium_sp._zg.B96 | NQ154_RS06000 | 561655..565215 | 2969409 | BREX-system-P-loop-protein-BrxC-[Microbacterium-sp.-zg.B96] |
| WP_257505179.1 | Type1.BREX | [illegible] | 1162 | Bacteria>Terrabacteria_group>Actinomycetota>Actinomycetes>Micrococcales>Microbacteriaceae | GCF_030246925.1 | Microbacterium_sp._zg.Y625 | NQ156_RS01310 | 276922..280410 | 2969411 | BREX-system-P-loop-protein-BrxC-[Microbacterium-sp.-zg.Y625] |
| WP_028495577.1 | Type1.BREX | [illegible] | 1160 | Bacteria>Terrabacteria_group>Actinomycetota>Actinomycetes>Micrococcales>Microbacteriaceae | GCF_900113885.1 | Microbacterium | BM226_RS05185 | 1039412..1042894 | 33882 | MULTISPECIES:-BREX-system-P-loop-protein-BrxC-[Microbacterium] |
| WP_201614919.1 | Type1.BREX | [illegible] | 1170 | Bacteria>Terrabacteria_group>Actinomycetota>Actinomycetes>Micrococcales>Microbacteriaceae | GCF_904848685.1 | Gulosibacter_hominis | JMX86_RS10045 | 19055..22567 | 2770504 | BREX-system-P-loop-protein-BrxC-[Gulosibacter-hominis] |
| WP_230672563.1 | Type1.BREX | [illegible] | 1176 | Bacteria>Terrabacteria_group>Actinomycetota>Actinomycetes>Micrococcales>Microbacteriaceae | GCF_920939495.1 | Rathayibacter_sp._Leaf248 | LRG51_RS10050 | 2075326..2078856 | 2876555 | BREX-system-P-loop-protein-BrxC-[Rathayibacter-sp.-Leaf248] |
| WP_222446684.1 | Type1.BREX | [illegible] | 1152 | Bacteria>Terrabacteria_group>Actinomycetota>Actinomycetes>Micrococcales>Microbacteriaceae | - | Microbacterium_esteraromaticum | KUV70_11015 | 269249..272707 | 57043 | BREX-system-P-loop-protein-BrxC-[Microbacterium-esteraromaticum] |
| WP_219885258.1 | Type1.BREX | [illegible] | 1210 | Bacteria>Terrabacteria_group>Actinomycetota>Actinomycetes>Micrococcales>Micrococcaceae | - | Arthrobacter_sp._PAMC25284 | KY499_09320 | 2016646..2020278 | 2861279 | BREX-system-P-loop-protein-BrxC-[Arthrobacter-sp.-PAMC25284] |
| AFR34584.1 | Type1.BREX | [illegible] | 1166 | Bacteria>Terrabacteria_group>Actinomycetota>Actinomycetes>Micrococcales>Micrococcaceae | GCA_000294695.2 | Arthrobacter_sp._Rue61a | ARUE_113p00760 | 81229..84729 | 1118963 | putative-ATP-binding-protein-[Arthrobacter-sp.-Rue61a] |
| ERI35655.1 | Type1.BREX | [illegible] | 1161 | Bacteria>Terrabacteria_group>Actinomycetota>Actinomycetes>Micrococcales>Micrococcaceae | GCA_000465895.2 | Arthrobacter_sp._AK-YN10 | M707_20590 | 5564..9049 | 1349820 | ATP-binding-protein-[Arthrobacter-sp.-AK-YN10] |
| ALV40633.1 | Type1.BREX | [illegible] | 1172 | Bacteria>Terrabacteria_group>Actinomycetota>Actinomycetes>Micrococcales>Micrococcaceae | GCA_001484605.1 | Pseudarthrobacter_sulfonivorans | AU252_05160 | 1179330..1182848 | 121292 | hypothetical-protein-AU252_05160-[Pseudarthrobacter-sulfonivorans] |
| AOY70771.1 | Type1.BREX | [illegible] | 1161 | Bacteria>Terrabacteria_group>Actinomycetota>Actinomycetes>Micrococcales>Micrococcaceae | GCA_001854365.1 | Arthrobacter_sp._ZXY-2 | ARZXY2_1218 | 1271114..1274599 | 1806905 | ATP-binding-protein-[Arthrobacter-sp.-ZXY-2] |
| ASN39993.1 | Type1.BREX | [illegible] | 1170 | Bacteria>Terrabacteria_group>Actinomycetota>Actinomycetes>Micrococcales>Micrococcaceae | GCA_002224445.1 | Arthrobacter_sp._7749 | CGQ24_13870 | 3050415..3053927 | 2020130 | ATP-binding-protein-[Arthrobacter-sp.-7749] |
| BBE21856.1 | Type1.BREX | [illegible] | 1144 | Bacteria>Terrabacteria_group>Actinomycetota>Actinomycetes>Micrococcales>Micrococcaceae | GCA_004001285.1 | Arthrobacter_sp._MN05-02 | MN0502_07390 | 781393..784827 | 1571833 | hypothetical-protein-MN0502_07390-[Arthrobacter-sp.-MN05-02] |
| THE19344.1 | Type1.BREX | [illegible] | 1166 | Bacteria>Terrabacteria_group>Actinomycetota>Actinomycetes>Micrococcales>Micrococcaceae | GCA_004799725.1 | Kocuria_rosea | E1J17_01420 | 4081..7581 | 1275 | BREX-system-P-loop-protein-BrxC-[Kocuria-rosea] |
| KAA9394087.1 | Type1.BREX | [illegible] | 1176 | Bacteria>Terrabacteria_group>Actinomycetota>Actinomycetes>Micrococcales>Micrococcaceae | GCA_008831305.1 | Kocuria_coralli | FCK90_09275 | 131322..134852 | 1461025 | BREX-system-P-loop-protein-BrxC-[Kocuria-coralli] |

| Protein-ID | System | Gene-Neighbourhoods | Length | Taxonomy | Assembly-ID | Strain | Locus-Tag | Genomic-coordinates | TaxID | Fasta-defline |
|---|---|---|---|---|---|---|---|---|---|---|
| NHQ58332.1 | Type1.BREX | ICLR-HTH-><-?<-?\|\|?-><-?\|\|?->BrxA-DNA-binding->BrxB-BrxF-iSTAND->DUF6079*->N6-METHYLASE->PglZ->SIGMA+OB-Fold+AAA-ATPase+HKD-DNase-BrxL-><-?\|\|?-><-RADICAL-SAM<-?<-ACET<-XIS-HTH | 1174 | Bacteria>Terrabacteria_group>Actinomycetota>Actinomycetes>Micrococcales>Micrococcaceae | GCA_011602345.1 | Micrococcus_luteus | G9B47_07855 | 17274..20798 | 1270 | BREX-system-P-loop-protein-BrxC-[Micrococcus-luteus] |
| MBB4882886.1 | Type1.BREX | 35exo->tRNA-><-Recombinase+ZnR\|\|ABhydrolase->XIS-HTH->BrxA-DNA-binding->BrxB-BrxF-iSTAND->DUF6079*->N6-METHYLASE->HTH+REase->PglZ->SIGMA+OB-Fold+AAA-ATPase+HKD-DNase-BrxL->SIR2->HIN-HTH->HTH_Tnp_1+HTH_21+rve+rve_3-> | 1173 | Bacteria>Terrabacteria_group>Actinomycetota>Actinomycetes>Micrococcales>Micrococcaceae | GCA_014204815.1 | Micrococcus_flavus | BJ976_001237 | 1342727..1346248 | 384602 | hypothetical-protein-[Micrococcus-flavus] |
| QOD04369.1 | Type1.BREX | NUDIX-><-URI-DNase<-MACRODOMAIN+SIR2<-REase<-?\|\|HKD-DNase+SF1-SF2-Helicase+HTH+HTH+SRA-><-ParB+HNH\|\|BrxA-DNA-binding->BrxB-BrxF-iSTAND->DUF6079*->N6-METHYLASE->N6-METHYLASE->PglZ->SIGMA+OB-Fold+AAA-ATPase+Lon-Protease-BrxL->SF1-SF2-Helicase->Sigma-HTH->Sigma-HTH->rve->HTH+rve-> | 1145 | Bacteria>Terrabacteria_group>Actinomycetota>Actinomycetes>Micrococcales>Micrococcaceae | GCA_014764445.1 | Pseudarthrobacter_sp._BIM_B-2242 | IDT60_04725 | 994235..997672 | 2772401 | BREX-system-P-loop-protein-BrxC-[Pseudarthrobacter-sp.-BIM-B-2242] |
| MBM7847604.1 | Type1.BREX | rve_3->?->?->?->BrxA-DNA-binding->BrxB-BrxF-iSTAND->DUF6079*->N6-METHYLASE->PglZ->SIGMA+OB-Fold+AAA-ATPase+Lon-Protease-BrxL-><-URI-DNase+RAMA | 1149 | Bacteria>Terrabacteria_group>Actinomycetota>Actinomycetes>Micrococcales>Micrococcaceae | GCA_016907875.1 | Arthrobacter_roseus | JOE65_000832 | 769391..772840 | 136274 | phage-tail-protein-X-[Arthrobacter-roseus] |
| MBR9643072.1 | Type1.BREX | <-DNA-glycosylase<-URI-DNase<-MACRODOMAIN+SIR2<-REase\|\|HKD-DNase+SF1-SF2-Helicase+HTH+HTH+SRA-><-SIGMA+OB-Fold+AAA-ATPase+HKD-DNase-BrxL<-PglZ<-N6-METHYLASE<-DUF6079*<-BrxB-BrxF-iSTAND<-BrxA-DNA-binding\|\|McrB-AAA->McrC-NTD+REase->ABhydrolase+ABhydrolase-> | 1173 | Bacteria>Terrabacteria_group>Actinomycetota>Actinomycetes>Micrococcales>Micrococcaceae | GCA_018215265.1 | Paenarthrobacter_sp._GOM3 | IRJ34_14045 | 199455..202976 | 2782567 | BREX-system-P-loop-protein-BrxC-[Paenarthrobacter-sp.-GOM3] |
| QYF88488.1 | Type1.BREX | HIN-HTH-><-SIGMA+OB-Fold+AAA-ATPase+HKD-DNase-BrxL<-PglZ<-N6-METHYLASE<-DUF6079*<-BrxB-BrxF-iSTAND<-XIS-HTH<-?<-?\|\|XIS-HTH->?->HNH-> | 1210 | Bacteria>Terrabacteria_group>Actinomycetota>Actinomycetes>Micrococcales>Micrococcaceae | GCA_019443425.1 | Arthrobacter_sp._PAMC25284 | KY499_09320 | 2016646..2020278 | 2861279 | BREX-system-P-loop-protein-BrxC-[Arthrobacter-sp.-PAMC25284] |
| MCO4261480.1 | Type1.BREX | <-SbcC<-JetB<-JetA<-HEPN<-SF1-SF2-Helicase<-REase<-SIGMA+OB-Fold+AAA-ATPase+Lon-Protease-BrxL<-PglZ<-N6-METHYLASE<-DUF6079*<-BrxB-BrxF-iSTAND<-BrxA-DNA-binding<-HKD-DNase+SF1-SF2-Helicase+HTH+HTH+SRA\|\|URI-DNase->MACRODOMAIN+SIR2->URI-DNase-><-?<-NUDIX | 1151 | Bacteria>Terrabacteria_group>Actinomycetota>Actinomycetes>Micrococcales>Micrococcaceae | GCA_023913835.1 | Pseudarthrobacter_sp._MDT3-26 | NG697_00745 | 166559..170014 | 2953651 | BREX-system-P-loop-protein-BrxC-[Pseudarthrobacter-sp.-MDT3-26] |
| WP_019309257.1 | Type1.BREX | <-MPTase<-HTH_3\|\|SMODS->?->?-><-SIGMA+OB-Fold+AAA-ATPase+HKD-DNase-BrxL<-PglZ<-N6-METHYLASE<-DUF6079*<-BrxB-BrxF-iSTAND<-BrxA-DNA-binding<-XIS-HTH<-HTH_3<-?<-?<-?<-?\|\|ABhydrolase-> | 1159 | Bacteria>Terrabacteria_group>Actinomycetota>Actinomycetes>Micrococcales>Micrococcaceae | GCF_000214115.1 | Kocuria_rhizophila | GGK_RS0101750 | 388045..391524 | 72000 | BREX-system-P-loop-protein-BrxC-[Kocuria-rhizophila] |
| WP_005267613.1 | Type1.BREX | <-DUF4011+SF1-SF2-Helicase+REase<-SIGMA+OB-Fold+AAA-ATPase+Lon-Protease-BrxL<-PglZ<-PglZ-p<-N6-METHYLASE<-DUF6079*<-BrxB-BrxF-iSTAND<-NUDIX\|\|?->ARG->?->?->RNase-> | 1151 | Bacteria>Terrabacteria_group>Actinomycetota>Actinomycetes>Micrococcales>Micrococcaceae | GCF_000328305.2 | Arthrobacter_crystallopoietes | D477_RS04285 | 11236..14691 | 37928 | BREX-system-P-loop-protein-BrxC-[Arthrobacter-crystallopoietes] |
| WP_017834907.1 | Type1.BREX | <-SIGMA+OB-Fold+AAA-ATPase+HKD-DNase-BrxL<-PglZ<-N6-METHYLASE<-DUF6079*<-BrxB-BrxF-iSTAND<-BrxA-DNA-binding<-XIS-HTH<-?<-McrC-NTD+REase<-McrB-AAA<-REase<-?\|\|<-Schlafen+HATPase+wHTH | 1190 | Bacteria>Terrabacteria_group>Actinomycetota>Actinomycetes>Micrococcales>Micrococcaceae | GCF_000349605.1 | Kocuria_sp._UCD-OTCP | H488_RS34905 | 37795..41367 | 1292021 | BREX-system-P-loop-protein-BrxC-[Kocuria-sp.-UCD-OTCP] |
| WP_155853802.1 | Type1.BREX | <-Primpol+PriCT1<-ArdCN+MPTase\|\|HTH->MPTase->?->ABhydrolase->XIS-HTH->BrxA-DNA-binding->BrxB-BrxF-iSTAND->DUF6079*->N6-METHYLASE->PglZ->SIGMA+OB-Fold+AAA-ATPase+HKD-DNase-BrxL->?->VirB11+MuB-ATPase-> | 1170 | Bacteria>Terrabacteria_group>Actinomycetota>Actinomycetes>Micrococcales>Micrococcaceae | GCF_000520595.1 | Arthrobacter_sp._H5 | Y708_RS0109785 | 51778..55290 | 1267973 | BREX-system-P-loop-protein-BrxC-[Arthrobacter-sp.-H5] |
| WP_062292549.1 | Type1.BREX | VirB11+MuB-ATPase->?-><-SIGMA+OB-Fold+AAA-ATPase+HKD-DNase-BrxL<-PglZ<-N6-METHYLASE-p<-Phage_int_SAM_3+Phage_integrase\|\|N6-METHYLASE-p-><-N6-METHYLASE<-DUF6079*<-BrxB-BrxF-iSTAND<-BrxA-DNA-binding<-XIS-HTH<-ArdCN+MPTase<-?<-VirD4-TraG+VirD4-FtsK<-nSTAND3 | 1169 | Bacteria>Terrabacteria_group>Actinomycetota>Actinomycetes>Micrococcales>Micrococcaceae | GCF_001445575.1 | Arthrobacter_alpinus | AS189_RS18910 | 4268679..4272188 | 656366 | BREX-system-P-loop-protein-BrxC-[Arthrobacter-alpinus] |
| WP_099093371.1 | Type1.BREX | <-HEPN+SF1-SF2-Helicase+REase<-?\|\|?-><-ParB+HNH<-SIGMA+OB-Fold+AAA-ATPase+HKD-DNase-BrxL<-PglZ<-N6-METHYLASE<-DUF6079*<-BrxB-BrxF-iSTAND<-BrxA-DNA-binding<-XIS-HTH<-HKD-DNase+SF1-SF2-Helicase+HTH+HTH+SRA\|\|?->?->McrB-AAA->McrC-NTD+REase-> | 1173 | Bacteria>Terrabacteria_group>Actinomycetota>Actinomycetes>Micrococcales>Micrococcaceae | GCF_001484605.1 | Pseudarthrobacter_sulfonivorans | AU252_RS05160 | 1179330..1182851 | 121292 | BREX-system-P-loop-protein-BrxC-[Pseudarthrobacter-sulfonivorans] |
| WP_078028155.1 | Type1.BREX | <-EVE+HNH\|\|?->?->CHTH->?->?-><-SIGMA+OB-Fold+AAA-ATPase+HKD-DNase-BrxL<-PglZ<-N6-METHYLASE<-DUF6079*<-BrxB-BrxF-iSTAND<-BrxA-DNA-binding<-XIS-HTH\|\|?-><-?\|\|?-><-Phage_integrase<-XIS-HTH<-AAA-ATPase | 1176 | Bacteria>Terrabacteria_group>Actinomycetota>Actinomycetes>Micrococcales>Micrococcaceae | GCF_002008315.2 | Arthrobacter_sp._KBS0703 | B1A87_RS03145 | 655510..659040 | 1955698 | BREX-system-P-loop-protein-BrxC-[Arthrobacter-sp.-KBS0703] |
| WP_104161527.1 | Type1.BREX | SF1-SF2-Helicase->HKD-DNase+SF1-SF2-Helicase+HTH+HTH+SRA->HNH-><-?<-REase+SF1-SF2-Helicase<-SIGMA+OB-Fold+AAA-ATPase+Lon-Protease-BrxL<-PglZ<-N6-METHYLASE<-DUF6079*<-BrxB-BrxF-iSTAND<-BrxA-DNA-binding-p\|\|REase->?->?-><-?\|\|ParB+HNH-> | 1151 | Bacteria>Terrabacteria_group>Actinomycetota>Actinomycetes>Micrococcales>Micrococcaceae | GCF_002929255.1 | Arthrobacter_sp._ZGTC212 | CXQ89_RS12670 | 85850..89305 | 2058899 | BREX-system-P-loop-protein-BrxC-[Arthrobacter-sp.-ZGTC212] |
| WP_112234372.1 | Type1.BREX | <-CHASE3+HISKIN<-?<-?<-REC+OmpR-HTH<-?<-?\|\|XIS-HTH->BrxA-DNA-binding->BrxB-BrxF-iSTAND->DUF6079*->N6-METHYLASE->PglZ->SIGMA+OB-Fold+AAA-ATPase+Lon-Protease-BrxL-><-HTH_3<-?<-?\|\|MarR-HTH-> | 1152 | Bacteria>Terrabacteria_group>Actinomycetota>Actinomycetes>Micrococcales>Micrococcaceae | GCF_003290245.1 | Kocuria_sp._BT304 | DNL27_RS00440 | 93123..96581 | 1702043 | BREX-system-P-loop-protein-BrxC-[Kocuria-sp.-BT304] |
| WP_242630351.1 | Type1.BREX | ParA-Soj-PloopNTPase->?-><-TIR\|\|XIS-HTH->BrxA-DNA-binding->BrxB-BrxF-iSTAND->DUF6079*->N6-METHYLASE->PglZ->SIGMA+OB-Fold+AAA-ATPase+HKD-DNase-BrxL-><-?\|\|REase->HTH+REase->URI-DNase->DUF6361->HKD-DNase-> | 1169 | Bacteria>Terrabacteria_group>Actinomycetota>Actinomycetes>Micrococcales>Micrococcaceae | GCF_004799725.1 | Kocuria_rosea | E1J17_RS01420 | 4072..7581 | 1275 | BREX-system-P-loop-protein-BrxC-[Kocuria-rosea] |
| WP_142936644.1 | Type1.BREX | <-NERD<-?<-PIN<-?<-NAD_binding_10<-SIGMA+OB-Fold+AAA-ATPase+HKD-DNase-BrxL<-PglZ<-N6-METHYLASE<-DUF6079*<-BrxB-BrxF-iSTAND<-BrxA-DNA-binding<-XIS-HTH\|\|?->XIS-HTH->Phage_int_SAM_3->Phage_integrase-> | 1161 | Bacteria>Terrabacteria_group>Actinomycetota>Actinomycetes>Micrococcales>Micrococcaceae | GCF_006964045.1 | Arthrobacter_sp._TS-15 | EU811_RS00880 | 63009..66494 | 2510797 | BREX-system-P-loop-protein-BrxC-[Arthrobacter-sp.-TS-15] |
| WP_181954261.1 | Type1.BREX | <-Primpol+PriCT1<-ArdCN+MPTase\|\|HTH->MPTase->?->ABhydrolase->XIS-HTH->BrxA-DNA-binding->BrxB-BrxF-iSTAND->DUF6079*->N6-METHYLASE->PglZ->SIGMA+OB-Fold+AAA-ATPase+Lon-Protease-BrxL->PHP+SbcC-><-DNA_METHYLASE\|\|HIN-HTH->HTH_Tnp_1+HTH_21+rve+rve_3->HIN-HTH+PAIREDC-HTH-> | 1152 | Bacteria>Terrabacteria_group>Actinomycetota>Actinomycetes>Micrococcales>Micrococcaceae | GCF_008831305.1 | Kocuria_coralli | FCK90_RS09280 | 131394..134852 | 1461025 | BREX-system-P-loop-protein-BrxC-[Kocuria-coralli] |
| WP_208011566.1 | Type1.BREX | ICLR-HTH-><-?<-?\|\|?-><-?\|\|?->BrxA-DNA-binding->BrxB-BrxF-iSTAND->DUF6079*->N6-METHYLASE->PglZ->SIGMA+OB-Fold+AAA-ATPase+HKD-DNase-BrxL-><-?\|\|?-><-RADICAL-SAM<-?<-ACET<-XIS-HTH | 1172 | Bacteria>Terrabacteria_group>Actinomycetota>Actinomycetes>Micrococcales>Micrococcaceae | GCF_011602345.1 | Micrococcus_luteus | G9B47_RS07850 | 17280..20798 | 1270 | BREX-system-P-loop-protein-BrxC-[Micrococcus-luteus] |
| WP_171585597.1 | Type1.BREX | Peptidase_M20-><-Peptidase_M20\|\|DNA_METHYLASE-><-?<-?<-?\|\|?-><-SIGMA+OB-Fold+AAA-ATPase+HKD-DNase-BrxL<-N6-METHYLASE<-DUF6079*<-BrxB-BrxF-iSTAND<-XIS-HTH<-?\|\|?-><-?<-HNH\|\|Peptidase-> | 1171 | Bacteria>Terrabacteria_group>Actinomycetota>Actinomycetes>Micrococcales>Micrococcaceae | GCF_013116745.1 | Arthrobacter_sp._260 | HMI57_RS08915 | 135623..139138 | 2735314 | BREX-system-P-loop-protein-BrxC-[Arthrobacter-sp.-260] |
| WP_135027971.1 | Type1.BREX | 35exo->tRNA-><-Recombinase+ZnR\|\|ABhydrolase->XIS-HTH->BrxA-DNA-binding->BrxB-BrxF-iSTAND->DUF6079*->N6-METHYLASE->HTH+REase->PglZ->SIGMA+OB-Fold+AAA-ATPase+HKD-DNase-BrxL->SIR2->HTH_Tnp_1+HTH_21+rve+rve_3-><-?\|\|?-><-HAD+HAD | 1170 | Bacteria>Terrabacteria_group>Actinomycetota>Actinomycetes>Micrococcales>Micrococcaceae | GCF_014644875.1 | Micrococcus_flavus | E4A47_RS01340 | 286387..289899 | 384602 | BREX-system-P-loop-protein-BrxC-[Micrococcus-flavus] |
| WP_190607902.1 | Type1.BREX | <-SIGMA-HTH<-ARG<-?\|\|NUDIX->ARPP2->HKD-DNase+SF1-SF2-Helicase+HTH+HTH+SRA->XIS-HTH->BrxA-DNA-binding->BrxB-BrxF-iSTAND->DUF6079*->N6-METHYLASE->PglZ->?->?->?-><-?<-GNTR-HTH | 1171 | Bacteria>Terrabacteria_group>Actinomycetota>Actinomycetes>Micrococcales>Micrococcaceae | GCF_014694315.1 | Arthrobacter_ipis | G9E11_RS17130 | 470424..473939 | 2716202 | BREX-system-P-loop-protein-BrxC-[Arthrobacter-ipis] |
| WP_223883883.1 | Type1.BREX | NUDIX-><-URI-DNase<-MACRODOMAIN+SIR2<-REase<-?\|\|HKD-DNase+SF1-SF2-Helicase+HTH+HTH+SRA-><-ParB+HNH\|\|BrxA-DNA-binding->BrxB-BrxF-iSTAND->DUF6079*->N6-METHYLASE->PglZ->SIGMA+OB-Fold+AAA-ATPase+Lon-Protease-BrxL->SF1-SF2-Helicase->Sigma-HTH->HTH+rve->AAA-ATPase-> | 1144 | Bacteria>Terrabacteria_group>Actinomycetota>Actinomycetes>Micrococcales>Micrococcaceae | GCF_014764445.1 | Pseudarthrobacter_sp._BIM_B-2242 | IDT60_RS04725 | 994238..997672 | 2772401 | BREX-system-P-loop-protein-BrxC-[Pseudarthrobacter-sp.-BIM-B-2242] |
| WP_193058359.1 | Type1.BREX | N6-METHYLASE-p->zf-ISL3+HTH_Tnp_ISL3->N6-METHYLASE-p-><-N6-METHYLASE<-DUF6079*<-BrxB-BrxF-iSTAND<-BrxA-DNA-binding\|\|?->?->?->?-><-?<-?\|\|LuxR-HTH-> | 1160 | Bacteria>Terrabacteria_group>Actinomycetota>Actinomycetes>Micrococcales>Micrococcaceae | GCF_014897395.1 | Glutamicibacter_arilaitensis | INT15_RS12685 | 5692..9174 | 256701 | BREX-system-P-loop-protein-BrxC-[Glutamicibacter-arilaitensis] |
| WP_193340403.1 | Type1.BREX | <-SIGMA+OB-Fold+AAA-ATPase+HKD-DNase-BrxL<-PglZ<-REase<-N6-METHYLASE<-DUF6079*<-BrxB-BrxF-iSTAND<-BrxA-DNA-binding<-XIS-HTH\|\|?-><-EVE+REase | 1153 | Bacteria>Terrabacteria_group>Actinomycetota>Actinomycetes>Micrococcales>Micrococcaceae | GCF_014926485.1 | Pseudarthrobacter_sp._AB1 | DAD99_RS03725 | 12045..15506 | 2138309 | BREX-system-P-loop-protein-BrxC-[Pseudarthrobacter-sp.-AB1] |
| WP_258190683.1 | Type1.BREX | HIN-HTH-><-SIGMA+OB-Fold+AAA-ATPase+HKD-DNase-BrxL<-PglZ<-N6-METHYLASE<-DUF6079*<-BrxA-DNA-binding<-XIS-HTH<-?<-?\|\|?->HNH-> | 1170 | Bacteria>Terrabacteria_group>Actinomycetota>Actinomycetes>Micrococcales>Micrococcaceae | GCF_019443425.1 | Arthrobacter_sp._PAMC25284 | KY499_RS09320 | 2016646..2020158 | 2861279 | BREX-system-P-loop-protein-BrxC-[Arthrobacter-sp.-PAMC25284] |
| WP_142938938.1 | Type1.BREX | ParB+HNH-p->ParB+HNH-><-?<-?<-URI-DNase\|\|?-><-SIGMA+OB-Fold+AAA-ATPase+HKD-DNase-BrxL<-PglZ<-N6-METHYLASE<-DUF6079*<-BrxB-BrxF-iSTAND<-BrxA-DNA-binding<-XIS-HTH\|\|?->Radical-SAM-><-METHYLASE<-Phage_integrase+Phage_integrase\|\|Phage_integrase-><-HTH_3 | 1162 | Bacteria>Terrabacteria_group>Actinomycetota>Actinomycetes>Micrococcales>Micrococcaceae | GCF_019976975.1 | unclassified_Arthrobacter | LDN90_RS14640 | 3248673..3252161 | 235627 | MULTISPECIES:-BREX-system-P-loop-protein-BrxC-[unclassified-Arthrobacter] |
| WP_224048794.1 | Type1.BREX | NUDIX->wHTH(SBD)+HNH-><-?\|\|?-><-SF1-SF2-Helicase\|\|XIS-HTH->BrxA-DNA-binding->BrxB-BrxF-iSTAND->DUF6079*->N6-METHYLASE->PglZ->N6-METHYLASE->SIGMA+OB-Fold+AAA-ATPase+HKD-DNase-BrxL-><-ParB+HNH\|\|HEPN+SF1-SF2-Helicase+REase-> | 1173 | Bacteria>Terrabacteria_group>Actinomycetota>Actinomycetes>Micrococcales>Micrococcaceae | GCF_019977335.1 | Arthrobacter_sp._NicSoilB4 | LDO13_RS04105 | 882114..885635 | 2830997 | BREX-system-P-loop-protein-BrxC-[Arthrobacter-sp.-NicSoilB4] |
| WP_226761965.1 | Type1.BREX | LexA-protease->?->?->?->XIS-HTH->BrxA-DNA-binding->BrxB-BrxF-iSTAND->DUF6079*->N6-METHYLASE->PglZ->SIGMA+OB-Fold+AAA-ATPase+HKD-DNase-BrxL-><-?<-?<-?\|\|HEPN->GDResolvase+HIN-HTH-> | 1161 | Bacteria>Terrabacteria_group>Actinomycetota>Actinomycetes>Micrococcales>Micrococcaceae | GCF_020532825.1 | Arthrobacter_sp._SO3 | BJQ90_RS20165 | 19380..22865 | 1897057 | BREX-system-P-loop-protein-BrxC-[Arthrobacter-sp.-SO3] |
| WP_241470046.1 | Type1.BREX | DNA_glycosylase->DNA_glycosylase->?->?-><-SIGMA+OB-Fold+AAA-ATPase+HKD-DNase-BrxL<-PglZ<-N6-METHYLASE<-DUF6079*<-BrxB-BrxF-iSTAND<-BrxA-DNA-binding\|\|?->HTH_26->?-><-ABC-ATPase+SbcC | 1168 | Bacteria>Terrabacteria_group>Actinomycetota>Actinomycetes>Micrococcales>Micrococcaceae | GCF_022565835.1 | Nesterenkonia_sp._DZ6 | LTH93_RS03075 | 681136..684642 | 2901229 | BREX-system-P-loop-protein-BrxC-[Nesterenkonia-sp.-DZ6] |
| WP_241464163.1 | Type1.BREX | <-REase\|\|DNA-glycosylase->?->DEDD_Tnp_IS110+Transposase_20-><-DEDD_Tnp_IS110+Transposase_20<-SIGMA+OB-Fold+AAA-ATPase+Lon-Protease-BrxL<-PglZ<-N6-METHYLASE<-DUF6079*<-BrxB-BrxF-iSTAND<-BrxA-DNA-binding<-XIS-HTH<-?<-?\|\|?-><-DNA_glycosylase | 1151 | Bacteria>Terrabacteria_group>Actinomycetota>Actinomycetes>Micrococcales>Micrococcaceae | GCF_022565915.1 | Nesterenkonia_sp._LB17 | LTH96_02575 | 576573..580028 | 2901230 | BREX-system-P-loop-protein-BrxC-[Nesterenkonia-sp.-LB17] |
| WP_241707532.1 | Type1.BREX | REase->ChlI+Lon_C+AAA->?->?->?->?->XIS-HTH->BrxA-DNA-binding->BrxB-BrxF-iSTAND->DUF6079*->N6-METHYLASE->PglZ->SIGMA+OB-Fold+AAA-ATPase+HKD-DNase-BrxL-><-?<-DNA-glycosylase\|\|?->NACHT-> | 1171 | Bacteria>Terrabacteria_group>Actinomycetota>Actinomycetes>Micrococcales>Micrococcaceae | GCF_022606295.1 | Arthrobacter_bambusae | KNN17_RS03605 | 221894..225409 | 1338426 | BREX-system-P-loop-protein-BrxC-[Arthrobacter-bambusae] |
| WP_247040359.1 | Type1.BREX | HTH_3-><-Phage_integrase\|\|Phage_integrase+Phage_integrase->METHYLASE-><-Radical-SAM<-?\|\|XIS-HTH->BrxA-DNA-binding->BrxB-BrxF-iSTAND->DUF6079*->N6-METHYLASE->PglZ->SIGMA+OB-Fold+AAA-ATPase+HKD-DNase-BrxL-><-?\|\|URI-DNase->?->?-><-ParB+HNH | 1168 | Bacteria>Terrabacteria_group>Actinomycetota>Actinomycetes>Micrococcales>Micrococcaceae | GCF_023062595.1 | Arthrobacter_rhizosphaerae | MU551_RS01475 | 284158..287664 | 2855490 | BREX-system-P-loop-protein-BrxC-[Arthrobacter-rhizosphaerae] |
| WP_248193800.1 | Type1.BREX | <-SIGMA+OB-Fold+AAA-ATPase+HKD-DNase-BrxL<-PglZ<-?<-N6-METHYLASE<-DUF6079*<-BrxB-BrxF-iSTAND<-BrxA-DNA-binding<-XIS-HTH\|\|?-><-HAD+HAD | 1173 | Bacteria>Terrabacteria_group>Actinomycetota>Actinomycetes>Micrococcales>Micrococcaceae | GCF_023147525.1 | Micrococcus_endophyticus | KZX39_RS06505 | 8911..12432 | 455343 | BREX-system-P-loop-protein-BrxC-[Micrococcus-endophyticus] |
| WP_091699215.1 | Type1.BREX | <-ABhydrolase\|\|?-><-?<-Calcineurin<-SIGMA+OB-Fold+AAA-ATPase+Lon-Protease-BrxL<-PglZ<-N6-METHYLASE<-DUF6079*<-BrxB-BrxF-iSTAND<-BrxA-DNA-binding<-XIS-HTH<-?\|\|?-><-ACET<-?<-ParB+HNH | 1151 | Bacteria>Terrabacteria_group>Actinomycetota>Actinomycetes>Micrococcales>Micrococcaceae | GCF_900116905.1 | Micrococcus_terreus | BM355_RS12655 | 54420..57875 | 574650 | BREX-system-P-loop-protein-BrxC-[Micrococcus-terreus] |
| WP_073116219.1 | Type1.BREX | ICLR-HTH-><-?<-?\|\|?-><-?\|\|?->BrxA-DNA-binding->BrxB-BrxF-iSTAND->DUF6079*->N6-METHYLASE->PglZ->SIGMA+OB-Fold+AAA-ATPase+HKD-DNase-BrxL-><-?\|\|?-><-RADICAL-SAM<-?<-ACET<-XIS-HTH | 1174 | Bacteria>Terrabacteria_group>Actinomycetota>Actinomycetes>Micrococcales>Micrococcaceae | GCF_900142795.1 | Micrococcus | BUB74_RS03185 | 94466..97990 | 1269 | MULTISPECIES:-BREX-system-P-loop-protein-BrxC-[Micrococcus] |
| WP_237187176.1 | Type1.BREX | <-HHH_5+SNF2_N+SF1-SF2-Helicase\|\|SWACOS-><-?<-SIGMA+OB-Fold+AAA-ATPase+HKD-DNase-BrxL<-PglZ<-Schlafen+HATPase+wHTH<-N6-METHYLASE<-DUF6079*<-BrxB-BrxF-iSTAND<-BrxA-DNA-binding\|\|?-><-Aminotran_4<-?<-ABC-ATPase+SbcC+ABC-ATPase+SbcC | 1190 | Bacteria>Terrabacteria_group>Actinomycetota>Actinomycetes>Micrococcales>Micrococcaceae | GCF_928381705.1 | Rothia_nasimurium | L6762_RS10435 | 40591..44163 | 85336 | BREX-system-P-loop-protein-BrxC-[Rothia-nasimurium] |
| WP_211713005.1 | Type1.BREX | <-DNA-glycosylase<-URI-DNase<-MACRODOMAIN+SIR2<-REase\|\|HKD-DNase+SF1-SF2-Helicase+HTH+HTH+SRA-><-SIGMA+OB-Fold+AAA-ATPase+HKD-DNase-BrxL<-PglZ<-N6-METHYLASE<-DUF6079*<-BrxB-BrxF-iSTAND<-BrxA-DNA-binding\|\|McrB-AAA->McrC-NTD+REase->ABhydrolase+ABhydrolase-> | 1173 | Bacteria>Terrabacteria_group>Actinomycetota>Actinomycetes>Micrococcales>Micrococcaceae | - | Paenarthrobacter_sp._GOM3 | IRJ34_14045 | 199455..202976 | 2782567 | BREX-system-P-loop-protein-BrxC-[Paenarthrobacter-sp.-GOM3] |
| WP_252613231.1 | Type1.BREX | <-SbcC<-JetB<-JetA<-HEPN<-SF1-SF2-Helicase<-REase<-SIGMA+OB-Fold+AAA-ATPase+Lon-Protease-BrxL<-PglZ<-N6-METHYLASE<-DUF6079*<-BrxB-BrxF-iSTAND<-BrxA-DNA-binding<-HKD-DNase+SF1-SF2-Helicase+HTH+HTH+SRA\|\|URI-DNase->MACRODOMAIN+SIR2->URI-DNase-><-?<-NUDIX | 1151 | Bacteria>Terrabacteria_group>Actinomycetota>Actinomycetes>Micrococcales>Micrococcaceae | - | Pseudarthrobacter_sp._MDT3-26 | NG697_00745 | 166559..170014 | 2953651 | BREX-system-P-loop-protein-BrxC-[Pseudarthrobacter-sp.-MDT3-26] |
| WP_100350249.1 | Type1.BREX | HEPN+SF1-SF2-Helicase+REase->?-><-Pkinase+CR-Kinase<-SIGMA+OB-Fold+AAA-ATPase+HKD-DNase-BrxL<-PglZ<-N6-METHYLASE-p<-Phage_int_SAM_3+Phage_integrase\|\|N6-METHYLASE-p-><-N6-METHYLASE<-DUF6079*<-BrxB-BrxF-iSTAND<-BrxA-DNA-binding\|\|?->ZNR+RmuC-> | 1179 | Bacteria>Terrabacteria_group>Actinomycetota>Actinomycetes>Micrococcales>Micrococcales_incertae_sedis | GCF_002798015.1 | Luteimicrobium_subarcticum | CLV34_RS10655 | 413171..416710 | 620910 | BREX-system-P-loop-protein-BrxC-[Luteimicrobium-subarcticum] |
| WP_075958588.1 | Type1.BREX | <-MazG<-nSTAND1+HNH<-REase+SF1-SF2-Helicase-p<-REase\|\|XIS-HTH->BrxA-DNA-binding->BrxB-BrxF-iSTAND->DUF6079-p*->DUF6079-p->N6-METHYLASE->SIGMA+OB-Fold+AAA-ATPase+Lon-Protease-BrxL->ParB+HNH-><-REase<-SF1-SF2-Helicase+REase<-REase<-REase<-REase | 295 | Bacteria>Terrabacteria_group>Actinomycetota>Actinomycetes>Micrococcales>Ornithinimicrobiaceae | GCF_001942405.1 | Ornithinimicrobium_sp._CNJ-824 | BJF81_RS12330 | 99001..99888 | 1904966 | hypothetical-protein-[Ornithinimicrobium-sp.-CNJ-824] |
| WP_075958589.1 | Type1.BREX | <-MazG<-nSTAND1+HNH<-REase+SF1-SF2-Helicase-p<-REase\|\|XIS-HTH->BrxA-DNA-binding->BrxB-BrxF-iSTAND->DUF6079-p->DUF6079-p*->N6-METHYLASE->SIGMA+OB-Fold+AAA-ATPase+Lon-Protease-BrxL->ParB+HNH-><-REase<-SF1-SF2-Helicase+REase<-REase<-REase<-REase | 882 | Bacteria>Terrabacteria_group>Actinomycetota>Actinomycetes>Micrococcales>Ornithinimicrobiaceae | GCF_001942405.1 | Ornithinimicrobium_sp._CNJ-824 | BJF81_RS12335 | 99810..102458 | 1904966 | BREX-system-P-loop-protein-BrxC-[Ornithinimicrobium-sp.-CNJ-824] |
| SDF46422.1 | Type1.BREX | ABC-ATPase+SbcC->Cysteine-Desulfurase->?->?->?->BrxA-DNA-binding->BrxB-BrxF-iSTAND->DUF6079*->N6-METHYLASE->PglZ->SIGMA+OB-Fold+AAA-ATPase+HKD-DNase-BrxL->REase->PAIREDC-HTH-> | 1166 | Bacteria>Terrabacteria_group>Actinomycetota>Actinomycetes>Micrococcales>Promicromonosporaceae | GCA_900102445.1 | Cellulosimicrobium_cellulans | SAMN04487781_1443 | 34344..37844 | 1710 | hypothetical-protein-SAMN04487781_1443-[Cellulosimicrobium-cellulans] |
| WP_162290006.1 | Type1.BREX | PIWI-><-HerA/FtsK<-SIR2+SIR2<-SIGMA+OB-Fold+AAA-ATPase+HKD-DNase-BrxL<-PglZ<-N6-METHYLASE<-DUF6079*<-BrxB-BrxF-iSTAND<-BrxA-DNA-binding<-XIS-HTH<-tRNA<-?<-GNTR-HTH\|\|DSBH+AraC-HTH+AraC-HTH-><-HTH_42<-ABhydrolase | 1167 | Bacteria>Terrabacteria_group>Actinomycetota>Actinomycetes>Micrococcales>Promicromonosporaceae | GCF_010287905.1 | Cellulosimicrobium_fucosivorans | GYH36_RS12555 | 12661..16164 | 2850388 | BREX-system-P-loop-protein-BrxC-[Cellulosimicrobium-fucosivorans] |
| WP_253873967.1 | Type1.BREX | <-URI-DNase\|\|?->SIR2->EVE-><-SIGMA+OB-Fold+AAA-ATPase+HKD-DNase-BrxL<-PglZ<-N6-METHYLASE<-DUF6079*<-BrxB-BrxF-iSTAND<-BrxA-DNA-binding\|\|?-><-CHTH\|\|?-><-REase<-REase<-ANK+ANK+ANK<-REase | 1157 | Bacteria>Terrabacteria_group>Actinomycetota>Actinomycetes>Micrococcales>Promicromonosporaceae | GCF_024171995.1 | Promicromonospora_umidemergens | APR04_RS20365 | 11407..14880 | 629679 | BREX-system-P-loop-protein-BrxC-[Promicromonospora-umidemergens] |
| WP_142121348.1 | Type1.BREX | CR-ATPase5+REase-><-REase+SF1-SF2-Helicase-p\|\|BrxA-DNA-binding->BrxB-BrxF-iSTAND->DUF6079*->N6-METHYLASE->?->PglZ->RVT->SIGMA+OB-Fold+AAA-ATPase+HKD-DNase-BrxL-><-?\|\|SF1-SF2-Helicase->wHTH-><-CR-ATPase5+REase | 1170 | Bacteria>Terrabacteria_group>Actinomycetota>Actinomycetes>Micrococcales>Rarobacteraceae | GCF_006716265.1 | Rarobacter_faecitabidus | FB461_RS08150 | 23162..26674 | 13243 | BREX-system-P-loop-protein-BrxC-[Rarobacter-faecitabidus] |
| WP_142113745.1 | Type1.BREX | <-VirB11+MuB-ATPase\|\|DNA_METHYLASE-><-HerA/FtsK<-SIR2<-SIGMA+OB-Fold+AAA-ATPase+HKD-DNase-BrxL<-PglZ<-Schlafen+HATPase+wHTH<-?<-N6-METHYLASE<-DUF6079*<-BrxB-BrxF-iSTAND<-BrxA-DNA-binding<-XIS-HTH<-ABhydrolase<-ABhydrolase<-?<-MPTase<-HTH\|\|ArdCN+MPTase-> | 1173 | Bacteria>Terrabacteria_group>Actinomycetota>Actinomycetes>Micrococcales>Rarobacteraceae | - | Rarobacter_incanus | FB389_RS10090 | 2435944..2439465 | 153494 | BREX-system-P-loop-protein-BrxC-[Rarobacter-incanus] |
| SNU02231.1 | Type1.BREX | <-SbcC<-?<-SIGMA+OB-Fold+AAA-ATPase+HKD-DNase-BrxL<-PglZ<-N6-METHYLASE-p<-Phage_int_SAM_3+Phage_integrase\|\|?->N6-METHYLASE-p-><-N6-METHYLASE<-DUF6079*<-BrxB-BrxF-iSTAND<-BrxA-DNA-binding\|\|SF1-SF2-Helicase+PhospholipaseD-><-?<-?\|\|Peptidase_M23-><-?\|\|TraM-> | 1159 | Bacteria>Terrabacteria_group>Actinomycetota>Actinomycetes>Micrococcales>Ruaniaceae | GCA_900199505.1 | Ruaniaceae_bacterium_KH17 | SAMN06298212_12910 | 14626..18105 | 1945888 | hypothetical-protein-SAMN06298212_12910-[Ruaniaceae-bacterium-KH17] |
| WP_147917199.1 | Type1.BREX | HicA-><-?\|\|?->DNA_glycosylase-><-?<-?<-?\|\|BrxA-DNA-binding->BrxB-BrxF-iSTAND->DUF6079*->N6-METHYLASE->PglZ->SIGMA+OB-Fold+AAA-ATPase+HKD-DNase-BrxL->HTH_3-><-?\|\|?->?->?->ABC-ATPase+SbcC-> | 1152 | Bacteria>Terrabacteria_group>Actinomycetota>Actinomycetes>Micrococcales>Ruaniaceae | GCF_008000995.1 | Ruania_zhangjianzhongii | FU260_RS11560 | 2487086..2490544 | 2603206 | BREX-system-P-loop-protein-BrxC-[Ruania-zhangjianzhongii] |
| WP_163542003.1 | Type1.BREX | <-HIN-HTH\|\|SbcC->?-><-?<-SIGMA+OB-Fold+AAA-ATPase+HKD-DNase-BrxL<-PglZ<-REase<-N6-METHYLASE<-DUF6079*<-BrxB-BrxF-iSTAND<-BrxA-DNA-binding<-XIS-HTH<-?<-?\|\|CN_hydrolase+NAD_synthase->?-><-Pkinase<-RelE-ParE | 1154 | Bacteria>Terrabacteria_group>Actinomycetota>Actinomycetes>Micrococcales>Ruaniaceae | GCF_010994285.1 | Occultella_kanbiaonis | G7B41_RS06055 | 118944..122408 | 2675754 | BREX-system-P-loop-protein-BrxC-[Occultella-kanbiaonis] |
| OJX67210.1 | Type1.BREX | <-MULE-TRANSPOSASE\|\|NTP_transf->?->Phage_holin->XIS-HTH->BrxA-DNA-binding->BrxB-BrxF-iSTAND->DUF6079*-> | 1179 | Bacteria>Terrabacteria_group>Actinomycetota>Actinomycetes>Micrococcales>unclassified_Micrococcales | GCA_001898855.1 | Micrococcales_bacterium_73-13 | BGO95_06600 | 6462..10001 | 1895791 | ATP-binding-protein-[Micrococcales-bacterium-73-13] |
| MCL2802837.1 | Type1.BREX | <-RADICAL-SAM<-sGTP<-?\|\|SWACOS->TPR->?-><-SIGMA+OB-Fold+AAA-ATPase+HKD-DNase-BrxL<-PglZ<-N6-METHYLASE<-DUF6079*<-BrxB-BrxF-iSTAND<-BrxA-DNA-binding\|\|?-><-?\|\|?-><-PhdYeFM_antitox | 1162 | Bacteria>Terrabacteria_group>Actinomycetota>Actinomycetes>Micrococcales>unclassified_Micrococcales | GCA_009784895.1 | Micrococcales_bacterium | FWD29_02600 | 43259..46747 | 2026762 | BREX-system-P-loop-protein-BrxC-[Micrococcales-bacterium] |
| WP_183221872.1 | Type1.BREX | <-STYKIN+Pkinase<-?<-?\|\|?-><-?\|\|BrxA-DNA-binding->BrxB-BrxF-iSTAND->DUF6079*->N6-METHYLASE->PglZ->SIGMA+OB-Fold+AAA-ATPase+HKD-DNase-BrxL-><-GGDEF+EAL<-?\|\|ABhydrolase->?->?->METHYLASE-> | 1164 | Bacteria>Terrabacteria_group>Actinomycetota>Actinomycetes>Micromonosporales>Micromonosporaceae | GCF_014647795.1 | Actinoplanes_campanulatus | IEZ13_RS15515 | 158515..162009 | 113559 | BREX-system-P-loop-protein-BrxC-[Actinoplanes-campanulatus] |
| WP_203833901.1 | Type1.BREX | <-METHYLASE<-?<-?<-ABhydrolase\|\|?->GGDEF+EAL-><-SIGMA+OB-Fold+AAA-ATPase+HKD-DNase-BrxL<-PglZ<-N6-METHYLASE<-DUF6079*<-BrxB-BrxF-iSTAND<-BrxA-DNA-binding\|\|?-><-?\|\|?->?->STYKIN+Pkinase-> | 1164 | Bacteria>Terrabacteria_group>Actinomycetota>Actinomycetes>Micromonosporales>Micromonosporaceae | GCF_016862035.1 | Actinoplanes_campanulatus | Aca09nite_RS24855 | 101506..105000 | 113559 | BREX-system-P-loop-protein-BrxC-[Actinoplanes-campanulatus] |
| WP_188126500.1 | Type1.BREX | <-METHYLASE<-?<-?<-ABhydrolase\|\|GGDEF+EAL-><-SIGMA+OB-Fold+AAA-ATPase+HKD-DNase-BrxL<-PglZ<-N6-METHYLASE<-DUF6079*<-BrxB-BrxF-iSTAND<-BrxA-DNA-binding\|\|?-><-?\|\|?->?->STYKIN+Pkinase-> | 1164 | Bacteria>Terrabacteria_group>Actinomycetota>Actinomycetes>Micromonosporales>Micromonosporaceae | GCF_016862255.1 | Actinoplanes_lobatus | IEZ29_RS29630 | 68673..72167 | 113568 | BREX-system-P-loop-protein-BrxC-[Actinoplanes-lobatus] |
| WP_203928003.1 | Type1.BREX | TPR-><-SIGMA+OB-Fold+AAA-ATPase+HKD-DNase-BrxL<-PglZ<-N6-METHYLASE<-DUF6079*<-BrxB-BrxF-iSTAND<-BrxA-DNA-binding<-APATPase+TPR+TPR+TPR+TPR+TPR+TPR+TPR\|\|NAD_binding_10->?-><-NACHT\|\|SIGMA-HTH-> | 1154 | Bacteria>Terrabacteria_group>Actinomycetota>Actinomycetes>Micromonosporales>Micromonosporaceae | GCF_016863655.1 | Virgisporangium_ochraceum | Voc01_RS14900 | 8921..12385 | 65505 | BREX-system-P-loop-protein-BrxC-[Virgisporangium-ochraceum] |
| WP_260726425.1 | Type1.BREX | <-SIGMA-HTH<-?<-SAM-METHYLASE\|\|?->?->?-><-SIGMA+OB-Fold+AAA-ATPase+HKD-DNase-BrxL<-PglZ<-N6-METHYLASE<-DUF6079*<-BrxB-BrxF-iSTAND<-BrxA-DNA-binding<-XIS-HTH\|\|HAMP+MA+MA->?->SIGMA-HTH-><-Glycos_transf_1+Glyco_trans_1_2<-Glycos_transf_1+Glyco_trans_1_2<-SF1-SF2-Helicase | 1154 | Bacteria>Terrabacteria_group>Actinomycetota>Actinomycetes>Micromonosporales>Micromonosporaceae | GCF_025264685.1 | Dactylosporangium_roseum | Drose_RS01775 | 392440..395904 | 47989 | BREX-system-P-loop-protein-BrxC-[Dactylosporangium-roseum] |
| WP_265248531.1 | Type1.BREX | HTH+MPTase-><-EVE+REase<-DNA-glycosylase<-?\|\|Phage_int_SAM_3+Phage_integrase->?->BrxA-DNA-binding->BrxB-BrxF-iSTAND->DUF6079*->N6-METHYLASE->PglZ->SIGMA+OB-Fold+AAA-ATPase+Lon-Protease-BrxL->?->?->RADICAL-SAM->MPTase-><-?<-MoxR-AAA | 1155 | Bacteria>Terrabacteria_group>Actinomycetota>Actinomycetes>Micromonosporales>Micromonosporaceae | GCF_026130045.1 | Micromonospora_sp._CPCC_205371 | K1W54_RS08550 | 13084..16551 | 2865942 | BREX-system-P-loop-protein-BrxC-[Micromonospora-sp.-CPCC-205371] |
| WP_091673594.1 | Type1.BREX | RADICAL-SAM->?->?-><-HAD\|\|XIS-HTH->Phage_int_SAM_3+Phage_integrase->XIS-HTH->BrxA-DNA-binding->BrxB-BrxF-iSTAND->DUF6079*->N6-METHYLASE->PglZ->SIGMA+OB-Fold+AAA-ATPase+HKD-DNase-BrxL->DNA_glycosylase-><-?<-SF1-SF2-Helicase<-HKD-DNase<-DUF6361<-RNase | 1156 | Bacteria>Terrabacteria_group>Actinomycetota>Actinomycetes>Micromonosporales>Micromonosporaceae | GCF_900089595.1 | Micromonospora_auratinigra | GA0070611_RS28700 | 6259973..6263443 | 261654 | BREX-system-P-loop-protein-BrxC-[Micromonospora-auratinigra] |
| WP_091348932.1 | Type1.BREX | XIS-HTH->BrxA-DNA-binding->BrxB-BrxF-iSTAND->DUF6079*->N6-METHYLASE->PglZ->SIGMA+OB-Fold+AAA-ATPase+HKD-DNase-BrxL->HIN-HTH-><-HTH_32<-HTH_32<-SF1-SF2-Helicase<-HKD-DNase+HKD-Rease | 1153 | Bacteria>Terrabacteria_group>Actinomycetota>Actinomycetes>Micromonosporales>Micromonosporaceae | GCF_900091465.1 | Micromonospora_rhizosphaerae | GA0070624_RS15965 | 3332590..3336051 | 568872 | BREX-system-P-loop-protein-BrxC-[Micromonospora-rhizosphaerae] |
| HJG42479.1 | Type1.BREX | BrxA-DNA-binding->BrxB-BrxF-iSTAND->DUF6079*->N6-METHYLASE->PglZ->SIGMA+OB-Fold+AAA-ATPase+HKD-DNase-BrxL-> | 1176 | Bacteria>Terrabacteria_group>Actinomycetota>Actinomycetes>Mycobacteriales>Corynebacteriaceae | GCA_020743465.1 | Corynebacterium_phoceense | K8V34_00330 | 34206..37736 | 1686286 | BREX-system-P-loop-protein-BrxC-[Corynebacterium-phoceense] |
| WP_102213741.1 | Type1.BREX | XIS-HTH->BrxA-DNA-binding->BrxB-BrxF-iSTAND->DUF6079*->N6-METHYLASE->HTH+REase->PglZ->SIGMA+OB-Fold+AAA-ATPase+HKD-DNase-BrxL-><-ParB+HNH+METHYLASE | 1173 | Bacteria>Terrabacteria_group>Actinomycetota>Actinomycetes>Mycobacteriales>Corynebacteriaceae | GCF_002871715.1 | Corynebacterium_xerosis | CJ204_09375 | 1693..5214 | 1725 | BREX-system-P-loop-protein-BrxC-[Corynebacterium-xerosis] |
| WP_220313057.1 | Type1.BREX | <-REC+OmpR-HTH<-MerR-HTH+TipAS<-?<-SIR2\|\|?->TetR-HTH->?->BrxA-DNA-binding->BrxB-BrxF-iSTAND->DUF6079*->N6-METHYLASE-><-N6-METHYLASE-p\|\|Phage_int_SAM_3+Phage_integrase->N6-METHYLASE-p->PglZ->SIGMA+OB-Fold+AAA-ATPase+HKD-DNase-BrxL-><-?<-?<-ADH_N+ADH_zinc_N | 1160 | Bacteria>Terrabacteria_group>Actinomycetota>Actinomycetes>Mycobacteriales>Corynebacteriaceae | GCF_019551735.1 | Corynebacterium_glutamicum | JJQ73_RS12320 | 2669346..2672828 | 1718 | BREX-system-P-loop-protein-BrxC-[Corynebacterium-glutamicum] |
| WP_259898411.1 | Type1.BREX | <-HIN-HTH<-ParB+HNH<-SIGMA+OB-Fold+AAA-ATPase+HKD-DNase-BrxL<-PglZ<-N6-METHYLASE<-DUF6079*<-BrxB-BrxF-iSTAND<-BrxA-DNA-binding<-XIS-HTH | 1160 | Bacteria>Terrabacteria_group>Actinomycetota>Actinomycetes>Mycobacteriales>Corynebacteriaceae | GCF_025152505.1 | Corynebacterium_sanguinis | M3G46_RS10600 | 10062..13544 | 2594913 | BREX-system-P-loop-protein-BrxC-[Corynebacterium-sanguinis] |
| WP_271034470.1 | Type1.BREX | <-HTH_23+rve+rve_3\|\|?->BrxA-DNA-binding->BrxB-BrxF-iSTAND->DUF6079*->N6-METHYLASE-><-N6-METHYLASE-p\|\|Phage_int_SAM_3+Phage_integrase->N6-METHYLASE-p->PglZ->SIGMA+OB-Fold+AAA-ATPase+HKD-DNase-BrxL-> | 1159 | Bacteria>Terrabacteria_group>Actinomycetota>Actinomycetes>Mycobacteriales>Corynebacteriaceae | GCF_027886475.1 | Corynebacterium_sp._SCR221107 | PAB09_RS02255 | 466456..469935 | 3017361 | BREX-system-P-loop-protein-BrxC-[Corynebacterium-sp.-SCR221107] |
| WP_245802167.1 | Type1.BREX | <-ARG<-SIR2<-SIGMA+OB-Fold+AAA-ATPase+HKD-DNase-BrxL<-PglZ<-Schlafen+HATPase+wHTH<-N6-METHYLASE<-DUF6079*<-BrxB-BrxF-iSTAND<-BrxA-DNA-binding | 1179 | Bacteria>Terrabacteria_group>Actinomycetota>Actinomycetes>Mycobacteriales>Corynebacteriaceae | GCF_900169525.1 | Corynebacterium_pacaense | B7E02_RS05025 | 1051363..1054902 | 1816684 | BREX-system-P-loop-protein-BrxC-[Corynebacterium-pacaense] |
| WP_047036580.1 | Type1.BREX | <-Phage_integrase\|\|ACET-><-HerA/FtsK<-SIR2+SIR2<-SIGMA+OB-Fold+AAA-ATPase+HKD-DNase-BrxL<-PglZ<-N6-METHYLASE<-DUF6079*<-BrxB-BrxF-iSTAND<-BrxA-DNA-binding<-XIS-HTH<-TetR+HTH<-Phage_integrase+Phage_integrase<-Phage_integrase+Phage_integrase<-Phage_int_SAM_1+Phage_integrase<-DNA_glycosylase | 1156 | Bacteria>Terrabacteria_group>Actinomycetota>Actinomycetes>Mycobacteriales | GCF_900141005.1 | Mycobacteriaceae | ABW05_RS31210 | 104572..108042 | 1762 | MULTISPECIES:-BREX-system-P-loop-protein-BrxC-[Mycobacteriaceae] |
| PZT90276.1 | Type1.BREX | HAD+HAD-><-?\|\|XIS-HTH->BrxA-DNA-binding->BrxB-BrxF-iSTAND->DUF6079*->N6-METHYLASE->HTH+REase->PglZ->SIGMA+OB-Fold+AAA-ATPase+HKD-DNase-BrxL->REase-> | 1137 | Bacteria>Terrabacteria_group>Actinomycetota>Actinomycetes>Mycobacteriales>Gordoniaceae | GCA_003248405.1 | Gordonia_sp._(in:_high_G+C_Gram-positive_bacteria) | DI630_31230 | 6915..10328 | 84139 | MAG:-BREX-system-P-loop-protein-BrxC-[Gordonia-sp.-(in:-high-G+C-Gram-positive-bacteria)] |
| RUP38447.1 | Type1.BREX | <-N6-METHYLASE-p<-DUF6079*<-BrxB-BrxF-iSTAND<-BrxA-DNA-binding\|\|URI-DNase+RAMA->?->ABhydrolase->\|\|?->DndD->HEPN-> | 1160 | Bacteria>Terrabacteria_group>Actinomycetota>Actinomycetes>Mycobacteriales>Gordoniaceae | GCA_003987705.1 | Gordonia_sp._(in:_high_G+C_Gram-positive_bacteria) | EKK60_09740 | 1345..4827 | 84139 | MAG:-BREX-system-P-loop-protein-BrxC-[Gordonia-sp.-(in:-high-G+C-Gram-positive-bacteria)] |
| WP_042377474.1 | Type1.BREX | TetR-HTH->?-><-?<-?<-SIGMA+OB-Fold+AAA-ATPase+HKD-DNase-BrxL<-PglZ<-N6-METHYLASE<-DUF6079*<-BrxB-BrxF-iSTAND<-BrxA-DNA-binding\|\|Exonuc_V_gamma+REase->SF1-SF2-Helicase+REase-><-?<-DNA_glycosylase<-HNH+HNH | 1161 | Bacteria>Terrabacteria_group>Actinomycetota>Actinomycetes>Mycobacteriales>Gordoniaceae | GCF_000529675.1 | Gordonia_alkanivorans | TY17_RS15305 | 47194..50679 | 84096 | BREX-system-P-loop-protein-BrxC-[Gordonia-alkanivorans] |
| WP_159370828.1 | Type1.BREX | <-DNA_glycosylase\|\|DUF6361->HKD-DNase->SF1-SF2-Helicase-><-SIGMA+OB-Fold+AAA-ATPase+HKD-DNase-BrxL<-PglZ<-N6-METHYLASE<-DUF6079*<-BrxB-BrxF-iSTAND<-BrxA-DNA-binding<-?<-?<-ARG<-?\|\|?-><-SAM-METHYLASE | 1174 | Bacteria>Terrabacteria_group>Actinomycetota>Actinomycetes>Mycobacteriales>Gordoniaceae | GCF_009856645.1 | Gordonia_sp._JH63 | GR168_RS12095 | 2701909..2705433 | 2698900 | BREX-system-P-loop-protein-BrxC-[Gordonia-sp.-JH63] |
| WP_213244950.1 | Type1.BREX | <-SIGMA+OB-Fold+AAA-ATPase+HKD-DNase-BrxL<-PglZ<-HTH+REase<-N6-METHYLASE-p<-?<-Phage_int_SAM_3+Phage_integrase<-Phage_int_SAM_3+Phage_integrase\|\|N6-METHYLASE-p-><-N6-METHYLASE<-DUF6079*<-BrxB-BrxF-iSTAND<-BrxA-DNA-binding | 1174 | Bacteria>Terrabacteria_group>Actinomycetota>Actinomycetes>Mycobacteriales>Gordoniaceae | GCF_009914535.1 | Gordonia_pseudamarae | GII33_RS19265 | 4440551..4444075 | 2831662 | BREX-system-P-loop-protein-BrxC-[Gordonia-pseudamarae] |
| WP_223374200.1 | Type1.BREX | REase->sGTP+sGTP+GTPase-AIG->sGTP-><-PHP+SbcC<-?<-SIGMA+OB-Fold+AAA-ATPase+HKD-DNase-BrxL<-PglZ<-N6-METHYLASE<-DUF6079*<-BrxB-BrxF-iSTAND<-BrxA-DNA-binding<-XIS-HTH<-?<-TnsC-ATPase\|\|PAIREDC-HTH-><-rve\|\|HTH_29+HTH_32+DDE_3-><-REase+wHTH | 1154 | Bacteria>Terrabacteria_group>Actinomycetota>Actinomycetes>Mycobacteriales>Gordoniaceae | GCF_019930645.1 | Gordonia_bronchialis | K8O93_RS05575 | 1152614..1156078 | 2054 | BREX-system-P-loop-protein-BrxC-[Gordonia-bronchialis] |
| WP_226512485.1 | Type1.BREX | MACRODOMAIN+SIR2-><-?<-?\|\|DUF6361->HKD-DNase->SF1-SF2-Helicase->URI-DNase->BrxA-DNA-binding->BrxB-BrxF-iSTAND->DUF6079-p*->DUF6079-p->XIS-HTH->PIN->N6-METHYLASE-p->N6-METHYLASE->PglZ->?->AbiEi_4+AbiEi_1->SIGMA+OB-Fold+AAA-ATPase+HKD-DNase-BrxL-> | 435 | Bacteria>Terrabacteria_group>Actinomycetota>Actinomycetes>Mycobacteriales>Gordoniaceae | GCF_020520305.1 | Gordonia_sp._WA4-43 | LEL84_RS18910 | 4162672..4163979 | 2878678 | BREX-system-P-loop-protein-BrxC-[Gordonia-sp.-WA4-43] |
| WP_226512486.1 | Type1.BREX | DUF6361->HKD-DNase->SF1-SF2-Helicase->URI-DNase->BrxA-DNA-binding->BrxB-BrxF-iSTAND->DUF6079-p->DUF6079-p*->XIS-HTH->PIN->N6-METHYLASE-p->N6-METHYLASE->PglZ->?->AbiEi_4+AbiEi_1->SIGMA+OB-Fold+AAA-ATPase+HKD-DNase-BrxL-><-ABhydrolase+ABhydrolase | 785 | Bacteria>Terrabacteria_group>Actinomycetota>Actinomycetes>Mycobacteriales>Gordoniaceae | GCF_020520305.1 | Gordonia_sp._WA4-43 | LEL84_RS18915 | 4163880..4166237 | 2878678 | hypothetical-protein-[Gordonia-sp.-WA4-43] |
| WP_246991513.1 | Type1.BREX | <-SIGMA+OB-Fold+AAA-ATPase+HKD-DNase-BrxL<-PglZ<-?<-?<-N6-METHYLASE-p<-Phage_int_SAM_3+Phage_integrase\|\|N6-METHYLASE-p-><-N6-METHYLASE<-DUF6079*<-BrxB-BrxF-iSTAND<-BrxA-DNA-binding<-sGTP<-?\|\|?->O6MGT-HTH-> | 1175 | Bacteria>Terrabacteria_group>Actinomycetota>Actinomycetes>Mycobacteriales>Gordoniaceae | GCF_023017165.1 | Gordonia_alkaliphila | MUG78_01460 | 307013..310540 | 1053547 | BREX-system-P-loop-protein-BrxC-[Gordonia-alkaliphila] |
| WP_247863885.1 | Type1.BREX | <-ParA-Soj-PloopNTPase\|\|SopB+HNH-><-SIGMA+OB-Fold+AAA-ATPase+Lon-Protease-BrxL<-PglZ<-N6-METHYLASE<-DUF6079*<-BrxB-BrxF-iSTAND<-BrxA-DNA-binding\|\|HTH_3->?-><-?<-Phage_int_SAM_1+Phage_integrase<-NUDIX | 1151 | Bacteria>Terrabacteria_group>Actinomycetota>Actinomycetes>Mycobacteriales>Gordoniaceae | GCF_023100845.1 | Gordonia_sp._PP30 | MYK68_RS11760 | 2542969..2546424 | 2935861 | BREX-system-P-loop-protein-BrxC-[Gordonia-sp.-PP30] |
| OBI43944.1 | Type1.BREX | HNH+HNH-><-SIR2\|\|HEPN->Classical-AAA->Peptidase_S8-><-SIGMA+OB-Fold+AAA-ATPase+Lon-Protease-BrxL<-PglZ<-N6-METHYLASE<-DUF6079*<-BrxB-BrxF-iSTAND<-BrxA-DNA-binding<-?\|\|DNA_glycosylase->?-><-?<-?<-TOPRIM+Primase<-XIS-HTH | 1152 | Bacteria>Terrabacteria_group>Actinomycetota>Actinomycetes>Mycobacteriales>Mycobacteriaceae | GCA_001672775.1 | Mycobacterium_kyorinense | A5707_03925 | 29624..33082 | 487514 | hypothetical-protein-A5707_03925-[Mycobacterium-kyorinense] |
| OKH76423.1 | Type1.BREX | <-NUDIX\|\|ARG-><-SbcC<-?<-SIGMA+OB-Fold+AAA-ATPase+HKD-DNase-BrxL<-PglZ<-N6-METHYLASE<-DUF6079*<-BrxB-BrxF-iSTAND<-BrxA-DNA-binding<-XIS-HTH | 1156 | Bacteria>Terrabacteria_group>Actinomycetota>Actinomycetes>Mycobacteriales>Mycobacteriaceae | GCA_001905565.1 | Mycobacterium_sp._SWH-M1 | EB72_01455 | 44646..48116 | 1490485 | ATP-binding-protein-[Mycobacterium-sp.-SWH-M1] |
| OKH79002.1 | Type1.BREX | XIS-HTH->BrxA-DNA-binding->BrxB-BrxF-iSTAND->DUF6079*->N6-METHYLASE->PglZ->SIGMA+OB-Fold+AAA-ATPase+HKD-DNase-BrxL->?->SbcC-><-MPTase<-?<-?<-Phage_integrase | 1156 | Bacteria>Terrabacteria_group>Actinomycetota>Actinomycetes>Mycobacteriales>Mycobacteriaceae | GCA_001905655.1 | Mycobacterium_sp._ST-F2 | EB75_25595 | 1846..5316 | 1490484 | ATP-binding-protein-[Mycobacterium-sp.-ST-F2] |
| RIT13053.1 | Type1.BREX | <-HTH_26<-Phage_int_SAM_4+Phage_integrase\|\|REase+wHTH->HTH_32+Mu-transpos_C->TnsC-ATPase->?->XIS-HTH->BrxA-DNA-binding->BrxB-BrxF-iSTAND->DUF6079*->N6-METHYLASE->PglZ->SIGMA+OB-Fold+AAA-ATPase+Lon-Protease-BrxL->PHP+SbcC->?->REase->?->REase+SF1-SF2-Helicase->N6-METHYLASE-> | 1175 | Bacteria>Terrabacteria_group>Actinomycetota>Actinomycetes>Mycobacteriales>Mycobacteriaceae | GCA_003582485.1 | Mycobacteroides_abscessus | D2E72_00925 | 191578..195105 | 36809 | BREX-system-P-loop-protein-BrxC-[Mycobacteroides-abscessus] |
| TXH24025.1 | Type1.BREX | <-N6-METHYLASE-p<-DUF6079*<-BrxB-BrxF-iSTAND<-BrxA-DNA-binding<-XIS-HTH<-?<-TnsC-ATPase<-HTH_32+Mu-transpos_C<-REase+wHTH | 1176 | Bacteria>Terrabacteria_group>Actinomycetota>Actinomycetes>Mycobacteriales>Mycobacteriaceae | GCA_008012155.1 | Mycobacterium_sp. | E6R06_13520 | 506..4036 | 1785 | MAG:-BREX-system-P-loop-protein-BrxC-[Mycobacterium-sp.] |
| TXI54389.1 | Type1.BREX | <-N6-METHYLASE<-DUF6079*<-BrxB-BrxF-iSTAND<-BrxA-DNA-binding<-XIS-HTH<-DNA_glycosylase<-TetR-HTH<-Phage_integrase<-Phage_integrase<-SF1-SF2-Helicase+REase | 1156 | Bacteria>Terrabacteria_group>Actinomycetota>Actinomycetes>Mycobacteriales>Mycobacteriaceae | GCA_008015735.1 | Mycolicibacterium_mageritense | E6Q55_32810 | 2674..6144 | 53462 | BREX-system-P-loop-protein-BrxC-[Mycolicibacterium-mageritense] |
| GFM17528.1 | Type1.BREX | <-N6-METHYLASE-p<-DUF6079*<-BrxB-BrxF-iSTAND<-BrxA-DNA-binding<-McrC-NTD+REase<-McrB-AAA<-HTH+REase<-HEPN+SF1-SF2-Helicase+REase<-?<-Preader7 | 1155 | Bacteria>Terrabacteria_group>Actinomycetota>Actinomycetes>Mycobacteriales>Mycobacteriaceae | GCA_017312405.1 | Mycobacterium_sp._PO1 | PO1_contig-018-2 | 841..4308 | 1882219 | putative-P-loop-containing-nucleoside-triphospha-te-hydrolase-[Mycobacterium-sp.-PO1] |
| MCA4756838.1 | Type1.BREX | <-N6-METHYLASE<-DUF6079*<-BrxB-BrxF-iSTAND<-BrxA-DNA-binding<-XIS-HTH<-TetR-HTH<-Phage_integrase<-Phage_integrase<-XIS-HTH | 1154 | Bacteria>Terrabacteria_group>Actinomycetota>Actinomycetes>Mycobacteriales>Mycobacteriaceae | GCA_020275685.1 | Mycolicibacterium_fortuitum | H8Z60_30510 | 3659..7123 | 1766 | BREX-system-P-loop-protein-BrxC-[Mycolicibacterium-fortuitum] |
| WP_018600838.1 | Type1.BREX | <-ABhydrolase\|\|?-><-?<-?\|\|?-><-HTH_3\|\|BrxA-DNA-binding->BrxB-BrxF-iSTAND->DUF6079*->N6-METHYLASE->PglZ->SIGMA+OB-Fold+AAA-ATPase+HKD-DNase-BrxL-><-?<-?\|\|?-><-?<-ACT<-SIR2 | 1152 | Bacteria>Terrabacteria_group>Actinomycetota>Actinomycetes>Mycobacteriales>Mycobacteriaceae | GCF_000373905.1 | Mycobacterium_sp._155 | B133_RS0110195 | 2156875..2160333 | 1157943 | BREX-system-P-loop-protein-BrxC-[Mycobacterium-sp.-155] |
| WP_020822248.1 | Type1.BREX | Pkinase+TPR+TPR-><-?\|\|TetR-HTH-><-?\|\|Glyoxalase-><-SIGMA+OB-Fold+AAA-ATPase+HKD-DNase-BrxL<-RVT<-PglZ<-N6-METHYLASE<-DUF6079*<-BrxB-BrxF-iSTAND<-BrxA-DNA-binding<-XIS-HTH<-zf-ISL3+HTH_Tnp_ISL3+DDE_Tnp_ISL3 | 1152 | Bacteria>Terrabacteria_group>Actinomycetota>Actinomycetes>Mycobacteriales>Mycobacteriaceae | GCF_000418535.1 | Mycobacterium_intracellulare | OEM_RS09395 | 2016549..2020007 | 1767 | BREX-system-P-loop-protein-BrxC-[Mycobacterium-intracellulare] |
| WP_052621181.1 | Type1.BREX | <-HerA/FtsK<-?<-?<-SF1-SF2-Helicase+REase<-HerA/FtsK<-SIR2<-SIGMA+OB-Fold+AAA-ATPase+HKD-DNase-BrxL<-PglZ<-N6-METHYLASE<-DUF6079*<-BrxB-BrxF-iSTAND<-BrxA-DNA-binding<-XIS-HTH<-TetR+HTH<-Phage_integrase+Phage_integrase<-Phage_integrase+Phage_integrase<-Phage_int_SAM_1+Phage_integrase\|\|HEPN-> | 1156 | Bacteria>Terrabacteria_group>Actinomycetota>Actinomycetes>Mycobacteriales>Mycobacteriaceae | GCF_001214625.1 | Mycobacteroides_abscessus | B5319_RS07405 | 1475995..1479465 | 36809 | BREX-system-P-loop-protein-BrxC-[Mycobacteroides-abscessus] |
| WP_064441930.1 | Type1.BREX | <-PglZ-p<-N6-METHYLASE<-DUF6079*<-BrxB-BrxF-iSTAND<-BrxA-DNA-binding\|\|ParB+HNH->ParB+HNH->?->DndD->HEPN->?->REase-> | 1148 | Bacteria>Terrabacteria_group>Actinomycetota>Actinomycetes>Mycobacteriales>Mycobacteriaceae | GCF_001653095.1 | Hoyosella_altamirensis | HAL02_RS18845 | 3788..7234 | 616997 | BREX-system-P-loop-protein-BrxC-[Hoyosella-altamirensis] |
| WP_067214457.1 | Type1.BREX | Calcineurin->SbcC+SbcC->?-><-REase\|\|MACRODOMAIN+SIR2-><-?<-SIGMA+OB-Fold+AAA-ATPase+Lon-Protease-BrxL<-PglZ<-N6-METHYLASE<-DUF6079*<-BrxB-BrxF-iSTAND<-BrxA-DNA-binding\|\|?->HKD-DNase+HKD-DNase-><-Peptidase_M48+TerB<-HIRAN<-?<-HEPN+SF1-SF2-Helicase+REase<-Peptidase_M23+SLT | 1155 | Bacteria>Terrabacteria_group>Actinomycetota>Actinomycetes>Mycobacteriales>Mycobacteriaceae | GCF_001667065.1 | Mycolicibacterium_celeriflavum | A5765_RS01610 | 25848..29315 | 1249101 | BREX-system-P-loop-protein-BrxC-[Mycolicibacterium-celeriflavum] |
| WP_082969488.1 | Type1.BREX | HNH+HNH-><-SIR2\|\|HEPN->Classical-AAA->Peptidase_S8-><-SIGMA+OB-Fold+AAA-ATPase+Lon-Protease-BrxL<-PglZ<-N6-METHYLASE<-DUF6079*<-BrxB-BrxF-iSTAND<-BrxA-DNA-binding\|\|DNA_glycosylase->?-><-?<-TOPRIM+Primase<-XIS-HTH\|\|<-tRNA<-?<-Calcineurin | 1154 | Bacteria>Terrabacteria_group>Actinomycetota>Actinomycetes>Mycobacteriales>Mycobacteriaceae | GCF_001672775.1 | Mycobacterium_kyorinense | A5707_RS15595 | 29624..33088 | 487514 | BREX-system-P-loop-protein-BrxC-[Mycobacterium-kyorinense] |
| WP_065022017.1 | Type1.BREX | <-N6-METHYLASE<-DUF6079*<-BrxB-BrxF-iSTAND<-BrxA-DNA-binding<-XIS-HTH<-?<-?<-HKD-DNase+SF1-SF2-Helicase+HTH+HTH+SRA<-HTH+REase\|\|DUF4352-> | 1156 | Bacteria>Terrabacteria_group>Actinomycetota>Actinomycetes>Mycobacteriales>Mycobacteriaceae | GCF_001672855.1 | Mycolicibacterium_fortuitum | A5667_RS26930 | 2805..6275 | 1766 | BREX-system-P-loop-protein-BrxC-[Mycolicibacterium-fortuitum] |
| WP_065289569.1 | Type1.BREX | DUF5086-><-ANK+ANK<-ZnR\|\|XIS-HTH->BrxA-DNA-binding->BrxB-BrxF-iSTAND->DUF6079*->N6-METHYLASE->PglZ->SIGMA+OB-Fold+AAA-ATPase+HKD-DNase-BrxL-> | 1152 | Bacteria>Terrabacteria_group>Actinomycetota>Actinomycetes>Mycobacteriales>Mycobacteriaceae | GCF_001679965.1 | Mycolicibacter_kumamotonensis | ACT18_RS22010 | 4536..7994 | 354243 | BREX-system-P-loop-protein-BrxC-[Mycolicibacter-kumamotonensis] |
| WP_070953149.1 | Type1.BREX | SAM-METHYLASE-><-SopB+HNH<-?<-?\|\|XIS-HTH->BrxA-DNA-binding->BrxB-BrxF-iSTAND->DUF6079*->N6-METHYLASE->PglZ->SIGMA+OB-Fold+AAA-ATPase+HKD-DNase-BrxL->Acyl_transf_3->?->?-><-ACET | 1156 | Bacteria>Terrabacteria_group>Actinomycetota>Actinomycetes>Mycobacteriales>Mycobacteriaceae | GCF_001853775.1 | Mycobacteroides_chelonae | BKG71_RS10405 | 2100910..2104380 | 1774 | BREX-system-P-loop-protein-BrxC-[Mycobacteroides-chelonae] |
| WP_070917649.1 | Type1.BREX | <-SKI<-ParD_antitoxin\|\|REase+wHTH->HTH_32+Mu-transpos_C->TnsC-ATPase->?->XIS-HTH->BrxA-DNA-binding->BrxB-BrxF-iSTAND->DUF6079*->N6-METHYLASE->PglZ->SIGMA+OB-Fold+AAA-ATPase+Lon-Protease-BrxL->PHP+SbcC->?->Glyoxalase-> | 1152 | Bacteria>Terrabacteria_group>Actinomycetota>Actinomycetes>Mycobacteriales>Mycobacteriaceae | GCF_001853985.1 | Mycobacteroides_chelonae | BKG83_RS22680 | 427497..430955 | 1774 | BREX-system-P-loop-protein-BrxC-[Mycobacteroides-chelonae] |
| WP_234813880.1 | Type1.BREX | <-ParB+HNH<-SIGMA+OB-Fold+AAA-ATPase+HKD-DNase-BrxL<-PglZ<-N6-METHYLASE<-DUF6079-p<-DUF6079-p*<-BrxB-BrxF-iSTAND<-BrxA-DNA-binding | 682 | Bacteria>Terrabacteria_group>Actinomycetota>Actinomycetes>Mycobacteriales>Mycobacteriaceae | GCF_002101705.1 | Mycolicibacterium_iranicum | AWC12_RS31570 | 96008..98056 | 912594 | BREX-system-P-loop-protein-BrxC-[Mycolicibacterium-iranicum] |
| WP_182875480.1 | Type1.BREX | <-HTH_26<-Phage_int_SAM_4+Phage_integrase\|\|REase+wHTH->HTH_32+Mu-transpos_C->TnsC-ATPase->?->XIS-HTH->BrxA-DNA-binding->BrxB-BrxF-iSTAND->DUF6079*->N6-METHYLASE->PglZ->SIGMA+OB-Fold+AAA-ATPase+Lon-Protease-BrxL->PHP+SbcC->REase->REase+SF1-SF2-Helicase->N6-METHYLASE->TRD->KAP-NTPase-> | 1151 | Bacteria>Terrabacteria_group>Actinomycetota>Actinomycetes>Mycobacteriales>Mycobacteriaceae | GCF_003582485.1 | Mycobacteroides_abscessus | D2E72_RS00925 | 191650..195105 | 36809 | BREX-system-P-loop-protein-BrxC-[Mycobacteroides-abscessus] |
| WP_119635996.1 | Type1.BREX | <-XIS-HTH<-?<-?<-SF1-SF2-Helicase+REase<-HerA/FtsK<-SIR2+SIR2<-SIGMA+OB-Fold+AAA-ATPase+HKD-DNase-BrxL<-PglZ<-N6-METHYLASE<-DUF6079*<-BrxB-BrxF-iSTAND<-BrxA-DNA-binding<-XIS-HTH<-TetR+HTH<-Phage_integrase+Phage_integrase<-Phage_integrase+Phage_integrase<-Phage_int_SAM_1+Phage_integrase\|\|HEPN-> | 1156 | Bacteria>Terrabacteria_group>Actinomycetota>Actinomycetes>Mycobacteriales>Mycobacteriaceae | GCF_003582985.1 | Mycobacteroides_abscessus | D2E92_RS23075 | 16528..19998 | 36809 | BREX-system-P-loop-protein-BrxC-[Mycobacteroides-abscessus] |
| WP_134206037.1 | Type1.BREX | <-REase<-MORC<-HEPN+DUF499<-METHYLASE-p<-HerA/FtsK<-SIR2+SIR2<-SIGMA+OB-Fold+AAA-ATPase+HKD-DNase-BrxL<-PglZ<-N6-METHYLASE<-DUF6079*<-BrxB-BrxF-iSTAND<-BrxA-DNA-binding<-XIS-HTH<-TetR+HTH<-Phage_integrase+Phage_integrase<-Phage_integrase+Phage_integrase<-Phage_int_SAM_1+Phage_integrase<-REase\|\|DNA_METHYLASE-> | 1155 | Bacteria>Terrabacteria_group>Actinomycetota>Actinomycetes>Mycobacteriales>Mycobacteriaceae | GCF_004366875.1 | Mycobacteroides_salmoniphilum | CCUG62472_RS09145 | 418307..421774 | 404941 | BREX-system-P-loop-protein-BrxC-[Mycobacteroides-salmoniphilum] |
| WP_085074024.1 | Type1.BREX | <-sGTP<-?<-sGTP<-HerA/FtsK<-SIR2+SIR2<-SIGMA+OB-Fold+AAA-ATPase+HKD-DNase-BrxL<-PglZ<-N6-METHYLASE<-DUF6079*<-BrxB-BrxF-iSTAND<-BrxA-DNA-binding<-XIS-HTH | 1154 | Bacteria>Terrabacteria_group>Actinomycetota>Actinomycetes>Mycobacteriales>Mycobacteriaceae | GCF_010723135.1 | Mycobacterium_kubicae | H7H99_RS20140 | 22484..25948 | 120959 | BREX-system-P-loop-protein-BrxC-[Mycobacterium-kubicae] |
| WP_163778779.1 | Type1.BREX | GGDEF+EAL->?-><-Glyco_tran_28_C<-?\|\|?-><-?<-tRNA\|\|XIS-HTH->BrxA-DNA-binding->BrxB-BrxF-iSTAND->DUF6079*->N6-METHYLASE->PglZ->SIGMA+OB-Fold+AAA-ATPase+HKD-DNase-BrxL->ParB+HNH->?->AAA-ATPase->ZnR->?->Phage_integrase-> | 1156 | Bacteria>Terrabacteria_group>Actinomycetota>Actinomycetes>Mycobacteriales>Mycobacteriaceae | GCF_010727945.1 | Mycobacterium_cookii | G6N27_RS18550 | 3937772..3941242 | 1775 | BREX-system-P-loop-protein-BrxC-[Mycobacterium-cookii] |
| WP_085152401.1 | Type1.BREX | <-ABhydrolase+ABhydrolase<-Preader7<-HEPN+SF1-SF2-Helicase+REase<-?<-SIGMA+OB-Fold+AAA-ATPase+HKD-DNase-BrxL<-PglZ<-N6-METHYLASE<-DUF6079*<-BrxB-BrxF-iSTAND<-BrxA-DNA-binding<-XIS-HTH<-HKD-DNase+SF1-SF2-Helicase+HTH+HTH+SRA\|\|35exo+SF1-SF2-Helicase->SF1-SF2-Helicase-> | 1156 | Bacteria>Terrabacteria_group>Actinomycetota>Actinomycetes>Mycobacteriales>Mycobacteriaceae | GCF_010729895.1 | Mycolicibacterium_confluentis | H7K00_RS24370 | 129281..132751 | 28047 | BREX-system-P-loop-protein-BrxC-[Mycolicibacterium-confluentis] |
| WP_196326342.1 | Type1.BREX | ABhydrolase->?-><-tRNA\|\|?-><-?\|\|XIS-HTH->XIS-HTH->BrxA-DNA-binding->BrxB-BrxF-iSTAND->DUF6079*->N6-METHYLASE->PglZ->SIGMA+OB-Fold+AAA-ATPase+HKD-DNase-BrxL->SIR2->HerA/FtsK-><-NUDIX\|\|?->?->HerA/FtsK-> | 1156 | Bacteria>Terrabacteria_group>Actinomycetota>Actinomycetes>Mycobacteriales>Mycobacteriaceae | GCF_015700725.1 | Mycobacteroides_chelonae | HA135_RS11130 | 2312216..2315686 | 1774 | BREX-system-P-loop-protein-BrxC-[Mycobacteroides-chelonae] |
| WP_074292199.1 | Type1.BREX | <-HerA/FtsK<-?<-?\|\|NUDIX-><-HerA/FtsK<-SIR2<-SIGMA+OB-Fold+AAA-ATPase+HKD-DNase-BrxL<-PglZ<-N6-METHYLASE<-DUF6079*<-BrxB-BrxF-iSTAND<-BrxA-DNA-binding<-XIS-HTH<-PIN<-XIS-HTH | 1156 | Bacteria>Terrabacteria_group>Actinomycetota>Actinomycetes>Mycobacteriales>Mycobacteriaceae | GCF_017183935.1 | Mycobacteroides_abscessus | I3U31_RS04170 | 833809..837279 | 36809 | BREX-system-P-loop-protein-BrxC-[Mycobacteroides-abscessus] |
| WP_222877735.1 | Type1.BREX | <-NUDIX\|\|<-HTH+REase<-HEPN+SF1-SF2-Helicase+REase<-?<-?\|\|BrxA-DNA-binding->BrxB-BrxF-iSTAND->DUF6079*->N6-METHYLASE->PglZ->SIGMA+OB-Fold+AAA-ATPase+Lon-Protease-BrxL->HIN-HTH->Phage_int_SAM_1+Phage_integrase->Phage_int_SAM_4+Phage_integrase->HIN-HTH->AAA-ATPase-> | 1150 | Bacteria>Terrabacteria_group>Actinomycetota>Actinomycetes>Mycobacteriales>Mycobacteriaceae | GCF_019856655.1 | Mycolicibacterium_austroafricanum | JN085_RS14455 | 3063577..3067029 | 39687 | BREX-system-P-loop-protein-BrxC-[Mycolicibacterium-austroafricanum] |
| WP_241204715.1 | Type1.BREX | <-DnaK<-?\|\|?->?-><-HerA/FtsK<-SIR2<-SIGMA+OB-Fold+AAA-ATPase+HKD-DNase-BrxL<-PglZ<-N6-METHYLASE<-DUF6079*<-BrxB-BrxF-iSTAND<-BrxA-DNA-binding<-XIS-HTH\|\|HEPN-><-AAA-ATPase<-HTH_7+rve+rve_3\|\|HEPN-> | 1156 | Bacteria>Terrabacteria_group>Actinomycetota>Actinomycetes>Mycobacteriales>Mycobacteriaceae | GCF_022348085.1 | Mycobacterium_sp._PSTR-4-N | MHN84_RS11260 | 180742..184212 | 2917745 | BREX-system-P-loop-protein-BrxC-[Mycobacterium-sp.-PSTR-4-N] |
| WP_260940163.1 | Type1.BREX | XIS-HTH->BrxA-DNA-binding->BrxB-BrxF-iSTAND->DUF6079*->N6-METHYLASE->PglZ->SIGMA+OB-Fold+AAA-ATPase+HKD-DNase-BrxL->?->ParB+HNH-p->HIN-HTH-><-wHTH\|\|MORC+DUF3684+DUF3883-><-REase+HTH+35exo+Small-B-barrel+SF1-SF2-Helicase | 1156 | Bacteria>Terrabacteria_group>Actinomycetota>Actinomycetes>Mycobacteriales>Mycobacteriaceae | GCF_025331195.1 | Mycolicibacterium_llatzerense | A7G45_RS21390 | 50451..53921 | 280871 | BREX-system-P-loop-protein-BrxC-[Mycolicibacterium-llatzerense] |
| WP_066911619.1 | Type1.BREX | <-UDG-conflict<-HerA/FtsK<-SIR2+SIR2<-SIGMA+OB-Fold+AAA-ATPase+HKD-DNase-BrxL<-PglZ<-N6-METHYLASE<-DUF6079*<-BrxB-BrxF-iSTAND<-BrxA-DNA-binding<-XIS-HTH<-Classical-AAA<-Peptidase_S8 | 1161 | Bacteria>Terrabacteria_group>Actinomycetota>Actinomycetes>Mycobacteriales>Mycobacteriaceae | GCF_025821415.1 | Mycobacterium_interjectum | H7H80_RS25830 | 12521..16006 | 33895 | BREX-system-P-loop-protein-BrxC-[Mycobacterium-interjectum] |
| WP_264004160.1 | Type1.BREX | <-SF1-SF2-Helicase<-HKD-DNase<-DUF6361<-SIGMA+OB-Fold+AAA-ATPase+Lon-Protease-BrxL<-PglZ<-N6-METHYLASE<-DUF6079*<-BrxB-BrxF-iSTAND<-BrxA-DNA-binding<-SF1-SF2-Helicase+DUF1998<-viral-protein\|\|GNTR-HTH-> | 1146 | Bacteria>Terrabacteria_group>Actinomycetota>Actinomycetes>Mycobacteriales>Mycobacteriaceae | GCF_025821515.1 | Mycolicibacterium_sphagni | H7H98_RS10450 | 77826..81266 | 1786 | BREX-system-P-loop-protein-BrxC-[Mycolicibacterium-sphagni] |
| WP_264033192.1 | Type1.BREX | Classical-AAA->HTH_3-><-SF1-SF2-Helicase+REase<-SF1-SF2-Helicase+REase<-HerA/FtsK<-SIR2<-SIGMA+OB-Fold+AAA-ATPase+HKD-DNase-BrxL<-PglZ<-N6-METHYLASE<-DUF6079*<-BrxB-BrxF-iSTAND | 1156 | Bacteria>Terrabacteria_group>Actinomycetota>Actinomycetes>Mycobacteriales>Mycobacteriaceae | GCF_025821885.1 | Mycolicibacterium_gilvum | H7I56_RS29610 | 57941..61411 | 1804 | BREX-system-P-loop-protein-BrxC-[Mycolicibacterium-gilvum] |
| WP_267319448.1 | Type1.BREX | wHTH(SBD)+HNH-><-HEPN<-DndD\|\|RAMA-><-ParB+HNH<-SIGMA+OB-Fold+AAA-ATPase+Lon-Protease-BrxL<-PglZ<-N6-METHYLASE<-DUF6079*<-BrxB-BrxF-iSTAND<-BrxA-DNA-binding\|\|DEDD_Tnp_IS110+Transposase_20-> | 1149 | Bacteria>Terrabacteria_group>Actinomycetota>Actinomycetes>Mycobacteriales>Mycobacteriaceae | GCF_026501225.1 | Mycolicibacterium_mucogenicum | OS122_RS24315 | 502363..505812 | 56689 | BREX-system-P-loop-protein-BrxC-[Mycolicibacterium-mucogenicum] |
| WP_074325944.1 | Type1.BREX | <-HEPN\|\|Phage_int_SAM_1+Phage_integrase->Phage_integrase+Phage_integrase->Phage_integrase+Phage_integrase->TetR+HTH->XIS-HTH->BrxA-DNA-binding->BrxB-BrxF-iSTAND->DUF6079*->N6-METHYLASE->PglZ->SIGMA+OB-Fold+AAA-ATPase+HKD-DNase-BrxL->SIR2+SIR2->HerA/FtsK->?->?->XIS-HTH->Phage_int_SAM_3+Phage_integrase-> | 1152 | Bacteria>Terrabacteria_group>Actinomycetota>Actinomycetes>Mycobacteriales>Mycobacteriaceae | GCF_900133325.1 | Mycobacteroides_abscessus | BUU36_RS03865 | 789941..793399 | 36809 | BREX-system-P-loop-protein-BrxC-[Mycobacteroides-abscessus] |
| WP_074304478.1 | Type1.BREX | <-HTH_26<-Phage_int_SAM_4+Phage_integrase\|\|REase+wHTH->HTH_32+Mu-transpos_C->TnsC-ATPase->?->XIS-HTH->BrxA-DNA-binding->BrxB-BrxF-iSTAND->DUF6079*->N6-METHYLASE->PglZ->SIGMA+OB-Fold+AAA-ATPase+Lon-Protease-BrxL-><-?\|\|PHP+SbcC->Phage_integrase->HTH_26->Phage_integrase+Phage_integrase-> | 1152 | Bacteria>Terrabacteria_group>Actinomycetota>Actinomycetes>Mycobacteriales>Mycobacteriaceae | GCF_900135295.1 | Mycobacteroides_abscessus | BUT45_RS11730 | 347735..351193 | 36809 | BREX-system-P-loop-protein-BrxC-[Mycobacteroides-abscessus] |
| WP_225596454.1 | Type1.BREX | <-N6-METHYLASE<-DUF6079*<-BrxB-BrxF-iSTAND<-BrxA-DNA-binding<-XIS-HTH<-TetR-HTH<-Phage_integrase<-Phage_integrase<-XIS-HTH | 1154 | Bacteria>Terrabacteria_group>Actinomycetota>Actinomycetes>Mycobacteriales>Mycobacteriaceae | - | Mycolicibacterium_fortuitum | H8Z60_30510 | 3659..7123 | 1766 | BREX-system-P-loop-protein-BrxC-[Mycolicibacterium-fortuitum] |
| ERB52864.1 | Type1.BREX | <-MIT_C<-?<-SIGMA+OB-Fold+AAA-ATPase+Lon-Protease-BrxL<-PglZ<-N6-METHYLASE-p<-N6-METHYLASE<-DUF6079-p<-DUF6079-p<-DUF6079-p*<-BrxB-BrxF-iSTAND<-BrxA-DNA-binding<-XIS-HTH<-REase\|\|?-><-?<-PIN | 456 | Bacteria>Terrabacteria_group>Actinomycetota>Actinomycetes>Mycobacteriales>Nocardiaceae | GCA_000454285.1 | Rhodococcus_sp._P27 | N806_17080 | 18800..20170 | 1384060 | hypothetical-protein-N806_17080-[Rhodococcus-sp.-P27] |
| ERB55725.1 | Type1.BREX | <-DUF6079*<-BrxB-BrxF-iSTAND<-BrxA-DNA-binding<-XIS-HTH<-REase\|\|?-><-?<-PIN | 1016 | Bacteria>Terrabacteria_group>Actinomycetota>Actinomycetes>Mycobacteriales>Nocardiaceae | GCA_000454285.1 | Rhodococcus_sp._P27 | N806_31935 | 1..3049 | 1384060 | ATP-binding-protein,-partial-[Rhodococcus-sp.-P27] |
| CDZ92533.1 | Type1.BREX | <-HTH_36<-?\|\|35exo+BRCT-><-SIGMA+OB-Fold+AAA-ATPase+Lon-Protease-BrxL<-PglZ\|\|?->?-><-N6-METHYLASE<-DUF6079*<-BrxB-BrxF-iSTAND<-BrxA-DNA-binding\|\|LACI-HTH+PBPI->?->CN_hydrolase-> | 1158 | Bacteria>Terrabacteria_group>Actinomycetota>Actinomycetes>Mycobacteriales>Nocardiaceae | GCA_000824945.2 | Rhodococcus_ruber | RHRU231_960062 | 73339..76815 | 1830 | putative-P-loop-containing-nucleoside-triphosphate-hydrolase-[Rhodococcus-ruber] |
| KSZ56505.1 | Type1.BREX | HDGYP+HDGYP-><-Glyoxalase<-ABhydrolase<-?\|\|?->?->Trypsin+REase-><-HTH_21+rve+rve_3<-HIN-HTH<-DUF6079* | 1039 | Bacteria>Terrabacteria_group>Actinomycetota>Actinomycetes>Mycobacteriales>Nocardiaceae | GCA_001465325.1 | Rhodococcus_pyridinivorans_KG-16 | Z045_23210 | 104692..107812 | 1441730 | ATP-binding-protein,-partial-[Rhodococcus-pyridinivorans-KG-16] |
| OFE08162.1 | Type1.BREX | <-Pkinase+TPR+TPR<-?<-?<-?<-?<-?\|\|tRNA-><-?\|\|BrxA-DNA-binding->BrxB-BrxF-iSTAND->DUF6079*->PglZ->SIGMA+OB-Fold+AAA-ATPase+HKD-DNase-BrxL->?->?->?-><-ParB+HNH\|\|?->?->LZ_Tnp_IS481+HTH_32+rve+rve_3-> | 1175 | Bacteria>Terrabacteria_group>Actinomycetota>Actinomycetes>Mycobacteriales>Nocardiaceae | GCA_001757665.1 | Rhodococcus_sp._1139 | A5N83_14165 | 58033..61560 | 1833762 | ATP-binding-protein-[Rhodococcus-sp.-1139] |
| ORL42035.1 | Type1.BREX | <-ANK+ANK<-SIGMA+OB-Fold+AAA-ATPase+HKD-DNase-BrxL<-AbiEii+ZnR<-AbiEi_4+AbiEi_1<-sGTP<-sGTP+sGTP+GTPase-AIG<-PglZ<-N6-METHYLASE<-DUF6079*<-BrxB-BrxF-iSTAND<-BrxA-DNA-binding | 1188 | Bacteria>Terrabacteria_group>Actinomycetota>Actinomycetes>Mycobacteriales>Nocardiaceae | GCA_002094315.1 | Prescottella_equi | A6F59_07450 | 52967..56533 | 43767 | hypothetical-protein-A6F59_07450-[Prescottella-equi] |
| RGP48319.1 | Type1.BREX | <-GDResolvase+HIN-HTH<-SIGMA+OB-Fold+AAA-ATPase+HKD-DNase-BrxL<-PglZ<-N6-METHYLASE<-DUF6079*<-BrxB-BrxF-iSTAND-p | 1175 | Bacteria>Terrabacteria_group>Actinomycetota>Actinomycetes>Mycobacteriales>Nocardiaceae | GCA_003444715.1 | Rhodococcus_erythropolis | AWH04_08905 | 69911..73438 | 1833 | ATP-binding-protein-[Rhodococcus-erythropolis] |
| MBM4589636.1 | Type1.BREX | <-ANK+ANK<-SIGMA+OB-Fold+AAA-ATPase+HKD-DNase-BrxL<-AbiEii+ZnR<-AbiEi_4+AbiEi_1<-sGTP<-sGTP<-ZnR<-sGTP<-PglZ<-N6-METHYLASE<-DUF6079*<-BrxB-BrxF-iSTAND<-BrxA-DNA-binding<-?\|\|?-><-Glyoxalase\|\|?->?->Calcineurin->MalT+LuxR-HTH-> | 1188 | Bacteria>Terrabacteria_group>Actinomycetota>Actinomycetes>Mycobacteriales>Nocardiaceae | GCA_016879295.1 | Prescottella_equi | GS507_15715 | 26092..29658 | 43767 | BREX-system-P-loop-protein-BrxC-[Prescottella-equi] |
| MBM4523030.1 | Type1.BREX | <-MalT+LuxR-HTH<-Calcineurin<-?<-?\|\|Glyoxalase-><-?\|\|?->BrxA-DNA-binding->BrxB-BrxF-iSTAND->DUF6079*->N6-METHYLASE->PglZ->sGTP+sGTP+GTPase-AIG->sGTP->AbiEi_4+AbiEi_1->AbiEii+ZnR->SIGMA+OB-Fold+AAA-ATPase+HKD-DNase-BrxL->ANK+ANK-> | 1188 | Bacteria>Terrabacteria_group>Actinomycetota>Actinomycetes>Mycobacteriales>Nocardiaceae | GCA_016879495.1 | Prescottella_equi | GS442_03230 | 599834..603400 | 43767 | BREX-system-P-loop-protein-BrxC-[Prescottella-equi] |
| MBP1050938.1 | Type1.BREX | <-REase<-?<-?<-ARG\|\|NUDIX->BrxA-DNA-binding->BrxB-BrxF-iSTAND->DUF6079*->N6-METHYLASE->PglZ->SIGMA+OB-Fold+AAA-ATPase+HKD-DNase-BrxL-><-?<-?\|\|GDResolvase+HIN-HTH-><-ZnR\|\|HISKIN-> | 1171 | Bacteria>Terrabacteria_group>Actinomycetota>Actinomycetes>Mycobacteriales>Nocardiaceae | GCA_017831275.1 | Rhodococcus_qingshengii | J6397_12385 | 68679..72194 | 334542 | BREX-system-P-loop-protein-BrxC-[Rhodococcus-qingshengii] |
| MBT2268266.1 | Type1.BREX | <-SIGMA+OB-Fold+AAA-ATPase+HKD-DNase-BrxL<-PglZ<-N6-METHYLASE-p<-N6-METHYLASE-p<-DUF6079*<-BrxB-BrxF-iSTAND<-BrxA-DNA-binding<-?<-TIR-like | 1170 | Bacteria>Terrabacteria_group>Actinomycetota>Actinomycetes>Mycobacteriales>Nocardiaceae | GCA_018615075.1 | Rhodococcus_erythropolis | J7E74_25040 | 11077..14589 | 1833 | BREX-system-P-loop-protein-BrxC-[Rhodococcus-erythropolis] |
| MBY6706229.1 | Type1.BREX | <-SF1-SF2-Helicase+DUF1998<-wHTH<-LON-AAA<-REase\|\|?-><-ABhydrolase+ABhydrolase<-HNH\|\|BrxA-DNA-binding->BrxB-BrxF-iSTAND->DUF6079*->N6-METHYLASE->PglZ->SIGMA+OB-Fold+AAA-ATPase+Lon-Protease-BrxL->ParB+HNH->\|\|Exonuc_V_gamma+REase->SF1-SF2-Helicase+REase-><-SF1-SF2-Helicase+Classical-AAA<-HD | 1147 | Bacteria>Terrabacteria_group>Actinomycetota>Actinomycetes>Mycobacteriales>Nocardiaceae | GCA_019836145.1 | Rhodococcus_sp._BP-241 | HQ308_05385 | 442671..446114 | 2739441 | BREX-system-P-loop-protein-BrxC-[Rhodococcus-sp.-BP-241] |
| BDU00126.1 | Type1.BREX | <-ABhydrolase\|\|Recombinase+ZnR->BrxA-DNA-binding->BrxB-BrxF-iSTAND->DUF6079*->N6-METHYLASE->?->PglZ->SIGMA+OB-Fold+AAA-ATPase+HKD-DNase-BrxL-><-HTH_21+rve+rve_3\|\|HIN-HTH->HTH_Tnp_1+HTH_21+rve+rve_3-><-HTH_21+rve+rve_3<-HIN-HTH | 1173 | Bacteria>Terrabacteria_group>Actinomycetota>Actinomycetes>Mycobacteriales>Nocardiaceae | GCA_027924405.1 | Nocardia_sp._ON39_IFM12276 | IFM12276_31540 | 3396523..3400044 | 2984338 | hypothetical-protein-IFM12276_31540-[Nocardia-sp.-ON39_IFM12276] |
| WP_020907622.1 | Type1.BREX | <-GDResolvase\|\|?-><-PIN<-?<-GDResolvase+HIN-HTH<-?<-SIGMA+OB-Fold+AAA-ATPase+HKD-DNase-BrxL<-PglZ<-N6-METHYLASE<-DUF6079*<-BrxB-BrxF-iSTAND<-BrxA-DNA-binding<-XIS-HTH<-Phage_integrase | 1168 | Bacteria>Terrabacteria_group>Actinomycetota>Actinomycetes>Mycobacteriales>Nocardiaceae | GCF_000010105.1 | Rhodococcus_erythropolis | RER_RS16440 | 3107254..3110760 | 1833 | BREX-system-P-loop-protein-BrxC-[Rhodococcus-erythropolis] |
| WP_039798161.1 | Type1.BREX | HTH+REase->?->?-><-DDE_Tnp_ISL3<-DDE_Tnp_ISL3<-DEDD_Tnp_IS110+Transposase_20\|\|BrxA-DNA-binding->BrxB-BrxF-iSTAND->DUF6079*->N6-METHYLASE->?->PglZ->SIGMA+OB-Fold+AAA-ATPase+HKD-DNase-BrxL-><-?<-?\|\|NUDIX-><-?<-RADICAL-SAM | 1173 | Bacteria>Terrabacteria_group>Actinomycetota>Actinomycetes>Mycobacteriales>Nocardiaceae | GCF_000308435.1 | Nocardia_araoensis | ON21_RS13315 | 16578..20099 | 228600 | BREX-system-P-loop-protein-BrxC-[Nocardia-araoensis] |
| WP_029256831.1 | Type1.BREX | ParB+HNH-><-?<-?<-SIGMA+OB-Fold+AAA-ATPase+HKD-DNase-BrxL<-PglZ<-N6-METHYLASE<-DUF6079*<-BrxB-BrxF-iSTAND<-BrxA-DNA-binding\|\|?-><-tRNA\|\|?->?->?->?->Pkinase+TPR+TPR-> | 1173 | Bacteria>Terrabacteria_group>Actinomycetota>Actinomycetes>Mycobacteriales>Nocardiaceae | GCF_000696675.2 | Rhodococcus_erythropolis | H351_RS02855 | 635383..638904 | 1833 | BREX-system-P-loop-protein-BrxC-[Rhodococcus-erythropolis] |
| WP_032384849.1 | Type1.BREX | <-EVE+REase<-ParB+HNH\|\|SF1-SF2-Helicase+REase->SF1-SF2-Helicase+HTH-><-HerA/FtsK<-SIR2<-SIGMA+OB-Fold+AAA-ATPase+HKD-DNase-BrxL<-PglZ<-N6-METHYLASE<-DUF6079*<-BrxB-BrxF-iSTAND<-BrxA-DNA-binding<-XIS-HTH<-REC+OmpR-HTH<-HisKA_3+HATPase_c\|\|DUF4352->?->URI-DNase-> | 1155 | Bacteria>Terrabacteria_group>Actinomycetota>Actinomycetes>Mycobacteriales>Nocardiaceae | GCF_000760875.1 | Rhodococcus_fascians | BH91_RS0103225 | 150682..154149 | 1828 | BREX-system-P-loop-protein-BrxC-[Rhodococcus-fascians] |
| WP_045066308.1 | Type1.BREX | IG+IG-><-?\|\|Acyltransferase->?-><-MORC+DUF3684+REase<-?\|\|BrxA-DNA-binding->BrxB-BrxF-iSTAND->DUF6079*->N6-METHYLASE->PglZ->SIGMA+OB-Fold+AAA-ATPase+HKD-DNase-BrxL->?-><-?<-?<-?\|\|HAD+HAD-> | 1173 | Bacteria>Terrabacteria_group>Actinomycetota>Actinomycetes>Mycobacteriales>Nocardiaceae | GCF_000949305.1 | unclassified_Rhodococcus_(in:_high_G+C_Gram-positive_bacteria) | SZ00_RS11305 | 2127210..2130731 | 192944 | MULTISPECIES:-BREX-system-P-loop-protein-BrxC-[unclassified-Rhodococcus-(in:-high-G+C-Gram-positive-bacteria)] |
| WP_060654441.1 | Type1.BREX | Trypsin+REase->?->?-><-?\|\|HTH_36-><-HIN-HTH<-N6-METHYLASE<-DUF6079* | 1039 | Bacteria>Terrabacteria_group>Actinomycetota>Actinomycetes>Mycobacteriales>Nocardiaceae | GCF_001465325.1 | Rhodococcus_pyridinivorans | Z045_RS23165 | 104692..107812 | 103816 | BREX-system-P-loop-protein-BrxC,-partial-[Rhodococcus-pyridinivorans] |
| WP_062991956.1 | Type1.BREX | <-MACRODOMAIN<-?\|\|HTH_3->ACET->SWACOS->Cysteine-Desulfurase->Pkinase->BrxA-DNA-binding->BrxB-BrxF-iSTAND->DUF6079*->N6-METHYLASE->?->PglZ->SIGMA+OB-Fold+AAA-ATPase+HKD-DNase-BrxL-><-URI-DNase+RAMA<-?<-AbiEii | 1173 | Bacteria>Terrabacteria_group>Actinomycetota>Actinomycetes>Mycobacteriales>Nocardiaceae | GCF_001613085.1 | Nocardia_salmonicida | NS1_RS36345 | 21477..24998 | 53431 | BREX-system-P-loop-protein-BrxC-[Nocardia-salmonicida] |
| WP_154100274.1 | Type1.BREX | BrxA-DNA-binding->BrxB-BrxF-iSTAND->DUF6079*->N6-METHYLASE->PglZ->SIGMA+OB-Fold+AAA-ATPase+HKD-DNase-BrxL->?-><-?<-RAMA<-?<-?\|\|GDResolvase+HIN-HTH-> | 1172 | Bacteria>Terrabacteria_group>Actinomycetota>Actinomycetes>Mycobacteriales>Nocardiaceae | GCF_001653035.1 | Rhodococcus_sp._YH3-3 | YH3_RS02030 | 410612..414130 | 1803579 | BREX-system-P-loop-protein-BrxC-[Rhodococcus-sp.-YH3-3] |
| WP_065351167.1 | Type1.BREX | XIS-HTH-><-?<-MarR-HTH<-?<-?<-?\|\|?->BrxA-DNA-binding->BrxB-BrxF-iSTAND->DUF6079*->N6-METHYLASE->PglZ->SIGMA+OB-Fold+AAA-ATPase+Lon-Protease-BrxL-><-HHH_4<-ParA-Soj-PloopNTPase<-?<-MORC<-SIGMA-HTH | 1149 | Bacteria>Terrabacteria_group>Actinomycetota>Actinomycetes>Mycobacteriales>Nocardiaceae | GCF_001682295.1 | Rhodococcus_sp._008 | AOT96_RS00395 | 76782..80231 | 1723645 | BREX-system-P-loop-protein-BrxC-[Rhodococcus-sp.-008] |
| WP_070161347.1 | Type1.BREX | <-Pkinase+TPR+TPR<-?<-?<-?<-?\|\|tRNA-><-?\|\|BrxA-DNA-binding->BrxB-BrxF-iSTAND->DUF6079*->N6-METHYLASE->N6-METHYLASE-p->PglZ->SIGMA+OB-Fold+AAA-ATPase+HKD-DNase-BrxL->?->?-><-ParB+HNH | 1175 | Bacteria>Terrabacteria_group>Actinomycetota>Actinomycetes>Mycobacteriales>Nocardiaceae | GCF_001757665.1 | Rhodococcus_sp._1139 | A5N83_RS14165 | 58033..61560 | 1833762 | BREX-system-P-loop-protein-BrxC-[Rhodococcus-sp.-1139] |
| WP_032407000.1 | Type1.BREX | <-DUF4352\|\|HisKA_3+HATPase_c->REC+OmpR-HTH->REase->?->XIS-HTH->BrxA-DNA-binding->BrxB-BrxF-iSTAND->DUF6079*->N6-METHYLASE->PglZ->SIGMA+OB-Fold+AAA-ATPase+HKD-DNase-BrxL->SIR2->HerA/FtsK-><-SF1-SF2-Helicase+HTH<-SF1-SF2-Helicase+REase\|\|ParB+HNH->EVE+REase-> | 1155 | Bacteria>Terrabacteria_group>Actinomycetota>Actinomycetes>Mycobacteriales>Nocardiaceae | GCF_001894785.1 | Rhodococcus_fascians | BI05_RS0109965 | 97728..101195 | 1828 | BREX-system-P-loop-protein-BrxC-[Rhodococcus-fascians] |
| WP_080909365.1 | Type1.BREX | <-GDResolvase+HIN-HTH\|\|?->?->RAMA->?->?-><-SIGMA+OB-Fold+AAA-ATPase+HKD-DNase-BrxL<-PglZ<-N6-METHYLASE<-DUF6079*<-BrxB-BrxF-iSTAND<-BrxA-DNA-binding\|\|?-><-tRNA\|\|?->?->?->?->Pkinase+TPR+TPR-> | 1173 | Bacteria>Terrabacteria_group>Actinomycetota>Actinomycetes>Mycobacteriales>Nocardiaceae | GCF_002076325.1 | Rhodococcus_sp._66b | B0E55_RS07060 | 288553..292074 | 1945511 | BREX-system-P-loop-protein-BrxC-[Rhodococcus-sp.-66b] |
| WP_081557823.1 | Type1.BREX | <-PIN<-?<-GDResolvase+HIN-HTH<-?<-SIGMA+OB-Fold+AAA-ATPase+HKD-DNase-BrxL<-PglZ<-N6-METHYLASE-p<-N6-METHYLASE<-DUF6079*<-BrxB-BrxF-iSTAND<-BrxA-DNA-binding<-XIS-HTH<-Phage_integrase | 1168 | Bacteria>Terrabacteria_group>Actinomycetota>Actinomycetes>Mycobacteriales>Nocardiaceae | GCF_002079265.1 | Rhodococcus_sp._BH4 | A0W34_RS13850 | 2965873..2969379 | 1807790 | BREX-system-P-loop-protein-BrxC-[Rhodococcus-sp.-BH4] |
| WP_088896782.1 | Type1.BREX | ParB+HNH->nSTAND1+HNH-><-?\|\|XIS-HTH->BrxA-DNA-binding->BrxB-BrxF-iSTAND->DUF6079*->N6-METHYLASE->PglZ->SIGMA+OB-Fold+AAA-ATPase+HKD-DNase-BrxL->?-><-HTH_21+rve+rve_3 | 1154 | Bacteria>Terrabacteria_group>Actinomycetota>Actinomycetes>Mycobacteriales>Nocardiaceae | GCF_002215235.1 | Rhodococcus_sp._BUPNP1 | B9C99_RS01850 | 433431..436895 | 1432786 | BREX-system-P-loop-protein-BrxC-[Rhodococcus-sp.-BUPNP1] |
| WP_094685397.1 | Type1.BREX | <-CASPASE\|\|SF1-SF2-Helicase+REase+HTH->SF1-SF2-Helicase+HTH-><-HARE-HTH<-SIGMA+OB-Fold+AAA-ATPase+Lon-Protease-BrxL<-PglZ<-N6-METHYLASE<-DUF6079*<-BrxB-BrxF-iSTAND<-BrxA-DNA-binding\|\|?->Exonuc_V_gamma+REase->SF1-SF2-Helicase+REase-><-McrC-NTD+REase<-HTH+McrB-AAA<-REC+OmpR-HTH<-HisKA_3+HATPase_c | 1150 | Bacteria>Terrabacteria_group>Actinomycetota>Actinomycetes>Mycobacteriales>Nocardiaceae | GCF_002259365.1 | Rhodococcus_sp._14-2496-1d | CH296_RS18070 | 16030..19482 | 2023146 | BREX-system-P-loop-protein-BrxC-[Rhodococcus-sp.-14-2496-1d] |

| Protein-ID | System | Gene-Neighbourhoods | Length | Taxonomy | Assembly-ID | Strain | Locus-Tag | Genomic-coordinates | TaxID | Fasta-defline |
|---|---|---|---|---|---|---|---|---|---|---|
| WP_094721084.1 | Type1.BREX | [illegible] | 1174 | Bacteria>Terrabacteria_group>Actinomycetota>Actinomycetes>Mycobacteriales>Nocardiaceae | GCF_002259415.1 | Rhodococcus_sp._14-2470-1b | CH293_RS18620 | 24333..27857 | 2023149 | BREX-system-P-loop-protein-BrxC-[Rhodococcus-sp.-14-2470-1b] |
| WP_198387782.1 | Type1.BREX | [illegible] | 1173 | Bacteria>Terrabacteria_group>Actinomycetota>Actinomycetes>Mycobacteriales>Nocardiaceae | GCF_002269365.1 | Rhodococcus_pyridinivorans | CJU69_RS21710 | 4620785..4624306 | 103816 | BREX-system-P-loop-protein-BrxC-[Rhodococcus-pyridinivorans] |
| WP_122191117.1 | Type1.BREX | [illegible] | 1173 | Bacteria>Terrabacteria_group>Actinomycetota>Actinomycetes>Mycobacteriales>Nocardiaceae | GCF_003696265.1 | Nocardia_stercoris | EBN03_RS27825 | 64799..68320 | 2483361 | BREX-system-P-loop-protein-BrxC-[Nocardia-stercoris] |
| WP_149431764.1 | Type1.BREX | [illegible] | 1184 | Bacteria>Terrabacteria_group>Actinomycetota>Actinomycetes>Mycobacteriales>Nocardiaceae | GCF_008297975.1 | Rhodococcus_cavernicola | FOY51_RS18630 | 121957..125511 | 2495913 | BREX-system-P-loop-protein-BrxC-[Rhodococcus-cavernicola] |
| WP_225320218.1 | Type1.BREX | [illegible] | 435 | Bacteria>Terrabacteria_group>Actinomycetota>Actinomycetes>Mycobacteriales>Nocardiaceae | GCF_011611865.1 | Rhodococcus_erythropolis | G9444_RS32380 | 46132..47439 | 1833 | BREX-system-P-loop-protein-BrxC-[Rhodococcus-erythropolis] |
| WP_191902974.1 | Type1.BREX | [illegible] | 769 | Bacteria>Terrabacteria_group>Actinomycetota>Actinomycetes>Mycobacteriales>Nocardiaceae | GCF_014837135.1 | Rhodococcus_ruber | ICV35_RS24625 | 37953..40259 | 1830 | BREX-system-P-loop-protein-BrxC,-partial-[Rhodococcus-ruber] |
| WP_194984549.1 | Type1.BREX | [illegible] | 1159 | Bacteria>Terrabacteria_group>Actinomycetota>Actinomycetes>Mycobacteriales>Nocardiaceae | GCF_015477645.1 | Nocardia_cyriacigeorgica | IU457_RS27110 | 9259..12738 | 135487 | BREX-system-P-loop-protein-BrxC-[Nocardia-cyriacigeorgica] |
| WP_195040827.1 | Type1.BREX | [illegible] | 1159 | Bacteria>Terrabacteria_group>Actinomycetota>Actinomycetes>Mycobacteriales>Nocardiaceae | GCF_015477975.1 | Nocardia_cyriacigeorgica | IU440_RS19520 | 298293..301772 | 135487 | BREX-system-P-loop-protein-BrxC-[Nocardia-cyriacigeorgica] |
| WP_238424144.1 | Type1.BREX | [illegible] | 1175 | Bacteria>Terrabacteria_group>Actinomycetota>Actinomycetes>Mycobacteriales>Nocardiaceae | GCF_016804345.1 | Rhodococcus_ruber | F1734_RS10025 | 2133745..2137272 | 1830 | BREX-system-P-loop-protein-BrxC-[Rhodococcus-ruber] |
| WP_203329368.1 | Type1.BREX | [illegible] | 1149 | Bacteria>Terrabacteria_group>Actinomycetota>Actinomycetes>Mycobacteriales>Nocardiaceae | GCF_016839185.1 | Rhodococcus | JYA75_RS22365 | 4928975..4932424 | 1827 | MULTISPECIES:-BREX-system-P-loop-protein-BrxC-[Rhodococcus] |
| WP_245219430.1 | Type1.BREX | [illegible] | 1173 | Bacteria>Terrabacteria_group>Actinomycetota>Actinomycetes>Mycobacteriales>Nocardiaceae | GCF_017831275.1 | Rhodococcus_qingshengii | J6397_RS12385 | 68673..72194 | 334542 | BREX-system-P-loop-protein-BrxC-[Rhodococcus-qingshengii] |
| WP_222635591.1 | Type1.BREX | [illegible] | 1188 | Bacteria>Terrabacteria_group>Actinomycetota>Actinomycetes>Mycobacteriales>Nocardiaceae | GCF_019834785.1 | unclassified_Rhodococcus_(in:_high_G+C_Gram-positive_bacteria) | HQ363_RS22195 | 12044..15610 | 192944 | MULTISPECIES:-BREX-system-P-loop-protein-BrxC-[unclassified-Rhodococcus-(in:-high-G+C-Gram-positive-bacteria)] |
| WP_261775798.1 | Type1.BREX | [illegible] | 1142 | Bacteria>Terrabacteria_group>Actinomycetota>Actinomycetes>Mycobacteriales>Nocardiaceae | GCF_019836145.1 | Rhodococcus_sp._BP-241 | HQ308_RS05380 | 442686..446114 | 2739441 | BREX-system-P-loop-protein-BrxC-[Rhodococcus-sp.-BP-241] |
| WP_230831750.1 | Type1.BREX | [illegible] | 1158 | Bacteria>Terrabacteria_group>Actinomycetota>Actinomycetes>Mycobacteriales>Nocardiaceae | GCF_021025995.1 | Rhodococcus_ruber | O4273_RS26160 | 16301..19777 | 1830 | BREX-system-P-loop-protein-BrxC-[Rhodococcus-ruber] |
| WP_230793610.1 | Type1.BREX | [illegible] | 1173 | Bacteria>Terrabacteria_group>Actinomycetota>Actinomycetes>Mycobacteriales>Nocardiaceae | GCF_021026075.1 | Rhodococcus_qingshengii | LQ424_RS30730 | 10749..14270 | 334542 | BREX-system-P-loop-protein-BrxC-[Rhodococcus-qingshengii] |
| WP_231782753.1 | Type1.BREX | [illegible] | 1173 | Bacteria>Terrabacteria_group>Actinomycetota>Actinomycetes>Mycobacteriales>Nocardiaceae | GCF_021165975.1 | Rhodococcus_qingshengii | LRL17_RS24605 | 5269427..5272948 | 334542 | BREX-system-P-loop-protein-BrxC-[Rhodococcus-qingshengii] |
| WP_233062445.1 | Type1.BREX | [illegible] | 1173 | Bacteria>Terrabacteria_group>Actinomycetota>Actinomycetes>Mycobacteriales>Nocardiaceae | GCF_021183645.1 | Nocardia_gipuzkoensis | LTT66_RS33135 | 7242905..7246426 | 2749991 | BREX-system-P-loop-protein-BrxC-[Nocardia-gipuzkoensis] |
| WP_242324362.1 | Type1.BREX | [illegible] | 1188 | Bacteria>Terrabacteria_group>Actinomycetota>Actinomycetes>Mycobacteriales>Nocardiaceae | GCF_022655485.1 | Rhodococcus_opacus | MOO23_RS40495 | 105196..108762 | 37919 | BREX-system-P-loop-protein-BrxC-[Rhodococcus-opacus] |
| WP_064077541.1 | Type1.BREX | [illegible] | 1188 | Bacteria>Terrabacteria_group>Actinomycetota>Actinomycetes>Mycobacteriales>Nocardiaceae | GCF_022691665.1 | Prescottella_equi | MPC39_RS12500 | 2698129..2701695 | 43767 | BREX-system-P-loop-protein-BrxC-[Prescottella-equi] |
| WP_242421152.1 | Type1.BREX | [illegible] | 1172 | Bacteria>Terrabacteria_group>Actinomycetota>Actinomycetes>Mycobacteriales>Nocardiaceae | GCF_022691685.1 | Prescottella_equi | MPC38_RS08825 | 1859263..1862781 | 43767 | BREX-system-P-loop-protein-BrxC-[Prescottella-equi] |
| WP_156525037.1 | Type1.BREX | [illegible] | 1173 | Bacteria>Terrabacteria_group>Actinomycetota>Actinomycetes>Mycobacteriales>Nocardiaceae | GCF_023221595.1 | Rhodococcus | M0639_RS29930 | 74142..77663 | 1827 | MULTISPECIES:-BREX-system-P-loop-protein-BrxC-[Rhodococcus] |
| WP_253659814.1 | Type1.BREX | [illegible] | 1156 | Bacteria>Terrabacteria_group>Actinomycetota>Actinomycetes>Mycobacteriales>Nocardiaceae | GCF_024171815.1 | Williamsia_maris | LX13_RS03015 | 647894..651364 | 72806 | BREX-system-P-loop-protein-BrxC-[Williamsia-maris] |
| WP_155277929.1 | Type1.BREX | [illegible] | 1173 | Bacteria>Terrabacteria_group>Actinomycetota>Actinomycetes>Mycobacteriales>Nocardiaceae | GCF_024349805.1 | Rhodococcus | RE2895_RS00215 | 46904..50425 | 1827 | MULTISPECIES:-BREX-system-P-loop-protein-BrxC-[Rhodococcus] |
| WP_256520207.1 | Type1.BREX | [illegible] | 1174 | Bacteria>Terrabacteria_group>Actinomycetota>Actinomycetes>Mycobacteriales>Nocardiaceae | GCF_024498635.1 | Rhodococcus_qingshengii | LRQ08_RS00215 | 46197..49721 | 334542 | BREX-system-P-loop-protein-BrxC-[Rhodococcus-qingshengii] |
| WP_269590671.1 | Type1.BREX | [illegible] | 379 | Bacteria>Terrabacteria_group>Actinomycetota>Actinomycetes>Mycobacteriales>Nocardiaceae | GCF_027270695.1 | Rhodococcus_ruber | O4272_RS12415 | 333881..335022 | 1830 | hypothetical-protein,-partial-[Rhodococcus-ruber] |
| WP_269590680.1 | Type1.BREX | [illegible] | 812 | Bacteria>Terrabacteria_group>Actinomycetota>Actinomycetes>Mycobacteriales>Nocardiaceae | GCF_027270695.1 | Rhodococcus_ruber | O4272_RS25860 | 12..2450 | 1830 | BREX-system-P-loop-protein-BrxC-[Rhodococcus-ruber] |
| WP_269577378.1 | Type1.BREX | [illegible] | 1173 | Bacteria>Terrabacteria_group>Actinomycetota>Actinomycetes>Mycobacteriales>Nocardiaceae | GCF_027270845.1 | Rhodococcus_erythropolis | O4216_RS13595 | 117955..121476 | 1833 | BREX-system-P-loop-protein-BrxC-[Rhodococcus-erythropolis] |
| WP_118604627.1 | Type1.BREX | [illegible] | 1173 | Bacteria>Terrabacteria_group>Actinomycetota>Actinomycetes>Mycobacteriales>Nocardiaceae | - | Rhodococcus_erythropolis | AWH04_RS14655 | 69911..73432 | 1833 | BREX-system-P-loop-protein-BrxC-[Rhodococcus-erythropolis] |
| WP_214887135.1 | Type1.BREX | [illegible] | 1170 | Bacteria>Terrabacteria_group>Actinomycetota>Actinomycetes>Mycobacteriales>Nocardiaceae | - | Rhodococcus_erythropolis | J7E74_25040 | 11077..14589 | 1833 | BREX-system-P-loop-protein-BrxC-[Rhodococcus-erythropolis] |
| WP_222645759.1 | Type1.BREX | [illegible] | 1147 | Bacteria>Terrabacteria_group>Actinomycetota>Actinomycetes>Mycobacteriales>Nocardiaceae | - | Rhodococcus_sp._BP-241 | HQ308_05385 | 442671..446114 | 2739441 | BREX-system-P-loop-protein-BrxC-[Rhodococcus-sp.-BP-241] |
| TWS24839.1 | Type1.BREX | [illegible] | 1298 | Bacteria>Terrabacteria_group>Actinomycetota>Actinomycetes>Mycobacteriales>Tsukamurellaceae | GCA_007858445.1 | Tsukamurella_sputi | FK268_06235 | 209185..213081 | 2591848 | BREX-system-P-loop-protein-BrxC-[Tsukamurella-sputi] |
| NKY16937.1 | Type1.BREX | [illegible] | 1182 | Bacteria>Terrabacteria_group>Actinomycetota>Actinomycetes>Mycobacteriales>Tsukamurellaceae | GCA_012396015.1 | Tsukamurella_spumae | HF999_00885 | 181603..185151 | 44753 | BREX-system-P-loop-protein-BrxC-[Tsukamurella-spumae] |
| WP_068569480.1 | Type1.BREX | [illegible] | 1182 | Bacteria>Terrabacteria_group>Actinomycetota>Actinomycetes>Mycobacteriales>Tsukamurellaceae | GCF_001575195.1 | Tsukamurella_pseudospumae | AXK60_RS03070 | 654047..657595 | 239498 | BREX-system-P-loop-protein-BrxC-[Tsukamurella-pseudospumae] |
| WP_068743364.1 | Type1.BREX | [illegible] | 1180 | Bacteria>Terrabacteria_group>Actinomycetota>Actinomycetes>Mycobacteriales>Tsukamurellaceae | GCF_001575205.1 | Tsukamurella_pseudospumae | AXK61_RS01225 | 254752..258294 | 239498 | BREX-system-P-loop-protein-BrxC-[Tsukamurella-pseudospumae] |
| WP_158636061.1 | Type1.BREX | [illegible] | 1180 | Bacteria>Terrabacteria_group>Actinomycetota>Actinomycetes>Mycobacteriales>Tsukamurellaceae | GCF_007858445.1 | Tsukamurella_sputi | FK268_RS06235 | 209185..212727 | 2591848 | BREX-system-P-loop-protein-BrxC-[Tsukamurella-sputi] |
| WP_246207064.1 | Type1.BREX | [illegible] | 1179 | Bacteria>Terrabacteria_group>Actinomycetota>Actinomycetes>Mycobacteriales>Tsukamurellaceae | GCF_012396015.1 | Tsukamurella_spumae | HF999_RS00885 | 181612..185151 | 44753 | BREX-system-P-loop-protein-BrxC-[Tsukamurella-spumae] |
| WP_248491899.1 | Type1.BREX | [illegible] | 1180 | Bacteria>Terrabacteria_group>Actinomycetota>Actinomycetes>Mycobacteriales>Tsukamurellaceae | GCF_023168325.1 | Tsukamurella_sp._PLM1 | MTP03_RS10785 | 2075466..2079008 | 2929795 | BREX-system-P-loop-protein-BrxC-[Tsukamurella-sp.-PLM1] |
| WP_211131543.1 | Type1.BREX | [illegible] | 1144 | Bacteria>Terrabacteria_group>Actinomycetota>Actinomycetes>Mycobacteriales>Tsukamurellaceae | - | Tsukamurella_tyrosinosolvens | JVY00_15335 | 3136296..3139730 | 57704 | BREX-system-P-loop-protein-BrxC-[Tsukamurella-tyrosinosolvens] |
| TCC44056.1 | Type1.BREX | [illegible] | 1159 | Bacteria>Terrabacteria_group>Actinomycetota>Actinomycetes>Propionibacteriales>Kribbellaceae | GCA_004331485.1 | Kribbella_capetownensis | E0H75_37810 | 267915..271394 | 1572659 | BREX-system-P-loop-protein-BrxC-[Kribbella-capetownensis] |
| WP_238163516.1 | Type1.BREX | [illegible] | 1162 | Bacteria>Terrabacteria_group>Actinomycetota>Actinomycetes>Propionibacteriales>Kribbellaceae | GCF_004331485.1 | Kribbella_capetownensis | E0H75_RS37805 | 267906..271394 | 1572659 | BREX-system-P-loop-protein-BrxC-[Kribbella-capetownensis] |
| GHE18922.1 | Type1.BREX | [illegible] | 886 | Bacteria>Terrabacteria_group>Actinomycetota>Actinomycetes>Propionibacteriales>Nocardioidaceae | GCA_014653115.1 | Nocardioides_flavus_Wang_et_al._2016 | GCM10011376_35320 | 22373..25033 | 2058780 | hypothetical-protein-GCM10011376_35320-[Nocardioides-flavus-Wang-et-al.-2016] |
| MBD8606020.1 | Type1.BREX | [illegible] | 1148 | Bacteria>Terrabacteria_group>Actinomycetota>Actinomycetes>Propionibacteriales>Nocardioidaceae | GCA_014841755.1 | Aeromicrobium_sp._CFBP_8757 | IFT73_04070 | 832068..835514 | 2775288 | BREX-system-P-loop-protein-BrxC-[Aeromicrobium-sp.-CFBP-8757] |
| WP_010833228.1 | Type1.BREX | [illegible] | 1160 | Bacteria>Terrabacteria_group>Actinomycetota>Actinomycetes>Propionibacteriales>Nocardioidaceae | GCF_000389985.1 | Nocardioides_sp._CF8 | CF8_RS11160 | 2288637..2292119 | 110319 | BREX-system-P-loop-protein-BrxC-[Nocardioides-sp.-CF8] |
| WP_028707702.1 | Type1.BREX | [illegible] | 1159 | Bacteria>Terrabacteria_group>Actinomycetota>Actinomycetes>Propionibacteriales>Nocardioidaceae | GCF_000423465.1 | Propionicicella_superfundia | G555_RS0102905 | 100593..104072 | 348582 | BREX-system-P-loop-protein-BrxC-[Propionicicella-superfundia] |
| WP_068111667.1 | Type1.BREX | [illegible] | 1154 | Bacteria>Terrabacteria_group>Actinomycetota>Actinomycetes>Propionibacteriales>Nocardioidaceae | GCF_001653335.1 | Nocardioides_dokdonensis | I601_RS15600 | 3293021..3296485 | 450734 | BREX-system-P-loop-protein-BrxC-[Nocardioides-dokdonensis] |
| WP_107767053.1 | Type1.BREX | [illegible] | 1154 | Bacteria>Terrabacteria_group>Actinomycetota>Actinomycetes>Propionibacteriales>Nocardioidaceae | GCF_003047265.1 | Nocardioides_terrigena | DAE25_RS15680 | 407524..410988 | 424797 | BREX-system-P-loop-protein-BrxC-[Nocardioides-terrigena] |
| WP_116571233.1 | Type1.BREX | [illegible] | 1152 | Bacteria>Terrabacteria_group>Actinomycetota>Actinomycetes>Propionibacteriales>Nocardioidaceae | GCF_003076135.1 | Nocardioides_gansuensis | DDE18_RS05460 | 245415..248873 | 2138300 | BREX-system-P-loop-protein-BrxC-[Nocardioides-gansuensis] |
| WP_147685627.1 | Type1.BREX | [illegible] | 1154 | Bacteria>Terrabacteria_group>Actinomycetota>Actinomycetes>Propionibacteriales>Nocardioidaceae | GCF_008039565.1 | Aeromicrobium_terrae | FHP06_RS08160 | 302941..306405 | 2498846 | BREX-system-P-loop-protein-BrxC-[Aeromicrobium-terrae] |
| WP_148615130.1 | Type1.BREX | [illegible] | 1154 | Bacteria>Terrabacteria_group>Actinomycetota>Actinomycetes>Propionibacteriales>Nocardioidaceae | GCF_008087085.1 | Nocardioides_rubriscoriae | FZC61_RS15785 | 254232..257696 | 642762 | BREX-system-P-loop-protein-BrxC-[Nocardioides-rubriscoriae] |
| WP_166189200.1 | Type1.BREX | [illegible] | 1154 | Bacteria>Terrabacteria_group>Actinomycetota>Actinomycetes>Propionibacteriales>Nocardioidaceae | GCF_011250565.1 | Nocardioides_sp._IC4_145 | G6553_RS03445 | 61321..64785 | 2714037 | BREX-system-P-loop-protein-BrxC-[Nocardioides-sp.-IC4_145] |
| WP_179652935.1 | Type1.BREX | [illegible] | 1154 | Bacteria>Terrabacteria_group>Actinomycetota>Actinomycetes>Propionibacteriales>Nocardioidaceae | GCF_013408525.1 | Nocardioides_aromaticivorans | BJ993_RS25170 | 157779..161243 | 200618 | BREX-system-P-loop-protein-BrxC-[Nocardioides-aromaticivorans] |
| WP_191280810.1 | Type1.BREX | [illegible] | 943 | Bacteria>Terrabacteria_group>Actinomycetota>Actinomycetes>Propionibacteriales>Nocardioidaceae | GCF_014653115.1 | Nocardioides_flavus_Wang_et_al._2016 | IE200_RS17700 | 22202..25033 | 2058780 | BREX-system-P-loop-protein-BrxC-[Nocardioides-flavus-Wang-et-al.-2016] |
| WP_256435434.1 | Type1.BREX | [illegible] | 1142 | Bacteria>Terrabacteria_group>Actinomycetota>Actinomycetes>Propionibacteriales>Nocardioidaceae | GCF_014841755.1 | Aeromicrobium_sp._CFBP_8757 | IFT73_RS04070 | 832086..835514 | 2775288 | BREX-system-P-loop-protein-BrxC-[Aeromicrobium-sp.-CFBP-8757] |
| BAK36822.1 | Type1.BREX | [illegible] | 1176 | Bacteria>Terrabacteria_group>Actinomycetota>Actinomycetes>Propionibacteriales>Propionibacteriaceae | GCA_000270245.1 | Microlunatus_phosphovorus_NM-1 | MLP_38080 | 4107257..4110787 | 1032480 | hypothetical-protein-MLP_38080-[Microlunatus-phosphovorus-NM-1] |
| AQP50316.1 | Type1.BREX | [illegible] | 1159 | Bacteria>Terrabacteria_group>Actinomycetota>Actinomycetes>Propionibacteriales>Propionibacteriaceae | GCA_001998865.1 | Tessaracoccus_flavescens | BW733_05195 | 1055640..1059119 | 399497 | ATP-binding-protein-[Tessaracoccus-flavescens] |
| NLI84511.1 | Type1.BREX | [illegible] | 1160 | Bacteria>Terrabacteria_group>Actinomycetota>Actinomycetes>Propionibacteriales>Propionibacteriaceae | GCA_012517405.1 | Propionibacterium_sp. | GX440_03735 | 112337..115819 | 1977903 | BREX-system-P-loop-protein-BrxC-[Propionibacterium-sp.] |
| MBK7820961.1 | Type1.BREX | [illegible] | 1176 | Bacteria>Terrabacteria_group>Actinomycetota>Actinomycetes>Propionibacteriales>Propionibacteriaceae | GCA_016714275.1 | Tessaracoccus_sp. | IPJ61_07760 | 1693709..1697239 | 1971211 | BREX-system-P-loop-protein-BrxC-[Tessaracoccus-sp.] |
| MBO1030854.1 | Type1.BREX | [illegible] | 1152 | Bacteria>Terrabacteria_group>Actinomycetota>Actinomycetes>Propionibacteriales>Propionibacteriaceae | GCA_017355265.1 | Tessaracoccus_sp._SD287 | IPV09_05845 | 30832..34290 | 2782008 | BREX-system-P-loop-protein-BrxC-[Tessaracoccus-sp.-SD287] |
| MCI1746855.1 | Type1.BREX | [illegible] | 1159 | Bacteria>Terrabacteria_group>Actinomycetota>Actinomycetes>Propionibacteriales>Propionibacteriaceae | GCA_022647765.1 | Acidipropionibacterium_sp. | LKI24_01350 | 263130..266609 | 1912218 | BREX-system-P-loop-protein-BrxC-[Acidipropionibacterium-sp.] |
| WP_041792941.1 | Type1.BREX | [illegible] | 1173 | Bacteria>Terrabacteria_group>Actinomycetota>Actinomycetes>Propionibacteriales>Propionibacteriaceae | GCF_000270245.1 | Microlunatus_phosphovorus | MLP_RS18400 | 4107266..4110787 | 29405 | BREX-system-P-loop-protein-BrxC-[Microlunatus-phosphovorus] |
| WP_028706416.1 | Type1.BREX | [illegible] | 1171 | Bacteria>Terrabacteria_group>Actinomycetota>Actinomycetes>Propionibacteriales>Propionibacteriaceae | GCF_000426605.1 | Propionibacterium_acidifaciens | K336_RS0104645 | 45562..49077 | 556499 | BREX-system-P-loop-protein-BrxC-[Propionibacterium-acidifaciens] |
| WP_021796710.1 | Type1.BREX | [illegible] | 1171 | Bacteria>Terrabacteria_group>Actinomycetota>Actinomycetes>Propionibacteriales>Propionibacteriaceae | GCF_000478805.1 | Propionibacterium_acidifaciens | HMPREF0682_RS02920 | 11275..14790 | 556499 | BREX-system-P-loop-protein-BrxC-[Propionibacterium-acidifaciens] |
| WP_048768895.1 | Type1.BREX | [illegible] | 1170 | Bacteria>Terrabacteria_group>Actinomycetota>Actinomycetes>Propionibacteriales>Propionibacteriaceae | GCF_001050955.1 | Propionibacterium_freudenreichii | PFCIRM1025_RS01850 | 33525..37037 | 1744 | BREX-system-P-loop-protein-BrxC-[Propionibacterium-freudenreichii] |
| WP_055346094.1 | Type1.BREX | [illegible] | 1170 | Bacteria>Terrabacteria_group>Actinomycetota>Actinomycetes>Propionibacteriales>Propionibacteriaceae | GCF_001369055.1 | Propionibacterium_freudenreichii | PFCIRM516_RS05500 | 48312..51824 | 1744 | BREX-system-P-loop-protein-BrxC-[Propionibacterium-freudenreichii] |
| WP_237268307.1 | Type1.BREX | [illegible] | 1194 | Bacteria>Terrabacteria_group>Actinomycetota>Actinomycetes>Propionibacteriales>Propionibacteriaceae | GCF_001998865.1 | Tessaracoccus_flavescens | BW733_RS05155 | 1055640..1059224 | 399497 | BREX-system-P-loop-protein-BrxC-[Tessaracoccus-flavescens] |
| WP_094364310.1 | Type1.BREX | [illegible] | 1173 | Bacteria>Terrabacteria_group>Actinomycetota>Actinomycetes>Propionibacteriales>Propionibacteriaceae | GCF_002250625.1 | Enemella_dayhoffiae | CGZ93_RS11755 | 175760..179281 | 2016507 | BREX-system-P-loop-protein-BrxC-[Enemella-dayhoffiae] |
| WP_097828824.1 | Type1.BREX | [illegible] | 1173 | Bacteria>Terrabacteria_group>Actinomycetota>Actinomycetes>Propionibacteriales>Propionibacteriaceae | GCF_004011075.1 | Acidipropionibacterium_jensenii | C0Z11_RS09870 | 2230980..2234501 | 1749 | BREX-system-P-loop-protein-BrxC-[Acidipropionibacterium-jensenii] |
| WP_182782739.1 | Type1.BREX | [illegible] | 1159 | Bacteria>Terrabacteria_group>Actinomycetota>Actinomycetes>Propionibacteriales>Propionibacteriaceae | GCF_014164725.1 | Tessaracoccus_sp._MC1627 | H5399_RS09110 | 126520..129999 | 2760312 | BREX-system-P-loop-protein-BrxC-[Tessaracoccus-sp.-MC1627] |
| WP_182799522.1 | Type1.BREX | [illegible] | 1173 | Bacteria>Terrabacteria_group>Actinomycetota>Actinomycetes>Propionibacteriales>Propionibacteriaceae | GCF_014164775.1 | Tessaracoccus_sp._MC1679 | H5398_RS07140 | 376087..379608 | 2760313 | BREX-system-P-loop-protein-BrxC-[Tessaracoccus-sp.-MC1679] |
| WP_232547416.1 | Type1.BREX | [illegible] | 1159 | Bacteria>Terrabacteria_group>Actinomycetota>Actinomycetes>Propionibacteriales>Propionibacteriaceae | GCF_014858005.1 | Propioniciclava_soli | IFY26_RS02480 | 540114..543593 | 2775081 | BREX-system-P-loop-protein-BrxC-[Propioniciclava-soli] |
| WP_204918880.1 | Type1.BREX | [illegible] | 1173 | Bacteria>Terrabacteria_group>Actinomycetota>Actinomycetes>Propionibacteriales>Propionibacteriaceae | GCF_016907535.1 | Microlunatus_panaciterrae | JOE57_RS13940 | 3049076..3052597 | 400768 | BREX-system-P-loop-protein-BrxC-[Microlunatus-panaciterrae] |
| WP_255521255.1 | Type1.BREX | [illegible] | 1146 | Bacteria>Terrabacteria_group>Actinomycetota>Actinomycetes>Propionibacteriales>Propionibacteriaceae | GCF_017355265.1 | Tessaracoccus_sp._SD287 | IPV09_RS05845 | 30832..34272 | 2782008 | BREX-system-P-loop-protein-BrxC-[Tessaracoccus-sp.-SD287] |
| WP_073185198.1 | Type1.BREX | [illegible] | 1159 | Bacteria>Terrabacteria_group>Actinomycetota>Actinomycetes>Propionibacteriales>Propionibacteriaceae | GCF_900141915.1 | Tessaracoccus_bendigoensis | BUC15_RS00075 | 21322..24801 | 72764 | BREX-system-P-loop-protein-BrxC-[Tessaracoccus-bendigoensis] |
| WP_073189906.1 | Type1.BREX | [illegible] | 1176 | Bacteria>Terrabacteria_group>Actinomycetota>Actinomycetes>Propionibacteriales>Propionibacteriaceae | GCF_900141915.1 | Tessaracoccus_bendigoensis | BUC15_RS15345 | 1766..5296 | 72764 | BREX-system-P-loop-protein-BrxC-[Tessaracoccus-bendigoensis] |
| WP_130875595.1 | Type1.BREX | [illegible] | 1171 | Bacteria>Terrabacteria_group>Actinomycetota>Actinomycetes>Propionibacteriales>Propionibacteriaceae | GCF_900607225.1 | [Pseudopropionibacterium]_massiliense | E0E39_RS01895 | 450669..454184 | 2220000 | BREX-system-P-loop-protein-BrxC-[[Pseudopropionibacterium] |
| CRK59369.1 | Type1.BREX | [illegible] | 1156 | Bacteria>Terrabacteria_group>Actinomycetota>Actinomycetes>Pseudonocardiales>Pseudonocardiaceae | GCA_900070365.1 | Alloactinosynnema_sp._L-07 | . | 4548794..4552264 | 1653480 | ATPase-like-protein-[Alloactinosynnema-sp.-L-07] |
| WP_027928180.1 | Type1.BREX | [illegible] | 1164 | Bacteria>Terrabacteria_group>Actinomycetota>Actinomycetes>Pseudonocardiales>Pseudonocardiaceae | GCF_000383915.1 | Amycolatopsis_benzoatilytica | AMYBE_RS0130435 | 6563032..6566526 | 346045 | BREX-system-P-loop-protein-BrxC-[Amycolatopsis-benzoatilytica] |
| WP_142096818.1 | Type1.BREX | [illegible] | 1155 | Bacteria>Terrabacteria_group>Actinomycetota>Actinomycetes>Pseudonocardiales>Pseudonocardiaceae | GCF_006717045.1 | Pseudonocardia_cypriaca | FB388_RS03550 | 743160..746627 | 882449 | BREX-system-P-loop-protein-BrxC-[Pseudonocardia-cypriaca] |
| WP_116112927.1 | Type1.BREX | [illegible] | 1159 | Bacteria>Terrabacteria_group>Actinomycetota>Actinomycetes>Pseudonocardiales>Pseudonocardiaceae | GCF_023376025.1 | Amycolatopsis | L1857_RS08640 | 1752725..1756204 | 1813 | MULTISPECIES:-BREX-system-P-loop-protein-BrxC-[Amycolatopsis] |
| WP_259630250.1 | Type1.BREX | [illegible] | 1159 | Bacteria>Terrabacteria_group>Actinomycetota>Actinomycetes>Pseudonocardiales>Pseudonocardiaceae | GCF_025061645.1 | Umezawaea_endophytica | NZH93_RS48935 | 8817..12296 | 1654476 | BREX-system-P-loop-protein-BrxC-[Umezawaea-endophytica] |
| WP_231949657.1 | Type1.BREX | [illegible] | 1159 | Bacteria>Terrabacteria_group>Actinomycetota>Actinomycetes>Pseudonocardiales>Pseudonocardiaceae | GCF_900070365.1 | Alloactinosynnema_sp._L-07 | BN1701_RS20310 | 4548794..4552273 | 1653480 | BREX-system-P-loop-protein-BrxC-[Alloactinosynnema-sp.-L-07] |
| NBT25028.1 | Type1.BREX | [illegible] | 1142 | Bacteria>Terrabacteria_group>Actinomycetota>Actinomycetes>unclassified_Actinomycetes | GCA_009922265.1 | Actinomycetia_bacterium | EBT09_00420 | 11532..14960 | 1883427 | BREX-system-P-loop-protein-BrxC-[Actinomycetia-bacterium] |
| MBU1800642.1 | Type1.BREX | [illegible] | 728 | Bacteria>Terrabacteria_group>Actinomycetota>Actinomycetes>unclassified_Actinomycetes | GCA_018823685.1 | Actinomycetia_bacterium | KKA97_00020 | 1..2189 | 1883427 | BREX-system-P-loop-protein-BrxC-[Actinomycetia-bacterium] |
| MBW3578771.1 | Type1.BREX | [illegible] | 1145 | Bacteria>Terrabacteria_group>Actinomycetota>Actinomycetes>unclassified_Actinomycetes | GCA_019347675.1 | Actinomycetia_bacterium | KY462_13720 | 21398..24835 | 1883427 | BREX-system-P-loop-protein-BrxC-[Actinomycetia-bacterium] |
| MCA0378127.1 | Type1.BREX | [illegible] | 1183 | Bacteria>Terrabacteria_group>Actinomycetota>Actinomycetes>unclassified_Actinomycetes | GCA_020161315.1 | Actinomycetia_bacterium | LCH36_01625 | 63135..66686 | 1883427 | BREX-system-P-loop-protein-BrxC-[Actinomycetia-bacterium] |
| WP_251163982.1 | Type1.BREX | [illegible] | 1193 | Bacteria>Terrabacteria_group>Actinomycetota>Coriobacteriia>Coriobacteriales>Atopobiaceae | - | Caniella_muris | IM763_04355 | 91958..95539 | 2941502 | BREX-system-P-loop-protein-BrxC-[Caniella-muris] |
| MBF0599289.1 | Type1.BREX | [illegible] | 1193 | Bacteria>Terrabacteria_group>Actinomycetota>Coriobacteriia>Coriobacteriales>Atopobiaceae | GCA_015234195.1 | Atopobiaceae_bacterium_FL090493 | IM763_04355 | 91958..95539 | 2661918 | BREX-system-P-loop-protein-BrxC-[Atopobiaceae-bacterium-FL090493] |
| MBM6989329.1 | Type1.BREX | [illegible] | 1040 | Bacteria>Terrabacteria_group>Actinomycetota>Coriobacteriia>Coriobacteriales>Atopobiaceae | GCA_016901295.1 | Parafannyhessea_umbonata | I3I99_06995 | 31435..34557 | 604330 | BREX-system-P-loop-protein-BrxC-[Parafannyhessea-umbonata] |
| MBR1828335.1 | Type1.BREX | [illegible] | 965 | Bacteria>Terrabacteria_group>Actinomycetota>Coriobacteriia>Coriobacteriales>Atopobiaceae | GCA_017479665.1 | Atopobiaceae_bacterium | IJ781_02320 | 5917..8812 | 2485926 | BREX-system-P-loop-protein-BrxC-[Atopobiaceae-bacterium] |
| MBR3314870.1 | Type1.BREX | [illegible] | 1189 | Bacteria>Terrabacteria_group>Actinomycetota>Coriobacteriia>Coriobacteriales>Atopobiaceae | GCA_017510745.1 | Atopobiaceae_bacterium | IKG18_12110 | 33041..36610 | 2485926 | BREX-system-P-loop-protein-BrxC-[Atopobiaceae-bacterium] |
| MBR3327180.1 | Type1.BREX | [illegible] | 1188 | Bacteria>Terrabacteria_group>Actinomycetota>Coriobacteriia>Coriobacteriales>Atopobiaceae | GCA_017513965.1 | Atopobiaceae_bacterium | IKG22_07650 | 7166..10732 | 2485926 | BREX-system-P-loop-protein-BrxC-[Atopobiaceae-bacterium] |
| MBR3160180.1 | Type1.BREX | [illegible] | 1186 | Bacteria>Terrabacteria_group>Actinomycetota>Coriobacteriia>Coriobacteriales>Atopobiaceae | GCA_017533825.1 | Atopobiaceae_bacterium | IKF14_13935 | 9053..12613 | 2485926 | BREX-system-P-loop-protein-BrxC-[Atopobiaceae-bacterium] |
| HIZ18751.1 | Type1.BREX | [illegible] | 1177 | Bacteria>Terrabacteria_group>Actinomycetota>Coriobacteriia>Coriobacteriales>Atopobiaceae | GCA_019116265.1 | Candidatus_Olsenella_stercoravium | IAA22_06560 | 20591..24124 | 2838713 | BREX-system-P-loop-protein-BrxC-[Candidatus-Olsenella-stercoravium] |
| MCC6106578.1 | Type1.BREX | [illegible] | 1195 | Bacteria>Terrabacteria_group>Actinomycetota>Coriobacteriia>Coriobacteriales>Atopobiaceae | GCA_020844585.1 | Atopobium_sp. | LIV26_02205 | 3285..6872 | 1872650 | BREX-system-P-loop-protein-BrxC-[Atopobium-sp.] |
| MCC6107793.1 | Type1.BREX | [illegible] | 1194 | Bacteria>Terrabacteria_group>Actinomycetota>Coriobacteriia>Coriobacteriales>Atopobiaceae | GCA_020844585.1 | Atopobium_sp. | LIV26_08545 | 381..3965 | 1872650 | BREX-system-P-loop-protein-BrxC-[Atopobium-sp.] |
| MCC6107938.1 | Type1.BREX | [illegible] | 1196 | Bacteria>Terrabacteria_group>Actinomycetota>Coriobacteriia>Coriobacteriales>Atopobiaceae | GCA_020844585.1 | Atopobium_sp. | LIV26_09295 | 8478..12068 | 1872650 | BREX-system-P-loop-protein-BrxC-[Atopobium-sp.] |
| MCH4081490.1 | Type1.BREX | [illegible] | 1194 | Bacteria>Terrabacteria_group>Actinomycetota>Coriobacteriia>Coriobacteriales>Atopobiaceae | GCA_022483965.1 | Atopobiaceae_bacterium | LKE92_05045 | 1068955..1072539 | 2485926 | BREX-system-P-loop-protein-BrxC-[Atopobiaceae-bacterium] |
| MCI1665141.1 | Type1.BREX | [illegible] | 1166 | Bacteria>Terrabacteria_group>Actinomycetota>Coriobacteriia>Coriobacteriales>Atopobiaceae | GCA_022648095.1 | Atopobiaceae_bacterium | LKI25_02075 | 510347..513847 | 2485926 | BREX-system-P-loop-protein-BrxC-[Atopobiaceae-bacterium] |
| SFW99921.1 | Type1.BREX | [illegible] | 1199 | Bacteria>Terrabacteria_group>Actinomycetota>Coriobacteriia>Coriobacteriales>Atopobiaceae | GCA_900119625.1 | Olsenella_sp._kh2p3 | SAMN04487823_101110 | 126088..129687 | 1797112 | hypothetical-protein-SAMN04487823_101110-[Olsenella-sp.-kh2p3] |
| WP_154435944.1 | Type1.BREX | [illegible] | 1196 | Bacteria>Terrabacteria_group>Actinomycetota>Coriobacteriia>Coriobacteriales>Atopobiaceae | GCF_009695875.1 | Olsenella_porci | FYJ68_RS09105 | 10364..13954 | 2652279 | BREX-system-P-loop-protein-BrxC-[Olsenella-porci] |
| WP_204829513.1 | Type1.BREX | [illegible] | 1193 | Bacteria>Terrabacteria_group>Actinomycetota>Coriobacteriia>Coriobacteriales>Atopobiaceae | GCF_905219715.1 | Olsenella_sp._CU969 | JX283_RS04440 | 12671..16252 | 2780101 | BREX-system-P-loop-protein-BrxC-[Olsenella-sp.-CU969] |
| WP_226802729.1 | Type1.BREX | [illegible] | 1218 | Bacteria>Terrabacteria_group>Actinomycetota>Coriobacteriia>Coriobacteriales>Coriobacteriaceae | - | Collinsella_aerofaciens | LIP24_01870 | 443737..447393 | 74426 | BREX-system-P-loop-protein-BrxC-[Collinsella-aerofaciens] |
| WP_226790532.1 | Type1.BREX | [illegible] | 829 | Bacteria>Terrabacteria_group>Actinomycetota>Coriobacteriia>Coriobacteriales>Coriobacteriaceae | - | Collinsella_aerofaciens | LIP30_03585 | 83156..85645 | 74426 | BREX-system-P-loop-protein-BrxC-[Collinsella-aerofaciens] |
| WP_204537521.1 | Type1.BREX | [illegible] | 1171 | Bacteria>Terrabacteria_group>Actinomycetota>Coriobacteriia>Coriobacteriales>Coriobacteriaceae | - | Collinsella_tanakaei | H6A33_RS00145 | 33492..37007 | 626935 | BREX-system-P-loop-protein-BrxC-[Collinsella-tanakaei] |
| CDA36127.1 | Type1.BREX | [illegible] | 496 | Bacteria>Terrabacteria_group>Actinomycetota>Coriobacteriia>Coriobacteriales>Coriobacteriaceae | GCA_000434535.1 | Collinsella_sp._CAG:166 | BN511_01363 | 33711..35201 | 1262850 | uncharacterized-protein-BN511_01363-[Collinsella-sp.-CAG:166] |
| OUO19305.1 | Type1.BREX | [illegible] | 1189 | Bacteria>Terrabacteria_group>Actinomycetota>Coriobacteriia>Coriobacteriales>Coriobacteriaceae | GCA_002161805.1 | Collinsella_sp._An307 | B5F89_08645 | 123238..126807 | 1965630 | hypothetical-protein-B5F89_08645-[Collinsella-sp.-An307] |
| RHB16634.1 | Type1.BREX | [illegible] | 1276 | Bacteria>Terrabacteria_group>Actinomycetota>Coriobacteriia>Coriobacteriales>Coriobacteriaceae | GCA_003468125.1 | Collinsella_sp._AM40-7AC | DW892_08460 | 19101..22931 | 2292319 | BREX-system-P-loop-protein-BrxC-[Collinsella-sp.-AM40-7AC] |
| RRF95308.1 | Type1.BREX | [illegible] | 1196 | Bacteria>Terrabacteria_group>Actinomycetota>Coriobacteriia>Coriobacteriales>Coriobacteriaceae | GCA_003862165.1 | Coriobacteriaceae_bacterium | DUD31_01665 | 3961..7551 | 2011094 | BREX-system-P-loop-protein-BrxC-[Coriobacteriaceae-bacterium] |
| MBD9244247.1 | Type1.BREX | [illegible] | 1191 | Bacteria>Terrabacteria_group>Actinomycetota>Coriobacteriia>Coriobacteriales>Coriobacteriaceae | GCA_014847015.1 | Coriobacteriaceae_bacterium | EGQ66_00120 | 16164..19739 | 2011094 | BREX-system-P-loop-protein-BrxC-[Coriobacteriaceae-bacterium] |
| MBN2945151.1 | Type1.BREX | [illegible] | 1220 | Bacteria>Terrabacteria_group>Actinomycetota>Coriobacteriia>Coriobacteriales>Coriobacteriaceae | GCA_016939135.1 | Collinsella_sp. | JTJ24_07700 | 2990..6652 | 1965294 | BREX-system-P-loop-protein-BrxC-[Collinsella-sp.] |
| MBN2939679.1 | Type1.BREX | [illegible] | 1196 | Bacteria>Terrabacteria_group>Actinomycetota>Coriobacteriia>Coriobacteriales>Coriobacteriaceae | GCA_016939175.1 | Collinsella_sp. | JTJ11_06140 | 12590..16180 | 1965294 | BREX-system-P-loop-protein-BrxC-[Collinsella-sp.] |
| MBS4780573.1 | Type1.BREX | [illegible] | 1220 | Bacteria>Terrabacteria_group>Actinomycetota>Coriobacteriia>Coriobacteriales>Coriobacteriaceae | GCA_018363345.1 | Collinsella_sp. | KH012_09635 | 5937..9599 | 1965294 | BREX-system-P-loop-protein-BrxC-[Collinsella-sp.] |
| MBS5146167.1 | Type1.BREX | [illegible] | 1157 | Bacteria>Terrabacteria_group>Actinomycetota>Coriobacteriia>Coriobacteriales>Coriobacteriaceae | GCA_018365915.1 | Collinsella_intestinalis | KHY67_00445 | 107883..111356 | 147207 | BREX-system-P-loop-protein-BrxC-[Collinsella-intestinalis] |
| MBS5295771.1 | Type1.BREX | [illegible] | 1237 | Bacteria>Terrabacteria_group>Actinomycetota>Coriobacteriia>Coriobacteriales>Coriobacteriaceae | GCA_018367145.1 | Collinsella_sp. | KHY16_04805 | 17070..20783 | 1965294 | BREX-system-P-loop-protein-BrxC-[Collinsella-sp.] |
| MBS5500285.1 | Type1.BREX | [illegible] | 1184 | Bacteria>Terrabacteria_group>Actinomycetota>Coriobacteriia>Coriobacteriales>Coriobacteriaceae | GCA_018368855.1 | Collinsella_stercoris | KHX02_04115 | 75105..78659 | 147206 | BREX-system-P-loop-protein-BrxC-[Collinsella-stercoris] |
| MBS6611921.1 | Type1.BREX | [illegible] | 1187 | Bacteria>Terrabacteria_group>Actinomycetota>Coriobacteriia>Coriobacteriales>Coriobacteriaceae | GCA_018377935.1 | Collinsella_intestinalis | KH279_01485 | 57380..60943 | 147207 | BREX-system-P-loop-protein-BrxC-[Collinsella-intestinalis] |
| MBS6608517.1 | Type1.BREX | [illegible] | 1220 | Bacteria>Terrabacteria_group>Actinomycetota>Coriobacteriia>Coriobacteriales>Coriobacteriaceae | GCA_018377985.1 | Collinsella_sp. | KH291_01665 | 76425..80087 | 1965294 | BREX-system-P-loop-protein-BrxC-[Collinsella-sp.] |
| MBS6695654.1 | Type1.BREX | [illegible] | 1220 | Bacteria>Terrabacteria_group>Actinomycetota>Coriobacteriia>Coriobacteriales>Coriobacteriaceae | GCA_018378775.1 | Collinsella_sp. | KH246_00985 | 77204..80866 | 1965294 | BREX-system-P-loop-protein-BrxC-[Collinsella-sp.] |
| MBS6858450.1 | Type1.BREX | [illegible] | 1220 | Bacteria>Terrabacteria_group>Actinomycetota>Coriobacteriia>Coriobacteriales>Coriobacteriaceae | GCA_018380015.1 | Collinsella_sp. | KH187_04800 | 313170..316832 | 1965294 | BREX-system-P-loop-protein-BrxC-[Collinsella-sp.] |
| MBS7081308.1 | Type1.BREX | [illegible] | 1220 | Bacteria>Terrabacteria_group>Actinomycetota>Coriobacteriia>Coriobacteriales>Coriobacteriaceae | GCA_018382125.1 | Collinsella_sp. | KH077_07095 | 14975..18637 | 1965294 | BREX-system-P-loop-protein-BrxC-[Collinsella-sp.] |
| HIU24258.1 | Type1.BREX | [illegible] | 553 | Bacteria>Terrabacteria_group>Actinomycetota>Coriobacteriia>Coriobacteriales>Coriobacteriaceae | GCA_018714895.1 | Candidatus_Coprovicinus_avistercoris | IAD17_04995 | 94..1755 | 2840754 | BREX-system-P-loop-protein-BrxC-[Candidatus-Coprovicinus-avistercoris] |
| HJG30547.1 | Type1.BREX | [illegible] | 1187 | Bacteria>Terrabacteria_group>Actinomycetota>Coriobacteriia>Coriobacteriales>Coriobacteriaceae | GCA_020743295.1 | Collinsella_ihumii | K8U80_04025 | 25860..29423 | 1720204 | BREX-system-P-loop-protein-BrxC-[Collinsella-ihumii] |
| HJG31856.1 | Type1.BREX | [illegible] | 1194 | Bacteria>Terrabacteria_group>Actinomycetota>Coriobacteriia>Coriobacteriales>Coriobacteriaceae | GCA_020743295.1 | Collinsella_ihumii | K8U80_10755 | 12811..16395 | 1720204 | BREX-system-P-loop-protein-BrxC-[Collinsella-ihumii] |
| MCI5950399.1 | Type1.BREX | [illegible] | 1191 | Bacteria>Terrabacteria_group>Actinomycetota>Coriobacteriia>Coriobacteriales>Coriobacteriaceae | GCA_022770045.1 | Coriobacteriaceae_bacterium | MRZ21_06930 | 3508..7083 | 2011094 | BREX-system-P-loop-protein-BrxC-[Coriobacteriaceae-bacterium] |
| MCI6404793.1 | Type1.BREX | [illegible] | 1219 | Bacteria>Terrabacteria_group>Actinomycetota>Coriobacteriia>Coriobacteriales>Coriobacteriaceae | GCA_022777095.1 | Collinsella_sp. | MR800_00245 | 28856..32515 | 1965294 | BREX-system-P-loop-protein-BrxC-[Collinsella-sp.] |
| VWL93952.1 | Type1.BREX | [illegible] | 1220 | Bacteria>Terrabacteria_group>Actinomycetota>Coriobacteriia>Coriobacteriales>Coriobacteriaceae | GCA_902501405.1 | Collinsella_aerofaciens | OJIJPFFD_00715 | 258553..262215 | 74426 | Uncharacterised-protein-[Collinsella-aerofaciens] |
| VWL63783.1 | Type1.BREX | [illegible] | 1196 | Bacteria>Terrabacteria_group>Actinomycetota>Coriobacteriia>Coriobacteriales>Coriobacteriaceae | GCA_902501535.1 | Collinsella_aerofaciens | AOBCEHNH_00485 | 524521..528111 | 74426 | Uncharacterised-protein-[Collinsella-aerofaciens] |
| WP_006722236.1 | Type1.BREX | [illegible] | 1184 | Bacteria>Terrabacteria_group>Actinomycetota>Coriobacteriia>Coriobacteriales>Coriobacteriaceae | GCF_000156175.1 | Collinsella_intestinalis | COLINT_01988 | 37923..41477 | 147207 | BREX-system-P-loop-protein-BrxC-[Collinsella-intestinalis] |
| WP_006235452.1 | Type1.BREX | [illegible] | 1220 | Bacteria>Terrabacteria_group>Actinomycetota>Coriobacteriia>Coriobacteriales>Coriobacteriaceae | GCF_000169035.1 | Collinsella_aerofaciens | COLAER_RS06810 | 43874..47536 | 74426 | BREX-system-P-loop-protein-BrxC-[Collinsella-aerofaciens] |
| WP_087204139.1 | Type1.BREX | [illegible] | 1199 | Bacteria>Terrabacteria_group>Actinomycetota>Coriobacteriia>Coriobacteriales>Coriobacteriaceae | GCF_002159765.1 | Collinsella_sp._An268 | B5F70_RS03250 | 39703..43302 | 1965612 | BREX-system-P-loop-protein-BrxC-[Collinsella-sp.-An268] |
| WP_094336026.1 | Type1.BREX | [illegible] | 1201 | Bacteria>Terrabacteria_group>Actinomycetota>Coriobacteriia>Coriobacteriales>Coriobacteriaceae | GCF_002161605.1 | [Collinsella]_massiliensis | B5G02_RS09410 | 12439..16044 | 1232426 | BREX-system-P-loop-protein-BrxC-[[Collinsella] |
| WP_158098455.1 | Type1.BREX | [illegible] | 1186 | Bacteria>Terrabacteria_group>Actinomycetota>Coriobacteriia>Coriobacteriales>Coriobacteriaceae | GCF_002161805.1 | Collinsella_sp._An307 | B5F89_RS08645 | 123238..126798 | 1965630 | BREX-system-P-loop-protein-BrxC-[Collinsella-sp.-An307] |
| WP_099431905.1 | Type1.BREX | [illegible] | 1219 | Bacteria>Terrabacteria_group>Actinomycetota>Coriobacteriia>Coriobacteriales>Coriobacteriaceae | GCF_002736145.1 | Collinsella_aerofaciens | FX111_RS04100 | 928827..932486 | 74426 | BREX-system-P-loop-protein-BrxC-[Collinsella-aerofaciens] |
| WP_117639687.1 | Type1.BREX | [illegible] | 1220 | Bacteria>Terrabacteria_group>Actinomycetota>Coriobacteriia>Coriobacteriales>Coriobacteriaceae | GCF_003436395.1 | Collinsella_sp._TM06-3 | DXD56_RS01895 | 14278..17940 | 2292342 | BREX-system-P-loop-protein-BrxC-[Collinsella-sp.-TM06-3] |
| WP_117735584.1 | Type1.BREX | [illegible] | 1196 | Bacteria>Terrabacteria_group>Actinomycetota>Coriobacteriia>Coriobacteriales>Coriobacteriaceae | GCF_003436895.1 | Collinsella_sp._TF09-1AT | DXC90_RS08415 | 29935..33525 | 2292334 | BREX-system-P-loop-protein-BrxC-[Collinsella-sp.-TF09-1AT] |
| WP_117716812.1 | Type1.BREX | [illegible] | 1220 | Bacteria>Terrabacteria_group>Actinomycetota>Coriobacteriia>Coriobacteriales>Coriobacteriaceae | GCF_003438115.1 | Collinsella_sp._OM08-14AT | DXC18_RS07380 | 14092..17754 | 2292329 | BREX-system-P-loop-protein-BrxC-[Collinsella-sp.-OM08-14AT] |
| WP_117743051.1 | Type1.BREX | [illegible] | 1189 | Bacteria>Terrabacteria_group>Actinomycetota>Coriobacteriia>Coriobacteriales>Coriobacteriaceae | GCF_003438275.1 | Collinsella_sp._OM07-12 | DXB98_RS02540 | 4181..7750 | 2292328 | BREX-system-P-loop-protein-BrxC-[Collinsella-sp.-OM07-12] |
| WP_117746709.1 | Type1.BREX | [illegible] | 1220 | Bacteria>Terrabacteria_group>Actinomycetota>Coriobacteriia>Coriobacteriales>Coriobacteriaceae | GCF_003438495.1 | Collinsella_sp._TF05-9AC | DXC52_RS07155 | 15430..19092 | 2292330 | BREX-system-P-loop-protein-BrxC-[Collinsella-sp.-TF05-9AC] |
| WP_117776129.1 | Type1.BREX | [illegible] | 1220 | Bacteria>Terrabacteria_group>Actinomycetota>Coriobacteriia>Coriobacteriales>Coriobacteriaceae | GCF_003465485.1 | Collinsella_sp._AF11-11 | DWV58_RS02900 | 1647..5309 | 2292212 | BREX-system-P-loop-protein-BrxC-[Collinsella-sp.-AF11-11] |
| WP_117817622.1 | Type1.BREX | [illegible] | 1220 | Bacteria>Terrabacteria_group>Actinomycetota>Coriobacteriia>Coriobacteriales>Coriobacteriaceae | GCF_003465595.1 | unclassified_Collinsella | DWV43_RS07190 | 19104..22766 | 2637548 | MULTISPECIES:-BREX-system-P-loop-protein-BrxC-[unclassified-Collinsella] |
| WP_117763119.1 | Type1.BREX | [illegible] | 1220 | Bacteria>Terrabacteria_group>Actinomycetota>Coriobacteriia>Coriobacteriales>Coriobacteriaceae | GCF_003466125.1 | Collinsella_sp._TM10-22 | DXD94_RS02350 | 80369..84031 | 2292344 | BREX-system-P-loop-protein-BrxC-[Collinsella-sp.-TM10-22] |
| WP_117786732.1 | Type1.BREX | [illegible] | 1224 | Bacteria>Terrabacteria_group>Actinomycetota>Coriobacteriia>Coriobacteriales>Coriobacteriaceae | GCF_003467795.1 | Collinsella_sp._AM36-4AA | DW850_RS07585 | 35525..39199 | 2292317 | BREX-system-P-loop-protein-BrxC-[Collinsella-sp.-AM36-4AA] |
| WP_118079631.1 | Type1.BREX | [illegible] | 1220 | Bacteria>Terrabacteria_group>Actinomycetota>Coriobacteriia>Coriobacteriales>Coriobacteriaceae | GCF_003469185.1 | Collinsella_sp._AM34-10 | DW823_RS01190 | 281874..285536 | 2292316 | BREX-system-P-loop-protein-BrxC-[Collinsella-sp.-AM34-10] |

| Protein-ID | System | Gene-Neighbourhoods | Length | Taxonomy | Assembly-ID | Strain | Locus-Tag | Genomic-coordinates | TaxID | Fasta-define |
|---|---|---|---|---|---|---|---|---|---|---|
| WP_117637789.1 | Type1.BREX | [illegible] | 1225 | [illegible] | GCF_003469205.1 | unclassified_Collinsella | DW821_RS01045 | 231685..235362 | 2637548 | [illegible] |
| WP_118226697.1 | Type1.BREX | [illegible] | 1224 | [illegible] | GCF_003470945.1 | Collinsella_sp._AM23-17 | DW645_RS05360 | 15377..19051 | 2292030 | [illegible] |
| WP_118259355.1 | Type1.BREX | [illegible] | 1217 | [illegible] | GCF_003471585.1 | Collinsella_sp._AM18-10 | DW211_RS03970 | 212680..216333 | 2292028 | [illegible] |
| WP_117583764.1 | Type1.BREX | [illegible] | 1220 | [illegible] | GCF_003472325.1 | unclassified_Collinsella | DW136_RS06205 | 18574..22236 | 2637548 | [illegible] |
| WP_118273143.1 | Type1.BREX | [illegible] | 1220 | [illegible] | GCF_003472465.1 | unclassified_Collinsella | DW129_RS04200 | 27076..30738 | 2637548 | [illegible] |
| WP_117629077.1 | Type1.BREX | [illegible] | 1220 | [illegible] | GCF_003474585.1 | unclassified_Collinsella | DW037_RS01545 | 67055..70717 | 2637548 | [illegible] |
| WP_118384652.1 | Type1.BREX | [illegible] | 1220 | [illegible] | GCF_003475325.1 | Collinsella_sp._AF33-16 | DWZ52_RS00745 | 185923..189585 | 2292012 | [illegible] |
| WP_124786083.1 | Type1.BREX | [illegible] | 1220 | [illegible] | GCF_003856815.1 | Collinsella_aerofaciens | CV096_RS03705 | 854606..858268 | 74426 | [illegible] |
| WP_129423162.1 | Type1.BREX | [illegible] | 1205 | [illegible] | GCF_004135645.1 | Senegalimassilia_faecalis | ET524_RS02380 | 534170..537787 | 2509433 | [illegible] |
| WP_154515691.1 | Type1.BREX | [illegible] | 1219 | [illegible] | GCF_009696325.1 | unclassified_Collinsella | FYJ40_RS02790 | 33433..37092 | 2637548 | [illegible] |
| WP_195443573.1 | Type1.BREX | [illegible] | 1199 | [illegible] | GCF_015555045.1 | Collinsella_aerofaciens | I2E45_RS00495 | 74218..77817 | 74426 | [illegible] |
| WP_204559883.1 | Type1.BREX | [illegible] | 1171 | [illegible] | GCF_016902615.1 | Collinsella_tanakaei | H6A21_RS01790 | 55502..59017 | 626935 | [illegible] |
| WP_215694813.1 | Type1.BREX | [illegible] | 1220 | [illegible] | GCF_018785265.1 | Collinsella_aerofaciens | GPK42_RS08810 | 321305..324967 | 74426 | [illegible] |
| WP_217107944.1 | Type1.BREX | [illegible] | 1218 | [illegible] | GCF_019041595.1 | Collinsella | KTF50_RS05115 | 99865..103521 | 102106 | [illegible] |
| WP_161156402.1 | Type1.BREX | [illegible] | 1220 | [illegible] | GCF_024125125.1 | Collinsella_aerofaciens | NH935_RS00365 | 80395..84057 | 74426 | [illegible] |
| WP_260699436.1 | Type1.BREX | [illegible] | 1220 | [illegible] | GCF_025289055.1 | Collinsella_aerofaciens | JMJ97_RS05540 | 45542..49204 | 74426 | [illegible] |
| WP_270849201.1 | Type1.BREX | [illegible] | 1216 | [illegible] | GCF_027669965.1 | Candidatus_Collinsella_stercoripullorum | PHA29_RS08595 | 12169..15819 | 2838522 | [illegible] |
| WP_270255955.1 | Type1.BREX | [illegible] | 1184 | [illegible] | GCF_027670685.1 | Collinsella_intestinalis | PHA22_RS06165 | 15600..19154 | 147207 | [illegible] |
| WP_270471419.1 | Type1.BREX | [illegible] | 1184 | [illegible] | GCF_027683425.1 | Collinsella_intestinalis | PGX73_RS02685 | 38070..41624 | 147207 | [illegible] |
| WP_271743294.1 | Type1.BREX | [illegible] | 1220 | [illegible] | GCF_028206795.1 | Collinsella_aerofaciens | PMX70_RS07965 | 7863..11525 | 74426 | [illegible] |
| WP_271729196.1 | Type1.BREX | [illegible] | 1220 | [illegible] | GCF_028206955.1 | Collinsella_aerofaciens | PMX73_RS03570 | 36978..40640 | 74426 | [illegible] |
| WP_196032768.1 | Type1.BREX | [illegible] | 1196 | [illegible] | GCF_028209425.1 | Collinsella_aerofaciens | PMX18_RS06205 | 35659..39249 | 74426 | [illegible] |
| WP_271743676.1 | Type1.BREX | [illegible] | 1220 | [illegible] | GCF_028209515.1 | Collinsella_aerofaciens | PMX12_RS01585 | 140989..144651 | 74426 | [illegible] |
| WP_138340596.1 | Type1.BREX | [illegible] | 1220 | [illegible] | GCF_028210055.1 | Collinsella_aerofaciens | PMX24_RS06035 | 8884..12546 | 74426 | [illegible] |
| WP_152076869.1 | Type1.BREX | [illegible] | 1220 | [illegible] | GCF_901212465.1 | Collinsella_aerofaciens | FGK86_RS10675 | 25896..29558 | 74426 | [illegible] |
| WP_261981442.1 | Type1.BREX | [illegible] | 1191 | [illegible] | GCF_901212495.1 | Collinsella_aerofaciens | SFLMC1_RS10875 | 26184..29759 | 74426 | [illegible] |
| WP_144744499.1 | Type1.BREX | [illegible] | 1188 | [illegible] | GCF_902150035.1 | Enorma_burkinafasonensis | FS885_RS07810 | 13812..17378 | 2590867 | [illegible] |
| WP_055252035.1 | Type1.BREX | [illegible] | 1220 | [illegible] | GCF_902362815.1 | Collinsella_aerofaciens | FY253_RS06375 | 17204..20866 | 74426 | [illegible] |
| WP_118094641.1 | Type1.BREX | [illegible] | 1220 | [illegible] | GCF_902364885.1 | Collinsella | FY305_RS06610 | 68794..72456 | 102106 | [illegible] |
| WP_158567048.1 | Type1.BREX | [illegible] | 1221 | [illegible] | GCF_902364945.1 | Collinsella | FY295_RS08455 | 19266..22931 | 102106 | [illegible] |
| WP_006722026.1 | Type1.BREX | [illegible] | 1184 | [illegible] | GCF_902377675.1 | Collinsella_stercoris | COLSTE_RS09805 | 32347..35901 | 147206 | [illegible] |
| WP_226814226.1 | Type1.BREX | [illegible] | 1191 | [illegible] | GCF_902501535.1 | Collinsella_aerofaciens | GDK10_RS03455 | 524536..528111 | 74426 | [illegible] |
| WP_156063073.1 | Type1.BREX | [illegible] | 1220 | [illegible] | GCF_902501595.1 | Collinsella_aerofaciens | CKJAJONC_RS02725 | 127137..130799 | 74426 | [illegible] |
| WP_273384065.1 | Type1.BREX | [illegible] | 1183 | [illegible] | GCF_944320695.1 | Enorma_phocaeensis | PUQ31_RS06660 | 14198..17749 | 1871019 | [illegible] |
| WP_136846141.1 | Type1.BREX | [illegible] | 1192 | [illegible] | GCF_948466405.1 | Parvibacter_caecicola | E5982_RS08750 | 41748..45326 | 747645 | [illegible] |
| MBR2835999.1 | Type1.BREX | [illegible] | 1194 | [illegible] | GCA_017515595.1 | Coriobacteriales_bacterium | IKE43_09885 | 4830..8414 | 2478480 | [illegible] |
| HIT45392.1 | Type1.BREX | [illegible] | 1225 | [illegible] | GCA_018715935.1 | Candidatus_Aphodovivens_excrementavium | IAC28_04775 | 5578..9255 | 2840682 | [illegible] |
| CDB32945.1 | Type1.BREX | [illegible] | 819 | [illegible] | GCA_000436075.1 | Eggerthella_sp._CAG:209 | BN534_00740 | 18377..20836 | 1262875 | [illegible] |
| MCI8424991.1 | Type1.BREX | [illegible] | 1237 | [illegible] | GCA_022791935.1 | Adlercreutzia_sp. | HFJ72_04945 | 8870..12583 | 1872387 | [illegible] |
| MCI9261625.1 | Type1.BREX | [illegible] | 1223 | [illegible] | GCA_022798975.1 | Eggerthellaceae_bacterium | HFJ71_04015 | 1625..5296 | 1972561 | [illegible] |
| MCI9494931.1 | Type1.BREX | [illegible] | 1225 | [illegible] | GCA_022800735.1 | Adlercreutzia_mucosicola | HFJ67_04665 | 45500..49177 | 580026 | [illegible] |
| WP_012797473.1 | Type1.BREX | [illegible] | 1180 | [illegible] | GCF_000023885.1 | Slackia_heliotrinireducens | EL225_RS01045 | 248773..252315 | 84110 | [illegible] |
| WP_028027182.1 | Type1.BREX | [illegible] | 1225 | [illegible] | GCF_000422625.1 | Adlercreutzia_mucosicola | G503_RS0108070 | 76011..79688 | 580026 | [illegible] |
| WP_114615436.1 | Type1.BREX | [illegible] | 1205 | [illegible] | GCF_003340315.1 | Slackia_isoflavoniconvertens | C1881_RS05050 | 87697..91314 | 572010 | [illegible] |
| WP_123219636.1 | Type1.BREX | [illegible] | 1205 | [illegible] | GCF_003725955.1 | Slackia_isoflavoniconvertens | DMP05_RS06270 | 85657..89274 | 572010 | [illegible] |
| WP_158049617.1 | Type1.BREX | [illegible] | 1253 | [illegible] | GCF_008831035.1 | Ellagibacter_isourolithinifaciens | F8C90_06285 | 49918..53679 | 2137581 | [illegible] |
| WP_160709085.1 | Type1.BREX | [illegible] | 1222 | [illegible] | GCF_009911695.1 | Enterorhabdus_sp._P55 | D1643_RS04550 | 2737..6405 | 2304571 | [illegible] |
| WP_165048948.1 | Type1.BREX | [illegible] | 1188 | [illegible] | GCF_011038975.1 | unclassified_Adlercreutzia | G7Y50_RS02135 | 57415..60981 | 2636013 | [illegible] |
| WP_260162736.1 | Type1.BREX | [illegible] | 1200 | [illegible] | GCF_014192995.1 | Slackia_isoflavoniconvertens | FHX70_RS04860 | 1129426..1133028 | 572010 | [illegible] |
| WP_195347157.1 | Type1.BREX | [illegible] | 1201 | [illegible] | GCF_028209285.1 | Eggerthella_lenta | PMV52_RS07270 | 251002..254607 | 84112 | [illegible] |
| WP_251198431.1 | Type1.BREX | [illegible] | 1200 | [illegible] | GCF_943192915.1 | Anaerotardibacter_muris | NFR25_RS06470 | 38827..42429 | 2941505 | [illegible] |
| WP_251178638.1 | Type1.BREX | [illegible] | 1185 | [illegible] | GCF_943193125.1 | Adlercreutzia_agrestimuris | NFR28_RS04520 | 318927..322484 | 2941324 | [illegible] |
| WP_251231669.1 | Type1.BREX | [illegible] | 1213 | [illegible] | GCF_943193145.1 | Adlercreutzia_aquisgranensis | NFR41_RS09570 | 19774..23415 | 2941323 | [illegible] |
| WP_227070741.1 | Type1.BREX | [illegible] | 1199 | [illegible] | GCF_015551405.1 | Eggerthellaceae | I2D50_RS06370 | 44195..47794 | 1643826 | [illegible] |
| MBS5450116.1 | Type1.BREX | [illegible] | 1261 | [illegible] | GCA_018368455.1 | Coriobacteriia_bacterium | KHZ24_02715 | 94274..98059 | 2052159 | [illegible] |
| QIN77519.1 | Type1.BREX | [illegible] | 1167 | [illegible] | GCA_011492965.1 | Rubrobacter_marinus | GBA65_02255 | 450674..454177 | 2653852 | [illegible] |
| MBA2375851.1 | Type1.BREX | [illegible] | 1057 | [illegible] | GCA_013695475.1 | Rubrobacter_sp. | HOV75_06930 | 3189..6362 | 2036010 | [illegible] |
| MBA3426223.1 | Type1.BREX | [illegible] | 773 | [illegible] | GCA_013812245.1 | Rubrobacter_sp. | HOU04_17185 | 1..2321 | 2036010 | [illegible] |
| MBA3474041.1 | Type1.BREX | [illegible] | 1219 | [illegible] | GCA_013813465.1 | Rubrobacter_sp. | HOT57_12585 | 3865..7524 | 2036010 | [illegible] |
| WP_166395201.1 | Type1.BREX | [illegible] | 1151 | [illegible] | GCF_011492965.1 | Rubrobacter_marinus | GBA65_RS02320 | 450722..454177 | 2653852 | [illegible] |
| WP_217922216.1 | Type1.BREX | [illegible] | 1154 | [illegible] | GCF_018141545.1 | Miltoncostaea_oceani | IU369_RS17235 | 3377817..3381281 | 2843216 | [illegible] |
| NCO65434.1 | Type1.BREX | [illegible] | 1161 | [illegible] | GCA_009993245.1 | Solirubrobacter_sp. | GW874_02325 | 6745..10230 | 1932788 | [illegible] |
| MBJ7470365.1 | Type1.BREX | [illegible] | 1157 | [illegible] | GCA_016463515.1 | Solirubrobacteraceae_bacterium | JHD16_03650 | 3440..6913 | 2586742 | [illegible] |
| MCC6224163.1 | Type1.BREX | [illegible] | 1147 | [illegible] | GCA_020845695.1 | Thermoleophilia_bacterium | IT201_11820 | 14236..17679 | 2026888 | [illegible] |
| MBX3097309.1 | Type1.BREX | [illegible] | 1160 | [illegible] | GCA_019638835.1 | Fimbriimonadaceae_bacterium | KF812_10640 | 838..4320 | 2682144 | [illegible] |
| MBX7135695.1 | Type1.BREX | [illegible] | 1157 | [illegible] | GCA_019694835.1 | Fimbriimonadaceae_bacterium | K1X67_23755 | 7706..11179 | 2682144 | [illegible] |
| OIO89397.1 | Type1.BREX | [illegible] | 1150 | [illegible] | GCA_001872605.1 | Armatimonadetes_bacterium_CG2_30_59_28 | AUJ92_21575 | 2496..5948 | 1805027 | [illegible] |
| PIX47499.1 | Type1.BREX | [illegible] | 296 | [illegible] | GCA_002782725.1 | Armatimonadetes_bacterium_CG_4_8_14_3_um_filter_66_20 | COZ57_08300 | 1787..2673 | 1973912 | [illegible] |
| PIX47500.1 | Type1.BREX | [illegible] | 827 | [illegible] | GCA_002782725.1 | Armatimonadetes_bacterium_CG_4_8_14_3_um_filter_66_20 | COZ57_08305 | 2749..5234 | 1973912 | [illegible] |
| PIX45092.1 | Type1.BREX | [illegible] | 1150 | [illegible] | GCA_002782765.1 | Armatimonadetes_bacterium_CG_4_8_14_3_um_filter_58_9 | COZ56_02690 | 2386..5838 | 1973911 | [illegible] |
| HCU38259.1 | Type1.BREX | [illegible] | 696 | [illegible] | GCA_003542955.1 | Armatimonadota_bacterium | DGT21_23410 | 1803..3893 | 2033014 | [illegible] |
| MBI3929085.1 | Type1.BREX | [illegible] | 807 | [illegible] | GCA_016199555.1 | Armatimonadota_bacterium | HY319_26310 | 4685..7108 | 2033014 | [illegible] |
| MBN1458231.1 | Type1.BREX | [illegible] | 1222 | [illegible] | GCA_016928475.1 | Armatimonadota_bacterium | JXA57_01760 | 13903..17571 | 2033014 | [illegible] |
| MCC6484060.1 | Type1.BREX | [illegible] | 1150 | [illegible] | GCA_020846715.1 | Armatimonadota_bacterium | IT209_04360 | 66495..69947 | 2033014 | [illegible] |
| MCL6447260.1 | Type1.BREX | [illegible] | 1148 | [illegible] | GCA_023511535.1 | Armatimonadota_bacterium | K6U04_03765 | 21161..24607 | 2033014 | [illegible] |
| MCL6447596.1 | Type1.BREX | [illegible] | 1222 | [illegible] | GCA_023511535.1 | Armatimonadota_bacterium | K6U04_05490 | 13164..16832 | 2033014 | [illegible] |
| QSO48381.1 | Type1.BREX | [illegible] | 1175 | [illegible] | GCA_017298635.1 | Alicyclobacillus_mengziensis | JZ786_05180 | 1119199..1122726 | 2931921 | [illegible] |
| MCL6598118.1 | Type1.BREX | [illegible] | 1205 | [illegible] | GCA_023512275.1 | Alicyclobacillus_macrosporangiidus | K6T81_05195 | 736..4353 | 392015 | [illegible] |
| WP_021297234.1 | Type1.BREX | [illegible] | 1177 | [illegible] | GCF_000444055.1 | Alicyclobacillus_acidoterrestris | N007_RS35530 | 2450..5983 | 1450 | [illegible] |
| WP_067624659.1 | Type1.BREX | [illegible] | 1178 | [illegible] | GCF_001544355.1 | Alicyclobacillus_acidiphilus | AAC01S_RS15875 | 2314..5850 | 182455 | [illegible] |
| WP_067929227.1 | Type1.BREX | [illegible] | 1178 | [illegible] | GCF_001552255.1 | Alicyclobacillus_shizuokensis | ASH01S_RS14820 | 6987..10523 | 392014 | [illegible] |
| WP_206657716.1 | Type1.BREX | [illegible] | 1175 | [illegible] | GCF_017298635.1 | Alicyclobacillus_mengziensis | JZ786_RS05180 | 1119199..1122726 | 2931921 | [illegible] |
| WP_241716776.1 | Type1.BREX | [illegible] | 1042 | [illegible] | GCF_022606465.1 | Sulfoacidibacillus_ferrooxidans | MM817_RS15405 | 72..3200 | 2005001 | [illegible] |
| WP_075389023.1 | Type1.BREX | [illegible] | 1189 | [illegible] | - | Alkalihalophilus_pseudofirmus | BTR23_RS23260 | 11771..15340 | 79885 | [illegible] |
| WP_130776765.1 | Type1.BREX | [illegible] | 1187 | [illegible] | - | Bacillus_cereus | BC2903_22190 | 87962..91525 | 1396 | [illegible] |
| WP_001993018.1 | Type1.BREX | [illegible] | 1192 | [illegible] | - | Bacillus_cereus | BCH308197_0965 | 98223..101801 | 1396 | [illegible] |
| WP_076855694.1 | Type1.BREX | [illegible] | 1194 | [illegible] | - | Bacillus_cereus | BKK42_RS28150 | 15538..19122 | 1396 | [illegible] |
| WP_016131320.1 | Type1.BREX | [illegible] | 1190 | [illegible] | - | Bacillus_cereus | KOW_RS09165 | 326636..330208 | 1396 | [illegible] |
| WP_249727160.1 | Type1.BREX | [illegible] | 1195 | [illegible] | - | Bacillus_paralicheniformis | M9A47_RS18470 | 136341..139928 | 1648923 | [illegible] |
| EOP54853.1 | Type1.BREX | [illegible] | 1193 | [illegible] | GCA_000399485.1 | Bacillus_cereus_VD136 | IIW_00987 | 109394..112975 | 1053234 | [illegible] |
| KEK19923.1 | Type1.BREX | [illegible] | 1190 | [illegible] | GCA_000712595.1 | Bacillus_manliponensis | BAMA_19340 | 136868..140440 | 574376 | [illegible] |
| OPA28160.1 | Type1.BREX | [illegible] | 1190 | [illegible] | GCA_002021465.1 | Bacillus_cereus | BHL47_19745 | 10940..14512 | 1396 | [illegible] |
| PAK40595.1 | Type1.BREX | [illegible] | 1191 | [illegible] | GCA_002276345.1 | Peribacillus_simplex | CHI08_14590 | 21852..25427 | 1478 | [illegible] |
| ATI52289.1 | Type1.BREX | [illegible] | 1190 | [illegible] | GCA_002504225.1 | Bacillus_cereus | CPZ32_18870 | 3450635..3454207 | 1396 | [illegible] |
| PDY65967.1 | Type1.BREX | [illegible] | 1192 | [illegible] | GCA_002550315.1 | Bacillus_cereus | COM93_22635 | 9795..13373 | 1396 | [illegible] |
| PFQ26669.1 | Type1.BREX | [illegible] | 1195 | [illegible] | GCA_002567415.1 | Bacillus_cereus | COK13_01020 | 52170..55757 | 1396 | [illegible] |
| PGK17286.1 | Type1.BREX | [illegible] | 1192 | [illegible] | GCA_002574715.1 | Bacillus_cereus | CN901_21555 | 30762..34340 | 1396 | [illegible] |
| PKF86734.1 | Type1.BREX | [illegible] | 1193 | [illegible] | GCA_002835675.1 | Bacillus_sp._BA3 | CW306_21620 | 276915..280496 | 2057910 | [illegible] |
| KAA6462805.1 | Type1.BREX | [illegible] | 1195 | [illegible] | GCA_008682215.1 | Bacillus_cereus | DX932_17685 | 80940..84527 | 1396 | [illegible] |
| MBB2481351.1 | Type1.BREX | [illegible] | 1187 | [illegible] | GCA_014174455.1 | Bacillus_sp._APMAM | H5P36_14260 | 24010..27573 | 2761103 | [illegible] |
| GGJ55411.1 | Type1.BREX | [illegible] | 1365 | [illegible] | GCA_014647615.1 | Virgibacillus_kapii | GCM10007111_17110 | 601764..605861 | 1638645 | [illegible] |
| MBE2917754.1 | Type1.BREX | [illegible] | 1179 | [illegible] | GCA_014890465.1 | Anoxybacillus_flavithermus | HPJ95_05660 | 21197..24736 | 33934 | [illegible] |
| MBE2912384.1 | Type1.BREX | [illegible] | 1179 | [illegible] | GCA_014890515.1 | Anoxybacillus_flavithermus | HPK00_02735 | 1719..5258 | 33934 | [illegible] |
| MBO1513528.1 | Type1.BREX | [illegible] | 1187 | [illegible] | GCA_017497975.1 | Metabacillus_bambusae | I7822_17955 | 21833..25396 | 2795218 | [illegible] |
| MBT2214695.1 | Type1.BREX | [illegible] | 1340 | [illegible] | GCA_018603665.1 | Virgibacillus_dakarensis | KK120_02535 | 179341..183363 | 1917889 | [illegible] |
| MBW7642206.1 | Type1.BREX | [illegible] | 1188 | [illegible] | GCA_019448595.1 | Geobacillus_thermoleovorans | K1I45_03795 | 253845..257411 | 33941 | [illegible] |
| MCD1158981.1 | Type1.BREX | [illegible] | 1194 | [illegible] | GCA_021012855.1 | Peribacillus_castrilensis | LQF35_01355 | 268240..271824 | 2897690 | [illegible] |
| MCD8502089.1 | Type1.BREX | [illegible] | 597 | [illegible] | GCA_021208445.1 | Bacillaceae_bacterium | LRY71_10940 | 4605..6398 | 1889774 | [illegible] |
| MCG1034434.1 | Type1.BREX | [illegible] | 1204 | [illegible] | GCA_021991685.1 | Bacillus_amyloliquefaciens | J5S76_20860 | 92651..96265 | 1390 | [illegible] |
| MCI1589808.1 | Type1.BREX | [illegible] | 1190 | [illegible] | GCA_022646685.1 | Heyndrickxia_oleronia | LKH79_04555 | 33937..37509 | 38875 | [illegible] |
| MCI1592662.1 | Type1.BREX | [illegible] | 1190 | [illegible] | GCA_022646685.1 | Heyndrickxia_oleronia | LKH79_19270 | 20507..24079 | 38875 | [illegible] |
| MCI3146523.1 | Type1.BREX | [illegible] | 1195 | [illegible] | GCA_022696175.1 | Bacillus_cereus | C5137_09385 | 1769234..1772821 | 1396 | [illegible] |
| UNT71505.1 | Type1.BREX | [illegible] | 1190 | [illegible] | GCA_022701035.1 | Bacillus_sp._N447-1 | IQ781_06225 | 1264486..1268058 | 2789208 | [illegible] |
| UQX55099.1 | Type1.BREX | [illegible] | 1206 | [illegible] | GCA_023516215.1 | Cytobacillus_oceanisediminis | M5V91_04885 | 909305..912925 | 665099 | [illegible] |
| MCM3314633.1 | Type1.BREX | [illegible] | 1194 | [illegible] | GCA_023713535.1 | Psychrobacillus_sp._MER_TA_17 | M3626_19750 | 42859..46443 | 2939576 | [illegible] |
| UUE89892.1 | Type1.BREX | [illegible] | 1193 | [illegible] | GCA_024498915.1 | Bacillus_cereus | L2I54_04675 | 914094..917675 | 1396 | [illegible] |
| SFA50349.1 | Type1.BREX | [illegible] | 1174 | [illegible] | GCA_900111795.1 | Anoxybacillus_pushchinoensis | SAMN05216169_10246 | 6894..10418 | 150248 | [illegible] |
| WP_238533070.1 | Type1.BREX | [illegible] | 1086 | [illegible] | GCF_000262755.1 | Bacillus_methanolicus | PB1_RS00020 | 9029..12289 | 1471 | [illegible] |
| WP_001166856.1 | Type1.BREX | [illegible] | 1195 | [illegible] | GCF_000293745.1 | Bacillus_toyonensis | IGI_RS16405 | 3258372..3261959 | 155322 | [illegible] |
| WP_233275777.1 | Type1.BREX | [illegible] | 1184 | [illegible] | GCF_000429685.1 | Salibacterium_aidingense | G475_RS0113415 | 15614..19168 | 384933 | [illegible] |
| WP_026584641.1 | Type1.BREX | [illegible] | 1194 | [illegible] | GCF_000518885.1 | Bacillus_sp._J33 | L634_RS0124025 | 6414..9998 | 935836 | [illegible] |
| WP_042147861.1 | Type1.BREX | [illegible] | 1194 | [illegible] | GCF_000577245.1 | Paucisalibacillus_sp._EB02 | TY11_RS14970 | 163346..166930 | 1347087 | [illegible] |
| WP_026673780.1 | Type1.BREX | [illegible] | 1188 | [illegible] | GCF_000621445.1 | Alkalihalobacillus_bogoriensis | T323_RS0113555 | 41147..44713 | 246272 | [illegible] |
| WP_031537547.1 | Type1.BREX | [illegible] | 1187 | [illegible] | GCF_000701605.1 | Bacillus_sp._MB2021 | T349_RS0112475 | 16440..20003 | 1408303 | [illegible] |
| WP_235184434.1 | Type1.BREX | [illegible] | 1193 | [illegible] | GCF_000712595.1 | Bacillus_manliponensis | BAMA_RS08125 | 136859..140440 | 574376 | [illegible] |
| WP_042224724.1 | Type1.BREX | [illegible] | 1180 | [illegible] | GCF_000724085.1 | Oceanobacillus_manasiensis | BN985_RS16170 | 528907..532449 | 586413 | [illegible] |
| WP_048824801.1 | Type1.BREX | [illegible] | 1190 | [illegible] | GCF_000821085.1 | Bacillus_sp._B-jedd | BN1002_01993 | 1962591..1966163 | 1476857 | [illegible] |
| WP_041847547.1 | Type1.BREX | [illegible] | 1194 | [illegible] | GCF_000832165.1 | Caldibacillus_thermoamylovorans | B4065_RS04110 | 1312..4896 | 35841 | [illegible] |
| WP_219913306.1 | Type1.BREX | [illegible] | 1195 | [illegible] | GCF_001117785.1 | Bacillus_paranthracis | AMO22_RS12700 | 404..3991 | 2026186 | [illegible] |
| WP_049624985.1 | Type1.BREX | [illegible] | 1183 | [illegible] | GCF_001183895.1 | Geobacillus_stearothermophilus | AA904_RS09790 | 5825..9376 | 1422 | [illegible] |
| WP_053997117.1 | Type1.BREX | [illegible] | 1190 | [illegible] | GCF_001281525.1 | Lysinibacillus_macroides | ADM90_RS22630 | 144729..148301 | 33935 | [illegible] |
| WP_072180298.1 | Type1.BREX | [illegible] | 1195 | [illegible] | GCF_001286825.1 | Bacillus_cereus | AOJ89_RS12960 | 22364..25951 | 1396 | [illegible] |
| WP_062353349.1 | Type1.BREX | [illegible] | 1190 | [illegible] | GCF_001375515.1 | Bacillus_kwashiorkori | BN1242_RS07000 | 23447..27019 | 1522318 | [illegible] |
| WP_062105395.1 | Type1.BREX | [illegible] | 1188 | [illegible] | GCF_001375535.1 | Bacillus_niameyensis | BN1243_RS06445 | 1312932..1316498 | 1522308 | [illegible] |
| WP_058950067.1 | Type1.BREX | [illegible] | 1366 | [illegible] | GCF_001485235.1 | Oceanobacillus_picturae | OPHB3_RS08850 | 6149..10249 | 171693 | [illegible] |
| WP_061859498.1 | Type1.BREX | [illegible] | 1192 | [illegible] | GCF_001593805.1 | Priestia_megaterium | BMBG_RS08635 | 1672480..1676058 | 1404 | [illegible] |
| WP_061860130.1 | Type1.BREX | [illegible] | 1192 | [illegible] | GCF_001593805.1 | Priestia_megaterium | BMBG_RS25870 | 30472..34050 | 1404 | [illegible] |
| WP_062756374.1 | Type1.BREX | [illegible] | 1189 | [illegible] | GCF_001598935.1 | Parageobacillus_toebii | GT3_RS15670 | 17547..21116 | 153151 | [illegible] |
| WP_064472570.1 | Type1.BREX | [illegible] | 1206 | [illegible] | GCF_001645405.1 | Bacillus_wiedmannii | A6278_RS13065 | 45022..48642 | 1890302 | [illegible] |
| WP_070141359.1 | Type1.BREX | [illegible] | 1189 | [illegible] | GCF_001757835.1 | Bacillus_mycoides | BWG0E8_RS06340 | 263356..266925 | 1405 | [illegible] |
| WP_071743491.1 | Type1.BREX | [illegible] | 1195 | [illegible] | GCF_001884125.1 | Bacillus | BAU22_RS23055 | 7102..10689 | 1386 | [illegible] |
| WP_071716603.1 | Type1.BREX | [illegible] | 1192 | [illegible] | GCF_001884225.1 | Bacillus_sp._N35-10-4 | BAU26_RS20735 | 58685..62263 | 1866315 | [illegible] |
| WP_075688961.1 | Type1.BREX | [illegible] | 1194 | [illegible] | GCF_001940785.1 | Bacillus_sp._MRMR6 | BTR25_RS15520 | 15107..18691 | 1928617 | [illegible] |

| Protein-ID | System | Gene-Neighbourhoods | Length | Taxonomy | Assembly-ID | Strain | Locus-Tag | Genomic-coordinates | TaxID | Fasta-define |
|---|---|---|---|---|---|---|---|---|---|---|
| WP_077362405.1 | Type1.BREX | <-McrC-NTD+REase<-AAA-ATPase<-?<-?<-SIGMA+OB-Fold+AAA-ATPase+Lon-Protease-BrxL<-PglZ<-N6-METHYLASE<-N6-METHYLASE<-DUF6079*<-BrxB-BrxF-iSTAND<-BrxA-DNA-binding | 1185 | Bacteria>Terrabacteria_group>Bacillota>Bacilli>Bacillales>Bacillaceae | GCF_001999465.1 | Fictibacillus_arsenicus | UN64_RS10450 | 648714..652271 | 255247 | BREX-system-P-loop-protein-BrxC-[Fictibacillus-arsenicus] |
| WP_078203472.1 | Type1.BREX | IONCHANNEL+PAS+HISKIN+MORC->REC+CITB-HTH+ZDNA-HTH->?->BrxA-DNA-binding->BrxB-BrxF-iSTAND->DUF6079*->N6-METHYLASE->PglZ->SIGMA+OB-Fold+AAA-ATPase+Lon-Protease-BrxL-> | 1197 | Bacteria>Terrabacteria_group>Bacillota>Bacilli>Bacillales>Bacillaceae | GCF_002014505.1 | Bacillus_mycoides | BW899_RS14845 | 198879..202472 | 1405 | BREX-system-P-loop-protein-BrxC-[Bacillus-mycoides] |
| WP_078427952.1 | Type1.BREX | HTH_6+SIS-><-?<-PAS+GAF+PAS+PAS+PAS+HISKIN<-N6-METHYLASE<-PglZ<-N6-METHYLASE<-N6-METHYLASE<-DUF6079*<-BrxB-BrxF-iSTAND<-BrxA-DNA-binding\|\|?->HISKIN->HISKIN-><-?<-MarR-HTH<-HAD+HAD\|\|HE[illegible] | 1189 | Bacteria>Terrabacteria_group>Bacillota>Bacilli>Bacillales>Bacillaceae | GCF_002019605.1 | Alkalihalobacterium_alkalinitrilicum | BK574_RS06200 | 1314904..1318473 | 427920 | BREX-system-P-loop-protein-BrxC-[Alkalihalobacterium-alkalinitrilicum] |
| WP_078386197.1 | Type1.BREX | PIN->REase->OLD-ABC->HNH->?->BrxA-DNA-binding->BrxB-BrxF-iSTAND->DUF6079*->N6-METHYLASE->PglZ->SIGMA+OB-Fold+AAA-ATPase+Lon-Protease-BrxL->MarR-HTH->?->?->BZIP+SF1-SF2-Helicase-> | 1190 | Bacteria>Terrabacteria_group>Bacillota>Bacilli>Bacillales>Bacillaceae | GCF_002021465.1 | Bacillus_cereus | BHL47_19745 | 10940..14512 | 1396 | BREX-system-P-loop-protein-BrxC-[Bacillus-cereus] |
| WP_095320972.1 | Type1.BREX | <-HISKIN<-REC+OmpR-HTH\|\|?-><-?<-OLD-ABC<-ParB+HNH<-SIGMA+OB-Fold+AAA-ATPase+Lon-Protease-BrxL<-PglZ<-N6-METHYLASE<-DUF6079*<-BrxB-BrxF-iSTAND<-BrxA-DNA-binding | 1194 | Bacteria>Terrabacteria_group>Bacillota>Bacilli>Bacillales>Bacillaceae | GCF_002271995.1 | Niallia_circulans | CHH55_RS22000 | 53589..57173 | 1397 | BREX-system-P-loop-protein-BrxC-[Niallia-circulans] |
| WP_097907522.1 | Type1.BREX | SF1-SF2-Helicase->BrxA-DNA-binding->BrxB-BrxF-iSTAND->DUF6079*->N6-METHYLASE->PglZ-> | 1186 | Bacteria>Terrabacteria_group>Bacillota>Bacilli>Bacillales>Bacillaceae | GCF_002553125.1 | Bacillus_wiedmannii | CON72_RS14770 | 3890..7450 | 1890302 | BREX-system-P-loop-protein-BrxC-[Bacillus-wiedmannii] |
| WP_098124645.1 | Type1.BREX | Phage_integrase->XIS-HTH+XIS-HTH+XIS-HTH+XIS-HTH+XIS-HTH->Phage_integrase->HORMA-><-?<-?<-SIGMA+OB-Fold+AAA-ATPase+Lon-Protease-BrxL<-PglZ<-N6-METHYLASE<-DUF6079*<-BrxB-BrxF-iSTAND<-B[illegible] | 1195 | Bacteria>Terrabacteria_group>Bacillota>Bacilli>Bacillales>Bacillaceae | GCF_002555245.1 | Bacillus_pseudomycoides | CN634_RS03625 | 84463..88050 | 64104 | BREX-system-P-loop-protein-BrxC-[Bacillus-pseudomycoides] |
| WP_098085629.1 | Type1.BREX | IONCHANNEL+PAS+HISKIN+MORC->REC+CITB-HTH+ZDNA-HTH->?->BrxA-DNA-binding->BrxB-BrxF-iSTAND->DUF6079*->N6-METHYLASE->PglZ->SIGMA+OB-Fold+AAA-ATPase+Lon-Protease-BrxL-><-35exo+wHTH+SF1-SF[illegible] | 1192 | Bacteria>Terrabacteria_group>Bacillota>Bacilli>Bacillales>Bacillaceae | GCF_002555855.1 | Bacillus_wiedmannii | CN905_RS18745 | 34066..37644 | 1890302 | BREX-system-P-loop-protein-BrxC-[Bacillus-wiedmannii] |
| WP_098154948.1 | Type1.BREX | SF1-SF2-Helicase-><-?<-DndB<-?\|\|?->\|\|\|\|BrxA-DNA-binding->BrxB-BrxF-iSTAND->DUF6079*->N6-METHYLASE->PglZ->SIGMA+OB-Fold+AAA-ATPase+Lon-Protease-BrxL->?->?->?-><-HNH<-?<-EVE | 1195 | Bacteria>Terrabacteria_group>Bacillota>Bacilli>Bacillales>Bacillaceae | GCF_002556885.1 | Bacillus_toyonensis | CN544_RS21920 | 45516..49103 | 155322 | BREX-system-P-loop-protein-BrxC-[Bacillus-toyonensis] |
| WP_098217368.1 | Type1.BREX | rve->?->HTH_Tnp_1+HTH_21+rve+rve_3-><-?\|\|?->Phage_int_SAM_1+Phage_integrase->TOPRIM+Topoisom_bac+TOPRIM_Crpt->BrxA-DNA-binding->BrxB-BrxF-iSTAND->DUF6079*->N6-METHYLASE->PglZ->SIGMA+O[illegible] | 1195 | Bacteria>Terrabacteria_group>Bacillota>Bacilli>Bacillales>Bacillaceae | GCF_002557585.1 | Bacillus_pseudomycoides | CN584_RS13740 | 37733..41320 | 64104 | BREX-system-P-loop-protein-BrxC-[Bacillus-pseudomycoides] |
| WP_097863054.1 | Type1.BREX | <-HTH_Tnp_1+HTH_21+rve+rve_3\|\|<-SIGMA+OB-Fold+AAA-ATPase+Lon-Protease-BrxL<-?<-PglZ<-N6-METHYLASE<-DUF6079*<-BrxB-BrxF-iSTAND<-BrxA-DNA-binding<-?<-REC+CITB-HTH+ZDNA-HTH<-IONCHANNEL+P[illegible] | 1192 | Bacteria>Terrabacteria_group>Bacillota>Bacilli>Bacillales>Bacillaceae | GCF_002561635.1 | Bacillus_cereus | COC44_RS19850 | 13392..16970 | 1396 | BREX-system-P-loop-protein-BrxC-[Bacillus-cereus] |
| WP_098557664.1 | Type1.BREX | TOPRIM+Topoisom_bac->?->?->SbcC->Phage_int_SAM_1+Phage_integrase->TOPRIM+Topoisom_bac+TOPRIM_Crpt->BrxA-DNA-binding->BrxB-BrxF-iSTAND->DUF6079*->N6-METHYLASE->PglZ->SIGMA+OB-Fold+AAA-[illegible] | 1195 | Bacteria>Terrabacteria_group>Bacillota>Bacilli>Bacillales>Bacillaceae | GCF_002566845.1 | Bacillus_cereus | COJ77_RS08250 | 133787..137374 | 1396 | BREX-system-P-loop-protein-BrxC-[Bacillus-cereus] |
| WP_098762758.1 | Type1.BREX | <-HTH_Tnp_1+HTH_21+rve+rve_3\|\|<-SIGMA+OB-Fold+AAA-ATPase+Lon-Protease-BrxL<-?<-PglZ<-N6-METHYLASE<-DUF6079*<-BrxB-BrxF-iSTAND<-BrxA-DNA-binding<-?<-REC+CITB-HTH+ZDNA-HTH<-IONCHANNEL+P[illegible] | 1192 | Bacteria>Terrabacteria_group>Bacillota>Bacilli>Bacillales>Bacillaceae | GCF_002574715.1 | Bacillus_cereus | CN901_RS21555 | 30762..34340 | 1396 | BREX-system-P-loop-protein-BrxC-[Bacillus-cereus] |
| WP_098584860.1 | Type1.BREX | IONCHANNEL+PAS+HISKIN+MORC->REC+CITB-HTH+ZDNA-HTH->?->BrxA-DNA-binding->BrxB-BrxF-iSTAND->DUF6079*->N6-METHYLASE->PglZ->SIGMA+OB-Fold+AAA-ATPase+Lon-Protease-BrxL-><-35exo+wHTH+SF1-SF[illegible] | 1195 | Bacteria>Terrabacteria_group>Bacillota>Bacilli>Bacillales>Bacillaceae | GCF_002575795.1 | Bacillus_cereus | CN985_RS11285 | 16461..20048 | 1396 | BREX-system-P-loop-protein-BrxC-[Bacillus-cereus] |
| WP_098882708.1 | Type1.BREX | <-ZnR\|\|?->?-><-?\|\|IONCHANNEL+PAS+HISKIN+MORC->REC+CITB-HTH+ZDNA-HTH->?->BrxA-DNA-binding->BrxB-BrxF-iSTAND->DUF6079*->N6-METHYLASE->PglZ->SIGMA+OB-Fold+AAA-ATPase+Lon-Protease-BrxL->N[illegible] | 1194 | Bacteria>Terrabacteria_group>Bacillota>Bacilli>Bacillales>Bacillaceae | GCF_002577745.1 | Bacillus_cereus_group | FS786_RS24760 | 17629..21213 | 86661 | MULTISPECIES:-BREX-system-P-loop-protein-BrxC-[Bacillus-cereus-group] |
| WP_098854104.1 | Type1.BREX | SLATT->Phage_integrase-><-REase+DUF5655\|\|EVE+HNH->BrxA-DNA-binding->BrxB-BrxF-iSTAND->DUF6079*->N6-METHYLASE->PglZ->SIGMA+OB-Fold+AAA-ATPase+Lon-Protease-BrxL->?->?->RVT+HNH->?->?->LA[illegible] | 1195 | Bacteria>Terrabacteria_group>Bacillota>Bacilli>Bacillales>Bacillaceae | GCF_002579245.1 | Bacillus_toyonensis | COE72_RS00880 | 66293..69880 | 155322 | BREX-system-P-loop-protein-BrxC-[Bacillus-toyonensis] |
| WP_098933724.1 | Type1.BREX | [illegible] | 1195 | Bacteria>Terrabacteria_group>Bacillota>Bacilli>Bacillales>Bacillaceae | GCF_002584635.1 | Bacillus_sp._AFS031507 | COE25_RS18930 | 113248..116835 | 2033496 | BREX-system-P-loop-protein-BrxC-[Bacillus-sp.-AFS031507] |
| WP_208738702.1 | Type1.BREX | [illegible] | 1194 | Bacteria>Terrabacteria_group>Bacillota>Bacilli>Bacillales>Bacillaceae | GCF_002953475.1 | Bacillus_licheniformis | C1T27_RS03565 | 718289..721873 | 1402 | BREX-system-P-loop-protein-BrxC-[Bacillus-licheniformis] |
| WP_106071134.1 | Type1.BREX | [illegible] | 1191 | Bacteria>Terrabacteria_group>Bacillota>Bacilli>Bacillales>Bacillaceae | GCF_002998255.1 | Bacillus_paralicheniformis | C6W27_RS19030 | 56980..60555 | 1648923 | BREX-system-P-loop-protein-BrxC-[Bacillus-paralicheniformis] |
| WP_108713788.1 | Type1.BREX | [illegible] | 1191 | Bacteria>Terrabacteria_group>Bacillota>Bacilli>Bacillales>Bacillaceae | GCF_003073375.1 | unclassified_Lysinibacillus | DCE79_RS14920 | 3050129..3053704 | 2636778 | MULTISPECIES:-BREX-system-P-loop-protein-BrxC-[unclassified-Lysinibacillus] |
| WP_116555157.1 | Type1.BREX | [illegible] | 1181 | Bacteria>Terrabacteria_group>Bacillota>Bacilli>Bacillales>Bacillaceae | GCF_003097615.1 | Pueribacillus_theae | DCC39_RS12060 | 24428..27973 | 2171751 | BREX-system-P-loop-protein-BrxC-[Pueribacillus-theae] |
| WP_114371647.1 | Type1.BREX | [illegible] | 1205 | Bacteria>Terrabacteria_group>Bacillota>Bacilli>Bacillales>Bacillaceae | GCF_003336365.1 | Salicibibacter_kimchii | DT065_05890 | 1163447..1167064 | 2099786 | BREX-system-P-loop-protein-BrxC-[Salicibibacter-kimchii] |
| WP_115749618.1 | Type1.BREX | [illegible] | 1190 | Bacteria>Terrabacteria_group>Bacillota>Bacilli>Bacillales>Bacillaceae | GCF_003369565.1 | Oceanobacillus_chungangensis | CWR45_RS09390 | 32723..36295 | 1229152 | BREX-system-P-loop-protein-BrxC-[Oceanobacillus-chungangensis] |
| WP_115773755.1 | Type1.BREX | [illegible] | 1182 | Bacteria>Terrabacteria_group>Bacillota>Bacilli>Bacillales>Bacillaceae | GCF_003369575.1 | Oceanobacillus_arenosus | CWR48_RS13395 | 42631..46179 | 1229153 | BREX-system-P-loop-protein-BrxC-[Oceanobacillus-arenosus] |
| WP_116279377.1 | Type1.BREX | [illegible] | 1184 | Bacteria>Terrabacteria_group>Bacillota>Bacilli>Bacillales>Bacillaceae | GCF_003399705.1 | Virgibacillus_dokdonensis | CAI16_RS17315 | 10248..13802 | 302167 | BREX-system-P-loop-protein-BrxC-[Virgibacillus-dokdonensis] |
| WP_119547238.1 | Type1.BREX | [illegible] | 1193 | Bacteria>Terrabacteria_group>Bacillota>Bacilli>Bacillales>Bacillaceae | GCF_003581585.1 | Bacillus_salacetis | D3H55_RS12405 | 3906..7487 | 2315464 | BREX-system-P-loop-protein-BrxC-[Bacillus-salacetis] |
| WP_121614394.1 | Type1.BREX | [illegible] | 1192 | Bacteria>Terrabacteria_group>Bacillota>Bacilli>Bacillales>Bacillaceae | GCF_003667765.1 | Mesobacillus_foraminis | D9829_RS27750 | 386716..390294 | 279826 | BREX-system-P-loop-protein-BrxC-[Mesobacillus-foraminis] |
| WP_121661845.1 | Type1.BREX | [illegible] | 1365 | Bacteria>Terrabacteria_group>Bacillota>Bacilli>Bacillales>Bacillaceae | GCF_003667825.1 | Metabacillus_litoralis | D9842_RS06665 | 1400368..1404465 | 152268 | hypothetical-protein-[Metabacillus-litoralis] |
| WP_126293910.1 | Type1.BREX | [illegible] | 1190 | Bacteria>Terrabacteria_group>Bacillota>Bacilli>Bacillales>Bacillaceae | GCF_003966145.1 | Lysinibacillus_telephonicus | EKG35_RS07955 | 47066..50638 | 1714840 | BREX-system-P-loop-protein-BrxC-[Lysinibacillus-telephonicus] |
| WP_126410856.1 | Type1.BREX | [illegible] | 1194 | Bacteria>Terrabacteria_group>Bacillota>Bacilli>Bacillales>Bacillaceae | GCF_003966255.1 | Bacillus_yapensis | EKG37_RS21695 | 35397..38981 | 2492960 | BREX-system-P-loop-protein-BrxC-[Bacillus-yapensis] |
| WP_134188914.1 | Type1.BREX | [illegible] | 1195 | Bacteria>Terrabacteria_group>Bacillota>Bacilli>Bacillales>Bacillaceae | GCF_004367815.1 | Bacillus_sp._BH2 | EZE46_RS16445 | 318521..322108 | 2528958 | BREX-system-P-loop-protein-BrxC-[Bacillus-sp.-BH2] |
| WP_149870410.1 | Type1.BREX | [illegible] | 1182 | Bacteria>Terrabacteria_group>Bacillota>Bacilli>Bacillales>Bacillaceae | GCF_005154805.1 | Bacillus_sp._S3 | FAY30_RS13735 | 2868936..2872484 | 486398 | BREX-system-P-loop-protein-BrxC-[Bacillus-sp.-S3] |
| WP_136999292.1 | Type1.BREX | [illegible] | 1195 | Bacteria>Terrabacteria_group>Bacillota>Bacilli>Bacillales>Bacillaceae | GCF_005217675.1 | Bacillus_cereus | FC683_RS05590 | 24419..28006 | 1396 | BREX-system-P-loop-protein-BrxC-[Bacillus-cereus] |
| WP_137744598.1 | Type1.BREX | [illegible] | 1190 | Bacteria>Terrabacteria_group>Bacillota>Bacilli>Bacillales>Bacillaceae | GCF_005502275.1 | Robertmurraya_siralis | FCJ93_RS20760 | 22138..25710 | 77777 | BREX-system-P-loop-protein-BrxC-[Robertmurraya-siralis] |
| WP_140968666.1 | Type1.BREX | [illegible] | 1195 | Bacteria>Terrabacteria_group>Bacillota>Bacilli>Bacillales>Bacillaceae | GCF_006494425.1 | Bacillus_sp._(in:_firmicutes) | FJ659_RS01030 | 197909..201496 | 1409 | BREX-system-P-loop-protein-BrxC-[Bacillus-sp.-(in:-firmicutes)] |
| WP_141604544.1 | Type1.BREX | [illegible] | 1191 | Bacteria>Terrabacteria_group>Bacillota>Bacilli>Bacillales>Bacillaceae | GCF_006553035.1 | Terrilactibacillus_laevilacticus | FKM94_RS13485 | 2906..6481 | 1380157 | BREX-system-P-loop-protein-BrxC-[Terrilactibacillus-laevilacticus] |
| WP_144533672.1 | Type1.BREX | [illegible] | 1190 | Bacteria>Terrabacteria_group>Bacillota>Bacilli>Bacillales>Bacillaceae | GCF_007665325.1 | Bacillus_pumilus | FSV19_RS14530 | 168165..171737 | 1408 | BREX-system-P-loop-protein-BrxC-[Bacillus-pumilus] |
| WP_144463068.1 | Type1.BREX | [illegible] | 1186 | Bacteria>Terrabacteria_group>Bacillota>Bacilli>Bacillales>Bacillaceae | GCF_007671835.1 | Siminovitchia_fortis | FS422_RS10505 | 2167..5727 | 254758 | BREX-system-P-loop-protein-BrxC-[Siminovitchia-fortis] |
| WP_145524022.1 | Type1.BREX | [illegible] | 1194 | Bacteria>Terrabacteria_group>Bacillota>Bacilli>Bacillales>Bacillaceae | GCF_007673165.1 | Virgibacillus_sp._SK37 | FS404_RS12660 | 5290..8874 | 403957 | BREX-system-P-loop-protein-BrxC-[Virgibacillus-sp.-SK37] |
| WP_144538762.1 | Type1.BREX | [illegible] | 1191 | Bacteria>Terrabacteria_group>Bacillota>Bacilli>Bacillales>Bacillaceae | GCF_007677425.1 | Bacillus_sp._FJAT-22090 | FS503_RS11600 | 136183..139758 | 1581038 | BREX-system-P-loop-protein-BrxC-[Bacillus-sp.-FJAT-22090] |
| WP_144557887.1 | Type1.BREX | [illegible] | 1191 | Bacteria>Terrabacteria_group>Bacillota>Bacilli>Bacillales>Bacillaceae | GCF_007725405.1 | Alkalihalobacillus_miscanthi | FPV17_RS02925 | 558061..561636 | 2598861 | BREX-system-P-loop-protein-BrxC-[Alkalihalobacillus-miscanthi] |
| WP_147667282.1 | Type1.BREX | [illegible] | 1192 | Bacteria>Terrabacteria_group>Bacillota>Bacilli>Bacillales>Bacillaceae | GCF_008039555.1 | Cerasibacillus_terrae | FHP05_RS09090 | 130074..133652 | 2498845 | BREX-system-P-loop-protein-BrxC-[Cerasibacillus-terrae] |
| WP_150158491.1 | Type1.BREX | [illegible] | 1195 | Bacteria>Terrabacteria_group>Bacillota>Bacilli>Bacillales>Bacillaceae | GCF_008682215.1 | Bacillus_cereus | DX932_RS17660 | 80940..84527 | 1396 | BREX-system-P-loop-protein-BrxC-[Bacillus-cereus] |
| WP_151681679.1 | Type1.BREX | [illegible] | 1192 | Bacteria>Terrabacteria_group>Bacillota>Bacilli>Bacillales>Bacillaceae | GCF_008974225.1 | Weizmannia_acidilactici | GBH70_RS14395 | 13994..17572 | 2607726 | BREX-system-P-loop-protein-BrxC-[Weizmannia-acidilactici] |
| WP_156645486.1 | Type1.BREX | [illegible] | 1185 | Bacteria>Terrabacteria_group>Bacillota>Bacilli>Bacillales>Bacillaceae | GCF_009741735.1 | Lentibacillus_sp._JNUCC-1 | JNUCC1_RS10425 | 5243..8800 | 2654513 | BREX-system-P-loop-protein-BrxC-[Lentibacillus-sp.-JNUCC-1] |
| WP_159347017.1 | Type1.BREX | [illegible] | 1187 | Bacteria>Terrabacteria_group>Bacillota>Bacilli>Bacillales>Bacillaceae | GCF_009849605.1 | Cytobacillus_firmus | KIS1582_RS21365 | 6662..10225 | 1399 | BREX-system-P-loop-protein-BrxC-[Cytobacillus-firmus] |
| WP_159376681.1 | Type1.BREX | [illegible] | 1194 | Bacteria>Terrabacteria_group>Bacillota>Bacilli>Bacillales>Bacillaceae | GCF_009866865.1 | Bacillus_subtilis | Bateq7PJ16_RS04040 | 771317..774901 | 1423 | BREX-system-P-loop-protein-BrxC-[Bacillus-subtilis] |
| WP_163174462.1 | Type1.BREX | [illegible] | 1189 | Bacteria>Terrabacteria_group>Bacillota>Bacilli>Bacillales>Bacillaceae | GCF_011008565.1 | Weizmannia_ginsengihumi | G4D61_RS16490 | 22544..26113 | 363870 | BREX-system-P-loop-protein-BrxC-[Weizmannia-ginsengihumi] |
| WP_163243257.1 | Type1.BREX | [illegible] | 1190 | Bacteria>Terrabacteria_group>Bacillota>Bacilli>Bacillales>Bacillaceae | GCF_011008855.1 | Bacillus_aquiflavi | H1Z61_RS15490 | 1670..5242 | 2672567 | BREX-system-P-loop-protein-BrxC-[Bacillus-aquiflavi] |
| WP_166025629.1 | Type1.BREX | [illegible] | 1191 | Bacteria>Terrabacteria_group>Bacillota>Bacilli>Bacillales>Bacillaceae | GCF_011393025.1 | Neobacillus_terrae | G8761_RS08595 | 43782..47357 | 3034837 | BREX-system-P-loop-protein-BrxC-[Neobacillus-terrae] |
| WP_171510262.1 | Type1.BREX | [illegible] | 1366 | Bacteria>Terrabacteria_group>Bacillota>Bacilli>Bacillales>Bacillaceae | GCF_013138395.1 | Niallia_circulans | HLK66_RS25080 | 5165831..5169931 | 1397 | hypothetical-protein-[Niallia-circulans] |
| WP_188632237.1 | Type1.BREX | [illegible] | 1181 | Bacteria>Terrabacteria_group>Bacillota>Bacilli>Bacillales>Bacillaceae | GCF_014646635.1 | Lentibacillus_kapialis | GCM10007063_12550 | 56169..59714 | 340214 | BREX-system-P-loop-protein-BrxC-[Lentibacillus-kapialis] |
| WP_191813942.1 | Type1.BREX | [illegible] | 1194 | Bacteria>Terrabacteria_group>Bacillota>Bacilli>Bacillales>Bacillaceae | GCF_014836495.1 | Cytobacillus_stercorigallinarum | H9655_RS11295 | 265009..268593 | 2762240 | BREX-system-P-loop-protein-BrxC-[Cytobacillus-stercorigallinarum] |
| WP_194950107.1 | Type1.BREX | [illegible] | 1196 | Bacteria>Terrabacteria_group>Bacillota>Bacilli>Bacillales>Bacillaceae | GCF_014890465.1 | Anoxybacillus_flavithermus | HPJ95_RS05650 | 21146..24736 | 33934 | BREX-system-P-loop-protein-BrxC-[Anoxybacillus-flavithermus] |
| WP_194950212.1 | Type1.BREX | [illegible] | 1196 | Bacteria>Terrabacteria_group>Bacillota>Bacilli>Bacillales>Bacillaceae | GCF_014890515.1 | Anoxybacillus_flavithermus | HPK00_RS02735 | 1719..5309 | 33934 | BREX-system-P-loop-protein-BrxC-[Anoxybacillus-flavithermus] |
| WP_193647953.1 | Type1.BREX | [illegible] | 1192 | Bacteria>Terrabacteria_group>Bacillota>Bacilli>Bacillales>Bacillaceae | GCF_015071065.1 | Bacillus_cereus | FT637_RS21110 | 102498..106076 | 1396 | BREX-system-P-loop-protein-BrxC-[Bacillus-cereus] |
| WP_196616266.1 | Type1.BREX | [illegible] | 1194 | Bacteria>Terrabacteria_group>Bacillota>Bacilli>Bacillales>Bacillaceae | GCF_015732475.1 | Bacillus_velezensis | IEN91_RS03495 | 706047..709631 | 492670 | BREX-system-P-loop-protein-BrxC-[Bacillus-velezensis] |
| WP_197207598.1 | Type1.BREX | [illegible] | 1194 | Bacteria>Terrabacteria_group>Bacillota>Bacilli>Bacillales>Bacillaceae | GCF_015844875.1 | Cytobacillus_firmus | ABE26_RS22015 | 1321572..1325156 | 1399 | BREX-system-P-loop-protein-BrxC-[Cytobacillus-firmus] |
| WP_197218912.1 | Type1.BREX | [illegible] | 1194 | Bacteria>Terrabacteria_group>Bacillota>Bacilli>Bacillales>Bacillaceae | GCF_015845055.1 | Bacillus_mycoides | ABE17_RS06425 | 419706..423290 | 1405 | BREX-system-P-loop-protein-BrxC-[Bacillus-mycoides] |
| WP_199696150.1 | Type1.BREX | [illegible] | 1195 | Bacteria>Terrabacteria_group>Bacillota>Bacilli>Bacillales>Bacillaceae | GCF_016483705.1 | Bacillus_cereus_group_sp._N6 | JDS99_RS27915 | 2655..6242 | 2794583 | BREX-system-P-loop-protein-BrxC-[Bacillus-cereus-group-sp.-N6] |
| WP_074543742.1 | Type1.BREX | [illegible] | 1186 | Bacteria>Terrabacteria_group>Bacillota>Bacilli>Bacillales>Bacillaceae | GCF_016757935.1 | Bacillus_cereus_group | PDN14_RS24715 | 18623..22183 | 86661 | MULTISPECIES:-BREX-system-P-loop-protein-BrxC-[Bacillus-cereus-group] |
| WP_071742543.1 | Type1.BREX | [illegible] | 1206 | Bacteria>Terrabacteria_group>Bacillota>Bacilli>Bacillales>Bacillaceae | GCF_016758535.1 | Bacillus | IRV24_RS20405 | 8695..12315 | 1386 | MULTISPECIES:-BREX-system-P-loop-protein-BrxC-[Bacillus] |
| WP_204955633.1 | Type1.BREX | [illegible] | 1192 | Bacteria>Terrabacteria_group>Bacillota>Bacilli>Bacillales>Bacillaceae | GCF_016908395.1 | Metabacillus_crassostreae | JOC75_RS19370 | 135240..138818 | 929098 | BREX-system-P-loop-protein-BrxC-[Metabacillus-crassostreae] |
| WP_243459544.1 | Type1.BREX | [illegible] | 1182 | Bacteria>Terrabacteria_group>Bacillota>Bacilli>Bacillales>Bacillaceae | GCF_017497975.1 | Metabacillus_bambusae | I7822_RS17965 | 21848..25396 | 2795218 | BREX-system-P-loop-protein-BrxC-[Metabacillus-bambusae] |
| WP_209482153.1 | Type1.BREX | [illegible] | 1187 | Bacteria>Terrabacteria_group>Bacillota>Bacilli>Bacillales>Bacillaceae | GCF_017873675.1 | Virgibacillus_litoralis | J2Z82_RS17835 | 25252..28815 | 578221 | BREX-system-P-loop-protein-BrxC-[Virgibacillus-litoralis] |
| WP_210367870.1 | Type1.BREX | [illegible] | 1190 | Bacteria>Terrabacteria_group>Bacillota>Bacilli>Bacillales>Bacillaceae | GCF_017912555.1 | Bacillus_sp._REN3 | JJQ88_RS20340 | 2531..6103 | 2802440 | BREX-system-P-loop-protein-BrxC-[Bacillus-sp.-REN3] |
| WP_213108269.1 | Type1.BREX | [illegible] | 1190 | Bacteria>Terrabacteria_group>Bacillota>Bacilli>Bacillales>Bacillaceae | GCF_018343555.1 | Bacillus_sp._FJAT-49731 | KHA89_RS14830 | 208852..212424 | 2833584 | BREX-system-P-loop-protein-BrxC-[Bacillus-sp.-FJAT-49731] |
| WP_214800485.1 | Type1.BREX | [illegible] | 1194 | Bacteria>Terrabacteria_group>Bacillota>Bacilli>Bacillales>Bacillaceae | GCF_018613535.1 | unclassified_Bacillus_(in:_firmicutes) | J7E26_RS00915 | 158819..162403 | 185979 | MULTISPECIES:-BREX-system-P-loop-protein-BrxC-[unclassified-Bacillus-(in:-firmicutes)] |
| WP_070170510.1 | Type1.BREX | [illegible] | 1193 | Bacteria>Terrabacteria_group>Bacillota>Bacilli>Bacillales>Bacillaceae | GCF_018736765.1 | Bacillus_mycoides | BWGOE6_RS18280 | 9483..13064 | 1405 | BREX-system-P-loop-protein-BrxC-[Bacillus-mycoides] |
| WP_216486703.1 | Type1.BREX | [illegible] | 1194 | Bacteria>Terrabacteria_group>Bacillota>Bacilli>Bacillales>Bacillaceae | GCF_018918075.1 | Virgibacillus_sp._MSJ-26 | KQI49_RS03110 | 240564..244148 | 2841522 | BREX-system-P-loop-protein-BrxC-[Virgibacillus-sp.-MSJ-26] |
| WP_216828510.1 | Type1.BREX | [illegible] | 1192 | Bacteria>Terrabacteria_group>Bacillota>Bacilli>Bacillales>Bacillaceae | GCF_019024285.1 | Alkalihalobacterium_elongatum | GKP30_RS05985 | 60975..64553 | 2675466 | BREX-system-P-loop-protein-BrxC-[Alkalihalobacterium-elongatum] |
| WP_216889200.1 | Type1.BREX | [illegible] | 1197 | Bacteria>Terrabacteria_group>Bacillota>Bacilli>Bacillales>Bacillaceae | GCF_019036865.1 | Bacillus_pumilus | KM878_RS13750 | 41263..44856 | 1408 | BREX-system-P-loop-protein-BrxC-[Bacillus-pumilus] |
| WP_216871762.1 | Type1.BREX | [illegible] | 1185 | Bacteria>Terrabacteria_group>Bacillota>Bacilli>Bacillales>Bacillaceae | GCF_019037265.1 | Virgibacillus_pantothenticus | KM909_RS12220 | 65907..69464 | 1473 | BREX-system-P-loop-protein-BrxC-[Virgibacillus-pantothenticus] |
| WP_220158113.1 | Type1.BREX | [illegible] | 1190 | Bacteria>Terrabacteria_group>Bacillota>Bacilli>Bacillales>Bacillaceae | GCF_019459225.1 | Bacillus_sp._IITD106 | KOH71_20660 | 11529..15101 | 2864214 | BREX-system-P-loop-protein-BrxC-[Bacillus-sp.-IITD106] |
| WP_221827426.1 | Type1.BREX | [illegible] | 1143 | Bacteria>Terrabacteria_group>Bacillota>Bacilli>Bacillales>Bacillaceae | GCF_019749295.1 | Neobacillus_niacini | HO184_RS18025 | 24955..28386 | 86668 | BREX-system-P-loop-protein-BrxC-[Neobacillus-niacini] |
| WP_222483047.1 | Type1.BREX | [illegible] | 1194 | Bacteria>Terrabacteria_group>Bacillota>Bacilli>Bacillales>Bacillaceae | GCF_019801725.1 | Priestia_flexa | KUV92_RS18295 | 34663..38247 | 86664 | BREX-system-P-loop-protein-BrxC-[Priestia-flexa] |
| WP_223087052.1 | Type1.BREX | [illegible] | 1190 | Bacteria>Terrabacteria_group>Bacillota>Bacilli>Bacillales>Bacillaceae | GCF_019915265.1 | Bacillus_aquiflavi | K6959_RS16340 | 3377801..3381373 | 2672567 | BREX-system-P-loop-protein-BrxC-[Bacillus-aquiflavi] |
| WP_226646168.1 | Type1.BREX | [illegible] | 1191 | Bacteria>Terrabacteria_group>Bacillota>Bacilli>Bacillales>Bacillaceae | GCF_020524415.1 | Mesobacillus_subterraneus | LC051_RS17725 | 565894..569469 | 285983 | BREX-system-P-loop-protein-BrxC-[Mesobacillus-subterraneus] |
| WP_226536426.1 | Type1.BREX | [illegible] | 1197 | Bacteria>Terrabacteria_group>Bacillota>Bacilli>Bacillales>Bacillaceae | GCF_020524645.1 | Fictibacillus_halophilus | LC044_RS10690 | 18428..22021 | 1610490 | BREX-system-P-loop-protein-BrxC-[Fictibacillus-halophilus] |
| WP_229142420.1 | Type1.BREX | [illegible] | 1192 | Bacteria>Terrabacteria_group>Bacillota>Bacilli>Bacillales>Bacillaceae | GCF_020731355.1 | Bacillus_sp._REN16 | LKW28_RS21250 | 18532..22110 | 2887296 | BREX-system-P-loop-protein-BrxC-[Bacillus-sp.-REN16] |
| WP_236787490.1 | Type1.BREX | [illegible] | 1364 | Bacteria>Terrabacteria_group>Bacillota>Bacilli>Bacillales>Bacillaceae | GCF_021654895.1 | Bacillus_cereus | L6489_RS02850 | 541168..545262 | 1396 | hypothetical-protein-[Bacillus-cereus] |
| WP_237075536.1 | Type1.BREX | [illegible] | 1192 | Bacteria>Terrabacteria_group>Bacillota>Bacilli>Bacillales>Bacillaceae | GCF_021864615.1 | Priestia_megaterium | L4R70_RS26975 | 37253..40831 | 1404 | BREX-system-P-loop-protein-BrxC-[Priestia-megaterium] |
| WP_240257149.1 | Type1.BREX | [illegible] | 1195 | Bacteria>Terrabacteria_group>Bacillota>Bacilli>Bacillales>Bacillaceae | GCF_022427965.1 | Fredinandcohnia_sp._SECRCQ15 | MJG50_RS17985 | 30135..33722 | 2918902 | BREX-system-P-loop-protein-BrxC-[Fredinandcohnia-sp.-SECRCQ15] |
| WP_241817599.1 | Type1.BREX | [illegible] | 1186 | Bacteria>Terrabacteria_group>Bacillota>Bacilli>Bacillales>Bacillaceae | GCF_022625245.1 | Bacillus_sp._TL12 | LOK41_RS17400 | 27409..30969 | 2894756 | BREX-system-P-loop-protein-BrxC-[Bacillus-sp.-TL12] |
| WP_242196467.1 | Type1.BREX | [illegible] | 1191 | Bacteria>Terrabacteria_group>Bacillota>Bacilli>Bacillales>Bacillaceae | GCF_022662815.1 | Bacillus_cereus_group_sp._BfR-BA-01492 | MR063_RS08665 | 67486..71061 | 2920361 | BREX-system-P-loop-protein-BrxC-[Bacillus-cereus-group-sp.-BfR-BA-01492] |
| WP_242554826.1 | Type1.BREX | [illegible] | 1195 | Bacteria>Terrabacteria_group>Bacillota>Bacilli>Bacillales>Bacillaceae | GCF_022696175.1 | Bacillus_cereus | C5137_09385 | 1769234..1772821 | 1396 | BREX-system-P-loop-protein-BrxC-[Bacillus-cereus] |
| WP_243824972.1 | Type1.BREX | [illegible] | 1194 | Bacteria>Terrabacteria_group>Bacillota>Bacilli>Bacillales>Bacillaceae | GCF_022819205.1 | Bacillus_altitudinis | MTX65_RS03395 | 674724..678308 | 293387 | BREX-system-P-loop-protein-BrxC-[Bacillus-altitudinis] |
| WP_248717077.1 | Type1.BREX | [illegible] | 1194 | Bacteria>Terrabacteria_group>Bacillota>Bacilli>Bacillales>Bacillaceae | GCF_023242215.1 | Lysinibacillus_sp._Ag94 | MY533_RS18340 | 3907339..3910923 | 2936682 | BREX-system-P-loop-protein-BrxC-[Lysinibacillus-sp.-Ag94] |
| WP_251685493.1 | Type1.BREX | [illegible] | 1194 | Bacteria>Terrabacteria_group>Bacillota>Bacilli>Bacillales>Bacillaceae | GCF_023713225.1 | Metabacillus_halosaccharovorans | M3683_RS22960 | 50474..54058 | 930124 | BREX-system-P-loop-protein-BrxC-[Metabacillus-halosaccharovorans] |
| WP_251628816.1 | Type1.BREX | [illegible] | 1194 | Bacteria>Terrabacteria_group>Bacillota>Bacilli>Bacillales>Bacillaceae | GCF_023713275.1 | Niallia_sp._MER_TA_168 | M3659_RS13820 | 26120..29704 | 2939568 | BREX-system-P-loop-protein-BrxC-[Niallia-sp.-MER-TA-168] |
| WP_251462564.1 | Type1.BREX | [illegible] | 1190 | Bacteria>Terrabacteria_group>Bacillota>Bacilli>Bacillales>Bacillaceae | GCF_023713865.1 | Priestia_aryabhattai | M3635_RS25970 | 27912..31484 | 412384 | BREX-system-P-loop-protein-BrxC-[Priestia-aryabhattai] |
| WP_251181366.1 | Type1.BREX | [illegible] | 1194 | Bacteria>Terrabacteria_group>Bacillota>Bacilli>Bacillales>Bacillaceae | GCF_023714765.1 | Bacillus_safensis | M3573_RS16070 | 195672..199256 | 561879 | BREX-system-P-loop-protein-BrxC-[Bacillus-safensis] |
| WP_252209316.1 | Type1.BREX | [illegible] | 1193 | Bacteria>Terrabacteria_group>Bacillota>Bacilli>Bacillales>Bacillaceae | GCF_023824095.1 | Bacillus_thuringiensis | LIT28_RS29000 | 287070..290651 | 1428 | BREX-system-P-loop-protein-BrxC-[Bacillus-thuringiensis] |
| WP_256226107.1 | Type1.BREX | [illegible] | 1193 | Bacteria>Terrabacteria_group>Bacillota>Bacilli>Bacillales>Bacillaceae | GCF_024494405.1 | Fictibacillus_sp._WQ_8-8 | M1K46_RS11725 | 422268..425849 | 2938788 | BREX-system-P-loop-protein-BrxC-[Fictibacillus-sp.-WQ-8-8] |
| WP_256240922.1 | Type1.BREX | [illegible] | 1192 | Bacteria>Terrabacteria_group>Bacillota>Bacilli>Bacillales>Bacillaceae | GCF_024494465.1 | Bacillus_sp._V3B | JMM81_RS07410 | 65568..69146 | 2804915 | BREX-system-P-loop-protein-BrxC-[Bacillus-sp.-V3B] |
| WP_061456853.1 | Type1.BREX | [illegible] | 1193 | Bacteria>Terrabacteria_group>Bacillota>Bacilli>Bacillales>Bacillaceae | GCF_024498915.1 | Bacillus_cereus | L2I54_04675 | 914094..917675 | 1396 | BREX-system-P-loop-protein-BrxC-[Bacillus-cereus] |
| WP_257351442.1 | Type1.BREX | [illegible] | 1188 | Bacteria>Terrabacteria_group>Bacillota>Bacilli>Bacillales>Bacillaceae | GCF_024609785.1 | Pseudalkalibacillus_decolorationis | NON16_RS21340 | 32931..36497 | 163879 | BREX-system-P-loop-protein-BrxC-[Pseudalkalibacillus-decolorationis] |
| WP_103102742.1 | Type1.BREX | [illegible] | 1186 | Bacteria>Terrabacteria_group>Bacillota>Bacilli>Bacillales>Bacillaceae | GCF_024712785.1 | Bacillus | NHG47_RS26605 | 5089487..5093047 | 1386 | MULTISPECIES:-BREX-system-P-loop-protein-BrxC-[Bacillus] |
| WP_259420301.1 | Type1.BREX | [illegible] | 1191 | Bacteria>Terrabacteria_group>Bacillota>Bacilli>Bacillales>Bacillaceae | GCF_024919535.1 | Lysinibacillus_sp._A4 | NY607_RS20580 | 136984..140559 | 2976269 | BREX-system-P-loop-protein-BrxC-[Lysinibacillus-sp.-A4] |
| WP_259548277.1 | Type1.BREX | [illegible] | 1187 | Bacteria>Terrabacteria_group>Bacillota>Bacilli>Bacillales>Bacillaceae | GCF_024971915.1 | Heyndrickxia_oleronia | KXF32_RS03880 | 741080..744643 | 38875 | BREX-system-P-loop-protein-BrxC-[Heyndrickxia-oleronia] |
| WP_265627628.1 | Type1.BREX | [illegible] | 1191 | Bacteria>Terrabacteria_group>Bacillota>Bacilli>Bacillales>Bacillaceae | GCF_026210435.1 | Bacillus_paralicheniformis | OPU65_RS10200 | 1974820..1978395 | 1648923 | BREX-system-P-loop-protein-BrxC-[Bacillus-paralicheniformis] |
| WP_268350015.1 | Type1.BREX | [illegible] | 1194 | Bacteria>Terrabacteria_group>Bacillota>Bacilli>Bacillales>Bacillaceae | GCF_026790155.1 | Bacillus_haynesii | MX159_RS10935 | 51273..54857 | 1925021 | BREX-system-P-loop-protein-BrxC-[Bacillus-haynesii] |
| WP_268348715.1 | Type1.BREX | [illegible] | 1194 | Bacteria>Terrabacteria_group>Bacillota>Bacilli>Bacillales>Bacillaceae | GCF_026796415.1 | Bacillus_haynesii | MOF36_RS21390 | 75295..78879 | 1925021 | BREX-system-P-loop-protein-BrxC-[Bacillus-haynesii] |
| WP_270721503.1 | Type1.BREX | [illegible] | 1193 | Bacteria>Terrabacteria_group>Bacillota>Bacilli>Bacillales>Bacillaceae | GCF_027713805.1 | unclassified_Bacillus_cereus_group | PDM96_RS19895 | 249516..253097 | 2750818 | MULTISPECIES:-BREX-system-P-loop-protein-BrxC-[unclassified-Bacillus-cereus-group] |
| WP_270755744.1 | Type1.BREX | [illegible] | 1190 | Bacteria>Terrabacteria_group>Bacillota>Bacilli>Bacillales>Bacillaceae | GCF_027716165.1 | unclassified_Bacillus_cereus_group | PDQ71_RS11875 | 42041..45613 | 2750818 | MULTISPECIES:-BREX-system-P-loop-protein-BrxC-[unclassified-Bacillus-cereus-group] |
| WP_275118136.1 | Type1.BREX | [illegible] | 1188 | Bacteria>Terrabacteria_group>Bacillota>Bacilli>Bacillales>Bacillaceae | GCF_029028005.1 | Alkalihalobacillus_sp._MEB203 | N7Z68_RS08935 | 51886..55452 | 2980103 | BREX-system-P-loop-protein-BrxC-[Alkalihalobacillus-sp.-MEB203] |
| WP_061531636.1 | Type1.BREX | [illegible] | 1195 | Bacteria>Terrabacteria_group>Bacillota>Bacilli>Bacillales>Bacillaceae | GCF_029504955.1 | Bacillus_cereus_group | M2246_RS18875 | 193407..196994 | 86661 | MULTISPECIES:-BREX-system-P-loop-protein-BrxC-[Bacillus] |
| WP_120075792.1 | Type1.BREX | [illegible] | 1186 | Bacteria>Terrabacteria_group>Bacillota>Bacilli>Bacillales>Bacillaceae | GCF_030123425.1 | Siminovitchia_fortis | D4N35_016900 | 7617..11177 | 254758 | BREX-system-P-loop-protein-BrxC-[Siminovitchia-fortis] |
| WP_110928088.1 | Type1.BREX | [illegible] | 1197 | Bacteria>Terrabacteria_group>Bacillota>Bacilli>Bacillales>Bacillaceae | GCF_900098925.1 | Bacillus_massiliglaciei | BQ4305_RS12315 | 67463..71056 | 1816693 | BREX-system-P-loop-protein-BrxC-[Bacillus-massiliglaciei] |
| WP_093194832.1 | Type1.BREX | [illegible] | 1182 | Bacteria>Terrabacteria_group>Bacillota>Bacilli>Bacillales>Bacillaceae | GCF_900100295.1 | Salimicrobium_halophilum | BLQ44_RS15655 | 20875..24423 | 86666 | BREX-system-P-loop-protein-BrxC-[Salimicrobium-halophilum] |
| WP_244149217.1 | Type1.BREX | [illegible] | 1191 | Bacteria>Terrabacteria_group>Bacillota>Bacilli>Bacillales>Bacillaceae | GCF_900111795.1 | Anoxybacillus_pushchinoensis | BM213_RS08155 | 6843..10418 | 150248 | BREX-system-P-loop-protein-BrxC-[Anoxybacillus-pushchinoensis] |
| WP_088049429.1 | Type1.BREX | [illegible] | 1191 | Bacteria>Terrabacteria_group>Bacillota>Bacilli>Bacillales>Bacillaceae | GCF_900155625.1 | Virgibacillus_dakarensis | CZ382_RS01190 | 203858..207433 | 1917889 | BREX-system-P-loop-protein-BrxC-[Virgibacillus-dakarensis] |
| WP_088051628.1 | Type1.BREX | [illegible] | 1366 | Bacteria>Terrabacteria_group>Bacillota>Bacilli>Bacillales>Bacillaceae | GCF_900155625.1 | Virgibacillus_dakarensis | CZ382_RS12920 | 682519..686619 | 1917889 | hypothetical-protein-[Virgibacillus-dakarensis] |
| WP_042357731.1 | Type1.BREX | [illegible] | 1190 | Bacteria>Terrabacteria_group>Bacillota>Bacilli>Bacillales>Bacillaceae | GCF_902375395.1 | Bacillus_rubiinfantis | FY453_RS20345 | 135883..139455 | 1499680 | BREX-system-P-loop-protein-BrxC-[Bacillus-rubiinfantis] |
| WP_033827164.1 | Type1.BREX | [illegible] | 1186 | Bacteria>Terrabacteria_group>Bacillota>Bacilli>Bacillales>Bacillaceae | GCF_902375605.1 | Bacillus_andreraoultii | FY465_RS04670 | 342053..345613 | 1499685 | BREX-system-P-loop-protein-BrxC-[Bacillus-andreraoultii] |
| WP_147675231.1 | Type1.BREX | [illegible] | 1260 | Bacteria>Terrabacteria_group>Bacillota>Bacilli>Bacillales>Bacillaceae | GCF_902375755.1 | Numidum_massiliense | BN1247_RS10940 | 2280999..2284781 | 1522315 | BREX-system-P-loop-protein-BrxC-[Numidum-massiliense] |
| WP_175638347.1 | Type1.BREX | [illegible] | 1193 | Bacteria>Terrabacteria_group>Bacillota>Bacilli>Bacillales>Bacillaceae | GCF_903166545.1 | Metabacillus_schmidteae | HWV59_RS06330 | 1297576..1301157 | 2730405 | BREX-system-P-loop-protein-BrxC-[Metabacillus-schmidteae] |
| WP_230497381.1 | Type1.BREX | [illegible] | 1193 | Bacteria>Terrabacteria_group>Bacillota>Bacilli>Bacillales>Bacillaceae | GCF_917563885.1 | Pseudoneobacillus_rhizosphaerae | NEOCIP111885_02886 | 60307..63888 | 2880968 | BREX-system-P-loop-protein-BrxC-[Pseudoneobacillus-rhizosphaerae] |
| WP_251541482.1 | Type1.BREX | [illegible] | 1194 | Bacteria>Terrabacteria_group>Bacillota>Bacilli>Bacillales>Bacillaceae | - | unclassified_Mesobacillus | M3593_RS10825 | 897349..900933 | 2675270 | MULTISPECIES:-BREX-system-P-loop-protein-BrxC-[unclassified-Mesobacillus] |
| WP_102415646.1 | Type1.BREX | [illegible] | 1340 | Bacteria>Terrabacteria_group>Bacillota>Bacilli>Bacillales>Bacillaceae | - | Virgibacillus_dakarensis | KK120_02535 | 179341..183363 | 1917889 | MULTISPECIES:-BREX-system-P-loop-protein-BrxC-[Terrabacteria-group] |
| WP_250889608.1 | Type1.BREX | [illegible] | 1335 | Bacteria>Terrabacteria_group>Bacillota>Bacilli>Bacillales>Bacillaceae | - | Virgibacillus_dakarensis | KK120_RS02530 | 179356..183363 | 1917889 | BREX-system-P-loop-protein-BrxC-[Virgibacillus-dakarensis] |
| MBR3215270.1 | Type1.BREX | [illegible] | 915 | Bacteria>Terrabacteria_group>Bacillota>Bacilli>Bacillales>Bacillales_incertae_sedis | GCA_017514325.1 | Exiguobacterium_sp. | IKF69_02280 | 1..2744 | 44751 | BREX-system-P-loop-protein-BrxC-[Exiguobacterium-sp.] |
| WP_012369225.1 | Type1.BREX | [illegible] | 1192 | Bacteria>Terrabacteria_group>Bacillota>Bacilli>Bacillales>Bacillales_incertae_sedis | GCF_000019905.1 | Exiguobacterium_sibiricum | Exig_0314 | 327084..330662 | 332410 | BREX-system-P-loop-protein-BrxC-[Exiguobacterium-sibiricum] |
| WP_028107090.1 | Type1.BREX | [illegible] | 1192 | Bacteria>Terrabacteria_group>Bacillota>Bacilli>Bacillales>Bacillales_incertae_sedis | GCF_000620805.1 | Exiguobacterium_undae | P397_RS0115435 | 2856472..2860050 | 169177 | BREX-system-P-loop-protein-BrxC-[Exiguobacterium-undae] |
| WP_131470342.1 | Type1.BREX | [illegible] | 1188 | Bacteria>Terrabacteria_group>Bacillota>Bacilli>Bacillales>Bacillales_incertae_sedis | GCF_004337105.1 | Exiguobacterium_sp._SH3S1 | EVJ26_RS09655 | 119198..122764 | 2510955 | BREX-system-P-loop-protein-BrxC-[Exiguobacterium-sp.-SH3S1] |
| WP_184405123.1 | Type1.BREX | [illegible] | 1206 | Bacteria>Terrabacteria_group>Bacillota>Bacilli>Bacillales>Bacillales_incertae_sedis | GCF_014207655.1 | Geomicrobium_halophilum | HNR44_RS14850 | 20482..24102 | 549000 | BREX-system-P-loop-protein-BrxC-[Geomicrobium-halophilum] |
| WP_195865812.1 | Type1.BREX | [illegible] | 1192 | Bacteria>Terrabacteria_group>Bacillota>Bacilli>Bacillales>Bacillales_incertae_sedis | GCF_015645365.1 | Exiguobacterium_sp._TBG-PICH-001 | ITJ88_RS11985 | 155888..159466 | 2785532 | BREX-system-P-loop-protein-BrxC-[Exiguobacterium-sp.-TBG-PICH-001] |
| WP_214711985.1 | Type1.BREX | [illegible] | 1188 | Bacteria>Terrabacteria_group>Bacillota>Bacilli>Bacillales>Bacillales_incertae_sedis | GCF_018617985.1 | Exiguobacterium_sp._s55 | KNP09_RS10355 | 38338..41904 | 2751245 | BREX-system-P-loop-protein-BrxC-[Exiguobacterium-sp.-s55] |
| WP_214837746.1 | Type1.BREX | [illegible] | 1188 | Bacteria>Terrabacteria_group>Bacillota>Bacilli>Bacillales>Bacillales_incertae_sedis | GCF_018618345.1 | Exiguobacterium_sp._s36 | KNP95_RS13400 | 83896..87462 | 2751227 | BREX-system-P-loop-protein-BrxC-[Exiguobacterium-sp.-s36] |
| WP_214844978.1 | Type1.BREX | [illegible] | 1192 | Bacteria>Terrabacteria_group>Bacillota>Bacilli>Bacillales>Bacillales_incertae_sedis | GCF_018618435.1 | Exiguobacterium_sp._s150 | KNP76_RS15010 | 3016..6594 | 2751221 | BREX-system-P-loop-protein-BrxC-[Exiguobacterium-sp.-s150] |
| WP_214855538.1 | Type1.BREX | [illegible] | 1189 | Bacteria>Terrabacteria_group>Bacillota>Bacilli>Bacillales>Bacillales_incertae_sedis | GCF_018618855.1 | Exiguobacterium_sp._s166 | KNP93_RS12475 | 27028..30597 | 2751204 | BREX-system-P-loop-protein-BrxC-[Exiguobacterium-sp.-s166] |
| WP_088074350.1 | Type1.BREX | [illegible] | 1205 | Bacteria>Terrabacteria_group>Bacillota>Bacilli>Bacillales | GCF_019042735.1 | Bacillaceae | KS407_RS09130 | 3322..6939 | 186817 | MULTISPECIES:-BREX-system-P-loop-protein-BrxC-[Bacillaceae] |
| PZF87824.1 | Type1.BREX | [illegible] | 709 | Bacteria>Terrabacteria_group>Bacillota>Bacilli>Bacillales>Listeriaceae | GCA_003236525.1 | Listeria_ivanovii | C1905_11805 | 25473..27602 | 1638 | BREX-system-P-loop-protein-BrxC-[Listeria-ivanovii] |
| WP_038408557.1 | Type1.BREX | [illegible] | 1193 | Bacteria>Terrabacteria_group>Bacillota>Bacilli>Bacillales>Listeriaceae | GCF_000763475.1 | Listeria_ivanovii | JL58_RS02435 | 518979..522560 | 1638 | BREX-system-P-loop-protein-BrxC-[Listeria-ivanovii] |
| WP_253255047.1 | Type1.BREX | [illegible] | 696 | Bacteria>Terrabacteria_group>Bacillota>Bacilli>Bacillales>Listeriaceae | GCF_003316755.1 | Listeria_ivanovii | C1899_RS11615 | 25473..27563 | 1638 | BREX-system-P-loop-protein-BrxC-[Listeria-ivanovii] |
| WP_185368496.1 | Type1.BREX | [illegible] | 1193 | Bacteria>Terrabacteria_group>Bacillota>Bacilli>Bacillales>Listeriaceae | GCF_014222345.1 | Listeria_booriae | HB817_RS11180 | 30544..34125 | 1552123 | BREX-system-P-loop-protein-BrxC-[Listeria-booriae] |
| WP_185362295.1 | Type1.BREX | [illegible] | 1193 | Bacteria>Terrabacteria_group>Bacillota>Bacilli>Bacillales>Listeriaceae | GCF_014222445.1 | Listeria_booriae | HB814_RS11985 | 175648..179229 | 1552123 | BREX-system-P-loop-protein-BrxC-[Listeria-booriae] |
| WP_185366528.1 | Type1.BREX | [illegible] | 1193 | Bacteria>Terrabacteria_group>Bacillota>Bacilli>Bacillales>Listeriaceae | GCF_014222665.1 | Listeria_booriae | HB820_RS07225 | 29407..32988 | 1552123 | BREX-system-P-loop-protein-BrxC-[Listeria-booriae] |
| WP_185386338.1 | Type1.BREX | [illegible] | 1193 | Bacteria>Terrabacteria_group>Bacillota>Bacilli>Bacillales>Listeriaceae | GCF_014222895.1 | Listeria_booriae | HB856_13205 | 326299..329880 | 1552123 | BREX-system-P-loop-protein-BrxC-[Listeria-booriae] |
| WP_185336408.1 | Type1.BREX | [illegible] | 1193 | Bacteria>Terrabacteria_group>Bacillota>Bacilli>Bacillales>Listeriaceae | GCF_014223285.1 | Listeria_seeligeri | HB832_RS11630 | 11793..15374 | 1640 | BREX-system-P-loop-protein-BrxC-[Listeria-seeligeri] |
| WP_185337616.1 | Type1.BREX | [illegible] | 1193 | Bacteria>Terrabacteria_group>Bacillota>Bacilli>Bacillales>Listeriaceae | GCF_014223645.1 | Listeria_seeligeri | HB843_RS07175 | 20354..23935 | 1640 | BREX-system-P-loop-protein-BrxC-[Listeria-seeligeri] |
| WP_185468191.1 | Type1.BREX | [illegible] | 1193 | Bacteria>Terrabacteria_group>Bacillota>Bacilli>Bacillales>Listeriaceae | GCF_014225465.1 | Listeria_seeligeri | HCA03_RS04485 | 873458..877039 | 1640 | BREX-system-P-loop-protein-BrxC-[Listeria-seeligeri] |
| WP_185614077.1 | Type1.BREX | [illegible] | 1193 | Bacteria>Terrabacteria_group>Bacillota>Bacilli>Bacillales>Listeriaceae | GCF_014228485.1 | Listeria_booriae | HCB15_RS12165 | 128944..132525 | 1552123 | BREX-system-P-loop-protein-BrxC-[Listeria-booriae] |

| Protein-ID | System | Gene-Neighbourhoods | Length | Taxonomy | Assembly-ID | Strain | Locus-Tag | Genomic-coordinates | TaxID | Fasta-defline |
|---|---|---|---|---|---|---|---|---|---|---|
| WP_185579015.1 | Type1.BREX | ZDNA-HTH-><-ABhydrolase\|\|?-><-?<-?\|\|GDResolvase+HIN-HTH->BrxB-BrxF-iSTAND->DUF6079*->N6-METHYLASE->PglZ->Cas9+HNH+RuvC+CRISPR_Cas9->Cas_Cas1->CRISPR_Cas2->Cas_Csn2->HTH_6+SIS-> | 1193 | Bacteria>Terrabacteria_group>Bacillota>Bacilli>Bacillales>Listeriaceae | GCF_014228555.1 | Listeria_booriae | HCB21_RS10060 | 340641..344222 | 1552123 | BREX-system-P-loop-protein-BrxC-[Listeria-booriae] |
| WP_185522090.1 | Type1.BREX | <-HTH_6+SIS<-Cas_Csn2<-CRISPR_Cas2<-Cas_Cas1<-Cas9+HNH+RuvC+CRISPR_Cas9<-PglZ<-N6-METHYLASE<-DUF6079*<-BrxB-BrxF-iSTAND<-GDResolvase+HIN-HTH\|\|?->?->ABhydrolase-><-ZDNA-HTH | 1193 | Bacteria>Terrabacteria_group>Bacillota>Bacilli>Bacillales>Listeriaceae | GCF_014228595.1 | Listeria_booriae | HCB29_RS12135 | 34445..38026 | 1552123 | BREX-system-P-loop-protein-BrxC-[Listeria-booriae] |
| WP_185628908.1 | Type1.BREX | <-HTH_6+SIS<-Cas_Csn2<-CRISPR_Cas2<-Cas_Cas1<-Cas9+HNH+RuvC+CRISPR_Cas9<-PglZ<-N6-METHYLASE<-DUF6079*<-BrxB-BrxF-iSTAND<-GDResolvase+HIN-HTH\|\|?->?-><-?\|\|ABhydrolase-><-ZDNA-HTH | 1193 | Bacteria>Terrabacteria_group>Bacillota>Bacilli>Bacillales>Listeriaceae | GCF_014229405.1 | Listeria_booriae | HCB69_RS07940 | 28391..31972 | 1552123 | BREX-system-P-loop-protein-BrxC-[Listeria-booriae] |
| WP_185642206.1 | Type1.BREX | <-HTH_6+SIS<-CRISPR_Cas2<-Cas_Cas1<-Cas9+HNH+RuvC+CRISPR_Cas9<-PglZ<-N6-METHYLASE<-DUF6079*<-BrxB-BrxF-iSTAND<-GDResolvase+HIN-HTH\|\|?->?-><-?\|\|ABhydrolase-><-ZDNA-HTH | 1193 | Bacteria>Terrabacteria_group>Bacillota>Bacilli>Bacillales>Listeriaceae | GCF_014229465.1 | Listeria_booriae | HCJ81_RS12860 | 26874..30455 | 1552123 | BREX-system-P-loop-protein-BrxC-[Listeria-booriae] |
| WP_185648199.1 | Type1.BREX | <-ABhydrolase<-?<-?\|\|GDResolvase+HIN-HTH->BrxB-BrxF-iSTAND->DUF6079*->N6-METHYLASE->PglZ->Cas9+HNH+RuvC+CRISPR_Cas9->Cas_Cas1->CRISPR_Cas2->Cas_Csn2->HTH_6+SIS-> | 1193 | Bacteria>Terrabacteria_group>Bacillota>Bacilli>Bacillales>Listeriaceae | GCF_014229625.1 | Listeria_booriae | HCX63_RS17175 | 5084..8665 | 1552123 | BREX-system-P-loop-protein-BrxC-[Listeria-booriae] |
| WP_185622114.1 | Type1.BREX | <-HTH_6+SIS<-Cas_Csn2<-CRISPR_Cas2<-Cas_Cas1<-Cas9+HNH+RuvC+CRISPR_Cas9<-PglZ<-N6-METHYLASE<-DUF6079*<-BrxB-BrxF-iSTAND<-GDResolvase+HIN-HTH\|\|?->?-><-?\|\|ABhydrolase-><-ZDNA-HTH | 1193 | Bacteria>Terrabacteria_group>Bacillota>Bacilli>Bacillales>Listeriaceae | GCF_014229925.1 | Listeria_booriae | HBP99_RS14050 | 28201..31782 | 1552123 | BREX-system-P-loop-protein-BrxC-[Listeria-booriae] |
| WP_203575133.1 | Type1.BREX | <-Glyco_hydro_38<-?<-?<-?<-Cas_Csn2<-CRISPR_Cas2<-Cas_Cas1<-Cas9+HNH+RuvC+CRISPR_Cas9<-PglZ<-DUF6079*<-BrxB-BrxF-iSTAND<-BrxA-DNA-binding<-GDResolvase+HIN-HTH\|\|His_Phos_1-><-?<-Imm-LR[illegible] | 1193 | Bacteria>Terrabacteria_group>Bacillota>Bacilli>Bacillales>Listeriaceae | GCF_016881625.1 | Listeria_seeligeri | D8X85_RS06090 | 1242417..1245998 | 1640 | BREX-system-P-loop-protein-BrxC-[Listeria-seeligeri] |
| WP_203576520.1 | Type1.BREX | <-PglZ<-HTH_6+SIS<-?<-ROK-HTH<-Cas_Csn2<-CRISPR_Cas2<-Cas_Cas1<-Cas9+HNH+RuvC+CRISPR_Cas9<-PglZ<-DUF6079*<-BrxB-BrxF-iSTAND<-BrxA-DNA-binding<-GDResolvase+HIN-HTH\|\|His_Phos_1-><-?<-I[illegible] | 1193 | Bacteria>Terrabacteria_group>Bacillota>Bacilli>Bacillales>Listeriaceae | GCF_016881645.1 | Listeria_seeligeri | D8X91_RS03000 | 624726..628307 | 1640 | BREX-system-P-loop-protein-BrxC-[Listeria-seeligeri] |
| WP_185463641.1 | Type1.BREX | <-Cas_Csn2<-CRISPR_Cas2<-Cas_Cas1<-Cas9+HNH+RuvC+CRISPR_Cas9<-PglZ<-N6-METHYLASE<-DUF6079*<-BrxB-BrxF-iSTAND<-BrxA-DNA-binding<-GDResolvase+HIN-HTH\|\|His_Phos_1-><-?<-Imm-LRR+IG\|\|MarR-[illegible] | 1193 | Bacteria>Terrabacteria_group>Bacillota>Bacilli>Bacillales>Listeriaceae | GCF_016881645.1 | Listeria_seeligeri | [illegible] | [illegible] | 1640 | BREX-system-P-loop-protein-BrxC-[Listeria-seeligeri] |
| WP_099222167.1 | Type1.BREX | <-ACET<-TetR-HTH\|\|?-><-?\|\|?->?->?->GDResolvase+HIN-HTH->BrxB-BrxF-iSTAND->DUF6079*->N6-METHYLASE->PglZ->Cas9+HNH+RuvC+CRISPR_Cas9->Cas_Cas1->CRISPR_Cas2->Cas_Csn2-> | 1193 | Bacteria>Terrabacteria_group>Bacillota>Bacilli>Bacillales>Listeriaceae | GCF_018475685.1 | Listeria_seeligeri | PH491_RS03775 | 52253..55834 | 1640 | BREX-system-P-loop-protein-BrxC-[Listeria-seeligeri] |
| WP_111130888.1 | Type1.BREX | <-HTH_6+SIS\|\|?-><-HTH_24+ROK-HTH<-?\|\|?-><-PglZ<-N6-METHYLASE-p<-N6-METHYLASE<-?<-DUF6079-p*<-DUF6079-p<-BrxB-BrxF-iSTAND<-BrxA-DNA-binding<-GDResolvase+HIN-HTH\|\|His_Phos_1-><-?\|\|MarR-[illegible] | 1195 | Bacteria>Terrabacteria_group>Bacillota>Bacilli>Bacillales>Listeriaceae | GCF_900183385.1 | Listeria_costaricensis | CS921_RS06065 | 258667..262254 | 2026604 | BREX-system-P-loop-protein-BrxC-[Listeria-costaricensis] |
| WP_238502397.1 | Type1.BREX | ParB+HNH->REase->?->REase->HTH_31->REase+DUF4268->?->BrxA-DNA-binding->BrxB-BrxF-iSTAND->DUF6079*->N6-METHYLASE->PglZ->SIGMA+OB-Fold+AAA-ATPase+Lon-Protease-BrxL-><-RHH<-?\|\|MarR-HTH->[illegible] | 709 | Bacteria>Terrabacteria_group>Bacillota>Bacilli>Bacillales>Listeriaceae | - | Listeria_ivanovii | C1905_11805 | 25473..27602 | 1638 | BREX-system-P-loop-protein-BrxC-[Listeria-ivanovii] |
| MXO77664.1 | Type1.BREX | <-SIGMA+OB-Fold+AAA-ATPase+Lon-Protease-BrxL-p<-PglZ<-N6-METHYLASE<-DUF6079*<-BrxB-BrxF-iSTAND<-BrxA-DNA-binding<-?<-REase+DUF4268<-?<-TRAM+METHYLASE<-?<-?<-TRPR-HTH | 1184 | Bacteria>Terrabacteria_group>Bacillota>Bacilli>Bacillales>Paenibacillaceae | - | Brevibacillus_agri | MCW62_14325 | 38506..42060 | 51101 | BREX-system-P-loop-protein-BrxC-[Brevibacillus-agri] |
| NBD25621.1 | Type1.BREX | MerR-HTH->wHTH+REase->SF1-SF2-Helicase->NUDIX->REase->HTH_31->?->BrxA-DNA-binding->BrxB-BrxF-iSTAND->DUF6079*->N6-METHYLASE->PglZ->SIGMA+OB-Fold+AAA-ATPase+Lon-Protease-BrxL->?-><-RHH[illegible] | 1188 | Bacteria>Terrabacteria_group>Bacillota>Bacilli>Bacillales>Paenibacillaceae | GCA_009827695.1 | Paenibacillus_sp._OT2-17 | GRP75_07340 | 6853..10419 | 2691605 | BREX-system-P-loop-protein-BrxC-[Paenibacillus-sp.-OT2-17] |
| NUU75083.1 | Type1.BREX | ParB+HNH->REase->HTH_31->REase+DUF4268->?->BrxA-DNA-binding->BrxB-BrxF-iSTAND->DUF6079*->N6-METHYLASE->PglZ->SIGMA+OB-Fold+AAA-ATPase+Lon-Protease-BrxL-><-RHH<-?\|\|?->VirB4-FtsK-> | [illegible] | [illegible] | [illegible] | [illegible] | [illegible] | [illegible] | [illegible] | [illegible] |
| MBN6185688.1 | Type1.BREX | <-VirB4-FtsK\|\|ZnR+ZnR->?-><-HEPN<-SIGMA+OB-Fold+AAA-ATPase+Lon-Protease-BrxL<-PglZ<-N6-METHYLASE<-DUF6079*<-BrxB-BrxF-iSTAND<-BrxA-DNA-binding<-?<-?<-MerR-HTH<-XIS-HTH<-ZnR+RHH_7\|\|?->[illegible] | [illegible] | [illegible] | [illegible] | [illegible] | [illegible] | [illegible] | [illegible] | [illegible] |
| MBO2945633.1 | Type1.BREX | [illegible] | [illegible] | [illegible] | [illegible] | [illegible] | [illegible] | [illegible] | [illegible] | [illegible] |
| MBP1153944.1 | Type1.BREX | [illegible] | [illegible] | [illegible] | [illegible] | [illegible] | [illegible] | [illegible] | [illegible] | [illegible] |
| MBT2292692.1 | Type1.BREX | [illegible] | [illegible] | [illegible] | [illegible] | [illegible] | [illegible] | [illegible] | [illegible] | [illegible] |
| MCG5252483.1 | Type1.BREX | [illegible] | [illegible] | [illegible] | [illegible] | [illegible] | [illegible] | [illegible] | [illegible] | [illegible] |
| MCG7380059.1 | Type1.BREX | [illegible] | [illegible] | [illegible] | [illegible] | [illegible] | [illegible] | [illegible] | [illegible] | [illegible] |
| MCM3342062.1 | Type1.BREX | [illegible] | [illegible] | [illegible] | [illegible] | [illegible] | [illegible] | [illegible] | [illegible] | [illegible] |
| SDT09037.1 | Type1.BREX | [illegible] | [illegible] | [illegible] | [illegible] | [illegible] | [illegible] | [illegible] | [illegible] | [illegible] |
| WP_010494807.1 | Type1.BREX | [illegible] | [illegible] | [illegible] | [illegible] | [illegible] | [illegible] | [illegible] | [illegible] | [illegible] |
| WP_018751260.1 | Type1.BREX | [illegible] | [illegible] | [illegible] | [illegible] | [illegible] | [illegible] | [illegible] | [illegible] | [illegible] |
| WP_027093447.1 | Type1.BREX | [illegible] | [illegible] | [illegible] | [illegible] | [illegible] | [illegible] | [illegible] | [illegible] | [illegible] |
| WP_036689483.1 | Type1.BREX | [illegible] | [illegible] | [illegible] | [illegible] | [illegible] | [illegible] | [illegible] | [illegible] | [illegible] |
| WP_037284690.1 | Type1.BREX | [illegible] | [illegible] | [illegible] | [illegible] | [illegible] | [illegible] | [illegible] | [illegible] | [illegible] |
| WP_047914199.1 | Type1.BREX | [illegible] | [illegible] | [illegible] | [illegible] | [illegible] | [illegible] | [illegible] | [illegible] | [illegible] |
| WP_038596865.1 | Type1.BREX | [illegible] | [illegible] | [illegible] | [illegible] | [illegible] | [illegible] | [illegible] | [illegible] | [illegible] |
| WP_038693632.1 | Type1.BREX | [illegible] | [illegible] | [illegible] | [illegible] | [illegible] | [illegible] | [illegible] | [illegible] | [illegible] |
| WP_056697878.1 | Type1.BREX | [illegible] | [illegible] | [illegible] | [illegible] | [illegible] | [illegible] | [illegible] | [illegible] | [illegible] |
| WP_060645783.1 | Type1.BREX | [illegible] | [illegible] | [illegible] | [illegible] | [illegible] | [illegible] | [illegible] | [illegible] | [illegible] |
| WP_068693300.1 | Type1.BREX | [illegible] | [illegible] | [illegible] | [illegible] | [illegible] | [illegible] | [illegible] | [illegible] | [illegible] |
| WP_079914137.1 | Type1.BREX | [illegible] | [illegible] | [illegible] | [illegible] | [illegible] | [illegible] | [illegible] | [illegible] | [illegible] |
| WP_095288526.1 | Type1.BREX | [illegible] | [illegible] | [illegible] | [illegible] | [illegible] | [illegible] | [illegible] | [illegible] | [illegible] |
| WP_096467390.1 | Type1.BREX | [illegible] | [illegible] | [illegible] | [illegible] | [illegible] | [illegible] | [illegible] | [illegible] | [illegible] |
| WP_099517056.1 | Type1.BREX | [illegible] | [illegible] | [illegible] | [illegible] | [illegible] | [illegible] | [illegible] | [illegible] | [illegible] |
| WP_126425785.1 | Type1.BREX | [illegible] | [illegible] | [illegible] | [illegible] | [illegible] | [illegible] | [illegible] | [illegible] | [illegible] |
| WP_127459520.1 | Type1.BREX | [illegible] | [illegible] | [illegible] | [illegible] | [illegible] | [illegible] | [illegible] | [illegible] | [illegible] |
| WP_134684889.1 | Type1.BREX | [illegible] | [illegible] | [illegible] | [illegible] | [illegible] | [illegible] | [illegible] | [illegible] | [illegible] |
| WP_062409619.1 | Type1.BREX | [illegible] | [illegible] | [illegible] | [illegible] | [illegible] | [illegible] | [illegible] | [illegible] | [illegible] |
| WP_148456431.1 | Type1.BREX | [illegible] | [illegible] | [illegible] | [illegible] | [illegible] | [illegible] | [illegible] | [illegible] | [illegible] |
| WP_153976927.1 | Type1.BREX | [illegible] | [illegible] | [illegible] | [illegible] | [illegible] | [illegible] | [illegible] | [illegible] | [illegible] |
| WP_202404033.1 | Type1.BREX | [illegible] | [illegible] | [illegible] | [illegible] | [illegible] | [illegible] | [illegible] | [illegible] | [illegible] |
| WP_201755430.1 | Type1.BREX | [illegible] | [illegible] | [illegible] | [illegible] | [illegible] | [illegible] | [illegible] | [illegible] | [illegible] |
| WP_163976037.1 | Type1.BREX | [illegible] | [illegible] | [illegible] | [illegible] | [illegible] | [illegible] | [illegible] | [illegible] | [illegible] |
| WP_172197184.1 | Type1.BREX | [illegible] | [illegible] | [illegible] | [illegible] | [illegible] | [illegible] | [illegible] | [illegible] | [illegible] |
| WP_217506352.1 | Type1.BREX | [illegible] | [illegible] | [illegible] | [illegible] | [illegible] | [illegible] | [illegible] | [illegible] | [illegible] |
| WP_243896166.1 | Type1.BREX | [illegible] | [illegible] | [illegible] | [illegible] | [illegible] | [illegible] | [illegible] | [illegible] | [illegible] |
| WP_245247302.1 | Type1.BREX | [illegible] | [illegible] | [illegible] | [illegible] | [illegible] | [illegible] | [illegible] | [illegible] | [illegible] |
| WP_209844448.1 | Type1.BREX | [illegible] | [illegible] | [illegible] | [illegible] | [illegible] | [illegible] | [illegible] | [illegible] | [illegible] |
| WP_212949696.1 | Type1.BREX | [illegible] | [illegible] | [illegible] | [illegible] | [illegible] | [illegible] | [illegible] | [illegible] | [illegible] |
| WP_155620894.1 | Type1.BREX | [illegible] | [illegible] | [illegible] | [illegible] | [illegible] | [illegible] | [illegible] | [illegible] | [illegible] |
| WP_213618121.1 | Type1.BREX | [illegible] | [illegible] | [illegible] | [illegible] | [illegible] | [illegible] | [illegible] | [illegible] | [illegible] |
| WP_251038338.1 | Type1.BREX | [illegible] | [illegible] | [illegible] | [illegible] | [illegible] | [illegible] | [illegible] | [illegible] | [illegible] |
| WP_222215805.1 | Type1.BREX | [illegible] | [illegible] | [illegible] | [illegible] | [illegible] | [illegible] | [illegible] | [illegible] | [illegible] |
| WP_224725205.1 | Type1.BREX | [illegible] | [illegible] | [illegible] | [illegible] | [illegible] | [illegible] | [illegible] | [illegible] | [illegible] |
| WP_274388573.1 | Type1.BREX | [illegible] | [illegible] | [illegible] | [illegible] | [illegible] | [illegible] | [illegible] | [illegible] | [illegible] |
| WP_275446318.1 | Type1.BREX | [illegible] | [illegible] | [illegible] | [illegible] | [illegible] | [illegible] | [illegible] | [illegible] | [illegible] |
| WP_251579171.1 | Type1.BREX | [illegible] | [illegible] | [illegible] | [illegible] | [illegible] | [illegible] | [illegible] | [illegible] | [illegible] |
| WP_258278098.1 | Type1.BREX | [illegible] | [illegible] | [illegible] | [illegible] | [illegible] | [illegible] | [illegible] | [illegible] | [illegible] |
| WP_262687656.1 | Type1.BREX | [illegible] | [illegible] | [illegible] | [illegible] | [illegible] | [illegible] | [illegible] | [illegible] | [illegible] |
| WP_272047030.1 | Type1.BREX | [illegible] | [illegible] | [illegible] | [illegible] | [illegible] | [illegible] | [illegible] | [illegible] | [illegible] |
| WP_274359496.1 | Type1.BREX | [illegible] | [illegible] | [illegible] | [illegible] | [illegible] | [illegible] | [illegible] | [illegible] | [illegible] |
| WP_091013323.1 | Type1.BREX | [illegible] | [illegible] | [illegible] | [illegible] | [illegible] | [illegible] | [illegible] | [illegible] | [illegible] |
| WP_110933554.1 | Type1.BREX | [illegible] | [illegible] | [illegible] | [illegible] | [illegible] | [illegible] | [illegible] | [illegible] | [illegible] |
| WP_208915620.1 | Type1.BREX | [illegible] | [illegible] | [illegible] | [illegible] | [illegible] | [illegible] | [illegible] | [illegible] | [illegible] |
| WP_215184617.1 | Type1.BREX | [illegible] | [illegible] | [illegible] | [illegible] | [illegible] | [illegible] | [illegible] | [illegible] | [illegible] |
| WP_212938911.1 | Type1.BREX | [illegible] | [illegible] | [illegible] | [illegible] | [illegible] | [illegible] | [illegible] | [illegible] | [illegible] |
| WP_239293644.1 | Type1.BREX | [illegible] | [illegible] | [illegible] | [illegible] | [illegible] | [illegible] | [illegible] | [illegible] | [illegible] |
| WP_251582630.1 | Type1.BREX | [illegible] | [illegible] | [illegible] | [illegible] | [illegible] | [illegible] | [illegible] | [illegible] | [illegible] |
| WP_155606600.1 | Type1.BREX | [illegible] | [illegible] | [illegible] | [illegible] | [illegible] | [illegible] | [illegible] | [illegible] | [illegible] |
| KGR89570.1 | Type1.BREX | [illegible] | [illegible] | [illegible] | [illegible] | [illegible] | [illegible] | [illegible] | [illegible] | [illegible] |
| PIC94624.1 | Type1.BREX | [illegible] | [illegible] | [illegible] | [illegible] | [illegible] | [illegible] | [illegible] | [illegible] | [illegible] |
| MBO0589396.1 | Type1.BREX | [illegible] | [illegible] | [illegible] | [illegible] | [illegible] | [illegible] | [illegible] | [illegible] | [illegible] |
| WP_252990012.1 | Type1.BREX | [illegible] | [illegible] | [illegible] | [illegible] | [illegible] | [illegible] | [illegible] | [illegible] | [illegible] |
| WP_053168317.1 | Type1.BREX | [illegible] | [illegible] | [illegible] | [illegible] | [illegible] | [illegible] | [illegible] | [illegible] | [illegible] |
| WP_058385280.1 | Type1.BREX | [illegible] | [illegible] | [illegible] | [illegible] | [illegible] | [illegible] | [illegible] | [illegible] | [illegible] |
| WP_068488209.1 | Type1.BREX | [illegible] | [illegible] | [illegible] | [illegible] | [illegible] | [illegible] | [illegible] | [illegible] | [illegible] |
| WP_077588117.1 | Type1.BREX | [illegible] | [illegible] | [illegible] | [illegible] | [illegible] | [illegible] | [illegible] | [illegible] | [illegible] |
| WP_083033314.1 | Type1.BREX | [illegible] | [illegible] | [illegible] | [illegible] | [illegible] | [illegible] | [illegible] | [illegible] | [illegible] |
| WP_036178904.1 | Type1.BREX | [illegible] | [illegible] | [illegible] | [illegible] | [illegible] | [illegible] | [illegible] | [illegible] | [illegible] |
| WP_233192413.1 | Type1.BREX | [illegible] | [illegible] | [illegible] | [illegible] | [illegible] | [illegible] | [illegible] | [illegible] | [illegible] |
| WP_107841263.1 | Type1.BREX | [illegible] | [illegible] | [illegible] | [illegible] | [illegible] | [illegible] | [illegible] | [illegible] | [illegible] |
| WP_108306485.1 | Type1.BREX | [illegible] | [illegible] | [illegible] | [illegible] | [illegible] | [illegible] | [illegible] | [illegible] | [illegible] |
| WP_125903398.1 | Type1.BREX | [illegible] | [illegible] | [illegible] | [illegible] | [illegible] | [illegible] | [illegible] | [illegible] | [illegible] |
| WP_208650578.1 | Type1.BREX | [illegible] | [illegible] | [illegible] | [illegible] | [illegible] | [illegible] | [illegible] | [illegible] | [illegible] |
| WP_215101348.1 | Type1.BREX | [illegible] | [illegible] | [illegible] | [illegible] | [illegible] | [illegible] | [illegible] | [illegible] | [illegible] |
| WP_216971036.1 | Type1.BREX | [illegible] | [illegible] | [illegible] | [illegible] | [illegible] | [illegible] | [illegible] | [illegible] | [illegible] |
| WP_223643511.1 | Type1.BREX | [illegible] | [illegible] | [illegible] | [illegible] | [illegible] | [illegible] | [illegible] | [illegible] | [illegible] |
| WP_124068694.1 | Type1.BREX | [illegible] | [illegible] | [illegible] | [illegible] | [illegible] | [illegible] | [illegible] | [illegible] | [illegible] |
| MCI1881447.1 | Type1.BREX | [illegible] | [illegible] | [illegible] | [illegible] | [illegible] | [illegible] | [illegible] | [illegible] | [illegible] |
| WP_010024517.1 | Type1.BREX | [illegible] | [illegible] | [illegible] | [illegible] | [illegible] | [illegible] | [illegible] | [illegible] | [illegible] |
| WP_010632733.1 | Type1.BREX | [illegible] | [illegible] | [illegible] | [illegible] | [illegible] | [illegible] | [illegible] | [illegible] | [illegible] |
| WP_027725313.1 | Type1.BREX | [illegible] | [illegible] | [illegible] | [illegible] | [illegible] | [illegible] | [illegible] | [illegible] | [illegible] |
| WP_152080341.1 | Type1.BREX | [illegible] | [illegible] | [illegible] | [illegible] | [illegible] | [illegible] | [illegible] | [illegible] | [illegible] |
| WP_205007179.1 | Type1.BREX | [illegible] | [illegible] | [illegible] | [illegible] | [illegible] | [illegible] | [illegible] | [illegible] | [illegible] |
| WP_093671255.1 | Type1.BREX | [illegible] | [illegible] | [illegible] | [illegible] | [illegible] | [illegible] | [illegible] | [illegible] | [illegible] |
| WP_130003775.1 | Type1.BREX | [illegible] | [illegible] | [illegible] | [illegible] | [illegible] | [illegible] | [illegible] | [illegible] | [illegible] |
| OZT77510.1 | Type1.BREX | [illegible] | [illegible] | [illegible] | [illegible] | [illegible] | [illegible] | [illegible] | [illegible] | [illegible] |
| HJF20832.1 | Type1.BREX | [illegible] | [illegible] | [illegible] | [illegible] | [illegible] | [illegible] | [illegible] | [illegible] | [illegible] |
| WP_256974727.1 | Type1.BREX | [illegible] | [illegible] | [illegible] | [illegible] | [illegible] | [illegible] | [illegible] | [illegible] | [illegible] |
| WP_115884287.1 | Type1.BREX | [illegible] | [illegible] | [illegible] | [illegible] | [illegible] | [illegible] | [illegible] | [illegible] | [illegible] |
| WP_094906236.1 | Type1.BREX | [illegible] | [illegible] | [illegible] | [illegible] | [illegible] | [illegible] | [illegible] | [illegible] | [illegible] |
| WP_094264422.1 | Type1.BREX | [illegible] | [illegible] | [illegible] | [illegible] | [illegible] | [illegible] | [illegible] | [illegible] | [illegible] |
| WP_108025903.1 | Type1.BREX | [illegible] | [illegible] | [illegible] | [illegible] | [illegible] | [illegible] | [illegible] | [illegible] | [illegible] |
| WP_212773156.1 | Type1.BREX | [illegible] | [illegible] | [illegible] | [illegible] | [illegible] | [illegible] | [illegible] | [illegible] | [illegible] |
| WP_091835632.1 | Type1.BREX | [illegible] | [illegible] | [illegible] | [illegible] | [illegible] | [illegible] | [illegible] | [illegible] | [illegible] |
| HHW38998.1 | Type1.BREX | [illegible] | [illegible] | [illegible] | [illegible] | [illegible] | [illegible] | [illegible] | [illegible] | [illegible] |
| MBK0347250.1 | Type1.BREX | [illegible] | [illegible] | [illegible] | [illegible] | [illegible] | [illegible] | [illegible] | [illegible] | [illegible] |
| WP_069286048.1 | Type1.BREX | [illegible] | [illegible] | [illegible] | [illegible] | [illegible] | [illegible] | [illegible] | [illegible] | [illegible] |
| WP_153832550.1 | Type1.BREX | [illegible] | [illegible] | [illegible] | [illegible] | [illegible] | [illegible] | [illegible] | [illegible] | [illegible] |
| WP_153861778.1 | Type1.BREX | [illegible] | [illegible] | [illegible] | [illegible] | [illegible] | [illegible] | [illegible] | [illegible] | [illegible] |
| WP_197559022.1 | Type1.BREX | [illegible] | [illegible] | [illegible] | [illegible] | [illegible] | [illegible] | [illegible] | [illegible] | [illegible] |
| HAJ70155.1 | Type1.BREX | [illegible] | [illegible] | [illegible] | [illegible] | [illegible] | [illegible] | [illegible] | [illegible] | [illegible] |
| KAF3303185.1 | Type1.BREX | [illegible] | [illegible] | [illegible] | [illegible] | [illegible] | [illegible] | [illegible] | [illegible] | [illegible] |
| HJA89654.1 | Type1.BREX | [illegible] | [illegible] | [illegible] | [illegible] | [illegible] | [illegible] | [illegible] | [illegible] | [illegible] |
| WP_027109249.1 | Type1.BREX | [illegible] | [illegible] | [illegible] | [illegible] | [illegible] | [illegible] | [illegible] | [illegible] | [illegible] |
| WP_051258577.1 | Type1.BREX | [illegible] | [illegible] | [illegible] | [illegible] | [illegible] | [illegible] | [illegible] | [illegible] | [illegible] |
| WP_034551880.1 | Type1.BREX | [illegible] | [illegible] | [illegible] | [illegible] | [illegible] | [illegible] | [illegible] | [illegible] | [illegible] |
| WP_157464084.1 | Type1.BREX | [illegible] | [illegible] | [illegible] | [illegible] | [illegible] | [illegible] | [illegible] | [illegible] | [illegible] |
| WP_076767471.1 | Type1.BREX | [illegible] | [illegible] | [illegible] | [illegible] | [illegible] | [illegible] | [illegible] | [illegible] | [illegible] |
| WP_106190321.1 | Type1.BREX | [illegible] | [illegible] | [illegible] | [illegible] | [illegible] | [illegible] | [illegible] | [illegible] | [illegible] |
| WP_108033171.1 | Type1.BREX | [illegible] | [illegible] | [illegible] | [illegible] | [illegible] | [illegible] | [illegible] | [illegible] | [illegible] |
| WP_091487342.1 | Type1.BREX | [illegible] | [illegible] | [illegible] | [illegible] | [illegible] | [illegible] | [illegible] | [illegible] | [illegible] |
| WP_146924061.1 | Type1.BREX | [illegible] | [illegible] | [illegible] | [illegible] | [illegible] | [illegible] | [illegible] | [illegible] | [illegible] |
| WP_188204587.1 | Type1.BREX | [illegible] | [illegible] | [illegible] | [illegible] | [illegible] | [illegible] | [illegible] | [illegible] | [illegible] |
| WP_220727330.1 | Type1.BREX | [illegible] | [illegible] | [illegible] | [illegible] | [illegible] | [illegible] | [illegible] | [illegible] | [illegible] |
| WP_230523906.1 | Type1.BREX | [illegible] | [illegible] | [illegible] | [illegible] | [illegible] | [illegible] | [illegible] | [illegible] | [illegible] |
| WP_272161471.1 | Type1.BREX | [illegible] | [illegible] | [illegible] | [illegible] | [illegible] | [illegible] | [illegible] | [illegible] | [illegible] |
| WP_274731382.1 | Type1.BREX | [illegible] | [illegible] | [illegible] | [illegible] | [illegible] | [illegible] | [illegible] | [illegible] | [illegible] |
| WP_274731953.1 | Type1.BREX | [illegible] | [illegible] | [illegible] | [illegible] | [illegible] | [illegible] | [illegible] | [illegible] | [illegible] |
| WP_068559440.1 | Type1.BREX | [illegible] | [illegible] | [illegible] | [illegible] | [illegible] | [illegible] | [illegible] | [illegible] | [illegible] |
| WP_091633361.1 | Type1.BREX | [illegible] | [illegible] | [illegible] | [illegible] | [illegible] | [illegible] | [illegible] | [illegible] | [illegible] |
| WP_085559185.1 | Type1.BREX | [illegible] | [illegible] | [illegible] | [illegible] | [illegible] | [illegible] | [illegible] | [illegible] | [illegible] |
| WP_085558466.1 | Type1.BREX | [illegible] | [illegible] | [illegible] | [illegible] | [illegible] | [illegible] | [illegible] | [illegible] | [illegible] |
| WP_092651006.1 | Type1.BREX | [illegible] | [illegible] | [illegible] | [illegible] | [illegible] | [illegible] | [illegible] | [illegible] | [illegible] |
| WP_091531492.1 | Type1.BREX | [illegible] | [illegible] | [illegible] | [illegible] | [illegible] | [illegible] | [illegible] | [illegible] | [illegible] |
| WP_092090777.1 | Type1.BREX | [illegible] | [illegible] | [illegible] | [illegible] | [illegible] | [illegible] | [illegible] | [illegible] | [illegible] |
| WP_119093786.1 | Type1.BREX | [illegible] | [illegible] | [illegible] | [illegible] | [illegible] | [illegible] | [illegible] | [illegible] | [illegible] |
| WP_201729912.1 | Type1.BREX | [illegible] | [illegible] | [illegible] | [illegible] | [illegible] | [illegible] | [illegible] | [illegible] | [illegible] |
| WP_225744536.1 | Type1.BREX | [illegible] | [illegible] | [illegible] | [illegible] | [illegible] | [illegible] | [illegible] | [illegible] | [illegible] |

| Protein-ID | System | Gene-Neighbourhoods | Length | Taxonomy | Assembly-ID | Strain | Locus-Tag | Genomic-coordinates | TaxID | Fasta-define |
|---|---|---|---|---|---|---|---|---|---|---|
| WP_240393577.1 | Type1.BREX | <-ABC-ATPase+SbcC\|\|HTH_3-><-SIGMA+OB-Fold+AAA-ATPase+Lon-Protease-BrxL<-PglZ<-N6-METHYLASE<-DUF6079*<-BrxB-BrxF-iSTAND<-BrxA-DNA-binding | 1223 | Bacteria>Terrabacteria_group>Bacillota>Bacilli>Lactobacillales>Enterococcaceae | - | Enterococcus_faecium | LC935_10425 | 13809..17480 | 1352 | BREX-system-P-loop-protein-BrxC-[Enterococcus-faecium] |
| HCT96458.1 | Type1.BREX | ABC-ATPase+SbcC->?->?->?->?->?->?->BrxA-DNA-binding->BrxB-BrxF-iSTAND->DUF6079*->N6-METHYLASE-p-> | 1186 | Bacteria>Terrabacteria_group>Bacillota>Bacilli>Lactobacillales>Enterococcaceae | GCA_003542775.1 | Vagococcus_sp. | DIC43_05855 | 39044..42604 | 1933889 | BREX-system-P-loop-protein-BrxC-[Vagococcus-sp.] |
| NTK19117.1 | Type1.BREX | <-ABC-ATPase+SbcC<-?\|\|HTH_3-><-SIGMA+OB-Fold+AAA-ATPase+Lon-Protease-BrxL<-PglZ<-N6-METHYLASE<-DUF6079*<-BrxB-BrxF-iSTAND<-BrxA-DNA-binding<-?<-?<-?<-?\|\|?-><-DNA_METHYLASE<-HTH_3 | 1223 | Bacteria>Terrabacteria_group>Bacillota>Bacilli>Lactobacillales>Enterococcaceae | GCA_013319725.1 | Enterococcus_faecium | HQ640_03130 | 37512..41183 | 1352 | BREX-system-P-loop-protein-BrxC-[Enterococcus-faecium] |
| NTK45186.1 | Type1.BREX | HTH_3-><-SIGMA+OB-Fold+AAA-ATPase+Lon-Protease-BrxL<-PglZ<-N6-METHYLASE<-DUF6079*<-BrxB-BrxF-iSTAND<-BrxA-DNA-binding | 1223 | Bacteria>Terrabacteria_group>Bacillota>Bacilli>Lactobacillales>Enterococcaceae | GCA_013319965.1 | Enterococcus_faecium | HQ652_10115 | 10379..14050 | 1352 | BREX-system-P-loop-protein-BrxC-[Enterococcus-faecium] |
| NTK69333.1 | Type1.BREX | <-ABC-ATPase+SbcC\|\|HTH_3-><-SIGMA+OB-Fold+AAA-ATPase+Lon-Protease-BrxL<-PglZ<-N6-METHYLASE<-DUF6079*<-BrxB-BrxF-iSTAND<-BrxA-DNA-binding<-?<-?<-?\|\|?-><-DNA_METHYLASE<-?<-HTH_3 | 1223 | Bacteria>Terrabacteria_group>Bacillota>Bacilli>Lactobacillales>Enterococcaceae | GCA_013320115.1 | Enterococcus_faecium | HQ665_07235 | 31996..35667 | 1352 | BREX-system-P-loop-protein-BrxC-[Enterococcus-faecium] |
| NTM85167.1 | Type1.BREX | DDE_Tnp_1_3->BrxA-DNA-binding->BrxB-BrxF-iSTAND->DUF6079*->N6-METHYLASE->PglZ->SIGMA+OB-Fold+AAA-ATPase+Lon-Protease-BrxL-><-HTH_3\|\|?->?->ABC-ATPase+SbcC-> | 1223 | Bacteria>Terrabacteria_group>Bacillota>Bacilli>Lactobacillales>Enterococcaceae | GCA_013323295.1 | Enterococcus_faecium | HQ718_09930 | 1613..5284 | 1352 | BREX-system-P-loop-protein-BrxC-[Enterococcus-faecium] |
| NTO21783.1 | Type1.BREX | <-ABC-ATPase+SbcC<-?<-?\|\|HTH_3-><-SIGMA+OB-Fold+AAA-ATPase+Lon-Protease-BrxL<-PglZ<-N6-METHYLASE<-DUF6079*<-BrxB-BrxF-iSTAND<-BrxA-DNA-binding<-DDE_Tnp_1_3 | 1223 | Bacteria>Terrabacteria_group>Bacillota>Bacilli>Lactobacillales>Enterococcaceae | GCA_013324745.1 | Enterococcus_faecium | HQ913_09345 | 18191..21862 | 1352 | BREX-system-P-loop-protein-BrxC-[Enterococcus-faecium] |
| EGO7696340.1 | Type1.BREX | <-DEOR-HTH<-?<-?<-?<-?\|\|Hpr-Kinase-N->?->BrxA-DNA-binding->BrxB-BrxF-iSTAND->DUF6079*->N6-METHYLASE-><-N6-METHYLASE-p<-Phage_int_SAM_5+Phage_integrase\|\|N6-METHYLASE-p->PglZ->SIGMA+OB-Fold+AAA-ATPase+Lon-Protease-BrxL->?-><-MerR-HTH+HCR+BAF\|\|PTSEIIC-> | 1221 | Bacteria>Terrabacteria_group>Bacillota>Bacilli>Lactobacillales>Enterococcaceae | GCA_015327805.1 | Enterococcus_faecalis | FP314_001561 | 23157..26822 | 1351 | BREX-system-P-loop-protein-BrxC-[Enterococcus-faecalis] |
| EGO8062761.1 | Type1.BREX | <-N6-METHYLASE<-DUF6079*<-BrxB-BrxF-iSTAND<-BrxA-DNA-binding<-?<-Hpr-Kinase-N\|\|?->?->?->?->DEOR-HTH-> | 1221 | Bacteria>Terrabacteria_group>Bacillota>Bacilli>Lactobacillales>Enterococcaceae | GCA_015330365.1 | Enterococcus_faecalis | FBR83_000486 | 3057..6722 | 1351 | BREX-system-P-loop-protein-BrxC-[Enterococcus-faecalis] |
| HAP7983067.1 | Type1.BREX | <-ABC-ATPase+SbcC<-?<-?\|\|HTH_3-><-SIGMA+OB-Fold+AAA-ATPase+Lon-Protease-BrxL<-PglZ<-N6-METHYLASE<-DUF6079*<-BrxB-BrxF-iSTAND<-BrxA-DNA-binding | 1223 | Bacteria>Terrabacteria_group>Bacillota>Bacilli>Lactobacillales>Enterococcaceae | GCA_015364525.1 | Enterococcus_faecium | IYY73_002092 | 18058..21729 | 1352 | BREX-system-P-loop-protein-BrxC-[Enterococcus-faecium] |
| HAQ6848321.1 | Type1.BREX | <-ABC-ATPase+SbcC<-?<-?\|\|HTH_3-><-SIGMA+OB-Fold+AAA-ATPase+Lon-Protease-BrxL<-PglZ<-N6-METHYLASE<-DUF6079*<-BrxB-BrxF-iSTAND<-BrxA-DNA-binding | 1223 | Bacteria>Terrabacteria_group>Bacillota>Bacilli>Lactobacillales>Enterococcaceae | GCA_015437545.1 | Enterococcus_faecium | IX463_002172 | 13730..17401 | 1352 | BREX-system-P-loop-protein-BrxC-[Enterococcus-faecium] |
| HAQ7306339.1 | Type1.BREX | <-ABC-ATPase+SbcC<-?<-?\|\|HTH_3-><-SIGMA+OB-Fold+AAA-ATPase+Lon-Protease-BrxL<-PglZ<-N6-METHYLASE<-DUF6079* | 1176 | Bacteria>Terrabacteria_group>Bacillota>Bacilli>Lactobacillales>Enterococcaceae | GCA_015446545.1 | Enterococcus_faecium | IX650_002001 | 13732..17263 | 1352 | BREX-system-P-loop-protein-BrxC-[Enterococcus-faecium] |
| HAQ8237001.1 | Type1.BREX | <-ABC-ATPase+SbcC<-?<-?\|\|HTH_3-><-SIGMA+OB-Fold+AAA-ATPase+Lon-Protease-BrxL<-PglZ<-N6-METHYLASE<-DUF6079*<-BrxB-BrxF-iSTAND<-BrxA-DNA-binding | 1223 | Bacteria>Terrabacteria_group>Bacillota>Bacilli>Lactobacillales>Enterococcaceae | GCA_015452345.1 | Enterococcus_faecium | IX970_001551 | 18057..21728 | 1352 | BREX-system-P-loop-protein-BrxC-[Enterococcus-faecium] |
| HAQ8954439.1 | Type1.BREX | <-ABC-ATPase+SbcC<-?<-?\|\|HTH_3-><-SIGMA+OB-Fold+AAA-ATPase+Lon-Protease-BrxL<-PglZ<-N6-METHYLASE<-DUF6079-p* | 939 | Bacteria>Terrabacteria_group>Bacillota>Bacilli>Lactobacillales>Enterococcaceae | GCA_015457805.1 | Enterococcus_faecium | IYC31_002159 | 18057..20878 | 1352 | BREX-system-P-loop-protein-BrxC-[Enterococcus-faecium] |
| HAQ9538118.1 | Type1.BREX | <-ABC-ATPase+SbcC<-?<-?\|\|HTH_3-><-SIGMA+OB-Fold+AAA-ATPase+Lon-Protease-BrxL<-PglZ<-N6-METHYLASE<-DUF6079*<-BrxB-BrxF-iSTAND<-BrxA-DNA-binding | 1223 | Bacteria>Terrabacteria_group>Bacillota>Bacilli>Lactobacillales>Enterococcaceae | GCA_015461205.1 | Enterococcus_faecium | IYE17_001922 | 18059..21730 | 1352 | BREX-system-P-loop-protein-BrxC-[Enterococcus-faecium] |
| HAR0082175.1 | Type1.BREX | <-ABC-ATPase+SbcC<-?<-?\|\|HTH_3-><-SIGMA+OB-Fold+AAA-ATPase+Lon-Protease-BrxL<-PglZ<-N6-METHYLASE<-DUF6079* | 1071 | Bacteria>Terrabacteria_group>Bacillota>Bacilli>Lactobacillales>Enterococcaceae | GCA_015465145.1 | Enterococcus_faecium | IYG34_001995 | 18058..21274 | 1352 | BREX-system-P-loop-protein-BrxC-[Enterococcus-faecium] |
| EGP4757178.1 | Type1.BREX | HTH_3-><-SIGMA+OB-Fold+AAA-ATPase+Lon-Protease-BrxL<-PglZ<-N6-METHYLASE<-DUF6079*<-BrxB-BrxF-iSTAND<-BrxA-DNA-binding | 1223 | Bacteria>Terrabacteria_group>Bacillota>Bacilli>Lactobacillales>Enterococcaceae | GCA_015501165.1 | Enterococcus_faecium | F6336_002291 | 15372..19043 | 1352 | BREX-system-P-loop-protein-BrxC-[Enterococcus-faecium] |
| EGP5122183.1 | Type1.BREX | <-ABC-ATPase+SbcC<-?\|\|HTH_3-><-SIGMA+OB-Fold+AAA-ATPase+Lon-Protease-BrxL<-PglZ<-N6-METHYLASE<-DUF6079*<-BrxB-BrxF-iSTAND<-BrxA-DNA-binding | 1223 | Bacteria>Terrabacteria_group>Bacillota>Bacilli>Lactobacillales>Enterococcaceae | GCA_015505205.1 | Enterococcus_faecium | EL427_11505 | 15481..19152 | 1352 | BREX-system-P-loop-protein-BrxC-[Enterococcus-faecium] |
| EGS7978425.1 | Type1.BREX | <-N6-METHYLASE<-DUF6079*<-BrxB-BrxF-iSTAND<-BrxA-DNA-binding<-?<-Hpr-Kinase-N\|\|?->?->?->?->DEOR-HTH-> | 1221 | Bacteria>Terrabacteria_group>Bacillota>Bacilli>Lactobacillales>Enterococcaceae | GCA_015920245.1 | Enterococcus_faecalis | I4L79_000250 | 3156..6821 | 1351 | BREX-system-P-loop-protein-BrxC-[Enterococcus-faecalis] |
| MBJ1581502.1 | Type1.BREX | <-ABC-ATPase+SbcC<-?<-?\|\|HTH_3-><-SIGMA+OB-Fold+AAA-ATPase+Lon-Protease-BrxL<-PglZ<-N6-METHYLASE<-DUF6079*<-BrxB-BrxF-iSTAND<-BrxA-DNA-binding | 1223 | Bacteria>Terrabacteria_group>Bacillota>Bacilli>Lactobacillales>Enterococcaceae | GCA_016394005.1 | Enterococcus_faecium | JFE65_09230 | 18064..21735 | 1352 | BREX-system-P-loop-protein-BrxC-[Enterococcus-faecium] |
| EHA7758737.1 | Type1.BREX | <-DEOR-HTH<-?<-?<-?<-?\|\|Hpr-Kinase-N->?->BrxA-DNA-binding->BrxB-BrxF-iSTAND->DUF6079-p*-> | 611 | Bacteria>Terrabacteria_group>Bacillota>Bacilli>Lactobacillales>Enterococcaceae | GCA_016733805.1 | Enterococcus_faecalis | JMJ06_002105 | 23231..25064 | 1351 | BREX-system-P-loop-protein-BrxC-[Enterococcus-faecalis] |
| HBB6805772.1 | Type1.BREX | <-ABC-ATPase+SbcC<-?\|\|HTH_3-><-SIGMA+OB-Fold+AAA-ATPase+Lon-Protease-BrxL<-PglZ<-N6-METHYLASE<-DUF6079*<-BrxB-BrxF-iSTAND<-BrxA-DNA-binding | 1223 | Bacteria>Terrabacteria_group>Bacillota>Bacilli>Lactobacillales>Enterococcaceae | GCA_017966085.1 | Enterococcus_faecium | KB900_002068 | 13733..17404 | 1352 | BREX-system-P-loop-protein-BrxC-[Enterococcus-faecium] |
| EHM3054557.1 | Type1.BREX | <-PglZ<-Y1_Tnp<-N6-METHYLASE<-DUF6079*<-BrxB-BrxF-iSTAND<-BrxA-DNA-binding | 1223 | Bacteria>Terrabacteria_group>Bacillota>Bacilli>Lactobacillales>Enterococcaceae | GCA_018220435.1 | Enterococcus_faecium | KGA03_002421 | 7403..11074 | 1352 | BREX-system-P-loop-protein-BrxC-[Enterococcus-faecium] |
| MCH3341469.1 | Type1.BREX | <-ABC-ATPase+SbcC\|\|HTH_3-><-SIGMA+OB-Fold+AAA-ATPase+Lon-Protease-BrxL<-PglZ<-N6-METHYLASE<-DUF6079*<-BrxB-BrxF-iSTAND<-BrxA-DNA-binding | 1223 | Bacteria>Terrabacteria_group>Bacillota>Bacilli>Lactobacillales>Enterococcaceae | GCA_022463855.1 | Enterococcus_faecium | LC935_10425 | 13809..17480 | 1352 | BREX-system-P-loop-protein-BrxC-[Enterococcus-faecium] |
| HCA4699593.1 | Type1.BREX | <-ABC-ATPase+SbcC<-?<-?\|\|HTH_3-><-SIGMA+OB-Fold+AAA-ATPase+Lon-Protease-BrxL<-PglZ<-N6-METHYLASE<-DUF6079*<-BrxB-BrxF-iSTAND | 1223 | Bacteria>Terrabacteria_group>Bacillota>Bacilli>Lactobacillales>Enterococcaceae | GCA_022922335.1 | Enterococcus_faecium | MWZ72_002010 | 18064..21735 | 1352 | BREX-system-P-loop-protein-BrxC-[Enterococcus-faecium] |
| HCA4993928.1 | Type1.BREX | <-ABC-ATPase+SbcC<-?<-?\|\|HTH_3-><-SIGMA+OB-Fold+AAA-ATPase+Lon-Protease-BrxL<-PglZ<-N6-METHYLASE<-DUF6079*<-BrxB-BrxF-iSTAND<-BrxA-DNA-binding | 1223 | Bacteria>Terrabacteria_group>Bacillota>Bacilli>Lactobacillales>Enterococcaceae | GCA_022932465.1 | Enterococcus_faecium | MW143_001926 | 18056..21727 | 1352 | BREX-system-P-loop-protein-BrxC-[Enterococcus-faecium] |
| EJB2765985.1 | Type1.BREX | <-DUF6079-p*<-BrxB-BrxF-iSTAND<-BrxA-DNA-binding<-?<-Hpr-Kinase-N\|\|?->?->?->?->DEOR-HTH-> | 524 | Bacteria>Terrabacteria_group>Bacillota>Bacilli>Lactobacillales>Enterococcaceae | GCA_022936025.1 | Enterococcus_faecalis | MT958_000792 | 1..1573 | 1351 | BREX-system-P-loop-protein-BrxC-[Enterococcus-faecalis] |
| HCI0787385.1 | Type1.BREX | <-ABC-ATPase+SbcC<-?<-?\|\|HTH_3-><-SIGMA+OB-Fold+AAA-ATPase+Lon-Protease-BrxL<-PglZ<-N6-METHYLASE<-DUF6079-p* | 881 | Bacteria>Terrabacteria_group>Bacillota>Bacilli>Lactobacillales>Enterococcaceae | GCA_024313135.1 | Enterococcus_faecium | NOA45_002008 | 18063..20710 | 1352 | BREX-system-P-loop-protein-BrxC-[Enterococcus-faecium] |
| HCK3992712.1 | Type1.BREX | HTH_3-><-SIGMA+OB-Fold+AAA-ATPase+Lon-Protease-BrxL<-PglZ<-N6-METHYLASE<-DUF6079*<-BrxB-BrxF-iSTAND<-BrxA-DNA-binding | 1223 | Bacteria>Terrabacteria_group>Bacillota>Bacilli>Lactobacillales>Enterococcaceae | GCA_024823995.1 | Enterococcus_faecium | NOA28_002014 | 9602..13273 | 1352 | BREX-system-P-loop-protein-BrxC-[Enterococcus-faecium] |
| BDP56409.1 | Type1.BREX | <-HIN-HTH\|\|?->BrxB-BrxF-iSTAND->?->MULE-TRANSPOSASE->BrxA-DNA-binding->BrxB-BrxF-iSTAND->DUF6079-p-><-zf-ISL3+HTH_Tnp_ISL3+DDE_Tnp_ISL3\|\|DUF6079-p*->N6-METHYLASE-p->N6-METHYLASE-p->N6-METHYLASE-p->PglZ->SIGMA+OB-Fold+AAA-ATPase+Lon-Protease-BrxL-><-HTH_3\|\|?->?->ABC-ATPase+SbcC-> | 949 | Bacteria>Terrabacteria_group>Bacillota>Bacilli>Lactobacillales>Enterococcaceae | GCA_026073375.1 | Enterococcus_faecium | EfmJHP36_08880 | 842990..845839 | 1352 | hypothetical-protein-EfmJHP36_08880-[Enterococcus-faecium] |
| BDP59934.1 | Type1.BREX | <-HIN-HTH\|\|?->MULE-TRANSPOSASE->MULE-TRANSPOSASE->BrxA-DNA-binding-p->BrxB-BrxF-iSTAND->DUF6079-p-><-zf-ISL3+HTH_Tnp_ISL3+DDE_Tnp_ISL3\|\|DUF6079-p*->?->N6-METHYLASE->PglZ->SIGMA+OB-Fold+AAA-ATPase+Lon-Protease-BrxL-><-HTH_3\|\|?->?->ABC-ATPase+SbcC-> | 739 | Bacteria>Terrabacteria_group>Bacillota>Bacilli>Lactobacillales>Enterococcaceae | GCA_026073395.1 | Enterococcus_faecium | EfmJHP38_08720 | 846169..848388 | 1352 | hypothetical-protein-EfmJHP38_08720-[Enterococcus-faecium] |
| WP_002358702.1 | Type1.BREX | <-ABC-ATPase+SbcC\|\|HTH_3-><-SIGMA+OB-Fold+AAA-ATPase+Lon-Protease-BrxL<-PglZ<-N6-METHYLASE<-DUF6079*<-BrxB-BrxF-iSTAND<-BrxA-DNA-binding | 1223 | Bacteria>Terrabacteria_group>Bacillota>Bacilli>Lactobacillales>Enterococcaceae | GCF_000295315.2 | Enterococcus_faecium | HMPREF1356_RS07125 | 13767..17438 | 1352 | BREX-system-P-loop-protein-BrxC-[Enterococcus-faecium] |
| WP_002340335.1 | Type1.BREX | <-ABC-ATPase+SbcC<-?\|\|CNMP+CRP-HTH->?->\|\|MULE-TRANSPOSASE->BrxA-DNA-binding->BrxB-BrxF-iSTAND->DUF6079*->N6-METHYLASE->PglZ->SIGMA+OB-Fold+AAA-ATPase+Lon-Protease-BrxL-><-HTH_3\|\|ABC-ATPase+SbcC->?-><-zf-ISL3+HTH_Tnp_ISL3+DDE_Tnp_ISL3 | 1223 | Bacteria>Terrabacteria_group>Bacillota>Bacilli>Lactobacillales>Enterococcaceae | GCF_000322065.1 | Enterococcus_faecium | OK3_RS07275 | 142504..146175 | 1352 | BREX-system-P-loop-protein-BrxC-[Enterococcus-faecium] |
| WP_028790423.1 | Type1.BREX | BrxA-DNA-binding->BrxB-BrxF-iSTAND->DUF6079*->N6-METHYLASE->PglZ-> | 1220 | Bacteria>Terrabacteria_group>Bacillota>Bacilli>Lactobacillales>Enterococcaceae | GCF_000423785.1 | Tetragenococcus_muriaticus | G577_RS0105945 | 2131..5793 | 64642 | BREX-system-P-loop-protein-BrxC-[Tetragenococcus-muriaticus] |
| WP_002397586.1 | Type1.BREX | <-DEOR-HTH<-?<-?<-?<-?\|\|Hpr-Kinase-N->?->BrxA-DNA-binding->BrxB-BrxF-iSTAND->DUF6079*->N6-METHYLASE-><-Phage_int_SAM_5+Phage_integrase\|\|N6-METHYLASE-p->PglZ->SIGMA+OB-Fold+AAA-ATPase+Lon-Protease-BrxL->?-><-MerR-HTH+HCR+BAF\|\|PTSEIIC->Glyco_hydro_1-> | 1221 | Bacteria>Terrabacteria_group>Bacillota>Bacilli>Lactobacillales>Enterococcaceae | GCF_000647995.1 | Enterococcus_faecalis | NNC36_RS11895 | 57506..61171 | 1351 | BREX-system-P-loop-protein-BrxC-[Enterococcus-faecalis] |
| WP_010830678.1 | Type1.BREX | <-SIGMA+OB-Fold+AAA-ATPase+Lon-Protease-BrxL<-PglZ<-N6-METHYLASE-p\|\|Phage_int_SAM_5+Phage_integrase->N6-METHYLASE-p->?-><-N6-METHYLASE<-DUF6079*<-BrxB-BrxF-iSTAND<-BrxA-DNA-binding<-?<-Hpr-Kinase-N\|\|?->?->?->DEOR-HTH-> | 1221 | Bacteria>Terrabacteria_group>Bacillota>Bacilli>Lactobacillales>Enterococcaceae | GCF_000648055.1 | Enterococcus_faecalis | P784_RS05735 | 43923..47588 | 1351 | BREX-system-P-loop-protein-BrxC-[Enterococcus-faecalis] |
| WP_061086322.1 | Type1.BREX | LZ_Tnp_IS66+zf-IS66+DDE_Tnp_IS66_C-><-?<-?<-?<-ABC-ATPase+SbcC\|\|HTH_3-><-SIGMA+OB-Fold+AAA-ATPase+Lon-Protease-BrxL<-PglZ<-N6-METHYLASE<-DUF6079-p* | 941 | Bacteria>Terrabacteria_group>Bacillota>Bacilli>Lactobacillales>Enterococcaceae | GCF_001547265.1 | Enterococcus_faecium | AS262_RS06445 | 37117..39942 | 1352 | BREX-system-P-loop-protein-BrxC,-partial-[Enterococcus-faecium] |
| WP_061100912.1 | Type1.BREX | <-DEOR-HTH<-?<-?<-?<-?\|\|Hpr-Kinase-N->?->BrxA-DNA-binding->BrxB-BrxF-iSTAND->DUF6079*->N6-METHYLASE-><-N6-METHYLASE-p<-?<-ParB+HNH<-Resolvase+Recombinase+ZnR<-Resolvase+Recombinase+ZnR<-HIN-HTH | 1221 | Bacteria>Terrabacteria_group>Bacillota>Bacilli>Lactobacillales>Enterococcaceae | GCF_001563075.1 | Enterococcus_faecalis | AW733_RS05605 | 279027..282692 | 1351 | BREX-system-P-loop-protein-BrxC-[Enterococcus-faecalis] |
| WP_002321316.1 | Type1.BREX | <-ABC-ATPase+SbcC<-?\|\|CNMP+CRP-HTH->?-><-HTH_Tnp_1+HTH_21+rve+rve_3\|\|MULE-TRANSPOSASE->BrxA-DNA-binding->BrxB-BrxF-iSTAND->DUF6079*->N6-METHYLASE->PglZ->SIGMA+OB-Fold+AAA-ATPase+Lon-Protease-BrxL-><-HTH_3\|\|ABC-ATPase+SbcC->?->?->?-><-LZ_Tnp_IS66+zf-IS66+DDE_Tnp_IS66+DDE_Tnp_IS66_C | 1223 | Bacteria>Terrabacteria_group>Bacillota>Bacilli>Lactobacillales>Enterococcaceae | GCF_001721005.1 | Enterococcus_faecium | AL017_RS02660 | 520733..524404 | 1352 | BREX-system-P-loop-protein-BrxC-[Enterococcus-faecium] |
| WP_086318169.1 | Type1.BREX | <-DEOR-HTH<-?<-?<-?<-?\|\|?->BrxA-DNA-binding->BrxB-BrxF-iSTAND->DUF6079*->N6-METHYLASE-><-N6-METHYLASE-p<-Phage_int_SAM_5+Phage_integrase\|\|N6-METHYLASE-p->PglZ->SIGMA+OB-Fold+AAA-ATPase+Lon-Protease-BrxL->?-><-?<-?\|\|Sulfate_transp+STAS-> | 1221 | Bacteria>Terrabacteria_group>Bacillota>Bacilli>Lactobacillales>Enterococcaceae | GCF_002141205.1 | Enterococcus_faecalis | A5830_RS12845 | 257941..261606 | 1351 | BREX-system-P-loop-protein-BrxC-[Enterococcus-faecalis] |
| WP_126762085.1 | Type1.BREX | Peptidase-><-KAP-NTPase<-HTH+REase<-?<-SIGMA+OB-Fold+AAA-ATPase+Lon-Protease-BrxL<-PglZ<-N6-METHYLASE<-DUF6079*<-BrxB-BrxF-iSTAND<-BrxA-DNA-binding<-?<-?<-?<-ABC-ATPase+SbcC<-ABC-ATPase+SbcC | 1186 | Bacteria>Terrabacteria_group>Bacillota>Bacilli>Lactobacillales>Enterococcaceae | GCF_003987555.1 | Vagococcus_lutrae | CBF33_RS04670 | 25387..28947 | 81947 | BREX-system-P-loop-protein-BrxC-[Vagococcus-lutrae] |
| WP_126809277.1 | Type1.BREX | TraJ-RHH->TraI-VirD2-RCRE-><-?<-Polbeta<-SIGMA+OB-Fold+AAA-ATPase+Lon-Protease-BrxL<-PglZ<-N6-METHYLASE<-DUF6079*<-BrxB-BrxF-iSTAND<-BrxA-DNA-binding<-HTH_26<-JAB<-ArdCN+LPD32+LPD36+N6_Mtase+SF1-SF2-Helicase<-DNA_METHYLASE<-TOPRIM+Topoisom_bac<-VWA+SIGMA-HTH | 1199 | Bacteria>Terrabacteria_group>Bacillota>Bacilli>Lactobacillales>Enterococcaceae | GCF_003987645.1 | Vagococcus_elongatus | CBF29_08285 | 56572..60171 | 180344 | BREX-system-P-loop-protein-BrxC-[Vagococcus-elongatus] |
| WP_130647944.1 | Type1.BREX | <-ABC-ATPase+SbcC\|\|HTH_3-><-SIGMA+OB-Fold+AAA-ATPase+Lon-Protease-BrxL<-PglZ<-N6-METHYLASE<-DUF6079*<-BrxB-BrxF-iSTAND<-BrxA-DNA-binding | 1223 | Bacteria>Terrabacteria_group>Bacillota>Bacilli>Lactobacillales>Enterococcaceae | GCF_004300425.1 | Enterococcus_faecium | EWU48_RS10220 | 18141..21812 | 1352 | BREX-system-P-loop-protein-BrxC-[Enterococcus-faecium] |
| WP_139896804.1 | Type1.BREX | MULE-TRANSPOSASE->?->?->?-><-?\|\|?->BrxA-DNA-binding->BrxB-BrxF-iSTAND->DUF6079*->N6-METHYLASE->PglZ->SIGMA+OB-Fold+AAA-ATPase+Lon-Protease-BrxL-><-HTH_3\|\|?->ABC-ATPase+SbcC-> | 1223 | Bacteria>Terrabacteria_group>Bacillota>Bacilli>Lactobacillales>Enterococcaceae | GCF_006351845.1 | Enterococcus_faecium | FHK65_RS02695 | 522260..525931 | 1352 | BREX-system-P-loop-protein-BrxC-[Enterococcus-faecium] |
| WP_141426751.1 | Type1.BREX | <-ABC-ATPase+SbcC<-?\|\|HTH_3-><-SIGMA+OB-Fold+AAA-ATPase+Lon-Protease-BrxL<-PglZ<-N6-METHYLASE<-DUF6079*<-BrxB-BrxF-iSTAND<-BrxA-DNA-binding<-?<-?<-?<-?\|\|?-><-DNA_METHYLASE<-HTH_3 | 1223 | Bacteria>Terrabacteria_group>Bacillota>Bacilli>Lactobacillales>Enterococcaceae | GCF_006541535.1 | Enterococcus_faecium | FKZ07_RS08010 | 37532..41203 | 1352 | BREX-system-P-loop-protein-BrxC-[Enterococcus-faecium] |
| WP_144771940.1 | Type1.BREX | <-MerR-HTH+SHS2+SHS2<-Tox-ABhydrolase<-PTSEIIC<-?<-MazG<-ParB+HNH<-SIGMA+OB-Fold+AAA-ATPase+Lon-Protease-BrxL<-PglZ<-N6-METHYLASE<-DUF6079*<-BrxB-BrxF-iSTAND<-BrxA-DNA-binding | 1223 | Bacteria>Terrabacteria_group>Bacillota>Bacilli>Lactobacillales>Enterococcaceae | GCF_007682015.1 | Enterococcus_casseliflavus | FS854_RS14505 | 69459..73130 | 37734 | BREX-system-P-loop-protein-BrxC-[Enterococcus-casseliflavus] |
| WP_156237125.1 | Type1.BREX | BrxA-DNA-binding->BrxB-BrxF-iSTAND->DUF6079*->N6-METHYLASE->PglZ->SIGMA+OB-Fold+AAA-ATPase+Lon-Protease-BrxL->ParB+HNH->MazG->SF1-SF2-Helicase->?->SF1-SF2-Helicase->CR-ATPase5+REase-> | 1223 | Bacteria>Terrabacteria_group>Bacillota>Bacilli>Lactobacillales>Enterococcaceae | GCF_009734725.1 | Enterococcus_casseliflavus | EZ072_RS09880 | 26969..30640 | 37734 | BREX-system-P-loop-protein-BrxC-[Enterococcus-casseliflavus] |
| WP_242459300.1 | Type1.BREX | <-ABC-ATPase+SbcC\|\|HTH_3-><-SIGMA+OB-Fold+AAA-ATPase+Lon-Protease-BrxL<-PglZ<-N6-METHYLASE<-DUF6079*<-DUF6079-p<-BrxB-BrxF-iSTAND<-BrxA-DNA-binding | 1001 | Bacteria>Terrabacteria_group>Bacillota>Bacilli>Lactobacillales>Enterococcaceae | GCF_009846515.2 | Enterococcus_faecium | X541_RS14255 | 19469..22474 | 1352 | BREX-system-P-loop-protein-BrxC-[Enterococcus-faecium] |
| WP_172875133.1 | Type1.BREX | <-ABC-ATPase+SbcC\|\|HTH_3-><-PglZ<-N6-METHYLASE<-DUF6079*<-BrxB-BrxF-iSTAND<-BrxA-DNA-binding | 1223 | Bacteria>Terrabacteria_group>Bacillota>Bacilli>Lactobacillales>Enterococcaceae | GCF_013266315.1 | Enterococcus_faecium | HR106_RS06795 | 13728..17399 | 1352 | BREX-system-P-loop-protein-BrxC-[Enterococcus-faecium] |
| WP_127820617.1 | Type1.BREX | BrxA-DNA-binding->BrxB-BrxF-iSTAND->DUF6079*->N6-METHYLASE->PglZ-><-HTH_3\|\|ABC-ATPase+SbcC-> | 1223 | Bacteria>Terrabacteria_group>Bacillota>Bacilli>Lactobacillales>Enterococcaceae | GCF_013266375.1 | Enterococcus_faecium | HR111_RS07860 | 1435..5106 | 1352 | BREX-system-P-loop-protein-BrxC-[Enterococcus-faecium] |
| WP_194194888.1 | Type1.BREX | <-ACET<-?<-TPR<-?<-MazG<-ParB+HNH<-SIGMA+OB-Fold+AAA-ATPase+Lon-Protease-BrxL<-PglZ<-N6-METHYLASE<-DUF6079*<-BrxB-BrxF-iSTAND<-BrxA-DNA-binding<-?<-?<-?\|\|HTH_3-> | 1223 | Bacteria>Terrabacteria_group>Bacillota>Bacilli>Lactobacillales>Enterococcaceae | GCF_015230945.1 | Enterococcus_casseliflavus | HAX40_RS05295 | 101566..105237 | 37734 | BREX-system-P-loop-protein-BrxC-[Enterococcus-casseliflavus] |
| WP_202585039.1 | Type1.BREX | <-TIR<-?<-TIR<-HTH+REase<-?<-ZnR<-SIGMA+OB-Fold+AAA-ATPase+Lon-Protease-BrxL<-PglZ<-N6-METHYLASE<-DUF6079*<-BrxB-BrxF-iSTAND<-BrxA-DNA-binding<-?<-?<-?<-?<-ABC-ATPase+SbcC<-ABC-ATPase+SbcC | 1186 | Bacteria>Terrabacteria_group>Bacillota>Bacilli>Lactobacillales>Enterococcaceae | GCF_016760265.1 | Vagococcus_lutrae | VL4N_RS03475 | 40566..44126 | 81947 | BREX-system-P-loop-protein-BrxC-[Vagococcus-lutrae] |
| WP_204654268.1 | Type1.BREX | His_biosynth->?->?->Aminotran_1_2-><-Glyoxalase\|\|YTCD-HTH->?->BrxA-DNA-binding->BrxB-BrxF-iSTAND->DUF6079*->N6-METHYLASE-><-Phage_integrase\|\|N6-METHYLASE-p->PglZ->SIGMA+OB-Fold+AAA-ATPase+Lon-Protease-BrxL->DOC+HTH-> | 1207 | Bacteria>Terrabacteria_group>Bacillota>Bacilli>Lactobacillales>Enterococcaceae | GCF_016909125.1 | Enterococcus_lemanii | JOD14_RS07795 | 27391..31014 | 1159752 | BREX-system-P-loop-protein-BrxC-[Enterococcus-lemanii] |
| WP_116593245.1 | Type1.BREX | <-DEOR-HTH<-?<-?<-?<-?\|\|Hpr-Kinase-N->?->BrxA-DNA-binding->BrxB-BrxF-iSTAND->DUF6079*->N6-METHYLASE-> | 1221 | Bacteria>Terrabacteria_group>Bacillota>Bacilli>Lactobacillales>Enterococcaceae | GCF_017642205.1 | Enterococcus_faecalis | EY652_RS11110 | 85892..89557 | 1351 | BREX-system-P-loop-protein-BrxC-[Enterococcus-faecalis] |
| WP_229444528.1 | Type1.BREX | <-HTH_Tnp_1+HTH_21+rve+rve_3<-SIGMA+OB-Fold+AAA-ATPase+Lon-Protease-BrxL<-PglZ\|\|Phage_integrase-><-N6-METHYLASE<-?<-?<-DUF6079*<-BrxB-BrxF-iSTAND<-BrxA-DNA-binding<-sGTP\|\|ACET->?->?->?->Transposase_20-> | 1206 | Bacteria>Terrabacteria_group>Bacillota>Bacilli>Lactobacillales>Enterococcaceae | GCF_020567645.1 | Enterococcus_sp._CWB-B31 | LI954_RS11380 | 72260..75880 | 2885159 | BREX-system-P-loop-protein-BrxC-[Enterococcus-sp.-CWB-B31] |
| WP_002327397.1 | Type1.BREX | <-ABC-ATPase+SbcC<-?\|\|HTH_3-><-PglZ<-Y1_Tnp<-N6-METHYLASE<-DUF6079*<-BrxB-BrxF-iSTAND<-BrxA-DNA-binding | 1223 | Bacteria>Terrabacteria_group>Bacillota>Bacilli>Lactobacillales>Enterococcaceae | GCF_021123205.1 | Enterococcus_faecium | IV423_RS10650 | 22640..26311 | 1352 | BREX-system-P-loop-protein-BrxC-[Enterococcus-faecium] |
| WP_231420058.1 | Type1.BREX | <-Glyco_hydro_1<-PTSEIIC\|\|MerR-HTH+HCR+BAF-><-?<-SIGMA+OB-Fold+AAA-ATPase+Lon-Protease-BrxL<-PglZ<-N6-METHYLASE-p\|\|Phage_int_SAM_5+Phage_integrase-><-N6-METHYLASE<-DUF6079*<-BrxB-BrxF-iSTAND<-BrxA-DNA-binding<-?<-Hpr-Kinase-N\|\|?->?->?->?->DEOR-HTH-> | 1221 | Bacteria>Terrabacteria_group>Bacillota>Bacilli>Lactobacillales>Enterococcaceae | GCF_021124535.1 | Enterococcus_faecalis | IVZ09_RS12045 | 46778..50443 | 1351 | BREX-system-P-loop-protein-BrxC-[Enterococcus-faecalis] |
| WP_234769699.1 | Type1.BREX | HTH_6+SIS->BrxA-DNA-binding->BrxB-BrxF-iSTAND->DUF6079*->PglZ->?->?->?->?->?->ZDNA-HTH-> | 1220 | Bacteria>Terrabacteria_group>Bacillota>Bacilli>Lactobacillales>Enterococcaceae | GCF_021474625.1 | Tetragenococcus_halophilus | LQF56_RS09850 | 6175..9837 | 51669 | BREX-system-P-loop-protein-BrxC-[Tetragenococcus-halophilus] |
| WP_243197397.1 | Type1.BREX | HTH_26->BrxA-DNA-binding->BrxB-BrxF-iSTAND->DUF6079*->wHTH+ZnR->Four-helical-protein->N6-METHYLASE->?->PglZ->SIGMA+OB-Fold+AAA-ATPase+Lon-Protease-BrxL-> | 1201 | Bacteria>Terrabacteria_group>Bacillota>Bacilli>Lactobacillales>Enterococcaceae | GCF_022807075.1 | Enterococcus_cecorum | MMJ55_RS10180 | 2645..6250 | 44008 | BREX-system-P-loop-protein-BrxC-[Enterococcus-cecorum] |
| WP_116495172.1 | Type1.BREX | <-PTSEIIC\|\|MerR-HTH+HCR+BAF-><-?<-SIGMA+OB-Fold+AAA-ATPase+Lon-Protease-BrxL<-PglZ<-N6-METHYLASE-p\|\|Phage_int_SAM_5+Phage_integrase->N6-METHYLASE-p-><-N6-METHYLASE<-DUF6079*<-BrxB-BrxF-iSTAND<-BrxA-DNA-binding<-?<-Hpr-Kinase-N\|\|?->?->?->?->DEOR-HTH-> | 1221 | Bacteria>Terrabacteria_group>Bacillota>Bacilli>Lactobacillales>Enterococcaceae | GCF_023300025.1 | Enterococcus_faecalis | M2914_RS11820 | 2452223..2455888 | 1351 | BREX-system-P-loop-protein-BrxC-[Enterococcus-faecalis] |
| WP_251851849.1 | Type1.BREX | DSBH+AraC-HTH-><-ParB+HNH<-PglZ<-N6-METHYLASE-p\|\|Phage_int_SAM_5+Phage_integrase->N6-METHYLASE-p-><-CR-ATPase5+REase<-N6-METHYLASE<-DUF6079*<-BrxB-BrxF-iSTAND<-BrxA-DNA-binding\|\|MULE-TRANSPOSASE->?->?-><-DDE_Tnp<-?\|\|DDE_Tnp_IS240-> | 1202 | Bacteria>Terrabacteria_group>Bacillota>Bacilli>Lactobacillales>Enterococcaceae | GCF_023744155.1 | Enterococcus_italicus | NE261_RS07355 | 549628..553236 | 246144 | BREX-system-P-loop-protein-BrxC-[Enterococcus-italicus] |
| WP_002344971.1 | Type1.BREX | BrxA-DNA-binding->BrxB-BrxF-iSTAND->DUF6079*->N6-METHYLASE->PglZ->SIGMA+OB-Fold+AAA-ATPase+Lon-Protease-BrxL-><-HTH_3\|\|ABC-ATPase+SbcC-> | 1223 | Bacteria>Terrabacteria_group>Bacillota>Bacilli>Lactobacillales>Enterococcaceae | GCF_026073455.1 | Enterococcus_faecium | OGY46_RS09105 | 1308..4979 | 1352 | BREX-system-P-loop-protein-BrxC-[Enterococcus-faecium] |
| WP_096573229.1 | Type1.BREX | HTH_3->?->DNA_METHYLASE-><-?\|\|?->?->?->BrxA-DNA-binding->BrxB-BrxF-iSTAND->DUF6079*->N6-METHYLASE->PglZ-><-HTH_3\|\|?->ABC-ATPase+SbcC-> | 1223 | Bacteria>Terrabacteria_group>Bacillota>Bacilli>Lactobacillales>Enterococcaceae | GCF_026621805.1 | Enterococcus_faecium | OY807_RS08695 | 133028..136699 | 1352 | BREX-system-P-loop-protein-BrxC-[Enterococcus-faecium] |
| WP_104843808.1 | Type1.BREX | NtrC-AAA+FIS-HTH->HAD+HAD->BrxA-DNA-binding->BrxB-BrxF-iSTAND->DUF6079*->N6-METHYLASE->Y1_Tnp->PglZ->SIGMA+OB-Fold+AAA-ATPase+Lon-Protease-BrxL-><-HTH_3\|\|?->ABC-ATPase+SbcC-> | 1223 | Bacteria>Terrabacteria_group>Bacillota>Bacilli>Lactobacillales>Enterococcaceae | GCF_027674745.1 | Enterococcus_faecium | PGU87_RS03230 | 14252..17923 | 1352 | BREX-system-P-loop-protein-BrxC-[Enterococcus-faecium] |
| WP_061605122.1 | Type1.BREX | HTH_3->?->DNA_METHYLASE-><-?\|\|?->?->?->BrxA-DNA-binding->BrxB-BrxF-iSTAND->DUF6079*->N6-METHYLASE->Y1_Tnp->PglZ->SIGMA+OB-Fold+AAA-ATPase+Lon-Protease-BrxL-><-HTH_3\|\|?->ABC-ATPase+SbcC-> | 1223 | Bacteria>Terrabacteria_group>Bacillota>Bacilli>Lactobacillales>Enterococcaceae | GCF_900066025.1 | Enterococcus_faecium | EFE1002_RS02625 | 566193..569864 | 1352 | BREX-system-P-loop-protein-BrxC-[Enterococcus-faecium] |
| WP_131769129.1 | Type1.BREX | <-PTSEIIC\|\|MerR-HTH+HCR+BAF-><-?<-SIGMA+OB-Fold+AAA-ATPase+Lon-Protease-BrxL<-PglZ<-N6-METHYLASE-p\|\|Phage_int_SAM_5+Phage_integrase->N6-METHYLASE-p-><-N6-METHYLASE<-DUF6079*<-BrxB-BrxF-iSTAND<-BrxA-DNA-binding<-?<-Hpr-Kinase-N\|\|?->?->?->?->DEOR-HTH-> | 1221 | Bacteria>Terrabacteria_group>Bacillota>Bacilli>Lactobacillales>Enterococcaceae | GCF_900683585.1 | Enterococcus_faecalis | E1058_RS12355 | 2454613..2458278 | 1351 | BREX-system-P-loop-protein-BrxC-[Enterococcus-faecalis] |
| WP_258233427.1 | Type1.BREX | <-CHTH<-?<-?<-?<-HTH+REase<-SIGMA+OB-Fold+AAA-ATPase+Lon-Protease-BrxL<-PglZ<-N6-METHYLASE<-DUF6079*<-BrxB-BrxF-iSTAND<-BrxA-DNA-binding<-?<-?<-?<-PTSEIIC | 1227 | Bacteria>Terrabacteria_group>Bacillota>Bacilli>Lactobacillales>Enterococcaceae | GCF_901212505.1 | Enterococcus_avium | EAVMC1_RS06665 | 43716..47399 | 33945 | BREX-system-P-loop-protein-BrxC-[Enterococcus-avium] |
| WP_142431712.1 | Type1.BREX | DNA_METHYLASE-><-?\|\|?->?->?->BrxA-DNA-binding->BrxB-BrxF-iSTAND->DUF6079*->N6-METHYLASE->PglZ->SIGMA+OB-Fold+AAA-ATPase+Lon-Protease-BrxL-><-HTH_3\|\|?->ABC-ATPase+SbcC-> | 1223 | Bacteria>Terrabacteria_group>Bacillota>Bacilli>Lactobacillales>Enterococcaceae | GCF_902159355.1 | Enterococcus_faecium | FN050_RS00635 | 125427..129098 | 1352 | BREX-system-P-loop-protein-BrxC-[Enterococcus-faecium] |
| WP_118249569.1 | Type1.BREX | Mrr_N+ACET-><-?\|\|BrxA-DNA-binding->BrxB-BrxF-iSTAND->DUF6079*->N6-METHYLASE->PglZ->SIGMA+OB-Fold+AAA-ATPase+Lon-Protease-BrxL-><-?<-?<-?<-?\|\|MarR-HTH->PhoH-> | 1209 | Bacteria>Terrabacteria_group>Bacillota>Bacilli>Lactobacillales>Enterococcaceae | GCF_902364615.1 | Vagococcus | FY248_RS00660 | 114586..118215 | 2737 | MULTISPECIES:-BREX-system-P-loop-protein-BrxC-[Vagococcus] |
| WP_078122584.1 | Type1.BREX | <-SIGMA+OB-Fold+AAA-ATPase+Lon-Protease-BrxL<-PglZ<-N6-METHYLASE-p\|\|Phage_int_SAM_5+Phage_integrase->N6-METHYLASE-p-><-N6-METHYLASE<-DUF6079*<-BrxB-BrxF-iSTAND<-BrxA-DNA-binding<-?<-Hpr-Kinase-N\|\|?->?->?->?->DEOR-HTH-> | 1221 | Bacteria>Terrabacteria_group>Bacillota>Bacilli>Lactobacillales>Enterococcaceae | GCF_945957195.1 | Enterococcus_faecalis | OC374_RS04755 | 43942..47607 | 1351 | BREX-system-P-loop-protein-BrxC-[Enterococcus-faecalis] |
| EQM54927.1 | Type1.BREX | LPAM_1+DUF4767->?->?->BrxA-DNA-binding->BrxB-BrxF-iSTAND->DUF6079*->N6-METHYLASE-> | 1223 | Bacteria>Terrabacteria_group>Bacillota>Bacilli>Lactobacillales>Lactobacillaceae | GCA_000463075.2 | Lactiplantibacillus_plantarum_EGD-AQ4 | N692_10815 | 71429..75100 | 1382301 | hypothetical-protein-N692_10815-[Lactiplantibacillus-plantarum-EGD-AQ4] |
| KGB15027.1 | Type1.BREX | S2P+PDZ->?->MazG->BrxA-DNA-binding->BrxB-BrxF-iSTAND->DUF6079*->N6-METHYLASE-p->PglZ->SIGMA+OB-Fold+AAA-ATPase+Lon-Protease-BrxL-><-wHTH | 1215 | Bacteria>Terrabacteria_group>Bacillota>Bacilli>Lactobacillales>Lactobacillaceae | GCA_000755625.1 | Latilactobacillus_sakei | KY41_03530 | 43765..47412 | 1599 | hypothetical-protein-KY41_03530-[Latilactobacillus-sakei] |
| ALJ23836.1 | Type1.BREX | HAD+HAD->?->?->CNMP+CRP-HTH->ParB+HNH->BrxA-DNA-binding->BrxB-BrxF-iSTAND->DUF6079*->N6-METHYLASE->nSTAND3->PglZ->TerBN+TPR-><-?<-Peptidase_M13_N+Peptidase_M13\|\|?->?->Peptidase_M20-> | 1242 | Bacteria>Terrabacteria_group>Bacillota>Bacilli>Lactobacillales>Lactobacillaceae | GCA_001314245.2 | Lactobacillus_gallinarum | AO203_08530 | 1678493..1682221 | 52242 | ATPase-[Lactobacillus-gallinarum] |
| KWU03693.1 | Type1.BREX | <-SIR2<-N6-METHYLASE<-DUF6079*<-BrxB-BrxF-iSTAND<-BrxA-DNA-binding<-HTH_3<-?<-PolB<-?\|\|?->?-><-HTH_26 | 1243 | Bacteria>Terrabacteria_group>Bacillota>Bacilli>Lactobacillales>Lactobacillaceae | GCA_001541385.1 | Lactobacillus_crispatus | AEL95_06275 | 4945..8676 | 47770 | ATPase-[Lactobacillus-crispatus] |
| PEH09492.1 | Type1.BREX | <-SIGMA+OB-Fold+AAA-ATPase+Lon-Protease-BrxL<-PglZ<-N6-METHYLASE<-DUF6079*<-BrxB-BrxF-iSTAND<-BrxA-DNA-binding<-HTH_26<-?<-N6-METHYLASE-p | 1183 | Bacteria>Terrabacteria_group>Bacillota>Bacilli>Lactobacillales>Lactobacillaceae | GCA_002554075.1 | Lactobacillus_sp._UMNPBX2 | CP353_08275 | 15088..18639 | 2042045 | ATPase-[Lactobacillus-sp.-UMNPBX2] |
| AUI77142.1 | Type1.BREX | <-Peptidase_M20\|\|<-?<-?\|\|Peptidase_M13_N+Peptidase_M13-><-HTH_Tnp_4\|\|?->\|\|?-><-MULE-TRANSPOSASE<-DUF6079*<-BrxB-BrxF-iSTAND<-BrxA-DNA-binding<-ParB+HNH<-CNMP+CRP-HTH<-?<-?<-HAD+HAD | 1240 | Bacteria>Terrabacteria_group>Bacillota>Bacilli>Lactobacillales>Lactobacillaceae | GCA_002849955.1 | Lactobacillus_helveticus | Lh22155_05455 | 1074208..1077930 | 1587 | ATPase-[Lactobacillus-helveticus] |
| AUX12764.1 | Type1.BREX | ADH_N+ADH_zinc_N-><-?<-ParB<-ParB<-CRISPR_Cas2<-Cas_Cas1<-Cas9+HNH+RuvC<-PglZ<-N6-METHYLASE<-DUF6079*<-BrxB-BrxF-iSTAND<-BrxA-DNA-binding<-rRNA<-rRNA<-rRNA<-rRNA<-rRNA<-rRNA<-ABC-ATPase+SbcC+ABC-ATPase+SbcC<-?<-DnaB+sGTP<-?<-Peptidase_M3_N+Peptidase_M3 | 1215 | Bacteria>Terrabacteria_group>Bacillota>Bacilli>Lactobacillales>Lactobacillaceae | GCA_002953655.1 | Latilactobacillus_sakei | CO213_10250 | 2077540..2081187 | 1599 | BREX-system-P-loop-protein-BrxC-[Latilactobacillus-sakei] |
| AVK61135.1 | Type1.BREX | <-ABhydrolase\|\|GNTR-HTH+UTRA-><-?<-?<-S1+GTPase-AIG<-PglZ\|\|N6-METHYLASE->Arm-DNA-bind_4+Phage_int_SAM_5+Phage_integrase-><-N6-METHYLASE<-DUF6079*<-BrxB-BrxF-iSTAND<-BrxA-DNA-binding<-ABhydrolase<-?<-?<-?\|\|?-><-GGDEF<-EAL | 1232 | Bacteria>Terrabacteria_group>Bacillota>Bacilli>Lactobacillales>Lactobacillaceae | GCA_002970915.1 | Lactobacillus_sp._CBA3605 | C5Z25_04875 | 984247..987945 | 2099788 | BREX-system-P-loop-protein-BrxC-[Lactobacillus-sp.-CBA3605] |
| AVK63726.1 | Type1.BREX | <-ABhydrolase\|\|GNTR-HTH+UTRA-><-?<-?<-S1+GTPase-AIG<-PglZ\|\|N6-METHYLASE->Arm-DNA-bind_4+Phage_int_SAM_5+Phage_integrase-><-N6-METHYLASE<-DUF6079*<-BrxB-BrxF-iSTAND<-BrxA-DNA-binding<-ROK-HTH<-HTH_OrfB_IS605+OrfB_IS605+OrfB_Zn_ribbon<-?<-?\|\|?-><-GGDEF<-EAL | 1232 | Bacteria>Terrabacteria_group>Bacillota>Bacilli>Lactobacillales>Lactobacillaceae | GCA_002970935.1 | Lactobacillus_sp._CBA3606 | C5Z26_06230 | 1252250..1255948 | 2099789 | BREX-system-P-loop-protein-BrxC-[Lactobacillus-sp.-CBA3606] |
| HBO56057.1 | Type1.BREX | <-SIGMA+OB-Fold+AAA-ATPase+Lon-Protease-BrxL<-PglZ\|\|N6-METHYLASE->Arm-DNA-bind_4+Phage_int_SAM_5+Phage_integrase-><-N6-METHYLASE<-DUF6079*<-BrxB-BrxF-iSTAND<-BrxA-DNA-binding<-?\|\|?->?-><-McrC-NTD+REase | 1194 | Bacteria>Terrabacteria_group>Bacillota>Bacilli>Lactobacillales>Lactobacillaceae | GCA_003533085.1 | Leuconostoc_mesenteroides | DD625_03795 | 14290..17874 | 1245 | BREX-system-P-loop-protein-BrxC-[Leuconostoc-mesenteroides] |
| RND53299.1 | Type1.BREX | <-SIGMA+OB-Fold+AAA-ATPase+Lon-Protease-BrxL<-rve<-PglZ\|\|N6-METHYLASE-p->HIN-HTH->HTH_21+rve+rve_3-><-DUF6079*<-BrxB-BrxF-iSTAND<-BrxA-DNA-binding\|\|MazE_antitoxin-><-?<-?\|\|PAIREDC-HTH+HIN-HTH-><-rve<-LACI-HTH+PBPI | 1062 | Bacteria>Terrabacteria_group>Bacillota>Bacilli>Lactobacillales>Lactobacillaceae | GCA_003712345.1 | Lacticaseibacillus_paracasei | FAM18121_02066 | 531834..535022 | 1597 | hypothetical-protein-FAM18121_02066-[Lacticaseibacillus-paracasei] |
| TXJ87453.1 | Type1.BREX | HDRnase->CHC2+TOPRIM-DNAG->SIGMA-HTH+SIGMA-HTH->SIGMA-HTH+SIGMA-HTH->\|\|HAD+HAD->\|\|?->BrxA-DNA-binding->BrxB-BrxF-iSTAND->\|\|DUF6079-p*->N6-METHYLASE->SIGMA+OB-Fold+AAA-ATPase+Lon-Protease-BrxL->HEPN-> | 917 | Bacteria>Terrabacteria_group>Bacillota>Bacilli>Lactobacillales>Lactobacillaceae | GCA_008016355.1 | Lactobacillus_delbrueckii | FU655_07470 | 24596..27349 | 1584 | BREX-system-P-loop-protein-BrxC-[Lactobacillus-delbrueckii] |
| MSB67176.1 | Type1.BREX | <-N6-METHYLASE-p<-DUF6079*<-BrxB-BrxF-iSTAND<-BrxA-DNA-binding<-?<-SAM-METHYLASE<-?\|\|MarR-HTH->HAD+HAD-><-ZnR+SLATT | 1220 | Bacteria>Terrabacteria_group>Bacillota>Bacilli>Lactobacillales>Lactobacillaceae | GCA_009678855.1 | Leuconostoc_lactis | GKC72_08220 | 557..4219 | 1246 | BREX-system-P-loop-protein-BrxC-[Leuconostoc-lactis] |
| MBA1433800.1 | Type1.BREX | <-Arm-DNA-bind_4+Phage_int_SAM_5+Phage_integrase<-N6-METHYLASE<-DUF6079*<-BrxB-BrxF-iSTAND<-BrxA-DNA-binding\|\|ZnR->?->?->Phage_integrase-> | 1222 | Bacteria>Terrabacteria_group>Bacillota>Bacilli>Lactobacillales>Lactobacillaceae | GCA_013607485.1 | Bombilactobacillus_bombi | EQ827_00800 | 5020..8688 | 1303590 | BREX-system-P-loop-protein-BrxC-[Bombilactobacillus-bombi] |
| MBD8998786.1 | Type1.BREX | <-wHTH+WYL+WCX<-TIR<-?<-SIGMA+OB-Fold+AAA-ATPase+Lon-Protease-BrxL<-?<-PglZ<-OLD-ABC+TOPRIM<-N6-METHYLASE<-TIR<-DUF6079*<-BrxB-BrxF-iSTAND<-BrxA-DNA-binding<-HTH_26<-?<-?<-?\|\|<-Classical-AAA<-PRTase | 1203 | Bacteria>Terrabacteria_group>Bacillota>Bacilli>Lactobacillales>Lactobacillaceae | GCA_014847545.1 | Ligilactobacillus_ruminis | EGQ92_00800 | 15462..19073 | 1623 | BREX-system-P-loop-protein-BrxC-[Ligilactobacillus-ruminis] |
| MBS0953503.1 | Type1.BREX | N6-METHYLASE-p->Arm-DNA-bind_4+Phage_int_SAM_5+Phage_integrase-><-N6-METHYLASE<-DUF6079* | 1057 | Bacteria>Terrabacteria_group>Bacillota>Bacilli>Lactobacillales>Lactobacillaceae | GCA_018257215.1 | Lactiplantibacillus_plantarum | JK218_15530 | 5561..8734 | 1590 | BREX-system-P-loop-protein-BrxC-[Lactiplantibacillus-plantarum] |
| MBT8803912.1 | Type1.BREX | ParB+HNH->BrxA-DNA-binding->BrxB-BrxF-iSTAND->DUF6079*->N6-METHYLASE->PglZ->SIGMA+OB-Fold+AAA-ATPase+Lon-Protease-BrxL->HEPN->?->D6MGT-HTH-><-?<-NUDIX\|\|METHYLASE-> | 1178 | Bacteria>Terrabacteria_group>Bacillota>Bacilli>Lactobacillales>Lactobacillaceae | GCA_018719835.1 | Lactobacillus_delbrueckii_subsp._bulgaricus | BTJ20_08000 | 3864..7400 | 1585 | ATPase-[Lactobacillus-delbrueckii-subsp.-bulgaricus] |
| MBT8900174.1 | Type1.BREX | <-SIGMA+OB-Fold+AAA-ATPase+Lon-Protease-BrxL<-PglZ<-N6-METHYLASE<-DUF6079*<-BrxB-BrxF-iSTAND<-BrxA-DNA-binding<-ParB+HNH\|\|<-HAD+HAD | 1178 | Bacteria>Terrabacteria_group>Bacillota>Bacilli>Lactobacillales>Lactobacillaceae | GCA_018720625.1 | Lactobacillus_delbrueckii_subsp._bulgaricus | BTI57_06430 | 8367..11903 | 1585 | ATPase-[Lactobacillus-delbrueckii-subsp.-bulgaricus] |
| MBT8901771.1 | Type1.BREX | <-METHYLASE\|\|NUDIX->?->\|\|<-AbiGi<-HEPN<-SIGMA+OB-Fold+AAA-ATPase+Lon-Protease-BrxL<-PglZ<-N6-METHYLASE<-DUF6079*<-BrxB-BrxF-iSTAND<-BrxA-DNA-binding<-ParB+HNH<-?\|\|<-HAD+HAD<-SIGMA-HTH+SIGMA-HTH<-SIGMA-HTH+SIGMA-HTH | 1178 | Bacteria>Terrabacteria_group>Bacillota>Bacilli>Lactobacillales>Lactobacillaceae | GCA_018720695.1 | Lactobacillus_delbrueckii_subsp._bulgaricus | BTI52_06840 | 21471..25007 | 1585 | ATPase-[Lactobacillus-delbrueckii-subsp.-bulgaricus] |
| MBT8914213.1 | Type1.BREX | ParB+HNH->BrxA-DNA-binding->BrxB-BrxF-iSTAND->DUF6079*->N6-METHYLASE->PglZ->SIGMA+OB-Fold+AAA-ATPase+Lon-Protease-BrxL->HEPN-> | 1178 | Bacteria>Terrabacteria_group>Bacillota>Bacilli>Lactobacillales>Lactobacillaceae | GCA_018720815.1 | Lactobacillus_delbrueckii_subsp._bulgaricus | BTI61_07605 | 4087..7623 | 1585 | ATPase-[Lactobacillus-delbrueckii-subsp.-bulgaricus] |
| MBU3830350.1 | Type1.BREX | SF1-SF2-Helicase-><-?<-?\|\|Sulfatase+Phosphodiest->?->?->?-><-?\|\|BrxA-DNA-binding->BrxB-BrxF-iSTAND->DUF6079*->N6-METHYLASE->MazE_antitoxin->DOC->PglZ-><-?\|\|?->?->TraI-VirD2-RCRE-> | 1221 | Bacteria>Terrabacteria_group>Bacillota>Bacilli>Lactobacillales>Lactobacillaceae | GCA_018883805.1 | Candidatus_Limosilactobacillus_merdavium | H9843_05610 | 43121..46786 | 2838651 | BREX-system-P-loop-protein-BrxC-[Candidatus-Limosilactobacillus-merdavium] |
| HJF09584.1 | Type1.BREX | <-Glyco_hydro_36N+Melibiase+Glyco_hydro_36C<-ABhydrolase\|\|?->?->?->?->Glyco_hydro_1->BrxA-DNA-binding->BrxB-BrxF-iSTAND->DUF6079*-> | 1241 | Bacteria>Terrabacteria_group>Bacillota>Bacilli>Lactobacillales>Lactobacillaceae | GCA_020742315.1 | Lactobacillus_crispatus | K8V23_02095 | 14734..18459 | 47770 | BREX-system-P-loop-protein-BrxC-[Lactobacillus-crispatus] |
| HJG15197.1 | Type1.BREX | <-PglZ<-?<-?<-N6-METHYLASE<-DUF6079*<-BrxB-BrxF-iSTAND<-BrxA-DNA-binding\|\|<-?<-?\|\|MODE-HTH+PBPII-><-?\|\|MODE-HTH+PBPII-><-REase | 1241 | Bacteria>Terrabacteria_group>Bacillota>Bacilli>Lactobacillales>Lactobacillaceae | GCA_020743205.1 | Ligilactobacillus_salivarius | K8V06_03525 | 11658..15383 | 1624 | BREX-system-P-loop-protein-BrxC-[Ligilactobacillus-salivarius] |
| MCE3043097.1 | Type1.BREX | <-SIGMA+OB-Fold+AAA-ATPase+Lon-Protease-BrxL<-PglZ\|\|N6-METHYLASE->Arm-DNA-bind_4+Phage_int_SAM_5+Phage_integrase-><-N6-METHYLASE<-DUF6079*<-BrxB-BrxF-iSTAND<-BrxA-DNA-binding | 1235 | Bacteria>Terrabacteria_group>Bacillota>Bacilli>Lactobacillales>Lactobacillaceae | GCA_021300595.1 | Lacticaseibacillus_rhamnosus | LW144_13175 | 12071..15778 | 47715 | BREX-system-P-loop-protein-BrxC-[Lacticaseibacillus-rhamnosus] |
| MCH3902414.1 | Type1.BREX | OrfB_IS605+OrfB_Zn_ribbon-><-tRNA\|\|RADICAL-SAM->?->?->Y1_Tnp->HTH_OrfB_IS605+OrfB_IS605+OrfB_Zn_ribbon->BrxA-DNA-binding->BrxB-BrxF-iSTAND->DUF6079*->N6-METHYLASE->PglZ->SIGMA+OB-Fold+AAA-ATPase+Lon-Protease-BrxL->HTH_Tnp_1->HTH_Tnp_1+HTH_21+rve+rve_3->?-><-MULE-TRANSPOSASE\|\|?-><-LZ_Tnp_IS66+zf-IS66+DDE_Tnp_IS66+DDE_Tnp_IS66_C | 1205 | Bacteria>Terrabacteria_group>Bacillota>Bacilli>Lactobacillales>Lactobacillaceae | GCA_022482265.1 | Limosilactobacillus_oris | LKE19_02070 | 430919..434536 | 1632 | BREX-system-P-loop-protein-BrxC-[Limosilactobacillus-oris] |
| MCH3921366.1 | Type1.BREX | <-TraI-VirD2-RCRE<-?<-?\|\|?-><-PglZ<-DOC<-MazE_antitoxin<-N6-METHYLASE<-DUF6079*\|\|HIN-HTH+PAIREDC-HTH-><-BrxB-BrxF-iSTAND<-BrxA-DNA-binding<-HTH_38\|\|?-><-?<-His_Phos_1 | 1221 | Bacteria>Terrabacteria_group>Bacillota>Bacilli>Lactobacillales>Lactobacillaceae | GCA_022482305.1 | Limosilactobacillus_sp. | LKE14_00865 | 208799..212464 | 2773925 | BREX-system-P-loop-protein-BrxC-[Limosilactobacillus-sp.] |
| MCH3996392.1 | Type1.BREX | Peptidase_M13_N+Peptidase_M13->HIN-HTH+PAIREDC-HTH->Acetate_kinase-><-TerBN+TPR<-?<-PglZ<-N6-METHYLASE<-DUF6079*<-BrxB-BrxF-iSTAND<-BrxA-DNA-binding<-ParB+HNH<-?<-?<-?<-AAA_33 | 1244 | Bacteria>Terrabacteria_group>Bacillota>Bacilli>Lactobacillales>Lactobacillaceae | GCA_022483225.1 | Lactobacillus_amylovorus | LKE67_02050 | 413910..417644 | 1604 | BREX-system-P-loop-protein-BrxC-[Lactobacillus-amylovorus] |
| MCI1849257.1 | Type1.BREX | <-SIGMA+OB-Fold+AAA-ATPase+Lon-Protease-BrxL<-PglZ\|\|N6-METHYLASE->Arm-DNA-bind_4+Phage_int_SAM_5+Phage_integrase-><-N6-METHYLASE<-DUF6079*<-BrxB-BrxF-iSTAND<-BrxA-DNA-binding\|\|?-><-AIPR | 1224 | Bacteria>Terrabacteria_group>Bacillota>Bacilli>Lactobacillales>Lactobacillaceae | GCA_022649255.1 | Schleiferilactobacillus_harbinensis | LKI81_00230 | 45023..48697 | 304207 | BREX-system-P-loop-protein-BrxC-[Schleiferilactobacillus-harbinensis] |
| MCI7336507.1 | Type1.BREX | <-DEDD_Tnp_IS110<-?<-?\|\|MODE-HTH+PBPII-><-N6-METHYLASE<-DUF6079*<-BrxB-BrxF-iSTAND-p | 1227 | Bacteria>Terrabacteria_group>Bacillota>Bacilli>Lactobacillales>Lactobacillaceae | GCA_022783965.1 | Lactobacillus_amylovorus | MSH35_07235 | 6941..10624 | 1604 | BREX-system-P-loop-protein-BrxC-[Lactobacillus-amylovorus] |
| MCI7713737.1 | Type1.BREX | AAA_33->?->?->ParB+HNH->BrxA-DNA-binding->BrxB-BrxF-iSTAND->DUF6079*->N6-METHYLASE-> | 1244 | Bacteria>Terrabacteria_group>Bacillota>Bacilli>Lactobacillales>Lactobacillaceae | GCA_022786405.1 | Lactobacillus_johnsonii | MSL49_00240 | 14783..18517 | 33959 | BREX-system-P-loop-protein-BrxC-[Lactobacillus-johnsonii] |
| MCO6542239.1 | Type1.BREX | <-HTH_26<-HTH_26\|\|BrxA-DNA-binding->BrxB-BrxF-iSTAND->DUF6079*->N6-METHYLASE-><-Arm-DNA-bind_4+Phage_int_SAM_5+Phage_integrase<-N6-METHYLASE\|\|PglZ->SIGMA+OB-Fold+AAA-ATPase+Lon-Protease-BrxL->SIR2->HerA/FtsK->McrB-AAA->McrC-NTD+REase-> | 1209 | Bacteria>Terrabacteria_group>Bacillota>Bacilli>Lactobacillales>Lactobacillaceae | GCA_024104005.1 | Lactobacillus_sp. | J6573_08070 | 13480..17109 | 1591 | BREX-system-P-loop-protein-BrxC-[Lactobacillus-sp.] |
| MCO6543132.1 | Type1.BREX | <-McrC-NTD+REase<-McrB-AAA<-HerA/FtsK<-SIR2<-SIGMA+OB-Fold+AAA-ATPase+Lon-Protease-BrxL<-PglZ\|\|N6-METHYLASE->Arm-DNA-bind_4+Phage_int_SAM_5+Phage_integrase-><-N6-METHYLASE<-DUF6079*<-BrxB-BrxF-iSTAND<-BrxA-DNA-binding\|\|HTH_26->HTH_26-><-Glyco_hydro_1<-?<-PTSEIIC<-?\|\|BirA-HTH-> | 1206 | Bacteria>Terrabacteria_group>Bacillota>Bacilli>Lactobacillales>Lactobacillaceae | GCA_024104055.1 | Lactobacillus_sp. | J6584_04075 | 172447..176067 | 1591 | BREX-system-P-loop-protein-BrxC-[Lactobacillus-sp.] |
| MCP9304657.1 | Type1.BREX | <-PglZ\|\|N6-METHYLASE->Arm-DNA-bind_4+Phage_int_SAM_5+Phage_integrase-><-N6-METHYLASE<-DUF6079*<-BrxB-BrxF-iSTAND<-BrxA-DNA-binding\|\|MazE_antitoxin-><-?<-?<-LACI-HTH+PBPI<-Glyco_hydro_32N+Glyco_hydro_32C<-?<-SAM-METHYLASE | 1217 | Bacteria>Terrabacteria_group>Bacillota>Bacilli>Lactobacillales>Lactobacillaceae | GCA_024329455.1 | Lacticaseibacillus_paracasei | KBX55_07815 | 9810..13463 | 1597 | BREX-system-P-loop-protein-BrxC-[Lacticaseibacillus-paracasei] |
| MCT6901727.1 | Type1.BREX | <-ZnR<-RVT<-?<-?<-SIGMA+OB-Fold+AAA-ATPase+Lon-Protease-BrxL<-PglZ<-N6-METHYLASE<-DUF6079*<-BrxB-BrxF-iSTAND<-BrxA-DNA-binding | 1195 | Bacteria>Terrabacteria_group>Bacillota>Bacilli>Lactobacillales>Lactobacillaceae | GCA_025291565.1 | Lactobacillus_sp. | M3Z48_00675 | 20204..23791 | 1591 | BREX-system-P-loop-protein-BrxC-[Lactobacillus-sp.] |
| MCT7705168.1 | Type1.BREX | <-ACET+ACET<-?\|\|?-><-PglZ<-?<-?<-?<-N6-METHYLASE<-DUF6079*<-BrxB-BrxF-iSTAND<-BrxA-DNA-binding<-?<-?<-?<-?<-Cysteine-Desulfurase<-?<-ABC-ATPase+SbcC | 1243 | Bacteria>Terrabacteria_group>Bacillota>Bacilli>Lactobacillales>Lactobacillaceae | GCA_025346185.1 | Lactobacillus_gasseri | N5846_06715 | 16441..20172 | 1596 | BREX-system-P-loop-protein-BrxC-[Lactobacillus-gasseri] |
| MCT7861392.1 | Type1.BREX | <-PglZ<-HEPN<-?<-?<-N6-METHYLASE<-?<-DUF6079*<-BrxB-BrxF-iSTAND<-BrxA-DNA-binding | 1165 | Bacteria>Terrabacteria_group>Bacillota>Bacilli>Lactobacillales>Lactobacillaceae | GCA_025347225.1 | Lactobacillus_crispatus | N4Q24_00945 | 8507..12000 | 47770 | BREX-system-P-loop-protein-BrxC-[Lactobacillus-crispatus] |
| UXI83507.1 | Type1.BREX | zf-ISL3+HTH_Tnp_ISL3-><-SIGMA+OB-Fold+AAA-ATPase+Lon-Protease-BrxL<-PglZ<-N6-METHYLASE<-DUF6079*<-BrxB-BrxF-iSTAND<-BrxA-DNA-binding<-HTH_26<-?<-N6-METHYLASE<-TRD\|\|Phage_int_SAM_5+Phage_integrase-> | 1183 | Bacteria>Terrabacteria_group>Bacillota>Bacilli>Lactobacillales>Lactobacillaceae | GCA_025398675.1 | Ligilactobacillus_salivarius | NYZ94_01970 | 392306..395857 | 1624 | BREX-system-P-loop-protein-BrxC-[Ligilactobacillus-salivarius] |
| WP_013245951.1 | Type1.BREX | <-SIGMA+OB-Fold+AAA-ATPase+Lon-Protease-BrxL<-PglZ\|\|N6-METHYLASE->Arm-DNA-bind_4+Phage_int_SAM_5+Phage_integrase-><-N6-METHYLASE<-DUF6079*<-BrxB-BrxF-iSTAND<-BrxA-DNA-binding\|\|MazE_antitoxin-><-?<-?<-LACI-HTH+PBPI<-Glyco_hydro_32N+Glyco_hydro_32C<-?<-SAM-METHYLASE | 1212 | Bacteria>Terrabacteria_group>Bacillota>Bacilli>Lactobacillales>Lactobacillaceae | GCF_000019245.4 | Lacticaseibacillus_paracasei | LCAZH_RS10350 | 2020392..2024030 | 1597 | BREX-system-P-loop-protein-BrxC-[Lacticaseibacillus-paracasei] |
| WP_014569899.1 | Type1.BREX | <-ParB+HNH<-SIGMA+OB-Fold+AAA-ATPase+Lon-Protease-BrxL<-PglZ\|\|N6-METHYLASE->Arm-DNA-bind_4+Phage_int_SAM_5+Phage_integrase-><-N6-METHYLASE<-DUF6079*<-BrxB-BrxF-iSTAND<-BrxA-DNA-binding\|\|MazE_antitoxin-><-?<-?<-LACI-HTH+PBPI<-Glyco_hydro_32N+Glyco_hydro_32C<-?<-SAM-METHYLASE | 1210 | Bacteria>Terrabacteria_group>Bacillota>Bacilli>Lactobacillales>Lactobacillaceae | GCF_000026505.1 | Lacticaseibacillus_rhamnosus | LGG_RS10100 | 2165561..2169193 | 47715 | BREX-system-P-loop-protein-BrxC-[Lacticaseibacillus-rhamnosus] |
| WP_012846188.1 | Type1.BREX | <-RFC-AAA<-PglZ<-?<-?\|\|MODE-HTH+PBPII-><-N6-METHYLASE<-DUF6079*<-BrxB-BrxF-iSTAND<-BrxA-DNA-binding<-?<-?<-ACET+ACET<-?<-?\|\|<-?<-REase+SF1-SF2-Helicase | 1243 | Bacteria>Terrabacteria_group>Bacillota>Bacilli>Lactobacillales>Lactobacillaceae | GCF_000091405.1 | Lactobacillus_johnsonii | FI9785_RS04615 | 948220..951951 | 33959 | BREX-system-P-loop-protein-BrxC-[Lactobacillus-johnsonii] |
| WP_013102196.1 | Type1.BREX | <-SIGMA+OB-Fold+AAA-ATPase+Lon-Protease-BrxL<-PglZ\|\|N6-METHYLASE->Arm-DNA-bind_4+Phage_int_SAM_5+Phage_integrase-><-N6-METHYLASE<-DUF6079*<-BrxB-BrxF-iSTAND<-BrxA-DNA-binding\|\|?->?-><-McrC-NTD+REase<-McrC-NTD+REase<-?<-ABC-ATPase+SbcC | 1194 | Bacteria>Terrabacteria_group>Bacillota>Bacilli>Lactobacillales>Lactobacillaceae | GCF_000092505.1 | Leuconostoc_kimchii | LKI_RS00700 | 46053..49637 | 136609 | BREX-system-P-loop-protein-BrxC-[Leuconostoc-kimchii] |
| WP_003671433.1 | Type1.BREX | His_Phos_1->?->?->rRNA->rRNA->rRNA->tRNA->BrxA-DNA-binding->BrxB-BrxF-iSTAND->DUF6079*->N6-METHYLASE->MazE_antitoxin->DOC->PglZ->?->HTH_Tnp_1+HTH_21+rve+rve_3-><-HTH_3 | 1221 | Bacteria>Terrabacteria_group>Bacillota>Bacilli>Lactobacillales>Lactobacillaceae | GCF_000159455.2 | Limosilactobacillus_reuteri | HMPREF0538_RS06415 | 1311284..1314949 | 1598 | BREX-system-P-loop-protein-BrxC-[Limosilactobacillus-reuteri] |
| WP_005684782.1 | Type1.BREX | Glyco_hydro_32N+Glyco_hydro_32C->LACI-HTH+PBPI->?->?->?-><-MazE_antitoxin\|\|BrxA-DNA-binding->BrxB-BrxF-iSTAND->DUF6079*->N6-METHYLASE-p-> | 1210 | Bacteria>Terrabacteria_group>Bacillota>Bacilli>Lactobacillales>Lactobacillaceae | GCF_000173255.4 | Lacticaseibacillus_rhamnosus | LRH_RS10375 | 137182..140814 | 47715 | BREX-system-P-loop-protein-BrxC-[Lacticaseibacillus-rhamnosus] |
| WP_010014398.1 | Type1.BREX | <-PglZ<-N6-METHYLASE<-DUF6079*<-BrxB-BrxF-iSTAND<-BrxA-DNA-binding | 1202 | Bacteria>Terrabacteria_group>Bacillota>Bacilli>Lactobacillales>Lactobacillaceae | GCF_000184285.1 | Loigolactobacillus_coryniformis | FC16_RS08220 | 7125..10733 | 1610 | BREX-system-P-loop-protein-BrxC-[Loigolactobacillus-coryniformis] |
| WP_014563593.1 | Type1.BREX | <-ParB+HNH<-HTH_21+rve<-HTH_23+HTH_28\|\|Calcineurin-><-TerBN+TPR<-PglZ<-RVT<-N6-METHYLASE<-DUF6079*<-BrxB-BrxF-iSTAND<-BrxA-DNA-binding<-ParB+HNH<-ParB+HNH<-?<-ACET+ACET<-CNMP+CRP-HTH | 1238 | Bacteria>Terrabacteria_group>Bacillota>Bacilli>Lactobacillales>Lactobacillaceae | GCF_000189515.1 | Lactobacillus_helveticus | LBHH_RS05435 | 1072223..1075939 | 1587 | BREX-system-P-loop-protein-BrxC-[Lactobacillus-helveticus] |
| WP_014565829.1 | Type1.BREX | AAA_33->?->?->BrxA-DNA-binding->BrxB-BrxF-iSTAND->DUF6079*->N6-METHYLASE-><-MODE-HTH+PBPII\|\|?->?->PglZ->TerBN+TPR-> | 1233 | Bacteria>Terrabacteria_group>Bacillota>Bacilli>Lactobacillales>Lactobacillaceae | GCF_000194115.1 | Lactobacillus_amylovorus | LAB52_RS05240 | 1027360..1031061 | 1604 | BREX-system-P-loop-protein-BrxC-[Lactobacillus-amylovorus] |
| WP_004903747.1 | Type1.BREX | Arm-DNA-bind_4+Phage_int_SAM_5+Phage_integrase-><-N6-METHYLASE<-DUF6079*<-BrxB-BrxF-iSTAND<-BrxA-DNA-binding<-HTH_Tnp_1+HTH_21+rve+rve_3<-HIN-HTH | 1194 | Bacteria>Terrabacteria_group>Bacillota>Bacilli>Lactobacillales>Lactobacillaceae | GCF_000239895.1 | Leuconostoc_citreum | LEUCOC10_RS09130 | 5196..8780 | 33964 | BREX-system-P-loop-protein-BrxC-[Leuconostoc-citreum] |
| WP_034992311.1 | Type1.BREX | GTPase-AIG->?->HTH+REase->BrxA-DNA-binding->BrxB-BrxF-iSTAND->DUF6079*->N6-METHYLASE-> | 1228 | Bacteria>Terrabacteria_group>Bacillota>Bacilli>Lactobacillales>Lactobacillaceae | GCF_000255515.1 | Liquorilactobacillus_vini | IC26_RS02660 | 4237..7923 | 238015 | BREX-system-P-loop-protein-BrxC-[Liquorilactobacillus-vini] |
| WP_003595619.1 | Type1.BREX | <-Arm-DNA-bind_4+Phage_int_SAM_5+Phage_integrase<-DUF6079*<-BrxB-BrxF-iSTAND<-BrxA-DNA-binding\|\|MazE_antitoxin-><-?<-?<-LACI-HTH+PBPI<-Glyco_hydro_32N+Glyco_hydro_32C<-?<-SAM-METHYLASE | 1217 | Bacteria>Terrabacteria_group>Bacillota>Bacilli>Lactobacillales>Lactobacillaceae | GCF_000309705.1 | Lacticaseibacillus_paracasei | LCAUCD174_RS03785 | 5913..9566 | 1597 | BREX-system-P-loop-protein-BrxC-[Lacticaseibacillus-paracasei] |
| WP_003606939.1 | Type1.BREX | <-SIGMA+OB-Fold+AAA-ATPase+Lon-Protease-BrxL<-PglZ\|\|N6-METHYLASE->Arm-DNA-bind_4+Phage_int_SAM_5+Phage_integrase-><-N6-METHYLASE<-DUF6079*<-BrxB-BrxF-iSTAND<-BrxA-DNA-binding\|\|MazE_antitoxin-><-?<-?<-LACI-HTH+PBPI<-Glyco_hydro_32N+Glyco_hydro_32C<-?<-SAM-METHYLASE | 1215 | Bacteria>Terrabacteria_group>Bacillota>Bacilli>Lactobacillales>Lactobacillaceae | GCF_000309785.1 | Lacticaseibacillus_paracasei | LCALPC37_RS08645 | 12743..16390 | 1597 | BREX-system-P-loop-protein-BrxC-[Lacticaseibacillus-paracasei] |
| WP_005715305.1 | Type1.BREX | <-ParB+HNH<-SIGMA+OB-Fold+AAA-ATPase+Lon-Protease-BrxL<-PglZ\|\|N6-METHYLASE->Arm-DNA-bind_4+Phage_int_SAM_5+Phage_integrase-><-N6-METHYLASE<-DUF6079*<-BrxB-BrxF-iSTAND<-BrxA-DNA-binding\|\|MazE_antitoxin-><-?<-?<-?<-LACI-HTH+PBPI<-Glyco_hydro_32N+Glyco_hydro_32C | 1210 | Bacteria>Terrabacteria_group>Bacillota>Bacilli>Lactobacillales>Lactobacillaceae | GCF_000311945.1 | Lacticaseibacillus_rhamnosus | LRHMDP2_RS10615 | 154290..157922 | 47715 | BREX-system-P-loop-protein-BrxC-[Lacticaseibacillus-rhamnosus] |
| WP_009555629.1 | Type1.BREX | HAD+S6PP->?-><-REase<-?<-HTH+REase<-PglZ<-N6-METHYLASE<-DUF6079*<-BrxB-BrxF-iSTAND<-BrxA-DNA-binding<-?<-?<-?<-HKD-DNase+SF1-SF2-Helicase+HTH+HTH+SRA | 1218 | Bacteria>Terrabacteria_group>Bacillota>Bacilli>Lactobacillales>Lactobacillaceae | GCF_000317165.1 | Ligilactobacillus_saerimneri | D271_RS07755 | 76703..80359 | 228229 | BREX-system-P-loop-protein-BrxC-[Ligilactobacillus-saerimneri] |
| WP_003602979.1 | Type1.BREX | <-GDResolvase+HIN-HTH<-?<-ParB+HNH<-SIGMA+OB-Fold+AAA-ATPase+Lon-Protease-BrxL<-PglZ\|\|N6-METHYLASE->Arm-DNA-bind_4+Phage_int_SAM_5+Phage_integrase-><-N6-METHYLASE<-DUF6079*<-BrxB-BrxF-iSTAND<-BrxA-DNA-binding<-?\|\|?->?->?->HTH_Tnp_1-> | 1241 | Bacteria>Terrabacteria_group>Bacillota>Bacilli>Lactobacillales>Lactobacillaceae | GCF_000376145.1 | Lacticaseibacillus | LCAUW4_RS03725 | 15865..19590 | 2759736 | MULTISPECIES:-BREX-system-P-loop-protein-BrxC-[Lacticaseibacillus] |
| WP_015638640.1 | Type1.BREX | ABC-ATPase+SbcC+SWACOS+SbcC->35exo->?->?->?-><-?<-?\|\|tRNA->tRNA->BrxA-DNA-binding->BrxB-BrxF-iSTAND->DUF6079*->N6-METHYLASE-><-Arm-DNA-bind_4+Phage_int_SAM_5+Phage_integrase<-N6-METHYLASE\|\|PglZ->SIGMA+OB-Fold+AAA-ATPase+Lon-Protease-BrxL->ParB+HNH->?->HTH_28-> | 1230 | Bacteria>Terrabacteria_group>Bacillota>Bacilli>Lactobacillales>Lactobacillaceae | GCF_000397165.1 | Limosilactobacillus_fermentum | LBFF_RS02010 | 427124..430816 | 1613 | BREX-system-P-loop-protein-BrxC-[Limosilactobacillus-fermentum] |
| WP_016383162.1 | Type1.BREX | <-HTH+REase<-SIGMA+OB-Fold+AAA-ATPase+Lon-Protease-BrxL<-PglZ\|\|N6-METHYLASE->Arm-DNA-bind_4+Phage_int_SAM_5+Phage_integrase->zf-ISL3+HTH_Tnp_ISL3+DDE_Tnp_ISL3-><-N6-METHYLASE<-DUF6079*<-BrxB-BrxF-iSTAND<-BrxA-DNA-binding\|\|MazE_antitoxin-><-?<-?<-LACI-HTH+PBPI<-Glyco_hydro_32N+Glyco_hydro_32C<-?<-SAM-METHYLASE | 1215 | Bacteria>Terrabacteria_group>Bacillota>Bacilli>Lactobacillales>Lactobacillaceae | GCF_000409835.1 | Lacticaseibacillus_paracasei | J3H89_RS10660 | 2208273..2211920 | 1597 | BREX-system-P-loop-protein-BrxC-[Lacticaseibacillus-paracasei] |
| WP_016373336.1 | Type1.BREX | <-DUF6079*<-BrxB-BrxF-iSTAND<-BrxA-DNA-binding\|\|MazE_antitoxin-><-PTSEIIA1 | 1012 | Bacteria>Terrabacteria_group>Bacillota>Bacilli>Lactobacillales>Lactobacillaceae | GCF_000410335.1 | Lacticaseibacillus_paracasei | LPL14_RS01095 | 1..3038 | 1597 | BREX-system-P-loop-protein-BrxC,-partial-[Lacticaseibacillus-paracasei] |
| WP_027828098.1 | Type1.BREX | SF1-SF2-Helicase-><-?\|\|BrxA-DNA-binding->BrxB-BrxF-iSTAND->DUF6079*->N6-METHYLASE-><-Arm-DNA-bind_4+Phage_int_SAM_5+Phage_integrase<-N6-METHYLASE\|\|PglZ->SIGMA+OB-Fold+AAA-ATPase+Lon-Protease-BrxL->ParB+HNH-> | 1241 | Bacteria>Terrabacteria_group>Bacillota>Bacilli>Lactobacillales>Lactobacillaceae | GCF_000425885.1 | Schleiferilactobacillus_harbinensis | FC91_RS07095 | 17902..21627 | 304207 | BREX-system-P-loop-protein-BrxC-[Schleiferilactobacillus-harbinensis] |
| WP_020843329.1 | Type1.BREX | <-SPOUT<-Phage_integrase<-PglZ<-DOC<-MazE_antitoxin\|\|PAIREDC-HTH->zf-ISL3+HTH_Tnp_ISL3-><-N6-METHYLASE<-DUF6079*<-BrxB-BrxF-iSTAND<-BrxA-DNA-binding<-?\|\|CR-ATPase5+REase-><-?<-Sulfatase+Phosphodiest | 1221 | Bacteria>Terrabacteria_group>Bacillota>Bacilli>Lactobacillales>Lactobacillaceae | GCF_000439275.1 | Limosilactobacillus_reuteri | N134_RS08695 | 1857524..1861189 | 1598 | BREX-system-P-loop-protein-BrxC-[Limosilactobacillus-reuteri] |
| WP_023192364.1 | Type1.BREX | ASCH->Glyoxalase->Glyoxalase->ParB+HNH->ParB+HNH->BrxA-DNA-binding->BrxB-BrxF-iSTAND->DUF6079-p->DUF6079*->PglZ->TerBN+TPR->DDE_Tnp_1-><-DUF772+DDE_Tnp_1+DDE_Tnp_1_6<-?<-?\|\|Peptidase->?->Glycos_transf-> | 1179 | Bacteria>Terrabacteria_group>Bacillota>Bacilli>Lactobacillales>Lactobacillaceae | GCF_000493335.1 | Lactobacillus_helveticus | LHE_RS17595 | 1223279..1226818 | 1587 | BREX-system-P-loop-protein-BrxC-[Lactobacillus-helveticus] |
| WP_023192226.1 | Type1.BREX | ASCH->Glyoxalase->ParB+HNH->?->ParB+HNH->BrxA-DNA-binding->BrxB-BrxF-iSTAND->DUF6079*->N6-METHYLASE->nSTAND3-> | 1243 | Bacteria>Terrabacteria_group>Bacillota>Bacilli>Lactobacillales>Lactobacillaceae | GCF_000493555.1 | Lactobacillus_helveticus | LHCIRMBIA104_RS09710 | 11167..14898 | 1587 | BREX-system-P-loop-protein-BrxC-[Lactobacillus-helveticus] |
| WP_025283578.1 | Type1.BREX | ASCH->Glyoxalase->Glyoxalase->ParB+HNH->BrxA-DNA-binding->BrxB-BrxF-iSTAND->DUF6079*->N6-METHYLASE->HEPN->HEPN->PglZ-><-?\|\|DDE_Tnp_1-><-Calcineurin<-Peptidase_M13_N+Peptidase_M13 | 1243 | Bacteria>Terrabacteria_group>Bacillota>Bacilli>Lactobacillales>Lactobacillaceae | GCF_000525715.1 | Lactobacillus_helveticus | LBH_0939 | 971853..975584 | 1587 | BREX-system-P-loop-protein-BrxC-[Lactobacillus-helveticus] |
| WP_047672904.1 | Type1.BREX | LPAM_1+DUF4767->?->?->BrxA-DNA-binding->BrxB-BrxF-iSTAND->DUF6079*->N6-METHYLASE-><-Arm-DNA-bind_4+Phage_int_SAM_5+Phage_integrase<-N6-METHYLASE\|\|PglZ->?->URI-DNase->HTH_28+HTH_32-> | 1216 | Bacteria>Terrabacteria_group>Bacillota>Bacilli>Lactobacillales>Lactobacillaceae | GCF_000762955.1 | Lactiplantibacillus_plantarum | CMPG5300_RS03655 | 792142..795792 | 1590 | BREX-system-P-loop-protein-BrxC-[Lactiplantibacillus-plantarum] |
| WP_047678144.1 | Type1.BREX | DUF6079*->N6-METHYLASE->N6-METHYLASE-p->Arm-DNA-bind_4+Phage_int_SAM_5+Phage_integrase-><-N6-METHYLASE\|\|PglZ->PglZ->SIGMA+OB-Fold+AAA-ATPase+Lon-Protease-BrxL->ParB+HNH-> | 1121 | Bacteria>Terrabacteria_group>Bacillota>Bacilli>Lactobacillales>Lactobacillaceae | GCF_000801045.1 | Lacticaseibacillus_rhamnosus | LaR116_RS11810 | 1..3366 | 47715 | BREX-system-P-loop-protein-BrxC,-partial-[Lacticaseibacillus-rhamnosus] |
| WP_025012991.1 | Type1.BREX | HTH_6+SIS-><-HTH+REase<-SIGMA+OB-Fold+AAA-ATPase+Lon-Protease-BrxL<-PglZ\|\|N6-METHYLASE->Phage_int_SAM_5+Phage_integrase-><-N6-METHYLASE<-DUF6079*<-BrxB-BrxF-iSTAND<-BrxA-DNA-binding<-?\|\|MazE_antitoxin-><-?<-?<-?<-LACI-HTH+PBPI<-Glyco_hydro_32N+Glyco_hydro_32C | 1214 | Bacteria>Terrabacteria_group>Bacillota>Bacilli>Lactobacillales>Lactobacillaceae | GCF_000829055.1 | Lacticaseibacillus | LBCZ_RS09700 | 1993700..1997344 | 2759736 | MULTISPECIES:-BREX-system-P-loop-protein-BrxC-[Lacticaseibacillus] |
| WP_043941308.1 | Type1.BREX | <-SIGMA+OB-Fold+AAA-ATPase+Lon-Protease-BrxL<-PglZ\|\|N6-METHYLASE->Arm-DNA-bind_4+Phage_int_SAM_5+Phage_integrase-><-N6-METHYLASE<-DUF6079*<-BrxB-BrxF-iSTAND<-BrxA-DNA-binding<-ADH_N+ADH_zinc_N | 1194 | Bacteria>Terrabacteria_group>Bacillota>Bacilli>Lactobacillales>Lactobacillaceae | GCF_000878185.1 | Weissella_cibaria | ab3b_RS06420 | 11942..15526 | 137591 | BREX-system-P-loop-protein-BrxC-[Weissella-cibaria] |
| WP_237751028.1 | Type1.BREX | <-SLATT\|\|HKD-DNase+HKD-DNase->?->ADH_N+ADH_zinc_N->ADH_N+ADH_zinc_N->BrxA-DNA-binding->BrxB-BrxF-iSTAND->DUF6079-p*-> | 520 | Bacteria>Terrabacteria_group>Bacillota>Bacilli>Lactobacillales>Lactobacillaceae | GCF_000878205.1 | Weissella_cibaria | QX99_RS03780 | 24305..25867 | 137591 | BREX-system-P-loop-protein-BrxC-[Weissella-cibaria] |
| WP_237750494.1 | Type1.BREX | <-DUF6079-p*<-BrxB-BrxF-iSTAND<-BrxA-DNA-binding<-ADH_N+ADH_zinc_N<-ADH_N+ADH_zinc_N<-?<-HKD-DNase+HKD-DNase<-?\|\|SLATT->SLATT-> | 520 | Bacteria>Terrabacteria_group>Bacillota>Bacilli>Lactobacillales>Lactobacillaceae | GCF_000878215.1 | Weissella_cibaria | ff3pr_RS09465 | 86..1648 | 137591 | BREX-system-P-loop-protein-BrxC-[Weissella-cibaria] |
| WP_048704318.1 | Type1.BREX | <-SIGMA+OB-Fold+AAA-ATPase+Lon-Protease-BrxL<-PglZ\|\|N6-METHYLASE->Arm-DNA-bind_4+Phage_int_SAM_5+Phage_integrase-><-N6-METHYLASE<-DUF6079*<-BrxB-BrxF-iSTAND<-BrxA-DNA-binding\|\|MarR-HTH-><-?<-?<-?<-Glycos_transf | 1231 | Bacteria>Terrabacteria_group>Bacillota>Bacilli>Lactobacillales>Lactobacillaceae | GCF_001050475.1 | Companilactobacillus_ginsenosidimutans | ABM34_RS06250 | 1191208..1194903 | 1007676 | BREX-system-P-loop-protein-BrxC-[Companilactobacillus-ginsenosidimutans] |
| WP_049181746.1 | Type1.BREX | <-N6-METHYLASE<-DUF6079*<-BrxB-BrxF-iSTAND<-BrxA-DNA-binding\|\|MazE_antitoxin-><-?<-?<-LACI-HTH+PBPI<-Glyco_hydro_32N+Glyco_hydro_32C<-?<-SAM-METHYLASE | 1215 | Bacteria>Terrabacteria_group>Bacillota>Bacilli>Lactobacillales>Lactobacillaceae | GCF_001076935.1 | Lacticaseibacillus_paracasei | ADJ78_RS08430 | 3680..7327 | 1597 | BREX-system-P-loop-protein-BrxC-[Lacticaseibacillus-paracasei] |
| WP_057137321.1 | Type1.BREX | AA_permease-><-?\|\|ParA-Soj-PloopNTPase->?->\|\|BrxA-DNA-binding->BrxB-BrxF-iSTAND->DUF6079*->N6-METHYLASE->HTH_Tnp_1->HTH_Tnp_1+HTH_21+rve+rve_3->MazE_antitoxin->DOC->PglZ-> | 1221 | Bacteria>Terrabacteria_group>Bacillota>Bacilli>Lactobacillales>Lactobacillaceae | GCF_001368775.1 | Lactiplantibacillus_plantarum | LP80_RS11410 | 36651..40316 | 1590 | BREX-system-P-loop-protein-BrxC-[Lactiplantibacillus-plantarum] |
| WP_010491697.1 | Type1.BREX | Glyco_hydro_32N+Glyco_hydro_32C->LACI-HTH+PBPI->?->?->?-><-MazE_antitoxin\|\|?->BrxA-DNA-binding->BrxB-BrxF-iSTAND->DUF6079*->N6-METHYLASE-><-Phage_int_SAM_5+Phage_integrase<-N6-METHYLASE\|\|PglZ->SIGMA+OB-Fold+AAA-ATPase+Lon-Protease-BrxL->HTH+REase->HTH+REase-><-HTH_6+SIS | 1214 | Bacteria>Terrabacteria_group>Bacillota>Bacilli>Lactobacillales>Lactobacillaceae | GCF_001433745.1 | Lacticaseibacillus_zeae | FD51_RS09320 | 70770..74414 | 57037 | BREX-system-P-loop-protein-BrxC-[Lacticaseibacillus-zeae] |
| WP_004894779.1 | Type1.BREX | MarR-HTH+DTXR-HTH->ABC-ATPase+SbcC->?-><-MODE-HTH+PBPII\|\|RFC-AAA->?->\|\|?->BrxA-DNA-binding->BrxB-BrxF-iSTAND->DUF6079*->N6-METHYLASE->nSTAND3->HEPN-p->REase->PglZ->TetR-HTH->?->?->ABC-ATPase-> | 1243 | Bacteria>Terrabacteria_group>Bacillota>Bacilli>Lactobacillales>Lactobacillaceae | GCF_001433975.1 | Lactobacillus_johnsonii | FC22_RS00455 | 98654..102385 | 33959 | BREX-system-P-loop-protein-BrxC-[Lactobacillus-johnsonii] |
| WP_020089143.1 | Type1.BREX | <-MarR-HTH\|\|?->tRNA-><-GGDEF<-HicB_lk_antitox\|\|?-><-35exo+BRCT\|\|BrxA-DNA-binding->BrxB-BrxF-iSTAND->DUF6079*->N6-METHYLASE->N6-METHYLASE-p-><-Arm-DNA-bind_4+Phage_int_SAM_5+Phage_integrase\|\|PglZ->?->?->?->HTH_IclR+ROK-HTH-> | 1223 | Bacteria>Terrabacteria_group>Bacillota>Bacilli>Lactobacillales>Lactobacillaceae | GCF_001434055.1 | Levilactobacillus_parabrevis | IV61_RS02185 | 59074..62745 | 357278 | BREX-system-P-loop-protein-BrxC-[Levilactobacillus-parabrevis] |

| Protein-ID | System | Gene-Neighbourhoods | Length | Taxonomy | Assembly-ID | Strain | Locus-Tag | Genomic-coordinates | TaxID | Fasta-define |
|---|---|---|---|---|---|---|---|---|---|---|
| WP_057734072.1 | Type1.BREX | <-HTH_IclR+ROK-HTH<-?<-?<-?<-PglZ\|\|N6-METHYLASE-p->N6-METHYLASE-p->Arm-DNA-bind_4+Phage_int_SAM_5+Phage_integrase-><-N6-METHYLASE<-DUF6079*<-BrxB-BrxF-iSTAND<-BrxA-DNA-binding\|\|35exo+[illegible] | 1222 | Bacteria>Terrabacteria_group>Bacillota>Bacilli>Lactobacillales>Lactobacillaceae | GCF_001434395.1 | Levilactobacillus_hammesii | FD28_RS07215 | 115937..119605 | 267633 | BREX-system-P-loop-protein-BrxC-[Levilactobacillus-hammesii] |
| WP_057739064.1 | Type1.BREX | <-PglZ<-PglZ\|\|N6-METHYLASE-p->N6-METHYLASE-p-><-Phage_int_SAM_5+Phage_integrase<-N6-METHYLASE<-?<-DUF6079*<-BrxB-BrxF-iSTAND<-BrxA-DNA-binding<-?<-?<-?<-?\|\|Abi-> | 1207 | Bacteria>Terrabacteria_group>Bacillota>Bacilli>Lactobacillales>Lactobacillaceae | GCF_001434745.1 | Companilactobacillus_alimentarius | FC67_RS06745 | 50927..54550 | 1602 | BREX-system-P-loop-protein-BrxC-[Companilactobacillus-alimentarius] |
| WP_056966680.1 | Type1.BREX | wHTH-><-HTH+REase<-SIGMA+OB-Fold+AAA-ATPase+Lon-Protease-BrxL<-PglZ<-N6-METHYLASE-p\|\|Phage_int_SAM_5+Phage_integrase->N6-METHYLASE-p->N6-METHYLASE-p-><-N6-METHYLASE<-DUF6079*<-BrxB-Br[illegible] | 1215 | Bacteria>Terrabacteria_group>Bacillota>Bacilli>Lactobacillales>Lactobacillaceae | GCF_001435495.1 | Latilactobacillus_curvatus | LCU_RS04405 | 828464..832111 | 28038 | BREX-system-P-loop-protein-BrxC-[Latilactobacillus-curvatus] |
| WP_048734157.1 | Type1.BREX | BrxA-DNA-binding->BrxB-BrxF-iSTAND->DUF6079*->N6-METHYLASE-><-Arm-DNA-bind_4+Phage_int_SAM_5+Phage_integrase<-N6-METHYLASE\|\|PglZ->?->DOC+HTH-> | 1216 | Bacteria>Terrabacteria_group>Bacillota>Bacilli>Lactobacillales>Lactobacillaceae | GCF_001435525.1 | Levilactobacillus_koreensis | FC99_RS10810 | 1494..5144 | 637971 | BREX-system-P-loop-protein-BrxC-[Levilactobacillus-koreensis] |
| WP_057821153.1 | Type1.BREX | <-Arm-DNA-bind_4+Phage_int_SAM_3+Phage_integrase<-HTH<-?<-SIGMA-HTH<-?\|\|\|\|?->BrxA-DNA-binding->BrxB-BrxF-iSTAND->DUF6079*->N6-METHYLASE->?->?->LZ_Tnp_IS66+zf-IS66+DDE_Tnp_IS66+DDE_Tnp[illegible] | 1227 | Bacteria>Terrabacteria_group>Bacillota>Bacilli>Lactobacillales>Lactobacillaceae | GCF_001435585.1 | Schleiferilactobacillus_perolens | FD09_RS10555 | 7707..11390 | 100468 | BREX-system-P-loop-protein-BrxC-[Schleiferilactobacillus-perolens] |
| WP_027107209.1 | Type1.BREX | ABhydrolase->?->BrxA-DNA-binding->BrxB-BrxF-iSTAND->DUF6079*->N6-METHYLASE->PglZ->NUDIX-> | 1195 | Bacteria>Terrabacteria_group>Bacillota>Bacilli>Lactobacillales>Lactobacillaceae | GCF_001438785.1 | Ligilactobacillus_ceti | G540_RS0105170 | 464815..468402 | 395085 | BREX-system-P-loop-protein-BrxC-[Ligilactobacillus-ceti] |
| WP_057880764.1 | Type1.BREX | <-N6-METHYLASE<-DUF6079*<-BrxB-BrxF-iSTAND<-BrxA-DNA-binding<-?<-MerR-HTH+SHS2+SHS2 | 1206 | Bacteria>Terrabacteria_group>Bacillota>Bacilli>Lactobacillales>Lactobacillaceae | GCF_001438805.1 | Companilactobacillus_kimchiensis | IV57_RS06820 | 1933..5553 | 993692 | BREX-system-P-loop-protein-BrxC-[Companilactobacillus-kimchiensis] |
| WP_060598044.1 | Type1.BREX | <-METHYLASE\|\|NUDIX->?-><-AraC-HTH+O6MGT-HTH<-?<-HEPN<-SIGMA+OB-Fold+AAA-ATPase+Lon-Protease-BrxL<-PglZ<-N6-METHYLASE<-DUF6079*<-BrxB-BrxF-iSTAND<-BrxA-DNA-binding<-ParB+HNH<-?<-?\|\|<-[illegible] | 1178 | Bacteria>Terrabacteria_group>Bacillota>Bacilli>Lactobacillales>Lactobacillaceae | GCF_001469775.1 | Lactobacillus_delbrueckii | AT236_RS05345 | 1068532..1072068 | 1584 | BREX-system-P-loop-protein-BrxC-[Lactobacillus-delbrueckii] |
| WP_060416898.1 | Type1.BREX | ParB+HNH->?->BrxA-DNA-binding->BrxB-BrxF-iSTAND->DUF6079*->N6-METHYLASE->\|\|PglZ->SIGMA+OB-Fold+AAA-ATPase+Lon-Protease-BrxL-><-GDResolvase+HIN-HTH<-?\|\|?->PAIREDC-HTH-> | 1226 | Bacteria>Terrabacteria_group>Bacillota>Bacilli>Lactobacillales>Lactobacillaceae | GCF_001540905.1 | Levilactobacillus_brevis | ABB39_RS11500 | 6540..10220 | 1580 | BREX-system-P-loop-protein-BrxC-[Levilactobacillus-brevis] |
| WP_061400584.1 | Type1.BREX | MarR-HTH+DTXR-HTH->ABC-ATPase+SbcC->?-><-LysR-HTH\|\|RFC-AAA->BrxA-DNA-binding->BrxB-BrxF-iSTAND->DUF6079*->N6-METHYLASE->nSTAND3->PglZ->TetR-HTH->TetR-HTH->?-><-ACET\|\|Amidohydro_2-> | 1243 | Bacteria>Terrabacteria_group>Bacillota>Bacilli>Lactobacillales>Lactobacillaceae | GCF_001572665.1 | Lactobacillus_johnsonii | AYJ53_03505 | 85204..88935 | 33959 | BREX-system-P-loop-protein-BrxC-[Lactobacillus-johnsonii] |
| WP_063515698.1 | Type1.BREX | <-N6-METHYLASE-p<-DUF6079*<-BrxB-BrxF-iSTAND<-BrxA-DNA-binding\|\|?-><-SF1-SF2-Helicase | 1241 | Bacteria>Terrabacteria_group>Bacillota>Bacilli>Lactobacillales>Lactobacillaceae | GCF_001636145.1 | Schleiferilactobacillus_harbinensis | AWV73_RS12760 | 2097..5822 | 304207 | BREX-system-P-loop-protein-BrxC-[Schleiferilactobacillus-harbinensis] |
| WP_046782810.1 | Type1.BREX | <-GDResolvase+HIN-HTH<-?<-ParB+HNH<-SIGMA+OB-Fold+AAA-ATPase+Lon-Protease-BrxL<-PglZ\|\|N6-METHYLASE->Arm-DNA-bind_4+Phage_int_SAM_5+Phage_integrase-><-N6-METHYLASE<-DUF6079*<-BrxB-BrxF[illegible] | 1241 | Bacteria>Terrabacteria_group>Bacillota>Bacilli>Lactobacillales>Lactobacillaceae | GCF_001656585.1 | Lacticaseibacillus_rhamnosus | PY74_RS00235 | 14729..18454 | 47715 | BREX-system-P-loop-protein-BrxC-[Lacticaseibacillus-rhamnosus] |
| WP_068224099.1 | Type1.BREX | Cas9+HNH+RuvC->Cas_Cas1->CRISPR_Cas2-><-?<-?<-?\|\|?->BrxA-DNA-binding->BrxB-BrxF-iSTAND->DUF6079*->?->N6-METHYLASE->PglZ->?->MazG->?-><-?\|\|DapB_N+DapB_C->NTP_transf+HDRnase-> | 1217 | Bacteria>Terrabacteria_group>Bacillota>Bacilli>Lactobacillales>Lactobacillaceae | GCF_001663655.1 | Loigolactobacillus_backii | AYR52_RS03885 | 851506..855159 | 375175 | BREX-system-P-loop-protein-BrxC-[Loigolactobacillus-backii] |
| WP_068376961.1 | Type1.BREX | Cas9+HNH+RuvC->Cas_Cas1->CRISPR_Cas2-><-?<-?<-?\|\|?->BrxA-DNA-binding->BrxB-BrxF-iSTAND->DUF6079*->N6-METHYLASE->PglZ->?->MazG->?-><-?\|\|DapB_N+DapB_C->NTP_transf+HDRnase->ABC-ATPase+Sb[illegible] | 1217 | Bacteria>Terrabacteria_group>Bacillota>Bacilli>Lactobacillales>Lactobacillaceae | GCF_001663715.1 | Loigolactobacillus_backii | AYR54_RS03895 | 856571..860224 | 375175 | BREX-system-P-loop-protein-BrxC-[Loigolactobacillus-backii] |
| WP_068851359.1 | Type1.BREX | ZnR+SLATT-><-HAD+HAD<-MarR-HTH\|\|?->SAM-METHYLASE->?->BrxA-DNA-binding->BrxB-BrxF-iSTAND->DUF6079*->N6-METHYLASE-><-Phage_integrase<-N6-METHYLASE\|\|PglZ->SIGMA+OB-Fold+AAA-ATPase+Lon-Pr[illegible] | 1216 | Bacteria>Terrabacteria_group>Bacillota>Bacilli>Lactobacillales>Lactobacillaceae | GCF_001698145.1 | Leuconostoc_lactis | BCR17_RS04240 | 864786..868436 | 1246 | BREX-system-P-loop-protein-BrxC-[Leuconostoc-lactis] |
| WP_069168466.1 | Type1.BREX | LZ_Tnp_IS66+zf-IS66+DDE_Tnp_IS66+DDE_Tnp_IS66_C-><-TetR-HTH<-TetR-HTH<-PglZ<-?<-?<-?<-N6-METHYLASE<-DUF6079*<-BrxB-BrxF-iSTAND<-BrxA-DNA-binding\|\|<-?<-DDE_5+DDE_Tnp_1\|\|LysR-HTH-><-ABC[illegible] | 1243 | Bacteria>Terrabacteria_group>Bacillota>Bacilli>Lactobacillales>Lactobacillaceae | GCF_001714745.1 | Lactobacillus_johnsonii | BBP16_01700 | 360123..363854 | 33959 | BREX-system-P-loop-protein-BrxC-[Lactobacillus-johnsonii] |
| WP_069469583.1 | Type1.BREX | [illegible] | 1183 | Bacteria>Terrabacteria_group>Bacillota>Bacilli>Lactobacillales>Lactobacillaceae | GCF_001723525.1 | Ligilactobacillus_salivarius | M3M92_RS08920 | 79825..83376 | 1624 | BREX-system-P-loop-protein-BrxC-[Ligilactobacillus-salivarius] |
| WP_069468096.1 | Type1.BREX | [illegible] | 1215 | Bacteria>Terrabacteria_group>Bacillota>Bacilli>Lactobacillales>Lactobacillaceae | GCF_001723545.1 | Latilactobacillus_curvatus | LCW_03905 | 751343..754990 | 28038 | BREX-system-P-loop-protein-BrxC-[Latilactobacillus-curvatus] |
| WP_070065450.1 | Type1.BREX | [illegible] | 1241 | Bacteria>Terrabacteria_group>Bacillota>Bacilli>Lactobacillales>Lactobacillaceae | GCF_001752545.1 | Oenococcus_kitaharae | NT95_RS03550 | 13261..16986 | 336988 | BREX-system-P-loop-protein-BrxC-[Oenococcus-kitaharae] |
| WP_070489741.1 | Type1.BREX | [illegible] | 1188 | Bacteria>Terrabacteria_group>Bacillota>Bacilli>Lactobacillales>Lactobacillaceae | GCF_001808415.1 | Lactobacillus_sp._HMSC08B12 | HMPREF3168_RS09245 | 1333..4899 | 1581136 | BREX-system-P-loop-protein-BrxC-[Lactobacillus-sp.-HMSC08B12] |
| WP_070792499.1 | Type1.BREX | [illegible] | 1227 | Bacteria>Terrabacteria_group>Bacillota>Bacilli>Lactobacillales>Lactobacillaceae | GCF_001831225.1 | Lacticaseibacillus_rhamnosus | BKP38_RS08735 | 14861..18544 | 47715 | BREX-system-P-loop-protein-BrxC-[Lacticaseibacillus-rhamnosus] |
| WP_071253015.1 | Type1.BREX | [illegible] | 1241 | Bacteria>Terrabacteria_group>Bacillota>Bacilli>Lactobacillales>Lactobacillaceae | GCF_001858275.1 | Lacticaseibacillus_paracasei | BBX46_RS14350 | 14873..18598 | 1597 | BREX-system-P-loop-protein-BrxC-[Lacticaseibacillus-paracasei] |
| WP_081371217.1 | Type1.BREX | [illegible] | 1243 | Bacteria>Terrabacteria_group>Bacillota>Bacilli>Lactobacillales>Lactobacillaceae | GCF_001895375.1 | Lactobacillus_helveticus | BEL10_RS03665 | 22378..26109 | 1587 | BREX-system-P-loop-protein-BrxC-[Lactobacillus-helveticus] |
| WP_081510105.1 | Type1.BREX | [illegible] | 1183 | Bacteria>Terrabacteria_group>Bacillota>Bacilli>Lactobacillales>Lactobacillaceae | GCF_002079325.1 | Ligilactobacillus_salivarius | B6U61_RS08785 | 65481..69032 | 1624 | BREX-system-P-loop-protein-BrxC-[Ligilactobacillus-salivarius] |
| WP_081535666.1 | Type1.BREX | [illegible] | 1183 | Bacteria>Terrabacteria_group>Bacillota>Bacilli>Lactobacillales>Lactobacillaceae | GCF_002079425.1 | Ligilactobacillus_salivarius | B6U56_RS10470 | 187776..191327 | 1624 | BREX-system-P-loop-protein-BrxC-[Ligilactobacillus-salivarius] |
| WP_081512952.1 | Type1.BREX | [illegible] | 1183 | Bacteria>Terrabacteria_group>Bacillota>Bacilli>Lactobacillales>Lactobacillaceae | GCF_002079645.1 | Ligilactobacillus_salivarius | B6U45_RS09365 | 86817..90368 | 1624 | BREX-system-P-loop-protein-BrxC-[Ligilactobacillus-salivarius] |
| WP_081538386.1 | Type1.BREX | [illegible] | 1183 | Bacteria>Terrabacteria_group>Bacillota>Bacilli>Lactobacillales>Lactobacillaceae | GCF_002079705.1 | Ligilactobacillus_salivarius | B6U43_RS08310 | 4936..8487 | 1624 | BREX-system-P-loop-protein-BrxC-[Ligilactobacillus-salivarius] |
| WP_085391474.1 | Type1.BREX | [illegible] | 1219 | Bacteria>Terrabacteria_group>Bacillota>Bacilli>Lactobacillales>Lactobacillaceae | GCF_002105575.1 | Latilactobacillus_sakei | LP065_RS02285 | 462739..466398 | 1599 | BREX-system-P-loop-protein-BrxC-[Latilactobacillus-sakei] |
| WP_085721967.1 | Type1.BREX | [illegible] | 1221 | Bacteria>Terrabacteria_group>Bacillota>Bacilli>Lactobacillales>Lactobacillaceae | GCF_002112805.1 | Limosilactobacillus_reuteri | B6J76_RS07075 | 2565..6230 | 1598 | BREX-system-P-loop-protein-BrxC-[Limosilactobacillus-reuteri] |
| WP_085769594.1 | Type1.BREX | [illegible] | 1241 | Bacteria>Terrabacteria_group>Bacillota>Bacilli>Lactobacillales>Lactobacillaceae | GCF_002117345.1 | Levilactobacillus_brevis | AZI11_RS13305 | 1..3726 | 1580 | BREX-system-P-loop-protein-BrxC-[Levilactobacillus-brevis] |
| WP_086131490.1 | Type1.BREX | [illegible] | 1221 | Bacteria>Terrabacteria_group>Bacillota>Bacilli>Lactobacillales>Lactobacillaceae | GCF_002128755.1 | Limosilactobacillus_reuteri | BHL75_RS03855 | 10373..14038 | 1598 | BREX-system-P-loop-protein-BrxC-[Limosilactobacillus-reuteri] |
| WP_086355469.1 | Type1.BREX | [illegible] | 1183 | Bacteria>Terrabacteria_group>Bacillota>Bacilli>Lactobacillales>Lactobacillaceae | GCF_002142575.1 | Lactobacillus_delbrueckii | B9N98_RS00590 | 112113..115664 | 1584 | BREX-system-P-loop-protein-BrxC-[Lactobacillus-delbrueckii] |
| WP_087236056.1 | Type1.BREX | [illegible] | 1244 | Bacteria>Terrabacteria_group>Bacillota>Bacilli>Lactobacillales>Lactobacillaceae | GCF_002160635.1 | Lactobacillus_gallinarum | B5E95_07165 | 9727..13461 | 52242 | BREX-system-P-loop-protein-BrxC-[Lactobacillus-gallinarum] |
| WP_087363710.1 | Type1.BREX | [illegible] | 1244 | Bacteria>Terrabacteria_group>Bacillota>Bacilli>Lactobacillales>Lactobacillaceae | GCF_002160975.1 | Lactobacillus_gallinarum | B5E63_RS08425 | 13791..17525 | 52242 | BREX-system-P-loop-protein-BrxC-[Lactobacillus-gallinarum] |
| WP_087716717.1 | Type1.BREX | [illegible] | 1221 | Bacteria>Terrabacteria_group>Bacillota>Bacilli>Lactobacillales>Lactobacillaceae | GCF_002179515.1 | Levilactobacillus_brevis | CA846_RS04700 | 148535..152200 | 1580 | BREX-system-P-loop-protein-BrxC-[Levilactobacillus-brevis] |
| WP_053267336.1 | Type1.BREX | [illegible] | 1216 | Bacteria>Terrabacteria_group>Bacillota>Bacilli>Lactobacillales>Lactobacillaceae | GCF_002205775.2 | Lactiplantibacillus_plantarum | CEF05_RS11175 | 2286410..2290060 | 1590 | BREX-system-P-loop-protein-BrxC-[Lactiplantibacillus-plantarum] |
| WP_089145461.1 | Type1.BREX | [illegible] | 1241 | Bacteria>Terrabacteria_group>Bacillota>Bacilli>Lactobacillales>Lactobacillaceae | GCF_002218565.1 | Lactobacillus_crispatus | AYP86_RS08235 | 29930..33655 | 47770 | BREX-system-P-loop-protein-BrxC-[Lactobacillus-crispatus] |
| WP_089150965.1 | Type1.BREX | [illegible] | 1241 | Bacteria>Terrabacteria_group>Bacillota>Bacilli>Lactobacillales>Lactobacillaceae | GCF_002218615.1 | Lactobacillus_crispatus | AYP77_RS06615 | 10349..14074 | 47770 | BREX-system-P-loop-protein-BrxC-[Lactobacillus-crispatus] |
| WP_089143244.1 | Type1.BREX | [illegible] | 1241 | Bacteria>Terrabacteria_group>Bacillota>Bacilli>Lactobacillales>Lactobacillaceae | GCF_002218685.1 | Lactobacillus_crispatus | AYP84_RS05225 | 10126..13851 | 47770 | BREX-system-P-loop-protein-BrxC-[Lactobacillus-crispatus] |
| WP_089147332.1 | Type1.BREX | [illegible] | 1241 | Bacteria>Terrabacteria_group>Bacillota>Bacilli>Lactobacillales>Lactobacillaceae | GCF_002218765.1 | Lactobacillus_crispatus | AYP81_RS05215 | 20138..23863 | 47770 | BREX-system-P-loop-protein-BrxC-[Lactobacillus-crispatus] |
| WP_089150406.1 | Type1.BREX | [illegible] | 1241 | Bacteria>Terrabacteria_group>Bacillota>Bacilli>Lactobacillales>Lactobacillaceae | GCF_002218775.1 | Lactobacillus_crispatus | AYP83_05520 | 19732..23457 | 47770 | BREX-system-P-loop-protein-BrxC-[Lactobacillus-crispatus] |
| WP_089151214.1 | Type1.BREX | [illegible] | 1241 | Bacteria>Terrabacteria_group>Bacillota>Bacilli>Lactobacillales>Lactobacillaceae | GCF_002218855.1 | Lactobacillus | AYP88_RS05225 | 6192..9917 | 1578 | MULTISPECIES:-BREX-system-P-loop-protein-BrxC-[Lactobacillus] |
| WP_089147455.1 | Type1.BREX | [illegible] | 1241 | Bacteria>Terrabacteria_group>Bacillota>Bacilli>Lactobacillales>Lactobacillaceae | GCF_002218895.1 | Lactobacillus_crispatus | AYP91_RS03130 | 36647..40372 | 47770 | BREX-system-P-loop-protein-BrxC-[Lactobacillus-crispatus] |
| WP_089146191.1 | Type1.BREX | [illegible] | 1241 | Bacteria>Terrabacteria_group>Bacillota>Bacilli>Lactobacillales>Lactobacillaceae | GCF_002218945.1 | Lactobacillus | AYP94_RS08355 | 41770..45495 | 1578 | MULTISPECIES:-BREX-system-P-loop-protein-BrxC-[Lactobacillus] |
| WP_089147834.1 | Type1.BREX | [illegible] | 1241 | Bacteria>Terrabacteria_group>Bacillota>Bacilli>Lactobacillales>Lactobacillaceae | GCF_002218975.1 | Lactobacillus | AYP96_RS04755 | 10061..13786 | 1578 | MULTISPECIES:-BREX-system-P-loop-protein-BrxC-[Lactobacillus] |
| WP_089197894.1 | Type1.BREX | [illegible] | 470 | Bacteria>Terrabacteria_group>Bacillota>Bacilli>Lactobacillales>Lactobacillaceae | GCF_002220815.1 | Lactiplantibacillus_plantarum | GBLP1_RS03710 | 792781..794193 | 1590 | BREX-system-P-loop-protein-BrxC-[Lactiplantibacillus-plantarum] |
| WP_094503029.1 | Type1.BREX | [illegible] | 1243 | Bacteria>Terrabacteria_group>Bacillota>Bacilli>Lactobacillales>Lactobacillaceae | GCF_002253385.1 | Lactobacillus_taiwanensis | CBF76_RS02150 | 6459..10190 | 508451 | BREX-system-P-loop-protein-BrxC-[Lactobacillus-taiwanensis] |
| WP_095342391.1 | Type1.BREX | [illegible] | 1246 | Bacteria>Terrabacteria_group>Bacillota>Bacilli>Lactobacillales>Lactobacillaceae | GCF_002276565.1 | Lactobacillus_kefiranofaciens | B8W86_RS09305 | 2234..5974 | 267818 | BREX-system-P-loop-protein-BrxC-[Lactobacillus-kefiranofaciens] |
| WP_049170967.1 | Type1.BREX | [illegible] | 1210 | Bacteria>Terrabacteria_group>Bacillota>Bacilli>Lactobacillales>Lactobacillaceae | GCF_002286235.1 | Lacticaseibacillus_rhamnosus | BGK71_RS10420 | 2042751..2046383 | 47715 | BREX-system-P-loop-protein-BrxC-[Lacticaseibacillus-rhamnosus] |
| WP_049166573.1 | Type1.BREX | [illegible] | 1210 | Bacteria>Terrabacteria_group>Bacillota>Bacilli>Lactobacillales>Lactobacillaceae | GCF_002287945.1 | Lacticaseibacillus_rhamnosus | N507_RS04265 | 896062..899694 | 47715 | BREX-system-P-loop-protein-BrxC-[Lacticaseibacillus-rhamnosus] |
| WP_095759131.1 | Type1.BREX | [illegible] | 1183 | Bacteria>Terrabacteria_group>Bacillota>Bacilli>Lactobacillales>Lactobacillaceae | GCF_002289875.1 | Ligilactobacillus_salivarius | A8C45_RS04465 | 18491..22042 | 1624 | BREX-system-P-loop-protein-BrxC-[Ligilactobacillus-salivarius] |
| WP_096584693.1 | Type1.BREX | [illegible] | 1215 | Bacteria>Terrabacteria_group>Bacillota>Bacilli>Lactobacillales>Lactobacillaceae | GCF_002370375.1 | Latilactobacillus_sakei | CCX78_RS03480 | 665145..668792 | 1599 | BREX-system-P-loop-protein-BrxC-[Latilactobacillus-sakei] |
| WP_097550471.1 | Type1.BREX | [illegible] | 1243 | Bacteria>Terrabacteria_group>Bacillota>Bacilli>Lactobacillales>Lactobacillaceae | GCF_002532085.1 | Lactobacillus_helveticus | CR094_RS03640 | 2555..6286 | 1587 | BREX-system-P-loop-protein-BrxC-[Lactobacillus-helveticus] |
| WP_098035805.1 | Type1.BREX | [illegible] | 1241 | Bacteria>Terrabacteria_group>Bacillota>Bacilli>Lactobacillales>Lactobacillaceae | GCF_002553955.1 | unclassified_Lactobacillus | CP362_RS02950 | 53500..57225 | 2620435 | MULTISPECIES:-BREX-system-P-loop-protein-BrxC-[unclassified-Lactobacillus] |
| WP_100665544.1 | Type1.BREX | [illegible] | 1194 | Bacteria>Terrabacteria_group>Bacillota>Bacilli>Lactobacillales>Lactobacillaceae | GCF_002804045.1 | Leuconostoc | CU067_RS06120 | 1080918..1084502 | 1243 | MULTISPECIES:-BREX-system-P-loop-protein-BrxC-[Leuconostoc] |
| WP_100908581.1 | Type1.BREX | [illegible] | 1212 | Bacteria>Terrabacteria_group>Bacillota>Bacilli>Lactobacillales>Lactobacillaceae | GCF_002813615.1 | Lacticaseibacillus_paracasei | BBD24_RS11115 | 2207874..2211512 | 1597 | BREX-system-P-loop-protein-BrxC-[Lacticaseibacillus-paracasei] |
| WP_101812672.1 | Type1.BREX | [illegible] | 1241 | Bacteria>Terrabacteria_group>Bacillota>Bacilli>Lactobacillales>Lactobacillaceae | GCF_002849955.1 | Lactobacillus_helveticus | Lh22155_RS05635 | 1074208..1077933 | 1587 | BREX-system-P-loop-protein-BrxC-[Lactobacillus-helveticus] |
| WP_102116084.1 | Type1.BREX | [illegible] | 1230 | Bacteria>Terrabacteria_group>Bacillota>Bacilli>Lactobacillales>Lactobacillaceae | GCF_002869825.2 | Limosilactobacillus_fermentum | C0965_RS02310 | 470388..474080 | 1613 | BREX-system-P-loop-protein-BrxC-[Limosilactobacillus-fermentum] |
| WP_097558087.1 | Type1.BREX | [illegible] | 1216 | Bacteria>Terrabacteria_group>Bacillota>Bacilli>Lactobacillales>Lactobacillaceae | GCF_002943545.1 | Lactiplantibacillus | GTP03_RS14070 | 6103..9753 | 2767842 | MULTISPECIES:-BREX-system-P-loop-protein-BrxC-[Lactiplantibacillus] |
| WP_158666994.1 | Type1.BREX | [illegible] | 1195 | Bacteria>Terrabacteria_group>Bacillota>Bacilli>Lactobacillales>Lactobacillaceae | GCF_002953655.1 | Latilactobacillus_sakei | C0213_RS10250 | 2077540..2081127 | 1599 | BREX-system-P-loop-protein-BrxC-[Latilactobacillus-sakei] |
| WP_158682902.1 | Type1.BREX | [illegible] | 1227 | Bacteria>Terrabacteria_group>Bacillota>Bacilli>Lactobacillales>Lactobacillaceae | GCF_002970915.1 | Lactobacillus_sp._CBA3605 | C5Z25_RS04875 | 984247..987930 | 2099788 | BREX-system-P-loop-protein-BrxC-[Lactobacillus-sp.-CBA3605] |
| WP_158682814.1 | Type1.BREX | [illegible] | 1227 | Bacteria>Terrabacteria_group>Bacillota>Bacilli>Lactobacillales>Lactobacillaceae | GCF_002970935.1 | Lactobacillus_sp._CBA3606 | C5Z26_RS06230 | 1252250..1255933 | 2099789 | BREX-system-P-loop-protein-BrxC-[Lactobacillus-sp.-CBA3606] |
| WP_105920579.1 | Type1.BREX | [illegible] | 1223 | Bacteria>Terrabacteria_group>Bacillota>Bacilli>Lactobacillales>Lactobacillaceae | GCF_002993485.1 | Lactiplantibacillus_pentosus | C6Y12_RS16130 | 244136..247807 | 1589 | BREX-system-P-loop-protein-BrxC-[Lactiplantibacillus-pentosus] |
| WP_106904923.1 | Type1.BREX | [illegible] | 1241 | Bacteria>Terrabacteria_group>Bacillota>Bacilli>Lactobacillales>Lactobacillaceae | GCF_003020005.1 | Lactiplantibacillus_plantarum | B5G53_RS16135 | 18984..22709 | 1590 | BREX-system-P-loop-protein-BrxC-[Lactiplantibacillus-plantarum] |
| WP_225367119.1 | Type1.BREX | [illegible] | 1182 | Bacteria>Terrabacteria_group>Bacillota>Bacilli>Lactobacillales>Lactobacillaceae | GCF_003046115.1 | Lacticaseibacillus_rhamnosus | DA801_RS04680 | 68737..72285 | 47715 | BREX-system-P-loop-protein-BrxC-[Lacticaseibacillus-rhamnosus] |
| WP_114788341.1 | Type1.BREX | [illegible] | 1241 | Bacteria>Terrabacteria_group>Bacillota>Bacilli>Lactobacillales>Lactobacillaceae | GCF_003347455.1 | Lactiplantibacillus_plantarum | DB726_RS11995 | 25864..29589 | 1590 | BREX-system-P-loop-protein-BrxC-[Lactiplantibacillus-plantarum] |
| WP_117285422.1 | Type1.BREX | [illegible] | 1243 | Bacteria>Terrabacteria_group>Bacillota>Bacilli>Lactobacillales>Lactobacillaceae | GCF_003428395.1 | Lactobacillus_johnsonii | D0Y49_RS01485 | 293356..297087 | 33959 | BREX-system-P-loop-protein-BrxC-[Lactobacillus-johnsonii] |
| WP_118272847.1 | Type1.BREX | [illegible] | 1183 | Bacteria>Terrabacteria_group>Bacillota>Bacilli>Lactobacillales>Lactobacillaceae | GCF_003472615.1 | Ligilactobacillus_salivarius | DW111_RS08715 | 20935..24486 | 1624 | BREX-system-P-loop-protein-BrxC-[Ligilactobacillus-salivarius] |
| WP_118908009.1 | Type1.BREX | [illegible] | 1206 | Bacteria>Terrabacteria_group>Bacillota>Bacilli>Lactobacillales>Lactobacillaceae | GCF_003522965.1 | Bombilactobacillus_bombi | DS830_RS01685 | 308484..312104 | 1303590 | BREX-system-P-loop-protein-BrxC-[Bombilactobacillus-bombi] |
| WP_113532397.1 | Type1.BREX | [illegible] | 1239 | Bacteria>Terrabacteria_group>Bacillota>Bacilli>Lactobacillales>Lactobacillaceae | GCF_003584685.1 | Lactobacillus_paragasseri | LpgJCM5343_RS05125 | 1058950..1062669 | 2107999 | BREX-system-P-loop-protein-BrxC-[Lactobacillus-paragasseri] |
| WP_129531714.1 | Type1.BREX | [illegible] | 1216 | Bacteria>Terrabacteria_group>Bacillota>Bacilli>Lactobacillales>Lactobacillaceae | GCF_003597635.1 | Lactiplantibacillus_plantarum | D5289_RS00270 | 52296..55946 | 1590 | BREX-system-P-loop-protein-BrxC-[Lactiplantibacillus-plantarum] |
| WP_123020673.1 | Type1.BREX | [illegible] | 1212 | Bacteria>Terrabacteria_group>Bacillota>Bacilli>Lactobacillales>Lactobacillaceae | GCF_003712245.1 | Lacticaseibacillus_paracasei | FAM18108_RS11170 | 41969..45607 | 1597 | BREX-system-P-loop-protein-BrxC-[Lacticaseibacillus-paracasei] |
| WP_123156030.1 | Type1.BREX | [illegible] | 1212 | Bacteria>Terrabacteria_group>Bacillota>Bacilli>Lactobacillales>Lactobacillaceae | GCF_003712395.1 | Lacticaseibacillus_paracasei | FAM18113_RS08130 | 717..4355 | 1597 | BREX-system-P-loop-protein-BrxC-[Lacticaseibacillus-paracasei] |
| WP_123156952.1 | Type1.BREX | [illegible] | 1215 | Bacteria>Terrabacteria_group>Bacillota>Bacilli>Lactobacillales>Lactobacillaceae | GCF_003712625.1 | Lacticaseibacillus_paracasei | FAM19404_RS10240 | 4290..7937 | 1597 | BREX-system-P-loop-protein-BrxC-[Lacticaseibacillus-paracasei] |
| WP_125765408.1 | Type1.BREX | [illegible] | 1216 | Bacteria>Terrabacteria_group>Bacillota>Bacilli>Lactobacillales>Lactobacillaceae | GCF_003946045.1 | Levilactobacillus_mulengensis | EKM84_RS08865 | 97572..101222 | 2486025 | BREX-system-P-loop-protein-BrxC-[Levilactobacillus-mulengensis] |
| WP_125573155.1 | Type1.BREX | [illegible] | 1207 | Bacteria>Terrabacteria_group>Bacillota>Bacilli>Lactobacillales>Lactobacillaceae | GCF_003946145.1 | Levilactobacillus_huananensis | EKM71_RS02790 | 155319..158942 | 2486019 | BREX-system-P-loop-protein-BrxC-[Levilactobacillus-huananensis] |
| WP_125583865.1 | Type1.BREX | [illegible] | 1220 | Bacteria>Terrabacteria_group>Bacillota>Bacilli>Lactobacillales>Lactobacillaceae | GCF_003946245.1 | Levilactobacillus_cerevisiae | EKM99_RS13110 | 767..4429 | 1704076 | BREX-system-P-loop-protein-BrxC-[Levilactobacillus-cerevisiae] |
| WP_125682913.1 | Type1.BREX | [illegible] | 1223 | Bacteria>Terrabacteria_group>Bacillota>Bacilli>Lactobacillales>Lactobacillaceae | GCF_003946265.1 | Levilactobacillus_yonginensis | EKM87_RS06125 | 5427..9098 | 1054041 | BREX-system-P-loop-protein-BrxC-[Levilactobacillus-yonginensis] |
| WP_125770444.1 | Type1.BREX | [illegible] | 1225 | Bacteria>Terrabacteria_group>Bacillota>Bacilli>Lactobacillales>Lactobacillaceae | GCF_003946325.1 | Companilactobacillus_furfuricola | EKM78_RS06175 | 4477..8154 | 1462575 | BREX-system-P-loop-protein-BrxC-[Companilactobacillus-furfuricola] |
| WP_125550128.1 | Type1.BREX | [illegible] | 1208 | Bacteria>Terrabacteria_group>Bacillota>Bacilli>Lactobacillales>Lactobacillaceae | GCF_003946385.1 | Loigolactobacillus_zhaoyuanensis | EKM76_RS07800 | 7684..11310 | 2486017 | BREX-system-P-loop-protein-BrxC-[Loigolactobacillus-zhaoyuanensis] |
| WP_125551973.1 | Type1.BREX | [illegible] | 1206 | Bacteria>Terrabacteria_group>Bacillota>Bacilli>Lactobacillales>Lactobacillaceae | GCF_003946445.1 | Loigolactobacillus_jiayinensis | EKM80_RS04790 | 68578..72198 | 2486016 | BREX-system-P-loop-protein-BrxC-[Loigolactobacillus-jiayinensis] |
| WP_125758091.1 | Type1.BREX | [illegible] | 1212 | Bacteria>Terrabacteria_group>Bacillota>Bacilli>Lactobacillales>Lactobacillaceae | GCF_003946465.1 | Lacticaseibacillus_hegangensis | EKM86_RS11340 | 12431..16069 | 2486010 | BREX-system-P-loop-protein-BrxC-[Lacticaseibacillus-hegangensis] |
| WP_125584965.1 | Type1.BREX | [illegible] | 1214 | Bacteria>Terrabacteria_group>Bacillota>Bacilli>Lactobacillales>Lactobacillaceae | GCF_003946485.1 | Lacticaseibacillus_jixianensis | EKM85_RS03935 | 121172..124816 | 2486012 | BREX-system-P-loop-protein-BrxC-[Lacticaseibacillus-jixianensis] |
| WP_125762723.1 | Type1.BREX | [illegible] | 1225 | Bacteria>Terrabacteria_group>Bacillota>Bacilli>Lactobacillales>Lactobacillaceae | GCF_003946555.1 | Companilactobacillus_hulinensis | EKM66_RS06280 | 16382..20059 | 2486007 | BREX-system-P-loop-protein-BrxC-[Companilactobacillus-hulinensis] |
| WP_125647605.1 | Type1.BREX | [illegible] | 1230 | Bacteria>Terrabacteria_group>Bacillota>Bacilli>Lactobacillales>Lactobacillaceae | GCF_003946675.1 | Lapidilactobacillus_gannanensis | EKM81_RS01890 | 14683..18375 | 2486002 | BREX-system-P-loop-protein-BrxC-[Lapidilactobacillus-gannanensis] |
| WP_125606901.1 | Type1.BREX | [illegible] | 1207 | Bacteria>Terrabacteria_group>Bacillota>Bacilli>Lactobacillales>Lactobacillaceae | GCF_003946695.1 | Lapidilactobacillus_bayanensis | EKM65_RS06650 | 19317..22940 | 2485998 | BREX-system-P-loop-protein-BrxC-[Lapidilactobacillus-bayanensis] |
| WP_125978511.1 | Type1.BREX | [illegible] | 1244 | Bacteria>Terrabacteria_group>Bacillota>Bacilli>Lactobacillales>Lactobacillaceae | GCF_003952845.1 | Lactobacillus_acidophilus | CXB72_RS05650 | 1103564..1107298 | 1579 | BREX-system-P-loop-protein-BrxC-[Lactobacillus-acidophilus] |
| WP_014918808.1 | Type1.BREX | [illegible] | 1244 | Bacteria>Terrabacteria_group>Bacillota>Bacilli>Lactobacillales>Lactobacillaceae | GCF_003955865.1 | Lactobacillus_helveticus | LH5_RS02000 | 414775..418509 | 1587 | BREX-system-P-loop-protein-BrxC-[Lactobacillus-helveticus] |
| WP_126133408.1 | Type1.BREX | [illegible] | 1219 | Bacteria>Terrabacteria_group>Bacillota>Bacilli>Lactobacillales>Lactobacillaceae | GCF_003957415.1 | Latilactobacillus_curvatus | CYK59_RS06580 | 1253905..1257564 | 28038 | BREX-system-P-loop-protein-BrxC-[Latilactobacillus-curvatus] |
| WP_126353447.1 | Type1.BREX | [illegible] | 1212 | Bacteria>Terrabacteria_group>Bacillota>Bacilli>Lactobacillales>Lactobacillaceae | GCF_003966835.1 | Lacticaseibacillus_paracasei | EL355_RS11595 | 2248100..2251738 | 1597 | BREX-system-P-loop-protein-BrxC-[Lacticaseibacillus-paracasei] |
| WP_127526369.1 | Type1.BREX | [illegible] | 1216 | Bacteria>Terrabacteria_group>Bacillota>Bacilli>Lactobacillales>Lactobacillaceae | GCF_004000705.1 | Lactiplantibacillus_plantarum | KB1253_31180 | 16316..19966 | 1590 | BREX-system-P-loop-protein-BrxC-[Lactiplantibacillus-plantarum] |
| WP_127848553.1 | Type1.BREX | [illegible] | 1214 | Bacteria>Terrabacteria_group>Bacillota>Bacilli>Lactobacillales>Lactobacillaceae | GCF_004010095.1 | Lacticaseibacillus_hulanensis | EKN00_RS02415 | 61527..65171 | 2493111 | BREX-system-P-loop-protein-BrxC-[Lacticaseibacillus-hulanensis] |
| WP_128537187.1 | Type1.BREX | [illegible] | 1241 | Bacteria>Terrabacteria_group>Bacillota>Bacilli>Lactobacillales>Lactobacillaceae | GCF_004055415.1 | Lactiplantibacillus_plantarum | EQG58_RS16175 | 99358..103083 | 1590 | BREX-system-P-loop-protein-BrxC-[Lactiplantibacillus-plantarum] |
| WP_128529840.1 | Type1.BREX | [illegible] | 1241 | Bacteria>Terrabacteria_group>Bacillota>Bacilli>Lactobacillales>Lactobacillaceae | GCF_004078645.1 | Lactiplantibacillus_plantarum | EQJ06_RS15675 | 35919..39644 | 1590 | BREX-system-P-loop-protein-BrxC-[Lactiplantibacillus-plantarum] |
| WP_128485902.1 | Type1.BREX | [illegible] | 1241 | Bacteria>Terrabacteria_group>Bacillota>Bacilli>Lactobacillales>Lactobacillaceae | GCF_004101645.1 | Lactiplantibacillus_plantarum | EQK45_RS16695 | 37408..41133 | 1590 | BREX-system-P-loop-protein-BrxC-[Lactiplantibacillus-plantarum] |
| WP_128739645.1 | Type1.BREX | [illegible] | 1194 | Bacteria>Terrabacteria_group>Bacillota>Bacilli>Lactobacillales>Lactobacillaceae | GCF_004103675.1 | Leuconostoc_mesenteroides | EQZ98_RS05410 | 1059689..1063273 | 1245 | BREX-system-P-loop-protein-BrxC-[Leuconostoc-mesenteroides] |
| WP_129335566.1 | Type1.BREX | [illegible] | 1188 | Bacteria>Terrabacteria_group>Bacillota>Bacilli>Lactobacillales>Lactobacillaceae | GCF_004123065.1 | Lactobacillus_delbrueckii | EST31_RS04460 | 19918..23484 | 1584 | BREX-system-P-loop-protein-BrxC-[Lactobacillus-delbrueckii] |
| WP_134086962.1 | Type1.BREX | [illegible] | 1194 | Bacteria>Terrabacteria_group>Bacillota>Bacilli>Lactobacillales>Lactobacillaceae | GCF_004368765.1 | Leuconostoc_mesenteroides | C7818_RS08810 | 4933..8517 | 1245 | BREX-system-P-loop-protein-BrxC-[Leuconostoc-mesenteroides] |
| WP_134833540.1 | Type1.BREX | [illegible] | 1194 | Bacteria>Terrabacteria_group>Bacillota>Bacilli>Lactobacillales>Lactobacillaceae | GCF_004551615.1 | Leuconostoc_kimchii | EW139_RS05650 | 1051849..1055433 | 136609 | BREX-system-P-loop-protein-BrxC-[Leuconostoc-kimchii] |
| WP_135517412.1 | Type1.BREX | [illegible] | 839 | Bacteria>Terrabacteria_group>Bacillota>Bacilli>Lactobacillales>Lactobacillaceae | GCF_004730965.1 | Lactiplantibacillus_plantarum | DVH03_RS13095 | 2657264..2659783 | 1590 | BREX-system-P-loop-protein-BrxC-[Lactiplantibacillus-plantarum] |
| WP_137641366.1 | Type1.BREX | [illegible] | 1213 | Bacteria>Terrabacteria_group>Bacillota>Bacilli>Lactobacillales>Lactobacillaceae | GCF_005405105.1 | Lactiplantibacillus_dongliensis | FFM80_RS13995 | 1986..5627 | 2559919 | BREX-system-P-loop-protein-BrxC-[Lactiplantibacillus-dongliensis] |
| WP_137638544.1 | Type1.BREX | [illegible] | 1204 | Bacteria>Terrabacteria_group>Bacillota>Bacilli>Lactobacillales>Lactobacillaceae | GCF_005405165.1 | Loigolactobacillus_binensis | FFM78_RS11390 | 22588..26202 | 2559922 | BREX-system-P-loop-protein-BrxC-[Loigolactobacillus-binensis] |
| WP_064613173.1 | Type1.BREX | [illegible] | 1219 | Bacteria>Terrabacteria_group>Bacillota>Bacilli>Lactobacillales>Lactobacillaceae | GCF_006151905.1 | Lacticaseibacillus_rhamnosus | FG342_RS13010 | 2567642..2571301 | 47715 | BREX-system-P-loop-protein-BrxC-[Lacticaseibacillus-rhamnosus] |
| WP_145463941.1 | Type1.BREX | [illegible] | 1194 | Bacteria>Terrabacteria_group>Bacillota>Bacilli>Lactobacillales>Lactobacillaceae | GCF_007785685.1 | Weissella_cibaria | F0435_RS06180 | 1309248..1312832 | 137591 | BREX-system-P-loop-protein-BrxC-[Weissella-cibaria] |
| WP_146988326.1 | Type1.BREX | [illegible] | 1204 | Bacteria>Terrabacteria_group>Bacillota>Bacilli>Lactobacillales>Lactobacillaceae | GCF_007954685.1 | Loigolactobacillus_coryniformis | FGL77_RS03560 | 710842..714456 | 1610 | BREX-system-P-loop-protein-BrxC-[Loigolactobacillus-coryniformis] |
| WP_146995455.1 | Type1.BREX | [illegible] | 1241 | Bacteria>Terrabacteria_group>Bacillota>Bacilli>Lactobacillales>Lactobacillaceae | GCF_007989965.1 | Schleiferilactobacillus_harbinensis | LHA01_RS13380 | 4622..8347 | 304207 | BREX-system-P-loop-protein-BrxC-[Schleiferilactobacillus-harbinensis] |
| WP_147768479.1 | Type1.BREX | [illegible] | 1183 | Bacteria>Terrabacteria_group>Bacillota>Bacilli>Lactobacillales>Lactobacillaceae | GCF_008016915.1 | Ligilactobacillus_salivarius | FG086_RS06640 | 61876..65427 | 1624 | BREX-system-P-loop-protein-BrxC-[Ligilactobacillus-salivarius] |
| WP_149350175.1 | Type1.BREX | [illegible] | 1212 | Bacteria>Terrabacteria_group>Bacillota>Bacilli>Lactobacillales>Lactobacillaceae | GCF_008329845.1 | Lacticaseibacillus_paracasei | D0638_RS11245 | 2196110..2199748 | 1597 | BREX-system-P-loop-protein-BrxC-[Lacticaseibacillus-paracasei] |
| WP_150288904.1 | Type1.BREX | [illegible] | 1194 | Bacteria>Terrabacteria_group>Bacillota>Bacilli>Lactobacillales>Lactobacillaceae | GCF_008689695.1 | Leuconostoc_carnosum | FE409_RS04225 | 31528..35112 | 1252 | BREX-system-P-loop-protein-BrxC-[Leuconostoc-carnosum] |
| WP_150289839.1 | Type1.BREX | [illegible] | 1194 | Bacteria>Terrabacteria_group>Bacillota>Bacilli>Lactobacillales>Lactobacillaceae | GCF_008690985.1 | Leuconostoc_carnosum | FD956_RS04085 | 690368..693952 | 1252 | BREX-system-P-loop-protein-BrxC-[Leuconostoc-carnosum] |
| WP_150392678.1 | Type1.BREX | [illegible] | 1241 | Bacteria>Terrabacteria_group>Bacillota>Bacilli>Lactobacillales>Lactobacillaceae | GCF_008694105.1 | Schleiferilactobacillus_harbinensis | FMM01_RS11950 | 2413874..2417599 | 304207 | BREX-system-P-loop-protein-BrxC-[Schleiferilactobacillus-harbinensis] |
| WP_152261659.1 | Type1.BREX | [illegible] | 1241 | Bacteria>Terrabacteria_group>Bacillota>Bacilli>Lactobacillales>Lactobacillaceae | GCF_009217765.1 | Schleiferilactobacillus_harbinensis | D1010_RS17280 | 3443993..3447718 | 304207 | BREX-system-P-loop-protein-BrxC-[Schleiferilactobacillus-harbinensis] |
| WP_153110820.1 | Type1.BREX | [illegible] | 1241 | Bacteria>Terrabacteria_group>Bacillota>Bacilli>Lactobacillales>Lactobacillaceae | GCF_009495575.1 | Lentilactobacillus_buchneri | FIU75_RS13270 | 20367..24092 | 1581 | BREX-system-P-loop-protein-BrxC-[Lentilactobacillus-buchneri] |
| WP_196246253.1 | Type1.BREX | [illegible] | 1193 | Bacteria>Terrabacteria_group>Bacillota>Bacilli>Lactobacillales>Lactobacillaceae | GCF_009687905.1 | Lacticaseibacillus_zhaodongensis | GN107_RS04555 | 97021..100602 | 2668065 | BREX-system-P-loop-protein-BrxC-[Lacticaseibacillus-zhaodongensis] |
| WP_160528668.1 | Type1.BREX | [illegible] | 1217 | Bacteria>Terrabacteria_group>Bacillota>Bacilli>Lactobacillales>Lactobacillaceae | GCF_009823605.1 | Lacticaseibacillus_paracasei | GRZ59_RS05150 | 1513..5166 | 1597 | BREX-system-P-loop-protein-BrxC-[Lacticaseibacillus-paracasei] |
| WP_206672604.1 | Type1.BREX | [illegible] | 998 | Bacteria>Terrabacteria_group>Bacillota>Bacilli>Lactobacillales>Lactobacillaceae | GCF_009863825.1 | Ligilactobacillus_salivarius | FYM86_RS06705 | 34358..37354 | 1624 | BREX-system-P-loop-protein-BrxC-[Ligilactobacillus-salivarius] |
| WP_160995430.1 | Type1.BREX | [illegible] | 1183 | Bacteria>Terrabacteria_group>Bacillota>Bacilli>Lactobacillales>Lactobacillaceae | GCF_009865755.1 | Ligilactobacillus_salivarius | FYL08_RS07580 | 23122..26673 | 1624 | BREX-system-P-loop-protein-BrxC-[Ligilactobacillus-salivarius] |
| WP_161023346.1 | Type1.BREX | [illegible] | 1183 | Bacteria>Terrabacteria_group>Bacillota>Bacilli>Lactobacillales>Lactobacillaceae | GCF_009866105.1 | Ligilactobacillus_salivarius | FYL18_RS08455 | 14755..18306 | 1624 | BREX-system-P-loop-protein-BrxC-[Ligilactobacillus-salivarius] |
| WP_065989573.1 | Type1.BREX | [illegible] | 1241 | Bacteria>Terrabacteria_group>Bacillota>Bacilli>Lactobacillales>Lactobacillaceae | GCF_009933525.1 | Lactobacillus_crispatus | GSR61_RS03550 | 679020..682745 | 47770 | BREX-system-P-loop-protein-BrxC-[Lactobacillus-crispatus] |
| WP_162100826.1 | Type1.BREX | [illegible] | 1219 | Bacteria>Terrabacteria_group>Bacillota>Bacilli>Lactobacillales>Lactobacillaceae | GCF_010092945.1 | Latilactobacillus_sakei | GSF71_RS01930 | 394668..398327 | 1599 | BREX-system-P-loop-protein-BrxC-[Latilactobacillus-sakei] |
| WP_163588066.1 | Type1.BREX | [illegible] | 1219 | Bacteria>Terrabacteria_group>Bacillota>Bacilli>Lactobacillales>Lactobacillaceae | GCF_010586905.1 | Ligilactobacillus_murinus | FEE40_RS11090 | 2232079..2235738 | 1622 | BREX-system-P-loop-protein-BrxC-[Ligilactobacillus-murinus] |
| WP_095662164.1 | Type1.BREX | [illegible] | 1243 | Bacteria>Terrabacteria_group>Bacillota>Bacilli>Lactobacillales>Lactobacillaceae | GCF_011392295.1 | Lactobacillus_helveticus | ERJ65_RS00195 | 7386..11117 | 1587 | BREX-system-P-loop-protein-BrxC-[Lactobacillus-helveticus] |
| WP_246220218.1 | Type1.BREX | [illegible] | 660 | Bacteria>Terrabacteria_group>Bacillota>Bacilli>Lactobacillales>Lactobacillaceae | GCF_012396505.1 | Periweissella_fabalis | HF964_RS09930 | 200728..202710 | 1070421 | BREX-system-P-loop-protein-BrxC-[Periweissella-fabalis] |
| WP_168899781.1 | Type1.BREX | [illegible] | 1222 | Bacteria>Terrabacteria_group>Bacillota>Bacilli>Lactobacillales>Lactobacillaceae | GCF_012641095.1 | Lactobacillus_sp._HBUAS51381 | HEQ50_RS06865 | 2003..5671 | 2722743 | BREX-system-P-loop-protein-BrxC-[Lactobacillus-sp.-HBUAS51381] |
| WP_168786060.1 | Type1.BREX | [illegible] | 1216 | Bacteria>Terrabacteria_group>Bacillota>Bacilli>Lactobacillales>Lactobacillaceae | GCF_012689225.1 | Lactiplantibacillus_plantarum | GJ674_RS13810 | 14796..18446 | 1590 | BREX-system-P-loop-protein-BrxC-[Lactiplantibacillus-plantarum] |
| WP_170080380.1 | Type1.BREX | [illegible] | 1241 | Bacteria>Terrabacteria_group>Bacillota>Bacilli>Lactobacillales>Lactobacillaceae | GCF_012843585.1 | Lactobacillus_crispatus | HF868_RS03010 | 100979..104704 | 47770 | BREX-system-P-loop-protein-BrxC-[Lactobacillus-crispatus] |
| WP_170102413.1 | Type1.BREX | [illegible] | 1183 | Bacteria>Terrabacteria_group>Bacillota>Bacilli>Lactobacillales>Lactobacillaceae | GCF_012843615.1 | Ligilactobacillus_salivarius | HF867_RS02660 | 30638..34189 | 1624 | BREX-system-P-loop-protein-BrxC-[Ligilactobacillus-salivarius] |
| WP_169461453.1 | Type1.BREX | [illegible] | 1243 | Bacteria>Terrabacteria_group>Bacillota>Bacilli>Lactobacillales>Lactobacillaceae | GCF_012843675.1 | Lactobacillus_sp._MRS-253-APC-2B | HF864_RS02000 | 82846..86577 | 2725305 | BREX-system-P-loop-protein-BrxC-[Lactobacillus-sp.-MRS-253-APC-2B] |
| WP_172993980.1 | Type1.BREX | [illegible] | 1243 | Bacteria>Terrabacteria_group>Bacillota>Bacilli>Lactobacillales>Lactobacillaceae | GCF_013280765.1 | Lactobacillus_helveticus | IMAU60206_RS01575 | 25352..29083 | 1587 | BREX-system-P-loop-protein-BrxC-[Lactobacillus-helveticus] |
| WP_172993225.1 | Type1.BREX | [illegible] | 597 | Bacteria>Terrabacteria_group>Bacillota>Bacilli>Lactobacillales>Lactobacillaceae | GCF_013280775.1 | Lactobacillus_helveticus | IMAU60211_RS07170 | 7274..9066 | 1587 | BREX-system-P-loop-protein-BrxC,-partial-[Lactobacillus-helveticus] |
| WP_172993587.1 | Type1.BREX | [illegible] | 1243 | Bacteria>Terrabacteria_group>Bacillota>Bacilli>Lactobacillales>Lactobacillaceae | GCF_013280875.1 | Lactobacillus_helveticus | IMAU60201_RS07155 | 4819..8550 | 1587 | BREX-system-P-loop-protein-BrxC-[Lactobacillus-helveticus] |
| WP_172987666.1 | Type1.BREX | [illegible] | 1243 | Bacteria>Terrabacteria_group>Bacillota>Bacilli>Lactobacillales>Lactobacillaceae | GCF_013280955.1 | Lactobacillus_helveticus | IMAU50022_RS01295 | 3445..7176 | 1587 | BREX-system-P-loop-protein-BrxC-[Lactobacillus-helveticus] |
| WP_172982106.1 | Type1.BREX | [illegible] | 1243 | Bacteria>Terrabacteria_group>Bacillota>Bacilli>Lactobacillales>Lactobacillaceae | GCF_013281265.1 | Lactobacillus_helveticus | IMAU30078_RS05865 | 2723..6454 | 1587 | BREX-system-P-loop-protein-BrxC-[Lactobacillus-helveticus] |
| WP_173006741.1 | Type1.BREX | [illegible] | 1241 | Bacteria>Terrabacteria_group>Bacillota>Bacilli>Lactobacillales>Lactobacillaceae | GCF_013281475.1 | Lactobacillus_helveticus | IMAU30014_RS01855 | 25052..28777 | 1587 | BREX-system-P-loop-protein-BrxC-[Lactobacillus-helveticus] |
| WP_173001654.1 | Type1.BREX | [illegible] | 1241 | Bacteria>Terrabacteria_group>Bacillota>Bacilli>Lactobacillales>Lactobacillaceae | GCF_013281675.1 | Lactobacillus_helveticus | IMAU10149_RS03700 | 20557..24282 | 1587 | BREX-system-P-loop-protein-BrxC-[Lactobacillus-helveticus] |
| WP_173819159.1 | Type1.BREX | [illegible] | 1216 | Bacteria>Terrabacteria_group>Bacillota>Bacilli>Lactobacillales>Lactobacillaceae | GCF_013305265.1 | Lactiplantibacillus_plantarum | HRW02_RS12740 | 2738517..2742167 | 1590 | BREX-system-P-loop-protein-BrxC-[Lactiplantibacillus-plantarum] |
| WP_173959264.1 | Type1.BREX | [illegible] | 1212 | Bacteria>Terrabacteria_group>Bacillota>Bacilli>Lactobacillales>Lactobacillaceae | GCF_013307125.1 | Lacticaseibacillus_paracasei | FBD73_RS12340 | 2603842..2607480 | 1597 | BREX-system-P-loop-protein-BrxC-[Lacticaseibacillus-paracasei] |
| WP_179394507.1 | Type1.BREX | [illegible] | 1200 | Bacteria>Terrabacteria_group>Bacillota>Bacilli>Lactobacillales>Lactobacillaceae | GCF_013407595.1 | Lacticaseibacillus_absianus | HII33_RS04075 | 947851..951453 | 2729623 | BREX-system-P-loop-protein-BrxC-[Lacticaseibacillus-absianus] |
| WP_180808196.1 | Type1.BREX | [illegible] | 1244 | Bacteria>Terrabacteria_group>Bacillota>Bacilli>Lactobacillales>Lactobacillaceae | GCF_013456995.1 | Lactobacillus_crispatus | HOG71_RS07705 | 1537548..1541282 | 47770 | BREX-system-P-loop-protein-BrxC-[Lactobacillus-crispatus] |
| WP_180872733.1 | Type1.BREX | [illegible] | 1239 | Bacteria>Terrabacteria_group>Bacillota>Bacilli>Lactobacillales>Lactobacillaceae | GCF_013487865.1 | Lactobacillus_johnsonii | GT082_04895 | 1041555..1045274 | 33959 | BREX-system-P-loop-protein-BrxC-[Lactobacillus-johnsonii] |
| WP_181189091.1 | Type1.BREX | [illegible] | 1207 | Bacteria>Terrabacteria_group>Bacillota>Bacilli>Lactobacillales>Lactobacillaceae | GCF_013607485.1 | Bombilactobacillus_bombi | EQ827_RS00810 | 5020..8643 | 1303590 | BREX-system-P-loop-protein-BrxC-[Bombilactobacillus-bombi] |
| WP_182611612.1 | Type1.BREX | [illegible] | 1241 | Bacteria>Terrabacteria_group>Bacillota>Bacilli>Lactobacillales>Lactobacillaceae | GCF_014155845.1 | Lacticaseibacillus_rhamnosus | FOS38_RS12550 | 14856..18581 | 47715 | BREX-system-P-loop-protein-BrxC-[Lacticaseibacillus-rhamnosus] |
| WP_187397365.1 | Type1.BREX | [illegible] | 1183 | Bacteria>Terrabacteria_group>Bacillota>Bacilli>Lactobacillales>Lactobacillaceae | GCF_014336755.1 | Ligilactobacillus_salivarius | DTL01_RS01875 | 9580..13131 | 1624 | BREX-system-P-loop-protein-BrxC-[Ligilactobacillus-salivarius] |

| Protein-ID | System | Gene-Neighbourhoods | Length | Taxonomy | Assembly-ID | Strain | Locus-Tag | Genomic-coordinates | TaxID | Fasta-defline |
|---|---|---|---|---|---|---|---|---|---|---|
| WP_188891961.1 | Type1.BREX | ParB+HNH->REase->MULE-TRANSPOSASE->ParB+HNH->BrxA-DNA-binding->BrxB-BrxF-iSTAND->DUF6079*->N6-METHYLASE->nSTAND3->PglZ->?->HTH+REase->TPR+TPR-> | 1241 | Bacteria>Terrabacteria_group>Bacillota>Bacilli>Lactobacillales>Lactobacillaceae | GCF_014656585.1 | Lactobacillus_kefiranofaciens | ICI50_RS03490 | 670073..673798 | 267818 | BREX-system-P-loop-protein-BrxC-[Lactobacillus-kefiranofaciens] |
| WP_194177171.1 | Type1.BREX | Resolvase+HTH_28-><-?<-PglZ<-DOC<-MazE_antitoxin<-N6-METHYLASE<-DUF6079*<-BrxB-BrxF-iSTAND<-BrxA-DNA-binding | 1221 | Bacteria>Terrabacteria_group>Bacillota>Bacilli>Lactobacillales>Lactobacillaceae | GCF_015234165.1 | Limosilactobacillus_oris | IRZ64_RS09035 | 9438..13103 | 1632 | BREX-system-P-loop-protein-BrxC-[Limosilactobacillus-oris] |
| WP_194321065.1 | Type1.BREX | [illegible] | 1194 | Bacteria>Terrabacteria_group>Bacillota>Bacilli>Lactobacillales>Lactobacillaceae | GCF_015277835.1 | Leuconostoc_citreum | IRM63_RS04335 | 861917..865501 | 33964 | BREX-system-P-loop-protein-BrxC-[Leuconostoc-citreum] |
| WP_195317824.1 | Type1.BREX | [illegible] | 1049 | Bacteria>Terrabacteria_group>Bacillota>Bacilli>Lactobacillales>Lactobacillaceae | GCF_015551705.1 | Weissella_cibaria | I2C84_RS00170 | 1..3151 | 137591 | BREX-system-P-loop-protein-BrxC,-partial-[Weissella-cibaria] |
| WP_064657275.1 | Type1.BREX | [illegible] | 1210 | Bacteria>Terrabacteria_group>Bacillota>Bacilli>Lactobacillales>Lactobacillaceae | GCF_015558325.1 | Lacticaseibacillus_rhamnosus | LW144_RS03870 | 70650..74282 | 47715 | BREX-system-P-loop-protein-BrxC-[Lacticaseibacillus-rhamnosus] |
| WP_108298644.1 | Type1.BREX | [illegible] | 1215 | Bacteria>Terrabacteria_group>Bacillota>Bacilli>Lactobacillales>Lactobacillaceae | GCF_015558835.1 | Lacticaseibacillus_paracasei | I2G54_RS02180 | 48912..52559 | 1597 | BREX-system-P-loop-protein-BrxC-[Lacticaseibacillus-paracasei] |
| WP_195558263.1 | Type1.BREX | [illegible] | 1183 | Bacteria>Terrabacteria_group>Bacillota>Bacilli>Lactobacillales>Lactobacillaceae | GCF_015560055.1 | Ligilactobacillus_salivarius | I2G60_RS06320 | 4891..8442 | 1624 | BREX-system-P-loop-protein-BrxC-[Ligilactobacillus-salivarius] |
| WP_195631091.1 | Type1.BREX | [illegible] | 1241 | Bacteria>Terrabacteria_group>Bacillota>Bacilli>Lactobacillales>Lactobacillaceae | GCF_015561225.1 | Lactiplantibacillus_plantarum | I2H95_RS14900 | 2281..6006 | 1590 | BREX-system-P-loop-protein-BrxC-[Lactiplantibacillus-plantarum] |
| WP_199412090.1 | Type1.BREX | <-DUF6079*<-BrxB-BrxF-iSTAND<-BrxA-DNA-binding<-?<-SIGMA-HTH<-?<-?<-HNH+ADA-DBD<-?<-PspA | 1194 | Bacteria>Terrabacteria_group>Bacillota>Bacilli>Lactobacillales>Lactobacillaceae | GCF_016465895.1 | Weissella_confusa | HAU13_RS10875 | 757..4341 | 1583 | BREX-system-P-loop-protein-BrxC-[Weissella-confusa] |
| WP_201308105.1 | Type1.BREX | <-RuvB-AAA+RuvB-HTH<-?<-MORC<-?<-?<-N6-METHYLASE<-DUF6079*<-BrxB-BrxF-iSTAND<-BrxA-DNA-binding<-?<-?<-MerR-HTH | 1210 | Bacteria>Terrabacteria_group>Bacillota>Bacilli>Lactobacillales>Lactobacillaceae | GCF_016586205.1 | Companilactobacillus_farciminis | JMF75_RS09670 | 25470..29102 | 1612 | BREX-system-P-loop-protein-BrxC-[Companilactobacillus-farciminis] |
| WP_200136379.1 | Type1.BREX | [illegible] | 1212 | Bacteria>Terrabacteria_group>Bacillota>Bacilli>Lactobacillales>Lactobacillaceae | GCF_016624925.1 | unclassified_Leuconostoc | IAE53_RS00085 | 21588..25226 | 2685106 | MULTISPECIES:-BREX-system-P-loop-protein-BrxC-[unclassified-Leuconostoc] |
| WP_193820257.1 | Type1.BREX | [illegible] | 1219 | Bacteria>Terrabacteria_group>Bacillota>Bacilli>Lactobacillales>Lactobacillaceae | GCF_016653515.1 | Lacticaseibacillus_rhamnosus | JIO02_RS10990 | 2305265..2308924 | 47715 | BREX-system-P-loop-protein-BrxC-[Lacticaseibacillus-rhamnosus] |
| WP_202861291.1 | Type1.BREX | [illegible] | 1220 | Bacteria>Terrabacteria_group>Bacillota>Bacilli>Lactobacillales>Lactobacillaceae | GCF_016775685.1 | Lactiplantibacillus_plantarum | JMO19_RS04040 | 797969..801631 | 1590 | BREX-system-P-loop-protein-BrxC-[Lactiplantibacillus-plantarum] |
| WP_204120188.1 | Type1.BREX | [illegible] | 1216 | Bacteria>Terrabacteria_group>Bacillota>Bacilli>Lactobacillales>Lactobacillaceae | GCF_016861585.1 | Levilactobacillus_wangkuiensis | JUF14_RS11710 | 5195..8845 | 2799566 | BREX-system-P-loop-protein-BrxC-[Levilactobacillus-wangkuiensis] |
| WP_203640134.1 | Type1.BREX | <-ROK-HTH<-?<-?<-PglZ\|\|N6-METHYLASE->Arm-DNA-bind_4+Phage_int_SAM_5+Phage_integrase-><-N6-METHYLASE<-DUF6079*<-BrxB-BrxF-iSTAND<-BrxA-DNA-binding | 1216 | Bacteria>Terrabacteria_group>Bacillota>Bacilli>Lactobacillales>Lactobacillaceae | GCF_016861605.1 | Levilactobacillus_wangkuiensis | JUF13_RS13680 | 14876..18526 | 2799566 | BREX-system-P-loop-protein-BrxC-[Levilactobacillus-wangkuiensis] |
| WP_204121707.1 | Type1.BREX | [illegible] | 1220 | Bacteria>Terrabacteria_group>Bacillota>Bacilli>Lactobacillales>Lactobacillaceae | GCF_016861675.1 | Levilactobacillus_sp._244-2 | JUF12_RS09870 | 83412..87074 | 2799569 | BREX-system-P-loop-protein-BrxC-[Levilactobacillus-sp.-244-2] |
| WP_203626028.1 | Type1.BREX | [illegible] | 1229 | Bacteria>Terrabacteria_group>Bacillota>Bacilli>Lactobacillales>Lactobacillaceae | GCF_016861715.1 | Lacticaseibacillus_mingshuiensis | JUE68_RS01590 | 378582..382271 | 2799574 | BREX-system-P-loop-protein-BrxC-[Lacticaseibacillus-mingshuiensis] |
| WP_204122461.1 | Type1.BREX | [illegible] | 1229 | Bacteria>Terrabacteria_group>Bacillota>Bacilli>Lactobacillales>Lactobacillaceae | GCF_016861765.1 | Lacticaseibacillus_mingshuiensis | JUF03_RS02150 | 248342..252031 | 2799574 | BREX-system-P-loop-protein-BrxC-[Lacticaseibacillus-mingshuiensis] |
| WP_203636934.1 | Type1.BREX | [illegible] | 1229 | Bacteria>Terrabacteria_group>Bacillota>Bacilli>Lactobacillales>Lactobacillaceae | GCF_016861785.1 | Lacticaseibacillus_mingshuiensis | JUE67_RS01480 | 348937..352626 | 2799574 | BREX-system-P-loop-protein-BrxC-[Lacticaseibacillus-mingshuiensis] |
| WP_064533504.1 | Type1.BREX | [illegible] | 1210 | Bacteria>Terrabacteria_group>Bacillota>Bacilli>Lactobacillales>Lactobacillaceae | GCF_016887865.1 | Lacticaseibacillus_rhamnosus | JNW95_RS08680 | 79348..82980 | 47715 | BREX-system-P-loop-protein-BrxC-[Lacticaseibacillus-rhamnosus] |
| WP_204759736.1 | Type1.BREX | [illegible] | 1183 | Bacteria>Terrabacteria_group>Bacillota>Bacilli>Lactobacillales>Lactobacillaceae | GCF_016902115.1 | Ligilactobacillus_salivarius | H5972_05415 | 6514..10065 | 1624 | BREX-system-P-loop-protein-BrxC-[Ligilactobacillus-salivarius] |
| WP_103658548.1 | Type1.BREX | [illegible] | 1243 | Bacteria>Terrabacteria_group>Bacillota>Bacilli>Lactobacillales>Lactobacillaceae | GCF_017565805.1 | Lactobacillus_helveticus | MVU88_RS02210 | 7396..11127 | 1587 | BREX-system-P-loop-protein-BrxC-[Lactobacillus-helveticus] |
| WP_208199753.1 | Type1.BREX | [illegible] | 1239 | Bacteria>Terrabacteria_group>Bacillota>Bacilli>Lactobacillales>Lactobacillaceae | GCF_017581995.1 | Schleiferilactobacillus_harbinensis | J5F27_RS12780 | 6081..9800 | 304207 | BREX-system-P-loop-protein-BrxC-[Schleiferilactobacillus-harbinensis] |
| WP_159257171.1 | Type1.BREX | [illegible] | 1223 | Bacteria>Terrabacteria_group>Bacillota>Bacilli>Lactobacillales>Lactobacillaceae | GCF_018069575.1 | Lactiplantibacillus_pentosus | FOM15_RS08440 | 491141..494812 | 1589 | BREX-system-P-loop-protein-BrxC-[Lactiplantibacillus-pentosus] |
| WP_213950276.1 | Type1.BREX | [illegible] | 1210 | Bacteria>Terrabacteria_group>Bacillota>Bacilli>Lactobacillales>Lactobacillaceae | GCF_018458775.1 | Lacticaseibacillus_rhamnosus | LHV47_RS13585 | 177771..181403 | 47715 | BREX-system-P-loop-protein-BrxC-[Lacticaseibacillus-rhamnosus] |
| WP_215485506.1 | Type1.BREX | [illegible] | 1239 | Bacteria>Terrabacteria_group>Bacillota>Bacilli>Lactobacillales>Lactobacillaceae | GCF_018728605.1 | Lactobacillus_gasseri | E6O77_RS03170 | 608983..612702 | 1596 | BREX-system-P-loop-protein-BrxC-[Lactobacillus-gasseri] |
| WP_216814454.1 | Type1.BREX | [illegible] | 1194 | Bacteria>Terrabacteria_group>Bacillota>Bacilli>Lactobacillales>Lactobacillaceae | GCF_018991665.1 | Weissella_paramesenteroides | KGP42_RS08110 | 64614..68198 | 1249 | BREX-system-P-loop-protein-BrxC-[Weissella-paramesenteroides] |
| WP_216746496.1 | Type1.BREX | [illegible] | 1223 | Bacteria>Terrabacteria_group>Bacillota>Bacilli>Lactobacillales>Lactobacillaceae | GCF_018993275.1 | Lactiplantibacillus_pentosus | KGF67_RS02355 | 5738..9409 | 1589 | BREX-system-P-loop-protein-BrxC-[Lactiplantibacillus-pentosus] |
| WP_218674449.1 | Type1.BREX | [illegible] | 1244 | Bacteria>Terrabacteria_group>Bacillota>Bacilli>Lactobacillales>Lactobacillaceae | GCF_019278055.1 | Lactobacillus_crispatus | KDZ21_RS07630 | 12982..16716 | 47770 | BREX-system-P-loop-protein-BrxC-[Lactobacillus-crispatus] |
| WP_220028697.1 | Type1.BREX | [illegible] | 1244 | Bacteria>Terrabacteria_group>Bacillota>Bacilli>Lactobacillales>Lactobacillaceae | GCF_019455965.1 | Lactobacillus_helveticus | E0712_RS03715 | 165606..169340 | 1587 | BREX-system-P-loop-protein-BrxC-[Lactobacillus-helveticus] |
| WP_220023636.1 | Type1.BREX | [illegible] | 1241 | Bacteria>Terrabacteria_group>Bacillota>Bacilli>Lactobacillales>Lactobacillaceae | GCF_019455975.1 | Lactobacillus_helveticus | E0715_RS08290 | 19434..23159 | 1587 | BREX-system-P-loop-protein-BrxC-[Lactobacillus-helveticus] |
| WP_220735234.1 | Type1.BREX | [illegible] | 1212 | Bacteria>Terrabacteria_group>Bacillota>Bacilli>Lactobacillales>Lactobacillaceae | GCF_019656015.1 | Leuconostoc_inhae | K5Q14_RS07795 | 32108..35746 | 178001 | BREX-system-P-loop-protein-BrxC-[Leuconostoc-inhae] |
| WP_222258122.1 | Type1.BREX | [illegible] | 1221 | Bacteria>Terrabacteria_group>Bacillota>Bacilli>Lactobacillales>Lactobacillaceae | GCF_019797825.1 | Limosilactobacillus_panis | KZE55_RS08430 | 1755887..1759552 | 47493 | BREX-system-P-loop-protein-BrxC-[Limosilactobacillus-panis] |
| WP_223320767.1 | Type1.BREX | [illegible] | 1194 | Bacteria>Terrabacteria_group>Bacillota>Bacilli>Lactobacillales>Lactobacillaceae | GCF_019927025.1 | Leuconostoc_mesenteroides | J3329_RS06880 | 12023..15607 | 1245 | BREX-system-P-loop-protein-BrxC-[Leuconostoc-mesenteroides] |
| WP_223660616.1 | Type1.BREX | [illegible] | 1239 | Bacteria>Terrabacteria_group>Bacillota>Bacilli>Lactobacillales>Lactobacillaceae | GCF_020026535.1 | Lactobacillus_johnsonii | K7D69_RS01700 | 3964..7683 | 33959 | BREX-system-P-loop-protein-BrxC-[Lactobacillus-johnsonii] |
| WP_060391297.1 | Type1.BREX | [illegible] | 1211 | Bacteria>Terrabacteria_group>Bacillota>Bacilli>Lactobacillales>Lactobacillaceae | GCF_020088925.1 | Leuconostoc_gasicomitatum | KII98_RS08285 | 305123..308758 | 115778 | BREX-system-P-loop-protein-BrxC-[Leuconostoc-gasicomitatum] |
| WP_225047865.1 | Type1.BREX | [illegible] | 1221 | Bacteria>Terrabacteria_group>Bacillota>Bacilli>Lactobacillales>Lactobacillaceae | GCF_020180945.1 | Lacticaseibacillus_kribbianus | LE416_RS07415 | 6539..10204 | 2926292 | BREX-system-P-loop-protein-BrxC-[Lacticaseibacillus-kribbianus] |
| WP_032957101.1 | Type1.BREX | [illegible] | 1210 | Bacteria>Terrabacteria_group>Bacillota>Bacilli>Lactobacillales>Lactobacillaceae | GCF_020826335.1 | Lacticaseibacillus_rhamnosus | LL671_RS03770 | 826686..830318 | 47715 | BREX-system-P-loop-protein-BrxC-[Lacticaseibacillus-rhamnosus] |
| WP_060462133.1 | Type1.BREX | [illegible] | 1244 | Bacteria>Terrabacteria_group>Bacillota>Bacilli>Lactobacillales>Lactobacillaceae | GCF_020887095.1 | Lactobacillus_crispatus | LPC26_RS08700 | 1768105..1771839 | 47770 | BREX-system-P-loop-protein-BrxC-[Lactobacillus-crispatus] |
| WP_231545262.1 | Type1.BREX | [illegible] | 1171 | Bacteria>Terrabacteria_group>Bacillota>Bacilli>Lactobacillales>Lactobacillaceae | GCF_021134235.1 | Lactobacillus_delbrueckii | LOB65_RS04175 | 29230..32745 | 1584 | BREX-system-P-loop-protein-BrxC-[Lactobacillus-delbrueckii] |
| WP_231523637.1 | Type1.BREX | [illegible] | 1183 | Bacteria>Terrabacteria_group>Bacillota>Bacilli>Lactobacillales>Lactobacillaceae | GCF_021134645.1 | Lactobacillus_delbrueckii | LOB39_RS07020 | 33020..36571 | 1584 | BREX-system-P-loop-protein-BrxC-[Lactobacillus-delbrueckii] |
| WP_231524175.1 | Type1.BREX | [illegible] | 1188 | Bacteria>Terrabacteria_group>Bacillota>Bacilli>Lactobacillales>Lactobacillaceae | GCF_021134795.1 | Lactobacillus_delbrueckii | LOB43_RS04555 | 15910..19476 | 1584 | BREX-system-P-loop-protein-BrxC-[Lactobacillus-delbrueckii] |
| WP_231529866.1 | Type1.BREX | [illegible] | 1188 | Bacteria>Terrabacteria_group>Bacillota>Bacilli>Lactobacillales>Lactobacillaceae | GCF_021135055.1 | Lactobacillus_delbrueckii | LOB49_05200 | 20821..24387 | 1584 | BREX-system-P-loop-protein-BrxC-[Lactobacillus-delbrueckii] |
| WP_231535729.1 | Type1.BREX | [illegible] | 1188 | Bacteria>Terrabacteria_group>Bacillota>Bacilli>Lactobacillales>Lactobacillaceae | GCF_021135595.1 | Lactobacillus_delbrueckii | LOC07_RS06205 | 4796..8362 | 1584 | BREX-system-P-loop-protein-BrxC-[Lactobacillus-delbrueckii] |
| WP_191671322.1 | Type1.BREX | [illegible] | 1230 | Bacteria>Terrabacteria_group>Bacillota>Bacilli>Lactobacillales>Lactobacillaceae | GCF_021278125.1 | Limosilactobacillus_fermentum | LR965_RS05320 | 12718..16410 | 1613 | BREX-system-P-loop-protein-BrxC-[Limosilactobacillus-fermentum] |
| WP_269843153.1 | Type1.BREX | [illegible] | 1235 | Bacteria>Terrabacteria_group>Bacillota>Bacilli>Lactobacillales>Lactobacillaceae | GCF_021300595.1 | Lacticaseibacillus_rhamnosus | LW144_RS13115 | 12071..15778 | 47715 | BREX-system-P-loop-protein-BrxC-[Lacticaseibacillus-rhamnosus] |
| WP_236150042.1 | Type1.BREX | [illegible] | 1171 | Bacteria>Terrabacteria_group>Bacillota>Bacilli>Lactobacillales>Lactobacillaceae | GCF_021600205.1 | Lactobacillus_delbrueckii | L5979_RS00560 | 22704..26219 | 1584 | BREX-system-P-loop-protein-BrxC-[Lactobacillus-delbrueckii] |
| WP_236153484.1 | Type1.BREX | [illegible] | 1183 | Bacteria>Terrabacteria_group>Bacillota>Bacilli>Lactobacillales>Lactobacillaceae | GCF_021600265.1 | Lactobacillus_delbrueckii | ME802_14840 | 21460..25011 | 1584 | BREX-system-P-loop-protein-BrxC-[Lactobacillus-delbrueckii] |
| WP_236149793.1 | Type1.BREX | [illegible] | 1188 | Bacteria>Terrabacteria_group>Bacillota>Bacilli>Lactobacillales>Lactobacillaceae | GCF_021600925.1 | Lactobacillus_delbrueckii | L5981_RS02870 | 31328..34894 | 1584 | BREX-system-P-loop-protein-BrxC-[Lactobacillus-delbrueckii] |
| WP_050952512.1 | Type1.BREX | [illegible] | 1188 | Bacteria>Terrabacteria_group>Bacillota>Bacilli>Lactobacillales>Lactobacillaceae | GCF_021601025.1 | Lactobacillus_delbrueckii | L5988_RS00790 | 158168..161734 | 1584 | BREX-system-P-loop-protein-BrxC-[Lactobacillus-delbrueckii] |
| WP_238059865.1 | Type1.BREX | [illegible] | 1203 | Bacteria>Terrabacteria_group>Bacillota>Bacilli>Lactobacillales>Lactobacillaceae | GCF_022131005.1 | Lacticaseibacillus_saniviri | K9861_RS11735 | 11859..15470 | 931533 | BREX-system-P-loop-protein-BrxC-[Lacticaseibacillus-saniviri] |
| WP_238064446.1 | Type1.BREX | [illegible] | 1241 | Bacteria>Terrabacteria_group>Bacillota>Bacilli>Lactobacillales>Lactobacillaceae | GCF_022131015.1 | Lacticaseibacillus_paracasei | K9853_RS00445 | 1326..5051 | 1597 | BREX-system-P-loop-protein-BrxC-[Lacticaseibacillus-paracasei] |
| WP_241479268.1 | Type1.BREX | [illegible] | 1216 | Bacteria>Terrabacteria_group>Bacillota>Bacilli>Lactobacillales>Lactobacillaceae | GCF_022568855.1 | Lactiplantibacillus_plantarum | MNV61_RS08805 | 14671..18321 | 1590 | BREX-system-P-loop-protein-BrxC-[Lactiplantibacillus-plantarum] |
| WP_241467420.1 | Type1.BREX | [illegible] | 809 | Bacteria>Terrabacteria_group>Bacillota>Bacilli>Lactobacillales>Lactobacillaceae | GCF_022568865.1 | Lactiplantibacillus_plantarum | MNV59_RS04265 | 204654..207083 | 1590 | BREX-system-P-loop-protein-BrxC-[Lactiplantibacillus-plantarum] |
| WP_242073729.1 | Type1.BREX | [illegible] | 1188 | Bacteria>Terrabacteria_group>Bacillota>Bacilli>Lactobacillales>Lactobacillaceae | GCF_022642945.1 | Lactobacillus_delbrueckii | G8B25_RS07935 | 1514865..1518431 | 1584 | BREX-system-P-loop-protein-BrxC-[Lactobacillus-delbrueckii] |
| WP_244056757.1 | Type1.BREX | [illegible] | 1221 | Bacteria>Terrabacteria_group>Bacillota>Bacilli>Lactobacillales>Lactobacillaceae | GCF_022836475.1 | Ligilactobacillus_pabuli | LPAF129_RS09900 | 19808..23473 | 2886039 | BREX-system-P-loop-protein-BrxC-[Ligilactobacillus-pabuli] |
| WP_248558916.1 | Type1.BREX | [illegible] | 1216 | Bacteria>Terrabacteria_group>Bacillota>Bacilli>Lactobacillales>Lactobacillaceae | GCF_023195585.1 | Lactiplantibacillus_plantarum | M0857_RS07205 | 62931..66581 | 1590 | BREX-system-P-loop-protein-BrxC-[Lactiplantibacillus-plantarum] |
| WP_249905369.1 | Type1.BREX | [illegible] | 1244 | Bacteria>Terrabacteria_group>Bacillota>Bacilli>Lactobacillales>Lactobacillaceae | GCF_023523445.1 | Lactobacillus_amylovorus | LOB74_RS05625 | 1115195..1118929 | 1604 | BREX-system-P-loop-protein-BrxC-[Lactobacillus-amylovorus] |
| WP_250825484.1 | Type1.BREX | [illegible] | 1247 | Bacteria>Terrabacteria_group>Bacillota>Bacilli>Lactobacillales>Lactobacillaceae | GCF_023674385.1 | Lactobacillus_kefiranofaciens | MU859_RS01790 | 311336..315079 | 267818 | BREX-system-P-loop-protein-BrxC-[Lactobacillus-kefiranofaciens] |
| WP_251945007.1 | Type1.BREX | [illegible] | 1195 | Bacteria>Terrabacteria_group>Bacillota>Bacilli>Lactobacillales>Lactobacillaceae | GCF_023734315.1 | Latilactobacillus_sakei | A4W83_RS09725 | 1916378..1919965 | 1599 | BREX-system-P-loop-protein-BrxC-[Latilactobacillus-sakei] |
| WP_251908041.1 | Type1.BREX | [illegible] | 1219 | Bacteria>Terrabacteria_group>Bacillota>Bacilli>Lactobacillales>Lactobacillaceae | GCF_023734335.1 | Latilactobacillus_sakei | A4W84_RS06360 | 1251250..1254909 | 1599 | BREX-system-P-loop-protein-BrxC-[Latilactobacillus-sakei] |
| WP_105300054.1 | Type1.BREX | [illegible] | 1215 | Bacteria>Terrabacteria_group>Bacillota>Bacilli>Lactobacillales>Lactobacillaceae | GCF_024022895.1 | Latilactobacillus_sakei | LAS9267_RS03030 | 155179..158826 | 1599 | BREX-system-P-loop-protein-BrxC-[Latilactobacillus-sakei] |
| WP_253613261.1 | Type1.BREX | [illegible] | 941 | Bacteria>Terrabacteria_group>Bacillota>Bacilli>Lactobacillales>Lactobacillaceae | GCF_024158965.2 | Lacticaseibacillus_paracasei | NJN40_RS04035 | 830871..833696 | 1597 | BREX-system-P-loop-protein-BrxC-[Lacticaseibacillus-paracasei] |
| WP_254681527.1 | Type1.BREX | [illegible] | 1219 | Bacteria>Terrabacteria_group>Bacillota>Bacilli>Lactobacillales>Lactobacillaceae | GCF_024273085.1 | Latilactobacillus_sakei | LOX63_RS01755 | 341729..345388 | 1599 | BREX-system-P-loop-protein-BrxC-[Latilactobacillus-sakei] |
| WP_261913355.1 | Type1.BREX | [illegible] | 1217 | Bacteria>Terrabacteria_group>Bacillota>Bacilli>Lactobacillales>Lactobacillaceae | GCF_024329455.1 | Lacticaseibacillus_paracasei | KBX55_RS07855 | 9810..13463 | 1597 | BREX-system-P-loop-protein-BrxC-[Lacticaseibacillus-paracasei] |
| WP_255033756.1 | Type1.BREX | [illegible] | 1241 | Bacteria>Terrabacteria_group>Bacillota>Bacilli>Lactobacillales>Lactobacillaceae | GCF_024330045.1 | Levilactobacillus_brevis | NMO39_RS12030 | 10442..14167 | 1580 | BREX-system-P-loop-protein-BrxC-[Levilactobacillus-brevis] |
| WP_261809685.1 | Type1.BREX | [illegible] | 1211 | Bacteria>Terrabacteria_group>Bacillota>Bacilli>Lactobacillales>Lactobacillaceae | GCF_024345185.1 | Levilactobacillus_humaensis | OCU14_RS02680 | 59918..63553 | 2950375 | BREX-system-P-loop-protein-BrxC-[Levilactobacillus-humaensis] |
| WP_135942480.1 | Type1.BREX | [illegible] | 1219 | Bacteria>Terrabacteria_group>Bacillota>Bacilli>Lactobacillales>Lactobacillaceae | GCF_024622325.1 | Ligilactobacillus_murinus | NSA12_RS07380 | 1361..5020 | 1622 | BREX-system-P-loop-protein-BrxC-[Ligilactobacillus-murinus] |
| WP_249391181.1 | Type1.BREX | [illegible] | 1241 | Bacteria>Terrabacteria_group>Bacillota>Bacilli>Lactobacillales>Lactobacillaceae | GCF_025122345.1 | Lactiplantibacillus_plantarum | M4S29_RS14340 | 1936..5661 | 1590 | BREX-system-P-loop-protein-BrxC-[Lactiplantibacillus-plantarum] |
| WP_046025494.1 | Type1.BREX | [illegible] | 1230 | Bacteria>Terrabacteria_group>Bacillota>Bacilli>Lactobacillales>Lactobacillaceae | GCF_025122475.1 | Limosilactobacillus_fermentum | EFE30_RS03130 | 13799..17491 | 1613 | BREX-system-P-loop-protein-BrxC-[Limosilactobacillus-fermentum] |
| WP_259713892.1 | Type1.BREX | [illegible] | 1194 | Bacteria>Terrabacteria_group>Bacillota>Bacilli>Lactobacillales>Lactobacillaceae | GCF_025127215.1 | Weissella_cibaria | EFL83_RS06745 | 62783..66367 | 137591 | BREX-system-P-loop-protein-BrxC-[Weissella-cibaria] |
| WP_259682117.1 | Type1.BREX | [illegible] | 1223 | Bacteria>Terrabacteria_group>Bacillota>Bacilli>Lactobacillales>Lactobacillaceae | GCF_025129205.1 | Lactiplantibacillus_pentosus | EFM55_RS05940 | 21301..24972 | 1589 | BREX-system-P-loop-protein-BrxC-[Lactiplantibacillus-pentosus] |
| WP_259693422.1 | Type1.BREX | [illegible] | 1242 | Bacteria>Terrabacteria_group>Bacillota>Bacilli>Lactobacillales>Lactobacillaceae | GCF_025129275.1 | Lactobacillus_helveticus | EFM10_RS03595 | 12781..16509 | 1587 | BREX-system-P-loop-protein-BrxC-[Lactobacillus-helveticus] |
| WP_259685554.1 | Type1.BREX | [illegible] | 1240 | Bacteria>Terrabacteria_group>Bacillota>Bacilli>Lactobacillales>Lactobacillaceae | GCF_025129335.1 | Lactobacillus_helveticus | EFM06_RS11120 | 12796..16518 | 1587 | BREX-system-P-loop-protein-BrxC-[Lactobacillus-helveticus] |
| WP_260265100.1 | Type1.BREX | [illegible] | 1171 | Bacteria>Terrabacteria_group>Bacillota>Bacilli>Lactobacillales>Lactobacillaceae | GCF_025186005.1 | Lactobacillus_delbrueckii | EFR42_RS08010 | 1230..4745 | 1584 | BREX-system-P-loop-protein-BrxC-[Lactobacillus-delbrueckii] |
| WP_260301636.1 | Type1.BREX | [illegible] | 1241 | Bacteria>Terrabacteria_group>Bacillota>Bacilli>Lactobacillales>Lactobacillaceae | GCF_025186395.1 | Schleiferilactobacillus_harbinensis | EFT87_RS04585 | 5213..8938 | 304207 | BREX-system-P-loop-protein-BrxC-[Schleiferilactobacillus-harbinensis] |
| WP_260155113.1 | Type1.BREX | [illegible] | 1194 | Bacteria>Terrabacteria_group>Bacillota>Bacilli>Lactobacillales>Lactobacillaceae | GCF_025188365.1 | Leuconostoc_mesenteroides | EFN50_RS00350 | 1772..5356 | 1245 | BREX-system-P-loop-protein-BrxC-[Leuconostoc-mesenteroides] |
| WP_260182661.1 | Type1.BREX | [illegible] | 1227 | Bacteria>Terrabacteria_group>Bacillota>Bacilli>Lactobacillales>Lactobacillaceae | GCF_025189415.1 | Lacticaseibacillus_rhamnosus | EFO73_RS10425 | 40181..43864 | 47715 | BREX-system-P-loop-protein-BrxC-[Lacticaseibacillus-rhamnosus] |
| WP_260198753.1 | Type1.BREX | [illegible] | 1212 | Bacteria>Terrabacteria_group>Bacillota>Bacilli>Lactobacillales>Lactobacillaceae | GCF_025190685.1 | Lacticaseibacillus_paracasei | EFP50_RS14565 | 1957..5595 | 1597 | BREX-system-P-loop-protein-BrxC-[Lacticaseibacillus-paracasei] |
| WP_260308143.1 | Type1.BREX | [illegible] | 1245 | Bacteria>Terrabacteria_group>Bacillota>Bacilli>Lactobacillales>Lactobacillaceae | GCF_025190905.1 | Lactobacillus_johnsonii | EFP49_06745 | 38785..42522 | 33959 | BREX-system-P-loop-protein-BrxC-[Lactobacillus-johnsonii] |
| WP_260274304.1 | Type1.BREX | [illegible] | 1243 | Bacteria>Terrabacteria_group>Bacillota>Bacilli>Lactobacillales>Lactobacillaceae | GCF_025191325.1 | Lactobacillus_helveticus | EFP83_RS06575 | 9420..13151 | 1587 | BREX-system-P-loop-protein-BrxC-[Lactobacillus-helveticus] |
| WP_260307210.1 | Type1.BREX | [illegible] | 1243 | Bacteria>Terrabacteria_group>Bacillota>Bacilli>Lactobacillales>Lactobacillaceae | GCF_025191425.1 | Lactobacillus_helveticus | EFP97_RS06690 | 9418..13149 | 1587 | BREX-system-P-loop-protein-BrxC-[Lactobacillus-helveticus] |
| WP_260212031.1 | Type1.BREX | [illegible] | 1244 | Bacteria>Terrabacteria_group>Bacillota>Bacilli>Lactobacillales>Lactobacillaceae | GCF_025194565.1 | Lactobacillus_amylovorus | EFR95_RS04430 | 29331..33065 | 1604 | BREX-system-P-loop-protein-BrxC-[Lactobacillus-amylovorus] |
| WP_260355673.1 | Type1.BREX | [illegible] | 1194 | Bacteria>Terrabacteria_group>Bacillota>Bacilli>Lactobacillales>Lactobacillaceae | GCF_025211155.1 | Leuconostoc_suionicum | EFM34_RS08570 | 2844..6428 | 1511761 | BREX-system-P-loop-protein-BrxC-[Leuconostoc-suionicum] |
| WP_260565630.1 | Type1.BREX | [illegible] | 1227 | Bacteria>Terrabacteria_group>Bacillota>Bacilli>Lactobacillales>Lactobacillaceae | GCF_025215665.1 | Lacticaseibacillus_paracasei | EOK76_RS04770 | 1022969..1026652 | 1597 | BREX-system-P-loop-protein-BrxC-[Lacticaseibacillus-paracasei] |
| WP_265000178.1 | Type1.BREX | [illegible] | 1218 | Bacteria>Terrabacteria_group>Bacillota>Bacilli>Lactobacillales>Lactobacillaceae | GCF_026013585.1 | Levilactobacillus_namurensis | OM945_RS10460 | 2686..6342 | 380393 | BREX-system-P-loop-protein-BrxC-[Levilactobacillus-namurensis] |
| WP_265678613.1 | Type1.BREX | [illegible] | 1215 | Bacteria>Terrabacteria_group>Bacillota>Bacilli>Lactobacillales>Lactobacillaceae | GCF_026193275.1 | Latilactobacillus_curvatus | OP752_RS00225 | 50217..53864 | 28038 | BREX-system-P-loop-protein-BrxC-[Latilactobacillus-curvatus] |
| WP_267160700.1 | Type1.BREX | [illegible] | 1241 | Bacteria>Terrabacteria_group>Bacillota>Bacilli>Lactobacillales>Lactobacillaceae | GCF_026410225.1 | Levilactobacillus_brevis | OSJ94_RS09260 | 5042..8767 | 1580 | BREX-system-P-loop-protein-BrxC-[Levilactobacillus-brevis] |
| WP_267667066.1 | Type1.BREX | [illegible] | 1194 | Bacteria>Terrabacteria_group>Bacillota>Bacilli>Lactobacillales>Lactobacillaceae | GCF_026616915.1 | Leuconostoc_gelidum | NZD49_RS09275 | 14139..17723 | 1244 | BREX-system-P-loop-protein-BrxC-[Leuconostoc-gelidum] |
| WP_268886389.1 | Type1.BREX | [illegible] | 1183 | Bacteria>Terrabacteria_group>Bacillota>Bacilli>Lactobacillales>Lactobacillaceae | GCF_026891475.1 | Lactobacillus_delbrueckii | OO549_RS05810 | 14866..18417 | 1584 | BREX-system-P-loop-protein-BrxC-[Lactobacillus-delbrueckii] |
| WP_269903045.1 | Type1.BREX | [illegible] | 1216 | Bacteria>Terrabacteria_group>Bacillota>Bacilli>Lactobacillales>Lactobacillaceae | GCF_027568335.1 | Lactiplantibacillus_plantarum | NUG30_RS00235 | 55667..59317 | 1590 | BREX-system-P-loop-protein-BrxC-[Lactiplantibacillus-plantarum] |
| WP_270411955.1 | Type1.BREX | [illegible] | 1184 | Bacteria>Terrabacteria_group>Bacillota>Bacilli>Lactobacillales>Lactobacillaceae | GCF_027672425.1 | Ligilactobacillus_ruminis | PGT38_RS00055 | 14514..18068 | 1623 | BREX-system-P-loop-protein-BrxC-[Ligilactobacillus-ruminis] |
| WP_270450877.1 | Type1.BREX | [illegible] | 1244 | Bacteria>Terrabacteria_group>Bacillota>Bacilli>Lactobacillales>Lactobacillaceae | GCF_027680585.1 | Lactobacillus_crispatus | PGX14_RS06760 | 16446..20180 | 47770 | BREX-system-P-loop-protein-BrxC-[Lactobacillus-crispatus] |
| WP_270447891.1 | Type1.BREX | [illegible] | 1188 | Bacteria>Terrabacteria_group>Bacillota>Bacilli>Lactobacillales>Lactobacillaceae | GCF_027681365.1 | Lactobacillus_delbrueckii | PGX26_RS03415 | 69740..73306 | 1584 | BREX-system-P-loop-protein-BrxC-[Lactobacillus-delbrueckii] |
| WP_270482187.1 | Type1.BREX | [illegible] | 1241 | Bacteria>Terrabacteria_group>Bacillota>Bacilli>Lactobacillales>Lactobacillaceae | GCF_027682305.1 | Lactobacillus_crispatus | PGX51_RS02195 | 50434..54159 | 47770 | BREX-system-P-loop-protein-BrxC-[Lactobacillus-crispatus] |
| WP_081510717.1 | Type1.BREX | [illegible] | 1183 | Bacteria>Terrabacteria_group>Bacillota>Bacilli>Lactobacillales>Lactobacillaceae | GCF_027686005.1 | Ligilactobacillus_salivarius | PGY81_RS07685 | 22761..26312 | 1624 | BREX-system-P-loop-protein-BrxC-[Ligilactobacillus-salivarius] |
| WP_270641829.1 | Type1.BREX | [illegible] | 1183 | Bacteria>Terrabacteria_group>Bacillota>Bacilli>Lactobacillales>Lactobacillaceae | GCF_027692205.1 | Ligilactobacillus_salivarius | PG139_RS08665 | 2046..5597 | 1624 | BREX-system-P-loop-protein-BrxC-[Ligilactobacillus-salivarius] |
| WP_270643022.1 | Type1.BREX | [illegible] | 1243 | Bacteria>Terrabacteria_group>Bacillota>Bacilli>Lactobacillales>Lactobacillaceae | GCF_027694665.1 | Limosilactobacillus_mucosae | PG747_RS07320 | 14640..18371 | 97478 | BREX-system-P-loop-protein-BrxC-[Limosilactobacillus-mucosae] |
| WP_271018243.1 | Type1.BREX | [illegible] | 1183 | Bacteria>Terrabacteria_group>Bacillota>Bacilli>Lactobacillales>Lactobacillaceae | GCF_027858035.1 | Lactobacillus_delbrueckii | PF599_RS07175 | 1642..5193 | 1584 | BREX-system-P-loop-protein-BrxC-[Lactobacillus-delbrueckii] |
| WP_271876485.1 | Type1.BREX | [illegible] | 1233 | Bacteria>Terrabacteria_group>Bacillota>Bacilli>Lactobacillales>Lactobacillaceae | GCF_028308365.1 | Lactobacillus_amylovorus | ODV29_RS04110 | 28769..32470 | 1604 | BREX-system-P-loop-protein-BrxC-[Lactobacillus-amylovorus] |
| WP_271882935.1 | Type1.BREX | [illegible] | 1227 | Bacteria>Terrabacteria_group>Bacillota>Bacilli>Lactobacillales>Lactobacillaceae | GCF_028308535.1 | Lactobacillus_amylovorus | ODV81_RS08115 | 8502..12185 | 1604 | BREX-system-P-loop-protein-BrxC-[Lactobacillus-amylovorus] |
| WP_271875350.1 | Type1.BREX | [illegible] | 1233 | Bacteria>Terrabacteria_group>Bacillota>Bacilli>Lactobacillales>Lactobacillaceae | GCF_028308735.1 | Lactobacillus_amylovorus | ODV82_RS03950 | 34428..38129 | 1604 | BREX-system-P-loop-protein-BrxC-[Lactobacillus-amylovorus] |
| WP_271873858.1 | Type1.BREX | [illegible] | 1244 | Bacteria>Terrabacteria_group>Bacillota>Bacilli>Lactobacillales>Lactobacillaceae | GCF_028308865.1 | Lactobacillus_amylovorus | ODV08_RS03455 | 54390..58124 | 1604 | BREX-system-P-loop-protein-BrxC-[Lactobacillus-amylovorus] |
| WP_271897403.1 | Type1.BREX | [illegible] | 1233 | Bacteria>Terrabacteria_group>Bacillota>Bacilli>Lactobacillales>Lactobacillaceae | GCF_028308945.1 | Lactobacillus_amylovorus | ODV12_RS07245 | 16097..19798 | 1604 | BREX-system-P-loop-protein-BrxC-[Lactobacillus-amylovorus] |
| WP_271862779.1 | Type1.BREX | [illegible] | 1244 | Bacteria>Terrabacteria_group>Bacillota>Bacilli>Lactobacillales>Lactobacillaceae | GCF_028309095.1 | Lactobacillus_amylovorus | ODU71_RS08740 | 22834..26568 | 1604 | BREX-system-P-loop-protein-BrxC-[Lactobacillus-amylovorus] |
| WP_271901215.1 | Type1.BREX | [illegible] | 1221 | Bacteria>Terrabacteria_group>Bacillota>Bacilli>Lactobacillales>Lactobacillaceae | GCF_028369155.1 | Limosilactobacillus_reuteri | PQK88_RS04815 | 2601..6266 | 1598 | BREX-system-P-loop-protein-BrxC-[Limosilactobacillus-reuteri] |
| WP_273645655.1 | Type1.BREX | [illegible] | 1221 | Bacteria>Terrabacteria_group>Bacillota>Bacilli>Lactobacillales>Lactobacillaceae | GCF_028607005.1 | Limosilactobacillus_vaginalis | PRK59_RS03875 | 829452..833117 | 1633 | BREX-system-P-loop-protein-BrxC-[Limosilactobacillus-vaginalis] |
| WP_273932798.1 | Type1.BREX | [illegible] | 1212 | Bacteria>Terrabacteria_group>Bacillota>Bacilli>Lactobacillales>Lactobacillaceae | GCF_028613785.1 | Lacticaseibacillus_paracasei | OP490_RS13270 | 2984..6622 | 1597 | BREX-system-P-loop-protein-BrxC-[Lacticaseibacillus-paracasei] |
| WP_273725609.1 | Type1.BREX | [illegible] | 1194 | Bacteria>Terrabacteria_group>Bacillota>Bacilli>Lactobacillales>Lactobacillaceae | GCF_028618945.1 | Leuconostoc_mesenteroides | PNN51_RS08050 | 12149..15733 | 1245 | BREX-system-P-loop-protein-BrxC-[Leuconostoc-mesenteroides] |
| WP_273709328.1 | Type1.BREX | [illegible] | 1194 | Bacteria>Terrabacteria_group>Bacillota>Bacilli>Lactobacillales>Lactobacillaceae | GCF_028619695.1 | Leuconostoc_mesenteroides | PNN69_RS09405 | 12528..16112 | 1245 | BREX-system-P-loop-protein-BrxC-[Leuconostoc-mesenteroides] |
| WP_273710227.1 | Type1.BREX | [illegible] | 1194 | Bacteria>Terrabacteria_group>Bacillota>Bacilli>Lactobacillales>Lactobacillaceae | GCF_028619795.1 | Leuconostoc_mesenteroides | PNK31_RS09760 | 1790..5374 | 1245 | BREX-system-P-loop-protein-BrxC-[Leuconostoc-mesenteroides] |
| WP_273711741.1 | Type1.BREX | [illegible] | 1194 | Bacteria>Terrabacteria_group>Bacillota>Bacilli>Lactobacillales>Lactobacillaceae | GCF_028620115.1 | Leuconostoc_mesenteroides | PNN68_RS09110 | 1573..5157 | 1245 | BREX-system-P-loop-protein-BrxC-[Leuconostoc-mesenteroides] |
| WP_273960123.1 | Type1.BREX | [illegible] | 1215 | Bacteria>Terrabacteria_group>Bacillota>Bacilli>Lactobacillales>Lactobacillaceae | GCF_028622855.1 | Latilactobacillus_curvatus | PSR33_RS06690 | 1305883..1309530 | 28038 | BREX-system-P-loop-protein-BrxC-[Latilactobacillus-curvatus] |
| WP_274784940.1 | Type1.BREX | [illegible] | 1239 | Bacteria>Terrabacteria_group>Bacillota>Bacilli>Lactobacillales>Lactobacillaceae | GCF_028878205.1 | Lacticaseibacillus_casei | LTZ17_RS09610 | 16306..20025 | 1582 | BREX-system-P-loop-protein-BrxC-[Lacticaseibacillus-casei] |
| WP_275091722.1 | Type1.BREX | [illegible] | 1217 | Bacteria>Terrabacteria_group>Bacillota>Bacilli>Lactobacillales>Lactobacillaceae | GCF_029025825.1 | Lactiplantibacillus_paraplantarum | PWO93_RS03745 | 824201..827854 | 60520 | BREX-system-P-loop-protein-BrxC-[Lactiplantibacillus-paraplantarum] |
| WP_099739799.1 | Type1.BREX | [illegible] | 1216 | Bacteria>Terrabacteria_group>Bacillota>Bacilli>Lactobacillales>Lactobacillaceae | GCF_029854335.1 | Lactiplantibacillus_plantarum | QC766_RS03755 | 808098..811748 | 1590 | BREX-system-P-loop-protein-BrxC-[Lactiplantibacillus-plantarum] |
| WP_160811299.1 | Type1.BREX | [illegible] | 1244 | Bacteria>Terrabacteria_group>Bacillota>Bacilli>Lactobacillales>Lactobacillaceae | GCF_030214445.1 | Lactobacillus_crispatus | EFS45_RS04625 | 2234..5968 | 47770 | BREX-system-P-loop-protein-BrxC-[Lactobacillus-crispatus] |
| WP_260165795.1 | Type1.BREX | [illegible] | 1212 | Bacteria>Terrabacteria_group>Bacillota>Bacilli>Lactobacillales>Lactobacillaceae | GCF_030361075.1 | Leuconostoc_citreum | EFN72_RS03350 | 147093..150731 | 33964 | BREX-system-P-loop-protein-BrxC-[Leuconostoc-citreum] |
| WP_061827034.1 | Type1.BREX | [illegible] | 1219 | Bacteria>Terrabacteria_group>Bacillota>Bacilli>Lactobacillales>Lactobacillaceae | GCF_030378265.1 | Latilactobacillus_sakei | A2G83_RS01845 | 86379..90038 | 1599 | BREX-system-P-loop-protein-BrxC-[Latilactobacillus-sakei] |
| WP_105961058.1 | Type1.BREX | [illegible] | 1223 | Bacteria>Terrabacteria_group>Bacillota>Bacilli>Lactobacillales>Lactobacillaceae | GCF_030549395.1 | Lactiplantibacillus | D6U18_RS15980 | 71104..74775 | 2767842 | MULTISPECIES:-BREX-system-P-loop-protein-BrxC-[Lactiplantibacillus] |
| WP_260390513.1 | Type1.BREX | [illegible] | 1216 | Bacteria>Terrabacteria_group>Bacillota>Bacilli>Lactobacillales>Lactobacillaceae | GCF_030710155.1 | Lactiplantibacillus_plantarum | EFS29_RS01035 | 1052..4702 | 1590 | BREX-system-P-loop-protein-BrxC-[Lactiplantibacillus-plantarum] |
| WP_074161956.1 | Type1.BREX | [illegible] | 1216 | Bacteria>Terrabacteria_group>Bacillota>Bacilli>Lactobacillales>Lactobacillaceae | GCF_900078525.1 | Lactiplantibacillus_plantarum | ERK20_RS15090 | 19950..23600 | 1590 | BREX-system-P-loop-protein-BrxC-[Lactiplantibacillus-plantarum] |
| WP_198033633.1 | Type1.BREX | [illegible] | 1195 | Bacteria>Terrabacteria_group>Bacillota>Bacilli>Lactobacillales>Lactobacillaceae | GCF_900215905.1 | Latilactobacillus_sakei | LSAJ18_RS08865 | 1804156..1807743 | 1599 | BREX-system-P-loop-protein-BrxC-[Latilactobacillus-sakei] |
| WP_099947387.1 | Type1.BREX | [illegible] | 1219 | Bacteria>Terrabacteria_group>Bacillota>Bacilli>Lactobacillales>Lactobacillaceae | GCF_900234395.1 | Latilactobacillus_sakei | LSAJ54_RS06515 | 1291898..1295557 | 1599 | BREX-system-P-loop-protein-BrxC-[Latilactobacillus-sakei] |
| WP_157771040.1 | Type1.BREX | [illegible] | 641 | Bacteria>Terrabacteria_group>Bacillota>Bacilli>Lactobacillales>Lactobacillaceae | GCF_900234395.1 | Latilactobacillus_sakei | LSAJ54_RS09745 | 1939205..1941130 | 1599 | BREX-system-P-loop-protein-BrxC-[Latilactobacillus-sakei] |
| WP_101511690.1 | Type1.BREX | [illegible] | 1243 | Bacteria>Terrabacteria_group>Bacillota>Bacilli>Lactobacillales>Lactobacillaceae | GCF_900248185.1 | Lactobacillus_helveticus | JM182_RS07250 | 23706..27437 | 1587 | BREX-system-P-loop-protein-BrxC-[Lactobacillus-helveticus] |
| WP_172436372.1 | Type1.BREX | [illegible] | 970 | Bacteria>Terrabacteria_group>Bacillota>Bacilli>Lactobacillales>Lactobacillaceae | GCF_901830435.1 | Lactiplantibacillus_plantarum | LKM53_RS04900 | 104356..107268 | 1590 | BREX-system-P-loop-protein-BrxC-[Lactiplantibacillus-plantarum] |
| WP_142500354.1 | Type1.BREX | [illegible] | 1217 | Bacteria>Terrabacteria_group>Bacillota>Bacilli>Lactobacillales>Lactobacillaceae | GCF_901933195.1 | Lacticaseibacillus_paracasei | LPCP272_RS05120 | 828..4481 | 1597 | BREX-system-P-loop-protein-BrxC-[Lacticaseibacillus-paracasei] |
| WP_143113415.1 | Type1.BREX | [illegible] | 1243 | Bacteria>Terrabacteria_group>Bacillota>Bacilli>Lactobacillales>Lactobacillaceae | GCF_901971775.1 | Limosilactobacillus_mucosae | LMUP508_RS08745 | 3931..7662 | 97478 | BREX-system-P-loop-protein-BrxC-[Limosilactobacillus-mucosae] |
| WP_048488336.1 | Type1.BREX | [illegible] | 1227 | Bacteria>Terrabacteria_group>Bacillota>Bacilli>Lactobacillales>Lactobacillaceae | GCF_901971785.1 | Lacticaseibacillus_rhamnosus | LRHP344_RS03410 | 15330..19013 | 47715 | BREX-system-P-loop-protein-BrxC-[Lacticaseibacillus-rhamnosus] |
| WP_118027524.1 | Type1.BREX | [illegible] | 1254 | Bacteria>Terrabacteria_group>Bacillota>Bacilli>Lactobacillales>Lactobacillaceae | GCF_902363955.1 | Lactobacillus_amylovorus | FY235_RS07460 | 2640..6404 | 1604 | BREX-system-P-loop-protein-BrxC-[Lactobacillus-amylovorus] |
| WP_048345183.1 | Type1.BREX | [illegible] | 1243 | Bacteria>Terrabacteria_group>Bacillota>Bacilli>Lactobacillales>Lactobacillaceae | GCF_902385785.1 | Limosilactobacillus_mucosae | FXZ18_RS04780 | 197559..201290 | 97478 | BREX-system-P-loop-protein-BrxC-[Limosilactobacillus-mucosae] |
| WP_047036116.1 | Type1.BREX | [illegible] | 1183 | Bacteria>Terrabacteria_group>Bacillota>Bacilli>Lactobacillales>Lactobacillaceae | GCF_902385835.1 | Ligilactobacillus_salivarius | FYL10_RS06665 | 51760..55311 | 1624 | BREX-system-P-loop-protein-BrxC-[Ligilactobacillus-salivarius] |
| WP_117689516.1 | Type1.BREX | [illegible] | 1243 | Bacteria>Terrabacteria_group>Bacillota>Bacilli>Lactobacillales>Lactobacillaceae | GCF_902399865.1 | Lactobacillus_gasseri | PGU81_RS09215 | 16213..19944 | 1596 | BREX-system-P-loop-protein-BrxC-[Lactobacillus-gasseri] |
| WP_021350006.1 | Type1.BREX | [illegible] | 1230 | Bacteria>Terrabacteria_group>Bacillota>Bacilli>Lactobacillales>Lactobacillaceae | GCF_925284415.1 | Limosilactobacillus_fermentum | QFK68_RS03250 | 14373..18065 | 1613 | BREX-system-P-loop-protein-BrxC-[Limosilactobacillus-fermentum] |
| WP_251920454.1 | Type1.BREX | [illegible] | 1183 | Bacteria>Terrabacteria_group>Bacillota>Bacilli>Lactobacillales>Lactobacillaceae | GCF_925291425.1 | Ligilactobacillus_salivarius | NGI32_RS06440 | 116014..119565 | 1624 | BREX-system-P-loop-protein-BrxC-[Ligilactobacillus-salivarius] |
| WP_064081740.1 | Type1.BREX | [illegible] | 1178 | Bacteria>Terrabacteria_group>Bacillota>Bacilli>Lactobacillales>Lactobacillaceae | GCF_925300675.1 | Lactobacillus_delbrueckii | NGL01_RS07850 | 3981..7517 | 1584 | BREX-system-P-loop-protein-BrxC-[Lactobacillus-delbrueckii] |
| MCG0730673.1 | Type1.BREX | [illegible] | 1213 | Bacteria>Terrabacteria_group>Bacillota>Bacilli>Lactobacillales>Lactobacillaceae | - | Lactiplantibacillus_plantarum | IMAU60171_02928 | 2439..6080 | 1590 | BREX-system-P-loop-protein-BrxC-[Lactiplantibacillus-plantarum] |
| WP_211758382.1 | Type1.BREX | [illegible] | 1057 | Bacteria>Terrabacteria_group>Bacillota>Bacilli>Lactobacillales>Lactobacillaceae | - | Lactiplantibacillus_plantarum | JK218_15530 | 5561..8734 | 1590 | BREX-system-P-loop-protein-BrxC-[Lactiplantibacillus-plantarum] |
| WP_249220179.1 | Type1.BREX | [illegible] | 1047 | Bacteria>Terrabacteria_group>Bacillota>Bacilli>Lactobacillales>Lactobacillaceae | - | Lactiplantibacillus_plantarum | JK218_RS15545 | 5561..8704 | 1590 | BREX-system-P-loop-protein-BrxC-[Lactiplantibacillus-plantarum] |
| WP_021336769.1 | Type1.BREX | [illegible] | 1223 | Bacteria>Terrabacteria_group>Bacillota>Bacilli>Lactobacillales>Lactobacillaceae | - | Lactiplantibacillus_plantarum | N692_10815 | 71429..75100 | 1590 | BREX-system-P-loop-protein-BrxC-[Lactiplantibacillus-plantarum] |
| WP_060472123.1 | Type1.BREX | [illegible] | 1242 | Bacteria>Terrabacteria_group>Bacillota>Bacilli>Lactobacillales>Lactobacillaceae | - | Lactobacillus_gallinarum | AO203_08530 | 1678493..1682221 | 52242 | BREX-system-P-loop-protein-BrxC-[Lactobacillus-gallinarum] |
| WP_236656790.1 | Type1.BREX | [illegible] | 1159 | Bacteria>Terrabacteria_group>Bacillota>Bacilli>Lactobacillales>Lactobacillaceae | - | Lactobacillus_helveticus | LHEW6_RS10210 | 11901..15380 | 1587 | BREX-system-P-loop-protein-BrxC-[Lactobacillus-helveticus] |

| Protein-ID | System | Gene-Neighbourhoods | Length | Taxonomy | Assembly-ID | Strain | Locus-Tag | Genomic-coordinates | TaxID | Fasta-define |
|---|---|---|---|---|---|---|---|---|---|---|
| WP_255262987.1 | Type1.BREX | <-N6-METHYLASE-p<-DUF6079*<-DUF6079-p<-BrxB-BrxF-iSTAND<-BrxA-DNA-binding | 1145 | Bacteria>Terrabacteria_group>[illegible] | [illegible] | [illegible] | LEUCOC11_RS06135 | 461..3898 | 33964 | [illegible] |
| WP_089998518.1 | Type1.BREX | HIN-HTH->HTH_Tnp_1+HTH_21+rve+rve_3->BrxA-DNA-binding->BrxB-BrxF-iSTAND->DUF6079*->N6-METHYLASE-> [illegible] | 1194 | Bacteria>Terrabacteria_group>[illegible] | [illegible] | [illegible] | PB1A_RS01525 | 5425..9009 | 178001 | [illegible] |
| WP_168924054.1 | Type1.BREX | McrB-AAA->McrC-NTD+REase->?->BrxA-DNA-binding->BrxB-BrxF-iSTAND->DUF6079*-><-N6-METHYLASE-p<-?\|\|PglZ-> [illegible] | 1191 | Bacteria>Terrabacteria_group>[illegible] | [illegible] | [illegible] | L3V66_04100 | 164372..167947 | 2722706 | [illegible] |
| COD97965.1 | Type1.BREX | IONCHANNEL+PAS+HISKIN+MORC->REC+CITB-HTH+ZDNA-HTH->?->BrxA-DNA-binding->BrxB-BrxF-iSTAND->DUF6079*-> [illegible] | 1193 | Bacteria>Terrabacteria_group>[illegible] | [illegible] | [illegible] | ERS020515_00205 | 198134..201715 | 1313 | [illegible] |
| WP_058211096.1 | Type1.BREX | <-PglZ<-TOPRIM<-OLD-ABC<-?<-?<-N6-METHYLASE<-DUF6079*<-BrxB-BrxF-iSTAND<-BrxA-DNA-binding<-XIS-HTH<-?<-?<-?<- [illegible] | 1179 | Bacteria>Terrabacteria_group>[illegible] | [illegible] | [illegible] | KF134_RS08810 | 324861..328400 | 1358 | [illegible] |
| WP_191672696.1 | Type1.BREX | Glyoxalase-><-?<-?\|\|?->XIS-HTH->BrxA-DNA-binding->BrxB-BrxF-iSTAND->DUF6079*->N6-METHYLASE->?->?->OLD-ABC-> [illegible] | 1179 | Bacteria>Terrabacteria_group>[illegible] | [illegible] | [illegible] | DXX96_RS03180 | 80359..83898 | 1940789 | [illegible] |
| WP_259747683.1 | Type1.BREX | Phage_integrase->N6-METHYLASE-p-><-?<-?\|\|?->TIR->N6-METHYLASE-p-><-N6-METHYLASE-p<-N6-METHYLASE<-DUF6079*<- [illegible] | 1216 | Bacteria>Terrabacteria_group>[illegible] | [illegible] | [illegible] | EFK39_RS07840 | 16583..20233 | 1358 | [illegible] |
| WP_153973612.1 | Type1.BREX | <-DUF6079*<-BrxB-BrxF-iSTAND<-BrxA-DNA-binding\|\|DEDD_Tnp_IS110->DEDD_Tnp_IS110->Transposase_20-><-?<-SSB<-?<- [illegible] | 1216 | Bacteria>Terrabacteria_group>[illegible] | [illegible] | [illegible] | GKM45_RS02085 | 1086..4736 | 1358 | [illegible] |
| WP_251420892.1 | Type1.BREX | <-PglZ<-TOPRIM<-OLD-ABC<-?<-N6-METHYLASE<-DUF6079*<-BrxB-BrxF-iSTAND<-BrxA-DNA-binding<-XIS-HTH<-?<-?<-?<- [illegible] | 1179 | Bacteria>Terrabacteria_group>[illegible] | [illegible] | [illegible] | NFR64_RS02430 | 42877..46416 | 1357 | [illegible] |
| MCH4057848.1 | Type1.BREX | <-HTH_28+HTH_32\|\|<-?<-ParB+HNH<-SIGMA+OB-Fold+AAA-ATPase+Lon-Protease-BrxL<-PglZ\|\|N6-METHYLASE-> [illegible] | 1214 | Bacteria>Terrabacteria_group>[illegible] | [illegible] | [illegible] | LKE89_08070 | 16697..20341 | 2291323 | [illegible] |
| HHU20134.1 | Type1.BREX | BrxA-DNA-binding->BrxB-BrxF-iSTAND->DUF6079*->N6-METHYLASE->PglZ->?->?->?-><-?<-HAD+HAD<-?\|\|NUDIX-> | 1183 | Bacteria>Terrabacteria_group>[illegible] | [illegible] | [illegible] | GXZ58_07985 | 5866..9417 | 1903720 | [illegible] |
| MBQ8901518.1 | Type1.BREX | <-SIGMA+OB-Fold+AAA-ATPase+Lon-Protease-BrxL<-PglZ<-N6-METHYLASE<-DUF6079* | 1176 | Bacteria>Terrabacteria_group>[illegible] | [illegible] | [illegible] | IJY87_00435 | 8544..12074 | 1903720 | [illegible] |
| MBR6516599.1 | Type1.BREX | N6-METHYLASE-p->BrxA-DNA-binding->BrxB-BrxF-iSTAND->DUF6079*->?->PglZ->SIGMA+OB-Fold+AAA-ATPase+Lon-Protease- [illegible] | 1196 | Bacteria>Terrabacteria_group>[illegible] | [illegible] | [illegible] | IKT40_07050 | 1703..5293 | 1903720 | [illegible] |
| HIQ90970.1 | Type1.BREX | <-ZnR+ZnR<-DUF6079*<-BrxB-BrxF-iSTAND<-BrxA-DNA-binding<-VbhA+DOC+HTH<-GDResolvase+Recombinase+ZnR<-?<-tRNA<- [illegible] | 1175 | Bacteria>Terrabacteria_group>[illegible] | [illegible] | [illegible] | IAB27_05045 | 1414..4941 | 2840752 | [illegible] |
| WP_076917462.1 | Type1.BREX | AT-HTH-HYD2->Tox-ART-HYD2->AT-HTH-HYD2-><-SbcC<-ParB+HNH<-SIGMA+OB-Fold+AAA-ATPase+Lon-Protease-BrxL<-PglZ<- [illegible] | 1178 | Bacteria>Terrabacteria_group>[illegible] | [illegible] | [illegible] | BQ5375_RS04300 | 340320..343856 | 1871035 | [illegible] |
| WP_076917491.1 | Type1.BREX | TraJ-RHH->TraI-VirD2-RCRE->HTH_26->BrxA-DNA-binding->BrxB-BrxF-iSTAND->DUF6079*->N6-METHYLASE->?->PglZ-> [illegible] | 1181 | Bacteria>Terrabacteria_group>[illegible] | [illegible] | [illegible] | BQ5375_RS04570 | 391811..395356 | 1871035 | [illegible] |
| HIZ73445.1 | Type1.BREX | [illegible] | 1196 | Bacteria>Terrabacteria_group>[illegible] | [illegible] | [illegible] | H9964_07670 | 4171..7761 | 2838603 | [illegible] |
| WP_231160542.1 | Type1.BREX | [illegible] | 1215 | Bacteria>Terrabacteria_group>[illegible] | [illegible] | [illegible] | G8S21_07290 | 73653..77300 | 1491 | [illegible] |
| WP_118538325.1 | Type1.BREX | [illegible] | 1205 | Bacteria>Terrabacteria_group>[illegible] | [illegible] | [illegible] | DWX91_03850 | 21550..25167 | 2293004 | [illegible] |
| WP_118669593.1 | Type1.BREX | [illegible] | 1196 | Bacteria>Terrabacteria_group>[illegible] | [illegible] | [illegible] | DXC82_15795 | 32814..36404 | 2293052 | [illegible] |
| WP_152887213.1 | Type1.BREX | [illegible] | 995 | Bacteria>Terrabacteria_group>[illegible] | [illegible] | [illegible] | GBZ86_01880 | 24585..27572 | 39493 | [illegible] |
| CCZ42093.1 | Type1.BREX | [illegible] | 1203 | Bacteria>Terrabacteria_group>[illegible] | [illegible] | [illegible] | BN479_02002 | 2349..5960 | 1262773 | [illegible] |
| CCZ63532.1 | Type1.BREX | [illegible] | 1215 | Bacteria>Terrabacteria_group>[illegible] | [illegible] | [illegible] | BN544_01435 | 55563..59210 | 1263067 | [illegible] |
| CDA58916.1 | Type1.BREX | [illegible] | 1178 | Bacteria>Terrabacteria_group>[illegible] | [illegible] | [illegible] | BN559_00263 | 10316..13852 | 1262784 | [illegible] |
| CDD75188.1 | Type1.BREX | [illegible] | 1200 | Bacteria>Terrabacteria_group>[illegible] | [illegible] | [illegible] | BN737_01375 | 69547..73149 | 1262828 | [illegible] |
| KJS18733.1 | Type1.BREX | [illegible] | 1210 | Bacteria>Terrabacteria_group>[illegible] | [illegible] | [illegible] | VR72_21175 | 10275..13907 | 1629719 | [illegible] |
| ODM26401.1 | Type1.BREX | [illegible] | 1139 | Bacteria>Terrabacteria_group>[illegible] | [illegible] | [illegible] | A7W90_09300 | 2213078..2216497 | 1848158 | [illegible] |
| ODM26889.1 | Type1.BREX | [illegible] | 1186 | Bacteria>Terrabacteria_group>[illegible] | [illegible] | [illegible] | A7W90_12080 | 2831354..2834914 | 1848158 | [illegible] |
| PHS35613.1 | Type1.BREX | [illegible] | 1156 | Bacteria>Terrabacteria_group>[illegible] | [illegible] | [illegible] | COA82_04040 | 61598..65068 | 1872455 | [illegible] |
| RGF28780.1 | Type1.BREX | [illegible] | 1196 | Bacteria>Terrabacteria_group>[illegible] | [illegible] | [illegible] | DW081_14990 | 34449..38039 | 2293020 | [illegible] |
| RHO49165.1 | Type1.BREX | [illegible] | 1205 | Bacteria>Terrabacteria_group>[illegible] | [illegible] | [illegible] | DW115_05780 | 132608..136225 | 2293022 | [illegible] |
| HAR85098.1 | Type1.BREX | [illegible] | 406 | Bacteria>Terrabacteria_group>[illegible] | [illegible] | [illegible] | DCR69_05160 | 4723..5940 | 1506 | [illegible] |
| RHQ86816.1 | Type1.BREX | [illegible] | 1205 | Bacteria>Terrabacteria_group>[illegible] | [illegible] | [illegible] | DWX91_03850 | 21550..25167 | 2293004 | [illegible] |
| RHU61449.1 | Type1.BREX | [illegible] | 1196 | Bacteria>Terrabacteria_group>[illegible] | [illegible] | [illegible] | DXC82_15795 | 32814..36404 | 2293052 | [illegible] |
| RHW04487.1 | Type1.BREX | [illegible] | 1182 | Bacteria>Terrabacteria_group>[illegible] | [illegible] | [illegible] | DXA90_04730 | 56515..60063 | 2292276 | [illegible] |
| HCO74350.1 | Type1.BREX | [illegible] | 1176 | Bacteria>Terrabacteria_group>[illegible] | [illegible] | [illegible] | DIT16_05785 | 6287..9817 | 1506 | [illegible] |
| MPQ42518.1 | Type1.BREX | [illegible] | 995 | Bacteria>Terrabacteria_group>[illegible] | [illegible] | [illegible] | GBZ86_01880 | 24585..27572 | 39493 | [illegible] |
| NBG87676.1 | Type1.BREX | [illegible] | 1195 | Bacteria>Terrabacteria_group>[illegible] | [illegible] | [illegible] | ISALK_04100 | 132779..136366 | 2565777 | [illegible] |
| NBH35981.1 | Type1.BREX | [illegible] | 1182 | Bacteria>Terrabacteria_group>[illegible] | [illegible] | [illegible] | D3Z58_21210 | 19493..23041 | 1898204 | [illegible] |
| NBH17641.1 | Type1.BREX | [illegible] | 1182 | Bacteria>Terrabacteria_group>[illegible] | [illegible] | [illegible] | D3Z55_09200 | 40728..44276 | 1898204 | [illegible] |
| NBH73079.1 | Type1.BREX | [illegible] | 1212 | Bacteria>Terrabacteria_group>[illegible] | [illegible] | [illegible] | D3Z51_13830 | 3930..7568 | 1898204 | [illegible] |
| NBI69485.1 | Type1.BREX | [illegible] | 1212 | Bacteria>Terrabacteria_group>[illegible] | [illegible] | [illegible] | D3Z50_00070 | 19913..23551 | 1898204 | [illegible] |
| NFG40378.1 | Type1.BREX | [illegible] | 1215 | Bacteria>Terrabacteria_group>[illegible] | [illegible] | [illegible] | FC789_04035 | 879792..883439 | 1491 | [illegible] |
| NFI95403.1 | Type1.BREX | [illegible] | 1215 | Bacteria>Terrabacteria_group>[illegible] | [illegible] | [illegible] | FC961_13560 | 15228..18875 | 1491 | [illegible] |
| NFH89561.1 | Type1.BREX | [illegible] | 1195 | Bacteria>Terrabacteria_group>[illegible] | [illegible] | [illegible] | FDA33_04900 | 24561..28148 | 1491 | [illegible] |
| NFO12999.1 | Type1.BREX | [illegible] | 1215 | Bacteria>Terrabacteria_group>[illegible] | [illegible] | [illegible] | FDB34_02025 | 196714..200361 | 1491 | [illegible] |
| NLE27040.1 | Type1.BREX | [illegible] | 1147 | Bacteria>Terrabacteria_group>[illegible] | [illegible] | [illegible] | GX625_17200 | 20680..24123 | 1898204 | [illegible] |
| NLN64679.1 | Type1.BREX | [illegible] | 1174 | Bacteria>Terrabacteria_group>[illegible] | [illegible] | [illegible] | GX144_04605 | 4627..8151 | 1898204 | [illegible] |
| NMA18734.1 | Type1.BREX | [illegible] | 1198 | Bacteria>Terrabacteria_group>[illegible] | [illegible] | [illegible] | GX939_08350 | 9727..13323 | 1898204 | [illegible] |
| HHU54035.1 | Type1.BREX | [illegible] | 1204 | Bacteria>Terrabacteria_group>[illegible] | [illegible] | [illegible] | GXZ43_08205 | 1499..5113 | 1898204 | [illegible] |
| HHW31257.1 | Type1.BREX | [illegible] | 1186 | Bacteria>Terrabacteria_group>[illegible] | [illegible] | [illegible] | GXX20_06235 | 1900..5460 | 1898204 | [illegible] |
| MBE5721948.1 | Type1.BREX | [illegible] | 1182 | Bacteria>Terrabacteria_group>[illegible] | [illegible] | [illegible] | EGR31_00720 | 13034..16582 | 1506 | [illegible] |
| MBE6059117.1 | Type1.BREX | [illegible] | 1176 | Bacteria>Terrabacteria_group>[illegible] | [illegible] | [illegible] | E7215_02940 | 4923..8453 | 318464 | [illegible] |
| HAT4275166.1 | Type1.BREX | [illegible] | 1221 | Bacteria>Terrabacteria_group>[illegible] | [illegible] | [illegible] | I9084_000700 | 10095..13760 | 1502 | [illegible] |
| EGT3599215.1 | Type1.BREX | [illegible] | 1221 | Bacteria>Terrabacteria_group>[illegible] | [illegible] | [illegible] | FT888_03085 | 92913..96578 | 1502 | [illegible] |
| MBN1067552.1 | Type1.BREX | [illegible] | 1215 | Bacteria>Terrabacteria_group>[illegible] | [illegible] | [illegible] | DVW02_06040 | 97050..100697 | 1491 | [illegible] |
| EHK2362606.1 | Type1.BREX | [illegible] | 1221 | Bacteria>Terrabacteria_group>[illegible] | [illegible] | [illegible] | KCL45_000419 | 449510..453175 | 1502 | [illegible] |
| QUO21490.1 | Type1.BREX | [illegible] | 1182 | Bacteria>Terrabacteria_group>[illegible] | [illegible] | [illegible] | KFE18_13085 | 2693065..2696613 | 2830657 | [illegible] |
| MBS5468798.1 | Type1.BREX | [illegible] | 1178 | Bacteria>Terrabacteria_group>[illegible] | [illegible] | [illegible] | KHX69_08270 | 35146..38682 | 1506 | [illegible] |
| MBS5781623.1 | Type1.BREX | [illegible] | 1171 | Bacteria>Terrabacteria_group>[illegible] | [illegible] | [illegible] | KID04_01935 | 432250..435765 | 1506 | [illegible] |
| MBS5995886.1 | Type1.BREX | [illegible] | 1221 | Bacteria>Terrabacteria_group>[illegible] | [illegible] | [illegible] | KIB05_13835 | 52036..55701 | 1502 | [illegible] |
| MBS6503618.1 | Type1.BREX | [illegible] | 1172 | Bacteria>Terrabacteria_group>[illegible] | [illegible] | [illegible] | KH415_18730 | 1781..5299 | 1506 | [illegible] |
| MBS6600456.1 | Type1.BREX | [illegible] | 1198 | Bacteria>Terrabacteria_group>[illegible] | [illegible] | [illegible] | KH333_05500 | 22181..25777 | 1506 | [illegible] |
| MBS7187675.1 | Type1.BREX | [illegible] | 994 | Bacteria>Terrabacteria_group>[illegible] | [illegible] | [illegible] | KH070_11040 | 23892..26873 | 1506 | [illegible] |
| MCC2165352.1 | Type1.BREX | [illegible] | 1217 | Bacteria>Terrabacteria_group>[illegible] | [illegible] | [illegible] | LKD32_10815 | 1974..5627 | 2885361 | [illegible] |
| MCD3245748.1 | Type1.BREX | [illegible] | 1215 | Bacteria>Terrabacteria_group>[illegible] | [illegible] | [illegible] | G8S21_07290 | 73653..77300 | 36828 | [illegible] |
| MCI6655647.1 | Type1.BREX | [illegible] | 1220 | Bacteria>Terrabacteria_group>[illegible] | [illegible] | [illegible] | MSH21_10420 | 3289..6951 | 1506 | [illegible] |
| MCI7419995.1 | Type1.BREX | [illegible] | 482 | Bacteria>Terrabacteria_group>[illegible] | [illegible] | [illegible] | MSS65_10400 | 1..1448 | 1506 | [illegible] |
| MCI7503208.1 | Type1.BREX | [illegible] | 1182 | Bacteria>Terrabacteria_group>[illegible] | [illegible] | [illegible] | MSA76_06060 | 12904..16452 | 1506 | [illegible] |
| MCI8726399.1 | Type1.BREX | [illegible] | 1179 | Bacteria>Terrabacteria_group>[illegible] | [illegible] | [illegible] | HFG66_09835 | 51517..55056 | 2613924 | [illegible] |
| MCI8983480.1 | Type1.BREX | [illegible] | 1182 | Bacteria>Terrabacteria_group>[illegible] | [illegible] | [illegible] | HFG78_13950 | 16181..19729 | 2613924 | [illegible] |
| MCI9069232.1 | Type1.BREX | [illegible] | 1204 | Bacteria>Terrabacteria_group>[illegible] | [illegible] | [illegible] | HFJ14_01010 | 2819..6433 | 1506 | [illegible] |
| MCI9636406.1 | Type1.BREX | [illegible] | 1182 | Bacteria>Terrabacteria_group>[illegible] | [illegible] | [illegible] | HFG77_08430 | 22407..25955 | 2613924 | [illegible] |
| HCG3022573.1 | Type1.BREX | [illegible] | 1221 | Bacteria>Terrabacteria_group>[illegible] | [illegible] | [illegible] | NJU98_002507 | 52406..56071 | 1502 | [illegible] |
| EJO5349026.1 | Type1.BREX | [illegible] | 1214 | Bacteria>Terrabacteria_group>[illegible] | [illegible] | [illegible] | NRP93_003178 | 1627..5271 | 1491 | [illegible] |
| UUM12003.1 | Type1.BREX | [illegible] | 1192 | Bacteria>Terrabacteria_group>[illegible] | [illegible] | [illegible] | NQU17_00105 | 23667..27245 | 2961947 | [illegible] |
| EJT5935287.1 | Type1.BREX | [illegible] | 1221 | Bacteria>Terrabacteria_group>[illegible] | [illegible] | [illegible] | N2W52_000012 | 19602..23267 | 1502 | [illegible] |
| EJT6164547.1 | Type1.BREX | [illegible] | 1221 | Bacteria>Terrabacteria_group>[illegible] | [illegible] | [illegible] | N2W22_000999 | 1099443..1103108 | 1502 | [illegible] |
| EJT6169494.1 | Type1.BREX | [illegible] | 1221 | Bacteria>Terrabacteria_group>[illegible] | [illegible] | [illegible] | N2W53_000036 | 52483..56148 | 1502 | [illegible] |
| EJT6540217.1 | Type1.BREX | [illegible] | 1221 | Bacteria>Terrabacteria_group>[illegible] | [illegible] | [illegible] | N2W31_000036 | 52406..56071 | 1502 | [illegible] |
| WP_208078150.1 | Type1.BREX | [illegible] | 1216 | Bacteria>Terrabacteria_group>[illegible] | [illegible] | [illegible] | HJU46_RS14910 | 6641..10291 | 1492 | [illegible] |
| SCG90365.1 | Type1.BREX | [illegible] | 527 | Bacteria>Terrabacteria_group>[illegible] | [illegible] | [illegible] | SAMEA3545239_03586 | 14803..16386 | 59620 | [illegible] |
| SCH67534.1 | Type1.BREX | [illegible] | 946 | Bacteria>Terrabacteria_group>[illegible] | [illegible] | [illegible] | SAMEA3545303_02047 | 234904..237744 | 59620 | [illegible] |
| VDG73855.1 | Type1.BREX | [illegible] | 967 | Bacteria>Terrabacteria_group>[illegible] | [illegible] | [illegible] | NCTC10913_04233 | 180980..183883 | 1530 | [illegible] |
| CAB1245762.1 | Type1.BREX | [illegible] | 1188 | Bacteria>Terrabacteria_group>[illegible] | [illegible] | [illegible] | CLOSBL3_11249 | 1215362..1218928 | 2799563 | [illegible] |
| WP_013239900.1 | Type1.BREX | [illegible] | 1188 | Bacteria>Terrabacteria_group>[illegible] | [illegible] | [illegible] | CLJU_RS16115 | 3575485..3579051 | 1538 | [illegible] |
| WP_003411890.1 | Type1.BREX | [illegible] | 1215 | Bacteria>Terrabacteria_group>[illegible] | [illegible] | [illegible] | CBY_RS17320 | 4352..7999 | 1492 | [illegible] |
| WP_003452047.1 | Type1.BREX | [illegible] | 1221 | Bacteria>Terrabacteria_group>[illegible] | [illegible] | [illegible] | CPC_RS09775 | 28796..32461 | 1502 | [illegible] |
| WP_003473497.1 | Type1.BREX | [illegible] | 1221 | Bacteria>Terrabacteria_group>[illegible] | [illegible] | [illegible] | CJD_RS01695 | 75268..78933 | 1502 | [illegible] |
| WP_013977767.1 | Type1.BREX | [illegible] | 1214 | Bacteria>Terrabacteria_group>[illegible] | [illegible] | [illegible] | CXIVA_RS09290 | 1976603..1980247 | 1042156 | [illegible] |
| WP_002598576.1 | Type1.BREX | [illegible] | 1208 | Bacteria>Terrabacteria_group>[illegible] | [illegible] | [illegible] | HMPREF1092_RS10115 | 2173864..2177490 | 29372 | [illegible] |
| WP_002599393.1 | Type1.BREX | [illegible] | 1218 | Bacteria>Terrabacteria_group>[illegible] | [illegible] | [illegible] | HMPREF1092_RS14350 | 173524..177180 | 29372 | [illegible] |
| WP_023388008.1 | Type1.BREX | [illegible] | 1166 | Bacteria>Terrabacteria_group>[illegible] | [illegible] | [illegible] | T472_RS11260 | 54891..58391 | 1408819 | [illegible] |
| WP_044037496.1 | Type1.BREX | [illegible] | 1185 | Bacteria>Terrabacteria_group>[illegible] | [illegible] | [illegible] | CM240_RS06275 | 1424237..1427794 | 1216932 | [illegible] |
| WP_035381181.1 | Type1.BREX | [illegible] | 1193 | Bacteria>Terrabacteria_group>[illegible] | [illegible] | [illegible] | Q428_RS11710 | 3478..7059 | 655338 | [illegible] |
| WP_029450765.1 | Type1.BREX | [illegible] | 1139 | Bacteria>Terrabacteria_group>[illegible] | [illegible] | [illegible] | BV55_RS0100675 | 144014..147433 | 37659 | [illegible] |
| WP_076044950.1 | Type1.BREX | [illegible] | 1215 | Bacteria>Terrabacteria_group>[illegible] | [illegible] | [illegible] | B2H88_RS17345 | 150245..153892 | 1491 | [illegible] |
| WP_035131431.1 | Type1.BREX | [illegible] | 1176 | Bacteria>Terrabacteria_group>[illegible] | [illegible] | [illegible] | IO99_RS06165 | 55628..59158 | 318464 | [illegible] |
| WP_039258635.1 | Type1.BREX | [illegible] | 1214 | Bacteria>Terrabacteria_group>[illegible] | [illegible] | [illegible] | ADU88_RS01575 | 72725..76369 | 1491 | [illegible] |
| WP_043662653.1 | Type1.BREX | [illegible] | 1216 | Bacteria>Terrabacteria_group>[illegible] | [illegible] | [illegible] | AWN73_RS08010 | 18592..22242 | 1492 | [illegible] |
| WP_152648993.1 | Type1.BREX | [illegible] | 946 | Bacteria>Terrabacteria_group>[illegible] | [illegible] | [illegible] | CLFS41_RS01500 | 248814..251654 | 1609975 | [illegible] |
| WP_060670433.1 | Type1.BREX | [illegible] | 1221 | Bacteria>Terrabacteria_group>[illegible] | [illegible] | [illegible] | FORC3_RS07365 | 1728066..1731731 | 1502 | [illegible] |
| WP_058953242.1 | Type1.BREX | [illegible] | 1206 | Bacteria>Terrabacteria_group>[illegible] | [illegible] | [illegible] | BN3431_RS06425 | 17547..21167 | 1519 | [illegible] |
| WP_089984090.1 | Type1.BREX | [illegible] | 1183 | Bacteria>Terrabacteria_group>[illegible] | [illegible] | [illegible] | BN3456_RS10470 | 25187..28738 | 1776045 | [illegible] |
| WP_061995750.1 | Type1.BREX | [illegible] | 1190 | Bacteria>Terrabacteria_group>[illegible] | [illegible] | [illegible] | AZI32_RS10280 | 12953..16525 | 1676991 | [illegible] |
| WP_065077880.1 | Type1.BREX | [illegible] | 1188 | Bacteria>Terrabacteria_group>[illegible] | [illegible] | [illegible] | CLRAG_RS07695 | 80363..83929 | 217158 | [illegible] |
| WP_070965028.1 | Type1.BREX | [illegible] | 1193 | Bacteria>Terrabacteria_group>[illegible] | [illegible] | [illegible] | BJL90_RS05380 | 1160755..1164336 | 1497 | [illegible] |
| WP_128609922.1 | Type1.BREX | [illegible] | 1190 | Bacteria>Terrabacteria_group>[illegible] | [illegible] | [illegible] | AXF41_RS14000 | 8813..12385 | 84025 | [illegible] |
| WP_077851369.1 | Type1.BREX | [illegible] | 1216 | Bacteria>Terrabacteria_group>[illegible] | [illegible] | [illegible] | CLAUR_RS09645 | 2191072..2194722 | 36839 | [illegible] |
| WP_078025149.1 | Type1.BREX | [illegible] | 1380 | Bacteria>Terrabacteria_group>[illegible] | [illegible] | [illegible] | BS637_RS12490 | 20116..24258 | 1962263 | [illegible] |
| WP_078054348.1 | Type1.BREX | [illegible] | 1197 | Bacteria>Terrabacteria_group>[illegible] | [illegible] | [illegible] | BS638_RS00485 | 111870..115463 | 1962263 | [illegible] |
| WP_087022170.1 | Type1.BREX | [illegible] | 1214 | Bacteria>Terrabacteria_group>[illegible] | [illegible] | [illegible] | CBW42_RS12580 | 21770..25414 | 1945634 | [illegible] |
| WP_087414865.1 | Type1.BREX | [illegible] | 1211 | Bacteria>Terrabacteria_group>[illegible] | [illegible] | [illegible] | B5F15_RS07050 | 102809..106444 | 501571 | [illegible] |
| WP_103227376.1 | Type1.BREX | [illegible] | 1215 | Bacteria>Terrabacteria_group>[illegible] | [illegible] | [illegible] | C0033_RS08180 | 119935..123582 | 2067550 | [illegible] |
| WP_106010897.1 | Type1.BREX | [illegible] | 1203 | Bacteria>Terrabacteria_group>[illegible] | [illegible] | [illegible] | CLLU_RS16660 | 4389..8000 | 1691940 | [illegible] |
| WP_117552964.1 | Type1.BREX | [illegible] | 1217 | Bacteria>Terrabacteria_group>[illegible] | [illegible] | [illegible] | DW826_RS13215 | 39134..42787 | 2305242 | [illegible] |
| WP_118320912.1 | Type1.BREX | [illegible] | 1183 | Bacteria>Terrabacteria_group>[illegible] | [illegible] | [illegible] | DW260_RS00800 | 177692..181243 | 2292009 | [illegible] |
| WP_118442699.1 | Type1.BREX | [illegible] | 1182 | Bacteria>Terrabacteria_group>[illegible] | [illegible] | [illegible] | DW019_RS04810 | 72641..76189 | 2293016 | [illegible] |
| WP_118513467.1 | Type1.BREX | [illegible] | 1183 | Bacteria>Terrabacteria_group>[illegible] | [illegible] | [illegible] | DW020_RS00895 | 209396..212947 | 2292007 | [illegible] |
| WP_118467570.1 | Type1.BREX | [illegible] | 1182 | Bacteria>Terrabacteria_group>[illegible] | [illegible] | [illegible] | DWZ45_RS03230 | 701093..704641 | 2614128 | [illegible] |
| WP_118532764.1 | Type1.BREX | [illegible] | 1217 | Bacteria>Terrabacteria_group>[illegible] | [illegible] | [illegible] | DWY84_RS09695 | 65896..69549 | 2292206 | [illegible] |
| WP_118728541.1 | Type1.BREX | [illegible] | 1217 | Bacteria>Terrabacteria_group>[illegible] | [illegible] | [illegible] | DW954_RS15865 | 16382..20035 | 2292306 | [illegible] |
| WP_118731364.1 | Type1.BREX | [illegible] | 1196 | Bacteria>Terrabacteria_group>[illegible] | [illegible] | [illegible] | DW835_RS14805 | 20266..23856 | 2293030 | [illegible] |
| WP_118740431.1 | Type1.BREX | [illegible] | 1182 | Bacteria>Terrabacteria_group>[illegible] | [illegible] | [illegible] | DW807_RS02880 | 180384..183932 | 2293029 | [illegible] |
| WP_118740660.1 | Type1.BREX | [illegible] | 1182 | Bacteria>Terrabacteria_group>[illegible] | [illegible] | [illegible] | DW807_RS07265 | 17813..21361 | 2293029 | [illegible] |
| WP_118742502.1 | Type1.BREX | [illegible] | 1182 | Bacteria>Terrabacteria_group>[illegible] | [illegible] | [illegible] | DW739_RS01385 | 291245..294793 | 2293027 | [illegible] |
| WP_118744885.1 | Type1.BREX | [illegible] | 1203 | Bacteria>Terrabacteria_group>[illegible] | [illegible] | [illegible] | DW721_RS03885 | 61324..64935 | 2293026 | [illegible] |
| WP_118670884.1 | Type1.BREX | [illegible] | 1196 | Bacteria>Terrabacteria_group>[illegible] | [illegible] | [illegible] | DXC24_RS05235 | 319406..322996 | 2293049 | [illegible] |
| WP_118650255.1 | Type1.BREX | [illegible] | 1182 | Bacteria>Terrabacteria_group>[illegible] | [illegible] | [illegible] | DXA91_RS01310 | 45088..48636 | 2292309 | [illegible] |
| WP_119970230.1 | Type1.BREX | [illegible] | 1207 | Bacteria>Terrabacteria_group>[illegible] | [illegible] | [illegible] | D4Z93_RS02740 | 562843..566466 | 2068654 | [illegible] |
| WP_122060107.1 | Type1.BREX | [illegible] | 1206 | Bacteria>Terrabacteria_group>[illegible] | [illegible] | [illegible] | D9040_RS18360 | 46156..49776 | 84023 | [illegible] |
| WP_129032313.1 | Type1.BREX | [illegible] | 1211 | Bacteria>Terrabacteria_group>[illegible] | [illegible] | [illegible] | DP122_RS12500 | 4965..8600 | 1513 | [illegible] |
| WP_129133083.1 | Type1.BREX | [illegible] | 1211 | Bacteria>Terrabacteria_group>[illegible] | [illegible] | [illegible] | DP154_01650 | 337342..340977 | 1513 | [illegible] |
| WP_129029509.1 | Type1.BREX | [illegible] | 1211 | Bacteria>Terrabacteria_group>[illegible] | [illegible] | [illegible] | DP139_RS14050 | 25791..29426 | 1513 | [illegible] |
| WP_130100563.1 | Type1.BREX | [illegible] | 1211 | Bacteria>Terrabacteria_group>[illegible] | [illegible] | [illegible] | DP144_RS01650 | 337398..341033 | 1513 | [illegible] |
| WP_132241425.1 | Type1.BREX | [illegible] | 1215 | Bacteria>Terrabacteria_group>[illegible] | [illegible] | [illegible] | EV214_RS00010 | 458..4105 | 2018667 | [illegible] |
| WP_035765204.1 | Type1.BREX | [illegible] | 1216 | Bacteria>Terrabacteria_group>[illegible] | [illegible] | [illegible] | FF104_RS11880 | 2636649..2640299 | 1492 | [illegible] |
| WP_156211829.1 | Type1.BREX | [illegible] | 1216 | Bacteria>Terrabacteria_group>[illegible] | [illegible] | [illegible] | GND98_RS05145 | 54704..58354 | 1492 | [illegible] |
| WP_160686263.1 | Type1.BREX | [illegible] | 1197 | Bacteria>Terrabacteria_group>[illegible] | [illegible] | [illegible] | GXK56_RS06185 | 118154..121747 | 2698832 | [illegible] |
| WP_164956997.1 | Type1.BREX | [illegible] | 1189 | Bacteria>Terrabacteria_group>[illegible] | [illegible] | [illegible] | G9F72_RS01180 | 248731..252300 | 238834 | [illegible] |
| WP_164958329.1 | Type1.BREX | [illegible] | 1363 | Bacteria>Terrabacteria_group>[illegible] | [illegible] | [illegible] | G9F72_RS19190 | 4031148..4035239 | 238834 | [illegible] |
| WP_164949146.1 | Type1.BREX | [illegible] | 1215 | Bacteria>Terrabacteria_group>[illegible] | [illegible] | [illegible] | G9F71_RS17965 | 3736090..3739737 | 2724481 | [illegible] |
| WP_096635663.1 | Type1.BREX | [illegible] | 1196 | Bacteria>Terrabacteria_group>[illegible] | [illegible] | [illegible] | HF847_RS11310 | 7987..11577 | 1494 | [illegible] |

| Protein-ID | System | Gene-Neighbourhoods | Length | Taxonomy | Assembly-ID | Strain | Locus-Tag | Genomic-coordinates | TaxID | Fasta-defline |
|---|---|---|---|---|---|---|---|---|---|---|
| WP_027639925.1 | Type1.BREX | [illegible] | 1214 | Bacteria>Terrabacteria_group>Bacillota>Clostridia>Eubacteriales>Clostridiaceae | GCF_012844035.1 | Clostridium_cadaveris | HF846_RS03710 | 105265..108909 | 1529 | BREX-system-P-loop-protein-BrxC-[Clostridium-cadaveris] |
| WP_169298298.1 | Type1.BREX | [illegible] | 1215 | Bacteria>Terrabacteria_group>Bacillota>Clostridia>Eubacteriales>Clostridiaceae | GCF_012926525.1 | Clostridium_muellerianum | HBE96_RS13730 | 164710..168357 | 2716538 | BREX-system-P-loop-protein-BrxC-[Clostridium-muellerianum] |
| WP_171786495.1 | Type1.BREX | [illegible] | 1216 | Bacteria>Terrabacteria_group>Bacillota>Clostridia>Eubacteriales>Clostridiaceae | GCF_013149405.1 | Clostridium_beijerinckii | B0H39_RS13645 | 3014066..3017716 | 1520 | BREX-system-P-loop-protein-BrxC-[Clostridium-beijerinckii] |
| WP_077854163.1 | Type1.BREX | [illegible] | 1216 | Bacteria>Terrabacteria_group>Bacillota>Clostridia>Eubacteriales>Clostridiaceae | GCF_013297045.1 | Clostridium_beijerinckii | BCD95_RS19475 | 4461966..4465616 | 1520 | BREX-system-P-loop-protein-BrxC-[Clostridium-beijerinckii] |
| WP_178312104.1 | Type1.BREX | [illegible] | 1218 | Bacteria>Terrabacteria_group>Bacillota>Clostridia>Eubacteriales>Clostridiaceae | GCF_013390975.1 | Clostridium_cadaveris | HXZ52_RS14490 | 15645..19301 | 1529 | BREX-system-P-loop-protein-BrxC-[Clostridium-cadaveris] |
| WP_182089446.1 | Type1.BREX | [illegible] | 1192 | Bacteria>Terrabacteria_group>Bacillota>Clostridia>Eubacteriales>Clostridiaceae | GCF_014050525.1 | Clostridium_sp._cel8 | H2684_RS10970 | 21279..24857 | 2663123 | BREX-system-P-loop-protein-BrxC-[Clostridium-sp.-cel8] |
| WP_185162822.1 | Type1.BREX | [illegible] | 1215 | Bacteria>Terrabacteria_group>Bacillota>Clostridia>Eubacteriales>Clostridiaceae | GCF_014212415.1 | Clostridium_algidicarnis | H7D93_RS12595 | 17014..20661 | 37659 | BREX-system-P-loop-protein-BrxC-[Clostridium-algidicarnis] |
| WP_173680574.1 | Type1.BREX | [illegible] | 1216 | Bacteria>Terrabacteria_group>Bacillota>Clostridia>Eubacteriales>Clostridiaceae | GCF_014230485.1 | Clostridium_tetanomorphum | FHX96_RS20165 | 4177507..4181157 | 1553 | BREX-system-P-loop-protein-BrxC-[Clostridium-tetanomorphum] |
| WP_171780276.1 | Type1.BREX | [illegible] | 1216 | Bacteria>Terrabacteria_group>Bacillota>Clostridia>Eubacteriales>Clostridiaceae | GCF_014230655.1 | Clostridium_beijerinckii | HGI39_RS06505 | 40889..44539 | 1520 | BREX-system-P-loop-protein-BrxC-[Clostridium-beijerinckii] |
| WP_191597330.1 | Type1.BREX | [illegible] | 1215 | Bacteria>Terrabacteria_group>Bacillota>Clostridia>Eubacteriales>Clostridiaceae | GCF_014805305.1 | Clostridium_botulinum | HYI18_RS06240 | 53962..57609 | 1491 | BREX-system-P-loop-protein-BrxC-[Clostridium-botulinum] |
| WP_191748096.1 | Type1.BREX | [illegible] | 1195 | Bacteria>Terrabacteria_group>Bacillota>Clostridia>Eubacteriales>Clostridiaceae | GCF_014836325.1 | Clostridium_gallinarum | H9660_RS02165 | 39157..42744 | 2762246 | BREX-system-P-loop-protein-BrxC-[Clostridium-gallinarum] |
| WP_195271030.1 | Type1.BREX | [illegible] | 1217 | Bacteria>Terrabacteria_group>Bacillota>Clostridia>Eubacteriales>Clostridiaceae | GCF_015549545.1 | Clostridium_paraputrificum | I2B30_RS05530 | 47168..50821 | 29363 | BREX-system-P-loop-protein-BrxC-[Clostridium-paraputrificum] |
| WP_195433362.1 | Type1.BREX | [illegible] | 1196 | Bacteria>Terrabacteria_group>Bacillota>Clostridia>Eubacteriales>Clostridiaceae | GCF_015555735.1 | Clostridium_perfringens | I2E72_RS09195 | 116528..120118 | 1502 | BREX-system-P-loop-protein-BrxC-[Clostridium-perfringens] |
| WP_195540766.1 | Type1.BREX | [illegible] | 1182 | Bacteria>Terrabacteria_group>Bacillota>Clostridia>Eubacteriales>Clostridiaceae | GCF_015558125.1 | Clostridium_sp._D43t1_170807_D5 | I2G39_RS02195 | 465793..469341 | 2787150 | BREX-system-P-loop-protein-BrxC-[Clostridium-sp.-D43t1_170807_D5] |
| WP_195958585.1 | Type1.BREX | [illegible] | 1216 | Bacteria>Terrabacteria_group>Bacillota>Clostridia>Eubacteriales>Clostridiaceae | GCF_015667735.1 | Clostridium_butyricum | I4V18_RS19095 | 6839..10489 | 1492 | BREX-system-P-loop-protein-BrxC-[Clostridium-butyricum] |
| WP_115613438.1 | Type1.BREX | [illegible] | 1221 | Bacteria>Terrabacteria_group>Bacillota>Clostridia>Eubacteriales>Clostridiaceae | GCF_016027155.1 | Clostridium_perfringens | I6G61_RS10175 | 2178205..2181870 | 1502 | BREX-system-P-loop-protein-BrxC-[Clostridium-perfringens] |
| WP_198605367.1 | Type1.BREX | [illegible] | 1221 | Bacteria>Terrabacteria_group>Bacillota>Clostridia>Eubacteriales>Clostridiaceae | GCF_016236125.1 | Clostridium_perfringens | H8J62_RS02875 | 34894..38559 | 1502 | BREX-system-P-loop-protein-BrxC-[Clostridium-perfringens] |
| WP_198621144.1 | Type1.BREX | [illegible] | 1221 | Bacteria>Terrabacteria_group>Bacillota>Clostridia>Eubacteriales>Clostridiaceae | GCF_016236545.1 | Clostridium_perfringens | H8J92_RS02870 | 79478..83143 | 1502 | BREX-system-P-loop-protein-BrxC-[Clostridium-perfringens] |
| WP_198616415.1 | Type1.BREX | [illegible] | 1221 | Bacteria>Terrabacteria_group>Bacillota>Clostridia>Eubacteriales>Clostridiaceae | GCF_016236635.1 | Clostridium_perfringens | H8J95_RS08945 | 20074..23739 | 1502 | BREX-system-P-loop-protein-BrxC-[Clostridium-perfringens] |
| WP_085295441.1 | Type1.BREX | [illegible] | 1195 | Bacteria>Terrabacteria_group>Bacillota>Clostridia>Eubacteriales>Clostridiaceae | GCF_016925995.1 | Clostridium_botulinum | DVV95_11085 | 64949..68536 | 1491 | BREX-system-P-loop-protein-BrxC-[Clostridium-botulinum] |
| WP_207130137.1 | Type1.BREX | [illegible] | 1215 | Bacteria>Terrabacteria_group>Bacillota>Clostridia>Eubacteriales>Clostridiaceae | GCF_017330945.1 | Clostridium_botulinum | EXQ34_RS04235 | 14589..18236 | 1491 | BREX-system-P-loop-protein-BrxC-[Clostridium-botulinum] |
| WP_208337082.1 | Type1.BREX | [illegible] | 1196 | Bacteria>Terrabacteria_group>Bacillota>Clostridia>Eubacteriales>Clostridiaceae | GCF_017590635.1 | Clostridium_perfringens | JJB73_RS09180 | 122575..126165 | 1502 | BREX-system-P-loop-protein-BrxC-[Clostridium-perfringens] |
| WP_209702448.1 | Type1.BREX | [illegible] | 1377 | Bacteria>Terrabacteria_group>Bacillota>Clostridia>Eubacteriales>Clostridiaceae | GCF_017874415.1 | Clostridium_algifaecis | J2Z42_RS09735 | 1889..6022 | 1472040 | hypothetical-protein-[Clostridium-algifaecis] |
| WP_164804802.1 | Type1.BREX | [illegible] | 1221 | Bacteria>Terrabacteria_group>Bacillota>Clostridia>Eubacteriales>Clostridiaceae | GCF_018135545.1 | Clostridium_perfringens | G6Z22_RS06045 | 21931..25596 | 1502 | BREX-system-P-loop-protein-BrxC-[Clostridium-perfringens] |
| WP_216131592.1 | Type1.BREX | [illegible] | 1215 | Bacteria>Terrabacteria_group>Bacillota>Clostridia>Eubacteriales>Clostridiaceae | GCF_018861395.1 | Clostridium_algidicarnis | KPL27_RS05405 | 140711..144358 | 37659 | BREX-system-P-loop-protein-BrxC-[Clostridium-algidicarnis] |
| WP_216114827.1 | Type1.BREX | [illegible] | 1215 | Bacteria>Terrabacteria_group>Bacillota>Clostridia>Eubacteriales>Clostridiaceae | GCF_018861505.1 | Clostridium_algidicarnis | KPL28_RS10550 | 250266..253913 | 37659 | BREX-system-P-loop-protein-BrxC-[Clostridium-algidicarnis] |
| WP_216309297.1 | Type1.BREX | [illegible] | 1362 | Bacteria>Terrabacteria_group>Bacillota>Clostridia>Eubacteriales>Clostridiaceae | GCF_018861735.1 | Clostridium_lacusfryxellense | KPL55_RS24035 | 5522..9610 | 205328 | hypothetical-protein-[Clostridium-lacusfryxellense] |
| WP_216137994.1 | Type1.BREX | [illegible] | 1203 | Bacteria>Terrabacteria_group>Bacillota>Clostridia>Eubacteriales>Clostridiaceae | GCF_018861855.1 | Clostridium_estertheticum | KPL49_RS19700 | 51196..54807 | 238834 | BREX-system-P-loop-protein-BrxC-[Clostridium-estertheticum] |
| WP_216464187.1 | Type1.BREX | [illegible] | 1210 | Bacteria>Terrabacteria_group>Bacillota>Clostridia>Eubacteriales>Clostridiaceae | GCF_018917145.1 | Clostridium_bornimense | KQI30_RS15155 | 1337..4969 | 1216932 | BREX-system-P-loop-protein-BrxC-[Clostridium-bornimense] |
| WP_216574650.1 | Type1.BREX | [illegible] | 1182 | Bacteria>Terrabacteria_group>Bacillota>Clostridia>Eubacteriales>Clostridiaceae | GCF_018918265.1 | Falcatimonas_sp._MSJ-15 | KQI85_RS03175 | 188313..191861 | 2841515 | BREX-system-P-loop-protein-BrxC-[Falcatimonas-sp.-MSJ-15] |
| WP_217303321.1 | Type1.BREX | [illegible] | 1188 | Bacteria>Terrabacteria_group>Bacillota>Clostridia>Eubacteriales>Clostridiaceae | GCF_019056555.1 | Clostridium_sp._001 | B5S50_RS05665 | 1268220..1271786 | 1970093 | BREX-system-P-loop-protein-BrxC-[Clostridium-sp.-001] |
| WP_221260245.1 | Type1.BREX | [illegible] | 1218 | Bacteria>Terrabacteria_group>Bacillota>Clostridia>Eubacteriales>Clostridiaceae | GCF_019705275.1 | Clostridium_chauvoei | K4H38_08460 | 40416..44072 | 46867 | BREX-system-P-loop-protein-BrxC-[Clostridium-chauvoei] |
| WP_053340835.1 | Type1.BREX | [illegible] | 1215 | Bacteria>Terrabacteria_group>Bacillota>Clostridia>Eubacteriales>Clostridiaceae | GCF_019838245.1 | Clostridium_botulinum | HYH70_RS05150 | 1157679..1161326 | 1491 | BREX-system-P-loop-protein-BrxC-[Clostridium-botulinum] |
| WP_061313931.1 | Type1.BREX | [illegible] | 1195 | Bacteria>Terrabacteria_group>Bacillota>Clostridia>Eubacteriales>Clostridiaceae | GCF_019838465.1 | Clostridium_botulinum | HYH32_RS08655 | 29233..32820 | 1491 | BREX-system-P-loop-protein-BrxC-[Clostridium-botulinum] |
| WP_223078572.1 | Type1.BREX | [illegible] | 1372 | Bacteria>Terrabacteria_group>Bacillota>Clostridia>Eubacteriales>Clostridiaceae | GCF_019913025.1 | Clostridium_butyricum | K7185_RS07970 | 91502..95620 | 1492 | hypothetical-protein-[Clostridium-butyricum] |
| WP_198609318.1 | Type1.BREX | [illegible] | 1221 | Bacteria>Terrabacteria_group>Bacillota>Clostridia>Eubacteriales>Clostridiaceae | GCF_020139275.1 | Clostridium_perfringens | KLF33_RS07605 | 1732833..1736498 | 1502 | BREX-system-P-loop-protein-BrxC-[Clostridium-perfringens] |
| WP_226090463.1 | Type1.BREX | [illegible] | 1215 | Bacteria>Terrabacteria_group>Bacillota>Clostridia>Eubacteriales>Clostridiaceae | GCF_020443435.1 | Clostridium_estertheticum | LGK96_RS12410 | 367687..371334 | 238834 | BREX-system-P-loop-protein-BrxC-[Clostridium-estertheticum] |
| WP_226260379.1 | Type1.BREX | [illegible] | 1189 | Bacteria>Terrabacteria_group>Bacillota>Clostridia>Eubacteriales>Clostridiaceae | GCF_020443565.1 | Clostridium_sp._CS001 | LGK97_RS14015 | 46160..49729 | 2880648 | BREX-system-P-loop-protein-BrxC-[Clostridium-sp.-CS001] |
| WP_226176799.1 | Type1.BREX | [illegible] | 1188 | Bacteria>Terrabacteria_group>Bacillota>Clostridia>Eubacteriales>Clostridiaceae | GCF_020443625.1 | Clostridium_tagluense | LGL05_RS19065 | 71229..74795 | 360422 | BREX-system-P-loop-protein-BrxC-[Clostridium-tagluense] |
| WP_226104726.1 | Type1.BREX | [illegible] | 1215 | Bacteria>Terrabacteria_group>Bacillota>Clostridia>Eubacteriales>Clostridiaceae | GCF_020443655.1 | Clostridium_algidicarnis | LGK99_RS11195 | 54237..57884 | 37659 | BREX-system-P-loop-protein-BrxC-[Clostridium-algidicarnis] |
| WP_118555970.1 | Type1.BREX | [illegible] | 1182 | Bacteria>Terrabacteria_group>Bacillota>Clostridia>Eubacteriales>Clostridiaceae | GCF_020686925.1 | Clostridium | LKD25_RS05345 | 60772..64320 | 1485 | MULTISPECIES:-BREX-system-P-loop-protein-BrxC-[Clostridium] |
| WP_228604542.1 | Type1.BREX | [illegible] | 1215 | Bacteria>Terrabacteria_group>Bacillota>Clostridia>Eubacteriales>Clostridiaceae | GCF_020819825.1 | Clostridium_botulinum | LMP43_RS01960 | 20879..24526 | 1491 | BREX-system-P-loop-protein-BrxC-[Clostridium-botulinum] |
| WP_228572337.1 | Type1.BREX | [illegible] | 1221 | Bacteria>Terrabacteria_group>Bacillota>Clostridia>Eubacteriales>Clostridiaceae | GCF_020819925.1 | Clostridium_perfringens | LMS36_RS03155 | 130857..134522 | 1502 | BREX-system-P-loop-protein-BrxC-[Clostridium-perfringens] |
| WP_229981609.1 | Type1.BREX | [illegible] | 1188 | Bacteria>Terrabacteria_group>Bacillota>Clostridia>Eubacteriales>Clostridiaceae | GCF_020908015.1 | Clostridium_sp._WLY-B-L2 | LN736_RS12085 | 83226..86792 | 2836848 | BREX-system-P-loop-protein-BrxC-[Clostridium-sp.-WLY-B-L2] |
| WP_231016614.1 | Type1.BREX | [illegible] | 1189 | Bacteria>Terrabacteria_group>Bacillota>Clostridia>Eubacteriales>Clostridiaceae | GCF_021043395.1 | Clostridium_guangxiense | LQK79_RS11195 | 7642..11211 | 1662055 | BREX-system-P-loop-protein-BrxC-[Clostridium-guangxiense] |
| WP_269205839.1 | Type1.BREX | [illegible] | 1208 | Bacteria>Terrabacteria_group>Bacillota>Clostridia>Eubacteriales>Clostridiaceae | GCF_021077705.1 | Clostridium_botulinum | G8S21_RS07265 | 73653..77279 | 1491 | BREX-system-P-loop-protein-BrxC-[Clostridium-botulinum] |
| WP_236911583.1 | Type1.BREX | [illegible] | 1193 | Bacteria>Terrabacteria_group>Bacillota>Clostridia>Eubacteriales>Clostridiaceae | GCF_021655095.1 | Clostridium_sp._Cult1 | C3E89_RS06110 | 1240047..1243628 | 2079002 | BREX-system-P-loop-protein-BrxC-[Clostridium-sp.-Cult1] |
| WP_238021483.1 | Type1.BREX | [illegible] | 1195 | Bacteria>Terrabacteria_group>Bacillota>Clostridia>Eubacteriales>Clostridiaceae | GCF_022133285.1 | Clostridium_cochlearium | L0O71_RS07755 | 6541..10128 | 1494 | BREX-system-P-loop-protein-BrxC-[Clostridium-cochlearium] |
| WP_173796807.1 | Type1.BREX | [illegible] | 1196 | Bacteria>Terrabacteria_group>Bacillota>Clostridia>Eubacteriales>Clostridiaceae | GCF_022133285.1 | Clostridium_cochlearium | L0O71_RS11235 | 1853..5443 | 1494 | BREX-system-P-loop-protein-BrxC-[Clostridium-cochlearium] |
| WP_252235883.1 | Type1.BREX | [illegible] | 1215 | Bacteria>Terrabacteria_group>Bacillota>Clostridia>Eubacteriales>Clostridiaceae | GCF_023846115.1 | Clostridium_sp._ZS1 | NG923_RS15480 | 30630..34277 | 2949989 | BREX-system-P-loop-protein-BrxC-[Clostridium-sp.-ZS1] |
| WP_252217802.1 | Type1.BREX | [illegible] | 1218 | Bacteria>Terrabacteria_group>Bacillota>Clostridia>Eubacteriales>Clostridiaceae | GCF_023846215.1 | Clostridium_chauvoei | J9878_RS07560 | 18587..22243 | 46867 | BREX-system-P-loop-protein-BrxC-[Clostridium-chauvoei] |
| WP_257415167.1 | Type1.BREX | [illegible] | 1197 | Bacteria>Terrabacteria_group>Bacillota>Clostridia>Eubacteriales>Clostridiaceae | GCF_024622825.1 | Clostridium_tepidum | NSA27_RS10235 | 93515..97108 | 1962263 | BREX-system-P-loop-protein-BrxC-[Clostridium-tepidum] |
| WP_257411542.1 | Type1.BREX | [illegible] | 1195 | Bacteria>Terrabacteria_group>Bacillota>Clostridia>Eubacteriales>Clostridiaceae | GCF_024622885.1 | Clostridium_sporogenes | NSA52_RS02830 | 6440..10027 | 1509 | BREX-system-P-loop-protein-BrxC-[Clostridium-sporogenes] |
| WP_058008254.1 | Type1.BREX | [illegible] | 1215 | Bacteria>Terrabacteria_group>Bacillota>Clostridia>Eubacteriales>Clostridiaceae | GCF_026130305.1 | Clostridium | LAV42_RS05410 | 123760..127407 | 1485 | MULTISPECIES:-BREX-system-P-loop-protein-BrxC-[Clostridium] |
| WP_265874436.1 | Type1.BREX | [illegible] | 1206 | Bacteria>Terrabacteria_group>Bacillota>Clostridia>Eubacteriales>Clostridiaceae | GCF_026240615.1 | Clostridium_kluyveri | OP486_RS05685 | 1250288..1253908 | 1534 | BREX-system-P-loop-protein-BrxC-[Clostridium-kluyveri] |
| WP_216171751.1 | Type1.BREX | [illegible] | 1215 | Bacteria>Terrabacteria_group>Bacillota>Clostridia>Eubacteriales>Clostridiaceae | GCF_026650905.1 | Clostridium_estertheticum | LL037_RS21145 | 4418214..4421861 | 238834 | BREX-system-P-loop-protein-BrxC-[Clostridium-estertheticum] |
| WP_270474687.1 | Type1.BREX | [illegible] | 1196 | Bacteria>Terrabacteria_group>Bacillota>Clostridia>Eubacteriales>Clostridiaceae | GCF_027681225.1 | Clostridium_cochlearium | PGX36_RS05095 | 39059..42649 | 1494 | BREX-system-P-loop-protein-BrxC-[Clostridium-cochlearium] |
| WP_270487117.1 | Type1.BREX | [illegible] | 1201 | Bacteria>Terrabacteria_group>Bacillota>Clostridia>Eubacteriales>Clostridiaceae | GCF_027681635.1 | Clostridium_fessum | PGX92_RS02760 | 620693..624298 | 2126740 | BREX-system-P-loop-protein-BrxC-[Clostridium-fessum] |
| WP_270659444.1 | Type1.BREX | [illegible] | 1185 | Bacteria>Terrabacteria_group>Bacillota>Clostridia>Eubacteriales>Clostridiaceae | GCF_027687525.1 | Paraclostridium_bifermentans | PGZ35_RS09255 | 5212..8769 | 1490 | BREX-system-P-loop-protein-BrxC-[Paraclostridium-bifermentans] |
| WP_270807575.1 | Type1.BREX | [illegible] | 1201 | Bacteria>Terrabacteria_group>Bacillota>Clostridia>Eubacteriales>Clostridiaceae | GCF_027723425.1 | Clostridium_fessum | PG965_RS01510 | 334129..337734 | 2126740 | BREX-system-P-loop-protein-BrxC-[Clostridium-fessum] |
| WP_111693597.1 | Type1.BREX | [illegible] | 1196 | Bacteria>Terrabacteria_group>Bacillota>Clostridia>Eubacteriales>Clostridiaceae | GCF_028867175.1 | Clostridium_perfringens | PVA22_RS02075 | 481632..485222 | 1502 | BREX-system-P-loop-protein-BrxC-[Clostridium-perfringens] |
| WP_220707229.1 | Type1.BREX | [illegible] | 1215 | Bacteria>Terrabacteria_group>Bacillota>Clostridia>Eubacteriales>Clostridiaceae | GCF_030585505.1 | Clostridium_estertheticum | K2182_RS19255 | 29060..32707 | 238834 | BREX-system-P-loop-protein-BrxC-[Clostridium-estertheticum] |
| WP_021876221.1 | Type1.BREX | [illegible] | 1218 | Bacteria>Terrabacteria_group>Bacillota>Clostridia>Eubacteriales>Clostridiaceae | GCF_900037355.1 | Clostridium_chauvoei | C6H62_RS09310 | 1987009..1990665 | 46867 | BREX-system-P-loop-protein-BrxC-[Clostridium-chauvoei] |
| WP_066893479.1 | Type1.BREX | [illegible] | 1215 | Bacteria>Terrabacteria_group>Bacillota>Clostridia>Eubacteriales>Clostridiaceae | GCF_900086595.1 | Clostridium_nigeriense | BN6774_RS10780 | 2270723..2274370 | 1805470 | BREX-system-P-loop-protein-BrxC-[Clostridium-nigeriense] |
| WP_089867191.1 | Type1.BREX | [illegible] | 1196 | Bacteria>Terrabacteria_group>Bacillota>Clostridia>Eubacteriales>Clostridiaceae | GCF_900103025.1 | Clostridium_cochlearium | BLS04_RS09710 | 7640..11230 | 1494 | BREX-system-P-loop-protein-BrxC-[Clostridium-cochlearium] |
| WP_089867383.1 | Type1.BREX | [illegible] | 1184 | Bacteria>Terrabacteria_group>Bacillota>Clostridia>Eubacteriales>Clostridiaceae | GCF_900103025.1 | Clostridium_cochlearium | BLS04_RS10080 | 14668..18222 | 1494 | BREX-system-P-loop-protein-BrxC-[Clostridium-cochlearium] |
| WP_052222505.1 | Type1.BREX | [illegible] | 1366 | Bacteria>Terrabacteria_group>Bacillota>Clostridia>Eubacteriales>Clostridiaceae | GCF_900113195.1 | Clostridium_homopropionicum | BM106_RS12185 | 166200..170300 | 36844 | hypothetical-protein-[Clostridium-homopropionicum] |
| WP_074913308.1 | Type1.BREX | [illegible] | 1195 | Bacteria>Terrabacteria_group>Bacillota>Clostridia>Eubacteriales>Clostridiaceae | GCF_900115135.1 | Proteiniclasticum_ruminis | BM243_RS14610 | 3534..7121 | 398199 | BREX-system-P-loop-protein-BrxC-[Proteiniclasticum-ruminis] |
| WP_072525015.1 | Type1.BREX | [illegible] | 1199 | Bacteria>Terrabacteria_group>Bacillota>Clostridia>Eubacteriales>Clostridiaceae | GCF_900120155.1 | Clostridium_sp._Marseille-P3244 | BQ7367_RS08330 | 1368431..1372030 | 1871020 | BREX-system-P-loop-protein-BrxC-[Clostridium-sp.-Marseille-P3244] |
| WP_072897339.1 | Type1.BREX | [illegible] | 1218 | Bacteria>Terrabacteria_group>Bacillota>Clostridia>Eubacteriales>Clostridiaceae | GCF_900129365.1 | Clostridium_fallax | BUA70_RS11140 | 23714..27370 | 1533 | BREX-system-P-loop-protein-BrxC-[Clostridium-fallax] |
| WP_073007111.1 | Type1.BREX | [illegible] | 1214 | Bacteria>Terrabacteria_group>Bacillota>Clostridia>Eubacteriales>Clostridiaceae | GCF_900142075.1 | Clostridium_amylolyticum | BUA82_RS09870 | 579511..583155 | 1121298 | BREX-system-P-loop-protein-BrxC-[Clostridium-amylolyticum] |
| WP_143148632.1 | Type1.BREX | [illegible] | 1377 | Bacteria>Terrabacteria_group>Bacillota>Clostridia>Eubacteriales>Clostridiaceae | GCF_900142075.1 | Clostridium_amylolyticum | BUA82_RS13560 | 29831..33964 | 1121298 | hypothetical-protein-[Clostridium-amylolyticum] |
| WP_073026825.1 | Type1.BREX | [illegible] | 1381 | Bacteria>Terrabacteria_group>Bacillota>Clostridia>Eubacteriales>Clostridiaceae | GCF_900142105.1 | Lutispora_thermophila | BUB48_RS13880 | 40274..44419 | 288966 | hypothetical-protein-[Lutispora-thermophila] |
| WP_073026962.1 | Type1.BREX | [illegible] | 1186 | Bacteria>Terrabacteria_group>Bacillota>Clostridia>Eubacteriales>Clostridiaceae | GCF_900142105.1 | Lutispora_thermophila | BUB48_RS14275 | 12515..16075 | 288966 | BREX-system-P-loop-protein-BrxC-[Lutispora-thermophila] |
| WP_072965766.1 | Type1.BREX | [illegible] | 1187 | Bacteria>Terrabacteria_group>Bacillota>Clostridia>Eubacteriales>Clostridiaceae | GCF_900142285.1 | Caminicella_sporogenes | BUA90_RS02195 | 9238..12801 | 166485 | BREX-system-P-loop-protein-BrxC-[Caminicella-sporogenes] |
| WP_102401523.1 | Type1.BREX | [illegible] | 1215 | Bacteria>Terrabacteria_group>Bacillota>Clostridia>Eubacteriales>Clostridiaceae | GCF_900184255.1 | Haloimpatiens_massiliensis | C1715_RS16875 | 1834111..1837758 | 1658110 | BREX-system-P-loop-protein-BrxC-[Haloimpatiens-massiliensis] |
| WP_150356414.1 | Type1.BREX | [illegible] | 1203 | Bacteria>Terrabacteria_group>Bacillota>Clostridia>Eubacteriales>Clostridiaceae | GCF_900184915.1 | Clostridium_sp._HV4-5-A1G | F3063_RS05640 | 35144..38755 | 2004595 | BREX-system-P-loop-protein-BrxC-[Clostridium-sp.-HV4-5-A1G] |
| WP_150355659.1 | Type1.BREX | [illegible] | 1188 | Bacteria>Terrabacteria_group>Bacillota>Clostridia>Eubacteriales>Clostridiaceae | GCF_900184915.1 | Clostridium_sp._HV4-5-A1G | J5G60_RS01905 | 31046..34612 | 2004595 | BREX-system-P-loop-protein-BrxC-[Clostridium-sp.-HV4-5-A1G] |
| WP_115641994.1 | Type1.BREX | [illegible] | 1215 | Bacteria>Terrabacteria_group>Bacillota>Clostridia>Eubacteriales>Clostridiaceae | GCF_900461105.1 | Clostridium_putrefaciens | DY168_RS12220 | 2676627..2680274 | 99675 | BREX-system-P-loop-protein-BrxC-[Clostridium-putrefaciens] |
| WP_111744218.1 | Type1.BREX | [illegible] | 1221 | Bacteria>Terrabacteria_group>Bacillota>Clostridia>Eubacteriales>Clostridiaceae | GCF_900475545.1 | Clostridium_perfringens | Cp4442_RS12820 | 4585..8250 | 1502 | BREX-system-P-loop-protein-BrxC-[Clostridium-perfringens] |
| WP_142710734.1 | Type1.BREX | [illegible] | 1221 | Bacteria>Terrabacteria_group>Bacillota>Clostridia>Eubacteriales>Clostridiaceae | GCF_902166005.1 | Clostridium_perfringens | FN719_RS05040 | 228419..232084 | 1502 | BREX-system-P-loop-protein-BrxC-[Clostridium-perfringens] |
| WP_142975592.1 | Type1.BREX | [illegible] | 1221 | Bacteria>Terrabacteria_group>Bacillota>Clostridia>Eubacteriales>Clostridiaceae | GCF_902166025.1 | Clostridium_perfringens | FN715_RS03295 | 751975..755640 | 1502 | BREX-system-P-loop-protein-BrxC-[Clostridium-perfringens] |
| WP_142420582.1 | Type1.BREX | [illegible] | 1196 | Bacteria>Terrabacteria_group>Bacillota>Clostridia>Eubacteriales>Clostridiaceae | GCF_902166135.1 | Clostridium_perfringens | FN714_RS08835 | 22939..26529 | 1502 | BREX-system-P-loop-protein-BrxC-[Clostridium-perfringens] |
| WP_055656468.1 | Type1.BREX | [illegible] | 1220 | Bacteria>Terrabacteria_group>Bacillota>Clostridia>Eubacteriales>Clostridiaceae | GCF_902362405.1 | Hungatella_hathewayi | FY393_RS15415 | 235272..238934 | 154046 | BREX-system-P-loop-protein-BrxC-[Hungatella-hathewayi] |
| WP_147564561.1 | Type1.BREX | [illegible] | 1206 | Bacteria>Terrabacteria_group>Bacillota>Clostridia>Eubacteriales>Clostridiaceae | GCF_902363545.1 | Clostridium_tyrobutyricum | FXW39_RS01670 | 332277..335897 | 1519 | BREX-system-P-loop-protein-BrxC-[Clostridium-tyrobutyricum] |
| WP_106999750.1 | Type1.BREX | [illegible] | 1182 | Bacteria>Terrabacteria_group>Bacillota>Clostridia>Eubacteriales>Clostridiaceae | GCF_937998705.1 | Clostridium | PGT52_RS02970 | 663128..666676 | 1485 | MULTISPECIES:-BREX-system-P-loop-protein-BrxC-[Clostridium] |
| WP_195381616.1 | Type1.BREX | [illegible] | 1221 | Bacteria>Terrabacteria_group>Bacillota>Clostridia>Eubacteriales>Clostridiaceae | GCF_951336195.1 | Clostridium_perfringens | H8J86_RS06960 | 66718..70383 | 1502 | BREX-system-P-loop-protein-BrxC-[Clostridium-perfringens] |
| WP_160719334.1 | Type1.BREX | [illegible] | 1195 | Bacteria>Terrabacteria_group>Bacillota>Clostridia>Eubacteriales>Clostridiaceae | - | Isachenkonia_alkalipeptolytica | ISALK_04100 | 132779..136366 | 2565777 | BREX-system-P-loop-protein-BrxC-[Isachenkonia-alkalipeptolytica] |
| WP_117466987.1 | Type1.BREX | [illegible] | 1196 | Bacteria>Terrabacteria_group>Bacillota>Clostridia>Eubacteriales>Clostridiaceae | - | unclassified_Clostridium | DW081_14990 | 34449..38039 | 2614128 | MULTISPECIES:-BREX-system-P-loop-protein-BrxC-[unclassified-Clostridium] |
| WP_117821980.1 | Type1.BREX | [illegible] | 1196 | Bacteria>Terrabacteria_group>Bacillota>Clostridia>Eubacteriales>Clostridiaceae | - | unclassified_Clostridium | DW959_RS12365 | 44943..48533 | 2614128 | MULTISPECIES:-BREX-system-P-loop-protein-BrxC-[unclassified-Clostridium] |
| MCL2361204.1 | Type1.BREX | [illegible] | 1185 | Bacteria>Terrabacteria_group>Bacillota>Clostridia>Eubacteriales>Defluviitaleaceae | GCA_009781725.1 | Defluviitaleaceae_bacterium | FWC73_05265 | 21378..24935 | 2660712 | BREX-system-P-loop-protein-BrxC-[Defluviitaleaceae-bacterium] |
| WP_006523745.1 | Type1.BREX | [illegible] | 1204 | Bacteria>Terrabacteria_group>Bacillota>Clostridia>Eubacteriales>Desulfallaceae | GCF_000233715.2 | Desulfoscipio_gibsoniae | Desgi_1207 | 1236850..1240464 | 102134 | BREX-system-P-loop-protein-BrxC-[Desulfoscipio-gibsoniae] |
| WP_195712192.1 | Type1.BREX | [illegible] | 1185 | Bacteria>Terrabacteria_group>Bacillota>Clostridia>Eubacteriales>Desulfallaceae | GCF_015594625.1 | Desulfallas_sp._Bu1-1 | IT084_RS07280 | 138582..142139 | 2787620 | BREX-system-P-loop-protein-BrxC-[Desulfallas-sp.-Bu1-1] |
| KGP77156.1 | Type1.BREX | [illegible] | 1012 | Bacteria>Terrabacteria_group>Bacillota>Clostridia>Eubacteriales>Desulfitobacteriaceae | GCA_000770645.1 | Desulfosporosinus_sp._Tol-M | JT05_00745 | 1..3040 | 1536651 | ATPase,-partial-[Desulfosporosinus-sp.-Tol-M] |
| KGP76494.1 | Type1.BREX | [illegible] | 1189 | Bacteria>Terrabacteria_group>Bacillota>Clostridia>Eubacteriales>Desulfitobacteriaceae | GCA_000770645.1 | Desulfosporosinus_sp._Tol-M | JT05_04435 | 4696..8265 | 1536651 | hypothetical-protein-JT05_04435-[Desulfosporosinus-sp.-Tol-M] |
| KUO77762.1 | Type1.BREX | [illegible] | 1193 | Bacteria>Terrabacteria_group>Bacillota>Clostridia>Eubacteriales>Desulfitobacteriaceae | GCA_001516045.1 | Desulfosporosinus_sp._BRH_c37 | APF81_01440 | 9315..12896 | 1734396 | MAG:-ATPase-[Desulfosporosinus-sp.-BRH_c37] |
| UWG97285.1 | Type1.BREX | [illegible] | 1185 | Bacteria>Terrabacteria_group>Bacillota>Clostridia>Eubacteriales>Desulfitobacteriaceae | GCA_024972775.1 | Dehalobacter_sp._DCM | LPY66_00125 | 21149..24706 | 2907827 | BREX-system-P-loop-protein-BrxC-[Dehalobacter-sp.-DCM] |
| UWG96530.1 | Type1.BREX | [illegible] | 1172 | Bacteria>Terrabacteria_group>Bacillota>Clostridia>Eubacteriales>Desulfitobacteriaceae | GCA_024972775.1 | Dehalobacter_sp._DCM | LPY66_16790 | 3636490..3640008 | 2907827 | BREX-system-P-loop-protein-BrxC-[Dehalobacter-sp.-DCM] |
| SPF40064.1 | Type1.BREX | [illegible] | 1194 | Bacteria>Terrabacteria_group>Bacillota>Clostridia>Eubacteriales>Desulfitobacteriaceae | GCA_900290375.1 | Candidatus_Desulfosporosinus_infrequens | SBF1_2170004 | 1616..5200 | 2043169 | conserved-hypothetical-protein-[Candidatus-Desulfosporosinus-infrequens] |
| WP_011459229.1 | Type1.BREX | [illegible] | 1172 | Bacteria>Terrabacteria_group>Bacillota>Clostridia>Eubacteriales>Desulfitobacteriaceae | GCF_000010045.1 | Desulfitobacterium_hafniense | AT727_RS10680 | 26709..30227 | 49338 | BREX-system-P-loop-protein-BrxC-[Desulfitobacterium-hafniense] |
| WP_014901355.1 | Type1.BREX | [illegible] | 1203 | Bacteria>Terrabacteria_group>Bacillota>Clostridia>Eubacteriales>Desulfitobacteriaceae | GCF_000231385.2 | Desulfosporosinus_meridiei | Desmer_0377 | 362087..365698 | 79209 | BREX-system-P-loop-protein-BrxC-[Desulfosporosinus-meridiei] |
| WP_018305995.1 | Type1.BREX | [illegible] | 1172 | Bacteria>Terrabacteria_group>Bacillota>Clostridia>Eubacteriales>Desulfitobacteriaceae | GCF_000379505.1 | Desulfitobacterium_hafniense | A37Y_RS0110035 | 20132..23650 | 49338 | BREX-system-P-loop-protein-BrxC-[Desulfitobacterium-hafniense] |
| WP_139314555.1 | Type1.BREX | [illegible] | 764 | Bacteria>Terrabacteria_group>Bacillota>Clostridia>Eubacteriales>Desulfitobacteriaceae | GCF_001936615.1 | Desulfosporosinus_metallidurans | DSOL_RS20630 | 27661..29954 | 1888891 | BREX-system-P-loop-protein-BrxC,-partial-[Desulfosporosinus-metallidurans] |
| KUK53483.1 | Type1.BREX | [illegible] | 782 | Bacteria>Terrabacteria_group>Bacillota>Clostridia>Eubacteriales>Desulfotomaculaceae | GCA_001508375.1 | Desulfotomaculum_sp._46_296 | DCW46_09075 | 1..2347 | 1635262 | MAG:-Putative-P-loop-containing-nucleoside-triphosphate-hydrolase,-partial-[Desulfotomaculum-sp.-46_296] |
| KUK52655.1 | Type1.BREX | [illegible] | 1148 | Bacteria>Terrabacteria_group>Bacillota>Clostridia>Eubacteriales>Desulfotomaculaceae | GCA_001508375.1 | Desulfotomaculum_sp._46_296 | XD78_1828 | 5958..9404 | 1635262 | MAG:-hypothetical-protein-XD78_1828-[Desulfotomaculum-sp.-46_296] |
| OPX91799.1 | Type1.BREX | [illegible] | 1181 | Bacteria>Terrabacteria_group>Bacillota>Clostridia>Eubacteriales>Desulfotomaculaceae | GCA_002067515.1 | Pelotomaculum_sp._PtaB.Bin013 | A4E52_00325 | 25..3570 | 1811692 | MAG:-hypothetical-protein-A4E52_00325-[Pelotomaculum-sp.-PtaB.Bin013] |
| OPX91092.1 | Type1.BREX | [illegible] | 1195 | Bacteria>Terrabacteria_group>Bacillota>Clostridia>Eubacteriales>Desulfotomaculaceae | GCA_002067515.1 | Pelotomaculum_sp._PtaB.Bin013 | A4E52_00617 | 7955..11542 | 1811692 | MAG:-hypothetical-protein-A4E52_00617-[Pelotomaculum-sp.-PtaB.Bin013] |
| HAG09972.1 | Type1.BREX | [illegible] | 1148 | Bacteria>Terrabacteria_group>Bacillota>Clostridia>Eubacteriales>Desulfotomaculaceae | GCA_003445665.1 | Desulfotomaculum_sp. | DCK76_00890 | 5882..9328 | 41211 | BREX-system-P-loop-protein-BrxC-[Desulfotomaculum-sp.] |
| WP_134214351.1 | Type1.BREX | [illegible] | 1198 | Bacteria>Terrabacteria_group>Bacillota>Clostridia>Eubacteriales>Desulfotomaculaceae | GCF_004369225.1 | Pelotomaculum_propionicicum | Pmgp_RS12670 | 23567..27163 | 258475 | BREX-system-P-loop-protein-BrxC-[Pelotomaculum-propionicicum] |
| WP_240084449.1 | Type1.BREX | [illegible] | 1192 | Bacteria>Terrabacteria_group>Bacillota>Clostridia>Eubacteriales>Desulfotomaculaceae | GCF_022410535.1 | Pelotomaculum_terephthalicicum | L9W92_RS14450 | 18120..21698 | 206393 | BREX-system-P-loop-protein-BrxC-[Pelotomaculum-terephthalicicum] |
| WP_117571353.1 | Type1.BREX | [illegible] | 1177 | Bacteria>Terrabacteria_group>Bacillota>Clostridia>Eubacteriales>Eubacteriaceae | - | Eubacterium_sp._TM05-53 | DXD51_03480 | 17212..20745 | 2292353 | BREX-system-P-loop-protein-BrxC-[Eubacterium-sp.-TM05-53] |
| KAI4451658.1 | Type1.BREX | [illegible] | 932 | Bacteria>Terrabacteria_group>Bacillota>Clostridia>Eubacteriales>Eubacteriaceae | GCA_000364225.2 | Eubacterium_plexicaudatum_ASF492 | C823_006218 | 5247159..5249957 | 1235802 | hypothetical-protein-C823_006218-[Eubacterium-plexicaudatum-ASF492] |
| OLA08999.1 | Type1.BREX | [illegible] | 1184 | Bacteria>Terrabacteria_group>Bacillota>Clostridia>Eubacteriales>Eubacteriaceae | GCA_001916415.1 | Eubacterium_sp._45_250 | BHW21_06160 | 165792..169346 | 1897004 | MAG:-ATPase-[Eubacterium-sp.-45_250] |
| RGJ41450.1 | Type1.BREX | [illegible] | 1182 | Bacteria>Terrabacteria_group>Bacillota>Clostridia>Eubacteriales>Eubacteriaceae | GCA_003466385.1 | Eubacterium_sp._TM06-47 | DXD61_10770 | 2336..5884 | 2292354 | BREX-system-P-loop-protein-BrxC-[Eubacterium-sp.-TM06-47] |
| RHE43175.1 | Type1.BREX | [illegible] | 1182 | Bacteria>Terrabacteria_group>Bacillota>Clostridia>Eubacteriales>Eubacteriaceae | GCA_003469815.1 | Eubacterium_sp._AM28-29 | DW744_00515 | 115473..119021 | 2292349 | BREX-system-P-loop-protein-BrxC-[Eubacterium-sp.-AM28-29] |
| NTW71836.1 | Type1.BREX | [illegible] | 914 | Bacteria>Terrabacteria_group>Bacillota>Clostridia>Eubacteriales>Eubacteriaceae | GCA_013335935.1 | Eubacteriaceae_bacterium | HGA49_06295 | 18702..21443 | 2049045 | BREX-system-P-loop-protein-BrxC-[Eubacteriaceae-bacterium] |
| MBR6404277.1 | Type1.BREX | [illegible] | 1160 | Bacteria>Terrabacteria_group>Bacillota>Clostridia>Eubacteriales>Eubacteriaceae | GCA_017539165.1 | Eubacterium_sp. | IKS48_12920 | 16986..20468 | 142586 | BREX-system-P-loop-protein-BrxC-[Eubacterium-sp.] |
| MBS6586983.1 | Type1.BREX | [illegible] | 1209 | Bacteria>Terrabacteria_group>Bacillota>Clostridia>Eubacteriales>Eubacteriaceae | GCA_018377805.1 | Eubacterium_sp. | KH345_12625 | 21110..24739 | 142586 | BREX-system-P-loop-protein-BrxC-[Eubacterium-sp.] |
| MCH3955225.1 | Type1.BREX | [illegible] | 1171 | Bacteria>Terrabacteria_group>Bacillota>Clostridia>Eubacteriales>Eubacteriaceae | GCA_022482865.1 | Eubacterium_sp. | LKE44_08530 | 1944805..1948320 | 142586 | BREX-system-P-loop-protein-BrxC-[Eubacterium-sp.] |
| MCH4007762.1 | Type1.BREX | [illegible] | 1173 | Bacteria>Terrabacteria_group>Bacillota>Clostridia>Eubacteriales>Eubacteriaceae | GCA_022483305.1 | Eubacterium_sp. | LKF03_02665 | 579207..582728 | 142586 | BREX-system-P-loop-protein-BrxC-[Eubacterium-sp.] |
| MCH4073094.1 | Type1.BREX | [illegible] | 1171 | Bacteria>Terrabacteria_group>Bacillota>Clostridia>Eubacteriales>Eubacteriaceae | GCA_022483875.1 | Pseudoramibacter_sp. | LKF02_09135 | 1854333..1857848 | 2034862 | BREX-system-P-loop-protein-BrxC-[Pseudoramibacter-sp.] |
| MCI5510134.1 | Type1.BREX | [illegible] | 1182 | Bacteria>Terrabacteria_group>Bacillota>Clostridia>Eubacteriales>Eubacteriaceae | GCA_022764165.1 | Eubacterium_sp. | MR436_06790 | 9461..13009 | 142586 | BREX-system-P-loop-protein-BrxC-[Eubacterium-sp.] |
| MCI9378046.1 | Type1.BREX | [illegible] | 1218 | Bacteria>Terrabacteria_group>Bacillota>Clostridia>Eubacteriales>Eubacteriaceae | GCA_022800015.1 | Eubacterium_sp. | HFI06_05840 | 4725..8381 | 142586 | BREX-system-P-loop-protein-BrxC-[Eubacterium-sp.] |
| MCI9420874.1 | Type1.BREX | [illegible] | 1213 | Bacteria>Terrabacteria_group>Bacillota>Clostridia>Eubacteriales>Eubacteriaceae | GCA_022800355.1 | Eubacterium_sp. | HFG32_12910 | 593..4234 | 142586 | BREX-system-P-loop-protein-BrxC-[Eubacterium-sp.] |
| CVI69463.1 | Type1.BREX | [illegible] | 1179 | Bacteria>Terrabacteria_group>Bacillota>Clostridia>Eubacteriales>Eubacteriaceae | GCA_900016875.1 | Eubacteriaceae_bacterium_CHKCI004 | BN3660_01565 | 6017..9556 | 1780380 | hypothetical-protein-BN3660_01565-[Eubacteriaceae-bacterium-CHKCI004] |
| SEG33478.1 | Type1.BREX | [illegible] | 1157 | Bacteria>Terrabacteria_group>Bacillota>Clostridia>Eubacteriales>Eubacteriaceae | GCA_900108205.1 | Eubacterium_ruminantium | SAMN04487934_11616 | 23559..27032 | 42322 | hypothetical-protein-SAMN04487934_11616-[Eubacterium-ruminantium] |
| WP_026396241.1 | Type1.BREX | [illegible] | 1187 | Bacteria>Terrabacteria_group>Bacillota>Clostridia>Eubacteriales>Eubacteriaceae | GCF_000472665.1 | Acetobacterium_dehalogenans | A3KS_RS0118990 | 31738..35301 | 82116 | BREX-system-P-loop-protein-BrxC-[Acetobacterium-dehalogenans] |
| WP_242834784.1 | Type1.BREX | [illegible] | 1017 | Bacteria>Terrabacteria_group>Bacillota>Clostridia>Eubacteriales>Eubacteriaceae | GCF_000472665.1 | Acetobacterium_dehalogenans | A3KS_RS21665 | 9901..12954 | 82116 | BREX-system-P-loop-protein-BrxC-[Acetobacterium-dehalogenans] |
| WP_055289382.1 | Type1.BREX | [illegible] | 1183 | Bacteria>Terrabacteria_group>Bacillota>Clostridia>Eubacteriales>Eubacteriaceae | GCF_001406295.1 | Eubacterium_ramulus | ARA08_RS03325 | 204677..208228 | 39490 | BREX-system-P-loop-protein-BrxC-[Eubacterium-ramulus] |
| WP_111888010.1 | Type1.BREX | [illegible] | 1189 | Bacteria>Terrabacteria_group>Bacillota>Clostridia>Eubacteriales>Eubacteriaceae | GCF_003260995.1 | Acetobacterium_sp._KB-1 | DOZ58_RS09185 | 1965484..1969053 | 2184575 | BREX-system-P-loop-protein-BrxC-[Acetobacterium-sp.-KB-1] |
| WP_186842510.1 | Type1.BREX | [illegible] | 1222 | Bacteria>Terrabacteria_group>Bacillota>Clostridia>Eubacteriales>Eubacteriaceae | GCF_014284475.1 | Acetobacterium_fimetarium | GH808_RS09300 | 27272..30940 | 52691 | BREX-system-P-loop-protein-BrxC-[Acetobacterium-fimetarium] |
| WP_186894780.1 | Type1.BREX | [illegible] | 1187 | Bacteria>Terrabacteria_group>Bacillota>Clostridia>Eubacteriales>Eubacteriaceae | GCF_014284495.1 | Acetobacterium_malicum | GH811_RS13415 | 62919..66482 | 52692 | BREX-system-P-loop-protein-BrxC-[Acetobacterium-malicum] |
| WP_250934564.1 | Type1.BREX | [illegible] | 1187 | Bacteria>Terrabacteria_group>Bacillota>Clostridia>Eubacteriales>Eubacteriaceae | GCF_023700225.1 | Acetobacterium_wieringae | CHL1_RS02680 | 557303..560866 | 52694 | BREX-system-P-loop-protein-BrxC-[Acetobacterium-wieringae] |
| WP_250934580.1 | Type1.BREX | [illegible] | 656 | Bacteria>Terrabacteria_group>Bacillota>Clostridia>Eubacteriales>Eubacteriaceae | GCF_023700225.1 | Acetobacterium_wieringae | CHL1_RS02785 | 587968..589938 | 52694 | BREX-system-P-loop-protein-BrxC-[Acetobacterium-wieringae] |
| WP_270367180.1 | Type1.BREX | [illegible] | 1182 | Bacteria>Terrabacteria_group>Bacillota>Clostridia>Eubacteriales>Eubacteriaceae | GCF_027667005.1 | Eubacterium_ramulus | PGS15_RS04515 | 102111..105659 | 39490 | BREX-system-P-loop-protein-BrxC-[Eubacterium-ramulus] |
| WP_078766837.1 | Type1.BREX | [illegible] | 1203 | Bacteria>Terrabacteria_group>Bacillota>Clostridia>Eubacteriales>Eubacteriaceae | GCF_900167115.1 | Eubacterium_uniforme | B5D16_RS09945 | 14532..18143 | 39495 | BREX-system-P-loop-protein-BrxC-[Eubacterium-uniforme] |
| WP_133969317.1 | Type1.BREX | [illegible] | 1184 | Bacteria>Terrabacteria_group>Bacillota>Clostridia>Eubacteriales>Eubacteriaceae | GCF_900683775.1 | Eubacterium_limosum | E4L76_RS20940 | 4393668..4397222 | 1736 | BREX-system-P-loop-protein-BrxC-[Eubacterium-limosum] |
| AVX30039.1 | Type1.BREX | [illegible] | 1148 | Bacteria>Terrabacteria_group>Bacillota>Clostridia>Eubacteriales>Eubacteriales_incertae_sedis | GCA_003054495.1 | Carboxydocella_thermautotrophica | CTH_0434 | 429289..432735 | 178899 | hypothetical-protein-CTH_0434-[Carboxydocella-thermautotrophica] |
| MCI6150868.1 | Type1.BREX | [illegible] | 1179 | Bacteria>Terrabacteria_group>Bacillota>Clostridia>Eubacteriales>Eubacteriales_incertae_sedis | GCA_022772165.1 | Flintibacter_sp. | MR705_10640 | 43131..46670 | 1918624 | BREX-system-P-loop-protein-BrxC-[Flintibacter-sp.] |
| MCI9638683.1 | Type1.BREX | [illegible] | 1159 | Bacteria>Terrabacteria_group>Bacillota>Clostridia>Eubacteriales>Eubacteriales_incertae_sedis | GCA_022802005.1 | Emergencia_sp. | HFE72_00905 | 28674..32153 | 1926557 | BREX-system-P-loop-protein-BrxC-[Emergencia-sp.] |
| WP_087184149.1 | Type1.BREX | [illegible] | 1178 | Bacteria>Terrabacteria_group>Bacillota>Clostridia>Eubacteriales>Eubacteriales_incertae_sedis | GCF_002159185.1 | Gemmiger_sp._An87 | B5G38_RS07650 | 33911..37447 | 1965662 | BREX-system-P-loop-protein-BrxC-[Gemmiger-sp.-An87] |
| WP_193503082.1 | Type1.BREX | [illegible] | 1176 | Bacteria>Terrabacteria_group>Bacillota>Clostridia>Eubacteriales>Eubacteriales_incertae_sedis | GCF_014982805.1 | Gemmiger_gallinarum | INF35_RS12765 | 36645..40175 | 2779354 | BREX-system-P-loop-protein-BrxC-[Gemmiger-gallinarum] |
| WP_270462427.1 | Type1.BREX | [illegible] | 1205 | Bacteria>Terrabacteria_group>Bacillota>Clostridia>Eubacteriales>Eubacteriales_incertae_sedis | GCF_027662265.1 | Gemmiger_formicilis | PGQ24_RS15845 | 6404..10021 | 745368 | BREX-system-P-loop-protein-BrxC-[Gemmiger-formicilis] |

| Protein-ID | System | Gene-Neighbourhoods | Length | Taxonomy | Assembly-ID | Strain | Locus-Tag | Genomic-coordinates | TaxID | Fasta-define |
|---|---|---|---|---|---|---|---|---|---|---|
| WP_272041140.1 | Type1.BREX | [illegible] | 1181 | [illegible] | GCF_028321865.1 | Intestinimonas_butyriciproducens | PND64_RS14385 | 23490..27035 | 1297617 | [illegible] |
| WP_091728844.1 | Type1.BREX | [illegible] | 1193 | [illegible] | GCF_900107485.1 | Proteiniborus_ethanoligenes | BLV37_RS06315 | 24858..28439 | 415015 | [illegible] |
| WP_130869775.1 | Type1.BREX | [illegible] | 1181 | [illegible] | GCF_900626285.1 | Intestinimonas_massiliensis | E0E41_RS10000 | 1985167..1988712 | 1673721 | [illegible] |
| WP_055219000.1 | Type1.BREX | [illegible] | 1182 | [illegible] | GCF_013300755.1 | Lachnospiraceae | G5A93_RS10915 | 4301..7849 | 186803 | [illegible] |
| WP_117739785.1 | Type1.BREX | [illegible] | 1183 | [illegible] | GCF_013301605.1 | Lachnospiraceae | G4434_RS12220 | 38820..42371 | 186803 | [illegible] |
| WP_138373666.1 | Type1.BREX | [illegible] | 1199 | [illegible] | GCF_015554965.1 | Oscillospiraceae | I2E95_RS13170 | 36042..39641 | 216572 | [illegible] |
| WP_138373965.1 | Type1.BREX | [illegible] | 1179 | [illegible] | GCF_015554965.1 | Oscillospiraceae | I2E95_RS16325 | 14974..18513 | 216572 | [illegible] |
| WP_118699091.1 | Type1.BREX | [illegible] | 1182 | [illegible] | GCF_020560115.1 | Lachnospiraceae | LI126_RS00410 | 82443..85991 | 186803 | [illegible] |
| WP_118447467.1 | Type1.BREX | [illegible] | 1182 | [illegible] | GCF_022900215.1 | Lachnospiraceae | MUY38_RS08320 | 63562..67110 | 186803 | [illegible] |
| WP_101871400.1 | Type1.BREX | [illegible] | 1217 | [illegible] | GCF_024463595.1 | Lachnospiraceae | LI221_RS06895 | 48954..52607 | 186803 | [illegible] |
| WP_118637679.1 | Type1.BREX | [illegible] | 1215 | [illegible] | GCF_027924705.1 | Oscillospiraceae | H6A32_RS13625 | 30472..34119 | 216572 | [illegible] |
| WP_117761890.1 | Type1.BREX | [illegible] | 1182 | [illegible] | GCF_902364755.1 | Lachnospiraceae | FXX70_RS12975 | 76873..80421 | 186803 | [illegible] |
| WP_256171760.1 | Type1.BREX | [illegible] | 1182 | [illegible] | - | Dorea_longicatena | NE607_07100 | 21358..24906 | 88431 | [illegible] |
| WP_186856038.1 | Type1.BREX | [illegible] | 1203 | [illegible] | - | Dorea_sp._NSJ-36 | H8S07_RS11320 | 56679..60290 | 2763040 | [illegible] |
| WP_227161425.1 | Type1.BREX | [illegible] | 946 | [illegible] | - | Enterocloster_citroniae | LI031_09475 | 245890..248730 | 358743 | [illegible] |
| EFU77068.1 | Type1.BREX | [illegible] | 1222 | [illegible] | GCA_000185385.1 | Lachnoanaerobaculum_saburreum_DSM_3986 | HMPREF0381_1067 | 36430..40098 | 887325 | [illegible] |
| EFV19659.1 | Type1.BREX | [illegible] | 1177 | [illegible] | GCA_000185545.1 | Lachnospiraceae_bacterium_8_1_57FAA | HMPREF1026_01169 | 31500..35033 | 665951 | [illegible] |
| EGG92088.1 | Type1.BREX | [illegible] | 1217 | [illegible] | GCA_000209465.1 | Lachnospiraceae_oral_taxon_107_str._F0167 | HMPREF0491_00115 | 36108..39761 | 575593 | [illegible] |
| EGN43144.1 | Type1.BREX | [illegible] | 1182 | [illegible] | GCA_000218445.1 | Lachnospiraceae_bacterium_1_1_57FAA | HMPREF0990_02349 | 46029..49577 | 658081 | [illegible] |
| EHI56636.1 | Type1.BREX | [illegible] | 1222 | [illegible] | GCA_000235445.1 | Johnsonella_ignava_ATCC_51276 | HMPREF9333_00083 | 47745..51413 | 679200 | [illegible] |
| EHO18585.1 | Type1.BREX | [illegible] | 1217 | [illegible] | GCA_000242235.1 | Stomatobaculum_longum | HMPREF9623_00053 | 10076..13729 | 796942 | [illegible] |
| EHO16994.1 | Type1.BREX | [illegible] | 1180 | [illegible] | GCA_000242235.1 | Stomatobaculum_longum | HMPREF9623_00593 | 29898..33440 | 796942 | [illegible] |
| EHO53370.1 | Type1.BREX | [illegible] | 1186 | [illegible] | GCA_000242315.1 | Lachnospiraceae_bacterium_oral_taxon_082_str._F0431 | HMPREF9099_00896 | 82439..85999 | 861454 | [illegible] |
| EOS24741.1 | Type1.BREX | [illegible] | 1175 | [illegible] | GCA_000403215.2 | Lachnospiraceae_bacterium_A4 | C804_05106 | 19579..23106 | 397291 | [illegible] |
| EOS28360.1 | Type1.BREX | [illegible] | 1224 | [illegible] | GCA_000403275.2 | Lachnospiraceae_bacterium_28-4 | C807_03898 | 219781..223455 | 397287 | [illegible] |
| EOS37361.1 | Type1.BREX | [illegible] | 1182 | [illegible] | GCA_000403295.2 | Lachnospiraceae_bacterium_M18-1 | C808_03339 | 122087..125635 | 1235792 | [illegible] |
| EOS37735.1 | Type1.BREX | [illegible] | 1218 | [illegible] | GCA_000403295.2 | Lachnospiraceae_bacterium_M18-1 | C808_03715 | 724875..728531 | 1235792 | [illegible] |
| EOS42584.1 | Type1.BREX | [illegible] | 1182 | [illegible] | GCA_000403315.2 | Lachnospiraceae_bacterium_A2 | C810_04454 | 133846..137394 | 397290 | [illegible] |
| CCY77707.1 | Type1.BREX | [illegible] | 1196 | [illegible] | GCA_000432395.1 | Butyrivibrio_crossotus_CAG:259 | BN569_00900 | 88927..92517 | 1263062 | [illegible] |
| CUO05580.1 | Type1.BREX | [illegible] | 1182 | [illegible] | GCA_001405875.1 | Roseburia_hominis | ERS852391_01284 | 73560..77108 | 301301 | [illegible] |
| CUM83721.1 | Type1.BREX | [illegible] | 1182 | [illegible] | GCA_001406375.1 | [Eubacterium]_rectale | ERS852580_00755 | 13381..16929 | 39491 | [illegible] |
| OLA13308.1 | Type1.BREX | [illegible] | 1222 | [illegible] | GCA_001916315.1 | Coprococcus_sp._CAG:131-related_45_246 | BHW16_02690 | 449..4117 | 1896998 | [illegible] |
| ONI39098.1 | Type1.BREX | [illegible] | 1197 | [illegible] | GCA_001983455.1 | Epulopiscium_sp._SCG-B11WGA-EpuloA1 | AN396_09505 | 82489..86082 | 1710701 | [illegible] |
| OON94626.1 | Type1.BREX | [illegible] | 1173 | [illegible] | GCA_002007295.1 | Epulopiscium_sp._AS2M-Bin001 | ATN31_00325 | 9781..13302 | 1764958 | [illegible] |
| OOO00080.1 | Type1.BREX | [illegible] | 1209 | [illegible] | GCA_002007315.1 | Epulopiscium_sp._Nele67-Bin004 | ATN35_09340 | 17804..21433 | 1764962 | [illegible] |
| PHV69589.1 | Type1.BREX | [illegible] | 1212 | [illegible] | GCA_002735925.1 | Lachnospiraceae_bacterium | CS063_14865 | 2119..5757 | 1898203 | [illegible] |
| PLT70230.1 | Type1.BREX | [illegible] | 1222 | [illegible] | GCA_002865325.1 | [Ruminococcus]_gnavus | CDL26_14520 | 14609..18277 | 33038 | [illegible] |
| RGC74026.1 | Type1.BREX | [illegible] | 1203 | [illegible] | GCA_003434385.1 | Lachnospiraceae_bacterium_AM23-2LB | DW655_01895 | 389201..392812 | 2302964 | [illegible] |
| RGI65884.1 | Type1.BREX | [illegible] | 1196 | [illegible] | GCA_003436035.1 | [Eubacterium]_rectale | DXD95_13285 | 1619..5209 | 39491 | [illegible] |
| RGO23301.1 | Type1.BREX | [illegible] | 1201 | [illegible] | GCA_003439775.1 | Dorea_sp._OM02-2LB | DXB23_13015 | 77589..81194 | 2292347 | [illegible] |
| HAB87400.1 | Type1.BREX | [illegible] | 1196 | [illegible] | GCA_003448155.1 | Coprococcus_sp. | DCE60_00030 | 3490..7080 | 2049024 | [illegible] |
| RGS69619.1 | Type1.BREX | [illegible] | 1182 | [illegible] | GCA_003459375.1 | Blautia_obeum | DWX77_14595 | 469..4017 | 40520 | [illegible] |
| RGW41485.1 | Type1.BREX | [illegible] | 1182 | [illegible] | GCA_003464935.1 | [Eubacterium]_rectale | DWV78_01400 | 115095..118643 | 39491 | [illegible] |
| RHA07177.1 | Type1.BREX | [illegible] | 1182 | [illegible] | GCA_003466745.1 | [Eubacterium]_rectale | DW951_01530 | 328230..331778 | 39491 | [illegible] |
| RGZ95621.1 | Type1.BREX | [illegible] | 1182 | [illegible] | GCA_003467085.1 | [Eubacterium]_rectale | DW967_00020 | 2042..5590 | 39491 | [illegible] |
| RGZ88026.1 | Type1.BREX | [illegible] | 1208 | [illegible] | GCA_003467085.1 | [Eubacterium]_rectale | DW967_16245 | 9325..12951 | 39491 | [illegible] |
| RHC12241.1 | Type1.BREX | [illegible] | 1044 | [illegible] | GCA_003467725.1 | Roseburia_intestinalis | DW856_19080 | 43..3177 | 166486 | [illegible] |
| RHD97433.1 | Type1.BREX | [illegible] | 1182 | [illegible] | GCA_003468965.1 | [Eubacterium]_rectale | DW775_02150 | 1680..5228 | 39491 | [illegible] |
| RHL80687.1 | Type1.BREX | [illegible] | 1205 | [illegible] | GCA_003474775.1 | [Eubacterium]_rectale | DW001_05755 | 128905..132522 | 39491 | [illegible] |
| HAX07244.1 | Type1.BREX | [illegible] | 1196 | [illegible] | GCA_003487615.1 | Butyrivibrio_sp. | DCX93_04145 | 4022..7612 | 28121 | [illegible] |
| HBA06127.1 | Type1.BREX | [illegible] | 1182 | [illegible] | GCA_003491515.1 | Roseburia_sp. | DC452_03320 | 12130..15678 | 2049040 | [illegible] |
| HBD78540.1 | Type1.BREX | [illegible] | 1138 | [illegible] | GCA_003497645.1 | Roseburia_sp. | DC044_10265 | 1..3413 | 2049040 | [illegible] |
| HBV83857.1 | Type1.BREX | [illegible] | 711 | [illegible] | GCA_003513705.1 | Lachnospiraceae_bacterium | DEB74_13945 | 1..2133 | 1898203 | [illegible] |
| HCI18472.1 | Type1.BREX | [illegible] | 799 | [illegible] | GCA_003521585.1 | Lachnospiraceae_bacterium | DFH32_07645 | 14441..16840 | 1898203 | [illegible] |
| HCR84102.1 | Type1.BREX | [illegible] | 437 | [illegible] | GCA_003524895.1 | Lachnospiraceae_bacterium | DIW07_11990 | 6618..7927 | 1898203 | [illegible] |
| HCI84254.1 | Type1.BREX | [illegible] | 782 | [illegible] | GCA_003526665.1 | Lachnospiraceae_bacterium | DHV79_06645 | 6068..8416 | 1898203 | [illegible] |
| HCS67503.1 | Type1.BREX | [illegible] | 1189 | [illegible] | GCA_003531095.1 | Oribacterium_sp. | DIW34_04725 | 7705..11274 | 1969407 | [illegible] |
| HBP24697.1 | Type1.BREX | [illegible] | 629 | [illegible] | GCA_003531935.1 | Oribacterium_sp. | DD632_04640 | 9060..10945 | 1969407 | [illegible] |
| HCX41706.1 | Type1.BREX | [illegible] | 1196 | [illegible] | GCA_003539645.1 | Lachnospiraceae_bacterium | DHH50_05105 | 179559..183149 | 1898203 | [illegible] |
| HCT91519.1 | Type1.BREX | [illegible] | 656 | [illegible] | GCA_003542935.1 | Lachnospiraceae_bacterium | DF613_09115 | 1..1968 | 1898203 | [illegible] |
| HCU04414.1 | Type1.BREX | [illegible] | 1182 | [illegible] | GCA_003543095.1 | Roseburia_sp. | DF667_12405 | 43487..47035 | 2049040 | [illegible] |
| RKW57361.1 | Type1.BREX | [illegible] | 1217 | [illegible] | GCA_003640285.1 | Lachnospiraceae_bacterium | D8H95_03165 | 3356..7009 | 1898203 | [illegible] |
| RKW51253.1 | Type1.BREX | [illegible] | 1217 | [illegible] | GCA_003640285.1 | Lachnospiraceae_bacterium | D8H95_17195 | 4877..8530 | 1898203 | [illegible] |
| TGY98248.1 | Type1.BREX | [illegible] | 1218 | [illegible] | GCA_004793545.1 | Petralouisia_muris | E5329_00220 | 47888..51544 | 3032872 | [illegible] |
| TGX97927.1 | Type1.BREX | [illegible] | 1201 | [illegible] | GCA_004793585.1 | Lachnospiraceae_bacterium | E5357_10825 | 35909..39514 | 1898203 | [illegible] |
| TYL58976.1 | Type1.BREX | [illegible] | 1182 | [illegible] | GCA_008122485.1 | [Eubacterium]_rectale | FYL37_04475 | 43133..46681 | 39491 | [illegible] |
| TYL56933.1 | Type1.BREX | [illegible] | 1196 | [illegible] | GCA_008123415.1 | [Eubacterium]_rectale | FYL31_13935 | 1744..5334 | 39491 | [illegible] |
| MSS15025.1 | Type1.BREX | [illegible] | 821 | [illegible] | GCA_009696445.1 | Porcincola_intestinalis | FYJ35_08215 | 8308..10773 | 2606632 | [illegible] |
| NBJ03627.1 | Type1.BREX | [illegible] | 1205 | [illegible] | GCA_009911175.1 | Lachnospiraceae_bacterium | D3Z62_26960 | 10883..14500 | 1898203 | [illegible] |
| NBI58577.1 | Type1.BREX | [illegible] | 1182 | [illegible] | GCA_009911275.1 | Lachnospiraceae_bacterium | D3Z53_11015 | 107826..111374 | 1898203 | [illegible] |
| NBI87409.1 | Type1.BREX | [illegible] | 1218 | [illegible] | GCA_009911505.1 | Lachnospiraceae_bacterium | D3Z47_14860 | 26627..30283 | 1898203 | [illegible] |
| GFI03050.1 | Type1.BREX | [illegible] | 1224 | [illegible] | GCA_011959285.1 | Lachnospiraceae_bacterium | IMSAGC005_01882 | 2895..6569 | 1898203 | [illegible] |
| GFI22895.1 | Type1.BREX | [illegible] | 1209 | [illegible] | GCA_011959805.1 | Lachnospiraceae_bacterium | IMSAGC011_01672 | 23760..27389 | 1898203 | [illegible] |
| GFI23757.1 | Type1.BREX | [illegible] | 1176 | [illegible] | GCA_011959805.1 | Lachnospiraceae_bacterium | IMSAGC011_02547 | 4111..7641 | 1898203 | [illegible] |
| GFI37407.1 | Type1.BREX | [illegible] | 1182 | [illegible] | GCA_011960265.1 | Lachnospiraceae_bacterium | IMSAGC015_01594 | 3920..7468 | 1898203 | [illegible] |
| NMA84127.1 | Type1.BREX | [illegible] | 1196 | [illegible] | GCA_012799415.1 | Epulopiscium_sp. | GX962_09740 | 6009..9599 | 2384 | [illegible] |
| NSE64224.1 | Type1.BREX | [illegible] | 626 | [illegible] | GCA_013301175.1 | Faecalicatena_fissicatena | HFM88_08960 | 28662..30542 | 290055 | [illegible] |
| NSJ91243.1 | Type1.BREX | [illegible] | 1139 | [illegible] | GCA_013304585.1 | Coprococcus_sp._MSK.21.13 | G7A79_19115 | 7512..10931 | 2709719 | [illegible] |
| QNM03383.1 | Type1.BREX | [illegible] | 1215 | [illegible] | GCA_014337175.1 | Simiaoa_sunii | H9Q77_04470 | 956158..959805 | 2763672 | [illegible] |
| MBD5550863.1 | Type1.BREX | [illegible] | 901 | [illegible] | GCA_014804795.1 | Lachnospiraceae_bacterium | HDQ96_06750 | 4317..7018 | 1898203 | [illegible] |
| MBD9030270.1 | Type1.BREX | [illegible] | 1205 | [illegible] | GCA_014847615.1 | Butyrivibrio_crossotus | EGR17_09315 | 97277..100894 | 45851 | [illegible] |
| MBD9288174.1 | Type1.BREX | [illegible] | 1182 | [illegible] | GCA_014848915.1 | Agathobacter_sp. | EGS40_13260 | 26721..30269 | 2021311 | [illegible] |
| MBE5871717.1 | Type1.BREX | [illegible] | 1196 | [illegible] | GCA_015056815.1 | Lachnospiraceae_bacterium | E7294_10725 | 54734..58324 | 1898203 | [illegible] |
| MBN2952027.1 | Type1.BREX | [illegible] | 1182 | [illegible] | GCA_016939215.1 | Fusicatenibacter_saccharivorans | JTJ23_00185 | 3299..6847 | 1150298 | [illegible] |
| MBN2955416.1 | Type1.BREX | [illegible] | 1217 | [illegible] | GCA_016939255.1 | Blautia_massiliensis | JTJ22_02515 | 22599..26252 | 1737424 | [illegible] |
| MBQ3163820.1 | Type1.BREX | [illegible] | 1358 | [illegible] | GCA_017397345.1 | Lachnospiraceae_bacterium | IJC02_04660 | 79026..83102 | 1898203 | [illegible] |
| MBQ8913711.1 | Type1.BREX | [illegible] | 1206 | [illegible] | GCA_017460945.1 | Lachnospiraceae_bacterium | IJ054_06675 | 2560..6180 | 1898203 | [illegible] |
| MBQ9134971.1 | Type1.BREX | [illegible] | 1213 | [illegible] | GCA_017463605.1 | Lachnospiraceae_bacterium | IJX66_02610 | 29830..33471 | 1898203 | [illegible] |
| MBQ8665324.1 | Type1.BREX | [illegible] | 1213 | [illegible] | GCA_017467715.1 | Lachnospiraceae_bacterium | IJ526_00325 | 437..4078 | 1898203 | [illegible] |
| MBR2123171.1 | Type1.BREX | [illegible] | 1174 | [illegible] | GCA_017474455.1 | Lachnospiraceae_bacterium | IJ930_09665 | 172252..175776 | 1898203 | [illegible] |
| MBR2401863.1 | Type1.BREX | [illegible] | 701 | [illegible] | GCA_017503825.1 | Lachnospiraceae_bacterium | IKB01_03695 | 1..2105 | 1898203 | [illegible] |
| MBR3242658.1 | Type1.BREX | [illegible] | 719 | [illegible] | GCA_017511245.1 | Parasporobacterium_sp. | IKF90_08180 | 2891..5047 | 2740533 | [illegible] |
| MBR3244379.1 | Type1.BREX | [illegible] | 1207 | [illegible] | GCA_017511245.1 | Parasporobacterium_sp. | IKF90_17090 | 4939..8562 | 2740533 | [illegible] |
| MBR3171576.1 | Type1.BREX | [illegible] | 1178 | [illegible] | GCA_017514845.1 | Lachnospiraceae_bacterium | IKF22_10080 | 15258..18794 | 1898203 | [illegible] |
| MBR4084227.1 | Type1.BREX | [illegible] | 1213 | [illegible] | GCA_017521605.1 | Lachnospiraceae_bacterium | IKK33_08100 | 377..4018 | 1898203 | [illegible] |
| MBR4606065.1 | Type1.BREX | [illegible] | 1191 | [illegible] | GCA_017536085.1 | Lachnospiraceae_bacterium | IKO41_07545 | 5884..9459 | 1898203 | [illegible] |
| MBR4608207.1 | Type1.BREX | [illegible] | 1155 | [illegible] | GCA_017536085.1 | Lachnospiraceae_bacterium | IKO41_18520 | 18020..21487 | 1898203 | [illegible] |
| MBR4345194.1 | Type1.BREX | [illegible] | 1173 | [illegible] | GCA_017542245.1 | Lachnospiraceae_bacterium | IKP88_21230 | 56947..60468 | 1898203 | [illegible] |
| MBO4458248.1 | Type1.BREX | [illegible] | 1216 | [illegible] | GCA_017615275.1 | Butyrivibrio_sp. | J5802_11055 | 10594..14244 | 28121 | [illegible] |
| MBP3802955.1 | Type1.BREX | [illegible] | 1204 | [illegible] | GCA_017940955.1 | Oribacterium_sp. | J6I76_03520 | 9410..13024 | 1969407 | [illegible] |
| MBP3803747.1 | Type1.BREX | [illegible] | 1173 | [illegible] | GCA_017940955.1 | Oribacterium_sp. | J6I76_07595 | 13930..17451 | 1969407 | [illegible] |
| MBP3882711.1 | Type1.BREX | [illegible] | 1195 | [illegible] | GCA_017941625.1 | Lachnospiraceae_bacterium | J6E40_11195 | 14786..18373 | 1898203 | [illegible] |
| MBS5544368.1 | Type1.BREX | [illegible] | 1211 | [illegible] | GCA_018369095.1 | Lachnospiraceae_bacterium | KHX49_00170 | 28664..32299 | 1898203 | [illegible] |
| MBS5764136.1 | Type1.BREX | [illegible] | 1205 | [illegible] | GCA_018371095.1 | Lachnospiraceae_bacterium | KHW68_11675 | 2915..6532 | 1898203 | [illegible] |
| MBS5881823.1 | Type1.BREX | [illegible] | 1214 | [illegible] | GCA_018372015.1 | Lachnoanaerobaculum_sp. | KIC55_05385 | 26019..29663 | 2049030 | [illegible] |
| HIS62962.1 | Type1.BREX | [illegible] | 1178 | [illegible] | GCA_018710905.1 | Candidatus_Scybalomonas_excrementigallinarum | IAC14_12110 | 84..3620 | 2840944 | [illegible] |
| MBU5248973.1 | Type1.BREX | [illegible] | 1201 | [illegible] | GCA_018916865.1 | Coprococcus_comes | KQH93_04110 | 138468..142073 | 410072 | [illegible] |
| HIV42030.1 | Type1.BREX | [illegible] | 1173 | [illegible] | GCA_019112735.1 | Candidatus_Mediterraneibacter_guildfordensis | H9904_08590 | 1629..5150 | 2838672 | [illegible] |
| HIV92250.1 | Type1.BREX | [illegible] | 1180 | [illegible] | GCA_019113175.1 | Candidatus_Eisenbergiella_stercoravium | H9744_11330 | 86529..90071 | 2838556 | [illegible] |
| HJB00421.1 | Type1.BREX | [illegible] | 484 | [illegible] | GCA_019118545.1 | Candidatus_Mediterraneibacter_merdavium | H9780_04575 | 1..1452 | 2838676 | [illegible] |
| HJC65843.1 | Type1.BREX | [illegible] | 1180 | [illegible] | GCA_019119955.1 | Candidatus_Enterocloster_excrementigallinarum | H9931_03865 | 69183..72725 | 2838558 | [illegible] |
| MCB7506159.1 | Type1.BREX | [illegible] | 1182 | [illegible] | GCA_020553125.1 | Blautia_sp._MSK20_18 | LI091_01875 | 117354..120902 | 2883186 | [illegible] |
| MCB7110231.1 | Type1.BREX | [illegible] | 1182 | [illegible] | GCA_020554685.1 | [Eubacterium]_rectale | LON19_09275 | 43133..46681 | 39491 | [illegible] |
| MCC2126187.1 | Type1.BREX | [illegible] | 1182 | [illegible] | GCA_020686965.1 | Lachnospiraceae_bacterium_CLA-AA-H276 | LKD36_08340 | 179380..182928 | 2885352 | [illegible] |
| MCC2167282.1 | Type1.BREX | [illegible] | 1217 | [illegible] | GCA_020687225.1 | Lachnospiraceae_bacterium_CLA-AA-H244 | LKD45_06170 | 37511..41164 | 2885350 | [illegible] |
| MCC2229908.1 | Type1.BREX | [illegible] | 1182 | [illegible] | GCA_020687485.1 | Lachnospiraceae_bacterium_CLA-AA-H215 | LKD81_02670 | 26793..30341 | 2885348 | [illegible] |
| MCC2230802.1 | Type1.BREX | [illegible] | 1217 | [illegible] | GCA_020687485.1 | Lachnospiraceae_bacterium_CLA-AA-H215 | LKD81_07295 | 25894..29547 | 2885348 | [illegible] |
| MCD7743820.1 | Type1.BREX | [illegible] | 1187 | [illegible] | GCA_021199475.1 | Lachnospiraceae_bacterium | LUI13_00760 | 15415..18978 | 1898203 | [illegible] |
| MCD7763957.1 | Type1.BREX | [illegible] | 1215 | [illegible] | GCA_021199655.1 | Lachnospiraceae_bacterium | LUI14_12340 | 45356..49003 | 1898203 | [illegible] |
| MCD7882907.1 | Type1.BREX | [illegible] | 1235 | [illegible] | GCA_021201095.1 | Lachnospiraceae_bacterium | LUI87_04245 | 13956..17663 | 1898203 | [illegible] |
| MCD7956582.1 | Type1.BREX | [illegible] | 1211 | [illegible] | GCA_021201665.1 | Lachnospiraceae_bacterium | LUG93_12740 | 57762..61397 | 1898203 | [illegible] |
| MCD8011880.1 | Type1.BREX | [illegible] | 1182 | [illegible] | GCA_021202075.1 | Lachnospiraceae_bacterium | LUG99_01630 | 126..3674 | 1898203 | [illegible] |
| MCD8362902.1 | Type1.BREX | [illegible] | 1212 | [illegible] | GCA_021205385.1 | Lachnospiraceae_bacterium | LUC98_08085 | 19577..23215 | 1898203 | [illegible] |
| MCH3986384.1 | Type1.BREX | [illegible] | 1223 | [illegible] | GCA_022483185.1 | Lachnospiraceae_bacterium | LKE69_00320 | 66250..69921 | 1898203 | [illegible] |
| MCH4064994.1 | Type1.BREX | [illegible] | 1223 | [illegible] | GCA_022483745.1 | Lachnospiraceae_bacterium | LKF12_06105 | 333794..337465 | 1898203 | [illegible] |
| MCI5856701.1 | Type1.BREX | [illegible] | 218 | [illegible] | GCA_022769125.1 | Agathobacter_sp. | MR016_04965 | 1..655 | 2021311 | [illegible] |
| MCI7121458.1 | Type1.BREX | [illegible] | 1201 | [illegible] | GCA_022775325.1 | [Ruminococcus]_gnavus | MR941_06720 | 10105..13710 | 33038 | [illegible] |
| MCI6977806.1 | Type1.BREX | [illegible] | 1182 | [illegible] | GCA_022775645.1 | Lachnospiraceae_bacterium | MR930_04145 | 182006..185554 | 1898203 | [illegible] |
| MCI6915425.1 | Type1.BREX | [illegible] | 1217 | [illegible] | GCA_022777965.1 | Lachnospiraceae_bacterium | MR799_08105 | 55214..58867 | 1898203 | [illegible] |
| MCI6533897.1 | Type1.BREX | [illegible] | 1173 | [illegible] | GCA_022782135.1 | Lachnospiraceae_bacterium | MR444_03700 | 25882..29403 | 1898203 | [illegible] |
| MCI6498729.1 | Type1.BREX | [illegible] | 1196 | [illegible] | GCA_022782285.1 | Lachnospiraceae_bacterium | MSA21_07600 | 105165..108755 | 1898203 | [illegible] |
| MCI7493454.1 | Type1.BREX | [illegible] | 1196 | [illegible] | GCA_022785005.1 | Lachnobacterium_sp. | MSA91_11815 | 73793..77383 | 2774292 | [illegible] |
| MCI8269180.1 | Type1.BREX | [illegible] | 1182 | [illegible] | GCA_022790625.1 | Lachnospiraceae_bacterium | HFG55_05920 | 9727..13275 | 1898203 | [illegible] |
| MCI8292339.1 | Type1.BREX | [illegible] | 1208 | [illegible] | GCA_022790755.1 | Hespellia_sp. | HFH53_02225 | 79420..83046 | 2714646 | [illegible] |
| MCI8490100.1 | Type1.BREX | [illegible] | 1198 | [illegible] | GCA_022792655.1 | Lachnospiraceae_bacterium | HFJ04_07660 | 19316..22912 | 1898203 | [illegible] |
| MCI8692001.1 | Type1.BREX | [illegible] | 1202 | [illegible] | GCA_022794455.1 | Lachnospiraceae_bacterium | HFH91_04535 | 2210..5818 | 1898203 | [illegible] |
| MCI8732218.1 | Type1.BREX | [illegible] | 1210 | [illegible] | GCA_022794815.1 | Lachnospiraceae_bacterium | HFH57_14060 | 7970..11602 | 1898203 | [illegible] |
| MCI8749656.1 | Type1.BREX | [illegible] | 1213 | [illegible] | GCA_022794875.1 | Lachnospiraceae_bacterium | HFH67_17680 | 8915..12556 | 1898203 | [illegible] |
| MCI8860127.1 | Type1.BREX | [illegible] | 1213 | [illegible] | GCA_022795635.1 | Lachnospiraceae_bacterium | HFI71_11555 | 14645..18286 | 1898203 | [illegible] |
| MCI8995241.1 | Type1.BREX | [illegible] | 1204 | [illegible] | GCA_022796795.1 | Lachnospiraceae_bacterium | HFI30_06070 | 7009..10623 | 1898203 | [illegible] |
| MCI9133533.1 | Type1.BREX | [illegible] | 1204 | [illegible] | GCA_022797975.1 | Lachnospiraceae_bacterium | HFI31_05000 | 7501..11115 | 1898203 | [illegible] |
| MCI9239442.1 | Type1.BREX | [illegible] | 1182 | [illegible] | GCA_022798835.1 | Dorea_sp. | HFH07_13900 | 2922..6470 | 2040332 | [illegible] |
| MCI9358063.1 | Type1.BREX | [illegible] | 1213 | [illegible] | GCA_022799775.1 | Lachnospiraceae_bacterium | HFH59_11085 | 6660..10301 | 1898203 | [illegible] |

| Protein-ID | System | Gene-Neighbourhoods | Length | Taxonomy | Assembly-ID | Strain | Locus-Tag | Genomic-coordinates | TaxID | Fasta-define |
|---|---|---|---|---|---|---|---|---|---|---|
| MCI9399935.1 | Type1.BREX | <-TetR-HTH\|\|?->BrxA-DNA-binding->BrxB-BrxF-iSTAND->DUF6079*->N6-METHYLASE->LRP-HTH->Pkinase+CR-Kinase->EAL->PglZ->SIGMA+OB-Fold+AAA-ATPase+Lon-Protease-BrxL->?->DUF1016+REase-> | 1176 | Bacteria>Terrabacteria_group>Bacillota>Clostridia>Eubacteriales>Lachnospiraceae | GCA_022800085.1 | Lachnospiraceae_bacterium | HFJ07_08990 | 28102..31632 | 1898203 | BREX-system-P-loop-protein-BrxC-[Lachnospiraceae-bacterium] |
| MCI9389752.1 | Type1.BREX | <-REC+OmpR-HTH<-?\|\|Amidohydro_1+Adenine_deam_C->Carboxypeptidase+METHYLASE->?->MODE-HTH+PBPII->?->BrxA-DNA-binding->BrxB-BrxF-iSTAND->DUF6079*->N6-METHYLASE-p-> | 1197 | Bacteria>Terrabacteria_group>Bacillota>Clostridia>Eubacteriales>Lachnospiraceae | GCA_022800095.1 | Lachnospiraceae_bacterium | HFH54_08745 | 31751..35344 | 1898203 | BREX-system-P-loop-protein-BrxC-[Lachnospiraceae-bacterium] |
| MCI9471722.1 | Type1.BREX | <-REC+OmpR-HTH<-?\|\|Amidohydro_1+Adenine_deam_C->Carboxypeptidase+METHYLASE->?->MODE-HTH+PBPII->?->BrxA-DNA-binding->BrxB-BrxF-iSTAND->DUF6079*->N6-METHYLASE-> | 1197 | Bacteria>Terrabacteria_group>Bacillota>Clostridia>Eubacteriales>Lachnospiraceae | GCA_022800715.1 | Lachnospiraceae_bacterium | HFH56_11080 | 31533..35126 | 1898203 | BREX-system-P-loop-protein-BrxC-[Lachnospiraceae-bacterium] |
| MCI9453449.1 | Type1.BREX | <-PglZ<-?<-?<-Classical-AAA<-DUF6079*<-BrxB-BrxF-iSTAND<-BrxA-DNA-binding<-Calcineurin\|\|?->Tox-RelE->PadR-HTH-> | 1222 | Bacteria>Terrabacteria_group>Bacillota>Clostridia>Eubacteriales>Lachnospiraceae | GCA_022800995.1 | Dorea_sp. | HFG83_06420 | 26015..29683 | 2040332 | BREX-system-P-loop-protein-BrxC-[Dorea-sp.] |
| MCI9595477.1 | Type1.BREX | <-PglZ<-N6-METHYLASE<-DUF6079*<-BrxB-BrxF-iSTAND<-BrxA-DNA-binding<-HTH_26 | 1176 | Bacteria>Terrabacteria_group>Bacillota>Clostridia>Eubacteriales>Lachnospiraceae | GCA_022801655.1 | Lachnospiraceae_bacterium | HFG51_15410 | 5386..8916 | 1898203 | BREX-system-P-loop-protein-BrxC-[Lachnospiraceae-bacterium] |
| GKH56658.1 | Type1.BREX | <-Tox-RelA<-SMODS<-SLATT<-SIGMA+OB-Fold+AAA-ATPase+Lon-Protease-BrxL<-PglZ<-AT-HTH-HYD2-2<-?<-Tox-ART-HYD2<-N6-METHYLASE<-DUF6079*<-BrxB-BrxF-iSTAND<-BrxA-DNA-binding<-XIS-HTH<-?<-?<-TraM<-TraJ-RHH<-ArdCN+LPD32+LPD36+N6_Mtase+SF1-SF2-Helicase | 1217 | Bacteria>Terrabacteria_group>Bacillota>Clostridia>Eubacteriales>Lachnospiraceae | GCA_022835195.1 | Lachnospiraceae_bacterium | CE91St58_40430 | 4462022..4465675 | 1898203 | hypothetical-protein-CE91St58_40430-[Lachnospiraceae-bacterium] |
| BDF25269.1 | Type1.BREX | <-REase<-?<-?<-?<-HHH_4+HTH+wHTH+SF1-SF2-Helicase+REase<-MORC+DUF3684+HNH<-HEPN+SH3-like+SF1-SF2-Helicase+wHTH<-PglZ<-N6-METHYLASE<-DUF6079*<-BrxB-BrxF-iSTAND<-BrxA-DNA-binding<-HTH_26<-?<-wHTH+WYL+WCX<-SSB<-?<-TraJ-RHH<-LPD2 | 1209 | Bacteria>Terrabacteria_group>Bacillota>Clostridia>Eubacteriales>Lachnospiraceae | GCA_022845855.1 | [Clostridium]_symbiosum | CE91St65_31490 | 3519815..3523444 | 1512 | hypothetical-protein-CE91St65_31490-[[Clostridium] |
| MCM1125306.1 | Type1.BREX | MoxR-AAA->LRR_5->Glyco_hydro_3_C+Fn3-like+Glyco_hydro_3->Peptidase_C26->tRNA->tRNA-><-?<-MerR-HTH\|\|HTH_26->BrxA-DNA-binding->BrxB-BrxF-iSTAND->DUF6079*->ParB+HNH->HEPN->N6-METHYLASE-p->?-><-tRNA\|\|?->HAD+HAD-> | 1205 | Bacteria>Terrabacteria_group>Bacillota>Clostridia>Eubacteriales>Lachnospiraceae | GCA_023663945.1 | Lachnospiraceae_bacterium | NC429_02425 | 28301..31918 | 1898203 | BREX-system-P-loop-protein-BrxC-[Lachnospiraceae-bacterium] |
| MCQ4892848.1 | Type1.BREX | HTH_36->?->CopG/RHH->?->TraJ-RHH->TraI-VirD2-RCRE->HTH_26->BrxA-DNA-binding->BrxB-BrxF-iSTAND->DUF6079*->?->N6-METHYLASE->HEPN->Schlafen+HATPase+wHTH->PglZ->SIGMA+OB-Fold+AAA-ATPase+Lon-Protease-BrxL->OLD-ABC+TOPRIM->SF1-SF2-Helicase-><-HTH | 1182 | Bacteria>Terrabacteria_group>Bacillota>Clostridia>Eubacteriales>Lachnospiraceae | GCA_024461355.1 | Dorea_longicatena | NE607_07100 | 21358..24906 | 88431 | BREX-system-P-loop-protein-BrxC-[Dorea-longicatena] |
| MCQ5166118.1 | Type1.BREX | Peptidase_M18+Peptidase_M42->?->TRAM+METHYLASE->ZnR+Phage_GP20->HTH_26->BrxA-DNA-binding->BrxB-BrxF-iSTAND->DUF6079*->N6-METHYLASE->PglZ->SIGMA+OB-Fold+AAA-ATPase+Lon-Protease-BrxL->?->?->ZnR+NACHT-> | 1208 | Bacteria>Terrabacteria_group>Bacillota>Clostridia>Eubacteriales>Lachnospiraceae | GCA_024462985.1 | Roseburia_hominis | NE645_08885 | 23142..26768 | 301301 | BREX-system-P-loop-protein-BrxC-[Roseburia-hominis] |
| MCR5205351.1 | Type1.BREX | ParA-Soj-PloopNTPase->ParD_antitoxin->HTH_26->BrxA-DNA-binding->BrxB-BrxF-iSTAND->DUF6079-p*-> | 821 | Bacteria>Terrabacteria_group>Bacillota>Clostridia>Eubacteriales>Lachnospiraceae | GCA_024696945.1 | Lachnospiraceae_bacterium | K6E47_09860 | 3021..5484 | 1898203 | BREX-system-P-loop-protein-BrxC-[Lachnospiraceae-bacterium] |
| MCR5776618.1 | Type1.BREX | wHTH+WYL+WCX->?->HTH_26->BrxA-DNA-binding->BrxB-BrxF-iSTAND->DUF6079*->N6-METHYLASE->?->Tox-ART-HYD2-> | 1211 | Bacteria>Terrabacteria_group>Bacillota>Clostridia>Eubacteriales>Lachnospiraceae | GCA_024698945.1 | Lachnospiraceae_bacterium | K6G84_04260 | 3400..7035 | 1898203 | BREX-system-P-loop-protein-BrxC-[Lachnospiraceae-bacterium] |
| MCR5777562.1 | Type1.BREX | Phage_integrase->Phage_int_SAM_1+Phage_integrase->LPD2->TraJ-RHH->?->?->HTH_26->BrxA-DNA-binding->BrxB-BrxF-iSTAND->DUF6079*->N6-METHYLASE-> | 1208 | Bacteria>Terrabacteria_group>Bacillota>Clostridia>Eubacteriales>Lachnospiraceae | GCA_024698945.1 | Lachnospiraceae_bacterium | K6G84_09145 | 10893..14519 | 1898203 | BREX-system-P-loop-protein-BrxC-[Lachnospiraceae-bacterium] |
| MCR5727513.1 | Type1.BREX | BrxA-DNA-binding->BrxB-BrxF-iSTAND->DUF6079*->VbhA+DOC+HTH->N6-METHYLASE->HEPN->SbcC+SbcC->?->PglZ->SIGMA+OB-Fold+AAA-ATPase+Lon-Protease-BrxL->?->SF1-SF2-Helicase+REase-> | 1173 | Bacteria>Terrabacteria_group>Bacillota>Clostridia>Eubacteriales>Lachnospiraceae | GCA_024699435.1 | Lachnospiraceae_bacterium | K6G24_08610 | 2669..6190 | 1898203 | BREX-system-P-loop-protein-BrxC-[Lachnospiraceae-bacterium] |
| SCG99927.1 | Type1.BREX | <-PglZ<-Schlafen+HATPase+wHTH<-?<-?<-AT-HTH-HYD2<-RNase<-RNaseH<-N6-METHYLASE-p<-N6-METHYLASE<-DUF6079*<-BrxB-BrxF-iSTAND<-BrxA-DNA-binding<-HTH_26<-?<-?<-?<-?<-REC+OmpR-HTH | 1182 | Bacteria>Terrabacteria_group>Bacillota>Clostridia>Eubacteriales>Lachnospiraceae | GCA_900066205.1 | uncultured_Blautia_sp. | SAMEA3545256_00122 | 140025..143573 | 765821 | Uncharacterised-protein-[uncultured-Blautia-sp.] |
| SCW50280.1 | Type1.BREX | Arm-DNA-bind_4+Phage_int_SAM_3+Phage_integrase->TraJ-RHH->ParA-Soj-PloopNTPase->ParB->?->?->HTH_26->BrxA-DNA-binding->BrxB-BrxF-iSTAND->DUF6079*->N6-METHYLASE->?->PglZ->HEPN->SIGMA+OB-Fold+AAA-ATPase+Lon-Protease-BrxL->Peptidase_C14-> | 1181 | Bacteria>Terrabacteria_group>Bacillota>Clostridia>Eubacteriales>Lachnospiraceae | GCA_900100095.1 | Lachnospiraceae_bacterium_C10 | SAMN02910400_01008 | 22627..26172 | 1520822 | hypothetical-protein-SAMN02910400_01008-[Lachnospiraceae-bacterium-C10] |
| SEQ84346.1 | Type1.BREX | Classical-AAA->MACRODOMAIN->?->HTH_26->BrxA-DNA-binding->BrxB-BrxF-iSTAND->DUF6079*->N6-METHYLASE->N6-METHYLASE->PglZ->?->?->Schlafen+HATPase+wHTH->SIGMA+OB-Fold+AAA-ATPase+Lon-Protease-BrxL->LRR_5+LRR_8-> | 1212 | Bacteria>Terrabacteria_group>Bacillota>Clostridia>Eubacteriales>Lachnospiraceae | GCA_900110635.1 | Lachnospiraceae_bacterium_NE2001 | SAMN02910369_02521 | 19203..22841 | 1520823 | hypothetical-protein-SAMN02910369_02521-[Lachnospiraceae-bacterium-NE2001] |
| SEP57791.1 | Type1.BREX | <-REase+HIN-HTH<-DUF6079-p*<-BrxB-BrxF-iSTAND<-BrxA-DNA-binding<-HTH_26<-RelE-ParE<-PhdYeFM_antitox<-RNase+DSRBD | 439 | Bacteria>Terrabacteria_group>Bacillota>Clostridia>Eubacteriales>Lachnospiraceae | GCA_900110975.1 | Lachnospiraceae_bacterium_RM5 | SAMN02910289_00002 | 294..1613 | 1200721 | hypothetical-protein-SAMN02910289_00002-[Lachnospiraceae-bacterium-RM5] |
| SEP83303.1 | Type1.BREX | DUF6079-p*->N6-METHYLASE->PglZ->Schlafen+HATPase+wHTH->SIGMA+OB-Fold+AAA-ATPase+Lon-Protease-BrxL->CR-Kinase->?->?->REase-> | 728 | Bacteria>Terrabacteria_group>Bacillota>Clostridia>Eubacteriales>Lachnospiraceae | GCA_900110975.1 | Lachnospiraceae_bacterium_RM5 | SAMN02910289_00625 | 328..2514 | 1200721 | hypothetical-protein-SAMN02910289_00625-[Lachnospiraceae-bacterium-RM5] |
| SFS09257.1 | Type1.BREX | <-HIN-HTH\|\|DUF6079-p*->N6-METHYLASE->PglZ->SIGMA+OB-Fold+AAA-ATPase+Lon-Protease-BrxL-><-rve_3 | 786 | Bacteria>Terrabacteria_group>Bacillota>Clostridia>Eubacteriales>Lachnospiraceae | GCA_900112775.1 | Anaeromicropila_populeti | SAMN05661086_03721 | 491..2851 | 37658 | hypothetical-protein-SAMN05661086_03721-[Anaeromicropila-populeti] |
| SFG07145.1 | Type1.BREX | ArdCN+MPTase-inact+LPD19->TOPRIM+Topoisom_bac->LPD2->TraJ-RHH->?->HTH_26->BrxA-DNA-binding->BrxB-BrxF-iSTAND->DUF6079*->N6-METHYLASE->?->PglZ->AbiGi->SIGMA+OB-Fold+AAA-ATPase+Lon-Protease-BrxL->ParB+HNH->HEPN-> | 1182 | Bacteria>Terrabacteria_group>Bacillota>Clostridia>Eubacteriales>Lachnospiraceae | GCA_900113385.1 | Lachnospiraceae_bacterium_C7 | SAMN04487761_10376 | 78580..82128 | 1792306 | hypothetical-protein-SAMN04487761_10376-[Lachnospiraceae-bacterium-C7] |
| SFG31088.1 | Type1.BREX | <-SIGMA+OB-Fold+AAA-ATPase+Lon-Protease-BrxL<-PglZ<-HTH+REase<-HNH+ADA-DBD<-?<-N6-METHYLASE<-DUF6079-p*<-BrxB-BrxF-iSTAND<-BrxA-DNA-binding<-HTH_26<-?<-?<-TraI-VirD2-RCRE | 783 | Bacteria>Terrabacteria_group>Bacillota>Clostridia>Eubacteriales>Lachnospiraceae | GCA_900113415.1 | Oribacterium_sp._WCC10 | SAMN05216356_105117 | 149481..151832 | 1855343 | hypothetical-protein-SAMN05216356_105117-[Oribacterium-sp.-WCC10] |
| SFG31099.1 | Type1.BREX | <-SIGMA+OB-Fold+AAA-ATPase+Lon-Protease-BrxL<-PglZ<-HTH+REase<-HNH+ADA-DBD<-?<-N6-METHYLASE<-DUF6079-p*<-BrxB-BrxF-iSTAND<-BrxA-DNA-binding<-HTH_26<-?<-?<-TraI-VirD2-RCRE | 380 | Bacteria>Terrabacteria_group>Bacillota>Clostridia>Eubacteriales>Lachnospiraceae | GCA_900113415.1 | Oribacterium_sp._WCC10 | SAMN05216356_105118 | 151957..153099 | 1855343 | hypothetical-protein-SAMN05216356_105118-[Oribacterium-sp.-WCC10] |
| SFL32270.1 | Type1.BREX | DDE_Tnp->?->HTH_26->BrxA-DNA-binding->BrxB-BrxF-iSTAND->DUF6079*->N6-METHYLASE->HTH+REase->?->PglZ->SIGMA+OB-Fold+AAA-ATPase+Lon-Protease-BrxL-><-DSBH+AraC-HTH | 1207 | Bacteria>Terrabacteria_group>Bacillota>Clostridia>Eubacteriales>Lachnospiraceae | GCA_900114405.1 | Lachnospiraceae_bacterium_KH1T2 | SAMN05216390_1196 | 2450..6073 | 1855374 | hypothetical-protein-SAMN05216390_1196-[Lachnospiraceae-bacterium-KH1T2] |
| SNU04610.1 | Type1.BREX | HTH_3->?->?->35exo->SbcC->?->wHTH+WYL+WCX->BrxA-DNA-binding->BrxB-BrxF-iSTAND->DUF6079*->KAP-NTPase->N6-METHYLASE->HEPN->PglZ->HEPN+SH3-like+SF1-SF2-Helicase+wHTH->MORC+DUF3684+HNH->MORC+DUF3684+HNH->SF1-SF2-Helicase->SF1-SF2-Helicase+REase-> | 1206 | Bacteria>Terrabacteria_group>Bacillota>Clostridia>Eubacteriales>Lachnospiraceae | GCA_900199595.1 | Lachnospiraceae_bacterium | SAMN06297422_101152 | 246659..250279 | 1898203 | hypothetical-protein-SAMN06297422_101152-[Lachnospiraceae-bacterium] |
| WP_013273420.1 | Type1.BREX | <-REase+iREase<-HEPN+PAS+DUF4011+SF1-SF2-Helicase+REase<-SIGMA+OB-Fold+AAA-ATPase+Lon-Protease-BrxL<-PglZ<-?<-TIR<-N6-METHYLASE<-?<-DUF6079*<-BrxB-BrxF-iSTAND<-BrxA-DNA-binding<-HTH_26<-Phage_int_SAM_1+Phage_integrase<-Phage_integrase<-Phage_int_SAM_4+Phage_integrase<-HTH_36<-VirD4-TraG+VirD4-FtsK<-ZnR+TOPRIM | 1207 | Bacteria>Terrabacteria_group>Bacillota>Clostridia>Eubacteriales>Lachnospiraceae | GCF_000144625.1 | Lacrimispora_saccharolytica | CLOSA_RS14020 | 3024764..3028387 | 84030 | BREX-system-P-loop-protein-BrxC-[Lacrimispora-saccharolytica] |
| WP_013274277.1 | Type1.BREX | <-SMODS<-SLATT<-SIGMA+OB-Fold+AAA-ATPase+Lon-Protease-BrxL<-PglZ<-AT-HTH-HYD2-2<-?<-Tox-ART-HYD2<-N6-METHYLASE<-DUF6079*<-BrxB-BrxF-iSTAND<-BrxA-DNA-binding<-XIS-HTH<-?<-MACRODOMAIN<-?<-?<-TraM<-TraJ-RHH | 1217 | Bacteria>Terrabacteria_group>Bacillota>Clostridia>Eubacteriales>Lachnospiraceae | GCF_000144625.1 | Lacrimispora_saccharolytica | CLOSA_RS18595 | 4003035..4006688 | 84030 | BREX-system-P-loop-protein-BrxC-[Lacrimispora-saccharolytica] |
| WP_154647080.1 | Type1.BREX | ABC-ATPase->?->?->MarR-HTH->XIS-HTH->BrxA-DNA-binding->BrxB-BrxF-iSTAND->DUF6079*->N6-METHYLASE->Tox-ART-HYD2->?->AT-HTH-HYD2-2->PglZ-> | 1217 | Bacteria>Terrabacteria_group>Bacillota>Clostridia>Eubacteriales>Lachnospiraceae | GCF_000185385.1 | Lachnoanaerobaculum_saburreum | HMPREF0381_RS05105 | 1086193..1089846 | 467210 | BREX-system-P-loop-protein-BrxC-[Lachnoanaerobaculum-saburreum] |
| WP_208913720.1 | Type1.BREX | HicA_toxin->ZnR+ZnR+ZnR->HTH_26->BrxA-DNA-binding->DUF6079-p*->N6-METHYLASE->OLD-ABC->NERD->PglZ->SIGMA+OB-Fold+AAA-ATPase+Lon-Protease-BrxL->LRR_5+LRR_8->?->STYKIN-> | 644 | Bacteria>Terrabacteria_group>Bacillota>Clostridia>Eubacteriales>Lachnospiraceae | GCF_000210655.1 | Roseburia_intestinalis | ROI_RS12900 | 2840710..2842643 | 166486 | BREX-system-P-loop-protein-BrxC,-partial-[Roseburia-intestinalis] |
| WP_007865588.1 | Type1.BREX | <-MODE-HTH+PBPII\|\|?->?->?->ABhydrolase-><-?\|\|?->?->TatD_DNase->DUF6079-p*->OLD-ABC->REase->METHYLASE->ZnR+ZnR->DUF4404->PglZ->SbcC-> | 946 | Bacteria>Terrabacteria_group>Bacillota>Clostridia>Eubacteriales>Lachnospiraceae | GCF_000233455.1 | Enterocloster_citroniae | HMPREF9469_RS19440 | 127107..129947 | 358743 | BREX-system-P-loop-protein-BrxC-[Enterocloster-citroniae] |
| WP_156797221.1 | Type1.BREX | <-RNase\|\|?-><-?<-?<-SIGMA+OB-Fold+AAA-ATPase+Lon-Protease-BrxL<-PglZ<-AT-HTH-HYD2-2<-?<-Tox-ART-HYD2<-N6-METHYLASE<-DUF6079*<-BrxB-BrxF-iSTAND<-BrxA-DNA-binding<-XIS-HTH<-?<-?<-?<-RADICAL-SAM | 1217 | Bacteria>Terrabacteria_group>Bacillota>Clostridia>Eubacteriales>Lachnospiraceae | GCF_000235445.1 | Johnsonella_ignava | HMPREF9333_RS00440 | 95732..99385 | 43995 | BREX-system-P-loop-protein-BrxC-[Johnsonella-ignava] |
| WP_009536434.1 | Type1.BREX | <-SF1-SF2-Helicase<-ParB<-SbcC<-SIGMA+OB-Fold+AAA-ATPase+Lon-Protease-BrxL<-PglZ<-Schlafen+HATPase+wHTH<-HEPN<-N6-METHYLASE-p<-N6-METHYLASE<-DUF6079*<-BrxB-BrxF-iSTAND<-BrxA-DNA-binding<-HTH_26<-HTH+rve<-Sigma-HTH<-TIR<-Peptidase_C14<-TIR\|\|DEDD_Tnp_IS110-> | 1217 | Bacteria>Terrabacteria_group>Bacillota>Clostridia>Eubacteriales>Lachnospiraceae | GCF_000238075.1 | Oribacterium_asaccharolyticum | HMPREF9624_00544 | 316049..319702 | 1501332 | BREX-system-P-loop-protein-BrxC-[Oribacterium-asaccharolyticum] |
| WP_008755223.1 | Type1.BREX | MarR-HTH->XIS-HTH->BrxA-DNA-binding->BrxB-BrxF-iSTAND->DUF6079*->TIR->N6-METHYLASE->Tox-ART-HYD2->?->AT-HTH-HYD2-2->PglZ->SIGMA+OB-Fold+AAA-ATPase+Lon-Protease-BrxL->?->ClpABC-AAA-> | 1217 | Bacteria>Terrabacteria_group>Bacillota>Clostridia>Eubacteriales>Lachnospiraceae | GCF_000257705.1 | Lachnoanaerobaculum_saburreum | HMPREF9970_RS14240 | 4033..7686 | 467210 | BREX-system-P-loop-protein-BrxC-[Lachnoanaerobaculum-saburreum] |
| WP_009534426.1 | Type1.BREX | <-XIS-HTH<-SF1-SF2-Helicase<-?<-?<-SIGMA+OB-Fold+AAA-ATPase+Lon-Protease-BrxL<-PglZ<-DOC+HTH<-N6-METHYLASE<-DUF6079*<-BrxB-BrxF-iSTAND<-BrxA-DNA-binding<-SbcC<-DNA_METHYLASE\|\|DEDD_Tnp_IS110->Transposase_20-> | 1214 | Bacteria>Terrabacteria_group>Bacillota>Clostridia>Eubacteriales>Lachnospiraceae | GCF_000277505.1 | Oribacterium_parvum | HMPREF9625_RS09235 | 2079969..2083613 | 1501329 | BREX-system-P-loop-protein-BrxC-[Oribacterium-parvum] |
| WP_002576384.1 | Type1.BREX | <-SMODS<-SLATT<-SIGMA+OB-Fold+AAA-ATPase+Lon-Protease-BrxL<-PglZ<-?<-N6-METHYLASE<-TIR<-DUF6079*<-BrxB-BrxF-iSTAND<-BrxA-DNA-binding<-XIS-HTH<-?<-?<-?<-TraM<-TraJ-RHH<-LPD32+LPD36+N6_Mtase+SF1-SF2-Helicase | 1217 | Bacteria>Terrabacteria_group>Bacillota>Clostridia>Eubacteriales>Lachnospiraceae | GCF_000371705.1 | Enterocloster_bolteae | HMPREF1085_RS16605 | 3667388..3671041 | 208479 | BREX-system-P-loop-protein-BrxC-[Enterocloster-bolteae] |
| WP_002578404.1 | Type1.BREX | HTH-><-SF1-SF2-Helicase<-OLD-ABC+TOPRIM<-SIGMA+OB-Fold+AAA-ATPase+Lon-Protease-BrxL<-PglZ<-Schlafen+HATPase+wHTH<-HEPN<-N6-METHYLASE<-?<-DUF6079*<-BrxB-BrxF-iSTAND<-BrxA-DNA-binding<-HTH_26<-HTH_36<-TraI-VirD2-RCRE<-TraJ-RHH<-CopG/RHH<-?<-HTH_36 | 1182 | Bacteria>Terrabacteria_group>Bacillota>Clostridia>Eubacteriales>Lachnospiraceae | GCF_000371725.1 | Enterocloster_bolteae | HMPREF1095_RS19895 | 1377442..1380990 | 208479 | BREX-system-P-loop-protein-BrxC-[Enterocloster-bolteae] |
| WP_016216871.1 | Type1.BREX | <-HTH\|\|\|\|HTH->\|HTH_26->BrxA-DNA-binding->BrxB-BrxF-iSTAND->DUF6079*-><-DEDD_Tnp_IS110+Transposase_20\|\|HEPN->N6-METHYLASE->HTH+REase->Schlafen+HATPase+wHTH->PglZ-> | 1182 | Bacteria>Terrabacteria_group>Bacillota>Clostridia>Eubacteriales>Lachnospiraceae | GCF_000403455.2 | Dorea_sp._5-2 | C817_RS00440 | 102069..105617 | 1235798 | BREX-system-P-loop-protein-BrxC-[Dorea-sp.-5-2] |
| WP_022775135.1 | Type1.BREX | HIN-HTH->?-><-?<-Peptidase_C14<-SIGMA+OB-Fold+AAA-ATPase+Lon-Protease-BrxL<-?<-PglZ<-RAMA<-N6-METHYLASE<-DUF6079*<-BrxB-BrxF-iSTAND<-BrxA-DNA-binding<-HTH_26 | 1182 | Bacteria>Terrabacteria_group>Bacillota>Clostridia>Eubacteriales>Lachnospiraceae | GCF_000420825.1 | Butyrivibrio_sp._AE2015 | G585_RS0111460 | 49364..52912 | 1280663 | BREX-system-P-loop-protein-BrxC-[Butyrivibrio-sp.-AE2015] |
| WP_022757456.1 | Type1.BREX | HKD-DNase+SF1-SF2-Helicase+HTH+HTH+SRA->?->?->?->HTH_26->BrxA-DNA-binding->BrxB-BrxF-iSTAND->DUF6079*->N6-METHYLASE->?->PglZ->HEPN+SH3-like+SF1-SF2-Helicase+wHTH->MORC+DUF3684+HNH->SIGMA-HTH+SIGMA-HTH->35exo+wHTH+SF1-SF2-Helicase+HTH-><-wHTH+WYL+WCX<-wHTH+WYL+WCX | 1210 | Bacteria>Terrabacteria_group>Bacillota>Clostridia>Eubacteriales>Lachnospiraceae | GCF_000420985.1 | Butyrivibrio_fibrisolvens | G616_RS0107275 | 19485..23117 | 831 | BREX-system-P-loop-protein-BrxC-[Butyrivibrio-fibrisolvens] |
| WP_022757624.1 | Type1.BREX | <-HNH+NERD+ZnR\|\|GDResolvase-><-?<-?<-SIGMA+OB-Fold+AAA-ATPase+Lon-Protease-BrxL<-HNH<-PglZ<-N6-METHYLASE<-DUF6079*<-BrxB-BrxF-iSTAND<-BrxA-DNA-binding<-HTH_26<-?<-TraI-VirD2-RCRE<-TraJ-RHH<-?<-HTH_3\|\|ParA-Soj-PloopNTPase-> | 1181 | Bacteria>Terrabacteria_group>Bacillota>Clostridia>Eubacteriales>Lachnospiraceae | GCF_000420985.1 | Butyrivibrio_fibrisolvens | G616_RS0108215 | 23876..27421 | 831 | BREX-system-P-loop-protein-BrxC-[Butyrivibrio-fibrisolvens] |
| WP_022770173.1 | Type1.BREX | <-SF1-SF2-Helicase+REase<-SF1-SF2-Helicase<-URI-DNase<-HEPN+REase<-SIGMA+OB-Fold+AAA-ATPase+Lon-Protease-BrxL<-PglZ<-N6-METHYLASE-p<-N6-METHYLASE<-DUF6079*<-BrxB-BrxF-iSTAND<-BrxA-DNA-binding | 1202 | Bacteria>Terrabacteria_group>Bacillota>Clostridia>Eubacteriales>Lachnospiraceae | GCF_000421405.1 | Butyrivibrio_sp._NC2007 | G632_RS0114760 | 79363..82971 | 1280683 | BREX-system-P-loop-protein-BrxC-[Butyrivibrio-sp.-NC2007] |
| WP_027207205.1 | Type1.BREX | XIS-HTH-><-?<-ParB+HNH<-SIGMA+OB-Fold+AAA-ATPase+Lon-Protease-BrxL<-HEPN<-PglZ<-N6-METHYLASE<-DUF6079*<-BrxB-BrxF-iSTAND<-BrxA-DNA-binding<-HTH_26<-SF1-SF2-Helicase<-?<-HEPN+PAS+DUF4011+SF1-SF2-Helicase+REase<-ParB<-DDE_Tnp_1_4 | 1196 | Bacteria>Terrabacteria_group>Bacillota>Clostridia>Eubacteriales>Lachnospiraceae | GCF_000423985.1 | Butyrivibrio_fibrisolvens | G591_RS0114720 | 29938..33528 | 831 | BREX-system-P-loop-protein-BrxC-[Butyrivibrio-fibrisolvens] |
| WP_026495918.1 | Type1.BREX | HTH_3->Resolvase+Recombinase->?->?->ParB->ParB->HTH_26->BrxA-DNA-binding->BrxB-BrxF-iSTAND->DUF6079*->N6-METHYLASE->?->METHYLASE->PglZ->SIGMA+OB-Fold+AAA-ATPase+Lon-Protease-BrxL->OLD-ABC+TOPRIM->SF1-SF2-Helicase-> | 1181 | Bacteria>Terrabacteria_group>Bacillota>Clostridia>Eubacteriales>Lachnospiraceae | GCF_000424265.1 | Butyrivibrio_sp._WCD3002 | G608_RS0110170 | 160583..164128 | 1280676 | BREX-system-P-loop-protein-BrxC-[Butyrivibrio-sp.-WCD3002] |
| WP_027204781.1 | Type1.BREX | ArdCN+MPTase-inact+LPD19->CHTH->TOPRIM+Topoisom_bac->LPD2->TraJ-RHH->?->HTH_26->BrxA-DNA-binding->BrxB-BrxF-iSTAND->DUF6079*->N6-METHYLASE->RAMA->?->PglZ->SIGMA+OB-Fold+AAA-ATPase+Lon-Protease-BrxL->?->?->ParB+HNH->HNH-> | 1181 | Bacteria>Terrabacteria_group>Bacillota>Clostridia>Eubacteriales>Lachnospiraceae | GCF_000424425.1 | Butyrivibrio_fibrisolvens | G624_RS0113140 | 43921..47466 | 831 | BREX-system-P-loop-protein-BrxC-[Butyrivibrio-fibrisolvens] |
| WP_026519395.1 | Type1.BREX | <-LRR_5<-BrxL<-SIGMA+OB-Fold+AAA-ATPase+Lon-Protease-BrxL<-PHP+SbcC<-PglZ<-HEPN<-N6-METHYLASE<-DUF6079*<-BrxB-BrxF-iSTAND<-BrxA-DNA-binding<-HTH_26<-?<-?<-?<-TraJ-RHH<-LPD2<-ZnR+TOPRIM | 1194 | Bacteria>Terrabacteria_group>Bacillota>Clostridia>Eubacteriales>Lachnospiraceae | GCF_000424525.1 | Butyrivibrio_sp._FCS006 | G633_RS0106250 | 43812..47396 | 1280684 | BREX-system-P-loop-protein-BrxC-[Butyrivibrio-sp.-FCS006] |
| WP_026508807.1 | Type1.BREX | <-LexA-protease<-?<-PIN<-?<-HHH_4+HTH+wHTH+SF1-SF2-Helicase+REase<-MORC+DUF3684+HNH<-HEPN+SH3-like+SF1-SF2-Helicase+wHTH<-PglZ<-N6-METHYLASE<-DUF6079*<-BrxB-BrxF-iSTAND<-BrxA-DNA-binding<-HTH_26<-?<-TraJ-RHH<-LPD2<-TOPRIM+Topoisom_bac<-ArdCN+MPTase-inact+LPD19 | 1210 | Bacteria>Terrabacteria_group>Bacillota>Clostridia>Eubacteriales>Lachnospiraceae | GCF_000424585.1 | Butyrivibrio_sp._MC2013 | G636_RS0115515 | 259305..262937 | 1280686 | BREX-system-P-loop-protein-BrxC-[Butyrivibrio-sp.-MC2013] |
| WP_029067206.1 | Type1.BREX | HTH_26->BrxA-DNA-binding->BrxB-BrxF-iSTAND->DUF6079*->N6-METHYLASE->PglZ->ParB+HNH->SIGMA+OB-Fold+AAA-ATPase+Lon-Protease-BrxL->RVT+HEPN->HTH_38->HTH_26->BrxA-DNA-binding->BrxB-BrxF-iSTAND-> | 1182 | Bacteria>Terrabacteria_group>Bacillota>Clostridia>Eubacteriales>Lachnospiraceae | GCF_000687515.1 | Lachnobacterium_bovis | CC91_RS0108435 | 378442..381990 | 140626 | BREX-system-P-loop-protein-BrxC-[Lachnobacterium-bovis] |
| WP_029067213.1 | Type1.BREX | N6-METHYLASE->PglZ->ParB+HNH->SIGMA+OB-Fold+AAA-ATPase+Lon-Protease-BrxL->RVT+HEPN->HTH_38->HTH_26->BrxA-DNA-binding->BrxB-BrxF-iSTAND->DUF6079*->N6-METHYLASE->RAMA->PglZ->?->SIGMA+OB-Fold+AAA-ATPase+Lon-Protease-BrxL->KAP-NTPase->KAP-glycosylase->?->TatD_DNase-> | 1182 | Bacteria>Terrabacteria_group>Bacillota>Clostridia>Eubacteriales>Lachnospiraceae | GCF_000687515.1 | Lachnobacterium_bovis | CC91_RS0108490 | 397340..400888 | 140626 | BREX-system-P-loop-protein-BrxC-[Lachnobacterium-bovis] |
| WP_055216182.1 | Type1.BREX | <-PIN<-ClpABC-AAA<-?<-SIGMA+OB-Fold+AAA-ATPase+Lon-Protease-BrxL<-PglZ<-HEPN<-N6-METHYLASE<-DUF6079*<-BrxB-BrxF-iSTAND<-BrxA-DNA-binding<-HTH_26<-?<-?<-?<-?<-ArdCN+LPD32+LPD36+N6_Mtase+SF1-SF2-Helicase | 1186 | Bacteria>Terrabacteria_group>Bacillota>Clostridia>Eubacteriales>Lachnospiraceae | GCF_001404435.1 | Lachnospira_eligens | ERS852490_02366 | 200381..203941 | 39485 | BREX-system-P-loop-protein-BrxC-[Lachnospira-eligens] |
| WP_055268215.1 | Type1.BREX | ZnR->HTH_26->BrxA-DNA-binding->BrxB-BrxF-iSTAND->DUF6079*->N6-METHYLASE->AT-HTH-HYD2->Schlafen+HATPase+wHTH->PglZ->SIGMA+OB-Fold+AAA-ATPase+Lon-Protease-BrxL->OLD-ABC+TOPRIM->SF1-SF2-Helicase->Phage_integrase-> | 1182 | Bacteria>Terrabacteria_group>Bacillota>Clostridia>Eubacteriales>Lachnospiraceae | GCF_001406335.1 | Fusicatenibacter_saccharivorans | ARA47_RS16350 | 4112..7660 | 1150298 | BREX-system-P-loop-protein-BrxC-[Fusicatenibacter-saccharivorans] |
| WP_059085682.1 | Type1.BREX | MACRODOMAIN->?->ZnR+ZnR+ZnR->HTH_26->BrxA-DNA-binding->BrxB-BrxF-iSTAND->DUF6079*->N6-METHYLASE->ParB+HNH->OLD-ABC->?->OLD-ABC->HNH->PglZ-> | 1196 | Bacteria>Terrabacteria_group>Bacillota>Clostridia>Eubacteriales>Lachnospiraceae | GCF_001487165.1 | Blautia_massiliensis | BN3261_RS05710 | 1028364..1031954 | 1737424 | BREX-system-P-loop-protein-BrxC-[Blautia-massiliensis] |
| WP_060932093.1 | Type1.BREX | Peptidase_M32->TPR->BrxA-DNA-binding->BrxB-BrxF-iSTAND->DUF6079*->Schlafen+HATPase+wHTH->N6-METHYLASE->PglZ->DOC+HTH->?->?-><-?<-?<-ABhydrolase | 1215 | Bacteria>Terrabacteria_group>Bacillota>Clostridia>Eubacteriales>Lachnospiraceae | GCF_001552975.1 | Lachnoanaerobaculum_saburreum | HMPREF1866_RS12095 | 13224..16871 | 467210 | BREX-system-P-loop-protein-BrxC-[Lachnoanaerobaculum-saburreum] |
| WP_075720910.1 | Type1.BREX | <-LRR_5+LRR_8<-?<-SIGMA+OB-Fold+AAA-ATPase+Lon-Protease-BrxL<-?<-PglZ<-Schlafen+HATPase+wHTH<-REase<-N6-METHYLASE<-DUF6079*<-BrxB-BrxF-iSTAND<-BrxA-DNA-binding<-HTH_26<-ZnR+ZnR+ZnR<-HicA_toxin<-?<-?<-MACRODOMAIN<-35exo+ADA-DBD | 1196 | Bacteria>Terrabacteria_group>Bacillota>Clostridia>Eubacteriales>Lachnospiraceae | GCF_001940225.1 | Roseburia_sp._499 | BIV20_RS10280 | 367523..371113 | 1261634 | BREX-system-P-loop-protein-BrxC-[Roseburia-sp.-499] |
| WP_075672058.1 | Type1.BREX | N-OB+HDRnase->TRAM+METHYLASE->HEPN->HEPN->?->?->HTH_26->BrxA-DNA-binding->BrxB-BrxF-iSTAND->DUF6079*->?->N6-METHYLASE->SIR2->AT-HTH-HYD2->Schlafen+HATPase+wHTH->PglZ->SIGMA+OB-Fold+AAA-ATPase+Lon-Protease-BrxL->PHP+SbcC-> | 1182 | Bacteria>Terrabacteria_group>Bacillota>Clostridia>Eubacteriales>Lachnospiraceae | GCF_001940245.1 | Anaerostipes_sp._992a | BHF69_RS12275 | 110494..114042 | 1261637 | BREX-system-P-loop-protein-BrxC-[Anaerostipes-sp.-992a] |
| WP_087163144.1 | Type1.BREX | <-HTH_Tnp_1<-SF1-SF2-Helicase<-OLD-ABC+TOPRIM<-SIGMA+OB-Fold+AAA-ATPase+Lon-Protease-BrxL<-PglZ<-Schlafen+HATPase+wHTH<-SIR2<-N6-METHYLASE<-?<-DUF6079*<-BrxB-BrxF-iSTAND<-BrxA-DNA-binding<-HTH_26\|\|HIN-HTH->?->LZ_Tnp_IS66+zf-IS66+DDE_Tnp_IS66+DDE_Tnp_IS66_C-> | 1179 | Bacteria>Terrabacteria_group>Bacillota>Clostridia>Eubacteriales>Lachnospiraceae | GCF_002160525.1 | Lachnoclostridium_sp._An169 | B5F07_RS20950 | 16171..19710 | 1965569 | BREX-system-P-loop-protein-BrxC-[Lachnoclostridium-sp.-An169] |
| WP_087175247.1 | Type1.BREX | <-RepA_N+HTH<-?<-TraJ-RHH<-?<-SIGMA+OB-Fold+AAA-ATPase+Lon-Protease-BrxL<-?<-OLD-ABC<-PglZ<-N6-METHYLASE<-DUF6079*<-BrxB-BrxF-iSTAND<-BrxA-DNA-binding<-HTH_26<-?<-?<-?<-wHTH+HNH<-wHTH+HNH<-?<-MazG | 1201 | Bacteria>Terrabacteria_group>Bacillota>Clostridia>Eubacteriales>Lachnospiraceae | GCF_002160985.1 | Lachnoclostridium_sp._An118 | B5E62_RS14545 | 43604..47209 | 1965547 | BREX-system-P-loop-protein-BrxC-[Lachnoclostridium-sp.-An118] |
| WP_087273661.1 | Type1.BREX | DNA_METHYLASE->?->ParB+HNH->HEPN-><-SbcC<-SIGMA+OB-Fold+AAA-ATPase+Lon-Protease-BrxL<-PglZ<-HEPN<-N6-METHYLASE<-DUF6079*<-BrxB-BrxF-iSTAND<-BrxA-DNA-binding<-HTH_26<-SIR2<-AbiEii\|\|transposase-> | 1212 | Bacteria>Terrabacteria_group>Bacillota>Clostridia>Eubacteriales>Lachnospiraceae | GCF_002161645.1 | Blautia_sp._An46 | B5G00_RS13700 | 83455..87093 | 1965636 | BREX-system-P-loop-protein-BrxC-[Blautia-sp.-An46] |
| WP_103203921.1 | Type1.BREX | RHH_8-><-?<-ZnR+TOPRIM\|\|?->SymE_toxin->?->wHTH(SBD)+HNH->BrxA-DNA-binding->BrxB-BrxF-iSTAND->DUF6079*->N6-METHYLASE->PglZ->?->SIGMA+OB-Fold+AAA-ATPase+Lon-Protease-BrxL->RAMA->ARG-> | 1186 | Bacteria>Terrabacteria_group>Bacillota>Clostridia>Eubacteriales>Lachnospiraceae | GCF_002904165.1 | Herbinix_hemicellulosilytica | HHT355_RS13290 | 82989..86549 | 1564487 | BREX-system-P-loop-protein-BrxC-[Herbinix-hemicellulosilytica] |
| WP_114002767.1 | Type1.BREX | <-PglZ<-HNH<-OLD-ABC<-?<-OLD-ABC<-ParB+HNH<-N6-METHYLASE<-DUF6079*<-BrxB-BrxF-iSTAND<-BrxA-DNA-binding<-HTH_26<-ZnR+ZnR+ZnR<-?<-MACRODOMAIN | 1196 | Bacteria>Terrabacteria_group>Bacillota>Clostridia>Eubacteriales>Lachnospiraceae | GCF_003324155.1 | Blautia_obeum | C4886_RS16030 | 22477..26067 | 40520 | BREX-system-P-loop-protein-BrxC-[Blautia-obeum] |
| WP_117638895.1 | Type1.BREX | wHTH+WYL+WCX->?->?->?->HTH_26->BrxA-DNA-binding->BrxB-BrxF-iSTAND->DUF6079*->N6-METHYLASE->?->PglZ->HEPN+SH3-like+SF1-SF2-Helicase+wHTH->MORC+DUF3684+HNH->SF1-SF2-Helicase->PIN->?->PIN-> | 1203 | Bacteria>Terrabacteria_group>Bacillota>Clostridia>Eubacteriales>Lachnospiraceae | GCF_003436075.1 | Blautia_obeum | DXD81_RS08455 | 28744..32355 | 40520 | BREX-system-P-loop-protein-BrxC-[Blautia-obeum] |
| WP_118348289.1 | Type1.BREX | <-SIGMA+OB-Fold+AAA-ATPase+Lon-Protease-BrxL<-PglZ<-Schlafen+HATPase+wHTH<-HEPN<-N6-METHYLASE<-?<-DUF6079*<-BrxB-BrxF-iSTAND<-BrxA-DNA-binding<-HTH_26<-?<-TraI-VirD2-RCRE<-TraJ-RHH<-?<-CopG/RHH<-HTH_36 | 1182 | Bacteria>Terrabacteria_group>Bacillota>Clostridia>Eubacteriales>Lachnospiraceae | GCF_003458885.1 | Coprococcus_sp._AF21-14LB | DWX73_RS07815 | 11153..14701 | 2292231 | BREX-system-P-loop-protein-BrxC-[Coprococcus-sp.-AF21-14LB] |
| WP_117765413.1 | Type1.BREX | <-PHP+SbcC<-SIGMA+OB-Fold+AAA-ATPase+Lon-Protease-BrxL<-PglZ<-AT-HTH-HYD2-2<-?<-Tox-ART-HYD2<-N6-METHYLASE<-DUF6079*<-BrxB-BrxF-iSTAND<-BrxA-DNA-binding<-XIS-HTH<-?<-MACRODOMAIN<-?\|\|YTCD-HTH-><-?<-SIR2 | 1217 | Bacteria>Terrabacteria_group>Bacillota>Clostridia>Eubacteriales>Lachnospiraceae | GCF_003463065.1 | Blautia_sp._OF03-15BH | DXA57_RS05510 | 273958..277611 | 2292287 | BREX-system-P-loop-protein-BrxC-[Blautia-sp.-OF03-15BH] |
| WP_118051545.1 | Type1.BREX | <-Phage_int_SAM_5+Phage_integrase\|\|TraJ-RHH->?-><-?\|\|HTH_26->BrxA-DNA-binding->BrxB-BrxF-iSTAND->DUF6079*->?->N6-METHYLASE->HEPN->Schlafen+HATPase+wHTH->PglZ->PHP+SbcC-> | 1183 | Bacteria>Terrabacteria_group>Bacillota>Clostridia>Eubacteriales>Lachnospiraceae | GCF_003464645.1 | Blautia_obeum | DWW07_RS08215 | 33645..37196 | 40520 | BREX-system-P-loop-protein-BrxC-[Blautia-obeum] |
| WP_118592376.1 | Type1.BREX | <-Schlafen+HATPase+wHTH-p<-ClpABC-AAA<-?<-SIGMA+OB-Fold+AAA-ATPase+Lon-Protease-BrxL<-PglZ<-HEPN<-N6-METHYLASE<-DUF6079*<-BrxB-BrxF-iSTAND<-BrxA-DNA-binding<-HTH_26<-?<-?<-?<-?<-SF1-SF2-Helicase<-N6-N4-METHYLASE | 1186 | Bacteria>Terrabacteria_group>Bacillota>Clostridia>Eubacteriales>Lachnospiraceae | GCF_003467035.1 | Roseburia_intestinalis | DW927_RS18160 | 14462..18022 | 166486 | BREX-system-P-loop-protein-BrxC-[Roseburia-intestinalis] |
| WP_118012961.1 | Type1.BREX | <-Phage_int_SAM_5+Phage_integrase\|\|TraJ-RHH->?-><-?\|\|HTH_26->BrxA-DNA-binding->BrxB-BrxF-iSTAND->DUF6079*->TIR->N6-METHYLASE->ZnR+ZnR->DUF4404->Schlafen+HATPase+wHTH->PglZ->SIGMA+OB-Fold+AAA-ATPase+Lon-Protease-BrxL-> | 1183 | Bacteria>Terrabacteria_group>Bacillota>Clostridia>Eubacteriales>Lachnospiraceae | GCF_003467455.1 | Blautia_obeum | DW880_RS12300 | 36915..40466 | 40520 | BREX-system-P-loop-protein-BrxC-[Blautia-obeum] |
| WP_118198114.1 | Type1.BREX | <-SIGMA+OB-Fold+AAA-ATPase+Lon-Protease-BrxL<-?<-PHP<-PglZ<-Schlafen+HATPase+wHTH<-HEPN<-N6-METHYLASE<-?<-DUF6079*<-BrxB-BrxF-iSTAND<-BrxA-DNA-binding<-HTH_26\|\|?-><-?<-TraJ-RHH\|\|Phage_int_SAM_5+Phage_integrase-> | 1183 | Bacteria>Terrabacteria_group>Bacillota>Clostridia>Eubacteriales>Lachnospiraceae | GCF_003471125.1 | Blautia_obeum | DW272_RS15845 | 24197..27748 | 40520 | BREX-system-P-loop-protein-BrxC-[Blautia-obeum] |
| WP_118390632.1 | Type1.BREX | wHTH+WYL+WCX->HTH_26->BrxA-DNA-binding->BrxB-BrxF-iSTAND->DUF6079*->N6-METHYLASE->REase->Schlafen+HATPase+wHTH->PglZ->Pkinase->SIGMA+OB-Fold+AAA-ATPase+Lon-Protease-BrxL->?->SF1-SF2-Helicase+TUDOR->MPTase-> | 1196 | Bacteria>Terrabacteria_group>Bacillota>Clostridia>Eubacteriales>Lachnospiraceae | GCF_003474935.1 | Lachnospira_eligens | DWZ79_RS02870 | 260992..264582 | 39485 | BREX-system-P-loop-protein-BrxC-[Lachnospira-eligens] |
| WP_118601699.1 | Type1.BREX | HTH_Tnp_1->?->ZnR->?->HTH_26->BrxA-DNA-binding->BrxB-BrxF-iSTAND->DUF6079*->N6-METHYLASE->AT-HTH-HYD2->Schlafen+HATPase+wHTH->PglZ->SIGMA+OB-Fold+AAA-ATPase+Lon-Protease-BrxL->OLD-ABC+TOPRIM->SF1-SF2-Helicase->Phage_integrase-> | 1182 | Bacteria>Terrabacteria_group>Bacillota>Clostridia>Eubacteriales>Lachnospiraceae | GCF_003479155.1 | Blautia_sp._AF19-1 | DWX26_RS14805 | 4595..8143 | 2292960 | BREX-system-P-loop-protein-BrxC-[Blautia-sp.-AF19-1] |
| WP_118742111.1 | Type1.BREX | ZnR->?->HTH_26->BrxA-DNA-binding->BrxB-BrxF-iSTAND->DUF6079*->N6-METHYLASE->RAMA->HEPN->Schlafen+HATPase+wHTH->PglZ->SIGMA+OB-Fold+AAA-ATPase+Lon-Protease-BrxL->OLD-ABC+TOPRIM->SF1-SF2-Helicase-> | 1182 | Bacteria>Terrabacteria_group>Bacillota>Clostridia>Eubacteriales>Lachnospiraceae | GCF_003480245.1 | Blautia_sp._AM42-2 | DW915_10705 | 4262..7810 | 2292976 | BREX-system-P-loop-protein-BrxC-[Blautia-sp.-AM42-2] |
| WP_111525527.1 | Type1.BREX | VirD4-TraG+VirD4-FtsK->?->?->Peptidase_M32->BrxA-DNA-binding->BrxB-BrxF-iSTAND->DUF6079*->N6-METHYLASE->AbiEi_4->SF1-SF2-Helicase->ATPase_c->MORC+REase->RelB->Tox-RelE->RADICAL-SAM-> | 1217 | Bacteria>Terrabacteria_group>Bacillota>Clostridia>Eubacteriales>Lachnospiraceae | GCF_003589745.1 | Lachnoanaerobaculum_umeaense | D4A81_RS04980 | 1085086..1088739 | 617123 | BREX-system-P-loop-protein-BrxC-[Lachnoanaerobaculum-umeaense] |
| WP_124952173.1 | Type1.BREX | ABhydrolase->?->?->?-><-?<-DOC+HTH<-PglZ<-N6-METHYLASE<-Schlafen+HATPase+wHTH<-DUF6079*<-BrxB-BrxF-iSTAND<-BrxA-DNA-binding<-TPR<-Peptidase_M32 | 1215 | Bacteria>Terrabacteria_group>Bacillota>Clostridia>Eubacteriales>Lachnospiraceae | GCF_003862485.1 | Lachnoanaerobaculum_orale | EHW90_RS07145 | 1507422..1511069 | 979627 | BREX-system-P-loop-protein-BrxC-[Lachnoanaerobaculum-orale] |
| WP_118582742.1 | Type1.BREX | <-MULE-TRANSPOSASE<-HTH+HTH+CHTH<-SIGMA+OB-Fold+AAA-ATPase+Lon-Protease-BrxL<-?<-PglZ<-?<-NERD<-OLD-ABC<-N6-METHYLASE<-DUF6079*<-BrxB-BrxF-iSTAND<-BrxA-DNA-binding<-HTH_26\|\|?->?-><-MODE-HTH+PBPII\|\|?->?-><-MULE-TRANSPOSASE | 1194 | Bacteria>Terrabacteria_group>Bacillota>Clostridia>Eubacteriales>Lachnospiraceae | GCF_003865035.1 | Anaerotignum_faecicola | EJE48_RS08260 | 191884..195468 | 2358141 | BREX-system-P-loop-protein-BrxC-[Anaerotignum-faecicola] |
| WP_135987576.1 | Type1.BREX | <-MarR-HTH<-?<-RhoN+RhoN+RNase+S1<-?<-?<-PglZ<-REase<-OLD-ABC<-N6-METHYLASE<-DUF6079*<-BrxB-BrxF-iSTAND<-BrxA-DNA-binding<-?<-PRTase | 1218 | Bacteria>Terrabacteria_group>Bacillota>Clostridia>Eubacteriales>Lachnospiraceae | GCF_004793545.1 | Petralouisia_muris | E5329_RS00220 | 47888..51544 | 3032872 | BREX-system-P-loop-protein-BrxC-[Petralouisia-muris] |
| WP_154525460.1 | Type1.BREX | CN_hydrolase->MerR-HTH->ABhydrolase->?->?->HTH_26->BrxA-DNA-binding->BrxB-BrxF-iSTAND->DUF6079-p*->METHYLASE-><-MobA_MobL<-?<-?<-GDResolvase+Recombinase+ZnR\|\|DUF6079-p->N6-METHYLASE-> | 821 | Bacteria>Terrabacteria_group>Bacillota>Clostridia>Eubacteriales>Lachnospiraceae | GCF_009696445.1 | Porcincola_intestinalis | FYJ35_RS08285 | 8308..10773 | 2606632 | BREX-system-P-loop-protein-BrxC-[Porcincola-intestinalis] |
| WP_154780312.1 | Type1.BREX | Peptidase->?->?->MACRODOMAIN->?->ZnR+ZnR+ZnR->HTH_26->BrxA-DNA-binding->BrxB-BrxF-iSTAND->DUF6079*->N6-METHYLASE->RVT->?->OLD-ABC->HNH->?->PglZ->HEPN->SIGMA+OB-Fold+AAA-ATPase+Lon-Protease-BrxL-> | 1201 | Bacteria>Terrabacteria_group>Bacillota>Clostridia>Eubacteriales>Lachnospiraceae | GCF_009707925.1 | Blautia_luti | GKZ57_RS08760 | 74022..77627 | 89014 | BREX-system-P-loop-protein-BrxC-[Blautia-luti] |
| WP_157350284.1 | Type1.BREX | <-HEPN+HEPN<-ParB+HNH<-AT-HTH-HYD2<-Tox-ART-HYD2<-AT-HTH-HYD2<-KAP-NTPase<-N6-METHYLASE<-TIR<-?<-DUF6079*<-BrxB-BrxF-iSTAND<-BrxA-DNA-binding<-HTH_26<-HTH_36<-TraI-VirD2-RCRE<-TraJ-RHH<-CopG/RHH<-?<-HTH_36 | 1182 | Bacteria>Terrabacteria_group>Bacillota>Clostridia>Eubacteriales>Lachnospiraceae | GCF_009759525.1 | Roseburia_intestinalis | GCK47_RS06015 | 115947..119495 | 166486 | BREX-system-P-loop-protein-BrxC-[Roseburia-intestinalis] |
| WP_161171751.1 | Type1.BREX | ZnR+ZnR+ZnR->HTH_26->BrxA-DNA-binding->BrxB-BrxF-iSTAND->DUF6079*->N6-METHYLASE-> | 1196 | Bacteria>Terrabacteria_group>Bacillota>Clostridia>Eubacteriales>Lachnospiraceae | GCF_009875795.1 | unclassified_Coprococcus | GT568_RS06940 | 430338..433928 | 2684943 | MULTISPECIES:-BREX-system-P-loop-protein-BrxC-[unclassified-Coprococcus] |
| WP_168932969.1 | Type1.BREX | <-OLD-ABC+TOPRIM<-SIGMA+OB-Fold+AAA-ATPase+Lon-Protease-BrxL<-PglZ<-Schlafen+HATPase+wHTH<-DUF4404<-ZnR+ZnR<-N6-METHYLASE<-TIR<-?<-DUF6079*<-BrxB-BrxF-iSTAND<-BrxA-DNA-binding<-HTH_26<-HTH_36<-TraI-VirD2-RCRE<-TraJ-RHH<-?<-?\|\|HTH_3-> | 1182 | Bacteria>Terrabacteria_group>Bacillota>Clostridia>Eubacteriales>Lachnospiraceae | GCF_012843825.1 | Dorea_formicigenerans | HF855_RS01350 | 86312..89860 | 39486 | BREX-system-P-loop-protein-BrxC-[Dorea-formicigenerans] |
| WP_173900038.1 | Type1.BREX | <-SIGMA+OB-Fold+AAA-ATPase+Lon-Protease-BrxL<-PglZ<-Schlafen+HATPase+wHTH<-DUF4404<-ZnR+ZnR<-N6-METHYLASE<-TIR<-DUF6079*<-BrxB-BrxF-iSTAND<-BrxA-DNA-binding<-HTH_26\|\|?-><-?<-TraJ-RHH\|\|Phage_int_SAM_5+Phage_integrase-> | 1183 | Bacteria>Terrabacteria_group>Bacillota>Clostridia>Eubacteriales>Lachnospiraceae | GCF_013299585.1 | Blautia_obeum | G4467_RS13410 | 30685..34236 | 40520 | BREX-system-P-loop-protein-BrxC-[Blautia-obeum] |
| WP_173865463.1 | Type1.BREX | <-STYKIN<-?<-LRR_5+LRR_8<-SIGMA+OB-Fold+AAA-ATPase+Lon-Protease-BrxL<-Schlafen+HATPase+wHTH<-PglZ<-N6-METHYLASE<-PIN<-OLD-ABC<-DUF6079*<-BrxB-BrxF-iSTAND<-BrxA-DNA-binding<-HTH_26<-ZnR+ZnR+ZnR<-35exo+ADA-DBD<-?<-MACRODOMAIN<-?<-DOC+zhth | 1201 | Bacteria>Terrabacteria_group>Bacillota>Clostridia>Eubacteriales>Lachnospiraceae | GCF_013300965.1 | Faecalicatena_fissicatena | HFN12_RS00390 | 75728..79333 | 290055 | BREX-system-P-loop-protein-BrxC-[Faecalicatena-fissicatena] |
| WP_173866293.1 | Type1.BREX | <-Polbeta<-MarR-HTH<-VbhA+DOC+HTH<-wHTH+WYL+WCX<-PglZ<-?<-N6-METHYLASE-p<-N6-METHYLASE<-DUF6079*<-BrxB-BrxF-iSTAND<-BrxA-DNA-binding<-HTH_26<-nSTAND3 | 1100 | Bacteria>Terrabacteria_group>Bacillota>Clostridia>Eubacteriales>Lachnospiraceae | GCF_013300965.1 | Faecalicatena_fissicatena | HFN12_RS07560 | 16187..19489 | 290055 | BREX-system-P-loop-protein-BrxC-[Faecalicatena-fissicatena] |
| WP_173813806.1 | Type1.BREX | ZnR->?->HTH_26->BrxA-DNA-binding->BrxB-BrxF-iSTAND->DUF6079*->N6-METHYLASE->AbiGi+seqA+RAMA->Schlafen+HATPase+wHTH->PglZ->SIGMA+OB-Fold+AAA-ATPase+Lon-Protease-BrxL->OLD-ABC+TOPRIM->SF1-SF2-Helicase->Phage_integrase->Phage_integrase-> | 1182 | Bacteria>Terrabacteria_group>Bacillota>Clostridia>Eubacteriales>Lachnospiraceae | GCF_013301105.1 | Fusicatenibacter_saccharivorans | G5B05_RS14750 | 4301..7849 | 1150298 | BREX-system-P-loop-protein-BrxC-[Fusicatenibacter-saccharivorans] |
| WP_173793922.1 | Type1.BREX | <-MazG<-SIGMA-HTH\|\|HNH+3C-ABC->?->?->HTH_26->BrxA-DNA-binding->BrxB-BrxF-iSTAND->DUF6079*->N6-METHYLASE->TIR->PglZ->SIGMA+OB-Fold+AAA-ATPase+Lon-Protease-BrxL->HTH+REase->?->CHTH-> | 1184 | Bacteria>Terrabacteria_group>Bacillota>Clostridia>Eubacteriales>Lachnospiraceae | GCF_013301385.1 | Dorea_formicigenerans | G4398_RS10070 | 34100..37654 | 39486 | BREX-system-P-loop-protein-BrxC-[Dorea-formicigenerans] |
| WP_173794457.1 | Type1.BREX | MACRODOMAIN->?->HicA_toxin->ZnR+ZnR+ZnR->HTH_26->BrxA-DNA-binding->BrxB-BrxF-iSTAND->DUF6079*->N6-METHYLASE->OLD-ABC->NERD->PglZ->Pkinase->SIGMA+OB-Fold+AAA-ATPase+Lon-Protease-BrxL->LRR_5+LRR_8->?->STYKIN-> | 1196 | Bacteria>Terrabacteria_group>Bacillota>Clostridia>Eubacteriales>Lachnospiraceae | GCF_013301435.1 | Coprococcus_eutactus | G4458_RS01680 | 390743..394333 | 33043 | BREX-system-P-loop-protein-BrxC-[Coprococcus-eutactus] |
| WP_173794823.1 | Type1.BREX | CopG/RHH->?->TraJ-RHH->TraI-VirD2-RCRE->?->HTH_26->BrxA-DNA-binding->BrxB-BrxF-iSTAND->DUF6079*->?->N6-METHYLASE->?->?->Tox-ART-HYD2->HEPN->Schlafen+HATPase+wHTH->PglZ->SIGMA+OB-Fold+AAA-ATPase+Lon-Protease-BrxL-> | 1183 | Bacteria>Terrabacteria_group>Bacillota>Clostridia>Eubacteriales>Lachnospiraceae | GCF_013301435.1 | Coprococcus_eutactus | G4458_RS07820 | 91077..94628 | 33043 | BREX-system-P-loop-protein-BrxC-[Coprococcus-eutactus] |
| WP_173753518.1 | Type1.BREX | wHTH+WYL+WCX->?->?->?->HTH_26->BrxA-DNA-binding->BrxB-BrxF-iSTAND->DUF6079*->N6-METHYLASE->DUF4365+REase->PglZ->HEPN+SH3-like+SF1-SF2-Helicase+wHTH->MORC+DUF3684+HNH->SF1-SF2-Helicase->PIN-> | 1203 | Bacteria>Terrabacteria_group>Bacillota>Clostridia>Eubacteriales>Lachnospiraceae | GCF_013301805.1 | Blautia | G5A92_RS05405 | 158405..162016 | 572511 | MULTISPECIES:-BREX-system-P-loop-protein-BrxC-[Blautia] |
| WP_173770663.1 | Type1.BREX | HTH-><-PHP+SbcC<-SIGMA+OB-Fold+AAA-ATPase+Lon-Protease-BrxL<-PglZ<-Schlafen+HATPase+wHTH<-?<-AT-HTH-HYD2<-N6-METHYLASE<-?<-DUF6079*<-BrxB-BrxF-iSTAND<-BrxA-DNA-binding<-HTH_26<-HTH_36<-?<-?<-ABhydrolase<-?<-SAM-METHYLASE | 1182 | Bacteria>Terrabacteria_group>Bacillota>Clostridia>Eubacteriales>Lachnospiraceae | GCF_013302375.1 | Blautia_faecis | G5A01_RS01800 | 135200..138748 | 871665 | BREX-system-P-loop-protein-BrxC-[Blautia-faecis] |
| WP_173904654.1 | Type1.BREX | DUF6079*->?->N6-METHYLASE->ZnR+ZnR->DUF4404->PglZ->SIGMA+OB-Fold+AAA-ATPase+Lon-Protease-BrxL-> | 1179 | Bacteria>Terrabacteria_group>Bacillota>Clostridia>Eubacteriales>Lachnospiraceae | GCF_013304225.1 | Enterocloster_clostridioformis | G5B20_RS15345 | 66..3605 | 1531 | BREX-system-P-loop-protein-BrxC-[Enterocloster-clostridioformis] |
| WP_002585535.1 | Type1.BREX | <-ClpABC-AAA<-?<-SIGMA+OB-Fold+AAA-ATPase+Lon-Protease-BrxL<-PglZ<-Schlafen+HATPase+wHTH<-SbcC+APATPase+SbcC<-N6-METHYLASE<-?<-DUF6079*<-BrxB-BrxF-iSTAND<-BrxA-DNA-binding<-HTH_26<-HTH\|\|?-><-?<-?<-TraJ-RHH<-RVT | 1178 | Bacteria>Terrabacteria_group>Bacillota>Clostridia>Eubacteriales>Lachnospiraceae | GCF_013304275.1 | Enterocloster_clostridioformis | HMPREF1088_RS04980 | 1098149..1101685 | 1531 | BREX-system-P-loop-protein-BrxC-[Enterocloster-clostridioformis] |
| WP_173727293.1 | Type1.BREX | GTPase-AIG->?->?->SH3->?->XIS-HTH->BrxA-DNA-binding->BrxB-BrxF-iSTAND->DUF6079*->TIR->N6-METHYLASE->Tox-ART-HYD2->?->AT-HTH-HYD2-2->PglZ->SIGMA+OB-Fold+AAA-ATPase+Lon-Protease-BrxL->Cap4_nuclease-> | 1217 | Bacteria>Terrabacteria_group>Bacillota>Clostridia>Eubacteriales>Lachnospiraceae | GCF_013304385.1 | Blautia_glucerasea | G5B11_06545 | 11040..14693 | 536633 | BREX-system-P-loop-protein-BrxC-[Blautia-glucerasea] |
| WP_173769033.1 | Type1.BREX | <-MORC+DUF3684+HNH<-?<-?<-?<-N6-METHYLASE<-DUF6079*<-BrxB-BrxF-iSTAND<-BrxA-DNA-binding<-HTH_26<-ZnR+ZnR+ZnR<-?<-MACRODOMAIN | 1201 | Bacteria>Terrabacteria_group>Bacillota>Clostridia>Eubacteriales>Lachnospiraceae | GCF_013305155.1 | Blautia_massiliensis | HQK10_RS13240 | 15504..19109 | 1737424 | BREX-system-P-loop-protein-BrxC-[Blautia-massiliensis] |
| WP_173741754.1 | Type1.BREX | <-MORC+DUF3684+HNH<-?<-?<-?<-STYKIN<-?<-N6-METHYLASE<-DUF6079*<-BrxB-BrxF-iSTAND<-BrxA-DNA-binding<-HTH_26<-ZnR+ZnR+ZnR<-?<-MACRODOMAIN | 1201 | Bacteria>Terrabacteria_group>Bacillota>Clostridia>Eubacteriales>Lachnospiraceae | GCF_013305165.1 | Blautia_massiliensis | G5B03_RS13685 | 17067..20672 | 1737424 | BREX-system-P-loop-protein-BrxC-[Blautia-massiliensis] |
| WP_181475440.1 | Type1.BREX | REase->?->Phage_integrase->?->HTH_36->HTH_36->HTH_26->BrxA-DNA-binding->BrxB-BrxF-iSTAND->DUF6079*->N6-METHYLASE->ZnR+ZnR->DUF4404->Schlafen+HATPase+wHTH->PglZ->SIGMA+OB-Fold+AAA-ATPase+Lon-Protease-BrxL->PHP+SbcC-><-HTH | 1182 | Bacteria>Terrabacteria_group>Bacillota>Clostridia>Eubacteriales>Lachnospiraceae | GCF_013694215.1 | Sellimonas_intestinalis | H1A12_RS00250 | 51099..54647 | 1653434 | BREX-system-P-loop-protein-BrxC-[Sellimonas-intestinalis] |
| WP_249326641.1 | Type1.BREX | GNTR-HTH+Aminotran_1_2->?->HTH_26->BrxA-DNA-binding->BrxB-BrxF-iSTAND->DUF6079*->Tox-ART-HYD2->AT-HTH-HYD2-1->N6-METHYLASE->HEPN->Schlafen+HATPase+wHTH->PglZ->SIGMA+OB-Fold+AAA-ATPase+Lon-Protease-BrxL->?->RNase-> | 1215 | Bacteria>Terrabacteria_group>Bacillota>Clostridia>Eubacteriales>Lachnospiraceae | GCF_014337175.1 | Simiaoa_sunii | H9Q77_RS04465 | 956158..959805 | 2763672 | BREX-system-P-loop-protein-BrxC-[Simiaoa-sunii] |
| WP_128132381.1 | Type1.BREX | MerR-HTH-><-?<-REase<-?<-SIGMA+OB-Fold+AAA-ATPase+Lon-Protease-BrxL<-PglZ<-HEPN<-N6-METHYLASE<-DUF6079*<-BrxB-BrxF-iSTAND<-BrxA-DNA-binding<-HTH_26<-HTH\|\|?-><-?<-TraJ-RHH<-SF1-SF2-Helicase | 1182 | Bacteria>Terrabacteria_group>Bacillota>Clostridia>Eubacteriales>Lachnospiraceae | GCF_015547745.1 | Enterocloster_bolteae | I2D45_RS23920 | 73790..77338 | 208479 | BREX-system-P-loop-protein-BrxC-[Enterocloster-bolteae] |
| WP_195415774.1 | Type1.BREX | <-MODE-HTH+PBPII\|\|?->?->?->ABhydrolase-><-?\|\|?->?->TatD_DNase->DUF6079-p*->OLD-ABC->REase-><-GDResolvase+Recombinase+ZnR<-?\|\|HTH_3->?-><-HTH_3 | 946 | Bacteria>Terrabacteria_group>Bacillota>Clostridia>Eubacteriales>Lachnospiraceae | GCF_015548445.1 | Enterocloster_citroniae | I2E22_RS12115 | 210359..213199 | 358743 | BREX-system-P-loop-protein-BrxC-[Enterocloster-citroniae] |
| WP_195230488.1 | Type1.BREX | <-SMODS<-SLATT<-SIGMA+OB-Fold+AAA-ATPase+Lon-Protease-BrxL<-PglZ<-AT-HTH-HYD2-2<-?<-Tox-ART-HYD2<-N6-METHYLASE<-DUF6079*<-BrxB-BrxF-iSTAND<-BrxA-DNA-binding<-XIS-HTH<-?<-?<-TraM<-TraJ-RHH<-LPD32+LPD36+N6_Mtase+SF1-SF2-Helicase | 1217 | Bacteria>Terrabacteria_group>Bacillota>Clostridia>Eubacteriales>Lachnospiraceae | GCF_015549285.1 | Coprococcus_comes | I1955_RS03720 | 51058..54711 | 410072 | BREX-system-P-loop-protein-BrxC-[Coprococcus-comes] |
| WP_195251421.1 | Type1.BREX | MACRODOMAIN->?->35exo+ADA-DBD->HTH_26->BrxA-DNA-binding->BrxB-BrxF-iSTAND->DUF6079*->N6-METHYLASE->ParB+HNH->HEPN-> | 1201 | Bacteria>Terrabacteria_group>Bacillota>Clostridia>Eubacteriales>Lachnospiraceae | GCF_015550485.1 | Coprococcus_comes | I2A60_RS13430 | 4839..8444 | 410072 | BREX-system-P-loop-protein-BrxC-[Coprococcus-comes] |
| WP_195441337.1 | Type1.BREX | CopG/RHH->?->TraJ-RHH->TraI-VirD2-RCRE->?->HTH_26->BrxA-DNA-binding->BrxB-BrxF-iSTAND->DUF6079*->N6-METHYLASE->AT-HTH-HYD2->?->AbiGi->Schlafen+HATPase+wHTH->PglZ->SIGMA+OB-Fold+AAA-ATPase+Lon-Protease-BrxL->TIR-><-HTH | 1182 | Bacteria>Terrabacteria_group>Bacillota>Clostridia>Eubacteriales>Lachnospiraceae | GCF_015554785.1 | Roseburia_intestinalis | I2F09_RS06795 | 71540..75088 | 166486 | BREX-system-P-loop-protein-BrxC-[Roseburia-intestinalis] |
| WP_195565783.1 | Type1.BREX | <-PglZ<-?<-?<-OLD-ABC<-TatD_DNase<-?<-KAP-glycosylase<-KAP-NTPase<-N6-METHYLASE<-DUF6079*<-BrxB-BrxF-iSTAND<-BrxA-DNA-binding<-HTH_26<-ZnR+ZnR+ZnR<-?<-MACRODOMAIN<-?<-NUDIX<-HKD-DNase+SF1-SF2-Helicase+HTH+HTH+SRA | 1201 | Bacteria>Terrabacteria_group>Bacillota>Clostridia>Eubacteriales>Lachnospiraceae | GCF_015556425.1 | Blautia_wexlerae | I2F87_RS05895 | 190232..193837 | 418240 | BREX-system-P-loop-protein-BrxC-[Blautia-wexlerae] |
| WP_195609804.1 | Type1.BREX | Peptidase_M20->MACRODOMAIN->?->?->?->?->HTH_26->BrxA-DNA-binding->BrxB-BrxF-iSTAND->DUF6079*->N6-METHYLASE->PglZ->OLD-ABC+TOPRIM->SIGMA+OB-Fold+AAA-ATPase+Lon-Protease-BrxL->LRR_5+LRR_8->Cas_Cas1->CRISPR_Cas2-> | 1203 | Bacteria>Terrabacteria_group>Bacillota>Clostridia>Eubacteriales>Lachnospiraceae | GCF_015560805.1 | Mediterraneibacter_glycyrrhizinilyticus | I2H87_RS09315 | 27635..31246 | 342942 | BREX-system-P-loop-protein-BrxC-[Mediterraneibacter-glycyrrhizinilyticus] |
| WP_138345918.1 | Type1.BREX | <-SIGMA-HTH+SIGMA-HTH+SIGMA-HTH<-AT-HTH-HYD2<-SIGMA+OB-Fold+AAA-ATPase+Lon-Protease-BrxL<-PglZ<-Schlafen+HATPase+wHTH<-?<-AT-HTH-HYD2<-N6-METHYLASE<-DUF6079*<-BrxB-BrxF-iSTAND<-BrxA-DNA-binding<-HTH_26 | 1182 | Bacteria>Terrabacteria_group>Bacillota>Clostridia>Eubacteriales>Lachnospiraceae | GCF_015669935.1 | Roseburia | FGQ84_RS06860 | 104625..108173 | 841 | MULTISPECIES:-BREX-system-P-loop-protein-BrxC-[Roseburia] |
| WP_208100468.1 | Type1.BREX | <-RADICAL-SAM<-PglZ<-Tox-RelE<-RelB<-N6-METHYLASE<-McrB-AAA<-DUF6079*<-BrxB-BrxF-iSTAND<-BrxA-DNA-binding\|\|ABhydrolase->?-><-?<-?<-?<-DOC+HTH<-Peptidase_M32 | 1217 | Bacteria>Terrabacteria_group>Bacillota>Clostridia>Eubacteriales>Lachnospiraceae | GCF_017565785.1 | Lachnoanaerobaculum_sp._Marseille-Q4761 | J4O15_RS08620 | 306409..310062 | 2819511 | BREX-system-P-loop-protein-BrxC-[Lachnoanaerobaculum-sp.-Marseille-Q4761] |
| WP_215726767.1 | Type1.BREX | ParB->?-><-?\|\|HTH_3->HTH_26->BrxA-DNA-binding->BrxB-BrxF-iSTAND->DUF6079*->?->N6-METHYLASE->HEPN+ZnR->HEPN->PglZ->?->SIGMA+OB-Fold+AAA-ATPase+Lon-Protease-BrxL->wHTH+WYL+WCX->HTH+REase-> | 1205 | Bacteria>Terrabacteria_group>Bacillota>Clostridia>Eubacteriales>Lachnospiraceae | GCF_018784865.1 | Dorea_longicatena | GPK48_RS02600 | 18956..22573 | 88431 | BREX-system-P-loop-protein-BrxC-[Dorea-longicatena] |
| WP_215631370.1 | Type1.BREX | GTPase-AIG->?->MACRODOMAIN->?->35exo+ADA-DBD->HTH_26->BrxB-BrxF-iSTAND->DUF6079*->Schlafen+HATPase+wHTH->N6-METHYLASE-><-DEDD_Tnp_IS110+Transposase_20\|\|?->OLD-ABC->NERD->OLD-ABC->PglZ->OLD-ABC-> | 1201 | Bacteria>Terrabacteria_group>Bacillota>Clostridia>Eubacteriales>Lachnospiraceae | GCF_018785175.1 | Coprococcus_catus | GPK82_RS00920 | 42842..46447 | 116085 | BREX-system-P-loop-protein-BrxC-[Coprococcus-catus] |
| WP_215631393.1 | Type1.BREX | <-HTH+REase<-wHTH+WYL+WCX<-SIGMA+OB-Fold+AAA-ATPase+Lon-Protease-BrxL<-?<-PglZ<-HNH+CXC3<-N6-METHYLASE<-Four-helical-protein<-wHTH+ZnR<-DUF6079*<-BrxB-BrxF-iSTAND<-BrxA-DNA-binding<-HTH_26<-HTH_3\|\|?->?<-Lysozyme+NLPC<-?<-?<-VirB4-FtsK | 1205 | Bacteria>Terrabacteria_group>Bacillota>Clostridia>Eubacteriales>Lachnospiraceae | GCF_018785175.1 | Coprococcus_catus | GPK82_RS01090 | 19804..23421 | 116085 | BREX-system-P-loop-protein-BrxC-[Coprococcus-catus] |
| WP_215689424.1 | Type1.BREX | GTPase-AIG->?->MACRODOMAIN->?->35exo+ADA-DBD->HTH_26->BrxA-DNA-binding->BrxB-BrxF-iSTAND->DUF6079*->N6-METHYLASE->OLD-ABC->NERD->?->PglZ->?->SIGMA+OB-Fold+AAA-ATPase+Lon-Protease-BrxL-> | 1203 | Bacteria>Terrabacteria_group>Bacillota>Clostridia>Eubacteriales>Lachnospiraceae | GCF_018785195.1 | Coprococcus_catus | GPK74_RS04355 | 145087..148698 | 116085 | BREX-system-P-loop-protein-BrxC-[Coprococcus-catus] |
| WP_252210442.1 | Type1.BREX | <-TIR<-SIGMA+OB-Fold+AAA-ATPase+Lon-Protease-BrxL<-PglZ<-Schlafen+HATPase+wHTH<-HEPN<-AT-HTH-HYD2<-Tox-ART-HYD2<-AT-HTH-HYD2<-SIR2<-DUF6079*<-BrxB-BrxF-iSTAND<-BrxA-DNA-binding<-HTH_26<-HTH_36<-ABhydrolase<-?<-SAM-METHYLASE | 1099 | Bacteria>Terrabacteria_group>Bacillota>Clostridia>Eubacteriales>Lachnospiraceae | GCF_018785625.1 | Blautia_faecis | GPK86_RS03045 | 13256..16555 | 871665 | BREX-system-P-loop-protein-BrxC-[Blautia-faecis] |
| WP_216418339.1 | Type1.BREX | <-STYKIN<-?<-LRR_5+LRR_8<-SIGMA+OB-Fold+AAA-ATPase+Lon-Protease-BrxL<-PglZ<-N6-METHYLASE<-DUF6079*<-BrxB-BrxF-iSTAND<-BrxA-DNA-binding<-HTH_26<-ZnR+ZnR+ZnR<-?<-MACRODOMAIN | 1203 | Bacteria>Terrabacteria_group>Bacillota>Clostridia>Eubacteriales>Lachnospiraceae | GCF_018917985.1 | Blautia_sp._MSJ-36 | KQI39_RS10905 | 43877..47488 | 2841530 | BREX-system-P-loop-protein-BrxC-[Blautia-sp.-MSJ-36] |
| WP_216429335.1 | Type1.BREX | HTH_26->BrxA-DNA-binding->BrxB-BrxF-iSTAND->DUF6079*->N6-METHYLASE->RAMA->HEPN->PglZ->SIGMA+OB-Fold+AAA-ATPase+Lon-Protease-BrxL->?->REase->?<-HTH_3 | 1182 | Bacteria>Terrabacteria_group>Bacillota>Clostridia>Eubacteriales>Lachnospiraceae | GCF_018918145.1 | Anaerotignum_sp._MSJ-24 | KQI66_RS10650 | 65878..69426 | 2841521 | BREX-system-P-loop-protein-BrxC-[Anaerotignum-sp.-MSJ-24] |
| WP_216523175.1 | Type1.BREX | <-wHTH+WYL+WCX<-?\|\|GDResolvase-><-?<-?<-SIGMA+OB-Fold+AAA-ATPase+Lon-Protease-BrxL<-PglZ<-HEPN<-N6-METHYLASE<-DUF6079*<-BrxB-BrxF-iSTAND<-BrxA-DNA-binding<-?<-DEOR-HTH<-?\|\|?->?->Phage_int_SAM_3->Phage_integrase-> | 1206 | Bacteria>Terrabacteria_group>Bacillota>Clostridia>Eubacteriales>Lachnospiraceae | GCF_018918205.1 | Lachnoclostridium_sp._MSJ-17 | KQI64_RS02435 | 23130..26750 | 2841516 | BREX-system-P-loop-protein-BrxC-[Lachnoclostridium-sp.-MSJ-17] |
| WP_274939928.1 | Type1.BREX | rve->AAA-ATPase->?->?->HTH_26->BrxA-DNA-binding->BrxB-BrxF-iSTAND->DUF6079*->SymE_toxin->?<-SIGMA-HTH<-?\|\|?->STAND->McrB-AAA->SymE_toxin->DNA_METHYLASE\|\|HTH_3-><-?<-TTRNase-> | 1214 | Bacteria>Terrabacteria_group>Bacillota>Clostridia>Eubacteriales>Lachnospiraceae | GCF_019355395.1 | Chordicoccus_furentiruminis | G4G92_RS11200 | 2473676..2477320 | 2709410 | BREX-system-P-loop-protein-BrxC-[Chordicoccus-furentiruminis] |
| WP_226870686.1 | Type1.BREX | HTH+HisKin->?->?->?->ABC-ATPase+ABC-ATPase->\|\|?->HTH_26->BrxA-DNA-binding->BrxB-BrxF-iSTAND->DUF6079*->?->N6-METHYLASE->?->?->AT-HTH-HYD2->Tox-ART-HYD2->AT-HTH-HYD2->HEPN->Schlafen+HATPase+wHTH-> | 1182 | Bacteria>Terrabacteria_group>Bacillota>Clostridia>Eubacteriales>Lachnospiraceae | GCF_020539685.1 | Mediterraneibacter_faecis | LIR06_RS00765 | 153448..156996 | 592978 | BREX-system-P-loop-protein-BrxC-[Mediterraneibacter-faecis] |
| WP_265190723.1 | Type1.BREX | <-ClpABN-AAA+ClpABC-AAA\|\|HTH-><-MULE-TRANSPOSASE<-PHP+SbcC<-SIGMA+OB-Fold+AAA-ATPase+Lon-Protease-BrxL<-PglZ<-Schlafen+HATPase+wHTH<-HEPN<-N6-METHYLASE<-DUF6079*<-BrxB-BrxF-iSTAND<-BrxA-DNA-binding<-HTH_26<-HTH_36 | 1182 | Bacteria>Terrabacteria_group>Bacillota>Clostridia>Eubacteriales>Lachnospiraceae | GCF_020553125.1 | Blautia_sp._MSK20_18 | LIO91_RS01865 | 117354..120902 | 2883186 | BREX-system-P-loop-protein-BrxC-[Blautia-sp.-MSK20_18] |
| WP_227147306.1 | Type1.BREX | LPD32+LPD36+N6_Mtase+SF1-SF2-Helicase->TraJ-RHH->TraM->?->?->XIS-HTH->BrxA-DNA-binding->BrxB-BrxF-iSTAND->DUF6079*->N6-METHYLASE->Tox-ART-HYD2->?->AT-HTH-HYD2-2->PglZ->SIGMA+OB-Fold+AAA-ATPase+Lon-Protease-BrxL->Cap4_nuclease-> | 1217 | Bacteria>Terrabacteria_group>Bacillota>Clostridia>Eubacteriales>Lachnospiraceae | GCF_020554525.1 | Enterocloster_bolteae | LIO19_RS21260 | 26213..29866 | 208479 | BREX-system-P-loop-protein-BrxC-[Enterocloster-bolteae] |
| WP_227148103.1 | Type1.BREX | LPD32+LPD36+N6_Mtase+SF1-SF2-Helicase->TraJ-RHH->TraM->?->?->XIS-HTH->BrxA-DNA-binding->BrxB-BrxF-iSTAND->DUF6079*->N6-METHYLASE->?->PglZ->SIGMA+OB-Fold+AAA-ATPase+Lon-Protease-BrxL->Tox-RelA->HKD-DNase->HKD-DNase->HKD-DNase->REC+OmpR-HTH->HAMP+HISKIN-> | 1217 | Bacteria>Terrabacteria_group>Bacillota>Clostridia>Eubacteriales>Lachnospiraceae | GCF_020554865.1 | Lacrimispora_sp._210928-DFI.3.58 | LIZ87_RS14625 | 23221..26874 | 2883214 | BREX-system-P-loop-protein-BrxC-[Lacrimispora-sp.-210928-DFI.3.58] |
| WP_227178521.1 | Type1.BREX | RADICAL-SAM->?->?-><-Peptidase<-?\|\|HTH->HTH_26->BrxA-DNA-binding->BrxB-BrxF-iSTAND->DUF6079*->N6-METHYLASE->Schlafen+HATPase+wHTH->PglZ->?->SIGMA+OB-Fold+AAA-ATPase+Lon-Protease-BrxL->?->REase->?-><-MPTase | 1182 | Bacteria>Terrabacteria_group>Bacillota>Clostridia>Eubacteriales>Lachnospiraceae | GCF_020555685.1 | Enterocloster_bolteae | LIO18_RS10100 | 522012..525560 | 208479 | BREX-system-P-loop-protein-BrxC-[Enterocloster-bolteae] |
| WP_227249221.1 | Type1.BREX | MNS+REase->TOPRIM+Topoisom_bac+TOPRIM_Crpt+TOPRIM_Crpt+TOPRIM_Crpt->?->?->?->HTH_26->BrxA-DNA-binding->BrxB-BrxF-iSTAND->DUF6079*->?->N6-METHYLASE->AT-HTH-HYD2->?->AbiGi->Schlafen+HATPase+wHTH->PglZ->SIGMA+OB-Fold+AAA-ATPase+Lon-Protease-BrxL->AT-HTH-HYD2-> | 1182 | Bacteria>Terrabacteria_group>Bacillota>Clostridia>Eubacteriales>Lachnospiraceae | GCF_020557615.1 | Roseburia_faecis | LIZ84_RS08395 | 26678..30226 | 301302 | BREX-system-P-loop-protein-BrxC-[Roseburia-faecis] |
| WP_227165162.1 | Type1.BREX | MACRODOMAIN->?->35exo+ADA-DBD->ZnR+ZnR+ZnR->HTH_26->BrxA-DNA-binding->BrxB-BrxF-iSTAND->DUF6079*->N6-METHYLASE->HEPN->PglZ->OLD-ABC+TOPRIM->SIGMA+OB-Fold+AAA-ATPase+Lon-Protease-BrxL->LRR_5+LRR_8->?->STYKIN-> | 1201 | Bacteria>Terrabacteria_group>Bacillota>Clostridia>Eubacteriales>Lachnospiraceae | GCF_020560115.1 | Mediterraneibacter_sp._210702-DFI.5.30 | LI126_RS08100 | 75769..79374 | 2883232 | BREX-system-P-loop-protein-BrxC-[Mediterraneibacter-sp.-210702-DFI.5.30] |
| WP_227083256.1 | Type1.BREX | REase+SPF-cons->wHTH+WYL+WCX->?->?->?->HTH_26->BrxA-DNA-binding->BrxB-BrxF-iSTAND->DUF6079*->N6-METHYLASE->DUF4365+REase->PglZ->HEPN+SH3-like+SF1-SF2-Helicase+wHTH->MORC+DUF3684+HNH-> | 1203 | Bacteria>Terrabacteria_group>Bacillota>Clostridia>Eubacteriales>Lachnospiraceae | GCF_020560805.1 | Coprococcus_catus | LI172_RS16280 | 10893..14504 | 116085 | BREX-system-P-loop-protein-BrxC-[Coprococcus-catus] |
| WP_227184330.1 | Type1.BREX | Peptidase_M20->MACRODOMAIN->?->?->HTH_26->BrxA-DNA-binding->BrxB-BrxF-iSTAND->DUF6079*->N6-METHYLASE-> | 1203 | Bacteria>Terrabacteria_group>Bacillota>Clostridia>Eubacteriales>Lachnospiraceae | GCF_020562365.1 | Mediterraneibacter_glycyrrhizinilyticus | LI216_RS12770 | 28291..31902 | 342942 | BREX-system-P-loop-protein-BrxC-[Mediterraneibacter-glycyrrhizinilyticus] |
| WP_227239829.1 | Type1.BREX | <-ABC-ATPase+SbcC<-REC+OmpR-HTH\|\|SIGMA-HTH->?->XIS-HTH->Integrase_DNA+Phage_integrase-><-N6-METHYLASE<-Schlafen+HATPase+wHTH<-DUF6079*<-BrxB-BrxF-iSTAND<-BrxA-DNA-binding<-HTH_26<-NUDIX<-?<-HATPase_c<-?<-SAM-METHYLASE<-STAS | 1203 | Bacteria>Terrabacteria_group>Bacillota>Clostridia>Eubacteriales>Lachnospiraceae | GCF_020563965.1 | Blautia_glucerasea | LI273_RS06155 | 27379..30990 | 536633 | BREX-system-P-loop-protein-BrxC-[Blautia-glucerasea] |
| WP_021651779.1 | Type1.BREX | <-PAS+HISKIN+REC<-?<-ParB<-ParB<-SymE_toxin<-HTH+REase<-SIGMA+OB-Fold+AAA-ATPase+Lon-Protease-BrxL<-PglZ<-N6-METHYLASE<-DUF6079*<-BrxB-BrxF-iSTAND<-BrxA-DNA-binding<-HTH_26<-?<-HNH+3C-ABC\|\|SIGMA-HTH-> | 1184 | Bacteria>Terrabacteria_group>Bacillota>Clostridia>Eubacteriales>Lachnospiraceae | GCF_020735365.1 | Blautia | LK422_RS11525 | 2410424..2413978 | 572511 | MULTISPECIES:-BREX-system-P-loop-protein-BrxC-[Blautia] |
| WP_238033441.1 | Type1.BREX | <-Phage_integrase<-Phage_integrase<-SF1-SF2-Helicase<-OLD-ABC+TOPRIM<-SIGMA+OB-Fold+AAA-ATPase+Lon-Protease-BrxL<-PglZ<-Schlafen+HATPase+wHTH<-AbiGi+seqA+RAMA<-N6-METHYLASE<-DUF6079*<-BrxB-BrxF-iSTAND<-BrxA-DNA-binding<-HTH_26<-?<-ZnR | 1182 | Bacteria>Terrabacteria_group>Bacillota>Clostridia>Eubacteriales>Lachnospiraceae | GCF_022136435.1 | Fusicatenibacter_saccharivorans | LON21_RS13045 | 78196..81744 | 1150298 | BREX-system-P-loop-protein-BrxC-[Fusicatenibacter-saccharivorans] |
| WP_246945856.1 | Type1.BREX | <-PglZ<-?<-Peptidase_M6<-MORC<-OLD-ABC<-OLD-ABC<-NERD<-OLD-ABC<-N6-METHYLASE<-DUF6079*<-BrxB-BrxF-iSTAND<-BrxA-DNA-binding<-HTH_26<-PIN<-?<-?<-MACRODOMAIN\|\|Phage_integrase-> | 1201 | Bacteria>Terrabacteria_group>Bacillota>Clostridia>Eubacteriales>Lachnospiraceae | GCF_023008345.1 | Dorea_longicatena | K5I24_RS06140 | 1316081..1319686 | 88431 | BREX-system-P-loop-protein-BrxC-[Dorea-longicatena] |
| WP_109627603.1 | Type1.BREX | HTH_26->BrxA-DNA-binding->BrxB-BrxF-iSTAND->DUF6079*->N6-METHYLASE->OLD-ABC->?->PglZ->Schlafen+HATPase+wHTH->SIGMA+OB-Fold+AAA-ATPase+Lon-Protease-BrxL->?-><-ACET | 1201 | Bacteria>Terrabacteria_group>Bacillota>Clostridia>Eubacteriales>Lachnospiraceae | GCF_024622195.1 | Murimonas_intestini | NSB30_RS07505 | 2822..6427 | 1337051 | BREX-system-P-loop-protein-BrxC-[Murimonas-intestini] |
| WP_257465570.1 | Type1.BREX | XIS-HTH->BrxA-DNA-binding->BrxB-BrxF-iSTAND->DUF6079*->N6-METHYLASE->Tox-ART-HYD2->?->AT-HTH-HYD2-2->PglZ->SIGMA+OB-Fold+AAA-ATPase+Lon-Protease-BrxL->SLATT->SMODS-> | 1217 | Bacteria>Terrabacteria_group>Bacillota>Clostridia>Eubacteriales>Lachnospiraceae | GCF_024622975.1 | Blautia_caecimuris | NSA57_RS17205 | 4109..7762 | 1796615 | BREX-system-P-loop-protein-BrxC-[Blautia-caecimuris] |
| WP_257465733.1 | Type1.BREX | <-RNase<-RNaseH<-N6-METHYLASE<-DUF6079*<-BrxB-BrxF-iSTAND<-BrxA-DNA-binding<-HTH_26<-?<-TraI-VirD2-RCRE<-TraJ-RHH | 1183 | Bacteria>Terrabacteria_group>Bacillota>Clostridia>Eubacteriales>Lachnospiraceae | GCF_024622975.1 | Blautia_caecimuris | NSA57_RS18375 | 5099..8650 | 1796615 | BREX-system-P-loop-protein-BrxC-[Blautia-caecimuris] |
| WP_158358547.1 | Type1.BREX | <-SIGMA+OB-Fold+AAA-ATPase+Lon-Protease-BrxL<-PglZ<-Schlafen+HATPase+wHTH<-HEPN+HEPN<-ParB+HNH<-AT-HTH-HYD2<-Tox-ART-HYD2<-AT-HTH-HYD2<-N6-METHYLASE<-DUF6079*<-BrxB-BrxF-iSTAND<-BrxA-DNA-binding<-HTH_26<-?<-?<-?<-DOC+HTH<-HTH_3<-PDZ_6+Peptidase_S55 | 1182 | Bacteria>Terrabacteria_group>Bacillota>Clostridia>Eubacteriales>Lachnospiraceae | GCF_025567195.1 | Altiscatomonas_aceti | OCV69_RS08300 | 70137..73685 | 2981724 | BREX-system-P-loop-protein-BrxC-[Altiscatomonas-aceti] |
| WP_270422341.1 | Type1.BREX | CopG/RHH->?->TraJ-RHH->TraI-VirD2-RCRE->?->HTH_26->BrxA-DNA-binding->BrxB-BrxF-iSTAND->DUF6079*->?->N6-METHYLASE->HEPN->Schlafen+HATPase+wHTH->PglZ->SIGMA+OB-Fold+AAA-ATPase+Lon-Protease-BrxL->OLD-ABC+TOPRIM->SF1-SF2-Helicase-><-HTH | 1182 | Bacteria>Terrabacteria_group>Bacillota>Clostridia>Eubacteriales>Lachnospiraceae | GCF_027661365.1 | Fusicatenibacter_saccharivorans | PGQ16_RS03035 | 254369..257917 | 1150298 | BREX-system-P-loop-protein-BrxC-[Fusicatenibacter-saccharivorans] |
| WP_270389656.1 | Type1.BREX | MACRODOMAIN->?->?->HTH_26->BrxA-DNA-binding->BrxB-BrxF-iSTAND->DUF6079*->N6-METHYLASE->AbiGi+seqA+RAMA->OLD-ABC->?->PglZ->OLD-ABC+TOPRIM->SIGMA+OB-Fold+AAA-ATPase+Lon-Protease-BrxL->LRR_5+LRR_8-> | 1201 | Bacteria>Terrabacteria_group>Bacillota>Clostridia>Eubacteriales>Lachnospiraceae | GCF_027661925.1 | Blautia_massiliensis | PGQ23_RS05910 | 13798..17403 | 1737424 | BREX-system-P-loop-protein-BrxC-[Blautia-massiliensis] |
| WP_118278843.1 | Type1.BREX | <-ParB+HNH<-?<-SIGMA+OB-Fold+AAA-ATPase+Lon-Protease-BrxL<-?<-PglZ<-?<-N6-METHYLASE<-Four-helical-protein<-wHTH+ZnR<-DUF6079*<-BrxB-BrxF-iSTAND<-BrxA-DNA-binding<-HTH_26<-HTH_3 | 1205 | Bacteria>Terrabacteria_group>Bacillota>Clostridia>Eubacteriales>Lachnospiraceae | GCF_027662525.1 | [Ruminococcus]_lactaris | DW116_RS04365 | 20004..23621 | 46228 | BREX-system-P-loop-protein-BrxC-[[Ruminococcus] |
| WP_270414325.1 | Type1.BREX | <-Cap4_nuclease<-SIGMA+OB-Fold+AAA-ATPase+Lon-Protease-BrxL<-PglZ<-?<-N6-METHYLASE<-DUF6079*<-BrxB-BrxF-iSTAND<-BrxA-DNA-binding<-XIS-HTH<-?<-?<-?<-?<-Phage_int_SAM_3+Phage_integrase<-Phage_int_SAM_3+Phage_integrase | 1217 | Bacteria>Terrabacteria_group>Bacillota>Clostridia>Eubacteriales>Lachnospiraceae | GCF_027663385.1 | Blautia_faecis | PGR36_RS14930 | 64613..68266 | 871665 | BREX-system-P-loop-protein-BrxC-[Blautia-faecis] |
| WP_270395874.1 | Type1.BREX | <-SIGMA+OB-Fold+AAA-ATPase+Lon-Protease-BrxL<-PglZ<-Schlafen+HATPase+wHTH<-AT-HTH-HYD2<-RNaseH<-N6-METHYLASE<-DUF6079*<-BrxB-BrxF-iSTAND<-BrxA-DNA-binding<-HTH_26<-?<-HTH_36<-?<-Cysteine-Desulfurase<-Peptidase_M20 | 1182 | Bacteria>Terrabacteria_group>Bacillota>Clostridia>Eubacteriales>Lachnospiraceae | GCF_027666095.1 | Mediterraneibacter_massiliensis | PGR65_RS11790 | 30843..34391 | 1720300 | BREX-system-P-loop-protein-BrxC-[Mediterraneibacter-massiliensis] |
| WP_119247667.1 | Type1.BREX | HTH_36->?->CopG/RHH->TraJ-RHH->TraI-VirD2-RCRE->HTH_26->BrxA-DNA-binding->BrxB-BrxF-iSTAND->DUF6079*->?->N6-METHYLASE->HEPN->Schlafen+HATPase+wHTH->PglZ->SIGMA+OB-Fold+AAA-ATPase+Lon-Protease-BrxL->OLD-ABC+TOPRIM->SF1-SF2-Helicase-><-HTH | 1182 | Bacteria>Terrabacteria_group>Bacillota>Clostridia>Eubacteriales>Lachnospiraceae | GCF_027674485.1 | Dorea_longicatena | PGV47_RS07575 | 41898..45446 | 88431 | BREX-system-P-loop-protein-BrxC-[Dorea-longicatena] |
| WP_270645287.1 | Type1.BREX | HTH_19-><-SIGMA+OB-Fold+AAA-ATPase+Lon-Protease-BrxL<-?<-PglZ<-PIN<-OLD-ABC<-OLD-ABC+TOPRIM<-N6-METHYLASE<-DUF6079*<-BrxB-BrxF-iSTAND<-BrxA-DNA-binding<-HTH_26<-?<-?<-?<-ArdCN+MPTase-inact+LPD19<-?<-TOPRIM+Topoisom_bac | 1205 | Bacteria>Terrabacteria_group>Bacillota>Clostridia>Eubacteriales>Lachnospiraceae | GCF_027692785.1 | Merdimonas_faecis | PG729_RS15130 | 28205..31822 | 1653435 | BREX-system-P-loop-protein-BrxC-[Merdimonas-faecis] |
| WP_244339353.1 | Type1.BREX | <-REase<-?<-?<-?<-HHH_4+HTH+wHTH+SF1-SF2-Helicase+REase<-MORC+DUF3684+HNH<-HEPN+SH3-like+SF1-SF2-Helicase+wHTH<-PglZ<-N6-METHYLASE<-DUF6079*<-BrxB-BrxF-iSTAND<-BrxA-DNA-binding<-HTH_26<-?<-?<-wHTH+WYL+WCX | 1209 | Bacteria>Terrabacteria_group>Bacillota>Clostridia>Eubacteriales>Lachnospiraceae | GCF_028207415.1 | [Clostridium]_symbiosum | PMZ71_RS08780 | 52022..55651 | 1512 | BREX-system-P-loop-protein-BrxC-[[Clostridium] |
| WP_271824191.1 | Type1.BREX | MPTase->AT-HTH-HYD2->Tox-ART-HYD2->AT-HTH-HYD2-><-SbcC<-ParB+HNH<-SIGMA+OB-Fold+AAA-ATPase+Lon-Protease-BrxL<-PglZ<-N6-METHYLASE<-DUF6079*<-BrxB-BrxF-iSTAND<-BrxA-DNA-binding<-HTH_26<-HTH\|\|?->Peptidase-><-?<-?<-TraJ-RHH | 1178 | Bacteria>Terrabacteria_group>Bacillota>Clostridia>Eubacteriales>Lachnospiraceae | GCF_028210655.1 | Enterocloster_clostridioformis | PMX39_RS12925 | 18239..21775 | 1531 | BREX-system-P-loop-protein-BrxC-[Enterocloster-clostridioformis] |
| WP_099385222.1 | Type1.BREX | <-Tox-RES<-?<-URI-DNase<-SIGMA+OB-Fold+AAA-ATPase+Lon-Protease-BrxL<-?<-PglZ<-HEPN<-N6-METHYLASE<-N6-METHYLASE<-DUF6079*<-BrxB-BrxF-iSTAND<-BrxA-DNA-binding<-HTH_26<-ACET | 1207 | Bacteria>Terrabacteria_group>Bacillota>Clostridia>Eubacteriales>Lachnospiraceae | GCF_028539255.1 | Agathobacter_ruminis | CSX02_RS00500 | 90797..94420 | 1712665 | BREX-system-P-loop-protein-BrxC-[Agathobacter-ruminis] |
| WP_226815507.1 | Type1.BREX | <-HNH<-?<-OLD-ABC<-SF1-SF2-Helicase<-?<-TIR<-N6-METHYLASE<-DUF6079*<-BrxB-BrxF-iSTAND<-BrxA-DNA-binding<-HTH_26<-35exo+ADA-DBD<-?<-MACRODOMAIN | 1203 | Bacteria>Terrabacteria_group>Bacillota>Clostridia>Eubacteriales>Lachnospiraceae | GCF_900066165.1 | Blautia_glucerasea | LIQ25_RS14525 | 37765..41376 | 536633 | BREX-system-P-loop-protein-BrxC-[Blautia-glucerasea] |
| WP_090167800.1 | Type1.BREX | <-HKD-DNase<-?<-SIGMA+OB-Fold+AAA-ATPase+Lon-Protease-BrxL<-BrxD-AAA<-?<-PglZ<-HEPN<-N6-METHYLASE<-DUF6079*<-BrxB-BrxF-iSTAND<-BrxA-DNA-binding<-HTH_26<-?<-?<-wHTH+WYL+WCX<-HNH<-?<-ARG | 1201 | Bacteria>Terrabacteria_group>Bacillota>Clostridia>Eubacteriales>Lachnospiraceae | GCF_900100545.1 | Pseudobutyrivibrio_sp._49 | BLP54_RS06100 | 185622..189227 | 1855344 | BREX-system-P-loop-protein-BrxC-[Pseudobutyrivibrio-sp.-49] |
| WP_090815514.1 | Type1.BREX | <-SF1-SF2-Helicase<-SF1-SF2-Helicase<-SF1-SF2-Helicase+TOPC<-ParB+HNH<-?<-?<-SIGMA+OB-Fold+AAA-ATPase+Lon-Protease-BrxL<-PglZ<-N6-METHYLASE<-DUF6079*<-BrxB-BrxF-iSTAND<-BrxA-DNA-binding<-HTH_26<-?<-?<-?<-TraJ-RHH<-ArdCN+MPTase-inact+LPD19 | 1182 | Bacteria>Terrabacteria_group>Bacillota>Clostridia>Eubacteriales>Lachnospiraceae | GCF_900107835.1 | Oribacterium_sp._KHPX15 | BLV66_RS05660 | 48724..52272 | 1855342 | BREX-system-P-loop-protein-BrxC-[Oribacterium-sp.-KHPX15] |
| WP_091825279.1 | Type1.BREX | AbiEii->?->?->TraJ-RHH->Integrase_DNA+Phage_integrase->ZnR+TOPRIM->XIS-HTH->BrxA-DNA-binding->BrxB-BrxF-iSTAND->DUF6079*->N6-METHYLASE->PglZ->SIGMA+OB-Fold+AAA-ATPase+Lon-Protease-BrxL->?->ParB+HNH->UDG-conflict->?->?->HTH_28-> | 1145 | Bacteria>Terrabacteria_group>Bacillota>Clostridia>Eubacteriales>Lachnospiraceae | GCF_900109475.1 | Butyrivibrio_sp._ob235 | BMW61_RS19080 | 17910..21347 | 1761780 | BREX-system-P-loop-protein-BrxC-[Butyrivibrio-sp.-ob235] |

| Protein-ID | System | Gene-Neighbourhoods | Length | Taxonomy | Assembly-ID | Strain | Locus-Tag | Genomic-coordinates | TaxID | Fasta-defline |
|---|---|---|---|---|---|---|---|---|---|---|
| WP_090393802.1 | Type1.BREX | wHTH+WYL+WCX->?->?->HKD-DNase->HTH_26->BrxA-DNA-binding->BrxB-BrxF-iSTAND->DUF6079*->N6-METHYLASE->?->PglZ->SIGMA+OB-Fold+AAA-ATPase+Lon-Protease-BrxL->HEPN->?->HKD-DNase->SF1-SF2-Helicase->REase-> | 1182 | Bacteria>Terrabacteria_group>Bacillota>Clostridia>Eubacteriales>Lachnospiraceae | GCF_900111635.1 | Pseudobutyrivibrio_sp._C4 | BMY66_RS00580 | 126012..129560 | 1520803 | BREX-system-P-loop-protein-BrxC-[Pseudobutyrivibrio-sp.-C4] |
| WP_242940588.1 | Type1.BREX | <-DUF6079-p*<-BrxB-BrxF-iSTAND<-BrxA-DNA-binding<-HTH_26<-SbcC<-?<-HNH\|\|?->SIGMA-HTH-> | 365 | Bacteria>Terrabacteria_group>Bacillota>Clostridia>Eubacteriales>Lachnospiraceae | GCF_900112775.1 | Anaeromicropila_populeti | BM305_RS18095 | 17..1114 | 37658 | BREX-system-P-loop-protein-BrxC-[Anaeromicropila-populeti] |
| WP_090545210.1 | Type1.BREX | <-HKD-DNase<-?<-SIGMA+OB-Fold+AAA-ATPase+Lon-Protease-BrxL<-BrxD-AAA<-?<-PglZ<-HEPN<-N6-METHYLASE<-N6-METHYLASE<-DUF6079*<-BrxB-BrxF-iSTAND<-BrxA-DNA-binding<-HTH_26<-?<-?<-wHTH+WYL+WCX<-HNH<-?<-ARG | 1201 | Bacteria>Terrabacteria_group>Bacillota>Clostridia>Eubacteriales>Lachnospiraceae | GCF_900113655.1 | Pseudobutyrivibrio_sp._OR37 | BM097_RS01535 | 134498..138103 | 1798186 | BREX-system-P-loop-protein-BrxC-[Pseudobutyrivibrio-sp.-OR37] |
| WP_091687629.1 | Type1.BREX | HTH_3->?-><-REase+N-glycosylase<-SF1-SF2-Helicase<-SbcC<-?<-SIGMA+OB-Fold+AAA-ATPase+Lon-Protease-BrxL<-PglZ<-N6-METHYLASE<-DUF6079*<-BrxB-BrxF-iSTAND<-BrxA-DNA-binding<-HTH_26<-?<-TraJ-RHH<-LPD2<-TOPRIM+Topoisom_bac<-?<-ArdCN+MPTase-inact+LPD19 | 1208 | Bacteria>Terrabacteria_group>Bacillota>Clostridia>Eubacteriales>Lachnospiraceae | GCF_900115365.1 | Anaerocolumna_aminovalerica | BM183_RS14685 | 24667..28293 | 1527 | BREX-system-P-loop-protein-BrxC-[Anaerocolumna-aminovalerica] |
| WP_143111148.1 | Type1.BREX | <-MODE-HTH+PBPII\|\|?->?->?->ABhydrolase-><-?\|\|?->?->TatD_DNase->DUF6079-p*->OLD-ABC->REase->METHYLASE->?->ZnR+ZnR->ABC-ATPase+SbcC-> | 946 | Bacteria>Terrabacteria_group>Bacillota>Clostridia>Eubacteriales>Lachnospiraceae | GCF_900115855.1 | Enterocloster_citroniae | BM366_RS01240 | 259291..262131 | 358743 | BREX-system-P-loop-protein-BrxC-[Enterocloster-citroniae] |
| WP_073386276.1 | Type1.BREX | OLD-ABC+TOPRIM->HNH->?->SF1-SF2-Helicase->?->SF1-SF2-Helicase->HTH_26->BrxA-DNA-binding->BrxB-BrxF-iSTAND->DUF6079*->N6-METHYLASE->?->N6-METHYLASE->PglZ->?->SIGMA+OB-Fold+AAA-ATPase+Lon-Protease-BrxL->?->SIGMA-HTH->HEPN+ParB-> | 1206 | Bacteria>Terrabacteria_group>Bacillota>Clostridia>Eubacteriales>Lachnospiraceae | GCF_900129945.1 | Butyrivibrio_fibrisolvens | BUC27_RS06005 | 31476..35096 | 831 | BREX-system-P-loop-protein-BrxC-[Butyrivibrio-fibrisolvens] |
| WP_072849732.1 | Type1.BREX | YTCD-HTH-><-SF1-SF2-Helicase<-SIGMA+OB-Fold+AAA-ATPase+Lon-Protease-BrxL<-PglZ<-Schlafen+HATPase+wHTH<-?<-N6-METHYLASE<-Four-helical-protein<-wHTH+ZnR<-DUF6079*<-BrxB-BrxF-iSTAND<-BrxA-DNA-binding<-HTH_26<-?<-?\|\|HTH_3-><-TraI-VirD2-RCRE<-TraJ-RHH | 1211 | Bacteria>Terrabacteria_group>Bacillota>Clostridia>Eubacteriales>Lachnospiraceae | GCF_900142265.1 | Anaerotignum_lactatifermentans | BUB35_RS04915 | 36734..40369 | 160404 | BREX-system-P-loop-protein-BrxC-[Anaerotignum-lactatifermentans] |
| WP_072699981.1 | Type1.BREX | <-HEPN<-?<-SIGMA+OB-Fold+AAA-ATPase+Lon-Protease-BrxL<-TIR<-PglZ<-RAMA<-N6-METHYLASE<-DUF6079*<-BrxB-BrxF-iSTAND<-BrxA-DNA-binding<-HTH_26<-?<-TraJ-RHH<-LPD2<-TOPRIM+Topoisom_bac<-?<-ArdCN+MPTase-inact+LPD19 | 1185 | Bacteria>Terrabacteria_group>Bacillota>Clostridia>Eubacteriales>Lachnospiraceae | GCF_900143205.1 | Butyrivibrio_hungatei | BUA02_RS00135 | 23142..26699 | 185008 | BREX-system-P-loop-protein-BrxC-[Butyrivibrio-hungatei] |
| WP_102289727.1 | Type1.BREX | <-TnpB_IS66\|\|?-><-SIGMA+OB-Fold+AAA-ATPase+Lon-Protease-BrxL<-Schlafen+HATPase+wHTH<-PglZ<-HEPN<-N6-METHYLASE<-DUF6079*<-BrxB-BrxF-iSTAND<-BrxA-DNA-binding<-HTH_26<-?<-?<-?<-?<-?\|\|HAMP+HisKin+HATPase_c-> | 1215 | Bacteria>Terrabacteria_group>Bacillota>Clostridia>Eubacteriales>Lachnospiraceae | GCF_900243045.1 | Eisenbergiella_massiliensis | C1733_RS26410 | 2142071..2145718 | 1720294 | BREX-system-P-loop-protein-BrxC-[Eisenbergiella-massiliensis] |
| WP_144362776.1 | Type1.BREX | <-Schlafen+HATPase+wHTH<-HEPN<-AT-HTH-HYD2<-Tox-ART-HYD2<-AT-HTH-HYD2<-?<-?<-N6-METHYLASE<-?<-DUF6079*<-BrxB-BrxF-iSTAND<-BrxA-DNA-binding<-HTH_26<-TraI-VirD2-RCRE<-TraJ-RHH<-CopG/RHH<-?<-HTH_36 | 1182 | Bacteria>Terrabacteria_group>Bacillota>Clostridia>Eubacteriales>Lachnospiraceae | GCF_902167785.1 | [Ruminococcus]_gnavus | FQ315_RS12895 | 14727..18275 | 33038 | BREX-system-P-loop-protein-BrxC-[[Ruminococcus] |
| WP_055164649.1 | Type1.BREX | <-ACET\|\|?->?->?-><-?<-?\|\|HTH_26->BrxA-DNA-binding->BrxB-BrxF-iSTAND->DUF6079*->N6-METHYLASE->OLD-ABC->NERD->PglZ->SIGMA+OB-Fold+AAA-ATPase+Lon-Protease-BrxL->?->CHTH-> | 1203 | Bacteria>Terrabacteria_group>Bacillota>Clostridia>Eubacteriales>Lachnospiraceae | GCF_902362465.1 | Blautia_hydrogenotrophica | FY213_RS07140 | 172247..175858 | 53443 | BREX-system-P-loop-protein-BrxC-[Blautia-hydrogenotrophica] |
| WP_055287913.1 | Type1.BREX | <-SF1-SF2-Helicase+TUDOR<-?<-SIGMA+OB-Fold+AAA-ATPase+Lon-Protease-BrxL<-OLD-ABC+TOPRIM<-PglZ<-Schlafen+HATPase+wHTH<-REase<-N6-METHYLASE<-DUF6079*<-BrxB-BrxF-iSTAND<-BrxA-DNA-binding<-HTH_26<-wHTH+WYL+WCX | 1196 | Bacteria>Terrabacteria_group>Bacillota>Clostridia>Eubacteriales>Lachnospiraceae | GCF_902362785.1 | Lachnospira_eligens | FXW70_RS12185 | 167515..171105 | 39485 | BREX-system-P-loop-protein-BrxC-[Lachnospira-eligens] |
| WP_117450344.1 | Type1.BREX | <-MODE-HTH+PBPII\|\|?->?->?->ABhydrolase-><-?\|\|?->?->TatD_DNase->DUF6079-p*->OLD-ABC->REase->METHYLASE->?->ZnR+ZnR->ABC-ATPase+SbcC-> | 946 | Bacteria>Terrabacteria_group>Bacillota>Clostridia>Eubacteriales>Lachnospiraceae | GCF_902364255.1 | Enterocloster_citroniae | FY458_RS04080 | 886658..889498 | 358743 | BREX-system-P-loop-protein-BrxC-[Enterocloster-citroniae] |
| WP_251353723.1 | Type1.BREX | <-TOPRIM<-?\|\|HTH_3->?->ParB->?->HTH_26->BrxA-DNA-binding->BrxB-BrxF-iSTAND->DUF6079*->N6-METHYLASE->?->PglZ-> | 1197 | Bacteria>Terrabacteria_group>Bacillota>Clostridia>Eubacteriales>Lachnospiraceae | GCF_943193015.1 | Hominisplanchenecus_sp._DSM_111139 | NFR99_RS11950 | 14312..17905 | 2941517 | BREX-system-P-loop-protein-BrxC-[Hominisplanchenecus-murintestinalis] |
| WP_251490385.1 | Type1.BREX | <-SIR2<-?<-PglZ<-N6-METHYLASE<-DUF6079*<-BrxB-BrxF-iSTAND<-BrxA-DNA-binding<-HTH_26\|\|?-><-HATPase+wHTH<-Schlafen+HATPase+wHTH-p<-ABC-ATPase+SbcC | 1182 | Bacteria>Terrabacteria_group>Bacillota>Clostridia>Eubacteriales>Lachnospiraceae | GCF_943193115.1 | Otoolea_muris | NFS31_RS00400 | 84330..87878 | 2941515 | BREX-system-P-loop-protein-BrxC-[Otoolea-muris] |
| WP_251499804.1 | Type1.BREX | <-ABC-ATPase+SbcC<-?<-REC+OmpR-HTH<-XIS-HTH\|\|HTH_26->BrxA-DNA-binding->DUF6079-p*->SIGMA+OB-Fold+AAA-ATPase+Lon-Protease-BrxL-p-><-MODE-HTH+PBPII\|\|?->?-><-AlcRN+AraC-HTH+AraC-HTH<-ArsR-HTH\|\|?->?->ArsR-HTH-> | 704 | Bacteria>Terrabacteria_group>Bacillota>Clostridia>Eubacteriales>Lachnospiraceae | GCF_943193115.1 | Otoolea_muris | NFS31_RS21535 | 7780..9894 | 2941515 | BREX-system-P-loop-protein-BrxC-[Otoolea-muris] |
| WP_065540591.1 | Type1.BREX | CHTH->HTH_26->BrxA-DNA-binding->BrxB-BrxF-iSTAND->DUF6079*->N6-METHYLASE->PglZ->Schlafen+HATPase+wHTH->?-><-MODE-HTH+PBPII\|\|?->?-><-HISKIN | 1209 | Bacteria>Terrabacteria_group>Bacillota>Clostridia>Eubacteriales>Lachnospiraceae | GCF_947644355.1 | Blautia_pseudococcoides | A4V09_RS00325 | 73624..77253 | 1796616 | BREX-system-P-loop-protein-BrxC-[Blautia-pseudococcoides] |
| WP_009531888.1 | Type1.BREX | XIS-HTH->BrxA-DNA-binding->BrxB-BrxF-iSTAND->DUF6079*->N6-METHYLASE->PglZ->SIGMA+OB-Fold+AAA-ATPase+Lon-Protease-BrxL->?->PADR-HTH->?->?-><-HicB | 1217 | Bacteria>Terrabacteria_group>Bacillota>Clostridia>Eubacteriales>Lachnospiraceae | - | Stomatobaculum_longum | HMPREF9623_00053 | 10076..13729 | 796942 | BREX-system-P-loop-protein-BrxC-[Stomatobaculum-longum] |
| WP_009532426.1 | Type1.BREX | <-PglZ<-HATPase+wHTH<-Schlafen+HATPase+wHTH<-DUF4404<-ZnR+ZnR<-N6-METHYLASE<-DUF6079*<-BrxB-BrxF-iSTAND<-BrxA-DNA-binding<-HTH_26<-?<-?<-?<-TrkH | 1180 | Bacteria>Terrabacteria_group>Bacillota>Clostridia>Eubacteriales>Lachnospiraceae | - | Stomatobaculum_longum | HMPREF9623_00593 | 29898..33440 | 796942 | BREX-system-P-loop-protein-BrxC-[Stomatobaculum-longum] |
| WP_256167163.1 | Type1.BREX | <-Cap4_nuclease<-SIGMA+OB-Fold+AAA-ATPase+Lon-Protease-BrxL<-PglZ<-AT-HTH-HYD2-2<-?<-Tox-ART-HYD2<-N6-METHYLASE<-TIR<-DUF6079*<-BrxB-BrxF-iSTAND<-BrxA-DNA-binding<-XIS-HTH<-?<-MACRODOMAIN<-REase+N-glycosylase | 1217 | Bacteria>Terrabacteria_group>Bacillota>Clostridia>Eubacteriales>Oscillospiraceae | - | Agathobaculum_butyriciproducens | NE606_05805 | 173642..177295 | 1628085 | BREX-system-P-loop-protein-BrxC-[Agathobaculum-butyriciproducens] |
| WP_120485415.1 | Type1.BREX | MACRODOMAIN->?->35exo+ADA-DBD->ZnR+ZnR+ZnR->HTH_26->BrxA-DNA-binding->BrxB-BrxF-iSTAND->DUF6079*->N6-METHYLASE->OLD-ABC->NERD->PglZ->OLD-ABC+TOPRIM->SIGMA+OB-Fold+AAA-ATPase+Lon-Protease-BrxL->LRR_5+LRR_8->?->STYKIN-> | 1201 | Bacteria>Terrabacteria_group>Bacillota>Clostridia>Eubacteriales>Oscillospiraceae | - | Anaerotruncus_sp._1XD22-93 | D3Z44_12640 | 10997..14602 | 2320082 | BREX-system-P-loop-protein-BrxC-[Anaerotruncus-sp.-1XD22-93] |
| WP_097772844.1 | Type1.BREX | <-HAMP+HISKIN<-REC+OmpR-HTH<-HKD-DNase+HKD-DNase<-HKD-DNase+HKD-DNase<-Tox-RelA<-SIGMA+OB-Fold+AAA-ATPase+Lon-Protease-BrxL<-PglZ<-?<-N6-METHYLASE<-DUF6079*<-BrxB-BrxF-iSTAND<-BrxA-DNA-binding<-XIS-HTH<-?<-?<-TraM<-TraJ-RHH<-LPD32+LPD36+N6_Mtase+SF1-SF2-Helicase | 1217 | Bacteria>Terrabacteria_group>Bacillota>Clostridia>Eubacteriales>Oscillospiraceae | - | Faecalibacterium_prausnitzii | CGS57_RS07390 | 76016..79669 | 853 | BREX-system-P-loop-protein-BrxC-[Faecalibacterium-prausnitzii] |
| WP_055192172.1 | Type1.BREX | <-MORC+DUF3684+HNH<-HEPN+SH3-like+SF1-SF2-Helicase+wHTH<-PglZ<-Schlafen+HATPase+wHTH<-HEPN<-?<-?<-N6-METHYLASE<-DUF6079*<-BrxB-BrxF-iSTAND<-BrxA-DNA-binding<-?<-35exo+wHTH+SF1-SF2-Helicase+HTH<-?<-35exo | 1204 | Bacteria>Terrabacteria_group>Bacillota>Clostridia>Eubacteriales>Oscillospiraceae | - | Faecalibacterium_prausnitzii | FXW87_RS13010 | 15894..19508 | 853 | BREX-system-P-loop-protein-BrxC-[Faecalibacterium-prausnitzii] |
| WP_156068326.1 | Type1.BREX | LPD32+LPD36+N6_Mtase+SF1-SF2-Helicase->TraJ-RHH->TraM->?->?->XIS-HTH->BrxA-DNA-binding->BrxB-BrxF-iSTAND->DUF6079*->N6-METHYLASE->Tox-ART-HYD2->?->AT-HTH-HYD2-2->PglZ->SIGMA+OB-Fold+AAA-ATPase+Lon-Protease-BrxL->SLATT->SMODS->Tox-RelA-> | 1217 | Bacteria>Terrabacteria_group>Bacillota>Clostridia>Eubacteriales>Oscillospiraceae | - | Faecalibacterium_prausnitzii | GO812_RS01965 | 48616..52269 | 853 | BREX-system-P-loop-protein-BrxC-[Faecalibacterium-prausnitzii] |
| EGJ45320.2 | Type1.BREX | <-HTH_3\|\|CHTH-><-?<-SIGMA+OB-Fold+AAA-ATPase+Lon-Protease-BrxL<-PglZ<-?<-N6-METHYLASE<-DUF6079*<-BrxB-BrxF-iSTAND<-BrxA-DNA-binding<-HTH_26<-TraI-VirD2-RCRE<-TraJ-RHH | 1181 | Bacteria>Terrabacteria_group>Bacillota>Clostridia>Eubacteriales>Oscillospiraceae | GCA_000177015.3 | Ruminococcaceae_bacterium_D16 | HMPREF0866_03146 | 49329..52874 | 552398 | hypothetical-protein-HMPREF0866_03146-[Ruminococcaceae-bacterium-D16] |
| CBL33286.1 | Type1.BREX | <-MoxR-AAA<-?<-LRR_5+LRR_5+LRR_5+LRR_5\|\|?->?->HTH_26->BrxA-DNA-binding->BrxB-BrxF-iSTAND-><-?\|\|DUF6079*->N6-METHYLASE->STYKinase-like->N6-METHYLASE-p->PglZ->SIGMA+OB-Fold+AAA-ATPase+Lon-Protease-BrxL-> | 1103 | Bacteria>Terrabacteria_group>Bacillota>Clostridia>Eubacteriales>Oscillospiraceae | GCA_000210635.1 | [Eubacterium]_siraeum_V10Sc8a | ES1_00670 | 50729..54040 | 717961 | hypothetical-protein-ES1_00670-[[Eubacterium] |
| CDA72644.1 | Type1.BREX | ABC-ATPase+SbcC->ABC-ATPase+SbcC->?->Phage_int_SAM_3+Phage_integrase->BrxA-DNA-binding->BrxB-BrxF-iSTAND->DUF6079*->N6-METHYLASE->?->N6-METHYLASE-p->PglZ->SIGMA+OB-Fold+AAA-ATPase+Lon-Protease-BrxL-> | 1163 | Bacteria>Terrabacteria_group>Bacillota>Clostridia>Eubacteriales>Oscillospiraceae | GCA_000434695.1 | Ruminococcus_sp._CAG:579 | BN718_01320 | 5394..8885 | 1262963 | putative-ATPase-involved-in-DNA-repair-[Ruminococcus-sp.-CAG:579] |
| CUQ84432.1 | Type1.BREX | <-SIGMA+OB-Fold+AAA-ATPase+Lon-Protease-BrxL<-PglZ<-N6-METHYLASE<-DUF6079*<-BrxB-BrxF-iSTAND<-BrxA-DNA-binding<-HTH_26<-?<-HNH+3C-ABC\|\|GDResolvase->AraC-HTH+AraC-HTH->LRR_5+LRR_5+LRR_5+LRR_5-> | 1184 | Bacteria>Terrabacteria_group>Bacillota>Clostridia>Eubacteriales>Oscillospiraceae | GCA_001404415.1 | [Eubacterium]_siraeum | ERS852540_00874 | 91022..94576 | 39492 | Uncharacterised-protein-[[Eubacterium] |
| ARP51069.1 | Type1.BREX | <-HTH_26<-?\|\|<-SIGMA-HTH\|\|HNH+3C-ABC->?->?->HTH_26->BrxA-DNA-binding->BrxB-BrxF-iSTAND->DUF6079*->N6-METHYLASE->PglZ->SIGMA+OB-Fold+AAA-ATPase+Lon-Protease-BrxL->STAS+HisKin-><-?<-?<-REase | 1171 | Bacteria>Terrabacteria_group>Bacillota>Clostridia>Eubacteriales>Oscillospiraceae | GCA_002119605.1 | Ruminococcaceae_bacterium_CPB6 | B6259_09405 | 1901650..1905165 | 1572656 | ATPase-[Ruminococcaceae-bacterium-CPB6] |
| OUP64017.1 | Type1.BREX | <-ClpABC-AAA<-?<-SIGMA+OB-Fold+AAA-ATPase+Lon-Protease-BrxL<-PglZ<-Schlafen+HATPase+wHTH<-SbcC+APATPase+SbcC<-N6-METHYLASE<-?<-DUF6079*<-BrxB-BrxF-iSTAND<-BrxA-DNA-binding<-HTH_26<-?<-SF1-SF2-Helicase<-?\|\|?-><-?<-HAD+HAD | 1178 | Bacteria>Terrabacteria_group>Bacillota>Clostridia>Eubacteriales>Oscillospiraceae | GCA_002160415.1 | Drancourtella_sp._An177 | B5F13_08895 | 15658..19194 | 1965573 | ATPase-[Drancourtella-sp.-An177] |
| PDX66641.1 | Type1.BREX | <-HAMP+HISKIN<-REC+OmpR-HTH<-HKD-DNase+HKD-DNase<-HKD-DNase+HKD-DNase<-Tox-RelA<-SIGMA+OB-Fold+AAA-ATPase+Lon-Protease-BrxL<-PglZ<-N6-METHYLASE<-DUF6079*<-BrxB-BrxF-iSTAND<-BrxA-DNA-binding<-XIS-HTH<-?<-?<-?<-TraM<-TraJ-RHH<-ArdCN+LPD32+LPD36+N6_Mtase+SF1-SF2-Helicase | 1222 | Bacteria>Terrabacteria_group>Bacillota>Clostridia>Eubacteriales>Oscillospiraceae | GCA_002549855.1 | Faecalibacterium_prausnitzii | CGS53_07635 | 76016..79684 | 853 | ATPase-[Faecalibacterium-prausnitzii] |
| ATO99674.1 | Type1.BREX | <-HAMP+HISKIN<-REC+OmpR-HTH<-HKD-DNase+HKD-DNase<-HKD-DNase+HKD-DNase<-Tox-RelA<-SIGMA+OB-Fold+AAA-ATPase+Lon-Protease-BrxL<-PglZ<-?<-N6-METHYLASE<-DUF6079*<-BrxB-BrxF-iSTAND<-BrxA-DNA-binding<-XIS-HTH<-?<-?<-?<-TraM<-TraJ-RHH<-ArdCN+LPD32+LPD36+N6_Mtase+SF1-SF2-Helicase | 1222 | Bacteria>Terrabacteria_group>Bacillota>Clostridia>Eubacteriales>Oscillospiraceae | GCA_002734145.1 | Faecalibacterium_prausnitzii | CG447_07050 | 1383970..1387638 | 853 | ATPase-[Faecalibacterium-prausnitzii] |
| PWJ70542.1 | Type1.BREX | HTH_3->Resolvase+Recombinase->?->?->ParB->ParB-><-?\|\|BrxA-DNA-binding->BrxB-BrxF-iSTAND->DUF6079*->N6-METHYLASE->PglZ->SIGMA+OB-Fold+AAA-ATPase+Lon-Protease-BrxL->URI-DNase-><-?<-HTH_36<-VirD4-TraG+VirD4-FtsK<-ZnR+TOPRIM<-MobL | 1194 | Bacteria>Terrabacteria_group>Bacillota>Clostridia>Eubacteriales>Oscillospiraceae | GCA_003149065.1 | Ruminococcaceae_bacterium_R-25 | SAMN06297423_0384 | 443223..446807 | 2202136 | hypothetical-protein-BO040_0384-[Ruminococcaceae-bacterium-R-25] |
| RGD21358.1 | Type1.BREX | <-STYKIN<-?<-LRR_5+LRR_8<-SIGMA+OB-Fold+AAA-ATPase+Lon-Protease-BrxL<-PglZ<-?<-?<-N6-METHYLASE<-DUF6079*<-BrxB-BrxF-iSTAND<-BrxA-DNA-binding\|\|HTH-> | 1214 | Bacteria>Terrabacteria_group>Bacillota>Clostridia>Eubacteriales>Oscillospiraceae | GCA_003434585.1 | Subdoligranulum_sp._AM23-21AC | DW651_08385 | 184245..187889 | 2302961 | BREX-system-P-loop-protein-BrxC-[Subdoligranulum-sp.-AM23-21AC] |
| RGU34205.1 | Type1.BREX | TetR-HTH->?->?->?->?->MACRODOMAIN->HTH_26->BrxA-DNA-binding->BrxB-BrxF-iSTAND->DUF6079-p*->N6-METHYLASE-p->?->SIR2->?->?->?->?->?->RADICAL-SAM-> | 703 | Bacteria>Terrabacteria_group>Bacillota>Clostridia>Eubacteriales>Oscillospiraceae | GCA_003460065.1 | Ruminococcus_sp._AF17-22AC | DWW86_04440 | 258277..260388 | 2292248 | BREX-system-P-loop-protein-BrxC-[Ruminococcus-sp.-AF17-22AC] |
| RGG69738.1 | Type1.BREX | SPOUT-><-AraC-HTH+AraC-HTH\|\|ABhydrolase-><-?<-SIGMA+OB-Fold+AAA-ATPase+Lon-Protease-BrxL<-PglZ-p<-?<-N6-METHYLASE<-DUF6079*<-BrxB-BrxF-iSTAND<-BrxA-DNA-binding<-Phage_int_SAM_3+Phage_integrase\|\|PhdYeFM_antitox-> | 1161 | Bacteria>Terrabacteria_group>Bacillota>Clostridia>Eubacteriales>Oscillospiraceae | GCA_003461215.1 | Ruminococcus_sp._AF19-15 | DWX34_02860 | 160576..164061 | 2293157 | BREX-system-P-loop-protein-BrxC-[Ruminococcus-sp.-AF19-15] |
| RGG58361.1 | Type1.BREX | DSBH+AraC-HTH+AraC-HTH-><-IONCHANNEL+HISKIN+REC<-HTH+REase<-SIGMA+OB-Fold+AAA-ATPase+Lon-Protease-BrxL<-PglZ<-?<-N6-METHYLASE<-DUF6079*<-BrxB-BrxF-iSTAND<-BrxA-DNA-binding<-HTH_26 | 1184 | Bacteria>Terrabacteria_group>Bacillota>Clostridia>Eubacteriales>Oscillospiraceae | GCA_003461215.1 | Ruminococcus_sp._AF19-15 | DWX34_06625 | 16920..20474 | 2293157 | BREX-system-P-loop-protein-BrxC-[Ruminococcus-sp.-AF19-15] |
| RGH96185.1 | Type1.BREX | HTH_19-><-tRNA\|\|PBPI+GGDEF->?-><-PhdYeFM_antitox\|\|Phage_int_SAM_3+Phage_integrase->BrxA-DNA-binding->BrxB-BrxF-iSTAND->DUF6079*->N6-METHYLASE->?->?->N6-METHYLASE-p->PglZ->SIGMA+OB-Fold+AAA-ATPase+Lon-Protease-BrxL->?-><-ABhydrolase | 1161 | Bacteria>Terrabacteria_group>Bacillota>Clostridia>Eubacteriales>Oscillospiraceae | GCA_003462205.1 | Ruminococcus_sp._AM28-13 | DW733_00275 | 57797..61282 | 2293194 | BREX-system-P-loop-protein-BrxC-[Ruminococcus-sp.-AM28-13] |
| RHO73087.1 | Type1.BREX | <-HTH_38\|\|?->HTH_36->CopG/RHH->?->HTH_26->BrxB-BrxF-iSTAND->DUF6079*-> | 1182 | Bacteria>Terrabacteria_group>Bacillota>Clostridia>Eubacteriales>Oscillospiraceae | GCA_003477475.1 | Ruminococcus_sp._AF45-4BH | DW073_15885 | 7917..11465 | 2292071 | BREX-system-P-loop-protein-BrxC-[Ruminococcus-sp.-AF45-4BH] |
| RJV95618.1 | Type1.BREX | TraJ-RHH->?->?-><-Peptidase<-?\|\|HTH->HTH_26->BrxA-DNA-binding->BrxB-BrxF-iSTAND->DUF6079*->TIR->N6-METHYLASE->Schlafen+HATPase+wHTH->PglZ->SIGMA+OB-Fold+AAA-ATPase+Lon-Protease-BrxL->?->ClpABC-AAA-> | 1178 | Bacteria>Terrabacteria_group>Bacillota>Clostridia>Eubacteriales>Oscillospiraceae | GCA_003603335.1 | Subdoligranulum_sp._AF14-43 | DWW15_14705 | 24178..27714 | 2302959 | BREX-system-P-loop-protein-BrxC-[Subdoligranulum-sp.-AF14-43] |
| MCL2124490.1 | Type1.BREX | <-sGTP<-?\|\|?->?->?->BrxA-DNA-binding->BrxB-BrxF-iSTAND->DUF6079*->N6-METHYLASE->NERD+SF1-SF2-Helicase->OLD-ABC->HNH->PolB->HEPN->PglZ->SIGMA+OB-Fold+AAA-ATPase+Lon-Protease-BrxL-> | 1207 | Bacteria>Terrabacteria_group>Bacillota>Clostridia>Eubacteriales>Oscillospiraceae | GCA_009776905.1 | Oscillospiraceae_bacterium | FWH33_00685 | 36555..40178 | 2485925 | BREX-system-P-loop-protein-BrxC-[Oscillospiraceae-bacterium] |
| MCL2053364.1 | Type1.BREX | TPR+TPR+TPR+TPR->?->TPR+TPR+TPR+TPR+TPR->HTH_26->BrxA-DNA-binding->BrxB-BrxF-iSTAND->DUF6079-p*->?->SLOG+BirA-HTH->N6-METHYLASE-p->?->HTH+REase->PglZ->SIGMA+OB-Fold+AAA-ATPase+Lon-Protease-BrxL-> | 377 | Bacteria>Terrabacteria_group>Bacillota>Clostridia>Eubacteriales>Oscillospiraceae | GCA_009777725.1 | Oscillospiraceae_bacterium | FWG90_02835 | 151410..152543 | 2485925 | BREX-system-P-loop-protein-BrxC-[Oscillospiraceae-bacterium] |
| MCL2045254.1 | Type1.BREX | SPOUT->SF1-SF2-Helicase+PRTase-p-><-?<-tRNA\|\|?-><-?<-?\|\|?->BrxA-DNA-binding->BrxB-BrxF-iSTAND->DUF6079*-><-CR-ATPase5+REase\|\|N6-METHYLASE->SbcC->Tox-ART-HYD2->?->?->PolB->Ntox45->HEPN-> | 1178 | Bacteria>Terrabacteria_group>Bacillota>Clostridia>Eubacteriales>Oscillospiraceae | GCA_009777775.1 | Oscillospiraceae_bacterium | FWG88_02570 | 99920..103456 | 2485925 | BREX-system-P-loop-protein-BrxC-[Oscillospiraceae-bacterium] |
| MBD5149892.1 | Type1.BREX | ArsR-HTH->HTH_3-><-?<-?<-NTP_transf_3\|\|?->HTH_26->BrxA-DNA-binding->BrxB-BrxF-iSTAND->DUF6079*->N6-METHYLASE->OLD-ABC->REase->PglZ->DDE_Tnp_1_assoc->?->?->ABC-ATPase+SbcC+ABC-ATPase+SbcC->HTH-> | 1183 | Bacteria>Terrabacteria_group>Bacillota>Clostridia>Eubacteriales>Oscillospiraceae | GCA_014799845.1 | Oscillibacter_sp. | HDT18_05880 | 26304..29855 | 1945593 | BREX-system-P-loop-protein-BrxC-[Oscillibacter-sp.] |
| MBM6924220.1 | Type1.BREX | HTH_26->BrxA-DNA-binding->BrxB-BrxF-iSTAND->DUF6079*->N6-METHYLASE->PglZ->NERD->SIGMA+OB-Fold+AAA-ATPase+Lon-Protease-BrxL->35exo+wHTH+SF1-SF2-Helicase+HTH->?-><-?<-?<-?<-ZnR+ZnR | 1178 | Bacteria>Terrabacteria_group>Bacillota>Clostridia>Eubacteriales>Oscillospiraceae | GCA_016901815.1 | Hydrogenoanaerobacterium_saccharovorans | H9X81_11035 | 2147..5683 | 474960 | BREX-system-P-loop-protein-BrxC-[Hydrogenoanaerobacterium-saccharovorans] |
| MBQ6611347.1 | Type1.BREX | <-SIGMA+OB-Fold+AAA-ATPase+Lon-Protease-BrxL<-PglZ<-HTH+REase<-N6-METHYLASE<-DUF6079*<-BrxB-BrxF-iSTAND<-BrxA-DNA-binding\|\|SIGMA-HTH->?->MqsA_antitoxin+DUF4065-><-?<-?<-N6-METHYLASE | 1182 | Bacteria>Terrabacteria_group>Bacillota>Clostridia>Eubacteriales>Oscillospiraceae | GCA_017434435.1 | Oscillospiraceae_bacterium | IJH70_13030 | 9925..13473 | 2485925 | BREX-system-P-loop-protein-BrxC-[Oscillospiraceae-bacterium] |
| MBQ7681865.1 | Type1.BREX | <-LRR_5+LRR_5+LRR_5+LRR_5\|\|HISKIN+REC+REC->?->BrxA-DNA-binding->BrxB-BrxF-iSTAND->DUF6079*->N6-METHYLASE-> | 1182 | Bacteria>Terrabacteria_group>Bacillota>Clostridia>Eubacteriales>Oscillospiraceae | GCA_017443745.1 | Oscillibacter_sp. | IJT31_06615 | 5929..9477 | 1945593 | BREX-system-P-loop-protein-BrxC-[Oscillibacter-sp.] |
| MBQ9664285.1 | Type1.BREX | REase-><-SIGMA+OB-Fold+AAA-ATPase+Lon-Protease-BrxL<-PglZ<-?<-N6-METHYLASE<-DUF6079*<-BrxB-BrxF-iSTAND<-BrxA-DNA-binding | 1198 | Bacteria>Terrabacteria_group>Bacillota>Clostridia>Eubacteriales>Oscillospiraceae | GCA_017458065.1 | Oscillospiraceae_bacterium | IJV40_14155 | 25902..29498 | 2485925 | BREX-system-P-loop-protein-BrxC-[Oscillospiraceae-bacterium] |
| MBR7037761.1 | Type1.BREX | DUF6079-p*->N6-METHYLASE-p-><-?<-Glyco_hydro_3<-?<-?<-SIGMA-HTH | 735 | Bacteria>Terrabacteria_group>Bacillota>Clostridia>Eubacteriales>Oscillospiraceae | GCA_017518725.1 | Oscillospiraceae_bacterium | IKI21_00715 | 1..2210 | 2485925 | BREX-system-P-loop-protein-BrxC-[Oscillospiraceae-bacterium] |
| MBR7008376.1 | Type1.BREX | BrxA-DNA-binding->BrxB-BrxF-iSTAND->DUF6079*->N6-METHYLASE->?->N6-METHYLASE-p->PglZ->SIGMA+OB-Fold+AAA-ATPase+Lon-Protease-BrxL-> | 1166 | Bacteria>Terrabacteria_group>Bacillota>Clostridia>Eubacteriales>Oscillospiraceae | GCA_017519245.1 | Ruminococcus_sp. | IKH90_07090 | 1298..4798 | 41978 | BREX-system-P-loop-protein-BrxC-[Ruminococcus-sp.] |
| MBR6617436.1 | Type1.BREX | <-PglZ<-N6-METHYLASE-p<-?<-N6-METHYLASE<-DUF6079*<-LRR_5<-BrxB-BrxF-iSTAND<-BrxA-DNA-binding\|\|Phage_integrase-><-IG+IG<-SF1-SF2-Helicase<-XIS-HTH | 1161 | Bacteria>Terrabacteria_group>Bacillota>Clostridia>Eubacteriales>Oscillospiraceae | GCA_017520925.1 | Oscillospiraceae_bacterium | IKL00_06140 | 5040..8525 | 2485925 | BREX-system-P-loop-protein-BrxC-[Oscillospiraceae-bacterium] |
| MBO5318820.1 | Type1.BREX | LPD32+LPD36+N6_Mtase+SF1-SF2-Helicase->ZnR+TOPRIM->BrxA-DNA-binding->BrxB-BrxF-iSTAND->DUF6079*->?->?->N6-METHYLASE->PglZ->SIGMA+OB-Fold+AAA-ATPase+Lon-Protease-BrxL->?-><-?\|\|HTH-><-TraI-VirD2-RCRE | 1206 | Bacteria>Terrabacteria_group>Bacillota>Clostridia>Eubacteriales>Oscillospiraceae | GCA_017624635.1 | Ruminococcus_sp. | J6B01_03305 | 52295..55915 | 41978 | BREX-system-P-loop-protein-BrxC-[Ruminococcus-sp.] |
| MBP0983334.1 | Type1.BREX | AAA-ATPase->?->?-><-?\|\|?->GDResolvase+Recombinase+Zn_ribbon_recom-><-HTH_3\|\|BrxA-DNA-binding->BrxB-BrxF-iSTAND->DUF6079*->N6-METHYLASE->PglZ->SIGMA+OB-Fold+AAA-ATPase+Lon-Protease-BrxL->35exo+wHTH+SF1-SF2-Helicase+HTH-><-?\|\|?->METHYLASE-> | 1218 | Bacteria>Terrabacteria_group>Bacillota>Clostridia>Eubacteriales>Oscillospiraceae | GCA_017829075.1 | Oscillospiraceae_bacterium | J6A19_06370 | 36629..40285 | 2485925 | BREX-system-P-loop-protein-BrxC-[Oscillospiraceae-bacterium] |
| MBP1566405.1 | Type1.BREX | <-CHTH\|\|?-><-CR-ATPase8+REase<-MNS+REase<-SIGMA+OB-Fold+AAA-ATPase+Lon-Protease-BrxL<-?<-PglZ<-OLD-ABC<-N6-METHYLASE<-DUF6079*<-BrxB-BrxF-iSTAND<-BrxA-DNA-binding<-SMODS<-?<-?<-CR-ATPase8+REase | 1178 | Bacteria>Terrabacteria_group>Bacillota>Clostridia>Eubacteriales>Oscillospiraceae | GCA_017848115.1 | Oscillospiraceae_bacterium | J6A58_11685 | 59197..62733 | 2485925 | BREX-system-P-loop-protein-BrxC-[Oscillospiraceae-bacterium] |
| MBP1547887.1 | Type1.BREX | <-DDE_Tnp\|\|BrxA-DNA-binding->BrxB-BrxF-iSTAND->DUF6079*->?->N6-METHYLASE->PglZ->SIGMA+OB-Fold+AAA-ATPase+Lon-Protease-BrxL->?->AbiEii-> | 1210 | Bacteria>Terrabacteria_group>Bacillota>Clostridia>Eubacteriales>Oscillospiraceae | GCA_017848125.1 | Oscillospiraceae_bacterium | J6A37_14990 | 4952..8584 | 2485925 | BREX-system-P-loop-protein-BrxC-[Oscillospiraceae-bacterium] |
| MBP3248829.1 | Type1.BREX | DUF6079*->N6-METHYLASE-p-><-?<-HTH_3<-?<-?\|\|?->?->?-><-XIS-HTH+GDResolvase | 1133 | Bacteria>Terrabacteria_group>Bacillota>Clostridia>Eubacteriales>Oscillospiraceae | GCA_017934345.1 | Ruminococcus_sp. | J6M48_00560 | 495..3896 | 41978 | BREX-system-P-loop-protein-BrxC-[Ruminococcus-sp.] |
| MBP8594426.1 | Type1.BREX | HTH_3->?-><-N6-METHYLASE-p<-DUF6079*<-?\|\|?-><-HISKIN<-REC+OmpR-HTH\|\|?-><-NTP_transf_3+Hexapep+Hexapep<-ABhydrolase\|\|?->MarR-HTH-> | 1133 | Bacteria>Terrabacteria_group>Bacillota>Clostridia>Eubacteriales>Oscillospiraceae | GCA_018052525.1 | Ruminococcus_sp. | KBI35_08440 | 128893..132294 | 41978 | BREX-system-P-loop-protein-BrxC-[Ruminococcus-sp.] |
| MBP9989226.1 | Type1.BREX | Phage_int_SAM_3+Phage_integrase->BrxA-DNA-binding->BrxB-BrxF-iSTAND->DUF6079*->N6-METHYLASE-p-> | 1160 | Bacteria>Terrabacteria_group>Bacillota>Clostridia>Eubacteriales>Oscillospiraceae | GCA_018064255.1 | Ruminococcus_sp. | KBT46_06980 | 3336..6818 | 41978 | BREX-system-P-loop-protein-BrxC-[Ruminococcus-sp.] |
| MBS1435198.1 | Type1.BREX | Phage_int_SAM_1+Phage_integrase-><-?<-TraJ-RHH\|\|HTH_26->BrxA-DNA-binding->BrxB-BrxF-iSTAND->DUF6079*->?->N6-METHYLASE->PglZ->SIGMA+OB-Fold+AAA-ATPase+Lon-Protease-BrxL->SMODS-> | 1178 | Bacteria>Terrabacteria_group>Bacillota>Clostridia>Eubacteriales>Oscillospiraceae | GCA_018265115.1 | Oscillospiraceae_bacterium | HPO15_04040 | 34076..37612 | 2485925 | BREX-system-P-loop-protein-BrxC-[Oscillospiraceae-bacterium] |
| MBS4908679.1 | Type1.BREX | DDE_Tnp_1-><-DUF6079*<-BrxB-BrxF-iSTAND<-BrxA-DNA-binding<-HTH_26<-?<-TraI-VirD2-RCRE | 1182 | Bacteria>Terrabacteria_group>Bacillota>Clostridia>Eubacteriales>Oscillospiraceae | GCA_018364055.1 | Ruminococcus_sp. | KH021_12110 | 782..4330 | 41978 | BREX-system-P-loop-protein-BrxC-[Ruminococcus-sp.] |
| MBS6624534.1 | Type1.BREX | XIS-HTH->BrxA-DNA-binding->BrxB-BrxF-iSTAND->DUF6079*->N6-METHYLASE->?->PglZ->SIGMA+OB-Fold+AAA-ATPase+Lon-Protease-BrxL->?->ClpABC-AAA-> | 1217 | Bacteria>Terrabacteria_group>Bacillota>Clostridia>Eubacteriales>Oscillospiraceae | GCA_018378145.1 | Ruminococcus_sp. | KH309_05615 | 1641..5294 | 41978 | BREX-system-P-loop-protein-BrxC-[Ruminococcus-sp.] |
| MBS7028836.1 | Type1.BREX | <-RVT<-SIGMA+OB-Fold+AAA-ATPase+Lon-Protease-BrxL<-PglZ<-HEPN<-N6-METHYLASE<-?<-?<-DUF6079*<-BrxB-BrxF-iSTAND<-BrxA-DNA-binding<-DOC+HTH\|\|DUF932->?->?->CR-ATPase5+REase->?->CR-ATPase8+REase-> | 1182 | Bacteria>Terrabacteria_group>Bacillota>Clostridia>Eubacteriales>Oscillospiraceae | GCA_018381555.1 | Faecalibacterium_prausnitzii | KH132_10095 | 50879..54427 | 853 | BREX-system-P-loop-protein-BrxC-[Faecalibacterium-prausnitzii] |
| MBS7188423.1 | Type1.BREX | ABhydrolase-><-?<-SIGMA+OB-Fold+AAA-ATPase+Lon-Protease-BrxL<-PglZ<-N6-METHYLASE-p<-PRTase<-RAMA<-N6-METHYLASE<-DUF6079*<-BrxB-BrxF-iSTAND<-BrxA-DNA-binding<-Phage_int_SAM_3+Phage_integrase<-?<-PBPI+GGDEF | 1161 | Bacteria>Terrabacteria_group>Bacillota>Clostridia>Eubacteriales>Oscillospiraceae | GCA_018382835.1 | Ruminococcus_bicirculans | KH055_02500 | 41675..45160 | 1160721 | BREX-system-P-loop-protein-BrxC-[Ruminococcus-bicirculans] |
| HIS77364.1 | Type1.BREX | <-SIGMA+OB-Fold+AAA-ATPase+Lon-Protease-BrxL<-PglZ<-N6-METHYLASE-p<-?<-N6-METHYLASE<-DUF6079*<-BrxB-BrxF-iSTAND<-BrxA-DNA-binding<-GDResolvase+HIN-HTH\|\|CR-ATPase5+REase->?-><-XIS-HTH+XIS-HTH | 1169 | Bacteria>Terrabacteria_group>Bacillota>Clostridia>Eubacteriales>Oscillospiraceae | GCA_018716725.1 | Candidatus_Merdivicinus_excrementipullorum | IAB51_11260 | 8193..11702 | 2840867 | BREX-system-P-loop-protein-BrxC-[Candidatus-Merdivicinus-excrementipullorum] |
| HJB38307.1 | Type1.BREX | <-PglZ-p<-SbcC+APATPase+SbcC<-N6-METHYLASE<-DUF6079*<-BrxB-BrxF-iSTAND<-BrxA-DNA-binding<-HTH_26 | 1180 | Bacteria>Terrabacteria_group>Bacillota>Clostridia>Eubacteriales>Oscillospiraceae | GCA_019118965.1 | Candidatus_Acutalibacter_ornithocaccae | H9942_09620 | 6171..9713 | 2838416 | BREX-system-P-loop-protein-BrxC-[Candidatus-Acutalibacter-ornithocaccae] |
| MCC3659975.1 | Type1.BREX | AraC-HTH+AraC-HTH-><-IONCHANNEL+HISKIN+REC<-SAD+HNH<-?<-PglZ<-?<-N6-METHYLASE<-DUF6079*<-BrxB-BrxF-iSTAND<-BrxA-DNA-binding<-HTH_26<-?<-?<-?<-?\|\|?-><-68TM-wHTH+CACHE+GGDEF | 1184 | Bacteria>Terrabacteria_group>Bacillota>Clostridia>Eubacteriales>Oscillospiraceae | GCA_020740265.1 | Ruminococcus_albus | LLP35_08975 | 12774..16328 | 1264 | BREX-system-P-loop-protein-BrxC-[Ruminococcus-albus] |
| MCC8130482.1 | Type1.BREX | <-AAA+<-RepA_N+HTH<-TraI-VirD2-RCRE<-TraJ-RHH\|\|BrxA-DNA-binding->BrxB-BrxF-iSTAND->DUF6079*->N6-METHYLASE->PglZ->SIGMA+OB-Fold+AAA-ATPase+Lon-Protease-BrxL-> | 1169 | Bacteria>Terrabacteria_group>Bacillota>Clostridia>Eubacteriales>Oscillospiraceae | GCA_020860605.1 | Ruminococcus_sp. | LIO72_01830 | 7361..10870 | 41978 | BREX-system-P-loop-protein-BrxC-[Ruminococcus-sp.] |
| MCD7769618.1 | Type1.BREX | RelB->PIN->HTH_26->BrxA-DNA-binding->BrxB-BrxF-iSTAND->DUF6079*->TPR+TPR-><-?\|\|N6-METHYLASE-p-> | 1211 | Bacteria>Terrabacteria_group>Bacillota>Clostridia>Eubacteriales>Oscillospiraceae | GCA_021199715.1 | Oscillospiraceae_bacterium | LUH36_05850 | 2620..6255 | 2485925 | BREX-system-P-loop-protein-BrxC-[Oscillospiraceae-bacterium] |
| MCI6015526.1 | Type1.BREX | ABC-ATPase+SbcC-><-?<-RADICAL-SAM\|\|?->?->HTH_26->BrxA-DNA-binding->BrxB-BrxF-iSTAND->DUF6079*->?->N6-METHYLASE->OLD-ABC->NERD->PglZ->SIGMA+OB-Fold+AAA-ATPase+Lon-Protease-BrxL-><-?<-?\|\|ZnR+TOPRIM-> | 1211 | Bacteria>Terrabacteria_group>Bacillota>Clostridia>Eubacteriales>Oscillospiraceae | GCA_022770595.1 | Dysosmobacter_sp. | MRZ31_02380 | 21851..25486 | 2591382 | BREX-system-P-loop-protein-BrxC-[Dysosmobacter-sp.] |
| MCI6194160.1 | Type1.BREX | HTH+LexA-protease->HTH_3-><-?<-Glyco_hydro_4+Glyco_hydro_4C<-?\|\|MODE-HTH+PBPII-><-MORC+DUF3684+HNH<-HEPN+SH3-like+SF1-SF2-Helicase+wHTH<-PglZ<-N6-METHYLASE<-DUF6079*<-BrxB-BrxF-iSTAND<-BrxA-DNA-binding<-?<-35exo+wHTH+SF1-SF2-Helicase+HTH<-?<-35exo<-?<-?<-DEOR-HTH | 1204 | Bacteria>Terrabacteria_group>Bacillota>Clostridia>Eubacteriales>Oscillospiraceae | GCA_022772255.1 | Faecalibacterium_sp. | MR651_04080 | 20634..24248 | 1971605 | BREX-system-P-loop-protein-BrxC-[Faecalibacterium-sp.] |
| MCI6921905.1 | Type1.BREX | ABC-ATPase+SbcC+ABC-ATPase+SbcC->?->?->?->LRR_5->HEPN->BrxA-DNA-binding->BrxB-BrxF-iSTAND->DUF6079*->N6-METHYLASE-p-> | 1161 | Bacteria>Terrabacteria_group>Bacillota>Clostridia>Eubacteriales>Oscillospiraceae | GCA_022778185.1 | Oscillospiraceae_bacterium | MR756_00805 | 25286..28771 | 2485925 | BREX-system-P-loop-protein-BrxC-[Oscillospiraceae-bacterium] |
| MCI6596832.1 | Type1.BREX | <-STYKIN<-?<-LRR_5+LRR_8<-SIGMA+OB-Fold+AAA-ATPase+Lon-Protease-BrxL<-PglZ<-?<-?<-N6-METHYLASE<-DUF6079*<-BrxB-BrxF-iSTAND<-BrxA-DNA-binding\|\|HTH-><-DUF772+DDE_Tnp_1 | 1214 | Bacteria>Terrabacteria_group>Bacillota>Clostridia>Eubacteriales>Oscillospiraceae | GCA_022779505.1 | Ruthenibacterium_lactatiformans | MSH17_05650 | 69035..72679 | 1550024 | BREX-system-P-loop-protein-BrxC-[Ruthenibacterium-lactatiformans] |
| MCI6580384.1 | Type1.BREX | <-SIGMA+OB-Fold+AAA-ATPase+Lon-Protease-BrxL-p<-PglZ<-OLD-ABC<-N6-METHYLASE<-DUF6079*<-BrxB-BrxF-iSTAND<-BrxA-DNA-binding<-SF1-SF2-Helicase | 1176 | Bacteria>Terrabacteria_group>Bacillota>Clostridia>Eubacteriales>Oscillospiraceae | GCA_022779535.1 | Oscillospiraceae_bacterium | MSH15_00090 | 8217..11747 | 2485925 | BREX-system-P-loop-protein-BrxC-[Oscillospiraceae-bacterium] |
| MCI7422253.1 | Type1.BREX | Glyco_hydro_3+Glyco_hydro_3_C+Fn3-like->?->?->?-><-PglZ<-?<-N6-METHYLASE<-DUF6079*<-BrxB-BrxF-iSTAND | 1161 | Bacteria>Terrabacteria_group>Bacillota>Clostridia>Eubacteriales>Oscillospiraceae | GCA_022785355.1 | Ruminococcus_sp. | MSA57_03345 | 21833..25318 | 41978 | BREX-system-P-loop-protein-BrxC-[Ruminococcus-sp.] |
| MCI7629773.1 | Type1.BREX | Glyco_hydro_3+Glyco_hydro_3_C+Fn3-like->?->?->?-><-PglZ<-HEPN<-N6-METHYLASE<-DUF6079-p* | 942 | Bacteria>Terrabacteria_group>Bacillota>Clostridia>Eubacteriales>Oscillospiraceae | GCA_022787645.1 | Ruminococcus_sp. | MSS55_04245 | 22217..25045 | 41978 | BREX-system-P-loop-protein-BrxC-[Ruminococcus-sp.] |
| MCI8801905.1 | Type1.BREX | <-DapB_N+DapB_C<-?\|\|Peptidase_M20-><-PglZ<-N6-METHYLASE<-DUF6079*<-BrxB-BrxF-iSTAND<-BrxA-DNA-binding<-HTH_26<-Peptidase_M29<-Peptidase_M20<-?<-ABC-ATPase+SbcC | 1213 | Bacteria>Terrabacteria_group>Bacillota>Clostridia>Eubacteriales>Oscillospiraceae | GCA_022795315.1 | Oscillibacter_sp. | HFG01_00990 | 12249..15890 | 1945593 | BREX-system-P-loop-protein-BrxC-[Oscillibacter-sp.] |
| MCI8835830.1 | Type1.BREX | ABC-ATPase->ABC-ATPase+SbcC->BrxA-DNA-binding->BrxB-BrxF-iSTAND->DUF6079*->Schlafen+HATPase+wHTH->N6-METHYLASE->OLD-ABC->PIN->PglZ-> | 1205 | Bacteria>Terrabacteria_group>Bacillota>Clostridia>Eubacteriales>Oscillospiraceae | GCA_022795995.1 | Ruminococcus_sp. | HFH24_10725 | 31305..34922 | 41978 | BREX-system-P-loop-protein-BrxC-[Ruminococcus-sp.] |
| MCI8909814.1 | Type1.BREX | <-PglZ<-N6-METHYLASE-p<-?<-N6-METHYLASE<-DUF6079*<-BrxB-BrxF-iSTAND<-BrxA-DNA-binding | 1175 | Bacteria>Terrabacteria_group>Bacillota>Clostridia>Eubacteriales>Oscillospiraceae | GCA_022796315.1 | Oscillibacter_sp. | HFG09_03955 | 6847..10374 | 1945593 | BREX-system-P-loop-protein-BrxC-[Oscillibacter-sp.] |
| MCI9433298.1 | Type1.BREX | <-SIGMA+OB-Fold+AAA-ATPase+Lon-Protease-BrxL<-PglZ<-DUF6079*<-BrxB-BrxF-iSTAND<-BrxA-DNA-binding<-HTH_26<-JetC<-?<-HNH\|\|HTH_3->PemK_toxin->ParB-> | 1178 | Bacteria>Terrabacteria_group>Bacillota>Clostridia>Eubacteriales>Oscillospiraceae | GCA_022800455.1 | Oscillospiraceae_bacterium | HFF70_12895 | 9735..13271 | 2485925 | BREX-system-P-loop-protein-BrxC-[Oscillospiraceae-bacterium] |
| MCI9506677.1 | Type1.BREX | REC+OmpR-HTH->?->HTH_3->?->ABC-ATPase+SbcC->BrxA-DNA-binding->BrxB-BrxF-iSTAND->DUF6079*->N6-METHYLASE->PglZ-><-?<-?<-?<-JetA | 1210 | Bacteria>Terrabacteria_group>Bacillota>Clostridia>Eubacteriales>Oscillospiraceae | GCA_022800665.1 | Oscillospiraceae_bacterium | HFF62_08790 | 7365..10997 | 2485925 | BREX-system-P-loop-protein-BrxC-[Oscillospiraceae-bacterium] |
| MCI9648503.1 | Type1.BREX | GDSL-><-?<-HTH_6+SIS<-?\|\|BrxA-DNA-binding->BrxB-BrxF-iSTAND->DUF6079*->PglZ->?-><-REC<-?<-?\|\|PBPI->ABC-ATPase+SbcC+ABC-ATPase+SbcC-> | 1198 | Bacteria>Terrabacteria_group>Bacillota>Clostridia>Eubacteriales>Oscillospiraceae | GCA_022804785.1 | Oscillibacter_sp. | HFG11_03585 | 14926..18522 | 1945593 | BREX-system-P-loop-protein-BrxC-[Oscillibacter-sp.] |
| MCM1231671.1 | Type1.BREX | <-Schlafen+HATPase+wHTH\|\|SF1-SF2-Helicase->XIS-HTH->\|\|Phage_int_SAM_3+Phage_integrase->BrxA-DNA-binding->BrxB-BrxF-iSTAND->DUF6079*->N6-METHYLASE->PglZ->SIGMA+OB-Fold+AAA-ATPase+Lon-Protease-BrxL->?->wHTH+WYL+WCX->?->RADICAL-SAM-> | 1205 | Bacteria>Terrabacteria_group>Bacillota>Clostridia>Eubacteriales>Oscillospiraceae | GCA_023664505.1 | Ruminococcus_flavefaciens | NC489_16260 | 22171..25788 | 1265 | BREX-system-P-loop-protein-BrxC-[Ruminococcus-flavefaciens] |
| MCQ4886543.1 | Type1.BREX | <-MORC+REase<-STYKinase-like<-SF1-SF2-Helicase<-OLD-ABC+TOPRIM<-SIGMA+OB-Fold+AAA-ATPase+Lon-Protease-BrxL<-PglZ<-NERD<-N6-METHYLASE<-?<-DUF6079*<-BrxB-BrxF-iSTAND<-BrxA-DNA-binding<-XIS-HTH\|\|HKD-DNase+SF1-SF2-Helicase+HTH+HTH+SRA->?->NUDIX->DUF932-> | 1218 | Bacteria>Terrabacteria_group>Bacillota>Clostridia>Eubacteriales>Oscillospiraceae | GCA_024460915.1 | Faecalibacterium_prausnitzii | NE589_05495 | 127412..131068 | 853 | BREX-system-P-loop-protein-BrxC-[Faecalibacterium-prausnitzii] |
| MCQ5054180.1 | Type1.BREX | <-Cap4_nuclease<-SIGMA+OB-Fold+AAA-ATPase+Lon-Protease-BrxL<-PglZ<-AT-HTH-HYD2-2<-?<-Tox-ART-HYD2<-N6-METHYLASE<-TIR<-DUF6079*<-BrxB-BrxF-iSTAND<-BrxA-DNA-binding<-XIS-HTH<-?<-MACRODOMAIN<-REase+N-glycosylase | 1217 | Bacteria>Terrabacteria_group>Bacillota>Clostridia>Eubacteriales>Oscillospiraceae | GCA_024462395.1 | Agathobaculum_butyriciproducens | NE606_05805 | 173642..177295 | 1628085 | BREX-system-P-loop-protein-BrxC-[Agathobaculum-butyriciproducens] |
| MCR4638957.1 | Type1.BREX | <-HTH_3<-SIGMA+OB-Fold+AAA-ATPase+Lon-Protease-BrxL<-PglZ<-AbiGi<-N6-METHYLASE<-ParB+HNH<-DUF6079*<-BrxB-BrxF-iSTAND<-BrxA-DNA-binding<-?<-?<-?<-?<-?<-HIRAN<-DnaJ | 1175 | Bacteria>Terrabacteria_group>Bacillota>Clostridia>Eubacteriales>Oscillospiraceae | GCA_024684275.1 | Ruminococcus_sp. | K5677_07000 | 11947..15474 | 41978 | BREX-system-P-loop-protein-BrxC-[Ruminococcus-sp.] |
| UYJ00845.1 | Type1.BREX | <-NUDIX<-?<-HKD-DNase+SF1-SF2-Helicase+HTH+HTH+SRA\|\|XIS-HTH->BrxA-DNA-binding->BrxB-BrxF-iSTAND->DUF6079*->HTH+REase->N6-METHYLASE->TIR->PglZ->SIGMA+OB-Fold+AAA-ATPase+Lon-Protease-BrxL->ParB+HNH->STYKinase-like-> | 1218 | Bacteria>Terrabacteria_group>Bacillota>Clostridia>Eubacteriales>Oscillospiraceae | GCA_025757765.1 | Faecalibacterium_prausnitzii | OGM65_10405 | 2149952..2153608 | 853 | MAG:-BREX-system-P-loop-protein-BrxC-[Faecalibacterium-prausnitzii] |
| UYJ05877.1 | Type1.BREX | <-HEPN+SH3-like+SF1-SF2-Helicase+wHTH<-PglZ<-Schlafen+HATPase+wHTH<-HEPN<-?<-?<-?<-?<-N6-METHYLASE<-DUF6079*<-BrxB-BrxF-iSTAND<-BrxA-DNA-binding<-?<-35exo+wHTH+SF1-SF2-Helicase+HTH<-?<-35exo<-?<-?\|\|HTH-> | 1204 | Bacteria>Terrabacteria_group>Bacillota>Clostridia>Eubacteriales>Oscillospiraceae | GCA_025757805.1 | Oscillospiraceae_bacterium | OGM79_08450 | 1764021..1767635 | 2485925 | MAG:-BREX-system-P-loop-protein-BrxC-[Oscillospiraceae-bacterium] |
| UYJ11714.1 | Type1.BREX | <-CR-ATPase8+REase<-?<-CR-ATPase5+REase<-?<-?\|\|DOC+HTH->BrxA-DNA-binding->BrxB-BrxF-iSTAND->DUF6079*->?->?->N6-METHYLASE->OLD-ABC->REase->PglZ->SIGMA+OB-Fold+AAA-ATPase+Lon-Protease-BrxL->ParB+HNH->REase+SF1-SF2-Helicase-> | 1182 | Bacteria>Terrabacteria_group>Bacillota>Clostridia>Eubacteriales>Oscillospiraceae | GCA_025757845.1 | Oscillospiraceae_bacterium | OGM78_02700 | 567882..571430 | 2485925 | MAG:-BREX-system-P-loop-protein-BrxC-[Oscillospiraceae-bacterium] |
| UYJ36126.1 | Type1.BREX | TraJ-RHH-><-CR-ATPase5+REase\|\|TraI-VirD2-RCRE->?->?->?->BrxA-DNA-binding->BrxB-BrxF-iSTAND->DUF6079*->N6-METHYLASE->PglZ->HEPN+SH3-like+SF1-SF2-Helicase+wHTH->MORC+DUF3684+HNH->?->Tox-RelA->HKD-DNase+HKD-DNase->HKD-DNase+HKD-DNase->REC+OmpR-HTH-> | 1205 | Bacteria>Terrabacteria_group>Bacillota>Clostridia>Eubacteriales>Oscillospiraceae | GCA_025757985.1 | Oscillospiraceae_bacterium | OGM67_07425 | 1479517..1483134 | 2485925 | MAG:-BREX-system-P-loop-protein-BrxC-[Oscillospiraceae-bacterium] |
| UYJ36609.1 | Type1.BREX | HTH-><-TIR<-SIGMA+OB-Fold+AAA-ATPase+Lon-Protease-BrxL<-PglZ<-Schlafen+HATPase+wHTH<-HEPN<-N6-METHYLASE<-?<-DUF6079*<-BrxB-BrxF-iSTAND<-BrxA-DNA-binding<-HTH_26<-TraI-VirD2-RCRE<-TraJ-RHH<-?<-CopG/RHH<-?<-HTH_36 | 1182 | Bacteria>Terrabacteria_group>Bacillota>Clostridia>Eubacteriales>Oscillospiraceae | GCA_025758005.1 | Oscillospiraceae_bacterium | OGM10_10415 | 2214157..2217705 | 2485925 | MAG:-BREX-system-P-loop-protein-BrxC-[Oscillospiraceae-bacterium] |
| WEG34878.1 | Type1.BREX | PTSEIIA1->?->?->?->?->HTH_26->BrxA-DNA-binding->BrxB-BrxF-iSTAND->DUF6079*->N6-METHYLASE->PglZ->SIGMA+OB-Fold+AAA-ATPase+Lon-Protease-BrxL->?->RNA_METHYLASE->?->?->?->SPOUT-> | 1216 | Bacteria>Terrabacteria_group>Bacillota>Clostridia>Eubacteriales>Oscillospiraceae | GCA_029101545.1 | Oscillospiraceae_bacterium_UPII_610-J | PYS60_02935 | 702599..706249 | 3029272 | BREX-system-P-loop-protein-BrxC-[Oscillospiraceae-bacterium-UPII-610-J] |
| WEG35252.1 | Type1.BREX | 35exo+BRCT->ASCH->?->HTH_26->BrxA-DNA-binding->BrxB-BrxF-iSTAND->DUF6079*->N6-METHYLASE->?->TIR->PglZ->SIGMA+OB-Fold+AAA-ATPase+Lon-Protease-BrxL->?->RNA_METHYLASE-> | 1184 | Bacteria>Terrabacteria_group>Bacillota>Clostridia>Eubacteriales>Oscillospiraceae | GCA_029101565.1 | Oscillospiraceae_bacterium_CHIC02_1186E3-8 | PYS61_04790 | 1135899..1139453 | 3029273 | BREX-system-P-loop-protein-BrxC-[Oscillospiraceae-bacterium-CHIC02-1186E3-8] |
| SCI31347.1 | Type1.BREX | 68TM-wHTH+CACHE+GGDEF->?->?->?->?->?->HTH_26->BrxA-DNA-binding->BrxB-BrxF-iSTAND->DUF6079*->N6-METHYLASE->?->PglZ->SIGMA+OB-Fold+AAA-ATPase+Lon-Protease-BrxL->?->SAD+HNH->IONCHANNEL+HISKIN+REC-><-DSBH+AraC-HTH+AraC-HTH | 1184 | Bacteria>Terrabacteria_group>Bacillota>Clostridia>Eubacteriales>Oscillospiraceae | GCA_900066635.1 | uncultured_Ruminococcus_sp. | SAMEA3545357_00481 | 41099..44653 | 165186 | Uncharacterised-protein-[uncultured-Ruminococcus-sp.] |
| SCH98752.1 | Type1.BREX | 68TM-wHTH+CACHE+GGDEF->?->?->?->?->?->HTH_26->BrxA-DNA-binding->BrxB-BrxF-iSTAND->DUF6079*->N6-METHYLASE->?->PglZ->SIGMA+OB-Fold+AAA-ATPase+Lon-Protease-BrxL->HTH+REase->IONCHANNEL+HISKIN+REC->?-><-DSBH+AraC-HTH+AraC-HTH | 1184 | Bacteria>Terrabacteria_group>Bacillota>Clostridia>Eubacteriales>Oscillospiraceae | GCA_900067025.1 | uncultured_Ruminococcus_sp. | SAMEA3545436_02036 | 58203..61757 | 165186 | Uncharacterised-protein-[uncultured-Ruminococcus-sp.] |
| SFX61047.1 | Type1.BREX | HTH+LexA-protease->?->?->?->?->?->BrxA-DNA-binding->BrxB-BrxF-iSTAND->DUF6079*->?->N6-METHYLASE->?->N6-METHYLASE-p->PglZ->SIGMA+OB-Fold+AAA-ATPase+Lon-Protease-BrxL->?->?->RADICAL-SAM-> | 1161 | Bacteria>Terrabacteria_group>Bacillota>Clostridia>Eubacteriales>Oscillospiraceae | GCA_900119535.1 | Ruminococcus_flavefaciens | SAMN04487832_10833 | 29488..32973 | 1265 | hypothetical-protein-SAMN04487832_10833-[Ruminococcus-flavefaciens] |
| WP_010247739.1 | Type1.BREX | <-Herpes_LMP1+REase<-SIGMA+OB-Fold+AAA-ATPase+Lon-Protease-BrxL<-PglZ<-REase<-OLD-ABC<-N6-METHYLASE<-DUF6079*<-BrxB-BrxF-iSTAND<-BrxA-DNA-binding<-?<-NUDIX+DUF429<-?<-DDE_Tnp_IS66+DDE_Tnp_IS66_C<-?<-HIN-HTH<-ParB+HNH | 1183 | Bacteria>Terrabacteria_group>Bacillota>Clostridia>Eubacteriales>Oscillospiraceae | GCF_000179595.2 | Acetivibrio_cellulolyticus | ACECE_RS0212925 | 179923..183474 | 35830 | BREX-system-P-loop-protein-BrxC-[Acetivibrio-cellulolyticus] |
| WP_044960386.1 | Type1.BREX | <-CR-ATPase8+REase<-?<-?<-DUF932\|\|DOC+HTH->BrxA-DNA-binding->BrxB-BrxF-iSTAND->DUF6079*->N6-METHYLASE->HEPN->PglZ->SIGMA+OB-Fold+AAA-ATPase+Lon-Protease-BrxL->RVT->RVT-> | 1182 | Bacteria>Terrabacteria_group>Bacillota>Clostridia>Eubacteriales>Oscillospiraceae | GCF_000209855.1 | Faecalibacterium_prausnitzii | FPR_RS00845 | 148257..151805 | 853 | BREX-system-P-loop-protein-BrxC-[Faecalibacterium-prausnitzii] |
| WP_014253628.1 | Type1.BREX | <-ZnR+TOPRIM\|\|?->?->DndB->?->BrxA-DNA-binding->BrxB-BrxF-iSTAND->DUF6079*->N6-METHYLASE->PglZ->?->SIGMA+OB-Fold+AAA-ATPase+Lon-Protease-BrxL->RAMA->ARG->ZnR-> | 1186 | Bacteria>Terrabacteria_group>Bacillota>Clostridia>Eubacteriales>Oscillospiraceae | GCF_000237085.1 | Acetivibrio_clariflavus | CLOCL_RS01215 | 254231..257791 | 288965 | BREX-system-P-loop-protein-BrxC-[Acetivibrio-clariflavus] |
| WP_014256029.1 | Type1.BREX | <-Glyoxalase<-N6-METHYLASE<-Peptidase<-?<-?<-SIGMA+OB-Fold+AAA-ATPase+Lon-Protease-BrxL<-PglZ<-Schlafen+HATPase+wHTH<-N6-METHYLASE<-DUF6079*<-BrxB-BrxF-iSTAND<-BrxA-DNA-binding<-wHTH(SBD)+HNH<-?<-?<-N6-METHYLASE | 1186 | Bacteria>Terrabacteria_group>Bacillota>Clostridia>Eubacteriales>Oscillospiraceae | GCF_000237085.1 | Acetivibrio_clariflavus | CLOCL_RS14360 | 3415236..3418796 | 288965 | BREX-system-P-loop-protein-BrxC-[Acetivibrio-clariflavus] |
| WP_040660344.1 | Type1.BREX | <-SAD+REase<-?<-SIGMA+OB-Fold+AAA-ATPase+Lon-Protease-BrxL<-PglZ<-?<-?<-DUF6079*<-BrxB-BrxF-iSTAND<-BrxA-DNA-binding<-HTH_26<-?<-?\|\|HTH-><-HTH_36<-VirD4-TraG+VirD4-FtsK<-ZnR+TOPRIM | 1197 | Bacteria>Terrabacteria_group>Bacillota>Clostridia>Eubacteriales>Oscillospiraceae | GCF_000307265.1 | Oscillibacter_ruminantium | ON16_RS06050 | 315197..318790 | 1263547 | BREX-system-P-loop-protein-BrxC-[Oscillibacter-ruminantium] |
| WP_027621802.1 | Type1.BREX | <-Glyoxalase<-N6-METHYLASE<-Peptidase<-?<-?<-SIGMA+OB-Fold+AAA-ATPase+Lon-Protease-BrxL<-PglZ<-Schlafen+HATPase+wHTH<-N6-METHYLASE<-DUF6079*<-BrxB-BrxF-iSTAND<-BrxA-DNA-binding<-wHTH(SBD)+HNH\|\|DDE_5+DDE_Tnp_1-><-?<-?<-N6-METHYLASE | 1186 | Bacteria>Terrabacteria_group>Bacillota>Clostridia>Eubacteriales>Oscillospiraceae | GCF_000519985.1 | Acetivibrio_clariflavus | CLOCLA_RS0107445 | 1195913..1199473 | 288965 | BREX-system-P-loop-protein-BrxC-[Acetivibrio-clariflavus] |
| WP_024861712.1 | Type1.BREX | Phage_int_SAM_3->?->GDResolvase+HIN-HTH->?->ParB+HNH->BrxA-DNA-binding->BrxB-BrxF-iSTAND->DUF6079*->N6-METHYLASE-p->N6-METHYLASE-p->PglZ->SIGMA+OB-Fold+AAA-ATPase+Lon-Protease-BrxL-><-?<-?<-tRNA<-?\|\|SF1-SF2-Helicase+TUDOR-> | 1171 | Bacteria>Terrabacteria_group>Bacillota>Clostridia>Eubacteriales>Oscillospiraceae | GCF_000526795.1 | Ruminococcus_flavefaciens | N774_RS0113310 | 3002422..3005937 | 1265 | BREX-system-P-loop-protein-BrxC-[Ruminococcus-flavefaciens] |
| WP_028506203.1 | Type1.BREX | BrxA-DNA-binding->BrxB-BrxF-iSTAND->DUF6079*->N6-METHYLASE->?->N6-METHYLASE-p->PglZ->SIGMA+OB-Fold+AAA-ATPase+Lon-Protease-BrxL->?->LRR_5+LRR_5+LRR_5->LRR_5+LRR_5+LRR_5+LRR_5-> | 1166 | Bacteria>Terrabacteria_group>Bacillota>Clostridia>Eubacteriales>Oscillospiraceae | GCF_000621805.1 | Ruminococcus_sp._FC2018 | T494_RS0110140 | 1729..5229 | 1410617 | BREX-system-P-loop-protein-BrxC-[Ruminococcus-sp.-FC2018] |
| WP_050006674.1 | Type1.BREX | <-HTH_26\|\|?->?->?-><-?<-TraI-VirD2-RCRE<-TraJ-RHH\|\|HTH_26->BrxA-DNA-binding->BrxB-BrxF-iSTAND->DUF6079*-> | 1214 | Bacteria>Terrabacteria_group>Bacillota>Clostridia>Eubacteriales>Oscillospiraceae | GCF_000949455.1 | Ruthenibacterium_lactatiformans | TQ39_RS18820 | 7667..11311 | 1550024 | BREX-system-P-loop-protein-BrxC-[Ruthenibacterium-lactatiformans] |
| WP_087287659.1 | Type1.BREX | Peptidase_M20-><-HTH+REase<-SIGMA+OB-Fold+AAA-ATPase+Lon-Protease-BrxL<-PglZ<-SbcC<-N6-METHYLASE<-DUF6079*<-BrxB-BrxF-iSTAND<-BrxA-DNA-binding<-HTH_26<-AbiEii<-AbiEi_4\|\|TraJ-RHH->RepA_N+HTH->AAA+->XIS-HTH-> | 1180 | Bacteria>Terrabacteria_group>Bacillota>Clostridia>Eubacteriales>Oscillospiraceae | GCF_002160305.1 | Pseudoflavonifractor_sp._An184 | B5F19_RS02685 | 31518..35060 | 1965576 | BREX-system-P-loop-protein-BrxC-[Pseudoflavonifractor-sp.-An184] |
| WP_087168431.1 | Type1.BREX | XIS-HTH-><-?<-?<-SIGMA+OB-Fold+AAA-ATPase+Lon-Protease-BrxL<-PglZ<-RNase<-RNaseH<-N6-METHYLASE<-DUF6079*<-BrxB-BrxF-iSTAND<-BrxA-DNA-binding<-HTH_26<-?<-ParB<-ParA-Soj-PloopNTPase<-SymE_toxin<-TRAM+METHYLASE | 1199 | Bacteria>Terrabacteria_group>Bacillota>Clostridia>Eubacteriales>Oscillospiraceae | GCF_002160935.1 | Drancourtella_sp._An12 | B5E64_RS10280 | 62703..66302 | 1965548 | BREX-system-P-loop-protein-BrxC-[Drancourtella-sp.-An12] |
| WP_087369154.1 | Type1.BREX | <-GspA-ATPase<-SIR2<-?<-?<-?<-?\|\|BrxA-DNA-binding->BrxB-BrxF-iSTAND->DUF6079*->N6-METHYLASE->?->HEPN->PglZ->SIGMA+OB-Fold+AAA-ATPase+Lon-Protease-BrxL-><-Lysozyme+NLPC<-VirB4-FtsK | 1178 | Bacteria>Terrabacteria_group>Bacillota>Clostridia>Eubacteriales>Oscillospiraceae | GCF_002161675.1 | Pseudoflavonifractor_sp._An44 | B5F98_RS00420 | 121430..124966 | 1965635 | BREX-system-P-loop-protein-BrxC-[Pseudoflavonifractor-sp.-An44] |
| WP_094549793.1 | Type1.BREX | <-ZnR<-ARG<-?<-TIR<-?<-HNH<-SF1-SF2-Helicase+wHTH+Small-B-barrel+DUF1998<-BrxD-AAA<-PglZ<-DUF6079*<-BrxB-BrxF-iSTAND<-BrxA-DNA-binding<-N6-METHYLASE<-REase<-?<-?<-?<-?<-DndB | 1381 | Bacteria>Terrabacteria_group>Bacillota>Clostridia>Eubacteriales>Oscillospiraceae | GCF_002252565.1 | Petroclostridium_xylanilyticum | CIB29_RS11425 | 318791..322936 | 1792311 | hypothetical-protein-[Petroclostridium-xylanilyticum] |
| WP_097800070.1 | Type1.BREX | <-HTH\|\|?->?->35exo->?->35exo+wHTH+SF1-SF2-Helicase+HTH->?->BrxA-DNA-binding->BrxB-BrxF-iSTAND->DUF6079*->N6-METHYLASE->?->SIR2->?->Schlafen+HATPase+wHTH->PglZ->?->HEPN+SH3-like+SF1-SF2-Helicase+wHTH->MORC+DUF3684+HNH-> | 1204 | Bacteria>Terrabacteria_group>Bacillota>Clostridia>Eubacteriales>Oscillospiraceae | GCF_002549775.1 | Faecalibacterium_prausnitzii | CGS49_RS00505 | 101133..104747 | 853 | BREX-system-P-loop-protein-BrxC-[Faecalibacterium-prausnitzii] |
| WP_097782652.1 | Type1.BREX | <-DUF932<-NUDIX<-HKD-DNase+SF1-SF2-Helicase+HTH+HTH+SRA\|\|XIS-HTH->BrxA-DNA-binding->BrxB-BrxF-iSTAND->DUF6079*->Trypsin->N6-METHYLASE->TIR->PglZ->SIGMA+OB-Fold+AAA-ATPase+Lon-Protease-BrxL->ParB+HNH->HKD-DNase+SF1-SF2-Helicase+HTH+HTH+SRA->STYKinase-like->HNH-> | 1220 | Bacteria>Terrabacteria_group>Bacillota>Clostridia>Eubacteriales>Oscillospiraceae | GCF_002549935.1 | Faecalibacterium_prausnitzii | CGS55_RS03365 | 35885..39547 | 853 | BREX-system-P-loop-protein-BrxC-[Faecalibacterium-prausnitzii] |
| WP_097839702.1 | Type1.BREX | <-RVT<-?<-SIGMA+OB-Fold+AAA-ATPase+Lon-Protease-BrxL<-PglZ<-NERD<-OLD-ABC<-N6-METHYLASE<-Trypsin<-DUF6079*<-BrxB-BrxF-iSTAND<-BrxA-DNA-binding<-XIS-HTH\|\|HKD-DNase+SF1-SF2-Helicase+HTH+HTH+SRA->NUDIX->DUF932->?->?->CR-ATPase5+REase-> | 1218 | Bacteria>Terrabacteria_group>Bacillota>Clostridia>Eubacteriales>Oscillospiraceae | GCF_002549975.1 | Faecalibacterium_prausnitzii | CGS58_RS09655 | 46670..50326 | 853 | BREX-system-P-loop-protein-BrxC-[Faecalibacterium-prausnitzii] |
| WP_101299018.1 | Type1.BREX | REase+wHTH-><-?<-AAA-ATPase<-HTH+rve\|\|BrxA-DNA-binding->BrxB-BrxF-iSTAND->DUF6079*->N6-METHYLASE->PglZ->SIGMA+OB-Fold+AAA-ATPase+Lon-Protease-BrxL->?->?->Phage_int_SAM_4+Phage_integrase->?->?->SF1-SF2-Helicase-> | 1184 | Bacteria>Terrabacteria_group>Bacillota>Clostridia>Eubacteriales>Oscillospiraceae | GCF_002844395.1 | Acetivibrio_saccincola | HVS_RS02805 | 628053..631607 | 1677857 | BREX-system-P-loop-protein-BrxC-[Acetivibrio-saccincola] |
| WP_114000050.1 | Type1.BREX | <-SIGMA+OB-Fold+AAA-ATPase+Lon-Protease-BrxL<-?<-PglZ<-?<-N6-METHYLASE<-?<-DUF6079*<-BrxB-BrxF-iSTAND<-BrxA-DNA-binding<-HTH_26<-?<-?\|\|?->?->?-><-ACET | 1202 | Bacteria>Terrabacteria_group>Bacillota>Clostridia>Eubacteriales>Oscillospiraceae | GCF_003324125.1 | Subdoligranulum_sp._APC924/74 | C4885_RS09230 | 15307..18915 | 2086273 | BREX-system-P-loop-protein-BrxC-[Subdoligranulum-sp.-APC924/74] |
| WP_113992229.1 | Type1.BREX | <-CR-ATPase5+REase<-?<-?<-DUF932<-NUDIX<-HKD-DNase+SF1-SF2-Helicase+HTH+HTH+SRA\|\|XIS-HTH->BrxA-DNA-binding->BrxB-BrxF-iSTAND->DUF6079*->wHTH+ZnR->Four-helical-protein->N6-METHYLASE->HEPN->PglZ->SIGMA+OB-Fold+AAA-ATPase+Lon-Protease-BrxL->ParB+HNH->HKD-DNase+SF1-SF2-Helicase+HTH+HTH+SRA->nSTAND-> | 1218 | Bacteria>Terrabacteria_group>Bacillota>Clostridia>Eubacteriales>Oscillospiraceae | GCF_003324185.1 | Faecalibacterium_prausnitzii | C7J97_RS04695 | 32898..36554 | 853 | BREX-system-P-loop-protein-BrxC-[Faecalibacterium-prausnitzii] |
| WP_117554516.1 | Type1.BREX | <-PIN<-HTH+MPTase<-STYKinase-like<-ParB+HNH<-SIGMA+OB-Fold+AAA-ATPase+Lon-Protease-BrxL<-PglZ<-?<-N6-METHYLASE<-Trypsin<-DUF6079*<-BrxB-BrxF-iSTAND<-BrxA-DNA-binding<-XIS-HTH\|\|HKD-DNase+SF1-SF2-Helicase+HTH+HTH+SRA->NUDIX->DUF932->?->?->CR-ATPase5+REase-> | 1217 | Bacteria>Terrabacteria_group>Bacillota>Clostridia>Eubacteriales>Oscillospiraceae | GCF_003434135.1 | Faecalibacterium_prausnitzii | DW855_RS10430 | 56056..59709 | 853 | BREX-system-P-loop-protein-BrxC-[Faecalibacterium-prausnitzii] |
| WP_117891999.1 | Type1.BREX | DSBH+AraC-HTH+AraC-HTH-><-IONCHANNEL+HISKIN+REC<-HTH+REase<-SIGMA+OB-Fold+AAA-ATPase+Lon-Protease-BrxL<-PglZ<-?<-N6-METHYLASE<-DUF6079*<-BrxB-BrxF-iSTAND<-BrxA-DNA-binding<-HTH_26 | 1184 | Bacteria>Terrabacteria_group>Bacillota>Clostridia>Eubacteriales>Oscillospiraceae | GCF_003461215.1 | Ruminococcus_sp._AF19-15 | DWX34_06625 | 16920..20474 | 2293157 | BREX-system-P-loop-protein-BrxC-[Ruminococcus-sp.-AF19-15] |
| WP_128705604.1 | Type1.BREX | Resolvase+Recombinase->SAM-METHYLASE->Resolvase+Recombinase->ABC-ATPase+SbcC->?->CHTH->BrxA-DNA-binding->BrxB-BrxF-iSTAND->DUF6079*->N6-METHYLASE->PglZ->SIGMA+OB-Fold+AAA-ATPase+Lon-Protease-BrxL->rve->DnaB+nSTAND3->ZnR+Phage_GP20->HTH_3->ParB+HNH-> | 1139 | Bacteria>Terrabacteria_group>Bacillota>Clostridia>Eubacteriales>Oscillospiraceae | GCF_004102745.1 | Acetivibrio_mesophilus | EFD62_RS00955 | 186850..190269 | 2487273 | BREX-system-P-loop-protein-BrxC-[Acetivibrio-mesophilus] |
| WP_128706494.1 | Type1.BREX | BrxA-DNA-binding->BrxB-BrxF-iSTAND->DUF6079*->N6-METHYLASE->PglZ->SIGMA+OB-Fold+AAA-ATPase+Lon-Protease-BrxL->?-><-?\|\|SF1-SF2-Helicase-> | 1191 | Bacteria>Terrabacteria_group>Bacillota>Clostridia>Eubacteriales>Oscillospiraceae | GCF_004102745.1 | Acetivibrio_mesophilus | EFD62_RS16120 | 3642..7217 | 2487273 | BREX-system-P-loop-protein-BrxC-[Acetivibrio-mesophilus] |
| WP_137698823.1 | Type1.BREX | <-PhdYeFM_antitox<-?<-?<-?<-?<-REase<-SIGMA+OB-Fold+AAA-ATPase+Lon-Protease-BrxL<-PglZ<-N6-METHYLASE<-DUF6079*<-BrxB-BrxF-iSTAND<-BrxA-DNA-binding<-?<-?<-?<-?<-?<-?<-Phage_int_SAM_5+Phage_integrase | 1191 | Bacteria>Terrabacteria_group>Bacillota>Clostridia>Eubacteriales>Oscillospiraceae | GCF_005473905.2 | Ruminiclostridium_herbifermentans | EHE19_RS13505 | 3333193..3336768 | 2488810 | BREX-system-P-loop-protein-BrxC-[Ruminiclostridium-herbifermentans] |
| WP_120485839.1 | Type1.BREX | <-SIGMA+OB-Fold+AAA-ATPase+Lon-Protease-BrxL<-PglZ<-REase<-OLD-ABC<-N6-METHYLASE<-DUF6079*<-BrxB-BrxF-iSTAND<-BrxA-DNA-binding<-HTH_26<-JetC<-?<-HNH\|\|HTH_3->PemK_toxin->ParB-> | 1178 | Bacteria>Terrabacteria_group>Bacillota>Clostridia>Eubacteriales>Oscillospiraceae | GCF_009917605.1 | unclassified_Anaerotruncus | D5269_RS12205 | 11128..14664 | 2641626 | MULTISPECIES:-BREX-system-P-loop-protein-BrxC-[unclassified-Anaerotruncus] |
| WP_195249802.1 | Type1.BREX | SPOUT-><-?\|\|ABhydrolase-><-?<-SIGMA+OB-Fold+AAA-ATPase+Lon-Protease-BrxL<-PglZ<-?<-?<-N6-METHYLASE<-DUF6079*<-BrxB-BrxF-iSTAND<-BrxA-DNA-binding | 1161 | Bacteria>Terrabacteria_group>Bacillota>Clostridia>Eubacteriales>Oscillospiraceae | GCF_015549395.1 | Ruminococcus_bicirculans | I2A70_RS11350 | 17655..21140 | 1160721 | BREX-system-P-loop-protein-BrxC-[Ruminococcus-bicirculans] |
| WP_195254595.1 | Type1.BREX | <-PIN<-SF1-SF2-Helicase<-TERNS3+HEPN<-SIGMA+OB-Fold+AAA-ATPase+Lon-Protease-BrxL<-PglZ<-N6-METHYLASE<-DUF6079*<-BrxB-BrxF-iSTAND<-BrxA-DNA-binding<-HTH_26<-TraI-VirD2-RCRE<-TraJ-RHH | 1211 | Bacteria>Terrabacteria_group>Bacillota>Clostridia>Eubacteriales>Oscillospiraceae | GCF_015550675.1 | Merdimmobilis_hominis | I2A64_RS11910 | 31886..35521 | 2897707 | BREX-system-P-loop-protein-BrxC-[Merdimmobilis-hominis] |
| WP_195494801.1 | Type1.BREX | <-HTH+sGTP\|\|HKD-DNase-><-?<-SF1-SF2-Helicase<-?<-GDSL<-SIGMA+OB-Fold+AAA-ATPase+Lon-Protease-BrxL<-PglZ<-?<-N6-METHYLASE<-DUF6079*<-BrxB-BrxF-iSTAND<-BrxA-DNA-binding<-HTH_26<-HNH+ADA-DBD<-?\|\|HTH_3-><-HTH_36<-VirD4-TraG+VirD4-FtsK<-ZnR+TOPRIM | 1211 | Bacteria>Terrabacteria_group>Bacillota>Clostridia>Eubacteriales>Oscillospiraceae | GCF_015554425.1 | Flavonifractor_plautii | I2F92_RS05225 | 188245..191880 | 292800 | BREX-system-P-loop-protein-BrxC-[Flavonifractor-plautii] |
| WP_207739085.1 | Type1.BREX | METHYLASE->?->?->?->HTH_26->BrxA-DNA-binding->BrxB-BrxF-iSTAND->DUF6079*->N6-METHYLASE->OLD-ABC->HNH->PglZ->SIGMA+OB-Fold+AAA-ATPase+Lon-Protease-BrxL->?->Arm-DNA-bind_4+Phage_int_SAM_3+Phage_integrase->?->XIS-HTH+XIS-HTH-> | 1173 | Bacteria>Terrabacteria_group>Bacillota>Clostridia>Eubacteriales>Oscillospiraceae | GCF_015558275.1 | Massiliimalia_timonensis | I2G46_RS07880 | 51772..55293 | 1987501 | BREX-system-P-loop-protein-BrxC-[Massiliimalia-timonensis] |
| WP_148337313.1 | Type1.BREX | LPD32+LPD36+N6_Mtase+SF1-SF2-Helicase->TraJ-RHH->TraM->?->?->XIS-HTH->BrxA-DNA-binding->BrxB-BrxF-iSTAND->DUF6079*->N6-METHYLASE->Tox-ART-HYD2->?->AT-HTH-HYD2-2->PglZ->SIGMA+OB-Fold+AAA-ATPase+Lon-Protease-BrxL->Cap4_nuclease-> | 1217 | Bacteria>Terrabacteria_group>Bacillota>Clostridia>Eubacteriales>Oscillospiraceae | GCF_015558925.1 | Flavonifractor_plautii | I2I15_RS11330 | 38217..41870 | 292800 | BREX-system-P-loop-protein-BrxC-[Flavonifractor-plautii] |
| WP_195991333.1 | Type1.BREX | <-KAP-glycosylase<-KAP-NTPase<-SIGMA+OB-Fold+AAA-ATPase+Lon-Protease-BrxL<-PglZ<-AT-HTH-HYD2-2<-?<-Tox-ART-HYD2<-N6-METHYLASE<-TIR<-DUF6079*<-BrxB-BrxF-iSTAND<-BrxA-DNA-binding<-XIS-HTH | 1217 | Bacteria>Terrabacteria_group>Bacillota>Clostridia>Eubacteriales>Oscillospiraceae | GCF_015668065.1 | unclassified_Ruminococcus | I4V84_RS04405 | 84805..88458 | 2608920 | MULTISPECIES:-BREX-system-P-loop-protein-BrxC-[unclassified-Ruminococcus] |
| WP_195982636.1 | Type1.BREX | SAM-METHYLASE->?->ABhydrolase->?->?->HTH_36->HTH_26->BrxA-DNA-binding->BrxB-BrxF-iSTAND->DUF6079*->TIR->N6-METHYLASE->AT-HTH-HYD2->?->Schlafen+HATPase+wHTH->PglZ->SIGMA+OB-Fold+AAA-ATPase+Lon-Protease-BrxL->TIR-><-HTH | 1182 | Bacteria>Terrabacteria_group>Bacillota>Clostridia>Eubacteriales>Oscillospiraceae | GCF_015669355.1 | Ruminococcus_sp._1001275B_160808_F8 | I4V83_RS01630 | 22407..25955 | 2787131 | BREX-system-P-loop-protein-BrxC-[Ruminococcus-sp.-1001275B_160808_F8] |
| WP_207751064.1 | Type1.BREX | HTH_26->BrxA-DNA-binding->BrxB-BrxF-iSTAND->DUF6079*->N6-METHYLASE->NERD->SIGMA+OB-Fold+AAA-ATPase+Lon-Protease-BrxL->35exo+wHTH+SF1-SF2-Helicase+HTH->?-><-?<-ZnR+ZnR<-HTH_26 | 1173 | Bacteria>Terrabacteria_group>Bacillota>Clostridia>Eubacteriales>Oscillospiraceae | GCF_016901815.1 | Hydrogenoanaerobacterium_saccharovorans | H9X81_RS11030 | 2162..5683 | 474960 | BREX-system-P-loop-protein-BrxC-[Hydrogenoanaerobacterium-saccharovorans] |
| WP_204881984.1 | Type1.BREX | VirD4-TraG+VirD4-FtsK->TraM->TrwI-><-?\|\|?->VirB4-FtsK->HMG+NLPC+NLPC->BrxA-DNA-binding->BrxB-BrxF-iSTAND->DUF6079*->N6-METHYLASE->Tox-ART-HYD3->AbiGi->PglZ->SIGMA+OB-Fold+AAA-ATPase+Lon-Protease-BrxL->ZnR->?->wHTH+WYL->SF1-SF2-Helicase-> | 1179 | Bacteria>Terrabacteria_group>Bacillota>Clostridia>Eubacteriales>Oscillospiraceae | GCF_016901955.1 | Pseudoflavonifractor_phocaeensis | H6B51_RS01515 | 67307..70846 | 1870988 | BREX-system-P-loop-protein-BrxC-[Pseudoflavonifractor-phocaeensis] |
| WP_204901129.1 | Type1.BREX | SF1-SF2-Helicase-><-SIGMA+OB-Fold+AAA-ATPase+Lon-Protease-BrxL<-PglZ<-REase<-OLD-ABC<-?<-Tox-RES<-N6-METHYLASE<-SIR2<-DUF6079*<-BrxB-BrxF-iSTAND<-BrxA-DNA-binding<-SIGMA-HTH | 1179 | Bacteria>Terrabacteria_group>Bacillota>Clostridia>Eubacteriales>Oscillospiraceae | GCF_016902575.1 | Pseudoflavonifractor_capillosus | H9X86_RS08600 | 19493..23032 | 106588 | BREX-system-P-loop-protein-BrxC-[Pseudoflavonifractor-capillosus] |
| WP_207700511.1 | Type1.BREX | <-Tox-RelA<-SMODS<-SLATT<-SIGMA+OB-Fold+AAA-ATPase+Lon-Protease-BrxL<-PglZ<-AT-HTH-HYD2-2<-?<-Tox-ART-HYD2<-N6-METHYLASE<-DUF6079*<-BrxB-BrxF-iSTAND<-BrxA-DNA-binding<-XIS-HTH<-?<-?<-TraM<-TraJ-RHH<-LPD32+LPD36+N6_Mtase+SF1-SF2-Helicase | 1217 | Bacteria>Terrabacteria_group>Bacillota>Clostridia>Eubacteriales>Oscillospiraceae | GCF_017377545.1 | Faecalibacterium_sp._Marseille-Q4164 | J3352_RS11645 | 12124..15777 | 2817020 | BREX-system-P-loop-protein-BrxC-[Faecalibacterium-sp.-Marseille-Q4164] |
| WP_212792861.1 | Type1.BREX | <-CR-ATPase8+REase<-?<-?<-?<-DUF932\|\|DOC+HTH->BrxA-DNA-binding->BrxB-BrxF-iSTAND->DUF6079*->?->?->N6-METHYLASE-> | 1182 | Bacteria>Terrabacteria_group>Bacillota>Clostridia>Eubacteriales>Oscillospiraceae | GCF_018327745.1 | Faecalibacterium_prausnitzii | MCC02041_25000 | 16287..19835 | 853 | BREX-system-P-loop-protein-BrxC-[Faecalibacterium-prausnitzii] |
| WP_215659170.1 | Type1.BREX | <-TIR<-SIGMA+OB-Fold+AAA-ATPase+Lon-Protease-BrxL<-PglZ<-DUF4404<-ZnR+ZnR<-N6-METHYLASE<-?<-HNH<-OLD-ABC<-DUF6079*<-BrxB-BrxF-iSTAND<-BrxA-DNA-binding<-HTH_26<-MazE_antitoxin+SpoVT_C<-?\|\|?-><-?<-RADICAL-SAM | 1184 | Bacteria>Terrabacteria_group>Bacillota>Clostridia>Eubacteriales>Oscillospiraceae | GCF_018783905.1 | Ruminococcus_sp._MCC718 | GPK60_RS07265 | 26956..30510 | 2592649 | BREX-system-P-loop-protein-BrxC-[Ruminococcus-sp.-MCC718] |
| WP_215696711.1 | Type1.BREX | LPD32+LPD36+N6_Mtase+SF1-SF2-Helicase->TraJ-RHH->TraM->?->?->XIS-HTH->BrxA-DNA-binding->BrxB-BrxF-iSTAND->DUF6079*->N6-METHYLASE->?->PglZ->SIGMA+OB-Fold+AAA-ATPase+Lon-Protease-BrxL->?->ClpABC-AAA-> | 1217 | Bacteria>Terrabacteria_group>Bacillota>Clostridia>Eubacteriales>Oscillospiraceae | GCF_018784555.1 | Faecalibacterium_prausnitzii | GPK63_RS06415 | 101308..104961 | 853 | BREX-system-P-loop-protein-BrxC-[Faecalibacterium-prausnitzii] |
| WP_215675194.1 | Type1.BREX | <-CR-ATPase8+REase<-?<-CR-ATPase5+REase<-?<-DUF932\|\|DOC+HTH->BrxA-DNA-binding->BrxB-BrxF-iSTAND->DUF6079*->TIR->N6-METHYLASE->?->PglZ->SIGMA+OB-Fold+AAA-ATPase+Lon-Protease-BrxL->ParB+HNH->HKD-DNase+SF1-SF2-Helicase+HTH+HTH+SRA->SF1-SF2-Helicase+TOPC->STYKinase-like-> | 1182 | Bacteria>Terrabacteria_group>Bacillota>Clostridia>Eubacteriales>Oscillospiraceae | GCF_018784645.1 | Faecalibacterium_prausnitzii | GPK62_RS07855 | 35115..38663 | 853 | BREX-system-P-loop-protein-BrxC-[Faecalibacterium-prausnitzii] |
| WP_216553535.1 | Type1.BREX | <-UDG-conflict<-?<-?\|\|HTH_19-><-tRNA\|\|PBPI+GGDEF->?->\|\|Phage_int_SAM_3+Phage_integrase->BrxA-DNA-binding->BrxB-BrxF-iSTAND->DUF6079*->N6-METHYLASE->?->PglZ->SIGMA+OB-Fold+AAA-ATPase+Lon-Protease-BrxL->?-><-ABhydrolase<-SPOUT<-?<-HAD | 1161 | Bacteria>Terrabacteria_group>Bacillota>Clostridia>Eubacteriales>Oscillospiraceae | GCF_018917825.1 | Ruminococcus_sp._MSJ-25 | KQH97_RS06365 | 89589..93074 | 2841536 | BREX-system-P-loop-protein-BrxC-[Ruminococcus-sp.-MSJ-25] |
| WP_223386332.1 | Type1.BREX | <-Tox-RelA<-SMODS<-SLATT<-SIGMA+OB-Fold+AAA-ATPase+Lon-Protease-BrxL<-PglZ<-AT-HTH-HYD2-2<-?<-Tox-ART-HYD2<-N6-METHYLASE<-DUF6079*<-BrxB-BrxF-iSTAND<-BrxA-DNA-binding<-XIS-HTH<-?<-?<-TraM<-TraJ-RHH | 1217 | Bacteria>Terrabacteria_group>Bacillota>Clostridia>Eubacteriales>Oscillospiraceae | GCF_019967975.1 | Faecalibacterium_prausnitzii | I5P95_RS12425 | 2568697..2572350 | 853 | BREX-system-P-loop-protein-BrxC-[Faecalibacterium-prausnitzii] |
| WP_223450814.1 | Type1.BREX | REase+N-glycosylase->MACRODOMAIN->?->?->XIS-HTH->BrxA-DNA-binding->BrxB-BrxF-iSTAND->DUF6079*->N6-METHYLASE->Tox-ART-HYD2->?->AT-HTH-HYD2-2->PglZ->SIGMA+OB-Fold+AAA-ATPase+Lon-Protease-BrxL->Tox-RelA->HKD-DNase+HKD-DNase->HKD-DNase+HKD-DNase-> | 1217 | Bacteria>Terrabacteria_group>Bacillota>Clostridia>Eubacteriales>Oscillospiraceae | GCF_019969405.1 | Faecalibacterium_prausnitzii | LC355_RS02555 | 495112..498765 | 853 | BREX-system-P-loop-protein-BrxC-[Faecalibacterium-prausnitzii] |
| WP_223452176.1 | Type1.BREX | <-RVT<-SIGMA+OB-Fold+AAA-ATPase+Lon-Protease-BrxL<-PglZ<-?<-N6-METHYLASE<-?<-DUF6079*<-BrxB-BrxF-iSTAND<-BrxA-DNA-binding<-DOC+HTH\|\|DUF932->?->?->?->CR-ATPase8+REase-> | 1178 | Bacteria>Terrabacteria_group>Bacillota>Clostridia>Eubacteriales>Oscillospiraceae | GCF_019969405.1 | Faecalibacterium_prausnitzii | LC355_RS13060 | 2672298..2675834 | 853 | BREX-system-P-loop-protein-BrxC-[Faecalibacterium-prausnitzii] |
| WP_223452849.1 | Type1.BREX | Cysteine-Desulfurase->?->?-><-RVT<-SIGMA+OB-Fold+AAA-ATPase+Lon-Protease-BrxL<-PglZ<-?<-N6-METHYLASE<-?<-DUF6079*<-BrxB-BrxF-iSTAND<-BrxA-DNA-binding<-DOC+HTH\|\|DUF932->?->?->?->CR-ATPase8+REase-> | 1183 | Bacteria>Terrabacteria_group>Bacillota>Clostridia>Eubacteriales>Oscillospiraceae | GCF_019969415.1 | Faecalibacterium_prausnitzii | LC358_RS12990 | 2672306..2675857 | 853 | BREX-system-P-loop-protein-BrxC-[Faecalibacterium-prausnitzii] |
| WP_005924150.1 | Type1.BREX | <-HTH_3\|\|REC+OmpR-HTH->HISKIN->ABC-ATPase+SbcC->?->SIGMA-HTH->?->GDResolvase+Recombinase+ZnR->DUF6079*->?->?->N6-METHYLASE->?->PglZ->SIGMA+OB-Fold+AAA-ATPase+Lon-Protease-BrxL->RVT-> | 1177 | Bacteria>Terrabacteria_group>Bacillota>Clostridia>Eubacteriales>Oscillospiraceae | GCF_019969455.1 | Faecalibacterium_prausnitzii | LC357_RS02670 | 540740..544273 | 853 | BREX-system-P-loop-protein-BrxC-[Faecalibacterium-prausnitzii] |
| WP_227612042.1 | Type1.BREX | BrxA-DNA-binding->BrxB-BrxF-iSTAND->DUF6079*->N6-METHYLASE->?->?->AbiGi->Schlafen+HATPase+wHTH->PglZ->HEPN+SH3-like+SF1-SF2-Helicase-> | 1204 | Bacteria>Terrabacteria_group>Bacillota>Clostridia>Eubacteriales>Oscillospiraceae | GCF_020686905.1 | Faecalibacterium_sp._CLA-AA-H283 | LKD27_RS12770 | 2733..6347 | 2881278 | BREX-system-P-loop-protein-BrxC-[Faecalibacterium-sp.-CLA-AA-H283] |
| WP_227612689.1 | Type1.BREX | LPD32+LPD36+N6_Mtase+SF1-SF2-Helicase->TraJ-RHH->TraM->?->?->XIS-HTH->BrxA-DNA-binding->BrxB-BrxF-iSTAND->DUF6079*->N6-METHYLASE->Tox-ART-HYD2->?->AT-HTH-HYD2-2->PglZ->SIGMA+OB-Fold+AAA-ATPase+Lon-Protease-BrxL->SLATT->SMODS->Tox-RelA-> | 1217 | Bacteria>Terrabacteria_group>Bacillota>Clostridia>Eubacteriales>Oscillospiraceae | GCF_020687145.1 | Faecalibacterium_sp._CLA-AA-H254 | LKD26_RS04550 | 119321..122974 | 2881273 | BREX-system-P-loop-protein-BrxC-[Faecalibacterium-sp.-CLA-AA-H254] |

| Protein-ID | System | Gene-Neighbourhoods | Length | Taxonomy | Assembly-ID | Strain | Locus-Tag | Genomic-coordinates | TaxID | Fasta-defline |
|---|---|---|---|---|---|---|---|---|---|---|
| WP_235255096.1 | Type1.BREX | <-SbcC+SbcC<-?<-JetA\|\|XIS-HTH->BrxA-DNA-binding->BrxB-BrxF-iSTAND->DUF6079*->HNH+SAVED->SMODS->?->N6-METHYLASE->ParB+HNH->sGTP->?->HerA/FtsK->HNH-> | 1199 | Bacteria>Terrabacteria_group>Bacillota>Clostridia>Eubacteriales>Oscillospiraceae | GCF_021531255.1 | Fournierella_massiliensis | JQM64_RS00320 | 59255..62854 | 1650663 | BREX-system-P-loop-protein-BrxC-[Fournierella-massiliensis] |
| WP_249257849.1 | Type1.BREX | [illegible] | 1217 | Bacteria>Terrabacteria_group>Bacillota>Clostridia>Eubacteriales>Oscillospiraceae | GCF_023347315.1 | Faecalibacterium_sp._I2-3-92 | MTP42_RS02040 | 437412..441065 | 2929490 | BREX-system-P-loop-protein-BrxC-[Faecalibacterium-sp.-I2-3-92] |
| WP_249234870.1 | Type1.BREX | [illegible] | 1217 | Bacteria>Terrabacteria_group>Bacillota>Clostridia>Eubacteriales>Oscillospiraceae | GCF_023347355.1 | Faecalibacterium_sp._IP-1-18 | MTP43_RS12455 | 2619017..2622670 | 2929488 | BREX-system-P-loop-protein-BrxC-[Faecalibacterium-sp.-IP-1-18] |
| WP_256266343.1 | Type1.BREX | [illegible] | 1218 | Bacteria>Terrabacteria_group>Bacillota>Clostridia>Eubacteriales>Oscillospiraceae | GCF_024460915.1 | Faecalibacterium_prausnitzii | NE589_RS05495 | 127412..131068 | 853 | BREX-system-P-loop-protein-BrxC-[Faecalibacterium-prausnitzii] |
| WP_267300356.1 | Type1.BREX | [illegible] | 1184 | Bacteria>Terrabacteria_group>Bacillota>Clostridia>Eubacteriales>Oscillospiraceae | GCF_025567495.1 | Hominimerdicola_aceti | OCV57_RS02360 | 41099..44653 | 2981726 | BREX-system-P-loop-protein-BrxC-[Hominimerdicola-aceti] |
| WP_263041866.1 | Type1.BREX | [illegible] | 1211 | Bacteria>Terrabacteria_group>Bacillota>Clostridia>Eubacteriales>Oscillospiraceae | GCF_025630755.1 | Acutalibacter_sp._LFL-21 | ODJ80_RS09375 | 227875..231510 | 2983399 | BREX-system-P-loop-protein-BrxC-[Acutalibacter-sp.-LFL-21] |
| WP_117473063.1 | Type1.BREX | [illegible] | 1214 | Bacteria>Terrabacteria_group>Bacillota>Clostridia>Eubacteriales>Oscillospiraceae | GCF_027690895.1 | Ruthenibacterium_lactatiformans | DW651_08385 | 184245..187889 | 1550024 | BREX-system-P-loop-protein-BrxC-[Ruthenibacterium-lactatiformans] |
| WP_275252236.1 | Type1.BREX | [illegible] | 1218 | Bacteria>Terrabacteria_group>Bacillota>Clostridia>Eubacteriales>Oscillospiraceae | GCF_029076575.1 | Faecalibacterium_sp._DFI.5.82 | POG09_RS06330 | 132484..136140 | 3031725 | BREX-system-P-loop-protein-BrxC-[Faecalibacterium-sp.-DFI.5.82] |
| WP_117891516.1 | Type1.BREX | [illegible] | 1161 | Bacteria>Terrabacteria_group>Bacillota>Clostridia>Eubacteriales>Oscillospiraceae | GCF_900067005.1 | Ruminococcus_sp._AF19-15 | DWX34_02860 | 160576..164061 | 2293157 | BREX-system-P-loop-protein-BrxC-[Ruminococcus-sp.-AF19-15] |
| WP_021658826.1 | Type1.BREX | [illegible] | 1211 | Bacteria>Terrabacteria_group>Bacillota>Clostridia>Eubacteriales>Oscillospiraceae | GCF_900129515.1 | Bittarella_massiliensis | BUB64_RS00425 | 83459..87094 | 1720313 | BREX-system-P-loop-protein-BrxC-[Bittarella-massiliensis] |
| WP_101908637.1 | Type1.BREX | [illegible] | 1192 | Bacteria>Terrabacteria_group>Bacillota>Clostridia>Eubacteriales>Oscillospiraceae | GCF_900186535.1 | Marasmitruncus_massiliensis | C1A10_RS02055 | 174899..178477 | 1944642 | BREX-system-P-loop-protein-BrxC-[Marasmitruncus-massiliensis] |
| WP_158398728.1 | Type1.BREX | [illegible] | 1182 | Bacteria>Terrabacteria_group>Bacillota>Clostridia>Eubacteriales>Oscillospiraceae | GCF_902167845.1 | Faecalibacterium_prausnitzii | FQ260_RS07280 | 12413..15961 | 853 | BREX-system-P-loop-protein-BrxC-[Faecalibacterium-prausnitzii] |
| WP_058352689.1 | Type1.BREX | <-REase<-?<-TIR<-?<-DUF6079*<-BrxB-BrxF-iSTAND<-BrxA-DNA-binding<-HTH_26<-HTH\|\|?->?->?-><-?<-GDResolvase+Recombinase+ZnR | 1179 | Bacteria>Terrabacteria_group>Bacillota>Clostridia>Eubacteriales>Oscillospiraceae | GCF_902387445.1 | Acetivibrio_ethanolgignens | FX109_RS09105 | 46912..50451 | 290052 | BREX-system-P-loop-protein-BrxC-[Acetivibrio-ethanolgignens] |
| WP_117922984.1 | Type1.BREX | [illegible] | 1161 | Bacteria>Terrabacteria_group>Bacillota>Clostridia>Eubacteriales>Oscillospiraceae | GCF_958404065.1 | Ruminococcus | DW733_00275 | 57797..61282 | 1263 | MULTISPECIES:-BREX-system-P-loop-protein-BrxC-[Ruminococcus] |
| WP_021683521.1 | Type1.BREX | <-SIGMA+OB-Fold+AAA-ATPase+Lon-Protease-BrxL<-PglZ<-PRTase<-N6-METHYLASE<-DUF6079*<-BrxB-BrxF-iSTAND<-BrxA-DNA-binding | 1162 | Bacteria>Terrabacteria_group>Bacillota>Clostridia>Eubacteriales>Oscillospiraceae | GCF_958428055.1 | Ruminococcus_callidus | RUMCAL_RS13370 | 9405..12893 | 40519 | BREX-system-P-loop-protein-BrxC-[Ruminococcus-callidus] |
| WP_235256054.1 | Type1.BREX | DOC+HTH-><-SIGMA+OB-Fold+AAA-ATPase+Lon-Protease-BrxL<-PglZ<-?<-?<-N6-METHYLASE<-DUF6079*<-BrxB-BrxF-iSTAND<-BrxA-DNA-binding | 1174 | Bacteria>Terrabacteria_group>Bacillota>Clostridia>Eubacteriales>Oscillospiraceae | - | Oscillibacter_valericigenes | JQM63_RS11255 | 11460..14984 | 351091 | BREX-system-P-loop-protein-BrxC-[Oscillibacter-valericigenes] |
| WP_118142807.1 | Type1.BREX | [illegible] | 1203 | Bacteria>Terrabacteria_group>Bacillota>Clostridia>Eubacteriales>Oscillospiraceae | - | Ruminococcus_sp._OM07-7 | DXC06_07755 | 161101..164712 | 2293234 | BREX-system-P-loop-protein-BrxC-[Ruminococcus-sp.-OM07-7] |
| WP_118330216.1 | Type1.BREX | [illegible] | 1205 | Bacteria>Terrabacteria_group>Bacillota>Clostridia>Eubacteriales>Oscillospiraceae | - | unclassified_Ruminococcus | DW225_05645 | 153142..156759 | 2608920 | MULTISPECIES:-BREX-system-P-loop-protein-BrxC-[unclassified-Ruminococcus] |
| WP_117958368.1 | Type1.BREX | [illegible] | 1184 | Bacteria>Terrabacteria_group>Bacillota>Clostridia>Eubacteriales>Oscillospiraceae | - | unclassified_Ruminococcus | LLP35_08975 | 12774..16328 | 2608920 | MULTISPECIES:-BREX-system-P-loop-protein-BrxC-[Ruminococcus] |
| WP_066638545.1 | Type1.BREX | [illegible] | 1184 | Bacteria>Terrabacteria_group>Bacillota>Clostridia>Eubacteriales>Peptococcaceae | - | Desulfolucianica_intricata | DIN01_RS10945 | 45737..49291 | 1285191 | BREX-system-P-loop-protein-BrxC-[Desulfolucianica-intricata] |
| KJS02126.1 | Type1.BREX | <-SIGMA+OB-Fold+AAA-ATPase+Lon-Protease-BrxL<-PglZ<-RAMPs<-?<-RAMPs<-HD<-RAMPs<-RAMPs<-N6-METHYLASE<-DUF6079*<-BrxB-BrxF-iSTAND<-BrxA-DNA-binding<-HTH_26 | 1198 | Bacteria>Terrabacteria_group>Bacillota>Clostridia>Eubacteriales>Peptococcaceae | GCA_000961585.1 | Peptococcaceae_bacterium_BRH_c4a | VR68_03640 | 83699..87295 | 1629716 | hypothetical-protein-VR68_03640-[Peptococcaceae-bacterium-BRH_c4a] |
| KLU38952.1 | Type1.BREX | [illegible] | 1148 | Bacteria>Terrabacteria_group>Bacillota>Clostridia>Eubacteriales>Peptococcaceae | GCA_001028815.1 | Peptococcaceae_bacterium_1109 | AA931_11510 | 14607..18053 | 1655638 | nucleoside-triphosphate-hydrolase-[Peptococcaceae-bacterium-1109] |
| NLW39193.1 | Type1.BREX | [illegible] | 1174 | Bacteria>Terrabacteria_group>Bacillota>Clostridia>Eubacteriales>Peptococcaceae | GCA_012728715.1 | Peptococcaceae_bacterium | GXY90_11620 | 6816..10340 | 2052179 | BREX-system-P-loop-protein-BrxC-[Peptococcaceae-bacterium] |
| MBE6111698.1 | Type1.BREX | [illegible] | 1223 | Bacteria>Terrabacteria_group>Bacillota>Clostridia>Eubacteriales>Peptococcaceae | GCA_015058785.1 | Peptococcaceae_bacterium | E7195_01600 | 73755..77426 | 2052179 | BREX-system-P-loop-protein-BrxC-[Peptococcaceae-bacterium] |
| MBS3950095.1 | Type1.BREX | [illegible] | 1183 | Bacteria>Terrabacteria_group>Bacillota>Clostridia>Eubacteriales>Peptococcaceae | GCA_018333435.1 | Peptococcaceae_bacterium | KGZ53_05495 | 2469..6020 | 2052179 | BREX-system-P-loop-protein-BrxC-[Peptococcaceae-bacterium] |
| WP_031517445.1 | Type1.BREX | [illegible] | 1139 | Bacteria>Terrabacteria_group>Bacillota>Clostridia>Eubacteriales>Peptococcaceae | GCF_000711975.1 | Desulfofalx_alkaliphila | BR02_RS0112060 | 1531..4950 | 105483 | BREX-system-P-loop-protein-BrxC-[Desulfofalx-alkaliphila] |
| WP_089612556.1 | Type1.BREX | [illegible] | 1191 | Bacteria>Terrabacteria_group>Bacillota>Clostridia>Eubacteriales>Peptococcaceae | GCF_002224645.1 | Dehalobacterium_formicoaceticum | CEQ75_RS08150 | 1656670..1660245 | 51515 | BREX-system-P-loop-protein-BrxC-[Dehalobacterium-formicoaceticum] |
| WP_152948294.1 | Type1.BREX | [illegible] | 1148 | Bacteria>Terrabacteria_group>Bacillota>Clostridia>Eubacteriales>Peptococcaceae | GCF_009428905.1 | Desulfofundulus_thermobenzoicus | GFC01_RS16720 | 1980..5426 | 29376 | BREX-system-P-loop-protein-BrxC-[Desulfofundulus-thermobenzoicus] |
| MBF1049794.1 | Type1.BREX | [illegible] | 1204 | Bacteria>Terrabacteria_group>Bacillota>Clostridia>Eubacteriales>Peptostreptococcaceae | GCA_015257205.1 | Peptostreptococcus_sp. | HXL23_03750 | 40553..44167 | 1262 | BREX-system-P-loop-protein-BrxC-[Peptostreptococcus-sp.] |
| MCA0486762.1 | Type1.BREX | [illegible] | 1204 | Bacteria>Terrabacteria_group>Bacillota>Clostridia>Eubacteriales>Peptostreptococcaceae | GCA_019168885.2 | Clostridioides_difficile | KOO17_000025 | 22445..26059 | 1496 | BREX-system-P-loop-protein-BrxC-[Clostridioides-difficile] |
| WP_013361249.1 | Type1.BREX | [illegible] | 1212 | Bacteria>Terrabacteria_group>Bacillota>Clostridia>Eubacteriales>Peptostreptococcaceae | GCF_000196455.1 | Acetoanaerobium_sticklandii | CLOST_RS05375 | 1173972..1177610 | 1511 | BREX-system-P-loop-protein-BrxC-[Acetoanaerobium-sticklandii] |
| WP_028829862.1 | Type1.BREX | [illegible] | 1139 | Bacteria>Terrabacteria_group>Bacillota>Clostridia>Eubacteriales>Peptostreptococcaceae | GCF_000423525.1 | Proteocatella_sphenisci | G558_RS0108875 | 8541..11960 | 181070 | BREX-system-P-loop-protein-BrxC-[Proteocatella-sphenisci] |
| WP_025434966.1 | Type1.BREX | [illegible] | 1207 | Bacteria>Terrabacteria_group>Bacillota>Clostridia>Eubacteriales>Peptostreptococcaceae | GCF_000597865.1 | Peptoclostridium_acidaminophilum | EAL2_RS03125 | 592925..596548 | 1731 | BREX-system-P-loop-protein-BrxC-[Peptoclostridium-acidaminophilum] |
| WP_057535321.1 | Type1.BREX | [illegible] | 1185 | Bacteria>Terrabacteria_group>Bacillota>Clostridia>Eubacteriales>Peptostreptococcaceae | GCF_001372455.1 | Paeniclostridium_sordellii | R26833_RS12520 | 91086..94643 | 1505 | BREX-system-P-loop-protein-BrxC-[Paeniclostridium-sordellii] |
| WP_057552934.1 | Type1.BREX | [illegible] | 1185 | Bacteria>Terrabacteria_group>Bacillota>Clostridia>Eubacteriales>Peptostreptococcaceae | GCF_001372915.1 | Paeniclostridium_sordellii | SSCC32135_RS03600 | 682084..685641 | 1505 | BREX-system-P-loop-protein-BrxC-[Paeniclostridium-sordellii] |
| WP_055087863.1 | Type1.BREX | [illegible] | 1190 | Bacteria>Terrabacteria_group>Bacillota>Clostridia>Eubacteriales>Peptostreptococcaceae | GCF_001406775.1 | Intestinibacter_bartlettii | ARA80_RS04190 | 939339..942911 | 261299 | BREX-system-P-loop-protein-BrxC-[Intestinibacter-bartlettii] |
| WP_099191952.1 | Type1.BREX | [illegible] | 1190 | Bacteria>Terrabacteria_group>Bacillota>Clostridia>Eubacteriales>Peptostreptococcaceae | GCF_002724055.1 | Tepidibacter_mesophilus | Tmeso_RS13165 | 34071..37643 | 655607 | BREX-system-P-loop-protein-BrxC-[Tepidibacter-mesophilus] |
| WP_106776089.1 | Type1.BREX | [illegible] | 1197 | Bacteria>Terrabacteria_group>Bacillota>Clostridia>Eubacteriales>Peptostreptococcaceae | GCF_003012055.1 | Peptostreptococcus_russellii | UF10_RS01565 | 145981..149574 | 215200 | BREX-system-P-loop-protein-BrxC-[Peptostreptococcus-russellii] |
| WP_138342246.1 | Type1.BREX | [illegible] | 1190 | Bacteria>Terrabacteria_group>Bacillota>Clostridia>Eubacteriales>Peptostreptococcaceae | GCF_005845095.1 | Intestinibacter_bartlettii | FGQ59_RS01165 | 255162..258734 | 261299 | BREX-system-P-loop-protein-BrxC-[Intestinibacter-bartlettii] |
| WP_226924212.1 | Type1.BREX | [illegible] | 1190 | Bacteria>Terrabacteria_group>Bacillota>Clostridia>Eubacteriales>Peptostreptococcaceae | GCF_020537215.1 | Intestinibacter_bartlettii | LIP50_RS08300 | 25949..29521 | 261299 | BREX-system-P-loop-protein-BrxC-[Intestinibacter-bartlettii] |
| WP_270839456.1 | Type1.BREX | [illegible] | 1215 | Bacteria>Terrabacteria_group>Bacillota>Clostridia>Eubacteriales>Peptostreptococcaceae | GCF_027668035.1 | Peptacetobacter_hiranonis | PHA06_RS05540 | 93069..96716 | 89152 | BREX-system-P-loop-protein-BrxC-[Peptacetobacter-hiranonis] |
| WP_271847095.1 | Type1.BREX | [illegible] | 1184 | Bacteria>Terrabacteria_group>Bacillota>Clostridia>Eubacteriales>Peptostreptococcaceae | GCF_028326405.1 | Peptostreptococcus_anaerobius | PN283_RS04660 | 12553..16107 | 1261 | BREX-system-P-loop-protein-BrxC-[Peptostreptococcus-anaerobius] |
| WP_073126806.1 | Type1.BREX | [illegible] | 1197 | Bacteria>Terrabacteria_group>Bacillota>Clostridia>Eubacteriales>Peptostreptococcaceae | GCF_900129815.1 | Asaccharospora_irregularis | BUB34_RS09565 | 11025..14618 | 29359 | BREX-system-P-loop-protein-BrxC-[Asaccharospora-irregularis] |
| WP_147616385.1 | Type1.BREX | [illegible] | 1190 | Bacteria>Terrabacteria_group>Bacillota>Clostridia>Eubacteriales>Peptostreptococcaceae | GCF_902362795.1 | Intestinibacter_bartlettii | FXW45_RS03040 | 649695..653267 | 261299 | BREX-system-P-loop-protein-BrxC-[Intestinibacter-bartlettii] |
| WP_153971752.1 | Type1.BREX | [illegible] | 1187 | Bacteria>Terrabacteria_group>Bacillota>Clostridia>Eubacteriales>Peptostreptococcaceae | - | Romboutsia_ilealis | EAI30_RS08825 | 173166..176729 | 1115758 | BREX-system-P-loop-protein-BrxC-[Romboutsia-ilealis] |
| WP_251617110.1 | Type1.BREX | [illegible] | 1178 | Bacteria>Terrabacteria_group>Bacillota>Clostridia>Eubacteriales>Pumilibacteraceae | GCF_943193045.1 | Pumilibacter_muris | NFR62_RS07810 | 48561..52097 | 2941510 | BREX-system-P-loop-protein-BrxC-[Pumilibacter-muris] |
| WP_245302581.1 | Type1.BREX | [illegible] | 1198 | Bacteria>Terrabacteria_group>Bacillota>Clostridia>Eubacteriales>Symbiobacteriaceae | GCF_017874315.1 | Symbiobacterium_terraclitae | J2Z79_RS10300 | 10592..14188 | 557451 | BREX-system-P-loop-protein-BrxC-[Symbiobacterium-terraclitae] |
| TCW39537.1 | Type1.BREX | [illegible] | 1173 | Bacteria>Terrabacteria_group>Bacillota>Clostridia>Eubacteriales>Syntrophomonadaceae | GCA_004343465.1 | Thermohydrogenium_kirishiense | EDC21_10596 | 99998..103519 | 937254 | hypothetical-protein-EDC21_10596-[Thermohydrogenium-kirishiense] |
| NLB52378.1 | Type1.BREX | [illegible] | 1201 | Bacteria>Terrabacteria_group>Bacillota>Clostridia>Eubacteriales>Syntrophomonadaceae | GCA_012511795.1 | Syntrophomonadaceae_bacterium | GX808_05495 | 6370..9975 | 2093811 | BREX-system-P-loop-protein-BrxC-[Syntrophomonadaceae-bacterium] |
| NLX88392.1 | Type1.BREX | [illegible] | 1190 | Bacteria>Terrabacteria_group>Bacillota>Clostridia>Eubacteriales>Syntrophomonadaceae | GCA_012728345.1 | Syntrophomonadaceae_bacterium | GXZ09_03155 | 18986..22558 | 2093811 | BREX-system-P-loop-protein-BrxC-[Syntrophomonadaceae-bacterium] |
| HHW42067.1 | Type1.BREX | [illegible] | 520 | Bacteria>Terrabacteria_group>Bacillota>Clostridia>Eubacteriales>Syntrophomonadaceae | GCA_012840385.1 | Syntrophomonadaceae_bacterium | GXX19_13180 | 195021..196579 | 2093811 | BREX-system-P-loop-protein-BrxC-[Syntrophomonadaceae-bacterium] |
| WP_011640409.1 | Type1.BREX | [illegible] | 1199 | Bacteria>Terrabacteria_group>Bacillota>Clostridia>Eubacteriales>Syntrophomonadaceae | GCF_000014725.1 | Syntrophomonas_wolfei | SWOL_RS05025 | 1120940..1124539 | 863 | BREX-system-P-loop-protein-BrxC-[Syntrophomonas-wolfei] |
| WP_011641865.1 | Type1.BREX | [illegible] | 1185 | Bacteria>Terrabacteria_group>Bacillota>Clostridia>Eubacteriales>Syntrophomonadaceae | GCF_000014725.1 | Syntrophomonas_wolfei | SWOL_RS12915 | 2833709..2837266 | 863 | BREX-system-P-loop-protein-BrxC-[Syntrophomonas-wolfei] |
| WP_061213686.1 | Type1.BREX | [illegible] | 1199 | Bacteria>Terrabacteria_group>Bacillota>Clostridia>Eubacteriales>Syntrophomonadaceae | GCF_001570625.1 | Syntrophomonas_wolfei | AWB54_RS05890 | 82988..86587 | 863 | BREX-system-P-loop-protein-BrxC-[Syntrophomonas-wolfei] |
| WP_061215054.1 | Type1.BREX | [illegible] | 1184 | Bacteria>Terrabacteria_group>Bacillota>Clostridia>Eubacteriales>Syntrophomonadaceae | GCF_001570625.1 | Syntrophomonas_wolfei | AWB54_RS13115 | 26224..29778 | 863 | BREX-system-P-loop-protein-BrxC-[Syntrophomonas-wolfei] |
| OAD88367.1 | Type1.BREX | [illegible] | 1209 | Bacteria>Terrabacteria_group>Bacillota>Clostridia>Eubacteriales>unclassified_Eubacteriales | GCA_001641065.1 | Clostridiales_bacterium_KLE1615 | HMPREF2738_01633 | 60233..63862 | 1715004 | hypothetical-protein-HMPREF2738_01633-[Clostridiales-bacterium-KLE1615] |
| OGO82624.1 | Type1.BREX | [illegible] | 1198 | Bacteria>Terrabacteria_group>Bacillota>Clostridia>Eubacteriales>unclassified_Eubacteriales | GCA_001795385.1 | Clostridiales_bacterium_GWD2_32_19 | A2Y18_05155 | 14818..18414 | 1797679 | ATPase-[Clostridiales-bacterium-GWD2_32_19] |
| OGO91750.1 | Type1.BREX | [illegible] | 1191 | Bacteria>Terrabacteria_group>Bacillota>Clostridia>Eubacteriales>unclassified_Eubacteriales | GCA_001797275.1 | Clostridiales_bacterium_GWF2_38_85 | A2Y17_13400 | 59822..63397 | 1797683 | hypothetical-protein-A2Y17_13400-[Clostridiales-bacterium-GWF2_38_85] |
| OKZ97591.1 | Type1.BREX | [illegible] | 1204 | Bacteria>Terrabacteria_group>Bacillota>Clostridia>Eubacteriales>unclassified_Eubacteriales | GCA_001915795.1 | Clostridiales_bacterium_42_27 | BHV90_19800 | 17930..21544 | 1897045 | hypothetical-protein-BHV90_19800-[Clostridiales-bacterium-42_27] |
| OKZ58849.1 | Type1.BREX | [illegible] | 1181 | Bacteria>Terrabacteria_group>Bacillota>Clostridia>Eubacteriales>unclassified_Eubacteriales | GCA_001915845.1 | Clostridiales_bacterium_45_37 | BHV92_02900 | 11586..15131 | 1897047 | ATPase-[Clostridiales-bacterium-45_37] |
| PWM66029.1 | Type1.BREX | [illegible] | 1180 | Bacteria>Terrabacteria_group>Bacillota>Clostridia>Eubacteriales>unclassified_Eubacteriales | GCA_003343885.1 | Clostridiales_bacterium | DBX61_08855 | 48363..51905 | 1898207 | BREX-system-P-loop-protein-BrxC-[Clostridiales-bacterium] |
| HAQ40519.1 | Type1.BREX | [illegible] | 1183 | Bacteria>Terrabacteria_group>Bacillota>Clostridia>Eubacteriales>unclassified_Eubacteriales | GCA_003457065.1 | Clostridiales_bacterium | DCM73_06580 | 77939..81490 | 1898207 | BREX-system-P-loop-protein-BrxC-[Clostridiales-bacterium] |
| RJW84978.1 | Type1.BREX | [illegible] | 1183 | Bacteria>Terrabacteria_group>Bacillota>Clostridia>Eubacteriales>unclassified_Eubacteriales | GCA_003604275.1 | Clostridiales_bacterium_AF36-10 | DWZ86_13085 | 1260..4811 | 2292883 | BREX-system-P-loop-protein-BrxC-[Clostridiales-bacterium-AF36-10] |
| MCL1808846.1 | Type1.BREX | [illegible] | 1193 | Bacteria>Terrabacteria_group>Bacillota>Clostridia>Eubacteriales>unclassified_Eubacteriales | GCA_009782285.1 | Clostridiales_bacterium | FWG42_03645 | 31775..35356 | 1898207 | BREX-system-P-loop-protein-BrxC-[Clostridiales-bacterium] |
| NBI63881.1 | Type1.BREX | [illegible] | 1226 | Bacteria>Terrabacteria_group>Bacillota>Clostridia>Eubacteriales>unclassified_Eubacteriales | GCA_009911365.1 | Clostridiales_bacterium | D3Z38_12635 | 51240..54920 | 1898207 | BREX-system-P-loop-protein-BrxC-[Clostridiales-bacterium] |
| NLC74531.1 | Type1.BREX | [illegible] | 1161 | Bacteria>Terrabacteria_group>Bacillota>Clostridia>Eubacteriales>unclassified_Eubacteriales | GCA_012512995.1 | Clostridiales_bacterium | GX685_04750 | 28468..31953 | 1898207 | BREX-system-P-loop-protein-BrxC-[Clostridiales-bacterium] |
| NLK72540.1 | Type1.BREX | [illegible] | 1375 | Bacteria>Terrabacteria_group>Bacillota>Clostridia>Eubacteriales>unclassified_Eubacteriales | GCA_012519645.1 | Clostridiales_bacterium | GX285_05945 | 4916..9043 | 1898207 | hypothetical-protein-[Clostridiales-bacterium] |
| NLO81519.1 | Type1.BREX | [illegible] | 1186 | Bacteria>Terrabacteria_group>Bacillota>Clostridia>Eubacteriales>unclassified_Eubacteriales | GCA_012523455.1 | Clostridiales_bacterium | GX094_00420 | 4845..8405 | 1898207 | BREX-system-P-loop-protein-BrxC-[Clostridiales-bacterium] |
| NLZ36694.1 | Type1.BREX | [illegible] | 1174 | Bacteria>Terrabacteria_group>Bacillota>Clostridia>Eubacteriales>unclassified_Eubacteriales | GCA_012800735.1 | Clostridiales_bacterium | GX897_04365 | 3762..7286 | 1898207 | BREX-system-P-loop-protein-BrxC-[Clostridiales-bacterium] |
| MBD5099706.1 | Type1.BREX | [illegible] | 1178 | Bacteria>Terrabacteria_group>Bacillota>Clostridia>Eubacteriales>unclassified_Eubacteriales | GCA_014799645.1 | Clostridiales_bacterium | HDT29_00135 | 31691..35227 | 1898207 | BREX-system-P-loop-protein-BrxC-[Clostridiales-bacterium] |
| MBD9283724.1 | Type1.BREX | [illegible] | 1202 | Bacteria>Terrabacteria_group>Bacillota>Clostridia>Eubacteriales>unclassified_Eubacteriales | GCA_014848695.1 | Clostridiales_bacterium | EGR96_02215 | 1987..5595 | 1898207 | BREX-system-P-loop-protein-BrxC-[Clostridiales-bacterium] |
| MBE6030170.1 | Type1.BREX | [illegible] | 1212 | Bacteria>Terrabacteria_group>Bacillota>Clostridia>Eubacteriales>unclassified_Eubacteriales | GCA_015058155.1 | Clostridiales_bacterium | E7225_01045 | 104723..108361 | 1898207 | BREX-system-P-loop-protein-BrxC-[Clostridiales-bacterium] |
| MBS5027730.1 | Type1.BREX | [illegible] | 1182 | Bacteria>Terrabacteria_group>Bacillota>Clostridia>Eubacteriales>unclassified_Eubacteriales | GCA_018365035.1 | Clostridiales_bacterium | KHZ67_05250 | 47682..51230 | 1898207 | BREX-system-P-loop-protein-BrxC-[Clostridiales-bacterium] |
| MBS5052649.1 | Type1.BREX | [illegible] | 1182 | Bacteria>Terrabacteria_group>Bacillota>Clostridia>Eubacteriales>unclassified_Eubacteriales | GCA_018365235.1 | Clostridiales_bacterium | KHZ49_10850 | 8851..12399 | 1898207 | BREX-system-P-loop-protein-BrxC-[Clostridiales-bacterium] |
| MBS5198011.1 | Type1.BREX | [illegible] | 1199 | Bacteria>Terrabacteria_group>Bacillota>Clostridia>Eubacteriales>unclassified_Eubacteriales | GCA_018366395.1 | Clostridiales_bacterium | KHY93_12815 | 21497..25096 | 1898207 | BREX-system-P-loop-protein-BrxC-[Clostridiales-bacterium] |
| MBS5313230.1 | Type1.BREX | [illegible] | 1205 | Bacteria>Terrabacteria_group>Bacillota>Clostridia>Eubacteriales>unclassified_Eubacteriales | GCA_018367215.1 | Clostridiales_bacterium | KHY53_10185 | 5342..8959 | 1898207 | BREX-system-P-loop-protein-BrxC-[Clostridiales-bacterium] |
| MCC8075264.1 | Type1.BREX | [illegible] | 1181 | Bacteria>Terrabacteria_group>Bacillota>Clostridia>Eubacteriales>unclassified_Eubacteriales | GCA_020859945.1 | Clostridiales_bacterium | LIO95_04860 | 22835..26380 | 1898207 | BREX-system-P-loop-protein-BrxC-[Clostridiales-bacterium] |
| MCC8066947.1 | Type1.BREX | [illegible] | 1187 | Bacteria>Terrabacteria_group>Bacillota>Clostridia>Eubacteriales>unclassified_Eubacteriales | GCA_020859965.1 | Clostridiales_bacterium | LIO94_07585 | 9778..13341 | 1898207 | BREX-system-P-loop-protein-BrxC-[Clostridiales-bacterium] |
| MCD4713411.1 | Type1.BREX | [illegible] | 980 | Bacteria>Terrabacteria_group>Bacillota>Clostridia>Eubacteriales>unclassified_Eubacteriales | GCA_021107455.1 | Clostridiales_bacterium | K8R73_09035 | 9509..12451 | 1898207 | BREX-system-P-loop-protein-BrxC-[Clostridiales-bacterium] |
| MCD5414444.1 | Type1.BREX | [illegible] | 1202 | Bacteria>Terrabacteria_group>Bacillota>Clostridia>Eubacteriales>unclassified_Eubacteriales | GCA_021129775.1 | Clostridiales_bacterium | LR001_05510 | 10994..14602 | 1898207 | BREX-system-P-loop-protein-BrxC-[Clostridiales-bacterium] |
| MCD5411055.1 | Type1.BREX | [illegible] | 1213 | Bacteria>Terrabacteria_group>Bacillota>Clostridia>Eubacteriales>unclassified_Eubacteriales | GCA_021129785.1 | Clostridiales_bacterium | LRZ93_05310 | 22732..26373 | 1898207 | BREX-system-P-loop-protein-BrxC-[Clostridiales-bacterium] |
| MCD7920553.1 | Type1.BREX | [illegible] | 1211 | Bacteria>Terrabacteria_group>Bacillota>Clostridia>Eubacteriales>unclassified_Eubacteriales | GCA_021201475.1 | Clostridiales_bacterium | LUG45_10925 | 23806..27441 | 1898207 | BREX-system-P-loop-protein-BrxC-[Clostridiales-bacterium] |
| MCD8018067.1 | Type1.BREX | [illegible] | 1215 | Bacteria>Terrabacteria_group>Bacillota>Clostridia>Eubacteriales>unclassified_Eubacteriales | GCA_021202175.1 | Clostridiales_bacterium | LUF92_00310 | 7758..11405 | 1898207 | BREX-system-P-loop-protein-BrxC-[Clostridiales-bacterium] |
| MCD8148190.1 | Type1.BREX | [illegible] | 773 | Bacteria>Terrabacteria_group>Bacillota>Clostridia>Eubacteriales>unclassified_Eubacteriales | GCA_021203005.1 | Clostridiales_bacterium | LUE92_01205 | 1..2322 | 1898207 | BREX-system-P-loop-protein-BrxC-[Clostridiales-bacterium] |
| MCD8221484.1 | Type1.BREX | [illegible] | 1183 | Bacteria>Terrabacteria_group>Bacillota>Clostridia>Eubacteriales>unclassified_Eubacteriales | GCA_021204025.1 | Clostridiales_bacterium | LUD07_04705 | 24313..27864 | 1898207 | BREX-system-P-loop-protein-BrxC-[Clostridiales-bacterium] |
| MCD8370861.1 | Type1.BREX | [illegible] | 1183 | Bacteria>Terrabacteria_group>Bacillota>Clostridia>Eubacteriales>unclassified_Eubacteriales | GCA_021205425.1 | Clostridiales_bacterium | LUC94_11190 | 2149..5700 | 1898207 | BREX-system-P-loop-protein-BrxC-[Clostridiales-bacterium] |
| MCH5164332.1 | Type1.BREX | [illegible] | 1174 | Bacteria>Terrabacteria_group>Bacillota>Clostridia>Eubacteriales>unclassified_Eubacteriales | GCA_022508465.1 | Clostridiales_bacterium | J1F36_04875 | 11791..15315 | 1898207 | BREX-system-P-loop-protein-BrxC-[Clostridiales-bacterium] |
| MCI2147015.1 | Type1.BREX | [illegible] | 1257 | Bacteria>Terrabacteria_group>Bacillota>Clostridia>Eubacteriales>unclassified_Eubacteriales | GCA_022654135.1 | Clostridiales_bacterium | LKK18_03325 | 16623..20396 | 1898207 | BREX-system-P-loop-protein-BrxC-[Clostridiales-bacterium] |
| MCI5810179.1 | Type1.BREX | [illegible] | 391 | Bacteria>Terrabacteria_group>Bacillota>Clostridia>Eubacteriales>unclassified_Eubacteriales | GCA_022769895.1 | Clostridiales_bacterium | MR018_01245 | 116842..118014 | 1898207 | BREX-system-P-loop-protein-BrxC-[Clostridiales-bacterium] |
| MCI6290154.1 | Type1.BREX | [illegible] | 1178 | Bacteria>Terrabacteria_group>Bacillota>Clostridia>Eubacteriales>unclassified_Eubacteriales | GCA_022773115.1 | Clostridiales_bacterium | MR620_02485 | 557..4093 | 1898207 | BREX-system-P-loop-protein-BrxC-[Clostridiales-bacterium] |
| MCI6325701.1 | Type1.BREX | [illegible] | 852 | Bacteria>Terrabacteria_group>Bacillota>Clostridia>Eubacteriales>unclassified_Eubacteriales | GCA_022777505.1 | Clostridiales_bacterium | MR790_03620 | 102230..104785 | 1898207 | BREX-system-P-loop-protein-BrxC-[Clostridiales-bacterium] |
| MCR4615485.1 | Type1.BREX | [illegible] | 1161 | Bacteria>Terrabacteria_group>Bacillota>Clostridia>Eubacteriales>unclassified_Eubacteriales | GCA_024684435.1 | Clostridiales_bacterium | K5756_04990 | 82244..85729 | 1898207 | BREX-system-P-loop-protein-BrxC-[Clostridiales-bacterium] |
| WP_129723506.1 | Type1.BREX | [illegible] | 1381 | Bacteria>Terrabacteria_group>Bacillota>Clostridia>Eubacteriales>Xylanivirgaceae | GCF_004138105.1 | Xylanivirga_thermophila | EJN67_RS06315 | 20747..24892 | 2496273 | hypothetical-protein-[Xylanivirga-thermophila] |
| WP_129724329.1 | Type1.BREX | [illegible] | 1183 | Bacteria>Terrabacteria_group>Bacillota>Clostridia>Eubacteriales>Xylanivirgaceae | GCF_004138105.1 | Xylanivirga_thermophila | EJN67_RS10845 | 9801..13352 | 2496273 | BREX-system-P-loop-protein-BrxC-[Xylanivirga-thermophila] |
| WP_099045440.1 | Type1.BREX | [illegible] | 1217 | Bacteria>Terrabacteria_group>Bacillota>Clostridia | GCF_000174195.1 | Eubacteriales | ROSEINA2194_RS00605 | 10712..14365 | 186802 | MULTISPECIES:-BREX-system-P-loop-protein-BrxC-[Eubacteriales] |
| WP_117965654.1 | Type1.BREX | [illegible] | 1217 | Bacteria>Terrabacteria_group>Bacillota>Clostridia | GCF_003461485.1 | Eubacteriales | DWW66_RS00315 | 65325..68978 | 186802 | MULTISPECIES:-BREX-system-P-loop-protein-BrxC-[Eubacteriales] |
| WP_173887209.1 | Type1.BREX | [illegible] | 1205 | Bacteria>Terrabacteria_group>Bacillota>Clostridia | GCF_013301225.1 | Eubacteriales | GPK60_RS01920 | 35129..38746 | 186802 | MULTISPECIES:-BREX-system-P-loop-protein-BrxC-[Eubacteriales] |
| WP_173717967.1 | Type1.BREX | [illegible] | 1179 | Bacteria>Terrabacteria_group>Bacillota>Clostridia | GCF_013302415.1 | Eubacteriales | G5A19_RS02670 | 21026..24565 | 186802 | MULTISPECIES:-BREX-system-P-loop-protein-BrxC-[Eubacteriales] |
| WP_118373617.1 | Type1.BREX | [illegible] | 1196 | Bacteria>Terrabacteria_group>Bacillota>Clostridia | GCF_014337155.1 | Eubacteriales | H9Q76_RS05865 | 1235551..1239141 | 186802 | MULTISPECIES:-BREX-system-P-loop-protein-BrxC-[Eubacteriales] |
| WP_195379248.1 | Type1.BREX | [illegible] | 1182 | Bacteria>Terrabacteria_group>Bacillota>Clostridia | GCF_015551645.1 | Eubacteriales | I2D69_RS01030 | 22703..26251 | 186802 | MULTISPECIES:-BREX-system-P-loop-protein-BrxC-[Eubacteriales] |
| WP_117886551.1 | Type1.BREX | [illegible] | 1205 | Bacteria>Terrabacteria_group>Bacillota>Clostridia | GCF_027670865.1 | Eubacteriales | DXA20_RS00560 | 123704..127321 | 186802 | MULTISPECIES:-BREX-system-P-loop-protein-BrxC-[Eubacteriales] |
| WP_005488038.1 | Type1.BREX | [illegible] | 1306 | Bacteria>Terrabacteria_group>Bacillota>Clostridia>Halanaerobiales>Halanaerobiaceae | GCF_000350165.1 | Halanaerobium_saccharolyticum | HSACCH_RS04005 | 779901..783821 | 43595 | DUF6079-family-protein-[Halanaerobium-saccharolyticum] |
| WP_110301119.1 | Type1.BREX | [illegible] | 1304 | Bacteria>Terrabacteria_group>Bacillota>Clostridia>Halanaerobiales>Halanaerobiaceae | GCF_003201135.1 | Halanaerobium_congolense | C8C78_RS09810 | 20697..24611 | 54121 | DUF6079-family-protein-[Halanaerobium-congolense] |
| WP_114458231.1 | Type1.BREX | [illegible] | 1191 | Bacteria>Terrabacteria_group>Bacillota>Clostridia>Halanaerobiales>Halanaerobiaceae | GCF_003337245.1 | Halanaerobium_sp._MA284_MarDTE_T2 | DFR78_RS07540 | 27891..31466 | 2183913 | BREX-system-P-loop-protein-BrxC-[Halanaerobium-sp.-MA284_MarDTE_T2] |
| WP_133529650.1 | Type1.BREX | [illegible] | 1191 | Bacteria>Terrabacteria_group>Bacillota>Clostridia>Halanaerobiales>Halanaerobiaceae | GCF_004362725.1 | Halanaerobium_saccharolyticum | C7957_RS01790 | 171598..175173 | 43595 | BREX-system-P-loop-protein-BrxC-[Halanaerobium-saccharolyticum] |
| WP_134059654.1 | Type1.BREX | [illegible] | 1304 | Bacteria>Terrabacteria_group>Bacillota>Clostridia>Halanaerobiales>Halanaerobiaceae | GCF_004366375.1 | Halanaerobium_congolense | C7954_RS09050 | 8410..12324 | 54121 | DUF6079-family-protein-[Halanaerobium-congolense] |
| WP_089716529.1 | Type1.BREX | [illegible] | 1190 | Bacteria>Terrabacteria_group>Bacillota>Clostridia>Halanaerobiales>Halanaerobiaceae | GCF_900100055.1 | Halanaerobium_congolense | BMY48_RS04305 | 58895..62467 | 54121 | BREX-system-P-loop-protein-BrxC-[Halanaerobium-congolense] |
| WP_149796815.1 | Type1.BREX | [illegible] | 1304 | Bacteria>Terrabacteria_group>Bacillota>Clostridia>Halanaerobiales>Halanaerobiaceae | GCF_900102605.1 | Halanaerobium_congolense | F3G65_RS08880 | 2073..5987 | 54121 | DUF6079-family-protein-[Halanaerobium-congolense] |
| WP_073159284.1 | Type1.BREX | [illegible] | 1304 | Bacteria>Terrabacteria_group>Bacillota>Clostridia>Halanaerobiales>Halanaerobiaceae | GCF_900143045.1 | Halanaerobium_congolense | BUC14_RS07890 | 22601..26515 | 54121 | DUF6079-family-protein-[Halanaerobium-congolense] |
| WP_127015284.1 | Type1.BREX | [illegible] | 1182 | Bacteria>Terrabacteria_group>Bacillota>Clostridia>Halanaerobiales>Halanaerobiales_incertae_sedis | GCF_003991135.1 | Anoxybacter_fermentans | BBF96_RS00140 | 30482..34030 | 1323375 | BREX-system-P-loop-protein-BrxC-[Anoxybacter-fermentans] |
| WP_015326777.1 | Type1.BREX | [illegible] | 1197 | Bacteria>Terrabacteria_group>Bacillota>Clostridia>Halanaerobiales>Halobacteroidaceae | GCF_000328625.1 | Halobacteroides_halobius | HALHA_RS05395 | 1087361..1090954 | 42422 | BREX-system-P-loop-protein-BrxC-[Halobacteroides-halobius] |
| WP_018248942.1 | Type1.BREX | [illegible] | 1197 | Bacteria>Terrabacteria_group>Bacillota>Clostridia>Halanaerobiales>Halobacteroidaceae | GCF_000379025.1 | Orenia_marismortui | OREMA_RS0108995 | 158007..161600 | 46469 | BREX-system-P-loop-protein-BrxC-[Orenia-marismortui] |
| WP_248626875.1 | Type1.BREX | [illegible] | 1201 | Bacteria>Terrabacteria_group>Bacillota>Clostridia>Halanaerobiales>Halobacteroidaceae | GCF_023227745.1 | Natroniella_acetigena | MWH25_RS06690 | 32057..35662 | 52004 | BREX-system-P-loop-protein-BrxC-[Natroniella-acetigena] |
| WP_089751015.1 | Type1.BREX | [illegible] | 1202 | Bacteria>Terrabacteria_group>Bacillota>Clostridia>Halanaerobiales>Halobacteroidaceae | GCF_900101375.1 | unclassified_Candidatus_Frackibacter | BM077_RS12865 | 2648..6256 | 2648818 | MULTISPECIES:-BREX-system-P-loop-protein-BrxC-[unclassified-Candidatus-Frackibacter] |
| WP_097018851.1 | Type1.BREX | [illegible] | 1197 | Bacteria>Terrabacteria_group>Bacillota>Clostridia>Halanaerobiales>Halobacteroidaceae | GCF_900215265.1 | Orenia_metallireducens | BX659_RS14840 | 51636..55229 | 1413210 | BREX-system-P-loop-protein-BrxC-[Orenia-metallireducens] |
| MCK4258501.1 | Type1.BREX | [illegible] | 1185 | Bacteria>Terrabacteria_group>Bacillota>Clostridia>Halanaerobiales>unclassified_Halanaerobiales | GCA_023128035.1 | Halanaerobiales_bacterium | KAX49_05955 | 28461..32018 | 2094029 | BREX-system-P-loop-protein-BrxC-[Halanaerobiales-bacterium] |
| QSQ09315.1 | Type1.BREX | [illegible] | 1187 | Bacteria>Terrabacteria_group>Bacillota>Clostridia>Koleobacterales>Koleobacteraceae | GCA_017301615.1 | Koleobacter_methoxysyntrophicus | H0A61_01676 | 1748127..1751690 | 2751313 | hypothetical-protein-H0A61_01676-[Koleobacter-methoxysyntrophicus] |
| WP_206706674.1 | Type1.BREX | [illegible] | 1191 | Bacteria>Terrabacteria_group>Bacillota>Clostridia>Koleobacterales>Koleobacteraceae | GCA_017301615.1 | Koleobacter_methoxysyntrophicus | H0A61_RS08405 | 1748115..1751690 | 2751313 | BREX-system-P-loop-protein-BrxC-[Koleobacter-methoxysyntrophicus] |
| RJX18832.1 | Type1.BREX | [illegible] | 1147 | Bacteria>Terrabacteria_group>Bacillota>Clostridia>Thermoanaerobacterales>Thermoanaerobacteraceae | GCA_003604985.1 | Ammonifex_sp. | C4570_06155 | 13420..16863 | 1872493 | MAG:-BREX-system-P-loop-protein-BrxC-[Ammonifex-sp.] |
| HHV16015.1 | Type1.BREX | [illegible] | 1184 | Bacteria>Terrabacteria_group>Bacillota>Clostridia>Thermoanaerobacterales>Thermoanaerobacteraceae | GCA_012839665.1 | Gelria_sp. | GXX58_05515 | 1705..5259 | 2699831 | BREX-system-P-loop-protein-BrxC-[Gelria-sp.] |
| WP_011345517.1 | Type1.BREX | [illegible] | 1187 | Bacteria>Terrabacteria_group>Bacillota>Clostridia>Thermoanaerobacterales>Thermoanaerobacteraceae | GCF_000012865.1 | Carboxydothermus_hydrogenoformans | CHY_RS12235 | 2357097..2360660 | 129958 | BREX-system-P-loop-protein-BrxC-[Carboxydothermus-hydrogenoformans] |
| WP_015051267.1 | Type1.BREX | [illegible] | 1148 | Bacteria>Terrabacteria_group>Bacillota>Clostridia>Thermoanaerobacterales>Thermoanaerobacteraceae | GCF_000305935.1 | Thermacetogenium_phaeum | TPH_RS10940 | 2236275..2239721 | 85874 | BREX-system-P-loop-protein-BrxC-[Thermacetogenium-phaeum] |
| WP_069591144.1 | Type1.BREX | [illegible] | 1148 | Bacteria>Terrabacteria_group>Bacillota>Clostridia>Thermoanaerobacterales>Thermoanaerobacteraceae | GCF_001729945.1 | Moorella_thermoacetica | Maut_RS13535 | 2689241..2692687 | 1525 | BREX-system-P-loop-protein-BrxC-[Moorella-thermoacetica] |
| WP_071522643.1 | Type1.BREX | [illegible] | 1148 | Bacteria>Terrabacteria_group>Bacillota>Clostridia>Thermoanaerobacterales>Thermoanaerobacteraceae | GCF_001874085.1 | Moorella_thermoacetica | MOOTH_RS12485 | 21888..25334 | 1525 | BREX-system-P-loop-protein-BrxC-[Moorella-thermoacetica] |
| WP_258359412.1 | Type1.BREX | [illegible] | 1148 | Bacteria>Terrabacteria_group>Bacillota>Clostridia>Thermoanaerobacterales>Thermoanaerobacteraceae | GCF_024733845.1 | Moorella_sp._SLA38 | NWF31_RS05005 | 117585..121031 | 2972948 | BREX-system-P-loop-protein-BrxC-[Moorella-sp.-SLA38] |
| WP_011393725.1 | Type1.BREX | [illegible] | 1148 | Bacteria>Terrabacteria_group>Bacillota>Clostridia>Thermoanaerobacterales>Thermoanaerobacteraceae | GCF_027925125.1 | Moorella_thermoacetica | QJ070_RS08400 | 457989..461435 | 1525 | BREX-system-P-loop-protein-BrxC-[Moorella-thermoacetica] |
| WP_013297042.1 | Type1.BREX | [illegible] | 1186 | Bacteria>Terrabacteria_group>Bacillota>Clostridia>Thermoanaerobacterales>Thermoanaerobacterales_Family_III._Incertae_Sedis | GCF_000145615.1 | Thermoanaerobacterium_thermosaccharolyticum | TTHE_RS02540 | 484510..488070 | 1517 | BREX-system-P-loop-protein-BrxC-[Thermoanaerobacterium-thermosaccharolyticum] |
| WP_094396730.1 | Type1.BREX | [illegible] | 1173 | Bacteria>Terrabacteria_group>Bacillota>Clostridia>Thermoanaerobacterales>Thermoanaerobacterales_Family_III._Incertae_Sedis | GCF_002250075.1 | Thermoanaerobacterium_thermosaccharolyticum | Thert_RS01160 | 231193..234714 | 1517 | BREX-system-P-loop-protein-BrxC-[Thermoanaerobacterium-thermosaccharolyticum] |
| WP_094397497.1 | Type1.BREX | [illegible] | 1381 | Bacteria>Terrabacteria_group>Bacillota>Clostridia>Thermoanaerobacterales>Thermoanaerobacterales_Family_III._Incertae_Sedis | GCF_002250075.1 | Thermoanaerobacterium_thermosaccharolyticum | Thert_RS08605 | 1721072..1725217 | 1517 | hypothetical-protein-[Thermoanaerobacterium-thermosaccharolyticum] |
| WP_099252528.1 | Type1.BREX | [illegible] | 1173 | Bacteria>Terrabacteria_group>Bacillota>Clostridia>Thermoanaerobacterales>Thermoanaerobacterales_Family_III._Incertae_Sedis | GCF_002701205.1 | Thermoanaerobacterium_thermosaccharolyticum | BFT35_RS01005 | 197977..201498 | 1517 | BREX-system-P-loop-protein-BrxC-[Thermoanaerobacterium-thermosaccharolyticum] |
| WP_132775540.1 | Type1.BREX | [illegible] | 1186 | Bacteria>Terrabacteria_group>Bacillota>Clostridia>Thermoanaerobacterales>Thermoanaerobacterales_Family_III._Incertae_Sedis | GCF_008632855.1 | Thermoanaerobacterium_thermosaccharolyticum | F1655_RS11615 | 3021..6581 | 1517 | BREX-system-P-loop-protein-BrxC-[Thermoanaerobacterium-thermosaccharolyticum] |
| WP_192404070.1 | Type1.BREX | [illegible] | 1186 | Bacteria>Terrabacteria_group>Bacillota>Clostridia>Thermoanaerobacterales>Thermoanaerobacterales_Family_III._Incertae_Sedis | GCF_014856575.1 | Thermoanaerobacterium_thermosaccharolyticum | F3K45_RS12105 | 13117..16677 | 1517 | BREX-system-P-loop-protein-BrxC-[Thermoanaerobacterium-thermosaccharolyticum] |
| WP_192404334.1 | Type1.BREX | [illegible] | 1173 | Bacteria>Terrabacteria_group>Bacillota>Clostridia>Thermoanaerobacterales>Thermoanaerobacterales_Family_III._Incertae_Sedis | GCF_014856575.1 | Thermoanaerobacterium_thermosaccharolyticum | F3K45_RS13085 | 29192..32713 | 1517 | BREX-system-P-loop-protein-BrxC-[Thermoanaerobacterium-thermosaccharolyticum] |
| WP_192453407.1 | Type1.BREX | [illegible] | 1173 | Bacteria>Terrabacteria_group>Bacillota>Clostridia>Thermoanaerobacterales>Thermoanaerobacterales_Family_III._Incertae_Sedis | GCF_014856605.1 | Thermoanaerobacterium_thermosaccharolyticum | F3K46_12200 | 29192..32713 | 1517 | BREX-system-P-loop-protein-BrxC-[Thermoanaerobacterium-thermosaccharolyticum] |
| WP_209454287.1 | Type1.BREX | [illegible] | 1186 | Bacteria>Terrabacteria_group>Bacillota>Clostridia>Thermoanaerobacterales>Thermoanaerobacterales_Family_III._Incertae_Sedis | GCF_017874545.1 | Thermoanaerobacterium_butyriciformans | J2Z80_RS10135 | 14658..18218 | 1702242 | BREX-system-P-loop-protein-BrxC-[Thermoanaerobacterium-butyriciformans] |
| WP_253666097.1 | Type1.BREX | [illegible] | 1173 | Bacteria>Terrabacteria_group>Bacillota>Clostridia>Thermoanaerobacterales>Thermoanaerobacterales_Family_III._Incertae_Sedis | GCF_024171655.1 | Thermoanaerobacterium_thermosaccharolyticum | BD412_RS01225 | 67583..71104 | 1517 | BREX-system-P-loop-protein-BrxC-[Thermoanaerobacterium-thermosaccharolyticum] |
| WP_271628813.1 | Type1.BREX | [illegible] | 1186 | Bacteria>Terrabacteria_group>Bacillota>Clostridia>Thermoanaerobacterales>Thermoanaerobacterales_Family_III._Incertae_Sedis | GCF_028198085.1 | Caldicellulosiruptor_sp._DIB_104C | PGC14_RS09960 | 2010036..2013596 | 3019889 | BREX-system-P-loop-protein-BrxC-[Caldicellulosiruptor-sp.-DIB-104C] |
| WP_096230883.1 | Type1.BREX | [illegible] | 1173 | Bacteria>Terrabacteria_group>Bacillota>Clostridia>Thermoanaerobacterales>Thermoanaerobacterales_Family_III._Incertae_Sedis | GCF_900205865.1 | Thermoanaerobacterium_sp._RBIITD | CPG45_RS04855 | 1042474..1045995 | 1550240 | BREX-system-P-loop-protein-BrxC-[Thermoanaerobacterium-sp.-RBIITD] |
| HHW04166.1 | Type1.BREX | [illegible] | 1186 | Bacteria>Terrabacteria_group>Bacillota>Clostridia>Thermoanaerobacterales>unclassified_Thermoanaerobacterales | GCA_012840135.1 | Thermoanaerobacterales_bacterium | GXX35_15425 | 56222..59782 | 2304039 | BREX-system-P-loop-protein-BrxC-[Thermoanaerobacterales-bacterium] |
| WP_013777644.1 | Type1.BREX | [illegible] | 1188 | Bacteria>Terrabacteria_group>Bacillota>Clostridia>Thermosediminibacterales>Tepidanaerobacteraceae | GCF_000213235.1 | Tepidanaerobacter_acetatoxydans | TEPRE1_RS02650 | 565756..569322 | 499229 | BREX-system-P-loop-protein-BrxC-[Tepidanaerobacter-acetatoxydans] |
| WP_059032619.1 | Type1.BREX | [illegible] | 1187 | Bacteria>Terrabacteria_group>Bacillota>Clostridia>Thermosediminibacterales>Tepidanaerobacteraceae | GCF_027925185.1 | Tepidanaerobacter_syntrophicus | TSYNT_RS06145 | 249546..253109 | 224999 | BREX-system-P-loop-protein-BrxC-[Tepidanaerobacter-syntrophicus] |
| NCB93717.1 | Type1.BREX | [illegible] | 1211 | Bacteria>Terrabacteria_group>Bacillota>Clostridia>unclassified_Clostridia | GCA_009929525.1 | Clostridia_bacterium | EOM40_14340 | 4870..8505 | 2044939 | BREX-system-P-loop-protein-BrxC-[Clostridia-bacterium] |
| NCC01269.1 | Type1.BREX | [illegible] | 1202 | Bacteria>Terrabacteria_group>Bacillota>Clostridia>unclassified_Clostridia | GCA_009929615.1 | Clostridia_bacterium | EOM34_11440 | 9159..12767 | 2044939 | BREX-system-P-loop-protein-BrxC-[Clostridia-bacterium] |

| Protein-ID | System | Gene-Neighbourhoods | Length | Taxonomy | Assembly-ID | Strain | Locus-Tag | Genomic-coordinates | TaxID | Fasta-define |
|---|---|---|---|---|---|---|---|---|---|---|
| NCC15938.1 | Type1.BREX | <-SIGMA+OB-Fold+AAA-ATPase+Lon-Protease-BrxL<-PglZ<-?<-N6-METHYLASE<-Four-helical-protein<-wHTH+ZnR<-DUF6079*<-BrxB-BrxF-iSTAND<-BrxA-DNA-binding<-HTH_26<-?<-TraJ-RHH<-?<-LPD32+LPD36+N6_Mtase+SF1-SF2-Helicase | 1207 | Bacteria>Terrabacteria_group>Bacillota>Clostridia>unclassified_Clostridia | GCA_009929735.1 | Clostridia_bacterium | EOM28_06255 | 34440..38063 | 2044939 | BREX-system-P-loop-protein-BrxC-[Clostridia-bacterium] |
| QRN85655.1 | Type1.BREX | APATPase+FtsK+VirB4->?->SMS-KaiC->HTH_36->HTH_26->BrxA-DNA-binding->BrxB-BrxF-iSTAND->DUF6079*->?->N6-METHYLASE->PglZ->SIGMA+OB-Fold+AAA-ATPase+Lon-Protease-BrxL->?->?->OLD-ABC+TOPRIM->SF1-SF2-Helicase-><-MPTase | 1213 | Bacteria>Terrabacteria_group>Bacillota>Clostridia>unclassified_Clostridia | GCA_016887505.1 | Clostridia_bacterium | JR334_12015 | 2538815..2542456 | 2044939 | BREX-system-P-loop-protein-BrxC-[Clostridia-bacterium] |
| MBQ5562930.1 | Type1.BREX | DEOR-HTH->?->?->BrxA-DNA-binding->BrxB-BrxF-iSTAND->DUF6079-p*-><-RADICAL-SAM | 670 | Bacteria>Terrabacteria_group>Bacillota>Clostridia>unclassified_Clostridia | GCA_017386515.1 | Clostridia_bacterium | IIT39_06055 | 5344..7356 | 2044939 | BREX-system-P-loop-protein-BrxC-[Clostridia-bacterium] |
| MBQ3415902.1 | Type1.BREX | <-SIGMA+OB-Fold+AAA-ATPase+Lon-Protease-BrxL<-PglZ<-?<-PIN<-OLD-ABC<-N6-METHYLASE<-?<-DUF6079*<-BrxB-BrxF-iSTAND<-BrxA-DNA-binding<-?<-?<-RHH_8 | 1216 | Bacteria>Terrabacteria_group>Bacillota>Clostridia>unclassified_Clostridia | GCA_017404025.1 | Clostridia_bacterium | IJH39_11340 | 11283..14933 | 2044939 | BREX-system-P-loop-protein-BrxC-[Clostridia-bacterium] |
| MBQ6380318.1 | Type1.BREX | N6-METHYLASE->TRD->SbcC->?->STYKinase-like-><-?<-SAM-METHYLASE\|\|BrxA-DNA-binding->BrxB-BrxF-iSTAND->DUF6079*->CR-ATPase5+REase->N6-METHYLASE-p-> | 1174 | Bacteria>Terrabacteria_group>Bacillota>Clostridia>unclassified_Clostridia | GCA_017430965.1 | Clostridia_bacterium | IJJ41_01780 | 27986..31510 | 2044939 | BREX-system-P-loop-protein-BrxC-[Clostridia-bacterium] |
| MBQ9210206.1 | Type1.BREX | XIS-HTH->XIS-HTH->Arm-DNA-bind_4+Phage_int_SAM_3+Phage_integrase-><-?<-Calcineurin<-SIGMA+OB-Fold+AAA-ATPase+Lon-Protease-BrxL<-N6-METHYLASE<-PglZ<-N6-METHYLASE<-DUF6079*<-BrxB-BrxF-iSTAND<-BrxA-DNA-binding<-HTH_26<-Zn_Tnp_IS1+DDE_Tnp_IS1595<-tRNA<-?<-HAD\|\|SIGMA-HTH-> | 1195 | Bacteria>Terrabacteria_group>Bacillota>Clostridia>unclassified_Clostridia | GCA_017464305.1 | Clostridia_bacterium | IJI53_00755 | 191732..195319 | 2044939 | BREX-system-P-loop-protein-BrxC-[Clostridia-bacterium] |
| MBQ9252308.1 | Type1.BREX | TOPRIM->AbiEi->?->AbiEii->REase->BrxA-DNA-binding->BrxB-BrxF-iSTAND->DUF6079*->N6-METHYLASE->Tox-ART-HYD2->PglZ->SIGMA+OB-Fold+AAA-ATPase+Lon-Protease-BrxL->?-><-Amidohydro_1\|\|Glyco_hydro_4+Glyco_hydro_4C->?->URI-DNase+UVR+UvrC_HhH_N+HHH-> | 1218 | Bacteria>Terrabacteria_group>Bacillota>Clostridia>unclassified_Clostridia | GCA_017470145.1 | Clostridia_bacterium | IJI88_06715 | 56969..60625 | 2044939 | BREX-system-P-loop-protein-BrxC-[Clostridia-bacterium] |
| MBQ8616934.1 | Type1.BREX | <-N6-METHYLASE<-ParB+HNH<-DUF6079*<-BrxB-BrxF-iSTAND<-BrxA-DNA-binding<-HTH_26<-?<-ARG<-MACRODOMAIN | 1186 | Bacteria>Terrabacteria_group>Bacillota>Clostridia>unclassified_Clostridia | GCA_017472845.1 | Clostridia_bacterium | IJ418_05400 | 5468..9028 | 2044939 | BREX-system-P-loop-protein-BrxC-[Clostridia-bacterium] |
| MBR1711355.1 | Type1.BREX | TraJ-RHH->?->?->?->MACRODOMAIN->?->HTH_26->BrxA-DNA-binding->BrxB-BrxF-iSTAND->DUF6079*->N6-METHYLASE->?->?->PglZ->OLD-ABC-> | 1196 | Bacteria>Terrabacteria_group>Bacillota>Clostridia>unclassified_Clostridia | GCA_017476605.1 | Clostridia_bacterium | IJ719_21465 | 11467..15057 | 2044939 | BREX-system-P-loop-protein-BrxC-[Clostridia-bacterium] |
| MBR1685959.1 | Type1.BREX | <-DUF6079*<-BrxB-BrxF-iSTAND<-BrxA-DNA-binding<-?\|\|ABhydrolase+ABhydrolase-> | 1211 | Bacteria>Terrabacteria_group>Bacillota>Clostridia>unclassified_Clostridia | GCA_017476845.1 | Clostridia_bacterium | IJ708_12570 | 1..3633 | 2044939 | BREX-system-P-loop-protein-BrxC-[Clostridia-bacterium] |
| MBR4003949.1 | Type1.BREX | <-HTH_26\|\|?->Response_reg-><-HTH_3\|\|BrxA-DNA-binding->BrxB-BrxF-iSTAND->?->DUF6079*->N6-METHYLASE->PglZ->SIGMA+OB-Fold+AAA-ATPase+Lon-Protease-BrxL-> | 1175 | Bacteria>Terrabacteria_group>Bacillota>Clostridia>unclassified_Clostridia | GCA_017523305.1 | Clostridia_bacterium | IKI95_07895 | 6107..9634 | 2044939 | BREX-system-P-loop-protein-BrxC-[Clostridia-bacterium] |
| MBR6253006.1 | Type1.BREX | <-NUDIX<-HEPN<-?<-REase<-SbcC<-ParB+HNH<-PglZ<-N6-METHYLASE<-DUF6079*<-BrxB-BrxF-iSTAND<-BrxA-DNA-binding<-FGase<-?<-?<-?<-HTH_3 | 1184 | Bacteria>Terrabacteria_group>Bacillota>Clostridia>unclassified_Clostridia | GCA_017539905.1 | Clostridia_bacterium | IKR04_04115 | 193581..197135 | 2044939 | BREX-system-P-loop-protein-BrxC-[Clostridia-bacterium] |
| MCI5963041.1 | Type1.BREX | HTH_26->BrxA-DNA-binding->BrxB-BrxF-iSTAND->DUF6079*->N6-METHYLASE->SbcC->PglZ->HEPN+SH3-like+SF1-SF2-Helicase+wHTH->MORC+DUF3684+HNH->SF1-SF2-Helicase->PIN->SF1-SF2-Helicase->SF1-SF2-Helicase+REase-> | 1205 | Bacteria>Terrabacteria_group>Bacillota>Clostridia>unclassified_Clostridia | GCA_022770105.1 | Clostridia_bacterium | MR061_06045 | 4040..7657 | 2044939 | BREX-system-P-loop-protein-BrxC-[Clostridia-bacterium] |
| MCR4418710.1 | Type1.BREX | REC+OmpR-HTH->SF1-SF2-Helicase->PolB->HEPN->ncRNA->Phage_int_SAM_4+Phage_integrase->?->MqsA_antitoxin->BrxB-BrxF-iSTAND->DUF6079*->N6-METHYLASE->PglZ->SIGMA+OB-Fold+AAA-ATPase+Lon-Protease-BrxL->PHP+SbcC->REase+EVE-><-?\|\|Calcineurin->SbcC-> | 1149 | Bacteria>Terrabacteria_group>Bacillota>Clostridia>unclassified_Clostridia | GCA_024653205.1 | Clostridia_bacterium | NUV99_00960 | 189881..193330 | 2044939 | BREX-system-P-loop-protein-BrxC-[Clostridia-bacterium] |
| MCR5783583.1 | Type1.BREX | <-ABhydrolase+ABhydrolase<-?<-GDResolvase+Recombinase<-ParB+HNH<-SIGMA+OB-Fold+AAA-ATPase+Lon-Protease-BrxL<-PglZ<-Schlafen+HATPase+wHTH<-Schlafen+HATPase+wHTH<-N6-METHYLASE<-DUF6079*<-BrxB-BrxF-iSTAND<-BrxA-DNA-binding<-HTH_26\|\|Cysteine-Desulfurase->HIN-HTH->DNA_METHYLASE->?->VirD4-TraG+VirD4-FtsK->ArdCN+MPTase-inact+LPD19-> | 1177 | Bacteria>Terrabacteria_group>Bacillota>Clostridia>unclassified_Clostridia | GCA_024699005.1 | Clostridia_bacterium | K6G90_12720 | 47894..51427 | 2044939 | BREX-system-P-loop-protein-BrxC-[Clostridia-bacterium] |
| CCX63523.1 | Type1.BREX | Phage_integrase-><-RelE-ParE<-MazE_antitoxin<-SIGMA+OB-Fold+AAA-ATPase+Lon-Protease-BrxL<-PglZ<-HNH<-OLD-ABC<-N6-METHYLASE<-DUF6079*<-BrxB-BrxF-iSTAND<-BrxA-DNA-binding<-XIS-HTH\|\|VirD4-TraG+VirD4-FtsK->?->?->ArdCN+MPTase-inact+LPD19->TraJ-RHH-> | 1217 | Bacteria>Terrabacteria_group>Bacillota>environmental_samples | GCA_000431495.1 | Firmicutes_bacterium_CAG:791 | BN785_00943 | 24533..28186 | 1262993 | uncharacterized-protein-BN785_00943-[Firmicutes-bacterium-CAG:791] |
| CDC28969.1 | Type1.BREX | MobL->?->GDResolvase+Recombinase+ZnR->?->?->?->HTH_26->BrxA-DNA-binding->BrxB-BrxF-iSTAND->DUF6079*->N6-METHYLASE->ZnR+ZnR->DUF4404->PglZ->SIGMA+OB-Fold+AAA-ATPase+Lon-Protease-BrxL->?->ClpABC-AAA-><-Peptidase_S66<-Peptidase_S66 | 1178 | Bacteria>Terrabacteria_group>Bacillota>environmental_samples | GCA_000432875.1 | Firmicutes_bacterium_CAG:466 | BN668_00732 | 29859..33395 | 1263025 | putative-uncharacterized-protein-[Firmicutes-bacterium-CAG:466] |
| EHM91311.1 | Type1.BREX | <-HTH_3\|\|HTH->Arm-DNA-bind_4+Phage_int_SAM_3+Phage_integrase->TraJ-RHH->?->?->GDResolvase+HIN-HTH->BrxA-DNA-binding-p->BrxB-BrxF-iSTAND->DUF6079-p*-><-Integrase_DNA+Phage_integrase<-?<-ABC-ATPase+SbcC<-?\|\|DUF6079-p->REase->N6-METHYLASE->METHYLASE->RADICAL-SAM-> | 917 | Bacteria>Terrabacteria_group>Bacillota>Erysipelotrichia>Erysipelotrichales>Coprobacillaceae | GCA_000239735.1 | Coprobacillus_sp._3_3_56FAA | HMPREF1021_02117 | 35070..37823 | 665941 | hypothetical-protein-HMPREF1021_02117-[Coprobacillus-sp.-3_3_56FAA] |
| MBN2931490.1 | Type1.BREX | HEPN->?->MqsA_antitoxin+DUF4065->?->?->HEPN->BrxA-DNA-binding->BrxB-BrxF-iSTAND->DUF6079*->N6-METHYLASE->TIR->PglZ->SbcC->SIGMA+OB-Fold+AAA-ATPase+Lon-Protease-BrxL->?->MazG-><-TetR-HTH | 1177 | Bacteria>Terrabacteria_group>Bacillota>Erysipelotrichia>Erysipelotrichales>Coprobacillaceae | GCA_016939095.1 | Catenibacterium_mitsuokai | JTJ30_05795 | 72892..76425 | 100886 | BREX-system-P-loop-protein-BrxC-[Catenibacterium-mitsuokai] |
| MCI6014853.1 | Type1.BREX | BrxA-DNA-binding->BrxB-BrxF-iSTAND->DUF6079*->N6-METHYLASE->N6-METHYLASE-p->N6-METHYLASE-p->N6-METHYLASE-p->PglZ->SIGMA+OB-Fold+AAA-ATPase+Lon-Protease-BrxL->SAVED-> | 1185 | Bacteria>Terrabacteria_group>Bacillota>Erysipelotrichia>Erysipelotrichales>Coprobacillaceae | GCA_022770505.1 | Coprobacillus_sp. | MRZ09_07695 | 2041..5598 | 2137881 | BREX-system-P-loop-protein-BrxC-[Coprobacillus-sp.] |
| MCR0160131.1 | Type1.BREX | <-XIS-HTH<-?<-HTH+REase<-SIGMA+OB-Fold+AAA-ATPase+Lon-Protease-BrxL<-PglZ<-N6-METHYLASE<-DUF6079*<-BrxB-BrxF-iSTAND<-BrxA-DNA-binding<-HTH_26<-?<-HNH+3C-ABC\|\|SIGMA-HTH->MazG->?->REase-> | 1184 | Bacteria>Terrabacteria_group>Bacillota>Erysipelotrichia>Erysipelotrichales>Coprobacillaceae | GCA_024590065.1 | [Clostridium]_innocuum | MKC37_06145 | 87759..91313 | 1522 | BREX-system-P-loop-protein-BrxC-[[Clostridium]] |
| MCR0222787.1 | Type1.BREX | TraJ-RHH->TraM->?->?->?->XIS-HTH->BrxA-DNA-binding->BrxB-BrxF-iSTAND->DUF6079*->N6-METHYLASE->?->PglZ->SIGMA+OB-Fold+AAA-ATPase+Lon-Protease-BrxL->SLATT->SMODS-> | 1217 | Bacteria>Terrabacteria_group>Bacillota>Erysipelotrichia>Erysipelotrichales>Coprobacillaceae | GCA_024590565.1 | [Clostridium]_innocuum | MKA57_06120 | 83560..87213 | 1522 | BREX-system-P-loop-protein-BrxC-[[Clostridium]] |
| MCR0365675.1 | Type1.BREX | TraJ-RHH->TraM->?->?->?->XIS-HTH->BrxA-DNA-binding->BrxB-BrxF-iSTAND->DUF6079*->N6-METHYLASE->?->PglZ->SIGMA+OB-Fold+AAA-ATPase+Lon-Protease-BrxL->SLATT-> | 1217 | Bacteria>Terrabacteria_group>Bacillota>Erysipelotrichia>Erysipelotrichales>Coprobacillaceae | GCA_024591215.1 | [Clostridium]_innocuum | MKD15_12735 | 65751..69404 | 1522 | BREX-system-P-loop-protein-BrxC-[[Clostridium]] |
| MCR0404518.1 | Type1.BREX | <-Tox-RelA<-SMODS<-SLATT<-SIGMA+OB-Fold+AAA-ATPase+Lon-Protease-BrxL<-PglZ<-AT-HTH-HYD2-2<-?<-Tox-ART-HYD2<-N6-METHYLASE<-DUF6079*<-BrxB-BrxF-iSTAND<-BrxA-DNA-binding<-XIS-HTH<-?<-?<-?<-TraM<-TraJ-RHH<-SF1-SF2-Helicase | 1217 | Bacteria>Terrabacteria_group>Bacillota>Erysipelotrichia>Erysipelotrichales>Coprobacillaceae | GCA_024591505.1 | [Clostridium]_innocuum | MKA56_06965 | 75017..78670 | 1522 | BREX-system-P-loop-protein-BrxC-[[Clostridium]] |
| WP_233109295.1 | Type1.BREX | <-REase<-?<-MazG<-SIGMA-HTH\|\|HNH+3C-ABC->?->HTH_26->BrxA-DNA-binding->BrxB-BrxF-iSTAND->DUF6079-p*->DUF6079-p->N6-METHYLASE-p->?->ParA-Soj-PloopNTPase->ParB-> | 818 | Bacteria>Terrabacteria_group>Bacillota>Erysipelotrichia>Erysipelotrichales>Coprobacillaceae | GCF_000763955.1 | [Clostridium]_innocuum | CIAN88_RS13715 | 34742..37198 | 1522 | BREX-system-P-loop-protein-BrxC-[[Clostridium]] |
| WP_118263816.1 | Type1.BREX | <-MODE-HTH+PBPIII\|\|?-><-HTH_3\|\|HTH->TraJ-RHH->?->?->GDResolvase+HIN-HTH->BrxB-BrxF-iSTAND->DUF6079*->N6-METHYLASE->TIR->SIGMA+OB-Fold+AAA-ATPase+Lon-Protease-BrxL-> | 1173 | Bacteria>Terrabacteria_group>Bacillota>Erysipelotrichia>Erysipelotrichales>Coprobacillaceae | GCF_015670235.1 | Thomasclavelia_ramosa | I4V48_RS05865 | 68656..72177 | 1547 | BREX-system-P-loop-protein-BrxC-[Thomasclavelia-ramosa] |
| WP_254151261.1 | Type1.BREX | HEPN->BrxA-DNA-binding->BrxB-BrxF-iSTAND->DUF6079-p->DUF6079*->N6-METHYLASE-p->?->PglZ->SbcC->SIGMA+OB-Fold+AAA-ATPase+Lon-Protease-BrxL->?->MazG-><-TetR-HTH | 1084 | Bacteria>Terrabacteria_group>Bacillota>Erysipelotrichia>Erysipelotrichales>Coprobacillaceae | GCF_019041565.1 | Catenibacterium_mitsuokai | KTF58_RS01735 | 17255..20509 | 100886 | BREX-system-P-loop-protein-BrxC-[Catenibacterium-mitsuokai] |
| WP_008791974.1 | Type1.BREX | <-Peptidase<-?\|\|HTH-><-HTH_38\|\|HTH_26->BrxA-DNA-binding->BrxB-BrxF-iSTAND->DUF6079*->N6-METHYLASE->AbiGi->PglZ->SIGMA+OB-Fold+AAA-ATPase+Lon-Protease-BrxL->?->REase-> | 1182 | Bacteria>Terrabacteria_group>Bacillota>Erysipelotrichia>Erysipelotrichales>Coprobacillaceae | GCF_020097255.1 | Thomasclavelia_ramosa | HMPREF0978_01713 | 101921..105469 | 1547 | BREX-system-P-loop-protein-BrxC-[Thomasclavelia-ramosa] |
| WP_226793210.1 | Type1.BREX | TetR-HTH-><-MazG<-?<-SIGMA+OB-Fold+AAA-ATPase+Lon-Protease-BrxL<-SbcC<-PglZ<-?<-N6-METHYLASE<-DUF6079*<-DUF6079-p<-BrxB-BrxF-iSTAND<-BrxA-DNA-binding<-HEPN | 1084 | Bacteria>Terrabacteria_group>Bacillota>Erysipelotrichia>Erysipelotrichales>Coprobacillaceae | GCF_020537355.1 | Catenibacterium_mitsuokai | LIP38_RS01520 | 50676..53930 | 100886 | BREX-system-P-loop-protein-BrxC-[Catenibacterium-mitsuokai] |
| WP_242410996.1 | Type1.BREX | REase+N-glycosylase->MACRODOMAIN->?->?->?->?->XIS-HTH->BrxA-DNA-binding->BrxB-BrxF-iSTAND->DUF6079*->N6-METHYLASE->?->PglZ->SIGMA+OB-Fold+AAA-ATPase+Lon-Protease-BrxL->Tox-RelA->HKD-DNase+HKD-DNase->HKD-DNase+HKD-DNase->REC+OmpR-HTH->HAMP+HISKIN-> | 1217 | Bacteria>Terrabacteria_group>Bacillota>Erysipelotrichia>Erysipelotrichales>Coprobacillaceae | GCF_022693605.1 | [Clostridium]_innocuum | LV810_RS05930 | 195675..199328 | 1522 | BREX-system-P-loop-protein-BrxC-[[Clostridium]] |
| WP_270396150.1 | Type1.BREX | BrxA-DNA-binding->BrxB-BrxF-iSTAND->DUF6079*->N6-METHYLASE->ParB+HNH->HEPN+HEPN->PglZ->SbcC->SIGMA+OB-Fold+AAA-ATPase+Lon-Protease-BrxL->REase-> | 1177 | Bacteria>Terrabacteria_group>Bacillota>Erysipelotrichia>Erysipelotrichales>Coprobacillaceae | GCF_027659655.1 | Catenibacterium_mitsuokai | PGO85_RS08450 | 1355..4888 | 100886 | BREX-system-P-loop-protein-BrxC-[Catenibacterium-mitsuokai] |
| WP_074685139.1 | Type1.BREX | <-REase<-?<-?<-Peptidase_C14<-SIGMA+OB-Fold+AAA-ATPase+Lon-Protease-BrxL<-TIR<-PglZ<-N6-METHYLASE<-DUF6079*<-BrxB-BrxF-iSTAND<-BrxA-DNA-binding<-HTH_26\|\|?->rve+rve_3-><-?<-MarR-HTH<-?<-RADICAL-SAM | 1182 | Bacteria>Terrabacteria_group>Bacillota>Erysipelotrichia>Erysipelotrichales>Coprobacillaceae | GCF_900106815.1 | Kandleria_vitulina | BLV52_RS00280 | 58615..62163 | 1630 | BREX-system-P-loop-protein-BrxC-[Kandleria-vitulina] |
| HAM30535.1 | Type1.BREX | <-REC\|\|HTH_3->?-><-?<-?\|\|Peptidase_M20->HTH_26->BrxA-DNA-binding->BrxB-BrxF-iSTAND->DUF6079*->?->PglZ-p-> | 1196 | Bacteria>Terrabacteria_group>Bacillota>Erysipelotrichia>Erysipelotrichales>Erysipelotrichaceae | GCA_003453155.1 | Erysipelotrichaceae_bacterium | DCP49_03800 | 6251..9841 | 2049044 | BREX-system-P-loop-protein-BrxC-[Erysipelotrichaceae-bacterium] |
| HCR67829.1 | Type1.BREX | DUF6079*->N6-METHYLASE->?->Schlafen+HATPase+wHTH->PglZ->HTH+REase->SIGMA+OB-Fold+AAA-ATPase+Lon-Protease-BrxL->DUF4365+REase-> | 1177 | Bacteria>Terrabacteria_group>Bacillota>Erysipelotrichia>Erysipelotrichales>Erysipelotrichaceae | GCA_003533525.1 | Erysipelotrichaceae_bacterium | DIV43_01625 | 18..3551 | 2049044 | BREX-system-P-loop-protein-BrxC-[Erysipelotrichaceae-bacterium] |
| TXT19373.1 | Type1.BREX | <-NERD+RNA-Helicase\|\|DUF488-1->DUF488-s->DUF4326->SF1-SF2-Helicase->SIGMA-HTH+SIGMA-HTH->35exo+wHTH+SF1-SF2-Helicase+HTH->BrxA-DNA-binding->BrxB-BrxF-iSTAND->DUF6079*->N6-METHYLASE->PglZ->?->SIGMA+OB-Fold+AAA-ATPase+Lon-Protease-BrxL->DUF3644-> | 1174 | Bacteria>Terrabacteria_group>Bacillota>Erysipelotrichia>Erysipelotrichales>Erysipelotrichaceae | GCA_008080535.1 | Erysipelotrichaceae_bacterium | FD133_371 | 19305..22829 | 2049044 | hypothetical-protein-FD133_371-[Erysipelotrichaceae-bacterium] |
| MBD8958802.1 | Type1.BREX | <-HEPN<-SIGMA+OB-Fold+AAA-ATPase+Lon-Protease-BrxL<-REase+DUF4268<-PglZ<-SbcC<-?<-?<-N6-METHYLASE<-?<-DUF6079*<-BrxB-BrxF-iSTAND<-BrxA-DNA-binding<-HTH_26 | 1182 | Bacteria>Terrabacteria_group>Bacillota>Erysipelotrichia>Erysipelotrichales>Erysipelotrichaceae | GCA_014847315.1 | Holdemanella_biformis | EGQ83_10165 | 17990..21538 | 1735 | BREX-system-P-loop-protein-BrxC-[Holdemanella-biformis] |
| MBQ6490577.1 | Type1.BREX | Resolvase+Recombinase->?->?->ParB->ParB->HTH_26->BrxA-DNA-binding->BrxB-BrxF-iSTAND->DUF6079*->TIR->N6-METHYLASE->PglZ->SIGMA+OB-Fold+AAA-ATPase+Lon-Protease-BrxL->?->TIR->?->ParB+HNH-> | 1214 | Bacteria>Terrabacteria_group>Bacillota>Erysipelotrichia>Erysipelotrichales>Erysipelotrichaceae | GCA_017432325.1 | Solobacterium_sp. | IJI75_15280 | 48018..51662 | 2060878 | BREX-system-P-loop-protein-BrxC-[Solobacterium-sp.] |
| MBR0419736.1 | Type1.BREX | <-SIGMA+OB-Fold+AAA-ATPase+Lon-Protease-BrxL<-N6-METHYLASE<-PglZ<-DUF6079*<-BrxB-BrxF-iSTAND<-BrxA-DNA-binding | 1173 | Bacteria>Terrabacteria_group>Bacillota>Erysipelotrichia>Erysipelotrichales>Erysipelotrichaceae | GCA_017432545.1 | Erysipelotrichaceae_bacterium | IJI66_11255 | 10561..14082 | 2049044 | BREX-system-P-loop-protein-BrxC-[Erysipelotrichaceae-bacterium] |
| MBP3871947.1 | Type1.BREX | <-N6-METHYLASE-p<-PglZ<-DUF6079*<-BrxB-BrxF-iSTAND<-BrxA-DNA-binding<-?<-HNH<-?<-HNH | 1175 | Bacteria>Terrabacteria_group>Bacillota>Erysipelotrichia>Erysipelotrichales>Erysipelotrichaceae | GCA_017941505.1 | Faecalicoccus_sp. | J6E46_13345 | 3304..6831 | 1971758 | BREX-system-P-loop-protein-BrxC-[Faecalicoccus-sp.] |
| MBS3987747.1 | Type1.BREX | BrxA-DNA-binding->BrxB-BrxF-iSTAND->DUF6079*->N6-METHYLASE-><-MULE-TRANSPOSASE | 1171 | Bacteria>Terrabacteria_group>Bacillota>Erysipelotrichia>Erysipelotrichales>Erysipelotrichaceae | GCA_018335705.1 | Erysipelothrix_sp. | KGZ38_07285 | 3111..6626 | 38403 | BREX-system-P-loop-protein-BrxC-[Erysipelothrix-sp.] |
| MCD7892352.1 | Type1.BREX | <-XIS-HTH<-PemK_toxin<-?<-SIGMA-HTH<-?\|\|?-><-SbcC\|\|BrxA-DNA-binding->BrxB-BrxF-iSTAND->DUF6079*->N6-METHYLASE->ACET->PglZ->?->SIGMA+OB-Fold+AAA-ATPase+Lon-Protease-BrxL->Tox-RES->ZnR+ZnR->ZnR+ZnR-> | 1186 | Bacteria>Terrabacteria_group>Bacillota>Erysipelotrichia>Erysipelotrichales>Erysipelotrichaceae | GCA_021201315.1 | Erysipelotrichaceae_bacterium | LUG60_01500 | 53355..56915 | 2049044 | BREX-system-P-loop-protein-BrxC-[Erysipelotrichaceae-bacterium] |
| MCF0260592.1 | Type1.BREX | BrxA-DNA-binding->BrxB-BrxF-iSTAND->DUF6079-p*->DUF6079-p->N6-METHYLASE-p->PglZ-p->SIGMA+OB-Fold+AAA-ATPase+Lon-Protease-BrxL-> | 497 | Bacteria>Terrabacteria_group>Bacillota>Erysipelotrichia>Erysipelotrichales>Erysipelotrichaceae | GCA_021443985.1 | Erysipelotrichaceae_bacterium | HUJ54_12110 | 1218..2710 | 2049044 | BREX-system-P-loop-protein-BrxC-[Erysipelotrichaceae-bacterium] |
| MCH4019936.1 | Type1.BREX | Schlafen+HATPase+wHTH->Schlafen+HATPase+wHTH->?->?->GDResolvase+HIN-HTH->HTH_26->BrxA-DNA-binding->BrxB-BrxF-iSTAND->?->DUF6079*->N6-METHYLASE->Schlafen+HATPase+wHTH->PglZ->SIGMA+OB-Fold+AAA-ATPase+Lon-Protease-BrxL->OLD-ABC+TOPRIM->SF1-SF2-Helicase-> | 1223 | Bacteria>Terrabacteria_group>Bacillota>Erysipelotrichia>Erysipelotrichales>Erysipelotrichaceae | GCA_022483385.1 | Erysipelotrichaceae_bacterium | LKE61_03575 | 742879..746550 | 2049044 | BREX-system-P-loop-protein-BrxC-[Erysipelotrichaceae-bacterium] |
| MCI6089875.1 | Type1.BREX | SF1-SF2-Helicase+DUF1998->?->BrxA-DNA-binding->BrxB-BrxF-iSTAND->DUF6079-p*-> | 870 | Bacteria>Terrabacteria_group>Bacillota>Erysipelotrichia>Erysipelotrichales>Erysipelotrichaceae | GCA_022771005.1 | Solobacterium_sp. | MR695_04825 | 12247..14856 | 2060878 | BREX-system-P-loop-protein-BrxC-[Solobacterium-sp.] |
| GJM58076.1 | Type1.BREX | SIGMA+OB-Fold+AAA-ATPase+Lon-Protease-BrxL->CR-ATPase5+REase-><-?<-SIGMA+OB-Fold+AAA-ATPase+Lon-Protease-BrxL<-PglZ<-HTH+REase<-HTH+REase<-N6-METHYLASE<-?<-DUF6079*<-BrxB-BrxF-iSTAND<-BrxA-DNA-binding<-HTH_26<-Peptidase_M20\|\|?-><-?<-GspA-ATPase<-HTH_32+rve | 1176 | Bacteria>Terrabacteria_group>Bacillota>Erysipelotrichia>Erysipelotrichales>Erysipelotrichaceae | GCA_022834735.1 | Erysipelotrichaceae_bacterium_OPF54 | EROP_17690 | 1809630..1813160 | 2894072 | hypothetical-protein-EROP_17690-[Erysipelotrichaceae-bacterium-OPF54] |
| MCQ4803584.1 | Type1.BREX | Phage_int_SAM_3+Phage_integrase->tRNA-><-Arm-DNA-bind_4+Phage_int_SAM_3+Phage_integrase<-Phage_integrase+Phage_integrase<-XIS-HTH<-PemK_toxin<-SIGMA-HTH\|\|HTH_26->BrxA-DNA-binding->BrxB-BrxF-iSTAND->DUF6079*->N6-METHYLASE->?->Schlafen+HATPase+wHTH->PglZ->HTH+REase->SIGMA+OB-Fold+AAA-ATPase+Lon-Protease-BrxL->DUF4365+REase-> | 1177 | Bacteria>Terrabacteria_group>Bacillota>Erysipelotrichia>Erysipelotrichales>Erysipelotrichaceae | GCA_024460475.1 | Holdemanella_sp._MSK.7.32 | NE544_03830 | 12806..16339 | 2965273 | BREX-system-P-loop-protein-BrxC-[Holdemanella-sp.-MSK.7.32] |
| WP_006060901.1 | Type1.BREX | <-BirA-HTH<-Schlafen+HATPase+wHTH<-?<-?<-HTH+REase<-?<-SIGMA+OB-Fold+AAA-ATPase+Lon-Protease-BrxL<-PglZ<-N6-METHYLASE<-DUF6079*<-BrxB-BrxF-iSTAND<-BrxA-DNA-binding<-HTH_26<-?<-HNH+3C-ABC\|\|LuxR-HTH->?->SF1-SF2-Helicase-> | 1180 | Bacteria>Terrabacteria_group>Bacillota>Erysipelotrichia>Erysipelotrichales>Erysipelotrichaceae | GCF_000157995.1 | Holdemania_filiformis | HOLDEFILI_RS15965 | 13193..16735 | 61171 | BREX-system-P-loop-protein-BrxC-[Holdemania-filiformis] |
| WP_019892646.1 | Type1.BREX | ArsR-HTH->SAM-METHYLASE->HKD-DNase+SF1-SF2-Helicase+HTH+HTH+SRA->REase+DUF4268->BrxA-DNA-binding->BrxB-BrxF-iSTAND->DUF6079*->?->N6-METHYLASE->PglZ->SIGMA+OB-Fold+AAA-ATPase+Lon-Protease-BrxL-><-AAA-ATPase<-HTH_7+rve+rve_3\|\|MACRODOMAIN->MACRODOMAIN-> | 1175 | Bacteria>Terrabacteria_group>Bacillota>Erysipelotrichia>Erysipelotrichales>Erysipelotrichaceae | GCF_000384195.1 | Allobaculum_stercoricanis | ALLST_RS0103345 | 705410..708937 | 174709 | BREX-system-P-loop-protein-BrxC-[Allobaculum-stercoricanis] |
| WP_075884671.1 | Type1.BREX | <-SIGMA+OB-Fold+AAA-ATPase+Lon-Protease-BrxL<-PglZ<-N6-METHYLASE<-DUF6079*<-BrxB-BrxF-iSTAND<-BrxA-DNA-binding | 1175 | Bacteria>Terrabacteria_group>Bacillota>Erysipelotrichia>Erysipelotrichales>Erysipelotrichaceae | GCF_001945535.1 | Faecalibaculum_rodentium | BO223_RS01545 | 8627..12154 | 1702221 | BREX-system-P-loop-protein-BrxC-[Faecalibaculum-rodentium] |
| WP_118011256.1 | Type1.BREX | <-N6-METHYLASE-p<-N6-METHYLASE-p<-N6-METHYLASE-p<-?<-DUF6079*<-BrxB-BrxF-iSTAND<-BrxA-DNA-binding<-HTH_26<-?<-?<-?<-REase<-PBPB+PBPB+HISKIN+MORC+REC | 1182 | Bacteria>Terrabacteria_group>Bacillota>Erysipelotrichia>Erysipelotrichales>Erysipelotrichaceae | GCF_003468025.1 | Holdemanella_biformis | DW907_RS05670 | 5357..8905 | 1735 | BREX-system-P-loop-protein-BrxC-[Holdemanella-biformis] |
| WP_118365305.1 | Type1.BREX | <-GspA-ATPase<-N6-METHYLASE<-AbiGi<-?<-DUF6079*<-BrxB-BrxF-iSTAND<-BrxA-DNA-binding\|\|<-?<-SIGMA+OB-Fold+AAA-ATPase+Lon-Protease-BrxL<-?<-?<-?<-?<-RADICAL-SAM | 1172 | Bacteria>Terrabacteria_group>Bacillota>Erysipelotrichia>Erysipelotrichales>Erysipelotrichaceae | GCF_003474925.1 | Amedibacillus_dolichus | DWZ83_02410 | 4709..8227 | 31971 | BREX-system-P-loop-protein-BrxC-[Amedibacillus-dolichus] |
| WP_166083468.1 | Type1.BREX | <-SF1-SF2-Helicase+REase-p<-OLD-ABC+TOPRIM<-SIGMA+OB-Fold+AAA-ATPase+Lon-Protease-BrxL<-PglZ<-N6-METHYLASE<-DUF6079*<-BrxB-BrxF-iSTAND<-BrxA-DNA-binding<-HTH_26<-?<-SF1-SF2-Helicase\|\|?-><-Recombinase<-VirD4-TraG+VirD4-FtsK<-MobL | 1192 | Bacteria>Terrabacteria_group>Bacillota>Erysipelotrichia>Erysipelotrichales>Erysipelotrichaceae | GCF_011421495.1 | Erysipelothrix_anatis | GDN62_RS10715 | 11860..15438 | 2683713 | BREX-system-P-loop-protein-BrxC-[Erysipelothrix-anatis] |
| WP_186999074.1 | Type1.BREX | HTH_26->BrxA-DNA-binding->BrxB-BrxF-iSTAND->DUF6079*->?->N6-METHYLASE->?->?->?->PglZ->REase+DUF4268->REase+DUF4268-> | 1182 | Bacteria>Terrabacteria_group>Bacillota>Erysipelotrichia>Erysipelotrichales>Erysipelotrichaceae | GCF_014306155.1 | Holdemanella_hominis | H8911_RS06490 | 3688..7236 | 2764327 | BREX-system-P-loop-protein-BrxC-[Holdemanella-hominis] |
| WP_195233263.1 | Type1.BREX | <-SIR2<-SIGMA+OB-Fold+AAA-ATPase+Lon-Protease-BrxL<-tRNA<-PglZ<-Schlafen+HATPase+wHTH<-?<-N6-METHYLASE<-DUF6079*<-BrxB-BrxF-iSTAND<-BrxA-DNA-binding<-HTH_26\|\|SIGMA-HTH->PemK_toxin->XIS-HTH->Phage_integrase+Phage_integrase->Arm-DNA-bind_4+Phage_int_SAM_3+Phage_integrase-> | 1177 | Bacteria>Terrabacteria_group>Bacillota>Erysipelotrichia>Erysipelotrichales>Erysipelotrichaceae | GCF_015547685.1 | Holdemanella_biformis | I1941_RS05785 | 11708..15241 | 1735 | BREX-system-P-loop-protein-BrxC-[Holdemanella-biformis] |
| WP_256268391.1 | Type1.BREX | Phage_int_SAM_3+Phage_integrase->tRNA-><-Arm-DNA-bind_4+Phage_int_SAM_3+Phage_integrase<-Phage_integrase+Phage_integrase<-XIS-HTH<-PemK_toxin<-SIGMA-HTH\|\|HTH_26->BrxA-DNA-binding->BrxB-BrxF-iSTAND->DUF6079*->N6-METHYLASE->?->Schlafen+HATPase+wHTH->PglZ->HTH+REase->SIGMA+OB-Fold+AAA-ATPase+Lon-Protease-BrxL->DUF4365+REase-> | 1177 | Bacteria>Terrabacteria_group>Bacillota>Erysipelotrichia>Erysipelotrichales>Erysipelotrichaceae | GCF_024460475.1 | Holdemanella_sp._MSK.7.32 | NE544_RS03830 | 12806..16339 | 2965273 | BREX-system-P-loop-protein-BrxC-[Holdemanella-sp.-MSK.7.32] |
| WP_270474052.1 | Type1.BREX | PBPB+PBPB+HISKIN+MORC+REC->REase->?->HTH_26->BrxA-DNA-binding->BrxB-BrxF-iSTAND->DUF6079*->?->N6-METHYLASE-p-> | 1182 | Bacteria>Terrabacteria_group>Bacillota>Erysipelotrichia>Erysipelotrichales>Erysipelotrichaceae | GCF_027662285.1 | Holdemanella_porci | PGP84_RS10070 | 8164..11712 | 2652276 | BREX-system-P-loop-protein-BrxC-[Holdemanella-porci] |
| WP_107029377.1 | Type1.BREX | <-wHTH+WYL+WCX\|\|PIN->?->?->HTH_26->BrxA-DNA-binding->BrxB-BrxF-iSTAND->DUF6079*->N6-METHYLASE->HEPN->PglZ->SIGMA+OB-Fold+AAA-ATPase+Lon-Protease-BrxL-> | 1178 | Bacteria>Terrabacteria_group>Bacillota>Erysipelotrichia>Erysipelotrichales>Erysipelotrichaceae | GCF_027669645.1 | Faecalibacillus_intestinalis | C7U54_RS03870 | 158603..162139 | 1982626 | BREX-system-P-loop-protein-BrxC-[Faecalibacillus-intestinalis] |
| WP_102342725.1 | Type1.BREX | Peptidase-><-?\|\|?-><-?\|\|RHH->BrxA-DNA-binding->BrxB-BrxF-iSTAND->DUF6079*->N6-METHYLASE->Schlafen+HATPase+wHTH->PglZ->SIGMA+OB-Fold+AAA-ATPase+Lon-Protease-BrxL-><-?\|\|?->HTH+rve->AAA+->OrfB_Zn_ribbon-> | 1203 | Bacteria>Terrabacteria_group>Bacillota>Erysipelotrichia>Erysipelotrichales>Erysipelotrichaceae | GCF_900240265.1 | Galactobacillus_timonensis | C1714_RS08210 | 1722051..1725662 | 2041840 | BREX-system-P-loop-protein-BrxC-[Galactobacillus-timonensis] |
| WP_102343499.1 | Type1.BREX | <-HTH_3<-wHTH+TPR<-HTH_3<-HerA/FtsK<-SIR2<-SIGMA+OB-Fold+AAA-ATPase+Lon-Protease-BrxL<-PglZ<-Schlafen+HATPase+wHTH<-N6-METHYLASE<-DUF6079*<-BrxB-BrxF-iSTAND<-BrxA-DNA-binding<-HTH_26\|\|Response_reg-><-?<-TraM<-TraJ-RHH | 1208 | Bacteria>Terrabacteria_group>Bacillota>Erysipelotrichia>Erysipelotrichales>Erysipelotrichaceae | GCF_900240265.1 | Galactobacillus_timonensis | C1714_RS12180 | 363466..367092 | 2041840 | BREX-system-P-loop-protein-BrxC-[Galactobacillus-timonensis] |
| WP_108775121.1 | Type1.BREX | <-RHH<-LZ_Tnp_IS66+DDE_Tnp_IS66+DDE_Tnp_IS66_C<-?<-Mob_Pre<-SIGMA+OB-Fold+AAA-ATPase+Lon-Protease-BrxL<-PglZ<-?<-N6-METHYLASE<-DUF6079*<-BrxB-BrxF-iSTAND<-BrxA-DNA-binding<-HTH_26<-TraJ-RHH<-Arm-DNA-bind_4+Phage_int_SAM_3+Phage_integrase<-HTH | 1204 | Bacteria>Terrabacteria_group>Bacillota>Erysipelotrichia>Erysipelotrichales>Erysipelotrichaceae | GCF_900343155.1 | Lactimicrobium_massiliense | DHU66_RS06845 | 223785..227399 | 2161814 | BREX-system-P-loop-protein-BrxC-[Lactimicrobium-massiliense] |
| WP_273235218.1 | Type1.BREX | <-SIGMA+OB-Fold+AAA-ATPase+Lon-Protease-BrxL<-PglZ<-N6-METHYLASE<-DUF6079*<-BrxB-BrxF-iSTAND-p<-BrxA-DNA-binding | 1174 | Bacteria>Terrabacteria_group>Bacillota>Erysipelotrichia>Erysipelotrichales>Erysipelotrichaceae | GCF_910589625.1 | Ileibacterium_valens | NW265_RS06600 | 8355..11879 | 1862668 | BREX-system-P-loop-protein-BrxC-[Ileibacterium-valens] |
| WP_212725106.1 | Type1.BREX | <-Phage_integrase\|\|N6-METHYLASE->?-><-SIGMA+OB-Fold+AAA-ATPase+Lon-Protease-BrxL<-PglZ<-N6-METHYLASE<-DUF6079*<-BrxB-BrxF-iSTAND<-BrxA-DNA-binding<-?<-?<-HORMA<-Phage_integrase<-XIS-HTH<-Phage_integrase<-AAA-ATPase-p | 1205 | Bacteria>Terrabacteria_group>Bacillota>Erysipelotrichia>Erysipelotrichales>Turicibacteraceae | GCF_024499075.1 | Turicibacter_bilis | J0J70_RS00075 | 21552..25169 | 2735723 | BREX-system-P-loop-protein-BrxC-[Turicibacter-bilis] |
| WP_262953490.1 | Type1.BREX | <-N6-METHYLASE<-DUF6079*<-BrxB-BrxF-iSTAND<-BrxA-DNA-binding<-SF1-SF2-Helicase<-ASCH<-tRNA<-?<-RNase+DSRBD<-?<-SF1-SF2-Helicase | 1175 | Bacteria>Terrabacteria_group>Bacillota>Erysipelotrichia>Erysipelotrichales>Turicibacteraceae | GCF_025595085.1 | Turicibacter_sp._1E2 | NEH72_RS01665 | 3622..7149 | 2951143 | BREX-system-P-loop-protein-BrxC-[Turicibacter-sp.-1E2] |
| NBK98923.1 | Type1.BREX | wHTH+WYL+WCX->?->SPOUT->PolB->HEPN->?->BrxA-DNA-binding->BrxB-BrxF-iSTAND->DUF6079-p*-> | 377 | Bacteria>Terrabacteria_group>Bacillota>Erysipelotrichia>unclassified_Erysipelotrichia | GCA_009916335.1 | Erysipelotrichia_bacterium | EOM50_13050 | 4855..5984 | 2184014 | BREX-system-P-loop-protein-BrxC-[Erysipelotrichia-bacterium] |
| NCB38812.1 | Type1.BREX | <-Schlafen+HATPase+wHTH-p<-DUF6079*<-BrxB-BrxF-iSTAND<-BrxA-DNA-binding<-?<-?<-?<-?<-Arm-DNA-bind_3+Phage_integrase | 1193 | Bacteria>Terrabacteria_group>Bacillota>Erysipelotrichia>unclassified_Erysipelotrichia | GCA_009928855.1 | Erysipelotrichia_bacterium | EOM80_08585 | 655..4236 | 2184014 | BREX-system-P-loop-protein-BrxC-[Erysipelotrichia-bacterium] |
| NCB40064.1 | Type1.BREX | <-Phage_integrase+Phage_integrase<-?\|\|BrxA-DNA-binding->BrxB-BrxF-iSTAND->DUF6079-p*-> | 385 | Bacteria>Terrabacteria_group>Bacillota>Erysipelotrichia>unclassified_Erysipelotrichia | GCA_009928855.1 | Erysipelotrichia_bacterium | EOM80_14980 | 3043..4197 | 2184014 | BREX-system-P-loop-protein-BrxC-[Erysipelotrichia-bacterium] |
| HAA34479.1 | Type1.BREX | DOC-><-HicB_lk_antitox\|\|?->BrxA-DNA-binding->BrxB-BrxF-iSTAND->BrxB-BrxF-iSTAND-p->DUF6079*->N6-METHYLASE->PolB->HEPN->PglZ->SIGMA+OB-Fold+AAA-ATPase+Lon-Protease-BrxL-p-> | 1003 | Bacteria>Terrabacteria_group>Bacillota | GCA_003444595.1 | Bacillota_bacterium | DCD97_04120 | 2822..5830 | 1879010 | BREX-system-P-loop-protein-BrxC-[Bacillota-bacterium] |
| HCT36708.1 | Type1.BREX | <-HEPN<-Ntox45\|\|ACET-><-ABC-ATPase+SbcC<-ABC-ATPase+SbcC\|\|RADICAL-SAM-><-NERD\|\|BrxA-DNA-binding->BrxB-BrxF-iSTAND->DUF6079*->N6-METHYLASE->Schlafen+HATPase+wHTH->PglZ->SIGMA+OB-Fold+AAA-ATPase+Lon-Protease-BrxL-><-?\|\|?-><-?\|\|?-><-HTH_Tnp_1+HTH_21+rve+rve_3 | 1172 | Bacteria>Terrabacteria_group>Bacillota | GCA_003543125.1 | Bacillota_bacterium | DF292_06730 | 25271..28789 | 1879010 | BREX-system-P-loop-protein-BrxC-[Bacillota-bacterium] |
| MTI67135.1 | Type1.BREX | BrxA-DNA-binding->BrxB-BrxF-iSTAND->DUF6079*->N6-METHYLASE->PglZ->SIGMA+OB-Fold+AAA-ATPase+Lon-Protease-BrxL->HEPN->?-><-Phage_int_SAM_4+Phage_integrase | 1184 | Bacteria>Terrabacteria_group>Bacillota | GCA_009711765.1 | Bacillota_bacterium | FH753_11140 | 12161..15715 | 1879010 | BREX-system-P-loop-protein-BrxC-[Bacillota-bacterium] |
| MCL2301622.1 | Type1.BREX | GDResolvase-><-GDResolvase\|\|?-><-?\|\|?->XIS-HTH->BrxA-DNA-binding->BrxB-BrxF-iSTAND->DUF6079*->N6-METHYLASE->DOC+HTH->PglZ->SIGMA+OB-Fold+AAA-ATPase+Lon-Protease-BrxL-> | 1174 | Bacteria>Terrabacteria_group>Bacillota | GCA_009786275.1 | Bacillota_bacterium | FWC27_15905 | 14080..17604 | 1879010 | BREX-system-P-loop-protein-BrxC-[Bacillota-bacterium] |
| NLC46665.1 | Type1.BREX | BrxA-DNA-binding->BrxB-BrxF-iSTAND->DUF6079*->N6-METHYLASE->PglZ->SIGMA+OB-Fold+AAA-ATPase+Lon-Protease-BrxL-> | 1172 | Bacteria>Terrabacteria_group>Bacillota | GCA_012512655.1 | Bacillota_bacterium | GX766_05910 | 1502..5020 | 1879010 | BREX-system-P-loop-protein-BrxC-[Bacillota-bacterium] |
| NLG84492.1 | Type1.BREX | <-Peptidase_M50<-S4+PSYN<-?<-?<-?<-SIGMA+OB-Fold+AAA-ATPase+Lon-Protease-BrxL<-PglZ<-PHP+SbcC<-DUF6079*<-BrxB-BrxF-iSTAND<-BrxA-DNA-binding | 1146 | Bacteria>Terrabacteria_group>Bacillota | GCA_012516115.1 | Bacillota_bacterium | GX493_07810 | 18483..21923 | 1879010 | BREX-system-P-loop-protein-BrxC-[Bacillota-bacterium] |
| NLG80270.1 | Type1.BREX | ABC-ATPase+SbcC->ABC-ATPase+SbcC-><-REase+NACHT<-PglZ<-N6-METHYLASE<-N6-METHYLASE<-DUF6079*<-BrxB-BrxF-iSTAND<-BrxA-DNA-binding<-SIGMA+OB-Fold+AAA-ATPase+Lon-Protease-BrxL<-ParB+HNH<-AAA-ATPase-p<-DEDD_Tnp_IS110+Transposase_20\|\|?->DDE_Tnp_ISL3-> | 1185 | Bacteria>Terrabacteria_group>Bacillota | GCA_012516125.1 | Bacillota_bacterium | GX492_10705 | 20631..24188 | 1879010 | BREX-system-P-loop-protein-BrxC-[Bacillota-bacterium] |
| NLJ79432.1 | Type1.BREX | <-N6-METHYLASE-p<-DUF6079*<-BrxB-BrxF-iSTAND<-BrxA-DNA-binding\|\|?-><-?<-?<-MarR-HTH | 1187 | Bacteria>Terrabacteria_group>Bacillota | GCA_012518645.1 | Bacillota_bacterium | GX335_00185 | 1101..4664 | 1879010 | BREX-system-P-loop-protein-BrxC-[Bacillota-bacterium] |
| NLJ85301.1 | Type1.BREX | OrfB_Zn_ribbon->BrxA-DNA-binding->BrxB-BrxF-iSTAND->DUF6079*->N6-METHYLASE-p-> | 1183 | Bacteria>Terrabacteria_group>Bacillota | GCA_012518905.1 | Bacillota_bacterium | GX322_02580 | 1602..5153 | 1879010 | BREX-system-P-loop-protein-BrxC-[Bacillota-bacterium] |
| NLX91917.1 | Type1.BREX | Exonuc_V_gamma+REase->SF1-SF2-Helicase+REase->TetR-HTH->?->?->?->BrxA-DNA-binding->BrxB-BrxF-iSTAND->DUF6079*->PolB->HEPN->PglZ->SIGMA+OB-Fold+AAA-ATPase+Lon-Protease-BrxL->EVE+McrB-AAA->McrC-NTD+REase->ARG->HEPN->RelE-ParE-> | 1176 | Bacteria>Terrabacteria_group>Bacillota | GCA_012728375.1 | Bacillota_bacterium | GXZ07_10120 | 36614..40144 | 1879010 | BREX-system-P-loop-protein-BrxC-[Bacillota-bacterium] |
| HHV57184.1 | Type1.BREX | <-SIGMA+OB-Fold+AAA-ATPase+Lon-Protease-BrxL<-PglZ<-N6-METHYLASE<-DUF6079*<-BrxB-BrxF-iSTAND<-BrxA-DNA-binding | 1147 | Bacteria>Terrabacteria_group>Bacillota | GCA_012839755.1 | Bacillota_bacterium | GXX50_05400 | 6921..10364 | 1879010 | BREX-system-P-loop-protein-BrxC-[Bacillota-bacterium] |
| NPV28413.1 | Type1.BREX | NAD_synthase->?->?->?->REase->BrxA-DNA-binding->BrxB-BrxF-iSTAND->DUF6079*->N6-METHYLASE->PHP+SbcC->PglZ->SIGMA+OB-Fold+AAA-ATPase+Lon-Protease-BrxL->?->35exo+T7DNApol-> | 1208 | Bacteria>Terrabacteria_group>Bacillota | GCA_013177945.1 | Bacillota_bacterium | HPY58_01910 | 11153..14779 | 1879010 | BREX-system-P-loop-protein-BrxC-[Bacillota-bacterium] |
| MBA1335117.1 | Type1.BREX | 35exo+T7DNApol->?->?->?->?->tRNA-><-SIGMA+OB-Fold+AAA-ATPase+Lon-Protease-BrxL<-PglZ<-N6-METHYLASE<-DUF6079*<-BrxB-BrxF-iSTAND<-BrxA-DNA-binding<-?<-DDE_3<-HTH_23+HTH_32+HTH_33 | 1190 | Bacteria>Terrabacteria_group>Bacillota | GCA_013540605.1 | Bacillota_bacterium | HPY66_0739 | 77367..80939 | 1879010 | BREX-system-P-loop-protein-BrxC-[Bacillota-bacterium] |
| MBQ7264807.1 | Type1.BREX | BrxA-DNA-binding->BrxB-BrxF-iSTAND->DUF6079*->N6-METHYLASE->?->?->PglZ-p-> | 1181 | Bacteria>Terrabacteria_group>Bacillota | GCA_017444985.1 | Bacillota_bacterium | IJS61_01765 | 4436..7981 | 1879010 | BREX-system-P-loop-protein-BrxC-[Bacillota-bacterium] |
| MBP2637786.1 | Type1.BREX | HTH_26->BrxA-DNA-binding->BrxB-BrxF-iSTAND->DUF6079*->N6-METHYLASE->?->?->PglZ->SIGMA+OB-Fold+AAA-ATPase+Lon-Protease-BrxL->REase+DUF5655-> | 1214 | Bacteria>Terrabacteria_group>Bacillota | GCA_017889705.1 | Bacillota_bacterium | H6Q72_3693 | 16334..19978 | 1879010 | hypothetical-protein-[Bacillota-bacterium] |
| MBP7331845.1 | Type1.BREX | SIGMA-HTH->?->?->?->BrxA-DNA-binding->BrxB-BrxF-iSTAND->DUF6079*->N6-METHYLASE-p-> | 1182 | Bacteria>Terrabacteria_group>Bacillota | GCA_017996655.1 | Bacillota_bacterium | KBA08_06005 | 6742..10290 | 1879010 | BREX-system-P-loop-protein-BrxC-[Bacillota-bacterium] |
| MBS4881094.1 | Type1.BREX | TraJ-RHH->TraJ-RHH->BrxA-DNA-binding->BrxB-BrxF-iSTAND->DUF6079*->N6-METHYLASE-p->N6-METHYLASE-p-> | 1195 | Bacteria>Terrabacteria_group>Bacillota | GCA_018363755.1 | Bacillota_bacterium | KH138_12510 | 2919..6506 | 1879010 | BREX-system-P-loop-protein-BrxC-[Bacillota-bacterium] |
| MBU0997904.1 | Type1.BREX | N6-METHYLASE-p->ABC-ATPase+SbcC->Glyco_transf_9->?-><-?\|\|?-><-?\|\|tRNA->BrxA-DNA-binding->BrxB-BrxF-iSTAND->DUF6079-p*-> | 911 | Bacteria>Terrabacteria_group>Bacillota | GCA_018816905.1 | Bacillota_bacterium | KKE16_07690 | 9752..12487 | 1879010 | BREX-system-P-loop-protein-BrxC-[Bacillota-bacterium] |
| MBU1143925.1 | Type1.BREX | <-MORC+DUF3684+REase<-?<-?<-REase+SF1-SF2-Helicase-p<-REase+SF1-SF2-Helicase<-?<-SIGMA+OB-Fold+AAA-ATPase+Lon-Protease-BrxL<-PglZ<-N6-METHYLASE<-DUF6079*<-BrxB-BrxF-iSTAND<-BrxA-DNA-binding<-MarR-HTH<-XIS-HTH\|\|Resolvase+Recombinase+ZnR->GDResolvase-> | 1173 | Bacteria>Terrabacteria_group>Bacillota | GCA_018818645.1 | Bacillota_bacterium | KKH92_09840 | 28547..32068 | 1879010 | BREX-system-P-loop-protein-BrxC-[Bacillota-bacterium] |
| MBU4541895.1 | Type1.BREX | <-PglZ-p<-TOPRIM<-OLD-ABC<-AbiEi_4<-N6-METHYLASE<-DUF6079*<-BrxB-BrxF-iSTAND<-BrxA-DNA-binding<-DOC+HTH | 1187 | Bacteria>Terrabacteria_group>Bacillota | GCA_018903255.1 | Bacillota_bacterium | KJ774_11820 | 7195..10758 | 1879010 | BREX-system-P-loop-protein-BrxC-[Bacillota-bacterium] |
| MCD8180314.1 | Type1.BREX | <-PglZ<-SIR2<-N6-METHYLASE<-?<-DUF6079*<-BrxB-BrxF-iSTAND<-BrxA-DNA-binding<-TraJ-RHH<-?<-HTH_36<-Integrase_DNA+Phage_integrase<-XIS-HTH<-ZnR | 1187 | Bacteria>Terrabacteria_group>Bacillota | GCA_021203665.1 | Bacillota_bacterium | LUF26_02365 | 8189..11752 | 1879010 | BREX-system-P-loop-protein-BrxC-[Bacillota-bacterium] |
| MCI8284119.1 | Type1.BREX | <-PHP+35exo+PHP+ICAde<-?<-?<-?<-SIGMA+OB-Fold+AAA-ATPase+Lon-Protease-BrxL<-PglZ<-Schlafen+HATPase+wHTH<-N6-METHYLASE<-DUF6079*<-BrxB-BrxF-iSTAND<-BrxA-DNA-binding<-S2P+PDZ | 1174 | Bacteria>Terrabacteria_group>Bacillota | GCA_022790955.1 | Bacillota_bacterium | HFE90_02465 | 72208..75732 | 1879010 | BREX-system-P-loop-protein-BrxC-[Bacillota-bacterium] |
| MCI8646068.1 | Type1.BREX | DUF6079-p*->Schlafen+HATPase+wHTH->N6-METHYLASE->?->PglZ->\|\|?->RNase+DSRBD->SbcC->SbcC->?->MMR_HSR1+sGTP+sGTP-> | 714 | Bacteria>Terrabacteria_group>Bacillota | GCA_022794115.1 | Bacillota_bacterium | HFE76_04520 | 340..2484 | 1879010 | BREX-system-P-loop-protein-BrxC-[Bacillota-bacterium] |
| MCI9596746.1 | Type1.BREX | <-METHYLASE<-?<-?\|\|?->?-><-?\|\|MerR-HTH-><-PglZ<-N6-METHYLASE<-DUF6079*<-BrxB-BrxF-iSTAND<-BrxA-DNA-binding<-ACET\|\|?-><-?<-MarR-HTH | 1226 | Bacteria>Terrabacteria_group>Bacillota | GCA_022801695.1 | Bacillota_bacterium | HFE75_05510 | 74881..78561 | 1879010 | BREX-system-P-loop-protein-BrxC-[Bacillota-bacterium] |
| MCL5289452.1 | Type1.BREX | REase->UDG-conflict->?->HKD-DNase->SF1-SF2-Helicase->BrxA-DNA-binding->BrxB-BrxF-iSTAND->DUF6079*->N6-METHYLASE->OLD-ABC-> | 1181 | Bacteria>Terrabacteria_group>Bacillota | GCA_023481335.1 | Bacillota_bacterium | M1489_00180 | 10460..14005 | 1879010 | BREX-system-P-loop-protein-BrxC-[Bacillota-bacterium] |
| MCL5289723.1 | Type1.BREX | RADICAL-SAM->NUDIX->?->?->BrxA-DNA-binding->BrxB-BrxF-iSTAND->DUF6079-p*-> | 904 | Bacteria>Terrabacteria_group>Bacillota | GCA_023481335.1 | Bacillota_bacterium | M1489_01590 | 22488..25200 | 1879010 | BREX-system-P-loop-protein-BrxC-[Bacillota-bacterium] |
| OLA37697.1 | Type1.BREX | <-HEPN<-SIGMA+OB-Fold+AAA-ATPase+Lon-Protease-BrxL<-PglZ<-RADICAL-SAM<-?<-REase<-OLD-ABC<-DUF6079*<-BrxB-BrxF-iSTAND<-BrxA-DNA-binding | 1207 | Bacteria>Terrabacteria_group>Bacillota>Negativicutes>Acidaminococcales>Acidaminococcaceae | GCA_001916865.1 | Phascolarctobacterium_succinatutens | BHW43_05680 | 71469..75092 | 626940 | hypothetical-protein-BHW43_05680-[Phascolarctobacterium-succinatutens] |
| MBS5426410.1 | Type1.BREX | <-Schlafen+HATPase+wHTH<-?\|\|?-><-SIGMA+OB-Fold+AAA-ATPase+Lon-Protease-BrxL<-PglZ<-REase<-OLD-ABC<-N6-METHYLASE<-Schlafen+HATPase+wHTH<-DUF6079*<-BrxB-BrxF-iSTAND<-BrxA-DNA-binding<-zf-ISL3+HTH_Tnp_ISL3+DDE_Tnp_ISL3 | 1207 | Bacteria>Terrabacteria_group>Bacillota>Negativicutes>Acidaminococcales>Acidaminococcaceae | GCA_018368575.1 | Phascolarctobacterium_succinatutens | KHX39_04785 | 71877..75500 | 626940 | BREX-system-P-loop-protein-BrxC-[Phascolarctobacterium-succinatutens] |
| MCD8174817.1 | Type1.BREX | <-SIGMA+OB-Fold+AAA-ATPase+Lon-Protease-BrxL<-PglZ<-N6-METHYLASE<-N6-METHYLASE-p<-DUF6079* | 1211 | Bacteria>Terrabacteria_group>Bacillota>Negativicutes>Acidaminococcales>Acidaminococcaceae | GCA_021203645.1 | Phascolarctobacterium_sp. | LUD41_02585 | 9418..13053 | 2049039 | BREX-system-P-loop-protein-BrxC-[Phascolarctobacterium-sp.] |
| WP_187771775.1 | Type1.BREX | BrxA-DNA-binding->BrxB-BrxF-iSTAND->DUF6079-p*->?->N6-METHYLASE->N6-METHYLASE-p->PglZ->PglZ-> | 626 | Bacteria>Terrabacteria_group>Bacillota>Negativicutes>Acidaminococcales>Acidaminococcaceae | GCF_014489955.1 | Phascolarctobacterium_faecium | H9Y09_RS05325 | 1118272..1120152 | 33025 | BREX-system-P-loop-protein-BrxC-[Phascolarctobacterium-faecium] |
| WP_270440215.1 | Type1.BREX | <-HTH_Tnp_1+HTH_21+rve+rve_3\|\|PAIREDC-HTH-><-HTH+REase<-SIGMA+OB-Fold+AAA-ATPase+Lon-Protease-BrxL<-PglZ<-N6-METHYLASE<-DUF6079*<-BrxB-BrxF-iSTAND<-BrxA-DNA-binding<-HTH_26<-?<-HNH+3C-ABC\|\|SIGMA-HTH-> | 1171 | Bacteria>Terrabacteria_group>Bacillota>Negativicutes>Acidaminococcales>Acidaminococcaceae | GCF_027680825.1 | Acidaminococcus_provencensis | PGX33_RS00725 | 148887..152402 | 2058289 | BREX-system-P-loop-protein-BrxC-[Acidaminococcus-provencensis] |
| WP_160101902.1 | Type1.BREX | <-SIGMA-HTH<-?<-N6-METHYLASE-p<-DUF6079*<-BrxB-BrxF-iSTAND<-BrxA-DNA-binding<-RVT | 1225 | Bacteria>Terrabacteria_group>Bacillota>Negativicutes>Acidaminococcales>Acidaminococcaceae | GCF_937976085.1 | Acidaminococcus_provencensis | C7E10_RS11555 | 4486..8163 | 2058289 | BREX-system-P-loop-protein-BrxC-[Acidaminococcus-provencensis] |
| MBE8950075.1 | Type1.BREX | <-sGTP-MglA+sGTP<-PglZ<-N6-METHYLASE<-DUF6079*<-BrxB-BrxF-iSTAND<-BrxA-DNA-binding<-His_biosynth<-?<-?<-?<-?<-?<-SIS_2+SIS | 1169 | Bacteria>Terrabacteria_group>Bacillota>Negativicutes>Negativicutes_incertae_sedis>Quinella | GCA_015206825.1 | Quinella_sp._3Q1 | SR3Q1_05680 | 79017..82526 | 1958791 | BREX-system-P-loop-protein-BrxC-[Quinella-sp.-3Q1] |
| EEX76839.1 | Type1.BREX | <-SIGMA+OB-Fold+AAA-ATPase+Lon-Protease-BrxL-p<-?<-PglZ<-Schlafen+BirA-HTH<-N6-METHYLASE-p<-?<-N6-METHYLASE<-DUF6079*<-BrxB-BrxF-iSTAND<-BrxA-DNA-binding<-CBS<-?<-?\|\|?-><-?\|\|REase-> | 1183 | Bacteria>Terrabacteria_group>Bacillota>Negativicutes>Selenomonadales>Selenomonadaceae | GCA_000160495.1 | Selenomonas_sputigena_ATCC_35185 | SELSPUOL_01796 | 555362..558913 | 546271 | hypothetical-protein-SELSPUOL_01796-[Selenomonas-sputigena-ATCC-35185] |
| MBE6098022.1 | Type1.BREX | <-SIGMA+OB-Fold+AAA-ATPase+Lon-Protease-BrxL<-PglZ<-REase<-OLD-ABC<-N6-METHYLASE<-DUF6079*<-BrxB-BrxF-iSTAND<-BrxA-DNA-binding<-PIN<-?<-Peptidase_M15 | 1220 | Bacteria>Terrabacteria_group>Bacillota>Negativicutes>Selenomonadales>Selenomonadaceae | GCA_015058685.1 | Schwartzia_succinivorans | E7198_09540 | 51689..55351 | 55507 | BREX-system-P-loop-protein-BrxC-[Schwartzia-succinivorans] |
| MBQ4404900.1 | Type1.BREX | <-sGTP-MglA+sGTP<-PglZ<-N6-METHYLASE<-DUF6079*<-BrxB-BrxF-iSTAND<-BrxA-DNA-binding<-His_biosynth<-?<-?<-?<-?<-?<-SIS_2+SIS | 1169 | Bacteria>Terrabacteria_group>Bacillota>Negativicutes>Selenomonadales>Selenomonadaceae | GCA_017412925.1 | Selenomonadaceae_bacterium | II857_10910 | 28330..31839 | 2652289 | BREX-system-P-loop-protein-BrxC-[Selenomonadaceae-bacterium] |
| MBQ1876908.1 | Type1.BREX | MORC->N6-METHYLASE->?->?->BrxA-DNA-binding->BrxB-BrxF-iSTAND->DUF6079*->N6-METHYLASE-> | 1191 | Bacteria>Terrabacteria_group>Bacillota>Negativicutes>Selenomonadales>Selenomonadaceae | GCA_017426625.1 | Selenomonas_sp. | II145_09455 | 12087..15662 | 2053611 | BREX-system-P-loop-protein-BrxC-[Selenomonas-sp.] |
| MBR3457117.1 | Type1.BREX | <-PglZ-p<-HNH<-OLD-ABC<-N6-METHYLASE-p<-?<-PIN<-N6-METHYLASE<-DUF6079*<-BrxB-BrxF-iSTAND-p | 1215 | Bacteria>Terrabacteria_group>Bacillota>Negativicutes>Selenomonadales>Selenomonadaceae | GCA_017509225.1 | Selenomonadaceae_bacterium | IKH16_03040 | 6412..10059 | 2652289 | BREX-system-P-loop-protein-BrxC-[Selenomonadaceae-bacterium] |
| MBR6889065.1 | Type1.BREX | <-PglZ<-N6-METHYLASE<-DUF6079*<-BrxB-BrxF-iSTAND<-BrxA-DNA-binding<-His_biosynth<-?<-?<-?<-?<-?<-SIS_2+SIS | 1169 | Bacteria>Terrabacteria_group>Bacillota>Negativicutes>Selenomonadales>Selenomonadaceae | GCA_017529935.1 | Selenomonadaceae_bacterium | IKN16_11585 | 6417..9926 | 2652289 | BREX-system-P-loop-protein-BrxC-[Selenomonadaceae-bacterium] |
| MBD6179950.1 | Type1.BREX | <-DUF6079*<-BrxB-BrxF-iSTAND<-BrxA-DNA-binding<-?<-DDE_Tnp_4<-Transposase_31\|\|?->MACRODOMAIN->?->STYKIN-> | 1182 | Bacteria>Terrabacteria_group>Bacillota>Negativicutes>Selenomonadales>Selenomonadaceae | GCA_017634675.1 | Selenomonadaceae_bacterium | J6004_12320 | 370..3918 | 2652289 | BREX-system-P-loop-protein-BrxC-[Selenomonadaceae-bacterium] |
| MBP8598624.1 | Type1.BREX | <-McrC-NTD+REase<-McrB-AAA<-PglZ<-HTH+REase<-N6-METHYLASE<-DUF6079*<-BrxB-BrxF-iSTAND<-BrxA-DNA-binding<-HTH_26<-?<-ACET<-?<-HAMP+HISKIN<-REC+OmpR-HTH<-BirA-HTH | 1194 | Bacteria>Terrabacteria_group>Bacillota>Negativicutes>Selenomonadales>Selenomonadaceae | GCA_018052705.1 | Selenomonas_sp. | KBI24_05335 | 21839..25423 | 2053611 | BREX-system-P-loop-protein-BrxC-[Selenomonas-sp.] |
| SEH31389.1 | Type1.BREX | <-PglZ<-PIN<-?<-N6-METHYLASE<-DUF6079*<-BrxB-BrxF-iSTAND<-BrxA-DNA-binding<-OLD-ABC<-ParB<-ParA-Soj-PloopNTPase<-?<-HTH_3\|\|?->ParA-Soj-PloopNTPase-> | 1188 | Bacteria>Terrabacteria_group>Bacillota>Negativicutes>Selenomonadales>Selenomonadaceae | GCA_900108415.1 | Selenomonas_ruminantium | SAMN05216583_11711 | 10616..14182 | 971 | hypothetical-protein-SAMN05216583_11711-[Selenomonas-ruminantium] |
| WP_013740677.1 | Type1.BREX | SAM-METHYLASE->?->?-><-REase<-?\|\|?->CBS->BrxA-DNA-binding->BrxB-BrxF-iSTAND->DUF6079*->N6-METHYLASE->Schlafen+BirA-HTH->PglZ->?->SIGMA+OB-Fold+AAA-ATPase+Lon-Protease-BrxL-p->?->?->?->sGTP-> | 1180 | Bacteria>Terrabacteria_group>Bacillota>Negativicutes>Selenomonadales>Selenomonadaceae | GCF_000208405.1 | Selenomonas_sputigena | SELSP_RS03620 | 821879..825421 | 69823 | BREX-system-P-loop-protein-BrxC-[Selenomonas-sputigena] |
| WP_037354735.1 | Type1.BREX | DDE_Tnp_1-><-?<-?<-?\|\|?->BrxA-DNA-binding->BrxB-BrxF-iSTAND->DUF6079*->N6-METHYLASE->OLD-ABC->HNH->?->PglZ->SIGMA+OB-Fold+AAA-ATPase+Lon-Protease-BrxL->HTH_3-> | 1182 | Bacteria>Terrabacteria_group>Bacillota>Negativicutes>Selenomonadales>Selenomonadaceae | GCF_000701565.1 | Selenomonas_sp._FC4001 | T320_RS08130 | 15634..19182 | 1408313 | BREX-system-P-loop-protein-BrxC-[Selenomonas-sp.-FC4001] |
| WP_149189168.1 | Type1.BREX | <-PglZ<-?<-N6-METHYLASE<-DUF6079*<-BrxB-BrxF-iSTAND<-BrxA-DNA-binding<-HAD+HAD<-REC+GGDEF+EAL<-?\|\|CNMP+CRP-HTH-> | 1189 | Bacteria>Terrabacteria_group>Bacillota>Negativicutes>Selenomonadales>Selenomonadaceae | GCF_008271345.1 | Selenomonas_sp._mPRGC8 | FZ041_RS07805 | 13049..16618 | 2606905 | BREX-system-P-loop-protein-BrxC-[Selenomonas-sp.-mPRGC8] |
| WP_273038291.1 | Type1.BREX | <-SIGMA+OB-Fold+AAA-ATPase+Lon-Protease-BrxL<-PglZ<-REase<-OLD-ABC<-N6-METHYLASE<-DUF6079*<-BrxB-BrxF-iSTAND<-BrxA-DNA-binding<-PIN<-?<-Peptidase_M15 | 1220 | Bacteria>Terrabacteria_group>Bacillota>Negativicutes>Selenomonadales>Selenomonadaceae | GCF_015058685.1 | Schwartzia_succinivorans | E7198_RS09640 | 51689..55351 | 55507 | BREX-system-P-loop-protein-BrxC-[Schwartzia-succinivorans] |
| WP_273698075.1 | Type1.BREX | <-REase<-PglZ<-N6-METHYLASE<-PIN<-OLD-ABC<-HNH<-OLD-ABC<-DUF6079*<-BrxB-BrxF-iSTAND<-BrxA-DNA-binding<-?<-?<-?<-?<-SKI<-?\|\|O6MGT-HTH-> | 1181 | Bacteria>Terrabacteria_group>Bacillota>Negativicutes>Selenomonadales>Selenomonadaceae | GCF_905371815.1 | Selenomonas_noxia | NW174_RS07850 | 36953..40498 | 135083 | BREX-system-P-loop-protein-BrxC-[Selenomonas-noxia] |
| WP_273079788.1 | Type1.BREX | BrxA-DNA-binding->BrxB-BrxF-iSTAND->DUF6079*->OLD-ABC->HNH->OLD-ABC->PIN->N6-METHYLASE->PglZ->SIR2->HerA/FtsK-> | 1187 | Bacteria>Terrabacteria_group>Bacillota>Negativicutes>Selenomonadales>Selenomonadaceae | GCF_905371945.1 | Selenomonas_felix | NW177_RS10645 | 2029..5592 | 1944634 | BREX-system-P-loop-protein-BrxC-[Selenomonas-felix] |
| WP_018704563.1 | Type1.BREX | <-REase<-?<-?<-SIGMA-HTH\|\|NACHT+SNaCT->XIS-HTH->BrxA-DNA-binding->BrxB-BrxF-iSTAND->DUF6079*->N6-METHYLASE->PglZ->SIGMA+OB-Fold+AAA-ATPase+Lon-Protease-BrxL->?->Resolvase+Recombinase+ZnR->GDResolvase-> | 1174 | Bacteria>Terrabacteria_group>Bacillota>Negativicutes>Selenomonadales>Sporomusaceae | GCF_000374545.1 | Anaeromusa_acidaminophila | C508_RS0115895 | 22597..26121 | 81464 | BREX-system-P-loop-protein-BrxC-[Anaeromusa-acidaminophila] |
| WP_145101507.1 | Type1.BREX | <-SF1-SF2-Helicase+PRTase<-RADICAL-SAM<-?<-ABC-ATPase+SbcC+ABC-ATPase+SbcC\|\|SF1-SF2-Helicase-><-PglZ<-?<-N6-METHYLASE<-N6-METHYLASE-p<-DUF6079*<-BrxB-BrxF-iSTAND<-BrxA-DNA-binding<-HTH_26<-TOPRIM+Topoisom_bac+TOPRIM_Crpt<-?<-SF1-SF2-Helicase\|\|Acyltransferase-> | 1211 | Bacteria>Terrabacteria_group>Bacillota>Negativicutes>Selenomonadales>Sporomusaceae | GCF_007830295.1 | Sporomusa_sp._KB1 | SALPA_RS27425 | 5094234..5097869 | 943346 | BREX-system-P-loop-protein-BrxC-[Sporomusa-sp.-KB1] |
| WP_250726535.1 | Type1.BREX | CR-ATPase5+REase-><-DOC<-?<-?<-SIGMA+OB-Fold+AAA-ATPase+Lon-Protease-BrxL<-PglZ<-N6-METHYLASE<-DUF6079*<-BrxB-BrxF-iSTAND<-BrxA-DNA-binding<-XIS-HTH<-?<-tRNA<-?<-?\|\|PAS+HISKIN->ACET-> | 1193 | Bacteria>Terrabacteria_group>Bacillota>Negativicutes>Selenomonadales>Sporomusaceae | GCF_023660125.1 | Sporomusa_sphaeroides | M7775_RS08455 | 32998..36579 | 47679 | BREX-system-P-loop-protein-BrxC-[Sporomusa-sphaeroides] |
| NCB77375.1 | Type1.BREX | <-OLD-ABC<-SIGMA+OB-Fold+AAA-ATPase+Lon-Protease-BrxL<-PglZ<-NERD<-OLD-ABC<-N6-METHYLASE<-Four-helical-protein<-DUF6079*<-BrxB-BrxF-iSTAND<-BrxA-DNA-binding<-HTH_26\|\|XIS-HTH->XIS-HTH-><-Schlafen+HATPase+wHTH<-?<-?<-MarR-HTH | 1210 | Bacteria>Terrabacteria_group>Bacillota>Negativicutes>unclassified_Negativicutes | GCA_009929485.1 | Negativicutes_bacterium | EOM42_09855 | 20012..23644 | 2499140 | BREX-system-P-loop-protein-BrxC-[Negativicutes-bacterium] |
| NCD09159.1 | Type1.BREX | <-DUF3644<-?<-HNH<-OLD-ABC<-N6-METHYLASE<-?<-DUF6079*<-BrxB-BrxF-iSTAND<-BrxA-DNA-binding<-?<-Phage_integrase | 1142 | Bacteria>Terrabacteria_group>Bacillota>Negativicutes>unclassified_Negativicutes | GCA_009930275.1 | Negativicutes_bacterium | EOL98_06980 | 8743..12171 | 2499140 | BREX-system-P-loop-protein-BrxC-[Negativicutes-bacterium] |
| ETI97421.1 | Type1.BREX | HTH_26->BrxA-DNA-binding->BrxB-BrxF-iSTAND->DUF6079*->N6-METHYLASE->?->?->PglZ->SIGMA+OB-Fold+AAA-ATPase+Lon-Protease-BrxL-p->SIGMA+OB-Fold+AAA-ATPase+Lon-Protease-BrxL->?-><-?<-HTH_Tnp_1+HTH_21+rve+rve_3 | 1184 | Bacteria>Terrabacteria_group>Bacillota>Negativicutes>Veillonellales>Veillonellaceae | GCA_000508645.1 | Veillonella_sp._DORA_B_18_19_23 | Q621_VSBC00051G0004 | 2127..5681 | 1403933 | MAG:-hypothetical-protein-Q621_VSBC00051G0004-[Veillonella-sp.-DORA_B_18_19_23] |
| HJE83014.1 | Type1.BREX | <-SIGMA+OB-Fold+AAA-ATPase+Lon-Protease-BrxL<-PglZ<-Schlafen+HATPase+wHTH<-N6-METHYLASE<-DUF6079*<-BrxB-BrxF-iSTAND<-BrxA-DNA-binding<-GDResolvase\|\|?->GDResolvase+HIN-HTH->?-><-HTH_3 | 1205 | Bacteria>Terrabacteria_group>Bacillota>Negativicutes>Veillonellales>Veillonellaceae | GCA_020742115.1 | Megasphaera_stantonii | K8V17_06240 | 9916..13533 | 2144175 | BREX-system-P-loop-protein-BrxC-[Megasphaera-stantonii] |
| MCH4217192.1 | Type1.BREX | BrxA-DNA-binding->BrxB-BrxF-iSTAND->DUF6079*->N6-METHYLASE->HTH+REase->SbcC->Schlafen+BirA-HTH->PglZ->SIGMA+OB-Fold+AAA-ATPase+Lon-Protease-BrxL-><-tRNA<-HHH+DNA_pol_B+RNase\|\|Calcineurin-> | 1206 | Bacteria>Terrabacteria_group>Bacillota>Negativicutes>Veillonellales>Veillonellaceae | GCA_022486845.1 | Megasphaera_sp. | LKF74_01360 | 2380..6000 | 2023260 | BREX-system-P-loop-protein-BrxC-[Megasphaera-sp.] |
| MCI1750263.1 | Type1.BREX | <-Cas_Cas1<-?<-HEPN\|\|DDE_5+DDE_Tnp_1-><-SIGMA+OB-Fold+AAA-ATPase+Lon-Protease-BrxL<-PglZ<-?<-?<-N6-METHYLASE<-DUF6079*<-BrxB-BrxF-iSTAND<-BrxA-DNA-binding<-HTH_26<-?<-HNH+3C-ABC\|\|D5N+D5ATPase->Vrr-REase->SF1-SF2-Helicase-p->SIGMA-HTH-> | 1166 | Bacteria>Terrabacteria_group>Bacillota>Negativicutes>Veillonellales>Veillonellaceae | GCA_022647805.1 | Megasphaera_cerevisiae | LKI17_02755 | 578892..582392 | 39029 | BREX-system-P-loop-protein-BrxC-[Megasphaera-cerevisiae] |
| UYJ17329.1 | Type1.BREX | <-Glyoxalase<-?<-HEPN<-SIGMA+OB-Fold+AAA-ATPase+Lon-Protease-BrxL<-PglZ<-?<-?<-N6-METHYLASE<-DUF6079*<-BrxB-BrxF-iSTAND<-BrxA-DNA-binding<-HTH_26<-?<-HNH-p<-HNH+3C-ABC\|\|SIGMA-HTH->MazG->D5N+D5ATPase-> | 1171 | Bacteria>Terrabacteria_group>Bacillota>Negativicutes>Veillonellales>Veillonellaceae | GCA_025757885.1 | Veillonellaceae_bacterium | OGM58_02475 | 522598..526113 | 2049049 | MAG:-BREX-system-P-loop-protein-BrxC-[Veillonellaceae-bacterium] |
| WP_020310321.1 | Type1.BREX | BrxA-DNA-binding->BrxB-BrxF-iSTAND->DUF6079*->N6-METHYLASE->PglZ->SIGMA+OB-Fold+AAA-ATPase+Lon-Protease-BrxL->?->ArdCN+MPTase->ABC-ATPase+ABC-ATPase+ABC-ATPase+SbcC-> | 1205 | Bacteria>Terrabacteria_group>Bacillota>Negativicutes>Veillonellales>Veillonellaceae | GCF_000417505.1 | Megasphaera_sp._NM10 | NM10_RS03990 | 1866..5483 | 1273103 | BREX-system-P-loop-protein-BrxC-[Megasphaera-sp.-NM10] |
| WP_020723763.1 | Type1.BREX | OrfB_Zn_ribbon-><-ABC-ATPase+ABC-ATPase+ABC-ATPase+SbcC<-ArdCN+MPTase<-?<-SIGMA+OB-Fold+AAA-ATPase+Lon-Protease-BrxL<-PglZ<-PglZ-p<-N6-METHYLASE<-DUF6079*<-BrxB-BrxF-iSTAND<-BrxA-DNA-binding | 1205 | Bacteria>Terrabacteria_group>Bacillota>Negativicutes>Veillonellales>Veillonellaceae | GCF_000417525.1 | Megasphaera_sp._BL7 | G153_RS02755 | 15363..18980 | 1285585 | BREX-system-P-loop-protein-BrxC-[Megasphaera-sp.-BL7] |
| WP_036202192.1 | Type1.BREX | OrfB_Zn_ribbon->Calcineurin->?->HicB_lk_antitox-><-HicB_lk_antitox<-?\|\|?->BrxA-DNA-binding->BrxB-BrxF-iSTAND->DUF6079*->N6-METHYLASE->PglZ->SIGMA+OB-Fold+AAA-ATPase+Lon-Protease-BrxL->?->ArdCN+MPTase-><-?\|\|?-><-HAD+HAD+NUDIX | 1205 | Bacteria>Terrabacteria_group>Bacillota>Negativicutes>Veillonellales>Veillonellaceae | GCF_000763195.1 | Megasphaera_elsdenii | JY94_RS00850 | 17323..20940 | 907 | BREX-system-P-loop-protein-BrxC-[Megasphaera-elsdenii] |
| WP_107195318.1 | Type1.BREX | Calcineurin->OrfB_Zn_ribbon-><-SIGMA+OB-Fold+AAA-ATPase+Lon-Protease-BrxL<-PglZ<-N6-METHYLASE<-DUF6079*<-BrxB-BrxF-iSTAND<-BrxA-DNA-binding<-GDResolvase\|\|?-><-tRNA<-?\|\|?->?->?->RADICAL-SAM | 1205 | Bacteria>Terrabacteria_group>Bacillota>Negativicutes>Veillonellales>Veillonellaceae | GCF_003367905.1 | Megasphaera_stantonii | DKB62_RS07720 | 1651247..1654864 | 2144175 | BREX-system-P-loop-protein-BrxC-[Megasphaera-stantonii] |
| WP_118021977.1 | Type1.BREX | ABhydrolase-><-HTH_3<-ArdCN<-SIGMA+OB-Fold+AAA-ATPase+Lon-Protease-BrxL<-PglZ<-DOC+HTH<-N6-METHYLASE<-DUF6079*<-BrxB-BrxF-iSTAND<-BrxA-DNA-binding<-tRNA<-TPR+TPR\|\|tRNA->?->?->?->?-><-?<-Classical-AAA | 1205 | Bacteria>Terrabacteria_group>Bacillota>Negativicutes>Veillonellales>Veillonellaceae | GCF_003467125.1 | Megasphaera_sp._AM44-1BH | DW949_RS04090 | 102789..106406 | 2292358 | BREX-system-P-loop-protein-BrxC-[Megasphaera-sp.-AM44-1BH] |
| WP_169013945.1 | Type1.BREX | <-ArdCN+MPTase<-?<-SIGMA+OB-Fold+AAA-ATPase+Lon-Protease-BrxL<-PglZ<-Schlafen+HATPase+wHTH<-N6-METHYLASE<-DUF6079*<-BrxB-BrxF-iSTAND<-BrxA-DNA-binding<-?\|\|?->HicB_lk_antitox-><-HicB_lk_antitox<-?<-Calcineurin | 1205 | Bacteria>Terrabacteria_group>Bacillota>Negativicutes>Veillonellales>Veillonellaceae | GCF_012928525.1 | Megasphaera_elsdenii | HG933_RS10950 | 13141..16758 | 907 | BREX-system-P-loop-protein-BrxC-[Megasphaera-elsdenii] |
| WP_195394664.1 | Type1.BREX | BrxA-DNA-binding->BrxB-BrxF-iSTAND->DUF6079*->PglZ->SWACOS->REase-><-?\|\|?-><-HAD+HAD | 1203 | Bacteria>Terrabacteria_group>Bacillota>Negativicutes>Veillonellales>Veillonellaceae | GCF_015554925.1 | Veillonella_parvula | I2E05_RS02830 | 246695..250306 | 29466 | BREX-system-P-loop-protein-BrxC-[Veillonella-parvula] |
| WP_221741438.1 | Type1.BREX | <-SIGMA+OB-Fold+AAA-ATPase+Lon-Protease-BrxL<-PglZ<-?<-?<-N6-METHYLASE<-DUF6079*<-BrxB-BrxF-iSTAND<-BrxA-DNA-binding<-HTH_26<-?<-?<-HAD+S6PP<-Peptidase_M20 | 1184 | Bacteria>Terrabacteria_group>Bacillota>Negativicutes>Veillonellales>Veillonellaceae | GCF_019731115.1 | Veillonella_parvula | HCG55_RS08060 | 301540..305094 | 29466 | BREX-system-P-loop-protein-BrxC-[Veillonella-parvula] |
| WP_245675156.1 | Type1.BREX | DDE_Tnp_IS66+DDE_Tnp_IS66_C-><-?<-SIGMA+OB-Fold+AAA-ATPase+Lon-Protease-BrxL<-PglZ<-Schlafen+HATPase+wHTH<-HerA/FtsK<-SIR2<-N6-METHYLASE<-DUF6079-p<-DUF6079*<-BrxB-BrxF-iSTAND<-BrxA-DNA-binding<-Y1_Tnp<-tRNA\|\|FN3+Calcineurin-> | 1013 | Bacteria>Terrabacteria_group>Bacillota>Negativicutes>Veillonellales>Veillonellaceae | GCF_900103535.1 | Megasphaera_paucivorans | BLR11_RS11785 | 16117..19158 | 349095 | BREX-system-P-loop-protein-BrxC-[Megasphaera-paucivorans] |
| WP_108850313.1 | Type1.BREX | <-Glyoxalase<-?<-HEPN<-SIGMA+OB-Fold+AAA-ATPase+Lon-Protease-BrxL<-PglZ<-?<-?<-N6-METHYLASE<-DUF6079*<-BrxB-BrxF-iSTAND<-BrxA-DNA-binding<-HTH_26<-?<-HNH+3C-ABC\|\|D5N+D5ATPase->Vrr-REase->SF1-SF2-Helicase->SIGMA-HTH-> | 1171 | Bacteria>Terrabacteria_group>Bacillota>Negativicutes>Veillonellales>Veillonellaceae | GCF_900343095.1 | Dialister_massiliensis | DHT67_RS06445 | 1390627..1394142 | 2161821 | BREX-system-P-loop-protein-BrxC-[Dialister-massiliensis] |
| WP_273096524.1 | Type1.BREX | <-N6-METHYLASE-p<-N6-METHYLASE<-DUF6079*<-BrxB-BrxF-iSTAND<-BrxA-DNA-binding<-?<-?<-?<-?<-?<-LuxR-HTH | 1218 | Bacteria>Terrabacteria_group>Bacillota>Negativicutes>Veillonellales>Veillonellaceae | GCF_905188205.1 | Dialister_succinatiphilus | NW100_RS05055 | 3776..7432 | 487173 | BREX-system-P-loop-protein-BrxC-[Dialister-succinatiphilus] |
| WP_128751700.1 | Type1.BREX | VWA+SIGMA-HTH->TOPRIM+Topoisom_bac->DNA_METHYLASE->ArdCN+LPD32+LPD36+N6_Mtase+SF1-SF2-Helicase->?->JAB->HTH_26->BrxA-DNA-binding->BrxB-BrxF-iSTAND->DUF6079*->N6-METHYLASE->?->PglZ->SIGMA+OB-Fold+AAA-ATPase+Lon-Protease-BrxL->Polbeta->?-><-TraI-VirD2-RCRE<-TraJ-RHH | 1197 | Bacteria>Terrabacteria_group>Bacillota>Tissierellia>Tissierellales>Acidilutibacteraceae | GCF_004103715.1 | Acidilutibacter_cellobiosedens | EQM13_RS01230 | 244231..247824 | 2507161 | BREX-system-P-loop-protein-BrxC-[Acidilutibacter-cellobiosedens] |
| WP_128752230.1 | Type1.BREX | Radical_SAM->?->?->?->?->?-><-?\|\|HTH_26->BrxA-DNA-binding->BrxB-BrxF-iSTAND->DUF6079*->N6-METHYLASE->?->?->?->?->RAMPs->?->RAMPs->PglZ-> | 1195 | Bacteria>Terrabacteria_group>Bacillota>Tissierellia>Tissierellales>Acidilutibacteraceae | GCF_004103715.1 | Acidilutibacter_cellobiosedens | EQM13_RS05940 | 1255764..1259351 | 2507161 | BREX-system-P-loop-protein-BrxC-[Acidilutibacter-cellobiosedens] |
| HHX69703.1 | Type1.BREX | <-N6-METHYLASE-p<-DUF6079*<-BrxB-BrxF-iSTAND<-BrxA-DNA-binding<-HTH_26\|\|PAIREDC-HTH->PAIREDC-HTH-> | 1209 | Bacteria>Terrabacteria_group>Bacillota>Tissierellia>Tissierellales>Peptoniphilaceae | GCA_012841535.1 | Gallicola_sp. | GX708_16865 | 1137..4766 | 2683910 | BREX-system-P-loop-protein-BrxC-[Gallicola-sp.] |
| MBS6525206.1 | Type1.BREX | HATPase_c_4->DEOR-HTH-><-ABC-ATPase+SbcC<-ABC-ATPase+SbcC<-?<-?<-?\|\|BrxA-DNA-binding->BrxB-BrxF-iSTAND->DUF6079*->N6-METHYLASE->PglZ->SIGMA+OB-Fold+AAA-ATPase+Lon-Protease-BrxL-> | 1183 | Bacteria>Terrabacteria_group>Bacillota>Tissierellia>Tissierellales>Peptoniphilaceae | GCA_018377275.1 | Peptoniphilaceae_bacterium | KH322_04220 | 38899..42450 | 1891242 | BREX-system-P-loop-protein-BrxC-[Peptoniphilaceae-bacterium] |
| WP_094203897.1 | Type1.BREX | <-PglZ<-N6-METHYLASE<-DUF6079*<-BrxB-BrxF-iSTAND<-BrxA-DNA-binding<-?<-?<-?\|\|?-><-ABC-ATPase+SbcC | 1183 | Bacteria>Terrabacteria_group>Bacillota>Tissierellia>Tissierellales>Peptoniphilaceae | GCF_002243175.1 | Finegoldia_magna | B9N53_RS04675 | 18982..22533 | 1260 | BREX-system-P-loop-protein-BrxC-[Finegoldia-magna] |
| WP_243659990.1 | Type1.BREX | <-ABC-ATPase+SbcC<-?<-?\|\|CHTH->Schlafen+HATPase+wHTH->RVT+HEPN->HTH_26->BrxA-DNA-binding->BrxB-BrxF-iSTAND->DUF6079*->N6-METHYLASE->PglZ->SIGMA+OB-Fold+AAA-ATPase+Lon-Protease-BrxL->?->?->?-><-?\|\|?->ABC-ATPase+SbcC-> | 1219 | Bacteria>Terrabacteria_group>Bacillota>Tissierellia>Tissierellales>Peptoniphilaceae | GCF_016458825.1 | Miniphocaeibacter_halophilus | JFY71_RS06390 | 1303116..1306775 | 2931922 | BREX-system-P-loop-protein-BrxC-[Miniphocaeibacter-halophilus] |
| WP_055078308.1 | Type1.BREX | ABC-ATPase+SbcC+ABC-ATPase+SbcC->?->BPD_transp_2->Amidohydro_1->?->BrxA-DNA-binding->BrxB-BrxF-iSTAND->DUF6079*->N6-METHYLASE->PglZ->?->?-><-Peptidase_M42+Peptidase_M28\|\|AA_kinase+ACT_7->?->NAD_binding_3+Homoserine_dh->Thr_synth_N+PALP-> | 1186 | Bacteria>Terrabacteria_group>Bacillota>Tissierellia>Tissierellales>Peptoniphilaceae | GCF_902375645.1 | Lagierella_massiliensis | FY373_RS02695 | 569444..573004 | 1689303 | BREX-system-P-loop-protein-BrxC-[Lagierella-massiliensis] |
| WP_226808069.1 | Type1.BREX | <-RNase<-PglZ<-N6-METHYLASE<-DUF6079*<-BrxB-BrxF-iSTAND<-BrxA-DNA-binding | 1185 | Bacteria>Terrabacteria_group>Bacillota>Tissierellia>Tissierellales>Sporanaerobacteraceae | - | Anaerosalibacter_bizertensis | KUA25_13485 | 7035..10592 | 932217 | BREX-system-P-loop-protein-BrxC-[Anaerosalibacter-bizertensis] |

| Protein-ID | System | Gene-Neighbourhoods | Length | Taxonomy | Assembly-ID | Strain | Locus-Tag | Genomic-coordinates | TaxID | Fasta-define |
|---|---|---|---|---|---|---|---|---|---|---|
| WP_072743893.1 | Type1.BREX | HTH_26->?->DOC+HTH->HAD+HAD->ParB+HNH->BrxA-DNA-binding->BrxB-BrxF-iSTAND->DUF6079*->N6-METHYLASE->OLD-ABC->TOPRIM->PglZ->SIGMA+OB-Fold+AAA-ATPase+Lon-Protease-BrxL->?->?->HDRnase-> | 1192 | Bacteria>Terrabacteria_group>Bacillota>Tissierellia>Tissierellales>Sporanaerobacteraceae | GCF_900130025.1 | Sporanaerobacter_acetigenes | BUA21_RS05940 | 73351..76929 | 165813 | BREX-system-P-loop-protein-BrxC-[Sporanaerobacter-acetigenes] |
| WP_077369895.1 | Type1.BREX | DNA_METHYLASE->?->?->RVT+HNH->?->JAB->HTH_26->BrxA-DNA-binding->BrxB-BrxF-iSTAND->DUF6079*->N6-METHYLASE->PglZ->SIGMA+OB-Fold+AAA-ATPase+Lon-Protease-BrxL->Polbeta->?-><-TraI-VirD2-RCRE<-TraJ-RHH\|\|?->Preader+URI-DNase-> | 1197 | Bacteria>Terrabacteria_group>Bacillota>Tissierellia>Tissierellales>Sporanaerobacteraceae | GCF_900155565.1 | Anaerosalibacter_sp._Marseille-P3206 | BQ9840_RS11430 | 2085841..2089434 | 1871005 | BREX-system-P-loop-protein-BrxC-[Anaerosalibacter-sp.-Marseille-P3206] |
| MBZ2174150.1 | Type1.BREX | <-Cysteine-Desulfurase<-ICLR-HTH<-?<-?<-?<-?<-SIGMA+OB-Fold+AAA-ATPase+Lon-Protease-BrxL<-PglZ<-N6-METHYLASE<-DUF6079*<-BrxB-BrxF-iSTAND<-BrxA-DNA-binding | 1184 | Bacteria>Terrabacteria_group>Bacillota>Tissierellia>Tissierellales>Tepidimicrobiaceae | GCA_019931165.1 | Schnuerera_sp._xch1 | K8M07_02700 | 74326..77880 | 2874283 | BREX-system-P-loop-protein-BrxC-[Schnuerera-sp.-xch1] |
| WP_132026817.1 | Type1.BREX | <-EVE+McrB-AAA<-?\|\|?-><-HKD-DNase<-ParB+HNH<-SIGMA+OB-Fold+AAA-ATPase+Lon-Protease-BrxL<-PglZ<-N6-METHYLASE<-N6-METHYLASE<-DUF6079*<-BrxB-BrxF-iSTAND<-BrxA-DNA-binding<-?<-?<-?<-CRISPR_Cas2<-Cas_Cas1<-Cas9+HNH+RuvC_III | 1197 | Bacteria>Terrabacteria_group>Bacillota>Tissierellia>Tissierellales>Tepidimicrobiaceae | GCF_016767175.1 | Keratinibaculum_paraultunense | JL105_RS06905 | 1394896..1398489 | 1278232 | BREX-system-P-loop-protein-BrxC-[Keratinibaculum-paraultunense] |
| WP_263282436.1 | Type1.BREX | <-Cysteine-Desulfurase<-ICLR-HTH<-?<-?<-?<-?<-SIGMA+OB-Fold+AAA-ATPase+Lon-Protease-BrxL<-PglZ<-N6-METHYLASE<-DUF6079*<-BrxB-BrxF-iSTAND<-BrxA-DNA-binding | 1179 | Bacteria>Terrabacteria_group>Bacillota>Tissierellia>Tissierellales>Tepidimicrobiaceae | GCF_019931165.1 | Schnuerera_sp._xch1 | K8M07_RS02695 | 74326..77865 | 2874283 | BREX-system-P-loop-protein-BrxC-[Schnuerera-sp.-xch1] |
| WP_223376349.1 | Type1.BREX | <-Cysteine-Desulfurase<-ICLR-HTH<-?<-?<-?<-?<-SIGMA+OB-Fold+AAA-ATPase+Lon-Protease-BrxL<-PglZ<-N6-METHYLASE<-DUF6079*<-BrxB-BrxF-iSTAND<-BrxA-DNA-binding | 1184 | Bacteria>Terrabacteria_group>Bacillota>Tissierellia>Tissierellales>Tepidimicrobiaceae | - | Schnuerera_sp._xch1 | K8M07_02700 | 74326..77880 | 2874283 | BREX-system-P-loop-protein-BrxC-[Schnuerera-sp.-xch1] |
| WP_054870024.1 | Type1.BREX | SIGMA-HTH+SIGMA-HTH+SIGMA-HTH-><-?\|\|METHYLASE->?->?->RNaseH->ncRNA->?->BrxA-DNA-binding->BrxB-BrxF-iSTAND->DUF6079*->N6-METHYLASE->OLD-ABC->TOPRIM->PglZ->SIGMA+OB-Fold+AAA-ATPase+Lon-Protease-BrxL->SbcC->HNH->?->ARG-> | 1187 | Bacteria>Terrabacteria_group>Bacillota>Tissierellia>Tissierellales>Thermohalobacteraceae | GCF_001316435.1 | Caloranaerobacter_sp._TR13 | TR13x_RS01045 | 223070..226633 | 1302151 | BREX-system-P-loop-protein-BrxC-[Caloranaerobacter-sp.-TR13] |
| MCK9445088.1 | Type1.BREX | ABC-ATPase+SbcC->ABC-ATPase+SbcC->?->?->?->?->BrxA-DNA-binding->BrxB-BrxF-iSTAND->DUF6079*->N6-METHYLASE-p-> | 1184 | Bacteria>Terrabacteria_group>Bacillota>Tissierellia>Tissierellales>Tissierellaceae | GCA_023229175.1 | Tissierellaceae_bacterium | MOQ14_11240 | 8982..12536 | 2562714 | BREX-system-P-loop-protein-BrxC-[Tissierellaceae-bacterium] |
| WP_154440136.1 | Type1.BREX | TetR-HTH->?->?->?->?->BrxA-DNA-binding->BrxB-BrxF-iSTAND->DUF6079*->PglZ->SIGMA+OB-Fold+AAA-ATPase+Lon-Protease-BrxL->?-><-ZnR | 1195 | Bacteria>Terrabacteria_group>Bacillota>Tissierellia>Tissierellales>Tissierellaceae | GCF_009695605.1 | Tissierella_pigra | FYJ83_RS09760 | 55642..59229 | 2607614 | BREX-system-P-loop-protein-BrxC-[Tissierella-pigra] |
| WP_216584698.1 | Type1.BREX | TetR-HTH->?->?->?->?->BrxA-DNA-binding->BrxB-BrxF-iSTAND->DUF6079*->PglZ->SIGMA+OB-Fold+AAA-ATPase+Lon-Protease-BrxL->?-><-ZnR<-?<-?\|\|HTH_Tnp_1+HTH_21+rve+rve_3-><-ZnR | 1195 | Bacteria>Terrabacteria_group>Bacillota>Tissierellia>Tissierellales>Tissierellaceae | GCF_018917845.1 | Tissierella_pigra | KQI41_RS01725 | 332242..335829 | 2607614 | BREX-system-P-loop-protein-BrxC-[Tissierella-pigra] |
| WP_154440327.1 | Type1.BREX | primpol+D5N+D5ATPase-><-?\|\|?->?->RHH_8->REase->N6-METHYLASE->BrxA-DNA-binding->BrxB-BrxF-iSTAND->DUF6079*->PglZ->BrxD-AAA->SF1-SF2-Helicase+wHTH+Small-B-barrel+DUF1998->?->CarD+HTH->ZnR+HARE-HTH-> | 1380 | Bacteria>Terrabacteria_group>Bacillota>Tissierellia>Tissierellales>Tissierellaceae | GCF_018917845.1 | Tissierella_pigra | KQI41_RS04690 | 949565..953707 | 2607614 | hypothetical-protein-[Tissierella-pigra] |
| MBV1758099.1 | Type1.BREX | HEPN+RES->APATPase+FtsK+VirB4->?->SMS-KaiC->HTH_36->HTH_26->BrxA-DNA-binding->BrxB-BrxF-iSTAND->DUF6079*->ZnR+ZnR->DUF4404->HAD+HAD-> | 1213 | Bacteria>Terrabacteria_group>Bacillota>Tissierellia>Tissierellia_incertae_sedis>Dethiosulfatibacter | GCA_019084435.1 | Dethiosulfatibacter_sp. | KMY55_09680 | 27127..30768 | 2812851 | BREX-system-P-loop-protein-BrxC-[Dethiosulfatibacter-sp.] |
| MBV1758946.1 | Type1.BREX | <-SIGMA+OB-Fold+AAA-ATPase-BrxL<-PglZ<-DUF6079*<-BrxB-BrxF-iSTAND<-DUF4007<-N6-METHYLASE-p | 1366 | Bacteria>Terrabacteria_group>Bacillota>Tissierellia>Tissierellia_incertae_sedis>Dethiosulfatibacter | GCA_019084435.1 | Dethiosulfatibacter_sp. | KMY55_14030 | 2857..6957 | 2812851 | hypothetical-protein-[Dethiosulfatibacter-sp.] |
| WP_073050525.1 | Type1.BREX | <-RNase<-NERD<-OLD-ABC<-REase<-SIGMA+OB-Fold+AAA-ATPase+Lon-Protease-BrxL<-PglZ<-HEPN<-N6-METHYLASE<-?<-DUF6079*<-BrxB-BrxF-iSTAND<-BrxA-DNA-binding<-HTH_26 | 1214 | Bacteria>Terrabacteria_group>Bacillota>Tissierellia>Tissierellia_incertae_sedis>Dethiosulfatibacter | GCF_900142005.1 | Dethiosulfatibacter_aminovorans | BUB05_RS15460 | 41174..44818 | 332095 | BREX-system-P-loop-protein-BrxC-[Dethiosulfatibacter-aminovorans] |
| OUL09467.1 | Type1.BREX | <-SF1-SF2-Helicase+REase<-SF1-SF2-Helicase+REase<-?<-SIGMA+OB-Fold+AAA-ATPase+Lon-Protease-BrxL<-PglZ<-REase<-N6-METHYLASE<-DUF6079*<-BrxB-BrxF-iSTAND<-BrxA-DNA-binding<-HTH_26<-?<-?<-HTH+REase<-?<-?<-Glyco_hydro_1 | 1207 | Bacteria>Terrabacteria_group>Bacillota>Tissierellia>Tissierellia_incertae_sedis>Sedimentibacter | GCA_002155105.1 | Sedimentibacter_sp._SX930 | B0533_04695 | 278529..282152 | 1955060 | ATPase-[Sedimentibacter-sp.-SX930] |
| MBP7221715.1 | Type1.BREX | <-McrB-NTD+McrB-AAA<-?<-?<-REase<-?<-SbcC<-SIGMA+OB-Fold+AAA-ATPase+Lon-Protease-BrxL<-PglZ<-N6-METHYLASE<-DUF6079*<-BrxB-BrxF-iSTAND<-BrxA-DNA-binding<-?<-?<-35exo<-?<-?<-REase+RNase<-ARG | 1201 | Bacteria>Terrabacteria_group>Bacillota>Tissierellia>Tissierellia_incertae_sedis>Sedimentibacter | GCA_017995755.1 | Sedimentibacter_sp. | KBA50_00585 | 18309..21914 | 1960295 | BREX-system-P-loop-protein-BrxC-[Sedimentibacter-sp.] |
| NLW23292.1 | Type1.BREX | TetR-HTH->HD+PspB->ACET->Peptidase_S58->?->BrxB-BrxF-iSTAND->DnaJ->BrxA-DNA-binding->BrxB-BrxF-iSTAND->DUF6079*->N6-METHYLASE->PglZ->?->N6-METHYLASE-> | 1176 | Bacteria>Terrabacteria_group>Bacillota>Tissierellia>unclassified_Tissierellia | GCA_012728725.1 | Tissierellia_bacterium | GXY88_08600 | 13444..16974 | 2049431 | BREX-system-P-loop-protein-BrxC-[Tissierellia-bacterium] |
| NMB27654.1 | Type1.BREX | BrxA-DNA-binding->BrxB-BrxF-iSTAND->DUF6079*->N6-METHYLASE->PglZ->?-><-PHP\|\|?->?->ABC-ATPase+SbcC->ACET-> | 1184 | Bacteria>Terrabacteria_group>Bacillota>Tissierellia>unclassified_Tissierellia | GCA_012798935.1 | Tissierellia_bacterium | GX987_06350 | 6707..10261 | 2049431 | BREX-system-P-loop-protein-BrxC-[Tissierellia-bacterium] |
| HHV26669.1 | Type1.BREX | <-Cysteine-Desulfurase<-RAMA<-?<-?<-?<-RNase<-PglZ<-N6-METHYLASE<-DUF6079*<-BrxB-BrxF-iSTAND<-BrxA-DNA-binding<-?<-?<-?<-?<-ABC-ATPase+SbcC\|\|REC+OmpR-HTH->HISKIN-> | 1185 | Bacteria>Terrabacteria_group>Bacillota>Tissierellia>unclassified_Tissierellia | GCA_012839575.1 | Tissierellia_bacterium | GXX63_05675 | 30640..34197 | 2049431 | BREX-system-P-loop-protein-BrxC-[Tissierellia-bacterium] |
| PKM61864.1 | Type1.BREX | HTH_26->BrxA-DNA-binding->BrxB-BrxF-iSTAND->DUF6079*->N6-METHYLASE->?->N6-METHYLASE->PglZ->SIGMA+OB-Fold+AAA-ATPase+Lon-Protease-BrxL-><-DOC<-Phd-YefM | 1189 | Bacteria>Terrabacteria_group>Bacillota>unclassified_Bacillota | GCA_002841405.1 | Firmicutes_bacterium_HGW-Firmicutes-4 | CVU99_01555 | 23012..26581 | 2013785 | BREX-system-P-loop-protein-BrxC-[Firmicutes-bacterium-HGW-Firmicutes-4] |
| PKM86293.1 | Type1.BREX | <-DUF6079-p*<-BrxB-BrxF-iSTAND<-BrxA-DNA-binding<-?<-?\|\|MACRODOMAIN-><-SpoOE-like | 806 | Bacteria>Terrabacteria_group>Bacillota>unclassified_Bacillota | GCA_002841595.1 | Firmicutes_bacterium_HGW-Firmicutes-12 | CVU87_12500 | 1..2418 | 2013773 | BREX-system-P-loop-protein-BrxC,-partial-[Firmicutes-bacterium-HGW-Firmicutes-12] |
| RGH23082.1 | Type1.BREX | PemK_toxin->?->?->BrxA-DNA-binding->BrxB-BrxF-iSTAND->DUF6079*->N6-METHYLASE->OLD-ABC->HNH->PglZ-><-?\|\|?->?-><-ZnR | 1180 | Bacteria>Terrabacteria_group>Bacillota>unclassified_Bacillota | GCA_003461705.1 | Firmicutes_bacterium_AF12-30 | DWV72_07055 | 48691..52233 | 2292884 | BREX-system-P-loop-protein-BrxC-[Firmicutes-bacterium-AF12-30] |
| RHP99067.1 | Type1.BREX | GNTR-HTH+Aminotran_1_2->?->HTH_26->BrxA-DNA-binding->BrxB-BrxF-iSTAND->DUF6079*->Tox-ART-HYD2->N6-METHYLASE->HEPN->Schlafen+HATPase+wHTH->PglZ->SIGMA+OB-Fold+AAA-ATPase+Lon-Protease-BrxL->?->RNase->SKI-> | 1215 | Bacteria>Terrabacteria_group>Bacillota>unclassified_Bacillota | GCA_003478205.1 | Firmicutes_bacterium_AM59-13 | DXA19_01120 | 43130..46777 | 2292897 | BREX-system-P-loop-protein-BrxC-[Firmicutes-bacterium-AM59-13] |
| RHT37458.1 | Type1.BREX | <-ClpABC-AAA<-?<-SIGMA+OB-Fold+AAA-ATPase+Lon-Protease-BrxL<-PglZ<-Schlafen+HATPase+wHTH<-HEPN<-N6-METHYLASE<-?<-DUF6079*<-BrxB-BrxF-iSTAND<-HTH_26<-HTH_36<-?<-?\|\|DSBH-><-ABhydrolase | 1182 | Bacteria>Terrabacteria_group>Bacillota>unclassified_Bacillota | GCA_003480425.1 | Firmicutes_bacterium_AM31-12AC | DW790_07855 | 25743..29291 | 2292892 | BREX-system-P-loop-protein-BrxC-[Firmicutes-bacterium-AM31-12AC] |
| MBJ7595965.1 | Type1.BREX | <-SIGMA+OB-Fold+AAA-ATPase+HKD-DNase-BrxL<-PglZ<-N6-METHYLASE<-DUF6079*<-BrxB-BrxF-iSTAND<-BrxA-DNA-binding | 1162 | Bacteria>Terrabacteria_group>Candidatus_Dormibacteraeota>unclassified_Candidatus_Dormibacteraeota | GCA_016464765.1 | Candidatus_Dormibacteraeota_bacterium | JF886_14125 | 8709..12197 | 2052315 | BREX-system-P-loop-protein-BrxC-[Candidatus-Dormibacteraeota-bacterium] |
| MAT43664.1 | Type1.BREX | <-AAA-ATPase<-TPR+TPR+TPR+TPR<-?<-?<-?<-?<-SIGMA+OB-Fold+AAA-ATPase+HKD-DNase-BrxL<-PglZ<-N6-METHYLASE<-DUF6079*<-BrxB-BrxF-iSTAND<-BrxA-DNA-binding<-CBS+HEPN\|\|?->SF1-SF2-Helicase+DUF1998->ParB<-?<-rve | 1169 | Bacteria>Terrabacteria_group>Chloroflexota>Anaerolineae>Anaerolineales>Anaerolineaceae | GCA_002702705.1 | Anaerolineaceae_bacterium | CL609_15105 | 30474..33983 | 2024896 | hypothetical-protein-[Anaerolineaceae-bacterium] |
| MAT45245.1 | Type1.BREX | <-SIGMA+OB-Fold+AAA-ATPase+HKD-DNase-BrxL<-PglZ<-HTH+MPTase<-HigB_toxin<-RADICAL-SAM<-?<-?<-NACHT<-N6-METHYLASE<-DUF6079*<-BrxB-BrxF-iSTAND<-BrxA-DNA-binding<-XIS-HTH<-?<-TniQ<-TnsC-ATPase<-rve+Mu-transpos_C<-REase | 1172 | Bacteria>Terrabacteria_group>Chloroflexota>Anaerolineae>Anaerolineales>Anaerolineaceae | GCA_002702705.1 | Anaerolineaceae_bacterium | CL609_23195 | 14888..18406 | 2024896 | hypothetical-protein-[Anaerolineaceae-bacterium] |
| MBE2233048.1 | Type1.BREX | <-SIGMA+OB-Fold+AAA-ATPase+HKD-DNase-BrxL<-PglZ<-HTH+MPTase<-HigB_toxin<-N6-METHYLASE<-SF1-SF2-Helicase<-DUF6079* | 1166 | Bacteria>Terrabacteria_group>Chloroflexota>Anaerolineae>Anaerolineales>Anaerolineaceae | GCA_014879765.1 | Anaerolinea_sp. | IAE85_06095 | 14605..18105 | 1872519 | BREX-system-P-loop-protein-BrxC-[Anaerolinea-sp.] |
| MBQ6502300.1 | Type1.BREX | DOC+HTH-><-?<-?\|\|?->?->BrxA-DNA-binding->BrxB-BrxF-iSTAND->DUF6079*->OLD-ABC->TOPRIM->?->?->N6-METHYLASE->PglZ->SIGMA+OB-Fold+AAA-ATPase+HKD-DNase-BrxL->ARG->ParB+HNH-> | 1194 | Bacteria>Terrabacteria_group>Chloroflexota>Anaerolineae>Anaerolineales>Anaerolineaceae | GCA_017432685.1 | Flexilinea_sp. | IJI57_00125 | 8965..12549 | 2017390 | BREX-system-P-loop-protein-BrxC-[Flexilinea-sp.] |
| MBQ6343447.1 | Type1.BREX | BrxA-DNA-binding->BrxB-BrxF-iSTAND->DUF6079*-><-?<-?\|\|PhdYeFM_antitox->PIN->?->?->?->N6-METHYLASE-> | 1175 | Bacteria>Terrabacteria_group>Chloroflexota>Anaerolineae>Anaerolineales>Anaerolineaceae | GCA_017432985.1 | Anaerolineaceae_bacterium | IJI41_10010 | 1911..5438 | 2024896 | BREX-system-P-loop-protein-BrxC-[Anaerolineaceae-bacterium] |
| MBQ6518172.1 | Type1.BREX | Recombinase->HTH_26->BrxA-DNA-binding->BrxB-BrxF-iSTAND->DUF6079*->ParB+HNH->?->N6-METHYLASE->MORC+REase->Schlafen+HATPase+wHTH+HTH->PglZ->TIR->SIGMA+OB-Fold+AAA-ATPase+Lon-Protease-BrxL->KAP-NTPase-> | 1176 | Bacteria>Terrabacteria_group>Chloroflexota>Anaerolineae>Anaerolineales>Anaerolineaceae | GCA_017433515.1 | Anaerolineaceae_bacterium | IJI14_05585 | 5340..8870 | 2024896 | BREX-system-P-loop-protein-BrxC-[Anaerolineaceae-bacterium] |
| MBP9040834.1 | Type1.BREX | SF1-SF2-Helicase-><-SIGMA+OB-Fold+AAA-ATPase+Lon-Protease-BrxL<-PglZ<-N6-METHYLASE<-HEPN<-DUF6079* | 1023 | Bacteria>Terrabacteria_group>Chloroflexota>Anaerolineae>Anaerolineales>Anaerolineaceae | GCA_018057145.1 | Anaerolineaceae_bacterium | KBF64_03565 | 8301..11374 | 2024896 | BREX-system-P-loop-protein-BrxC-[Anaerolineaceae-bacterium] |
| MCD4671323.1 | Type1.BREX | <-DUF6079-p*<-BrxB-BrxF-iSTAND<-BrxA-DNA-binding<-XIS-HTH<-?<-HTH_3\|\|XIS-HTH->Phage_integrase-><-tRNA<-REase | 728 | Bacteria>Terrabacteria_group>Chloroflexota>Anaerolineae>Anaerolineales>Anaerolineaceae | GCA_021107195.1 | Anaerolineaceae_bacterium | K8R77_01555 | 1..2186 | 2024896 | BREX-system-P-loop-protein-BrxC-[Anaerolineaceae-bacterium] |
| WP_062190148.1 | Type1.BREX | <-sGTP<-DDE_Tnp<-SIGMA+OB-Fold+AAA-ATPase+Lon-Protease-BrxL<-PglZ<-N6-METHYLASE<-HEPN<-DUF6079*<-?<-?<-N6-METHYLASE<-BrxB-BrxF-iSTAND<-BrxA-DNA-binding<-HTH_26<-CHTH | 1245 | Bacteria>Terrabacteria_group>Chloroflexota>Anaerolineae>Anaerolineales>Anaerolineaceae | GCF_001050195.2 | Anaerolinea_thermolimosa | ATHL_RS04285 | 935293..939030 | 229919 | BREX-system-P-loop-protein-BrxC-[Anaerolinea-thermolimosa] |
| WP_062419207.1 | Type1.BREX | ACET->?->MULE-TRANSPOSASE->DDE_Tnp-><-N6-METHYLASE-p<-SIGMA+OB-Fold+AAA-ATPase-BrxL<-PglZ<-?<-N6-METHYLASE<-DUF6079*<-LON-AAA<-DUF4007\|\|DnaB+SMS-KaiC->XIS-HTH->Phage_integrase->?-><-Phage_int_SAM_4+Phage_integrase\|\|?->ParA-Soj-PloopNTPase-> | 1371 | Bacteria>Terrabacteria_group>Chloroflexota>Anaerolineae>Anaerolineales>Anaerolineaceae | GCF_001050255.2 | Levilinea_saccharolytica | LSAC_RS17710 | 3184296..3188411 | 229921 | chitobiase/beta-hexosaminidase-C-terminal-domain-containing-protein-[Levilinea-saccharolytica] |
| WP_062421183.1 | Type1.BREX | <-Phage_integrase<-XIS-HTH<-?\|\|LuxR-HTH->CHTH->XIS-HTH->?->DUF4007->LON-AAA->DUF6079*->N6-METHYLASE->PglZ->SIGMA+OB-Fold+AAA-ATPase-BrxL->N6-METHYLASE-p->MarR-HTH->?->pPIWI-><-METHYLASE+SF1-SF2-Helicase | 1371 | Bacteria>Terrabacteria_group>Chloroflexota>Anaerolineae>Anaerolineales>Anaerolineaceae | GCF_001050275.1 | Leptolinea_tardivitalis | ADM99_RS14205 | 186230..190345 | 229920 | chitobiase/beta-hexosaminidase-C-terminal-domain-containing-protein-[Leptolinea-tardivitalis] |
| WP_116223389.1 | Type1.BREX | <-SAM-METHYLASE\|\|?->?->?->?-><-?\|\|DUF4007->LON-AAA->DUF6079*->N6-METHYLASE->NERD->PglZ->SIGMA+OB-Fold+AAA-ATPase-BrxL->N6-METHYLASE-p-><-HIN-HTH<-DUF488\|\|?->RNasePH+RNasePH+KH+S1-> | 1371 | Bacteria>Terrabacteria_group>Chloroflexota>Anaerolineae>Anaerolineales>Anaerolineaceae | GCF_003966975.1 | Pelolinea_submarina | Pelsub_RS12620 | 2863892..2868007 | 913107 | chitobiase/beta-hexosaminidase-C-terminal-domain-containing-protein-[Pelolinea-submarina] |
| MBN1316703.1 | Type1.BREX | XIS-HTH->?-><-URI-DNase<-?\|\|HTH_3->MPTase->BrxA-DNA-binding->BrxB-BrxF-iSTAND->DUF6079-p*-> | 471 | Bacteria>Terrabacteria_group>Chloroflexota>Anaerolineae>Anaerolineales>unclassified_Anaerolineales | - | Anaerolineales_bacterium | JXA42_14590 | 5464..6877 | 2073117 | hypothetical-protein-[Anaerolineales-bacterium] |
| TEU18421.1 | Type1.BREX | <-ARG<-?<-Lon_2<-?<-DUF433<-SIGMA+OB-Fold+AAA-ATPase-BrxL<-PglZ<-DUF6079*<-BrxB-BrxF-iSTAND<-?<-DUF4007<-N6-METHYLASE<-NTP_transf_5 | 1365 | Bacteria>Terrabacteria_group>Chloroflexota>Anaerolineae>Anaerolineales>unclassified_Anaerolineales | GCA_004377365.1 | Anaerolineales_bacterium | E3J21_06050 | 13257..17354 | 2073117 | ATP-binding-protein-[Anaerolineales-bacterium] |
| NLF02163.1 | Type1.BREX | N6-METHYLASE-p->DUF6079*->HEPN->PglZ->SIGMA+OB-Fold+AAA-ATPase+Lon-Protease-BrxL->HIN-HTH-> | 1237 | Bacteria>Terrabacteria_group>Chloroflexota>Anaerolineae>Anaerolineales>unclassified_Anaerolineales | GCA_012514695.1 | Anaerolineales_bacterium | GX601_14420 | 1151..4864 | 2073117 | BREX-system-P-loop-protein-BrxC-[Anaerolineales-bacterium] |
| NLF02862.1 | Type1.BREX | Phage_integrase->LuxR-HTH->Phage_integrase->?->BrxA-DNA-binding->BrxB-BrxF-iSTAND->DUF6079*->N6-METHYLASE->ParB+HNH->PglZ->SIGMA+OB-Fold+AAA-ATPase+Lon-Protease-BrxL-> | 1157 | Bacteria>Terrabacteria_group>Chloroflexota>Anaerolineae>Anaerolineales>unclassified_Anaerolineales | GCA_012514695.1 | Anaerolineales_bacterium | GX601_18015 | 11185..14658 | 2073117 | BREX-system-P-loop-protein-BrxC-[Anaerolineales-bacterium] |
| MBN1450028.1 | Type1.BREX | <-SIGMA+OB-Fold+AAA-ATPase+Lon-Protease-BrxL<-PglZ<-REase+DUF5655<-N6-METHYLASE<-HTH+HKD-DNase<-Pkinase+SF1-SF2-Helicase<-?<-NERD+Pkinase<-DUF6079*<-BrxB-BrxF-iSTAND<-BrxA-DNA-binding | 1209 | Bacteria>Terrabacteria_group>Chloroflexota>Anaerolineae>Anaerolineales>unclassified_Anaerolineales | GCA_016928525.1 | Anaerolineales_bacterium | JW963_03360 | 18534..22163 | 2073117 | BREX-system-P-loop-protein-BrxC-[Anaerolineales-bacterium] |
| MBN1451020.1 | Type1.BREX | <-DUF6079*<-BrxB-BrxF-iSTAND<-BrxA-DNA-binding<-XIS-HTH\|\|HEPN-><-?<-?<-?<-Phage_integrase | 1079 | Bacteria>Terrabacteria_group>Chloroflexota>Anaerolineae>Anaerolineales>unclassified_Anaerolineales | GCA_016928525.1 | Anaerolineales_bacterium | JW963_08400 | 1..3239 | 2073117 | BREX-system-P-loop-protein-BrxC-[Anaerolineales-bacterium] |
| MCA9933330.1 | Type1.BREX | HTH_28+rve->XIS-HTH-><-?\|\|BrxA-DNA-binding->BrxB-BrxF-iSTAND->DUF6079*->N6-METHYLASE->HEPN->PglZ->SIGMA+OB-Fold+AAA-ATPase+HKD-DNase-BrxL-><-CopG_antitoxin<-BrnT_toxin<-GDResolvase+HIN-HTH\|\|?->TIR-> | 1174 | Bacteria>Terrabacteria_group>Chloroflexota>Anaerolineae>Anaerolineales>unclassified_Anaerolineales | GCA_020432585.1 | Anaerolineales_bacterium | KC415_05370 | 4959..8483 | 2073117 | BREX-system-P-loop-protein-BrxC-[Anaerolineales-bacterium] |
| MCB0100628.1 | Type1.BREX | PTSEIIA1->MarR-HTH->?->BrxA-DNA-binding->BrxB-BrxF-iSTAND->DUF6079*->HEPN->N6-METHYLASE->PglZ->SIGMA+OB-Fold+AAA-ATPase+Lon-Protease-BrxL->ParB+HNH->?->HTH_3-> | 1152 | Bacteria>Terrabacteria_group>Chloroflexota>Anaerolineae>Anaerolineales>unclassified_Anaerolineales | GCA_020433725.1 | Anaerolineales_bacterium | KDD74_01185 | 15066..18524 | 2073117 | BREX-system-P-loop-protein-BrxC-[Anaerolineales-bacterium] |
| MCC6192471.1 | Type1.BREX | <-SIGMA+OB-Fold+AAA-ATPase+Lon-Protease-BrxL<-PglZ<-Schlafen+HATPase+wHTH<-N6-METHYLASE<-DUF6079*<-BrxB-BrxF-iSTAND<-BrxA-DNA-binding\|\|?->REase-> | 1152 | Bacteria>Terrabacteria_group>Chloroflexota>Anaerolineae>Anaerolineales>unclassified_Anaerolineales | GCA_020845955.1 | Anaerolineales_bacterium | IT318_25875 | 21631..25089 | 2073117 | BREX-system-P-loop-protein-BrxC-[Anaerolineales-bacterium] |
| MCJ7529874.1 | Type1.BREX | rve->?->?->PIN->?->PemK_toxin->BrxA-DNA-binding->BrxB-BrxF-iSTAND->DUF6079*->?->Schlafen->N6-METHYLASE->HigB_toxin->HTH+MPTase->PglZ->SIGMA+OB-Fold+AAA-ATPase+HKD-DNase-BrxL-><-HTH_37 | 1170 | Bacteria>Terrabacteria_group>Chloroflexota>Anaerolineae>Anaerolineales>unclassified_Anaerolineales | GCA_022865645.1 | Anaerolineales_bacterium | MUO64_02430 | 4866..8378 | 2073117 | BREX-system-P-loop-protein-BrxC-[Anaerolineales-bacterium] |
| MCU0485175.1 | Type1.BREX | <-SbcC<-?<-Sulfate_transp+STAS\|\|?->?->DOC+HTH->BrxA-DNA-binding->BrxB-BrxF-iSTAND->DUF6079*->N6-METHYLASE->?->SF1-SF2-Helicase->PglZ->SIGMA+OB-Fold+AAA-ATPase+Lon-Protease-BrxL-> | 1163 | Bacteria>Terrabacteria_group>Chloroflexota>Anaerolineae>Anaerolineales>unclassified_Anaerolineales | GCA_025454595.1 | Anaerolineales_bacterium | MUC85_03595 | 6459..9950 | 2073117 | BREX-system-P-loop-protein-BrxC-[Anaerolineales-bacterium] |
| OIO89050.1 | Type1.BREX | ParA-Soj-PloopNTPase->?-><-?\|\|?->?->?->BrxA-DNA-binding->BrxB-BrxF-iSTAND->DUF6079*-> | 1228 | Bacteria>Terrabacteria_group>Chloroflexota>Anaerolineae>unclassified_Anaerolineae | GCA_001871755.1 | Anaerolineae_bacterium_CG2_30_64_16 | AUK03_15190 | 14542..18228 | 1805005 | MAG:-hypothetical-protein-AUK03_15190-[Anaerolineae-bacterium-CG2_30_64_16] |
| HEC36340.1 | Type1.BREX | <-SIGMA+OB-Fold+AAA-ATPase+Lon-Protease-BrxL<-PglZ<-N6-METHYLASE<-TPR<-HicB<-HicA_toxin<-N6-METHYLASE<-DUF6079*<-HEPN<-N6-METHYLASE<-?<-MarR-HTH<-Schlafen+HATPase+wHTH+HTH\|\|RNase-> | 1243 | Bacteria>Terrabacteria_group>Chloroflexota>Anaerolineae>unclassified_Anaerolineae | GCA_011052725.1 | Anaerolineae_bacterium | ENI39_07385 | 20646..24377 | 2052143 | BREX-system-P-loop-protein-BrxC-[Anaerolineae-bacterium] |
| MBK9093475.1 | Type1.BREX | RNase->?->?->?->BrxA-DNA-binding->BrxB-BrxF-iSTAND->DUF6079*->OLD-ABC+TOPRIM->N6-METHYLASE->APATPase->TPR+TPR+TPR+TPR->TPR+TPR+TPR+TPR->PglZ-p->OLD-ABC->?->PglZ-> | 1151 | Bacteria>Terrabacteria_group>Chloroflexota>Anaerolineae>unclassified_Anaerolineae | GCA_016719785.1 | Anaerolineae_bacterium | IPM84_12015 | 171966..175421 | 2052143 | BREX-system-P-loop-protein-BrxC-[Anaerolineae-bacterium] |
| MBK9096490.1 | Type1.BREX | <-SIGMA+OB-Fold+AAA-ATPase+Lon-Protease-BrxL<-PglZ<-ParB+HNH<-HEPN+REase<-?<-N6-METHYLASE<-TOPRIM<-OLD-ABC<-DUF6079-p*<-DUF6079-p<-BrxB-BrxF-iSTAND<-BrxA-DNA-binding<-?\|\|?->N6-METHYLASE-p->RVT-> | 900 | Bacteria>Terrabacteria_group>Chloroflexota>Anaerolineae>unclassified_Anaerolineae | GCA_016719785.1 | Anaerolineae_bacterium | IPM84_27780 | 244922..247624 | 2052143 | hypothetical-protein-[Anaerolineae-bacterium] |
| MBL7202371.1 | Type1.BREX | <-N6-METHYLASE-p<-DUF6079*<-BrxB-BrxF-iSTAND<-BrxA-DNA-binding<-?<-PolB<-PolB | 1168 | Bacteria>Terrabacteria_group>Chloroflexota>Anaerolineae>unclassified_Anaerolineae | GCA_016783145.1 | Anaerolineae_bacterium | ISS56_19725 | 1222..4728 | 2052143 | BREX-system-P-loop-protein-BrxC-[Anaerolineae-bacterium] |
| MBN1139877.1 | Type1.BREX | <-HTH+REase<-NACHT+GUN4<-?<-SIGMA+OB-Fold+AAA-ATPase+Lon-Protease-BrxL<-PglZ<-N6-METHYLASE<-?<-DUF6079*<-BrxB-BrxF-iSTAND<-BrxA-DNA-binding | 1204 | Bacteria>Terrabacteria_group>Chloroflexota>Anaerolineae>unclassified_Anaerolineae | GCA_016926595.1 | Anaerolineae_bacterium | JXM73_25115 | 15012..18626 | 2052143 | BREX-system-P-loop-protein-BrxC-[Anaerolineae-bacterium] |
| MBN1933573.1 | Type1.BREX | DUF6079*->PglZ->SIGMA+OB-Fold+AAA-ATPase-BrxL->?->SAM-METHYLASE->?->Glyoxalase->Beta-lactamase-> | 1063 | Bacteria>Terrabacteria_group>Chloroflexota>Anaerolineae>unclassified_Anaerolineae | GCA_016931365.1 | Anaerolineae_bacterium | JW934_02850 | 1..3193 | 2052143 | hypothetical-protein-[Anaerolineae-bacterium] |
| UCC65222.1 | Type1.BREX | <-PIN<-Phd-YefM\|\|?->MarR-HTH-><-?<-?<-HHH+DNA_pol_B+RNase<-DUF6079-p*<-BrxB-BrxF-iSTAND\|\|NTP_transferase+MannoseP_isomer->?-><-?<-SWACOS<-NTP_transferase\|\|PSBP->Peptidase_M22-> | 700 | Bacteria>Terrabacteria_group>Chloroflexota>Anaerolineae>unclassified_Anaerolineae | GCA_020344275.1 | Anaerolineae_bacterium | JSV36_09375 | 1994468..1996567 | 2052143 | MAG:-BREX-system-P-loop-protein-BrxC,-partial-[Anaerolineae-bacterium] |
| MCL4299606.1 | Type1.BREX | <-DnaJ<-?<-N6-METHYLASE<-N6-METHYLASE<-?<-?<-?<-REase<-DUF6079*<-BrxB-BrxF-iSTAND<-BrxA-DNA-binding<-MPTase<-HTH_3\|\|?->?->SMODS->?->JAB-> | 1231 | Bacteria>Terrabacteria_group>Chloroflexota>Anaerolineae>unclassified_Anaerolineae | GCA_023382535.1 | Anaerolineae_bacterium | KJ077_27970 | 96973..100668 | 2052143 | BREX-system-P-loop-protein-BrxC-[Anaerolineae-bacterium] |
| MBK7780587.1 | Type1.BREX | <-PglZ-p\|\|TOPRIM->?->NUDIX-><-N6-METHYLASE\|\|MqsA_antitoxin+DUF4065->?->NACHT-><-DUF6079-p<-DUF6079-p*<-BrxB-BrxF-iSTAND<-BrxA-DNA-binding\|\|?->Peptidase->TPR+TPR-> | 931 | Bacteria>Terrabacteria_group>Chloroflexota>Ardenticatenia>unclassified_Ardenticatenia | GCA_016709185.1 | Ardenticatenia_bacterium | IPJ58_07315 | 1935822..1938617 | 2099665 | BREX-system-P-loop-protein-BrxC-[Ardenticatenia-bacterium] |
| MBK7781823.1 | Type1.BREX | HAD+HAD->?->REase->?->Peptidase->?->?->BrxA-DNA-binding->BrxB-BrxF-iSTAND->DUF6079*-><-DOC+zhth<-MazE_antitoxin\|\|?->?-><-?<-?<-PIN<-PhdYeFM_antitox | 1168 | Bacteria>Terrabacteria_group>Chloroflexota>Ardenticatenia>unclassified_Ardenticatenia | GCA_016709185.1 | Ardenticatenia_bacterium | IPJ58_13850 | 157782..161288 | 2099665 | BREX-system-P-loop-protein-BrxC-[Ardenticatenia-bacterium] |
| MXZ22476.1 | Type1.BREX | <-PAPS-reduct\|\|BrxA-DNA-binding->BrxB-BrxF-iSTAND->DUF6079*->N6-METHYLASE->PglZ->SIGMA+OB-Fold+AAA-ATPase+Lon-Protease-BrxL-> | 1172 | Bacteria>Terrabacteria_group>Chloroflexota>Caldilineae>Caldilineales>Caldilineaceae | GCA_009837805.1 | Caldilineaceae_bacterium_SB0665_bin_25 | F4Y84_18080 | 7617..11135 | 2605262 | BREX-system-P-loop-protein-BrxC-[Caldilineaceae-bacterium-SB0665_bin_25] |
| MBW7883550.1 | Type1.BREX | DUF6079*->N6-METHYLASE-p->DOC+HTH->PglZ->SIGMA+OB-Fold+AAA-ATPase+Lon-Protease-BrxL-> | 1157 | Bacteria>Terrabacteria_group>Chloroflexota>Caldilineae>Caldilineales>Caldilineaceae | GCA_019455155.1 | Caldilineaceae_bacterium | H3C34_13100 | 829..4302 | 2614250 | BREX-system-P-loop-protein-BrxC-[Caldilineaceae-bacterium] |
| MBW7885561.1 | Type1.BREX | Phage_int_SAM_4+Phage_integrase->?->BrxA-DNA-binding->BrxB-BrxF-iSTAND->DUF6079-p*-> | 720 | Bacteria>Terrabacteria_group>Chloroflexota>Caldilineae>Caldilineales>Caldilineaceae | GCA_019455155.1 | Caldilineaceae_bacterium | H3C34_23620 | 3376..5536 | 2614250 | hypothetical-protein-[Caldilineaceae-bacterium] |
| MBX3053885.1 | Type1.BREX | ABC-ATPase+SbcC-><-MerR-HTH<-?\|\|BrxA-DNA-binding->BrxB-BrxF-iSTAND->DUF6079*->N6-METHYLASE->PglZ->SIGMA+OB-Fold+AAA-ATPase+Lon-Protease-BrxL->McrB-AAA-> | 1153 | Bacteria>Terrabacteria_group>Chloroflexota>Caldilineae>Caldilineales>Caldilineaceae | GCA_019637685.1 | Caldilineaceae_bacterium | KF753_20575 | 58506..61967 | 2614250 | BREX-system-P-loop-protein-BrxC-[Caldilineaceae-bacterium] |
| MCB0024047.1 | Type1.BREX | BrxA-DNA-binding->BrxB-BrxF-iSTAND->DUF6079-p*->DUF6079->N6-METHYLASE-p-> | 86 | Bacteria>Terrabacteria_group>Chloroflexota>Caldilineae>Caldilineales>Caldilineaceae | GCA_020433165.1 | Caldilinea_sp. | KDD91_13525 | 1905..2161 | 2293560 | hypothetical-protein-[Caldilinea-sp.] |
| MCB0024048.1 | Type1.BREX | BrxA-DNA-binding->BrxB-BrxF-iSTAND->DUF6079-p->DUF6079*->N6-METHYLASE-p-> | 1006 | Bacteria>Terrabacteria_group>Chloroflexota>Caldilineae>Caldilineales>Caldilineaceae | GCA_020433165.1 | Caldilinea_sp. | KDD91_13530 | 2333..5353 | 2293560 | BREX-system-P-loop-protein-BrxC-[Caldilinea-sp.] |
| MCB0149305.1 | Type1.BREX | <-DUF6079*<-DUF6079-p<-BrxB-BrxF-iSTAND<-BrxA-DNA-binding<-SIGMA-HTH<-SIGMA-HTH | 1029 | Bacteria>Terrabacteria_group>Chloroflexota>Caldilineae>Caldilineales>Caldilineaceae | GCA_020433585.1 | Caldilineaceae_bacterium | KDE01_16850 | 104..3193 | 2614250 | BREX-system-P-loop-protein-BrxC-[Caldilineaceae-bacterium] |
| MCB9117629.1 | Type1.BREX | <-DUF6079*<-BrxB-BrxF-iSTAND<-BrxA-DNA-binding<-SIGMA-HTH<-SIGMA-HTH\|\|Peptidase-> | 1145 | Bacteria>Terrabacteria_group>Chloroflexota>Caldilineae>Caldilineales>Caldilineaceae | GCA_020634815.1 | Caldilineaceae_bacterium | H6639_21995 | 310..3747 | 2614250 | BREX-system-P-loop-protein-BrxC-[Caldilineaceae-bacterium] |
| MCL4835316.1 | Type1.BREX | <-N6-METHYLASE-p<-?<-URI-DNase<-DUF6079*<-BrxB-BrxF-iSTAND<-BrxA-DNA-binding<-RADICAL-SAM | 1206 | Bacteria>Terrabacteria_group>Chloroflexota>Caldilineae>Caldilineales>Caldilineaceae | GCA_023384455.1 | Caldilineaceae_bacterium | KJZ86_22950 | 2317..5937 | 2614250 | BREX-system-P-loop-protein-BrxC-[Caldilineaceae-bacterium] |
| NUQ36643.1 | Type1.BREX | <-N6-METHYLASE-p<-OLD-ABC+TOPRIM<-DUF6079*<-BrxB-BrxF-iSTAND<-BrxA-DNA-binding<-?\|\|?->?->N6-N4-METHYLASE-p->AbiEi_4+AbiEi_1->AbiEii+ZnR->PglZ-p-> | 1152 | Bacteria>Terrabacteria_group>Chloroflexota>Caldilineae>Caldilineales>unclassified_Caldilineales | GCA_013360575.1 | Caldilineales_bacterium | HUU23_02375 | 3098..6556 | 2306055 | BREX-system-P-loop-protein-BrxC-[Caldilineales-bacterium] |
| MBX7236823.1 | Type1.BREX | CNMP->MNS+REase->?->BrxA-DNA-binding->BrxB-BrxF-iSTAND->DUF6079*->RNase->N6-METHYLASE-p-> | 1151 | Bacteria>Terrabacteria_group>Chloroflexota>Caldilineae>Caldilineales>unclassified_Caldilineales | GCA_019695115.1 | Caldilineales_bacterium | K1X65_20755 | 25151..28606 | 2306055 | BREX-system-P-loop-protein-BrxC-[Caldilineales-bacterium] |
| RRR74017.1 | Type1.BREX | HTH_3->ParA-Soj-PloopNTPase->ParB->35exo->?-><-SIGMA+OB-Fold+AAA-ATPase+Lon-Protease-BrxL<-PglZ<-?<-N6-METHYLASE<-DUF6079*<-BrxB-BrxF-iSTAND<-BrxA-DNA-binding\|\|?->?->?->APATPase+TPR+TPR+TPR+TPR+TPR+TPR+TPR->APATPase+TPR+TPR+TPR+TPR+TPR+TPR+TPR-> | 1326 | Bacteria>Terrabacteria_group>Chloroflexota>Chloroflexia>Chloroflexales>Chloroflexineae | GCA_003934145.1 | Candidatus_Viridilinea_halotolerans | EI684_07910 | 16787..20767 | 2491704 | BREX-system-P-loop-protein-BrxC-[Candidatus-Viridilinea-halotolerans] |
| NJN66674.1 | Type1.BREX | <-N6-METHYLASE-p<-PIN<-CopG/RHH<-N6-METHYLASE-p<-DUF6079-p<-DUF6079-p*<-DUF6079-p<-BrxB-BrxF-iSTAND<-BrxA-DNA-binding<-HTH_3\|\|?->?->?-><-DDE_Tnp\|\|DUF4158-> | 607 | Bacteria>Terrabacteria_group>Chloroflexota>Chloroflexia>Chloroflexales>Chloroflexineae | GCA_012032475.1 | Chloroflexaceae_bacterium | HC884_08105 | 4075..5898 | 2562117 | hypothetical-protein-[Chloroflexaceae-bacterium] |
| WP_097651212.1 | Type1.BREX | <-PglZ<-PIN<-?<-N6-METHYLASE-p<-PIN<-CopG/RHH<-?<-PolB<-N6-METHYLASE<-DUF6079*<-BrxB-BrxF-iSTAND<-BrxA-DNA-binding<-?\|\|?-><-?<-?<-ParA-Soj-PloopNTPase+ParA-Soj-PloopNTPase | 1204 | Bacteria>Terrabacteria_group>Chloroflexota>Chloroflexia>Chloroflexales>Chloroflexineae | GCF_002532075.1 | Candidatus_Chloroploca_asiatica | A9Q02_RS04750 | 75845..79459 | 1506545 | BREX-system-P-loop-protein-BrxC-[Candidatus-Chloroploca-asiatica] |
| WP_205712750.1 | Type1.BREX | <-N6-METHYLASE-p\|\|REase+N6_Mtase->SbcC->?-><-DUF6079*<-Four-helical-protein | 1238 | Bacteria>Terrabacteria_group>Chloroflexota>Chloroflexia>Chloroflexales>Chloroflexineae | GCF_004762035.2 | Candidatus_Chloroploca_sp._M50-1 | EYB53_RS23925 | 6374..10090 | 2528176 | BREX-system-P-loop-protein-BrxC-[Candidatus-Chloroploca-sp.-M50-1] |
| WP_220593725.1 | Type1.BREX | <-DDE_Tnp_ISAZ013<-?<-?<-RRXRR+HNH\|\|HTH_3->XIS-HTH->BrxA-DNA-binding->BrxB-BrxF-iSTAND->N6-METHYLASE->DUF6079*-><-NTP_transferase\|\|His_Phos_1->?->NTP_transf->?->ACET-> | 1101 | Bacteria>Terrabacteria_group>Chloroflexota>Chloroflexia>Chloroflexales>Chloroflexineae | GCF_019599295.1 | Oscillochloris_sp._ZM17-4 | K2Z83_RS03180 | 298417..301719 | 2866714 | BREX-system-P-loop-protein-BrxC,-partial-[Oscillochloris-sp.-ZM17-4] |
| KPV54581.1 | Type1.BREX | <-N6-METHYLASE<-REase<-DUF6079-p*<-DUF6079-p<-HEPN<-BrxB-BrxF-iSTAND<-BrxA-DNA-binding<-XIS-HTH | 802 | Bacteria>Terrabacteria_group>Chloroflexota>Chloroflexia>Chloroflexales>Roseiflexineae | GCA_001399705.1 | Kouleothrix_aurantiaca | SE17_02870 | 3869..6277 | 186479 | hypothetical-protein-SE17_02870-[Kouleothrix-aurantiaca] |
| MCG8347006.1 | Type1.BREX | <-PolB<-CopG/RHH<-?<-wHTH<-REase+HTH<-N6-METHYLASE<-Y1_Tnp<-DUF6079*<-BrxB-BrxF-iSTAND | 1203 | Bacteria>Terrabacteria_group>Chloroflexota>Chloroflexia>unclassified_Chloroflexales | GCA_022360155.1 | Chloroflexales_bacterium | MI924_04420 | 5729..9340 | 2725990 | BREX-system-P-loop-protein-BrxC-[Chloroflexales-bacterium] |
| WP_041330520.1 | Type1.BREX | SIGMA-HTH->DNA_pol3_beta_2+DNA_pol3_beta_3->?->?->?->HTH->BrxA-DNA-binding->BrxB-BrxF-iSTAND->DUF6079*->N6-METHYLASE->PglZ->SIGMA+OB-Fold+AAA-ATPase+Lon-Protease-BrxL->ABhydrolase->SMODS->?->JAB->HNH->MerR-HTH-> | 1147 | Bacteria>Terrabacteria_group>Chloroflexota>Dehalococcoidia>Dehalococcoidales>Dehalococcoidaceae | GCF_000830925.1 | Dehalococcoides_mccartyi | X792_RS01140 | 211516..214959 | 61435 | BREX-system-P-loop-protein-BrxC-[Dehalococcoides-mccartyi] |
| ADJ26904.1 | Type1.BREX | <-MULE-TRANSPOSASE<-SF1-SF2-Helicase<-?<-SLOG+BirA-HTH<-?<-SIGMA+OB-Fold+AAA-ATPase+HKD-DNase-BrxL<-PglZ<-N6-METHYLASE<-DUF6079*<-BrxB-BrxF-iSTAND<-BrxA-DNA-binding<-XIS-HTH<-?<-?<-?<-?<-wHTH | 1208 | Bacteria>Terrabacteria_group>Chloroflexota>Dehalococcoidia>Dehalogenimonas>Dehalogenimonas_lykanthroporepellens | GCA_000143165.1 | Dehalogenimonas_lykanthroporepellens_BL-DC-9 | Dehly_1626 | 1598643..1602269 | 552811 | conserved-hypothetical-protein-[Dehalogenimonas-lykanthroporepellens-BL-DC-9] |
| AKG53298.1 | Type1.BREX | TPR+TPR+TPR->?->BrxA-DNA-binding->BrxB-BrxF-iSTAND-p->DUF6079*->N6-METHYLASE->Calcineurin-SbcD->SbcC+CCDC158->PglZ->SIGMA+OB-Fold+AAA-ATPase+Lon-Protease-BrxL->RNase-><-?\|\|RNase->MORC+REase-> | 1138 | Bacteria>Terrabacteria_group>Chloroflexota>Dehalococcoidia>Dehalogenimonas>unclassified_Dehalogenimonas | GCA_001005265.1 | Dehalogenimonas_sp._WBC-2 | DGWBC_0620 | 571606..575022 | 943347 | hypothetical-protein-DGWBC_0620-[Dehalogenimonas-sp.-WBC-2] |
| MBE0430956.1 | Type1.BREX | XIS-HTH->BrxA-DNA-binding->BrxB-BrxF-iSTAND->DUF6079*->Schlafen+HATPase+wHTH+HTH->OLD-ABC->REase->N6-METHYLASE->PglZ-p-> | 1193 | Bacteria>Terrabacteria_group>Chloroflexota>Dehalococcoidia>unclassified_Dehalococcoidia | GCA_014859205.1 | Dehalococcoidia_bacterium | IBX67_03920 | 8033..11614 | 2026734 | BREX-system-P-loop-protein-BrxC-[Dehalococcoidia-bacterium] |
| MCC7365290.1 | Type1.BREX | <-VirB6<-?<-SIGMA+OB-Fold+AAA-ATPase+Lon-Protease-BrxL<-PglZ<-?<-N6-METHYLASE<-DUF6079*<-BrxB-BrxF-iSTAND<-LysM+wHTH+TPR | 1151 | Bacteria>Terrabacteria_group>Chloroflexota>Dehalococcoidia>unclassified_Dehalococcoidia | GCA_020850615.1 | Dehalococcoidia_bacterium | IT303_13065 | 99884..103339 | 2026734 | BREX-system-P-loop-protein-BrxC-[Dehalococcoidia-bacterium] |
| MCD6599573.1 | Type1.BREX | <-ACET<-?<-?\|\|MerR-HTH->GspA-ATPase->?->BrxA-DNA-binding->BrxB-BrxF-iSTAND->DUF6079-p*-> | 612 | Bacteria>Terrabacteria_group>Chloroflexota>Dehalococcoidia>unclassified_Dehalococcoidia | GCA_021162245.1 | Dehalococcoidia_bacterium | J7L19_03270 | 8143..9979 | 2026734 | BREX-system-P-loop-protein-BrxC-[Dehalococcoidia-bacterium] |
| WP_201375473.1 | Type1.BREX | <-SIGMA+OB-Fold+AAA-ATPase+Lon-Protease-BrxL<-PglZ<-?<-N6-METHYLASE<-DUF6079*<-BrxB-BrxF-iSTAND<-BrxA-DNA-binding<-?<-?\|\|REase-><-?\|\|HKD-DNase->?->SF1-SF2-Helicase-> | 1216 | Bacteria>Terrabacteria_group>Chloroflexota>Ktedonobacteria>Ktedonobacterales>Ktedonobacteraceae | GCF_016587375.1 | Ktedonobacter_robiniae | KSB_77480 | 1130085..1133735 | 2778365 | BREX-system-P-loop-protein-BrxC-[Ktedonobacter-robiniae] |
| WP_201380155.1 | Type1.BREX | <-DndH<-DndG<-NERD+Pkinase+Pkinase+DndF<-SIGMA+OB-Fold+AAA-ATPase+Lon-Protease-BrxL<-PglZ<-N6-METHYLASE<-DUF6079*<-BrxB-BrxF-iSTAND<-BrxA-DNA-binding<-?<-HTH_3\|\|?->?->?->?->REase-> | 1154 | Bacteria>Terrabacteria_group>Chloroflexota>Ktedonobacteria>Ktedonobacterales>Ktedonobacteraceae | GCF_016587415.1 | Ktedonobacter_sp._SOSP1-85 | KTE_RS08295 | 1959841..1963305 | 2778367 | BREX-system-P-loop-protein-BrxC-[Ktedonobacter-sp.-SOSP1-85] |
| WP_129893875.1 | Type1.BREX | HIT-><-?<-?<-?\|\|?->BrxA-DNA-binding->BrxB-BrxF-iSTAND->DUF6079*->N6-METHYLASE->PglZ->SIGMA+OB-Fold+AAA-ATPase+Lon-Protease-BrxL->?-><-?<-REC\|\|TPR-><-HTH+HKD-DNase<-SF1-SF2-Helicase | 1205 | Bacteria>Terrabacteria_group>Chloroflexota>Ktedonobacteria>Ktedonobacterales>Ktedonosporobacteraceae | GCF_004208415.1 | Ktedonosporobacter_rubrisoli | EPA93_RS45330 | 10598722..10602339 | 2509675 | BREX-system-P-loop-protein-BrxC-[Ktedonosporobacter-rubrisoli] |
| GHP00803.1 | Type1.BREX | NTP_transf-><-DDE_Tnp_ISL3<-?\|\|HIN-HTH->HTH_21+rve+rve_3->?->DUF4007->LON-AAA->DUF6079*->PglZ->SIGMA+OB-Fold+AAA-ATPase-BrxL->?-><-?<-?<-?<-LexA-protease<-NACHT+68TM-wHTH+NACHT<-CR-Kinase+CR-Kinase | 1383 | Bacteria>Terrabacteria_group>Chloroflexota>Ktedonobacteria>Ktedonobacterales>Reticulibacteraceae | GCA_016587455.1 | Reticulibacter_mediterranei | KSF_108500 | 208272..212423 | 2778369 | hypothetical-protein-KSF_108500-[Reticulibacter-mediterranei] |
| WP_220211383.1 | Type1.BREX | NACHT->?->NTP_transf-><-zf-ISL3+HTH_Tnp_ISL3\|\|HIN-HTH->HTH_21+rve+rve_3->?->DUF4007->LON-AAA->DUF6079*->PglZ->SIGMA+OB-Fold+AAA-ATPase-BrxL->?-><-?<-?<-?<-LexA-protease<-NACHT+68TM-wHTH+NACHT<-CR-Kinase+CR-Kinase | 1378 | Bacteria>Terrabacteria_group>Chloroflexota>Ktedonobacteria>Ktedonobacterales>Reticulibacteraceae | GCF_016587455.1 | Reticulibacter_mediterranei | K3T83_RS53825 | 208287..212423 | 2778369 | hypothetical-protein-[Reticulibacter-mediterranei] |
| KAA3643964.1 | Type1.BREX | <-ParA-Soj-PloopNTPase\|\|GspA-ATPase->?->?->?->?->BrxA-DNA-binding->BrxB-BrxF-iSTAND->DUF6079*->N6-METHYLASE->PglZ->SIGMA+OB-Fold+AAA-ATPase+Lon-Protease-BrxL->REase-><-?<-NTP_transf_5<-REC | 1152 | Bacteria>Terrabacteria_group>Chloroflexota>unclassified_Chloroflexota | GCA_008501685.1 | Chloroflexota_bacterium | DWQ07_17765 | 184634..188092 | 2026724 | BREX-system-P-loop-protein-BrxC-[Chloroflexota-bacterium] |
| MXZ45726.1 | Type1.BREX | <-DOC+HTH<-?\|\|Recombinase+ZnR->AbiEi_4-><-?<-SIGMA+OB-Fold+AAA-ATPase+Lon-Protease-BrxL<-PglZ<-N6-METHYLASE<-DUF6079*<-BrxB-BrxF-iSTAND<-BrxA-DNA-binding\|\|PAPS-reduct-><-PIN<-HTH+MPTase\|\|ParB+HNH-><-PIN\|\|GDResolvase+Recombinase+ZnR-><-DNA_pol3_beta+DNA_pol3_beta_2+DNA_pol3_beta_3 | 1155 | Bacteria>Terrabacteria_group>Chloroflexota>unclassified_Chloroflexota | GCA_009837285.1 | Chloroflexota_bacterium | F4Z08_01820 | 13972..17439 | 2026724 | BREX-system-P-loop-protein-BrxC-[Chloroflexota-bacterium] |
| MYB75653.1 | Type1.BREX | <-ParB+HNH\|\|Peptidase_M78-><-SIGMA+OB-Fold+AAA-ATPase+Lon-Protease-BrxL<-PglZ<-N6-METHYLASE<-DUF6079*<-BrxB-BrxF-iSTAND<-BrxA-DNA-binding\|\|PAPS-reduct-><-URI-DNase | 1154 | Bacteria>Terrabacteria_group>Chloroflexota>unclassified_Chloroflexota | GCA_009839775.1 | Chloroflexota_bacterium | F4X83_00835 | 12467..15931 | 2026724 | BREX-system-P-loop-protein-BrxC-[Chloroflexota-bacterium] |
| MYF79701.1 | Type1.BREX | <-N6-METHYLASE-p<-DUF6079*<-BrxB-BrxF-iSTAND<-BrxA-DNA-binding\|\|?-><-ArdC<-MPTase<-REase<-PIWI | 1253 | Bacteria>Terrabacteria_group>Chloroflexota>unclassified_Chloroflexota | GCA_009843025.1 | Chloroflexota_bacterium | F4180_07600 | 1797..5558 | 2026724 | BREX-system-P-loop-protein-BrxC-[Chloroflexota-bacterium] |
| HDQ70548.1 | Type1.BREX | <-SIGMA+OB-Fold+AAA-ATPase+HKD-DNase-BrxL<-PglZ<-N6-METHYLASE<-?<-DUF6079*<-BrxB-BrxF-iSTAND<-BrxA-DNA-binding | 1168 | Bacteria>Terrabacteria_group>Chloroflexota>unclassified_Chloroflexota | GCA_011046405.1 | Chloroflexota_bacterium | ENN19_00445 | 9344..12850 | 2026724 | BREX-system-P-loop-protein-BrxC-[Chloroflexota-bacterium] |
| NLW72147.1 | Type1.BREX | <-HTH_3<-?<-SIGMA+OB-Fold+AAA-ATPase+Lon-Protease-BrxL<-PglZ<-HEPN<-N6-METHYLASE<-?<-DUF6079*<-BrxB-BrxF-iSTAND<-BrxA-DNA-binding<-HTH_26\|\|Calcineurin-> | 1224 | Bacteria>Terrabacteria_group>Chloroflexota>unclassified_Chloroflexota | GCA_012729755.1 | Chloroflexota_bacterium | GXY37_04595 | 11724..15398 | 2026724 | BREX-system-P-loop-protein-BrxC-[Chloroflexota-bacterium] |
| HHY55161.1 | Type1.BREX | <-ACET<-PglZ<-?<-OLD-ABC<-?<-N6-METHYLASE<-OLD-ABC+TOPRIM<-DUF6079*<-BrxB-BrxF-iSTAND<-BrxA-DNA-binding<-?<-?<-NTP_transferase | 1149 | Bacteria>Terrabacteria_group>Chloroflexota>unclassified_Chloroflexota | GCA_012842785.1 | Chloroflexota_bacterium | GYA08_06960 | 29927..33376 | 2026724 | BREX-system-P-loop-protein-BrxC-[Chloroflexota-bacterium] |
| NWF82102.1 | Type1.BREX | <-N6-METHYLASE<-?<-?<-DUF6079*<-BrxB-BrxF-iSTAND<-BrxA-DNA-binding<-?\|\|?->?->SIGMA-HTH+SIGMA-HTH-><-?\|\|LPD36+METHYLASE+SF1-SF2-Helicase-> | 1158 | Bacteria>Terrabacteria_group>Chloroflexota>unclassified_Chloroflexota | GCA_013388805.1 | Chloroflexota_bacterium | HXY37_18960 | 3120..6596 | 2026724 | BREX-system-P-loop-protein-BrxC-[Chloroflexota-bacterium] |
| NWJ94851.1 | Type1.BREX | <-SIGMA+OB-Fold+AAA-ATPase+Lon-Protease-BrxL<-PglZ<-?<-REase<-?<-N6-METHYLASE<-?<-?<-DUF6079*<-BrxB-BrxF-iSTAND<-BrxA-DNA-binding<-XIS-HTH<-?<-XIS-HTH\|\|?->MoxR-AAA->VWA+vWA-L+CR-ATPase9->CR-ATPase-> | 1218 | Bacteria>Terrabacteria_group>Chloroflexota>unclassified_Chloroflexota | GCA_013390945.1 | Chloroflexota_bacterium | HXX20_03630 | 11914..15570 | 2026724 | BREX-system-P-loop-protein-BrxC-[Chloroflexota-bacterium] |
| NWJ96922.1 | Type1.BREX | HTH_3->BrxA-DNA-binding->BrxB-BrxF-iSTAND->DUF6079*->?->HEPN->ParA-Soj-PloopNTPase+ORC-CDC6-like->?->?->N6-METHYLASE-> | 1214 | Bacteria>Terrabacteria_group>Chloroflexota>unclassified_Chloroflexota | GCA_013390945.1 | Chloroflexota_bacterium | HXX20_14155 | 2064..5708 | 2026724 | BREX-system-P-loop-protein-BrxC-[Chloroflexota-bacterium] |
| NWJ97618.1 | Type1.BREX | BrxA-DNA-binding->BrxB-BrxF-iSTAND->DUF6079*->MetJarc-HTH->?->REase->N6-METHYLASE-> | 1157 | Bacteria>Terrabacteria_group>Chloroflexota>unclassified_Chloroflexota | GCA_013390945.1 | Chloroflexota_bacterium | HXX20_17665 | 1496..4969 | 2026724 | BREX-system-P-loop-protein-BrxC-[Chloroflexota-bacterium] |
| MBC8504021.1 | Type1.BREX | HTH_26->HTH_26->BrxA-DNA-binding->BrxB-BrxF-iSTAND->DUF6079*->N6-METHYLASE-p-> | 1174 | Bacteria>Terrabacteria_group>Chloroflexota>unclassified_Chloroflexota | GCA_014384465.1 | Chloroflexota_bacterium | H8D34_04145 | 3789..7313 | 2026724 | BREX-system-P-loop-protein-BrxC-[Chloroflexota-bacterium] |
| MBC8508906.1 | Type1.BREX | wHTH+WYL+WCX->BrxA-DNA-binding->BrxB-BrxF-iSTAND->DUF6079*->N6-METHYLASE->PglZ->SIGMA+OB-Fold+AAA-ATPase+Lon-Protease-BrxL-><-Phage_lysozyme<-Phage_holin | 1220 | Bacteria>Terrabacteria_group>Chloroflexota>unclassified_Chloroflexota | GCA_014384465.1 | Chloroflexota_bacterium | H8D34_28965 | 2565..6227 | 2026724 | BREX-system-P-loop-protein-BrxC-[Chloroflexota-bacterium] |
| MBE9475406.1 | Type1.BREX | BrxB-BrxF-iSTAND-p->BrxA-DNA-binding->BrxB-BrxF-iSTAND->DUF6079*->N6-METHYLASE->REase+AIPR->OLD-ABC->HNH->PglZ->SIGMA+OB-Fold+AAA-ATPase+Lon-Protease-BrxL-> | 1145 | Bacteria>Terrabacteria_group>Chloroflexota>unclassified_Chloroflexota | GCA_015222395.1 | Chloroflexota_bacterium | IMY85_10995 | 2370..5807 | 2026724 | BREX-system-P-loop-protein-BrxC-[Chloroflexota-bacterium] |
| MBI2864205.1 | Type1.BREX | <-ABC-ATPase+SbcC<-ABC-ATPase+SbcC\|\|PolB-><-PhdYeFM_antitox\|\|?->PIN->BrxA-DNA-binding->BrxB-BrxF-iSTAND->DUF6079-p*-> | 773 | Bacteria>Terrabacteria_group>Chloroflexota>unclassified_Chloroflexota | GCA_016190625.1 | Chloroflexota_bacterium | HYX94_06540 | 21104..23423 | 2026724 | BREX-system-P-loop-protein-BrxC-[Chloroflexota-bacterium] |
| MBI2765415.1 | Type1.BREX | SF1-SF2-Helicase-><-?<-?<-?<-VirB6\|\|BrxA-DNA-binding->BrxB-BrxF-iSTAND->DUF6079-p->DUF6079*->HEPN-><-?\|\|N6-METHYLASE->N6-METHYLASE-p->PglZ->SIGMA+OB-Fold+AAA-ATPase+Lon-Protease-BrxL-><-LysM+wHTH+TPR | 1100 | Bacteria>Terrabacteria_group>Chloroflexota>unclassified_Chloroflexota | GCA_016191435.1 | Chloroflexota_bacterium | HYX53_05820 | 46909..50211 | 2026724 | BREX-system-P-loop-protein-BrxC-[Chloroflexota-bacterium] |
| MBI4317923.1 | Type1.BREX | Radical_SAM->?->?->?->BrxA-DNA-binding->BrxB-BrxF-iSTAND->DUF6079*->N6-METHYLASE->OLD-ABC->HNH->PglZ->SIGMA+OB-Fold+AAA-ATPase+Lon-Protease-BrxL-> | 1145 | Bacteria>Terrabacteria_group>Chloroflexota>unclassified_Chloroflexota | GCA_016210495.1 | Chloroflexota_bacterium | HY675_05490 | 15525..18962 | 2026724 | BREX-system-P-loop-protein-BrxC-[Chloroflexota-bacterium] |
| MBK7180413.1 | Type1.BREX | <-REase<-SF1-SF2-Helicase+DUF1998<-?<-?<-wHTH+WYL+WCX<-?\|\|?->BrxA-DNA-binding->BrxB-BrxF-iSTAND->DUF6079*->N6-METHYLASE->?->?-><-PIN<-RHH\|\|PglZ-> | 1173 | Bacteria>Terrabacteria_group>Chloroflexota>unclassified_Chloroflexota | GCA_016707865.1 | Chloroflexota_bacterium | IPH82_25035 | 72782..76303 | 2026724 | BREX-system-P-loop-protein-BrxC-[Chloroflexota-bacterium] |
| MBK8989553.1 | Type1.BREX | PIN->CRISPR_Cas6->HTH_Tnp_1+HTH_21+rve+rve_3-><-?<-SIGMA+OB-Fold+AAA-ATPase+HKD-DNase-BrxL<-PglZ\|\|RHH-><-?<-N6-METHYLASE<-DUF6079*<-BrxB-BrxF-iSTAND<-BrxA-DNA-binding<-?\|\|wHTH+WYL+WCX->RNase_T+35exo->?->ABhydrolase+ABhydrolase->?->BrxB-BrxF-iSTAND-> | 1173 | Bacteria>Terrabacteria_group>Chloroflexota>unclassified_Chloroflexota | GCA_016716065.1 | Chloroflexota_bacterium | IPM39_26430 | 84319..87840 | 2026724 | BREX-system-P-loop-protein-BrxC-[Chloroflexota-bacterium] |
| MBK9123540.1 | Type1.BREX | HerA/FtsK->BirA-HTH->BirA-HTH->?->?->CR-ATPase5->HTH->BrxA-DNA-binding->BrxB-BrxF-iSTAND->DUF6079*->N6-METHYLASE->N6-METHYLASE->PHP+SbcC->PglZ->SIGMA+OB-Fold+AAA-ATPase+Lon-Protease-BrxL-><-GDResolvase+ZnR | 1144 | Bacteria>Terrabacteria_group>Chloroflexota>unclassified_Chloroflexota | GCA_016716525.1 | Chloroflexota_bacterium | IPM16_10545 | 313837..317271 | 2026724 | BREX-system-P-loop-protein-BrxC-[Chloroflexota-bacterium] |
| MBK9750791.1 | Type1.BREX | <-Phage_holin<-Relaxase<-MetJarc-HTH<-SIGMA+OB-Fold+AAA-ATPase+Lon-Protease-BrxL<-PglZ<-PglZ<-PemK_toxin<-HTH+MPTase<-N6-METHYLASE<-DUF6079*<-BrxB-BrxF-iSTAND<-BrxA-DNA-binding<-?<-?<-?\|\|TIR-> | 1206 | Bacteria>Terrabacteria_group>Chloroflexota>unclassified_Chloroflexota | GCA_016717205.1 | Chloroflexota_bacterium | IPO91_29070 | 213079..216699 | 2026724 | BREX-system-P-loop-protein-BrxC-[Chloroflexota-bacterium] |
| MBM3127996.1 | Type1.BREX | XIS-HTH-><-SIGMA+OB-Fold+AAA-ATPase+Lon-Protease-BrxL<-PglZ<-TOPRIM<-OLD-ABC<-N6-METHYLASE<-?<-?<-?<-DUF6079*<-BrxB-BrxF-iSTAND<-BrxA-DNA-binding\|\|?->REase->HTH_3->?->?->HerA/FtsK-><-FGS | 1153 | Bacteria>Terrabacteria_group>Chloroflexota>unclassified_Chloroflexota | GCA_016865935.1 | Chloroflexota_bacterium | FJ009_05090 | 25630..29091 | 2026724 | BREX-system-P-loop-protein-BrxC-[Chloroflexota-bacterium] |
| MCC6629583.1 | Type1.BREX | HIN-HTH->BrxA-DNA-binding->BrxB-BrxF-iSTAND->DUF6079*->?->PIN->N6-METHYLASE->PolB->HEPN->?->PglZ->SIGMA+OB-Fold+AAA-ATPase+Lon-Protease-BrxL-> | 1204 | Bacteria>Terrabacteria_group>Chloroflexota>unclassified_Chloroflexota | GCA_020849725.1 | Chloroflexota_bacterium | IT340_19565 | 31214..34828 | 2026724 | BREX-system-P-loop-protein-BrxC-[Chloroflexota-bacterium] |
| MCL4487040.1 | Type1.BREX | BrxA-DNA-binding->BrxB-BrxF-iSTAND->Peptidase_M91->N6-METHYLASE->HKD-DNase->DUF6079*->HEPN->PhdYeFM_antitox->PIN->PglZ->?->?->?->MerR-HTH+TipAS-> | 1221 | Bacteria>Terrabacteria_group>Chloroflexota>unclassified_Chloroflexota | GCA_023381515.1 | Chloroflexota_bacterium | M1570_02770 | 10289..13954 | 2026724 | BREX-system-P-loop-protein-BrxC-[Chloroflexota-bacterium] |
| MCL4509264.1 | Type1.BREX | SPOUT->Glycos_transf_1+Glycos_trans_1_2-><-?<-?\|\|BrxA-DNA-binding->BrxB-BrxF-iSTAND->DUF6079*->N6-METHYLASE->RAMPs->RAMPs->?->?->RAMPs-> | 1205 | Bacteria>Terrabacteria_group>Chloroflexota>unclassified_Chloroflexota | GCA_023381555.1 | Chloroflexota_bacterium | M1296_07040 | 60116..63733 | 2026724 | BREX-system-P-loop-protein-BrxC-[Chloroflexota-bacterium] |
| MCL5996402.1 | Type1.BREX | <-N6-METHYLASE<-HicA_toxin<-?<-DUF6079*<-BrxB-BrxF-iSTAND<-BrxA-DNA-binding | 1149 | Bacteria>Terrabacteria_group>Chloroflexota>unclassified_Chloroflexota | GCA_023488685.1 | Chloroflexota_bacterium | M1546_10155 | 4457..7906 | 2026724 | BREX-system-P-loop-protein-BrxC-[Chloroflexota-bacterium] |
| MCP4164415.1 | Type1.BREX | <-rve<-rve<-PAIREDC-HTH\|\|BrxA-DNA-binding->BrxB-BrxF-iSTAND->DUF6079*->Y1_Tnp-> | 1177 | Bacteria>Terrabacteria_group>Chloroflexota>unclassified_Chloroflexota | GCA_024224665.1 | Chloroflexota_bacterium | GY759_00800 | 4074..7607 | 2026724 | BREX-system-P-loop-protein-BrxC-[Chloroflexota-bacterium] |
| OGH97200.1 | Type1.BREX | ANK-><-?\|\|BrxA-DNA-binding->BrxB-BrxF-iSTAND->DUF6079*->REase->N6-METHYLASE->HTH+REase->PglZ->SIGMA+OB-Fold+AAA-ATPase+Lon-Protease-BrxL->Y2_Tnp-> | 1198 | Bacteria>Terrabacteria_group>Cyanobacteriota/Melainabacteria_group>Candidatus_Margulisbacteria>Candidatus_Riflemargulisbacteria | GCA_001783265.1 | Candidatus_Margulisbacteria_bacterium_GWD2_39_127 | A2X43_08520 | 54910..58506 | 1817871 | hypothetical-protein-A2X43_08520-[Candidatus-Margulisbacteria-bacterium-GWD2_39_127] |
| OGI09205.1 | Type1.BREX | <-Y2_Tnp<-SIGMA+OB-Fold+AAA-ATPase+Lon-Protease-BrxL<-PglZ<-HTH+REase<-N6-METHYLASE<-REase<-DUF6079*<-BrxB-BrxF-iSTAND<-BrxA-DNA-binding\|\|?-><-ANK | 1198 | Bacteria>Terrabacteria_group>Cyanobacteriota/Melainabacteria_group>Candidatus_Margulisbacteria>unclassified_Candidatus_Margulisbacteria | GCA_001784485.1 | Candidatus_Margulisbacteria_bacterium_GWE2_39_32 | A2X41_01355 | 17633..21229 | 1817872 | hypothetical-protein-A2X41_01355-[Candidatus-Margulisbacteria-bacterium-GWE2_39_32] |
| PZM83737.1 | Type1.BREX | <-Y2_Tnp<-SIGMA+OB-Fold+AAA-ATPase+Lon-Protease-BrxL<-PglZ<-HTH+REase<-N6-METHYLASE<-REase<-DUF6079*<-BrxB-BrxF-iSTAND<-BrxA-DNA-binding\|\|?-><-ANK | 1198 | Bacteria>Terrabacteria_group>Cyanobacteriota/Melainabacteria_group>Candidatus_Margulisbacteria>unclassified_Candidatus_Margulisbacteria | GCA_003242895.1 | Candidatus_Margulisbacteria_bacterium | DKM50_01295 | 12966..16562 | 2053573 | BREX-system-P-loop-protein-BrxC-[Candidatus-Margulisbacteria-bacterium] |
| MBX9724057.1 | Type1.BREX | <-SIGMA+OB-Fold+AAA-ATPase+HKD-DNase-BrxL\|\|BrxA-DNA-binding->BrxB-BrxF-iSTAND->DUF6079*->OLD-ABC->REase->N6-METHYLASE->PglZ-p-> | 1175 | Bacteria>Terrabacteria_group>Cyanobacteriota/Melainabacteria_group>Candidatus_Melainabacteria>Candidatus_Obscuribacterales>unclassified_Candidatus_Obscuribacterales | GCA_019747345.1 | Candidatus_Obscuribacterales_bacterium | K2X81_21805 | 3793..7320 | 2364082 | BREX-system-P-loop-protein-BrxC-[Candidatus-Obscuribacterales-bacterium] |
| MBX9573698.1 | Type1.BREX | <-N6-METHYLASE-p<-N6-METHYLASE<-DUF6079*<-BrxB-BrxF-iSTAND<-BrxA-DNA-binding<-TPR<-TPR<-Sulfate_transp+STAS\|\|?->SIGMA-HTH+SIGMA-HTH+SIGMA-HTH-> | 1207 | Bacteria>Terrabacteria_group>Cyanobacteriota/Melainabacteria_group>Candidatus_Melainabacteria>Candidatus_Obscuribacterales>unclassified_Candidatus_Obscuribacterales | GCA_019747875.1 | Candidatus_Obscuribacterales_bacterium | K2X77_32675 | 4402..8025 | 2364082 | BREX-system-P-loop-protein-BrxC-[Candidatus-Obscuribacterales-bacterium] |
| MBT9546703.1 | Type1.BREX | Peptidase_M2->BrxA-DNA-binding->BrxB-BrxF-iSTAND->DUF6079*->N6-METHYLASE->RVT->N6-METHYLASE-p->OLD-ABC->HNH->PglZ->SIGMA+OB-Fold+AAA-ATPase+HKD-DNase-BrxL-> | 1179 | Bacteria>Terrabacteria_group>Cyanobacteriota/Melainabacteria_group>Candidatus_Sericytochromatia>unclassified_Candidatus_Sericytochromatia | GCA_018780035.1 | Candidatus_Sericytochromatia_bacterium | IV090_15030 | 94995..98534 | 2762020 | BREX-system-P-loop-protein-BrxC-[Candidatus-Sericytochromatia-bacterium] |
| WAS06096.1 | Type1.BREX | <-PglZ<-PIN<-PhdYeFM_antitox<-?<-PIN<-PhdYeFM_antitox<-N6-METHYLASE<-PIN<-PhdYeFM_antitox<-DUF6079*<-BrxB-BrxF-iSTAND<-BrxA-DNA-binding\|\|?-><-NTP_transferase+MannoseP_isomer<-?<-TP_METHYLASE<-NTP_transf_3+Hexapep+Hexapep<-?<-RNase+DSRBD | 1221 | Bacteria>Terrabacteria_group>Cyanobacteriota/Melainabacteria_group>Cyanobacteriota>Cyanophyceae>Gloeomargaritales | GCA_022206215.1 | Gloeomargaritales_cyanobacterium_VI4D9 | LQF76_04175 | 903259..906924 | 2898145 | BREX-system-P-loop-protein-BrxC-[Gloeomargaritales-cyanobacterium-VI4D9] |
| WP_071453316.1 | Type1.BREX | <-N6-METHYLASE<-RelE-ParE<-?<-REase+HTH\|\|?->PIN-><-HEPN<-DUF6079*<-BrxB-BrxF-iSTAND<-BrxA-DNA-binding<-?\|\|?->?->?->tRNA-><-?\|\|MPTase-> | 1210 | Bacteria>Terrabacteria_group>Cyanobacteriota/Melainabacteria_group>Cyanobacteriota>Cyanophyceae>Gloeomargaritales | GCF_001870225.1 | Gloeomargarita_lithophora | GlitD10_RS01485 | 315738..319370 | 1188228 | BREX-system-P-loop-protein-BrxC-[Gloeomargarita-lithophora] |
| WP_220184431.1 | Type1.BREX | <-SIGMA+OB-Fold+AAA-ATPase+Lon-Protease-BrxL-p<-PglZ<-N6-METHYLASE-p<-N6-METHYLASE<-OLD-ABC<-PIN<-DUF433<-DUF6079-p<-DUF6079-p*<-BrxB-BrxF-iSTAND\|\|<-?<-?<-ParA-Soj-PloopNTPase<-?\|\|ACT-><-?<-Peptidase_M17_N+Peptidase_M17<-ABhydrolase | 838 | Bacteria>Terrabacteria_group>Cyanobacteriota/Melainabacteria_group>Cyanobacteriota>Cyanophyceae>Nostocales | - | Cylindrospermopsis_raciborskii | FHK99_RS13050 | 13287..15803 | 77022 | BREX-system-P-loop-protein-BrxC-[Cylindrospermopsis-raciborskii] |
| MBG0744678.1 | Type1.BREX | <-SIGMA-HTH<-TPR+TPR\|\|?->TPR+TPR->ARG-><-RADICAL-SAM\|\|?->BrxB-BrxF-iSTAND->DUF6079*->RNaseT2->N6-METHYLASE-> | 1184 | Bacteria>Terrabacteria_group>Cyanobacteriota/Melainabacteria_group>Cyanobacteriota>Cyanophyceae>Nostocales | - | Cylindrospermopsis_raciborskii_KL1 | IV298_14570 | 6960..10514 | 2787621 | BREX-system-P-loop-protein-BrxC-[Cylindrospermopsis-raciborskii-KL1] |
| EFA74474.1 | Type1.BREX | <-N6-METHYLASE-p<-N6-METHYLASE<-RNaseT2<-DUF6079*<-BrxB-BrxF-iSTAND\|\|RADICAL-SAM->REase->Schlafen+HATPase+wHTH->DUF29->?-><-?<-RFC-AAA+RuvB-AAA | 1176 | Bacteria>Terrabacteria_group>Cyanobacteriota/Melainabacteria_group>Cyanobacteriota>Cyanophyceae>Nostocales | GCA_000175855.1 | Raphidiopsis_brookii_D9 | CRD_00004 | 5426..8956 | 533247 | hypothetical-protein-CRD_00004-[Raphidiopsis-brookii-D9] |
| BAZ90345.1 | Type1.BREX | <-SIGMA+OB-Fold+AAA-ATPase+HKD-DNase-BrxL<-PglZ<-HTH+REase<-N6-METHYLASE<-REase<-OLD-ABC<-PIN<-DUF433<-DUF6079*<-BrxB-BrxF-iSTAND\|\|?->?-><-?<-?\|\|?-><-?<-ABC-ATPase+ABC-ATPase<-NTP_transferase+MannoseP_isomer | 1166 | Bacteria>Terrabacteria_group>Cyanobacteriota/Melainabacteria_group>Cyanobacteriota>Cyanophyceae>Nostocales | GCA_002368135.1 | Raphidiopsis_curvata_NIES-932 | NIES932_18440 | 1957862..1961362 | 1973482 | hypothetical-protein-NIES932_18440-[Raphidiopsis-curvata-NIES-932] |
| MBU7587299.1 | Type1.BREX | ParA-Soj-PloopNTPase->?->BrxA-DNA-binding->BrxB-BrxF-iSTAND->DUF6079*->REase->N6-METHYLASE-> | 1221 | Bacteria>Terrabacteria_group>Cyanobacteriota/Melainabacteria_group>Cyanobacteriota>Cyanophyceae>Nostocales | GCA_018993265.1 | Nostoc_sp._TH1S01 | KAF91_31390 | 3365..7030 | 442893 | BREX-system-P-loop-protein-BrxC-[Nostoc-sp.-TH1S01] |
| MBW4478483.1 | Type1.BREX | REase+HTH->REase+HTH-><-N6-METHYLASE<-DUF6079*<-BrxB-BrxF-iSTAND<-?<-RelE-ParE<-MetJarc-HTH | 1184 | Bacteria>Terrabacteria_group>Cyanobacteriota/Melainabacteria_group>Cyanobacteriota>Cyanophyceae>Nostocales | GCA_019358985.1 | Tolypothrix_brevis_GSE-NOS-MK-07-07A | KME54_16860 | 5705..9259 | 2835095 | BREX-system-P-loop-protein-BrxC-[Tolypothrix-brevis-GSE-NOS-MK-07-07A] |
| MBW4511575.1 | Type1.BREX | <-DDE_Tnp_ISAZ013\|\|?->?->?->?->Phage_int_SAM_4-><-?<-OLD-ABC\|\|BrxB-BrxF-iSTAND->DUF6079-p*->ACET->MerR-HTH+TipAS->?->REase+HTH-><-?\|\|DSBH+AraC-HTH+AraC-HTH->ABhydrolase-> | 463 | Bacteria>Terrabacteria_group>Cyanobacteriota/Melainabacteria_group>Cyanobacteriota>Cyanophyceae>Nostocales | GCA_019359035.1 | Scytonematopsis_contorta_HA4267-MV1 | KME64_34500 | 69099..70490 | 1049028 | hypothetical-protein-[Scytonematopsis-contorta-HA4267-MV1] |
| MCC5669622.1 | Type1.BREX | DDE_Tnp->BrxA-DNA-binding-><-SIGMA+OB-Fold+AAA-ATPase+Lon-Protease-BrxL<-PglZ+Phosphodiest<-?<-ART-DarT<-HTH+ART-DarT<-SF1-SF2-Helicase<-N6-METHYLASE<-DUF6079-p* | 898 | Bacteria>Terrabacteria_group>Cyanobacteriota/Melainabacteria_group>Cyanobacteriota>Cyanophyceae>Nostocales | GCA_020829645.1 | Nostoc_mirabile_CHAB5784 | LC653_38870 | 16322..19018 | 2780401 | BREX-system-P-loop-protein-BrxC-[Nostoc-mirabile-CHAB5784] |
| WP_052304589.1 | Type1.BREX | Glyoxalase->?-><-?<-SF1-SF2-Helicase+wHTH+Small-B-barrel+DUF1998<-SF1-SF2-Helicase+wHTH+Small-B-barrel+DUF1998-p<-BrxD-AAA<-PglZ<-N6-METHYLASE<-DUF6079-p*<-?<-?\|\|?->?-><-?<-Thioredoxin | 745 | Bacteria>Terrabacteria_group>Cyanobacteriota/Melainabacteria_group>Cyanobacteriota>Cyanophyceae>Nostocales | GCF_000020025.1 | Nostoc_punctiforme | NPUN_RS11740 | 2881762..2883999 | 272131 | hypothetical-protein-[Nostoc-punctiforme] |
| WP_012411904.1 | Type1.BREX | REase-><-?<-?<-?\|\|?-><-?\|\|tRNA-><-Cas12k\|\|MerR-HTH->BrxB-BrxF-iSTAND->DUF6079*->N6-METHYLASE-><-?<-REase\|\|PglZ+Phosphodiest->SIGMA+OB-Fold+AAA-ATPase+Lon-Protease-BrxL-><-BrxA-DNA-binding<-RHH\|\|?->CHTH-> | 1179 | Bacteria>Terrabacteria_group>Cyanobacteriota/Melainabacteria_group>Cyanobacteriota>Cyanophyceae>Nostocales | GCF_000020025.1 | Nostoc_punctiforme | NPUN_RS28630 | 6987385..6990924 | 272131 | BREX-system-P-loop-protein-BrxC-[Nostoc-punctiforme] |
| WP_006277878.1 | Type1.BREX | <-REase\|\|?-><-SIGMA+OB-Fold+AAA-ATPase+HKD-DNase-BrxL<-PglZ<-HTH+REase<-N6-METHYLASE-p<-OLD-ABC<-PIN<-DUF433<-DUF6079*<-BrxB-BrxF-iSTAND\|\|?->?-><-?<-?<-ParA-Soj-PloopNTPase<-?\|\|ACT-> | 1166 | Bacteria>Terrabacteria_group>Cyanobacteriota/Melainabacteria_group>Cyanobacteriota>Cyanophyceae>Nostocales | GCF_000175835.1 | Cylindrospermopsis_raciborskii | CRC_RS10895 | 11801..15301 | 77022 | BREX-system-P-loop-protein-BrxC-[Cylindrospermopsis-raciborskii] |
| WP_057176814.1 | Type1.BREX | <-ACT\|\|?->ParA-Soj-PloopNTPase->?->?->?-><-?<-?\|\|BrxB-BrxF-iSTAND->DUF6079*->DUF433->PIN->OLD-ABC->REase->REase->N6-METHYLASE->HTH+REase->PglZ->SIGMA+OB-Fold+AAA-ATPase+HKD-DNase-BrxL-> | 1166 | Bacteria>Terrabacteria_group>Cyanobacteriota/Melainabacteria_group>Cyanobacteriota>Cyanophyceae>Nostocales | GCF_001432185.1 | Cylindrospermopsis_sp._CR12 | ASL19_RS00210 | 43411..46911 | 1747196 | BREX-system-P-loop-protein-BrxC-[Cylindrospermopsis-sp.-CR12] |
| WP_057176920.1 | Type1.BREX | Schlafen+HATPase+wHTH-><-SIGMA+OB-Fold+AAA-ATPase+Lon-Protease-BrxL<-PglZ+Phosphodiest<-N6-METHYLASE<-RNaseT2<-DUF6079*<-BrxB-BrxF-iSTAND<-?\|\|RADICAL-SAM-> | 1184 | Bacteria>Terrabacteria_group>Cyanobacteriota/Melainabacteria_group>Cyanobacteriota>Cyanophyceae>Nostocales | GCF_001432185.1 | Cylindrospermopsis_sp._CR12 | ASL19_RS00905 | 19893..23447 | 1747196 | BREX-system-P-loop-protein-BrxC-[Cylindrospermopsis-sp.-CR12] |
| WP_061546830.1 | Type1.BREX | RFC-AAA+RuvB-AAA->?-><-?<-DUF29<-Schlafen+HATPase+wHTH<-REase<-RADICAL-SAM\|\|?->BrxB-BrxF-iSTAND->DUF6079*->RNaseT2->N6-METHYLASE-> | 1177 | Bacteria>Terrabacteria_group>Cyanobacteriota/Melainabacteria_group>Cyanobacteriota>Cyanophyceae>Nostocales | GCF_001586755.1 | Cylindrospermopsis_raciborskii | A0J49_RS12215 | 40600..44133 | 77022 | BREX-system-P-loop-protein-BrxC-[Cylindrospermopsis-raciborskii] |
| WP_069067979.1 | Type1.BREX | CHTH->?->?->BrxA-DNA-binding->BrxB-BrxF-iSTAND->DUF6079*->?->N6-METHYLASE->OLD-ABC->HNH->PglZ->SIGMA+OB-Fold+AAA-ATPase+Lon-Protease-BrxL-><-?<-HTH_23 | 1220 | Bacteria>Terrabacteria_group>Cyanobacteriota/Melainabacteria_group>Cyanobacteriota>Cyanophyceae>Nostocales | GCF_001712795.1 | Nostoc_sp._KVJ20 | A4S05_RS00980 | 24778..28440 | 457944 | BREX-system-P-loop-protein-BrxC-[Nostoc-sp.-KVJ20] |
| WP_071242349.1 | Type1.BREX | <-ACT\|\|?->ParA-Soj-PloopNTPase->?->?->?-><-?<-?\|\|BrxB-BrxF-iSTAND->DUF6079*->DUF433->PIN->OLD-ABC->REase->N6-METHYLASE->HTH+REase->PglZ->SIGMA+OB-Fold+AAA-ATPase+HKD-DNase-BrxL->REase-> | 1166 | Bacteria>Terrabacteria_group>Cyanobacteriota/Melainabacteria_group>Cyanobacteriota>Cyanophyceae>Nostocales | GCF_001858115.1 | Cylindrospermopsis_raciborskii | BCV63_RS11865 | 45142..48642 | 77022 | BREX-system-P-loop-protein-BrxC-[Cylindrospermopsis-raciborskii] |
| WP_071249054.1 | Type1.BREX | RFC-AAA+RuvB-AAA->?-><-?<-DUF29<-Schlafen+HATPase+wHTH<-RADICAL-SAM\|\|?->BrxB-BrxF-iSTAND->DUF6079*->RNaseT2->N6-METHYLASE-><-AbiEii<-AbiEi_4\|\|PglZ+Phosphodiest->SIGMA+OB-Fold+AAA-ATPase+Lon-Protease-BrxL-><-?<-?<-RAMPs | 1176 | Bacteria>Terrabacteria_group>Cyanobacteriota/Melainabacteria_group>Cyanobacteriota>Cyanophyceae>Nostocales | GCF_001858125.1 | Cylindrospermopsis_raciborskii | BCV64_RS05180 | 29216..32746 | 77022 | BREX-system-P-loop-protein-BrxC-[Cylindrospermopsis-raciborskii] |
| WP_071248143.1 | Type1.BREX | <-REase<-REase<-REase<-REase\|\|?->BrxA-DNA-binding->BrxB-BrxF-iSTAND->DUF6079*->OLD-ABC->REase->N6-METHYLASE->HTH+REase->PglZ->SIGMA+OB-Fold+AAA-ATPase+HKD-DNase-BrxL-><-DsbD<-Thioredoxin | 1172 | Bacteria>Terrabacteria_group>Cyanobacteriota/Melainabacteria_group>Cyanobacteriota>Cyanophyceae>Nostocales | GCF_001858125.1 | Cylindrospermopsis_raciborskii | CENA302_RS09780 | 17155..20673 | 77022 | BREX-system-P-loop-protein-BrxC-[Cylindrospermopsis-raciborskii] |
| WP_079290320.1 | Type1.BREX | RAMPs->?-><-SIGMA+OB-Fold+AAA-ATPase+Lon-Protease-BrxL<-PglZ+Phosphodiest\|\|AbiEi_4->AbiEii-><-N6-METHYLASE<-RNaseT2<-DUF6079*<-BrxB-BrxF-iSTAND<-?\|\|RADICAL-SAM->Schlafen+HATPase+wHTH->DUF29->?-><-?<-RFC-AAA+RuvB-AAA | 1177 | Bacteria>Terrabacteria_group>Cyanobacteriota/Melainabacteria_group>Cyanobacteriota>Cyanophyceae>Nostocales | GCF_002027345.1 | Cylindrospermopsis_raciborskii | CENA302_RS00080 | 28001..31534 | 77022 | BREX-system-P-loop-protein-BrxC-[Cylindrospermopsis-raciborskii] |
| WP_085727554.1 | Type1.BREX | ParB+HNH->URI-DNase-><-SIGMA+OB-Fold+AAA-ATPase+Lon-Protease-BrxL<-PglZ+Phosphodiest<-HTH_28+HTH_32+DDE_3<-?<-?<-N6-METHYLASE<-RNaseT2<-DUF6079*<-BrxB-BrxF-iSTAND<-?\|\|RADICAL-SAM->Schlafen+HATPase+wHTH->DUF29->?-><-?<-RFC-AAA+RuvB-AAA | 1177 | Bacteria>Terrabacteria_group>Cyanobacteriota/Melainabacteria_group>Cyanobacteriota>Cyanophyceae>Nostocales | GCF_002114155.1 | Cylindrospermopsis_raciborskii | B7087_RS05495 | 18208..21741 | 77022 | BREX-system-P-loop-protein-BrxC-[Cylindrospermopsis-raciborskii] |

<table>
<tr><th>Protein-ID</th><th>System</th><th>Gene-Neighbourhoods</th><th>Length</th><th>Taxonomy</th><th>Assembly-ID</th><th>Strain</th><th>Locus-Tag</th><th>Genomic-coordinates</th><th>TaxID</th><th>Fasta-define</th></tr>
<tr><td>WP_094331741.1</td><td>Type1.BREX</td><td>BrxA-DNA-binding-><-SIGMA+OB-Fold+AAA-ATPase+Lon-Protease-BrxL<-PglZ+Phosphodiest<-?<-HTH+ART-DarT<-SF1-SF2-Helicase<-N6-METHYLASE<-DUF6079*<-BrxB-BrxF-iSTAND</td><td>1180</td><td>Bacteria>Terrabacteria_group>Cyanobacteriota/Melainabacteria_group>Cyanobacteriota>Cyanophyceae>Nostocales</td><td>GCF_002245975.1</td><td>Nostoc_sp._'Peltigera_membranacea_cyanobiont'_213</td><td>CDG77_RS23320</td><td>21997..25539</td><td>2014530</td><td>BREX-system-P-loop-protein-BrxC-[Nostoc-sp.-'Peltigera-membranacea-cyanobiont'-213]</td></tr>
<tr><td>WP_102947438.1</td><td>Type1.BREX</td><td><-SIGMA+OB-Fold+AAA-ATPase+HKD-DNase-BrxL<-PglZ<-HTH+REase<-N6-METHYLASE<-REase<-OLD-ABC<-PIN<-DUF433<-DUF6079*<-BrxB-BrxF-iSTAND||?->?-><-?<-?<-?<-ParA-Soj-PloopNTPase<-?||ACT-></td><td>1166</td><td>Bacteria>Terrabacteria_group>Cyanobacteriota/Melainabacteria_group>Cyanobacteriota>Cyanophyceae>Nostocales</td><td>GCF_002893285.1</td><td>Cylindrospermopsis_raciborskii</td><td>CEP07_RS06090</td><td>15682..19182</td><td>77022</td><td>BREX-system-P-loop-protein-BrxC-[Cylindrospermopsis-raciborskii]</td></tr>
<tr><td>WP_115538370.1</td><td>Type1.BREX</td><td><-HTH+REase<-N6-METHYLASE-p<-DDE_3<-HTH_28+HTH_32+DDE_3<-N6-METHYLASE<-REase<-OLD-ABC<-PIN<-DUF433<-DUF6079*<-BrxB-BrxF-iSTAND||?->?-><-?<-?<-?<-ParA-Soj-PloopNTPase<-?||ACT-></td><td>1166</td><td>Bacteria>Terrabacteria_group>Cyanobacteriota/Melainabacteria_group>Cyanobacteriota>Cyanophyceae>Nostocales</td><td>GCF_003367075.2</td><td>Cylindrospermopsis_raciborskii</td><td>C6N34_008515</td><td>1862557..1866057</td><td>77022</td><td>BREX-system-P-loop-protein-BrxC-[Cylindrospermopsis-raciborskii]</td></tr>
<tr><td>WP_115538435.1</td><td>Type1.BREX</td><td><-RADICAL-SAM||?->BrxB-BrxF-iSTAND->DUF6079*->RNaseT2->N6-METHYLASE->PglZ+Phosphodiest->SIGMA+OB-Fold+AAA-ATPase+Lon-Protease-BrxL-><-Schlafen+HATPase+wHTH</td><td>1184</td><td>Bacteria>Terrabacteria_group>Cyanobacteriota/Melainabacteria_group>Cyanobacteriota>Cyanophyceae>Nostocales</td><td>GCF_003367075.2</td><td>Cylindrospermopsis_raciborskii</td><td>C6N34_RS02565</td><td>554978..558532</td><td>77022</td><td>BREX-system-P-loop-protein-BrxC-[Cylindrospermopsis-raciborskii]</td></tr>
<tr><td>WP_127084959.1</td><td>Type1.BREX</td><td>CASPASE->?-><-?<-TOPRIM<-OLD-ABC<-Schlafen+HATPase+wHTH||BrxA-DNA-binding->BrxB-BrxF-iSTAND->DUF6079-p*->?->?->PIN->CopG/RHH->N6-METHYLASE-p->OLD-ABC->HNH-></td><td>725</td><td>Bacteria>Terrabacteria_group>Cyanobacteriota/Melainabacteria_group>Cyanobacteriota>Cyanophyceae>Nostocales</td><td>GCF_003991905.1</td><td>Dulcicalothrix_desertica</td><td>DSM106972_RS34665</td><td>70206..72383</td><td>32056</td><td>BREX-system-P-loop-protein-BrxC-[Dulcicalothrix-desertica]</td></tr>
<tr><td>WP_141302954.1</td><td>Type1.BREX</td><td><-PglZ||DDE_Tnp_IS1-><-N6-METHYLASE-p<-N6-METHYLASE-p<-REase<-OLD-ABC<-PIN<-DUF433<-DUF6079*<-BrxB-BrxF-iSTAND||?->?-><-?<-?<-?<-ParA-Soj-PloopNTPase<-?||ACT-></td><td>1166</td><td>Bacteria>Terrabacteria_group>Cyanobacteriota/Melainabacteria_group>Cyanobacteriota>Cyanophyceae>Nostocales</td><td>GCF_006523545.1</td><td>Cylindrospermopsis_raciborskii</td><td>FIV49_RS01515</td><td>324659..328159</td><td>77022</td><td>BREX-system-P-loop-protein-BrxC-[Cylindrospermopsis-raciborskii]</td></tr>
<tr><td>WP_152591724.1</td><td>Type1.BREX</td><td>SAM-METHYLASE-><-AAA-ATPase<-EVE<-?<-?<-?<-?<-REase+SF1-SF2-Helicase||BrxB-BrxF-iSTAND->DUF6079*->N6-METHYLASE->SF1-SF2-Helicase->HTH+ART-DarT->?->PglZ+Phosphodiest->SIGMA+OB-Fold+[illegible]</td><td>1180</td><td>Bacteria>Terrabacteria_group>Cyanobacteriota/Melainabacteria_group>Cyanobacteriota>Cyanophyceae>Nostocales</td><td>GCF_009372195.1</td><td>Nostoc_sphaeroides</td><td>GXM_RS34850</td><td>1375509..1379051</td><td>446679</td><td>BREX-system-P-loop-protein-BrxC-[Nostoc-sphaeroides]</td></tr>
<tr><td>WP_114083280.1</td><td>Type1.BREX</td><td>NACHT->?->BrxA-DNA-binding-><-SIGMA+OB-Fold+AAA-ATPase+Lon-Protease-BrxL<-PglZ+Phosphodiest<-SbcC+ABC-ATPase<-N6-METHYLASE<-HEPN<-DUF6079*<-BrxB-BrxF-iSTAND||?->?->REase->Transposase-[illegible]</td><td>1180</td><td>Bacteria>Terrabacteria_group>Cyanobacteriota/Melainabacteria_group>Cyanobacteriota>Cyanophyceae>Nostocales</td><td>GCF_009873495.1</td><td>Nostoc_sp._ATCC_53789</td><td>GJB62_RS33695</td><td>77191..80733</td><td>76335</td><td>BREX-system-P-loop-protein-BrxC-[Nostoc-sp.-ATCC-53789]</td></tr>
<tr><td>WP_179070115.1</td><td>Type1.BREX</td><td>RADICAL-SAM-><-?||BrxA-DNA-binding-><-SIGMA+OB-Fold+AAA-ATPase+Lon-Protease-BrxL<-PglZ+Phosphodiest<-?<-HTH+ART-DarT<-SF1-SF2-Helicase<-N6-METHYLASE<-DUF6079*<-BrxB-BrxF-iSTAND<-TPR+[illegible]</td><td>1180</td><td>Bacteria>Terrabacteria_group>Cyanobacteriota/Melainabacteria_group>Cyanobacteriota>Cyanophyceae>Nostocales</td><td>GCF_013393905.1</td><td>Nostoc_sp._C052</td><td>FD723_RS36665</td><td>99740..103282</td><td>2576902</td><td>BREX-system-P-loop-protein-BrxC-[Nostoc-sp.-C052]</td></tr>
<tr><td>WP_187707239.1</td><td>Type1.BREX</td><td>Schlafen+HATPase+wHTH-><-SIGMA+OB-Fold+AAA-ATPase+Lon-Protease-BrxL<-PglZ+Phosphodiest<-N6-METHYLASE<-RNaseT2<-DUF6079*<-BrxB-BrxF-iSTAND<-?||RADICAL-SAM-></td><td>1189</td><td>Bacteria>Terrabacteria_group>Cyanobacteriota/Melainabacteria_group>Cyanobacteriota>Cyanophyceae>Nostocales</td><td>GCF_014489415.1</td><td>Cylindrospermopsis_curvispora</td><td>IAR63_RS08410</td><td>1849314..1852883</td><td>747548</td><td>BREX-system-P-loop-protein-BrxC-[Cylindrospermopsis-curvispora]</td></tr>
<tr><td>WP_187705577.1</td><td>Type1.BREX</td><td>NTP_transferase+MannoseP_isomer->ABC-ATPase+ABC-ATPase->?-><-?||?->?-><-?<-?||BrxB-BrxF-iSTAND->DUF6079*->DUF433->PIN->OLD-ABC->REase->N6-METHYLASE->HTH+REase->PglZ->SIGMA+OB-Fold+AAA[illegible]</td><td>1166</td><td>Bacteria>Terrabacteria_group>Cyanobacteriota/Melainabacteria_group>Cyanobacteriota>Cyanophyceae>Nostocales</td><td>GCF_014489415.1</td><td>Cylindrospermopsis_curvispora</td><td>IAR63_RS13115</td><td>2945663..2949163</td><td>747548</td><td>BREX-system-P-loop-protein-BrxC-[Cylindrospermopsis-curvispora]</td></tr>
<tr><td>WP_190970522.1</td><td>Type1.BREX</td><td><-MerR-HTH||BrxB-BrxF-iSTAND->DUF6079*->N6-METHYLASE->SbcC+ABC-ATPase->PglZ+Phosphodiest->SIGMA+OB-Fold+AAA-ATPase+Lon-Protease-BrxL-><-BrxA-DNA-binding||SF1-SF2-Helicase-><-?<-CHTH</td><td>1178</td><td>Bacteria>Terrabacteria_group>Cyanobacteriota/Melainabacteria_group>Cyanobacteriota>Cyanophyceae>Nostocales</td><td>GCF_014698475.1</td><td>Nostoc_spongiaeforme</td><td>H6G74_RS26715</td><td>22051..25587</td><td>502487</td><td>BREX-system-P-loop-protein-BrxC-[Nostoc-spongiaeforme]</td></tr>
<tr><td>WP_194144969.1</td><td>Type1.BREX</td><td>BrxA-DNA-binding-><-SIGMA+OB-Fold+AAA-ATPase+Lon-Protease-BrxL<-PglZ+Phosphodiest||REase->?-><-N6-METHYLASE<-DUF6079*<-BrxB-BrxF-iSTAND<-?||?->CHTH-></td><td>1179</td><td>Bacteria>Terrabacteria_group>Cyanobacteriota/Melainabacteria_group>Cyanobacteriota>Cyanophyceae>Nostocales</td><td>GCF_015206985.1</td><td>Nostoc_sp._LEGE_12447</td><td>IQ274_RS34165</td><td>10352..13891</td><td>1828640</td><td>BREX-system-P-loop-protein-BrxC-[Nostoc-sp.-LEGE-12447]</td></tr>
<tr><td>WP_235116163.1</td><td>Type1.BREX</td><td><-Glyco_hydro_63||BrxA-DNA-binding-><-SIGMA+OB-Fold+AAA-ATPase+Lon-Protease-BrxL<-PglZ+Phosphodiest||REase->?-><-N6-METHYLASE||MetJarc-HTH->PemK_toxin-><-DUF6079*<-BrxB-BrxF-iSTAND<-[illegible]</td><td>1180</td><td>Bacteria>Terrabacteria_group>Cyanobacteriota/Melainabacteria_group>Cyanobacteriota>Cyanophyceae>Nostocales</td><td>GCF_015207005.2</td><td>Desmonostoc_muscorum</td><td>IQ276_RS32480</td><td>7902408..7905950</td><td>1179</td><td>BREX-system-P-loop-protein-BrxC-[Desmonostoc-muscorum]</td></tr>
<tr><td>WP_200987085.1</td><td>Type1.BREX</td><td><-Acyltransferase<-?<-?<-?<-Peptidase_M17_N+Peptidase_M17||BrxB-BrxF-iSTAND->DUF6079*->N6-METHYLASE->SF1-SF2-Helicase->HTH+ART-DarT->?-><-?||PglZ+Phosphodiest->?->SIGMA+OB-Fold+AAA-A[illegible]</td><td>1185</td><td>Bacteria>Terrabacteria_group>Cyanobacteriota/Melainabacteria_group>Cyanobacteriota>Cyanophyceae>Nostocales</td><td>GCF_015245355.1</td><td>Anabaenopsis_elenkinii</td><td>IM676_RS11755</td><td>2659649..2663206</td><td>156213</td><td>BREX-system-P-loop-protein-BrxC-[Anabaenopsis-elenkinii]</td></tr>
<tr><td>WP_196520822.1</td><td>Type1.BREX</td><td><-Cas12k||MerR-HTH->BrxB-BrxF-iSTAND->DUF6079*->N6-METHYLASE-><-AIPR||PglZ+Phosphodiest-></td><td>1180</td><td>Bacteria>Terrabacteria_group>Cyanobacteriota/Melainabacteria_group>Cyanobacteriota>Cyanophyceae>Nostocales</td><td>GCF_015714725.1</td><td>Nostoc_sp._WHI</td><td>F8S12_RS34870</td><td>51345..54887</td><td>2650611</td><td>BREX-system-P-loop-protein-BrxC-[Nostoc-sp.-WHI]</td></tr>
<tr><td>WP_211942103.1</td><td>Type1.BREX</td><td>[illegible]</td><td>1166</td><td>Bacteria>Terrabacteria_group>Cyanobacteriota/Melainabacteria_group>Cyanobacteriota>Cyanophyceae>Nostocales</td><td>GCF_018139025.1</td><td>Cylindrospermopsis_raciborskii</td><td>J9B92_RS02395</td><td>526727..530227</td><td>77022</td><td>BREX-system-P-loop-protein-BrxC-[Cylindrospermopsis-raciborskii]</td></tr>
<tr><td>WP_211941867.1</td><td>Type1.BREX</td><td>[illegible]</td><td>1189</td><td>Bacteria>Terrabacteria_group>Cyanobacteriota/Melainabacteria_group>Cyanobacteriota>Cyanophyceae>Nostocales</td><td>GCF_018139025.1</td><td>Cylindrospermopsis_raciborskii</td><td>J9B92_RS13845</td><td>3094080..3097649</td><td>77022</td><td>BREX-system-P-loop-protein-BrxC-[Cylindrospermopsis-raciborskii]</td></tr>
<tr><td>WP_224095538.1</td><td>Type1.BREX</td><td>[illegible]</td><td>1178</td><td>Bacteria>Terrabacteria_group>Cyanobacteriota/Melainabacteria_group>Cyanobacteriota>Cyanophyceae>Nostocales</td><td>GCF_019976755.1</td><td>Nostoc_sp._MS1</td><td>NSMS1_RS32170</td><td>256782..260318</td><td>2764711</td><td>BREX-system-P-loop-protein-BrxC-[Nostoc-sp.-MS1]</td></tr>
<tr><td>WP_227788897.1</td><td>Type1.BREX</td><td>[illegible]</td><td>1169</td><td>Bacteria>Terrabacteria_group>Cyanobacteriota/Melainabacteria_group>Cyanobacteriota>Cyanophyceae>Nostocales</td><td>GCF_020702635.1</td><td>Nodularia_sp._LEGE_04288</td><td>IQ240_RS09380</td><td>506..4015</td><td>1828639</td><td>BREX-system-P-loop-protein-BrxC-[Nodularia-sp.-LEGE-04288]</td></tr>
<tr><td>WP_229553232.1</td><td>Type1.BREX</td><td>[illegible]</td><td>1231</td><td>Bacteria>Terrabacteria_group>Cyanobacteriota/Melainabacteria_group>Cyanobacteriota>Cyanophyceae>Nostocales</td><td>GCF_020829625.1</td><td>Nostoc_sp._XA010</td><td>LC608_RS09180</td><td>421109..424804</td><td>2780407</td><td>BREX-system-P-loop-protein-BrxC-[Nostoc-sp.-XA010]</td></tr>
<tr><td>WP_236308453.1</td><td>Type1.BREX</td><td>[illegible]</td><td>1166</td><td>Bacteria>Terrabacteria_group>Cyanobacteriota/Melainabacteria_group>Cyanobacteriota>Cyanophyceae>Nostocales</td><td>GCF_021650815.1</td><td>Cylindrospermopsis_raciborskii</td><td>L3I90_RS00125</td><td>24709..28209</td><td>77022</td><td>BREX-system-P-loop-protein-BrxC-[Cylindrospermopsis-raciborskii]</td></tr>
<tr><td>WP_236308178.1</td><td>Type1.BREX</td><td>[illegible]</td><td>1184</td><td>Bacteria>Terrabacteria_group>Cyanobacteriota/Melainabacteria_group>Cyanobacteriota>Cyanophyceae>Nostocales</td><td>GCF_021650815.1</td><td>Cylindrospermopsis_raciborskii</td><td>L3I90_RS11055</td><td>2527770..2531324</td><td>77022</td><td>BREX-system-P-loop-protein-BrxC-[Cylindrospermopsis-raciborskii]</td></tr>
<tr><td>WP_240869109.1</td><td>Type1.BREX</td><td>[illegible]</td><td>1166</td><td>Bacteria>Terrabacteria_group>Cyanobacteriota/Melainabacteria_group>Cyanobacteriota>Cyanophyceae>Nostocales</td><td>GCF_022496815.1</td><td>Cylindrospermopsis_raciborskii</td><td>GSN00_13735</td><td>14003..17503</td><td>77022</td><td>BREX-system-P-loop-protein-BrxC-[Cylindrospermopsis-raciborskii]</td></tr>
<tr><td>WP_240869635.1</td><td>Type1.BREX</td><td>[illegible]</td><td>1184</td><td>Bacteria>Terrabacteria_group>Cyanobacteriota/Melainabacteria_group>Cyanobacteriota>Cyanophyceae>Nostocales</td><td>GCF_022496815.1</td><td>Cylindrospermopsis_raciborskii</td><td>GSN00_RS16395</td><td>100185..103739</td><td>77022</td><td>BREX-system-P-loop-protein-BrxC-[Cylindrospermopsis-raciborskii]</td></tr>
<tr><td>WP_271763145.1</td><td>Type1.BREX</td><td>[illegible]</td><td>1182</td><td>Bacteria>Terrabacteria_group>Cyanobacteriota/Melainabacteria_group>Cyanobacteriota>Cyanophyceae>Nostocales</td><td>GCF_028331085.1</td><td>Anabaena_sp._CS-542/02</td><td>PN447_RS11410</td><td>43296..46844</td><td>3021719</td><td>BREX-system-P-loop-protein-BrxC-[Anabaena-sp.-CS-542/02]</td></tr>
<tr><td>WP_162398350.1</td><td>Type1.BREX</td><td>[illegible]</td><td>1178</td><td>Bacteria>Terrabacteria_group>Cyanobacteriota/Melainabacteria_group>Cyanobacteriota>Cyanophyceae>Nostocales</td><td>-</td><td>Nostoc_sp._B(2019)</td><td>GS682_RS23175</td><td>25673..29209</td><td>2690446</td><td>BREX-system-P-loop-protein-BrxC-[Nostoc-sp.-B(2019)]</td></tr>
<tr><td>CCI01315.1</td><td>Type1.BREX</td><td>[illegible]</td><td>1188</td><td>Bacteria>Terrabacteria_group>Cyanobacteriota/Melainabacteria_group>Cyanobacteriota>Cyanophyceae>Oscillatoriophycideae</td><td>GCA_000312185.1</td><td>Microcystis_aeruginosa_PCC_9443</td><td>MICAC_1830011</td><td>77203..80769</td><td>1160281</td><td>conserved-hypothetical-protein-[Microcystis-aeruginosa-PCC-9443]</td></tr>
<tr><td>REJ45948.1</td><td>Type1.BREX</td><td>[illegible]</td><td>1168</td><td>Bacteria>Terrabacteria_group>Cyanobacteriota/Melainabacteria_group>Cyanobacteriota>Cyanophyceae>Oscillatoriophycideae</td><td>GCA_003389435.1</td><td>Microcystis_flos-aquae_DF17</td><td>DWQ53_12245</td><td>7047..10553</td><td>2060470</td><td>BREX-system-P-loop-protein-BrxC-[Microcystis-flos-aquae-DF17]</td></tr>
<tr><td>BBD57221.1</td><td>Type1.BREX</td><td>[illegible]</td><td>1176</td><td>Bacteria>Terrabacteria_group>Cyanobacteriota/Melainabacteria_group>Cyanobacteriota>Cyanophyceae>Oscillatoriophycideae</td><td>GCA_003609755.1</td><td>Planktothrix_agardhii_NIES-204</td><td>NIES204_45570</td><td>14633..18163</td><td>282423</td><td>hypothetical-protein-NIES204_45570-[Planktothrix-agardhii-NIES-204]</td></tr>
<tr><td>TRU20104.1</td><td>Type1.BREX</td><td>[illegible]</td><td>1168</td><td>Bacteria>Terrabacteria_group>Cyanobacteriota/Melainabacteria_group>Cyanobacteriota>Cyanophyceae>Oscillatoriophycideae</td><td>GCA_007095795.1</td><td>Microcystis_aeruginosa_Ma_MB_S_20031200_S102D</td><td>EWV79_18805</td><td>31642..35148</td><td>2486255</td><td>BREX-system-P-loop-protein-BrxC-[Microcystis-aeruginosa-Ma_MB_S_20031200_S102D]</td></tr>
<tr><td>NCR78979.1</td><td>Type1.BREX</td><td>[illegible]</td><td>1168</td><td>Bacteria>Terrabacteria_group>Cyanobacteriota/Melainabacteria_group>Cyanobacteriota>Cyanophyceae>Oscillatoriophycideae</td><td>GCA_009996105.1</td><td>Microcystis_aeruginosa_K13-10</td><td>GPI91_02970</td><td>22657..26163</td><td>2685970</td><td>BREX-system-P-loop-protein-BrxC-[Microcystis-aeruginosa-K13-10]</td></tr>
<tr><td>NCS28722.1</td><td>Type1.BREX</td><td>[illegible]</td><td>1168</td><td>Bacteria>Terrabacteria_group>Cyanobacteriota/Melainabacteria_group>Cyanobacteriota>Cyanophyceae>Oscillatoriophycideae</td><td>GCA_009996455.1</td><td>Microcystis_aeruginosa_F13-15</td><td>GPJ18_09070</td><td>7084..10590</td><td>2685954</td><td>BREX-system-P-loop-protein-BrxC-[Microcystis-aeruginosa-F13-15]</td></tr>
<tr><td>NCS23184.1</td><td>Type1.BREX</td><td>[illegible]</td><td>1168</td><td>Bacteria>Terrabacteria_group>Cyanobacteriota/Melainabacteria_group>Cyanobacteriota>Cyanophyceae>Oscillatoriophycideae</td><td>GCA_009996485.1</td><td>Microcystis_aeruginosa_BS13-02</td><td>GPI96_01080</td><td>43062..46568</td><td>2685952</td><td>BREX-system-P-loop-protein-BrxC-[Microcystis-aeruginosa-BS13-02]</td></tr>
<tr><td>MBD2550293.1</td><td>Type1.BREX</td><td>[illegible]</td><td>1071</td><td>Bacteria>Terrabacteria_group>Cyanobacteriota/Melainabacteria_group>Cyanobacteriota>Cyanophyceae>Oscillatoriophycideae</td><td>GCA_014698445.1</td><td>Microcystis_elabens_FACHB-917</td><td>H6G65_12015</td><td>37472..40687</td><td>2692898</td><td>BREX-system-P-loop-protein-BrxC-[Microcystis-elabens-FACHB-917]</td></tr>
<tr><td>MBE5230557.1</td><td>Type1.BREX</td><td>[illegible]</td><td>1168</td><td>Bacteria>Terrabacteria_group>Cyanobacteriota/Melainabacteria_group>Cyanobacteriota>Cyanophyceae>Oscillatoriophycideae</td><td>GCA_014946735.1</td><td>Microcystis_aeruginosa_PMC_728.11</td><td>INR81_16200</td><td>47803..51309</td><td>2316423</td><td>BREX-system-P-loop-protein-BrxC-[Microcystis-aeruginosa-PMC-728.11]</td></tr>
<tr><td>MCA2504292.1</td><td>Type1.BREX</td><td>[illegible]</td><td>1168</td><td>Bacteria>Terrabacteria_group>Cyanobacteriota/Melainabacteria_group>Cyanobacteriota>Cyanophyceae>Oscillatoriophycideae</td><td>GCA_020244565.1</td><td>Microcystis_sp._M62BS1</td><td>IM561_02895</td><td>20537..24043</td><td>2771204</td><td>BREX-system-P-loop-protein-BrxC-[Microcystis-sp.-M62BS1]</td></tr>
<tr><td>WP_235758970.1</td><td>Type1.BREX</td><td>[illegible]</td><td>1176</td><td>Bacteria>Terrabacteria_group>Cyanobacteriota/Melainabacteria_group>Cyanobacteriota>Cyanophyceae>Oscillatoriophycideae</td><td>GCA_030820635.1</td><td>Planktothrix_agardhii</td><td>L2E81_RS22490</td><td>2176..5706</td><td>1160</td><td>BREX-system-P-loop-protein-BrxC-[Planktothrix-agardhii]</td></tr>
<tr><td>WP_007306283.1</td><td>Type1.BREX</td><td>[illegible]</td><td>655</td><td>Bacteria>Terrabacteria_group>Cyanobacteriota/Melainabacteria_group>Cyanobacteriota>Cyanophyceae>Oscillatoriophycideae</td><td>GCF_000167195.1</td><td>Crocosphaera_watsonii</td><td>CWATDRAFT_RS17495</td><td>8394..10361</td><td>263511</td><td>BREX-system-P-loop-protein-BrxC-[Crocosphaera-watsonii]</td></tr>
<tr><td>WP_007306520.1</td><td>Type1.BREX</td><td>[illegible]</td><td>1164</td><td>Bacteria>Terrabacteria_group>Cyanobacteriota/Melainabacteria_group>Cyanobacteriota>Cyanophyceae>Oscillatoriophycideae</td><td>GCF_000167195.1</td><td>Crocosphaera_watsonii</td><td>CWATDRAFT_RS18760</td><td>19121..22612</td><td>263511</td><td>BREX-system-P-loop-protein-BrxC,-partial-[Crocosphaera-watsonii]</td></tr>
<tr><td>WP_004266780.1</td><td>Type1.BREX</td><td>[illegible]</td><td>1168</td><td>Bacteria>Terrabacteria_group>Cyanobacteriota/Melainabacteria_group>Cyanobacteriota>Cyanophyceae>Oscillatoriophycideae</td><td>GCF_000312165.1</td><td>Microcystis_aeruginosa</td><td>MICAB_RS21670</td><td>36173..39679</td><td>1126</td><td>BREX-system-P-loop-protein-BrxC-[Microcystis-aeruginosa]</td></tr>
<tr><td>WP_043996480.1</td><td>Type1.BREX</td><td>[illegible]</td><td>1168</td><td>Bacteria>Terrabacteria_group>Cyanobacteriota/Melainabacteria_group>Cyanobacteriota>Cyanophyceae>Oscillatoriophycideae</td><td>GCF_000312185.1</td><td>Microcystis_aeruginosa</td><td>MICAC_RS06610</td><td>77263..80769</td><td>1126</td><td>BREX-system-P-loop-protein-BrxC-[Microcystis-aeruginosa]</td></tr>
<tr><td>WP_002790046.1</td><td>Type1.BREX</td><td>[illegible]</td><td>532</td><td>Bacteria>Terrabacteria_group>Cyanobacteriota/Melainabacteria_group>Cyanobacteriota>Cyanophyceae>Oscillatoriophycideae</td><td>GCF_000312225.1</td><td>Microcystis_aeruginosa</td><td>MICAF_RS30295</td><td>136943..138541</td><td>1126</td><td>BREX-system-P-loop-protein-BrxC-[Microcystis-aeruginosa]</td></tr>
<tr><td>WP_027255660.1</td><td>Type1.BREX</td><td>[illegible]</td><td>1176</td><td>Bacteria>Terrabacteria_group>Cyanobacteriota/Melainabacteria_group>Cyanobacteriota>Cyanophyceae>Oscillatoriophycideae</td><td>GCF_000464825.1</td><td>Planktothrix_agardhii</td><td>N316_RS0124710</td><td>8429..11959</td><td>1160</td><td>BREX-system-P-loop-protein-BrxC-[Planktothrix-agardhii]</td></tr>
<tr><td>WP_159293306.1</td><td>Type1.BREX</td><td>[illegible]</td><td>1168</td><td>Bacteria>Terrabacteria_group>Cyanobacteriota/Melainabacteria_group>Cyanobacteriota>Cyanophyceae>Oscillatoriophycideae</td><td>GCF_009811835.1</td><td>Microcystis_aeruginosa</td><td>GVO68_RS08615</td><td>57243..60749</td><td>1126</td><td>BREX-system-P-loop-protein-BrxC-[Microcystis-aeruginosa]</td></tr>
<tr><td>WP_190429845.1</td><td>Type1.BREX</td><td>[illegible]</td><td>1222</td><td>Bacteria>Terrabacteria_group>Cyanobacteriota/Melainabacteria_group>Cyanobacteriota>Cyanophyceae>Oscillatoriophycideae</td><td>GCF_014695635.1</td><td>Coleofasciculus_sp._FACHB-712</td><td>H6F50_RS18615</td><td>18748..22416</td><td>2692789</td><td>BREX-system-P-loop-protein-BrxC-[Coleofasciculus-sp.-FACHB-712]</td></tr>
<tr><td>WP_190817045.1</td><td>Type1.BREX</td><td>[illegible]</td><td>1234</td><td>Bacteria>Terrabacteria_group>Cyanobacteriota/Melainabacteria_group>Cyanobacteriota>Cyanophyceae>Oscillatoriophycideae</td><td>GCF_014698845.1</td><td>Coleofasciculus_sp._FACHB-1120</td><td>H6H02_RS09830</td><td>171769..175473</td><td>2692783</td><td>BREX-system-P-loop-protein-BrxC-[Coleofasciculus-sp.-FACHB-1120]</td></tr>
<tr><td>WP_193959346.1</td><td>Type1.BREX</td><td>[illegible]</td><td>1168</td><td>Bacteria>Terrabacteria_group>Cyanobacteriota/Melainabacteria_group>Cyanobacteriota>Cyanophyceae>Oscillatoriophycideae</td><td>GCF_015207185.1</td><td>Microcystis_sp._LEGE_08355</td><td>IQ242_RS05955</td><td>8581..12087</td><td>1828687</td><td>BREX-system-P-loop-protein-BrxC-[Microcystis-sp.-LEGE-08355]</td></tr>
<tr><td>WP_271989206.1</td><td>Type1.BREX</td><td>[illegible]</td><td>1168</td><td>Bacteria>Terrabacteria_group>Cyanobacteriota/Melainabacteria_group>Cyanobacteriota>Cyanophyceae>Oscillatoriophycideae</td><td>GCF_028330905.1</td><td>Microcystis_aeruginosa</td><td>PN502_RS07790</td><td>14255..17761</td><td>1126</td><td>BREX-system-P-loop-protein-BrxC-[Microcystis-aeruginosa]</td></tr>
<tr><td>WP_272076547.1</td><td>Type1.BREX</td><td>[illegible]</td><td>1168</td><td>Bacteria>Terrabacteria_group>Cyanobacteriota/Melainabacteria_group>Cyanobacteriota>Cyanophyceae>Oscillatoriophycideae</td><td>GCF_028331105.1</td><td>Microcystis_aeruginosa</td><td>PN487_RS04630</td><td>38433..41939</td><td>1126</td><td>BREX-system-P-loop-protein-BrxC-[Microcystis-aeruginosa]</td></tr>
<tr><td>WP_275522152.1</td><td>Type1.BREX</td><td>[illegible]</td><td>1181</td><td>Bacteria>Terrabacteria_group>Cyanobacteriota/Melainabacteria_group>Cyanobacteriota>Cyanophyceae>Oscillatoriophycideae</td><td>GCF_029167235.1</td><td>Kamptonema_sp._UHCC_0994</td><td>PY364_RS09785</td><td>12149..15694</td><td>3031329</td><td>BREX-system-P-loop-protein-BrxC-[Kamptonema-sp.-UHCC-0994]</td></tr>
<tr><td>WP_254034082.1</td><td>Type1.BREX</td><td>[illegible]</td><td>1176</td><td>Bacteria>Terrabacteria_group>Cyanobacteriota/Melainabacteria_group>Cyanobacteriota>Cyanophyceae>Oscillatoriophycideae</td><td>GCF_904830845.1</td><td>Planktothrix_agardhii</td><td>NMG87_RS21740</td><td>24173..27703</td><td>1160</td><td>BREX-system-P-loop-protein-BrxC-[Planktothrix-agardhii]</td></tr>
<tr><td>WP_190546543.1</td><td>Type1.BREX</td><td>[illegible]</td><td>1071</td><td>Bacteria>Terrabacteria_group>Cyanobacteriota/Melainabacteria_group>Cyanobacteriota>Cyanophyceae>Oscillatoriophycideae</td><td>-</td><td>Microcystis_elabens</td><td>H6G65_12015</td><td>37472..40687</td><td>44824</td><td>BREX-system-P-loop-protein-BrxC-[Microcystis-elabens]</td></tr>
<tr><td>WP_156114218.1</td><td>Type1.BREX</td><td>[illegible]</td><td>835</td><td>Bacteria>Terrabacteria_group>Cyanobacteriota/Melainabacteria_group>Cyanobacteriota>Cyanophyceae>Pleurocapsales</td><td>GCF_000756305.1</td><td>Myxosarcina_sp._GI1</td><td>KV40_RS27950</td><td>18823..21330</td><td>1541065</td><td>BREX-system-P-loop-protein-BrxC-[Myxosarcina-sp.-GI1]</td></tr>
<tr><td>KPQ32454.1</td><td>Type1.BREX</td><td>[illegible]</td><td>1200</td><td>Bacteria>Terrabacteria_group>Cyanobacteriota/Melainabacteria_group>Cyanobacteriota>Cyanophyceae>Pseudanabaenales</td><td>GCA_001314865.1</td><td>Phormidesmis_priestleyi_Ana</td><td>HLUCCA11_21420</td><td>1901..5503</td><td>1666911</td><td>Brex-phage-defense-system-ATPase-BrxC-[Phormidesmis-priestleyi-Ana]</td></tr>
<tr><td>PZO46844.1</td><td>Type1.BREX</td><td>[illegible]</td><td>1224</td><td>Bacteria>Terrabacteria_group>Cyanobacteriota/Melainabacteria_group>Cyanobacteriota>Cyanophyceae>Pseudanabaenales</td><td>GCA_003242115.1</td><td>Phormidesmis_priestleyi</td><td>DCF15_19745</td><td>1652..5326</td><td>268141</td><td>BREX-system-P-loop-protein-BrxC-[Phormidesmis-priestleyi]</td></tr>
<tr><td>PZV12558.1</td><td>Type1.BREX</td><td>[illegible]</td><td>1216</td><td>Bacteria>Terrabacteria_group>Cyanobacteriota/Melainabacteria_group>Cyanobacteriota>Cyanophyceae>Pseudanabaenales</td><td>GCA_003249025.1</td><td>Leptolyngbya_sp.</td><td>DCF22_12140</td><td>5388..9038</td><td>47254</td><td>BREX-system-P-loop-protein-BrxC-[Leptolyngbya-sp.]</td></tr>
<tr><td>MBC6953093.1</td><td>Type1.BREX</td><td>[illegible]</td><td>1150</td><td>Bacteria>Terrabacteria_group>Cyanobacteriota/Melainabacteria_group>Cyanobacteriota>Cyanophyceae>Pseudanabaenales</td><td>GCA_014338235.1</td><td>Leptolyngbya_sp.</td><td>DWB46_02725</td><td>400090..403542</td><td>47254</td><td>BREX-system-P-loop-protein-BrxC-[Leptolyngbya-sp.]</td></tr>
<tr><td>MBW4655645.1</td><td>Type1.BREX</td><td>[illegible]</td><td>1141</td><td>Bacteria>Terrabacteria_group>Cyanobacteriota/Melainabacteria_group>Cyanobacteriota>Cyanophyceae>Pseudanabaenales</td><td>GCA_019359435.1</td><td>Kaiparowitsia_implicata_GSE-PSE-MK54-09C</td><td>KME20_21790</td><td>10175..13600</td><td>2057771</td><td>BREX-system-P-loop-protein-BrxC-[Kaiparowitsia-implicata-GSE-PSE-MK54-09C]</td></tr>
<tr><td>MCA6503076.1</td><td>Type1.BREX</td><td>[illegible]</td><td>1202</td><td>Bacteria>Terrabacteria_group>Cyanobacteriota/Melainabacteria_group>Cyanobacteriota>Cyanophyceae>Pseudanabaenales</td><td>GCA_020403835.1</td><td>Pseudanabaena_sp._M090S1SP2A07QC</td><td>IM580_10820</td><td>20780..24388</td><td>2771254</td><td>BREX-system-P-loop-protein-BrxC-[Pseudanabaena-sp.-M090S1SP2A07QC]</td></tr>
<tr><td>MCA6523516.1</td><td>Type1.BREX</td><td>[illegible]</td><td>1202</td><td>Bacteria>Terrabacteria_group>Cyanobacteriota/Melainabacteria_group>Cyanobacteriota>Cyanophyceae>Pseudanabaenales</td><td>GCA_020403865.1</td><td>Pseudanabaena_sp._M051S1SP2A07QC</td><td>IM596_17155</td><td>23967..27575</td><td>2771253</td><td>BREX-system-P-loop-protein-BrxC-[Pseudanabaena-sp.-M051S1SP2A07QC]</td></tr>
<tr><td>MCA6571981.1</td><td>Type1.BREX</td><td>[illegible]</td><td>1202</td><td>Bacteria>Terrabacteria_group>Cyanobacteriota/Melainabacteria_group>Cyanobacteriota>Cyanophyceae>Pseudanabaenales</td><td>GCA_020404105.1</td><td>Pseudanabaena_sp._M53BS1SP1A06MG</td><td>IM536_21850</td><td>2730..6338</td><td>2771239</td><td>BREX-system-P-loop-protein-BrxC-[Pseudanabaena-sp.-M53BS1SP1A06MG]</td></tr>
<tr><td>MCA6593937.1</td><td>Type1.BREX</td><td>[illegible]</td><td>813</td><td>Bacteria>Terrabacteria_group>Cyanobacteriota/Melainabacteria_group>Cyanobacteriota>Cyanophyceae>Pseudanabaenales</td><td>GCA_020404945.1</td><td>Pseudanabaena_sp._M38BS1SP1A06MG</td><td>IM535_17950</td><td>20684..23126</td><td>2771238</td><td>BREX-system-P-loop-protein-BrxC-[Pseudanabaena-sp.-M38BS1SP1A06MG]</td></tr>
<tr><td>MCA6597699.1</td><td>Type1.BREX</td><td>[illegible]</td><td>1202</td><td>Bacteria>Terrabacteria_group>Cyanobacteriota/Melainabacteria_group>Cyanobacteriota>Cyanophyceae>Pseudanabaenales</td><td>GCA_020405005.1</td><td>Pseudanabaena_sp._M046S1SP1A06QC</td><td>IM539_15710</td><td>2728..6336</td><td>2771236</td><td>BREX-system-P-loop-protein-BrxC-[Pseudanabaena-sp.-M046S1SP1A06QC]</td></tr>
<tr><td>MCE2975831.1</td><td>Type1.BREX</td><td>[illegible]</td><td>1202</td><td>Bacteria>Terrabacteria_group>Cyanobacteriota/Melainabacteria_group>Cyanobacteriota>Cyanophyceae>Pseudanabaenales</td><td>GCA_021298775.1</td><td>Pseudanabaena_sp._CoA8_M7</td><td>LW851_05905</td><td>10182..13790</td><td>2904992</td><td>BREX-system-P-loop-protein-BrxC-[Pseudanabaena-sp.-CoA8_M7]</td></tr>
<tr><td>MCL1489692.1</td><td>Type1.BREX</td><td>[illegible]</td><td>1200</td><td>Bacteria>Terrabacteria_group>Cyanobacteriota/Melainabacteria_group>Cyanobacteriota>Cyanophyceae>Pseudanabaenales</td><td>GCA_023336205.1</td><td>Pseudanabaena_sp._Salubria-1</td><td>M1G31_02900</td><td>157927..161529</td><td>2938918</td><td>BREX-system-P-loop-protein-BrxC-[Pseudanabaena-sp.-Salubria-1]</td></tr>
<tr><td>WP_009625661.1</td><td>Type1.BREX</td><td>[illegible]</td><td>1178</td><td>Bacteria>Terrabacteria_group>Cyanobacteriota/Melainabacteria_group>Cyanobacteriota>Cyanophyceae>Pseudanabaenales</td><td>GCF_000332215.1</td><td>Pseudanabaena_biceps</td><td>PSE7429_RS03435</td><td>33637..37173</td><td>927669</td><td>MULTISPECIES:-BREX-system-P-loop-protein-BrxC-[Pseudanabaena]</td></tr>
<tr><td>WP_068818822.1</td><td>Type1.BREX</td><td>[illegible]</td><td>1182</td><td>Bacteria>Terrabacteria_group>Cyanobacteriota/Melainabacteria_group>Cyanobacteriota>Cyanophyceae>Pseudanabaenales</td><td>GCF_001650195.1</td><td>Phormidesmis_priestleyi</td><td>A8707_RS19220</td><td>3171..6719</td><td>268141</td><td>BREX-system-P-loop-protein-BrxC-[Phormidesmis-priestleyi]</td></tr>
<tr><td>WP_094532951.1</td><td>Type1.BREX</td><td>[illegible]</td><td>1181</td><td>Bacteria>Terrabacteria_group>Cyanobacteriota/Melainabacteria_group>Cyanobacteriota>Cyanophyceae>Pseudanabaenales</td><td>GCF_002251945.1</td><td>Pseudanabaena_sp._SR411</td><td>B9G53_RS13005</td><td>14836..18381</td><td>1980935</td><td>BREX-system-P-loop-protein-BrxC-[Pseudanabaena-sp.-SR411]</td></tr>
<tr><td>WP_169613736.1</td><td>Type1.BREX</td><td>[illegible]</td><td>1150</td><td>Bacteria>Terrabacteria_group>Cyanobacteriota/Melainabacteria_group>Cyanobacteriota>Cyanophyceae>Pseudanabaenales</td><td>GCF_012911975.1</td><td>Nodosilinea_sp._P-1105</td><td>E1H13_RS12425</td><td>20381..23833</td><td>2546229</td><td>BREX-system-P-loop-protein-BrxC-[Nodosilinea-sp.-P-1105]</td></tr>
<tr><td>WP_190797467.1</td><td>Type1.BREX</td><td>[illegible]</td><td>1186</td><td>Bacteria>Terrabacteria_group>Cyanobacteriota/Melainabacteria_group>Cyanobacteriota>Cyanophyceae>Pseudanabaenales</td><td>GCF_014695875.1</td><td>Leptolyngbya_sp._FACHB-541</td><td>H6G00_RS15685</td><td>19029..22589</td><td>2692810</td><td>BREX-system-P-loop-protein-BrxC-[Leptolyngbya-sp.-FACHB-541]</td></tr>
<tr><td>WP_193855895.1</td><td>Type1.BREX</td><td>[illegible]</td><td>1185</td><td>Bacteria>Terrabacteria_group>Cyanobacteriota/Melainabacteria_group>Cyanobacteriota>Cyanophyceae>Pseudanabaenales</td><td>GCF_015207105.1</td><td>cf._Phormidesmis_sp._LEGE_11477</td><td>IQ256_RS12585</td><td>42701..46258</td><td>1828680</td><td>BREX-system-P-loop-protein-BrxC-[cf.-Phormidesmis-sp.-LEGE-11477]</td></tr>
<tr><td>AFY73453.1</td><td>Type1.BREX</td><td>[illegible]</td><td>1120</td><td>Bacteria>Terrabacteria_group>Cyanobacteriota/Melainabacteria_group>Cyanobacteriota>Cyanophyceae>Synechococcales</td><td>GCA_000317085.1</td><td>Synechococcus_sp._PCC_7502</td><td>Syn7502_01364</td><td>1387283..1390645</td><td>1173263</td><td>hypothetical-protein-Syn7502_01364-[Synechococcus-sp.-PCC-7502]</td></tr>
<tr><td>PZV04575.1</td><td>Type1.BREX</td><td>[illegible]</td><td>1162</td><td>Bacteria>Terrabacteria_group>Cyanobacteriota/Melainabacteria_group>Cyanobacteriota>Cyanophyceae>Synechococcales</td><td>GCA_003248995.1</td><td>Cyanobium_sp.</td><td>DCF23_05760</td><td>6472..9960</td><td>2164130</td><td>MAG:-BREX-system-P-loop-protein-BrxC-[Cyanobium-sp.]</td></tr>
<tr><td>HBH74800.1</td><td>Type1.BREX</td><td>[illegible]</td><td>1163</td><td>Bacteria>Terrabacteria_group>Cyanobacteriota/Melainabacteria_group>Cyanobacteriota>Cyanophyceae>Synechococcales</td><td>GCA_003508015.1</td><td>Synechococcales_bacterium_UBA10510</td><td>DDY43_15550</td><td>2736..6227</td><td>2055795</td><td>BREX-system-P-loop-protein-BrxC-[Synechococcales-bacterium-UBA10510]</td></tr>
<tr><td>NJL99823.1</td><td>Type1.BREX</td><td>[illegible]</td><td>1145</td><td>Bacteria>Terrabacteria_group>Cyanobacteriota/Melainabacteria_group>Cyanobacteriota>Cyanophyceae>Synechococcales</td><td>GCA_012031565.1</td><td>Synechococcaceae_cyanobacterium_SM2_3_2</td><td>HC924_13960</td><td>2310..5747</td><td>2720471</td><td>BREX-system-P-loop-protein-BrxC-[Synechococcaceae-cyanobacterium-SM2_3_2]</td></tr>
<tr><td>MCE2837751.1</td><td>Type1.BREX</td><td>[illegible]</td><td>600</td><td>Bacteria>Terrabacteria_group>Cyanobacteriota/Melainabacteria_group>Cyanobacteriota>Cyanophyceae>Synechococcales</td><td>GCA_021298515.1</td><td>Cyanobium_sp._49614_E6</td><td>LW834_12445</td><td>8564..10363</td><td>2904989</td><td>BREX-system-P-loop-protein-BrxC-[Cyanobium-sp.-49614_E6]</td></tr>
<tr><td>WP_007100811.1</td><td>Type1.BREX</td><td>[illegible]</td><td>1161</td><td>Bacteria>Terrabacteria_group>Cyanobacteriota/Melainabacteria_group>Cyanobacteriota>Cyanophyceae>Synechococcales</td><td>GCF_000153065.1</td><td>unclassified_Synechococcus</td><td>RS9917_RS03965</td><td>839955..843440</td><td>2626047</td><td>MULTISPECIES:-BREX-system-P-loop-protein-BrxC-[unclassified-Synechococcus]</td></tr>
<tr><td>WP_015123655.1</td><td>Type1.BREX</td><td>[illegible]</td><td>1221</td><td>Bacteria>Terrabacteria_group>Cyanobacteriota/Melainabacteria_group>Cyanobacteriota>Cyanophyceae>Synechococcales</td><td>GCF_000316685.1</td><td>Synechococcus_sp._PCC_6312</td><td>SYN6312_RS04385</td><td>886176..889841</td><td>195253</td><td>BREX-system-P-loop-protein-BrxC-[Synechococcus-sp.-PCC-6312]</td></tr>
<tr><td>WP_028946553.1</td><td>Type1.BREX</td><td>[illegible]</td><td>1191</td><td>Bacteria>Terrabacteria_group>Cyanobacteriota/Melainabacteria_group>Cyanobacteriota>Cyanophyceae>Synechococcales</td><td>GCF_000478825.2</td><td>Synechocystis_sp._PCC_6714</td><td>D082_RS16385</td><td>87258..90833</td><td>1147</td><td>BREX-system-P-loop-protein-BrxC-[Synechocystis-sp.-PCC-6714]</td></tr>
<tr><td>WP_071799667.1</td><td>Type1.BREX</td><td>[illegible]</td><td>1161</td><td>Bacteria>Terrabacteria_group>Cyanobacteriota/Melainabacteria_group>Cyanobacteriota>Cyanophyceae>Synechococcales</td><td>GCF_001885215.1</td><td>unclassified_Synechococcus</td><td>BM449_RS00610</td><td>111734..115219</td><td>2626047</td><td>MULTISPECIES:-BREX-system-P-loop-protein-BrxC-[unclassified-Synechococcus]</td></tr>
<tr><td>WP_094560560.1</td><td>Type1.BREX</td><td>[illegible]</td><td>1161</td><td>Bacteria>Terrabacteria_group>Cyanobacteriota/Melainabacteria_group>Cyanobacteriota>Cyanophyceae>Synechococcales</td><td>GCF_002252665.1</td><td>Synechococcus_sp._8F6</td><td>CJZ78_RS08320</td><td>43413..46898</td><td>2025606</td><td>BREX-system-P-loop-protein-BrxC-[Synechococcus-sp.-8F6]</td></tr>
<tr><td>WP_188020741.1</td><td>Type1.BREX</td><td>[illegible]</td><td>1163</td><td>Bacteria>Terrabacteria_group>Cyanobacteriota/Melainabacteria_group>Cyanobacteriota>Cyanophyceae>Synechococcales</td><td>GCF_014279615.1</td><td>Synechococcus_sp._RS9907</td><td>SynRS9907_RS08975</td><td>1616845..1620336</td><td>221350</td><td>BREX-system-P-loop-protein-BrxC-[Synechococcus-sp.-RS9907]</td></tr>
<tr><td>WP_186471764.1</td><td>Type1.BREX</td><td>[illegible]</td><td>1163</td><td>Bacteria>Terrabacteria_group>Cyanobacteriota/Melainabacteria_group>Cyanobacteriota>Cyanophyceae>Synechococcales</td><td>GCF_014279955.1</td><td>Synechococcus_sp._BMK-MC-1</td><td>SynBMKMC1_RS11005</td><td>1938101..1941592</td><td>1442551</td><td>BREX-system-P-loop-protein-BrxC-[Synechococcus-sp.-BMK-MC-1]</td></tr>
<tr><td>WP_186513675.1</td><td>Type1.BREX</td><td>[illegible]</td><td>1163</td><td>Bacteria>Terrabacteria_group>Cyanobacteriota/Melainabacteria_group>Cyanobacteriota>Cyanophyceae>Synechococcales</td><td>GCF_014280115.1</td><td>Synechococcus_sp._A15-44</td><td>SynA1544_02490</td><td>2002733..2006224</td><td>1050646</td><td>BREX-system-P-loop-protein-BrxC-[Synechococcus-sp.-A15-44]</td></tr>
<tr><td>WP_186696561.1</td><td>Type1.BREX</td><td>[illegible]</td><td>1164</td><td>Bacteria>Terrabacteria_group>Cyanobacteriota/Melainabacteria_group>Cyanobacteriota>Cyanophyceae>Synechococcales</td><td>GCF_014280235.1</td><td>Cyanobium_sp._NS01</td><td>CyaNS01_RS07595</td><td>1454207..1457701</td><td>261284</td><td>BREX-system-P-loop-protein-BrxC-[Cyanobium-sp.-NS01]</td></tr>
<tr><td>WP_194019596.1</td><td>Type1.BREX</td><td>[illegible]</td><td>1205</td><td>Bacteria>Terrabacteria_group>Cyanobacteriota/Melainabacteria_group>Cyanobacteriota>Cyanophyceae>Synechococcales</td><td>GCF_015207985.1</td><td>Synechocystis_salina</td><td>IQ217_RS08375</td><td>30847..34464</td><td>945780</td><td>BREX-system-P-loop-protein-BrxC-[Synechocystis-salina]</td></tr>
<tr><td>WP_197158751.1</td><td>Type1.BREX</td><td>[illegible]</td><td>1161</td><td>Bacteria>Terrabacteria_group>Cyanobacteriota/Melainabacteria_group>Cyanobacteriota>Cyanophyceae>Synechococcales</td><td>GCF_015840525.1</td><td>Synechococcus_sp._CBW1006</td><td>H8F26_RS08495</td><td>1771673..1775158</td><td>1353138</td><td>BREX-system-P-loop-protein-BrxC-[Synechococcus-sp.-CBW1006]</td></tr>
<tr><td>WP_197170384.1</td><td>Type1.BREX</td><td>[illegible]</td><td>1161</td><td>Bacteria>Terrabacteria_group>Cyanobacteriota/Melainabacteria_group>Cyanobacteriota>Cyanophyceae>Synechococcales</td><td>GCF_015840915.1</td><td>Synechococcus_sp._CBW1002</td><td>H8F24_RS17545</td><td>3558842..3562327</td><td>1353134</td><td>BREX-system-P-loop-protein-BrxC-[Synechococcus-sp.-CBW1002]</td></tr>
<tr><td>WP_247215746.1</td><td>Type1.BREX</td><td>[illegible]</td><td>1221</td><td>Bacteria>Terrabacteria_group>Cyanobacteriota/Melainabacteria_group>Cyanobacteriota>Cyanophyceae>Synechococcales</td><td>GCF_022984075.1</td><td>Synechococcus_sp._C9</td><td>MLD66_RS04495</td><td>907547..911212</td><td>102119</td><td>BREX-system-P-loop-protein-BrxC-[Synechococcus-sp.-C9]</td></tr>
<tr><td>WP_250398658.1</td><td>Type1.BREX</td><td>[illegible]</td><td>1163</td><td>Bacteria>Terrabacteria_group>Cyanobacteriota/Melainabacteria_group>Cyanobacteriota>Cyanophyceae>Synechococcales</td><td>GCF_023630615.1</td><td>Synechococcus_sp._MU1648</td><td>NCN06_RS03450</td><td>583154..586645</td><td>2508351</td><td>BREX-system-P-loop-protein-BrxC-[Synechococcus-sp.-MU1648]</td></tr>
<tr><td>WP_255087766.1</td><td>Type1.BREX</td><td>[illegible]</td><td>1161</td><td>Bacteria>Terrabacteria_group>Cyanobacteriota/Melainabacteria_group>Cyanobacteriota>Cyanophyceae>Synechococcales</td><td>GCF_024345415.1</td><td>unclassified_Synechococcus</td><td>KBY92_RS03480</td><td>114966..118451</td><td>2626047</td><td>MULTISPECIES:-BREX-system-P-loop-protein-BrxC-[unclassified-Synechococcus]</td></tr>
<tr><td>WP_255021231.1</td><td>Type1.BREX</td><td>[illegible]</td><td>1166</td><td>Bacteria>Terrabacteria_group>Cyanobacteriota/Melainabacteria_group>Cyanobacteriota>Cyanophyceae>Synechococcales</td><td>GCF_024345755.1</td><td>Cyanobium_sp._L1E-Cus</td><td>KBZ22_RS04695</td><td>26198..29698</td><td>2823714</td><td>BREX-system-P-loop-protein-BrxC-[Cyanobium-sp.-L1E-Cus]</td></tr>
<tr><td>WP_255142981.1</td><td>Type1.BREX</td><td>[illegible]</td><td>1163</td><td>Bacteria>Terrabacteria_group>Cyanobacteriota/Melainabacteria_group>Cyanobacteriota>Cyanophyceae>Synechococcales</td><td>GCF_024345875.1</td><td>Synechococcus_sp._EJ6-Ellesmere</td><td>KBY78_RS06980</td><td>8981..12472</td><td>2823734</td><td>BREX-system-P-loop-protein-BrxC-[Synechococcus-sp.-EJ6-Ellesmere]</td></tr>
<tr><td>WP_254955669.1</td><td>Type1.BREX</td><td>[illegible]</td><td>1161</td><td>Bacteria>Terrabacteria_group>Cyanobacteriota/Melainabacteria_group>Cyanobacteriota>Cyanophyceae>Synechococcales</td><td>GCF_024346055.1</td><td>unclassified_Cyanobium</td><td>KBZ12_RS00475</td><td>78178..81663</td><td>2627006</td><td>MULTISPECIES:-BREX-system-P-loop-protein-BrxC-[unclassified-Cyanobium]</td></tr>
<tr><td>WP_254967828.1</td><td>Type1.BREX</td><td>[illegible]</td><td>1161</td><td>Bacteria>Terrabacteria_group>Cyanobacteriota/Melainabacteria_group>Cyanobacteriota>Cyanophyceae>Synechococcales</td><td>GCF_024346135.1</td><td>Cyanobium_sp._CH-040</td><td>KBY90_RS07090</td><td>13099..16584</td><td>2823708</td><td>BREX-system-P-loop-protein-BrxC-[Cyanobium-sp.-CH-040]</td></tr>
<tr><td>WP_254970971.1</td><td>Type1.BREX</td><td>[illegible]</td><td>1161</td><td>Bacteria>Terrabacteria_group>Cyanobacteriota/Melainabacteria_group>Cyanobacteriota>Cyanophyceae>Synechococcales</td><td>GCF_024346155.1</td><td>Cyanobium_sp._Candia_9D4</td><td>KBZ08_RS02055</td><td>407526..411011</td><td>2823707</td><td>BREX-system-P-loop-protein-BrxC-[Cyanobium-sp.-Candia-9D4]</td></tr>
<tr><td>WP_255125140.1</td><td>Type1.BREX</td><td>[illegible]</td><td>1162</td><td>Bacteria>Terrabacteria_group>Cyanobacteriota/Melainabacteria_group>Cyanobacteriota>Cyanophyceae>Synechococcales</td><td>GCF_024346215.1</td><td>Synechococcus_lacustris</td><td>KBY79_RS11620</td><td>2627..6115</td><td>2116544</td><td>BREX-system-P-loop-protein-BrxC-[Synechococcus-lacustris]</td></tr>
<tr><td>WP_255125212.1</td><td>Type1.BREX</td><td>[illegible]</td><td>1165</td><td>Bacteria>Terrabacteria_group>Cyanobacteriota/Melainabacteria_group>Cyanobacteriota>Cyanophyceae>Synechococcales</td><td>GCF_024346215.1</td><td>Synechococcus_lacustris</td><td>KBY79_RS12110</td><td>7241..10738</td><td>2116544</td><td>BREX-system-P-loop-protein-BrxC-[Synechococcus-lacustris]</td></tr>
<tr><td>WP_255015050.1</td><td>Type1.BREX</td><td>[illegible]</td><td>986</td><td>Bacteria>Terrabacteria_group>Cyanobacteriota/Melainabacteria_group>Cyanobacteriota>Cyanophyceae>Synechococcales</td><td>GCF_024346275.1</td><td>Cyanobium_sp._BA20m-p-22</td><td>KBZ15_RS09925</td><td>8060..11022</td><td>2823704</td><td>BREX-system-P-loop-protein-BrxC,-partial-[Cyanobium-sp.-BA20m-p-22]</td></tr>
<tr><td>WP_255005473.1</td><td>Type1.BREX</td><td>[illegible]</td><td>1161</td><td>Bacteria>Terrabacteria_group>Cyanobacteriota/Melainabacteria_group>Cyanobacteriota>Cyanophyceae>Synechococcales</td><td>GCF_024346315.1</td><td>Cyanobium_sp._ATX_6F1</td><td>KBZ13_RS02100</td><td>369855..373340</td><td>2823702</td><td>BREX-system-P-loop-protein-BrxC-[Cyanobium-sp.-ATX-6F1]</td></tr>
<tr><td>WP_254945611.1</td><td>Type1.BREX</td><td>[illegible]</td><td>1244</td><td>Bacteria>Terrabacteria_group>Cyanobacteriota/Melainabacteria_group>Cyanobacteriota>Cyanophyceae>Synechococcales</td><td>GCF_024346395.1</td><td>Cyanobium_sp._AMD-g</td><td>KBY82_RS12625</td><td>126184..129918</td><td>2823699</td><td>BREX-system-P-loop-protein-BrxC-[Cyanobium-sp.-AMD-g]</td></tr>
<tr><td>WP_254995750.1</td><td>Type1.BREX</td><td>[illegible]</td><td>1161</td><td>Bacteria>Terrabacteria_group>Cyanobacteriota/Melainabacteria_group>Cyanobacteriota>Cyanophyceae>Synechococcales</td><td>GCF_024346415.1</td><td>Cyanobium_sp._Aljojuca_7D2</td><td>KBY57_RS10265</td><td>32496..35981</td><td>2823698</td><td>BREX-system-P-loop-protein-BrxC-[Cyanobium-sp.-Aljojuca-7D2]</td></tr>
<tr><td>WP_255017269.1</td><td>Type1.BREX</td><td>[illegible]</td><td>1162</td><td>Bacteria>Terrabacteria_group>Cyanobacteriota/Melainabacteria_group>Cyanobacteriota>Cyanophyceae>Synechococcales</td><td>GCF_024346475.1</td><td>Cyanobium_sp._A2C-AMD</td><td>KBZ17_RS12415</td><td>5448..8936</td><td>2823695</td><td>BREX-system-P-loop-protein-BrxC-[Cyanobium-sp.-A2C-AMD]</td></tr>
<tr><td>WP_259727792.1</td><td>Type1.BREX</td><td>[illegible]</td><td>1162</td><td>Bacteria>Terrabacteria_group>Cyanobacteriota/Melainabacteria_group>Cyanobacteriota>Cyanophyceae>Synechococcales</td><td>GCF_025128115.1</td><td>Synechococcus_sp._CS-1326</td><td>KQ307_RS09240</td><td>37418..40906</td><td>2847978</td><td>BREX-system-P-loop-protein-BrxC-[Synechococcus-sp.-CS-1326]</td></tr>
<tr><td>WP_210391379.1</td><td>Type1.BREX</td><td>[illegible]</td><td>1155</td><td>Bacteria>Terrabacteria_group>Cyanobacteriota/Melainabacteria_group>Cyanobacteriota>Cyanophyceae>Synechococcales</td><td>-</td><td>Synechococcus_sp._PCC_7502</td><td>SYN7502_RS06685</td><td>1387178..1390645</td><td>1173263</td><td>BREX-system-P-loop-protein-BrxC-[Synechococcus-sp.-PCC-7502]</td></tr>
<tr><td>MBD1853918.1</td><td>Type1.BREX</td><td>[illegible]</td><td>1186</td><td>Bacteria>Terrabacteria_group>Cyanobacteriota/Melainabacteria_group>Cyanobacteriota>unclassified_Cyanobacteriota</td><td>GCA_014695325.1</td><td>Cyanobacteria_bacterium_FACHB-502</td><td>H6F87_28810</td><td>26444..30004</td><td>2692765</td><td>BREX-system-P-loop-protein-BrxC-[Cyanobacteria-bacterium-FACHB-502]</td></tr>
<tr><td>MBU6345615.1</td><td>Type1.BREX</td><td>[illegible]</td><td>1166</td><td>Bacteria>Terrabacteria_group>Cyanobacteriota/Melainabacteria_group>Cyanobacteriota>unclassified_Cyanobacteriota</td><td>GCA_018969285.1</td><td>Cyanobacteria_bacterium_REEB494</td><td>KGR70_09985</td><td>4368..7868</td><td>2800107</td><td>BREX-system-P-loop-protein-BrxC-[Cyanobacteria-bacterium-REEB494]</td></tr>
<tr><td>MBU6197215.1</td><td>Type1.BREX</td><td>[illegible]</td><td>1179</td><td>Bacteria>Terrabacteria_group>Cyanobacteriota/Melainabacteria_group>Cyanobacteriota>unclassified_Cyanobacteriota</td><td>GCA_018970325.1</td><td>Cyanobacteria_bacterium_REEB446</td><td>KGO93_06600</td><td>1417..4956</td><td>2800105</td><td>BREX-system-P-loop-protein-BrxC-[Cyanobacteria-bacterium-REEB446]</td></tr>
<tr><td>WP_221587161.1</td><td>Type1.BREX</td><td>[illegible]</td><td>1145</td><td>Bacteria>Terrabacteria_group>Deinococcota>Deinococci>Deinococcales>Deinococcaceae</td><td>-</td><td>Deinococcus_sp._RIT780</td><td>K5D85_00055</td><td>17975..21412</td><td>2870472</td><td>BREX-system-P-loop-protein-BrxC-[Deinococcus-sp.-RIT780]</td></tr>
<tr><td>MBB6099350.1</td><td>Type1.BREX</td><td>[illegible]</td><td>1151</td><td>Bacteria>Terrabacteria_group>Deinococcota>Deinococci>Deinococcales>Deinococcaceae</td><td>GCA_014202645.1</td><td>Deinobacterium_chartae</td><td>HNR42_002791</td><td>17098..20553</td><td>521158</td><td>hypothetical-protein-[Deinobacterium-chartae]</td></tr>
<tr><td>GGM57141.1</td><td>Type1.BREX</td><td>[illegible]</td><td>1153</td><td>Bacteria>Terrabacteria_group>Deinococcota>Deinococci>Deinococcales>Deinococcaceae</td><td>GCA_014647995.1</td><td>Deinococcus_arenae</td><td>GCM10008956_36020</td><td>13025..16486</td><td>1452751</td><td>hypothetical-protein-GCM10008956_36020-[Deinococcus-arenae]</td></tr>
<tr><td>MBX8463533.1</td><td>Type1.BREX</td><td>[illegible]</td><td>1145</td><td>Bacteria>Terrabacteria_group>Deinococcota>Deinococci>Deinococcales>Deinococcaceae</td><td>GCA_019718715.1</td><td>Deinococcus_sp._RIT780</td><td>K5D85_00055</td><td>17975..21412</td><td>2870472</td><td>BREX-system-P-loop-protein-BrxC-[Deinococcus-sp.-RIT780]</td></tr>
<tr><td>WP_017869746.1</td><td>Type1.BREX</td><td>[illegible]</td><td>1176</td><td>Bacteria>Terrabacteria_group>Deinococcota>Deinococci>Deinococcales>Deinococcaceae</td><td>GCF_000348665.1</td><td>Deinococcus_wulumuqiensis</td><td>G6R31_RS14545</td><td>153090..156620</td><td>980427</td><td>BREX-system-P-loop-protein-BrxC-[Deinococcus-wulumuqiensis]</td></tr>
<tr><td>WP_027460516.1</td><td>Type1.BREX</td><td>[illegible]</td><td>1140</td><td>Bacteria>Terrabacteria_group>Deinococcota>Deinococci>Deinococcales>Deinococcaceae</td><td>GCF_000482805.1</td><td>Deinococcus_murrayi</td><td>H565_RS0107590</td><td>4623..8045</td><td>68910</td><td>BREX-system-P-loop-protein-BrxC-[Deinococcus-murrayi]</td></tr>
<tr><td>WP_046843320.1</td><td>Type1.BREX</td><td>[illegible]</td><td>1151</td><td>Bacteria>Terrabacteria_group>Deinococcota>Deinococci>Deinococcales>Deinococcaceae</td><td>GCF_001007995.1</td><td>Deinococcus_soli_(ex_Cha_et_al._2016)</td><td>SY84_RS06385</td><td>1288437..1291892</td><td>1309411</td><td>BREX-system-P-loop-protein-BrxC-[Deinococcus-soli-(ex-Cha-et-al.-2016)]</td></tr>
<tr><td>WP_104992429.1</td><td>Type1.BREX</td><td>[illegible]</td><td>1140</td><td>Bacteria>Terrabacteria_group>Deinococcota>Deinococci>Deinococcales>Deinococcaceae</td><td>GCF_002953415.1</td><td>Deinococcus_sp._NW-56</td><td>C3K08_RS15840</td><td>30048..33470</td><td>2080419</td><td>BREX-system-P-loop-protein-BrxC-[Deinococcus-sp.-NW-56]</td></tr>
<tr><td>WP_110887791.1</td><td>Type1.BREX</td><td>[illegible]</td><td>1192</td><td>Bacteria>Terrabacteria_group>Deinococcota>Deinococci>Deinococcales>Deinococcaceae</td><td>GCF_003217515.1</td><td>Deinococcus_yavapaiensis</td><td>DES52_RS15755</td><td>92720..96298</td><td>309889</td><td>BREX-system-P-loop-protein-BrxC-[Deinococcus-yavapaiensis]</td></tr>
<tr><td>WP_135229998.1</td><td>Type1.BREX</td><td>[illegible]</td><td>1141</td><td>Bacteria>Terrabacteria_group>Deinococcota>Deinococci>Deinococcales>Deinococcaceae</td><td>GCF_004634215.1</td><td>Deinococcus_fonticola</td><td>E5Z01_RS14400</td><td>28954..32379</td><td>2528713</td><td>BREX-system-P-loop-protein-BrxC-[Deinococcus-fonticola]</td></tr>
<tr><td>WP_143721449.1</td><td>Type1.BREX</td><td>[illegible]</td><td>1148</td><td>Bacteria>Terrabacteria_group>Deinococcota>Deinococci>Deinococcales>Deinococcaceae</td><td>GCF_007280555.1</td><td>Deinococcus_detaillensis</td><td>FNU79_RS14110</td><td>42957..46403</td><td>2592048</td><td>BREX-system-P-loop-protein-BrxC-[Deinococcus-detaillensis]</td></tr>
<tr><td>WP_152872063.1</td><td>Type1.BREX</td><td>[illegible]</td><td>1188</td><td>Bacteria>Terrabacteria_group>Deinococcota>Deinococci>Deinococcales>Deinococcaceae</td><td>GCF_009377345.1</td><td>Deinococcus_terrestris</td><td>F8S09_RS13770</td><td>69508..73074</td><td>2651870</td><td>BREX-system-P-loop-protein-BrxC-[Deinococcus-terrestris]</td></tr>
<tr><td>WP_189007839.1</td><td>Type1.BREX</td><td>[illegible]</td><td>1186</td><td>Bacteria>Terrabacteria_group>Deinococcota>Deinococci>Deinococcales>Deinococcaceae</td><td>GCF_014646895.1</td><td>Deinococcus_roseus</td><td>IEY52_RS23290</td><td>14191..17751</td><td>392414</td><td>BREX-system-P-loop-protein-BrxC-[Deinococcus-roseus]</td></tr>
<tr><td>WP_216328012.1</td><td>Type1.BREX</td><td>[illegible]</td><td>1198</td><td>Bacteria>Terrabacteria_group>Deinococcota>Deinococci>Deinococcales>Deinococcaceae</td><td>GCF_018863415.1</td><td>Deinococcus_aestuarii</td><td>IC605_RS19250</td><td>17503..21099</td><td>2774531</td><td>BREX-system-P-loop-protein-BrxC-[Deinococcus-aestuarii]</td></tr>
<tr><td>WP_221091410.1</td><td>Type1.BREX</td><td>[illegible]</td><td>1200</td><td>Bacteria>Terrabacteria_group>Deinococcota>Deinococci>Deinococcales>Deinococcaceae</td><td>GCF_019693445.1</td><td>Deinococcus_aquaedulcis</td><td>KMW22_RS17995</td><td>15477..19079</td><td>2840455</td><td>BREX-system-P-loop-protein-BrxC-[Deinococcus-aquaedulcis]</td></tr>
<tr><td>WP_255578705.1</td><td>Type1.BREX</td><td>[illegible]</td><td>1153</td><td>Bacteria>Terrabacteria_group>Deinococcota>Deinococci>Deinococcales>Deinococcaceae</td><td>GCF_019718715.1</td><td>Deinococcus_sp._RIT780</td><td>K5D85_RS00055</td><td>17975..21436</td><td>2870472</td><td>BREX-system-P-loop-protein-BrxC-[Deinococcus-sp.-RIT780]</td></tr>
<tr><td>WP_249395097.1</td><td>Type1.BREX</td><td>[illegible]</td><td>1149</td><td>Bacteria>Terrabacteria_group>Deinococcota>Deinococci>Deinococcales>Deinococcaceae</td><td>GCF_023370075.1</td><td>Deinococcus_sp._QL22</td><td>M1R55_RS19505</td><td>18739..22188</td><td>2939437</td><td>BREX-system-P-loop-protein-BrxC-[Deinococcus-sp.-QL22]</td></tr>
<tr><td>WP_092265648.1</td><td>Type1.BREX</td><td>[illegible]</td><td>1160</td><td>Bacteria>Terrabacteria_group>Deinococcota>Deinococci>Deinococcales>Deinococcaceae</td><td>GCF_900109185.1</td><td>Deinococcus_reticulitermitis</td><td>BMY43_RS15290</td><td>11757..15239</td><td>856736</td><td>BREX-system-P-loop-protein-BrxC-[Deinococcus-reticulitermitis]</td></tr>
<tr><td>HHU80770.1</td><td>Type1.BREX</td><td>[illegible]</td><td>1173</td><td>Bacteria>Terrabacteria_group>Mycoplasmatota>Mollicutes>Acholeplasmatales>Acholeplasmataceae</td><td>GCA_012838945.1</td><td>Acholeplasmataceae_bacterium</td><td>GXZ35_05495</td><td>1540..5061</td><td>1898209</td><td>BREX-system-P-loop-protein-BrxC-[Acholeplasmataceae-bacterium]</td></tr>
<tr><td>MCK9492972.1</td><td>Type1.BREX</td><td>[illegible]</td><td>1198</td><td>Bacteria>Terrabacteria_group>Mycoplasmatota>Mollicutes>Acholeplasmatales>Acholeplasmataceae</td><td>GCA_023228755.1</td><td>Acholeplasmataceae_bacterium</td><td>M0Q00_01245</td><td>13412..17008</td><td>1898209</td><td>BREX-system-P-loop-protein-BrxC-[Acholeplasmataceae-bacterium]</td></tr>
<tr><td>HIS91144.1</td><td>Type1.BREX</td><td>[illegible]</td><td>1175</td><td>Bacteria>Terrabacteria_group>Mycoplasmatota>Mycoplasmatota_incertae_sedis>Candidatus_Faecisoma</td><td>GCA_018716405.1</td><td>Candidatus_Faecisoma_merdavium</td><td>IAC20_05750</td><td>4406..7933</td><td>2840801</td><td>BREX-system-P-loop-protein-BrxC-[Candidatus-Faecisoma-merdavium]</td></tr>
<tr><td>KPA14790.1</td><td>Type1.BREX</td><td>[illegible]</td><td>1187</td><td>Bacteria>Thermodesulfobacteriota>Desulfobacteria>Desulfobacterales>Candidatus_Magnetomoraceae>Candidatus_Magnetomorum</td><td>GCA_001292585.1</td><td>Candidatus_Magnetomorum_sp._HK-1</td><td>MHK_004993</td><td>4620..8183</td><td>1509431</td><td>DNA-repair-ATPase-[Candidatus-Magnetomorum-sp.-HK-1]</td></tr>
<tr><td>WP_073476316.1</td><td>Type1.BREX</td><td>[illegible]</td><td>1215</td><td>Bacteria>Thermodesulfobacteriota>Desulfobacteria>Desulfobacterales>Desulfatibacillaceae>Desulfatibacillum</td><td>GCF_900142135.1</td><td>Desulfatibacillum_alkenivorans</td><td>BUE18_RS12840</td><td>56211..59858</td><td>259354</td><td>BREX-system-P-loop-protein-BrxC-[Desulfatibacillum-alkenivorans]</td></tr>
<tr><td>WP_028324416.1</td><td>Type1.BREX</td><td>[illegible]</td><td>1145</td><td>Bacteria>Thermodesulfobacteriota>Desulfobacteria>Desulfobacterales>Desulfatirhabdiaceae>Desulfatirhabdium</td><td>GCF_000429925.1</td><td>Desulfatirhabdium_butyrativorans</td><td>G492_RS0109380</td><td>84085..87522</td><td>340467</td><td>BREX-system-P-loop-protein-BrxC-[Desulfatirhabdium-butyrativorans]</td></tr>
<tr><td>MBF0232702.1</td><td>Type1.BREX</td><td>[illegible]</td><td>1220</td><td>Bacteria>Thermodesulfobacteriota>Desulfobacteria>Desulfobacterales>Desulfobacteraceae>Desulfamplus</td><td>GCA_015231525.1</td><td>Desulfamplus_sp.</td><td>HQK65_06645</td><td>4455..8117</td><td>2006184</td><td>BREX-system-P-loop-protein-BrxC-[Desulfamplus-sp.]</td></tr>
<tr><td>MBF0232743.1</td><td>Type1.BREX</td><td>[illegible]</td><td>466</td><td>Bacteria>Thermodesulfobacteriota>Desulfobacteria>Desulfobacterales>Desulfobacteraceae>Desulfamplus</td><td>GCA_015231525.1</td><td>Desulfamplus_sp.</td><td>HQK65_06860</td><td>6630..8029</td><td>2006184</td><td>BREX-system-P-loop-protein-BrxC-[Desulfamplus-sp.]</td></tr>
<tr><td>MBF0468020.1</td><td>Type1.BREX</td><td>[illegible]</td><td>1187</td><td>Bacteria>Thermodesulfobacteriota>Desulfobacteria>Desulfobacterales>Desulfobacteraceae>Desulfamplus</td><td>GCA_015232935.1</td><td>Desulfamplus_sp.</td><td>HQK61_03910</td><td>4881..8444</td><td>2006184</td><td>BREX-system-P-loop-protein-BrxC-[Desulfamplus-sp.]</td></tr>
<tr><td>WP_080799293.1</td><td>Type1.BREX</td><td>[illegible]</td><td>1193</td><td>Bacteria>Thermodesulfobacteriota>Desulfobacteria>Desulfobacterales>Desulfobacteraceae>Desulfamplus</td><td>GCF_900170035.1</td><td>Desulfamplus_magnetovallimortis</td><td>MTBBW1_RS03710</td><td>949012..952593</td><td>1246637</td><td>BREX-system-P-loop-protein-BrxC-[Desulfamplus-magnetovallimortis]</td></tr>
<tr><td>WP_163354568.1</td><td>Type1.BREX</td><td>[illegible]</td><td>1187</td><td>Bacteria>Thermodesulfobacteriota>Desulfobacteria>Desulfobacterales>Desulfobacteraceae>Desulfobacter</td><td>-</td><td>Desulfobacter_hydrogenophilus</td><td>G3N28_17830</td><td>17749..21312</td><td>2291</td><td>BREX-system-P-loop-protein-BrxC-[Desulfobacter-hydrogenophilus]</td></tr>
<tr><td>HBT88578.1</td><td>Type1.BREX</td><td>[illegible]</td><td>1186</td><td>Bacteria>Thermodesulfobacteriota>Desulfobacteria>Desulfobacterales>Desulfobacteraceae>Desulfobacter</td><td>GCA_003510365.1</td><td>Desulfobacter_sp.</td><td>DEB50_08960</td><td>38159..41719</td><td>2294</td><td>BREX-system-P-loop-protein-BrxC-[Desulfobacter-sp.]</td></tr>
<tr><td>HBT89391.1</td><td>Type1.BREX</td><td>[illegible]</td><td>1187</td><td>Bacteria>Thermodesulfobacteriota>Desulfobacteria>Desulfobacterales>Desulfobacteraceae>Desulfobacter</td><td>GCA_003510365.1</td><td>Desulfobacter_sp.</td><td>DEB50_13295</td><td>35068..38631</td><td>2294</td><td>BREX-system-P-loop-protein-BrxC-[Desulfobacter-sp.]</td></tr>
<tr><td>NDY73873.1</td><td>Type1.BREX</td><td>[illegible]</td><td>1187</td><td>Bacteria>Thermodesulfobacteriota>Desulfobacteria>Desulfobacterales>Desulfobacteraceae>Desulfobacter</td><td>GCA_010499545.1</td><td>Desulfobacter_hydrogenophilus</td><td>G3N28_17830</td><td>17749..21312</td><td>2291</td><td>BREX-system-P-loop-protein-BrxC-[Desulfobacter-hydrogenophilus]</td></tr>
<tr><td>MBP8828964.1</td><td>Type1.BREX</td><td>[illegible]</td><td>1186</td><td>Bacteria>Thermodesulfobacteriota>Desulfobacteria>Desulfobacterales>Desulfobacteraceae>Desulfobacter</td><td>GCA_018055085.1</td><td>Desulfobacter_sp.</td><td>KBG98_04840</td><td>1249..4809</td><td>2294</td><td>BREX-system-P-loop-protein-BrxC-[Desulfobacter-sp.]</td></tr>
<tr><td>WP_035236115.1</td><td>Type1.BREX</td><td>[illegible]</td><td>1189</td><td>Bacteria>Thermodesulfobacteriota>Desulfobacteria>Desulfobacterales>Desulfobacteraceae>Desulfobacter</td><td>GCF_000745975.1</td><td>Desulfobacter_vibrioformis</td><td>Q366_RS02915</td><td>253933..257502</td><td>34031</td><td>BREX-system-P-loop-protein-BrxC-[Desulfobacter-vibrioformis]</td></tr>
</table>

| Protein-ID | System | Gene-Neighbourhoods | Length | Taxonomy | Assembly-ID | Strain | Locus-Tag | Genomic-coordinates | TaxID | Fasta-define |
|---|---|---|---|---|---|---|---|---|---|---|
| WP_111959628.1 | Type1.BREX | Zn_Tnp_IS91+Y2_Tnp-><-SIGMA+OB-Fold+AAA-ATPase+HKD-DNase-BrxL<-PglZ<-N6-METHYLASE<-REase<-OLD-ABC<-DUF6079*<-BrxB-BrxF-iSTAND<-BrxA-DNA-binding<-XIS-HTH\|\|HHH_5+SF1-SF2-Helicase->?-><- [illegible] | 1188 | Bacteria>Thermodesulfobacteriota>Desulfobacteria>Desulfobacterales>Desulfobacteraceae>Desulfobacter | GCF_004319545.1 | Desulfobacter_hydrogenophilus | DO021_RS19095 | 11943..15509 | 2291 | BREX-system-P-loop-protein-BrxC-[Desulfobacter-hydrogenophilus] |
| WP_178365089.1 | Type1.BREX | <-DDE_Tnp\|\|?->?->PRTase->RADICAL-SAM->BrxA-DNA-binding->BrxB-BrxF-iSTAND->DUF6079*->N6-METHYLASE->OLD-ABC->HNH->PglZ->SIGMA+OB-Fold+AAA-ATPase+HKD-DNase-BrxL->tRNA-><-?<-PAS+NtrC-AAA+ [illegible] | 1189 | Bacteria>Thermodesulfobacteriota>Desulfobacteria>Desulfobacterales>Desulfobacteraceae>Desulfobacter | GCF_013389365.1 | Desulfobacter_latus | HXW94_RS01255 | 9015..12584 | 2292 | BREX-system-P-loop-protein-BrxC-[Desulfobacter-latus] |
| MBL6994747.1 | Type1.BREX | BrxA-DNA-binding->BrxB-BrxF-iSTAND->DUF6079*->N6-METHYLASE->RVT->N6-METHYLASE-p->?->PglZ->SIGMA+OB-Fold+AAA-ATPase+Lon-Protease-BrxL->REase+nSTAND3->?->SbcC-> | 1186 | Bacteria>Thermodesulfobacteriota>Desulfobacteria>Desulfobacterales>Desulfobacteraceae>Desulfobacula | GCA_016785005.1 | Desulfobacula_sp. | ISR66_12690 | 4271..7831 | 2593537 | BREX-system-P-loop-protein-BrxC-[Desulfobacula-sp.] |
| MBT3178936.1 | Type1.BREX | <-N6-METHYLASE-p<-DUF6079*<-BrxB-BrxF-iSTAND<-BrxA-DNA-binding<-Phage_holin | 1196 | Bacteria>Thermodesulfobacteriota>Desulfobacteria>Desulfobacterales>Desulfobacteraceae>Desulfobacula | GCA_018645325.1 | Desulfobacula_sp. | HN339_18650 | 1350..4940 | 2593537 | BREX-system-P-loop-protein-BrxC-[Desulfobacula-sp.] |
| MBT3484108.1 | Type1.BREX | Phage_stabilise->?->?->?->?->?->Phage_holin->BrxA-DNA-binding->BrxB-BrxF-iSTAND->DUF6079*-> | 1084 | Bacteria>Thermodesulfobacteriota>Desulfobacteria>Desulfobacterales>Desulfobacteraceae>Desulfobacula | GCA_018646965.1 | Desulfobacula_sp. | HN468_02475 | 117211..120462 | 2593537 | BREX-system-P-loop-protein-BrxC-[Desulfobacula-sp.] |
| MBT4024573.1 | Type1.BREX | Phage_stabilise->?->?->?->?->?->Phage_holin->BrxA-DNA-binding->BrxB-BrxF-iSTAND->DUF6079-p*-> | 640 | Bacteria>Thermodesulfobacteriota>Desulfobacteria>Desulfobacterales>Desulfobacteraceae>Desulfobacula | GCA_018650485.1 | Desulfobacula_sp. | HOE60_06830 | 94483..96404 | 2593537 | BREX-system-P-loop-protein-BrxC-[Desulfobacula-sp.] |
| MBT4199889.1 | Type1.BREX | Phage_stabilise->?->?->?->?->?->Phage_holin->BrxA-DNA-binding->BrxB-BrxF-iSTAND->DUF6079*-> | 1130 | Bacteria>Thermodesulfobacteriota>Desulfobacteria>Desulfobacterales>Desulfobacteraceae>Desulfobacula | GCA_018651405.1 | Desulfobacula_sp. | HOE13_14760 | 42577..45967 | 2593537 | BREX-system-P-loop-protein-BrxC-[Desulfobacula-sp.] |
| MBT3805240.1 | Type1.BREX | Phage_stabilise->?->?->?->?->?->Phage_holin->BrxA-DNA-binding->BrxB-BrxF-iSTAND->DUF6079*-> | 1104 | Bacteria>Thermodesulfobacteriota>Desulfobacteria>Desulfobacterales>Desulfobacteraceae>Desulfobacula | GCA_018654565.1 | Desulfobacula_sp. | HOG03_11660 | 51501..54811 | 2593537 | BREX-system-P-loop-protein-BrxC-[Desulfobacula-sp.] |
| MBT4875636.1 | Type1.BREX | <-DUF6079-p*<-BrxB-BrxF-iSTAND<-BrxA-DNA-binding<-Phage_holin<-?<-?<-?<-?<-?<-Phage_stabilise | 600 | Bacteria>Thermodesulfobacteriota>Desulfobacteria>Desulfobacterales>Desulfobacteraceae>Desulfobacula | GCA_018664585.1 | Desulfobacula_sp. | HON48_10555 | 1..1800 | 2593537 | BREX-system-P-loop-protein-BrxC-[Desulfobacula-sp.] |
| MBT5544616.1 | Type1.BREX | <-DUF6079*<-BrxB-BrxF-iSTAND<-BrxA-DNA-binding<-Phage_holin<-?<-?<-?<-?<-?<-Phage_stabilise | 1148 | Bacteria>Thermodesulfobacteriota>Desulfobacteria>Desulfobacterales>Desulfobacteraceae>Desulfobacula | GCA_018665755.1 | Desulfobacula_sp. | HOK24_09040 | 1..3445 | 2593537 | BREX-system-P-loop-protein-BrxC-[Desulfobacula-sp.] |
| MBT7628744.1 | Type1.BREX | Phage_holin->BrxA-DNA-binding->BrxB-BrxF-iSTAND->DUF6079*->N6-METHYLASE->RAMA->PglZ->SIGMA+OB-Fold+AAA-ATPase+HKD-DNase-BrxL-> | 1196 | Bacteria>Thermodesulfobacteriota>Desulfobacteria>Desulfobacterales>Desulfobacteraceae>Desulfobacula | GCA_018675165.1 | Desulfobacula_sp. | HN597_03445 | 17169..20759 | 2593537 | BREX-system-P-loop-protein-BrxC-[Desulfobacula-sp.] |
| MBT7792445.1 | Type1.BREX | <-DUF6079-p*<-BrxB-BrxF-iSTAND<-BrxA-DNA-binding<-Phage_holin<-?<-?<-?<-?<-?<-Phage_stabilise | 744 | Bacteria>Thermodesulfobacteriota>Desulfobacteria>Desulfobacterales>Desulfobacteraceae>Desulfobacula | GCA_018676235.1 | Desulfobacula_sp. | HN767_07870 | 1..2233 | 2593537 | BREX-system-P-loop-protein-BrxC-[Desulfobacula-sp.] |
| MBT7050920.1 | Type1.BREX | <-DUF6079-p*<-BrxB-BrxF-iSTAND<-BrxA-DNA-binding<-Phage_holin<-?<-?<-?<-?<-?<-Phage_stabilise | 777 | Bacteria>Thermodesulfobacteriota>Desulfobacteria>Desulfobacterales>Desulfobacteraceae>Desulfobacula | GCA_018694265.1 | Desulfobacula_sp. | HN978_14790 | 1..2331 | 2593537 | BREX-system-P-loop-protein-BrxC-[Desulfobacula-sp.] |
| MBT6750309.1 | Type1.BREX | Phage_stabilise->?->?->?->?->?->Phage_holin->BrxA-DNA-binding->BrxB-BrxF-iSTAND->DUF6079*-> | 1102 | Bacteria>Thermodesulfobacteriota>Desulfobacteria>Desulfobacterales>Desulfobacteraceae>Desulfobacula | GCA_018696655.1 | Desulfobacula_sp. | HOA72_15355 | 44795..48100 | 2593537 | BREX-system-P-loop-protein-BrxC-[Desulfobacula-sp.] |
| MBT5971202.1 | Type1.BREX | <-DUF6079-p*<-BrxB-BrxF-iSTAND<-BrxA-DNA-binding<-Phage_holin<-?<-?<-?<-?<-?<-Phage_stabilise | 606 | Bacteria>Thermodesulfobacteriota>Desulfobacteria>Desulfobacterales>Desulfobacteraceae>Desulfobacula | GCA_018697915.1 | Desulfobacula_sp. | HOG59_08615 | 1..1820 | 2593537 | BREX-system-P-loop-protein-BrxC-[Desulfobacula-sp.] |
| WP_014958234.1 | Type1.BREX | <-SIGMA+OB-Fold+AAA-ATPase+HKD-DNase-BrxL<-PglZ<-?<-OLD-ABC<-ParB<-N6-METHYLASE<-DUF6079*<-BrxB-BrxF-iSTAND<-BrxA-DNA-binding\|\|?->GspA-ATPase-> | 1193 | Bacteria>Thermodesulfobacteriota>Desulfobacteria>Desulfobacterales>Desulfobacteraceae>Desulfobacula | GCF_000307105.1 | Desulfobacula_toluolica | TOL2_RS15330 | 3369625..3373206 | 28223 | BREX-system-P-loop-protein-BrxC-[Desulfobacula-toluolica] |
| WP_092229947.1 | Type1.BREX | <-Phage_integrase\|\|?->DSBH->?-><-?\|\|BrxA-DNA-binding->BrxB-BrxF-iSTAND->DUF6079*->N6-METHYLASE->ParB->OLD-ABC->?->PglZ->SIGMA+OB-Fold+AAA-ATPase+HKD-DNase-BrxL-> | 1193 | Bacteria>Thermodesulfobacteriota>Desulfobacteria>Desulfobacterales>Desulfobacteraceae>Desulfobacula | GCF_900105645.1 | Desulfobacula_phenolica | BLT94_RS02150 | 503008..506589 | 90732 | BREX-system-P-loop-protein-BrxC-[Desulfobacula-phenolica] |
| WP_139446530.1 | Type1.BREX | <-SIGMA+OB-Fold+AAA-ATPase+Lon-Protease-BrxL\|\|DiS_P_DiS+Peptidase_A24-><-?\|\|?-><-SIGMA+OB-Fold+AAA-ATPase+HKD-DNase-BrxL<-PglZ<-HNH<-OLD-ABC<-N6-METHYLASE<-DUF6079*<-BrxB-BrxF-iSTAND< [illegible] | 1199 | Bacteria>Thermodesulfobacteriota>Desulfobacteria>Desulfobacterales>Desulfobacteraceae>Desulfobotulus | GCF_006175995.1 | Desulfobotulus_mexicanus | FIM25_RS03945 | 130812..134411 | 2586642 | BREX-system-P-loop-protein-BrxC-[Desulfobotulus-mexicanus] |
| MCF8089785.1 | Type1.BREX | OLD-ABC->REase->DUF4007->LON-AAA->N6-METHYLASE->OLD-ABC->TOPRIM->DUF6079*->PglZ-> | 1350 | Bacteria>Thermodesulfobacteriota>Desulfobacteria>Desulfobacterales>Desulfobacteraceae>Desulfotignum | GCA_021737465.1 | Desulfotignum_sp. | K9L23_17210 | 8544..12596 | 2049434 | hypothetical-protein-[Desulfotignum-sp.] |
| MCF8090824.1 | Type1.BREX | <-SIGMA+OB-Fold+AAA-ATPase+Lon-Protease-BrxL-p<-PglZ<-OLD-ABC<-N6-METHYLASE<-MqsA_antitoxin<-DUF6079-p* | 425 | Bacteria>Thermodesulfobacteriota>Desulfobacteria>Desulfobacterales>Desulfobacteraceae>Desulfotignum | GCA_021737465.1 | Desulfotignum_sp. | K9L23_22535 | 8128..9407 | 2049434 | BREX-system-P-loop-protein-BrxC-[Desulfotignum-sp.] |
| MCF8138751.1 | Type1.BREX | BrxA-DNA-binding->BrxB-BrxF-iSTAND->DUF6079*->MqsA_antitoxin->N6-METHYLASE->OLD-ABC->PglZ->SIGMA+OB-Fold+AAA-ATPase+Lon-Protease-BrxL->ParB+N6_N4_METHYLASE-> | 1149 | Bacteria>Thermodesulfobacteriota>Desulfobacteria>Desulfobacterales>Desulfobacteraceae>Desulfotignum | GCA_021737695.1 | Desulfotignum_sp. | K9K63_15720 | 2047..5496 | 2049434 | BREX-system-P-loop-protein-BrxC-[Desulfotignum-sp.] |
| WP_024334643.1 | Type1.BREX | <-DDE_Tnp_ISAZ013<-DDE_Tnp_ISAZ013\|\|AIPR->BrxA-DNA-binding->BrxB-BrxF-iSTAND->DUF6079*->Schlafen+HATPase+wHTH->N6-METHYLASE->PHP+SbcC->ParB+HNH->DUF932->OLD-ABC->HNH->PglZ->SIGMA+OB-F [illegible] | 1144 | Bacteria>Thermodesulfobacteriota>Desulfobacteria>Desulfobacterales>Desulfobacteraceae>Desulfotignum | GCF_000421285.1 | Desulfotignum_balticum | K365_RS0111330 | 2237205..2240639 | 115781 | BREX-system-P-loop-protein-BrxC-[Desulfotignum-balticum] |
| OQY14226.1 | Type1.BREX | <-Y1_Tnp<-DUF6079*<-BrxB-BrxF-iSTAND<-BrxA-DNA-binding<-O6MGT-HTH<-?\|\|?->ABC-ATPase+ABC-ATPase+ABC-ATPase+SbcC-><-?<-?<-GGDEF | 1181 | Bacteria>Thermodesulfobacteriota>Desulfobacteria>Desulfobacterales>Desulfobacteraceae>unclassified_Desulfobacteraceae | GCA_002084375.1 | Desulfobacteraceae_bacterium_4572_187 | B6I30_01085 | 4692..8237 | 1971620 | ATPase-[Desulfobacteraceae-bacterium-4572_187] |
| OQY59120.1 | Type1.BREX | BrxA-DNA-binding->BrxB-BrxF-iSTAND->DUF6079*->DUF6079-p->?->PemK_toxin->N6-METHYLASE->PglZ->SIGMA+OB-Fold+AAA-ATPase+Lon-Protease-BrxL->?->?->PTSEIIA1-> | 1028 | Bacteria>Thermodesulfobacteriota>Desulfobacteria>Desulfobacterales>Desulfobacteraceae>unclassified_Desulfobacteraceae | GCA_002085455.1 | Desulfobacteraceae_bacterium_4572_88 | B6245_08220 | 2969..6055 | 1972444 | hypothetical-protein-B6245_08220-[Desulfobacteraceae-bacterium-4572_88] |
| RZB38310.1 | Type1.BREX | <-TPR+TPR<-?\|\|?->?->?-><-?\|\|?->BrxA-DNA-binding->BrxB-BrxF-iSTAND->DUF6079*->N6-METHYLASE-p->N6-METHYLASE->OLD-ABC->HNH->PglZ->SIGMA+OB-Fold+AAA-ATPase+HKD-DNase-BrxL-><-?<-?<-ABC-ATP [illegible] | 1173 | Bacteria>Thermodesulfobacteriota>Desulfobacteria>Desulfobacterales>Desulfobacteraceae>unclassified_Desulfobacteraceae | GCA_004193595.1 | Desulfobacteraceae_bacterium_Eth-SRB2 | SRB2_00058 | 60586..64107 | 2528438 | hypothetical-protein-SRB2_00058-[Desulfobacteraceae-bacterium-Eth-SRB2] |
| RZB36557.1 | Type1.BREX | <-SIGMA+OB-Fold+AAA-ATPase+Lon-Protease-BrxL<-PglZ<-Schlafen+HATPase+wHTH<-N6-METHYLASE<-CMP-hydrolase<-?<-DUF6079*<-BrxB-BrxF-iSTAND<-BrxA-DNA-binding\|\|?->?-><-?\|\|?->?->HTH_23+HTH_32 [illegible] | 1148 | Bacteria>Thermodesulfobacteriota>Desulfobacteria>Desulfobacterales>Desulfobacteraceae>unclassified_Desulfobacteraceae | GCA_004193595.1 | Desulfobacteraceae_bacterium_Eth-SRB2 | SRB2_02053 | 72440..75886 | 2528438 | hypothetical-protein-SRB2_02053-[Desulfobacteraceae-bacterium-Eth-SRB2] |
| HDR14882.1 | Type1.BREX | <-SIGMA+OB-Fold+AAA-ATPase+Lon-Protease-BrxL<-PglZ<-Schlafen+HATPase+wHTH<-N6-METHYLASE<-DUF6079*<-BrxB-BrxF-iSTAND<-BrxA-DNA-binding | 1147 | Bacteria>Thermodesulfobacteriota>Desulfobacteria>Desulfobacterales>Desulfobacteraceae>unclassified_Desulfobacteraceae | GCA_011048855.1 | Desulfobacteraceae_bacterium | ENN79_05225 | 10108..13551 | 2049433 | BREX-system-P-loop-protein-BrxC-[Desulfobacteraceae-bacterium] |
| NQT69312.1 | Type1.BREX | wHTH+WYL+WCX-p->RapA+SF1-SF2-Helicase+APATPase->?->BrxD-AAA->BrxD->SF1-SF2-Helicase->BrxA-DNA-binding->BrxB-BrxF-iSTAND->DUF6079*->N6-METHYLASE-p-> | 1145 | Bacteria>Thermodesulfobacteriota>Desulfobacteria>Desulfobacterales>Desulfobacteraceae>unclassified_Desulfobacteraceae | GCA_013203195.1 | Desulfobacteraceae_bacterium | HQ552_06990 | 10372..13809 | 2049433 | BREX-system-P-loop-protein-BrxC-[Desulfobacteraceae-bacterium] |
| MBL6967423.1 | Type1.BREX | BrxA-DNA-binding->BrxB-BrxF-iSTAND->DUF6079*->PhdYeFM_antitox->RelE-ParE->RNase->OLD-ABC-> | 1196 | Bacteria>Thermodesulfobacteriota>Desulfobacteria>Desulfobacterales>Desulfobacteraceae>unclassified_Desulfobacteraceae | GCA_016776415.1 | Desulfobacteraceae_bacterium | ISR62_03265 | 4310..7900 | 2049433 | BREX-system-P-loop-protein-BrxC-[Desulfobacteraceae-bacterium] |
| MBL7173047.1 | Type1.BREX | <-wHTH+WYL+WCX<-AbiEi_4\|\|?->BrxA-DNA-binding->BrxB-BrxF-iSTAND->DUF6079*->OLD-ABC->REase->N6-METHYLASE-p-> | 1172 | Bacteria>Thermodesulfobacteriota>Desulfobacteria>Desulfobacterales>Desulfobacteraceae>unclassified_Desulfobacteraceae | GCA_016782905.1 | Desulfobacteraceae_bacterium | ISS68_09440 | 5839..9357 | 2049433 | BREX-system-P-loop-protein-BrxC-[Desulfobacteraceae-bacterium] |
| MBL7102042.1 | Type1.BREX | <-OLD-ABC<-DUF6079*<-BrxB-BrxF-iSTAND<-BrxA-DNA-binding<-?\|\|AbiEi_4->AbiEii->REase->wHTH+WYL+WCX-> | 1172 | Bacteria>Thermodesulfobacteriota>Desulfobacteria>Desulfobacterales>Desulfobacteraceae>unclassified_Desulfobacteraceae | GCA_016783045.1 | Desulfobacteraceae_bacterium | ISS60_04735 | 1060..4578 | 2049433 | BREX-system-P-loop-protein-BrxC-[Desulfobacteraceae-bacterium] |
| MBL7224733.1 | Type1.BREX | <-CR-Kinase+pi3KINASE<-?<-Radical-SAM<-RelE-ParE<-?<-?<-?<-RNase<-DUF6079*<-BrxB-BrxF-iSTAND<-BrxA-DNA-binding\|\|wHTH+WYL+WCX-> | 1196 | Bacteria>Thermodesulfobacteriota>Desulfobacteria>Desulfobacterales>Desulfobacteraceae>unclassified_Desulfobacteraceae | GCA_016783055.1 | Desulfobacteraceae_bacterium | ISS59_01250 | 5905..9495 | 2049433 | BREX-system-P-loop-protein-BrxC-[Desulfobacteraceae-bacterium] |
| MCB9481003.1 | Type1.BREX | HAMP+MA+MA->wHTH+WYL+WCX->?->?->?->?->SF1-SF2-Helicase+DUF1998->BrxA-DNA-binding->BrxB-BrxF-iSTAND->DUF6079*->N6-METHYLASE->PglZ-p->PglZ->SIGMA+OB-Fold+AAA-ATPase+HKD-DNase-BrxL->?->? [illegible] | 1181 | Bacteria>Thermodesulfobacteriota>Desulfobacteria>Desulfobacterales>Desulfobacteraceae>unclassified_Desulfobacteraceae | GCA_020633955.1 | Desulfobacteraceae_bacterium | H6680_04155 | 92278..95823 | 2049433 | BREX-system-P-loop-protein-BrxC-[Desulfobacteraceae-bacterium] |
| MCF8035328.1 | Type1.BREX | <-Thermonuclease<-Phage_integrase<-?<-SIGMA+OB-Fold+AAA-ATPase+Lon-Protease-BrxL<-PglZ<-RelE-ParE<-Unstab_antitox<-N6-METHYLASE<-DUF6079*<-BrxB-BrxF-iSTAND<-BrxA-DNA-binding<-?<-?<-?\| [illegible] | 1185 | Bacteria>Thermodesulfobacteriota>Desulfobacteria>Desulfobacterales>Desulfobacteraceae>unclassified_Desulfobacteraceae | GCA_021737225.1 | Desulfobacteraceae_bacterium | K9K62_00505 | 14563..18120 | 2049433 | BREX-system-P-loop-protein-BrxC-[Desulfobacteraceae-bacterium] |
| MCP3943397.1 | Type1.BREX | <-SIGMA+OB-Fold+AAA-ATPase+Lon-Protease-BrxL<-PglZ<-Schlafen+HATPase+wHTH<-PHP+SbcC<-N6-METHYLASE<-DUF6079*<-BrxB-BrxF-iSTAND<-BrxA-DNA-binding<-XIS-HTH\|\|GDResolvase-> | 1213 | Bacteria>Thermodesulfobacteriota>Desulfobacteria>Desulfobacterales>Desulfobacteraceae>unclassified_Desulfobacteraceae | GCA_024226935.1 | Desulfobacteraceae_bacterium | GY710_18195 | 13498..17139 | 2049433 | BREX-system-P-loop-protein-BrxC-[Desulfobacteraceae-bacterium] |
| MCP4747910.1 | Type1.BREX | <-SIGMA+OB-Fold+AAA-ATPase+Lon-Protease-BrxL<-PglZ<-N6-METHYLASE<-DUF6079*<-BrxB-BrxF-iSTAND<-BrxA-DNA-binding | 1187 | Bacteria>Thermodesulfobacteriota>Desulfobacteria>Desulfobacterales>Desulfobacteraceae>unclassified_Desulfobacteraceae | GCA_024230435.1 | Desulfobacteraceae_bacterium | GY874_17495 | 47064..50627 | 2049433 | BREX-system-P-loop-protein-BrxC-[Desulfobacteraceae-bacterium] |
| WP_207688112.1 | Type1.BREX | <-SF1-SF2-Helicase<-?<-TPR+TPR+TPR+TPR<-SIGMA-HTH+SIGMA-HTH\|\|tRNA->XIS-HTH->BrxA-DNA-binding->BrxB-BrxF-iSTAND->DUF6079*->?->Schlafen+HATPase+wHTH->N6-METHYLASE->HTH+REase->SbcC->PglZ [illegible] | 1185 | Bacteria>Thermodesulfobacteriota>Desulfobacteria>Desulfobacterales>Desulfococcaceae>Desulfonema | GCF_017377355.1 | Desulfonema_limicola | dnl_RS20625 | 4834354..4837911 | 45656 | BREX-system-P-loop-protein-BrxC-[Desulfonema-limicola] |
| WP_207689226.1 | Type1.BREX | <-PglZ<-?<-?<-?<-N6-METHYLASE-p<-N6-METHYLASE\|\|PolB->HEPN-><-DUF6079*<-BrxB-BrxF-iSTAND<-BrxA-DNA-binding\|\|?->?->REase-><-?<-?<-PAS+NtrC-AAA+FIS-HTH | 1199 | Bacteria>Thermodesulfobacteriota>Desulfobacteria>Desulfobacterales>Desulfococcaceae>Desulfonema | GCF_017377355.1 | Desulfonema_limicola | dnl_RS26385 | 6146847..6150446 | 45656 | BREX-system-P-loop-protein-BrxC-[Desulfonema-limicola] |
| WP_236888848.1 | Type1.BREX | DUF3670+SF1-SF2-Helicase->?->?->?->?->ZDNA-HTH->BrxA-DNA-binding->BrxB-BrxF-iSTAND->DUF6079*->N6-METHYLASE->PglZ->SIGMA+OB-Fold+AAA-ATPase+HKD-DNase-BrxL->GTPase-AIG->sGTP->?->JetB [illegible] | 1177 | Bacteria>Thermodesulfobacteriota>Desulfobacteria>Desulfobacterales>Desulfolunaceae>Desulfoluna | GCF_021654575.1 | Desulfoluna_limicola | L6434_RS14765 | 3845772..3849305 | 2810562 | BREX-system-P-loop-protein-BrxC-[Desulfoluna-limicola] |
| WP_181552438.1 | Type1.BREX | <-N6-METHYLASE<-HEPN<-DUF6079*<-BrxB-BrxF-iSTAND<-BrxA-DNA-binding<-?\|\|HTH->Phage_int_SAM_3+Phage_integrase-> | 1209 | Bacteria>Thermodesulfobacteriota>Desulfobacteria>Desulfobacterales>Desulfosalsimonadaceae>Desulfosalsimonas | GCF_013761005.1 | Desulfosalsimonas_propionicica | HNR65_RS15495 | 3160..6789 | 332175 | BREX-system-P-loop-protein-BrxC-[Desulfosalsimonas-propionicica] |
| KJS33128.1 | Type1.BREX | XIS-HTH->BrxA-DNA-binding->BrxB-BrxF-iSTAND->DUF6079*->N6-METHYLASE->REase+AIPR->ParB+HNH->DUF3644->PglZ->SIGMA+OB-Fold+AAA-ATPase+Lon-Protease-BrxL->MULE-TRANSPOSASE->?->OLD-ABC+TOPR [illegible] | 1145 | Bacteria>Thermodesulfobacteriota>Desulfobacteria>Desulfobacterales>Desulfosarcinaceae>Desulfatitalea | GCA_000961655.1 | Desulfatitalea_sp._BRH_c12 | VR64_03780 | 109216..112653 | 1629708 | MAG:-hypothetical-protein-VR64_03780-[Desulfatitalea-sp.-BRH_c12] |
| WP_155308948.1 | Type1.BREX | VirB10->?->DnaB+HerA/FtsK->TraH->TraG_N-><-?\|\|XIS-HTH->BrxA-DNA-binding->BrxB-BrxF-iSTAND->DUF6079*->OLD-ABC->HNH->N6-METHYLASE->N6-METHYLASE->REase+AIPR->PglZ->SIGMA+OB-Fold+AAA-ATPa [illegible] | 1145 | Bacteria>Thermodesulfobacteriota>Desulfobacteria>Desulfobacterales>Desulfosarcinaceae>Desulfosarcina | GCF_009689005.1 | Desulfosarcina_ovata | GN112_RS03415 | 763655..767092 | 83564 | BREX-system-P-loop-protein-BrxC-[Desulfosarcina-ovata] |
| PIP37810.1 | Type1.BREX | <-DUF6079*<-BrxB-BrxF-iSTAND<-BrxA-DNA-binding<-ParB+HNH<-?<-?<-REase+HTH+35exo+Small-B-barrel+SF1-SF2-Helicase<-URI-DNase | 1174 | Bacteria>Thermodesulfobacteriota>Desulfobacteria>Desulfobacterales>unclassified_Desulfobacterales | GCA_002772195.1 | Desulfobacterales_bacterium_CG23_combo_of_CG06-09_8_20_14_all_51_8 | COX19_15630 | 1..3524 | 1973983 | hypothetical-protein-COX19_15630,-partial-[Desulfobacterales-bacterium-CG23_combo_of_CG06-09_8_20_14_all_51_8] |
| PIU49967.1 | Type1.BREX | <-DUF6079-p*<-BrxB-BrxF-iSTAND<-BrxA-DNA-binding<-AbiEii<-AbiEi_3_N+AbiEi_3 | 217 | Bacteria>Thermodesulfobacteriota>Desulfobacteria>Desulfobacterales>unclassified_Desulfobacterales | GCA_002779465.1 | Desulfobacterales_bacterium_CG07_land_8_20_14_0_80_52_14 | COS92_03800 | 1..651 | 1973982 | BREX-system-P-loop-protein-BrxC,-partial-[Desulfobacterales-bacterium-CG07_land_8_20_14_0_80_52_14] |
| MBC8421197.1 | Type1.BREX | <-AbiEii<-AbiEi_4\|\|?->BrxA-DNA-binding->BrxB-BrxF-iSTAND->DUF6079*->OLD-ABC->REase->N6-METHYLASE-> | 1172 | Bacteria>Thermodesulfobacteriota>Desulfobacteria>Desulfobacterales>unclassified_Desulfobacterales | GCA_014382905.1 | Desulfobacterales_bacterium | H8E10_21680 | 4515..8033 | 2044940 | BREX-system-P-loop-protein-BrxC-[Desulfobacterales-bacterium] |
| MBC8430784.1 | Type1.BREX | <-N6-METHYLASE<-DUF6079*<-BrxB-BrxF-iSTAND<-BrxA-DNA-binding<-?<-?\|\|?->?->RADICAL-SAM-> | 1187 | Bacteria>Thermodesulfobacteriota>Desulfobacteria>Desulfobacterales>unclassified_Desulfobacterales | GCA_014383215.1 | Desulfobacterales_bacterium | H8D96_02585 | 3776..7339 | 2044940 | BREX-system-P-loop-protein-BrxC-[Desulfobacterales-bacterium] |
| MBF0101465.1 | Type1.BREX | <-Glyco_hydro_19<-SIGMA+OB-Fold+AAA-ATPase+Lon-Protease-BrxL<-PglZ<-TOPRIM<-OLD-ABC<-HNH<-OLD-ABC<-N6-METHYLASE<-DUF6079*<-BrxB-BrxF-iSTAND<-BrxA-DNA-binding<-XIS-HTH | 1145 | Bacteria>Thermodesulfobacteriota>Desulfobacteria>Desulfobacterales>unclassified_Desulfobacterales | GCA_015228675.1 | Desulfobacterales_bacterium | HQK77_11205 | 12812..16249 | 2044940 | BREX-system-P-loop-protein-BrxC-[Desulfobacterales-bacterium] |
| MBF0120841.1 | Type1.BREX | <-SIGMA+OB-Fold+AAA-ATPase+Lon-Protease-BrxL<-PglZ<-N6-METHYLASE<-CR-ATPase8+REase<-DUF6079*<-BrxB-BrxF-iSTAND<-BrxA-DNA-binding<-HEPN | 1201 | Bacteria>Thermodesulfobacteriota>Desulfobacteria>Desulfobacterales>unclassified_Desulfobacterales | GCA_015228685.1 | Desulfobacterales_bacterium | HQK79_18595 | 27077..30682 | 2044940 | BREX-system-P-loop-protein-BrxC-[Desulfobacterales-bacterium] |
| MBF0121242.1 | Type1.BREX | <-PglZ<-RelE-ParE<-?<-?<-N6-METHYLASE-p<-HTH+REase<-Schlafen+HATPase+wHTH<-N6-METHYLASE<-Four-helical-protein<-DUF6079*<-BrxB-BrxF-iSTAND<-BrxA-DNA-binding | 1189 | Bacteria>Thermodesulfobacteriota>Desulfobacteria>Desulfobacterales>unclassified_Desulfobacterales | GCA_015228685.1 | Desulfobacterales_bacterium | HQK79_20605 | 12595..16164 | 2044940 | BREX-system-P-loop-protein-BrxC-[Desulfobacterales-bacterium] |
| MBF0225339.1 | Type1.BREX | <-ROK-HTH<-?<-REC+NtrC-AAA+FIS-HTH<-CHASE3+HISKIN<-Peptidase_S49+Peptidase_S49<-?\|\|tRNA->BrxA-DNA-binding->BrxB-BrxF-iSTAND->DUF6079*->Y1_Tnp->N6-METHYLASE->?->PglZ->SIGMA+OB-Fold+AAA [illegible] | 1198 | Bacteria>Thermodesulfobacteriota>Desulfobacteria>Desulfobacterales>unclassified_Desulfobacterales | GCA_015231595.1 | Desulfobacterales_bacterium | HQK76_07780 | 43716..47312 | 2044940 | BREX-system-P-loop-protein-BrxC-[Desulfobacterales-bacterium] |
| MCF6289521.1 | Type1.BREX | <-Peptidase_M20<-?\|\|?->?->tRNA-><-SIGMA+OB-Fold+AAA-ATPase+Lon-Protease-BrxL<-PglZ<-N6-METHYLASE<-Schlafen+HATPase+wHTH<-DUF6079*<-BrxB-BrxF-iSTAND<-BrxA-DNA-binding\|\|TRPR-HTH->?->Hip [illegible] | 1147 | Bacteria>Thermodesulfobacteriota>Desulfobacteria>Desulfobacterales>unclassified_Desulfobacterales | GCA_021647905.1 | Desulfobacterales_bacterium | L3J03_00740 | 80901..84344 | 2044940 | BREX-system-P-loop-protein-BrxC-[Desulfobacterales-bacterium] |
| MCG2778948.1 | Type1.BREX | DUF6079-p*->HEPN->N6-METHYLASE->OLD-ABC->TOPRIM->PglZ->SIGMA+OB-Fold+AAA-ATPase+Lon-Protease-BrxL-> | 757 | Bacteria>Thermodesulfobacteriota>Desulfobacteria>Desulfobacterales>unclassified_Desulfobacterales | GCA_022013615.1 | Desulfobacterales_bacterium | L6406_25005 | 1..2276 | 2044940 | hypothetical-protein-[Desulfobacterales-bacterium] |
| KGO34922.1 | Type1.BREX | <-PglZ<-ParB+HNH<-N6-METHYLASE<-DUF6079*<-BrxB-BrxF-iSTAND<-BrxA-DNA-binding | 1193 | Bacteria>Thermodesulfobacteriota>Desulfobulbia>Desulfobulbales>Desulfobulbaceae>Desulfobulbus | GCA_000769715.1 | Desulfobulbus_sp._Tol-SR | JT06_05465 | 8780..12361 | 1536652 | hypothetical-protein-JT06_05465-[Desulfobulbus-sp.-Tol-SR] |
| KGO34873.1 | Type1.BREX | MlaE->ABC-ATPase+SbcC->?-><-?\|\|XIS-HTH->BrxA-DNA-binding->BrxB-BrxF-iSTAND->DUF6079*->OLD-ABC->REase->N6-METHYLASE->?->OLD-ABC->HNH->PglZ->SIGMA+OB-Fold+AAA-ATPase+Lon-Protease-BrxL-> | 1146 | Bacteria>Thermodesulfobacteriota>Desulfobulbia>Desulfobulbales>Desulfobulbaceae>Desulfobulbus | GCA_000769715.1 | Desulfobulbus_sp._Tol-SR | JT06_05895 | 26559..29999 | 1536652 | hypothetical-protein-JT06_05895-[Desulfobulbus-sp.-Tol-SR] |
| WP_205241479.1 | Type1.BREX | <-Arm-DNA-bind_3+Phage_integrase\|\|?->AAA-ATPase->?-><-?<-SIGMA+OB-Fold+AAA-ATPase+Lon-Protease-BrxL<-PglZ<-HTH+REase<-N6-METHYLASE<-DUF6079*<-BrxB-BrxF-iSTAND<-BrxA-DNA-binding | 1148 | Bacteria>Thermodesulfobacteriota>Desulfobulbia>Desulfobulbales>Desulfobulbaceae>Desulfobulbus | GCF_016918545.1 | Desulfobulbus_alkaliphilus | JWG43_RS03775 | 164535..167981 | 869814 | BREX-system-P-loop-protein-BrxC-[Desulfobulbus-alkaliphilus] |
| WP_205243275.1 | Type1.BREX | <-SIGMA+OB-Fold+AAA-ATPase+Lon-Protease-BrxL<-REase<-PglZ<-Schlafen+HATPase+wHTH<-N6-METHYLASE<-?<-DUF6079*<-REase<-BrxB-BrxF-iSTAND<-BrxA-DNA-binding<-?\|\|wHTH+WYL+WCX-> | 1162 | Bacteria>Thermodesulfobacteriota>Desulfobulbia>Desulfobulbales>Desulfobulbaceae>Desulfobulbus | GCF_016918545.1 | Desulfobulbus_alkaliphilus | JWG43_RS12510 | 49790..53278 | 869814 | BREX-system-P-loop-protein-BrxC-[Desulfobulbus-alkaliphilus] |
| WP_205228200.1 | Type1.BREX | <-ParB+HNH<-SbcC<-Calcineurin+Calcineurin-SbcD<-SbcC+SbcC<-SIGMA+OB-Fold+AAA-ATPase+HKD-DNase-BrxL<-PglZ<-N6-METHYLASE<-REase<-ABC-ATPase<-DUF6079*<-BrxB-BrxF-iSTAND<-BrxA-DNA-binding [illegible] | 1182 | Bacteria>Thermodesulfobacteriota>Desulfobulbia>Desulfobulbales>Desulfobulbaceae>Desulfobulbus | GCF_016919065.1 | Desulfobulbus_rhabdoformis | JWJ90_RS15890 | 30219..33767 | 34032 | BREX-system-P-loop-protein-BrxC-[Desulfobulbus-rhabdoformis] |
| WP_028584971.1 | Type1.BREX | <-ZnR<-SIGMA+OB-Fold+AAA-ATPase+HKD-DNase-BrxL<-PglZ<-PadR-HTH<-Schlafen+HATPase+wHTH<-N6-METHYLASE<-REase<-OLD-ABC<-DUF6079*<-BrxB-BrxF-iSTAND<-BrxA-DNA-binding<-?<-VirB11<-?<-PCMT+M [illegible] | 1201 | Bacteria>Thermodesulfobacteriota>Desulfobulbia>Desulfobulbales>Desulfobulbaceae>Desulfogranum | GCF_000429965.1 | Desulfogranum_mediterraneum | G494_RS0113505 | 11738..15343 | 160661 | BREX-system-P-loop-protein-BrxC-[Desulfogranum-mediterraneum] |
| RJX32575.1 | Type1.BREX | Phage_integrase->BrxA-DNA-binding->BrxB-BrxF-iSTAND->DUF6079*->?->N6-METHYLASE->Schlafen+HATPase+wHTH->PglZ->SIGMA+OB-Fold+AAA-ATPase+Lon-Protease-BrxL->HD->SLATT-> | 1147 | Bacteria>Thermodesulfobacteriota>Desulfobulbia>Desulfobulbales>Desulfobulbaceae>Desulfurivibrio | GCA_003605035.1 | Desulfurivibrio_sp. | C4531_06340 | 2310..5753 | 2093369 | MAG:-BREX-system-P-loop-protein-BrxC-[Desulfurivibrio-sp.] |
| WP_013164351.1 | Type1.BREX | <-SIGMA+OB-Fold+AAA-ATPase+Lon-Protease-BrxL<-PglZ<-TOPRIM<-OLD-ABC<-N6-METHYLASE<-Schlafen+HATPase+wHTH<-DUF6079*<-BrxB-BrxF-iSTAND<-BrxA-DNA-binding<-XIS-HTH<-RNase<-?\|\|MULE-TRANSPO [illegible] | 1145 | Bacteria>Thermodesulfobacteriota>Desulfobulbia>Desulfobulbales>Desulfobulbaceae>Desulfurivibrio | GCF_000092205.1 | Desulfurivibrio_alkaliphilus | DAAHT2_RS10985 | 2524530..2527967 | 427923 | BREX-system-P-loop-protein-BrxC-[Desulfurivibrio-alkaliphilus] |
| HBI14530.1 | Type1.BREX | Arm-DNA-bind_3+Phage_integrase->XIS-HTH->?->HTH_58->Phage_integrase->?->SWIM->BrxA-DNA-binding->BrxB-BrxF-iSTAND->DUF6079*->N6-METHYLASE->REase+AIPR->PglZ->SIGMA+OB-Fold+AAA-ATPase+Lo [illegible] | 1145 | Bacteria>Thermodesulfobacteriota>Desulfobulbia>Desulfobulbales>Desulfobulbaceae>unclassified_Desulfobulbaceae | GCF_003508005.1 | Desulfobulbaceae_bacterium | DDY20_03275 | 148942..152379 | 2053307 | BREX-system-P-loop-protein-BrxC-[Desulfobulbaceae-bacterium] |
| MBC8316504.1 | Type1.BREX | <-N6-METHYLASE-p<-?<-DUF6079*<-BrxB-BrxF-iSTAND<-BrxA-DNA-binding<-?<-?<-?<-?<-?<-?<-XIS-HTH | 1176 | Bacteria>Thermodesulfobacteriota>Desulfobulbia>Desulfobulbales>Desulfobulbaceae>unclassified_Desulfobulbaceae | GCA_014382295.1 | Desulfobulbaceae_bacterium | H8E41_01265 | 2255..5785 | 2053307 | BREX-system-P-loop-protein-BrxC-[Desulfobulbaceae-bacterium] |
| MCJ7602132.1 | Type1.BREX | GDResolvase+HIN-HTH->BrxA-DNA-binding->BrxB-BrxF-iSTAND->DUF6079*->N6-METHYLASE->Schlafen+HATPase+wHTH->PglZ->SIGMA+OB-Fold+AAA-ATPase+Lon-Protease-BrxL->SbcC->SF1-SF2-Helicase-><-Tra [illegible] | 1147 | Bacteria>Thermodesulfobacteriota>Desulfobulbia>Desulfobulbales>Desulfobulbaceae>unclassified_Desulfobulbaceae | GCA_022866315.1 | Desulfobulbaceae_bacterium | MUO63_11635 | 3420..6863 | 2053307 | BREX-system-P-loop-protein-BrxC-[Desulfobulbaceae-bacterium] |
| MCJ7602405.1 | Type1.BREX | <-SIGMA+OB-Fold+AAA-ATPase+Lon-Protease-BrxL<-PglZ<-N6-METHYLASE<-DUF6079*<-BrxB-BrxF-iSTAND<-BrxA-DNA-binding<-CopG_antitoxin<-BrnT_toxin | 1186 | Bacteria>Thermodesulfobacteriota>Desulfobulbia>Desulfobulbales>Desulfobulbaceae>unclassified_Desulfobulbaceae | GCA_022866315.1 | Desulfobulbaceae_bacterium | MUO63_13025 | 17464..21024 | 2053307 | BREX-system-P-loop-protein-BrxC-[Desulfobulbaceae-bacterium] |
| MBA3007526.1 | Type1.BREX | Phage_integrase->XIS-HTH->BrxA-DNA-binding->BrxB-BrxF-iSTAND->DUF6079*->Schlafen+HATPase+wHTH->OLD-ABC->REase->N6-METHYLASE->Schlafen+HATPase+wHTH->PglZ->SIGMA+OB-Fold+AAA-ATPase+Lon- [illegible] | 1147 | Bacteria>Thermodesulfobacteriota>Desulfobulbia>Desulfobulbales>Desulfocapsaceae>Desulfocapsa | GCA_013792135.1 | Desulfocapsa_sp. | FP810_13320 | 53855..57298 | 2066021 | BREX-system-P-loop-protein-BrxC-[Desulfocapsa-sp.] |
| TKB07808.1 | Type1.BREX | <-HEPN<-SbcC<-Calcineurin+Calcineurin-SbcD<-SbcC+SbcC<-SIGMA+OB-Fold+AAA-ATPase+HKD-DNase-BrxL<-PglZ<-N6-METHYLASE<-REase<-OLD-ABC<-DUF6079*<-BrxB-BrxF-iSTAND<-BrxA-DNA-binding<-XIS-H [illegible] | 1183 | Bacteria>Thermodesulfobacteriota>Desulfobulbia>Desulfobulbales>Desulfocapsaceae>Desulforhopalus | GCA_005116575.1 | Desulforhopalus_sp._IMCC35007 | FCL48_15775 | 80281..83832 | 2569543 | BREX-system-P-loop-protein-BrxC-[Desulforhopalus-sp.-IMCC35007] |
| MCK9175761.1 | Type1.BREX | HicA_toxin->HicB-><-N6-METHYLASE<-DUF6079*<-BrxB-BrxF-iSTAND<-BrxA-DNA-binding | 1186 | Bacteria>Thermodesulfobacteriota>Desulfobulbia>Desulfobulbales>Desulfocapsaceae>Desulforhopalus | GCA_023231425.1 | Desulforhopalus_sp. | MOO96_10850 | 5461..9021 | 2099376 | BREX-system-P-loop-protein-BrxC-[Desulforhopalus-sp.] |
| WP_169309248.1 | Type1.BREX | <-HEPN<-SbcC<-Calcineurin+Calcineurin-SbcD<-SbcC+SbcC<-SIGMA+OB-Fold+AAA-ATPase+HKD-DNase-BrxL<-PglZ<-N6-METHYLASE<-REase<-OLD-ABC<-DUF6079*<-BrxB-BrxF-iSTAND<-BrxA-DNA-binding<-XIS-H [illegible] | 1182 | Bacteria>Thermodesulfobacteriota>Desulfobulbia>Desulfobulbales>Desulfocapsaceae>Desulforhopalus | GCF_005116575.1 | Desulforhopalus_sp._IMCC35007 | FCL48_RS15780 | 80281..83829 | 2569543 | BREX-system-P-loop-protein-BrxC-[Desulforhopalus-sp.-IMCC35007] |
| WP_205217989.1 | Type1.BREX | BrxA-DNA-binding->BrxB-BrxF-iSTAND->?->?->DUF6079*->N6-METHYLASE-><-?<-?<-Y1_Tnp<-?<-?<-?\|\|Phage_int_SAM_4-><-Y1_Tnp | 1186 | Bacteria>Thermodesulfobacteriota>Desulfobulbia>Desulfobulbales>Desulfocapsaceae>Desulforhopalus | GCF_016918505.1 | Desulforhopalus_vacuolatus | JWG39_RS03600 | 214603..218163 | 40414 | BREX-system-P-loop-protein-BrxC-[Desulforhopalus-vacuolatus] |
| WP_027370868.1 | Type1.BREX | <-PIN<-MazE_antitoxin\|\|?->RHH->PIN->?->GGDEF+EAL->tRNA->BrxA-DNA-binding->BrxB-BrxF-iSTAND->DUF6079*->N6-METHYLASE->?->Schlafen+HATPase+wHTH->PglZ->?->HicB_lk_antitox->REase->SIGMA+OB [illegible] | 1190 | Bacteria>Thermodesulfobacteriota>Desulfovibrionia>Desulfovibrionales>Desulfohalobiaceae>Desulfovermiculus | GCF_000620765.1 | Desulfovermiculus_halophilus | N902_RS0110295 | 13006..16578 | 339722 | BREX-system-P-loop-protein-BrxC-[Desulfovermiculus-halophilus] |
| MCF8038487.1 | Type1.BREX | BrxB-BrxF-iSTAND->DUF6079*->N6-METHYLASE->?->PglZ->SIGMA+OB-Fold+AAA-ATPase+Lon-Protease-BrxL-> | 1188 | Bacteria>Thermodesulfobacteriota>Desulfovibrionia>Desulfovibrionales>Desulfohalobiaceae>unclassified_Desulfohalobiaceae | GCA_021737115.1 | Desulfohalobiaceae_bacterium | K9K79_04150 | 650..4216 | 2683721 | BREX-system-P-loop-protein-BrxC-[Desulfohalobiaceae-bacterium] |
| WP_015773519.1 | Type1.BREX | <-SMODS<-?<-?<-MIT_C<-SIGMA+OB-Fold+AAA-ATPase+Lon-Protease-BrxL<-PglZ<-REase+AIPR<-N6-METHYLASE<-DUF6079*<-BrxB-BrxF-iSTAND<-BrxA-DNA-binding<-NERD+ZnR<-?<-Calcineurin-SbcD<-Calcineu [illegible] | 1142 | Bacteria>Thermodesulfobacteriota>Desulfovibrionia>Desulfovibrionales>Desulfomicrobiaceae>Desulfomicrobium | GCF_000023225.1 | Desulfomicrobium_baculatum | DBAC_RS06650 | 1466575..1470003 | 899 | BREX-system-P-loop-protein-BrxC-[Desulfomicrobium-baculatum] |
| WP_066605667.1 | Type1.BREX | HDRnase->?-><-?<-tRNA<-?\|\|?->?->?->BrxA-DNA-binding->BrxB-BrxF-iSTAND->DUF6079*->N6-METHYLASE->OLD-ABC->HNH->PglZ->SIGMA+OB-Fold+AAA-ATPase+Lon-Protease-BrxL-><-RNA_METHYLASE | 1145 | Bacteria>Thermodesulfobacteriota>Desulfovibrionia>Desulfovibrionales>Desulfomicrobiaceae>Desulfomicrobium | GCF_001553625.1 | Desulfomicrobium_orale | AXF15_RS07915 | 1725501..1728938 | 132132 | BREX-system-P-loop-protein-BrxC-[Desulfomicrobium-orale] |
| WP_192624162.1 | Type1.BREX | AbrB-HTH->BrxA-DNA-binding->BrxB-BrxF-iSTAND->DUF6079*->HEPN->N6-METHYLASE->PHP+SbcC->Schlafen+HATPase+wHTH->PglZ->SIGMA+OB-Fold+AAA-ATPase+Lon-Protease-BrxL->HEPN-> | 1142 | Bacteria>Thermodesulfobacteriota>Desulfovibrionia>Desulfovibrionales>Desulfomicrobiaceae>Desulfomicrobium | GCF_014873765.1 | Desulfomicrobium_macestii | H4684_RS13755 | 68182..71610 | 90731 | BREX-system-P-loop-protein-BrxC-[Desulfomicrobium-macestii] |
| WP_092376845.1 | Type1.BREX | Phage_integrase->TOPRIM->?-><-TRPR-HTH\|\|?-><-HTH_3\|\|AbrB-HTH->BrxA-DNA-binding->BrxB-BrxF-iSTAND->DUF6079*->ParB+HNH->N6-METHYLASE->?->?->PglZ->HEPN+SH3-like+SF1-SF2-Helicase+wHTH->MO [illegible] | 1183 | Bacteria>Thermodesulfobacteriota>Desulfovibrionia>Desulfovibrionales>Desulfomicrobiaceae>Desulfomicrobium | GCF_900114115.1 | Desulfomicrobium_apsheronum | BMZ40_RS13600 | 74396..77947 | 52560 | BREX-system-P-loop-protein-BrxC-[Desulfomicrobium-apsheronum] |
| WP_008869235.1 | Type1.BREX | <-Y1_Tnp<-ARG<-?<-SIGMA+OB-Fold+AAA-ATPase+Lon-Protease-BrxL<-PglZ<-TOPRIM<-OLD-ABC<-N6-METHYLASE<-DUF6079*<-BrxB-BrxF-iSTAND<-BrxA-DNA-binding<-?<-ARG<-HTH_3<-HTH_3<-Tox-RelE<-HTH_3< [illegible] | 1147 | Bacteria>Thermodesulfobacteriota>Desulfovibrionia>Desulfovibrionales>Desulfonatronovibrionaceae>Desulfonatronospira | GCF_000174435.1 | Desulfonatronospira_thiodismutans | DTHIO_RS04855 | 1069759..1073202 | 488939 | BREX-system-P-loop-protein-BrxC-[Desulfonatronospira-thiodismutans] |
| WP_035067990.1 | Type1.BREX | <-35exo<-?<-SbcC+SbcC<-SIGMA+OB-Fold+AAA-ATPase+HKD-DNase-BrxL<-PglZ<-N6-METHYLASE<-REase<-OLD-ABC<-DUF6079*<-BrxB-BrxF-iSTAND<-BrxA-DNA-binding<-wHTH+WYL+WCX\|\|RHH_1->RelE-ParE-><-?\|\| [illegible] | 1182 | Bacteria>Thermodesulfobacteriota>Desulfovibrionia>Desulfovibrionales>Desulfovibrionaceae>Cupidesulfovibrio | GCF_000504305.1 | Cupidesulfovibrio_termitidis | DESTE_RS12450 | 3095808..3099356 | 42252 | BREX-system-P-loop-protein-BrxC-[Cupidesulfovibrio-termitidis] |
| AAS96500.1 | Type1.BREX | <-GGDEF<-ZnR+HTH_Tnp_ISL3+DDE_Tnp_ISL3<-SIGMA+OB-Fold+AAA-ATPase+HKD-DNase-BrxL<-PglZ<-REase<-N6-METHYLASE<-REase<-OLD-ABC<-DUF6079*<-BrxB-BrxF-iSTAND\|\|?-><-wHTH+WYL+WCX-p\|\|HIN-HTH->H [illegible] | 1183 | Bacteria>Thermodesulfobacteriota>Desulfovibrionia>Desulfovibrionales>Desulfovibrionaceae>Desulfovibrio | GCA_000195755.1 | Desulfovibrio_vulgaris_str._Hildenborough | DVU_2025 | 2106300..2109851 | 882 | conserved-hypothetical-protein-[Desulfovibrio-vulgaris-str.-Hildenborough] |
| ATD82493.1 | Type1.BREX | zf-ISL3+HTH_Tnp_ISL3+DDE_Tnp_ISL3->?->?->?->BrxB-BrxF-iSTAND->BrxA-DNA-binding->BrxB-BrxF-iSTAND->DUF6079*->?->N6-METHYLASE->PglZ->SIGMA+OB-Fold+AAA-ATPase+HKD-DNase-BrxL->SbcC+SbcC-> [illegible] | 1183 | Bacteria>Thermodesulfobacteriota>Desulfovibrionia>Desulfovibrionales>Desulfovibrionaceae>Desulfovibrio | GCA_002334685.1 | Desulfovibrio_sp._G11 | CNY67_14735 | 3238216..3241767 | 631220 | hypothetical-protein-CNY67_14735-[Desulfovibrio-sp.-G11] |
| MBI4803787.1 | Type1.BREX | <-LuxR-HTH<-SIGMA+OB-Fold+AAA-ATPase+Lon-Protease-BrxL<-PglZ<-ParB+HNH<-PglZ<-HTH_21+rve<-PIN<-OLD-ABC<-DUF6079*<-BrxB-BrxF-iSTAND<-BrxA-DNA-binding<-XIS-HTH<-?\|\|TraJ-RHH->TraI-Vi [illegible] | 1145 | Bacteria>Thermodesulfobacteriota>Desulfovibrionia>Desulfovibrionales>Desulfovibrionaceae>Desulfovibrio | GCA_016208105.1 | Desulfovibrio_sp. | HY795_00975 | 45852..49289 | 885 | BREX-system-P-loop-protein-BrxC-[Desulfovibrio-sp.] |
| HJA77858.1 | Type1.BREX | BrxB-BrxF-iSTAND-p-><-?\|\|ACET->BrxA-DNA-binding->BrxB-BrxF-iSTAND->DUF6079*->OLD-ABC->PIN->N6-METHYLASE->PHP+SbcC->PglZ->SIGMA+OB-Fold+AAA-ATPase+Lon-Protease-BrxL->SIR2-> | 1145 | Bacteria>Thermodesulfobacteriota>Desulfovibrionia>Desulfovibrionales>Desulfovibrionaceae>Desulfovibrio | GCA_019118745.1 | Candidatus_Desulfovibrio_gallistercoris | H9774_13535 | 7700..11137 | 2838533 | BREX-system-P-loop-protein-BrxC-[Candidatus-Desulfovibrio-gallistercoris] |
| MCI7405014.1 | Type1.BREX | <-35exo<-?\|\|?->?->?-><-SIGMA+OB-Fold+AAA-ATPase+Lon-Protease-BrxL<-PglZ<-TOPRIM<-OLD-ABC<-N6-METHYLASE<-DUF6079*<-BrxB-BrxF-iSTAND<-BrxA-DNA-binding | 1199 | Bacteria>Thermodesulfobacteriota>Desulfovibrionia>Desulfovibrionales>Desulfovibrionaceae>Desulfovibrio | GCA_022783735.1 | Desulfovibrio_piger | MSS64_01040 | 35161..38760 | 901 | BREX-system-P-loop-protein-BrxC-[Desulfovibrio-piger] |
| MCQ4861473.1 | Type1.BREX | PAS+PAS+GGDEF-><-?<-?\|\|PhdYeFM_antitox->BrxA-DNA-binding->BrxB-BrxF-iSTAND->DUF6079*->?->N6-METHYLASE->PglZ->SIGMA+OB-Fold+AAA-ATPase+HKD-DNase-BrxL->SbcC+SbcC->Calcineurin+Calcineuri [illegible] | 1179 | Bacteria>Thermodesulfobacteriota>Desulfovibrionia>Desulfovibrionales>Desulfovibrionaceae>Desulfovibrio | GCA_024460775.1 | Desulfovibrio_desulfuricans | NE637_09955 | 25210..28749 | 876 | BREX-system-P-loop-protein-BrxC-[Desulfovibrio-desulfuricans] |
| SBW06947.1 | Type1.BREX | Arm-DNA-bind_3+Phage_int_SAM_3+Phage_integrase-><-?<-?<-?\|\|HIN-HTH->HTH_Tnp_1+HTH_21+rve+rve_3-><-N6-METHYLASE-p<-?<-DUF6079*<-BrxB-BrxF-iSTAND<-BrxA-DNA-binding<-PhdYeFM_antitox\|\|?-> [illegible] | 1179 | Bacteria>Thermodesulfobacteriota>Desulfovibrionia>Desulfovibrionales>Desulfovibrionaceae>Desulfovibrio | GCA_900079715.1 | uncultured_Desulfovibrio_sp. | KM92DES2_12263 | 2357980..2361519 | 167968 | conserved-hypothetical-protein-[uncultured-Desulfovibrio-sp.] |
| WP_014524353.1 | Type1.BREX | <-GGDEF<-ZnR+HTH_Tnp_ISL3+DDE_Tnp_ISL3<-SIGMA+OB-Fold+AAA-ATPase+HKD-DNase-BrxL<-PglZ<-REase<-N6-METHYLASE<-REase<-OLD-ABC<-DUF6079*<-BrxB-BrxF-iSTAND<-BrxA-DNA-binding<-wHTH+WY [illegible] | 1182 | Bacteria>Thermodesulfobacteriota>Desulfovibrionia>Desulfovibrionales>Desulfovibrionaceae>Desulfovibrio | GCF_000195755.1 | Desulfovibrio_vulgaris | DVU_RS09565 | 2106300..2109848 | 881 | BREX-system-P-loop-protein-BrxC-[Desulfovibrio-vulgaris] |
| WP_043638458.1 | Type1.BREX | NTP_transferase-><-?\|\|GAF+GAF+HISKIN-><-?<-?\|\|ParB+N6_N4_METHYLASE->?->BrxA-DNA-binding->BrxB-BrxF-iSTAND->DUF6079*->OLD-ABC->REase->N6-METHYLASE->PglZ->SIGMA+OB-Fold+AAA-ATPase+HKD- [illegible] | 1182 | Bacteria>Thermodesulfobacteriota>Desulfovibrionia>Desulfovibrionales>Desulfovibrionaceae>Desulfovibrio | GCF_000801335.2 | Desulfovibrio_sp._TomC | NY78_RS16925 | 95476..99024 | 1562888 | BREX-system-P-loop-protein-BrxC-[Desulfovibrio-sp.-TomC] |
| WP_256267712.1 | Type1.BREX | PAS+PAS+GGDEF-><-?<-?\|\|PhdYeFM_antitox->BrxA-DNA-binding->BrxB-BrxF-iSTAND->DUF6079*->?->N6-METHYLASE->PglZ->SIGMA+OB-Fold+AAA-ATPase+HKD-DNase-BrxL->SbcC+SbcC->Calcineurin+Calcineuri [illegible] | 1179 | Bacteria>Thermodesulfobacteriota>Desulfovibrionia>Desulfovibrionales>Desulfovibrionaceae>Desulfovibrio | GCF_900079715.1 | Desulfovibrio_desulfuricans | NE637_09955 | 25210..28749 | 876 | BREX-system-P-loop-protein-BrxC-[Desulfovibrio-desulfuricans] |
| WP_187008409.1 | Type1.BREX | <-HIN-HTH\|\|zf-ISL3+HTH_Tnp_ISL3+DDE_Tnp_ISL3->?->?->?->BrxB-BrxF-iSTAND->DUF6079-p*->XIS-HTH->BrxA-DNA-binding->BrxB-BrxF-iSTAND->DUF6079->?->N6-METHYLASE->PglZ->SIGMA+OB-Fold+AAA-ATP [illegible] | 191 | Bacteria>Thermodesulfobacteriota>Desulfovibrionia>Desulfovibrionales>Desulfovibrionaceae>Desulfovibrio | GCF_900243745.1 | Desulfovibrio_sp._G11 | DSVG11_RS05805 | 1310698..1311273 | 631220 | DUF6079-family-protein-[Desulfovibrio-sp.-G11] |
| WP_072312019.1 | Type1.BREX | <-HIN-HTH\|\|zf-ISL3+HTH_Tnp_ISL3+DDE_Tnp_ISL3->?->?->?->BrxB-BrxF-iSTAND->XIS-HTH->BrxA-DNA-binding->BrxB-BrxF-iSTAND->DUF6079*->?->N6-METHYLASE->PglZ->SIGMA+OB-Fold+AAA-ATPase+HKD-DNa [illegible] | 1182 | Bacteria>Thermodesulfobacteriota>Desulfovibrionia>Desulfovibrionales>Desulfovibrionaceae | GCF_002334685.1 | Desulfovibrio | CNY67_RS14735 | 3238219..3241767 | 872 | MULTISPECIES:-BREX-system-P-loop-protein-BrxC-[Desulfovibrio] |
| WP_160714268.1 | Type1.BREX | Phage_integrase->?->?->?-><-URI-DNase+RAMA\|\|<-SIGMA+OB-Fold+AAA-ATPase+Lon-Protease-BrxL<-PglZ<-N6-METHYLASE<-DUF6079*<-BrxB-BrxF-iSTAND<-BrxA-DNA-binding<-XIS-HTH | 1153 | Bacteria>Thermodesulfobacteriota>Desulfovibrionia>Desulfovibrionales>Desulfovibrionaceae>Halodesulfovibrio | GCF_009914765.1 | Halodesulfovibrio_sp._MK-HDV | MKHDV_RS08455 | 165116..168577 | 2599925 | BREX-system-P-loop-protein-BrxC-[Halodesulfovibrio-sp.-MK-HDV] |
| MBG0773462.1 | Type1.BREX | <-SIGMA+OB-Fold+AAA-ATPase+Lon-Protease-BrxL<-PglZ<-HEPN<-N6-METHYLASE<-DUF6079*<-BrxB-BrxF-iSTAND<-BrxA-DNA-binding<-Phage_integrase<-tRNA\|\|?->SIGMA-HTH->ncRNA->?->?->?->REC+PP2C-> | 1190 | Bacteria>Thermodesulfobacteriota>Desulfovibrionia>Desulfovibrionales>Desulfovibrionaceae>Oleidesulfovibrio | GCA_015709285.1 | Oleidesulfovibrio_alaskensis | HOS83_07675 | 9061..12633 | 58180 | BREX-system-P-loop-protein-BrxC-[Oleidesulfovibrio-alaskensis] |
| MBL3581670.1 | Type1.BREX | <-KAP-glycosylase<-KAP-NTPase<-HARE-HTH+REase<-SIGMA+OB-Fold+AAA-ATPase+Lon-Protease-BrxL<-PglZ<-ParB+HNH<-PHP+SbcC<-N6-METHYLASE<-DUF6079*<-BrxB-BrxF-iSTAND<-BrxA-DNA-binding<-XIS-HT [illegible] | 1145 | Bacteria>Thermodesulfobacteriota>Desulfovibrionia>Desulfovibrionales>Desulfovibrionaceae>Oleidesulfovibrio | GCA_016756815.1 | Oleidesulfovibrio_alaskensis | JMJ96_04565 | 108279..111716 | 58180 | BREX-system-P-loop-protein-BrxC-[Oleidesulfovibrio-alaskensis] |
| WP_273238273.1 | Type1.BREX | <-SIGMA+OB-Fold+AAA-ATPase+Lon-Protease-BrxL<-PglZ<-HEPN<-N6-METHYLASE<-DUF6079*<-BrxB-BrxF-iSTAND<-BrxA-DNA-binding<-Phage_integrase<-tRNA\|\|?->SIGMA-HTH->ncRNA->?->?->?->REC+PP2C-> | 1191 | Bacteria>Thermodesulfobacteriota>Desulfovibrionia>Desulfovibrionales>Desulfovibrionaceae>Oleidesulfovibrio | GCF_015709285.1 | Oleidesulfovibrio_alaskensis | HOS83_RS07680 | 9061..12636 | 58180 | BREX-system-P-loop-protein-BrxC-[Oleidesulfovibrio-alaskensis] |
| WP_273192722.1 | Type1.BREX | <-KAP-glycosylase<-KAP-NTPase<-HARE-HTH+REase<-SIGMA+OB-Fold+AAA-ATPase+Lon-Protease-BrxL<-PglZ<-ParB+HNH<-PHP+SbcC<-N6-METHYLASE<-DUF6079*<-BrxB-BrxF-iSTAND<-BrxA-DNA-binding<-XIS-HT [illegible] | 1146 | Bacteria>Thermodesulfobacteriota>Desulfovibrionia>Desulfovibrionales>Desulfovibrionaceae>Oleidesulfovibrio | GCF_016756815.1 | Oleidesulfovibrio_alaskensis | JMJ96_RS04575 | 108279..111719 | 58180 | BREX-system-P-loop-protein-BrxC-[Oleidesulfovibrio-alaskensis] |
| SOB57200.1 | Type1.BREX | <-SIGMA+OB-Fold+AAA-ATPase+Lon-Protease-BrxL<-PglZ<-HEPN<-N6-METHYLASE-p\|\|HTH_Tnp_1->HTH_Tnp_1+HTH_21+rve+rve_3-><-N6-METHYLASE-p<-N6-METHYLASE-p<-DUF6079*<-BrxB-BrxF-iSTAND<-BrxA-DNA [illegible] | 1190 | Bacteria>Thermodesulfobacteriota>Desulfovibrionia>Desulfovibrionales>Desulfovibrionaceae>Pseudodesulfovibrio | GCA_900217235.1 | Pseudodesulfovibrio_profundus | DPRO_0321 | 331811..335383 | 57320 | conserved-protein-of-unknown-function-[Pseudodesulfovibrio-profundus] |
| WP_232005669.1 | Type1.BREX | <-SIGMA+OB-Fold+AAA-ATPase+Lon-Protease-BrxL<-PglZ<-HEPN<-N6-METHYLASE-p\|\|HTH_Tnp_1+HTH_21+rve+rve_3-><-N6-METHYLASE-p<-DUF6079*<-BrxB-BrxF-iSTAND<-BrxA-DNA-binding<-P [illegible] | 1191 | Bacteria>Thermodesulfobacteriota>Desulfovibrionia>Desulfovibrionales>Desulfovibrionaceae>Pseudodesulfovibrio | GCF_900217235.1 | Pseudodesulfovibrio_profundus | DPRO_RS01600 | 331811..335386 | 57320 | BREX-system-P-loop-protein-BrxC-[Pseudodesulfovibrio-profundus] |
| WP_015862248.1 | Type1.BREX | <-Phage_integrase<-?<-?\|\|ParB+N6_N4_METHYLASE->PHP+SbcC-><-SIGMA+OB-Fold+AAA-ATPase+Lon-Protease-BrxL<-PglZ<-HEPN<-N6-METHYLASE<-DUF6079*<-BrxB-BrxF-iSTAND<-BrxA-DNA-binding<-HKD-DNas [illegible] | 1188 | Bacteria>Thermodesulfobacteriota>Desulfovibrionia>Desulfovibrionales>Desulfovibrionaceae>Solidesulfovibrio | GCF_000010665.1 | Solidesulfovibrio_magneticus | DMR_RS17110 | 4099313..4102879 | 184917 | BREX-system-P-loop-protein-BrxC-[Solidesulfovibrio-magneticus] |
| MCH5276204.1 | Type1.BREX | <-SIGMA+OB-Fold+AAA-ATPase+Lon-Protease-BrxL<-PglZ<-?<-N6-METHYLASE-p<-HTH+REase<-N6-METHYLASE<-REase<-OLD-ABC<-DUF6079*<-BrxB-BrxF-iSTAND<-BrxA-DNA-binding<-Arm-DNA-bind_3+Phage_inte [illegible] | 1145 | Bacteria>Thermodesulfobacteriota>Desulfovibrionia>Desulfovibrionales>Desulfovibrionaceae>unclassified_Desulfovibrionaceae | GCA_022510785.1 | Desulfovibrionaceae_bacterium | J1E80_00045 | 14777..18214 | 2049043 | BREX-system-P-loop-protein-BrxC-[Desulfovibrionaceae-bacterium] |
| NLW81341.1 | Type1.BREX | <-TetR-HTH\|\|BrxA-DNA-binding->BrxB-BrxF-iSTAND->DUF6079*->DOC+HTH->N6-METHYLASE-> | 1184 | Bacteria>Thermodesulfobacteriota>Desulfovibrionia>Desulfovibrionales>unclassified_Desulfovibrionales | GCA_012729655.1 | Desulfovibrionales_bacterium | GXY42_06675 | 10723..14277 | 2448050 | BREX-system-P-loop-protein-BrxC-[Desulfovibrionales-bacterium] |
| PLX83270.1 | Type1.BREX | <-STYKinase-like\|\|Thermonuclease-><-SIGMA+OB-Fold+AAA-ATPase+HKD-DNase-BrxL<-ZnR<-PglZ<-TOPRIM<-OLD-ABC<-N6-METHYLASE<-DUF6079*<-BrxB-BrxF-iSTAND<-BrxA-DNA-binding\|\|?->?->?-><-?\|\|?->G [illegible] | 1175 | Bacteria>Thermodesulfobacteriota>Desulfuromonadia>Desulfuromonadales>Desulfuromonadaceae>Desulfuromonas | GCA_002868845.1 | Desulfuromonas_sp. | C0617_12115 | 157475..161002 | 892 | MAG:-BREX-system-P-loop-protein-BrxC-[Desulfuromonas-sp.] |
| WP_011736152.1 | Type1.BREX | <-SIGMA+OB-Fold+AAA-ATPase+Lon-Protease-BrxL<-PglZ<-DOC+HTH<-Tox-RelA<-ParA-Soj-PloopNTPase<-N6-METHYLASE<-DUF6079*<-BrxB-BrxF-iSTAND<-BrxA-DNA-binding<-RHH_5 | 1192 | Bacteria>Thermodesulfobacteriota>Desulfuromonadia>Desulfuromonadales>Desulfuromonadaceae>Pelobacter | GCF_000015045.1 | Pelobacter_propionicus | PPRO_RS11345 | 2470743..2474321 | 29543 | BREX-system-P-loop-protein-BrxC-[Pelobacter-propionicus] |
| MCF6266282.1 | Type1.BREX | <-ZnR+HTH_Tnp_ISL3+DDE_Tnp_ISL3\|\|?-><-NACHT\|\|ZnR+TOPRIM+AAA-helicase+wHTH->?->CHTH->BrxA-DNA-binding->BrxB-BrxF-iSTAND->DUF6079*->SF1-SF2-Helicase+PRTase->SLOG+BirA-HTH->N6-METHYLASE- [illegible] | 1145 | Bacteria>Thermodesulfobacteriota>Desulfuromonadia>Desulfuromonadales>Geopsychrobacteraceae>Desulfuromusa | GCA_021647765.1 | Desulfuromusa_sp. | L3J57_07025 | 24640..28077 | 2614228 | BREX-system-P-loop-protein-BrxC-[Desulfuromusa-sp.] |
| MBE0574417.1 | Type1.BREX | <-VirD4-TraG+VirD4-FtsK<-?<-?\|\|XIS-HTH->BrxA-DNA-binding->BrxB-BrxF-iSTAND->DUF6079*->OLD-ABC->REase->N6-METHYLASE->REase->PglZ->SIGMA+OB-Fold+AAA-ATPase+HKD-DNase-BrxL->SbcC+APATPase [illegible] | 1183 | Bacteria>Thermodesulfobacteriota>Desulfuromonadia>Desulfuromonadales>unclassified_Desulfuromonadales | GCA_014860225.1 | Desulfuromonadales_bacterium | IH613_00815 | 71489..75040 | 2099678 | BREX-system-P-loop-protein-BrxC-[Desulfuromonadales-bacterium] |
| MBW6509585.1 | Type1.BREX | Amidohydro_1->?->?-><-Four-helical-protein\|\|XIS-HTH->BrxA-DNA-binding->BrxB-BrxF-iSTAND->DUF6079*->Schlafen+HATPase+wHTH+HTH->OLD-ABC->REase->N6-METHYLASE->REase-> | 1178 | Bacteria>Thermodesulfobacteriota>Desulfuromonadia>Desulfuromonadales>unclassified_Desulfuromonadales | GCA_019429255.1 | Desulfuromonadales_bacterium | K0A94_08605 | 43514..47050 | 2099678 | BREX-system-P-loop-protein-BrxC-[Desulfuromonadales-bacterium] |
| AJY71182.1 | Type1.BREX | NTP_transf_8->Calcineurin+Calcineurin-SbcD->SbcC+CCDC158->PIN-><-DUF4372+DDE_Tnp_1\|\|HTH_Tnp_1->HTH_Tnp_1+HTH_21+rve+rve_3->BrxA-DNA-binding->BrxB-BrxF-iSTAND->DUF6079*->?->N6-METHYLAS [illegible] | 1183 | Bacteria>Thermodesulfobacteriota>Desulfuromonadia>Geobacterales>Geobacteraceae>Geobacter | GCA_000961035.1 | Geobacter_sulfurreducens | RW64_17280 | 3775110..3778661 | 35554 | hypothetical-protein-RW64_17280-[Geobacter-sulfurreducens] |
| MBI1919696.1 | Type1.BREX | <-REase+SF1-SF2-Helicase<-MORC+DUF3684+REase<-SbcC+CCDC158<-Calcineurin-SbcD<-SLATT<-SMODS<-SIGMA+OB-Fold+AAA-ATPase+Lon-Protease-BrxL<-PglZ<-N6-METHYLASE<-DUF6079*<-BrxB-BrxF-iSTAND< [illegible] | 1150 | Bacteria>Thermodesulfobacteriota>Desulfuromonadia>Geobacterales>Geobacteraceae>Geobacter | GCA_016177775.1 | Geobacter_sp. | HYS23_01305 | 177481..180933 | 46610 | BREX-system-P-loop-protein-BrxC-[Geobacter-sp.] |
| WP_010942753.1 | Type1.BREX | <-Ferredoxin-RRM+Radical_SAM<-?<-SIGMA+OB-Fold+AAA-ATPase+Lon-Protease-BrxL<-PglZ<-N6-METHYLASE<-HEPN<-DUF6079*<-BrxB-BrxF-iSTAND<-BrxA-DNA-binding<-?<-?\|\|CHTH-><-?<-?\|\|Phage_integras [illegible] | 1214 | Bacteria>Thermodesulfobacteriota>Desulfuromonadia>Geobacterales>Geobacteraceae>Geobacter | GCF_000007985.2 | Geobacter_sulfurreducens | GS_RS10585 | 2325892..2329536 | 35554 | BREX-system-P-loop-protein-BrxC-[Geobacter-sulfurreducens] |
| WP_207162791.1 | Type1.BREX | <-Peptidase_C14<-TIR<-SIGMA+OB-Fold+AAA-ATPase+HKD-DNase-BrxL<-PglZ<-TOPRIM<-OLD-ABC<-N6-METHYLASE<-REase<-OLD-ABC<-DUF6079*<-BrxB-BrxF-iSTAND<-BrxA-DNA-binding<-AbiEi_4+AbiEi_1<-ParB [illegible] | 1175 | Bacteria>Thermodesulfobacteriota>Desulfuromonadia>Geobacterales>Geobacteraceae>Geobacter | GCF_017338855.1 | Geobacter_benzoatilyticus | JZM60_RS12610 | 2741421..2744948 | 2815309 | BREX-system-P-loop-protein-BrxC-[Geobacter-benzoatilyticus] |
| MBJ6802709.1 | Type1.BREX | BrxA-DNA-binding->BrxB-BrxF-iSTAND->DUF6079*->OLD-ABC->REase->N6-METHYLASE->PglZ->SIGMA+OB-Fold+AAA-ATPase+HKD-DNase-BrxL->?-><-?<-NERD+ZnR<-wHTH+SAD+HNH | 1183 | Bacteria>Thermodesulfobacteriota>Desulfuromonadia>Geobacterales>Geobacteraceae>Geomonas | GCA_016458235.1 | Geomonas_propionica | JFN90_21475 | 2703..6254 | 2798582 | BREX-system-P-loop-protein-BrxC-[Geomonas-propionica] |
| WP_236019113.1 | Type1.BREX | BrxA-DNA-binding->BrxB-BrxF-iSTAND->DUF6079*->OLD-ABC->REase->N6-METHYLASE->PglZ->SIGMA+OB-Fold+AAA-ATPase+HKD-DNase-BrxL->?-><-?<-NERD+ZnR<-wHTH+SAD+HNH | 1191 | Bacteria>Thermodesulfobacteriota>Desulfuromonadia>Geobacterales>Geobacteraceae>Geomonas | GCF_016458235.1 | Geomonas_propionica | JFN90_RS21475 | 2679..6254 | 2798582 | BREX-system-P-loop-protein-BrxC-[Geomonas-propionica] |
| OPY81706.1 | Type1.BREX | HIN-HTH->AAA-ATPase-><-DnaB+TnsC-ATPase<-?<-?<-XIS-HTH<-?\|\|BrxA-DNA-binding->BrxB-BrxF-iSTAND->DUF6079*->Schlafen+HATPase+wHTH+HTH->N6-METHYLASE->OLD-ABC->TOPRIM->PglZ->SIGMA+OB-Fold+ [illegible] | 1200 | Bacteria>Thermodesulfobacteriota>Syntrophia>Syntrophales>Syntrophaceae>Syntrophus | GCA_002067745.1 | Syntrophus_sp._PtaU1.Bin005 | A4E70_01072 | 12725..16327 | 1811712 | MAG:-hypothetical-protein-A4E70_01072-[Syntrophus-sp.-PtaU1.Bin005] |
| OPY81323.1 | Type1.BREX | <-ART-DarT<-SF1-SF2-Helicase<-HARE-HTH<-SIGMA+OB-Fold+AAA-ATPase+Lon-Protease-BrxL<-PglZ<-ParB+HNH<-HTH+REase<-N6-METHYLASE<-HATPase+wHTH<-DUF6079*<-BrxB-BrxF-iSTAND<-BrxA-DNA-binding [illegible] | 1144 | Bacteria>Thermodesulfobacteriota>Syntrophia>Syntrophales>Syntrophaceae>Syntrophus | GCA_002067745.1 | Syntrophus_sp._PtaU1.Bin005 | A4E70_01341 | 26340..29774 | 1811712 | MAG:-hypothetical-protein-A4E70_01341-[Syntrophus-sp.-PtaU1.Bin005] |
| WP_011416818.1 | Type1.BREX | <-SIGMA+OB-Fold+AAA-ATPase+Lon-Protease-BrxL<-PglZ<-Schlafen+HATPase+wHTH<-SbcC<-DUF932<-N6-METHYLASE<-?<-TIR<-TIR<-DUF6079*<-BrxB-BrxF-iSTAND<-BrxA-DNA-binding-p\|\|?-><-DEDD_Tnp_IS110 [illegible] | 1165 | Bacteria>Thermodesulfobacteriota>Syntrophia>Syntrophales>Syntrophaceae>Syntrophus | GCF_000013405.1 | Syntrophus_aciditrophicus | SYN_RS04305 | 947033..950530 | 316277 | BREX-system-P-loop-protein-BrxC-[Syntrophus-aciditrophicus] |
| HCS77930.1 | Type1.BREX | <-TOPRIM+Topoisom_bac<-?\|\|?->tRNA->tRNA->tRNA->tRNA->BrxA-DNA-binding->BrxB-BrxF-iSTAND->DUF6079*-><-HicB<-HicA_toxin\|\|N6-METHYLASE->PglZ->SIGMA+OB-Fold+AAA-ATPase+Lon-Protease-BrxL-> [illegible] | 1147 | Bacteria>Thermodesulfobacteriota>Syntrophia>Syntrophales>Syntrophaceae>unclassified_Syntrophaceae | GCA_003531355.1 | Syntrophaceae_bacterium | DIW05_09685 | 10653..14096 | 2052184 | BREX-system-P-loop-protein-BrxC-[Syntrophaceae-bacterium] |
| KAF0159518.1 | Type1.BREX | <-REC+NtrC-AAA+FIS-HTH<-PAS+SHELIX+PAS+PAS+PAS<-REC+GAF+HDGYP<-?\|\|?-><-RNase<-?<-SIGMA+OB-Fold+AAA-ATPase-BrxL<-PglZ<-DUF6079*<-AbiEii+ZnR<-AbiEi_4+AbiEi_1<-PIN<-HTH+MPTase<-?<-N6-MET [illegible] | 1347 | Bacteria>Thermodesulfobacteriota>Syntrophia>Syntrophales>Syntrophaceae>unclassified_Syntrophaceae | GCA_009773315.1 | Syntrophaceae_bacterium | FD159_459 | 44035..48078 | 2052184 | Uncharacterized-protein-FD159_459-[Syntrophaceae-bacterium] |
| MBP7766239.1 | Type1.BREX | <-TOPRIM<-XIS-HTH<-?<-DUF932\|\|MetJarc-HTH->?->BrxA-DNA-binding->BrxB-BrxF-iSTAND->DUF6079*->N6-METHYLASE->AbiEi_4+AbiEi_1->AbiEii+ZnR->PglZ->SIGMA+OB-Fold+AAA-ATPase+Lon-Protease-BrxL [illegible] | 1184 | Bacteria>Thermodesulfobacteriota>Syntrophia>Syntrophales>Syntrophaceae>unclassified_Syntrophaceae | GCA_018000275.1 | Syntrophaceae_bacterium | KA113_13715 | 59522..63076 | 2052184 | BREX-system-P-loop-protein-BrxC-[Syntrophaceae-bacterium] |
| MCG2741590.1 | Type1.BREX | REase+RNase->REase+N-glycosylase->BrxA-DNA-binding->BrxB-BrxF-iSTAND->DUF6079*-> | 1115 | Bacteria>Thermodesulfobacteriota>Syntrophia>Syntrophales>Syntrophaceae>unclassified_Syntrophaceae | GCA_022013455.1 | Syntrophaceae_bacterium | L6300_15350 | 5284..8629 | 2052184 | BREX-system-P-loop-protein-BrxC-[Syntrophaceae-bacterium] |
| WP_258930562.1 | Type1.BREX | <-PRTase<-SIGMA+OB-Fold+AAA-ATPase+Lon-Protease-BrxL<-PglZ<-Tox-RelE<-?<-DUF6079-p*<-BrxB-BrxF-iSTAND<-BrxA-DNA-binding\|\|?-><-SPOUT<-SAM-METHYLASE | 198 | Bacteria>Thermodesulfobacteriota>Syntrophobacteria>Syntrophobacterales>Syntrophobacteraceae>Desulfoferrobacter | GCF_022603395.1 | Desulfoferrobacter_suflitae | K1V27_RS13550 | 105134..105730 | 2865782 | hypothetical-protein-[Desulfoferrobacter-suflitae] |
| PIV05610.1 | Type1.BREX | HerA/FtsK->Schlafen+HATPase+wHTH-><-CR-ATPase5+REase\|\|ParB->PolB->?->?->OLD-ABC->TOPRIM->DUF6079*->PglZ->SIGMA+OB-Fold+AAA-ATPase-BrxL->?->?-><-?<-?<-DsbD | 1346 | Bacteria>Thermodesulfobacteriota>Syntrophobacteria>Syntrophobacterales>unclassified_Syntrophobacterales | GCA_002779905.1 | Syntrophobacterales_bacterium_CG03_land_8_20_14_0_80_58_14 | COS57_07000 | 12627..16667 | 1974100 | hypothetical-protein-COS57_07000-[Syntrophobacterales-bacterium-CG03_land_8_20_14_0_80_58_14] |
| MBW6486409.1 | Type1.BREX | <-SIR2<-?<-SSB<-RADICAL-SAM\|\|CR-ATPase5+REase->Beta-lactamase->BrxA-DNA-binding->BrxB-BrxF-iSTAND->DUF6079*->?->N6-METHYLASE-><-?<-?\|\|DOC+HTH->Calcineurin-SbcD->SbcC+CCDC158->DUF932-> [illegible] | 1049 | Bacteria>Thermodesulfobacteriota>Syntrophobacteria>Syntrophobacterales>unclassified_Syntrophobacterales | GCA_019429105.1 | Syntrophobacterales_bacterium | K0B01_09695 | 13495..16644 | 2282155 | BREX-system-P-loop-protein-BrxC-[Syntrophobacterales-bacterium] |
| CCU85336.1 | Type1.BREX | <-DnaB<-HTH_36<-?\|\|BrxA-DNA-binding->BrxB-BrxF-iSTAND->DUF6079*->N6-METHYLASE->PglZ->SIGMA+OB-Fold+AAA-ATPase+Lon-Protease-BrxL->?->REase+DUF4268->McrB-AAA->McrC-NTD+REase->REase->Abi [illegible] | 1166 | Bacteria>Thermotogota>Thermotogae>Kosmotogales>Kosmotogaceae>Mesotoga | GCA_000367705.1 | Mesotoga_infera | PHOSAC3_150336 | 356643..360143 | 1236046 | hypothetical-protein-PHOSAC3_150336-[Mesotoga-infera] |
| WP_103134081.1 | Type1.BREX | <-SIGMA+OB-Fold+AAA-ATPase+Lon-Protease-BrxL<-PglZ<-ParB+HNH<-N6-METHYLASE<-DUF6079*<-BrxB-BrxF-iSTAND<-BrxA-DNA-binding\|\|?->HTH_36->HTH_36->AAA+-> | 1167 | Bacteria>Thermotogota>Thermotogae>Kosmotogales>Kosmotogaceae>Mesotoga | GCF_002896535.1 | Mesotoga_sp._B105.6.4 | RJ60_RS04180 | 254841..258344 | 1582224 | BREX-system-P-loop-protein-BrxC-[Mesotoga-sp.-B105.6.4] |
| WP_110990429.1 | Type1.BREX | <-CASPASE<-?<-SIGMA+OB-Fold+AAA-ATPase+Lon-Protease-BrxL<-PglZ<-ParB+HNH<-N6-METHYLASE<-DUF6079*<-BrxB-BrxF-iSTAND<-BrxA-DNA-binding<-PAS+NtrC-AAA+FIS-HTH<-METHYLASE<-tRNA\|\|?->nSTAND1 [illegible] | 1163 | Bacteria>Thermotogota>Thermotogae>Kosmotogales>Kosmotogaceae>Mesotoga | GCF_003228205.1 | Mesotoga_sp._TolDC | LH53_RS05220 | 29090..32581 | 1389250 | BREX-system-P-loop-protein-BrxC-[Mesotoga-sp.-TolDC] |
| WP_121509576.1 | Type1.BREX | <-DnaB<-HTH_36<-?\|\|BrxA-DNA-binding->BrxB-BrxF-iSTAND->DUF6079*->N6-METHYLASE->PglZ->SIGMA+OB-Fold+AAA-ATPase+Lon-Protease-BrxL->HNH->?->ParB+HNH->REase->AbiEi-> | 1162 | Bacteria>Thermotogota>Thermotogae>Kosmotogales>Kosmotogaceae>Mesotoga | GCF_003664935.1 | Mesotoga_sp._H07pep.5.4 | Y696_RS00810 | 21677..25165 | 1463664 | BREX-system-P-loop-protein-BrxC-[Mesotoga-sp.-H07pep.5.4] |
| WP_121550387.1 | Type1.BREX | <-AbiEii<-AbiEi<-REase<-McrC-NTD+REase<-REase+DUF4268<-?<-SIGMA+OB-Fold+AAA-ATPase+Lon-Protease-BrxL<-PglZ<-N6-METHYLASE<-DUF6079*<-BrxB-BrxF-iSTAND<-BrxA-DNA-binding\|\|?->HTH_36->DnaB [illegible] | 1166 | Bacteria>Thermotogota>Thermotogae>Kosmotogales>Kosmotogaceae>Mesotoga | GCF_003664995.1 | Mesotoga_sp._BH458_6_3_2_1 | Y697_RS03850 | 31631..35131 | 1437446 | BREX-system-P-loop-protein-BrxC-[Mesotoga-sp.-BH458_6_3_2_1] |
| WP_169698881.1 | Type1.BREX | Phage_integrase-><-McrC-NTD+REase<-McrB-AAA<-REase+DUF4268<-SIGMA+OB-Fold+AAA-ATPase+Lon-Protease-BrxL<-PglZ<-N6-METHYLASE<-DUF6079*<-BrxB-BrxF-iSTAND<-BrxA-DNA-binding\|\|?->RVT-><-HAD [illegible] | 1164 | Bacteria>Thermotogota>Thermotogae>Kosmotogales>Kosmotogaceae>Mesotoga | GCF_900157305.1 | Mesotoga_infera | MESINF_RS05385 | 1186645..1190139 | 1236046 | BREX-system-P-loop-protein-BrxC-[Mesotoga-infera] |
| WP_091403827.1 | Type1.BREX | HIN-HTH->rve-><-?<-XIS-HTH<-DOC+zhth\|\|BrxA-DNA-binding->BrxB-BrxF-iSTAND->DUF6079*->N6-METHYLASE->PglZ->SIGMA+OB-Fold+AAA-ATPase+Lon-Protease-BrxL-><-?<-?<-PolB<-McrC-NTD+REase<-EVE+M [illegible] | 1185 | Bacteria>Thermotogota>Thermotogae>Petrotogales>Petrotogaceae>Geotoga | GCF_900102615.1 | Geotoga_petraea | BLS00_RS06420 | 5189..8746 | 28234 | BREX-system-P-loop-protein-BrxC-[Geotoga-petraea] |
| WP_165973936.1 | Type1.BREX | <-RADICAL-SAM<-?\|\|?->?->BrxA-DNA-binding->BrxB-BrxF-iSTAND->DUF6079-p*->DUF6079-p->?->PglZ->SIGMA+OB-Fold+AAA-ATPase+Lon-Protease-BrxL->REase->HTH+REase->?->DDE_Tnp_1->SF1-SF2-Helicas [illegible] | 535 | Bacteria>Thermotogota>Thermotogae>Petrotogales>Petrotogaceae>Marinitoga | GCF_004135675.1 | Marinitoga_lauensis | DJ007_RS01225 | 233414..235021 | 2201189 | BREX-system-P-loop-protein-BrxC-[Marinitoga-lauensis] |
| WP_254867722.1 | Type1.BREX | <-HIN-HTH\|\|BrxA-DNA-binding->BrxB-BrxF-iSTAND->DUF6079*->N6-METHYLASE->PglZ->SIGMA+OB-Fold+AAA-ATPase+Lon-Protease-BrxL->HNH->CR-ATPase-><-?<-?\|\|ABhydrolase-> | 1163 | Bacteria>Thermotogota>Thermotogae>Petrotogales>Petrotogaceae>Marinitoga | GCF_013363285.1 | Marinitoga_sp._1154 | XO12_RS09410 | 1913..5404 | 1643335 | BREX-system-P-loop-protein-BrxC-[Marinitoga-sp.-1154] |
| NLI08170.1 | Type1.BREX | DUF6079-p*->N6-METHYLASE->?->PglZ->SIGMA+OB-Fold+AAA-ATPase+Lon-Protease-BrxL-p-> | 421 | Bacteria>Thermotogota>Thermotogae>Thermotogales>Thermotogaceae>unclassified_Thermotogaceae | GCA_012517055.1 | Thermotogaceae_bacterium | GX457_13930 | 1..1268 | 2268206 | BREX-system-P-loop-protein-BrxC-[Thermotogaceae-bacterium] |
| KUK92978.1 | Type1.BREX | <-N6-METHYLASE-p<-DUF6079*<-BrxB-BrxF-iSTAND<-BrxA-DNA-binding\|\|REase+iREase-> | 1163 | Bacteria>Thermotogota>Thermotogae>Thermotogales>unclassified_Thermotogales | GCA_001509135.1 | Thermotogales_bacterium_46_20 | XE05_1214 | 393..3884 | 1635293 | Uncharacterized-protein-XE05_1214-[Thermotogales-bacterium-46_20] |
| HCZ06893.1 | Type1.BREX | <-N6-METHYLASE-p<-DUF6079*<-BrxB-BrxF-iSTAND<-BrxA-DNA-binding<-RAMPs<-?<-RAMPs<-?<-RAMPs<-?<-CARF+HEPN | 1168 | Bacteria>Thermotogota>unclassified_Thermotogota | GCA_003538775.1 | Thermotogota_bacterium | DHV12_07180 | 952..4458 | 2053689 | BREX-system-P-loop-protein-BrxC-[Thermotogota-bacterium] |
| KUO61378.1 | Type1.BREX | <-Schlafen+HATPase+wHTH<-SIGMA+OB-Fold+AAA-ATPase+HKD-DNase-BrxL<-PglZ<-HTH+REase<-?<-N6-METHYLASE<-DUF6079*<-BrxB-BrxF-iSTAND<-BrxA-DNA-binding<-HTH_Tnp_1+HTH_21+rve+rve_3 | 1157 | Bacteria>unclassified_Bacteria | GCA_001516025.1 | bacterium_BRH_c32 | APF79_11995 | 11772..15245 | 1734400 | hypothetical-protein-APF79_11995-[bacterium-BRH_c32] |
| OQA19974.1 | Type1.BREX | BrxA-DNA-binding->BrxB-BrxF-iSTAND->DUF6079*->?->RelE-ParE->VapB_antitoxin->PIN->N6-METHYLASE->REase->PglZ->SIGMA+OB-Fold+AAA-ATPase+Lon-Protease-BrxL-> | 1164 | Bacteria>unclassified_Bacteria | GCA_002069435.1 | bacterium_ADurb.Bin363 | BWY64_00492 | 2230..5724 | 1866927 | hypothetical-protein-BWY64_00492-[bacterium-ADurb.Bin363] |
| GBE48762.1 | Type1.BREX | METHYLASE->REC+OmpR-HTH->REC+OmpR-HTH->?->URI-DNase+UVR+RNase+HHH->?->tRNA->tRNA->tRNA->?->BrxB-BrxF-iSTAND->DUF6079*->PIN->N6-METHYLASE->PHP+SbcC->Schlafen+HATPase+wHTH->PglZ->SIGMA+ [illegible] | 1155 | Bacteria>unclassified_Bacteria | GCA_002897795.1 | bacterium_BMS3Bbin12 | BMS3Bbin12_01950 | 17892..21359 | 2005735 | hypothetical-protein-BMS3Bbin12_01950-[bacterium-BMS3Bbin12] |
| RKJ05437.1 | Type1.BREX | <-REC<-PglZ<-Schlafen+HATPase+wHTH<-N6-METHYLASE<-DUF6079*<-BrxB-BrxF-iSTAND<-BrxA-DNA-binding<-PDZ_6+Peptidase_S55<-SbcC+SbcC<-wHTH+WYL<-?<-?<-SAM-METHYLASE | 1221 | Bacteria>unclassified_Bacteria | GCA_003611905.1 | bacterium_D16-54 | D7X87_08505 | 80521..84186 | 2320104 | BREX-system-P-loop-protein-BrxC-[bacterium-D16-54] |
| RKJ48602.1 | Type1.BREX | HTH-><-PHP+SbcC<-SIGMA+OB-Fold+AAA-ATPase+Lon-Protease-BrxL<-PglZ<-Schlafen+HATPase+wHTH<-HEPN<-N6-METHYLASE<-TIR<-?<-DUF6079*<-BrxB-BrxF-iSTAND<-BrxA-DNA-binding<-HTH_26<-?<-TraI-Vir [illegible] | 1182 | Bacteria>unclassified_Bacteria | GCA_003612585.1 | bacterium_1XD42-54 | D7Y05_12400 | 29825..33373 | 2320110 | BREX-system-P-loop-protein-BrxC-[bacterium-1XD42-54] |
| RLJ22005.1 | Type1.BREX | <-N6-METHYLASE<-DUF6079*<-BrxB-BrxF-iSTAND<-BrxA-DNA-binding<-HTH_26<-?<-HTH<-?\|\|?-><-DSBH<-N6-METHYLASE-p | 1188 | Bacteria>unclassified_Bacteria | GCA_003660225.1 | bacterium_endosymbiont_of_Escarpia_laminata | DJ030_02690 | 2258..5824 | 1221601 | BREX-system-P-loop-protein-BrxC-[bacterium-endosymbiont-of-Escarpia-laminata] |
| TGU46786.1 | Type1.BREX | <-SIGMA+OB-Fold+AAA-ATPase+HKD-DNase-BrxL<-PglZ<-N6-METHYLASE<-DUF6079*<-BrxB-BrxF-iSTAND<-BrxA-DNA-binding<-wHTH+WYL+WCX\|\|35Exo+REase->SF1-SF2-Helicase+REase->Small-B-barrel+SF1-SF2- [illegible] | 1179 | Bacteria>unclassified_Bacteria | GCA_004791775.1 | bacterium_M00.F.Ca.ET.152.01.1.1 | EN795_30760 | 8346..11885 | 2563900 | BREX-system-P-loop-protein-BrxC-[bacterium-M00.F.Ca.ET.152.01.1.1] |
| TRZ93676.1 | Type1.BREX | <-REase<-HerA/FtsK<-SIR2<-REase<-SIGMA+OB-Fold+AAA-ATPase+Lon-Protease-BrxL<-PglZ<-N6-METHYLASE<-DUF6079*<-BrxB-BrxF-iSTAND<-BrxA-DNA-binding<-XIS-HTH<-?<-REase<-?<-REase+35exo | 1181 | Bacteria>unclassified_Bacteria | GCA_007280235.1 | bacterium | D4R78_07780 | 14482..18027 | 1869227 | BREX-system-P-loop-protein-BrxC-[bacterium] |
| NBJ82986.1 | Type1.BREX | RADICAL-SAM-><-?<-?\|\|REC+SHELIX+HISKIN->REase->?->HTH_26->BrxA-DNA-binding->BrxB-BrxF-iSTAND->DUF6079*->N6-METHYLASE->HTH+REase->PglZ->?->?->ZDNA-HTH->AAA+->?->PAS+NtrC-AAA+FIS-HTH-> | 1182 | Bacteria>unclassified_Bacteria | GCA_009917545.1 | bacterium_1XD42-76 | D5271_13855 | 76500..80048 | 2320881 | BREX-system-P-loop-protein-BrxC-[bacterium-1XD42-76] |
| NBK91895.1 | Type1.BREX | MoxR-AAA->TraB-><-HTH_3\|\|?->?->HTH_26->BrxA-DNA-binding->BrxB-BrxF-iSTAND->DUF6079*->N6-METHYLASE->Schlafen+HATPase+wHTH->PglZ->?->?->?->?->?->?->RNasePH+RNasePH+KH+S1-> | 1180 | Bacteria>unclassified_Bacteria | GCA_009917555.1 | bacterium_1XD21-13 | D5278_07865 | 171021..174563 | 2320888 | BREX-system-P-loop-protein-BrxC-[bacterium-1XD21-13] |
| NCP08807.1 | Type1.BREX | HTH_3-><-PIN<-wHTH<-?\|\|CR-ATPase5+REase->BrxA-DNA-binding->BrxB-BrxF-iSTAND->DUF6079*->Schlafen+HATPase+wHTH+HTH->?->?->HTH+REase->PglZ->?->SIGMA+OB-Fold+AAA-ATPase+HKD-DNase-BrxL-> | 1186 | Bacteria>unclassified_Bacteria | GCA_009993525.1 | bacterium | GW860_07900 | 18171..21731 | 1869227 | BREX-system-P-loop-protein-BrxC-[bacterium] |

| Protein-ID | System | Gene-Neighbourhoods | Length | Taxonomy | Assembly-ID | Strain | Locus-Tag | Genomic-coordinates | TaxID | Fasta-defline |
|---|---|---|---|---|---|---|---|---|---|---|
| NDC75310.1 | Type1.BREX | <-PglZ-p<-N6-METHYLASE<-DUF6079*<-BrxA-DNA-binding<-XIS-HTH | 1146 | Bacteria>unclassified_Bacteria | GCA_010027235.1 | bacterium | EBZ70_08510 | 4161..7601 | 1869227 | BREX-system-P-loop-protein-BrxC-[bacterium] |
| NLB11218.1 | Type1.BREX | <-DUF6079-p*<-BrxB-BrxF-iSTAND<-BrxA-DNA-binding<-wHTH+WYL+WCX\|\|HTH_3->SF1-SF2-Helicase-> | 978 | Bacteria>unclassified_Bacteria | GCA_012511355.1 | bacterium | GX831_02050 | 1..2935 | 1869227 | BREX-system-P-loop-protein-BrxC-[bacterium] |
| MBD5384945.1 | Type1.BREX | <-Acyltransferase<-?<-?\|\|HISKIN+MORC->REC+OmpR-HTH->BrxA-DNA-binding->BrxB-BrxF-iSTAND->DUF6079-p*->N6-METHYLASE->OLD-ABC->TOPRIM->PglZ->SIGMA+OB-Fold+AAA-ATPase+HKD-DNase-BrxL-> | 843 | Bacteria>unclassified_Bacteria | GCA_014802625.1 | bacterium | HDR69_00765 | 93850..96381 | 1869227 | BREX-system-P-loop-protein-BrxC-[bacterium] |
| MBI1293571.1 | Type1.BREX | HHH_5+SF1-SF2-Helicase->?->?-><-Glyco_transf_28+UDPGT<-TetR-HTH<-PhdYeFM_antitox\|\|BrxA-DNA-binding->BrxB-BrxF-iSTAND->DUF6079*->Y1_Tnp->N6-METHYLASE->REase->?->HTH_3->?->PglZ->SIGMA+OB-Fold+AAA-ATPase+HKD-DNase-BrxL-><-SF1-SF2-Helicase+REase | 1173 | Bacteria>unclassified_Bacteria | GCA_016124945.1 | bacterium | GC175_01265 | 40562..44083 | 1869227 | BREX-system-P-loop-protein-BrxC-[bacterium] |
| MBI1292832.1 | Type1.BREX | DDE_Tnp->?-><-D5N+viral-AAA+CDC6-HTH<-?<-?<-?\|\|BrxA-DNA-binding->BrxB-BrxF-iSTAND->DUF6079*->?->N6-METHYLASE->DOC+HTH->ParB+HNH->PglZ->SIGMA+OB-Fold+AAA-ATPase+Lon-Protease-BrxL-><-?\|\|?->Recombinase+ZnR-> | 1147 | Bacteria>unclassified_Bacteria | GCA_016125005.1 | bacterium | GC173_16580 | 8915..12358 | 1869227 | BREX-system-P-loop-protein-BrxC-[bacterium] |
| MBI9030472.1 | Type1.BREX | REase+DUF5655-><-SF1-SF2-Helicase+PhospholipaseD<-?<-SIGMA+OB-Fold+AAA-ATPase+HKD-DNase-BrxL<-PglZ<-Schlafen+HATPase+wHTH<-N6-METHYLASE<-DUF6079*<-BrxB-BrxF-iSTAND<-BrxA-DNA-binding | 1162 | Bacteria>unclassified_Bacteria | GCA_016342615.1 | bacterium | JEZ13_00510 | 21679..25167 | 1869227 | BREX-system-P-loop-protein-BrxC-[bacterium] |
| MBK6734349.1 | Type1.BREX | Peptidase_M48+TPR_9->?->?->HKD-DNase->?->?->?->BrxA-DNA-binding->DUF6079-p->DUF6079-p*->RVT->PglZ->PglZ-p->PglZ->?->SIGMA+OB-Fold+AAA-ATPase+Lon-Protease-BrxL-p-> | 924 | Bacteria>unclassified_Bacteria | GCA_016703265.1 | bacterium | IPG61_09705 | 532459..535233 | 1869227 | BREX-system-P-loop-protein-BrxC-[bacterium] |
| MBN2373590.1 | Type1.BREX | BrxA-DNA-binding->BrxB-BrxF-iSTAND->DUF6079*->N6-METHYLASE->Schlafen+HATPase+wHTH+HTH->Peptidase_M29+DUF2971-> | 1198 | Bacteria>unclassified_Bacteria | GCA_016934595.1 | bacterium | JXL19_07380 | 2083..5679 | 1869227 | BREX-system-P-loop-protein-BrxC-[bacterium] |
| MBP7862612.1 | Type1.BREX | HTH+MPTase->SLOG+BirA-HTH->?->Glycos_transf-><-?\|\|?-><-SIGMA+OB-Fold+AAA-ATPase+HKD-DNase-BrxL<-PglZ<-N6-METHYLASE<-DUF6079*<-BrxB-BrxF-iSTAND<-BrxA-DNA-binding<-?<-ABhydrolase\|\|?->?-><-TPR+TPR+TPR+TPR+TPR<-Trypsin+PDZ\|\|URI-DNase-> | 1181 | Bacteria>unclassified_Bacteria | GCA_018001945.1 | bacterium | KA183_13090 | 24829..28374 | 1869227 | BREX-system-P-loop-protein-BrxC-[bacterium] |
| MBQ0129310.1 | Type1.BREX | <-SIGMA+OB-Fold+AAA-ATPase+HKD-DNase-BrxL<-Schlafen+HATPase+wHTH<-PglZ<-?<-?<-N6-METHYLASE-p<-Schlafen+HATPase+wHTH<-SIR2<-N6-METHYLASE<-DUF6079*<-BrxB-BrxF-iSTAND<-BrxA-DNA-binding | 1169 | Bacteria>unclassified_Bacteria | GCA_018065885.1 | bacterium | KBT57_05675 | 15436..18945 | 1869227 | BREX-system-P-loop-protein-BrxC-[bacterium] |
| MBS5372264.1 | Type1.BREX | <-ABC-ATPase+SbcC<-ABC-ATPase+SbcC<-MarR-HTH\|\|HTH_26->BrxA-DNA-binding->BrxB-BrxF-iSTAND->DUF6079-p*-> | 454 | Bacteria>unclassified_Bacteria | GCA_018367885.1 | butyrate-producing_bacterium | KHY84_03545 | 45706..47067 | 2811711 | BREX-system-P-loop-protein-BrxC-[butyrate-producing-bacterium] |
| MBS5529915.1 | Type1.BREX | <-SIGMA+OB-Fold+AAA-ATPase+HKD-DNase-BrxL<-PglZ<-?<-N6-METHYLASE<-HD<-N6-METHYLASE-p<-DUF6079*<-BrxB-BrxF-iSTAND<-BrxA-DNA-binding<-XIS-HTH | 1169 | Bacteria>unclassified_Bacteria | GCA_018369055.1 | bacterium | KH017_03920 | 10993..14502 | 1869227 | BREX-system-P-loop-protein-BrxC-[bacterium] |
| MBS5642190.1 | Type1.BREX | <-ABC-ATPase+SbcC<-MarR-HTH\|\|?->HTH_26->BrxA-DNA-binding->BrxB-BrxF-iSTAND->DUF6079-p*-> | 723 | Bacteria>unclassified_Bacteria | GCA_018369965.1 | butyrate-producing_bacterium | KHX12_12210 | 5332..7500 | 2811711 | BREX-system-P-loop-protein-BrxC-[butyrate-producing-bacterium] |
| MBU4348826.1 | Type1.BREX | BrxB-BrxF-iSTAND->DUF6079*->N6-METHYLASE->N6-METHYLASE-p->PglZ->SIGMA+OB-Fold+AAA-ATPase+Lon-Protease-BrxL->HNH->?->SF1-SF2-Helicase->?->REase-> | 1193 | Bacteria>unclassified_Bacteria | GCA_018896635.1 | bacterium | KJ599_00710 | 356..3937 | 1869227 | BREX-system-P-loop-protein-BrxC-[bacterium] |
| MCB6991564.1 | Type1.BREX | <-ACET<-?<-ABC-ATPase+SbcC<-ABC-ATPase+SbcC<-MarR-HTH\|\|?->HTH_26->BrxA-DNA-binding->BrxB-BrxF-iSTAND->DUF6079*->OLD-ABC->HNH->N6-METHYLASE->KAP-NTPase-> | 1183 | Bacteria>unclassified_Bacteria | GCA_020558425.1 | bacterium_210820-DFI.6.38 | LI187_13870 | 12055..15606 | 2883293 | BREX-system-P-loop-protein-BrxC-[bacterium-210820-DFI.6.38] |
| MCE5247410.1 | Type1.BREX | <-REase<-?<-SIGMA+OB-Fold+AAA-ATPase+Lon-Protease-BrxL<-PglZ<-N6-METHYLASE\|\|?-><-DUF6079*<-BrxB-BrxF-iSTAND<-BrxA-DNA-binding | 1148 | Bacteria>unclassified_Bacteria | GCA_021372775.1 | bacterium | LLG88_10905 | 14182..17628 | 1869227 | BREX-system-P-loop-protein-BrxC-[bacterium] |
| MCI0601822.1 | Type1.BREX | <-MORC<-HTH+MPTase<-?<-SIGMA+OB-Fold+AAA-ATPase+Lon-Protease-BrxL<-PglZ<-N6-METHYLASE<-HEPN<-DUF6079*<-BrxB-BrxF-iSTAND<-BrxA-DNA-binding | 1220 | Bacteria>unclassified_Bacteria | GCA_022616075.1 | bacterium | L0156_02305 | 63267..66929 | 1869227 | BREX-system-P-loop-protein-BrxC-[bacterium] |
| MCP4571821.1 | Type1.BREX | Arm-DNA-bind_3+Phage_integrase->XIS-HTH->AAA-ATPase->?->BrxA-DNA-binding->BrxB-BrxF-iSTAND->DUF6079*-> | 1152 | Bacteria>unclassified_Bacteria | GCA_024231905.1 | bacterium | GY838_05660 | 49684..53142 | 1869227 | BREX-system-P-loop-protein-BrxC-[bacterium] |
| MCP4550320.1 | Type1.BREX | <-IG+IG\|\|METHYLASE->SAM-METHYLASE-><-Aminotran_1_2\|\|?->tRNA->tRNA-><-Phage_int_SAM_5+Phage_integrase\|\|BrxA-DNA-binding->BrxB-BrxF-iSTAND->DUF6079*->N6-METHYLASE->PglZ->SIGMA+OB-Fold+AAA-ATPase+Lon-Protease-BrxL->?-><-?<-XIS-HTH<-?<-HTH_3 | 1152 | Bacteria>unclassified_Bacteria | GCA_024231955.1 | bacterium | GY835_28000 | 114570..118028 | 1869227 | BREX-system-P-loop-protein-BrxC-[bacterium] |
| MCR9098968.1 | Type1.BREX | <-PglZ\|\|?->ParB+OLD-ABC->TOPRIM-><-HTH_3<-SbcC<-REase<-SH3<-N6-METHYLASE<-DUF6079*<-BrxB-BrxF-iSTAND<-BrxA-DNA-binding<-RNase<-SF1-SF2-Helicase+REase\|\|NtrC-AAA+FIS-HTH-><-ZnR<-ZnR | 1186 | Bacteria>unclassified_Bacteria | GCA_024742755.1 | bacterium | NXI25_03390 | 27983..31543 | 1869227 | BREX-system-P-loop-protein-BrxC-[bacterium] |
| EGR4851343.1 | Type1.BREX | <-ZnR<-?\|\|?->SF1-SF2-Helicase->AbiEi->viral-protein->wHTH+WYL+WCX->BrxA-DNA-binding->BrxB-BrxF-iSTAND->DUF6079*->N6-METHYLASE->ParB+HNH->GrpE->?->PglZ->SIGMA+OB-Fold+AAA-ATPase+Lon-Protease-BrxL-><-wHTH\|\|DUF4113-> | 1225 | - | - | - | C4G37_RS13070 | 147789..151466 | - | DNA-repair-protein-[Vibrio-parahaemolyticus] |
| EGR4833273.1 | Type1.BREX | <-ZnR<-?\|\|?->SF1-SF2-Helicase->AbiEi->viral-protein->wHTH+WYL+WCX->BrxA-DNA-binding->BrxB-BrxF-iSTAND->DUF6079*->N6-METHYLASE->?->?->PglZ->SIGMA+OB-Fold+AAA-ATPase+Lon-Protease-BrxL-><-wHTH\|\|DUF4113-><-?<-CR-ATPase4 | 1225 | - | - | - | C4G52_RS17670 | 119644..123321 | - | DNA-repair-protein-[Vibrio-parahaemolyticus] |
| EGR4808858.1 | Type1.BREX | <-ZnR<-?\|\|?->SF1-SF2-Helicase->AbiEi->viral-protein->wHTH+WYL+WCX->BrxA-DNA-binding->BrxB-BrxF-iSTAND->DUF6079*->N6-METHYLASE->ParB+HNH->GrpE->?->PglZ->SIGMA+OB-Fold+AAA-ATPase+Lon-Protease-BrxL-><-wHTH\|\|DUF4113-> | 1225 | - | - | - | C4G65_RS20915 | 24307..27984 | - | DNA-repair-protein-[Vibrio-parahaemolyticus] |
| MCH8487209.1 | Type1.BREX | BrxA-DNA-binding->BrxB-BrxF-iSTAND->DUF6079*->N6-METHYLASE->AbiGi+seqA+RAMA->?->SF1-SF2-Helicase->?->PglZ->SIGMA+OB-Fold+AAA-ATPase+HKD-DNase-BrxL->HigB_toxin->HTH_3-> | 1177 | - | GCA_022565315.1 | - | LAT75_10090 | 1579..5112 | 2875933 | BREX-system-P-loop-protein-BrxC-[Cyclonatronum-sp.] |
| WP_086519900.1 | Type1.BREX | <-PIN\|\|nSTAND1+HNH->RNaseH->NACHT+SF1-SF2-Helicase+HKD-DNase->REase-><-HKD-DNase+OGG-like<-SIGMA+OB-Fold+AAA-ATPase+Lon-Protease-BrxL<-PglZ<-?<-N6-METHYLASE<-DUF6079*<-BrxB-BrxF-iSTAND<-BrxA-DNA-binding<-?<-ColE7+HNH<-ACET\|\|NUDIX->?-><-DUF2754 | 1213 | - | GCF_900775425.1 | Bacteria | E7W97_RS22710 | 357533..361174 | 2 | MULTISPECIES:-BREX-system-P-loop-protein-BrxC-[Bacteria] |
| WP_142449058.1 | Type1.BREX | BrxA-DNA-binding->BrxB-BrxF-iSTAND->DUF6079*->N6-METHYLASE->PglZ->SIGMA+OB-Fold+AAA-ATPase+Lon-Protease-BrxL-> | 1223 | - | GCF_902164105.1 | Bacteria | FN517_RS25265 | 1603..5274 | 2 | MULTISPECIES:-BREX-system-P-loop-protein-BrxC-[Bacteria] |
| EGR5004813.1 | Type1.BREX | CR-ATPase4->?-><-DUF4113\|\|wHTH-><-SIGMA+OB-Fold+AAA-ATPase+Lon-Protease-BrxL<-PglZ<-?<-?<-N6-METHYLASE<-DUF6079*<-BrxB-BrxF-iSTAND<-BrxA-DNA-binding<-wHTH+WYL+WCX<-viral-protein<-AbiEi<-SF1-SF2-Helicase<-?\|\|?->ZnR-> | 1225 | - | - | - | L5169_000073 | 92916..96593 | - | DNA-repair-protein-[Vibrio-parahaemolyticus] |
| EIL2714655.1 | Type1.BREX | <-wHTH+WYL<-?<-?\|\|?->XIS-HTH->ZnR->wHTH+WYL+WCX->BrxA-DNA-binding->BrxB-BrxF-iSTAND->DUF6079*->ParB+HNH->N6-METHYLASE->PglZ->SIGMA+OB-Fold+AAA-ATPase+Lon-Protease-BrxL->CARF+NtrC-AAA+FIS-HTH-> | 1225 | - | - | - | M0K80_RS10540 | 473799..477476 | - | BREX-system-P-loop-protein-BrxC-[Providencia-rettgeri] |
| -- | | | | | | | | | | |
| --Type-5-BREX | | | | | | | | | | |
| --Duplication-of-BrxC/BrxX | | | | | | | | | | |
| --Canonical-BREX | | | | | | | | | | |
| -- | | | | | | | | | | |
| MUV48482.1 | Type5.BREX | BrxA-DNA-binding->DUF6079*->BrxB-BrxF-iSTAND->DUF6079->N6-METHYLASE->PglZ->N6-METHYLASE->TUDOR+SF1-SF2-Helicase+iREase+CTD-><-?<-?<-ABC-ATPase+SbcC | 1227 | Archaea>Euryarchaeota>Stenosarchaea_group>Halobacteria>Halobacteriales>Haloarculaceae | GCA_009741785.1 | Haloarcula_sp._CBA1122 | GJ630_01725 | 3031..6714 | 2668069 | hypothetical-protein-[Haloarcula-sp.-CBA1122] |
| WP_014030941.1 | Type5.BREX | <-TUDOR+SF1-SF2-Helicase+iREase+CTD<-N6-METHYLASE<-PglZ<-?<-N6-METHYLASE\|\|viral-protein-><-REase<-DUF6079*<-BrxB-BrxF-iSTAND<-DUF6079*<-BrxA-DNA-binding<-ParA-Soj-PloopNTPase | 1280 | Archaea>Euryarchaeota>Stenosarchaea_group>Halobacteria>Halobacteriales>Haloarculaceae | GCF_000223905.1 | Haloarcula_hispanica | HISP_RS16540 | 291914..295756 | 51589 | hypothetical-protein-[Haloarcula-hispanica] |
| WP_014030943.1 | Type5.BREX | <-TUDOR+SF1-SF2-Helicase+iREase+CTD<-N6-METHYLASE<-PglZ<-?<-N6-METHYLASE\|\|viral-protein-><-REase<-DUF6079<-BrxB-BrxF-iSTAND<-DUF6079*<-BrxA-DNA-binding<-ParA-Soj-PloopNTPase | 1227 | Archaea>Euryarchaeota>Stenosarchaea_group>Halobacteria>Halobacteriales>Haloarculaceae | GCF_000223905.1 | Haloarcula_hispanica | HISP_RS16550 | 296331..300014 | 51589 | hypothetical-protein-[Haloarcula-hispanica] |
| WP_007190712.1 | Type5.BREX | <-REC+PAS+HISKIN<-GDResolvase\|\|BrxA-DNA-binding->DUF6079*->BrxB-BrxF-iSTAND->DUF6079-><-?\|\|N6-METHYLASE->?->PglZ->TUDOR+SF1-SF2-Helicase+iREase+CTD-> | 1248 | Archaea>Euryarchaeota>Stenosarchaea_group>Halobacteria>Halobacteriales>Haloarculaceae | GCF_000337755.1 | Haloarcula_californiae | C435_RS20625 | 4933..8679 | 244363 | hypothetical-protein-[Haloarcula-californiae] |
| WP_007190714.1 | Type5.BREX | <-REC+PAS+HISKIN<-GDResolvase\|\|BrxA-DNA-binding->DUF6079->BrxB-BrxF-iSTAND->DUF6079*-><-?\|\|N6-METHYLASE->?->PglZ->TUDOR+SF1-SF2-Helicase+iREase+CTD-><-?<-?\|\|Peptidase_M29+DUF2971->HTH_28-> | 1280 | Archaea>Euryarchaeota>Stenosarchaea_group>Halobacteria>Halobacteriales>Haloarculaceae | GCF_000337755.1 | Haloarcula_californiae | C435_RS20635 | 9293..13135 | 244363 | hypothetical-protein-[Haloarcula-californiae] |
| WP_050036525.1 | Type5.BREX | ParA-Soj-PloopNTPase->BrxA-DNA-binding->DUF6079*->BrxB-BrxF-iSTAND->DUF6079-><-viral-protein\|\|N6-METHYLASE->?->PglZ->N6-METHYLASE->TUDOR+SF1-SF2-Helicase+iREase+CTD-> | 1227 | Archaea>Euryarchaeota>Stenosarchaea_group>Halobacteria>Halobacteriales>Haloarculaceae | GCF_001485555.1 | unclassified_Haloarcula | SG26_00325 | 77874..81557 | 2624677 | MULTISPECIES:-hypothetical-protein-[unclassified-Haloarcula] |
| WP_050036527.1 | Type5.BREX | ParA-Soj-PloopNTPase->BrxA-DNA-binding->DUF6079->BrxB-BrxF-iSTAND->DUF6079*-><-viral-protein\|\|N6-METHYLASE->?->PglZ->N6-METHYLASE->TUDOR+SF1-SF2-Helicase+iREase+CTD-><-?<-?<-ABC-ATPase+SbcC | 1280 | Archaea>Euryarchaeota>Stenosarchaea_group>Halobacteria>Halobacteriales>Haloarculaceae | GCF_001485555.1 | unclassified_Haloarcula | SG26_00335 | 82132..85974 | 2624677 | MULTISPECIES:-hypothetical-protein-[unclassified-Haloarcula] |
| WP_064287185.1 | Type5.BREX | <-TUDOR+SF1-SF2-Helicase+iREase+CTD<-N6-METHYLASE<-PglZ<-?<-N6-METHYLASE<-DUF6079<-BrxB-BrxF-iSTAND<-DUF6079*<-BrxA-DNA-binding<-ParA-Soj-PloopNTPase | 1227 | Archaea>Euryarchaeota>Stenosarchaea_group>Halobacteria>Halobacteriales>Haloarculaceae | GCF_001647155.1 | Haloarcula_sp._K1 | AV929_RS01580 | 168563..172246 | 1622207 | hypothetical-protein-[Haloarcula-sp.-K1] |
| WP_064287183.1 | Type5.BREX | ABC-ATPase+SbcC->?->?-><-TUDOR+SF1-SF2-Helicase+iREase+CTD<-N6-METHYLASE<-PglZ<-?<-N6-METHYLASE<-DUF6079*<-BrxB-BrxF-iSTAND<-DUF6079<-BrxA-DNA-binding<-ParA-Soj-PloopNTPase | 1280 | Archaea>Euryarchaeota>Stenosarchaea_group>Halobacteria>Halobacteriales>Haloarculaceae | GCF_001647155.1 | unclassified_Haloarcula | AV929_RS01570 | 164146..167988 | 2624677 | MULTISPECIES:-hypothetical-protein-[unclassified-Haloarcula] |
| WP_121538347.1 | Type5.BREX | <-HTH_20\|\|?->?->ParA-Soj-PloopNTPase->?-><-TUDOR+SF1-SF2-Helicase+iREase+CTD<-PglZ<-N6-METHYLASE\|\|?-><-DUF6079*<-BrxB-BrxF-iSTAND<-DUF6079<-BrxA-DNA-binding\|\|Exonuc_V_gamma+REase->SF1-SF2-Helicase+REase-><-REase\|\|ORC-CDC6+HTH-><-?<-McrC-NTD+REase | 1272 | Archaea>Euryarchaeota>Stenosarchaea_group>Halobacteria>Halobacteriales>Haloarculaceae | GCF_003665885.1 | Haloarcula_sp._Atlit-120R | DVK01_RS19340 | 38493..42311 | 2282135 | hypothetical-protein-[Haloarcula-sp.-Atlit-120R] |
| WP_121538351.1 | Type5.BREX | ParA-Soj-PloopNTPase->?-><-TUDOR+SF1-SF2-Helicase+iREase+CTD<-PglZ<-N6-METHYLASE\|\|?-><-DUF6079<-BrxB-BrxF-iSTAND<-DUF6079*<-BrxA-DNA-binding\|\|Exonuc_V_gamma+REase->SF1-SF2-Helicase+REase-><-REase\|\|ORC-CDC6+HTH-><-?<-McrC-NTD+REase<-McrB-AAA | 1251 | Archaea>Euryarchaeota>Stenosarchaea_group>Halobacteria>Halobacteriales>Haloarculaceae | GCF_003665885.1 | Haloarcula_sp._Atlit-120R | DVK01_RS19350 | 42895..46650 | 2282135 | hypothetical-protein-[Haloarcula-sp.-Atlit-120R] |
| WP_135304847.1 | Type5.BREX | <-REase\|\|?-><-MORC<-TUDOR+SF1-SF2-Helicase+iREase+CTD<-N6-METHYLASE<-PglZ<-N6-METHYLASE<-PIN<-?<-DUF6079*<-BrxB-BrxF-iSTAND<-DUF6079<-BrxA-DNA-binding\|\|?->?->?-><-?\|\|McrB-AAA->McrC-NTD+REase-> | 1273 | Archaea>Euryarchaeota>Stenosarchaea_group>Halobacteria>Halobacteriales>Haloarculaceae | GCF_004681185.1 | Halomicroarcula_amylolytica | E4S61_RS15390 | 199089..202910 | 2562280 | hypothetical-protein-[Halomicroarcula-amylolytica] |
| WP_135304849.1 | Type5.BREX | <-MORC<-TUDOR+SF1-SF2-Helicase+iREase+CTD<-N6-METHYLASE<-PglZ<-N6-METHYLASE<-PIN<-?<-DUF6079<-BrxB-BrxF-iSTAND<-DUF6079*<-BrxA-DNA-binding\|\|?->?->?-><-?\|\|McrB-AAA->McrC-NTD+REase-><-SF1-SF2-Helicase | 1250 | Archaea>Euryarchaeota>Stenosarchaea_group>Halobacteria>Halobacteriales>Haloarculaceae | GCF_004681185.1 | Halomicroarcula_amylolytica | E4S61_RS15400 | 203494..207246 | 2562280 | hypothetical-protein-[Halomicroarcula-amylolytica] |
| WP_151148304.1 | Type5.BREX | ParA-Soj-PloopNTPase->BrxA-DNA-binding->DUF6079*->BrxB-BrxF-iSTAND->DUF6079->?-><-viral-protein\|\|N6-METHYLASE->?->PglZ->N6-METHYLASE->TUDOR+SF1-SF2-Helicase+iREase+CTD-> | 1227 | Archaea>Euryarchaeota>Stenosarchaea_group>Halobacteria>Halobacteriales>Haloarculaceae | GCF_008728975.1 | Haloarcula_sp._CBA1131 | Har1131_RS19690 | 405415..409098 | 1853686 | hypothetical-protein-[Haloarcula-sp.-CBA1131] |
| WP_225308203.1 | Type5.BREX | ParA-Soj-PloopNTPase->BrxA-DNA-binding->DUF6079->BrxB-BrxF-iSTAND->DUF6079*->?-><-viral-protein\|\|N6-METHYLASE->?->PglZ->N6-METHYLASE->TUDOR+SF1-SF2-Helicase+iREase+CTD-> | 1005 | Archaea>Euryarchaeota>Stenosarchaea_group>Halobacteria>Halobacteriales>Haloarculaceae | GCF_008728975.1 | Haloarcula_sp._CBA1131 | Har1131_RS19700 | 409673..412690 | 1853686 | hypothetical-protein-[Haloarcula-sp.-CBA1131] |
| WP_151104049.1 | Type5.BREX | ABC-ATPase+SbcC->?->?-><-TUDOR+SF1-SF2-Helicase+iREase+CTD<-N6-METHYLASE<-PglZ<-?<-N6-METHYLASE\|\|viral-protein-><-DUF6079*<-BrxB-BrxF-iSTAND<-DUF6079<-BrxA-DNA-binding<-ParA-Soj-PloopNTPase | 1280 | Archaea>Euryarchaeota>Stenosarchaea_group>Halobacteria>Halobacteriales>Haloarculaceae | GCF_008729095.1 | Haloarcula_hispanica | EGD51_RS16380 | 224591..228433 | 51589 | hypothetical-protein-[Haloarcula-hispanica] |
| WP_151104050.1 | Type5.BREX | <-TUDOR+SF1-SF2-Helicase+iREase+CTD<-N6-METHYLASE<-PglZ<-?<-N6-METHYLASE\|\|viral-protein-><-DUF6079<-BrxB-BrxF-iSTAND<-DUF6079*<-BrxA-DNA-binding<-ParA-Soj-PloopNTPase | 1227 | Archaea>Euryarchaeota>Stenosarchaea_group>Halobacteria>Halobacteriales>Haloarculaceae | GCF_008729095.1 | Haloarcula_hispanica | EGD51_RS16390 | 229008..232691 | 51589 | hypothetical-protein-[Haloarcula-hispanica] |
| WP_165896131.1 | Type5.BREX | <-REase\|\|?-><-TUDOR+SF1-SF2-Helicase+iREase+CTD<-?<-N6-METHYLASE\|\|BrxA-DNA-binding->DUF6079*->BrxB-BrxF-iSTAND->DUF6079->PglZ->DOC+HTH-><-SF1-SF2-Helicase+REase | 1233 | Archaea>Euryarchaeota>Stenosarchaea_group>Halobacteria>Halobacteriales>Haloarculaceae | GCF_011440255.1 | Haloarcula_sp._JP-Z28 | G9463_RS04925 | 80940..84641 | 2716715 | hypothetical-protein-[Haloarcula-sp.-JP-Z28] |
| WP_165896132.1 | Type5.BREX | <-REase\|\|?-><-TUDOR+SF1-SF2-Helicase+iREase+CTD<-?<-N6-METHYLASE\|\|BrxA-DNA-binding->DUF6079->BrxB-BrxF-iSTAND->DUF6079*->PglZ->DOC+HTH-><-SF1-SF2-Helicase+REase | 1231 | Archaea>Euryarchaeota>Stenosarchaea_group>Halobacteria>Halobacteriales>Haloarculaceae | GCF_011440255.1 | Haloarcula_sp._JP-Z28 | G9463_RS04935 | 85257..88952 | 2716715 | hypothetical-protein-[Haloarcula-sp.-JP-Z28] |
| WP_127015186.1 | Type5.BREX | HTH_29+HTH_32+DDE_3-><-?<-TUDOR+SF1-SF2-Helicase+iREase+CTD<-PglZ<-?<-RelE<-?<-N6-METHYLASE\|\|?-><-DUF6079*<-BrxB-BrxF-iSTAND<-DUF6079<-BrxA-DNA-binding | 1280 | Archaea>Euryarchaeota>Stenosarchaea_group>Halobacteria>Halobacteriales>Haloarculaceae | GCF_011627385.1 | Haloarcula | G9C82_RS20125 | 32096..35938 | 2237 | MULTISPECIES:-hypothetical-protein-[Haloarcula] |
| WP_127015187.1 | Type5.BREX | <-TUDOR+SF1-SF2-Helicase+iREase+CTD<-PglZ<-?<-RelE<-?<-N6-METHYLASE\|\|?-><-DUF6079<-BrxB-BrxF-iSTAND<-DUF6079*<-BrxA-DNA-binding | 1248 | Archaea>Euryarchaeota>Stenosarchaea_group>Halobacteria>Halobacteriales>Haloarculaceae | GCF_011627385.1 | Haloarcula | G9C82_RS20135 | 36552..40298 | 2237 | MULTISPECIES:-hypothetical-protein-[Haloarcula] |
| WP_255153072.1 | Type5.BREX | RNase-><-SF1-SF2-Helicase<-REase<-SF1-SF2-Helicase\|\|CR-ATPase9+REase-><-REase<-TBP+TBP<-PglZ<-DUF6079*<-BrxB-BrxF-iSTAND<-DUF6079<-BrxA-DNA-binding\|\|N6-METHYLASE->?->TUDOR+SF1-SF2-Helicase+iREase+CTD-><-?\|\|REase-> | 1230 | Archaea>Euryarchaeota>Stenosarchaea_group>Halobacteria>Halobacteriales>Haloarculaceae | GCF_024298865.1 | Halosegnis_sp._ZY10 | NL112_RS22055 | 49161..52853 | 2962671 | hypothetical-protein-[Halosegnis-sp.-ZY10] |
| WP_255153074.1 | Type5.BREX | <-SF1-SF2-Helicase<-REase<-SF1-SF2-Helicase\|\|CR-ATPase9+REase-><-REase<-TBP+TBP<-PglZ<-DUF6079<-BrxB-BrxF-iSTAND<-DUF6079*<-BrxA-DNA-binding\|\|N6-METHYLASE->?->TUDOR+SF1-SF2-Helicase+iREase+CTD-><-?\|\|REase-><-?\|\|MarR-HTH-> | 1233 | Archaea>Euryarchaeota>Stenosarchaea_group>Halobacteria>Halobacteriales>Haloarculaceae | GCF_024298865.1 | Halosegnis_sp._ZY10 | NL112_RS22065 | 53469..57170 | 2962671 | hypothetical-protein-[Halosegnis-sp.-ZY10] |
| WP_092662309.1 | Type5.BREX | PIWI-><-TUDOR+SF1-SF2-Helicase+iREase+CTD<-PglZ<-N6-METHYLASE<-DUF6079*<-BrxB-BrxF-iSTAND<-DUF6079<-BrxA-DNA-binding<-REase<-?\|\|GDResolvase-><-SF1-SF2-Helicase | 1279 | Archaea>Euryarchaeota>Stenosarchaea_group>Halobacteria>Halobacteriales>Haloarculaceae | GCF_900110215.1 | Halorientalis_persicus | BMX17_RS17055 | 56359..60198 | 1367881 | hypothetical-protein-[Halorientalis-persicus] |
| WP_092662313.1 | Type5.BREX | PIWI-><-TUDOR+SF1-SF2-Helicase+iREase+CTD<-PglZ<-N6-METHYLASE<-DUF6079<-BrxB-BrxF-iSTAND<-DUF6079*<-BrxA-DNA-binding<-REase<-?\|\|GDResolvase-><-SF1-SF2-Helicase<-?\|\|?->?->NAD_binding_10-> | 1248 | Archaea>Euryarchaeota>Stenosarchaea_group>Halobacteria>Halobacteriales>Haloarculaceae | GCF_900110215.1 | Halorientalis_persicus | BMX17_RS17065 | 60812..64558 | 1367881 | hypothetical-protein-[Halorientalis-persicus] |
| WP_050049402.1 | Type5.BREX | BrxA-DNA-binding->DUF6079*->BrxB-BrxF-iSTAND->DUF6079->?-><-?<-TBP+TBP\|\|N6-METHYLASE->PglZ->TUDOR+SF1-SF2-Helicase+iREase+CTD-> | 1248 | Archaea>Euryarchaeota>Stenosarchaea_group>Halobacteria>Halobacteriales>Halobacteriaceae | GCF_001011115.1 | Halanaeroarchaeum_sulfurireducens | HLASF_RS10545 | 1321..5067 | 1604004 | hypothetical-protein-[Halanaeroarchaeum-sulfurireducens] |
| WP_050049404.1 | Type5.BREX | BrxA-DNA-binding->DUF6079->BrxB-BrxF-iSTAND->DUF6079*->?-><-?<-TBP+TBP\|\|N6-METHYLASE->PglZ->TUDOR+SF1-SF2-Helicase+iREase+CTD-><-?\|\|Phage_integrase-><-RelE-ParE | 1275 | Archaea>Euryarchaeota>Stenosarchaea_group>Halobacteria>Halobacteriales>Halobacteriaceae | GCF_001011115.1 | Halanaeroarchaeum_sulfurireducens | HLASF_RS10555 | 5681..9508 | 1604004 | hypothetical-protein-[Halanaeroarchaeum-sulfurireducens] |
| WP_054520042.1 | Type5.BREX | BrxA-DNA-binding->DUF6079*->BrxB-BrxF-iSTAND->DUF6079->?-><-?<-TBP+TBP\|\|N6-METHYLASE->PglZ->TUDOR+SF1-SF2-Helicase+iREase+CTD-> | 1248 | Archaea>Euryarchaeota>Stenosarchaea_group>Halobacteria>Halobacteriales>Halobacteriaceae | GCF_001305655.1 | Halanaeroarchaeum_sulfurireducens | HLASA_RS10775 | 1321..5067 | 1604004 | hypothetical-protein-[Halanaeroarchaeum-sulfurireducens] |
| WP_058984838.1 | Type5.BREX | <-SF1-SF2-Helicase+REase<-Exonuc_V_gamma+REase<-TUDOR+SF1-SF2-Helicase+iREase+CTD<-REase+METHYLASE<-PglZ<-?<-N6-METHYLASE\|\|viral-protein-><-REase<-DUF6079*<-BrxB-BrxF-iSTAND<-DUF6079<-BrxA-DNA-binding\|\|ORC-CDC6->HTH_28->NTP_transf_5->?->?->35exo-> | 1277 | Archaea>Euryarchaeota>Stenosarchaea_group>Halobacteria>Halobacteriales>Halobacteriaceae | GCF_001485535.1 | Halobacterium | AV266_RS14305 | 153821..157654 | 2239 | MULTISPECIES:-hypothetical-protein-[Halobacterium] |
| WP_058984840.1 | Type5.BREX | <-TUDOR+SF1-SF2-Helicase+iREase+CTD<-REase+METHYLASE<-PglZ<-?<-N6-METHYLASE\|\|viral-protein-><-REase<-DUF6079<-BrxB-BrxF-iSTAND<-DUF6079*<-BrxA-DNA-binding\|\|ORC-CDC6->HTH_28->NTP_transf_5->?->?->35exo-><-?<-HTH_20 | 1227 | Archaea>Euryarchaeota>Stenosarchaea_group>Halobacteria>Halobacteriales>Halobacteriaceae | GCF_001485535.1 | Halobacterium | AV266_RS14315 | 158229..161912 | 2239 | MULTISPECIES:-hypothetical-protein-[Halobacterium] |
| WP_124956320.1 | Type5.BREX | <-PADR-HTH<-?<-OrfB_Zn_ribbon\|\|?->?->?->BrxA-DNA-binding->DUF6079*->BrxB-BrxF-iSTAND->DUF6079->Peptidase_M1->N6-METHYLASE->PglZ->TUDOR+SF1-SF2-Helicase+iREase+CTD-> | 1239 | Archaea>Euryarchaeota>Stenosarchaea_group>Halobacteria>Halobacteriales>Halobacteriaceae | GCF_003862495.1 | Halocatena_pleomorpha | EIK79_RS15545 | 19257..22976 | 1785090 | hypothetical-protein-[Halocatena-pleomorpha] |
| WP_124956324.1 | Type5.BREX | <-PADR-HTH<-?<-OrfB_Zn_ribbon\|\|?->?->?->BrxA-DNA-binding->DUF6079->BrxB-BrxF-iSTAND->DUF6079*->Peptidase_M1->N6-METHYLASE->PglZ->TUDOR+SF1-SF2-Helicase+iREase+CTD-> | 1271 | Archaea>Euryarchaeota>Stenosarchaea_group>Halobacteria>Halobacteriales>Halobacteriaceae | GCF_003862495.1 | Halocatena_pleomorpha | EIK79_RS15555 | 23560..27375 | 1785090 | hypothetical-protein-[Halocatena-pleomorpha] |
| WP_232700617.1 | Type5.BREX | <-TUDOR+SF1-SF2-Helicase+iREase+CTD\|\|TBP+TBP-><-PglZ<-?<-N6-METHYLASE<-?<-DUF6079*<-BrxB-BrxF-iSTAND<-DUF6079<-BrxA-DNA-binding<-SF1-SF2-Helicase+REase<-SF1-SF2-Helicase+REase\|\|?->?->?->ABC-ATPase+SbcC-> | 1277 | Archaea>Euryarchaeota>Stenosarchaea_group>Halobacteria>Halobacteriales>Halobacteriaceae | GCF_021249345.1 | Halobacterium_sp._Gai3-17 | LT965_RS09790 | 1831133..1834966 | 2902623 | hypothetical-protein-[Halobacterium-sp.-Gai3-17] |
| WP_232700619.1 | Type5.BREX | <-TUDOR+SF1-SF2-Helicase+iREase+CTD\|\|TBP+TBP-><-PglZ<-?<-N6-METHYLASE<-?<-DUF6079<-BrxB-BrxF-iSTAND<-DUF6079*<-BrxA-DNA-binding<-SF1-SF2-Helicase+REase<-SF1-SF2-Helicase+REase\|\|?->?->?->ABC-ATPase+SbcC->ABC-ATPase+SbcC-> | 1249 | Archaea>Euryarchaeota>Stenosarchaea_group>Halobacteria>Halobacteriales>Halobacteriaceae | GCF_021249345.1 | Halobacterium_sp._Gai3-17 | LT965_RS09800 | 1835580..1839329 | 2902623 | hypothetical-protein-[Halobacterium-sp.-Gai3-17] |
| WP_247420977.1 | Type5.BREX | McrC-NTD+REase->SF1-SF2-Helicase->REase->SF1-SF2-Helicase-><-TUDOR+SF1-SF2-Helicase+iREase+CTD\|\|?->TBP+TBP-><-PglZ<-N6-METHYLASE<-DUF6079*<-BrxB-BrxF-iSTAND<-DUF6079<-BrxA-DNA-binding\|\|REase+METHYLASE-><-PIN<-CARF<-?<-ZnR+REase<-MarR-HTH | 1270 | Archaea>Euryarchaeota>Stenosarchaea_group>Halobacteria>Halobacteriales>Halobacteriaceae | GCF_023074835.1 | Halomarina_salina | MX571_RS20530 | 26025..29837 | 1872699 | hypothetical-protein-[Halomarina-salina] |
| WP_247420981.1 | Type5.BREX | REase->SF1-SF2-Helicase-><-TUDOR+SF1-SF2-Helicase+iREase+CTD\|\|?->TBP+TBP-><-PglZ<-N6-METHYLASE<-DUF6079<-BrxB-BrxF-iSTAND<-DUF6079*<-BrxA-DNA-binding\|\|REase+METHYLASE-><-PIN<-CARF<-?<-ZnR+REase<-MarR-HTH\|\|?-><-McrC-NTD+REase | 1239 | Archaea>Euryarchaeota>Stenosarchaea_group>Halobacteria>Halobacteriales>Halobacteriaceae | GCF_023074835.1 | Halomarina_salina | MX571_RS20540 | 30421..34140 | 1872699 | hypothetical-protein-[Halomarina-salina] |
| WP_247994824.1 | Type5.BREX | ORC-CDC6+HTH-><-?<-?<-ABC-ATPase+SbcC\|\|SF1-SF2-Helicase+REase->SF1-SF2-Helicase+REase-><-TUDOR+SF1-SF2-Helicase+iREase+CTD<-PglZ<-N6-METHYLASE<-DUF6079*<-BrxB-BrxF-iSTAND<-DUF6079<-BrxA-DNA-binding<-?<-?\|\|?-><-PIWI | 1274 | Archaea>Euryarchaeota>Stenosarchaea_group>Halobacteria>Halobacteriales>Halobacteriaceae | GCF_023115355.1 | Halocatena_salina | MW046_RS14210 | 287042..290866 | 2934340 | hypothetical-protein-[Halocatena-salina] |
| WP_247994826.1 | Type5.BREX | <-ABC-ATPase+SbcC\|\|SF1-SF2-Helicase+REase->SF1-SF2-Helicase+REase-><-TUDOR+SF1-SF2-Helicase+iREase+CTD<-PglZ<-N6-METHYLASE<-DUF6079<-BrxB-BrxF-iSTAND<-DUF6079*<-BrxA-DNA-binding<-?<-?\|\|?-><-PIWI | 1247 | Archaea>Euryarchaeota>Stenosarchaea_group>Halobacteria>Halobacteriales>Halobacteriaceae | GCF_023115355.1 | Halocatena_salina | MW046_RS14220 | 291483..295226 | 2934340 | hypothetical-protein-[Halocatena-salina] |
| WP_247995979.1 | Type5.BREX | <-ARG<-McrC-NTD+REase<-McrB-AAA<-SF1-SF2-Helicase+REase<-Exonuc_V_gamma+REase\|\|BrxA-DNA-binding->DUF6079*->BrxB-BrxF-iSTAND->DUF6079->REase->N6-METHYLASE-><-OLD-ABC+TOPRIM\|\|PglZ->TUDOR+SF1-SF2-Helicase+iREase+CTD->?->BrxD-AAA-> | 1225 | Archaea>Euryarchaeota>Stenosarchaea_group>Halobacteria>Halobacteriales>Halobacteriaceae | GCF_023115355.1 | Halocatena_salina | MW046_RS18845 | 37719..41396 | 2934340 | hypothetical-protein-[Halocatena-salina] |
| WP_247995981.1 | Type5.BREX | <-ARG<-McrC-NTD+REase<-McrB-AAA<-SF1-SF2-Helicase+REase<-Exonuc_V_gamma+REase\|\|BrxA-DNA-binding->DUF6079->BrxB-BrxF-iSTAND->DUF6079*->REase->N6-METHYLASE-><-OLD-ABC+TOPRIM\|\|PglZ->TUDOR+SF1-SF2-Helicase+iREase+CTD->?->BrxD-AAA->SF1-SF2-Helicase+wHTH+Small-B-barrel+DUF1998-><-CARF+HTH | 1278 | Archaea>Euryarchaeota>Stenosarchaea_group>Halobacteria>Halobacteriales>Halobacteriaceae | GCF_023115355.1 | Halocatena_salina | MW046_RS18855 | 41971..45807 | 2934340 | hypothetical-protein-[Halocatena-salina] |
| WP_253227448.1 | Type5.BREX | <-Peptidase_M29+DUF2971<-SF1-SF2-Helicase<-TUDOR+SF1-SF2-Helicase+iREase+CTD<-PglZ<-N6-METHYLASE<-DUF6079*<-BrxB-BrxF-iSTAND<-DUF6079<-BrxA-DNA-binding<-REase\|\|REase+METHYLASE->DOC+HTH-> | 1268 | Archaea>Euryarchaeota>Stenosarchaea_group>Halobacteria>Halobacteriales>Halobacteriaceae | GCF_024136975.1 | Haladaptatus_sp._AB643 | NKF25_RS14930 | 104157..107963 | 2934174 | hypothetical-protein-[Haladaptatus-sp.-AB643] |
| WP_253227450.1 | Type5.BREX | <-Peptidase_M29+DUF2971<-SF1-SF2-Helicase<-TUDOR+SF1-SF2-Helicase+iREase+CTD<-PglZ<-N6-METHYLASE<-DUF6079<-BrxB-BrxF-iSTAND<-DUF6079*<-BrxA-DNA-binding<-REase\|\|REase+METHYLASE->DOC+HTH->?->?->?-><-?\|\|OrfB_IS605+OrfB_Zn_ribbon-> | 1239 | Archaea>Euryarchaeota>Stenosarchaea_group>Halobacteria>Halobacteriales>Halobacteriaceae | GCF_024136975.1 | Haladaptatus_sp._AB643 | NKF25_RS14940 | 108547..112266 | 2934174 | hypothetical-protein-[Haladaptatus-sp.-AB643] |
| WP_254538609.1 | Type5.BREX | <-SF1-SF2-Helicase+REase<-SF1-SF2-Helicase+REase\|\|?->GspA-ATPase->?-><-ORC-CDC6<-OrfB_Zn_ribbon-><-DOC+HTH\|\|BrxA-DNA-binding->DUF6079*->BrxB-BrxF-iSTAND->DUF6079->N6-METHYLASE->PglZ->TUDOR+SF1-SF2-Helicase+iREase+CTD-><-?\|\|?->?->PADR-HTH | 1258 | Archaea>Euryarchaeota>Stenosarchaea_group>Halobacteria>Halobacteriales>Halobacteriaceae | GCF_024227715.1 | Halomarina_sp._BCD28 | NKG96_RS19415 | 93398..97174 | 2961595 | hypothetical-protein-[Halomarina-sp.-BCD28] |
| WP_254538611.1 | Type5.BREX | GspA-ATPase->?-><-ORC-CDC6-><-OrfB_Zn_ribbon-><-DOC+HTH\|\|BrxA-DNA-binding->DUF6079->BrxB-BrxF-iSTAND->DUF6079*->N6-METHYLASE->PglZ->TUDOR+SF1-SF2-Helicase+iREase+CTD-><-?\|\|?->?->PADR-HTH | 1269 | Archaea>Euryarchaeota>Stenosarchaea_group>Halobacteria>Halobacteriales>Halobacteriaceae | GCF_024227715.1 | Halomarina_sp._BCD28 | NKG96_RS19425 | 97758..101567 | 2961595 | hypothetical-protein-[Halomarina-sp.-BCD28] |
| WP_257301028.1 | Type5.BREX | CARF->PIN-><-CR-ATPase9+REase\|\|?-><-SF1-SF2-Helicase<-TUDOR+SF1-SF2-Helicase+iREase+CTD<-N6-METHYLASE\|\|BrxA-DNA-binding->DUF6079->BrxB-BrxF-iSTAND->DUF6079->PglZ->SF1-SF2-Helicase->REase->SF1-SF2-Helicase-><-DrmB<-SF1-SF2-Helicase | 1246 | Archaea>Euryarchaeota>Stenosarchaea_group>Halobacteria>Halobacteriales>Halobacteriaceae | GCF_024609245.1 | Haloarchaeobius_sp._FL176 | N0978_RS18575 | 32605..36345 | 2967129 | hypothetical-protein-[Haloarchaeobius-sp.-FL176] |
| WP_257301030.1 | Type5.BREX | PIN-><-CR-ATPase9+REase\|\|?-><-SF1-SF2-Helicase<-TUDOR+SF1-SF2-Helicase+iREase+CTD<-N6-METHYLASE\|\|BrxA-DNA-binding->DUF6079->BrxB-BrxF-iSTAND->DUF6079*->PglZ->SF1-SF2-Helicase->REase->SF1-SF2-Helicase-><-DrmB<-SF1-SF2-Helicase<-?<-N6-METHYLASE+DAM\|\|REase-> | 1228 | Archaea>Euryarchaeota>Stenosarchaea_group>Halobacteria>Halobacteriales>Halobacteriaceae | GCF_024609245.1 | Haloarchaeobius_sp._FL176 | N0978_RS18585 | 36973..40659 | 2967129 | hypothetical-protein-[Haloarchaeobius-sp.-FL176] |
| WP_264783322.1 | Type5.BREX | DOC+HTH-><-?<-REase\|\|?-><-TUDOR+SF1-SF2-Helicase+iREase+CTD<-?<-N6-METHYLASE\|\|BrxA-DNA-binding->DUF6079*->BrxB-BrxF-iSTAND->DUF6079->PglZ->TBP+TBP->REase->MORC+DUF3684+REase-><-ORC-CDC6<-?<-GspA-ATPase | 1233 | Archaea>Euryarchaeota>Stenosarchaea_group>Halobacteria>Halobacteriales>Halobacteriaceae | GCF_026008575.1 | Haladaptatus_sp._T7 | OPC11_RS21275 | 54807..58508 | 2029368 | hypothetical-protein-[Haladaptatus-sp.-T7] |
| WP_264783324.1 | Type5.BREX | <-REase\|\|?-><-TUDOR+SF1-SF2-Helicase+iREase+CTD<-?<-N6-METHYLASE\|\|BrxA-DNA-binding->DUF6079->BrxB-BrxF-iSTAND->DUF6079*->PglZ->TBP+TBP->REase->MORC+DUF3684+REase-><-ORC-CDC6<-?<-GspA-ATPase<-?\|\|SF1-SF2-Helicase+REase-> | 1230 | Archaea>Euryarchaeota>Stenosarchaea_group>Halobacteria>Halobacteriales>Halobacteriaceae | GCF_026008575.1 | Haladaptatus_sp._T7 | OPC11_RS21285 | 59124..62816 | 2029368 | hypothetical-protein-[Haladaptatus-sp.-T7] |
| WP_267638585.1 | Type5.BREX | Phage_integrase-><-?<-?<-ARG\|\|?-><-?<-Preader+URI-DNase\|\|BrxA-DNA-binding->DUF6079*->BrxB-BrxF-iSTAND->DUF6079->N6-METHYLASE->PglZ->TUDOR+SF1-SF2-Helicase+iREase+CTD-><-SF1-SF2-Helicase+REase<-Exonuc_V_gamma+REase\|\|HAD->HKD-DNase+HKD-Rease-> | 1251 | Archaea>Euryarchaeota>Stenosarchaea_group>Halobacteria>Halobacteriales>Halobacteriaceae | GCF_026586665.1 | Halovenus_rubra | N5098_RS14425 | 45600..49355 | 869890 | hypothetical-protein-[Halovenus-rubra] |
| WP_089670450.1 | Type5.BREX | <-TUDOR+SF1-SF2-Helicase+iREase+CTD<-N6-METHYLASE\|\|BrxA-DNA-binding->DUF6079*->BrxB-BrxF-iSTAND->DUF6079->PglZ-><-N6-METHYLASE+DAM\|\|REase->N6-METHYLASE+DAM-><-N6-METHYLASE+DAM\|\|SF1-SF2-Helicase+REase->SF1-SF2-Helicase+REase-> | 1247 | Archaea>Euryarchaeota>Stenosarchaea_group>Halobacteria>Halobacteriales>Halobacteriaceae | GCF_900110535.1 | Halobacterium_jilantaiense | BMW35_RS14745 | 7544..11287 | 355548 | hypothetical-protein-[Halobacterium-jilantaiense] |
| WP_089670452.1 | Type5.BREX | <-TUDOR+SF1-SF2-Helicase+iREase+CTD<-N6-METHYLASE\|\|BrxA-DNA-binding->DUF6079->BrxB-BrxF-iSTAND->DUF6079*->PglZ-><-N6-METHYLASE+DAM\|\|REase->N6-METHYLASE+DAM-><-N6-METHYLASE+DAM\|\|SF1-SF2-Helicase+REase->SF1-SF2-Helicase+REase-><-Calcineurin-SbcD | 1227 | Archaea>Euryarchaeota>Stenosarchaea_group>Halobacteria>Halobacteriales>Halobacteriaceae | GCF_900110535.1 | Halobacterium_jilantaiense | BMW35_RS14755 | 11918..15601 | 355548 | hypothetical-protein-[Halobacterium-jilantaiense] |
| WP_244705671.1 | Type5.BREX | <-SF1-SF2-Helicase+REase<-SF1-SF2-Helicase+REase<-TUDOR+SF1-SF2-Helicase+iREase+CTD<-PglZ<-N6-METHYLASE\|\|TBP+TBP-><-DUF6079*<-BrxB-BrxF-iSTAND<-DUF6079-p\|\|?-><-BrxA-DNA-binding\|\|?-><-?<-Glyoxalase | 1269 | Archaea>Euryarchaeota>Stenosarchaea_group>Halobacteria>Halobacteriales>Halococcaceae | GCF_022870485.1 | Halococcus_dombrowskii | MUK72_RS16110 | 375497..379306 | 179637 | hypothetical-protein-[Halococcus-dombrowskii] |
| WP_244705675.1 | Type5.BREX | <-SF1-SF2-Helicase+REase<-SF1-SF2-Helicase+REase<-TUDOR+SF1-SF2-Helicase+iREase+CTD<-PglZ<-N6-METHYLASE\|\|TBP+TBP-><-DUF6079<-BrxB-BrxF-iSTAND<-DUF6079-p*\|\|?-><-BrxA-DNA-binding\|\|?-><-?<-Glyoxalase\|\|?->?->AbiEi_4+AbiEi_1->AbiEii+ZnR-> | 848 | Archaea>Euryarchaeota>Stenosarchaea_group>Halobacteria>Halobacteriales>Halococcaceae | GCF_022870485.1 | Halococcus_dombrowskii | MUK72_RS16120 | 379890..382436 | 179637 | hypothetical-protein-[Halococcus-dombrowskii] |
| WP_020446108.1 | Type5.BREX | <-SF1-SF2-Helicase+REase\|\|?-><-SF1-SF2-Helicase+REase\|\|RHH->Phage_integrase-><-?<-DOC+HTH<-Phage_integrase\|\|BrxA-DNA-binding->DUF6079*->BrxB-BrxF-iSTAND->DUF6079-><-?\|\|N6-METHYLASE->PglZ->N6-METHYLASE->REase->TUDOR+SF1-SF2-Helicase+iREase+CTD-> | 1264 | Archaea>Euryarchaeota>Stenosarchaea_group>Halobacteria>Halobacteriales>Natronoarchaeaceae | GCF_000403645.1 | Salinarchaeum_sp._Harcht-Bsk1 | L593_RS06325 | 1375537..1379331 | 1333523 | hypothetical-protein-[Salinarchaeum-sp.-Harcht-Bsk1] |
| WP_020446110.1 | Type5.BREX | <-SF1-SF2-Helicase+REase\|\|RHH->Phage_integrase-><-?<-DOC+HTH<-Phage_integrase\|\|BrxA-DNA-binding->DUF6079->BrxB-BrxF-iSTAND->DUF6079*-><-?\|\|N6-METHYLASE->PglZ->N6-METHYLASE->REase->TUDOR+SF1-SF2-Helicase+iREase+CTD->?-><-REC+PAS+PAS+HISKIN | 1276 | Archaea>Euryarchaeota>Stenosarchaea_group>Halobacteria>Halobacteriales>Natronoarchaeaceae | GCF_000403645.1 | Salinarchaeum_sp._Harcht-Bsk1 | L593_RS06335 | 1379948..1383778 | 1333523 | hypothetical-protein-[Salinarchaeum-sp.-Harcht-Bsk1] |
| WP_114604755.1 | Type5.BREX | HTH+McrB-AAA->McrC-NTD+REase-><-TUDOR+SF1-SF2-Helicase+iREase+CTD<-N6-METHYLASE\|\|BrxA-DNA-binding->DUF6079*->BrxB-BrxF-iSTAND->DUF6079->PglZ-><-N6-METHYLASE+DAM<-N6-METHYLASE-p\|\|REase->N6-METHYLASE+DAM-><-N6-METHYLASE+DAM<-Calcineurin-SbcD | 1250 | Archaea>Euryarchaeota>Stenosarchaea_group>Halobacteria>Haloferacales>Haloferacaceae | GCF_003342675.1 | Haloplanus_rubicundus | DU484_RS00410 | 102411..106163 | 1547898 | hypothetical-protein-[Haloplanus-rubicundus] |
| WP_114604757.1 | Type5.BREX | HTH+McrB-AAA->McrC-NTD+REase-><-TUDOR+SF1-SF2-Helicase+iREase+CTD<-N6-METHYLASE\|\|BrxA-DNA-binding->DUF6079->BrxB-BrxF-iSTAND->DUF6079*->PglZ-><-N6-METHYLASE+DAM<-N6-METHYLASE-p\|\|REase->N6-METHYLASE+DAM-><-N6-METHYLASE+DAM<-Calcineurin-SbcD\|\|SF1-SF2-Helicase->REase-> | 1223 | Archaea>Euryarchaeota>Stenosarchaea_group>Halobacteria>Haloferacales>Haloferacaceae | GCF_003342675.1 | Haloplanus_rubicundus | DU484_RS00420 | 106791..110462 | 1547898 | hypothetical-protein-[Haloplanus-rubicundus] |
| WP_121920987.1 | Type5.BREX | PIN->?->?->?->?-><-?\|\|?->BrxA-DNA-binding->DUF6079*->BrxB-BrxF-iSTAND->DUF6079->N6-METHYLASE->PglZ->TUDOR+SF1-SF2-Helicase+iREase+CTD-><-tRNA<-DTXR-HTH | 1258 | Archaea>Euryarchaeota>Stenosarchaea_group>Halobacteria>Haloferacales>Haloferacaceae | GCF_003856835.1 | Haloplanus_aerogenes | DU502_RS05450 | 1062216..1065992 | 660522 | hypothetical-protein-[Haloplanus-aerogenes] |
| WP_121920985.1 | Type5.BREX | PIN->?->?->?->?-><-?\|\|?->BrxA-DNA-binding->DUF6079->BrxB-BrxF-iSTAND->DUF6079*->N6-METHYLASE->PglZ->TUDOR+SF1-SF2-Helicase+iREase+CTD-><-tRNA<-DTXR-HTH<-?<-?<-?<-ABC-ATPase+SbcC | 1264 | Archaea>Euryarchaeota>Stenosarchaea_group>Halobacteria>Haloferacales>Haloferacaceae | GCF_003856835.1 | Haloplanus_aerogenes | DU502_RS05460 | 1066576..1070370 | 660522 | hypothetical-protein-[Haloplanus-aerogenes] |
| WP_103992123.1 | Type5.BREX | <-Calcineurin<-HKD-DNase\|\|CARF+HTH->?-><-?<-?<-?\|\|ParB+HNH->BrxA-DNA-binding->DUF6079*->BrxB-BrxF-iSTAND->DUF6079-><-OLD-ABC+TOPRIM\|\|N6-METHYLASE-><-REase\|\|PglZ->REase+METHYLASE->TUDOR+SF1-SF2-Helicase+iREase+CTD-> | 1227 | Archaea>Euryarchaeota>Stenosarchaea_group>Halobacteria>Haloferacales>Haloferacaceae | GCF_004799685.1 | Halobellus_limi | DV707_RS08430 | 1701258..1704941 | 699433 | hypothetical-protein-[Halobellus-limi] |
| WP_103992125.1 | Type5.BREX | CARF+HTH->?-><-?<-?<-?\|\|ParB+HNH->BrxA-DNA-binding->DUF6079->BrxB-BrxF-iSTAND->DUF6079*-><-OLD-ABC+TOPRIM\|\|N6-METHYLASE-><-REase\|\|PglZ->REase+METHYLASE->TUDOR+SF1-SF2-Helicase+iREase+CTD-> | 1284 | Archaea>Euryarchaeota>Stenosarchaea_group>Halobacteria>Haloferacales>Haloferacaceae | GCF_004799685.1 | Halobellus_limi | DV707_RS08440 | 1705516..1709370 | 699433 | hypothetical-protein-[Halobellus-limi] |
| WP_191452134.1 | Type5.BREX | BrxA-DNA-binding->DUF6079*->BrxB-BrxF-iSTAND->DUF6079->N6-METHYLASE->N6-METHYLASE-p->PglZ->?->TUDOR+SF1-SF2-Helicase+iREase+CTD-><-BirA-HTH | 1239 | Archaea>Euryarchaeota>Stenosarchaea_group>Halobacteria>Haloferacales>Haloferacaceae | GCF_009741955.1 | Halogeometricum_sp._CBA1124 | GJ632_RS03790 | 1006..4725 | 2668071 | hypothetical-protein-[Halogeometricum-sp.-CBA1124] |
| WP_191452136.1 | Type5.BREX | BrxA-DNA-binding->DUF6079->BrxB-BrxF-iSTAND->DUF6079*->?->N6-METHYLASE->N6-METHYLASE-p->PglZ->?->TUDOR+SF1-SF2-Helicase+iREase+CTD-><-BirA-HTH | 1270 | Archaea>Euryarchaeota>Stenosarchaea_group>Halobacteria>Haloferacales>Haloferacaceae | GCF_009741955.1 | Halogeometricum_sp._CBA1124 | GJ632_RS03800 | 5309..9121 | 2668071 | hypothetical-protein-[Halogeometricum-sp.-CBA1124] |
| WP_188786134.1 | Type5.BREX | <-ABC-ATPase+SbcC<-?<-?\|\|tRNA->tRNA-><-TUDOR+SF1-SF2-Helicase+iREase+CTD<-PglZ<-N6-METHYLASE<-PIN<-?<-DUF6079*<-BrxB-BrxF-iSTAND<-DUF6079<-BrxA-DNA-binding<-Phage_integrase<-REase<-PAS\|\|?-><-DSBH | 1265 | Archaea>Euryarchaeota>Stenosarchaea_group>Halobacteria>Haloferacales>Haloferacaceae | GCF_014646875.1 | Halobellus_salinus | IEX72_RS04225 | 173567..177364 | 931585 | hypothetical-protein-[Halobellus-salinus] |
| WP_188786136.1 | Type5.BREX | <-TUDOR+SF1-SF2-Helicase+iREase+CTD<-PglZ<-N6-METHYLASE<-PIN<-?<-DUF6079<-BrxB-BrxF-iSTAND<-DUF6079*<-BrxA-DNA-binding<-Phage_integrase<-REase<-PAS\|\|?-><-DSBH<-?<-Sulfatase+Phosphodiest+DUF4976<-NUDIX+DUF429 | 1239 | Archaea>Euryarchaeota>Stenosarchaea_group>Halobacteria>Haloferacales>Haloferacaceae | GCF_014646875.1 | Halobellus_salinus | IEX72_RS04235 | 177948..181667 | 931585 | hypothetical-protein-[Halobellus-salinus] |
| WP_251330659.1 | Type5.BREX | <-TUDOR+SF1-SF2-Helicase+iREase+CTD<-REase<-PglZ<-N6-METHYLASE\|\|?-><-?<-DUF6079*<-BrxB-BrxF-iSTAND<-DUF6079<-BrxA-DNA-binding<-?\|\|?-><-DnaB+DnaB | 1274 | Archaea>Euryarchaeota>Stenosarchaea_group>Halobacteria>Haloferacales>Haloferacaceae | GCF_023703795.1 | Haloplanus_sp._HW8-1 | NBT82_RS06060 | 1119588..1123412 | 2949995 | hypothetical-protein-[Haloplanus-sp.-HW8-1] |
| WP_251330661.1 | Type5.BREX | <-TUDOR+SF1-SF2-Helicase+iREase+CTD<-REase<-PglZ<-N6-METHYLASE\|\|?-><-?<-DUF6079<-BrxB-BrxF-iSTAND<-DUF6079*<-BrxA-DNA-binding<-?\|\|?-><-DnaB+DnaB\|\|?-><-?<-?\|\|?->TrkH-> | 1248 | Archaea>Euryarchaeota>Stenosarchaea_group>Halobacteria>Haloferacales>Haloferacaceae | GCF_023703795.1 | Haloplanus_sp._HW8-1 | NBT82_RS06070 | 1124026..1127772 | 2949995 | hypothetical-protein-[Haloplanus-sp.-HW8-1] |
| WP_256309267.1 | Type5.BREX | <-DDE_Tnp_1<-?<-REase<-?<-MetJarc-HTH\|\|?-><-SF1-SF2-Helicase<-TBP+TBP<-PglZ<-DUF6079*<-BrxB-BrxF-iSTAND<-DUF6079<-BrxA-DNA-binding\|\|N6-METHYLASE->?->TUDOR+SF1-SF2-Helicase+iREase+CTD-> | 1231 | Archaea>Euryarchaeota>Stenosarchaea_group>Halobacteria>Haloferacales>Haloferacaceae | GCF_024464595.1 | Halobellus_litoreus | N0360_RS18265 | 57150..60845 | 755310 | hypothetical-protein-[Halobellus-litoreus] |
| WP_256309268.1 | Type5.BREX | <-REase<-?<-MetJarc-HTH\|\|?-><-SF1-SF2-Helicase<-TBP+TBP<-PglZ<-DUF6079<-BrxB-BrxF-iSTAND<-DUF6079*<-BrxA-DNA-binding\|\|N6-METHYLASE->?->TUDOR+SF1-SF2-Helicase+iREase+CTD-> | 1233 | Archaea>Euryarchaeota>Stenosarchaea_group>Halobacteria>Haloferacales>Haloferacaceae | GCF_024464595.1 | Halobellus_litoreus | N0360_RS18275 | 61461..65162 | 755310 | hypothetical-protein-[Halobellus-litoreus] |
| WP_089872534.1 | Type5.BREX | PADR-HTH->DOC+HTH-><-?<-REase\|\|?-><-TUDOR+SF1-SF2-Helicase+iREase+CTD<-N6-METHYLASE<-TBP+TBP\|\|BrxA-DNA-binding->DUF6079*->BrxB-BrxF-iSTAND->DUF6079->PglZ-><-REase\|\|ORC-CDC6->SMODS-> | 1233 | Archaea>Euryarchaeota>Stenosarchaea_group>Halobacteria>Haloferacales>Haloferacaceae | GCF_900114455.1 | Halogranum_rubrum | BM083_RS21560 | 31014..34715 | 553466 | hypothetical-protein-[Halogranum-rubrum] |
| WP_089872538.1 | Type5.BREX | <-REase\|\|?-><-TUDOR+SF1-SF2-Helicase+iREase+CTD<-N6-METHYLASE<-TBP+TBP\|\|BrxA-DNA-binding->DUF6079->BrxB-BrxF-iSTAND->DUF6079*->PglZ-><-REase\|\|ORC-CDC6->SMODS-> | 1230 | Archaea>Euryarchaeota>Stenosarchaea_group>Halobacteria>Haloferacales>Haloferacaceae | GCF_900114455.1 | Halogranum_rubrum | BM083_RS21570 | 35331..39023 | 553466 | hypothetical-protein-[Halogranum-rubrum] |
| TQQ78814.1 | Type5.BREX | <-Polbeta<-HTH_20<-?<-ParA-Soj-PloopNTPase<-SF1-SF2-Helicase+REase<-Exonuc_V_gamma+REase<-PglZ<-DUF6079*<-BrxB-BrxF-iSTAND<-DUF6079<-BrxA-DNA-binding\|\|N6-METHYLASE->TUDOR+SF1-SF2-Helicase+iREase+CTD->ORC-CDC6->?-><-REase | 1233 | Archaea>Euryarchaeota>Stenosarchaea_group>Halobacteria>Haloferacales>Halorubraceae | GCA_006861665.1 | Halonotius_terrestris | EGH24_12695 | 42218..45919 | 2487750 | hypothetical-protein-EGH24_12695-[Halonotius-terrestris] |
| WP_004047662.1 | Type5.BREX | Peptidase_M29-><-?<-BirA-HTH\|\|?-><-?<-ParA-Soj-PloopNTPase<-?\|\|?->BrxA-DNA-binding->DUF6079*->BrxB-BrxF-iSTAND->DUF6079->?->HEPN->N6-METHYLASE->PglZ->N6-METHYLASE->TUDOR+SF1-SF2-Helicase+iREase+CTD-> | 1249 | Archaea>Euryarchaeota>Stenosarchaea_group>Halobacteria>Haloferacales>Halorubraceae | GCF_000337915.1 | Halorubrum_saccharovorum | C471_RS07440 | 145399..149148 | 2248 | hypothetical-protein-[Halorubrum-saccharovorum] |
| WP_004047666.1 | Type5.BREX | <-BirA-HTH\|\|?-><-?<-ParA-Soj-PloopNTPase<-?\|\|?->BrxA-DNA-binding->DUF6079->BrxB-BrxF-iSTAND->DUF6079*->?->HEPN->N6-METHYLASE->PglZ->N6-METHYLASE->TUDOR+SF1-SF2-Helicase+iREase+CTD->?->?-><-DDE_Tnp_IS240 | 1276 | Archaea>Euryarchaeota>Stenosarchaea_group>Halobacteria>Haloferacales>Halorubraceae | GCF_000337915.1 | Halorubrum_saccharovorum | C471_RS07450 | 149762..153592 | 2248 | hypothetical-protein-[Halorubrum-saccharovorum] |
| WP_129452643.1 | Type5.BREX | <-GDResolvase\|\|?-><-DDE_Tnp_IS240<-TUDOR+SF1-SF2-Helicase+iREase+CTD<-REase+METHYLASE<-PglZ<-?<-N6-METHYLASE\|\|viral-protein-><-DUF6079*<-BrxB-BrxF-iSTAND<-DUF6079<-BrxA-DNA-binding<-ORC-CDC6<-zf-IS66+DDE_Tnp_IS66\|\|SbcC-><-DDE_Tnp_1\|\|OLD-ABC+TOPRIM-> | 1280 | Archaea>Euryarchaeota>Stenosarchaea_group>Halobacteria>Haloferacales>Halorubraceae | GCF_004126515.1 | Halorubrum_ezzemoulense | E0776_RS18050 | 513790..517632 | 337243 | hypothetical-protein-[Halorubrum-ezzemoulense] |
| WP_129452645.1 | Type5.BREX | <-DDE_Tnp_IS240<-TUDOR+SF1-SF2-Helicase+iREase+CTD<-REase+METHYLASE<-PglZ<-?<-N6-METHYLASE\|\|viral-protein-><-DUF6079<-BrxB-BrxF-iSTAND<-DUF6079*<-BrxA-DNA-binding<-ORC-CDC6<-zf-IS66+DDE_Tnp_IS66\|\|SbcC-><-DDE_Tnp_1\|\|OLD-ABC+TOPRIM->?->?->SbcC-> | 1227 | Archaea>Euryarchaeota>Stenosarchaea_group>Halobacteria>Haloferacales>Halorubraceae | GCF_004126515.1 | Halorubrum_ezzemoulense | E0776_RS18060 | 518207..521890 | 337243 | hypothetical-protein-[Halorubrum-ezzemoulense] |
| WP_136591377.1 | Type5.BREX | <-HTH_28\|\|ParA-Soj-PloopNTPase->?-><-Phage_integrase\|\|SF1-SF2-Helicase->SF1-SF2-Helicase+REase->SF1-SF2-Helicase+REase->BrxA-DNA-binding->DUF6079*->BrxB-BrxF-iSTAND->DUF6079-><-?\|\|N6-METHYLASE->?->PglZ->TUDOR+SF1-SF2-Helicase+iREase+CTD->?-><-OrfB_IS605+OrfB_Zn_ribbon | 1258 | Archaea>Euryarchaeota>Stenosarchaea_group>Halobacteria>Haloferacales>Halorubraceae | GCF_004916935.1 | Salinigranum_halophilum | E6C73_RS15450 | 221772..225548 | 2565931 | hypothetical-protein-[Salinigranum-halophilum] |
| WP_136591379.1 | Type5.BREX | ParA-Soj-PloopNTPase->?-><-Phage_integrase\|\|SF1-SF2-Helicase->SF1-SF2-Helicase+REase->SF1-SF2-Helicase+REase->BrxA-DNA-binding->DUF6079->BrxB-BrxF-iSTAND->DUF6079*-><-?\|\|N6-METHYLASE->?->PglZ->TUDOR+SF1-SF2-Helicase+iREase+CTD->?-><-OrfB_IS605+OrfB_Zn_ribbon | 1269 | Archaea>Euryarchaeota>Stenosarchaea_group>Halobacteria>Haloferacales>Halorubraceae | GCF_004916935.1 | Salinigranum_halophilum | E6C73_RS15460 | 226132..229941 | 2565931 | hypothetical-protein-[Salinigranum-halophilum] |
| WP_138006592.1 | Type5.BREX | <-TUDOR+SF1-SF2-Helicase+iREase+CTD<-PglZ<-N6-METHYLASE<-?<-HEPN<-Peptidase_M1<-DUF6079*<-BrxB-BrxF-iSTAND<-DUF6079<-BrxA-DNA-binding | 1272 | Archaea>Euryarchaeota>Stenosarchaea_group>Halobacteria>Haloferacales>Halorubraceae | GCF_005543295.1 | Halalkalirubrum_salinum | E5K03_RS09380 | 11807..15625 | 2563889 | hypothetical-protein-[Halalkalirubrum-salinum] |
| WP_138006594.1 | Type5.BREX | <-TUDOR+SF1-SF2-Helicase+iREase+CTD<-PglZ<-N6-METHYLASE<-?<-HEPN<-Peptidase_M1<-DUF6079<-BrxB-BrxF-iSTAND<-DUF6079*<-BrxA-DNA-binding | 1258 | Archaea>Euryarchaeota>Stenosarchaea_group>Halobacteria>Haloferacales>Halorubraceae | GCF_005543295.1 | Halalkalirubrum_salinum | E5K03_RS09390 | 16209..19985 | 2563889 | hypothetical-protein-[Halalkalirubrum-salinum] |
| WP_142980503.1 | Type5.BREX | <-Phage_int_SAM_5<-?<-Polbeta<-HTH_20<-?<-ParA-Soj-PloopNTPase<-SF1-SF2-Helicase+REase<-Exonuc_V_gamma+REase<-PglZ<-DUF6079*<-BrxB-BrxF-iSTAND<-DUF6079<-BrxA-DNA-binding\|\|N6-METHYLASE->TUDOR+SF1-SF2-Helicase+iREase+CTD->ORC-CDC6->?-><-REase | 1230 | Archaea>Euryarchaeota>Stenosarchaea_group>Halobacteria>Haloferacales>Halorubraceae | GCF_006861665.1 | Halonotius_terrestris | EGH24_RS12690 | 37910..41602 | 2487750 | hypothetical-protein-[Halonotius-terrestris] |
| WP_221886744.1 | Type5.BREX | <-Polbeta<-HTH_20<-?<-ParA-Soj-PloopNTPase<-SF1-SF2-Helicase+REase<-Exonuc_V_gamma+REase<-PglZ<-DUF6079<-BrxB-BrxF-iSTAND<-DUF6079*<-BrxA-DNA-binding\|\|N6-METHYLASE->TUDOR+SF1-SF2-Helicase+iREase+CTD->ORC-CDC6->?-><-REase | 1230 | Archaea>Euryarchaeota>Stenosarchaea_group>Halobacteria>Haloferacales>Halorubraceae | GCF_006861665.1 | Halonotius_terrestris | EGH24_RS12700 | 42218..45910 | 2487750 | hypothetical-protein-[Halonotius-terrestris] |
| WP_144927821.1 | Type5.BREX | <-REase<-TUDOR+SF1-SF2-Helicase+iREase+CTD<-N6-METHYLASE\|\|BrxA-DNA-binding->DUF6079*->BrxB-BrxF-iSTAND->DUF6079->PglZ-><-N6-METHYLASE+DAM\|\|REase->N6-METHYLASE+DAM-><-N6-METHYLASE+DAM<-McrC-NTD+REase | 1250 | Archaea>Euryarchaeota>Stenosarchaea_group>Halobacteria>Haloferacales>Halorubraceae | GCF_007671685.1 | Halorubrum_salsamenti | FGL94_RS17080 | 8993..12745 | 2583990 | hypothetical-protein-[Halorubrum-salsamenti] |
| WP_144927825.1 | Type5.BREX | <-REase<-TUDOR+SF1-SF2-Helicase+iREase+CTD<-N6-METHYLASE\|\|BrxA-DNA-binding->DUF6079->BrxB-BrxF-iSTAND->DUF6079*->PglZ-><-N6-METHYLASE+DAM\|\|REase->N6-METHYLASE+DAM-><-N6-METHYLASE+DAM<-McrC-NTD+REase | 1228 | Archaea>Euryarchaeota>Stenosarchaea_group>Halobacteria>Haloferacales>Halorubraceae | GCF_007671685.1 | Halorubrum_salsamenti | FGL94_RS17090 | 13373..17059 | 2583990 | hypothetical-protein-[Halorubrum-salsamenti] |
| WP_159369387.1 | Type5.BREX | <-TUDOR+SF1-SF2-Helicase+iREase+CTD<-REase+METHYLASE<-PglZ<-N6-METHYLASE\|\|?->?-><-REase<-DUF6079*<-BrxB-BrxF-iSTAND<-DUF6079<-BrxA-DNA-binding\|\|?-><-?<-?<-?\|\|ZDNA-HTH-> | 1277 | Archaea>Euryarchaeota>Stenosarchaea_group>Halobacteria>Haloferacales>Halorubraceae | GCF_009856205.1 | Halorubrum_terrestre | GLW36_RS12720 | 29017..32850 | 267368 | hypothetical-protein-[Halorubrum-terrestre] |
| WP_159369389.1 | Type5.BREX | <-TUDOR+SF1-SF2-Helicase+iREase+CTD<-REase+METHYLASE<-PglZ<-N6-METHYLASE\|\|?->?-><-REase<-DUF6079<-BrxB-BrxF-iSTAND<-DUF6079*<-BrxA-DNA-binding\|\|?-><-?<-?<-?\|\|ZDNA-HTH-> | 1227 | Archaea>Euryarchaeota>Stenosarchaea_group>Halobacteria>Haloferacales>Halorubraceae | GCF_009856205.1 | Halorubrum_terrestre | GLW36_RS12730 | 33425..37108 | 267368 | hypothetical-protein-[Halorubrum-terrestre] |
| WP_256418943.1 | Type5.BREX | <-ORC-CDC6+HTH\|\|?-><-TUDOR+SF1-SF2-Helicase+iREase+CTD<-PglZ<-?<-N6-METHYLASE\|\|PIN-><-DUF6079<-BrxB-BrxF-iSTAND<-DUF6079*<-BrxA-DNA-binding\|\|Exonuc_V_gamma+REase->SF1-SF2-Helicase+REase->ParA-Soj-PloopNTPase-> | 1248 | Archaea>Euryarchaeota>Stenosarchaea_group>Halobacteria>Haloferacales>Halorubraceae | GCF_024494655.1 | Halorubrum_laminariae | N0887_RS14550 | 91131..94877 | 1433523 | hypothetical-protein-[Halorubrum-laminariae] |
| WP_256419350.1 | Type5.BREX | <-HTH_3<-ZnR\|\|?-><-?\|\|?->BrxA-DNA-binding->DUF6079*->BrxB-BrxF-iSTAND->DUF6079-><-ORC-CDC6+TIP49-AAA+CDC6-HTH\|\|N6-METHYLASE->SbcC->PglZ-><-REase\|\|?->TUDOR+SF1-SF2-Helicase+iREase+CTD-> | 1226 | Archaea>Euryarchaeota>Stenosarchaea_group>Halobacteria>Haloferacales>Halorubraceae | GCF_024494655.1 | Halorubrum_laminariae | N0887_RS16900 | 14820..18500 | 1433523 | hypothetical-protein-[Halorubrum-laminariae] |
| AEN07515.1 | Type5.BREX | MazE_antitoxin->?-><-DDE_Tnp_1\|\|?->SF1-SF2-Helicase+REase->SF1-SF2-Helicase+REase->?->BrxA-DNA-binding->DUF6079*->BrxB-BrxF-iSTAND->DUF6079-><-PIN<-MazE_antitoxin\|\|N6-METHYLASE->?->PglZ->TUDOR+SF1-SF2-Helicase+iREase+CTD-> | 1239 | Archaea>Euryarchaeota>Stenosarchaea_group>Halobacteria>Haloferacales>unclassified_Haloferacales | GCA_000224475.1 | halophilic_archaeon_DL31 | Halar_0255 | 246678..250397 | 756883 | hypothetical-protein-Halar_0255-[halophilic-archaeon-DL31] |
| AEN07517.1 | Type5.BREX | <-DDE_Tnp_1\|\|?->SF1-SF2-Helicase+REase->SF1-SF2-Helicase+REase->?->BrxA-DNA-binding->DUF6079->BrxB-BrxF-iSTAND->DUF6079*-><-PIN<-MazE_antitoxin\|\|N6-METHYLASE->?->PglZ->TUDOR+SF1-SF2-Helicase+iREase+CTD-><-?<-SF1-SF2-Helicase | 1270 | Archaea>Euryarchaeota>Stenosarchaea_group>Halobacteria>Haloferacales>unclassified_Haloferacales | GCA_000224475.1 | halophilic_archaeon_DL31 | Halar_0257 | 250981..254793 | 756883 | hypothetical-protein-Halar_0257-[halophilic-archaeon-DL31] |
| ELY62517.1 | Type5.BREX | <-SF1-SF2-Helicase+REase<-SF1-SF2-Helicase+REase\|\|?->DDE_Tnp-><-N6-METHYLASE+DAM\|\|HTH->N6-METHYLASE+DAM->BrxA-DNA-binding-><-TUDOR+SF1-SF2-Helicase+iREase+CTD\|\|DUF6079*-><-?\|\|DUF6079->PglZ-><-?<-?<-?<-?\|\|DDE_Tnp_IS66->HTH_Tnp_1-> | 1276 | Archaea>Euryarchaeota>Stenosarchaea_group>Halobacteria>Natrialbales>Natrialbaceae | GCA_000337695.1 | Natronococcus_jeotgali_DSM_18795 | C492_07975 | 41020..44850 | 1227498 | hypothetical-protein-C492_07975-[Natronococcus-jeotgali-DSM-18795] |
| ELY62519.1 | Type5.BREX | DDE_Tnp-><-N6-METHYLASE+DAM\|\|HTH->N6-METHYLASE+DAM->BrxA-DNA-binding-><-TUDOR+SF1-SF2-Helicase+iREase+CTD\|\|DUF6079-><-?\|\|DUF6079*->PglZ-><-?<-?<-?<-?\|\|DDE_Tnp_IS66->HTH_Tnp_1-> | 1264 | Archaea>Euryarchaeota>Stenosarchaea_group>Halobacteria>Natrialbales>Natrialbaceae | GCA_000337695.1 | Natronococcus_jeotgali_DSM_18795 | C492_07985 | 45481..49275 | 1227498 | hypothetical-protein-C492_07985-[Natronococcus-jeotgali-DSM-18795] |
| WP_006182470.1 | Type5.BREX | <-ParA-Soj-PloopNTPase<-SF1-SF2-Helicase+REase<-Exonuc_V_gamma+REase\|\|BrxA-DNA-binding->DUF6079*->BrxB-BrxF-iSTAND->DUF6079->?->N6-METHYLASE->N6-METHYLASE-p->?->PglZ->TUDOR+SF1-SF2-Helicase+iREase+CTD-> | 1248 | Archaea>Euryarchaeota>Stenosarchaea_group>Halobacteria>Natrialbales>Natrialbaceae | GCF_000230735.2 | Natrinema_pellirubrum | NATPE_RS20815 | 248294..252040 | 69525 | hypothetical-protein-[Natrinema-pellirubrum] |
| WP_006182472.1 | Type5.BREX | <-ParA-Soj-PloopNTPase<-SF1-SF2-Helicase+REase<-Exonuc_V_gamma+REase\|\|BrxA-DNA-binding->DUF6079->BrxB-BrxF-iSTAND->DUF6079*->?->N6-METHYLASE->N6-METHYLASE-p->?->PglZ->TUDOR+SF1-SF2-Helicase+iREase+CTD-><-?<-DDE_Tnp_1 | 1277 | Archaea>Euryarchaeota>Stenosarchaea_group>Halobacteria>Natrialbales>Natrialbaceae | GCF_000230735.2 | Natrinema_pellirubrum | NATPE_RS20825 | 252654..256487 | 69525 | hypothetical-protein-[Natrinema-pellirubrum] |
| WP_014865882.1 | Type5.BREX | <-ParB+HNH\|\|?-><-ORC-CDC6\|\|?-><-TUDOR+SF1-SF2-Helicase+iREase+CTD<-PglZ<-N6-METHYLASE\|\|?-><-DUF6079*<-BrxB-BrxF-iSTAND<-DUF6079<-BrxA-DNA-binding<-?\|\|Exonuc_V_gamma+REase->SF1-SF2-Helicase+REase->ParA-Soj-PloopNTPase->?->Phage_int_SAM_5-> | 1280 | Archaea>Euryarchaeota>Stenosarchaea_group>Halobacteria>Natrialbales>Natrialbaceae | GCF_000281695.1 | Natrinema_sp._J7-2 | NJ7G_RS18310 | 64410..68252 | 406552 | hypothetical-protein-[Natrinema-sp.-J7-2] |
| WP_014865884.1 | Type5.BREX | <-ORC-CDC6\|\|?-><-TUDOR+SF1-SF2-Helicase+iREase+CTD<-PglZ<-N6-METHYLASE\|\|?-><-DUF6079<-BrxB-BrxF-iSTAND<-DUF6079*<-BrxA-DNA-binding<-?\|\|Exonuc_V_gamma+REase->SF1-SF2-Helicase+REase->ParA-Soj-PloopNTPase->?->Phage_int_SAM_5->HNH->REase-> | 1248 | Archaea>Euryarchaeota>Stenosarchaea_group>Halobacteria>Natrialbales>Natrialbaceae | GCF_000281695.1 | Natrinema_sp._J7-2 | NJ7G_RS18320 | 68866..72612 | 406552 | hypothetical-protein-[Natrinema-sp.-J7-2] |
| WP_006093018.1 | Type5.BREX | <-REase\|\|?-><-TUDOR+SF1-SF2-Helicase+iREase+CTD<-?<-N6-METHYLASE\|\|BrxA-DNA-binding->DUF6079*->BrxB-BrxF-iSTAND->DUF6079->PglZ->DOC+HTH-><-SF1-SF2-Helicase+REase\|\|Polbeta-><-?<-Polbeta | 1233 | Archaea>Euryarchaeota>Stenosarchaea_group>Halobacteria>Natrialbales>Natrialbaceae | GCF_000337235.1 | Natronorubrum_tibetense | NATTI_RS0124815 | 143759..147460 | 63128 | hypothetical-protein-[Natronorubrum-tibetense] |
| WP_006093016.1 | Type5.BREX | <-REase\|\|?-><-TUDOR+SF1-SF2-Helicase+iREase+CTD<-?<-N6-METHYLASE\|\|BrxA-DNA-binding->DUF6079->BrxB-BrxF-iSTAND->DUF6079*->PglZ->DOC+HTH-><-SF1-SF2-Helicase+REase<-SF1-SF2-Helicase+REase\|\|Polbeta-><-?<-Polbeta\|\|Phage_integrase-> | 1231 | Archaea>Euryarchaeota>Stenosarchaea_group>Halobacteria>Natrialbales>Natrialbaceae | GCF_000337235.1 | Natronorubrum_tibetense | NATTI_RS0124825 | 148076..151771 | 63128 | hypothetical-protein-[Natronorubrum-tibetense] |
| WP_049967097.1 | Type5.BREX | <-TUDOR+SF1-SF2-Helicase+iREase+CTD<-PglZ\|\|ZnR+REase->OLD-ABC+TOPRIM-><-N6-METHYLASE\|\|?-><-DUF6079*<-BrxB-BrxF-iSTAND<-DUF6079<-BrxA-DNA-binding\|\|ORC-CDC6-><-?\|\|?->?->?->35exo-><-?<-HTH_20 | 1227 | Archaea>Euryarchaeota>Stenosarchaea_group>Halobacteria>Natrialbales>Natrialbaceae | GCF_000710605.1 | Natrinema_thermotolerans | HL44_RS19275 | 148133..151816 | 121872 | hypothetical-protein-[Natrinema-thermotolerans] |
| WP_120246840.1 | Type5.BREX | ParB-><-?<-MORC+REase-p<-McrC-NTD+REase<-McrB-AAA\|\|?-><-?\|\|BrxA-DNA-binding->DUF6079*->BrxB-BrxF-iSTAND->DUF6079-><-TBP+TBP\|\|N6-METHYLASE-><-?<-ZnR+REase\|\|PglZ->TUDOR+SF1-SF2-Helicase+iREase+CTD-> | 1227 | Archaea>Euryarchaeota>Stenosarchaea_group>Halobacteria>Natrialbales>Natrialbaceae | GCF_003610195.1 | Halopiger_aswanensis | ATJ93_RS22195 | 27105..30788 | 148449 | hypothetical-protein-[Halopiger-aswanensis] |
| WP_120246842.1 | Type5.BREX | <-MORC+REase-p<-McrC-NTD+REase<-McrB-AAA\|\|?-><-?\|\|BrxA-DNA-binding->DUF6079->BrxB-BrxF-iSTAND->DUF6079*-><-TBP+TBP\|\|N6-METHYLASE-><-?<-ZnR+REase\|\|PglZ->TUDOR+SF1-SF2-Helicase+iREase+CTD->?-><-REase\|\|AAA-ATPase-p-> | 1280 | Archaea>Euryarchaeota>Stenosarchaea_group>Halobacteria>Natrialbales>Natrialbaceae | GCF_003610195.1 | Halopiger_aswanensis | ATJ93_RS22205 | 31363..35205 | 148449 | hypothetical-protein-[Halopiger-aswanensis] |
| WP_160047006.1 | Type5.BREX | <-TUDOR+SF1-SF2-Helicase+iREase+CTD<-N6-METHYLASE\|\|BrxA-DNA-binding->DUF6079*->BrxB-BrxF-iSTAND->DUF6079->PglZ->DOC+HTH->HTH_23+HTH_32+HTH_33->BirA-HTH->PIN-> | 1233 | Archaea>Euryarchaeota>Stenosarchaea_group>Halobacteria>Natrialbales>Natrialbaceae | GCF_009807105.1 | Natrialba_sp._INN-245 | GRS80_RS07395 | 8592..12293 | 2690967 | hypothetical-protein-[Natrialba-sp.-INN-245] |
| WP_160047007.1 | Type5.BREX | <-TUDOR+SF1-SF2-Helicase+iREase+CTD<-N6-METHYLASE\|\|BrxA-DNA-binding->DUF6079->BrxB-BrxF-iSTAND->DUF6079*->PglZ->DOC+HTH->HTH_23+HTH_32+HTH_33->BirA-HTH->PIN-> | 1232 | Archaea>Euryarchaeota>Stenosarchaea_group>Halobacteria>Natrialbales>Natrialbaceae | GCF_009807105.1 | Natrialba_sp._INN-245 | GRS80_RS07405 | 12909..16607 | 2690967 | hypothetical-protein-[Natrialba-sp.-INN-245] |
| WP_254526165.1 | Type5.BREX | ParA-Soj-PloopNTPase->?-><-TUDOR+SF1-SF2-Helicase+iREase+CTD<-PglZ<-N6-METHYLASE\|\|?-><-?<-REase+METHYLASE<-DUF6079*<-BrxB-BrxF-iSTAND<-DUF6079<-BrxA-DNA-binding\|\|Exonuc_V_gamma+REase->SF1-SF2-Helicase+REase-><-REase\|\|ORC-CDC6+HTH-><-?<-McrC-NTD+REase | 1270 | Archaea>Euryarchaeota>Stenosarchaea_group>Halobacteria>Natrialbales>Natrialbaceae | GCF_024227435.1 | Haloterrigena_sp._ZJ2 | NJT13_RS22765 | 13272..17084 | 2961570 | hypothetical-protein-[Natrinema-caseinilyticum] |
| WP_254526167.1 | Type5.BREX | <-TUDOR+SF1-SF2-Helicase+iREase+CTD<-PglZ<-N6-METHYLASE\|\|?-><-?<-REase+METHYLASE<-DUF6079<-BrxB-BrxF-iSTAND<-DUF6079*<-BrxA-DNA-binding\|\|Exonuc_V_gamma+REase->SF1-SF2-Helicase+REase-><-REase\|\|ORC-CDC6+HTH-><-?<-McrC-NTD+REase<-McrB-AAA<-Classical-AAA | 1251 | Archaea>Euryarchaeota>Stenosarchaea_group>Halobacteria>Natrialbales>Natrialbaceae | GCF_024227435.1 | Haloterrigena_sp._ZJ2 | NJT13_RS22775 | 17668..21423 | 2961570 | hypothetical-protein-[Natrinema-caseinilyticum] |
| WP_141198542.1 | Type5.BREX | STYKIN+CR-Kinase->SF1-SF2-Helicase->HTH+HEPN->McrB-AAA->REase-><-CR-ATPase<-TUDOR+SF1-SF2-Helicase+iREase+CTD<-PglZ<-N6-METHYLASE<-DUF6079*<-BrxB-BrxF-iSTAND<-DUF6079<-BrxA-DNA-binding\|\|?->?->?-><-?\|\|?-><-TIR | 1147 | Bacteria>delta/epsilon_subdivisions>Deltaproteobacteria>Bradymonadales>Bradymonadaceae>Persicimonas | GCF_006517175.1 | Persicimonas_caeni | FIV42_RS15335 | 4159002..4162445 | 2292766 | hypothetical-protein-[Persicimonas-caeni] |
| WP_103016193.1 | Type5.BREX | <-SbcC<-Calcineurin-SbcD<-SF1-SF2-Helicase<-OLD-ABC+TOPRIM<-TUDOR+SF1-SF2-Helicase+iREase+CTD<-PglZ<-SIR2<-N6-METHYLASE<-DUF6079*<-BrxB-BrxF-iSTAND<-DUF6079<-BrxA-DNA-binding<-REase+RNase<-viral-protein<-?<-Phage_holin_2_4<-RHH<-RHH | 1108 | Bacteria>FCB_group>Bacteroidetes/Chlorobi_group>Bacteroidota>Bacteroidetes_incertae_sedis>Rhodothermaceae | GCF_002894575.1 | Salinibacter_ruber | B7R92_RS05385 | 33133..36459 | 146919 | hypothetical-protein-[Salinibacter-ruber] |
| WP_146031917.1 | Type5.BREX | <-Calcineurin-SbcD<-SF1-SF2-Helicase<-OLD-ABC+TOPRIM<-TUDOR+SF1-SF2-Helicase+iREase+CTD<-PglZ<-SIR2<-N6-METHYLASE<-DUF6079<-BrxB-BrxF-iSTAND<-DUF6079*<-BrxA-DNA-binding<-REase+RNase<-viral-protein<-?<-Phage_holin_2_4<-RHH<-RHH<-TraJ-RHH\|\|HTH_3+Peptidase_S24-> | 1245 | Bacteria>FCB_group>Bacteroidetes/Chlorobi_group>Bacteroidota>Bacteroidetes_incertae_sedis>Rhodothermaceae | GCF_002894575.1 | Salinibacter_ruber | B7R92_RS05395 | 37102..40839 | 146919 | hypothetical-protein-[Salinibacter-ruber] |
| WP_259165690.1 | Type5.BREX | <-SbcC<-Calcineurin-SbcD<-OLD-ABC+TOPRIM<-TUDOR+SF1-SF2-Helicase+iREase+CTD<-PglZ<-N6-METHYLASE<-DUF6079*<-BrxB-BrxF-iSTAND<-DUF6079*<-BrxA-DNA-binding<-REase+RNase<-viral-protein<-?<-ZnR | 1245 | Bacteria>FCB_group>Bacteroidetes/Chlorobi_group>Bacteroidota>Bacteroidetes_incertae_sedis>Rhodothermaceae | GCF_024806515.1 | Salinibacter_ruber | GGP90_RS03930 | 969591..973328 | 146919 | hypothetical-protein-[Salinibacter-ruber] |
| WP_259182568.1 | Type5.BREX | <-PIN\|\|METHYLASE+SF1-SF2-Helicase->ArdCN+MPTase->RHH->CMP-hydrolase->?->HKD-DNase+SF1-SF2-Helicase->REase+RNase->BrxA-DNA-binding->DUF6079*->BrxB-BrxF-iSTAND->DUF6079->N6-METHYLASE->PglZ->TUDOR+SF1-SF2-Helicase+iREase+CTD->OLD-ABC+TOPRIM->SF1-SF2-Helicase->Calcineurin-SbcD->SbcC-> | 1248 | Bacteria>FCB_group>Bacteroidetes/Chlorobi_group>Bacteroidota>Bacteroidetes_incertae_sedis>Rhodothermaceae | GCF_024806795.1 | Salinibacter_ruber | GGQ19_RS12625 | 57748..61494 | 146919 | hypothetical-protein-[Salinibacter-ruber] |

| Protein-ID | System | Gene-Neighbourhoods | Length | Taxonomy | Assembly-ID | Strain | Locus-Tag | Genomic-coordinates | TaxID | Fasta-define |
|---|---|---|---|---|---|---|---|---|---|---|
| WP_259125819.1 | Type5.BREX | <-SbcC<-Calcineurin-SbcD<-?<-TUDOR+SF1-SF2-Helicase+iREase+CTD<-?<-PglZ<-N6-METHYLASE<-DUF6079<-BrxB-BrxF-iSTAND<-DUF6079*<-BrxA-DNA-binding<-REase+RNase<-?\|\|?->HORMA-> | 1245 | Bacteria>FCB_group>Bacteroidetes/Chlorobi_group>Bacteroidota>Bacteroidetes_incertae_sedis>Rhodothermaceae | GCF_024806955.1 | Salinibacter_ruber | GGQ05_001713 | 83114..86851 | 146919 | hypothetical-protein-[Salinibacter-ruber] |
| WP_259220531.1 | Type5.BREX | [illegible] | 1245 | Bacteria>FCB_group>Bacteroidetes/Chlorobi_group>Bacteroidota>Bacteroidetes_incertae_sedis>Rhodothermaceae | GCF_024807065.1 | Salinibacter_ruber | GGP94_RS13020 | 49040..52777 | 146919 | hypothetical-protein-[Salinibacter-ruber] |
| WP_259253366.1 | Type5.BREX | [illegible] | 1245 | Bacteria>FCB_group>Bacteroidetes/Chlorobi_group>Bacteroidota>Bacteroidetes_incertae_sedis>Rhodothermaceae | GCF_024807415.1 | Salinibacter_ruber | GGP63_RS14335 | 56454..60191 | 146919 | hypothetical-protein-[Salinibacter-ruber] |
| WP_259129819.1 | Type5.BREX | [illegible] | 1245 | Bacteria>FCB_group>Bacteroidetes/Chlorobi_group>Bacteroidota>Bacteroidetes_incertae_sedis>Rhodothermaceae | GCF_024807455.1 | Salinibacter_ruber | GGQ04_RS13395 | 34758..38495 | 146919 | hypothetical-protein-[Salinibacter-ruber] |
| WP_259113946.1 | Type5.BREX | [illegible] | 1245 | Bacteria>FCB_group>Bacteroidetes/Chlorobi_group>Bacteroidota>Bacteroidetes_incertae_sedis>Rhodothermaceae | GCF_024807465.1 | Salinibacter_ruber | GGP53_RS11770 | 37254..40991 | 146919 | hypothetical-protein-[Salinibacter-ruber] |
| WP_259127211.1 | Type5.BREX | [illegible] | 1255 | Bacteria>FCB_group>Bacteroidetes/Chlorobi_group>Bacteroidota>Bacteroidetes_incertae_sedis>Rhodothermaceae | GCF_024807525.1 | Salinibacter_ruber | GGP44_RS04645 | 1121352..1125119 | 146919 | hypothetical-protein-[Salinibacter-ruber] |
| WP_259123903.1 | Type5.BREX | <-HORMA<-?\|\|REase+RNase->BrxA-DNA-binding->DUF6079*->BrxB-BrxF-iSTAND->DUF6079->N6-METHYLASE->PglZ->?->TUDOR+SF1-SF2-Helicase+iREase+CTD->?->Calcineurin-SbcD->SbcC-> | 1245 | Bacteria>FCB_group>Bacteroidetes/Chlorobi_group>Bacteroidota>Bacteroidetes_incertae_sedis>Rhodothermaceae | GCF_024808095.1 | Salinibacter_ruber | GGP68_RS06295 | 37891..41628 | 146919 | hypothetical-protein-[Salinibacter-ruber] |
| WP_259086338.1 | Type5.BREX | REase+RNase->BrxA-DNA-binding->DUF6079*->BrxB-BrxF-iSTAND->DUF6079->N6-METHYLASE->PglZ->?->TUDOR+SF1-SF2-Helicase+iREase+CTD->OLD-ABC+TOPRIM->Calcineurin-SbcD->SbcC-> | 1245 | Bacteria>FCB_group>Bacteroidetes/Chlorobi_group>Bacteroidota>Bacteroidetes_incertae_sedis>Rhodothermaceae | GCF_024808115.1 | Salinibacter_ruber | GGP80_RS14460 | 1984..5721 | 146919 | hypothetical-protein-[Salinibacter-ruber] |
| WP_259107773.1 | Type5.BREX | [illegible] | 1245 | Bacteria>FCB_group>Bacteroidetes/Chlorobi_group>Bacteroidota>Bacteroidetes_incertae_sedis>Rhodothermaceae | GCF_024808295.1 | Salinibacter_ruber | GGP92_RS13390 | 54298..58035 | 146919 | hypothetical-protein-[Salinibacter-ruber] |
| WP_259048220.1 | Type5.BREX | [illegible] | 1245 | Bacteria>FCB_group>Bacteroidetes/Chlorobi_group>Bacteroidota>Bacteroidetes_incertae_sedis>Rhodothermaceae | GCF_024808395.1 | Salinibacter_ruber | GGP52_RS13845 | 3637..7374 | 146919 | hypothetical-protein-[Salinibacter-ruber] |
| WP_259120983.1 | Type5.BREX | [illegible] | 1245 | Bacteria>FCB_group>Bacteroidetes/Chlorobi_group>Bacteroidota>Bacteroidetes_incertae_sedis>Rhodothermaceae | GCF_024808535.1 | Salinibacter_ruber | GGP96_RS14190 | 56803..60540 | 146919 | hypothetical-protein-[Salinibacter-ruber] |
| WP_259122105.1 | Type5.BREX | [illegible] | 1245 | Bacteria>FCB_group>Bacteroidetes/Chlorobi_group>Bacteroidota>Bacteroidetes_incertae_sedis>Rhodothermaceae | GCF_024809285.1 | Salinibacter_ruber | GGQ10_RS14855 | 72088..75825 | 146919 | hypothetical-protein-[Salinibacter-ruber] |
| WP_259134094.1 | Type5.BREX | [illegible] | 1245 | Bacteria>FCB_group>Bacteroidetes/Chlorobi_group>Bacteroidota>Bacteroidetes_incertae_sedis>Rhodothermaceae | GCF_024810035.1 | Salinibacter_ruber | GGP48_RS11375 | 70595..74332 | 146919 | hypothetical-protein-[Salinibacter-ruber] |
| WP_251951637.1 | Type5.BREX | [illegible] | 1108 | Bacteria>FCB_group>Bacteroidetes/Chlorobi_group>Bacteroidota>Bacteroidetes_incertae_sedis>Rhodothermaceae | GCF_943908255.1 | Salinibacter_ruber | NGL47_RS11295 | 22718..26044 | 146919 | hypothetical-protein-[Salinibacter-ruber] |
| WP_251951639.1 | Type5.BREX | [illegible] | 1245 | Bacteria>FCB_group>Bacteroidetes/Chlorobi_group>Bacteroidota>Bacteroidetes_incertae_sedis>Rhodothermaceae | GCF_943908255.1 | Salinibacter_ruber | NGL47_RS11305 | 26687..30424 | 146919 | hypothetical-protein-[Salinibacter-ruber] |
| -- | | | | | | | | | | |
| --Type-5-BREX | | | | | | | | | | |
| --No-identified-Duplication | | | | | | | | | | |
| --Canonical-BREX | | | | | | | | | | |
| -- | | | | | | | | | | |
| WP_254821928.1 | Type5.BREX | HKD-DNase->?->MORC+REase-><-TUDOR+SF1-SF2-Helicase+iREase+CTD<-N6-METHYLASE\|\|BrxA-DNA-binding->BrxB-BrxF-iSTAND->DUF6079*->PglZ-><-REase\|\|CARF->PIN-> | 1231 | Archaea>Euryarchaeota>Stenosarchaea_group>Halobacteria>Halobacteriales>Haloarculaceae | GCF_024298885.1 | Haloglomus_sp._ZY41 | NL114_RS16070 | 3304298..3307993 | 2962672 | [illegible] |
| WP_158204981.1 | Type5.BREX | BrxB-BrxF-iSTAND->DUF6079*->N6-METHYLASE->PglZ->REase->TUDOR+SF1-SF2-Helicase+iREase+CTD-> | 1269 | Archaea>Euryarchaeota>Stenosarchaea_group>Halobacteria>Halobacteriales>Halobacteriaceae | GCF_009791395.1 | Halomarina_oriensis | GQS65_RS12490 | 799..4608 | 671145 | [illegible] |
| WP_157746008.1 | Type5.BREX | [illegible] | 1280 | Archaea>Euryarchaeota>Stenosarchaea_group>Halobacteria>Haloferacales>Halorubraceae | GCF_002355655.1 | Halorubrum_trapanicum | CPZ01_RS15580 | 1415231..3842 | 29284 | [illegible] |
| WP_049967095.1 | Type5.BREX | [illegible] | 1277 | Archaea>Euryarchaeota>Stenosarchaea_group>Halobacteria>Natrialbales>Natrialbaceae | GCF_000710605.1 | Natrinema_thermotolerans | DVR14_24745 | 889875..893708 | 121872 | [illegible] |
| WP_048047670.1 | Type5.BREX | [illegible] | 1212 | Archaea>Euryarchaeota>Stenosarchaea_group>Methanomicrobia>Methanosarcinales>Methanosarcinaceae | GCF_000979995.1 | Methanosarcina_mazei | DU74_RS10165 | 54975..58613 | 2209 | [illegible] |
| WP_233675470.1 | Type5.BREX | [illegible] | 1173 | Bacteria>Pseudomonadota>Alphaproteobacteria>Rhodobacterales>Paracoccaceae>Sinirhodobacter | - | Sinirhodobacter_sp._WL0062 | LZA78_02955 | 53668..57189 | 2907145 | [illegible] |
| OYZ13690.1 | Type5.BREX | [illegible] | 1197 | Bacteria>Pseudomonadota>Alphaproteobacteria>Sphingomonadales>unclassified_Sphingomonadales | GCA_002281615.1 | Sphingomonadales_bacterium_28-64-96 | B7Y35_07475 | 14573..18166 | 1970591 | [illegible] |
| MBX3681311.1 | Type5.BREX | [illegible] | 1203 | Bacteria>Pseudomonadota>Betaproteobacteria>Rhodocyclales>Zoogloeaceae>Thauera | GCA_019635105.1 | Thauera_sp. | KF731_01205 | 3180..6791 | 1905334 | [illegible] |
| WP_012584348.1 | Type5.BREX | [illegible] | 1203 | Bacteria>Pseudomonadota>Betaproteobacteria>Rhodocyclales>Zoogloeaceae>Thauera | GCF_000021765.1 | Thauera_sp._MZ1T | TMZ1T_RS01390 | 330563..334174 | 85643 | [illegible] |
| WP_004357336.1 | Type5.BREX | [illegible] | 1203 | Bacteria>Pseudomonadota>Betaproteobacteria>Rhodocyclales>Zoogloeaceae>Thauera | GCF_000310225.1 | Thauera_phenylacetica | C667_RS03740 | 10784..14395 | 164400 | [illegible] |
| WP_247733862.1 | Type5.BREX | [illegible] | 1203 | Bacteria>Pseudomonadota>Betaproteobacteria>Rhodocyclales>Zoogloeaceae>Thauera | GCF_023093695.1 | Thauera_aromatica | MX652_RS11970 | 18297..21908 | 59405 | [illegible] |
| WP_139442435.1 | Type5.BREX | [illegible] | 1231 | Bacteria>Pseudomonadota>Gammaproteobacteria>Aeromonadales>Aeromonadaceae>Aeromonas | GCF_006246325.1 | Aeromonas_veronii | CF108_RS03265 | 124178..127873 | 654 | [illegible] |
| WP_007349691.1 | Type5.BREX | [illegible] | 1194 | Bacteria>Pseudomonadota>Gammaproteobacteria>Pseudomonadales>Marinobacteraceae>Marinobacter | GCF_000169375.1 | Marinobacter_sp._ELB17 | MELB17_RS08075 | 35418..39002 | 270374 | [illegible] |
| MQT58298.1 | Type5.BREX | [illegible] | 1223 | Bacteria>Pseudomonadota>Gammaproteobacteria>Pseudomonadales>Pseudomonadaceae>Pseudomonas | GCA_009600165.1 | Pseudomonas_sp._FSL_R10-0399 | GHO41_13210 | 14571..18242 | 2662194 | [illegible] |
| MBJ2206056.1 | Type5.BREX | [illegible] | 1223 | Bacteria>Pseudomonadota>Gammaproteobacteria>Pseudomonadales>Pseudomonadaceae>Pseudomonas | GCA_016405205.1 | Pseudomonas_carnis | JFT33_05595 | 489982..493653 | 2487355 | [illegible] |
| MBJ2272566.1 | Type5.BREX | [illegible] | 1224 | [illegible] | GCA_016405625.1 | Pseudomonas_haemolytica | JFT41_06185 | 268773..272447 | 2600065 | [illegible] |
| MCF5056337.1 | Type5.BREX | [illegible] | 1223 | [illegible] | GCA_021605255.1 | Pseudomonas_proteolytica | GIW75_05045 | 72064..75735 | 219574 | [illegible] |
| VVQ23891.1 | Type5.BREX | [illegible] | 1225 | [illegible] | GCA_902498245.1 | Pseudomonas_fluorescens | PS928_05643 | 91300..94977 | 294 | [illegible] |
| WP_138763800.1 | Type5.BREX | [illegible] | 1223 | [illegible] | GCF_005930675.1 | Pseudomonas_lactis | FHL14_RS16420 | 47911..51582 | 1615674 | [illegible] |
| WP_198848263.1 | Type5.BREX | [illegible] | 1223 | [illegible] | GCF_029960145.1 | Pseudomonas_sp._GBPI_506 | NIM88_RS17440 | 41945..45616 | 1735795 | [illegible] |
| WP_150787878.1 | Type5.BREX | [illegible] | 1222 | [illegible] | GCF_902498245.1 | Pseudomonas_fluorescens | F8N86_RS29045 | 91300..94968 | 294 | [illegible] |
| WP_236299165.1 | Type5.BREX | [illegible] | 1223 | [illegible] | - | Pseudomonas_proteolytica | GIW75_05045 | 72064..75735 | 219574 | [illegible] |
| WP_198809634.1 | Type5.BREX | [illegible] | 1224 | [illegible] | - | Pseudomonas_shahriarae | NEE03_RS00560 | 109508..113182 | 2745512 | [illegible] |
| WP_096305156.1 | Type5.BREX | [illegible] | 1161 | [illegible] | GCF_900230215.1 | Jatrophihabitans_sp._GAS493 | CPH63_RS13795 | 2948299..2951784 | 1907575 | [illegible] |
| WP_104138811.1 | Type5.BREX | [illegible] | 1173 | [illegible] | GCF_002929275.1 | Arthrobacter_sp._ZGTC131 | CXQ90_RS09900 | 131121..134642 | 2058898 | [illegible] |
| WP_235519134.1 | Type5.BREX | [illegible] | 1162 | [illegible] | GCF_021538675.1 | Paenarthrobacter_sp._AR_02 | LRQ04_RS07575 | 297017..300505 | 2899821 | [illegible] |
| WP_036372476.1 | Type5.BREX | [illegible] | 1152 | [illegible] | GCF_000612725.1 | Mycolicibacterium_austroafricanum | BN976_RS17355 | 59223..62681 | 39687 | [illegible] |
| WP_036470026.1 | Type5.BREX | [illegible] | 1155 | [illegible] | GCF_000689255.1 | Mycobacterium_triplex | BN973_RS18575 | 3994507..3997974 | 47839 | [illegible] |
| WP_265123865.1 | Type5.BREX | [illegible] | 1159 | [illegible] | GCF_026041215.1 | Hoyosella_sp._YIM_151337 | ONR57_RS12750 | 17833..21312 | 2992742 | [illegible] |
| OUS97312.1 | Type5.BREX | [illegible] | 1175 | [illegible] | GCA_002165735.1 | Rhodococcus_sp._NCIMB_12038 | CA951_02905 | 126594..130121 | 933800 | [illegible] |
| WP_256973250.1 | Type5.BREX | [illegible] | 1213 | [illegible] | GCF_002165735.1 | Rhodococcus_sp._NCIMB_12038 | CA951_RS02920 | 126480..130121 | 933800 | [illegible] |
| WP_153653111.1 | Type5.BREX | [illegible] | 1160 | [illegible] | GCF_009649075.1 | Aeromicrobium_yanjiei | GEV26_RS10980 | 2228820..2232302 | 2662028 | [illegible] |
| WP_154708534.1 | Type5.BREX | [illegible] | 1187 | [illegible] | - | Cutibacterium_acnes | CP883_05580 | 10794..14357 | 1747 | [illegible] |
| REB77450.1 | Type5.BREX | [illegible] | 1187 | [illegible] | GCA_003384685.1 | Cutibacterium_acnes | CP883_05580 | 10794..14357 | 1747 | [illegible] |
| WP_029421470.1 | Type5.BREX | [illegible] | 1205 | [illegible] | GCF_000702485.1 | Alicyclobacillus_macrosporangiidus | N687_RS0108610 | 1729200..1732817 | 392015 | [illegible] |
| WP_002116626.1 | Type5.BREX | [illegible] | 1188 | [illegible] | - | Bacillus_cereus | IEQ_RS03950 | 791262..794828 | 1396 | [illegible] |
| OHR66989.1 | Type5.BREX | [illegible] | 1195 | [illegible] | GCA_001838545.1 | Bacillus_sp._HMSC76G11 | HMPREF3291_01045 | 70062..73649 | 1608910 | [illegible] |
| ASF41331.1 | Type5.BREX | [illegible] | 1188 | [illegible] | GCA_002209425.1 | Halobacillus_halophilus | CEH05_20035 | 4059294..4062860 | 1570 | [illegible] |
| ASS85704.1 | Type5.BREX | [illegible] | 1183 | [illegible] | GCA_002243605.1 | Geobacillus_lituanicus | GLN3_00240 | 44558..48109 | 169283 | [illegible] |
| PFG07777.1 | Type5.BREX | [illegible] | 1195 | [illegible] | GCA_002563655.1 | Bacillus_sp._es.034 | ATG71_4686 | 4473829..4477416 | 1761763 | [illegible] |
| PGR68857.1 | Type5.BREX | [illegible] | 1191 | [illegible] | GCA_002576445.1 | Bacillus_cereus | COC49_05330 | 39030..42605 | 1396 | [illegible] |
| PRT22691.1 | Type5.BREX | [illegible] | 1188 | [illegible] | GCA_002998815.1 | Bacillus_thuringiensis | C6351_31305 | 13182..16748 | 1428 | [illegible] |
| MBU9710538.1 | Type5.BREX | [illegible] | 1188 | [illegible] | GCA_019042215.1 | Evansella_tamaricis | KS419_02125 | 46685..50251 | 2069301 | [illegible] |
| WP_019720353.1 | Type5.BREX | [illegible] | 1187 | [illegible] | GCF_000223155.1 | Weizmannia_coagulans | IOE_RS0101945 | 1303..4866 | 1398 | [illegible] |
| WP_003250543.1 | Type5.BREX | [illegible] | 1194 | [illegible] | GCF_000258725.1 | Parageobacillus_thermoglucosidasius | GT20_RS09290 | 1817833..1821417 | 1426 | [illegible] |
| WP_223254242.1 | Type5.BREX | [illegible] | 1203 | [illegible] | GCF_000284515.1 | Halobacillus_halophilus | HBHAL_RS19750 | 4059450..4063061 | 1570 | [illegible] |
| WP_026590665.1 | Type5.BREX | [illegible] | 1185 | [illegible] | GCF_000482345.1 | Bacillus_sp._UNC437CL72CviS29 | MO14_RS0104560 | 312641..316198 | 1340430 | [illegible] |
| WP_026908399.1 | Type5.BREX | [illegible] | 1187 | [illegible] | GCF_000482485.1 | Paucisalibacillus_globulus | K332_RS0116125 | 154598..158161 | 351095 | [illegible] |
| WP_042385359.1 | Type5.BREX | [illegible] | 1194 | [illegible] | GCF_000648295.1 | Parageobacillus_thermoglucosidasius | GT2_RS14370 | 66511..70095 | 1426 | [illegible] |
| WP_048718293.1 | Type5.BREX | [illegible] | 1187 | [illegible] | GCF_001076885.1 | Bacillus_sp._522_BSPC | ADJ83_RS15690 | 17567..21130 | 1579338 | [illegible] |
| WP_061570112.1 | Type5.BREX | [illegible] | 1201 | [illegible] | GCF_001587535.1 | Caldibacillus_debilis | B4135_RS15850 | 22498..26103 | 301148 | [illegible] |
| WP_066159884.1 | Type5.BREX | [illegible] | 1205 | [illegible] | GCF_001591945.1 | Halalkalibacter_krulwichiae | BKR01S_RS20150 | 13948..17565 | 199441 | [illegible] |
| WP_077428558.1 | Type5.BREX | [illegible] | 1179 | [illegible] | GCF_001996285.1 | Anoxybacillus_karvacharensis | BO219_RS04435 | 413742..417281 | 1919277 | [illegible] |
| WP_047753295.1 | Type5.BREX | [illegible] | 1183 | [illegible] | GCF_002243665.1 | Geobacillus | GT3921_RS13000 | 2562536..2566087 | 129337 | [illegible] |
| WP_096225439.1 | Type5.BREX | [illegible] | 1183 | [illegible] | GCF_002335725.1 | Geobacillus_sp._FJAT-46040 | CD000_RS08375 | 1580958..1584509 | 2011017 | [illegible] |
| WP_098359294.1 | Type5.BREX | [illegible] | 1186 | [illegible] | GCF_002561295.1 | Bacillus_cereus | CN373_RS10800 | 80983..84543 | 1396 | [illegible] |
| WP_098613368.1 | Type5.BREX | [illegible] | 1188 | [illegible] | GCF_002567495.1 | Bacillus_cereus | COK05_RS16060 | 61384..64950 | 1396 | [illegible] |
| WP_098794042.1 | Type5.BREX | [illegible] | 1188 | [illegible] | GCF_002576235.1 | Priestia_megaterium | COA18_RS28020 | 1940977..1944543 | 1404 | [illegible] |
| WP_098878446.1 | Type5.BREX | [illegible] | 1191 | [illegible] | GCF_002576445.1 | Bacillus_cereus | COC49_RS05330 | 39030..42605 | 1396 | [illegible] |
| WP_100332651.1 | Type5.BREX | [illegible] | 1187 | [illegible] | GCF_002797355.1 | Bacillus_xiapuensis | CEF20_RS14915 | 671484..675047 | 2014075 | [illegible] |
| WP_066188676.1 | Type5.BREX | [illegible] | 1187 | [illegible] | GCF_002835735.1 | Cytobacillus_horneckiae | CWS20_RS08115 | 27127..30690 | 549687 | [illegible] |
| WP_106082566.1 | Type5.BREX | [illegible] | 1188 | [illegible] | GCF_002998815.1 | Bacillus_thuringiensis | C6351_31305 | 13182..16748 | 1428 | [illegible] |
| WP_117327463.1 | Type5.BREX | [illegible] | 1195 | [illegible] | GCF_003429095.1 | Peribacillus_saganii | D0469_RS14520 | 34367..37954 | 2303992 | [illegible] |
| WP_132948756.1 | Type5.BREX | [illegible] | 1183 | [illegible] | GCF_004341205.1 | Thermolongibacillus_altinsuensis | EDD69_RS09635 | 59871..63422 | 575256 | [illegible] |
| WP_136831108.1 | Type5.BREX | [illegible] | 1195 | [illegible] | GCF_005116465.1 | Robertmurraya_kyonggiensis | FA727_RS11510 | 51719..55306 | 1037680 | [illegible] |
| WP_144565381.1 | Type5.BREX | [illegible] | 1186 | [illegible] | GCF_007681185.1 | Bacillus_mobilis | FS794_RS23165 | 89317..92877 | 2026190 | [illegible] |
| WP_144543246.1 | Type5.BREX | [illegible] | 1192 | [illegible] | GCF_007830235.1 | Cytobacillus_oceanisediminis | IQ19_RS15315 | 231804..235382 | 665099 | [illegible] |
| WP_093855906.1 | Type5.BREX | [illegible] | 1187 | [illegible] | GCF_007989445.1 | Tenuibacillus_multivorans | BLP77_RS04470 | 353994..357557 | 237069 | [illegible] |
| WP_155112638.1 | Type5.BREX | [illegible] | 1186 | [illegible] | GCF_009711125.1 | Metabacillus_mangrovi | GKZ89_RS11910 | 46455..50015 | 1491830 | [illegible] |
| WP_167560822.1 | Type5.BREX | [illegible] | 1192 | [illegible] | GCF_012030395.1 | Bacillus_subtilis | HC742_RS20760 | 3449689..3453267 | 1423 | [illegible] |
| WP_184528635.1 | Type5.BREX | [illegible] | 1192 | [illegible] | GCF_014207535.1 | Bacillus_benzoevorans | HNR53_RS18175 | 67020..70598 | 1456 | [illegible] |
| WP_188383791.1 | Type5.BREX | [illegible] | 1187 | [illegible] | GCF_014637405.1 | Ornithinibacillus_halotolerans | IEV35_RS05945 | 32497..36060 | 1274357 | [illegible] |
| WP_188386543.1 | Type5.BREX | [illegible] | 1187 | [illegible] | GCF_014638355.1 | Priestia_taiwanensis | IEV41_RS00670 | 120045..123608 | 1347902 | [illegible] |
| WP_199714161.1 | Type5.BREX | [illegible] | 1191 | [illegible] | GCF_016483555.1 | Bacillus_cereus_group | JDS82_RS26190 | 13772..17347 | 86661 | [illegible] |
| WP_202778095.1 | Type5.BREX | [illegible] | 1192 | [illegible] | GCF_016772275.1 | Heyndrickxia_vini | I5776_RS19095 | 3841015..3844593 | 1476025 | [illegible] |
| WP_205168141.1 | Type5.BREX | [illegible] | 1189 | [illegible] | GCF_016908495.1 | Bacillus_pakistanensis | JOC86_RS02215 | 441948..445517 | 992288 | [illegible] |
| WP_212956143.1 | Type5.BREX | [illegible] | 1188 | [illegible] | GCF_018332475.1 | Bacillus | KI369_RS12465 | 38598..42164 | 1386 | [illegible] |
| WP_214723414.1 | Type5.BREX | [illegible] | 1189 | [illegible] | GCF_018613175.1 | Bacillus_sp._ISL-47 | J7I93_RS03705 | 56511..60080 | 2819130 | [illegible] |
| WP_215562746.1 | Type5.BREX | [illegible] | 1185 | [illegible] | GCF_018741585.1 | Bacillus_wiedmannii | EXW48_RS05020 | 1017548..1021105 | 1890302 | [illegible] |
| WP_221874921.1 | Type5.BREX | [illegible] | 1192 | [illegible] | GCF_019749255.1 | Mesobacillus_maritimus | H0185_RS18235 | 45712..49290 | 1643336 | [illegible] |
| WP_224936976.1 | Type5.BREX | [illegible] | 1187 | [illegible] | GCF_020171845.1 | Cytobacillus_kochii | LCM23_RS10855 | 44257..47820 | 859143 | [illegible] |
| WP_226516206.1 | Type5.BREX | [illegible] | 1187 | [illegible] | GCF_020519665.1 | Bacillus_shivajii | LGQ02_RS20940 | 4386157..4389720 | 1983719 | [illegible] |
| WP_229143299.1 | Type5.BREX | [illegible] | 1188 | [illegible] | GCF_020686245.1 | Bacillus_paranthracis | LKL84_RS22410 | 92907..96473 | 2026186 | [illegible] |
| WP_048558547.1 | Type5.BREX | [illegible] | 1185 | [illegible] | GCF_021655055.1 | Bacillus_cereus | L6497_RS04770 | 943584..947141 | 1396 | [illegible] |
| WP_200152411.1 | Type5.BREX | [illegible] | 1185 | [illegible] | GCF_021655215.1 | Bacillus_cereus | L6484_RS04895 | 983694..987251 | 1396 | [illegible] |
| WP_242261892.1 | Type5.BREX | [illegible] | 1188 | [illegible] | GCF_022662915.1 | Bacillus_cereus_group_sp._BfR-BA-01453 | MR070_RS20010 | 124720..128286 | 2920355 | [illegible] |
| WP_242225824.1 | Type5.BREX | [illegible] | 1185 | [illegible] | GCF_022663655.1 | Bacillus_cereus_group_sp._BfR-BA-01358 | MR078_RS05890 | 38308..41865 | 2920320 | [illegible] |
| WP_242320708.1 | Type5.BREX | [illegible] | 1186 | [illegible] | GCF_022663735.1 | Bacillus_cereus_group_sp._BfR-BA-01354 | MR162_RS10425 | 194779..198339 | 2920317 | [illegible] |
| WP_243571747.1 | Type5.BREX | [illegible] | 1192 | [illegible] | GCF_022814905.1 | Bacillus_stercoris | MSS26_RS04990 | 1003415..1006993 | 2054641 | [illegible] |
| WP_247084294.1 | Type5.BREX | [illegible] | 1184 | [illegible] | GCF_023022965.1 | Bacillus_sp._JY-23 | MUG87_RS19160 | 3790720..3794274 | 2933872 | [illegible] |
| WP_251625212.1 | Type5.BREX | [illegible] | 1192 | [illegible] | GCF_023712125.1 | Metabacillus_idriensis | M4D55_RS07610 | 46091..49669 | 324768 | [illegible] |
| WP_251646840.1 | Type5.BREX | [illegible] | 1195 | [illegible] | GCF_023714395.1 | Niallia_sp._MER_6 | M3600_17125 | 94088..97675 | 2939567 | [illegible] |
| WP_251339432.1 | Type5.BREX | [illegible] | 1192 | [illegible] | GCF_023714665.1 | Heyndrickxia_oleronia | M3589_RS18030 | 387742..391320 | 38875 | [illegible] |
| WP_252289705.1 | Type5.BREX | [illegible] | 1182 | [illegible] | GCF_023823935.1 | Peribacillus_asahii | LIS76_RS13265 | 2733994..2737542 | 228899 | [illegible] |
| WP_258922098.1 | Type5.BREX | [illegible] | 1187 | [illegible] | GCF_024758645.1 | Weizmannia_coagulans | L1879_RS05770 | 1225557..1229120 | 1398 | [illegible] |
| WP_260358530.1 | Type5.BREX | [illegible] | 1177 | [illegible] | GCF_025209795.1 | Peribacillus_frigoritolerans | NOU24_RS18785 | 747765..751298 | 450367 | [illegible] |
| WP_268298773.1 | Type5.BREX | [illegible] | 1192 | [illegible] | GCF_026789485.1 | Bacillus_inaquosorum | MOB75_RS13790 | 96491..100069 | 483913 | [illegible] |
| WP_269897765.1 | Type5.BREX | [illegible] | 1196 | [illegible] | GCF_027557635.1 | Lysinibacillus_sp._JK80 | HXV90_RS18610 | 3707833..3711423 | 2749809 | [illegible] |
| WP_231419114.1 | Type5.BREX | [illegible] | 1188 | [illegible] | GCF_030166655.1 | Pontibacillus_sp._HN14 | LQ226_RS16640 | 43379..46945 | 2898421 | [illegible] |
| WP_098147710.1 | Type5.BREX | [illegible] | 1186 | [illegible] | GCF_030676405.1 | Bacillus_wiedmannii | CN685_RS18690 | 6510..10070 | 1890302 | [illegible] |
| WP_073518919.1 | Type5.BREX | [illegible] | 1185 | [illegible] | GCF_900094815.1 | Bacillus_cereus_group | CCY53_RS04725 | 210220..213777 | 86661 | [illegible] |
| WP_099159038.1 | Type5.BREX | [illegible] | 1187 | [illegible] | GCF_900187325.1 | Virgibacillus_ndiopensis | CS937_RS13675 | 175761..179324 | 2004408 | [illegible] |
| WP_163530739.1 | Type5.BREX | [illegible] | 1188 | [illegible] | GCF_902726645.1 | Halobacillus_ihumii | G6R08_RS20435 | 4081488..4085054 | 2686092 | [illegible] |
| WP_026834104.1 | Type5.BREX | [illegible] | 1192 | [illegible] | GCF_000620825.1 | Exiguobacterium_undae | P399_RS0115220 | 2864676..2868254 | 169177 | [illegible] |
| WP_026826561.1 | Type5.BREX | [illegible] | 1191 | [illegible] | GCF_000620845.1 | Exiguobacterium_marinum | P400_RS0112700 | 2404557..2408132 | 273528 | [illegible] |
| WP_078145973.1 | Type5.BREX | [illegible] | 1187 | [illegible] | GCF_002009685.1 | Exiguobacterium_sp._HVEsp1 | BV910_RS00630 | 122796..126359 | 1934003 | [illegible] |
| WP_131508621.1 | Type5.BREX | [illegible] | 1200 | [illegible] | GCF_004337165.1 | Exiguobacterium_sp._SH0S1 | EVJ20_RS13610 | 61897..65499 | 2510949 | [illegible] |
| WP_089525998.1 | Type5.BREX | [illegible] | 1208 | [illegible] | GCF_002233675.1 | Paenibacillus_herberti | CGZ75_RS19630 | 18692..22318 | 1619309 | [illegible] |
| WP_138881991.1 | Type5.BREX | [illegible] | 1205 | [illegible] | GCF_005938385.1 | Paenibacillus_mesophilus | FE783_RS24340 | 175558..179175 | 2582849 | [illegible] |
| WP_272441147.1 | Type5.BREX | [illegible] | 1201 | [illegible] | GCF_028421645.1 | Aneurinibacillus_sp._Ricciae_BoGa-3 | PP175_RS01545 | 299012..302617 | 3022697 | [illegible] |
| WP_066463066.1 | Type5.BREX | [illegible] | 1200 | [illegible] | GCF_001700325.1 | Caryophanon_latum | A6K76_RS05925 | 1324..4926 | 33977 | [illegible] |
| WP_107933601.1 | Type5.BREX | [illegible] | 1192 | [illegible] | GCF_003049615.1 | Ureibacillus_chungkukjangi | C9J33_RS05745 | 69413..72991 | 1202712 | [illegible] |
| WP_108587666.1 | Type5.BREX | [illegible] | 1191 | [illegible] | GCF_003058165.1 | Paenisporosarcina_sp._OV554 | C8K15_RS17245 | 15921..19496 | 2135694 | [illegible] |
| WP_274307983.1 | Type5.BREX | [illegible] | 1196 | [illegible] | GCF_028747805.1 | Solibacillus_sp._ZS111008 | O7776_RS16115 | 3287543..3291133 | 2912187 | [illegible] |
| WP_092984062.1 | Type5.BREX | [illegible] | 1184 | [illegible] | GCF_900101075.1 | Salinicoccus_qingdaonensis | BLR15_RS01740 | 53033..56587 | 576118 | [illegible] |
| WP_271398742.1 | Type5.BREX | [illegible] | 1184 | [illegible] | GCF_947502135.1 | Salinicoccus_roseus | POR62_RS08110 | 202088..205642 | 45670 | [illegible] |
| KKE80057.1 | Type5.BREX | [illegible] | 1187 | [illegible] | GCA_000974285.1 | Bacilli_bacterium_VT-13-104 | WH51_03830 | 66407..69970 | 1637974 | [illegible] |
| WP_164950923.1 | Type5.BREX | [illegible] | 1191 | [illegible] | GCF_011065955.2 | Clostridium_sp._FP2 | G9F71_RS16320 | 3398773..3402348 | 2724481 | [illegible] |
| WP_185649263.1 | Type5.BREX | [illegible] | 1189 | [illegible] | GCF_014231265.1 | Clostridium_sp._DJ247 | HGI79_RS03445 | 40571..44140 | 2726188 | [illegible] |
| WP_207649636.1 | Type5.BREX | [illegible] | 1190 | [illegible] | GCF_001936615.1 | Desulfosporosinus_metallidurans | DSOL_RS15020 | 40493..44065 | 1888891 | [illegible] |
| KJS50653.1 | Type5.BREX | [illegible] | 1187 | [illegible] | GCA_000961805.1 | Peptococcaceae_bacterium_BRH_c23 | VR66_01655 | 19236..22799 | 1629714 | [illegible] |
| WP_146889206.1 | Type5.BREX | [illegible] | 1186 | [illegible] | GCF_007990775.1 | Deinococcus_cellulosilyticus | DC3_RS23585 | 23663..27223 | 401558 | [illegible] |
| WP_189062792.1 | Type5.BREX | [illegible] | 1145 | [illegible] | GCF_014647995.1 | Deinococcus_arenae | IEY66_RS18050 | 13025..16462 | 1452751 | [illegible] |

| Protein-ID | System | Gene-Neighbourhoods | Length | Taxonomy | Assembly-ID | Strain | Locus-Tag | Genomic-coordinates | TaxID | Fasta-define |
|---|---|---|---|---|---|---|---|---|---|---|
| MCP3686167.1 | Type5.BREX | SF1-SF2-Helicase->?-><-SIGMA+OB-Fold+AAA-ATPase+HKD-DNase-BrxL<-PglZ<-N6-METHYLASE<-DUF6079*<-BrxB-BrxF-iSTAND<-BrxA-DNA-binding<-ANK+ANK<-?<-TUDOR+SF1-SF2-Helicase+iREase+CTD\|\|?->SIR [illegible] | 1183 | Bacteria>unclassified_Bacteria | GCA_024206195.1 | bacterium | GY861_26265 | 21131..24682 | 1869227 | BREX-system-P-loop-protein-BrxC-[bacterium] |
| -- | | | | | | | | | | |
| --Type-6-BREX | | | | | | | | | | |
| --Neighbourhoods-containing-BrxE | | | | | | | | | | |
| --Canonical-BREX | | | | | | | | | | |
| -- | | | | | | | | | | |
| VVB72030.1 | Type6.BREX | <-DUF6079*<-BrxB-BrxF-iSTAND<-BrxE-p<-BrxA-DNA-binding<-?<-MULE-TRANSPOSASE | 1215 | Archaea>environmental_samples | GCA_902385025.1 | uncultured_archaeon | LFW1512_01783 | 2594..6241 | 115547 | Uncharacterised-protein-[uncultured-archaeon] |
| MCQ4334814.1 | Type6.BREX | NUDIX+DUF429-><-SLATT<-SMODS<-?\|\|?-><-Calcineurin-SbcD\|\|BrxE->BrxA-DNA-binding->BrxB-BrxF-iSTAND->DUF6079*->N6-METHYLASE-p-> | 1214 | Archaea>Euryarchaeota>Stenosarchaea_group>Halobacteria>Halobacteriales>Haloarculaceae | GCA_024449025.1 | Natronomonas_aquatica | KM295_15270 | 10999..14643 | 2841590 | BREX-system-P-loop-protein-BrxC-[Natronomonas-aquatica] |
| WP_004594182.1 | Type6.BREX | <-Cas7\|\|BirA-HTH-><-SIGMA+OB-Fold+AAA-ATPase+Lon-Protease-BrxL<-PglZ<-?<-N6-METHYLASE\|\|TBP+TBP-><-DUF6079*<-BrxB-BrxF-iSTAND<-BrxA-DNA-binding<-BrxE\|\|Calcineurin-SbcD->SF1-SF2-Helicas [illegible] | 1214 | Archaea>Euryarchaeota>Stenosarchaea_group>Halobacteria>Halobacteriales>Haloarculaceae | GCF_000336635.1 | Haloarcula_japonica | C444_RS16625 | 447725..451369 | 29282 | BREX-system-P-loop-protein-BrxC-[Haloarcula-japonica] |
| WP_226023962.1 | Type6.BREX | <-Polbeta\|\|EVE+McrB-AAA->McrC-NTD+REase-><-?<-Calcineurin-SbcD\|\|BrxE->BrxA-DNA-binding->BrxB-BrxF-iSTAND->DUF6079*-><-REase\|\|N6-METHYLASE->PglZ-><-?\|\|SIGMA+OB-Fold+AAA-ATPase+Lon-Prot [illegible] | 1213 | Archaea>Euryarchaeota>Stenosarchaea_group>Halobacteria>Halobacteriales>Haloarculaceae | GCF_020405245.1 | Halomicrobium_salinisoli | LE161_RS18755 | 257437..261078 | 2878391 | BREX-system-P-loop-protein-BrxC-[Halomicrobium-salinisoli] |
| WP_256030949.1 | Type6.BREX | NUDIX+DUF429-><-SLATT<-SMODS<-?\|\|?-><-Calcineurin-SbcD\|\|BrxE->BrxA-DNA-binding->BrxB-BrxF-iSTAND->DUF6079*->N6-METHYLASE-p-> | 1214 | Archaea>Euryarchaeota>Stenosarchaea_group>Halobacteria>Halobacteriales>Haloarculaceae | GCF_024449025.1 | Natronomonas_aquatica | KM295_RS15415 | 10999..14643 | 2841590 | BREX-system-P-loop-protein-BrxC-[Natronomonas-aquatica] |
| WP_262178340.1 | Type6.BREX | <-RNase<-HNH<-35exo<-?<-Cas7<-SIGMA+OB-Fold+AAA-ATPase+Lon-Protease-BrxL<-PglZ\|\|?-><-N6-METHYLASE<-DUF6079*<-BrxB-BrxF-iSTAND<-BrxA-DNA-binding<-BrxE\|\|Calcineurin-SbcD->?->?-><-SF1-SF [illegible] | 1214 | Archaea>Euryarchaeota>Stenosarchaea_group>Halobacteria>Halobacteriales>Haloarculaceae | GCF_025457585.1 | unclassified_Halomicroarcula | NJQ96_RS10490 | 2043246..2046890 | 2630810 | MULTISPECIES:-BREX-system-P-loop-protein-BrxC-[unclassified-Halomicroarcula] |
| WP_145847379.1 | Type6.BREX | <-Cas7<-ZnR\|\|HTH-><-SIGMA+OB-Fold+AAA-ATPase+Lon-Protease-BrxL<-PglZ<-N6-METHYLASE\|\|REase-><-DUF6079*<-BrxB-BrxF-iSTAND<-BrxA-DNA-binding<-BrxE\|\|Calcineurin-SbcD->?-><-?<-?\|\|McrA+HTH+ [illegible] | 1213 | Archaea>Euryarchaeota>Stenosarchaea_group>Halobacteria>Halobacteriales>Halobacteriaceae | GCF_007833275.1 | Salarchaeum_sp._JOR-1 | FQU85_RS09825 | 1692629..1696270 | 2599399 | BREX-system-P-loop-protein-BrxC-[Salarchaeum-sp.-JOR-1] |
| WP_115795723.1 | Type6.BREX | <-Calcineurin<-McrA+HTH+HNH\|\|REase->REase->REase+METHYLASE-><-Calcineurin-SbcD\|\|BrxE->BrxA-DNA-binding->BrxB-BrxF-iSTAND->DUF6079*->N6-METHYLASE->PglZ->SIGMA+OB-Fold+AAA-ATPase+Lon-Pr [illegible] | 1214 | Archaea>Euryarchaeota>Stenosarchaea_group>Halobacteria>Halobacteriales>Halobacteriaceae | GCF_011440385.1 | Halorussus | G9466_RS05520 | 1085078..1088722 | 1070314 | MULTISPECIES:-BREX-system-P-loop-protein-BrxC-[Halorussus] |
| WP_188979080.1 | Type6.BREX | <-35exo<-?<-?<-Cas7\|\|BirA-HTH-><-SIGMA+OB-Fold+AAA-ATPase+Lon-Protease-BrxL\|\|?-><-PglZ<-N6-METHYLASE<-DUF6079*<-BrxB-BrxF-iSTAND<-BrxA-DNA-binding<-BrxE\|\|Calcineurin-SbcD->?->DDE_Tnp_ [illegible] | 1214 | Archaea>Euryarchaeota>Stenosarchaea_group>Halobacteria>Halobacteriales>Halobacteriaceae | GCF_014647475.1 | Halocalculus_aciditolerans | IEY26_RS11485 | 81184..84828 | 1383812 | BREX-system-P-loop-protein-BrxC-[Halocalculus-aciditolerans] |
| WP_254547347.1 | Type6.BREX | <-URI-DNase\|\|?->ArdCN+MPTase-inact->?->BrxE->BrxA-DNA-binding->BrxB-BrxF-iSTAND->DUF6079*->N6-METHYLASE-><-Zn_Tnp_IS1595+DDE_Tnp_IS1595<-?\|\|PglZ->SIGMA+OB-Fold+AAA-ATPase+Lon-Protease [illegible] | 1206 | Archaea>Euryarchaeota>Stenosarchaea_group>Halobacteria>Halobacteriales>Halobacteriaceae | GCF_024228315.1 | Halomarina_sp._BND7 | NKI68_RS22870 | 57828..61448 | 2961599 | BREX-system-P-loop-protein-BrxC-[Halomarina-sp.-BND7] |
| WP_256401541.1 | Type6.BREX | PIN->?->ORC-CDC6+HTH->?-><-SIGMA+OB-Fold+AAA-ATPase+Lon-Protease-BrxL<-PglZ<-?<-N6-METHYLASE\|\|?-><-DUF6079*<-BrxB-BrxF-iSTAND<-BrxA-DNA-binding<-BrxE\|\|Calcineurin-SbcD->SF1-SF2-Helica [illegible] | 1214 | Archaea>Euryarchaeota>Stenosarchaea_group>Halobacteria>Halobacteriales>Halobacteriaceae | GCF_024495425.1 | Haloarchaeobius_litoreus | NOW55_RS18175 | 109801..113445 | 755306 | BREX-system-P-loop-protein-BrxC-[Haloarchaeobius-litoreus] |
| WP_007738385.1 | Type6.BREX | BrxE->BrxA-DNA-binding->BrxB-BrxF-iSTAND->DUF6079*->DDE_Tnp->N6-METHYLASE-p-> | 1213 | Archaea>Euryarchaeota>Stenosarchaea_group>Halobacteria>Halobacteriales>Halococcaceae | GCF_000336715.1 | Halococcus_thailandensis | C451_RS04955 | 2197..5838 | 335952 | BREX-system-P-loop-protein-BrxC-[Halococcus-thailandensis] |
| WP_049936156.1 | Type6.BREX | HTH+BirA-HTH-><-?<-ABC-ATPase+SbcC<-?<-?\|\|tRNA-><-SIGMA+OB-Fold+AAA-ATPase+Lon-Protease-BrxL<-PglZ<-N6-METHYLASE<-REase<-DUF6079*<-BrxB-BrxF-iSTAND<-BrxA-DNA-binding<-BrxE\|\|Calcineuri [illegible] | 1215 | Archaea>Euryarchaeota>Stenosarchaea_group>Halobacteria>Haloferacales>Haloferacaceae | GCF_000427685.1 | Haloplanus_natans | HALNA_RS09570 | 1440330..1443977 | 376171 | BREX-system-P-loop-protein-BrxC-[Haloplanus-natans] |
| WP_122105591.1 | Type6.BREX | <-PAS+HISKIN+ST-Hiskin\|\|?-><-SF1-SF2-Helicase<-Calcineurin-SbcD\|\|BrxE->BrxA-DNA-binding->BrxB-BrxF-iSTAND->DUF6079*->N6-METHYLASE->REase->PglZ->SIGMA+OB-Fold+AAA-ATPase+Lon-Protease-B [illegible] | 1214 | Archaea>Euryarchaeota>Stenosarchaea_group>Halobacteria>Haloferacales>Haloferacaceae | GCF_003697845.1 | Halobellus_captivus | EA078_RS09395 | 31813..35457 | 2592614 | BREX-system-P-loop-protein-BrxC-[Halobellus-captivus] |
| WP_170076589.1 | Type6.BREX | <-SbcC<-Calcineurin<-REase<-?<-Calcineurin-SbcD\|\|BrxE->BrxA-DNA-binding->BrxB-BrxF-iSTAND->DUF6079*->N6-METHYLASE->?->SbcC->PglZ->SIGMA+OB-Fold+AAA-ATPase+Lon-Protease-BrxL->PIN->ARG- [illegible] | 1214 | Archaea>Euryarchaeota>Stenosarchaea_group>Halobacteria>Haloferacales>Haloferacaceae | GCF_012726105.1 | Haloferax_alexandrinus | GOC85_RS10615 | 2063512..2067156 | 114529 | BREX-system-P-loop-protein-BrxC-[Haloferax-alexandrinus] |
| WP_089880634.1 | Type6.BREX | <-Cas7\|\|BirA-HTH-><-SIGMA+OB-Fold+AAA-ATPase+Lon-Protease-BrxL<-PglZ<-TBP+TBP<-N6-METHYLASE<-?<-DUF6079*<-BrxB-BrxF-iSTAND<-BrxA-DNA-binding<-BrxE\|\|Calcineurin-SbcD->McrA+HTH+HNH->Cal [illegible] | 1214 | Archaea>Euryarchaeota>Stenosarchaea_group>Halobacteria>Haloferacales>Haloferacaceae | GCF_900115785.1 | Halogeometricum_limi | BM282_RS10840 | 420011..423655 | 555875 | BREX-system-P-loop-protein-BrxC-[Halogeometricum-limi] |
| WP_092566453.1 | Type6.BREX | MazE_antitoxin->GDResolvase+HIN-HTH-><-?<-?<- [illegible] | 1214 | Archaea>Euryarchaeota>Stenosarchaea_group>Halobacteria>Haloferacales>Halorubraceae | GCF_001261915.1 | Halorubrum_sp._SD626R | SD626R_RS11260 | 44358..48002 | 1419722 | BREX-system-P-loop-protein-BrxC-[Halorubrum-sp.-SD626R] |
| WP_096390020.1 | Type6.BREX | <-Calcineurin<-GDResolvase+HIN-HTH<-REase<-?<- [illegible] | 1214 | Archaea>Euryarchaeota>Stenosarchaea_group>Halobacteria>Haloferacales>Halorubraceae | GCF_002355635.1 | Halopenitus_persicus | CPZ00_RS05685 | 1169858..1173502 | 1048396 | BREX-system-P-loop-protein-BrxC-[Halopenitus-persicus] |
| WP_152043548.1 | Type6.BREX | <-SbcC<-Calcineurin\|\|?->\|\|BrxE->BrxA-DNA-binding [illegible] | 1204 | Archaea>Euryarchaeota>Stenosarchaea_group>Halobacteria>Haloferacales>Halorubraceae | GCF_009176545.1 | Salinigranum_salinum | FZP69_RS21845 | 9948..13562 | 1364937 | BREX-system-P-loop-protein-BrxC-[Salinigranum-salinum] |
| WP_256395827.1 | Type6.BREX | REase-><-Calcineurin-SbcD\|\|BrxE->BrxA-DNA-binding [illegible] | 1214 | Archaea>Euryarchaeota>Stenosarchaea_group>Halobacteria>Haloferacales>Halorubraceae | GCF_024494645.1 | Halohasta_litorea | NO877_RS08165 | 663760..667404 | 869891 | BREX-system-P-loop-protein-BrxC-[Halohasta-litorea] |
| WP_271945465.1 | Type6.BREX | <-SLATT<-SMODS<-?\|\|?-><-Calcineurin-SbcD\|\|BrxE-> [illegible] | 1214 | Archaea>Euryarchaeota>Stenosarchaea_group>Halobacteria>Haloferacales>Halorubraceae | GCF_028211275.1 | Halorubrum_ezzemoulense | PM036_RS03905 | 12589..16233 | 337243 | BREX-system-P-loop-protein-BrxC-[Halorubrum-ezzemoulense] |
| WP_076580859.1 | Type6.BREX | REase->BrxE->BrxA-DNA-binding->BrxB-BrxF-iSTAND- [illegible] | 1206 | Archaea>Euryarchaeota>Stenosarchaea_group>Halobacteria>Natrialbales>Natrialbaceae | GCF_001971705.1 | Natronorubrum_daqingense | BW960_RS07735 | 340245..343865 | 588898 | BREX-system-P-loop-protein-BrxC-[Natronorubrum-daqingense] |
| WP_121744813.1 | Type6.BREX | Thermonuclease->?->?->McrC-NTD+REase->SF1-SF2-He [illegible] | 1206 | Archaea>Euryarchaeota>Stenosarchaea_group>Halobacteria>Natrialbales>Natrialbaceae | GCF_003670115.1 | Natronorubrum_halophilum | DWB23_RS21360 | 42652..46272 | 1702106 | BREX-system-P-loop-protein-BrxC-[Natronorubrum-halophilum] |
| WP_179261303.1 | Type6.BREX | <-HD<-?\|\|ABhydrolase->?-><-SIGMA+OB-Fold+AAA-ATP [illegible] | 1214 | Archaea>Euryarchaeota>Stenosarchaea_group>Halobacteria>Natrialbales>Natrialbaceae | GCF_013402815.2 | Natrinema_halophilum | HYG82_RS33105 | 2537554..2541198 | 1699371 | BREX-system-P-loop-protein-BrxC-[Natrinema-halophilum] |
| WP_253430473.1 | Type6.BREX | Phage_integrase+Phage_integr_3-><-?\|\|?-><-REase\| [illegible] | 1206 | Archaea>Euryarchaeota>Stenosarchaea_group>Halobacteria>Natrialbales>Natrialbaceae | GCF_024138145.1 | Halovivax_sp._AGai3-5 | NGM15_RS10380 | 2104102..2107722 | 2953746 | BREX-system-P-loop-protein-BrxC-[Halovivax-sp.-AGai3-5] |
| WP_273738559.1 | Type6.BREX | McrB-AAA->McrC-NTD+REase->BrxE->BrxA-DNA-binding [illegible] | 1214 | Archaea>Euryarchaeota>Stenosarchaea_group>Halobacteria>Natrialbales>Natrialbaceae | GCF_028618695.1 | Natrinema_soli | PGX04_RS11075 | 6409..10053 | 1930624 | BREX-system-P-loop-protein-BrxC-[Natrinema-soli] |
| MBX7223615.1 | Type6.BREX | HIRAN->?->TUDOR+SF1-SF2-Helicase+iREase+CTD->?-> [illegible] | 1210 | Bacteria>Acidobacteriota>Blastocatellia>unclassified_Blastocatellia | GCA_019695185.1 | Blastocatellia_bacterium | K1Y36_27120 | 9976..13608 | 2052146 | BREX-system-P-loop-protein-BrxC-[Blastocatellia-bacterium] |
| MCI0487530.1 | Type6.BREX | <-N6-METHYLASE<-DUF6079*<-BrxB-BrxF-iSTAND<- [illegible] | 1208 | Bacteria>Acidobacteriota>Blastocatellia>unclassified_Blastocatellia | GCA_022615215.1 | Blastocatellia_bacterium | L0229_13130 | 4420..8046 | 2052146 | BREX-system-P-loop-protein-BrxC-[Blastocatellia-bacterium] |
| MBK9963544.1 | Type6.BREX | HTH+MPTase-><-SF1-SF2-Helicase+wHTH+Small-B-barre [illegible] | 1207 | Bacteria>Acidobacteriota>Holophagae>Holophagales>unclassified_Holophagales | GCA_016722225.1 | Holophagales_bacterium | IPP07_01030 | 234150..237773 | 2478486 | BREX-system-P-loop-protein-BrxC-[Holophagales-bacterium] |
| BDC48932.1 | Type6.BREX | <-HKD-DNase+HKD-Rease<-?<-SIGMA+OB-Fold+AAA-ATPas [illegible] | 1202 | Bacteria>Acidobacteriota>Terriglobia>Bryobacterales>unclassified_Bryobacterales | GCA_016865505.2 | Bryobacterales_bacterium_F-183 | F183_A12480 | 1423409..1427017 | 2802972 | hypothetical-protein-F183_A12480-[Bryobacterales-bacterium-F-183] |
| MBZ5596030.1 | Type6.BREX | <-SIGMA+OB-Fold+AAA-ATPase+Lon-Protease-BrxL<- [illegible] | 1208 | Bacteria>Acidobacteriota>Terriglobia>unclassified_Terriglobia | GCA_020073305.1 | Acidobacteriia_bacterium | LAP39_27625 | 24377..28003 | 2081523 | BREX-system-P-loop-protein-BrxC-[Acidobacteriia-bacterium] |
| MBP7797364.1 | Type6.BREX | <-RNase<-ANK+ANK<-REase<-SbcC<-?<-HNH<-SIGMA+OB- [illegible] | 1177 | Bacteria>Acidobacteriota>Thermoanaerobaculia>Thermoanaerobaculales>Thermoanaerobaculaceae>unclassified_Thermoanaerobaculaceae | GCA_018000955.1 | Thermoanaerobaculaceae_bacterium | KA072_01920 | 40724..44257 | 2699758 | BREX-system-P-loop-protein-BrxC-[Thermoanaerobaculaceae-bacterium] |
| MCG3192298.1 | Type6.BREX | <-DUF932<-ParB+HNH<-?<-SIGMA+OB-Fold+AAA-ATPase+ [illegible] | 1183 | Bacteria>Acidobacteriota>Thermoanaerobaculia>unclassified_Thermoanaerobaculia | GCA_022071945.1 | Thermoanaerobaculia_bacterium | DIJKHBIC_01536 | 22288..25839 | 2651171 | hypothetical-protein-[Thermoanaerobaculia-bacterium] |
| OQC41890.1 | Type6.BREX | CASPASE->?->?->?-><-SIGMA+OB-Fold+AAA-ATPase+Lon- [illegible] | 1181 | Bacteria>Acidobacteriota>unclassified_Acidobacteriota | GCA_002071135.1 | Acidobacteria_bacterium_ADurb.Bin051 | BWX64_00587 | 14644..18189 | 1852786 | hypothetical-protein-BWX64_00587-[Acidobacteria-bacterium-ADurb.Bin051] |
| MBS1808657.1 | Type6.BREX | <-SF1-SF2-Helicase+DUF1998<-?<-35exo\|\|NtrC-AAA+ [illegible] | 1212 | Bacteria>Acidobacteriota>unclassified_Acidobacteriota | GCA_018268995.1 | Acidobacteriota_bacterium | JST84_10740 | 129628..133266 | 1978231 | BREX-system-P-loop-protein-BrxC-[Acidobacteriota-bacterium] |
| MCC6130024.1 | Type6.BREX | HTH+LexA-protease->?->?->?->BrxE->BrxA-DNA-bindin [illegible] | 1183 | Bacteria>Acidobacteriota>unclassified_Acidobacteriota | GCA_020845555.1 | Acidobacteriota_bacterium | IT186_08845 | 15940..19491 | 1978231 | BREX-system-P-loop-protein-BrxC-[Acidobacteriota-bacterium] |
| MCI0390260.1 | Type6.BREX | MPTase-><-NTP_transf_5<-?<-RHH<-?\|\|BrxE->BrxA-DN [illegible] | 1215 | Bacteria>Acidobacteriota>unclassified_Acidobacteriota | GCA_022614935.1 | Acidobacteriota_bacterium | MOB07_16040 | 5937..9584 | 1978231 | BREX-system-P-loop-protein-BrxC-[Acidobacteriota-bacterium] |
| MCD6541517.1 | Type6.BREX | BrxE->BrxA-DNA-binding->BrxB-BrxF-iSTAND->DUF607 [illegible] | 1162 | Bacteria>Bacteria_incertae_sedis>Bacteria_candidate_phyla>Candidatus_Bipolaricaulota>unclassified_Candidatus_Bipolaricaulota | GCA_021163465.1 | Candidatus_Bipolaricaulota_bacterium | J7K76_06780 | 2223..5711 | 2306012 | BREX-system-P-loop-protein-BrxC-[Candidatus-Bipolaricaulota-bacterium] |
| MBP7775371.1 | Type6.BREX | HTH_3->?->?->XIS-HTH->BrxE->BrxA-DNA-binding->Br [illegible] | 472 | Bacteria>Bacteria_incertae_sedis>Bacteria_candidate_phyla>Candidatus_Saccharibacteria>Candidatus_Saccharimonadia>Candidatus_Saccharimonadales | GCA_018000475.1 | Candidatus_Saccharimonas_sp. | KA069_02090 | 5805..7222 | 1917697 | BREX-system-P-loop-protein-BrxC-[Candidatus-Saccharimonas-sp.] |
| MCB0345181.1 | Type6.BREX | wHTH+HNH->SAM-METHYLASE->SF1-SF2-Helicase->AbiGi [illegible] | 811 | Bacteria>Bdellovibrionota>Bdellovibrionia>Bdellovibrionales>unclassified_Bdellovibrionales | GCA_020434585.1 | Bdellovibrionales_bacterium | KDD66_08680 | 10999..13432 | 2053517 | BREX-system-P-loop-protein-BrxC-[Bdellovibrionales-bacterium] |
| NOY28205.1 | Type6.BREX | <-DUF6079-p*<-BrxB-BrxF-iSTAND<-BrxA-DNA-binding [illegible] | 408 | Bacteria>Bdellovibrionota>Oligoflexia>unclassified_Oligoflexia | GCA_013152105.1 | Oligoflexia_bacterium | GXP62_20270 | 1..1225 | 2599941 | BREX-system-P-loop-protein-BrxC-[Oligoflexia-bacterium] |
| WP_146981109.1 | Type6.BREX | Peptidase_S8-><-SF1-SF2-Helicase+wHTH+Small-B-ba [illegible] | 1216 | Bacteria>delta/epsilon_subdivisions>Deltaproteobacteria>Bradymonadales>Lujinxingiaceae>Lujinxingia | GCF_007997015.1 | Lujinxingia_vulgaris | FRC98_RS09600 | 57836..61486 | 2600176 | BREX-system-P-loop-protein-BrxC-[Lujinxingia-vulgaris] |
| QED28443.1 | Type6.BREX | <-N6-METHYLASE\|\|?->?->?->?->?-><-DUF6079*<-BrxB- [illegible] | 1202 | Bacteria>delta/epsilon_subdivisions>Deltaproteobacteria>Bradymonadales>Microvenatoraceae>Microvenator | GCA_007993755.1 | Microvenator_marinus | FRD01_14605 | 3560726..3564334 | 2600177 | BREX-system-P-loop-protein-BrxC-[Microvenator-marinus] |
| WP_146960870.1 | Type6.BREX | <-PglZ<-?\|\|?->?-><-N6-METHYLASE\|\|?->?->?->?-><- [illegible] | 1202 | Bacteria>delta/epsilon_subdivisions>Deltaproteobacteria>Bradymonadales>Microvenatoraceae>Microvenator | GCF_007993755.1 | Microvenator_marinus | FRD01_RS14605 | 3560726..3564334 | 2600177 | BREX-system-P-loop-protein-BrxC-[Microvenator-marinus] |
| WP_163297755.1 | Type6.BREX | <-DUF4236<-?<-REase<-SF1-SF2-Helicase+REase<- [illegible] | 1226 | Bacteria>delta/epsilon_subdivisions>Deltaproteobacteria>Deltaproteobacteria_incertae_sedis>Dissulfurirhabdaceae>Dissulfurirhabdus | GCF_010499435.1 | Dissulfurirhabdus_thermomarina | G3N55_RS01855 | 13905..17585 | 1765737 | BREX-system-P-loop-protein-BrxC-[Dissulfurirhabdus-thermomarina] |
| PJB45699.1 | Type6.BREX | <-SF1-SF2-Helicase+wHTH+Small-B-barrel+DUF1998-p [illegible] | 1207 | Bacteria>delta/epsilon_subdivisions>Deltaproteobacteria>unclassified_Deltaproteobacteria | GCA_002789955.1 | Deltaproteobacteria_bacterium_CG_4_9_14_3_um_filter_63_12 | C0108_06840 | 12981..16604 | 1973967 | BREX-system-P-loop-protein-BrxC-[Deltaproteobacteria-bacterium-CG_4_9_14_3_um_filter_63_12] |
| RME21859.1 | Type6.BREX | <-OLD-ABC<-N6-METHYLASE<-DUF6079*<-BrxB-BrxF- [illegible] | 1180 | Bacteria>delta/epsilon_subdivisions>Deltaproteobacteria>unclassified_Deltaproteobacteria | GCA_003694325.1 | Deltaproteobacteria_bacterium | D6798_17180 | 2990..6532 | 2026735 | BREX-system-P-loop-protein-BrxC-[Deltaproteobacteria-bacterium] |
| TNF28011.1 | Type6.BREX | HEPN+PAS+DUF4011+SF1-SF2-Helicase+REase->Classica [illegible] | 1215 | Bacteria>delta/epsilon_subdivisions>Deltaproteobacteria>unclassified_Deltaproteobacteria | GCA_006226565.1 | Deltaproteobacteria_bacterium | EP329_18685 | 31419..35066 | 2026735 | BREX-system-P-loop-protein-BrxC-[Deltaproteobacteria-bacterium] |
| TVQ94624.1 | Type6.BREX | Schlafen+PIWI-><-N6-METHYLASE<-DUF6079*<-BrxB-Br [illegible] | 1211 | Bacteria>delta/epsilon_subdivisions>Deltaproteobacteria>unclassified_Deltaproteobacteria | GCA_007693595.1 | Deltaproteobacteria_bacterium | EA397_01285 | 5402..9037 | 2026735 | BREX-system-P-loop-protein-BrxC-[Deltaproteobacteria-bacterium] |
| MBI1948544.1 | Type6.BREX | <-HTH+MPTase<-?\|\|?-><-?<-APATPase<-SF1-SF2-Helic [illegible] | 1213 | Bacteria>delta/epsilon_subdivisions>Deltaproteobacteria>unclassified_Deltaproteobacteria | GCA_016177705.1 | Deltaproteobacteria_bacterium | HYS27_22850 | 64502..68143 | 2026735 | BREX-system-P-loop-protein-BrxC-[Deltaproteobacteria-bacterium] |
| MBI2378669.1 | Type6.BREX | PIN-><-?<-PglZ<-Schlafen+HATPase+wHTH<-PglZ-p<-? [illegible] | 1207 | Bacteria>delta/epsilon_subdivisions>Deltaproteobacteria>unclassified_Deltaproteobacteria | GCA_016183035.1 | Deltaproteobacteria_bacterium | HYV07_32040 | 22898..26521 | 2026735 | BREX-system-P-loop-protein-BrxC-[Deltaproteobacteria-bacterium] |
| MBI4818899.1 | Type6.BREX | <-PglZ-p<-TOPRIM<-OLD-ABC<-Schlafen+HATPase+wHTH [illegible] | 1207 | Bacteria>delta/epsilon_subdivisions>Deltaproteobacteria>unclassified_Deltaproteobacteria | GCA_016208175.1 | Deltaproteobacteria_bacterium | HY791_21690 | 12157..15780 | 2026735 | BREX-system-P-loop-protein-BrxC-[Deltaproteobacteria-bacterium] |
| MBI4706185.1 | Type6.BREX | XIS-HTH->BrxE->BrxA-DNA-binding->BrxB-BrxF-iSTAN [illegible] | 1236 | Bacteria>delta/epsilon_subdivisions>Deltaproteobacteria>unclassified_Deltaproteobacteria | GCA_016209145.1 | Deltaproteobacteria_bacterium | HY744_34245 | 2867..6577 | 2026735 | BREX-system-P-loop-protein-BrxC-[Deltaproteobacteria-bacterium] |
| MBI4519050.1 | Type6.BREX | <-PIN<-DUF433<-N6-METHYLASE-p\|\|?->?->?-><-N6-MET [illegible] | 1218 | Bacteria>delta/epsilon_subdivisions>Deltaproteobacteria>unclassified_Deltaproteobacteria | GCA_016210005.1 | Deltaproteobacteria_bacterium | HY699_24940 | 63374..67030 | 2026735 | BREX-system-P-loop-protein-BrxC-[Deltaproteobacteria-bacterium] |
| MBI5543786.1 | Type6.BREX | XIS-HTH->Classical-AAA->Peptidase_S8-><-SF1-SF2- [illegible] | 1206 | Bacteria>delta/epsilon_subdivisions>Deltaproteobacteria>unclassified_Deltaproteobacteria | GCA_016218885.1 | Deltaproteobacteria_bacterium | HY901_07870 | 25804..29424 | 2026735 | BREX-system-P-loop-protein-BrxC-[Deltaproteobacteria-bacterium] |
| MBI5547484.1 | Type6.BREX | BrxE->BrxA-DNA-binding->BrxB-BrxF-iSTAND->DUF607 [illegible] | 1206 | Bacteria>delta/epsilon_subdivisions>Deltaproteobacteria>unclassified_Deltaproteobacteria | GCA_016218885.1 | Deltaproteobacteria_bacterium | HY901_26680 | 1772..5392 | 2026735 | BREX-system-P-loop-protein-BrxC-[Deltaproteobacteria-bacterium] |
| MBK8011741.1 | Type6.BREX | <-TIR<-REase+HTH+35exo+Small-B-barrel+SF1-SF2-He [illegible] | 1179 | Bacteria>delta/epsilon_subdivisions>Deltaproteobacteria>unclassified_Deltaproteobacteria | GCA_016713425.1 | Deltaproteobacteria_bacterium | IPK13_10335 | 2729207..2732746 | 2026735 | BREX-system-P-loop-protein-BrxC-[Deltaproteobacteria-bacterium] |
| MBK9643799.1 | Type6.BREX | ParA-Soj-PloopNTPase->?-><-?<-?\|\|BrxE->BrxA-DNA- [illegible] | 437 | Bacteria>delta/epsilon_subdivisions>Deltaproteobacteria>unclassified_Deltaproteobacteria | GCA_016717595.1 | Deltaproteobacteria_bacterium | IPO67_01340 | 265315..266626 | 2026735 | BREX-system-P-loop-protein-BrxC-[Deltaproteobacteria-bacterium] |
| MBK9368670.1 | Type6.BREX | HNH->?->?->?-><-?\|\|DNA-glycosylase->BrxE->BrxA-D [illegible] | 1184 | Bacteria>delta/epsilon_subdivisions>Deltaproteobacteria>unclassified_Deltaproteobacteria | GCA_016719515.1 | Deltaproteobacteria_bacterium | IPN01_20575 | 101681..105235 | 2026735 | BREX-system-P-loop-protein-BrxC-[Deltaproteobacteria-bacterium] |
| MCB9735989.1 | Type6.BREX | <-HEPN<-Polbeta\|\|BrxE->BrxA-DNA-binding->BrxB-Br [illegible] | 1207 | Bacteria>delta/epsilon_subdivisions>Deltaproteobacteria>unclassified_Deltaproteobacteria | GCA_020632715.1 | Deltaproteobacteria_bacterium | H6745_25655 | 182878..186501 | 2026735 | BREX-system-P-loop-protein-BrxC-[Deltaproteobacteria-bacterium] |
| MCB9489065.1 | Type6.BREX | <-DUF6079-p*<-BrxB-BrxF-iSTAND<-BrxA-DNA-binding [illegible] | 892 | Bacteria>delta/epsilon_subdivisions>Deltaproteobacteria>unclassified_Deltaproteobacteria | GCA_020633895.1 | Deltaproteobacteria_bacterium | H6684_10070 | 1..2676 | 2026735 | BREX-system-P-loop-protein-BrxC-[Deltaproteobacteria-bacterium] |
| MCC6811534.1 | Type6.BREX | STYKinase-like->?->HTH_32+DDE_3->HTH_32+DDE_3-> [illegible] | 1210 | Bacteria>delta/epsilon_subdivisions>Deltaproteobacteria>unclassified_Deltaproteobacteria | GCA_020848545.1 | Deltaproteobacteria_bacterium | IT381_29155 | 76632..80264 | 2026735 | BREX-system-P-loop-protein-BrxC-[Deltaproteobacteria-bacterium] |
| MCC7070407.1 | Type6.BREX | VirB6->?->?->CR-Kinase+pi3KINASE->?->?->BrxE->Br [illegible] | 1209 | Bacteria>delta/epsilon_subdivisions>Deltaproteobacteria>unclassified_Deltaproteobacteria | GCA_020850995.1 | Deltaproteobacteria_bacterium | IT383_03735 | 19295..22924 | 2026735 | BREX-system-P-loop-protein-BrxC-[Deltaproteobacteria-bacterium] |
| MCC6848734.1 | Type6.BREX | <-Sulfatase+DUF4976\|\|?->Peptidase_S15+PepX_C-><- [illegible] | 1181 | Bacteria>delta/epsilon_subdivisions>Deltaproteobacteria>unclassified_Deltaproteobacteria | GCA_020852955.1 | Deltaproteobacteria_bacterium | IT294_09545 | 30719..34264 | 2026735 | BREX-system-P-loop-protein-BrxC-[Deltaproteobacteria-bacterium] |
| MCH7912323.1 | Type6.BREX | <-CHTH<-XIS-HTH<-?<-Primpol+PriCT<-?<-?\|\|BrxE->B [illegible] | 1217 | Bacteria>delta/epsilon_subdivisions>Deltaproteobacteria>unclassified_Deltaproteobacteria | GCA_022563955.1 | Deltaproteobacteria_bacterium | IH856_04695 | 13112..16765 | 2026735 | BREX-system-P-loop-protein-BrxC-[Deltaproteobacteria-bacterium] |
| BAH89618.1 | Type6.BREX | CHTH-><-?\|\|?->ZnR+TOPRIM+AAA-helicase+wHTH-><-?\| [illegible] | 1211 | Bacteria>environmental_samples | - | uncultured_bacterium | . | 12703..16338 | 77133 | ATPase-[uncultured-bacterium] |
| AGC72568.1 | Type6.BREX | METHYLASE-><-SIGMA+OB-Fold+AAA-ATPase+Lon-Protea [illegible] | 1181 | Bacteria>environmental_samples | - | uncultured_bacterium_A1Q1_fos_1870 | . | 12880..16425 | 1256554 | hypothetical-protein-[uncultured-bacterium-A1Q1_fos_1870] |
| WP_259091014.1 | Type6.BREX | SF1-SF2-Helicase+REase-><-HORMA\|\|BrxE->BrxA-DNA- [illegible] | 1192 | Bacteria>FCB_group>Bacteroidetes/Chlorobi_group>Bacteroidota>Bacteroidetes_incertae_sedis>Rhodothermaceae | GCF_024807395.1 | Salinibacter_ruber | GGP54_RS13575 | 8828..12406 | 146919 | BREX-system-P-loop-protein-BrxC-[Salinibacter-ruber] |
| WP_259130005.1 | Type6.BREX | <-SbcC+SbcC<-Calcineurin-SbcD<-HORMA<-SIGMA+OB-F [illegible] | 1199 | Bacteria>FCB_group>Bacteroidetes/Chlorobi_group>Bacteroidota>Bacteroidetes_incertae_sedis>Rhodothermaceae | GCF_024807455.1 | Salinibacter_ruber | GGQ04_003012 | 22826..26425 | 146919 | BREX-system-P-loop-protein-BrxC-[Salinibacter-ruber] |
| WP_259228204.1 | Type6.BREX | <-REC+OmpR-HTH<-PAS+PAS+PAS+GAF+HISKIN<-Phage_in [illegible] | 1201 | Bacteria>FCB_group>Bacteroidetes/Chlorobi_group>Bacteroidota>Bacteroidetes_incertae_sedis>Rhodothermaceae | GCF_024809755.1 | Salinibacter_ruber | GGP93_RS14280 | 71185..74790 | 146919 | BREX-system-P-loop-protein-BrxC-[Salinibacter-ruber] |
| WP_251961450.1 | Type6.BREX | SLOG+BirA-HTH-><-?\|\|?-><-HD<-SbcC+SbcC<-Calcineu [illegible] | 1188 | Bacteria>FCB_group>Bacteroidetes/Chlorobi_group>Bacteroidota>Bacteroidetes_incertae_sedis>Rhodothermaceae | GCF_943908195.1 | Salinibacter_ruber | NGL62_RS09035 | 20957..24523 | 146919 | BREX-system-P-loop-protein-BrxC-[Salinibacter-ruber] |
| WP_117964426.1 | Type6.BREX | <-TRAM+METHYLASE\|\|?-><-UDG<-?<-ARG\|\|RHH+MobC->?- [illegible] | 938 | Bacteria>FCB_group>Bacteroidetes/Chlorobi_group>Bacteroidota>Bacteroidia>Bacteroidales | - | Bacteroides_uniformis | IYV59_08305 | 23505..26321 | 820 | MULTISPECIES:-BREX-system-P-loop-protein-BrxC-[Bacteroides] |
| EFG14417.1 | Type6.BREX | PRTase->?->?->tRNA->tRNA-><-?<-SIGMA+OB-Fold+AAA- [illegible] | 762 | Bacteria>FCB_group>Bacteroidetes/Chlorobi_group>Bacteroidota>Bacteroidia>Bacteroidales | GCA_000178215.1 | Bacteroides_xylanisolvens_SD_CC_1b | CW3_0497 | 100900..103188 | 702447 | hypothetical-protein-CW3_0497-[Bacteroides-xylanisolvens-SD-CC-1b] |
| EFF57551.1 | Type6.BREX | CR-ATPase4->ZnR+TOPRIM->TraJ-RHH->?->HTH_3->BrxE [illegible] | 706 | Bacteria>FCB_group>Bacteroidetes/Chlorobi_group>Bacteroidota>Bacteroidia>Bacteroidales | GCA_000178295.1 | Bacteroides_xylanisolvens_SD_CC_2a | CW1_2295 | 25339..27459 | 702444 | hypothetical-protein-CW1_2295-[Bacteroides-xylanisolvens-SD-CC-2a] |
| CCX63114.1 | Type6.BREX | VirD4-TraG+VirD4-FtsK->Phage_lysozyme-><-?\|\|HTH_ [illegible] | 1224 | Bacteria>FCB_group>Bacteroidetes/Chlorobi_group>Bacteroidota>Bacteroidia>Bacteroidales | GCA_000431055.1 | Bacteroides_sp._CAG:598 | BN727_01191 | 30404..34078 | 1262743 | predicted-protein-[Bacteroides-sp.-CAG:598] |
| CDM00531.1 | Type6.BREX | PRTase->?->?->tRNA->tRNA-><-?<-SIGMA+OB-Fold+AAA- [illegible] | 689 | Bacteria>FCB_group>Bacteroidetes/Chlorobi_group>Bacteroidota>Bacteroidia>Bacteroidales | GCA_000577955.1 | Bacteroides_xylanisolvens_SD_CC_2a | BN891_34560 | 156968..159037 | 702444 | hypothetical-protein-BN891_34560-[Bacteroides-xylanisolvens-SD-CC-2a] |
| RGJ49912.1 | Type6.BREX | <-TRAM+METHYLASE\|\|<-UDG<-?<-ARG\|\|RHH+MobC->?-><- [illegible] | 938 | Bacteria>FCB_group>Bacteroidetes/Chlorobi_group>Bacteroidota>Bacteroidia>Bacteroidales | GCA_003436345.1 | Bacteroides_sp._D20 | DXD58_12440 | 30883..33699 | 585543 | BREX-system-P-loop-protein-BrxC-[Bacteroides-sp.-D20] |
| HAM28416.1 | Type6.BREX | ChlI+Lon_C+AAA-><-RNase<-AraC-HTH\|\|?-><-?\|\|HTH_3 [illegible] | 1218 | Bacteria>FCB_group>Bacteroidetes/Chlorobi_group>Bacteroidota>Bacteroidia>Bacteroidales | GCA_003453655.1 | Prevotella_sp. | DCP79_00675 | 66224..69880 | 59823 | BREX-system-P-loop-protein-BrxC-[Prevotella-sp.] |
| RHK71295.1 | Type6.BREX | <-ARG\|\|RHH+MobC->?-><-?<-SIGMA+OB-Fold+AAA-ATPas [illegible] | 145 | Bacteria>FCB_group>Bacteroidetes/Chlorobi_group>Bacteroidota>Bacteroidia>Bacteroidales | GCA_003474125.1 | Bacteroides_uniformis | DW051_06915 | 97313..97750 | 820 | hypothetical-protein-DW051_06915-[Bacteroides-uniformis] |
| MBF7062545.1 | Type6.BREX | <-TRAM+METHYLASE\|\|?-><-UDG<-?<-ARG\|\|RHH+MobC->?- [illegible] | 938 | Bacteria>FCB_group>Bacteroidetes/Chlorobi_group>Bacteroidota>Bacteroidia>Bacteroidales | GCA_015594615.1 | Bacteroides_sp._HF-5613 | ITJ87_11475 | 2819894..2822710 | 2785794 | BREX-system-P-loop-protein-BrxC-[Bacteroides-sp.-HF-5613] |
| MBM6671973.1 | Type6.BREX | <-SIGMA+OB-Fold+AAA-ATPase+HKD-DNase-BrxL<-PglZ< [illegible] | 1224 | Bacteria>FCB_group>Bacteroidetes/Chlorobi_group>Bacteroidota>Bacteroidia>Bacteroidales | GCA_016900015.1 | Phocaeicola_coprophilus | H6B14_14560 | 8845..12519 | 387090 | BREX-system-P-loop-protein-BrxC-[Phocaeicola-coprophilus] |
| MBP8934558.1 | Type6.BREX | HTH_3->BrxE->BrxA-DNA-binding->BrxB-BrxF-iSTAND- [illegible] | 1232 | Bacteria>FCB_group>Bacteroidetes/Chlorobi_group>Bacteroidota>Bacteroidia>Bacteroidales | GCA_018056085.1 | Prevotella_sp. | KBG68_02510 | 3234..6932 | 59823 | BREX-system-P-loop-protein-BrxC-[Prevotella-sp.] |
| MBV3827782.1 | Type6.BREX | <-UDG<-?<-ARG\|\|RHH+MobC->?-><-?<-SIGMA+OB-Fold+A [illegible] | 938 | Bacteria>FCB_group>Bacteroidetes/Chlorobi_group>Bacteroidota>Bacteroidia>Bacteroidales | GCA_019129685.1 | Bacteroides_uniformis | DW051_06905 | 94105..96921 | 820 | BREX-system-P-loop-protein-BrxC-[Bacteroides-uniformis] |
| MCI6342424.1 | Type6.BREX | DnaB+TnsC-ATPase->?-><-SIGMA+OB-Fold+AAA-ATPase+ [illegible] | 1217 | Bacteria>FCB_group>Bacteroidetes/Chlorobi_group>Bacteroidota>Bacteroidia>Bacteroidales | GCA_022777695.1 | Prevotella_sp. | MR788_07720 | 61686..65339 | 59823 | BREX-system-P-loop-protein-BrxC-[Prevotella-sp.] |
| MCI7387531.1 | Type6.BREX | Arm-DNA-bind_5-><-UDG<-?<-ARG\|\|?-><-SIGMA+OB-Fol [illegible] | 1215 | Bacteria>FCB_group>Bacteroidetes/Chlorobi_group>Bacteroidota>Bacteroidia>Bacteroidales | GCA_022784345.1 | Bacteroides_uniformis | MSH64_12855 | 232214..235861 | 820 | BREX-system-P-loop-protein-BrxC-[Bacteroides-uniformis] |
| MCR5643213.1 | Type6.BREX | <-N6-METHYLASE<-DUF6079*<-BrxB-BrxF-iSTAND<-BrxA [illegible] | 1209 | Bacteria>FCB_group>Bacteroidetes/Chlorobi_group>Bacteroidota>Bacteroidia>Bacteroidales | GCA_024690905.1 | Prevotella_sp. | K6G32_07765 | 4331..7960 | 59823 | BREX-system-P-loop-protein-BrxC-[Prevotella-sp.] |
| WP_232506819.1 | Type6.BREX | PRTase->?->?->tRNA->tRNA-><-?<-SIGMA+OB-Fold+AAA- [illegible] | 794 | Bacteria>FCB_group>Bacteroidetes/Chlorobi_group>Bacteroidota>Bacteroidia>Bacteroidales | GCF_000178215.1 | Bacteroides_xylanisolvens | CW3_RS0123000 | 100900..103284 | 371601 | hypothetical-protein-[Bacteroides-xylanisolvens] |
| WP_008016782.1 | Type6.BREX | <-SIGMA+OB-Fold+AAA-ATPase+HKD-DNase-BrxL<-PglZ< [illegible] | 425 | Bacteria>FCB_group>Bacteroidetes/Chlorobi_group>Bacteroidota>Bacteroidia>Bacteroidales | GCF_000178215.1 | Bacteroides_xylanisolvens | CW3_RS26225 | 103281..104558 | 371601 | DUF6079-family-protein-[Bacteroides-xylanisolvens] |
| WP_004314649.1 | Type6.BREX | XIS-HTH->CR-ATPase4->ZnR+TOPRIM->TraJ-RHH->?->?- [illegible] | 515 | Bacteria>FCB_group>Bacteroidetes/Chlorobi_group>Bacteroidota>Bacteroidia>Bacteroidales | GCF_000178295.1 | Bacteroides_xylanisolvens | CW1_RS26610 | 23799..25346 | 371601 | BREX-system-P-loop-protein-BrxC-[Bacteroides-xylanisolvens] |
| WP_232505074.1 | Type6.BREX | CR-ATPase4->ZnR+TOPRIM->TraJ-RHH->?->?->BrxE->Br [illegible] | 711 | Bacteria>FCB_group>Bacteroidetes/Chlorobi_group>Bacteroidota>Bacteroidia>Bacteroidales | GCF_000178295.1 | Bacteroides_xylanisolvens | CW1_RS26615 | 25324..27459 | 371601 | hypothetical-protein-[Bacteroides-xylanisolvens] |
| WP_155941928.1 | Type6.BREX | Glyoxalase-><-?<-?\|\|HD->wHTH+WYL+WCX->HTH_3->Brx [illegible] | 1198 | Bacteria>FCB_group>Bacteroidetes/Chlorobi_group>Bacteroidota>Bacteroidia>Bacteroidales | GCF_000424185.1 | Prevotella_sp._AGR2160 | G604_RS15270 | 9168..12764 | 1280674 | BREX-system-P-loop-protein-BrxC-[Prevotella-sp.-AGR2160] |
| WP_044064719.1 | Type6.BREX | <-REase<-?<-SIGMA+OB-Fold+AAA-ATPase+HKD-DNase-B [illegible] | 1217 | Bacteria>FCB_group>Bacteroidetes/Chlorobi_group>Bacteroidota>Bacteroidia>Bacteroidales | GCF_000833995.1 | Prevotella_pectinovora | ST45_RS02505 | 69518..73171 | 1602169 | BREX-system-P-loop-protein-BrxC-[Prevotella-pectinovora] |
| WP_008139697.1 | Type6.BREX | HTH_3->BrxE->BrxA-DNA-binding-p->BrxB-BrxF-iSTAN [illegible] | 1215 | Bacteria>FCB_group>Bacteroidetes/Chlorobi_group>Bacteroidota>Bacteroidia>Bacteroidales | GCF_001315785.1 | Phocaeicola_coprophilus | JCM13818_RS17975 | 17242..20889 | 387090 | BREX-system-P-loop-protein-BrxC-[Phocaeicola-coprophilus] |
| WP_140392864.1 | Type6.BREX | <-TatD_DNase<-?<-?<-KAP-NTPase<-SIGMA+OB-Fold+AA [illegible] | 1211 | Bacteria>FCB_group>Bacteroidetes/Chlorobi_group>Bacteroidota>Bacteroidia>Bacteroidales | GCF_002159645.1 | Parabacteroides_sp._An277 | B5F77_RS15595 | 54457..58092 | 1965619 | BREX-system-P-loop-protein-BrxC-[Parabacteroides-sp.-An277] |
| WP_131226118.1 | Type6.BREX | <-N6-METHYLASE-p<-DUF6079*<-BrxB-BrxF-iSTAND<-Br [illegible] | 1213 | Bacteria>FCB_group>Bacteroidetes/Chlorobi_group>Bacteroidota>Bacteroidia>Bacteroidales | GCF_004328515.1 | Bacteroides | L0M94_RS22830 | 2375..6016 | 816 | MULTISPECIES:-BREX-system-P-loop-protein-BrxC-[Bacteroides] |
| WP_181757796.1 | Type6.BREX | <-DUF772+DDE_Tnp_1+DDE_Tnp_1_6\|\|Arm-DNA-bind_5-> [illegible] | 722 | Bacteria>FCB_group>Bacteroidetes/Chlorobi_group>Bacteroidota>Bacteroidia>Bacteroidales | GCF_013874375.1 | Bacteroides_sp._CACC_737 | H1A11_RS04980 | 1170653..1172821 | 2755405 | hypothetical-protein-[Bacteroides-sp.-CACC-737] |
| WP_181757797.1 | Type6.BREX | Arm-DNA-bind_5-><-UDG<-?<-ARG\|\|?-><-SIGMA+OB-Fol [illegible] | 496 | Bacteria>FCB_group>Bacteroidetes/Chlorobi_group>Bacteroidota>Bacteroidia>Bacteroidales | GCF_013874375.1 | Bacteroides_sp._CACC_737 | H1A11_RS04985 | 1172743..1174233 | 2755405 | DUF6079-family-protein-[Bacteroides-sp.-CACC-737] |
| WP_186905779.1 | Type6.BREX | <-TRAM+METHYLASE\|\|?->Arm-DNA-bind_5-><-UDG<-?<-A [illegible] | 1213 | Bacteria>FCB_group>Bacteroidetes/Chlorobi_group>Bacteroidota>Bacteroidia>Bacteroidales | GCF_014287575.1 | Bacteroides_sp._NSJ-21 | H8S53_RS11040 | 2778988..2782629 | 2763025 | BREX-system-P-loop-protein-BrxC-[Bacteroides-sp.-NSJ-21] |
| WP_195289444.1 | Type6.BREX | CR-ATPase4->ZnR+TOPRIM->TraJ-RHH->?->GNAT->HTH_3 [illegible] | 1224 | Bacteria>FCB_group>Bacteroidetes/Chlorobi_group>Bacteroidota>Bacteroidia>Bacteroidales | GCF_015549385.1 | Phocaeicola_massiliensis | I2C40_RS17240 | 9600..13274 | 204516 | BREX-system-P-loop-protein-BrxC-[Phocaeicola-massiliensis] |
| WP_195366148.1 | Type6.BREX | <-TatD_DNase<-?<-KAP-glycosylase<-KAP-NTPase<-SI [illegible] | 921 | Bacteria>FCB_group>Bacteroidetes/Chlorobi_group>Bacteroidota>Bacteroidia>Bacteroidales | GCF_015550085.1 | Bacteroides_ovatus | PO368_RS20360 | 46610..49375 | 28116 | BREX-system-P-loop-protein-BrxC-[Bacteroides-ovatus] |
| WP_117964425.1 | Type6.BREX | Arm-DNA-bind_5-><-UDG<-?<-ARG\|\|RHH+MobC->?-><-SI [illegible] | 265 | Bacteria>FCB_group>Bacteroidetes/Chlorobi_group>Bacteroidota>Bacteroidia>Bacteroidales | GCF_015679285.1 | Bacteroides | ITJ87_RS11480 | 2822744..2823541 | 816 | MULTISPECIES:-DUF6079-family-protein-[Bacteroides] |
| WP_203049164.1 | Type6.BREX | <-TOPRIM<-NUDIX<-?<-DUF4369+Thioredoxin_8\|\|RNase [illegible] | 1216 | Bacteria>FCB_group>Bacteroidetes/Chlorobi_group>Bacteroidota>Bacteroidia>Bacteroidales | GCF_016803255.1 | Prevotella_copri | F8Q89_RS01960 | 71962..75612 | 165179 | BREX-system-P-loop-protein-BrxC-[Prevotella-copri] |
| WP_215758559.1 | Type6.BREX | Phage_int_SAM_5+Phage_integrase->?->XIS-HTH->?-> [illegible] | 1217 | Bacteria>FCB_group>Bacteroidetes/Chlorobi_group>Bacteroidota>Bacteroidia>Bacteroidales | GCF_018789675.1 | Hallella_faecis | KM934_00340 | 78197..81850 | 2841596 | BREX-system-P-loop-protein-BrxC-[Hallella-faecis] |
| WP_138333738.1 | Type6.BREX | STYKinase-like->Pkinase->SIR2+SIR2->?->?-><-REas [illegible] | 1224 | Bacteria>FCB_group>Bacteroidetes/Chlorobi_group>Bacteroidota>Bacteroidia>Bacteroidales | GCF_019412725.1 | Bacteroides | GPL18_RS15350 | 126249..129923 | 816 | MULTISPECIES:-BREX-system-P-loop-protein-BrxC-[Bacteroides] |
| WP_237798701.1 | Type6.BREX | DSBH+AraC-HTH+AraC-HTH->\|\|?->?-><-?<-SF1-SF2-Hel [illegible] | 1216 | Bacteria>FCB_group>Bacteroidetes/Chlorobi_group>Bacteroidota>Bacteroidia>Bacteroidales | GCF_022024195.1 | Prevotella_bryantii | L6467_12860 | 2115705..2119355 | 77095 | BREX-system-P-loop-protein-BrxC-[Prevotella-bryantii] |
| WP_272706625.1 | Type6.BREX | XIS-HTH-><-?<-?\|\|?->?->HTH_3->BrxE->BrxA-DNA-bin [illegible] | 191 | Bacteria>FCB_group>Bacteroidetes/Chlorobi_group>Bacteroidota>Bacteroidia>Bacteroidales | GCF_028528065.1 | Bacteroides_faecis | PQH00_RS23500 | 20443..21015 | 674529 | DUF6079-family-protein,-partial-[Bacteroides-faecis] |
| WP_272706631.1 | Type6.BREX | HTH_3->BrxE->BrxA-DNA-binding->BrxB-BrxF-iSTAND- [illegible] | 862 | Bacteria>FCB_group>Bacteroidetes/Chlorobi_group>Bacteroidota>Bacteroidia>Bacteroidales | GCF_028528065.1 | Bacteroides_faecis | PQH00_RS23505 | 21026..23611 | 674529 | BREX-system-P-loop-protein-BrxC,-partial-[Bacteroides-faecis] |
| WP_274990748.1 | Type6.BREX | Arm-DNA-bind_5-><-UDG<-?<-ARG\|\|?-><-SIGMA+OB-Fol [illegible] | 772 | Bacteria>FCB_group>Bacteroidetes/Chlorobi_group>Bacteroidota>Bacteroidia>Bacteroidales | GCF_028993715.1 | Bacteroides_uniformis | PX688_RS13675 | 79793..82111 | 820 | hypothetical-protein-[Bacteroides-uniformis] |
| WP_274990749.1 | Type6.BREX | Arm-DNA-bind_5-><-UDG<-?<-ARG\|\|?-><-SIGMA+OB-Fol [illegible] | 411 | Bacteria>FCB_group>Bacteroidetes/Chlorobi_group>Bacteroidota>Bacteroidia>Bacteroidales | GCF_028993715.1 | Bacteroides_uniformis | PX688_RS13680 | 82199..83434 | 820 | DUF6079-family-protein-[Bacteroides-uniformis] |
| WP_008770354.1 | Type6.BREX | DDE_Tnp-><-?<-SIGMA+OB-Fold+AAA-ATPase+HKD-DNase- [illegible] | 1219 | Bacteria>FCB_group>Bacteroidetes/Chlorobi_group>Bacteroidota>Bacteroidia>Bacteroidales | GCF_902374095.1 | Bacteroides | BSAG_RS19905 | 423901..427560 | 816 | MULTISPECIES:-BREX-system-P-loop-protein-BrxC-[Bacteroides] |
| WP_174646033.1 | Type6.BREX | AbiEii->XIS-HTH-><-?\|\|?->HTH_3->BrxE->BrxA-DNA-b [illegible] | 1215 | Bacteria>FCB_group>Bacteroidetes/Chlorobi_group>Bacteroidota>Bacteroidia>Bacteroidales | GCF_902825465.1 | Bacteroides_sp._Marseille-P8574 | HUX74_RS00900 | 196439..200086 | 2697504 | BREX-system-P-loop-protein-BrxC-[Bacteroides-sp.-Marseille-P8574] |
| WP_046149424.1 | Type6.BREX | AbiEii->?-><-?<-?\|\|XIS-HTH->HTH_3->BrxE->BrxA-DN [illegible] | 1217 | Bacteria>FCB_group>Bacteroidetes/Chlorobi_group>Bacteroidota>Bacteroidia>Bacteroidales | - | Parabacteroides_sp._HGS0025 | HMPREF1212_RS26675 | 2885204..2888857 | 1078087 | BREX-system-P-loop-protein-BrxC-[Parabacteroides-sp.-HGS0025] |
| WP_117695280.1 | Type6.BREX | <-SLATT_4<-SIGMA+OB-Fold+AAA-ATPase+HKD-DNase-Br [illegible] | 1216 | Bacteria>FCB_group>Bacteroidetes/Chlorobi_group>Bacteroidota>Bacteroidia>Bacteroidales | - | Prevotella_copri | DXC61_RS12020 | 26690..30340 | 165179 | BREX-system-P-loop-protein-BrxC-[Prevotella-copri] |
| WP_233339205.1 | Type6.BREX | <-SLATT_4<-SIGMA+OB-Fold+AAA-ATPase+HKD-DNase-Br [illegible] | 1216 | Bacteria>FCB_group>Bacteroidetes/Chlorobi_group>Bacteroidota>Bacteroidia>Bacteroidales | - | Prevotella_copri | LYY06_RS08410 | 34942..38592 | 165179 | BREX-system-P-loop-protein-BrxC-[Prevotella-copri] |
| PLW98642.1 | Type6.BREX | MarR-HTH->?->REase->BrxE->BrxA-DNA-binding->BrxB [illegible] | 249 | Bacteria>FCB_group>Bacteroidetes/Chlorobi_group>Bacteroidota>Bacteroidia>Marinilabiliales | GCA_002869405.1 | Marinilabiliales_bacterium | C0593_05050 | 18447..19193 | 2053303 | MAG:-BREX-system-P-loop-protein-BrxC,-partial-[Marinilabiliales-bacterium] |
| WP_173247258.1 | Type6.BREX | <-METHYLASE<-?<-?<-N6-METHYLASE-p<-?<-?<-DUF6079 [illegible] | 240 | Bacteria>FCB_group>Bacteroidetes/Chlorobi_group>Bacteroidota>Cytophagia>Cytophagales | GCF_013256425.1 | Hymenobacter_sp._BRD67 | GKZ67_RS20920 | 36982..37704 | 2675877 | hypothetical-protein-[Hymenobacter-sp.-BRD67] |
| WP_191006191.1 | Type6.BREX | HEPN->Phage_integrase+Phage_integrase-><-?<-HEPN [illegible] | 1204 | Bacteria>FCB_group>Bacteroidetes/Chlorobi_group>Bacteroidota>Cytophagia>Cytophagales | GCF_014699115.1 | Hymenobacter_montanus | IC235_RS15945 | 31815..35429 | 2771359 | BREX-system-P-loop-protein-BrxC-[Hymenobacter-montanus] |
| WP_204660265.1 | Type6.BREX | <-HTH_3\|\|?->?->CopG/RHH->CR-ATPase5+REase->AbiEi [illegible] | 1216 | Bacteria>FCB_group>Bacteroidetes/Chlorobi_group>Bacteroidota>Cytophagia>Cytophagales | GCF_016894445.1 | Dyadobacter_sandarakinus | HWI92_RS00525 | 139788..143438 | 2747268 | BREX-system-P-loop-protein-BrxC-[Dyadobacter-sandarakinus] |
| WP_243520943.1 | Type6.BREX | Glyoxalase-><-?<-HEPN<-?\|\|?->?->BrxE->BrxA-DNA-b [illegible] | 1204 | Bacteria>FCB_group>Bacteroidetes/Chlorobi_group>Bacteroidota>Cytophagia>Cytophagales | GCF_022811645.1 | Hymenobacter_monticola | MTP16_RS25545 | 68480..72094 | 1705399 | BREX-system-P-loop-protein-BrxC-[Hymenobacter-monticola] |
| WP_235164443.1 | Type6.BREX | <-STYKIN<-?<-?<-?<-McrC-NTD+REase<-HTH+McrB-AAA< [illegible] | 1216 | Bacteria>FCB_group>Bacteroidetes/Chlorobi_group>Bacteroidota>Cytophagia>Cytophagales | GCF_023821985.2 | Dyadobacter_chenhuakuii | NFI80_RS01415 | 331734..335384 | 2909339 | BREX-system-P-loop-protein-BrxC-[Dyadobacter-chenhuakuii] |
| WP_027137224.1 | Type6.BREX | DarT+Peptidase->MACRODOMAIN->TPR+TPR->?-><-SIGMA [illegible] | 1196 | Bacteria>FCB_group>Bacteroidetes/Chlorobi_group>Bacteroidota>Flavobacteriia>Flavobacteriales | GCF_000425645.1 | Gaetbulibacter_saemankumensis | H611_RS0104475 | 476638..480228 | 311208 | BREX-system-P-loop-protein-BrxC-[Gaetbulibacter-saemankumensis] |
| WP_166386804.1 | Type6.BREX | Arm-DNA-bind_5+Phage_int_SAM_5+Phage_integrase-> [illegible] | 1195 | Bacteria>FCB_group>Bacteroidetes/Chlorobi_group>Bacteroidota>Flavobacteriia>Flavobacteriales | GCF_011379025.1 | Polaribacter_sp._11A2H | GQS58_RS13590 | 79017..82604 | 2687290 | BREX-system-P-loop-protein-BrxC-[Polaribacter-sp.-11A2H] |
| WP_193813001.1 | Type6.BREX | <-wHTH+WYL+WCX\|\|ZnR+CC+REase->?-><-?<-SIGMA+OB-F [illegible] | 1217 | Bacteria>FCB_group>Bacteroidetes/Chlorobi_group>Bacteroidota>Flavobacteriia>Flavobacteriales | GCF_015191005.1 | Kaistella_flava_(ex_Peng_et_al._2021) | Q73A0000_05090 | 1135416..1139069 | 2038776 | BREX-system-P-loop-protein-BrxC-[Kaistella-flava-(ex-Peng-et-al.-2021)] |
| MBK7566196.1 | Type6.BREX | <-SF1-SF2-Helicase<-MORC+DUF3684+REase<-?<-ParB< [illegible] | 1193 | Bacteria>FCB_group>Bacteroidetes/Chlorobi_group>Bacteroidota>unclassified_Bacteroidota | GCA_016706865.1 | Bacteroidota_bacterium | IPI31_00060 | 18734..22315 | 1898104 | BREX-system-P-loop-protein-BrxC-[Bacteroidota-bacterium] |
| NMW22669.1 | Type6.BREX | <-SIGMA+OB-Fold+AAA-ATPase+HKD-DNase-BrxL<-PglZ< [illegible] | 1210 | Bacteria>FCB_group>Bacteroidetes/Chlorobi_group>Chlorobiota>Chlorobiia>Chlorobiales | GCA_012972835.1 | Chlorobiaceae_bacterium | HKK67_13830 | 33011..36643 | 2067991 | BREX-system-P-loop-protein-BrxC-[Chlorobiaceae-bacterium] |
| MBO6574434.1 | Type6.BREX | ABC-ATPase+SbcC+ABC-ATPase+SbcC->?->?-><-HKD-DNa [illegible] | 1180 | Bacteria>FCB_group>Bacteroidetes/Chlorobi_group>Rhodothermota>Rhodothermia>Rhodothermales | GCA_017642665.1 | Rhodothermales_bacterium | JJ976_03365 | 859179..862721 | 2306054 | BREX-system-P-loop-protein-BrxC-[Rhodothermales-bacterium] |
| MCB5285398.1 | Type6.BREX | <-DNA_METHYLASE<-SIGMA+OB-Fold+AAA-ATPase+Lon-Pr [illegible] | 1227 | Bacteria>FCB_group>Candidatus_Cloacimonetes>unclassified_Candidatus_Cloacimonetes | GCA_020532085.1 | Candidatus_Cloacimonetes_bacterium | LHW45_07405 | 38085..41768 | 2030808 | BREX-system-P-loop-protein-BrxC-[Candidatus-Cloacimonadota-bacterium] |
| NQV48959.1 | Type6.BREX | ParB+HNH-p->XIS-HTH->BrxE-p->BrxE-p->BrxA-DNA-bi [illegible] | 1210 | Bacteria>FCB_group>Candidatus_Marinimicrobia | GCA_013202385.1 | Candidatus_Marinimicrobia_bacterium | HQ507_00570 | 3171..6803 | 2026760 | BREX-system-P-loop-protein-BrxC-[Candidatus-Marinimicrobia-bacterium] |
| ODT04869.1 | Type6.BREX | <-SIGMA+OB-Fold+AAA-ATPase+Lon-Protease-BrxL<-Pg [illegible] | 1181 | Bacteria>FCB_group>Gemmatimonadota>unclassified_Gemmatimonadota | GCA_001724275.1 | Gemmatimonadetes_bacterium_SCN_70-22 | ABS52_01570 | 17993..21538 | 1660163 | hypothetical-protein-ABS52_01570-[Gemmatimonadetes-bacterium-SCN-70-22] |
| MBK6845930.1 | Type6.BREX | Integrase_DNA+Phage_integrase-><-?<-?<-TPR+TPR+T [illegible] | 1212 | Bacteria>FCB_group>Gemmatimonadota>unclassified_Gemmatimonadota | GCA_016702715.1 | Gemmatimonadota_bacterium | IPG88_27295 | 59207..62845 | 2026742 | BREX-system-P-loop-protein-BrxC-[Gemmatimonadota-bacterium] |
| MBX6362419.1 | Type6.BREX | Peptidase-><-?\|\|BrxE->BrxA-DNA-binding->BrxB-Brx [illegible] | 1201 | Bacteria>FCB_group>Gemmatimonadota>unclassified_Gemmatimonadota | GCA_019689775.1 | Gemmatimonadota_bacterium | IRZ00_01015 | 11407..15012 | 2026742 | BREX-system-P-loop-protein-BrxC-[Gemmatimonadota-bacterium] |
| WP_240631539.1 | Type6.BREX | <-Tox-RES<-XIS-HTH\|\|HDRnase-><-SIGMA+OB-Fold+AAA [illegible] | 1234 | Bacteria | GCF_003671915.1 | Pseudomonadota | D3M96_RS03425 | 754727..758431 | 1224 | MULTISPECIES:-BREX-system-P-loop-protein-BrxC-[Pseudomonadota] |
| WP_203328719.1 | Type6.BREX | <-TraM<-DUF3262<-?<-?<-SMODS<-?\|\|wHTH+WYL+WCX->B [illegible] | 1213 | Bacteria | GCF_016836415.1 | Pseudomonadota | JQ482_RS25705 | 5512214..5515855 | 1224 | MULTISPECIES:-BREX-system-P-loop-protein-BrxC-[Pseudomonadota] |
| WP_240685338.1 | Type6.BREX | SMODS->?->JAB-><-SIGMA+OB-Fold+AAA-ATPase+HKD-DN [illegible] | 1213 | Bacteria | GCF_025461165.1 | Pseudomonadota | JAK52_RS02505 | 287897..291538 | 1224 | MULTISPECIES:-BREX-system-P-loop-protein-BrxC-[Pseudomonadota] |
| WP_012632492.1 | Type6.BREX | <-REase+35exo<-?\|\|BrxE->BrxA-DNA-binding->BrxB-B [illegible] | 1206 | Bacteria>Myxococcota>Myxococcia>Myxococcales>Cystobacterineae>Anaeromyxobacteraceae | GCF_000022145.1 | Anaeromyxobacter_dehalogenans | A2CP1_RS05765 | 1286335..1289955 | 161493 | BREX-system-P-loop-protein-BrxC-[Anaeromyxobacter-dehalogenans] |
| WP_242370713.1 | Type6.BREX | HTH_3-><-?\|\|BrxE->BrxA-DNA-binding->BrxB-BrxF-iS [illegible] | 1210 | Bacteria>Myxococcota>Myxococcia>Myxococcales>Cystobacterineae>Anaeromyxobacteraceae | GCF_022690725.1 | Anaeromyxobacter_sp._SG26 | MNY63_RS01430 | 49071..52703 | 2925407 | BREX-system-P-loop-protein-BrxC-[Anaeromyxobacter-sp.-SG26] |
| WP_242352153.1 | Type6.BREX | <-RVT+Trypsin\|\|REase->REase-><-DNA_METHYLASE<-?\| [illegible] | 1179 | Bacteria>Myxococcota>Myxococcia>Myxococcales>Cystobacterineae>Anaeromyxobacteraceae | GCF_022690745.1 | Anaeromyxobacter_sp._SG64 | MNZ29_RS00510 | 126977..130516 | 2925409 | BREX-system-P-loop-protein-BrxC-[Anaeromyxobacter-sp.-SG64] |

| Protein-ID | System | Gene-Neighbourhoods | Length | Taxonomy | Assembly-ID | Strain | Locus-Tag | Genomic-coordinates | TaxID | Fasta-define |
|---|---|---|---|---|---|---|---|---|---|---|
| WP_242352647.1 | Type6.BREX | <-N6-METHYLASE<-?<-DUF6079*<-BrxB-BrxF-iSTAND<-BrxA-DNA-binding<-BrxE<-?<-?<-?<-ZnR+TOPRIM+TOPRIM | 1179 | Bacteria>Myxococcota>Myxococcia>Myxococcales>Cystobacterineae>Anaeromyxobacteraceae | GCF_022690745.1 | Anaeromyxobacter_sp._SG64 | MNZ29_RS02505 | 2253..5792 | 2925409 | BREX-system-P-loop-protein-BrxC-[Anaeromyxobacter-sp.-SG64] |
| QRK08524.1 | Type6.BREX | <-Peptidase_S66+Peptidase_S66C<-DNA_METHYLASE\|\|tRNA-><-glycosylase<-SIGMA+OB-Fold+AAA-ATPase+Lon-Protease-BrxL<-PglZ<-N6-METHYLASE<-DUF6079*<-BrxB-BrxF-iSTAND<-BrxA-DNA-binding<-BrxE<-XIS-HTH<-?<-GDResolvase+ZnR<-?<-RAMA | 1181 | Bacteria>Myxococcota>Myxococcia>Myxococcales>Cystobacterineae>Archangiaceae | GCA_016859125.1 | Archangium_violaceum | JQX13_52995 | 12541464..12545009 | 83451 | BREX-system-P-loop-protein-BrxC-[Archangium-violaceum] |
| WP_239014411.1 | Type6.BREX | <-Peptidase_S66+Peptidase_S66C<-DNA_METHYLASE\|\|tRNA-><-glycosylase<-SIGMA+OB-Fold+AAA-ATPase+Lon-Protease-BrxL<-PglZ<-N6-METHYLASE<-DUF6079*<-BrxB-BrxF-iSTAND<-BrxA-DNA-binding<-BrxE<-XIS-HTH<-?<-GDResolvase+ZnR<-?<-RAMA | 1179 | Bacteria>Myxococcota>Myxococcia>Myxococcales>Cystobacterineae>Archangiaceae | GCF_016859125.1 | Archangium_violaceum | JQX13_RS52990 | 12541464..12545003 | 83451 | BREX-system-P-loop-protein-BrxC-[Archangium-violaceum] |
| WP_095958445.1 | Type6.BREX | <-Phage_integrase<-DDE_Tnp\|\|<-URI-DNase<-?\|\|HNH-><-SF1-SF2-Helicase+wHTH+Small-B-barrel+DUF1998<-BrxD-AAA<-PglZ<-N6-METHYLASE<-DUF6079*<-BrxB-BrxF-iSTAND<-BrxA-DNA-binding<-BrxE<-?<-?\|\|N6-METHYLASE-> | 1203 | Bacteria>Myxococcota>Myxococcia>Myxococcales>Cystobacterineae>Myxococcaceae | GCF_002305895.1 | Corallococcus_macrosporus | MYMAC_RS14000 | 3308377..3311988 | 35 | BREX-system-P-loop-protein-BrxC-[Corallococcus-macrosporus] |
| WP_120625208.1 | Type6.BREX | <-SF1-SF2-Helicase+wHTH+Small-B-barrel+DUF1998<-BrxD-AAA<-PglZ<-N6-METHYLASE<-DUF6079*<-BrxB-BrxF-iSTAND<-BrxA-DNA-binding<-BrxE<-wHTH+REase+ZnR<-McrB-AAA<-HTH+HEPN<-SF1-SF2-Helicase<-STYKIN | 1203 | Bacteria>Myxococcota>Myxococcia>Myxococcales>Cystobacterineae>Myxococcaceae | GCF_003611735.1 | Corallococcus_sicarius | D7X12_RS10925 | 19643..23254 | 2316726 | BREX-system-P-loop-protein-BrxC-[Corallococcus-sicarius] |
| WP_120577073.1 | Type6.BREX | PGK->?->?->tRNA->XIS-HTH->BrxE->BrxA-DNA-binding->BrxB-BrxF-iSTAND->DUF6079*->Tox-RelA->N6-METHYLASE->PglZ->SIGMA+OB-Fold+AAA-ATPase+Lon-Protease-BrxL->glycosylase-> | 1188 | Bacteria>Myxococcota>Myxococcia>Myxococcales>Cystobacterineae>Myxococcaceae | GCF_003612075.1 | Corallococcus_sp._CA041A | D7V77_RS15540 | 7194..10760 | 2316727 | BREX-system-P-loop-protein-BrxC-[Corallococcus-sp.-CA041A] |
| WP_140874522.1 | Type6.BREX | rve->AAA-ATPase->?->HTH_58->?-><-SF1-SF2-Helicase+wHTH+Small-B-barrel+DUF1998<-BrxD-AAA<-PglZ<-N6-METHYLASE<-DUF6079*<-BrxB-BrxF-iSTAND<-BrxA-DNA-binding<-BrxE<-Phage_int_SAM_3+Phage_integrase<-XIS-HTH<-?<-?\|\|?->Pkinase-> | 1201 | Bacteria>Myxococcota>Myxococcia>Myxococcales>Cystobacterineae>Myxococcaceae | GCF_006402015.1 | Myxococcus_xanthus | BHS07_RS13795 | 3252653..3256258 | 34 | BREX-system-P-loop-protein-BrxC-[Myxococcus-xanthus] |
| WP_169348680.1 | Type6.BREX | ACET->?->?->?->BrxE->BrxA-DNA-binding->BrxB-BrxF-iSTAND->DUF6079*->N6-METHYLASE->PglZ->Pkinase-> | 1203 | Bacteria>Myxococcota>Myxococcia>Myxococcales>Cystobacterineae>Myxococcaceae | GCF_012933655.1 | Pyxidicoccus_fallax | HPC49_RS25470 | 6020..9631 | 394095 | BREX-system-P-loop-protein-BrxC-[Pyxidicoccus-fallax] |
| WP_174479070.1 | Type6.BREX | <-SIGMA+OB-Fold+AAA-ATPase+Lon-Protease-BrxL<-PglZ<-N6-METHYLASE<-N6-METHYLASE<-DUF6079*<-BrxB-BrxF-iSTAND<-BrxA-DNA-binding<-BrxE<-XIS-HTH<-DDE_Tnp_1<-?<-?<-?<-rve_3 | 1179 | Bacteria>Myxococcota>Myxococcia>Myxococcales>Cystobacterineae>Myxococcaceae | GCF_013336625.1 | Myxococcus_sp._CA033 | HUA78_RS45185 | 10597..14136 | 2741516 | BREX-system-P-loop-protein-BrxC-[Myxococcus-sp.-CA033] |
| WP_174464828.1 | Type6.BREX | <-Glycos_transf<-Glycos_transf\|\|?->XIS-HTH->MORC+DUF3684+REase->SF1-SF2-Helicase->BrxE->BrxA-DNA-binding->BrxB-BrxF-iSTAND->DUF6079*->N6-METHYLASE->PglZ-> | 1203 | Bacteria>Myxococcota>Myxococcia>Myxococcales>Cystobacterineae>Myxococcaceae | GCF_013336645.1 | Myxococcus_sp._CA039A | HR086_RS03110 | 56984..60595 | 2741737 | BREX-system-P-loop-protein-BrxC-[Myxococcus-sp.-CA039A] |
| WP_176433375.1 | Type6.BREX | ABC-ATPase+SbcC->ADH_N+ADH_zinc_N->?->?->SF1-SF2-Helicase->BrxE->BrxA-DNA-binding->BrxB-BrxF-iSTAND->DUF6079*->N6-METHYLASE->PglZ-> | 1203 | Bacteria>Myxococcota>Myxococcia>Myxococcales>Cystobacterineae>Myxococcaceae | GCF_013372595.1 | Myxococcus_sp._AM011 | HUW62_RS01785 | 94774..98385 | 2745200 | BREX-system-P-loop-protein-BrxC-[Myxococcus-sp.-AM011] |
| WP_201422032.1 | Type6.BREX | <-glycosylase<-SIGMA+OB-Fold+AAA-ATPase+Lon-Protease-BrxL<-PglZ<-HNH<-OLD-ABC<-N6-METHYLASE<-N6-METHYLASE<-Tox-RelA<-DUF6079*<-BrxB-BrxF-iSTAND<-BrxA-DNA-binding<-BrxE<-XIS-HTH\|\|<-ACET\|\|?->?->REC+NtrC-AAA+FIS-HTH-> | 1188 | Bacteria>Myxococcota>Myxococcia>Myxococcales>Cystobacterineae>Myxococcaceae | GCF_016698685.1 | Myxococcus_xanthus | JXA73_RS29390 | 7346078..7349644 | 34 | BREX-system-P-loop-protein-BrxC-[Myxococcus-xanthus] |
| WP_267556343.1 | Type6.BREX | <-rve_3<-SIGMA+OB-Fold+AAA-ATPase+Lon-Protease-BrxL<-PglZ<-HNH<-OLD-ABC<-N6-METHYLASE<-Tox-RelA<-DUF6079*<-BrxB-BrxF-iSTAND<-BrxA-DNA-binding<-BrxE<-XIS-HTH\|\|?-><-?<-wHTH(SBD)+HNH<-?<-rve_3 | 1189 | Bacteria>Myxococcota>Myxococcia>Myxococcales>Cystobacterineae>Myxococcaceae | GCF_026626625.1 | Corallococcus_sp._BB11-1 | OV207_RS17695 | 240885..244454 | 2996783 | BREX-system-P-loop-protein-BrxC-[Corallococcus-sp.-BB11-1] |
| AKU90922.1 | Type6.BREX | APATPase+TPR+TPR+TPR+TPR+TPR+TPR+TPR-><-?\|\|?->?->YjhX_toxin->BrxE->BrxA-DNA-binding->BrxB-BrxF-iSTAND->DUF6079*->N6-METHYLASE->PglZ->BrxD-AAA->?->SF1-SF2-Helicase+wHTH+Small-B-barrel+DUF1998->HTH+REase-> | 1203 | Bacteria>Myxococcota>Myxococcia>Myxococcales>Cystobacterineae>Vulgatibacteraceae | GCA_001263175.1 | Vulgatibacter_incomptus | AKJ08_1309 | 1574430..1578041 | 1391653 | hypothetical-protein-AKJ08_1309-[Vulgatibacter-incomptus] |
| WP_050725309.1 | Type6.BREX | APATPase+TPR+TPR+TPR+TPR+TPR+TPR+TPR->?-><-?\|\|?->YjhX_toxin->BrxE->BrxA-DNA-binding->BrxB-BrxF-iSTAND->DUF6079*->N6-METHYLASE->PglZ->BrxD-AAA->Calcineurin->ABC-3C->?->SF1-SF2-Helicase+wHTH+Small-B-barrel+DUF1998->HTH+REase-> | 1202 | Bacteria>Myxococcota>Myxococcia>Myxococcales>Cystobacterineae>Vulgatibacteraceae | GCF_001263175.1 | Vulgatibacter_incomptus | AKJ08_RS06425 | 1574433..1578041 | 1391653 | BREX-system-P-loop-protein-BrxC-[Vulgatibacter-incomptus] |
| WP_106091681.1 | Type6.BREX | <-HTH_3\|\|?-><-?\|\|?->?-><-?\|\|BrxE->BrxA-DNA-binding->BrxB-BrxF-iSTAND->DUF6079*->N6-METHYLASE->?->PglZ->BrxD-AAA->SF1-SF2-Helicase+wHTH+Small-B-barrel+35exo->?->HKD-DNase+HKD-Rease-> | 1214 | Bacteria>Myxococcota>Myxococcia>Myxococcales>Myxococcales_incertae_sedis>Enhygromyxa | GCF_002994635.1 | Enhygromyxa_salina | ENSA7_RS24060 | 62772..66416 | 215803 | BREX-system-P-loop-protein-BrxC-[Enhygromyxa-salina] |
| HCF56393.1 | Type6.BREX | <-Schlafen+HATPase+wHTH<-N6-METHYLASE-p\|\|?->PIN-><-TOPRIM<-OLD-ABC<-DUF6079-p<-APATPase<-DUF6079-p*<-DUF6079-p<-HNH<-BrxB-BrxF-iSTAND<-BrxA-DNA-binding<-BrxA-DNA-binding<-BrxE | 508 | Bacteria>Myxococcota>Myxococcia>Myxococcales>unclassified_Myxococcales | GCA_003534605.1 | Myxococcales_bacterium | DFS52_00150 | 7726..9251 | 2026763 | BREX-system-P-loop-protein-BrxC-[Myxococcales-bacterium] |
| HCF56394.1 | Type6.BREX | <-Schlafen+HATPase+wHTH<-N6-METHYLASE-p\|\|?->PIN-><-TOPRIM<-OLD-ABC<-DUF6079-p<-APATPase<-DUF6079-p<-DUF6079-p*<-HNH<-BrxB-BrxF-iSTAND<-BrxA-DNA-binding<-BrxA-DNA-binding<-BrxE | 363 | Bacteria>Myxococcota>Myxococcia>Myxococcales>unclassified_Myxococcales | GCA_003534605.1 | Myxococcales_bacterium | DFS52_00155 | 9470..10559 | 2026763 | BREX-system-P-loop-protein-BrxC-[Myxococcales-bacterium] |
| NUN14061.1 | Type6.BREX | HTH+REase->BrxE->BrxA-DNA-binding->BrxB-BrxF-iSTAND->DUF6079*->?->N6-METHYLASE->PglZ->BrxD-AAA->SF1-SF2-Helicase+wHTH+Small-B-barrel+DUF1998->Schlafen+HATPase+wHTH-> | 1213 | Bacteria>Myxococcota>Myxococcia>Myxococcales>unclassified_Myxococcales | GCA_013360285.1 | Myxococcales_bacterium | HUU55_10555 | 3445..7086 | 2026763 | BREX-system-P-loop-protein-BrxC-[Myxococcales-bacterium] |
| MBK6463374.1 | Type6.BREX | <-Sulfatase+DUF4976\|\|?->?-><-?\|\|?->?->BrxE->BrxA-DNA-binding->BrxB-BrxF-iSTAND->DUF6079*->SbcC->N6-METHYLASE-p->?->N6-METHYLASE-p->N6-METHYLASE-p->REase->N6-METHYLASE-p->HTH+REase-> | 1235 | Bacteria>Myxococcota>Myxococcia>Myxococcales>unclassified_Myxococcales | GCA_016704525.1 | Myxococcales_bacterium | IPF92_20555 | 135329..139036 | 2026763 | BREX-system-P-loop-protein-BrxC-[Myxococcales-bacterium] |
| MBL9099429.1 | Type6.BREX | MerR-HTH->BrxE->BrxA-DNA-binding->BrxB-BrxF-iSTAND->DUF6079*->N6-METHYLASE->PglZ->BrxD-AAA->?->SF1-SF2-Helicase+wHTH+Small-B-barrel+35exo->?->GT4+RADICAL-SAM->Pkinase-> | 1209 | Bacteria>Myxococcota>Myxococcia>Myxococcales>unclassified_Myxococcales | GCA_016793465.1 | Myxococcales_bacterium | JNL82_00645 | 4641..8270 | 2026763 | BREX-system-P-loop-protein-BrxC-[Myxococcales-bacterium] |
| MBT9554535.1 | Type6.BREX | STAS+HisKin-><-SF1-SF2-Helicase+wHTH+Small-B-barrel+DUF1998<-HNH<-OLD-ABC<-BrxD-AAA<-PglZ<-N6-METHYLASE<-DUF6079*<-BrxB-BrxF-iSTAND<-BrxA-DNA-binding<-BrxE<-?<-?<-RADICAL-SAM<-?\|\|SWACOS-><-SF1-SF2-Helicase+DUF3418 | 1211 | Bacteria>Myxococcota>Myxococcia>Myxococcales>unclassified_Myxococcales | GCA_018780435.1 | Myxococcales_bacterium | IV100_00845 | 29582..33217 | 2026763 | BREX-system-P-loop-protein-BrxC-[Myxococcales-bacterium] |
| MBX3245755.1 | Type6.BREX | RAMA->?->GDResolvase+ZnR->?->XIS-HTH->BrxE->BrxA-DNA-binding->BrxB-BrxF-iSTAND->DUF6079*->N6-METHYLASE->PglZ->SIGMA+OB-Fold+AAA-ATPase+Lon-Protease-BrxL->N4-N6-METHYLASE->ParB+HNH->HEPN-><-HerA/FtsK<-?\|\|glycosylase-> | 1181 | Bacteria>Myxococcota>Myxococcia>Myxococcales>unclassified_Myxococcales | GCA_019637115.1 | Myxococcales_bacterium | KF901_01090 | 38728..42273 | 2026763 | BREX-system-P-loop-protein-BrxC-[Myxococcales-bacterium] |
| MBX3247840.1 | Type6.BREX | FGS->?->?->RADICAL-SAM->?->BrxE->BrxA-DNA-binding->BrxB-BrxF-iSTAND->DUF6079*->N6-METHYLASE-><-AbiEii<-wHTH+AbiEi_2\|\|HTH+MPTase->Peptidase->PglZ->BrxD-AAA->SF1-SF2-Helicase+wHTH+Small-B-barrel+35exo->HTH+REase-> | 1203 | Bacteria>Myxococcota>Myxococcia>Myxococcales>unclassified_Myxococcales | GCA_019637115.1 | Myxococcales_bacterium | KF901_11730 | 14699..18310 | 2026763 | BREX-system-P-loop-protein-BrxC-[Myxococcales-bacterium] |
| MCA9627282.1 | Type6.BREX | <-DUF6079*<-BrxB-BrxF-iSTAND<-BrxA-DNA-binding<-BrxE\|\|XIS-HTH->TIR+PNTD2-><-?\|\|PIWI-> | 1208 | Bacteria>Myxococcota>Myxococcia>Myxococcales>unclassified_Myxococcales | GCA_020430865.1 | Myxococcales_bacterium | KC766_06435 | 393..4019 | 2026763 | BREX-system-P-loop-protein-BrxC-[Myxococcales-bacterium] |
| MCA9678853.1 | Type6.BREX | BrxE->BrxA-DNA-binding->BrxB-BrxF-iSTAND->DUF6079*->N6-METHYLASE->PglZ->?->BrxD-AAA->SF1-SF2-Helicase+wHTH+Small-B-barrel+35exo-> | 1205 | Bacteria>Myxococcota>Myxococcia>Myxococcales>unclassified_Myxococcales | GCA_020431345.1 | Myxococcales_bacterium | KC464_27740 | 2206..5823 | 2026763 | BREX-system-P-loop-protein-BrxC-[Myxococcales-bacterium] |
| WP_012825864.1 | Type6.BREX | RAMA->GDResolvase+ZnR->?->?->BrxE->BrxA-DNA-binding->BrxB-BrxF-iSTAND->DUF6079*->N6-METHYLASE->PglZ->BrxD-AAA->SF1-SF2-Helicase+wHTH+Small-B-barrel+35exo-> | 1207 | Bacteria>Myxococcota>Polyangia>Haliangiales>Kofleriaceae>Haliangium | GCF_000024805.1 | Haliangium_ochraceum | HOCH_RS03020 | 800520..804143 | 80816 | BREX-system-P-loop-protein-BrxC-[Haliangium-ochraceum] |
| WP_012826405.1 | Type6.BREX | TPR-><-?<-?<-?\|\|?->BrxE->BrxA-DNA-binding->BrxB-BrxF-iSTAND->DUF6079*->N6-METHYLASE->PglZ->BrxD-AAA->SF1-SF2-Helicase+wHTH+Small-B-barrel+35exo-> | 1207 | Bacteria>Myxococcota>Polyangia>Haliangiales>Kofleriaceae>Haliangium | GCF_000024805.1 | Haliangium_ochraceum | HOCH_RS05940 | 1613340..1616963 | 80816 | BREX-system-P-loop-protein-BrxC-[Haliangium-ochraceum] |
| MBP9202688.1 | Type6.BREX | <-ChlI+Lon_C+AAA<-HEPN<-SF1-SF2-Helicase+wHTH+Small-B-barrel+35exo<-BrxD-AAA<-PglZ<-HTH+REase<-N6-METHYLASE<-?<-DUF6079*<-BrxB-BrxF-iSTAND<-BrxA-DNA-binding<-BrxE | 1203 | Bacteria>Myxococcota>Polyangia>Haliangiales>Kofleriaceae>unclassified_Kofleriaceae | GCA_018058095.1 | Kofleriaceae_bacterium | KBG28_01800 | 90937..94548 | 2212474 | BREX-system-P-loop-protein-BrxC-[Kofleriaceae-bacterium] |
| MBK8262770.1 | Type6.BREX | <-DnaJ\|\|?-><-?\|\|BrxE->BrxA-DNA-binding->BrxB-BrxF-iSTAND->DUF6079*->OLD-ABC->TOPRIM->N6-METHYLASE->PglZ->BrxD-AAA->ParB+HNH->SF1-SF2-Helicase+wHTH+Small-B-barrel+35exo-><-HEPN | 1208 | Bacteria>Myxococcota>Polyangia>Nannocystales>Nannocystaceae>Nannocystis | GCA_016712315.1 | Nannocystis_sp. | IPK80_15725 | 176860..180486 | 1962667 | BREX-system-P-loop-protein-BrxC-[Nannocystis-sp.] |
| WP_129580216.1 | Type6.BREX | <-ACET\|\|?->MPTase->tRNA-><-SF1-SF2-Helicase+wHTH+Small-B-barrel+DUF1998<-BrxD-AAA<-PglZ<-N6-METHYLASE<-DUF6079*<-BrxB-BrxF-iSTAND<-BrxA-DNA-binding<-BrxE\|\|REase-> | 1207 | Bacteria>Myxococcota>Polyangia>Polyangiales>Polyangiaceae | GCF_004135755.1 | Sorangium | NQZ70_RS48040 | 13122794..13126417 | 39643 | MULTISPECIES:-BREX-system-P-loop-protein-BrxC-[Sorangium] |
| WP_272422163.1 | Type6.BREX | <-RADICAL-SAM<-?\|\|BrxE->BrxA-DNA-binding->BrxB-BrxF-iSTAND->DUF6079*->N6-METHYLASE->Tox-RelA->PglZ->SIGMA+OB-Fold+AAA-ATPase+Lon-Protease-BrxL-><-MORC+DUF3684+REase\|\|KAP-NTPase->?->REase->SIGMA-HTH-> | 1182 | Bacteria>Myxococcota>Polyangia>Polyangiales>Polyangiaceae>Polyangium | GCF_028435265.1 | Polyangium_jinanense | KEG38_RS23925 | 101001..104549 | 2829994 | BREX-system-P-loop-protein-BrxC-[Polyangium-jinanense] |
| KYF76807.1 | Type6.BREX | <-Pkinase<-GT4+RADICAL-SAM<-?<-SIGMA+OB-Fold+AAA-ATPase+Lon-Protease-BrxL<-?\|\|BrxE->BrxA-DNA-binding->BrxB-BrxF-iSTAND->DUF6079*->N6-METHYLASE-><-?<-?<-?\|\|PglZ->BrxD-AAA->SF1-SF2-Helicase+wHTH+Small-B-barrel+DUF1998-><-?<-REase | 1203 | Bacteria>Myxococcota>Polyangia>Polyangiales>Polyangiaceae>Sorangium | GCA_001589275.1 | Sorangium_cellulosum | BE11_22350 | 12334..15945 | 56 | hypothetical-protein-BE11_22350-[Sorangium-cellulosum] |
| WP_020733584.1 | Type6.BREX | ABC-ATPase+SbcC-><-SIGMA+OB-Fold+AAA-ATPase+Lon-Protease-BrxL<-PglZ<-HNH<-OLD-ABC<-N6-METHYLASE<-N6-METHYLASE<-Schlafen+HATPase+wHTH<-DUF6079*<-BrxB-BrxF-iSTAND<-BrxA-DNA-binding<-BrxE<-XIS-HTH\|\|?->?->ACET-> | 1182 | Bacteria>Myxococcota>Polyangia>Polyangiales>Polyangiaceae>Sorangium | GCF_000418325.1 | Sorangium_cellulosum | SCE1572_RS08150 | 2251441..2254989 | 56 | BREX-system-P-loop-protein-BrxC-[Sorangium-cellulosum] |
| WP_061609317.1 | Type6.BREX | <-REase<-URI-DNase+RAMA\|\|BrxE->BrxA-DNA-binding->BrxB-BrxF-iSTAND->DUF6079*->N6-METHYLASE->PglZ->BrxD-AAA->AIPR->SF1-SF2-Helicase+wHTH+Small-B-barrel+DUF1998-><-?<-?\|\|REase-> | 1208 | Bacteria>Myxococcota>Polyangia>Polyangiales>Polyangiaceae>Sorangium | GCF_001589285.1 | Sorangium_cellulosum | BE15_RS23340 | 23274..26900 | 56 | BREX-system-P-loop-protein-BrxC-[Sorangium-cellulosum] |
| WP_129347587.1 | Type6.BREX | ParA-Soj-PloopNTPase->?->?-><-N6-METHYLASE<-?<-?<-DUF6079*<-BrxB-BrxF-iSTAND<-BrxA-DNA-binding<-BrxE<-?\|\|?-><-?<-?\|\|PAS+STAS-> | 1203 | Bacteria>Myxococcota>Polyangia>Polyangiales>Polyangiaceae>Sorangium | GCF_004135735.1 | Sorangium_cellulosum | SOCEGT47_RS14535 | 3823991..3827602 | 56 | BREX-system-P-loop-protein-BrxC-[Sorangium-cellulosum] |
| WP_129354457.1 | Type6.BREX | <-Glyco_hydro_15<-?\|\|?-><-SIGMA+OB-Fold+AAA-ATPase+Lon-Protease-BrxL<-PglZ<-N6-METHYLASE<-Schlafen+HATPase+wHTH<-DUF6079*<-BrxB-BrxF-iSTAND<-BrxA-DNA-binding<-BrxE<-REC+FIS-HTH<-HISKIN\|\|Methyltransf_33->FGS-><-SAM-METHYLASE | 1180 | Bacteria>Myxococcota>Polyangia>Polyangiales>Polyangiaceae>Sorangium | GCF_004135735.1 | Sorangium_cellulosum | SOCEGT47_RS35865 | 9457770..9461312 | 56 | BREX-system-P-loop-protein-BrxC-[Sorangium-cellulosum] |
| MCL2723928.1 | Type6.BREX | Glyoxalase->?->?-><-SIGMA+OB-Fold+AAA-ATPase+Lon-Protease-BrxL<-PglZ<-Schlafen+HATPase+wHTH<-SbcC<-N6-METHYLASE<-N6-METHYLASE<-DUF6079*<-BrxB-BrxF-iSTAND<-BrxA-DNA-binding<-BrxE<-?<-?\|\|?-><-?<-?\|\|Pkinase+Pkinase-> | 1179 | Bacteria>Myxococcota>Polyangia>Polyangiales>Polyangiaceae>unclassified_Polyangiaceae | GCA_009781005.1 | Polyangiaceae_bacterium | FWD69_05765 | 18977..22516 | 2268199 | BREX-system-P-loop-protein-BrxC-[Polyangiaceae-bacterium] |
| NOU34914.1 | Type6.BREX | <-DUF3670+SF1-SF2-Helicase<-ASCH<-SF1-SF2-Helicase+wHTH+Small-B-barrel+DUF1998<-BrxD-AAA<-PglZ<-N6-METHYLASE<-N6-METHYLASE<-?<-DUF6079*<-BrxB-BrxF-iSTAND<-BrxA-DNA-binding<-BrxE | 1207 | Bacteria>Myxococcota>Polyangia>Polyangiales>Polyangiaceae>unclassified_Polyangiaceae | GCA_013141295.1 | Polyangiaceae_bacterium | HDO96_44040 | 28048..31671 | 2268199 | BREX-system-P-loop-protein-BrxC-[Polyangiaceae-bacterium] |
| NRA34737.1 | Type6.BREX | <-DUF6079-p*<-BrxB-BrxF-iSTAND<-BrxA-DNA-binding<-BrxE\|\|?->?->zf-IS66+DDE_Tnp_IS66-> | 526 | Bacteria>Myxococcota>Polyangia>Polyangiales>Polyangiaceae>unclassified_Polyangiaceae | GCA_013214025.1 | Polyangiaceae_bacterium | HRU17_15500 | 1..1578 | 2268199 | BREX-system-P-loop-protein-BrxC-[Polyangiaceae-bacterium] |
| MBK6517532.1 | Type6.BREX | Exonuc_V_gamma+REase->SF1-SF2-Helicase+REase->REase-><-TOPRIM<-OLD-ABC<-?\|\|BrxE->BrxA-DNA-binding->BrxB-BrxF-iSTAND->DUF6079*-><-PemK_toxin<-?\|\|N6-METHYLASE->N6-METHYLASE-p->?->?->PIN->PglZ->BrxD-AAA-> | 1209 | Bacteria>Myxococcota>Polyangia>Polyangiales>Polyangiaceae>unclassified_Polyangiaceae | GCA_016704335.1 | Polyangiaceae_bacterium | IPG04_26305 | 282797..286426 | 2268199 | BREX-system-P-loop-protein-BrxC-[Polyangiaceae-bacterium] |
| MBK8256776.1 | Type6.BREX | <-HKD-DNase<-CASPASE<-?<-?<-SF1-SF2-Helicase+wHTH+Small-B-barrel+DUF1998<-BrxD-AAA<-PglZ<-N6-METHYLASE<-N6-METHYLASE<-DUF6079*<-BrxB-BrxF-iSTAND<-BrxA-DNA-binding<-BrxE\|\|?-><-SIGMA-HTH<-CR-ATPase8+REase | 1209 | Bacteria>Myxococcota>Polyangia>Polyangiales>Polyangiaceae>unclassified_Polyangiaceae | GCA_016712345.1 | Polyangiaceae_bacterium | IPK82_29405 | 1486518..1490147 | 2268199 | BREX-system-P-loop-protein-BrxC-[Polyangiaceae-bacterium] |
| MBK8937968.1 | Type6.BREX | <-BrxD-AAA<-PglZ<-PIN<-?<-?<-N6-METHYLASE-p<-N6-METHYLASE\|\|?->PemK_toxin-><-DUF6079*<-BrxB-BrxF-iSTAND<-BrxA-DNA-binding<-BrxE\|\|OLD-ABC->TOPRIM-><-REase<-SF1-SF2-Helicase+REase<-SF1-SF2-Helicase<-SF1-SF2-Helicase+REase-p | 1016 | Bacteria>Myxococcota>Polyangia>Polyangiales>Polyangiaceae>unclassified_Polyangiaceae | GCA_016719945.1 | Polyangiaceae_bacterium | IPM79_10070 | 425737..428787 | 2268199 | BREX-system-P-loop-protein-BrxC-[Polyangiaceae-bacterium] |
| MCB9613493.1 | Type6.BREX | <-SF1-SF2-Helicase+wHTH+Small-B-barrel+35exo<-BrxD-AAA<-PglZ\|\|wHTH+AbiEi_2->AbiEii-><-?<-N6-METHYLASE<-TraT<-Tox-RelA<-DUF6079*<-BrxB-BrxF-iSTAND<-BrxA-DNA-binding<-BrxE\|\|?->?->XIS-HTH->?-><-?<-TPR | 1203 | Bacteria>Myxococcota>Polyangia>Polyangiales>Sandaracinaceae>Sandaracinus | GCA_020633155.1 | Sandaracinus_sp. | H6722_13670 | 29710..33321 | 2024858 | BREX-system-P-loop-protein-BrxC-[Sandaracinus-sp.] |
| MCB9599931.1 | Type6.BREX | HTH_3-><-XIS-HTH<-?\|\|?-><-?<-?\|\|BrxE->BrxA-DNA-binding->BrxB-BrxF-iSTAND->DUF6079*->N6-METHYLASE->?-><-AbiEii<-wHTH+AbiEi_2\|\|PglZ->BrxD-AAA->SF1-SF2-Helicase+wHTH+Small-B-barrel+35exo-> | 1205 | Bacteria>Myxococcota>Polyangia>Polyangiales>Sandaracinaceae>Sandaracinus | GCA_020633175.1 | Sandaracinus_sp. | H6720_06200 | 129083..132700 | 2024858 | BREX-system-P-loop-protein-BrxC-[Sandaracinus-sp.] |
| MCK6514475.1 | Type6.BREX | <-SF1-SF2-Helicase<-HamA<-SIGMA+OB-Fold+AAA-ATPase+Lon-Protease-BrxL<-PglZ<-OLD-ABC<-REase<-N6-METHYLASE-p<-N6-METHYLASE<-Schlafen+HATPase+wHTH<-DUF6079*<-DUF6079-p<-BrxB-BrxF-iSTAND<-BrxA-DNA-binding<-BrxE<-DNA-glycosylase\|\|RADICAL-SAM-> | 1083 | Bacteria>Myxococcota>unclassified_Myxococcota | GCA_023150775.1 | Myxococcota_bacterium | L6R46_05405 | 17071..20323 | 2818507 | BREX-system-P-loop-protein-BrxC-[Myxococcota-bacterium] |
| MCK6527600.1 | Type6.BREX | RADICAL-SAM->?-><-SIGMA+OB-Fold+AAA-ATPase+Lon-Protease-BrxL<-?<-?<-?<-HNH<-OLD-ABC<-N6-METHYLASE<-DUF6079*<-BrxB-BrxF-iSTAND<-BrxA-DNA-binding<-BrxE\|\|?->GDResolvase+Recombinase+ZnR-> | 1184 | Bacteria>Myxococcota>unclassified_Myxococcota | GCA_023150875.1 | Myxococcota_bacterium | L6R50_08565 | 57228..60782 | 2818507 | BREX-system-P-loop-protein-BrxC-[Myxococcota-bacterium] |
| MCK6575226.1 | Type6.BREX | <-PglZ-p<-TOPRIM<-OLD-ABC<-N6-METHYLASE<-?<-OLD-ABC<-DUF6079*<-BrxB-BrxF-iSTAND<-BrxA-DNA-binding<-BrxE\|\|MPTase->?->?-><-REase | 1247 | Bacteria>Myxococcota>unclassified_Myxococcota | GCA_023150895.1 | Myxococcota_bacterium | L6V77_29520 | 8524..12267 | 2818507 | BREX-system-P-loop-protein-BrxC-[Myxococcota-bacterium] |
| MCU0671310.1 | Type6.BREX | <-TOPRIM<-OLD-ABC<-DUF6079*<-BrxB-BrxF-iSTAND<-BrxA-DNA-binding<-BrxE\|\|?-><-?\|\|?->XIS-HTH-><-HTH_3 | 1203 | Bacteria>Myxococcota>unclassified_Myxococcota | GCA_025453555.1 | Myxococcota_bacterium | MUE69_00810 | 1555..5166 | 2818507 | BREX-system-P-loop-protein-BrxC-[Myxococcota-bacterium] |
| WP_123104890.1 | Type6.BREX | SIGMA-HTH->tRNA-><-SIGMA+OB-Fold+AAA-ATPase+Lon-Protease-BrxL<-PglZ<-ParB+HNH<-TOPRIM<-OLD-ABC<-N6-METHYLASE<-N6-METHYLASE<-DUF6079*<-BrxB-BrxF-iSTAND<-BrxA-DNA-binding<-BrxE\|\|<-?<-XIS-HTH\|\|?->?->TOPRIM-> | 1210 | Bacteria>Pseudomonadota>Acidithiobacillia>Acidithiobacillales>Acidithiobacillaceae>Acidithiobacillus | GCF_003721225.1 | Acidithiobacillus_sulfuriphilus | EC580_RS10670 | 12374..16006 | 1867749 | BREX-system-P-loop-protein-BrxC-[Acidithiobacillus-sulfuriphilus] |
| WP_215855738.1 | Type6.BREX | <-SF1-SF2-Helicase\|\|tRNA->Phage_integrase->BrxE->BrxA-DNA-binding->BrxB-BrxF-iSTAND->DUF6079*->?->N6-METHYLASE->ParB+HNH->PglZ->SIGMA+OB-Fold+AAA-ATPase+Lon-Protease-BrxL->?-><-DDE_Tnp_IS1595 | 1210 | Bacteria>Pseudomonadota>Acidithiobacillia>Acidithiobacillales>Acidithiobacillaceae>Acidithiobacillus | GCF_018854635.1 | Acidithiobacillus_ferridurans | HF641_RS00655 | 27295..30927 | 1232575 | BREX-system-P-loop-protein-BrxC-[Acidithiobacillus-ferridurans] |
| WP_029415251.1 | Type6.BREX | <-Exonuc_V_gamma+REase<-wHTH+WYL+WCX\|\|?->DUF6361->HKD-DNase->SF1-SF2-Helicase-><-SIGMA+OB-Fold+AAA-ATPase+Lon-Protease-BrxL<-PglZ<-N6-METHYLASE<-DUF6079*<-BrxB-BrxF-iSTAND<-BrxA-DNA-binding<-BrxE | 1176 | Bacteria>Pseudomonadota>Alphaproteobacteria>Caulobacterales>Caulobacteraceae>Brevundimonas | GCF_000701445.1 | Brevundimonas_bacteroides | Q333_RS0102925 | 140583..144113 | 74311 | BREX-system-P-loop-protein-BrxC-[Brevundimonas-bacteroides] |
| WP_056505736.1 | Type6.BREX | <-Classical-AAA<-REase+wHTH<-?<-?<-Calcineurin\|\|BrxE->BrxA-DNA-binding->BrxB-BrxF-iSTAND->DUF6079*->N6-METHYLASE-><-?\|\|N6-METHYLASE-p->PglZ->SIGMA+OB-Fold+AAA-ATPase+Lon-Protease-BrxL->?-><-?\|\|REase+wHTH->rve_3+Mu-transpos_C-> | 1180 | Bacteria>Pseudomonadota>Alphaproteobacteria>Hyphomicrobiales>Aurantimonadaceae>Aureimonas | GCF_001424685.1 | Aureimonas_sp._Leaf454 | ASG43_09150 | 425473..429015 | 1736381 | BREX-system-P-loop-protein-BrxC-[Aureimonas-sp.-Leaf454] |
| WP_204157651.1 | Type6.BREX | <-TUDOR+SF1-SF2-Helicase+iREase+CTD<-SIGMA+OB-Fold+AAA-ATPase+HKD-DNase-BrxL<-PglZ<-N6-METHYLASE-p<-AAA-ATPase<-rve<-N6-METHYLASE-p<-N6-METHYLASE<-DUF6079*<-BrxB-BrxF-iSTAND<-BrxE<-PhdYeFM_antitox<-TraI-VirD2-RCRE<-TraJ-RHH\|\|?-><-DOC+HTH | 1208 | Bacteria>Pseudomonadota>Alphaproteobacteria>Hyphomicrobiales>Brucellaceae>Brucella/Ochrobactrum_group | GCF_016887905.1 | Brucella_anthropi | JQC79_RS05930 | 1236117..1239743 | 529 | BREX-system-P-loop-protein-BrxC-[Brucella-anthropi] |
| WP_046141798.1 | Type6.BREX | <-XIS-HTH<-?\|\|RAMA->GDResolvase+ZnR->ZnR->?->BrxE->BrxA-DNA-binding->BrxB-BrxF-iSTAND->DUF6079*->N6-METHYLASE->N6-METHYLASE->PglZ->SIGMA+OB-Fold+AAA-ATPase+HKD-DNase-BrxL-><-tRNA<-ADH_N+ADH_zinc_N | 1208 | Bacteria>Pseudomonadota>Alphaproteobacteria>Hyphomicrobiales>Devosiaceae>Devosia | - | Devosia_soli | VW35_RS04250 | 28883..32509 | 361041 | BREX-system-P-loop-protein-BrxC-[Devosia-soli] |
| WP_090228177.1 | Type6.BREX | PhdYeFM_antitox->BrxE->BrxA-DNA-binding->BrxB-BrxF-iSTAND->DUF6079*->N6-METHYLASE->PglZ->SIGMA+OB-Fold+AAA-ATPase+Lon-Protease-BrxL-><-SF1-SF2-Helicase<-?\|\|?-><-?<-XIS-HTH<-HIN-HTH | 1176 | Bacteria>Pseudomonadota>Alphaproteobacteria>Hyphomicrobiales>Hyphomicrobiaceae>Filomicrobium | GCF_900104305.1 | Filomicrobium_insigne | BLS30_RS09065 | 671961..675491 | 418854 | BREX-system-P-loop-protein-BrxC-[Filomicrobium-insigne] |
| WP_160533296.1 | Type6.BREX | <-FA_desaturase\|\|MORC-><-?\|\|?->BrxE->BrxA-DNA-binding->BrxB-BrxF-iSTAND->DUF6079*->N6-METHYLASE->?->N6-METHYLASE-p->PglZ->SIGMA+OB-Fold+AAA-ATPase+Lon-Protease-BrxL-><-MORC<-REase<-DNA_METHYLASE | 1180 | Bacteria>Pseudomonadota>Alphaproteobacteria>Hyphomicrobiales>Methylobacteriaceae>Methylobacterium | GCF_009806555.1 | Methylobacterium_sp._2A | FVE89_RS06905 | 159464..163006 | 2603816 | BREX-system-P-loop-protein-BrxC-[Methylobacterium-sp.-2A] |
| KFG67132.1 | Type6.BREX | <-AbiGi+seqA+RAMA\|\|BrxE->BrxA-DNA-binding->BrxB-BrxF-iSTAND->DUF6079*->N6-METHYLASE-p->PglZ->SIGMA+OB-Fold+AAA-ATPase+HKD-DNase-BrxL->HNH->HNH->HNH->?-><-HEPN<-Polbeta | 1210 | Bacteria>Pseudomonadota>Alphaproteobacteria>Hyphomicrobiales>Methylobacteriaceae>Microvirga | GCA_000739015.1 | Microvirga_sp._BSC39 | JH26_23850 | 105316..108948 | 1549810 | ATPase-[Microvirga-sp.-BSC39] |
| WP_036357316.1 | Type6.BREX | <-AbiGi+seqA+RAMA<-?\|\|PhdYeFM_antitox->BrxE->BrxA-DNA-binding->BrxB-BrxF-iSTAND->DUF6079*->N6-METHYLASE->PglZ->SIGMA+OB-Fold+AAA-ATPase+HKD-DNase-BrxL->HNH->HNH->HNH->?-><-HEPN<-Polbeta | 1208 | Bacteria>Pseudomonadota>Alphaproteobacteria>Hyphomicrobiales>Methylobacteriaceae>Microvirga | GCF_000739015.1 | Microvirga_sp._BSC39 | MA07_RS22775 | 105322..108948 | 1549810 | BREX-system-P-loop-protein-BrxC-[Microvirga-sp.-BSC39] |
| WP_152708987.1 | Type6.BREX | Immank+ANK+ANK-><-?<-?\|\|?-><-SIGMA+OB-Fold+AAA-ATPase+Lon-Protease-BrxL<-PglZ<-N6-METHYLASE<-N6-METHYLASE<-DUF6079*<-BrxB-BrxF-iSTAND<-BrxA-DNA-binding<-BrxE<-CopG/RHH\|\|Phage_AlpA->Ntox49->UDG->HTH_3->MPTase-> | 1183 | Bacteria>Pseudomonadota>Alphaproteobacteria>Hyphomicrobiales>Methylobacteriaceae>Microvirga | GCF_009296195.1 | Microvirga_tunisiensis | FS317_RS04420 | 59212..62763 | 2108360 | BREX-system-P-loop-protein-BrxC-[Microvirga-tunisiensis] |
| WP_199050625.1 | Type6.BREX | <-ParB+HNH<-ARG\|\|?-><-?<-SF1-SF2-Helicase+DUF1998<-?\|\|BrxE->BrxA-DNA-binding->BrxB-BrxF-iSTAND->DUF6079*->N6-METHYLASE->N6-METHYLASE-p->PglZ->SIGMA+OB-Fold+AAA-ATPase+HKD-DNase-BrxL-><-Phage_integrase | 1206 | Bacteria>Pseudomonadota>Alphaproteobacteria>Hyphomicrobiales>Methylobacteriaceae>Microvirga | GCF_016427565.1 | Microvirga_splendida | JAO75_RS18515 | 48792..52412 | 2795727 | BREX-system-P-loop-protein-BrxC-[Microvirga-splendida] |
| WP_253589372.1 | Type6.BREX | <-HIRAN<-BirA-HTH<-SF1-SF2-Helicase+REase<-35Exo+REase<-?\|\|wHTH+WYL+WCX->BrxE->BrxA-DNA-binding->BrxB-BrxF-iSTAND->DUF6079*->N6-METHYLASE->N6-METHYLASE->PglZ->SIGMA+OB-Fold+AAA-ATPase+HKD-DNase-BrxL->?-><-?<-?\|\|?->?->HNH-> | 1208 | Bacteria>Pseudomonadota>Alphaproteobacteria>Hyphomicrobiales>Nitrobacteraceae>Bradyrhizobium | - | Bradyrhizobium_elkanii | J2R96_RS28580 | 6233220..6236846 | 29448 | BREX-system-P-loop-protein-BrxC-[Bradyrhizobium-elkanii] |
| WP_212207485.1 | Type6.BREX | <-PIWI<-SIGMA+OB-Fold+AAA-ATPase+HKD-DNase-BrxL<-PglZ<-ParB+HNH<-N6-METHYLASE-p<-N6-METHYLASE<-DUF6079*<-BrxB-BrxF-iSTAND<-BrxA-DNA-binding<-BrxE<-wHTH+WYL+WCX\|\|35Exo+REase->SF1-SF2-Helicase+REase-> | 1208 | Bacteria>Pseudomonadota>Alphaproteobacteria>Hyphomicrobiales>Nitrobacteraceae>Bradyrhizobium | - | Bradyrhizobium_sp._AUGA_SZCCT0274 | JQ620_09165 | 74241..77867 | 2807670 | BREX-system-P-loop-protein-BrxC-[Bradyrhizobium-sp.-AUGA-SZCCT0274] |
| QAU48836.1 | Type6.BREX | HEPN-><-?<-PIWI<-REase+PNTD1+PNTD2<-SIGMA+OB-Fold+AAA-ATPase+Lon-Protease-BrxL<-PglZ<-N6-METHYLASE<-N6-METHYLASE<-DUF6079*<-BrxB-BrxF-iSTAND<-BrxA-DNA-binding<-BrxE<-CopG/RHH\|\|?->?->Calcineurin->REase+wHTH-> | 1179 | Bacteria>Pseudomonadota>Alphaproteobacteria>Hyphomicrobiales>Nitrobacteraceae>Bradyrhizobium | GCA_004114955.1 | Bradyrhizobium_guangzhouense | XH91_28100 | 5872801..5876340 | 1325095 | BREX-system-P-loop-protein-BrxC-[Bradyrhizobium-guangzhouense] |
| NEU96935.1 | Type6.BREX | <-ANK+ANK+ANK<-SbcC<-REase<-SIR2<-?<-SIGMA+OB-Fold+AAA-ATPase+Lon-Protease-BrxL<-PglZ<-N6-METHYLASE<-N6-METHYLASE<-DUF6079*<-BrxB-BrxF-iSTAND<-BrxA-DNA-binding<-BrxE<-GDResolvase+HIN-HTH\|\|?->?-><-ACYC+TPR+TPR<-ACYC+TPR+TPR<-HISKIN | 1176 | Bacteria>Pseudomonadota>Alphaproteobacteria>Hyphomicrobiales>Nitrobacteraceae>Bradyrhizobium | GCA_010811875.1 | Bradyrhizobium_uaiense | FNJ47_14065 | 18759..22289 | 2594946 | BREX-system-P-loop-protein-BrxC-[Bradyrhizobium-uaiense] |
| MBR1240293.1 | Type6.BREX | <-PIWI<-SIGMA+OB-Fold+AAA-ATPase+HKD-DNase-BrxL<-PglZ<-ParB+HNH<-N6-METHYLASE-p<-N6-METHYLASE<-DUF6079*<-BrxB-BrxF-iSTAND<-BrxA-DNA-binding<-BrxE<-wHTH+WYL+WCX\|\|35Exo+REase->SF1-SF2-Helicase+REase-> | 1208 | Bacteria>Pseudomonadota>Alphaproteobacteria>Hyphomicrobiales>Nitrobacteraceae>Bradyrhizobium | GCA_018130325.1 | Bradyrhizobium_sp._AUGA_SZCCT0274 | JQ620_09165 | 74241..77867 | 2807670 | BREX-system-P-loop-protein-BrxC-[Bradyrhizobium-sp.-AUGA-SZCCT0274] |
| WP_244433425.1 | Type6.BREX | AAA-ATPase-><-SIGMA+OB-Fold+AAA-ATPase+Lon-Protease-BrxL<-PglZ<-N6-METHYLASE<-DUF6079*<-BrxB-BrxF-iSTAND<-BrxA-DNA-binding<-BrxE<-?\|\|Phage_AlpA-> | 1176 | Bacteria>Pseudomonadota>Alphaproteobacteria>Hyphomicrobiales>Nitrobacteraceae>Bradyrhizobium | GCF_000261565.1 | Bradyrhizobium_japonicum | IC67_RS0140285 | 14408..17938 | 375 | BREX-system-P-loop-protein-BrxC-[Bradyrhizobium-japonicum] |
| WP_036025094.1 | Type6.BREX | <-BirA-HTH<-MetJarc-HTH<-UDG<-Ntox49<-Phage_AlpA\|\|BrxE->BrxA-DNA-binding->BrxB-BrxF-iSTAND->DUF6079*->N6-METHYLASE->N6-METHYLASE->PglZ->SIGMA+OB-Fold+AAA-ATPase+Lon-Protease-BrxL->HKD-DNase-><-?<-?\|\|?->CR-Trypsin1+HKD-DNase-> | 1179 | Bacteria>Pseudomonadota>Alphaproteobacteria>Hyphomicrobiales>Nitrobacteraceae>Bradyrhizobium | GCF_000261805.1 | Bradyrhizobium_yuanmingense | IC68_RS15665 | 22684..26223 | 108015 | BREX-system-P-loop-protein-BrxC-[Bradyrhizobium-yuanmingense] |
| WP_245477239.1 | Type6.BREX | HEPN-><-?<-PIWI<-REase+PNTD1+PNTD2<-SIGMA+OB-Fold+AAA-ATPase+Lon-Protease-BrxL<-PglZ<-N6-METHYLASE<-N6-METHYLASE<-DUF6079*<-BrxB-BrxF-iSTAND<-BrxA-DNA-binding<-BrxE<-CopG/RHH\|\|?->?->Calcineurin->REase+wHTH-> | 1176 | Bacteria>Pseudomonadota>Alphaproteobacteria>Hyphomicrobiales>Nitrobacteraceae>Bradyrhizobium | GCF_004114955.1 | Bradyrhizobium_guangzhouense | XH91_RS28040 | 5872801..5876331 | 1325095 | BREX-system-P-loop-protein-BrxC-[Bradyrhizobium-guangzhouense] |
| WP_225048696.1 | Type6.BREX | <-UDG<-Ntox49<-Phage_AlpA\|\|BrxE->BrxA-DNA-binding->BrxB-BrxF-iSTAND-><-N6-METHYLASE<-HTH_Tnp_IS630<-N6-METHYLASE<-DUF6079-p*\|\|PglZ->SIGMA+OB-Fold+AAA-ATPase+Lon-Protease-BrxL->HKD-DNase-><-?<-?<-?\|\|HTH_29+HTH_32+DDE_3->?->CR-Trypsin1+HKD-DNase-> | 404 | Bacteria>Pseudomonadota>Alphaproteobacteria>Hyphomicrobiales>Nitrobacteraceae>Bradyrhizobium | GCF_017565685.3 | Bradyrhizobium_barranii | J4G48_RS15880 | 3299543..3300757 | 2992140 | BREX-system-P-loop-protein-BrxC-[Bradyrhizobium-barranii] |
| WP_249144787.1 | Type6.BREX | <-PIWI<-SIGMA+OB-Fold+AAA-ATPase+HKD-DNase-BrxL<-PglZ<-ParB+HNH<-N6-METHYLASE-p<-N6-METHYLASE<-DUF6079*<-BrxB-BrxF-iSTAND<-BrxA-DNA-binding<-BrxE<-wHTH+WYL+WCX\|\|35Exo+REase->SF1-SF2-Helicase+REase-> | 1203 | Bacteria>Pseudomonadota>Alphaproteobacteria>Hyphomicrobiales>Nitrobacteraceae>Bradyrhizobium | GCF_018130325.1 | Bradyrhizobium_sp._AUGA_SZCCT0274 | JQ620_RS09170 | 74241..77852 | 2807670 | BREX-system-P-loop-protein-BrxC-[Bradyrhizobium-sp.-AUGA-SZCCT0274] |
| WP_213286691.1 | Type6.BREX | Phage_integrase->MarR-HTH->AbiEii->?->?->BrxE->BrxA-DNA-binding->BrxB-BrxF-iSTAND->DUF6079*->N6-METHYLASE->N6-METHYLASE->ParB+HNH->PglZ->SIGMA+OB-Fold+AAA-ATPase+HKD-DNase-BrxL-><-TPR<-?<-?\|\|wHTH+WYL+WCX-> | 1208 | Bacteria>Pseudomonadota>Alphaproteobacteria>Hyphomicrobiales>Nitrobacteraceae>Bradyrhizobium | GCF_018398875.1 | Bradyrhizobium_sp._sGM-13 | KJQ96_RS11300 | 145012..148638 | 2831781 | BREX-system-P-loop-protein-BrxC-[Bradyrhizobium-sp.-sGM-13] |
| WP_247477718.1 | Type6.BREX | BrxE->BrxA-DNA-binding->BrxB-BrxF-iSTAND->DUF6079*->N6-METHYLASE->PglZ->SIGMA+OB-Fold+AAA-ATPase+HKD-DNase-BrxL->SbcC->HNH-><-?<-?\|\|TPR+TPR+TPR+TPR-> | 1206 | Bacteria>Pseudomonadota>Alphaproteobacteria>Hyphomicrobiales>Nitrobacteraceae>Bradyrhizobium | GCF_023072855.1 | Bradyrhizobium_sp._CW9 | IVB57_RS09580 | 204980..208600 | 2782689 | BREX-system-P-loop-protein-BrxC-[Bradyrhizobium-sp.-CW9] |
| WP_247461860.1 | Type6.BREX | <-REase+wHTH<-?<-?<-REase+wHTH<-Calcineurin<-?\|\|CopG/RHH->BrxE->BrxB-BrxF-iSTAND->DUF6079*->N6-METHYLASE->PglZ->SIGMA+OB-Fold+AAA-ATPase+Lon-Protease-BrxL-><-ABhydrolase<-SMODS<-SAVED | 1180 | Bacteria>Pseudomonadota>Alphaproteobacteria>Hyphomicrobiales>Nitrobacteraceae>Bradyrhizobium | GCF_023073035.1 | Bradyrhizobium_sp._62 | IVB47_RS12420 | 413436..416978 | 1043588 | BREX-system-P-loop-protein-BrxC-[Bradyrhizobium-sp.-62] |
| WP_247475826.1 | Type6.BREX | HTH_3->BrxE->BrxA-DNA-binding->BrxB-BrxF-iSTAND->DUF6079*->N6-METHYLASE->N6-METHYLASE->Tox-RelA->PglZ->SIGMA+OB-Fold+AAA-ATPase+Lon-Protease-BrxL->GDResolvase+HIN-HTH-><-LuxR-HTH | 1180 | Bacteria>Pseudomonadota>Alphaproteobacteria>Hyphomicrobiales>Nitrobacteraceae>Bradyrhizobium | GCF_023073675.1 | Bradyrhizobium_sp._169 | IVB04_RS03585 | 13561..17103 | 2782640 | BREX-system-P-loop-protein-BrxC-[Bradyrhizobium-sp.-169] |
| WP_259215308.1 | Type6.BREX | DSBH->?->tRNA->BrxE->BrxA-DNA-binding->BrxB-BrxF-iSTAND->DUF6079*->N6-METHYLASE->PglZ->SIGMA+OB-Fold+AAA-ATPase+HKD-DNase-BrxL->SbcC->HNH-><-?<-?<-?<-SIGMA-HTH+HIN-HTH | 1207 | Bacteria>Pseudomonadota>Alphaproteobacteria>Hyphomicrobiales>Nitrobacteraceae>Bradyrhizobium | GCF_024809395.1 | Bradyrhizobium_centrosematis | GGE24_RS17830 | 223880..227503 | 1300039 | BREX-system-P-loop-protein-BrxC-[Bradyrhizobium-centrosematis] |
| WP_271557646.1 | Type6.BREX | HKD-DNase+HKD-Rease->BrxE->BrxA-DNA-binding->BrxB-BrxF-iSTAND->DUF6079*->N6-METHYLASE->N6-METHYLASE->PglZ->SIGMA+OB-Fold+AAA-ATPase+Lon-Protease-BrxL-> | 1176 | Bacteria>Pseudomonadota>Alphaproteobacteria>Hyphomicrobiales>Nitrobacteraceae>Bradyrhizobium | GCF_028128655.1 | Bradyrhizobium_sp._CCBAU_45394 | WN73_RS39375 | 411495..415025 | 1325087 | BREX-system-P-loop-protein-BrxC-[Bradyrhizobium-sp.-CCBAU-45394] |
| WP_245288697.1 | Type6.BREX | <-REase+wHTH<-Calcineurin<-?<-?<-?\|\|CopG/RHH->BrxE->BrxA-DNA-binding->BrxB-BrxF-iSTAND->DUF6079*-><-?\|\|N6-METHYLASE->PglZ->SIGMA+OB-Fold+AAA-ATPase+Lon-Protease-BrxL-> | 1176 | Bacteria>Pseudomonadota>Alphaproteobacteria>Hyphomicrobiales>Nitrobacteraceae>Bradyrhizobium | GCF_030584845.1 | Bradyrhizobium_elkanii | A3AK_RS0111990 | 95945..99475 | 29448 | BREX-system-P-loop-protein-BrxC-[Bradyrhizobium-elkanii] |
| WP_012494683.1 | Type6.BREX | <-TnpB_IS66<-?<-?<-?\|\|?->CopG/RHH->BrxE->BrxA-DNA-binding->BrxB-BrxF-iSTAND->DUF6079*->N6-METHYLASE->PglZ->SIGMA+OB-Fold+AAA-ATPase+Lon-Protease-BrxL->HKD-DNase-><-HisKin<-?<-HIRAN<-BirA-HTH | 1177 | Bacteria>Pseudomonadota>Alphaproteobacteria>Hyphomicrobiales>Nitrobacteraceae | GCF_000020445.1 | Rhodopseudomonas | RPAL_RS05615 | 1188010..1191543 | 1073 | MULTISPECIES:-BREX-system-P-loop-protein-BrxC-[Rhodopseudomonas] |
| RWC69660.1 | Type6.BREX | <-TraJ-RHH\|\|ParA-Soj-PloopNTPase->CopG/RHH->HTH_3->BrxE->BrxA-DNA-binding->BrxB-BrxF-iSTAND->DUF6079*->N6-METHYLASE->Tox-RelA->PglZ-> | 1180 | Bacteria>Pseudomonadota>Alphaproteobacteria>Hyphomicrobiales>Phyllobacteriaceae>Mesorhizobium | GCA_004017165.1 | Mesorhizobium_sp. | EOS71_27685 | 41950..45492 | 1871066 | BREX-system-P-loop-protein-BrxC-[Mesorhizobium-sp.] |
| RWD70309.1 | Type6.BREX | Glycos_transf->?-><-SAM-METHYLASE<-RADICAL-SAM<-SIGMA+OB-Fold+AAA-ATPase+Lon-Protease-BrxL<-PglZ<-N6-METHYLASE<-N6-METHYLASE<-Tox-RelA<-DUF6079*<-BrxB-BrxF-iSTAND<-BrxA-DNA-binding<-BrxE\|\|ParB+HNH->?->GDResolvase+HIN-HTH-><-?<-Primpol | 1181 | Bacteria>Pseudomonadota>Alphaproteobacteria>Hyphomicrobiales>Phyllobacteriaceae>Mesorhizobium | GCA_004018245.1 | Mesorhizobium_sp. | EOS37_15285 | 26400..29945 | 1871066 | BREX-system-P-loop-protein-BrxC-[Mesorhizobium-sp.] |
| RWG10409.1 | Type6.BREX | Peptidase_S11->ZnR+SLATT->tRNA->XIS-HTH->BrxE->BrxA-DNA-binding->BrxB-BrxF-iSTAND->DUF6079-p*-> | 694 | Bacteria>Pseudomonadota>Alphaproteobacteria>Hyphomicrobiales>Phyllobacteriaceae>Mesorhizobium | GCA_004018325.1 | Mesorhizobium_sp. | EOQ58_27065 | 13724..15805 | 1871066 | BREX-system-P-loop-protein-BrxC,-partial-[Mesorhizobium-sp.] |
| RWG25813.1 | Type6.BREX | <-DUF6079-p*<-BrxB-BrxF-iSTAND<-BrxA-DNA-binding<-BrxE<-XIS-HTH<-tRNA<-ZnR+SLATT<-Peptidase_S11 | 574 | Bacteria>Pseudomonadota>Alphaproteobacteria>Hyphomicrobiales>Phyllobacteriaceae>Mesorhizobium | GCA_004018725.1 | Mesorhizobium_sp. | EOQ61_27945 | 1..1722 | 1871066 | BREX-system-P-loop-protein-BrxC,-partial-[Mesorhizobium-sp.] |
| RWH18602.1 | Type6.BREX | Peptidase_S11->ZnR+SLATT->tRNA->XIS-HTH->BrxE->BrxA-DNA-binding->BrxB-BrxF-iSTAND->DUF6079-p*-> | 542 | Bacteria>Pseudomonadota>Alphaproteobacteria>Hyphomicrobiales>Phyllobacteriaceae>Mesorhizobium | GCA_004019225.1 | Mesorhizobium_sp. | EOQ77_29380 | 12025..13651 | 1871066 | BREX-system-P-loop-protein-BrxC,-partial-[Mesorhizobium-sp.] |
| RWM12913.1 | Type6.BREX | <-McrB-NTD+HNH<-SIGMA+OB-Fold+AAA-ATPase+Lon-Protease-BrxL<-PglZ<-Tox-RelA<-N6-METHYLASE<-N6-METHYLASE<-DUF6079*<-BrxB-BrxF-iSTAND<-BrxA-DNA-binding<-BrxE | 1180 | Bacteria>Pseudomonadota>Alphaproteobacteria>Hyphomicrobiales>Phyllobacteriaceae>Mesorhizobium | GCA_004020315.1 | Mesorhizobium_sp. | EOR73_30525 | 23274..26816 | 1871066 | BREX-system-P-loop-protein-BrxC-[Mesorhizobium-sp.] |
| RWM88364.1 | Type6.BREX | <-N6-METHYLASE<-DUF6079*<-BrxB-BrxF-iSTAND<-BrxA-DNA-binding<-BrxE<-?<-HKD-DNase+HKD-Rease<-Peptidase-M15<-REC+NtrC-AAA+FIS-HTH\|\|Peptidase_M3_N+Peptidase_M3-> | 1211 | Bacteria>Pseudomonadota>Alphaproteobacteria>Hyphomicrobiales>Phyllobacteriaceae>Mesorhizobium | GCA_004020365.1 | Mesorhizobium_sp. | EOR84_28155 | 2131..5766 | 1871066 | BREX-system-P-loop-protein-BrxC-[Mesorhizobium-sp.] |
| RWO43571.1 | Type6.BREX | CopG/RHH->BrxE->BrxA-DNA-binding->BrxB-BrxF-iSTAND->DUF6079*->N6-METHYLASE->N6-METHYLASE->PglZ->SIGMA+OB-Fold+AAA-ATPase+Lon-Protease-BrxL-> | 1180 | Bacteria>Pseudomonadota>Alphaproteobacteria>Hyphomicrobiales>Phyllobacteriaceae>Mesorhizobium | GCA_004021685.1 | Mesorhizobium_sp. | EOS13_30605 | 42300..45842 | 1871066 | BREX-system-P-loop-protein-BrxC-[Mesorhizobium-sp.] |
| TIL20465.1 | Type6.BREX | BrxE->BrxA-DNA-binding->BrxB-BrxF-iSTAND->DUF6079*->N6-METHYLASE->N6-METHYLASE->ParB+HNH->PglZ->SIGMA+OB-Fold+AAA-ATPase+HKD-DNase-BrxL-> | 1208 | Bacteria>Pseudomonadota>Alphaproteobacteria>Hyphomicrobiales>Phyllobacteriaceae>Mesorhizobium | GCA_004961195.1 | Mesorhizobium_sp. | E5Y88_32415 | 5019..8645 | 1871066 | BREX-system-P-loop-protein-BrxC-[Mesorhizobium-sp.] |
| TIS34142.1 | Type6.BREX | Zn_Tnp_IS91+Y2_Tnp->ABhydrolase-><-?\|\|GDResolvase+HIN-HTH->?->CopG/RHH->BrxE->BrxA-DNA-binding->BrxB-BrxF-iSTAND->DUF6079*->N6-METHYLASE->PglZ->SIGMA+OB-Fold+AAA-ATPase+Lon-Protease-BrxL-><-?<-?<-?<-?<-HTH_3 | 1178 | Bacteria>Pseudomonadota>Alphaproteobacteria>Hyphomicrobiales>Phyllobacteriaceae>Mesorhizobium | GCA_004964075.1 | Mesorhizobium_sp. | E5W95_31550 | 60338..63874 | 1871066 | BREX-system-P-loop-protein-BrxC-[Mesorhizobium-sp.] |
| TIR32688.1 | Type6.BREX | <-DUF6079-p*<-BrxB-BrxF-iSTAND<-BrxA-DNA-binding<-BrxE<-XIS-HTH<-tRNA<-ZnR+SLATT<-Peptidase_S11 | 592 | Bacteria>Pseudomonadota>Alphaproteobacteria>Hyphomicrobiales>Phyllobacteriaceae>Mesorhizobium | GCA_004964495.1 | Mesorhizobium_sp. | E5X27_28680 | 1..1777 | 1871066 | BREX-system-P-loop-protein-BrxC,-partial-[Mesorhizobium-sp.] |
| WP_023760219.1 | Type6.BREX | <-SIGMA+OB-Fold+AAA-ATPase+Lon-Protease-BrxL<-PglZ<-Tox-RelA<-N6-METHYLASE<-N6-METHYLASE<-DUF6079*<-BrxB-BrxF-iSTAND<-BrxA-DNA-binding<-BrxE<-XIS-HTH<-tRNA<-?\|\|SF1-SF2-Helicase->?-><-ZnR+SLATT<-DDE_Tnp_1_assoc+DDE_Tnp_1 | 1180 | Bacteria>Pseudomonadota>Alphaproteobacteria>Hyphomicrobiales>Phyllobacteriaceae>Mesorhizobium | GCF_000502955.1 | Mesorhizobium_sp._LNHC252B00 | X744_RS152620 | 73471..77013 | 1287252 | BREX-system-P-loop-protein-BrxC-[Mesorhizobium-sp.-LNHC252B00] |
| WP_206520489.1 | Type6.BREX | <-SF1-SF2-Helicase\|\|?->\|\|?->?->AbiEi_4->RVT->RVT->BrxE->DUF6079-p*->SF1-SF2-Helicase-><-HTH_3<-STYKIN<-?\|\|?-><-?<-Phage_integrase<-Phage_int_SAM_1+Phage_integrase\|\|<-?<-HTH_3 | 286 | Bacteria>Pseudomonadota>Alphaproteobacteria>Hyphomicrobiales>Phyllobacteriaceae>Mesorhizobium | GCF_003997555.1 | Mesorhizobium_sp._M1A.T.Ca.IN.004.03.1.1 | EOD29_RS24250 | 126802..127662 | 2496795 | hypothetical-protein-[Mesorhizobium-sp.-M1A.T.Ca.IN.004.03.1.1] |
| WP_140781888.1 | Type6.BREX | <-D5ATPase<-XIS-HTH\|\|MerR-HTH->?->HKD-DNase+HKD-Rease->?->BrxE->BrxA-DNA-binding->BrxB-BrxF-iSTAND->DUF6079*->N6-METHYLASE->PglZ->SIGMA+OB-Fold+AAA-ATPase+HKD-DNase-BrxL-><-?<-?<-CHTH | 1211 | Bacteria>Pseudomonadota>Alphaproteobacteria>Hyphomicrobiales>Phyllobacteriaceae>Mesorhizobium | GCF_006442585.1 | Mesorhizobium_sp._B2-4-8 | FJ947_RS26560 | 105800..109435 | 2589941 | BREX-system-P-loop-protein-BrxC-[Mesorhizobium-sp.-B2-4-8] |
| WP_224719979.1 | Type6.BREX | <-Ntox49<-REase+HTH+35exo+Small-B-barrel+SF1-SF2-Helicase\|\|wHTH+WYL->wHTH+WYL+WCX->?->GreA_GreB+SF1-SF2-Helicase-p->BrxE->BrxA-DNA-binding->BrxB-BrxF-iSTAND->DUF6079*->N6-METHYLASE->N6-METHYLASE->PglZ->SIGMA+OB-Fold+AAA-ATPase+HKD-DNase-BrxL-> | 1208 | Bacteria>Pseudomonadota>Alphaproteobacteria>Hyphomicrobiales>Phyllobacteriaceae>Mesorhizobium | GCF_020164885.1 | Mesorhizobium_sp._C01-1-7 | LB516_RS16265 | 26347..29973 | 2876632 | BREX-system-P-loop-protein-BrxC-[Mesorhizobium-sp.-C01-1-7] |
| WP_183807573.1 | Type6.BREX | <-SIGMA+OB-Fold+AAA-ATPase+HKD-DNase-BrxL<-PglZ<-N6-METHYLASE<-N6-METHYLASE<-DUF6079*<-BrxB-BrxF-iSTAND<-BrxE<-PhdYeFM_antitox<-PIN+OST-HTH | 1206 | Bacteria>Pseudomonadota>Alphaproteobacteria>Hyphomicrobiales>Rhizobiaceae>Mycoplana | GCF_014196765.1 | Mycoplana_azooxidifex | GGQ64_RS22460 | 165035..168655 | 1636188 | BREX-system-P-loop-protein-BrxC-[Mycoplana-azooxidifex] |
| TBF64186.1 | Type6.BREX | <-XIS-HTH\|\|HTH_19->?->Couple_hipA+HipA_C->CHTH-><-SIGMA+OB-Fold+AAA-ATPase+HKD-DNase-BrxL<-PglZ<-N6-METHYLASE<-N6-METHYLASE<-DUF6079*<-BrxB-BrxF-iSTAND<-BrxA-DNA-binding<-BrxE<-PhdYeFM_antitox<-VirD4-TraG+VirD4-FtsK<-TraI-VirD2-RCRE<-TraJ-RHH<-HTH_36\|\|HIN-HTH-> | 1208 | Bacteria>Pseudomonadota>Alphaproteobacteria>Hyphomicrobiales>Rhizobiaceae>Rhizobium/Agrobacterium_group | GCA_004306065.1 | Rhizobium_leguminosarum | ELG90_15310 | 3002859..3006485 | 384 | BREX-system-P-loop-protein-BrxC-[Rhizobium-leguminosarum] |
| MBA8801280.1 | Type6.BREX | <-SIGMA+OB-Fold+AAA-ATPase+HKD-DNase-BrxL<-PglZ<-N6-METHYLASE<-N6-METHYLASE<-DUF6079*<-BrxB-BrxF-iSTAND<-BrxA-DNA-binding<-BrxE\|\|?-><-wHTH+WYL+WCX\|\|?->35Exo+REase->SMODS-> | 1208 | Bacteria>Pseudomonadota>Alphaproteobacteria>Hyphomicrobiales>Rhizobiaceae>Rhizobium/Agrobacterium_group | GCA_014138475.1 | Agrobacterium_sp._RC10-4-1 | FHW77_005035 | 9105..12731 | 2587039 | hypothetical-protein-[Agrobacterium-sp.-RC10-4-1] |
| MBX5048875.1 | Type6.BREX | TetR-HTH->?-><-SIGMA+OB-Fold+AAA-ATPase+HKD-DNase-BrxL<-PglZ<-ParB+HNH<-N6-METHYLASE<-N6-METHYLASE<-DUF6079*<-BrxB-BrxF-iSTAND<-BrxA-DNA-binding<-BrxE<-tRNA\|\|?->Thioredoxin-> | 1213 | Bacteria>Pseudomonadota>Alphaproteobacteria>Hyphomicrobiales>Rhizobiaceae>Rhizobium/Agrobacterium_group | GCA_019684595.1 | Rhizobium_lentis | HJB64_19910 | 192083..195724 | 1138194 | BREX-system-P-loop-protein-BrxC-[Rhizobium-lentis] |
| WP_130750054.1 | Type6.BREX | <-XIS-HTH\|\|HTH_19->?->Couple_hipA+HipA_C->CHTH-><-SIGMA+OB-Fold+AAA-ATPase+HKD-DNase-BrxL<-PglZ<-N6-METHYLASE<-N6-METHYLASE<-DUF6079*<-BrxB-BrxF-iSTAND<-BrxA-DNA-binding<-BrxE<-PhdYeFM_antitox<-VirD4-TraG+VirD4-FtsK<-?<-?<-TraJ-RHH<-HTH_36 | 1208 | Bacteria>Pseudomonadota>Alphaproteobacteria>Hyphomicrobiales>Rhizobiaceae>Rhizobium/Agrobacterium_group | GCF_004304245.1 | Rhizobium_leguminosarum | ELI03_RS14940 | 2921080..2924706 | 384 | BREX-system-P-loop-protein-BrxC-[Rhizobium-leguminosarum] |
| WP_197729861.1 | Type6.BREX | <-XIS-HTH\|\|HTH_19->?->Couple_hipA+HipA_C->CHTH-><-SIGMA+OB-Fold+AAA-ATPase+HKD-DNase-BrxL<-PglZ<-N6-METHYLASE<-N6-METHYLASE<-DUF6079*<-BrxB-BrxF-iSTAND<-BrxA-DNA-binding-p<-BrxE<-PhdYeFM_antitox<-VirD4-TraG+VirD4-FtsK<-?<-?<-TraJ-RHH<-HTH_36 | 1210 | Bacteria>Pseudomonadota>Alphaproteobacteria>Hyphomicrobiales>Rhizobiaceae>Rhizobium/Agrobacterium_group | GCF_004306065.1 | Rhizobium_leguminosarum | ELG90_RS15310 | 3002859..3006491 | 384 | BREX-system-P-loop-protein-BrxC-[Rhizobium-leguminosarum] |
| WP_132661711.1 | Type6.BREX | Glyoxalase+ZnR->HipA_C->?->HKD-DNase+HKD-Rease-><-SIGMA+OB-Fold+AAA-ATPase+HKD-DNase-BrxL<-PglZ<-N6-METHYLASE<-N6-METHYLASE<-DUF6079*<-BrxB-BrxF-iSTAND<-BrxA-DNA-binding<-BrxE<-PhdYeFM_antitox<-wHTH+WYL+WCX\|\|SMODS->?->Classical-AAA->Cap4_nuclease+SAVED-> | 1208 | Bacteria>Pseudomonadota>Alphaproteobacteria>Hyphomicrobiales>Rhizobiaceae>Rhizobium/Agrobacterium_group | GCF_004342325.1 | Rhizobium_azibense | EV130_RS24200 | 158902..162528 | 1136135 | BREX-system-P-loop-protein-BrxC-[Rhizobium-azibense] |
| WP_259669791.1 | Type6.BREX | HTH_35->N6-METHYLASE->REase-><-SIGMA+OB-Fold+AAA-ATPase+Lon-Protease-BrxL<-PglZ<-N6-METHYLASE<-N6-METHYLASE<-DUF6079*<-BrxB-BrxF-iSTAND<-BrxA-DNA-binding<-BrxE<-XIS-HTH\|\|<-?\|\|DDE_Tnp_IS66->Phage_integrase->ParA-Soj-PloopNTPase->ParB+RepB-> | 1181 | Bacteria>Pseudomonadota>Alphaproteobacteria>Hyphomicrobiales>Rhizobiaceae>Rhizobium/Agrobacterium_group | GCF_019684975.1 | Rhizobium_sp._NZLR5 | HJB88_RS30900 | 14141..17686 | 2731103 | BREX-system-P-loop-protein-BrxC-[Rhizobium-sp.-NZLR5] |
| WP_183856131.1 | Type6.BREX | <-ParB+HNH\|\|<-HIN-HTH\|\|HTH_35->N6-METHYLASE->REase-><-SIGMA+OB-Fold+AAA-ATPase+Lon-Protease-BrxL<-PglZ<-N6-METHYLASE<-N6-METHYLASE<-DUF6079*<-BrxB-BrxF-iSTAND<-BrxA-DNA-binding<-BrxE<-XIS-HTH\|\|DDE_Tnp_IS66->Phage_integrase-><-?<-?<-ZDNA-HTH+HTH | 1180 | Bacteria>Pseudomonadota>Alphaproteobacteria>Hyphomicrobiales>Rhizobiaceae>Rhizobium/Agrobacterium_group | GCF_019685155.1 | unclassified_Rhizobium | FHX03_RS25930 | 236740..240282 | 2613769 | MULTISPECIES:-BREX-system-P-loop-protein-BrxC-[unclassified-Rhizobium] |
| WP_168274407.1 | Type6.BREX | <-Polbeta+HEPN\|\|PhdYeFM_antitox->PIN-><-Phage_integrase<-DDE_Tnp_IS66\|\|XIS-HTH->BrxE->BrxA-DNA-binding->BrxB-BrxF-iSTAND->DUF6079*->N6-METHYLASE->N6-METHYLASE->Tox-RelA->PglZ->SIGMA+OB-Fold+AAA-ATPase+Lon-Protease-BrxL-><-REase<-N6-METHYLASE<-HTH_35\|\|ParB+HNH-> | 1180 | Bacteria>Pseudomonadota>Alphaproteobacteria>Hyphomicrobiales>Rhizobiaceae>Rhizobium/Agrobacterium_group | GCF_019778525.1 | Rhizobium | HFD68_RS26235 | 204618..208160 | 379 | MULTISPECIES:-BREX-system-P-loop-protein-BrxC-[Rhizobium] |
| WP_222037895.1 | Type6.BREX | VirB11+VirB4-FtsK-><-?<-Polbeta+HEPN\|\|PhdYeFM_antitox->PIN->\|\|DOC+HTH->\|\|BrxE->BrxA-DNA-binding->BrxB-BrxF-iSTAND->DUF6079*->N6-METHYLASE->N6-METHYLASE->PglZ->SIGMA+OB-Fold+AAA-ATPase+Lon-Protease-BrxL-><-McrC-NTD+REase<-RAMA+McrB-AAA<-SF1-SF2-Helicase+REase<-SF1-SF2-Helicase+REase\|\|wHTH+WYL+WCX-> | 1178 | Bacteria>Pseudomonadota>Alphaproteobacteria>Hyphomicrobiales>Rhizobiaceae>Rhizobium/Agrobacterium_group | GCF_019779625.1 | Rhizobium_laguerreae | HFN80_16895 | 150024..153560 | 1076926 | BREX-system-P-loop-protein-BrxC-[Rhizobium-laguerreae] |
| WP_250810295.1 | Type6.BREX | <-GreA_GreB+SF1-SF2-Helicase<-PIWI<-REase+TIR<-?\|\|wHTH+WYL+WCX->BrxE->BrxA-DNA-binding->BrxB-BrxF-iSTAND->DUF6079*->N6-METHYLASE->PglZ->SIGMA+OB-Fold+AAA-ATPase+HKD-DNase-BrxL-><-?<-ParB+HNH\|\|GDResolvase+HIN-HTH-><-HTH_3 | 1208 | Bacteria>Pseudomonadota>Alphaproteobacteria>Hyphomicrobiales>Rhizobiaceae>Rhizobium/Agrobacterium_group | GCF_023667605.1 | Neorhizobium_tomejilense | L1082_RS18485 | 51181..54807 | 2093828 | BREX-system-P-loop-protein-BrxC-[Neorhizobium-tomejilense] |
| WP_164064485.1 | Type6.BREX | <-TIR<-wHTH+WYL+WCX\|\|REase+HTH+35exo+Small-B-barrel+SF1-SF2-Helicase->McrB-AAA->McrC-NTD+REase-><-SIGMA+OB-Fold+AAA-ATPase+Lon-Protease-BrxL<-PglZ<-N6-METHYLASE<-N6-METHYLASE<-DUF6079*<-BrxB-BrxF-iSTAND<-BrxA-DNA-binding<-BrxE\|\|<-DOC+HTH\|\|PhdYeFM_antitox-><-ZDNA-HTH+HTH<-ParB+RepB<-ParA-Soj-PloopNTPase | 1178 | Bacteria>Pseudomonadota>Alphaproteobacteria>Hyphomicrobiales>Rhizobiaceae>Rhizobium/Agrobacterium_group | - | Rhizobium_leguminosarum | GR223_RS33875 | 57482..61018 | 384 | BREX-system-P-loop-protein-BrxC-[Rhizobium-leguminosarum] |
| WP_221145756.1 | Type6.BREX | TetR-HTH->?-><-SIGMA+OB-Fold+AAA-ATPase+HKD-DNase-BrxL<-PglZ<-ParB+HNH<-N6-METHYLASE<-N6-METHYLASE<-DUF6079*<-BrxB-BrxF-iSTAND<-BrxA-DNA-binding<-BrxE<-tRNA\|\|?->Thioredoxin-> | 1213 | Bacteria>Pseudomonadota>Alphaproteobacteria>Hyphomicrobiales>Rhizobiaceae>Rhizobium/Agrobacterium_group | - | Rhizobium_lentis | HJB64_19910 | 192083..195724 | 1138194 | BREX-system-P-loop-protein-BrxC-[Rhizobium-lentis] |
| WP_192067374.1 | Type6.BREX | <-SIGMA+OB-Fold+AAA-ATPase+HKD-DNase-BrxL<-PglZ<-N6-METHYLASE<-N6-METHYLASE<-DUF6079*<-BrxB-BrxF-iSTAND<-BrxA-DNA-binding<-BrxE<-Phage_integrase<-XIS-HTH<-Phage_integrase\|\|REase->wHTH+REase->MPTase-> | 1208 | Bacteria>Pseudomonadota>Alphaproteobacteria>Hyphomicrobiales>Rhizobiaceae>Rhizobium/Agrobacterium_group | - | unclassified_Rhizobium | IFT53_20370 | 8816..12442 | 2613769 | MULTISPECIES:-BREX-system-P-loop-protein-BrxC-[unclassified-Rhizobium] |
| KSV61619.1 | Type6.BREX | <-SIGMA+OB-Fold+AAA-ATPase+HKD-DNase-BrxL<-PglZ<-N6-METHYLASE<-DUF6079*<-BrxB-BrxF-iSTAND<-BrxA-DNA-binding<-BrxE<-tRNA\|\|?->?->ACET->ACET-><-?\|\|sGTP-> | 1203 | Bacteria>Pseudomonadota>Alphaproteobacteria>Hyphomicrobiales>Rhizobiaceae>Sinorhizobium/Ensifer_group | GCA_001461715.1 | Sinorhizobium_sp._GL2 | N182_13660 | 115788..119399 | 1358416 | hypothetical-protein-N182_13660-[Sinorhizobium-sp.-GL2] |
| WP_042775876.1 | Type6.BREX | <-HTH_Tnp<-?<-TUDOR+SF1-SF2-Helicase+iREase+CTD<-SIGMA+OB-Fold+AAA-ATPase+HKD-DNase-BrxL<-PglZ<-N6-METHYLASE<-N6-METHYLASE<-DUF6079*<-BrxB-BrxF-iSTAND<-BrxA-DNA-binding<-BrxE<-?\|\|?-><-HIN-HTH | 1208 | Bacteria>Pseudomonadota>Alphaproteobacteria>Hyphomicrobiales>Rhizobiaceae>Sinorhizobium/Ensifer_group | GCF_000261925.1 | Sinorhizobium_fredii | IC54_RS06660 | 41741..45367 | 380 | BREX-system-P-loop-protein-BrxC-[Sinorhizobium-fredii] |
| WP_209439290.1 | Type6.BREX | Calcineurin->SbcC+SbcC->TUDOR+SF1-SF2-Helicase+iREase+CTD->?->?-><-SIGMA+OB-Fold+AAA-ATPase+HKD-DNase-BrxL<-PglZ<-N6-METHYLASE<-N6-METHYLASE<-DUF6079*<-BrxB-BrxF-iSTAND<-BrxA-DNA-binding<-BrxE<-PhdYeFM_antitox\|\|?->PhdYeFM_antitox->PIN-><-PhdYeFM_antitox\|\|Phage_integrase-> | 1210 | Bacteria>Pseudomonadota>Alphaproteobacteria>Hyphomicrobiales>Rhizobiaceae>Sinorhizobium/Ensifer_group | GCF_000427465.1 | Sinorhizobium_arboris | SINAR_RS0100665 | 116577..120209 | 76745 | BREX-system-P-loop-protein-BrxC-[Sinorhizobium-arboris] |
| WP_113887228.1 | Type6.BREX | SIR2-><-tRNA\|\|?->?->tRNA-><-?<-URI-DNase<-SIGMA+OB-Fold+AAA-ATPase+Lon-Protease-BrxL<-PglZ<-N6-METHYLASE<-DUF6079*<-BrxB-BrxF-iSTAND<-BrxA-DNA-binding<-BrxE\|\|GDResolvase-><-?<-DnaB | 1177 | Bacteria>Pseudomonadota>Alphaproteobacteria>Hyphomicrobiales>Roseiarcaceae>Roseiarcus | GCF_003315135.1 | Roseiarcus_fermentans | DFR50_RS00495 | 107260..110793 | 1473586 | BREX-system-P-loop-protein-BrxC-[Roseiarcus-fermentans] |
| MCP4384519.1 | Type6.BREX | <-Phage_int_SAM_3+Phage_integrase<-XIS-HTH\|\|?->?->BrxE->BrxA-DNA-binding->BrxB-BrxF-iSTAND->DUF6079*->N6-METHYLASE->N6-METHYLASE-p-> | 1179 | Bacteria>Pseudomonadota>Alphaproteobacteria>Hyphomicrobiales>unclassified_Hyphomicrobiales | GCA_024222895.1 | Hyphomicrobiales_bacterium | GY798_24430 | 4683..8222 | 1909294 | BREX-system-P-loop-protein-BrxC-[Hyphomicrobiales-bacterium] |
| GLS17293.1 | Type6.BREX | HTH_3-><-?<-SbcC\|\|XIS-HTH->BrxE->BrxA-DNA-binding->BrxB-BrxF-iSTAND->DUF6079*->N6-METHYLASE->Tox-RelA->PglZ->SIGMA+OB-Fold+AAA-ATPase+Lon-Protease-BrxL-> | 1180 | Bacteria>Pseudomonadota>Alphaproteobacteria>Hyphomicrobiales>Xanthobacteraceae>Labrys | GCA_030160835.1 | Labrys_miyagiensis | GCM10007874_03080 | 168594..172136 | 346912 | hypothetical-protein-GCM10007874_03080-[Labrys-miyagiensis] |
| WP_066770455.1 | Type6.BREX | McrB-AAA->McrC-NTD+REase-><-SIGMA+OB-Fold+AAA-ATPase+Lon-Protease-BrxL<-PglZ<-N6-METHYLASE<-DUF6079*<-BrxB-BrxF-iSTAND<-BrxA-DNA-binding<-BrxE<-APATPase<-tRNA<-REC+OmpR-HTH\|\|FGase-> | 1176 | Bacteria>Pseudomonadota>Alphaproteobacteria>Hyphomonadales>Hyphomonadaceae>Candidatus_Viadribacter | GCF_001679665.1 | Candidatus_Viadribacter_manganicus | ATE48_RS09195 | 1777065..1780595 | 1759059 | BREX-system-P-loop-protein-BrxC-[Candidatus-Viadribacter-manganicus] |
| WP_212538477.1 | Type6.BREX | <-TetR-HTH\|\|?->wHTH+WYL+WCX->BrxE->BrxA-DNA-binding->BrxB-BrxF-iSTAND->DUF6079*->N6-METHYLASE->N6-METHYLASE-p->ParB+HNH->PglZ->SIGMA+OB-Fold+AAA-ATPase+HKD-DNase-BrxL-><-XIS-HTH<-?\|\|HTH+LexA-protease->UDG-> | 1208 | Bacteria>Pseudomonadota>Alphaproteobacteria>Rhodobacterales>Paracoccaceae>Aestuariicoccus | GCF_018224955.1 | Aestuariicoccus_sp._KMU-90 | KB874_RS20730 | 16568..20194 | 2827239 | BREX-system-P-loop-protein-BrxC-[Aestuariicoccus-sp.-KMU-90] |
| WP_263337868.1 | Type6.BREX | Peptidase-M15->?-><-SIGMA+OB-Fold+AAA-ATPase+HKD-DNase-BrxL<-PglZ<-N6-METHYLASE<-DUF6079*<-BrxB-BrxF-iSTAND<-BrxA-DNA-binding<-BrxE<-XIS-HTH<-Arm-DNA-bind_3+Phage_int_SAM_3+Phage_integrase\|\|MerR-HTH-> | 1203 | Bacteria>Pseudomonadota>Alphaproteobacteria>Rhodobacterales>Paracoccaceae>Defluviimonas | GCF_025673475.1 | Defluviimonas_sp._WL0024 | OEZ60_RS15020 | 60055..63666 | 2984153 | BREX-system-P-loop-protein-BrxC-[Defluviimonas-sp.-WL0024] |
| WP_114283705.1 | Type6.BREX | <-SIGMA+OB-Fold+AAA-ATPase+Lon-Protease-BrxL<-PglZ<-N6-METHYLASE<-DUF6079*<-BrxB-BrxF-iSTAND<-BrxA-DNA-binding<-BrxE<-HORMA\|\|?-><-McrC-NTD+REase<-McrB-AAA<-REase\|\|Exonuc_V_gamma+REase-> | 1180 | Bacteria>Pseudomonadota>Alphaproteobacteria>Rhodobacterales>Paracoccaceae>Halocynthiibacter | GCF_003333265.1 | Candidatus_Halocynthiibacter_alkanivorans | DWA01_RS00685 | 21438..24980 | 2267619 | BREX-system-P-loop-protein-BrxC-[Candidatus-Halocynthiibacter-alkanivorans] |
| WP_249060201.1 | Type6.BREX | RelE-ParE->TPR->RNase_T+35exo->XIS-HTH->BrxE->BrxA-DNA-binding->BrxB-BrxF-iSTAND->DUF6079*->N6-METHYLASE->N6-METHYLASE->PglZ->SIGMA+OB-Fold+AAA-ATPase+HKD-DNase-BrxL->?->DUF4236+TPR->McrB-AAA->McrC-NTD+REase-> | 1206 | Bacteria>Pseudomonadota>Alphaproteobacteria>Rhodobacterales>Paracoccaceae>Roseibaca | GCF_023336755.1 | Roseibaca_sp._V10 | M3N55_RS14200 | 5935..9555 | 2940293 | BREX-system-P-loop-protein-BrxC-[Roseibaca-sp.-V10] |
| NTT88404.1 | Type6.BREX | <-SWIM\|\|Arm-DNA-bind_3+Phage_int_SAM_3+Phage_integrase->wHTH+WYL+WCX->BrxE->BrxA-DNA-binding->BrxB-BrxF-iSTAND->DUF6079*->N6-METHYLASE->PglZ->SIGMA+OB-Fold+AAA-ATPase+HKD-DNase-BrxL-> | 1208 | Bacteria>Pseudomonadota>Alphaproteobacteria>Rhodobacterales>Paracoccaceae>Tabrizicola | GCA_013328815.1 | Tabrizicola_sp._SY72 | HS053_21140 | 11855..15481 | 2741673 | BREX-system-P-loop-protein-BrxC-[Tabrizicola-sp.-SY72] |
| WP_194245628.1 | Type6.BREX | <-SWIM\|\|Arm-DNA-bind_3+Phage_int_SAM_3+Phage_integrase->wHTH+WYL+WCX->BrxE->BrxA-DNA-binding->BrxB-BrxF-iSTAND->DUF6079*->N6-METHYLASE->N6-METHYLASE-p->PglZ->SIGMA+OB-Fold+AAA-ATPase+HKD-DNase-BrxL-> | 1213 | Bacteria>Pseudomonadota>Alphaproteobacteria>Rhodobacterales>Paracoccaceae>Tabrizicola | GCA_013328815.1 | Tabrizicola_sp._SY72 | HS053_RS21145 | 11840..15481 | 2741673 | BREX-system-P-loop-protein-BrxC-[Tabrizicola-sp.-SY72] |
| WP_107845004.1 | Type6.BREX | <-TraI-VirD2-RCRE<-HIN-HTH+TraJ-RHH<-?<-FurR-HTH<-ArdCN+MPTase\|\|wHTH+WYL+WCX->BrxE->BrxA-DNA-binding->BrxB-BrxF-iSTAND->DUF6079*->N6-METHYLASE->N6-METHYLASE-p->PglZ->SIGMA+OB-Fold+AAA-ATPase+HKD-DNase-BrxL-><-?<-Couple_hipA+HipA_C<-HTH_3<-HTH_36 | 1208 | Bacteria>Pseudomonadota>Alphaproteobacteria>Rhodobacterales>Roseobacteraceae>Litoreibacter | GCF_003054285.1 | Litoreibacter_ponti | C8N43_RS07515 | 1489253..1492879 | 1510457 | BREX-system-P-loop-protein-BrxC-[Litoreibacter-ponti] |
| WP_188483502.1 | Type6.BREX | Glyco_hydro_53+Phage-tail_3->?->?->Peptidase-M15->?-><-SIGMA+OB-Fold+AAA-ATPase+HKD-DNase-BrxL<-PglZ<-N6-METHYLASE<-DUF6079*<-BrxB-BrxF-iSTAND<-BrxA-DNA-binding<-BrxE<-wHTH+WYL+WCX<-McrC-NTD+REase<-EVE+McrB-AAA<-?<-ParB<-ParB | 1208 | Bacteria>Pseudomonadota>Alphaproteobacteria>Rhodobacterales>Roseobacteraceae>Marivita | GCF_014637725.1 | Marivita_lacus | IEV82_RS18225 | 47405..51031 | 1323742 | BREX-system-P-loop-protein-BrxC-[Marivita-lacus] |
| WP_109670829.1 | Type6.BREX | XIS-HTH->BrxE->BrxA-DNA-binding->BrxB-BrxF-iSTAND->DUF6079*->N6-METHYLASE->Tox-RelA->PglZ->SIGMA+OB-Fold+AAA-ATPase+Lon-Protease-BrxL-><-?<-?<-Peptidase-M15 | 1180 | Bacteria>Pseudomonadota>Alphaproteobacteria>Rhodobacterales>Roseobacteraceae>Roseicyclus | GCF_003148775.1 | Roseicyclus_mahoneyensis | C7455_RS18625 | 49433..52975 | 164332 | BREX-system-P-loop-protein-BrxC-[Roseicyclus-mahoneyensis] |
| WP_076451079.1 | Type6.BREX | ABhydrolase-><-?<-Phage_integrase\|\|XIS-HTH->wHTH+WYL+WCX->BrxE->BrxA-DNA-binding->BrxB-BrxF-iSTAND->DUF6079*->N6-METHYLASE->N6-METHYLASE->PglZ->SIGMA+OB-Fold+AAA-ATPase+HKD-DNase-BrxL-><-?<-?<-Peptidase-M15 | 1208 | Bacteria>Pseudomonadota>Alphaproteobacteria>Rhodobacterales>Roseobacteraceae>Roseivivax | GCF_900156805.1 | Roseivivax_lentus | BW990_RS19885 | 19994..23620 | 633194 | BREX-system-P-loop-protein-BrxC-[Roseivivax-lentus] |
| WP_268956598.1 | Type6.BREX | <-SIGMA+OB-Fold+AAA-ATPase+HKD-DNase-BrxL<-PglZ<-N6-METHYLASE<-DUF6079*<-BrxB-BrxF-iSTAND<-BrxA-DNA-binding<-BrxE<-XIS-HTH<-HTH_3\|\|?->?->wHTH+HNH-> | 1203 | Bacteria>Pseudomonadota>Alphaproteobacteria>Rhodobacterales>Roseobacteraceae>Roseovarius | - | Roseovarius_sp._EGI_FJ00037 | N5A93_RS18625 | 8608..12219 | 2978479 | BREX-system-P-loop-protein-BrxC-[Roseovarius-sp.-EGI-FJ00037] |
| PSL17302.1 | Type6.BREX | <-HAMP+HISKIN\|\|?->tRNA-><-SIR2<-SIGMA+OB-Fold+AAA-ATPase+HKD-DNase-BrxL<-PglZ<-N6-METHYLASE<-N6-METHYLASE<-DUF6079*<-BrxB-BrxF-iSTAND<-BrxA-DNA-binding<-BrxE<-XIS-HTH<-HEPN<-Polbeta<-wHTH+AbiEi_2\|\|REase-> | 1216 | Bacteria>Pseudomonadota>Alphaproteobacteria>Rhodobacterales>Roseobacteraceae>Shimia | GCA_003014475.1 | Shimia_abyssi | CLV88_11975 | 80664..84314 | 1662395 | hypothetical-protein-CLV88_11975-[Shimia-abyssi] |
| WP_207797113.1 | Type6.BREX | <-HAMP+HISKIN\|\|?->tRNA-><-SIR2<-SIGMA+OB-Fold+AAA-ATPase+HKD-DNase-BrxL<-PglZ<-N6-METHYLASE<-N6-METHYLASE<-DUF6079*<-BrxB-BrxF-iSTAND<-BrxA-DNA-binding<-BrxE<-XIS-HTH<-HEPN<-Polbeta<-wHTH+AbiEi_2\|\|REase-> | 1207 | Bacteria>Pseudomonadota>Alphaproteobacteria>Rhodobacterales>Roseobacteraceae>Shimia | GCF_003014475.1 | Shimia_abyssi | CLV88_RS20200 | 80664..84287 | 1662395 | BREX-system-P-loop-protein-BrxC-[Shimia-abyssi] |
| WP_220429122.1 | Type6.BREX | <-DDE_Tnp_1_6\|\|OLD-ABC->HNH-><-?<-HIN-HTH\|\|XIS-HTH->BrxE->BrxA-DNA-binding->BrxB-BrxF-iSTAND->DUF6079*->N6-METHYLASE->PglZ->SIGMA+OB-Fold+AAA-ATPase+HKD-DNase-BrxL->tRNA->ABC-ATPase+SbcC->?->?->ABhydrolase-><-Phosphodiest | 1207 | Bacteria>Pseudomonadota>Alphaproteobacteria>Rhodobacterales>Roseobacteraceae>Sulfitobacter | GCF_900313045.1 | Sulfitobacter_mediterraneus | D9A10_RS06915 | 36169..39792 | 83219 | BREX-system-P-loop-protein-BrxC-[Sulfitobacter-mediterraneus] |
| WP_013639411.1 | Type6.BREX | VirB10->RHH->TraJ-RHH->ACET+ACET-><-SIGMA+OB-Fold+AAA-ATPase+Lon-Protease-BrxL<-PglZ<-Tox-RelA<-N6-METHYLASE<-N6-METHYLASE<-DUF6079*<-BrxB-BrxF-iSTAND<-BrxA-DNA-binding<-BrxE<-XIS-HTH<-Arm-DNA-bind_3+Phage_int_SAM_3+Phage_integrase | 1180 | Bacteria>Pseudomonadota>Alphaproteobacteria>Rhodospirillales>Acetobacteraceae>Acidiphilium | GCF_000202835.1 | Acidiphilium_multivorum | ACMV_RS02190 | 507130..510672 | 62140 | BREX-system-P-loop-protein-BrxC-[Acidiphilium-multivorum] |
| WP_265637943.1 | Type6.BREX | <-TetR-HTH\|\|?->XIS-HTH->BrxE->BrxA-DNA-binding->BrxB-BrxF-iSTAND->DDE_Tnp_1_4->BrxB-BrxF-iSTAND->DUF6079-p->DUF6079*->N6-METHYLASE-p->HIN-HTH->AAA-ATPase-><-TnsC-ATPase<-HTH_28+rve+Mu-transpos_C\|\|VirB6->VirB8->VirB9->VirB10-> | 1221 | Bacteria>Pseudomonadota>Alphaproteobacteria>Rhodospirillales>Acetobacteraceae>Acidiphilium | GCF_026191915.1 | Acidiphilium_sp._PA | AruPA_RS16835 | 28443..32108 | 2871705 | BREX-system-P-loop-protein-BrxC-[Acidiphilium-sp.-PA] |
| WP_201727547.1 | Type6.BREX | VirB8->VirB9->VirB10->RHH->TraJ-RHH->ACET+ACET->BrxE->BrxA-DNA-binding->BrxB-BrxF-iSTAND->DUF6079-p->HIN-HTH->?->LZ_Tnp_IS66+zf-IS66+DDE_Tnp_IS66+DDE_Tnp_IS66_C->?->DUF6079-p->N6-METHYLASE->N6-METHYLASE-p->ParB+HNH->HTH_29+HTH_32+HTH_33+DDE_3-> | 650 | Bacteria>Pseudomonadota>Alphaproteobacteria>Rhodospirillales>Acetobacteraceae>Acidocella | GCF_907164655.1 | Acidocella_sp._C78 | ACIDOCELLA78_RS32075 | 228724..230676 | 1671486 | BREX-system-P-loop-protein-BrxC-[Acidocella-sp.-C78] |
| WP_201727548.1 | Type6.BREX | ACET+ACET->BrxE->BrxA-DNA-binding->BrxB-BrxF-iSTAND->DUF6079-p->HIN-HTH->?->LZ_Tnp_IS66+zf-IS66+DDE_Tnp_IS66+DDE_Tnp_IS66_C->?->DUF6079-p*->N6-METHYLASE->N6-METHYLASE-p->ParB+HNH->HTH_29+HTH_32+HTH_33+DDE_3->ParB+HNH->PglZ->SIGMA+OB-Fold+AAA-ATPase+HKD-DNase-BrxL-><-Arm-DNA-bind_3+Phage_int_SAM_3+Phage_integrase | 495 | Bacteria>Pseudomonadota>Alphaproteobacteria>Rhodospirillales>Acetobacteraceae>Acidocella | GCF_907164655.1 | Acidocella_sp._C78 | ACIDOCELLA78_RS32100 | 233672..235159 | 1671486 | hypothetical-protein-[Acidocella-sp.-C78] |
| WP_213670186.1 | Type6.BREX | TPR-><-?\|\|?-><-MODE-HTH+PBPII\|\|?->BrxE->BrxA-DNA-binding->BrxB-BrxF-iSTAND->DUF6079*->N6-METHYLASE->PglZ->SIGMA+OB-Fold+AAA-ATPase+Lon-Protease-BrxL-><-tRNA<-?\|\|?->?->Peptidase_S11+PBP5_C->Thymidylate_kin-> | 1178 | Bacteria>Pseudomonadota>Alphaproteobacteria>Rhodospirillales>Acetobacteraceae>Roseococcus | GCF_018413645.1 | Roseococcus_pinisoli | KHU32_RS11245 | 519199..522735 | 2835040 | BREX-system-P-loop-protein-BrxC-[Roseococcus-pinisoli] |
| KAA0676972.1 | Type6.BREX | <-DDE_Tnp_IS240<-DDE_Tnp_1_5+DDE_Tnp_1<-Phage_integrase+Phage_integrase\|\|?-><-REC+LuxR-HTH<-?<-SIGMA+OB-Fold+AAA-ATPase+HKD-DNase-BrxL<-PglZ<-N6-METHYLASE-p<-DUF6079*<-BrxB-BrxF-iSTAND<-BrxA-DNA-binding<-BrxE<-?<-HHH_4+HTH+wHTH+SF1-SF2-Helicase+REase<-SF1-SF2-Helicase<-MORC+DUF3684+REase\|\|XIS-HTH->NtrC-AAA-> | 1207 | Bacteria>Pseudomonadota>Alphaproteobacteria>Rhodospirillales>Acetobacteraceae>Roseomonas | GCA_008365375.1 | Roseomonas_genomospecies_6 | DS843_25180 | 43725..47348 | 214106 | BREX-system-P-loop-protein-BrxC-[Roseomonas-genomospecies-6] |
| MBI1207569.1 | Type6.BREX | ACT->?-><-?\|\|?->BrxE->BrxA-DNA-binding->BrxB-BrxF-iSTAND->DUF6079*->N6-METHYLASE->PglZ->SIGMA+OB-Fold+AAA-ATPase+HKD-DNase-BrxL-><-?<-SAM-METHYLASE<-?<-NUDIX | 1222 | Bacteria>Pseudomonadota>Alphaproteobacteria>Rhodospirillales>Azospirillaceae>Azospirillum | GCA_016124535.1 | Azospirillum_sp. | GC191_09820 | 259214..262882 | 34012 | BREX-system-P-loop-protein-BrxC-[Azospirillum-sp.] |
| WP_109072015.1 | Type6.BREX | <-McrC-NTD+REase<-McrB-AAA<-wHTH+WYL+WCX<-SF1-SF2-Helicase+wHTH+Small-B-barrel+DUF1998<-BrxD-AAA<-PglZ<-?<-N6-METHYLASE<-?<-DUF6079*<-BrxB-BrxF-iSTAND<-BrxA-DNA-binding<-BrxE<-?<-?<-?\|\|METHYLASE-> | 1208 | Bacteria>Pseudomonadota>Alphaproteobacteria>Rhodospirillales>Azospirillaceae>Azospirillum | GCF_003119115.1 | Azospirillum_sp._TSH58 | TSH58p_RS25495 | 880428..884054 | 664962 | BREX-system-P-loop-protein-BrxC-[Azospirillum-sp.-TSH58] |
| WP_149270940.1 | Type6.BREX | <-McrC-NTD+REase<-wHTH+McrB-AAA<-SF1-SF2-Helicase+wHTH+Small-B-barrel+DUF1998<-BrxD-AAA<-PglZ<-?<-N6-METHYLASE<-?<-DUF6079*<-BrxB-BrxF-iSTAND<-BrxA-DNA-binding<-BrxE<-GDResolvase+Recombinase+ZnR<-Phage_integrase<-Phage_integrase<-XIS-HTH | 1208 | Bacteria>Pseudomonadota>Alphaproteobacteria>Rhodospirillales>Azospirillaceae>Azospirillum | GCF_008365135.1 | Azospirillum_sp._Sh1 | FZ029_RS18855 | 113384..117010 | 2607285 | BREX-system-P-loop-protein-BrxC-[Azospirillum-sp.-Sh1] |
| WP_200477164.1 | Type6.BREX | Phage_integrase-><-HHH_4+HTH+wHTH+SF1-SF2-Helicase+REase<-SF1-SF2-Helicase<-MORC+DUF3684+REase<-SIGMA+OB-Fold+AAA-ATPase+HKD-DNase-BrxL<-PglZ<-N6-METHYLASE<-N6-METHYLASE<-DUF6079*<-BrxB-BrxF-iSTAND<-BrxA-DNA-binding<-BrxE | 1208 | Bacteria>Pseudomonadota>Alphaproteobacteria>Rhodospirillales>Azospirillaceae>Azospirillum | GCF_016622105.1 | Azospirillum_brasilense | GAY29_RS06575 | 31945..35571 | 192 | BREX-system-P-loop-protein-BrxC-[Azospirillum-brasilense] |
| MCA1971289.1 | Type6.BREX | TetR-HTH-><-Trypsin<-Phage_integrase<-?\|\|BrxE->BrxA-DNA-binding->BrxB-BrxF-iSTAND->DUF6079*->?->N6-METHYLASE-> | 1211 | Bacteria>Pseudomonadota>Alphaproteobacteria>Rhodospirillales>Rhodospirillaceae>Caenispirillum | GCA_020217205.1 | Caenispirillum_sp. | LDL44_00475 | 10512..14147 | 1954482 | BREX-system-P-loop-protein-BrxC-[Caenispirillum-sp.] |
| KJS39375.1 | Type6.BREX | <-N6-METHYLASE-p<-?<-DUF6079*<-BrxB-BrxF-iSTAND<-BrxA-DNA-binding<-BrxE<-HTH_Tnp_1+HTH_21+rve+rve_3<-HTH_Tnp_1\|\|?->HEPN->Arm-DNA-bind_3+Phage_int_SAM_3+Phage_integrase-> | 1208 | Bacteria>Pseudomonadota>Alphaproteobacteria>Rhodospirillales>Rhodospirillaceae>unclassified_Rhodospirillaceae | GCA_000961745.1 | Rhodospirillaceae_bacterium_BRH_c57 | VR70_08020 | 665..4291 | 1629718 | hypothetical-protein-VR70_08020-[Rhodospirillaceae-bacterium-BRH_c57] |
| HCW69913.1 | Type6.BREX | VirB11->VirD4-TraG+VirD4-FtsK->?->ArdCN+MPTase+TOPRIM->TraJ-RHH->TraI-VirD2-RCRE-><-SIGMA+OB-Fold+AAA-ATPase+HKD-DNase-BrxL<-PglZ<-ParB+HNH<-DUF6079*<-BrxB-BrxF-iSTAND<-BrxA-DNA-binding<-BrxE\|\|GDResolvase+Recombinase->?->?-><-?<-HTH_36 | 1208 | Bacteria>Pseudomonadota>Alphaproteobacteria>Rhodospirillales>Thalassospiraceae>Thalassospira | GCA_003539675.1 | Thalassospira_lucentensis | DHR80_22445 | 45804..49430 | 168935 | BREX-system-P-loop-protein-BrxC-[Thalassospira-lucentensis] |

| Protein-ID | System | Gene-Neighbourhoods | Length | Taxonomy | Assembly-ID | Strain | Locus-Tag | Genomic-coordinates | TaxID | Fasta-define |
|---|---|---|---|---|---|---|---|---|---|---|
| TVR84541.1 | Type6.BREX | [illegible] | 1206 | [illegible] | GCA_007695045.1 | Rhodospirillales_bacterium | EA405_01435 | 12860..16480 | 2026786 | [illegible] |
| WP_193715186.1 | Type6.BREX | [illegible] | 1215 | [illegible] | GCF_015143595.1 | Sneathiella_sp._P13V-1 | GUA87_RS03840 | 775657..779304 | 2697366 | [illegible] |
| WP_069309464.1 | Type6.BREX | [illegible] | 1208 | [illegible] | GCF_001720465.1 | Porphyrobacter_sp._LM_6 | BG023_RS04915 | 1019092..1022718 | 1896196 | [illegible] |
| MBB3862674.1 | Type6.BREX | [illegible] | 1208 | [illegible] | GCA_014196055.1 | Novosphingobium_hassiacum | GGQ88_003976 | 14858..18484 | 173676 | [illegible] |
| WP_054434404.1 | Type6.BREX | [illegible] | 1208 | [illegible] | GCF_001298105.1 | Novosphingobium_sp._ST904 | EDF59_RS28295 | 5043..8669 | 1684385 | [illegible] |
| WP_246386386.1 | Type6.BREX | [illegible] | 1213 | [illegible] | GCF_014196055.1 | Novosphingobium_hassiacum | GGQ88_RS19485 | 14843..18484 | 173676 | [illegible] |
| WP_230189894.1 | Type6.BREX | [illegible] | 1141 | [illegible] | GCF_918741285.1 | Sphingobium_sp._CECT_9361 | LQ220_RS19990 | 41177..44602 | 2845384 | [illegible] |
| NJC05753.1 | Type6.BREX | [illegible] | 1209 | [illegible] | GCA_011927725.1 | Sphingomonas_kaistensis | GGQ97_001546 | 1560422..1564051 | 298708 | [illegible] |
| RYG90169.1 | Type6.BREX | [illegible] | 1208 | [illegible] | GCA_004145145.1 | Alphaproteobacteria_bacterium | EON59_00500 | 24262..27888 | 1913988 | [illegible] |
| NCC22319.1 | Type6.BREX | [illegible] | 1212 | [illegible] | GCA_009929775.1 | Alphaproteobacteria_bacterium | EOM26_07635 | 14398..18036 | 1913988 | [illegible] |
| MBF0372584.1 | Type6.BREX | [illegible] | 1175 | [illegible] | GCA_015232435.1 | Alphaproteobacteria_bacterium | HQL39_04115 | 4738..8265 | 1913988 | [illegible] |
| MCB9697628.1 | Type6.BREX | [illegible] | 719 | [illegible] | GCA_020632835.1 | Alphaproteobacteria_bacterium | H6738_12675 | 1..2156 | 1913988 | [illegible] |
| QKS29579.1 | Type6.BREX | [illegible] | 1180 | [illegible] | GCA_013347225.1 | Candidatus_Accumulibacter_similis | HT579_12075 | 2666587..2670129 | 2954388 | [illegible] |
| MBN8451986.1 | Type6.BREX | [illegible] | 1213 | [illegible] | GCA_017302415.1 | Candidatus_Accumulibacter_sp. | J0M31_02680 | 99081..102722 | 2053492 | [illegible] |
| WP_042797405.1 | Type6.BREX | [illegible] | 1207 | [illegible] | GCF_000783435.2 | Achromobacter_insolitus | MC81_20460 | 4405602..4409225 | 217204 | [illegible] |
| WP_076471941.1 | Type6.BREX | [illegible] | 1207 | [illegible] | GCF_001957055.1 | Achromobacter_xylosoxidans | BIZ53_RS00410 | 81330..84953 | 85698 | [illegible] |
| WP_212312045.1 | Type6.BREX | [illegible] | 1217 | [illegible] | GCF_018195695.1 | Achromobacter_sp._Marseille-Q0513 | KDH83_RS00260 | 59261..62914 | 2829161 | [illegible] |
| WP_233436978.1 | Type6.BREX | [illegible] | 1234 | [illegible] | GCF_022976495.1 | Achromobacter_xylosoxidans | DBA06_RS11680 | 120350..124054 | 85698 | [illegible] |
| WP_259250627.1 | Type6.BREX | [illegible] | 1213 | [illegible] | GCF_024806975.1 | Achromobacter_sp._JUb104 | M2361_RS16380 | 989592..993233 | 2940590 | [illegible] |
| WP_180097564.1 | Type6.BREX | [illegible] | 1213 | [illegible] | GCF_902859995.1 | Achromobacter_mucicolens | H0520_RS02915 | 616832..620473 | 1389922 | [illegible] |
| AYN22647.1 | Type6.BREX | [illegible] | 1213 | [illegible] | GCA_003671915.1 | Alcaligenes_aquatilis | D3M96_03425 | 754727..758368 | 323284 | [illegible] |
| WP_199265315.1 | Type6.BREX | [illegible] | 1213 | [illegible] | GCF_016446305.1 | Alcaligenes_faecalis | GTU60_RS02720 | 606582..610223 | 511 | [illegible] |
| WP_032972587.1 | Type6.BREX | [illegible] | 1213 | [illegible] | GCF_006770345.1 | Bordetella_hinzii | CBR67_RS06420 | 1279803..1283444 | 103855 | [illegible] |
| MAO51251.1 | Type6.BREX | [illegible] | 1209 | [illegible] | GCA_002693665.1 | Pusillimonas_sp._(ex_Stolz_et_al._2005) | CML16_10275 | 63709..67338 | 1979962 | [illegible] |
| HBT31937.1 | Type6.BREX | [illegible] | 210 | [illegible] | GCA_003509985.1 | Pusillimonas_sp._(ex_Stolz_et_al._2005) | DEB15_03430 | 7486..8117 | 1979962 | [illegible] |
| HBT31938.1 | Type6.BREX | [illegible] | 153 | [illegible] | GCA_003509985.1 | Pusillimonas_sp._(ex_Stolz_et_al._2005) | DEB15_03435 | 8203..8663 | 1979962 | [illegible] |
| HCP76921.1 | Type6.BREX | [illegible] | 618 | [illegible] | GCA_003525685.1 | Pusillimonas_sp._(ex_Stolz_et_al._2005) | DIU11_04070 | 51387..53240 | 1979962 | [illegible] |
| NYT45622.1 | Type6.BREX | [illegible] | 1214 | [illegible] | GCA_013416345.1 | Alcaligenaceae_bacterium | H0A64_12395 | 218321..221965 | 2021368 | [illegible] |
| OQC82779.1 | Type6.BREX | [illegible] | 1181 | [illegible] | - | Burkholderia_cenocepacia | UE98_23645 | 24667..28212 | 95486 | [illegible] |
| OQD19343.1 | Type6.BREX | [illegible] | 1181 | [illegible] | GCA_002071935.1 | Burkholderia_cenocepacia | UE98_23645 | 24667..28212 | 95486 | [illegible] |
| MCA8264434.1 | Type6.BREX | [illegible] | 1210 | [illegible] | GCA_020420005.1 | Burkholderia_multivorans | LGM79_28445 | 10767..14399 | 87883 | [illegible] |
| WP_039318161.1 | Type6.BREX | [illegible] | 1181 | [illegible] | GCF_000801975.1 | Burkholderia_sp._A9 | PI86_RS01910 | 215778..219323 | 1365108 | [illegible] |
| WP_059944780.1 | Type6.BREX | [illegible] | 1213 | [illegible] | GCF_001527205.1 | Burkholderia_territorii | WT00_RS19595 | 122446..126087 | 1503055 | [illegible] |
| WP_059936887.1 | Type6.BREX | [illegible] | 1213 | [illegible] | GCF_001528605.1 | Burkholderia_multivorans | WK33_RS03230 | 12913..16554 | 87883 | [illegible] |
| WP_218673170.1 | Type6.BREX | [illegible] | 1182 | [illegible] | GCF_002071875.2 | Burkholderia_cenocepacia | UE97_RS31760 | 3013..6561 | 95486 | [illegible] |
| WP_124667309.1 | Type6.BREX | [illegible] | 1181 | [illegible] | GCF_003854225.1 | Burkholderia_sp._Bp8986 | DID99_RS19395 | 111864..115409 | 2184550 | [illegible] |
| WP_124661146.1 | Type6.BREX | [illegible] | 1210 | [illegible] | GCF_003854235.1 | Burkholderia_sp._Bp8998 | DIE06_RS09175 | 82371..86003 | 2184557 | [illegible] |
| WP_226255239.1 | Type6.BREX | [illegible] | 1208 | [illegible] | GCF_020419005.1 | Burkholderia_cepacia | LGM39_RS24930 | 366713..370339 | 292 | [illegible] |
| WP_175041065.1 | Type6.BREX | [illegible] | 1215 | [illegible] | GCF_902499225.1 | Burkholderia_contaminans | BLA3211_RS35460 | 10621..14268 | 488447 | [illegible] |
| WP_175704189.1 | Type6.BREX | [illegible] | 1215 | [illegible] | GCF_902829775.1 | Burkholderia_ambifaria | HWV86_RS26205 | 192524..196171 | 152480 | [illegible] |
| WP_175725598.1 | Type6.BREX | [illegible] | 1181 | [illegible] | GCF_902830165.1 | Burkholderia_ambifaria | HWV94_RS18190 | 262570..266115 | 152480 | [illegible] |
| WP_186043215.1 | Type6.BREX | [illegible] | 1213 | [illegible] | GCF_902830235.1 | Burkholderia_gladioli | H9C96_RS14825 | 15196..18837 | 28095 | [illegible] |
| WP_186236015.1 | Type6.BREX | [illegible] | 1215 | [illegible] | GCF_902832475.1 | Burkholderia_gladioli | H9G27_RS00755 | 169921..173568 | 28095 | [illegible] |
| WP_075360437.1 | Type6.BREX | [illegible] | 1210 | [illegible] | GCF_001935975.1 | Caballeronia_sordidicola | BSU03_RS34875 | 87497..91129 | 196367 | [illegible] |
| WP_187608173.1 | Type6.BREX | [illegible] | 1215 | [illegible] | GCF_014384845.1 | Caballeronia_sp._EK | IAG25_RS30620 | 154529..158176 | 2767469 | [illegible] |
| WP_250462416.1 | Type6.BREX | [illegible] | 1181 | [illegible] | GCF_023631485.1 | Caballeronia_sp._GAFFF2 | NCT04_RS00125 | 29887..33432 | 2921741 | [illegible] |
| WP_091908362.1 | Type6.BREX | [illegible] | 1213 | [illegible] | GCF_900095735.1 | Chitinasiproducens_palmae | BLU54_RS11860 | 213511..217152 | 1770053 | [illegible] |
| WP_013958181.1 | Type6.BREX | [illegible] | 1214 | [illegible] | GCF_000219215.1 | Cupriavidus_necator | CNE_1c33280 | 3480419..3484063 | 106590 | [illegible] |
| WP_174424431.1 | Type6.BREX | [illegible] | 1215 | [illegible] | GCF_013337645.1 | Cupriavidus_basilensis | FCJ59_RS11575 | 146108..149755 | 68895 | [illegible] |
| WP_059038859.1 | Type6.BREX | [illegible] | 1215 | [illegible] | GCF_001485395.1 | Ralstonia | SAMD00023378_RS10640 | 9198..12845 | 48736 | [illegible] |
| WP_220363411.1 | Type6.BREX | [illegible] | 1215 | [illegible] | GCF_019511605.1 | Paraburkholderia_phenoliruptrix | JNB89_RS20360 | 10910..14557 | 252970 | [illegible] |
| WP_230560751.1 | Type6.BREX | [illegible] | 1215 | [illegible] | GCF_020881035.1 | Paraburkholderia_sp._MMS20-SJTN17 | LJ655_RS08170 | 67515..71162 | 2886945 | [illegible] |
| WP_217272796.1 | Type6.BREX | [illegible] | 1182 | [illegible] | GCF_900141745.1 | Paraburkholderia_phenazinium | BUS06_RS04315 | 916329..919877 | 60549 | [illegible] |
| WP_183964658.1 | Type6.BREX | [illegible] | 1210 | [illegible] | GCF_014202215.1 | Quisquiliibacterium_transsilvanicum | HNQ70_RS04390 | 209103..212735 | 1549638 | [illegible] |
| WP_152905885.1 | Type6.BREX | [illegible] | 1102 | [illegible] | - | Ralstonia_solanacearum | ADE39_RS16210 | 148615..151923 | 305 | [illegible] |
| TXH34387.1 | Type6.BREX | [illegible] | 1189 | [illegible] | GCA_008012415.1 | Burkholderiaceae_bacterium | E6Q92_12965 | 10756..14325 | 2030806 | [illegible] |
| NUN59723.1 | Type6.BREX | [illegible] | 1210 | [illegible] | GCA_013360785.1 | Burkholderiaceae_bacterium | HUU13_01210 | 9533..13165 | 2030806 | [illegible] |
| MBP6902893.1 | Type6.BREX | [illegible] | 1219 | [illegible] | GCA_017992055.1 | Burkholderiaceae_bacterium | KBC73_22560 | 2680..6339 | 2030806 | [illegible] |
| MCP5271536.1 | Type6.BREX | [illegible] | 1219 | [illegible] | GCA_024235855.1 | Burkholderiaceae_bacterium | H6932_09900 | 152250..155909 | 2030806 | [illegible] |
| WP_143968626.1 | Type6.BREX | [illegible] | 1210 | [illegible] | GCF_007556755.1 | Tepidimonas_fonticaldi | Tfont_RS05070 | 95231..98863 | 1101373 | [illegible] |
| WP_054257718.1 | Type6.BREX | [illegible] | 1215 | [illegible] | GCF_001298675.1 | Acidovorax_caeni | BN2503_RS16450 | 13891..17538 | 343013 | [illegible] |
| WP_057202647.1 | Type6.BREX | [illegible] | 1207 | [illegible] | GCF_001428645.1 | Acidovorax_sp._Root217 | ASE31_RS15170 | 91296..94919 | 1736492 | [illegible] |
| GAO21740.1 | Type6.BREX | [illegible] | 1213 | [illegible] | GCA_000974645.1 | Alicycliphilus_sp._B1 | ALISP_1560 | 91569..95210 | 1603291 | [illegible] |
| WP_034379112.1 | Type6.BREX | [illegible] | 1210 | [illegible] | GCF_000761335.1 | Comamonas_thiooxydans | P245_RS10570 | 85855..89487 | 363952 | [illegible] |
| WP_096915618.1 | Type6.BREX | [illegible] | 1224 | [illegible] | GCF_002443055.1 | Delftia_acidovorans | CHL79_RS18790 | 4226898..4230572 | 80866 | [illegible] |
| WP_234638190.1 | Type6.BREX | [illegible] | 1219 | [illegible] | GCF_903684635.1 | Delftia_tsuruhatensis | MJ845_RS20380 | 81444..85103 | 180282 | [illegible] |
| WP_047349524.1 | Type6.BREX | [illegible] | 1213 | [illegible] | GCF_001023515.1 | Diaphorobacter_sp._J5-51 | OX89_RS03130 | 14085..17726 | 680496 | [illegible] |
| WP_106685100.1 | Type6.BREX | [illegible] | 1213 | [illegible] | GCF_003008575.1 | Melaminivora_suipulveris | C6568_RS16475 | 3505730..3509371 | 2109913 | [illegible] |
| CDS49862.1 | Type6.BREX | [illegible] | 1210 | [illegible] | - | Polaromonas_sp._CG9_12 | . | 485322..488954 | 1504672 | [illegible] |
| MCB1996567.1 | Type6.BREX | [illegible] | 373 | [illegible] | GCA_020446165.1 | Rhodoferax_sp. | KDG57_11955 | 8309..9428 | 50421 | [illegible] |
| MBW7834374.1 | Type6.BREX | [illegible] | 1210 | [illegible] | GCA_019455265.1 | Simplicispira_suum | H3C29_14305 | 45952..49584 | 2109915 | [illegible] |
| WP_273364231.1 | Type6.BREX | [illegible] | 1210 | [illegible] | GCF_019455265.1 | Simplicispira_suum | H3C29_RS14325 | 45952..49584 | 2109915 | [illegible] |
| WP_068673039.1 | Type6.BREX | [illegible] | 1212 | [illegible] | GCF_001541225.1 | unclassified_Variovorax | APY03_RS01000 | 202384..206022 | 663243 | [illegible] |
| WP_218509735.1 | Type6.BREX | [illegible] | 1213 | [illegible] | GCF_019207085.1 | Variovorax_sp._dw_308 | HCU76_RS14255 | 36638..40279 | 2721546 | [illegible] |
| WP_093162784.1 | Type6.BREX | [illegible] | 1212 | [illegible] | GCF_900107745.1 | Variovorax_sp._YR216 | BLW05_RS12080 | 465668..469306 | 1882828 | [illegible] |
| WP_141835430.1 | Type6.BREX | [illegible] | 1209 | [illegible] | - | Herbaspirillum_sp._SJZ107 | FBX97_RS19505 | 315565..319194 | 2572881 | [illegible] |
| WP_152879050.1 | Type6.BREX | [illegible] | 1215 | [illegible] | GCF_009295655.1 | Duganella_sp._FT27W | F8197_RS00385 | 74619..78266 | 2654636 | [illegible] |
| OGB35307.1 | Type6.BREX | [illegible] | 1215 | [illegible] | GCA_001770935.1 | Burkholderiales_bacterium_RIFCSPLOWO2_12_FULL_61_40 | A3F78_11695 | 46897..50544 | 1797566 | [illegible] |
| NMA29242.1 | Type6.BREX | [illegible] | 1208 | [illegible] | GCA_012799975.1 | Burkholderiales_bacterium | GX934_15915 | 6559..10185 | 1891238 | [illegible] |
| MBL8528754.1 | Type6.BREX | [illegible] | 1180 | [illegible] | GCA_016791285.1 | Burkholderiales_bacterium | JNL68_13790 | 29505..33047 | 1891238 | [illegible] |
| WP_194117489.1 | Type6.BREX | [illegible] | 1213 | [illegible] | GCF_015223195.1 | Chitinilyticum_piscinae | INR99_RS16780 | 39673..43314 | 2866724 | [illegible] |
| WP_088965422.1 | Type6.BREX | [illegible] | 1215 | [illegible] | GCF_900090205.1 | Vogesella_sp._LIG4 | PSELUDRAFT_RS02905 | 629858..633505 | 1192162 | [illegible] |
| WP_153715435.1 | Type6.BREX | [illegible] | 1213 | [illegible] | GCF_009650905.1 | Eikenella_corrodens | GJ661_RS08680 | 18367..22008 | 539 | [illegible] |
| MBA0014163.1 | Type6.BREX | [illegible] | 1210 | [illegible] | GCA_013450215.1 | Nitrosospira_sp. | H0Z53_00860 | 54117..57749 | 38080 | [illegible] |
| WP_090458040.1 | Type6.BREX | [illegible] | 1213 | [illegible] | GCF_900100985.1 | Nitrosospira_sp._Nsp1 | BLP78_RS07395 | 18423..22064 | 136547 | [illegible] |
| SFW23276.1 | Type6.BREX | [illegible] | 1213 | [illegible] | GCA_900119085.1 | Nitrosovibrio_sp._Nv17 | SAMN05216414_10762 | 70156..73797 | 1855339 | [illegible] |
| WP_218184829.1 | Type6.BREX | [illegible] | 1234 | [illegible] | GCF_900119085.1 | Nitrosovibrio_sp._Nv17 | BT108_RS05915 | 70156..73860 | 1855339 | [illegible] |
| WP_121241046.1 | Type6.BREX | [illegible] | 1202 | [illegible] | GCF_003865015.1 | Sulfurisoma_sediminicola | EJE94_RS09880 | 239721..243329 | 1381557 | [illegible] |
| MBK7235806.1 | Type6.BREX | [illegible] | 1180 | [illegible] | GCA_016707455.1 | Sterolibacteriaceae_bacterium | IPI02_09355 | 160184..163726 | 2682146 | [illegible] |
| MBK6972675.1 | Type6.BREX | [illegible] | 1180 | [illegible] | GCA_016708905.1 | Sterolibacteriaceae_bacterium | IPH26_06870 | 229623..233165 | 2682146 | [illegible] |
| ODU51178.1 | Type6.BREX | [illegible] | 1213 | [illegible] | GCA_001725135.1 | Thiobacillus_sp._SCN_63-374 | ABS92_00645 | 9244..12885 | 1660149 | [illegible] |
| MBW8470844.1 | Type6.BREX | [illegible] | 1215 | [illegible] | GCA_019459125.1 | Thiobacillus_sp. | K0M67_21445 | 16263..19910 | 924 | [illegible] |
| WP_246167890.1 | Type6.BREX | [illegible] | 243 | [illegible] | GCF_009469695.1 | Propionivibrio_limicola | GH659_RS14290 | 2233..2964 | 167645 | [illegible] |
| TRZ91915.1 | Type6.BREX | [illegible] | 1212 | [illegible] | GCA_007280105.1 | Rhodocyclaceae_bacterium | D4R84_13335 | 10594..14232 | 1898103 | [illegible] |
| MBK6907454.1 | Type6.BREX | [illegible] | 1208 | [illegible] | GCA_016702335.1 | Rhodocyclaceae_bacterium | IPH08_10320 | 1000303..1003929 | 1898103 | [illegible] |
| MBK6907823.1 | Type6.BREX | [illegible] | 1202 | [illegible] | GCA_016702335.1 | Rhodocyclaceae_bacterium | IPH08_12425 | 301349..304957 | 1898103 | [illegible] |
| MCA3121411.1 | Type6.BREX | [illegible] | 1210 | [illegible] | GCA_020248305.1 | Rhodocyclaceae_bacterium | ING90_02100 | 367371..371003 | 1898103 | [illegible] |
| TVT60730.1 | Type6.BREX | [illegible] | 1213 | [illegible] | GCA_007713545.1 | Azoarcus_sp._PHD | FHK80_01725 | 174831..178472 | 1628740 | [illegible] |
| WP_141017182.1 | Type6.BREX | [illegible] | 1207 | [illegible] | GCF_006496635.1 | Azoarcus_sp._DD4 | CJ010_RS05935 | 1245819..1249442 | 2027405 | [illegible] |
| MBL8465274.1 | Type6.BREX | [illegible] | 1209 | [illegible] | GCA_016790365.1 | Thauera_sp. | JNL37_10520 | 3981..7610 | 1905334 | [illegible] |
| MCK6408296.1 | Type6.BREX | [illegible] | 1034 | [illegible] | GCA_023150215.1 | Thauera_sp. | L6Q70_03770 | 1..3103 | 1905334 | [illegible] |
| CAH1745218.1 | Type6.BREX | [illegible] | 1209 | [illegible] | GCA_932126575.1 | Thauera_sp._PIV-1 | C4PIVTH_0053 | 44546..48175 | 204461 | [illegible] |
| WP_096450985.1 | Type6.BREX | [illegible] | 1180 | [illegible] | GCF_002354895.1 | Thauera_sp._K11 | CCZ27_RS19540 | 4450815..4454357 | 2005884 | [illegible] |
| WP_247725420.1 | Type6.BREX | [illegible] | 1209 | [illegible] | GCF_023093595.1 | Thauera_aromatica | MXC99_RS10610 | 105350..108979 | 59405 | [illegible] |
| GHT91750.1 | Type6.BREX | [illegible] | 1209 | [illegible] | GCA_026000515.1 | Betaproteobacteria_bacterium | FACS1894116_00820 | 42503..46132 | 1891241 | [illegible] |
| WP_100653505.1 | Type6.BREX | [illegible] | 1216 | [illegible] | GCF_002803925.1 | Aeromonas_veronii | CVS41_08490 | 1813618..1817268 | 654 | [illegible] |
| WP_103244178.1 | Type6.BREX | [illegible] | 1210 | [illegible] | GCF_002902865.1 | Aeromonas_sp._ASNIH5 | C2U30_RS17315 | 3671715..3675347 | 1758179 | [illegible] |
| WP_216948840.1 | Type6.BREX | [illegible] | 1216 | [illegible] | GCF_019046985.1 | Aeromonas_sp._FDAARGOS_1417 | I6L47_RS05350 | 1228193..1231843 | 2778066 | [illegible] |
| WP_241417643.1 | Type6.BREX | [illegible] | 1216 | [illegible] | GCF_022557195.1 | Aeromonas_hydrophila | MKW86_RS08070 | 1734303..1737953 | 644 | [illegible] |
| WP_217358045.1 | Type6.BREX | [illegible] | 951 | [illegible] | GCF_002499975.2 | Alteromonas_pelagimontana | CA267_RS01545 | 356718..359573 | 1858656 | [illegible] |
| WP_220104111.1 | Type6.BREX | [illegible] | 1208 | [illegible] | GCF_019457915.1 | Neiella_holothuriorum | K0504_RS10305 | 24423..28049 | 2870530 | [illegible] |
| MCD8559955.1 | Type6.BREX | [illegible] | 181 | [illegible] | GCA_021209305.1 | Shewanella_xiamenensis | LRY75_14310 | 8249..8794 | 332186 | [illegible] |
| MCD8559956.1 | Type6.BREX | [illegible] | 1026 | [illegible] | GCA_021209305.1 | Shewanella_xiamenensis | LRY75_14315 | 8805..11885 | 332186 | [illegible] |
| WP_173496635.1 | Type6.BREX | [illegible] | 1211 | [illegible] | GCF_013283205.1 | Shewanella_sp._ISTPL2 | HUB64_RS06505 | 1443171..1446806 | 2699425 | [illegible] |
| WP_188922770.1 | Type6.BREX | [illegible] | 1210 | [illegible] | GCF_014648295.1 | Shewanella_saliphila | IEY48_RS17840 | 13004..16636 | 2282698 | [illegible] |
| WP_219012197.1 | Type6.BREX | [illegible] | 1211 | [illegible] | GCF_019303235.1 | Shewanella_algae | K0E59_RS06910 | 134402..138037 | 38313 | [illegible] |
| MAT50029.1 | Type6.BREX | [illegible] | 1215 | [illegible] | GCA_002702725.1 | Porticoccaceae_bacterium | CMK32_02455 | 2974..6621 | 2026782 | [illegible] |
| WP_020507127.1 | Type6.BREX | [illegible] | 1179 | [illegible] | GCF_000379525.1 | Lamprocystis_purpurea | A390_RS0123535 | 17517..21056 | 61598 | [illegible] |
| WP_132582321.1 | Type6.BREX | [illegible] | 1208 | [illegible] | GCF_004345025.1 | Rheinheimera_sp._D18 | E0Z06_RS04605 | 954479..958105 | 2545632 | [illegible] |
| CRI67889.1 | Type6.BREX | [illegible] | 1209 | [illegible] | GCA_001499735.1 | Thiocapsa_sp._KS1 | THIOKS1890022 | 27242..30871 | 610332 | [illegible] |
| QVL47244.1 | Type6.BREX | [illegible] | 1179 | [illegible] | GCA_018399035.1 | Thiocapsa_sp. | KFB96_16145 | 3553103..3556642 | 2024551 | [illegible] |
| WP_093191699.1 | Type6.BREX | [illegible] | 1203 | [illegible] | GCF_001499735.1 | Thiocapsa_sp._KS1 | THIOKS1_RS05340 | 1192857..1196468 | 610332 | [illegible] |
| WP_014777950.1 | Type6.BREX | [illegible] | 1206 | [illegible] | GCF_000227745.2 | Thiocystis_violascens | THIVI_RS07160 | 1619300..1622920 | 73141 | [illegible] |
| WP_200373981.1 | Type6.BREX | [illegible] | 1206 | [illegible] | GCF_016583575.1 | Thiocystis_violacea | CKO27_RS06060 | 39556..43176 | 13725 | [illegible] |
| WP_100920354.1 | Type6.BREX | [illegible] | 1178 | [illegible] | GCF_002813775.1 | Candidatus_Thiodictyon_syntrophicum | THSYN_17920 | 4302311..4305847 | 1166950 | [illegible] |
| WP_015282553.1 | Type6.BREX | [illegible] | 1209 | [illegible] | GCF_000327045.1 | Thioflavicoccus_mobilis | THIMO_RS17980 | 83065..86694 | 80679 | [illegible] |
| WP_058554449.1 | Type6.BREX | [illegible] | 1214 | [illegible] | GCF_001469165.1 | Thiohalocapsa_sp._ML1 | ML01_RS05745 | 155025..158669 | 1431688 | [illegible] |
| WP_164452822.1 | Type6.BREX | [illegible] | 1214 | [illegible] | GCF_010820565.1 | Thiorhodococcus_minor | G3446_RS10685 | 55754..59398 | 57489 | [illegible] |
| WP_009149681.1 | Type6.BREX | [illegible] | 1208 | [illegible] | GCF_000228725.2 | Thiorhodovibrio_sp._970 | THI970DRAFT_RS14125 | 1743457..1747083 | 631362 | [illegible] |
| WP_201065314.1 | Type6.BREX | [illegible] | 1212 | [illegible] | GCF_016653415.1 | Thiorhodovibrio_winogradskyi | CCR91_RS06145 | 56528..60166 | 77007 | [illegible] |
| TVQ85671.1 | Type6.BREX | [illegible] | 1212 | [illegible] | GCA_007693535.1 | Chromatiaceae_bacterium | EA400_15015 | 6841..10479 | 2049432 | [illegible] |
| MBK8639641.1 | Type6.BREX | [illegible] | 1178 | [illegible] | GCA_016714645.1 | Chromatiaceae_bacterium | IPN92_15725 | 846355..849891 | 2049432 | [illegible] |
| MBP6733683.1 | Type6.BREX | [illegible] | 1181 | [illegible] | GCA_017990755.1 | Chromatiaceae_bacterium | KA142_03150 | 2562..6107 | 2049432 | [illegible] |
| MCC7280699.1 | Type6.BREX | [illegible] | 1181 | [illegible] | GCA_020851955.1 | Chromatiaceae_bacterium | IT487_20620 | 13992..17537 | 2049432 | [illegible] |
| MCF7977306.1 | Type6.BREX | [illegible] | 1182 | [illegible] | GCA_021736685.1 | Chromatiaceae_bacterium | K9L82_04715 | 1611..5159 | 2049432 | [illegible] |
| WP_239679777.1 | Type6.BREX | [illegible] | 1223 | [illegible] | GCF_022227625.1 | Ectothiorhodospira_variabilis | LPW30_RS00090 | 25011..28682 | 505694 | [illegible] |
| WP_109962896.1 | Type6.BREX | [illegible] | 985 | [illegible] | GCF_002356555.2 | Halorhodospira_halochloris | HH1059_RS10060 | 2230108..2233065 | 1052 | [illegible] |
| WP_238639758.1 | Type6.BREX | [illegible] | 1207 | [illegible] | GCF_022229325.1 | Halorhodospira_halochloris | LRD17_RS10705 | 4342..7965 | 1052 | [illegible] |
| WP_253479345.1 | Type6.BREX | [illegible] | 1216 | [illegible] | GCF_024170285.1 | Natronocella_acetinitrilica | J2T57_RS14030 | 326743..330393 | 414046 | [illegible] |
| WP_019611713.1 | Type6.BREX | [illegible] | 1216 | [illegible] | GCF_000381705.1 | Thioalkalivibrio_sp._AKL7 | G315_RS0100950 | 222198..225848 | 1158155 | [illegible] |
| WP_132538615.1 | Type6.BREX | [illegible] | 1179 | [illegible] | GCF_004341245.1 | Plasticicumulans_lactativorans | EV699_RS02475 | 34554..38093 | 1133106 | [illegible] |
| MBK8754669.1 | Type6.BREX | [illegible] | 1209 | [illegible] | GCA_016710535.1 | Candidatus_Competibacteraceae_bacterium | IPL99_24865 | 32033..35662 | 2053538 | [illegible] |
| MCB1919725.1 | Type6.BREX | [illegible] | 1177 | [illegible] | GCA_020444905.1 | Candidatus_Competibacteraceae_bacterium | KDJ28_07085 | 14941..18474 | 2053538 | [illegible] |

| Protein-ID | System | Gene-Neighbourhoods | Length | Taxonomy | Assembly-ID | Strain | Locus-Tag | Genomic-coordinates | TaxID | Fasta-define |
|---|---|---|---|---|---|---|---|---|---|---|
| WP_068809899.1 | Type6.BREX | [illegible] | 1214 | Bacteria>Pseudomonadota>Gammaproteobacteria>Gammaproteobacteria_incertae_sedis>Pseudohongiella | GCF_014642795.1 | Pseudohongiella_nitratireducens | IEW51_RS03970 | 206813..210457 | 1768907 | BREX-system-P-loop-protein-BrxC-[Pseudohongiella-nitratireducens] |
| MCG8006852.1 | Type6.BREX | [illegible] | 1182 | Bacteria>Pseudomonadota>Gammaproteobacteria>Gammaproteobacteria_incertae_sedis>sulfur-oxidizing_symbionts>Candidatus_Thiodiazotropha | GCA_016842225.1 | Candidatus_Thiodiazotropha_lotti | JAY76_04265 | 12912..16460 | 2792787 | BREX-system-P-loop-protein-BrxC-[Candidatus-Thiodiazotropha-lotti] |
| MCG7876672.1 | Type6.BREX | [illegible] | 1215 | Bacteria>Pseudomonadota>Gammaproteobacteria>Gammaproteobacteria_incertae_sedis>sulfur-oxidizing_symbionts>Candidatus_Thiodiazotropha | GCA_016842655.1 | Candidatus_Thiodiazotropha_taylori | JAY78_10080 | 4978..8625 | 2792791 | BREX-system-P-loop-protein-BrxC-[Candidatus-Thiodiazotropha-taylori] |
| MCG7891938.1 | Type6.BREX | TraG_N-><-?<-?\|\|wHTH+WYL+WCX->BrxE->BrxA-DNA-binding->BrxB-BrxF-iSTAND->DUF6079*-> | 1184 | Bacteria>Pseudomonadota>Gammaproteobacteria>Gammaproteobacteria_incertae_sedis>sulfur-oxidizing_symbionts>Candidatus_Thiodiazotropha | GCA_016842745.1 | Candidatus_Thiodiazotropha_taylori | JAY79_16965 | 5079..8630 | 2792791 | BREX-system-P-loop-protein-BrxC-[Candidatus-Thiodiazotropha-taylori] |
| MCG7951159.1 | Type6.BREX | [illegible] | 1215 | Bacteria>Pseudomonadota>Gammaproteobacteria>Gammaproteobacteria_incertae_sedis>sulfur-oxidizing_symbionts>Candidatus_Thiodiazotropha | GCA_016842765.1 | Candidatus_Thiodiazotropha_taylori | JAZ09_08050 | 22496..26143 | 2792791 | BREX-system-P-loop-protein-BrxC-[Candidatus-Thiodiazotropha-taylori] |
| MCG8102611.1 | Type6.BREX | [illegible] | 1215 | Bacteria>Pseudomonadota>Gammaproteobacteria>Gammaproteobacteria_incertae_sedis>sulfur-oxidizing_symbionts>Candidatus_Thiodiazotropha | GCA_016842845.1 | Candidatus_Thiodiazotropha_taylori | JAY70_09490 | 22526..26173 | 2792791 | BREX-system-P-loop-protein-BrxC-[Candidatus-Thiodiazotropha-taylori] |
| MCG8058214.1 | Type6.BREX | [illegible] | 1215 | Bacteria>Pseudomonadota>Gammaproteobacteria>Gammaproteobacteria_incertae_sedis>sulfur-oxidizing_symbionts>Candidatus_Thiodiazotropha | GCA_016843165.1 | Candidatus_Thiodiazotropha_taylori | JAZ21_00070 | 27190..30837 | 2792791 | BREX-system-P-loop-protein-BrxC-[Candidatus-Thiodiazotropha-taylori] |
| WP_077729128.1 | Type6.BREX | <-N6-METHYLASE<-GreA_GreB+SF1-SF2-Helicase<-DUF6079*<-BrxB-BrxF-iSTAND<-BrxA-DNA-binding<-BrxE<-wHTH+WYL+WCX\|\|JetA->JetB->SbcC-> | 1213 | Bacteria>Pseudomonadota>Gammaproteobacteria>Methylococcales>Methylococcaceae>Methylocaldum | GCF_002005105.1 | Methylocaldum_sp._14B | B2A78_RS06605 | 5139..8780 | 1912213 | BREX-system-P-loop-protein-BrxC-[Methylocaldum-sp.-14B] |
| UOH75125.1 | Type6.BREX | [illegible] | 1219 | Bacteria>Pseudomonadota>Gammaproteobacteria>Moraxellales>Moraxellaceae>Acinetobacter | GCA_022832955.1 | Acinetobacter_schindleri | MOW08_00685 | 158728..162387 | 108981 | BREX-system-P-loop-protein-BrxC-[Acinetobacter-schindleri] |
| WP_005162499.1 | Type6.BREX | [illegible] | 1063 | Bacteria>Pseudomonadota>Gammaproteobacteria>Moraxellales>Moraxellaceae>Acinetobacter | GCF_000369425.1 | Acinetobacter_sp._NIPH_284 | F908_RS12580 | 1400419..1403610 | 1217704 | BREX-system-P-loop-protein-BrxC-[Acinetobacter-sp.-NIPH-284] |
| WP_140963539.1 | Type6.BREX | [illegible] | 1214 | Bacteria>Pseudomonadota>Gammaproteobacteria>Moraxellales>Moraxellaceae>Acinetobacter | GCF_006494575.1 | Acinetobacter_baumannii | FJU84_RS09090 | 25284..28928 | 470 | BREX-system-P-loop-protein-BrxC-[Acinetobacter-baumannii] |
| WP_032068282.1 | Type6.BREX | [illegible] | 1214 | Bacteria>Pseudomonadota>Gammaproteobacteria>Moraxellales>Moraxellaceae>Acinetobacter | GCF_014559915.1 | Acinetobacter_baumannii | HPJ29_RS06280 | 177170..180814 | 470 | BREX-system-P-loop-protein-BrxC-[Acinetobacter-baumannii] |
| WP_264453044.1 | Type6.BREX | DUF932->?->?->XIS-HTH->SAM-METHYLASE->wHTH(SBD)+HNH->BrxE->BrxA-DNA-binding->BrxB-BrxF-iSTAND->DUF6079*->N6-METHYLASE->ParB+HNH->PglZ->SIGMA+OB-Fold+AAA-ATPase+HKD-DNase-BrxL-> | 1214 | Bacteria>Pseudomonadota>Gammaproteobacteria>Moraxellales>Moraxellaceae>Acinetobacter | GCF_025938795.1 | Acinetobacter_baumannii | OLN40_RS03910 | 58425..62069 | 470 | BREX-system-P-loop-protein-BrxC-[Acinetobacter-baumannii] |
| WP_134244656.1 | Type6.BREX | <-SIGMA+OB-Fold+AAA-ATPase+HKD-DNase-BrxL<-PglZ<-?<-N6-METHYLASE<-N6-METHYLASE<-DUF6079*<-BrxB-BrxF-iSTAND<-BrxA-DNA-binding<-BrxE<-DDE_Tnp_4 | 1217 | Bacteria>Pseudomonadota>Gammaproteobacteria>Moraxellales>Moraxellaceae>Alkanindiges | GCF_004377485.1 | Alkanindiges_illinoisensis | E2B99_RS09020 | 10631..14284 | 197183 | BREX-system-P-loop-protein-BrxC-[Alkanindiges-illinoisensis] |
| WP_151794804.1 | Type6.BREX | [illegible] | 1214 | Bacteria>Pseudomonadota>Gammaproteobacteria>Moraxellales>Moraxellaceae | GCF_008997035.1 | Acinetobacter | GBP95_RS04055 | 337470..341114 | 469 | MULTISPECIES:-BREX-system-P-loop-protein-BrxC-[Acinetobacter] |
| WP_102077888.1 | Type6.BREX | [illegible] | 1219 | Bacteria>Pseudomonadota>Gammaproteobacteria>Moraxellales>Moraxellaceae>Psychrobacter | GCF_001742065.1 | unclassified_Psychrobacter | CXF69_RS01065 | 108247..111906 | 196806 | MULTISPECIES:-BREX-system-P-loop-protein-BrxC-[unclassified-Psychrobacter] |
| WP_218690652.1 | Type6.BREX | [illegible] | 1213 | Bacteria>Pseudomonadota>Gammaproteobacteria>Moraxellales>Moraxellaceae>Psychrobacter | GCF_019279065.1 | Psychrobacter_sp._BF1 | KZA12_RS14005 | 8359..12000 | 2821147 | BREX-system-P-loop-protein-BrxC-[Psychrobacter-sp.-BF1] |
| WP_238051284.1 | Type6.BREX | [illegible] | 1213 | Bacteria>Pseudomonadota>Gammaproteobacteria>Moraxellales>Moraxellaceae>Psychrobacter | GCF_022129405.1 | Psychrobacter_sp._Ps2 | I3254_RS03805 | 13291..16932 | 2790956 | BREX-system-P-loop-protein-BrxC-[Psychrobacter-sp.-Ps2] |
| WP_201561160.1 | Type6.BREX | [illegible] | 1219 | Bacteria>Pseudomonadota>Gammaproteobacteria>Moraxellales>Moraxellaceae>Psychrobacter | GCF_904846215.1 | Psychrobacter_glacincola | JMW90_RS00425 | 116435..120094 | 56810 | BREX-system-P-loop-protein-BrxC-[Psychrobacter-glacincola] |
| WP_206606031.1 | Type6.BREX | [illegible] | 1016 | Bacteria>Pseudomonadota>Gammaproteobacteria>Nevskiales>Steroidobacteraceae>Steroidobacter | GCF_003104955.1 | Steroidobacter_cummioxidans | D0550_RS01455 | 17780..20830 | 1803913 | BREX-system-P-loop-protein-BrxC-[Steroidobacter-cummioxidans] |
| WP_153499070.1 | Type6.BREX | [illegible] | 1214 | Bacteria>Pseudomonadota>Gammaproteobacteria>Oceanospirillales>Alcanivoracaceae>Alcanivorax | GCF_009601165.1 | Alcanivorax_sediminis | GFN93_RS03795 | 861896..865540 | 2663008 | BREX-system-P-loop-protein-BrxC-[Alcanivorax-sediminis] |
| WP_159558342.1 | Type6.BREX | [illegible] | 1213 | Bacteria>Pseudomonadota>Gammaproteobacteria>Oceanospirillales>Alcanivoracaceae>Alcanivorax | GCF_009846755.1 | Alcanivorax_sp._S71-1-4 | A167_RS05685 | 90732..94373 | 1177159 | BREX-system-P-loop-protein-BrxC-[Alcanivorax-sp.-S71-1-4] |
| MBR2514422.1 | Type6.BREX | [illegible] | 1224 | Bacteria>Pseudomonadota>Gammaproteobacteria>Oceanospirillales>Halomonadaceae>Halomonas | GCA_017515565.1 | Halomonas_sp. | IKE45_10465 | 23089..26763 | 1486246 | BREX-system-P-loop-protein-BrxC-[Halomonas-sp.] |
| WP_161432258.1 | Type6.BREX | [illegible] | 1223 | Bacteria>Pseudomonadota>Gammaproteobacteria>Oceanospirillales>Halomonadaceae>Halomonas | GCF_009902005.1 | Halomonas_alimentaria | GRB96_RS11410 | 2397586..2401257 | 147248 | BREX-system-P-loop-protein-BrxC-[Halomonas-alimentaria] |
| WP_110669866.1 | Type6.BREX | [illegible] | 1225 | Bacteria>Pseudomonadota>Gammaproteobacteria>Oceanospirillales>Halomonadaceae>Salinicola | GCF_003206155.1 | Salinicola_halophilus | DOM33_RS13680 | 401271..404948 | 184065 | BREX-system-P-loop-protein-BrxC-[Salinicola-halophilus] |
| WP_254579406.1 | Type6.BREX | [illegible] | 1212 | Bacteria>Pseudomonadota>Gammaproteobacteria>Oceanospirillales>Oceanospirillaceae>Marinobacterium | GCF_024267675.1 | Marinobacterium_sedimentorum | MRS74_RS06180 | 150386..154024 | 2927804 | BREX-system-P-loop-protein-BrxC-[Marinobacterium-sedimentorum] |
| QEW05875.1 | Type6.BREX | [illegible] | 1212 | Bacteria>Pseudomonadota>Gammaproteobacteria>Oceanospirillales>Oceanospirillaceae>Nitrincola | GCA_008727795.1 | Nitrincola_iocasae | F5I99_04885 | 1032629..1036267 | 2614693 | BREX-system-P-loop-protein-BrxC-[Nitrincola-iocasae] |
| WP_225307559.1 | Type6.BREX | [illegible] | 1214 | Bacteria>Pseudomonadota>Gammaproteobacteria>Oceanospirillales>Oceanospirillaceae>Nitrincola | GCF_008727795.1 | Nitrincola_iocasae | F5I99_RS04885 | 1032623..1036267 | 2614693 | BREX-system-P-loop-protein-BrxC-[Nitrincola-iocasae] |
| WP_058011517.1 | Type6.BREX | [illegible] | 1213 | Bacteria>Pseudomonadota>Gammaproteobacteria>Pseudomonadales | GCF_901482535.1 | Pseudomonadaceae | FQU16_RS01885 | 279271..282912 | 135621 | MULTISPECIES:-BREX-system-P-loop-protein-BrxC-[Pseudomonadaceae] |
| WP_269819144.1 | Type6.BREX | [illegible] | 1219 | Bacteria>Pseudomonadota>Gammaproteobacteria>Pseudomonadales>Pseudomonadaceae>Denitrificimonas | GCF_027498055.1 | Denitrificimonas_caeni | O6P33_RS05150 | 1067699..1071358 | 521720 | BREX-system-P-loop-protein-BrxC-[Denitrificimonas-caeni] |
| WP_019470415.1 | Type6.BREX | [illegible] | 1212 | Bacteria>Pseudomonadota>Gammaproteobacteria>Pseudomonadales>Pseudomonadaceae | GCF_000292285.1 | Pseudomonas | N805_RS25690 | 5809538..5813176 | 286 | MULTISPECIES:-BREX-system-P-loop-protein-BrxC-[Pseudomonas] |
| WP_033986129.1 | Type6.BREX | [illegible] | 1212 | Bacteria>Pseudomonadota>Gammaproteobacteria>Pseudomonadales>Pseudomonadaceae | GCF_020180035.1 | Pseudomonas | A0894_RS16050 | 13512..17150 | 286 | MULTISPECIES:-BREX-system-P-loop-protein-BrxC-[Pseudomonas] |
| EQL43544.1 | Type6.BREX | [illegible] | 1213 | Bacteria>Pseudomonadota>Gammaproteobacteria>Pseudomonadales>Pseudomonadaceae>Pseudomonas | GCA_000467675.1 | Pseudomonas_aeruginosa_VRFPA03 | M770_31820 | 18788..22429 | 1350465 | ATPase-[Pseudomonas-aeruginosa-VRFPA03] |
| MBG3983980.1 | Type6.BREX | [illegible] | 1212 | Bacteria>Pseudomonadota>Gammaproteobacteria>Pseudomonadales>Pseudomonadaceae>Pseudomonas | GCA_015733145.1 | Pseudomonas_aeruginosa | I4919_20885 | 86674..90312 | 287 | BREX-system-P-loop-protein-BrxC-[Pseudomonas-aeruginosa] |
| MBG4647655.1 | Type6.BREX | [illegible] | 1213 | Bacteria>Pseudomonadota>Gammaproteobacteria>Pseudomonadales>Pseudomonadaceae>Pseudomonas | GCA_015735165.1 | Pseudomonas_aeruginosa | I5F68_27785 | 4447..8088 | 287 | BREX-system-P-loop-protein-BrxC-[Pseudomonas-aeruginosa] |
| MBG6382130.1 | Type6.BREX | [illegible] | 1210 | Bacteria>Pseudomonadota>Gammaproteobacteria>Pseudomonadales>Pseudomonadaceae>Pseudomonas | GCA_015762815.1 | Pseudomonas_aeruginosa | I5I12_28760 | 1691..5323 | 287 | BREX-system-P-loop-protein-BrxC-[Pseudomonas-aeruginosa] |
| MBH3980671.1 | Type6.BREX | [illegible] | 1213 | Bacteria>Pseudomonadota>Gammaproteobacteria>Pseudomonadales>Pseudomonadaceae>Pseudomonas | GCA_016011075.1 | Pseudomonas_aeruginosa | I5Q70_02570 | 503331..506972 | 287 | BREX-system-P-loop-protein-BrxC-[Pseudomonas-aeruginosa] |
| MBL7230767.1 | Type6.BREX | [illegible] | 1214 | Bacteria>Pseudomonadota>Gammaproteobacteria>Pseudomonadales>Pseudomonadaceae>Pseudomonas | GCA_016785205.1 | Pseudomonas_sp. | JJK56_22660 | 12252..15896 | 306 | BREX-system-P-loop-protein-BrxC-[Pseudomonas-sp.] |
| MBN0299744.1 | Type6.BREX | [illegible] | 1213 | Bacteria>Pseudomonadota>Gammaproteobacteria>Pseudomonadales>Pseudomonadaceae>Pseudomonas | GCA_016922145.1 | Pseudomonas_aeruginosa | JTM78_25120 | 125675..129316 | 287 | BREX-system-P-loop-protein-BrxC-[Pseudomonas-aeruginosa] |
| MBN0536373.1 | Type6.BREX | [illegible] | 1206 | Bacteria>Pseudomonadota>Gammaproteobacteria>Pseudomonadales>Pseudomonadaceae>Pseudomonas | GCA_016922535.1 | Pseudomonas_aeruginosa | JTM59_25840 | 125679..129299 | 287 | BREX-system-P-loop-protein-BrxC-[Pseudomonas-aeruginosa] |
| HBN8119635.1 | Type6.BREX | [illegible] | 1213 | Bacteria>Pseudomonadota>Gammaproteobacteria>Pseudomonadales>Pseudomonadaceae>Pseudomonas | GCA_021586615.1 | Pseudomonas_aeruginosa | L3655_000128 | 111220..114861 | 287 | BREX-system-P-loop-protein-BrxC-[Pseudomonas-aeruginosa] |
| HBO3309314.1 | Type6.BREX | [illegible] | 1188 | Bacteria>Pseudomonadota>Gammaproteobacteria>Pseudomonadales>Pseudomonadaceae>Pseudomonas | GCA_021652015.1 | Pseudomonas_aeruginosa | L4R58_004351 | 30228..33794 | 287 | BREX-system-P-loop-protein-BrxC-[Pseudomonas-aeruginosa] |
| HBO4843314.1 | Type6.BREX | [illegible] | 1207 | Bacteria>Pseudomonadota>Gammaproteobacteria>Pseudomonadales>Pseudomonadaceae>Pseudomonas | GCA_021660595.1 | Pseudomonas_aeruginosa | L4V37_002405 | 61513..65136 | 287 | BREX-system-P-loop-protein-BrxC-[Pseudomonas-aeruginosa] |
| HBP1093904.1 | Type6.BREX | [illegible] | 1213 | Bacteria>Pseudomonadota>Gammaproteobacteria>Pseudomonadales>Pseudomonadaceae>Pseudomonas | GCA_021687815.1 | Pseudomonas_aeruginosa | L5S32_002755 | 180884..184525 | 287 | BREX-system-P-loop-protein-BrxC-[Pseudomonas-aeruginosa] |
| HBP5527123.1 | Type6.BREX | [illegible] | 1212 | Bacteria>Pseudomonadota>Gammaproteobacteria>Pseudomonadales>Pseudomonadaceae>Pseudomonas | GCA_021712095.1 | Pseudomonas_aeruginosa | L6B52_09510 | 5147..8785 | 287 | BREX-system-P-loop-protein-BrxC-[Pseudomonas-aeruginosa] |
| HCE6004670.1 | Type6.BREX | [illegible] | 1213 | Bacteria>Pseudomonadota>Gammaproteobacteria>Pseudomonadales>Pseudomonadaceae>Pseudomonas | GCA_023904815.1 | Pseudomonas_aeruginosa | NHK94_000107 | 121075..124716 | 287 | BREX-system-P-loop-protein-BrxC-[Pseudomonas-aeruginosa] |
| HCF1683404.1 | Type6.BREX | [illegible] | 1213 | Bacteria>Pseudomonadota>Gammaproteobacteria>Pseudomonadales>Pseudomonadaceae>Pseudomonas | GCA_023941775.1 | Pseudomonas_aeruginosa | NH867_002024 | 49404..53045 | 287 | BREX-system-P-loop-protein-BrxC-[Pseudomonas-aeruginosa] |
| MCO7642712.1 | Type6.BREX | [illegible] | 1204 | Bacteria>Pseudomonadota>Gammaproteobacteria>Pseudomonadales>Pseudomonadaceae>Pseudomonas | GCA_024128275.1 | Pseudomonas_sp._S_311-6 | NJI34_38735 | 21195..24809 | 2956727 | BREX-system-P-loop-protein-BrxC-[Pseudomonas-sp.-S-311-6] |
| HCL4225641.1 | Type6.BREX | [illegible] | 1213 | Bacteria>Pseudomonadota>Gammaproteobacteria>Pseudomonadales>Pseudomonadaceae>Pseudomonas | GCA_025005345.1 | Pseudomonas_aeruginosa | N2C25_003868 | 36520..40161 | 287 | BREX-system-P-loop-protein-BrxC-[Pseudomonas-aeruginosa] |
| WCY64468.1 | Type6.BREX | [illegible] | 1213 | Bacteria>Pseudomonadota>Gammaproteobacteria>Pseudomonadales>Pseudomonadaceae>Pseudomonas | GCA_028610105.1 | Pseudomonas_aeruginosa | NHK94_000107 | 121075..124716 | 287 | BREX-system-P-loop-protein-BrxC-[Pseudomonas-aeruginosa] |
| WP_013692225.1 | Type6.BREX | [illegible] | 1213 | Bacteria>Pseudomonadota>Gammaproteobacteria>Pseudomonadales>Pseudomonadaceae>Pseudomonas | GCF_000194805.1 | Pseudomonas_brassicacearum | PSEBR_RS03955 | 941705..945346 | 930166 | BREX-system-P-loop-protein-BrxC-[Pseudomonas-brassicacearum] |
| WP_007159175.1 | Type6.BREX | [illegible] | 1212 | Bacteria>Pseudomonadota>Gammaproteobacteria>Pseudomonadales>Pseudomonadaceae>Pseudomonas | GCF_000236825.1 | Pseudomonas_psychrotolerans | PPL19_RS03910 | 152166..155804 | 237610 | BREX-system-P-loop-protein-BrxC-[Pseudomonas-psychrotolerans] |
| WP_017337797.1 | Type6.BREX | [illegible] | 1213 | Bacteria>Pseudomonadota>Gammaproteobacteria>Pseudomonadales>Pseudomonadaceae>Pseudomonas | GCF_000293885.2 | Pseudomonas_fluorescens | B723_RS16820 | 3631858..3635499 | 294 | BREX-system-P-loop-protein-BrxC-[Pseudomonas-fluorescens] |
| WP_044305280.1 | Type6.BREX | [illegible] | 1213 | Bacteria>Pseudomonadota>Gammaproteobacteria>Pseudomonadales>Pseudomonadaceae>Pseudomonas | GCF_000793505.1 | Pseudomonas_aeruginosa | NT87_RS20770 | 137629..141270 | 287 | BREX-system-P-loop-protein-BrxC-[Pseudomonas-aeruginosa] |
| WP_058181876.1 | Type6.BREX | [illegible] | 1213 | Bacteria>Pseudomonadota>Gammaproteobacteria>Pseudomonadales>Pseudomonadaceae>Pseudomonas | GCF_001450675.1 | Pseudomonas_aeruginosa | AO933_RS04770 | 131954..135595 | 287 | BREX-system-P-loop-protein-BrxC-[Pseudomonas-aeruginosa] |
| WP_077575502.1 | Type6.BREX | [illegible] | 1211 | Bacteria>Pseudomonadota>Gammaproteobacteria>Pseudomonadales>Pseudomonadaceae>Pseudomonas | GCF_002003595.1 | Pseudomonas_aeruginosa | BWR00_RS05555 | 37630..41265 | 287 | BREX-system-P-loop-protein-BrxC-[Pseudomonas-aeruginosa] |
| WP_101152285.1 | Type6.BREX | [illegible] | 1211 | Bacteria>Pseudomonadota>Gammaproteobacteria>Pseudomonadales>Pseudomonadaceae>Pseudomonas | GCF_002836195.1 | Pseudomonas_sp._Choline-3u-10 | CXF92_RS00360 | 72818..76453 | 2058311 | BREX-system-P-loop-protein-BrxC-[Pseudomonas-sp.-Choline-3u-10] |
| WP_124126664.1 | Type6.BREX | [illegible] | 1210 | Bacteria>Pseudomonadota>Gammaproteobacteria>Pseudomonadales>Pseudomonadaceae>Pseudomonas | GCF_003836525.1 | Pseudomonas_aeruginosa | IPC776_RS06145 | 34793..38425 | 287 | BREX-system-P-loop-protein-BrxC-[Pseudomonas-aeruginosa] |
| WP_126632346.1 | Type6.BREX | [illegible] | 1213 | Bacteria>Pseudomonadota>Gammaproteobacteria>Pseudomonadales>Pseudomonadaceae>Pseudomonas | GCF_003975645.1 | Pseudomonas_aeruginosa | IPC447_RS09565 | 370705..374346 | 287 | BREX-system-P-loop-protein-BrxC-[Pseudomonas-aeruginosa] |
| WP_023094202.1 | Type6.BREX | [illegible] | 1213 | Bacteria>Pseudomonadota>Gammaproteobacteria>Pseudomonadales>Pseudomonadaceae>Pseudomonas | GCF_004346985.1 | Pseudomonas_aeruginosa | C9I70_RS03025 | 613762..617403 | 287 | BREX-system-P-loop-protein-BrxC-[Pseudomonas-aeruginosa] |
| WP_134249510.1 | Type6.BREX | [illegible] | 1213 | Bacteria>Pseudomonadota>Gammaproteobacteria>Pseudomonadales>Pseudomonadaceae>Pseudomonas | GCF_004374205.1 | Pseudomonas_aeruginosa | IPC115_RS16040 | 120938..124579 | 287 | BREX-system-P-loop-protein-BrxC-[Pseudomonas-aeruginosa] |
| WP_134295820.1 | Type6.BREX | [illegible] | 1213 | Bacteria>Pseudomonadota>Gammaproteobacteria>Pseudomonadales>Pseudomonadaceae>Pseudomonas | GCF_004374605.1 | Pseudomonas_aeruginosa | IPC112_RS05880 | 126222..129863 | 287 | BREX-system-P-loop-protein-BrxC-[Pseudomonas-aeruginosa] |
| WP_143505426.1 | Type6.BREX | [illegible] | 1212 | Bacteria>Pseudomonadota>Gammaproteobacteria>Pseudomonadales>Pseudomonadaceae>Pseudomonas | GCF_007049795.1 | Pseudomonas_mendocina | EQ826_RS09455 | 197604..201242 | 300 | BREX-system-P-loop-protein-BrxC-[Pseudomonas-mendocina] |
| WP_161928857.1 | Type6.BREX | [illegible] | 1212 | Bacteria>Pseudomonadota>Gammaproteobacteria>Pseudomonadales>Pseudomonadaceae>Pseudomonas | GCF_009932815.1 | Pseudomonas_yangonensis | GYH25_RS19705 | 87173..90811 | 2579922 | BREX-system-P-loop-protein-BrxC-[Pseudomonas-yangonensis] |
| WP_169883492.1 | Type6.BREX | [illegible] | 1213 | Bacteria>Pseudomonadota>Gammaproteobacteria>Pseudomonadales>Pseudomonadaceae>Pseudomonas | GCF_012986805.1 | Pseudomonas_fragi | HBN91_RS17750 | 10864..14505 | 296 | BREX-system-P-loop-protein-BrxC-[Pseudomonas-fragi] |
| WP_180216146.1 | Type6.BREX | [illegible] | 1213 | Bacteria>Pseudomonadota>Gammaproteobacteria>Pseudomonadales>Pseudomonadaceae>Pseudomonas | GCF_013421145.1 | Pseudomonas_aeruginosa | H0B52_RS17845 | 402112..405753 | 287 | BREX-system-P-loop-protein-BrxC-[Pseudomonas-aeruginosa] |
| WP_181088903.1 | Type6.BREX | [illegible] | 1214 | Bacteria>Pseudomonadota>Gammaproteobacteria>Pseudomonadales>Pseudomonadaceae>Pseudomonas | GCF_013523035.1 | Pseudomonas_plecoglossicida | G7011_RS14160 | 93028..96672 | 70775 | BREX-system-P-loop-protein-BrxC-[Pseudomonas-plecoglossicida] |
| WP_185818333.1 | Type6.BREX | [illegible] | 1214 | Bacteria>Pseudomonadota>Gammaproteobacteria>Pseudomonadales>Pseudomonadaceae>Pseudomonas | GCF_014236655.1 | Pseudomonas_kielensis | H7995_RS10625 | 125852..129496 | 2762577 | BREX-system-P-loop-protein-BrxC-[Pseudomonas-kielensis] |
| WP_186598283.1 | Type6.BREX | [illegible] | 1212 | Bacteria>Pseudomonadota>Gammaproteobacteria>Pseudomonadales>Pseudomonadaceae>Pseudomonas | GCF_014269085.1 | Pseudomonas_taiwanensis | HU747_RS04800 | 431376..435014 | 470150 | BREX-system-P-loop-protein-BrxC-[Pseudomonas-taiwanensis] |
| WP_273507518.1 | Type6.BREX | [illegible] | 1214 | Bacteria>Pseudomonadota>Gammaproteobacteria>Pseudomonadales>Pseudomonadaceae>Pseudomonas | GCF_016785205.1 | Pseudomonas_sp. | JJK56_22660 | 12252..15896 | 306 | BREX-system-P-loop-protein-BrxC-[Pseudomonas-sp.] |
| WP_203328783.1 | Type6.BREX | [illegible] | 1213 | Bacteria>Pseudomonadota>Gammaproteobacteria>Pseudomonadales>Pseudomonadaceae>Pseudomonas | GCF_016836545.1 | Pseudomonas_aeruginosa | JQ483_RS07485 | 1620720..1624361 | 287 | BREX-system-P-loop-protein-BrxC-[Pseudomonas-aeruginosa] |
| WP_209939193.1 | Type6.BREX | [illegible] | 1213 | Bacteria>Pseudomonadota>Gammaproteobacteria>Pseudomonadales>Pseudomonadaceae>Pseudomonas | GCF_017838075.1 | Pseudomonas_aeruginosa | J0A89_RS01675 | 341330..344971 | 287 | BREX-system-P-loop-protein-BrxC-[Pseudomonas-aeruginosa] |
| WP_223717957.1 | Type6.BREX | [illegible] | 1213 | Bacteria>Pseudomonadota>Gammaproteobacteria>Pseudomonadales>Pseudomonadaceae>Pseudomonas | GCF_020036535.1 | Pseudomonas_aeruginosa | LC626_RS08340 | 482522..486163 | 287 | BREX-system-P-loop-protein-BrxC-[Pseudomonas-aeruginosa] |
| WP_227364598.1 | Type6.BREX | [illegible] | 1213 | Bacteria>Pseudomonadota>Gammaproteobacteria>Pseudomonadales>Pseudomonadaceae>Pseudomonas | GCF_020639995.1 | Pseudomonas_aeruginosa | LJG07_RS13375 | 107262..110903 | 287 | BREX-system-P-loop-protein-BrxC-[Pseudomonas-aeruginosa] |
| WP_229605965.1 | Type6.BREX | [illegible] | 1212 | Bacteria>Pseudomonadota>Gammaproteobacteria>Pseudomonadales>Pseudomonadaceae>Pseudomonas | GCF_020831405.1 | Pseudomonas_oryzagri | LMJ29_RS13110 | 91421..95059 | 2892443 | BREX-system-P-loop-protein-BrxC-[Pseudomonas-oryzagri] |
| WP_033990403.1 | Type6.BREX | [illegible] | 1213 | Bacteria>Pseudomonadota>Gammaproteobacteria>Pseudomonadales>Pseudomonadaceae>Pseudomonas | GCF_020907765.1 | Pseudomonas_aeruginosa | LOH47_RS10865 | 120991..124632 | 287 | BREX-system-P-loop-protein-BrxC-[Pseudomonas-aeruginosa] |
| WP_236165768.1 | Type6.BREX | [illegible] | 1212 | Bacteria>Pseudomonadota>Gammaproteobacteria>Pseudomonadales>Pseudomonadaceae>Pseudomonas | GCF_021603865.1 | Pseudomonas_fulva | L6D10_RS06765 | 41152..44790 | 47880 | BREX-system-P-loop-protein-BrxC-[Pseudomonas-fulva] |
| WP_247022253.1 | Type6.BREX | [illegible] | 1213 | Bacteria>Pseudomonadota>Gammaproteobacteria>Pseudomonadales>Pseudomonadaceae>Pseudomonas | GCF_023060065.1 | Pseudomonas_aeruginosa | M9Y84_RS04355 | 402804..406445 | 287 | BREX-system-P-loop-protein-BrxC-[Pseudomonas-aeruginosa] |
| WP_247325109.1 | Type6.BREX | [illegible] | 371 | Bacteria>Pseudomonadota>Gammaproteobacteria>Pseudomonadales>Pseudomonadaceae>Pseudomonas | GCF_023101305.1 | Pseudomonas_aeruginosa | MX621_RS22735 | 4826578..4827693 | 287 | DUF6079-family-protein-[Pseudomonas-aeruginosa] |
| WP_262867297.1 | Type6.BREX | [illegible] | 796 | Bacteria>Pseudomonadota>Gammaproteobacteria>Pseudomonadales>Pseudomonadaceae>Pseudomonas | GCF_025515535.1 | Pseudomonas_aeruginosa | ODQ25_RS15790 | 61608..63994 | 287 | BREX-system-P-loop-protein-BrxC,-partial-[Pseudomonas-aeruginosa] |
| WP_010792139.1 | Type6.BREX | [illegible] | 1210 | Bacteria>Pseudomonadota>Gammaproteobacteria>Pseudomonadales>Pseudomonadaceae>Pseudomonas | GCF_026126725.1 | Pseudomonas_aeruginosa | K3Z93_RS02355 | 20406..24038 | 287 | BREX-system-P-loop-protein-BrxC-[Pseudomonas-aeruginosa] |
| WP_057380122.1 | Type6.BREX | [illegible] | 1213 | Bacteria>Pseudomonadota>Gammaproteobacteria>Pseudomonadales>Pseudomonadaceae>Pseudomonas | GCF_026727755.1 | Pseudomonas_aeruginosa | PAC42_RS26725 | 5539866..5543507 | 287 | BREX-system-P-loop-protein-BrxC-[Pseudomonas-aeruginosa] |
| WP_058409982.1 | Type6.BREX | [illegible] | 1212 | Bacteria>Pseudomonadota>Gammaproteobacteria>Pseudomonadales>Pseudomonadaceae>Pseudomonas | GCF_029959515.1 | Pseudomonas_sp._ICMP_19500 | AO239_RS04345 | 8658..12296 | 1208102 | BREX-system-P-loop-protein-BrxC-[Pseudomonas-sp.-ICMP-19500] |
| WP_034038617.1 | Type6.BREX | [illegible] | 1213 | Bacteria>Pseudomonadota>Gammaproteobacteria>Pseudomonadales>Pseudomonadaceae>Pseudomonas | GCF_900147165.1 | Pseudomonas_aeruginosa | K3Z85_RS02500 | 4652..8293 | 287 | BREX-system-P-loop-protein-BrxC-[Pseudomonas-aeruginosa] |
| WP_150756947.1 | Type6.BREX | [illegible] | 1210 | Bacteria>Pseudomonadota>Gammaproteobacteria>Pseudomonadales>Pseudomonadaceae>Pseudomonas | GCF_902498135.1 | Pseudomonas_fluorescens | F8N72_RS03340 | 145208..148840 | 294 | BREX-system-P-loop-protein-BrxC-[Pseudomonas-fluorescens] |
| WP_151213669.1 | Type6.BREX | [illegible] | 1210 | Bacteria>Pseudomonadota>Gammaproteobacteria>Pseudomonadales>Pseudomonadaceae>Pseudomonas | GCF_911283405.1 | Pseudomonas_fluorescens | F9263_RS07600 | 503497..507129 | 294 | BREX-system-P-loop-protein-BrxC-[Pseudomonas-fluorescens] |
| WP_095027056.1 | Type6.BREX | [illegible] | 1213 | Bacteria>Pseudomonadota>Gammaproteobacteria>Pseudomonadales>Pseudomonadaceae | - | Pseudomonas | JFT63_RS16080 | 178886..182527 | 286 | MULTISPECIES:-BREX-system-P-loop-protein-BrxC-[Pseudomonas] |
| WP_121740032.1 | Type6.BREX | [illegible] | 1213 | Bacteria>Pseudomonadota>Gammaproteobacteria>Pseudomonadales>Pseudomonadaceae>Pseudomonas | - | Pseudomonas_aeruginosa | D3M96_03425 | 754727..758368 | 287 | MULTISPECIES:-BREX-system-P-loop-protein-BrxC-[Proteobacteria] |
| WP_150298006.1 | Type6.BREX | [illegible] | 1212 | Bacteria>Pseudomonadota>Gammaproteobacteria>Pseudomonadales>Pseudomonadaceae>Pseudomonas | - | Pseudomonas_aeruginosa | F3G60_11255 | 510071..513709 | 287 | BREX-system-P-loop-protein-BrxC-[Pseudomonas-aeruginosa] |
| WP_223843585.1 | Type6.BREX | [illegible] | 858 | Bacteria>Pseudomonadota>Gammaproteobacteria>Pseudomonadales>Pseudomonadaceae>Pseudomonas | - | Pseudomonas_sp._CFA | F7661_RS08255 | 1849794..1852370 | 2651048 | BREX-system-P-loop-protein-BrxC-[Pseudomonas-sp.-CFA] |
| WP_223843584.1 | Type6.BREX | [illegible] | 372 | Bacteria>Pseudomonadota>Gammaproteobacteria>Pseudomonadales>Pseudomonadaceae>Pseudomonas | - | Pseudomonas_sp._CFA | F7661_RS27875 | 1848730..1849848 | 2651048 | hypothetical-protein-[Pseudomonas-sp.-CFA] |
| WP_023443891.1 | Type6.BREX | [illegible] | 1213 | Bacteria>Pseudomonadota>Gammaproteobacteria>Pseudomonadales>Pseudomonadaceae>Stutzerimonas | GCF_000495915.1 | Stutzerimonas_stutzeri_group | F753_RS26950 | 57885..61526 | 136846 | MULTISPECIES:-BREX-system-P-loop-protein-BrxC-[Stutzerimonas-stutzeri-group] |
| WP_120997717.1 | Type6.BREX | [illegible] | 1212 | Bacteria>Pseudomonadota>Gammaproteobacteria>Pseudomonadales>Pseudomonadaceae>Stutzerimonas | GCF_003640395.1 | [Pseudomonas]_urumqiensis | D7325_RS17370 | 36612..40250 | 638269 | BREX-system-P-loop-protein-BrxC-[[Pseudomonas] |
| WP_125866641.1 | Type6.BREX | [illegible] | 1213 | Bacteria>Pseudomonadota>Gammaproteobacteria>Pseudomonadales>Pseudomonadaceae>Stutzerimonas | GCF_003935625.1 | Stutzerimonas_stutzeri | EGJ12_RS17010 | 223453..227094 | 316 | BREX-system-P-loop-protein-BrxC-[Stutzerimonas-stutzeri] |
| WP_212739636.1 | Type6.BREX | [illegible] | 1207 | Bacteria>Pseudomonadota>Gammaproteobacteria>Pseudomonadales>Pseudomonadaceae>Stutzerimonas | GCF_018316915.2 | Stutzerimonas_balearica | D3P44_RS15315 | 3232724..3236347 | 74829 | BREX-system-P-loop-protein-BrxC-[Stutzerimonas-balearica] |
| WP_041106841.1 | Type6.BREX | [illegible] | 1213 | Bacteria>Pseudomonadota>Gammaproteobacteria>Pseudomonadales>Pseudomonadaceae>Stutzerimonas | - | Stutzerimonas_stutzeri | QX25_RS10015 | 148024..151665 | 316 | BREX-system-P-loop-protein-BrxC-[Stutzerimonas-stutzeri] |
| WP_210226002.1 | Type6.BREX | [illegible] | 1210 | Bacteria>Pseudomonadota>Gammaproteobacteria>Thiotrichales>Thiotrichaceae>Thiothrix | GCF_017901155.1 | Candidatus_Thiothrix_anitrata | J8380_RS12830 | 2543544..2547176 | 2823902 | BREX-system-P-loop-protein-BrxC-[Candidatus-Thiothrix-anitrata] |
| TAK52340.1 | Type6.BREX | [illegible] | 1210 | Bacteria>Pseudomonadota>Gammaproteobacteria>unclassified_Gammaproteobacteria | GCA_004297335.1 | Gammaproteobacteria_bacterium | EPO25_13990 | 58344..61976 | 1913989 | BREX-system-P-loop-protein-BrxC-[Gammaproteobacteria-bacterium] |
| NPU94904.1 | Type6.BREX | [illegible] | 1213 | Bacteria>Pseudomonadota>Gammaproteobacteria>unclassified_Gammaproteobacteria | GCA_013178375.1 | Gammaproteobacteria_bacterium | HPY82_23615 | 18761..22402 | 1913989 | BREX-system-P-loop-protein-BrxC-[Gammaproteobacteria-bacterium] |
| WP_058119139.1 | Type6.BREX | [illegible] | 1208 | Bacteria>Pseudomonadota>Gammaproteobacteria>Vibrionales>Vibrionaceae>Photobacterium | GCF_001455895.1 | Photobacterium_kishitanii | AT442_RS02840 | 630344..633970 | 318456 | BREX-system-P-loop-protein-BrxC-[Photobacterium-kishitanii] |
| GIB72257.1 | Type6.BREX | [illegible] | 1211 | Bacteria>Pseudomonadota>Gammaproteobacteria>Vibrionales>Vibrionaceae>Vibrio | GCA_023164365.1 | Vibrio_cholerae | VCSRO19_0643 | 268485..272120 | 666 | ATPase-[Vibrio-cholerae] |
| WP_228097414.1 | Type6.BREX | [illegible] | 1208 | Bacteria>Pseudomonadota>Gammaproteobacteria>Vibrionales>Vibrionaceae>Vibrio | GCF_020741135.1 | Vibrio_parahaemolyticus | IB300_RS08390 | 552360..555986 | 670 | BREX-system-P-loop-protein-BrxC-[Vibrio-parahaemolyticus] |
| WP_017462916.1 | Type6.BREX | [illegible] | 1213 | Bacteria>Pseudomonadota>Gammaproteobacteria>Xanthomonadales>Rhodanobacteraceae>Dyella | GCF_000334915.1 | Dyella_ginsengisoli | C693_RS0114050 | 37588..41229 | 363848 | BREX-system-P-loop-protein-BrxC-[Dyella-ginsengisoli] |
| WP_049622247.1 | Type6.BREX | [illegible] | 1210 | Bacteria>Pseudomonadota>Gammaproteobacteria>Xanthomonadales>Rhodanobacteraceae>Frateuria | GCF_001182895.1 | Frateuria_defendens | ACH65_RS08940 | 10628..14260 | 2219559 | BREX-system-P-loop-protein-BrxC-[Frateuria-defendens] |
| WP_024958028.1 | Type6.BREX | [illegible] | 1210 | Bacteria>Pseudomonadota>Gammaproteobacteria>Xanthomonadales>Xanthomonadaceae | GCF_001591205.1 | Stenotrophomonas | SMA02S_RS05105 | 16595..20227 | 40323 | MULTISPECIES:-BREX-system-P-loop-protein-BrxC-[Stenotrophomonas] |
| WP_070426722.1 | Type6.BREX | [illegible] | 1209 | Bacteria>Pseudomonadota>Gammaproteobacteria>Xanthomonadales>Xanthomonadaceae | GCF_900766255.1 | Stenotrophomonas | BIZ42_RS13545 | 3079736..3083365 | 40323 | MULTISPECIES:-BREX-system-P-loop-protein-BrxC-[Stenotrophomonas] |
| WP_130516891.1 | Type6.BREX | [illegible] | 1209 | Bacteria>Pseudomonadota>Gammaproteobacteria>Xanthomonadales>Xanthomonadaceae>Pseudoxanthomonas | GCF_004284745.1 | Pseudoxanthomonas_winnipegensis | EYC46_RS06610 | 172143..175772 | 2480810 | BREX-system-P-loop-protein-BrxC-[Pseudoxanthomonas-winnipegensis] |
| WP_162203842.1 | Type6.BREX | [illegible] | 1213 | Bacteria>Pseudomonadota>Gammaproteobacteria>Xanthomonadales>Xanthomonadaceae>Pseudoxanthomonas | GCF_010093235.1 | Pseudoxanthomonas_suwonensis | CSC68_RS12285 | 54892..58533 | 314722 | BREX-system-P-loop-protein-BrxC-[Pseudoxanthomonas-suwonensis] |
| MBH1753233.1 | Type6.BREX | [illegible] | 1209 | Bacteria>Pseudomonadota>Gammaproteobacteria>Xanthomonadales>Xanthomonadaceae>Stenotrophomonas | GCA_015998385.1 | Stenotrophomonas_maltophilia | I5V37_01935 | 25775..29404 | 40324 | BREX-system-P-loop-protein-BrxC-[Stenotrophomonas-maltophilia] |
| MBN5094965.1 | Type6.BREX | [illegible] | 1213 | Bacteria>Pseudomonadota>Gammaproteobacteria>Xanthomonadales>Xanthomonadaceae>Stenotrophomonas | GCA_017150075.1 | Stenotrophomonas_maltophilia | JY450_00935 | 90079..93720 | 40324 | BREX-system-P-loop-protein-BrxC-[Stenotrophomonas-maltophilia] |
| MBN5083090.1 | Type6.BREX | [illegible] | 1207 | Bacteria>Pseudomonadota>Gammaproteobacteria>Xanthomonadales>Xanthomonadaceae>Stenotrophomonas | GCA_017150495.1 | Stenotrophomonas_maltophilia | JY467_05250 | 61681..65304 | 40324 | BREX-system-P-loop-protein-BrxC-[Stenotrophomonas-maltophilia] |
| WP_004146422.1 | Type6.BREX | [illegible] | 1207 | Bacteria>Pseudomonadota>Gammaproteobacteria>Xanthomonadales>Xanthomonadaceae>Stenotrophomonas | GCF_000355725.1 | Stenotrophomonas_maltophilia | G5A34_RS07915 | 81973..85596 | 40324 | BREX-system-P-loop-protein-BrxC-[Stenotrophomonas-maltophilia] |
| WP_088610961.1 | Type6.BREX | [illegible] | 1209 | Bacteria>Pseudomonadota>Gammaproteobacteria>Xanthomonadales>Xanthomonadaceae>Stenotrophomonas | GCF_002208885.2 | Stenotrophomonas_maltophilia | CEQ03_RS09960 | 2076489..2080118 | 40324 | BREX-system-P-loop-protein-BrxC-[Stenotrophomonas-maltophilia] |
| WP_099491466.1 | Type6.BREX | [illegible] | 1209 | Bacteria>Pseudomonadota>Gammaproteobacteria>Xanthomonadales>Xanthomonadaceae>Stenotrophomonas | GCF_002739275.1 | Stenotrophomonas_maltophilia | BRB91_RS07170 | 53995..57624 | 40324 | BREX-system-P-loop-protein-BrxC-[Stenotrophomonas-maltophilia] |
| WP_099562737.1 | Type6.BREX | [illegible] | 1209 | Bacteria>Pseudomonadota>Gammaproteobacteria>Xanthomonadales>Xanthomonadaceae>Stenotrophomonas | GCF_002740395.1 | Stenotrophomonas_maltophilia | BRC18_RS18230 | 18705..22334 | 40324 | BREX-system-P-loop-protein-BrxC-[Stenotrophomonas-maltophilia] |
| WP_164085131.1 | Type6.BREX | [illegible] | 1209 | Bacteria>Pseudomonadota>Gammaproteobacteria>Xanthomonadales>Xanthomonadaceae>Stenotrophomonas | GCF_010605045.1 | Stenotrophomonas_maltophilia | G5A32_RS08645 | 68360..71989 | 40324 | BREX-system-P-loop-protein-BrxC-[Stenotrophomonas-maltophilia] |
| WP_180875343.1 | Type6.BREX | [illegible] | 1209 | Bacteria>Pseudomonadota>Gammaproteobacteria>Xanthomonadales>Xanthomonadaceae>Stenotrophomonas | GCF_013463485.1 | Stenotrophomonas_maltophilia | D7U74_RS02160 | 10322..13951 | 40324 | BREX-system-P-loop-protein-BrxC-[Stenotrophomonas-maltophilia] |
| WP_049450109.1 | Type6.BREX | [illegible] | 1209 | Bacteria>Pseudomonadota>Gammaproteobacteria>Xanthomonadales>Xanthomonadaceae>Stenotrophomonas | GCF_030549525.1 | Stenotrophomonas_maltophilia | MT454_RS13320 | 40759..44388 | 40324 | BREX-system-P-loop-protein-BrxC-[Stenotrophomonas-maltophilia] |
| WP_111722876.1 | Type6.BREX | [illegible] | 1209 | Bacteria>Pseudomonadota>Gammaproteobacteria>Xanthomonadales>Xanthomonadaceae>Stenotrophomonas | GCF_900475685.1 | Stenotrophomonas_maltophilia | DQN96_RS04535 | 962467..966096 | 40324 | BREX-system-P-loop-protein-BrxC-[Stenotrophomonas-maltophilia] |
| WP_180834305.1 | Type6.BREX | [illegible] | 1209 | Bacteria>Pseudomonadota>Gammaproteobacteria>Xanthomonadales>Xanthomonadaceae>Stenotrophomonas | - | Stenotrophomonas_maltophilia | D7U93_RS19145 | 16028..19657 | 40324 | BREX-system-P-loop-protein-BrxC-[Stenotrophomonas-maltophilia] |
| WP_262214630.1 | Type6.BREX | [illegible] | 1213 | Bacteria>Pseudomonadota>Gammaproteobacteria>Xanthomonadales>Xanthomonadaceae>Stenotrophomonas | - | Stenotrophomonas_maltophilia | JAK45_RS19240 | 75512..79153 | 40324 | BREX-system-P-loop-protein-BrxC-[Stenotrophomonas-maltophilia] |
| VUJ46804.1 | Type6.BREX | [illegible] | 1207 | Bacteria>Pseudomonadota>Gammaproteobacteria>Xanthomonadales>Xanthomonadaceae>Stenotrophomonas | - | Stenotrophomonas_maltophilia | N5D45_RS14200 | 24687..28310 | 40324 | hypothetical-protein-NGKLMNPD_00401-[Stenotrophomonas-maltophilia] |
| MBS3958360.1 | Type6.BREX | [illegible] | 1213 | Bacteria>Pseudomonadota>Gammaproteobacteria>Xanthomonadales>Xanthomonadaceae>unclassified_Xanthomonadaceae | GCA_018333835.1 | Xanthomonadaceae_bacterium | KGZ52_03015 | 3241..6882 | 1926873 | BREX-system-P-loop-protein-BrxC-[Xanthomonadaceae-bacterium] |
| WP_078567675.1 | Type6.BREX | [illegible] | 1213 | Bacteria>Pseudomonadota>Gammaproteobacteria>Xanthomonadales>Xanthomonadaceae>Xanthomonas | GCF_002019295.1 | Xanthomonas_campestris | Xvtf_RS11430 | 4159..7800 | 339 | BREX-system-P-loop-protein-BrxC-[Xanthomonas-campestris] |
| WP_142804161.1 | Type6.BREX | [illegible] | 1210 | Bacteria>Pseudomonadota>Hydrogenophilia>Hydrogenophilales>Hydrogenophilaceae>Tepidiphilus | GCF_006980785.1 | Tepidiphilus_sp._J10 | EPD81_RS04405 | 897239..900871 | 2502185 | BREX-system-P-loop-protein-BrxC-[Tepidiphilus-sp.-J10] |
| WP_169116165.1 | Type6.BREX | [illegible] | 1210 | Bacteria>Pseudomonadota>Hydrogenophilia>Hydrogenophilales>Hydrogenophilaceae>Tepidiphilus | GCF_012911495.1 | Tepidiphilus_baoligensis | GV368_RS08140 | 10579..14211 | 2698687 | BREX-system-P-loop-protein-BrxC-[Tepidiphilus-baoligensis] |
| NCC04627.1 | Type6.BREX | [illegible] | 1206 | Bacteria>Pseudomonadota>unclassified_Pseudomonadota | GCA_009929575.1 | Pseudomonadota_bacterium | EOM37_11440 | 1963..5583 | 1977087 | BREX-system-P-loop-protein-BrxC-[Pseudomonadota-bacterium] |
| MBS0339220.1 | Type6.BREX | [illegible] | 1180 | Bacteria>Pseudomonadota>unclassified_Pseudomonadota | GCA_018241905.1 | Pseudomonadota_bacterium | JSS56_01750 | 21347..24889 | 1977087 | BREX-system-P-loop-protein-BrxC-[Pseudomonadota-bacterium] |
| MBU0973505.1 | Type6.BREX | [illegible] | 1209 | Bacteria>Pseudomonadota>unclassified_Pseudomonadota | GCA_018816605.1 | Pseudomonadota_bacterium | KKC20_22890 | 2683..6312 | 1977087 | BREX-system-P-loop-protein-BrxC-[Pseudomonadota-bacterium] |
| PIX55556.1 | Type6.BREX | [illegible] | 1215 | Bacteria>Pseudomonadota>Zetaproteobacteria>unclassified_Zetaproteobacteria | GCA_002782825.1 | Zetaproteobacteria_bacterium_CG_4_10_14_3_um_filter_54_28 | COZ50_02095 | 2235..5882 | 1974114 | BREX-system-P-loop-protein-BrxC-[Zetaproteobacteria-bacterium-CG_4_10_14_3_um_filter_54_28] |
| NPU98919.1 | Type6.BREX | [illegible] | 1224 | Bacteria>PVC_group>Candidatus_Omnitrophica>unclassified_Candidatus_Omnitrophica | GCA_013177805.1 | Candidatus_Omnitrophica_bacterium | HPY51_17110 | 7762..11436 | 2035772 | BREX-system-P-loop-protein-BrxC-[Candidatus-Omnitrophota-bacterium] |
| MBX3380229.1 | Type6.BREX | [illegible] | 1207 | Bacteria>PVC_group>Planctomycetota>Phycisphaerae>Phycisphaerales>Phycisphaeraceae | GCA_019636565.1 | Phycisphaeraceae_bacterium | KF805_09030 | 72021..75644 | 2026777 | BREX-system-P-loop-protein-BrxC-[Phycisphaeraceae-bacterium] |
| HIB67895.1 | Type6.BREX | [illegible] | 1116 | Bacteria>PVC_group>Planctomycetota>Phycisphaerae>Phycisphaerales>unclassified_Phycisphaerales | GCA_012964585.1 | Phycisphaerales_bacterium | EYO33_23065 | 26327..29673 | 2052180 | BREX-system-P-loop-protein-BrxC-[Phycisphaerales-bacterium] |
| MBY0113214.1 | Type6.BREX | [illegible] | 1179 | Bacteria>PVC_group>Planctomycetota>Phycisphaerae>Phycisphaerales>unclassified_Phycisphaerales | GCA_019744815.1 | Phycisphaerales_bacterium | K2Y21_10355 | 73058..76597 | 2052180 | BREX-system-P-loop-protein-BrxC-[Phycisphaerales-bacterium] |
| MCG8407728.1 | Type6.BREX | [illegible] | 1181 | Bacteria>PVC_group>Planctomycetota>Phycisphaerae>Phycisphaerales>unclassified_Phycisphaerales | GCA_022360095.1 | Phycisphaerales_bacterium | MI923_21220 | 28816..32361 | 2052180 | BREX-system-P-loop-protein-BrxC-[Phycisphaerales-bacterium] |
| WP_009097835.1 | Type6.BREX | [illegible] | 1207 | Bacteria>PVC_group>Planctomycetota>Planctomycetia>Pirellulales>Pirellulaceae | GCF_000346425.1 | Rhodopirellula_sp._SWK7 | RRSWK_RS13975 | 105510..109133 | 595460 | BREX-system-P-loop-protein-BrxC-[Rhodopirellula-sp.-SWK7] |
| WP_145167965.1 | Type6.BREX | [illegible] | 1219 | Bacteria>PVC_group>Planctomycetota>Planctomycetia>Pirellulales>Pirellulaceae | GCF_007741535.1 | Rubripirellula_lacrimiformis | K227x_RS02725 | 780099..783758 | 1930273 | BREX-system-P-loop-protein-BrxC-[Rubripirellula-lacrimiformis] |
| WP_146394139.1 | Type6.BREX | [illegible] | 1207 | Bacteria>PVC_group>Planctomycetota>Planctomycetia>Pirellulales>Pirellulaceae | GCF_007859855.1 | Rhodopirellula_solitaria | CA85_RS23950 | 9786..13409 | 2527987 | BREX-system-P-loop-protein-BrxC-[Rhodopirellula-solitaria] |
| TVP98542.1 | Type6.BREX | [illegible] | 1207 | Bacteria>PVC_group>Planctomycetota>Planctomycetia>Planctomycetales>Planctomycetaceae | GCA_007694185.1 | Planctomycetaceae_bacterium | EA381_12175 | 2603..6226 | 2026779 | BREX-system-P-loop-protein-BrxC-[Planctomycetaceae-bacterium] |
| MCA9087267.1 | Type6.BREX | [illegible] | 1180 | Bacteria>PVC_group>Planctomycetota>Planctomycetia>Planctomycetales>Planctomycetaceae | GCA_020426515.1 | Planctomycetaceae_bacterium | KDA90_01390 | 104120..107662 | 2026779 | BREX-system-P-loop-protein-BrxC-[Planctomycetaceae-bacterium] |
| WP_013111330.1 | Type6.BREX | [illegible] | 1208 | Bacteria>PVC_group>Planctomycetota>Planctomycetia>Planctomycetales>Planctomycetaceae | GCF_000092105.1 | Planctopirus_limnophila | PLIM_RS15820 | 3989938..3993564 | 120 | BREX-system-P-loop-protein-BrxC-[Planctopirus-limnophila] |
| WP_145294603.1 | Type6.BREX | [illegible] | 1181 | Bacteria>PVC_group>Planctomycetota>Planctomycetia>Planctomycetales>Planctomycetaceae | GCF_007752935.1 | Crateriforma_conspicua | Mal65_RS05525 | 1500908..1504453 | 2527996 | BREX-system-P-loop-protein-BrxC-[Crateriforma-conspicua] |
| MCA9173122.1 | Type6.BREX | [illegible] | 1180 | Bacteria>PVC_group>Planctomycetota>Planctomycetia>Planctomycetales>unclassified_Planctomycetales | GCA_020426845.1 | Planctomycetales_bacterium | KDB14_01345 | 5365..8907 | 2053591 | BREX-system-P-loop-protein-BrxC-[Planctomycetales-bacterium] |

| Protein-ID | System | Gene-Neighbourhoods | Length | Taxonomy | Assembly-ID | Strain | Locus-Tag | Genomic-coordinates | TaxID | Fasta-define |
|---|---|---|---|---|---|---|---|---|---|---|
| MCA9231627.1 | Type6.BREX | <-REase<-SIGMA+OB-Fold+AAA-ATPase+Lon-Protease-BrxL<-PglZ<-N6-METHYLASE<-DUF6079*<-BrxB-BrxF-iSTAND<-BrxA-DNA-binding<-BrxE<-?\|\|REase-> | 1178 | Bacteria>PVC_group>Planctomycetota>Planctomycetia>Planctomycetales>unclassified_Planctomycetales | GCA_020427155.1 | Planctomycetales_bacterium | KDA57_13330 | 9427..12963 | 2053591 | BREX-system-P-loop-protein-BrxC-[Planctomycetales-bacterium] |
| NQW48456.1 | Type6.BREX | <-MarR-HTH<-DUF6079*<-BrxB-BrxF-iSTAND<-BrxA-DNA-binding<-BrxE\|\|Peptidase_M48+TerB->?->?->?->35exo+T7DNApol-> | 1207 | Bacteria>PVC_group>Planctomycetota>unclassified_Planctomycetota | GCA_013205375.1 | Planctomycetota_bacterium | HQ464_11920 | 495..4118 | 2026780 | BREX-system-P-loop-protein-BrxC-[Planctomycetota-bacterium] |
| MBL8841980.1 | Type6.BREX | HTH_3->?->?->ParB+N6_N4_METHYLASE-><-?\|\|?->BrxE->BrxA-DNA-binding->BrxB-BrxF-iSTAND->DUF6079*->N6-METHYLASE->N6-METHYLASE->N6-METHYLASE->PglZ->SIGMA+OB-Fold+AAA-ATPase+Lon-Protease-BrxL-> | 1181 | Bacteria>PVC_group>Planctomycetota>unclassified_Planctomycetota | GCA_016795285.1 | Planctomycetota_bacterium | JNL90_10705 | 176280..179825 | 2026780 | BREX-system-P-loop-protein-BrxC-[Planctomycetota-bacterium] |
| MBM4012727.1 | Type6.BREX | <-Peptidase_M48+TerB\|\|BrxE->BrxA-DNA-binding->BrxB-BrxF-iSTAND->DUF6079-p*-> | 378 | Bacteria>PVC_group>Planctomycetota>unclassified_Planctomycetota | GCA_016872925.1 | Planctomycetota_bacterium | FJ286_15375 | 4393..5526 | 2026780 | BREX-system-P-loop-protein-BrxC-[Planctomycetota-bacterium] |
| MBM4057033.1 | Type6.BREX | <-Peptidase_M48+TerB\|\|BrxE->BrxA-DNA-binding->BrxB-BrxF-iSTAND->DUF6079-p*-> | 352 | Bacteria>PVC_group>Planctomycetota>unclassified_Planctomycetota | GCA_016873005.1 | Planctomycetota_bacterium | FJ275_02185 | 3104..4159 | 2026780 | BREX-system-P-loop-protein-BrxC-[Planctomycetota-bacterium] |
| RRK00196.1 | Type6.BREX | <-DNA_METHYLASE<-PglZ-p<-PglZ<-PglZ<-?<-OLD-ABC<-ParB<-DUF6079*<-BrxB-BrxF-iSTAND<-BrxA-DNA-binding<-BrxE<-REase<-?<-HEPN | 1207 | Bacteria>PVC_group>Verrucomicrobiota>Opitutae>Opitutales>Opitutaceae | GCA_001653815.2 | Opitutaceae_bacterium_TAV3 | Ga0100230_019740 | 2814873..2818496 | 278958 | BREX-system-P-loop-protein-BrxC-[Opitutaceae-bacterium-TAV3] |
| RRJ96051.1 | Type6.BREX | <-TrwC+SF1-SF2-Helicase\|\|?-><-DNA_METHYLASE<-SIGMA+OB-Fold+AAA-ATPase+HKD-DNase-BrxL<-PglZ<-?<-OLD-ABC<-ParB<-DUF6079*<-BrxB-BrxF-iSTAND<-BrxA-DNA-binding<-BrxE<-REase<-?<-Phage_int_SAM_1+Phage_integrase<-HEPN<-ZnR+ATPcone | 1206 | Bacteria>PVC_group>Verrucomicrobiota>Opitutae>Opitutales>Opitutaceae | GCA_001653825.2 | Opitutaceae_bacterium_TAV4 | Ga0100231_019065 | 2732930..2736550 | 278959 | BREX-system-P-loop-protein-BrxC-[Opitutaceae-bacterium-TAV4] |
| TAF24165.1 | Type6.BREX | <-ACYC<-?<-SIGMA+OB-Fold+AAA-ATPase+HKD-DNase-BrxL<-PglZ<-N6-METHYLASE<-DUF6079*<-BrxB-BrxF-iSTAND<-BrxA-DNA-binding<-BrxE<-?<-MPTase<-HTH_3\|\|?->REase-> | 1215 | Bacteria>PVC_group>Verrucomicrobiota>unclassified_Verrucomicrobiota | GCA_004293445.1 | Verrucomicrobiota_bacterium | EAZ71_11350 | 54847..58494 | 2026799 | BREX-system-P-loop-protein-BrxC-[Verrucomicrobiota-bacterium] |
| GDY21840.1 | Type6.BREX | BrxE->BrxA-DNA-binding->BrxB-BrxF-iSTAND->DUF6079*->N6-METHYLASE->HTH+REase-> | 1208 | Bacteria>PVC_group>Verrucomicrobiota>unclassified_Verrucomicrobiota | GCA_014193395.1 | Verrucomicrobiota_bacterium | LBMAG56_31870 | 2824..6450 | 2026799 | hypothetical-protein-LBMAG56_31870-[Verrucomicrobiota-bacterium] |
| MBV8228025.1 | Type6.BREX | <-N6-METHYLASE-p<-DUF6079*<-BrxB-BrxF-iSTAND<-BrxA-DNA-binding<-BrxE\|\|?->METHYLASE->Radical-SAM-> | 1212 | Bacteria>PVC_group>Verrucomicrobiota>unclassified_Verrucomicrobiota | GCA_019236535.1 | Verrucomicrobiota_bacterium | JO232_22865 | 1592..5230 | 2026799 | BREX-system-P-loop-protein-BrxC-[Verrucomicrobiota-bacterium] |
| MCP5535434.1 | Type6.BREX | <-XIS-HTH\|\|?->Phage_integrase->?->?->?->BrxE->BrxA-DNA-binding->BrxB-BrxF-iSTAND->DUF6079*->N6-METHYLASE->DrHyd->Schlafen+HATPase+wHTH->PglZ->SIGMA+OB-Fold+AAA-ATPase+HKD-DNase-BrxL->?->?->?->RAMPs-> | 1186 | Bacteria>PVC_group>Verrucomicrobiota>Verrucomicrobiae>Verrucomicrobiales>Akkermansiaceae | GCA_024233115.1 | Akkermansiaceae_bacterium | H7A51_04270 | 18569..22129 | 2562705 | BREX-system-P-loop-protein-BrxC-[Akkermansiaceae-bacterium] |
| MBE7502027.1 | Type6.BREX | HTH_3-><-?<-?<-?<-GNAT<-NERD+ZnR\|\|BrxE->BrxA-DNA-binding->BrxB-BrxF-iSTAND->DUF6079*->N6-METHYLASE-><-DDE_Tnp_1\|\|N6-METHYLASE->ParB+HNH->OLD-ABC->TOPRIM->DDE_Tnp_1->PglZ->SIGMA+OB-Fold+AAA-ATPase+HKD-DNase-BrxL-> | 1231 | Bacteria>PVC_group>Verrucomicrobiota>Verrucomicrobiae>Verrucomicrobiales>unclassified_Verrucomicrobiales | GCA_015075445.1 | Verrucomicrobiales_bacterium | HS113_17385 | 108161..111856 | 2026801 | BREX-system-P-loop-protein-BrxC-[Verrucomicrobiales-bacterium] |
| WP_200392181.1 | Type6.BREX | <-DUF6079-p*<-BrxB-BrxF-iSTAND<-BrxA-DNA-binding<-BrxE<-?<-GNTR-HTH<-HTH_3<-tRNA<-METHYLASE | 500 | Bacteria>PVC_group>Verrucomicrobiota>Verrucomicrobiae>Verrucomicrobiales>Verrucomicrobiaceae | GCF_016595415.1 | Roseibacillus_ishigakijimensis | JIN78_RS11805 | 43..1545 | 454146 | hypothetical-protein-[Roseibacillus-ishigakijimensis] |
| WP_015675681.1 | Type6.BREX | <-SF1-SF2-Helicase+wHTH+Small-B-barrel+DUF1998<-BrxD-AAA<-PglZ<-N6-METHYLASE-p<-AbiEii<-AbiEi_3_N+AbiEi_3<-ParB+HNH<-N6-METHYLASE<-DUF6079*<-BrxB-BrxF-iSTAND<-BrxA-DNA-binding<-BrxE<-TPR+TPR<-TPR<-MarR-HTH<-ParB<-ParA-Soj-PloopNTPase | 1203 | Bacteria>Spirochaetota>Spirochaetia>Leptospirales>Leptospiraceae>Leptospira | GCF_000332475.2 | Leptospira_yanagawae | LEP1GSC202_0423 | 26532..30143 | 293069 | BREX-system-P-loop-protein-BrxC-[Leptospira-yanagawae] |
| WP_135697923.1 | Type6.BREX | <-DUF6079-p*<-BrxB-BrxF-iSTAND<-BrxA-DNA-binding<-BrxE<-TPR+TPR<-MarR-HTH<-ParB<-ParA-Soj-PloopNTPase | 907 | Bacteria>Spirochaetota>Spirochaetia>Leptospirales>Leptospiraceae>Leptospira | GCF_004770525.1 | Leptospira_meyeri | EHQ94_RS11050 | 1..2721 | 29508 | BREX-system-P-loop-protein-BrxC,-partial-[Leptospira-meyeri] |
| WP_135754988.1 | Type6.BREX | <-PglZ<-N6-METHYLASE-p<-?<-N6-METHYLASE<-DUF6079*<-BrxB-BrxF-iSTAND<-BrxA-DNA-binding<-BrxE<-TPR+TPR<-MarR-HTH<-ParB<-ParA-Soj-PloopNTPase | 1201 | Bacteria>Spirochaetota>Spirochaetia>Leptospirales>Leptospiraceae>Leptospira | GCF_004770625.1 | Leptospira_bouyouniensis | EHQ10_RS18560 | 6082..9687 | 2484911 | BREX-system-P-loop-protein-BrxC-[Leptospira-bouyouniensis] |
| WP_238754851.1 | Type6.BREX | ANK->ParA-Soj-PloopNTPase->ParB->MarR-HTH->TPR+TPR->BrxE->BrxA-DNA-binding->BrxB-BrxF-iSTAND->DUF6079*->N6-METHYLASE->?->?->?->ParB+HNH->AbiEi_3_N+AbiEi_3->DOC+HTH->N6-METHYLASE-p->PglZ-> | 1201 | Bacteria>Spirochaetota>Spirochaetia>Leptospirales>Leptospiraceae>Leptospira | GCF_022267385.1 | Leptospira_bandrabouensis | LFX13_RS16565 | 34041..37646 | 2484903 | BREX-system-P-loop-protein-BrxC-[Leptospira-bandrabouensis] |
| MBK6607326.1 | Type6.BREX | BrxE->BrxA-DNA-binding->BrxB-BrxF-iSTAND->DUF6079-p*->DUF6079-p*->N6-METHYLASE-p->N6-METHYLASE-p->URI-DNase->HTH+REase->N6-METHYLASE-p->?->N6-METHYLASE-p->PglZ-p-> | 320 | Bacteria>Spirochaetota>Spirochaetia>Leptospirales>Leptospiraceae>unclassified_Leptospiraceae | GCA_016703885.1 | Leptospiraceae_bacterium | IPG24_18070 | 61452..62414 | 2026757 | hypothetical-protein-[Leptospiraceae-bacterium] |
| MBK6607327.1 | Type6.BREX | BrxE->BrxA-DNA-binding->BrxB-BrxF-iSTAND->DUF6079-p->DUF6079-p*->N6-METHYLASE-p->N6-METHYLASE-p->URI-DNase->HTH+REase->N6-METHYLASE-p->?->N6-METHYLASE-p->PglZ-p-> | 877 | Bacteria>Spirochaetota>Spirochaetia>Leptospirales>Leptospiraceae>unclassified_Leptospiraceae | GCA_016703885.1 | Leptospiraceae_bacterium | IPG24_18075 | 62425..65058 | 2026757 | hypothetical-protein-[Leptospiraceae-bacterium] |
| OQB98767.1 | Type6.BREX | ANK+ANK-><-?<-?<-SIGMA+OB-Fold+AAA-ATPase+HKD-DNase-BrxL<-PglZ<-N6-METHYLASE<-?<-?<-N6-METHYLASE<-DUF6079*<-BrxB-BrxF-iSTAND<-BrxA-DNA-binding<-BrxE\|\|?->SF1-SF2-Helicase+DUF1998-><-REase<-Peptidase_C14 | 1205 | Bacteria>Spirochaetota>unclassified_Spirochaetota | GCA_002070755.1 | Spirochaetes_bacterium_ADurb.Bin110 | BWX81_00245 | 23321..26938 | 1852908 | hypothetical-protein-BWX81_00245-[Spirochaetes-bacterium-ADurb.Bin110] |
| QVL35508.1 | Type6.BREX | Arm-DNA-bind_3+Phage_integrase->XIS-HTH->TOPRIM->?-><-SF1-SF2-Helicase+REase<-DNA_METHYLASE<-SIGMA+OB-Fold+AAA-ATPase+Lon-Protease-BrxL<-PglZ<-N6-METHYLASE<-DUF6079*<-BrxB-BrxF-iSTAND<-BrxA-DNA-binding<-BrxE<-CHTH<-?\|\|HIN-HTH->HARE-HTH+REase-><-ABC-ATPase+SbcC | 1227 | Bacteria>Synergistota>unclassified_Synergistota | GCA_018398915.1 | Synergistota_bacterium | KIH16_09955 | 2165249..2168932 | 2268202 | BREX-system-P-loop-protein-BrxC-[Synergistota-bacterium] |
| WP_208027651.1 | Type6.BREX | ACET-><-?<-REC+EAL\|\|HISKIN->REC+OmpR-HTH->?-><-SIGMA+OB-Fold+AAA-ATPase+Lon-Protease-BrxL<-PglZ<-N6-METHYLASE<-DUF6079*<-BrxB-BrxF-iSTAND<-BrxA-DNA-binding<-BrxE\|\|DUF433->?->Sulfatase-> | 1163 | Bacteria>Terrabacteria_group>Actinomycetota>Acidimicrobiia>Acidimicrobiales>Iamiaceae | GCF_014805525.1 | Rhabdothermincola_sediminis | HZF19_RS04995 | 196224..199715 | 2751370 | BREX-system-P-loop-protein-BrxC-[Rhabdothermincola-sediminis] |
| MCB1001743.1 | Type6.BREX | <-ARPP2<-SF1-SF2-Helicase<-35exo+TUDOR+SF1-SF2-Helicase+iREase+CTD<-TrwC+SF1-SF2-Helicase+TOPRIM<-?<-SIGMA-HTH\|\|BrxE->BrxA-DNA-binding->BrxB-BrxF-iSTAND->DUF6079*->N6-METHYLASE->PglZ->SIGMA+OB-Fold+AAA-ATPase+Lon-Protease-BrxL-><-?<-?<-REase<-SbcC<-JetB<-JetA | 1151 | Bacteria>Terrabacteria_group>Actinomycetota>Acidimicrobiia>Acidimicrobiales>unclassified_Acidimicrobiales | GCA_020439435.1 | Acidimicrobiales_bacterium | KDB40_20800 | 40059..43514 | 2201156 | BREX-system-P-loop-protein-BrxC-[Acidimicrobiales-bacterium] |
| SBW20208.1 | Type6.BREX | BrxE->BrxA-DNA-binding->BrxB-BrxF-iSTAND->DUF6079*->Schlafen+HATPase+wHTH->N6-METHYLASE->N6-METHYLASE-> | 1180 | Bacteria>Terrabacteria_group>Actinomycetota>Actinomycetes>Frankiales>Frankiaceae | - | Candidatus_Frankia_californiensis | FDG2_1623 | 2594..6136 | 1839754 | hypothetical-protein-FDG2_1623-[Candidatus-Frankia-californiensis] |
| KQC36858.1 | Type6.BREX | REase->?->HicB_lk_antitox-><-?<-SF1-SF2-Helicase+wHTH+Small-B-barrel+DUF1998<-BrxD-AAA<-PglZ<-N6-METHYLASE<-DUF6079*<-BrxB-BrxF-iSTAND<-BrxA-DNA-binding<-BrxE<-?<-HTH_32+DDE_3 | 1205 | Bacteria>Terrabacteria_group>Actinomycetota>Actinomycetes>Frankiales>Frankiaceae | GCA_001414035.1 | Frankia_sp._ACN1ag | UK82_18990 | 119450..123067 | 102891 | hypothetical-protein-UK82_18990-[Frankia-sp.-ACN1ag] |
| PGH49734.1 | Type6.BREX | <-LysM+Peptidase_M23<-?<-?\|\|NTP_transf_5->\|\|?-><-SF1-SF2-Helicase+wHTH+Small-B-barrel+DUF1998<-BrxD-AAA<-PglZ<-N6-METHYLASE<-DUF6079*<-BrxB-BrxF-iSTAND<-BrxA-DNA-binding<-BrxE<-pPIWI+MID+RNaseH-p<-pPIWI+SF1-SF2-Helicase<-pPIWI+pPIWI<-HTH_3+HTH_3 | 1206 | Bacteria>Terrabacteria_group>Actinomycetota>Actinomycetes>Kitasatosporales>Streptomycetaceae | GCA_002573955.1 | Streptomyces_sp._Ru87 | CRI70_15985 | 31482..35102 | 2044307 | hypothetical-protein-CRI70_15985-[Streptomyces-sp.-Ru87] |
| UJV46968.1 | Type6.BREX | <-VirB6<-?<-TraM<-?\|\|N6-N4-METHYLASE->\|\|BrxE->BrxA-DNA-binding->BrxB-BrxF-iSTAND->DUF6079*->N6-METHYLASE->PglZ->BrxD-AAA->SF1-SF2-Helicase+wHTH+Small-B-barrel+35exo->XIS-HTH->Phage_integrase->N6-METHYLASE-><-SF1-SF2-Helicase<-McrC-NTD+REase | 1206 | Bacteria>Terrabacteria_group>Actinomycetota>Actinomycetes>Kitasatosporales>Streptomycetaceae | GCA_021665875.1 | Streptomyces_sp._AMCC400023 | CVT30_32940 | 7360114..7363734 | 2056258 | BREX-system-P-loop-protein-BrxC-[Streptomyces-sp.-AMCC400023] |
| WP_218940480.1 | Type6.BREX | <-LysM+Peptidase_M23<-LysM+Peptidase_M23<-?\|\|NTP_transf_5->?-><-SF1-SF2-Helicase+wHTH+Small-B-barrel+DUF1998<-BrxD-AAA<-PglZ<-N6-METHYLASE<-DUF6079*<-BrxB-BrxF-iSTAND<-BrxA-DNA-binding<-BrxE<-pPIWI+MID+RNaseH<-pPIWI+SF1-SF2-Helicase<-pPIWI+pPIWI<-HTH_3+HTH_3 | 1210 | Bacteria>Terrabacteria_group>Actinomycetota>Actinomycetes>Kitasatosporales>Streptomycetaceae | GCF_002573955.1 | Streptomyces_sp._Ru87 | CRI70_RS15985 | 31482..35114 | 2044307 | BREX-system-P-loop-protein-BrxC-[Streptomyces-sp.-Ru87] |
| WP_128375281.1 | Type6.BREX | BrxA-DNA-binding->?->?->BrxE->BrxA-DNA-binding->BrxB-BrxF-iSTAND->DUF6079*->N6-METHYLASE->PglZ->BrxD-AAA->SF1-SF2-Helicase+wHTH+Small-B-barrel+35exo->HTH_3+HTH_3->pPIWI+pPIWI->pPIWI+SF1-SF2-Helicase->pPIWI+MID+RNaseH->OLD-ABC+TOPRIM-> | 1206 | Bacteria>Terrabacteria_group>Actinomycetota>Actinomycetes>Kitasatosporales>Streptomycetaceae | GCF_004023625.1 | Streptomyces_cavernae | EF387_RS05155 | 27680..31300 | 2259034 | BREX-system-P-loop-protein-BrxC-[Streptomyces-cavernae] |
| WP_237310553.1 | Type6.BREX | <-VirB6<-?<-TraM<-?\|\|N6-N4-METHYLASE->\|\|BrxE->BrxA-DNA-binding->BrxB-BrxF-iSTAND->DUF6079*->N6-METHYLASE->PglZ->BrxD-AAA->SF1-SF2-Helicase+wHTH+Small-B-barrel+35exo->XIS-HTH->Phage_integrase->N6-METHYLASE-><-SF1-SF2-Helicase<-McrC-NTD+REase | 1210 | Bacteria>Terrabacteria_group>Actinomycetota>Actinomycetes>Kitasatosporales>Streptomycetaceae | GCF_021665875.1 | Streptomyces_sp._AMCC400023 | CVT30_RS32680 | 7360102..7363734 | 2056258 | BREX-system-P-loop-protein-BrxC-[Streptomyces-sp.-AMCC400023] |
| WP_252545223.1 | Type6.BREX | ABhydrolase-><-?\|\|?->?-><-?\|\|SWACOS->BrxE->BrxA-DNA-binding->BrxB-BrxF-iSTAND->DUF6079*->N6-METHYLASE->PglZ->BrxD-AAA->SF1-SF2-Helicase+wHTH+Small-B-barrel+35exo->HTH_3+HTH_3->pPIWI+pPIWI->pPIWI+SF1-SF2-Helicase->pPIWI+MID+RNaseH-> | 1210 | Bacteria>Terrabacteria_group>Actinomycetota>Actinomycetes>Kitasatosporales>Streptomycetaceae | GCF_023923245.1 | Streptomyces_phaeoluteigriseus | NFX46_RS01495 | 327017..330649 | 114686 | BREX-system-P-loop-protein-BrxC-[Streptomyces-phaeoluteigriseus] |
| KKK07472.1 | Type6.BREX | HTH_3->?-><-?\|\|XIS-HTH->BrxE->BrxA-DNA-binding->BrxB-BrxF-iSTAND->DUF6079*->PglZ->BrxD-AAA->pPIWI+pPIWI->pPIWI+SF1-SF2-Helicase->pPIWI+MID+RNaseH-><-tRNA<-?\|\|?-><-?\|\|AraC-HTH+AraC-HTH-> | 1206 | Bacteria>Terrabacteria_group>Actinomycetota>Actinomycetes>Micromonosporales>Micromonosporaceae | GCA_000982955.1 | Micromonospora_sp._HK10 | LQ51_02370 | 30923..34543 | 1538294 | hypothetical-protein-LQ51_02370-[Micromonospora-sp.-HK10] |
| WP_030441358.1 | Type6.BREX | <-APATPase+TPR+TPR+TPR+TPR<-?<-?\|\|?-><-SIGMA+OB-Fold+AAA-ATPase+Lon-Protease-BrxL<-PglZ<-N6-METHYLASE<-N6-METHYLASE-p<-DUF6079*<-BrxB-BrxF-iSTAND<-BrxA-DNA-binding<-BrxE<-?<-wHTH | 1180 | Bacteria>Terrabacteria_group>Actinomycetota>Actinomycetes>Micromonosporales>Micromonosporaceae | GCF_000721705.1 | Actinoplanes_subtropicus | IH26_RS0139195 | 17752..21294 | 543632 | BREX-system-P-loop-protein-BrxC-[Actinoplanes-subtropicus] |
| WP_197087687.1 | Type6.BREX | HTH_3->?->XIS-HTH->BrxE->BrxA-DNA-binding->BrxB-BrxF-iSTAND->DUF6079*->N6-METHYLASE->PglZ->BrxD-AAA->SF1-SF2-Helicase+wHTH+Small-B-barrel+35exo->pPIWI+pPIWI->pPIWI+SF1-SF2-Helicase->pPIWI+MID+RNaseH-> | 1210 | Bacteria>Terrabacteria_group>Actinomycetota>Actinomycetes>Micromonosporales>Micromonosporaceae | GCF_000982955.1 | Micromonospora_sp._HK10 | LQ51_RS12845 | 30911..34543 | 1538294 | BREX-system-P-loop-protein-BrxC-[Micromonospora-sp.-HK10] |
| WP_233625589.1 | Type6.BREX | <-DUF6079-p*<-BrxB-BrxF-iSTAND<-BrxA-DNA-binding<-BrxE\|\|McrC-NTD+REase->McrC-NTD+REase-><-DDE_Tnp | 383 | Bacteria>Terrabacteria_group>Actinomycetota>Actinomycetes>Micromonosporales>Micromonosporaceae | GCF_003947245.1 | Actinoplanes_sp._ATCC_53533 | DMB66_RS58195 | 1..1150 | 1288362 | hypothetical-protein,-partial-[Actinoplanes-sp.-ATCC-53533] |
| WP_208468634.1 | Type6.BREX | XIS-HTH->URI-DNase->REase->BrxE->BrxA-DNA-binding->BrxB-BrxF-iSTAND->DUF6079*->Schlafen+HATPase+wHTH->N6-METHYLASE->N6-METHYLASE->PglZ->SIGMA+OB-Fold+AAA-ATPase+Lon-Protease-BrxL-><-DDE_Tnp_1_assoc+DDE_Tnp_1 | 1183 | Bacteria>Terrabacteria_group>Actinomycetota>Actinomycetes>Micromonosporales>Micromonosporaceae | GCF_017592555.1 | Actinoplanes_flavus | J5X75_RS18430 | 129103..132654 | 2820290 | BREX-system-P-loop-protein-BrxC-[Actinoplanes-flavus] |
| WP_250791051.1 | Type6.BREX | <-DUF6307<-?<-?\|\|HTH_3->STYKIN-><-SF1-SF2-Helicase+wHTH+Small-B-barrel+35exo<-BrxD-AAA<-PglZ<-N6-METHYLASE<-DUF6079*<-BrxB-BrxF-iSTAND<-BrxA-DNA-binding<-BrxE<-XIS-HTH<-SF1-SF2-Helicase<-SF1-SF2-Helicase<-ParB+HNH<-?<-REase | 1210 | Bacteria>Terrabacteria_group>Actinomycetota>Actinomycetes>Micromonosporales>Micromonosporaceae | GCF_023656545.1 | Micromonospora_phytophila | NCC78_RS00575 | 40120..43752 | 709888 | BREX-system-P-loop-protein-BrxC-[Micromonospora-phytophila] |
| WP_269739547.1 | Type6.BREX | <-RADICAL-SAM<-?\|\|HTH_3->?->REase-><-SF1-SF2-Helicase+wHTH+Small-B-barrel+35exo<-BrxD-AAA<-PglZ<-N6-METHYLASE<-DUF6079*<-BrxB-BrxF-iSTAND<-BrxA-DNA-binding<-BrxE<-XIS-HTH<-Phage_int_SAM_3+Phage_integrase<-XIS-HTH<-?<-BirA-HTH\|\|NUDIX-> | 1209 | Bacteria>Terrabacteria_group>Actinomycetota>Actinomycetes>Micromonosporales>Micromonosporaceae | GCF_027460185.1 | Verrucosispora_sp._WMMA2121 | O7605_RS13740 | 1051345..1054974 | 3015164 | BREX-system-P-loop-protein-BrxC-[Verrucosispora-sp.-WMMA2121] |
| OBF48784.1 | Type6.BREX | HEPN+SF1-SF2-Helicase<-?<-HIRAN-><-HKD-DNase<-?\|\|BrxE->REase->BrxE->BrxA-DNA-binding->BrxB-BrxF-iSTAND->DUF6079*->N6-METHYLASE->PglZ->SIGMA+OB-Fold+AAA-ATPase+Lon-Protease-BrxL-><-SF1-SF2-Helicase+REase\|\|?-><-SF1-SF2-Helicase+REase | 1143 | Bacteria>Terrabacteria_group>Actinomycetota>Actinomycetes>Mycobacteriales>Mycobacteriaceae | GCA_001667465.1 | Mycolicibacterium_monacense | A5778_22445 | 215738..219169 | 85693 | hypothetical-protein-A5778_22445-[Mycolicibacterium-monacense] |
| MBP2454134.1 | Type6.BREX | <-McrB-AAA<-Preader7<-HEPN+SF1-SF2-Helicase+REase<-?\|\|BrxE-p->BrxA-DNA-binding->BrxB-BrxF-iSTAND->DUF6079-p->zf-ISL3+HTH_Tnp_ISL3+DDE_Tnp_ISL3->DUF6079-p*->N6-METHYLASE-p-><-REase<-N6-METHYLASE<-?<-TetR+HTH<-Phage_integrase+Phage_integrase<-Phage_integrase+Phage_integrase<-Phage_int_SAM_1+Phage_integrase<-DNA_glycosylase | 870 | Bacteria>Terrabacteria_group>Actinomycetota>Actinomycetes>Mycobacteriales>Mycobacteriaceae | GCA_017876775.1 | Mycolicibacterium_lutetiense | J0F57_004047 | 2230868..2233480 | 1641992 | hypothetical-protein-[Mycolicibacterium-lutetiense] |
| WP_120314218.1 | Type6.BREX | HEPN+SF1-SF2-Helicase+REase->HIRAN-><-HKD-DNase+HKD-DNase<-?\|\|BrxE->BrxA-DNA-binding->BrxB-BrxF-iSTAND->DUF6079*->N6-METHYLASE->PglZ->SIGMA+OB-Fold+AAA-ATPase+Lon-Protease-BrxL-><-SF1-SF2-Helicase+REase<-SF1-SF2-Helicase<-SF1-SF2-Helicase+REase<-?<-?<-SbcC<-SbcC | 1163 | Bacteria>Terrabacteria_group>Actinomycetota>Actinomycetes>Mycobacteriales>Mycobacteriaceae | GCF_001667465.1 | Mycolicibacterium_monacense | A5778_RS22240 | 215678..219169 | 85693 | BREX-system-P-loop-protein-BrxC-[Mycolicibacterium-monacense] |
| WP_068042721.1 | Type6.BREX | <-Phage_integrase<-?<-ZnR<-AAA-ATPase<-ParB+HNH\|\|?-><-SIGMA+OB-Fold+AAA-ATPase+Lon-Protease-BrxL<-PglZ<-N6-METHYLASE<-DUF6079*<-BrxB-BrxF-iSTAND<-BrxA-DNA-binding<-BrxE\|\|ABhydrolase+ACYC-> | 1161 | Bacteria>Terrabacteria_group>Actinomycetota>Actinomycetes>Mycobacteriales>Mycobacteriaceae | GCF_001667585.1 | Mycobacterium_sp._E2733 | A5678_RS12510 | 65198..68683 | 1834138 | BREX-system-P-loop-protein-BrxC-[Mycobacterium-sp.-E2733] |
| WP_166680331.1 | Type6.BREX | <-REase<-?<-?<-?\|\|?-><-SIGMA+OB-Fold+AAA-ATPase+Lon-Protease-BrxL<-PglZ<-N6-METHYLASE<-DUF6079*<-BrxB-BrxF-iSTAND<-BrxA-DNA-binding<-BrxE\|\|RAMA-> | 1161 | Bacteria>Terrabacteria_group>Actinomycetota>Actinomycetes>Propionibacteriales>Kribbellaceae | GCF_004366005.1 | Kribbella_sp._VKM_Ac-2566 | EV647_RS14620 | 664432..667917 | 2512218 | BREX-system-P-loop-protein-BrxC-[Kribbella-sp.-VKM-Ac-2566] |
| WP_228644184.1 | Type6.BREX | HEPN+APATPase+SlCOLD->?-><-SF1-SF2-Helicase+wHTH+Small-B-barrel+35exo<-BrxD-AAA<-PglZ<-N6-METHYLASE<-DUF6079*<-BrxB-BrxF-iSTAND<-BrxA-DNA-binding<-BrxE<-HTH_Tnp_1+HTH_21+rve+rve_3\|\|HIN-HTH-> | 1205 | Bacteria>Terrabacteria_group>Actinomycetota>Actinomycetes>Streptosporangiales>Streptosporangiaceae | GCF_020826795.1 | Microtetraspora_sp._AC03309 | IMZ11_RS21685 | 78339..81956 | 2779376 | BREX-system-P-loop-protein-BrxC-[Microtetraspora-sp.-AC03309] |
| WP_091094284.1 | Type6.BREX | <-MULE-TRANSPOSASE\|\|TPR-><-?\|\|SF1-SF2-Helicase->BrxE->BrxA-DNA-binding->BrxB-BrxF-iSTAND->DUF6079*->N6-METHYLASE->N6-METHYLASE->PglZ->SIGMA+OB-Fold+AAA-ATPase+Lon-Protease-BrxL-><-DDE_Tnp\|\|<-RVT\|\|REase-><-ParB+HNH\|\|DDE_Tnp_ISL3-> | 1179 | Bacteria>Terrabacteria_group>Actinomycetota>Actinomycetes>Streptosporangiales>Streptosporangiaceae | GCF_900111685.1 | Nonomuraea_wenchangensis | BMW85_RS47095 | 18289..21828 | 568860 | BREX-system-P-loop-protein-BrxC-[Nonomuraea-wenchangensis] |
| WP_111865227.1 | Type6.BREX | <-LuxR-HTH<-?<-BetaPropeller+ABhydrolase<-LuxR-HTH\|\|?->DUF4096+DDE_Tnp_1+DDE_Tnp_1_2->BrxE->BrxA-DNA-binding->BrxB-BrxF-iSTAND->DUF6079*->N6-METHYLASE->PglZ->BrxD-AAA->SF1-SF2-Helicase+wHTH+Small-B-barrel+35exo->?-><-MPTase<-HTH_3 | 1205 | Bacteria>Terrabacteria_group>Actinomycetota>Actinomycetes>Streptosporangiales>Thermomonosporaceae | GCF_003289645.1 | Actinomadura_craniellae | DPM19_RS09740 | 52224..55841 | 2231787 | BREX-system-P-loop-protein-BrxC-[Actinomadura-craniellae] |
| WP_131935960.1 | Type6.BREX | Calcineurin->SAM-METHYLASE->tRNA-><-?<-SF1-SF2-Helicase+wHTH+Small-B-barrel+35exo<-BrxD-AAA<-PglZ<-N6-METHYLASE<-DUF6079*<-BrxB-BrxF-iSTAND<-BrxA-DNA-binding<-BrxE<-XIS-HTH<-?<-Relaxase<-?<-HISKIN\|\|HTH_31+DUF5753-> | 1205 | Bacteria>Terrabacteria_group>Actinomycetota>Actinomycetes>Streptosporangiales>Thermomonosporaceae | GCF_004348335.1 | Actinomadura_bangladeshensis | E1284_RS00900 | 46072..49689 | 453573 | BREX-system-P-loop-protein-BrxC-[Actinomadura-bangladeshensis] |
| KUK48423.1 | Type6.BREX | CHTH->BrxE->BrxA-DNA-binding->BrxB-BrxF-iSTAND->DUF6079*->N6-METHYLASE->OLD-ABC+TOPRIM->PglZ->SIGMA+OB-Fold+AAA-ATPase+Lon-Protease-BrxL->DNA_METHYLASE->SF1-SF2-Helicase+REase->REase-> | 1227 | Bacteria>Terrabacteria_group>Actinomycetota>Actinomycetes>unclassified_Actinomycetes | GCA_001508275.1 | Actinobacteria_bacterium_66_15 | XD74_0976 | 7706..11389 | 1635289 | Uncharacterized-protein-XD74_0976-[Actinobacteria-bacterium-66_15] |
| MBX7132279.1 | Type6.BREX | <-REase<-?<-ARG\|\|NUDIX->?->?->BrxE->BrxA-DNA-binding->BrxB-BrxF-iSTAND->DUF6079*->N6-METHYLASE->PglZ->SIGMA+OB-Fold+AAA-ATPase+Lon-Protease-BrxL->McrB-AAA-> | 1161 | Bacteria>Terrabacteria_group>Armatimonadota>Fimbriimonadia>Fimbriimonadales>Fimbriimonadaceae | GCA_019694835.1 | Fimbriimonadaceae_bacterium | K1X67_06340 | 6954..10439 | 2682144 | BREX-system-P-loop-protein-BrxC-[Fimbriimonadaceae-bacterium] |
| WP_075749471.1 | Type6.BREX | <-TetR-HTH\|\|?->?->TRAM+METHYLASE->BrxE->BrxA-DNA-binding->BrxB-BrxF-iSTAND->DUF6079*->N6-METHYLASE->PglZ->BrxD-AAA->?-><-?<-?<-WXG+PT+HNH\|\|?->REase+DUF5655-> | 1209 | Bacteria>Terrabacteria_group>Bacillota>Bacilli>Bacillales>Bacillaceae | - | Bacillus_licheniformis | BHT94_10545 | 89762..93391 | 1402 | BREX-system-P-loop-protein-BrxC-[Bacillus-licheniformis] |
| APC47779.1 | Type6.BREX | <-TetR-HTH\|\|ABC-ATPase+SbcC->?->HEPN+PAS+DUF4011+SF1-SF2-Helicase+REase->REase->BrxE->BrxA-DNA-binding->BrxB-BrxF-iSTAND->DUF6079*->N6-METHYLASE->PglZ->BrxD-AAA->SF1-SF2-Helicase+wHTH+Small-B-barrel+DUF1998->ParB+HNH->NUDIX->?-><-HKD-DNase+SF1-SF2-Helicase+HTH+HTH+SRA | 1225 | Bacteria>Terrabacteria_group>Bacillota>Bacilli>Bacillales>Bacillaceae | GCA_001878675.1 | Virgibacillus_halodenitrificans | BME96_06155 | 1179738..1183415 | 1482 | hypothetical-protein-BME96_06155-[Virgibacillus-halodenitrificans] |
| PJW20776.1 | Type6.BREX | SF1-SF2-Helicase->REase->?-><-?<-?\|\|HTH->BrxE->BrxA-DNA-binding->BrxB-BrxF-iSTAND->DUF6079*->N6-METHYLASE->PglZ->BrxD-AAA->SF1-SF2-Helicase+wHTH+Small-B-barrel+DUF1998->HTH_Tnp+HTH_Tnp->Phage_integrase-><-Phage_int_SAM_1+Phage_integrase | 1210 | Bacteria>Terrabacteria_group>Bacillota>Bacilli>Bacillales>Bacillaceae | GCA_002810005.1 | Geobacillus_thermodenitrificans | CV632_09710 | 419847..423479 | 33940 | hypothetical-protein-CV632_09710-[Geobacillus-thermodenitrificans] |
| WP_028784627.1 | Type6.BREX | ArdCN+MPTase-inact->TOPRIM->BrxE->BrxA-DNA-binding->BrxB-BrxF-iSTAND->DUF6079*->N6-METHYLASE->PglZ->ArsR-HTH->?->?->PNPase-> | 1207 | Bacteria>Terrabacteria_group>Bacillota>Bacilli>Bacillales>Bacillaceae | GCF_000496835.1 | Thalassobacillus_devorans | P855_RS0116640 | 86998..90621 | 279813 | BREX-system-P-loop-protein-BrxC-[Thalassobacillus-devorans] |
| WP_032871746.1 | Type6.BREX | WXG+PT+HNH->?->?-><-BrxD-AAA<-PglZ<-N6-METHYLASE<-DUF6079*<-BrxB-BrxF-iSTAND<-BrxA-DNA-binding<-BrxE<-TRAM+METHYLASE<-?<-?\|\|TetR-HTH-> | 1208 | Bacteria>Terrabacteria_group>Bacillota>Bacilli>Bacillales>Bacillaceae | GCF_000746035.1 | Bacillus_sp._UNC69MF | BR62_RS0118440 | 55963..59589 | 1449047 | BREX-system-P-loop-protein-BrxC-[Bacillus-sp.-UNC69MF] |
| WP_197035844.1 | Type6.BREX | <-TetR-HTH\|\|ABC-ATPase+SbcC->?->HEPN+PAS+DUF4011+SF1-SF2-Helicase+REase->BrxE->BrxA-DNA-binding->BrxB-BrxF-iSTAND->DUF6079*->N6-METHYLASE->PglZ->BrxD-AAA->SF1-SF2-Helicase+wHTH+Small-B-barrel+DUF1998->ParB+HNH->NUDIX-><-HKD-DNase+SF1-SF2-Helicase+HTH+HTH+SRA<-?<-zf-ISL3+HTH_Tnp_ISL3+DDE_Tnp_ISL3 | 1207 | Bacteria>Terrabacteria_group>Bacillota>Bacilli>Bacillales>Bacillaceae | GCF_001878675.1 | Virgibacillus_halodenitrificans | BME96_RS06225 | 1179792..1183415 | 1482 | BREX-system-P-loop-protein-BrxC-[Virgibacillus-halodenitrificans] |
| WP_080529022.1 | Type6.BREX | TRAM+METHYLASE->BrxE->BrxA-DNA-binding->BrxB-BrxF-iSTAND->DUF6079*->N6-METHYLASE->PglZ->BrxD-AAA->?-><-?<-WXG+PT+HNH\|\|?->REase+DUF5655->?->TPR+TPR+TPR-> | 1209 | Bacteria>Terrabacteria_group>Bacillota>Bacilli>Bacillales>Bacillaceae | GCF_002072735.1 | Bacillus_subtilis | FC605_RS03715 | 716511..720140 | 1423 | BREX-system-P-loop-protein-BrxC-[Bacillus-subtilis] |
| WP_094244383.1 | Type6.BREX | TRAM+METHYLASE->BrxE->?->BrxA-DNA-binding->BrxB-BrxF-iSTAND->DUF6079*->N6-METHYLASE->p-><-DEDD_Tnp_IS110+Transposase_20\|\|N6-METHYLASE->PglZ->BrxD-AAA->SF1-SF2-Helicase+wHTH+Small-B-barrel+DUF1998-p-><-LZ_Tnp_IS66+zf-IS66+DDE_Tnp_IS66+DDE_Tnp_IS66_C<-?<-HIN-HTH | 1205 | Bacteria>Terrabacteria_group>Bacillota>Bacilli>Bacillales>Bacillaceae | GCF_002243625.1 | Aeribacillus_pallidus | AP3564_00890 | 191752..195369 | 33936 | BREX-system-P-loop-protein-BrxC-[Aeribacillus-pallidus] |
| WP_157806984.1 | Type6.BREX | REase->?-><-?<-?<-?\|\|HTH->BrxE->BrxA-DNA-binding->BrxB-BrxF-iSTAND->DUF6079*->N6-METHYLASE->PglZ->BrxD-AAA->SF1-SF2-Helicase+wHTH+Small-B-barrel+DUF1998->HTH_Tnp+HTH_Tnp->Phage_integrase-><-Phage_int_SAM_1+Phage_integrase | 1207 | Bacteria>Terrabacteria_group>Bacillota>Bacilli>Bacillales>Bacillaceae | GCF_002810005.1 | Geobacillus_thermodenitrificans | CV632_RS09695 | 419856..423479 | 33940 | BREX-system-P-loop-protein-BrxC-[Geobacillus-thermodenitrificans] |
| WP_106034770.1 | Type6.BREX | <-TetR-HTH\|\|?->?->TRAM+METHYLASE->BrxE->BrxA-DNA-binding->BrxB-BrxF-iSTAND->DUF6079*->N6-METHYLASE->PglZ->BrxD-AAA->SF1-SF2-Helicase-p-><-?<-?<-WXG+PT+HNH\|\|TPR+TPR+TPR-> | 1208 | Bacteria>Terrabacteria_group>Bacillota>Bacilli>Bacillales>Bacillaceae | GCF_002998075.1 | Bacillus_atrophaeus | C6W23_RS17370 | 106732..110358 | 1452 | BREX-system-P-loop-protein-BrxC-[Bacillus-atrophaeus] |
| WP_151706155.1 | Type6.BREX | Resolvase+Recombinase+ZnR->Recombinase-><-?<-?<-SF1-SF2-Helicase+wHTH+Small-B-barrel+DUF1998<-BrxD-AAA<-PglZ<-N6-METHYLASE<-DUF6079*<-BrxB-BrxF-iSTAND<-BrxA-DNA-binding<-BrxE | 1205 | Bacteria>Terrabacteria_group>Bacillota>Bacilli>Bacillales>Bacillaceae | GCF_008974205.1 | Weizmannia_acidilactici | GBI00_RS13195 | 20055..23672 | 2607726 | BREX-system-P-loop-protein-BrxC-[Weizmannia-acidilactici] |
| WP_174532678.1 | Type6.BREX | WXG+PT+HNH->?->?-><-BrxD-AAA<-PglZ<-N6-METHYLASE-p<-DUF6079*<-BrxB-BrxF-iSTAND<-BrxA-DNA-binding<-BrxE<-TRAM+METHYLASE<-?\|\|TetR-HTH-> | 1208 | Bacteria>Terrabacteria_group>Bacillota>Bacilli>Bacillales>Bacillaceae | GCF_013341255.1 | Bacillus_amyloliquefaciens | HT132_RS17105 | 3489605..3493231 | 1390 | BREX-system-P-loop-protein-BrxC-[Bacillus-amyloliquefaciens] |
| WP_215020742.1 | Type6.BREX | Phage_int_SAM_1+Phage_integrase->?->HTH_29+rve+rve_3->?->GspA-ATPase-><-SF1-SF2-Helicase+wHTH+Small-B-barrel+DUF1998<-BrxD-AAA<-PglZ<-N6-METHYLASE<-DUF6079*<-BrxB-BrxF-iSTAND<-BrxA-DNA-binding<-BrxE\|\|GDResolvase+HIN-HTH-><-?<-ABhydrolase | 1207 | Bacteria>Terrabacteria_group>Bacillota>Bacilli>Bacillales>Bacillaceae | GCF_018613245.1 | Bacillus_sp._ISL-40 | J7E79_RS03690 | 15057..18680 | 2819126 | BREX-system-P-loop-protein-BrxC-[Bacillus-sp.-ISL-40] |
| WP_251188026.1 | Type6.BREX | <-REase+DUF5655<-?\|\|WXG+PT+HNH->?->?-><-?<-BrxD-AAA<-PglZ<-N6-METHYLASE<-DUF6079*<-BrxB-BrxF-iSTAND<-BrxA-DNA-binding<-BrxE<-TRAM+METHYLASE | 1209 | Bacteria>Terrabacteria_group>Bacillota>Bacilli>Bacillales>Bacillaceae | GCF_026795275.1 | Bacillus_subtilis | MOE40_RS05525 | 198177..201806 | 1423 | BREX-system-P-loop-protein-BrxC-[Bacillus-subtilis] |
| WP_263326060.1 | Type6.BREX | GDResolvase+Recombinase->AAA-ATPase->REase->BrxE->BrxA-DNA-binding->BrxB-BrxF-iSTAND->DUF6079*->N6-METHYLASE->PglZ->SIGMA+OB-Fold+AAA-ATPase+HKD-DNase-BrxL->Trypsin->GDResolvase+HIN-HTH->ParB+HNH->HEPN-> | 1207 | Bacteria>Terrabacteria_group>Bacillota>Bacilli>Bacillales>Bacillaceae | GCF_946151075.1 | Neobacillus_sp._Marseille-Q6967 | OGZ30_RS15230 | 466591..470214 | 2963105 | BREX-system-P-loop-protein-BrxC-[Neobacillus-sp.-Marseille-Q6967] |
| ETI91650.1 | Type6.BREX | LZ_Tnp_IS66+zf-IS66+DDE_Tnp_IS66+DDE_Tnp_IS66_C->?-><-URI-DNase<-SF1-SF2-Helicase+wHTH+Small-B-barrel+DUF1998<-BrxD-AAA<-PglZ<-N6-METHYLASE<-DUF6079*<-BrxB-BrxF-iSTAND<-BrxA-DNA-binding<-BrxE<-RHH_8<-?<-SF1-SF2-Helicase<-?<-?<-TnsC-ATPase | 1215 | Bacteria>Terrabacteria_group>Bacillota>Clostridia>Eubacteriales>Clostridiaceae | GCA_000508465.1 | Clostridium_butyricum_DORA_1 | Q607_CBUC00021G0092 | 106260..109907 | 1403941 | hypothetical-protein-Q607_CBUC00021G0092-[Clostridium-butyricum-DORA_1] |
| WP_055669755.1 | Type6.BREX | 35exo+BRCT->?->wHTH+HNH->?->?-><-REase\|\|BrxE->BrxA-DNA-binding->BrxB-BrxF-iSTAND->DUF6079*->N6-METHYLASE->PglZ->SIGMA+OB-Fold+AAA-ATPase+Lon-Protease-BrxL->DOC+HTH->RNase->?->?->?->transposase-> | 1232 | Bacteria>Terrabacteria_group>Bacillota>Clostridia>Eubacteriales>Clostridiaceae | GCF_001403615.1 | Desnuesiella_massiliensis | BN2665_RS24780 | 1439782..1443480 | 1650662 | BREX-system-P-loop-protein-BrxC-[Desnuesiella-massiliensis] |
| WP_103082719.1 | Type6.BREX | HTH_28+HTH_32+rve+rve_3+Mu-transpos_C->GspA-ATPase-><-?<-SF1-SF2-Helicase+wHTH+Small-B-barrel+DUF1998<-BrxD-AAA<-PglZ<-N6-METHYLASE<-DUF6079*<-BrxB-BrxF-iSTAND<-BrxA-DNA-binding<-BrxE<-?<-RHH_8<-?\|\|?-><-SF1-SF2-Helicase | 1207 | Bacteria>Terrabacteria_group>Bacillota>Clostridia>Eubacteriales>Oscillospiraceae | GCF_002896855.1 | Pseudoclostridium_thermosuccinogenes | CDO33_RS17620 | 4030030..4033653 | 84032 | BREX-system-P-loop-protein-BrxC-[Pseudoclostridium-thermosuccinogenes] |
| MCL6478738.1 | Type6.BREX | <-SIGMA+OB-Fold+AAA-ATPase+Lon-Protease-BrxL<-PglZ<-N6-METHYLASE<-DUF6079*<-BrxB-BrxF-iSTAND<-BrxA-DNA-binding<-BrxE\|\|Calcineurin+Calcineurin-SbcD-> | 1193 | Bacteria>Terrabacteria_group>Bacillota>Clostridia>Eubacteriales>Peptococcaceae | GCA_023511305.1 | Peptococcaceae_bacterium | K6T65_10015 | 10799..14380 | 2052179 | BREX-system-P-loop-protein-BrxC-[Peptococcaceae-bacterium] |
| WP_072866936.1 | Type6.BREX | <-MNS+REase\|\|Polbeta-><-SIGMA+OB-Fold+AAA-ATPase+Lon-Protease-BrxL<-PglZ\|\|STAS+HisKin->?-><-N6-METHYLASE<-DUF6079*<-BrxB-BrxF-iSTAND<-BrxA-DNA-binding<-BrxE | 1188 | Bacteria>Terrabacteria_group>Bacillota>Clostridia>Eubacteriales>Peptococcaceae | GCF_900142025.1 | Desulfofundulus_thermosubterraneus | BUA65_RS01250 | 259510..263076 | 348840 | BREX-system-P-loop-protein-BrxC-[Desulfofundulus-thermosubterraneus] |
| WP_206924346.1 | Type6.BREX | <-SF1-SF2-Helicase+wHTH+Small-B-barrel+DUF1998-p<-SF1-SF2-Helicase+wHTH+Small-B-barrel+DUF1998-p<-BrxD-AAA<-PglZ<-N6-METHYLASE<-DUF6079*<-BrxB-BrxF-iSTAND<-BrxA-DNA-binding<-BrxE\|\|HIN-HTH->?-><-?<-?<-?<-HEPN | 1206 | Bacteria>Terrabacteria_group>Bacillota>Clostridia>Eubacteriales>Peptostreptococcaceae | - | Terrisporobacter_glycolicus | JOL31_RS08125 | 1701033..1704653 | 36841 | BREX-system-P-loop-protein-BrxC-[Terrisporobacter-glycolicus] |
| MBP2018669.1 | Type6.BREX | <-REase<-SIGMA+OB-Fold+AAA-ATPase+Lon-Protease-BrxL<-PglZ<-N6-METHYLASE<-DUF6079*<-BrxB-BrxF-iSTAND<-BrxA-DNA-binding<-BrxE\|\|McrC-NTD+REase->McrB-AAA-><-SF1-SF2-Helicase+wHTH+Small-B-barrel+DUF1998<-?\|\|SF1-SF2-Helicase-> | 1199 | Bacteria>Terrabacteria_group>Bacillota>Clostridia>Eubacteriales>Symbiobacteriaceae | GCA_017874315.1 | Symbiobacterium_terraclitae | J2Z79_002084 | 10592..14191 | 557451 | hypothetical-protein-[Symbiobacterium-terraclitae] |
| MBY6275820.1 | Type6.BREX | <-HEPN<-SIGMA+OB-Fold+AAA-ATPase+Lon-Protease-BrxL<-PglZ<-N6-METHYLASE<-N6-METHYLASE<-DUF6079*<-BrxB-BrxF-iSTAND<-BrxA-DNA-binding<-BrxE\|\|McrC-NTD+REase->AAA-ATPase-> | 1196 | Bacteria>Terrabacteria_group>Bacillota>Clostridia>Eubacteriales>Symbiobacteriaceae | GCA_019806665.1 | Symbiobacterium_thermophilum | CWE10_06285 | 9595..13185 | 2734 | BREX-system-P-loop-protein-BrxC-[Symbiobacterium-thermophilum] |
| WP_273378700.1 | Type6.BREX | <-HEPN<-?<-SIGMA+OB-Fold+AAA-ATPase+Lon-Protease-BrxL<-PglZ<-N6-METHYLASE<-N6-METHYLASE<-DUF6079*<-BrxB-BrxF-iSTAND<-BrxA-DNA-binding<-BrxE\|\|McrC-NTD+REase-> | 1196 | Bacteria>Terrabacteria_group>Bacillota>Clostridia>Eubacteriales>Symbiobacteriaceae | GCF_019806665.1 | Symbiobacterium_thermophilum | CWE10_RS06335 | 9595..13185 | 2734 | BREX-system-P-loop-protein-BrxC-[Symbiobacterium-thermophilum] |
| NLO22061.1 | Type6.BREX | <-DUF6079-p*<-BrxB-BrxF-iSTAND<-BrxA-DNA-binding<-BrxE<-GDResolvase+HIN-HTH | 579 | Bacteria>Terrabacteria_group>Bacillota>Clostridia>Eubacteriales>Syntrophomonadaceae | GCA_012522955.1 | Syntrophomonadaceae_bacterium | GX119_08680 | 1..1736 | 2093811 | BREX-system-P-loop-protein-BrxC-[Syntrophomonadaceae-bacterium] |
| KXM11538.1 | Type6.BREX | <-PAS+NtrC-AAA+FIS-HTH<-DDE_Tnp_ISAZ013\|\|MPTase->PIN-><-SIGMA+OB-Fold+AAA-ATPase+HKD-DNase-BrxL<-PglZ<-N6-METHYLASE<-DUF6079*<-BrxB-BrxF-iSTAND<-BrxA-DNA-binding<-BrxE<-TPR<-TIR<-?<-?<-?\|\|DDE_Tnp_1-> | 1207 | Bacteria>Terrabacteria_group>Bacillota>Clostridia>Eubacteriales>unclassified_Eubacteriales | GCA_000987045.1 | Clostridiales_bacterium_PH28_bin88 | SY88_07480 | 24699..28322 | 1605376 | ATPase-[Clostridiales-bacterium-PH28_bin88] |
| NLN06909.1 | Type6.BREX | BrxE->BrxA-DNA-binding->BrxB-BrxF-iSTAND->DUF6079*->OLD-ABC+TOPRIM->PglZ->SIGMA+OB-Fold+AAA-ATPase+Lon-Protease-BrxL->DNA_METHYLASE->SF1-SF2-Helicase+REase->REase-> | 1222 | Bacteria>Terrabacteria_group>Bacillota | GCA_012522075.1 | Bacillota_bacterium | GX167_04740 | 2708..6376 | 1879010 | BREX-system-P-loop-protein-BrxC-[Bacillota-bacterium] |
| WP_025641432.1 | Type6.BREX | GspA-ATPase-><-Phage_int_SAM_4<-DndB<-DndB<-SF1-SF2-Helicase+wHTH+Small-B-barrel+DUF1998<-BrxD-AAA<-PglZ<-PglZ<-N6-METHYLASE<-DUF6079*<-BrxB-BrxF-iSTAND<-BrxA-DNA-binding<-BrxE<-?<-?<-?<-?<-AAA-ATPase | 1207 | Bacteria>Terrabacteria_group>Bacillota>Tissierellia>Tissierellales>Tepidimicrobiaceae | GCF_000511955.1 | Schnuerera_ultunensis | CUL_RS0109785 | 516055..519678 | 45497 | BREX-system-P-loop-protein-BrxC-[Schnuerera-ultunensis] |
| WP_062419291.1 | Type6.BREX | METHYLASE+SF1-SF2-Helicase-><-?<-SIGMA+OB-Fold+AAA-ATPase+HKD-DNase-BrxL<-PglZ<-TOPRIM<-OLD-ABC<-N6-METHYLASE<-N6-METHYLASE<-DUF6079*<-BrxB-BrxF-iSTAND<-BrxA-DNA-binding<-BrxE<-XIS-HTH<-CHTH<-LuxR-HTH\|\|?->XIS-HTH->Phage_integrase-> | 1206 | Bacteria>Terrabacteria_group>Chloroflexota>Anaerolineae>Anaerolineales>Anaerolineaceae | GCF_001050255.2 | Levilinea_saccharolytica | LSAC_RS18205 | 3303565..3307185 | 229921 | BREX-system-P-loop-protein-BrxC-[Levilinea-saccharolytica] |
| WP_062280889.1 | Type6.BREX | <-Phage_int_SAM_1+Phage_integrase<-XIS-HTH<-?\|\|LuxR-HTH->CHTH->XIS-HTH->BrxE->BrxA-DNA-binding->BrxB-BrxF-iSTAND->DUF6079*->N6-METHYLASE->PglZ->SIGMA+OB-Fold+AAA-ATPase+HKD-DNase-BrxL->?->HerA/FtsK-><-METHYLASE+SF1-SF2-Helicase<-?\|\|SIGMA-HTH-> | 1206 | Bacteria>Terrabacteria_group>Chloroflexota>Anaerolineae>Anaerolineales>Anaerolineaceae | GCF_001192795.1 | Flexilinea_flocculi | ATC1_RS09365 | 1036215..1039835 | 1678840 | BREX-system-P-loop-protein-BrxC-[Flexilinea-flocculi] |
| MCC6190561.1 | Type6.BREX | MPTase->?->HerA/FtsK->HEPN->REase->?->BrxE->BrxA-DNA-binding->BrxB-BrxF-iSTAND->DUF6079*->N6-METHYLASE->Lipase->HTH+HEPN->McrB-AAA->REase->PglZ->SIGMA+OB-Fold+AAA-ATPase+Lon-Protease-BrxL->?->Phage_int+DNA-glycosylase-> | 1208 | Bacteria>Terrabacteria_group>Chloroflexota>Anaerolineae>Anaerolineales>unclassified_Anaerolineales | - | Anaerolineales_bacterium | IT318_16150 | 14120..17746 | 2073117 | BREX-system-P-loop-protein-BrxC-[Anaerolineales-bacterium] |
| NPV77432.1 | Type6.BREX | LuxR-HTH->CHTH->XIS-HTH->BrxE->BrxA-DNA-binding->BrxB-BrxF-iSTAND->DUF6079*->N6-METHYLASE->SIGMA+OB-Fold+AAA-ATPase+HKD-DNase-BrxL->?-><-METHYLASE+SF1-SF2-Helicase<-?\|\|wHTH+AbiEi_2->AbiEii-><-DNA_pol3_beta+DNA_pol3_beta_2+DNA_pol3_beta_3 | 1206 | Bacteria>Terrabacteria_group>Chloroflexota>Anaerolineae>unclassified_Anaerolineae | GCA_013178185.1 | Anaerolineae_bacterium | HPY59_13805 | 3635..7255 | 2052143 | BREX-system-P-loop-protein-BrxC-[Anaerolineae-bacterium] |
| MCL4264723.1 | Type6.BREX | <-SIGMA-HTH<-SIGMA-HTH\|\|?->?->TPR+TPR-><-SIGMA+OB-Fold+AAA-ATPase+HKD-DNase-BrxL<-PglZ<-N6-METHYLASE<-DUF6079*<-BrxB-BrxF-iSTAND<-BrxA-DNA-binding<-BrxE\|\|?->TPR+TPR+TPR->?-><-TPR+TPR+TPR+TPR<-CASPASE | 1209 | Bacteria>Terrabacteria_group>Chloroflexota>Anaerolineae>unclassified_Anaerolineae | GCA_023382655.1 | Anaerolineae_bacterium | KJ069_15985 | 16020..19649 | 2052143 | BREX-system-P-loop-protein-BrxC-[Anaerolineae-bacterium] |
| MBE2236346.1 | Type6.BREX | <-MarR-HTH<-NAD_binding_10\|\|?-><-?<-RADICAL-SAM\|\|BrxE->BrxA-DNA-binding->BrxB-BrxF-iSTAND->DUF6079*-> | 1204 | Bacteria>Terrabacteria_group>Chloroflexota>Caldilineae>Caldilineales>Caldilineaceae | GCA_014879855.1 | Caldilineaceae_bacterium | IAE81_01055 | 8862..12476 | 2614250 | BREX-system-P-loop-protein-BrxC-[Caldilineaceae-bacterium] |
| MCB0069888.1 | Type6.BREX | METHYLASE+SF1-SF2-Helicase-><-?\|\|GDResolvase+HIN-HTH->BrxE->BrxA-DNA-binding->BrxB-BrxF-iSTAND->DUF6079*->N6-METHYLASE->?->?->PglZ->SIGMA+OB-Fold+AAA-ATPase+Lon-Protease-BrxL-p-> | 1235 | Bacteria>Terrabacteria_group>Chloroflexota>Caldilineae>Caldilineales>Caldilineaceae | GCA_020433325.1 | Caldilineaceae_bacterium | KDE20_00460 | 85387..89094 | 2614250 | BREX-system-P-loop-protein-BrxC-[Caldilineaceae-bacterium] |
| WP_012881599.1 | Type6.BREX | HTH->?-><-REase<-SF1-SF2-Helicase+REase<-DNA_METHYLASE<-SIGMA+OB-Fold+AAA-ATPase+Lon-Protease-BrxL<-PglZ<-N6-METHYLASE<-DUF6079*<-BrxB-BrxF-iSTAND<-BrxA-DNA-binding<-BrxE<-CHTH<-tRNA<-?<-ACET | 1228 | Bacteria>Terrabacteria_group>Chloroflexota>Dehalococcoidia>Dehalococcoidales>Dehalococcoidaceae | GCF_000025025.1 | Dehalococcoides_mccartyi | DHCVS_RS01280 | 252867..256553 | 61435 | BREX-system-P-loop-protein-BrxC-[Dehalococcoides-mccartyi] |
| WP_015407539.1 | Type6.BREX | HTH->?-><-REase<-REase<-SF1-SF2-Helicase+REase<-DNA_METHYLASE<-SIGMA+OB-Fold+AAA-ATPase+Lon-Protease-BrxL<-PglZ<-N6-METHYLASE<-DUF6079*<-BrxB-BrxF-iSTAND<-BrxA-DNA-binding<-BrxE<-CHTH<-tRNA<-?<-ACET | 1226 | Bacteria>Terrabacteria_group>Chloroflexota>Dehalococcoidia>Dehalococcoidales>Dehalococcoidaceae | GCF_000341695.1 | Dehalococcoides_mccartyi | BTF_RS01445 | 279971..283651 | 61435 | BREX-system-P-loop-protein-BrxC-[Dehalococcoides-mccartyi] |
| OJX40202.1 | Type6.BREX | <-MerR-HTH\|\|?->METHYLASE+SF1-SF2-Helicase-><-?<-SIGMA+OB-Fold+AAA-ATPase+HKD-DNase-BrxL<-PglZ<-N6-METHYLASE<-DUF6079*<-BrxB-BrxF-iSTAND<-BrxA-DNA-binding<-BrxE<-XIS-HTH<-CHTH<-LuxR-HTH\|\|?->XIS-HTH->Phage_integrase-> | 1199 | Bacteria>Terrabacteria_group>Chloroflexota>unclassified_Chloroflexota | GCA_001899005.1 | Chloroflexi_bacterium_44-23 | BG078_14595 | 64174..67773 | 1895927 | ATPase-[Chloroflexi-bacterium-44-23] |
| MBI3971803.1 | Type6.BREX | <-N6-METHYLASE-p<-DUF6079*<-BrxB-BrxF-iSTAND<-BrxA-DNA-binding<-BrxE<-?\|\|DSBH+AraC-HTH+AraC-HTH->?->?->Sulfatase+DUF229+DUF4976-> | 1224 | Bacteria>Terrabacteria_group>Chloroflexota>unclassified_Chloroflexota | GCA_016199265.1 | Chloroflexota_bacterium | HY332_10990 | 386..4060 | 2026724 | BREX-system-P-loop-protein-BrxC-[Chloroflexota-bacterium] |
| MBI3748365.1 | Type6.BREX | <-DUF6079-p*<-BrxB-BrxF-iSTAND<-BrxA-DNA-binding<-BrxE<-MORC+DUF3684+DUF3883<-SF1-SF2-Helicase+REase<-Preader17<-?\|\|HEPN-><-HIN-HTH | 659 | Bacteria>Terrabacteria_group>Chloroflexota>unclassified_Chloroflexota | GCA_016200745.1 | Chloroflexota_bacterium | HY262_05925 | 1..1978 | 2026724 | BREX-system-P-loop-protein-BrxC-[Chloroflexota-bacterium] |
| MBI3732787.1 | Type6.BREX | <-McrB-AAA<-?<-SIGMA+OB-Fold+AAA-ATPase+Lon-Protease-BrxL<-PglZ\|\|<-N6-METHYLASE<-HEPN<-DUF6079*<-BrxB-BrxF-iSTAND<-BrxA-DNA-binding<-BrxE<-?<-?<-?<-HerA/FtsK | 1243 | Bacteria>Terrabacteria_group>Chloroflexota>unclassified_Chloroflexota | GCA_016200765.1 | Chloroflexota_bacterium | HY259_04935 | 12602..16333 | 2026724 | BREX-system-P-loop-protein-BrxC-[Chloroflexota-bacterium] |
| MBI4322558.1 | Type6.BREX | <-DUF6079-p*<-BrxB-BrxF-iSTAND<-BrxA-DNA-binding<-BrxE<-Phage_integrase<-?\|\|?->HKD-DNase+HKD-Rease->REase-> | 427 | Bacteria>Terrabacteria_group>Chloroflexota>unclassified_Chloroflexota | GCA_016210495.1 | Chloroflexota_bacterium | HY675_29045 | 1..1283 | 2026724 | BREX-system-P-loop-protein-BrxC-[Chloroflexota-bacterium] |
| MBK8030403.1 | Type6.BREX | <-Phage_holin<-Relaxase<-MetJarc-HTH<-?\|\|BrxE->BrxA-DNA-binding->BrxB-BrxF-iSTAND->N6-METHYLASE->DUF6079*->HEPN->N6-METHYLASE->N6-METHYLASE-p->PglZ->PglZ->SIGMA+OB-Fold+AAA-ATPase+Lon-Protease-BrxL-> | 1258 | Bacteria>Terrabacteria_group>Chloroflexota>unclassified_Chloroflexota | GCA_016713355.1 | Chloroflexota_bacterium | IPK17_13050 | 320822..324598 | 2026724 | BREX-system-P-loop-protein-BrxC-[Chloroflexota-bacterium] |
| MBK8935331.1 | Type6.BREX | <-Peptidase<-HerA/FtsK<-?<-?<-?\|\|BrxE->BrxA-DNA-binding->BrxB-BrxF-iSTAND->DUF6079-p*-> | 259 | Bacteria>Terrabacteria_group>Chloroflexota>unclassified_Chloroflexota | GCA_016720015.1 | Chloroflexota_bacterium | IPM76_24040 | 25896..26672 | 2026724 | BREX-system-P-loop-protein-BrxC-[Chloroflexota-bacterium] |
| MCL4488347.1 | Type6.BREX | HerA/FtsK->MarR-HTH->?->ParB->?-><-?<-SIGMA+OB-Fold+AAA-ATPase+Lon-Protease-BrxL<-PglZ<-N6-METHYLASE<-DUF6079*<-BrxB-BrxF-iSTAND<-BrxA-DNA-binding<-BrxE<-SF1-SF2-Helicase+REase<-SF1-SF2-Helicase+REase\|\|?->?->DDE_Tnp_ISAZ013-> | 1220 | Bacteria>Terrabacteria_group>Chloroflexota>unclassified_Chloroflexota | GCA_023381515.1 | Chloroflexota_bacterium | M1570_09495 | 91471..95133 | 2026724 | BREX-system-P-loop-protein-BrxC-[Chloroflexota-bacterium] |
| MCL5952139.1 | Type6.BREX | <-REC<-?<-HKD-DNase+Preader14<-REase<-SIGMA+OB-Fold+AAA-ATPase+Lon-Protease-BrxL<-PglZ<-N6-METHYLASE<-DUF6079*<-BrxB-BrxF-iSTAND<-BrxA-DNA-binding<-BrxE<-SF1-SF2-Helicase+REase<-SF1-SF2-Helicase+REase\|\|?->?->REase->SbcC+SbcC-> | 1220 | Bacteria>Terrabacteria_group>Chloroflexota>unclassified_Chloroflexota | GCA_023489485.1 | Chloroflexota_bacterium | M1132_10550 | 55663..59325 | 2026724 | BREX-system-P-loop-protein-BrxC-[Chloroflexota-bacterium] |
| WP_190710345.1 | Type6.BREX | <-EVE\|\|BrxE->BrxA-DNA-binding->BrxB-BrxF-iSTAND->DUF6079*->N6-METHYLASE-> | 1205 | Bacteria>Terrabacteria_group>Cyanobacteriota/Melainabacteria_group>Cyanobacteriota>Cyanophyceae>Nostocales | - | Anabaena_minutissima | H6G36_RS29955 | 4664..8281 | 2303152 | BREX-system-P-loop-protein-BrxC-[Anabaena-minutissima] |
| WP_190963244.1 | Type6.BREX | HTH_26->?->?->?->HEPN->BrxE->BrxA-DNA-binding->BrxB-BrxF-iSTAND->DUF6079*->N6-METHYLASE->MORC+DUF3684->MORC+DUF3684+REase->PglZ->BrxD-AAA->SF1-SF2-Helicase+wHTH+Small-B-barrel+DUF1998-> | 1207 | Bacteria>Terrabacteria_group>Cyanobacteriota/Melainabacteria_group>Cyanobacteriota>Cyanophyceae>Nostocales | - | Desmonostoc_muscorum | H6G91_22885 | 8366..11989 | 1179 | BREX-system-P-loop-protein-BrxC-[Desmonostoc-muscorum] |
| OYE05053.1 | Type6.BREX | <-DNA_METHYLASE\|\|?-><-?<-SIGMA+OB-Fold+AAA-ATPase+Lon-Protease-BrxL<-PglZ<-DUF6079*<-BrxB-BrxF-iSTAND<-BrxA-DNA-binding<-BrxE\|\|HTH_26-> | 1239 | Bacteria>Terrabacteria_group>Cyanobacteriota/Melainabacteria_group>Cyanobacteriota>Cyanophyceae>Nostocales | GCA_002245985.1 | Nostoc_sp._'Peltigera_membranacea_cyanobiont'_232 | CDG79_09605 | 16194..19913 | 2014531 | hypothetical-protein-CDG79_09605-[Nostoc-sp.-'Peltigera-membranacea-cyanobiont'-232] |
| BAY35694.1 | Type6.BREX | TPR-><-Peptidase_C14<-?<-?<-SIGMA+OB-Fold+AAA-ATPase+Lon-Protease-BrxL<-PglZ<-PglZ<-N6-METHYLASE<-DUF6079*<-BrxB-BrxF-iSTAND<-BrxA-DNA-binding<-BrxE\|\|HTH_26-> | 1238 | Bacteria>Terrabacteria_group>Cyanobacteriota/Melainabacteria_group>Cyanobacteriota>Cyanophyceae>Nostocales | GCA_002368215.1 | Nostoc_sp._NIES-2111 | NIES2111_00100 | 19397..23113 | 1973475 | hypothetical-protein-NIES2111_00100-[Nostoc-sp.-NIES-2111] |
| TVP63092.1 | Type6.BREX | OrfB_IS605+OrfB_Zn_ribbon->?-><-ABhydrolase\|\|BrxE->BrxA-DNA-binding->BrxB-BrxF-iSTAND->DUF6079*->N6-METHYLASE->?-><-?\|\|PglZ->BrxD-AAA->SF1-SF2-Helicase+wHTH+Small-B-barrel+DUF1998->ParA-Soj-PloopNTPase->ParB+HTH-> | 1206 | Bacteria>Terrabacteria_group>Cyanobacteriota/Melainabacteria_group>Cyanobacteriota>Cyanophyceae>Nostocales | GCA_007692755.1 | Nodularia_sp._(in:_cyanobacteria) | EA343_08975 | 5740..9360 | 2047085 | BREX-system-P-loop-protein-BrxC-[Nodularia-sp.-(in:-cyanobacteria)] |
| MBR8839459.1 | Type6.BREX | NACHT-><-?\|\|?-><-CBS+CBS\|\|?->BrxE->BrxA-DNA-binding->BrxB-BrxF-iSTAND->DUF6079*->N6-METHYLASE->PglZ-p-> | 1206 | Bacteria>Terrabacteria_group>Cyanobacteriota/Melainabacteria_group>Cyanobacteriota>Cyanophyceae>Nostocales | GCA_016903675.1 | Stigonema_ocellatum_SAG_48.90_=_DSM_106950 | DSM106950_37010 | 12253..15873 | 221294 | BREX-system-P-loop-protein-BrxC-[Stigonema-ocellatum-SAG-48.90-=-DSM-106950] |
| MBN3948789.1 | Type6.BREX | <-DUF6079-p*<-BrxB-BrxF-iSTAND<-BrxA-DNA-binding<-BrxE\|\|?-><-wHTH+REase+ZnR<-McrB-AAA<-HTH+HEPN<-SF1-SF2-Helicase<-STYKIN | 200 | Bacteria>Terrabacteria_group>Cyanobacteriota/Melainabacteria_group>Cyanobacteriota>Cyanophyceae>Nostocales | GCA_017115085.1 | Nostoc_sp._NMS7 | HWQ38_20950 | 1..601 | 2815391 | BREX-system-P-loop-protein-BrxC-[Nostoc-sp.-NMS7] |
| MBN3923061.1 | Type6.BREX | <-N6-N4-METHYLASE<-?\|\|?-><-?\|\|BrxE->BrxA-DNA-binding->BrxB-BrxF-iSTAND->DUF6079*->N6-METHYLASE->\|\|DDE_Tnp_1_assoc->DDE_Tnp_1_assoc->PglZ->BrxD-AAA->SF1-SF2-Helicase+wHTH+Small-B-barrel+DUF1998-> | 1206 | Bacteria>Terrabacteria_group>Cyanobacteriota/Melainabacteria_group>Cyanobacteriota>Cyanophyceae>Nostocales | GCA_017115095.1 | Nostoc_sp._NMS4 | HWQ37_07850 | 5829..9449 | 2815390 | BREX-system-P-loop-protein-BrxC-[Nostoc-sp.-NMS4] |
| MBW4670333.1 | Type6.BREX | <-Glycos_transf<-TPR\|\|REase-><-Acetate_kinase\|\|BrxE->BrxA-DNA-binding->BrxB-BrxF-iSTAND->DUF6079*->?->PIN->?-><-REase<-?\|\|N6-METHYLASE-> | 1211 | Bacteria>Terrabacteria_group>Cyanobacteriota/Melainabacteria_group>Cyanobacteriota>Cyanophyceae>Nostocales | GCA_019359595.1 | Cyanomargarita_calcarea_GSE-NOS-MK-12-04C | KME60_23685 | 99014..102649 | 2839659 | BREX-system-P-loop-protein-BrxC-[Cyanomargarita-calcarea-GSE-NOS-MK-12-04C] |
| WP_223278078.1 | Type6.BREX | <-DNA_METHYLASE\|\|?-><-SIGMA+OB-Fold+AAA-ATPase+Lon-Protease-BrxL<-PglZ<-N6-METHYLASE<-DUF6079*<-BrxB-BrxF-iSTAND<-BrxA-DNA-binding<-BrxE\|\|HTH_26-> | 1238 | Bacteria>Terrabacteria_group>Cyanobacteriota/Melainabacteria_group>Cyanobacteriota>Cyanophyceae>Nostocales | GCF_002245985.1 | Nostoc_sp._'Peltigera_membranacea_cyanobiont'_232 | CDG79_RS09650 | 16194..19910 | 2014531 | BREX-system-P-loop-protein-BrxC-[Nostoc-sp.-'Peltigera-membranacea-cyanobiont'-232] |
| WP_094351990.1 | Type6.BREX | <-DNA_METHYLASE<-?<-?\|\|?->?->METHYLASE->BrxE->BrxA-DNA-binding->BrxB-BrxF-iSTAND->DUF6079*->N6-METHYLASE->PglZ->BrxD-AAA->SF1-SF2-Helicase+wHTH+Small-B-barrel+DUF1998-><-?<-Polbeta | 1206 | Bacteria>Terrabacteria_group>Cyanobacteriota/Melainabacteria_group>Cyanobacteriota>Cyanophyceae>Nostocales | GCF_002246015.1 | Nostoc_sp._'Peltigera_membranacea_cyanobiont'_210A | CDG76_RS27200 | 197984..201604 | 2014529 | BREX-system-P-loop-protein-BrxC-[Nostoc-sp.-'Peltigera-membranacea-cyanobiont'-210A] |
| WP_096585923.1 | Type6.BREX | METHYLASE->?->SF1-SF2-Helicase+PRTase-><-SF1-SF2-Helicase\|\|BrxE->BrxA-DNA-binding->BrxB-BrxF-iSTAND->DUF6079*->N6-METHYLASE->PglZ->BrxD-AAA->SF1-SF2-Helicase+wHTH+Small-B-barrel+DUF1998->?->?->ACET-><-?<-Glyoxalase | 1206 | Bacteria>Terrabacteria_group>Cyanobacteriota/Melainabacteria_group>Cyanobacteriota>Cyanophyceae>Nostocales | GCF_002367975.1 | Anabaenopsis_circularis | CA723_RS28450 | 110380..114000 | 109266 | BREX-system-P-loop-protein-BrxC-[Anabaenopsis-circularis] |
| WP_096637339.1 | Type6.BREX | <-Transposase<-?<-?<-HTH_26\|\|BrxE->BrxA-DNA-binding->BrxB-BrxF-iSTAND-><-Y1_Tnp\|\|OrfB_IS605+OrfB_Zn_ribbon->DUF6079*->N6-METHYLASE->PglZ->BrxD-AAA->SF1-SF2-Helicase+wHTH+Small-B-barrel+DUF1998-><-HEPN<-HNH<-?<-?\|\|Peptidase_C14-> | 1204 | Bacteria>Terrabacteria_group>Cyanobacteriota/Melainabacteria_group>Cyanobacteriota>Cyanophyceae>Nostocales | GCF_002368015.1 | Trichormus_variabilis | CA725_RS30025 | 10233..13847 | 264691 | BREX-system-P-loop-protein-BrxC-[Trichormus-variabilis] |
| WP_096678048.1 | Type6.BREX | TPR-><-?<-?<-SIGMA+OB-Fold+AAA-ATPase+Lon-Protease-BrxL<-PglZ<-PglZ<-DUF6079*<-BrxB-BrxF-iSTAND<-BrxA-DNA-binding<-BrxE\|\|HTH_26-> | 1239 | Bacteria>Terrabacteria_group>Cyanobacteriota/Melainabacteria_group>Cyanobacteriota>Cyanophyceae>Nostocales | GCF_002368215.1 | Nostoc_sp._NIES-2111 | CA737_RS00055 | 19397..23116 | 1973475 | BREX-system-P-loop-protein-BrxC-[Nostoc-sp.-NIES-2111] |
| WP_096585011.1 | Type6.BREX | <-MerR-HTH<-SF1-SF2-Helicase+wHTH+Small-B-barrel+DUF1998<-BrxD-AAA<-BrxD-AAA<-?<-SF1-SF2-Helicase+wHTH+Small-B-barrel+DUF1998<-BrxD-AAA<-PglZ<-N6-METHYLASE<-DUF6079*<-BrxB-BrxF-iSTAND<-BrxA-DNA-binding<-BrxE\|\|SF1-SF2-Helicase-><-SF1-SF2-Helicase+PRTase<-?<-OrfB_IS605+OrfB_Zn_ribbon | 1206 | Bacteria>Terrabacteria_group>Cyanobacteriota/Melainabacteria_group>Cyanobacteriota>Cyanophyceae>Nostocales | GCF_002368295.1 | Tolypothrix_tenuis | CDC28_RS35840 | 19688..23308 | 457083 | BREX-system-P-loop-protein-BrxC-[Tolypothrix-tenuis] |
| WP_168499564.1 | Type6.BREX | <-N6-METHYLASE<-?<-?<-DUF6079*<-BrxB-BrxF-iSTAND<-BrxA-DNA-binding<-BrxE<-?\|\|TraJ-RHH-> | 1207 | Bacteria>Terrabacteria_group>Cyanobacteriota/Melainabacteria_group>Cyanobacteriota>Cyanophyceae>Nostocales | GCF_009711975.1 | Anabaena_sp._UHCC_0204 | FJR04_RS23910 | 3610..7233 | 2590009 | BREX-system-P-loop-protein-BrxC-[Anabaena-sp.-UHCC-0204] |
| WP_190755500.1 | Type6.BREX | METHYLASE->?->SF1-SF2-Helicase+PRTase-><-SF1-SF2-Helicase\|\|BrxE->BrxA-DNA-binding->BrxB-BrxF-iSTAND->DUF6079*->N6-METHYLASE->PglZ->BrxD-AAA->SF1-SF2-Helicase+wHTH+Small-B-barrel+DUF1998->MerR-HTH-> | 1206 | Bacteria>Terrabacteria_group>Cyanobacteriota/Melainabacteria_group>Cyanobacteriota>Cyanophyceae>Nostocales | GCF_014697455.1 | Nostoc_sp._FACHB-145 | H6G90_RS25450 | 31450..35070 | 2692836 | BREX-system-P-loop-protein-BrxC-[Nostoc-sp.-FACHB-145] |
| WP_190951034.1 | Type6.BREX | MoxR-AAA->VWA+vWA-L+CR-ATPase9->?-><-SF1-SF2-Helicase+wHTH+Small-B-barrel+DUF1998<-BrxD-AAA<-PglZ<-MORC+DUF3684+REase<-MORC+DUF3684<-N6-METHYLASE<-DUF6079*<-BrxB-BrxF-iSTAND<-BrxA-DNA-binding<-BrxE<-CASPASE<-?\|\|Phage_int_SAM_1+Phage_integrase->RHH_9->?->ACET-> | 1206 | Bacteria>Terrabacteria_group>Cyanobacteriota/Melainabacteria_group>Cyanobacteriota>Cyanophyceae>Nostocales | GCF_014698495.1 | Nostoc_punctiforme | H6G94_RS21770 | 51100..54720 | 272131 | BREX-system-P-loop-protein-BrxC-[Nostoc-punctiforme] |
| WP_229485753.1 | Type6.BREX | <-REase+URI-DNase<-?<-?<-?<-SF1-SF2-Helicase+wHTH+Small-B-barrel+DUF1998-p<-BrxD-AAA<-PglZ<-N6-METHYLASE<-DUF6079*<-BrxB-BrxF-iSTAND<-BrxA-DNA-binding<-BrxE<-METHYLASE<-?<-?\|\|?->DNA_METHYLASE-> | 1206 | Bacteria>Terrabacteria_group>Cyanobacteriota/Melainabacteria_group>Cyanobacteriota>Cyanophyceae>Nostocales | GCF_020829495.1 | Nostoc_favosum | LC586_RS15955 | 88159..91779 | 2907819 | BREX-system-P-loop-protein-BrxC-[Nostoc-favosum] |
| WP_190881152.1 | Type6.BREX | <-HTH_26\|\|?->?->HEPN-><-SF1-SF2-Helicase+wHTH+Small-B-barrel+DUF1998<-BrxD-AAA<-PglZ<-N6-METHYLASE<-DUF6079*<-BrxB-BrxF-iSTAND<-BrxA-DNA-binding<-BrxE\|\|?-><-METHYLASE\|\|RADICAL-SAM-> | 1206 | Bacteria>Terrabacteria_group>Cyanobacteriota/Melainabacteria_group>Cyanobacteriota>Cyanophyceae>Nostocales | - | Nostoc_sp._FACHB-973 | H6G93_19595 | 51403..55023 | 2692844 | BREX-system-P-loop-protein-BrxC-[Nostoc-sp.-FACHB-973] |
| MBD2549124.1 | Type6.BREX | BrxE->BrxA-DNA-binding->BrxB-BrxF-iSTAND->DUF6079*->N6-METHYLASE->Schlafen+HATPase+wHTH->OLD-ABC->?->Peptidase_M78->PglZ->Schlafen+HATPase+wHTH-p->Schlafen+HATPase+wHTH+HTH->SIGMA+OB-Fold+AAA-ATPase+Lon-Protease-BrxL-> | 1176 | Bacteria>Terrabacteria_group>Cyanobacteriota/Melainabacteria_group>Cyanobacteriota>Cyanophyceae>Oscillatoriophycideae | GCA_014698445.1 | Microcystis_elabens_FACHB-917 | H6G65_05915 | 2261..5791 | 2692898 | BREX-system-P-loop-protein-BrxC-[Microcystis-elabens-FACHB-917] |
| MBW4696203.1 | Type6.BREX | Phage_int_SAM_4+Phage_integrase->ParA-Soj-PloopNTPase->ParB+HTH->BrxE->BrxA-DNA-binding->BrxB-BrxF-iSTAND->DUF6079*-><-?\|\|?->N6-METHYLASE-p->?->PIN-> | 1212 | Bacteria>Terrabacteria_group>Cyanobacteriota/Melainabacteria_group>Cyanobacteriota>Cyanophyceae>Oscillatoriophycideae | GCA_019359375.1 | Lyngbya_sp._HA4199-MV5 | KME27_31195 | 7651..11289 | 2835090 | BREX-system-P-loop-protein-BrxC-[Lyngbya-sp.-HA4199-MV5] |
| WP_194061351.1 | Type6.BREX | <-REase+URI-DNase\|\|Phage_integrase->BrxE->BrxA-DNA-binding->BrxB-BrxF-iSTAND->DUF6079*->N6-METHYLASE->PglZ->BrxD-AAA->SF1-SF2-Helicase+wHTH+Small-B-barrel+DUF1998->?-><-?\|\|HEPN->Phage_integrase-> | 1206 | Bacteria>Terrabacteria_group>Cyanobacteriota/Melainabacteria_group>Cyanobacteriota>Cyanophyceae>Oscillatoriophycideae | GCF_015207285.1 | Tychonema_sp._LEGE_07203 | IQ252_RS06740 | 5314..8934 | 1828671 | BREX-system-P-loop-protein-BrxC-[Tychonema-sp.-LEGE-07203] |
| WP_194008286.1 | Type6.BREX | <-HEPN<-?<-?<-?<-REase+URI-DNase\|\|Phage_integrase->BrxE->BrxA-DNA-binding->BrxB-BrxF-iSTAND->DUF6079*->N6-METHYLASE->PglZ->BrxD-AAA->SF1-SF2-Helicase+wHTH+Small-B-barrel+DUF1998-><-REase<-?\|\|?->\|\|HTH_23+HTH_32+HTH_33-> | 1208 | Bacteria>Terrabacteria_group>Cyanobacteriota/Melainabacteria_group>Cyanobacteriota>Cyanophyceae>Oscillatoriophycideae | GCF_015207345.1 | unclassified_Tychonema | IQ245_RS02235 | 107207..110833 | 2642144 | MULTISPECIES:-BREX-system-P-loop-protein-BrxC-[unclassified-Tychonema] |
| WP_190543924.1 | Type6.BREX | BrxE->BrxA-DNA-binding->BrxB-BrxF-iSTAND->DUF6079*->N6-METHYLASE->Schlafen+HATPase+wHTH->OLD-ABC->?->Peptidase_M78->PglZ->Schlafen+HATPase+wHTH+HTH->SIGMA+OB-Fold+AAA-ATPase+Lon-Protease-BrxL-> | 1176 | Bacteria>Terrabacteria_group>Cyanobacteriota/Melainabacteria_group>Cyanobacteriota>Cyanophyceae>Oscillatoriophycideae | - | Microcystis_elabens | H6G65_05915 | 2261..5791 | 44824 | BREX-system-P-loop-protein-BrxC-[Microcystis-elabens] |
| KPQ33031.1 | Type6.BREX | <-SF1-SF2-Helicase+wHTH+Small-B-barrel+DUF1998<-BrxD-AAA<-PglZ<-PIN<-?<-N6-METHYLASE<-DUF433<-DUF6079*<-BrxB-BrxF-iSTAND<-BrxA-DNA-binding<-BrxE<-?<-AAA-ATPase | 1209 | Bacteria>Terrabacteria_group>Cyanobacteriota/Melainabacteria_group>Cyanobacteriota>Cyanophyceae>Pseudanabaenales | GCA_001314865.1 | Phormidesmis_priestleyi_Ana | HLUCCA11_19925 | 26535..30164 | 1666911 | hypothetical-protein-HLUCCA11_19925-[Phormidesmis-priestleyi-Ana] |
| PZO41937.1 | Type6.BREX | TnsC-ATPase->?-><-?\|\|RHH->HEPN+SPOCS-><-SF1-SF2-Helicase+wHTH+Small-B-barrel+DUF1998<-BrxD-AAA<-PglZ<-N6-METHYLASE<-DUF6079*<-BrxB-BrxF-iSTAND<-BrxA-DNA-binding<-BrxE<-MerR-HTH\|\|Cas12k-> | 1213 | Bacteria>Terrabacteria_group>Cyanobacteriota/Melainabacteria_group>Cyanobacteriota>Cyanophyceae>Pseudanabaenales | GCA_003242085.1 | Pseudanabaena_frigida | DCF19_08710 | 105719..109360 | 945775 | BREX-system-P-loop-protein-BrxC-[Pseudanabaena-frigida] |
| NJM99703.1 | Type6.BREX | BrxE->BrxA-DNA-binding->BrxB-BrxF-iSTAND->DUF6079-p*->DUF6079-p->DUF6079-p-> | 167 | Bacteria>Terrabacteria_group>Cyanobacteriota/Melainabacteria_group>Cyanobacteriota>Cyanophyceae>Pseudanabaenales | GCA_012033015.1 | Phormidesmis_sp._RL_2_1 | HC800_23455 | 2510..3013 | 2720422 | hypothetical-protein-[Phormidesmis-sp.-RL_2_1] |
| NJM99705.1 | Type6.BREX | BrxE->BrxA-DNA-binding->BrxB-BrxF-iSTAND->DUF6079-p->DUF6079-p->DUF6079-p*-> | 365 | Bacteria>Terrabacteria_group>Cyanobacteriota/Melainabacteria_group>Cyanobacteriota>Cyanophyceae>Pseudanabaenales | GCA_012033015.1 | Phormidesmis_sp._RL_2_1 | HC800_23465 | 3377..4474 | 2720422 | hypothetical-protein-[Phormidesmis-sp.-RL_2_1] |
| MBW4580237.1 | Type6.BREX | <-ABC-ATPase+SbcC\|\|?->?-><-SAM-METHYLASE<-?\|\|CHTH->BrxE->BrxA-DNA-binding->BrxB-BrxF-iSTAND->DUF6079*->?->REase->DUF29->AbrB-HTH->PIN->PglZ->BrxD-AAA->SF1-SF2-Helicase+wHTH+Small-B-barrel+DUF1998-> | 1217 | Bacteria>Terrabacteria_group>Cyanobacteriota/Melainabacteria_group>Cyanobacteriota>Cyanophyceae>Pseudanabaenales | GCA_019358895.1 | Tildeniella_nuda_ZEHNDER_1965/U140 | KME42_11780 | 246570..250223 | 2136405 | BREX-system-P-loop-protein-BrxC-[Tildeniella-nuda-ZEHNDER-1965/U140] |
| MBW4653775.1 | Type6.BREX | <-Trypsin<-?<-?\|\|CopG/RHH->TrwC+SF1-SF2-Helicase->BrxE->BrxA-DNA-binding->BrxB-BrxF-iSTAND->DUF6079*->N6-METHYLASE->HicB_lk_antitox->HicA_toxin->?->?->?->N6-METHYLASE-p-> | 1212 | Bacteria>Terrabacteria_group>Cyanobacteriota/Melainabacteria_group>Cyanobacteriota>Cyanophyceae>Pseudanabaenales | GCA_019359435.1 | Kaiparowitsia_implicata_GSE-PSE-MK54-09C | KME20_12170 | 106200..109838 | 2057771 | BREX-system-P-loop-protein-BrxC-[Kaiparowitsia-implicata-GSE-PSE-MK54-09C] |
| MCL1489589.1 | Type6.BREX | HTH_32+DDE_3->?->?-><-N6-METHYLASE-p<-DDE_Tnp_1_assoc+DDE_Tnp_1<-N6-METHYLASE-p<-?<-BrxB-BrxF-iSTAND<-REase<-DUF6079*<-BrxB-BrxF-iSTAND<-BrxA-DNA-binding<-BrxE\|\|ParA-Soj-PloopNTPase->ParB-><-NACHT<-PrimPol | 1227 | Bacteria>Terrabacteria_group>Cyanobacteriota/Melainabacteria_group>Cyanobacteriota>Cyanophyceae>Pseudanabaenales | GCA_023336205.1 | Pseudanabaena_sp._Salubria-1 | M1G31_02380 | 45126..48809 | 2938918 | BREX-system-P-loop-protein-BrxC-[Pseudanabaena-sp.-Salubria-1] |
| MCU0565455.1 | Type6.BREX | <-SF1-SF2-Helicase+wHTH+Small-B-barrel+DUF1998<-BrxD-AAA<-PglZ\|\|?->HNH-><-N6-METHYLASE<-?<-SKI<-DUF6079*<-BrxB-BrxF-iSTAND<-BrxA-DNA-binding<-BrxE | 1209 | Bacteria>Terrabacteria_group>Cyanobacteriota/Melainabacteria_group>Cyanobacteriota>Cyanophyceae>Pseudanabaenales | GCA_025454315.1 | Oculatellaceae_cyanobacterium_Prado106 | MUF49_02535 | 23046..26675 | 2929566 | BREX-system-P-loop-protein-BrxC-[Oculatellaceae-cyanobacterium-Prado106] |
| WP_009625711.1 | Type6.BREX | <-SF1-SF2-Helicase+wHTH+Small-B-barrel+DUF1998<-BrxD-AAA<-PglZ\|\|PhdYeFM_antitox->REase->?-><-N6-METHYLASE<-ACET<-BrxB-BrxF-iSTAND<-DUF6079*<-BrxB-BrxF-iSTAND<-BrxA-DNA-binding<-BrxE\|\|ParA-Soj-PloopNTPase->ParB-> | 1222 | Bacteria>Terrabacteria_group>Cyanobacteriota/Melainabacteria_group>Cyanobacteriota>Cyanophyceae>Pseudanabaenales | GCF_000332215.1 | Pseudanabaena_biceps | PSE7429_RS03680 | 23473..27141 | 927669 | MULTISPECIES:-BREX-system-P-loop-protein-BrxC-[Pseudanabaena] |
| WP_023076661.1 | Type6.BREX | <-REase+iREase\|\|?->?->?->REase+SPF-cons->BrxE->BrxA-DNA-binding->BrxB-BrxF-iSTAND->DUF6079*->N6-METHYLASE->PglZ->BrxD-AAA->SF1-SF2-Helicase+wHTH+Small-B-barrel+DUF1998-><-REase\|\|?-><-REase<-HerA/FtsK<-TrwC+SF1-SF2-Helicase+TOPRIM | 1206 | Bacteria>Terrabacteria_group>Cyanobacteriota/Melainabacteria_group>Cyanobacteriota>Cyanophyceae>Pseudanabaenales | GCF_000482245.1 | Leptolyngbya_sp._Heron_Island_J | N836_RS70195 | 33447..37067 | 1385935 | BREX-system-P-loop-protein-BrxC-[Leptolyngbya-sp.-Heron-Island-J] |
| WP_094536005.1 | Type6.BREX | TOPRIM+SF1-SF2-Helicase->NACHT-><-ParB<-ParA-Soj-PloopNTPase\|\|BrxE->BrxA-DNA-binding->BrxB-BrxF-iSTAND->DUF6079*->REase->REase->N6-METHYLASE-> | 1222 | Bacteria>Terrabacteria_group>Cyanobacteriota/Melainabacteria_group>Cyanobacteriota>Cyanophyceae>Pseudanabaenales | GCF_002251945.1 | Pseudanabaena_sp._SR411 | B9G53_RS24225 | 12197..15865 | 1980935 | BREX-system-P-loop-protein-BrxC-[Pseudanabaena-sp.-SR411] |
| WP_121969249.1 | Type6.BREX | CHTH->MazG->BrxE->BrxA-DNA-binding->BrxB-BrxF-iSTAND->DUF6079*->N6-METHYLASE->?->PIN->N6-METHYLASE-p->?->?->PglZ->SIGMA+OB-Fold+AAA-ATPase+HKD-DNase-BrxL-> | 1213 | Bacteria>Terrabacteria_group>Cyanobacteriota/Melainabacteria_group>Cyanobacteriota>Cyanophyceae>Pseudanabaenales | GCF_002286735.1 | Leptolyngbya_sp._BC1307 | CKL79_RS08920 | 112124..115765 | 2029589 | BREX-system-P-loop-protein-BrxC-[Leptolyngbya-sp.-BC1307] |
| WP_194058630.1 | Type6.BREX | pPIWI+pPIWI->pPIWI+SF1-SF2-Helicase->pPIWI-><-SF1-SF2-Helicase+wHTH+Small-B-barrel+DUF1998<-BrxD-AAA<-PglZ<-?<-REase<-N6-METHYLASE<-DUF6079*<-BrxB-BrxF-iSTAND<-BrxA-DNA-binding<-BrxE<-CHTH\|\|?-><-pPIWI<-?<-N6-METHYLASE | 1220 | Bacteria>Terrabacteria_group>Cyanobacteriota/Melainabacteria_group>Cyanobacteriota>Cyanophyceae>Pseudanabaenales | GCF_015207525.1 | Nodosilinea_sp._LEGE_06152 | IQ265_RS24920 | 20188..23850 | 2777966 | BREX-system-P-loop-protein-BrxC-[Nodosilinea-sp.-LEGE-06152] |
| WP_275335484.1 | Type6.BREX | <-DUF3970<-?<-?\|\|CopG/RHH->TrwC+SF1-SF2-Helicase->BrxE->BrxA-DNA-binding->BrxB-BrxF-iSTAND->DUF6079*->N6-METHYLASE->MetJarc-HTH->RelE-ParE->N6-METHYLASE-p->?-><-?\|\|?->PIN->Schlafen+PIWI-> | 1212 | Bacteria>Terrabacteria_group>Cyanobacteriota/Melainabacteria_group>Cyanobacteriota>Cyanophyceae>Pseudanabaenales | GCF_029109325.1 | Nodosilinea_sp._TSF1-S3 | PGN35_RS19040 | 478733..482371 | 3020489 | BREX-system-P-loop-protein-BrxC-[Nodosilinea-sp.-TSF1-S3] |
| PZV04977.1 | Type6.BREX | GAF->?-><-?\|\|<-SIGMA+OB-Fold+AAA-ATPase+Lon-Protease-BrxL<-HTH_3<-?<-PglZ<-?<-N6-METHYLASE<-DUF6079*<-BrxB-BrxF-iSTAND<-BrxA-DNA-binding<-BrxE<-?<-Peptidase\|\|PhdYeFM_antitox->PIN->DUF4236-> | 1178 | Bacteria>Terrabacteria_group>Cyanobacteriota/Melainabacteria_group>Cyanobacteriota>Cyanophyceae>Synechococcales | GCA_003248995.1 | Cyanobium_sp. | DCF23_04805 | 51461..54997 | 2164130 | MAG:-BREX-system-P-loop-protein-BrxC-[Cyanobium-sp.] |
| MCF8139566.1 | Type6.BREX | Trypsin+PDZ-><-SIGMA-HTH\|\|REase->HNH-><-SIGMA+OB-Fold+AAA-ATPase+Lon-Protease-BrxL<-PglZ<-Peptidase_M78<-?<-N6-METHYLASE<-DUF6079*<-BrxB-BrxF-iSTAND<-BrxA-DNA-binding<-BrxE<-?\|\|AAA+wHTH+WYL->?->PemK_toxin-><-?<-Thermonuclease | 1178 | Bacteria>Terrabacteria_group>Cyanobacteriota/Melainabacteria_group>Cyanobacteriota>Cyanophyceae>Synechococcales | GCA_021737825.1 | Cyanobium_usitatum_Tobar12.5m-G36 | K9J75_00470 | 22428..25964 | 2874848 | BREX-system-P-loop-protein-BrxC-[Cyanobium-usitatum-Tobar12.5m-G36] |
| MCF8140154.1 | Type6.BREX | <-AAA+wHTH+WYL\|\|?-><-SF1-SF2-Helicase+wHTH+Small-B-barrel+DUF1998<-BrxD-AAA<-PglZ<-TOPRIM<-OLD-ABC<-N6-METHYLASE<-DUF6079*<-BrxB-BrxF-iSTAND<-BrxA-DNA-binding<-BrxE\|\|HEPN-> | 1215 | Bacteria>Terrabacteria_group>Cyanobacteriota/Melainabacteria_group>Cyanobacteriota>Cyanophyceae>Synechococcales | GCA_021737825.1 | Cyanobium_usitatum_Tobar12.5m-G36 | K9J75_03480 | 41417..45064 | 2874848 | BREX-system-P-loop-protein-BrxC-[Cyanobium-usitatum-Tobar12.5m-G36] |
| WP_051039072.1 | Type6.BREX | <-PglZ<-XIS-HTH\|\|NTP_transf_3->HTH-><-N6-METHYLASE-p<-PglZ\|\|?->DDE_Tnp_1-><-DUF6079-p\|\|<-DUF6079-p*<-BrxB-BrxF-iSTAND<-BrxA-DNA-binding<-BrxE<-TPR+TPR<-RNase | 595 | Bacteria>Terrabacteria_group>Cyanobacteriota/Melainabacteria_group>Cyanobacteriota>Cyanophyceae>Synechococcales | GCF_000317145.1 | Chamaesiphon_minutus | CHA6605_RS30405 | 314871..316658 | 1173032 | BREX-system-P-loop-protein-BrxC-[Chamaesiphon-minutus] |
| WP_186514738.1 | Type6.BREX | <-DUF4236+TPR<-?\|\|Thermonuclease->BrxE->BrxA-DNA-binding->BrxB-BrxF-iSTAND->DUF6079*->N6-METHYLASE->PglZ->Schlafen+HATPase+wHTH+HTH->SIGMA+OB-Fold+AAA-ATPase+Lon-Protease-BrxL->?-><-?\|\|SIGMA-HTH-> | 1176 | Bacteria>Terrabacteria_group>Cyanobacteriota/Melainabacteria_group>Cyanobacteriota>Cyanophyceae>Synechococcales | GCF_014280115.1 | Synechococcus_sp._A15-44 | SynA1544_RS02755 | 498877..502407 | 1050646 | BREX-system-P-loop-protein-BrxC-[Synechococcus-sp.-A15-44] |
| WP_193814304.1 | Type6.BREX | <-SIGMA+OB-Fold+AAA-ATPase+Lon-Protease-BrxL<-PglZ<-N6-METHYLASE<-ORC-CDC6<-ParA-Soj-PloopNTPase<-HEPN<-N6-METHYLASE<-DUF6079*<-BrxB-BrxF-iSTAND<-BrxA-DNA-binding<-BrxE | 1182 | Bacteria>Terrabacteria_group>Cyanobacteriota/Melainabacteria_group>Cyanobacteriota>Cyanophyceae>Synechococcales | GCF_015207535.1 | Cyanobium_sp._LEGE_06143 | IQ216_RS03215 | 12673..16221 | 945727 | BREX-system-P-loop-protein-BrxC-[Cyanobium-sp.-LEGE-06143] |
| WP_197211783.1 | Type6.BREX | XIS-HTH-><-Arm-DNA-bind_3+Phage_integrase<-tRNA<-SIGMA+OB-Fold+AAA-ATPase+Lon-Protease-BrxL<-HTH_3<-?<-PglZ<-Schlafen+HATPase+wHTH+HTH<-N6-METHYLASE<-DUF6079*<-BrxB-BrxF-iSTAND<-BrxA-DNA-binding<-BrxE<-?<-FA_desaturase | 1176 | Bacteria>Terrabacteria_group>Cyanobacteriota/Melainabacteria_group>Cyanobacteriota>Cyanophyceae>Synechococcales | GCF_015840715.1 | Synechococcus_sp._CBW1004 | H8F25_RS01965 | 424832..428362 | 1353136 | BREX-system-P-loop-protein-BrxC-[Synechococcus-sp.-CBW1004] |
| WP_254937144.1 | Type6.BREX | <-SIGMA-HTH\|\|REase->HNH-><-SIGMA+OB-Fold+AAA-ATPase+Lon-Protease-BrxL<-PglZ<-N6-METHYLASE<-DUF6079*<-BrxB-BrxF-iSTAND<-BrxA-DNA-binding<-BrxE<-?\|\|Sak4+WYL->?->PemK_toxin-><-?<-SAM-METHYLASE | 1178 | Bacteria>Terrabacteria_group>Cyanobacteriota/Melainabacteria_group>Cyanobacteriota>Cyanophyceae>Synechococcales | GCF_024345395.1 | Cyanobium_sp._Tous-M-B4 | KBY73_RS11030 | 158493..162029 | 2823724 | BREX-system-P-loop-protein-BrxC-[Cyanobium-sp.-Tous-M-B4] |
| WP_255021816.1 | Type6.BREX | <-SIGMA-HTH<-HEPN<-SIGMA+OB-Fold+AAA-ATPase+Lon-Protease-BrxL<-?<-Schlafen+HATPase+wHTH<-PglZ<-N6-METHYLASE<-DUF6079*<-BrxB-BrxF-iSTAND<-BrxA-DNA-binding<-BrxE<-?\|\|Sak4+WYL-><-?<-Thermonuclease\|\|?->DUF4236+TPR-> | 1178 | Bacteria>Terrabacteria_group>Cyanobacteriota/Melainabacteria_group>Cyanobacteriota>Cyanophyceae>Synechococcales | GCF_024345755.1 | Cyanobium_sp._L1E-Cus | KBZ22_RS09750 | 55554..59090 | 2823714 | BREX-system-P-loop-protein-BrxC-[Cyanobium-sp.-L1E-Cus] |

| Protein-ID | System | Gene-Neighbourhoods | Length | Taxonomy | Assembly-ID | Strain | Locus-Tag | Genomic-coordinates | TaxID | Fasta-defline |
|---|---|---|---|---|---|---|---|---|---|---|
| WP_255118717.1 | Type6.BREX | HNH-><-SIGMA+OB-Fold+AAA-ATPase+Lon-Protease-BrxL<-PglZ<-Schlafen+HATPase+wHTH<-N6-METHYLASE<-DUF6079*<-BrxB-BrxF-iSTAND<-BrxA-DNA-binding<-BrxE<-Pkinase+S1COLD<-CR-Kinase+pi3KINASE<-?<-Thermonuclease | 1176 | Bacteria>Terrabacteria_group>Cyanobacteriota/Melainabacteria_group>Cyanobacteriota>Cyanophyceae>Synechococcales | GCF_024345775.1 | unclassified_Synechococcus | KBY80_RS12355 | 16408..19938 | 2626047 | MULTISPECIES:-BREX-system-P-loop-protein-BrxC-[unclassified-Synechococcus] |
| WP_254954829.1 | Type6.BREX | <-SIGMA-HTH<-?<-HEPN\|\|BrxE->BrxA-DNA-binding->BrxB-BrxF-iSTAND->DUF6079*-> | 1215 | Bacteria>Terrabacteria_group>Cyanobacteriota/Melainabacteria_group>Cyanobacteriota>Cyanophyceae>Synechococcales | GCF_024345955.1 | unclassified_Cyanobium | KBZ34_RS18230 | 14804..18451 | 2627006 | MULTISPECIES:-BREX-system-P-loop-protein-BrxC-[unclassified-Cyanobium] |
| WP_254988813.1 | Type6.BREX | wHTH+HNH+HIT->EVE+HNH-><-SIGMA+OB-Fold+AAA-ATPase+Lon-Protease-BrxL<-DOC+HTH<-PglZ<-N6-METHYLASE<-DUF6079*<-BrxB-BrxF-iSTAND<-BrxA-DNA-binding<-BrxE<-?<-ArdCN+MPTase | 1178 | Bacteria>Terrabacteria_group>Cyanobacteriota/Melainabacteria_group>Cyanobacteriota>Cyanophyceae>Synechococcales | GCF_024346335.1 | Cyanobium_sp._ATX_6E8 | KBY70_RS13205 | 12848..16384 | 2823701 | BREX-system-P-loop-protein-BrxC-[Cyanobium-sp.-ATX-6E8] |
| WP_255017387.1 | Type6.BREX | LexA-protease->tRNA->?-><-HTH_Tnp_1+HTH_21+rve+rve_3<-HIN-HTH<-DUF6079-p*<-BrxB-BrxF-iSTAND<-BrxA-DNA-binding<-BrxE-p | 818 | Bacteria>Terrabacteria_group>Cyanobacteriota/Melainabacteria_group>Cyanobacteriota>Cyanophyceae>Synechococcales | GCF_024346475.1 | Cyanobium_sp._A2C-AMD | KBZ17_RS14945 | 4219..6675 | 2823695 | BREX-system-P-loop-protein-BrxC-[Cyanobium-sp.-A2C-AMD] |
| ETR72221.1 | Type6.BREX | SAM-METHYLASE->BrxE->BrxA-DNA-binding->BrxB-BrxF-iSTAND->DUF6079-p*->DUF6079-p->DOC+HTH->PIN->Phage_int_SAM_4+Phage_integrase-> | 719 | Bacteria>Thermodesulfobacteriota>Desulfobacteria>Desulfobacterales>Desulfobacteraceae>Candidatus_Magnetoglobus | GCA_000516475.1 | Candidatus_Magnetoglobus_multicellularis_str._Araruama | OMM_01887 | 4829..6988 | 890399 | hypothetical-protein-OMM_01887-[Candidatus-Magnetoglobus-multicellularis-str.-Araruama] |
| MBF0231918.1 | Type6.BREX | <-DUF6079*<-BrxB-BrxF-iSTAND<-BrxA-DNA-binding<-BrxE\|\|?->Calcineurin-SbcD->SbcC+CCDC158-> | 1187 | Bacteria>Thermodesulfobacteriota>Desulfobacteria>Desulfobacterales>Desulfobacteraceae>Desulfamplus | GCA_015231525.1 | Desulfamplus_sp. | HQK65_02630 | 1..3563 | 2006184 | BREX-system-P-loop-protein-BrxC-[Desulfamplus-sp.] |
| MBF0258414.1 | Type6.BREX | <-Phage_integrase<-?\|\|BrxE->BrxA-DNA-binding->BrxB-BrxF-iSTAND->DUF6079*-><-?<-BrnT_toxin\|\|N6-METHYLASE->PglZ->BrxD-AAA->SF1-SF2-Helicase+wHTH+Small-B-barrel+DUF1998-> | 1217 | Bacteria>Thermodesulfobacteriota>Desulfobacteria>Desulfobacterales>Desulfobacteraceae>Desulfamplus | GCA_015231705.1 | Desulfamplus_sp. | HQK62_06190 | 6144..9797 | 2006184 | BREX-system-P-loop-protein-BrxC-[Desulfamplus-sp.] |
| WP_080803995.1 | Type6.BREX | MerR-HTH->?-><-SIGMA+OB-Fold+AAA-ATPase+HKD-DNase-BrxL<-HTH_3<-?<-PglZ<-N6-METHYLASE<-?<-BrnT_toxin<-DUF6079*<-BrxB-BrxF-iSTAND<-BrxA-DNA-binding<-BrxE\|\|?-><-?<-REC+PAS+HISKIN<-Shikimate_dh_N+SDH_C<-REC+OmpR-HTH<-BirA-HTH | 1205 | Bacteria>Thermodesulfobacteriota>Desulfobacteria>Desulfobacterales>Desulfobacteraceae>Desulfamplus | GCF_900170035.1 | Desulfamplus_magnetovallimortis | MTBBW1_RS12175 | 100549..104166 | 1246637 | BREX-system-P-loop-protein-BrxC-[Desulfamplus-magnetovallimortis] |
| MBT5546118.1 | Type6.BREX | <-Schlafen+HATPase+wHTH<-ParB+HNH<-NERD<-?<-N6-METHYLASE-p<-N6-METHYLASE<-DUF6079*<-BrxB-BrxF-iSTAND<-BrxA-DNA-binding<-BrxE<-SF1-SF2-Helicase<-BrxD<-BrxD-AAA<-?<-RapA+SF1-SF2-Helicase+APATPase | 1206 | Bacteria>Thermodesulfobacteriota>Desulfobacteria>Desulfobacterales>Desulfobacteraceae>Desulfobacula | GCA_018665755.1 | Desulfobacula_sp. | HOK24_16785 | 13994..17614 | 2593537 | BREX-system-P-loop-protein-BrxC-[Desulfobacula-sp.] |
| MCD4719518.1 | Type6.BREX | BrxE->BrxA-DNA-binding->BrxB-BrxF-iSTAND->DUF6079*->OLD-ABC->REase->N6-METHYLASE->PglZ-> | 1208 | Bacteria>Thermodesulfobacteriota>Desulfobacteria>Desulfobacterales>Desulfobacteraceae>Desulfobacula | GCA_021107415.1 | Desulfobacula_sp. | K8S13_06615 | 2783..6409 | 2593537 | BREX-system-P-loop-protein-BrxC-[Desulfobacula-sp.] |
| MCF8075502.1 | Type6.BREX | BrxE->BrxA-DNA-binding->BrxB-BrxF-iSTAND->DUF6079*->N6-METHYLASE->OLD-ABC->PglZ->SIGMA+OB-Fold+AAA-ATPase+Lon-Protease-BrxL-p-> | 1219 | Bacteria>Thermodesulfobacteriota>Desulfobacteria>Desulfobacterales>Desulfobacteraceae>Desulfotignum | GCA_021737305.1 | Desulfotignum_sp. | K9K87_04985 | 1803..5462 | 2049434 | BREX-system-P-loop-protein-BrxC-[Desulfotignum-sp.] |
| MCF8089162.1 | Type6.BREX | ZetaToxin->?->?->BrxE->BrxA-DNA-binding->BrxB-BrxF-iSTAND->DUF6079*->N6-METHYLASE->OLD-ABC->PglZ->SIGMA+OB-Fold+AAA-ATPase+HKD-DNase-BrxL->?->SbcC-> | 1219 | Bacteria>Thermodesulfobacteriota>Desulfobacteria>Desulfobacterales>Desulfobacteraceae>Desulfotignum | GCA_021737465.1 | Desulfotignum_sp. | K9L23_14045 | 5260..8919 | 2049434 | BREX-system-P-loop-protein-BrxC-[Desulfotignum-sp.] |
| RJP85376.1 | Type6.BREX | URI-DNase->?->REase-><-?<-SF1-SF2-Helicase+wHTH+Small-B-barrel+DUF1998<-BrxD-AAA<-PglZ<-N6-METHYLASE<-DUF6079*<-BrxB-BrxF-iSTAND<-BrxA-DNA-binding<-BrxE\|\|NACHT->Classical-AAA-> | 1206 | Bacteria>Thermodesulfobacteriota>Desulfobacteria>Desulfobacterales>Desulfobacteraceae>unclassified_Desulfobacteraceae | GCA_003599885.1 | Desulfobacteraceae_bacterium | C4518_16165 | 18537..22157 | 2049433 | BREX-system-P-loop-protein-BrxC-[Desulfobacteraceae-bacterium] |
| MBT3312293.1 | Type6.BREX | Phage_integrase-><-?<-Schlafen+HATPase+wHTH<-ParB+HNH<-NERD<-?<-N6-METHYLASE-p<-N6-METHYLASE<-DUF6079*<-BrxB-BrxF-iSTAND<-BrxA-DNA-binding<-BrxE<-SF1-SF2-Helicase<-BrxD<-BrxD-AAA<-?<-RapA+SF1-SF2-Helicase+APATPase | 1206 | Bacteria>Thermodesulfobacteriota>Desulfobacteria>Desulfobacterales>Desulfobacteraceae>unclassified_Desulfobacteraceae | GCA_018648885.1 | Desulfobacteraceae_bacterium | HN379_09900 | 17655..21275 | 2049433 | BREX-system-P-loop-protein-BrxC-[Desulfobacteraceae-bacterium] |
| CBX26795.1 | Type6.BREX | <-rve<-SIGMA+OB-Fold+AAA-ATPase+HKD-DNase-BrxL<-PglZ<-Schlafen+HATPase+wHTH<-DrHyd<-N6-METHYLASE<-REase<-REase<-OLD-ABC<-DUF6079*<-BrxB-BrxF-iSTAND<-BrxA-DNA-binding<-BrxE\|\|URI-DNase-><-tRNA<-?\|\|?-><-PRTase<-DnaJ | 1208 | Bacteria>Thermodesulfobacteriota>Desulfobacteria>Desulfobacterales>Desulfobacteriaceae>Desulfobacterium | - | uncultured_Desulfobacterium_sp. | N47_A08240 | 98686..102312 | 201089 | hypothetical-protein-N47_A08240-[uncultured-Desulfobacterium-sp.] |
| MBC8432684.1 | Type6.BREX | <-REase<-SF1-SF2-Helicase+REase<-DNA_METHYLASE<-SIGMA+OB-Fold+AAA-ATPase+Lon-Protease-BrxL<-TOPRIM<-OLD-ABC<-PglZ<-ParB+HNH+Schlafen<-N6-METHYLASE<-DUF6079*<-BrxB-BrxF-iSTAND<-BrxA-DNA-binding<-BrxE<-XIS-HTH\|\|TatD_DNase-><-?<-HEPN<-PolB | 1227 | Bacteria>Thermodesulfobacteriota>Desulfobacteria>Desulfobacterales>unclassified_Desulfobacterales | GCA_014383215.1 | Desulfobacterales_bacterium | H8D96_12300 | 20226..23909 | 2044940 | BREX-system-P-loop-protein-BrxC-[Desulfobacterales-bacterium] |
| MBL7179931.1 | Type6.BREX | <-CHTH\|\|AbiEi_4+AbiEi_1->wHTH+WYL->HTH+HTH->N6-N4-METHYLASE-p->XIS-HTH->BrxE->BrxA-DNA-binding->BrxB-BrxF-iSTAND->DUF6079*->N6-METHYLASE-> | 1209 | Bacteria>Thermodesulfobacteriota>Desulfobacteria>Desulfobacterales>unclassified_Desulfobacterales | GCA_016782965.1 | Desulfobacterales_bacterium | ISS65_06935 | 17457..21086 | 2044940 | BREX-system-P-loop-protein-BrxC-[Desulfobacterales-bacterium] |
| KGO33733.1 | Type6.BREX | <-DUF6079*<-BrxB-BrxF-iSTAND<-BrxA-DNA-binding<-BrxE-p\|\|?-><-?<-?<-?\|\|wHTH+WYL+WCX-><-tRNA<-S1+RNase | 1223 | Bacteria>Thermodesulfobacteriota>Desulfobulbia>Desulfobulbales>Desulfobulbaceae>Desulfobulbus | GCA_000769715.1 | Desulfobulbus_sp._Tol-SR | JT06_12305 | 116..3787 | 1536652 | hypothetical-protein-JT06_12305-[Desulfobulbus-sp.-Tol-SR] |
| WP_205243962.1 | Type6.BREX | <-N6-METHYLASE-p<-HATPase+wHTH<-DUF6079*<-BrxB-BrxF-iSTAND<-BrxA-DNA-binding<-BrxE<-tRNA\|\|DNA_METHYLASE->?->?->?->?->METHYLASE-> | 1223 | Bacteria>Thermodesulfobacteriota>Desulfobulbia>Desulfobulbales>Desulfobulbaceae>Desulfobulbus | GCF_016918545.1 | Desulfobulbus_alkaliphilus | JWG43_RS16050 | 2786..6457 | 869814 | BREX-system-P-loop-protein-BrxC-[Desulfobulbus-alkaliphilus] |
| OQX18971.1 | Type6.BREX | ABC-ATPase+SbcC-><-?\|\|?->PadR-HTH-><-SIGMA+OB-Fold+AAA-ATPase+HKD-DNase-BrxL<-PglZ<-N6-METHYLASE<-HATPase+wHTH<-DUF6079*<-BrxB-BrxF-iSTAND<-BrxA-DNA-binding<-BrxE-p | 1229 | Bacteria>Thermodesulfobacteriota>Desulfobulbia>Desulfobulbales>Desulfobulbaceae>unclassified_Desulfobulbaceae | GCA_002083975.1 | Desulfobulbaceae_bacterium_A2 | BWK76_05990 | 20716..24405 | 1934250 | hypothetical-protein-BWK76_05990-[Desulfobulbaceae-bacterium-A2] |
| BDV01246.1 | Type6.BREX | <-DTXR-HTH\|\|?->BrxE->BrxA-DNA-binding->BrxB-BrxF-iSTAND->DUF6079*->N6-METHYLASE->N6-METHYLASE->OLD-ABC->TOPRIM->PglZ->SIGMA+OB-Fold+AAA-ATPase+HKD-DNase-BrxL->?-><-?\|\|HAMP+MA+MA-> | 1210 | Bacteria>Thermodesulfobacteriota>Desulfovibrionia>Desulfovibrionales>Desulfomicrobiaceae>Thermodesulfomicrobium | GCA_027925145.1 | Thermodesulfomicrobium_sp._WS | TDMWS_13310 | 1427384..1431016 | 3004129 | hypothetical-protein-TDMWS_13310-[Thermodesulfomicrobium-sp.-WS] |
| WP_176632187.1 | Type6.BREX | <-HNH<-ParB+HNH<-REase<-SF1-SF2-Helicase+REase<-DNA_METHYLASE<-SIGMA+OB-Fold+AAA-ATPase+Lon-Protease-BrxL<-PglZ<-N6-METHYLASE<-DUF6079*<-BrxB-BrxF-iSTAND<-BrxA-DNA-binding<-BrxE<-CHTH<-?<-?<-?<-?<-Peptidase_S78 | 1228 | Bacteria>Thermodesulfobacteriota>Desulfovibrionia>Desulfovibrionales>Desulfovibrionaceae>Desulfolutivibrio | GCF_013376495.1 | Desulfolutivibrio_sulfoxidireducens | GD605_RS15160 | 3521039..3524725 | 2773299 | BREX-system-P-loop-protein-BrxC-[Desulfolutivibrio-sulfoxidireducens] |
| GFH62752.1 | Type6.BREX | HISKIN->?->?->SIGMA-HTH->ncRNA->?->?->BrxE->BrxA-DNA-binding->BrxB-BrxF-iSTAND->DUF6079*->N6-METHYLASE->OLD-ABC->NERD->PglZ->SIGMA+OB-Fold+AAA-ATPase+HKD-DNase-BrxL-> | 1209 | Bacteria>Thermodesulfobacteriota>Desulfovibrionia>Desulfovibrionales>Desulfovibrionaceae>Desulfovibrio | GCA_013374015.1 | Candidatus_Desulfovibrio_kirbyi | ZNDK_0523 | 67387..71016 | 2696086 | BREX-system-P-loop-protein-[Candidatus-Desulfovibrio-kirbyi] |
| KIH76252.1 | Type6.BREX | <-REase+wHTH\|\|?-><-SIGMA+OB-Fold+AAA-ATPase+HKD-DNase-BrxL<-PglZ<-N6-METHYLASE<-DUF6079*<-BrxB-BrxF-iSTAND<-BrxA-DNA-binding<-BrxE<-ANK+ANK<-SIR2 | 1206 | Bacteria>Thermodesulfobacteriota>Desulfuromonadia>Desulfuromonadales>Geoalkalibacteraceae>Geoalkalibacter | GCA_000820505.1 | Geoalkalibacter_ferrihydriticus_DSM_17813 | BLS07_RS00630 | 142762..146382 | 1121915 | hypothetical-protein-GFER_11575-[Geoalkalibacter-ferrihydriticus-DSM-17813] |
| WP_040099782.1 | Type6.BREX | <-REase+wHTH\|\|?-><-SIGMA+OB-Fold+AAA-ATPase+HKD-DNase-BrxL<-PglZ<-N6-METHYLASE<-DUF6079*<-BrxB-BrxF-iSTAND<-BrxA-DNA-binding<-BrxE<-ANK+ANK<-SIR2 | 1206 | Bacteria>Thermodesulfobacteriota>Desulfuromonadia>Desulfuromonadales>Geoalkalibacteraceae>Geoalkalibacter | GCF_900103065.1 | Geoalkalibacter_ferrihydriticus | BLS07_RS00630 | 142762..146382 | 392333 | BREX-system-P-loop-protein-BrxC-[Geoalkalibacter-ferrihydriticus] |
| UPU37481.1 | Type6.BREX | <-XIS-HTH<-Arm-DNA-bind_3+Phage_integrase<-Arm-DNA-bind_2+Phage_integrase<-?<-?<-?<-?\|\|BrxE->BrxB-BrxF-iSTAND->DUF6079*->N6-METHYLASE->PglZ->BrxD-AAA->SF1-SF2-Helicase+wHTH+Small-B-barrel+DUF1998->McrB-AAA->?->?->McrB-AAA->McrC-NTD+REase-> | 1210 | Bacteria>Thermodesulfobacteriota>Desulfuromonadia>Geobacterales>Geobacteraceae>Geomonas | GCA_023221575.1 | Geomonas_paludis | M1B72_07185 | 1682397..1686029 | 2740185 | BREX-system-P-loop-protein-BrxC-[Geomonas-paludis] |
| WP_199394243.1 | Type6.BREX | ParB+HTH->XIS-HTH-><-SIGMA+OB-Fold+AAA-ATPase+Lon-Protease-BrxL<-PglZ<-AbiGi+seqA+RAMA<-N6-METHYLASE<-DUF6079*<-BrxB-BrxF-iSTAND<-BrxA-DNA-binding<-BrxE\|\|?->?->DUF2787-> | 1180 | Bacteria>Thermodesulfobacteriota>Desulfuromonadia>Geobacterales>Geobacteraceae>Geomonas | GCF_016458235.1 | Geomonas_propionica | JFN90_RS06275 | 364633..368175 | 2798582 | BREX-system-P-loop-protein-BrxC-[Geomonas-propionica] |
| WP_216500608.1 | Type6.BREX | Phage_integrase-><-?<-SIGMA+OB-Fold+AAA-ATPase+HKD-DNase-BrxL<-PglZ<-N6-METHYLASE<-DUF6079*<-BrxB-BrxF-iSTAND<-BrxA-DNA-binding<-BrxE<-MORC+REase<-wHTH+REase+ZnR<-McrB-AAA<-HTH+HEPN<-SF1-SF2-Helicase<-STYKIN+CR-Kinase | 1206 | Bacteria>Thermodesulfobacteriota>Desulfuromonadia>Geobacterales>Geobacteraceae>Geomonas | GCF_018919385.1 | Geomonas_diazotrophica | KOM00_RS19070 | 11745..15365 | 2843197 | BREX-system-P-loop-protein-BrxC-[Geomonas-diazotrophica] |
| WP_248647070.1 | Type6.BREX | <-XIS-HTH<-Arm-DNA-bind_3+Phage_integrase<-Arm-DNA-bind_2+Phage_integrase<-?<-?<-?<-?\|\|BrxE->BrxB-BrxF-iSTAND->DUF6079*->N6-METHYLASE->PglZ->BrxD-AAA->SF1-SF2-Helicase+wHTH+Small-B-barrel+DUF1998->McrB-AAA->?->?->McrB-AAA->McrC-NTD+REase-> | 1210 | Bacteria>Thermodesulfobacteriota>Desulfuromonadia>Geobacterales>Geobacteraceae>Geomonas | GCF_023221575.1 | Geomonas_paludis | M1B72_RS07185 | 1682397..1686029 | 2740185 | BREX-system-P-loop-protein-BrxC-[Geomonas-paludis] |
| WP_073040092.1 | Type6.BREX | <-HEPN<-PolB<-?<-?\|\|CHTH->BrxE->BrxA-DNA-binding->BrxB-BrxF-iSTAND->DUF6079*->N6-METHYLASE-> | 1226 | Bacteria>Thermodesulfobacteriota>Syntrophobacteria>Syntrophobacterales>Syntrophobacteraceae>Desulfacinum | GCF_900129305.1 | Desulfacinum_infernum | BUB04_RS12725 | 58235..61915 | 35837 | BREX-system-P-loop-protein-BrxC-[Desulfacinum-infernum] |
| OPY73251.1 | Type6.BREX | Phage_integrase-><-REase<-Phage_int_SAM_1+Phage_integrase<-AbiEii<-AbiEi_4+AbiEi_1\|\|CHTH->BrxE->BrxA-DNA-binding->BrxB-BrxF-iSTAND->DUF6079*->N6-METHYLASE->PglZ->OLD-ABC->TOPRIM->SIGMA+OB-Fold+AAA-ATPase+Lon-Protease-BrxL->DNA_METHYLASE->REase-> | 1224 | Bacteria>Thermodesulfobacteriota>Syntrophorhabdia>Syntrophorhabdales>Syntrophorhabdaceae>Syntrophorhabdus | GCA_002067415.1 | Syntrophorhabdus_sp._PtaU1.Bin002 | A4E62_00567 | 11466..15140 | 1811702 | MAG:-hypothetical-protein-A4E62_00567-[Syntrophorhabdus-sp.-PtaU1.Bin002] |
| TGQ70838.1 | Type6.BREX | HKD-DNase+HKD-Rease->?->BrxE->BrxA-DNA-binding->BrxB-BrxF-iSTAND->DUF6079*->N6-METHYLASE->N6-METHYLASE->PglZ->SIGMA+OB-Fold+AAA-ATPase+HKD-DNase-BrxL-><-?<-?<-CHTH | 1211 | Bacteria>unclassified_Bacteria | GCA_004791255.1 | bacterium_M00.F.Ca.ET.205.01.1.1 | EN848_08645 | 8202..11837 | 2563914 | BREX-system-P-loop-protein-BrxC-[bacterium-M00.F.Ca.ET.205.01.1.1] |
| NQX01743.1 | Type6.BREX | BrxE->BrxA-DNA-binding->BrxB-BrxF-iSTAND->DUF6079*->CR-ATPase5+REase->DOC+HTH->Four-helical-protein->N6-METHYLASE-p-> | 1210 | Bacteria>unclassified_Bacteria | GCA_013205625.1 | bacterium | HQ447_13895 | 2028..5660 | 1869227 | BREX-system-P-loop-protein-BrxC-[bacterium] |
| MBS2037451.1 | Type6.BREX | BrxE->BrxA-DNA-binding->BrxB-BrxF-iSTAND->DUF6079*-><-REase\|\|<-REase<-REase\|\|?-><-?<-?<-?\|\|MerR-HTH-><-MODE-HTH+PBPII | 1202 | Bacteria>unclassified_Bacteria | GCA_018266105.1 | bacterium | JST97_20850 | 5122..8730 | 1869227 | BREX-system-P-loop-protein-BrxC-[bacterium] |
| MBT9585258.1 | Type6.BREX | <-DUF6079-p*<-BrxB-BrxF-iSTAND<-BrxA-DNA-binding<-BrxE<-TPR+TPR+TPR+TPR<-Classical-AAA | 553 | Bacteria>unclassified_Bacteria | GCA_018780265.1 | bacterium | IV102_18080 | 1..1660 | 1869227 | BREX-system-P-loop-protein-BrxC-[bacterium] |
| MCE5200791.1 | Type6.BREX | <-REase<-SbcC+CCDC158<-Calcineurin-SbcD<-SF1-SF2-Helicase<-MORC+DUF3684+REase<-SIGMA+OB-Fold+AAA-ATPase+Lon-Protease-BrxL<-PglZ<-?<-N6-METHYLASE<-DUF6079*<-BrxB-BrxF-iSTAND<-BrxA-DNA-binding<-BrxE<-XIS-HTH<-?<-GDResolvase+Recombinase+Zn_ribbon_recom<-GDResolvase+Recombinase+Zn_ribbon_recom | 1217 | Bacteria>unclassified_Bacteria | GCA_021373255.1 | bacterium | LLG39_17655 | 101989..105642 | 1869227 | BREX-system-P-loop-protein-BrxC-[bacterium] |
| MCP4549289.1 | Type6.BREX | <-XIS-HTH<-?\|\|BrxE->BrxA-DNA-binding->BrxB-BrxF-iSTAND->N6-METHYLASE->?->REase->DUF6079*->HEPN->N6-METHYLASE-><-?\|\|?->N6-METHYLASE->AbiEi_4+AbiEi_1->AbiEii+ZnR-> | 1242 | Bacteria>unclassified_Bacteria | GCA_024231955.1 | bacterium | GY835_22785 | 11465..15193 | 1869227 | BREX-system-P-loop-protein-BrxC-[bacterium] |

| Protein-ID | System | Gene-Neighbourhoods | Length | Taxonomy | Assembly-ID | Strain | Locus-Tag | Genomic-coordinates | TaxID | Fasta-defline |
|---|---|---|---|---|---|---|---|---|---|---|
| -- | | | | | | | | | | |
| --Type2.Pgl.like.BREX | | | | | | | | | | |
| -- | | | | | | | | | | |
| MBX7222602.1 | Type2.Pgl.like.BREX | <-Sigma70_r2+Sigma70_r4_2<-PglZ<-DUF6079*<-?<-ParA-Soj-PloopNTPase<-N6-METHYLASE<-Y1_Tnp<-PglW<-?<-XIS-HTH | 1255 | Bacteria>Acidobacteriota>Blastocatellia>unclassified_Blastocatellia | GCA_019695185.1 | Blastocatellia_bacterium | K1Y36_21830 | 5409..9176 | 2052146 | phage-resistance-protein-[Blastocatellia-bacterium] |
| WP_179587197.1 | Type2.Pgl.like.BREX | <-HNH<-?\|\|TPR+TPR->TPR+TPR+TPR+TPR-><-SF1-SF2-Helicase+wHTH+Small-B-barrel+DUF1998<-BrxD-AAA<-PglZ<-DUF6079*<-N6-METHYLASE<-PglW<-?<-?\|\|ParB-><-?\|\|MODE-HTH+PBPII-> | 1224 | Bacteria>Acidobacteriota>Terriglobia>Terriglobales>Acidobacteriaceae>Edaphobacter | GCF_013410115.1 | Edaphobacter_lichenicola | HDF12_RS21445 | 9884..13558 | 2051959 | phage-resistance-protein-[Edaphobacter-lichenicola] |
| MBZ5495689.1 | Type2.Pgl.like.BREX | rve_3-><-SF1-SF2-Helicase+wHTH+Small-B-barrel+DUF1998-p<-BrxD-AAA<-PglZ<-DUF6079*<-Schlafen+HATPase+wHTH<-N6-METHYLASE<-PglW<-?<-?<-?<-DEOR-HTH+ISOCOT<-?<-SSB | 1230 | Bacteria>Acidobacteriota>Terriglobia>unclassified_Terriglobia | GCA_020072565.1 | Acidobacteriia_bacterium | LAP85_04755 | 203598..207290 | 2081523 | phage-resistance-protein-[Acidobacteriia-bacterium] |
| MBZ5659872.1 | Type2.Pgl.like.BREX | <-SF1-SF2-Helicase<-HKD-DNase<-?<-MORC+REase\|\|XIS-HTH->SF1-SF2-Helicase+REase->SF1-SF2-Helicase+REase->PglW->N6-METHYLASE->DUF6079*->PglZ->BrxD-AAA->SF1-SF2-Helicase+wHTH+Small-B-barrel+DUF1998-> | 1221 | Bacteria>Acidobacteriota>Terriglobia>unclassified_Terriglobia | GCA_020073485.1 | Acidobacteriia_bacterium | LA008_05635 | 166999..170664 | 2081523 | phage-resistance-protein-[Acidobacteriia-bacterium] |
| KAA0252888.1 | Type2.Pgl.like.BREX | <-SF1-SF2-Helicase+wHTH+Small-B-barrel+DUF1998<-BrxD-AAA<-PglZ<-DUF6079*<-CR-ATPase5+REase<-PIN<-RHH_6<-?<-AIPR<-N6-METHYLASE<-MazE_antitoxin<-?<-PglW | 1212 | Bacteria>Acidobacteriota>unclassified_Acidobacteriota | GCA_008363385.1 | Acidobacteriota_bacterium | EDX89_16490 | 7189..10827 | 1978231 | phage-resistance-protein-[Acidobacteriota-bacterium] |
| MBS1807695.1 | Type2.Pgl.like.BREX | ABC-ATPase+SbcC->?->Thioredoxin->?->?-><-?<-?<-PAS+HISKIN+REC\|\|tRNA->DUF6079-p*->?->?->PhdYeFM_antitox->RelE-ParE->?->?->Peptidase-> | 181 | Bacteria>Acidobacteriota>unclassified_Acidobacteriota | GCA_018268995.1 | Acidobacteriota_bacterium | JST84_05850 | 44896..45441 | 1978231 | hypothetical-protein-[Acidobacteriota-bacterium] |
| MCB1051856.1 | Type2.Pgl.like.BREX | N6-METHYLASE-p->DUF6079*->PglZ-><-RelE-ParE<-HEPN\|\|BrxD-AAA->SF1-SF2-Helicase+wHTH+Small-B-barrel+DUF1998->SF1-SF2-Helicase+HKD-DNase-> | 1210 | Bacteria>Acidobacteriota>unclassified_Acidobacteriota | GCA_020439765.1 | Acidobacteriota_bacterium | KDC71_14760 | 1153..4785 | 1978231 | phage-resistance-protein-[Acidobacteriota-bacterium] |
| MCB9398403.1 | Type2.Pgl.like.BREX | SLOG+BirA-HTH->?-><-SF1-SF2-Helicase+HKD-DNase<-SF1-SF2-Helicase+wHTH+Small-B-barrel+DUF1998<-BrxD-AAA\|\|HEPN->RelE-ParE-><-PglZ<-DUF6079*<-N6-METHYLASE<-PglW<-?\|\|<-?<-?\|\|?->Y1_Tnp-> | 1210 | Bacteria>Acidobacteriota>unclassified_Acidobacteriota | GCA_020637335.1 | Acidobacteriota_bacterium | H6510_11360 | 13924..17556 | 1978231 | phage-resistance-protein-[Acidobacteriota-bacterium] |
| MCI0393078.1 | Type2.Pgl.like.BREX | <-SF1-SF2-Helicase<-?<-SF1-SF2-Helicase+wHTH+Small-B-barrel+DUF1998<-BrxD-AAA<-PglZ<-DUF6079*<-PIN<-RHH<-N6-METHYLASE-p | 1229 | Bacteria>Acidobacteriota>unclassified_Acidobacteriota | GCA_022614935.1 | Acidobacteriota_bacterium | MOB07_30470 | 15158..18847 | 1978231 | phage-resistance-protein-[Acidobacteriota-bacterium] |
| WP_239491482.1 | Type2.Pgl.like.BREX | <-REase<-Vrr-REase\|\|PglW->TPR+TPR->N6-METHYLASE->SF1-SF2-Helicase+REase+HTH->wHTH->?->RelE-ParE->DUF6079*->PglZ->BrxD-AAA->SF1-SF2-Helicase+wHTH+Small-B-barrel+DUF1998-><-ParB<-Classical-AAA<-?<-?<-TPR\|\|SF1-SF2-Helicase+wHTH+Small-B-barrel+DUF1998-> | 1207 | Bacteria>Acidobacteriota>Vicinamibacteria>Vicinamibacterales>Vicinamibacteraceae>Luteitalea | GCF_016865485.1 | Luteitalea_sp._TBR-22 | TBR22_RS03065 | 752096..755719 | 2802971 | hypothetical-protein-[Luteitalea-sp.-TBR-22] |
| MCK6556375.1 | Type2.Pgl.like.BREX | <-URI-DNase<-PglZ<-DUF6079*<-PglW<-Aminotran_1_2<-tRNA<-tRNA\|\|?-><-?<-?<-?<-Glyco_transf_9<-?\|\|CR-ATPase5+REase-> | 1216 | Bacteria>Bacteria_incertae_sedis>Bacteria_candidate_phyla>Candidatus_Binatota>Candidatus_Binatia>unclassified_Candidatus_Binatia | GCA_023150935.1 | Candidatus_Binatia_bacterium | L6Q96_17615 | 3747..7397 | 2838779 | phage-resistance-protein-[Candidatus-Binatia-bacterium] |
| MBK8233303.1 | Type2.Pgl.like.BREX | N6-METHYLASE-><-DDE_Tnp_ISL3<-HTH_Tnp_ISL3+DDE_Tnp_ISL3<-DDE_Tnp_ISL3<-?<-?<-?<-PglZ<-?<-DUF6079*<-N6-METHYLASE-p\|\|zf-ISL3+HTH_Tnp_ISL3+DDE_Tnp_ISL3->DDE_Tnp_ISL3-><-SF1-SF2-Helicase+wHTH+Small-B-barrel+DUF1998<-BrxD-AAA\|\|?-><-Pkinase<-SIGMA-HTH | 1188 | Bacteria>Bacteria_incertae_sedis>Bacteria_candidate_phyla>Candidatus_Eisenbacteria | GCA_016712505.1 | Candidatus_Eisenbacteria_bacterium | IPK72_22705 | 37540..41106 | 2212470 | hypothetical-protein-[Candidatus-Eisenbacteria-bacterium] |
| MBK8233500.1 | Type2.Pgl.like.BREX | <-CR-ATPase5+REase<-?\|\|?->?->?->?-><-SF1-SF2-Helicase+wHTH+Small-B-barrel+DUF1998<-BrxD-AAA<-DUF6079*\|\|?->?->KAP-NTPase->?-><-METHYLASE<-?<-DUF433<-N6-METHYLASE | 1224 | Bacteria>Bacteria_incertae_sedis>Bacteria_candidate_phyla>Candidatus_Eisenbacteria | GCA_016712505.1 | Candidatus_Eisenbacteria_bacterium | IPK72_23715 | 273119..276793 | 2212470 | phage-resistance-protein-[Candidatus-Eisenbacteria-bacterium] |
| MBI4865891.1 | Type2.Pgl.like.BREX | <-ABhydrolase\|\|?->?->?-><-?\|\|NTP_transf_3-><-SF1-SF2-Helicase+wHTH+Small-B-barrel+DUF1998<-BrxD-AAA<-PglZ<-DUF6079*<-PIN<-wHTH<-N6-METHYLASE<-PglW<-?\|\|?->HMA+HAD+HAD->?-><-Peptidase_S58 | 1245 | Bacteria>Bacteria_incertae_sedis>Bacteria_candidate_phyla>Candidatus_Wallbacteria | GCA_016213845.1 | Candidatus_Wallbacteria_bacterium | HY816_02970 | 106548..110285 | 2053640 | phage-resistance-protein-[Candidatus-Wallbacteria-bacterium] |
| MBI2373943.1 | Type2.Pgl.like.BREX | PglW-><-PIN<-PhdYeFM_antitox\|\|?->\|\|N6-METHYLASE->?->Tox-RelE->HTH_3->?->DUF6079*->PglZ->BrxD-AAA->SF1-SF2-Helicase+wHTH+Small-B-barrel+35exo-> | 1216 | Bacteria>delta/epsilon_subdivisions>Deltaproteobacteria>unclassified_Deltaproteobacteria | GCA_016183035.1 | Deltaproteobacteria_bacterium | HYV07_08095 | 51496..55146 | 2026735 | hypothetical-protein-[Deltaproteobacteria-bacterium] |
| MBI5446424.1 | Type2.Pgl.like.BREX | PglW->N6-METHYLASE->NTP_transf_5->?->DUF6079*->PglZ->BrxD-AAA->SF1-SF2-Helicase+wHTH+Small-B-barrel+DUF1998->REase-> | 1225 | Bacteria>delta/epsilon_subdivisions>Deltaproteobacteria>unclassified_Deltaproteobacteria | GCA_016218905.1 | Deltaproteobacteria_bacterium | HY900_35070 | 8054..11731 | 2026735 | phage-resistance-protein-[Deltaproteobacteria-bacterium] |
| MBK7069433.1 | Type2.Pgl.like.BREX | SF1-SF2-Helicase+REase->?-><-SF1-SF2-Helicase+wHTH+Small-B-barrel+DUF1998<-BrxD-AAA<-PglZ<-DUF6079*<-?<-?<-?\|\|McrB-AAA->McrC-NTD+REase-> | 1234 | Bacteria>delta/epsilon_subdivisions>Deltaproteobacteria>unclassified_Deltaproteobacteria | GCA_016708475.1 | Deltaproteobacteria_bacterium | IPH47_31710 | 9535..13239 | 2026735 | hypothetical-protein-[Deltaproteobacteria-bacterium] |
| MBK8240682.1 | Type2.Pgl.like.BREX | AbiEi_4->AbiEii->ACET-><-HEPN<-SF1-SF2-Helicase+wHTH+Small-B-barrel+DUF1998<-BrxD-AAA<-PglZ<-DUF6079*<-N6-METHYLASE-p<-METHYLASE-p<-N6-METHYLASE | 1233 | Bacteria>delta/epsilon_subdivisions>Deltaproteobacteria>unclassified_Deltaproteobacteria | GCA_016712475.1 | Deltaproteobacteria_bacterium | IPK74_34620 | 583423..587124 | 2026735 | hypothetical-protein-[Deltaproteobacteria-bacterium] |
| MBO6936777.1 | Type2.Pgl.like.BREX | HIN-HTH->?->?->?->PglW->wHTH+REase->N6-METHYLASE->OLD-ABC->TOPRIM->DUF6079*->PglZ->Tox-RelA->BrxD-AAA->SF1-SF2-Helicase+wHTH+Small-B-barrel+35exo->NACHT->?->DUF932-> | 1220 | Bacteria>delta/epsilon_subdivisions>Deltaproteobacteria>unclassified_Deltaproteobacteria | GCA_017644045.1 | Deltaproteobacteria_bacterium | JJ863_17540 | 237441..241103 | 2026735 | hypothetical-protein-[Deltaproteobacteria-bacterium] |
| MCC6750664.1 | Type2.Pgl.like.BREX | TPR+REase-><-SF1-SF2-Helicase\|\|?->PglW->SbcC->?->N6-METHYLASE->DUF6079*->PglZ->BrxD-AAA->SF1-SF2-Helicase+wHTH+Small-B-barrel+DUF1998-><-?<-HAD+PRTase+PRTase+REase+SF1-SF2-Helicase | 1239 | Bacteria>delta/epsilon_subdivisions>Deltaproteobacteria>unclassified_Deltaproteobacteria | GCA_020848905.1 | Deltaproteobacteria_bacterium | IT371_23605 | 21330..25049 | 2026735 | hypothetical-protein-[Deltaproteobacteria-bacterium] |
| MCC7380504.1 | Type2.Pgl.like.BREX | N6-METHYLASE-p->DUF6079*->PglZ->BrxD-AAA->SF1-SF2-Helicase+wHTH+Small-B-barrel+DUF1998->?->SF1-SF2-Helicase+REase->TrwC+SF1-SF2-Helicase-> | 1227 | Bacteria>delta/epsilon_subdivisions>Deltaproteobacteria>unclassified_Deltaproteobacteria | GCA_020850435.1 | Deltaproteobacteria_bacterium | IT384_01640 | 825..4508 | 2026735 | hypothetical-protein-[Deltaproteobacteria-bacterium] |
| MCE9575760.1 | Type2.Pgl.like.BREX | <-Glycos_transf<-ASCH<-?<-SF1-SF2-Helicase+wHTH+Small-B-barrel+DUF1998<-BrxD-AAA<-PglZ<-DUF6079*<-?<-?<-ParB+HNH<-N6-METHYLASE<-PglW<-?<-wHTH+WYL+WCX\|\|SF1-SF2-Helicase->SF1-SF2-Helicase-> | 1225 | Bacteria>delta/epsilon_subdivisions>Deltaproteobacteria>unclassified_Deltaproteobacteria | GCA_021413675.1 | Deltaproteobacteria_bacterium | K8W52_21600 | 58772..62449 | 2026735 | hypothetical-protein-[Deltaproteobacteria-bacterium] |
| MVZ89494.1 | Type2.Pgl.like.BREX | ABC-ATPase-><-ParB+HNH<-?<-SF1-SF2-Helicase+wHTH+Small-B-barrel+DUF1998<-BrxD-AAA<-PglZ<-DUF6079*<-N6-METHYLASE\|\|HTH-><-PglW\|\|?->?->HNH-><-Cysteine-Desulfurase | 1285 | Bacteria>FCB_group>Bacteroidetes/Chlorobi_group>Bacteroidota>Bacteroidia>Bacteroidales | GCA_009768845.1 | Microbacter_sp._ANSKLAB05 | GQF49_03195 | 691555..695412 | 2684407 | phage-resistance-protein-[Microbacter-sp.-ANSKLAB05] |
| MBK9578953.1 | Type2.Pgl.like.BREX | wHTH+HNH->nSTAND1+HNH->REase-><-SF1-SF2-Helicase+wHTH+Small-B-barrel+DUF1998<-BrxD-AAA<-PglZ<-DUF6079*<-AbiEii<-AbiEi_3_N+AbiEi_3<-REase<-N6-METHYLASE<-PglW<-?<-wHTH+WYL+WCX<-PVC-AAA+Classical-AAA<-tRNA<-tRNA\|\|tRNA->TPR+TPR-> | 1216 | Bacteria>FCB_group>Fibrobacterota>unclassified_Fibrobacterota | GCA_016718115.1 | Fibrobacterota_bacterium | IPO40_17925 | 4308238..4311888 | 2052160 | hypothetical-protein-[Fibrobacterota-bacterium] |
| MBK9978474.1 | Type2.Pgl.like.BREX | <-DnaJ<-?<-BrxD-AAA<-PglZ<-PglZ-p<-DUF6079-p*<-DUF6079-p<-DUF6079-p<-N6-METHYLASE-p<-HIN-HTH<-N6-METHYLASE<-N6-METHYLASE-p<-?<-PglW | 771 | Bacteria>FCB_group>Gemmatimonadota>unclassified_Gemmatimonadota | GCA_016721935.1 | Gemmatimonadota_bacterium | IPP20_11200 | 618963..621278 | 2026742 | hypothetical-protein-[Gemmatimonadota-bacterium] |
| MBK9978475.1 | Type2.Pgl.like.BREX | <-DnaJ<-?<-BrxD-AAA<-PglZ<-PglZ-p<-DUF6079-p<-DUF6079-p*<-DUF6079-p<-N6-METHYLASE-p<-HIN-HTH<-N6-METHYLASE<-N6-METHYLASE-p<-?<-PglW | 297 | Bacteria>FCB_group>Gemmatimonadota>unclassified_Gemmatimonadota | GCA_016721935.1 | Gemmatimonadota_bacterium | IPP20_11205 | 621191..622084 | 2026742 | hypothetical-protein-[Gemmatimonadota-bacterium] |
| MBK9978476.1 | Type2.Pgl.like.BREX | <-DnaJ<-?<-BrxD-AAA<-PglZ<-PglZ-p<-DUF6079-p<-DUF6079-p<-DUF6079-p*<-N6-METHYLASE-p<-HIN-HTH<-N6-METHYLASE<-N6-METHYLASE-p<-?<-PglW\|\|?->?->REase-> | 204 | Bacteria>FCB_group>Gemmatimonadota>unclassified_Gemmatimonadota | GCA_016721935.1 | Gemmatimonadota_bacterium | IPP20_11210 | 621991..622605 | 2026742 | hypothetical-protein-[Gemmatimonadota-bacterium] |
| WP_204489965.1 | Type2.Pgl.like.BREX | XIS-HTH->?->PglW->N6-METHYLASE->NACHT->?->?->DUF6079*->PglZ->BrxD-AAA->SF1-SF2-Helicase+wHTH+Small-B-barrel+DUF1998-><-MORC+DUF3684+DUF3883<-NtrC-AAA<-?\|\|AAA-ATPase->REase->SF1-SF2-Helicase-> | 1232 | Bacteria>Myxococcota>Myxococcia>Myxococcales>Cystobacterineae>Archangiaceae | GCF_016904885.1 | Archangium_primigenium | I3V78_RS19475 | 4682490..4686188 | 2792470 | hypothetical-protein-[Archangium-primigenium] |
| WP_169825674.1 | Type2.Pgl.like.BREX | Pkinase->?->?-><-?\|\|XIS-HTH->?->PglW->N6-METHYLASE->DUF6079*->PglZ->BrxD-AAA->SF1-SF2-Helicase+wHTH+Small-B-barrel+DUF1998-><-REase<-DDE_Tnp_1 | 1232 | Bacteria>Myxococcota>Myxococcia>Myxococcales>Cystobacterineae>Myxococcaceae | - | Corallococcus_exiguus | HJC10_12660 | 18357..22055 | 83462 | hypothetical-protein-[Corallococcus-exiguus] |
| WP_147444643.1 | Type2.Pgl.like.BREX | HTH_58->?->zf-IS66+DDE_Tnp_IS66->?-><-?<-SF1-SF2-Helicase+wHTH+Small-B-barrel+DUF1998<-BrxD-AAA<-PglZ<-DUF6079*<-N6-METHYLASE<-?<-?<-PglW<-?\|\|?-><-tRNA<-?<-?<-PGK | 1209 | Bacteria>Myxococcota>Myxococcia>Myxococcales>Cystobacterineae>Myxococcaceae | GCF_003611675.1 | Corallococcus_sp._CA053C | D7V97_RS01525 | 32146..35775 | 2316732 | hypothetical-protein-[Corallococcus-sp.-CA053C] |
| WP_169345285.1 | Type2.Pgl.like.BREX | <-nSTAND1+HNH\|\|REase->Pkinase-><-NERD+SF1-SF2-Helicase\|\|SIR2-><-SF1-SF2-Helicase+wHTH+Small-B-barrel+DUF1998<-BrxD-AAA<-PglZ<-DUF6079*<-N6-METHYLASE<-PglW<-?<-tRNA<-?<-?<-PGK | 1233 | Bacteria>Myxococcota>Myxococcia>Myxococcales>Cystobacterineae>Myxococcaceae | GCF_012933655.1 | Pyxidicoccus_fallax | HG543_RS14220 | 30199..33900 | 394095 | hypothetical-protein-[Pyxidicoccus-fallax] |
| WP_193346317.1 | Type2.Pgl.like.BREX | REC->?->?->?->?-><-SF1-SF2-Helicase+wHTH+Small-B-barrel+DUF1998<-BrxD-AAA<-PglZ<-DUF6079*<-N6-METHYLASE<-PglW<-?<-XIS-HTH\|\|?-><-?\|\|Aminotran_1_2-> | 1231 | Bacteria>Myxococcota>Myxococcia>Myxococcales>Cystobacterineae>Myxococcaceae | GCF_014930455.1 | Corallococcus_soli | G4177_RS01770 | 428654..432349 | 2710757 | hypothetical-protein-[Corallococcus-soli] |
| WP_052548156.1 | Type2.Pgl.like.BREX | <-CARF+NtrC-AAA+FIS-HTH<-HKD-DNase<-?<-SF1-SF2-Helicase\|\|?->PglW->N6-METHYLASE->DUF6079*->PglZ->BrxD-AAA->SF1-SF2-Helicase+wHTH+Small-B-barrel+DUF1998->HTH+HTH+SF1-SF2-Helicase-> | 1232 | Bacteria>Myxococcota>Myxococcia>Myxococcales>Myxococcales_incertae_sedis>Enhygromyxa | GCF_000737335.2 | Enhygromyxa_salina | DB30_RS10015 | 151565..155263 | 215803 | hypothetical-protein-[Enhygromyxa-salina] |
| MBK6462400.1 | Type2.Pgl.like.BREX | PglW->N6-METHYLASE-p->N6-METHYLASE-p->N6-METHYLASE-p->SbcC->PIN->?-><-?\|\|DUF6079*->PglZ->BrxD-AAA->SF1-SF2-Helicase+wHTH+Small-B-barrel+DUF1998-> | 1239 | Bacteria>Myxococcota>Myxococcia>Myxococcales>unclassified_Myxococcales | GCA_016704525.1 | Myxococcales_bacterium | IPF92_15495 | 609713..613432 | 2026763 | hypothetical-protein-[Myxococcales-bacterium] |
| MBK7581086.1 | Type2.Pgl.like.BREX | HEPN-><-REase<-REase<-?<-Pkinase+TPR+TPR<-SF1-SF2-Helicase+wHTH+Small-B-barrel+DUF1998<-BrxD-AAA<-PglZ<-DUF6079*\|\|ZnR+HARE-HTH->?-><-?<-?\|\|HKD-DNase->REase-> | 1215 | Bacteria>Myxococcota>Myxococcia>Myxococcales>unclassified_Myxococcales | GCA_016706225.1 | Myxococcales_bacterium | IPI67_12840 | 1168179..1171826 | 2026763 | hypothetical-protein-[Myxococcales-bacterium] |
| MBK7402882.1 | Type2.Pgl.like.BREX | <-Phage_integrase\|\|?->Phage_integrase-><-SF1-SF2-Helicase-p<-SF1-SF2-Helicase+wHTH+Small-B-barrel+DUF1998-p<-PglZ<-DUF6079-p<-DUF6079-p*<-?<-OLD-ABC<-SbcC | 496 | Bacteria>Myxococcota>Myxococcia>Myxococcales>unclassified_Myxococcales | GCA_016709545.1 | Myxococcales_bacterium | IPJ34_43325 | 17328..18818 | 2026763 | hypothetical-protein-[Myxococcales-bacterium] |
| MBK8994477.1 | Type2.Pgl.like.BREX | <-XIS-HTH<-?\|\|?->?-><-Pkinase+TPR+TPR<-?<-SF1-SF2-Helicase+wHTH+Small-B-barrel+DUF1998<-BrxD-AAA<-PglZ<-DUF6079*<-?\|\|TPR->?->?-><-?<-?\|\|HKD-DNase-> | 1215 | Bacteria>Myxococcota>Myxococcia>Myxococcales>unclassified_Myxococcales | GCA_016716175.1 | Myxococcales_bacterium | IPM35_01815 | 439325..442972 | 2026763 | hypothetical-protein-[Myxococcales-bacterium] |
| MBK9034632.1 | Type2.Pgl.like.BREX | N6-METHYLASE-p->?->?->?->?-><-HTH_3\|\|?->MarR-HTH->DUF6079*->PglZ->BrxD-AAA->SF1-SF2-Helicase+wHTH+Small-B-barrel+DUF1998-><-?<-?<-?<-?<-?<-ADH_N+ADH_zinc_N | 1229 | Bacteria>Myxococcota>Myxococcia>Myxococcales>unclassified_Myxococcales | GCA_016717005.1 | Myxococcales_bacterium | IPL61_25765 | 505253..508942 | 2026763 | phage-resistance-protein-[Myxococcales-bacterium] |
| MBL8678385.1 | Type2.Pgl.like.BREX | HTH_Tnp_1+HTH_21+rve+rve_3->?->?->?-><-?\|\|?->?->DUF6079*->PglZ->BrxD-AAA->SF1-SF2-Helicase+wHTH+Small-B-barrel+DUF1998-><-SF1-SF2-Helicase+REase<-?\|\|CR-Kinase+CR-Kinase->?->NtrC-AAA+FIS-HTH-> | 1238 | Bacteria>Myxococcota>Myxococcia>Myxococcales>unclassified_Myxococcales | GCA_016793325.1 | Myxococcales_bacterium | JNK05_04435 | 11346..15062 | 2026763 | hypothetical-protein-[Myxococcales-bacterium] |
| MBL9101052.1 | Type2.Pgl.like.BREX | Pkinase->?-><-?\|\|?->?-><-SF1-SF2-Helicase+wHTH+Small-B-barrel+DUF1998<-?<-BrxD-AAA<-PglZ<-DUF6079*<-N6-METHYLASE<-PglW<-?\|\|SMODS->?-><-SAVED | 1239 | Bacteria>Myxococcota>Myxococcia>Myxococcales>unclassified_Myxococcales | GCA_016793465.1 | Myxococcales_bacterium | JNL82_08850 | 16272..19991 | 2026763 | hypothetical-protein-[Myxococcales-bacterium] |
| MBL9042574.1 | Type2.Pgl.like.BREX | <-SF1-SF2-Helicase+wHTH+Small-B-barrel+DUF1998<-BrxD-AAA<-PglZ<-DUF6079-p<-DUF6079-p*<-DUF6079-p\|\|REase->?-><-N6-METHYLASE-p<-nSTAND1+HNH | 741 | Bacteria>Myxococcota>Myxococcia>Myxococcales>unclassified_Myxococcales | GCA_016794465.1 | Myxococcales_bacterium | JNM83_13290 | 13478..15703 | 2026763 | hypothetical-protein-[Myxococcales-bacterium] |
| MBL9042575.1 | Type2.Pgl.like.BREX | <-SF1-SF2-Helicase+wHTH+Small-B-barrel+DUF1998<-BrxD-AAA<-PglZ<-DUF6079-p<-DUF6079-p<-DUF6079-p*\|\|REase->?-><-N6-METHYLASE-p<-nSTAND1+HNH | 196 | Bacteria>Myxococcota>Myxococcia>Myxococcales>unclassified_Myxococcales | GCA_016794465.1 | Myxococcales_bacterium | JNM83_13295 | 15767..16357 | 2026763 | hypothetical-protein-[Myxococcales-bacterium] |
| MBL9044563.1 | Type2.Pgl.like.BREX | <-SF1-SF2-Helicase+wHTH+Small-B-barrel+DUF1998<-BrxD-AAA<-PglZ<-DUF6079*<-?<-N6-METHYLASE<-PglW | 1228 | Bacteria>Myxococcota>Myxococcia>Myxococcales>unclassified_Myxococcales | GCA_016794465.1 | Myxococcales_bacterium | JNM83_23325 | 9267..12953 | 2026763 | hypothetical-protein-[Myxococcales-bacterium] |
| MCA9574057.1 | Type2.Pgl.like.BREX | PglW-><-ACET<-?\|\|N6-METHYLASE->?->Trypsin->Trypsin->?->DUF6079*->PglZ->BrxD-AAA->SF1-SF2-Helicase+wHTH+Small-B-barrel+DUF1998->?-><-NERD<-OLD-ABC\|\|?->OLD-ABC->HNH-> | 1219 | Bacteria>Myxococcota>Myxococcia>Myxococcales>unclassified_Myxococcales | GCA_020430805.1 | Myxococcales_bacterium | KC668_01430 | 17634..21293 | 2026763 | hypothetical-protein-[Myxococcales-bacterium] |
| MCB9544268.1 | Type2.Pgl.like.BREX | <-HTH_3<-SF1-SF2-Helicase+wHTH+Small-B-barrel+DUF1998<-BrxD-AAA<-PglZ<-DUF6079*<-N6-METHYLASE<-TOPRIM<-OLD-ABC<-PglW | 1210 | Bacteria>Myxococcota>Myxococcia>Myxococcales>unclassified_Myxococcales | GCA_020633435.1 | Myxococcales_bacterium | H6706_00065 | 12002..15634 | 2026763 | phage-resistance-protein-[Myxococcales-bacterium] |
| MCG5052490.1 | Type2.Pgl.like.BREX | <-HTH+HTH+SF1-SF2-Helicase\|\|?->PglW->N6-METHYLASE->N6-METHYLASE-p->OLD-ABC->TOPRIM->?->?->DUF6079*->PglZ->BrxD-AAA->SF1-SF2-Helicase+wHTH+Small-B-barrel+DUF1998->?->?-><-?\|\|?->REase-> | 1212 | Bacteria>Myxococcota>Myxococcia>Myxococcales>unclassified_Myxococcales | GCA_022184915.1 | Myxococcales_bacterium | KA712_05985 | 1448480..1452118 | 2026763 | hypothetical-protein-[Myxococcales-bacterium] |
| WP_224191848.1 | Type2.Pgl.like.BREX | SIGMA-HTH->?->?->CASPASE->?->?->PglW->N6-METHYLASE->DUF6079*->PglZ->BrxD-AAA->?->SF1-SF2-Helicase+wHTH+Small-B-barrel+DUF1998-> | 1247 | Bacteria>Myxococcota>Polyangia>Nannocystales>Nannocystaceae>Nannocystis | GCF_020073745.1 | Nannocystis_pusilla | K7C98_RS12500 | 226910..230653 | 889268 | hypothetical-protein-[Nannocystis-pusilla] |
| WP_129573695.1 | Type2.Pgl.like.BREX | ABC-ATPase+SbcC-><-SIGMA+OB-Fold+AAA-ATPase+Lon-Protease-BrxL<-?<-?<-RVT<-SF1-SF2-Helicase+wHTH+Small-B-barrel+DUF1998<-BrxD-AAA<-PglZ<-DUF6079*<-?<-?<-NACHT<-N6-METHYLASE<-TOPRIM<-OLD-ABC<-PglW<-?<-XIS-HTH | 1233 | Bacteria>Myxococcota>Polyangia>Polyangiales>Polyangiaceae | GCF_004135755.1 | Sorangium | SOCE836_RS08180 | 2224817..2228518 | 39643 | MULTISPECIES:-hypothetical-protein-[Sorangium] |
| WP_012240290.1 | Type2.Pgl.like.BREX | PglW->N6-METHYLASE-><-?<-?\|\|?-><-MerR-HTH<-?\|\|DUF6079*->OLD-ABC->HNH->PglZ->BrxD-AAA->SF1-SF2-Helicase+wHTH+Small-B-barrel+DUF1998-><-?<-?<-?<-ZnR | 1215 | Bacteria>Myxococcota>Polyangia>Polyangiales>Polyangiaceae>Sorangium | GCF_000067165.1 | Sorangium_cellulosum | SCE_RS49865 | 10694583..10698230 | 56 | hypothetical-protein-[Sorangium-cellulosum] |
| WP_104981119.1 | Type2.Pgl.like.BREX | MODE-HTH+PBPII->?->STYKIN->?-><-ACET\|\|MODE-HTH+PBPII-><-SF1-SF2-Helicase+wHTH+Small-B-barrel+DUF1998<-BrxD-AAA<-PglZ<-DUF6079*<-?<-GspA-ATPase<-?<-?<-Phage_GPD+Phage_base_V+Sarcoglycan_1<-?<-SbcC<-N6-METHYLASE<-PglW | 1205 | Bacteria>Myxococcota>Polyangia>Polyangiales>Polyangiaceae>Sorangium | GCF_002950945.1 | Sorangium_cellulosum | SOCE26_RS18530 | 5002376..5005993 | 56 | hypothetical-protein-[Sorangium-cellulosum] |
| WP_129345937.1 | Type2.Pgl.like.BREX | <-ParB+HNH\|\|?->Peptidase_M90->Classical-AAA->Peptidase_S8-><-SF1-SF2-Helicase+wHTH+Small-B-barrel+DUF1998<-BrxD-AAA<-PglZ<-DUF6079*<-N6-METHYLASE<-TOPRIM<-OLD-ABC<-PglW<-?<-XIS-HTH | 1232 | Bacteria>Myxococcota>Polyangia>Polyangiales>Polyangiaceae>Sorangium | GCF_004135735.1 | Sorangium_cellulosum | SOCEGT47_RS05500 | 1447833..1451531 | 56 | hypothetical-protein-[Sorangium-cellulosum] |
| WP_129349661.1 | Type2.Pgl.like.BREX | Glyco_hydro_19->?->?-><-?<-DUF6079-p*\|\|?->DDE_Tnp_1_assoc+DDE_Tnp_1->?->?->\|\|DDE_Tnp->LuxR-HTH->?->RHH_6->HNH-> | 190 | Bacteria>Myxococcota>Polyangia>Polyangiales>Polyangiaceae>Sorangium | GCF_004135735.1 | Sorangium_cellulosum | SOCEGT47_RS22655 | 6035496..6036068 | 56 | hypothetical-protein-[Sorangium-cellulosum] |
| NUP07975.1 | Type2.Pgl.like.BREX | AIPR->?->REase->SF1-SF2-Helicase+REase-><-?<-TPR\|\|?->PglW->N6-METHYLASE->DUF6079*->PglZ->BrxD-AAA->SF1-SF2-Helicase+wHTH+Small-B-barrel+DUF1998-><-SF1-SF2-Helicase<-REase<-EVE+HNH<-?<-?<-McrB-AAA | 1219 | Bacteria>Myxococcota>Polyangia>Polyangiales>Polyangiaceae>unclassified_Polyangiaceae | GCA_013361525.1 | Polyangiaceae_bacterium | HOW73_18165 | 31951..35610 | 2268199 | hypothetical-protein-[Polyangiaceae-bacterium] |
| MBK6514240.1 | Type2.Pgl.like.BREX | Amidohydro_2->?->?->?->TPR+TPR->?-><-SF1-SF2-Helicase+wHTH+Small-B-barrel+DUF1998<-BrxD-AAA<-PglZ<-DUF6079*\|\|?-><-?<-?<-N6-METHYLASE<-PglW | 1208 | Bacteria>Myxococcota>Polyangia>Polyangiales>Polyangiaceae>unclassified_Polyangiaceae | GCA_016704335.1 | Polyangiaceae_bacterium | IPG04_08985 | 133320..136946 | 2268199 | hypothetical-protein-[Polyangiaceae-bacterium] |
| MBK8255146.1 | Type2.Pgl.like.BREX | CNMP->Phd-YefM->PIN-><-SF1-SF2-Helicase+wHTH+Small-B-barrel+DUF1998<-BrxD-AAA<-PglZ<-?<-DUF6079*<-N6-METHYLASE<-PglW<-?<-RHS+RHS+HNH | 1238 | Bacteria>Myxococcota>Polyangia>Polyangiales>Polyangiaceae>unclassified_Polyangiaceae | GCA_016712345.1 | Polyangiaceae_bacterium | IPK82_21100 | 190917..194633 | 2268199 | hypothetical-protein-[Polyangiaceae-bacterium] |
| MBK8939661.1 | Type2.Pgl.like.BREX | TPR+TPR->?->?-><-?<-SF1-SF2-Helicase+wHTH+Small-B-barrel+DUF1998<-BrxD-AAA<-PglZ<-DUF6079*\|\|?-><-?<-?<-N6-METHYLASE-p<-N6-METHYLASE<-PglW | 1208 | Bacteria>Myxococcota>Polyangia>Polyangiales>Polyangiaceae>unclassified_Polyangiaceae | GCA_016719945.1 | Polyangiaceae_bacterium | IPM79_19085 | 122396..126022 | 2268199 | hypothetical-protein-[Polyangiaceae-bacterium] |
| MAT26109.1 | Type2.Pgl.like.BREX | URI-DNase-><-DUF932<-REase<-KAP-NTPase<-SF1-SF2-Helicase+wHTH+Small-B-barrel+35exo<-BrxD-AAA<-TraT<-Tox-RelA<-PglZ<-DUF6079*<-N6-METHYLASE<-wHTH+REase<-PglW<-?\|\|?->XIS-HTH->?->35exo-> | 1225 | Bacteria>Myxococcota>Polyangia>Polyangiales>Sandaracinaceae>Sandaracinus | GCA_002699025.1 | Sandaracinus_sp. | CMN29_14285 | 38263..41940 | 2024858 | hypothetical-protein-[Sandaracinus-sp.] |
| MBJ71781.1 | Type2.Pgl.like.BREX | <-35exo<-?<-XIS-HTH<-?\|\|?->PglW->wHTH+REase->N6-METHYLASE->DUF6079*->PglZ->Tox-RelA->TraT->BrxD-AAA->SF1-SF2-Helicase+wHTH+Small-B-barrel+35exo->KAP-NTPase->REase->DUF932-><-URI-DNase | 1225 | Bacteria>Myxococcota>Polyangia>Polyangiales>Sandaracinaceae>Sandaracinus | GCA_002720945.1 | Sandaracinus_sp. | CMN31_10645 | 12624..16301 | 2024858 | hypothetical-protein-[Sandaracinus-sp.] |
| MCB9616422.1 | Type2.Pgl.like.BREX | <-HKD-DNase+HKD-Rease<-?<-SF1-SF2-Helicase+wHTH+Small-B-barrel+35exo<-BrxD-AAA<-PglZ<-DUF6079*\|\|TPR-><-?\|\|?-><-HicA\|\|?->CHTH->?-><-N6-METHYLASE-p<-N6-METHYLASE | 1219 | Bacteria>Myxococcota>Polyangia>Polyangiales>Sandaracinaceae>Sandaracinus | GCA_020633155.1 | Sandaracinus_sp. | H6722_28635 | 10274..13933 | 2024858 | hypothetical-protein-[Sandaracinus-sp.] |
| MBK8409896.1 | Type2.Pgl.like.BREX | DUF6079-p*->DUF6079-p->PglZ->SF1-SF2-Helicase+wHTH+Small-B-barrel+DUF1998-p->?->SF1-SF2-Helicase+wHTH+Small-B-barrel+DUF1998-><-URI-DNase\|\|?->?->GDResolvase+Recombinase+ZnR-> | 400 | Bacteria>Myxococcota>Polyangia>Polyangiales>Sandaracinaceae>unclassified_Sandaracinaceae | GCA_016711615.1 | Sandaracinaceae_bacterium | IPL19_18190 | 1..1204 | 2510337 | phage-resistance-protein-[Sandaracinaceae-bacterium] |
| MCB9660515.1 | Type2.Pgl.like.BREX | LysM+PTSEIIA1-><-Pkinase+TPR+TPR\|\|tRNA->sGTP->?->sGTP->sGTP->?->PglW->N6-METHYLASE->DUF6079*->PglZ->BrxD-AAA->SF1-SF2-Helicase+wHTH+Small-B-barrel+DUF1998->McrB-NTD+HNH->Peptidase_M20->?->?->Acyltransferase-> | 1251 | Bacteria>Myxococcota>Polyangia>Polyangiales>Sandaracinaceae>unclassified_Sandaracinaceae | GCA_020633055.1 | Sandaracinaceae_bacterium | H6726_22920 | 102836..106591 | 2510337 | hypothetical-protein-[Sandaracinaceae-bacterium] |
| MCB9596957.1 | Type2.Pgl.like.BREX | HKD-DNase-><-?\|\|?->STYKinase-like->?->Peptidase-><-SF1-SF2-Helicase+wHTH+Small-B-barrel+35exo<-BrxD-AAA<-PglZ<-DUF6079*\|\|?->?->?-><-NERD<-OLD-ABC<-N6-METHYLASE<-PglW | 1224 | Bacteria>Myxococcota>Polyangia>Polyangiales>Sandaracinaceae>unclassified_Sandaracinaceae | GCA_020633195.1 | Sandaracinaceae_bacterium | H6719_29815 | 49488..53162 | 2510337 | hypothetical-protein-[Sandaracinaceae-bacterium] |
| WP_134500750.1 | Type2.Pgl.like.BREX | ARG-><-ARG\|\|?-><-?<-SF1-SF2-Helicase+DUF1998<-?<-SF1-SF2-Helicase+wHTH+Small-B-barrel+DUF1998<-BrxD-AAA<-PglZ<-DUF6079*<-N6-METHYLASE<-PglW<-NACHT\|\|tRNA-><-Phage_integrase | 1241 | Bacteria>Pseudomonadota>Alphaproteobacteria>Hyphomicrobiales>Methylobacteriaceae>Microvirga | GCF_004458765.1 | Microvirga_pakistanensis | E4K09_RS27275 | 130220..133945 | 1682650 | phage-resistance-protein-[Microvirga-pakistanensis] |
| CCD90940.1 | Type2.Pgl.like.BREX | MORC+DUF3684+REase-><-?\|\|SNF+SF1-SF2-Helicase->?-><-SF1-SF2-Helicase+wHTH+Small-B-barrel+DUF1998<-BrxD-AAA<-PglZ<-DUF6079*<-N6-METHYLASE<-PglW<-?<-XIS-HTH\|\|HTH-> | 1229 | Bacteria>Pseudomonadota>Alphaproteobacteria>Hyphomicrobiales>Nitrobacteraceae>Bradyrhizobium | GCA_000239775.2 | Bradyrhizobium_sp._ORS_375 | BRAO375_1270017 | 25541..29230 | 566679 | Bacteriophage-phiC31-resistance-gene-[Bradyrhizobium-sp.-ORS-375] |
| WP_139031288.1 | Type2.Pgl.like.BREX | CASPASE->?->?->?->\|\|MORC+DUF3684+REase->SF1-SF2-Helicase-><-SF1-SF2-Helicase+wHTH+Small-B-barrel+DUF1998<-BrxD-AAA<-PglZ<-DUF6079*<-N6-METHYLASE<-PglW<-?\|\|HTH->?-><-HIN-HTH<-DnaB | 1233 | Bacteria>Pseudomonadota>Alphaproteobacteria>Hyphomicrobiales>Nitrobacteraceae>Bradyrhizobium | GCF_000239775.1 | Bradyrhizobium_sp._ORS_375 | BRAO375_RS02955 | 25541..29242 | 566679 | hypothetical-protein-[Bradyrhizobium-sp.-ORS-375] |
| WP_247539401.1 | Type2.Pgl.like.BREX | CASPASE-><-PIWI<-REase+TIR\|\|?->GreA_GreB+SF1-SF2-Helicase->PglW->N6-METHYLASE->N6-METHYLASE-p->DUF6079*->PglZ->BrxD-AAA->SF1-SF2-Helicase+wHTH+Small-B-barrel+DUF1998-><-?<-?\|\|GDResolvase+HIN-HTH-> | 1261 | Bacteria>Pseudomonadota>Alphaproteobacteria>Hyphomicrobiales>Nitrobacteraceae>Bradyrhizobium | GCF_023073655.1 | Bradyrhizobium_sp._168 | IVB03_RS03610 | 137059..140844 | 2782639 | hypothetical-protein-[Bradyrhizobium-sp.-168] |
| RWO03902.1 | Type2.Pgl.like.BREX | <-GDResolvase+HIN-HTH\|\|DOC+HTH->?->?->?->RADICAL-SAM->MetJarc-HTH->PglW->N6-METHYLASE->DUF6079*->PglZ->BrxD-AAA->SF1-SF2-Helicase+wHTH+Small-B-barrel+DUF1998-> | 1254 | Bacteria>Pseudomonadota>Alphaproteobacteria>Hyphomicrobiales>Phyllobacteriaceae>Mesorhizobium | GCA_004021405.1 | Mesorhizobium_sp. | EOS07_31950 | 27266..31030 | 1871066 | phage-resistance-protein-[Mesorhizobium-sp.] |
| TIL48218.1 | Type2.Pgl.like.BREX | ParA-Soj-PloopNTPase->CopG/RHH->?->HTH-><-SF1-SF2-Helicase+wHTH+Small-B-barrel+DUF1998<-BrxD-AAA<-PglZ<-DUF6079*<-N6-METHYLASE<-PglW | 1233 | Bacteria>Pseudomonadota>Alphaproteobacteria>Hyphomicrobiales>Phyllobacteriaceae>Mesorhizobium | GCA_004961135.1 | Mesorhizobium_sp. | E5Y83_32340 | 14902..18603 | 1871066 | phage-resistance-protein-[Mesorhizobium-sp.] |
| TIR29941.1 | Type2.Pgl.like.BREX | ParA-Soj-PloopNTPase->CopG/RHH->?->HTH_3-><-SF1-SF2-Helicase+wHTH+Small-B-barrel+DUF1998<-BrxD-AAA<-PglZ<-DUF6079*<-N6-METHYLASE<-PglW\|\|XIS-HTH->Tox-RES->?->?-><-REase | 1233 | Bacteria>Pseudomonadota>Alphaproteobacteria>Hyphomicrobiales>Phyllobacteriaceae>Mesorhizobium | GCA_005063535.1 | Mesorhizobium_sp. | E5X35_24510 | 10898..14599 | 1871066 | phage-resistance-protein-[Mesorhizobium-sp.] |
| WP_259979852.1 | Type2.Pgl.like.BREX | <-SF1-SF2-Helicase+wHTH+Small-B-barrel+DUF1998<-BrxD-AAA<-PglZ<-DUF6079*<-N6-METHYLASE<-PglW | 1230 | Bacteria>Pseudomonadota>Alphaproteobacteria>Hyphomicrobiales>Phyllobacteriaceae>Mesorhizobium | GCF_025153915.1 | unclassified_Mesorhizobium | P3C30_RS29480 | 7340..11032 | 325217 | MULTISPECIES:-hypothetical-protein-[unclassified-Mesorhizobium] |
| SDR29277.1 | Type2.Pgl.like.BREX | Primpol+D5ATPase->?->ACYC+TPR+TPR->?-><-AAA-ATPase<-HIN-HTH<-BrxD-AAA<-PglZ<-DUF6079*<-N6-METHYLASE-p<-HTH_29+HTH_32+HTH_33+DDE_3<-N6-METHYLASE<-PglW\|\|HNH-> | 1254 | Bacteria>Pseudomonadota>Alphaproteobacteria>Hyphomicrobiales>unclassified_Hyphomicrobiales | GCA_900103895.1 | Rhizobiales_bacterium_GAS113 | SAMN05519103_01783 | 1902144..1905908 | 1884352 | hypothetical-protein-SAMN05519103_01783-[Rhizobiales-bacterium-GAS113] |
| WP_085440420.1 | Type2.Pgl.like.BREX | ParA-Soj-PloopNTPase->?->PglW->?->N6-METHYLASE->HEPN->Polbeta->DUF6079*->PglZ->BrxD-AAA->SF1-SF2-Helicase+wHTH+Small-B-barrel+DUF1998->\|\|AAA-ATPase->Phage_integrase->RelE-ParE->?->MPTase->NUDIX-> | 1216 | Bacteria>Pseudomonadota>Alphaproteobacteria>Magnetococcales>Magnetococcaceae>Magnetofaba | GCF_002109495.1 | Magnetofaba_australis | MAIT1_RS02375 | 14899..18549 | 1472297 | hypothetical-protein-[Magnetofaba-australis] |
| MBF0145158.1 | Type2.Pgl.like.BREX | <-SF1-SF2-Helicase<-?<-SF1-SF2-Helicase+wHTH+Small-B-barrel+DUF1998<-BrxD-AAA<-PglZ<-DUF6079*<-N6-METHYLASE<-?<-PglW\|\|ArsR-HTH->?->?->?->?->PolB-> | 1221 | Bacteria>Pseudomonadota>Alphaproteobacteria>Magnetococcales>unclassified_Magnetococcales | GCA_015229035.1 | Magnetococcales_bacterium | HQL85_03105 | 34578..38243 | 2026759 | phage-resistance-protein-[Magnetococcales-bacterium] |
| MBF0182076.1 | Type2.Pgl.like.BREX | PglW->?->N6-METHYLASE->DUF6079*->PglZ->BrxD-AAA->SF1-SF2-Helicase+wHTH+Small-B-barrel+DUF1998-> | 1221 | Bacteria>Pseudomonadota>Alphaproteobacteria>Magnetococcales>unclassified_Magnetococcales | GCA_015231235.1 | Magnetococcales_bacterium | HQM03_18820 | 9121..12786 | 2026759 | phage-resistance-protein-[Magnetococcales-bacterium] |
| MBF0261195.1 | Type2.Pgl.like.BREX | ParB->AAA-ATPase->PglW->N6-METHYLASE->HEPN->Polbeta->DUF6079*->PglZ->BrxD-AAA->SF1-SF2-Helicase+wHTH+Small-B-barrel+DUF1998-><-tRNA<-PAS+HISKIN+REC | 1224 | Bacteria>Pseudomonadota>Alphaproteobacteria>Magnetococcales>unclassified_Magnetococcales | GCA_015231775.1 | Magnetococcales_bacterium | HQL97_05040 | 14025..17699 | 2026759 | phage-resistance-protein-[Magnetococcales-bacterium] |
| MBF0427535.1 | Type2.Pgl.like.BREX | SF1-SF2-Helicase->DUF6079-p*->?->RADICAL-SAM-><-SF1-SF2-Helicase+DUF3418<-?<-CHASE2+ACYC | 115 | Bacteria>Pseudomonadota>Alphaproteobacteria>Magnetococcales>unclassified_Magnetococcales | GCA_015232765.1 | Magnetococcales_bacterium | HQL94_01320 | 2715..3062 | 2026759 | hypothetical-protein-[Magnetococcales-bacterium] |
| WP_090664771.1 | Type2.Pgl.like.BREX | <-GDResolvase<-KAP-NTPase\|\|?->HTH+sGTP->?->?-><-SF1-SF2-Helicase+wHTH+Small-B-barrel+DUF1998<-BrxD-AAA<-PglZ<-DUF6079*<-N6-METHYLASE<-HTH_3<-Tox-RelE<-PglW<-SF1-SF2-Helicase<-SF1-SF2-Helicase+DUF1998<-SIGMA-HTH<-SF1-SF2-Helicase<-sGTP+sGTP | 1235 | Bacteria>Pseudomonadota>Alphaproteobacteria>Rhodospirillales>Acetobacteraceae>Belnapia | GCF_900101615.1 | Belnapia_rosea | BLR02_RS22905 | 34881..38588 | 938405 | hypothetical-protein-[Belnapia-rosea] |
| WP_133290013.1 | Type2.Pgl.like.BREX | HNH+SAVED-><-SF1-SF2-Helicase+wHTH+Small-B-barrel+DUF1998<-BrxD-AAA<-PglZ<-DUF6079*<-N6-METHYLASE<-HTH_3<-Tox-RelE<-PglW | 1111 | Bacteria>Pseudomonadota>Alphaproteobacteria>Rhodospirillales>Acetobacteraceae>Dankookia | GCF_004355005.1 | Dankookia_rubra | E2C06_RS18055 | 7876..11211 | 1442381 | phage-resistance-protein-[Dankookia-rubra] |
| WP_160939352.1 | Type2.Pgl.like.BREX | <-HNH\|\|HTH_3-><-?<-SAM-METHYLASE<-HEPN<-SF1-SF2-Helicase+wHTH+Small-B-barrel+DUF1998<-BrxD-AAA<-PglZ<-DUF6079*<-N6-METHYLASE<-HTH_3<-PglW\|\|?->?-><-TraJ-RHH | 1234 | Bacteria>Pseudomonadota>Alphaproteobacteria>Rhodospirillales>Acetobacteraceae>Roseomonas | GCF_009829925.1 | Roseomonas_coralli | E0493_RS21655 | 10402..14106 | 2545983 | phage-resistance-protein-[Roseomonas-coralli] |
| WP_029008651.1 | Type2.Pgl.like.BREX | <-Phage_integrase<-HTH_3\|\|PglW->?->?-><-DOC+HTH\|\|N6-METHYLASE->?->?->DUF6079*->PglZ->BrxD-AAA->SF1-SF2-Helicase+wHTH+Small-B-barrel+DUF1998-><-?<-?<-TraJ-RHH<-?\|\|MODE-HTH+PBPII-> | 1248 | Bacteria>Pseudomonadota>Alphaproteobacteria>Rhodospirillales>Azospirillaceae>Azospirillum | GCF_000429625.1 | Azospirillum_halopraeferens | G472_RS0112395 | 159669..163415 | 34010 | hypothetical-protein-[Azospirillum-halopraeferens] |
| WP_119508925.1 | Type2.Pgl.like.BREX | Phage_integrase->Phage_integrase->PglW->REase->MACRODOMAIN-><-?\|\|N6-METHYLASE->?->?->DUF6079*->PglZ->BrxD-AAA->SF1-SF2-Helicase+wHTH+Small-B-barrel+DUF1998-><-ACET<-tRNA\|\|?->?->SWIM+DUF3670+SF1-SF2-Helicase-> | 1234 | Bacteria>Pseudomonadota>Alphaproteobacteria>Rhodospirillales>Azospirillaceae>Azospirillum | GCF_003584185.1 | Azospirillum_brasilense | D2T81_RS18095 | 28359..32063 | 192 | phage-resistance-protein-[Azospirillum-brasilense] |
| WP_119830362.1 | Type2.Pgl.like.BREX | ABC-ATPase+SbcC->?-><-?\|\|?->?-><-REase\|\|PglW->?->N6-METHYLASE->DUF6079*->PglZ->PglZ-p->BrxD-AAA->SF1-SF2-Helicase+wHTH+Small-B-barrel+DUF1998-><-XIS-HTH<-?\|\|?->Phage_integrase-> | 1240 | Bacteria>Pseudomonadota>Alphaproteobacteria>Rhodospirillales>Azospirillaceae>Azospirillum | GCF_003590795.1 | Azospirillum_cavernae | D3877_RS09520 | 2073388..2077110 | 2320860 | phage-resistance-protein-[Azospirillum-cavernae] |
| WP_149226934.1 | Type2.Pgl.like.BREX | PglW->N6-METHYLASE->?->?->DUF6079*->PglZ->BrxD-AAA->SF1-SF2-Helicase+wHTH+Small-B-barrel+DUF1998-><-rve+rve_3\|\|?->ZnR-><-?<-TetR-HTH | 1248 | Bacteria>Pseudomonadota>Alphaproteobacteria>Rhodospirillales>Azospirillaceae>Azospirillum | GCF_008364825.1 | Azospirillum_sp._B21 | FZ983_RS26270 | 90469..94215 | 2607496 | phage-resistance-protein-[Azospirillum-sp.-B21] |
| WP_149273804.1 | Type2.Pgl.like.BREX | <-DUF3670+SF1-SF2-Helicase<-?<-?<-?\|\|?-><-SF1-SF2-Helicase+wHTH+Small-B-barrel+DUF1998<-BrxD-AAA<-PglZ<-DUF6079*<-?<-?<-N6-METHYLASE<-PglW\|\|ParB+HNH->HTH_3-> | 1253 | Bacteria>Pseudomonadota>Alphaproteobacteria>Rhodospirillales>Azospirillaceae>Azospirillum | GCF_008365135.1 | Azospirillum_sp._Sh1 | FZ029_RS30830 | 15392..19153 | 2607285 | phage-resistance-protein-[Azospirillum-sp.-Sh1] |
| WP_149201507.1 | Type2.Pgl.like.BREX | TOPRIM-><-SWIM+DUF3670+SF1-SF2-Helicase<-?<-?<-?\|\|?-><-SF1-SF2-Helicase+wHTH+Small-B-barrel+DUF1998<-BrxD-AAA<-PglZ<-DUF6079*<-?<-?<-N6-METHYLASE<-PglW\|\|ParB+HNH->HTH_3-> | 1249 | Bacteria>Pseudomonadota>Alphaproteobacteria>Rhodospirillales>Azospirillaceae>Azospirillum | GCF_030160695.1 | Azospirillum_oryzae | FZ938_RS27750 | 31384..35133 | 286727 | phage-resistance-protein-[Azospirillum-oryzae] |
| WP_002730571.1 | Type2.Pgl.like.BREX | VirB10->RHH->PglW->PhdYeFM_antitox->PIN->N6-METHYLASE-><-RVT<-?\|\|DUF6079*->PglZ->BrxD-AAA->SF1-SF2-Helicase+wHTH+Small-B-barrel+DUF1998-><-ACET+ACET<-TraJ-RHH<-Arm-DNA-bind_3+Phage_int_SAM_3+Phage_integrase<-HTH_37<-Tox-RelE | 1226 | Bacteria>Pseudomonadota>Alphaproteobacteria>Rhodospirillales>Rhodospirillaceae>Magnetospirillum | GCF_000294655.1 | Magnetospirillum_molischianum | PHAMO_RS15140 | 20012..23692 | 1083 | hypothetical-protein-[Magnetospirillum-molischianum] |
| OQC18060.1 | Type2.Pgl.like.BREX | <-SF1-SF2-Helicase+wHTH+Small-B-barrel+DUF1998<-BrxD-AAA<-PglZ<-DUF6079*<-SecB<-MqsA_antitoxin+HTH_3<-MqsR_toxin<-HEPN<-N6-METHYLASE<-PglW<-HTH_26<-TraG_N<-TraH | 1239 | Bacteria>Pseudomonadota>Alphaproteobacteria>unclassified_Alphaproteobacteria | GCA_002070795.1 | Alphaproteobacteria_bacterium_ADurb.Bin100 | BWX79_00004 | 6983..10702 | 1852791 | hypothetical-protein-BWX79_00004-[Alphaproteobacteria-bacterium-ADurb.Bin100] |
| WP_248595568.1 | Type2.Pgl.like.BREX | HTH_26->PglW->N6-METHYLASE->PIN->N6-METHYLASE-p->AbrB-HTH->PIN->DUF6079*->DUF6079-p->PglZ->BirA-HTH->?->SF1-SF2-Helicase+wHTH+Small-B-barrel+DUF1998-><-?<-MNS+REase<-N6-N4-METHYLASE<-SF1-SF2-Helicase+REase | 1101 | Bacteria>Pseudomonadota>Betaproteobacteria>Betaproteobacteria_incertae_sedis>Candidatus_Accumulibacter | GCF_012939955.1 | Candidatus_Accumulibacter_phosphatis | E4Q23_RS18415 | 9811..13116 | 327160 | phage-resistance-protein-[Candidatus-Accumulibacter-phosphatis] |
| WP_186407482.1 | Type2.Pgl.like.BREX | <-PIN<-PrlF_antitoxin<-PglZ<-DUF6079*<-ORC-CDC6<-ParA-Soj-PloopNTPase<-HEPN<-N6-METHYLASE-p<-AbrB-HTH<-N6-METHYLASE<-REase<-PglW<-HTH_26 | 1230 | Bacteria>Pseudomonadota>Betaproteobacteria>Betaproteobacteria_incertae_sedis>Candidatus_Accumulibacter | GCF_900089955.1 | Candidatus_Accumulibacter_aalborgensis | ACCAA_40006 | 4735..8427 | 1860102 | phage-resistance-protein-[Candidatus-Accumulibacter-aalborgensis] |
| EXI73732.1 | Type2.Pgl.like.BREX | <-REase\|\|?->?->?-><-?<-SF1-SF2-Helicase+wHTH+Small-B-barrel+DUF1998<-BrxD-AAA<-PglZ<-DUF6079-p<-DUF6079*<-ORC-CDC6<-ParA-Soj-PloopNTPase<-HEPN<-N6-METHYLASE<-PglW | 1111 | Bacteria>Pseudomonadota>Betaproteobacteria>Betaproteobacteria_incertae_sedis>Candidatus_Accumulibacter>unclassified_Candidatus_Accumulibacter | GCA_000584995.1 | Candidatus_Accumulibacter_sp._SK-11 | AW07_02418 | 96349..99684 | 1454000 | MAG:-hypothetical-protein-AW07_02418-[Candidatus-Accumulibacter-sp.-SK-11] |
| HAY27851.1 | Type2.Pgl.like.BREX | <-SF1-SF2-Helicase+wHTH+Small-B-barrel+DUF1998<-BrxD-AAA<-PglZ<-DUF6079*\|\|?->?-><-ParA-Soj-PloopNTPase<-N6-METHYLASE<-PglW\|\|ParA-Soj-PloopNTPase->ParA-Soj-PloopNTPase-> | 1226 | Bacteria>Pseudomonadota>Betaproteobacteria>Betaproteobacteria_incertae_sedis>Candidatus_Accumulibacter>unclassified_Candidatus_Accumulibacter | GCA_003487685.1 | Candidatus_Accumulibacter_sp. | DCY47_10225 | 9448..13128 | 2053492 | phage-resistance-protein-[Candidatus-Accumulibacter-sp.] |
| MBK8115206.1 | Type2.Pgl.like.BREX | HTH_26->PglW-p->AbrB-HTH->N6-METHYLASE-p->OLD-ABC->NERD->AbrB-HTH->DUF6079*->PglZ->PrlF_antitoxin->PIN->SIR2->REase->SbcC->GNAT-> | 1245 | Bacteria>Pseudomonadota>Betaproteobacteria>Betaproteobacteria_incertae_sedis>Candidatus_Accumulibacter>unclassified_Candidatus_Accumulibacter | GCA_016712935.1 | Candidatus_Accumulibacter_sp. | IPK44_12010 | 14320..18057 | 2053492 | phage-resistance-protein-[Candidatus-Accumulibacter-sp.] |
| MBK7955387.1 | Type2.Pgl.like.BREX | ABC-ATPase+ABC-ATPase->?->?->PglW->N6-METHYLASE->HEPN->ParA-Soj-PloopNTPase->ORC-CDC6->DUF6079*->PglZ->BrxD-AAA-><-tRNA\|\|<-?<-?\|\|Aminotran_1_2->Aminotran_1_2->NAD_binding_3+Homoserine_dh+ACT-> | 1240 | Bacteria>Pseudomonadota>Betaproteobacteria>Betaproteobacteria_incertae_sedis>Candidatus_Accumulibacter>unclassified_Candidatus_Accumulibacter | GCA_016713625.1 | Candidatus_Accumulibacter_sp. | IPK02_16365 | 365185..368907 | 2053492 | phage-resistance-protein-[Candidatus-Accumulibacter-sp.] |
| MBN8498947.1 | Type2.Pgl.like.BREX | PglW->N6-METHYLASE->OLD-ABC->HNH->?->AbrB-HTH->DUF6079*->PglZ->BrxD-AAA->SF1-SF2-Helicase+wHTH+Small-B-barrel+DUF1998-> | 1215 | Bacteria>Pseudomonadota>Betaproteobacteria>Betaproteobacteria_incertae_sedis>Candidatus_Accumulibacter>unclassified_Candidatus_Accumulibacter | GCA_017302455.1 | Candidatus_Accumulibacter_sp. | JOM32_19155 | 11909..15556 | 2053492 | phage-resistance-protein-[Candidatus-Accumulibacter-sp.] |
| MBN8437012.1 | Type2.Pgl.like.BREX | HTH_26->PglW->N6-METHYLASE->OLD-ABC->HNH->N6-METHYLASE-p->OLD-ABC->NERD->AbrB-HTH->DUF6079*->PglZ->PrlF_antitoxin->PIN->CR-Kinase->TPR+TPR+TPR+TPR+TPR+TPR+TPR->?->HTH->DOC+HTH->BrxD-AAA-> | 1242 | Bacteria>Pseudomonadota>Betaproteobacteria>Betaproteobacteria_incertae_sedis>Candidatus_Accumulibacter>unclassified_Candidatus_Accumulibacter | GCA_017302555.1 | Candidatus_Accumulibacter_sp. | JOL95_03060 | 58448..62176 | 2053492 | phage-resistance-protein-[Candidatus-Accumulibacter-sp.] |
| MCM8596143.1 | Type2.Pgl.like.BREX | Four-helical-protein->N6-METHYLASE->AbrB-HTH->PIN->N6-METHYLASE-p->HEPN->ParA-Soj-PloopNTPase->ORC-CDC6->DOC+HTH->DUF6079-p*->DOC+HTH->DUF6079->PglZ->BrxD-AAA->SF1-SF2-Helicase+wHTH+Small-B-barrel+DUF1998->?->?->?->ABhydrolase-> | 167 | Bacteria>Pseudomonadota>Betaproteobacteria>Betaproteobacteria_incertae_sedis>Candidatus_Accumulibacter>unclassified_Candidatus_Accumulibacter | GCA_023806595.1 | Candidatus_Accumulibacter_sp. | NFV71_12460 | 22138..22641 | 2053492 | hypothetical-protein-[Candidatus-Accumulibacter-sp.] |
| MCM8596145.1 | Type2.Pgl.like.BREX | AbrB-HTH->PIN->N6-METHYLASE-p->HEPN->ParA-Soj-PloopNTPase->ORC-CDC6->DOC+HTH->DUF6079-p->DOC+HTH->DUF6079*->PglZ->BrxD-AAA->SF1-SF2-Helicase+wHTH+Small-B-barrel+DUF1998->?->?->?->ABhydrolase-><-ABC-ATPase+SbcC | 1210 | Bacteria>Pseudomonadota>Betaproteobacteria>Betaproteobacteria_incertae_sedis>Candidatus_Accumulibacter>unclassified_Candidatus_Accumulibacter | GCA_023806595.1 | Candidatus_Accumulibacter_sp. | NFV71_12470 | 24231..27863 | 2053492 | phage-resistance-protein-[Candidatus-Accumulibacter-sp.] |
| MCM8625579.1 | Type2.Pgl.like.BREX | AbrB-HTH->PIN->N6-METHYLASE-p->HEPN->ParA-Soj-PloopNTPase->ORC-CDC6->DOC+HTH->DUF6079-p->DOC+HTH->DUF6079*->PglZ->BrxD-AAA->SF1-SF2-Helicase+wHTH+Small-B-barrel+DUF1998->?->?->?->ABhydrolase-><-ABC-ATPase+SbcC | 1210 | Bacteria>Pseudomonadota>Betaproteobacteria>Betaproteobacteria_incertae_sedis>Candidatus_Accumulibacter>unclassified_Candidatus_Accumulibacter | GCA_023806625.1 | Candidatus_Accumulibacter_sp. | NFW05_06375 | 11612..15244 | 2053492 | phage-resistance-protein-[Candidatus-Accumulibacter-sp.] |
| WP_034949743.1 | Type2.Pgl.like.BREX | PglW->N6-METHYLASE->GrpE->?->MqsR_toxin->MqsA_antitoxin+HTH->SecB->DUF6079*->PglZ->BrxD-AAA->SF1-SF2-Helicase+wHTH+Small-B-barrel+DUF1998-><-MobA_MobL\|\|?->?-><-VirD4-TraG+VirD4-FtsK | 1232 | Bacteria>Pseudomonadota>Betaproteobacteria>Betaproteobacteria_incertae_sedis | GCF_000584975.1 | Candidatus_Accumulibacter | ACCUM_RS01425 | 33268..36966 | 327159 | phage-resistance-protein-[Candidatus-Accumulibacter-phosphatis] |
| WP_097389166.1 | Type2.Pgl.like.BREX | <-Integrase_DNA+Phage_integrase\|\|REase->?->REase+iREase-><-?<-HNH+Polbeta<-SF1-SF2-Helicase+wHTH+Small-B-barrel+DUF1998<-BrxD-AAA<-PglZ<-DUF6079*<-N6-METHYLASE<-PglW<-HTH_26 | 1232 | Bacteria>Pseudomonadota>Betaproteobacteria>Burkholderiales>Burkholderiaceae>Burkholderia | - | Burkholderia_sp._OK233 | SAMN05446635_0228 | 218410..222108 | 1882795 | phage-resistance-protein-[Burkholderia-sp.-OK233] |
| WP_009893665.1 | Type2.Pgl.like.BREX | <-DUF932<-?\|\|wHTH+WYL+WCX->PglW->N6-METHYLASE->DUF6079*->PglZ-><-?<-Vrr-REase<-?<-tRNA\|\|?->?-><-?\|\|MODE-HTH+PBPII-><-TOPRIM+Topoisom_bac | 1227 | Bacteria>Pseudomonadota>Betaproteobacteria>Burkholderiales>Burkholderiaceae>Burkholderia | GCF_000012365.1 | Burkholderia_thailandensis | BTH_RS12790 | 139845..143528 | 57975 | hypothetical-protein-[Burkholderia-thailandensis] |
| WP_043294106.1 | Type2.Pgl.like.BREX | <-TPR+TPR\|\|?->?->?->?->\|\|AA_kinase+ACT+ACT_7->tRNA->PglW->N6-METHYLASE->DUF6079*->PglZ-><-HDRnase\|\|?-><-?<-AbiEii<-AbiEi_3_N+AbiEi_3\|\|?-><-TraG_N | 1230 | Bacteria>Pseudomonadota>Betaproteobacteria>Burkholderiales>Burkholderiaceae>Burkholderia | GCF_000775085.1 | Burkholderia_pseudomallei | Y026_RS18545 | 31385..35077 | 28450 | hypothetical-protein-[Burkholderia-pseudomallei] |
| WP_066579362.1 | Type2.Pgl.like.BREX | HDRnase-><-PglZ<-DUF6079*<-N6-METHYLASE<-PglW<-tRNA<-AA_kinase+ACT+ACT_7\|\|<-?<-?<-?<-?\|\|TPR+TPR-> | 1230 | Bacteria>Pseudomonadota>Betaproteobacteria>Burkholderiales>Burkholderiaceae>Burkholderia | GCF_001523835.1 | Burkholderia_sp._ABCPW_14 | WS62_RS31385 | 6102..9794 | 1637860 | hypothetical-protein-[Burkholderia-sp.-ABCPW-14] |
| WP_076853767.1 | Type2.Pgl.like.BREX | <-PIN<-HTH_3<-Phage_integrase\|\|?->CHTH->Phage_integrase-><-PglZ<-DUF6079*<-N6-METHYLASE<-PglW<-wHTH+WYL+WCX<-?<-?<-?<-?<-DUF932 | 1232 | Bacteria>Pseudomonadota>Betaproteobacteria>Burkholderiales>Burkholderiaceae>Burkholderia | GCF_001976185.1 | Burkholderia_pseudomallei | AQ710_RS14945 | 7213..10911 | 28450 | hypothetical-protein-[Burkholderia-pseudomallei] |
| WP_089468784.1 | Type2.Pgl.like.BREX | <-RNaseP<-?\|\|RFC-AAA+DNAA-HTH->?->MORC+DNA_gyrase+TOPRIM->PglW->N6-METHYLASE->DUF6079*->PglZ->SymE_toxin-> | 1231 | Bacteria>Pseudomonadota>Betaproteobacteria>Burkholderiales>Burkholderiaceae>Burkholderia | GCF_002223345.1 | Burkholderia_sp._AU16741 | CFB89_RS16680 | 753667..757362 | 2015347 | hypothetical-protein-[Burkholderia-sp.-AU16741] |
| WP_201762955.1 | Type2.Pgl.like.BREX | TOPRIM+Topoisom_bac-><-MODE-HTH+PBPII\|\|?-><-?<-?\|\|tRNA->?->Vrr-REase->?-><-PglZ<-DUF6079*<-N6-METHYLASE<-PglW<-wHTH+WYL+WCX\|\|?->DUF932-> | 1218 | Bacteria>Pseudomonadota>Betaproteobacteria>Burkholderiales>Burkholderiaceae>Burkholderia | GCF_009909285.1 | Burkholderia_thailandensis | GT630_RS08725 | 154943..158599 | 57975 | hypothetical-protein-[Burkholderia-thailandensis] |
| WP_165987030.1 | Type2.Pgl.like.BREX | <-rve+rve_3<-?<-CASPASE\|\|?->TPR-><-?\|\|HTH_26->PglW->N6-METHYLASE->DUF6079*->PglZ->BrxD-AAA->SF1-SF2-Helicase+wHTH+Small-B-barrel+DUF1998->ANK+ANK-><-REase<-SbcC\|\|SIGMA-HTH+HIN-HTH->AAA-ATPase-> | 1232 | Bacteria>Pseudomonadota>Betaproteobacteria>Burkholderiales>Burkholderiaceae>Caballeronia | GCF_011493005.1 | Caballeronia_sp._SBC1 | SBC1_RS01400 | 309404..313102 | 2705548 | phage-resistance-protein-[Caballeronia-sp.-SBC1] |
| WP_109581345.1 | Type2.Pgl.like.BREX | Phage_integrase-><-UDG<-SF1-SF2-Helicase+wHTH+Small-B-barrel+DUF1998<-BrxD-AAA<-PglZ<-DUF6079*<-N6-METHYLASE<-PglW<-HTH_26<-HKD-DNase+DUF3150<-?<-SF1-SF2-Helicase+REase<-SF1-SF2-Helicase<-Exonuc_V_gamma+REase<-wHTH+WYL+WCX | 1238 | Bacteria>Pseudomonadota>Betaproteobacteria>Burkholderiales>Burkholderiaceae>Cupriavidus | GCF_003148725.1 | Cupriavidus_plantarum | C7419_RS07730 | 1681669..1685385 | 942865 | phage-resistance-protein-[Cupriavidus-plantarum] |
| WP_224079890.1 | Type2.Pgl.like.BREX | Phage_int_SAM_2+Integrase_1->?-><-SWIM+DUF3670+SF1-SF2-Helicase<-?<-?<-?\|\|HTH_26->PglW->N6-METHYLASE->DUF6079*->PglZ->BrxD-AAA->SF1-SF2-Helicase+wHTH+Small-B-barrel+DUF1998-> | 1238 | Bacteria>Pseudomonadota>Betaproteobacteria>Burkholderiales>Burkholderiaceae>Cupriavidus | GCF_914271435.1 | Cupriavidus_laharis | LDO29_RS11410 | 343301..347017 | 151654 | phage-resistance-protein-[Cupriavidus-laharis] |
| ODS98668.1 | Type2.Pgl.like.BREX | <-DUF772+DDE_Tnp_1\|\|Phage_integrase->HTH_26->PglW->N6-METHYLASE-><-?\|\|?->DOC+HTH->DUF6079*->PglZ->BrxD-AAA->SF1-SF2-Helicase+wHTH+Small-B-barrel+DUF1998-><-?<-?<-?<-?<-?<-ABhydrolase | 1232 | Bacteria>Pseudomonadota>Betaproteobacteria>Burkholderiales>Burkholderiaceae>Lautropia | GCA_001724855.1 | Lautropia_sp._SCN_69-89 | ABS56_03945 | 16954..20652 | 1660104 | MAG:-phage-resistance-protein-[Lautropia-sp.-SCN-69-89] |
| WP_028211990.1 | Type2.Pgl.like.BREX | DUF932->?->?->?->?->wHTH+WYL+WCX->PglW->N6-METHYLASE->DUF6079*->PglZ-><-Phage_integrase<-?\|\|Arm-DNA-bind_3+Phage_integrase-> | 1232 | Bacteria>Pseudomonadota>Betaproteobacteria>Burkholderiales>Burkholderiaceae>Paraburkholderia | GCF_000472845.1 | Paraburkholderia_mimosarum | A19S_RS0116540 | 25178..28876 | 312026 | hypothetical-protein-[Paraburkholderia-mimosarum] |
| WP_085482634.1 | Type2.Pgl.like.BREX | <-SF1-SF2-Helicase+wHTH+Small-B-barrel+DUF1998<-BrxD-AAA<-PglZ<-DUF6079*<-N6-METHYLASE<-PglW<-HTH_26<-tRNA<-?<-?<-SIS_2+SIS<-REase\|\|TP_METHYLASE-> | 1232 | Bacteria>Pseudomonadota>Betaproteobacteria>Burkholderiales>Burkholderiaceae>Paraburkholderia | GCF_900177725.1 | Paraburkholderia_susongensis | B9Z01_RS10495 | 309671..313369 | 1515439 | phage-resistance-protein-[Paraburkholderia-susongensis] |
| WP_200621125.1 | Type2.Pgl.like.BREX | <-ParB+HNH<-AAA-ATPase<-HTH+rve\|\|?->PhdYeFM_antitox->RelE-ParE-><-SF1-SF2-Helicase+wHTH+Small-B-barrel+DUF1998<-BrxD-AAA<-PglZ<-DUF6079*<-N6-METHYLASE<-PglW<-HTH_26 | 1233 | Bacteria>Pseudomonadota>Betaproteobacteria>Burkholderiales>Burkholderiaceae>Paraburkholderia | GCF_905220685.1 | Paraburkholderia_aspalathi | KHO25_RS29095 | 31330..35031 | 1324617 | phage-resistance-protein-[Paraburkholderia-aspalathi] |
| NKA78072.1 | Type2.Pgl.like.BREX | RFC-AAA+DNAA-HTH->?->MORC+DNA_gyrase+TOPRIM-><-PglZ<-DUF6079*<-N6-METHYLASE<-PglW<-?\|\|METHYLASE->LexA-protease->Radical-SAM->METHYLASE+DsrD-> | 1232 | Bacteria>Pseudomonadota>Betaproteobacteria>Burkholderiales>Burkholderiaceae>Ralstonia | GCA_012062495.1 | Ralstonia_solanacearum | GO291_00005 | 8535..12233 | 305 | hypothetical-protein-[Ralstonia-solanacearum] |
| WP_231408557.1 | Type2.Pgl.like.BREX | RFC-AAA+DNAA-HTH->?->MORC+DNA_gyrase+TOPRIM-><-HTH_29+HTH_32+DDE_3\|\|SbcC+SbcC->\|\|HIN-HTH->AAA-ATPase-><-PglZ<-DUF6079*<-N6-METHYLASE<-PglW<-?\|\|DUF4096+DDE_Tnp_1+DDE_Tnp_1_2->METHYLASE->LexA-protease->Radical-SAM->METHYLASE+DsrD-> | 1240 | Bacteria>Pseudomonadota>Betaproteobacteria>Burkholderiales>Burkholderiaceae>Ralstonia | GCA_021117135.1 | Ralstonia_solanacearum | LRP28_RS08930 | 1882275..1885997 | 305 | hypothetical-protein-[Ralstonia-solanacearum] |
| WP_245209875.1 | Type2.Pgl.like.BREX | RFC-AAA+DNAA-HTH->?->MORC+DNA_gyrase+TOPRIM-><-PglZ<-DUF6079*<-N6-METHYLASE<-PglW<-?\|\|METHYLASE->LexA-protease->Radical-SAM->METHYLASE+DsrD-><-?<-35exo | 1265 | Bacteria>Pseudomonadota>Betaproteobacteria>Burkholderiales>Burkholderiaceae>Ralstonia | - | Ralstonia_solanacearum | GO291_RS00025 | 8535..12332 | 305 | hypothetical-protein-[Ralstonia-solanacearum] |
| MBC7213753.1 | Type2.Pgl.like.BREX | <-rve\|\|?->HTH_26->PglW->N6-METHYLASE->OLD-ABC->NERD->DUF6079*->PglZ->BrxD-AAA->SF1-SF2-Helicase+wHTH+Small-B-barrel+DUF1998->Classical-AAA->Peptidase-S8+Subtilisin->SIGMA-HTH->HTH_3-> | 1231 | Bacteria>Pseudomonadota>Betaproteobacteria>Burkholderiales>Burkholderiaceae>unclassified_Burkholderiaceae | GCA_014359545.1 | Burkholderiaceae_bacterium | H5U28_00855 | 13751..17446 | 2030806 | phage-resistance-protein-[Burkholderiaceae-bacterium] |
| MCL4738012.1 | Type2.Pgl.like.BREX | <-SF1-SF2-Helicase+wHTH+Small-B-barrel+DUF1998-p<-BrxD-AAA<-PglZ<-DUF6079*<-DOC+HTH<-DUF6079-p<-HEPN<-N6-METHYLASE<-PglW<-HTH_26<-TPR+TPR<-Peptidase | 1233 | Bacteria>Pseudomonadota>Betaproteobacteria>Burkholderiales>Burkholderiaceae>unclassified_Burkholderiaceae | GCA_023383975.1 | Burkholderiaceae_bacterium | KJZ81_07380 | 5499..9200 | 2030806 | phage-resistance-protein-[Burkholderiaceae-bacterium] |
| WP_182661252.1 | Type2.Pgl.like.BREX | <-Peptidase\|\|?-><-?<-SF1-SF2-Helicase<-OLD-ABC+TOPRIM\|\|HTH_26->PglW->N6-METHYLASE->DOC+HTH->DUF6079*->PglZ->SF1-SF2-Helicase+wHTH+Small-B-barrel+DUF1998-><-AbiEii<-AbiEi_4<-REase<-?<-?<-VirD4-TraG+VirD4-FtsK<-VirB11+GTPase-AIG | 1238 | Bacteria>Pseudomonadota>Betaproteobacteria>Burkholderiales>Burkholderiales_genera_incertae_sedis>Aquariibacter | GCF_014145335.1 | Aquariibacter_albus | H4F90_RS02755 | 606925..610641 | 2759899 | DUF6079-family-protein-[Aquariibacter-albus] |
| WP_026434334.1 | Type2.Pgl.like.BREX | MULE-TRANSPOSASE->\|\|RHH_5->PglW->N6-METHYLASE->DUF6079*->PglZ->ParB+HNH-><-tRNA<-?<-?<-?<-?<-ACET | 1230 | Bacteria>Pseudomonadota>Betaproteobacteria>Burkholderiales>Comamonadaceae>Acidovorax | GCF_000687165.1 | Acidovorax_oryzae | T336_RS0123470 | 234027..237719 | 862720 | hypothetical-protein-[Acidovorax-oryzae] |
| WP_087745225.1 | Type2.Pgl.like.BREX | <-REase<-?<-?<-?<-?<-?<-SF1-SF2-Helicase+wHTH+Small-B-barrel+DUF1998<-BrxD-AAA<-PglZ<-DUF6079*<-?<-N6-METHYLASE<-PglW<-HTH_26\|\|?-><-TraG_N<-TraH | 1238 | Bacteria>Pseudomonadota>Betaproteobacteria>Burkholderiales>Comamonadaceae>Acidovorax | GCF_002176815.1 | unclassified_Acidovorax | CCX87_RS07735 | 1627243..1630959 | 2684926 | MULTISPECIES:-phage-resistance-protein-[unclassified-Acidovorax] |
| WP_272568621.1 | Type2.Pgl.like.BREX | HTH_29+HTH_32+DDE_3-><-?\|\|HTH_29+HTH_32+DDE_3->HTH_26->PglW->N6-METHYLASE->OLD-ABC->TOPRIM->DUF6079*->PglZ->BrxD-AAA->SF1-SF2-Helicase+wHTH+Small-B-barrel+DUF1998->?->?->?->BZIP+SF1-SF2-Helicase-><-GDResolvase\|\|RADICAL-SAM+TRAM-> | 1231 | Bacteria>Pseudomonadota>Betaproteobacteria>Burkholderiales>Comamonadaceae>Acidovorax | GCF_030159035.1 | Acidovorax_sp._GBBC_1281 | M5C96_RS08970 | 1970971..1974666 | 2940492 | phage-resistance-protein-[Acidovorax-sp.-GBBC-1281] |
| WP_172219748.1 | Type2.Pgl.like.BREX | <-35exo<-LRR+LRR\|\|?-><-Peptidase_M48+TPR_9<-ParB+HNH<-?\|\|HNH->DUF6079-p*->PglZ->BrxD-AAA->SF1-SF2-Helicase+wHTH+Small-B-barrel+DUF1998-><-?<-wHTH+WYL+WCX<-?<-Ntox49 | 872 | Bacteria>Pseudomonadota>Betaproteobacteria>Burkholderiales>Comamonadaceae>Caenimonas | GCF_013155545.1 | Caenimonas_soli | HNE04_RS13600 | 18189..20807 | 2735555 | hypothetical-protein-[Caenimonas-soli] |
| WP_045266902.1 | Type2.Pgl.like.BREX | <-ParA-Soj-PloopNTPase+ParA-Soj-PloopNTPase<-?<-?<-ArsR-HTH\|\|?-><-SF1-SF2-Helicase+wHTH+Small-B-barrel+DUF1998<-BrxD-AAA<-PglZ<-DUF6079*<-NERD<-OLD-ABC<-SecB<-MqsA_antitoxin+HTH_3<-MqsR_toxin<-ORC-CDC6<-ParA-Soj-PloopNTPase<-HEPN<-N6-METHYLASE | 1229 | Bacteria>Pseudomonadota>Betaproteobacteria>Burkholderiales>Comamonadaceae>Comamonas | GCA_030820635.1 | Comamonas_aquatica | MA05_RS11355 | 2505937..2509626 | 225991 | hypothetical-protein-[Comamonas-aquatica] |
| WP_066704901.1 | Type2.Pgl.like.BREX | <-HTH_3<-SIGMA-HTH<-Peptidase-S8+Subtilisin<-Classical-AAA<-SF1-SF2-Helicase+wHTH+Small-B-barrel+DUF1998<-BrxD-AAA<-PglZ<-DUF6079*<-N6-METHYLASE<-PglW<-HTH_26\|\|RHH_6->TraI-VirD2-RCRE-> | 1231 | Bacteria>Pseudomonadota>Betaproteobacteria>Burkholderiales>Comamonadaceae>Curvibacter | GCF_001592265.1 | Curvibacter_delicatus | CDE01S_RS05840 | 75691..79386 | 80879 | phage-resistance-protein-[Curvibacter-delicatus] |
| WP_047351120.1 | Type2.Pgl.like.BREX | <-TraI-VirD2-RCRE<-RHH_6<-?<-MqsA_antitoxin+DUF4065\|\|HTH_26->PglW->N6-METHYLASE->OLD-ABC->NERD->DUF6079*->PglZ->BrxD-AAA->SF1-SF2-Helicase+wHTH+Small-B-barrel+DUF1998->?->?->TUDOR+SF1-SF2-Helicase+iREase+CTD-> | 1231 | Bacteria>Pseudomonadota>Betaproteobacteria>Burkholderiales>Comamonadaceae>Diaphorobacter | GCF_001023515.1 | Diaphorobacter_sp._J5-51 | OX89_RS14220 | 15400..19095 | 680496 | hypothetical-protein-[Diaphorobacter-sp.-J5-51] |
| WP_144731145.1 | Type2.Pgl.like.BREX | <-TraI-VirD2-RCRE<-RHH_6\|\|HTH_26->PglW->N6-METHYLASE->DUF6079*->PglZ->BrxD-AAA->SF1-SF2-Helicase+wHTH+Small-B-barrel+DUF1998->?->?->TUDOR+SF1-SF2-Helicase+iREase+CTD->?-><-?<-wHTH+WYL+WCX | 1231 | Bacteria>Pseudomonadota>Betaproteobacteria>Burkholderiales>Comamonadaceae>Extensimonas | GCF_007655255.1 | Extensimonas_perlucida | FK527_RS06160 | 10962..14657 | 2590786 | phage-resistance-protein-[Extensimonas-perlucida] |
| WP_105731121.1 | Type2.Pgl.like.BREX | TraJ-RHH->HerA/FtsK->?->TrwC+SF1-SF2-Helicase->?->?-><-SF1-SF2-Helicase+wHTH+Small-B-barrel+DUF1998<-BrxD-AAA<-PglZ<-DUF6079*<-ORC-CDC6<-ParA-Soj-PloopNTPase<-HEPN<-NERD<-OLD-ABC<-N6-METHYLASE<-PglW<-HTH_26 | 1236 | Bacteria>Pseudomonadota>Betaproteobacteria>Burkholderiales>Comamonadaceae>Malikia | GCF_002980625.1 | Malikia_spinosa | C6P61_RS17195 | 22160..25870 | 86180 | phage-resistance-protein-[Malikia-spinosa] |
| WP_077032751.1 | Type2.Pgl.like.BREX | <-MarR-HTH<-?\|\|?->HTH_3->?->RHH_5->BrxA-DNA-binding->BrxB-BrxF-iSTAND->DUF6079*->?->N6-METHYLASE->PHP+SbcC->ParB+HNH->NERD+iPkinase+Pkinase+SF1-SF2-Helicase->?->PglZ->SIGMA+OB-Fold+AAA-ATPase+Lon-Protease-BrxL-> | 1153 | Bacteria>Pseudomonadota>Betaproteobacteria>Burkholderiales>Comamonadaceae>Pelomonas | GCF_001984095.1 | Pelomonas_sp._KK5 | B0W88_RS00865 | 20538..23999 | 1855730 | BREX-system-P-loop-protein-BrxC-[Pelomonas-sp.-KK5] |
| HAL37809.1 | Type2.Pgl.like.BREX | ABC-ATPase+SbcC->?->CN_hydrolase-><-SF1-SF2-Helicase+wHTH+Small-B-barrel+DUF1998<-BrxD-AAA<-PglZ<-DUF6079*<-SecB<-MqsA_antitoxin+HTH_3<-MqsR_toxin<-N6-METHYLASE<-PglW<-HTH_26\|\|?->DNA_METHYLASE-><-Schlafen+HATPase+wHTH | 1243 | Bacteria>Pseudomonadota>Betaproteobacteria>Burkholderiales>Comamonadaceae>Polaromonas | GCA_003454315.1 | Polaromonas_sp. | DCPO3_06690 | 19090..22821 | 1869339 | phage-resistance-protein-[Polaromonas-sp.] |
| WP_011797935.1 | Type2.Pgl.like.BREX | Classical-AAA->?-><-HTH_Tnp_1+HTH_21+rve+rve_3<-?<-SF1-SF2-Helicase<-OLD-ABC+TOPRIM<-SF1-SF2-Helicase+wHTH+Small-B-barrel+DUF1998<-BrxD-AAA<-PglZ<-DUF6079*<-NERD<-OLD-ABC<-N6-METHYLASE<-CR-ATPase<-PglW<-HTH_26\|\|?-><-AAA-ATPase<-HTH+rve | 1240 | Bacteria>Pseudomonadota>Betaproteobacteria>Burkholderiales>Comamonadaceae>Polaromonas | GCF_000015505.1 | Polaromonas_naphthalenivorans | PNAP_RS28015 | 176847..180569 | 216465 | hypothetical-protein-[Polaromonas-naphthalenivorans] |
| OYX60388.1 | Type2.Pgl.like.BREX | N6-METHYLASE-p->ParA-Soj-PloopNTPase->ORC-CDC6->DUF6079*->PglZ->BrxD-AAA->SF1-SF2-Helicase+wHTH+Small-B-barrel+DUF1998->HisKin-><-RADICAL-SAM+TRAM<-?\|\|ACET-> | 1229 | Bacteria>Pseudomonadota>Betaproteobacteria>Burkholderiales>Comamonadaceae>unclassified_Comamonadaceae | GCA_002280595.1 | Comamonadaceae_bacterium_32-67-11 | B7Y96_02165 | 4072..7761 | 1970505 | phage-resistance-protein-[Comamonadaceae-bacterium-32-67-11] |
| RJP65479.1 | Type2.Pgl.like.BREX | MqsR_toxin->MqsA_antitoxin+HTH_3->SecB->DUF6079*->PglZ->BrxD-AAA->SF1-SF2-Helicase+wHTH+Small-B-barrel+DUF1998->Phage_int_SAM_1+Phage_integrase->?->XIS-HTH->Tox-RES->wHTH+AbiEi_2->Polbeta-> | 1229 | Bacteria>Pseudomonadota>Betaproteobacteria>Burkholderiales>Comamonadaceae>unclassified_Comamonadaceae | GCA_003599455.1 | Comamonadaceae_bacterium | C4535_14920 | 1829..5518 | 1871071 | phage-resistance-protein-[Comamonadaceae-bacterium] |
| MBH2042929.1 | Type2.Pgl.like.BREX | HTH_26->PglW->N6-METHYLASE->MqsR_toxin->MqsA_antitoxin+HTH_3->SecB->DOC+HTH->DUF6079*->PglZ->BrxD-AAA->SF1-SF2-Helicase+wHTH+Small-B-barrel+DUF1998-> | 1240 | Bacteria>Pseudomonadota>Betaproteobacteria>Burkholderiales>Comamonadaceae>unclassified_Comamonadaceae | GCA_016000285.1 | Comamonadaceae_bacterium | I8H87_05185 | 12695..16417 | 1871071 | phage-resistance-protein-[Comamonadaceae-bacterium] |
| WP_215246545.1 | Type2.Pgl.like.BREX | <-CHTH<-Calcineurin<-Calcineurin<-HAD+CHTH<-Calcineurin-SbcD\|\|HTH_26->PglW->N6-METHYLASE->?->DUF6079*->PglZ->BrxD-AAA->SF1-SF2-Helicase+wHTH+Small-B-barrel+DUF1998-><-GreA_GreB+SF1-SF2-Helicase-p<-PIN+OST-HTH<-HTH_3\|\|REase-><-?<-RADICAL-SAM | 1231 | Bacteria>Pseudomonadota>Betaproteobacteria>Burkholderiales>Comamonadaceae>Variovorax | GCF_018614875.1 | Variovorax_paradoxus | J7E49_RS09220 | 16942..20637 | 34073 | phage-resistance-protein-[Variovorax-paradoxus] |
| WP_265288960.1 | Type2.Pgl.like.BREX | <-HIN-HTH<-?<-?<-?<-SF1-SF2-Helicase+wHTH+Small-B-barrel+DUF1998<-BrxD-AAA<-PglZ<-DUF6079*<-TOPRIM<-OLD-ABC<-SecB<-MqsA_antitoxin+HTH_3<-MqsR_toxin<-?<-N6-METHYLASE-p\|\|?-><-HTH_23+HTH_32 | 1232 | Bacteria>Pseudomonadota>Betaproteobacteria>Burkholderiales>Comamonadaceae>Verminephrobacter | GCF_026127525.1 | Verminephrobacter_eiseniae | D5047_RS18225 | 3902767..3906465 | 364317 | phage-resistance-protein-[Verminephrobacter-eiseniae] |
| WP_070303454.1 | Type2.Pgl.like.BREX | <-ArsR-HTH\|\|?->tRNA->Phage_integrase->Radical-SAM->METHYLASE+DsrD->METHYLASE-><-SF1-SF2-Helicase+wHTH+Small-B-barrel+DUF1998<-BrxD-AAA<-PglZ<-DUF6079*<-N6-METHYLASE<-PglW<-HTH_26\|\|?->?->?->HIN-HTH-><-HTH_3 | 1230 | Bacteria>Pseudomonadota>Betaproteobacteria>Burkholderiales>Oxalobacteraceae>Janthinobacterium | GCF_001758715.1 | Janthinobacterium_sp._HH104 | JAB6_RS14510 | 225223..228915 | 1537276 | phage-resistance-protein-[Janthinobacterium-sp.-HH104] |
| WP_139089367.1 | Type2.Pgl.like.BREX | Phage_integrase-><-DDE_Tnp_1\|\|PHP+SbcC->HTH_26->PglW->N6-METHYLASE->DUF6079*->PglZ->BrxD-AAA->SF1-SF2-Helicase+wHTH+Small-B-barrel+DUF1998-><-SF1-SF2-Helicase\|\|HTH_3->VirB11->?->?->VirD4-TraG+VirD4-FtsK-> | 1228 | Bacteria>Pseudomonadota>Betaproteobacteria>Burkholderiales>Oxalobacteraceae>Janthinobacterium | GCF_006152155.1 | Janthinobacterium_lividum | FHI69_RS02570 | 572923..576609 | 29581 | phage-resistance-protein-[Janthinobacterium-lividum] |
| HCY61923.1 | Type2.Pgl.like.BREX | XIS-HTH->?->HKD-DNase+HKD-DNase->SF1-SF2-Helicase->HTH_26->PglW->N6-METHYLASE->?->DUF6079*->PglZ->BrxD-AAA->SF1-SF2-Helicase+wHTH+Small-B-barrel+DUF1998-><-PRTase<-Arm-DNA-bind_3+Phage_integrase<-HAD+HAD<-SAM-METHYLASE | 1231 | Bacteria>Pseudomonadota>Betaproteobacteria>Burkholderiales>Oxalobacteraceae>unclassified_Oxalobacteraceae | GCA_003537575.1 | Oxalobacteraceae_bacterium | DHV59_03605 | 66758..70453 | 1871070 | phage-resistance-protein-[Oxalobacteraceae-bacterium] |
| WP_028998074.1 | Type2.Pgl.like.BREX | DEDD_Tnp_IS110+Transposase_20->?->?-><-?<-?<-SF1-SF2-Helicase+wHTH+Small-B-barrel+DUF1998<-BrxD-AAA<-PglZ<-DUF6079*<-SecB<-MqsA_antitoxin+HTH_3<-MqsR_toxin<-DOC+HTH<-HNH<-?<-ANK<-N6-METHYLASE<-PglW | 1231 | Bacteria>Pseudomonadota>Betaproteobacteria>Burkholderiales>Sphaerotilaceae>Azohydromonas | GCF_000430725.1 | Azohydromonas_australica | H537_RS0111665 | 33180..36875 | 364039 | hypothetical-protein-[Azohydromonas-australica] |

| Protein-ID | System | Gene-Neighbourhoods | Length | Taxonomy | Assembly-ID | Strain | Locus-Tag | Genomic-coordinates | TaxID | Fasta-define |
|---|---|---|---|---|---|---|---|---|---|---|
| WP_241924402.1 | Type2.Pgl.like.BREX | REC+OmpR-HTH-><-NUDIX\|\|?->?->?->?-><-SF1-SF2-Helicase+wHTH+Small-B-barrel+DUF1998<-BrxD-AAA<-PglZ<-DUF6079*<-DOC+HTH<-?<-NERD<-OLD-ABC<-N6-METHYLASE<-PglW<-HTH_26 | 1226 | Bacteria>Pseudomonadota>Betaproteobacteria>Burkholderiales>Sphaerotilaceae>Azohydromonas | GCF_022631395.1 | Calidifontimicrobium_sp._SYSU_G02091 | MOJ79_RS12180 | 67829..71509 | 2926421 | DUF6079-family-protein-[Calidifontimicrobium-sp.-SYSU-G02091] |
| MBL0085872.1 | Type2.Pgl.like.BREX | HD->HD+PspB-><-?\|\|HTH_26->PglW-><-?\|\|N6-METHYLASE-p->HEPN->DUF6079*->PglZ->BrxD-AAA->SF1-SF2-Helicase+wHTH+Small-B-barrel+DUF1998-> | 1236 | Bacteria>Pseudomonadota>Betaproteobacteria>Burkholderiales>Sphaerotilaceae>Ideonella | GCA_016721465.1 | Ideonella_sp. | IPP44_03995 | 47182..50892 | 1929293 | phage-resistance-protein-[Ideonella-sp.] |
| MBL0086259.1 | Type2.Pgl.like.BREX | HDRnase->VirD4-TraG+VirB4-FtsK->?-><-HNH<-ABC-ATPase<-SF1-SF2-Helicase+wHTH+Small-B-barrel+DUF1998<-BrxD-AAA<-PglZ<-DUF6079*<-N6-METHYLASE<-PglW<-HTH_26\|\|OLD-ABC+TOPRIM->SF1-SF2-Helicase-> | 1243 | Bacteria>Pseudomonadota>Betaproteobacteria>Burkholderiales>Sphaerotilaceae>Ideonella | GCA_016721465.1 | Ideonella_sp. | IPP44_06015 | 500695..504426 | 1929293 | phage-resistance-protein-[Ideonella-sp.] |
| WP_234414047.1 | Type2.Pgl.like.BREX | <-Phage_integrase<-?\|\|HTH_26->PglW->N6-METHYLASE->DOC+HTH->DUF6079*->PglZ->BrxD-AAA->SF1-SF2-Helicase+wHTH+Small-B-barrel+DUF1998->OLD-ABC->HNH-><-?<-?\|\|Phage_int_SAM_1+Phage_integrase-> | 1226 | Bacteria>Pseudomonadota>Betaproteobacteria>Burkholderiales>Sphaerotilaceae>Ideonella | GCF_002198735.1 | Ideonella_sp._A_288 | EOINJALH_RS25855 | 77971..81651 | 1962181 | DUF6079-family-protein-[Ideonella-sp.-A-288] |
| WP_130480155.1 | Type2.Pgl.like.BREX | <-Shikimate_dh_N+SDH_C\|\|ICLR-HTH->?->?->HTH_26->PglW->N6-METHYLASE->?->DOC+HTH->DUF6079*->PglZ->BrxD-AAA->SF1-SF2-Helicase+wHTH+Small-B-barrel+DUF1998-><-ParB+HNH\|\|?-><-GTPase-AIG<-GTPase-AIG | 1244 | Bacteria>Pseudomonadota>Betaproteobacteria>Burkholderiales>Sphaerotilaceae>Leptothrix | GCF_004216565.1 | Leptothrix_mobilis | EV685_RS01090 | 181615..185349 | 47994 | DUF6079-family-protein-[Leptothrix-mobilis] |
| MCE2659973.1 | Type2.Pgl.like.BREX | Tox-RelE-><-SF1-SF2-Helicase+wHTH+Small-B-barrel+DUF1998<-BrxD-AAA<-PglZ<-DUF6079*<-SecB<-MqsA_antitoxin+HTH_3<-MqsR_toxin | 1219 | Bacteria>Pseudomonadota>Betaproteobacteria>Burkholderiales>Sphaerotilaceae>Rubrivivax | GCA_021298085.1 | Rubrivivax_sp. | LW854_17330 | 6458..10117 | 50259 | phage-resistance-protein-[Rubrivivax-sp.] |
| WP_164962560.1 | Type2.Pgl.like.BREX | <-SIGMA+OB-Fold+AAA-ATPase+Lon-Protease-BrxL<-PglZ<-?<-NERD+iPkinase+Pkinase+SF1-SF2-Helicase<-ParB+HNH<-SF1-SF2-Helicase<-OLD-ABC+TOPRIM<-N6-METHYLASE<-DUF6079*<-BrxB-BrxF-iSTAND<-BrxA-DNA-binding<-RHH_5<-HTH_26\|\|Phage_AlpA->Arm-DNA-bind_2+Phage_int_SAM_3+Phage_integrase->?->?->URI-DNase-> | 1153 | Bacteria>Pseudomonadota>Betaproteobacteria>Burkholderiales>Sphaerotilaceae>Rubrivivax | GCF_011065555.1 | Rubrivivax_sp._JA1026 | G5B56_RS05565 | 402578..406039 | 2710888 | BREX-system-P-loop-protein-BrxC-[Rubrivivax-sp.-JA1026] |
| WP_244954261.1 | Type2.Pgl.like.BREX | HTH_26->PglW-p->PglW-p->N6-METHYLASE-p->N6-METHYLASE->HEPN->ParA-Soj-PloopNTPase->ORC-CDC6->DUF6079-p*->PglZ->BrxD-AAA->\|\|wHTH+WYL+WCX->PVC-AAA+Classical-AAA->SIR2-> | 909 | Bacteria>Pseudomonadota>Betaproteobacteria>Burkholderiales>Sphaerotilaceae>Sphaerotilus | GCF_008329925.1 | Sphaerotilus_sulfidivorans | EWH46_RS00085 | 20429..23158 | 639200 | DUF6079-family-protein-[Sphaerotilus-sulfidivorans] |
| WP_179636391.1 | Type2.Pgl.like.BREX | N6-METHYLASE-p->?->DOC->DUF6079*->PglZ->BrxD-AAA->SF1-SF2-Helicase+wHTH+Small-B-barrel+DUF1998->OLD-ABC+TOPRIM->SF1-SF2-Helicase-> | 1246 | Bacteria>Pseudomonadota>Betaproteobacteria>Burkholderiales>Sphaerotilaceae>Sphaerotilus | GCF_013410775.1 | Sphaerotilus_montanus | HZU83_RS20715 | 3769..7509 | 522889 | phage-resistance-protein-[Sphaerotilus-montanus] |
| WP_180693050.1 | Type2.Pgl.like.BREX | N6-METHYLASE->HEPN->ParA-Soj-PloopNTPase->ORC-CDC6->DUF6079*->PglZ->BrxD-AAA->SF1-SF2-Helicase+wHTH+Small-B-barrel+DUF1998->JetB->SbcC->?-><-ParB<-ParA-Soj-PloopNTPase | 1248 | Bacteria>Pseudomonadota>Betaproteobacteria>Burkholderiales>Sphaerotilaceae>Sphaerotilus | GCF_013426975.1 | Sphaerotilus_sulfidivorans | HZU84_RS18045 | 6467..10213 | 639200 | phage-resistance-protein-[Sphaerotilus-sulfidivorans] |
| WP_251969637.1 | Type2.Pgl.like.BREX | URI-DNase->DDE_Tnp_1+DDE_Tnp_1->N6-METHYLASE-p->HEPN->ParA-Soj-PloopNTPase->ORC-CDC6->DOC+HTH->OLD-ABC->TOPRIM->DUF6079*->PglZ->BrxD-AAA->SF1-SF2-Helicase+wHTH+Small-B-barrel+DUF1998-><-tRNA\|\|GGDEF-><-?<-?<-SAM-METHYLASE | 1228 | Bacteria>Pseudomonadota>Betaproteobacteria>Burkholderiales>Sphaerotilaceae>Sphaerotilus | GCF_023734135.1 | Sphaerotilus_sp._FB-5 | NGK70_RS16750 | 3933187..3936873 | 2914710 | phage-resistance-protein-[Sphaerotilus-sp.-FB-5] |
| WP_251970424.1 | Type2.Pgl.like.BREX | <-DEDD_Tnp_IS110+Transposase_20<-?\|\|?-><-?\|\|?->?->RHH_5->BrxA-DNA-binding->BrxB-BrxF-iSTAND->DUF6079*->N6-METHYLASE->PHP+SbcC->ParB+HNH->NERD+iPkinase+Pkinase+SF1-SF2-Helicase->?->PglZ->SIGMA+OB-Fold+AAA-ATPase+Lon-Protease-BrxL-><-SIGMA-HTH\|\|Tox-RelE-> | 1153 | Bacteria>Pseudomonadota>Betaproteobacteria>Burkholderiales>Sphaerotilaceae>Sphaerotilus | GCF_023734135.1 | Sphaerotilus_sp._FB-5 | NGK70_RS21040 | 4913419..4916880 | 2914710 | BREX-system-P-loop-protein-BrxC-[Sphaerotilus-sp.-FB-5] |
| MBI3156220.1 | Type2.Pgl.like.BREX | <-SIR2<-?\|\|DSBA+Thioredoxin->BrnA_antitoxin-><-?<-SF1-SF2-Helicase+wHTH+Small-B-barrel+DUF1998<-BrxD-AAA<-PglZ<-DUF6079*<-DOC+HTH<-DDE_Tnp_ISL3 | 1236 | Bacteria>Pseudomonadota>Betaproteobacteria>Burkholderiales>unclassified_Burkholderiales | GCA_016191995.1 | Burkholderiales_bacterium | HYZ20_12520 | 19795..23505 | 1891238 | phage-resistance-protein-[Burkholderiales-bacterium] |
| MBK6715972.1 | Type2.Pgl.like.BREX | DSBH+AraC-HTH+AraC-HTH-><-?<-?\|\|Peptidase_M24->HD-><-?<-SF1-SF2-Helicase+wHTH+Small-B-barrel+DUF1998<-BrxD-AAA<-PglZ<-DUF6079*<-TOPRIM<-OLD-ABC<-DOC+HTH<-SecB<-MqsA_antitoxin+HTH_3<-MqsR_toxin<-ORC-CDC6<-ParA-Soj-PloopNTPase<-HEPN | 1235 | Bacteria>Pseudomonadota>Betaproteobacteria>Burkholderiales>unclassified_Burkholderiales | GCA_016703325.1 | Burkholderiales_bacterium | IPG57_12970 | 74426..78133 | 1891238 | phage-resistance-protein-[Burkholderiales-bacterium] |
| MBK7615719.1 | Type2.Pgl.like.BREX | <-SF1-SF2-Helicase+wHTH+Small-B-barrel+DUF1998<-BrxD-AAA<-PglZ<-DUF6079*<-N6-METHYLASE-p<-?<-METHYLASE<-N6-METHYLASE<-PglW<-HTH_26<-?<-?\|\|wHTH+AbiEi_2-> | 1193 | Bacteria>Pseudomonadota>Betaproteobacteria>Burkholderiales>unclassified_Burkholderiales | GCA_016710065.1 | Burkholderiales_bacterium | IPJ08_15120 | 15240..18821 | 1891238 | phage-resistance-protein-[Burkholderiales-bacterium] |
| VWX60042.1 | Type2.Pgl.like.BREX | XIS-HTH->?->TOPRIM->SMS-KaiC->?->BrxE->BrxA-DNA-binding->BrxB-BrxF-iSTAND->DUF6079*->?->Tox-RelA->N6-METHYLASE->N6-METHYLASE->PglZ->SIGMA+OB-Fold+AAA-ATPase+Lon-Protease-BrxL->NERD+iPkinase+Pkinase+SF1-SF2-Helicase->?->McrB-AAA-> | 1181 | Bacteria>Pseudomonadota>Betaproteobacteria>Burkholderiales>unclassified_Burkholderiales | GCA_902506565.1 | Burkholderiales_bacterium_8X | VARIO8X_20150 | 134929..138474 | 2653183 | conserved-hypothetical-protein-[Burkholderiales-bacterium-8X] |
| MBA4142467.1 | Type2.Pgl.like.BREX | <-ABhydrolase\|\|?->?->?->?->tRNA-><-PglZ<-DUF6079*<-N6-METHYLASE\|\|GDResolvase+HIN-HTH->?-><-?<-AraC-HTH\|\|?->LexA-protease->Glyoxalase-><-MerR-HTH | 1232 | Bacteria>Pseudomonadota>Betaproteobacteria>Nitrosomonadales>Nitrosomonadaceae>Nitrosospira | GCA_013821425.1 | Nitrosospira_sp. | HOX43_05540 | 80286..83984 | 38080 | hypothetical-protein-[Nitrosospira-sp.] |
| WP_074775093.1 | Type2.Pgl.like.BREX | <-OLD-ABC+TOPRIM<-?\|\|?->?->?->HARE-HTH->?->PglW->N6-METHYLASE->DUF6079*->PglZ->BrxD-AAA->SF1-SF2-Helicase+wHTH+Small-B-barrel+DUF1998->?->?->?-><-?<-Peptidase_M32 | 1234 | Bacteria>Pseudomonadota>Betaproteobacteria>Nitrosomonadales>Nitrosomonadaceae>Nitrosospira | GCF_900107715.1 | Nitrosospira_multiformis | BLV78_RS11110 | 23359..27063 | 1231 | phage-resistance-protein-[Nitrosospira-multiformis] |
| WP_169259375.1 | Type2.Pgl.like.BREX | REC+GGDEF+EAL-><-PIN+HTH<-SF1-SF2-Helicase+wHTH+Small-B-barrel+DUF1998<-?<-?<-?<-PglZ<-DUF6079*<-N6-METHYLASE<-PglW<-HTH_26<-SF1-SF2-Helicase<-?<-?<-?<-METHYLASE-p | 1232 | Bacteria>Pseudomonadota>Betaproteobacteria>Rhodocyclales>Rhodocyclaceae>Aromatoleum | GCF_012910955.1 | Aromatoleum_diolicum | GPA25_RS05620 | 140631..144329 | 75796 | phage-resistance-protein-[Aromatoleum-diolicum] |
| KAF0162607.1 | Type2.Pgl.like.BREX | <-VirD4-TraG+VirB4-FtsK<-?<-?<-?\|\|HTH_26->PglW->N6-METHYLASE->?->DOC+HTH->DUF6079*->PglZ->BrxD-AAA->SF1-SF2-Helicase+wHTH+Small-B-barrel+DUF1998->?->?->?->?->HTH_3->Couple_hipA+HipA_C-> | 1246 | Bacteria>Pseudomonadota>Betaproteobacteria>Rhodocyclales>Rhodocyclaceae>unclassified_Rhodocyclaceae | GCA_009773275.1 | Rhodocyclaceae_bacterium | FD157_3838 | 25776..29516 | 1898103 | putative-phage-resistance-protein-[Rhodocyclaceae-bacterium] |
| MBK9311591.1 | Type2.Pgl.like.BREX | <-TraE<-?<-?<-?<-?<-?\|\|HTH_26->PglW->N6-METHYLASE->DUF6079*->PglZ->?->?->BrxD-AAA-><-?<-HD | 1234 | Bacteria>Pseudomonadota>Betaproteobacteria>Rhodocyclales>Rhodocyclaceae>unclassified_Rhodocyclaceae | GCA_016719325.1 | Rhodocyclaceae_bacterium | IPN12_12840 | 2522887..2526591 | 1898103 | phage-resistance-protein-[Rhodocyclaceae-bacterium] |
| MBI3093458.1 | Type2.Pgl.like.BREX | <-HTH_26\|\|Phage_AlpA->?->?->NERD+Pkinase+Pkinase+SF1-SF2-Helicase->?-><-ParB+HNH<-PglZ<-DUF6079*<-N6-METHYLASE<-PglW<-XIS-HTH<-HTH_3\|\|?->Peptidase_M90->?-><-ABhydrolase<-ADH_N+ADH_zinc_N+METHYLASE | 1234 | Bacteria>Pseudomonadota>Betaproteobacteria>Rhodocyclales>unclassified_Rhodocyclales | GCA_016192435.1 | Rhodocyclales_bacterium | HYY97_01150 | 54238..57942 | 2072421 | hypothetical-protein-[Rhodocyclales-bacterium] |
| WP_015434799.1 | Type2.Pgl.like.BREX | <-AAA-ATPase<-HTH+rve<-?<-?\|\|HTH_26->PglW->N6-METHYLASE->SbcC->Schlafen+HATPase+wHTH->DUF6079*->PglZ->BrxD-AAA->SF1-SF2-Helicase+wHTH+Small-B-barrel+DUF1998->HAD+CHTH-><-XIS-HTH\|\|HAD+CHTH-><-?\|\|HicA_toxin-> | 1232 | Bacteria>Pseudomonadota>Betaproteobacteria>Rhodocyclales>Zoogloeaceae>Azoarcus | GCF_000349945.1 | Azoarcus_sp._KH32C | AZKH_RS05730 | 1276072..1279770 | 748247 | phage-resistance-protein-[Azoarcus-sp.-KH32C] |
| WP_065339872.1 | Type2.Pgl.like.BREX | <-Phage_integrase<-?<-wHTH+WYL+WCX<-ARG<-Ntox49<-?<-?\|\|BrxA-DNA-binding->BrxB-BrxF-iSTAND->DUF6079*->?->N6-METHYLASE->NERD+iPkinase+Pkinase+SF1-SF2-Helicase->?->PglZ->SIGMA+OB-Fold+AAA-ATPase+Lon-Protease-BrxL-><-XIS-HTH | 1153 | Bacteria>Pseudomonadota>Betaproteobacteria>Rhodocyclales>Zoogloeaceae>Azoarcus | GCF_001682385.1 | Azoarcus_olearius | dqs_RS05275 | 1112648..1116109 | 418699 | BREX-system-P-loop-protein-BrxC-[Azoarcus-olearius] |
| WP_133588700.1 | Type2.Pgl.like.BREX | <-SbcC+SidE<-JetB<-?\|\|XIS-HTH->PglW->N6-METHYLASE->DUF6079*->PglZ->ParB+HNH-><-?<-VirD4-TraG+VirB4-FtsK<-HDRnase<-?<-HD<-DUF932<-TOPRIM | 1233 | Bacteria>Pseudomonadota>Betaproteobacteria>Rhodocyclales>Zoogloeaceae>Azoarcus | GCF_004361735.1 | Azoarcus_indigens | C7389_RS04285 | 316481..320182 | 29545 | hypothetical-protein-[Azoarcus-indigens] |
| WP_021249969.1 | Type2.Pgl.like.BREX | <-VirB5<-?<-?<-?<-VirB6<-SF1-SF2-Helicase+wHTH+Small-B-barrel+DUF1998<-BrxD-AAA<-PglZ<-DUF6079*<-?<-SecB<-MqsA_antitoxin+HTH_3<-MqsR_toxin<-N6-METHYLASE\|\|Y1_Tnp-> | 1237 | Bacteria>Pseudomonadota>Betaproteobacteria>Rhodocyclales>Zoogloeaceae>Thauera | GCF_000443165.1 | Thauera_terpenica | M622_RS33100 | 17284..20997 | 76113 | hypothetical-protein-[Thauera-terpenica] |
| WP_096448011.1 | Type2.Pgl.like.BREX | TraN->?->?->TraH->TraG_N-><-?\|\|HTH_26->PglW->N6-METHYLASE->DUF6079*->PglZ->BrxD-AAA->SF1-SF2-Helicase+wHTH+Small-B-barrel+DUF1998->OLD-ABC+TOPRIM->SF1-SF2-Helicase->Phage_int_SAM_2+Integrase_1->XIS-HTH->SAM-METHYLASE-><-SF1-SF2-Helicase | 1231 | Bacteria>Pseudomonadota>Betaproteobacteria>Rhodocyclales>Zoogloeaceae>Thauera | GCF_002354895.1 | Thauera_sp._K11 | CCZ27_RS10595 | 2428363..2432058 | 2005884 | phage-resistance-protein-[Thauera-sp.-K11] |
| PKO27557.1 | Type2.Pgl.like.BREX | <-SIGMA+OB-Fold+AAA-ATPase+Lon-Protease-BrxL<-PglZ<-?<-NERD+iPkinase+Pkinase+SF1-SF2-Helicase<-ParB+HNH<-SF1-SF2-Helicase<-OLD-ABC+TOPRIM<-N6-METHYLASE<-DUF6079*<-BrxB-BrxF-iSTAND<-BrxA-DNA-binding<-RHH_5<-?<-DnaB<-XIS-HTH<-?<-?\|\|MarR-HTH-> | 1153 | Bacteria>Pseudomonadota>Betaproteobacteria>unclassified_Betaproteobacteria | GCA_002842215.1 | Betaproteobacteria_bacterium_HGW-Betaproteobacteria-9 | CVU36_19700 | 65227..68688 | 2013721 | BREX-system-P-loop-protein-BrxC-[Betaproteobacteria-bacterium-HGW-Betaproteobacteria-9] |
| MBI3147358.1 | Type2.Pgl.like.BREX | <-TraE<-?<-?<-?<-?<-?\|\|HTH_26->PglW->N6-METHYLASE->DUF6079*->PglZ->RVT->BrxD-AAA->SF1-SF2-Helicase+wHTH+Small-B-barrel+DUF1998-><-?<-VirD4-TraG+VirB4-FtsK | 1234 | Bacteria>Pseudomonadota>Betaproteobacteria>unclassified_Betaproteobacteria | GCA_016192065.1 | Betaproteobacteria_bacterium | HYZ17_02460 | 12898..16602 | 1891241 | phage-resistance-protein-[Betaproteobacteria-bacterium] |
| MBK7517443.1 | Type2.Pgl.like.BREX | HD->DDE_5+DDE_Tnp_1->HD+PspB-><-?\|\|HTH_26->PglW->N6-METHYLASE->SWACOS->?->DUF6079*->PglZ->BrxD-AAA->SF1-SF2-Helicase+wHTH+Small-B-barrel+DUF1998-> | 1237 | Bacteria>Pseudomonadota>Betaproteobacteria>unclassified_Betaproteobacteria | GCA_016706525.1 | Betaproteobacteria_bacterium | IPI51_17995 | 205572..209285 | 1891241 | phage-resistance-protein-[Betaproteobacteria-bacterium] |
| MCL5969864.1 | Type2.Pgl.like.BREX | PAS+HISKIN->REC+NtrC-AAA+FIS-HTH-><-?\|\|PglW->N6-METHYLASE->HEPN->ParA-Soj-PloopNTPase->ORC-CDC6->DUF6079-p*-> | 120 | Bacteria>Pseudomonadota>Betaproteobacteria>unclassified_Betaproteobacteria | GCA_023488365.1 | Betaproteobacteria_bacterium | M1359_11710 | 32177..32537 | 1891241 | hypothetical-protein-[Betaproteobacteria-bacterium] |
| GHU42418.1 | Type2.Pgl.like.BREX | HIN-HTH->?-><-Schlafen+HATPase+wHTH+HTH<-PglZ<-DUF6079*<-N6-METHYLASE<-PglW<-XIS-HTH<-AbiEi_4\|\|HTH_Tnp_1+HTH_21+rve+rve_3-> | 1231 | Bacteria>Pseudomonadota>Betaproteobacteria>unclassified_Betaproteobacteria | GCA_025997095.1 | Betaproteobacteria_bacterium | AGMMS50289_07090 | 5308..9003 | 1891241 | hypothetical-protein-AGMMS50289_07090-[Betaproteobacteria-bacterium] |
| WP_265048421.1 | Type2.Pgl.like.BREX | AbiEi_4->AbiEii-><-?<-SF1-SF2-Helicase<-?\|\|HTH_26->PglW->N6-METHYLASE->DUF6079*->PglZ->BrxD-AAA->SF1-SF2-Helicase+wHTH+Small-B-barrel+DUF1998->?-><-?\|\|?->TOPRIM->AAA+HTH-> | 1223 | Bacteria>Pseudomonadota>Gammaproteobacteria>Alteromonadales>Alteromonadaceae>Alkalimarinus | GCF_026016465.1 | Alkalimarinus_sp._A2M4 | NKI27_RS04095 | 933028..936699 | 2961619 | hypothetical-protein-[Alkalimarinus-sp.-A2M4] |
| WP_121637688.1 | Type2.Pgl.like.BREX | <-NERD+iPkinase+Pkinase+SF1-SF2-Helicase<-SIGMA+OB-Fold+AAA-ATPase+Lon-Protease-BrxL<-PglZ<-REase+NACHT<-?<-RADICAL-SAM<-N6-METHYLASE<-ParB+HNH<-DUF6079*<-BrxB-BrxF-iSTAND<-BrxA-DNA-binding<-wHTH+WYL+WCX\|\|?-><-?\|\|?-><-SF1-SF2-Helicase<-PadR-HTH\|\|HTH_3-> | 1229 | Bacteria>Pseudomonadota>Gammaproteobacteria>Alteromonadales>Pseudoalteromonadaceae>Pseudoalteromonas | GCF_003668795.1 | Pseudoalteromonas_agarivorans | D9T18_RS10095 | 2262491..2266180 | 176102 | BREX-system-P-loop-protein-BrxC-[Pseudoalteromonas-agarivorans] |
| MCF1457977.1 | Type2.Pgl.like.BREX | <-AbiEi_4<-AbiGi+seqA+RAMA<-SF1-SF2-Helicase+wHTH+Small-B-barrel+DUF1998<-BrxD-AAA<-PglZ<-DUF6079*<-N6-METHYLASE<-PglW\|\|?-><-RelE-ParE<-PhdYeFM_antitox\|\|XIS-HTH->Phage_integrase->Phage_integrase->Phage_integrase-> | 1220 | Bacteria>Pseudomonadota>Gammaproteobacteria>Alteromonadales>Shewanellaceae>Shewanella | GCA_021462595.1 | Shewanella_sp. | LPH21_10625 | 15615..19277 | 50422 | hypothetical-protein-[Shewanella-sp.] |
| WP_202960803.1 | Type2.Pgl.like.BREX | HTH_3-><-?<-?<-?<-DUF932<-?\|\|XIS-HTH->PglW->N6-METHYLASE->DUF6079*->PglZ->BrxD-AAA->SF1-SF2-Helicase+wHTH+Small-B-barrel+DUF1998->?-><-Arm-DNA-bind_3+Phage_integrase<-?<-?<-?<-? | 1217 | Bacteria>Pseudomonadota>Gammaproteobacteria>Cellvibrionales>Cellvibrionaceae>Marinimicrobium | GCF_000463525.1 | Marinimicrobium_sp._LS-A18 | N645_RS0114975 | 897740..901393 | 1381596 | hypothetical-protein-[Marinimicrobium-sp.-LS-A18] |
| WP_207002182.1 | Type2.Pgl.like.BREX | SF1-SF2-Helicase-><-HEPN+RES<-?<-?<-HEPN\|\|HTH_26->PglW->N6-METHYLASE->DUF6079*->PglZ->BrxD-AAA->SF1-SF2-Helicase+wHTH+Small-B-barrel+DUF1998->ABC-ATPase->HNH-> | 1223 | Bacteria>Pseudomonadota>Gammaproteobacteria>Cellvibrionales>Microbulbiferaceae>Microbulbifer | GCF_017303155.1 | Microbulbifer_salipaludis | JF535_RS11380 | 1357642..1361313 | 187980 | hypothetical-protein-[Microbulbifer-salipaludis] |
| WP_155450778.1 | Type2.Pgl.like.BREX | <-SF1-SF2-Helicase+wHTH+Small-B-barrel+DUF1998<-BrxD-AAA<-PglZ<-DUF6079*<-REase<-N6-METHYLASE<-PIN<-ParD_antitoxin<-PglW\|\|AbiEi_4->AbiEii-> | 1221 | Bacteria>Pseudomonadota>Gammaproteobacteria>Chromatiales>Chromatiaceae>Allochromatium | GCF_009720725.1 | Allochromatium_palmeri | GJ668_RS14065 | 7650..11315 | 231048 | phage-resistance-protein-[Allochromatium-palmeri] |
| WP_200346143.1 | Type2.Pgl.like.BREX | PglW->SbcC->TOPRIM->?->N6-METHYLASE->Unstab_antitox->DUF6079*->RHH_6->PIN->PglZ->BrxD-AAA->MazE_antitoxin->DOC->SF1-SF2-Helicase+wHTH+Small-B-barrel+DUF1998->Cysteine-Desulfurase-> | 1272 | Bacteria>Pseudomonadota>Gammaproteobacteria>Chromatiales>Chromatiaceae>Halochromatium | GCF_016584085.1 | Halochromatium_glycolicum | CKO40_RS10425 | 44898..48716 | 85075 | phage-resistance-protein-[Halochromatium-glycolicum] |
| WP_201217441.1 | Type2.Pgl.like.BREX | <-ParB+HNH<-?<-?<-SF1-SF2-Helicase+wHTH+Small-B-barrel+DUF1998<-BrxD-AAA<-PglZ<-PIN<-RHH_6<-DUF6079*<-RelE-ParE<-Unstab_antitox<-N6-METHYLASE<-PIN<-ParD_antitoxin<-PglW<-?<-PrlF_antitoxin\|\|HigB_toxin-> | 1227 | Bacteria>Pseudomonadota>Gammaproteobacteria>Chromatiales>Chromatiaceae>Halochromatium | GCF_016653315.1 | Halochromatium_roseum | CCR96_RS03585 | 10744..14427 | 391920 | phage-resistance-protein-[Halochromatium-roseum] |
| WP_020503528.1 | Type2.Pgl.like.BREX | TraH->?->?->?->PglW->N6-METHYLASE->PolB->N6-METHYLASE-p->DUF6079*->PglZ->BrxD-AAA->SF1-SF2-Helicase+wHTH+Small-B-barrel+DUF1998-><-DOC+HTH<-?<-HTH+MPTase | 1224 | Bacteria>Pseudomonadota>Gammaproteobacteria>Chromatiales>Chromatiaceae>Lamprocystis | GCF_000379525.1 | Lamprocystis_purpurea | A390_RS0104590 | 31846..35520 | 61598 | hypothetical-protein-[Lamprocystis-purpurea] |
| WP_020506075.1 | Type2.Pgl.like.BREX | <-SF1-SF2-Helicase+wHTH+Small-B-barrel+DUF1998<-BrxD-AAA<-PglZ<-DUF6079*<-Unstab_antitox<-PIN<-?<-N6-METHYLASE<-HTH_37<-ParD_antitoxin<-REase<-PglW | 1221 | Bacteria>Pseudomonadota>Gammaproteobacteria>Chromatiales>Chromatiaceae>Lamprocystis | GCF_000379525.1 | Lamprocystis_purpurea | A390_RS0118090 | 24140..27805 | 61598 | DUF6079-family-protein-[Lamprocystis-purpurea] |
| QVL47101.1 | Type2.Pgl.like.BREX | PglW->N6-METHYLASE->PolB->HEPN->?->N6-METHYLASE-p->DUF6079*->PglZ->BrxD-AAA->SF1-SF2-Helicase+wHTH+Small-B-barrel+DUF1998-><-?<-NUDIX\|\|HAD-><-SF1-SF2-Helicase | 1225 | Bacteria>Pseudomonadota>Gammaproteobacteria>Chromatiales>Chromatiaceae>Thiocapsa | GCA_018399035.1 | Thiocapsa_sp. | KFB96_15350 | 3389147..3392824 | 2024551 | MAG:-phage-resistance-protein-[Thiocapsa-sp.] |
| WP_120798742.1 | Type2.Pgl.like.BREX | <-SbcC<-NtrC-AAA+FIS-HTH<-RHH_1\|\|?-><-?<-SF1-SF2-Helicase+wHTH+Small-B-barrel+DUF1998<-BrxD-AAA<-PglZ<-DUF6079*<-REase<-N6-METHYLASE<-PIN<-ParD_antitoxin<-HicB_lk_antitox<-PglW | 1232 | Bacteria>Pseudomonadota>Gammaproteobacteria>Chromatiales>Chromatiaceae>Thiocapsa | GCF_003634315.1 | Thiocapsa_rosea | BDD21_RS20530 | 4622209..4625907 | 69360 | phage-resistance-protein-[Thiocapsa-rosea] |
| WP_200389032.1 | Type2.Pgl.like.BREX | <-PAS+PAS+PAS+GGDEF<-ParB<-MerR-HTH+ParA-Soj-PloopNTPase\|\|?->LexA-protease-><-?<-SF1-SF2-Helicase+wHTH+Small-B-barrel+DUF1998<-BrxD-AAA<-PglZ<-DUF6079* | 1033 | Bacteria>Pseudomonadota>Gammaproteobacteria>Chromatiales>Chromatiaceae>Thiocapsa | GCF_016583985.1 | Thiocapsa_imhoffii | CKO25_RS16430 | 71688..74791 | 382777 | phage-resistance-protein,-partial-[Thiocapsa-imhoffii] |
| QVL48761.1 | Type2.Pgl.like.BREX | HicA_toxin->ParD_antitoxin->HTH_3->N6-METHYLASE-p->METHYLASE-p->N6-METHYLASE->REase->ParD_antitoxin->RelE-ParE->DUF6079*->RHH_6->PIN->PglZ->SF1-SF2-Helicase+wHTH+Small-B-barrel+DUF1998-><-?<-D5N+D5ATPase\|\|?-><-PVC-AAA | 1186 | Bacteria>Pseudomonadota>Gammaproteobacteria>Chromatiales>Chromatiaceae>Thiocapsa | GCA_018399035.1 | Thiocapsa_sp. | KFB96_24975 | 5465265..5468825 | 2024551 | MAG:-phage-resistance-protein-[Thiocapsa-sp.] |
| WP_200333081.1 | Type2.Pgl.like.BREX | <-SIGMA-HTH+SIGMA-HTH+SIGMA-HTH\|\|PglW->REase->ParD_antitoxin->N6-METHYLASE->?->Unstab_antitox->RelE-ParE->DUF6079*->RHH_6->PIN->PglZ->SF1-SF2-Helicase+wHTH+Small-B-barrel+DUF1998-><-DDE_Tnp_1_assoc | 1216 | Bacteria>Pseudomonadota>Gammaproteobacteria>Chromatiales>Chromatiaceae>Thiocystis | GCF_016583565.1 | Thiocystis_violacea | CKO23_RS17345 | 19925..23575 | 13725 | DUF6079-family-protein-[Thiocystis-violacea] |
| WP_100922885.1 | Type2.Pgl.like.BREX | <-SF1-SF2-Helicase+wHTH+Small-B-barrel+DUF1998<-BrxD-AAA<-PglZ<-DUF6079*<-ORC-CDC6<-ParA-Soj-PloopNTPase<-HEPN<-VapB_antitoxin<-?<-N6-METHYLASE-p<-AbrB-HTH<-N6-METHYLASE<-PemK_toxin | 1239 | Bacteria>Pseudomonadota>Gammaproteobacteria>Chromatiales>Chromatiaceae>Thiodictyon | GCF_002813775.1 | Candidatus_Thiodictyon_syntrophicum | THSYN_RS30800 | 328512..332231 | 1166950 | phage-resistance-protein-[Candidatus-Thiodictyon-syntrophicum] |
| WP_058553885.1 | Type2.Pgl.like.BREX | N6-METHYLASE->?->?->PIN->HEPN->ParA-Soj-PloopNTPase->ORC-CDC6-><-MPTase<-RelE\|\|DUF6079*->PglZ->BrxD-AAA->SF1-SF2-Helicase+wHTH+Small-B-barrel+DUF1998->?->?-><-VirB11<-?<-Glycos_transf | 1215 | Bacteria>Pseudomonadota>Gammaproteobacteria>Chromatiales>Chromatiaceae>Thiohalocapsa | GCF_001469165.1 | Thiohalocapsa_sp._ML1 | ML01_RS02835 | 105338..108985 | 1431688 | hypothetical-protein-[Thiohalocapsa-sp.-ML1] |
| WP_150094150.1 | Type2.Pgl.like.BREX | PglW->N6-METHYLASE->Unstab_antitox->RelE-ParE->DUF6079*->PglZ->BrxD-AAA->SF1-SF2-Helicase+wHTH+Small-B-barrel+DUF1998-> | 1221 | Bacteria>Pseudomonadota>Gammaproteobacteria>Chromatiales>Chromatiaceae>Thiohalocapsa | GCF_008632335.1 | Thiohalocapsa_marina | F2Q65_RS14605 | 20024..23689 | 424902 | phage-resistance-protein-[Thiohalocapsa-marina] |
| WP_200236884.1 | Type2.Pgl.like.BREX | PhdYeFM_antitox->PIN->?->N6-METHYLASE->TPR+TPR+TPR+TPR->?->N6-METHYLASE-p->?-><-MPTase\|\|DUF6079*-><-PhdYeFM_antitox\|\|?->RHH_6-><-?\|\|PglZ->BrxD-AAA-><-PIN<-RelB\|\|SF1-SF2-Helicase+wHTH+Small-B-barrel+DUF1998-> | 1236 | Bacteria>Pseudomonadota>Gammaproteobacteria>Chromatiales>Chromatiaceae>Thiohalocapsa | GCF_016583825.1 | Thiohalocapsa_halophila | CKO31_RS10635 | 25991..29701 | 69359 | phage-resistance-protein-[Thiohalocapsa-halophila] |
| WP_164653515.1 | Type2.Pgl.like.BREX | <-SF1-SF2-Helicase+wHTH+Small-B-barrel+DUF1998<-BrxD-AAA\|\|?-><-PglZ<-DUF6079*\|\|CopG/RHH->PIN-><-?<-REase<-REase<-Unstab_antitox\|\|Tox-RelE->HTH_3-><-N6-METHYLASE | 1220 | Bacteria>Pseudomonadota>Gammaproteobacteria>Chromatiales>Chromatiaceae>Thiorhodococcus | GCF_010915725.1 | Thiorhodococcus_mannitoliphagus | G3480_RS08855 | 7645..11307 | 329406 | phage-resistance-protein-[Thiorhodococcus-mannitoliphagus] |
| WP_201065793.1 | Type2.Pgl.like.BREX | NAD_binding_10->?-><-SF1-SF2-Helicase+wHTH+Small-B-barrel+DUF1998<-BrxD-AAA<-PglZ<-DUF6079*<-RelE-ParE<-Unstab_antitox<-N6-METHYLASE<-?<-RNase<-PglW | 1260 | Bacteria>Pseudomonadota>Gammaproteobacteria>Chromatiales>Chromatiaceae>Thiorhodovibrio | GCF_016653415.1 | Thiorhodovibrio_winogradskyi | CCR91_RS07050 | 10750..14532 | 77007 | phage-resistance-protein-[Thiorhodovibrio-winogradskyi] |
| MCF7993507.1 | Type2.Pgl.like.BREX | <-SAM-METHYLASE<-?<-SAM-METHYLASE<-REase\|\|<-?\|\|?-><-REase<-PglZ<-DUF6079*<-N6-METHYLASE<-PglW<-?<-REase<-REase | 1237 | Bacteria>Pseudomonadota>Gammaproteobacteria>Chromatiales>Chromatiaceae>unclassified_Chromatiaceae | GCA_021736825.1 | Chromatiaceae_bacterium | K9L88_01525 | 9566..13279 | 2049432 | phage-resistance-protein-[Chromatiaceae-bacterium] |
| MCP5414111.1 | Type2.Pgl.like.BREX | <-Phage_integrase<-?\|\|HTH_26->PglW->N6-METHYLASE->DUF6079*->PglZ->BrxD-AAA->SF1-SF2-Helicase+wHTH+Small-B-barrel+DUF1998->?->?->?->SbcC-> | 1216 | Bacteria>Pseudomonadota>Gammaproteobacteria>Chromatiales>Chromatiaceae>unclassified_Chromatiaceae | GCA_024235295.1 | Chromatiaceae_bacterium | H6961_05835 | 63898..67548 | 2049432 | phage-resistance-protein-[Chromatiaceae-bacterium] |
| WP_210386919.1 | Type2.Pgl.like.BREX | <-HNH<-?<-?<-SF1-SF2-Helicase+wHTH+Small-B-barrel+DUF1998<-BrxD-AAA<-PglZ<-DUF6079*<-DUF6079-p<-PIN<-?<-N6-METHYLASE-p | 1060 | Bacteria>Pseudomonadota>Gammaproteobacteria>Chromatiales>Ectothiorhodospiraceae>Ectothiorhodospira | GCF_000225005.1 | Ectothiorhodospira_sp._PHS-1 | ECTPHS_RS11290 | 30529..33711 | 519989 | phage-resistance-protein-[Ectothiorhodospira-sp.-PHS-1] |
| WP_210386920.1 | Type2.Pgl.like.BREX | <-HNH<-?<-?<-SF1-SF2-Helicase+wHTH+Small-B-barrel+DUF1998<-BrxD-AAA<-PglZ<-DUF6079<-DUF6079-p*<-PIN<-?<-N6-METHYLASE-p | 162 | Bacteria>Pseudomonadota>Gammaproteobacteria>Chromatiales>Ectothiorhodospiraceae>Ectothiorhodospira | GCF_000225005.1 | Ectothiorhodospira_sp._PHS-1 | ECTPHS_RS14965 | 33781..34269 | 519989 | hypothetical-protein-[Ectothiorhodospira-sp.-PHS-1] |
| WP_238617951.1 | Type2.Pgl.like.BREX | <-HNH<-?<-?<-SF1-SF2-Helicase+wHTH+Small-B-barrel+DUF1998<-BrxD-AAA<-PglZ<-DUF6079*<-TOPRIM<-OLD-ABC<-N6-METHYLASE<-CopG_antitoxin<-BrnT_toxin<-PglW<-?<-Peptidase_M48+TPR_9\|\|ABhydrolase-> | 1228 | Bacteria>Pseudomonadota>Gammaproteobacteria>Chromatiales>Ectothiorhodospiraceae>Ectothiorhodospira | GCF_022227765.1 | Ectothiorhodospira_shaposhnikovii | LQD11_RS13150 | 102159..105845 | 1054 | phage-resistance-protein-[Ectothiorhodospira-shaposhnikovii] |
| WP_018719626.1 | Type2.Pgl.like.BREX | HAD+HAD-><-HTH_Tnp_1+HTH_21+rve+rve_3\|\|PglW->N6-METHYLASE->RHH_1->RelE-ParE->?->DUF6079*->PglZ->BrxD-AAA->SF1-SF2-Helicase+wHTH+Small-B-barrel+DUF1998->?-><-HTH_21+rve+rve_3 | 1219 | Bacteria>Pseudomonadota>Gammaproteobacteria>Chromatiales>Ectothiorhodospiraceae | GCF_000374645.1 | Arhodomonas | C516_RS0117010 | 26950..30609 | 2368 | MULTISPECIES:-hypothetical-protein-[Arhodomonas] |
| WP_238634085.1 | Type2.Pgl.like.BREX | <-SF1-SF2-Helicase+wHTH+Small-B-barrel+DUF1998<-BrxD-AAA<-PglZ<-DUF6079*<-?<-N6-METHYLASE<-PglW | 1221 | Bacteria>Pseudomonadota>Gammaproteobacteria>Chromatiales>Ectothiorhodospiraceae>Halorhodospira | GCF_022227985.1 | unclassified_Halorhodospira | LRA98_RS11675 | 6347..10012 | 2626748 | MULTISPECIES:-phage-resistance-protein-[unclassified-Halorhodospira] |
| WP_005004488.1 | Type2.Pgl.like.BREX | <-DDE_Tnp_1+DDE_Tnp_1\|\|PglW->N6-METHYLASE->MazE_antitoxin->PIN->DUF6079*->PglZ->BrxD-AAA->SF1-SF2-Helicase+wHTH+Small-B-barrel+DUF1998-><-SAM-METHYLASE\|\|PhdYeFM_antitox->PIN-><-tRNA\|\|RhoN+RNase+S1->SPOUT-> | 1222 | Bacteria>Pseudomonadota>Gammaproteobacteria>Chromatiales>Ectothiorhodospiraceae>Nitrococcus | GCF_000153205.1 | Nitrococcus_mobilis | NB231_RS19260 | 3295318..3298986 | 35797 | hypothetical-protein-[Nitrococcus-mobilis] |
| MCK7580665.1 | Type2.Pgl.like.BREX | N6-METHYLASE-p->N6-METHYLASE-p->N6-METHYLASE-p-><-REase\|\|DDE_Tnp_1-><-DDE_Tnp_1\|\|HTH_Tnp_4+DDE_Tnp_4->\|\|DUF6079-p*->DUF6079-p-><-?\|\|?->?-><-?\|\|PglZ-p-><-PIN | 892 | Bacteria>Pseudomonadota>Gammaproteobacteria>Chromatiales>unclassified_Chromatiales | GCA_023156155.1 | Chromatiales_bacterium | MZV65_36870 | 34509..37187 | 2026725 | phage-resistance-protein-[Chromatiales-bacterium] |
| WP_048671262.1 | Type2.Pgl.like.BREX | <-DSBH\|\|?->?->PglW-><-PIN<-PhdYeFM_antitox\|\|N6-METHYLASE->?->?->DUF6079*->PglZ->Amidohydro_1-><-?\|\|?->?->ABC-ATPase+SbcC-> | 1239 | Bacteria>Pseudomonadota>Gammaproteobacteria>Gammaproteobacteria_incertae_sedis>Candidatus_Competibacteraceae>Candidatus_Competibacter | GCF_001051235.1 | Candidatus_Competibacter_denitrificans | BN873_RS04980 | 19269..22988 | 1400862 | hypothetical-protein-[Candidatus-Competibacter-denitrificans] |
| MCG7935437.1 | Type2.Pgl.like.BREX | <-AbiEi_4<-AbiGi+seqA+RAMA<-SF1-SF2-Helicase+wHTH+Small-B-barrel+DUF1998<-BrxD-AAA<-PglZ<-DUF6079*<-N6-METHYLASE<-PglW\|\|?-><-RelE-ParE<-PhdYeFM_antitox\|\|XIS-HTH->Phage_integrase-> | 1215 | Bacteria>Pseudomonadota>Gammaproteobacteria>Gammaproteobacteria_incertae_sedis>sulfur-oxidizing_symbionts>Candidatus_Thiodiazotropha | GCA_016842345.1 | Candidatus_Thiodiazotropha_taylori | JAY68_12350 | 15615..19262 | 2792791 | phage-resistance-protein-[Candidatus-Thiodiazotropha-taylori] |
| MCG8094254.1 | Type2.Pgl.like.BREX | PglW->N6-METHYLASE->DUF6079*->PglZ->BrxD-AAA->SF1-SF2-Helicase+wHTH+Small-B-barrel+DUF1998->AbiGi+seqA+RAMA->\|\|?->?->?->?->ABC-ATPase+SbcC-> | 1215 | Bacteria>Pseudomonadota>Gammaproteobacteria>Gammaproteobacteria_incertae_sedis>sulfur-oxidizing_symbionts>Candidatus_Thiodiazotropha | GCA_016842965.1 | Candidatus_Thiodiazotropha_endolucinida | JAZ17_11620 | 8627..12274 | 1655433 | phage-resistance-protein-[Candidatus-Thiodiazotropha-endolucinida] |
| WP_162851400.1 | Type2.Pgl.like.BREX | NTP_transf_3+Hexapep+Hexapep-><-NTP_transf_3<-?<-?<-?\|\|?->HTH_26->PglW->N6-METHYLASE->DUF6079*->PglZ->BrxD-AAA->SF1-SF2-Helicase+wHTH+Small-B-barrel+DUF1998->?->TniQ-><-TnsC-ATPase<-rve+Mu-transpos_C<-REase\|\|SIR2-> | 1212 | Bacteria>Pseudomonadota>Gammaproteobacteria>Nevskiales>Sinobacteraceae>Panacagrimonas | GCF_004364935.1 | Panacagrimonas_perspica | DFR24_RS24595 | 97223..100861 | 381431 | DUF6079-family-protein-[Panacagrimonas-perspica] |
| WP_136249411.1 | Type2.Pgl.like.BREX | NTP_transf_3+Hexapep+Hexapep-><-NTP_transf_3<-?<-?<-?\|\|?->HTH_26->PglW->N6-METHYLASE->DUF6079*-> | 1152 | Bacteria>Pseudomonadota>Gammaproteobacteria>Nevskiales>Sinobacteraceae>Panacagrimonas | GCF_004799025.1 | Panacagrimonas_perspica | B1810_RS21740 | 24184..27638 | 381431 | DUF6079-family-protein,-partial-[Panacagrimonas-perspica] |
| WP_252180111.1 | Type2.Pgl.like.BREX | <-HIN-HTH\|\|?->?-><-sGTP<-?<-SF1-SF2-Helicase+wHTH+Small-B-barrel+DUF1998<-BrxD-AAA<-PglZ<-DUF6079*<-?<-N6-METHYLASE<-PglW<-HTH_26\|\|PIN->RADICAL-SAM->?->PRTase->ParB+HNH->ParB+HNH-> | 1224 | Bacteria>Pseudomonadota>Gammaproteobacteria>Oceanospirillales>Endozoicomonadaceae>Endozoicomonas | GCF_023822025.1 | Endozoicomonas_sp._4G | K7B67_RS09560 | 2457320..2460994 | 2872754 | DUF6079-family-protein-[Endozoicomonas-sp.-4G] |
| WP_011397281.1 | Type2.Pgl.like.BREX | ABC-ATPase+SbcC->?->?-><-?<-HAMP+MA+MA<-SF1-SF2-Helicase+wHTH+Small-B-barrel+DUF1998<-BrxD-AAA<-PglZ<-DUF6079*<-N6-METHYLASE<-TOPRIM<-OLD-ABC<-PglW<-XIS-HTH | 1227 | Bacteria>Pseudomonadota>Gammaproteobacteria>Oceanospirillales>Hahellaceae>Hahella | GCF_000012985.1 | Hahella_chejuensis | HCH_RS15580 | 3593467..3597150 | 158327 | hypothetical-protein-[Hahella-chejuensis] |
| WP_138801530.1 | Type2.Pgl.like.BREX | Phage_integrase->Arm-DNA-bind_3+Phage_integrase-><-?<-TOPRIM<-OLD-ABC<-SF1-SF2-Helicase+wHTH+Small-B-barrel+DUF1998<-BrxD-AAA<-PglZ<-DUF6079*<-N6-METHYLASE<-PglW<-XIS-HTH\|\|?-><-ParB+HNH<-ParB+HNH\|\|tRNA->?-><-XIS-HTH<-ZnR+TOPRIM+AAA-helicase+wHTH | 1236 | Bacteria>Pseudomonadota>Gammaproteobacteria>Oceanospirillales>Halomonadaceae>Halomonas | GCF_005937175.1 | Halomonas_sp._ATBC28 | E0L35_RS23015 | 16131..19841 | 2545264 | DUF6079-family-protein-[Halomonas-sp.-ATBC28] |
| WP_175072159.1 | Type2.Pgl.like.BREX | XIS-HTH-><-?\|\|?-><-?\|\|XIS-HTH->PglW->N6-METHYLASE->OLD-ABC->HNH->DUF6079*->PglZ->BrxD-AAA->SF1-SF2-Helicase+wHTH+Small-B-barrel+DUF1998-><-?\|\|REase-><-?<-Arm-DNA-bind_3+Phage_integrase\|\|HTH+LexA-protease->N6-N4-METHYLASE-> | 1237 | Bacteria>Pseudomonadota>Gammaproteobacteria>Oceanospirillales>Halomonadaceae>Halomonas | GCF_013349955.1 | Halomonas_taeanensis | HUS66_RS15805 | 34399..38112 | 284577 | DUF6079-family-protein-[Halomonas-taeanensis] |
| WP_233973947.1 | Type2.Pgl.like.BREX | XIS-HTH-><-?<-?\|\|XIS-HTH->PglW->N6-METHYLASE->DUF6079*->PglZ->BrxD-AAA->SF1-SF2-Helicase+wHTH+Small-B-barrel+DUF1998->OLD-ABC->?-><-Arm-DNA-bind_3+Phage_integrase | 1236 | Bacteria>Pseudomonadota>Gammaproteobacteria>Oceanospirillales>Halomonadaceae>Halomonas | GCF_021395175.1 | Halomonas_titanicae | LZG37_RS03195 | 12394..16104 | 664683 | DUF6079-family-protein-[Halomonas-titanicae] |
| WP_240569311.1 | Type2.Pgl.like.BREX | XIS-HTH->PglW->OLD-ABC->HNH->N6-METHYLASE->DUF6079*->PglZ->BrxD-AAA->SF1-SF2-Helicase+wHTH+Small-B-barrel+DUF1998->?->SF1-SF2-Helicase+wHTH+Small-B-barrel+DUF1998-><-?<-GspA-ATPase | 1239 | Bacteria>Pseudomonadota>Gammaproteobacteria>Oceanospirillales>Halomonadaceae>Halomonas | GCF_022487945.1 | Halomonas_sp._EGI_63088 | MKP05_RS16485 | 45674..49393 | 2920385 | DUF6079-family-protein-[Halomonas-sp.-EGI-63088] |
| WP_251308249.1 | Type2.Pgl.like.BREX | <-EAL<-?<-GreAB-C\|\|?-><-?<-SF1-SF2-Helicase+wHTH+Small-B-barrel+DUF1998<-BrxD-AAA<-PglZ<-DUF6079*<-N6-METHYLASE<-HNH<-OLD-ABC<-PglW<-XIS-HTH<-?\|\|?->wHTH+WYL+WCX->ParB+HNH-> | 1235 | Bacteria>Pseudomonadota>Gammaproteobacteria>Oceanospirillales>Halomonadaceae>Halomonas | GCF_023708045.1 | Halomonas_sp._NCCP-2165 | NFU04_RS04185 | 82862..86569 | 2942298 | DUF6079-family-protein-[Halomonas-sp.-NCCP-2165] |
| WP_072820441.1 | Type2.Pgl.like.BREX | XIS-HTH->?->?-><-?\|\|XIS-HTH->PglW->OLD-ABC->HNH->N6-METHYLASE->DUF6079*->PglZ->BrxD-AAA->SF1-SF2-Helicase+wHTH+Small-B-barrel+DUF1998->HARE-HTH+REase-><-?\|\|?->?->?-><-HD | 1236 | Bacteria>Pseudomonadota>Gammaproteobacteria>Oceanospirillales>Halomonadaceae>Halomonas | GCF_900128925.1 | Halomonas_ilicicola | BUA41_RS05240 | 148075..151785 | 480814 | phage-resistance-protein-[Halomonas-ilicicola] |
| GLQ31491.1 | Type2.Pgl.like.BREX | <-Thioredoxin_4\|\|wHTH+WYL+WCX->BrxA-DNA-binding->BrxB-BrxF-iSTAND->DUF6079*->N6-METHYLASE->ParB+HNH->PglZ->SIGMA+OB-Fold+AAA-ATPase+Lon-Protease-BrxL->NERD+iPkinase+Pkinase+SF1-SF2-Helicase->?->REase+iREase->REase+HTH+35exo+Small-B-barrel+SF1-SF2-Helicase->PHP+SbcC-> | 1219 | Bacteria>Pseudomonadota>Gammaproteobacteria>Oceanospirillales>Oceanospirillaceae>Litoribrevibacter | GCA_030159995.1 | Litoribrevibacter_albus | GCM10007876_19700 | 85769..89428 | 1473156 | hypothetical-protein-GCM10007876_19700-[Litoribrevibacter-albus] |
| WP_230588680.1 | Type2.Pgl.like.BREX | ACET-><-?<-NERD+iPkinase+Pkinase+SF1-SF2-Helicase\|\|BrxA-DNA-binding->BrxB-BrxF-iSTAND->DUF6079-p*->DUF6079-p->N6-METHYLASE->HTH+REase->PglZ->SIGMA+OB-Fold+AAA-ATPase+Lon-Protease-BrxL-> | 790 | Bacteria>Pseudomonadota>Gammaproteobacteria>Pasteurellales>Pasteurellaceae>Gallibacterium | GCF_000771855.1 | Gallibacterium_anatis | JP27_RS00190 | 44436..46808 | 750 | BREX-system-P-loop-protein-BrxC-[Gallibacterium-anatis] |
| WP_230588682.1 | Type2.Pgl.like.BREX | ACET-><-?<-NERD+iPkinase+Pkinase+SF1-SF2-Helicase\|\|BrxA-DNA-binding->BrxB-BrxF-iSTAND->DUF6079-p->DUF6079-p*->N6-METHYLASE->HTH+REase->PglZ->SIGMA+OB-Fold+AAA-ATPase+Lon-Protease-BrxL-> | 366 | Bacteria>Pseudomonadota>Gammaproteobacteria>Pasteurellales>Pasteurellaceae>Gallibacterium | GCF_000771855.1 | Gallibacterium_anatis | JP27_RS12010 | 46763..47863 | 750 | hypothetical-protein-[Gallibacterium-anatis] |
| WP_039087873.1 | Type2.Pgl.like.BREX | ACET-><-?<-NERD+iPkinase+Pkinase+SF1-SF2-Helicase\|\|BrxA-DNA-binding->BrxB-BrxF-iSTAND->DUF6079*->N6-METHYLASE->HTH+REase->PglZ->SIGMA+OB-Fold+AAA-ATPase+Lon-Protease-BrxL-> | 1142 | Bacteria>Pseudomonadota>Gammaproteobacteria>Pasteurellales>Pasteurellaceae>Gallibacterium | GCF_000772345.1 | Gallibacterium_anatis | IO45_RS09495 | 41290..44718 | 750 | BREX-system-P-loop-protein-BrxC-[Gallibacterium-anatis] |
| WP_039158978.1 | Type2.Pgl.like.BREX | ACET-><-?<-NERD+iPkinase+Pkinase+SF1-SF2-Helicase\|\|BrxA-DNA-binding->BrxB-BrxF-iSTAND->DUF6079*->N6-METHYLASE->HTH+REase->PglZ->SIGMA+OB-Fold+AAA-ATPase+Lon-Protease-BrxL-> | 1142 | Bacteria>Pseudomonadota>Gammaproteobacteria>Pasteurellales>Pasteurellaceae>Gallibacterium | GCF_000772365.1 | Gallibacterium_anatis | IO49_RS08695 | 41238..44666 | 750 | BREX-system-P-loop-protein-BrxC-[Gallibacterium-anatis] |
| WP_196767790.1 | Type2.Pgl.like.BREX | ACET-><-?<-NERD+iPkinase+Pkinase+SF1-SF2-Helicase\|\|BrxA-DNA-binding->BrxB-BrxF-iSTAND->DUF6079-p*->DUF6079-p->N6-METHYLASE->?->PglZ->SIGMA+OB-Fold+AAA-ATPase+Lon-Protease-BrxL-><-RuvB-AAA+RuvB-HTH<-?\|\|?->O6MGT-HTH-> | 782 | Bacteria>Pseudomonadota>Gammaproteobacteria>Pasteurellales>Pasteurellaceae>Gallibacterium | GCF_001678565.1 | Gallibacterium_anatis | QVO3_RS05900 | 31345..33693 | 750 | BREX-system-P-loop-protein-BrxC-[Gallibacterium-anatis] |
| WP_196767791.1 | Type2.Pgl.like.BREX | ACET-><-?<-NERD+iPkinase+Pkinase+SF1-SF2-Helicase\|\|BrxA-DNA-binding->BrxB-BrxF-iSTAND->DUF6079-p->DUF6079-p*->N6-METHYLASE->?->PglZ->SIGMA+OB-Fold+AAA-ATPase+Lon-Protease-BrxL-><-RuvB-AAA+RuvB-HTH<-?\|\|?->O6MGT-HTH-> | 325 | Bacteria>Pseudomonadota>Gammaproteobacteria>Pasteurellales>Pasteurellaceae>Gallibacterium | GCF_001678565.1 | Gallibacterium_anatis | QVO3_RS12370 | 33801..34778 | 750 | hypothetical-protein-[Gallibacterium-anatis] |
| WP_094873724.1 | Type2.Pgl.like.BREX | <-MULE-TRANSPOSASE<-SIGMA+OB-Fold+AAA-ATPase+Lon-Protease-BrxL<-PglZ<-HNH<-OLD-ABC<-N6-METHYLASE<-DUF6079*<-BrxB-BrxF-iSTAND<-BrxA-DNA-binding\|\|NERD+iPkinase+Pkinase+SF1-SF2-Helicase->?-><-MULE-TRANSPOSASE<-ACET | 1142 | Bacteria>Pseudomonadota>Gammaproteobacteria>Pasteurellales>Pasteurellaceae>Gallibacterium | GCF_002263255.2 | Gallibacterium_anatis | CF557_RS03210 | 664755..668183 | 750 | BREX-system-P-loop-protein-BrxC-[Gallibacterium-anatis] |
| WP_007350230.1 | Type2.Pgl.like.BREX | <-NACHT\|\|?->?->?-><-?<-SF1-SF2-Helicase+wHTH+Small-B-barrel+DUF1998<-BrxD-AAA<-PglZ<-DUF6079*<-N6-METHYLASE-p\|\|HIN-HTH->?->?->DDE_Tnp_IS66-><-?\|\|?->?-><-N6-METHYLASE<-TOPRIM | 1224 | Bacteria>Pseudomonadota>Gammaproteobacteria>Pseudomonadales>Marinobacteraceae>Marinobacter | GCF_000169375.1 | Marinobacter_sp._ELB17 | MELB17_RS10590 | 56251..59925 | 270374 | DUF6079-family-protein-[Marinobacter-sp.-ELB17] |
| WP_091642160.1 | Type2.Pgl.like.BREX | <-ABC-ATPase+SbcC<-MlaE\|\|tRNA->tRNA->Calcineurin->SbcC+CCDC158->NERD+iPkinase+Pkinase+SF1-SF2-Helicase->?->BrxE->BrxA-DNA-binding->BrxB-BrxF-iSTAND->DUF6079*->N6-METHYLASE->N6-METHYLASE->ParB+HNH->PglZ->SIGMA+OB-Fold+AAA-ATPase+HKD-DNase-BrxL-><-HTH_Tnp_1+HTH_21+rve+rve_3 | 1213 | Bacteria>Pseudomonadota>Gammaproteobacteria>Pseudomonadales>Marinobacteraceae>Marinobacter | GCF_900114695.1 | Marinobacter_salarius | BMZ08_RS07625 | 233412..237053 | 1420917 | BREX-system-P-loop-protein-BrxC-[Marinobacter-salarius] |
| TXH88799.1 | Type2.Pgl.like.BREX | PglW->N6-METHYLASE->DUF6079*->PglZ->BrxD-AAA->SF1-SF2-Helicase+wHTH+Small-B-barrel+DUF1998-> | 1224 | Bacteria>Pseudomonadota>Gammaproteobacteria>Pseudomonadales>Pseudomonadaceae>Pseudomonas | GCA_008015405.1 | Pseudomonas_sp. | E6Q71_04395 | 8189..11863 | 306 | phage-resistance-protein-[Pseudomonas-sp.] |
| MBG4805295.1 | Type2.Pgl.like.BREX | MerR-HTH->tRNA->PglW->N6-METHYLASE->DUF6079*->PglZ->BrxD-AAA->SF1-SF2-Helicase+wHTH+Small-B-barrel+DUF1998->XIS-HTH->RNaseH+ArgR-HTH->ABC-ATPase->HNH-> | 1225 | Bacteria>Pseudomonadota>Gammaproteobacteria>Pseudomonadales>Pseudomonadaceae>Pseudomonas | GCA_015735905.1 | Pseudomonas_aeruginosa | I5F80_09645 | 33397..37074 | 287 | phage-resistance-protein-[Pseudomonas-aeruginosa] |
| MBH3530270.1 | Type2.Pgl.like.BREX | <-HNH<-ABC-ATPase<-RNaseH+ArgR-HTH<-XIS-HTH<-SF1-SF2-Helicase+wHTH+Small-B-barrel+DUF1998<-BrxD-AAA<-PglZ<-DUF6079*<-N6-METHYLASE<-PglW<-tRNA<-MerR-HTH | 1225 | Bacteria>Pseudomonadota>Gammaproteobacteria>Pseudomonadales>Pseudomonadaceae>Pseudomonas | GCA_016009595.1 | Pseudomonas_aeruginosa | I5L62_06380 | 108500..112177 | 287 | phage-resistance-protein-[Pseudomonas-aeruginosa] |
| HBO3398854.1 | Type2.Pgl.like.BREX | MerR-HTH->tRNA->PglW->N6-METHYLASE->DUF6079*->PglZ->BrxD-AAA->SF1-SF2-Helicase+wHTH+Small-B-barrel+DUF1998-> | 1224 | Bacteria>Pseudomonadota>Gammaproteobacteria>Pseudomonadales>Pseudomonadaceae>Pseudomonas | GCA_021652755.1 | Pseudomonas_aeruginosa | LZL90_23935 | 48172..51846 | 287 | phage-resistance-protein-[Pseudomonas-aeruginosa] |
| HBP0156834.1 | Type2.Pgl.like.BREX | MerR-HTH->tRNA->PglW->N6-METHYLASE->DUF6079*->PglZ->BrxD-AAA->SF1-SF2-Helicase+wHTH+Small-B-barrel+DUF1998-><-RNase<-HTH+MPTase<-?<-?\|\|XIS-HTH-> | 1225 | Bacteria>Pseudomonadota>Gammaproteobacteria>Pseudomonadales>Pseudomonadaceae>Pseudomonas | GCA_021684135.1 | Pseudomonas_aeruginosa | L5P47_004332 | 89020..92697 | 287 | phage-resistance-protein-[Pseudomonas-aeruginosa] |
| HBP6640957.1 | Type2.Pgl.like.BREX | <-HNH<-OLD-ABC<-SF1-SF2-Helicase<-XIS-HTH<-SF1-SF2-Helicase+wHTH+Small-B-barrel+DUF1998<-BrxD-AAA<-PglZ<-DUF6079*<-N6-METHYLASE<-PglW<-tRNA<-MerR-HTH | 1225 | Bacteria>Pseudomonadota>Gammaproteobacteria>Pseudomonadales>Pseudomonadaceae>Pseudomonas | GCA_021711635.1 | Pseudomonas_aeruginosa | L6B20_01530 | 336625..340302 | 287 | phage-resistance-protein-[Pseudomonas-aeruginosa] |
| HCF2670340.1 | Type2.Pgl.like.BREX | MerR-HTH->tRNA->PglW->N6-METHYLASE->DUF6079*->PglZ->BrxD-AAA->SF1-SF2-Helicase+wHTH+Small-B-barrel+DUF1998-> | 1224 | Bacteria>Pseudomonadota>Gammaproteobacteria>Pseudomonadales>Pseudomonadaceae>Pseudomonas | GCA_023944855.1 | Pseudomonas_aeruginosa | NIB23_005013 | 45223..48897 | 287 | phage-resistance-protein-[Pseudomonas-aeruginosa] |
| HCI3598031.1 | Type2.Pgl.like.BREX | MerR-HTH->tRNA->PglW->N6-METHYLASE->DUF6079*->PglZ->BrxD-AAA->SF1-SF2-Helicase+wHTH+Small-B-barrel+DUF1998->XIS-HTH-><-?<-KAP_NTPase<-DUF4969 | 1225 | Bacteria>Pseudomonadota>Gammaproteobacteria>Pseudomonadales>Pseudomonadaceae>Pseudomonas | GCA_024369735.1 | Pseudomonas_aeruginosa | NQI16_004638 | 48075..51752 | 287 | phage-resistance-protein-[Pseudomonas-aeruginosa] |
| UTX28290.1 | Type2.Pgl.like.BREX | MerR-HTH->tRNA->PglW->N6-METHYLASE->DUF6079*->PglZ->BrxD-AAA->SF1-SF2-Helicase+wHTH+Small-B-barrel+DUF1998-><-HIN-HTH+PAIREDC-HTH<-SymE_toxin<-FUSC<-ZnR+ZnR+Pkinase+SF1-SF2-Helicase | 1224 | Bacteria>Pseudomonadota>Gammaproteobacteria>Pseudomonadales>Pseudomonadaceae>Pseudomonas | GCA_024397355.1 | Pseudomonas_aeruginosa | NNL25_11420 | 2392777..2396451 | 287 | phage-resistance-protein-[Pseudomonas-aeruginosa] |
| MCT5096227.1 | Type2.Pgl.like.BREX | MerR-HTH->tRNA->PglW->N6-METHYLASE->DUF6079*->PglZ->BrxD-AAA->SF1-SF2-Helicase+wHTH+Small-B-barrel+DUF1998-> | 1224 | Bacteria>Pseudomonadota>Gammaproteobacteria>Pseudomonadales>Pseudomonadaceae>Pseudomonas | GCA_025235805.1 | Pseudomonas_aeruginosa | LZL90_23935 | 48172..51846 | 287 | phage-resistance-protein-[Pseudomonas-aeruginosa] |
| WP_132789291.1 | Type2.Pgl.like.BREX | MerR-HTH->tRNA->PglW->N6-METHYLASE->DUF6079*->PglZ->BrxD-AAA->SF1-SF2-Helicase+wHTH+Small-B-barrel+DUF1998->XIS-HTH->RNaseH+ArgR-HTH->ABC-ATPase->HNH-> | 1225 | Bacteria>Pseudomonadota>Gammaproteobacteria>Pseudomonadales>Pseudomonadaceae>Pseudomonas | GCA_030820635.1 | Pseudomonas_aeruginosa | EQ600_RS01155 | 9673..13350 | 287 | phage-resistance-protein-[Pseudomonas-aeruginosa] |
| WP_055137475.1 | Type2.Pgl.like.BREX | Exonuc_V_gamma+REase->SF1-SF2-Helicase+REase->?->?->HNH-><-SF1-SF2-Helicase+wHTH+Small-B-barrel+DUF1998<-BrxD-AAA<-PglZ<-DUF6079*<-N6-METHYLASE<-PglW<-MerR-HTH | 1225 | Bacteria>Pseudomonadota>Gammaproteobacteria>Pseudomonadales>Pseudomonadaceae>Pseudomonas | GCF_001412065.1 | Pseudomonas_corrugata | ADY50_RS22470 | 36819..40496 | 47879 | phage-resistance-protein-[Pseudomonas-corrugata] |
| WP_023091210.1 | Type2.Pgl.like.BREX | MerR-HTH->tRNA->PglW->N6-METHYLASE->DUF6079*->PglZ->BrxD-AAA->SF1-SF2-Helicase+wHTH+Small-B-barrel+DUF1998->XIS-HTH->?->HIN-HTH-> | 1224 | Bacteria>Pseudomonadota>Gammaproteobacteria>Pseudomonadales>Pseudomonadaceae>Pseudomonas | GCF_001542795.2 | Pseudomonas_aeruginosa | IDX03_RS11970 | 482235..485909 | 287 | hypothetical-protein-[Pseudomonas-aeruginosa] |
| WP_069864079.1 | Type2.Pgl.like.BREX | MerR-HTH->tRNA->PglW->N6-METHYLASE->DUF6079*->PglZ->BrxD-AAA->SF1-SF2-Helicase+wHTH+Small-B-barrel+DUF1998->XIS-HTH->ACET->ASCH->?-><-?<-GTPase-AIG+IIGP-C | 1224 | Bacteria>Pseudomonadota>Gammaproteobacteria>Pseudomonadales>Pseudomonadaceae>Pseudomonas | GCF_001748265.1 | Pseudomonas_humi | BJI51_RS19505 | 4380954..4384628 | 1826671 | phage-resistance-protein-[Pseudomonas-humi] |
| WP_070087178.1 | Type2.Pgl.like.BREX | URI-DNase-><-SIR2<-?<-?<-SF1-SF2-Helicase+REase<-?<-SF1-SF2-Helicase+wHTH+Small-B-barrel+DUF1998<-BrxD-AAA<-PglZ<-DUF6079*<-N6-METHYLASE<-PglW<-tRNA<-MerR-HTH | 1224 | Bacteria>Pseudomonadota>Gammaproteobacteria>Pseudomonadales>Pseudomonadaceae>Pseudomonas | GCF_001753915.1 | Pseudomonas_sp._NBRC_111122 | BJQ15_RS15375 | 44649..48323 | 1661037 | phage-resistance-protein-[Pseudomonas-sp.-NBRC-111122] |
| WP_088513981.1 | Type2.Pgl.like.BREX | N6-METHYLASE-p->DUF6079*->PglZ->BrxD-AAA->SF1-SF2-Helicase+wHTH+Small-B-barrel+DUF1998->ParB+HNH-><-?<-?<-?<-Arm-DNA-bind_3+Phage_integrase | 1224 | Bacteria>Pseudomonadota>Gammaproteobacteria>Pseudomonadales>Pseudomonadaceae>Pseudomonas | GCF_002157495.1 | Pseudomonas_sp._1239 | B8W70_RS03315 | 1000..4674 | 1985343 | phage-resistance-protein-[Pseudomonas-sp.-1239] |
| WP_094776636.1 | Type2.Pgl.like.BREX | SymE_toxin->HTH_Tnp_1+HTH_21+rve+rve_3-><-HNH<-ABC-ATPase<-RNaseH+ArgR-HTH<-XIS-HTH<-SF1-SF2-Helicase+wHTH+Small-B-barrel+DUF1998<-BrxD-AAA<-PglZ<-DUF6079*<-N6-METHYLASE<-TOPRIM<-OLD-ABC<-PglW<-tRNA<-MerR-HTH | 1225 | Bacteria>Pseudomonadota>Gammaproteobacteria>Pseudomonadales>Pseudomonadaceae>Pseudomonas | GCF_002239565.1 | Pseudomonas_aeruginosa | CF337_RS09100 | 62592..66269 | 287 | phage-resistance-protein-[Pseudomonas-aeruginosa] |
| WP_095557984.1 | Type2.Pgl.like.BREX | <-RNaseH<-XIS-HTH<-SF1-SF2-Helicase+wHTH+Small-B-barrel+DUF1998<-BrxD-AAA<-PglZ<-DUF6079*<-N6-METHYLASE<-PglW<-tRNA<-MerR-HTH | 1224 | Bacteria>Pseudomonadota>Gammaproteobacteria>Pseudomonadales>Pseudomonadaceae>Pseudomonas | GCF_002329925.1 | Pseudomonas_aeruginosa | CJU41_RS00035 | 8156..11830 | 287 | phage-resistance-protein-[Pseudomonas-aeruginosa] |
| WP_010794554.1 | Type2.Pgl.like.BREX | <-sGTP<-wHTH+WYL+WCX<-ZnR<-?<-NERD+iPkinase+Pkinase+SF1-SF2-Helicase<-XIS-HTH<-SF1-SF2-Helicase+wHTH+Small-B-barrel+DUF1998<-BrxD-AAA<-PglZ<-DUF6079*<-N6-METHYLASE<-PglW<-tRNA<-MerR-HTH | 1225 | Bacteria>Pseudomonadota>Gammaproteobacteria>Pseudomonadales>Pseudomonadaceae>Pseudomonas | GCF_002473025.1 | Pseudomonas_sp._HPB0071 | HMPREF1487_RS00125 | 33141..36818 | 1203578 | hypothetical-protein-[Pseudomonas-sp.-HPB0071] |
| WP_121335244.1 | Type2.Pgl.like.BREX | HTH+MPTase->RNase->HIN-HTH->?->LZ_Tnp_IS66+zf-IS66+DDE_Tnp_IS66+DDE_Tnp_IS66_C-><-SF1-SF2-Helicase+wHTH+Small-B-barrel+DUF1998<-BrxD-AAA<-PglZ<-DUF6079*<-N6-METHYLASE<-PglW<-tRNA<-MerR-HTH | 1224 | Bacteria>Pseudomonadota>Gammaproteobacteria>Pseudomonadales>Pseudomonadaceae>Pseudomonas | GCF_003633185.1 | Pseudomonas_aeruginosa | DQG94_RS17545 | 26628..30302 | 287 | phage-resistance-protein-[Pseudomonas-aeruginosa] |
| WP_124175829.1 | Type2.Pgl.like.BREX | DUF4969->?->KAP_NTPase->?-><-XIS-HTH<-SF1-SF2-Helicase+wHTH+Small-B-barrel+DUF1998<-BrxD-AAA<-PglZ<-DUF6079*<-N6-METHYLASE<-PglW<-tRNA<-MerR-HTH | 1225 | Bacteria>Pseudomonadota>Gammaproteobacteria>Pseudomonadales>Pseudomonadaceae>Pseudomonas | GCF_003839665.1 | Pseudomonas_aeruginosa | IPC273_RS12760 | 16437..20114 | 287 | phage-resistance-protein-[Pseudomonas-aeruginosa] |
| WP_124182862.1 | Type2.Pgl.like.BREX | <-XIS-HTH\|\|?->?->HTH+MPTase->RNase-><-SF1-SF2-Helicase+wHTH+Small-B-barrel+DUF1998<-BrxD-AAA<-PglZ<-DUF6079*<-N6-METHYLASE<-PglW<-tRNA<-MerR-HTH | 1224 | Bacteria>Pseudomonadota>Gammaproteobacteria>Pseudomonadales>Pseudomonadaceae>Pseudomonas | GCF_003839985.1 | Pseudomonas_aeruginosa | PAPB9_RS00045 | 10359..14033 | 287 | phage-resistance-protein-[Pseudomonas-aeruginosa] |
| WP_169919065.1 | Type2.Pgl.like.BREX | <-OLD-ABC+TOPRIM<-MAD-NTD+REase<-35exo+SF1-SF2-Helicase<-?<-NERD+iPkinase+Pkinase+SF1-SF2-Helicase<-XIS-HTH<-SF1-SF2-Helicase+wHTH+Small-B-barrel+DUF1998<-BrxD-AAA<-PglZ<-DUF6079*<-HNH<-OLD-ABC<-N6-METHYLASE<-PglW<-tRNA<-MerR-HTH | 1231 | Bacteria>Pseudomonadota>Gammaproteobacteria>Pseudomonadales>Pseudomonadaceae>Pseudomonas | GCF_012985845.1 | Pseudomonas_sp._WS_5013 | HBN76_RS00225 | 58998..62693 | 2717475 | phage-resistance-protein-[Pseudomonas-sp.-WS-5013] |
| WP_151896782.1 | Type2.Pgl.like.BREX | <-HNH<-?<-?<-XIS-HTH<-?<-BrxD-AAA<-PglZ<-DUF6079*<-N6-METHYLASE<-PglW<-tRNA<-MerR-HTH | 1225 | Bacteria>Pseudomonadota>Gammaproteobacteria>Pseudomonadales>Pseudomonadaceae>Pseudomonas | GCF_018026635.1 | Pseudomonas_aeruginosa | PPJ03_RS07590 | 16865..20542 | 287 | phage-resistance-protein-[Pseudomonas-aeruginosa] |
| WP_070870403.1 | Type2.Pgl.like.BREX | MerR-HTH->tRNA->PglW->N6-METHYLASE->DUF6079*->PglZ->BrxD-AAA->SF1-SF2-Helicase+wHTH+Small-B-barrel+DUF1998->XIS-HTH-><-?<-KAP_NTPase<-?<-DUF4969 | 1225 | Bacteria>Pseudomonadota>Gammaproteobacteria>Pseudomonadales>Pseudomonadaceae>Pseudomonas | GCF_021497425.1 | Pseudomonas_aeruginosa | EJ594_RS10920 | 33430..37107 | 287 | phage-resistance-protein-[Pseudomonas-aeruginosa] |
| WP_247248395.1 | Type2.Pgl.like.BREX | MerR-HTH->tRNA->PglW->N6-METHYLASE->DUF6079*->PglZ->BrxD-AAA->SF1-SF2-Helicase+wHTH+Small-B-barrel+DUF1998->XIS-HTH->SF1-SF2-Helicase->OLD-ABC->HNH->HHH_4+HTH+wHTH+SF1-SF2-Helicase+REase-> | 1224 | Bacteria>Pseudomonadota>Gammaproteobacteria>Pseudomonadales>Pseudomonadaceae>Pseudomonas | GCF_023060395.1 | Pseudomonas_aeruginosa | MX103_RS25025 | 62098..65772 | 287 | phage-resistance-protein-[Pseudomonas-aeruginosa] |
| WP_255869924.1 | Type2.Pgl.like.BREX | MerR-HTH->tRNA->PglW->N6-METHYLASE->DUF6079*->PglZ->BrxD-AAA->SF1-SF2-Helicase+wHTH+Small-B-barrel+DUF1998-><-HIN-HTH+PAIREDC-HTH<-SymE_toxin<-FUSC<-ZnR+ZnR+Pkinase+SF1-SF2-Helicase<-?<-ClpABC-AAA | 1224 | Bacteria>Pseudomonadota>Gammaproteobacteria>Pseudomonadales>Pseudomonadaceae>Pseudomonas | GCF_024397355.1 | Pseudomonas_aeruginosa | NNL25_RS11425 | 2392777..2396451 | 287 | phage-resistance-protein-[Pseudomonas-aeruginosa] |
| WP_265969626.1 | Type2.Pgl.like.BREX | <-NERD+iPkinase+Pkinase+SF1-SF2-Helicase\|\|?->?->?-><-SIGMA+OB-Fold+AAA-ATPase+Lon-Protease-BrxL<-PglZ<-ParB+HNH<-N6-METHYLASE<-DUF6079*<-BrxB-BrxF-iSTAND<-BrxA-DNA-binding<-wHTH+WYL+WCX<-?<-?<-XIS-HTH\|\|CopG/RHH-><-GNTR-HTH+Aminotran_1_2\|\|ACET-> | 1146 | Bacteria>Pseudomonadota>Gammaproteobacteria>Pseudomonadales>Pseudomonadaceae>Pseudomonas | GCF_026240895.1 | Pseudomonas_sp._DCB_AW | OO306_RS19225 | 23705..27145 | 2993596 | BREX-system-P-loop-protein-BrxC-[Pseudomonas-sp.-DCB_AW] |
| WP_269974942.1 | Type2.Pgl.like.BREX | MerR-HTH->tRNA->PglW->N6-METHYLASE->DUF6079*->PglZ->BrxD-AAA->SF1-SF2-Helicase+wHTH+Small-B-barrel+DUF1998->SF1-SF2-Helicase->OLD-ABC->HNH->HHH_4+HTH+wHTH+SF1-SF2-Helicase+REase-><-LZ_Tnp_IS66+zf-IS66+DDE_Tnp_IS66+DDE_Tnp_IS66_C | 1224 | Bacteria>Pseudomonadota>Gammaproteobacteria>Pseudomonadales>Pseudomonadaceae>Pseudomonas | GCF_027571095.1 | Pseudomonas_aeruginosa | PALA35_RS12150 | 2526857..2530531 | 287 | phage-resistance-protein-[Pseudomonas-aeruginosa] |
| WP_090241711.1 | Type2.Pgl.like.BREX | <-NERD+iPkinase+Pkinase+SF1-SF2-Helicase<-35exo+SF1-SF2-Helicase\|\|?->wHTH+WYL+WCX->BrxE->BrxA-DNA-binding->BrxB-BrxF-iSTAND->DUF6079*->N6-METHYLASE->ParB+HNH->PglZ->SIGMA+OB-Fold+AAA-ATPase+HKD-DNase-BrxL-><-HNH<-DUF4969 | 1214 | Bacteria>Pseudomonadota>Gammaproteobacteria>Pseudomonadales>Pseudomonadaceae>Pseudomonas | GCF_900113745.1 | Pseudomonas_guineae | BMZ32_RS09345 | 81047..84691 | 425504 | BREX-system-P-loop-protein-BrxC-[Pseudomonas-guineae] |
| WP_058013165.1 | Type2.Pgl.like.BREX | <-HNH<-?<-RNaseH+ArgR-HTH<-XIS-HTH<-SF1-SF2-Helicase+wHTH+Small-B-barrel+DUF1998<-BrxD-AAA<-PglZ<-DUF6079*<-N6-METHYLASE<-PglW<-tRNA<-MerR-HTH | 1225 | Bacteria>Pseudomonadota>Gammaproteobacteria>Pseudomonadales>Pseudomonadaceae>Pseudomonas | GCF_900707485.1 | Pseudomonas_aeruginosa | H2N61_RS04240 | 152347..156024 | 287 | hypothetical-protein-[Pseudomonas-aeruginosa] |
| WP_040259325.1 | Type2.Pgl.like.BREX | <-HTH_Tnp_1+HTH_21+rve+rve_3<-DUF932<-?\|\|?->XIS-HTH->?->PglW->N6-METHYLASE->DUF6079*->PglZ->BrxD-AAA->SF1-SF2-Helicase+wHTH+Small-B-barrel+DUF1998->?->NUDIX+NAAR1+NAAR2->NUDIX+NAAR1+NAAR2-><-URI-DNase\|\|?-><-HicB | 1224 | Bacteria>Pseudomonadota>Gammaproteobacteria>Pseudomonadales>Pseudomonadaceae>Pseudomonas | GCF_902375475.1 | Pseudomonas_massiliensis | BN1188_RS01065 | 92541..96215 | 522492 | hypothetical-protein-[Pseudomonas-massiliensis] |
| WP_064590780.1 | Type2.Pgl.like.BREX | <-SF1-SF2-Helicase<-?<-SF1-SF2-Helicase+wHTH+Small-B-barrel+DUF1998<-BrxD-AAA<-PglZ<-DUF6079*<-N6-METHYLASE<-PglW<-tRNA<-MerR-HTH | 1224 | Bacteria>Pseudomonadota>Gammaproteobacteria>Pseudomonadales>Pseudomonadaceae>Pseudomonas | GCF_947311875.1 | Pseudomonas_putida | QME24_RS02585 | 81199..84873 | 303 | hypothetical-protein-[Pseudomonas-putida] |
| WP_150761534.1 | Type2.Pgl.like.BREX | MerR-HTH->tRNA->PglW->N6-METHYLASE->DUF6079*->PglZ->BrxD-AAA->SF1-SF2-Helicase+wHTH+Small-B-barrel+DUF1998-><-HNH<-?<-?<-SF1-SF2-Helicase+REase<-Exonuc_V_gamma+REase | 1225 | Bacteria>Pseudomonadota>Gammaproteobacteria>Pseudomonadales>Pseudomonadaceae>Pseudomonas | - | Pseudomonas_fluorescens | F8N70_RS18085 | 85409..89086 | 294 | phage-resistance-protein-[Pseudomonas-fluorescens] |
| WP_003298378.1 | Type2.Pgl.like.BREX | MerR-HTH->tRNA->PglW->N6-METHYLASE->N6-METHYLASE->DUF6079*->PglZ->BrxD-AAA->SF1-SF2-Helicase+wHTH+Small-B-barrel+DUF1998->XIS-HTH->NERD+iPkinase+Pkinase+SF1-SF2-Helicase-> | 1224 | Bacteria>Pseudomonadota>Gammaproteobacteria>Pseudomonadales>Pseudomonadaceae>Stutzerimonas | GCF_000341615.1 | Stutzerimonas_stutzeri | B381_RS02880 | 85171..88845 | 316 | hypothetical-protein-[Stutzerimonas-stutzeri] |
| WP_025242139.1 | Type2.Pgl.like.BREX | MerR-HTH->tRNA->PglW->N6-METHYLASE->DUF6079*->PglZ->BrxD-AAA->SF1-SF2-Helicase+wHTH+Small-B-barrel+DUF1998->XIS-HTH->ABC-ATPase->HNH-><-?<-DUF4969 | 1225 | Bacteria>Pseudomonadota>Gammaproteobacteria>Pseudomonadales>Pseudomonadaceae>Stutzerimonas | GCF_000590475.1 | Stutzerimonas_stutzeri | NAV34_RS15425 | 259406..263083 | 316 | hypothetical-protein-[Stutzerimonas-stutzeri] |
| WP_102829213.1 | Type2.Pgl.like.BREX | <-sGTP<-wHTH+WYL+WCX<-ZnR<-?<-NERD+iPkinase+Pkinase+SF1-SF2-Helicase<-XIS-HTH<-SF1-SF2-Helicase+wHTH+Small-B-barrel+DUF1998<-BrxD-AAA<-PglZ<-DUF6079*<-N6-METHYLASE<-PglW<-tRNA<-MerR-HTH | 1224 | Bacteria>Pseudomonadota>Gammaproteobacteria>Pseudomonadales>Pseudomonadaceae>Stutzerimonas | GCF_016028635.1 | Stutzerimonas_degradans | I6G33_RS06065 | 1309743..1313417 | 2968968 | phage-resistance-protein-[Stutzerimonas-degradans] |
| WP_065983933.1 | Type2.Pgl.like.BREX | MerR-HTH->tRNA->PglW->N6-METHYLASE->DUF6079*->PglZ->BrxD-AAA->SF1-SF2-Helicase+wHTH+Small-B-barrel+DUF1998-><-RNase<-HTH+MPTase<-?<-?\|\|XIS-HTH-> | 1224 | Bacteria>Pseudomonadota>Gammaproteobacteria>Pseudomonadales>Pseudomonadaceae>Stutzerimonas | - | Stutzerimonas_xanthomarina | BBI09_RS07365 | 172008..175682 | 271420 | phage-resistance-protein-[Stutzerimonas-xanthomarina] |
| MBA1445772.1 | Type2.Pgl.like.BREX | <-RepB_primase<-XIS-HTH\|\|HTH_3->Couple_hipA+HipA_C->HTH_26->PglW->N6-METHYLASE->PolB->HEPN->DUF6079*->PglZ-p-><-?\|\|PglZ->BrxD-AAA->SF1-SF2-Helicase+wHTH+Small-B-barrel+DUF1998-> | 1216 | Bacteria>Pseudomonadota>Gammaproteobacteria>unclassified_Gammaproteobacteria | GCA_013607755.1 | Gammaproteobacteria_bacterium | FE835_12900 | 11927..15577 | 1913989 | phage-resistance-protein-[Gammaproteobacteria-bacterium] |
| MBL4761306.1 | Type2.Pgl.like.BREX | <-Phage_int_SAM_2+Integrase_1<-PAIREDC-HTH<-SF1-SF2-Helicase+wHTH+Small-B-barrel+DUF1998<-BrxD-AAA<-PglZ<-DUF6079*<-N6-METHYLASE<-PglW<-HTH_26<-?<-REase<-HEPN+SPOCS<-?<-MqsA_antitoxin+HTH_3 | 1223 | Bacteria>Pseudomonadota>Gammaproteobacteria>unclassified_Gammaproteobacteria | GCA_016764495.1 | Gammaproteobacteria_bacterium | JKY93_01235 | 10400..14071 | 1913989 | phage-resistance-protein-[Gammaproteobacteria-bacterium] |
| MBR9910596.1 | Type2.Pgl.like.BREX | TraJ-RHH->?->?->?->HTH_26->PglW->N6-METHYLASE->DUF6079*->PglZ->BrxD-AAA->SF1-SF2-Helicase+wHTH+Small-B-barrel+DUF1998->MORC+DUF3684+REase->SNF+SF1-SF2-Helicase-> | 1223 | Bacteria>Pseudomonadota>Gammaproteobacteria>unclassified_Gammaproteobacteria | GCA_018222935.1 | Gammaproteobacteria_bacterium | GYB33_09625 | 163798..167469 | 1913989 | cell-division-protein-ZapE-[Gammaproteobacteria-bacterium] |
| WP_136411735.1 | Type2.Pgl.like.BREX | <-CHTH<-HDRnase<-Arm-DNA-bind_2+Phage_int_SAM_3+Phage_integrase<-Phage_AlpA\|\|HTH_26->PglW->N6-METHYLASE->REase+HTH->NTP_transf_8->DUF6079*->PglZ->BrxD-AAA->SF1-SF2-Helicase+wHTH+Small-B-barrel+DUF1998-><-tRNA\|\|?->?-><-HISKIN+REC+HPT<-REC+GGDEF+EAL | 1213 | Bacteria>Pseudomonadota>Gammaproteobacteria>Xanthomonadales>Xanthomonadaceae>Luteimonas | GCF_004803715.2 | Luteimonas_yindakuii | E5843_RS02575 | 574939..578580 | 2565782 | phage-resistance-protein-[Luteimonas-yindakuii] |
| WP_199374480.1 | Type2.Pgl.like.BREX | SF1-SF2-Helicase+PRTase->SLOG+BirA-HTH->TPR+REase->?->PglW->N6-METHYLASE->REase+HTH->NTP_transf_8->DUF6079*->PglZ->BrxD-AAA->SF1-SF2-Helicase+wHTH+Small-B-barrel+DUF1998-> | 1217 | Bacteria>Pseudomonadota>Gammaproteobacteria>Xanthomonadales>Xanthomonadaceae>Luteimonas | GCF_016615815.1 | Luteimonas_sp._MC1572 | JGR64_RS00505 | 110209..113862 | 2799325 | hypothetical-protein-[Luteimonas-sp.-MC1572] |
| WP_213604607.1 | Type2.Pgl.like.BREX | <-Arm-DNA-bind_2+Phage_int_SAM_3+Phage_integrase<-Phage_AlpA\|\|HTH_26->PglW->N6-METHYLASE->REase+HTH->NTP_transf_8->DUF6079*->PglZ->BrxD-AAA->SF1-SF2-Helicase+wHTH+Small-B-barrel+DUF1998-><-tRNA\|\|?->?-><-HISKIN+REC+HPT<-REC+GGDEF+EAL | 1212 | Bacteria>Pseudomonadota>Gammaproteobacteria>Xanthomonadales>Xanthomonadaceae>Pseudoxanthomonas | GCF_018403535.1 | Pseudoxanthomonas_japonensis | KJS38_RS14075 | 14510..18148 | 69284 | hypothetical-protein-[Pseudoxanthomonas-japonensis] |
| MBS0213657.1 | Type2.Pgl.like.BREX | <-DUF6079-p*<-NTP_transf_8<-REase+HTH<-SecB<-MqsA_antitoxin+HTH_3<-MqsR_toxin<-N6-METHYLASE-p\|\|?-><-AAA-ATPase<-HTH+rve | 68 | Bacteria>Pseudomonadota>unclassified_Pseudomonadota | GCA_018242445.1 | Pseudomonadota_bacterium | JSR26_10835 | 381..587 | 1977087 | hypothetical-protein-[Pseudomonadota-bacterium] |
| MBT3378823.1 | Type2.Pgl.like.BREX | <-Calcineurin<-nSTAND1+HNH\|\|?-><-?<-?<-SF1-SF2-Helicase+wHTH+Small-B-barrel+DUF1998<-BrxD-AAA<-PglZ<-DUF6079*<-N6-METHYLASE | 1222 | Bacteria>PVC_group>Lentisphaerota>unclassified_Lentisphaerota | GCA_018648355.1 | Lentisphaerota_bacterium | HN406_24690 | 58119..61787 | 1932692 | phage-resistance-protein-[Lentisphaerota-bacterium] |
| HAI12821.1 | Type2.Pgl.like.BREX | <-SF1-SF2-Helicase+wHTH+Small-B-barrel+DUF1998<-BrxD-AAA<-PglZ<-DUF6079*<-MPTase<-N6-METHYLASE<-?<-PglW-p | 1223 | Bacteria>PVC_group>Planctomycetota>Phycisphaerae>Phycisphaerales>unclassified_Phycisphaerales | GCA_003450355.1 | Phycisphaerales_bacterium | DCM28_14025 | 6274..9945 | 2052180 | phage-resistance-protein-[Phycisphaerales-bacterium] |
| MCO6436556.1 | Type2.Pgl.like.BREX | XIS-HTH->PglW->N6-METHYLASE->?->DUF6079*->PglZ->BrxD-AAA->SF1-SF2-Helicase+wHTH+Small-B-barrel+DUF1998-><-?<-?<-DDE_3 | 1230 | Bacteria>PVC_group>Planctomycetota>Phycisphaerae>unclassified_Phycisphaerae | GCA_024102815.1 | Phycisphaerae_bacterium | J5J06_05665 | 9563..13255 | 2026778 | phage-resistance-protein-[Phycisphaerae-bacterium] |
| WP_010033155.1 | Type2.Pgl.like.BREX | PglW->N6-METHYLASE->OLD-ABC->HNH->DUF6079*->PglZ->BrxD-AAA->SF1-SF2-Helicase+wHTH+Small-B-barrel+DUF1998-><-?<-?<-?<-?<-?<-HTH_Tnp | 1233 | Bacteria>PVC_group>Planctomycetota>Planctomycetia>Gemmatales>Gemmataceae | GCF_003149495.1 | Gemmata_obscuriglobus | GOBSN_RS25255 | 6052898..6056599 | 114 | hypothetical-protein-[Gemmata-obscuriglobus] |
| WP_149112382.1 | Type2.Pgl.like.BREX | <-ABhydrolase<-?\|\|XIS-HTH->PglW->N6-METHYLASE->DUF6079*->PglZ->BrxD-AAA->SF1-SF2-Helicase+wHTH+Small-B-barrel+DUF1998-> | 1238 | Bacteria>PVC_group>Planctomycetota>Planctomycetia>Gemmatales>Gemmataceae | GCF_008254045.1 | Limnoglobus_roseus | PX52LOC_RS23570 | 5575302..5579018 | 2598579 | hypothetical-protein-[Limnoglobus-roseus] |
| VIP11468.1 | Type2.Pgl.like.BREX | <-MoxR-AAA\|\|XIS-HTH->PglW->N6-METHYLASE->DUF6079*->PglZ->BrxD-AAA->SF1-SF2-Helicase+wHTH+Small-B-barrel+DUF1998->?-><-?<-Phage_integrase<-tRNA<-?\|\|ZnR->?->NTP_transferase-> | 1236 | Bacteria>PVC_group>Planctomycetota>Planctomycetia>Gemmatales>Gemmataceae | - | Gemmataceae_bacterium | GCJUQL4_50210 | 6797846..6801556 | 2052164 | Uncharacterized-protein-OS=Singulisphaera-acidiphila-[Gemmataceae-bacterium] |
| WP_015245482.1 | Type2.Pgl.like.BREX | <-NACHT<-NACHT<-NACHT\|\|PglW->N6-METHYLASE->?->PIN->DUF6079*->PglZ->BrxD-AAA->SF1-SF2-Helicase+wHTH+Small-B-barrel+DUF1998-><-?<-Phage_integrase<-tRNA\|\|?->SIGMA-HTH-> | 1235 | Bacteria>PVC_group>Planctomycetota>Planctomycetia>Isosphaerales>Isosphaeraceae | GCF_000242455.2 | Singulisphaera_acidiphila | OSM_RS0102255 | 267953..271660 | 466153 | DUF6079-family-protein-[Singulisphaera-acidiphila] |
| NLY00515.1 | Type2.Pgl.like.BREX | Schlafen+HATPase+wHTH->DUF6079-p*->DUF6079-p->PglZ->BrxD-AAA->SF1-SF2-Helicase+wHTH+Small-B-barrel+DUF1998-p-><-?<-Pkinase+pi3KINASE<-?\|\|?->Pkinase-> | 290 | Bacteria>PVC_group>Planctomycetota>Planctomycetia>Pirellulales>Pirellulaceae | GCA_012728855.1 | Rhodopirellula_sp. | GXY83_30840 | 2961..3833 | 2024855 | hypothetical-protein-[Rhodopirellula-sp.] |
| NLY00516.1 | Type2.Pgl.like.BREX | Schlafen+HATPase+wHTH->DUF6079-p->DUF6079-p*->PglZ->BrxD-AAA->SF1-SF2-Helicase+wHTH+Small-B-barrel+DUF1998-p-><-?<-Pkinase+pi3KINASE<-?\|\|?->Pkinase-> | 632 | Bacteria>PVC_group>Planctomycetota>Planctomycetia>Pirellulales>Pirellulaceae | GCA_012728855.1 | Rhodopirellula_sp. | GXY83_30845 | 3837..5735 | 2024855 | hypothetical-protein-[Rhodopirellula-sp.] |
| NQV23511.1 | Type2.Pgl.like.BREX | <-SF1-SF2-Helicase<-HKD-DNase+HKD-Rease<-?<-SF1-SF2-Helicase+wHTH+Small-B-barrel+DUF1998<-BrxD-AAA<-PglZ<-DUF6079*<-HTH+REase<-?<-?<-HDRnase<-N6-METHYLASE<-PemK_toxin<-?<-XIS-HTH | 1260 | Bacteria>PVC_group>Planctomycetota>Planctomycetia>Pirellulales>Pirellulaceae | GCA_013203845.1 | Rhodopirellula_sp. | HQ518_03995 | 13896..17648 | 2024855 | phage-resistance-protein-[Rhodopirellula-sp.] |
| WP_095741475.1 | Type2.Pgl.like.BREX | PglW->?->?->?->REase->N6-METHYLASE->?->?->ABhydrolase->DUF6079*->PglZ->BrxD-AAA->SF1-SF2-Helicase+wHTH+Small-B-barrel+DUF1998->?->?->?->?->ClpABC-AAA-> | 1227 | Bacteria>PVC_group>Planctomycetota>Planctomycetia>Pirellulales>Pirellulaceae | GCF_002289565.1 | Rhodopirellula_sp._SM50 | CKO51_RS27155 | 38510..42193 | 595453 | phage-resistance-protein-[Rhodopirellula-sp.-SM50] |
| WP_105358292.1 | Type2.Pgl.like.BREX | <-SF1-SF2-Helicase\|\|?->GDResolvase+PPP1-HTH->ParB->ParB->RAMA-><-SF1-SF2-Helicase+wHTH+Small-B-barrel+DUF1998<-BrxD-AAA<-PglZ<-DUF6079*<-HNH<-OLD-ABC<-N6-METHYLASE<-?<-?<-PglW<-XIS-HTH | 1220 | Bacteria>PVC_group>Planctomycetota>Planctomycetia>Pirellulales>Pirellulaceae | GCF_002967725.1 | Blastopirellula_marina | C5Y98_RS24525 | 392384..396046 | 124 | phage-resistance-protein-[Blastopirellula-marina] |
| WP_145058317.1 | Type2.Pgl.like.BREX | XIS-HTH->PglW->N6-METHYLASE->OLD-ABC->HNH->DUF6079*->PglZ->BrxD-AAA->SF1-SF2-Helicase+wHTH+Small-B-barrel+DUF1998-><-HTH+MPTase<-?\|\|?->N6-METHYLASE-> | 1225 | Bacteria>PVC_group>Planctomycetota>Planctomycetia>Pirellulales>Pirellulaceae | GCF_007751035.1 | Lignipirellula_cremea | Pla8534_RS33060 | 8881234..8884911 | 2528010 | phage-resistance-protein-[Lignipirellula-cremea] |

| Protein-ID | System | Gene-Neighbourhoods | Length | Taxonomy | Assembly-ID | Strain | Locus-Tag | Genomic-coordinates | TaxID | Fasta-defline |
|---|---|---|---|---|---|---|---|---|---|---|
| WP_075083674.1 | Type2.Pgl.like.BREX | [illegible] | 1225 | Bacteria>PVC_group>Planctomycetota>Planctomycetia>Pirellulales>Pirellulaceae | GCF_008087665.1 | Mariniblastus_fucicola | MFFC18_RS22280 | 5944763..5948440 | 980251 | DUF6079-family-protein-[Mariniblastus-fucicola] |
| WP_164104212.1 | Type2.Pgl.like.BREX | [illegible] | 1237 | Bacteria>PVC_group>Planctomycetota>Planctomycetia>Pirellulales>Pirellulaceae | GCF_016837145.1 | Candidatus_Laterigemmans_baculatus | GNK13_RS19370 | 34997..38710 | 2770505 | phage-resistance-protein-[Candidatus-Laterigemmans-baculatus] |
| WP_230218085.1 | Type2.Pgl.like.BREX | [illegible] | 1226 | Bacteria>PVC_group>Planctomycetota>Planctomycetia>Pirellulales>Pirellulaceae | GCF_020966755.1 | Blastopirellula_sediminis | LOC68_RS09615 | 1467290..1470970 | 2894196 | hypothetical-protein-[Blastopirellula-sediminis] |
| WP_250928722.1 | Type2.Pgl.like.BREX | [illegible] | 1244 | Bacteria>PVC_group>Planctomycetota>Planctomycetia>Pirellulales>Pirellulaceae | GCF_023701485.1 | Rhodopirellula_aestuarii | NB063_RS10685 | 26062..29796 | 2950107 | DUF6079-family-protein-[Rhodopirellula-aestuarii] |
| MAX35346.1 | Type2.Pgl.like.BREX | <-SF1-SF2-Helicase+wHTH+Small-B-barrel+DUF1998<-REase<-BrxD-AAA<-PglZ<-DUF6079*<-HNH<-OLD-ABC<-N6-METHYLASE<-HNH<-OLD-ABC<-PglW | 1230 | Bacteria>PVC_group>Planctomycetota>Planctomycetia>Planctomycetales>Planctomycetaceae | GCA_002706765.1 | Gimesia_sp. | CME33_02125 | 36301..39993 | 2024833 | phage-resistance-protein-[Gimesia-sp.] |
| MBB02275.1 | Type2.Pgl.like.BREX | PglW-p-><-?\|\|N6-METHYLASE->HDRnase->?->?->DUF6079*->PglZ->N6-METHYLASE->BrxD-AAA->REase->SF1-SF2-Helicase+wHTH+Small-B-barrel+DUF1998->?->RAMA-> | 1233 | Bacteria>PVC_group>Planctomycetota>Planctomycetia>Planctomycetales>Planctomycetaceae | GCA_002709545.1 | Planctomyces_sp. | CMJ47_06490 | 8150..11851 | 37635 | phage-resistance-protein-[Planctomyces-sp.] |
| HBL44677.1 | Type2.Pgl.like.BREX | [illegible] | 783 | Bacteria>PVC_group>Planctomycetota>Planctomycetia>Planctomycetales>Planctomycetaceae | GCA_003505085.1 | Planctomycetaceae_bacterium | DDZ90_14925 | 11270..13618 | 2026779 | phage-resistance-protein-[Planctomycetaceae-bacterium] |
| HBL44678.1 | Type2.Pgl.like.BREX | [illegible] | 347 | Bacteria>PVC_group>Planctomycetota>Planctomycetia>Planctomycetales>Planctomycetaceae | GCA_003505085.1 | Planctomycetaceae_bacterium | DDZ90_14930 | 13810..14849 | 2026779 | phage-resistance-protein-[Planctomycetaceae-bacterium] |
| MBL8888645.1 | Type2.Pgl.like.BREX | [illegible] | 1223 | Bacteria>PVC_group>Planctomycetota>Planctomycetia>Planctomycetales>Planctomycetaceae | GCA_016794985.1 | Planctomycetaceae_bacterium | JNL67_01620 | 144198..147869 | 2026779 | phage-resistance-protein-[Planctomycetaceae-bacterium] |
| WP_002646768.1 | Type2.Pgl.like.BREX | <-HIN-HTH<-?<-?<-REase+iREase\|\|PglW->N6-METHYLASE->DUF6079*->PglZ->BrxD-AAA->SF1-SF2-Helicase+wHTH+Small-B-barrel+DUF1998-> | 1234 | Bacteria>PVC_group>Planctomycetota>Planctomycetia>Planctomycetales>Planctomycetaceae | GCF_000181475.1 | Gimesia_maris | GmarT_RS19720 | 5354403..5358107 | 122 | hypothetical-protein-[Gimesia-maris] |
| WP_082846329.1 | Type2.Pgl.like.BREX | [illegible] | 1224 | Bacteria>PVC_group>Planctomycetota>Planctomycetia>Planctomycetales>Planctomycetaceae | GCF_001610835.1 | Planctomyces_sp._SH-PL14 | VT03_RS18120 | 4718094..4721768 | 1632864 | phage-resistance-protein-[Planctomyces-sp.-SH-PL14] |
| WP_145372504.1 | Type2.Pgl.like.BREX | <-N6-METHYLASE-p<-SF1-SF2-Helicase<-HKD-DNase+HKD-Rease<-?<-?<-SF1-SF2-Helicase+wHTH+Small-B-barrel+DUF1998<-BrxD-AAA<-?<-PglZ<-DUF6079*<-N6-METHYLASE<-PglW | 1250 | Bacteria>PVC_group>Planctomycetota>Planctomycetia>Planctomycetales>Planctomycetaceae | GCF_007747795.1 | Maioricimonas_rarisocia | Mal4_RS27290 | 7391498..7395250 | 2528026 | phage-resistance-protein-[Maioricimonas-rarisocia] |
| WP_144981417.1 | Type2.Pgl.like.BREX | [illegible] | 1247 | Bacteria>PVC_group>Planctomycetota>Planctomycetia>Planctomycetales>Planctomycetaceae | GCF_007748195.1 | Gimesia_aquarii | V144x_RS03450 | 831512..835255 | 2527964 | hypothetical-protein-[Gimesia-aquarii] |
| WP_206028721.1 | Type2.Pgl.like.BREX | [illegible] | 1227 | Bacteria>PVC_group>Planctomycetota>Planctomycetia>Planctomycetales>Planctomycetaceae | GCF_011634775.1 | Thalassoroseus_pseudoceratinae | G6R38_RS25200 | 222067..225750 | 2713176 | phage-resistance-protein-[Thalassoroseus-pseudoceratinae] |
| QGJ70865.1 | Type2.Pgl.like.BREX | [illegible] | 1225 | Bacteria>PVC_group>Planctomycetota>Planctomycetia>Planctomycetales>unclassified_Planctomycetales | GCA_009664345.1 | Planctomycetales_bacterium_10988 | PBC10988_25640 | 4110375..4114052 | 2603660 | Bacteriophage-resistance-protein-PglY-[Planctomycetales-bacterium-10988] |
| MCE9632143.1 | Type2.Pgl.like.BREX | <-SF1-SF2-Helicase+wHTH+Small-B-barrel+DUF1998<-?<-BrxD-AAA<-PglZ<-DUF6079*<-N6-METHYLASE<-PglW | 1219 | Bacteria>PVC_group>Planctomycetota>Planctomycetia>unclassified_Planctomycetia | GCA_021414115.1 | Planctomycetia_bacterium | K8S94_15685 | 70185..73844 | 2052181 | phage-resistance-protein-[Planctomycetia-bacterium] |
| MBK6942983.1 | Type2.Pgl.like.BREX | [illegible] | 1211 | Bacteria>PVC_group>Planctomycetota>unclassified_Planctomycetota | GCA_016709145.1 | Planctomycetota_bacterium | IPH13_22660 | 256601..260236 | 2026780 | phage-resistance-protein-[Planctomycetota-bacterium] |
| MBK9387524.1 | Type2.Pgl.like.BREX | REase+DUF5655-><-OLD-ABC+TOPRIM\|\|PglW->N6-METHYLASE->DUF6079*->?->PglZ->HTH+MPTase->?->BrxD-AAA->SF1-SF2-Helicase+wHTH+Small-B-barrel+DUF1998-> | 1227 | Bacteria>PVC_group>Planctomycetota>unclassified_Planctomycetota | GCA_016719295.1 | Planctomycetota_bacterium | IPN34_22145 | 5837658..5841341 | 2026780 | phage-resistance-protein-[Planctomycetota-bacterium] |
| MBL8842560.1 | Type2.Pgl.like.BREX | [illegible] | 1261 | Bacteria>PVC_group>Planctomycetota>unclassified_Planctomycetota | GCA_016795285.1 | Planctomycetota_bacterium | JNL90_13655 | 295259..299044 | 2026780 | phage-resistance-protein-[Planctomycetota-bacterium] |
| MCE9553883.1 | Type2.Pgl.like.BREX | XIS-HTH->PglW->?->N6-METHYLASE->HEPN->?->ParB+HNH->DUF6079*->PglZ->BrxD-AAA->SF1-SF2-Helicase+wHTH+Small-B-barrel+DUF1998-><-?<-Pkinase+pi3KINASE<-?<-?\|\|?->PIN-> | 1232 | Bacteria>PVC_group>Planctomycetota>unclassified_Planctomycetota | GCA_021413705.1 | Planctomycetota_bacterium | K8T91_10985 | 60988..64686 | 2026780 | phage-resistance-protein-[Planctomycetota-bacterium] |
| WED64126.1 | Type2.Pgl.like.BREX | [illegible] | 1260 | Bacteria>PVC_group>Verrucomicrobiota>Opitutae>Opitutales>Opitutaceae | GCA_029023725.1 | Opitutaceae_bacterium_LMO-M01 | PXH66_17450 | 4532197..4535979 | 2909285 | hypothetical-protein-PXH66_17450-[Opitutaceae-bacterium-LMO-M01] |
| TAF23831.1 | Type2.Pgl.like.BREX | [illegible] | 1139 | Bacteria>PVC_group>Verrucomicrobiota>unclassified_Verrucomicrobiota | GCA_004293445.1 | Verrucomicrobiota_bacterium | EAZ71_12135 | 30329..33748 | 2026799 | phage-resistance-protein-[Verrucomicrobiota-bacterium] |
| MBI4662620.1 | Type2.Pgl.like.BREX | [illegible] | 1322 | Bacteria>PVC_group>Verrucomicrobiota>unclassified_Verrucomicrobiota | GCA_016209525.1 | Verrucomicrobiota_bacterium | HY735_27725 | 8107..12075 | 2026799 | phage-resistance-protein-[Verrucomicrobiota-bacterium] |
| MBN9688997.1 | Type2.Pgl.like.BREX | [illegible] | 1324 | Bacteria>PVC_group>Verrucomicrobiota>unclassified_Verrucomicrobiota | GCA_017309115.1 | Verrucomicrobiota_bacterium | J0M24_02070 | 214686..218660 | 2026799 | phage-resistance-protein-[Verrucomicrobiota-bacterium] |
| MBX3732320.1 | Type2.Pgl.like.BREX | [illegible] | 1335 | Bacteria>PVC_group>Verrucomicrobiota>Verrucomicrobiae>unclassified_Verrucomicrobiae | GCA_019634635.1 | Verrucomicrobiae_bacterium | KF791_06965 | 147710..151717 | 2499141 | phage-resistance-protein-[Verrucomicrobiae-bacterium] |
| MCP5560212.1 | Type2.Pgl.like.BREX | [illegible] | 1270 | Bacteria>PVC_group>Verrucomicrobiota>Verrucomicrobiae>Verrucomicrobiales>Verrucomicrobiaceae | GCA_024232965.1 | Verrucomicrobiaceae_bacterium | H7A55_20880 | 48036..51848 | 2026800 | phage-resistance-protein-[Verrucomicrobiaceae-bacterium] |
| WP_153758471.1 | Type2.Pgl.like.BREX | [illegible] | 1198 | Bacteria>Terrabacteria_group>Actinomycetota>Acidimicrobiia>Acidimicrobiales>Iamiaceae | GCF_009650215.1 | Actinomarinicola_tropica | GH723_RS04185 | 829858..833454 | 2789776 | phage-resistance-protein-[Actinomarinicola-tropica] |
| MCB0954591.1 | Type2.Pgl.like.BREX | [illegible] | 1219 | Bacteria>Terrabacteria_group>Actinomycetota>Acidimicrobiia>Acidimicrobiales>Ilumatobacteraceae | GCA_020439425.1 | Ilumatobacter_sp. | KDB12_00345 | 10363..14022 | 1967498 | hypothetical-protein-[Ilumatobacter-sp.] |
| WP_015443641.1 | Type2.Pgl.like.BREX | [illegible] | 1203 | Bacteria>Terrabacteria_group>Actinomycetota>Acidimicrobiia>Acidimicrobiales>Ilumatobacteraceae | GCF_000348785.1 | Ilumatobacter_coccineus | YM304_RS20475 | 4610450..4614061 | 467094 | hypothetical-protein-[Ilumatobacter-coccineus] |
| WP_116997338.1 | Type2.Pgl.like.BREX | [illegible] | 1228 | Bacteria>Terrabacteria_group>Actinomycetota>Acidimicrobiia>Acidimicrobiales>Ilumatobacteraceae | GCF_027365655.1 | Desertimonas_flava | M6J74_RS15035 | 88246..91932 | 2064846 | phage-resistance-protein-[Desertimonas-flava] |
| MCO5304916.1 | Type2.Pgl.like.BREX | [illegible] | 674 | Bacteria>Terrabacteria_group>Actinomycetota>Acidimicrobiia>Acidimicrobiales>Microthrixaceae | GCA_023958015.1 | Microthrixaceae_bacterium | M9906_03105 | 1..2021 | 2497627 | hypothetical-protein-[Microthrixaceae-bacterium] |
| MBU6244326.1 | Type2.Pgl.like.BREX | [illegible] | 1239 | Bacteria>Terrabacteria_group>Actinomycetota>Actinomycetes>Actinomycetales>unclassified_Actinomycetales | GCA_018970025.1 | Actinomycetales_bacterium | KGP12_03835 | 142062..145781 | 1911520 | hypothetical-protein-[Actinomycetales-bacterium] |
| WP_165632998.1 | Type2.Pgl.like.BREX | [illegible] | 1256 | Bacteria>Terrabacteria_group>Actinomycetota>Actinomycetes>Actinopolysporales>Actinopolysporaceae | GCF_011291845.1 | unclassified_Actinopolyspora | G9Q41_RS17510 | 60223..63993 | 2639451 | MULTISPECIES:-hypothetical-protein-[unclassified-Actinopolyspora] |
| WP_179533807.1 | Type2.Pgl.like.BREX | [illegible] | 1268 | Bacteria>Terrabacteria_group>Actinomycetota>Actinomycetes>Actinopolysporales>Actinopolysporaceae | GCF_013408175.1 | Actinopolyspora_biskrensis | FHR84_RS02705 | 601142..604948 | 1470178 | DUF6079-family-protein-[Actinopolyspora-biskrensis] |
| WP_092520725.1 | Type2.Pgl.like.BREX | [illegible] | 1268 | Bacteria>Terrabacteria_group>Actinomycetota>Actinomycetes>Actinopolysporales>Actinopolysporaceae | GCF_900100925.1 | Actinopolyspora_saharensis | BLR67_RS02345 | 518029..521835 | 995062 | DUF6079-family-protein-[Actinopolyspora-saharensis] |
| NEA84080.1 | Type2.Pgl.like.BREX | [illegible] | 1295 | Bacteria>Terrabacteria_group>Actinomycetota>Actinomycetes>Catenulisporales>Actinospicaceae | GCA_010548365.1 | Actinospica_acidiphila | G3I30_34735 | 68217..72104 | 304899 | phage-resistance-protein-[Actinospica-acidiphila] |
| WP_034267098.1 | Type2.Pgl.like.BREX | [illegible] | 1299 | Bacteria>Terrabacteria_group>Actinomycetota>Actinomycetes>Catenulisporales>Actinospicaceae | GCF_000504285.1 | Actinospica_robiniae | ACTRO_RS25910 | 6084378..6088277 | 304901 | hypothetical-protein-[Actinospica-robiniae] |
| WP_034270629.1 | Type2.Pgl.like.BREX | [illegible] | 1285 | Bacteria>Terrabacteria_group>Actinomycetota>Actinomycetes>Catenulisporales>Actinospicaceae | GCF_000504285.1 | Actinospica_robiniae | ACTRO_RS36570 | 8499799..8503656 | 304901 | hypothetical-protein-[Actinospica-robiniae] |
| WP_212516926.1 | Type2.Pgl.like.BREX | [illegible] | 1317 | Bacteria>Terrabacteria_group>Actinomycetota>Actinomycetes>Catenulisporales>Actinospicaceae | GCF_018139625.1 | Actinospica_acidithermotolerans | KDK95_RS05575 | 11445..15398 | 2828514 | hypothetical-protein-[Actinospica-acidithermotolerans] |
| WP_165034952.1 | Type2.Pgl.like.BREX | [illegible] | 73 | Bacteria>Terrabacteria_group>Actinomycetota>Actinomycetes>Frankiales>Frankiaceae | - | Candidatus_Frankia_californiensis | DD1DGVC_RS27975 | 25606..25825 | 1839754 | hypothetical-protein,-partial-[Candidatus-Frankia-californiensis] |
| WP_165034953.1 | Type2.Pgl.like.BREX | [illegible] | 987 | Bacteria>Terrabacteria_group>Actinomycetota>Actinomycetes>Frankiales>Frankiaceae | - | Candidatus_Frankia_californiensis | DD1DGVC_RS27980 | 26047..29007 | 1839754 | phage-resistance-protein,-partial-[Candidatus-Frankia-californiensis] |
| WP_131626881.1 | Type2.Pgl.like.BREX | [illegible] | 1295 | Bacteria>Terrabacteria_group>Actinomycetota>Actinomycetes>Frankiales>Frankiaceae | - | Frankia_sp._BMG5.11 | E0504_39210 | 174282..178169 | 222540 | phage-resistance-protein-[Frankia-sp.-BMG5.11] |
| KEZ35700.1 | Type2.Pgl.like.BREX | [illegible] | 1227 | Bacteria>Terrabacteria_group>Actinomycetota>Actinomycetes>Frankiales>Frankiaceae | GCA_000732115.1 | Frankia_sp._CeD | CEDDRAFT_02913 | 8708..12391 | 258230 | hypothetical-protein-CEDDRAFT_02913-[Frankia-sp.-CeD] |
| KFB03010.1 | Type2.Pgl.like.BREX | [illegible] | 1227 | Bacteria>Terrabacteria_group>Actinomycetota>Actinomycetes>Frankiales>Frankiaceae | GCA_000733325.1 | Frankia_sp._Allo2 | ALLO2DRAFT_04243 | 9890..13573 | 981405 | hypothetical-protein-ALLO2DRAFT_04243-[Frankia-sp.-Allo2] |
| OAA24559.1 | Type2.Pgl.like.BREX | [illegible] | 1298 | Bacteria>Terrabacteria_group>Actinomycetota>Actinomycetes>Frankiales>Frankiaceae | GCA_001636565.1 | Frankia_sp._EI5c | UG55_102981 | 105558..109454 | 683316 | hypothetical-protein-UG55_102981-[Frankia-sp.-EI5c] |
| OAA19520.1 | Type2.Pgl.like.BREX | [illegible] | 1227 | Bacteria>Terrabacteria_group>Actinomycetota>Actinomycetes>Frankiales>Frankiaceae | GCA_001636575.1 | Frankia_casuarinae | AAY23_11079 | 8703..12386 | 106370 | hypothetical-protein-AAY23_11079-[Frankia-casuarinae] |
| TCJ33271.1 | Type2.Pgl.like.BREX | [illegible] | 1295 | Bacteria>Terrabacteria_group>Actinomycetota>Actinomycetes>Frankiales>Frankiaceae | GCA_004337455.1 | Frankia_sp._BMG5.11 | E0504_39210 | 174282..178169 | 222540 | phage-resistance-protein-[Frankia-sp.-BMG5.11] |
| WP_011437340.1 | Type2.Pgl.like.BREX | [illegible] | 1227 | Bacteria>Terrabacteria_group>Actinomycetota>Actinomycetes>Frankiales>Frankiaceae | GCF_000013345.1 | Frankia | FRANCCI3_RS27065 | 3497641..3501324 | 1854 | MULTISPECIES:-hypothetical-protein-[Frankia] |
| WP_013426769.1 | Type2.Pgl.like.BREX | [illegible] | 1313 | Bacteria>Terrabacteria_group>Actinomycetota>Actinomycetes>Frankiales>Frankiaceae | GCF_000166135.1 | Frankia_inefficax | FRAEUI1C_RS28165 | 6960453..6964394 | 298654 | hypothetical-protein-[Pseudofrankia-inefficax] |
| WP_006541917.1 | Type2.Pgl.like.BREX | [illegible] | 1380 | Bacteria>Terrabacteria_group>Actinomycetota>Actinomycetes>Frankiales>Frankiaceae | GCF_000177675.1 | Frankia_sp._EUN1f | FREUN1FDRAFT_RS19740 | 13376..17518 | 102897 | hypothetical-protein-[Frankia-sp.-EUN1f] |
| WP_018638825.1 | Type2.Pgl.like.BREX | [illegible] | 1294 | Bacteria>Terrabacteria_group>Actinomycetota>Actinomycetes>Frankiales>Frankiaceae | GCF_000374165.1 | Frankia_elaeagni | B207_RS0122310 | 25036..28920 | 222534 | hypothetical-protein-[Parafrankia-elaeagni] |
| WP_023841472.1 | Type2.Pgl.like.BREX | [illegible] | 1227 | Bacteria>Terrabacteria_group>Actinomycetota>Actinomycetes>Frankiales>Frankiaceae | GCF_000503735.2 | unclassified_Frankia | CCI6_RS26135 | 9664..13347 | 2632575 | MULTISPECIES:-hypothetical-protein-[unclassified-Frankia] |
| WP_054566588.1 | Type2.Pgl.like.BREX | [illegible] | 1365 | Bacteria>Terrabacteria_group>Actinomycetota>Actinomycetes>Frankiales>Frankiaceae | GCF_001306465.1 | Frankia_sp._R43 | ACG83_RS12130 | 992224..996321 | 269536 | hypothetical-protein-[Frankia-sp.-R43] |
| WP_242424514.1 | Type2.Pgl.like.BREX | [illegible] | 1099 | Bacteria>Terrabacteria_group>Actinomycetota>Actinomycetes>Frankiales>Frankiaceae | GCF_001636565.1 | Frankia_sp._EI5c | UG55_RS15565 | 105558..108854 | 683316 | phage-resistance-protein,-partial-[Frankia-sp.-EI5c] |
| WP_070131879.1 | Type2.Pgl.like.BREX | [illegible] | 1227 | Bacteria>Terrabacteria_group>Actinomycetota>Actinomycetes>Frankiales>Frankiaceae | GCF_001756285.1 | Frankia_sp._CgIM4 | Manayef4_RS25020 | 8942..12625 | 1742262 | hypothetical-protein-[Frankia-sp.-CgIM4] |
| WP_071086063.1 | Type2.Pgl.like.BREX | [illegible] | 1419 | Bacteria>Terrabacteria_group>Actinomycetota>Actinomycetes>Frankiales>Frankiaceae | GCF_001854655.1 | Frankia_colletiae | CC117_RS19055 | 10795..15054 | 573497 | phage-resistance-protein-[Parafrankia-colletiae] |
| WP_071065308.1 | Type2.Pgl.like.BREX | [illegible] | 1400 | Bacteria>Terrabacteria_group>Actinomycetota>Actinomycetes>Frankiales>Frankiaceae | GCF_001854695.1 | Frankia_soli | BBK14_RS22795 | 38185..42387 | 2599596 | phage-resistance-protein-[Parafrankia-soli] |
| WP_076807001.1 | Type2.Pgl.like.BREX | [illegible] | 1227 | Bacteria>Terrabacteria_group>Actinomycetota>Actinomycetes>Frankiales>Frankiaceae | GCF_001983015.1 | Frankia_sp._CcI156 | CcI156_RS25830 | 37320..41003 | 1745380 | hypothetical-protein-[Frankia-sp.-CcI156] |
| WP_076822254.1 | Type2.Pgl.like.BREX | [illegible] | 1233 | Bacteria>Terrabacteria_group>Actinomycetota>Actinomycetes>Frankiales>Frankiaceae | GCF_001983105.1 | Frankia_asymbiotica | BL253_RS35820 | 24017..27718 | 1834516 | hypothetical-protein-[Pseudofrankia-asymbiotica] |
| WP_085016236.1 | Type2.Pgl.like.BREX | [illegible] | 1227 | Bacteria>Terrabacteria_group>Actinomycetota>Actinomycetes>Frankiales>Frankiaceae | GCF_002099325.1 | Frankia_sp._KB5 | KBI5_RS26565 | 8418..12101 | 683318 | hypothetical-protein-[Frankia-sp.-KB5] |
| WP_085003686.1 | Type2.Pgl.like.BREX | [illegible] | 1227 | Bacteria>Terrabacteria_group>Actinomycetota>Actinomycetes>Frankiales>Frankiaceae | GCF_002099355.1 | Frankia_casuarinae | UK99_RS24715 | 1925..5608 | 106370 | hypothetical-protein-[Frankia-casuarinae] |
| WP_248824787.1 | Type2.Pgl.like.BREX | [illegible] | 1332 | Bacteria>Terrabacteria_group>Actinomycetota>Actinomycetes>Frankiales>Frankiaceae | GCF_023242945.1 | Frankia_sp._Ag45/Mut15 | MXD59_RS12285 | 88835..92833 | 573489 | phage-resistance-protein-[Frankia-sp.-Ag45/Mut15] |
| WP_248829680.1 | Type2.Pgl.like.BREX | [illegible] | 1285 | Bacteria>Terrabacteria_group>Actinomycetota>Actinomycetes>Frankiales>Frankiaceae | GCF_023242985.1 | Frankia_sp._Mgl5 | MXD62_RS13890 | 22992..26849 | 2933793 | phage-resistance-protein-[Frankia-sp.-Mgl5] |
| WP_248819225.1 | Type2.Pgl.like.BREX | [illegible] | 1317 | Bacteria>Terrabacteria_group>Actinomycetota>Actinomycetes>Frankiales>Frankiaceae | GCF_023243045.1 | Frankia_sp._AgB32 | MXD60_RS12600 | 19114..23067 | 631119 | phage-resistance-protein-[Frankia-sp.-AgB32] |
| WP_251735898.1 | Type2.Pgl.like.BREX | [illegible] | 1329 | Bacteria>Terrabacteria_group>Actinomycetota>Actinomycetes>Frankiales>Frankiaceae | GCF_023716925.1 | Frankia_sp._R82 | ND747_RS21805 | 21178..25167 | 2950553 | phage-resistance-protein-[Frankia-sp.-R82] |
| WP_261574303.1 | Type2.Pgl.like.BREX | [illegible] | 1359 | Bacteria>Terrabacteria_group>Actinomycetota>Actinomycetes>Frankiales>Frankiaceae | GCF_025403375.1 | Frankia_gtarii | NBZ83_RS03385 | 25374..29453 | 2950102 | phage-resistance-protein-[Frankia-gtarii] |
| WP_112106787.1 | Type2.Pgl.like.BREX | [illegible] | 1295 | Bacteria>Terrabacteria_group>Actinomycetota>Actinomycetes>Frankiales>Frankiaceae | GCF_900465275.1 | Frankia_sp._Ea1.12 | FMEAI12_RS16915 | 15369..19256 | 573499 | phage-resistance-protein-[Frankia-sp.-Ea1.12] |
| WP_131786019.1 | Type2.Pgl.like.BREX | [illegible] | 1210 | Bacteria>Terrabacteria_group>Actinomycetota>Actinomycetes>Frankiales>Frankiaceae | GCF_900634745.1 | Frankia_symbiont_of_Coriaria_ruscifolia | CV1CTNDD_RS11015 | 13120..16752 | 1306542 | phage-resistance-protein-[Frankia-symbiont-of-Coriaria-ruscifolia] |
| WP_131770045.1 | Type2.Pgl.like.BREX | [illegible] | 1339 | Bacteria>Terrabacteria_group>Actinomycetota>Actinomycetes>Frankiales>Frankiaceae | GCF_900634765.1 | Candidatus_Frankia_californiensis | DD1DGNDD_RS11025 | 25505..29524 | 1839754 | phage-resistance-protein-[Candidatus-Frankia-californiensis] |
| WP_239405408.1 | Type2.Pgl.like.BREX | [illegible] | 1293 | Bacteria>Terrabacteria_group>Actinomycetota>Actinomycetes>Frankiales>Frankiaceae | GCF_917627395.1 | Frankia_sp._Cj3 | MJ790_RS14755 | 2539..6420 | 2880976 | phage-resistance-protein-[Frankia-sp.-Cj3] |
| WP_239395557.1 | Type2.Pgl.like.BREX | [illegible] | 936 | Bacteria>Terrabacteria_group>Actinomycetota>Actinomycetes>Frankiales>Frankiaceae | GCF_917627415.1 | Frankia_sp._CiP3 | MJ788_RS15840 | 24306..27114 | 2880971 | phage-resistance-protein,-partial-[Frankia-sp.-CiP3] |
| WP_106281978.1 | Type2.Pgl.like.BREX | [illegible] | 1240 | Bacteria>Terrabacteria_group>Actinomycetota>Actinomycetes>Geodermatophilales>Geodermatophilaceae | GCF_003002915.1 | Geodermatophilus_tzadiensis | LY71_RS22350 | 37561..41283 | 1137988 | phage-resistance-protein-[Geodermatophilus-tzadiensis] |
| WP_269185209.1 | Type2.Pgl.like.BREX | [illegible] | 1235 | Bacteria>Terrabacteria_group>Actinomycetota>Actinomycetes>Geodermatophilales>Geodermatophilaceae | GCF_026967925.1 | Modestobacter_sp._VKM_Ac-2985 | O2V61_RS21000 | 1062673..1066380 | 3004139 | hypothetical-protein-[Modestobacter-sp.-VKM-Ac-2985] |
| WP_147142479.1 | Type2.Pgl.like.BREX | [illegible] | 1246 | Bacteria>Terrabacteria_group>Actinomycetota>Actinomycetes>Glycomycetales>Glycomycetaceae | GCF_007994225.1 | Stackebrandtia_albiflava | LX16_RS21785 | 161033..164773 | 406432 | phage-resistance-protein-[Stackebrandtia-albiflava] |
| WP_199044191.1 | Type2.Pgl.like.BREX | [illegible] | 1232 | Bacteria>Terrabacteria_group>Actinomycetota>Actinomycetes>Glycomycetales>Glycomycetaceae | GCF_016428645.1 | Glycomyces_salinus | JFB92_RS23465 | 10873..14571 | 980294 | hypothetical-protein-[Glycomyces-salinus] |
| WP_213171151.1 | Type2.Pgl.like.BREX | [illegible] | 1229 | Bacteria>Terrabacteria_group>Actinomycetota>Actinomycetes>Glycomycetales>Glycomycetaceae | GCF_016925535.1 | Natronoglycomyces_albus | JQS30_RS15565 | 3645498..3649187 | 2811108 | DUF6079-family-protein-[Natronoglycomyces-albus] |
| WP_162453106.1 | Type2.Pgl.like.BREX | [illegible] | 1252 | Bacteria>Terrabacteria_group>Actinomycetota>Actinomycetes>Jiangellales>Jiangellaceae | GCF_010122465.1 | Phytoactinopolyspora_mesophila | F7O44_RS25385 | 166247..170005 | 2650750 | DUF6079-family-protein-[Phytoactinopolyspora-mesophila] |
| MBK6874125.1 | Type2.Pgl.like.BREX | [illegible] | 1245 | Bacteria>Terrabacteria_group>Actinomycetota>Actinomycetes>Kineosporiales>Kineosporiaceae | GCA_016702615.1 | Kineosporiaceae_bacterium | IPG94_22930 | 45769..49506 | 2487580 | phage-resistance-protein-[Kineosporiaceae-bacterium] |
| MBK7624680.1 | Type2.Pgl.like.BREX | [illegible] | 1245 | Bacteria>Terrabacteria_group>Actinomycetota>Actinomycetes>Kineosporiales>Kineosporiaceae | GCA_016709945.1 | Kineosporiaceae_bacterium | IPJ14_18995 | 362278..366015 | 2487580 | phage-resistance-protein-[Kineosporiaceae-bacterium] |
| MBK8076944.1 | Type2.Pgl.like.BREX | [illegible] | 929 | Bacteria>Terrabacteria_group>Actinomycetota>Actinomycetes>Kineosporiales>Kineosporiaceae | GCA_016713225.1 | Kineosporiaceae_bacterium | IPK24_15565 | 998937..1001726 | 2487580 | phage-resistance-protein-[Kineosporiaceae-bacterium] |
| MBK8076946.1 | Type2.Pgl.like.BREX | [illegible] | 78 | Bacteria>Terrabacteria_group>Actinomycetota>Actinomycetes>Kineosporiales>Kineosporiaceae | GCA_016713225.1 | Kineosporiaceae_bacterium | IPK24_15575 | 1002439..1002675 | 2487580 | hypothetical-protein-[Kineosporiaceae-bacterium] |
| MBL8929410.1 | Type2.Pgl.like.BREX | [illegible] | 1036 | Bacteria>Terrabacteria_group>Actinomycetota>Actinomycetes>Kineosporiales>Kineosporiaceae | GCA_016794595.1 | Kineosporiaceae_bacterium | JNL54_04730 | 81721..84833 | 2487580 | hypothetical-protein-[Kineosporiaceae-bacterium] |
| MBL8931661.1 | Type2.Pgl.like.BREX | [illegible] | 255 | Bacteria>Terrabacteria_group>Actinomycetota>Actinomycetes>Kineosporiales>Kineosporiaceae | GCA_016794595.1 | Kineosporiaceae_bacterium | JNL54_16190 | 1..766 | 2487580 | hypothetical-protein-[Kineosporiaceae-bacterium] |
| UER54528.1 | Type2.Pgl.like.BREX | [illegible] | 1216 | Bacteria>Terrabacteria_group>Actinomycetota>Actinomycetes>Kineosporiales>Kineosporiaceae | GCA_020881835.1 | Kineosporiaceae_bacterium_SCSIO_59966 | HJG43_08230 | 1761516..1765166 | 2730919 | phage-resistance-protein-[Kineosporiaceae-bacterium-SCSIO-59966] |
| WP_109776034.1 | Type2.Pgl.like.BREX | [illegible] | 1268 | Bacteria>Terrabacteria_group>Actinomycetota>Actinomycetes>Kineosporiales>Kineosporiaceae | GCF_003149145.1 | Quadrisphaera_granulorum | BXY45_RS18605 | 6830..10636 | 317664 | phage-resistance-protein-[Quadrisphaera-granulorum] |
| WP_267593973.1 | Type2.Pgl.like.BREX | [illegible] | 1229 | Bacteria>Terrabacteria_group>Actinomycetota>Actinomycetes>Kitasatosporales>Carbonactinosporaceae | GCF_026614415.1 | Carbonactinospora_thermoautotrophica | C3Y87_RS08180 | 69147..72836 | 1469144 | hypothetical-protein-[Carbonactinospora-thermoautotrophica] |
| EST37408.1 | Type2.Pgl.like.BREX | [illegible] | 1295 | Bacteria>Terrabacteria_group>Actinomycetota>Actinomycetes>Kitasatosporales>Streptomycetaceae | GCA_000497405.1 | Streptomycetaceae_bacterium_MP113-05 | N566_13090 | 5298..9185 | 1380770 | hypothetical-protein-N566_13090-[Streptomycetaceae-bacterium-MP113-05] |
| KMS82322.1 | Type2.Pgl.like.BREX | [illegible] | 1321 | Bacteria>Terrabacteria_group>Actinomycetota>Actinomycetes>Kitasatosporales>Streptomycetaceae | GCA_001047335.1 | Streptomyces_regensis | ACZ91_59210 | 21770..25735 | 68263 | PglY-protein-[Streptomyces-regensis] |
| KPC88168.1 | Type2.Pgl.like.BREX | [illegible] | 1295 | Bacteria>Terrabacteria_group>Actinomycetota>Actinomycetes>Kitasatosporales>Streptomycetaceae | GCA_001294395.1 | Streptomyces_sp._NRRL_WC-3753 | ADL35_06445 | 142151..146038 | 1609104 | bacteriophage-resistance-protein-[Streptomyces-sp.-NRRL-WC-3753] |
| ODA74657.1 | Type2.Pgl.like.BREX | [illegible] | 610 | Bacteria>Terrabacteria_group>Actinomycetota>Actinomycetes>Kitasatosporales>Streptomycetaceae | GCA_001469455.2 | Streptomyces_sp._AVP053U2 | APS67_001319 | 261591..263423 | 1737066 | hypothetical-protein-APS67_001319-[Streptomyces-sp.-AVP053U2] |
| KUN89644.1 | Type2.Pgl.like.BREX | [illegible] | 139 | Bacteria>Terrabacteria_group>Actinomycetota>Actinomycetes>Kitasatosporales>Streptomycetaceae | GCA_001514215.1 | Streptomyces_bungoensis | AQJ66_04195 | 996583..997002 | 285568 | hypothetical-protein-AQJ66_04195-[Streptomyces-bungoensis] |
| AMM07779.1 | Type2.Pgl.like.BREX | [illegible] | 1295 | Bacteria>Terrabacteria_group>Actinomycetota>Actinomycetes>Kitasatosporales>Streptomycetaceae | GCA_001577385.1 | Streptomyces_albidoflavus | Salbus254_1235 | 1445801..1449688 | 1886 | bacteriophage-resistance-protein-[Streptomyces-albidoflavus] |
| ARF59786.1 | Type2.Pgl.like.BREX | [illegible] | 1299 | Bacteria>Terrabacteria_group>Actinomycetota>Actinomycetes>Kitasatosporales>Streptomycetaceae | GCA_002082195.1 | Streptomyces_gilvosporeus | B1H19_30120 | 6750768..6754667 | 553510 | phage-resistance-protein-[Streptomyces-gilvosporeus] |
| ROR38155.1 | Type2.Pgl.like.BREX | [illegible] | 1320 | Bacteria>Terrabacteria_group>Actinomycetota>Actinomycetes>Kitasatosporales>Streptomycetaceae | GCA_003751605.1 | Kitasatospora_cineracea | EDD39_6327 | 1175437..1179399 | 88074 | hypothetical-protein-EDD39_6327-[Kitasatospora-cineracea] |
| RPK54894.1 | Type2.Pgl.like.BREX | [illegible] | 1295 | Bacteria>Terrabacteria_group>Actinomycetota>Actinomycetes>Kitasatosporales>Streptomycetaceae | GCA_003846265.1 | Streptomyces_sp._ADI96-15 | EES44_29045 | 154353..158240 | 1522761 | hypothetical-protein-EES44_29045-[Streptomyces-sp.-ADI96-15] |
| TKA04731.1 | Type2.Pgl.like.BREX | [illegible] | 1316 | Bacteria>Terrabacteria_group>Actinomycetota>Actinomycetes>Kitasatosporales>Streptomycetaceae | GCA_005047355.1 | Actinacidiphila_oryziradicis | FCI23_34795 | 22737..26687 | 2571141 | PglY-protein-[Actinacidiphila-oryziradicis] |
| GDY52272.1 | Type2.Pgl.like.BREX | [illegible] | 606 | Bacteria>Terrabacteria_group>Actinomycetota>Actinomycetes>Kitasatosporales>Streptomycetaceae | GCA_005405945.1 | Streptomyces_violaceusniger | SVIO_028950 | 3096806..3098626 | 68280 | hypothetical-protein-SVIO_028950-[Streptomyces-violaceusniger] |
| GDY52273.1 | Type2.Pgl.like.BREX | [illegible] | 222 | Bacteria>Terrabacteria_group>Actinomycetota>Actinomycetes>Kitasatosporales>Streptomycetaceae | GCA_005405945.1 | Streptomyces_violaceusniger | SVIO_028960 | 3098551..3099219 | 68280 | hypothetical-protein-SVIO_028960-[Streptomyces-violaceusniger] |
| GDY56597.1 | Type2.Pgl.like.BREX | [illegible] | 292 | Bacteria>Terrabacteria_group>Actinomycetota>Actinomycetes>Kitasatosporales>Streptomycetaceae | GCA_005405945.1 | Streptomyces_violaceusniger | SVIO_072200 | 7502674..7503552 | 68280 | hypothetical-protein-SVIO_072200-[Streptomyces-violaceusniger] |
| GDY56598.1 | Type2.Pgl.like.BREX | [illegible] | 1032 | Bacteria>Terrabacteria_group>Actinomycetota>Actinomycetes>Kitasatosporales>Streptomycetaceae | GCA_005405945.1 | Streptomyces_violaceusniger | SVIO_072210 | 7503470..7506568 | 68280 | hypothetical-protein-SVIO_072210-[Streptomyces-violaceusniger] |
| BCK67994.1 | Type2.Pgl.like.BREX | [illegible] | 1298 | Bacteria>Terrabacteria_group>Actinomycetota>Actinomycetes>Kitasatosporales>Streptomycetaceae | GCA_009811615.2 | Streptomyces_libani_subsp._rufus | Srufu_019470 | 2037295..2041191 | 249582 | hypothetical-protein-Srufu_019470-[Streptomyces-libani-subsp.-rufus] |
| NEE18804.1 | Type2.Pgl.like.BREX | [illegible] | 1307 | Bacteria>Terrabacteria_group>Actinomycetota>Actinomycetes>Kitasatosporales>Streptomycetaceae | GCA_010550285.1 | Streptomyces_sp._SID7499 | G3M58_71575 | 24154..28077 | 2706086 | type-II-toxin-antitoxin-system-VapC-family-toxin-[Streptomyces-sp.-SID7499] |
| NED01599.1 | Type2.Pgl.like.BREX | [illegible] | 1299 | Bacteria>Terrabacteria_group>Actinomycetota>Actinomycetes>Kitasatosporales>Streptomycetaceae | GCA_010550305.1 | Streptomyces_sp._SID6648 | G3I55_08000 | 34752..38651 | 2706084 | phage-resistance-protein-[Streptomyces-sp.-SID6648] |
| NEE38448.1 | Type2.Pgl.like.BREX | [illegible] | 1298 | Bacteria>Terrabacteria_group>Actinomycetota>Actinomycetes>Kitasatosporales>Streptomycetaceae | GCA_010550345.1 | Streptomyces_sp._SID7982 | G3M53_75280 | 35734..39630 | 2706094 | phage-resistance-protein-[Streptomyces-sp.-SID7982] |
| NUP22768.1 | Type2.Pgl.like.BREX | [illegible] | 1306 | Bacteria>Terrabacteria_group>Actinomycetota>Actinomycetes>Kitasatosporales>Streptomycetaceae | GCA_013361465.1 | Streptomyces_sp. | HOZ81_43260 | 19809..23729 | 1931 | PglY-protein-[Streptomyces-sp.] |
| NUV53534.1 | Type2.Pgl.like.BREX | [illegible] | 1294 | Bacteria>Terrabacteria_group>Actinomycetota>Actinomycetes>Kitasatosporales>Streptomycetaceae | GCA_013363625.1 | Streptomyces_coelicolor | G6W51_11545 | 76881..80765 | 1902 | phage-resistance-protein-[Streptomyces-coelicolor] |
| MBB5815397.1 | Type2.Pgl.like.BREX | [illegible] | 1298 | Bacteria>Terrabacteria_group>Actinomycetota>Actinomycetes>Kitasatosporales>Streptomycetaceae | GCA_014204745.1 | Streptomyces_collinus | HNR72_006425 | 7190212..7194108 | 42684 | hypothetical-protein-[Streptomyces-collinus] |
| MBD0735466.1 | Type2.Pgl.like.BREX | [illegible] | 1301 | Bacteria>Terrabacteria_group>Actinomycetota>Actinomycetes>Kitasatosporales>Streptomycetaceae | GCA_014596445.1 | Streptomyces_sp._CBMA29 | BGM09_02935 | 64585..68490 | 1896314 | PglY-protein-[Streptomyces-sp.-CBMA29] |
| GGW17154.1 | Type2.Pgl.like.BREX | [illegible] | 1307 | Bacteria>Terrabacteria_group>Actinomycetota>Actinomycetes>Kitasatosporales>Streptomycetaceae | GCA_014649295.1 | Streptomyces_libani_subsp._rufus | GCM10010501_36450 | 249806..253729 | 249582 | hypothetical-protein-GCM10010501_36450-[Streptomyces-libani-subsp.-rufus] |
| GGQ80746.1 | Type2.Pgl.like.BREX | [illegible] | 1294 | Bacteria>Terrabacteria_group>Actinomycetota>Actinomycetes>Kitasatosporales>Streptomycetaceae | GCA_014649355.1 | Streptomyces_althioticus | GCM10010250_60480 | 30959..34843 | 83380 | hypothetical-protein-GCM10010250_60480-[Streptomyces-althioticus] |
| GGX84370.1 | Type2.Pgl.like.BREX | [illegible] | 874 | Bacteria>Terrabacteria_group>Actinomycetota>Actinomycetes>Kitasatosporales>Streptomycetaceae | GCA_014650655.1 | Streptomyces_anandii_JCM_4720 | GCM10010510_31730 | 222719..225343 | 1306180 | hypothetical-protein-GCM10010510_31730-[Streptomyces-anandii-JCM-4720] |
| GHG37483.1 | Type2.Pgl.like.BREX | [illegible] | 1303 | Bacteria>Terrabacteria_group>Actinomycetota>Actinomycetes>Kitasatosporales>Streptomycetaceae | GCA_014654935.1 | Streptomyces_capoamus | GCM10018980_09390 | 255178..259089 | 68183 | hypothetical-protein-GCM10018980_09390-[Streptomyces-capoamus] |
| MBL0799495.1 | Type2.Pgl.like.BREX | [illegible] | 1298 | Bacteria>Terrabacteria_group>Actinomycetota>Actinomycetes>Kitasatosporales>Streptomycetaceae | GCA_016734865.1 | Streptomyces_albidoflavus | G6539_02235 | 12332..16228 | 1886 | phage-resistance-protein-[Streptomyces-albidoflavus] |
| MBT2380833.1 | Type2.Pgl.like.BREX | [illegible] | 1314 | Bacteria>Terrabacteria_group>Actinomycetota>Actinomycetes>Kitasatosporales>Streptomycetaceae | GCA_018614515.1 | Streptomyces_sp._ISL-111 | J7E90_26860 | 114837..118781 | 2819175 | hypothetical-protein-[Streptomyces-sp.-ISL-111] |
| MBW8800526.1 | Type2.Pgl.like.BREX | [illegible] | 1295 | Bacteria>Terrabacteria_group>Actinomycetota>Actinomycetes>Kitasatosporales>Streptomycetaceae | GCA_019510245.1 | Streptomyces_sp. | JF597_45235 | 8355..12242 | 1931 | phage-resistance-protein-[Streptomyces-sp.] |
| MCC8482294.1 | Type2.Pgl.like.BREX | [illegible] | 1299 | Bacteria>Terrabacteria_group>Actinomycetota>Actinomycetes>Kitasatosporales>Streptomycetaceae | GCA_020881045.1 | Streptomyces_globisporus | LMJ41_31040 | 42048..45947 | 1908 | phage-resistance-protein-[Streptomyces-globisporus] |
| URN12051.1 | Type2.Pgl.like.BREX | [illegible] | 732 | Bacteria>Terrabacteria_group>Actinomycetota>Actinomycetes>Kitasatosporales>Streptomycetaceae | GCA_023614295.1 | Streptomyces_radiopugnans | LUW77_10700 | 2146838..2149036 | 403935 | hypothetical-protein-LUW77_10700-[Streptomyces-radiopugnans] |
| MCP9962035.1 | Type2.Pgl.like.BREX | [illegible] | 1300 | Bacteria>Terrabacteria_group>Actinomycetota>Actinomycetes>Kitasatosporales>Streptomycetaceae | GCA_024349105.1 | Streptomyces_somaliensis | LUX05_12805 | 2765612..2769514 | 78355 | hypothetical-protein-[Streptomyces-somaliensis] |
| MCP9974853.1 | Type2.Pgl.like.BREX | [illegible] | 761 | Bacteria>Terrabacteria_group>Actinomycetota>Actinomycetes>Kitasatosporales>Streptomycetaceae | GCA_024349205.1 | Streptomyces_somaliensis | LUX02_14085 | 3014227..3016512 | 78355 | hypothetical-protein-[Streptomyces-somaliensis] |
| MCQ4043243.1 | Type2.Pgl.like.BREX | [illegible] | 1314 | Bacteria>Terrabacteria_group>Actinomycetota>Actinomycetes>Kitasatosporales>Streptomycetaceae | GCA_024436055.1 | Streptomyces_rubrisoli | NON19_14685 | 110508..114452 | 1387313 | phage-resistance-protein-[Streptomyces-rubrisoli] |
| MCQ4203515.1 | Type2.Pgl.like.BREX | [illegible] | 1318 | Bacteria>Terrabacteria_group>Actinomycetota>Actinomycetes>Kitasatosporales>Streptomycetaceae | GCA_024448155.1 | Streptomyces_coelicoflavus | M4J06_000980 | 19266..23222 | 285562 | hypothetical-protein-[Streptomyces-coelicoflavus] |
| MCQ4207771.1 | Type2.Pgl.like.BREX | [illegible] | 1296 | Bacteria>Terrabacteria_group>Actinomycetota>Actinomycetes>Kitasatosporales>Streptomycetaceae | GCA_024448165.1 | Streptomyces_longispororuber | M4J07_005562 | 11824..15714 | 68230 | DUF6079-family-protein-[Streptomyces-longispororuber] |
| WDV31004.1 | Type2.Pgl.like.BREX | [illegible] | 1272 | Bacteria>Terrabacteria_group>Actinomycetota>Actinomycetes>Kitasatosporales>Streptomycetaceae | GCA_028884895.1 | Streptomyces_sp._AD16 | OIM90_05400 | 1179426..1183244 | 1816220 | phage-resistance-protein-[Streptomyces-sp.-AD16] |
| WP_173872026.1 | Type2.Pgl.like.BREX | [illegible] | 1321 | Bacteria>Terrabacteria_group>Actinomycetota>Actinomycetes>Kitasatosporales>Streptomycetaceae | GCA_030820635.1 | Streptomyces_albus | FM076_18390 | 4166115..4170080 | 1888 | PglY-protein-[Streptomyces-albus] |
| WP_252932836.1 | Type2.Pgl.like.BREX | [illegible] | 1298 | Bacteria>Terrabacteria_group>Actinomycetota>Actinomycetes>Kitasatosporales>Streptomycetaceae | GCA_030820635.1 | Streptomyces_sp._Vc17.3-30 | KQI59_RS04665 | 4060..7956 | 2841672 | phage-resistance-protein-[Streptomyces-sp.-Vc17.3-30] |
| WP_012378658.1 | Type2.Pgl.like.BREX | [illegible] | 1299 | Bacteria>Terrabacteria_group>Actinomycetota>Actinomycetes>Kitasatosporales>Streptomycetaceae | GCF_000010605.1 | Streptomyces | SGR_RS07860 | 1889546..1893445 | 1883 | MULTISPECIES:-hypothetical-protein-[Streptomyces] |
| WP_095756750.1 | Type2.Pgl.like.BREX | [illegible] | 1295 | Bacteria>Terrabacteria_group>Actinomycetota>Actinomycetes>Kitasatosporales>Streptomycetaceae | GCF_000220705.2 | Streptomyces_xinghaiensis | SXIN_RS07035 | 1667275..1671162 | 1038928 | phage-resistance-protein-[Streptomyces-xinghaiensis] |
| WP_019359905.1 | Type2.Pgl.like.BREX | [illegible] | 1303 | Bacteria>Terrabacteria_group>Actinomycetota>Actinomycetes>Kitasatosporales>Streptomycetaceae | GCF_000280905.1 | Streptomyces_sp._AA1529 | A949_RS0130745 | 193487..197398 | 1203257 | hypothetical-protein-[Streptomyces-sp.-AA1529] |
| WP_006380121.1 | Type2.Pgl.like.BREX | [illegible] | 1295 | Bacteria>Terrabacteria_group>Actinomycetota>Actinomycetes>Kitasatosporales>Streptomycetaceae | GCF_000331005.1 | Streptomyces | STRTUCAR8_RS30870 | 20354..24241 | 1883 | MULTISPECIES:-hypothetical-protein-[Streptomyces] |
| WP_006130674.1 | Type2.Pgl.like.BREX | [illegible] | 1294 | Bacteria>Terrabacteria_group>Actinomycetota>Actinomycetes>Kitasatosporales>Streptomycetaceae | GCF_000342345.1 | Streptomyces_gancidicus | H114_RS05695 | 82720..86604 | 68204 | hypothetical-protein-[Streptomyces-gancidicus] |
| WP_015660833.1 | Type2.Pgl.like.BREX | [illegible] | 1294 | Bacteria>Terrabacteria_group>Actinomycetota>Actinomycetes>Kitasatosporales>Streptomycetaceae | GCF_000349325.1 | Streptomyces_davaonensis | BN159_RS30305 | 6862049..6865933 | 348043 | hypothetical-protein-[Streptomyces-davaonensis] |
| WP_015662322.1 | Type2.Pgl.like.BREX | [illegible] | 1300 | Bacteria>Terrabacteria_group>Actinomycetota>Actinomycetes>Kitasatosporales>Streptomycetaceae | GCF_000349325.1 | Streptomyces_davaonensis | BN159_RS37770 | 8561539..8565441 | 348043 | hypothetical-protein-[Streptomyces-davaonensis] |
| WP_018530191.1 | Type2.Pgl.like.BREX | [illegible] | 1308 | Bacteria>Terrabacteria_group>Actinomycetota>Actinomycetes>Kitasatosporales>Streptomycetaceae | GCF_000373565.1 | unclassified_Streptomyces | B072_RS0112005 | 248212..252138 | 2593676 | MULTISPECIES:-hypothetical-protein-[unclassified-Streptomyces] |
| WP_018383906.1 | Type2.Pgl.like.BREX | [illegible] | 1305 | Bacteria>Terrabacteria_group>Actinomycetota>Actinomycetes>Kitasatosporales>Streptomycetaceae | GCF_000380165.1 | Wenjunlia_vitaminophila | AQ490_RS15435 | 79089..83006 | 76728 | hypothetical-protein-[Wenjunlia-vitaminophila] |
| WP_020115998.1 | Type2.Pgl.like.BREX | [illegible] | 1301 | Bacteria>Terrabacteria_group>Actinomycetota>Actinomycetes>Kitasatosporales>Streptomycetaceae | GCF_000383615.1 | Streptomyces_canus | H293_RS0100895 | 199491..203396 | 58343 | hypothetical-protein-[Streptomyces-canus] |
| WP_020140010.1 | Type2.Pgl.like.BREX | [illegible] | 1301 | Bacteria>Terrabacteria_group>Actinomycetota>Actinomycetes>Kitasatosporales>Streptomycetaceae | GCF_000383655.1 | Streptomyces_sp._351MFTsu5.1 | H295_RS0134960 | 91744..95649 | 1172180 | hypothetical-protein-[Streptomyces-sp.-351MFTsu5.1] |
| WP_016431596.1 | Type2.Pgl.like.BREX | [illegible] | 1294 | Bacteria>Terrabacteria_group>Actinomycetota>Actinomycetes>Kitasatosporales>Streptomycetaceae | GCF_000411315.1 | Streptomyces_sp._HGB0020 | HMPREF1211_RS08040 | 1785199..1789083 | 1078086 | hypothetical-protein-[Streptomyces-sp.-HGB0020] |
| WP_020868709.1 | Type2.Pgl.like.BREX | [illegible] | 1306 | Bacteria>Terrabacteria_group>Actinomycetota>Actinomycetes>Kitasatosporales>Streptomycetaceae | GCF_000418455.1 | Streptomyces_rapamycinicus | M271_RS19160 | 4730013..4733933 | 1226757 | hypothetical-protein-[Streptomyces-rapamycinicus] |
| WP_023590424.1 | Type2.Pgl.like.BREX | [illegible] | 1296 | Bacteria>Terrabacteria_group>Actinomycetota>Actinomycetes>Kitasatosporales>Streptomycetaceae | GCF_000478605.2 | Streptomyces_thermolilacinus | J116_RS05090 | 1191219..1195109 | 285540 | hypothetical-protein-[Streptomyces-thermolilacinus] |
| WP_023416579.1 | Type2.Pgl.like.BREX | [illegible] | 1298 | Bacteria>Terrabacteria_group>Actinomycetota>Actinomycetes>Kitasatosporales>Streptomycetaceae | GCF_000495755.1 | unclassified_Streptomyces | B591_RS05565 | 1334058..1337954 | 2593676 | MULTISPECIES:-hypothetical-protein-[unclassified-Streptomyces] |
| WP_037951095.1 | Type2.Pgl.like.BREX | [illegible] | 1320 | Bacteria>Terrabacteria_group>Actinomycetota>Actinomycetes>Kitasatosporales>Streptomycetaceae | GCF_000590515.1 | Streptomyces_sp._PRh5 | Z951_RS11140 | 96158..100120 | 1158056 | hypothetical-protein-[Streptomyces-sp.-PRh5] |
| WP_033825778.1 | Type2.Pgl.like.BREX | [illegible] | 1312 | Bacteria>Terrabacteria_group>Actinomycetota>Actinomycetes>Kitasatosporales>Streptomycetaceae | GCF_000710405.2 | unclassified_Kitasatospora | BI06_RS11260 | 47992..51930 | 2633591 | MULTISPECIES:-DUF6079-family-protein-[unclassified-Kitasatospora] |
| WP_033203941.1 | Type2.Pgl.like.BREX | [illegible] | 1286 | Bacteria>Terrabacteria_group>Actinomycetota>Actinomycetes>Kitasatosporales>Streptomycetaceae | GCF_000716445.1 | Streptomyces | OO54_RS16745 | 55018..58878 | 1883 | MULTISPECIES:-PglY-protein-[Streptomyces] |

| Protein-ID | System | Gene-Neighbourhoods | Length | Taxonomy | Assembly-ID | Strain | Locus-Tag | Genomic-coordinates | TaxID | Fasta-defline |
|---|---|---|---|---|---|---|---|---|---|---|
| WP_030643413.1 | Type2.Pgl.like.BREX | [illegible] | 1297 | Bacteria>Terrabacteria_group>Actinomycetota>Actinomycetes>Kitasatosporales>Streptomycetaceae | GCF_000716535.1 | Streptomyces | OO52_RS12535 | 131539..135432 | 1883 | MULTISPECIES:-hypothetical-protein-[Streptomyces] |
| WP_033324489.1 | Type2.Pgl.like.BREX | [illegible] | 1307 | Bacteria>Terrabacteria_group>Actinomycetota>Actinomycetes>Kitasatosporales>Streptomycetaceae | GCF_000716805.1 | Streptomyces_yerevanensis | OP79_RS29515 | 12583..16506 | 66378 | PglY-protein-[Streptomyces-yerevanensis] |
| WP_030581836.1 | Type2.Pgl.like.BREX | [illegible] | 1295 | Bacteria>Terrabacteria_group>Actinomycetota>Actinomycetes>Kitasatosporales>Streptomycetaceae | GCF_000717105.1 | Streptomyces_anulatus | IG90_RS0111990 | 218972..222859 | 1892 | hypothetical-protein-[Streptomyces-anulatus] |
| WP_030321491.1 | Type2.Pgl.like.BREX | [illegible] | 1318 | Bacteria>Terrabacteria_group>Actinomycetota>Actinomycetes>Kitasatosporales>Streptomycetaceae | GCF_000717595.1 | Streptomyces_flavochromogenes | IF00_RS0124625 | 51178..55134 | 68199 | hypothetical-protein-[Streptomyces-flavochromogenes] |
| WP_030293901.1 | Type2.Pgl.like.BREX | [illegible] | 1300 | Bacteria>Terrabacteria_group>Actinomycetota>Actinomycetes>Kitasatosporales>Streptomycetaceae | GCF_000717655.1 | Streptomyces_katrae | IF09_RS0109400 | 71835..75737 | 68223 | hypothetical-protein-[Streptomyces-katrae] |
| WP_030145801.1 | Type2.Pgl.like.BREX | [illegible] | 1303 | Bacteria>Terrabacteria_group>Actinomycetota>Actinomycetes>Kitasatosporales>Streptomycetaceae | GCF_000717995.1 | Streptomyces_violaceorubidus | IE94_RS0103125 | 657978..661889 | 284042 | hypothetical-protein-[Streptomyces-violaceorubidus] |
| WP_030698798.1 | Type2.Pgl.like.BREX | [illegible] | 1324 | Bacteria>Terrabacteria_group>Actinomycetota>Actinomycetes>Kitasatosporales>Streptomycetaceae | GCF_000719495.1 | Streptomyces_albidoflavus | IF80_RS0123230 | 18564..22538 | 1886 | hypothetical-protein-[Streptomyces-albidoflavus] |
| WP_030709491.1 | Type2.Pgl.like.BREX | [illegible] | 1302 | Bacteria>Terrabacteria_group>Actinomycetota>Actinomycetes>Kitasatosporales>Streptomycetaceae | GCF_000720115.1 | Streptomyces_sp._NRRL_F-2580 | IF88_RS0110190 | 2870..6778 | 1463841 | hypothetical-protein-[Streptomyces-sp.-NRRL-F-2580] |
| WP_030881234.1 | Type2.Pgl.like.BREX | [illegible] | 1298 | Bacteria>Terrabacteria_group>Actinomycetota>Actinomycetes>Kitasatosporales>Streptomycetaceae | GCF_000720405.1 | Streptomyces_sp._NRRL_B-24051 | IG70_RS0127770 | 73848..77744 | 1463832 | hypothetical-protein-[Streptomyces-sp.-NRRL-B-24051] |
| WP_063767425.1 | Type2.Pgl.like.BREX | [illegible] | 1353 | Bacteria>Terrabacteria_group>Actinomycetota>Actinomycetes>Kitasatosporales>Streptomycetaceae | GCF_000721305.1 | Streptomyces_sp._NRRL_S-1868 | IG92_RS0121360 | 18400..22461 | 1463892 | hypothetical-protein-[Streptomyces-sp.-NRRL-S-1868] |
| WP_063759483.1 | Type2.Pgl.like.BREX | [illegible] | 1384 | Bacteria>Terrabacteria_group>Actinomycetota>Actinomycetes>Kitasatosporales>Streptomycetaceae | GCF_000721605.1 | Streptomyces_sp._NRRL_S-37 | IH50_RS0114185 | 274675..278829 | 1463903 | hypothetical-protein-[Streptomyces-sp.-NRRL-S-37] |
| WP_063836858.1 | Type2.Pgl.like.BREX | [illegible] | 1247 | Bacteria>Terrabacteria_group>Actinomycetota>Actinomycetes>Kitasatosporales>Streptomycetaceae | GCF_000722185.1 | Streptomyces_sp._NRRL_S-15 | IH15_RS0135825 | 5942..9683 | 1463886 | hypothetical-protein,-partial-[Streptomyces-sp.-NRRL-S-15] |
| WP_031093998.1 | Type2.Pgl.like.BREX | [illegible] | 494 | Bacteria>Terrabacteria_group>Actinomycetota>Actinomycetes>Kitasatosporales>Streptomycetaceae | GCF_000722185.1 | Streptomyces_sp._NRRL_S-15 | IH15_RS39765 | 56981..58463 | 1463886 | DUF6079-family-protein,-partial-[Streptomyces-sp.-NRRL-S-15] |
| WP_037930213.1 | Type2.Pgl.like.BREX | [illegible] | 1323 | Bacteria>Terrabacteria_group>Actinomycetota>Actinomycetes>Kitasatosporales>Streptomycetaceae | GCF_000725125.1 | Streptomyces_toyocaensis | BU52_RS07635 | 197363..201334 | 55952 | hypothetical-protein-[Streptomyces-toyocaensis] |
| WP_031178323.1 | Type2.Pgl.like.BREX | [illegible] | 1313 | Bacteria>Terrabacteria_group>Actinomycetota>Actinomycetes>Kitasatosporales>Streptomycetaceae | GCF_000725635.1 | Streptomyces_sp._NRRL_B-3253 | IO21_RS0123010 | 108342..112283 | 1463837 | hypothetical-protein-[Streptomyces-sp.-NRRL-B-3253] |
| WP_031126955.1 | Type2.Pgl.like.BREX | [illegible] | 1299 | Bacteria>Terrabacteria_group>Actinomycetota>Actinomycetes>Kitasatosporales>Streptomycetaceae | GCF_000725705.1 | Streptomyces | AOK13_RS15160 | 17218..21117 | 1883 | MULTISPECIES:-hypothetical-protein-[Streptomyces] |
| WP_043382132.1 | Type2.Pgl.like.BREX | [illegible] | 1316 | Bacteria>Terrabacteria_group>Actinomycetota>Actinomycetes>Kitasatosporales>Streptomycetaceae | GCF_000739045.1 | Streptomyces_mutabilis | FM21_RS27915 | 2768735..2772685 | 67332 | hypothetical-protein-[Streptomyces-mutabilis] |
| WP_043500398.1 | Type2.Pgl.like.BREX | [illegible] | 1294 | Bacteria>Terrabacteria_group>Actinomycetota>Actinomycetes>Kitasatosporales>Streptomycetaceae | GCF_000761215.1 | Streptomyces_glaucescens | SGLAU_RS10295 | 2386733..2390617 | 1907 | hypothetical-protein-[Streptomyces-glaucescens] |
| WP_042366886.1 | Type2.Pgl.like.BREX | [illegible] | 1311 | Bacteria>Terrabacteria_group>Actinomycetota>Actinomycetes>Kitasatosporales>Streptomycetaceae | GCF_000787815.1 | Streptacidiphilus_neutrinimicus | TR48_RS14550 | 88859..92794 | 105420 | hypothetical-protein-[Streptacidiphilus-neutrinimicus] |
| WP_052192753.1 | Type2.Pgl.like.BREX | [illegible] | 1298 | Bacteria>Terrabacteria_group>Actinomycetota>Actinomycetes>Kitasatosporales>Streptomycetaceae | GCF_000797385.1 | Streptomyces_sp._CNQ431 | PH39_RS06250 | 109132..113028 | 1571532 | hypothetical-protein-[Streptomyces-sp.-CNQ431] |
| WP_086083008.1 | Type2.Pgl.like.BREX | [illegible] | 1302 | Bacteria>Terrabacteria_group>Actinomycetota>Actinomycetes>Kitasatosporales>Streptomycetaceae | GCF_000802245.2 | Streptomyces_pluripotens | LK06_RS01830 | 426923..430831 | 1355015 | phage-resistance-protein-[Streptomyces-pluripotens] |
| WP_039653908.1 | Type2.Pgl.like.BREX | [illegible] | 1302 | Bacteria>Terrabacteria_group>Actinomycetota>Actinomycetes>Kitasatosporales>Streptomycetaceae | GCF_000816485.1 | Streptomyces_sp._MUSC_125 | LK08_RS17410 | 151118..155026 | 1428624 | hypothetical-protein-[Streptomyces-sp.-MUSC-125] |
| WP_042176278.1 | Type2.Pgl.like.BREX | [illegible] | 1297 | Bacteria>Terrabacteria_group>Actinomycetota>Actinomycetes>Kitasatosporales>Streptomycetaceae | GCF_000829695.1 | unclassified_Streptomyces | HUV60_RS03255 | 695180..699073 | 2593676 | MULTISPECIES:-hypothetical-protein-[unclassified-Streptomyces] |
| WP_044379847.1 | Type2.Pgl.like.BREX | [illegible] | 1298 | Bacteria>Terrabacteria_group>Actinomycetota>Actinomycetes>Kitasatosporales>Streptomycetaceae | GCF_000935185.3 | Streptomyces_noursei | SE95_RS29285 | 6702907..6706803 | 1971 | hypothetical-protein-[Streptomyces-noursei] |
| WP_047015461.1 | Type2.Pgl.like.BREX | [illegible] | 1287 | Bacteria>Terrabacteria_group>Actinomycetota>Actinomycetes>Kitasatosporales>Streptomycetaceae | GCF_001011035.1 | Streptomyces_sp._CNQ-509 | AA958_RS07555 | 1821426..1825289 | 444103 | DUF6079-family-protein-[Streptomyces-sp.-CNQ-509] |
| WP_047139866.1 | Type2.Pgl.like.BREX | [illegible] | 1298 | Bacteria>Terrabacteria_group>Actinomycetota>Actinomycetes>Kitasatosporales>Streptomycetaceae | GCF_001014595.1 | Streptomyces_sp._KE1 | WQ59_RS08655 | 50789..54685 | 1638939 | hypothetical-protein-[Streptomyces-sp.-KE1] |
| WP_234319629.1 | Type2.Pgl.like.BREX | [illegible] | 1343 | Bacteria>Terrabacteria_group>Actinomycetota>Actinomycetes>Kitasatosporales>Streptomycetaceae | GCF_001014595.1 | Streptomyces_sp._KE1 | WQ59_RS14830 | 95422..99453 | 1638939 | hypothetical-protein-[Streptomyces-sp.-KE1] |
| WP_048572280.1 | Type2.Pgl.like.BREX | [illegible] | 1294 | Bacteria>Terrabacteria_group>Actinomycetota>Actinomycetes>Kitasatosporales>Streptomycetaceae | GCF_001047315.1 | Streptomyces_leeuwenhoekii | ACH49_RS07740 | 40979..44863 | 1437453 | hypothetical-protein-[Streptomyces-leeuwenhoekii] |
| WP_053135729.1 | Type2.Pgl.like.BREX | [illegible] | 1305 | Bacteria>Terrabacteria_group>Actinomycetota>Actinomycetes>Kitasatosporales>Streptomycetaceae | GCF_001267885.1 | Streptomyces_ambofaciens | SAM23877_RS21640 | 4814542..4818459 | 1889 | hypothetical-protein-[Streptomyces-ambofaciens] |
| WP_053623542.1 | Type2.Pgl.like.BREX | [illegible] | 1308 | Bacteria>Terrabacteria_group>Actinomycetota>Actinomycetes>Kitasatosporales>Streptomycetaceae | GCF_001279425.1 | Streptomyces_sp._NRRL_F-2295 | ADK88_RS11800 | 69838..73764 | 1519477 | hypothetical-protein-[Streptomyces-sp.-NRRL-F-2295] |
| WP_053725463.1 | Type2.Pgl.like.BREX | [illegible] | 1298 | Bacteria>Terrabacteria_group>Actinomycetota>Actinomycetes>Kitasatosporales>Streptomycetaceae | GCF_001279695.1 | Streptomyces_sp._WM6378 | ADK54_RS11410 | 673..4569 | 1415557 | DUF6079-family-protein-[Streptomyces-sp.-WM6378] |
| WP_053763868.1 | Type2.Pgl.like.BREX | [illegible] | 1294 | Bacteria>Terrabacteria_group>Actinomycetota>Actinomycetes>Kitasatosporales>Streptomycetaceae | GCF_001279985.1 | Streptomyces_sp._AS58 | ADL00_RS43605 | 7603..11487 | 1519489 | hypothetical-protein-[Streptomyces-sp.-AS58] |
| WP_055507306.1 | Type2.Pgl.like.BREX | [illegible] | 1304 | Bacteria>Terrabacteria_group>Actinomycetota>Actinomycetes>Kitasatosporales>Streptomycetaceae | GCF_001418335.1 | Streptomyces_aurantiacus | AOK01_RS00950 | 70246..74160 | 47760 | hypothetical-protein-[Streptomyces-aurantiacus] |
| WP_055511936.1 | Type2.Pgl.like.BREX | [illegible] | 1295 | Bacteria>Terrabacteria_group>Actinomycetota>Actinomycetes>Kitasatosporales>Streptomycetaceae | GCF_001418475.1 | Streptomyces_ossamyceticus | AOK15_RS01705 | 11384..15271 | 249581 | hypothetical-protein-[Streptomyces-ossamyceticus] |
| WP_056798438.1 | Type2.Pgl.like.BREX | [illegible] | 1295 | Bacteria>Terrabacteria_group>Actinomycetota>Actinomycetes>Kitasatosporales>Streptomycetaceae | GCF_001426615.1 | unclassified_Streptomyces | ASD26_RS20305 | 149044..152931 | 2593676 | MULTISPECIES:-hypothetical-protein-[unclassified-Streptomyces] |
| WP_057583202.1 | Type2.Pgl.like.BREX | [illegible] | 1296 | Bacteria>Terrabacteria_group>Actinomycetota>Actinomycetes>Kitasatosporales>Streptomycetaceae | GCF_001428885.1 | Streptomyces_sp._Root264 | ASE41_RS32505 | 254370..258260 | 1736503 | hypothetical-protein-[Streptomyces-sp.-Root264] |
| WP_058042191.1 | Type2.Pgl.like.BREX | [illegible] | 1299 | Bacteria>Terrabacteria_group>Actinomycetota>Actinomycetes>Kitasatosporales>Streptomycetaceae | GCF_001445655.1 | Streptomyces_roseifaciens | AS857_RS06470 | 89914..93813 | 1488406 | hypothetical-protein-[Streptomyces-roseifaciens] |
| WP_059145331.1 | Type2.Pgl.like.BREX | [illegible] | 1306 | Bacteria>Terrabacteria_group>Actinomycetota>Actinomycetes>Kitasatosporales>Streptomycetaceae | GCF_001509775.1 | Streptomyces_violaceusniger | ADL28_RS21280 | 183244..187164 | 68280 | hypothetical-protein-[Streptomyces-violaceusniger] |
| WP_067237065.1 | Type2.Pgl.like.BREX | [illegible] | 1325 | Bacteria>Terrabacteria_group>Actinomycetota>Actinomycetes>Kitasatosporales>Streptomycetaceae | GCF_001514125.1 | Streptomyces_longwoodensis | AQJ30_RS22670 | 258338..262315 | 68231 | PglY-protein-[Streptomyces-longwoodensis] |
| WP_061915750.1 | Type2.Pgl.like.BREX | [illegible] | 1299 | Bacteria>Terrabacteria_group>Actinomycetota>Actinomycetes>Kitasatosporales>Streptomycetaceae | GCF_001514215.1 | Streptomyces_bungoensis | AQJ66_RS02630 | 666355..670254 | 285568 | hypothetical-protein-[Streptomyces-bungoensis] |
| WP_061920651.1 | Type2.Pgl.like.BREX | [illegible] | 1315 | Bacteria>Terrabacteria_group>Actinomycetota>Actinomycetes>Kitasatosporales>Streptomycetaceae | GCF_001514215.1 | Streptomyces_bungoensis | AQJ66_RS13685 | 278004..281951 | 285568 | hypothetical-protein-[Streptomyces-bungoensis] |
| WP_060944831.1 | Type2.Pgl.like.BREX | [illegible] | 1320 | Bacteria>Terrabacteria_group>Actinomycetota>Actinomycetes>Kitasatosporales>Streptomycetaceae | GCF_001553455.1 | Streptomyces_hygroscopicus | AXW62_RS14140 | 42881..46843 | 1912 | hypothetical-protein-[Streptomyces-hygroscopicus] |
| WP_062010201.1 | Type2.Pgl.like.BREX | [illegible] | 1320 | Bacteria>Terrabacteria_group>Actinomycetota>Actinomycetes>Kitasatosporales>Streptomycetaceae | GCF_001570645.1 | Streptomyces_hygroscopicus | AXW65_RS13440 | 81555..85517 | 1912 | hypothetical-protein-[Streptomyces-hygroscopicus] |
| WP_061404090.1 | Type2.Pgl.like.BREX | [illegible] | 1298 | Bacteria>Terrabacteria_group>Actinomycetota>Actinomycetes>Kitasatosporales>Streptomycetaceae | GCF_001577385.1 | Streptomyces_albidoflavus | Salbus254_RS06075 | 1445792..1449688 | 1886 | hypothetical-protein-[Streptomyces-albidoflavus] |
| WP_210565920.1 | Type2.Pgl.like.BREX | [illegible] | 1306 | Bacteria>Terrabacteria_group>Actinomycetota>Actinomycetes>Kitasatosporales>Streptomycetaceae | GCF_001598115.1 | Streptomyces_sp._AgN23 | AS97_RS28970 | 6649928..6653848 | 1188315 | hypothetical-protein-[Streptomyces-sp.-AgN23] |
| WP_062926533.1 | Type2.Pgl.like.BREX | [illegible] | 1309 | Bacteria>Terrabacteria_group>Actinomycetota>Actinomycetes>Kitasatosporales>Streptomycetaceae | GCF_001611795.1 | Streptomyces_qaidamensis | A4E84_RS11860 | 2715868..2719797 | 1783515 | hypothetical-protein-[Streptomyces-qaidamensis] |
| WP_067040782.1 | Type2.Pgl.like.BREX | [illegible] | 1295 | Bacteria>Terrabacteria_group>Actinomycetota>Actinomycetes>Kitasatosporales>Streptomycetaceae | GCF_001636945.2 | Streptomyces_sp._FXJ1.172 | A6P39_029160 | 6509387..6513274 | 710705 | phage-resistance-protein-[Streptomyces-sp.-FXJ1.172] |
| WP_063727790.1 | Type2.Pgl.like.BREX | [illegible] | 1299 | Bacteria>Terrabacteria_group>Actinomycetota>Actinomycetes>Kitasatosporales>Streptomycetaceae | GCF_001650215.1 | Streptomyces_sp._RTd22 | A7B19_RS30860 | 7245544..7249443 | 1841249 | hypothetical-protein-[Streptomyces-sp.-RTd22] |
| WP_067397487.1 | Type2.Pgl.like.BREX | [illegible] | 1298 | Bacteria>Terrabacteria_group>Actinomycetota>Actinomycetes>Kitasatosporales>Streptomycetaceae | GCF_001653115.1 | Streptomyces_sp._F-3 | STXM2123_RS19035 | 36552..40448 | 1840095 | phage-resistance-protein-[Streptomyces-sp.-F-3] |
| WP_065004487.1 | Type2.Pgl.like.BREX | [illegible] | 1315 | Bacteria>Terrabacteria_group>Actinomycetota>Actinomycetes>Kitasatosporales>Streptomycetaceae | GCF_001672315.1 | Streptomyces_sp._H-KF8 | A4U61_RS32740 | 4079733..4083680 | 1727216 | hypothetical-protein-[Streptomyces-sp.-H-KF8] |
| WP_067441133.1 | Type2.Pgl.like.BREX | [illegible] | 1297 | Bacteria>Terrabacteria_group>Actinomycetota>Actinomycetes>Kitasatosporales>Streptomycetaceae | GCF_001685355.1 | Streptomyces_lincolnensis | SLCG_RS11545 | 2586949..2590842 | 1915 | phage-resistance-protein-[Streptomyces-lincolnensis] |
| WP_069627934.1 | Type2.Pgl.like.BREX | [illegible] | 1306 | Bacteria>Terrabacteria_group>Actinomycetota>Actinomycetes>Kitasatosporales>Streptomycetaceae | GCF_001723055.1 | Streptomyces_niveus | BHU15_RS16810 | 32011..35931 | 193462 | PglY-protein-[Streptomyces-niveus] |
| WP_070388471.1 | Type2.Pgl.like.BREX | [illegible] | 1317 | Bacteria>Terrabacteria_group>Actinomycetota>Actinomycetes>Kitasatosporales>Streptomycetaceae | GCF_001767375.1 | Streptomyces_olivaceus | BC342_RS15735 | 3615707..3619660 | 47716 | hypothetical-protein-[Streptomyces-olivaceus] |
| WP_071373158.1 | Type2.Pgl.like.BREX | [illegible] | 1279 | Bacteria>Terrabacteria_group>Actinomycetota>Actinomycetes>Kitasatosporales>Streptomycetaceae | GCF_001866675.1 | Streptomyces_sp._MUSC_14 | BIV25_RS51055 | 161897..165736 | 1354889 | DUF6079-family-protein-[Streptomyces-sp.-MUSC-14] |
| WP_073727340.1 | Type2.Pgl.like.BREX | [illegible] | 1307 | Bacteria>Terrabacteria_group>Actinomycetota>Actinomycetes>Kitasatosporales>Streptomycetaceae | GCF_001905125.1 | Streptomyces_sp._TSRI0281 | A6A29_RS36835 | 505881..509804 | 1718998 | phage-resistance-protein-[Streptomyces-sp.-TSRI0281] |
| WP_073931399.1 | Type2.Pgl.like.BREX | [illegible] | 1293 | Bacteria>Terrabacteria_group>Actinomycetota>Actinomycetes>Kitasatosporales>Streptomycetaceae | GCF_001905725.1 | Streptomyces_sp._CB02400 | AMK33_04750 | 1152826..1156707 | 1703944 | phage-resistance-protein-[Streptomyces-sp.-CB02400] |
| WP_073897500.1 | Type2.Pgl.like.BREX | [illegible] | 1294 | Bacteria>Terrabacteria_group>Actinomycetota>Actinomycetes>Kitasatosporales>Streptomycetaceae | GCF_001905855.1 | Streptomyces_sp._CB01883 | AMK32_RS27770 | 364608..368492 | 1703943 | phage-resistance-protein-[Streptomyces-sp.-CB01883] |
| WP_073948333.1 | Type2.Pgl.like.BREX | [illegible] | 1294 | Bacteria>Terrabacteria_group>Actinomycetota>Actinomycetes>Kitasatosporales>Streptomycetaceae | GCF_001906585.1 | Streptomyces_kebangsaanensis | RH94_RS16410 | 41994..45878 | 864058 | hypothetical-protein-[Streptomyces-kebangsaanensis] |
| WP_077193671.1 | Type2.Pgl.like.BREX | [illegible] | 1301 | Bacteria>Terrabacteria_group>Actinomycetota>Actinomycetes>Kitasatosporales>Streptomycetaceae | GCF_001984445.1 | Streptomyces_lydicus | BXT88_RS28535 | 6380732..6384637 | 47763 | phage-resistance-protein-[Streptomyces-lydicus] |
| WP_078633283.1 | Type2.Pgl.like.BREX | [illegible] | 1295 | Bacteria>Terrabacteria_group>Actinomycetota>Actinomycetes>Kitasatosporales>Streptomycetaceae | GCF_002019855.1 | Streptomyces_antibioticus | HCX60_RS12060 | 2686146..2690033 | 1890 | phage-resistance-protein-[Streptomyces-antibioticus] |
| WP_081522440.1 | Type2.Pgl.like.BREX | [illegible] | 1312 | Bacteria>Terrabacteria_group>Actinomycetota>Actinomycetes>Kitasatosporales>Streptomycetaceae | GCF_002080455.1 | Streptomyces_sp._Sge12 | B6R96_RS11890 | 2759625..2763563 | 1972846 | phage-resistance-protein-[Streptomyces-sp.-Sge12] |
| WP_107426180.1 | Type2.Pgl.like.BREX | [illegible] | 1301 | Bacteria>Terrabacteria_group>Actinomycetota>Actinomycetes>Kitasatosporales>Streptomycetaceae | GCF_002082195.1 | Streptomyces_gilvosporeus | B1H19_RS30135 | 6750768..6754673 | 553510 | phage-resistance-protein-[Streptomyces-gilvosporeus] |
| WP_084898312.1 | Type2.Pgl.like.BREX | [illegible] | 1326 | Bacteria>Terrabacteria_group>Actinomycetota>Actinomycetes>Kitasatosporales>Streptomycetaceae | GCF_002099185.1 | Streptomyces_sp._CB03238 | BKD26_RS14940 | 390423..394403 | 1907777 | PglY-protein-[Streptomyces-sp.-CB03238] |
| WP_086823282.1 | Type2.Pgl.like.BREX | [illegible] | 1297 | Bacteria>Terrabacteria_group>Actinomycetota>Actinomycetes>Kitasatosporales>Streptomycetaceae | GCF_002156145.1 | Streptomyces_sp._NRRL_B-24572 | CCN56_RS03195 | 35384..39277 | 1962156 | phage-resistance-protein-[Streptomyces-sp.-NRRL-B-24572] |
| WP_087772686.1 | Type2.Pgl.like.BREX | [illegible] | 1298 | Bacteria>Terrabacteria_group>Actinomycetota>Actinomycetes>Kitasatosporales>Streptomycetaceae | GCF_002188355.1 | Streptomyces_sp._CS227 | B9W68_RS05745 | 1341091..1344987 | 1982763 | phage-resistance-protein-[Streptomyces-sp.-CS227] |
| WP_096622839.1 | Type2.Pgl.like.BREX | [illegible] | 1325 | Bacteria>Terrabacteria_group>Actinomycetota>Actinomycetes>Kitasatosporales>Streptomycetaceae | GCF_002382885.1 | Streptomyces_sp._WZ.A104 | CIB93_RS00460 | 105119..109096 | 2023771 | PglY-protein-[Streptomyces-sp.-WZ.A104] |
| WP_218923825.1 | Type2.Pgl.like.BREX | [illegible] | 1348 | Bacteria>Terrabacteria_group>Actinomycetota>Actinomycetes>Kitasatosporales>Streptomycetaceae | GCF_002551015.1 | unclassified_Streptomyces | CR491_RS01065 | 17513..21559 | 2593676 | MULTISPECIES:-hypothetical-protein-[unclassified-Streptomyces] |
| WP_097880367.1 | Type2.Pgl.like.BREX | [illegible] | 1299 | Bacteria>Terrabacteria_group>Actinomycetota>Actinomycetes>Kitasatosporales>Streptomycetaceae | GCF_002551135.1 | Streptomyces_sp._st140 | CR469_RS01835 | 68816..72715 | 1828052 | phage-resistance-protein-[Streptomyces-sp.-st140] |
| WP_098891635.1 | Type2.Pgl.like.BREX | [illegible] | 1299 | Bacteria>Terrabacteria_group>Actinomycetota>Actinomycetes>Kitasatosporales>Streptomycetaceae | GCF_002551295.1 | Streptomyces_sp._or3 | CR480_RS10855 | 74126..78025 | 1828020 | phage-resistance-protein-[Streptomyces-sp.-or3] |
| WP_097873672.1 | Type2.Pgl.like.BREX | [illegible] | 1300 | Bacteria>Terrabacteria_group>Actinomycetota>Actinomycetes>Kitasatosporales>Streptomycetaceae | GCF_002551335.1 | Streptomyces_sp._ms184 | CR468_RS02325 | 19903..23805 | 1827974 | phage-resistance-protein-[Streptomyces-sp.-ms184] |
| WP_098752344.1 | Type2.Pgl.like.BREX | [illegible] | 1295 | Bacteria>Terrabacteria_group>Actinomycetota>Actinomycetes>Kitasatosporales>Streptomycetaceae | GCF_002573955.1 | Streptomyces_sp._Ru87 | CRI70_RS14895 | 37032..40919 | 2044307 | hypothetical-protein-[Streptomyces-sp.-Ru87] |
| WP_099217838.1 | Type2.Pgl.like.BREX | [illegible] | 1299 | Bacteria>Terrabacteria_group>Actinomycetota>Actinomycetes>Kitasatosporales>Streptomycetaceae | GCF_002705725.1 | unclassified_Streptomyces | GTY74_RS33690 | 48529..52428 | 2593676 | MULTISPECIES:-phage-resistance-protein-[unclassified-Streptomyces] |
| WP_100203497.1 | Type2.Pgl.like.BREX | [illegible] | 1295 | Bacteria>Terrabacteria_group>Actinomycetota>Actinomycetes>Kitasatosporales>Streptomycetaceae | GCF_002794255.1 | Streptomyces_carminius | CUT44_RS21120 | 73830..77717 | 2665496 | phage-resistance-protein-[Streptomyces-carminius] |
| WP_100596575.1 | Type2.Pgl.like.BREX | [illegible] | 1309 | Bacteria>Terrabacteria_group>Actinomycetota>Actinomycetes>Kitasatosporales>Streptomycetaceae | GCF_002803065.1 | Streptomyces_sp._CB01635 | CG723_29500 | 36939..40868 | 2020326 | PglY-protein-[Streptomyces-sp.-CB01635] |
| WP_101279159.1 | Type2.Pgl.like.BREX | [illegible] | 1298 | Bacteria>Terrabacteria_group>Actinomycetota>Actinomycetes>Kitasatosporales>Streptomycetaceae | GCF_002844215.1 | Streptomyces_sp._EAG2 | CWE27_RS16835 | 41462..45358 | 2056495 | phage-resistance-protein-[Streptomyces-sp.-EAG2] |
| WP_102639909.1 | Type2.Pgl.like.BREX | [illegible] | 1302 | Bacteria>Terrabacteria_group>Actinomycetota>Actinomycetes>Kitasatosporales>Streptomycetaceae | GCF_002879675.3 | Streptomyces_sp._SMS_SU21 | C1C94_RS24615 | 1664504..1668412 | 2069440 | phage-resistance-protein-[Streptomyces-sp.-SMS_SU21] |
| WP_103503695.1 | Type2.Pgl.like.BREX | [illegible] | 1297 | Bacteria>Terrabacteria_group>Actinomycetota>Actinomycetes>Kitasatosporales>Streptomycetaceae | GCF_002910755.1 | Streptomyces_sp._SM14 | B9S65_RS21040 | 21301..25194 | 1736045 | phage-resistance-protein-[Streptomyces-sp.-SM14] |
| WP_103493996.1 | Type2.Pgl.like.BREX | [illegible] | 1295 | Bacteria>Terrabacteria_group>Actinomycetota>Actinomycetes>Kitasatosporales>Streptomycetaceae | GCF_002910775.2 | unclassified_Streptomyces | B9S64_RS11515 | 2609512..2613399 | 2593676 | MULTISPECIES:-phage-resistance-protein-[unclassified-Streptomyces] |
| WP_103506852.1 | Type2.Pgl.like.BREX | [illegible] | 1298 | Bacteria>Terrabacteria_group>Actinomycetota>Actinomycetes>Kitasatosporales>Streptomycetaceae | GCF_002910815.1 | Streptomyces_sp._FM008 | B9S63_RS32625 | 27016..30912 | 1983802 | phage-resistance-protein-[Streptomyces-sp.-FM008] |
| WP_258040143.1 | Type2.Pgl.like.BREX | [illegible] | 1387 | Bacteria>Terrabacteria_group>Actinomycetota>Actinomycetes>Kitasatosporales>Streptomycetaceae | GCF_002910825.1 | Streptomyces_sp._SM1 | B9S61_RS40115 | 11465..15626 | 402229 | hypothetical-protein,-partial-[Streptomyces-sp.-SM1] |
| WP_103529453.1 | Type2.Pgl.like.BREX | [illegible] | 1296 | Bacteria>Terrabacteria_group>Actinomycetota>Actinomycetes>Kitasatosporales>Streptomycetaceae | GCF_002910855.1 | Streptomyces_sp._SM12 | B9S59_RS23865 | 21242..25132 | 1071602 | phage-resistance-protein-[Streptomyces-sp.-SM12] |
| WP_103534295.1 | Type2.Pgl.like.BREX | [illegible] | 1307 | Bacteria>Terrabacteria_group>Actinomycetota>Actinomycetes>Kitasatosporales>Streptomycetaceae | GCF_002910905.1 | Streptomyces_sp._SM11 | B9S58_RS24885 | 17104..21027 | 565557 | phage-resistance-protein-[Streptomyces-sp.-SM11] |
| WP_099498686.1 | Type2.Pgl.like.BREX | [illegible] | 1297 | Bacteria>Terrabacteria_group>Actinomycetota>Actinomycetes>Kitasatosporales>Streptomycetaceae | GCF_002946835.1 | Streptomyces | C4B68_RS07930 | 1717869..1721762 | 1883 | MULTISPECIES:-phage-resistance-protein-[Streptomyces] |
| WP_099498944.1 | Type2.Pgl.like.BREX | [illegible] | 1303 | Bacteria>Terrabacteria_group>Actinomycetota>Actinomycetes>Kitasatosporales>Streptomycetaceae | GCF_002946835.1 | Streptomyces | C4B68_RS09525 | 2094134..2098045 | 1883 | MULTISPECIES:-phage-resistance-protein-[Streptomyces] |
| WP_099504556.1 | Type2.Pgl.like.BREX | [illegible] | 1312 | Bacteria>Terrabacteria_group>Actinomycetota>Actinomycetes>Kitasatosporales>Streptomycetaceae | GCF_002946835.1 | Streptomyces | C4B68_RS28490 | 6170305..6174243 | 1883 | MULTISPECIES:-PglY-protein-[Streptomyces] |
| WP_105973292.1 | Type2.Pgl.like.BREX | [illegible] | 1328 | Bacteria>Terrabacteria_group>Actinomycetota>Actinomycetes>Kitasatosporales>Streptomycetaceae | GCF_002954775.1 | Streptomyces_geranii | CW359_RS29840 | 34991..38977 | 2058923 | phage-resistance-protein-[Streptomyces-geranii] |
| WP_107018395.1 | Type2.Pgl.like.BREX | [illegible] | 1299 | Bacteria>Terrabacteria_group>Actinomycetota>Actinomycetes>Kitasatosporales>Streptomycetaceae | GCF_003024195.1 | Streptomyces_dioscori | C6Y14_RS21480 | 188736..192635 | 2109333 | phage-resistance-protein-[Streptomyces-dioscori] |
| WP_108006154.1 | Type2.Pgl.like.BREX | [illegible] | 1295 | Bacteria>Terrabacteria_group>Actinomycetota>Actinomycetes>Kitasatosporales>Streptomycetaceae | GCF_003046555.1 | Streptomyces_sp._VMFN-G11Ma | C7821_RS00505 | 113242..117129 | 2135609 | phage-resistance-protein-[Streptomyces-sp.-VMFN-G11Ma] |
| WP_109284035.1 | Type2.Pgl.like.BREX | [illegible] | 1304 | Bacteria>Terrabacteria_group>Actinomycetota>Actinomycetes>Kitasatosporales>Streptomycetaceae | GCF_003121295.1 | Streptomyces_orinoci | CV755_RS31530 | 49510..53424 | 67339 | phage-resistance-protein-[Streptomyces-orinoci] |
| WP_109293254.1 | Type2.Pgl.like.BREX | [illegible] | 1296 | Bacteria>Terrabacteria_group>Actinomycetota>Actinomycetes>Kitasatosporales>Streptomycetaceae | GCF_003122365.1 | Streptomyces_spongiicola | DDQ41_RS04155 | 979567..983457 | 1690221 | phage-resistance-protein-[Streptomyces-spongiicola] |
| WP_109378206.1 | Type2.Pgl.like.BREX | [illegible] | 1323 | Bacteria>Terrabacteria_group>Actinomycetota>Actinomycetes>Kitasatosporales>Streptomycetaceae | GCF_003143855.1 | Streptomyces_sp._NWU339 | DIZ27_RS11255 | 179446..183417 | 2185284 | PglY-protein-[Streptomyces-sp.-NWU339] |
| WP_111004202.1 | Type2.Pgl.like.BREX | [illegible] | 1309 | Bacteria>Terrabacteria_group>Actinomycetota>Actinomycetes>Kitasatosporales>Streptomycetaceae | GCF_003236365.1 | Streptomyces_sp._NTH33 | C1I97_RS17780 | 5348..9277 | 1735453 | hypothetical-protein-[Streptomyces-sp.-NTH33] |
| WP_111335439.1 | Type2.Pgl.like.BREX | [illegible] | 1293 | Bacteria>Terrabacteria_group>Actinomycetota>Actinomycetes>Kitasatosporales>Streptomycetaceae | GCF_003248335.1 | unclassified_Streptomyces | DNK55_RS27680 | 629182..633063 | 2593676 | MULTISPECIES:-hypothetical-protein-[unclassified-Streptomyces] |
| WP_112468596.1 | Type2.Pgl.like.BREX | [illegible] | 1292 | Bacteria>Terrabacteria_group>Actinomycetota>Actinomycetes>Kitasatosporales>Streptomycetaceae | GCF_003270085.1 | Streptomyces_triticisoli | DEP02_RS17985 | 1404..5282 | 2182797 | phage-resistance-protein-[Streptomyces-triticisoli] |
| WP_112469147.1 | Type2.Pgl.like.BREX | [illegible] | 1305 | Bacteria>Terrabacteria_group>Actinomycetota>Actinomycetes>Kitasatosporales>Streptomycetaceae | GCF_003270085.1 | Streptomyces_triticisoli | DEP02_RS20520 | 21209..25126 | 2182797 | hypothetical-protein-[Streptomyces-triticisoli] |
| WP_114054613.1 | Type2.Pgl.like.BREX | [illegible] | 1298 | Bacteria>Terrabacteria_group>Actinomycetota>Actinomycetes>Kitasatosporales>Streptomycetaceae | GCF_003325375.1 | Streptomyces_globosus | C0216_RS08100 | 1744779..1748675 | 68209 | phage-resistance-protein-[Streptomyces-globosus] |
| WP_114246647.1 | Type2.Pgl.like.BREX | [illegible] | 1313 | Bacteria>Terrabacteria_group>Actinomycetota>Actinomycetes>Kitasatosporales>Streptomycetaceae | GCF_003330865.1 | Streptomyces_atratus | C5746_RS28330 | 6153269..6157210 | 1893 | phage-resistance-protein-[Streptomyces-atratus] |
| WP_114626835.1 | Type2.Pgl.like.BREX | [illegible] | 1320 | Bacteria>Terrabacteria_group>Actinomycetota>Actinomycetes>Kitasatosporales>Streptomycetaceae | GCF_003346515.1 | Streptomyces_corynorhini | DVH02_RS29045 | 18264..22226 | 2282652 | phage-resistance-protein-[Streptomyces-corynorhini] |
| WP_128502442.1 | Type2.Pgl.like.BREX | [illegible] | 1304 | Bacteria>Terrabacteria_group>Actinomycetota>Actinomycetes>Kitasatosporales>Streptomycetaceae | GCF_003389455.1 | Streptomyces_inhibens | DY245_RS00990 | 15167..19081 | 2293571 | PglY-protein-[Streptomyces-inhibens] |
| WP_116428654.1 | Type2.Pgl.like.BREX | [illegible] | 1296 | Bacteria>Terrabacteria_group>Actinomycetota>Actinomycetes>Kitasatosporales>Streptomycetaceae | GCF_003402595.1 | Streptomyces_spongiicola | SSP531S_RS24190 | 6042..9932 | 1690221 | phage-resistance-protein-[Streptomyces-spongiicola] |
| WP_120685184.1 | Type2.Pgl.like.BREX | [illegible] | 1300 | Bacteria>Terrabacteria_group>Actinomycetota>Actinomycetes>Kitasatosporales>Streptomycetaceae | GCF_003626535.1 | Streptomyces_hoynatensis | D7294_RS31160 | 7577..11479 | 1141874 | phage-resistance-protein-[Streptomyces-hoynatensis] |
| WP_121719925.1 | Type2.Pgl.like.BREX | [illegible] | 1295 | Bacteria>Terrabacteria_group>Actinomycetota>Actinomycetes>Kitasatosporales>Streptomycetaceae | GCF_003671675.1 | Streptomyces_sp._E2N171 | EBF82_RS15280 | 2148..6035 | 1851914 | phage-resistance-protein-[Streptomyces-sp.-E2N171] |
| WP_121748720.1 | Type2.Pgl.like.BREX | [illegible] | 1300 | Bacteria>Terrabacteria_group>Actinomycetota>Actinomycetes>Kitasatosporales>Streptomycetaceae | GCF_003671715.1 | Streptomyces_sp._E2N166 | EBH09_RS22240 | 41286..45188 | 1851909 | hypothetical-protein-[Streptomyces-sp.-E2N166] |
| WP_121887751.1 | Type2.Pgl.like.BREX | [illegible] | 1295 | Bacteria>Terrabacteria_group>Actinomycetota>Actinomycetes>Kitasatosporales>Streptomycetaceae | GCF_003688995.1 | Streptomyces_shenzhenensis | CTZ28_RS03710 | 158817..162704 | 943815 | phage-resistance-protein-[Streptomyces-shenzhenensis] |
| WP_123562585.1 | Type2.Pgl.like.BREX | [illegible] | 1307 | Bacteria>Terrabacteria_group>Actinomycetota>Actinomycetes>Kitasatosporales>Streptomycetaceae | GCF_003751605.1 | Kitasatospora_cineracea | EDD39_RS31525 | 1175476..1179399 | 88074 | phage-resistance-protein-[Kitasatospora-cineracea] |
| WP_123459797.1 | Type2.Pgl.like.BREX | [illegible] | 1295 | Bacteria>Terrabacteria_group>Actinomycetota>Actinomycetes>Kitasatosporales>Streptomycetaceae | GCF_003752275.1 | Streptomyces_sp._CEV_2-1 | EDD95_RS00765 | 172440..176327 | 2485153 | phage-resistance-protein-[Streptomyces-sp.-CEV-2-1] |
| WP_124266060.1 | Type2.Pgl.like.BREX | [illegible] | 1295 | Bacteria>Terrabacteria_group>Actinomycetota>Actinomycetes>Kitasatosporales>Streptomycetaceae | GCF_003846255.1 | Streptomyces_sp._ADI98-10 | EES46_RS27970 | 1709341..1713228 | 1522763 | hypothetical-protein-[Streptomyces-sp.-ADI98-10] |
| WP_124273797.1 | Type2.Pgl.like.BREX | [illegible] | 1298 | Bacteria>Terrabacteria_group>Actinomycetota>Actinomycetes>Kitasatosporales>Streptomycetaceae | GCF_003846265.1 | Streptomyces_sp._ADI96-15 | EES44_RS29825 | 154353..158249 | 1522761 | phage-resistance-protein-[Streptomyces-sp.-ADI96-15] |
| WP_125502694.1 | Type2.Pgl.like.BREX | [illegible] | 1295 | Bacteria>Terrabacteria_group>Actinomycetota>Actinomycetes>Kitasatosporales>Streptomycetaceae | GCF_003945525.1 | unclassified_Streptomyces | DLM49_RS04285 | 990903..994790 | 2593676 | MULTISPECIES:-phage-resistance-protein-[unclassified-Streptomyces] |
| WP_125530906.1 | Type2.Pgl.like.BREX | [illegible] | 1298 | Bacteria>Terrabacteria_group>Actinomycetota>Actinomycetes>Kitasatosporales>Streptomycetaceae | GCF_003947585.1 | Streptomyces_sp._WAC_06725 | DMH15_RS04680 | 32673..36569 | 2203209 | phage-resistance-protein-[Streptomyces-sp.-WAC-06725] |
| WP_260614935.1 | Type2.Pgl.like.BREX | [illegible] | 1241 | Bacteria>Terrabacteria_group>Actinomycetota>Actinomycetes>Kitasatosporales>Streptomycetaceae | GCF_003949955.1 | Streptomyces_sp._WAC05292 | EF903_RS07655 | 26077..29800 | 2487418 | phage-resistance-protein,-partial-[Streptomyces-sp.-WAC05292] |
| WP_127468975.1 | Type2.Pgl.like.BREX | [illegible] | 1306 | Bacteria>Terrabacteria_group>Actinomycetota>Actinomycetes>Kitasatosporales>Streptomycetaceae | GCF_004000405.1 | Streptomyces_sp._B27 | EBQ22_RS28690 | 51055..54975 | 2485015 | phage-resistance-protein-[Streptomyces-sp.-B27] |
| WP_128462336.1 | Type2.Pgl.like.BREX | [illegible] | 1298 | Bacteria>Terrabacteria_group>Actinomycetota>Actinomycetes>Kitasatosporales>Streptomycetaceae | GCF_004078575.1 | Streptomyces_albidoflavus | EQK42_RS04975 | 17321..21217 | 1886 | phage-resistance-protein-[Streptomyces-albidoflavus] |
| WP_128801436.1 | Type2.Pgl.like.BREX | [illegible] | 1295 | Bacteria>Terrabacteria_group>Actinomycetota>Actinomycetes>Kitasatosporales>Streptomycetaceae | GCF_004104505.1 | unclassified_Streptomyces | DBP13_RS07075 | 1487876..1491763 | 2593676 | MULTISPECIES:-phage-resistance-protein-[unclassified-Streptomyces] |
| WP_136222946.1 | Type2.Pgl.like.BREX | [illegible] | 1301 | Bacteria>Terrabacteria_group>Actinomycetota>Actinomycetes>Kitasatosporales>Streptomycetaceae | GCF_004794975.1 | Streptomyces_sp._A1547 | E6W17_RS04085 | 285079..288984 | 2563105 | phage-resistance-protein-[Streptomyces-sp.-A1547] |
| WP_136324456.1 | Type2.Pgl.like.BREX | [illegible] | 1284 | Bacteria>Terrabacteria_group>Actinomycetota>Actinomycetes>Kitasatosporales>Streptomycetaceae | GCF_004795105.1 | Streptomyces_sp._A0642 | E6R60_RS01260 | 305327..309181 | 2563100 | PglY-protein-[Streptomyces-sp.-A0642] |
| WP_136181970.1 | Type2.Pgl.like.BREX | [illegible] | 1295 | Bacteria>Terrabacteria_group>Actinomycetota>Actinomycetes>Kitasatosporales>Streptomycetaceae | GCF_004795255.1 | Streptomyces_sp._RKND-216 | E4198_RS04240 | 941855..945742 | 2562581 | phage-resistance-protein-[Streptomyces-sp.-RKND-216] |
| WP_136728164.1 | Type2.Pgl.like.BREX | [illegible] | 1316 | Bacteria>Terrabacteria_group>Actinomycetota>Actinomycetes>Kitasatosporales>Streptomycetaceae | GCF_005047355.1 | Actinacidiphila_oryziradicis | FCI23_RS62555 | 22737..26687 | 2571141 | PglY-protein-[Actinacidiphila-oryziradicis] |
| WP_137230275.1 | Type2.Pgl.like.BREX | [illegible] | 1299 | Bacteria>Terrabacteria_group>Actinomycetota>Actinomycetes>Kitasatosporales>Streptomycetaceae | GCF_005234185.1 | Streptomyces_sp._BPSDS2 | FD916_RS06870 | 197367..201266 | 2571021 | phage-resistance-protein-[Streptomyces-sp.-BPSDS2] |
| WP_137992471.1 | Type2.Pgl.like.BREX | [illegible] | 1286 | Bacteria>Terrabacteria_group>Actinomycetota>Actinomycetes>Kitasatosporales>Streptomycetaceae | GCF_005517195.1 | Streptomyces_exfoliatus | FDM97_RS23450 | 5014702..5018562 | 1905 | PglY-protein-[Streptomyces-exfoliatus] |
| WP_142263956.1 | Type2.Pgl.like.BREX | [illegible] | 1295 | Bacteria>Terrabacteria_group>Actinomycetota>Actinomycetes>Kitasatosporales>Streptomycetaceae | GCF_006715185.1 | Streptomyces_sp._SLBN-115 | FBY34_RS01600 | 357905..361792 | 2768453 | phage-resistance-protein-[Streptomyces-sp.-SLBN-115] |
| WP_142151610.1 | Type2.Pgl.like.BREX | [illegible] | 1294 | Bacteria>Terrabacteria_group>Actinomycetota>Actinomycetes>Kitasatosporales>Streptomycetaceae | GCF_006715395.1 | Streptomyces_sp._SLBN-31 | FBY22_RS32840 | 2801489..2805373 | 2768444 | phage-resistance-protein-[Streptomyces-sp.-SLBN-31] |
| WP_144316319.1 | Type2.Pgl.like.BREX | [illegible] | 1300 | Bacteria>Terrabacteria_group>Actinomycetota>Actinomycetes>Kitasatosporales>Streptomycetaceae | GCF_007113425.2 | Streptomyces_sp._RLB3-17 | FNV62_RS53960 | 11911719..11915621 | 2594455 | PglY-protein-[Streptomyces-sp.-RLB3-17] |
| WP_143613014.1 | Type2.Pgl.like.BREX | [illegible] | 1280 | Bacteria>Terrabacteria_group>Actinomycetota>Actinomycetes>Kitasatosporales>Streptomycetaceae | GCF_007113465.1 | unclassified_Streptomyces | FNV65_RS16625 | 3594205..3598047 | 2593676 | MULTISPECIES:-PglY-protein-[unclassified-Streptomyces] |
| WP_143606190.1 | Type2.Pgl.like.BREX | [illegible] | 1300 | Bacteria>Terrabacteria_group>Actinomycetota>Actinomycetes>Kitasatosporales>Streptomycetaceae | GCF_007113545.2 | unclassified_Streptomyces | FNV68_RS00625 | 125740..129642 | 2593676 | MULTISPECIES:-PglY-protein-[unclassified-Streptomyces] |
| WP_143654853.1 | Type2.Pgl.like.BREX | [illegible] | 1280 | Bacteria>Terrabacteria_group>Actinomycetota>Actinomycetes>Kitasatosporales>Streptomycetaceae | GCF_007113565.2 | Streptomyces_sp._S1A1-7 | FNV64_RS18410 | 3725417..3729259 | 2594459 | PglY-protein-[Streptomyces-sp.-S1A1-7] |
| WP_147989639.1 | Type2.Pgl.like.BREX | [illegible] | 1296 | Bacteria>Terrabacteria_group>Actinomycetota>Actinomycetes>Kitasatosporales>Streptomycetaceae | GCF_008042045.1 | Streptomyces_sp._t39 | EAO77_RS19880 | 46286..50176 | 1828156 | phage-resistance-protein-[Streptomyces-sp.-t39] |
| WP_149182066.1 | Type2.Pgl.like.BREX | [illegible] | 1296 | Bacteria>Terrabacteria_group>Actinomycetota>Actinomycetes>Kitasatosporales>Streptomycetaceae | GCF_008271805.1 | Streptomyces_sp._TRM49041 | FVS66_RS17245 | 78613..82503 | 2603216 | phage-resistance-protein-[Streptomyces-sp.-TRM49041] |
| WP_150231687.1 | Type2.Pgl.like.BREX | [illegible] | 1310 | Bacteria>Terrabacteria_group>Actinomycetota>Actinomycetes>Kitasatosporales>Streptomycetaceae | GCF_008634015.1 | Streptomyces_filamentosus | CP979_RS24300 | 5261397..5265329 | 67294 | phage-resistance-protein-[Streptomyces-filamentosus] |
| WP_016826541.1 | Type2.Pgl.like.BREX | [illegible] | 1300 | Bacteria>Terrabacteria_group>Actinomycetota>Actinomycetes>Kitasatosporales>Streptomycetaceae | GCF_008704515.1 | Streptomyces_viridosporus | CP969_RS12545 | 2677846..2681748 | 67581 | hypothetical-protein-[Streptomyces-viridosporus] |
| WP_150501927.1 | Type2.Pgl.like.BREX | [illegible] | 1286 | Bacteria>Terrabacteria_group>Actinomycetota>Actinomycetes>Kitasatosporales>Streptomycetaceae | GCF_008705255.1 | Streptomyces_venezuelae | DEJ45_RS06855 | 1615979..1619839 | 54571 | PglY-protein-[Streptomyces-venezuelae] |
| WP_150950778.1 | Type2.Pgl.like.BREX | [illegible] | 1308 | Bacteria>Terrabacteria_group>Actinomycetota>Actinomycetes>Kitasatosporales>Streptomycetaceae | GCF_008806595.1 | Streptomyces_luteolifulvus | F7R91_RS21840 | 77489..81415 | 2615112 | phage-resistance-protein-[Streptomyces-luteolifulvus] |
| WP_134111864.1 | Type2.Pgl.like.BREX | [illegible] | 1294 | Bacteria>Terrabacteria_group>Actinomycetota>Actinomycetes>Kitasatosporales>Streptomycetaceae | GCF_008931305.1 | Streptomyces_coelicolor | FQ762_RS34260 | 7363057..7366941 | 1902 | phage-resistance-protein-[Streptomyces-coelicolor] |
| WP_152168305.1 | Type2.Pgl.like.BREX | [illegible] | 1293 | Bacteria>Terrabacteria_group>Actinomycetota>Actinomycetes>Kitasatosporales>Streptomycetaceae | GCF_009184865.1 | Streptomyces_phaeolivaceus | F9278_RS11920 | 2549005..2552886 | 2653200 | phage-resistance-protein-[Streptomyces-phaeolivaceus] |
| WP_077964595.1 | Type2.Pgl.like.BREX | [illegible] | 1296 | Bacteria>Terrabacteria_group>Actinomycetota>Actinomycetes>Kitasatosporales>Streptomycetaceae | GCF_009296025.1 | Streptomyces_tsukubensis | GBW32_RS05195 | 1168905..1172795 | 83656 | hypothetical-protein-[Streptomyces-tsukubensis] |
| WP_228126181.1 | Type2.Pgl.like.BREX | [illegible] | 1318 | Bacteria>Terrabacteria_group>Actinomycetota>Actinomycetes>Kitasatosporales>Streptomycetaceae | GCF_009495655.1 | Streptomyces | GEV49_RS20575 | 4651210..4655166 | 1883 | MULTISPECIES:-hypothetical-protein-[Streptomyces] |
| WP_153520868.1 | Type2.Pgl.like.BREX | [illegible] | 1297 | Bacteria>Terrabacteria_group>Actinomycetota>Actinomycetes>Kitasatosporales>Streptomycetaceae | GCF_009600885.1 | Streptomyces_jumonjinensis | FF041_RS02770 | 20638..24531 | 1945 | phage-resistance-protein-[Streptomyces-jumonjinensis] |
| WP_155072369.1 | Type2.Pgl.like.BREX | [illegible] | 1296 | Bacteria>Terrabacteria_group>Actinomycetota>Actinomycetes>Kitasatosporales>Streptomycetaceae | GCF_009709575.1 | Streptomyces_taklimakanensis | FOL17_RS21690 | 4932691..4936581 | 2569853 | phage-resistance-protein-[Streptomyces-taklimakanensis] |
| WP_156207776.1 | Type2.Pgl.like.BREX | [illegible] | 1295 | Bacteria>Terrabacteria_group>Actinomycetota>Actinomycetes>Kitasatosporales>Streptomycetaceae | GCF_009735685.1 | Streptomyces_lycii | GCU69_RS30080 | 21210..25097 | 2654337 | hypothetical-protein-[Streptomyces-lycii] |
| WP_156694667.1 | Type2.Pgl.like.BREX | [illegible] | 1295 | Bacteria>Terrabacteria_group>Actinomycetota>Actinomycetes>Kitasatosporales>Streptomycetaceae | GCF_009739905.1 | Streptomyces_ficellus | EIZ62_RS23460 | 5231851..5235738 | 1977088 | phage-resistance-protein-[Streptomyces-ficellus] |
| WP_190140957.1 | Type2.Pgl.like.BREX | [illegible] | 1298 | Bacteria>Terrabacteria_group>Actinomycetota>Actinomycetes>Kitasatosporales>Streptomycetaceae | GCF_009811575.1 | Streptomyces_glebosus | Sgleb_RS29750 | 6831668..6835564 | 249580 | PglY-protein-[Streptomyces-glebosus] |
| WP_159742973.1 | Type2.Pgl.like.BREX | [illegible] | 1299 | Bacteria>Terrabacteria_group>Actinomycetota>Actinomycetes>Kitasatosporales>Streptomycetaceae | GCF_009811635.1 | Streptomyces_tubercidicus | Stube_RS06795 | 1610444..1614343 | 47759 | phage-resistance-protein-[Streptomyces-tubercidicus] |
| WP_161069245.1 | Type2.Pgl.like.BREX | [illegible] | 1308 | Bacteria>Terrabacteria_group>Actinomycetota>Actinomycetes>Kitasatosporales>Streptomycetaceae | GCF_009827025.1 | Streptomyces_sp._HUCO-GS316 | GR925_RS33060 | 14814..18740 | 2692198 | phage-resistance-protein-[Streptomyces-sp.-HUCO-GS316] |

| Protein-ID | System | Gene-Neighbourhoods | Length | Taxonomy | Assembly-ID | Strain | Locus-Tag | Genomic-coordinates | TaxID | Fasta-defline |
|---|---|---|---|---|---|---|---|---|---|---|
| WP_031128307.1 | Type2.Pgl.like.BREX | [illegible] | 1299 | Bacteria>Terrabacteria_group>Actinomycetota>Actinomycetes>Kitasatosporales>Streptomycetaceae | GCF_009834665.1 | Streptomyces | K701_RS20870 | 114425..118324 | 1883 | MULTISPECIES:-hypothetical-protein-[Streptomyces] |
| WP_159767012.1 | Type2.Pgl.like.BREX | [illegible] | 1295 | Bacteria>Terrabacteria_group>Actinomycetota>Actinomycetes>Kitasatosporales>Streptomycetaceae | GCF_009857135.1 | Streptomyces_sp._HM190 | GSY65_RS10110 | 2414615..2418502 | 2695266 | phage-resistance-protein-[Streptomyces-sp.-HM190] |
| WP_111382238.1 | Type2.Pgl.like.BREX | [illegible] | 1300 | Bacteria>Terrabacteria_group>Actinomycetota>Actinomycetes>Kitasatosporales>Streptomycetaceae | GCF_009862055.1 | Streptomyces_sp._SID8358 | GTW52_RS08645 | 18687..22589 | 2690342 | phage-resistance-protein-[Streptomyces-sp.-SID8358] |
| WP_161168814.1 | Type2.Pgl.like.BREX | [illegible] | 1296 | Bacteria>Terrabacteria_group>Actinomycetota>Actinomycetes>Kitasatosporales>Streptomycetaceae | GCF_009862125.1 | Streptomyces_sp._SID5910 | GTW67_RS35430 | 5133..9023 | 2690312 | phage-resistance-protein-[Streptomyces-sp.-SID5910] |
| WP_011031061.1 | Type2.Pgl.like.BREX | [illegible] | 1294 | Bacteria>Terrabacteria_group>Actinomycetota>Actinomycetes>Kitasatosporales>Streptomycetaceae | GCF_009862445.1 | Streptomyces_sp._SID7813 | GTY87_RS33150 | 7362943..7366827 | 2690330 | hypothetical-protein-[Streptomyces-sp.-SID7813] |
| WP_052855836.1 | Type2.Pgl.like.BREX | [illegible] | 1280 | Bacteria>Terrabacteria_group>Actinomycetota>Actinomycetes>Kitasatosporales>Streptomycetaceae | GCF_009862505.1 | Streptomyces_sp._SID7805 | GTY53_RS24335 | 41453..45295 | 2690328 | DUF6079-family-protein-[Streptomyces-sp.-SID7805] |
| WP_161326401.1 | Type2.Pgl.like.BREX | [illegible] | 1296 | Bacteria>Terrabacteria_group>Actinomycetota>Actinomycetes>Kitasatosporales>Streptomycetaceae | GCF_009872535.1 | Streptomyces_sp._SID5789 | GTW93_RS03680 | 29135..33025 | 2690310 | phage-resistance-protein-[Streptomyces-sp.-SID5789] |
| WP_164183062.1 | Type2.Pgl.like.BREX | [illegible] | 1298 | Bacteria>Terrabacteria_group>Actinomycetota>Actinomycetes>Kitasatosporales>Streptomycetaceae | GCF_010523235.1 | Streptomyces_sp._604F | C3K23_RS25915 | 5696613..5700509 | 1476754 | phage-resistance-protein-[Streptomyces-sp.-604F] |
| WP_164407618.1 | Type2.Pgl.like.BREX | [illegible] | 1306 | Bacteria>Terrabacteria_group>Actinomycetota>Actinomycetes>Kitasatosporales>Streptomycetaceae | GCF_010548275.1 | Streptomyces_sp._SID12488 | G3I56_05390 | 276309..280229 | 2706040 | PglY-protein-[Streptomyces-sp.-SID12488] |
| WP_164615103.1 | Type2.Pgl.like.BREX | [illegible] | 1299 | Bacteria>Terrabacteria_group>Actinomycetota>Actinomycetes>Kitasatosporales>Streptomycetaceae | GCF_010548505.1 | Streptomyces_sp._SID14515 | G3I54_RS15510 | 49251..53150 | 2706074 | phage-resistance-protein-[Streptomyces-sp.-SID14515] |
| WP_164373105.1 | Type2.Pgl.like.BREX | [illegible] | 1322 | Bacteria>Terrabacteria_group>Actinomycetota>Actinomycetes>Kitasatosporales>Streptomycetaceae | GCF_010548515.1 | Streptomyces_sp._SID14446 | G3I62_RS24970 | 24029..27997 | 2706072 | phage-resistance-protein-[Streptomyces-sp.-SID14446] |
| WP_164382621.1 | Type2.Pgl.like.BREX | [illegible] | 1323 | Bacteria>Terrabacteria_group>Actinomycetota>Actinomycetes>Kitasatosporales>Streptomycetaceae | GCF_010993835.1 | Streptomyces_sp._OM5714 | STPH1_RS07315 | 1721493..1725464 | 2602736 | PglY-protein-[Streptomyces-sp.-OM5714] |
| WP_165234047.1 | Type2.Pgl.like.BREX | [illegible] | 1325 | Bacteria>Terrabacteria_group>Actinomycetota>Actinomycetes>Kitasatosporales>Streptomycetaceae | GCF_011044975.1 | Streptomyces_coryli | G5C51_RS08080 | 27992..31969 | 1128680 | PglY-protein-[Streptomyces-coryli] |
| WP_165263660.1 | Type2.Pgl.like.BREX | [illegible] | 1309 | Bacteria>Terrabacteria_group>Actinomycetota>Actinomycetes>Kitasatosporales>Streptomycetaceae | GCF_011045015.1 | Streptomyces_scabichelini | G5C60_RS27730 | 6556..10485 | 2711217 | PglY-protein-[Streptomyces-scabichelini] |
| WP_193454094.1 | Type2.Pgl.like.BREX | [illegible] | 1298 | Bacteria>Terrabacteria_group>Actinomycetota>Actinomycetes>Kitasatosporales>Streptomycetaceae | GCF_011170105.1 | Streptomyces_rutgersensis | Srut_RS22845 | 716645..720541 | 53451 | phage-resistance-protein-[Streptomyces-rutgersensis] |
| WP_166002742.1 | Type2.Pgl.like.BREX | [illegible] | 1295 | Bacteria>Terrabacteria_group>Actinomycetota>Actinomycetes>Kitasatosporales>Streptomycetaceae | GCF_011193355.1 | Streptomyces_sp._JB150 | G7Z13_RS26575 | 5772314..5776201 | 2714844 | phage-resistance-protein-[Streptomyces-sp.-JB150] |
| WP_165984583.1 | Type2.Pgl.like.BREX | [illegible] | 1356 | Bacteria>Terrabacteria_group>Actinomycetota>Actinomycetes>Kitasatosporales>Streptomycetaceae | GCF_011421595.1 | Streptomyces_sp._YIM_98790 | GQS09_RS04065 | 22949..27019 | 2689077 | hypothetical-protein-[Streptomyces-sp.-YIM-98790] |
| WP_173262696.1 | Type2.Pgl.like.BREX | [illegible] | 1296 | Bacteria>Terrabacteria_group>Actinomycetota>Actinomycetes>Kitasatosporales>Streptomycetaceae | GCF_011766325.1 | Streptomyces_pacificus | SCWH03_RS05415 | 43859..47749 | 2705029 | phage-resistance-protein-[Streptomyces-pacificus] |
| WP_167967748.1 | Type2.Pgl.like.BREX | [illegible] | 1298 | Bacteria>Terrabacteria_group>Actinomycetota>Actinomycetes>Kitasatosporales>Streptomycetaceae | GCF_012033735.1 | Streptomyces_lonarensis | HCN56_RS02350 | 22960..26856 | 700599 | phage-resistance-protein-[Streptomyces-lonarensis] |
| WP_167931758.1 | Type2.Pgl.like.BREX | [illegible] | 1298 | Bacteria>Terrabacteria_group>Actinomycetota>Actinomycetes>Kitasatosporales>Streptomycetaceae | GCF_012034385.1 | Streptomyces_spiramenti | HCJ92_RS02745 | 20478..24374 | 2720606 | phage-resistance-protein-[Streptomyces-spiramenti] |
| WP_169440217.1 | Type2.Pgl.like.BREX | [illegible] | 1300 | Bacteria>Terrabacteria_group>Actinomycetota>Actinomycetes>Kitasatosporales>Streptomycetaceae | GCF_012726235.1 | Streptomyces_sp._HNM0574 | HCC30_RS02200 | 531963..535865 | 2714954 | PglY-protein-[Streptomyces-sp.-HNM0574] |
| WP_169086808.1 | Type2.Pgl.like.BREX | [illegible] | 1309 | Bacteria>Terrabacteria_group>Actinomycetota>Actinomycetes>Kitasatosporales>Streptomycetaceae | GCF_012927225.1 | unclassified_Streptomyces | HHL18_RS29280 | 65110..69039 | 2593676 | MULTISPECIES:-phage-resistance-protein-[unclassified-Streptomyces] |
| WP_171109616.1 | Type2.Pgl.like.BREX | [illegible] | 1326 | Bacteria>Terrabacteria_group>Actinomycetota>Actinomycetes>Kitasatosporales>Streptomycetaceae | GCF_013009255.1 | Streptomyces | HJD23_RS18045 | 4041978..4045958 | 1883 | MULTISPECIES:-PglY-protein-[Streptomyces] |
| WP_171141282.1 | Type2.Pgl.like.BREX | [illegible] | 1320 | Bacteria>Terrabacteria_group>Actinomycetota>Actinomycetes>Kitasatosporales>Streptomycetaceae | GCF_013044505.1 | Streptomyces_sp._S3(2020) | HLK59_RS11655 | 32079..36041 | 2732044 | PglY-protein-[Streptomyces-sp.-S3(2020)] |
| WP_172387270.1 | Type2.Pgl.like.BREX | [illegible] | 1299 | Bacteria>Terrabacteria_group>Actinomycetota>Actinomycetes>Kitasatosporales>Streptomycetaceae | GCF_013177615.1 | Streptomyces_sp._MNP-20 | HCO73_RS37710 | 26479..30378 | 2721165 | phage-resistance-protein-[Streptomyces-sp.-MNP-20] |
| WP_175410415.1 | Type2.Pgl.like.BREX | [illegible] | 1323 | Bacteria>Terrabacteria_group>Actinomycetota>Actinomycetes>Kitasatosporales>Streptomycetaceae | GCF_013362995.1 | Streptomyces_sp._TRM64462 | HTZ82_RS17995 | 58952..62923 | 2741726 | phage-resistance-protein-[Streptomyces-sp.-TRM64462] |
| WP_175458172.1 | Type2.Pgl.like.BREX | [illegible] | 1298 | Bacteria>Terrabacteria_group>Actinomycetota>Actinomycetes>Kitasatosporales>Streptomycetaceae | GCF_013363465.1 | Streptomyces | G6W59_RS10675 | 242164..246060 | 1883 | MULTISPECIES:-phage-resistance-protein-[Streptomyces] |
| WP_176117073.1 | Type2.Pgl.like.BREX | [illegible] | 1309 | Bacteria>Terrabacteria_group>Actinomycetota>Actinomycetes>Kitasatosporales>Streptomycetaceae | GCF_013364175.1 | Streptomyces_sp._NA02536 | HUT14_RS10185 | 2278747..2282676 | 2742133 | phage-resistance-protein-[Streptomyces-sp.-NA02536] |
| WP_179084887.1 | Type2.Pgl.like.BREX | [illegible] | 1304 | Bacteria>Terrabacteria_group>Actinomycetota>Actinomycetes>Kitasatosporales>Streptomycetaceae | GCF_013394065.1 | Streptomyces_rectiverticillatus | FGW37_RS24660 | 5638152..5642066 | 173860 | DUF6079-family-protein-[Streptomyces-rectiverticillatus] |
| WP_179172839.1 | Type2.Pgl.like.BREX | [illegible] | 1298 | Bacteria>Terrabacteria_group>Actinomycetota>Actinomycetes>Kitasatosporales>Streptomycetaceae | GCF_013401435.1 | Streptomyces_sp._BR123 | HYE82_RS05840 | 43587..47483 | 2749828 | phage-resistance-protein-[Streptomyces-sp.-BR123] |
| WP_179181139.1 | Type2.Pgl.like.BREX | [illegible] | 1318 | Bacteria>Terrabacteria_group>Actinomycetota>Actinomycetes>Kitasatosporales>Streptomycetaceae | GCF_013401535.1 | Streptomyces_sp._CB04723 | HXS80_RS07185 | 1703905..1707861 | 2748873 | PglY-protein-[Streptomyces-sp.-CB04723] |
| WP_181356189.1 | Type2.Pgl.like.BREX | [illegible] | 1295 | Bacteria>Terrabacteria_group>Actinomycetota>Actinomycetes>Kitasatosporales>Streptomycetaceae | GCF_013618545.1 | Streptomyces_alkaliterrae | H3147_RS03280 | 30298..34185 | 2213162 | phage-resistance-protein-[Streptomyces-alkaliterrae] |
| WP_181660536.1 | Type2.Pgl.like.BREX | [illegible] | 1304 | Bacteria>Terrabacteria_group>Actinomycetota>Actinomycetes>Kitasatosporales>Streptomycetaceae | GCF_013778455.1 | Streptomyces_himalayensis | H1D24_RS28240 | 43695..47609 | 2820085 | PglY-protein-[Streptomyces-himalayensis] |
| WP_191852498.1 | Type2.Pgl.like.BREX | [illegible] | 1324 | Bacteria>Terrabacteria_group>Actinomycetota>Actinomycetes>Kitasatosporales>Streptomycetaceae | GCF_013913525.1 | Streptomyces_griseoaurantiacus | H1X69_RS09375 | 152668..156642 | 68213 | PglY-protein-[Streptomyces-griseoaurantiacus] |
| WP_161173771.1 | Type2.Pgl.like.BREX | [illegible] | 1294 | Bacteria>Terrabacteria_group>Actinomycetota>Actinomycetes>Kitasatosporales>Streptomycetaceae | GCF_014138545.1 | Streptomyces | BJ962_RS25945 | 2051930..2055814 | 1883 | MULTISPECIES:-phage-resistance-protein-[Streptomyces] |
| WP_184969721.1 | Type2.Pgl.like.BREX | [illegible] | 1319 | Bacteria>Terrabacteria_group>Actinomycetota>Actinomycetes>Kitasatosporales>Streptomycetaceae | GCF_014203595.1 | Streptomyces_echinatus | FHS34_RS26250 | 60873..64832 | 67293 | PglY-protein-[Streptomyces-echinatus] |
| WP_184574795.1 | Type2.Pgl.like.BREX | [illegible] | 1305 | Bacteria>Terrabacteria_group>Actinomycetota>Actinomycetes>Kitasatosporales>Streptomycetaceae | GCF_014203645.1 | Streptomyces_zagrosensis | FHS42_RS22770 | 242905..246822 | 1042984 | hypothetical-protein-[Streptomyces-zagrosensis] |
| WP_187063990.1 | Type2.Pgl.like.BREX | [illegible] | 1297 | Bacteria>Terrabacteria_group>Actinomycetota>Actinomycetes>Kitasatosporales>Streptomycetaceae | GCF_014295035.1 | Streptomyces_buecherae | H7H31_RS06855 | 1887405..1891298 | 2763006 | phage-resistance-protein-[Streptomyces-buecherae] |
| WP_223186495.1 | Type2.Pgl.like.BREX | [illegible] | 1324 | Bacteria>Terrabacteria_group>Actinomycetota>Actinomycetes>Kitasatosporales>Streptomycetaceae | GCF_014596445.1 | Streptomyces_sp._CBMA29 | BGM09_RS08230 | 64585..68559 | 1896314 | PglY-protein-[Streptomyces-sp.-CBMA29] |
| WP_188302216.1 | Type2.Pgl.like.BREX | [illegible] | 1312 | Bacteria>Terrabacteria_group>Actinomycetota>Actinomycetes>Kitasatosporales>Streptomycetaceae | GCF_014596465.1 | Streptomyces_sp._CBMA123 | BG452_RS01300 | 274066..278004 | 1896313 | DUF6079-family-protein-[Streptomyces-sp.-CBMA123] |
| WP_188337083.1 | Type2.Pgl.like.BREX | [illegible] | 1306 | Bacteria>Terrabacteria_group>Actinomycetota>Actinomycetes>Kitasatosporales>Streptomycetaceae | GCF_014621695.1 | Streptomyces_xanthii | IAG42_RS12405 | 2760683..2764603 | 2768069 | PglY-protein-[Streptomyces-xanthii] |
| WP_189191867.1 | Type2.Pgl.like.BREX | [illegible] | 1308 | Bacteria>Terrabacteria_group>Actinomycetota>Actinomycetes>Kitasatosporales>Streptomycetaceae | GCF_014645835.1 | Streptomyces_albiflavescens | IEY95_RS43770 | 44051..47977 | 1623582 | phage-resistance-protein-[Streptomyces-albiflavescens] |
| WP_189215567.1 | Type2.Pgl.like.BREX | [illegible] | 1309 | Bacteria>Terrabacteria_group>Actinomycetota>Actinomycetes>Kitasatosporales>Streptomycetaceae | GCF_014648075.1 | Streptomyces_ruber | IEZ04_RS05700 | 261607..265536 | 83378 | phage-resistance-protein-[Streptomyces-ruber] |
| WP_189273365.1 | Type2.Pgl.like.BREX | [illegible] | 1310 | Bacteria>Terrabacteria_group>Actinomycetota>Actinomycetes>Kitasatosporales>Streptomycetaceae | GCF_014648655.1 | Streptomyces_atratus | GCM10010207_01540 | 179323..183255 | 1893 | phage-resistance-protein-[Streptomyces-atratus] |
| WP_189233278.1 | Type2.Pgl.like.BREX | [illegible] | 1302 | Bacteria>Terrabacteria_group>Actinomycetota>Actinomycetes>Kitasatosporales>Streptomycetaceae | GCF_014648815.1 | Streptomyces_flaveolus | IEZ05_RS29325 | 142464..146372 | 67297 | phage-resistance-protein-[Streptomyces-flaveolus] |
| WP_189925600.1 | Type2.Pgl.like.BREX | [illegible] | 1299 | Bacteria>Terrabacteria_group>Actinomycetota>Actinomycetes>Kitasatosporales>Streptomycetaceae | GCF_014649075.1 | Streptomyces_asoensis | Saso_RS30340 | 1677261..1681160 | 249586 | phage-resistance-protein-[Streptomyces-asoensis] |
| WP_189531547.1 | Type2.Pgl.like.BREX | [illegible] | 1298 | Bacteria>Terrabacteria_group>Actinomycetota>Actinomycetes>Kitasatosporales>Streptomycetaceae | GCF_014649335.1 | Streptomyces_roseolilacinus | IE106_RS08455 | 240309..244205 | 66904 | phage-resistance-protein-[Streptomyces-roseolilacinus] |
| WP_190076975.1 | Type2.Pgl.like.BREX | [illegible] | 1321 | Bacteria>Terrabacteria_group>Actinomycetota>Actinomycetes>Kitasatosporales>Streptomycetaceae | GCF_014649475.1 | Streptomyces_daghestanicus | IE186_RS08935 | 129191..133156 | 66885 | PglY-protein-[Streptomyces-daghestanicus] |
| WP_189596065.1 | Type2.Pgl.like.BREX | [illegible] | 1294 | Bacteria>Terrabacteria_group>Actinomycetota>Actinomycetes>Kitasatosporales>Streptomycetaceae | GCF_014649515.1 | Streptomyces_pilosus | IE141_RS07070 | 323618..327502 | 28893 | phage-resistance-protein-[Streptomyces-pilosus] |
| WP_189553697.1 | Type2.Pgl.like.BREX | [illegible] | 1297 | Bacteria>Terrabacteria_group>Actinomycetota>Actinomycetes>Kitasatosporales>Streptomycetaceae | GCF_014649735.1 | Streptomyces_lavendofoliae | IE109_RS25595 | 27723..31616 | 67314 | phage-resistance-protein-[Streptomyces-lavendofoliae] |
| WP_189965530.1 | Type2.Pgl.like.BREX | [illegible] | 1299 | Bacteria>Terrabacteria_group>Actinomycetota>Actinomycetes>Kitasatosporales>Streptomycetaceae | GCF_014649955.1 | Streptomyces_violascens | IE206_RS24480 | 114519..118418 | 67381 | PglY-protein-[Streptomyces-violascens] |
| WP_189744979.1 | Type2.Pgl.like.BREX | [illegible] | 1295 | Bacteria>Terrabacteria_group>Actinomycetota>Actinomycetes>Kitasatosporales>Streptomycetaceae | GCF_014650435.1 | Streptomyces_tendae | IE258_RS18980 | 415118..419005 | 1932 | phage-resistance-protein-[Streptomyces-tendae] |
| WP_189823142.1 | Type2.Pgl.like.BREX | [illegible] | 1310 | Bacteria>Terrabacteria_group>Actinomycetota>Actinomycetes>Kitasatosporales>Streptomycetaceae | GCF_014650515.1 | Streptomyces_finlayi | IE282_RS09950 | 502304..506236 | 67296 | PglY-protein-[Streptomyces-finlayi] |
| WP_190179476.1 | Type2.Pgl.like.BREX | [illegible] | 1295 | Bacteria>Terrabacteria_group>Actinomycetota>Actinomycetes>Kitasatosporales>Streptomycetaceae | GCF_014650575.1 | Streptomyces_naganishii | IE217_RS21210 | 138490..142377 | 285447 | phage-resistance-protein-[Streptomyces-naganishii] |
| WP_193485689.1 | Type2.Pgl.like.BREX | [illegible] | 1295 | Bacteria>Terrabacteria_group>Actinomycetota>Actinomycetes>Kitasatosporales>Streptomycetaceae | GCF_014650795.1 | Streptomyces | KBZ75_RS22000 | 119443..123330 | 1883 | MULTISPECIES:-phage-resistance-protein-[Streptomyces] |
| WP_189852660.1 | Type2.Pgl.like.BREX | [illegible] | 1297 | Bacteria>Terrabacteria_group>Actinomycetota>Actinomycetes>Kitasatosporales>Streptomycetaceae | GCF_014650895.1 | Streptomyces_omiyaensis | IE287_RS31550 | 67641..71534 | 68247 | phage-resistance-protein-[Streptomyces-omiyaensis] |
| WP_189888567.1 | Type2.Pgl.like.BREX | [illegible] | 1299 | Bacteria>Terrabacteria_group>Actinomycetota>Actinomycetes>Kitasatosporales>Streptomycetaceae | GCF_014651015.1 | Streptomyces_xantholiticus | IE198_RS20240 | 155904..159803 | 68285 | phage-resistance-protein-[Streptomyces-xantholiticus] |
| WP_190048743.1 | Type2.Pgl.like.BREX | [illegible] | 1295 | Bacteria>Terrabacteria_group>Actinomycetota>Actinomycetes>Kitasatosporales>Streptomycetaceae | GCF_014651035.1 | Streptomyces_lomondensis | IE255_RS04370 | 1003302..1007189 | 68229 | phage-resistance-protein-[Streptomyces-lomondensis] |
| WP_189780601.1 | Type2.Pgl.like.BREX | [illegible] | 1322 | Bacteria>Terrabacteria_group>Actinomycetota>Actinomycetes>Kitasatosporales>Streptomycetaceae | GCF_014653855.1 | Streptomyces_capitiformicae | CES90_RS08320 | 1865463..1869431 | 2014920 | PglY-protein-[Streptomyces-capitiformicae] |
| WP_189784303.1 | Type2.Pgl.like.BREX | [illegible] | 1308 | Bacteria>Terrabacteria_group>Actinomycetota>Actinomycetes>Kitasatosporales>Streptomycetaceae | GCF_014653855.1 | Streptomyces_capitiformicae | IE327_RS22350 | 174729..178655 | 2014920 | phage-resistance-protein-[Streptomyces-capitiformicae] |
| WP_190127345.1 | Type2.Pgl.like.BREX | [illegible] | 1300 | Bacteria>Terrabacteria_group>Actinomycetota>Actinomycetes>Kitasatosporales>Streptomycetaceae | GCF_014654785.1 | Streptomyces_mashuensis | IE305_RS00650 | 166112..170014 | 33904 | phage-resistance-protein-[Streptomyces-mashuensis] |
| WP_190043128.1 | Type2.Pgl.like.BREX | [illegible] | 1297 | Bacteria>Terrabacteria_group>Actinomycetota>Actinomycetes>Kitasatosporales>Streptomycetaceae | GCF_014654895.1 | Streptomyces_filamentosus | IE326_RS25570 | 1478299..1482192 | 67294 | phage-resistance-protein-[Streptomyces-filamentosus] |
| WP_189978619.1 | Type2.Pgl.like.BREX | [illegible] | 1296 | Bacteria>Terrabacteria_group>Actinomycetota>Actinomycetes>Kitasatosporales>Streptomycetaceae | GCF_014654935.1 | Streptomyces_capoamus | IE312_RS04620 | 255199..259089 | 68183 | hypothetical-protein-[Streptomyces-capoamus] |
| WP_190135917.1 | Type2.Pgl.like.BREX | [illegible] | 1298 | Bacteria>Terrabacteria_group>Actinomycetota>Actinomycetes>Kitasatosporales>Streptomycetaceae | GCF_014656095.1 | Streptomyces_longispororuber | IE330_RS11995 | 43958..47854 | 68230 | phage-resistance-protein-[Streptomyces-longispororuber] |
| WP_150527865.1 | Type2.Pgl.like.BREX | [illegible] | 1301 | Bacteria>Terrabacteria_group>Actinomycetota>Actinomycetes>Kitasatosporales>Streptomycetaceae | GCF_014656155.1 | Streptomyces_vinaceus | IE310_RS19410 | 98676..102581 | 1960 | phage-resistance-protein-[Streptomyces-vinaceus] |
| WP_189832564.1 | Type2.Pgl.like.BREX | [illegible] | 1298 | Bacteria>Terrabacteria_group>Actinomycetota>Actinomycetes>Kitasatosporales>Streptomycetaceae | GCF_014656215.1 | Streptomyces_zaomyceticus | IE311_RS17210 | 206117..210013 | 68286 | phage-resistance-protein-[Streptomyces-zaomyceticus] |
| WP_189999228.1 | Type2.Pgl.like.BREX | [illegible] | 1299 | Bacteria>Terrabacteria_group>Actinomycetota>Actinomycetes>Kitasatosporales>Streptomycetaceae | GCF_014656255.1 | Streptomyces_rubradiris | IE323_RS35215 | 23790..27689 | 285531 | phage-resistance-protein-[Streptomyces-rubradiris] |
| WP_190849997.1 | Type2.Pgl.like.BREX | [illegible] | 1304 | Bacteria>Terrabacteria_group>Actinomycetota>Actinomycetes>Kitasatosporales>Streptomycetaceae | GCF_014701115.1 | Streptomyces_aurantiacus | IGS71_RS09715 | 2395233..2399147 | 47760 | PglY-protein-[Streptomyces-aurantiacus] |
| WP_192364278.1 | Type2.Pgl.like.BREX | [illegible] | 1294 | Bacteria>Terrabacteria_group>Actinomycetota>Actinomycetes>Kitasatosporales>Streptomycetaceae | GCF_014852565.2 | Streptomyces_caniscabiei | IHE65_RS32180 | 7252963..7256847 | 2746961 | phage-resistance-protein-[Streptomyces-caniscabiei] |
| WP_194046780.1 | Type2.Pgl.like.BREX | [illegible] | 1301 | Bacteria>Terrabacteria_group>Actinomycetota>Actinomycetes>Kitasatosporales>Streptomycetaceae | GCF_015160855.1 | Streptomyces_ferrugineus | IM697_RS10270 | 2254443..2258348 | 1413221 | phage-resistance-protein-[Streptomyces-ferrugineus] |
| WP_194277017.1 | Type2.Pgl.like.BREX | [illegible] | 1317 | Bacteria>Terrabacteria_group>Actinomycetota>Actinomycetes>Kitasatosporales>Streptomycetaceae | GCF_015244315.1 | Streptomyces_olivaceus | FYC74_RS20195 | 4691722..4695675 | 47716 | hypothetical-protein-[Streptomyces-olivaceus] |
| WP_196191504.1 | Type2.Pgl.like.BREX | [illegible] | 1324 | Bacteria>Terrabacteria_group>Actinomycetota>Actinomycetes>Kitasatosporales>Streptomycetaceae | GCF_015680775.2 | Streptomyces_sp._BRB081 | ICN76_RS15730 | 3658471..3662445 | 2769544 | PglY-protein-[Streptomyces-sp.-BRB081] |
| WP_197923435.1 | Type2.Pgl.like.BREX | [illegible] | 1235 | Bacteria>Terrabacteria_group>Actinomycetota>Actinomycetes>Kitasatosporales>Streptomycetaceae | GCF_016017415.1 | Streptomyces_sp._HB-N217 | I3J14_RS01390 | 16464..20171 | 2792016 | phage-resistance-protein-[Streptomyces-sp.-HB-N217] |
| WP_197988023.1 | Type2.Pgl.like.BREX | [illegible] | 1303 | Bacteria>Terrabacteria_group>Actinomycetota>Actinomycetes>Kitasatosporales>Streptomycetaceae | GCF_016031615.1 | Streptomyces_pactum | IHE55_RS05645 | 1413949..1417860 | 68249 | PglY-protein-[Streptomyces-pactum] |
| WP_200308438.1 | Type2.Pgl.like.BREX | [illegible] | 1297 | Bacteria>Terrabacteria_group>Actinomycetota>Actinomycetes>Kitasatosporales>Streptomycetaceae | GCF_016598475.1 | Streptomyces_adelaidensis | I8N67_RS35680 | 6729..10622 | 2796465 | phage-resistance-protein-[Streptomyces-adelaidensis] |
| WP_201851327.1 | Type2.Pgl.like.BREX | [illegible] | 1319 | Bacteria>Terrabacteria_group>Actinomycetota>Actinomycetes>Kitasatosporales>Streptomycetaceae | GCF_016741935.1 | Streptomyces_endocoffeicus | JK364_RS14425 | 428407..432366 | 2898945 | phage-resistance-protein-[Streptomyces-endocoffeicus] |
| WP_129245459.1 | Type2.Pgl.like.BREX | [illegible] | 1307 | Bacteria>Terrabacteria_group>Actinomycetota>Actinomycetes>Kitasatosporales>Streptomycetaceae | GCF_016876065.1 | Streptomyces_sioyaensis | EST54_RS05070 | 11707..15630 | 67364 | PglY-protein-[Streptomyces-sioyaensis] |
| WP_205359716.1 | Type2.Pgl.like.BREX | [illegible] | 1302 | Bacteria>Terrabacteria_group>Actinomycetota>Actinomycetes>Kitasatosporales>Streptomycetaceae | GCF_016918855.1 | Actinacidiphila_acididurans | ITX44_RS25570 | 260304..264212 | 2784346 | phage-resistance-protein-[Actinacidiphila-acididurans] |
| WP_205378071.1 | Type2.Pgl.like.BREX | [illegible] | 1297 | Bacteria>Terrabacteria_group>Actinomycetota>Actinomycetes>Kitasatosporales>Streptomycetaceae | GCF_016919245.1 | Streptomyces_zhihengii | JE024_RS36170 | 1980886..1984779 | 1818004 | phage-resistance-protein-[Streptomyces-zhihengii] |
| WP_206960808.1 | Type2.Pgl.like.BREX | [illegible] | 1306 | Bacteria>Terrabacteria_group>Actinomycetota>Actinomycetes>Kitasatosporales>Streptomycetaceae | GCF_017315755.1 | Streptomyces_beijiangensis | J0695_RS06135 | 11838..15758 | 163361 | hypothetical-protein-[Streptomyces-beijiangensis] |
| WP_207248053.1 | Type2.Pgl.like.BREX | [illegible] | 1302 | Bacteria>Terrabacteria_group>Actinomycetota>Actinomycetes>Kitasatosporales>Streptomycetaceae | GCF_017356985.1 | Streptomyces | J1792_RS22685 | 920556..924464 | 1883 | MULTISPECIES:-phage-resistance-protein-[Streptomyces] |
| WP_208779390.1 | Type2.Pgl.like.BREX | [illegible] | 1299 | Bacteria>Terrabacteria_group>Actinomycetota>Actinomycetes>Kitasatosporales>Streptomycetaceae | GCF_017614965.1 | Streptomyces_griseorubiginosus | GRC12_RS13030 | 31883..35782 | 67304 | hypothetical-protein-[Streptomyces-griseorubiginosus] |
| WP_209214544.1 | Type2.Pgl.like.BREX | [illegible] | 1314 | Bacteria>Terrabacteria_group>Actinomycetota>Actinomycetes>Kitasatosporales>Streptomycetaceae | GCF_017676345.1 | Streptomyces_smyrnaeus | JW613_RS33010 | 30286..34230 | 1387713 | phage-resistance-protein-[Streptomyces-smyrnaeus] |
| WP_209240964.1 | Type2.Pgl.like.BREX | [illegible] | 1229 | Bacteria>Terrabacteria_group>Actinomycetota>Actinomycetes>Kitasatosporales>Streptomycetaceae | GCF_017676365.1 | Streptomyces_oryzae | ITI46_RS19580 | 22640..26326 | 1434886 | phage-resistance-protein,-partial-[Streptomyces-oryzae] |
| WP_229376318.1 | Type2.Pgl.like.BREX | [illegible] | 1350 | Bacteria>Terrabacteria_group>Actinomycetota>Actinomycetes>Kitasatosporales>Streptomycetaceae | GCF_017676385.1 | Streptomyces_spirodelae | JW592_RS18225 | 58574..62626 | 2812904 | hypothetical-protein-[Streptomyces-spirodelae] |
| WP_209268533.1 | Type2.Pgl.like.BREX | [illegible] | 1303 | Bacteria>Terrabacteria_group>Actinomycetota>Actinomycetes>Kitasatosporales>Streptomycetaceae | GCF_017676385.1 | Streptomyces_spirodelae | JW592_RS30635 | 2117..6028 | 2812904 | phage-resistance-protein-[Streptomyces-spirodelae] |
| WP_209340089.1 | Type2.Pgl.like.BREX | [illegible] | 1324 | Bacteria>Terrabacteria_group>Actinomycetota>Actinomycetes>Kitasatosporales>Streptomycetaceae | GCF_017813245.1 | Streptomyces_montanisoli | JFN87_RS12560 | 26746..30720 | 2798581 | PglY-protein-[Streptomyces-montanisoli] |
| WP_247692466.1 | Type2.Pgl.like.BREX | [illegible] | 1364 | Bacteria>Terrabacteria_group>Actinomycetota>Actinomycetes>Kitasatosporales>Streptomycetaceae | GCF_018070225.1 | Streptomyces_sp._RK31 | KBZ00_RS21455 | 166561..170655 | 2824892 | hypothetical-protein-[Streptomyces-sp.-RK31] |
| WP_210988319.1 | Type2.Pgl.like.BREX | [illegible] | 1325 | Bacteria>Terrabacteria_group>Actinomycetota>Actinomycetes>Kitasatosporales>Streptomycetaceae | GCF_018070345.1 | Streptomyces_sp._F63 | KBZ10_RS16035 | 122891..126868 | 2824887 | PglY-protein-[Streptomyces-sp.-F63] |
| WP_210921437.1 | Type2.Pgl.like.BREX | [illegible] | 1296 | Bacteria>Terrabacteria_group>Actinomycetota>Actinomycetes>Kitasatosporales>Streptomycetaceae | GCF_018070565.1 | Streptomyces_sp._B93 | KBY47_RS13480 | 35274..39164 | 2824875 | hypothetical-protein-[Streptomyces-sp.-B93] |
| WP_210930997.1 | Type2.Pgl.like.BREX | [illegible] | 1299 | Bacteria>Terrabacteria_group>Actinomycetota>Actinomycetes>Kitasatosporales>Streptomycetaceae | GCF_018070665.1 | Streptomyces_sp._404i | KBY46_RS00835 | 204983..208882 | 2824902 | phage-resistance-protein-[Streptomyces-sp.-404i] |
| WP_211038996.1 | Type2.Pgl.like.BREX | [illegible] | 1297 | Bacteria>Terrabacteria_group>Actinomycetota>Actinomycetes>Kitasatosporales>Streptomycetaceae | GCF_018101125.2 | Streptomyces_tardus | JGS22_RS25655 | 96725..100618 | 2780544 | phage-resistance-protein-[Streptomyces-tardus] |
| WP_212672406.1 | Type2.Pgl.like.BREX | [illegible] | 1292 | Bacteria>Terrabacteria_group>Actinomycetota>Actinomycetes>Kitasatosporales>Streptomycetaceae | GCF_018128905.1 | Streptomyces_sp._A2-16 | IOD14_RS34895 | 7765786..7769664 | 2781734 | phage-resistance-protein-[Streptomyces-sp.-A2-16] |
| WP_212726653.1 | Type2.Pgl.like.BREX | [illegible] | 1299 | Bacteria>Terrabacteria_group>Actinomycetota>Actinomycetes>Kitasatosporales>Streptomycetaceae | GCF_018310535.1 | Streptomyces_mirabilis | SMIR_RS05040 | 1089234..1093133 | 68239 | phage-resistance-protein-[Streptomyces-mirabilis] |
| WP_212908905.1 | Type2.Pgl.like.BREX | [illegible] | 1296 | Bacteria>Terrabacteria_group>Actinomycetota>Actinomycetes>Kitasatosporales>Streptomycetaceae | GCF_018327685.1 | Streptomyces_sp._TS71-3 | Sm713_RS07750 | 1878907..1882797 | 2733862 | phage-resistance-protein-[Streptomyces-sp.-TS71-3] |
| WP_214331468.1 | Type2.Pgl.like.BREX | [illegible] | 1295 | Bacteria>Terrabacteria_group>Actinomycetota>Actinomycetes>Kitasatosporales>Streptomycetaceae | GCF_018530775.1 | Streptomyces_sp._Tu102 | KJZ88_RS09240 | 198289..202176 | 2838019 | phage-resistance-protein-[Streptomyces-sp.-Tu102] |
| WP_215129314.1 | Type2.Pgl.like.BREX | [illegible] | 1301 | Bacteria>Terrabacteria_group>Actinomycetota>Actinomycetes>Kitasatosporales>Streptomycetaceae | GCF_018614035.1 | Streptomyces_sp._ISL-96 | J7E96_RS09030 | 27274..31179 | 2819191 | PglY-protein-[Streptomyces-sp.-ISL-96] |
| WP_214957746.1 | Type2.Pgl.like.BREX | [illegible] | 1299 | Bacteria>Terrabacteria_group>Actinomycetota>Actinomycetes>Kitasatosporales>Streptomycetaceae | GCF_018614195.1 | Streptomyces_sp._ISL-43 | J7F03_RS38005 | 20163..24062 | 2819183 | DUF6079-family-protein-[Streptomyces-sp.-ISL-43] |
| WP_215167460.1 | Type2.Pgl.like.BREX | [illegible] | 1307 | Bacteria>Terrabacteria_group>Actinomycetota>Actinomycetes>Kitasatosporales>Streptomycetaceae | GCF_018614415.1 | Streptomyces_sp._ISL-12 | J7I94_RS14400 | 76255..80178 | 2819177 | hypothetical-protein-[Streptomyces-sp.-ISL-12] |
| WP_251074221.1 | Type2.Pgl.like.BREX | [illegible] | 1338 | Bacteria>Terrabacteria_group>Actinomycetota>Actinomycetes>Kitasatosporales>Streptomycetaceae | GCF_018614515.1 | Streptomyces_sp._ISL-111 | J7E90_RS26865 | 114837..118853 | 2819175 | hypothetical-protein-[Streptomyces-sp.-ISL-111] |
| WP_215084214.1 | Type2.Pgl.like.BREX | [illegible] | 1302 | Bacteria>Terrabacteria_group>Actinomycetota>Actinomycetes>Kitasatosporales>Streptomycetaceae | GCF_018614525.1 | Streptomyces_sp._ISL-11 | J7E86_RS24340 | 24343..28251 | 2819174 | phage-resistance-protein-[Streptomyces-sp.-ISL-11] |
| WP_214971885.1 | Type2.Pgl.like.BREX | [illegible] | 1309 | Bacteria>Terrabacteria_group>Actinomycetota>Actinomycetes>Kitasatosporales>Streptomycetaceae | GCF_018614615.1 | Streptomyces_sp._ISL-100 | J7E89_RS10735 | 38743..42672 | 2819173 | DUF6079-family-protein-[Streptomyces-sp.-ISL-100] |
| WP_215208284.1 | Type2.Pgl.like.BREX | [illegible] | 1294 | Bacteria>Terrabacteria_group>Actinomycetota>Actinomycetes>Kitasatosporales>Streptomycetaceae | GCF_018638295.1 | Streptomyces_sp._CHD11 | HTV45_RS25870 | 133410..137294 | 2741325 | phage-resistance-protein-[Streptomyces-sp.-CHD11] |
| WP_215455030.1 | Type2.Pgl.like.BREX | [illegible] | 1295 | Bacteria>Terrabacteria_group>Actinomycetota>Actinomycetes>Kitasatosporales>Streptomycetaceae | GCF_018728545.1 | Streptomyces_sp._ATCC_21386 | GWK25_RS31995 | 7442023..7445910 | 2699428 | phage-resistance-protein-[Streptomyces-sp.-ATCC-21386] |
| WP_216339501.1 | Type2.Pgl.like.BREX | [illegible] | 1320 | Bacteria>Terrabacteria_group>Actinomycetota>Actinomycetes>Kitasatosporales>Streptomycetaceae | GCF_018883605.1 | Streptomyces_sp._4503 | KN815_RS01825 | 18413..22375 | 2842201 | phage-resistance-protein-[Streptomyces-sp.-4503] |
| WP_217199264.1 | Type2.Pgl.like.BREX | [illegible] | 1297 | Bacteria>Terrabacteria_group>Actinomycetota>Actinomycetes>Kitasatosporales>Streptomycetaceae | GCF_019049085.1 | Streptomyces_buecherae | KUM04_RS25325 | 52290..56183 | 2763006 | phage-resistance-protein-[Streptomyces-buecherae] |
| WP_217245869.1 | Type2.Pgl.like.BREX | [illegible] | 1323 | Bacteria>Terrabacteria_group>Actinomycetota>Actinomycetes>Kitasatosporales>Streptomycetaceae | GCF_019049125.1 | Streptomyces_sp._AC602_WCS936 | KUM03_RS05570 | 51050..55021 | 2823685 | PglY-protein-[Streptomyces-sp.-AC602_WCS936] |
| WP_217240418.1 | Type2.Pgl.like.BREX | [illegible] | 1316 | Bacteria>Terrabacteria_group>Actinomycetota>Actinomycetes>Kitasatosporales>Streptomycetaceae | GCF_019049245.1 | Streptomyces_sp._AC555_RSS877 | KUM49_RS31700 | 25237..29187 | 2823688 | hypothetical-protein-[Streptomyces-sp.-AC555_RSS877] |
| WP_218197629.1 | Type2.Pgl.like.BREX | [illegible] | 1313 | Bacteria>Terrabacteria_group>Actinomycetota>Actinomycetes>Kitasatosporales>Streptomycetaceae | GCF_019192735.1 | Kitasatospora_aureofaciens | KV557_RS01080 | 237396..241337 | 1894 | DUF6079-family-protein-[Kitasatospora-aureofaciens] |
| WP_219205063.1 | Type2.Pgl.like.BREX | [illegible] | 1299 | Bacteria>Terrabacteria_group>Actinomycetota>Actinomycetes>Kitasatosporales>Streptomycetaceae | GCF_019350955.1 | Streptomyces_griseus | DTB58_RS07815 | 2762..6661 | 1911 | phage-resistance-protein-[Streptomyces-griseus] |
| WP_219664043.1 | Type2.Pgl.like.BREX | [illegible] | 1299 | Bacteria>Terrabacteria_group>Actinomycetota>Actinomycetes>Kitasatosporales>Streptomycetaceae | GCF_019400985.1 | Streptomyces_bambusae | GPJ59_RS00105 | 29061..32960 | 1550616 | DUF6079-family-protein-[Streptomyces-bambusae] |
| WP_220649266.1 | Type2.Pgl.like.BREX | [illegible] | 1295 | Bacteria>Terrabacteria_group>Actinomycetota>Actinomycetes>Kitasatosporales>Streptomycetaceae | GCF_019599145.1 | Streptomyces_akebiae | K1J60_RS32165 | 7429892..7433779 | 2865673 | phage-resistance-protein-[Streptomyces-akebiae] |
| WP_222995822.1 | Type2.Pgl.like.BREX | [illegible] | 1299 | Bacteria>Terrabacteria_group>Actinomycetota>Actinomycetes>Kitasatosporales>Streptomycetaceae | GCF_019890645.1 | Streptomyces_sp._SP2-10 | K7864_RS12155 | 58567..62466 | 2873385 | phage-resistance-protein-[Streptomyces-sp.-SP2-10] |
| WP_223735561.1 | Type2.Pgl.like.BREX | [illegible] | 1298 | Bacteria>Terrabacteria_group>Actinomycetota>Actinomycetes>Kitasatosporales>Streptomycetaceae | GCF_020024005.1 | Streptomyces_purpurogeneiscleroticus | CCS38_RS25010 | 26602..30498 | 68259 | phage-resistance-protein-[Streptomyces-purpurogeneiscleroticus] |
| WP_223780477.1 | Type2.Pgl.like.BREX | [illegible] | 1303 | Bacteria>Terrabacteria_group>Actinomycetota>Actinomycetes>Kitasatosporales>Streptomycetaceae | GCF_020034615.1 | Streptomyces_sp._A144 | K5X85_RS23165 | 5194666..5198577 | 2871487 | hypothetical-protein-[Streptomyces-sp.-A144] |
| WP_223770120.1 | Type2.Pgl.like.BREX | [illegible] | 1299 | Bacteria>Terrabacteria_group>Actinomycetota>Actinomycetes>Kitasatosporales>Streptomycetaceae | GCF_020037025.1 | Streptomyces_huiliensis | JNW98_RS22720 | 14304..18203 | 2876027 | phage-resistance-protein-[Streptomyces-huiliensis] |
| WP_224302004.1 | Type2.Pgl.like.BREX | [illegible] | 1298 | Bacteria>Terrabacteria_group>Actinomycetota>Actinomycetes>Kitasatosporales>Streptomycetaceae | GCF_020092605.1 | Streptomyces_olivaceus | KVH22_RS20560 | 335759..339655 | 47716 | phage-resistance-protein-[Streptomyces-olivaceus] |
| WP_226071739.1 | Type2.Pgl.like.BREX | [illegible] | 1302 | Bacteria>Terrabacteria_group>Actinomycetota>Actinomycetes>Kitasatosporales>Streptomycetaceae | GCF_020410765.1 | Streptomyces_marincola | LC193_RS04365 | 1027592..1031500 | 2878388 | phage-resistance-protein-[Streptomyces-marincola] |
| WP_229318956.1 | Type2.Pgl.like.BREX | [illegible] | 1299 | Bacteria>Terrabacteria_group>Actinomycetota>Actinomycetes>Kitasatosporales>Streptomycetaceae | GCF_020640855.1 | Streptomyces_californicus | LJC81_RS05680 | 41284..45183 | 67351 | phage-resistance-protein-[Streptomyces-californicus] |
| WP_229322137.1 | Type2.Pgl.like.BREX | [illegible] | 1299 | Bacteria>Terrabacteria_group>Actinomycetota>Actinomycetes>Kitasatosporales>Streptomycetaceae | GCF_020740835.1 | Streptomyces_sp._UNOC14_S4 | K6I34_RS01490 | 11397..15296 | 2872340 | phage-resistance-protein-[Streptomyces-sp.-UNOC14_S4] |
| WP_228951258.1 | Type2.Pgl.like.BREX | [illegible] | 1318 | Bacteria>Terrabacteria_group>Actinomycetota>Actinomycetes>Kitasatosporales>Streptomycetaceae | GCF_020861705.1 | Streptomyces_sp._R1 | LMJ38_RS02080 | 462940..466896 | 1509279 | hypothetical-protein-[Streptomyces-sp.-R1] |
| WP_230535305.1 | Type2.Pgl.like.BREX | [illegible] | 1298 | Bacteria>Terrabacteria_group>Actinomycetota>Actinomycetes>Kitasatosporales>Streptomycetaceae | GCF_020881855.1 | Streptomyces_sp._OUCMDZ-3434 | HN026_RS05010 | 1183168..1187064 | 1535304 | phage-resistance-protein-[Streptomyces-sp.-OUCMDZ-3434] |
| WP_230212047.1 | Type2.Pgl.like.BREX | [illegible] | 1295 | Bacteria>Terrabacteria_group>Actinomycetota>Actinomycetes>Kitasatosporales>Streptomycetaceae | GCF_020982545.1 | Streptomyces_sp._MNU76 | E4N62_RS04710 | 1087748..1091635 | 2560026 | phage-resistance-protein-[Streptomyces-sp.-MNU76] |
| WP_233868581.1 | Type2.Pgl.like.BREX | [illegible] | 1299 | Bacteria>Terrabacteria_group>Actinomycetota>Actinomycetes>Kitasatosporales>Streptomycetaceae | GCF_021390475.1 | Streptomyces_sp._ST2-7A | LZF96_RS09305 | 68853..72752 | 2907214 | phage-resistance-protein-[Streptomyces-sp.-ST2-7A] |
| WP_222810629.1 | Type2.Pgl.like.BREX | [illegible] | 1295 | Bacteria>Terrabacteria_group>Actinomycetota>Actinomycetes>Kitasatosporales>Streptomycetaceae | GCF_021441825.1 | Streptomyces | LXH13_RS32450 | 7097003..7100890 | 1883 | MULTISPECIES:-phage-resistance-protein-[Streptomyces] |
| WP_234526785.1 | Type2.Pgl.like.BREX | [illegible] | 1295 | Bacteria>Terrabacteria_group>Actinomycetota>Actinomycetes>Kitasatosporales>Streptomycetaceae | GCF_021442415.1 | unclassified_Streptomyces | B0E36_RS09890 | 28033..31920 | 2593676 | MULTISPECIES:-phage-resistance-protein-[unclassified-Streptomyces] |
| WP_235094310.1 | Type2.Pgl.like.BREX | [illegible] | 1280 | Bacteria>Terrabacteria_group>Actinomycetota>Actinomycetes>Kitasatosporales>Streptomycetaceae | GCF_021496785.1 | Streptomyces_sp._A1-5 | HRD51_RS26445 | 5961782..5965624 | 2738410 | DUF6079-family-protein-[Streptomyces-sp.-A1-5] |
| WP_235479559.1 | Type2.Pgl.like.BREX | [illegible] | 1327 | Bacteria>Terrabacteria_group>Actinomycetota>Actinomycetes>Kitasatosporales>Streptomycetaceae | GCF_021545335.1 | Streptomyces_sp._GQFP | LUX31_RS36385 | 8065984..8069967 | 2907545 | phage-resistance-protein-[Streptomyces-sp.-GQFP] |
| WP_237319137.1 | Type2.Pgl.like.BREX | [illegible] | 1296 | Bacteria>Terrabacteria_group>Actinomycetota>Actinomycetes>Kitasatosporales>Streptomycetaceae | GCF_021696945.1 | Streptomyces_sp._JJ36 | HOY81_RS06425 | 61898..65788 | 2736645 | DUF6079-family-protein-[Streptomyces-sp.-JJ36] |
| WP_237876962.1 | Type2.Pgl.like.BREX | [illegible] | 1295 | Bacteria>Terrabacteria_group>Actinomycetota>Actinomycetes>Kitasatosporales>Streptomycetaceae | GCF_022015835.1 | Streptomyces_sp._ICN903 | L7D48_RS02640 | 29310..33197 | 2964654 | phage-resistance-protein-[Streptomyces-sp.-ICN903] |
| WP_238862156.1 | Type2.Pgl.like.BREX | [illegible] | 1313 | Bacteria>Terrabacteria_group>Actinomycetota>Actinomycetes>Kitasatosporales>Streptomycetaceae | GCF_022288725.1 | Kitasatospora_sp._A2-31 | MCM47_RS15260 | 4435..8376 | 2916414 | DUF6079-family-protein-[Kitasatospora-sp.-A2-31] |
| WP_239764494.1 | Type2.Pgl.like.BREX | [illegible] | 1295 | Bacteria>Terrabacteria_group>Actinomycetota>Actinomycetes>Kitasatosporales>Streptomycetaceae | GCF_022392825.1 | Streptomyces_sp._CL12-4 | JRC61_RS06915 | 1575845..1579732 | 2810306 | phage-resistance-protein-[Streptomyces-sp.-CL12-4] |
| WP_043405293.1 | Type2.Pgl.like.BREX | [illegible] | 1302 | Bacteria>Terrabacteria_group>Actinomycetota>Actinomycetes>Kitasatosporales>Streptomycetaceae | GCF_022414615.1 | Streptomyces_sp._MUM_16J | I3F55_RS09130 | 112948..116856 | 2791988 | hypothetical-protein-[Streptomyces-sp.-MUM-16J] |
| WP_240868184.1 | Type2.Pgl.like.BREX | [illegible] | 1294 | Bacteria>Terrabacteria_group>Actinomycetota>Actinomycetes>Kitasatosporales>Streptomycetaceae | GCF_022513565.1 | Streptomyces_sp._CME_23 | MKW14_RS31030 | 68697..72581 | 2920937 | phage-resistance-protein-[Streptomyces-sp.-CME-23] |
| WP_049978563.1 | Type2.Pgl.like.BREX | [illegible] | 1298 | Bacteria>Terrabacteria_group>Actinomycetota>Actinomycetes>Kitasatosporales>Streptomycetaceae | GCF_022695635.1 | Streptomyces | ABZ411_RS16935 | 12358..16254 | 1883 | MULTISPECIES:-hypothetical-protein-[Streptomyces] |
| WP_242708185.1 | Type2.Pgl.like.BREX | [illegible] | 1294 | Bacteria>Terrabacteria_group>Actinomycetota>Actinomycetes>Kitasatosporales>Streptomycetaceae | GCF_022698145.1 | Streptomyces_sp._7R016 | MQN93_RS03240 | 755887..759771 | 2927582 | phage-resistance-protein-[Streptomyces-sp.-7R016] |
| WP_242752939.1 | Type2.Pgl.like.BREX | [illegible] | 1303 | Bacteria>Terrabacteria_group>Actinomycetota>Actinomycetes>Kitasatosporales>Streptomycetaceae | GCF_022699385.1 | Streptomyces_tubbatahanensis | MMF93_RS18290 | 4256977..4260888 | 2923272 | phage-resistance-protein-[Streptomyces-tubbatahanensis] |
| WP_243084264.1 | Type2.Pgl.like.BREX | [illegible] | 1350 | Bacteria>Terrabacteria_group>Actinomycetota>Actinomycetes>Kitasatosporales>Streptomycetaceae | GCF_022760275.1 | Streptomyces_sp._891-h | HC362_RS14135 | 3269091..3273143 | 2720714 | hypothetical-protein-[Streptomyces-sp.-891-h] |
| WP_243378512.1 | Type2.Pgl.like.BREX | [illegible] | 1299 | Bacteria>Terrabacteria_group>Actinomycetota>Actinomycetes>Kitasatosporales>Streptomycetaceae | GCF_022807935.1 | Streptomyces_sp._HP-A2021 | MQE23_RS06345 | 1370327..1374226 | 2927875 | phage-resistance-protein-[Streptomyces-sp.-HP-A2021] |
| WP_244419349.1 | Type2.Pgl.like.BREX | [illegible] | 1324 | Bacteria>Terrabacteria_group>Actinomycetota>Actinomycetes>Kitasatosporales>Streptomycetaceae | GCF_022846675.1 | Streptomyces_hygroscopicus | HOK021_RS33880 | 7777796..7781770 | 1912 | PglY-protein-[Streptomyces-hygroscopicus] |
| WP_247860072.1 | Type2.Pgl.like.BREX | [illegible] | 1298 | Bacteria>Terrabacteria_group>Actinomycetota>Actinomycetes>Kitasatosporales>Streptomycetaceae | GCF_023096395.1 | Streptomyces_sp._WAC00276 | MWG58_RS05435 | 1231905..1235801 | 2933778 | phage-resistance-protein-[Streptomyces-sp.-WAC00276] |
| WP_248631220.1 | Type2.Pgl.like.BREX | [illegible] | 1304 | Bacteria>Terrabacteria_group>Actinomycetota>Actinomycetes>Kitasatosporales>Streptomycetaceae | GCF_023218175.1 | Streptomyces_lichenis | M1O15_RS01075 | 264491..268405 | 2306967 | phage-resistance-protein-[Streptomyces-lichenis] |
| WP_248868244.1 | Type2.Pgl.like.BREX | [illegible] | 1309 | Bacteria>Terrabacteria_group>Actinomycetota>Actinomycetes>Kitasatosporales>Streptomycetaceae | GCF_023277745.1 | Streptomyces_sp._ST13-2-2 | K9S39_RS40370 | 8888555..8892484 | 2879846 | PglY-protein-[Streptomyces-sp.-ST13-2-2] |

| Protein-ID | System | Gene-Neighbourhoods | Length | Taxonomy | Assembly-ID | Strain | Locus-Tag | Genomic-coordinates | TaxID | Fasta-defline |
|---|---|---|---|---|---|---|---|---|---|---|
| WP_249635773.1 | Type2.Pgl.like.BREX | <-SF1-SF2-Helicase+wHTH+Small-B-barrel+DUF1998<-BrxD-AAA<-PglZ<-DUF6079*<-N6-METHYLASE<-PglW\|\|?->?-><-?\|\|?->?-><-Sigma70_r2+Sigma70_r4_2 | 1299 | Bacteria>Terrabacteria_group>Actinomycetota>Actinomycetes>Kitasatosporales>Streptomycetaceae | GCF_023502025.1 | Streptomyces_kronopolitis | M1201_RS11845 | 69758..73657 | 1612435 | PglY-protein-[Streptomyces-kronopolitis] |
| WP_249836752.1 | Type2.Pgl.like.BREX | RhoN+RNase+S1->REase->DSBH+AraC-HTH+AraC-HTH-><-?\|\|ncRNA-><-SF1-SF2-Helicase+wHTH+Small-B-barrel+DUF1998<-BrxD-AAA\|\|ParB+HNH->CBS+HEPN-><-PglZ<-DUF6079*<-MNS+REase<-DUF397<-HTH_31+DUF5753\|\|ST-Hiskin-><-N6-METHYLASE<-PglW\|\|?->N6-METHYLASE->TRD+TRD-> | 1308 | Bacteria>Terrabacteria_group>Actinomycetota>Actinomycetes>Kitasatosporales>Streptomycetaceae | GCF_023516615.1 | Streptomyces_panaciradicis | M8J74_RS14335 | 101006..104932 | 1470261 | phage-resistance-protein-[Streptomyces-panaciradicis] |
| WP_275557517.1 | Type2.Pgl.like.BREX | <-WXG+ABhydrolase<-?<-?\|\|?->RNaseH-><-SF1-SF2-Helicase+wHTH+Small-B-barrel+DUF1998<-BrxD-AAA<-PglZ<-DUF6079*\|\|?-><-?<-ST-Hiskin\|\|HTH_31+DUF5753->DUF397-><-N6-METHYLASE\|\|SMODS->Schlafen+HATPase+wHTH->Schlafen+HATPase+wHTH-> | 1298 | Bacteria>Terrabacteria_group>Actinomycetota>Actinomycetes>Kitasatosporales>Streptomycetaceae | GCF_023630275.1 | Streptomyces_sp._5-6(2022) | MY491_RS01205 | 285835..289731 | 2936510 | PglY-protein-[Streptomyces-sp.-5-6(2022)] |
| WP_250743033.1 | Type2.Pgl.like.BREX | <-REase<-SF1-SF2-Helicase+wHTH+Small-B-barrel+DUF1998<-BrxD-AAA\|\|?-><-PglZ<-DUF6079*<-N6-METHYLASE<-PglW\|\|REase+DUF5655-> | 1299 | Bacteria>Terrabacteria_group>Actinomycetota>Actinomycetes>Kitasatosporales>Streptomycetaceae | GCF_023669525.1 | Streptomyces_sp._G1 | NC490_RS24275 | 34238..38137 | 361572 | phage-resistance-protein-[Streptomyces-sp.-G1] |
| WP_251415650.1 | Type2.Pgl.like.BREX | PhdYeFM_antitox->PIN-><-?<-ACET+ACET<-?\|\|?-><-SF1-SF2-Helicase+wHTH+Small-B-barrel+DUF1998<-BrxD-AAA<-PglZ<-DUF6079*\|\|?->?-><-?\|\|?->?->HTH_31+DUF5753->DUF397-><-N6-METHYLASE | 1308 | Bacteria>Terrabacteria_group>Actinomycetota>Actinomycetes>Kitasatosporales>Streptomycetaceae | GCF_023703325.1 | Streptomyces_sp._MTZ3.1 | M1E25_RS15345 | 49476..53402 | 2938945 | PglY-protein-[Streptomyces-sp.-MTZ3.1] |
| WP_252540700.1 | Type2.Pgl.like.BREX | DUF4011+SF1-SF2-Helicase+REase-><-REase<-SF1-SF2-Helicase+wHTH+Small-B-barrel+DUF1998<-BrxD-AAA\|\|?->ParB+HNH->CBS+HEPN-><-PglZ<-DUF6079*\|\|?->?-><-?<-DUF397<-HTH_31+DUF5753\|\|ST-Hiskin-><-N6-METHYLASE<-PglW | 1295 | Bacteria>Terrabacteria_group>Actinomycetota>Actinomycetes>Kitasatosporales>Streptomycetaceae | GCF_023930305.1 | Streptomyces_sp._RO-S4 | LRR80_RS21265 | 34147..38034 | 2902486 | phage-resistance-protein-[Streptomyces-sp.-RO-S4] |
| WP_254377938.1 | Type2.Pgl.like.BREX | <-SF1-SF2-Helicase+wHTH+Small-B-barrel+DUF1998<-BrxD-AAA<-PglZ<-DUF6079*<-?<-?<-ST-Hiskin<-PhdYeFM_antitox<-?<-Phage_rep\|\|?->HTH_31+DUF5753->DUF397-> | 1314 | Bacteria>Terrabacteria_group>Actinomycetota>Actinomycetes>Kitasatosporales>Streptomycetaceae | GCF_024213005.1 | Streptomyces_sp._MAR25Y5 | NLY10_RS07385 | 289073..293017 | 2962028 | hypothetical-protein-[Streptomyces-sp.-MAR25Y5] |
| WP_255137544.1 | Type2.Pgl.like.BREX | NERD->Pkinase+CR-Kinase->PglW-p->HEPN->ParA-Soj-PloopNTPase->ORC-CDC6->N6-METHYLASE-><-DUF397\|\|ST-Hiskin->DUF6079*->?-><-?<-AraC-HTH | 1303 | Bacteria>Terrabacteria_group>Actinomycetota>Actinomycetes>Kitasatosporales>Streptomycetaceae | GCF_024349085.1 | Streptomyces_somaliensis | LUX12_RS08075 | 1807469..1811380 | 78355 | hypothetical-protein-[Streptomyces-somaliensis] |
| WP_255928156.1 | Type2.Pgl.like.BREX | <-CHTH<-?\|\|?->?-><-SF1-SF2-Helicase+wHTH+Small-B-barrel+DUF1998<-BrxD-AAA\|\|ParB+HNH->CBS+HEPN-><-PglZ<-DUF6079*<-ST-Hiskin\|\|HTH_31+DUF5753->?-><-N6-METHYLASE<-PglW<-REase+SF1-SF2-Helicase\|\|?->DUF2637+HTH->McrB-AAA-> | 1314 | Bacteria>Terrabacteria_group>Actinomycetota>Actinomycetes>Kitasatosporales>Streptomycetaceae | GCF_024436055.1 | Streptomyces_rubrisoli | NON19_RS14730 | 110508..114452 | 1387313 | phage-resistance-protein-[Streptomyces-rubrisoli] |
| WP_256111799.1 | Type2.Pgl.like.BREX | METHYLASE->?-><-?<-LysM+Peptidase_M23<-?<-tRNA<-?<-SF1-SF2-Helicase+wHTH+Small-B-barrel+DUF1998<-BrxD-AAA<-PglZ<-DUF6079* | 1318 | Bacteria>Terrabacteria_group>Actinomycetota>Actinomycetes>Kitasatosporales>Streptomycetaceae | GCF_024448155.1 | Streptomyces_coelicoflavus | M4J06_RS26830 | 19266..23222 | 285562 | hypothetical-protein-[Streptomyces-coelicoflavus] |
| WP_255970744.1 | Type2.Pgl.like.BREX | AA_permease_2->TRAM+SAM-methylase-><-SF1-SF2-Helicase+wHTH+Small-B-barrel+DUF1998<-BrxD-AAA<-DndB<-PglZ<-DUF6079*<-REase+SF1-SF2-Helicase<-DUF2171<-DUF1995\|\|DUF397-><-N6-METHYLASE<-PglW\|\|?->REase-> | 1296 | Bacteria>Terrabacteria_group>Actinomycetota>Actinomycetes>Kitasatosporales>Streptomycetaceae | GCF_024448165.1 | Streptomyces_longispororuber | M4J07_RS11740 | 11824..15714 | 68230 | DUF6079-family-protein-[Streptomyces-longispororuber] |
| WP_215220766.1 | Type2.Pgl.like.BREX | ADH_N+ADH_zinc_N->BrxA-DNA-binding->?->PglW->N6-METHYLASE-><-DUF397<-HTH_31+DUF5753\|\|DUF6079*->PglZ->BrxD-AAA->SF1-SF2-Helicase+wHTH+Small-B-barrel+DUF1998-><-HTH_Tnp_4+DDE_Tnp_4\|\|HTH_Tnp_4+DDE_Tnp_4-><-CR-ATPase9<-CR-ATPase9 | 1298 | Bacteria>Terrabacteria_group>Actinomycetota>Actinomycetes>Kitasatosporales>Streptomycetaceae | GCF_024499195.1 | unclassified_Streptomyces | KNZ81_RS05560 | 1357363..1361259 | 2593676 | MULTISPECIES:-phage-resistance-protein-[unclassified-Streptomyces] |
| WP_257203796.1 | Type2.Pgl.like.BREX | ACET-><-ABhydrolase<-?<-ABhydrolase\|\|HisKA_3+HATPase_c->REC+OmpR-HTH-><-SF1-SF2-Helicase+wHTH+Small-B-barrel+DUF1998<-BrxD-AAA<-PglZ<-DUF6079*<-?<-?\|\|GntR-HTH-><-N6-METHYLASE<-ORC-CDC6<-ParA-Soj-PloopNTPase<-?<-HEPN<-PglW | 1313 | Bacteria>Terrabacteria_group>Actinomycetota>Actinomycetes>Kitasatosporales>Streptomycetaceae | GCF_024586905.1 | Streptomyces_sp._BSP1 | M7752_RS13310 | 125622..129563 | 2944804 | hypothetical-protein-[Streptomyces-sp.-BSP1] |
| WP_257546101.1 | Type2.Pgl.like.BREX | <-MerR-HTH\|\|REase->DSBH+AraC-HTH+AraC-HTH->ncRNA-><-SF1-SF2-Helicase+wHTH+Small-B-barrel+DUF1998<-BrxD-AAA\|\|ParB+HNH->CBS+HEPN-><-PglZ<-DUF6079*<-?<-?<-?\|\|?->APATPase+TPR+TPR+TPR-><-DUF397<-HTH_31+DUF5753\|\|ST-Hiskin-><-N6-METHYLASE | 1308 | Bacteria>Terrabacteria_group>Actinomycetota>Actinomycetes>Kitasatosporales>Streptomycetaceae | GCF_024612015.1 | Streptomyces_sp._CA-210063 | JIX56_RS33355 | 7634726..7638652 | 2801029 | phage-resistance-protein-[Streptomyces-sp.-CA-210063] |
| WP_262037360.1 | Type2.Pgl.like.BREX | <-Peptidase_M22<-SF1-SF2-Helicase+wHTH+Small-B-barrel+DUF1998<-BrxD-AAA<-PglZ<-DUF6079*<-N6-METHYLASE<-PglW\|\|?->?->HTH_Tnp_1+HTH_21+rve+rve_3-> | 1307 | Bacteria>Terrabacteria_group>Actinomycetota>Actinomycetes>Kitasatosporales>Streptomycetaceae | GCF_024721345.1 | unclassified_Streptomyces | NE399_RS03500 | 98241..102164 | 2593676 | MULTISPECIES:-PglY-protein-[unclassified-Streptomyces] |
| WP_260636294.1 | Type2.Pgl.like.BREX | <-JetB<-PadR-wHTH\|\|PglW-><-ORC-CDC6<-ParA-Soj-PloopNTPase<-HEPN\|\|N6-METHYLASE-><-DUF397\|\|ST-Hiskin->DUF6079*->PglZ->BrxD-AAA->SF1-SF2-Helicase+wHTH+Small-B-barrel+DUF1998-><-?<-NUDIX | 1324 | Bacteria>Terrabacteria_group>Actinomycetota>Actinomycetes>Kitasatosporales>Streptomycetaceae | GCF_025252265.1 | Streptomyces_angustmyceticus | NW605_RS13680 | 3192049..3196023 | 285578 | PglY-protein-[Streptomyces-angustmyceticus] |
| WP_260826689.1 | Type2.Pgl.like.BREX | <-PhdYeFM_antitox\|\|PglW->HEPN->ParA-Soj-PloopNTPase->ORC-CDC6->N6-METHYLASE-><-DUF397<-HTH_31+DUF5753\|\|ST-Hiskin->DUF6079*-> | 1310 | Bacteria>Terrabacteria_group>Actinomycetota>Actinomycetes>Kitasatosporales>Streptomycetaceae | GCF_025290915.1 | Streptomyces_sp._15-116A | N4P33_RS13725 | 22419..26351 | 2259035 | hypothetical-protein-[Streptomyces-sp.-15-116A] |
| WP_261725450.1 | Type2.Pgl.like.BREX | PIN-><-SF1-SF2-Helicase+wHTH+Small-B-barrel+DUF1998<-BrxD-AAA<-PglZ<-DUF6079*<-N6-METHYLASE<-PglW\|\|DUF6731->?-><-REase+SF1-SF2-Helicase<-?<-HTH_3 | 1302 | Bacteria>Terrabacteria_group>Actinomycetota>Actinomycetes>Kitasatosporales>Streptomycetaceae | GCF_025399775.1 | Streptomyces_fulvoviolaceus | N4G65_RS27540 | 6857..10765 | 285535 | PglY-protein-[Streptomyces-fulvoviolaceus] |
| WP_261722751.1 | Type2.Pgl.like.BREX | REase+PNTD1+PNTD2->PIWI->Small-B-barrel+SF1-SF2-Helicase+RdgC+REase-><-?<-SF1-SF2-Helicase+wHTH+Small-B-barrel+DUF1998<-BrxD-AAA<-PglZ<-DUF6079*<-N6-METHYLASE<-PglW<-ncRNA<-?<-?\|\|?->?->SIR2-><-HAD | 1301 | Bacteria>Terrabacteria_group>Actinomycetota>Actinomycetes>Kitasatosporales>Streptomycetaceae | GCF_025399775.1 | Streptomyces_fulvoviolaceus | N4G65_RS01450 | 26727..30632 | 285535 | phage-resistance-protein-[Streptomyces-fulvoviolaceus] |
| WP_261719041.1 | Type2.Pgl.like.BREX | Peptidase_C14->PglW-><-ORC-CDC6<-ParA-Soj-PloopNTPase<-HEPN\|\|N6-METHYLASE-><-DUF397<-HTH_31+DUF5753\|\|ST-Hiskin->DUF6079*->PglZ->BrxD-AAA->SF1-SF2-Helicase+wHTH+Small-B-barrel+DUF1998->DDE_Tnp_1_4-><-tRNA<-?<-Calcineurin | 1324 | Bacteria>Terrabacteria_group>Actinomycetota>Actinomycetes>Kitasatosporales>Streptomycetaceae | GCF_025402955.1 | Streptomyces_lusitanus | N6J21_RS20815 | 4654985..4658959 | 68232 | PglY-protein-[Streptomyces-lusitanus] |
| WP_263277353.1 | Type2.Pgl.like.BREX | DCM->?->?->PglW->N6-METHYLASE->DUF6079*->PglZ->BrxD-AAA->SF1-SF2-Helicase+wHTH+Small-B-barrel+DUF1998-><-NUDIX<-REase+DUF5655\|\|<-PIWI<-HNH | 1299 | Bacteria>Terrabacteria_group>Actinomycetota>Actinomycetes>Kitasatosporales>Streptomycetaceae | GCF_025642175.1 | Streptomyces_sp._HUAS_14-6 | N8186_RS06515 | 1385275..1389174 | 2981135 | phage-resistance-protein-[Streptomyces-sp.-HUAS-14-6] |
| WP_264244287.1 | Type2.Pgl.like.BREX | ABhydrolase->?->?-><-?\|\|ACET->?-><-?<-DUF6079-p*<-?<-?\|\|GntR-HTH-><-N6-METHYLASE-p<-HTH_Tnp_4+DDE_Tnp_4 | 464 | Bacteria>Terrabacteria_group>Actinomycetota>Actinomycetes>Kitasatosporales>Streptomycetaceae | GCF_025854275.1 | Streptomyces_peucetius | OGH68_RS14745 | 3255631..3257025 | 1950 | hypothetical-protein-[Streptomyces-peucetius] |
| WP_265945318.1 | Type2.Pgl.like.BREX | TetR-HTH->Peptidase_S8-><-SF1-SF2-Helicase+wHTH+Small-B-barrel+DUF1998<-BrxD-AAA<-PglZ<-DUF6079*\|\|?->?->?->?->?->?->?->?->GNTR-HTH-> | 1298 | Bacteria>Terrabacteria_group>Actinomycetota>Actinomycetes>Kitasatosporales>Streptomycetaceae | GCF_026262575.1 | Streptomyces_sp._JHD_1 | OSB71_RS06795 | 60465..64361 | 2984864 | phage-resistance-protein-[Streptomyces-sp.-JHD-1] |
| WP_266396801.1 | Type2.Pgl.like.BREX | REase+DUF5655-><-pPIWI+SF1-SF2-Helicase<-pPIWI\|\|DUF4096+DDE_Tnp_1+DDE_Tnp_1_2->PglW->N6-METHYLASE->DUF6079*->PglZ->BrxD-AAA->SF1-SF2-Helicase+wHTH+Small-B-barrel+DUF1998->?->HTH+MPTase-> | 1297 | Bacteria>Terrabacteria_group>Actinomycetota>Actinomycetes>Kitasatosporales>Streptomycetaceae | GCF_026339775.1 | Streptomyces_anulatus | OOK53_RS00785 | 206425..210318 | 1892 | phage-resistance-protein-[Streptomyces-anulatus] |
| WP_266534767.1 | Type2.Pgl.like.BREX | <-PRTase\|\|?->?->?->tRNA-><-SF1-SF2-Helicase+wHTH+Small-B-barrel+DUF1998<-BrxD-AAA<-PglZ<-DUF6079*<-?<-?<-?<-?\|\|GntR-HTH-><-N6-METHYLASE\|\|?-><-ORC-CDC6<-ParA-Soj-PloopNTPase | 1327 | Bacteria>Terrabacteria_group>Actinomycetota>Actinomycetes>Kitasatosporales>Streptomycetaceae | GCF_026339795.1 | Streptomyces_anulatus | OG890_RS18040 | 4024087..4028070 | 1892 | hypothetical-protein-[Streptomyces-anulatus] |
| WP_266642610.1 | Type2.Pgl.like.BREX | <-HTH_31+DUF5753<-?\|\|?->?-><-ST-Hiskin->XIS-HTH+DUF433->PIN->?->?->DUF6079*->PglZ->BrxD-AAA->SF1-SF2-Helicase+wHTH+Small-B-barrel+DUF1998-><-ACET<-RADICAL-SAM<-?<-APATPase+TPR+TPR+TPR+TPR+TPR+TPR+TPR<-?<-BrxA-DNA-binding | 1296 | Bacteria>Terrabacteria_group>Actinomycetota>Actinomycetes>Kitasatosporales>Streptomycetaceae | GCF_026339975.1 | Streptomyces_sp._NBC_01500 | OG576_RS16220 | 3684738..3688628 | 2903886 | hypothetical-protein-[Streptomyces-sp.-NBC_01500] |
| WP_266487410.1 | Type2.Pgl.like.BREX | N6-METHYLASE-><-DUF397\|\|<-ABhydrolase\|\|ST-Hiskin->PhdYeFM_antitox->PIN-><-?<-?\|\|DUF6079*->PglZ->BrxD-AAA->SF1-SF2-Helicase+wHTH+Small-B-barrel+DUF1998-><-RNaseH<-REase+N6_Mtase<-TRD+TRD<-PIN<-XIS-HTH<-SbcC | 1299 | Bacteria>Terrabacteria_group>Actinomycetota>Actinomycetes>Kitasatosporales>Streptomycetaceae | GCF_026339995.1 | Streptomyces_sp._NBC_01481 | OOK40_RS29220 | 698872..702771 | 2975869 | PglY-protein-[Streptomyces-sp.-NBC_01481] |
| WP_266550550.1 | Type2.Pgl.like.BREX | N6-METHYLASE-><-ST-Hiskin\|\|HTH_31+DUF5753->DUF397-><-?<-?\|\|?->APATPase+TPR+TPR+TPR->DUF6079*->PglZ-><-CBS+HEPN<-ParB+HNH\|\|BrxD-AAA->SF1-SF2-Helicase+wHTH+Small-B-barrel+DUF1998->REase->DUF4258-> | 1295 | Bacteria>Terrabacteria_group>Actinomycetota>Actinomycetes>Kitasatosporales>Streptomycetaceae | GCF_026340025.1 | Streptomyces_viridodiastaticus | OG359_RS29015 | 6295887..6299774 | 67382 | phage-resistance-protein-[Streptomyces-viridodiastaticus] |
| WP_266662808.1 | Type2.Pgl.like.BREX | REase+SF1-SF2-Helicase->?->CASPASE+RNA-Helicase+REase->PglW->N6-METHYLASE-><-DUF397<-HTH_31+DUF5753\|\|DUF1995->DUF2171->DUF6079*->PglZ-><-CBS+HEPN<-ParB+HNH\|\|BrxD-AAA->SF1-SF2-Helicase+wHTH+Small-B-barrel+DUF1998->DUF4011+SF1-SF2-Helicase+REase-><-TRAM+SAM-methylase<-AA_permease_2 | 1302 | Bacteria>Terrabacteria_group>Actinomycetota>Actinomycetes>Kitasatosporales>Streptomycetaceae | GCF_026340075.1 | Streptomyces_sp._NBC_01443 | OG755_RS09945 | 2239790..2243698 | 2903868 | phage-resistance-protein-[Streptomyces-sp.-NBC_01443] |
| WP_267065183.1 | Type2.Pgl.like.BREX | <-REase\|\|?->HKD-DNase+HKD-DNase-><-REase\|\|?-><-SF1-SF2-Helicase+wHTH+Small-B-barrel+DUF1998<-BrxD-AAA<-PglZ<-DUF6079*\|\|?-><-PIN<-XIS-HTH+DUF433<-?<-ST-Hiskin<-N6-METHYLASE<-ORC-CDC6<-ParA-Soj-PloopNTPase | 1294 | Bacteria>Terrabacteria_group>Actinomycetota>Actinomycetes>Kitasatosporales>Streptomycetaceae | GCF_026340135.1 | Streptomyces_microflavus | OG249_RS13840 | 3118904..3122788 | 1919 | hypothetical-protein-[Streptomyces-microflavus] |
| WP_266738643.1 | Type2.Pgl.like.BREX | <-REase<-?\|\|PglW->N6-METHYLASE-><-DUF397<-HTH_31+DUF5753\|\|ST-Hiskin-><-PIN<-CopG/RHH\|\|DUF6079*->PglZ->BrxD-AAA->SF1-SF2-Helicase+wHTH+Small-B-barrel+DUF1998-><-TRAM+SAM-methylase<-AA_permease_2 | 1298 | Bacteria>Terrabacteria_group>Actinomycetota>Actinomycetes>Kitasatosporales>Streptomycetaceae | GCF_026340175.1 | Streptomyces_platensis | OG775_RS34870 | 29984..33880 | 58346 | phage-resistance-protein-[Streptomyces-platensis] |
| WP_266868158.1 | Type2.Pgl.like.BREX | SF1-SF2-Helicase+REase->N6-METHYLASE-><-DUF397<-HTH_31+DUF5753\|\|ST-Hiskin->OLD-ABC+TOPRIM-><-?\|\|PIN->?->DUF6079*->PglZ->BrxD-AAA->SF1-SF2-Helicase+wHTH+Small-B-barrel+DUF1998-><-?\|\|HTH+REase-><-AbiEii+ZnR<-AbiEi_4+AbiEi_1<-?<-APATPase+TPR+TPR+TPR+TPR+TPR+TPR+TPR | 1279 | Bacteria>Terrabacteria_group>Actinomycetota>Actinomycetes>Kitasatosporales>Streptomycetaceae | GCF_026340305.1 | Streptomyces_sp._NBC_01381 | OG453_RS15075 | 3225829..3229668 | 2903845 | PglY-protein-[Streptomyces-sp.-NBC_01381] |
| WP_266427954.1 | Type2.Pgl.like.BREX | <-CASPASE\|\|TPR+TPR+TPR-><-?\|\|?-><-SF1-SF2-Helicase+wHTH+Small-B-barrel+DUF1998<-BrxD-AAA<-DndB<-PglZ<-DUF6079*<-REase+SF1-SF2-Helicase<-DUF2171<-DUF1995\|\|HTH_31+DUF5753->DUF397-><-N6-METHYLASE<-PglW | 1306 | Bacteria>Terrabacteria_group>Actinomycetota>Actinomycetes>Kitasatosporales>Streptomycetaceae | GCF_026340595.1 | Streptomyces_sp._NBC_01363 | OG611_RS31550 | 814146..818066 | 2903840 | phage-resistance-protein-[Streptomyces-sp.-NBC_01363] |
| WP_266768081.1 | Type2.Pgl.like.BREX | HTH_Tnp->?->?-><-?<-MORC+DUF3684+REase\|\|<-?<-?\|\|PglW->N6-METHYLASE->DUF6079*->PglZ->BrxD-AAA->SF1-SF2-Helicase+wHTH+Small-B-barrel+DUF1998-><-HTH_3\|\|Phage_int_SAM_3+Phage_integrase->HTH_Tnp_4+DDE_Tnp_4-> | 1303 | Bacteria>Terrabacteria_group>Actinomycetota>Actinomycetes>Kitasatosporales>Streptomycetaceae | GCF_026340815.1 | Streptomyces_sp._NBC_01239 | OG601_RS08495 | 1832338..1836249 | 2903792 | PglY-protein-[Streptomyces-sp.-NBC_01239] |
| WP_266905180.1 | Type2.Pgl.like.BREX | METHYLASE->HEPN+VirB11+NACHT+FGS->?->SLATT->?-><-?\|\|DUF1995->DUF2171->REase+SF1-SF2-Helicase->DUF6079*->PglZ-><-RNaseH\|\|BrxD-AAA->SF1-SF2-Helicase+wHTH+Small-B-barrel+DUF1998-><-Peptidase_S15+PepX_C<-?<-ABhydrolase | 1298 | Bacteria>Terrabacteria_group>Actinomycetota>Actinomycetes>Kitasatosporales>Streptomycetaceae | GCF_026340885.1 | Streptomyces_sp._NBC_01242 | OG497_RS09405 | 2040067..2043963 | 2903795 | phage-resistance-protein-[Streptomyces-sp.-NBC_01242] |
| WP_266503384.1 | Type2.Pgl.like.BREX | <-AAA-ATPase\|\|?-><-?<-SF1-SF2-Helicase+wHTH+Small-B-barrel+DUF1998<-BrxD-AAA\|\|ParB+HNH->CBS+HEPN-><-PglZ<-DUF6079*<-MNS+REase<-DUF397<-HTH_31+DUF5753\|\|ST-Hiskin-><-N6-METHYLASE<-PglW<-?<-?\|\|HDRnase-> | 1314 | Bacteria>Terrabacteria_group>Actinomycetota>Actinomycetes>Kitasatosporales>Streptomycetaceae | GCF_026341515.1 | Streptomyces_sp._NBC_00878 | OHA11_RS35725 | 8238632..8242576 | 2975854 | phage-resistance-protein-[Streptomyces-sp.-NBC_00878] |
| WP_266688157.1 | Type2.Pgl.like.BREX | PglW->N6-METHYLASE-><-DUF397<-HTH_31+DUF5753\|\|DUF1995->DUF2171->REase+SF1-SF2-Helicase->DUF6079*->PglZ->DndB->BrxD-AAA->SF1-SF2-Helicase+wHTH+Small-B-barrel+DUF1998-><-TRAM+SAM-methylase<-AA_permease_2<-PALP<-GNTR-HTH | 1306 | Bacteria>Terrabacteria_group>Actinomycetota>Actinomycetes>Kitasatosporales>Streptomycetaceae | GCF_026341545.1 | unclassified_Streptomyces | OOK30_RS07470 | 1710511..1714431 | 2593676 | MULTISPECIES:-phage-resistance-protein-[unclassified-Streptomyces] |
| WP_266585102.1 | Type2.Pgl.like.BREX | DUF2171->?->SAM-METHYLASE->?-><-?<-?\|\|?->APATPase+TPR+TPR+TPR->DUF6079*->PglZ-><-CBS+CBS+HEPN<-ParB+HNH\|\|BrxD-AAA->SF1-SF2-Helicase+wHTH+Small-B-barrel+DUF1998-><-REC+OmpR-HTH<-HisKA_3+HATPase_c\|\|Phage_holin-> | 1294 | Bacteria>Terrabacteria_group>Actinomycetota>Actinomycetes>Kitasatosporales>Streptomycetaceae | GCF_026341995.1 | Streptomyces_sp._NBC_00452 | OOK12_RS10975 | 2378201..2382085 | 2975746 | phage-resistance-protein-[Streptomyces-sp.-NBC_00452] |
| WP_266728674.1 | Type2.Pgl.like.BREX | HISKIN->REC+OmpR-HTH-><-?\|\|NTP_transf_3->?-><-SF1-SF2-Helicase+wHTH+Small-B-barrel+DUF1998<-BrxD-AAA<-PglZ<-DUF6079*<-?<-PIN<-OLD-ABC+TOPRIM<-ST-Hiskin<-SF1-SF2-Helicase\|\|?->HTH_31+DUF5753->DUF397-><-N6-METHYLASE | 1285 | Bacteria>Terrabacteria_group>Actinomycetota>Actinomycetes>Kitasatosporales>Streptomycetaceae | GCF_026342115.1 | Streptomyces_sp._NBC_00378 | OOK13_RS07585 | 1669600..1673457 | 2975732 | PglY-protein-[Streptomyces-sp.-NBC_00378] |
| WP_266582205.1 | Type2.Pgl.like.BREX | <-TRD+TRD<-N6-METHYLASE<-?\|\|PglW->N6-METHYLASE-><-ST-Hiskin\|\|HTH_31+DUF5753->DUF397->MNS+REase->DUF6079*->PglZ-><-CBS+CBS+HEPN<-ParB+HNH\|\|BrxD-AAA->SF1-SF2-Helicase+wHTH+Small-B-barrel+DUF1998-><-ncRNA<-DSBH+AraC-HTH+AraC-HTH\|\|MerR-HTH->?-><-RhoN+RNase+S1 | 1308 | Bacteria>Terrabacteria_group>Actinomycetota>Actinomycetes>Kitasatosporales>Streptomycetaceae | GCF_026342135.1 | Streptomyces_sp._NBC_00365 | OOK36_RS13475 | 2959938..2963864 | 2975726 | phage-resistance-protein-[Streptomyces-sp.-NBC_00365] |
| WP_267050657.1 | Type2.Pgl.like.BREX | HTH_Tnp->?->?-><-?<-MORC+DUF3684+REase\|\|<-?<-?\|\|PglW->N6-METHYLASE->DUF6079*->PglZ->BrxD-AAA->SF1-SF2-Helicase+wHTH+Small-B-barrel+DUF1998-><-HTH_3\|\|Phage_int_SAM_3+Phage_integrase-><-?<-?\|\|APATPase+TPR+TPR+TPR+TPR+TPR+TPR+TPR-> | 1302 | Bacteria>Terrabacteria_group>Actinomycetota>Actinomycetes>Kitasatosporales>Streptomycetaceae | GCF_026342575.1 | Streptomyces_sp._NBC_00198 | OHA93_RS23215 | 1413305..1417213 | 2975677 | PglY-protein-[Streptomyces-sp.-NBC_00198] |
| WP_266923860.1 | Type2.Pgl.like.BREX | <-Phage_holin\|\|HisKA_3+HATPase_c->REC+OmpR-HTH-><-SF1-SF2-Helicase+wHTH+Small-B-barrel+DUF1998<-BrxD-AAA\|\|ParB+HNH->CBS+CBS+HEPN-><-PglZ<-DUF6079*<-APATPase+TPR+TPR+TPR<-?\|\|?->?-><-?<-SAM-METHYLASE<-?<-?\|\|TrmB-> | 1294 | Bacteria>Terrabacteria_group>Actinomycetota>Actinomycetes>Kitasatosporales>Streptomycetaceae | GCF_026342595.1 | Streptomyces_sp._NBC_00201 | OG895_RS10055 | 2209586..2213470 | 2975679 | phage-resistance-protein-[Streptomyces-sp.-NBC_00201] |
| WP_266840683.1 | Type2.Pgl.like.BREX | DUF2171->?->SAM-METHYLASE->?-><-?<-?\|\|?->APATPase+TPR+TPR+TPR->DUF6079*->PglZ-><-CBS+CBS+HEPN<-ParB+HNH\|\|BrxD-AAA->SF1-SF2-Helicase+wHTH+Small-B-barrel+DUF1998->REC+OmpR-HTH->HisKA_3+HATPase_c\|\|Phage_holin-> | 1294 | Bacteria>Terrabacteria_group>Actinomycetota>Actinomycetes>Kitasatosporales>Streptomycetaceae | GCF_026342695.1 | Streptomyces_sp._NBC_00183 | OG260_RS12155 | 2626615..2630499 | 2903633 | phage-resistance-protein-[Streptomyces-sp.-NBC_00183] |
| WP_267056864.1 | Type2.Pgl.like.BREX | AA_permease_2->TRAM+SAM-methylase-><-ANK+ANK<-SF1-SF2-Helicase+wHTH+Small-B-barrel+DUF1998<-BrxD-AAA<-PglZ<-DUF6079*<-N6-METHYLASE<-PglW\|\|?->?-><-PIN<-MetJarc-HTH | 1214 | Bacteria>Terrabacteria_group>Actinomycetota>Actinomycetes>Kitasatosporales>Streptomycetaceae | GCF_026343295.1 | Streptomyces_sp._NBC_00083 | OOJ98_RS23245 | 1548585..1552229 | 2975647 | PglY-protein-[Streptomyces-sp.-NBC_00083] |
| WP_266614476.1 | Type2.Pgl.like.BREX | REase+SF1-SF2-Helicase->?-><-REase+SF1-SF2-Helicase\|\|PglW->N6-METHYLASE-><-DUF397<-HTH_31+DUF5753\|\|DUF1995->DUF2171->DUF6079*->PglZ->DndB->BrxD-AAA->SF1-SF2-Helicase+wHTH+Small-B-barrel+DUF1998-><-TRAM+SAM-methylase<-AA_permease_2 | 1296 | Bacteria>Terrabacteria_group>Actinomycetota>Actinomycetes>Kitasatosporales>Streptomycetaceae | GCF_026343545.1 | Streptomyces_sp._NBC_00047 | OHB39_RS29685 | 1749641..1753531 | 2975627 | phage-resistance-protein-[Streptomyces-sp.-NBC_00047] |
| WP_267241498.1 | Type2.Pgl.like.BREX | <-DUF5086\|\|DUF4096+DDE_Tnp_1+DDE_Tnp_1_2-><-SF1-SF2-Helicase+wHTH+Small-B-barrel+DUF1998<-BrxD-AAA<-PglZ<-DUF6079*<-N6-METHYLASE\|\|HEPN->ParA-Soj-PloopNTPase->ORC-CDC6-><-PglW\|\|PadR-wHTH->JetB->?->DUF3323+TOPRIM-> | 1322 | Bacteria>Terrabacteria_group>Actinomycetota>Actinomycetes>Kitasatosporales>Streptomycetaceae | GCF_026420845.1 | Streptomyces_sp._PR69 | OF819_RS01825 | 9515..13483 | 2984950 | PglY-protein-[Streptomyces-sp.-PR69] |
| WP_267841177.1 | Type2.Pgl.like.BREX | SAM-METHYLASE->ACET-><-Pkinase\|\|REase-><-SF1-SF2-Helicase+wHTH+Small-B-barrel+DUF1998<-BrxD-AAA<-PglZ<-DUF6079*<-N6-METHYLASE<-PglW\|\|?-><-?\|\|BirA-HTH->XIS-HTH->Phage_integrase-><-tRNA\|\|GTPase-AIG->sGTP-> | 1299 | Bacteria>Terrabacteria_group>Actinomycetota>Actinomycetes>Kitasatosporales>Streptomycetaceae | GCF_026627185.1 | unclassified_Streptomyces | OTB23_RS20485 | 12477..16376 | 2593676 | MULTISPECIES:-phage-resistance-protein-[unclassified-Streptomyces] |
| WP_267842808.1 | Type2.Pgl.like.BREX | <-JetB<-?\|\|?->PglW->N6-METHYLASE-><-DUF397<-HTH_31+DUF5753\|\|DUF1995->DUF2171->DUF6079*->PglZ-><-CBS+HEPN<-ParB+HNH\|\|BrxD-AAA->SF1-SF2-Helicase+wHTH+Small-B-barrel+DUF1998-> | 1299 | Bacteria>Terrabacteria_group>Actinomycetota>Actinomycetes>Kitasatosporales>Streptomycetaceae | GCF_026627185.1 | unclassified_Streptomyces | OTB23_RS33960 | 52811..56710 | 2593676 | MULTISPECIES:-DUF6079-family-protein-[unclassified-Streptomyces] |
| WP_268974408.1 | Type2.Pgl.like.BREX | MarR-HTH->NTP_transf_9->DUF4011+SF1-SF2-Helicase+REase-><-SF1-SF2-Helicase+wHTH+Small-B-barrel+DUF1998<-BrxD-AAA\|\|ParB+HNH->CBS+HEPN-><-PglZ<-DUF6079*<-DUF397<-HTH_31+DUF5753\|\|ST-Hiskin-><-N6-METHYLASE<-PglW\|\|REase+SF1-SF2-Helicase->EVE+HNH->TOPRIM-><-HTH_3 | 1320 | Bacteria>Terrabacteria_group>Actinomycetota>Actinomycetes>Kitasatosporales>Streptomycetaceae | GCF_026898115.1 | Streptomyces_sp._S465 | N6H00_RS26955 | 6348483..6352445 | 2979468 | phage-resistance-protein-[Streptomyces-sp.-S465] |
| WP_269120498.1 | Type2.Pgl.like.BREX | <-HAD+HAD<-?<-?<-N6-METHYLASE<-SF1-SF2-Helicase+wHTH+Small-B-barrel+DUF1998<-BrxD-AAA<-PglZ<-DUF6079*<-ACET<-Pkinase<-?<-?\|\|HTH_3->HD-><-?<-?<-PSBP | 1324 | Bacteria>Terrabacteria_group>Actinomycetota>Actinomycetes>Kitasatosporales>Streptomycetaceae | GCF_026965885.1 | Streptomyces_sp._HB2AG | ODJ75_RS20345 | 11144..15118 | 2983400 | hypothetical-protein-[Streptomyces-sp.-HB2AG] |
| WP_269735388.1 | Type2.Pgl.like.BREX | <-Pkinase+pi3KINASE\|\|PglW->N6-METHYLASE-><-DUF397<-HTH_31+DUF5753\|\|ABhydrolase->DUF1995->DUF2171->DUF6079*->PglZ->BrxD-AAA->SF1-SF2-Helicase+wHTH+Small-B-barrel+DUF1998-><-?\|\|DEOR-HTH+ISOCOT-><-?\|\|?->?-><-GNTR-HTH+Aminotran_1_2 | 1301 | Bacteria>Terrabacteria_group>Actinomycetota>Actinomycetes>Kitasatosporales>Streptomycetaceae | GCF_027460205.1 | Streptomyces_sp._WMMC897 | O3441_RS16190 | 3323700..3327605 | 3014782 | phage-resistance-protein-[Streptomyces-sp.-WMMC897] |
| WP_269744057.1 | Type2.Pgl.like.BREX | GNTR-HTH+Aminotran_1_2-><-?<-?\|\|?-><-DEOR-HTH+ISOCOT\|\|?-><-SF1-SF2-Helicase+wHTH+Small-B-barrel+DUF1998<-BrxD-AAA<-PglZ<-DUF6079*<-DUF2171<-DUF1995<-ABhydrolase\|\|HTH_31+DUF5753->DUF397-><-N6-METHYLASE<-PglW\|\|Pkinase+pi3KINASE-> | 1301 | Bacteria>Terrabacteria_group>Actinomycetota>Actinomycetes>Kitasatosporales>Streptomycetaceae | GCF_027460245.1 | Streptomyces_sp._WMMC1477 | O7609_RS21380 | 4799252..4803157 | 3015155 | phage-resistance-protein-[Streptomyces-sp.-WMMC1477] |
| WP_271665636.1 | Type2.Pgl.like.BREX | <-ORC-CDC6<-ParA-Soj-PloopNTPase<-HEPN\|\|N6-METHYLASE-><-DUF397<-HTH_31+DUF5753\|\|ST-Hiskin-><-?<-BrxA-DNA-binding\|\|DUF6079*->PglZ->BrxD-AAA->SF1-SF2-Helicase+wHTH+Small-B-barrel+DUF1998->MNS+REase-><-?<-?\|\|SF1-SF2-Helicase-><-MODE-HTH+PBPII | 1309 | Bacteria>Terrabacteria_group>Actinomycetota>Actinomycetes>Kitasatosporales>Streptomycetaceae | GCF_028198125.1 | Streptomyces_sp._HUAS_31 | PGH47_RS27530 | 6141637..6145566 | 3020055 | PglY-protein-[Streptomyces-sp.-HUAS-31] |
| WP_271684120.1 | Type2.Pgl.like.BREX | <-HTH_31+DUF5753<-DUF397<-HTH_31+DUF5753<-PCMT+METHYLASE<-?\|\|HTH_3->HEPN->DDE_Tnp_ISL3->REase+SF1-SF2-Helicase->DUF6079*->PglZ->PhdYeFM_antitox->BrxD-AAA->SF1-SF2-Helicase+wHTH+Small-B-barrel+DUF1998-><-?<-DDE_Tnp\|\|CDP-OH_P_transf-><-SAM-METHYLASE | 1306 | Bacteria>Terrabacteria_group>Actinomycetota>Actinomycetes>Kitasatosporales>Streptomycetaceae | GCF_028200735.1 | Streptomyces_sp._ACA25 | PJ985_RS14290 | 102103..106023 | 3022596 | phage-resistance-protein-[Streptomyces-sp.-ACA25] |
| WP_271684148.1 | Type2.Pgl.like.BREX | <-FGS<-DUF397<-nSTAND1+SbcC+ABC-ATPase+SbcC\|\|?-><-?\|\|?-><-SF1-SF2-Helicase+wHTH+Small-B-barrel+DUF1998<-BrxD-AAA<-PglZ<-DUF6079*\|\|SF1-SF2-Helicase+HKD-DNase->?->DUF397-><-PCMT+METHYLASE<-ST-Hiskin\|\|HTH_31+DUF5753->DUF397-><-N6-METHYLASE\|\|HEPN-> | 1326 | Bacteria>Terrabacteria_group>Actinomycetota>Actinomycetes>Kitasatosporales>Streptomycetaceae | GCF_028200735.1 | Streptomyces_sp._ACA25 | PJ985_RS14445 | 141560..145540 | 3022596 | PglY-protein-[Streptomyces-sp.-ACA25] |
| WP_272168111.1 | Type2.Pgl.like.BREX | NACHT+68TM-wHTH-><-?\|\|?->?-><-PIN<-MetJarc-HTH\|\|DUF6079*->PglZ-><-CBS+CBS+HEPN<-ParB+HNH\|\|BrxD-AAA->SF1-SF2-Helicase+wHTH+Small-B-barrel+DUF1998-><-?\|\|?->?-><-REC+OmpR-HTH | 1295 | Bacteria>Terrabacteria_group>Actinomycetota>Actinomycetes>Kitasatosporales>Streptomycetaceae | GCF_028401405.1 | Streptomyces_heilongjiangensis | PO613_RS02325 | 86488..90375 | 945052 | phage-resistance-protein-[Streptomyces-heilongjiangensis] |
| WP_274043588.1 | Type2.Pgl.like.BREX | DOC-><-HISKIN<-?<-METHYLASE<-SF1-SF2-Helicase+wHTH+Small-B-barrel+DUF1998<-BrxD-AAA<-DndB<-PglZ<-DUF6079*<-REase+SF1-SF2-Helicase<-DUF2171\|\|HTH_31+DUF5753->DUF397-><-N6-METHYLASE<-PglW\|\|?->CASPASE+RNA-Helicase+REase-> | 1294 | Bacteria>Terrabacteria_group>Actinomycetota>Actinomycetes>Kitasatosporales>Streptomycetaceae | GCF_028657195.3 | Streptomyces_iconiensis | NMN56_RS24265 | 206305..210189 | 1384038 | phage-resistance-protein-[Streptomyces-iconiensis] |
| WP_275462590.1 | Type2.Pgl.like.BREX | <-SbcCD_C<-JetB<-?\|\|PglW->N6-METHYLASE-><-DUF397<-HTH_31+DUF5753\|\|DUF1995->DUF2171->DUF6079*->PglZ-><-CBS+HEPN<-ParB+HNH\|\|BrxD-AAA->SF1-SF2-Helicase+wHTH+Small-B-barrel+DUF1998->REase+SF1-SF2-Helicase->?-><-REase+SF1-SF2-Helicase<-TRAM+SAM-methylase | 1299 | Bacteria>Terrabacteria_group>Actinomycetota>Actinomycetes>Kitasatosporales>Streptomycetaceae | GCF_029167025.1 | Streptomyces_noursei | POD74_RS11305 | 2584662..2588561 | 1971 | phage-resistance-protein-[Streptomyces-noursei] |
| WP_051778415.1 | Type2.Pgl.like.BREX | <-ZnR+REase<-DUF3323+TOPRIM<-?<-?<-?<-?<-SF1-SF2-Helicase+wHTH+Small-B-barrel+DUF1998<-BrxD-AAA<-PglZ<-DUF6079*<-N6-METHYLASE<-PglW\|\|?->DDE_3-><-?\|\|NTP_transf_5-><-?<-?<-HTH_3 | 1313 | Bacteria>Terrabacteria_group>Actinomycetota>Actinomycetes>Kitasatosporales>Streptomycetaceae | GCF_030268825.1 | Kitasatospora_phosalacinea | OO57_RS29790 | 18332..22273 | 2065 | DUF6079-family-protein-[Kitasatospora-phosalacinea] |
| WP_055487734.1 | Type2.Pgl.like.BREX | SAM-METHYLASE-><-?\|\|PadR-HTH->?-><-?<-SF1-SF2-Helicase+wHTH+Small-B-barrel+DUF1998<-BrxD-AAA\|\|RNaseH-><-PglZ<-DUF6079*<-?<-?<-?<-?<-DUF2171<-DUF1995\|\|HTH_31+DUF5753->DUF397-><-N6-METHYLASE | 1296 | Bacteria>Terrabacteria_group>Actinomycetota>Actinomycetes>Kitasatosporales>Streptomycetaceae | GCF_900090145.1 | Streptomyces_sp._WMMB_322 | H180DRAFT_RS21380 | 15365..19255 | 1286821 | hypothetical-protein-[Streptomyces-sp.-WMMB-322] |
| WP_093749477.1 | Type2.Pgl.like.BREX | <-TPR<-SF1-SF2-Helicase+wHTH+Small-B-barrel+DUF1998<-BrxD-AAA\|\|ParB+HNH->CBS+HEPN-><-PglZ<-DUF6079*<-?<-?<-?<-DUF397<-DUF397<-HTH_31+DUF5753\|\|ST-Hiskin->PIN-><-PglW | 1301 | Bacteria>Terrabacteria_group>Actinomycetota>Actinomycetes>Kitasatosporales>Streptomycetaceae | GCF_900091865.1 | Streptomyces_sp._DvalAA-19 | GA0115244_RS01770 | 13027..16932 | 1839761 | phage-resistance-protein-[Streptomyces-sp.-DvalAA-19] |
| WP_093779314.1 | Type2.Pgl.like.BREX | <-DUF397<-HTH_31+DUF5753\|\|ST-Hiskin-><-?\|\|?-><-SF1-SF2-Helicase+wHTH+Small-B-barrel+DUF1998<-BrxD-AAA\|\|ParB+HNH->CBS+HEPN-><-PglZ<-DUF6079*<-N6-METHYLASE<-PglW<-DUF397<-?\|\|?-><-REase+SF1-SF2-Helicase\|\|REase+SF1-SF2-Helicase-> | 1297 | Bacteria>Terrabacteria_group>Actinomycetota>Actinomycetes>Kitasatosporales>Streptomycetaceae | GCF_900101975.1 | Streptomyces_sp._136MFCol5.1 | BLQ64_RS33290 | 55492..59385 | 1172182 | phage-resistance-protein-[Streptomyces-sp.-136MFCol5.1] |
| WP_093907967.1 | Type2.Pgl.like.BREX | PglW-><-ORC-CDC6<-ParA-Soj-PloopNTPase<-HEPN\|\|N6-METHYLASE-><-DUF397<-HTH_31+DUF5753\|\|ST-Hiskin-><-?\|\|DUF6079*->PglZ->BrxD-AAA->SF1-SF2-Helicase+wHTH+Small-B-barrel+DUF1998-><-?<-DOC+HTH<-PhdYeFM_antitox | 1306 | Bacteria>Terrabacteria_group>Actinomycetota>Actinomycetes>Kitasatosporales>Streptomycetaceae | GCF_900103585.1 | Streptomyces_sp._cf386 | BLU79_RS24200 | 224601..228521 | 1761904 | PglY-protein-[Streptomyces-sp.-cf386] |
| WP_074991486.1 | Type2.Pgl.like.BREX | SF1-SF2-Helicase->?-><-Phage_integrase\|\|?-><-REase+N6_METHYLASE\|\|PglW->N6-METHYLASE->DUF6079*->PglZ->BrxD-AAA->SF1-SF2-Helicase+wHTH+Small-B-barrel+DUF1998-><-REase\|\|?-><-?<-HTH_Tnp_4+DDE_Tnp_4<-Phage_holin | 1299 | Bacteria>Terrabacteria_group>Actinomycetota>Actinomycetes>Kitasatosporales>Streptomycetaceae | GCF_900104815.1 | Streptomyces_misionensis | BLW85_RS06965 | 1205526..1209425 | 67331 | phage-resistance-protein-[Streptomyces-misionensis] |
| WP_093484993.1 | Type2.Pgl.like.BREX | Phage_int_SAM_3->Phage_integrase->MetJarc-HTH->PIN-><-RNaseH<-SF1-SF2-Helicase+wHTH+Small-B-barrel+DUF1998<-BrxD-AAA\|\|PglW->N6-METHYLASE->DUF6079*->PglZ-><-?\|\|?-><-?<-?<-?\|\|?-><-?<-ICLR-HTH | 1281 | Bacteria>Terrabacteria_group>Actinomycetota>Actinomycetes>Kitasatosporales>Streptomycetaceae | GCF_900104865.1 | unclassified_Streptomyces | BX264_RS23020 | 5242558..5246403 | 2593676 | MULTISPECIES:-DUF6079-family-protein-[unclassified-Streptomyces] |
| WP_093460196.1 | Type2.Pgl.like.BREX | PIN-><-?<-?\|\|?->?-><-PglZ<-DUF6079*<-N6-METHYLASE<-PglW\|\|SF1-SF2-Helicase+wHTH+Small-B-barrel+DUF1998-><-?<-DUF2637+HTH\|\|<-?\|\|<-ADH_N+ADH_zinc_N<-DSBH | 1279 | Bacteria>Terrabacteria_group>Actinomycetota>Actinomycetes>Kitasatosporales>Streptomycetaceae | GCF_900105695.1 | Streptomyces_melanosporofaciens | BLV72_RS03380 | 155017..158856 | 67327 | DUF6079-family-protein-[Streptomyces-melanosporofaciens] |
| WP_093774790.1 | Type2.Pgl.like.BREX | SAM-METHYLASE-><-?<-?<-?\|\|?->APATPase+TPR+TPR+TPR-><-RelE-ParE<-PhdYeFM_antitox\|\|DUF6079*->PglZ-><-?\|\|?-><-CBS+HEPN<-ParB+HNH\|\|BrxD-AAA->SF1-SF2-Helicase+wHTH+Small-B-barrel+DUF1998-> | 1296 | Bacteria>Terrabacteria_group>Actinomycetota>Actinomycetes>Kitasatosporales>Streptomycetaceae | GCF_900110755.1 | Streptomyces_sp._yr375 | BMZ82_RS12295 | 40338..44228 | 1761906 | phage-resistance-protein-[Streptomyces-sp.-yr375] |
| WP_093840849.1 | Type2.Pgl.like.BREX | HTH_31+DUF5753->DUF397->N6-METHYLASE->RADICAL-SAM->HTH_42->NUDIX->DUF6079*->PglZ->BrxD-AAA->SF1-SF2-Helicase+wHTH+Small-B-barrel+DUF1998-><-DUF3323+TOPRIM | 1356 | Bacteria>Terrabacteria_group>Actinomycetota>Actinomycetes>Kitasatosporales>Streptomycetaceae | GCF_900112355.1 | Streptomyces_aidingensis | BM242_RS23160 | 100333..104403 | 910347 | hypothetical-protein-[Streptomyces-aidingensis] |
| WP_093931017.1 | Type2.Pgl.like.BREX | <-ABC-ATPase+SbcC<-?<-?<-DUF397<-HTH_31+DUF5753\|\|ST-Hiskin-><-?\|\|PglW->N6-METHYLASE->DUF6079*->PglZ->BrxD-AAA->SF1-SF2-Helicase+wHTH+Small-B-barrel+DUF1998->?-><-REase+SF1-SF2-Helicase\|\|REase+SF1-SF2-Helicase-><-?\|\|?-><-ABhydrolase | 1302 | Bacteria>Terrabacteria_group>Actinomycetota>Actinomycetes>Kitasatosporales>Streptomycetaceae | GCF_900116325.1 | Streptomyces_sp._ok210 | BM432_RS28965 | 56303..60211 | 1761905 | PglY-protein-[Streptomyces-sp.-ok210] |
| WP_072486156.1 | Type2.Pgl.like.BREX | ST-Hiskin->REC+OmpR-HTH-><-?\|\|NTP_transf_3->?-><-SF1-SF2-Helicase+wHTH+Small-B-barrel+DUF1998<-BrxD-AAA<-PglZ<-DUF6079*<-?<-PIN\|\|?-><-OLD-ABC+TOPRIM\|\|?-><-SF1-SF2-Helicase<-SF1-SF2-Helicase<-?\|\|HTH_31+DUF5753-> | 1285 | Bacteria>Terrabacteria_group>Actinomycetota>Actinomycetes>Kitasatosporales>Streptomycetaceae | GCF_900119365.1 | Streptomyces_atratus | BT364_RS17760 | 144134..147991 | 1893 | PglY-protein-[Streptomyces-atratus] |
| WP_097216299.1 | Type2.Pgl.like.BREX | <-SIGMA-HTH<-?<-McrC-NTD+REase<-RAMA+McrB-AAA<-URI-DNase\|\|?-><-SF1-SF2-Helicase+wHTH+Small-B-barrel+DUF1998<-BrxD-AAA<-PglZ<-DUF6079*\|\|?-><-ST-Hiskin\|\|HTH_31+DUF5753->DUF397-><-N6-METHYLASE\|\|HEPN->ParA-Soj-PloopNTPase->ORC-CDC6-><-PglW | 1306 | Bacteria>Terrabacteria_group>Actinomycetota>Actinomycetes>Kitasatosporales>Streptomycetaceae | GCF_900182715.1 | Streptomyces_sp._Ag82_G6-1 | CRP41_RS11255 | 2422725..2426645 | 1938853 | PglY-protein-[Streptomyces-sp.-Ag82_G6-1] |
| WP_205042538.1 | Type2.Pgl.like.BREX | PglW-><-PIN<-HTH+MPTase\|\|N6-METHYLASE-><-DUF397<-HTH_31+DUF5753\|\|ST-Hiskin->GNTR-HTH->?->DUF6079*->PglZ->BrxD-AAA->SF1-SF2-Helicase+wHTH+Small-B-barrel+DUF1998-><-?<-Peptidase_M4+Peptidase_M4_C\|\|?->?->?->HisKA_3+HATPase_c-> | 1306 | Bacteria>Terrabacteria_group>Actinomycetota>Actinomycetes>Kitasatosporales>Streptomycetaceae | GCF_910593825.1 | Actinacidiphila_bryophytorum | NFU06_RS12440 | 458824..462744 | 1436133 | PglY-protein-[Actinacidiphila-bryophytorum] |
| WP_250297835.1 | Type2.Pgl.like.BREX | <-SAD(SRA)+wHTH+HNH<-SF1-SF2-Helicase+HKD-DNase\|\|PglW->N6-METHYLASE-><-PIN<-PhdYeFM_antitox<-ST-Hiskin\|\|HTH_31+DUF5753->DUF397->DUF6079*->PglZ-><-CBS+HEPN<-ParB+HNH\|\|BrxD-AAA->SF1-SF2-Helicase+wHTH+Small-B-barrel+DUF1998-><-SWACOS+TPR+TPR+LuxR-HTH<-Peptidase_M4+Peptidase_M4_C\|\|?-><-PIN | 1306 | Bacteria>Terrabacteria_group>Actinomycetota>Actinomycetes>Kitasatosporales>Streptomycetaceae | GCF_940670765.1 | Streptomyces_sp._A_4/2 | NBY69_RS01990 | 291198..295118 | 2934314 | phage-resistance-protein-[Streptomyces-sp.-A-4/2] |
| WP_084751006.1 | Type2.Pgl.like.BREX | HMA+HAD+HAD->?-><-?\|\|ANK+ANK->BrxA-DNA-binding->?-><-SF1-SF2-Helicase+wHTH+Small-B-barrel+DUF1998<-BrxD-AAA<-PglZ<-DUF6079*<-N6-METHYLASE<-PglW<-?<-?\|\|XIS-HTH-><-?<-ABhydrolase | 1300 | Bacteria>Terrabacteria_group>Actinomycetota>Actinomycetes>Kitasatosporales>Streptomycetaceae | - | Kitasatospora_albolonga | B7C62_29555 | 6697658..6701560 | 68173 | phage-resistance-protein-[Kitasatospora-albolonga] |
| WP_189289187.1 | Type2.Pgl.like.BREX | <-wHTH+SAD+HNH<-SF1-SF2-Helicase+wHTH+Small-B-barrel+DUF1998<-BrxD-AAA\|\|RNaseH-><-PglZ<-DUF6079*\|\|?-><-DUF2171<-DUF1995\|\|HTH_31+DUF5753->DUF397-><-N6-METHYLASE<-PglW | 1304 | Bacteria>Terrabacteria_group>Actinomycetota>Actinomycetes>Kitasatosporales>Streptomycetaceae | - | Streptomyces_fumigatiscleroticus | IEZ18_RS17085 | 289621..293535 | 66371 | phage-resistance-protein-[Streptomyces-fumigatiscleroticus] |
| WP_058081575.1 | Type2.Pgl.like.BREX | SIGMA-HTH-><-METHYLASE\|\|?->?->?->?-><-SF1-SF2-Helicase+wHTH+Small-B-barrel+DUF1998<-BrxD-AAA<-PglZ<-DUF6079*<-ST-Hiskin\|\|HEPN->ParA-Soj-PloopNTPase-><-Phage_rep\|\|HTH_31+DUF5753->DUF397-><-N6-METHYLASE<-?<-TOPRIM | 1296 | Bacteria>Terrabacteria_group>Actinomycetota>Actinomycetes>Kitasatosporales>Streptomycetaceae | - | Streptomyces_hygroscopicus | SHL15_4501 | 5034773..5038663 | 1912 | hypothetical-protein-[Streptomyces-hygroscopicus] |
| WP_058082291.1 | Type2.Pgl.like.BREX | SAM-METHYLASE->?->?->?->REase-><-SF1-SF2-Helicase+wHTH+Small-B-barrel+DUF1998<-BrxD-AAA<-PglZ<-DUF6079*<-N6-METHYLASE<-PglW\|\|REase+N6_METHYLASE-><-pPIWI+MID+RNaseH-p<-pPIWI+SF1-SF2-Helicase<-pPIWI+pPIWI\|\|BirA-HTH->XIS-HTH->Phage_int_SAM_3+Phage_integrase-> | 1299 | Bacteria>Terrabacteria_group>Actinomycetota>Actinomycetes>Kitasatosporales>Streptomycetaceae | - | Streptomyces_hygroscopicus | SHL15_RS28210 | 6415785..6419684 | 1912 | hypothetical-protein-[Streptomyces-hygroscopicus] |
| WP_058082605.1 | Type2.Pgl.like.BREX | NUDIX->NUDIX->DUF4096+DDE_Tnp_1+DDE_Tnp_1_2->HTH->HEPN->URI-DNase-><-SF1-SF2-Helicase+wHTH+Small-B-barrel+DUF1998<-BrxD-AAA<-PglZ<-DUF6079*<-N6-METHYLASE<-PglW<-?<-DnaB+DnaB<-DNA_METHYLASE<-PAPS-reduct | 1299 | Bacteria>Terrabacteria_group>Actinomycetota>Actinomycetes>Kitasatosporales>Streptomycetaceae | - | Streptomyces_hygroscopicus | SHL15_RS30585 | 6999502..7003401 | 1912 | hypothetical-protein-[Streptomyces-hygroscopicus] |
| WP_100109645.1 | Type2.Pgl.like.BREX | <-STAS<-?\|\|MerR-HTH->?-><-?\|\|?->\|\|MORC+DUF3684+DUF3883-><-SF1-SF2-Helicase+wHTH+Small-B-barrel+DUF1998<-DndB<-DUF6079*<-REase+SF1-SF2-Helicase<-SAM-METHYLASE<-?<-DUF2171<-DUF1995\|\|HTH_31+DUF5753->DUF397-><-SAM-METHYLASE | 1298 | Bacteria>Terrabacteria_group>Actinomycetota>Actinomycetes>Kitasatosporales>Streptomycetaceae | - | Streptomyces_peucetius | CGZ69_RS28000 | 6102761..6106657 | 1950 | phage-resistance-protein-[Streptomyces-peucetius] |
| WP_254585481.1 | Type2.Pgl.like.BREX | <-REase<-SF1-SF2-Helicase+wHTH+Small-B-barrel+DUF1998\|\|?->ParB+HNH->CBS+HEPN-><-PglZ<-DUF6079*<-APATPase+TPR+TPR+TPR<-?\|\|?->?-><-?<-DUF397<-HTH_31+DUF5753\|\|ST-Hiskin-><-N6-METHYLASE | 1295 | Bacteria>Terrabacteria_group>Actinomycetota>Actinomycetes>Kitasatosporales>Streptomycetaceae | - | Streptomyces_sp._AC04842 | IGB19_RS03905 | 11329..15216 | 2775327 | phage-resistance-protein-[Streptomyces-sp.-AC04842] |
| WP_254588049.1 | Type2.Pgl.like.BREX | REase+REase->PhdYeFM_antitox->RelE-ParE->ST-Hiskin->DUF6079*->PglZ->BrxD-AAA->SF1-SF2-Helicase+wHTH+Small-B-barrel+DUF1998-> | 1323 | Bacteria>Terrabacteria_group>Actinomycetota>Actinomycetes>Kitasatosporales>Streptomycetaceae | - | Streptomyces_sp._AC04842 | IGB19_RS28940 | 3037..7008 | 2775327 | hypothetical-protein-[Streptomyces-sp.-AC04842] |
| WP_048458866.1 | Type2.Pgl.like.BREX | N6-METHYLASE-><-ST-Hiskin\|\|HTH_31+DUF5753->DUF397->?-><-?<-?\|\|?->APATPase+TPR+TPR+TPR->DUF6079*->PglZ-><-CBS+HEPN<-ParB+HNH<-?\|\|BrxD-AAA->SF1-SF2-Helicase+wHTH+Small-B-barrel+DUF1998->REase-> | 1295 | Bacteria>Terrabacteria_group>Actinomycetota>Actinomycetes>Kitasatosporales>Streptomycetaceae | - | Streptomyces_sp._HNS054 | AC003_RS21940 | 71998..75885 | 1662446 | bacteriophage-resistance-protein-[Streptomyces-sp.-HNS054] |
| WP_215187711.1 | Type2.Pgl.like.BREX | DUF4011+SF1-SF2-Helicase+REase-><-REase<-SF1-SF2-Helicase+wHTH+Small-B-barrel+DUF1998<-BrxD-AAA\|\|?->ParB+HNH->CBS+HEPN-><-PglZ<-DUF6079*\|\|?->?-><-?<-DUF397<-HTH_31+DUF5753\|\|ST-Hiskin-><-N6-METHYLASE<-PglW\|\|SAD(SRA)+wHTH+HNH-> | 1295 | Bacteria>Terrabacteria_group>Actinomycetota>Actinomycetes>Kitasatosporales>Streptomycetaceae | - | Streptomyces_sp._McG8 | HET68_RS00340 | 23201..27088 | 2725487 | phage-resistance-protein-[Streptomyces-sp.-McG8] |
| WP_266686669.1 | Type2.Pgl.like.BREX | <-N6-METHYLASE\|\|OLD-ABC->TOPRIM-><-?\|\|PglW->N6-METHYLASE-><-ST-Hiskin\|\|HTH_31+DUF5753->DUF397->DUF6079*->PglZ-><-CBS+CBS+HEPN<-ParB+HNH\|\|BrxD-AAA->SF1-SF2-Helicase+wHTH+Small-B-barrel+DUF1998-><-SAD(SRA)+wHTH+HNH<-?<-DUF2637+HTH | 1318 | Bacteria>Terrabacteria_group>Actinomycetota>Actinomycetes>Kitasatosporales>Streptomycetaceae | - | Streptomyces_sp._NBC_00268 | OOK47_RS16810 | 3624646..3628602 | 2975695 | phage-resistance-protein-[Streptomyces-sp.-NBC_00268] |
| WP_269962286.1 | Type2.Pgl.like.BREX | HISKIN->?-><-?<-XIS-HTH\|\|REase+DUF5655->REase-><-SF1-SF2-Helicase+wHTH+Small-B-barrel+DUF1998<-BrxD-AAA<-PglZ<-DUF6079*<-N6-METHYLASE<-PglW\|\|REase+N6_METHYLASE-><-DnaB+DnaB<-DNA_METHYLASE\|\|Pkinase-><-Phage_integrase<-Phage_integrase | 1299 | Bacteria>Terrabacteria_group>Actinomycetota>Actinomycetes>Kitasatosporales>Streptomycetaceae | - | Streptomyces_sp._NBC_00760 | OHA36_RS41800 | 8909353..8913252 | 2975823 | phage-resistance-protein-[Streptomyces-sp.-NBC_00760] |
| WP_167901571.1 | Type2.Pgl.like.BREX | PglW->?-><-?<-?\|\|?->DUF6079*-> | 1297 | Bacteria>Terrabacteria_group>Actinomycetota>Actinomycetes>Kitasatosporales>Streptomycetaceae | GCF_011882225.1 | Streptomyces_sp._NEAU-H3 | HC023_RS01900 | 16923..20816 | 2720636 | phage-resistance-protein-[Streptomyces-sp.-NEAU-H3] |
| WP_031082512.1 | Type2.Pgl.like.BREX | <-DUF6079*<-?\|\|?->?->?->HTH_31+DUF5753->DUF397-><-N6-METHYLASE\|\|HEPN->ParA-Soj-PloopNTPase-> | 1276 | Bacteria>Terrabacteria_group>Actinomycetota>Actinomycetes>Kitasatosporales>Streptomycetaceae | - | Streptomyces_sp._NRRL_WC-3549 | IH13_RS0108495 | 999..4827 | 1463925 | hypothetical-protein,-partial-[Streptomyces-sp.-NRRL-WC-3549] |
| WP_125621745.1 | Type2.Pgl.like.BREX | MerR-HTH->?->?-><-?\|\|?-><-SF1-SF2-Helicase+wHTH+Small-B-barrel+DUF1998<-BrxD-AAA<-PglZ<-DUF6079*<-?<-?<-?<-PhdYeFM_antitox<-DUF2171<-DUF1995<-SAM-METHYLASE<-HTH_3 | 1310 | Bacteria>Terrabacteria_group>Actinomycetota>Actinomycetes>Kitasatosporales>Streptomycetaceae | - | Streptomyces_sp._WAC_00631 | DMH02_RS23130 | 1121160..1125092 | 2203201 | phage-resistance-protein-[Streptomyces-sp.-WAC-00631] |
| WP_225627839.1 | Type2.Pgl.like.BREX | MoxR-AAA->?-><-?<-REase<-SF1-SF2-Helicase+wHTH+Small-B-barrel+DUF1998\|\|?->ParB+HNH->CBS+HEPN-><-PglZ<-DUF6079*<-?<-?<-?<-?<-DUF397<-HTH_31+DUF5753\|\|<-N6-METHYLASE<-PglW\|\|PIN-> | 1294 | Bacteria>Terrabacteria_group>Actinomycetota>Actinomycetes>Kitasatosporales>Streptomycetaceae | - | Streptomyces_werraensis | GN202_RS21145 | 54247..58131 | 68284 | phage-resistance-protein-[Streptomyces-werraensis] |
| WP_152196006.1 | Type2.Pgl.like.BREX | HisKA_3+HATPase_c->REC+OmpR-HTH-><-DNA_glycosylase\|\|REase+SF1-SF2-Helicase->nSTAND1+HNH-><-DUF932\|\|PglW-p->N6-METHYLASE->DUF6079*->PglZ->BrxD-AAA->SF1-SF2-Helicase+wHTH+Small-B-barrel+DUF1998->ParB+HNH->Tox-RelA-><-?\|\|REase-> | 1250 | Bacteria>Terrabacteria_group>Actinomycetota>Actinomycetes>Micrococcales>Bogoriellaceae | GCF_009193155.1 | Georgenia_subflava | GD748_RS14900 | 328674..332426 | 1622177 | hypothetical-protein-[Georgenia-subflava] |
| WP_236865296.1 | Type2.Pgl.like.BREX | <-Peptidase_S8<-Classical-AAA<-HKD-DNase+SF1-SF2-Helicase+HTH+HTH+SRA\|\|?->ParB+HNH->ParB+HNH->PglW->N6-METHYLASE->DUF6079*->PglZ->BrxD-AAA->SF1-SF2-Helicase+wHTH+Small-B-barrel+DUF1998-><-SF1-SF2-Helicase+HTH\|\|HEPN+PAS+DUF4011+SF1-SF2-Helicase+REase->CASPASE-> | 1273 | Bacteria>Terrabacteria_group>Actinomycetota>Actinomycetes>Micrococcales>Brevibacteriaceae | GCF_021729135.1 | Brevibacterium_daeguense | LXN01_RS11515 | 52930..56751 | 909936 | DUF6079-family-protein-[Brevibacterium-daeguense] |
| PJJ73862.1 | Type2.Pgl.like.BREX | D6+SF1-SF2-Helicase->?->MazG->URI-DNase+SF1-SF2-Helicase-><-DUF932\|\|PglW->N6-METHYLASE->DUF6079*->PglZ->BrxD-AAA->SF1-SF2-Helicase+wHTH+Small-B-barrel+DUF1998-><-?\|\|HIN-HTH->HTH_21+rve+rve_2-> | 1269 | Bacteria>Terrabacteria_group>Actinomycetota>Actinomycetes>Micrococcales>Cellulomonadaceae | GCA_002797595.1 | Sediminihabitans_luteus | CLV28_1344 | 144291..148100 | 1138585 | hypothetical-protein-CLV28_1344-[Sediminihabitans-luteus] |
| WP_051681727.1 | Type2.Pgl.like.BREX | <-ABhydrolase\|\|ClpABN-AAA+ClpABC-AAA->?-><-DndB<-SF1-SF2-Helicase+wHTH+Small-B-barrel+DUF1998<-AbiEii<-wHTH+AbiEi_2<-BrxD-AAA<-PglZ<-DUF6079*<-N6-METHYLASE<-PglW<-SIGMA-HTH<-SIGMA-HTH<-LACI-HTH+PBPI\|\|?->?->?->Glyco_hydro_3+Glyco_hydro_3_C+Fn3-like->DEOR-HTH+ISOCOT-> | 1207 | Bacteria>Terrabacteria_group>Actinomycetota>Actinomycetes>Micrococcales>Cellulomonadaceae | GCF_000708885.1 | Cellulomonas_sp._HZM | AW16_RS18595 | 49076..52699 | 1454010 | hypothetical-protein-[Cellulomonas-sp.-HZM] |
| WP_164983783.1 | Type2.Pgl.like.BREX | <-ParB<-SF1-SF2-Helicase<-SF1-SF2-Helicase\|\|HHH+wHTH-><-ANK+ANK\|\|PglW->N6-METHYLASE->DUF6079*->PglZ->BrxD-AAA->nSTAND1+HNH-><-DNA_METHYLASE\|\|SF1-SF2-Helicase+wHTH+Small-B-barrel+DUF1998->REase->SLATT-> | 1245 | Bacteria>Terrabacteria_group>Actinomycetota>Actinomycetes>Micrococcales>Cellulomonadaceae | GCF_004123805.1 | Cellulomonas_endophytica | EJA20_RS18530 | 69675..73412 | 2494735 | hypothetical-protein-[Cellulomonas-endophytica] |
| WP_169324635.1 | Type2.Pgl.like.BREX | <-SF1-SF2-Helicase+wHTH+Small-B-barrel+DUF1998<-BrxD-AAA<-PglZ<-DUF6079*<-N6-METHYLASE<-PglW<-?<-?\|\|?->?->SopB+HNH-> | 1222 | Bacteria>Terrabacteria_group>Actinomycetota>Actinomycetes>Micrococcales>Cellulomonadaceae | GCF_012952065.1 | Cellulomonas_fimi | HIR71_RS08545 | 82200..85868 | 1708 | DUF6079-family-protein-[Cellulomonas-fimi] |
| WP_175348384.1 | Type2.Pgl.like.BREX | <-REase\|\|35exo+TUDOR+SF1-SF2-Helicase+iREase+CTD-><-?<-?\|\|EVE+REase-><-REase+iREase\|\|PglW->N6-METHYLASE->N6-METHYLASE->DUF6079*->PglZ->BrxD-AAA->SF1-SF2-Helicase+wHTH+Small-B-barrel+DUF1998-><-HHH+wHTH<-URI-DNase+SF1-SF2-Helicase<-MazG\|\|sGTP+sGTP+GTPase-AIG->?->SF1-SF2-Helicase+REase-> | 1257 | Bacteria>Terrabacteria_group>Actinomycetota>Actinomycetes>Micrococcales>Cellulomonadaceae | GCF_013359775.1 | Cellulomonas_humilata | HP550_RS14505 | 196986..200759 | 144055 | hypothetical-protein-[Cellulomonas-humilata] |
| WP_157802544.1 | Type2.Pgl.like.BREX | HEPN-><-Phage_integrase<-NERD\|\|<-HTH_Tnp_1+HTH_21+rve+rve_3\|\|?-><-SF1-SF2-Helicase+wHTH+Small-B-barrel+DUF1998<-BrxD-AAA<-REase<-PglZ<-DUF6079*<-N6-METHYLASE<-PglW\|\|DUF932-><-URI-DNase+SF1-SF2-Helicase<-MazG<-?<-DNA-glycosylase<-SF1-SF2-Helicase<-HKD-DNase | 1279 | Bacteria>Terrabacteria_group>Actinomycetota>Actinomycetes>Micrococcales>Cellulomonadaceae | GCF_016863455.1 | Sediminihabitans_luteus | S1u03_RS03045 | 303914..307753 | 1138585 | hypothetical-protein-[Sediminihabitans-luteus] |
| WP_207377935.1 | Type2.Pgl.like.BREX | SLATT->ARG->?-><-?<-?<-?\|\|PglW->N6-METHYLASE->DUF6079*->PglZ->REase->SF1-SF2-Helicase+wHTH+Small-B-barrel+DUF1998-><-?<-SF1-SF2-Helicase<-?\|\|?->SF1-SF2-Helicase-p-> | 1280 | Bacteria>Terrabacteria_group>Actinomycetota>Actinomycetes>Micrococcales>Cellulomonadaceae | GCF_017353535.1 | Cellulomonas_sp._zg-ZUI22 | J1G43_RS02980 | 636874..640716 | 2816955 | DUF6079-family-protein-[Cellulomonas-sp.-zg-ZUI22] |
| WP_207339658.1 | Type2.Pgl.like.BREX | <-SF1-SF2-Helicase-p\|\|?->SF1-SF2-Helicase-><-?<-SF1-SF2-Helicase+wHTH+Small-B-barrel+DUF1998<-BrxD-AAA<-REase<-PglZ<-DUF6079*<-N6-METHYLASE<-PglW\|\|?->?->?-><-?<-ARG<-SLATT | 1280 | Bacteria>Terrabacteria_group>Actinomycetota>Actinomycetes>Micrococcales>Cellulomonadaceae | GCF_018388445.1 | Cellulomonas_wangleii | KG103_RS17020 | 3724016..3727858 | 2816956 | DUF6079-family-protein-[Cellulomonas-wangleii] |
| WP_227580442.1 | Type2.Pgl.like.BREX | HIN-HTH->?->?-><-PIN\|\|TPR+TPR+TPR+TPR->?-><-?\|\|PglW->N6-METHYLASE->DUF6079*->PglZ->REase->SF1-SF2-Helicase+wHTH+Small-B-barrel+DUF1998-><-?<-SF1-SF2-Helicase<-HKD-DNase<-DUF6361 | 1280 | Bacteria>Terrabacteria_group>Actinomycetota>Actinomycetes>Micrococcales>Cellulomonadaceae | GCF_020686105.1 | Cellulomonas_sp._zg-B12 | LJN51_RS08140 | 86713..90555 | 2886199 | DUF6079-family-protein-[Cellulomonas-sp.-zg-B12] |
| WP_227583990.1 | Type2.Pgl.like.BREX | ABhydrolase->Glyoxalase->PglW->N6-METHYLASE->DUF6079*->PglZ->REase->SF1-SF2-Helicase+wHTH+Small-B-barrel+DUF1998-><-?<-?\|\|SIGMA-HTH-><-?<-NAD_binding_10 | 1280 | Bacteria>Terrabacteria_group>Actinomycetota>Actinomycetes>Micrococcales>Cellulomonadaceae | GCF_020686725.1 | Cellulomonas_sp._zg-Y138 | LJN54_RS01240 | 102829..106671 | 2886201 | cell-envelope-integrity-protein-TolA-[Cellulomonas-sp.-zg-Y138] |
| WP_209893504.1 | Type2.Pgl.like.BREX | <-HTH_Tnp_1+HTH_21+rve+rve_3<-SbcC\|\|REase->?->NTP_transf->?->PglW->N6-METHYLASE->DUF6079*->PglZ->BrxD-AAA->SF1-SF2-Helicase+wHTH+Small-B-barrel+DUF1998->SbcC->?-><-URI-DNase+SF1-SF2-Helicase\|\|HTH+rve->AAA-ATPase->\|\|Phage_int_SAM_4+Phage_integrase-> | 1240 | Bacteria>Terrabacteria_group>Actinomycetota>Actinomycetes>Micrococcales>Dermabacteraceae | - | Brachybacterium_fresconis | JOF44_RS15670 | 3524448..3528170 | 173363 | DUF6079-family-protein-[Brachybacterium-fresconis] |
| WP_096196447.1 | Type2.Pgl.like.BREX | <-Peptidase<-HKD-DNase<-REase<-SbcC<-JetB<-JetA\|\|REase->PglW->N6-METHYLASE->DUF6079*->PglZ->BrxD-AAA->SF1-SF2-Helicase+wHTH+Small-B-barrel+DUF1998-><-URI-DNase+SF1-SF2-Helicase<-MazG<-?\|\|?-><-?\|\|KAP-NTPase-> | 1240 | Bacteria>Terrabacteria_group>Actinomycetota>Actinomycetes>Micrococcales>Dermabacteraceae | GCF_002332305.1 | Brachybacterium_alimentarium | CIK66_RS02685 | 237897..241619 | 47845 | DUF6079-family-protein-[Brachybacterium-alimentarium] |
| WP_259784964.1 | Type2.Pgl.like.BREX | MazG->URI-DNase+SF1-SF2-Helicase->?-><-SF1-SF2-Helicase+wHTH+Small-B-barrel+DUF1998<-BrxD-AAA<-PglZ<-DUF6079*<-PglW\|\|<-?<-SF1-SF2-Helicase<-HKD-DNase<-DUF6361\|\|REase->URI-DNase-> | 1240 | Bacteria>Terrabacteria_group>Actinomycetota>Actinomycetes>Micrococcales>Dermabacteraceae | GCF_025146185.1 | Brachybacterium_muris | M3B40_RS08420 | 23285..27007 | 219301 | DUF6079-family-protein-[Brachybacterium-muris] |
| WP_259849918.1 | Type2.Pgl.like.BREX | <-DNA-glycosylase\|\|?->?-><-REase\|\|?->?->McrB-AAA->McrC-NTD+REase->PglW->N6-METHYLASE->DUF6079*->PglZ->BrxD-AAA->SF1-SF2-Helicase+wHTH+Small-B-barrel+DUF1998-><-?<-URI-DNase+SF1-SF2-Helicase<-MazG | 1240 | Bacteria>Terrabacteria_group>Actinomycetota>Actinomycetes>Micrococcales>Dermabacteraceae | GCF_025151835.1 | Brachybacterium_muris | M3F57_RS06940 | 23850..27572 | 219301 | DUF6079-family-protein-[Brachybacterium-muris] |
| WP_259803636.1 | Type2.Pgl.like.BREX | ACET-><-?<-?\|\|ParB+HNH-><-DNA-glycosylase\|\|?->DUF6079*->PglZ->BrxD-AAA->SF1-SF2-Helicase+wHTH+Small-B-barrel+DUF1998-><-URI-DNase+SF1-SF2-Helicase<-MazG<-DNA_glycosylase\|\|?->?-><-ZnR | 1240 | Bacteria>Terrabacteria_group>Actinomycetota>Actinomycetes>Micrococcales>Dermabacteraceae | GCF_025151965.1 | Brachybacterium_muris | M3F59_RS02775 | 135293..139015 | 219301 | DUF6079-family-protein-[Brachybacterium-muris] |
| WP_038566393.1 | Type2.Pgl.like.BREX | <-REase\|\|?->PglW->N6-METHYLASE->DUF6079*->PglZ->BrxD-AAA->SF1-SF2-Helicase+wHTH+Small-B-barrel+DUF1998->?->wHTH+SAD+HNH->?->ABhydrolase-> | 1273 | Bacteria>Terrabacteria_group>Actinomycetota>Actinomycetes>Micrococcales>Dermacoccaceae | GCF_000725405.1 | Dermacoccus_nishinomiyaensis | HX89_RS01015 | 214037..217858 | 1274 | DUF6079-family-protein-[Dermacoccus-nishinomiyaensis] |
| WP_052589426.1 | Type2.Pgl.like.BREX | <-REase<-SbcC<-JetB<-?<-?<-?<-SF1-SF2-Helicase+wHTH+Small-B-barrel+DUF1998<-BrxD-AAA<-PglZ<-DUF6079*<-ParB+HNH<-N6-METHYLASE<-PglW<-?\|\|NERD-><-?<-?<-?<-PTSEIIA1 | 1246 | Bacteria>Terrabacteria_group>Actinomycetota>Actinomycetes>Micrococcales>Dermacoccaceae | GCF_001190945.1 | Luteipulveratus_mongoliensis | VV02_RS01635 | 337415..341155 | 571913 | DUF6079-family-protein-[Luteipulveratus-mongoliensis] |
| WP_141928904.1 | Type2.Pgl.like.BREX | <-zf-ISL3+HTH_Tnp_ISL3+DDE_Tnp_ISL3\|\|REase+SF1-SF2-Helicase->?->nSTAND1+HNH->GTPase-AIG->sGTP->PglW->N6-METHYLASE->ParB+HNH->DUF6079*->PglZ->BrxD-AAA->SF1-SF2-Helicase+wHTH+Small-B-barrel+DUF1998-p->\|\|<-MetJarc-HTH<-?\|\|?->DADA-Ligase-> | 1257 | Bacteria>Terrabacteria_group>Actinomycetota>Actinomycetes>Micrococcales>Dermacoccaceae | GCF_006715095.1 | Yimella_lutea | FB459_RS13790 | 2861920..2865693 | 587872 | phage-resistance-protein-[Yimella-lutea] |
| QQS02055.1 | Type2.Pgl.like.BREX | <-REC+OmpR-HTH<-ST-Hiskin\|\|Glyoxalase->YTCD-HTH->PhdYeFM_antitox->RelE-ParE->SF1-SF2-Helicase->PglW->N6-METHYLASE->DUF6079*->PglZ->BrxD-AAA->SF1-SF2-Helicase+wHTH+Small-B-barrel+DUF1998->DDE_Tnp_1->\|\|<-DnaB+TnsC-ATPase\|\|?->?->GNTR-HTH+UTRA-> | 1266 | Bacteria>Terrabacteria_group>Actinomycetota>Actinomycetes>Micrococcales>Dermatophilaceae | GCA_016699675.1 | Austwickia_sp. | IPK37_06725 | 1456783..1460583 | 1872664 | MAG:-phage-resistance-protein-[Austwickia-sp.] |
| WP_226347228.1 | Type2.Pgl.like.BREX | <-PhdYeFM_antitox<-?\|\|DOC+HTH->AIPR-><-SF1-SF2-Helicase+wHTH+Small-B-barrel+DUF1998<-PIN<-XIS-HTH<-PglZ<-DUF6079*<-N6-METHYLASE-p | 1239 | Bacteria>Terrabacteria_group>Actinomycetota>Actinomycetes>Micrococcales>Dermatophilaceae | GCF_014857905.1 | Agilicoccus_flavus | ICW24_RS18630 | 19284..23003 | 2775968 | hypothetical-protein-[Agilicoccus-flavus] |
| EWT07978.1 | Type2.Pgl.like.BREX | REase->SF1-SF2-Helicase-><-?<-?<-?<-SF1-SF2-Helicase+wHTH+Small-B-barrel+DUF1998<-BrxD-AAA<-PglZ<-DUF6079*<-N6-METHYLASE<-PglW\|\|REase+SF1-SF2-Helicase->?->SF1-SF2-Helicase+HKD-DNase-><-Peptidase_M1_N+Peptidase_M1 | 1241 | Bacteria>Terrabacteria_group>Actinomycetota>Actinomycetes>Micrococcales>Intrasporangiaceae | GCA_000576575.1 | Intrasporangium_chromatireducens_Q5-1 | N864_11720 | 48345..52070 | 584657 | phage-resistance-protein-[Intrasporangium-chromatireducens-Q5-1] |
| KRE61577.1 | Type2.Pgl.like.BREX | REase+SF1-SF2-Helicase->HNH-><-SF1-SF2-Helicase<-?\|\|PglW->N6-METHYLASE->DUF6079*->PglZ->BrxD-AAA->SF1-SF2-Helicase+wHTH+Small-B-barrel+DUF1998-><-?<-?<-Ribonuc_red_2_N+Ribonuc_red_lgC<-ZnR+ATPcone | 1256 | Bacteria>Terrabacteria_group>Actinomycetota>Actinomycetes>Micrococcales>Intrasporangiaceae | GCA_001428065.1 | Tetrasphaera_sp._Soil756 | ASG78_09445 | 108460..112230 | 1736399 | hypothetical-protein-ASG78_09445-[Tetrasphaera-sp.-Soil756] |
| WP_034711951.1 | Type2.Pgl.like.BREX | URI-DNase->REase->SF1-SF2-Helicase-><-?<-?<-?<-SF1-SF2-Helicase+wHTH+Small-B-barrel+DUF1998<-BrxD-AAA<-PglZ<-DUF6079*<-N6-METHYLASE<-PglW<-?<-sGTP\|\|REase+SF1-SF2-Helicase->?->VirB11+RNA-Helicase-><-Peptidase_M1_N+Peptidase_M1 | 1240 | Bacteria>Terrabacteria_group>Actinomycetota>Actinomycetes>Micrococcales>Intrasporangiaceae | GCF_000576575.1 | Intrasporangium_chromatireducens | N864_RS00195 | 48345..52067 | 1386088 | DUF6079-family-protein-[Intrasporangium-chromatireducens] |
| WP_128365313.1 | Type2.Pgl.like.BREX | <-nSTAND1+HNH<-MazG\|\|REase+SF1-SF2-Helicase->nSTAND1+HNH-><-HARE-HTH<-?\|\|PglW->ParB+HNH->N6-METHYLASE->DUF6079*->PglZ->BrxD-AAA->SF1-SF2-Helicase+wHTH+Small-B-barrel+DUF1998-> | 1247 | Bacteria>Terrabacteria_group>Actinomycetota>Actinomycetes>Micrococcales>Intrasporangiaceae | GCF_004025965.2 | Phycicoccus_flavus | EPD83_RS02120 | 100309..104052 | 2502783 | phage-resistance-protein-[Phycicoccus-flavus] |
| WP_159902715.1 | Type2.Pgl.like.BREX | Peptidase_M1_N+Peptidase_M1-><-?\|\|PALP-><-METHYLASE\|\|?-><-SF1-SF2-Helicase+wHTH+Small-B-barrel+DUF1998<-BrxD-AAA<-PglZ<-DUF6079*<-ParB+HNH<-N6-METHYLASE<-PglW\|\|HEPN+SF1-SF2-Helicase+REase->NTP_transf_3-> | 1250 | Bacteria>Terrabacteria_group>Actinomycetota>Actinomycetes>Micrococcales>Intrasporangiaceae | GCF_009858245.1 | Ornithinibacter_aureus | C8E84_RS12855 | 2701418..2705170 | 622664 | DUF6079-family-protein-[Ornithinibacter-aureus] |
| PPH05309.1 | Type2.Pgl.like.BREX | <-Glyco_hydro_130<-?<-?<-LACI-HTH+PBPI\|\|?->?-><-?\|\|PglW->N6-METHYLASE->DUF6079*->PglZ->BrxD-AAA->SF1-SF2-Helicase+wHTH+Small-B-barrel+DUF1998-><-?<-?\|\|ParB+HNH+METHYLASE-><-wHTH | 1198 | Bacteria>Terrabacteria_group>Actinomycetota>Actinomycetes>Micrococcales>Microbacteriaceae | GCA_002932035.1 | Rathayibacter_sp._AY1F6 | C5C44_04265 | 114455..118051 | 2080560 | hypothetical-protein-C5C44_04265-[Rathayibacter-sp.-AY1F6] |
| WP_022890259.1 | Type2.Pgl.like.BREX | <-LACI-HTH+PBPI<-?\|\|<-?<-SF1-SF2-Helicase+wHTH+Small-B-barrel+DUF1998<-BrxD-AAA<-PglZ<-DUF6079*<-N6-METHYLASE<-PglW\|\|?-><-?<-?<-HNH | 1241 | Bacteria>Terrabacteria_group>Actinomycetota>Actinomycetes>Micrococcales>Microbacteriaceae | GCF_000421545.1 | Agromyces_italicus | GQQ89_RS11740 | 22885..26610 | 279572 | hypothetical-protein-[Agromyces-italicus] |
| WP_025102575.1 | Type2.Pgl.like.BREX | ACYC-><-VirB11+MuB-ATPase\|\|PglW->N6-METHYLASE->DUF6079*->PglZ->BrxD-AAA->SF1-SF2-Helicase+wHTH+Small-B-barrel+DUF1998-><-ABhydrolase<-?<-MPTase<-HTH\|\|ArdCN+MPTase->Primpol+PriCT1-> | 1209 | Bacteria>Terrabacteria_group>Actinomycetota>Actinomycetes>Micrococcales>Microbacteriaceae | GCF_000633215.1 | Microbacterium_paraoxydans | U857_RS0101335 | 242027..245656 | 199592 | hypothetical-protein-[Microbacterium-paraoxydans] |
| WP_056125657.1 | Type2.Pgl.like.BREX | <-SF1-SF2-Helicase+wHTH+Small-B-barrel+DUF1998<-BrxD-AAA<-PglZ<-DUF6079*<-N6-METHYLASE<-PglW<-AraC-HTH\|\|?->ABC-ATPase+SbcC->?->HisKA_3+HATPase_c->REC+OmpR-HTH-> | 1194 | Bacteria>Terrabacteria_group>Actinomycetota>Actinomycetes>Micrococcales>Microbacteriaceae | GCF_001424385.1 | Curtobacterium_sp._Leaf183 | ASG04_RS14590 | 792626..796210 | 1736291 | hypothetical-protein-[Curtobacterium-sp.-Leaf183] |
| WP_067228573.1 | Type2.Pgl.like.BREX | <-URI-DNase\|\|HNH-><-?<-HTH_3<-?<-SF1-SF2-Helicase+wHTH+Small-B-barrel+DUF1998<-BrxD-AAA<-PglZ<-DUF6079*<-N6-METHYLASE<-PglW\|\|?-><-SF1-SF2-Helicase<-35exo+BRCT<-?<-ParB+HNH | 1240 | Bacteria>Terrabacteria_group>Actinomycetota>Actinomycetes>Micrococcales>Microbacteriaceae | GCF_001542775.1 | Microterricola_viridarii | AWU67_RS10280 | 2230743..2234465 | 412690 | hypothetical-protein-[Microterricola-viridarii] |
| WP_104090841.1 | Type2.Pgl.like.BREX | <-Primpol+PriCT1\|\|HTH->MPTase->?->ABhydrolase->?->PglW->N6-METHYLASE->DUF6079*->PglZ->BrxD-AAA->SF1-SF2-Helicase+wHTH+Small-B-barrel+DUF1998-><-MORC+DUF3684+HNH<-SH3-like+SF1-SF2-Helicase+wHTH\|\|SIR2->?->SbcC->VirB11+MuB-ATPase-> | 1240 | Bacteria>Terrabacteria_group>Actinomycetota>Actinomycetes>Micrococcales>Microbacteriaceae | GCF_002929595.1 | Cryobacterium_sp._N19 | CXY78_RS19830 | 367057..370779 | 2048288 | hypothetical-protein-[Cryobacterium-sp.-N19] |
| WP_104250216.1 | Type2.Pgl.like.BREX | <-Glyco_hydro_130<-?<-?<-LACI-HTH+PBPI\|\|?-><-?\|\|PglW->N6-METHYLASE->DUF6079*->PglZ->BrxD-AAA->SF1-SF2-Helicase+wHTH+Small-B-barrel+DUF1998->ParB+HNH+METHYLASE-><-wHTH | 1202 | Bacteria>Terrabacteria_group>Actinomycetota>Actinomycetes>Micrococcales>Microbacteriaceae | GCF_002931595.1 | Rathayibacter_sp._AY2B9 | C5C25_RS07400 | 22881..26489 | 2080572 | hypothetical-protein-[Rathayibacter-sp.-AY2B9] |
| WP_181132711.1 | Type2.Pgl.like.BREX | <-LACI-HTH+PBPI\|\|?->?-><-HEPN<-?\|\|PglW->N6-METHYLASE->DUF6079*->PglZ->BrxD-AAA->SF1-SF2-Helicase+wHTH+Small-B-barrel+DUF1998-><-?<-?\|\|ParB+HNH+METHYLASE-><-wHTH | 1173 | Bacteria>Terrabacteria_group>Actinomycetota>Actinomycetes>Micrococcales>Microbacteriaceae | GCF_002932035.1 | Rathayibacter_sp._AY1F6 | C5C44_RS04260 | 114530..118051 | 2080560 | hypothetical-protein-[Rathayibacter-sp.-AY1F6] |
| WP_104339419.1 | Type2.Pgl.like.BREX | PglW->N6-METHYLASE->DUF6079*->PglZ->BrxD-AAA->SF1-SF2-Helicase+wHTH+Small-B-barrel+DUF1998-><-?<-?\|\|ParB+HNH-> | 1198 | Bacteria>Terrabacteria_group>Actinomycetota>Actinomycetes>Micrococcales>Microbacteriaceae | GCF_002932195.1 | Rathayibacter_sp._AY1C4 | C5C99_RS00390 | 87157..90753 | 2080537 | hypothetical-protein-[Rathayibacter-sp.-AY1C4] |
| WP_104355140.1 | Type2.Pgl.like.BREX | wHTH-><-ParB+HNH\|\|?->?-><-SF1-SF2-Helicase+wHTH+Small-B-barrel+DUF1998<-BrxD-AAA<-PglZ<-DUF6079*<-N6-METHYLASE<-PglW | 1198 | Bacteria>Terrabacteria_group>Actinomycetota>Actinomycetes>Micrococcales>Microbacteriaceae | GCF_002932515.1 | Rathayibacter_sp._AY1D5 | C5C82_RS12010 | 27442..31038 | 2080546 | hypothetical-protein-[Rathayibacter-sp.-AY1D5] |
| WP_104319439.1 | Type2.Pgl.like.BREX | wHTH-><-ParB+HNH+METHYLASE\|\|?->?-><-SF1-SF2-Helicase+wHTH+Small-B-barrel+DUF1998<-BrxD-AAA<-PglZ<-DUF6079*<-N6-METHYLASE<-PglW\|\|?->?-><-?\|\|LACI-HTH+PBPI-> | 1198 | Bacteria>Terrabacteria_group>Actinomycetota>Actinomycetes>Micrococcales>Microbacteriaceae | GCF_002932635.1 | Rathayibacter_sp._AY1D1 | C5C56_RS09115 | 81440..85036 | 2080542 | hypothetical-protein-[Rathayibacter-sp.-AY1D1] |
| WP_136709617.1 | Type2.Pgl.like.BREX | <-HNH\|\|McrB-AAA->McrC-NTD+REase->?->HD-><-?<-?\|\|PglW->N6-METHYLASE->DUF6079*->PglZ->BrxD-AAA->SF1-SF2-Helicase+wHTH+Small-B-barrel+DUF1998-><-?\|\|REase-><-MULE-TRANSPOSASE<-?\|\|TraJ-RHH-> | 1233 | Bacteria>Terrabacteria_group>Actinomycetota>Actinomycetes>Micrococcales>Microbacteriaceae | GCF_004959775.1 | Agromyces_sp._H66 | EZM51_RS17590 | 40844..44545 | 2529859 | DUF6079-family-protein-[Agromyces-sp.-H66] |
| WP_147826323.1 | Type2.Pgl.like.BREX | ParB+HNH-><-?<-?\|\|HEPN+SF1-SF2-Helicase+REase->?->HTH_3->?->PglW->N6-METHYLASE->DUF6079*->PglZ->BrxD-AAA->SF1-SF2-Helicase+wHTH+Small-B-barrel+DUF1998->?->HNH-><-Acyl_transf_3 | 1213 | Bacteria>Terrabacteria_group>Actinomycetota>Actinomycetes>Micrococcales>Microbacteriaceae | GCF_008017445.1 | Microbacterium_mitrae | FVP60_RS10910 | 63863..67504 | 664640 | hypothetical-protein-[Microbacterium-mitrae] |
| WP_148733493.1 | Type2.Pgl.like.BREX | <-ACET\|\|?-><-HTH_3\|\|?->?->ParB+HNH->?->PglW->N6-METHYLASE->DUF6079*->PglZ->BrxD-AAA->SF1-SF2-Helicase+wHTH+Small-B-barrel+DUF1998->?->RVT->?->?->SopB+HNH-> | 1233 | Bacteria>Terrabacteria_group>Actinomycetota>Actinomycetes>Micrococcales>Microbacteriaceae | GCF_008122505.1 | Agromyces_mariniharenae | FYC51_RS10555 | 2282310..2286011 | 2604423 | DUF6079-family-protein-[Agromyces-mariniharenae] |
| WP_157001138.1 | Type2.Pgl.like.BREX | HIN-HTH+PAIREDC-HTH-><-rve+rve_3\|\|HEPN-><-MORC+DUF3684+HNH<-SH3-like+SF1-SF2-Helicase+wHTH<-SF1-SF2-Helicase+wHTH+Small-B-barrel+DUF1998<-BrxD-AAA<-?<-PglZ<-DUF6079*<-N6-METHYLASE<-PglW\|\|?-><-?\|\|?-><-HNH<-?<-?<-VirB11+MuB-ATPase | 1223 | Bacteria>Terrabacteria_group>Actinomycetota>Actinomycetes>Micrococcales>Microbacteriaceae | GCF_009749475.1 | Agromyces_laixinhei | GQQ90_RS03440 | 20902..24573 | 2585717 | hypothetical-protein-[Agromyces-laixinhei] |
| WP_161886743.1 | Type2.Pgl.like.BREX | <-Calcineurin\|\|DDE_Tnp_1+DDE_Tnp_1->?-><-REase<-SF1-SF2-Helicase+wHTH+Small-B-barrel+DUF1998<-BrxD-AAA<-PglZ<-DUF6079*<-N6-METHYLASE<-PglW<-?\|\|?->Phage_integrase->Phage_integrase->DDE_Tnp_1+DDE_Tnp_1-><-SbcC | 1227 | Bacteria>Terrabacteria_group>Actinomycetota>Actinomycetes>Micrococcales>Microbacteriaceae | GCF_009930795.1 | Marisediminicola_antarctica | BHD05_RS12615 | 2706873..2710556 | 674079 | DUF6079-family-protein-[Marisediminicola-antarctica] |
| WP_168625653.1 | Type2.Pgl.like.BREX | <-REase<-SbcC<-JetB<-JetA\|\|DOC+HTH->?->?-><-?<-SF1-SF2-Helicase+wHTH+Small-B-barrel+DUF1998<-PglZ<-DUF6079*<-N6-METHYLASE<-PglW\|\|ParB+HNH-><-SF1-SF2-Helicase+HTH<-SbcC+CCDC158<-Calcineurin<-HEPN+PAS+DUF4011+SF1-SF2-Helicase+REase<-ARG | 1190 | Bacteria>Terrabacteria_group>Actinomycetota>Actinomycetes>Micrococcales>Microbacteriaceae | GCF_012274275.1 | Cryobacterium_sp._BB307 | HGK58_RS00720 | 149088..152660 | 2716317 | hypothetical-protein-[Cryobacterium-sp.-BB307] |
| WP_206833857.1 | Type2.Pgl.like.BREX | PRTase-><-?\|\|PglW->DUF6079*->PglZ->BrxD-AAA->SF1-SF2-Helicase+wHTH+Small-B-barrel+DUF1998->ARPP2->JetA->JetB->SbcC->REase-><-HTH_26 | 1209 | Bacteria>Terrabacteria_group>Actinomycetota>Actinomycetes>Micrococcales>Microbacteriaceae | GCF_017303175.1 | Microbacterium_esteraromaticum | JF550_RS13935 | 10603..14232 | 57043 | DUF6079-family-protein-[Microbacterium-esteraromaticum] |
| WP_206824857.1 | Type2.Pgl.like.BREX | PRTase-><-?\|\|PglW->DUF6079-p*->DUF6079->PglZ->BrxD-AAA->SF1-SF2-Helicase+wHTH+Small-B-barrel+DUF1998->ARPP2->JetA->JetB->SbcC->REase-> | 159 | Bacteria>Terrabacteria_group>Actinomycetota>Actinomycetes>Micrococcales>Microbacteriaceae | GCF_017303215.1 | Microbacterium_esteraromaticum | JF543_RS13920 | 10603..11082 | 57043 | DUF6079-family-protein-[Microbacterium-esteraromaticum] |
| WP_206824858.1 | Type2.Pgl.like.BREX | PRTase-><-?\|\|PglW->DUF6079-p->DUF6079*->PglZ->BrxD-AAA->SF1-SF2-Helicase+wHTH+Small-B-barrel+DUF1998->ARPP2->JetA->JetB->SbcC->REase-><-HTH_26 | 1055 | Bacteria>Terrabacteria_group>Actinomycetota>Actinomycetes>Micrococcales>Microbacteriaceae | GCF_017303215.1 | Microbacterium_esteraromaticum | JF543_RS13925 | 11066..14233 | 57043 | hypothetical-protein-[Microbacterium-esteraromaticum] |
| WP_267659168.1 | Type2.Pgl.like.BREX | <-SMS-KaiC<-?\|\|tRNA->?-><-?<-TnsC-ATPase\|\|?->?->PglW->N6-METHYLASE->DUF6079*->PglZ->BrxD-AAA->SF1-SF2-Helicase+wHTH+Small-B-barrel+DUF1998-> | 1195 | Bacteria>Terrabacteria_group>Actinomycetota>Actinomycetes>Micrococcales>Microbacteriaceae | GCF_026625765.1 | Frigoribacterium_sp._SL97 | OVA02_RS02525 | 513826..517413 | 2994664 | hypothetical-protein-[Frigoribacterium-sp.-SL97] |
| WP_210007017.1 | Type2.Pgl.like.BREX | XIS-HTH->?->?->HNH->?-><-?\|\|PglW->N6-METHYLASE->DUF6079*->PglZ->BrxD-AAA->SF1-SF2-Helicase+wHTH+Small-B-barrel+DUF1998->?->TrwC+SF1-SF2-Helicase->MAD-NTD->ParB+HNH-><-SSB | 1213 | Bacteria>Terrabacteria_group>Actinomycetota>Actinomycetes>Micrococcales>Microbacteriaceae | GCF_027921865.1 | Microbacterium | PKE82_RS12790 | 698433..702074 | 33882 | MULTISPECIES:-hypothetical-protein-[Microbacterium] |
| WP_271637797.1 | Type2.Pgl.like.BREX | <-AbiEii<-AbiEi_4<-SF1-SF2-Helicase+wHTH+Small-B-barrel+DUF1998<-BrxD-AAA<-PglZ<-DUF6079*<-N6-METHYLASE<-PglW<-XIS-HTH<-TrwC+SF1-SF2-Helicase+TOPRIM<-Phage_integrase<-XIS-HTH<-tRNA<-?<-?<-SAM-METHYLASE | 1194 | Bacteria>Terrabacteria_group>Actinomycetota>Actinomycetes>Micrococcales>Microbacteriaceae | GCF_028198145.1 | Microbacterium_sp._nov._GSS16 | PGB26_RS11740 | 2458262..2461846 | 3019890 | hypothetical-protein-[Microbacterium-sp.-nov.-GSS16] |
| WP_068479726.1 | Type2.Pgl.like.BREX | <-SF1-SF2-Helicase+wHTH+Small-B-barrel+DUF1998<-BrxD-AAA<-PglZ<-DUF6079*<-N6-METHYLASE<-PglW | 1210 | Bacteria>Terrabacteria_group>Actinomycetota>Actinomycetes>Micrococcales>Microbacteriaceae | GCF_900016305.1 | Pseudoclavibacter_helvolus | BN2270_RS09190 | 6467..10099 | 255205 | hypothetical-protein-[Pseudoclavibacter-helvolus] |
| WP_092387762.1 | Type2.Pgl.like.BREX | REase-><-RNase<-ARG\|\|NUDIX->HTH_6+SIS->?->?-><-?<-PglZ<-DUF6079*<-N6-METHYLASE<-PglW<-ADH_N+ADH_zinc_N<-URI-DNase+SF1-SF2-Helicase<-MazG<-McrC-NTD+REase<-McrB-AAA<-ABhydrolase+ABhydrolase | 1223 | Bacteria>Terrabacteria_group>Actinomycetota>Actinomycetes>Micrococcales>Microbacteriaceae | GCF_900113095.1 | Curtobacterium_sp._YR515 | BM028_RS16065 | 396808..400479 | 1855316 | hypothetical-protein-[Curtobacterium-sp.-YR515] |
| WP_183357096.1 | Type2.Pgl.like.BREX | <-His_Phos_1\|\|?-><-?<-?\|\|?->\|\|ParB->?->PglW->N6-METHYLASE->DUF6079*->PglZ->?->?->SF1-SF2-Helicase+wHTH+Small-B-barrel+DUF1998->MazG->URI-DNase+SF1-SF2-Helicase->?-><-?\|\|HTH_29+HTH_32+DDE_3-> | 1252 | Bacteria>Terrabacteria_group>Actinomycetota>Actinomycetes>Micrococcales>Micrococcaceae | - | Garicola_koreensis | FHX47_RS01325 | 271475..275233 | 1262554 | phage-resistance-protein-[Garicola-koreensis] |
| PZO69535.1 | Type2.Pgl.like.BREX | N6-METHYLASE->DUF6079*->PglZ->BrxD-AAA->SF1-SF2-Helicase+wHTH+Small-B-barrel+DUF1998->?->Thioredoxin-><-?\|\|SF1-SF2-Helicase->HAMP+HISKIN->REC+OmpR-HTH-> | 1239 | Bacteria>Terrabacteria_group>Actinomycetota>Actinomycetes>Micrococcales>Micrococcaceae | GCA_003240705.1 | Kocuria_palustris | DI634_05665 | 1578..5297 | 71999 | phage-resistance-protein-[Kocuria-palustris] |

| Protein-ID | System | Gene-Neighbourhoods | Length | Taxonomy | Assembly-ID | Strain | Locus-Tag | Genomic-coordinates | TaxID | Fasta-defline |
|---|---|---|---|---|---|---|---|---|---|---|
| BCW18380.1 | Type2.Pgl.like.BREX | <-ParB+HNH\|\|HKD-DNase+SF1-SF2-Helicase->RVT->REase-><-SF1-SF2-Helicase+wHTH+Small-B-barrel+DUF1998<-BrxD-AAA<-PglZ<-DUF6079*<-N6-METHYLASE<-PglW<-?<-?<-ACET | 1246 | Bacteria>Terrabacteria_group>Actinomycetota>Actinomycetes>Micrococcales>Micrococcaceae | GCA_019977115.1 | Arthrobacter_sp._NtRootA9 | NtRootA9_10880 | 1149021..1152761 | 2830986 | hypothetical-protein-NtRootA9_10880-[Arthrobacter-sp.-NtRootA9] |
| WP_035923652.1 | Type2.Pgl.like.BREX | [illegible] | 1280 | Bacteria>Terrabacteria_group>Actinomycetota>Actinomycetes>Micrococcales>Micrococcaceae | GCF_000786195.1 | Kocuria_polaris | GY22_RS01830 | 406168..410010 | 136273 | DUF6079-family-protein-[Kocuria-polaris] |
| WP_052690527.1 | Type2.Pgl.like.BREX | [illegible] | 1260 | Bacteria>Terrabacteria_group>Actinomycetota>Actinomycetes>Micrococcales>Micrococcaceae | GCF_000962805.1 | Pseudarthrobacter_chlorophenolicus | UC32_RS05580 | 209469..213251 | 85085 | DUF6079-family-protein-[Pseudarthrobacter-chlorophenolicus] |
| WP_091607352.1 | Type2.Pgl.like.BREX | [illegible] | 1259 | Bacteria>Terrabacteria_group>Actinomycetota>Actinomycetes>Micrococcales>Micrococcaceae | GCF_001373495.1 | Arthrobacter_koreensis | BN2404_RS13470 | 181235..185014 | 199136 | DUF6079-family-protein-[Arthrobacter-koreensis] |
| WP_078026244.1 | Type2.Pgl.like.BREX | [illegible] | 1254 | Bacteria>Terrabacteria_group>Actinomycetota>Actinomycetes>Micrococcales>Micrococcaceae | GCF_002008305.4 | Micrococcus_sp._KBS0714 | B1A86_RS09745 | 2132395..2136159 | 1179670 | hypothetical-protein-[Micrococcus-sp.-KBS0714] |
| WP_089594056.1 | Type2.Pgl.like.BREX | [illegible] | 1258 | Bacteria>Terrabacteria_group>Actinomycetota>Actinomycetes>Micrococcales>Micrococcaceae | GCF_002224285.1 | Arthrobacter_sp._YN | CGK93_RS06110 | 1322963..1326739 | 2020486 | DUF6079-family-protein-[Arthrobacter-sp.-YN] |
| WP_095346967.1 | Type2.Pgl.like.BREX | [illegible] | 1258 | Bacteria>Terrabacteria_group>Actinomycetota>Actinomycetes>Micrococcales>Micrococcaceae | GCF_002276855.1 | Micrococcus_luteus | B8X03_RS04485 | 14066..17842 | 1270 | DUF6079-family-protein-[Micrococcus-luteus] |
| WP_115787466.1 | Type2.Pgl.like.BREX | [illegible] | 1280 | Bacteria>Terrabacteria_group>Actinomycetota>Actinomycetes>Micrococcales>Micrococcaceae | GCF_003369445.1 | Arthrobacter_silvisoli | DWV11_RS03295 | 77253..81095 | 2291022 | DUF6079-family-protein-[Arthrobacter-silvisoli] |
| WP_141161302.1 | Type2.Pgl.like.BREX | [illegible] | 1263 | Bacteria>Terrabacteria_group>Actinomycetota>Actinomycetes>Micrococcales>Micrococcaceae | GCF_006517215.1 | Pseudarthrobacter_sp._NIBRBAC000502771 | NIBR502771_RS20345 | 4407176..4410967 | 2590774 | DUF6079-family-protein-[Pseudarthrobacter-sp.-NIBRBAC000502771] |
| WP_142057738.1 | Type2.Pgl.like.BREX | [illegible] | 1252 | Bacteria>Terrabacteria_group>Actinomycetota>Actinomycetes>Micrococcales>Micrococcaceae | GCF_006680095.1 | Pseudarthrobacter_sp._B4EP4b | FNL14_RS03740 | 117771..121529 | 2590664 | hypothetical-protein-[Pseudarthrobacter-sp.-B4EP4b] |
| WP_144825788.1 | Type2.Pgl.like.BREX | <-ParB+HNH<-ParB+HNH\|\|PglW->N6-METHYLASE->DUF6079*->PglZ->BrxD-AAA->SF1-SF2-Helicase+wHTH+Small-B-barrel+DUF1998->MazG->URI-DNase+SF1-SF2-Helicase-> | 1254 | Bacteria>Terrabacteria_group>Actinomycetota>Actinomycetes>Micrococcales>Micrococcaceae | GCF_007677505.1 | Micrococcus_luteus | FS660_RS01700 | 358900..362664 | 1270 | phage-resistance-protein-[Micrococcus-luteus] |
| WP_058857753.1 | Type2.Pgl.like.BREX | [illegible] | 1240 | Bacteria>Terrabacteria_group>Actinomycetota>Actinomycetes>Micrococcales>Micrococcaceae | GCF_007992135.1 | Kocuria_flava | AS188_RS03960 | 864929..868651 | 446860 | DUF6079-family-protein-[Kocuria-flava] |
| WP_062735895.1 | Type2.Pgl.like.BREX | [illegible] | 1239 | Bacteria>Terrabacteria_group>Actinomycetota>Actinomycetes>Micrococcales>Micrococcaceae | GCF_007992155.1 | Kocuria_turfanensis | KTU01_RS15735 | 17791..21510 | 388357 | DUF6079-family-protein-[Kocuria-turfanensis] |
| WP_159803039.1 | Type2.Pgl.like.BREX | [illegible] | 1276 | Bacteria>Terrabacteria_group>Actinomycetota>Actinomycetes>Micrococcales>Micrococcaceae | GCF_009828595.1 | Arthrobacter_zhaoguopingii | GVV86_RS12955 | 137933..141763 | 2681491 | DUF6079-family-protein-[Arthrobacter-zhaoguopingii] |
| WP_176635883.1 | Type2.Pgl.like.BREX | [illegible] | 1261 | Bacteria>Terrabacteria_group>Actinomycetota>Actinomycetes>Micrococcales>Micrococcaceae | GCF_013376105.1 | Arthrobacter_wenxiniae | G6034_RS14835 | 48958..52743 | 2713570 | DUF6079-family-protein-[Arthrobacter-wenxiniae] |
| WP_185260728.1 | Type2.Pgl.like.BREX | <-ParB+HNH<-?<-TnsC-ATPase<-rve<-?<-REase+wHTH\|\|?->PglW->N6-METHYLASE->DUF6079*->PglZ->BrxD-AAA->SF1-SF2-Helicase+wHTH+Small-B-barrel+DUF1998-> | 1258 | Bacteria>Terrabacteria_group>Actinomycetota>Actinomycetes>Micrococcales>Micrococcaceae | GCF_014217545.1 | Pseudarthrobacter_sp._NBSH8 | FYJ92_RS10705 | 2331004..2334780 | 2596911 | DUF6079-family-protein-[Pseudarthrobacter-sp.-NBSH8] |
| WP_193591824.1 | Type2.Pgl.like.BREX | [illegible] | 1280 | Bacteria>Terrabacteria_group>Actinomycetota>Actinomycetes>Micrococcales>Micrococcaceae | GCF_015040095.1 | Paenarthrobacter_sp._YJN-5 | HMI59_RS15770 | 3390318..3394160 | 2735316 | DUF6079-family-protein-[Paenarthrobacter-sp.-YJN-5] |
| WP_205653537.1 | Type2.Pgl.like.BREX | [illegible] | 1256 | Bacteria>Terrabacteria_group>Actinomycetota>Actinomycetes>Micrococcales>Micrococcaceae | GCF_039526125.1 | Arthrobacter_pascens | JYA52_RS12660 | 115488..119258 | 1677 | hypothetical-protein-[Arthrobacter-pascens] |
| WP_236821494.1 | Type2.Pgl.like.BREX | [illegible] | 621 | Bacteria>Terrabacteria_group>Actinomycetota>Actinomycetes>Micrococcales>Micrococcaceae | GCF_021800405.1 | Arthrobacter_sp._FW306-07-I | LFT46_RS05330 | 1137913..1139778 | 2879622 | hypothetical-protein-[Arthrobacter-sp.-FW306-07-I] |
| WP_227909310.1 | Type2.Pgl.like.BREX | [illegible] | 1269 | Bacteria>Terrabacteria_group>Actinomycetota>Actinomycetes>Micrococcales>Micrococcaceae | GCF_023022985.1 | Arthrobacter_sp._zg-Y809 | MUG94_RS03215 | 693422..697231 | 2886940 | DUF6079-family-protein-[Arthrobacter-gengyunqii] |
| WP_251332900.1 | Type2.Pgl.like.BREX | [illegible] | 1279 | Bacteria>Terrabacteria_group>Actinomycetota>Actinomycetes>Micrococcales>Micrococcaceae | GCF_023715525.1 | Kocuria_rosea | M3185_RS06715 | 44856..48695 | 1275 | DUF6079-family-protein-[Kocuria-rosea] |
| WP_256246191.1 | Type2.Pgl.like.BREX | [illegible] | 1231 | Bacteria>Terrabacteria_group>Actinomycetota>Actinomycetes>Micrococcales>Micrococcaceae | GCF_024494415.1 | Pseudarthrobacter_sp._R1 | M8J71_RS01190 | 281938..285633 | 2944934 | hypothetical-protein-[Pseudarthrobacter-sp.-R1] |
| WP_144820012.1 | Type2.Pgl.like.BREX | [illegible] | 1258 | Bacteria>Terrabacteria_group>Actinomycetota>Actinomycetes>Micrococcales>Micrococcaceae | GCF_024655705.1 | Micrococcus_luteus | NUV33_RS00310 | 22108..25884 | 1270 | phage-resistance-protein-[Micrococcus-luteus] |
| WP_268158799.1 | Type2.Pgl.like.BREX | [illegible] | 1239 | Bacteria>Terrabacteria_group>Actinomycetota>Actinomycetes>Micrococcales>Micrococcaceae | GCF_026672455.1 | Pseudarthrobacter_sp._SL88 | OVA06_RS12985 | 2873680..2877399 | 2994666 | DUF6079-family-protein-[Pseudarthrobacter-sp.-SL88] |
| WP_091695611.1 | Type2.Pgl.like.BREX | [illegible] | 1242 | Bacteria>Terrabacteria_group>Actinomycetota>Actinomycetes>Micrococcales>Micrococcaceae | GCF_900116905.1 | Micrococcus_terreus | BM355_RS04880 | 97122..100850 | 574650 | DUF6079-family-protein-[Micrococcus-terreus] |
| WP_262103916.1 | Type2.Pgl.like.BREX | [illegible] | 1229 | Bacteria>Terrabacteria_group>Actinomycetota>Actinomycetes>Micrococcales>Micrococcaceae | GCF_946892675.1 | Arthrobacter_sp._Marseille-P9274 | OC550_RS03490 | 772318..776007 | 2866572 | DUF6079-family-protein-[Arthrobacter-sp.-Marseille-P9274] |
| WP_272026547.1 | Type2.Pgl.like.BREX | [illegible] | 1279 | Bacteria>Terrabacteria_group>Actinomycetota>Actinomycetes>Micrococcales>Micrococcaceae | GCF_947492165.1 | Kocuria_rosea | PQG49_RS13695 | 8067..11906 | 1275 | DUF6079-family-protein-[Kocuria-rosea] |
| WP_022924594.1 | Type2.Pgl.like.BREX | [illegible] | 1254 | Bacteria>Terrabacteria_group>Actinomycetota>Actinomycetes>Micrococcales>Ornithinimicrobiaceae | GCF_000421245.1 | Serinicoccus_marinus | FU792_RS06180 | 1326112..1329876 | 247333 | hypothetical-protein-[Serinicoccus-marinus] |
| WP_139720892.1 | Type2.Pgl.like.BREX | [illegible] | 1258 | Bacteria>Terrabacteria_group>Actinomycetota>Actinomycetes>Micrococcales>Ornithinimicrobiaceae | GCF_006337125.1 | Serinicoccus_chungangensis | FHD63_RS05785 | 1305359..1309135 | 767452 | DUF6079-family-protein-[Serinicoccus-chungangensis] |
| WP_097187146.1 | Type2.Pgl.like.BREX | [illegible] | 1251 | Bacteria>Terrabacteria_group>Actinomycetota>Actinomycetes>Micrococcales>Ornithinimicrobiaceae | GCF_900221015.1 | Ornithinimicrobium_cerasi | CRP33_RS04660 | 109157..112912 | 2248773 | phage-resistance-protein-[Ornithinimicrobium-cerasi] |
| MCK9795949.1 | Type2.Pgl.like.BREX | [illegible] | 1279 | Bacteria>Terrabacteria_group>Actinomycetota>Actinomycetes>Micrococcales>Promicromonosporaceae | GCA_023241745.1 | Isoptericola_sp._4D.3 | M1843_19560 | 49170..53009 | 2918523 | hypothetical-protein-[Isoptericola-sp.-4D.3] |
| WP_069982322.1 | Type2.Pgl.like.BREX | [illegible] | 1279 | Bacteria>Terrabacteria_group>Actinomycetota>Actinomycetes>Micrococcales>Promicromonosporaceae | GCF_001750215.1 | Isoptericola_variabilis | JZ7iCDA_RS01635 | 7887..11726 | 139208 | DUF6079-family-protein-[Isoptericola-variabilis] |
| WP_085387167.1 | Type2.Pgl.like.BREX | [illegible] | 1286 | Bacteria>Terrabacteria_group>Actinomycetota>Actinomycetes>Micrococcales>Promicromonosporaceae | GCF_002105775.1 | Cellulosimicrobium_sp._TH-20 | B8281_RS05525 | 1243526..1247386 | 1980001 | hypothetical-protein-[Cellulosimicrobium-sp.-TH-20] |
| WP_155100109.1 | Type2.Pgl.like.BREX | [illegible] | 1224 | Bacteria>Terrabacteria_group>Actinomycetota>Actinomycetes>Micrococcales>Promicromonosporaceae | GCF_009711085.1 | Cellulosimicrobium_composti | GJV82_RS17455 | 8365..12039 | 2672572 | hypothetical-protein-[Cellulosimicrobium-composti] |
| WP_223403307.1 | Type2.Pgl.like.BREX | [illegible] | 1225 | Bacteria>Terrabacteria_group>Actinomycetota>Actinomycetes>Micrococcales>Ruaniaceae | GCF_019947135.1 | Occultella_gossypii | KCQ71_RS04525 | 6371..10048 | 2800820 | hypothetical-protein-[Occultella-gossypii] |
| ADL44857.1 | Type2.Pgl.like.BREX | [illegible] | 1268 | Bacteria>Terrabacteria_group>Actinomycetota>Actinomycetes>Micromonosporales>Micromonosporaceae | GCA_000145235.1 | Micromonospora_aurantiaca_ATCC_27029 | Micau_1295 | 1340460..1344266 | 644283 | phage-resistance-protein-[Micromonospora-aurantiaca-ATCC-27029] |
| EEP73860.1 | Type2.Pgl.like.BREX | [illegible] | 1384 | Bacteria>Terrabacteria_group>Actinomycetota>Actinomycetes>Micromonosporales>Micromonosporaceae | GCA_000158815.1 | Micromonospora_sp._ATCC_39149 | MCAG_04187 | 4972490..4976644 | 219305 | phage-resistance-protein-[Micromonospora-sp.-ATCC-39149] |
| OWV13587.1 | Type2.Pgl.like.BREX | [illegible] | 1277 | Bacteria>Terrabacteria_group>Actinomycetota>Actinomycetes>Micromonosporales>Micromonosporaceae | GCA_002210435.1 | Micromonospora_wenchangensis | B5D80_00150 | 34626..38459 | 1185415 | phage-resistance-protein-[Micromonospora-wenchangensis] |
| PMR63113.1 | Type2.Pgl.like.BREX | [illegible] | 1257 | Bacteria>Terrabacteria_group>Actinomycetota>Actinomycetes>Micromonosporales>Micromonosporaceae | GCA_002879555.1 | Verrucosispora_sp._ts21 | C1A38_00595 | 137346..141119 | 2069341 | phage-resistance-protein-[Verrucosispora-sp.-ts21] |
| PWR15835.1 | Type2.Pgl.like.BREX | [illegible] | 1259 | Bacteria>Terrabacteria_group>Actinomycetota>Actinomycetes>Micromonosporales>Micromonosporaceae | GCA_003172935.1 | Micromonospora_sp._4G51 | DKT69_08935 | 26626..30405 | 2202420 | phage-resistance-protein-[Micromonospora-sp.-4G51] |
| RBJ11687.1 | Type2.Pgl.like.BREX | [illegible] | 1248 | Bacteria>Terrabacteria_group>Actinomycetota>Actinomycetes>Micromonosporales>Micromonosporaceae | GCA_003298855.1 | Micromonospora_chalcea | DRA43_00175 | 44960..48706 | 1874 | phage-resistance-protein-[Micromonospora-chalcea] |
| RGC69899.1 | Type2.Pgl.like.BREX | [illegible] | 1259 | Bacteria>Terrabacteria_group>Actinomycetota>Actinomycetes>Micromonosporales>Micromonosporaceae | GCA_003434305.1 | Micromonospora_sp._MW-13 | C5N14_05680 | 359766..363545 | 2094022 | hypothetical-protein-C5N14_05680-[Micromonospora-sp.-MW-13] |
| NIL64524.1 | Type2.Pgl.like.BREX | [illegible] | 1320 | Bacteria>Terrabacteria_group>Actinomycetota>Actinomycetes>Micromonosporales>Micromonosporaceae | GCA_011765705.1 | Salinispora_arenicola | HCB39_24800 | 6501..10463 | 168697 | phage-resistance-protein-[Salinispora-arenicola] |
| NIL64662.1 | Type2.Pgl.like.BREX | [illegible] | 136 | Bacteria>Terrabacteria_group>Actinomycetota>Actinomycetes>Micromonosporales>Micromonosporaceae | GCA_011765705.1 | Salinispora_arenicola | HCB39_25670 | 7737..8147 | 168697 | hypothetical-protein-[Salinispora-arenicola] |
| NMO57591.1 | Type2.Pgl.like.BREX | [illegible] | 1229 | Bacteria>Terrabacteria_group>Actinomycetota>Actinomycetes>Micromonosporales>Micromonosporaceae | GCA_012933445.1 | Actinoplanes_sp._TBRC_11911 | HH310_41300 | 30518..34207 | 2729386 | phage-resistance-protein-[Actinoplanes-sp.-TBRC-11911] |
| MBF9127685.1 | Type2.Pgl.like.BREX | [illegible] | 1251 | Bacteria>Terrabacteria_group>Actinomycetota>Actinomycetes>Micromonosporales>Micromonosporaceae | GCA_015690345.1 | Plantactinospora_alkalitolerans | IOC86_01545 | 11246..15001 | 2789879 | phage-resistance-protein-[Plantactinospora-alkalitolerans] |
| MBM0236142.1 | Type2.Pgl.like.BREX | [illegible] | 1104 | Bacteria>Terrabacteria_group>Actinomycetota>Actinomycetes>Micromonosporales>Micromonosporaceae | GCA_016802835.1 | Micromonospora_sp._ATA32 | JNW88_01625 | 16339..19653 | 2806103 | hypothetical-protein-[Micromonospora-sp.-ATA32] |
| GID80948.1 | Type2.Pgl.like.BREX | [illegible] | 1245 | Bacteria>Terrabacteria_group>Actinomycetota>Actinomycetes>Micromonosporales>Micromonosporaceae | GCA_016862115.1 | Actinoplanes_deccanensis | Ade02nite_95890 | 18490..22227 | 113561 | hypothetical-protein-Ade02nite_95890-[Actinoplanes-deccanensis] |
| GIF85896.1 | Type2.Pgl.like.BREX | [illegible] | 1243 | Bacteria>Terrabacteria_group>Actinomycetota>Actinomycetes>Micromonosporales>Micromonosporaceae | GCA_016862575.1 | Catellatospora_bangladeshensis | Cba03nite_72450 | 55149..58880 | 310355 | hypothetical-protein-Cba03nite_72450-[Catellatospora-bangladeshensis] |
| GIJ13476.1 | Type2.Pgl.like.BREX | [illegible] | 1278 | Bacteria>Terrabacteria_group>Actinomycetota>Actinomycetes>Micromonosporales>Micromonosporaceae | GCA_016863515.1 | Micromonospora_gifhornensis | Vgi01_01600 | 80178..84014 | 84594 | hypothetical-protein-Vgi01_01600-[Micromonospora-gifhornensis] |
| GIJ20358.1 | Type2.Pgl.like.BREX | [illegible] | 1248 | Bacteria>Terrabacteria_group>Actinomycetota>Actinomycetes>Micromonosporales>Micromonosporaceae | GCA_016863535.1 | Micromonospora_lutea | Vlu01_09820 | 13113..16859 | 419825 | hypothetical-protein-Vlu01_09820-[Micromonospora-lutea] |
| GIJ25135.1 | Type2.Pgl.like.BREX | [illegible] | 1265 | Bacteria>Terrabacteria_group>Actinomycetota>Actinomycetes>Micromonosporales>Micromonosporaceae | GCA_016863555.1 | Micromonospora_qiuiae | Vqi01_02970 | 72395..76192 | 502268 | hypothetical-protein-Vqi01_02970-[Micromonospora-qiuiae] |
| GIJ31530.1 | Type2.Pgl.like.BREX | [illegible] | 1249 | Bacteria>Terrabacteria_group>Actinomycetota>Actinomycetes>Micromonosporales>Micromonosporaceae | GCA_016863575.1 | Micromonospora_sediminimaris | Vse01_06780 | 106066..109815 | 547162 | hypothetical-protein-Vse01_06780-[Micromonospora-sediminimaris] |
| GIJ63077.1 | Type2.Pgl.like.BREX | [illegible] | 1235 | Bacteria>Terrabacteria_group>Actinomycetota>Actinomycetes>Micromonosporales>Micromonosporaceae | GCA_016863635.1 | Virgisporangium_aurantiacum | Vau01_105930 | 35466..39173 | 175570 | hypothetical-protein-Vau01_105930-[Virgisporangium-aurantiacum] |
| GIJ74035.1 | Type2.Pgl.like.BREX | [illegible] | 1225 | Bacteria>Terrabacteria_group>Actinomycetota>Actinomycetes>Micromonosporales>Micromonosporaceae | GCA_016863655.1 | Virgisporangium_ochraceum | Voc01_089520 | 17095..20772 | 65505 | hypothetical-protein-Voc01_089520-[Virgisporangium-ochraceum] |
| MBQ1057676.1 | Type2.Pgl.like.BREX | [illegible] | 1249 | Bacteria>Terrabacteria_group>Actinomycetota>Actinomycetes>Micromonosporales>Micromonosporaceae | GCA_018070525.1 | Micromonospora_sp._C32 | KBX35_23085 | 127089..130838 | 2824877 | hypothetical-protein-[Micromonospora-sp.-C32] |
| UWZ39780.1 | Type2.Pgl.like.BREX | [illegible] | 1237 | Bacteria>Terrabacteria_group>Actinomycetota>Actinomycetes>Micromonosporales>Micromonosporaceae | GCA_025264685.1 | Dactylosporangium_roseum | Drose_17080 | 3689537..3693250 | 47989 | phage-resistance-protein-[Dactylosporangium-roseum] |
| WBB47306.1 | Type2.Pgl.like.BREX | [illegible] | 1247 | Bacteria>Terrabacteria_group>Actinomycetota>Actinomycetes>Micromonosporales>Micromonosporaceae | GCA_027497155.1 | Verrucosispora_sp._WMMA2044 | O3597_19410 | 4271873..4275616 | 3016419 | hypothetical-protein-O3597_19410-[Verrucosispora-sp.-WMMA2044] |
| WBB91554.1 | Type2.Pgl.like.BREX | [illegible] | 1248 | Bacteria>Terrabacteria_group>Actinomycetota>Actinomycetes>Micromonosporales>Micromonosporaceae | GCA_027497295.1 | Verrucosispora_sp._WMMC514 | O7597_00415 | 88656..92402 | 3015156 | hypothetical-protein-O7597_00415-[Verrucosispora-sp.-WMMC514] |
| WBB97459.1 | Type2.Pgl.like.BREX | [illegible] | 1278 | Bacteria>Terrabacteria_group>Actinomycetota>Actinomycetes>Micromonosporales>Micromonosporaceae | GCA_027497315.1 | Solwaraspora_sp._WMMA2059 | O7553_00200 | 51311..55147 | 3015160 | phage-resistance-protein-[Solwaraspora-sp.-WMMA2059] |
| GHJ17063.1 | Type2.Pgl.like.BREX | [illegible] | 1250 | Bacteria>Terrabacteria_group>Actinomycetota>Actinomycetes>Micromonosporales>Micromonosporaceae | GCA_027925725.1 | Micromonospora_sp._AKA38 | TPA0908_50580 | 1808836..1812588 | 2733861 | hypothetical-protein-TPA0908_50580-[Micromonospora-sp.-AKA38] |
| SCG34899.1 | Type2.Pgl.like.BREX | [illegible] | 1235 | Bacteria>Terrabacteria_group>Actinomycetota>Actinomycetes>Micromonosporales>Micromonosporaceae | GCA_900090105.1 | Micromonospora_humi | GA0070213_101291 | 330964..334671 | 745366 | Protein-of-unknown-function-[Micromonospora-humi] |
| SCL45916.1 | Type2.Pgl.like.BREX | [illegible] | 1261 | Bacteria>Terrabacteria_group>Actinomycetota>Actinomycetes>Micromonosporales>Micromonosporaceae | GCA_900090315.1 | Micromonospora_citrea | GA0070606_0885 | 915217..919002 | 47855 | hypothetical-protein-GA0070606_0885-[Micromonospora-citrea] |
| SCF44468.1 | Type2.Pgl.like.BREX | [illegible] | 1262 | Bacteria>Terrabacteria_group>Actinomycetota>Actinomycetes>Micromonosporales>Micromonosporaceae | GCA_900091555.1 | Micromonospora_mirobrigensis | GA0070564_11094 | 104640..108428 | 262898 | hypothetical-protein-GA0070564_11094-[Micromonospora-mirobrigensis] |
| WP_244879734.1 | Type2.Pgl.like.BREX | [illegible] | 1312 | Bacteria>Terrabacteria_group>Actinomycetota>Actinomycetes>Micromonosporales>Micromonosporaceae | GCF_000145235.1 | Micromonospora_aurantiaca | MICAU_RS06490 | 1340328..1344266 | 47850 | phage-resistance-protein-[Micromonospora-aurantiaca] |
| WP_036374655.1 | Type2.Pgl.like.BREX | [illegible] | 1268 | Bacteria>Terrabacteria_group>Actinomycetota>Actinomycetes>Micromonosporales>Micromonosporaceae | GCF_000158815.1 | Micromonospora_sp._ATCC_39149 | MCAG_RS21715 | 4972838..4976644 | 219305 | hypothetical-protein-[Micromonospora-sp.-ATCC-39149] |
| WP_018791391.1 | Type2.Pgl.like.BREX | [illegible] | 1295 | Bacteria>Terrabacteria_group>Actinomycetota>Actinomycetes>Micromonosporales>Micromonosporaceae | GCF_000375005.1 | Salinispora_arenicola | C570_RS0124430 | 7009..10896 | 168697 | hypothetical-protein-[Salinispora-arenicola] |
| WP_023560516.1 | Type2.Pgl.like.BREX | [illegible] | 1242 | Bacteria>Terrabacteria_group>Actinomycetota>Actinomycetes>Micromonosporales>Micromonosporaceae | GCF_000494755.1 | Actinoplanes_friuliensis | AFR_RS29710 | 6440598..6444326 | 196914 | DUF499-domain-containing-protein-[Actinoplanes-friuliensis] |
| WP_018253482.1 | Type2.Pgl.like.BREX | [illegible] | 1300 | Bacteria>Terrabacteria_group>Actinomycetota>Actinomycetes>Micromonosporales>Micromonosporaceae | GCF_000514435.1 | Salinispora_mooreana | B174_RS0103450 | 111334..115236 | 999545 | hypothetical-protein-[Salinispora-mooreana] |
| WP_030270088.1 | Type2.Pgl.like.BREX | [illegible] | 1268 | Bacteria>Terrabacteria_group>Actinomycetota>Actinomycetes>Micromonosporales>Micromonosporaceae | GCF_000717585.1 | Micromonospora_globosa | IF06_RS0108605 | 37560..41366 | 47863 | phage-resistance-protein-[Micromonospora-globosa] |
| WP_030490600.1 | Type2.Pgl.like.BREX | [illegible] | 1237 | Bacteria>Terrabacteria_group>Actinomycetota>Actinomycetes>Micromonosporales>Micromonosporaceae | GCF_000718555.1 | Micromonospora_chokoriensis | IF46_RS0123835 | 124154..127867 | 356851 | hypothetical-protein-[Micromonospora-chokoriensis] |
| WP_046566816.1 | Type2.Pgl.like.BREX | [illegible] | 1247 | Bacteria>Terrabacteria_group>Actinomycetota>Actinomycetes>Micromonosporales>Micromonosporaceae | GCF_000982955.1 | Micromonospora_sp._HK10 | LQ51_RS16800 | 146901..150644 | 1538294 | DUF6079-family-protein-[Micromonospora-sp.-HK10] |
| WP_071094722.1 | Type2.Pgl.like.BREX | [illegible] | 1246 | Bacteria>Terrabacteria_group>Actinomycetota>Actinomycetes>Micromonosporales>Micromonosporaceae | GCF_001855515.1 | Micromonospora_sp._WMMB235 | BFV98_RS02480 | 625702..629442 | 1172030 | hypothetical-protein-[Micromonospora-sp.-WMMB235] |
| WP_071098590.1 | Type2.Pgl.like.BREX | [illegible] | 1246 | Bacteria>Terrabacteria_group>Actinomycetota>Actinomycetes>Micromonosporales>Micromonosporaceae | GCF_001855515.1 | Micromonospora_sp._WMMB235 | BFV98_RS29930 | 6717187..6720927 | 1172030 | DUF6079-family-protein-[Micromonospora-sp.-WMMB235] |
| WP_245854123.1 | Type2.Pgl.like.BREX | [illegible] | 1282 | Bacteria>Terrabacteria_group>Actinomycetota>Actinomycetes>Micromonosporales>Micromonosporaceae | GCF_002210435.1 | Micromonospora_wenchangensis | B5D80_RS00150 | 34626..38474 | 1185415 | phage-resistance-protein-[Micromonospora-wenchangensis] |
| WP_109817248.1 | Type2.Pgl.like.BREX | [illegible] | 1253 | Bacteria>Terrabacteria_group>Actinomycetota>Actinomycetes>Micromonosporales>Micromonosporaceae | GCF_003172955.1 | Micromonospora_acroterricola | DKT68_RS10675 | 45241..49002 | 2202421 | phage-resistance-protein-[Micromonospora-acroterricola] |
| WP_109902741.1 | Type2.Pgl.like.BREX | [illegible] | 1234 | Bacteria>Terrabacteria_group>Actinomycetota>Actinomycetes>Micromonosporales>Micromonosporaceae | GCF_003176715.1 | Micromonospora_sp._S4605 | DLJ47_RS23310 | 37628..41332 | 1420897 | phage-resistance-protein-[Micromonospora-sp.-S4605] |
| WP_109902821.1 | Type2.Pgl.like.BREX | [illegible] | 1275 | Bacteria>Terrabacteria_group>Actinomycetota>Actinomycetes>Micromonosporales>Micromonosporaceae | GCF_003176715.1 | Micromonospora_sp._S4605 | DLJ47_RS23535 | 18706..22533 | 1420897 | phage-resistance-protein-[Micromonospora-sp.-S4605] |
| WP_109965346.1 | Type2.Pgl.like.BREX | [illegible] | 1232 | Bacteria>Terrabacteria_group>Actinomycetota>Actinomycetes>Micromonosporales>Micromonosporaceae | GCF_003176745.1 | Micromonospora_globispora | DLE60_RS16080 | 8802..12500 | 1450148 | phage-resistance-protein-[Micromonospora-globispora] |
| WP_112699616.1 | Type2.Pgl.like.BREX | [illegible] | 1239 | Bacteria>Terrabacteria_group>Actinomycetota>Actinomycetes>Micromonosporales>Micromonosporaceae | GCF_003264365.1 | Micromonospora_noduli | GUI43_RS21395 | 33589..37308 | 709876 | phage-resistance-protein-[Micromonospora-noduli] |
| WP_117666184.1 | Type2.Pgl.like.BREX | [illegible] | 1272 | Bacteria>Terrabacteria_group>Actinomycetota>Actinomycetes>Micromonosporales>Micromonosporaceae | GCF_003434305.1 | Micromonospora_sp._MW-13 | C5N14_RS05745 | 359727..363545 | 2094022 | phage-resistance-protein-[Micromonospora-sp.-MW-13] |
| WP_120326728.1 | Type2.Pgl.like.BREX | [illegible] | 1252 | Bacteria>Terrabacteria_group>Actinomycetota>Actinomycetes>Micromonosporales>Micromonosporaceae | GCF_003610785.1 | Micromonospora_globbae | D7I43_RS02485 | 15181..18939 | 1894969 | phage-resistance-protein-[Micromonospora-globbae] |
| WP_120682047.1 | Type2.Pgl.like.BREX | [illegible] | 1249 | Bacteria>Terrabacteria_group>Actinomycetota>Actinomycetes>Micromonosporales>Micromonosporaceae | GCF_003626545.1 | Micromonospora_musae | D7147_RS25550 | 8535..12284 | 1894970 | phage-resistance-protein-[Micromonospora-musae] |
| WP_120781542.1 | Type2.Pgl.like.BREX | [illegible] | 1253 | Bacteria>Terrabacteria_group>Actinomycetota>Actinomycetes>Micromonosporales>Micromonosporaceae | GCF_003626655.1 | Micromonospora_costi | D7193_RS22530 | 149877..153638 | 1530042 | DUF499-domain-containing-protein-[Micromonospora-costi] |
| WP_121684609.1 | Type2.Pgl.like.BREX | [illegible] | 1247 | Bacteria>Terrabacteria_group>Actinomycetota>Actinomycetes>Micromonosporales>Micromonosporaceae | GCF_003667485.1 | Micromonospora_sp._BL1 | EAD96_RS14345 | 62472..66215 | 2478709 | phage-resistance-protein-[Micromonospora-sp.-BL1] |
| WP_123605937.1 | Type2.Pgl.like.BREX | [illegible] | 1277 | Bacteria>Terrabacteria_group>Actinomycetota>Actinomycetes>Micromonosporales>Micromonosporaceae | GCF_003751085.1 | Micromonospora_sp._Llam0 | EDC02_RS31830 | 993308..997141 | 2485143 | phage-resistance-protein-[Micromonospora-sp.-Llam0] |
| WP_123558026.1 | Type2.Pgl.like.BREX | [illegible] | 1266 | Bacteria>Terrabacteria_group>Actinomycetota>Actinomycetes>Micromonosporales>Micromonosporaceae | GCF_003762835.1 | Micromonospora_sp._HM5-17 | EF879_RS12655 | 725957..729757 | 2487710 | phage-resistance-protein-[Micromonospora-sp.-HM5-17] |
| WP_127508173.1 | Type2.Pgl.like.BREX | [illegible] | 1246 | Bacteria>Terrabacteria_group>Actinomycetota>Actinomycetes>Micromonosporales>Micromonosporaceae | GCF_003999975.1 | Actinoplanes_sp._LAM7112 | EER30_RS43470 | 52334..56074 | 2486853 | phage-resistance-protein-[Actinoplanes-sp.-LAM7112] |
| WP_130403424.1 | Type2.Pgl.like.BREX | [illegible] | 1276 | Bacteria>Terrabacteria_group>Actinomycetota>Actinomycetes>Micromonosporales>Micromonosporaceae | GCF_004217135.1 | Micromonospora_violae | EV382_RS18010 | 4090540..4094370 | 1278207 | phage-resistance-protein-[Micromonospora-violae] |
| WP_132338410.1 | Type2.Pgl.like.BREX | [illegible] | 1242 | Bacteria>Terrabacteria_group>Actinomycetota>Actinomycetes>Micromonosporales>Micromonosporaceae | GCF_004348325.1 | Micromonospora_sp._KC213 | E1166_RS11800 | 17015..20743 | 2530378 | phage-resistance-protein-[Micromonospora-sp.-KC213] |
| WP_139583841.1 | Type2.Pgl.like.BREX | [illegible] | 1276 | Bacteria>Terrabacteria_group>Actinomycetota>Actinomycetes>Micromonosporales>Micromonosporaceae | GCF_006228125.1 | Micromonospora_orduensis | FHG89_RS08630 | 13298..17128 | 1420891 | phage-resistance-protein-[Micromonospora-orduensis] |
| WP_141908168.1 | Type2.Pgl.like.BREX | [illegible] | 1254 | Bacteria>Terrabacteria_group>Actinomycetota>Actinomycetes>Micromonosporales>Micromonosporaceae | GCF_006715105.1 | Micromonospora_sp._A202 | FBZ33_RS05010 | 1059749..1063513 | 2572899 | phage-resistance-protein-[Micromonospora-sp.-A202] |
| WP_144127959.1 | Type2.Pgl.like.BREX | [illegible] | 1222 | Bacteria>Terrabacteria_group>Actinomycetota>Actinomycetes>Micromonosporales>Micromonosporaceae | GCF_007483665.1 | Catellatospora_sichuanensis | FLL47_RS40300 | 47041..50709 | 1969805 | phage-resistance-protein-[Catellatospora-sichuanensis] |
| WP_154942606.1 | Type2.Pgl.like.BREX | [illegible] | 1291 | Bacteria>Terrabacteria_group>Actinomycetota>Actinomycetes>Micromonosporales>Micromonosporaceae | GCF_007829925.1 | Micromonospora_palomenae | FHX75_RS24210 | 697879..701754 | 1461247 | phage-resistance-protein-[Micromonospora-palomenae] |
| WP_148425376.1 | Type2.Pgl.like.BREX | [illegible] | 1250 | Bacteria>Terrabacteria_group>Actinomycetota>Actinomycetes>Micromonosporales>Micromonosporaceae | GCF_008085935.1 | Micromonospora_sp._AP08 | FXF50_RS01520 | 333481..337233 | 2604467 | DUF499-domain-containing-protein-[Micromonospora-sp.-AP08] |
| WP_151490982.1 | Type2.Pgl.like.BREX | [illegible] | 1278 | Bacteria>Terrabacteria_group>Actinomycetota>Actinomycetes>Micromonosporales>Micromonosporaceae | GCF_008868155.1 | Micromonospora_sp._ALFpr18c | F8271_RS18295 | 14793..18629 | 1458665 | phage-resistance-protein-[Micromonospora-sp.-ALFpr18c] |
| WP_155331844.1 | Type2.Pgl.like.BREX | [illegible] | 1252 | Bacteria>Terrabacteria_group>Actinomycetota>Actinomycetes>Micromonosporales>Micromonosporaceae | GCF_009707425.1 | Micromonospora_sp._WMMC415 | GKC29_17435 | 3702765..3706523 | 2675222 | phage-resistance-protein-[Micromonospora-sp.-WMMC415] |
| WP_248294924.1 | Type2.Pgl.like.BREX | [illegible] | 1227 | Bacteria>Terrabacteria_group>Actinomycetota>Actinomycetes>Micromonosporales>Micromonosporaceae | GCF_012933445.1 | Actinoplanes_sp._TBRC_11911 | HH310_RS41300 | 30518..34201 | 2729386 | phage-resistance-protein-[Actinoplanes-sp.-TBRC-11911] |
| WP_178063746.1 | Type2.Pgl.like.BREX | [illegible] | 1268 | Bacteria>Terrabacteria_group>Actinomycetota>Actinomycetes>Micromonosporales>Micromonosporaceae | GCF_013389765.1 | Micromonospora_carbonacea | HXZ27_RS07685 | 1786517..1790323 | 47853 | phage-resistance-protein-[Micromonospora-carbonacea] |
| WP_179805403.1 | Type2.Pgl.like.BREX | [illegible] | 91 | Bacteria>Terrabacteria_group>Actinomycetota>Actinomycetes>Micromonosporales>Micromonosporaceae | GCF_013410185.1 | Micromonospora_purpureochromogenes | HDA35_RS28050 | 6023294..6023569 | 47872 | hypothetical-protein-[Micromonospora-purpureochromogenes] |
| WP_179779345.1 | Type2.Pgl.like.BREX | [illegible] | 1251 | Bacteria>Terrabacteria_group>Actinomycetota>Actinomycetes>Micromonosporales>Micromonosporaceae | GCF_013410645.1 | Micromonospora_jinlongensis | HNR22_RS05305 | 1251485..1255240 | 1287877 | phage-resistance-protein-[Micromonospora-jinlongensis] |
| WP_180200415.1 | Type2.Pgl.like.BREX | [illegible] | 1303 | Bacteria>Terrabacteria_group>Actinomycetota>Actinomycetes>Micromonosporales>Micromonosporaceae | GCF_013416675.1 | Salinispora_sp._H7-4 | H0524_RS22350 | 30513..34424 | 2748321 | phage-resistance-protein-[Salinispora-sp.-H7-4] |
| WP_181572023.1 | Type2.Pgl.like.BREX | [illegible] | 1249 | Bacteria>Terrabacteria_group>Actinomycetota>Actinomycetes>Micromonosporales>Micromonosporaceae | GCF_013694245.1 | Micromonospora_ferruginea | H1D33_RS12900 | 2682005..2685754 | 2749844 | DUF6079-family-protein-[Micromonospora-ferruginea] |
| WP_260322248.1 | Type2.Pgl.like.BREX | [illegible] | 1257 | Bacteria>Terrabacteria_group>Actinomycetota>Actinomycetes>Micromonosporales>Micromonosporaceae | GCF_014201145.1 | Micromonospora_parathelypteridis | HNR20_RS20430 | 4527676..4531449 | 1839617 | phage-resistance-protein-[Micromonospora-parathelypteridis] |
| WP_184686293.1 | Type2.Pgl.like.BREX | [illegible] | 1247 | Bacteria>Terrabacteria_group>Actinomycetota>Actinomycetes>Micromonosporales>Micromonosporaceae | GCF_014203425.1 | Micromonospora_echinospora | FHU28_RS21490 | 4613810..4617553 | 1877 | DUF499-domain-containing-protein-[Micromonospora-echinospora] |
| WP_221487643.1 | Type2.Pgl.like.BREX | [illegible] | 1266 | Bacteria>Terrabacteria_group>Actinomycetota>Actinomycetes>Micromonosporales>Micromonosporaceae | GCF_014205165.1 | Micromonospora_carbonacea | HDA31_RS22155 | 5299472..5303272 | 47853 | phage-resistance-protein-[Micromonospora-carbonacea] |
| WP_184182217.1 | Type2.Pgl.like.BREX | [illegible] | 1259 | Bacteria>Terrabacteria_group>Actinomycetota>Actinomycetes>Micromonosporales>Micromonosporaceae | GCF_014646315.1 | Micromonospora_parathelypteridis | IEY65_RS28755 | 129382..133161 | 1839617 | phage-resistance-protein-[Micromonospora-parathelypteridis] |
| WP_196923277.1 | Type2.Pgl.like.BREX | [illegible] | 1239 | Bacteria>Terrabacteria_group>Actinomycetota>Actinomycetes>Micromonosporales>Micromonosporaceae | GCF_015751785.1 | Micromonospora_vinacea | IW249_RS26745 | 5669487..5673206 | 709878 | hypothetical-protein-[Micromonospora-vinacea] |
| WP_146231013.1 | Type2.Pgl.like.BREX | [illegible] | 1252 | Bacteria>Terrabacteria_group>Actinomycetota>Actinomycetes>Micromonosporales>Micromonosporaceae | GCF_016802855.1 | unclassified_Micromonospora | JNW87_RS06290 | 11705..15463 | 2617518 | MULTISPECIES:-phage-resistance-protein-[unclassified-Micromonospora] |
| WP_238447619.1 | Type2.Pgl.like.BREX | [illegible] | 1256 | Bacteria>Terrabacteria_group>Actinomycetota>Actinomycetes>Micromonosporales>Micromonosporaceae | GCF_016803375.1 | Micromonospora_sp._4G55 | JNW89_RS03210 | 7127..10897 | 2806102 | hypothetical-protein-[Micromonospora-sp.-4G55] |
| WP_133876212.1 | Type2.Pgl.like.BREX | [illegible] | 1255 | Bacteria>Terrabacteria_group>Actinomycetota>Actinomycetes>Micromonosporales>Micromonosporaceae | GCF_016862015.1 | Actinoplanes_brasiliensis | C8E87_RS30125 | 6729671..6733438 | 52695 | phage-resistance-protein-[Actinoplanes-brasiliensis] |
| WP_203778155.1 | Type2.Pgl.like.BREX | [illegible] | 1218 | Bacteria>Terrabacteria_group>Actinomycetota>Actinomycetes>Micromonosporales>Micromonosporaceae | GCF_016862115.1 | Actinoplanes_deccanensis | Ade02nite_RS48075 | 18490..22146 | 113561 | hypothetical-protein-[Actinoplanes-deccanensis] |
| WP_092547197.1 | Type2.Pgl.like.BREX | [illegible] | 1243 | Bacteria>Terrabacteria_group>Actinomycetota>Actinomycetes>Micromonosporales>Micromonosporaceae | GCF_016862135.1 | Actinoplanes_derwentensis | Ade03nite_RS39185 | 21276..25007 | 113562 | hypothetical-protein-[Actinoplanes-derwentensis] |
| WP_203807939.1 | Type2.Pgl.like.BREX | [illegible] | 1220 | Bacteria>Terrabacteria_group>Actinomycetota>Actinomycetes>Micromonosporales>Micromonosporaceae | GCF_016862435.1 | Actinoplanes_tereljensis | Ate02nite_RS20625 | 60893..64555 | 571912 | phage-resistance-protein-[Actinoplanes-tereljensis] |
| WP_043526153.1 | Type2.Pgl.like.BREX | [illegible] | 1242 | Bacteria>Terrabacteria_group>Actinomycetota>Actinomycetes>Micromonosporales>Micromonosporaceae | GCF_016862455.1 | Actinoplanes_utahensis | Aut01nite_RS42125 | 11222..14950 | 1869 | hypothetical-protein-[Actinoplanes-utahensis] |
| WP_109591754.1 | Type2.Pgl.like.BREX | [illegible] | 1232 | Bacteria>Terrabacteria_group>Actinomycetota>Actinomycetes>Micromonosporales>Micromonosporaceae | GCF_016862475.1 | Actinoplanes_xinjiangensis | Axi01nite_RS08915 | 148861..152559 | 512350 | DUF6079-family-protein-[Actinoplanes-xinjiangensis] |
| WP_147315484.1 | Type2.Pgl.like.BREX | [illegible] | 1233 | Bacteria>Terrabacteria_group>Actinomycetota>Actinomycetes>Micromonosporales>Micromonosporaceae | GCF_016862495.1 | Asanoa_ferruginea | DFJ67_RS10715 | 2233471..2237172 | 53367 | phage-resistance-protein-[Asanoa-ferruginea] |
| WP_203702533.1 | Type2.Pgl.like.BREX | [illegible] | 1241 | Bacteria>Terrabacteria_group>Actinomycetota>Actinomycetes>Micromonosporales>Micromonosporaceae | GCF_016862515.1 | Asanoa_iriomotensis | Air01nite_RS13795 | 57096..60821 | 234613 | phage-resistance-protein-[Asanoa-iriomotensis] |
| WP_090804547.1 | Type2.Pgl.like.BREX | [illegible] | 1232 | Bacteria>Terrabacteria_group>Actinomycetota>Actinomycetes>Micromonosporales>Micromonosporaceae | GCF_016862535.1 | Asanoa_ishikariensis | BLV39_RS40650 | 257503..261201 | 137265 | phage-resistance-protein-[Asanoa-ishikariensis] |
| WP_203714740.1 | Type2.Pgl.like.BREX | [illegible] | 1213 | Bacteria>Terrabacteria_group>Actinomycetota>Actinomycetes>Micromonosporales>Micromonosporaceae | GCF_016862555.1 | Asanoa_siamensis | Asi02nite_RS19070 | 66429..70070 | 926357 | phage-resistance-protein-[Asanoa-siamensis] |
| WP_203756441.1 | Type2.Pgl.like.BREX | [illegible] | 1233 | Bacteria>Terrabacteria_group>Actinomycetota>Actinomycetes>Micromonosporales>Micromonosporaceae | GCF_016862575.1 | Catellatospora_bangladeshensis | Cba03nite_RS36035 | 55179..58880 | 310355 | hypothetical-protein-[Catellatospora-bangladeshensis] |
| WP_203846757.1 | Type2.Pgl.like.BREX | [illegible] | 1231 | Bacteria>Terrabacteria_group>Actinomycetota>Actinomycetes>Micromonosporales>Micromonosporaceae | GCF_016862795.1 | Dactylosporangium_siamense | Dsi01nite_RS15110 | 108084..111779 | 685454 | hypothetical-protein-[Dactylosporangium-siamense] |
| WP_168116349.1 | Type2.Pgl.like.BREX | [illegible] | 1227 | Bacteria>Terrabacteria_group>Actinomycetota>Actinomycetes>Micromonosporales>Micromonosporaceae | GCF_016863255.1 | Planosporangium_mesophilum | Pme01_RS16095 | 151646..155329 | 689768 | phage-resistance-protein-[Planosporangium-mesophilum] |
| WP_204289856.1 | Type2.Pgl.like.BREX | [illegible] | 1277 | Bacteria>Terrabacteria_group>Actinomycetota>Actinomycetes>Micromonosporales>Micromonosporaceae | GCF_016863515.1 | Micromonospora_gifhornensis | Vgi01_RS00810 | 80181..84014 | 84594 | phage-resistance-protein-[Micromonospora-gifhornensis] |
| WP_203994567.1 | Type2.Pgl.like.BREX | [illegible] | 1245 | Bacteria>Terrabacteria_group>Actinomycetota>Actinomycetes>Micromonosporales>Micromonosporaceae | GCF_016863535.1 | Micromonospora_lutea | Vlu01_RS04910 | 13122..16859 | 419825 | hypothetical-protein-[Micromonospora-lutea] |
| WP_204032214.1 | Type2.Pgl.like.BREX | [illegible] | 1247 | Bacteria>Terrabacteria_group>Actinomycetota>Actinomycetes>Micromonosporales>Micromonosporaceae | GCF_016863555.1 | Micromonospora_qiuiae | Vqi01_RS01475 | 72395..76138 | 502268 | hypothetical-protein-[Micromonospora-qiuiae] |
| WP_093402597.1 | Type2.Pgl.like.BREX | [illegible] | 1249 | Bacteria>Terrabacteria_group>Actinomycetota>Actinomycetes>Micromonosporales>Micromonosporaceae | GCF_016863575.1 | Micromonospora_sediminimaris | Vse01_RS03400 | 106066..109815 | 547162 | hypothetical-protein-[Micromonospora-sediminimaris] |
| WP_204009107.1 | Type2.Pgl.like.BREX | [illegible] | 1232 | Bacteria>Terrabacteria_group>Actinomycetota>Actinomycetes>Micromonosporales>Micromonosporaceae | GCF_016863635.1 | Virgisporangium_aurantiacum | Vau01_RS52800 | 35475..39173 | 175570 | phage-resistance-protein-[Virgisporangium-aurantiacum] |
| WP_239160936.1 | Type2.Pgl.like.BREX | [illegible] | 1222 | Bacteria>Terrabacteria_group>Actinomycetota>Actinomycetes>Micromonosporales>Micromonosporaceae | GCF_016863655.1 | Virgisporangium_ochraceum | Voc01_RS44730 | 17095..20763 | 65505 | phage-resistance-protein-[Virgisporangium-ochraceum] |
| WP_204944668.1 | Type2.Pgl.like.BREX | [illegible] | 1248 | Bacteria>Terrabacteria_group>Actinomycetota>Actinomycetes>Micromonosporales>Micromonosporaceae | GCF_016907275.1 | Micromonospora_luteifusca | JOD64_RS26115 | 5720487..5724233 | 709860 | hypothetical-protein-[Micromonospora-luteifusca] |
| WP_208577495.1 | Type2.Pgl.like.BREX | [illegible] | 1252 | Bacteria>Terrabacteria_group>Actinomycetota>Actinomycetes>Micromonosporales>Micromonosporaceae | GCF_017599015.1 | Micromonospora_tulbaghiae | J5U46_RS17340 | 18691..22449 | 479978 | phage-resistance-protein-[Micromonospora-tulbaghiae] |
| WP_210914512.1 | Type2.Pgl.like.BREX | [illegible] | 1286 | Bacteria>Terrabacteria_group>Actinomycetota>Actinomycetes>Micromonosporales>Micromonosporaceae | GCF_018070045.1 | Micromonospora_sp._U21 | KBX63_RS24020 | 115436..119296 | 2824899 | phage-resistance-protein-[Micromonospora-sp.-U21] |
| WP_210891808.1 | Type2.Pgl.like.BREX | [illegible] | 1279 | Bacteria>Terrabacteria_group>Actinomycetota>Actinomycetes>Micromonosporales>Micromonosporaceae | GCF_018070505.1 | Micromonospora_sp._C72 | KBX03_RS03155 | 128407..132246 | 2824880 | hypothetical-protein-[Micromonospora-sp.-C72] |
| WP_247688340.1 | Type2.Pgl.like.BREX | [illegible] | 1226 | Bacteria>Terrabacteria_group>Actinomycetota>Actinomycetes>Micromonosporales>Micromonosporaceae | GCF_018070525.1 | Micromonospora_sp._C32 | KBX35_RS23085 | 127158..130838 | 2824877 | hypothetical-protein-[Micromonospora-sp.-C32] |
| WP_212843868.1 | Type2.Pgl.like.BREX | [illegible] | 1245 | Bacteria>Terrabacteria_group>Actinomycetota>Actinomycetes>Micromonosporales>Micromonosporaceae | GCF_018326265.1 | Catellatospora_sp._IY07-71 | CS0771_57280 | 6381779..6385516 | 2728827 | phage-resistance-protein-[Catellatospora-sp.-IY07-71] |
| WP_215792561.1 | Type2.Pgl.like.BREX | [illegible] | 1237 | Bacteria>Terrabacteria_group>Actinomycetota>Actinomycetes>Micromonosporales>Micromonosporaceae | GCF_018829635.1 | Actinoplanes_bogorensis | KOI35_RS32765 | 171762..175475 | 1610840 | phage-resistance-protein-[Actinoplanes-bogorensis] |
| WP_230415349.1 | Type2.Pgl.like.BREX | [illegible] | 1330 | Bacteria>Terrabacteria_group>Actinomycetota>Actinomycetes>Micromonosporales>Micromonosporaceae | GCF_019697375.1 | Micromonospora_tarapacensis | KIF24_RS07250 | 261955..265947 | 2835305 | phage-resistance-protein-[Micromonospora-tarapacensis] |
| WP_229403999.1 | Type2.Pgl.like.BREX | [illegible] | 1242 | Bacteria>Terrabacteria_group>Actinomycetota>Actinomycetes>Micromonosporales>Micromonosporaceae | GCF_020884815.1 | Micromonospora_sp._NBRC_110038 | LPU61_RS02740 | 644366..648094 | 1550034 | phage-resistance-protein-[Micromonospora-sp.-NBRC-110038] |
| WP_259859975.1 | Type2.Pgl.like.BREX | [illegible] | 1224 | Bacteria>Terrabacteria_group>Actinomycetota>Actinomycetes>Micromonosporales>Micromonosporaceae | GCF_025137375.1 | Dactylosporangium_fulvum | Dfulv_RS45280 | 9836074..9839748 | 53359 | phage-resistance-protein-[Dactylosporangium-fulvum] |
| WP_260729733.1 | Type2.Pgl.like.BREX | [illegible] | 1235 | Bacteria>Terrabacteria_group>Actinomycetota>Actinomycetes>Micromonosporales>Micromonosporaceae | GCF_025264685.1 | Dactylosporangium_roseum | Drose_RS17225 | 3689543..3693250 | 47989 | phage-resistance-protein-[Dactylosporangium-roseum] |
| WP_052387983.1 | Type2.Pgl.like.BREX | [illegible] | 1223 | Bacteria>Terrabacteria_group>Actinomycetota>Actinomycetes>Micromonosporales>Micromonosporaceae | GCF_025264705.1 | Dactylosporangium_aurantiacum | Daura_RS48855 | 10914607..10918278 | 35754 | hypothetical-protein-[Dactylosporangium-aurantiacum] |
| WP_266267207.1 | Type2.Pgl.like.BREX | [illegible] | 1272 | Bacteria>Terrabacteria_group>Actinomycetota>Actinomycetes>Micromonosporales>Micromonosporaceae | GCF_026339755.1 | Micromonospora_sp._NBC_01655 | OOK41_RS27490 | 6244497..6248315 | 2975983 | phage-resistance-protein-[Micromonospora-sp.-NBC_01655] |
| WP_043326008.1 | Type2.Pgl.like.BREX | [illegible] | 1262 | Bacteria>Terrabacteria_group>Actinomycetota>Actinomycetes>Micromonosporales>Micromonosporaceae | GCF_027497275.1 | unclassified_Micromonospora | FJ969_RS15330 | 46960..50748 | 2617518 | MULTISPECIES:-phage-resistance-protein-[unclassified-Micromonospora] |

| Protein-ID | System | Gene-Neighbourhoods | Length | Taxonomy | Assembly-ID | Strain | Locus-Tag | Genomic-coordinates | TaxID | Fasta-defline |
|---|---|---|---|---|---|---|---|---|---|---|
| WP_047891313.1 | Type2.Pgl.like.BREX | <-HKD-DNase+HKD-Rease<-?<-?\|\|?-><-sGTP<-sGTP+sGTP+GTPase-AIG\|\|<-SF1-SF2-Helicase+wHTH+Small-B-barrel+DUF1998<-BrxD-AAA<-PglZ<-DUF6079*<-N6-METHYLASE<-PglW<-?<-?\|\|?->URI-DNase-> | 1248 | Bacteria>Terrabacteria_group>Actinomycetota>Actinomycetes>Micromonosporales>Micromonosporaceae | GCF_029626075.1 | Micromonospora_sp._RV43 | 07630_RS17935 | 4122924..4126670 | 1661387 | MULTISPECIES:-DUF6079-family-protein-[unclassified-Micromonospora] |
| WP_091056625.1 | Type2.Pgl.like.BREX | RADICAL-SAM->?->?->?->?-><-?<-AAA+HEPN<-ParB+HNH\|\|DUF6079*->PglZ->BrxD-AAA->SF1-SF2-Helicase+wHTH+Small-B-barrel+DUF1998-><-TetR-HTH\|\|?->ABhydrolase->?->Phage_holin->Peptidase_M36+CarboxypepD_reg-> | 1228 | Bacteria>Terrabacteria_group>Actinomycetota>Actinomycetes>Micromonosporales>Micromonosporaceae | GCF_900090105.1 | Micromonospora_humi | GA0070213_RS01445 | 330985..334671 | 745366 | DUF499-domain-containing-protein-[Micromonospora-humi] |
| WP_088971857.1 | Type2.Pgl.like.BREX | ABhydrolase->?->?->?->?->?-><-SF1-SF2-Helicase+wHTH+Small-B-barrel+DUF1998<-BrxD-AAA<-PglZ<-DUF6079*\|\|MULE-TRANSPOSASE-><-CASPASE\|\|DEDD_Tnp_IS110+Transposase_20->DNA_glycosylase-><-?<-?<-REase<-RNase<-N6-METHYLASE | 1257 | Bacteria>Terrabacteria_group>Actinomycetota>Actinomycetes>Micromonosporales>Micromonosporaceae | GCF_900090305.1 | Micromonospora_siamensis | GA0074704_RS19665 | 4245777..4249550 | 299152 | phage-resistance-protein-[Micromonospora-siamensis] |
| WP_091095298.1 | Type2.Pgl.like.BREX | LACI-HTH+PBPI->Abhydrolase_2->AraC-HTH+AraC-HTH->HD-><-SF1-SF2-Helicase+wHTH+Small-B-barrel+DUF1998<-BrxD-AAA<-PglZ<-DUF6079*<-CASPASE\|\|?->?-><-RNase\|\|DNA_glycosylase->?-><-N6-METHYLASE<-GntR-HTH | 1249 | Bacteria>Terrabacteria_group>Actinomycetota>Actinomycetes>Micromonosporales>Micromonosporaceae | GCF_900090315.1 | Micromonospora_citrea | GA0070606_RS04390 | 915217..918966 | 47855 | hypothetical-protein-[Micromonospora-citrea] |
| WP_091458195.1 | Type2.Pgl.like.BREX | PIN->?-><-DUF397<-HTH_31+DUF5753\|\|?->DUF6307->?-><-?<-?\|\|DUF6079*->PglZ->BrxD-AAA->SF1-SF2-Helicase+wHTH+Small-B-barrel+DUF1998->?-><-?<-?\|\|GGDEF+EAL->SIR2->ACT-> | 1252 | Bacteria>Terrabacteria_group>Actinomycetota>Actinomycetes>Micromonosporales>Micromonosporaceae | GCF_900091415.1 | Micromonospora_inyonensis | GA0074694_RS14325 | 3160410..3164168 | 47866 | phage-resistance-protein-[Micromonospora-inyonensis] |
| WP_088980550.1 | Type2.Pgl.like.BREX | Peptidase_S8-><-?<-SF1-SF2-Helicase+wHTH+Small-B-barrel+DUF1998<-BrxD-AAA<-PglZ<-DUF6079*\|\|?->?-><-DUF6307<-?\|\|HTH_31+DUF5753->DUF397-><-?<-N6-METHYLASE<-ParB+HNH | 1274 | Bacteria>Terrabacteria_group>Actinomycetota>Actinomycetes>Micromonosporales>Micromonosporaceae | GCF_900091495.1 | Micromonospora_echinospora | GA0070618_RS04800 | 1070135..1073959 | 1877 | phage-resistance-protein-[Micromonospora-echinospora] |
| WP_088962571.1 | Type2.Pgl.like.BREX | <-HTH_32<-?\|\|?->?->OLD-ABC+TOPRIM->SF1-SF2-Helicase-><-SF1-SF2-Helicase+wHTH+Small-B-barrel+DUF1998<-BrxD-AAA<-PglZ<-DUF6079*<-?<-N6-METHYLASE\|\|OLD-ABC->TOPRIM->ParB+HNH-><-PglW<-STYKIN<-HTH+TPR\|\|DEDD_Tnp_IS110+Transposase_20-> | 1239 | Bacteria>Terrabacteria_group>Actinomycetota>Actinomycetes>Micromonosporales>Micromonosporaceae | GCF_900091515.1 | Micromonospora_purpureochromogenes | GA0074696_RS20340 | 4398204..4401923 | 47872 | phage-resistance-protein-[Micromonospora-purpureochromogenes] |
| WP_245670258.1 | Type2.Pgl.like.BREX | <-REC+OmpR-HTH<-ST-Hiskin<-?<-ABC-ATPase+SbcC\|\|ABhydrolase-><-SF1-SF2-Helicase+wHTH+Small-B-barrel+DUF1998<-BrxD-AAA<-PglZ<-DUF6079*<-DUF6307<-?\|\|HTH_31->ACET->REase-><-N6-METHYLASE<-ParB+HNH<-TOPRIM<-OLD-ABC | 1266 | Bacteria>Terrabacteria_group>Actinomycetota>Actinomycetes>Micromonosporales>Micromonosporaceae | GCF_900091555.1 | Micromonospora_mirobrigensis | GA0070564_RS25905 | 104640..108440 | 262898 | phage-resistance-protein-[Micromonospora-mirobrigensis] |
| WP_091084811.1 | Type2.Pgl.like.BREX | <-DUF397<-HTH_31+DUF5753\|\|?->DUF6307->?-><-?<-?\|\|DUF6079*->PglZ->BrxD-AAA->SF1-SF2-Helicase+wHTH+Small-B-barrel+DUF1998-><-?\|\|GGDEF+EAL->SIR2->ACT-> | 1261 | Bacteria>Terrabacteria_group>Actinomycetota>Actinomycetes>Micromonosporales>Micromonosporaceae | GCF_900091585.1 | Micromonospora_nigra | GA0070616_RS19340 | 4441175..4444960 | 145857 | phage-resistance-protein-[Micromonospora-nigra] |
| WP_218106981.1 | Type2.Pgl.like.BREX | DDE_Tnp->N6-METHYLASE-p->RnaseH-><-SWIM<-DUF397<-HTH_31+DUF5753\|\|?->DUF6079*->PglZ->BrxD-AAA->SF1-SF2-Helicase+wHTH+Small-B-barrel+DUF1998-><-Peptidase_S8<-URI-DNase+UVR+UvrC_HhH_N+HHH<-?\|\|?-><-?<-nSTAND1+SbcC+ABC-ATPase+SbcC | 1252 | Bacteria>Terrabacteria_group>Actinomycetota>Actinomycetes>Micromonosporales>Micromonosporaceae | GCF_900091595.1 | Micromonospora_haikouensis | GA0070558_RS10795 | 5421..9179 | 686309 | DUF6079-family-protein-[Micromonospora-haikouensis] |
| WP_091562973.1 | Type2.Pgl.like.BREX | CASPASE->?->?->?->?-><-?<-?\|\|DUF6079*->PglZ->BrxD-AAA->Schlafen+HATPase+wHTH->ParA-Soj-PloopNTPase->SF1-SF2-Helicase+wHTH+Small-B-barrel+DUF1998->?->?->?-><-APATPase+ABC-ATPase+SbcC | 1244 | Bacteria>Terrabacteria_group>Actinomycetota>Actinomycetes>Micromonosporales>Micromonosporaceae | GCF_900107255.1 | Micromonospora_pattaloongensis | BLW47_RS24170 | 9662..13396 | 405436 | phage-resistance-protein-[Micromonospora-pattaloongensis] |
| WP_076473582.1 | Type2.Pgl.like.BREX | OLD-ABC->TOPRIM->ParB+HNH->N6-METHYLASE-><-?<-DUF397<-HTH_31+DUF5753\|\|?->DUF6307->DUF6079*->PglZ->BrxD-AAA->SF1-SF2-Helicase+wHTH+Small-B-barrel+DUF1998->?-><-HEPN | 1275 | Bacteria>Terrabacteria_group>Actinomycetota>Actinomycetes>Micromonosporales>Micromonosporaceae | GCF_900156065.1 | Micromonospora_avicenniae | BXA10_RS26535 | 37443..41270 | 1198245 | phage-resistance-protein-[Micromonospora-avicenniae] |
| WP_128136019.1 | Type2.Pgl.like.BREX | GNAT->DDE_5+DDE_Tnp_1->DDE_5+DDE_Tnp_1-><-?<-PIWI<-HNH+PNTD1+PNTD2\|\|DDE_Tnp->PglW->N6-METHYLASE->DUF6079*->BrxD-AAA->SF1-SF2-Helicase+wHTH+Small-B-barrel+DUF1998->\|\|MACRODOMAIN->sGTP+sGTP+GTPase-AIG->sGTP->\|\|ParB+HNH-> | 1248 | Bacteria>Terrabacteria_group>Actinomycetota>Actinomycetes>Micromonosporales>Micromonosporaceae | - | Micromonospora_chalcea | DRA43_RS00185 | 44960..48706 | 1874 | DUF6079-family-protein-[Micromonospora-chalcea] |
| WP_230393352.1 | Type2.Pgl.like.BREX | NACHT+68TM-wHTH-><-SF1-SF2-Helicase+wHTH+Small-B-barrel+DUF1998<-BrxD-AAA<-PglZ<-DUF6079*<-?\|\|HTH_31+DUF5753->DUF397->?->?->HTH-><-?<-N6-METHYLASE | 1249 | Bacteria>Terrabacteria_group>Actinomycetota>Actinomycetes>Micromonosporales>Micromonosporaceae | - | Plantactinospora_alkalitolerans | IOC86_RS01545 | 11246..14995 | 2789879 | phage-resistance-protein-[Plantactinospora-alkalitolerans] |
| WP_014010192.1 | Type2.Pgl.like.BREX | EVE+REase->REase+SF1-SF2-Helicase->HNH->REase+SF1-SF2-Helicase-><-HEPN\|\|PglW->?->N6-METHYLASE->DUF6079*->PglZ->BrxD-AAA->SIR2->SF1-SF2-Helicase+wHTH+Small-B-barrel+DUF1998-><-Aminotran_1_2\|\|zf-ISL3+HTH_Tnp_ISL3+DDE_Tnp_ISL3-><-Aminotran_3 | 1304 | Bacteria>Terrabacteria_group>Actinomycetota>Actinomycetes>Mycobacteriales>Corynebacteriaceae | GCF_000179395.2 | Corynebacterium_variabile | CVAR_RS08820 | 1922130..1926044 | 1727 | DUF6079-family-protein-[Corynebacterium-variabile] |
| WP_059289748.1 | Type2.Pgl.like.BREX | <-ICLR-HTH<-DDE_Tnp_1+DDE_Tnp_1_2\|\|PglW->N6-METHYLASE->DUF6079*->PglZ->BrxD-AAA->SF1-SF2-Helicase+wHTH+Small-B-barrel+DUF1998-><-?\|\|HEPN-> | 1277 | Bacteria>Terrabacteria_group>Actinomycetota>Actinomycetes>Mycobacteriales>Corynebacteriaceae | GCF_001518935.2 | Corynebacterium_glutamicum | APT58_RS12165 | 2569631..2573464 | 1718 | DUF6079-family-protein-[Corynebacterium-glutamicum] |
| WP_105324910.1 | Type2.Pgl.like.BREX | ABC-ATPase+SbcC->FAD_binding_9+SIP+Methyltransf_2->ABC-ATPase->ABC-ATPase->?->tRNA-><-SF1-SF2-Helicase+wHTH+Small-B-barrel+DUF1998<-BrxD-AAA<-PglZ<-DUF6079*<-N6-METHYLASE<-PglW<-?\|\|?->?-><-?<-?\|\|?->ABC-ATPase+SbcC-> | 1270 | Bacteria>Terrabacteria_group>Actinomycetota>Actinomycetes>Mycobacteriales>Corynebacteriaceae | GCF_002967075.1 | Corynebacterium_sp._J010B-136 | C5Y44_RS10645 | 148454..152266 | 2099401 | DUF6079-family-protein-[Corynebacterium-sp.-J010B-136] |
| WP_191733017.1 | Type2.Pgl.like.BREX | <-HEPN\|\|?-><-SF1-SF2-Helicase+wHTH+Small-B-barrel+DUF1998<-BrxD-AAA<-PglZ<-DUF6079*<-N6-METHYLASE<-PglW<-?<-?\|\|?->?->?->?-><-MODE-HTH+PBPII | 1285 | Bacteria>Terrabacteria_group>Actinomycetota>Actinomycetes>Mycobacteriales>Corynebacteriaceae | GCF_014837045.1 | Corynebacterium_gallinarum | H9627_RS05525 | 176693..180550 | 2762214 | DUF6079-family-protein-[Corynebacterium-gallinarum] |
| WP_244805207.1 | Type2.Pgl.like.BREX | <-SF1-SF2-Helicase+wHTH+Small-B-barrel+DUF1998\|\|?->HTH_3-><-BrxD-AAA<-PglZ<-DUF6079*<-N6-METHYLASE<-PglW<-PIN<-OLD-ABC<-URI-DNase+SF1-SF2-Helicase<-REase+SF1-SF2-Helicase\|\|35exo+TerB+TerB+BRCT+SIGMA-HTH->OLD-ABC->HNH-> | 1292 | Bacteria>Terrabacteria_group>Actinomycetota>Actinomycetes>Mycobacteriales>Corynebacteriaceae | GCF_022869005.1 | Corynebacterium_kalidii | MUN33_RS12285 | 327139..331017 | 2931982 | DUF6079-family-protein-[Corynebacterium-kalidii] |
| WP_073883613.1 | Type2.Pgl.like.BREX | <-SF1-SF2-Helicase+wHTH+Small-B-barrel+DUF1998<-SbcC<-BrxD-AAA<-PglZ<-DUF6079*<-N6-METHYLASE<-?<-PglW<-PIN<-OLD-ABC<-REase+SF1-SF2-Helicase<-REase+SF1-SF2-Helicase\|\|OLD-ABC->HNH-> | 1316 | Bacteria>Terrabacteria_group>Actinomycetota>Actinomycetes>Mycobacteriales>Corynebacteriaceae | GCF_900015285.1 | Corynebacterium_variabile | CVAR292_RS03495 | 9616..13566 | 1727 | DUF6079-family-protein-[Corynebacterium-variabile] |
| WP_158726429.1 | Type2.Pgl.like.BREX | REase+SF1-SF2-Helicase->nSTAND1+HNH->?->PglW->MPTase->PIN->N6-METHYLASE->ParB+HNH->DUF6079*->PglZ->?->?->?-><-?\|\|BrxD-AAA->SF1-SF2-Helicase+wHTH+Small-B-barrel+DUF1998->ParB+HNH-><-PAIREDC-HTH<-TetR-HTH | 1243 | Bacteria>Terrabacteria_group>Actinomycetota>Actinomycetes>Mycobacteriales>Corynebacteriales_incertae_sedis | GCF_009765205.1 | Tomitella_fengzijianii | GSQ71_RS04965 | 52130..55861 | 2597660 | DUF6079-family-protein-[Tomitella-fengzijianii] |
| WP_182347794.1 | Type2.Pgl.like.BREX | <-SF1-SF2-Helicase+wHTH+Small-B-barrel+DUF1998<-BrxD-AAA<-OLD-ABC+TOPRIM<-PglZ<-DUF6079*<-N6-METHYLASE<-PIN<-MPTase<-PglW<-HNH<-REase+SF1-SF2-Helicase | 1249 | Bacteria>Terrabacteria_group>Actinomycetota>Actinomycetes>Mycobacteriales>Corynebacteriales_incertae_sedis | GCF_014121175.1 | Tomitella_gaofuii | H7F09_RS10440 | 55689..59438 | 2760083 | DUF6079-family-protein-[Tomitella-gaofuii] |
| MBB0993316.1 | Type2.Pgl.like.BREX | <-HNH<-?<-?<-?<-?\|\|PglW->?-><-HTH\|\|N6-METHYLASE->DUF6079-p*->DUF6079->PglZ-> | 165 | Bacteria>Terrabacteria_group>Actinomycetota>Actinomycetes>Mycobacteriales>Dietziaceae | GCA_014144915.1 | Dietzia_sp._SLG510A3-40A3 | G6015_02995 | 12473..12966 | 1630646 | phage-resistance-protein-[Dietzia-sp.-SLG510A3-40A3] |
| MBB1010598.1 | Type2.Pgl.like.BREX | <-PglZ<-PglZ-p<-DUF6079-p*<-N6-METHYLASE\|\|HTH-><-?<-PglW\|\|?->?->?->?->HNH-> | 157 | Bacteria>Terrabacteria_group>Actinomycetota>Actinomycetes>Mycobacteriales>Dietziaceae | GCA_014144945.1 | Dietzia_sp._SLG510A3-3B2-2 | G6010_14190 | 5611..6082 | 1630645 | phage-resistance-protein-[Dietzia-sp.-SLG510A3-3B2-2] |
| WP_067717302.1 | Type2.Pgl.like.BREX | <-ParB+HNH<-ParB+HNH<-SF1-SF2-Helicase+wHTH+Small-B-barrel+DUF1998<-BrxD-AAA<-PglZ<-DUF6079*<-N6-METHYLASE\|\|HTH-><-PglW | 1284 | Bacteria>Terrabacteria_group>Actinomycetota>Actinomycetes>Mycobacteriales>Dietziaceae | GCF_001658045.1 | Dietzia_sp._111N12-1 | AYO52_RS11410 | 8543..12397 | 1785156 | DUF6079-family-protein-[Dietzia-sp.-111N12-1] |
| WP_096906343.1 | Type2.Pgl.like.BREX | <-URI-DNase<-MULE-TRANSPOSASE<-?<-SF1-SF2-Helicase+wHTH+Small-B-barrel+DUF1998<-BrxD-AAA<-PglZ<-DUF6079*<-N6-METHYLASE<-PglW<-DnaB+TnsC-ATPase<-?<-SSB<-?<-LuxR-HTH\|\|AAA-ATPase-> | 1253 | Bacteria>Terrabacteria_group>Actinomycetota>Actinomycetes>Mycobacteriales>Dietziaceae | GCF_002441635.1 | Dietzia_sp._WMMA184 | CK206_RS10365 | 2261537..2265298 | 2039808 | DUF6079-family-protein-[Dietzia-sp.-WMMA184] |
| WP_106386057.1 | Type2.Pgl.like.BREX | Cysteine-Desulfurase-><-?<-?\|\|PglW->N6-METHYLASE->DUF6079*->PglZ->BrxD-AAA->SF1-SF2-Helicase+wHTH+Small-B-barrel+DUF1998->ParB+HNH-><-ABC-ATPase | 1294 | Bacteria>Terrabacteria_group>Actinomycetota>Actinomycetes>Mycobacteriales>Dietziaceae | GCF_002999155.1 | Dietzia_sp._oral_taxon_368 | C3V38_RS13950 | 2983091..2986975 | 712270 | DUF6079-family-protein-[Dietzia-sp.-oral-taxon-368] |
| WP_182635011.1 | Type2.Pgl.like.BREX | PglW-><-HTH\|\|N6-METHYLASE->DUF6079*->BrxD-AAA->SF1-SF2-Helicase+wHTH+Small-B-barrel+DUF1998->ParB+HNH-> | 1286 | Bacteria>Terrabacteria_group>Actinomycetota>Actinomycetes>Mycobacteriales>Dietziaceae | GCF_014144995.1 | Dietzia_sp._DQ11-71 | G6023_RS08665 | 8723..12583 | 1630638 | DUF6079-family-protein-[Dietzia-sp.-DQ11-71] |
| WP_182642241.1 | Type2.Pgl.like.BREX | ABC-ATPase-><-ParB+HNH<-SF1-SF2-Helicase+wHTH+Small-B-barrel+DUF1998<-BrxD-AAA<-PglZ<-DUF6079*<-N6-METHYLASE\|\|HTH-><-PglW\|\|?->?->HNH-> | 1307 | Bacteria>Terrabacteria_group>Actinomycetota>Actinomycetes>Mycobacteriales>Dietziaceae | GCF_014145015.1 | Dietzia_sp._E1 | G6030_RS03660 | 11166..15089 | 328361 | DUF6079-family-protein-[Dietzia-sp.-E1] |
| WP_182609282.1 | Type2.Pgl.like.BREX | <-PglZ-p<-DUF6079*<-N6-METHYLASE\|\|HTH->?-><-PglW\|\|?-><-ABhydrolase+ABhydrolase\|\|?->HNH-> | 1277 | Bacteria>Terrabacteria_group>Actinomycetota>Actinomycetes>Mycobacteriales>Dietziaceae | GCF_014145155.1 | Dietzia_sp._B19 | G6020_RS00295 | 1041..4874 | 1630632 | DUF6079-family-protein-[Dietzia-sp.-B19] |
| WP_182640550.1 | Type2.Pgl.like.BREX | <-PglZ<-DUF6079*<-N6-METHYLASE\|\|HTH-><-PglW\|\|?-><-ABhydrolase+ABhydrolase\|\|?->HNH-><-Cysteine-Desulfurase | 1286 | Bacteria>Terrabacteria_group>Actinomycetota>Actinomycetes>Mycobacteriales>Dietziaceae | GCF_014145265.1 | Dietzia_sp._CW19 | G6021_RS01915 | 2156..6016 | 1630634 | DUF6079-family-protein-[Dietzia-sp.-CW19] |
| WP_213184396.1 | Type2.Pgl.like.BREX | Cysteine-Desulfurase-><-?<-?\|\|PglW->N6-METHYLASE->DUF6079*->PglZ->BrxD-AAA->SF1-SF2-Helicase+wHTH+Small-B-barrel+DUF1998->ParB+HNH-> | 1295 | Bacteria>Terrabacteria_group>Actinomycetota>Actinomycetes>Mycobacteriales>Dietziaceae | GCF_018390595.1 | Dietzia_massiliensis | KHA85_RS00290 | 66468..70355 | 2697499 | DUF6079-family-protein-[Dietzia-massiliensis] |
| WP_259881467.1 | Type2.Pgl.like.BREX | Cysteine-Desulfurase-><-?<-?\|\|PglW->N6-METHYLASE->DUF6079*->PglZ->BrxD-AAA->SF1-SF2-Helicase+wHTH+Small-B-barrel+DUF1998->?->ParB+HNH-><-ABC-ATPase | 1278 | Bacteria>Terrabacteria_group>Actinomycetota>Actinomycetes>Mycobacteriales>Dietziaceae | GCF_025145775.1 | Dietzia_cinnamea | M3B38_RS03585 | 36331..40167 | 321318 | DUF6079-family-protein-[Dietzia-cinnamea] |
| WP_259826368.1 | Type2.Pgl.like.BREX | ABC-ATPase-><-ParB+HNH<-SF1-SF2-Helicase+wHTH+Small-B-barrel+DUF1998<-BrxD-AAA<-PglZ<-DUF6079*<-N6-METHYLASE<-PglW\|\|?->?-><-Cysteine-Desulfurase<-?<-?<-TetR-HTH | 1278 | Bacteria>Terrabacteria_group>Actinomycetota>Actinomycetes>Mycobacteriales>Dietziaceae | GCF_025151765.1 | Dietzia_cinnamea | M3F32_RS00695 | 148483..152319 | 321318 | DUF6079-family-protein-[Dietzia-cinnamea] |
| WP_268092105.1 | Type2.Pgl.like.BREX | PglW-><-HTH\|\|N6-METHYLASE->DUF6079*->PglZ->BrxD-AAA->SF1-SF2-Helicase+wHTH+Small-B-barrel+DUF1998-><-HEPN<-ABC-ATPase | 1305 | Bacteria>Terrabacteria_group>Actinomycetota>Actinomycetes>Mycobacteriales>Dietziaceae | GCF_026672565.1 | Dietzia_sp._SL131 | OVA21_RS06975 | 1316811..1320728 | 2995149 | DUF6079-family-protein-[Dietzia-sp.-SL131] |
| WP_154827457.1 | Type2.Pgl.like.BREX | PglW-><-HTH\|\|N6-METHYLASE->DUF6079*->PglZ->BrxD-AAA->SF1-SF2-Helicase+wHTH+Small-B-barrel+DUF1998->?->ParB+HNH-><-ABC-ATPase | 1279 | Bacteria>Terrabacteria_group>Actinomycetota>Actinomycetes>Mycobacteriales>Dietziaceae | GCF_039531565.1 | Dietzia_kunjamensis | FN843_RS00140 | 36045..39884 | 322509 | DUF6079-family-protein-[Dietzia-kunjamensis] |
| NDK90272.1 | Type2.Pgl.like.BREX | <-HKD-DNase<-SAM-METHYLASE<-HKD-DNase<-EVE+REase\|\|PglW->N6-METHYLASE->DUF6079*->PglZ->BrxD-AAA->SF1-SF2-Helicase+wHTH+Small-B-barrel+DUF1998->?->?->?-><-?\|\|?->TetR-HTH-> | 1267 | Bacteria>Terrabacteria_group>Actinomycetota>Actinomycetes>Mycobacteriales>Gordoniaceae | GCA_010119475.1 | Gordonia_desulfuricans | GYA93_11860 | 13484..17287 | 89051 | phage-resistance-protein-[Gordonia-desulfuricans] |
| WP_020173080.1 | Type2.Pgl.like.BREX | METHYLASE-><-?<-EVE+REase<-REase\|\|PglW->N6-METHYLASE->DUF6079*->PglZ->BrxD-AAA->SF1-SF2-Helicase+wHTH+Small-B-barrel+DUF1998->sGTP+sGTP+GTPase-AIG->sGTP-><-HDRnase | 1265 | Bacteria>Terrabacteria_group>Actinomycetota>Actinomycetes>Mycobacteriales>Gordoniaceae | GCF_000241325.1 | Gordonia_polyisoprenivorans | GOPIP_RS06445 | 59683..63480 | 84595 | DUF6079-family-protein-[Gordonia-polyisoprenivorans] |
| WP_006438715.1 | Type2.Pgl.like.BREX | <-REase+SF1-SF2-Helicase\|\|PglW->N6-METHYLASE->DUF6079*->PglZ->BrxD-AAA->SF1-SF2-Helicase+wHTH+Small-B-barrel+DUF1998-> | 1243 | Bacteria>Terrabacteria_group>Actinomycetota>Actinomycetes>Mycobacteriales>Gordoniaceae | GCF_000332995.1 | Gordonia_amicalis | GOAMI_RS21405 | 11687..15418 | 89053 | DUF6079-family-protein-[Gordonia-amicalis] |
| WP_006368781.1 | Type2.Pgl.like.BREX | HDRnase-><-sGTP<-sGTP+sGTP+GTPase-AIG<-SF1-SF2-Helicase+wHTH+Small-B-barrel+DUF1998<-BrxD-AAA<-PglZ<-DUF6079*<-N6-METHYLASE<-PglW\|\|REase->EVE+REase->?-><-METHYLASE | 1264 | Bacteria>Terrabacteria_group>Actinomycetota>Actinomycetes>Mycobacteriales>Gordoniaceae | GCF_000385355.1 | Gordonia_polyisoprenivorans | A3OC_RS0127250 | 5988355..5992149 | 84595 | DUF6079-family-protein-[Gordonia-polyisoprenivorans] |
| WP_231632802.1 | Type2.Pgl.like.BREX | Glycos_transf->?->Glycos_transf-><-ARG<-?\|\|PglW->N6-METHYLASE->DUF6079*->PglZ->BrxD-AAA->SF1-SF2-Helicase+wHTH+Small-B-barrel+DUF1998->REase-> | 1261 | Bacteria>Terrabacteria_group>Actinomycetota>Actinomycetes>Mycobacteriales>Gordoniaceae | GCF_001418765.1 | Gordonia_sp._HS-NH1 | AMD30_RS20035 | 29569..33354 | 1435068 | DUF6079-family-protein-[Gordonia-sp.-HS-NH1] |
| WP_238403739.1 | Type2.Pgl.like.BREX | <-HKD-DNase<-SAM-METHYLASE<-HKD-DNase<-EVE+REase\|\|PglW->N6-METHYLASE->DUF6079*->PglZ->BrxD-AAA->SF1-SF2-Helicase+wHTH+Small-B-barrel+DUF1998-> | 1263 | Bacteria>Terrabacteria_group>Actinomycetota>Actinomycetes>Mycobacteriales>Gordoniaceae | GCF_001485495.1 | Gordonia_desulfuricans | GD1_RS13525 | 13275..17066 | 89051 | DUF6079-family-protein-[Gordonia-desulfuricans] |
| WP_101818926.1 | Type2.Pgl.like.BREX | <-ABhydrolase\|\|JetA->JetB->SbcC->?->REase->PglW-p->N6-METHYLASE->N6-METHYLASE->DUF6079*->PglZ->BrxD-AAA->SF1-SF2-Helicase+wHTH+Small-B-barrel+DUF1998->EVE+REase-><-URI-DNase<-ParB+HNH\|\|REase+SF1-SF2-Helicase-> | 1238 | Bacteria>Terrabacteria_group>Actinomycetota>Actinomycetes>Mycobacteriales>Gordoniaceae | GCF_002847865.1 | Gordonia_terrae | CYJ73_RS02695 | 593466..597182 | 2055 | phage-resistance-protein-[Gordonia-terrae] |
| WP_210738523.1 | Type2.Pgl.like.BREX | <-SF1-SF2-Helicase+wHTH+Small-B-barrel+DUF1998<-BrxD-AAA<-PglZ<-DUF6079*<-N6-METHYLASE<-PglW\|\|REase+SF1-SF2-Helicase-> | 1232 | Bacteria>Terrabacteria_group>Actinomycetota>Actinomycetes>Mycobacteriales>Gordoniaceae | GCF_012395955.1 | Gordonia_amicalis | HGA01_RS21730 | 6639..10337 | 89053 | DUF6079-family-protein-[Gordonia-amicalis] |
| WP_168702342.1 | Type2.Pgl.like.BREX | <-ABhydrolase\|\|JetA->JetB->?-><-HKD-DNase\|\|PglW->N6-METHYLASE->DUF6079*->PglZ->BrxD-AAA->SF1-SF2-Helicase+wHTH+Small-B-barrel+DUF1998->SF1-SF2-Helicase+REase->SF1-SF2-Helicase-> | 1270 | Bacteria>Terrabacteria_group>Actinomycetota>Actinomycetes>Mycobacteriales>Gordoniaceae | GCF_012498525.1 | Gordonia_paraffinivorans | HHO43_RS17825 | 31065..34877 | 175628 | DUF6079-family-protein-[Gordonia-paraffinivorans] |
| WP_191834363.1 | Type2.Pgl.like.BREX | HIN-HTH->AAA-ATPase-><-SF1-SF2-Helicase+REase<-AAA+<-HTH_7+rve+rve_3<-?<-REase+SF1-SF2-Helicase\|\|PglW->N6-METHYLASE->DUF6079*->PglZ->BrxD-AAA->SF1-SF2-Helicase+wHTH+Small-B-barrel+DUF1998->HTH_7+rve+rve_3->AAA+->?->McrB-AAA->McrC-NTD+REase-><-sGTP | 1243 | Bacteria>Terrabacteria_group>Actinomycetota>Actinomycetes>Mycobacteriales>Gordoniaceae | GCF_012974285.1 | Gordonia_ajococcus | GAA2_RS11805 | 2605571..2609302 | 1292359 | DUF6079-family-protein-[Gordonia-ajococcus] |
| WP_226511910.1 | Type2.Pgl.like.BREX | <-SF1-SF2-Helicase\|\|35exo+TUDOR+SF1-SF2-Helicase+iREase+CTD->SF1-SF2-Helicase-><-SF1-SF2-Helicase+wHTH+Small-B-barrel+DUF1998<-BrxD-AAA<-PglZ<-DUF6079*<-N6-METHYLASE<-PglW\|\|?->ARG->?->ParB+HNH-><-DnaB+TnsC-ATPase | 1266 | Bacteria>Terrabacteria_group>Actinomycetota>Actinomycetes>Mycobacteriales>Gordoniaceae | GCF_020520305.1 | Gordonia_sp._WA4-43 | LEL84_RS13660 | 3060068..3063868 | 2878678 | DUF6079-family-protein-[Gordonia-sp.-WA4-43] |
| WP_247864250.1 | Type2.Pgl.like.BREX | <-DNA_glycosylase\|\|HEPN+SF1-SF2-Helicase+REase->PglW->N6-METHYLASE->RAMA->?->HKD-DNase+HKD-DNase->DUF6079*->PglZ->BrxD-AAA->SF1-SF2-Helicase+wHTH+Small-B-barrel+DUF1998-><-?\|\|BetaPropeller+BetaPropeller-><-ABC-ATPase+SbcC | 1260 | Bacteria>Terrabacteria_group>Actinomycetota>Actinomycetes>Mycobacteriales>Gordoniaceae | GCF_023100845.1 | Gordonia_sp._PP30 | MYK68_RS13790 | 3003051..3006833 | 2935861 | DUF6079-family-protein-[Gordonia-sp.-PP30] |
| WP_075863498.1 | Type2.Pgl.like.BREX | <-HIN-HTH<-rve\|\|REase->PglW->N6-METHYLASE->DUF6079*->PglZ->BrxD-AAA->SF1-SF2-Helicase+wHTH+Small-B-barrel+DUF1998->EVE+REase-><-URI-DNase<-ParB+HNH\|\|SF1-SF2-Helicase+REase+HTH->SF1-SF2-Helicase+HTH-><-HKD-DNase | 1241 | Bacteria>Terrabacteria_group>Actinomycetota>Actinomycetes>Mycobacteriales>Gordoniaceae | GCF_029853985.1 | Gordonia_sp._CNJ-863 | QEN35_RS21110 | 9812..13537 | 1904963 | MULTISPECIES:-phage-resistance-protein-[Gordonia] |
| WP_240860430.1 | Type2.Pgl.like.BREX | PglW->N6-METHYLASE->DUF6079*->PglZ->BrxD-AAA->SF1-SF2-Helicase+wHTH+Small-B-barrel+DUF1998-> | 1275 | Bacteria>Terrabacteria_group>Actinomycetota>Actinomycetes>Mycobacteriales>Gordoniaceae | GCF_022513585.1 | Gordonia_sp._ABSL49_1 | MKW15_RS17550 | 54041..57868 | 2920941 | DUF6079-family-protein-[Gordonia-sp.-ABSL49_1] |
| EUA08687.1 | Type2.Pgl.like.BREX | <-SF1-SF2-Helicase+wHTH+Small-B-barrel+DUF1998<-SF1-SF2-Helicase+wHTH+Small-B-barrel+DUF1998-p<-BrxD-AAA-p<-BrxD-AAA<-BrxD-AAA-p<-PglZ-p<-PglZ-p<-PglZ-p<-DUF6079-p<-DUF6079-p*<-DUF6079-p<-N6-METHYLASE-p<-N6-METHYLASE-p<-?<-?<-METHYLASE-p<-N6-METHYLASE-p<-N6-METHYLASE-p | 523 | Bacteria>Terrabacteria_group>Actinomycetota>Actinomycetes>Mycobacteriales>Mycobacteriaceae | GCA_000523695.1 | Mycobacterium_xenopi_4042 | I553_9743 | 29875..31446 | 1299334 | phage-resistance-PglY-domain-protein-[Mycobacterium-xenopi-4042] |
| EUA08688.1 | Type2.Pgl.like.BREX | <-SF1-SF2-Helicase+wHTH+Small-B-barrel+DUF1998-p<-BrxD-AAA-p<-BrxD-AAA<-BrxD-AAA-p<-PglZ-p<-PglZ-p<-PglZ-p<-DUF6079-p<-DUF6079-p<-DUF6079-p*<-N6-METHYLASE-p<-N6-METHYLASE-p<-?<-?<-METHYLASE-p<-N6-METHYLASE-p<-N6-METHYLASE-p<-?<-RNase | 425 | Bacteria>Terrabacteria_group>Actinomycetota>Actinomycetes>Mycobacteriales>Mycobacteriaceae | GCA_000523695.1 | Mycobacterium_xenopi_4042 | I553_9744 | 31443..32720 | 1299334 | phage-resistance-PglY-domain-protein-[Mycobacterium-xenopi-4042] |
| OBG19347.1 | Type2.Pgl.like.BREX | Glyoxalase-><-?<-SF1-SF2-Helicase+wHTH+Small-B-barrel+DUF1998<-BrxD-AAA<-PglZ-p\|\|?-><-DUF6079*<-N6-METHYLASE<-PglW<-?<-?\|\|Phage_holin-><-?\|\|?-><-?<-ABhydrolase | 1274 | Bacteria>Terrabacteria_group>Actinomycetota>Actinomycetes>Mycobacteriales>Mycobacteriaceae | GCA_001667035.1 | Mycobacterium_sp._852002-51057_SCH5723018 | A5764_17090 | 132599..136423 | 1834094 | hypothetical-protein-A5764_17090-[Mycobacterium-sp.-852002-51057_SCH5723018] |
| OBJ70979.1 | Type2.Pgl.like.BREX | HKD-DNase+SF1-SF2-Helicase->URI-DNase->PglW-><-?<-MULE-TRANSPOSASE\|\|HTH_Tnp_ISL3+DDE_Tnp_ISL3->N6-METHYLASE-p->DUF6079*->PglZ->BrxD-AAA->SF1-SF2-Helicase+wHTH+Small-B-barrel+DUF1998->UDG-conflict-><-SF1-SF2-Helicase+REase<-?<-SNF+SF1-SF2-Helicase | 1234 | Bacteria>Terrabacteria_group>Actinomycetota>Actinomycetes>Mycobacteriales>Mycobacteriaceae | GCA_001668625.1 | Mycobacterium_sp._1274756.6 | A5643_08790 | 97079..100783 | 1834076 | phage-resistance-protein-[Mycobacterium-sp.-1274756.6] |
| OKH83306.1 | Type2.Pgl.like.BREX | <-HKD-DNase+HKD-DNase<-?\|\|PglW->N6-METHYLASE->DUF6079*->?<-?<-?\|\|PglZ-p->BrxD-AAA->SF1-SF2-Helicase+wHTH+Small-B-barrel+DUF1998-p-> | 1275 | Bacteria>Terrabacteria_group>Actinomycetota>Actinomycetes>Mycobacteriales>Mycobacteriaceae | GCA_001905655.1 | Mycobacterium_sp._ST-F2 | EB75_09970 | 75844..79671 | 1490484 | hypothetical-protein-EB75_09970-[Mycobacterium-sp.-ST-F2] |
| OLP01888.1 | Type2.Pgl.like.BREX | SF1-SF2-Helicase->?-><-SF1-SF2-Helicase+REase<-ParB+HNH<-SF1-SF2-Helicase+wHTH+Small-B-barrel+DUF1998<-BrxD-AAA<-PglZ<-DUF6079*<-N6-METHYLASE<-PglW\|\|Classical-AAA->?->GTPase-AIG+sGTP->MMR_HSR1-> | 1226 | Bacteria>Terrabacteria_group>Actinomycetota>Actinomycetes>Mycobacteriales>Mycobacteriaceae | GCA_001942045.1 | Mycolicibacterium_porcinum | BVU76_13180 | 83307..86987 | 39693 | phage-resistance-protein-[Mycolicibacterium-porcinum] |
| ORW04232.1 | Type2.Pgl.like.BREX | N6-METHYLASE-p->DUF6079*->PglZ->BrxD-AAA->SF1-SF2-Helicase+wHTH+Small-B-barrel+DUF1998-><-MACRODOMAIN+SIR2<-?<-SbcC+SbcC<-Calcineurin<-HerA/FtsK | 1274 | Bacteria>Terrabacteria_group>Actinomycetota>Actinomycetes>Mycobacteriales>Mycobacteriaceae | GCA_002101745.1 | Mycobacterium_kubicae | AWC13_00680 | 1814..5638 | 120959 | hypothetical-protein-AWC13_00680-[Mycobacterium-kubicae] |
| PND55336.1 | Type2.Pgl.like.BREX | DUF6079*->PglZ->BrxD-AAA->SF1-SF2-Helicase+wHTH+Small-B-barrel+DUF1998->McrB-AAA->McrC-NTD+REase-><-REase+SF1-SF2-Helicase<-Preader7<-SF1-SF2-Helicase+DUF1998<-viral-protein | 1227 | Bacteria>Terrabacteria_group>Actinomycetota>Actinomycetes>Mycobacteriales>Mycobacteriaceae | GCA_002887815.1 | Mycobacterium_sp._ENV421 | CRM90_22935 | 1271..4954 | 1213407 | phage-resistance-protein-[Mycobacterium-sp.-ENV421] |
| POY32161.1 | Type2.Pgl.like.BREX | <-DUF6079-p*<-N6-METHYLASE<-?<-RNase<-PglW<-PIN+35exo+T7DNApol\|\|?->?-><-REC+OmpR-HTH\|\|tRNA->DUF488-> | 189 | Bacteria>Terrabacteria_group>Actinomycetota>Actinomycetes>Mycobacteriales>Mycobacteriaceae | GCA_002920845.1 | Mycobacterium_kansasii | C3478_12880 | 1..568 | 1768 | phage-resistance-protein,-partial-[Mycobacterium-kansasii] |
| KAA0110610.1 | Type2.Pgl.like.BREX | <-DSBH<-?\|\|?-><-MMR_HSR1<-GTPase-AIG+sGTP\|\|PglW->N6-METHYLASE->DUF6079*->PglZ->BrxD-AAA->SF1-SF2-Helicase+wHTH+Small-B-barrel+DUF1998->McrB-AAA-ATPase->McrC-NTD+REase->\|\|35exo+SF1-SF2-Helicase->SNF+SF1-SF2-Helicase-> | 1226 | Bacteria>Terrabacteria_group>Actinomycetota>Actinomycetes>Mycobacteriales>Mycobacteriaceae | GCA_008329565.1 | Mycolicibacterium_sp._P1-5 | CIW47_06215 | 77198..80878 | 2024617 | phage-resistance-protein-[Mycolicibacterium-sp.-P1-5] |
| QEN12603.1 | Type2.Pgl.like.BREX | DNA-glycosylase-><-SNF+SF1-SF2-Helicase<-35exo+SF1-SF2-Helicase<-ParB+HNH<-SF1-SF2-Helicase+wHTH+Small-B-barrel+DUF1998<-BrxD-AAA<-PglZ<-DUF6079*<-N6-METHYLASE<-PglW<-?<-PIN<-Phd-YefM\|\|DSBH-> | 1226 | Bacteria>Terrabacteria_group>Actinomycetota>Actinomycetes>Mycobacteriales>Mycobacteriaceae | GCA_008329905.1 | Mycobacterium_sp._ELW1 | D3H54_04400 | 962839..966519 | 1547487 | phage-resistance-protein-[Mycobacterium-sp.-ELW1] |
| MBP3083666.1 | Type2.Pgl.like.BREX | ABhydrolase+ABhydrolase-><-MACRODOMAIN+SIR2<-HEPN+SF1-SF2-Helicase+REase<-SF1-SF2-Helicase+wHTH+Small-B-barrel+DUF1998<-BrxD-AAA<-PglZ<-DUF6079*<-N6-METHYLASE<-PglW<-?<-HKD-DNase+SF1-SF2-Helicase | 1275 | Bacteria>Terrabacteria_group>Actinomycetota>Actinomycetes>Mycobacteriales>Mycobacteriaceae | GCA_017915355.1 | Mycolicibacterium_fortuitum | JKJ01_10820 | 214262..218089 | 1766 | hypothetical-protein-[Mycolicibacterium-fortuitum] |
| MCB1288101.1 | Type2.Pgl.like.BREX | <-ARG<-?<-?\|\|HTH_3->?->ParB+HNH-><-SF1-SF2-Helicase+wHTH+Small-B-barrel+DUF1998<-BrxD-AAA<-PglZ<-DUF6079*<-N6-METHYLASE | 1218 | Bacteria>Terrabacteria_group>Actinomycetota>Actinomycetes>Mycobacteriales>Mycobacteriaceae | GCA_020439025.1 | Mycobacterium_sp. | KDB47_10475 | 12456..16112 | 1785 | phage-resistance-protein-[Mycobacterium-sp.] |
| MCB9410783.1 | Type2.Pgl.like.BREX | N6-METHYLASE->DUF6079*->PglZ->BrxD-AAA->SF1-SF2-Helicase+wHTH+Small-B-barrel+DUF1998->REase+iREase->?-><-?<-DNA_glycosylase\|\|XIS-HTH->Phage_int_SAM_3+Phage_integrase-> | 1214 | Bacteria>Terrabacteria_group>Actinomycetota>Actinomycetes>Mycobacteriales>Mycobacteriaceae | GCA_020637165.1 | Mycolicibacterium_sp. | H6521_15710 | 1891..5535 | 2320850 | phage-resistance-protein-[Mycolicibacterium-sp.] |
| SDF04848.1 | Type2.Pgl.like.BREX | viral-protein->SF1-SF2-Helicase->McrB-AAA-ATPase->McrC-NTD+REase->REase->PglW->N6-METHYLASE->DUF6079*->PglZ->BrxD-AAA->SF1-SF2-Helicase+wHTH+Small-B-barrel+DUF1998-><-HIN-HTH+PAIREDC-HTH<-AAA-ATPase-p | 1261 | Bacteria>Terrabacteria_group>Actinomycetota>Actinomycetes>Mycobacteriales>Mycobacteriaceae | GCA_900101555.1 | Mycolicibacterium_neoaurum | SAMN04488581_5267 | 158509..162294 | 1795 | hypothetical-protein-SAMN04488581_5267-[Mycolicibacterium-neoaurum] |
| WP_009955556.1 | Type2.Pgl.like.BREX | <-Arm-DNA-bind_4+Phage_int_SAM_3+Phage_integrase<-GDResolvase+Recombinase<-?<-ZnR<-Primpol<-Cox-HTH<-?\|\|PglW->N6-METHYLASE->DUF6079*->PglZ->BrxD-AAA->SF1-SF2-Helicase+wHTH+Small-B-barrel+DUF1998-><-BrxB-BrxF-iSTAND<-?\|\|NACHT+GTPase-AIG+sGTP-><-REase | 1246 | Bacteria>Terrabacteria_group>Actinomycetota>Actinomycetes>Mycobacteriales>Mycobacteriaceae | GCF_000172115.1 | Mycobacterium_intracellulare | JN195_RS24125 | 5066243..5069983 | 1767 | hypothetical-protein-[Mycobacterium-intracellulare] |
| WP_003920378.1 | Type2.Pgl.like.BREX | <-DUF488<-tRNA\|\|REC+LuxR-HTH-><-?<-?\|\|PIN+35exo+T7DNApol->PglW->RNase->?->N6-METHYLASE->DUF6079*->PglZ->BrxD-AAA->SF1-SF2-Helicase+wHTH+Small-B-barrel+DUF1998-> | 1231 | Bacteria>Terrabacteria_group>Actinomycetota>Actinomycetes>Mycobacteriales>Mycobacteriaceae | GCF_000257745.1 | Mycobacterium_xenopi | MXEN_RS07795 | 225316..229011 | 1789 | DUF6079-family-protein-[Mycobacterium-xenopi] |
| WP_044516882.1 | Type2.Pgl.like.BREX | <-Phage_integrase+Phage_integrase<-?<-Phage_int_SAM_1+Phage_integrase\|\|?->REase->SbcC->PglW->N6-METHYLASE->DUF6079*->PglZ->BrxD-AAA->SF1-SF2-Helicase+wHTH+Small-B-barrel+DUF1998-><-?<-?<-TetR-HTH<-?<-N6-METHYLASE | 1260 | Bacteria>Terrabacteria_group>Actinomycetota>Actinomycetes>Mycobacteriales>Mycobacteriaceae | GCF_000455325.1 | Mycolicibacterium_septicum | TX88_RS11525 | 2445100..2448882 | 98668 | DUF6079-family-protein-[Mycolicibacterium-septicum] |
| WP_242616555.1 | Type2.Pgl.like.BREX | <-McrB-AAA<-Preader7<-HEPN+SF1-SF2-Helicase+REase<-?<-?<-Peptidase_S8<-Classical-AAA\|\|PglW->N6-METHYLASE->DUF6079*->PglZ->BrxD-AAA->SF1-SF2-Helicase+wHTH+Small-B-barrel+DUF1998->ParB+HNH->ParB-><-HIN-HTH\|\|ParB+HNH-><-SF1-SF2-Helicase+DUF1998<-viral-protein | 1188 | Bacteria>Terrabacteria_group>Actinomycetota>Actinomycetes>Mycobacteriales>Mycobacteriaceae | GCF_000612825.1 | Mycolicibacterium_mageritense | BN978_RS05415 | 1124749..1128315 | 53462 | DUF6079-family-protein-[Mycolicibacterium-mageritense] |
| WP_047333090.1 | Type2.Pgl.like.BREX | DNA-glycosylase-><-SF1-SF2-Helicase<-35exo+SF1-SF2-Helicase<-ParB+HNH<-SF1-SF2-Helicase+wHTH+Small-B-barrel+DUF1998<-BrxD-AAA<-PglZ<-DUF6079*<-N6-METHYLASE<-PglW<-?\|\|NUDIX->?->DSBH-> | 1223 | Bacteria>Terrabacteria_group>Actinomycetota>Actinomycetes>Mycobacteriales>Mycobacteriaceae | GCF_001021385.1 | Mycobacterium_sp._EPa45 | AB431_RS04725 | 1023056..1026727 | 1545728 | phage-resistance-protein-[Mycobacterium-sp.-EPa45] |
| WP_075511384.1 | Type2.Pgl.like.BREX | <-DUF488<-tRNA\|\|REC+OmpR-HTH-><-?<-?\|\|PIN+35exo+T7DNApol->PglW->RNase->?->N6-METHYLASE->DUF6079*->PglZ->BrxD-AAA->SF1-SF2-Helicase+wHTH+Small-B-barrel+DUF1998->?->?-><-?<-?<-?\|\|SF1-SF2-Helicase-> | 1222 | Bacteria>Terrabacteria_group>Actinomycetota>Actinomycetes>Mycobacteriales>Mycobacteriaceae | GCF_001632895.1 | Mycobacterium_ostraviense | A4G28_RS13615 | 27338..31006 | 2738409 | phage-resistance-protein-[Mycobacterium-ostraviense] |
| WP_064929122.1 | Type2.Pgl.like.BREX | PglW->N6-METHYLASE->DUF6079*->PglZ->BrxD-AAA->SF1-SF2-Helicase+wHTH+Small-B-barrel+DUF1998->HTH+REase->?->?->REase->LuxR-HTH-> | 1233 | Bacteria>Terrabacteria_group>Actinomycetota>Actinomycetes>Mycobacteriales>Mycobacteriaceae | GCF_001665365.1 | Mycolicibacterium_elephantis | A5633_RS17090 | 6734..10435 | 81858 | hypothetical-protein-[Mycolicibacterium-elephantis] |
| WP_156664862.1 | Type2.Pgl.like.BREX | Glyoxalase-><-?<-?<-SF1-SF2-Helicase+wHTH+Small-B-barrel+DUF1998<-BrxD-AAA<-PglZ<-DUF6079*<-N6-METHYLASE<-PglW<-?<-?\|\|Phage_holin->\|\|?-><-?<-ABhydrolase | 1260 | Bacteria>Terrabacteria_group>Actinomycetota>Actinomycetes>Mycobacteriales>Mycobacteriaceae | GCF_001667035.1 | Mycobacterium_sp._852002-51057_SCH5723018 | A5764_RS25100 | 132599..136381 | 1834094 | DUF6079-family-protein-[Mycobacterium-sp.-852002-51057_SCH5723018] |
| WP_064959219.1 | Type2.Pgl.like.BREX | <-ParB+HNH<-HKD-DNase+HKD-DNase<-?\|\|PglW->N6-METHYLASE->DUF6079*->PglZ->BrxD-AAA->SF1-SF2-Helicase+wHTH+Small-B-barrel+DUF1998->?->sGTP+sGTP+GTPase-AIG->sGTP-><-REase+HTH+35exo+Small-B-barrel+SF1-SF2-Helicase | 1238 | Bacteria>Terrabacteria_group>Actinomycetota>Actinomycetes>Mycobacteriales>Mycobacteriaceae | GCF_001667205.1 | Mycolicibacterium_peregrinum | A5719_RS16195 | 14293..18009 | 43304 | DUF6079-family-protein-[Mycolicibacterium-peregrinum] |
| WP_065288057.1 | Type2.Pgl.like.BREX | <-CopG/RHH\|\|HTH_3-><-RVT\|\|PglW->HEPN->N6-METHYLASE->DUF6079*->PglZ->PglZ-p->BrxD-AAA->SF1-SF2-Helicase+wHTH+Small-B-barrel+DUF1998->McrB-AAA->McrC-NTD+REase-><-?<-?<-SF1-SF2-Helicase | 1226 | Bacteria>Terrabacteria_group>Actinomycetota>Actinomycetes>Mycobacteriales>Mycobacteriaceae | GCF_001679965.1 | Mycolicibacter_kumamotonensis | ACT18_RS09910 | 11886..15566 | 354243 | DUF6079-family-protein-[Mycolicibacter-kumamotonensis] |
| WP_065496726.1 | Type2.Pgl.like.BREX | Glyoxalase->?->CR-ATPase-><-HKD-DNase+HKD-DNase<-?\|\|PglW->N6-METHYLASE->DUF6079*->PglZ->BrxD-AAA->SF1-SF2-Helicase+wHTH+Small-B-barrel+DUF1998->?-><-?<-RAMA\|\|TetR-HTH-> | 1274 | Bacteria>Terrabacteria_group>Actinomycetota>Actinomycetes>Mycobacteriales>Mycobacteriaceae | GCF_001686735.1 | Mycobacterium_malmoense | A5676_RS12500 | 113280..117104 | 1780 | DUF6079-family-protein-[Mycobacterium-malmoense] |
| WP_071700409.1 | Type2.Pgl.like.BREX | <-SF1-SF2-Helicase+wHTH+Small-B-barrel+DUF1998<-BrxD-AAA<-PglZ<-DUF6079*<-N6-METHYLASE<-?<-RNase<-PglW<-PIN+35exo+T7DNApol\|\|?->?-><-REC+OmpR-HTH\|\|tRNA->DUF488-> | 1220 | Bacteria>Terrabacteria_group>Actinomycetota>Actinomycetes>Mycobacteriales>Mycobacteriaceae | GCF_001881585.2 | Mycobacterium_heckeshornense | BMW24_RS10415 | 61736..65398 | 110505 | DUF6079-family-protein-[Mycobacterium-heckeshornense] |
| WP_083082915.1 | Type2.Pgl.like.BREX | HTH_3-><-RVT\|\|PglW->HEPN->N6-METHYLASE->DUF6079*->PglZ->PglZ-p->BrxD-AAA->SF1-SF2-Helicase+wHTH+Small-B-barrel+DUF1998->McrB-AAA->McrC-NTD+REase-> | 1228 | Bacteria>Terrabacteria_group>Actinomycetota>Actinomycetes>Mycobacteriales>Mycobacteriaceae | GCF_002086285.1 | Mycolicibacter_kumamotonensis | BST28_RS21220 | 11421..15107 | 354243 | DUF6079-family-protein-[Mycolicibacter-kumamotonensis] |
| WP_245917586.1 | Type2.Pgl.like.BREX | <-DUF488<-tRNA\|\|REC+OmpR-HTH-><-?<-?\|\|PIN+35exo+T7DNApol->PglW->RNase->?->N6-METHYLASE->DUF6079-p*-> | 191 | Bacteria>Terrabacteria_group>Actinomycetota>Actinomycetes>Mycobacteriales>Mycobacteriaceae | GCF_002920805.1 | Mycobacterium_kansasii | C3479_RS20280 | 29080..29655 | 1768 | hypothetical-protein-[Mycobacterium-kansasii] |
| WP_103851013.1 | Type2.Pgl.like.BREX | <-SF1-SF2-Helicase+wHTH+Small-B-barrel+DUF1998<-BrxD-AAA<-PglZ<-DUF6079*<-N6-METHYLASE<-?<-RNase<-PglW<-PIN+35exo+T7DNApol\|\|?->?-><-REC+OmpR-HTH\|\|tRNA->DUF488-> | 1223 | Bacteria>Terrabacteria_group>Actinomycetota>Actinomycetes>Mycobacteriales>Mycobacteriaceae | GCF_002920875.1 | Mycobacterium_kansasii | C3469_RS01095 | 19547..23218 | 1768 | phage-resistance-protein-[Mycobacterium-kansasii] |
| WP_105365951.1 | Type2.Pgl.like.BREX | HKD-DNase+SF1-SF2-Helicase+HTH+HTH+SRA->viral-protein->SF1-SF2-Helicase+DUF1998->PglW->N6-METHYLASE->DUF6079*->PglZ->BrxD-AAA->SF1-SF2-Helicase+wHTH+Small-B-barrel+DUF1998->SF1-SF2-Helicase+REase->McrB-AAA->McrC-NTD+REase->REase->REase-> | 1274 | Bacteria>Terrabacteria_group>Actinomycetota>Actinomycetes>Mycobacteriales>Mycobacteriaceae | GCF_002968335.1 | Mycobacterium_sp._ITM-2016-00316 | C6A86_RS28955 | 32635..36459 | 2099695 | DUF6079-family-protein-[Mycobacterium-sp.-ITM-2016-00316] |
| WP_138743881.1 | Type2.Pgl.like.BREX | STYKIN-><-ParB+HNH<-HKD-DNase+HKD-DNase<-?\|\|PglW->N6-METHYLASE->DUF6079*->PglZ->BrxD-AAA->SF1-SF2-Helicase+wHTH+Small-B-barrel+DUF1998->sGTP+sGTP+GTPase-AIG->sGTP-><-SF1-SF2-Helicase\|\|DDE_Tnp_1-><-REase+HTH+35exo+Small-B-barrel+SF1-SF2-Helicase | 1261 | Bacteria>Terrabacteria_group>Actinomycetota>Actinomycetes>Mycobacteriales>Mycobacteriaceae | GCF_005924235.1 | Mycobacterium_sp._DBP42 | E0T84_RS04295 | 18614..22399 | 2545267 | DUF6079-family-protein-[Mycobacterium-sp.-DBP42] |
| WP_149480516.1 | Type2.Pgl.like.BREX | <-REase<-?\|\|SF1-SF2-Helicase+REase->SF1-SF2-Helicase+REase->MACRODOMAIN+SIR2-><-SF1-SF2-Helicase+wHTH+Small-B-barrel+DUF1998<-BrxD-AAA<-PglZ<-DUF6079*<-N6-METHYLASE<-PglW\|\|?->?->HKD-DNase+HKD-DNase->SF1-SF2-Helicase-><-HTH_3 | 1269 | Bacteria>Terrabacteria_group>Actinomycetota>Actinomycetes>Mycobacteriales>Mycobacteriaceae | GCF_008329535.1 | Mycolicibacterium_sp._P1-18 | CIW49_RS13615 | 579586..583395 | 2024615 | DUF6079-family-protein-[Mycolicibacterium-sp.-P1-18] |
| WP_070351272.1 | Type2.Pgl.like.BREX | <-McrC-NTD+REase<-McrB-AAA<-SF1-SF2-Helicase+wHTH+Small-B-barrel+DUF1998<-BrxD-AAA<-PglZ<-DUF6079*<-N6-METHYLASE<-PglW\|\|sGTP+sGTP+GTPase-AIG->sGTP-><-?<-REase<-TUDOR+SF1-SF2-Helicase+iREase+CTD+REase\|\|?->SbcC-> | 1222 | Bacteria>Terrabacteria_group>Actinomycetota>Actinomycetes>Mycobacteriales>Mycobacteriaceae | GCF_008329645.1 | Mycobacterium_grossiae | BEL07_RS01095 | 46083..49751 | 1552759 | DUF6079-family-protein-[Mycobacterium-grossiae] |
| WP_085198627.1 | Type2.Pgl.like.BREX | PglW->RNase->?->N6-METHYLASE->DUF6079*->PglZ->BrxD-AAA->SF1-SF2-Helicase+wHTH+Small-B-barrel+DUF1998-> | 1220 | Bacteria>Terrabacteria_group>Actinomycetota>Actinomycetes>Mycobacteriales>Mycobacteriaceae | GCF_009936235.1 | Mycobacterium_xenopi | PFJ02_RS01280 | 264585..268247 | 1789 | DUF6079-family-protein-[Mycobacterium-xenopi] |
| WP_193489332.1 | Type2.Pgl.like.BREX | <-SIR2<-SbcC<-?<-?\|\|PglW->N6-METHYLASE->DUF6079*->PglZ->BrxD-AAA->SF1-SF2-Helicase+wHTH+Small-B-barrel+DUF1998->ParB+HNH-> | 1222 | Bacteria>Terrabacteria_group>Actinomycetota>Actinomycetes>Mycobacteriales>Mycobacteriaceae | GCF_039528895.1 | Mycolicibacterium_murale | ABD253_RS31590 | 186413..190081 | 182220 | DUF6079-family-protein-[Mycolicibacterium-murale] |
| WP_139823017.1 | Type2.Pgl.like.BREX | HIRAN->PglW->N6-METHYLASE->\|\|<-Phage_integrase<-Phage_integrase<-Phage_integrase<-Phage_int_SAM_1\|\|\|N6-METHYLASE-p->DUF6079*->PglZ->BrxD-AAA->SF1-SF2-Helicase+wHTH+Small-B-barrel+DUF1998-><-MACRODOMAIN+SIR2<-?<-SbcC+SbcC<-Calcineurin<-HerA/FtsK | 1260 | Bacteria>Terrabacteria_group>Actinomycetota>Actinomycetes>Mycobacteriales>Mycobacteriaceae | GCF_010723135.1 | Mycobacterium_kubicae | G6N40_RS06420 | 72998..76780 | 120959 | DUF6079-family-protein-[Mycobacterium-kubicae] |
| WP_163642253.1 | Type2.Pgl.like.BREX | MACRODOMAIN->viral-protein->SF1-SF2-Helicase+DUF1998-><-ParB+HNH\|\|<-ParB<-ParB+HNH<-SF1-SF2-Helicase+wHTH+Small-B-barrel+DUF1998<-BrxD-AAA<-PglZ<-DUF6079*<-N6-METHYLASE<-PglW\|\|Classical-AAA->Peptidase_S8->?->?->HEPN+SF1-SF2-Helicase+REase->Preader7->McrB-AAA-> | 1226 | Bacteria>Terrabacteria_group>Actinomycetota>Actinomycetes>Mycobacteriales>Mycobacteriaceae | GCF_010727475.1 | Mycolicibacterium_mageritense | G6N67_RS20345 | 4225776..4229456 | 53462 | DUF6079-family-protein-[Mycolicibacterium-mageritense] |
| WP_138248772.1 | Type2.Pgl.like.BREX | JetB->SbcC->REase-><-?<-PIN<-?<-SPOUT<-SF1-SF2-Helicase+wHTH+Small-B-barrel+DUF1998<-BrxD-AAA<-PglZ<-DUF6079*<-HEPN<-N6-METHYLASE<-PglW\|\|HTH_3->nSTAND1+HNH-> | 1223 | Bacteria>Terrabacteria_group>Actinomycetota>Actinomycetes>Mycobacteriales>Mycobacteriaceae | GCF_010731115.1 | Mycolicibacterium_phocaicum | C1S79_RS09450 | 100122..103793 | 319706 | DUF6079-family-protein-[Mycolicibacterium-phocaicum] |
| WP_174400020.1 | Type2.Pgl.like.BREX | PhdYeFM_antitox->PIN->?->?-><-DSBH<-?\|\|N6-METHYLASE->DUF6079*->PglZ->BrxD-AAA->SF1-SF2-Helicase+wHTH+Small-B-barrel+DUF1998->ParB+HNH-><-?\|\|TPR+LuxR-HTH-> | 1224 | Bacteria>Terrabacteria_group>Actinomycetota>Actinomycetes>Mycobacteriales>Mycobacteriaceae | GCF_013337765.1 | Mycolicibacterium_sphagni | FEG63_RS22430 | 107447..111121 | 1786 | DUF6079-family-protein-[Mycolicibacterium-sphagni] |
| WP_193043845.1 | Type2.Pgl.like.BREX | HD-><-?<-?\|\|LACI-HTH+PBPI->?->?-><-SF1-SF2-Helicase+wHTH+Small-B-barrel+DUF1998<-BrxD-AAA<-PglZ<-DUF6079*<-N6-METHYLASE<-PglW<-SbcC<-REase<-SIR2\|\|RelE-ParE->HTH_3-> | 1260 | Bacteria>Terrabacteria_group>Actinomycetota>Actinomycetes>Mycobacteriales>Mycobacteriaceae | GCF_016313185.1 | Mycolicibacterium_baixiangningiae | I7X18_RS20525 | 4329743..4333525 | 2761578 | DUF6079-family-protein-[Mycolicibacterium-baixiangningiae] |
| WP_237161840.1 | Type2.Pgl.like.BREX | <-Preader7<-?<-?\|\|ABhydrolase+ABhydrolase-><-SIR2<-SF1-SF2-Helicase+wHTH+Small-B-barrel+DUF1998<-BrxD-AAA<-PglZ<-DUF6079*<-N6-METHYLASE<-PglW<-?<-HKD-DNase+SF1-SF2-Helicase+HTH+HTH+SRA\|\|?->?->?-><-?\|\|DUF4352-> | 1261 | Bacteria>Terrabacteria_group>Actinomycetota>Actinomycetes>Mycobacteriales>Mycobacteriaceae | GCF_017915355.1 | Mycolicibacterium_fortuitum | JKJ01_RS10815 | 214262..218047 | 1766 | DUF6079-family-protein-[Mycolicibacterium-fortuitum] |
| WP_210686514.1 | Type2.Pgl.like.BREX | <-MMR_HSR1<-GTPase-AIG+sGTP<-?<-Classical-AAA\|\|PglW->N6-METHYLASE->DUF6079*->PglZ->BrxD-AAA->SF1-SF2-Helicase+wHTH+Small-B-barrel+DUF1998->ParB+HNH->SF1-SF2-Helicase+REase-><-?<-SF1-SF2-Helicase | 1225 | Bacteria>Terrabacteria_group>Actinomycetota>Actinomycetes>Mycobacteriales>Mycobacteriaceae | GCF_017948465.1 | Mycolicibacterium_sp._GESEQ-9 | KCH01_RS04985 | 99902..103579 | 2812656 | DUF6079-family-protein-[Mycolicibacterium-sp.-GESEQ-9] |
| WP_011894071.1 | Type2.Pgl.like.BREX | <-McrC-NTD+REase<-McrB-AAA-ATPase<-SF1-SF2-Helicase+wHTH+Small-B-barrel+DUF1998<-BrxD-AAA<-PglZ<-DUF6079*<-N6-METHYLASE<-PglW<-ZnR<-?<-HTH<-?<-RelE-ParE | 1230 | Bacteria>Terrabacteria_group>Actinomycetota>Actinomycetes>Mycobacteriales>Mycobacteriaceae | GCF_019148985.1 | Mycolicibacterium_sp._PAM1 | KUF57_RS24765 | 9840..13532 | 2853535 | DUF6079-family-protein-[Mycolicibacterium-sp.-PAM1] |
| WP_221852024.1 | Type2.Pgl.like.BREX | <-DNA_glycosylase<-HEPN+SF1-SF2-Helicase+REase\|\|35exo+TUDOR+SF1-SF2-Helicase+iREase+CTD->SF1-SF2-Helicase-><-SF1-SF2-Helicase+wHTH+Small-B-barrel+DUF1998<-BrxD-AAA<-PglZ<-DUF6079*<-N6-METHYLASE<-PglW<-?<-HKD-DNase+SF1-SF2-Helicase+HTH+HTH+SRA\|\|?->?-><-?\|\|DUF4352-> | 1275 | Bacteria>Terrabacteria_group>Actinomycetota>Actinomycetes>Mycobacteriales>Mycobacteriaceae | GCF_019739075.1 | Mycolicibacterium_farcinogenes | K6L26_RS26085 | 5468799..5472626 | 1802 | DUF6079-family-protein-[Mycolicibacterium-farcinogenes] |
| WP_225342387.1 | Type2.Pgl.like.BREX | <-Arm-DNA-bind_4+Phage_int_SAM_3+Phage_integrase<-GDResolvase+Recombinase<-?<-Primpol<-Cox-HTH<-?\|\|PglW->N6-METHYLASE->DUF6079*->PglZ->BrxD-AAA->SF1-SF2-Helicase+wHTH+Small-B-barrel+DUF1998-><-HEPN<-ParA-Soj-PloopNTPase\|\|ParB+HNH->HEPN-> | 1246 | Bacteria>Terrabacteria_group>Actinomycetota>Actinomycetes>Mycobacteriales>Mycobacteriaceae | GCF_020217625.1 | Mycobacterium_intracellulare | JF729_RS19575 | 23932..27672 | 1767 | phage-resistance-protein-[Mycobacterium-intracellulare] |
| WP_099319304.1 | Type2.Pgl.like.BREX | <-SF1-SF2-Helicase\|\|?->?->?-><-?<-?<-SF1-SF2-Helicase+wHTH+Small-B-barrel+DUF1998<-BrxD-AAA<-PglZ<-DUF6079*<-N6-METHYLASE<-?<-RNase<-PglW<-PIN+35exo+T7DNApol\|\|?->?-><-REC+OmpR-HTH\|\|tRNA->DUF488-> | 1222 | Bacteria>Terrabacteria_group>Actinomycetota>Actinomycetes>Mycobacteriales>Mycobacteriaceae | GCF_021183725.1 | Mycobacterium_ostraviense | CDN38_RS00905 | 225380..229048 | 2738409 | phage-resistance-protein-[Mycobacterium-ostraviense] |
| WP_235881583.1 | Type2.Pgl.like.BREX | <-ABhydrolase\|\|?->?-><-ParB+HNH<-SF1-SF2-Helicase+wHTH+Small-B-barrel+DUF1998<-BrxD-AAA<-PglZ<-DUF6079*<-N6-METHYLASE\|\|Phage_int_SAM_1+Phage_integrase->Phage_integrase+Phage_integrase->Phage_integrase+Phage_integrase->TetR+HTH->XIS-HTH->SF1-SF2-Helicase->?->N6-N4-METHYLASE-> | 1229 | Bacteria>Terrabacteria_group>Actinomycetota>Actinomycetes>Mycobacteriales>Mycobacteriaceae | GCF_021559935.1 | Mycolicibacterium_vanbaalenii | HZU38_RS11485 | 2346374..2350063 | 110539 | phage-resistance-protein-[Mycolicibacterium-vanbaalenii] |
| WP_241205339.1 | Type2.Pgl.like.BREX | ParB+HNH->PglW->N6-METHYLASE->DUF6079*->PglZ->BrxD-AAA->SF1-SF2-Helicase+wHTH+Small-B-barrel+DUF1998->ParB+HNH+Schlafen-> | 1274 | Bacteria>Terrabacteria_group>Actinomycetota>Actinomycetes>Mycobacteriales>Mycobacteriaceae | GCF_022348085.1 | Mycobacterium_sp._PSTR-4-N | MHN84_RS15080 | 183100..186924 | 2917745 | DUF6079-family-protein-[Mycobacterium-sp.-PSTR-4-N] |
| WP_260991664.1 | Type2.Pgl.like.BREX | Phage_int_SAM_3+Phage_integrase->XIS-HTH->BirA-HTH->?->?-><-SF1-SF2-Helicase+wHTH+Small-B-barrel+DUF1998<-BrxD-AAA<-PglZ<-DUF6079*<-N6-METHYLASE<-PglW\|\|TIR->?->?->TIR->tRNA-><-HTH_3<-MarR-HTH | 1246 | Bacteria>Terrabacteria_group>Actinomycetota>Actinomycetes>Mycobacteriales>Mycobacteriaceae | GCF_025345615.1 | Mycobacterium_sp._CPCC_205710 | N4S67_RS04285 | 850216..853956 | 2978347 | phage-resistance-protein-[Mycobacterium-sp.-CPCC-205710] |
| WP_115328865.1 | Type2.Pgl.like.BREX | viral-protein->SF1-SF2-Helicase+DUF1998->?-><-ParB+HNH<-PglZ<-DUF6079*<-N6-METHYLASE<-PglW-p<-PglW-p | 1228 | Bacteria>Terrabacteria_group>Actinomycetota>Actinomycetes>Mycobacteriales>Mycobacteriaceae | GCF_025821885.1 | Mycolicibacterium_gilvum | H7I56_RS12860 | 22806..26492 | 1804 | phage-resistance-protein-[Mycolicibacterium-gilvum] |
| WP_264058319.1 | Type2.Pgl.like.BREX | SbcC+SbcC->?->?-><-MMR_HSR1<-GTPase-AIG+sGTP\|\|PglW->N6-METHYLASE->DUF6079*->PglZ->BrxD-AAA->SF1-SF2-Helicase+wHTH+Small-B-barrel+DUF1998->ParB+HNH->HIN-HTH->MULE-TRANSPOSASE-><-HKD-DNase+SF1-SF2-Helicase | 1224 | Bacteria>Terrabacteria_group>Actinomycetota>Actinomycetes>Mycobacteriales>Mycobacteriaceae | GCF_025822145.1 | Mycolicibacterium_frederiksbergense | H7JO8_RS19230 | 370396..374070 | 117567 | DUF6079-family-protein-[Mycolicibacterium-frederiksbergense] |
| WP_268785399.1 | Type2.Pgl.like.BREX | <-SF1-SF2-Helicase+wHTH+Small-B-barrel+DUF1998<-BrxD-AAA<-PglZ<-DUF6079*<-N6-METHYLASE<-PglW<-Peptidase_M48+TerB<-?<-HORMA<-SF1-SF2-Helicase+DUF1998<-viral-protein | 1274 | Bacteria>Terrabacteria_group>Actinomycetota>Actinomycetes>Mycobacteriales>Mycobacteriaceae | GCF_026805495.1 | Mycolicibacterium_iranicum | OY187_RS04300 | 6356..10180 | 912594 | DUF6079-family-protein-[Mycolicibacterium-iranicum] |
| WP_270923259.1 | Type2.Pgl.like.BREX | <-SF1-SF2-Helicase+wHTH+Small-B-barrel+DUF1998<-BrxD-AAA<-PglZ<-DUF6079*<-N6-METHYLASE<-?<-RNase<-PglW<-PIN+35exo+T7DNApol\|\|?->?-><-REC+OmpR-HTH\|\|tRNA->DUF488-> | 1220 | Bacteria>Terrabacteria_group>Actinomycetota>Actinomycetes>Mycobacteriales>Mycobacteriaceae | GCF_027854195.1 | Mycobacterium_xenopi | PFJ03_RS00260 | 62216..65878 | 1789 | phage-resistance-protein-[Mycobacterium-xenopi] |
| WP_036463067.1 | Type2.Pgl.like.BREX | viral-protein->SF1-SF2-Helicase+DUF1998->McrB-AAA->McrC-NTD+REase->REase->PglW->N6-METHYLASE->DUF6079*->PglZ->BrxD-AAA->SF1-SF2-Helicase+wHTH+Small-B-barrel+DUF1998-><-HIN-HTH+PAIREDC-HTH<-AAA-ATPase-p | 1275 | Bacteria>Terrabacteria_group>Actinomycetota>Actinomycetes>Mycobacteriales>Mycobacteriaceae | GCF_900101555.1 | Mycolicibacterium_neoaurum | BLS83_RS26015 | 158467..162294 | 1795 | DUF6079-family-protein-[Mycolicibacterium-neoaurum] |
| WP_066820955.1 | Type2.Pgl.like.BREX | <-ADH_N+ADH_zinc_N<-?\|\|MODE-HTH+PBPII-><-SopB+HNH<-?\|\|N6-METHYLASE-p->DUF6079*->PglZ->BrxD-AAA->SF1-SF2-Helicase+wHTH+Small-B-barrel+DUF1998-><-?<-Glyoxalase\|\|?-><-ABhydrolase+ACYC | 1230 | Bacteria>Terrabacteria_group>Actinomycetota>Actinomycetes>Mycobacteriales>Mycobacteriaceae | - | Mycobacterium_sp._1100029.7 | A5647_15270 | 11611..15303 | 1834067 | DUF6079-family-protein-[Mycobacterium-sp.-1100029.7] |
| WP_083060791.1 | Type2.Pgl.like.BREX | <-McrC-NTD+REase<-McrB-AAA-ATPase<-SF1-SF2-Helicase+wHTH+Small-B-barrel+DUF1998<-BrxD-AAA<-PglZ<-DUF6079*<-N6-METHYLASE<-PglW | 1222 | Bacteria>Terrabacteria_group>Actinomycetota>Actinomycetes>Mycobacteriales>Mycobacteriaceae | GCF_025823285.1 | Mycolicibacterium_bacteremicum | BST17_RS20760 | 42057..45725 | 564198 | DUF6079-family-protein-[Mycolicibacterium-bacteremicum] |
| EME22377.1 | Type2.Pgl.like.BREX | <-RAMA\|\|Thermonuclease->HKD-DNase->?->PglW->N6-METHYLASE->?->Tox-RelA->HEPN->DUF6079*->PglZ->BrxD-AAA->SF1-SF2-Helicase+wHTH+Small-B-barrel+DUF1998->REase+SF1-SF2-Helicase-><-?\|\|SF1-SF2-Helicase+REase->HKD-DNase+SF1-SF2-Helicase+HTH+HTH+SRA-> | 1256 | Bacteria>Terrabacteria_group>Actinomycetota>Actinomycetes>Mycobacteriales>Nocardiaceae | GCA_000341795.1 | Rhodococcus_triatomae_BKS_15-14 | G419_05577 | 69562..73332 | 1278078 | phage-resistance-protein-PglY-[Rhodococcus-triatomae-BKS-15-14] |
| KXF57185.1 | Type2.Pgl.like.BREX | <-SAM-METHYLASE<-?\|\|?->HHH_5+SNF2_N+SF1-SF2-Helicase-><-?<-SF1-SF2-Helicase+wHTH+Small-B-barrel+DUF1998<-BrxD-AAA<-PglZ<-DUF6079*<-N6-METHYLASE<-PglW<-?<-HTH_3\|\|?->?-><-?<-?<-ADH_N+ADH_zinc_N+METHYLASE | 1265 | Bacteria>Terrabacteria_group>Actinomycetota>Actinomycetes>Mycobacteriales>Nocardiaceae | GCA_001555475.1 | Rhodococcus_sp._SC4 | AXA44_30850 | 16533..20330 | 1793160 | phage-resistance-protein-[Rhodococcus-sp.-SC4] |
| AMY54745.1 | Type2.Pgl.like.BREX | HHH_5+SNF2_N+SF1-SF2-Helicase->RAMA->HATPase->REase->?->PglW->N6-METHYLASE->DUF6079-p*->N6-METHYLASE->DUF6079->PglZ->BrxD-AAA->SF1-SF2-Helicase+wHTH+Small-B-barrel+DUF1998-><-SF1-SF2-Helicase<-wHTH<-LON-AAA<-REase | 130 | Bacteria>Terrabacteria_group>Actinomycetota>Actinomycetes>Mycobacteriales>Nocardiaceae | GCA_001620305.1 | Rhodococcus_fascians_D188 | A3L23_03420 | 3540502..3540894 | 1051973 | hypothetical-protein-A3L23_03420-[Rhodococcus-fascians-D188] |
| OZD41665.1 | Type2.Pgl.like.BREX | <-CASPASE\|\|SF1-SF2-Helicase+REase->SF1-SF2-Helicase+HTH->wHTH+REase-><-?\|\|URI-DNase-><-SF1-SF2-Helicase+wHTH+Small-B-barrel+DUF1998<-BrxD-AAA<-PglZ<-DUF6079*<-N6-METHYLASE<-PglW<-CBS+HEPN+HEPN<-HEPN+SF1-SF2-Helicase+REase<-wHTH+HNH\|\|?-><-?\|\|MODE-HTH+PBPII-> | 1245 | Bacteria>Terrabacteria_group>Actinomycetota>Actinomycetes>Mycobacteriales>Nocardiaceae | GCA_002258825.1 | Rhodococcus_sp._06-1477-1B | CH252_27875 | 148678..152415 | 2022485 | phage-resistance-protein-[Rhodococcus-sp.-06-1477-1B] |
| PPJ35991.1 | Type2.Pgl.like.BREX | HTH->?-><-SF1-SF2-Helicase+wHTH+Small-B-barrel+DUF1998<-BrxD-AAA<-PglZ<-DUF6079*<-N6-METHYLASE<-PglW<-?<-?\|\|N6-METHYLASE->N6-METHYLASE->Pkinase+TPR-> | 1300 | Bacteria>Terrabacteria_group>Actinomycetota>Actinomycetes>Mycobacteriales>Nocardiaceae | GCA_002933535.1 | Nocardia_nova | C5E45_23110 | 140922..144824 | 37330 | phage-resistance-protein-[Nocardia-nova] |
| MQY22654.1 | Type2.Pgl.like.BREX | PglW->N6-METHYLASE->DUF6079*->PglZ->BrxD-AAA->SF1-SF2-Helicase+wHTH+Small-B-barrel+DUF1998-><-?\|\|?->?->APATPase+TPR+TPR+TPR+TPR->?->METHYLASE-> | 1295 | Bacteria>Terrabacteria_group>Actinomycetota>Actinomycetes>Mycobacteriales>Nocardiaceae | GCA_009604405.1 | Nocardia_macrotermitis | NRB20_57720 | 166823..170710 | 2585198 | hypothetical-protein-[Nocardia-macrotermitis] |
| MXQ77398.1 | Type2.Pgl.like.BREX | REase+SF1-SF2-Helicase->nSTAND1+HNH->HTH_36-><-?<-HNH<-?<-MORC\|\|PglW->N6-METHYLASE->DUF6079*->PglZ->BrxD-AAA->SF1-SF2-Helicase+wHTH+Small-B-barrel+DUF1998->URI-DNase+URI-DNase->MAD-NTD+EVE+McrB-AAA->McrC-NTD+REase->ANK+ANK->SF1-SF2-Helicase->REase-> | 1253 | Bacteria>Terrabacteria_group>Actinomycetota>Actinomycetes>Mycobacteriales>Nocardiaceae | GCA_009831075.1 | Rhodococcus_rhodochrous | GSM98_14630 | 113574..117335 | 1829 | phage-resistance-protein-[Rhodococcus-rhodochrous] |
| NKS26554.1 | Type2.Pgl.like.BREX | <-MalT+LuxR-HTH<-Calcineurin<-?<-?\|\|PglW->N6-METHYLASE->ParB+HNH-><-RNase+Kinase+Kinase\|\|DUF6079*->PglZ->BrxD-AAA->?->SF1-SF2-Helicase+wHTH+Small-B-barrel+DUF1998-><-?\|\|?->\|\|Glyoxalase-> | 1241 | Bacteria>Terrabacteria_group>Actinomycetota>Actinomycetes>Mycobacteriales>Nocardiaceae | GCA_012393535.1 | Prescottella_equi | GS505_12170 | 246707..250432 | 43767 | phage-resistance-protein-[Prescottella-equi] |
| NKS50173.1 | Type2.Pgl.like.BREX | <-HTH_3\|\|MORC+SF1-SF2-Helicase->SF1-SF2-Helicase->DrmB-><-?<-?<-SF1-SF2-Helicase+wHTH+Small-B-barrel+DUF1998<-BrxD-AAA<-PglZ<-DUF6079*\|\|?-><-?\|\|LZ_Tnp_IS481+HTH_32+rve+rve_3-><-MULE-TRANSPOSASE\|\|?->?-><-McrC-NTD+REase<-N6-METHYLASE<-PglW | 1264 | Bacteria>Terrabacteria_group>Actinomycetota>Actinomycetes>Mycobacteriales>Nocardiaceae | GCA_012393545.1 | Prescottella_equi | GS480_15720 | 3393685..3397479 | 43767 | phage-resistance-protein-[Prescottella-equi] |
| NKR55757.1 | Type2.Pgl.like.BREX | ParA-Soj-PloopNTPase->HHH_4->SIGMA-HTH+SIGMA-HTH-><-nSTAND1+HNH<-REase+SF1-SF2-Helicase<-ParB+HNH+Schlafen<-SF1-SF2-Helicase+wHTH+Small-B-barrel+DUF1998<-BrxD-AAA<-PglZ<-DUF6079*<-EVE+REase<-McrC-NTD+REase<-McrB-AAA<-N6-METHYLASE<-?<-MORC\|\|HHH_5+SNF2_N+SF1-SF2-Helicase-><-wHTH<-MAD-NTD | 1264 | Bacteria>Terrabacteria_group>Actinomycetota>Actinomycetes>Mycobacteriales>Nocardiaceae | GCA_012393885.1 | Prescottella_equi | GS481_17030 | 1591471..1595265 | 43767 | phage-resistance-protein-[Prescottella-equi] |
| NKR97090.1 | Type2.Pgl.like.BREX | PglW->N6-METHYLASE->HKD-DNase->RADICAL-SAM-><-HEPN\|\|DUF6079*->PglZ->BrxD-AAA->SF1-SF2-Helicase+wHTH+Small-B-barrel+DUF1998->REase+SF1-SF2-Helicase->nSTAND1+HNH-><-SIGMA-HTH+SIGMA-HTH<-HHH_4<-ParA-Soj-PloopNTPase<-SF1-SF2-Helicase+HTH | 1264 | Bacteria>Terrabacteria_group>Actinomycetota>Actinomycetes>Mycobacteriales>Nocardiaceae | GCA_012394005.1 | Prescottella_equi | GS473_09460 | 79040..82834 | 43767 | phage-resistance-protein-[Prescottella-equi] |
| NKW20297.1 | Type2.Pgl.like.BREX | <-McrB-AAA<-?\|\|?->?->?->DUF6361->HKD-DNase->SF1-SF2-Helicase->RVT->DUF6079*->SF1-SF2-Helicase+wHTH+Small-B-barrel+DUF1998-><-?<-nSTAND1+HNH | 1252 | Bacteria>Terrabacteria_group>Actinomycetota>Actinomycetes>Mycobacteriales>Nocardiaceae | GCA_012397345.1 | Prescottella_equi | GS940_23010 | 133803..137561 | 43767 | phage-resistance-protein-[Prescottella-equi] |
| NME80339.1 | Type2.Pgl.like.BREX | <-MAD-NTD+EVE+McrB-AAA<-URI-DNase<-SF1-SF2-Helicase+wHTH+Small-B-barrel+DUF1998<-BrxD-AAA<-PglZ<-DUF6079*<-N6-METHYLASE<-PglW<-REase+SF1-SF2-Helicase | 1239 | Bacteria>Terrabacteria_group>Actinomycetota>Actinomycetes>Mycobacteriales>Nocardiaceae | GCA_012844225.1 | Rhodococcus_sp._105337 | HF886_14710 | 10453..14172 | 2725310 | phage-resistance-protein-[Rhodococcus-sp.-105337] |
| MBF6175293.1 | Type2.Pgl.like.BREX | SF1-SF2-Helicase-><-SF1-SF2-Helicase+wHTH+Small-B-barrel+DUF1998<-BrxD-AAA<-PglZ<-DUF6079*<-N6-METHYLASE<-PglW | 1288 | Bacteria>Terrabacteria_group>Actinomycetota>Actinomycetes>Mycobacteriales>Nocardiaceae | GCA_015477355.1 | Nocardia_blacklockiae | IU476_27890 | 116199..120065 | 480036 | phage-resistance-protein-[Nocardia-blacklockiae] |
| MBF6225868.1 | Type2.Pgl.like.BREX | HTH_3->STYKIN->?->DUF397->PglW->N6-METHYLASE->DUF6079*->PglZ->BrxD-AAA->SF1-SF2-Helicase+wHTH+Small-B-barrel+DUF1998-><-DUF3323+TOPRIM<-?<-?<-DUF2397-<-?<-NTP_transf_9 | 1256 | Bacteria>Terrabacteria_group>Actinomycetota>Actinomycetes>Mycobacteriales>Nocardiaceae | GCA_015477435.1 | Nocardia_abscessus | IU470_12230 | 97362..101132 | 120957 | phage-resistance-protein-[Nocardia-abscessus] |
| MBF6301579.1 | Type2.Pgl.like.BREX | PadR-wHTH->?->?->DUF3323+TOPRIM-><-SF1-SF2-Helicase+wHTH+Small-B-barrel+DUF1998<-BrxD-AAA<-PglZ<-DUF6079*<-N6-METHYLASE<-PglW<-DUF397<-HTH_31+DUF5753\|\|?->?-><-?<-Abhydrolase_3<-SF1-SF2-Helicase | 1256 | Bacteria>Terrabacteria_group>Actinomycetota>Actinomycetes>Mycobacteriales>Nocardiaceae | GCA_015477605.1 | Nocardia_amamiensis | IU459_29170 | 22411..26181 | 404578 | phage-resistance-protein-[Nocardia-amamiensis] |
| MBF6346442.1 | Type2.Pgl.like.BREX | <-N6-METHYLASE<-?<-?\|\|PglW->N6-METHYLASE->DUF6079*->PglZ->BrxD-AAA->SF1-SF2-Helicase+wHTH+Small-B-barrel+DUF1998-><-?<-DUF397<-HTH_31+DUF5753 | 1251 | Bacteria>Terrabacteria_group>Actinomycetota>Actinomycetes>Mycobacteriales>Nocardiaceae | GCA_015477865.1 | Nocardia_cyriacigeorgica | IU409_23450 | 21396..25151 | 135487 | phage-resistance-protein-[Nocardia-cyriacigeorgica] |
| BCN48390.1 | Type2.Pgl.like.BREX | SIGMA-HTH+SIGMA-HTH-><-URI-DNase+URI-DNase<-?<-nSTAND1+HNH<-REase+SF1-SF2-Helicase<-ParB+HNH+Schlafen<-SF1-SF2-Helicase+wHTH+Small-B-barrel+DUF1998<-BrxD-AAA<-PglZ<-DUF6079*\|\|SopB+HNH-><-HKD-DNase\|\|?->?->?-><-MarR-HTH<-?<-N6-METHYLASE<-PglW | 1264 | Bacteria>Terrabacteria_group>Actinomycetota>Actinomycetes>Mycobacteriales>Nocardiaceae | GCA_019703935.1 | Prescottella_equi | RE9416_16910 | 1800636..1804430 | 43767 | hypothetical-protein-RE9416_16910-[Prescottella-equi] |
| WP_014350030.1 | Type2.Pgl.like.BREX | <-HEPN+REase<-DUF4352<-?<-?<-35exo<-?<-SF1-SF2-Helicase+wHTH+Small-B-barrel+DUF1998<-BrxD-AAA<-PglZ<-DUF6079*<-N6-METHYLASE<-PglW<-?\|\|N6-METHYLASE->TRD+TRD->Pkinase+TPR+TPR->HNH-><-SF1-SF2-Helicase | 1302 | Bacteria>Terrabacteria_group>Actinomycetota>Actinomycetes>Mycobacteriales>Nocardiaceae | GCF_000284035.1 | Nocardia_cyriacigeorgica | NOCYR_RS08920 | 1981623..1985531 | 135487 | hypothetical-protein-[Nocardia-cyriacigeorgica] |
| WP_040863307.1 | Type2.Pgl.like.BREX | REase+SF1-SF2-Helicase-><-CBS+CBS+HEPN<-ParB+HNH<-SF1-SF2-Helicase+wHTH+Small-B-barrel+DUF1998<-BrxD-AAA<-PglZ<-DUF6079*<-DndB\|\|?->?->HTH_31+DUF5753->DUF397-><-N6-METHYLASE<-PglW\|\|REase-> | 1308 | Bacteria>Terrabacteria_group>Actinomycetota>Actinomycetes>Mycobacteriales>Nocardiaceae | GCF_000308655.1 | Nocardia_niigatensis | ON33_RS34230 | 29321..33247 | 209249 | hypothetical-protein-[Nocardia-niigatensis] |
| WP_003934894.1 | Type2.Pgl.like.BREX | <-ZnR\|\|?-><-MULE-TRANSPOSASE<-SF1-SF2-Helicase+wHTH+Small-B-barrel+DUF1998\|\|HicA->HicB-><-?<-BrxD-AAA<-PglZ<-DUF6079*\|\|NTP_transf-><-DOC+HTH\|\|?-><-N6-METHYLASE<-PglW<-?<-STYKinase-like<-?<-His_Phos_1 | 1231 | Bacteria>Terrabacteria_group>Actinomycetota>Actinomycetes>Mycobacteriales>Nocardiaceae | GCF_000341965.1 | Rhodococcus_ruber | G352_RS03930 | 14064..17759 | 1830 | DUF6079-family-protein-[Rhodococcus-ruber] |
| WP_019927709.1 | Type2.Pgl.like.BREX | YTCD-HTH-><-SF1-SF2-Helicase+wHTH+Small-B-barrel+DUF1998<-BrxD-AAA<-PglZ<-DUF6079*<-ParB+HNH<-N6-METHYLASE<-PglW\|\|REase-><-?\|\|TetR-HTH-><-?<-Glyoxalase | 1247 | Bacteria>Terrabacteria_group>Actinomycetota>Actinomycetes>Mycobacteriales>Nocardiaceae | GCF_000381925.1 | Nocardia_sp._BMG111209 | G361_RS0113980 | 3029981..3033724 | 1160137 | DUF6079-family-protein-[Nocardia-sp.-BMG111209] |
| WP_019931234.1 | Type2.Pgl.like.BREX | MqsA_antitoxin+DUF4065->?->N6-METHYLASE-><-CBS+HEPN<-ParB+HNH<-SF1-SF2-Helicase+wHTH+Small-B-barrel+DUF1998<-BrxD-AAA<-PglZ<-DUF6079*\|\|DDE_Tnp_1_4-><-N6-METHYLASE<-DDE_Tnp_1<-N6-METHYLASE-p<-PglW\|\|?->HTH+MPTase->\|\|DUF772+DDE_Tnp_1_6-><-HTH_Tnp_1 | 1281 | Bacteria>Terrabacteria_group>Actinomycetota>Actinomycetes>Mycobacteriales>Nocardiaceae | GCF_000381925.1 | Nocardia_sp._BMG111209 | G361_RS0132065 | 198837..202682 | 1160137 | hypothetical-protein-[Nocardia-sp.-BMG111209] |
| WP_029543714.1 | Type2.Pgl.like.BREX | His_Phos_1->?->STYKinase-like->?->PglW->N6-METHYLASE->\|\|\|\|?->DDE_Tnp_1_assoc+DDE_Tnp_1->DUF6079*->PglZ->BrxD-AAA-><-?\|\|SF1-SF2-Helicase+wHTH+Small-B-barrel+DUF1998->?->ZnR-><-SbcC/MukB | 1238 | Bacteria>Terrabacteria_group>Actinomycetota>Actinomycetes>Mycobacteriales>Nocardiaceae | GCF_000470885.1 | Rhodococcus_aetherivorans | N505_RS10680 | 2237043..2240759 | 191292 | DUF6079-family-protein-[Rhodococcus-aetherivorans] |
| WP_032369004.1 | Type2.Pgl.like.BREX | ABC-ATPase+SbcC-><-?<-SbcC\|\|PglW->N6-METHYLASE->DUF6079*->PglZ->BrxD-AAA->SF1-SF2-Helicase+wHTH+Small-B-barrel+DUF1998-><-wHTH<-?<-SbcC | 1244 | Bacteria>Terrabacteria_group>Actinomycetota>Actinomycetes>Mycobacteriales>Nocardiaceae | GCF_000760995.1 | Rhodococcus_fascians | BIO2_RS0103085 | 151180..154914 | 1828 | DUF6079-family-protein-[Rhodococcus-fascians] |
| WP_230594698.1 | Type2.Pgl.like.BREX | HD-><-wHTH+REase<-HIRAN\|\|HEPN+SF1-SF2-Helicase+REase->CBS+HEPN+HEPN->?->PglW->N6-METHYLASE->DUF6079*->PglZ->BrxD-AAA->SF1-SF2-Helicase+wHTH+Small-B-barrel+DUF1998-><-wHTH<-ANK+ANK<-?<-SbcC | 1216 | Bacteria>Terrabacteria_group>Actinomycetota>Actinomycetes>Mycobacteriales>Nocardiaceae | GCF_000761075.1 | Rhodococcus_fascians | AA17_RS0121225 | 153530..157180 | 1828 | phage-resistance-protein-[Rhodococcus-fascians] |
| WP_033096655.1 | Type2.Pgl.like.BREX | His_Phos_1->?->STYKinase-like->?->PglW->N6-METHYLASE->DUF6079*->PglZ->BrxD-AAA->HHH_4+HTH+wHTH+SF1-SF2-Helicase+REase->SF1-SF2-Helicase+wHTH+Small-B-barrel+DUF1998->?->URI-DNase->?-><-Glyoxalase | 1233 | Bacteria>Terrabacteria_group>Actinomycetota>Actinomycetes>Mycobacteriales>Nocardiaceae | GCF_000763325.2 | Rhodococcus_sp._p52 | IM25_RS16950 | 3659539..3663240 | 935199 | DUF6079-family-protein-[Rhodococcus-sp.-p52] |
| WP_054373663.1 | Type2.Pgl.like.BREX | <-ZnR\|\|?-><-SF1-SF2-Helicase+wHTH+Small-B-barrel+DUF1998\|\|HicB-><-HHH_4+HTH+wHTH+SF1-SF2-Helicase+REase<-BrxD-AAA<-PglZ<-DUF6079*\|\|HEPN+REase-><-HTH_Tnp_1+HTH_21+rve+rve_3<-N6-METHYLASE<-PglW-p | 1238 | Bacteria>Terrabacteria_group>Actinomycetota>Actinomycetes>Mycobacteriales>Nocardiaceae | GCF_001278665.1 | Rhodococcus_rhodochrous | Z051_RS16935 | 26716..30432 | 1829 | DUF6079-family-protein-[Rhodococcus-rhodochrous] |
| WP_156399392.1 | Type2.Pgl.like.BREX | DndB->HIN-HTH->\|\|\|\|AAA-ATPase->TIR-><-?<-REase<-MORC\|\|PglW->N6-METHYLASE->DUF6079*->PglZ->BrxD-AAA->SF1-SF2-Helicase+wHTH+Small-B-barrel+DUF1998-><-wHTH<-ANK+ANK<-?<-SbcC | 1235 | Bacteria>Terrabacteria_group>Actinomycetota>Actinomycetes>Mycobacteriales>Nocardiaceae | GCF_001426085.1 | Rhodococcus_sp._Leaf278 | ASG84_RS04935 | 127966..131673 | 1736319 | DUF6079-family-protein-[Rhodococcus-sp.-Leaf278] |
| WP_057474819.1 | Type2.Pgl.like.BREX | HATPase->?->?->PglW->N6-METHYLASE->DUF6079*->PglZ->BrxD-AAA->SF1-SF2-Helicase+wHTH+Small-B-barrel+DUF1998-><-SF1-SF2-Helicase | 1290 | Bacteria>Terrabacteria_group>Actinomycetota>Actinomycetes>Mycobacteriales>Nocardiaceae | GCF_001426165.1 | Rhodococcus_sp._Leaf233 | ASH04_RS24720 | 139714..143586 | 1736302 | DUF6079-family-protein-[Rhodococcus-sp.-Leaf233] |
| WP_056816725.1 | Type2.Pgl.like.BREX | <-TetR-HTH\|\|?->?->?-><-Tox-RelA<-Pkinase+TPR+TPR\|\|PglW->N6-METHYLASE->DUF6079*->PglZ->BrxD-AAA->ParB+HNH->CBS+HEPN->SF1-SF2-Helicase+wHTH+Small-B-barrel+DUF1998-><-REase+SF1-SF2-Helicase<-?<-?<-SAM-METHYLASE | 1293 | Bacteria>Terrabacteria_group>Actinomycetota>Actinomycetes>Mycobacteriales>Nocardiaceae | GCF_001427185.1 | Nocardia_sp._Root136 | ASD42_RS14795 | 207384..211265 | 1736458 | hypothetical-protein-[Nocardia-sp.-Root136] |
| WP_238077028.1 | Type2.Pgl.like.BREX | SIGMA-HTH-><-MORC->?->ParA-Soj-PloopNTPase->HHH_4-><-RVT<-SF1-SF2-Helicase+wHTH+Small-B-barrel+DUF1998<-BrxD-AAA<-PglZ<-DUF6079*<-N6-METHYLASE<-PglW<-DNA_glycosylase\|\|HNH-><-nSTAND1+HNH<-REase+SF1-SF2-Helicase<-HEPN+SF1-SF2-Helicase+REase | 1303 | Bacteria>Terrabacteria_group>Actinomycetota>Actinomycetes>Mycobacteriales>Nocardiaceae | GCF_001552595.1 | Rhodococcus_erythropolis | RE4_RS16525 | 46882..50793 | 1833 | DUF6079-family-protein-[Rhodococcus-erythropolis] |
| WP_062795547.1 | Type2.Pgl.like.BREX | <-REase+SF1-SF2-Helicase\|\|sGTP+sGTP+GTPase-AIG->sGTP->ANK+ANK-><-McrC-NTD+REase<-McrB-AAA<-SF1-SF2-Helicase+wHTH+Small-B-barrel+DUF1998<-BrxD-AAA<-PglZ<-DUF6079*<-N6-METHYLASE<-PglW\|\|HNH->DUF6361->HKD-DNase->SF1-SF2-Helicase->Exonuc_V_gamma+REase->SF1-SF2-Helicase+REase-> | 1241 | Bacteria>Terrabacteria_group>Actinomycetota>Actinomycetes>Mycobacteriales>Nocardiaceae | GCF_001598915.1 | Williamsia_muralis | WIMU1_RS06335 | 1314541..1318266 | 85044 | DUF6079-family-protein-[Williamsia-muralis] |
| WP_063049769.1 | Type2.Pgl.like.BREX | HTH_3->STYKIN-><-DEDD_Tnp_IS110+Transposase_20<-?\|\|?-><-?\|\|PglW->N6-METHYLASE->DUF6079*->PglZ->BrxD-AAA->SF1-SF2-Helicase+wHTH+Small-B-barrel+DUF1998-> | 1256 | Bacteria>Terrabacteria_group>Actinomycetota>Actinomycetes>Mycobacteriales>Nocardiaceae | GCF_001612765.1 | Nocardia_arthritidis | NA8_RS08195 | 99706..103476 | 228602 | DUF6079-family-protein-[Nocardia-arthritidis] |
| WP_067804308.1 | Type2.Pgl.like.BREX | <-SF1-SF2-Helicase+wHTH+Small-B-barrel+DUF1998<-BrxD-AAA\|\|?->?-><-PglZ<-DUF6079*<-N6-METHYLASE<-PglW<-?<-?\|\|N6-METHYLASE->TRD+TRD->Pkinase+TPR+TPR-> | 1304 | Bacteria>Terrabacteria_group>Actinomycetota>Actinomycetes>Mycobacteriales>Nocardiaceae | GCF_001612785.1 | Nocardia_beijingensis | NB3_RS17840 | 47825..51739 | 95162 | phage-resistance-protein-[Nocardia-beijingensis] |
| WP_067496132.1 | Type2.Pgl.like.BREX | SF1-SF2-Helicase->McrB-AAA->McrC-NTD+REase-><-SF1-SF2-Helicase+HKD-DNase<-Pkinase+TPR+TPR\|\|?->PglW->N6-METHYLASE->DUF6079*->PglZ->BrxD-AAA->SF1-SF2-Helicase+wHTH+Small-B-barrel+DUF1998-><-REase+SF1-SF2-Helicase | 1299 | Bacteria>Terrabacteria_group>Actinomycetota>Actinomycetes>Mycobacteriales>Nocardiaceae | GCF_001612905.1 | Nocardia_ignorata | NI2_RS26665 | 97698..101597 | 145285 | phage-resistance-protein-[Nocardia-ignorata] |

| Protein-ID | System | Gene-Neighbourhoods | Length | Taxonomy | Assembly-ID | Strain | Locus-Tag | Genomic-coordinates | TaxID | Fasta-defline |
|---|---|---|---|---|---|---|---|---|---|---|
| WP_245660406.1 | Type2.Pgl.like.BREX | [illegible] | 1331 | Bacteria>Terrabacteria_group>Actinomycetota>Actinomycetes>Mycobacteriales>Nocardiaceae | GCF_001612965.1 | Nocardia_kruczakiae | NK1_RS09245 | 23385..27380 | 261477 | DUF6079-family-protein-[Nocardia-kruczakiae] |
| WP_068008803.1 | Type2.Pgl.like.BREX | [illegible] | 1230 | Bacteria>Terrabacteria_group>Actinomycetota>Actinomycetes>Mycobacteriales>Nocardiaceae | GCF_001613105.1 | Nocardia_pseudobrasiliensis | DFR76_RS37550 | 11394..15086 | 45979 | phage-resistance-protein-[Nocardia-pseudobrasiliensis] |
| WP_068057216.1 | Type2.Pgl.like.BREX | [illegible] | 1303 | Bacteria>Terrabacteria_group>Actinomycetota>Actinomycetes>Mycobacteriales>Nocardiaceae | GCF_001613365.1 | Nocardia_xishanensis | NX1_RS10700 | 31578..35489 | 238964 | DUF6079-family-protein-[Nocardia-xishanensis] |
| WP_245713296.1 | Type2.Pgl.like.BREX | [illegible] | 1271 | Bacteria>Terrabacteria_group>Actinomycetota>Actinomycetes>Mycobacteriales>Nocardiaceae | GCF_001613485.1 | Nocardia_rhamnosiphila | IG71_RS0111080 | 476296..480111 | 426716 | DUF6079-family-protein-[Nocardia-rhamnosiphila] |
| WP_145916598.1 | Type2.Pgl.like.BREX | [illegible] | 1290 | Bacteria>Terrabacteria_group>Actinomycetota>Actinomycetes>Mycobacteriales>Nocardiaceae | GCF_001620305.1 | Rhodococcus_fascians | A3L23_RS16890 | 3544457..3548329 | 1828 | DUF6079-family-protein-[Rhodococcus-fascians] |
| WP_067565329.1 | Type2.Pgl.like.BREX | [illegible] | 1307 | Bacteria>Terrabacteria_group>Actinomycetota>Actinomycetes>Mycobacteriales>Nocardiaceae | GCF_001625085.1 | Nocardia_acidivorans | NA10_RS08140 | 207417..211340 | 404580 | phage-resistance-protein-[Nocardia-acidivorans] |
| WP_064851885.1 | Type2.Pgl.like.BREX | [illegible] | 1235 | Bacteria>Terrabacteria_group>Actinomycetota>Actinomycetes>Mycobacteriales>Nocardiaceae | GCF_001665495.1 | Rhodococcus_sp._852002-51564_SCH6189132-a | A5767_RS05820 | 50780..54487 | 1834103 | DUF6079-family-protein-[Rhodococcus-sp.-852002-51564_SCH6189132-a] |
| WP_065922559.1 | Type2.Pgl.like.BREX | [illegible] | 1241 | Bacteria>Terrabacteria_group>Actinomycetota>Actinomycetes>Mycobacteriales>Nocardiaceae | GCF_001700945.1 | unclassified_Rhodococcus_(in:_high_G+C_Gram-positive_bacteria) | A4U64_RS11180 | 2546960..2550685 | 192944 | MULTISPECIES:-DUF6079-family-protein-[unclassified-Rhodococcus-(in:-high-G+C-Gram-positive-bacteria)] |
| WP_071934933.1 | Type2.Pgl.like.BREX | [illegible] | 1233 | Bacteria>Terrabacteria_group>Actinomycetota>Actinomycetes>Mycobacteriales>Nocardiaceae | GCF_001886355.1 | Rhodococcus_sp._2G | B0226_RS07740 | 1652822..1656523 | 1570939 | DUF6079-family-protein-[Rhodococcus-sp.-2G] |
| WP_071930295.1 | Type2.Pgl.like.BREX | [illegible] | 1251 | Bacteria>Terrabacteria_group>Actinomycetota>Actinomycetes>Mycobacteriales>Nocardiaceae | GCF_001886715.1 | Nocardia_mangyaensis | BOX37_RS28435 | 6245947..6249702 | 2213200 | phage-resistance-protein-[Nocardia-mangyaensis] |
| WP_073366415.1 | Type2.Pgl.like.BREX | [illegible] | 1278 | Bacteria>Terrabacteria_group>Actinomycetota>Actinomycetes>Mycobacteriales>Nocardiaceae | GCF_001894825.1 | Rhodococcus_jostii | RJ1_RS31445 | 27857..31693 | 132919 | DUF6079-family-protein-[Rhodococcus-jostii] |
| WP_088943629.1 | Type2.Pgl.like.BREX | [illegible] | 1250 | Bacteria>Terrabacteria_group>Actinomycetota>Actinomycetes>Mycobacteriales>Nocardiaceae | GCF_002091955.1 | Rhodococcus_sp._1168 | BJ147_RS10230 | 43432..47184 | 2018041 | DUF6079-family-protein-[Rhodococcus-sp.-1168] |
| WP_084960882.1 | Type2.Pgl.like.BREX | [illegible] | 1264 | Bacteria>Terrabacteria_group>Actinomycetota>Actinomycetes>Mycobacteriales>Nocardiaceae | GCF_002094405.3 | Prescottella_equi | A6I88_RS04030 | 891226..895020 | 43767 | DUF6079-family-protein-[Prescottella-equi] |
| WP_094714576.1 | Type2.Pgl.like.BREX | [illegible] | 1245 | Bacteria>Terrabacteria_group>Actinomycetota>Actinomycetes>Mycobacteriales>Nocardiaceae | GCF_002258885.1 | Rhodococcus_sp._06-1474-1B | CH266_RS02185 | 148879..152616 | 2022499 | DUF6079-family-protein-[Rhodococcus-sp.-06-1474-1B] |
| WP_098693855.1 | Type2.Pgl.like.BREX | [illegible] | 1269 | Bacteria>Terrabacteria_group>Actinomycetota>Actinomycetes>Mycobacteriales>Nocardiaceae | GCF_002568625.1 | Nocardia_terpenica | CRH09_RS11100 | 2476073..2479882 | 455432 | phage-resistance-protein-[Nocardia-terpenica] |
| WP_104375110.1 | Type2.Pgl.like.BREX | [illegible] | 1300 | Bacteria>Terrabacteria_group>Actinomycetota>Actinomycetes>Mycobacteriales>Nocardiaceae | GCF_002933465.1 | Nocardia_nova | C5E41_RS07640 | 327754..331656 | 37330 | DUF6079-family-protein-[Nocardia-nova] |
| WP_258081613.1 | Type2.Pgl.like.BREX | [illegible] | 1329 | Bacteria>Terrabacteria_group>Actinomycetota>Actinomycetes>Mycobacteriales>Nocardiaceae | GCF_002933535.1 | Nocardia_nova | C5E45_RS23115 | 140922..144911 | 37330 | DUF6079-family-protein-[Nocardia-nova] |
| WP_119699787.1 | Type2.Pgl.like.BREX | [illegible] | 1235 | Bacteria>Terrabacteria_group>Actinomycetota>Actinomycetes>Mycobacteriales>Nocardiaceae | GCF_003586525.1 | Rhodococcus_ruber | YT1_RS14630 | 2910213..2913920 | 1830 | DUF6079-family-protein-[Rhodococcus-ruber] |
| WP_260395788.1 | Type2.Pgl.like.BREX | [illegible] | 831 | Bacteria>Terrabacteria_group>Actinomycetota>Actinomycetes>Mycobacteriales>Nocardiaceae | GCF_003852455.1 | Rhodococcus_sp._KBW08 | DBV08_RS33050 | 139953..142448 | 2144188 | DUF6079-family-protein-[Rhodococcus-sp.-KBW08] |
| WP_127946806.1 | Type2.Pgl.like.BREX | [illegible] | 1232 | Bacteria>Terrabacteria_group>Actinomycetota>Actinomycetes>Mycobacteriales>Nocardiaceae | GCF_004011835.1 | Rhodococcus_spongiicola | EF834_RS08540 | 73116..76814 | 2487352 | DUF6079-family-protein-[Rhodococcus-spongiicola] |
| WP_242496869.1 | Type2.Pgl.like.BREX | [illegible] | 1242 | Bacteria>Terrabacteria_group>Actinomycetota>Actinomycetes>Mycobacteriales>Nocardiaceae | GCF_004153645.1 | Rhodococcus_sp._Q1 | EQ874_RS21100 | 15779..19507 | 2508718 | DUF6079-family-protein-[Rhodococcus-sp.-Q1] |
| WP_135046364.1 | Type2.Pgl.like.BREX | [illegible] | 1217 | Bacteria>Terrabacteria_group>Actinomycetota>Actinomycetes>Mycobacteriales>Nocardiaceae | GCF_004563845.1 | Rhodococcus_sp._1R11 | E4P29_RS16905 | 125769..129422 | 2559614 | phage-resistance-protein-[Rhodococcus-sp.-1R11] |
| WP_137797778.1 | Type2.Pgl.like.BREX | [illegible] | 1264 | Bacteria>Terrabacteria_group>Actinomycetota>Actinomycetes>Mycobacteriales>Nocardiaceae | GCF_005484805.1 | Rhodococcus_sp._SGAir0479 | E7742_RS04110 | 877804..881598 | 2567884 | DUF6079-family-protein-[Rhodococcus-sp.-SGAir0479] |
| WP_138997424.1 | Type2.Pgl.like.BREX | [illegible] | 1260 | Bacteria>Terrabacteria_group>Actinomycetota>Actinomycetes>Mycobacteriales>Nocardiaceae | GCF_006148895.1 | Rhodococcus_zopfii | FEF02_RS08580 | 134443..138225 | 43772 | DUF6079-family-protein-[Rhodococcus-zopfii] |
| WP_227833967.1 | Type2.Pgl.like.BREX | [illegible] | 1284 | Bacteria>Terrabacteria_group>Actinomycetota>Actinomycetes>Mycobacteriales>Nocardiaceae | GCF_009604405.1 | Nocardia_macrotermitis | NRB20_RS28450 | 166856..170710 | 2585198 | phage-resistance-protein-[Nocardia-macrotermitis] |
| WP_202400078.1 | Type2.Pgl.like.BREX | [illegible] | 1242 | Bacteria>Terrabacteria_group>Actinomycetota>Actinomycetes>Mycobacteriales>Nocardiaceae | GCF_009831075.1 | Rhodococcus_rhodochrous | GSM98_RS14610 | 113607..117335 | 1829 | DUF6079-family-protein-[Rhodococcus-rhodochrous] |
| WP_168579650.1 | Type2.Pgl.like.BREX | [illegible] | 1237 | Bacteria>Terrabacteria_group>Actinomycetota>Actinomycetes>Mycobacteriales>Nocardiaceae | GCF_012272955.1 | Rhodococcus_sp._DMU1 | HFP48_RS14070 | 3003998..3007711 | 2722825 | DUF6079-family-protein-[Rhodococcus-sp.-DMU1] |
| WP_039783856.1 | Type2.Pgl.like.BREX | [illegible] | 1295 | Bacteria>Terrabacteria_group>Actinomycetota>Actinomycetes>Mycobacteriales>Nocardiaceae | GCF_012396065.1 | Nocardia_cerradoensis | HGB45_RS12885 | 405357..409244 | 85688 | DUF6079-family-protein-[Nocardia-cerradoensis] |
| WP_206038827.1 | Type2.Pgl.like.BREX | [illegible] | 1228 | Bacteria>Terrabacteria_group>Actinomycetota>Actinomycetes>Mycobacteriales>Nocardiaceae | GCF_012844225.1 | Rhodococcus_sp._105337 | HF886_RS14710 | 10453..14139 | 2725310 | DUF6079-family-protein-[Rhodococcus-sp.-105337] |
| WP_249343668.1 | Type2.Pgl.like.BREX | [illegible] | 1271 | Bacteria>Terrabacteria_group>Actinomycetota>Actinomycetes>Mycobacteriales>Nocardiaceae | GCF_012971805.1 | Rhodococcus_opacus | HJ581_RS11030 | 86597..90412 | 37919 | DUF6079-family-protein-[Rhodococcus-opacus] |
| WP_188543681.1 | Type2.Pgl.like.BREX | [illegible] | 1212 | Bacteria>Terrabacteria_group>Actinomycetota>Actinomycetes>Mycobacteriales>Nocardiaceae | GCF_014635345.1 | Rhodococcus_trifolii | IEW02_RS05775 | 1232248..1235886 | 908250 | phage-resistance-protein-[Rhodococcus-trifolii] |
| WP_063000546.1 | Type2.Pgl.like.BREX | [illegible] | 1249 | Bacteria>Terrabacteria_group>Actinomycetota>Actinomycetes>Mycobacteriales>Nocardiaceae | GCF_014645735.1 | Nocardia_jinanensis | IEY68_RS31650 | 40816..44565 | 382504 | DUF6079-family-protein-[Nocardia-jinanensis] |
| WP_229684222.1 | Type2.Pgl.like.BREX | [illegible] | 1364 | Bacteria>Terrabacteria_group>Actinomycetota>Actinomycetes>Mycobacteriales>Nocardiaceae | GCF_014646135.1 | Nocardia_camponoti | IEX78_RS21925 | 193544..197638 | 1616106 | phage-resistance-protein-[Nocardia-camponoti] |
| WP_257233264.1 | Type2.Pgl.like.BREX | [illegible] | 1296 | Bacteria>Terrabacteria_group>Actinomycetota>Actinomycetes>Mycobacteriales>Nocardiaceae | GCF_015034605.1 | Rhodococcus_qingshengii | IM699_RS24685 | 5335189..5339079 | 334542 | DUF6079-family-protein-[Rhodococcus-qingshengii] |
| WP_228538510.1 | Type2.Pgl.like.BREX | [illegible] | 1278 | Bacteria>Terrabacteria_group>Actinomycetota>Actinomycetes>Mycobacteriales>Nocardiaceae | GCF_015355755.1 | Nocardia_sp._XZ_19_385 | IBX22_RS05375 | 1166777..1170613 | 2769488 | phage-resistance-protein-[Nocardia-sp.-XZ_19_385] |
| WP_228824462.1 | Type2.Pgl.like.BREX | [illegible] | 1278 | Bacteria>Terrabacteria_group>Actinomycetota>Actinomycetes>Mycobacteriales>Nocardiaceae | GCF_015477355.1 | Nocardia_blacklockiae | IU476_RS27875 | 116199..120035 | 480036 | phage-resistance-protein-[Nocardia-blacklockiae] |
| WP_195024438.1 | Type2.Pgl.like.BREX | [illegible] | 1300 | Bacteria>Terrabacteria_group>Actinomycetota>Actinomycetes>Mycobacteriales>Nocardiaceae | GCF_015477405.1 | Nocardia_farcinica | IU473_RS11255 | 584605..588507 | 37329 | DUF6079-family-protein-[Nocardia-farcinica] |
| WP_228782161.1 | Type2.Pgl.like.BREX | [illegible] | 1245 | Bacteria>Terrabacteria_group>Actinomycetota>Actinomycetes>Mycobacteriales>Nocardiaceae | GCF_015477435.1 | Nocardia_abscessus | IU470_RS12230 | 97395..101132 | 120957 | DUF6079-family-protein-[Nocardia-abscessus] |
| WP_228838699.1 | Type2.Pgl.like.BREX | [illegible] | 1245 | Bacteria>Terrabacteria_group>Actinomycetota>Actinomycetes>Mycobacteriales>Nocardiaceae | GCF_015477605.1 | Nocardia_amamiensis | IU459_RS29160 | 22411..26148 | 404578 | DUF6079-family-protein-[Nocardia-amamiensis] |
| WP_195000848.1 | Type2.Pgl.like.BREX | [illegible] | 1316 | Bacteria>Terrabacteria_group>Actinomycetota>Actinomycetes>Mycobacteriales>Nocardiaceae | GCF_015477835.1 | Nocardia_higoensis | IU449_RS05565 | 1242547..1246497 | 228599 | DUF6079-family-protein-[Nocardia-higoensis] |
| WP_228801228.1 | Type2.Pgl.like.BREX | [illegible] | 1271 | Bacteria>Terrabacteria_group>Actinomycetota>Actinomycetes>Mycobacteriales>Nocardiaceae | GCF_015477865.1 | Nocardia_cyriacigeorgica | IU409_RS23450 | 21336..25151 | 135487 | DUF6079-family-protein-[Nocardia-cyriacigeorgica] |
| WP_195065211.1 | Type2.Pgl.like.BREX | [illegible] | 1285 | Bacteria>Terrabacteria_group>Actinomycetota>Actinomycetes>Mycobacteriales>Nocardiaceae | GCF_015478245.1 | Nocardia_abscessus | IU428_RS07225 | 143380..147237 | 120957 | phage-resistance-protein-[Nocardia-abscessus] |
| WP_195027567.1 | Type2.Pgl.like.BREX | [illegible] | 1300 | Bacteria>Terrabacteria_group>Actinomycetota>Actinomycetes>Mycobacteriales>Nocardiaceae | GCF_015478425.1 | Nocardia_farcinica | AB8M53_RS07390 | 596656..600558 | 37329 | DUF6079-family-protein-[Nocardia-farcinica] |
| WP_072634863.1 | Type2.Pgl.like.BREX | [illegible] | 1243 | Bacteria>Terrabacteria_group>Actinomycetota>Actinomycetes>Mycobacteriales>Nocardiaceae | GCF_015654185.1 | Rhodococcus_sp._M8 | IS016_RS27035 | 5936618..5940349 | 1925550 | DUF6079-family-protein-[Rhodococcus-sp.-M8] |
| WP_203010703.1 | Type2.Pgl.like.BREX | [illegible] | 1235 | Bacteria>Terrabacteria_group>Actinomycetota>Actinomycetes>Mycobacteriales>Nocardiaceae | GCF_016804345.1 | Rhodococcus_ruber | F1734_RS14140 | 3038038..3041745 | 1830 | DUF6079-family-protein-[Rhodococcus-ruber] |
| WP_254206127.1 | Type2.Pgl.like.BREX | [illegible] | 1278 | Bacteria>Terrabacteria_group>Actinomycetota>Actinomycetes>Mycobacteriales>Nocardiaceae | GCF_019038435.1 | Nocardia_sp._ncl2 | JZ791_RS22515 | 9851..13687 | 2815723 | phage-resistance-protein-[Nocardia-sp.-ncl2] |
| WP_218474947.1 | Type2.Pgl.like.BREX | [illegible] | 1263 | Bacteria>Terrabacteria_group>Actinomycetota>Actinomycetes>Mycobacteriales>Nocardiaceae | GCF_039543325.1 | Nocardia_iowensis | ACGFNT_RS21195 | 377182..380973 | 204891 | phage-resistance-protein-[Nocardia-iowensis] |
| WP_219130349.1 | Type2.Pgl.like.BREX | [illegible] | 1235 | Bacteria>Terrabacteria_group>Actinomycetota>Actinomycetes>Mycobacteriales>Nocardiaceae | GCF_019317345.1 | Rhodococcus_sp._LW-XY12 | KXC42_RS11380 | 2418472..2422179 | 2856851 | DUF6079-family-protein-[Rhodococcus-sp.-LW-XY12] |
| WP_084848434.1 | Type2.Pgl.like.BREX | [illegible] | 1264 | Bacteria>Terrabacteria_group>Actinomycetota>Actinomycetes>Mycobacteriales>Nocardiaceae | GCF_019703975.1 | Prescottella_equi | A5N78_RS19000 | 135247..139041 | 43767 | DUF6079-family-protein-[Prescottella-equi] |
| WP_222677599.1 | Type2.Pgl.like.BREX | [illegible] | 1213 | Bacteria>Terrabacteria_group>Actinomycetota>Actinomycetes>Mycobacteriales>Nocardiaceae | GCF_019834505.1 | Rhodococcus_corynebacterioides | HQ609_RS09015 | 112218..115859 | 53972 | phage-resistance-protein-[Rhodococcus-corynebacterioides] |
| WP_222635286.1 | Type2.Pgl.like.BREX | [illegible] | 1237 | Bacteria>Terrabacteria_group>Actinomycetota>Actinomycetes>Mycobacteriales>Nocardiaceae | GCF_019834785.1 | unclassified_Rhodococcus_(in:_high_G+C_Gram-positive_bacteria) | HQ360_RS18085 | 40159..43872 | 192944 | MULTISPECIES:-DUF6079-family-protein-[unclassified-Rhodococcus-(in:-high-G+C-Gram-positive-bacteria)] |
| WP_222632854.1 | Type2.Pgl.like.BREX | [illegible] | 1293 | Bacteria>Terrabacteria_group>Actinomycetota>Actinomycetes>Mycobacteriales>Nocardiaceae | GCF_019836125.1 | unclassified_Rhodococcus_(in:_high_G+C_Gram-positive_bacteria) | HQ302_RS17065 | 242661..246542 | 192944 | MULTISPECIES:-DUF6079-family-protein-[unclassified-Rhodococcus-(in:-high-G+C-Gram-positive-bacteria)] |
| WP_227983186.1 | Type2.Pgl.like.BREX | [illegible] | 1295 | Bacteria>Terrabacteria_group>Actinomycetota>Actinomycetes>Mycobacteriales>Nocardiaceae | GCF_020733635.1 | Nocardia_sp._DSM_111726 | LKD76_RS21560 | 4819802..4823689 | 2887190 | DUF6079-family-protein-[Nocardia-sp.-DSM-111726] |
| WP_252173979.1 | Type2.Pgl.like.BREX | [illegible] | 1242 | Bacteria>Terrabacteria_group>Actinomycetota>Actinomycetes>Mycobacteriales>Nocardiaceae | GCF_023822085.1 | Rhodococcus_pyridinivorans | LLA01_RS07335 | 1548248..1551976 | 103816 | DUF6079-family-protein-[Rhodococcus-pyridinivorans] |
| WP_269419422.1 | Type2.Pgl.like.BREX | [illegible] | 1219 | Bacteria>Terrabacteria_group>Actinomycetota>Actinomycetes>Mycobacteriales>Nocardiaceae | GCF_027213855.1 | Rhodococcus_yunnanensis | O4H54_RS22340 | 14314..17973 | 278209 | phage-resistance-protein-[Rhodococcus-yunnanensis] |
| WP_274752681.1 | Type2.Pgl.like.BREX | [illegible] | 1269 | Bacteria>Terrabacteria_group>Actinomycetota>Actinomycetes>Mycobacteriales>Nocardiaceae | GCF_028863315.1 | Nocardia_gipuzkoensis | PWJ90_RS35990 | 6594..10403 | 2749991 | hypothetical-protein-[Nocardia-gipuzkoensis] |
| WP_068365410.1 | Type2.Pgl.like.BREX | [illegible] | 1246 | Bacteria>Terrabacteria_group>Actinomycetota>Actinomycetes>Mycobacteriales>Nocardiaceae | GCF_900111805.1 | Rhodococcus_kroppenstedtii | BMZ18_RS12920 | 141320..145060 | 293050 | DUF6079-family-protein-[Rhodococcus-kroppenstedtii] |
| WP_076482662.1 | Type2.Pgl.like.BREX | [illegible] | 1235 | Bacteria>Terrabacteria_group>Actinomycetota>Actinomycetes>Mycobacteriales>Nocardiaceae | GCF_900156495.1 | Williamsia_sterculiae | BW971_RS19020 | 25213..28920 | 1344003 | DUF6079-family-protein-[Williamsia-sterculiae] |
| WP_097245973.1 | Type2.Pgl.like.BREX | [illegible] | 1301 | Bacteria>Terrabacteria_group>Actinomycetota>Actinomycetes>Mycobacteriales>Nocardiaceae | GCF_900215415.1 | Nocardia_amikacinitolerans | CRP45_RS17645 | 723031..726936 | 756689 | phage-resistance-protein-[Nocardia-amikacinitolerans] |
| WP_104380333.1 | Type2.Pgl.like.BREX | [illegible] | 1300 | Bacteria>Terrabacteria_group>Actinomycetota>Actinomycetes>Mycobacteriales>Nocardiaceae | - | Nocardia_nova | C5E45_23110 | 140922..144824 | 37330 | DUF6079-family-protein-[Nocardia-nova] |
| KQU58166.1 | Type2.Pgl.like.BREX | [illegible] | 1248 | Bacteria>Terrabacteria_group>Actinomycetota>Actinomycetes>Mycobacteriales>Nocardiaceae | - | Rhodococcus_sp._Leaf278 | ASG84_16105 | 127927..131673 | 1736319 | phage-resistance-protein-[Rhodococcus-sp.-Leaf278] |
| WP_068569102.1 | Type2.Pgl.like.BREX | [illegible] | 1251 | Bacteria>Terrabacteria_group>Actinomycetota>Actinomycetes>Mycobacteriales>Tsukamurellaceae | GCF_001575195.1 | Tsukamurella_pseudospumae | AXK60_RS00825 | 181660..185415 | 239498 | DUF6079-family-protein-[Tsukamurella-pseudospumae] |
| WP_224646689.1 | Type2.Pgl.like.BREX | [illegible] | 1249 | Bacteria>Terrabacteria_group>Actinomycetota>Actinomycetes>Mycobacteriales>Tsukamurellaceae | GCF_020169415.1 | Tsukamurella_sp._M9C | LB823_RS09350 | 44778..48527 | 2877520 | DUF6079-family-protein-[Tsukamurella-sp.-M9C] |
| WP_188941325.1 | Type2.Pgl.like.BREX | [illegible] | 1250 | Bacteria>Terrabacteria_group>Actinomycetota>Actinomycetes>Nakamurellales>Nakamurellaceae | GCF_014646215.1 | Nakamurella_endophytica | IEY37_RS09760 | 571447..575199 | 1748367 | DUF6079-family-protein-[Nakamurella-endophytica] |
| WP_179789934.1 | Type2.Pgl.like.BREX | [illegible] | 1219 | Bacteria>Terrabacteria_group>Actinomycetota>Actinomycetes>Propionibacteriales>Actinopolymorphaceae | GCF_013407855.1 | Actinopolymorpha_rutila | F4554_RS25530 | 5730929..5734588 | 446787 | DUF6079-family-protein-[Actinopolymorpha-rutila] |
| WP_192748454.1 | Type2.Pgl.like.BREX | [illegible] | 1234 | Bacteria>Terrabacteria_group>Actinomycetota>Actinomycetes>Propionibacteriales>Actinopolymorphaceae | GCF_014873875.1 | Actinopolymorpha_pittospori | HEB94_RS02555 | 543266..546970 | 648752 | DUF6079-family-protein-[Actinopolymorpha-pittospori] |
| WP_092652883.1 | Type2.Pgl.like.BREX | [illegible] | 1221 | Bacteria>Terrabacteria_group>Actinomycetota>Actinomycetes>Propionibacteriales>Actinopolymorphaceae | GCF_900104745.1 | Actinopolymorpha_singaporensis | BLU27_RS10730 | 2355351..2359016 | 117157 | hypothetical-protein-[Actinopolymorpha-singaporensis] |
| WP_139239151.1 | Type2.Pgl.like.BREX | [illegible] | 1219 | Bacteria>Terrabacteria_group>Actinomycetota>Actinomycetes>Propionibacteriales>Actinopolymorphaceae | GCF_900113145.1 | Actinopolymorpha_cephalotaxi | BM325_RS25875 | 20933..24592 | 504797 | phage-resistance-protein-[Actinopolymorpha-cephalotaxi] |
| WP_131296141.1 | Type2.Pgl.like.BREX | [illegible] | 1261 | Bacteria>Terrabacteria_group>Actinomycetota>Actinomycetes>Propionibacteriales>Kribbellaceae | GCF_004331435.1 | Kribbella_sindirgiensis | E0H50_RS38770 | 82473..86258 | 1124744 | DUF6079-family-protein-[Kribbella-sindirgiensis] |
| WP_270133768.1 | Type2.Pgl.like.BREX | [illegible] | 1250 | Bacteria>Terrabacteria_group>Actinomycetota>Actinomycetes>Propionibacteriales>Kribbellaceae | GCF_027627575.1 | Kribbella_sp._CA-293567 | OX958_31410 | 6777047..6780799 | 3002436 | DUF6079-family-protein-[Kribbella-sp.-CA-293567] |
| GGD24638.1 | Type2.Pgl.like.BREX | [illegible] | 1227 | Bacteria>Terrabacteria_group>Actinomycetota>Actinomycetes>Propionibacteriales>Nocardioidaceae | GCA_014635145.1 | Nocardioides_daphniae | GCM10007231_24780 | 66469..70152 | 402297 | hypothetical-protein-GCM10007231_24780-[Nocardioides-daphniae] |
| MBY9075953.1 | Type2.Pgl.like.BREX | [illegible] | 1223 | Bacteria>Terrabacteria_group>Actinomycetota>Actinomycetes>Propionibacteriales>Nocardioidaceae | GCA_019679455.1 | Nocardioides_sp._WL0053 | K1X13_14055 | 1162868..1166539 | 2866161 | DUF6079-family-protein-[Nocardioides-sp.-WL0053] |
| WP_056862668.1 | Type2.Pgl.like.BREX | [illegible] | 1261 | Bacteria>Terrabacteria_group>Actinomycetota>Actinomycetes>Propionibacteriales>Nocardioidaceae | GCF_001422955.1 | Nocardioides_sp._Leaf285 | ASF47_RS04185 | 904317..908102 | 1736322 | DUF6079-family-protein-[Nocardioides-sp.-Leaf285] |
| WP_056707023.1 | Type2.Pgl.like.BREX | [illegible] | 1260 | Bacteria>Terrabacteria_group>Actinomycetota>Actinomycetes>Propionibacteriales>Nocardioidaceae | GCF_001427545.1 | unclassified_Nocardioides | ASD81_RS04170 | 894964..898746 | 2615069 | MULTISPECIES:-DUF6079-family-protein-[unclassified-Nocardioides] |
| WP_135832439.1 | Type2.Pgl.like.BREX | [illegible] | 1164 | Bacteria>Terrabacteria_group>Actinomycetota>Actinomycetes>Propionibacteriales>Nocardioidaceae | GCF_004777465.1 | Nocardioides_daphniae | E2C04_RS09760 | 1990153..1993647 | 402297 | hypothetical-protein-[Nocardioides-daphniae] |
| WP_168929655.1 | Type2.Pgl.like.BREX | [illegible] | 1236 | Bacteria>Terrabacteria_group>Actinomycetota>Actinomycetes>Propionibacteriales>Nocardioidaceae | GCF_004919085.1 | Nocardioides_sp._GY_10127 | E8D37_RS19755 | 682983..686693 | 2569762 | phage-resistance-protein-[Nocardioides-sp.-GY-10127] |
| WP_166390126.1 | Type2.Pgl.like.BREX | [illegible] | 1218 | Bacteria>Terrabacteria_group>Actinomycetota>Actinomycetes>Propionibacteriales>Nocardioidaceae | GCF_011421535.1 | Nocardioides_ochotonae | HBD44_RS08040 | 518625..522281 | 2685869 | hypothetical-protein-[Nocardioides-ochotonae] |
| WP_179517479.1 | Type2.Pgl.like.BREX | [illegible] | 1220 | Bacteria>Terrabacteria_group>Actinomycetota>Actinomycetes>Propionibacteriales>Nocardioidaceae | GCF_013409425.1 | Nocardioides_perillae | BJ989_RS06365 | 1361369..1365031 | 1119534 | phage-resistance-protein-[Nocardioides-perillae] |
| WP_188421964.1 | Type2.Pgl.like.BREX | [illegible] | 1223 | Bacteria>Terrabacteria_group>Actinomycetota>Actinomycetes>Propionibacteriales>Nocardioidaceae | GCF_014635145.1 | Nocardioides_daphniae | IEV66_RS12415 | 66469..70140 | 402297 | hypothetical-protein-[Nocardioides-daphniae] |
| WP_193638512.1 | Type2.Pgl.like.BREX | [illegible] | 1238 | Bacteria>Terrabacteria_group>Actinomycetota>Actinomycetes>Propionibacteriales>Nocardioidaceae | GCF_015070385.1 | Nocardioides_malaquae | IEQ44_RS10970 | 8267..11983 | 2773426 | DUF6079-family-protein-[Nocardioides-malaquae] |
| WP_225915232.1 | Type2.Pgl.like.BREX | [illegible] | 1221 | Bacteria>Terrabacteria_group>Actinomycetota>Actinomycetes>Propionibacteriales>Nocardioidaceae | GCF_019679455.1 | Nocardioides_sp._WL0053 | K1X13_RS14055 | 1162874..1166539 | 2866161 | DUF6079-family-protein-[Nocardioides-sp.-WL0053] |
| WP_264701740.1 | Type2.Pgl.like.BREX | [illegible] | 1240 | Bacteria>Terrabacteria_group>Actinomycetota>Actinomycetes>Propionibacteriales>Nocardioidaceae | GCF_025972045.1 | Micropruina_sp._KQZ13P-5 | OIE40_RS08295 | 24265..27987 | 2986940 | DUF6079-family-protein-[Micropruina-sp.-KQZ13P-5] |
| NLI49793.1 | Type2.Pgl.like.BREX | [illegible] | 1241 | Bacteria>Terrabacteria_group>Actinomycetota>Actinomycetes>Propionibacteriales>Propionibacteriaceae | GCA_012517965.1 | Propionibacterium_sp. | GX412_05150 | 15390..19115 | 1977903 | phage-resistance-protein-[Propionibacterium-sp.] |
| NNG21072.1 | Type2.Pgl.like.BREX | [illegible] | 1242 | Bacteria>Terrabacteria_group>Actinomycetota>Actinomycetes>Propionibacteriales>Propionibacteriaceae | GCA_013002685.1 | Naumannella_sp._ID2617S | HJ590_16175 | 146853..150581 | 1656893 | phage-resistance-protein-[Naumannella-sp.-ID2617S] |
| WP_013863744.1 | Type2.Pgl.like.BREX | [illegible] | 1239 | Bacteria>Terrabacteria_group>Actinomycetota>Actinomycetes>Propionibacteriales>Propionibacteriaceae | GCF_000270245.1 | Microlunatus_phosphovorus | MLP_RS13845 | 3081945..3085664 | 29405 | DUF6079-family-protein-[Microlunatus-phosphovorus] |
| WP_133626269.1 | Type2.Pgl.like.BREX | [illegible] | 1247 | Bacteria>Terrabacteria_group>Actinomycetota>Actinomycetes>Propionibacteriales>Propionibacteriaceae | GCF_004362375.1 | Enemella_evansiae | C8D81_RS13920 | 550694..554437 | 2016499 | phage-resistance-protein-[Enemella-evansiae] |
| WP_182817604.1 | Type2.Pgl.like.BREX | [illegible] | 1251 | Bacteria>Terrabacteria_group>Actinomycetota>Actinomycetes>Propionibacteriales>Propionibacteriaceae | GCF_014164675.1 | Propioniciclava_sp._MC1683 | H5397_RS11645 | 19536..23291 | 2760309 | phage-resistance-protein-[Propioniciclava-sp.-MC1683] |
| WP_182799709.1 | Type2.Pgl.like.BREX | [illegible] | 1247 | Bacteria>Terrabacteria_group>Actinomycetota>Actinomycetes>Propionibacteriales>Propionibacteriaceae | GCF_014164775.1 | Tessaracoccus_sp._MC1679 | H5398_RS08510 | 279146..282889 | 2760313 | DUF6079-family-protein-[Tessaracoccus-sp.-MC1679] |
| WP_188333772.1 | Type2.Pgl.like.BREX | [illegible] | 1233 | Bacteria>Terrabacteria_group>Actinomycetota>Actinomycetes>Propionibacteriales>Propionibacteriaceae | GCF_017569205.1 | Propioniciclava_sp._MC1595 | J4N02_RS00605 | 137322..141023 | 2760308 | phage-resistance-protein-[Propioniciclava-sp.-MC1595] |
| WP_257478499.1 | Type2.Pgl.like.BREX | [illegible] | 1236 | Bacteria>Terrabacteria_group>Actinomycetota>Actinomycetes>Propionibacteriales>Propionibacteriaceae | GCF_024623345.1 | Acidipropionibacterium_jensenii | NVR01_RS09600 | 147995..151705 | 1749 | DUF6079-family-protein-[Acidipropionibacterium-jensenii] |
| WP_093250694.1 | Type2.Pgl.like.BREX | [illegible] | 1240 | Bacteria>Terrabacteria_group>Actinomycetota>Actinomycetes>Propionibacteriales>Propionibacteriaceae | GCF_900103275.1 | Tessaracoccus_oleiagri | BLR46_RS07920 | 616502..620224 | 686624 | DUF6079-family-protein-[Tessaracoccus-oleiagri] |
| WP_073188979.1 | Type2.Pgl.like.BREX | [illegible] | 1235 | Bacteria>Terrabacteria_group>Actinomycetota>Actinomycetes>Propionibacteriales>Propionibacteriaceae | GCF_900141915.1 | Tessaracoccus_bendigoensis | BUC15_RS12935 | 18375..22082 | 72764 | DUF6079-family-protein-[Tessaracoccus-bendigoensis] |
| WP_030106731.1 | Type2.Pgl.like.BREX | [illegible] | 896 | Bacteria>Terrabacteria_group>Actinomycetota>Actinomycetes>Pseudonocardiales>Pseudonocardiaceae | - | Actinoalloteichus_caeruleus | HY74_RS31825 | 75989..78677 | 2893586 | hypothetical-protein,-partial-[Actinoalloteichus-caeruleus] |
| OKI30339.1 | Type2.Pgl.like.BREX | [illegible] | 1220 | Bacteria>Terrabacteria_group>Actinomycetota>Actinomycetes>Pseudonocardiales>Pseudonocardiaceae | GCA_001905065.1 | Saccharothrix_sp._CB00851 | A6A25_29210 | 108531..112193 | 1835005 | hypothetical-protein-A6A25_29210-[Saccharothrix-sp.-CB00851] |
| ONI70891.1 | Type2.Pgl.like.BREX | [illegible] | 1287 | Bacteria>Terrabacteria_group>Actinomycetota>Actinomycetes>Pseudonocardiales>Pseudonocardiaceae | GCA_001984155.1 | Actinosynnema_sp._ALI-1.44 | ALI144C_50850 | 853553..857416 | 1933779 | PglY-protein-[Actinosynnema-sp.-ALI-1.44] |
| RJQ78833.1 | Type2.Pgl.like.BREX | [illegible] | 1241 | Bacteria>Terrabacteria_group>Actinomycetota>Actinomycetes>Pseudonocardiales>Pseudonocardiaceae | GCA_003600225.1 | Pseudonocardiaceae_bacterium_YIM_PH_21723 | DSS17_12215 | 80514..84239 | 2340915 | PglY-protein-[Pseudonocardiaceae-bacterium-YIM-PH-21723] |
| RSM79384.1 | Type2.Pgl.like.BREX | [illegible] | 1140 | Bacteria>Terrabacteria_group>Actinomycetota>Actinomycetes>Pseudonocardiales>Pseudonocardiaceae | GCA_003947235.1 | Kibdelosporangium_aridum | DMH04_32060 | 79921..83343 | 2030 | phage-resistance-protein-[Kibdelosporangium-aridum] |
| MPZ65695.1 | Type2.Pgl.like.BREX | [illegible] | 1154 | Bacteria>Terrabacteria_group>Actinomycetota>Actinomycetes>Pseudonocardiales>Pseudonocardiaceae | GCA_009377865.1 | Pseudonocardiaceae_bacterium | GEU83_09335 | 36177..39641 | 1873463 | phage-resistance-protein-[Pseudonocardiaceae-bacterium] |
| MBB4967435.1 | Type2.Pgl.like.BREX | [illegible] | 1272 | Bacteria>Terrabacteria_group>Actinomycetota>Actinomycetes>Pseudonocardiales>Pseudonocardiaceae | GCA_014203755.1 | Saccharothrix_violaceirubra | F4559_004794 | 5188653..5192471 | 413306 | hypothetical-protein-[Saccharothrix-violaceirubra] |
| MCE7001655.1 | Type2.Pgl.like.BREX | [illegible] | 1125 | Bacteria>Terrabacteria_group>Actinomycetota>Actinomycetes>Pseudonocardiales>Pseudonocardiaceae | GCA_021390435.3 | Kibdelosporangium_philippinense | LWC34_02185 | 453095..456472 | 211113 | DUF6079-family-protein-[Kibdelosporangium-philippinense] |
| SDG04215.1 | Type2.Pgl.like.BREX | [illegible] | 1266 | Bacteria>Terrabacteria_group>Actinomycetota>Actinomycetes>Pseudonocardiales>Pseudonocardiaceae | GCA_900100955.1 | Lentzea_fradiae | SAMN05216553_10535 | 41765..45565 | 200378 | hypothetical-protein-SAMN05216553_10535-[Lentzea-fradiae] |
| WP_012796025.1 | Type2.Pgl.like.BREX | [illegible] | 1285 | Bacteria>Terrabacteria_group>Actinomycetota>Actinomycetes>Pseudonocardiales>Pseudonocardiaceae | GCF_000023865.1 | Saccharomonospora_viridis | SVIR_RS02555 | 520806..524663 | 1852 | hypothetical-protein-[Saccharomonospora-viridis] |
| WP_009942486.1 | Type2.Pgl.like.BREX | [illegible] | 1273 | Bacteria>Terrabacteria_group>Actinomycetota>Actinomycetes>Pseudonocardiales>Pseudonocardiaceae | GCF_000062885.1 | Saccharopolyspora_erythraea | HP400_RS18960 | 4310719..4314540 | 1836 | hypothetical-protein-[Saccharopolyspora-erythraea] |
| WP_005443481.1 | Type2.Pgl.like.BREX | [illegible] | 1295 | Bacteria>Terrabacteria_group>Actinomycetota>Actinomycetes>Pseudonocardiales>Pseudonocardiaceae | GCF_000231055.2 | Saccharomonospora_azurea | SACAZDRAFT_RS16200 | 3524892..3528779 | 40988 | hypothetical-protein-[Saccharomonospora-azurea] |
| WP_005461571.1 | Type2.Pgl.like.BREX | [illegible] | 1295 | Bacteria>Terrabacteria_group>Actinomycetota>Actinomycetes>Pseudonocardiales>Pseudonocardiaceae | GCF_000243395.2 | Saccharomonospora_glauca | SACGLDRAFT_RS02735 | 552674..556561 | 40990 | hypothetical-protein-[Saccharomonospora-glauca] |
| WP_005453484.1 | Type2.Pgl.like.BREX | [illegible] | 1293 | Bacteria>Terrabacteria_group>Actinomycetota>Actinomycetes>Pseudonocardiales>Pseudonocardiaceae | GCF_000244975.1 | Saccharomonospora_cyanea | SACCYDRAFT_RS03075 | 615748..619629 | 40989 | hypothetical-protein-[Saccharomonospora-cyanea] |
| WP_006237865.1 | Type2.Pgl.like.BREX | [illegible] | 1336 | Bacteria>Terrabacteria_group>Actinomycetota>Actinomycetes>Pseudonocardiales>Pseudonocardiaceae | GCF_000258175.1 | Saccharomonospora_xinjiangensis | SACXIDRAFT_RS07210 | 1702574..1706584 | 75294 | hypothetical-protein-[Saccharomonospora-xinjiangensis] |
| WP_015099262.1 | Type2.Pgl.like.BREX | [illegible] | 1218 | Bacteria>Terrabacteria_group>Actinomycetota>Actinomycetes>Pseudonocardiales>Pseudonocardiaceae | GCF_000328705.1 | Saccharothrix_espanaensis | BN6_RS08985 | 1889017..1892673 | 103731 | hypothetical-protein-[Saccharothrix-espanaensis] |
| WP_005155554.1 | Type2.Pgl.like.BREX | [illegible] | 1234 | Bacteria>Terrabacteria_group>Actinomycetota>Actinomycetes>Pseudonocardiales>Pseudonocardiaceae | GCF_000340415.1 | Amycolatopsis_azurea | C791_RS14085 | 36298..40002 | 36819 | hypothetical-protein-[Amycolatopsis-azurea] |
| WP_019811608.1 | Type2.Pgl.like.BREX | [illegible] | 1282 | Bacteria>Terrabacteria_group>Actinomycetota>Actinomycetes>Pseudonocardiales>Pseudonocardiaceae | GCF_000383775.1 | Saccharomonospora_halophila | SACHA_RS0116725 | 69160..73008 | 129922 | hypothetical-protein-[Saccharomonospora-halophila] |
| WP_037055275.1 | Type2.Pgl.like.BREX | [illegible] | 1224 | Bacteria>Terrabacteria_group>Actinomycetota>Actinomycetes>Pseudonocardiales>Pseudonocardiaceae | GCF_000423625.1 | Pseudonocardia_asaccharolytica | G567_RS26205 | 347067..350741 | 54010 | hypothetical-protein-[Pseudonocardia-asaccharolytica] |
| WP_026454408.1 | Type2.Pgl.like.BREX | [illegible] | 1256 | Bacteria>Terrabacteria_group>Actinomycetota>Actinomycetes>Pseudonocardiales>Pseudonocardiaceae | GCF_000430445.1 | Saccharomonospora_iraqiensis | ACTIR_RS0111530 | 106700..110470 | 52698 | DUF6079-family-protein-[Saccharomonospora-iraqiensis] |
| WP_026455414.1 | Type2.Pgl.like.BREX | [illegible] | 1286 | Bacteria>Terrabacteria_group>Actinomycetota>Actinomycetes>Pseudonocardiales>Pseudonocardiaceae | GCF_000430445.1 | Saccharomonospora_iraqiensis | ACTIR_RS0118640 | 37174..41034 | 52698 | hypothetical-protein-[Saccharomonospora-iraqiensis] |
| WP_027930282.1 | Type2.Pgl.like.BREX | [illegible] | 1265 | Bacteria>Terrabacteria_group>Actinomycetota>Actinomycetes>Pseudonocardiales>Pseudonocardiaceae | GCF_000473265.1 | Amycolatopsis_thermoflava | AMYTH_RS0110485 | 2108493..2112290 | 84480 | hypothetical-protein-[Amycolatopsis-thermoflava] |
| WP_030227514.1 | Type2.Pgl.like.BREX | [illegible] | 1275 | Bacteria>Terrabacteria_group>Actinomycetota>Actinomycetes>Pseudonocardiales>Pseudonocardiaceae | GCF_000718005.1 | Actinoalloteichus_caeruleus | IF15_RS0118090 | 70929..74756 | 2893586 | hypothetical-protein-[Actinoalloteichus-caeruleus] |
| WP_042190379.1 | Type2.Pgl.like.BREX | [illegible] | 1227 | Bacteria>Terrabacteria_group>Actinomycetota>Actinomycetes>Pseudonocardiales>Pseudonocardiaceae | GCF_000826545.1 | Kibdelosporangium_sp._MJ126-NF4 | BN1199_RS31465 | 645585..649268 | 703222 | DUF6079-family-protein-[Kibdelosporangium-sp.-MJ126-NF4] |
| WP_069849549.1 | Type2.Pgl.like.BREX | [illegible] | 1249 | Bacteria>Terrabacteria_group>Actinomycetota>Actinomycetes>Pseudonocardiales>Pseudonocardiaceae | GCF_001747425.1 | Actinoalloteichus_hymeniacidonis | TL08_RS14250 | 3139183..3142932 | 340345 | PglY-protein-[Actinoalloteichus-hymeniacidonis] |
| WP_073891944.1 | Type2.Pgl.like.BREX | [illegible] | 1228 | Bacteria>Terrabacteria_group>Actinomycetota>Actinomycetes>Pseudonocardiales>Pseudonocardiaceae | GCF_001905065.1 | Saccharothrix_sp._CB00851 | A6A25_RS19115 | 108507..112193 | 1835005 | hypothetical-protein-[Saccharothrix-sp.-CB00851] |
| WP_237774699.1 | Type2.Pgl.like.BREX | [illegible] | 1269 | Bacteria>Terrabacteria_group>Actinomycetota>Actinomycetes>Pseudonocardiales>Pseudonocardiaceae | GCF_001984155.1 | Actinosynnema_sp._ALI-1.44 | ALI144C_RS50860 | 853607..857416 | 1933779 | PglY-protein-[Actinosynnema-sp.-ALI-1.44] |
| WP_113691957.1 | Type2.Pgl.like.BREX | [illegible] | 1264 | Bacteria>Terrabacteria_group>Actinomycetota>Actinomycetes>Pseudonocardiales>Pseudonocardiaceae | GCF_003312875.1 | Amycolatopsis_albispora | A4R43_RS09335 | 2006879..2010673 | 1804986 | PglY-protein-[Amycolatopsis-albispora] |
| WP_123687428.1 | Type2.Pgl.like.BREX | [illegible] | 1265 | Bacteria>Terrabacteria_group>Actinomycetota>Actinomycetes>Pseudonocardiales>Pseudonocardiaceae | GCF_003752125.1 | Amycolatopsis_thermoflava | EDD35_RS38725 | 75705..79502 | 84480 | PglY-protein-[Amycolatopsis-thermoflava] |
| WP_051792781.1 | Type2.Pgl.like.BREX | [illegible] | 1228 | Bacteria>Terrabacteria_group>Actinomycetota>Actinomycetes>Pseudonocardiales>Pseudonocardiaceae | GCF_003947235.1 | Kibdelosporangium_aridum | DMH04_RS31890 | 79657..83343 | 2030 | hypothetical-protein-[Kibdelosporangium-aridum] |
| WP_130291362.1 | Type2.Pgl.like.BREX | [illegible] | 1241 | Bacteria>Terrabacteria_group>Actinomycetota>Actinomycetes>Pseudonocardiales>Pseudonocardiaceae | GCF_004217185.1 | Pseudonocardia_sediminis | EV383_RS20195 | 4368275..4372000 | 1397368 | PglY-protein-[Pseudonocardia-sediminis] |
| WP_133853968.1 | Type2.Pgl.like.BREX | [illegible] | 1236 | Bacteria>Terrabacteria_group>Actinomycetota>Actinomycetes>Pseudonocardiales>Pseudonocardiaceae | GCF_004362825.1 | Labedaea_rhizosphaerae | EV186_RS35890 | 197342..201052 | 598644 | DUF6079-family-protein-[Labedaea-rhizosphaerae] |
| WP_134712580.1 | Type2.Pgl.like.BREX | [illegible] | 1329 | Bacteria>Terrabacteria_group>Actinomycetota>Actinomycetes>Pseudonocardiales>Pseudonocardiaceae | GCF_004519465.1 | Saccharomonospora_xinjiangensis | EYD13_RS02955 | 603059..607048 | 75294 | PglY-protein-[Saccharomonospora-xinjiangensis] |
| WP_134731708.1 | Type2.Pgl.like.BREX | [illegible] | 1258 | Bacteria>Terrabacteria_group>Actinomycetota>Actinomycetes>Pseudonocardiales>Pseudonocardiaceae | GCF_004522235.1 | Amycolatopsis_nivea | EIZ44_RS25765 | 135092..138868 | 1644109 | PglY-protein-[Amycolatopsis-nivea] |
| WP_137816032.1 | Type2.Pgl.like.BREX | [illegible] | 1215 | Bacteria>Terrabacteria_group>Actinomycetota>Actinomycetes>Pseudonocardiales>Pseudonocardiaceae | GCF_005405885.1 | Gandjariella_thermophila | FFT21_RS23075 | 3826..7473 | 1931992 | phage-resistance-protein-[Gandjariella-thermophila] |
| WP_142002153.1 | Type2.Pgl.like.BREX | [illegible] | 1234 | Bacteria>Terrabacteria_group>Actinomycetota>Actinomycetes>Pseudonocardiales>Pseudonocardiaceae | GCF_006715045.1 | Amycolatopsis_cihanbeyliensis | FB471_RS19210 | 4232478..4236182 | 1128664 | PglY-protein-[Amycolatopsis-cihanbeyliensis] |
| WP_158887127.1 | Type2.Pgl.like.BREX | [illegible] | 1230 | Bacteria>Terrabacteria_group>Actinomycetota>Actinomycetes>Pseudonocardiales>Pseudonocardiaceae | GCF_009765355.1 | Amycolatopsis_anabasis | FXO23_RS21260 | 177715..181407 | 1840409 | DUF6079-family-protein-[Amycolatopsis-anabasis] |
| WP_167176974.1 | Type2.Pgl.like.BREX | [illegible] | 1270 | Bacteria>Terrabacteria_group>Actinomycetota>Actinomycetes>Pseudonocardiales>Pseudonocardiaceae | GCF_011761185.1 | Saccharomonospora_amisosensis | FHU38_RS24910 | 361613..365425 | 1128677 | PglY-protein-[Saccharomonospora-amisosensis] |
| WP_184860151.1 | Type2.Pgl.like.BREX | [illegible] | 1308 | Bacteria>Terrabacteria_group>Actinomycetota>Actinomycetes>Pseudonocardiales>Pseudonocardiaceae | GCF_014203355.1 | Kutzneria_kofuensis | BJ998_RS08830 | 1945314..1949240 | 103725 | phage-resistance-protein-[Kutzneria-kofuensis] |
| WP_190817006.1 | Type2.Pgl.like.BREX | [illegible] | 1322 | Bacteria>Terrabacteria_group>Actinomycetota>Actinomycetes>Pseudonocardiales>Pseudonocardiaceae | GCF_014697215.1 | Saccharopolyspora_pogona | DL519_RS19955 | 4381105..4385073 | 333966 | phage-resistance-protein-[Saccharopolyspora-pogona] |
| WP_229373475.1 | Type2.Pgl.like.BREX | [illegible] | 1283 | Bacteria>Terrabacteria_group>Actinomycetota>Actinomycetes>Pseudonocardiales>Pseudonocardiaceae | GCF_016464305.1 | Umezawaea_beigongshangensis | IMZ35_RS27315 | 28668..32519 | 2780383 | PglY-protein-[Umezawaea-beigongshangensis] |
| WP_201439211.1 | Type2.Pgl.like.BREX | [illegible] | 1227 | Bacteria>Terrabacteria_group>Actinomycetota>Actinomycetes>Pseudonocardiales>Pseudonocardiaceae | GCF_016695385.1 | Saccharothrix_sp._6-C | IOD16_RS05995 | 1267317..1271000 | 2781735 | hypothetical-protein-[Saccharothrix-sp.-6-C] |
| WP_206801084.1 | Type2.Pgl.like.BREX | [illegible] | 1230 | Bacteria>Terrabacteria_group>Actinomycetota>Actinomycetes>Pseudonocardiales>Pseudonocardiaceae | GCF_017309385.1 | Amycolatopsis_sp._195334CR | JYK18_RS05610 | 1136023..1139715 | 2814588 | DUF6079-family-protein-[Amycolatopsis-sp.-195334CR] |
| WP_209616138.1 | Type2.Pgl.like.BREX | [illegible] | 1229 | Bacteria>Terrabacteria_group>Actinomycetota>Actinomycetes>Pseudonocardiales>Pseudonocardiaceae | GCF_017876325.1 | Saccharothrix_coeruleofusca | JOF41_RS03780 | 767838..771527 | 33919 | hypothetical-protein-[Saccharothrix-coeruleofusca] |
| WP_211762971.1 | Type2.Pgl.like.BREX | [illegible] | 1256 | Bacteria>Terrabacteria_group>Actinomycetota>Actinomycetes>Pseudonocardiales>Pseudonocardiaceae | GCF_018263855.1 | Kutzneria_sp._CA-103260 | JJ691_RS30110 | 6839790..6843560 | 2802641 | PglY-protein-[Kutzneria-sp.-CA-103260] |
| WP_250557035.1 | Type2.Pgl.like.BREX | [illegible] | 1232 | Bacteria>Terrabacteria_group>Actinomycetota>Actinomycetes>Pseudonocardiales>Pseudonocardiaceae | GCF_018531445.1 | Pseudonocardia_sp._H11425 | KIK08_RS14850 | 18305..22003 | 2835865 | PglY-protein-[Pseudonocardia-sp.-H11425] |
| WP_219414043.1 | Type2.Pgl.like.BREX | [illegible] | 1222 | Bacteria>Terrabacteria_group>Actinomycetota>Actinomycetes>Pseudonocardiales>Pseudonocardiaceae | GCF_019396355.1 | Pseudonocardia_nigra | KME75_RS08690 | 8187..11855 | 1921578 | hypothetical-protein-[Pseudonocardia-nigra] |
| WP_224966475.1 | Type2.Pgl.like.BREX | [illegible] | 1270 | Bacteria>Terrabacteria_group>Actinomycetota>Actinomycetes>Pseudonocardiales>Pseudonocardiaceae | GCF_020177455.1 | unclassified_Saccharopolyspora | LCD33_RS01215 | 29823..33635 | 2646250 | MULTISPECIES:-PglY-protein-[unclassified-Saccharopolyspora] |
| WP_233722714.1 | Type2.Pgl.like.BREX | [illegible] | 1125 | Bacteria>Terrabacteria_group>Actinomycetota>Actinomycetes>Pseudonocardiales>Pseudonocardiaceae | GCF_021390435.2 | Kibdelosporangium_philippinense | ACFPQC_RS16680 | 2262963..2266340 | 211113 | DUF6079-family-protein-[Kibdelosporangium-philippinense] |
| WP_241868093.1 | Type2.Pgl.like.BREX | [illegible] | 1268 | Bacteria>Terrabacteria_group>Actinomycetota>Actinomycetes>Pseudonocardiales>Pseudonocardiaceae | GCF_022392385.1 | Saccharopolyspora_erythraea | HP400_RS18960 | 4310719..4314525 | 1836 | PglY-protein-[Saccharopolyspora-erythraea] |
| WP_242191032.1 | Type2.Pgl.like.BREX | [illegible] | 1228 | Bacteria>Terrabacteria_group>Actinomycetota>Actinomycetes>Pseudonocardiales>Pseudonocardiaceae | GCF_022678645.1 | Saccharopolyspora_soli | MOQ72_RS39180 | 5990..9676 | 2926618 | hypothetical-protein-[Saccharopolyspora-soli] |

| Protein-ID | System | Gene-Neighbourhoods | Length | Taxonomy | Assembly-ID | Strain | Locus-Tag | Genomic-coordinates | TaxID | Fasta-define |
|---|---|---|---|---|---|---|---|---|---|---|
| WP_253862673.1 | Type2.Pgl.like.BREX | <-SF1-SF2-Helicase+wHTH+Small-B-barrel+DUF1998<-BrxD-AAA<-PglZ\|\|XIS-HTH-><-SF1-SF2-Helicase+REase+HTH<-DUF6079*<-N6-METHYLASE<-PglW\|\|<-AbiEii<-wHTH+AbiEi_2<-HIN-HTH | 1258 | Bacteria>Terrabacteria_group>Actinomycetota>Actinomycetes>Pseudonocardiales>Pseudonocardiaceae | GCF_024171895.1 | Prauserella_halophila | LY41_RS05410 | 1127371..1131147 | 185641 | DUF6079-family-protein-[Prauserella-halophila] |
| WP_258953530.1 | Type2.Pgl.like.BREX | <-MazG\|\|NAD_binding_10-><-?<-?<-HTH+MPTase<-SF1-SF2-Helicase+wHTH+Small-B-barrel+DUF1998<-BrxD-AAA<-PglZ<-DUF6079*\|\|?-><-?<-?<-WXG<-?<-BirA-HTH<-N6-METHYLASE | 1200 | Bacteria>Terrabacteria_group>Actinomycetota>Actinomycetes>Pseudonocardiales>Pseudonocardiaceae | GCF_024648825.1 | Lentzea_californiensis | LX88_RS43215 | 11818..15420 | 438851 | hypothetical-protein-[Lentzea-californiensis] |
| WP_090048714.1 | Type2.Pgl.like.BREX | MODE-HTH+PBPII-><-SF1-SF2-Helicase+wHTH+Small-B-barrel+DUF1998<-BrxD-AAA<-PglZ<-DUF6079*<-N6-METHYLASE<-PglW<-ParB+HNH\|\|YbaB_DNA_bd->?->?->?-><-?\|\|ABhydrolase-> | 1271 | Bacteria>Terrabacteria_group>Actinomycetota>Actinomycetes>Pseudonocardiales>Pseudonocardiaceae | GCF_900100955.1 | Lentzea_fradiae | BLT23_RS12390 | 41765..45580 | 200378 | PglY-protein-[Lentzea-fradiae] |
| WP_034307025.1 | Type2.Pgl.like.BREX | <-SF1-SF2-Helicase\|\|wHTH+SAD+HNH->REase-><-?\|\|PglW->N6-METHYLASE->DUF6079*->PglZ->BrxD-AAA->SF1-SF2-Helicase+wHTH+Small-B-barrel+DUF1998->REase+SF1-SF2-Helicase->?->HKD-DNase+HKD-DNase-> | 1273 | Bacteria>Terrabacteria_group>Actinomycetota>Actinomycetes>Pseudonocardiales>Pseudonocardiaceae | GCF_900105055.1 | Amycolatopsis_lurida | BLW75_RS07195 | 425359..429180 | 31959 | hypothetical-protein-[Amycolatopsis-lurida] |
| WP_091629290.1 | Type2.Pgl.like.BREX | <-XIS-HTH\|\|HTH_26->NUDIX->NUDIX-><-?<-?<-SF1-SF2-Helicase+wHTH+Small-B-barrel+DUF1998<-BrxD-AAA<-PglZ<-DUF6079*<-N6-METHYLASE<-PglW<-?<-?<-?<-?<-?<-HTH_Tnp_1+HTH_21+rve+rve_3 | 1246 | Bacteria>Terrabacteria_group>Actinomycetota>Actinomycetes>Pseudonocardiales>Pseudonocardiaceae | GCF_900110575.1 | Amycolatopsis_saalfeldensis | BMY97_RS43650 | 24261..28001 | 394193 | DUF6079-family-protein-[Amycolatopsis-saalfeldensis] |
| WP_143086863.1 | Type2.Pgl.like.BREX | <-Pkinase+TPR+TPR\|\|REase->pPIWI+pPIWI->pPIWI+SF1-SF2-Helicase->pPIWI+MID+RNaseH-><-DEDD_Tnp_IS110+Transposase_20<-SF1-SF2-Helicase+wHTH+Small-B-barrel+DUF1998<-BrxD-AAA<-PglZ<-DUF6079*\|\|?-><-?<-?<-WXG<-?<-N6-METHYLASE<-HTH_31+DUF5753<-PglW<-ParB+HNH | 1225 | Bacteria>Terrabacteria_group>Actinomycetota>Actinomycetes>Pseudonocardiales>Pseudonocardiaceae | GCF_900110985.1 | Lentzea_flaviverrucosa | BMZ12_RS31225 | 187049..190726 | 200379 | phage-resistance-protein-[Lentzea-flaviverrucosa] |
| WP_037310056.1 | Type2.Pgl.like.BREX | FAP+Peptidase_S11-><-?<-?<-?<-?<-?<-SF1-SF2-Helicase+wHTH+Small-B-barrel+DUF1998<-BrxD-AAA<-PglZ<-DUF6079*<-N6-METHYLASE\|\|?->RHH-><-?<-?<-PglW<-?<-SH3 | 1285 | Bacteria>Terrabacteria_group>Actinomycetota>Actinomycetes>Pseudonocardiales>Pseudonocardiaceae | GCF_900115515.1 | Saccharomonospora_viridis | BM105_RS06060 | 39350..43207 | 1852 | hypothetical-protein-[Saccharomonospora-viridis] |
| WP_125727805.1 | Type2.Pgl.like.BREX | wHTH+SAD+HNH-><-REase+SF1-SF2-Helicase<-?\|\|PglW->N6-METHYLASE-><-CHTH\|\|DUF6079*->PglZ->BrxD-AAA->SF1-SF2-Helicase+wHTH+Small-B-barrel+DUF1998-><-tRNA\|\|REC+OmpR-HTH-> | 1140 | Bacteria>Terrabacteria_group>Actinomycetota>Actinomycetes>Pseudonocardiales>Pseudonocardiaceae | - | Kibdelosporangium_aridum | DMH04_32060 | 79921..83343 | 2030 | phage-resistance-protein-[Kibdelosporangium-aridum] |
| WP_251785089.1 | Type2.Pgl.like.BREX | <-ACET\|\|?-><-?<-?<-ParB+HNH<-SF1-SF2-Helicase+wHTH+Small-B-barrel+DUF1998<-BrxD-AAA<-PglZ<-DUF6079*<-N6-METHYLASE<-PglW<-ParB+HNH<-ParB+HNH\|\|?->Peptidase_C14+SF1-SF2-Helicase-> | 1229 | Bacteria>Terrabacteria_group>Actinomycetota>Actinomycetes>Pseudonocardiales>Pseudonocardiaceae | - | Pseudonocardia_sp._DR1-2 | ND486_RS16895 | 241560..245249 | 2951168 | PglY-protein-[Pseudonocardia-sp.-DR1-2] |
| MBO0801657.1 | Type2.Pgl.like.BREX | HNH-><-?<-?\|\|PglW->N6-METHYLASE-><-SAM-METHYLASE\|\|DUF6079*->PglZ->BrxD-AAA->SF1-SF2-Helicase+wHTH+Small-B-barrel+DUF1998->SAM-METHYLASE-><-?\|\|Tox-RelA->HTH+MPTase-><-OLD-ABC\|\|Phage_integrase-> | 1226 | Bacteria>Terrabacteria_group>Actinomycetota>Actinomycetes>Streptosporangiales>Nocardiopsaceae | GCA_017354375.1 | Nocardiopsaceae_bacterium | J2P25_01075 | 13072..16752 | 2100131 | phage-resistance-protein-[Nocardiopsaceae-bacterium] |
| MBO2529395.1 | Type2.Pgl.like.BREX | <-35exo<-?\|\|?-><-SF1-SF2-Helicase+wHTH+Small-B-barrel+DUF1998<-BrxD-AAA<-PglZ<-DUF6079*<-DUF397<-DUF397<-HTH_31+DUF5753\|\|?->ST-Hiskin-><-PglW<-35exo | 1261 | Bacteria>Terrabacteria_group>Actinomycetota>Actinomycetes>Streptosporangiales>Nocardiopsaceae | GCA_017577275.1 | Thermobifida_sp. | CW348_05995 | 216229..220014 | 2027107 | phage-resistance-protein-[Thermobifida-sp.] |
| SIO85591.1 | Type2.Pgl.like.BREX | ZnR->PglW->N6-METHYLASE->DUF6079*->PglZ->BrxD-AAA->SF1-SF2-Helicase+wHTH+Small-B-barrel+DUF1998-><-SbcC<-Cap4_nuclease | 1302 | Bacteria>Terrabacteria_group>Actinomycetota>Actinomycetes>Streptosporangiales>Nocardiopsaceae | GCA_900143625.1 | Nocardiopsis_sp._JB363 | BQ8420_07725 | 9588..13496 | 1434837 | Bacteriophage-(PhiC31)-resistance-gene-pglY-[Nocardiopsis-sp.-JB363] |
| WP_017604990.1 | Type2.Pgl.like.BREX | <-DnaA<-SF1-SF2-Helicase+wHTH+Small-B-barrel+DUF1998<-BrxD-AAA<-PglZ<-DUF6079*<-N6-METHYLASE<-ZnR<-ST-Hiskin\|\|HTH_31+DUF5753->DUF397->?->TrmB->HTH+DUF5753->PCMT+METHYLASE-> | 1278 | Bacteria>Terrabacteria_group>Actinomycetota>Actinomycetes>Streptosporangiales>Nocardiopsaceae | GCF_000341005.1 | Nocardiopsis_alkaliphila | D472_RS0113360 | 13776..17612 | 225762 | DUF6079-family-protein-[Nocardiopsis-alkaliphila] |
| WP_026127376.1 | Type2.Pgl.like.BREX | TrmB->HTH+DUF5753->HTH+HKD-DNase-><-DUF397<-HTH_31+DUF5753\|\|ST-Hiskin->ZnR->N6-METHYLASE->DUF6079*->PglZ->SF1-SF2-Helicase+wHTH+Small-B-barrel+DUF1998->?->ZnR-> | 1267 | Bacteria>Terrabacteria_group>Actinomycetota>Actinomycetes>Streptosporangiales>Nocardiopsaceae | GCF_000341125.1 | Nocardiopsis_lucentensis | D471_RS0103590 | 44350..48153 | 53441 | bacteriophage-resistance-gene-pglY-[Nocardiopsis-lucentensis] |
| WP_017608530.1 | Type2.Pgl.like.BREX | <-DDE_Tnp_1<-?\|\|APATPase+TPR+TPR+TPR+TPR->?->SWIM->?-><-SF1-SF2-Helicase+wHTH+Small-B-barrel+DUF1998<-BrxD-AAA<-PglZ<-DUF6079*<-N6-METHYLASE<-ZnR<-ST-Hiskin\|\|HTH_31+DUF5753->DUF397->?-><-?<-HNH | 1275 | Bacteria>Terrabacteria_group>Actinomycetota>Actinomycetes>Streptosporangiales>Nocardiopsaceae | GCF_000341145.1 | Nocardiopsis_xinjiangensis | D473_RS0107855 | 131266..135093 | 124285 | DUF6079-family-protein-[Nocardiopsis-xinjiangensis] |
| WP_017620543.1 | Type2.Pgl.like.BREX | <-PglZ-p<-DUF6079*<-N6-METHYLASE<-DUF397<-HTH_31+DUF5753\|\|?->?->?->MNS+REase->APATPase+TPR+TPR+TPR+TPR->REase-> | 1323 | Bacteria>Terrabacteria_group>Actinomycetota>Actinomycetes>Streptosporangiales>Nocardiopsaceae | GCF_000341165.1 | Nocardiopsis_gilva | D475_RS0121120 | 1143..5114 | 280236 | hypothetical-protein-[Nocardiopsis-gilva] |
| WP_016188301.1 | Type2.Pgl.like.BREX | 35exo+BRCT->PglW-><-?\|\|HTH_OrfB_IS605+OrfB_Zn_ribbon-><-ST-Hiskin<-?\|\|HTH_31+DUF5753->DUF397->DUF6079*->PglZ->BrxD-AAA->SF1-SF2-Helicase+wHTH+Small-B-barrel+DUF1998-><-?\|\|35exo+BRCT-> | 1261 | Bacteria>Terrabacteria_group>Actinomycetota>Actinomycetes>Streptosporangiales>Nocardiopsaceae | GCF_000401915.1 | Thermobifida_fusca | TM51_RS03845 | 129068..132853 | 2021 | hypothetical-protein-[Thermobifida-fusca] |
| WP_028648233.1 | Type2.Pgl.like.BREX | <-SbcC\|\|?->?-><-?<-DUF397<-HTH_31+DUF5753\|\|ST-Hiskin->ZnR->N6-METHYLASE->DUF6079*->PglZ->BrxD-AAA->SF1-SF2-Helicase+wHTH+Small-B-barrel+DUF1998->?->CASPASE+RNA-Helicase+REase->HEPN+PAS+DUF4011+SF1-SF2-Helicase+REase-><-LACI-HTH+PBPI | 1274 | Bacteria>Terrabacteria_group>Actinomycetota>Actinomycetes>Streptosporangiales>Nocardiopsaceae | GCF_000515115.1 | Nocardiopsis_sp._CNT312 | B042_RS0110365 | 88421..92245 | 1137268 | hypothetical-protein-[Nocardiopsis-sp.-CNT312] |
| WP_047867701.1 | Type2.Pgl.like.BREX | McrB-AAA->McrC-NTD+REase->Cap4_nuclease->?->SbcC-><-SF1-SF2-Helicase+wHTH+Small-B-barrel+DUF1998<-BrxD-AAA<-PglZ<-DUF6079*<-N6-METHYLASE<-ZnR<-ST-Hiskin\|\|HTH_31+DUF5753->DUF397->?->?->APATPase+TPR+TPR+TPR+TPR-><-TOPRIM | 1282 | Bacteria>Terrabacteria_group>Actinomycetota>Actinomycetes>Streptosporangiales>Nocardiopsaceae | GCF_001029525.1 | Nocardiopsis_sp._RV163 | ABD54_RS01945 | 50619..54467 | 1661388 | DUF6079-family-protein-[Nocardiopsis-sp.-RV163] |
| WP_068754110.1 | Type2.Pgl.like.BREX | ParA-Soj-PloopNTPase->?->PglW-><-?<-?\|\|HTH_31+DUF5753->DUF397->N6-METHYLASE->DUF6079*->PglZ->BrxD-AAA-><-?<-?<-?\|\|?-><-NUDIX<-NUDIX<-Thymidylate_kin | 1284 | Bacteria>Terrabacteria_group>Actinomycetota>Actinomycetes>Streptosporangiales>Nocardiopsaceae | GCF_001517975.1 | Thermobifida_cellulosilytica | AC529_RS05645 | 1315907..1319761 | 144786 | PglY-protein-[Thermobifida-cellulosilytica] |
| WP_011291156.1 | Type2.Pgl.like.BREX | <-35exo+BRCT\|\|?-><-SF1-SF2-Helicase+wHTH+Small-B-barrel+DUF1998<-BrxD-AAA<-PglZ<-DUF6079*<-DUF397<-HTH_31+DUF5753\|\|?->ST-Hiskin-> | 1261 | Bacteria>Terrabacteria_group>Actinomycetota>Actinomycetes>Streptosporangiales>Nocardiopsaceae | GCF_001592045.1 | Thermobifida_fusca | TF1_RS04110 | 284445..288230 | 2021 | hypothetical-protein-[Thermobifida-fusca] |
| WP_262391364.1 | Type2.Pgl.like.BREX | <-DUF397\|\|?->?->TrmB-><-DUF397<-HTH_31+DUF5753<-HTH_3\|\|N6-METHYLASE->DUF6079-p->DUF6079-p*->?->PglZ->PglZ-p->BrxD-AAA->McrB-AAA->McrC-NTD+REase-> | 986 | Bacteria>Terrabacteria_group>Actinomycetota>Actinomycetes>Streptosporangiales>Nocardiopsaceae | GCF_001942255.1 | Nocardiopsis_sp._CNR-923 | BJF83_RS06995 | 197288..200248 | 1904965 | phage-resistance-protein-[Nocardiopsis-sp.-CNR-923] |
| WP_262391363.1 | Type2.Pgl.like.BREX | HTH_31+DUF5753-><-DUF397\|\|?->?->TrmB-><-DUF397<-HTH_31+DUF5753<-HTH_3\|\|N6-METHYLASE->DUF6079-p->DUF6079-p*->?->PglZ->PglZ-p->BrxD-AAA->McrB-AAA->McrC-NTD+REase-> | 277 | Bacteria>Terrabacteria_group>Actinomycetota>Actinomycetes>Streptosporangiales>Nocardiopsaceae | GCF_001942255.1 | Nocardiopsis_sp._CNR-923 | BJF83_RS28590 | 196458..197291 | 1904965 | DUF6079-family-protein-[Nocardiopsis-sp.-CNR-923] |
| WP_110049005.1 | Type2.Pgl.like.BREX | PglW-><-?<-APATPase+TPR+TPR+TPR+TPR<-?<-?<-DUF397<-HTH_31+DUF5753\|\|ST-Hiskin->N6-METHYLASE->DUF6079*->PglZ->BrxD-AAA->SF1-SF2-Helicase+wHTH+Small-B-barrel+DUF1998->?-><-pPIWI+MID+RNaseH<-pPIWI+SF1-SF2-Helicase<-pPIWI+pPIWI<-REase\|\|Pkinase+TPR+TPR-> | 1273 | Bacteria>Terrabacteria_group>Actinomycetota>Actinomycetes>Streptosporangiales>Nocardiopsaceae | GCF_003182025.1 | Nocardiopsis_sp._L17-MgMaSL7 | BDW27_RS09285 | 338461..342282 | 1938893 | phage-resistance-protein-[Nocardiopsis-sp.-L17-MgMaSL7] |
| WP_116247802.1 | Type2.Pgl.like.BREX | SIGMA-HTH-><-?<-?<-SF1-SF2-Helicase+wHTH+Small-B-barrel+DUF1998<-BrxD-AAA<-PglZ<-DUF6079*<-N6-METHYLASE<-ZnR<-ST-Hiskin\|\|HTH_31+DUF5753->DUF397->?-><-?\|\|SbcC-><-SF1-SF2-Helicase | 1267 | Bacteria>Terrabacteria_group>Actinomycetota>Actinomycetes>Streptosporangiales>Nocardiopsaceae | GCF_003386285.1 | Nocardiopsis_sp._FIRDI_009 | DVR05_RS21965 | 42358..46161 | 714197 | phage-resistance-protein-[Nocardiopsis-sp.-FIRDI-009] |
| WP_068687705.1 | Type2.Pgl.like.BREX | <-SF1-SF2-Helicase+wHTH+Small-B-barrel+DUF1998<-BrxD-AAA<-PglZ<-DUF6079*<-PglW<-TetR-HTH\|\|?-><-?\|\|?-><-35exo | 1263 | Bacteria>Terrabacteria_group>Actinomycetota>Actinomycetes>Streptosporangiales>Nocardiopsaceae | GCF_003574835.2 | Thermobifida_halotolerans | ABH917_RS14650 | 3154400..3158191 | 483545 | phage-resistance-protein-[Thermobifida-halotolerans] |
| WP_068692654.1 | Type2.Pgl.like.BREX | RAMPs->?->?->RAMPs-><-CRISPR_Cas6\|\|RVT->?-><-BrxD-AAA<-PglZ<-DUF6079*<-N6-METHYLASE<-DUF397<-HTH_31+DUF5753\|\|?->?-><-PglW<-?<-ParA-Soj-PloopNTPase | 1282 | Bacteria>Terrabacteria_group>Actinomycetota>Actinomycetes>Streptosporangiales>Nocardiopsaceae | GCF_003574835.2 | Thermobifida_halotolerans | NI17_RS20570 | 4597638..4601486 | 483545 | PglY-protein-[Thermobifida-halotolerans] |
| WP_133740675.1 | Type2.Pgl.like.BREX | <-REase<-?<-ST-Hiskin<-?\|\|HTH_31+DUF5753->DUF397->N6-METHYLASE->DUF6079*->PglZ->BrxD-AAA->SF1-SF2-Helicase+wHTH+Small-B-barrel+DUF1998-><-?<-Tox-RelA\|\|?-><-?<-?<-CR-ATPase9 | 1252 | Bacteria>Terrabacteria_group>Actinomycetota>Actinomycetes>Streptosporangiales>Nocardiopsaceae | GCF_004363075.1 | Actinorugispora_endophytica | EV190_RS07190 | 236181..239939 | 1605990 | phage-resistance-protein-[Actinorugispora-endophytica] |
| WP_141923749.1 | Type2.Pgl.like.BREX | <-SF1-SF2-Helicase+wHTH+Small-B-barrel+DUF1998<-BrxD-AAA<-?<-?<-?\|\|HTH_31+DUF5753->DUF397\|\|?->?-><-PglZ<-DUF6079*\|\|?->?->TrmB->APATPase+TPR+TPR+TPR+TPR-><-DUF397<-HTH_31+DUF5753\|\|?-><-N6-METHYLASE<-PglW | 1291 | Bacteria>Terrabacteria_group>Actinomycetota>Actinomycetes>Streptosporangiales>Nocardiopsaceae | GCF_006717075.1 | Haloactinospora_alba | FHX37_RS10640 | 2355147..2359022 | 405555 | phage-resistance-protein-[Haloactinospora-alba] |
| WP_150241185.1 | Type2.Pgl.like.BREX | REase-><-APATPase+TPR+TPR+TPR+TPR<-?<-?<-DUF397<-HTH_31+DUF5753\|\|ST-Hiskin->ZnR->N6-METHYLASE->DUF6079*->PglZ->BrxD-AAA->SF1-SF2-Helicase+wHTH+Small-B-barrel+DUF1998->?-><-SbcC<-?<-Cap4_nuclease | 1276 | Bacteria>Terrabacteria_group>Actinomycetota>Actinomycetes>Streptosporangiales>Nocardiopsaceae | GCF_008638365.1 | Nocardiopsis_quinghaiensis | FYX01_RS13585 | 272770..276600 | 464995 | DUF6079-family-protein-[Nocardiopsis-quinghaiensis] |
| WP_161111646.1 | Type2.Pgl.like.BREX | HNH-><-HTH_Tnp_1+HTH_21+rve+rve_3<-HIN-HTH\|\|Schlafen+HATPase+wHTH->ParA-Soj-PloopNTPase->N6-METHYLASE->DUF6079*->PglZ->BrxD-AAA->SF1-SF2-Helicase+wHTH+Small-B-barrel+DUF1998->ParB+HNH->DnaA-><-SbcC<-?<-Cap4_nuclease<-McrC-NTD+REase | 1262 | Bacteria>Terrabacteria_group>Actinomycetota>Actinomycetes>Streptosporangiales>Nocardiopsaceae | GCF_009862005.1 | Nocardiopsis_alba | GTW20_RS21080 | 4844148..4847936 | 53437 | DUF6079-family-protein-[Nocardiopsis-alba] |
| WP_179642184.1 | Type2.Pgl.like.BREX | <-Phage_AlpA<-HicA<-ST-Hiskin<-?\|\|HTH_31+DUF5753-><-?<-OLD-ABC<-ParB\|\|N6-METHYLASE->DUF6079*->PglZ->BrxD-AAA->SF1-SF2-Helicase+wHTH+Small-B-barrel+DUF1998->?->?-><-SF1-SF2-Helicase+DUF1998<-viral-protein\|\|?->GrpE-> | 1270 | Bacteria>Terrabacteria_group>Actinomycetota>Actinomycetes>Streptosporangiales>Nocardiopsaceae | GCF_013408795.1 | Spinactinospora_alkalitolerans | HDA32_RS05605 | 1214349..1218161 | 687207 | phage-resistance-protein-[Spinactinospora-alkalitolerans] |
| WP_179811709.1 | Type2.Pgl.like.BREX | <-HTH+REase<-Pentapeptide+Pentapeptide+Pentapeptide+Pentapeptide<-?<-SF1-SF2-Helicase+wHTH+Small-B-barrel+DUF1998<-BrxD-AAA<-PglZ<-DUF6079*<-N6-METHYLASE<-ZnR<-ST-Hiskin\|\|HTH_31+DUF5753->DUF397->?->?->APATPase+TPR+TPR+TPR+TPR->SbcC-> | 1276 | Bacteria>Terrabacteria_group>Actinomycetota>Actinomycetes>Streptosporangiales>Nocardiopsaceae | GCF_013410445.1 | Nocardiopsis_sinuspersici | HNR06_RS26520 | 6072310..6076140 | 501010 | DUF6079-family-protein-[Nocardiopsis-sinuspersici] |
| WP_179821037.1 | Type2.Pgl.like.BREX | SF1-SF2-Helicase-><-?\|\|?->?->Cap4_nuclease->?->SbcC-><-SF1-SF2-Helicase+wHTH+Small-B-barrel+DUF1998<-PglZ<-DUF6079*<-N6-METHYLASE<-ZnR<-ST-Hiskin\|\|HTH_31+DUF5753->DUF397-><-NUDIX<-HTH+HKD-DNase | 1278 | Bacteria>Terrabacteria_group>Actinomycetota>Actinomycetes>Streptosporangiales>Nocardiopsaceae | GCF_013410755.1 | Nocardiopsis_aegyptia | HNR10_RS04420 | 953805..957641 | 220378 | DUF6079-family-protein-[Nocardiopsis-aegyptia] |
| WP_184578247.1 | Type2.Pgl.like.BREX | PglW->?->?-><-SIGMA-HTH<-?<-TetR-HTH<-ABhydrolase\|\|N6-METHYLASE->DUF6079*->PglZ->BrxD-AAA->SF1-SF2-Helicase+wHTH+Small-B-barrel+DUF1998-><-RNase\|\|TetR-HTH-><-AraC-HTH+AraC-HTH\|\|ADH_N+ADH_zinc_N-><-N6-METHYLASE<-ACET | 1253 | Bacteria>Terrabacteria_group>Actinomycetota>Actinomycetes>Streptosporangiales>Nocardiopsaceae | GCF_014203805.1 | Lipingzhangella_halophila | F4561_RS12250 | 2695388..2699149 | 1783352 | phage-resistance-protein-[Lipingzhangella-halophila] |
| WP_184584483.1 | Type2.Pgl.like.BREX | <-SAM-METHYLASE<-METHYLASE\|\|TRD+TRD-><-?<-?\|\|MerR-HTH-><-?<-BrxD-AAA<-PglZ<-DUF6079*<-N6-METHYLASE<-DUF397<-HTH_31+DUF5753\|\|?-><-Peptidase_M20<-HTH_3<-?\|\|?->N6-METHYLASE-> | 1258 | Bacteria>Terrabacteria_group>Actinomycetota>Actinomycetes>Streptosporangiales>Nocardiopsaceae | GCF_014203805.1 | Lipingzhangella_halophila | F4561_RS27640 | 134044..137820 | 1783352 | phage-resistance-protein-[Lipingzhangella-halophila] |
| WP_193120339.1 | Type2.Pgl.like.BREX | SWIM-><-?<-DSBH\|\|SWIM->SIGMA-HTH-><-SIGMA-HTH+SIGMA-HTH<-SF1-SF2-Helicase+wHTH+Small-B-barrel+DUF1998<-BrxD-AAA<-PglZ<-DUF6079*<-N6-METHYLASE<-ZnR<-ST-Hiskin\|\|HTH_31+DUF5753->DUF397->?->?->APATPase+TPR+TPR+TPR+TPR-><-PglW | 1270 | Bacteria>Terrabacteria_group>Actinomycetota>Actinomycetes>Streptosporangiales>Nocardiopsaceae | GCF_014892575.1 | Nocardiopsis_coralli | IDM40_RS03065 | 266608..270420 | 2772213 | DUF6079-family-protein-[Nocardiopsis-coralli] |
| WP_077690681.1 | Type2.Pgl.like.BREX | zf-ISL3+HTH_Tnp_ISL3-><-ParB<-ParB+HNH<-SF1-SF2-Helicase+wHTH+Small-B-barrel+DUF1998<-BrxD-AAA<-PglZ<-DUF6079*<-N6-METHYLASE<-ZnR\|\|HTH_31+DUF5753->DUF397-><-?\|\|SbcC-><-Peptidase_C14 | 1272 | Bacteria>Terrabacteria_group>Actinomycetota>Actinomycetes>Streptosporangiales>Nocardiopsaceae | GCF_014927145.1 | Nocardiopsis | NOSIN_RS11215 | 2608839..2612657 | 2013 | MULTISPECIES:-phage-resistance-protein-[Nocardiopsis] |
| WP_193587651.1 | Type2.Pgl.like.BREX | <-REase<-Phage_AlpA<-HicA\|\|35exo+BRCT->PglW->HTH_OrfB_IS605+OrfB_IS605+OrfB_Zn_ribbon->HTH_31+DUF5753->DUF397->DUF397->DUF6079*->PglZ->BrxD-AAA->SF1-SF2-Helicase+wHTH+Small-B-barrel+DUF1998-><-?\|\|35exo+BRCT-> | 1261 | Bacteria>Terrabacteria_group>Actinomycetota>Actinomycetes>Streptosporangiales>Nocardiopsaceae | GCF_015034585.1 | Thermobifida_fusca | IM867_RS07895 | 1754239..1758024 | 2021 | phage-resistance-protein-[Thermobifida-fusca] |
| WP_248591734.1 | Type2.Pgl.like.BREX | 35exo+BRCT-><-REase\|\|?->PglW-><-McrC-NTD+REase<-McrB-AAA<-?<-CR-ATPase9\|\|DUF6079*->PglZ->BrxD-AAA->SF1-SF2-Helicase+wHTH+Small-B-barrel+DUF1998->DUF4096+DDE_Tnp_1+DDE_Tnp_1_2-> | 1264 | Bacteria>Terrabacteria_group>Actinomycetota>Actinomycetes>Streptosporangiales>Nocardiopsaceae | GCF_023208015.1 | Thermobifida_alba | FOF52_RS21680 | 4853600..4857394 | 53522 | phage-resistance-protein-[Thermobifida-alba] |
| WP_270679261.1 | Type2.Pgl.like.BREX | <-MarR-HTH<-?<-Hiskin<-?<-SF1-SF2-Helicase+wHTH+Small-B-barrel+DUF1998<-BrxD-AAA<-PglZ<-DUF6079*\|\|?-><-ZnR<-ST-Hiskin\|\|HTH_31+DUF5753->DUF397-><-N6-METHYLASE<-TOPRIM<-OLD-ABC\|\|SopB+HNH-> | 1283 | Bacteria>Terrabacteria_group>Actinomycetota>Actinomycetes>Streptosporangiales>Nocardiopsaceae | GCF_027706745.1 | Nocardiopsis_sp._LSu2-4 | O4U47_RS19095 | 15792..19643 | 3018444 | phage-resistance-protein-[Nocardiopsis-sp.-LSu2-4] |
| WP_274811271.1 | Type2.Pgl.like.BREX | <-HTH_31+DUF5753\|\|ST-Hiskin->ZnR-><-SF1-SF2-Helicase<-DrmB<-SF1-SF2-Helicase\|\|SIGMA-HTH+SIGMA-HTH+SIGMA-HTH->PglW->N6-METHYLASE->DUF6079*->PglZ->BrxD-AAA->SF1-SF2-Helicase+wHTH+Small-B-barrel+DUF1998->?->?->?->HTH+REase-><-SWIM | 1304 | Bacteria>Terrabacteria_group>Actinomycetota>Actinomycetes>Streptosporangiales>Nocardiopsaceae | GCF_028882275.1 | Nocardiopsis_sp._N85 | PWG71_RS19280 | 25935..29849 | 3029400 | hypothetical-protein-[Nocardiopsis-sp.-N85] |
| WP_073377653.1 | Type2.Pgl.like.BREX | <-CASPASE+RNA-Helicase+REase<-?<-HTH+REase\|\|SWIM->McrB-AAA->McrC-NTD+REase-><-SF1-SF2-Helicase+wHTH+Small-B-barrel+DUF1998<-BrxD-AAA<-PglZ<-DUF6079*<-N6-METHYLASE<-PglW<-SIGMA-HTH+SIGMA-HTH\|\|SF1-SF2-Helicase->DrmB->SF1-SF2-Helicase-><-?<-REase+SF1-SF2-Helicase<-EVE+HNH | 1305 | Bacteria>Terrabacteria_group>Actinomycetota>Actinomycetes>Streptosporangiales>Nocardiopsaceae | GCF_900141985.1 | Nocardiopsis_flavescens | BUE45_RS09185 | 48689..52606 | 758803 | phage-resistance-protein-[Nocardiopsis-flavescens] |
| WP_087097129.1 | Type2.Pgl.like.BREX | ZnR->PglW->N6-METHYLASE->DUF6079*->PglZ->BrxD-AAA->SF1-SF2-Helicase+wHTH+Small-B-barrel+DUF1998-><-SbcC<-Cap4_nuclease | 1312 | Bacteria>Terrabacteria_group>Actinomycetota>Actinomycetes>Streptosporangiales>Nocardiopsaceae | GCF_900143625.1 | Nocardiopsis_sp._JB363 | BQ8420_RS07680 | 9558..13496 | 1434837 | phage-resistance-protein-[Nocardiopsis-sp.-JB363] |
| GES11111.1 | Type2.Pgl.like.BREX | PglW->N6-METHYLASE->DUF6079*->PglZ->BrxD-AAA->SF1-SF2-Helicase+wHTH+Small-B-barrel+DUF1998-><-MORC+REase<-HTH_31+DUF5753<-TrmB | 1297 | Bacteria>Terrabacteria_group>Actinomycetota>Actinomycetes>Streptosporangiales>Streptosporangiaceae | GCA_009687865.1 | Acrocarpospora_macrocephala | Amac_047080 | 9043..12936 | 150177 | hypothetical-protein-Amac_047080-[Acrocarpospora-macrocephala] |
| GII83983.1 | Type2.Pgl.like.BREX | <-HISKIN<-Abi\|\|?-><-OLD-ABC+TOPRIM\|\|PglW-><-?<-?\|\|HTH_31->N6-METHYLASE->DUF6079*->PglZ->BrxD-AAA->SF1-SF2-Helicase+wHTH+Small-B-barrel+DUF1998-><-REase+SF1-SF2-Helicase\|\|REase+SF1-SF2-Helicase->EVE+REase-> | 1270 | Bacteria>Terrabacteria_group>Actinomycetota>Actinomycetes>Streptosporangiales>Streptosporangiaceae | GCA_016863415.1 | Sphaerisporangium_siamense | Ssi03_19730 | 391245..395057 | 795645 | hypothetical-protein-Ssi03_19730-[Sphaerisporangium-siamense] |
| MBV9853174.1 | Type2.Pgl.like.BREX | <-HEPN<-Ntox45\|\|?-><-SF1-SF2-Helicase+wHTH+Small-B-barrel+DUF1998<-BrxD-AAA<-PglZ<-DUF6079*<-N6-METHYLASE-p<-N6-METHYLASE<-?<-PglW | 1226 | Bacteria>Terrabacteria_group>Actinomycetota>Actinomycetes>Streptosporangiales>Streptosporangiaceae | GCA_019239975.1 | Streptosporangiaceae_bacterium | JOY82_01435 | 99785..103465 | 2339230 | phage-resistance-protein-[Streptosporangiaceae-bacterium] |
| GLK08222.1 | Type2.Pgl.like.BREX | viral-protein->?->GrpE->HSP70+PIN->SF1-SF2-Helicase->PglW->N6-METHYLASE->DUF6079*->PglZ-><-PIN\|\|?->REase->pPIWI+pPIWI->pPIWI+SFII-RAD3->pPIWI+MID+RNaseH-> | 1230 | Bacteria>Terrabacteria_group>Actinomycetota>Actinomycetes>Streptosporangiales>Streptosporangiaceae | GCA_027922005.1 | Streptosporangium_carneum | GCM10017600_16270 | 450196..453888 | 47481 | hypothetical-protein-GCM10017600_16270-[Streptosporangium-carneum] |
| WP_013131794.1 | Type2.Pgl.like.BREX | <-SF1-SF2-Helicase+DUF1998<-HSP70+PIN<-GrpE<-?<-viral-protein\|\|PglW->N6-METHYLASE->DUF6079*->PglZ->?-><-CN_hydrolase<-?<-?<-?<-?\|\|LACI-HTH+PBPI-> | 1238 | Bacteria>Terrabacteria_group>Actinomycetota>Actinomycetes>Streptosporangiales>Streptosporangiaceae | GCF_000092645.1 | Thermobispora_bispora | TBIS_RS07700 | 1773045..1776761 | 2006 | hypothetical-protein-[Thermobispora-bispora] |
| WP_030506727.1 | Type2.Pgl.like.BREX | REase+PNTD1+PNTD2->PIWI-><-?<-ACET<-APATPase+TPR+TPR+TPR+TPR<-TrmB<-?\|\|PglW->N6-METHYLASE->DUF6079*->PglZ->BrxD-AAA->SF1-SF2-Helicase+wHTH+Small-B-barrel+DUF1998-><-?\|\|HEPN-><-ST-Hiskin\|\|TPR-> | 1277 | Bacteria>Terrabacteria_group>Actinomycetota>Actinomycetes>Streptosporangiales>Streptosporangiaceae | GCF_000718475.1 | Microbispora_rosea | IF49_RS0109560 | 22195..26028 | 58117 | hypothetical-protein-[Microbispora-rosea] |
| WP_052424042.1 | Type2.Pgl.like.BREX | <-CBS+HEPN+HEPN<-ParB+HNH\|\|PglW->N6-METHYLASE->DUF6079*->PglZ->BrxD-AAA->SF1-SF2-Helicase+wHTH+Small-B-barrel+DUF1998-> | 1293 | Bacteria>Terrabacteria_group>Actinomycetota>Actinomycetes>Streptosporangiales>Streptosporangiaceae | GCF_000725485.1 | Nonomuraea_candida | IO39_RS41140 | 11445..15326 | 359159 | hypothetical-protein-[Nonomuraea-candida] |
| WP_068899802.1 | Type2.Pgl.like.BREX | SAM-METHYLASE->?-><-?\|\|?->?->?-><-REase\|\|PglW->N6-METHYLASE->DUF6079*->PglZ->BrxD-AAA->SF1-SF2-Helicase+wHTH+Small-B-barrel+DUF1998->?->?->?-><-?\|\|?->Phage_integrase+Phage_integrase-> | 1312 | Bacteria>Terrabacteria_group>Actinomycetota>Actinomycetes>Streptosporangiales>Streptosporangiaceae | GCF_001653075.1 | Planomonospora_sphaerica | A9J22_RS22300 | 198932..202870 | 161355 | phage-resistance-protein-[Planomonospora-sphaerica] |
| WP_084008592.1 | Type2.Pgl.like.BREX | <-ST-Hiskin\|\|?->?->PCMT+METHYLASE->HTH+TPR-><-?<-SF1-SF2-Helicase+wHTH+Small-B-barrel+DUF1998<-BrxD-AAA<-PglZ<-DUF6079*<-N6-METHYLASE<-PglW<-SF1-SF2-Helicase+DUF1998<-HSP70+PIN<-GrpE<-?<-viral-protein\|\|Pkinase+TPR+TPR->SF1-SF2-Helicase-> | 1231 | Bacteria>Terrabacteria_group>Actinomycetota>Actinomycetes>Streptosporangiales>Streptosporangiaceae | GCF_001653075.1 | Planomonospora_sphaerica | A9J22_RS38830 | 78980..82675 | 161355 | hypothetical-protein-[Planomonospora-sphaerica] |
| WP_084958881.1 | Type2.Pgl.like.BREX | ArsR-HTH->?->?-><-?\|\|?->SWIM->PglW->N6-METHYLASE->?->DUF6079*->PglZ->BrxD-AAA->SF1-SF2-Helicase+wHTH+Small-B-barrel+DUF1998->?->JetB->SbcCD_C->DUF3323+TOPRIM-> | 1281 | Bacteria>Terrabacteria_group>Actinomycetota>Actinomycetes>Streptosporangiales>Streptosporangiaceae | GCF_002093975.1 | Thermoactinospora_rubra | BSB02_RS12220 | 40318..44163 | 1088767 | phage-resistance-protein-[Thermoactinospora-rubra] |
| WP_150933328.1 | Type2.Pgl.like.BREX | XIS-HTH-><-?<-REase<-?<-?<-?<-TPR+TPR+TPR+TPR<-DUF6079*<-N6-METHYLASE<-PglW<-SF1-SF2-Helicase+DUF1998<-HSP70+PIN<-GrpE<-?<-viral-protein\|\|N6-METHYLASE->Pkinase+TPR+TPR-> | 1216 | Bacteria>Terrabacteria_group>Actinomycetota>Actinomycetes>Streptosporangiales>Streptosporangiaceae | GCF_008728085.1 | Microbispora_cellulosiformans | F5972_RS10950 | 644784..648434 | 2614688 | phage-resistance-protein-[Microbispora-cellulosiformans] |
| WP_155353331.1 | Type2.Pgl.like.BREX | PglW->N6-METHYLASE->DUF6079*->PglZ->BrxD-AAA->SF1-SF2-Helicase+wHTH+Small-B-barrel+DUF1998-><-?<-?<-?<-REase+SF1-SF2-Helicase | 1266 | Bacteria>Terrabacteria_group>Actinomycetota>Actinomycetes>Streptosporangiales>Streptosporangiaceae | GCF_039525385.1 | Acrocarpospora_macrocephala | Amac_RS06090 | 52601..56401 | 150177 | phage-resistance-protein-[Acrocarpospora-macrocephala] |
| WP_155356502.1 | Type2.Pgl.like.BREX | HTH_3->PglW->N6-METHYLASE->DUF6079*->PglZ->BrxD-AAA->SF1-SF2-Helicase+wHTH+Small-B-barrel+DUF1998-><-MORC+REase<-HTH+DUF5753<-TrmB<-?<-REase<-Calcineurin-SbcD | 1290 | Bacteria>Terrabacteria_group>Actinomycetota>Actinomycetes>Streptosporangiales>Streptosporangiaceae | GCF_009687865.1 | Acrocarpospora_macrocephala | Amac_RS23220 | 9064..12936 | 150177 | phage-resistance-protein-[Acrocarpospora-macrocephala] |
| WP_155347967.1 | Type2.Pgl.like.BREX | Schlafen+HATPase+wHTH-><-?<-DDE_Tnp_1\|\|REase+SF1-SF2-Helicase->EVE+REase->REase-><-SF1-SF2-Helicase+wHTH+Small-B-barrel+DUF1998<-BrxD-AAA<-PglZ<-DUF6079*<-N6-METHYLASE<-PglW\|\|NACHT->?-><-DDE_Tnp_1_4\|\|REase+SF1-SF2-Helicase-> | 1311 | Bacteria>Terrabacteria_group>Actinomycetota>Actinomycetes>Streptosporangiales>Streptosporangiaceae | GCF_009687885.1 | Acrocarpospora_pleiomorpha | Aple_RS29485 | 96149..100084 | 90975 | phage-resistance-protein-[Acrocarpospora-pleiomorpha] |
| WP_182886644.1 | Type2.Pgl.like.BREX | REase+SF1-SF2-Helicase->?->?->?->SLATT->?->PglW->N6-METHYLASE->DUF6079*->PglZ->BrxD-AAA->SF1-SF2-Helicase+wHTH+Small-B-barrel+DUF1998-><-DDE_Tnp_ISL3<-zf-ISL3+HTH_Tnp_ISL3<-ST-Hiskin\|\|TPR->STYKIN-><-LysM+wHTH+TPR | 1288 | Bacteria>Terrabacteria_group>Actinomycetota>Actinomycetes>Streptosporangiales>Streptosporangiaceae | GCF_014161415.1 | Microbispora_sp._H10885 | HHX37_RS22465 | 251220..255086 | 2729110 | phage-resistance-protein-[Microbispora-sp.-H10885] |
| WP_185054172.1 | Type2.Pgl.like.BREX | HTH+REase->PglW->N6-METHYLASE->DUF6079*->PglZ->SF1-SF2-Helicase+wHTH+Small-B-barrel+DUF1998->HDRnase+SF1-SF2-Helicase->CRISPR_Cas6->CRISPR_Cas6-> | 1299 | Bacteria>Terrabacteria_group>Actinomycetota>Actinomycetes>Streptosporangiales>Streptosporangiaceae | GCF_014202315.1 | Thermocatellispora_tengchongensis | HNP84_RS34090 | 51634..55533 | 1073253 | phage-resistance-protein-[Thermocatellispora-tengchongensis] |
| WP_184941226.1 | Type2.Pgl.like.BREX | <-RelE-ParE<-PhdYeFM_antitox<-ST-Hiskin<-?\|\|HTH_31+DUF5753->DUF397->N6-METHYLASE->DUF6079*->PglZ->SF1-SF2-Helicase+wHTH+Small-B-barrel+DUF1998-><-REase+SF1-SF2-Helicase | 1243 | Bacteria>Terrabacteria_group>Actinomycetota>Actinomycetes>Streptosporangiales>Streptosporangiaceae | GCF_014203605.1 | Planomonospora_venezuelensis | FHS22_RS12470 | 313622..317353 | 1999 | phage-resistance-protein-[Planomonospora-venezuelensis] |
| WP_184985626.1 | Type2.Pgl.like.BREX | N6-METHYLASE->?->?->NUDIX-><-DUF397<-HTH_31+DUF5753\|\|ST-Hiskin-><-OLD-ABC<-OLD-ABC\|\|DUF6079*->PglZ->BrxD-AAA->SF1-SF2-Helicase+wHTH+Small-B-barrel+DUF1998->?->TPR+TPR+TPR+TPR+TPR+TPR->?-><-?\|\|?-><-wHTH+WYL+WCX | 1282 | Bacteria>Terrabacteria_group>Actinomycetota>Actinomycetes>Streptosporangiales>Streptosporangiaceae | GCF_014207705.1 | Sphaerisporangium_rubeum | BJ992_RS26770 | 6342516..6346364 | 321317 | phage-resistance-protein-[Sphaerisporangium-rubeum] |
| WP_189126795.1 | Type2.Pgl.like.BREX | STAS->ACET->?->?-><-GAF+PP2C<-SF1-SF2-Helicase+wHTH+Small-B-barrel+DUF1998<-BrxD-AAA<-PglZ<-DUF6079*<-NUDIX<-?<-?\|\|?->?-><-?\|\|HTH_23+HTH_32+HTH_33->DDE_3-> | 1335 | Bacteria>Terrabacteria_group>Actinomycetota>Actinomycetes>Streptosporangiales>Streptosporangiaceae | GCF_014646355.1 | Nonomuraea_cavernae | IEY82_RS25795 | 124541..128548 | 2045107 | phage-resistance-protein-[Nonomuraea-cavernae] |
| WP_189139358.1 | Type2.Pgl.like.BREX | GntR-HTH-><-ABhydrolase\|\|REase-><-?<-?<-SF1-SF2-Helicase+wHTH+Small-B-barrel+DUF1998<-BrxD-AAA<-PglZ<-DUF6079*<-NUDIX<-?\|\|?-><-?<-?<-?<-?\|\|SAM-METHYLASE-> | 1320 | Bacteria>Terrabacteria_group>Actinomycetota>Actinomycetes>Streptosporangiales>Streptosporangiaceae | GCF_014646515.1 | Nonomuraea_glycinis | IEY87_RS15745 | 488552..492514 | 2047744 | phage-resistance-protein-[Nonomuraea-glycinis] |
| WP_189142469.1 | Type2.Pgl.like.BREX | HHH_4+HTH+wHTH+SF1-SF2-Helicase+REase->?-><-REase+SF1-SF2-Helicase<-EVE+REase\|\|PglW->ParB+HNH+METHYLASE->N6-METHYLASE->DUF6079*->PglZ->BrxD-AAA->SF1-SF2-Helicase+wHTH+Small-B-barrel+DUF1998-><-?<-?\|\|HEPN-> | 1233 | Bacteria>Terrabacteria_group>Actinomycetota>Actinomycetes>Streptosporangiales>Streptosporangiaceae | GCF_014646515.1 | Nonomuraea_glycinis | IEY87_RS31590 | 255508..259209 | 2047744 | phage-resistance-protein-[Nonomuraea-glycinis] |
| WP_189236152.1 | Type2.Pgl.like.BREX | N6-METHYLASE->DUF6079*->PglZ->BrxD-AAA->SF1-SF2-Helicase+wHTH+Small-B-barrel+DUF1998-><-SF1-SF2-Helicase+DUF1998<-HSP70+PIN<-GrpE<-?<-viral-protein\|\|Pkinase+TPR+TPR-> | 1235 | Bacteria>Terrabacteria_group>Actinomycetota>Actinomycetes>Streptosporangiales>Streptosporangiaceae | GCF_014647955.1 | Planomonospora_parontospora | IEZ06_RS08615 | 2034..5741 | 58119 | phage-resistance-protein-[Planomonospora-parontospora] |
| WP_189653452.1 | Type2.Pgl.like.BREX | SF1-SF2-Helicase->NERD+Pkinase+Pkinase+DndF->DndG->DndH-><-?\|\|rve_3->PglW->N6-METHYLASE->DUF6079*->PglZ->BrxD-AAA->SF1-SF2-Helicase+wHTH+Small-B-barrel+DUF1998->HTH_3->rve-><-?\|\|?-><-?\|\|ST-Hiskin-> | 1270 | Bacteria>Terrabacteria_group>Actinomycetota>Actinomycetes>Streptosporangiales>Streptosporangiaceae | GCF_014648435.1 | Nonomuraea_spiralis | IEZ27_RS48575 | 48346..52158 | 46182 | PglY-protein-[Nonomuraea-spiralis] |
| WP_196469679.1 | Type2.Pgl.like.BREX | ABhydrolase->LACI-HTH+PBPI->?->MazG-><-REase+SF1-SF2-Helicase->REase+SF1-SF2-Helicase-><-SF1-SF2-Helicase+wHTH+Small-B-barrel+DUF1998<-BrxD-AAA<-PglZ<-DUF6079*<-N6-METHYLASE<-DUF397<-HTH_31+DUF5753\|\|?->ST-Hiskin-><-ParB+HNH+METHYLASE\|\|ParA-Soj-PloopNTPase+ORC-CDC6-like->?-><-PglW | 1201 | Bacteria>Terrabacteria_group>Actinomycetota>Actinomycetes>Streptosporangiales>Streptosporangiaceae | GCF_015711595.1 | Planomonospora_sp._ID91781 | HS048_RS31375 | 228950..232555 | 2738135 | hypothetical-protein-[Planomonospora-sp.-ID91781] |
| WP_196470002.1 | Type2.Pgl.like.BREX | ABC-ATPase+SbcC->SAM-METHYLASE->?-><-?<-REase\|\|PglW->N6-METHYLASE->DUF6079*->PglZ->SF1-SF2-Helicase+wHTH+Small-B-barrel+DUF1998-><-REase<-?<-Pkinase+Pkinase<-?<-?<-NUDIX | 1310 | Bacteria>Terrabacteria_group>Actinomycetota>Actinomycetes>Streptosporangiales>Streptosporangiaceae | GCF_015711595.1 | Planomonospora_sp._ID91781 | HS048_RS34225 | 62425..66357 | 2738135 | hypothetical-protein-[Planomonospora-sp.-ID91781] |
| WP_204288898.1 | Type2.Pgl.like.BREX | <-DDE_Tnp_ISL3<-zf-ISL3<-?<-TPR+TPR+TPR<-DndB\|\|Pkinase+TPR+TPR-><-SF1-SF2-Helicase+wHTH+Small-B-barrel+DUF1998<-BrxD-AAA<-PglZ<-DUF6079*<-N6-METHYLASE<-PglW<-SF1-SF2-Helicase+DUF1998<-HSP70+PIN<-GrpE<-?<-viral-protein\|\|N6-METHYLASE-> | 1232 | Bacteria>Terrabacteria_group>Actinomycetota>Actinomycetes>Streptosporangiales>Streptosporangiaceae | GCF_016863015.1 | Microbispora_amethystogenes | Mam01_RS31900 | 59581..63279 | 1427754 | hypothetical-protein-[Microbispora-amethystogenes] |
| WP_184886687.1 | Type2.Pgl.like.BREX | <-REC+OmpR-HTH<-HISKIN<-Abi<-OLD-ABC+TOPRIM\|\|PglW-><-?<-?\|\|HTH_31->N6-METHYLASE->DUF6079*->PglZ->BrxD-AAA->SF1-SF2-Helicase+wHTH+Small-B-barrel+DUF1998-><-REase+SF1-SF2-Helicase\|\|REase+SF1-SF2-Helicase->EVE+REase-> | 1272 | Bacteria>Terrabacteria_group>Actinomycetota>Actinomycetes>Streptosporangiales>Streptosporangiaceae | GCF_016863415.1 | Sphaerisporangium_siamense | Ssi03_RS09565 | 391239..395057 | 795645 | phage-resistance-protein-[Sphaerisporangium-siamense] |
| WP_206206219.1 | Type2.Pgl.like.BREX | <-CRISPR_Cas6<-HDRnase+SF1-SF2-Helicase<-SF1-SF2-Helicase+DUF1998<-HSP70+PIN<-GrpE<-?<-viral-protein\|\|PglW->N6-METHYLASE->DUF6079*->PglZ->?-><-CN_hydrolase<-?<-?<-?<-?\|\|LACI-HTH+PBPI-> | 1238 | Bacteria>Terrabacteria_group>Actinomycetota>Actinomycetes>Streptosporangiales>Streptosporangiaceae | GCF_017161365.1 | Thermobispora_bispora | CYL17_RS09670 | 2202892..2206608 | 2006 | hypothetical-protein-[Thermobispora-bispora] |
| WP_214643361.1 | Type2.Pgl.like.BREX | SAM-METHYLASE->?->His_Phos_1->?->PglW->N6-METHYLASE-p->N6-METHYLASE->DUF6079*->PglZ->BrxD-AAA->SF1-SF2-Helicase+wHTH+Small-B-barrel+DUF1998-><-?\|\|?-><-HerA/FtsK<-?<-DUF2637+HIN-HTH<-Phage_integrase | 1294 | Bacteria>Terrabacteria_group>Actinomycetota>Actinomycetes>Streptosporangiales>Streptosporangiaceae | GCF_018603905.1 | Nonomuraea_sp._NEAU-A123 | KLK06_RS52340 | 80125..84009 | 2839649 | hypothetical-protein-[Nonomuraea-sp.-NEAU-A123] |
| WP_219550272.1 | Type2.Pgl.like.BREX | <-Phage_integrase<-HTH_3\|\|XIS-HTH->ZnR+TOPRIM->ZnR+TOPRIM+AAA-helicase+wHTH->?-><-?<-?<-SF1-SF2-Helicase+wHTH+Small-B-barrel+DUF1998<-BrxD-AAA<-PglZ<-DUF6079*<-N6-METHYLASE<-PglW\|\|?->?->SF1-SF2-Helicase+DUF1998->?-><-METHYLASE | 1310 | Bacteria>Terrabacteria_group>Actinomycetota>Actinomycetes>Streptosporangiales>Streptosporangiaceae | GCF_019396385.1 | Nonomuraea_harbinensis | KME79_RS37710 | 11993..15925 | 1286938 | hypothetical-protein-[Nonomuraea-harbinensis] |
| WP_271277236.1 | Type2.Pgl.like.BREX | HIN-HTH->HTH_Tnp_1+HTH_21+rve+rve_3->?-><-?<-?\|\|METHYLASE->PglW->N6-METHYLASE->DUF6079*->PglZ->BrxD-AAA->SF1-SF2-Helicase+wHTH+Small-B-barrel+DUF1998->?-><-?<-?<-?<-ZnR+TOPRIM+AAA-helicase+wHTH<-ZnR+TOPRIM<-XIS-HTH | 1310 | Bacteria>Terrabacteria_group>Actinomycetota>Actinomycetes>Streptosporangiales>Streptosporangiaceae | GCF_027620355.1 | Nonomuraea_ferruginea | OUY24_RS19315 | 20830..24762 | 46174 | hypothetical-protein-[Nonomuraea-ferruginea] |
| WP_271216728.1 | Type2.Pgl.like.BREX | <-Pkinase+TPR+TPR<-N6-METHYLASE\|\|viral-protein->?->GrpE->HSP70+PIN->SF1-SF2-Helicase+DUF1998->PglW->N6-METHYLASE->DUF6079*->PglZ-><-PIN\|\|REase->pPIWI+pPIWI->pPIWI+SF1-SF2-Helicase->pPIWI+MID+RNaseH->?->JetB->SbcCD_C-> | 1232 | Bacteria>Terrabacteria_group>Actinomycetota>Actinomycetes>Streptosporangiales>Streptosporangiaceae | GCF_027922005.1 | Streptosporangium_carneum | PKF00_RS07965 | 450190..453888 | 47481 | phage-resistance-protein-[Streptosporangium-carneum] |
| WP_091094240.1 | Type2.Pgl.like.BREX | DDE_3-><-?<-AAA-ATPase<-rve+rve_3\|\|VirD4-TraG+VirD4-FtsK->?-><-SF1-SF2-Helicase+wHTH+Small-B-barrel+DUF1998<-BrxD-AAA<-PglZ<-DUF6079*<-N6-METHYLASE<-PglW\|\|?->?->?->ArsR-HTH-> | 1307 | Bacteria>Terrabacteria_group>Actinomycetota>Actinomycetes>Streptosporangiales>Streptosporangiaceae | GCF_900111685.1 | Nonomuraea_wenchangensis | BMW85_RS46850 | 12858..16781 | 568860 | phage-resistance-protein-[Nonomuraea-wenchangensis] |
| WP_189790886.1 | Type2.Pgl.like.BREX | <-TetR-HTH\|\|?-><-PemK_toxin<-ST-Hiskin\|\|HTH_31+DUF5753->DUF397-><-SF1-SF2-Helicase+wHTH+Small-B-barrel+DUF1998<-BrxD-AAA<-PglZ<-DUF6079*<-N6-METHYLASE<-DUF397<-HTH_31+DUF5753\|\|?->ST-Hiskin->HEPN+HEPN->?->ParA-Soj-PloopNTPase+ORC-CDC6-like-><-PglW | 1240 | Bacteria>Terrabacteria_group>Actinomycetota>Actinomycetes>Streptosporangiales>Streptosporangiaceae | - | Streptosporangium_violaceochromogenes | IE292_RS10545 | 46299..50021 | 46188 | DUF6079-family-protein-[Streptosporangium-violaceochromogenes] |
| NDU73743.1 | Type2.Pgl.like.BREX | HIRAN->?-><-?<-SF1-SF2-Helicase+REase<-SF1-SF2-Helicase+wHTH+Small-B-barrel+DUF1998<-BrxD-AAA<-PglZ<-DUF6079*<-N6-METHYLASE<-PglW\|\|<-?<-PCMT+METHYLASE<-ABhydrolase<-?<-?\|\|TrmB-><-HTH+DUF5753 | 1308 | Bacteria>Terrabacteria_group>Actinomycetota>Actinomycetes>Streptosporangiales>Thermomonosporaceae | GCA_010491635.1 | Actinomadura_lepetitiana | GWI34_14000 | 24527..28453 | 2126066 | phage-resistance-protein-[Actinomadura-lepetitiana] |
| WP_026414530.1 | Type2.Pgl.like.BREX | <-Pkinase+Pkinase<-?<-SF1-SF2-Helicase+wHTH+Small-B-barrel+DUF1998<-BrxD-AAA<-PglZ<-DUF6079*<-HTH_37<-ST-Hiskin\|\|HTH_31+DUF5753->DUF397->DUF397-><-N6-METHYLASE<-PglW\|\|?->JetB-> | 1269 | Bacteria>Terrabacteria_group>Actinomycetota>Actinomycetes>Streptosporangiales>Thermomonosporaceae | GCF_000518265.1 | Actinomadura_oligospora | P696_RS0125405 | 67406..71215 | 111804 | hypothetical-protein-[Actinomadura-oligospora] |
| WP_067795492.1 | Type2.Pgl.like.BREX | Calcineurin->tRNA-><-?<-ParB+HNH<-TOPRIM<-OLD-ABC<-SF1-SF2-Helicase+wHTH+Small-B-barrel+DUF1998<-BrxD-AAA<-PglZ<-DUF6079*<-N6-METHYLASE<-?<-?\|\|HTH_31+DUF5753->DUF397-> | 1307 | Bacteria>Terrabacteria_group>Actinomycetota>Actinomycetes>Streptosporangiales>Thermomonosporaceae | GCF_001552155.1 | Actinomadura_formosensis | AF1_RS09895 | 52263..56186 | 60706 | phage-resistance-protein-[Actinomadura-formosensis] |
| WP_067640687.1 | Type2.Pgl.like.BREX | <-SF1-SF2-Helicase+wHTH+Small-B-barrel+DUF1998<-BrxD-AAA<-PglZ<-DUF6079*<-?<-ST-Hiskin<-?\|\|?->HTH-><-SF1-SF2-Helicase<-?<-MORC+REase\|\|HTH_31+DUF5753-> | 1215 | Bacteria>Terrabacteria_group>Actinomycetota>Actinomycetes>Streptosporangiales>Thermomonosporaceae | GCF_001552195.1 | Actinomadura_latina | AL2_RS33015 | 71429..75076 | 163603 | hypothetical-protein-[Actinomadura-latina] |
| WP_067479645.1 | Type2.Pgl.like.BREX | <-SF1-SF2-Helicase\|\|?-><-?<-?<-OLD-ABC+TOPRIM<-SF1-SF2-Helicase+wHTH+Small-B-barrel+DUF1998<-BrxD-AAA<-PglZ<-DUF6079*<-HTH_37\|\|DUF4096+DDE_Tnp_1+DDE_Tnp_1_2->?->?->?->?->?-><-N6-METHYLASE<-PglW | 1281 | Bacteria>Terrabacteria_group>Actinomycetota>Actinomycetes>Streptosporangiales>Thermomonosporaceae | GCF_001552635.1 | Actinomadura_hibisca | AH6_RS14655 | 33213..37058 | 68565 | PglY-protein-[Actinomadura-hibisca] |
| WP_067482812.1 | Type2.Pgl.like.BREX | SbcC+SbcC->?->PglW-><-DUF397<-HTH_31+DUF5753\|\|?->N6-METHYLASE->DUF6079*->PglZ->BrxD-AAA->SF1-SF2-Helicase+wHTH+Small-B-barrel+DUF1998->OLD-ABC->TOPRIM->ParB+HNH-><-?<-N6-METHYLASE | 1293 | Bacteria>Terrabacteria_group>Actinomycetota>Actinomycetes>Streptosporangiales>Thermomonosporaceae | GCF_001552635.1 | Actinomadura_hibisca | AH6_RS22035 | 103171..107052 | 68565 | phage-resistance-protein-[Actinomadura-hibisca] |
| WP_103561564.1 | Type2.Pgl.like.BREX | <-HD<-?\|\|?->?->NTP_transferase+MannoseP_isomer-><-?<-?\|\|PglW->N6-METHYLASE->DUF6079*->PglZ->BrxD-AAA->SF1-SF2-Helicase+wHTH+Small-B-barrel+DUF1998-><-tRNA\|\|Peptidase_M20->?->?-><-?\|\|TOPRIM->Thymidylate_kin-> | 1302 | Bacteria>Terrabacteria_group>Actinomycetota>Actinomycetes>Streptosporangiales>Thermomonosporaceae | GCF_002911665.1 | Actinomadura_rubteroloni | BTM25_RS05045 | 1146873..1150781 | 1926885 | DUF6079-family-protein-[Actinomadura-rubteroloni] |
| WP_111864830.1 | Type2.Pgl.like.BREX | DnaA->?->?->STYKIN+S1+S1->?-><-SF1-SF2-Helicase+wHTH+Small-B-barrel+DUF1998<-BrxD-AAA<-PglZ<-DUF6079*<-ST-Hiskin\|\|?->?->?->HTH_31+DUF5753->DUF397-><-N6-METHYLASE<-PglW\|\|ParB+HNH-> | 1283 | Bacteria>Terrabacteria_group>Actinomycetota>Actinomycetes>Streptosporangiales>Thermomonosporaceae | GCF_003289645.1 | Actinomadura_craniellae | DPM19_RS09315 | 525187..529038 | 2231787 | phage-resistance-protein-[Actinomadura-craniellae] |
| WP_111870039.1 | Type2.Pgl.like.BREX | <-DOC<-?<-?<-ABC-ATPase+SbcC\|\|HisKA_3+HATPase_c->REC+LuxR-HTH-><-SF1-SF2-Helicase+wHTH+Small-B-barrel+DUF1998<-BrxD-AAA<-PglZ<-DUF6079*<-N6-METHYLASE<-PglW\|\|?->SIGMA-HTH->?->Abhydrolase->ADH_N+ADH_zinc_N-> | 1254 | Bacteria>Terrabacteria_group>Actinomycetota>Actinomycetes>Streptosporangiales>Thermomonosporaceae | GCF_003289645.1 | Actinomadura_craniellae | DPM19_RS22815 | 100013..103777 | 2231787 | DUF6079-family-protein-[Actinomadura-craniellae] |
| WP_119727666.1 | Type2.Pgl.like.BREX | <-SF1-SF2-Helicase+wHTH+Small-B-barrel+DUF1998<-BrxD-AAA<-PglZ<-DUF6079*<-N6-METHYLASE<-PglW\|\|Pkinase+TPR+TPR->REase->?-><-?\|\|?->LACI-HTH+PBPI-> | 1223 | Bacteria>Terrabacteria_group>Actinomycetota>Actinomycetes>Streptosporangiales>Thermomonosporaceae | GCF_003589885.1 | Thermomonospora_amylolytica | D3U04_RS08210 | 1778450..1782121 | 1411117 | phage-resistance-protein-[Thermomonospora-amylolytica] |
| WP_122192829.1 | Type2.Pgl.like.BREX | <-JetB<-?\|\|PglW->N6-METHYLASE-p-><-HTH_31+DUF5753\|\|ST-Hiskin->?->HTH_37->?->DUF6079-p->DEDD_Tnp_IS110->SF1-SF2-Helicase+wHTH+Small-B-barrel+DUF1998-><-NAD_binding_10<-NAD_binding_10<-?<-SIGMA-HTH<-MerR-HTH | 101 | Bacteria>Terrabacteria_group>Actinomycetota>Actinomycetes>Streptosporangiales>Thermomonosporaceae | GCF_003696215.1 | Actinomadura_harenae | EBO15_RS03505 | 98734..99039 | 2483351 | phage-resistance-protein-[Actinomadura-harenae] |
| WP_122192830.1 | Type2.Pgl.like.BREX | PglW->N6-METHYLASE-p-><-HTH_31+DUF5753\|\|ST-Hiskin->?->HTH_37->?->DUF6079-p->DUF6079-p*->DEDD_Tnp_IS110->SF1-SF2-Helicase+wHTH+Small-B-barrel+DUF1998-><-NAD_binding_10<-NAD_binding_10<-?<-SIGMA-HTH<-MerR-HTH\|\|?->Pkinase+Pkinase-> | 154 | Bacteria>Terrabacteria_group>Actinomycetota>Actinomycetes>Streptosporangiales>Thermomonosporaceae | GCF_003696215.1 | Actinomadura_harenae | EBO15_RS03510 | 99082..99546 | 2483351 | hypothetical-protein-[Actinomadura-harenae] |
| WP_123661993.1 | Type2.Pgl.like.BREX | <-DUF3323+TOPRIM<-?<-?<-?\|\|?-><-REase+SF1-SF2-Helicase<-SF1-SF2-Helicase+wHTH+Small-B-barrel+DUF1998<-BrxD-AAA<-PglZ<-DUF6079*<-N6-METHYLASE<-PglW<-DUF397<-HTH_31+DUF5753\|\|?-><-HTH_29+HTH_32+DDE_3<-HTH_28+HTH_32+DDE_3 | 1269 | Bacteria>Terrabacteria_group>Actinomycetota>Actinomycetes>Streptosporangiales>Thermomonosporaceae | GCF_003751225.1 | Actinocorallia_herbida | EDD29_RS02590 | 501794..505603 | 58109 | PglY-protein-[Actinocorallia-herbida] |
| WP_131962485.1 | Type2.Pgl.like.BREX | DnaA->?->STYKIN+S1+S1->?-><-SF1-SF2-Helicase+wHTH+Small-B-barrel+DUF1998<-BrxD-AAA<-PglZ<-DUF6079*<-HKD-DNase+HKD-DNase<-?<-?<-?\|\|?-><-REase<-?\|\|?-><-DUF397 | 1296 | Bacteria>Terrabacteria_group>Actinomycetota>Actinomycetes>Streptosporangiales>Thermomonosporaceae | GCF_004348735.1 | Actinomadura_sp._KC06 | E1287_RS00900 | 195026..198916 | 2530369 | PglY-protein-[Actinomadura-sp.-KC06] |
| WP_138633114.1 | Type2.Pgl.like.BREX | <-ParB+HNH\|\|PglW->N6-METHYLASE-><-?<-?\|\|HTH_31+DUF5753->DUF397-><-TOPRIM<-OLD-ABC\|\|DUF6079*->PglZ->BrxD-AAA->SF1-SF2-Helicase+wHTH+Small-B-barrel+DUF1998-><-LuxR-HTH | 1216 | Bacteria>Terrabacteria_group>Actinomycetota>Actinomycetes>Streptosporangiales>Thermomonosporaceae | GCF_005889745.1 | Actinomadura_geliboluensis | ETD96_02095 | 30714..34364 | 882440 | phage-resistance-protein-[Actinomadura-geliboluensis] |
| WP_141576271.1 | Type2.Pgl.like.BREX | <-PADR-HTH\|\|?-><-?<-SF1-SF2-Helicase+wHTH+Small-B-barrel+DUF1998<-BrxD-AAA<-PglZ<-DUF6079*<-N6-METHYLASE\|\|?-><-?<-?<-?<-?<-?\|\|?->HTH-> | 1210 | Bacteria>Terrabacteria_group>Actinomycetota>Actinomycetes>Streptosporangiales>Thermomonosporaceae | GCF_006547145.1 | Actinomadura_sp._WMMA1423 | FKD65_RS04415 | 852046..855678 | 2591108 | phage-resistance-protein-[Actinomadura-sp.-WMMA1423] |
| WP_148766853.1 | Type2.Pgl.like.BREX | HTH_31+DUF5753->DUF397-><-?\|\|?->REase+REase->REase+REase->?->HKD-DNase+HKD-DNase->DUF6079*->PglZ->BrxD-AAA->SF1-SF2-Helicase+wHTH+Small-B-barrel+DUF1998-><-?\|\|Pkinase+Pkinase->?->Pkinase+Pkinase-> | 1297 | Bacteria>Terrabacteria_group>Actinomycetota>Actinomycetes>Streptosporangiales>Thermomonosporaceae | GCF_008121305.1 | Actinomadura_decatromicini | FXF68_RS34725 | 406652..410545 | 2604572 | PglY-protein-[Actinomadura-decatromicini] |
| WP_148767668.1 | Type2.Pgl.like.BREX | PglW->N6-METHYLASE-><-DUF397<-DUF397<-HTH_31+DUF5753\|\|ST-Hiskin->?-><-?\|\|N6-METHYLASE->DUF6079*->PglZ->BrxD-AAA-><-?<-?\|\|?->?->?->?-><-TetR-HTH | 1210 | Bacteria>Terrabacteria_group>Actinomycetota>Actinomycetes>Streptosporangiales>Thermomonosporaceae | GCF_008121305.1 | Actinomadura_decatromicini | FXF68_RS38030 | 95114..98746 | 2604572 | phage-resistance-protein-[Actinomadura-decatromicini] |
| WP_161103223.1 | Type2.Pgl.like.BREX | DUF397-><-APATPase+TPR+TPR+TPR+TPR+TPR+TPR\|\|APATPase-><-?\|\|?->TPR+TPR+TPR+TPR+TPR-><-?<-SF1-SF2-Helicase+wHTH+Small-B-barrel+DUF1998<-BrxD-AAA<-PglZ<-DUF6079*<-N6-METHYLASE<-PglW\|\|?-><-?<-?\|\|?->HD->\|\|TetR-HTH-><-ABC-ATPase+SbcC+VirB11+ABC-ATPase+SbcC | 1419 | Bacteria>Terrabacteria_group>Actinomycetota>Actinomycetes>Streptosporangiales>Thermomonosporaceae | GCF_009831215.1 | Actinomadura_rayongensis | GQ466_RS13650 | 940291..944550 | 1429076 | phage-resistance-protein-[Actinomadura-rayongensis] |
| WP_182841320.1 | Type2.Pgl.like.BREX | Peptidase_M20->?->?-><-ParB+HNH<-TOPRIM<-OLD-ABC<-SF1-SF2-Helicase+wHTH+Small-B-barrel+DUF1998<-PglZ<-DUF6079*<-N6-METHYLASE<-?\|\|HTH_31+DUF5753->DUF397-><-PglW<-?<-SbcC+SbcC | 1290 | Bacteria>Terrabacteria_group>Actinomycetota>Actinomycetes>Streptosporangiales>Thermomonosporaceae | GCF_014138665.1 | Actinomadura_namibiensis | HNR61_RS01575 | 344427..348299 | 182080 | phage-resistance-protein-[Actinomadura-namibiensis] |
| WP_182706297.1 | Type2.Pgl.like.BREX | Peptidase_M24->?-><-?<-SF1-SF2-Helicase+wHTH+Small-B-barrel+DUF1998<-ParA-Soj-PloopNTPase<-Schlafen+HATPase+wHTH<-BrxD-AAA<-PglZ<-DUF6079*<-?<-NUDIX<-?<-?<-?<-Peptidase_S8<-?<-?<-DUF397 | 1207 | Bacteria>Terrabacteria_group>Actinomycetota>Actinomycetes>Streptosporangiales>Thermomonosporaceae | GCF_014138835.1 | Thermomonospora_cellulosilytica | HNR21_RS19325 | 4113272..4116895 | 1411118 | phage-resistance-protein-[Thermomonospora-cellulosilytica] |
| WP_184883128.1 | Type2.Pgl.like.BREX | N6-METHYLASE-p-><-HTH_31+DUF5753<-HTH<-HTH<-?\|\|?-><-NACHT+68TM-wHTH\|\|ST-Hiskin->?->DUF6079-p*->?-><-?\|\|?->?->?-><-ADH_N+ADH_zinc_N<-NAD_binding_10\|\|AraC-HTH+AraC-HTH-><-?\|\|SIGMA-HTH+PR-1-> | 912 | Bacteria>Terrabacteria_group>Actinomycetota>Actinomycetes>Streptosporangiales>Thermomonosporaceae | GCF_014648675.1 | Actinomadura | F4557_RS14345 | 3159444..3162182 | 1988 | MULTISPECIES:-DUF6079-family-protein-[Actinomadura] |
| WP_184883516.1 | Type2.Pgl.like.BREX | <-LuxR-HTH\|\|Glyoxalase->?-><-?<-SF1-SF2-Helicase+wHTH+Small-B-barrel+DUF1998<-BrxD-AAA<-PglZ<-DUF6079*\|\|?-><-?\|\|SWACOS->Pkinase-><-DUF397<-HTH_31+DUF5753 | 1282 | Bacteria>Terrabacteria_group>Actinomycetota>Actinomycetes>Streptosporangiales>Thermomonosporaceae | GCF_014648675.1 | Actinomadura | IEZ03_RS16650 | 331737..335585 | 1988 | MULTISPECIES:-PglY-protein-[Actinomadura] |
| WP_184890069.1 | Type2.Pgl.like.BREX | <-Relaxase<-TraJ-RHH<-SF1-SF2-Helicase+wHTH+Small-B-barrel+DUF1998<-BrxD-AAA<-PglZ<-DUF6079*<-N6-METHYLASE<-?<-?\|\|HTH_31+DUF5753->DUF397-> | 1296 | Bacteria>Terrabacteria_group>Actinomycetota>Actinomycetes>Streptosporangiales>Thermomonosporaceae | GCF_014648675.1 | Actinomadura | IEZ03_RS36965 | 8014..11904 | 1988 | MULTISPECIES:-phage-resistance-protein-[Actinomadura] |
| WP_192757840.1 | Type2.Pgl.like.BREX | <-YTCD-HTH\|\|?->NACHT+68TM-wHTH-><-TOPRIM<-OLD-ABC\|\|DUF6079*->PglZ->BrxD-AAA->SF1-SF2-Helicase+wHTH+Small-B-barrel+DUF1998-> | 1242 | Bacteria>Terrabacteria_group>Actinomycetota>Actinomycetes>Streptosporangiales>Thermomonosporaceae | GCF_014873935.1 | Actinomadura_algeriensis | H4W34_RS03575 | 878652..882380 | 1679523 | phage-resistance-protein-[Actinomadura-algeriensis] |
| WP_208265635.1 | Type2.Pgl.like.BREX | HTH_3->METHYLASE-><-XIS-HTH<-?<-Relaxase<-TraJ-RHH\|\|wHTH+SAD+HNH-><-SF1-SF2-Helicase+wHTH+Small-B-barrel+DUF1998<-PglZ<-DUF6079*<-N6-METHYLASE<-?\|\|HTH_31+DUF5753->DUF397-><-PglW\|\|?->HKD-DNase+HKD-Rease-> | 1303 | Bacteria>Terrabacteria_group>Actinomycetota>Actinomycetes>Streptosporangiales>Thermomonosporaceae | GCF_017573545.1 | Actinomadura_nitritigenes | J4557_RS07160 | 173118..177029 | 134602 | DUF6079-family-protein-[Actinomadura-nitritigenes] |
| WP_230618667.1 | Type2.Pgl.like.BREX | <-ADH_N+ADH_zinc_N\|\|?-><-?\|\|SAM-METHYLASE->?->?->?->PglW->N6-METHYLASE->DUF6079*->PglZ->BrxD-AAA->SF1-SF2-Helicase+wHTH+Small-B-barrel+DUF1998-> | 1296 | Bacteria>Terrabacteria_group>Actinomycetota>Actinomycetes>Streptosporangiales>Thermomonosporaceae | GCF_020991275.1 | Actinocorallia_sp._API_0066 | LO762_RS26150 | 235431..239321 | 2896846 | hypothetical-protein-[Actinocorallia-sp.-API-0066] |
| WP_242884440.1 | Type2.Pgl.like.BREX | HTH_31+DUF5753->DUF397->REase->MerR-HTH->?-><-TOPRIM<-OLD-ABC\|\|DUF6079*->PglZ->BrxD-AAA->SF1-SF2-Helicase+wHTH+Small-B-barrel+DUF1998-> | 1227 | Bacteria>Terrabacteria_group>Actinomycetota>Actinomycetes>Streptosporangiales>Thermomonosporaceae | GCF_022751195.1 | Actinomadura_litoris | K2F48_RS09350 | 533875..537558 | 2678616 | phage-resistance-protein-[Actinomadura-litoris] |
| WP_242911417.1 | Type2.Pgl.like.BREX | <-TPR+TPR<-?\|\|PglW->N6-METHYLASE-><-?\|\|HTH_31+DUF5753->DUF397->?->HKD-DNase+HKD-DNase->DUF6079*->PglZ->BrxD-AAA->SF1-SF2-Helicase+wHTH+Small-B-barrel+DUF1998-><-METHYLASE<-GNTR-HTH\|\|RADICAL-SAM->RADICAL-SAM->Peptidase-> | 1291 | Bacteria>Terrabacteria_group>Actinomycetota>Actinomycetes>Streptosporangiales>Thermomonosporaceae | GCF_022751215.1 | Actinomadura_terrae | K2F47_RS39680 | 90630..94505 | 604353 | PglY-protein-[Actinomadura-terrae] |
| WP_250355172.1 | Type2.Pgl.like.BREX | DSBH+AraC-HTH+AraC-HTH-><-?<-?\|\|?->?->?->ST-Hiskin->?->DUF6079-p*->?-><-?\|\|?->?->?->?-><-ABhydrolase | 65 | Bacteria>Terrabacteria_group>Actinomycetota>Actinomycetes>Streptosporangiales>Thermomonosporaceae | GCF_023614255.1 | Actinomadura_madurae | LUW76_RS00540 | 117273..117470 | 1993 | hypothetical-protein-[Actinomadura-madurae] |
| WP_250370435.1 | Type2.Pgl.like.BREX | DSBH+AraC-HTH+AraC-HTH-><-?<-?\|\|?->?->?->ST-Hiskin->?->DUF6079-p*->?-><-?\|\|?->?->?->?-><-ABhydrolase | 65 | Bacteria>Terrabacteria_group>Actinomycetota>Actinomycetes>Streptosporangiales>Thermomonosporaceae | GCF_023614275.1 | Actinomadura_madurae | LUW74_RS10305 | 2191601..2191798 | 1993 | hypothetical-protein-[Actinomadura-madurae] |
| WP_252780948.1 | Type2.Pgl.like.BREX | MqsA_antitoxin+DUF4065->?-><-ParB+HNH<-?\|\|DUF6079-p*->?->PglZ->BrxD-AAA->SF1-SF2-Helicase+wHTH+Small-B-barrel+DUF1998->SF1-SF2-Helicase+HKD-DNase->REase+SF1-SF2-Helicase-> | 428 | Bacteria>Terrabacteria_group>Actinomycetota>Actinomycetes>Streptosporangiales>Thermomonosporaceae | GCF_024055675.1 | Actinoallomurus_sp._WRP9H-5 | NE234_RS15470 | 379702..380988 | 2952536 | hypothetical-protein-[Actinoallomurus-sp.-WRP9H-5] |
| WP_255273511.1 | Type2.Pgl.like.BREX | PNPase->?-><-Peptidase_C14+SF1-SF2-Helicase<-?<-SF1-SF2-Helicase+wHTH+Small-B-barrel+DUF1998<-BrxD-AAA<-PglZ<-PglZ<-?<-DUF6079-p*<-HKD-DNase+HKD-DNase<-?<-?<-REase+REase\|\|SWACOS->Pkinase-><-DUF397<-HTH_31+DUF5753 | 945 | Bacteria>Terrabacteria_group>Actinomycetota>Actinomycetes>Streptosporangiales>Thermomonosporaceae | GCF_024348945.1 | Actinomadura_madurae | LUX33_RS08005 | 1455133..1457970 | 1993 | hypothetical-protein-[Actinomadura-madurae] |
| WP_255276709.1 | Type2.Pgl.like.BREX | PNPase->?->?-><-Peptidase_C14+SF1-SF2-Helicase<-?<-BrxD-AAA<-PglZ<-PglZ<-?<-DUF6079*<-HKD-DNase+HKD-DNase<-?<-?<-REase+REase\|\|SWACOS->Pkinase-><-DUF397<-HTH_31+DUF5753 | 1212 | Bacteria>Terrabacteria_group>Actinomycetota>Actinomycetes>Streptosporangiales>Thermomonosporaceae | GCF_024348965.1 | Actinomadura_madurae | LUX57_RS08355 | 1455061..1458699 | 1993 | PglY-protein-[Actinomadura-madurae] |
| WP_255282877.1 | Type2.Pgl.like.BREX | HTH_31+DUF5753->DUF397-><-Pkinase<-SWACOS\|\|REase+REase->?->?->HKD-DNase+HKD-DNase->DUF6079-p*->DUF6079-p->PglZ->?->Peptidase_C14+SF1-SF2-Helicase-><-?<-PNPase<-NUDIX | 920 | Bacteria>Terrabacteria_group>Actinomycetota>Actinomycetes>Streptosporangiales>Thermomonosporaceae | GCF_024348985.1 | Actinomadura_madurae | LUX73_RS44465 | 8619981..8622743 | 1993 | hypothetical-protein-[Actinomadura-madurae] |
| WP_255279493.1 | Type2.Pgl.like.BREX | <-Peptidase_C14+SF1-SF2-Helicase<-?<-SF1-SF2-Helicase+wHTH+Small-B-barrel+DUF1998<-BrxD-AAA<-PglZ-p<-PglZ-p<-PglZ<-DUF6079*<-HKD-DNase+HKD-DNase<-?<-?<-REase+REase\|\|SWACOS->Pkinase-><-DUF397<-HTH_31+DUF5753 | 1017 | Bacteria>Terrabacteria_group>Actinomycetota>Actinomycetes>Streptosporangiales>Thermomonosporaceae | GCF_024349005.1 | Actinomadura_madurae | LUX32_RS08265 | 1454911..1457964 | 1993 | PglY-protein-[Actinomadura-madurae] |
| WP_255267968.1 | Type2.Pgl.like.BREX | <-Peptidase_C14+SF1-SF2-Helicase<-?<-SF1-SF2-Helicase+wHTH+Small-B-barrel+DUF1998\|\|?-><-PglZ<-PglZ<-DUF6079-p<-DUF6079-p*<-?<-HKD-DNase+HKD-DNase<-?<-?<-REase+REase\|\|SWACOS->Pkinase-><-DUF397<-HTH_31+DUF5753 | 557 | Bacteria>Terrabacteria_group>Actinomycetota>Actinomycetes>Streptosporangiales>Thermomonosporaceae | GCF_024349025.1 | Actinomadura_madurae | LUX39_RS08485 | 1456946..1458619 | 1993 | hypothetical-protein-[Actinomadura-madurae] |
| MSW49533.1 | Type2.Pgl.like.BREX | PglW->N6-METHYLASE-><-OLD-ABC+TOPRIM\|\|DUF6079*->PglZ->BrxD-AAA->SF1-SF2-Helicase+wHTH+Small-B-barrel+DUF1998-><-?<-?\|\|?-><-?\|\|?->XIS-HTH-> | 1227 | Bacteria>Terrabacteria_group>Actinomycetota>Actinomycetes>unclassified_Actinomycetes | GCA_009699505.1 | Actinomycetia_bacterium | F2817_01410 | 14220..17903 | 1883427 | phage-resistance-protein-[Actinomycetia-bacterium] |
| MSW90813.1 | Type2.Pgl.like.BREX | N6-METHYLASE->Glycos_transf->?->?-><-?\|\|?->PglW->DUF6079*->PglZ->BrxD-AAA->ParB+HNH->SF1-SF2-Helicase+wHTH+Small-B-barrel+DUF1998-> | 1199 | Bacteria>Terrabacteria_group>Actinomycetota>Actinomycetes>unclassified_Actinomycetes | GCA_009699605.1 | Actinomycetia_bacterium | F2812_04345 | 18009..21608 | 1883427 | hypothetical-protein-[Actinomycetia-bacterium] |
| MBK9741407.1 | Type2.Pgl.like.BREX | PglW->N6-METHYLASE->DUF6079*->PglZ->BrxD-AAA->REase->SF1-SF2-Helicase+wHTH+Small-B-barrel+DUF1998-><-?\|\|?->?->McrB-AAA->McrC-NTD+REase-> | 1234 | Bacteria>Terrabacteria_group>Actinomycetota>Actinomycetes>unclassified_Actinomycetes | GCA_016717115.1 | Actinomycetia_bacterium | IP093_18460 | 25042..28746 | 1883427 | phage-resistance-protein-[Actinomycetia-bacterium] |
| WP_131153514.1 | Type2.Pgl.like.BREX | HTH_Tnp_1+HTH_21+rve+rve_3->?->PemK_toxin-><-SF1-SF2-Helicase+wHTH+Small-B-barrel+DUF1998<-BrxD-AAA<-PglZ<-DUF6079*<-NTP_transf<-HEPN<-N6-METHYLASE<-PglW\|\|?-><-?<-?<-MoxR-AAA | 1260 | Bacteria>Terrabacteria_group>Actinomycetota>Nitriliruptoria>Egibacterales>Egibacteraceae | GCF_004322855.1 | Egibacter_rhizosphaerae | ER308_RS02310 | 491517..495299 | 1670831 | phage-resistance-protein-[Egibacter-rhizosphaerae] |
| AXV04977.1 | Type2.Pgl.like.BREX | <-SF1-SF2-Helicase+wHTH+Small-B-barrel+DUF1998\|\|Schlafen+HATPase+wHTH-><-BrxD-AAA<-?\|\|EVE->?-><-PglZ<-REase<-?<-DUF6079*<-DUF4011+SF1-SF2-Helicase+REase\|\|MULE-TRANSPOSASE-><-NACHT+nSTAND4<-N6-METHYLASE<-PglW<-?\|\|?-><-STYKIN | 1213 | Bacteria>Terrabacteria_group>Actinomycetota>Nitriliruptoria>Euzebyales>Euzebyaceae | GCA_003344865.1 | Euzebya_pacifica | DVS28_a0270 | 300271..303912 | 1608957 | Bacteriophage-(PhiC31)-resistance-protein-PglY-[Euzebya-pacifica] |
| WP_164709775.1 | Type2.Pgl.like.BREX | <-SF1-SF2-Helicase+wHTH+Small-B-barrel+DUF1998\|\|Schlafen+HATPase+wHTH-><-BrxD-AAA\|\|EVE+REase->DNA-glycosylase->?-><-PglZ<-REase<-?<-DUF6079*<-DUF4011+SF1-SF2-Helicase+REase\|\|MULE-TRANSPOSASE-><-NACHT+nSTAND4<-N6-METHYLASE<-PglW<-?\|\|?-><-STYKIN | 1215 | Bacteria>Terrabacteria_group>Actinomycetota>Nitriliruptoria>Euzebyales>Euzebyaceae | GCF_003344865.1 | Euzebya_pacifica | DVS28_RS01320 | 300271..303918 | 1608957 | phage-resistance-protein-[Euzebya-pacifica] |
| MTV27839.1 | Type2.Pgl.like.BREX | PglW->OLD-ABC+TOPRIM->N6-METHYLASE->DUF6079*->PglZ->MPTase->PIN->BrxD-AAA->ParB+HNH->SF1-SF2-Helicase+wHTH+Small-B-barrel+DUF1998-> | 1204 | Bacteria>Terrabacteria_group>Actinomycetota>Nitriliruptoria>Nitriliruptorales>Nitriliruptoraceae | GCA_009720505.1 | Nitriliruptoraceae_bacterium_ZYF776 | FTX61_20835 | 10823..14437 | 2604685 | phage-resistance-protein-[Nitriliruptoraceae-bacterium-ZYF776] |
| MCP9485022.1 | Type2.Pgl.like.BREX | PglW-><-DEDD_Tnp_IS110+Transposase_20\|\|N6-METHYLASE->DUF6079*->PglZ->BrxD-AAA->SF1-SF2-Helicase+wHTH+Small-B-barrel+DUF1998->?->DUF932->ParB->TrwC+SF1-SF2-Helicase-> | 1214 | Bacteria>Terrabacteria_group>Actinomycetota>Rubrobacteria>Gaiellales>Gaiellaceae | GCA_024330215.1 | Candidatus_Gaiellasilicea_maunaloa | MSC30_04130 | 13693..17337 | 2928748 | hypothetical-protein-[Candidatus-Gaiellasilicea-maunaloa] |
| WP_270012295.1 | Type2.Pgl.like.BREX | <-Aminotran_3<-Calcineurin<-ACET\|\|?->?->tRNA->ParB+HNH->PglW->N6-METHYLASE->N6-METHYLASE->DUF6079*->PglZ->BrxD-AAA->SF1-SF2-Helicase+wHTH+Small-B-barrel+DUF1998->?->?->SF1-SF2-Helicase+DUF1998-><-nSTAND1+HNH<-?\|\|GNTR-HTH-> | 1201 | Bacteria>Terrabacteria_group>Actinomycetota>Thermoleophilia>Solirubrobacterales>Solirubrobacteraceae | - | Solirubrobacter_taibaiensis | OJ998_RS11960 | 133856..137461 | 1395955 | hypothetical-protein-[Solirubrobacter-taibaiensis] |
| MBX3168148.1 | Type2.Pgl.like.BREX | Calcineurin->?->Peptidase_S8->?-><-?\|\|ANK+ANK+ANK-><-MACRODOMAIN<-?<-PglZ<-DUF6079*\|\|Classical-AAA->Peptidase_S8-><-?<-N6-METHYLASE<-Phage_integrase\|\|N6-METHYLASE->ASCH-><-?<-PglW | 1198 | Bacteria>Terrabacteria_group>Candidatus_Eremiobacterota>unclassified_Candidatus_Eremiobacteraeota | GCA_019637395.1 | Candidatus_Eremiobacteraeota_bacterium | KF760_12090 | 289709..293305 | 2052317 | phage-resistance-protein-[Candidatus-Eremiobacteraeota-bacterium] |
| MCK6564971.1 | Type2.Pgl.like.BREX | DUF6079*->PglZ->BrxD-AAA->SF1-SF2-Helicase+wHTH+Small-B-barrel+DUF1998->N6-N4-METHYLASE->\|\|ParB+HNH->HEPN-><-wHTH+WYL+WCX | 1239 | Bacteria>Terrabacteria_group>Chloroflexota>Dehalococcoidia>unclassified_Dehalococcoidia | GCA_023151015.1 | Dehalococcoidia_bacterium | L6Q80_09490 | 238..3957 | 2026734 | hypothetical-protein-[Dehalococcoidia-bacterium] |
| WP_230840664.1 | Type2.Pgl.like.BREX | <-Thymidylate_kin\|\|?-><-?\|\|ABC-ATPase+SbcC->?-><-PglZ<-DUF6079*\|\|?->DDE_Tnp_1-><-DUF4326<-DUF488-s<-?<-DDE_Tnp_ISL3<-zf-ISL3+HTH_Tnp_ISL3<-N6-METHYLASE-p<-PglW | 1226 | Bacteria>Terrabacteria_group>Cyanobacteriota/Melainabacteria_group>Cyanobacteriota>Cyanophyceae>Gloeobacterales | GCF_021018745.1 | Gloeobacter_morelensis | ISF26_RS17800 | 3706277..3709957 | 2907343 | hypothetical-protein-[Gloeobacter-morelensis] |
| MCA6572687.1 | Type2.Pgl.like.BREX | TrkH->?->?-><-?\|\|?->?-><-?<-LexA-protease<-PglZ<-DUF6079*<-PIN<-wHTH<-Phd-YefM<-N6-METHYLASE\|\|BrnT_toxin->CopG_antitoxin-> | 1268 | Bacteria>Terrabacteria_group>Cyanobacteriota/Melainabacteria_group>Cyanobacteriota>Cyanophyceae>Pseudanabaenales | GCA_020404105.1 | Pseudanabaena_sp._M53BS1SP1A06MG | IM549_05450 | 57481..61287 | 2771239 | phage-resistance-protein-[Pseudanabaena-sp.-M53BS1SP1A06MG] |
| MCA6589826.1 | Type2.Pgl.like.BREX | <-CopG_antitoxin<-BrnT_toxin\|\|N6-METHYLASE->Phd-YefM->wHTH->PIN->DUF6079*->PglZ->LexA-protease-> | 1268 | Bacteria>Terrabacteria_group>Cyanobacteriota/Melainabacteria_group>Cyanobacteriota>Cyanophyceae>Pseudanabaenales | GCA_020404125.1 | Pseudanabaena_sp._M109S1SP1A06QC | IM522_11420 | 30251..34057 | 2771234 | phage-resistance-protein-[Pseudanabaena-sp.-M109S1SP1A06QC] |
| MCA6595660.1 | Type2.Pgl.like.BREX | TrkH->?->?-><-?<-LexA-protease<-PglZ<-DUF6079*<-PIN<-wHTH<-Phd-YefM<-N6-METHYLASE\|\|BrnT_toxin-> | 1271 | Bacteria>Terrabacteria_group>Cyanobacteriota/Melainabacteria_group>Cyanobacteriota>Cyanophyceae>Pseudanabaenales | GCA_020405005.1 | Pseudanabaena_sp._M046S1SP1A06QC | IM539_04925 | 45461..49276 | 2771236 | phage-resistance-protein-[Pseudanabaena-sp.-M046S1SP1A06QC] |
| WP_068818407.1 | Type2.Pgl.like.BREX | CHTH->?->?-><-ACET<-?<-?<-?\|\|?-><-PglZ<-DUF6079*<-REase<-HEPN<-PolB<-N6-METHYLASE<-?<-REase<-PglW\|\|HTH_Tnp_4+DDE_Tnp_4-> | 1257 | Bacteria>Terrabacteria_group>Cyanobacteriota/Melainabacteria_group>Cyanobacteriota>Cyanophyceae>Pseudanabaenales | GCF_001650195.1 | Phormidesmis_priestleyi | A8707_RS17055 | 19871..23644 | 268141 | DUF6079-family-protein-[Phormidesmis-priestleyi] |
| WP_121970611.1 | Type2.Pgl.like.BREX | Pkinase+STYKIN->?-><-TPR+TPR+TPR<-ABC-ATPase+SbcC<-?<-PglZ<-DUF6079*<-PemK_toxin<-?<-N6-METHYLASE<-REase<-PglW | 1229 | Bacteria>Terrabacteria_group>Cyanobacteriota/Melainabacteria_group>Cyanobacteriota>Cyanophyceae>Pseudanabaenales | GCF_002286735.1 | Leptolyngbya_sp._BC1307 | CKL79_RS16140 | 33982..37671 | 2029589 | phage-resistance-protein-[Leptolyngbya-sp.-BC1307] |
| WP_142653836.1 | Type2.Pgl.like.BREX | N6-METHYLASE-p-><-nSTAND1+HNH\|\|N6-METHYLASE-p->Phd-YefM->PIN-><-REase<-REase\|\|REase->DUF6079*->PglZ-><-VIT+VWA | 1259 | Bacteria>Terrabacteria_group>Cyanobacteriota/Melainabacteria_group>Cyanobacteriota>Cyanophyceae>Pseudanabaenales | GCF_006937785.1 | Pseudanabaena_sp._UWO311 | PseudUWO311_RS04160 | 162300..166079 | 2487337 | phage-resistance-protein-[Pseudanabaena-sp.-UWO311] |
| WP_017326151.1 | Type2.Pgl.like.BREX | MODE-HTH+PBPII->?->?-><-PglZ<-DUF6079*<-?<-HEPN<-PolB<-N6-METHYLASE<-PglW\|\|?-><-NUDIX\|\|?-><-SKI | 1246 | Bacteria>Terrabacteria_group>Cyanobacteriota/Melainabacteria_group>Cyanobacteriota>Cyanophyceae>Synechococcales | GCF_000332275.1 | Synechococcus_sp._PCC_7336 | SYN7336_RS11805 | 2475439..2479179 | 195250 | hypothetical-protein-[Synechococcus-sp.-PCC-7336] |
| WP_020876283.1 | Type2.Pgl.like.BREX | SIR2-><-?<-?\|\|?-><-?<-?<-DUF499<-DUF6079-p*<-?<-PolB<-N6-METHYLASE-p<-N6-METHYLASE-p<-REase\|\|ARG-><-tRNA<-?<-?<-RADICAL-SAM+TRAM | 103 | Bacteria>Thermodesulfobacteriota>Desulfobacteria>Desulfobacterales>Desulfococcaceae>Desulfococcus | GCF_000422185.1 | Desulfococcus_multivorans | DSMV_RS06230 | 93711..94022 | 897 | hypothetical-protein-[Desulfococcus-multivorans] |
| RLJ22034.1 | Type2.Pgl.like.BREX | <-RepB_primase<-XIS-HTH\|\|HTH_26->?->Couple_hipA+HipA_C->HTH_26->PglW->N6-METHYLASE->PolB->HEPN->DUF6079*->PglZ->BrxD-AAA->SF1-SF2-Helicase+wHTH+Small-B-barrel+DUF1998->ParB+HNH-> | 1215 | Bacteria>unclassified_Bacteria | GCA_003660225.1 | bacterium_endosymbiont_of_Escarpia_laminata | DJ030_02855 | 37781..41428 | 1221601 | phage-resistance-protein-[bacterium-endosymbiont-of-Escarpia-laminata] |
| MBN9519352.1 | Type2.Pgl.like.BREX | <-SF1-SF2-Helicase+wHTH+Small-B-barrel+DUF1998<-BrxD-AAA<-PglZ<-DUF6079*<-N6-METHYLASE<-PglW\|\|?->HKD-DNase+HKD-Rease->SF1-SF2-Helicase->?->?-><-MORC+DUF3684+REase\|\|ArdCN+MPTase-> | 1230 | Bacteria>unclassified_Bacteria | GCA_017308215.1 | bacterium | J0H58_12655 | 35569..39261 | 1869227 | hypothetical-protein-[bacterium] |
| MBO9539709.1 | Type2.Pgl.like.BREX | Peptidase->?->EVE-><-SF1-SF2-Helicase+wHTH+Small-B-barrel+DUF1998<-BrxD-AAA<-PglZ<-DUF6079*<-N6-METHYLASE<-PglW<-REase<-McrC-NTD+REase<-HTH+McrB-AAA<-ARG<-35exo<-SF1-SF2-Helicase+REase<-Exonuc_V_gamma+REase | 1211 | Bacteria>unclassified_Bacteria | GCA_017744355.1 | bacterium | J7643_03855 | 785728..789363 | 1869227 | phage-resistance-protein-[bacterium] |
| MCR9292939.1 | Type2.Pgl.like.BREX | <-SIGMA-HTH<-?<-SAM-METHYLASE<-?<-KAP-NTPase<-SF1-SF2-Helicase+wHTH+Small-B-barrel+DUF1998<-?<-BrxD-AAA<-PglZ<-DUF6079*<-N6-METHYLASE<-?<-PglW-p | 1230 | Bacteria>unclassified_Bacteria | GCA_024743355.1 | bacterium | NXI32_09495 | 14350..18042 | 1869227 | hypothetical-protein-[bacterium] |
| MBP0304453.1 | Type2.Pgl.like.BREX | <-HHH_4+HTH+wHTH+SF1-SF2-Helicase+REase-p<-HHH_4+HTH+wHTH+SF1-SF2-Helicase+REase-p<-HNH<-OLD-ABC<-SF1-SF2-Helicase<-XIS-HTH<-SF1-SF2-Helicase+wHTH+Small-B-barrel+DUF1998<-BrxD-AAA<-PglZ<-DUF6079*<-N6-METHYLASE<-PglW<-tRNA<-MerR-HTH | 1225 | - | - | - | P1J24_RS13185 | 18168..21845 | - | phage-resistance-protein-[Pseudomonas-aeruginosa] |
| ABP45679.1 | Type2.Pgl.like.BREX | HTH_21+rve+rve_2-><-?<-McrC-NTD+REase<-McrB-AAA-ATPase<-SF1-SF2-Helicase+wHTH+Small-B-barrel+DUF1998<-BrxD-AAA<-PglZ<-DUF6079*<-N6-METHYLASE<-PglW<-ZnR<-?<-HTH<-HigB<-RelE-ParE | 1230 | Bacteria>Terrabacteria_group>Actinomycetota>Actinomycetes>Mycobacteriales | - | Mycolicibacterium_gilvum_PYR-GCK | Mflv_3202 | 3392986..3396678 | 350054 | putative-phage-resistance-protein-[Mycolicibacterium-gilvum-PYR-GCK] |
| AEK37027.1 | Type2.Pgl.like.BREX | PglW->?->DUF6079*-> | 1304 | Bacteria>Terrabacteria_group>Actinomycetota>Actinomycetes>Mycobacteriales | GCA_000179395.2 | Corynebacterium_variabile_DSM_44702 | CVAR_1675 | 1922130..1926044 | 858619 | putative-phage-resistance-protein-[Corynebacterium-variabile-DSM-44702] |
| ALO97239.1 | Type2.Pgl.like.BREX | DUF4096+DDE_Tnp_1+DDE_Tnp_1_2->?->HEPN->?-><-SF1-SF2-Helicase+wHTH+Small-B-barrel+DUF1998<-BrxD-AAA<-PglZ<-DUF6079*<-N6-METHYLASE<-PglW<-?<-?<-DCM | 1299 | Bacteria>Terrabacteria_group>Actinomycetota>Actinomycetes>Kitasatosporales | GCA_001447075.1 | Streptomyces_hygroscopicus_subsp._limoneus | SHL15_6201 | 6999502..7003401 | 264445 | Bacteriophage-resistance-protein-[Streptomyces-hygroscopicus-subsp.-limoneus] |

| Protein-ID | System | Gene-Neighbourhoods | Length | Taxonomy | Assembly-ID | Strain | Locus-Tag | Genomic-coordinates | TaxID | Fasta-define |
|---|---|---|---|---|---|---|---|---|---|---|
| AMY54747.1 | Type2.Pgl.like.BREX | HATPase->REase->?->PglW->N6-METHYLASE->DUF6079-p->N6-METHYLASE->DUF6079*->PglZ->BrxD-AAA->SF1-SF2-Helicase+wHTH+Small-B-barrel+DUF1998-><-SF1-SF2-Helicase<-wHTH<-LON-AAA<-REase | 1290 | Bacteria>Terrabacteria_group>Actinomycetota>Actinomycetes>Mycobacteriales | GCA_001620305.1 | Rhodococcus_fascians_D188 | A3L23_03422 | 3544457..3548329 | 1051973 | Chromosome-partition-protein-Smc-[Rhodococcus-fascians-D188] |
| ASN19310.1 | Type2.Pgl.like.BREX | <-DUF6079*<-?<-?<-PglW | 1258 | Bacteria>Terrabacteria_group>Actinomycetota>Actinomycetes>Micrococcales | GCA_002224285.1 | Arthrobacter_sp._YN | CGK93_06110 | 1322963..1326739 | 2020486 | phage-resistance-protein-[Arthrobacter-sp.-YN] |
| AVH59039.1 | Type2.Pgl.like.BREX | HEPN+VirB11+NACHT+FGS->?->?-><-?<-DUF6731\|\|PglW->N6-METHYLASE->DUF6079*->PglZ->BrxD-AAA->SF1-SF2-Helicase+wHTH+Small-B-barrel+DUF1998-><-?\|\|?-><-Phage_int_SAM_3+Phage_integrase\|\|HTH-> | 1312 | Bacteria>Terrabacteria_group>Actinomycetota>Actinomycetes>Kitasatosporales | GCA_002946835.1 | Streptomyces_dengpaensis | C4B68_28490 | 6170305..6174243 | 2049881 | PglY-protein-[Streptomyces-dengpaensis] |
| AXY52317.1 | Type2.Pgl.like.BREX | STYKinase-like->?->PglW->N6-METHYLASE-><-?\|\|DUF6079*->PglZ->BrxD-AAA->SF1-SF2-Helicase-><-?\|\|SF1-SF2-Helicase+wHTH+Small-B-barrel+DUF1998-> | 1235 | Bacteria>Terrabacteria_group>Actinomycetota>Actinomycetes>Mycobacteriales | GCA_003586525.1 | Rhodococcus_ruber | YT1_2908 | 2910213..2913920 | 1830 | putative-phage-resistance-protein-[Rhodococcus-ruber] |
| BAN04394.1 | Type2.Pgl.like.BREX | PglW->N6-METHYLASE->DUF6079*->PglZ-><-PIN\|\|BrxD-AAA->SF1-SF2-Helicase+wHTH+Small-B-barrel+DUF1998->CASPASE+SF1-SF2-Helicase-><-SF1-SF2-Helicase | 1203 | Bacteria>Terrabacteria_group>Actinomycetota>Acidimicrobiia>Acidimicrobiales | GCA_000348785.1 | Ilumatobacter_coccineus_YM16-304 | YM304_40800 | 4610450..4614061 | 1313172 | putative-phage-resistance-protein-PglY-[Ilumatobacter-coccineus-YM16-304] |
| BCL27147.1 | Type2.Pgl.like.BREX | PglW->?->DUF6079*-> | 1304 | Bacteria>Terrabacteria_group>Actinomycetota>Actinomycetes>Kitasatosporales | GCA_014701115.1 | Streptomyces_aurantiacus | GCM10017557_20060 | 2395233..2399147 | 47760 | hypothetical-protein-GCM10017557_20060-[Streptomyces-aurantiacus] |
| BDH15728.1 | Type2.Pgl.like.BREX | PglW-><-BrxD-AAA<-ParA-Soj-PloopNTPase<-HEPN\|\|N6-METHYLASE-><-HTH_31+DUF5753\|\|ST-Hiskin->DUF6079*->PglZ->BrxD-AAA->SF1-SF2-Helicase+wHTH+Small-B-barrel+DUF1998-><-?<-NUDIX | 1324 | Bacteria>Terrabacteria_group>Actinomycetota>Actinomycetes>Kitasatosporales | GCA_022846675.1 | Streptomyces_hygroscopicus | HOK021_69070 | 7777796..7781770 | 1912 | hypothetical-protein-HOK021_69070-[Streptomyces-hygroscopicus] |
| BFE41879.1 | Type2.Pgl.like.BREX | PglW->N6-METHYLASE->DUF6079*->PglZ->BrxD-AAA->SF1-SF2-Helicase+wHTH+Small-B-barrel+DUF1998->?->SF1-SF2-Helicase->?->ParB+HNH-> | 1213 | Bacteria>Terrabacteria_group>Actinomycetota>Actinomycetes>Micrococcales | - | Microbacterium_imperiale | GCM10017544_28350 | 2932208..2935849 | 33884 | hypothetical-protein-GCM10017544_28350-[Microbacterium-imperiale] |
| BFE56536.1 | Type2.Pgl.like.BREX | SNF+SF1-SF2-Helicase->SF1-SF2-Helicase+REase->?->REase-><-SF1-SF2-Helicase+wHTH+Small-B-barrel+DUF1998<-BrxD-AAA<-PglZ<-DUF6079*<-N6-METHYLASE<-PglW\|\|?-><-?<-Phage_integrase | 1228 | Bacteria>Terrabacteria_group>Actinomycetota>Actinomycetes>Micromonosporales | - | Dactylosporangium_thailandense | GCM10020063_010620 | 1217275..1220961 | 47849 | hypothetical-protein-GCM10020063_010620-[Dactylosporangium-thailandense] |
| CAM04375.1 | Type2.Pgl.like.BREX | <-HTH_21+rve+rve_2<-HIN-HTH<-REase+SAVED\|\|rve_3-><-SF1-SF2-Helicase+wHTH+Small-B-barrel+DUF1998<-BrxD-AAA<-PglZ<-DUF6079*<-N6-METHYLASE<-PglW\|\|SF1-SF2-Helicase+REase->HTH_29+HTH_32+DDE_3-><-tRNA\|\|?-><-?<-ABhydrolase | 1273 | Bacteria>Terrabacteria_group>Actinomycetota>Actinomycetes>Pseudonocardiales | GCA_000062885.1 | Saccharopolyspora_erythraea_NRRL_2338 | SACE_5129 | 5720356..5724177 | 405948 | bacteriophage-(phiC31)-resistance-gene-PglY-[Saccharopolyspora-erythraea-NRRL-2338] |
| CTQ94910.1 | Type2.Pgl.like.BREX | <-SF1-SF2-Helicase+wHTH+Small-B-barrel+DUF1998<-BrxD-AAA<-PglZ<-DUF6079*<-N6-METHYLASE<-PglW<-REase | 1227 | Bacteria>Terrabacteria_group>Actinomycetota>Actinomycetes>Pseudonocardiales | - | Kibdelosporangium_sp._MJ126-NF4 | . | 7089226..7092909 | 703222 | Bacteriophage-(PhiC31)-resistance-gene-pglY-[Kibdelosporangium-sp.-MJ126-NF4] |
| EAR23329.1 | Type2.Pgl.like.BREX | REase-><-?\|\|PglW->N6-METHYLASE->?->PIN->DUF6079*->PglZ->BrxD-AAA->SF1-SF2-Helicase+wHTH+Small-B-barrel+DUF1998-><-SAM-methylase\|\|?->PIN-> | 1222 | Bacteria>Pseudomonadota>Gammaproteobacteria>Chromatiales>Ectothiorhodospiraceae | GCA_000153205.1 | Nitrococcus_mobilis_Nb-231 | NB231_15953 | 3295318..3298986 | 314278 | pglY-[Nitrococcus-mobilis-Nb-231] |
| EME66622.1 | Type2.Pgl.like.BREX | <-SF1-SF2-Helicase+wHTH+Small-B-barrel+DUF1998\|\|?-><-?<-BrxD-AAA<-PglZ<-DUF6079*\|\|?-><-DOC<-N6-METHYLASE<-PglW<-?<-STYKinase-like | 1231 | Bacteria>Terrabacteria_group>Actinomycetota>Actinomycetes>Mycobacteriales | GCA_000341965.1 | Rhodococcus_ruber_BKS_20-38 | G352_04031 | 14064..17759 | 1278076 | putative-phage-resistance-protein-[Rhodococcus-ruber-BKS-20-38] |
| EQD84160.1 | Type2.Pgl.like.BREX | <-REase+SAVED\|\|rve_3-><-SF1-SF2-Helicase+wHTH+Small-B-barrel+DUF1998<-BrxD-AAA<-PglZ<-DUF6079*<-N6-METHYLASE<-PglW\|\|SF1-SF2-Helicase+REase->HTH_29+HTH_32+DDE_3-> | 1273 | Bacteria>Terrabacteria_group>Actinomycetota>Actinomycetes>Pseudonocardiales | GCA_000448385.1 | Saccharopolyspora_erythraea_D | N599_21475 | 12158..15979 | 1382595 | hypothetical-protein-N599_21475-[Saccharopolyspora-erythraea-D] |
| ESQ06628.1 | Type2.Pgl.like.BREX | PglW->N6-METHYLASE-><-DUF397<-HTH_31+DUF5753\|\|ST-Hiskin->DUF6079*->PglZ->BrxD-AAA->SF1-SF2-Helicase+wHTH+Small-B-barrel+DUF1998-><-TRAM+SAM-methylase+METHYLASE<-AA_permease_2 | 1298 | Bacteria>Terrabacteria_group>Actinomycetota>Actinomycetes>Kitasatosporales | GCA_000495755.1 | Streptomyces_sp._PVA_94-07 | B590_05664 | 1375843..1379739 | 1225337 | bacteriophage-resistance-protein-[Streptomyces-sp.-PVA_94-07] |
| GAA0245427.1 | Type2.Pgl.like.BREX | <-TraJ-RHH<-?\|\|?-><-SF1-SF2-Helicase+wHTH+Small-B-barrel+DUF1998<-BrxD-AAA<-PglZ<-DUF6079*<-N6-METHYLASE<-?<-?\|\|HTH_31+DUF5753->DUF397-><-PglW | 1303 | Bacteria>Terrabacteria_group>Actinomycetota>Actinomycetes>Streptosporangiales | GCA_039524295.1 | Actinomadura_nitritigenes | GCM10009527_047640 | 51744..55655 | 134602 | hypothetical-protein-GCM10009527_047640-[Actinomadura-nitritigenes] |
| GAA0438063.1 | Type2.Pgl.like.BREX | REase-><-PglZ<-DUF6079*<-N6-METHYLASE<-PglW\|\|?->REase->REase-> | 1299 | Bacteria>Terrabacteria_group>Actinomycetota>Actinomycetes>Kitasatosporales | GCA_039521205.1 | Streptomyces_luteireticuli | GCM10010357_69310 | 4022..7921 | 173858 | hypothetical-protein-GCM10010357_69310-[Streptomyces-luteireticuli] |
| GAA0539957.1 | Type2.Pgl.like.BREX | <-SF1-SF2-Helicase+HKD-DNase\|\|PglW->N6-METHYLASE-><-ST-Hiskin\|\|HTH_31+DUF5753->DUF397->DUF6079*->PglZ-><-CBS+HEPN<-ParB+HNH\|\|BrxD-AAA->SF1-SF2-Helicase+wHTH+Small-B-barrel+DUF1998->SAD+REase-> | 1321 | Bacteria>Terrabacteria_group>Actinomycetota>Actinomycetes>Kitasatosporales | GCA_039521245.1 | Streptomyces_mordarskii | GCM10010390_47430 | 363832..367797 | 1226758 | hypothetical-protein-GCM10010390_47430-[Streptomyces-mordarskii] |
| GAA0739616.1 | Type2.Pgl.like.BREX | <-SF1-SF2-Helicase\|\|SNF+SF1-SF2-Helicase-><-?\|\|?-><-ParB\|\|PglW->SNF+SF1-SF2-Helicase->DUF6079*->PglZ->BrxD-AAA->SF1-SF2-Helicase+wHTH+Small-B-barrel+DUF1998-><-REase\|\|XIS-HTH-><-?<-Phage_integrase | 1237 | Bacteria>Terrabacteria_group>Actinomycetota>Actinomycetes>Micromonosporales | GCA_039521105.1 | Dactylosporangium_roseum | GCM10010199_63760 | 50168..53881 | 47989 | hypothetical-protein-GCM10010199_63760-[Dactylosporangium-roseum] |
| GAA0926818.1 | Type2.Pgl.like.BREX | <-HNH<-SF1-SF2-Helicase\|\|PglW->N6-METHYLASE->?->NACHT+REase-DUF3883->ParB+HNH->DUF6079*->PglZ->BrxD-AAA->SF1-SF2-Helicase+wHTH+Small-B-barrel+DUF1998-> | 1228 | Bacteria>Terrabacteria_group>Actinomycetota>Actinomycetes>Pseudonocardiales | GCA_039525585.1 | Pseudonocardia_zijingensis | GCM10009559_12190 | 45144..48830 | 153376 | hypothetical-protein-GCM10009559_12190-[Pseudonocardia-zijingensis] |
| GAA1025581.1 | Type2.Pgl.like.BREX | <-BrxD-AAA<-PglZ<-DUF6079*\|\|?->?->?-><-N6-METHYLASE-p<-N6-METHYLASE<-PglW<-Phage_int_SAM_3+Phage_integrase | 1222 | Bacteria>Terrabacteria_group>Actinomycetota>Actinomycetes>Micromonosporales | GCA_039525495.1 | Virgisporangium_ochraceum | GCM10009557_00030 | 5732..9400 | 65505 | hypothetical-protein-GCM10009557_00030-[Virgisporangium-ochraceum] |
| GAA1089909.1 | Type2.Pgl.like.BREX | REase->SF1-SF2-Helicase-><-?<-?<-SF1-SF2-Helicase+wHTH+Small-B-barrel+DUF1998<-BrxD-AAA<-PglZ<-DUF6079*<-N6-METHYLASE<-PglW<-?<-tRNA<-His_Phos_1 | 1261 | Bacteria>Terrabacteria_group>Actinomycetota>Actinomycetes>Propionibacteriales | GCA_039528155.1 | Nocardioides_dubius | GCM10009668_00430 | 62597..66382 | 317019 | hypothetical-protein-GCM10009668_00430-[Nocardioides-dubius] |
| GAA1254424.1 | Type2.Pgl.like.BREX | <-SF1-SF2-Helicase+HKD-DNase\|\|PglW->N6-METHYLASE-><-ST-Hiskin\|\|HTH_31+DUF5753->DUF397->DUF6079*->PglZ-><-CBS+HEPN<-ParB+HNH\|\|BrxD-AAA->SF1-SF2-Helicase+wHTH+Small-B-barrel+DUF1998-><-REase+SF1-SF2-Helicase\|\|REase+SF1-SF2-Helicase-> | 1320 | Bacteria>Terrabacteria_group>Actinomycetota>Actinomycetes>Kitasatosporales | GCA_039525965.1 | Streptomyces_yogyakartensis | GCM10009579_10490 | 273908..277870 | 114692 | hypothetical-protein-GCM10009579_10490-[Streptomyces-yogyakartensis] |
| GAA1393352.1 | Type2.Pgl.like.BREX | PglW->?->DUF6079*-> | 1247 | Bacteria>Terrabacteria_group>Actinomycetota>Actinomycetes>Propionibacteriales | GCA_039526385.1 | Luteococcus_peritonei | GCM10009599_16100 | 160074..163817 | 88874 | hypothetical-protein-GCM10009599_16100-[Luteococcus-peritonei] |
| GAA1555978.1 | Type2.Pgl.like.BREX | HIN-HTH-><-SF1-SF2-Helicase+wHTH+Small-B-barrel+DUF1998<-BrxD-AAA\|\|ParB+HNH->CBS+HEPN-><-PglZ<-DUF6079*<-?<-?\|\|HTH_31+DUF5753->DUF397-><-N6-METHYLASE<-PglW | 1298 | Bacteria>Terrabacteria_group>Actinomycetota>Actinomycetes>Kitasatosporales | GCA_039529415.1 | Streptomyces_globosus | GCM10009731_07930 | 81890..85786 | 68209 | hypothetical-protein-GCM10009731_07930-[Streptomyces-globosus] |
| GAA1575606.1 | Type2.Pgl.like.BREX | <-GNTR-HTH\|\|?->?->?->?->PglW->N6-METHYLASE->DUF6079*->PglZ->BrxD-AAA->SF1-SF2-Helicase+wHTH+Small-B-barrel+DUF1998-><-REase<-?<-Phage_int_SAM_3+Phage_integrase | 1246 | Bacteria>Terrabacteria_group>Actinomycetota>Actinomycetes>Micromonosporales | GCA_039531485.1 | Dactylosporangium_maewongense | GCM10009827_116810 | 18969..22709 | 634393 | hypothetical-protein-GCM10009827_116810-[Dactylosporangium-maewongense] |
| GAA1828696.1 | Type2.Pgl.like.BREX | PglW->?->DUF6079*-> | 1308 | Bacteria>Terrabacteria_group>Actinomycetota>Actinomycetes>Streptosporangiales | GCA_039529475.1 | Actinomadura_chokoriensis | GCM10009735_75380 | 10389..14315 | 454156 | hypothetical-protein-GCM10009735_75380-[Actinomadura-chokoriensis] |
| GAA1837104.1 | Type2.Pgl.like.BREX | <-ParB+HNH<-SF1-SF2-Helicase+wHTH+Small-B-barrel+DUF1998<-BrxD-AAA<-PglZ<-DUF6079*<-N6-METHYLASE<-PglW<-ParB+HNH\|\|?->CASPASE+SF1-SF2-Helicase-> | 1229 | Bacteria>Terrabacteria_group>Actinomycetota>Actinomycetes>Pseudonocardiales | GCA_039530145.1 | Pseudonocardia_carboxydivorans | GCM10009772_13250 | 35431..39120 | 415010 | hypothetical-protein-GCM10009772_13250-[Pseudonocardia-carboxydivorans] |
| GAA2269146.1 | Type2.Pgl.like.BREX | REase+SF1-SF2-Helicase->?->PglW->N6-METHYLASE->DUF6079*->PglZ->BrxD-AAA->SF1-SF2-Helicase+wHTH+Small-B-barrel+DUF1998-><-SF1-SF2-Helicase | 1234 | Bacteria>Terrabacteria_group>Actinomycetota>Actinomycetes>Glycomycetales | GCA_039531935.1 | Glycomyces_scopariae | GCM10009853_023280 | 496758..500462 | 480997 | hypothetical-protein-GCM10009853_023280-[Glycomyces-scopariae] |
| GAA2342654.1 | Type2.Pgl.like.BREX | <-SF1-SF2-Helicase+wHTH+Small-B-barrel+DUF1998<-BrxD-AAA\|\|ParB+HNH->CBS+HEPN-><-PglZ<-DUF6079*<-DUF397<-DUF397<-HTH_31+DUF5753\|\|ST-Hiskin-><-N6-METHYLASE<-PglW\|\|SF1-SF2-Helicase+HKD-DNase-> | 1313 | Bacteria>Terrabacteria_group>Actinomycetota>Actinomycetes>Kitasatosporales | GCA_039532705.1 | Streptomyces_cuspidosporus | GCM10010246_30030 | 66393..70334 | 66882 | hypothetical-protein-GCM10010246_30030-[Streptomyces-cuspidosporus] |
| GAA2354795.1 | Type2.Pgl.like.BREX | rve_3-><-PglZ<-DUF6079*<-N6-METHYLASE<-PglW\|\|BrxD-AAA->SF1-SF2-Helicase+wHTH+Small-B-barrel+DUF1998-> | 1279 | Bacteria>Terrabacteria_group>Actinomycetota>Actinomycetes>Kitasatosporales | GCA_039532925.1 | Streptomyces_violaceusniger | GCM10010376_91980 | 4742..8581 | 68280 | hypothetical-protein-GCM10010376_91980-[Streptomyces-violaceusniger] |
| GAA2369392.1 | Type2.Pgl.like.BREX | <-ParB\|\|PglW->N6-METHYLASE->DUF6079*->PglZ->BrxD-AAA->SF1-SF2-Helicase+wHTH+Small-B-barrel+DUF1998-><-REase\|\|XIS-HTH-> | 1237 | Bacteria>Terrabacteria_group>Actinomycetota>Actinomycetes>Micromonosporales | GCA_039532525.1 | Dactylosporangium_salmoneum | GCM10010170_069770 | 10412..14125 | 53361 | hypothetical-protein-GCM10010170_069770-[Dactylosporangium-salmoneum] |
| GAA2411559.1 | Type2.Pgl.like.BREX | <-DUF6079*<-?<-PglW | 1297 | Bacteria>Terrabacteria_group>Actinomycetota>Actinomycetes>Kitasatosporales | GCA_039533345.1 | Streptomyces_glaucosporus | GCM10010420_45640 | 9336..13229 | 284044 | hypothetical-protein-GCM10010420_45640-[Streptomyces-glaucosporus] |
| GAA2489816.1 | Type2.Pgl.like.BREX | PglW->?-><-?<-?\|\|?->?->?->DUF6079*-> | 1307 | Bacteria>Terrabacteria_group>Actinomycetota>Actinomycetes>Kitasatosporales | GCA_039532985.1 | Streptomyces_gobitricini | GCM10010393_21890 | 267541..271464 | 68211 | hypothetical-protein-GCM10010393_21890-[Streptomyces-gobitricini] |
| GAA2535801.1 | Type2.Pgl.like.BREX | <-SF1-SF2-Helicase+wHTH+Small-B-barrel+DUF1998<-BrxD-AAA<-PglZ<-DUF6079*<-N6-METHYLASE<-PglW<-?<-HTH | 1213 | Bacteria>Terrabacteria_group>Actinomycetota>Actinomycetes>Micrococcales | GCA_039532075.1 | Microbacterium_mitrae | GCM10009860_15300 | 1674826..1678467 | 664640 | hypothetical-protein-GCM10009860_15300-[Microbacterium-mitrae] |
| GAA2711449.1 | Type2.Pgl.like.BREX | CASPASE+SF1-SF2-Helicase->PglW->N6-METHYLASE->DUF6079*->PglZ-><-REase\|\|BrxD-AAA->SF1-SF2-Helicase+wHTH+Small-B-barrel+DUF1998-><-TRAM+SAM-methylase+METHYLASE<-AA_permease_2 | 1299 | Bacteria>Terrabacteria_group>Actinomycetota>Actinomycetes>Kitasatosporales | GCA_039532865.1 | Streptomyces_luteosporeus | GCM10010315_13380 | 224428..228327 | 173856 | hypothetical-protein-GCM10010315_13380-[Streptomyces-luteosporeus] |
| GAA2935464.1 | Type2.Pgl.like.BREX | GTPase-AIG+sGTP->MMR_HSR1->PglW->N6-METHYLASE->DUF6079*->PglZ->BrxD-AAA->SF1-SF2-Helicase+wHTH+Small-B-barrel+DUF1998->?-><-N6-METHYLASE | 1297 | Bacteria>Terrabacteria_group>Actinomycetota>Actinomycetes>Kitasatosporales | GCA_039534465.1 | Kitasatospora_cinereorecta | GCM10010518_23030 | 146197..150090 | 285560 | hypothetical-protein-GCM10010518_23030-[Kitasatospora-cinereorecta] |
| GAA3213060.1 | Type2.Pgl.like.BREX | <-SF1-SF2-Helicase\|\|?-><-SF1-SF2-Helicase+wHTH+Small-B-barrel+DUF1998<-BrxD-AAA<-PglZ<-DUF6079*<-N6-METHYLASE<-PglW | 1218 | Bacteria>Terrabacteria_group>Actinomycetota>Actinomycetes>Micrococcales | GCA_039534975.1 | Microbacterium_terregens | GCM10017690_25860 | 2793768..2797424 | 69363 | hypothetical-protein-GCM10017690_25860-[Microbacterium-terregens] |
| GAA3473660.1 | Type2.Pgl.like.BREX | DUF4096+DDE_Tnp_1+DDE_Tnp_1_2->DDE_Tnp_1_2-><-?\|\|N6-METHYLASE-><-SF1-SF2-Helicase+wHTH+Small-B-barrel+DUF1998<-BrxD-AAA<-PglZ<-DUF6079*<-N6-METHYLASE<-PglW\|\|ParB+HNH->CBS+HEPN+HEPN-> | 1214 | Bacteria>Terrabacteria_group>Actinomycetota>Actinomycetes>Streptosporangiales | GCA_039535395.1 | Nonomuraea_roseola | GCM10018965_082130 | 3620564..3624208 | 46179 | hypothetical-protein-GCM10018965_082130-[Nonomuraea-roseola] |
| GAA3523410.1 | Type2.Pgl.like.BREX | PglW->N6-METHYLASE-><-ST-Hiskin\|\|HTH_31+DUF5753->DUF397->?->DUF6079*->PglZ-><-CBS+HEPN<-ParB+HNH\|\|BrxD-AAA->SF1-SF2-Helicase+wHTH+Small-B-barrel+DUF1998-> | 1308 | Bacteria>Terrabacteria_group>Actinomycetota>Actinomycetes>Kitasatosporales | GCA_039538545.1 | Streptomyces_osmaniensis | GCM10022295_01140 | 136238..140164 | 593134 | hypothetical-protein-GCM10022295_01140-[Streptomyces-osmaniensis] |
| GAA3739320.1 | Type2.Pgl.like.BREX | <-SAM-methylase<-ST-Hiskin\|\|HTH_31+DUF5753->DUF397->PglW-><-?\|\|N6-METHYLASE->DUF6079*->PglZ->BrxD-AAA->SF1-SF2-Helicase+wHTH+Small-B-barrel+DUF1998-> | 1298 | Bacteria>Terrabacteria_group>Actinomycetota>Actinomycetes>Streptosporangiales | GCA_039539445.1 | Salinactinospora_qingdaonensis | GCM10022402_18870 | 16540..20436 | 702744 | hypothetical-protein-GCM10022402_18870-[Salinactinospora-qingdaonensis] |
| GAA3754331.1 | Type2.Pgl.like.BREX | <-HKD-DNase<-?<-URI-DNase+SF1-SF2-Helicase<-MazG\|\|?->PglW->N6-METHYLASE->DUF6079*->PglZ->BrxD-AAA->SF1-SF2-Helicase+wHTH+Small-B-barrel+DUF1998-><-HKD-DNase | 1211 | Bacteria>Terrabacteria_group>Actinomycetota>Actinomycetes>Micrococcales | GCA_039537285.1 | Microbacterium_kribbense | GCM10022240_04080 | 449466..453101 | 433645 | hypothetical-protein-GCM10022240_04080-[Microbacterium-kribbense] |
| GAA4256551.1 | Type2.Pgl.like.BREX | ACET->?->REase-><-SF1-SF2-Helicase+wHTH+Small-B-barrel+DUF1998<-BrxD-AAA<-PglZ<-DUF6079*<-N6-METHYLASE<-PglW\|\|?-><-?<-ACET<-HTH | 1240 | Bacteria>Terrabacteria_group>Actinomycetota>Actinomycetes>Micromonosporales | GCA_039537565.1 | Dactylosporangium_darangshiense | GCM10022255_069850 | 134567..138289 | 579108 | hypothetical-protein-GCM10022255_069850-[Dactylosporangium-darangshiense] |
| GAA4260842.1 | Type2.Pgl.like.BREX | <-DUF6079*<-?<-PglW | 1302 | Bacteria>Terrabacteria_group>Actinomycetota>Actinomycetes>Mycobacteriales | GCA_039538505.1 | Dietzia_aurantiaca | GCM10022294_06940 | 17390..21298 | 983873 | hypothetical-protein-GCM10022294_06940-[Dietzia-aurantiaca] |
| GAA4393356.1 | Type2.Pgl.like.BREX | <-DUF6079*<-?<-PglW | 1250 | Bacteria>Terrabacteria_group>Actinomycetota>Actinomycetes>Micrococcales | GCA_039541275.1 | Ornithinibacter_aureus | GCM10023153_13220 | 45478..49230 | 622664 | hypothetical-protein-GCM10023153_13220-[Ornithinibacter-aureus] |
| GAA4479485.1 | Type2.Pgl.like.BREX | D6+SF1-SF2-Helicase->?->PglW->N6-METHYLASE->?-><-HEPN<-ParB+HNH\|\|DUF6079*->PglZ->BrxD-AAA->SF1-SF2-Helicase+wHTH+Small-B-barrel+DUF1998-><-SIGMA-HTH+SIGMA-HTH<-?<-?<-SF1-SF2-Helicase | 1256 | Bacteria>Terrabacteria_group>Actinomycetota>Actinomycetes>Mycobacteriales | GCA_039540555.1 | Rhodococcus_olei | GCM10023094_24680 | 43453..47223 | 2161675 | hypothetical-protein-GCM10023094_24680-[Rhodococcus-olei] |
| GAA4535316.1 | Type2.Pgl.like.BREX | <-SF1-SF2-Helicase+wHTH+Small-B-barrel+DUF1998<-BrxD-AAA<-PglZ<-DUF6079*<-N6-METHYLASE<-PglW | 1204 | Bacteria>Terrabacteria_group>Actinomycetota>Actinomycetes>Pseudonocardiales | GCA_039541715.1 | Pseudonocardia_xishanensis | GCM10023175_00740 | 72067..75681 | 630995 | hypothetical-protein-GCM10023175_00740-[Pseudonocardia-xishanensis] |
| GAA4543128.1 | Type2.Pgl.like.BREX | <-Pkinase+TPR\|\|REase+SF1-SF2-Helicase->REase+SF1-SF2-Helicase-><-?\|\|?->PglW->N6-METHYLASE->DUF6079*->PglZ-><-DUF397<-HTH_31+DUF5753\|\|ST-Hiskin-><-CBS+HEPN<-ParB+HNH | 1298 | Bacteria>Terrabacteria_group>Actinomycetota>Actinomycetes>Kitasatosporales | GCA_039540655.1 | Streptomyces_collinus | GCM10023097_15710 | 621077..624973 | 42684 | hypothetical-protein-GCM10023097_15710-[Streptomyces-collinus] |
| GAA4742170.1 | Type2.Pgl.like.BREX | HNH->ABhydrolase-><-?<-SF1-SF2-Helicase+wHTH+Small-B-barrel+DUF1998<-BrxD-AAA<-PglZ<-DUF6079*<-N6-METHYLASE<-PglW<-?<-HEPN+SF1-SF2-Helicase+REase | 1192 | Bacteria>Terrabacteria_group>Actinomycetota>Actinomycetes>Micrococcales | GCA_039544415.1 | Amnibacterium_soli | GCM10025783_11850 | 395664..399242 | 1282736 | hypothetical-protein-GCM10025783_11850-[Amnibacterium-soli] |
| GAA4855310.1 | Type2.Pgl.like.BREX | <-SF1-SF2-Helicase+wHTH+Small-B-barrel+DUF1998<-BrxD-AAA<-PglZ<-DUF6079*<-N6-METHYLASE<-PglW | 1258 | Bacteria>Terrabacteria_group>Actinomycetota>Actinomycetes>Pseudonocardiales | GCA_039544495.1 | Saccharopolyspora_rosea | GCM10025787_44210 | 6730..10506 | 524884 | hypothetical-protein-GCM10025787_44210-[Saccharopolyspora-rosea] |
| GAA4873509.1 | Type2.Pgl.like.BREX | <-HNH<-REase+SF1-SF2-Helicase\|\|ParB+HNH-><-SF1-SF2-Helicase+wHTH+Small-B-barrel+DUF1998<-BrxD-AAA<-?<-PglZ<-DUF6079*<-ParB+HNH<-ParB+HNH<-N6-METHYLASE<-PglW | 1258 | Bacteria>Terrabacteria_group>Actinomycetota>Actinomycetes>Micrococcales | GCA_039544515.1 | Serinicoccus_chungangensis | GCM10025788_09860 | 605540..609316 | 767452 | hypothetical-protein-GCM10025788_09860-[Serinicoccus-chungangensis] |
| GAA4991782.1 | Type2.Pgl.like.BREX | ParB+HNH->PglW->DOC->N6-METHYLASE->DUF6079*->PglZ->BrxD-AAA->SF1-SF2-Helicase+wHTH+Small-B-barrel+DUF1998-><-SF1-SF2-Helicase+REase\|\|?-><-?\|\|ACET-> | 1247 | Bacteria>Terrabacteria_group>Actinomycetota>Actinomycetes>Pseudonocardiales | GCA_039544595.1 | Pseudonocardia_tropica | GCM10025792_45750 | 212546..216289 | 681289 | hypothetical-protein-GCM10025792_45750-[Pseudonocardia-tropica] |
| GAA5071929.1 | Type2.Pgl.like.BREX | <-Phage_integrase<-?<-SF1-SF2-Helicase+wHTH+Small-B-barrel+DUF1998<-BrxD-AAA<-PglZ<-DUF6079*<-N6-METHYLASE<-PglW\|\|HTH-><-tRNA\|\|?-><-tRNA<-35exo | 1263 | Bacteria>Terrabacteria_group>Actinomycetota>Actinomycetes>Mycobacteriales | GCA_039543325.1 | Nocardia_iowensis | GCM10023319_03500 | 372703..376494 | 204891 | hypothetical-protein-GCM10023319_03500-[Nocardia-iowensis] |
| GES07634.1 | Type2.Pgl.like.BREX | PglW->N6-METHYLASE->DUF6079*->PglZ->BrxD-AAA->SF1-SF2-Helicase+wHTH+Small-B-barrel+DUF1998-><-SF1-SF2-Helicase<-APATPase+TPR+TPR+TPR+TPR+TPR+TPR+TPR<-SF1-SF2-Helicase<-REase+SF1-SF2-Helicase | 1266 | Bacteria>Terrabacteria_group>Actinomycetota>Actinomycetes>Streptosporangiales | GCA_009687865.1 | Acrocarpospora_macrocephala | Amac_012290 | 52601..56401 | 150177 | hypothetical-protein-Amac_012290-[Acrocarpospora-macrocephala] |
| GGF99355.1 | Type2.Pgl.like.BREX | HKD-DNase->D6+SF1-SF2-Helicase->?-><-SF1-SF2-Helicase+wHTH+Small-B-barrel+DUF1998<-BrxD-AAA<-PglZ<-DUF6079*<-N6-METHYLASE<-PglW\|\|PglZ-><-BrxD-AAA<-HKD-DNase | 1212 | Bacteria>Terrabacteria_group>Actinomycetota>Actinomycetes>Mycobacteriales | GCA_014635345.1 | Rhodococcus_trifolii | GCM10007304_11520 | 1232248..1235886 | 908250 | hypothetical-protein-GCM10007304_11520-[Rhodococcus-trifolii] |
| GGL40855.1 | Type2.Pgl.like.BREX | PglW->?->DUF6079*-> | 1249 | Bacteria>Terrabacteria_group>Actinomycetota>Actinomycetes>Mycobacteriales | GCA_014645735.1 | Nocardia_jinanensis | GCM10011588_64570 | 40816..44565 | 382504 | hypothetical-protein-GCM10011588_64570-[Nocardia-jinanensis] |
| GGL87829.1 | Type2.Pgl.like.BREX | <-HNH<-SF1-SF2-Helicase+wHTH+Small-B-barrel+DUF1998<-BrxD-AAA\|\|RNaseH-><-PglZ<-DUF6079*\|\|?-><-?<-?\|\|HTH_31+DUF5753->DUF397-><-N6-METHYLASE<-PglW | 1304 | Bacteria>Terrabacteria_group>Actinomycetota>Actinomycetes>Kitasatosporales | GCA_014647975.1 | Streptomyces_fumigatiscleroticus | GCM10010129_34570 | 289621..293535 | 66371 | hypothetical-protein-GCM10010129_34570-[Streptomyces-fumigatiscleroticus] |
| GGL99858.1 | Type2.Pgl.like.BREX | <-DUF6079*<-?\|\|?->?-><-?<-PglW | 1250 | Bacteria>Terrabacteria_group>Actinomycetota>Actinomycetes>Nakamurellales | GCA_014646215.1 | Nakamurella_endophytica | GCM10011594_19780 | 571447..575199 | 1748367 | hypothetical-protein-GCM10011594_19780-[Nakamurella-endophytica] |
| GGQ83180.1 | Type2.Pgl.like.BREX | <-DUF6079*<-?<-PglW | 1299 | Bacteria>Terrabacteria_group>Actinomycetota>Actinomycetes>Kitasatosporales | GCA_014649075.1 | Streptomyces_asoensis | GCM10010496_53450 | 17626..21525 | 249586 | hypothetical-protein-GCM10010496_53450-[Streptomyces-asoensis] |
| GGU27936.1 | Type2.Pgl.like.BREX | <-SF1-SF2-Helicase+wHTH+Small-B-barrel+DUF1998<-BrxD-AAA<-PglZ<-DUF6079*<-ST-Hiskin<-N6-METHYLASE\|\|HEPN->ParA-Soj-PloopNTPase->BrxD-AAA-><-PglW\|\|DUF2397-> | 1321 | Bacteria>Terrabacteria_group>Actinomycetota>Actinomycetes>Kitasatosporales | GCA_014649475.1 | Streptomyces_daghestanicus | GCM10010259_18110 | 129191..133156 | 66885 | hypothetical-protein-GCM10010259_18110-[Streptomyces-daghestanicus] |
| GID29526.1 | Type2.Pgl.like.BREX | PIN->?-><-SF1-SF2-Helicase+wHTH+Small-B-barrel+DUF1998<-BrxD-AAA<-PglZ<-DUF6079*<-N6-METHYLASE<-PglW<-ACET+ACET | 1255 | Bacteria>Terrabacteria_group>Actinomycetota>Actinomycetes>Micromonosporales | GCA_016862015.1 | Actinoplanes_brasiliensis | Abr02nite_45090 | 34564..38331 | 52695 | hypothetical-protein-Abr02nite_45090-[Actinoplanes-brasiliensis] |
| GIF56652.1 | Type2.Pgl.like.BREX | <-DUF6079*<-?<-PglW | 1241 | Bacteria>Terrabacteria_group>Actinomycetota>Actinomycetes>Micromonosporales | GCA_016862515.1 | Asanoa_iriomotensis | Air01nite_27470 | 57096..60821 | 234613 | hypothetical-protein-Air01nite_27470-[Asanoa-iriomotensis] |
| GII23647.1 | Type2.Pgl.like.BREX | SF1-SF2-Helicase->PglW->N6-METHYLASE->DUF6079*->PglZ-><-PIN<-?\|\|BrxD-AAA->SF1-SF2-Helicase+wHTH+Small-B-barrel+DUF1998-> | 1227 | Bacteria>Terrabacteria_group>Actinomycetota>Actinomycetes>Micromonosporales | GCA_016863255.1 | Planosporangium_mesophilum | Pme01_32440 | 151646..155329 | 689768 | hypothetical-protein-Pme01_32440-[Planosporangium-mesophilum] |
| GKW48594.1 | Type2.Pgl.like.BREX | <-SF1-SF2-Helicase+wHTH+Small-B-barrel+DUF1998<-BrxD-AAA<-PglZ<-DUF6079*<-N6-METHYLASE<-HNH<-OLD-ABC<-PglW<-XIS-HTH | 1235 | Bacteria>Pseudomonadota>Gammaproteobacteria>Oceanospirillales>Halomonadaceae | GCA_023708045.1 | Halomonas_sp._NCCP-2165 | NCCP2165_08090 | 82862..86569 | 2942298 | hypothetical-protein-NCCP2165_08090-[Halomonas-sp.-NCCP-2165] |
| GLW58780.1 | Type2.Pgl.like.BREX | <-PAIREDC-HTH<-?\|\|PglW->N6-METHYLASE->DUF6079*->PglZ->BrxD-AAA->SF1-SF2-Helicase+wHTH+Small-B-barrel+DUF1998-> | 1313 | Bacteria>Terrabacteria_group>Actinomycetota>Actinomycetes>Kitasatosporales | GCA_030268825.1 | Kitasatospora_phosalacinea | Kpho01_67910 | 26328..30269 | 2065 | hypothetical-protein-Kpho01_67910-[Kitasatospora-phosalacinea] |
| GLZ41161.1 | Type2.Pgl.like.BREX | <-SF1-SF2-Helicase+wHTH+Small-B-barrel+DUF1998<-BrxD-AAA<-PglZ<-DUF6079*<-N6-METHYLASE<-PglW<-STYKinase-like<-ParB+HNH | 1262 | Bacteria>Terrabacteria_group>Actinomycetota>Actinomycetes>Pseudonocardiales | GCA_030269645.1 | Actinokineospora_sp._NBRC_105648 | Acsp05_47850 | 11746..15534 | 3032206 | hypothetical-protein-Acsp05_47850-[Actinokineospora-sp.-NBRC-105648] |
| HCD34860.1 | Type2.Pgl.like.BREX | XIS-HTH->PglW->?->N6-METHYLASE->?->DUF6079*->PglZ-> | 1223 | Bacteria>PVC_group>Planctomycetota>Phycisphaerae>Phycisphaerales | GCA_003517325.1 | Phycisphaerales_bacterium | DER01_20800 | 15809..19480 | 2052180 | phage-resistance-protein-[Phycisphaerales-bacterium] |
| HCF3662598.1 | Type2.Pgl.like.BREX | <-RNaseH+ArgR-HTH<-?<-SF1-SF2-Helicase+wHTH+Small-B-barrel+DUF1998<-BrxD-AAA<-PglZ<-DUF6079*<-N6-METHYLASE<-PglW | 1225 | Bacteria>Pseudomonadota>Gammaproteobacteria>Pseudomonadales>Pseudomonadaceae | GCA_023947915.1 | Pseudomonas_aeruginosa | NID08_005848 | 7719..11396 | 287 | TPA:-phage-resistance-protein-[Pseudomonas-aeruginosa] |
| HCJ0493611.1 | Type2.Pgl.like.BREX | MerR-HTH->tRNA->PglW->N6-METHYLASE->DUF6079*->PglZ->BrxD-AAA->SF1-SF2-Helicase+wHTH+Small-B-barrel+DUF1998->XIS-HTH-><-?<-KAP_NTPase<-DUF4969 | 1225 | Bacteria>Pseudomonadota>Gammaproteobacteria>Pseudomonadales>Pseudomonadaceae | GCA_024481025.1 | Pseudomonas_aeruginosa | NQI16_004638 | 48075..51752 | 287 | phage-resistance-protein-[Pseudomonas-aeruginosa] |
| HCL4045445.1 | Type2.Pgl.like.BREX | MerR-HTH->tRNA->PglW->N6-METHYLASE->DUF6079*->PglZ->BrxD-AAA->SF1-SF2-Helicase+wHTH+Small-B-barrel+DUF1998-> | 1225 | Bacteria>Pseudomonadota>Gammaproteobacteria>Pseudomonadales>Pseudomonadaceae | GCA_025004865.1 | Pseudomonas_aeruginosa | N2C00_003383 | 45038..48715 | 287 | TPA:-phage-resistance-protein-[Pseudomonas-aeruginosa] |
| HCR1633842.1 | Type2.Pgl.like.BREX | <-DUF6079*<-?<-PglW | 1225 | Bacteria>Pseudomonadota>Gammaproteobacteria>Pseudomonadales>Pseudomonadaceae | GCA_025922285.1 | Pseudomonas_aeruginosa | ONA63_005504 | 10905..14582 | 287 | TPA:-phage-resistance-protein-[Pseudomonas-aeruginosa] |
| HDY6530432.1 | Type2.Pgl.like.BREX | <-DUF6079*<-?<-PglW | 1225 | Bacteria>Pseudomonadota>Gammaproteobacteria>Pseudomonadales>Pseudomonadaceae | GCA_032039175.1 | Pseudomonas_aeruginosa | RRC64_001078 | 216194..219871 | 287 | TPA:-phage-resistance-protein-[Pseudomonas-aeruginosa] |
| HEU5155675.1 | Type2.Pgl.like.BREX | HTH_31+DUF5753->DUF397-><-?<-?<-?\|\|PglW->N6-METHYLASE->DUF6079*->PglZ->BrxD-AAA->SF1-SF2-Helicase+wHTH+Small-B-barrel+DUF1998->?->SAM-methylase-> | 1218 | Bacteria>Terrabacteria_group>Actinomycetota>Actinomycetes>Streptosporangiales | GCA_035765195.1 | Streptosporangiaceae_bacterium | VFU43_01665 | 40597..44253 | 2339230 | phage-resistance-protein-[Streptosporangiaceae-bacterium] |
| HEV2377799.1 | Type2.Pgl.like.BREX | <-SF1-SF2-Helicase<-BrxD-AAA<-PglZ<-DUF6079*<-N6-METHYLASE<-PglW<-?<-?<-?\|\|TraJ-RHH-> | 1253 | Bacteria>Terrabacteria_group>Actinomycetota>Actinomycetes>Streptosporangiales | GCA_035946955.1 | Streptosporangiaceae_bacterium | VGS19_37260 | 5187..8948 | 2339230 | hypothetical-protein-[Streptosporangiaceae-bacterium] |
| HEX6373915.1 | Type2.Pgl.like.BREX | <-McrC-NTD+REase<-McrB-AAA-ATPase<-?\|\|?-><-SF1-SF2-Helicase+wHTH+Small-B-barrel+DUF1998<-BrxD-AAA<-PglZ<-DUF6079*<-N6-METHYLASE<-PglW | 1223 | Bacteria>FCB_group>Gemmatimonadota>Longimicrobiia>Longimicrobiales | GCA_036379135.1 | Longimicrobium_sp. | VF006_33610 | 25618..29289 | 2029185 | DUF6079-family-protein-[Longimicrobium-sp.] |
| HEX7316680.1 | Type2.Pgl.like.BREX | PglW->N6-METHYLASE->DUF6079*->PglZ->HKD-DNase-> | 1211 | Bacteria>Acidobacteriota>Blastocatellia>Blastocatellales>Pyrinomonadaceae | GCA_036379655.1 | Pyrinomonadaceae_bacterium | VF297_22450 | 69697..73332 | 2283092 | hypothetical-protein-[Pyrinomonadaceae-bacterium] |
| HEY0513781.1 | Type2.Pgl.like.BREX | <-SF1-SF2-Helicase+wHTH+Small-B-barrel+DUF1998<-BrxD-AAA<-PglZ<-DUF6079*<-N6-METHYLASE<-PglW | 1244 | Bacteria>Acidobacteriota>Thermoanaerobaculia>unclassified_Thermoanaerobaculia | GCA_036402955.1 | Thermoanaerobaculia_bacterium | VGH73_17870 | 16504..20238 | 2651171 | phage-resistance-protein-[Thermoanaerobaculia-bacterium] |
| HEY1068995.1 | Type2.Pgl.like.BREX | PglW->N6-METHYLASE->?->MqsA_antitoxin+HTH->?->?->?->DUF6079*->PglZ->BrxD-AAA->SF1-SF2-Helicase+wHTH+Small-B-barrel+DUF1998-> | 1213 | Bacteria>Pseudomonadota>Gammaproteobacteria>Lysobacterales>Lysobacteraceae | GCA_036405455.1 | Thermomonas_sp. | VGE33_00460 | 53302..56943 | 1971895 | DUF6079-family-protein-[Thermomonas-sp.] |
| HEY5375208.1 | Type2.Pgl.like.BREX | PglW->N6-METHYLASE-><-?<-?\|\|DUF6079*-> | 1197 | Bacteria>Myxococcota>Polyangia>Polyangiales>Polyangiaceae | GCA_036509605.1 | Polyangiaceae_bacterium | VIK01_16125 | 12362..15955 | 2268199 | hypothetical-protein-[Polyangiaceae-bacterium] |
| HJW55338.1 | Type2.Pgl.like.BREX | HKD-DNase->D6+SF1-SF2-Helicase-><-SF1-SF2-Helicase+wHTH+Small-B-barrel+DUF1998<-BrxD-AAA<-PglZ<-DUF6079*<-N6-METHYLASE<-PglW<-SNF+SF1-SF2-Helicase<-?<-XIS-HTH | 1242 | Bacteria>Pseudomonadota>Betaproteobacteria>Burkholderiales>Burkholderiaceae | GCA_035255485.1 | Burkholderiaceae_bacterium | VJ577_08710 | 14043..17771 | 2030806 | phage-resistance-protein-[Burkholderiaceae-bacterium] |
| HNU10908.1 | Type2.Pgl.like.BREX | HTH_26->PglW->N6-METHYLASE->?->MqsA_antitoxin+HTH->?->?->DUF6079*->PglZ->BrxD-AAA->SF1-SF2-Helicase+wHTH+Small-B-barrel+DUF1998-> | 1233 | Bacteria>Pseudomonadota>Betaproteobacteria>Burkholderiales>Sphaerotilaceae | GCA_035340415.1 | Rubrivivax_sp. | PKJ45_06045 | 15543..19244 | 50259 | phage-resistance-protein-[Rubrivivax-sp.] |
| HOX59788.1 | Type2.Pgl.like.BREX | HTH->?->?->?-><-PglZ<-DUF6079*<-N6-METHYLASE<-PglW<-WYL | 1231 | Bacteria>PVC_group>Verrucomicrobiota>unclassified_Verrucomicrobiota | GCA_035389725.1 | Verrucomicrobiota_bacterium | PLC99_23140 | 6657..10352 | 2026799 | hypothetical-protein-[Verrucomicrobiota-bacterium] |
| HQA31350.1 | Type2.Pgl.like.BREX | PglW->ParB+HNH->N6-METHYLASE->DUF6079*->PglZ->BrxD-AAA->SF1-SF2-Helicase+wHTH+Small-B-barrel+DUF1998-> | 1248 | Bacteria>Terrabacteria_group>Actinomycetota>Actinomycetes>Propionibacteriales | GCA_035412375.1 | Propioniciclava_tarda | PKX10_08015 | 10082..13828 | 433330 | hypothetical-protein-[Propioniciclava-tarda] |
| HQC94916.1 | Type2.Pgl.like.BREX | HIN-HTH->?->?-><-SF1-SF2-Helicase+wHTH+Small-B-barrel+DUF1998<-BrxD-AAA<-PglZ<-DUF6079*<-?<-MqsA_antitoxin+HTH<-?<-N6-METHYLASE<-PglW | 1228 | Bacteria>Pseudomonadota>Betaproteobacteria>Burkholderiales>Burkholderiales_genera_incertae_sedis | GCA_035409685.1 | Aquabacterium_sp. | PK306_04330 | 15048..18734 | 1872578 | phage-resistance-protein-[Aquabacterium-sp.] |
| HQS63998.1 | Type2.Pgl.like.BREX | <-ParB+HNH<-PglZ<-DUF6079*<-N6-METHYLASE<-PglW<-WYL\|\|?->?->SbcC-> | 1232 | Bacteria>Pseudomonadota>Betaproteobacteria>Burkholderiales>Comamonadaceae | GCA_035428745.1 | Acidovorax_defluvii | PKV65_11570 | 11326..15024 | 86669 | hypothetical-protein-[Acidovorax-defluvii] |
| HRD10259.1 | Type2.Pgl.like.BREX | SF1-SF2-Helicase->ParB+HNH-><-SF1-SF2-Helicase+wHTH+Small-B-barrel+DUF1998<-BrxD-AAA<-PglZ<-DUF6079*<-N6-METHYLASE<-PglW | 1214 | Bacteria>Terrabacteria_group>Actinomycetota>Actinomycetes>Mycobacteriales | GCA_035441545.1 | Mycobacterium_sp. | PLI79_00200 | 19625..23269 | 1785 | DUF6079-family-protein-[Mycobacterium-sp.] |
| HRG94673.1 | Type2.Pgl.like.BREX | <-SF1-SF2-Helicase+wHTH+Small-B-barrel+DUF1998<-BrxD-AAA<-PglZ<-DUF6079*<-N6-METHYLASE<-REase<-PglW<-?\|\|REase->?->ParB+HNH-> | 1216 | Bacteria>Myxococcota>Polyangia>Polyangiales>Polyangiaceae | GCA_035439005.1 | Polyangiaceae_bacterium | PLR99_00400 | 14126..17776 | 2268199 | DUF6079-family-protein-[Polyangiaceae-bacterium] |
| HST50353.1 | Type2.Pgl.like.BREX | <-TetR-HTH\|\|PglW->N6-METHYLASE->DUF6079*->PglZ->BrxD-AAA->SF1-SF2-Helicase+wHTH+Small-B-barrel+DUF1998-> | 1240 | Bacteria>Terrabacteria_group>Actinomycetota>Actinomycetes>Jatrophihabitantales | GCA_035474675.1 | Jatrophihabitans_sp. | VLJ82_21095 | 43579..47301 | 1932789 | hypothetical-protein-[Jatrophihabitans-sp.] |
| HUP68595.1 | Type2.Pgl.like.BREX | HisKA_3+HATPase_c->REC+LuxR-HTH->?->?->PglW->N6-METHYLASE->DUF6079*->PglZ->BrxD-AAA->SF1-SF2-Helicase+wHTH+Small-B-barrel+DUF1998->SbcC->?->?-><-METHYLASE+DAM-Fungal | 1207 | Bacteria>Terrabacteria_group>Actinomycetota>Acidimicrobiia>Acidimicrobiales | GCA_035525365.1 | Acidimicrobiales_bacterium | VM142_02150 | 78808..82431 | 2201156 | hypothetical-protein-[Acidimicrobiales-bacterium] |
| HUY47908.1 | Type2.Pgl.like.BREX | ST-Hiskin->?->METHYLASE->?->?->PglW->N6-METHYLASE->DUF6079*->PglZ->BrxD-AAA->SF1-SF2-Helicase+wHTH+Small-B-barrel+DUF1998-><-REase | 1252 | Bacteria>Terrabacteria_group>Actinomycetota>Actinomycetes>Streptosporangiales | GCA_035531875.1 | Streptosporangiaceae_bacterium | VMV92_19625 | 26912..30670 | 2339230 | hypothetical-protein-[Streptosporangiaceae-bacterium] |
| HUY49759.1 | Type2.Pgl.like.BREX | His_Phos_1->tRNA->PglW->N6-METHYLASE->DUF6079*->PglZ->BrxD-AAA->SF1-SF2-Helicase+wHTH+Small-B-barrel+DUF1998-> | 1239 | Bacteria>Terrabacteria_group>Actinomycetota>Actinomycetes>Streptosporangiales | GCA_035531875.1 | Streptosporangiaceae_bacterium | VMV92_29260 | 80637..84356 | 2339230 | DUF6079-family-protein-[Streptosporangiaceae-bacterium] |
| HUZ27453.1 | Type2.Pgl.like.BREX | <-SF1-SF2-Helicase<-BrxD-AAA<-PglZ<-DUF6079*<-N6-METHYLASE<-PglW | 1244 | Bacteria>Terrabacteria_group>Actinomycetota>Actinomycetes>Streptosporangiales | GCA_035532295.1 | Streptosporangiaceae_bacterium | VMV07_27190 | 5131..8865 | 2339230 | phage-resistance-protein-[Streptosporangiaceae-bacterium] |
| HUZ54489.1 | Type2.Pgl.like.BREX | <-SF1-SF2-Helicase+wHTH+Small-B-barrel+DUF1998<-BrxD-AAA<-PglZ<-DUF6079*<-N6-METHYLASE<-PglW\|\|?->TraJ-RHH-> | 1255 | Bacteria>Terrabacteria_group>Actinomycetota>Actinomycetes>Streptosporangiales | GCA_035532775.1 | Streptosporangiaceae_bacterium | VMU94_18445 | 10901..14668 | 2339230 | DUF6079-family-protein-[Streptosporangiaceae-bacterium] |
| HUZ56112.1 | Type2.Pgl.like.BREX | <-SF1-SF2-Helicase+wHTH+Small-B-barrel+DUF1998<-BrxD-AAA<-PglZ<-DUF6079*<-N6-METHYLASE<-PglW<-SF1-SF2-Helicase<-?\|\|Pkinase+TPR-> | 1243 | Bacteria>Terrabacteria_group>Actinomycetota>Actinomycetes>Streptosporangiales | GCA_035532775.1 | Streptosporangiaceae_bacterium | VMU94_26755 | 8951..12682 | 2339230 | hypothetical-protein-[Streptosporangiaceae-bacterium] |
| HVV74600.1 | Type2.Pgl.like.BREX | PglW->N6-METHYLASE->DUF6079*->PglZ->BrxD-AAA->SF1-SF2-Helicase+wHTH+Small-B-barrel+DUF1998->?-><-?\|\|?-><-Phage_int_SAM_3+Phage_integrase | 1246 | Bacteria>Terrabacteria_group>Actinomycetota>Actinomycetes>Mycobacteriales | GCA_035547035.1 | Mycobacteriales_bacterium | VHC43_01085 | 11771..15511 | 1957017 | phage-resistance-protein-[Mycobacteriales-bacterium] |
| HWA24379.1 | Type2.Pgl.like.BREX | OLD-ABC+TOPRIM->?->?-><-SF1-SF2-Helicase+wHTH+Small-B-barrel+DUF1998<-BrxD-AAA<-PglZ<-DUF6079*<-N6-METHYLASE<-PglW | 1210 | Bacteria>PVC_group>Verrucomicrobiota>Opitutia>Opitutales | GCA_035549955.1 | Lacunisphaera_sp. | VG734_01800 | 43957..47589 | 1961806 | hypothetical-protein-[Lacunisphaera-sp.] |
| HXL95674.1 | Type2.Pgl.like.BREX | <-SF1-SF2-Helicase+wHTH+Small-B-barrel+DUF1998<-BrxD-AAA<-PglZ<-DUF6079*<-N6-METHYLASE<-PglW | 1262 | Bacteria>Terrabacteria_group>Actinomycetota>Actinomycetes>Streptosporangiales | GCA_035601975.1 | Streptosporangiaceae_bacterium | VN969_42685 | 38986..42774 | 2339230 | hypothetical-protein-[Streptosporangiaceae-bacterium] |
| HZR52088.1 | Type2.Pgl.like.BREX | <-YjhX_toxin<-YjhX_toxin<-SF1-SF2-Helicase+wHTH+Small-B-barrel+DUF1998<-BrxD-AAA<-PglZ<-DUF6079*<-N6-METHYLASE<-PglW | 1219 | Bacteria>Terrabacteria_group>Actinomycetota>Actinomycetes>Streptosporangiales | GCA_035651595.1 | Streptosporangiaceae_bacterium | VFB06_21565 | 41477..45136 | 2339230 | phage-resistance-protein-[Streptosporangiaceae-bacterium] |
| KAA0098913.1 | Type2.Pgl.like.BREX | REase->SF1-SF2-Helicase->?-><-SF1-SF2-Helicase+wHTH+Small-B-barrel+DUF1998<-BrxD-AAA<-PglZ<-DUF6079*<-N6-METHYLASE<-PglW | 1269 | Bacteria>Terrabacteria_group>Actinomycetota>Actinomycetes>Mycobacteriales | GCA_008329535.1 | Mycolicibacterium_sp._P1-18 | CIW49_13635 | 579586..583395 | 2024615 | hypothetical-protein-CIW49_13635-[Mycolicibacterium-sp.-P1-18] |
| KFU81956.1 | Type2.Pgl.like.BREX | <-SF1-SF2-Helicase\|\|wHTH+SAD+HNH->?->?->PglW->N6-METHYLASE->DUF6079*->PglZ->BrxD-AAA->SF1-SF2-Helicase+wHTH+Small-B-barrel+DUF1998->REase+SF1-SF2-Helicase->?->HKD-DNase-> | 1273 | Bacteria>Terrabacteria_group>Actinomycetota>Actinomycetes>Pseudonocardiales | GCA_000749465.3 | Amycolatopsis_lurida_NRRL_2430 | BB31_06335 | 65008..68829 | 1460371 | hypothetical-protein-BB31_06335-[Amycolatopsis-lurida-NRRL-2430] |
| KHD99079.1 | Type2.Pgl.like.BREX | <-DUF6079*<-?<-PglW | 1280 | Bacteria>Terrabacteria_group>Actinomycetota>Actinomycetes>Micrococcales | GCA_000786195.1 | Kocuria_polaris | GY22_01830 | 406168..410010 | 136273 | hypothetical-protein-GY22_01830-[Kocuria-polaris] |
| KOU07188.1 | Type2.Pgl.like.BREX | PglW-><-BrxD-AAA<-ParA-Soj-PloopNTPase\|\|N6-METHYLASE-><-DUF397<-?<-?\|\|DUF6079*->PglZ->BrxD-AAA->SF1-SF2-Helicase+wHTH+Small-B-barrel+DUF1998->Phd-YefM-> | 1308 | Bacteria>Terrabacteria_group>Actinomycetota>Actinomycetes>Kitasatosporales | GCA_001279425.1 | Streptomyces_sp._NRRL_F-2295 | ADK88_11870 | 69838..73764 | 1519477 | PglY-protein-[Streptomyces-sp.-NRRL-F-2295] |
| KQU28756.1 | Type2.Pgl.like.BREX | RAMA->HATPase->REase->?->PglW->N6-METHYLASE->DUF6079*->PglZ->BrxD-AAA->SF1-SF2-Helicase+wHTH+Small-B-barrel+DUF1998-><-SF1-SF2-Helicase<-wHTH<-LON-AAA | 1290 | Bacteria>Terrabacteria_group>Actinomycetota>Actinomycetes>Mycobacteriales | GCA_001426165.1 | Rhodococcus_sp._Leaf233 | ASH04_22300 | 139714..143586 | 1736302 | hypothetical-protein-ASH04_22300-[Rhodococcus-sp.-Leaf233] |
| KUN36385.1 | Type2.Pgl.like.BREX | <-SAD+REase\|\|PglW-><-BrxD-AAA<-ParA-Soj-PloopNTPase<-HEPN\|\|N6-METHYLASE->DUF6079*->PglZ->BrxD-AAA->SF1-SF2-Helicase+wHTH+Small-B-barrel+DUF1998-> | 1325 | Bacteria>Terrabacteria_group>Actinomycetota>Actinomycetes>Kitasatosporales | GCA_001514125.1 | Streptomyces_longwoodensis | AQJ30_22680 | 258338..262315 | 68231 | PglY-protein-[Streptomyces-longwoodensis] |
| KYF91859.1 | Type2.Pgl.like.BREX | <-SF1-SF2-Helicase+wHTH+Small-B-barrel+DUF1998<-PglZ<-DUF6079*\|\|ACET-><-N6-METHYLASE<-?<-?<-PglW | 1184 | Bacteria>Myxococcota>Polyangia>Polyangiales>Polyangiaceae | GCA_001589295.1 | Sorangium_cellulosum | BE20_13430 | 9126..12680 | 56 | hypothetical-protein-BE20_13430-[Sorangium-cellulosum] |
| MBB1018369.1 | Type2.Pgl.like.BREX | PglW-><-?\|\|N6-METHYLASE->DUF6079*->PglZ->BrxD-AAA->SF1-SF2-Helicase+wHTH+Small-B-barrel+DUF1998->ParB+HNH-> | 1286 | Bacteria>Terrabacteria_group>Actinomycetota>Actinomycetes>Mycobacteriales | GCA_014144995.1 | Dietzia_sp._DQ11-71 | G6023_08685 | 8723..12583 | 1630638 | phage-resistance-protein-[Dietzia-sp.-DQ11-71] |
| MBB1020394.1 | Type2.Pgl.like.BREX | <-ParB+AlbA_2<-SF1-SF2-Helicase+wHTH+Small-B-barrel+DUF1998<-BrxD-AAA<-PglZ<-DUF6079*<-N6-METHYLASE\|\|?-><-PglW\|\|?->?->HNH-> | 1307 | Bacteria>Terrabacteria_group>Actinomycetota>Actinomycetes>Mycobacteriales | GCA_014145015.1 | Dietzia_sp._E1 | G6030_03665 | 11166..15089 | 328361 | phage-resistance-protein-[Dietzia-sp.-E1] |
| MBB1049148.1 | Type2.Pgl.like.BREX | PglW-><-?\|\|?->DUF6079*-> | 1288 | Bacteria>Terrabacteria_group>Actinomycetota>Actinomycetes>Mycobacteriales | GCA_014145145.1 | Dietzia_cercidiphylli | G6028_15410 | 9663..13529 | 498199 | phage-resistance-protein-[Dietzia-cercidiphylli] |
| MBB1049897.1 | Type2.Pgl.like.BREX | <-PglZ<-DUF6079*<-N6-METHYLASE\|\|?-><-PglW\|\|?-><-ABhydrolase\|\|?->HNH-> | 1286 | Bacteria>Terrabacteria_group>Actinomycetota>Actinomycetes>Mycobacteriales | GCA_014145265.1 | Dietzia_sp._CW19 | G6021_01915 | 2156..6016 | 1630634 | phage-resistance-protein-[Dietzia-sp.-CW19] |
| MBB1055869.1 | Type2.Pgl.like.BREX | <-PglZ<-DUF6079*<-N6-METHYLASE\|\|?->?-><-PglW\|\|?->\|\|?->HNH-> | 1277 | Bacteria>Terrabacteria_group>Actinomycetota>Actinomycetes>Mycobacteriales | GCA_014145155.1 | Dietzia_sp._B19 | G6020_00295 | 1041..4874 | 1630632 | phage-resistance-protein-[Dietzia-sp.-B19] |
| MBB1516007.1 | Type2.Pgl.like.BREX | <-SF1-SF2-Helicase+wHTH+Small-B-barrel+DUF1998<-BrxD-AAA<-?<-HTH<-?<-PglZ<-DUF6079*<-N6-METHYLASE<-PglW<-?<-?\|\|?->?-><-ParB+HNH | 1247 | Bacteria>Terrabacteria_group>Actinomycetota>Actinomycetes>Propionibacteriales | GCA_014164775.1 | Tessaracoccus_sp._MC1679 | H5398_08510 | 279146..282889 | 2760313 | phage-resistance-protein-[Tessaracoccus-sp.-MC1679] |
| MBB5890596.1 | Type2.Pgl.like.BREX | PglW->N6-METHYLASE->DUF6079*->PglZ-><-CBS+HEPN<-ParB+HNH\|\|BrxD-AAA->SF1-SF2-Helicase+wHTH+Small-B-barrel+DUF1998-> | 1308 | Bacteria>Terrabacteria_group>Actinomycetota>Actinomycetes>Pseudonocardiales | GCA_014203355.1 | Kutzneria_kofuensis | BJ998_001792 | 1945314..1949240 | 103725 | hypothetical-protein-[Kutzneria-kofuensis] |
| MBE4746901.1 | Type2.Pgl.like.BREX | <-DUF6079*<-?<-PglW | 1231 | Bacteria>Myxococcota>Myxococcia>Myxococcales>Cystobacterineae | GCA_014930455.1 | Corallococcus_soli | G4177_01775 | 428654..432349 | 2710757 | hypothetical-protein-[Corallococcus-soli] |
| MBF0116836.1 | Type2.Pgl.like.BREX | PglW->?->?->?->DUF6079*-> | 1225 | Bacteria>Pseudomonadota>Alphaproteobacteria>Magnetococcales>unclassified_Magnetococcales | GCA_015228775.1 | Magnetococcales_bacterium | HQM04_17560 | 5951..9628 | 2026759 | phage-resistance-protein-[Magnetococcales-bacterium] |
| MBI9206403.1 | Type2.Pgl.like.BREX | MerR-HTH->tRNA->PglW->N6-METHYLASE->DUF6079*->PglZ->BrxD-AAA->SF1-SF2-Helicase+wHTH+Small-B-barrel+DUF1998-> | 1224 | Bacteria>Pseudomonadota>Gammaproteobacteria>Pseudomonadales>Pseudomonadaceae | GCA_016344045.1 | Pseudomonas_aeruginosa | JEW14_25110 | 33282..36956 | 287 | phage-resistance-protein-[Pseudomonas-aeruginosa] |
| MBM4487178.1 | Type2.Pgl.like.BREX | HNH->?-><-REase+SF1-SF2-Helicase<-SF1-SF2-Helicase+wHTH+Small-B-barrel+DUF1998<-BrxD-AAA<-PglZ<-DUF6079*<-?<-N6-METHYLASE<-PglW<-?<-D6+SF1-SF2-Helicase<-HKD-DNase | 1264 | Bacteria>Terrabacteria_group>Actinomycetota>Actinomycetes>Mycobacteriales | GCA_016874395.1 | Prescottella_equi | GS471_03215 | 674526..678320 | 43767 | phage-resistance-protein-[Prescottella-equi] |
| MBT2334085.1 | Type2.Pgl.like.BREX | HTH_26->PglW->?->DUF6079*->PglZ->BrxD-AAA->SF1-SF2-Helicase+wHTH+Small-B-barrel+DUF1998-><-SF1-SF2-Helicase<-?<-HTH | 1231 | Bacteria>Pseudomonadota>Betaproteobacteria>Burkholderiales>Comamonadaceae | GCA_018614875.1 | Variovorax_paradoxus | J7E49_09230 | 16942..20637 | 34073 | phage-resistance-protein-[Variovorax-paradoxus] |
| MBT2396441.1 | Type2.Pgl.like.BREX | <-DUF6079*<-?<-?<-?\|\|?->?-><-PglW | 1309 | Bacteria>Terrabacteria_group>Actinomycetota>Actinomycetes>Kitasatosporales | GCA_018614615.1 | Streptomyces_sp._ISL-100 | J7E89_10730 | 38743..42672 | 2819173 | phage-resistance-protein-[Streptomyces-sp.-ISL-100] |
| MBU2668296.1 | Type2.Pgl.like.BREX | N6-METHYLASE->Pkinase+TPR->PglW->N6-METHYLASE->DUF6079*->PglZ->BrxD-AAA->SF1-SF2-Helicase+wHTH+Small-B-barrel+DUF1998-><-REase+SF1-SF2-Helicase | 1237 | Bacteria>Terrabacteria_group>Actinomycetota>Actinomycetes>Micromonosporales | GCA_018829635.1 | Actinoplanes_bogorensis | KOI35_32770 | 171762..175475 | 1610840 | phage-resistance-protein-[Actinoplanes-bogorensis] |
| MBV6695717.1 | Type2.Pgl.like.BREX | HTH->XIS-HTH->Phage_integrase-><-?<-SF1-SF2-Helicase+wHTH+Small-B-barrel+DUF1998<-BrxD-AAA<-PglZ<-DUF6079*<-N6-METHYLASE<-PglW | 1313 | Bacteria>Terrabacteria_group>Actinomycetota>Actinomycetes>Kitasatosporales | GCA_019192735.1 | Kitasatospora_aureofaciens | KV557_01080 | 237396..241337 | 1894 | PglY-protein-[Kitasatospora-aureofaciens] |
| MBY6363249.1 | Type2.Pgl.like.BREX | <-SAD+REase\|\|SF1-SF2-Helicase+REase->SF1-SF2-Helicase-><-REase<-SF1-SF2-Helicase+wHTH+Small-B-barrel+DUF1998<-BrxD-AAA<-PglZ<-DUF6079*<-N6-METHYLASE<-PglW\|\|PglZ-><-BrxD-AAA\|\|McrB-AAA-ATPase->DUF2357+REase->HATPase-> | 1213 | Bacteria>Terrabacteria_group>Actinomycetota>Actinomycetes>Mycobacteriales | GCA_019834505.1 | Rhodococcus_corynebacterioides | HQ609_09010 | 112218..115859 | 53972 | phage-resistance-protein-[Rhodococcus-corynebacterioides] |
| MCH5644432.1 | Type2.Pgl.like.BREX | <-Phage_integrase\|\|PglW->N6-METHYLASE->DUF6079*->PglZ->BrxD-AAA->SF1-SF2-Helicase+wHTH+Small-B-barrel+DUF1998-> | 1275 | Bacteria>Terrabacteria_group>Actinomycetota>Actinomycetes>Mycobacteriales | GCA_022513585.1 | Gordonia_sp._ABSL49_1 | MKW15_17550 | 54041..57868 | 2920941 | DUF6079-family-protein-[Gordonia-sp.-ABSL49_1] |
| MCK1806707.1 | Type2.Pgl.like.BREX | <-DUF6079*<-?<-?<-?<-PglW | 1262 | Bacteria>Terrabacteria_group>Actinomycetota>Actinomycetes>Micromonosporales | GCA_023072705.1 | Micromonospora_sp._R42106 | MTQ09_12210 | 51303..55091 | 2929778 | phage-resistance-protein-[Micromonospora-sp.-R42106] |
| MCK6574439.1 | Type2.Pgl.like.BREX | XIS-HTH->?->PglW->N6-METHYLASE->OLD-ABC->?->DUF6079*->PglZ->BrxD-AAA->SF1-SF2-Helicase+wHTH+Small-B-barrel+DUF1998->N6-METHYLASE->\|\|ParB+HNH->HEPN-> | 1239 | Bacteria>Myxococcota>unclassified_Myxococcota | GCA_023150895.1 | Myxococcota_bacterium | L6V77_25430 | 11492..15211 | 2818507 | hypothetical-protein-[Myxococcota-bacterium] |
| MCK9876541.1 | Type2.Pgl.like.BREX | <-SF1-SF2-Helicase+wHTH+Small-B-barrel+DUF1998<-BrxD-AAA<-PglZ<-DUF6079*<-N6-METHYLASE<-PglW\|\|SbcC->?-><-?\|\|nSTAND3-> | 1332 | Bacteria>Terrabacteria_group>Actinomycetota>Actinomycetes>Frankiales | GCA_023242945.1 | Frankia_umida | MXD59_12270 | 88835..92833 | 573489 | phage-resistance-protein-[Frankia-umida] |
| MCK9895423.1 | Type2.Pgl.like.BREX | <-SF1-SF2-Helicase+wHTH+Small-B-barrel+DUF1998<-BrxD-AAA<-PglZ<-DUF6079*<-N6-METHYLASE<-PglW | 1317 | Bacteria>Terrabacteria_group>Actinomycetota>Actinomycetes>Frankiales | GCA_023243045.1 | Frankia_sp._AgB32 | MXD60_12600 | 19114..23067 | 631119 | phage-resistance-protein-[Frankia-sp.-AgB32] |
| MCM2371079.1 | Type2.Pgl.like.BREX | URI-DNase->?-><-?<-?\|\|XIS-HTH->PglW->N6-METHYLASE->DUF6079*->Schlafen+HATPase+wHTH->DUF1016_N->PglZ->BrxD-AAA->ParB+HNH->SF1-SF2-Helicase+wHTH+Small-B-barrel+DUF1998->NUDIX-> | 1244 | Bacteria>PVC_group>Planctomycetota>Planctomycetia>Pirellulales | GCA_023701485.1 | Aporhodopirellula_aestuarii | NB063_10705 | 26062..29796 | 2950107 | DUF6079-family-protein-[Aporhodopirellula-aestuarii] |
| MCM3886261.1 | Type2.Pgl.like.BREX | METHYLASE->?-><-?<-SF1-SF2-Helicase+wHTH+Small-B-barrel+DUF1998<-BrxD-AAA<-PglZ<-DUF6079*<-N6-METHYLASE<-PglW\|\|SbcC-><-?\|\|nSTAND3-> | 1329 | Bacteria>Terrabacteria_group>Actinomycetota>Actinomycetes>Frankiales | GCA_023716925.1 | Frankia_sp._R82 | ND747_21810 | 21178..25167 | 2950553 | phage-resistance-protein-[Frankia-sp.-R82] |
| MCP5439823.1 | Type2.Pgl.like.BREX | TraG_N-><-SbcC<-REase<-SIR2<-SF1-SF2-Helicase+wHTH+Small-B-barrel+DUF1998<-BrxD-AAA<-PglZ<-DUF6079*<-N6-METHYLASE<-PglW<-HTH_26\|\|HDRnase->Phage_integrase-> | 1216 | Bacteria>Pseudomonadota>Gammaproteobacteria>Chromatiales>Chromatiaceae | GCA_024235435.1 | Chromatiaceae_bacterium | H6958_04270 | 29187..32837 | 2049432 | phage-resistance-protein-[Chromatiaceae-bacterium] |
| MCR9166538.1 | Type2.Pgl.like.BREX | PglW->N6-METHYLASE->DUF6079*-> | 1229 | Bacteria>unclassified_Bacteria | GCA_024742475.1 | bacterium | NXI35_38075 | 10228..13917 | 1869227 | hypothetical-protein-[bacterium] |
| MCT2260556.1 | Type2.Pgl.like.BREX | ParB+HNH-><-DNA-glycosylase\|\|?->PglW->N6-METHYLASE->DUF6079*->PglZ->BrxD-AAA->SF1-SF2-Helicase+wHTH+Small-B-barrel+DUF1998-><-URI-DNase+SF1-SF2-Helicase<-MazG | 1240 | Bacteria>Terrabacteria_group>Actinomycetota>Actinomycetes>Micrococcales | GCA_025151965.1 | Brachybacterium_muris | M3F59_02755 | 135293..139015 | 219301 | DUF6079-family-protein-[Brachybacterium-muris] |
| MCT2273894.1 | Type2.Pgl.like.BREX | <-ParB+HNH<-?<-SF1-SF2-Helicase+wHTH+Small-B-barrel+DUF1998<-BrxD-AAA<-PglZ<-DUF6079*<-N6-METHYLASE<-PglW | 1278 | Bacteria>Terrabacteria_group>Actinomycetota>Actinomycetes>Mycobacteriales | GCA_025151825.1 | Dietzia_cinnamea | M3F76_05320 | 10881..14717 | 321318 | DUF6079-family-protein-[Dietzia-cinnamea] |
| MCT9080286.1 | Type2.Pgl.like.BREX | PIN-><-SF1-SF2-Helicase+wHTH+Small-B-barrel+DUF1998<-BrxD-AAA<-PglZ<-DUF6079*<-N6-METHYLASE<-PglW\|\|DUF6731->?-><-REase+SF1-SF2-Helicase<-HD<-HTH_3 | 1302 | Bacteria>Terrabacteria_group>Actinomycetota>Actinomycetes>Kitasatosporales | GCA_025399775.1 | Streptomyces_fulvoviolaceus | N4G65_27580 | 6857..10765 | 285535 | PglY-protein-[Streptomyces-fulvoviolaceus] |
| MCU7804185.1 | Type2.Pgl.like.BREX | <-HTH\|\|Phage_integrase->?->?->HTH->PglW->N6-METHYLASE->DUF6079*->PglZ->BrxD-AAA->SF1-SF2-Helicase+wHTH+Small-B-barrel+DUF1998-><-HTH_21+rve+rve_2<-PIN | 1215 | Bacteria>Pseudomonadota>Gammaproteobacteria>Chromatiales>Sedimenticolaceae | GCA_025655695.1 | Candidatus_Thiodiazotropha_sp._(ex_Lucinoma_borealis) | KZQ96_13390 | 42647..46294 | 2857020 | phage-resistance-protein-[Candidatus-Thiodiazotropha-sp.-(ex-Lucinoma-borealis)] |
| MCV7510960.1 | Type2.Pgl.like.BREX | <-URI-DNase+SF1-SF2-Helicase<-MazG<-SF1-SF2-Helicase+wHTH+Small-B-barrel+DUF1998<-BrxD-AAA<-PglZ<-DUF6079*<-N6-METHYLASE<-PglW\|\|ParB+HNH-> | 1254 | Bacteria>Terrabacteria_group>Actinomycetota>Actinomycetes>Micrococcales | GCA_025151565.2 | Micrococcus_luteus | M3E17_001990 | 70362..74126 | 1270 | hypothetical-protein-[Micrococcus-luteus] |
| MCV7623696.1 | Type2.Pgl.like.BREX | <-ParB+HNH\|\|PglW->N6-METHYLASE->DUF6079*->PglZ->BrxD-AAA->SF1-SF2-Helicase+wHTH+Small-B-barrel+DUF1998->MazG->URI-DNase+SF1-SF2-Helicase-> | 1254 | Bacteria>Terrabacteria_group>Actinomycetota>Actinomycetes>Micrococcales | GCA_025145605.2 | Micrococcus_luteus | M3B28_001605 | 64841..68605 | 1270 | hypothetical-protein-[Micrococcus-luteus] |
| MCX6543496.1 | Type2.Pgl.like.BREX | HEPN-><-?<-?<-?\|\|PglW->N6-METHYLASE->DUF6079*->PglZ->BrxD-AAA->SF1-SF2-Helicase+wHTH+Small-B-barrel+DUF1998-><-?<-?\|\|McrB-AAA-ATPase->McrC-NTD+REase-> | 1232 | Bacteria>Acidobacteriota>unclassified_Acidobacteriota | GCA_026393755.1 | Acidobacteriota_bacterium | NTV05_03680 | 26296..29994 | 1978231 | phage-resistance-protein-[Acidobacteriota-bacterium] |
| MCY3845511.1 | Type2.Pgl.like.BREX | <-SF1-SF2-Helicase+wHTH+Small-B-barrel+DUF1998<-BrxD-AAA<-PglZ<-DUF6079*<-?<-?<-?<-PglW | 1210 | Bacteria>Acidobacteriota>unclassified_Acidobacteriota | GCA_026706755.1 | Acidobacteriota_bacterium | OXH69_18455 | 9895..13527 | 1978231 | phage-resistance-protein-[Acidobacteriota-bacterium] |
| MCZ2341281.1 | Type2.Pgl.like.BREX | <-SF1-SF2-Helicase+wHTH+Small-B-barrel+DUF1998<-BrxD-AAA<-PglZ<-DUF6079*<-Schlafen+HATPase+wHTH<-N6-METHYLASE<-REase<-PglW | 1232 | Bacteria>FCB_group>Bacteroidota/Chlorobiota_group>Bacteroidota>Bacteroidia | GCA_026931725.1 | Bacteroidales_bacterium | LC104_05725 | 6258..9956 | 2030927 | DUF6079-family-protein-[Bacteroidales-bacterium] |
| MCZ2342607.1 | Type2.Pgl.like.BREX | <-SF1-SF2-Helicase+wHTH+Small-B-barrel+DUF1998<-BrxD-AAA<-PglZ<-DUF6079*<-N6-METHYLASE<-PglW<-XIS-HTH | 1226 | Bacteria>FCB_group>Bacteroidota/Chlorobiota_group>Bacteroidota>Bacteroidia | GCA_026931725.1 | Bacteroidales_bacterium | LC104_12575 | 25004..28684 | 2030927 | DUF6079-family-protein-[Bacteroidales-bacterium] |
| MCZ8257084.1 | Type2.Pgl.like.BREX | <-ParB+HNH<-PglZ<-DUF6079*<-N6-METHYLASE<-PglW<-WYL | 1232 | Bacteria>Pseudomonadota>Betaproteobacteria>Burkholderiales>Comamonadaceae | GCA_027532105.1 | Polaromonas_sp. | O9327_15650 | 4032..7730 | 1869339 | hypothetical-protein-[Polaromonas-sp.] |
| MDA0169812.1 | Type2.Pgl.like.BREX | <-ACET\|\|?->?->tRNA->ParB+HNH->PglW->N6-METHYLASE->N6-METHYLASE->DUF6079*->PglZ->BrxD-AAA->SF1-SF2-Helicase+wHTH+Small-B-barrel+DUF1998->?->?->SF1-SF2-Helicase+DUF1998-><-HNH | 1201 | Bacteria>Terrabacteria_group>Actinomycetota>Thermoleophilia>Solirubrobacterales | GCA_027587225.1 | Solirubrobacter_taibaiensis | OJ998_12015 | 133856..137461 | 1395955 | hypothetical-protein-[Solirubrobacter-taibaiensis] |
| MDB5350149.1 | Type2.Pgl.like.BREX | <-ArdCN+MPTase\|\|?->XIS-HTH->PglW->N6-METHYLASE->DUF6079*->PglZ->BrxD-AAA->SF1-SF2-Helicase+wHTH+Small-B-barrel+DUF1998-> | 1232 | Bacteria>PVC_group>Planctomycetota>unclassified_Planctomycetota | GCA_028291545.1 | Planctomycetota_bacterium | JWN86_1396 | 86192..89890 | 2026780 | hypothetical-protein-[Planctomycetota-bacterium] |

| Protein-ID | System | Gene-Neighbourhoods | Length | Taxonomy | Assembly-ID | Strain | Locus-Tag | Genomic-coordinates | TaxID | Fasta-defline |
|---|---|---|---|---|---|---|---|---|---|---|
| MDE1007015.1 | Type2.Pgl.like.BREX | <-SF1-SF2-Helicase<-BrxD-AAA<-PglZ<-DUF6079*<-N6-METHYLASE<-PglW<-HTH_26<-HEPN<-MORC+REase | 1234 | Bacteria>Pseudomonadota>Betaproteobacteria>Burkholderiales>Burkholderiaceae | GCA_028825355.1 | Paraburkholderia_fungorum | OSB38_15240 | 5113..8817 | 134537 | phage-resistance-protein-[Paraburkholderia-fungorum] |
| MDF2916037.1 | Type2.Pgl.like.BREX | <-HIN-HTH\|\|?->HIN-HTH->?-><-SF1-SF2-Helicase+wHTH+Small-B-barrel+DUF1998<-BrxD-AAA<-PglZ<-DUF6079*<-N6-METHYLASE<-PglW | 1203 | Bacteria>Terrabacteria_group>Actinomycetota>Actinomycetes>Micrococcales | GCA_029256205.1 | Microbacterium_sp. | K0S70_254 | 19597..23208 | 51671 | hypothetical-protein-[Microbacterium-sp.] |
| MDF9813068.1 | Type2.Pgl.like.BREX | <-DUF6079*<-?<-PglW | 1266 | Bacteria>Terrabacteria_group>Actinomycetota>Actinomycetes>Kitasatosporales | GCA_029504575.1 | Streptomyces_sp._SPB162 | M2266_002299 | 2470015..2473815 | 2940560 | hypothetical-protein-[Streptomyces-sp.-SPB162] |
| MDJ0841533.1 | Type2.Pgl.like.BREX | PglW->N6-METHYLASE->DUF6079*->PglZ-> | 1207 | Bacteria>Acidobacteriota>unclassified_Acidobacteriota | GCA_030066195.1 | Acidobacteriota_bacterium | QNK37_33810 | 8259..11882 | 1978231 | phage-resistance-protein-[Acidobacteriota-bacterium] |
| MDN5789362.1 | Type2.Pgl.like.BREX | <-PIN<-?<-REase\|\|?->?->PglW->N6-METHYLASE->DUF6079*->PglZ->BrxD-AAA->SF1-SF2-Helicase+wHTH+Small-B-barrel+DUF1998-><-XIS-HTH | 1246 | Bacteria>Terrabacteria_group>Actinomycetota>Actinomycetes>Micrococcales | GCA_030452775.1 | Micrococcales_bacterium | LOH25_00640 | 14544..18284 | 2026762 | DUF6079-family-protein-[Micrococcales-bacterium] |
| MDO9018002.1 | Type2.Pgl.like.BREX | PglW->?->?->?->N6-METHYLASE->?->ParB+HNH->DUF6079*->PglZ->BrxD-AAA->SF1-SF2-Helicase+wHTH+Small-B-barrel+DUF1998-> | 1198 | Bacteria>Myxococcota>Myxococcia>Myxococcales>unclassified_Myxococcales | GCA_030652255.1 | Myxococcales_bacterium | Q7V43_13890 | 24236..27832 | 2026763 | hypothetical-protein-[Myxococcales-bacterium] |
| MDO9020473.1 | Type2.Pgl.like.BREX | DOC->PglW->Pkinase+TPR->?->HEPN->DUF6079*->PglZ-> | 1226 | Bacteria>Myxococcota>Myxococcia>Myxococcales>unclassified_Myxococcales | GCA_030652255.1 | Myxococcales_bacterium | Q7V43_26340 | 12050..15730 | 2026763 | hypothetical-protein-[Myxococcales-bacterium] |
| MDQ2706432.1 | Type2.Pgl.like.BREX | <-SF1-SF2-Helicase+wHTH+Small-B-barrel+DUF1998<-PglZ<-DUF6079*<-?\|\|?-><-ParB+HNH<-N6-METHYLASE<-PglW | 1190 | Bacteria>Terrabacteria_group>Actinomycetota>unclassified_Actinomycetota | GCA_030852965.1 | Actinomycetota_bacterium | M3Z25_01800 | 5964..9536 | 2900548 | phage-resistance-protein-[Actinomycetota-bacterium] |
| MDQ3363903.1 | Type2.Pgl.like.BREX | PglW->N6-METHYLASE->?->DUF6079*->PglZ->BrxD-AAA->SF1-SF2-Helicase+wHTH+Small-B-barrel+DUF1998->?->?->REase-> | 1211 | Bacteria>Myxococcota>unclassified_Myxococcota | GCA_030857245.1 | Myxococcota_bacterium | M3680_00520 | 21741..25376 | 2818507 | DUF6079-family-protein-[Myxococcota-bacterium] |
| MDQ3647621.1 | Type2.Pgl.like.BREX | <-SF1-SF2-Helicase+wHTH+Small-B-barrel+DUF1998<-BrxD-AAA<-PglZ<-DUF6079*<-N6-METHYLASE<-PglW | 1212 | Bacteria>Terrabacteria_group>Actinomycetota>unclassified_Actinomycetota | GCA_030859845.1 | Actinomycetota_bacterium | M3433_03365 | 15908..19546 | 2900548 | DUF6079-family-protein-[Actinomycetota-bacterium] |
| MDR3405897.1 | Type2.Pgl.like.BREX | <-SF1-SF2-Helicase+wHTH+Small-B-barrel+DUF1998<-?<-Classical-AAA<-BrxD-AAA<-PglZ<-DUF6079*<-?<-?<-?<-N6-METHYLASE<-?<-REase<-PglW | 1266 | Bacteria>PVC_group>Verrucomicrobiota>Spartobacteria>Chthoniobacterales | GCA_031290995.1 | Chthoniobacter_sp. | P4L99_25655 | 42029..45829 | 2510640 | DUF6079-family-protein-[Chthoniobacter-sp.] |
| MDR6857864.1 | Type2.Pgl.like.BREX | <-HNH<-SNF+SF1-SF2-Helicase<-?<-SF1-SF2-Helicase<-SF1-SF2-Helicase+wHTH+Small-B-barrel+DUF1998<-BrxD-AAA<-PglZ<-DUF6079*<-N6-METHYLASE<-PglW<-?\|\|CR-ATPase5-><-ArdCN+MPTase<-GDResolvase+HIN-HTH | 1240 | Bacteria>Pseudomonadota>Betaproteobacteria>Burkholderiales>Comamonadaceae | GCA_031456095.1 | Variovorax_guangxiensis | J2W96_004189 | 402514..406236 | 1775474 | hypothetical-protein-[Variovorax-guangxiensis] |
| MDV2977864.1 | Type2.Pgl.like.BREX | <-McrC-NTD+REase<-McrB-AAA-ATPase<-McrB-AAA-ATPase<-HNH\|\|?->PglW->N6-METHYLASE->DUF6079*->PglZ->BrxD-AAA->SF1-SF2-Helicase+wHTH+Small-B-barrel+DUF1998-> | 1263 | Bacteria>Terrabacteria_group>Actinomycetota>Actinomycetes>unclassified_Actinomycetes | GCA_032849035.1 | Actinomycetes_bacterium_ARC8 | Q9R71_11840 | 409954..413745 | 3081831 | DUF6079-family-protein-[Actinomycetes-bacterium-ARC8] |
| MDW9414210.1 | Type2.Pgl.like.BREX | SbcC+SbcC->SNF+SF1-SF2-Helicase->?-><-SF1-SF2-Helicase+wHTH+Small-B-barrel+DUF1998<-BrxD-AAA<-PglZ<-DUF6079*<-N6-METHYLASE<-PglW<-MetJarc-HTH\|\|REase->SF1-SF2-Helicase-> | 1254 | Bacteria>Pseudomonadota>Alphaproteobacteria>Hyphomicrobiales>Rhizobiaceae | GCA_033712955.1 | Sinorhizobium_meliloti | GDB15_07010 | 21385..25149 | 382 | phage-resistance-protein-[Sinorhizobium-meliloti] |
| MDX0772250.1 | Type2.Pgl.like.BREX | <-SF1-SF2-Helicase\|\|MetJarc-HTH->PglW->N6-METHYLASE->DUF6079*->PglZ->BrxD-AAA->SF1-SF2-Helicase+wHTH+Small-B-barrel+DUF1998-><-?<-SNF+SF1-SF2-Helicase<-SbcC+SbcC | 1254 | Bacteria>Pseudomonadota>Alphaproteobacteria>Hyphomicrobiales>Rhizobiaceae | GCA_033718235.1 | Sinorhizobium_medicae | GDD63_28105 | 39129..42893 | 110321 | phage-resistance-protein-[Sinorhizobium-medicae] |
| MDX2267520.1 | Type2.Pgl.like.BREX | <-SF1-SF2-Helicase+wHTH+Small-B-barrel+DUF1998<-BrxD-AAA<-PglZ<-DUF6079*<-N6-METHYLASE<-PglW | 1238 | Bacteria>Acidobacteriota>Terriglobia>Bryobacterales>Bryobacteraceae | GCA_033765085.1 | Bryobacter_sp. | NW208_05400 | 12576..16292 | 1962911 | hypothetical-protein-[Bryobacter-sp.] |
| MDX3354815.1 | Type2.Pgl.like.BREX | <-SF1-SF2-Helicase+wHTH+Small-B-barrel+DUF1998<-BrxD-AAA<-PglZ<-DUF6079*<-N6-METHYLASE<-PglW<-?\|\|?->HKD-DNase-> | 1306 | Bacteria>Terrabacteria_group>Actinomycetota>Actinomycetes>Kitasatosporales | GCA_033794295.1 | Streptomyces_sp._ME01-24h | PV703_16185 | 17592..21512 | 3028670 | PglY-protein-[Streptomyces-sp.-ME01-24h] |
| MDY0058882.1 | Type2.Pgl.like.BREX | <-N6-METHYLASE\|\|ArdCN+MPTase->ABhydrolase->?->PglW->?->N6-METHYLASE->DUF6079*->PglZ->BrxD-AAA->SF1-SF2-Helicase+wHTH+Small-B-barrel+DUF1998->ABhydrolase-> | 1238 | Bacteria>Myxococcota>unclassified_Myxococcota | GCA_034005005.1 | Myxococcota_bacterium | RBU45_03650 | 24108..27824 | 2818507 | hypothetical-protein-[Myxococcota-bacterium] |
| MDZ4800307.1 | Type2.Pgl.like.BREX | <-Phage_integrase<-SF1-SF2-Helicase+wHTH+Small-B-barrel+DUF1998<-BrxD-AAA<-PglZ<-DUF6079*<-N6-METHYLASE<-PglW<-?\|\|?->?->?-><-SF1-SF2-Helicase | 1212 | Bacteria>Acidobacteriota>Terriglobia>Bryobacterales>Bryobacteraceae | GCA_034408745.1 | Bryobacteraceae_bacterium | SGI92_19285 | 38020..41658 | 2212468 | hypothetical-protein-[Bryobacteraceae-bacterium] |
| MEA5119840.1 | Type2.Pgl.like.BREX | <-SF1-SF2-Helicase+wHTH+Small-B-barrel+DUF1998<-BrxD-AAA<-?<-PglZ<-DUF6079*<-N6-METHYLASE<-PglW | 1265 | Bacteria>Terrabacteria_group>Actinomycetota>Actinomycetes>Propionibacteriales | GCA_034933485.1 | Propionibacterium_sp. | VB040_01040 | 50756..54553 | 1977903 | hypothetical-protein-[Propionibacterium-sp.] |
| MEB5061967.1 | Type2.Pgl.like.BREX | <-SF1-SF2-Helicase+wHTH+Small-B-barrel+DUF1998<-BrxD-AAA<-PglZ<-DUF6079*<-N6-METHYLASE<-PglW<-tRNA<-MerR-HTH | 1225 | Bacteria>Pseudomonadota>Gammaproteobacteria>Pseudomonadales>Pseudomonadaceae | GCA_035728105.1 | Pseudomonas_aeruginosa | RXR61_23145 | 16267..19944 | 287 | phage-resistance-protein-[Pseudomonas-aeruginosa] |
| MEC5193206.1 | Type2.Pgl.like.BREX | PglW->?->DUF6079*-> | 1279 | Bacteria>Terrabacteria_group>Actinomycetota>Actinomycetes>Micrococcales | GCA_035984855.1 | Arthrobacter_sp._MP_M4 | RCH15_003522 | 25882..29721 | 3071714 | hypothetical-protein-[Arthrobacter-sp.-MP_M4] |
| MEC5204627.1 | Type2.Pgl.like.BREX | <-SF1-SF2-Helicase+wHTH+Small-B-barrel+DUF1998<-BrxD-AAA<-PglZ<-DUF6079*<-N6-METHYLASE<-PglW\|\|?-><-?<-?<-Phage_int_SAM_3+Phage_integrase<-XIS-HTH | 1279 | Bacteria>Terrabacteria_group>Actinomycetota>Actinomycetes>Micrococcales | GCA_035984945.1 | Arthrobacter_sp._MP_M7 | RCH17_003457 | 9108..12947 | 3071716 | hypothetical-protein-[Arthrobacter-sp.-MP_M7] |
| MXQ65080.1 | Type2.Pgl.like.BREX | <-SF1-SF2-Helicase+wHTH+Small-B-barrel+DUF1998<-BrxD-AAA<-PglZ<-DUF6079*<-N6-METHYLASE<-PglW | 1419 | Bacteria>Terrabacteria_group>Actinomycetota>Actinomycetes>Streptosporangiales | GCA_009831215.1 | Actinomadura_rayongensis | GQ466_13650 | 940291..944550 | 1429076 | phage-resistance-protein-[Actinomadura-rayongensis] |
| NIL92568.1 | Type2.Pgl.like.BREX | <-ParB+HNH<-SF1-SF2-Helicase+wHTH+Small-B-barrel+DUF1998<-BrxD-AAA<-PglZ<-DUF6079*<-N6-METHYLASE<-PglW | 1216 | Bacteria>Terrabacteria_group>Actinomycetota>Actinomycetes>Mycobacteriales | GCA_011770045.1 | Rhodococcus_fascians | RhoFasSB10_04929 | 8256..11906 | 1828 | hypothetical-protein-[Rhodococcus-fascians] |
| NJA55277.1 | Type2.Pgl.like.BREX | <-PocR+AraC-HTH+AraC-HTH\|\|?->PglW->N6-METHYLASE-><-DUF397<-HTH_31+DUF5753\|\|ST-Hiskin->DUF6079*->PglZ->BrxD-AAA->SF1-SF2-Helicase+wHTH+Small-B-barrel+DUF1998-> | 1297 | Bacteria>Terrabacteria_group>Actinomycetota>Actinomycetes>Kitasatosporales | GCA_011882225.1 | Streptomyces_sp._NEAU-H3 | HC023_01900 | 16923..20816 | 2720636 | phage-resistance-protein-[Streptomyces-sp.-NEAU-H3] |
| NJC84888.1 | Type2.Pgl.like.BREX | REase+SF1-SF2-Helicase-><-?<-SF1-SF2-Helicase+wHTH+Small-B-barrel+DUF1998<-BrxD-AAA\|\|?->PIN-><-PglZ<-DUF6079*<-N6-METHYLASE<-PglW<-SF1-SF2-Helicase | 1227 | Bacteria>Terrabacteria_group>Actinomycetota>Actinomycetes>Micromonosporales | GCA_011947235.1 | Planosporangium_mesophilum | HC030_20345 | 11443..15126 | 689768 | phage-resistance-protein-[Planosporangium-mesophilum] |
| NKX80096.1 | Type2.Pgl.like.BREX | <-DUF6079*<-?<-PglW | 1243 | Bacteria>Terrabacteria_group>Actinomycetota>Actinomycetes>Mycobacteriales | GCA_012395955.1 | Gordonia_amicalis | HGA01_21735 | 6639..10370 | 89053 | phage-resistance-protein-[Gordonia-amicalis] |
| NKZ67066.1 | Type2.Pgl.like.BREX | <-DUF6079*<-?<-?<-?<-PglW | 1264 | Bacteria>Terrabacteria_group>Actinomycetota>Actinomycetes>Mycobacteriales | GCA_012443735.1 | Prescottella_equi | GTG23_22850 | 1396484..1400278 | 43767 | phage-resistance-protein-[Prescottella-equi] |
| NLU66093.1 | Type2.Pgl.like.BREX | <-HisKA_3+HATPase_c<-REC+LuxR-HTH\|\|tRNA-><-?<-?<-DUF6731\|\|PglW->N6-METHYLASE->DUF6079*->PglZ->BrxD-AAA->SF1-SF2-Helicase+wHTH+Small-B-barrel+DUF1998->?-><-?<-TetR-HTH | 1300 | Bacteria>Terrabacteria_group>Actinomycetota>Actinomycetes>Kitasatosporales | GCA_012726235.1 | Streptomyces_sp._HNM0574 | HCC30_02200 | 531963..535865 | 2714954 | PglY-protein-[Streptomyces-sp.-HNM0574] |
| NMO15997.1 | Type2.Pgl.like.BREX | Pkinase+TPR-><-REase\|\|?-><-SF1-SF2-Helicase+wHTH+Small-B-barrel+DUF1998<-BrxD-AAA<-PglZ<-DUF6079*<-N6-METHYLASE<-PglW | 1233 | Bacteria>Myxococcota>Myxococcia>Myxococcales>Cystobacterineae | GCA_012933655.1 | Pyxidicoccus_fallax | HG543_14225 | 30199..33900 | 394095 | hypothetical-protein-[Pyxidicoccus-fallax] |
| NSC23019.1 | Type2.Pgl.like.BREX | DUF6079*-> | 1321 | Bacteria>Terrabacteria_group>Actinomycetota>Actinomycetes>Kitasatosporales | GCA_013302895.1 | Streptomyces_albus_subsp._chlorinus | FM076_RS18165 | 4166115..4170080 | 337066 | PglY-protein-[Streptomyces-albus-subsp.-chlorinus] |
| NTY62303.1 | Type2.Pgl.like.BREX | Phd-YefM->PIN-><-?<-?\|\|PglW->N6-METHYLASE->DUF6079*->PglZ->BrxD-AAA->SF1-SF2-Helicase+wHTH+Small-B-barrel+DUF1998->ParB+HNH-> | 1224 | Bacteria>Terrabacteria_group>Actinomycetota>Actinomycetes>Mycobacteriales | GCA_013337765.1 | Mycolicibacterium_sphagni | FEG63_22450 | 107447..111121 | 1786 | phage-resistance-protein-[Mycolicibacterium-sphagni] |
| NUJ61074.1 | Type2.Pgl.like.BREX | PglW->?->?->?->DUF6079*-> | 1237 | Bacteria>Pseudomonadota>Gammaproteobacteria>Oceanospirillales>Halomonadaceae | GCA_013349955.1 | Onishia_taeanensis | HUS66_15815 | 34399..38112 | 284577 | phage-resistance-protein-[Halomonas-taeanensis] |
| NUU18463.1 | Type2.Pgl.like.BREX | SAD+REase-><-REase\|\|PglW->N6-METHYLASE->N6-METHYLASE->DUF6079*->PglZ->BrxD-AAA->SF1-SF2-Helicase+wHTH+Small-B-barrel+DUF1998-><-PglZ<-URI-DNase+SF1-SF2-Helicase<-MazG | 1257 | Bacteria>Terrabacteria_group>Actinomycetota>Actinomycetes>Micrococcales | GCA_013359775.1 | Cellulomonas_humilata | HP550_14495 | 196986..200759 | 144055 | hypothetical-protein-[Cellulomonas-humilata] |
| NVM96158.1 | Type2.Pgl.like.BREX | <-SF1-SF2-Helicase<-?\|\|McrB-AAA-ATPase->McrC-NTD+REase-><-SAD+REase\|\|PglW->N6-METHYLASE->DUF6079*->PglZ->BrxD-AAA->SF1-SF2-Helicase+wHTH+Small-B-barrel+DUF1998-> | 1261 | Bacteria>Terrabacteria_group>Actinomycetota>Actinomycetes>Micrococcales | GCA_013376105.1 | Arthrobacter_wenxiniae | G6034_14860 | 48958..52743 | 2713570 | phage-resistance-protein-[Arthrobacter-wenxiniae] |
| NYG54950.1 | Type2.Pgl.like.BREX | <-REase\|\|MORC->?->?->PglW->N6-METHYLASE->DUF6079*->PglZ->?->?->SF1-SF2-Helicase->?->SF1-SF2-Helicase+DUF1998-> | 1220 | Bacteria>Terrabacteria_group>Actinomycetota>Actinomycetes>Propionibacteriales | GCA_013409425.1 | Nocardioides_perillae | BJ989_001254 | 1361369..1365031 | 1119534 | hypothetical-protein-[Nocardioides-perillae] |
| NYH77234.1 | Type2.Pgl.like.BREX | PglW->?->DUF6079*-> | 1268 | Bacteria>Terrabacteria_group>Actinomycetota>Actinomycetes>Actinopolysporales | GCA_013408175.1 | Actinopolyspora_biskrensis | FHR84_000548 | 601142..604948 | 1470178 | hypothetical-protein-[Actinopolyspora-biskrensis] |
| OBF38943.1 | Type2.Pgl.like.BREX | <-ParB+HNH<-HKD-DNase<-?\|\|PglW->N6-METHYLASE->DUF6079*->PglZ->BrxD-AAA->SF1-SF2-Helicase+wHTH+Small-B-barrel+DUF1998->?->GTPase-AIG+sGTP->MMR_HSR1-><-REase | 1238 | Bacteria>Terrabacteria_group>Actinomycetota>Actinomycetes>Mycobacteriales | GCA_001667205.1 | Mycolicibacterium_peregrinum | A5719_17590 | 14293..18009 | 43304 | phage-resistance-protein-[Mycolicibacterium-peregrinum] |
| OCB36722.1 | Type2.Pgl.like.BREX | PglW->N6-METHYLASE->DUF6079*->PglZ->BrxD-AAA->SF1-SF2-Helicase+wHTH+Small-B-barrel+DUF1998->?-><-?<-?\|\|TetR-HTH-> | 1274 | Bacteria>Terrabacteria_group>Actinomycetota>Actinomycetes>Mycobacteriales | GCA_001686735.1 | Mycobacterium_malmoense | A5676_20310 | 113280..117104 | 1780 | hypothetical-protein-A5676_20310-[Mycobacterium-malmoense] |
| OCX20186.1 | Type2.Pgl.like.BREX | MerR-HTH->tRNA->PglW->N6-METHYLASE->DUF6079*->PglZ->BrxD-AAA->SF1-SF2-Helicase+wHTH+Small-B-barrel+DUF1998-> | 1224 | Bacteria>Pseudomonadota>Gammaproteobacteria>Pseudomonadales>Pseudomonadaceae | GCA_001705225.1 | Stutzerimonas_xanthomarina | BBI09_07390 | 172008..175682 | 271420 | phage-resistance-protein-[Stutzerimonas-xanthomarina] |
| ODS68498.1 | Type2.Pgl.like.BREX | WYL->PglW->N6-METHYLASE->DUF6079*-> | 1232 | Bacteria>Pseudomonadota>Betaproteobacteria>Burkholderiales>Alcaligenaceae>Bordetella | GCA_001724095.1 | Bordetella_sp._SCN_67-23 | ABS43_28105 | 8827..12525 | 1660091 | MAG:-hypothetical-protein-ABS43_28105-[Bordetella-sp.-SCN-67-23] |
| OHV25444.1 | Type2.Pgl.like.BREX | <-SF1-SF2-Helicase+wHTH+Small-B-barrel+DUF1998<-BrxD-AAA<-PglZ<-DUF6079*<-N6-METHYLASE<-PglW<-TetR-HTH\|\|?-><-Pkinase+TPR | 1400 | Bacteria>Terrabacteria_group>Actinomycetota>Actinomycetes>Frankiales | GCA_001854695.1 | Parafrankia_soli | BBK14_22215 | 38185..42387 | 2599596 | phage-resistance-protein-[Parafrankia-soli] |
| OHX06912.1 | Type2.Pgl.like.BREX | <-HNH\|\|PglW-><-?\|\|N6-METHYLASE-p->DUF6079*->PglZ->BrxD-AAA->SF1-SF2-Helicase+wHTH+Small-B-barrel+DUF1998->?->GTPase-AIG+sGTP->?->MMR_HSR1-> | 1246 | Bacteria>Terrabacteria_group>Actinomycetota>Actinomycetes>Micromonosporales | GCA_001855515.1 | Micromonospora_sp._WMMB235 | BFV98_30000 | 6717187..6720927 | 1172030 | hypothetical-protein-BFV98_30000-[Micromonospora-sp.-WMMB235] |
| OLL18356.1 | Type2.Pgl.like.BREX | <-HIN-HTH<-?\|\|<-?\|\|?-><-SF1-SF2-Helicase+wHTH+Small-B-barrel+DUF1998<-BrxD-AAA<-PglZ<-DUF6079*<-N6-METHYLASE<-PglW<-?\|\|?->\|\|DDE_Tnp_1-><-SNF+SF1-SF2-Helicase\|\|SF1-SF2-Helicase-> | 1243 | Bacteria>Terrabacteria_group>Actinomycetota>Actinomycetes>Mycobacteriales | GCA_001890475.2 | Rhodococcus_sp._M8 | BKE56_017695 | 433490..437221 | 1925550 | phage-resistance-protein-[Rhodococcus-sp.-M8] |
| OOC03654.1 | Type2.Pgl.like.BREX | <-REase+SF1-SF2-Helicase\|\|REase+SF1-SF2-Helicase-><-?<-SF1-SF2-Helicase+wHTH+Small-B-barrel+DUF1998<-BrxD-AAA<-PglZ<-DUF6079*<-N6-METHYLASE<-?<-Classical-AAA\|\|HIN-HTH-><-?<-?<-PglW | 1234 | Bacteria>Terrabacteria_group>Actinomycetota>Actinomycetes>Pseudonocardiales | GCA_001995215.1 | Amycolatopsis_azurea_DSM_43854 | B0293_25575 | 80450..84154 | 1238180 | PglY-protein-[Amycolatopsis-azurea-DSM-43854] |
| ORL86702.1 | Type2.Pgl.like.BREX | HEPN+SF1-SF2-Helicase+REase-><-?\|\|PglW->N6-METHYLASE->McrB-AAA-ATPase->McrC-NTD+REase->SAD+REase->DUF6079*->PglZ->BrxD-AAA->SF1-SF2-Helicase+wHTH+Small-B-barrel+DUF1998->?->DDE_Tnp_1_2->REase+SF1-SF2-Helicase->HNH-> | 1264 | Bacteria>Terrabacteria_group>Actinomycetota>Actinomycetes>Mycobacteriales | GCA_002095035.1 | Prescottella_equi | A5N78_18870 | 135247..139041 | 43767 | phage-resistance-protein-[Prescottella-equi] |
| ORT59286.1 | Type2.Pgl.like.BREX | <-SF1-SF2-Helicase+wHTH+Small-B-barrel+DUF1998<-BrxD-AAA<-PglZ<-DUF6079*<-N6-METHYLASE\|\|HEPN->ParA-Soj-PloopNTPase->BrxD-AAA-><-PglW | 1326 | Bacteria>Terrabacteria_group>Actinomycetota>Actinomycetes>Kitasatosporales | GCA_002099185.1 | Streptomyces_sp._CB03238 | BKD26_14935 | 390423..394403 | 1907777 | PglY-protein-[Streptomyces-sp.-CB03238] |
| PPG31569.1 | Type2.Pgl.like.BREX | PglW->N6-METHYLASE->DUF6079*->PglZ->BrxD-AAA->SF1-SF2-Helicase+wHTH+Small-B-barrel+DUF1998->ParB+HNH-> | 1202 | Bacteria>Terrabacteria_group>Actinomycetota>Actinomycetes>Micrococcales | GCA_002931595.1 | Rathayibacter_sp._AY2B9 | C5C25_07400 | 22881..26489 | 2080572 | hypothetical-protein-C5C25_07400-[Rathayibacter-sp.-AY2B9] |
| PPH99377.1 | Type2.Pgl.like.BREX | <-ParB+HNH\|\|?->?-><-SF1-SF2-Helicase+wHTH+Small-B-barrel+DUF1998<-BrxD-AAA<-PglZ<-DUF6079*<-N6-METHYLASE<-PglW | 1198 | Bacteria>Terrabacteria_group>Actinomycetota>Actinomycetes>Micrococcales | GCA_002932635.1 | Rathayibacter_sp._AY1D1 | C5C56_09115 | 81440..85036 | 2080542 | hypothetical-protein-C5C56_09115-[Rathayibacter-sp.-AY1D1] |
| PWK49465.1 | Type2.Pgl.like.BREX | <-MMR_HSR1<-GTPase-AIG+sGTP<-SF1-SF2-Helicase+wHTH+Small-B-barrel+DUF1998<-BrxD-AAA<-PglZ<-DUF6079*<-?<-Schlafen+HATPase+wHTH<-N6-METHYLASE<-PglW<-?\|\|?->N6-METHYLASE-> | 1232 | Bacteria>Terrabacteria_group>Actinomycetota>Actinomycetes>Micromonosporales | GCA_003148685.1 | Actinoplanes_xinjiangensis | BC793_104138 | 149029..152727 | 512350 | hypothetical-protein-BC793_104138-[Actinoplanes-xinjiangensis] |
| QBI18516.1 | Type2.Pgl.like.BREX | <-HIN-HTH\|\|HTH_21+rve+rve_2->?->?-><-SF1-SF2-Helicase+wHTH+Small-B-barrel+DUF1998<-BrxD-AAA<-PglZ<-DUF6079*<-?<-HEPN<-N6-METHYLASE<-PglW | 1260 | Bacteria>Terrabacteria_group>Actinomycetota>Nitriliruptoria>Egibacterales | GCA_004322855.1 | Egibacter_rhizosphaerae | ER308_02310 | 491517..495299 | 1670831 | phage-resistance-protein-[Egibacter-rhizosphaerae] |
| QCQ90489.1 | Type2.Pgl.like.BREX | <-HNH<-REase+SF1-SF2-Helicase<-ParB+HNH<-SF1-SF2-Helicase+wHTH+Small-B-barrel+DUF1998<-BrxD-AAA<-PglZ<-DUF6079*\|\|?-><-?<-N6-METHYLASE<-PglW<-?<-HATPase\|\|SF1-SF2-Helicase-> | 1264 | Bacteria>Terrabacteria_group>Actinomycetota>Actinomycetes>Mycobacteriales | GCA_005484805.1 | Rhodococcus_sp._SGAir0479 | E7742_04110 | 877804..881598 | 2567884 | phage-resistance-protein-[Rhodococcus-sp.-SGAir0479] |
| QDG64425.1 | Type2.Pgl.like.BREX | <-ParB+HNH<-SF1-SF2-Helicase+REase<-SF1-SF2-Helicase+REase<-HNH\|\|?->PglW->N6-METHYLASE->DUF6079*->PglZ->BrxD-AAA->SF1-SF2-Helicase+wHTH+Small-B-barrel+DUF1998-><-?<-OLD-ABC | 1263 | Bacteria>Terrabacteria_group>Actinomycetota>Actinomycetes>Micrococcales | GCA_006517215.1 | Pseudarthrobacter_sp._NIBRBAC000502771 | NIBR502771_20345 | 4407176..4410967 | 2590774 | phage-resistance-protein-[Pseudarthrobacter-sp.-NIBRBAC000502771] |
| QDU98792.1 | Type2.Pgl.like.BREX | XIS-HTH->PglW->N6-METHYLASE->OLD-ABC->HNH->DUF6079*->PglZ->BrxD-AAA->SF1-SF2-Helicase+wHTH+Small-B-barrel+DUF1998-> | 1225 | Bacteria>PVC_group>Planctomycetota>Planctomycetia>Pirellulales | GCA_007751035.1 | Lignipirellula_cremea | Pla8534_66660 | 8881234..8884911 | 2528010 | hypothetical-protein-Pla8534_66660-[Lignipirellula-cremea] |
| QEA05979.1 | Type2.Pgl.like.BREX | <-HTH_21+rve+rve_2<-HIN-HTH\|\|PglW->N6-METHYLASE->?->?->?->DUF6079*->PglZ->BrxD-AAA->SF1-SF2-Helicase+wHTH+Small-B-barrel+DUF1998-><-HIN-HTH | 1219 | unclassified_entries>unclassified_sequences>environmental_samples | - | uncultured_organism | KBTEX_02308 | 12215..15874 | 155900 | hypothetical-protein-KBTEX_02308-[uncultured-organism] |
| QGG94365.1 | Type2.Pgl.like.BREX | <-DNA_glycosylase<-D6+SF1-SF2-Helicase<-HKD-DNase<-DUF6361<-?\|\|PglW->N6-METHYLASE->DUF6079*->PglZ->BrxD-AAA->STYKinase-like->Couple_hipA+HIRAN->SF1-SF2-Helicase+wHTH+Small-B-barrel+DUF1998->RADICAL-SAM->SF1-SF2-Helicase-> | 1198 | Bacteria>Terrabacteria_group>Actinomycetota>Acidimicrobiia>Acidimicrobiales | GCA_009650215.1 | Actinomarinicola_tropica | GH723_04185 | 829858..833454 | 2789776 | phage-resistance-protein-[Actinomarinicola-tropica] |
| QKW00216.1 | Type2.Pgl.like.BREX | PglW->N6-METHYLASE-><-ST-Hiskin\|\|HTH_31+DUF5753->DUF397->HTH+MPTase->PIN->DUF6079*->PglZ-><-CBS+HEPN<-ParB+HNH\|\|BrxD-AAA->SF1-SF2-Helicase+wHTH+Small-B-barrel+DUF1998-> | 1309 | Bacteria>Terrabacteria_group>Actinomycetota>Actinomycetes>Kitasatosporales | GCA_013364175.1 | Streptomyces_sp._NA02536 | HUT14_10190 | 2278747..2282676 | 2742133 | phage-resistance-protein-[Streptomyces-sp.-NA02536] |
| QNI11044.1 | Type2.Pgl.like.BREX | PglW->N6-METHYLASE-><-?\|\|<-Phage_integrase<-?<-?\|\|\|\|N6-METHYLASE-p->DUF6079*->PglZ->BrxD-AAA->SF1-SF2-Helicase+wHTH+Small-B-barrel+DUF1998-><-?<-?<-SbcC+SbcC | 1260 | Bacteria>Terrabacteria_group>Actinomycetota>Actinomycetes>Mycobacteriales | GCA_014263335.1 | Mycobacterium_kubicae | GAN18_07330 | 1523162..1526944 | 120959 | hypothetical-protein-GAN18_07330-[Mycobacterium-kubicae] |
| QRK88134.1 | Type2.Pgl.like.BREX | <-REase+SAVED\|\|?->rve_3-><-SF1-SF2-Helicase+wHTH+Small-B-barrel+DUF1998<-BrxD-AAA<-PglZ<-DUF6079*<-N6-METHYLASE<-PglW\|\|SF1-SF2-Helicase+REase->?->HTH_29+HTH_32+DDE_3-> | 1273 | Bacteria>Terrabacteria_group>Actinomycetota>Actinomycetes>Pseudonocardiales | GCA_016859185.1 | Saccharopolyspora_erythraea | JQX30_25960 | 5815005..5818826 | 1836 | PglY-protein-[Saccharopolyspora-erythraea] |
| QXE29463.1 | Type2.Pgl.like.BREX | <-SF1-SF2-Helicase\|\|PglW->N6-METHYLASE->DUF6079*->PglZ->BrxD-AAA->SF1-SF2-Helicase+wHTH+Small-B-barrel+DUF1998-><-ABhydrolase | 1209 | Bacteria>Terrabacteria_group>Actinomycetota>Actinomycetes>Micrococcales | GCA_019056515.1 | Microbacterium_paraoxydans | IZR02_14000 | 2840251..2843880 | 199592 | hypothetical-protein-IZR02_14000-[Microbacterium-paraoxydans] |
| QXN93149.1 | Type2.Pgl.like.BREX | <-Phage_integrase<-AbrB<-SF1-SF2-Helicase+wHTH+Small-B-barrel+DUF1998<-BrxD-AAA<-PglZ<-DUF6079*<-N6-METHYLASE<-PglW\|\|HTH-><-tRNA\|\|?-><-tRNA<-35exo\|\|HerA/FtsK+A32-> | 1263 | Bacteria>Terrabacteria_group>Actinomycetota>Actinomycetes>Mycobacteriales | GCA_019222765.1 | Nocardia_iowensis | KV110_08615 | 1908465..1912256 | 204891 | phage-resistance-protein-[Nocardia-iowensis] |
| QXU55728.1 | Type2.Pgl.like.BREX | STYKinase-like->?-><-SF1-SF2-Helicase\|\|PglW->N6-METHYLASE-><-HEPN+REase\|\|DUF6079*->PglZ->BrxD-AAA->SF1-SF2-Helicase->SF1-SF2-Helicase+wHTH+Small-B-barrel+DUF1998->?-><-Glyoxalase | 1235 | Bacteria>Terrabacteria_group>Actinomycetota>Actinomycetes>Mycobacteriales | GCA_019317345.1 | Rhodococcus_sp._LW-XY12 | KXC42_11365 | 2418472..2422179 | 2856851 | phage-resistance-protein-[Rhodococcus-sp.-LW-XY12] |
| RAY12851.1 | Type2.Pgl.like.BREX | <-DUF6079*<-?<-PglW | 1254 | Bacteria>Terrabacteria_group>Actinomycetota>Actinomycetes>Streptosporangiales | GCA_003289645.1 | Actinomadura_craniellae | DPM19_22805 | 100013..103777 | 2231787 | PglY-protein-[Actinomadura-craniellae] |
| REK92078.1 | Type2.Pgl.like.BREX | <-SF1-SF2-Helicase+wHTH+Small-B-barrel+DUF1998<-BrxD-AAA<-PglZ<-DUF6079*<-N6-METHYLASE<-PglW\|\|DUF6731-><-?<-SAM-methylase | 1304 | Bacteria>Terrabacteria_group>Actinomycetota>Actinomycetes>Kitasatosporales | GCA_003389455.1 | Streptomyces_inhibens | DY245_00990 | 15167..19081 | 2293571 | PglY-protein-[Streptomyces-inhibens] |
| RKN35098.1 | Type2.Pgl.like.BREX | <-SF1-SF2-Helicase+wHTH+Small-B-barrel+DUF1998<-BrxD-AAA<-PglZ<-DUF6079*<-N6-METHYLASE<-PglW | 1300 | Bacteria>Terrabacteria_group>Actinomycetota>Actinomycetes>Kitasatosporales | GCA_003626535.1 | Streptomyces_hoynatensis | D7294_31150 | 7577..11479 | 1141874 | phage-resistance-protein-[Streptomyces-hoynatensis] |
| ROO83041.1 | Type2.Pgl.like.BREX | <-REase+SF1-SF2-Helicase<-SF1-SF2-Helicase+wHTH+Small-B-barrel+DUF1998<-BrxD-AAA<-PglZ<-DUF6079*<-N6-METHYLASE<-PglW<-?<-HTH_31+DUF5753\|\|?-><-HTH_29+HTH_32+DDE_3<-HTH_29+HTH_32+DDE_3 | 1269 | Bacteria>Terrabacteria_group>Actinomycetota>Actinomycetes>Streptosporangiales | GCA_003751225.1 | Actinocorallia_herbida | EDD29_0530 | 501794..505603 | 58109 | hypothetical-protein-EDD29_0530-[Actinocorallia-herbida] |
| ROQ80618.1 | Type2.Pgl.like.BREX | <-SF1-SF2-Helicase+wHTH+Small-B-barrel+DUF1998<-BrxD-AAA<-PglZ<-DUF6079*<-DUF397<-HTH_31+DUF5753\|\|?->?-><-PglW\|\|OLD-ABC-> | 1295 | Bacteria>Terrabacteria_group>Actinomycetota>Actinomycetes>Kitasatosporales | GCA_003752275.1 | Streptomyces_sp._CEV_2-1 | EDD95_0145 | 172440..176327 | 2485153 | hypothetical-protein-EDD95_0145-[Streptomyces-sp.-CEV-2-1] |
| SDL43696.1 | Type2.Pgl.like.BREX | PglW->N6-METHYLASE->DUF6079*->PglZ->?->HEPN+SNF+SF1-SF2-Helicase->?->BrxD-AAA->SF1-SF2-Helicase+wHTH+Small-B-barrel+DUF1998-> | 1240 | Bacteria>Terrabacteria_group>Actinomycetota>Actinomycetes>Propionibacteriales | GCA_900103275.1 | Tessaracoccus_oleiagri | SAMN04488242_1594 | 616502..620224 | 686624 | hypothetical-protein-SAMN04488242_1594-[Tessaracoccus-oleiagri] |
| SOC53729.1 | Type2.Pgl.like.BREX | PglW->N6-METHYLASE->?->?->?-><-ParB+HNH\|\|?->DUF6079*->PglZ->BrxD-AAA->SF1-SF2-Helicase+wHTH+Small-B-barrel+DUF1998->?->REase+SF1-SF2-Helicase->?-><-METHYLASE+DAM-Fungal | 1251 | Bacteria>Terrabacteria_group>Actinomycetota>Actinomycetes>Micrococcales | GCA_900221015.1 | Ornithinimicrobium_cerasi | SAMN05421879_10290 | 109157..112912 | 2248773 | hypothetical-protein-SAMN05421879_10290-[Ornithinimicrobium-cerasi] |
| TDO42299.1 | Type2.Pgl.like.BREX | PIN->?-><-SF1-SF2-Helicase+wHTH+Small-B-barrel+DUF1998<-BrxD-AAA<-PglZ<-DUF6079*<-N6-METHYLASE<-PglW<-ACET+ACET | 1255 | Bacteria>Terrabacteria_group>Actinomycetota>Actinomycetes>Micromonosporales | GCA_004362215.1 | Actinoplanes_brasiliensis | C8E87_6066 | 6729671..6733438 | 52695 | hypothetical-protein-C8E87_6066-[Actinoplanes-brasiliensis] |
| TMU14881.1 | Type2.Pgl.like.BREX | <-DUF6079*<-?<-PglW | 1236 | Bacteria>Pseudomonadota>Gammaproteobacteria>Oceanospirillales>Halomonadaceae | GCA_005937175.1 | Halomonas_sp._ATBC28 | E0L35_23020 | 16131..19841 | 2545264 | phage-resistance-protein-[Halomonas-sp.-ATBC28] |
| UOB08700.1 | Type2.Pgl.like.BREX | Phage_holin->ParB+HNH->CBS+HEPN-><-ST-Hiskin\|\|HTH_31+DUF5753->DUF397-><-PglZ<-DUF6079*<-N6-METHYLASE<-PglW<-?\|\|SF1-SF2-Helicase+HKD-DNase->?-><-REase+SF1-SF2-Helicase<-REase+SF1-SF2-Helicase | 1299 | Bacteria>Terrabacteria_group>Actinomycetota>Actinomycetes>Kitasatosporales | GCA_022807935.1 | Streptomyces_sp._HP-A2021 | MQE23_06345 | 1370327..1374226 | 2927875 | phage-resistance-protein-[Streptomyces-sp.-HP-A2021] |
| UXY34411.1 | Type2.Pgl.like.BREX | PAPS-reduct->DCM->DnaB+DnaB->?->PglW->N6-METHYLASE->DUF6079*->PglZ->BrxD-AAA->SF1-SF2-Helicase+wHTH+Small-B-barrel+DUF1998-><-NUDIX<-REase+DUF5655\|\|<-PIWI<-HNH | 1299 | Bacteria>Terrabacteria_group>Actinomycetota>Actinomycetes>Kitasatosporales | GCA_025642175.1 | Streptomyces_sp._HUAS_14-6 | N8I86_06515 | 1385275..1389174 | 2981135 | phage-resistance-protein-[Streptomyces-sp.-HUAS-14-6] |
| VVN60021.1 | Type2.Pgl.like.BREX | MerR-HTH->tRNA->PglW->N6-METHYLASE->DUF6079*->PglZ->BrxD-AAA->SF1-SF2-Helicase+wHTH+Small-B-barrel+DUF1998-><-?<-?<-?<-SF1-SF2-Helicase | 1225 | Bacteria>Pseudomonadota>Gammaproteobacteria>Pseudomonadales>Pseudomonadaceae | GCA_902497805.1 | Pseudomonas_fluorescens | PS687_03579 | 85409..89086 | 294 | hypothetical-protein-PS687_03579-[Pseudomonas-fluorescens] |
| WFG45383.1 | Type2.Pgl.like.BREX | ParB+HNH->PglW->DOC->N6-METHYLASE->DUF6079*->PglZ->BrxD-AAA->SF1-SF2-Helicase+wHTH+Small-B-barrel+DUF1998->ParB+HNH-> | 1247 | Bacteria>Terrabacteria_group>Actinomycetota>Actinomycetes>Pseudonocardiales | GCA_045159825.1 | Pseudonocardia_alni | PaSha_20215 | 4112617..4116360 | 33907 | PglY-protein-[Pseudonocardia-alni] |
| WP_015767813.1 | Type2.Pgl.like.BREX | <-SF1-SF2-Helicase+wHTH+Small-B-barrel+DUF1998<-BrxD-AAA<-PglZ<-DUF6079*<-?<-?<-?<-OLD-ABC<-N6-METHYLASE<-PglW | 1215 | Bacteria>Pseudomonadota>Betaproteobacteria>Betaproteobacteria_incertae_sedis>Candidatus_Accumulibacter | GCF_017302455.1 | Accumulibacter_sp. | J5X23_RS16850 | 3801310..3804957 | 2053492 | hypothetical-protein-[Accumulibacter-sp.] |
| WP_055470265.1 | Type2.Pgl.like.BREX | <-SF1-SF2-Helicase+wHTH+Small-B-barrel+DUF1998<-BrxD-AAA<-PglZ<-DUF6079*<-N6-METHYLASE<-PglW\|\|DUF6731-> | 1299 | Bacteria>Terrabacteria_group>Actinomycetota>Actinomycetes>Kitasatosporales>Streptomycetaceae | GCF_044387625.1 | Streptomyces_pathocidini | ACH429_RS00805 | 205406..209305 | 1650571 | hypothetical-protein-[Streptomyces-pathocidini] |
| WP_055814081.1 | Type2.Pgl.like.BREX | <-MazG\|\|REase+SF1-SF2-Helicase->HNH-><-SF1-SF2-Helicase<-?\|\|PglW->N6-METHYLASE->DUF6079*->PglZ->BrxD-AAA->SF1-SF2-Helicase+wHTH+Small-B-barrel+DUF1998-><-?<-Ribonuc_red_2_N+Ribonuc_red_lgC | 1256 | Bacteria>Terrabacteria_group>Actinomycetota>Actinomycetes>Micrococcales | GCF_001428065.1 | Nostocoides_sp._Soil756 | ASG78_RS09445 | 108460..112230 | 1736399 | hypothetical-protein-[Tetrasphaera-sp.-Soil756] |
| WP_061229212.1 | Type2.Pgl.like.BREX | PglW-><-?\|\|N6-METHYLASE->DUF6079*->PglZ->BrxD-AAA->SF1-SF2-Helicase+wHTH+Small-B-barrel+DUF1998-><-?<-ABC-ATPase | 1327 | Bacteria>Terrabacteria_group>Actinomycetota>Actinomycetes>Mycobacteriales>Dietziaceae | GCF_001571065.1 | Dietzia_cinnamea | AB6N35_RS14285 | 3001285..3005268 | 321318 | DUF6079-family-protein-[Dietzia-cinnamea] |
| WP_066853731.1 | Type2.Pgl.like.BREX | PglW->OLD-ABC+TOPRIM-><-?<-?\|\|?->N6-METHYLASE->DUF6079*->PglZ->BrxD-AAA->SF1-SF2-Helicase+wHTH+Small-B-barrel+DUF1998->?-><-SF1-SF2-Helicase+REase<-?<-SNF+SF1-SF2-Helicase | 1234 | Bacteria>Terrabacteria_group>Actinomycetota>Actinomycetes>Mycobacteriales | GCF_001668625.1 | Mycobacterium_sp._1274756.6 | A5643_RS08105 | 97079..100783 | 1834076 | DUF6079-family-protein-[Mycobacterium-sp.-1274756.6] |
| WP_073695752.1 | Type2.Pgl.like.BREX | <-HKD-DNase<-?\|\|PglW->N6-METHYLASE->DUF6079*->PglZ->BrxD-AAA-> | 1275 | Bacteria>Terrabacteria_group>Actinomycetota>Actinomycetes>Mycobacteriales | GCF_039668335.1 | unclassified_Mycobacterium | EB75_RS12775 | 75844..79671 | 2642494 | DUF6079-family-protein-[Mycobacterium-sp.-ST-F2] |
| WP_102141148.1 | Type2.Pgl.like.BREX | <-SF1-SF2-Helicase+wHTH+Small-B-barrel+DUF1998<-BrxD-AAA<-PglZ<-DUF6079*<-N6-METHYLASE<-HTH<-?<-?<-?<-PglW | 1220 | Bacteria>Terrabacteria_group>Actinomycetota>Actinomycetes>Mycobacteriales>Mycobacteriaceae | GCF_002838065.1 | Mycobacterium_sp._QGD_101 | CWS39_RS00170 | 36255..39917 | 1867256 | DUF6079-family-protein-[Mycobacterium-sp.-QGD-101] |
| WP_115328993.1 | Type2.Pgl.like.BREX | HTH->?->ZnR->PglW->?->PglW->N6-METHYLASE->DUF6079*->PglZ-> | 1229 | Bacteria>Terrabacteria_group>Actinomycetota>Actinomycetes>Mycobacteriales>Mycobacteriaceae | GCF_025821885.1 | Mycolicibacterium_gilvum | H7I56_RS00795 | 17644..21333 | 1804 | phage-resistance-protein-[Mycolicibacterium-gilvum] |
| WP_149378030.1 | Type2.Pgl.like.BREX | <-DUF6079*<-?<-PglW | 1226 | Bacteria>Terrabacteria_group>Actinomycetota>Actinomycetes>Mycobacteriales | GCF_008329905.1 | Mycobacterium_sp._ELW1 | D3H54_RS04400 | 962839..966519 | 1547487 | phage-resistance-protein-[Mycobacterium-sp.-ELW1] |
| WP_149438602.1 | Type2.Pgl.like.BREX | <-MMR_HSR1<-GTPase-AIG+sGTP\|\|PglW->N6-METHYLASE->DUF6079*->PglZ->BrxD-AAA->SF1-SF2-Helicase+wHTH+Small-B-barrel+DUF1998->McrB-AAA-ATPase->McrC-NTD+REase->\|\|35exo+SF1-SF2-Helicase->SNF+SF1-SF2-Helicase-> | 1226 | Bacteria>Terrabacteria_group>Actinomycetota>Actinomycetes>Mycobacteriales | GCF_008329565.1 | Mycolicibacterium_sp._P1-5 | CIW47_RS06200 | 77198..80878 | 2024617 | phage-resistance-protein-[Mycolicibacterium-sp.-P1-5] |
| WP_156045472.1 | Type2.Pgl.like.BREX | MMR_HSR1->?-><-McrC-NTD+REase<-McrB-AAA-ATPase<-SF1-SF2-Helicase+wHTH+Small-B-barrel+DUF1998<-BrxD-AAA<-PglZ<-DUF6079*<-N6-METHYLASE<-PglW\|\|HNH-> | 1195 | Bacteria>Terrabacteria_group>Actinomycetota | GCF_000717275.1 | Actinomycetes | OP91_RS11225 | 176978..180565 | 1760 | MULTISPECIES:-DUF6079-family-protein-[Actinomycetes] |
| WP_167733812.1 | Type2.Pgl.like.BREX | PglW->?->DUF6079*-> | 1246 | Bacteria>Terrabacteria_group>Actinomycetota>Actinomycetes>Micromonosporales | GCF_011765725.1 | Salinispora_arenicola | HCB17_RS05545 | 251906..255646 | 168697 | PglY-protein-[Salinispora-arenicola] |
| WP_169181230.1 | Type2.Pgl.like.BREX | <-GNTR-HTH\|\|MazG-><-REase+SF1-SF2-Helicase<-N6-METHYLASE<-SF1-SF2-Helicase+wHTH+Small-B-barrel+DUF1998<-BrxD-AAA<-PglZ<-DUF6079*<-N6-METHYLASE<-PglW<-Pkinase+TPR\|\|?->SbcC+SbcC->HEPN-><-DUF397 | 1229 | Bacteria>Terrabacteria_group>Actinomycetota>Actinomycetes>Micromonosporales | GCF_012933445.1 | Actinoplanes_sp._TBRC_11911 | HH310_RS41300 | 30518..34207 | 2729386 | phage-resistance-protein-[Actinoplanes-sp.-TBRC-11911] |
| WP_171810739.1 | Type2.Pgl.like.BREX | <-REase<-?<-?<-PglZ<-DUF6079*<-MACRODOMAIN<-DarT+Peptidase<-?<-PglW | 1208 | Bacteria>Myxococcota>Myxococcia>Myxococcales>Cystobacterineae>Myxococcaceae | GCF_013155575.1 | Corallococcus_exiguus | HRD50_RS12025 | 24201..27827 | 83462 | DUF6079-family-protein-[Corallococcus-exiguus] |
| WP_182870882.1 | Type2.Pgl.like.BREX | <-SF1-SF2-Helicase+wHTH+Small-B-barrel+DUF1998<-BrxD-AAA<-PglZ<-DUF6079*<-Schlafen+HATPase+wHTH<-N6-METHYLASE<-PglW<-XIS-HTH\|\|?-><-Phage_integrase | 1236 | Bacteria>PVC_group>Planctomycetota>Planctomycetia>Pirellulales | GCF_014154935.1 | Rhodopirellula_sp._JC639 | H1B37_RS29500 | 14885..18595 | 2755585 | hypothetical-protein-[Rhodopirellula-sp.-JC639] |
| WP_184672148.1 | Type2.Pgl.like.BREX | <-HTH<-?\|\|?-><-?\|\|PglW->N6-METHYLASE->DUF6079*->PglZ->BrxD-AAA->SF1-SF2-Helicase+wHTH+Small-B-barrel+DUF1998-><-SAM-methylase<-?\|\|PglZ->METHYLASE-> | 1272 | Bacteria>Terrabacteria_group>Actinomycetota>Actinomycetes>Pseudonocardiales | GCF_039537345.1 | Saccharothrix_violaceirubra | F4559_RS23655 | 5188653..5192471 | 413306 | PglY-protein-[Saccharothrix-violaceirubra] |
| WP_184852634.1 | Type2.Pgl.like.BREX | <-Pkinase+TPR\|\|<-?\|\|?->PglW->N6-METHYLASE->DUF6079*->PglZ-><-DUF397<-HTH_31+DUF5753\|\|ST-Hiskin-><-CBS+HEPN<-ParB+HNH | 1298 | Bacteria>Terrabacteria_group>Actinomycetota>Actinomycetes>Kitasatosporales | GCF_039540655.1 | Streptomyces_collinus | HNR72_RS32100 | 7190212..7194108 | 42684 | phage-resistance-protein-[Streptomyces-collinus] |
| WP_195033066.1 | Type2.Pgl.like.BREX | HTH_3->STYKIN->?->?->?->PglW->N6-METHYLASE->DUF6079*->?->BrxD-AAA->SF1-SF2-Helicase+wHTH+Small-B-barrel+DUF1998-><-?<-?<-?<-PadR-wHTH | 1256 | Bacteria>Terrabacteria_group>Actinomycetota>Actinomycetes>Mycobacteriales | GCF_015477435.1 | Nocardia_abscessus | IU470_RS12230 | 97362..101132 | 120957 | DUF6079-family-protein-[Nocardia-abscessus] |
| WP_195132798.1 | Type2.Pgl.like.BREX | DUF2397->?->?->?-><-SF1-SF2-Helicase+wHTH+Small-B-barrel+DUF1998<-BrxD-AAA<-PglZ<-DUF6079*<-N6-METHYLASE<-PglW<-DUF397<-HTH_31+DUF5753 | 1256 | Bacteria>Terrabacteria_group>Actinomycetota>Actinomycetes>Mycobacteriales | GCF_015477605.1 | Nocardia_amamiensis | IU459_RS29160 | 22411..26181 | 404578 | DUF6079-family-protein-[Nocardia-amamiensis] |
| WP_199444075.1 | Type2.Pgl.like.BREX | <-REase+SF1-SF2-Helicase<-wHTH+SAD+HNH<-REase+SF1-SF2-Helicase<-SF1-SF2-Helicase+wHTH+Small-B-barrel+DUF1998<-BrxD-AAA<-PglZ<-DUF6079*<-N6-METHYLASE<-PglW<-HTH_31+DUF5753\|\|CHTH+SAM-methylase-><-?<-CHTH+SAM-methylase<-HTH | 1288 | Bacteria>Terrabacteria_group>Actinomycetota>Actinomycetes>Pseudonocardiales | GCF_016464305.1 | Umezawaea_beigongshangensis | IMZ35_RS27315 | 28668..32534 | 2780383 | PglY-protein-[Umezawaea-beigongshangensis] |
| WP_210943087.1 | Type2.Pgl.like.BREX | <-SAD+REase\|\|REase->PIN-><-N6-METHYLASE\|\|?->PglW->N6-METHYLASE->DUF6079*->PglZ->BrxD-AAA->SF1-SF2-Helicase+wHTH+Small-B-barrel+DUF1998-><-Phage_int_SAM_3+Phage_integrase<-XIS-HTH-><-?<-BirA-HTH | 1249 | Bacteria>Terrabacteria_group>Actinomycetota>Actinomycetes>Micromonosporales | GCF_018070525.1 | Micromonospora_sp._C32 | KBX35_RS23085 | 127089..130838 | 2824877 | DUF6079-family-protein-[Micromonospora-sp.-C32] |
| WP_213457626.1 | Type2.Pgl.like.BREX | PglW->N6-METHYLASE->?->?->HEPN->?->N6-METHYLASE-p->DUF6079*->PglZ->BrxD-AAA->SF1-SF2-Helicase+wHTH+Small-B-barrel+DUF1998-><-?<-NUDIX\|\|?-><-SF1-SF2-Helicase | 1225 | Bacteria>Pseudomonadota>Gammaproteobacteria>Chromatiales>Chromatiaceae | GCF_018399035.1 | Thiocapsa_sp. | KFB96_RS15350 | 3389147..3392824 | 2024551 | phage-resistance-protein-[Thiocapsa-sp.] |
| WP_214403725.1 | Type2.Pgl.like.BREX | SF1-SF2-Helicase->ParB+HNH->ARPP2-><-SF1-SF2-Helicase+wHTH+Small-B-barrel+DUF1998<-BrxD-AAA<-PglZ<-DUF6079*<-N6-METHYLASE<-PglW\|\|XIS-HTH->Phage_int_SAM_3+Phage_integrase-><-ncRNA\|\|?-><-MGS+DUF1205+UDPGT | 1237 | Bacteria>Terrabacteria_group>Actinomycetota>Actinomycetes>Pseudonocardiales>Pseudonocardiaceae | GCF_018531445.1 | Pseudonocardia_sp._H11425 | KIK08_RS14850 | 18305..22018 | 2835865 | PglY-protein-[Pseudonocardia-sp.-H11425] |
| WP_217614697.1 | Type2.Pgl.like.BREX | SAD+REase-><-REase\|\|?->PglW->N6-METHYLASE->DUF6079*->PglZ->BrxD-AAA->SF1-SF2-Helicase+wHTH+Small-B-barrel+DUF1998-> | 1280 | Bacteria>Terrabacteria_group>Actinomycetota>Actinomycetes>Micrococcales>Cellulomonadaceae | GCF_019091725.1 | Cellulomonas_sp._GbtcB1 | KVT14_RS05210 | 47286..51128 | 2824746 | hypothetical-protein-[Cellulomonas-sp.-GbtcB1] |
| WP_221025642.1 | Type2.Pgl.like.BREX | <-REase\|\|MORC->SF1-SF2-Helicase->REase->PglW->N6-METHYLASE->DUF6079*->PglZ-><-?<-?<-ParB+HNH<-ParB+HNH | 1223 | Bacteria>Terrabacteria_group>Actinomycetota>Actinomycetes>Propionibacteriales | GCF_019679455.1 | Nocardioides_jiangsuensis | K1X13_RS14055 | 1162868..1166539 | 2866161 | DUF6079-family-protein-[Nocardioides-jiangsuensis] |
| WP_265868465.1 | Type2.Pgl.like.BREX | PglW->N6-METHYLASE-><-DUF397<-HTH_31+DUF5753\|\|?->?->REase+SF1-SF2-Helicase->DUF6079*->PglZ->AIPR->BrxD-AAA->SF1-SF2-Helicase+wHTH+Small-B-barrel+DUF1998->?-><-REase+SF1-SF2-Helicase | 1307 | Bacteria>Terrabacteria_group>Actinomycetota>Actinomycetes>Kitasatosporales>Streptomycetaceae | GCF_026236535.1 | Streptomyces_sp._SKN60 | KV205_RS32645 | 76897..80820 | 2855506 | DUF6079-family-protein-[Streptomyces-sp.-SKN60] |
| WP_267686051.1 | Type2.Pgl.like.BREX | PglW->?->DUF6079*-> | 1268 | Bacteria>Myxococcota>Polyangia>Nannocystales>Nannocystaceae | GCF_026626685.1 | Nannocystis_sp._SCPEA4 | OV142_RS31705 | 1094415..1098221 | 2996787 | hypothetical-protein-[Nannocystis-sp.-SCPEA4] |
| WP_268784737.1 | Type2.Pgl.like.BREX | PglW->N6-METHYLASE->DUF6079*->PglZ->BrxD-AAA->SF1-SF2-Helicase+wHTH+Small-B-barrel+DUF1998-><-SF1-SF2-Helicase<-SF1-SF2-Helicase<-?<-XIS-HTH | 1212 | Bacteria>Myxococcota>Myxococcia>Myxococcales>Cystobacterineae | GCF_026810205.1 | Myxococcus_sp._NMCA1 | OZ403_RS18695 | 4350343..4353981 | 2996785 | hypothetical-protein-[Myxococcus-sp.-NMCA1] |
| WP_269037386.1 | Type2.Pgl.like.BREX | PglW->?->?->?->?->DUF6079*-> | 1233 | Bacteria>Myxococcota>Polyangia>Nannocystales>Nannocystaceae | GCF_026965555.1 | Nannocystis_poenicansa | O0S08_RS02735 | 740509..744210 | 2995304 | hypothetical-protein-[Nannocystis-poenicansa] |
| WP_272090199.1 | Type2.Pgl.like.BREX | <-SF1-SF2-Helicase+wHTH+Small-B-barrel+DUF1998<-?<-BrxD-AAA<-PglZ<-DUF6079*<-N6-METHYLASE<-PglW | 1179 | Bacteria>Myxococcota>Polyangia>Nannocystales>Nannocystaceae | GCF_028368995.1 | Nannocystis_bainbridge | POL25_RS32610 | 1868845..1872384 | 2995303 | DUF6079-family-protein-[Nannocystis-bainbridge] |
| WP_272136732.1 | Type2.Pgl.like.BREX | HTH_Tnp_1+HTH_21+rve+rve_3->?-><-SF1-SF2-Helicase+wHTH+Small-B-barrel+DUF1998<-BrxD-AAA<-PglZ<-DUF6079*<-N6-METHYLASE<-PglW<-?<-XIS-HTH | 1234 | Bacteria>Myxococcota>Myxococcia>Myxococcales>Cystobacterineae | GCF_028368975.1 | Stigmatella_ashevillena | POL68_RS09770 | 1769957..1773661 | 2995309 | hypothetical-protein-[Stigmatella-ashevillena] |
| WP_276096852.1 | Type2.Pgl.like.BREX | rve_3-><-SF1-SF2-Helicase+wHTH+Small-B-barrel+DUF1998<-BrxD-AAA<-PglZ<-DUF6079*<-ST-Hiskin<-N6-METHYLASE\|\|HEPN->ParA-Soj-PloopNTPase->BrxD-AAA-><-PglW | 1328 | Bacteria>Terrabacteria_group>Actinomycetota>Actinomycetes>Kitasatosporales | GCF_029269905.1 | Streptomyces_silvisoli | P3G67_RS33410 | 7645..11631 | 3034235 | PglY-protein-[Streptomyces-silvisoli] |
| WP_276475786.1 | Type2.Pgl.like.BREX | <-SF1-SF2-Helicase+wHTH+Small-B-barrel+DUF1998<-BrxD-AAA<-PglZ<-DUF6079*<-N6-METHYLASE<-DUF397<-HTH_31+DUF5753\|\|?->?-><-PglW\|\|OLD-ABC-> | 1307 | Bacteria>Terrabacteria_group>Actinomycetota>Actinomycetes>Kitasatosporales | GCF_029351045.1 | Streptomyces_sp._WMMB303 | P2424_RS12255 | 2755258..2759181 | 3034154 | phage-resistance-protein-[Streptomyces-sp.-WMMB303] |
| WP_277210618.1 | Type2.Pgl.like.BREX | PglW->N6-METHYLASE->DUF6079*->PglZ->?->?->?->METHYLASE-> | 1271 | Bacteria>Terrabacteria_group>Actinomycetota>Actinomycetes>Micrococcales | GCF_029382255.1 | Isoptericola_croceus | PZC42_RS10170 | 342796..346611 | 3031406 | DUF6079-family-protein-[Isoptericola-croceus] |
| WP_277311123.1 | Type2.Pgl.like.BREX | PglW->N6-METHYLASE-p->N6-METHYLASE-p->DUF6079*->PglZ->BrxD-AAA->SF1-SF2-Helicase+wHTH+Small-B-barrel+DUF1998-> | 1294 | Bacteria>Terrabacteria_group>Actinomycetota>Actinomycetes>Streptosporangiales | GCF_029382225.1 | Spongiactinospora_sp._TRM90649 | P3X83_RS42450 | 14204..18088 | 3031114 | hypothetical-protein-[Spongiactinospora-sp.-TRM90649] |
| WP_277377410.1 | Type2.Pgl.like.BREX | REase+SF1-SF2-Helicase->HNH-><-HNH\|\|PglW->N6-METHYLASE->DUF6079*->PglZ->BrxD-AAA->SF1-SF2-Helicase+wHTH+Small-B-barrel+DUF1998->?->?-><-HHH_4<-ParA-Soj-PloopNTPase | 1303 | Bacteria>Terrabacteria_group>Actinomycetota>Actinomycetes>Mycobacteriales | GCF_029457415.1 | Rhodococcus_sp._RCBS9 | POM12_RS02265 | 466988..470899 | 3031999 | DUF6079-family-protein-[Rhodococcus-sp.-RCBS9] |
| WP_278145170.1 | Type2.Pgl.like.BREX | <-N6-METHYLASE\|\|?->PglW->N6-METHYLASE->DUF6079*->PglZ->BrxD-AAA->SF1-SF2-Helicase+wHTH+Small-B-barrel+DUF1998-><-?<-OLD-ABC+TOPRIM\|\|?-><-Phage_int_SAM_3+Phage_integrase | 1248 | Bacteria>Terrabacteria_group>Actinomycetota>Actinomycetes>Micromonosporales | GCF_029626165.1 | Micromonospora_sp._WMMD967 | O7631_RS17565 | 3776154..3779900 | 3016101 | hypothetical-protein-[Micromonospora-sp.-WMMD967] |
| WP_278468317.1 | Type2.Pgl.like.BREX | <-SF1-SF2-Helicase+wHTH+Small-B-barrel+DUF1998<-BrxD-AAA<-PglZ<-DUF6079*<-HNH<-OLD-ABC<-N6-METHYLASE<-PglW\|\|REase-> | 1234 | Bacteria>PVC_group>Planctomycetota>Planctomycetia>Planctomycetales | GCF_963971965.1 | Gimesia_maris | B4K77_RS12150 | 525086..528790 | 122 | phage-resistance-protein-[Gimesia-maris] |
| WP_279100766.1 | Type2.Pgl.like.BREX | SF1-SF2-Helicase+REase-><-?<-?<-SAD+REase<-SF1-SF2-Helicase+wHTH+Small-B-barrel+DUF1998<-BrxD-AAA<-PglZ<-DUF6079*<-N6-METHYLASE<-PglW\|\|URI-DNase-> | 1237 | Bacteria>Terrabacteria_group>Actinomycetota>Actinomycetes>Mycobacteriales | GCF_946223135.1 | Gordonia_bronchialis | QDO25_RS10180 | 21950..25663 | 2054 | DUF6079-family-protein-[Gordonia-bronchialis] |
| WP_279329418.1 | Type2.Pgl.like.BREX | <-SF1-SF2-Helicase+wHTH+Small-B-barrel+DUF1998<-BrxD-AAA\|\|ParB+HNH->CBS+HEPN-><-PglZ<-DUF6079*<-?<-?<-DUF397<-HTH_31+DUF5753\|\|ST-Hiskin-><-N6-METHYLASE<-PglW | 1309 | Bacteria>Terrabacteria_group>Actinomycetota>Actinomycetes>Kitasatosporales | GCF_029759795.1 | Streptomyces_sp._OS603R | P3T58_RS14585 | 166207..170136 | 3035287 | phage-resistance-protein-[Streptomyces-sp.-OS603R] |
| WP_279639433.1 | Type2.Pgl.like.BREX | <-wHTH+WYL+WCX<-?<-?<-?<-?<-?<-BrxD-AAA<-?<-DUF6079*<-?<-PglW | 1224 | Bacteria>Pseudomonadota>Gammaproteobacteria>Pseudomonadales>Pseudomonadaceae | GCF_029834775.1 | Stutzerimonas_stutzeri | N7329_RS03690 | 79252..82926 | 316 | phage-resistance-protein-[Stutzerimonas-stutzeri] |
| WP_279949630.1 | Type2.Pgl.like.BREX | REase+SF1-SF2-Helicase-><-?\|\|HTH_31+DUF5753->DUF397->?->PglW->N6-METHYLASE->DUF6079*->PglZ-><-?\|\|BrxD-AAA->SF1-SF2-Helicase+wHTH+Small-B-barrel+DUF1998-> | 1295 | Bacteria>Terrabacteria_group>Actinomycetota>Actinomycetes>Kitasatosporales | GCF_039540175.1 | Streptomyces_chitinivorans | ACG5V6_RS01050 | 76981..80868 | 1257027 | phage-resistance-protein-[Streptomyces-chitinivorans] |
| WP_280250352.1 | Type2.Pgl.like.BREX | PglW->N6-METHYLASE->DUF6079*->PglZ->BrxD-AAA->SF1-SF2-Helicase+wHTH+Small-B-barrel+DUF1998->?->?-><-HTH | 1285 | Bacteria>Terrabacteria_group>Actinomycetota>Actinomycetes>Mycobacteriales | GCF_029868845.1 | Nocardia_abscessus | QH188_RS09210 | 8986..12843 | 120957 | phage-resistance-protein-[Nocardia-abscessus] |
| WP_280274518.1 | Type2.Pgl.like.BREX | DUF2397->?->?->?->PglW->N6-METHYLASE->DUF6079*->PglZ->BrxD-AAA->SF1-SF2-Helicase+wHTH+Small-B-barrel+DUF1998-> | 1257 | Bacteria>Terrabacteria_group>Actinomycetota>Actinomycetes>Mycobacteriales | GCF_029868745.1 | Nocardia_wallacei | QH470_RS23465 | 21090..24863 | 480035 | DUF6079-family-protein-[Nocardia-wallacei] |
| WP_280288877.1 | Type2.Pgl.like.BREX | PglW->?->DUF6079*-> | 1224 | Bacteria>Pseudomonadota>Gammaproteobacteria>Pseudomonadales>Pseudomonadaceae | GCF_029871205.1 | Pseudomonas_sp._BN417 | E8F11_RS11045 | 33583..37257 | 2567890 | phage-resistance-protein-[Pseudomonas-sp.-BN417] |
| WP_280294506.1 | Type2.Pgl.like.BREX | PglW->N6-METHYLASE->DUF6079*->PglZ->BrxD-AAA->SF1-SF2-Helicase+wHTH+Small-B-barrel+DUF1998->?->?-><-HTH | 1285 | Bacteria>Terrabacteria_group>Actinomycetota>Actinomycetes>Mycobacteriales | GCF_029867355.1 | Nocardia_abscessus | QH274_RS12405 | 9124..12981 | 120957 | phage-resistance-protein-[Nocardia-abscessus] |
| WP_280310786.1 | Type2.Pgl.like.BREX | CASPASE+SF1-SF2-Helicase->?->?->?->PglW->N6-METHYLASE->DUF6079*->PglZ-> | 1285 | Bacteria>Terrabacteria_group>Actinomycetota>Actinomycetes>Mycobacteriales | GCF_029869045.1 | Nocardia_abscessus | QH193_RS31580 | 30327..34184 | 120957 | phage-resistance-protein-[Nocardia-abscessus] |
| WP_280369382.1 | Type2.Pgl.like.BREX | SF1-SF2-Helicase-><-SF1-SF2-Helicase+wHTH+Small-B-barrel+DUF1998<-BrxD-AAA<-PglZ<-DUF6079*<-N6-METHYLASE<-PglW<-?<-TetR-HTH\|\|ABhydrolase-> | 1288 | Bacteria>Terrabacteria_group>Actinomycetota>Actinomycetes>Mycobacteriales | GCF_029868875.1 | Nocardia_wallacei | QH200_RS30080 | 33237..37103 | 480035 | phage-resistance-protein-[Nocardia-wallacei] |
| WP_280447666.1 | Type2.Pgl.like.BREX | HTH_31+DUF5753->DUF397->?-><-SF1-SF2-Helicase+wHTH+Small-B-barrel+DUF1998<-BrxD-AAA<-PglZ<-DUF6079*<-N6-METHYLASE<-PglW\|\|?-><-?<-?<-?<-HEPN+SF1-SF2-Helicase+REase | 1250 | Bacteria>Terrabacteria_group>Actinomycetota>Actinomycetes>Mycobacteriales | GCF_029868825.1 | Nocardia_cyriacigeorgica | QH034_RS24375 | 157159..160911 | 135487 | DUF6079-family-protein-[Nocardia-cyriacigeorgica] |
| WP_280458879.1 | Type2.Pgl.like.BREX | XIS-HTH->?-><-?<-SF1-SF2-Helicase+wHTH+Small-B-barrel+DUF1998<-BrxD-AAA<-PglZ<-DUF6079*<-N6-METHYLASE<-PglW<-DUF397<-HTH_31+DUF5753 | 1251 | Bacteria>Terrabacteria_group>Actinomycetota>Actinomycetes>Mycobacteriales | GCF_029869845.1 | Nocardia_carnea | QH182_RS24940 | 91561..95316 | 37328 | DUF6079-family-protein-[Nocardia-carnea] |
| WP_280669740.1 | Type2.Pgl.like.BREX | PglW->N6-METHYLASE-><-DUF397<-HTH_31+DUF5753\|\|ST-Hiskin->?->Pkinase+TPR->DUF6079*->PglZ->BrxD-AAA->SF1-SF2-Helicase+wHTH+Small-B-barrel+DUF1998->SAD(SRA)+wHTH+HNH->?->?->\|\|BrxD-AAA-> | 1313 | Bacteria>Terrabacteria_group>Actinomycetota>Actinomycetes>Kitasatosporales | GCF_029892095.1 | unclassified_Kitasatospora | ABIA34_RS11505 | 265653..269594 | 2633591 | phage-resistance-protein-[Kitasatospora-sp.-MAP12-44] |
| WP_280879660.1 | Type2.Pgl.like.BREX | PglW->N6-METHYLASE->REase+SF1-SF2-Helicase->?->DUF6079*->PglZ->AIPR->BrxD-AAA->SF1-SF2-Helicase+wHTH+Small-B-barrel+DUF1998->YjhX_toxin-><-TRAM+SAM-methylase+METHYLASE<-AA_permease_2 | 1297 | Bacteria>Terrabacteria_group>Actinomycetota>Actinomycetes>Kitasatosporales | GCF_029892565.1 | Streptomyces_pseudovenezuelae | M2283_RS30540 | 263505..267398 | 67350 | DUF6079-family-protein-[Streptomyces-pseudovenezuelae] |
| WP_281145638.1 | Type2.Pgl.like.BREX | PglW->?->DUF6079*-> | 1282 | Bacteria>Terrabacteria_group>Actinomycetota>Actinomycetes>Propionibacteriales | GCF_029917005.1 | Tessaracoccus_sp._T21 | QH948_RS04195 | 925172..929020 | 3041766 | hypothetical-protein-[Tessaracoccus-sp.-T21] |
| WP_281601969.1 | Type2.Pgl.like.BREX | <-SAD+REase<-N6-METHYLASE\|\|?->PglW->N6-METHYLASE-p->HTH_Tnp_4+DDE_Tnp_4->N6-METHYLASE->DUF6079*->PglZ->BrxD-AAA->SF1-SF2-Helicase+wHTH+Small-B-barrel+DUF1998->DNA-glycosylase-><-Phage_int_SAM_3+Phage_integrase<-XIS-HTH | 1247 | Bacteria>Terrabacteria_group>Actinomycetota>Actinomycetes>Micromonosporales | GCF_027497155.1 | Verrucosispora_sp._WMMA2044 | O3597_RS19410 | 4271873..4275616 | 3016419 | hypothetical-protein-[Verrucosispora-sp.-WMMA2044] |
| WP_282203631.1 | Type2.Pgl.like.BREX | <-SF1-SF2-Helicase+wHTH+Small-B-barrel+DUF1998<-BrxD-AAA<-PglZ<-DUF6079*<-PglW<-?\|\|?->N6-METHYLASE-><-REase+SF1-SF2-Helicase | 1346 | Bacteria>Terrabacteria_group>Actinomycetota>Actinomycetes>Kitasatosporales | GCF_948472415.1 | Kitasatospora_fiedleri | QMQ26_RS31720 | 7019286..7023326 | 2991545 | phage-resistance-protein-[Kitasatospora-fiedleri] |
| WP_282697753.1 | Type2.Pgl.like.BREX | pPIWI+pPIWI->pPIWI+SFII-RAD3->pPIWI+MID+RNaseH->?->PglW->N6-METHYLASE->DUF6079*->PglZ->BrxD-AAA->SF1-SF2-Helicase+wHTH+Small-B-barrel+DUF1998-> | 1300 | Bacteria>Terrabacteria_group>Actinomycetota>Actinomycetes>Kitasatosporales | GCF_030010625.1 | Streptomyces_sp._CC208A | QK442_RS32750 | 16894..20796 | 3044573 | phage-resistance-protein-[Streptomyces-sp.-CC208A] |
| WP_282880752.1 | Type2.Pgl.like.BREX | <-REase\|\|PglW->N6-METHYLASE-><-DUF397<-HTH_31+DUF5753\|\|ST-Hiskin->DUF6079*->PglZ->BrxD-AAA->SF1-SF2-Helicase+wHTH+Small-B-barrel+DUF1998-> | 1298 | Bacteria>Terrabacteria_group>Actinomycetota>Actinomycetes>Kitasatosporales | GCF_030016275.1 | Streptomyces_coelicoflavus | QMA61_RS13290 | 283279..287175 | 285562 | phage-resistance-protein-[Streptomyces-coelicoflavus] |
| WP_283136706.1 | Type2.Pgl.like.BREX | <-TetR-HTH\|\|?->?->PglW->N6-METHYLASE->DUF6079*->PglZ->BrxD-AAA->SF1-SF2-Helicase+wHTH+Small-B-barrel+DUF1998-> | 1293 | Bacteria>Terrabacteria_group>Actinomycetota>Actinomycetes>Streptosporangiales | GCF_025996515.1 | Rhizohabitans_arisaemae | RUR_RS20470 | 1062407..1066288 | 2720610 | hypothetical-protein-[Rhizohabitans-arisaemae] |
| WP_283518500.1 | Type2.Pgl.like.BREX | HNH-><-PIN<-?<-?<-DUF6731\|\|PglW->N6-METHYLASE->DUF6079*->PglZ->BrxD-AAA->SF1-SF2-Helicase+wHTH+Small-B-barrel+DUF1998-><-PIN<-?\|\|<-N6-METHYLASE | 1296 | Bacteria>Terrabacteria_group>Actinomycetota>Actinomycetes>Kitasatosporales | GCF_030063415.1 | Streptomyces_sp._H27-C3 | QMK30_RS15520 | 162925..166815 | 3046305 | PglY-protein-[Streptomyces-sp.-H27-C3] |
| WP_283567920.1 | Type2.Pgl.like.BREX | HEPN+SF1-SF2-Helicase+REase->REase+SF1-SF2-Helicase->HNH-><-HNH\|\|?->PglW->N6-METHYLASE->DUF6079*->PglZ->BrxD-AAA->SF1-SF2-Helicase+wHTH+Small-B-barrel+DUF1998->McrB-AAA-ATPase->McrC-NTD+REase-> | 1303 | Bacteria>Terrabacteria_group>Actinomycetota>Actinomycetes>Mycobacteriales | GCF_030063525.1 | Rhodococcus_erythropolis | QNA23_RS16655 | 148115..152026 | 1833 | DUF6079-family-protein-[Rhodococcus-erythropolis] |
| WP_284918334.1 | Type2.Pgl.like.BREX | <-SF1-SF2-Helicase+wHTH+Small-B-barrel+DUF1998<-BrxD-AAA<-PglZ<-DUF6079*<-?<-N6-METHYLASE<-PglW<-CASPASE+SF1-SF2-Helicase<-XIS-HTH<-Phage_integrase<-Phage_int_SAM_3+Phage_integrase | 1232 | Bacteria>Terrabacteria_group>Actinomycetota>Actinomycetes>Micromonosporales | GCF_030252345.1 | Actinoplanes_oblitus | Actob_RS02425 | 536459..540157 | 3040509 | DUF6079-family-protein-[Actinoplanes-oblitus] |
| WP_285505841.1 | Type2.Pgl.like.BREX | <-SF1-SF2-Helicase+wHTH+Small-B-barrel+DUF1998<-BrxD-AAA<-PglZ<-DUF6079*<-N6-METHYLASE<-PglW<-STYKinase-like<-ParB+HNH | 1262 | Bacteria>Terrabacteria_group>Actinomycetota>Actinomycetes>Pseudonocardiales | GCF_030269645.1 | Actinokineospora_sp._NBRC_105648 | QSK30_RS23520 | 11746..15534 | 3032206 | PglY-protein-[Actinokineospora-sp.-NBRC-105648] |
| WP_285588125.1 | Type2.Pgl.like.BREX | ABhydrolase->HTH_29+HTH_32+DDE_3-><-?<-?<-SF1-SF2-Helicase+wHTH+Small-B-barrel+DUF1998<-BrxD-AAA<-PglZ<-DUF6079*<-?<-N6-METHYLASE<-PglW\|\|SNF+SF1-SF2-Helicase->SF1-SF2-Helicase-><-REase<-SF1-SF2-Helicase | 1221 | Bacteria>Terrabacteria_group>Actinomycetota>Actinomycetes>Pseudonocardiales | GCF_030269685.1 | Actinomycetospora_sp._NBRC_106378 | QSK00_RS04795 | 77910..81575 | 3032208 | hypothetical-protein-[Actinomycetospora-sp.-NBRC-106378] |
| WP_285664606.1 | Type2.Pgl.like.BREX | HisKA_3+HATPase_c->REC+LuxR-HTH-><-SF1-SF2-Helicase+wHTH+Small-B-barrel+DUF1998<-?<-Schlafen+HATPase+wHTH<-BrxD-AAA<-PglZ<-DUF6079*<-N6-METHYLASE<-PglW\|\|CASPASE+SF1-SF2-Helicase-><-?<-?<-?\|\|HTH_31+DUF5753-> | 1230 | Bacteria>Terrabacteria_group>Actinomycetota>Actinomycetes>Micromonosporales | GCF_030269765.1 | Actinorhabdospora_filicis | QSK10_RS20935 | 124926..128618 | 1785913 | hypothetical-protein-[Actinorhabdospora-filicis] |
| WP_285698961.1 | Type2.Pgl.like.BREX | HisKA_3+HATPase_c->REC+LuxR-HTH-><-?\|\|METHYLASE-><-SF1-SF2-Helicase+wHTH+Small-B-barrel+DUF1998<-BrxD-AAA<-PglZ<-DUF6079*<-N6-METHYLASE<-?<-?\|\|HTH_31+DUF5753->DUF397-><-PglW | 1298 | Bacteria>Terrabacteria_group>Actinomycetota>Actinomycetes>Streptosporangiales | GCF_030269515.1 | Actinomadura_sp._NBRC_104412 | QSK21_RS23565 | 61669..65565 | 3032203 | DUF6079-family-protein-[Actinomadura-sp.-NBRC-104412] |
| WP_285760465.1 | Type2.Pgl.like.BREX | McrB-AAA-ATPase->McrC-NTD+REase->PglW->N6-METHYLASE->DUF6079*->PglZ->BrxD-AAA->SF1-SF2-Helicase+wHTH+Small-B-barrel+DUF1998-> | 1296 | Bacteria>Terrabacteria_group>Actinomycetota>Actinomycetes>Streptosporangiales | GCF_030268225.1 | Nocardiopsis_ansamitocini | QSJ59_RS16540 | 381683..385573 | 1670832 | hypothetical-protein-[Nocardiopsis-ansamitocini] |
| WP_285777368.1 | Type2.Pgl.like.BREX | HTH->NUDIX-><-?\|\|PglW->N6-METHYLASE->DUF6079*->PglZ->BrxD-AAA->SF1-SF2-Helicase+wHTH+Small-B-barrel+DUF1998-> | 1299 | Bacteria>Terrabacteria_group>Actinomycetota>Actinomycetes>Streptosporangiales | GCF_030268685.1 | Microtetraspora_sp._NBRC_13810 | QSK26_RS23345 | 14983..18882 | 3030990 | hypothetical-protein-[Microtetraspora-sp.-NBRC-13810] |

| Protein-ID | System | Gene-Neighbourhoods | Length | Taxonomy | Assembly-ID | Strain | Locus-Tag | Genomic-coordinates | TaxID | Fasta-defline |
|---|---|---|---|---|---|---|---|---|---|---|
| WP_286267699.1 | Type2.Pgl.like.BREX | METHYLASE->?->?->tRNA->PglW->N6-METHYLASE->DUF6079*->PglZ->?->BrxD-AAA->SF1-SF2-Helicase+wHTH+Small-B-barrel+DUF1998-> | 1265 | Bacteria>Terrabacteria_group>Actinomycetota>Actinomycetes>Propionibacteriales | GCF_030297015.1 | Brooklawnia_propionicigenes | QUE25_RS03735 | 818664..822461 | 3041175 | hypothetical-protein-[Brooklawnia-sp.-SH051] |
| WP_287001672.1 | Type2.Pgl.like.BREX | <-REase<-SAD+REase<-SF1-SF2-Helicase+wHTH+Small-B-barrel+DUF1998<-BrxD-AAA<-PglZ<-DUF6079*<-N6-METHYLASE<-PglW\|\|URI-DNase-> | 1238 | Bacteria>Terrabacteria_group>Actinomycetota>Actinomycetes>Mycobacteriales | GCF_032084605.1 | Gordonia | RT393_RS15455 | 15359..19075 | 2053 | MULTISPECIES:-DUF6079-family-protein-[Gordonia] |
| WP_287381674.1 | Type2.Pgl.like.BREX | HTH-><-SF1-SF2-Helicase+wHTH+Small-B-barrel+DUF1998<-BrxD-AAA<-PglZ<-DUF6079*<-N6-METHYLASE<-PglW\|\|XIS-HTH-> | 1233 | Bacteria>Pseudomonadota>Alphaproteobacteria>Hyphomicrobiales>Phyllobacteriaceae | GCF_005063535.1 | Mesorhizobium_sp. | E5X35_RS24620 | 10898..14599 | 1871066 | hypothetical-protein-[Mesorhizobium-sp.] |
| WP_289218832.1 | Type2.Pgl.like.BREX | <-REase\|\|PglW->N6-METHYLASE-><-DUF397<-HTH_31+DUF5753\|\|ST-Hiskin->DUF6079*->PglZ->BrxD-AAA->SF1-SF2-Helicase+wHTH+Small-B-barrel+DUF1998-><-TRAM+SAM-methylase+METHYLASE<-AA_permease_2 | 1298 | Bacteria>Terrabacteria_group>Actinomycetota>Actinomycetes>Kitasatosporales | GCF_030345275.1 | Streptomyces_albidoflavus | QIA47_RS05575 | 1324085..1327981 | 1886 | phage-resistance-protein-[Streptomyces-albidoflavus] |
| WP_291980612.1 | Type2.Pgl.like.BREX | <-SF1-SF2-Helicase+wHTH+Small-B-barrel+DUF1998<-BrxD-AAA<-PglZ<-DUF6079*<-N6-METHYLASE<-PglW<-SF1-SF2-Helicase | 1208 | Bacteria>Acidobacteriota>Vicinamibacteria>Vicinamibacterales>Vicinamibacteraceae | GCF_022842525.1 | Luteitalea_sp. | Q0C41_RS01205 | 28219..31845 | 2004800 | hypothetical-protein-[Luteitalea-sp.] |
| WP_291996285.1 | Type2.Pgl.like.BREX | PglW->N6-METHYLASE->HEPN->ParA-Soj-PloopNTPase->ORC-CDC6->?->PIN->DUF6079*->PglZ->?->?->?->BrxD-AAA->SF1-SF2-Helicase+wHTH+Small-B-barrel+DUF1998-> | 1223 | Bacteria>Pseudomonadota>Betaproteobacteria>Betaproteobacteria_incertae_sedis>Candidatus_Accumulibacter | GCF_907163155.1 | Candidatus_Accumulibacter_sp._ACC012 | QZ648_RS04955 | 327559..331230 | 2823332 | phage-resistance-protein-[Candidatus-Accumulibacter-sp.-ACC012] |
| WP_293603323.1 | Type2.Pgl.like.BREX | HTH_26->PglW->N6-METHYLASE->N6-METHYLASE-p->?->MqsA_antitoxin+HTH->?->DUF6079*->PglZ->BrxD-AAA->SF1-SF2-Helicase+wHTH+Small-B-barrel+DUF1998-> | 1243 | Bacteria>Pseudomonadota>Betaproteobacteria>Burkholderiales>Comamonadaceae | GCF_002381615.1 | Polaromonas_sp._UBA4122 | B3F77_RS01950 | 22015..25746 | 1947074 | phage-resistance-protein-[Polaromonas-sp.-UBA4122] |
| WP_295584575.1 | Type2.Pgl.like.BREX | PglW->N6-METHYLASE->DUF6079*->PglZ->?->HNH-> | 1232 | Bacteria>Pseudomonadota>Gammaproteobacteria>Chromatiales>Chromatiaceae | GCF_903930065.1 | uncultured_Lamprocystis_sp. | QZY03_RS21840 | 14863..18561 | 543132 | phage-resistance-protein-[uncultured-Lamprocystis-sp.] |
| WP_295590672.1 | Type2.Pgl.like.BREX | PglW->N6-METHYLASE->DUF6079*->PglZ-> | 1232 | Bacteria>Pseudomonadota>Gammaproteobacteria>Chromatiales>Chromatiaceae | GCF_903960425.1 | uncultured_Lamprocystis_sp. | QZY11_RS13335 | 14923..18621 | 543132 | phage-resistance-protein-[uncultured-Lamprocystis-sp.] |
| WP_296554869.1 | Type2.Pgl.like.BREX | HEPN->?-><-SF1-SF2-Helicase+wHTH+Small-B-barrel+DUF1998<-BrxD-AAA<-PglZ<-DUF6079*<-N6-METHYLASE<-PglW<-?\|\|HNH-><-HNH<-REase+SF1-SF2-Helicase<-HEPN+SF1-SF2-Helicase+REase | 1303 | Bacteria>Terrabacteria_group>Actinomycetota>Actinomycetes>Mycobacteriales | GCF_016463425.1 | Rhodococcus_sp._(in:_high_G+C_Gram-positive_bacteria) | JHC75_RS20655 | 40983..44894 | 1831 | DUF6079-family-protein-[Rhodococcus-sp.-(in:-high-G+C-Gram-positive-bacteria)] |
| WP_298626467.1 | Type2.Pgl.like.BREX | HTH_26->PglW->N6-METHYLASE->HEPN->ParA-Soj-PloopNTPase->ORC-CDC6->OLD-ABC->DUF6079*->PglZ->BrxD-AAA->SF1-SF2-Helicase+wHTH+Small-B-barrel+DUF1998-><-?\|\|HTH-> | 1231 | Bacteria>Pseudomonadota>Betaproteobacteria>Rhodocyclales>Zoogloeaceae | GCF_937894855.1 | uncultured_Zoogloea_sp. | Q0672_RS22915 | 12500..16195 | 160237 | phage-resistance-protein-[uncultured-Zoogloea-sp.] |
| WP_300083179.1 | Type2.Pgl.like.BREX | HTH_26->PglW->N6-METHYLASE->?->MqsA_antitoxin+HTH->?->DUF6079*->PglZ->BrxD-AAA->SF1-SF2-Helicase+wHTH+Small-B-barrel+DUF1998-> | 1229 | Bacteria>Pseudomonadota>Betaproteobacteria>Burkholderiales>Comamonadaceae | GCF_028688095.1 | Rhodoferax_sp. | PHS22_RS04580 | 37371..41060 | 50421 | phage-resistance-protein-[Rhodoferax-sp.] |
| WP_300083660.1 | Type2.Pgl.like.BREX | <-PglZ<-DUF6079*<-PglW<-PglW<-SF1-SF2-Helicase | 1239 | Bacteria>Terrabacteria_group>Actinomycetota>Actinomycetes>Propionibacteriales | GCF_029977985.1 | Propioniciclava_sp. | Q1K57_RS13985 | 1803..5522 | 2038686 | hypothetical-protein-[Propioniciclava-sp.] |
| WP_300336619.1 | Type2.Pgl.like.BREX | HTH_26->PglW->N6-METHYLASE->?->N6-METHYLASE-p->?->DOC->DUF6079*->PglZ->?->?->SNF+SF1-SF2-Helicase-> | 1227 | Bacteria>Pseudomonadota>Betaproteobacteria>Betaproteobacteria_incertae_sedis>Candidatus_Accumulibacter | GCF_029255145.1 | Accumulibacter_sp. | Q1L50_RS05715 | 73126..76809 | 2053492 | DUF6079-family-protein-[Accumulibacter-sp.] |
| WP_300443210.1 | Type2.Pgl.like.BREX | PglW->N6-METHYLASE->HEPN->ParA-Soj-PloopNTPase->ORC-CDC6->OLD-ABC->?->DUF6079*->PglZ->BrxD-AAA->SF1-SF2-Helicase+wHTH+Small-B-barrel+DUF1998-> | 1229 | Bacteria>Pseudomonadota>Betaproteobacteria>Rhodocyclales>Zoogloeaceae | GCF_030157465.1 | Zoogloea_sp. | Q1W26_RS18740 | 24950..28639 | 49181 | phage-resistance-protein-[Zoogloea-sp.] |
| WP_300537122.1 | Type2.Pgl.like.BREX | HTH_26->PglW->N6-METHYLASE->?->DUF6079*->PglZ->BrxD-AAA->SF1-SF2-Helicase+wHTH+Small-B-barrel+DUF1998->?->?->?-><-HKD-DNase | 1233 | Bacteria>Pseudomonadota>Betaproteobacteria>Burkholderiales>Comamonadaceae | GCF_029945925.1 | Polaromonas_sp. | PSV40_RS10270 | 39659..43360 | 1869339 | phage-resistance-protein-[Polaromonas-sp.] |
| WP_301143627.1 | Type2.Pgl.like.BREX | HEPN+SF1-SF2-Helicase+REase->?-><-REase+SF1-SF2-Helicase\|\|MazG->PglW->N6-METHYLASE->DUF6079*->PglZ->BrxD-AAA->SF1-SF2-Helicase+wHTH+Small-B-barrel+DUF1998-><-?<-McrC-NTD+REase<-McrB-AAA-ATPase\|\|REase-> | 1232 | Bacteria>Terrabacteria_group>Actinomycetota>Actinomycetes>Micrococcales | GCF_030418295.1 | Demequina_sp._EGI_L300058 | QQX02_RS02065 | 443985..447683 | 3051664 | DUF6079-family-protein-[Demequina-sp.-EGI-L300058] |
| WP_301174165.1 | Type2.Pgl.like.BREX | PglW->N6-METHYLASE-><-?\|\|HTH_31+DUF5753->DUF397-><-?<-OLD-ABC\|\|DUF6079*->PglZ->BrxD-AAA->SF1-SF2-Helicase+wHTH+Small-B-barrel+DUF1998-> | 1216 | Bacteria>Terrabacteria_group>Actinomycetota>Actinomycetes>Streptosporangiales | GCF_030417495.1 | Actinomadura_geliboluensis | Q0Q90_RS04900 | 100085..103735 | 882440 | phage-resistance-protein-[Actinomadura-geliboluensis] |
| WP_301229242.1 | Type2.Pgl.like.BREX | PglW->N6-METHYLASE->DUF6079*->PglZ->BrxD-AAA->SF1-SF2-Helicase+wHTH+Small-B-barrel+DUF1998-><-?\|\|ParB+HNH->HKD-DNase+SF1-SF2-Helicase-> | 1234 | Bacteria>Terrabacteria_group>Actinomycetota>Actinomycetes>Micrococcales | GCF_030433645.1 | Arthrobacter_burdickii | P5G52_RS15705 | 232263..235967 | 3035920 | DUF6079-family-protein-[Arthrobacter-burdickii] |
| WP_301672196.1 | Type2.Pgl.like.BREX | PglW->N6-METHYLASE->DUF6079*->PglZ->BrxD-AAA->SF1-SF2-Helicase+wHTH+Small-B-barrel+DUF1998-> | 1214 | Bacteria>Pseudomonadota>Gammaproteobacteria>Methylococcales>Methylococcaceae | GCF_030490285.1 | Methylomonas_montana | QZJ86_RS19535 | 4233231..4236875 | 3058963 | DUF6079-family-protein-[Methylomonas-montana] |
| WP_301915527.1 | Type2.Pgl.like.BREX | PglW->?->DDE_Tnp_1_4->N6-METHYLASE->\|\|N6-METHYLASE-p->ParB+HNH-><-?\|\|DUF6079*->PglZ->BrxD-AAA->SF1-SF2-Helicase-><-HicB\|\|SF1-SF2-Helicase+wHTH+Small-B-barrel+DUF1998-> | 1234 | Bacteria>Terrabacteria_group>Actinomycetota>Actinomycetes>Mycobacteriales | GCF_030490385.1 | Rhodococcus_ruber | QYN14_RS07640 | 1601122..1604826 | 1830 | DUF6079-family-protein-[Rhodococcus-ruber] |
| WP_304042055.1 | Type2.Pgl.like.BREX | <-HTH<-?\|\|REase+SF1-SF2-Helicase->URI-DNase+SF1-SF2-Helicase->PglW->N6-METHYLASE->DUF6079*->PglZ->BrxD-AAA->?->?->SF1-SF2-Helicase+wHTH+Small-B-barrel+DUF1998-> | 1289 | Bacteria>Terrabacteria_group>Actinomycetota>Actinomycetes>Mycobacteriales | GCF_030461435.1 | Corynebacterium_glyciniphilum | L0L97_RS09505 | 19609..23478 | 1404244 | DUF6079-family-protein-[Corynebacterium-glyciniphilum] |
| WP_305683716.1 | Type2.Pgl.like.BREX | HTH_26->PglW->N6-METHYLASE->DUF6079*->PglZ->BrxD-AAA->SF1-SF2-Helicase+wHTH+Small-B-barrel+DUF1998-> | 1236 | Bacteria>Pseudomonadota>Betaproteobacteria>Burkholderiales>Comamonadaceae | GCF_030696875.1 | Hydrogenophaga_sp. | Q8S12_RS11645 | 19495..23205 | 1904254 | phage-resistance-protein-[Hydrogenophaga-sp.] |
| WP_306086581.1 | Type2.Pgl.like.BREX | PglW->N6-METHYLASE-><-DUF397<-HTH_31+DUF5753\|\|?->?->DUF6079*->PglZ->AIPR->BrxD-AAA->SF1-SF2-Helicase+wHTH+Small-B-barrel+DUF1998-> | 1306 | Bacteria>Terrabacteria_group>Actinomycetota>Actinomycetes>Kitasatosporales | GCF_030719275.1 | Streptomyces_laculatispora | P8A22_RS08810 | 2077617..2081537 | 887464 | phage-resistance-protein-[Streptomyces-laculatispora] |
| WP_306316462.1 | Type2.Pgl.like.BREX | GNTR-HTH-><-?<-SF1-SF2-Helicase+wHTH+Small-B-barrel+DUF1998<-BrxD-AAA\|\|PglW->N6-METHYLASE->DUF6079*->PglZ-><-DUF397<-HTH_31+DUF5753\|\|ST-Hiskin-><-?\|\|McrB-AAA-ATPase->McrC-NTD+REase-> | 1278 | Bacteria>Terrabacteria_group>Actinomycetota>Actinomycetes>Kitasatosporales | GCF_030766935.1 | unclassified_Streptomyces | RBJ65_RS07385 | 337477..341313 | 2593676 | MULTISPECIES:-DUF6079-family-protein-[unclassified-Streptomyces] |
| WP_306332893.1 | Type2.Pgl.like.BREX | <-DUF6731\|\|PglW->N6-METHYLASE->DUF6079*->PglZ->BrxD-AAA->SF1-SF2-Helicase+wHTH+Small-B-barrel+DUF1998-><-PIN | 1279 | Bacteria>Terrabacteria_group>Actinomycetota>Actinomycetes>Kitasatosporales | GCF_030766985.1 | Streptomyces_sp._KL118A | RBJ68_RS04450 | 56197..60036 | 3045153 | PglY-protein-[Streptomyces-sp.-KL118A] |
| WP_306948864.1 | Type2.Pgl.like.BREX | PglW->N6-METHYLASE-><-DUF397<-HTH_31+DUF5753\|\|?->?->REase+SF1-SF2-Helicase->DUF6079*->PglZ->AIPR->BrxD-AAA->SF1-SF2-Helicase+wHTH+Small-B-barrel+DUF1998-> | 1294 | Bacteria>Terrabacteria_group>Actinomycetota>Actinomycetes>Kitasatosporales | GCF_030817335.1 | Streptomyces_sp._B4I13 | QFZ66_RS12495 | 2616168..2620052 | 3042271 | phage-resistance-protein-[Streptomyces-sp.-B4I13] |
| WP_307099892.1 | Type2.Pgl.like.BREX | Phage_int_SAM_3+Phage_integrase->?->?->?->?->PglW->N6-METHYLASE->DUF6079*->PglZ->BrxD-AAA->SF1-SF2-Helicase+wHTH+Small-B-barrel+DUF1998-><-?<-?\|\|?-><-N6-METHYLASE | 1279 | Bacteria>Terrabacteria_group>Actinomycetota>Actinomycetes>Micrococcales | GCF_030817195.1 | Arthrobacter_globiformis | QF036_RS05365 | 1159174..1163013 | 1665 | hypothetical-protein-[Arthrobacter-globiformis] |
| WP_307240194.1 | Type2.Pgl.like.BREX | <-SF1-SF2-Helicase+wHTH+Small-B-barrel+DUF1998<-BrxD-AAA<-PglZ<-DUF6079*<-N6-METHYLASE\|\|?-><-?<-PglW | 1244 | Bacteria>Terrabacteria_group>Actinomycetota>Actinomycetes>Micromonosporales | GCF_030813715.1 | Catenuloplanes_indicus | J2S42_RS16660 | 3706448..3710182 | 137267 | hypothetical-protein-[Catenuloplanes-indicus] |
| WP_307361509.1 | Type2.Pgl.like.BREX | <-HIN-HTH<-?\|\|PglW->N6-METHYLASE->DUF6079*->PglZ->BrxD-AAA->SF1-SF2-Helicase+wHTH+Small-B-barrel+DUF1998-><-URI-DNase+SF1-SF2-Helicase<-MazG<-HKD-DNase+SF1-SF2-Helicase | 1263 | Bacteria>Terrabacteria_group>Actinomycetota>Actinomycetes>Micrococcales | GCF_030812615.1 | Pseudarthrobacter_niigatensis | J2T24_RS14755 | 9863..13654 | 369935 | DUF6079-family-protein-[Pseudarthrobacter-niigatensis] |
| WP_307610921.1 | Type2.Pgl.like.BREX | <-SF1-SF2-Helicase+wHTH+Small-B-barrel+DUF1998<-BrxD-AAA<-PglZ<-DUF6079*<-N6-METHYLASE<-N6-METHYLASE<-PglW\|\|?->HKD-DNase->D6+SF1-SF2-Helicase->SNF+SF1-SF2-Helicase-> | 1286 | Bacteria>Terrabacteria_group>Actinomycetota>Actinomycetes>Micrococcales | GCF_030817835.1 | Pseudarthrobacter_sp._W1I19 | QF038_RS15270 | 3284680..3288540 | 3042288 | DUF6079-family-protein-[Pseudarthrobacter-sp.-W1I19] |
| WP_308011205.1 | Type2.Pgl.like.BREX | Phage_integrase-><-Phage_integrase<-AbrB\|\|?->PglW->N6-METHYLASE->DUF6079*->PglZ->BrxD-AAA->SF1-SF2-Helicase+wHTH+Small-B-barrel+DUF1998-><-SF1-SF2-Helicase\|\|?-><-?<-MazG | 1267 | Bacteria>Terrabacteria_group>Actinomycetota>Actinomycetes>Pseudonocardiales | GCF_020177455.1 | unclassified_Saccharopolyspora | LCD33_RS01215 | 29832..33635 | 2646250 | MULTISPECIES:-PglY-protein-[unclassified-Saccharopolyspora] |
| WP_308676743.1 | Type2.Pgl.like.BREX | SF1-SF2-Helicase-><-SF1-SF2-Helicase+wHTH+Small-B-barrel+DUF1998<-BrxD-AAA<-PglZ<-DUF6079*<-N6-METHYLASE<-PglW<-?<-?\|\|REase+SF1-SF2-Helicase-> | 1262 | Bacteria>Terrabacteria_group>Actinomycetota>Actinomycetes>Pseudonocardiales | GCF_031082405.1 | Amycolatopsis_sp._A133 | Q5425_RS35455 | 68266..72054 | 3064472 | PglY-protein-[Amycolatopsis-sp.-A133] |
| WP_309182153.1 | Type2.Pgl.like.BREX | PglW->N6-METHYLASE->DUF6079*->PglZ->BrxD-AAA->SF1-SF2-Helicase+wHTH+Small-B-barrel+DUF1998-> | 1237 | Bacteria>Pseudomonadota>Alphaproteobacteria>Hyphomicrobiales>Nitrobacteraceae | GCF_031202265.1 | Bradyrhizobium_sp._Ash2021 | NL528_RS07945 | 1660498..1664211 | 2954771 | hypothetical-protein-[Bradyrhizobium-sp.-Ash2021] |
| WP_309961937.1 | Type2.Pgl.like.BREX | HTH_26->PglW->?->N6-METHYLASE->?->DUF6079*->HTH_Tnp_1+HTH_21+rve+rve_3->PglZ->BrxD-AAA->SF1-SF2-Helicase+wHTH+Small-B-barrel+DUF1998-><-?<-?<-N6-METHYLASE | 1229 | Bacteria>Pseudomonadota>Betaproteobacteria>Burkholderiales>Burkholderiaceae | GCF_031450875.1 | Caballeronia_sp._LZ001 | P9292_RS25930 | 3201527..3205216 | 3038553 | phage-resistance-protein-[Caballeronia-sp.-LZ001] |
| WP_309967004.1 | Type2.Pgl.like.BREX | <-HNH<-SNF+SF1-SF2-Helicase<-?<-SF1-SF2-Helicase<-SF1-SF2-Helicase+wHTH+Small-B-barrel+DUF1998<-BrxD-AAA<-PglZ<-DUF6079*<-N6-METHYLASE<-PglW\|\|CR-ATPase5-><-ArdCN+MPTase<-GDResolvase+HIN-HTH | 1240 | Bacteria>Pseudomonadota>Betaproteobacteria>Burkholderiales>Comamonadaceae | GCF_031456095.1 | Variovorax_guangxiensis | J2W96_RS20835 | 402514..406236 | 1775474 | DUF6079-family-protein-[Variovorax-guangxiensis] |
| WP_309976827.1 | Type2.Pgl.like.BREX | HNH->REase->PglW->N6-METHYLASE->DUF6079*->PglZ->BrxD-AAA->SF1-SF2-Helicase+wHTH+Small-B-barrel+DUF1998->ParB+HNH->ParB+HNH-> | 1223 | Bacteria>Terrabacteria_group>Actinomycetota>Actinomycetes>Micrococcales | GCF_031456545.1 | Agromyces_sp._3263 | J2X63_RS10530 | 2285618..2289289 | 2817750 | hypothetical-protein-[Agromyces-sp.-3263] |
| WP_310135047.1 | Type2.Pgl.like.BREX | Phage_integrase-><-?<-?<-?\|\|HTH_26->PglW->N6-METHYLASE->DUF6079*->PglZ->BrxD-AAA->SF1-SF2-Helicase+wHTH+Small-B-barrel+DUF1998->?->?->SbcC-> | 1232 | Bacteria>Pseudomonadota>Betaproteobacteria>Burkholderiales>Burkholderiaceae | GCF_031454725.1 | Paraburkholderia_terricola | J2797_RS27565 | 57993..61691 | 169427 | phage-resistance-protein-[Paraburkholderia-terricola] |
| WP_310769024.1 | Type2.Pgl.like.BREX | Pkinase+TPR->\|\|<-SF1-SF2-Helicase+wHTH+Small-B-barrel+DUF1998<-BrxD-AAA<-AIPR<-PglZ<-DUF6079*<-REase+SF1-SF2-Helicase<-?<-?\|\|HTH_31+DUF5753->DUF397-><-N6-METHYLASE<-PglW | 1295 | Bacteria>Terrabacteria_group>Actinomycetota>Actinomycetes>Kitasatosporales | GCF_031583065.1 | Streptomyces_sp._CGMCC_4.7035 | Q2K21_RS09895 | 2364072..2367959 | 3061628 | phage-resistance-protein-[Streptomyces-sp.-CGMCC-4.7035] |
| WP_310872976.1 | Type2.Pgl.like.BREX | PglW-><-?<-?\|\|?->?->?->DUF6079*-> | 1302 | Bacteria>Terrabacteria_group>Actinomycetota>Actinomycetes>Kitasatosporales | GCF_031597775.1 | Streptomyces_sp._BB1-1-1 | RI578_RS23835 | 5160580..5164488 | 3074430 | DUF6079-family-protein-[Streptomyces-sp.-BB1-1-1] |
| WP_311205221.1 | Type2.Pgl.like.BREX | <-SF1-SF2-Helicase<-SF1-SF2-Helicase\|\|PglW->N6-METHYLASE->DUF6079*->PglZ->BrxD-AAA->SF1-SF2-Helicase+wHTH+Small-B-barrel+DUF1998-><-ARPP2 | 1194 | Bacteria>Terrabacteria_group>Actinomycetota>Actinomycetes>Micrococcales | GCF_031844345.1 | Curtobacterium_sp._BRD11 | Q9R28_RS00190 | 48127..51711 | 2962581 | hypothetical-protein-[Curtobacterium-sp.-BRD11] |
| WP_311262937.1 | Type2.Pgl.like.BREX | <-HEPN+SF1-SF2-Helicase+REase<-?\|\|?-><-SF1-SF2-Helicase+wHTH+Small-B-barrel+DUF1998<-BrxD-AAA<-PglZ<-DUF6079*<-N6-METHYLASE<-PglW\|\|?->HKD-DNase->D6+SF1-SF2-Helicase->SNF+SF1-SF2-Helicase->SF1-SF2-Helicase+REase-> | 1286 | Bacteria>Terrabacteria_group>Actinomycetota>Actinomycetes>Micrococcales | GCF_031844305.1 | Pseudarthrobacter_sp._BRE9 | Q9R31_RS11915 | 178460..182320 | 2962582 | DUF6079-family-protein-[Pseudarthrobacter-sp.-BRE9] |
| WP_312777488.1 | Type2.Pgl.like.BREX | <-SF1-SF2-Helicase+wHTH+Small-B-barrel+DUF1998\|\|Phd-YefM->PIN-><-BrxD-AAA<-PglZ<-DUF6079*<-N6-METHYLASE\|\|OLD-ABC->HNH-><-PglW\|\|HEPN-> | 1302 | Bacteria>Terrabacteria_group>Actinomycetota>Actinomycetes>Mycobacteriales | GCF_032056685.1 | Corynebacterium_variabile | RT519_RS10550 | 105740..109648 | 1727 | DUF6079-family-protein-[Corynebacterium-variabile] |
| WP_312800196.1 | Type2.Pgl.like.BREX | REase+SF1-SF2-Helicase->REase+SF1-SF2-Helicase-><-?<-HEPN\|\|PglW->?->N6-METHYLASE->DUF6079*->PglZ->BrxD-AAA-><-PIN<-Phd-YefM\|\|SF1-SF2-Helicase+wHTH+Small-B-barrel+DUF1998-> | 1313 | Bacteria>Terrabacteria_group>Actinomycetota>Actinomycetes>Mycobacteriales | GCF_032053705.1 | Corynebacterium_variabile | RTY08_RS06030 | 395242..399183 | 1727 | DUF6079-family-protein-[Corynebacterium-variabile] |
| WP_312898988.1 | Type2.Pgl.like.BREX | DNA-glycosylase-><-DDE_Tnp_1<-SF1-SF2-Helicase+wHTH+Small-B-barrel+DUF1998\|\|Phd-YefM->PIN-><-BrxD-AAA<-PglZ<-DUF6079*<-N6-METHYLASE<-PglW\|\|HEPN->?-><-REase+SF1-SF2-Helicase<-HNH<-REase+SF1-SF2-Helicase | 1307 | Bacteria>Terrabacteria_group>Actinomycetota>Actinomycetes>Mycobacteriales | GCF_032057675.1 | Corynebacterium_variabile | RTZ97_RS05800 | 87842..91765 | 1727 | DUF6079-family-protein-[Corynebacterium-variabile] |
| WP_313326129.1 | Type2.Pgl.like.BREX | MerR-HTH->tRNA->PglW->N6-METHYLASE->DUF6079*->PglZ->BrxD-AAA->SF1-SF2-Helicase+wHTH+Small-B-barrel+DUF1998-><-?<-?\|\|?->HKD-DNase-> | 1224 | Bacteria>Pseudomonadota>Gammaproteobacteria>Pseudomonadales>Pseudomonadaceae | GCF_032080385.1 | Pseudomonas_qingdaonensis | RT286_RS06030 | 104602..108276 | 2056231 | phage-resistance-protein-[Pseudomonas-qingdaonensis] |
| WP_313566034.1 | Type2.Pgl.like.BREX | OLD-ABC->?->PglW->N6-METHYLASE->DUF6079*->PglZ->BrxD-AAA->XIS-HTH->PIN->SF1-SF2-Helicase+wHTH+Small-B-barrel+DUF1998-> | 1238 | Bacteria>Terrabacteria_group>Actinomycetota>Actinomycetes>Micrococcales | GCF_032084925.1 | Mobilicoccus_sp. | RT527_RS09215 | 10218..13934 | 2034349 | hypothetical-protein-[Mobilicoccus-sp.] |
| WP_315933119.1 | Type2.Pgl.like.BREX | HIN-HTH->?-><-PIN<-XIS-HTH\|\|?->PglW->N6-METHYLASE->DUF6079*->PglZ->BrxD-AAA->SF1-SF2-Helicase+wHTH+Small-B-barrel+DUF1998-><-?<-HTH | 1223 | Bacteria>Terrabacteria_group>Actinomycetota>Actinomycetes>Pseudonocardiales | GCF_032460545.1 | Umezawaea_sp._Da_62-37 | RM788_RS19500 | 4346679..4350350 | 3075927 | hypothetical-protein-[Umezawaea-sp.-Da-62-37] |
| WP_316738727.1 | Type2.Pgl.like.BREX | <-REase<-ParB+HNH<-SF1-SF2-Helicase+wHTH+Small-B-barrel+DUF1998<-BrxD-AAA<-PglZ<-DUF6079*<-N6-METHYLASE<-PglW\|\|?-><-REase+SF1-SF2-Helicase\|\|?->?-><-ACET | 1297 | Bacteria>Terrabacteria_group>Actinomycetota>Actinomycetes>Kitasatosporales | GCF_032595465.1 | Streptomyces_mirabilis | PU648_RS58380 | 85140..89033 | 68239 | PglY-protein-[Streptomyces-mirabilis] |
| WP_317550158.1 | Type2.Pgl.like.BREX | <-ACET\|\|?->HNH-><-?<-?\|\|PglW->N6-METHYLASE->DUF6079*->PglZ->BrxD-AAA->SF1-SF2-Helicase+wHTH+Small-B-barrel+DUF1998->ParB+HNH-><-HNH | 1238 | Bacteria>Terrabacteria_group>Actinomycetota>Actinomycetes>Mycobacteriales | GCF_033042295.1 | Rhodococcus_aetherivorans | R3Q02_RS04875 | 136468..140184 | 191292 | DUF6079-family-protein-[Rhodococcus-aetherivorans] |
| WP_317565877.1 | Type2.Pgl.like.BREX | <-SbcC\|\|PglW->N6-METHYLASE->DUF6079*->PglZ->BrxD-AAA->SF1-SF2-Helicase+wHTH+Small-B-barrel+DUF1998-><-PglZ<-?<-?<-SbcC | 1218 | Bacteria>Terrabacteria_group>Actinomycetota>Actinomycetes>Mycobacteriales | GCF_033042415.1 | Rhodococcus_yunnanensis | R3P96_RS20715 | 84839..88495 | 278209 | phage-resistance-protein-[Rhodococcus-yunnanensis] |
| WP_317798668.1 | Type2.Pgl.like.BREX | SF1-SF2-Helicase->?->HATPase->?->?->PglW->N6-METHYLASE->DUF6079*->PglZ->BrxD-AAA->SF1-SF2-Helicase+wHTH+Small-B-barrel+DUF1998-> | 1290 | Bacteria>Terrabacteria_group>Actinomycetota>Actinomycetes>Mycobacteriales | GCF_033142385.1 | Rhodococcus_sp._IEGM_1374 | R4P52_RS23455 | 54974..58846 | 3082221 | DUF6079-family-protein-[Rhodococcus-sp.-IEGM-1374] |
| WP_317859883.1 | Type2.Pgl.like.BREX | AA_permease_2->TRAM+SAM-methylase+METHYLASE-><-SF1-SF2-Helicase+wHTH+Small-B-barrel+DUF1998<-BrxD-AAA<-AIPR<-PglZ<-DUF6079*<-REase+SF1-SF2-Helicase<-?<-?\|\|HTH_31+DUF5753->DUF397-><-N6-METHYLASE<-PglW | 1306 | Bacteria>Terrabacteria_group>Actinomycetota>Actinomycetes>Kitasatosporales | GCF_033194775.1 | Streptomyces_sp._Wh19 | R6M67_RS01060 | 35919..39839 | 3076629 | phage-resistance-protein-[Streptomyces-sp.-Wh19] |
| WP_319111775.1 | Type2.Pgl.like.BREX | PglW->N6-METHYLASE-><-DUF397<-HTH_31+DUF5753\|\|?->?->REase+SF1-SF2-Helicase->DUF6079*-> | 1228 | Bacteria>Terrabacteria_group>Actinomycetota>Actinomycetes>Kitasatosporales | GCF_033792445.1 | Streptomyces_europaeiscabiei | PV386_RS20530 | 23122..26806 | 146819 | phage-resistance-protein,-partial-[Streptomyces-europaeiscabiei] |
| WP_319343133.1 | Type2.Pgl.like.BREX | DUF4096+DDE_Tnp_1+DDE_Tnp_1_2->REase-><-SF1-SF2-Helicase+wHTH+Small-B-barrel+DUF1998<-BrxD-AAA<-PglZ<-DUF6079*<-N6-METHYLASE<-PglW\|\|N6-METHYLASE-><-pPIWI+MID+RNaseH<-pPIWI+SFII-RAD3<-pPIWI+pPIWI\|\|<-DUF4096+DDE_Tnp_1+DDE_Tnp_1_2 | 1300 | Bacteria>Terrabacteria_group>Actinomycetota>Actinomycetes>Kitasatosporales | GCF_033790905.1 | Streptomyces_microflavus | NJD91_RS12255 | 51195..55097 | 1919 | phage-resistance-protein-[Streptomyces-microflavus] |
| WP_319723082.1 | Type2.Pgl.like.BREX | AA_permease_2->TRAM+SAM-methylase+METHYLASE-><-SF1-SF2-Helicase+wHTH+Small-B-barrel+DUF1998<-BrxD-AAA<-AIPR<-PglZ<-DUF6079*<-REase+SF1-SF2-Helicase<-?<-?\|\|HTH_31+DUF5753->DUF397-><-N6-METHYLASE<-PglW | 1301 | Bacteria>Terrabacteria_group>Actinomycetota>Actinomycetes>Kitasatosporales | GCF_033794505.1 | Streptomyces_sp._MB09-01 | PV721_RS11725 | 18486..22391 | 3028666 | phage-resistance-protein-[Streptomyces-sp.-MB09-01] |
| WP_320277584.1 | Type2.Pgl.like.BREX | <-Phage_integrase<-?<-?\|\|?-><-SF1-SF2-Helicase+wHTH+Small-B-barrel+DUF1998<-BrxD-AAA<-PglZ<-DUF6079*<-N6-METHYLASE<-PglW | 1231 | Bacteria>Pseudomonadota>Alphaproteobacteria>Hyphomicrobiales>Phyllobacteriaceae | GCF_033977165.1 | Mesorhizobium_sp._VK22E | RFM40_RS12370 | 144406..148101 | 3072319 | hypothetical-protein-[Mesorhizobium-sp.-VK22E] |
| WP_320291258.1 | Type2.Pgl.like.BREX | <-DUF6079*<-?<-PglW | 1231 | Bacteria>Pseudomonadota>Alphaproteobacteria>Hyphomicrobiales>Phyllobacteriaceae | GCF_033977205.1 | Mesorhizobium_sp._VK24D | RFM28_RS32540 | 10252..13947 | 3072314 | hypothetical-protein-[Mesorhizobium-sp.-VK24D] |
| WP_320536138.1 | Type2.Pgl.like.BREX | ParB+HNH->SF1-SF2-Helicase->?-><-?\|\|?->PglW->N6-METHYLASE->DUF6079*->PglZ->BrxD-AAA->SF1-SF2-Helicase+wHTH+Small-B-barrel+DUF1998-> | 1259 | Bacteria>Terrabacteria_group>Actinomycetota>Actinomycetes>Micrococcales | GCF_034048115.1 | Pseudarthrobacter_sp._IC2-21 | SBP01_RS13660 | 2977169..2980948 | 3092262 | DUF6079-family-protein-[Pseudarthrobacter-sp.-IC2-21] |
| WP_321213707.1 | Type2.Pgl.like.BREX | <-REase\|\|?->PglW->N6-METHYLASE->DUF6079*->PglZ->BrxD-AAA->SF1-SF2-Helicase+wHTH+Small-B-barrel+DUF1998-><-?<-?<-REase\|\|HTH-> | 1263 | Bacteria>Terrabacteria_group>Actinomycetota>Actinomycetes>Micrococcales | GCF_034258515.1 | Pseudarthrobacter_oxydans | SMD14_RS11375 | 2523525..2527316 | 1671 | DUF6079-family-protein-[Pseudarthrobacter-oxydans] |
| WP_322110049.1 | Type2.Pgl.like.BREX | <-DUF6079*<-?<-Phd-YefM | 1308 | Bacteria>Terrabacteria_group>Actinomycetota>Actinomycetes>Kitasatosporales | GCF_034300035.1 | Streptomyces_atratus | P6B95_RS31130 | 6727188..6731114 | 1893 | phage-resistance-protein-[Streptomyces-atratus] |
| WP_322753649.1 | Type2.Pgl.like.BREX | <-BrxD-AAA<-PglZ<-DUF6079*<-N6-METHYLASE<-PglW\|\|SF1-SF2-Helicase+wHTH+Small-B-barrel+DUF1998-><-CBS+HEPN<-ParB+HNH | 1320 | Bacteria>Terrabacteria_group>Actinomycetota>Actinomycetes>Frankiales | GCF_963555955.1 | Frankia_sp._Cas3 | U9983_RS09500 | 20534..24496 | 3073926 | DUF6079-family-protein-[Frankia-sp.-Cas3] |
| WP_322753661.1 | Type2.Pgl.like.BREX | PglW-><-RelE-ParE<-Phd-YefM\|\|N6-METHYLASE->DUF6079*->PglZ->BrxD-AAA->SF1-SF2-Helicase+wHTH+Small-B-barrel+DUF1998->HTH-> | 1214 | Bacteria>Terrabacteria_group>Actinomycetota>Actinomycetes>Frankiales | GCF_963555955.1 | Frankia_sp._Cas3 | U9983_RS09565 | 8568..12212 | 3073926 | hypothetical-protein-[Frankia-sp.-Cas3] |
| WP_323366507.1 | Type2.Pgl.like.BREX | SbcC->?-><-?\|\|?-><-PglZ<-DUF6079*<-?<-?<-N6-METHYLASE<-REase<-PglW | 1241 | Bacteria>Terrabacteria_group>Cyanobacteriota/Melainabacteria_group>Cyanobacteriota>Cyanophyceae | GCF_034932105.1 | Leptothoe_sp._PORK10_BA2 | VB741_RS17645 | 55512..59237 | 3110254 | hypothetical-protein-[Leptothoe-sp.-PORK10-BA2] |
| WP_324980990.1 | Type2.Pgl.like.BREX | PglW->?-><-?\|\|?->?->?->DUF6079*-> | 1308 | Bacteria>Terrabacteria_group>Actinomycetota>Actinomycetes>Kitasatosporales | GCF_035687565.1 | Streptomyces_sp. | VFG16_RS18990 | 19668..23594 | 1931 | phage-resistance-protein-[Streptomyces-sp.] |
| WP_325031808.1 | Type2.Pgl.like.BREX | PglW->N6-METHYLASE->DUF6079*->PglZ->BrxD-AAA->SF1-SF2-Helicase+wHTH+Small-B-barrel+DUF1998-><-?<-?<-ParB+HNH\|\|N6-METHYLASE-> | 1191 | Bacteria>Terrabacteria_group>Actinomycetota>Actinomycetes>Micrococcales | GCF_035687345.1 | Microbacterium_sp. | VFF97_RS01950 | 28446..32021 | 51671 | hypothetical-protein-[Microbacterium-sp.] |
| WP_326435021.1 | Type2.Pgl.like.BREX | PglW->?->?-><-?<-?\|\|?->DUF6079*-> | 1323 | Bacteria>Terrabacteria_group>Actinomycetota>Actinomycetes>Kitasatosporales | GCF_035828615.1 | Streptomyces_sp._H27-D2 | QMK28_RS01135 | 61413..65384 | 3046304 | PglY-protein-[Streptomyces-sp.-H27-D2] |
| WP_326661331.1 | Type2.Pgl.like.BREX | <-OLD-ABC\|\|PglW-><-?\|\|HTH_31+DUF5753->DUF397->N6-METHYLASE->DUF6079*->PglZ->BrxD-AAA->SF1-SF2-Helicase+wHTH+Small-B-barrel+DUF1998-> | 1295 | Bacteria>Terrabacteria_group>Actinomycetota>Actinomycetes>Kitasatosporales | GCF_035917155.1 | Streptomyces_sp._NBC_00385 | OG959_RS23215 | 5186083..5189970 | 2975733 | phage-resistance-protein-[Streptomyces-sp.-NBC_00385] |
| WP_326669072.1 | Type2.Pgl.like.BREX | PglW-><-ORC-CDC6\|\|N6-METHYLASE-><-DUF397<-HTH_31+DUF5753\|\|ST-Hiskin-><-?\|\|DUF6079*->PglZ->BrxD-AAA->SF1-SF2-Helicase+wHTH+Small-B-barrel+DUF1998-><-REase+SF1-SF2-Helicase<-ParB+HNH | 1301 | Bacteria>Terrabacteria_group>Actinomycetota>Actinomycetes>Kitasatosporales | GCF_035918175.1 | Streptomyces_canus | OG925_RS34790 | 7576793..7580698 | 58343 | PglY-protein-[Streptomyces-canus] |
| WP_326712555.1 | Type2.Pgl.like.BREX | HTH-><-?\|\|PglW->HEPN->ParA-Soj-PloopNTPase->BrxD-AAA->N6-METHYLASE->DUF6079*->PglZ->BrxD-AAA->SF1-SF2-Helicase+wHTH+Small-B-barrel+DUF1998->HTH-><-?\|\|?-><-ACET | 1292 | Bacteria>Terrabacteria_group>Actinomycetota>Actinomycetes>Kitasatosporales | GCF_036178905.1 | unclassified_Streptomyces | OHA28_RS28115 | 6167427..6171305 | 2593676 | MULTISPECIES:-hypothetical-protein-[unclassified-Streptomyces] |
| WP_326723507.1 | Type2.Pgl.like.BREX | REase->?->ncRNA-><-SF1-SF2-Helicase+wHTH+Small-B-barrel+DUF1998<-BrxD-AAA\|\|ParB+HNH->CBS+HEPN-><-PglZ<-DUF6079*<-?<-DUF397<-HTH_31+DUF5753\|\|ST-Hiskin-><-N6-METHYLASE<-PglW\|\|SAD(SRA)+wHTH+HNH-> | 1308 | Bacteria>Terrabacteria_group>Actinomycetota>Actinomycetes>Kitasatosporales | GCF_035917175.1 | Streptomyces_sp._NBC_00243 | OHT59_RS33045 | 7427338..7431244 | 2975688 | phage-resistance-protein-[Streptomyces-sp.-NBC_00243] |
| WP_326746002.1 | Type2.Pgl.like.BREX | <-pPIWI+SFII-RAD3<-?<-pPIWI+pPIWI<-SF1-SF2-Helicase+wHTH+Small-B-barrel+DUF1998<-BrxD-AAA<-PglZ<-DUF6079*<-N6-METHYLASE<-PglW\|\|REase+DUF5655-><-?<-?<-?<-DCM | 1297 | Bacteria>Terrabacteria_group>Actinomycetota>Actinomycetes>Kitasatosporales | GCF_045206205.1 | Streptomyces_sp._NBC_01760 | OG807_RS36730 | 8213139..8217032 | 2975931 | phage-resistance-protein-[Streptomyces-sp.-NBC_01760] |
| WP_326752420.1 | Type2.Pgl.like.BREX | PglW->N6-METHYLASE-><-ST-Hiskin\|\|HTH_31+DUF5753->DUF397->?->?->DUF6079*->PglZ-><-CBS+HEPN<-ParB+HNH\|\|BrxD-AAA->SF1-SF2-Helicase+wHTH+Small-B-barrel+DUF1998-><-REase | 1310 | Bacteria>Terrabacteria_group>Actinomycetota>Actinomycetes>Kitasatosporales | GCF_035918035.1 | Streptomyces_hirsutus | OIE73_RS11080 | 2415560..2419492 | 35620 | phage-resistance-protein-[Streptomyces-hirsutus] |
| WP_326795582.1 | Type2.Pgl.like.BREX | <-MerR-HTH<-?\|\|?-><-?\|\|ncRNA-><-PglZ<-DUF6079*<-N6-METHYLASE<-PglW\|\|BrxD-AAA->SF1-SF2-Helicase+wHTH+Small-B-barrel+DUF1998->N6-METHYLASE-> | 1284 | Bacteria>Terrabacteria_group>Actinomycetota>Actinomycetes>Kitasatosporales | GCF_035917395.1 | Streptomyces_sp._NBC_01808 | OG946_RS08875 | 2094020..2097874 | 2975947 | DUF6079-family-protein-[Streptomyces-sp.-NBC_01808] |
| WP_326818697.1 | Type2.Pgl.like.BREX | <-pPIWI+MID+RNaseH<-pPIWI+SFII-RAD3<-pPIWI+pPIWI<-SF1-SF2-Helicase+wHTH+Small-B-barrel+DUF1998<-BrxD-AAA<-PglZ<-DUF6079*<-N6-METHYLASE<-PglW\|\|REase+DUF5655-><-?<-?<-?<-DCM | 1297 | Bacteria>Terrabacteria_group>Actinomycetota>Actinomycetes>Kitasatosporales | GCF_035917875.1 | unclassified_Streptomyces | OHA26_RS39625 | 8793533..8797426 | 2593676 | MULTISPECIES:-phage-resistance-protein-[unclassified-Streptomyces] |
| WP_326854224.1 | Type2.Pgl.like.BREX | <-PIN<-?\|\|PglW->N6-METHYLASE->DUF6079*->PglZ->BrxD-AAA->SF1-SF2-Helicase+wHTH+Small-B-barrel+DUF1998-><-N6-METHYLASE\|\|REase-> | 1348 | Bacteria>Terrabacteria_group>Actinomycetota>Actinomycetes>Catenulisporales | GCF_035946395.1 | Actinocrinis_sp. | VGS97_RS15515 | 22413..26459 | 1920516 | hypothetical-protein-[Actinocrinis-sp.] |
| WP_328677725.1 | Type2.Pgl.like.BREX | <-REase<-?\|\|PglW->N6-METHYLASE-><-DUF397<-HTH_31+DUF5753\|\|REase+SF1-SF2-Helicase->DUF6079*->PglZ->AIPR->BrxD-AAA->SF1-SF2-Helicase+wHTH+Small-B-barrel+DUF1998-><-?<-TRAM+SAM-methylase+METHYLASE<-AA_permease_2 | 1295 | Bacteria>Terrabacteria_group>Actinomycetota>Actinomycetes>Kitasatosporales | GCF_036118775.1 | Streptomyces_sp._NBC_00343 | OHA62_RS11625 | 2752223..2756110 | 2975719 | phage-resistance-protein-[Streptomyces-sp.-NBC_00343] |
| WP_328712364.1 | Type2.Pgl.like.BREX | <-SF1-SF2-Helicase+wHTH+Small-B-barrel+DUF1998<-BrxD-AAA<-?\|\|REase+SF1-SF2-Helicase-><-PglZ<-DUF6079*<-N6-METHYLASE<-PglW\|\|Pkinase+TPR-> | 1293 | Bacteria>Terrabacteria_group>Actinomycetota>Actinomycetes>Mycobacteriales | GCF_036150455.1 | Nocardia_salmonicida | OHA85_RS00120 | 34364..38245 | 53431 | phage-resistance-protein-[Nocardia-salmonicida] |
| WP_328764025.1 | Type2.Pgl.like.BREX | <-GDResolvase\|\|?->HTH_29+HTH_32+DDE_3-><-SF1-SF2-Helicase+wHTH+Small-B-barrel+DUF1998<-BrxD-AAA<-PglZ<-DUF6079*<-N6-METHYLASE<-PglW\|\|DUF6731-> | 1295 | Bacteria>Terrabacteria_group>Actinomycetota>Actinomycetes>Kitasatosporales | GCF_036178565.1 | Streptomyces_sp._NBC_00272 | OG827_RS28355 | 6050650..6054537 | 2975698 | PglY-protein-[Streptomyces-sp.-NBC_00272] |
| WP_328775746.1 | Type2.Pgl.like.BREX | <-REase+SF1-SF2-Helicase\|\|PglW->N6-METHYLASE-><-DUF397<-HTH_31+DUF5753\|\|?->?->DUF6079*->PglZ-><-CBS+HEPN<-ParB+HNH\|\|BrxD-AAA->SF1-SF2-Helicase+wHTH+Small-B-barrel+DUF1998-><-TRAM+SAM-methylase+METHYLASE<-AA_permease_2 | 1299 | Bacteria>Terrabacteria_group>Actinomycetota>Actinomycetes>Kitasatosporales | GCF_036175285.1 | Streptomyces_goshikiensis | OHU17_RS09805 | 2191702..2195601 | 1942 | phage-resistance-protein-[Streptomyces-goshikiensis] |
| WP_328811366.1 | Type2.Pgl.like.BREX | SF1-SF2-Helicase+REase->SF1-SF2-Helicase->?->PglZ-><-SF1-SF2-Helicase+wHTH+Small-B-barrel+DUF1998<-BrxD-AAA<-PglZ<-DUF6079*<-N6-METHYLASE<-PglW<-HKD-DNase\|\|McrB-AAA-ATPase->DUF2357+REase-><-CBS+HEPN+HEPN<-HEPN+SF1-SF2-Helicase+REase | 1245 | Bacteria>Terrabacteria_group>Actinomycetota>Actinomycetes>Mycobacteriales | GCF_036163995.1 | Rhodococcus_sp._NBC_00294 | OHB23_RS09005 | 1915935..1919672 | 2976004 | DUF6079-family-protein-[Rhodococcus-sp.-NBC_00294] |
| WP_328846446.1 | Type2.Pgl.like.BREX | <-REase+SF1-SF2-Helicase<-?<-SF1-SF2-Helicase+wHTH+Small-B-barrel+DUF1998<-BrxD-AAA\|\|ParB+HNH->CBS+HEPN-><-PglZ<-DUF6079*<-?<-DUF397<-HTH_31+DUF5753\|\|ST-Hiskin-><-N6-METHYLASE<-PglW | 1326 | Bacteria>Terrabacteria_group>Actinomycetota>Actinomycetes>Kitasatosporales | GCF_036182465.1 | Streptomyces_sp._NBC_00258 | OG718_RS39415 | 8843524..8847504 | 2903642 | phage-resistance-protein-[Streptomyces-sp.-NBC_00258] |
| WP_328932270.1 | Type2.Pgl.like.BREX | <-REase<-?\|\|PglW->N6-METHYLASE-><-DUF397<-HTH_31+DUF5753\|\|REase+SF1-SF2-Helicase->DUF6079*->PglZ->AIPR->BrxD-AAA->SF1-SF2-Helicase+wHTH+Small-B-barrel+DUF1998-><-?<-TRAM+SAM-methylase+METHYLASE<-AA_permease_2 | 1295 | Bacteria>Terrabacteria_group>Actinomycetota>Actinomycetes>Kitasatosporales | GCF_036196325.1 | Streptomyces_sp._NBC_00228 | OG537_RS11480 | 2681580..2685467 | 2903637 | phage-resistance-protein-[Streptomyces-sp.-NBC_00228] |
| WP_328934070.1 | Type2.Pgl.like.BREX | <-DUF6079*<-?<-PglW | 1304 | Bacteria>Terrabacteria_group>Actinomycetota>Actinomycetes>Kitasatosporales | GCF_036196325.1 | Streptomyces_sp._NBC_00228 | OG537_RS29365 | 6628887..6632801 | 2903637 | PglY-protein-[Streptomyces-sp.-NBC_00228] |
| WP_328937459.1 | Type2.Pgl.like.BREX | PglW->N6-METHYLASE-><-DUF397<-HTH_31+DUF5753\|\|<-?<-?\|\|DUF6079*->PglZ->BrxD-AAA->SF1-SF2-Helicase+wHTH+Small-B-barrel+DUF1998-><-?\|\|?->HKD-DNase-> | 1298 | Bacteria>Terrabacteria_group>Actinomycetota>Actinomycetes>Kitasatosporales | GCF_036204065.1 | Streptomyces_tauricus | OG288_RS12455 | 3035680..3039576 | 68274 | phage-resistance-protein-[Streptomyces-tauricus] |
| WP_329024165.1 | Type2.Pgl.like.BREX | PglW->?-><-?<-?\|\|?->?->DUF6079*-> | 1309 | Bacteria>Terrabacteria_group>Actinomycetota>Actinomycetes>Kitasatosporales | GCF_036226685.1 | Streptomyces_sp._NBC_00690 | OID54_RS28780 | 6564231..6568160 | 2975808 | phage-resistance-protein-[Streptomyces-sp.-NBC_00690] |
| WP_329045578.1 | Type2.Pgl.like.BREX | <-DUF6079*<-?<-PglW | 1263 | Bacteria>Terrabacteria_group>Actinomycetota>Actinomycetes>Pseudonocardiales | GCF_036227105.1 | Amycolatopsis_sp._NBC_01488 | OG738_RS28905 | 6084944..6088735 | 2903563 | PglY-protein-[Amycolatopsis-sp.-NBC_01488] |
| WP_329144525.1 | Type2.Pgl.like.BREX | PglW->N6-METHYLASE-><-ST-Hiskin\|\|HTH_31+DUF5753->DUF397->DUF6079*->PglZ-><-CBS+HEPN<-ParB+HNH\|\|BrxD-AAA->SF1-SF2-Helicase+wHTH+Small-B-barrel+DUF1998-><-ncRNA\|\|MerR-HTH-> | 1326 | Bacteria>Terrabacteria_group>Actinomycetota>Actinomycetes>Kitasatosporales | GCF_036227585.1 | Streptomyces_niveus | OG275_RS09080 | 2069272..2073252 | 193462 | phage-resistance-protein-[Streptomyces-niveus] |
| WP_329173286.1 | Type2.Pgl.like.BREX | PglW->N6-METHYLASE->DUF6079*->PglZ->BrxD-AAA->SF1-SF2-Helicase+wHTH+Small-B-barrel+DUF1998-> | 1299 | Bacteria>Terrabacteria_group>Actinomycetota>Actinomycetes>Kitasatosporales | GCF_036227245.1 | Streptomyces_sp._NBC_01477 | OHA86_RS06555 | 1702839..1706738 | 2976015 | phage-resistance-protein-[Streptomyces-sp.-NBC_01477] |
| WP_329194497.1 | Type2.Pgl.like.BREX | PglW->N6-METHYLASE-><-ST-Hiskin\|\|HTH_31+DUF5753->DUF397->Phd-YefM->PIN->DUF6079*->PglZ-><-CBS+HEPN<-ParB+HNH\|\|BrxD-AAA->SF1-SF2-Helicase+wHTH+Small-B-barrel+DUF1998-><-SAD(SRA)+wHTH+HNH | 1309 | Bacteria>Terrabacteria_group>Actinomycetota>Actinomycetes>Kitasatosporales | GCF_036228565.1 | Streptomyces_sp._NBC_01435 | OG769_RS10315 | 2474520..2478449 | 2903865 | phage-resistance-protein-[Streptomyces-sp.-NBC_01435] |
| WP_329248535.1 | Type2.Pgl.like.BREX | Phd-YefM->RelE-ParE+Tox-RelE->PglW->N6-METHYLASE-><-DUF397<-HTH_31+DUF5753\|\|REase+SF1-SF2-Helicase->DUF6079*->PglZ->AIPR->BrxD-AAA->SF1-SF2-Helicase+wHTH+Small-B-barrel+DUF1998-><-ncRNA\|\|?-><-?<-TetR-HTH | 1296 | Bacteria>Terrabacteria_group>Actinomycetota>Actinomycetes>Kitasatosporales | GCF_036227225.1 | Streptomyces_sp._NBC_01478 | OG223_RS16420 | 3653394..3657284 | 2903882 | DUF6079-family-protein-[Streptomyces-sp.-NBC_01478] |
| WP_329283861.1 | Type2.Pgl.like.BREX | AA_permease_2->TRAM+SAM-methylase+METHYLASE-><-YjhX_toxin<-MORC+REase\|\|REase+DUF5655->PglW->N6-METHYLASE->DUF6079*->PglZ->BrxD-AAA->SF1-SF2-Helicase+wHTH+Small-B-barrel+DUF1998-><-?\|\|NUDIX-> | 1297 | Bacteria>Terrabacteria_group>Actinomycetota>Actinomycetes>Kitasatosporales | GCF_036226665.1 | Streptomyces_sp._NBC_00691 | OG392_RS27675 | 6136672..6140565 | 2903671 | phage-resistance-protein-[Streptomyces-sp.-NBC_00691] |
| WP_329299357.1 | Type2.Pgl.like.BREX | PglW->N6-METHYLASE-><-ST-Hiskin\|\|HTH_31+DUF5753->DUF397->DUF6079*->PglZ-><-CBS+HEPN<-ParB+HNH\|\|BrxD-AAA->SF1-SF2-Helicase+wHTH+Small-B-barrel+DUF1998-><-SAD(SRA)+wHTH+HNH | 1322 | Bacteria>Terrabacteria_group>Actinomycetota>Actinomycetes>Kitasatosporales | GCF_036226925.1 | Streptomyces_sp._NBC_00659 | OG410_RS13480 | 3111600..3115568 | 2903669 | phage-resistance-protein-[Streptomyces-sp.-NBC_00659] |
| WP_329359101.1 | Type2.Pgl.like.BREX | <-DUF4096+DDE_Tnp_1+DDE_Tnp_1_2<-?\|\|PglW->N6-METHYLASE->DUF6079*->PglZ->BrxD-AAA->SF1-SF2-Helicase+wHTH+Small-B-barrel+DUF1998-> | 1296 | Bacteria>Terrabacteria_group>Actinomycetota>Actinomycetes>Kitasatosporales | GCF_036227165.1 | Streptomyces_sp._NBC_01483 | OG318_RS52040 | 11321104..11324994 | 2903883 | PglY-protein-[Streptomyces-sp.-NBC_01483] |
| WP_329428925.1 | Type2.Pgl.like.BREX | <-HEPN\|\|PglW->N6-METHYLASE->DUF6079*->PglZ->BrxD-AAA->SF1-SF2-Helicase+wHTH+Small-B-barrel+DUF1998-><-DUF397<-HTH_31+DUF5753 | 1309 | Bacteria>Terrabacteria_group>Actinomycetota>Actinomycetes>Streptosporangiales | GCF_036250735.1 | Streptosporangium_sp._NBC_01495 | OG339_RS06920 | 1583981..1587910 | 2903899 | hypothetical-protein-[Streptosporangium-sp.-NBC_01495] |
| WP_329787392.1 | Type2.Pgl.like.BREX | SAM-methylase->HisKA_3+HATPase_c-><-?<-?<-SF1-SF2-Helicase+wHTH+Small-B-barrel+DUF1998<-BrxD-AAA<-PglZ<-DUF6079*<-N6-METHYLASE<-PglW\|\|REase+SF1-SF2-Helicase->REase+SF1-SF2-Helicase-> | 1250 | Bacteria>Terrabacteria_group>Actinomycetota>Actinomycetes>Pseudonocardiales | GCF_036321065.1 | Lentzea_sp._DG1S-22 | V1227_RS21575 | 4717613..4721365 | 3108822 | PglY-protein-[Lentzea-sp.-DG1S-22] |
| WP_330156324.1 | Type2.Pgl.like.BREX | <-HTH_31+DUF5753\|\|ST-Hiskin->?->PglW->N6-METHYLASE->DUF6079*->PglZ->BrxD-AAA->SF1-SF2-Helicase+wHTH+Small-B-barrel+DUF1998-><-McrC-NTD+REase<-McrB-AAA-ATPase | 1308 | Bacteria>Terrabacteria_group>Actinomycetota>Actinomycetes>Streptosporangiales | GCF_039526505.1 | Nocardiopsis_umidischolae | ABD221_RS06595 | 276268..280194 | 163605 | hypothetical-protein-[Nocardiopsis-umidischolae] |
| WP_330232093.1 | Type2.Pgl.like.BREX | PglW-><-?\|\|N6-METHYLASE->DUF6079*->PglZ->BrxD-AAA->SF1-SF2-Helicase+wHTH+Small-B-barrel+DUF1998-> | 1285 | Bacteria>Terrabacteria_group>Actinomycetota>Actinomycetes>Mycobacteriales | GCF_036346875.1 | Nocardia_sp._NBC_00508 | OHA40_RS06140 | 1395433..1399290 | 2975992 | phage-resistance-protein-[Nocardia-sp.-NBC_00508] |
| WP_330333133.1 | Type2.Pgl.like.BREX | AA_permease_2->TRAM+SAM-methylase+METHYLASE-><-SF1-SF2-Helicase+wHTH+Small-B-barrel+DUF1998<-BrxD-AAA<-AIPR<-PglZ<-DUF6079*<-?<-?\|\|HTH_31+DUF5753->DUF397-><-N6-METHYLASE<-PglW\|\|REase+SF1-SF2-Helicase-> | 1298 | Bacteria>Terrabacteria_group>Actinomycetota>Actinomycetes>Kitasatosporales | GCF_036346295.1 | Streptomyces_sp._NBC_00536 | OHS33_RS27775 | 6379336..6383232 | 2975769 | phage-resistance-protein-[Streptomyces-sp.-NBC_00536] |
| WP_330344465.1 | Type2.Pgl.like.BREX | PglW->N6-METHYLASE-><-ST-Hiskin\|\|HTH_31+DUF5753->DUF397->?->DUF6079*->PglZ-><-CBS+HEPN<-ParB+HNH\|\|BrxD-AAA->SF1-SF2-Helicase+wHTH+Small-B-barrel+DUF1998-> | 1308 | Bacteria>Terrabacteria_group>Actinomycetota>Actinomycetes>Kitasatosporales | GCF_036346085.1 | Streptomyces_longwoodensis | OHA09_RS12455 | 3070861..3074787 | 68231 | phage-resistance-protein-[Streptomyces-longwoodensis] |
| WP_330354472.1 | Type2.Pgl.like.BREX | PglW->HEPN->ParA-Soj-PloopNTPase->BrxD-AAA->N6-METHYLASE->DUF6079*->PglZ->BrxD-AAA->SF1-SF2-Helicase+wHTH+Small-B-barrel+DUF1998-><-?<-SAD+REase | 1305 | Bacteria>Terrabacteria_group>Actinomycetota>Actinomycetes>Kitasatosporales | GCF_036345055.1 | Streptomyces_chartreusis | OG997_RS17120 | 3862934..3866851 | 1969 | hypothetical-protein-[Streptomyces-chartreusis] |
| WP_330466073.1 | Type2.Pgl.like.BREX | PglW->N6-METHYLASE-><-ST-Hiskin\|\|HTH_31+DUF5753->DUF397->?->DUF6079*->PglZ-><-CBS+HEPN<-ParB+HNH\|\|BrxD-AAA->SF1-SF2-Helicase+wHTH+Small-B-barrel+DUF1998-> | 1308 | Bacteria>Terrabacteria_group>Actinomycetota>Actinomycetes>Kitasatosporales | GCF_045161325.1 | Streptomyces_longwoodensis | OG547_RS11280 | 2752021..2755947 | 68231 | phage-resistance-protein-[Streptomyces-longwoodensis] |
| WP_330811472.1 | Type2.Pgl.like.BREX | <-SAD+REase<-SF1-SF2-Helicase+wHTH+Small-B-barrel+DUF1998<-BrxD-AAA\|\|ParB+HNH->CBS+HEPN-><-PglZ<-DUF6079*<-DUF397<-HTH_31+DUF5753\|\|ST-Hiskin-><-N6-METHYLASE<-PglW\|\|REase+SF1-SF2-Helicase->HNH-> | 1320 | Bacteria>Terrabacteria_group>Actinomycetota>Actinomycetes>Kitasatosporales | GCF_036352935.1 | Streptomyces | V2J94_RS25890 | 112923..116885 | 1883 | phage-resistance-protein-[Streptomyces-sp.-DSM-41524] |
| WP_330928031.1 | Type2.Pgl.like.BREX | <-REC+LuxR-HTH\|\|?-><-?\|\|?->PglW->N6-METHYLASE->DUF6079*->PglZ->BrxD-AAA->SF1-SF2-Helicase+wHTH+Small-B-barrel+DUF1998-> | 1260 | Bacteria>PVC_group>Verrucomicrobiota>Opitutia>Opitutales | GCF_029023725.1 | Synoicihabitans_lomoniglobus | PXH66_RS17435 | 4532197..4535979 | 2909285 | hypothetical-protein-[Synoicihabitans-lomoniglobus] |
| WP_331447533.1 | Type2.Pgl.like.BREX | <-N6-METHYLASE\|\|PglW->N6-METHYLASE->DUF6079*->PglZ->PglZ->BrxD-AAA->SF1-SF2-Helicase+wHTH+Small-B-barrel+DUF1998-><-REase\|\|?->Phage_integrase-> | 1297 | Bacteria>Terrabacteria_group>Actinomycetota>Actinomycetes>Kitasatosporales | GCF_036541015.1 | Streptomyces_xanthochromogenes | V3060_RS08980 | 2066752..2070645 | 67384 | phage-resistance-protein-[Streptomyces-xanthochromogenes] |
| WP_332348677.1 | Type2.Pgl.like.BREX | PglW->N6-METHYLASE->OLD-ABC->?->?->?->DUF6079*->PglZ->BrxD-AAA->SF1-SF2-Helicase+wHTH+Small-B-barrel+DUF1998-><-?<-?<-rve | 1226 | Bacteria>Pseudomonadota>Betaproteobacteria>Betaproteobacteria_incertae_sedis>Candidatus_Accumulibacter | GCF_036625195.1 | Accumulibacter_sp. | V5B60_RS17530 | 4000160..4003840 | 2053492 | phage-resistance-protein-[Accumulibacter-sp.] |
| WP_334580654.1 | Type2.Pgl.like.BREX | HEPN+VirB11+NACHT+FGS->?->?-><-?<-DUF6731\|\|PglW->N6-METHYLASE->DUF6079*-><-DEDD_Tnp_IS110+Transposase_20\|\|PglZ->BrxD-AAA->SF1-SF2-Helicase+wHTH+Small-B-barrel+DUF1998-><-?\|\|?-><-Phage_int_SAM_3+Phage_integrase | 1312 | Bacteria>Terrabacteria_group>Actinomycetota>Actinomycetes>Kitasatosporales | GCF_036898445.1 | Streptomyces_sp._B21-108 | QBA54_RS35325 | 7740309..7744247 | 3039419 | PglY-protein-[Streptomyces-sp.-B21-108] |
| WP_334662749.1 | Type2.Pgl.like.BREX | HTH-><-?\|\|PglW->HEPN->ParA-Soj-PloopNTPase->BrxD-AAA->N6-METHYLASE->\|\|DUF6079*->PglZ->BrxD-AAA->SF1-SF2-Helicase+wHTH+Small-B-barrel+DUF1998-><-SF1-SF2-Helicase<-HTH_Tnp_1+HTH_21+rve+rve_3 | 1347 | Bacteria>Terrabacteria_group>Actinomycetota>Actinomycetes>Kitasatosporales | GCF_036919415.1 | Streptomyces_cyaneofuscatus | VVF06_RS00805 | 19993..24036 | 66883 | hypothetical-protein-[Streptomyces-cyaneofuscatus] |
| WP_336050190.1 | Type2.Pgl.like.BREX | <-SF1-SF2-Helicase+wHTH+Small-B-barrel+DUF1998<-BrxD-AAA\|\|ParB+HNH->CBS+HEPN-><-PglZ<-DUF6079*<-DUF397<-HTH_31+DUF5753\|\|ST-Hiskin-><-N6-METHYLASE<-PglW | 1317 | Bacteria>Terrabacteria_group>Actinomycetota>Actinomycetes>Kitasatosporales | GCF_037034225.1 | Streptomyces_sp._CA2R101 | WA779_RS15055 | 59886..63839 | 3120152 | phage-resistance-protein-[Streptomyces-sp.-CA2R101] |
| WP_336521091.1 | Type2.Pgl.like.BREX | <-SF1-SF2-Helicase+wHTH+Small-B-barrel+DUF1998<-BrxD-AAA<-PglZ<-DUF6079*<-ParB+HNH<-?<-HNH<-OLD-ABC<-N6-METHYLASE<-PglW | 1216 | Bacteria>PVC_group>Planctomycetota>Planctomycetia>Planctomycetales | GCF_030622045.1 | Planctoellipticum_variicoloris | SH412_RS26695 | 6848891..6852541 | 3064265 | phage-resistance-protein-[Planctoellipticum-variicoloris] |
| WP_336820102.1 | Type2.Pgl.like.BREX | <-SF1-SF2-Helicase+wHTH+Small-B-barrel+DUF1998<-BrxD-AAA<-PglZ<-DUF6079*<-N6-METHYLASE<-PglW<-SF1-SF2-Helicase<-?<-?\|\|REase+SF1-SF2-Helicase->URI-DNase+SF1-SF2-Helicase-> | 1255 | Bacteria>Terrabacteria_group>Actinomycetota>Actinomycetes>Mycobacteriales | GCF_037145975.1 | Gordonia_sp._MMO-8 | WFX31_RS17110 | 235945..239712 | 3127886 | phage-resistance-protein-[Gordonia-sp.-MMO-8] |
| WP_337179472.1 | Type2.Pgl.like.BREX | HTH_26->PglW->N6-METHYLASE->OLD-ABC->?->DUF6079*->PglZ->BrxD-AAA->SF1-SF2-Helicase+wHTH+Small-B-barrel+DUF1998-> | 1231 | Bacteria>Pseudomonadota>Betaproteobacteria>Burkholderiales>Comamonadaceae | GCF_037189435.1 | Hydrogenophaga_borbori | WG214_RS16370 | 26694..30389 | 2294117 | phage-resistance-protein-[Hydrogenophaga-borbori] |
| WP_337304542.1 | Type2.Pgl.like.BREX | DUF4096+DDE_Tnp_1+DDE_Tnp_1_2-><-?<-?\|\|?-><-SF1-SF2-Helicase+wHTH+Small-B-barrel+DUF1998<-BrxD-AAA<-PglZ<-DUF6079*<-ST-Hiskin\|\|HTH_31+DUF5753->DUF397-><-N6-METHYLASE<-PglW\|\|?-><-ACET | 1298 | Bacteria>Terrabacteria_group>Actinomycetota>Actinomycetes>Kitasatosporales | GCF_037204065.1 | unclassified_Streptomyces | V9L26_RS11245 | 111223..115119 | 2593676 | MULTISPECIES:-phage-resistance-protein-[unclassified-Streptomyces] |
| WP_337351099.1 | Type2.Pgl.like.BREX | <-SF1-SF2-Helicase+wHTH+Small-B-barrel+DUF1998<-BrxD-AAA<-PglZ<-DUF6079*<-N6-METHYLASE<-PglW<-?\|\|REase->?-><-?\|\|SF1-SF2-Helicase-> | 1263 | Bacteria>Terrabacteria_group>Actinomycetota>Actinomycetes>Micrococcales | GCF_037203945.1 | Pseudarthrobacter_sp._CCNWLW207 | V9L17_RS08305 | 83051..86842 | 3127468 | DUF6079-family-protein-[Pseudarthrobacter-sp.-CCNWLW207] |
| WRY82586.1 | Type2.Pgl.like.BREX | <-DUF6079*<-?\|\|?-><-?<-?<-?<-?<-PglW | 1352 | Bacteria>Terrabacteria_group>Actinomycetota>Actinomycetes>Kitasatosporales | GCA_041431075.1 | Streptomyces_clavifer | OG388_15840 | 3623075..3627133 | 68188 | hypothetical-protein-OG388_15840-[Streptomyces-clavifer] |
| WSB23153.1 | Type2.Pgl.like.BREX | HTH_Tnp_4+DDE_Tnp_4-><-SF1-SF2-Helicase+wHTH+Small-B-barrel+DUF1998<-BrxD-AAA<-PglZ<-DUF6079*\|\|HTH_31+DUF5753->DUF397-><-N6-METHYLASE<-PglW<-?<-?<-METHYLASE | 1298 | Bacteria>Terrabacteria_group>Actinomycetota>Actinomycetes>Kitasatosporales | GCA_041438605.1 | Streptomyces_albidoflavus | OHB02_24585 | 5605105..5609001 | 1886 | phage-resistance-protein-[Streptomyces-albidoflavus] |
| WSD55918.1 | Type2.Pgl.like.BREX | AA_permease_2->TRAM+SAM-methylase+METHYLASE-><-SF1-SF2-Helicase+wHTH+Small-B-barrel+DUF1998<-BrxD-AAA<-PglZ<-DUF6079*\|\|HTH_31+DUF5753->DUF397-><-N6-METHYLASE<-PglW\|\|REase->?->CASPASE+SF1-SF2-Helicase-> | 1298 | Bacteria>Terrabacteria_group>Actinomycetota>Actinomycetes>Kitasatosporales | GCA_041438845.1 | Streptomyces_albidoflavus | OHA76_25630 | 5812916..5816812 | 1886 | phage-resistance-protein-[Streptomyces-albidoflavus] |
| WSG45999.1 | Type2.Pgl.like.BREX | REase-><-SF1-SF2-Helicase+wHTH+Small-B-barrel+DUF1998<-BrxD-AAA<-PglZ<-DUF6079*<-N6-METHYLASE<-PglW<-?<-HTH | 1241 | Bacteria>Terrabacteria_group>Actinomycetota>Actinomycetes>Micromonosporales | GCA_035963215.1 | Dactylosporangium_sp._NBC_01737 | OHA72_22035 | 4831842..4835567 | 2975959 | phage-resistance-protein-[Dactylosporangium-sp.-NBC_01737] |
| WSL10657.1 | Type2.Pgl.like.BREX | <-Phage_int_SAM_3+Phage_integrase\|\|HTH-><-SF1-SF2-Helicase+wHTH+Small-B-barrel+DUF1998<-BrxD-AAA<-PglZ<-DUF6079*<-N6-METHYLASE<-PglW\|\|DUF6731->?-><-?\|\|REase-> | 1302 | Bacteria>Terrabacteria_group>Actinomycetota>Actinomycetes>Kitasatosporales | GCA_035982055.1 | Streptomyces_sp._NBC_01288 | OG194_33500 | 7410147..7414055 | 2903814 | PglY-protein-[Streptomyces-sp.-NBC_01288] |
| WSN43126.1 | Type2.Pgl.like.BREX | AA_permease_2->TRAM+SAM-methylase+METHYLASE-><-SF1-SF2-Helicase+wHTH+Small-B-barrel+DUF1998<-BrxD-AAA<-AIPR<-PglZ<-DUF6079*<-REase+SF1-SF2-Helicase<-?<-?\|\|HTH_31+DUF5753->DUF397-><-N6-METHYLASE<-PglW | 1296 | Bacteria>Terrabacteria_group>Actinomycetota>Actinomycetes>Kitasatosporales | GCA_035983595.1 | Streptomyces_sp._NBC_01334 | OG736_32455 | 7219943..7223833 | 2903827 | DUF6079-family-protein-[Streptomyces-sp.-NBC_01334] |
| WSN45804.1 | Type2.Pgl.like.BREX | <-DUF4096+DDE_Tnp_1+DDE_Tnp_1_2\|\|PglW->HEPN->ParA-Soj-PloopNTPase->BrxD-AAA->N6-METHYLASE->DUF6079*->PglZ->BrxD-AAA->SF1-SF2-Helicase+wHTH+Small-B-barrel+DUF1998-> | 1304 | Bacteria>Terrabacteria_group>Actinomycetota>Actinomycetes>Kitasatosporales | GCA_035983595.1 | Streptomyces_sp._NBC_01334 | OG736_44640 | 28052..31966 | 2903827 | hypothetical-protein-OG736_44640-[Streptomyces-sp.-NBC_01334] |
| WSP58015.1 | Type2.Pgl.like.BREX | PglW->?->DUF6079*-> | 1297 | Bacteria>Terrabacteria_group>Actinomycetota>Actinomycetes>Kitasatosporales | GCA_041435435.1 | Streptomyces_sp._NBC_01241 | OG306_29295 | 6528082..6531975 | 2903794 | phage-resistance-protein-[Streptomyces-sp.-NBC_01241] |
| WST78460.1 | Type2.Pgl.like.BREX | <-SF1-SF2-Helicase+wHTH+Small-B-barrel+DUF1998<-BrxD-AAA<-PglZ<-DUF6079*<-?<-N6-METHYLASE<-BrxD-AAA<-ParA-Soj-PloopNTPase<-HEPN<-PglW | 1312 | Bacteria>Terrabacteria_group>Actinomycetota>Actinomycetes>Kitasatosporales | GCA_045039715.1 | Streptomyces_sp._NBC_01136 | OG762_30240 | 6569338..6573276 | 2903754 | hypothetical-protein-OG762_30240-[Streptomyces-sp.-NBC_01136] |
| WSU11745.1 | Type2.Pgl.like.BREX | <-HTH_Tnp_1+HTH_21+rve+rve_3<-N6-METHYLASE\|\|?->\|\|YjhX_toxin-><-SF1-SF2-Helicase+wHTH+Small-B-barrel+DUF1998<-BrxD-AAA<-PglZ<-DUF6079*\|\|DUF397-><-N6-METHYLASE<-BrxD-AAA<-ParA-Soj-PloopNTPase<-HEPN<-PglW\|\|Phage_integrase-> | 1488 | Bacteria>Terrabacteria_group>Actinomycetota>Actinomycetes>Kitasatosporales | GCA_041435495.1 | Streptomyces_sp._NBC_01116 | OG245_30230 | 6895556..6900022 | 2903752 | hypothetical-protein-OG245_30230-[Streptomyces-sp.-NBC_01116] |
| WSU80365.1 | Type2.Pgl.like.BREX | PglW->N6-METHYLASE-><-DUF397<-HTH_31+DUF5753\|\|?->?->REase+SF1-SF2-Helicase->DUF6079*->PglZ->AIPR->BrxD-AAA->SF1-SF2-Helicase+wHTH+Small-B-barrel+DUF1998-><-TRAM+SAM-methylase+METHYLASE<-AA_permease_2 | 1308 | Bacteria>Terrabacteria_group>Actinomycetota>Actinomycetes>Kitasatosporales | GCA_045162865.1 | Streptomyces_globisporus | OG215_06810 | 1568760..1572686 | 1908 | phage-resistance-protein-[Streptomyces-globisporus] |
| WSU88910.1 | Type2.Pgl.like.BREX | PglW->N6-METHYLASE-><-DUF397<-HTH_31+DUF5753\|\|?->?->REase+SF1-SF2-Helicase->DUF6079*->PglZ-><-RNaseH\|\|BrxD-AAA->SF1-SF2-Helicase+wHTH+Small-B-barrel+DUF1998-><-GNTR-HTH | 1299 | Bacteria>Terrabacteria_group>Actinomycetota>Actinomycetes>Kitasatosporales | GCA_045162845.1 | Streptomyces_anulatus | OG575_09635 | 2228005..2231904 | 1892 | phage-resistance-protein-[Streptomyces-anulatus] |
| WSW28437.1 | Type2.Pgl.like.BREX | <-pPIWI+SFII-RAD3<-pPIWI+pPIWI<-PIWI<-?<-SF1-SF2-Helicase+wHTH+Small-B-barrel+DUF1998<-BrxD-AAA<-PglZ<-DUF6079*<-N6-METHYLASE<-PglW\|\|REase+DUF5655-> | 1297 | Bacteria>Terrabacteria_group>Actinomycetota>Actinomycetes>Kitasatosporales | GCA_045162695.1 | Streptomyces_sp._NBC_01003 | OG698_37525 | 8236620..8240513 | 2903714 | phage-resistance-protein-[Streptomyces-sp.-NBC_01003] |
| WSW86094.1 | Type2.Pgl.like.BREX | <-pPIWI+MID+RNaseH<-pPIWI+SFII-RAD3<-pPIWI+pPIWI<-SF1-SF2-Helicase+wHTH+Small-B-barrel+DUF1998<-BrxD-AAA<-PglZ<-DUF6079*<-N6-METHYLASE<-PglW\|\|Phage_integrase->XIS-HTH->Phage_integrase-><-YjhX_toxin | 1297 | Bacteria>Terrabacteria_group>Actinomycetota>Actinomycetes>Kitasatosporales | GCA_045162665.1 | Streptomyces_anulatus | OG536_29215 | 6647451..6651344 | 1892 | phage-resistance-protein-[Streptomyces-anulatus] |
| WSX42624.1 | Type2.Pgl.like.BREX | <-SF1-SF2-Helicase+wHTH+Small-B-barrel+DUF1998<-BrxD-AAA<-PglZ<-DUF6079*<-N6-METHYLASE<-DUF397<-HTH_31+DUF5753\|\|?-><-PglW\|\|OLD-ABC-> | 1296 | Bacteria>Terrabacteria_group>Actinomycetota>Actinomycetes>Kitasatosporales | GCA_045162215.1 | Streptomyces_sp._NBC_00963 | OG760_13435 | 3049147..3053037 | 2903697 | phage-resistance-protein-[Streptomyces-sp.-NBC_00963] |
| WSY41481.1 | Type2.Pgl.like.BREX | <-ST-Hiskin\|\|HTH_31+DUF5753->DUF397-><-SF1-SF2-Helicase+wHTH+Small-B-barrel+DUF1998<-BrxD-AAA<-PglZ<-DUF6079*<-N6-METHYLASE<-PglW<-tRNA\|\|REC+LuxR-HTH-> | 1295 | Bacteria>Terrabacteria_group>Actinomycetota>Actinomycetes>Kitasatosporales | GCA_045207655.1 | Embleya_sp._NBC_00888 | OG948_23980 | 5593191..5597078 | 2975960 | PglY-protein-[Embleya-sp.-NBC_00888] |
| WSY60842.1 | Type2.Pgl.like.BREX | <-SF1-SF2-Helicase+wHTH+Small-B-barrel+DUF1998<-BrxD-AAA<-PglZ<-DUF6079*<-N6-METHYLASE<-PglW\|\|?-><-DUF4096+DDE_Tnp_1+DDE_Tnp_1_2 | 1230 | Bacteria>Terrabacteria_group>Actinomycetota>Actinomycetes>Mycobacteriales | GCA_045207455.1 | Nocardia_sp._NBC_00881 | OH799_32575 | 7714027..7717719 | 2975995 | hypothetical-protein-OH799_32575-[Nocardia-sp.-NBC_00881] |
| WTA22046.1 | Type2.Pgl.like.BREX | PglW->N6-METHYLASE->DUF6079*->PglZ->BrxD-AAA->SF1-SF2-Helicase+wHTH+Small-B-barrel+DUF1998-><-pPIWI+MID+RNaseH<-pPIWI+SFII-RAD3<-pPIWI+pPIWI\|\|REase+DUF5655-> | 1297 | Bacteria>Terrabacteria_group>Actinomycetota>Actinomycetes>Kitasatosporales | GCA_045287305.1 | Streptomyces_sp._NBC_00853 | OG365_30705 | 6611587..6615480 | 2903681 | phage-resistance-protein-[Streptomyces-sp.-NBC_00853] |
| WTB16643.1 | Type2.Pgl.like.BREX | PIN-><-SF1-SF2-Helicase+wHTH+Small-B-barrel+DUF1998<-BrxD-AAA<-PglZ<-DUF6079*<-N6-METHYLASE<-PglW\|\|DUF6731-> | 1306 | Bacteria>Terrabacteria_group>Actinomycetota>Actinomycetes>Kitasatosporales | GCA_045161795.1 | Streptomyces_sp._NBC_00829 | OG293_25295 | 5698243..5702163 | 2903679 | PglY-protein-[Streptomyces-sp.-NBC_00829] |
| WTB22519.1 | Type2.Pgl.like.BREX | <-SF1-SF2-Helicase+REase<-SAD(SRA)+wHTH+HNH\|\|PglW->N6-METHYLASE-><-ST-Hiskin\|\|HTH_31+DUF5753->DUF397->DUF6079*->PglZ-><-CBS+HEPN<-ParB+HNH\|\|BrxD-AAA->SF1-SF2-Helicase+wHTH+Small-B-barrel+DUF1998-> | 1306 | Bacteria>Terrabacteria_group>Actinomycetota>Actinomycetes>Kitasatosporales | GCA_045161805.1 | Streptomyces_virginiae | OG253_14010 | 3171225..3175145 | 1961 | phage-resistance-protein-[Streptomyces-virginiae] |
| WTC44582.1 | Type2.Pgl.like.BREX | AA_permease_2->TRAM+SAM-methylase+METHYLASE-><-SF1-SF2-Helicase+wHTH+Small-B-barrel+DUF1998<-BrxD-AAA<-PglZ<-DUF6079*\|\|HTH_31+DUF5753->DUF397-><-N6-METHYLASE<-PglW\|\|REase->?->CASPASE+SF1-SF2-Helicase-> | 1298 | Bacteria>Terrabacteria_group>Actinomycetota>Actinomycetes>Kitasatosporales | - | Streptomyces_albidoflavus | OH810_24955 | 5684021..5687917 | 1886 | phage-resistance-protein-[Streptomyces-albidoflavus] |
| WTC74393.1 | Type2.Pgl.like.BREX | GNTR-HTH-><-SF1-SF2-Helicase+wHTH+Small-B-barrel+DUF1998<-BrxD-AAA\|\|RNaseH-><-PglZ<-DUF6079*<-REase+SF1-SF2-Helicase<-?<-?\|\|HTH_31+DUF5753->DUF397-><-N6-METHYLASE<-PglW | 1299 | Bacteria>Terrabacteria_group>Actinomycetota>Actinomycetes>Kitasatosporales | GCA_045209825.1 | Streptomyces_anulatus | OG882_30305 | 6746657..6750556 | 1892 | phage-resistance-protein-[Streptomyces-anulatus] |
| WTK31124.1 | Type2.Pgl.like.BREX | HTH->?-><-DUF4096+DDE_Tnp_1+DDE_Tnp_1_2\|\|XIS-HTH-><-SF1-SF2-Helicase+wHTH+Small-B-barrel+DUF1998<-BrxD-AAA<-PglZ<-DUF6079*<-N6-METHYLASE<-BrxD-AAA<-ParA-Soj-PloopNTPase<-HEPN<-PglW<-Pkinase+TPR<-OLD-ABC+TOPRIM | 1305 | Bacteria>Terrabacteria_group>Actinomycetota>Actinomycetes>Kitasatosporales | GCA_045040415.1 | Streptomyces_sp._NBC_01527 | OG612_29580 | 6536222..6540139 | 2903894 | hypothetical-protein-OG612_29580-[Streptomyces-sp.-NBC_01527] |
| WTK69774.1 | Type2.Pgl.like.BREX | <-REase\|\|?-><-?<-DUF6731\|\|PglW->N6-METHYLASE->DUF6079*->PglZ->BrxD-AAA->SF1-SF2-Helicase+wHTH+Small-B-barrel+DUF1998-><-?<-LuxR-HTH-><-?<-HTH | 1302 | Bacteria>Terrabacteria_group>Actinomycetota>Actinomycetes>Kitasatosporales | GCA_045040345.1 | Streptomyces_sp._NBC_01515 | OG407_16685 | 3775438..3779346 | 2903890 | PglY-protein-[Streptomyces-sp.-NBC_01515] |
| WTK77647.1 | Type2.Pgl.like.BREX | PglW->N6-METHYLASE-><-DUF397<-HTH_31+DUF5753\|\|?->?->REase+SF1-SF2-Helicase->DUF6079*->PglZ-><-RNaseH\|\|BrxD-AAA->SF1-SF2-Helicase+wHTH+Small-B-barrel+DUF1998->?-><-?<-ACET+ACET | 1303 | Bacteria>Terrabacteria_group>Actinomycetota>Actinomycetes>Kitasatosporales | GCA_045040335.1 | Streptomyces_sp._NBC_01511 | OG292_07480 | 1627933..1631844 | 2903889 | phage-resistance-protein-[Streptomyces-sp.-NBC_01511] |
| WTK93289.1 | Type2.Pgl.like.BREX | PglW->N6-METHYLASE-><-DUF397<-HTH_31+DUF5753\|\|?->?->REase+SF1-SF2-Helicase->DUF6079*->PglZ-><-RNaseH\|\|BrxD-AAA->SF1-SF2-Helicase+wHTH+Small-B-barrel+DUF1998->?-><-?<-ACET+ACET | 1303 | Bacteria>Terrabacteria_group>Actinomycetota>Actinomycetes>Kitasatosporales | GCA_045040315.1 | Streptomyces_sp._NBC_01508 | OG700_07120 | 1583357..1587268 | 2903888 | phage-resistance-protein-[Streptomyces-sp.-NBC_01508] |
| WTV30283.1 | Type2.Pgl.like.BREX | PglW->HEPN->ParA-Soj-PloopNTPase->BrxD-AAA->N6-METHYLASE-><-?<-?\|\|DUF6079*->PglZ->BrxD-AAA->SF1-SF2-Helicase+wHTH+Small-B-barrel+DUF1998->HTH-> | 1291 | Bacteria>Terrabacteria_group>Actinomycetota>Actinomycetes>Kitasatosporales | GCA_045041085.1 | [Kitasatospora]_papulosa | OG795_18585 | 4232384..4236259 | 1464011 | hypothetical-protein-OG795_18585-[[Kitasatospora] |
| WTW28784.1 | Type2.Pgl.like.BREX | PglW-><-BrxD-AAA<-ParA-Soj-PloopNTPase<-HEPN\|\|N6-METHYLASE-><-?\|\|DUF6079*->PglZ->BrxD-AAA->SF1-SF2-Helicase+wHTH+Small-B-barrel+DUF1998-><-?<-?<-?<-?\|\|REase-> | 1319 | Bacteria>Terrabacteria_group>Actinomycetota>Actinomycetes>Kitasatosporales | GCA_045040895.1 | Streptomyces_purpurascens | OHU35_23245 | 5102700..5106659 | 1924 | PglY-protein-[Streptomyces-purpurascens] |
| WTY85025.1 | Type2.Pgl.like.BREX | SF1-SF2-Helicase->PglW->N6-METHYLASE->DUF6079*->PglZ->BrxD-AAA->SF1-SF2-Helicase+wHTH+Small-B-barrel+DUF1998-><-Pkinase+TPR<-REase+SF1-SF2-Helicase\|\|REase+SF1-SF2-Helicase-><-HTH | 1234 | Bacteria>Terrabacteria_group>Actinomycetota>Actinomycetes>Streptosporangiales | GCA_045040145.1 | Sphaerisporangium_sp._NBC_01403 | OG589_24520 | 5528893..5532597 | 2903599 | hypothetical-protein-OG589_24520-[Sphaerisporangium-sp.-NBC_01403] |

<table>
<tr><th>Protein-ID</th><th>System</th><th>Gene-Neighbourhoods</th><th>Length</th><th>Taxonomy</th><th>Assembly-ID</th><th>Strain</th><th>Locus-Tag</th><th>Genomic-coordinates</th><th>TaxID</th><th>Fasta-defline</th></tr>
<tr><td>WTZ16473.1</td><td>Type2.Pgl.like.BREX</td><td>PglW->N6-METHYLASE-><-DUF397<-HTH_31+DUF5753||?->?->REase+SF1-SF2-Helicase->DUF6079*->PglZ-><-RNaseH||BrxD-AAA->SF1-SF2-Helicase+wHTH+Small-B-barrel+DUF1998-><-HKD-DNase+LuxR-HTH<-GNTR-HTH</td><td>1299</td><td>Bacteria>Terrabacteria_group>Actinomycetota>Actinomycetes>Kitasatosporales</td><td>GCA_045040115.1</td><td>Streptomyces_microflavus</td><td>OG270_07070</td><td>1564013..1567912</td><td>1919</td><td>phage-resistance-protein-[Streptomyces-microflavus]</td></tr>
<tr><td>WTZ27295.1</td><td>Type2.Pgl.like.BREX</td><td>HTH-><-NUDIX||?->?-><-?||PglW->N6-METHYLASE->DUF6079*->PglZ->BrxD-AAA->SF1-SF2-Helicase+wHTH+Small-B-barrel+DUF1998-></td><td>1231</td><td>Bacteria>Terrabacteria_group>Actinomycetota>Actinomycetes>Streptosporangiales</td><td>GCA_045040095.1</td><td>Microbispora_sp._NBC_01389</td><td>OG344_12180</td><td>2697446..2701141</td><td>2903584</td><td>hypothetical-protein-OG344_12180-[Microbispora-sp.-NBC_01389]</td></tr>
<tr><td>WTZ86643.1</td><td>Type2.Pgl.like.BREX</td><td><-HIN-HTH<-pPIWI+MID+RNaseH<-pPIWI+SFII-RAD3<-pPIWI+pPIWI<-SF1-SF2-Helicase+wHTH+Small-B-barrel+DUF1998<-BrxD-AAA<-PglZ<-DUF6079*<-N6-METHYLASE<-PglW||REase+DUF5655-><-TetR-HTH<-?||<-?<-DCM</td><td>1299</td><td>Bacteria>Terrabacteria_group>Actinomycetota>Actinomycetes>Kitasatosporales</td><td>GCA_045040035.1</td><td>Streptomyces_sp._NBC_01384</td><td>OG747_40680</td><td>8843149..8847048</td><td>2903847</td><td>phage-resistance-protein-[Streptomyces-sp.-NBC_01384]</td></tr>
<tr><td>WUA45283.1</td><td>Type2.Pgl.like.BREX</td><td>AA_permease_2->TRAM+SAM-methylase+METHYLASE-><-SF1-SF2-Helicase+wHTH+Small-B-barrel+DUF1998<-BrxD-AAA<-PglZ<-DUF6079*||DUF397-><-N6-METHYLASE<-BrxD-AAA<-ParA-Soj-PloopNTPase<-HEPN<-PglW<-DUF4096+DDE_Tnp_1+DDE_Tnp_1_2</td><td>1488</td><td>Bacteria>Terrabacteria_group>Actinomycetota>Actinomycetes>Kitasatosporales</td><td>GCA_045039935.1</td><td>Streptomyces_sp._NBC_01367</td><td>OG465_34035</td><td>7195206..7199672</td><td>2903841</td><td>hypothetical-protein-OG465_34035-[Streptomyces-sp.-NBC_01367]</td></tr>
<tr><td>WUE94592.1</td><td>Type2.Pgl.like.BREX</td><td><-DUF6079*<-?<-PglW</td><td>1248</td><td>Bacteria>Terrabacteria_group>Actinomycetota>Actinomycetes>Kitasatosporales</td><td>GCA_036013645.1</td><td>Streptomyces_sp._NBC_00490</td><td>OG381_36845</td><td>8096083..8099829</td><td>2903657</td><td>hypothetical-protein-OG381_36845-[Streptomyces-sp.-NBC_00490]</td></tr>
<tr><td>WUQ54585.1</td><td>Type2.Pgl.like.BREX</td><td><-GDResolvase||<-SF1-SF2-Helicase+wHTH+Small-B-barrel+DUF1998<-BrxD-AAA<-PglZ<-DUF6079*<-N6-METHYLASE||REase->?->?->?-><-N6-METHYLASE-p<-PglW</td><td>1294</td><td>Bacteria>Terrabacteria_group>Actinomycetota>Actinomycetes>Kitasatosporales</td><td>GCA_041435175.1</td><td>Streptomyces_virginiae</td><td>OG340_29295</td><td>6282308..6286192</td><td>1961</td><td>PglY-protein-[Streptomyces-virginiae]</td></tr>
<tr><td>WXC23206.1</td><td>Type2.Pgl.like.BREX</td><td>ACET+ACET->?->PIN-><-BrxD-AAA<-PglZ<-DUF6079*<-N6-METHYLASE<-PglW||DUF6731->?-><-?<-ACET+ACET</td><td>1306</td><td>Bacteria>Terrabacteria_group>Actinomycetota>Actinomycetes>Kitasatosporales</td><td>GCA_037201975.1</td><td>Streptomyces_sp._TG1A-60</td><td>WBG99_24810</td><td>5724208..5728128</td><td>3129111</td><td>PglY-protein-[Streptomyces-sp.-TG1A-60]</td></tr>
<tr><td>WXH56079.1</td><td>Type2.Pgl.like.BREX</td><td>REase+SF1-SF2-Helicase->HNH-><-HNH||?->PglW->N6-METHYLASE->DUF6079*->PglZ->BrxD-AAA->SF1-SF2-Helicase+wHTH+Small-B-barrel+DUF1998-></td><td>1303</td><td>Bacteria>Terrabacteria_group>Actinomycetota>Actinomycetes>Mycobacteriales</td><td>GCA_037463925.1</td><td>Rhodococcus_qingshengii</td><td>WDS21_19065</td><td>4152918..4156829</td><td>334542</td><td>DUF6079-family-protein-[Rhodococcus-qingshengii]</td></tr>
<tr><td>WXK63892.1</td><td>Type2.Pgl.like.BREX</td><td><-REC+LuxR-HTH<-HisKA_3+HATPase_c||tRNA-><-Phage_int_SAM_3+Phage_integrase||LuxR-HTH-><-SF1-SF2-Helicase+wHTH+Small-B-barrel+DUF1998<-BrxD-AAA<-PglZ<-DUF6079*<-?<-?<-N6-METHYLASE<-PglW||?->HTH_Tnp_4+DDE_Tnp_4-></td><td>1282</td><td>Bacteria>Terrabacteria_group>Actinomycetota>Actinomycetes>Streptosporangiales</td><td>GCA_045291505.1</td><td>Nonomuraea_sp._N2-4H</td><td>WBK31_35085</td><td>7621057..7624905</td><td>3128898</td><td>hypothetical-protein-WBK31_35085-[Nonomuraea-sp.-N2-4H]</td></tr>
</table>

| Protein-ID | System | Gene-Neighbourhoods | Length | Taxonomy | Assembly-ID | Strain | Locus-Tag | Genomic-coordinates | TaxID | Fasta-defline |
|---|---|---|---|---|---|---|---|---|---|---|
| -- | | | | | | | | | | |
| --Type3.BREX | | | | | | | | | | |
| -- | | | | | | | | | | |
| EMR73852.1 | Type3.MTaseII.containing.BREX | BrxB-BrxF-iSTAND->DUF6079*->?->N6-N4-METHYLASE->TUDOR+SF1-SF2-Helicase+iREase+CTD->PglZ->BrxA-DNA-binding-><-RNase<-?<-REase | 1209 | [illegible] | GCA_000349645.1 | Thermoplasmatales_archaeon_SCGC_AB-539-N05 | MBGDN05_00445 | 2273..5902 | 1198116 | hypothetical-protein-MBGDN05_00445-[Thermoplasmatales-archaeon-SCGC-AB-539-N05] |
| MBU0460852.1 | Type3.MTaseII.containing.BREX | REase+35exo->?->BrxB-BrxF-iSTAND->DUF6079*->Phage_holin->?->N6-N4-METHYLASE->?->TUDOR+SF1-SF2-Helicase+iREase+CTD->PglZ-> | 1220 | [illegible] | GCA_018812355.1 | Nanoarchaeota_archaeon | KJ771_08675 | 1826..5488 | 2026764 | hypothetical-protein-[Nanoarchaeota-archaeon] |
| WP_255152836.1 | Type3.MTaseII.containing.BREX | SF1-SF2-Helicase->?->HTH_28->NTP_transf_5-><-SF1-SF2-Helicase+REase<-Exonuc_V_gamma+REase\|\|N6-N4-METHYLASE->BrxA-DNA-binding->BrxB-BrxF-iSTAND->[illegible] | 1301 | [illegible] | GCF_024298865.1 | Halosegnis_sp._ZY10 | NL112_RS19810 | 17918..21823 | 2962671 | hypothetical-protein-[Halosegnis-sp.-ZY10] |
| WP_137694122.1 | Type3.MTaseII.containing.BREX | <-MetJarc-HTH<-?\|\|HNH->?->\|\|CopG/RHH-><-?\|\|N6-N4-METHYLASE->BrxA-DNA-binding->BrxB-BrxF-iSTAND->DUF6079*->PglZ->ORC-CDC6-><-?\|\|?->?->?-><-?\|\|?-><-[illegible] | 1291 | [illegible] | GCF_005435015.1 | Halorubrum_sp._SP9 | EXE45_RS09105 | 79102..82977 | 1537267 | hypothetical-protein-[Halorubrum-sp.-SP9] |
| WP_013878504.1 | Type3.MTaseII.containing.BREX | GDResolvase-><-?<-?<-?<-?\|\|?-><-ParB+N6_N4_METHYLASE\|\|BrxA-DNA-binding->DUF6079*->BrxB-BrxF-iSTAND->DUF6079->REase->HTH+ICLR-HTH->NTP_transf+wHTH-><-[illegible] | 1227 | [illegible] | GCF_000217715.1 | Halopiger_xanaduensis | HALXA_RS01365 | 276282..279965 | 387343 | hypothetical-protein-[Halopiger-xanaduensis] |
| WP_138651679.1 | Type3.MTaseII.containing.BREX | <-REase\|\|?-><-NTP_transf+wHTH<-HTH+ICLR-HTH<-REase<-DUF6079<-BrxB-BrxF-iSTAND<-DUF6079*<-BrxA-DNA-binding\|\|ParB+N6_N4_METHYLASE-><-?<-GDResolvase | 1227 | [illegible] | GCF_005890195.1 | Natrinema_pallidum | FGF80_RS00920 | 199338..203021 | 69527 | hypothetical-protein-[Natrinema-pallidum] |
| WP_250138967.1 | Type3.MTaseII.containing.BREX | McrC-NTD+REase->?-><-McrA+HTH+HNH<-ParB<-PglZ<-DUF6079*<-BrxB-BrxF-iSTAND<-BrxA-DNA-binding<-N6-N4-METHYLASE<-?\|\|ZnR->Cas7->?->?->35exo-> | 1300 | [illegible] | GCF_023566055.1 | Halosolutus_amylolyticus | MUN73_RS02990 | 634929..638831 | 2932267 | hypothetical-protein-[Halosolutus-amylolyticus] |
| HIH77326.1 | Type3.MTaseII.containing.BREX | <-SIGMA+OB-Fold+AAA-ATPase+Lon-Protease-BrxL-p<-PglZ<-PglZ<-?<-N6-N4-METHYLASE<-N6-N4-METHYLASE<-DUF6079*<-?<-BrxA-DNA-binding\|\|?-><-?<-RADICAL-SAM | 1247 | [illegible] | GCA_013329575.1 | Halobacteria_archaeon | HA341_00095 | 6086..9829 | 2670410 | hypothetical-protein-[Halobacteria-archaeon] |
| MBP7299139.1 | Type3.MTaseII.containing.BREX | <-BrxA-DNA-binding<-PglZ<-TUDOR+SF1-SF2-Helicase+iREase+CTD<-N6-N4-METHYLASE<-DUF6079-p* | 119 | [illegible] | GCA_017996475.1 | Methanoculleus_sp. | KA967_01905 | 8295..8655 | 90427 | hypothetical-protein-[Methanoculleus-sp.] |
| WP_062399977.1 | Type3.MTaseII.containing.BREX | <-TUDOR+SF1-SF2-Helicase+iREase+CTD-p<-?<-OLD-ABC<-N6-N4-METHYLASE<-N6-N4-METHYLASE<-N6-N4-METHYLASE-p<-DUF6079*<-BrxB-BrxF-iSTAND\|\|?->?-><-RelE-Par[illegible] | 1212 | [illegible] | GCF_001315945.1 | Methanogenium_cariaci | JCM10550_RS13830 | 38387..42025 | 2197 | DUF6079-family-protein-[Methanogenium-cariaci] |
| WP_066953691.1 | Type3.MTaseII.containing.BREX | BrxB-BrxF-iSTAND->DUF6079*->N6-N4-METHYLASE->TUDOR+SF1-SF2-Helicase+iREase+CTD->PglZ->BrxA-DNA-binding->?->MULE-TRANSPOSASE-> | 1226 | [illegible] | GCF_001571405.1 | Methanoculleus_thermophilus | MCUTH_RS00015 | 808..4488 | 2200 | DUF6079-family-protein-[Methanoculleus-thermophilus] |
| WP_209691006.1 | Type3.MTaseII.containing.BREX | <-RADICAL-SAM\|\|?-><-BrxA-DNA-binding<-PglZ<-TUDOR+SF1-SF2-Helicase+iREase+CTD<-MORC<-?<-N6-N4-METHYLASE<-Schlafen+HATPase+wHTH<-DUF6079*<-BrxB-BrxF-[illegible] | 1246 | [illegible] | GCF_017875315.1 | Methanomicrobium_sp._W14 | J2128_RS09600 | 65374..69114 | 2817839 | DUF6079-family-protein-[Methanomicrobium-sp.-W14] |
| WP_209629647.1 | Type3.MTaseII.containing.BREX | SF1-SF2-Helicase-><-?\|\|?->HEPN+SH3-like+SF1-SF2-Helicase+wHTH->MORC+REase-><-?<-JetB<-?<-RNase\|\|DUF6079*->N6-N4-METHYLASE->TUDOR+SF1-SF2-Helicase+iR[illegible] | 1225 | [illegible] | GCF_017875325.1 | Methanofollis_sp._W23 | J2129_RS04035 | 974119..977796 | 2817849 | DUF6079-family-protein-[Methanofollis-sp.-W23] |
| PWR75057.1 | Type3.MTaseII.containing.BREX | <-BrxA-DNA-binding<-PglZ<-TUDOR+SF1-SF2-Helicase+iREase+CTD<-Schlafen+HATPase+wHTH<-PHP+SbcC<-N6-N4-METHYLASE<-N6-N4-METHYLASE<-DUF6079-p<-DUF6079*<-[illegible] | 1073 | [illegible] | GCA_003173335.1 | Methanospirillum_stamsii | DLD82_07525 | 192047..195268 | 1277351 | ATPase-[Methanospirillum-stamsii] |
| NLV27395.1 | Type3.MTaseII.containing.BREX | DUF499->?-><-?<-HAD+PRTase+PRTase+REase+SF1-SF2-Helicase<-?<-?<-Glyco_hydro\|\|BrxB-BrxF-iSTAND->DUF6079*->N6-N4-METHYLASE->N6-N4-METHYLASE->?->TUDOR+[illegible] | 1230 | [illegible] | GCA_012729535.1 | Methanomicrobiales_archaeon | GXY48_09555 | 22039..25731 | 2184052 | ATP-binding-protein-[Methanomicrobiales-archaeon] |
| HHV24727.1 | Type3.MTaseII.containing.BREX | TOPRIM+Topoisom_bac+TOPRIM_Crpt-><-?<-?<-?<-TPR+TPR+TPR+TPR+TPR+TPR+TPR\|\|HEPN->BrxB-BrxF-iSTAND->DUF6079*->Schlafen+HATPase+wHTH->N6-N4-METHYLASE->HEPN-[illegible] | 1240 | [illegible] | GCA_012839505.1 | Methanosarcina_sp. | GXX65_09470 | 12369..16091 | 2213 | ATP-binding-protein-[Methanosarcina-sp.] |
| WP_013898594.1 | Type3.MTaseII.containing.BREX | [illegible] | 1246 | [illegible] | GCF_000217995.1 | Methanosalsum_zhilinae | MZHIL_RS06595 | 1417456..1421196 | 39669 | DUF6079-family-protein-[Methanosalsum-zhilinae] |
| WP_048159302.1 | Type3.MTaseII.containing.BREX | [illegible] | 1241 | [illegible] | GCF_000969965.1 | Methanosarcina_sp._WWM596 | MSWHS_RS14780 | 3345634..3349359 | 1434103 | DUF6079-family-protein-[Methanosarcina-sp.-WWM596] |
| WP_148705386.1 | Type3.MTaseII.containing.BREX | [illegible] | 1240 | [illegible] | GCF_000970125.1 | Methanosarcina_siciliae | MSSIH_RS10180 | 2386528..2390250 | 38027 | DUF6079-family-protein-[Methanosarcina-siciliae] |
| WP_048125636.1 | Type3.MTaseII.containing.BREX | [illegible] | 1246 | [illegible] | GCF_000970265.1 | Methanosarcina_lacustris | MSLAZ_1365 | 1708978..1712718 | 170861 | DUF6079-family-protein-[Methanosarcina-lacustris] |
| WP_048041739.1 | Type3.MTaseII.containing.BREX | [illegible] | 1241 | [illegible] | GCF_000979335.1 | Methanosarcina_mazei | DU31_RS16150 | 61828..65553 | 2209 | DUF6079-family-protein-[Methanosarcina-mazei] |
| WP_209616537.1 | Type3.MTaseII.containing.BREX | [illegible] | 1246 | [illegible] | GCF_017873415.1 | Methanolobus_bombayensis | J2746_RS03805 | 332561..336301 | 38023 | DUF6079-family-protein-[Methanolobus-bombayensis] |
| NYT10423.1 | Type3.MTaseII.containing.BREX | [illegible] | 245 | [illegible] | GCA_013415605.1 | Methanosarcinales_archaeon | GKC09_10925 | 8497..9234 | 2250255 | hypothetical-protein-[Methanosarcinales-archaeon] |
| OPX79392.1 | Type3.MTaseII.containing.BREX | [illegible] | 1243 | [illegible] | GCA_002067795.1 | Methanosaeta_sp._PtaB.Bin039 | A4E45_00671 | 40596..44327 | 1811685 | MAG:-hypothetical-protein-A4E45_00671-[Methanosaeta-sp.-PtaB.Bin039] |
| NLJ23755.1 | Type3.MTaseII.containing.BREX | [illegible] | 1243 | [illegible] | GCA_012517695.1 | Methanothrix_soehngenii | GX426_11730 | 9029..12760 | 2223 | ATP-binding-protein-[Methanothrix-soehngenii] |
| KYK31514.1 | Type3.MTaseII.containing.BREX | [illegible] | 1227 | [illegible] | GCA_001595815.1 | Theionarchaea_archaeon_DG-70-1 | AYK19_16750 | 12752..16435 | 1803814 | ATPase-[Theionarchaea-archaeon-DG-70-1] |
| MBE3050154.1 | Type3.MTaseII.containing.BREX | [illegible] | 1447 | [illegible] | GCA_014894645.1 | Candidatus_Bathyarchaeota_archaeon | IMZ48_48190 | 6822..11165 | 2026714 | hypothetical-protein-[Candidatus-Bathyarchaeota-archaeon] |
| MBT6646600.1 | Type3.MTaseII.containing.BREX | [illegible] | 1219 | [illegible] | GCA_018697615.1 | Nitrososphaerota_archaeon | HOB51_03670 | 4275..7934 | 2026795 | hypothetical-protein-[Nitrososphaerota-archaeon] |
| MCA2972183.1 | Type3.MTaseII.containing.BREX | [illegible] | 1239 | [illegible] | GCA_020247055.1 | Acidobacteriaceae_bacterium | INH43_27010 | 15323..19042 | 2052142 | ATP-binding-protein-[Acidobacteriaceae-bacterium] |
| MYN66483.1 | Type3.MTaseII.containing.BREX | [illegible] | 1235 | [illegible] | GCA_009861545.1 | Acidobacteriota_bacterium | F4X11_15860 | 71321..75028 | 1978231 | ATP-binding-protein-[Acidobacteriota-bacterium] |
| QQS46113.1 | Type3.MTaseII.containing.BREX | [illegible] | 1422 | [illegible] | GCA_016700195.1 | Acidobacteriota_bacterium | IPM66_19690 | 4950965..4955233 | 1978231 | hypothetical-protein-IPM66_19690-[Acidobacteriota-bacterium] |
| NLJ38119.1 | Type3.MTaseII.containing.BREX | [illegible] | 1243 | [illegible] | GCA_012517485.1 | Candidatus_Atribacteria_bacterium | GX432_05075 | 6404..10135 | 2053509 | ATP-binding-protein-[Candidatus-Atribacteria-bacterium] |
| MBI4537318.1 | Type3.MTaseII.containing.BREX | [illegible] | 208 | [illegible] | GCA_016209735.1 | candidate_division_NC10_bacterium | HY712_05115 | 15842..16468 | 2072417 | hypothetical-protein-[candidate-division-NC10-bacterium] |
| UCH96958.1 | Type3.MTaseII.containing.BREX | [illegible] | 1097 | [illegible] | GCA_020355945.1 | Candidatus_Aminicenantes_bacterium | JSV88_08940 | 2459667..2462960 | 2052149 | MAG:-ATP-binding-protein,-partial-[Candidatus-Aminicenantes-bacterium] |
| UCH92727.1 | Type3.MTaseII.containing.BREX | [illegible] | 465 | [illegible] | GCA_020355945.1 | Candidatus_Aminicenantes_bacterium | JSV88_20860 | 5485993..5487389 | 2052149 | MAG:-ATP-binding-protein,-partial-[Candidatus-Aminicenantes-bacterium] |
| MCD6192837.1 | Type3.MTaseII.containing.BREX | [illegible] | 1245 | [illegible] | GCA_021157535.1 | Candidatus_Aminicenantes_bacterium | J7L26_05130 | 23251..26988 | 2052149 | ATP-binding-protein-[Candidatus-Aminicenantes-bacterium] |
| MBD3236778.1 | Type3.MTaseII.containing.BREX | [illegible] | 431 | [illegible] | GCA_014729815.1 | Candidatus_Eisenbacteria_bacterium | GF330_08750 | 1770..3065 | 2212470 | BREX-3-system-P-loop-containing-protein-BrxF-[Candidatus-Eisenbacteria-bacterium] |
| MBK8230678.1 | Type3.MTaseII.containing.BREX | [illegible] | 1232 | [illegible] | GCA_016712505.1 | Candidatus_Eisenbacteria_bacterium | IPK72_08880 | 13493..17191 | 2212470 | ATP-binding-protein-[Candidatus-Eisenbacteria-bacterium] |
| MBQ3643693.1 | Type3.MTaseII.containing.BREX | [illegible] | 1227 | [illegible] | GCA_017417215.1 | Candidatus_Riflebacteria_bacterium | II961_03765 | 12317..16000 | 2053586 | hypothetical-protein-[Candidatus-Riflebacteria-bacterium] |
| OGK77274.1 | Type3.MTaseII.containing.BREX | [illegible] | 1235 | [illegible] | GCA_001788395.1 | Candidatus_Rokubacteria_bacterium_GWA2_70_23 | A2X53_04110 | 12396..16103 | 1802095 | MAG:-ATPase-[Candidatus-Rokubacteria-bacterium-GWA2_70_23] |
| MBC7752530.1 | Type3.MTaseII.containing.BREX | [illegible] | 1243 | [illegible] | GCA_014378805.1 | Candidatus_Saccharibacteria_bacterium | H7Z73_12610 | 10509..14240 | 2026720 | ATP-binding-protein-[Candidatus-Saccharibacteria-bacterium] |
| NLI41205.1 | Type3.MTaseII.containing.BREX | [illegible] | 1242 | [illegible] | GCA_012517765.1 | Caldisericales_bacterium | GX421_08515 | 1471..5199 | 2614251 | ATP-binding-protein-[Caldisericales-bacterium] |
| HIP19035.1 | Type3.MTaseII.containing.BREX | [illegible] | 1244 | [illegible] | GCA_015663065.1 | Sulfurovum_sp. | EYG78_06850 | 23973..27707 | 1969726 | ATP-binding-protein-[Sulfurovum-sp.] |
| MBN1944774.1 | Type3.MTaseII.containing.BREX | [illegible] | 1204 | [illegible] | GCA_016931535.1 | Bradymonadales_bacterium | JW797_03815 | 4633..8247 | 2099667 | hypothetical-protein-[Bradymonadales-bacterium] |
| WP_163298138.1 | Type3.MTaseII.containing.BREX | [illegible] | 1313 | [illegible] | GCF_010499435.1 | Dissulfurirhabdus_thermomarina | G3N55_RS03885 | 13781..17722 | 1765737 | DUF6079-family-protein-[Dissulfurirhabdus-thermomarina] |
| MBW1792990.1 | Type3.MTaseII.containing.BREX | [illegible] | 1213 | [illegible] | - | Deltaproteobacteria_bacterium | JRJ38_00920 | 3864..7505 | 2026735 | ATP-binding-protein-[Deltaproteobacteria-bacterium] |
| EFK08826.1 | Type3.MTaseII.containing.BREX | [illegible] | 1248 | [illegible] | GCA_000179315.1 | delta_proteobacterium_NaphS2 | NPH_0523 | 21640..25386 | 88274 | conserved-hypothetical-protein-[delta-proteobacterium-NaphS2] |
| OGP84719.1 | Type3.MTaseII.containing.BREX | [illegible] | 1186 | [illegible] | GCA_001797945.1 | Deltaproteobacteria_bacterium_RBG_16_58_17 | A2V87_08860 | 18905..22465 | 1797845 | ATPase-[Deltaproteobacteria-bacterium-RBG_16_58_17] |
| OGR04297.1 | Type3.MTaseII.containing.BREX | [illegible] | 1285 | [illegible] | GCA_001799195.1 | Deltaproteobacteria_bacterium_RIFOXYA12_FULL_61_11 | A2284_01915 | 10657..14514 | 1797896 | ATPase-[Deltaproteobacteria-bacterium-RIFOXYA12_FULL_61_11] |
| OGQ88897.1 | Type3.MTaseII.containing.BREX | [illegible] | 1252 | [illegible] | GCA_001799285.1 | Deltaproteobacteria_bacterium_RIFOXYD12_FULL_56_24 | A2512_08460 | 1712..5470 | 1797905 | ATPase-[Deltaproteobacteria-bacterium-RIFOXYD12_FULL_56_24] |
| OIP43034.1 | Type3.MTaseII.containing.BREX | [illegible] | 1041 | [illegible] | GCA_001873925.1 | Deltaproteobacteria_bacterium_CG2_30_63_29 | AUK47_03375 | 38912..42037 | 1805126 | hypothetical-protein-AUK47_03375-[Deltaproteobacteria-bacterium-CG2_30_63_29] |
| OQB76675.1 | Type3.MTaseII.containing.BREX | [illegible] | 1444 | [illegible] | GCA_002070575.1 | Deltaproteobacteria_bacterium_ADurb.Bin135 | BWX92_01600 | 30522..34856 | 1852867 | hypothetical-protein-BWX92_01600-[Deltaproteobacteria-bacterium-ADurb.Bin135] |
| PIV21312.1 | Type3.MTaseII.containing.BREX | [illegible] | 1235 | [illegible] | GCA_002780715.1 | Deltaproteobacteria_bacterium_CG03_land_8_20_14_0_80_45_14 | COS40_08020 | 9785..13492 | 1973969 | ATPase-[Deltaproteobacteria-bacterium-CG03_land_8_20_14_0_80_45_14] |
| PIX22666.1 | Type3.MTaseII.containing.BREX | [illegible] | 1233 | [illegible] | GCA_002783725.1 | Deltaproteobacteria_bacterium_CG_4_8_14_3_um_filter_45_9 | COZ69_10545 | 3046..6748 | 1973962 | ATPase,-partial-[Deltaproteobacteria-bacterium-CG_4_8_14_3_um_filter_45_9] |
| PKN28382.1 | Type3.MTaseII.containing.BREX | [illegible] | 1244 | [illegible] | GCA_002840495.1 | Deltaproteobacteria_bacterium_HGW-Deltaproteobacteria-21 | CVU64_13605 | 8246..11980 | 2013749 | ATPase-[Deltaproteobacteria-bacterium-HGW-Deltaproteobacteria-21] |
| PXF60115.1 | Type3.MTaseII.containing.BREX | [illegible] | 1283 | [illegible] | GCA_003194495.1 | Deltaproteobacteria_bacterium | C4B58_01095 | 57714..61565 | 2026735 | ATPase-[Deltaproteobacteria-bacterium] |
| TNF32509.1 | Type3.MTaseII.containing.BREX | [illegible] | 1236 | [illegible] | GCA_006226565.1 | Deltaproteobacteria_bacterium | EP329_09915 | 32814..36524 | 2026735 | ATP-binding-protein-[Deltaproteobacteria-bacterium] |
| NCO52936.1 | Type3.MTaseII.containing.BREX | [illegible] | 1042 | [illegible] | GCA_009993185.1 | Deltaproteobacteria_bacterium | GW875_12685 | 1..3131 | 2026735 | ATP-binding-protein-[Deltaproteobacteria-bacterium] |
| NCO56856.1 | Type3.MTaseII.containing.BREX | [illegible] | 1229 | [illegible] | GCA_009993275.1 | Deltaproteobacteria_bacterium | GW870_01520 | 18374..22063 | 2026735 | ATP-binding-protein-[Deltaproteobacteria-bacterium] |
| NCP02216.1 | Type3.MTaseII.containing.BREX | [illegible] | 132 | [illegible] | GCA_009993495.1 | Deltaproteobacteria_bacterium | GW861_02410 | 1..401 | 2026735 | ATP-binding-protein-[Deltaproteobacteria-bacterium] |
| NCS74602.1 | Type3.MTaseII.containing.BREX | [illegible] | 1229 | [illegible] | GCA_009994935.1 | Deltaproteobacteria_bacterium | GW783_10820 | 16268..19957 | 2026735 | ATP-binding-protein-[Deltaproteobacteria-bacterium] |
| NLF25483.1 | Type3.MTaseII.containing.BREX | [illegible] | 149 | [illegible] | GCA_012514925.1 | Deltaproteobacteria_bacterium | GX589_07485 | 21465..21914 | 2026735 | hypothetical-protein-[Deltaproteobacteria-bacterium] |
| MBI4774968.1 | Type3.MTaseII.containing.BREX | [illegible] | 1445 | [illegible] | GCA_016208165.1 | Deltaproteobacteria_bacterium | HY788_12455 | 28517..32854 | 2026735 | hypothetical-protein-[Deltaproteobacteria-bacterium] |
| MBI4764648.1 | Type3.MTaseII.containing.BREX | [illegible] | 1242 | [illegible] | GCA_016208255.1 | Deltaproteobacteria_bacterium | HY787_08590 | 9212..12940 | 2026735 | ATP-binding-protein-[Deltaproteobacteria-bacterium] |
| MBI5515241.1 | Type3.MTaseII.containing.BREX | [illegible] | 1233 | [illegible] | GCA_016218725.1 | Deltaproteobacteria_bacterium | HY909_15800 | 137952..141653 | 2026735 | ATP-binding-protein-[Deltaproteobacteria-bacterium] |
| MBI5489032.1 | Type3.MTaseII.containing.BREX | [illegible] | 1382 | [illegible] | GCA_016218815.1 | Deltaproteobacteria_bacterium | HY905_16985 | 60479..64627 | 2026735 | hypothetical-protein-[Deltaproteobacteria-bacterium] |
| MBK8716786.1 | Type3.MTaseII.containing.BREX | [illegible] | 1232 | [illegible] | GCA_016715415.1 | Deltaproteobacteria_bacterium | IPN32_18840 | 22806..26504 | 2026735 | ATP-binding-protein-[Deltaproteobacteria-bacterium] |
| MBS3918752.1 | Type3.MTaseII.containing.BREX | [illegible] | 1248 | [illegible] | GCA_018333535.1 | Deltaproteobacteria_bacterium | KG012_07670 | 20571..24317 | 2026735 | ATP-binding-protein-[Deltaproteobacteria-bacterium] |
| MBT4287260.1 | Type3.MTaseII.containing.BREX | [illegible] | 1398 | [illegible] | GCA_018651875.1 | Deltaproteobacteria_bacterium | HOD92_07955 | 4635..8831 | 2026735 | hypothetical-protein-[Deltaproteobacteria-bacterium] |
| MBW1982273.1 | Type3.MTaseII.containing.BREX | [illegible] | 1276 | [illegible] | GCA_019308175.1 | Deltaproteobacteria_bacterium | JRJ12_13740 | 13642..17472 | 2026735 | ATP-binding-protein-[Deltaproteobacteria-bacterium] |
| MBW2068238.1 | Type3.MTaseII.containing.BREX | [illegible] | 1245 | [illegible] | GCA_019309115.1 | Deltaproteobacteria_bacterium | JRI31_05095 | 91446..95183 | 2026735 | ATP-binding-protein-[Deltaproteobacteria-bacterium] |
| MCB9731980.1 | Type3.MTaseII.containing.BREX | [illegible] | 1233 | [illegible] | GCA_020632715.1 | Deltaproteobacteria_bacterium | H6745_05025 | 1400710..1404411 | 2026735 | ATP-binding-protein-[Deltaproteobacteria-bacterium] |
| MCD6261582.1 | Type3.MTaseII.containing.BREX | [illegible] | 1261 | [illegible] | GCA_021163815.1 | Deltaproteobacteria_bacterium | J7J52_00980 | 62923..66708 | 2026735 | ATP-binding-protein-[Deltaproteobacteria-bacterium] |
| CAB1056976.1 | Type3.MTaseII.containing.BREX | [illegible] | 1242 | [illegible] | GCA_902749685.1 | Olavius_sp._associated_proteobacterium_Delta_1 | D1BOALGB6SA_1715 | 1820..5548 | 698986 | hypothetical-protein-BVU_3741-[Olavius-sp.-associated-proteobacterium-Delta-1] |
| BBM69555.1 | Type3.MTaseII.containing.BREX | [illegible] | 1248 | [illegible] | GCA_009936255.1 | Rhodothermus_marinus | RmaAA213_14010 | 1619922..1623668 | 29549 | hypothetical-protein-RmaAA213_14010-[Rhodothermus-marinus] |
| WP_197738551.1 | Type3.MTaseII.containing.BREX | [illegible] | 1248 | [illegible] | GCF_009936255.1 | Rhodothermus_marinus | Rma1338_RS07045 | 1619922..1623668 | 29549 | DUF6079-family-protein-[Rhodothermus-marinus] |
| WP_250716305.1 | Type3.MTaseII.containing.BREX | [illegible] | 1221 | [illegible] | - | Bacteroides_fragilis | J8L04_RS14220 | 56443..60108 | 817 | DUF6079-family-protein-[Bacteroides-fragilis] |
| CCX79044.1 | Type3.MTaseII.containing.BREX | [illegible] | 1231 | [illegible] | GCA_000431255.1 | Parabacteroides_johnsonii_CAG:246 | BN560_00043 | 5288..8983 | 1263091 | putative-uncharacterized-protein-[Parabacteroides-johnsonii-CAG:246] |
| CCX55298.1 | Type3.MTaseII.containing.BREX | [illegible] | 1194 | [illegible] | GCA_000432655.1 | Bacteroides_sp._CAG:1060 | BN459_01197 | 8514..12098 | 1262734 | putative-uncharacterized-protein-[Bacteroides-sp.-CAG:1060] |
| CEA16533.1 | Type3.MTaseII.containing.BREX | [illegible] | 1222 | [illegible] | GCA_000953535.1 | Fermentimonas_caenicola | ING2E5B_1789 | 2009127..2012795 | 1562970 | hypothetical-protein-ING2E5B_1789-[Fermentimonas-caenicola] |
| PWL82904.1 | Type3.MTaseII.containing.BREX | [illegible] | 943 | [illegible] | GCA_003150315.1 | Prevotellaceae_bacterium | DBY24_02745 | 4283..7114 | 2049047 | hypothetical-protein-DBY24_02745-[Prevotellaceae-bacterium] |
| ROS80360.1 | Type3.MTaseII.containing.BREX | [illegible] | 1218 | [illegible] | GCA_003762605.1 | Muribaculaceae_bacterium_Isolate-042_(Harlan) | EEL35_09185 | 87209..90865 | 2486462 | hypothetical-protein-EEL35_09185-[Muribaculaceae-bacterium-Isolate-042-(Harlan)] |
| MBE6232697.1 | Type3.MTaseII.containing.BREX | [illegible] | 1221 | [illegible] | GCA_015060345.1 | Bacteroidales_bacterium | E7117_10120 | 3411..7076 | 2030927 | hypothetical-protein-[Bacteroidales-bacterium] |
| MBE6339841.1 | Type3.MTaseII.containing.BREX | [illegible] | 1225 | [illegible] | GCA_015061285.1 | Bacteroidales_bacterium | E7069_03750 | 60116..63793 | 2030927 | hypothetical-protein-[Bacteroidales-bacterium] |
| MBK8808662.1 | Type3.MTaseII.containing.BREX | [illegible] | 834 | [illegible] | GCA_016714605.1 | Bacteroidales_bacterium | IPO21_19315 | 57274..59778 | 2030927 | hypothetical-protein-[Bacteroidales-bacterium] |
| MBQ4549131.1 | Type3.MTaseII.containing.BREX | [illegible] | 1221 | [illegible] | GCA_017399385.1 | Bacteroidales_bacterium | IJA42_08135 | 153861..157526 | 2030927 | hypothetical-protein-[Bacteroidales-bacterium] |
| MBQ9466675.1 | Type3.MTaseII.containing.BREX | [illegible] | 1223 | [illegible] | GCA_017454355.1 | Muribaculaceae_bacterium | IJU62_06845 | 5125..8796 | 2498093 | hypothetical-protein-[Muribaculaceae-bacterium] |
| MBQ9636772.1 | Type3.MTaseII.containing.BREX | [illegible] | 1230 | [illegible] | GCA_017458165.1 | Prevotella_sp. | IJV36_02605 | 23790..27482 | 59823 | hypothetical-protein-[Prevotella-sp.] |
| MBQ8866661.1 | Type3.MTaseII.containing.BREX | [illegible] | 1221 | [illegible] | GCA_017464815.1 | Bacteroidaceae_bacterium | IJ013_02920 | 86455..90120 | 2212467 | hypothetical-protein-[Bacteroidaceae-bacterium] |
| MBQ7946199.1 | Type3.MTaseII.containing.BREX | [illegible] | 1221 | [illegible] | GCA_017468435.1 | Bacteroidales_bacterium | IJ280_05750 | 3462..7127 | 2030927 | hypothetical-protein-[Bacteroidales-bacterium] |
| MBQ8602359.1 | Type3.MTaseII.containing.BREX | [illegible] | 1221 | [illegible] | GCA_017472925.1 | Bacteroides_sp. | IJ413_07405 | 190228..193893 | 29523 | hypothetical-protein-[Bacteroides-sp.] |
| MBR2226364.1 | Type3.MTaseII.containing.BREX | [illegible] | 494 | [illegible] | GCA_017475135.1 | Bacteroidales_bacterium | IJ893_00685 | 26261..27742 | 2030927 | hypothetical-protein-[Bacteroidales-bacterium] |
| MBR2228193.1 | Type3.MTaseII.containing.BREX | [illegible] | 774 | [illegible] | GCA_017475135.1 | Bacteroidales_bacterium | IJ893_10045 | 1..2325 | 2030927 | hypothetical-protein-[Bacteroidales-bacterium] |
| MBR1790999.1 | Type3.MTaseII.containing.BREX | [illegible] | 1017 | [illegible] | GCA_017480355.1 | Bacteroidaceae_bacterium | IJ754_04495 | 101089..104142 | 2212467 | hypothetical-protein-[Bacteroidaceae-bacterium] |
| MBR2017127.1 | Type3.MTaseII.containing.BREX | [illegible] | 583 | [illegible] | GCA_017505865.1 | Prevotella_sp. | IKA00_07530 | 7585..9334 | 59823 | hypothetical-protein-[Prevotella-sp.] |
| MBR6863258.1 | Type3.MTaseII.containing.BREX | [illegible] | 1221 | [illegible] | GCA_017531205.1 | Bacteroidales_bacterium | IKM75_00205 | 26210..29875 | 2030927 | hypothetical-protein-[Bacteroidales-bacterium] |
| MBR4381371.1 | Type3.MTaseII.containing.BREX | [illegible] | 1226 | [illegible] | GCA_017534535.1 | Bacteroidaceae_bacterium | IKP48_08945 | 182570..186250 | 2212467 | hypothetical-protein-[Bacteroidaceae-bacterium] |
| MBR5729013.1 | Type3.MTaseII.containing.BREX | [illegible] | 1222 | [illegible] | GCA_017561895.1 | Prevotella_sp. | IKX61_02230 | 1187..4855 | 59823 | hypothetical-protein-[Prevotella-sp.] |
| MBO5155857.1 | Type3.MTaseII.containing.BREX | [illegible] | 1227 | [illegible] | GCA_017622935.1 | Prevotella_sp. | J6C05_01780 | 52026..55709 | 59823 | hypothetical-protein-[Prevotella-sp.] |
| MBO6222745.1 | Type3.MTaseII.containing.BREX | [illegible] | 940 | [illegible] | GCA_017635115.1 | Bacteroidales_bacterium | J6N56_03080 | 12019..14841 | 2030927 | hypothetical-protein-[Bacteroidales-bacterium] |
| MBS6225156.1 | Type3.MTaseII.containing.BREX | [illegible] | 1180 | [illegible] | GCA_018374715.1 | Parabacteroides_johnsonii | KH749_10645 | 947..4489 | 387661 | hypothetical-protein-[Parabacteroides-johnsonii] |
| MBV4244188.1 | Type3.MTaseII.containing.BREX | [illegible] | 584 | [illegible] | GCA_019132015.1 | Parabacteroides_johnsonii | KSZ16_09845 | 152717..154471 | 387661 | hypothetical-protein-[Parabacteroides-johnsonii] |
| MCI6627987.1 | Type3.MTaseII.containing.BREX | [illegible] | 1227 | [illegible] | GCA_022778295.1 | Bacteroidales_bacterium | MR722_01200 | 21727..25410 | 2030927 | DUF6079-family-protein-[Bacteroidales-bacterium] |
| MCI6526370.1 | Type3.MTaseII.containing.BREX | [illegible] | 1216 | [illegible] | GCA_022781865.1 | Bacteroidales_bacterium | MR441_04010 | 4737..8387 | 2030927 | DUF6079-family-protein-[Bacteroidales-bacterium] |
| MCK4344535.1 | Type3.MTaseII.containing.BREX | [illegible] | 1410 | [illegible] | GCA_023128755.1 | Bacteroidales_bacterium | KAX05_04555 | 6498..10730 | 2030927 | BREX-3-system-P-loop-containing-protein-BrxF-[Bacteroidales-bacterium] |
| MCK9303764.1 | Type3.MTaseII.containing.BREX | [illegible] | 1253 | [illegible] | GCA_023230265.1 | Bacteroidales_bacterium | MOP27_00015 | 2858..6619 | 2030927 | DUF6079-family-protein-[Bacteroidales-bacterium] |
| MCM1091477.1 | Type3.MTaseII.containing.BREX | [illegible] | 1226 | [illegible] | GCA_023663505.1 | Muribaculum_sp. | NC413_11600 | 23059..26739 | 1918611 | DUF6079-family-protein-[Muribaculum-sp.] |
| MCQ2258857.1 | Type3.MTaseII.containing.BREX | [illegible] | 1226 | [illegible] | GCA_024401215.1 | Bacteroidaceae_bacterium | MJZ41_12830 | 10847..14527 | 2212467 | DUF6079-family-protein-[Bacteroidaceae-bacterium] |
| MCS2643501.1 | Type3.MTaseII.containing.BREX | [illegible] | 1222 | [illegible] | GCA_024789685.1 | Bacteroides_fragilis | NXV97_11510 | 2860096..2863764 | 817 | DUF6079-family-protein-[Bacteroides-fragilis] |
| SCH13695.1 | Type3.MTaseII.containing.BREX | [illegible] | 1222 | [illegible] | GCA_900066445.1 | uncultured_Bacteroides_sp. | SAMEA3545309_00360 | 401489..405157 | 162156 | Uncharacterised-protein-[uncultured-Bacteroides-sp.] |
| SFU53504.1 | Type3.MTaseII.containing.BREX | [illegible] | 1229 | [illegible] | GCA_900116745.1 | Porphyromonadaceae_bacterium_KHP3R9 | SAMN05216364_102150 | 48696..52385 | 1855398 | hypothetical-protein-SAMN05216364_102150-[Porphyromonadaceae-bacterium-KHP3R9] |
| WP_012055867.1 | Type3.MTaseII.containing.BREX | [illegible] | 1222 | [illegible] | GCF_000012825.1 | Phocaeicola | I6I55_RS09890 | 2560138..2563806 | 909656 | MULTISPECIES:-DUF6079-family-protein-[Phocaeicola] |
| WP_008149081.1 | Type3.MTaseII.containing.BREX | [illegible] | 423 | [illegible] | GCF_000156495.1 | Parabacteroides_johnsonii | PRABACTJOHN_RS08640 | 13064..14332 | 387661 | DUF6079-family-protein,-partial-[Parabacteroides-johnsonii] |
| WP_039848176.1 | Type3.MTaseII.containing.BREX | [illegible] | 795 | [illegible] | GCF_000156495.1 | Parabacteroides_johnsonii | PRABACTJOHN_RS08645 | 14500..16889 | 387661 | DUF6079-family-protein,-partial-[Parabacteroides-johnsonii] |
| WP_018109372.1 | Type3.MTaseII.containing.BREX | [illegible] | 1220 | [illegible] | GCF_000375405.1 | Bacteroides_propionicifaciens | B091_RS0107385 | 7742..11404 | 392838 | DUF6079-family-protein-[Bacteroides-propionicifaciens] |
| WP_025069425.1 | Type3.MTaseII.containing.BREX | [illegible] | 831 | [illegible] | GCF_000613745.1 | Bacteroides_propionicifaciens | JCM14649_RS04025 | 73042..75537 | 392838 | DUF6079-family-protein-[Bacteroides-propionicifaciens] |
| WP_025069424.1 | Type3.MTaseII.containing.BREX | [illegible] | 400 | [illegible] | GCF_000613745.1 | Bacteroides_propionicifaciens | JCM14649_RS13845 | 71874..73076 | 392838 | DUF6079-family-protein-[Bacteroides-propionicifaciens] |
| WP_087375328.1 | Type3.MTaseII.containing.BREX | [illegible] | 1231 | [illegible] | GCF_002161725.1 | Parabacteroides | B5F96_RS06240 | 134438..138133 | 375288 | MULTISPECIES:-DUF6079-family-protein-[Parabacteroides] |
| WP_094386846.1 | Type3.MTaseII.containing.BREX | [illegible] | 1222 | [illegible] | GCF_002251285.1 | Prevotella_sp._P5-64 | CIK87_RS08690 | 37700..41368 | 2024226 | DUF6079-family-protein-[Prevotella-sp.-P5-64] |
| WP_094391617.1 | Type3.MTaseII.containing.BREX | [illegible] | 1230 | [illegible] | GCF_002251345.1 | Prevotella_sp._P4-119 | CIK89_RS12080 | 3914..7606 | 2024218 | DUF6079-family-protein-[Prevotella-sp.-P4-119] |
| WP_122342973.1 | Type3.MTaseII.containing.BREX | [illegible] | 1232 | [illegible] | GCF_003475265.1 | Parabacteroides_merdae | DWZ81_RS11700 | 138252..141950 | 46503 | DUF6079-family-protein-[Parabacteroides-merdae] |
| WP_135993298.1 | Type3.MTaseII.containing.BREX | [illegible] | 1218 | [illegible] | GCF_004793655.1 | Muribaculum_intestinale | E5333_RS08100 | 35638..39294 | 1796646 | DUF6079-family-protein-[Muribaculum-intestinale] |
| WP_172278700.1 | Type3.MTaseII.containing.BREX | [illegible] | 1220 | [illegible] | GCF_013166635.1 | Lentimicrobium_sp._S6 | HNS40_RS06010 | 45604..49266 | 2735872 | DUF6079-family-protein-[Lentimicrobium-sp.-S6] |
| WP_229034112.1 | Type3.MTaseII.containing.BREX | [illegible] | 1232 | [illegible] | GCF_015547825.1 | Parabacteroides_merdae | PN574_RS14080 | 4436..8134 | 46503 | DUF6079-family-protein-[Parabacteroides-merdae] |
| WP_204447185.1 | Type3.MTaseII.containing.BREX | [illegible] | 1222 | [illegible] | GCF_016902245.1 | Bacteroides_caecigallinarum | H6A66_RS14140 | 17963..21631 | 1411144 | DUF6079-family-protein-[Bacteroides-caecigallinarum] |
| WP_233471284.1 | Type3.MTaseII.containing.BREX | [illegible] | 626 | [illegible] | GCF_019132015.1 | Parabacteroides_johnsonii | KSZ16_RS09840 | 152717..154597 | 387661 | DUF6079-family-protein-[Parabacteroides-johnsonii] |
| WP_220368246.1 | Type3.MTaseII.containing.BREX | [illegible] | 1222 | [illegible] | GCF_019551475.1 | Prevotella_sp._Rep29 | GRF55_RS09890 | 2305278..2308946 | 2691580 | DUF6079-family-protein-[Prevotella-sp.-Rep29] |
| WP_221728838.1 | Type3.MTaseII.containing.BREX | [illegible] | 1231 | [illegible] | GCF_019733935.1 | Parabacteroides_johnsonii | HCG90_RS18570 | 811..4506 | 387661 | DUF6079-family-protein-[Parabacteroides-johnsonii] |
| WP_227963242.1 | Type3.MTaseII.containing.BREX | [illegible] | 1231 | [illegible] | GCF_020735865.1 | Parabacteroides_johnsonii | PGY44_RS11600 | 147925..151620 | 387661 | DUF6079-family-protein-[Parabacteroides-johnsonii] |
| WP_237797820.1 | Type3.MTaseII.containing.BREX | [illegible] | 1222 | [illegible] | GCF_022024195.1 | Prevotella_bryantii | L6467_RS02565 | 638923..642591 | 77095 | DUF6079-family-protein-[Prevotella-bryantii] |
| WP_259033665.1 | Type3.MTaseII.containing.BREX | [illegible] | 1231 | [illegible] | GCF_024792215.1 | Parabacteroides_johnsonii | NXV05_RS13750 | 3320605..3324300 | 387661 | DUF6079-family-protein-[Parabacteroides-johnsonii] |
| WP_270226497.1 | Type3.MTaseII.containing.BREX | [illegible] | 936 | [illegible] | GCF_027662465.1 | Phocaeicola_massiliensis | PGQ49_RS05935 | 189386..192196 | 204516 | DUF6079-family-protein-[Phocaeicola-massiliensis] |
| WP_154042813.1 | Type3.MTaseII.containing.BREX | [illegible] | 1222 | [illegible] | GCF_900066445.1 | Bacteroides_cellulolyticus | OCV82_RS01750 | 401489..405157 | 2981780 | DUF6079-family-protein-[Bacteroides-cellulolyticus] |
| WP_076930955.1 | Type3.MTaseII.containing.BREX | [illegible] | 1222 | [illegible] | GCF_900095135.1 | Proteiniphilum_saccharofermentans | PSM36_RS11115 | 2843030..2846698 | 1642647 | DUF6079-family-protein-[Proteiniphilum-saccharofermentans] |
| WP_074549953.1 | Type3.MTaseII.containing.BREX | [illegible] | 1228 | [illegible] | GCF_900103315.1 | Prevotella_bryantii | BLR39_RS11125 | 82929..86615 | 77095 | DUF6079-family-protein-[Prevotella-bryantii] |
| WP_073205024.1 | Type3.MTaseII.containing.BREX | [illegible] | 1222 | [illegible] | GCF_900142525.1 | Prevotella_ruminicola | BUB86_RS02805 | 117368..121036 | 839 | DUF6079-family-protein-[Prevotella-ruminicola] |
| WP_106830576.1 | Type3.MTaseII.containing.BREX | [illegible] | 1221 | [illegible] | GCF_900292045.1 | Parabacteroides_pacaensis | C9976_RS12075 | 555231..558896 | 2086575 | DUF6079-family-protein-[Parabacteroides-pacaensis] |
| WP_129650005.1 | Type3.MTaseII.containing.BREX | [illegible] | 1231 | [illegible] | GCF_900624885.1 | Parabacteroides_johnsonii | EXC77_RS08430 | 1966930..1970625 | 387661 | DUF6079-family-protein-[Parabacteroides-johnsonii] |
| WP_148480194.1 | Type3.MTaseII.containing.BREX | [illegible] | 1231 | [illegible] | GCF_902363645.1 | Parabacteroides_johnsonii | FY367_RS18035 | 86681..90376 | 387661 | DUF6079-family-protein-[Parabacteroides-johnsonii] |
| WP_195486431.1 | Type3.MTaseII.containing.BREX | [illegible] | 1231 | [illegible] | GCF_937876155.1 | Parabacteroides_johnsonii | PQG89_RS17770 | 5670..9365 | 387661 | DUF6079-family-protein-[Parabacteroides-johnsonii] |

| Protein-ID | System | Gene-Neighbourhoods | Length | Taxonomy | Assembly-ID | Strain | Locus-Tag | Genomic-coordinates | TaxID | Fasta-defline |
|---|---|---|---|---|---|---|---|---|---|---|
| MBP7508731.1 | Type3.MTaseII.containing.BREX | <-BrxA-DNA-binding<-PglZ<-TUDOR+SF1-SF2-Helicase+iREase+CTD<-N6-N4-METHYLASE<-DUF6079*<-BrxB-BrxF-iSTAND | 1221 | Bacteria>FCB_group>Bacteroidetes/Chlorobi_group>Bacteroidota>Bacteroidia>Marinilabiliales | GCA_017998095.1 | Prolixibacteraceae_bacterium | KA807_13005 | 8330..11995 | 2053594 | hypothetical-protein-[Prolixibacteraceae-bacterium] |
| NCB25597.1 | Type3.MTaseII.containing.BREX | <-DUF6079-p*<-BrxB-BrxF-iSTAND<-XIS-HTH<-NACHT+SNaCT\|\|SIGMA-HTH->MazG->?->REase-> | 589 | Bacteria>FCB_group>Bacteroidetes/Chlorobi_group>Bacteroidota>Bacteroidia>unclassified_Bacteroidia | GCA_009929165.1 | Bacteroidia_bacterium | EOM62_09015 | 1..1766 | 2044936 | exonuclease-SbcC-[Bacteroidia-bacterium] |
| MBP6412560.1 | Type3.MTaseII.containing.BREX | BrxB-BrxF-iSTAND->DUF6079*->?->N6-N4-METHYLASE->N6-N4-METHYLASE->?->TUDOR+SF1-SF2-Helicase+iREase+CTD->PglZ->BrxA-DNA-binding-><-McrC-NTD+REase+ZnR<-McrB-AAA | 1221 | Bacteria>FCB_group>Bacteroidetes/Chlorobi_group>Bacteroidota>Bacteroidia>unclassified_Bacteroidia | GCA_017988095.1 | Bacteroidia_bacterium | KA450_03870 | 2312..5977 | 2044936 | hypothetical-protein-[Bacteroidia-bacterium] |
| MBK6380106.1 | Type3.MTaseII.containing.BREX | <-TUDOR+SF1-SF2-Helicase+iREase+CTD-p<-N6-N4-METHYLASE-p<-?<-N6-N4-METHYLASE-p<-DUF6079-p<-DUF6079-p<-DUF6079-p<-?<-DUF6079-p<-DUF6079-p*<-DUF6079-p<-?<-?<-XIS-HTH<-ZetaToxin<-Phage_integrase<-Arm-DNA-bind_5+Phage_int_SAM_5+Phage_integrase | 115 | Bacteria>FCB_group>Bacteroidetes/Chlorobi_group>Bacteroidota>Chitinophagia>Chitinophagales | GCA_016704945.1 | Chitinophagaceae_bacterium | IPF72_10440 | 17726..18073 | 1869212 | hypothetical-protein-[Chitinophagaceae-bacterium] |
| MBK6991305.1 | Type3.MTaseII.containing.BREX | <-HTH+HKD-DNase+PIN<-SF1-SF2-Helicase<-PIWI<-BrxA-DNA-binding<-PglZ<-N6-N4-METHYLASE<-N6-N4-METHYLASE-p<-N6-N4-METHYLASE<-DUF6079*<-HTH<-?<-XIS-HTH<-?<-Arm-DNA-bind_5+Phage_int_SAM_5+Phage_integrase<-XIS-HTH<-TonBC+TonBC | 1407 | Bacteria>FCB_group>Bacteroidetes/Chlorobi_group>Bacteroidota>Chitinophagia>Chitinophagales | GCA_016708765.1 | Chitinophagaceae_bacterium | IPH34_05510 | 135970..140193 | 1869212 | BREX-3-system-P-loop-containing-protein-BrxF-[Chitinophagaceae-bacterium] |
| MBK9956994.1 | Type3.MTaseII.containing.BREX | <-HTH+HKD-DNase+PIN<-SF1-SF2-Helicase<-SF1-SF2-Helicase<-PIWI<-BrxA-DNA-binding<-PglZ<-N6-N4-METHYLASE<-DUF6079*<-HTH<-?<-XIS-HTH<-?<-Arm-DNA-bind_5+Phage_int_SAM_5+Phage_integrase<-XIS-HTH | 1407 | Bacteria>FCB_group>Bacteroidetes/Chlorobi_group>Bacteroidota>Chitinophagia>Chitinophagales | GCA_016722125.1 | Chitinophagaceae_bacterium | IPP11_00085 | 20536..24759 | 1869212 | BREX-3-system-P-loop-containing-protein-BrxF-[Chitinophagaceae-bacterium] |
| MBP6046015.1 | Type3.MTaseII.containing.BREX | TonBC+TonBC->sGTP->XIS-HTH->Arm-DNA-bind_5+Phage_int_SAM_5+Phage_integrase->?->XIS-HTH->?->HTH->BrxB-BrxF-iSTAND->DUF6079*->N6-N4-METHYLASE->HTH_3->Couple_hipA->HipA_C->TUDOR+SF1-SF2-Helicase+iREase+CTD->PglZ->BrxA-DNA-binding-><-?\|\|PIWI-> | 1221 | Bacteria>FCB_group>Bacteroidetes/Chlorobi_group>Bacteroidota>Chitinophagia>Chitinophagales | GCA_017984145.1 | Ferruginibacter_sp. | KA530_02925 | 110131..113796 | 1940288 | hypothetical-protein-[Ferruginibacter-sp.] |
| MBP6625017.1 | Type3.MTaseII.containing.BREX | <-PglZ-p<-N6-N4-METHYLASE<-DUF6079-p* | 676 | Bacteria>FCB_group>Bacteroidetes/Chlorobi_group>Bacteroidota>Chitinophagia>Chitinophagales | GCA_017989935.1 | Chitinophagaceae_bacterium | KA198_07580 | 2929..4959 | 1869212 | hypothetical-protein-[Chitinophagaceae-bacterium] |
| MCA6489951.1 | Type3.MTaseII.containing.BREX | <-METHYLASE<-?<-?\|\|?->?-><-?<-?\|\|Phage_int_SAM_5+Phage_integrase->HTH->DUF6079*->N6-N4-METHYLASE->PglZ->?->BrxA-DNA-binding-> | 1406 | Bacteria>FCB_group>Bacteroidetes/Chlorobi_group>Bacteroidota>Chitinophagia>Chitinophagales | GCA_020403635.1 | Chitinophagaceae_bacterium | IM551_07990 | 29442..33662 | 1869212 | BREX-3-system-P-loop-containing-protein-BrxF-[Chitinophagaceae-bacterium] |
| MCA6513727.1 | Type3.MTaseII.containing.BREX | <-METHYLASE<-?<-?\|\|?->?-><-?<-?\|\|Phage_int_SAM_5+Phage_integrase->BrxB-BrxF-iSTAND->DUF6079*->N6-N4-METHYLASE->?->TUDOR+SF1-SF2-Helicase+iREase+CTD->PglZ->BrxA-DNA-binding-> | 1221 | Bacteria>FCB_group>Bacteroidetes/Chlorobi_group>Bacteroidota>Chitinophagia>Chitinophagales | GCA_020403725.1 | Chitinophagaceae_bacterium | IM569_12790 | 33461..37126 | 1869212 | hypothetical-protein-[Chitinophagaceae-bacterium] |
| MCG9898371.1 | Type3.MTaseII.containing.BREX | <-BrxA-DNA-binding<-PglZ<-TUDOR+SF1-SF2-Helicase+iREase+CTD<-AbiEii<-wHTH+AbiEi_2<-?<-N6-N4-METHYLASE<-HHH_4+SF1-SF2-Helicase+REase<-DUF6079*<-BrxB-BrxF-iSTAND<-HTH_3<-?<-?<-CR-ATPase<-Classical-AAA<-HTH_36<-XIS-HTH<-HTH_19 | 1219 | Bacteria>FCB_group>Bacteroidetes/Chlorobi_group>Bacteroidota>Chitinophagia>Chitinophagales | GCA_022402565.1 | Hydrotalea_sp. | MH132_00055 | 16449..20108 | 2881279 | DUF6079-family-protein-[Hydrotalea-sp.] |
| WP_026751103.1 | Type3.MTaseII.containing.BREX | TonBC+TonBC->sGTP->XIS-HTH->Arm-DNA-bind_5+Phage_int_SAM_5+Phage_integrase->?->XIS-HTH->?->HTH->DUF6079*->N6-N4-METHYLASE->PglZ->BrxA-DNA-binding->PIWI->?->?->McrB-AAA->McrC-NTD+REase-> | 1407 | Bacteria>FCB_group>Bacteroidetes/Chlorobi_group>Bacteroidota>Chitinophagia>Chitinophagales | GCF_000508085.1 | Sediminibacterium_sp._C3 | F777_RS0105110 | 1077798..1082021 | 1267211 | BREX-3-system-P-loop-containing-protein-BrxF-[Sediminibacterium-sp.-C3] |
| KPQ15860.1 | Type3.MTaseII.containing.BREX | CR-ATPase5+REase->ABhydrolase->HTH_3->DUF6079*->N6-N4-METHYLASE->PglZ->BrxA-DNA-binding->?->?->Peptidase_M14->Glyoxalase-> | 1408 | Bacteria>FCB_group>Bacteroidetes/Chlorobi_group>Bacteroidota>Cytophagia>Cytophagales | GCA_001314815.1 | Algoriphagus_marincola_HL-49 | HLUCCX10_08570 | 4601..8827 | 1305737 | hypothetical-protein-HLUCCX10_08570-[Algoriphagus-marincola-HL-49] |
| WP_150904082.1 | Type3.MTaseII.containing.BREX | <-IG+IG\|\|?-><-SH3\|\|tRNA-><-?\|\|HTH_3-><-?\|\|?->BrxB-BrxF-iSTAND->DUF6079*->?->N6-N4-METHYLASE->TUDOR+SF1-SF2-Helicase+iREase+CTD->PglZ->BrxA-DNA-binding-><-HNH<-NUDIX+NUDIX<-HNH | 1221 | Bacteria>FCB_group>Bacteroidetes/Chlorobi_group>Bacteroidota>Cytophagia>Cytophagales | GCF_008727925.1 | Adhaeribacter_soli | F0P94_RS11720 | 112130..115795 | 2607655 | DUF6079-family-protein-[Adhaeribacter-soli] |
| WP_187176865.1 | Type3.MTaseII.containing.BREX | <-ROK-HTH<-?<-ANK+ANK\|\|TPR+TPR->MazG->McrB-AAA->McrC-NTD+REase->BrxB-BrxF-iSTAND->DUF6079*->HigB->HTH_3->?->N6-N4-METHYLASE->?->HEPN->HEPN->TUDOR+SF1-SF2-Helicase+iREase+CTD->PglZ-> | 1219 | Bacteria>FCB_group>Bacteroidetes/Chlorobi_group>Bacteroidota>Cytophagia>Cytophagales | GCF_014324365.1 | Algoriphagus_sp._AK58 | DDT91_RS14250 | 249398..253057 | 1406877 | DUF6079-family-protein-[Algoriphagus-sp.-AK58] |
| WP_254082868.1 | Type3.MTaseII.containing.BREX | <-BrxA-DNA-binding<-PglZ<-TUDOR+SF1-SF2-Helicase+iREase+CTD<-N6-N4-METHYLASE<-N6-N4-METHYLASE<-DUF6079*<-BrxB-BrxF-iSTAND<-HTH\|\|CNMP+CRP-HTH-><-?<-?\|\|N6-METHYLASE-p-> | 1221 | Bacteria>FCB_group>Bacteroidetes/Chlorobi_group>Bacteroidota>Cytophagia>Cytophagales | GCF_018598175.1 | Dawidia_cretensis | KK062_RS03625 | 834196..837861 | 2782350 | DUF6079-family-protein-[Dawidia-cretensis] |
| WP_270130055.1 | Type3.MTaseII.containing.BREX | <-ABhydrolase<-?\|\|?->?->PIN-><-BrxA-DNA-binding<-PglZ<-N6-N4-METHYLASE<-DUF6079*<-HTH<-ArdCN+MPTase<-?<-REase | 1408 | Bacteria>FCB_group>Bacteroidetes/Chlorobi_group>Bacteroidota>Cytophagia>Cytophagales | GCF_027594545.1 | Algoriphagus_sp._TR-M9 | PBT90_RS16355 | 3870472..3874698 | 3015169 | DUF6079-family-protein-[Algoriphagus-sp.-TR-M9] |
| WP_076420670.1 | Type3.MTaseII.containing.BREX | MORC+SF1-SF2-Helicase-><-BrxA-DNA-binding<-PglZ<-TUDOR+SF1-SF2-Helicase+iREase+CTD<-N6-N4-METHYLASE<-HHH_4+SF1-SF2-Helicase+REase<-DUF6079*<-BrxB-BrxF-iSTAND<-TPR+TPR+TPR+TPR<-?<-SIGMA-HTH | 1219 | Bacteria>FCB_group>Bacteroidetes/Chlorobi_group>Bacteroidota>Cytophagia>Cytophagales | GCF_900156415.1 | Pontibacter_lucknowensis | BW972_RS01755 | 406183..409842 | 1077936 | DUF6079-family-protein-[Pontibacter-lucknowensis] |
| WP_089242506.1 | Type3.MTaseII.containing.BREX | <-BrxA-DNA-binding<-PglZ<-TUDOR+SF1-SF2-Helicase+iREase+CTD<-ZnR+HEPN<-N6-N4-METHYLASE<-DUF6079*<-BrxB-BrxF-iSTAND<-HTH_3 | 1219 | Bacteria>FCB_group>Bacteroidetes/Chlorobi_group>Bacteroidota>Cytophagia>Cytophagales | GCF_900188245.1 | Belliella_buryatensis | CHB66_RS16440 | 9048..12707 | 1500549 | DUF6079-family-protein-[Belliella-buryatensis] |
| MBT5530412.1 | Type3.MTaseII.containing.BREX | <-wHTH+WYL+WCX<-?<-?<-BrxA-DNA-binding<-PglZ<-TUDOR+SF1-SF2-Helicase+iREase+CTD<-N6-N4-METHYLASE<-DUF6079*<-BrxB-BrxF-iSTAND<-XIS-HTH\|\|GDResolvase+HIN-HTH->ARG-> | 1235 | Bacteria>FCB_group>Bacteroidetes/Chlorobi_group>Bacteroidota>Cytophagia>unclassified_Cytophagia | GCA_018665725.1 | Cytophagia_bacterium | HOK35_14730 | 10077..13784 | 2026730 | ATP-binding-protein-[Cytophagia-bacterium] |
| KUO63976.1 | Type3.MTaseII.containing.BREX | TPR->BrxB-BrxF-iSTAND->DUF6079*->N6-N4-METHYLASE->N6-N4-METHYLASE->TUDOR+SF1-SF2-Helicase+iREase+CTD->PglZ->BrxA-DNA-binding-><-?<-PTSEIIA1 | 1219 | Bacteria>FCB_group>Bacteroidetes/Chlorobi_group>Bacteroidota>Flavobacteriia>Flavobacteriales | GCA_001516145.1 | Lutibacter_sp._BRH_c52 | APF83_08755 | 2841..6500 | 1734397 | MAG:-hypothetical-protein-APF83_08755-[Lutibacter-sp.-BRH_c52] |
| HAQ70461.1 | Type3.MTaseII.containing.BREX | wHTH+WYL+WCX-><-SF1-SF2-Helicase+PIN<-MORC+DUF3684+REase<-BrxA-DNA-binding<-PglZ<-N6-N4-METHYLASE<-DUF6079* | 1015 | Bacteria>FCB_group>Bacteroidetes/Chlorobi_group>Bacteroidota>Flavobacteriia>Flavobacteriales | GCA_003477125.1 | Flavobacteriales_bacterium | DCR48_05750 | 15379..18426 | 2021391 | hypothetical-protein-[Flavobacteriales-bacterium] |
| MCH8535025.1 | Type3.MTaseII.containing.BREX | TPR->BrxB-BrxF-iSTAND->DUF6079*->N6-N4-METHYLASE->SbcC->TUDOR+SF1-SF2-Helicase+iREase+CTD-> | 1219 | Bacteria>FCB_group>Bacteroidetes/Chlorobi_group>Bacteroidota>Flavobacteriia>Flavobacteriales | GCA_022565635.1 | Flavobacteriaceae_bacterium | LAT51_08160 | 2647..6306 | 1871037 | DUF6079-family-protein-[Flavobacteriaceae-bacterium] |
| WP_060524793.1 | Type3.MTaseII.containing.BREX | REase->?->XIS-HTH->HTH_36->Classical-AAA->HD->?->HTH_3->BrxB-BrxF-iSTAND->DUF6079*->HHH_4+SF1-SF2-Helicase+REase->N6-N4-METHYLASE->N6-N4-METHYLASE->?->TUDOR+SF1-SF2-Helicase+iREase+CTD->PglZ->BrxA-DNA-binding->NERD+SF1-SF2-Helicase-><-HTH | 1219 | Bacteria>FCB_group>Bacteroidetes/Chlorobi_group>Bacteroidota>Flavobacteriia>Flavobacteriales | GCF_001430865.1 | Nonlabens_sp._MIC269 | AAT17_RS12580 | 2784203..2787862 | 1476901 | DUF6079-family-protein-[Nonlabens-sp.-MIC269] |
| WP_133549725.1 | Type3.MTaseII.containing.BREX | Arm-DNA-bind_5+Phage_int_SAM_5+Phage_integrase-><-XIS-HTH<-BrxA-DNA-binding<-PglZ<-TUDOR+SF1-SF2-Helicase+iREase+CTD<-AbiEii<-wHTH+AbiEi_2<-N6-N4-METHYLASE<-DUF6079*<-BrxB-BrxF-iSTAND<-?<-?<-?<-?<-HTH_26\|\|SIGMA-HTH->OrfB_IS605+OrfB_Zn_ribbon->DOC+HTH-> | 1218 | Bacteria>FCB_group>Bacteroidetes/Chlorobi_group>Bacteroidota>Flavobacteriia>Flavobacteriales | GCF_004362395.1 | Salegentibacter_sp._24 | DET49_RS16540 | 17021..20677 | 2183986 | DUF6079-family-protein-[Salegentibacter-sp.-24] |
| WP_133532119.1 | Type3.MTaseII.containing.BREX | <-PVC-AAA<-ParB+HNH<-ParB+HNH<-35exo<-wHTH+WYL+WCX<-TRPR-HTH\|\|BrxB-BrxF-iSTAND->DUF6079*->HHH_4+SF1-SF2-Helicase+REase->N6-N4-METHYLASE->REase->N6-N4-METHYLASE->TUDOR+SF1-SF2-Helicase+iREase+CTD->PglZ->BrxA-DNA-binding->CR-Kinase-> | 1219 | Bacteria>FCB_group>Bacteroidetes/Chlorobi_group>Bacteroidota>Flavobacteriia>Flavobacteriales | GCF_004362665.1 | Flavobacterium_dankookense | BC748_RS03755 | 552688..556347 | 706186 | DUF6079-family-protein-[Flavobacterium-dankookense] |
| WP_181886021.1 | Type3.MTaseII.containing.BREX | D5ATPase+DUF3874->BrxB-BrxF-iSTAND->DUF6079*->HigB->HTH_3->?->N6-N4-METHYLASE->HTH_3->Couple_hipA+HipA_C->TUDOR+SF1-SF2-Helicase+iREase+CTD->PglZ->BrxA-DNA-binding-> | 1219 | Bacteria>FCB_group>Bacteroidetes/Chlorobi_group>Bacteroidota>Flavobacteriia>Flavobacteriales | GCF_014041915.1 | Chryseobacterium_sp._CX-624 | H1R16_RS01395 | 294504..298163 | 2758037 | DUF6079-family-protein-[Chryseobacterium-sp.-CX-624] |
| WP_188650283.1 | Type3.MTaseII.containing.BREX | Phage_int_SAM_5+Phage_integrase->?->?->XIS-HTH->XIS-HTH->?->DUF6079*->N6-N4-METHYLASE->PglZ->BrxA-DNA-binding->?->?->REase->XIS-HTH->METHYLASE-p-> | 1411 | Bacteria>FCB_group>Bacteroidetes/Chlorobi_group>Bacteroidota>Flavobacteriia>Flavobacteriales | GCF_014646655.1 | Yeosuana_aromativorans | IEW94_RS04035 | 151268..155503 | 288019 | DUF6079-family-protein-[Yeosuana-aromativorans] |
| WP_230475679.1 | Type3.MTaseII.containing.BREX | ArsR-HTH->?->?->?->?->?->XIS-HTH->D5ATPase+DUF3874->BrxB-BrxF-iSTAND-p->DUF6079*->HigB->HTH_3->?->N6-N4-METHYLASE->TUDOR+SF1-SF2-Helicase+iREase+CTD->PglZ->BrxA-DNA-binding-><-SLATT_1 | 1219 | Bacteria>FCB_group>Bacteroidetes/Chlorobi_group>Bacteroidota>Flavobacteriia>Flavobacteriales | GCF_017948305.1 | Faecalibacter_bovis | J9309_RS09625 | 1999568..2003227 | 2898187 | DUF6079-family-protein-[Faecalibacter-bovis] |
| WP_223443585.1 | Type3.MTaseII.containing.BREX | N6-METHYLASE->Peptidase_S66+Peptidase_S66C-><-?\|\|REase->\|\|Arm-DNA-bind_5+Phage_int_SAM_5+Phage_integrase->?->XIS-HTH->?->TPR+TPR->DUF6079*->N6-N4-METHYLASE->PglZ->BrxA-DNA-binding->?->MqsA_antitoxin+DUF4065->?->DOC+HTH-> | 1411 | Bacteria>FCB_group>Bacteroidetes/Chlorobi_group>Bacteroidota>Flavobacteriia>Flavobacteriales | GCF_019968605.1 | Polaribacter_litorisediminis | K8354_RS16920 | 3941599..3945834 | 1908341 | DUF6079-family-protein-[Polaribacter-litorisediminis] |
| MBK7606677.1 | Type3.MTaseII.containing.BREX | <-SF1-SF2-Helicase<-STYKIN<-?<-?<-PIWI<-BrxA-DNA-binding<-PglZ<-PglZ<-TUDOR+SF1-SF2-Helicase+iREase+CTD<-DUF6079*<-DUF6079-p<-BrxB-BrxF-iSTAND<-HTH<-?<-XIS-HTH<-?<-Arm-DNA-bind_5+Phage_int_SAM_5+Phage_integrase<-XIS-HTH<-sGTP | 1068 | Bacteria>FCB_group>Bacteroidetes/Chlorobi_group>Bacteroidota>Saprospiria>Saprospirales | GCA_016707125.1 | Saprospiraceae_bacterium | IPI18_05450 | 33724..36930 | 2202734 | hypothetical-protein-[Saprospiraceae-bacterium] |
| MBK7606678.1 | Type3.MTaseII.containing.BREX | <-STYKIN<-?<-?<-PIWI<-BrxA-DNA-binding<-PglZ<-PglZ<-TUDOR+SF1-SF2-Helicase+iREase+CTD<-DUF6079<-DUF6079-p*<-BrxB-BrxF-iSTAND<-HTH<-?<-XIS-HTH<-?<-Arm-DNA-bind_5+Phage_int_SAM_5+Phage_integrase<-XIS-HTH<-sGTP | 138 | Bacteria>FCB_group>Bacteroidetes/Chlorobi_group>Bacteroidota>Saprospiria>Saprospirales | GCA_016707125.1 | Saprospiraceae_bacterium | IPI18_05455 | 36971..37387 | 2202734 | hypothetical-protein-[Saprospiraceae-bacterium] |
| MBK7095332.1 | Type3.MTaseII.containing.BREX | <-Couple_hipA+HipA_C<-HTH_3<-N6-N4-METHYLASE<-?<-BrxD<-HHH_4+HTH+wHTH+SF1-SF2-Helicase+REase-p<-HHH_4+SF1-SF2-Helicase+REase<-DUF6079-p<-?<-DUF6079-p*<-BrxB-BrxF-iSTAND-p<-?<-TIR<-?<-?<-?<-?<-Phage_integrase<-Phage_int_SAM_5 | 791 | Bacteria>FCB_group>Bacteroidetes/Chlorobi_group>Bacteroidota>Saprospiria>Saprospirales | GCA_016708265.1 | Saprospiraceae_bacterium | IPH57_09915 | 176914..179289 | 2202734 | hypothetical-protein-[Saprospiraceae-bacterium] |
| MBP6236749.1 | Type3.MTaseII.containing.BREX | <-HEPN<-AbiGi+seqA+RAMA<-BrxA-DNA-binding<-PglZ<-TUDOR+SF1-SF2-Helicase+iREase+CTD<-HEPN<-N6-N4-METHYLASE<-DUF6079*<-BrxB-BrxF-iSTAND\|\|SF1-SF2-Helicase-><-NERD+ZnR | 1138 | Bacteria>FCB_group>Bacteroidetes/Chlorobi_group>Bacteroidota>Saprospiria>Saprospirales | GCA_017986015.1 | Saprospiraceae_bacterium | KA536_11425 | 32858..36274 | 2202734 | hypothetical-protein-[Saprospiraceae-bacterium] |
| NLJ08470.1 | Type3.MTaseII.containing.BREX | <-BrxA-DNA-binding<-PglZ<-TUDOR+SF1-SF2-Helicase+iREase+CTD<-Four-helical-protein<-N6-N4-METHYLASE<-N6-N4-METHYLASE<-?<-Schlafen+HATPase+wHTH<-DUF6079*<-BrxB-BrxF-iSTAND | 1223 | Bacteria>FCB_group>Bacteroidetes/Chlorobi_group>Bacteroidota>Sphingobacteriia>Sphingobacteriales | GCA_012517435.1 | Sphingobacteriales_bacterium | GX437_12475 | 14156..17827 | 2044944 | hypothetical-protein-[Sphingobacteriales-bacterium] |
| WP_118194776.1 | Type3.MTaseII.containing.BREX | sGTP->DOC+HTH-><-?\|\|REase->DOC+HTH->BrxB-BrxF-iSTAND->DUF6079*->HigB->HTH_3->?->N6-N4-METHYLASE->TUDOR+SF1-SF2-Helicase+iREase+CTD->PglZ->BrxA-DNA-binding-> | 1219 | Bacteria>FCB_group>Bacteroidetes/Chlorobi_group>Bacteroidota>Sphingobacteriia>Sphingobacteriales | GCF_003449035.1 | Pedobacter_indicus | D3P12_RS08990 | 1991719..1995378 | 2292082 | DUF6079-family-protein-[Pedobacter-indicus] |
| WP_207425049.1 | Type3.MTaseII.containing.BREX | <-35exo+T7DNApol<-?\|\|?-><-?<-?<-SIR2<-?<-?\|\|DUF6079*->N6-N4-METHYLASE->ParB+HNH->?->TUDOR+SF1-SF2-Helicase+iREase+CTD->PglZ->BrxA-DNA-binding->?->SIR2-><-HTH_3 | 1221 | Bacteria>FCB_group>Bacteroidetes/Chlorobi_group>Bacteroidota>Sphingobacteriia>Sphingobacteriales | GCF_017355855.1 | Pedobacter_sp._SYSU_D00535 | J4N23_RS00515 | 144649..148314 | 2810308 | DUF6079-family-protein-[Pedobacter-sp.-SYSU-D00535] |
| WP_091209299.1 | Type3.MTaseII.containing.BREX | AraC-HTH->?->?->CASPASE-><-HATPase_c<-Calcineurin<-MORC+REase\|\|?->BrxB-BrxF-iSTAND->DUF6079*->N6-N4-METHYLASE->TUDOR+SF1-SF2-Helicase+iREase+CTD->PglZ->BrxA-DNA-binding->Arm-DNA-bind_5+Phage_int_SAM_5+Phage_integrase-> | 1221 | Bacteria>FCB_group>Bacteroidetes/Chlorobi_group>Bacteroidota>Sphingobacteriia>Sphingobacteriales | GCF_900110105.1 | Mucilaginibacter_gossypiicola | SAMN05192574_102262 | 254385..258050 | 551995 | DUF6079-family-protein-[Mucilaginibacter-gossypiicola] |
| OQB31949.1 | Type3.MTaseII.containing.BREX | Arm-DNA-bind_5+Phage_int_SAM_5+Phage_integrase->?->XIS-HTH->?->HTH->DUF6079*->N6-N4-METHYLASE->PglZ->BrxA-DNA-binding->Peptidase_S8-><-Classical-AAA<-XIS-HTH\|\|METHYLASE-p-> | 1407 | Bacteria>FCB_group>Bacteroidetes/Chlorobi_group>Bacteroidota>unclassified_Bacteroidota | GCA_002070265.1 | Bacteroidetes_bacterium_ADurb.Bin174 | BWY08_00140 | 8925..13148 | 1852808 | hypothetical-protein-BWY08_00140-[Bacteroidetes-bacterium-ADurb.Bin174] |
| TNF50601.1 | Type3.MTaseII.containing.BREX | <-BrxA-DNA-binding<-PglZ<-TUDOR+SF1-SF2-Helicase+iREase+CTD<-N6-N4-METHYLASE<-DUF6079*<-BrxB-BrxF-iSTAND<-HTH_3<-Classical-AAA | 1219 | Bacteria>FCB_group>Bacteroidetes/Chlorobi_group>Bacteroidota>unclassified_Bacteroidota | GCA_006227105.1 | Bacteroidota_bacterium | EP305_00070 | 7893..11552 | 1898104 | hypothetical-protein-EP305_00070-[Bacteroidota-bacterium] |
| HDO65631.1 | Type3.MTaseII.containing.BREX | DUF6079*->Schlafen+HATPase+wHTH->DNA_METHYLASE->SbcC->N6-N4-METHYLASE->OLD-ABC+TOPRIM-> | 1130 | Bacteria>FCB_group>Bacteroidetes/Chlorobi_group>Bacteroidota>unclassified_Bacteroidota | GCA_011043795.1 | Bacteroidota_bacterium | ENH10_06315 | 1..3394 | 1898104 | ATP-binding-protein-[Bacteroidota-bacterium] |
| MBK7887715.1 | Type3.MTaseII.containing.BREX | <-Recombinase+ZnR<-GDResolvase\|\|?-><-?<-ZnR+CC+REase<-?\|\|HTH->BrxB-BrxF-iSTAND->DUF6079-p*->DUF6079-p->N6-N4-METHYLASE->N6-N4-METHYLASE->ParB+HNH->TUDOR+SF1-SF2-Helicase+iREase+CTD->PglZ-p->?->PglZ-p->BrxA-DNA-binding-> | 323 | Bacteria>FCB_group>Bacteroidetes/Chlorobi_group>Bacteroidota>unclassified_Bacteroidota | GCA_016713925.1 | Bacteroidota_bacterium | IPJ86_10610 | 600602..601573 | 1898104 | hypothetical-protein-[Bacteroidota-bacterium] |
| MBK7887716.1 | Type3.MTaseII.containing.BREX | <-Recombinase+ZnR<-GDResolvase\|\|?-><-?<-ZnR+CC+REase<-?\|\|HTH->BrxB-BrxF-iSTAND->DUF6079-p->DUF6079-p*->N6-N4-METHYLASE->N6-N4-METHYLASE->ParB+HNH->TUDOR+SF1-SF2-Helicase+iREase+CTD->PglZ-p->?->PglZ-p->BrxA-DNA-binding-> | 891 | Bacteria>FCB_group>Bacteroidetes/Chlorobi_group>Bacteroidota>unclassified_Bacteroidota | GCA_016713925.1 | Bacteroidota_bacterium | IPJ86_10615 | 601549..604224 | 1898104 | hypothetical-protein-[Bacteroidota-bacterium] |
| MBK9455370.1 | Type3.MTaseII.containing.BREX | XIS-HTH->?->?->HTH->BrxB-BrxF-iSTAND->DUF6079-p->DUF6079*->N6-N4-METHYLASE->N6-N4-METHYLASE-p->N6-N4-METHYLASE->PglZ->BrxA-DNA-binding->DrHyd->Schlafen+HATPase+wHTH->?->NUDIX+NUDIX-> | 1029 | Bacteria>FCB_group>Bacteroidetes/Chlorobi_group>Bacteroidota>unclassified_Bacteroidota | GCA_016718365.1 | Bacteroidota_bacterium | IPO24_07165 | 107735..110824 | 1898104 | hypothetical-protein-[Bacteroidota-bacterium] |
| MBK9503947.1 | Type3.MTaseII.containing.BREX | HNH-><-NUDIX+NUDIX<-?\|\|?-><-Schlafen+HATPase+wHTH<-DrHyd<-BrxA-DNA-binding<-PglZ<-N6-N4-METHYLASE<-DUF6079*<-HTH<-?<-?<-XIS-HTH<-?<-Arm-DNA-bind_5+Phage_int_SAM_5+Phage_integrase\|\|?-><-DnaJ | 1406 | Bacteria>FCB_group>Bacteroidetes/Chlorobi_group>Bacteroidota>unclassified_Bacteroidota | GCA_016718695.1 | Bacteroidota_bacterium | IPO03_00815 | 171188..175408 | 1898104 | BREX-3-system-P-loop-containing-protein-BrxF-[Bacteroidota-bacterium] |
| MBT3747344.1 | Type3.MTaseII.containing.BREX | <-ARG<-GDResolvase+HIN-HTH\|\|XIS-HTH->BrxB-BrxF-iSTAND->DUF6079*->N6-N4-METHYLASE->TUDOR+SF1-SF2-Helicase+iREase+CTD-p-> | 1262 | Bacteria>FCB_group>Bacteroidetes/Chlorobi_group>Bacteroidota>unclassified_Bacteroidota | GCA_018653945.1 | Bacteroidota_bacterium | HOG34_00060 | 2374..6162 | 1898104 | ATP-binding-protein-[Bacteroidota-bacterium] |
| MBU6204551.1 | Type3.MTaseII.containing.BREX | <-BrxA-DNA-binding<-PglZ<-TUDOR+SF1-SF2-Helicase+iREase+CTD<-N6-N4-METHYLASE<-DUF6079*<-BrxB-BrxF-iSTAND<-HTH_3 | 1230 | Bacteria>FCB_group>Bacteroidetes/Chlorobi_group>Bacteroidota>unclassified_Bacteroidota | GCA_018970345.1 | Bacteroidota_bacterium | KJS45_04130 | 7690..11382 | 1898104 | hypothetical-protein-[Bacteroidota-bacterium] |
| MCB0536139.1 | Type3.MTaseII.containing.BREX | XIS-HTH->XIS-HTH->?->?->?->HTH_3->DOC+HTH->HTH->DUF6079*->N6-N4-METHYLASE->PglZ->BrxA-DNA-binding->PIWI->MORC+DUF3684+REase->SF1-SF2-Helicase+PIN-> | 1412 | Bacteria>FCB_group>Bacteroidetes/Chlorobi_group>Bacteroidota>unclassified_Bacteroidota | GCA_020436205.1 | Bacteroidota_bacterium | KDE33_01305 | 6874..11112 | 1898104 | hypothetical-protein-[Bacteroidota-bacterium] |
| MCB0536152.1 | Type3.MTaseII.containing.BREX | <-BrxA-DNA-binding<-PglZ<-TUDOR+SF1-SF2-Helicase+iREase+CTD<-?<-N6-N4-METHYLASE<-HHH_4+SF1-SF2-Helicase+REase<-DUF6079*<-BrxB-BrxF-iSTAND<-HTH_3<-SLOG<-?<-HAD | 1222 | Bacteria>FCB_group>Bacteroidetes/Chlorobi_group>Bacteroidota>unclassified_Bacteroidota | GCA_020436205.1 | Bacteroidota_bacterium | KDE33_01370 | 11578..15246 | 1898104 | hypothetical-protein-[Bacteroidota-bacterium] |
| MCB0536182.1 | Type3.MTaseII.containing.BREX | HTH->DUF6079*->N6-N4-METHYLASE->PglZ->BrxA-DNA-binding->DrHyd->Schlafen+HATPase+wHTH->PIWI->SF1-SF2-Helicase->HTH+HKD-DNase+PIN-> | 1407 | Bacteria>FCB_group>Bacteroidetes/Chlorobi_group>Bacteroidota>unclassified_Bacteroidota | GCA_020436205.1 | Bacteroidota_bacterium | KDE33_01520 | 2623..6846 | 1898104 | BREX-3-system-P-loop-containing-protein-BrxF-[Bacteroidota-bacterium] |
| MCC5934986.1 | Type3.MTaseII.containing.BREX | <-BrxA-DNA-binding<-PglZ<-TUDOR+SF1-SF2-Helicase+iREase+CTD<-TOPRIM<-OLD-ABC<-REase+AIPR<-N6-N4-METHYLASE<-DUF6079*<-BrxB-BrxF-iSTAND | 1245 | Bacteria>FCB_group>Bacteroidetes/Chlorobi_group>Balneolota>Balneolia>Balneolales | GCA_020832405.1 | Balneolales_bacterium | JJU35_12140 | 14385..18122 | 2497613 | hypothetical-protein-[Balneolales-bacterium] |
| MCC5935599.1 | Type3.MTaseII.containing.BREX | <-BrxA-DNA-binding<-PglZ<-TUDOR+SF1-SF2-Helicase+iREase+CTD<-N6-N4-METHYLASE<-?<-HTH_3<-HigB<-DUF6079*\|\|?-><-HTH_3 | 1219 | Bacteria>FCB_group>Bacteroidetes/Chlorobi_group>Balneolota>Balneolia>Balneolales | GCA_020832405.1 | Balneolales_bacterium | JJU35_15230 | 11042..14701 | 2497613 | hypothetical-protein-[Balneolales-bacterium] |
| KZK74371.1 | Type3.MTaseII.containing.BREX | <-PARG+MACRODOMAIN<-BrxA-DNA-binding<-PglZ<-TUDOR+SF1-SF2-Helicase+iREase+CTD<-SF1-SF2-Helicase<-OLD-ABC+TOPRIM<-N6-N4-METHYLASE<-Schlafen+HATPase+wHTH<-DUF6079-p* | 824 | Bacteria>FCB_group>Bacteroidetes/Chlorobi_group>Chlorobiota>Chlorobiia>Chlorobiales | GCA_001622165.1 | Pelodictyon_luteolum | A3K90_02905 | 32401..34875 | 1100 | ATPase,-partial-[Pelodictyon-luteolum] |
| TNJ34106.1 | Type3.MTaseII.containing.BREX | Calcineurin+Calcineurin-SbcD->SbcC->REase+RNase->?->?->REase->?->XIS-HTH->BrxB-BrxF-iSTAND->DUF6079*->?->N6-N4-METHYLASE->PHP+SbcC->Schlafen+HATPase+wHTH->HTH_Tnp_1-> | 1255 | Bacteria>FCB_group>Bacteroidetes/Chlorobi_group>Chlorobiota>Chlorobiia>Chlorobiales | GCA_006265245.1 | Prosthecochloris_vibrioformis | FGF68_10360 | 10460..14227 | 1098 | ATP-binding-protein-[Prosthecochloris-vibrioformis] |
| NMW20448.1 | Type3.MTaseII.containing.BREX | <-N6-N4-METHYLASE<-HTH+REase<-DUF6079*<-BrxB-BrxF-iSTAND\|\|?-><-?<-?<-?<-ABhydrolase | 1245 | Bacteria>FCB_group>Bacteroidetes/Chlorobi_group>Chlorobiota>Chlorobiia>Chlorobiales | GCA_012972835.1 | Chlorobiaceae_bacterium | HKK67_02190 | 4149..7886 | 2067991 | ATP-binding-protein-[Chlorobiaceae-bacterium] |
| MBN1279366.1 | Type3.MTaseII.containing.BREX | <-DUF6079-p*<-BrxB-BrxF-iSTAND<-HTH+REase<-?<-?\|\|?-><-?\|\|?->?->REC+OmpR-HTH-> | 212 | Bacteria>FCB_group>Bacteroidetes/Chlorobi_group>Chlorobiota>Chlorobiia>Chlorobiales | GCA_016927285.1 | Chlorobiaceae_bacterium | JW989_06260 | 1..638 | 2067991 | hypothetical-protein-[Chlorobiaceae-bacterium] |
| MBV5329906.1 | Type3.MTaseII.containing.BREX | wHTH+WYL+WCX->BrxB-BrxF-iSTAND->DUF6079*->N6-N4-METHYLASE->Schlafen+HATPase+wHTH->TUDOR+SF1-SF2-Helicase+iREase+CTD->PglZ-p-> | 1245 | Bacteria>FCB_group>Bacteroidetes/Chlorobi_group>Chlorobiota>Chlorobiia>Chlorobiales | GCA_019162945.1 | Chlorobium_sp. | JZU65_20140 | 1991..5728 | 1095 | hypothetical-protein-[Chlorobium-sp.] |
| WP_076789317.1 | Type3.MTaseII.containing.BREX | <-ArdCN+MPTase<-?<-BrxA-DNA-binding<-PglZ<-TUDOR+SF1-SF2-Helicase+iREase+CTD<-REase+AIPR<-N6-N4-METHYLASE<-Schlafen+HATPase+wHTH<-DUF6079*<-BrxB-BrxF-iSTAND<-XIS-HTH<-?<-VirB10 | 1245 | Bacteria>FCB_group>Bacteroidetes/Chlorobi_group>Chlorobiota>Chlorobiia>Chlorobiales | GCF_001975465.1 | Chlorobium_sp._KB01 | BOO20_RS00745 | 13983..17720 | 1917528 | DUF6079-family-protein-[Chlorobium-sp.-KB01] |
| WP_211340434.1 | Type3.MTaseII.containing.BREX | SbcC->REase+RNase->?->?->REase->?->?->XIS-HTH->BrxB-BrxF-iSTAND->DUF6079*->?->N6-N4-METHYLASE->PHP+SbcC->Schlafen+HATPase+wHTH->TUDOR+SF1-SF2-Helicase+iREase+CTD-p-> | 1277 | Bacteria>FCB_group>Bacteroidetes/Chlorobi_group>Chlorobiota>Chlorobiia>Chlorobiales | GCF_006265245.1 | Prosthecochloris_vibrioformis | FGF68_RS10355 | 10394..14227 | 1098 | DUF6079-family-protein-[Prosthecochloris-vibrioformis] |
| MBK7265049.1 | Type3.MTaseII.containing.BREX | <-ABC-ATPase+ABC-ATPase<-?<-?<-?<-?<-ABC-ATPase+SbcC<-?<-PBPB\|\|BrxB-BrxF-iSTAND->DUF6079*->N6-N4-METHYLASE->N6-N4-METHYLASE->N6-N4-METHYLASE-p->OLD-ABC->TOPRIM->TUDOR+SF1-SF2-Helicase+iREase+CTD-p->TUDOR+SF1-SF2-Helicase+iREase+CTD->?->PglZ-p-> | 1222 | Bacteria>FCB_group>Bacteroidetes/Chlorobi_group>Ignavibacteriota>Ignavibacteria>Ignavibacteriales | GCA_016707285.1 | Ignavibacteriales_bacterium | IPI12_01680 | 352924..356592 | 2049428 | hypothetical-protein-[Ignavibacteriales-bacterium] |
| MBK7577375.1 | Type3.MTaseII.containing.BREX | <-HTH_Tnp_1+HTH_21+rve+rve_3<-HTH_21<-HTH_21<-HIN-HTH\|\|SIR2->SF1-SF2-Helicase->BrxB-BrxF-iSTAND->DUF6079*->N6-N4-METHYLASE->HHH_4+HTH+wHTH+SF1-SF2-Helicase+REase->?->N6-N4-METHYLASE->PglZ->BrxA-DNA-binding-><-XIS-HTH<-Phage_int_SAM_5+Phage_integrase<-tRNA\|\|TPR+TPR-> | 1236 | Bacteria>FCB_group>Bacteroidetes/Chlorobi_group>Ignavibacteriota>Ignavibacteria>unclassified_Ignavibacteria | GCA_016706965.1 | Ignavibacteria_bacterium | IPI24_08070 | 197510..201220 | 2053306 | ATP-binding-protein-[Ignavibacteria-bacterium] |
| MBK9183263.1 | Type3.MTaseII.containing.BREX | <-rve<-HTH_21<-HIN-HTH\|\|SIR2->?->SF1-SF2-Helicase->BrxB-BrxF-iSTAND->DUF6079*->?->PglZ-p->PglZ->BrxA-DNA-binding-><-XIS-HTH<-Phage_int_SAM_5+Phage_integrase<-tRNA\|\|TPR+TPR-> | 1159 | Bacteria>FCB_group>Bacteroidetes/Chlorobi_group>Ignavibacteriota>Ignavibacteria>unclassified_Ignavibacteria | GCA_016719845.1 | Ignavibacteria_bacterium | IPM83_09090 | 230869..234348 | 2053306 | ATP-binding-protein-[Ignavibacteria-bacterium] |
| NOQ23473.1 | Type3.MTaseII.containing.BREX | <-TUDOR+SF1-SF2-Helicase+iREase+CTD<-TOPRIM<-SbcC<-MORC<-N6-N4-METHYLASE<-?<-?<-TPR+HEPN<-HTH+REase<-DUF6079*<-BrxB-BrxF-iSTAND<-XIS-HTH\|\|?->OLD-ABC->OLD-ABC-> | 1242 | Bacteria>FCB_group>Candidatus_Aegiribacteria>unclassified_Candidatus_Aegiribacteria | GCA_013138965.1 | Candidatus_Aegiribacteria_sp. | GQ565_12615 | 17846..21574 | 2528271 | ATP-binding-protein-[Candidatus-Aegiribacteria-sp.] |
| NLO12465.1 | Type3.MTaseII.containing.BREX | <-PglZ-p<-N6-N4-METHYLASE<-Schlafen+HATPase+wHTH<-DUF6079-p* | 904 | Bacteria>FCB_group>Candidatus_Cloacimonetes>unclassified_Candidatus_Cloacimonetes | GCA_012522935.1 | Candidatus_Cloacimonetes_bacterium | GX122_08625 | 5538..8254 | 2030808 | ATP-binding-protein-[Candidatus-Cloacimonadota-bacterium] |
| MCB5228480.1 | Type3.MTaseII.containing.BREX | <-Peptidase_M1_N+Peptidase_M1<-?\|\|ACET+ACET->?-><-?<-SF1-SF2-Helicase<-OLD-ABC+TOPRIM\|\|BrxB-BrxF-iSTAND->DUF6079*->Schlafen+HATPase+wHTH->N6-N4-METHYLASE->PglZ->BrxA-DNA-binding->MORC+REase-> | 1245 | Bacteria>FCB_group>Candidatus_Cloacimonetes>unclassified_Candidatus_Cloacimonetes | GCA_020532325.1 | Candidatus_Cloacimonetes_bacterium | LHW44_00170 | 36950..40687 | 2030808 | DUF6079-family-protein-[Candidatus-Cloacimonadota-bacterium] |
| MCB5253472.1 | Type3.MTaseII.containing.BREX | DUF6079-p*->Schlafen+HATPase+wHTH->N6-N4-METHYLASE->PglZ->BrxA-DNA-binding->MORC+REase-> | 752 | Bacteria>FCB_group>Candidatus_Cloacimonetes>unclassified_Candidatus_Cloacimonetes | GCA_020532345.1 | Candidatus_Cloacimonetes_bacterium | LHW51_10915 | 1..2260 | 2030808 | DUF6079-family-protein-[Candidatus-Cloacimonadota-bacterium] |
| MCK9335092.1 | Type3.MTaseII.containing.BREX | <-PglZ-p<-N6-N4-METHYLASE<-Schlafen+HATPase+wHTH<-DUF6079-p* | 748 | Bacteria>FCB_group>Candidatus_Cloacimonetes>unclassified_Candidatus_Cloacimonetes | GCA_023230025.1 | Candidatus_Cloacimonetes_bacterium | MOQ99_07265 | 5735..7981 | 2030808 | DUF6079-family-protein-[Candidatus-Cloacimonadota-bacterium] |
| MCK9328444.1 | Type3.MTaseII.containing.BREX | HNH->?->SbcC->HTH_26->BrxB-BrxF-iSTAND->DUF6079*->N6-N4-METHYLASE->TUDOR+SF1-SF2-Helicase+iREase+CTD->PglZ->BrxA-DNA-binding-><-?<-?<-REase | 1244 | Bacteria>FCB_group>Candidatus_Cloacimonetes>unclassified_Candidatus_Cloacimonetes | GCA_023230175.1 | Candidatus_Cloacimonetes_bacterium | MOQ94_01075 | 3820..7554 | 2030808 | DUF6079-family-protein-[Candidatus-Cloacimonadota-bacterium] |
| MCK5036100.1 | Type3.MTaseII.containing.BREX | CR-ATPase5->?->DOC+HTH->?->BrxB-BrxF-iSTAND->DUF6079*->TPR+HEPN->?->?->N6-N4-METHYLASE-p-> | 1243 | Bacteria>FCB_group>Candidatus_Fermentibacteria>Candidatus_Sabulitectum>unclassified_Candidatus_Sabulitectum | GCA_023136985.1 | Candidatus_Sabulitectum_sp. | KAS73_09435 | 6070..9801 | 2822438 | ATP-binding-protein-[Candidatus-Sabulitectum-sp.] |
| MBW7865069.1 | Type3.MTaseII.containing.BREX | ZnR+TOPRIM+AAA-helicase+wHTH->?-><-?<-LexA-protease<-?<-HTH_3\|\|?->BrxB-BrxF-iSTAND->DUF6079*->Schlafen+HATPase+wHTH-> | 1242 | Bacteria>FCB_group>Candidatus_Hydrogenedentes>unclassified_Candidatus_Hydrogenedentes | GCA_019455225.1 | Candidatus_Hydrogenedentes_bacterium | H3C30_11745 | 23031..26759 | 2030809 | ATP-binding-protein-[Candidatus-Hydrogenedentes-bacterium] |
| KMQ51971.1 | Type3.MTaseII.containing.BREX | <-BrxA-DNA-binding<-PglZ<-?<-TUDOR+SF1-SF2-Helicase+iREase+CTD<-Schlafen+HATPase+wHTH<-N6-N4-METHYLASE<-Schlafen+HATPase+wHTH<-DUF6079*<-BrxB-BrxF-iSTAND | 1295 | Bacteria>FCB_group>Fibrobacterota>Chitinispirillia>Chitinispirillales>Chitinispirillaceae | GCA_001045525.1 | Chitinispirillum_alkaliphilum | CHISP_1226 | 137443..141330 | 1008392 | ATPase-[Chitinispirillum-alkaliphilum] |
| NLE00367.1 | Type3.MTaseII.containing.BREX | DUF6079*->RNaseH->?->N6-N4-METHYLASE->TUDOR+SF1-SF2-Helicase+iREase+CTD->PglZ->BrxA-DNA-binding-> | 1398 | Bacteria>FCB_group>Fibrobacterota>Fibrobacteria>Fibrobacterales>Fibrobacteraceae | GCA_012513935.1 | Fibrobacter_sp. | GX640_10875 | 880..5076 | 35828 | BREX-3-system-P-loop-containing-protein-BrxF-[Fibrobacter-sp.] |
| MBF0431034.1 | Type3.MTaseII.containing.BREX | <-Phage_integrase<-XIS-HTH<-?<-?<-?<-SF1-SF2-Helicase<-?\|\|?->BrxB-BrxF-iSTAND->DUF6079*->N6-N4-METHYLASE-> | 1244 | Bacteria>FCB_group>Fibrobacterota>Fibrobacteria>unclassified_Fibrobacteria | GCA_015232855.1 | Fibrobacteria_bacterium | HQK83_07130 | 30062..33796 | 2725311 | ATP-binding-protein-[Fibrobacteria-bacterium] |
| QQS04457.1 | Type3.MTaseII.containing.BREX | <-PhdYeFM_antitox<-HTH+REase\|\|?->HKD-DNase-><-BrxA-DNA-binding<-PglZ<-TUDOR+SF1-SF2-Helicase+iREase+CTD<-Schlafen+HATPase+wHTH<-DUF6079*<-BrxB-BrxF-iSTAND\|\|?-><-ParB+HNH<-UDG-conflict<-?\|\|HTH_33+DDE_3-><-?<-HTH_Tnp_1+HTH_21+rve+rve_3<-HIN-HTH | 1240 | Bacteria>FCB_group>Fibrobacterota>unclassified_Fibrobacterota | GCA_016699655.1 | Fibrobacterota_bacterium | IPK50_19530 | 4749323..4753045 | 2052160 | ATP-binding-protein-[Fibrobacterota-bacterium] |
| MYA65074.1 | Type3.MTaseII.containing.BREX | KAP-NTPase->KAP-glycosylase->?->TatD_DNase-><-BrxA-DNA-binding<-PglZ<-?<-N6-N4-METHYLASE<-Schlafen+HATPase+wHTH<-DUF6079*<-BrxB-BrxF-iSTAND<-?<-MORC+DUF3684+DUF3883 | 1242 | Bacteria>FCB_group>Gemmatimonadota>unclassified_Gemmatimonadota | GCA_009839035.1 | Gemmatimonadota_bacterium | F4139_01755 | 19843..23571 | 2026742 | ATP-binding-protein-[Gemmatimonadota-bacterium] |
| MBA4157055.1 | Type3.MTaseII.containing.BREX | <-N6-N4-METHYLASE<-?<-?<-?<-?<-DUF6079*<-BrxB-BrxF-iSTAND<-MORC+DUF3684+HNH<-MORC+DUF3684+HNH<-SH3-like+SF1-SF2-Helicase+wHTH\|\|HTH_23+HTH_32->HTH_Tnp_IS630-> | 1251 | Bacteria>FCB_group>Gemmatimonadota>unclassified_Gemmatimonadota | GCA_013820915.1 | Gemmatimonadota_bacterium | HOX65_06220 | 5513..9268 | 2026742 | ATP-binding-protein-[Gemmatimonadota-bacterium] |
| WP_235924374.1 | Type3.MTaseII.containing.BREX | <-MerR-HTH\|\|?->?->?->?->?->?->?-><-DDE_Tnp_1_5+DDE_Tnp_1<-DUF6079-p*<-BrxB-BrxF-iSTAND<-wHTH+WYL+WCX\|\|MODE-HTH+PBPII->HTH_3->Classical-AAA->Peptidase_S8-><-HTH_3\|\|LysR-HTH-> | 155 | Bacteria | GCF_003332705.2 | Pseudomonadota | PA34_RS24465 | 5081824..5082291 | 1224 | MULTISPECIES:-DUF6079-family-protein-[Pseudomonadota] |
| WP_204219988.1 | Type3.MTaseII.containing.BREX | <-Glyco_hydro_43+GH43_C2\|\|Glyco_hydro_32N+Glyco_hydro_43->?->ABC-ATPase+SbcC+ABC-ATPase+SbcC->?->?-><-DUF6079*<-N6-N4-METHYLASE<-BrxB-BrxF-iSTAND<-BrxA-DNA-binding<-SIGMA+OB-Fold+AAA-ATPase+Lon-Protease-BrxL<-PglZ<-?<-?\|\|HTH_3-><-CR-Kinase | 1378 | Bacteria>Myxococcota>Myxococcia>Myxococcales>Cystobacterineae>Archangiaceae | GCF_016887565.1 | Archangium_violaceum | JRI60_RS33460 | 8428766..8432902 | 83451 | hypothetical-protein-[Archangium-violaceum] |
| MBL8619839.1 | Type3.MTaseII.containing.BREX | BrxB-BrxF-iSTAND->DUF6079*->HEPN->N6-N4-METHYLASE->TUDOR+SF1-SF2-Helicase+iREase+CTD->PglZ->BrxA-DNA-binding->HHH_4+HTH+wHTH+SF1-SF2-Helicase+REase->TPR-> | 1236 | Bacteria>Myxococcota>Myxococcia>Myxococcales>unclassified_Myxococcales | GCA_016794525.1 | Myxococcales_bacterium | JNK64_01015 | 489..4199 | 2026763 | hypothetical-protein-[Myxococcales-bacterium] |
| MBE7447418.1 | Type3.MTaseII.containing.BREX | RAMA->?->GDResolvase+ZnR->?->XIS-HTH->DUF488->?->XIS-HTH->BrxB-BrxF-iSTAND->DUF6079*->N6-N4-METHYLASE->TUDOR+SF1-SF2-Helicase+iREase+CTD->PglZ->BrxA-DNA-binding->glycosylase-><-?<-ACET | 1229 | Bacteria>Myxococcota>Polyangia>Haliangiales>Kofleriaceae>unclassified_Kofleriaceae | GCA_015075455.1 | Kofleriaceae_bacterium | HS111_00575 | 104107..107796 | 2212474 | ATP-binding-protein-[Kofleriaceae-bacterium] |
| WP_271916064.1 | Type3.MTaseII.containing.BREX | SIGMA-HTH-><-?\|\|?->?->?-><-HTH_3<-?\|\|?->BrxB-BrxF-iSTAND->DUF6079*->N6-N4-METHYLASE->Schlafen+HATPase+wHTH-p->TUDOR+SF1-SF2-Helicase+iREase+CTD->PglZ->BrxA-DNA-binding->?->RADICAL-SAM-> | 1229 | Bacteria>Myxococcota>Polyangia>Polyangiales>Polyangiaceae>Polyangium | GCF_028369105.1 | Polyangium_sp._rjm3 | POL67_RS05840 | 1409765..1413454 | 2995306 | DUF6079-family-protein-[Polyangium-sp.-rjm3] |
| MBX3129303.1 | Type3.MTaseII.containing.BREX | DUF499-><-?<-ACET<-glycosylase<-BrxA-DNA-binding<-PglZ<-TUDOR+SF1-SF2-Helicase+iREase+CTD<-N6-N4-METHYLASE<-DUF6079*<-BrxB-BrxF-iSTAND<-XIS-HTH<-?<-GDResolvase+ZnR<-?<-RAMA\|\|?-><-Phage_int_SAM_1+Phage_integrase<-Phage_integrase | 1232 | Bacteria>Myxococcota>Polyangia>Polyangiales>Polyangiaceae>unclassified_Polyangiaceae | GCA_019637295.1 | Polyangiaceae_bacterium | KF718_21475 | 30512..34210 | 2268199 | hypothetical-protein-[Polyangiaceae-bacterium] |
| MCB9598269.1 | Type3.MTaseII.containing.BREX | <-DUF6079*<-BrxB-BrxF-iSTAND<-?\|\|?->SF1-SF2-Helicase->?->SF1-SF2-Helicase-> | 1231 | Bacteria>Myxococcota>Polyangia>Polyangiales>Sandaracinaceae>unclassified_Sandaracinaceae | GCA_020633195.1 | Sandaracinaceae_bacterium | H6719_36485 | 301..3996 | 2510337 | ATP-binding-protein-[Sandaracinaceae-bacterium] |
| MBT6716170.1 | Type3.MTaseII.containing.BREX | <-Arm-DNA-bind_3+Phage_integrase\|\|tRNA-><-Trypsin+PDZ<-CHTH\|\|?->wHTH+WYL+WCX->Calcineurin-SbcD->SbcC->BrxB-BrxF-iSTAND->DUF6079*->?->N6-N4-METHYLASE->HEPN->TUDOR+SF1-SF2-Helicase+iREase+CTD->PglZ->BrxA-DNA-binding-> | 1242 | Bacteria>Nitrospinae/Tectomicrobia_group>Nitrospinota>Nitrospinia>Nitrospinales>Nitrospinaceae | GCA_018669775.1 | Nitrospina_sp. | HOB18_00830 | 15038..18766 | 2024844 | ATP-binding-protein-[Nitrospina-sp.] |
| WP_168062758.1 | Type3.MTaseII.containing.BREX | ParB->LuxR-HTH-><-REase\|\|?-><-?\|\|Peptidase->?->BrxB-BrxF-iSTAND->DUF6079*->N6-N4-METHYLASE->HEPN->TUDOR+SF1-SF2-Helicase+iREase+CTD->PglZ->BrxA-DNA-binding->?-><-VirD4-TraG+VirD4-FtsK<-?<-HDRnase | 1242 | Bacteria>Nitrospirota>Nitrospiria>Candidatus_Troglogloeales>Candidatus_Manganitrophaceae>Candidatus_Manganitrophus | GCF_012184425.1 | Candidatus_Manganitrophus_noduliformans | MNODULE_RS19050 | 146292..150020 | 2606439 | DUF6079-family-protein-[Candidatus-Manganitrophus-noduliformans] |
| HBP90844.1 | Type3.MTaseII.containing.BREX | <-BrxA-DNA-binding<-PglZ<-TUDOR+SF1-SF2-Helicase+iREase+CTD<-?<-SbcC<-N6-N4-METHYLASE<-DUF6079*<-BrxB-BrxF-iSTAND\|\|GDResolvase+Recombinase+ZnR-><-Glycos_transf<-NUDIX<-?\|\|HD->ABC-3C-> | 1241 | Bacteria>Nitrospirota>Nitrospiria>Nitrospirales>Nitrospiraceae>unclassified_Nitrospiraceae | GCA_003523945.1 | Nitrospiraceae_bacterium | DD706_24520 | 35721..39446 | 2026770 | ATPase-[Nitrospiraceae-bacterium] |
| GIW56389.1 | Type3.MTaseII.containing.BREX | XIS-HTH->BrxB-BrxF-iSTAND-><-?<-?<-?\|\|DUF6079-p*->Schlafen+HATPase+wHTH-><-?\|\|Schlafen+HATPase+wHTH->N6-N4-METHYLASE-p->N6-N4-METHYLASE-p->?->?->?->N6-N4-METHYLASE-> | 386 | Bacteria>Nitrospirota>Nitrospiria>Nitrospirales>Nitrospiraceae>unclassified_Nitrospiraceae | GCA_026004315.1 | Nitrospiraceae_bacterium | KatS3mg082_2793 | 3690..4850 | 2026770 | MAG:-hypothetical-protein-KatS3mg082_2793-[Nitrospiraceae-bacterium] |
| WP_225487366.1 | Type3.MTaseII.containing.BREX | <-PBPI\|\|TP_METHYLASE->tRNA->Phage_integrase-><-AAA-ATPase<-HTH+rve<-BrxA-DNA-binding<-PglZ<-TUDOR+SF1-SF2-Helicase+iREase+CTD<-N6-N4-METHYLASE<-DUF6079*<-BrxB-BrxF-iSTAND<-REase\|\|TrwC+SF1-SF2-Helicase->HTH_58->?->?->?-><-ZnR+HTH_Tnp_ISL3+DDE_Tnp_ISL3 | 1250 | Bacteria>Pseudomonadota>Acidithiobacillia>Acidithiobacillales>Acidithiobacillaceae>Acidithiobacillus | - | Acidithiobacillus_ferrooxidans | LDB30_00720 | 146829..150581 | 920 | DUF6079-family-protein-[Acidithiobacillus-ferrooxidans] |
| MBU2804262.1 | Type3.MTaseII.containing.BREX | <-REase+NACHT<-REase\|\|wHTH+WYL+WCX->BrxB-BrxF-iSTAND->DUF6079*->HEPN->N6-N4-METHYLASE->?->TUDOR+SF1-SF2-Helicase+iREase+CTD->PglZ->BrxA-DNA-binding-><-?<-GspA-ATPase | 1242 | Bacteria>Pseudomonadota>Acidithiobacillia>Acidithiobacillales>Acidithiobacillaceae>Acidithiobacillus | GCA_018854695.1 | Acidithiobacillus_ferridurans | HF668_03640 | 12541..16269 | 1232575 | ATP-binding-protein-[Acidithiobacillus-ferridurans] |
| WP_035192522.1 | Type3.MTaseII.containing.BREX | Arm-DNA-bind_3+Phage_integrase->HTH_3->?-><-BrxA-DNA-binding<-PglZ<-TUDOR+SF1-SF2-Helicase+iREase+CTD<-HEPN<-N6-N4-METHYLASE<-Schlafen+HATPase+wHTH+HTH<-DUF6079*<-BrxB-BrxF-iSTAND<-XIS-HTH<-?\|\|?-><-?<-?\|\|Integrase_1->XIS-HTH-> | 1244 | Bacteria>Pseudomonadota>Acidithiobacillia>Acidithiobacillales>Acidithiobacillaceae>Acidithiobacillus | GCF_000750615.1 | Acidithiobacillus_ferrivorans | AFERRI_RS08540 | 128425..132159 | 160808 | DUF6079-family-protein-[Acidithiobacillus-ferrivorans] |
| WP_215854619.1 | Type3.MTaseII.containing.BREX | TnsC-ATPase->?->?->XIS-HTH->BrxB-BrxF-iSTAND->DUF6079*->Schlafen+HATPase+wHTH+HTH->N6-N4-METHYLASE->Classical-AAA->Peptidase_S8->TUDOR+SF1-SF2-Helicase+iREase+CTD->SIR2->PglZ->BrxA-DNA-binding-><-SF1-SF2-Helicase | 1244 | Bacteria>Pseudomonadota>Acidithiobacillia>Acidithiobacillales>Acidithiobacillaceae>Acidithiobacillus | GCF_018854475.1 | Acidithiobacillus_ferrooxidans | HF290_RS06270 | 6092..9826 | 920 | DUF6079-family-protein-[Acidithiobacillus-ferrooxidans] |
| WP_215878275.1 | Type3.MTaseII.containing.BREX | <-REase+NACHT<-REase\|\|wHTH+WYL+WCX->BrxB-BrxF-iSTAND->DUF6079*->HEPN->N6-N4-METHYLASE->?->TUDOR+SF1-SF2-Helicase+iREase+CTD->PglZ->BrxA-DNA-binding-><-?<-GspA-ATPase | 1246 | Bacteria>Pseudomonadota>Acidithiobacillia>Acidithiobacillales>Acidithiobacillaceae>Acidithiobacillus | GCF_018854695.1 | Acidithiobacillus_ferridurans | HF668_RS03540 | 12529..16269 | 1232575 | DUF6079-family-protein-[Acidithiobacillus-ferridurans] |
| WP_215861187.1 | Type3.MTaseII.containing.BREX | DnaB->?->?->XIS-HTH->BrxB-BrxF-iSTAND->DUF6079*->Schlafen+HATPase+wHTH+HTH->N6-N4-METHYLASE->SbcC->TOPRIM->OLD-ABC+TOPRIM->TUDOR+SF1-SF2-Helicase+iREase+CTD->SIR2->PglZ->BrxA-DNA-binding-> | 1244 | Bacteria>Pseudomonadota>Acidithiobacillia>Acidithiobacillales>Acidithiobacillaceae>Acidithiobacillus | GCF_018854875.1 | Acidithiobacillus_ferriphilus | HF928_RS01635 | 4629..8363 | 1689834 | DUF6079-family-protein-[Acidithiobacillus-ferriphilus] |
| WP_257245726.1 | Type3.MTaseII.containing.BREX | FURR-HTH->tRNA->?-><-BrxA-DNA-binding<-PglZ<-TUDOR+SF1-SF2-Helicase+iREase+CTD<-N6-N4-METHYLASE<-DUF6079*<-BrxB-BrxF-iSTAND<-MULE-TRANSPOSASE\|\|TrwC+SF1-SF2-Helicase->HTH_58->?->?-><-SIGMA-HTH<-NACHT<-TOPRIM | 1246 | Bacteria>Pseudomonadota>Acidithiobacillia>Acidithiobacillales>Acidithiobacillaceae>Acidithiobacillus | GCF_024606765.1 | Acidithiobacillus_ferrooxidans | NRY68_RS07345 | 31174..34914 | 920 | DUF6079-family-protein-[Acidithiobacillus-ferrooxidans] |
| KAF0228152.1 | Type3.MTaseII.containing.BREX | XIS-HTH-><-CHTH\|\|GDResolvase+PPP1-HTH->ParB->ParB->wHTH+WYL+WCX->BrxB-BrxF-iSTAND->DUF6079*->HEPN->N6-N4-METHYLASE->TUDOR+SF1-SF2-Helicase+iREase+CTD->PglZ->BrxA-DNA-binding-> | 1243 | Bacteria>Pseudomonadota>Alphaproteobacteria>Hyphomicrobiales>Beijerinckiaceae>unclassified_Beijerinckiaceae | GCA_009773635.1 | Beijerinckiaceae_bacterium | FD175_2258 | 41673..45404 | 1978229 | hypothetical-protein-FD175_2258-[Beijerinckiaceae-bacterium] |
| WP_236823724.1 | Type3.MTaseII.containing.BREX | <-BrxA-DNA-binding<-PglZ<-TUDOR+SF1-SF2-Helicase+iREase+CTD<-N6-N4-METHYLASE<-Schlafen+HATPase+wHTH<-DUF6079*<-BrxB-BrxF-iSTAND<-wHTH+WYL+WCX\|\|GNAT->?->?->SF1-SF2-Helicase+DUF1998-><-?<-GDResolvase+ZnR<-RAMA | 1265 | Bacteria>Pseudomonadota>Alphaproteobacteria>Hyphomicrobiales>Blastochloridaceae>Blastochloris | GCF_001548155.2 | Blastochloris_viridis | BV133_RS16120 | 3700078..3703875 | 1079 | DUF6079-family-protein-[Blastochloris-viridis] |
| NKE76483.1 | Type3.MTaseII.containing.BREX | <-35exo\|\|?->GGDEF->?->?->?-><-?\|\|XIS-HTH->wHTH+WYL+WCX->BrxB-BrxF-iSTAND->DUF6079*->N6-N4-METHYLASE->TUDOR+SF1-SF2-Helicase+iREase+CTD->PglZ->BrxA-DNA-binding-> | 1241 | Bacteria>Pseudomonadota>Alphaproteobacteria>Hyphomicrobiales>Brucellaceae>Brucella/Ochrobactrum_group | GCA_012241515.1 | Brucella_intermedia | HC237_13895 | 35495..39220 | 94625 | ATP-binding-protein-[Brucella-intermedia] |
| WP_207488589.1 | Type3.MTaseII.containing.BREX | VirB10->RHH-><-?<-ABhydrolase<-?<-TetR-HTH\|\|?->XIS-HTH->BrxB-BrxF-iSTAND->DUF6079*->Schlafen+HATPase+wHTH->MORC->N6-N4-METHYLASE->TUDOR+SF1-SF2-Helicase+iREase+CTD->PglZ->BrxA-DNA-binding-><-?<-Peptidase_M29 | 1246 | Bacteria>Pseudomonadota>Alphaproteobacteria>Hyphomicrobiales>Brucellaceae>Brucella/Ochrobactrum_group | GCF_017355465.1 | Brucella_pituitosa | IPV26_RS10115 | 74903..78643 | 571256 | DUF6079-family-protein-[Brucella-pituitosa] |
| WP_245514510.1 | Type3.MTaseII.containing.BREX | RHH->wHTH+WYL+WCX->BrxB-BrxF-iSTAND->DUF6079*->HEPN->N6-N4-METHYLASE->N6-N4-METHYLASE-p-> | 1243 | Bacteria>Pseudomonadota>Alphaproteobacteria>Hyphomicrobiales>Chelatococcaceae>Camelimonas | GCF_004342915.1 | Camelimonas_lactis | EV666_RS17715 | 2787..6518 | 659006 | DUF6079-family-protein-[Camelimonas-lactis] |
| WP_244643259.1 | Type3.MTaseII.containing.BREX | TrwJ->TrbK->VirB6->VirB8->VirB9->VirB10->RHH->wHTH+WYL+WCX->BrxB-BrxF-iSTAND->DUF6079*->HEPN->N6-N4-METHYLASE->TUDOR+SF1-SF2-Helicase+iREase+CTD->PglZ->BrxA-DNA-binding-><-HEPN | 1243 | Bacteria>Pseudomonadota>Alphaproteobacteria>Hyphomicrobiales>Chelatococcaceae>Camelimonas | GCF_014656355.1 | Camelimonas_fluminis | IE243_RS18850 | 12165..15896 | 1576911 | DUF6079-family-protein-[Camelimonas-fluminis] |
| CDP52011.1 | Type3.MTaseII.containing.BREX | Glyco_hydro_53+Phage-tail_3->?->?->?->?->?-><-PglZ<-TUDOR+SF1-SF2-Helicase+iREase+CTD<-N6-N4-METHYLASE<-DUF6079*<-wHTH+WYL+WCX<-XIS-HTH<-?<-?<-?<-?<-?<-?<-ABC-ATPase+SbcC | 1243 | Bacteria>Pseudomonadota>Alphaproteobacteria>Hyphomicrobiales>Devosiaceae>Devosia | GCA_000689495.1 | Devosia_sp._DBB001 | . | 17582..21313 | 1380412 | hypothetical-protein-BVU_3741-[Devosia-sp.-DBB001] |
| GLQ55357.1 | Type3.MTaseII.containing.BREX | <-TOPRIM\|\|?->?-><-GAF+PAS+GGDEF+EAL\|\|TPR->ACET-><-?\|\|BrxB-BrxF-iSTAND->DUF6079*->HEPN->N6-N4-METHYLASE->TUDOR+SF1-SF2-Helicase+iREase+CTD->PglZ->BrxA-DNA-binding-> | 1243 | Bacteria>Pseudomonadota>Alphaproteobacteria>Hyphomicrobiales>Devosiaceae>Devosia | GCA_030160115.1 | Devosia_nitrariae | GCM10010862_26160 | 527986..531717 | 2071872 | hypothetical-protein-GCM10010862_26160-[Devosia-nitrariae] |
| MBX9864944.1 | Type3.MTaseII.containing.BREX | <-PglZ<-TUDOR+SF1-SF2-Helicase+iREase+CTD<-N6-N4-METHYLASE<-DUF6079*<-BrxB-BrxF-iSTAND<-wHTH+WYL+WCX\|\|?->Preader+URI-DNase-> | 1229 | Bacteria>Pseudomonadota>Alphaproteobacteria>Hyphomicrobiales>Hyphomicrobiaceae>Hyphomicrobium | GCA_019744875.1 | Hyphomicrobium_sp. | K2Y42_19555 | 7941..11630 | 82 | ATP-binding-protein-[Hyphomicrobium-sp.] |
| MCC0001392.1 | Type3.MTaseII.containing.BREX | <-TOPRIM<-?<-?<-ParB<-DUF932\|\|wHTH+WYL+WCX->BrxB-BrxF-iSTAND->DUF6079*->?->N6-N4-METHYLASE->TUDOR+SF1-SF2-Helicase+iREase+CTD->PglZ->BrxA-DNA-binding->HEPN-> | 1241 | Bacteria>Pseudomonadota>Alphaproteobacteria>Hyphomicrobiales>Methylobacteriaceae>unclassified_Methylobacteriaceae | GCA_020638195.1 | Methylobacteriaceae_bacterium | H6870_20860 | 164083..167808 | 2021371 | ATP-binding-protein-[Methylobacteriaceae-bacterium] |
| GLI95561.1 | Type3.MTaseII.containing.BREX | TOPRIM-><-HKD-DNase\|\|XIS-HTH->BrxB-BrxF-iSTAND->DUF6079*->Schlafen+HATPase+wHTH->N6-N4-METHYLASE->TUDOR+SF1-SF2-Helicase+iREase+CTD->PglZ->BrxA-DNA-binding->?->ParB->?-><-METHYLASE+DsrD | 1229 | Bacteria>Pseudomonadota>Alphaproteobacteria>Hyphomicrobiales>Methylocystaceae>Methylocystis | GCA_027923385.1 | Methylocystis_echinoides | LMG27198_45530 | 58571..62260 | 29468 | hypothetical-protein-LMG27198_45530-[Methylocystis-echinoides] |
| WP_028350240.1 | Type3.MTaseII.containing.BREX | <-SIGMA-HTH+SIGMA-HTH<-S4+PSYN\|\|?->tRNA->tRNA->?-><-BrxA-DNA-binding<-PglZ<-TUDOR+SF1-SF2-Helicase+iREase+CTD<-N6-N4-METHYLASE<-HEPN<-DUF6079*<-BrxB-BrxF-iSTAND<-XIS-HTH<-?<-?<-?<-N6-METHYLASE<-?\|\|?-><-RADICAL-SAM | 1229 | Bacteria>Pseudomonadota>Alphaproteobacteria>Hyphomicrobiales>Nitrobacteraceae>Bradyrhizobium | GCF_000472965.1 | Bradyrhizobium_murdochi | YUO_RS0127990 | 107889..111578 | 1038859 | DUF6079-family-protein-[Bradyrhizobium-murdochi] |
| WP_085349327.1 | Type3.MTaseII.containing.BREX | ACET-><-Peptidase<-?<-?\|\|?->Pkinase+Pkinase->XIS-HTH->BrxB-BrxF-iSTAND->DUF6079-p*-><-tRNA\|\|?->?-><-?\|\|?->?->ADH_N+ADH_zinc_N-> | 454 | Bacteria>Pseudomonadota>Alphaproteobacteria>Hyphomicrobiales>Nitrobacteraceae>Bradyrhizobium | GCF_002108675.1 | Bradyrhizobium_canariense | BSZ20_RS14710 | 27527..28891 | 255045 | DUF6079-family-protein-[Bradyrhizobium-canariense] |
| WP_208738467.1 | Type3.MTaseII.containing.BREX | GDResolvase+PPP1-HTH->ParB->ParB-><-?\|\|?->?->McrB-AAA->XIS-HTH->BrxB-BrxF-iSTAND->DUF6079-p*-><-N6-N4-METHYLASE<-DUF6079-p\|\|N6-N4-METHYLASE->SbcC->?->TUDOR+SF1-SF2-Helicase+iREase+CTD->PglZ->BrxA-DNA-binding-> | 322 | Bacteria>Pseudomonadota>Alphaproteobacteria>Hyphomicrobiales>Nitrobacteraceae>Bradyrhizobium | GCF_004114425.1 | Bradyrhizobium_vignae | EAV90_RS04615 | 73969..74937 | 1549949 | DUF6079-family-protein-[Bradyrhizobium-vignae] |
| WP_129142724.1 | Type3.MTaseII.containing.BREX | ParB-><-?\|\|?->?->McrB-AAA->XIS-HTH->BrxB-BrxF-iSTAND->DUF6079-p-><-N6-N4-METHYLASE<-DUF6079-p*\|\|N6-N4-METHYLASE->SbcC->?->TUDOR+SF1-SF2-Helicase+iREase+CTD->PglZ->BrxA-DNA-binding-><-?\|\|Phage_integrase-> | 366 | Bacteria>Pseudomonadota>Alphaproteobacteria>Hyphomicrobiales>Nitrobacteraceae>Bradyrhizobium | GCF_004114425.1 | Bradyrhizobium_vignae | EAV90_RS04625 | 76559..77659 | 1549949 | DUF6079-family-protein-[Bradyrhizobium-vignae] |
| WP_141934535.1 | Type3.MTaseII.containing.BREX | <-wHTH+WYL+WCX\|\|REase+HTH+35exo+Small-B-barrel+SF1-SF2-Helicase-><-GDResolvase+HIN-HTH\|\|?-><-BrxA-DNA-binding<-PglZ<-TUDOR+SF1-SF2-Helicase+iREase+CTD<-N6-N4-METHYLASE<-DUF6079*<-BrxB-BrxF-iSTAND<-HHH_4+SF1-SF2-Helicase+REase<-?<-TUDOR+SF1-SF2-Helicase+iREase+CTD<-?\|\|HD+PspB->FGase-> | 1241 | Bacteria>Pseudomonadota>Alphaproteobacteria>Hyphomicrobiales>Nitrobacteraceae>Bradyrhizobium | GCF_006658635.1 | Bradyrhizobium_sp._UNPA324 | UNPA324_RS09470 | 1934333..1938058 | 1141174 | DUF6079-family-protein-[Bradyrhizobium-sp.-UNPA324] |
| WP_264623141.1 | Type3.MTaseII.containing.BREX | Arm-DNA-bind_3+Phage_integrase-><-Phage_AlpA<-BrxA-DNA-binding<-PglZ<-TUDOR+SF1-SF2-Helicase+iREase+CTD<-N6-N4-METHYLASE<-MORC<-Schlafen+HATPase+wHTH<-DUF6079*<-BrxB-BrxF-iSTAND<-XIS-HTH\|\|SIGMA-HTH->LuxR-HTH->?->MODE-HTH+PBPII-><-RHH<-VirB9<-VirB8 | 1246 | Bacteria>Pseudomonadota>Alphaproteobacteria>Hyphomicrobiales>Nitrobacteraceae>Bradyrhizobium | GCF_025960955.1 | Bradyrhizobium_elkanii | M2210_RS06800 | 1521940..1525680 | 29448 | DUF6079-family-protein-[Bradyrhizobium-elkanii] |
| WP_119855769.1 | Type3.MTaseII.containing.BREX | <-RADICAL-SAM\|\|?->SIGMA-HTH->ParA-Soj-PloopNTPase->?->LON-AAA->XIS-HTH->BrxB-BrxF-iSTAND->DUF6079*->HEPN->N6-N4-METHYLASE->TUDOR+SF1-SF2-Helicase+iREase+CTD->PglZ->BrxA-DNA-binding-><-Phage_int_SAM_1+Phage_integrase\|\|HTH_3->HipA_C-> | 1229 | Bacteria>Pseudomonadota>Alphaproteobacteria>Hyphomicrobiales>Nitrobacteraceae>Rhodopseudomonas | GCF_003591005.1 | Rhodopseudomonas_palustris | D4Q52_RS06700 | 131316..135005 | 1076 | DUF6079-family-protein-[Rhodopseudomonas-palustris] |
| WP_012110066.1 | Type3.MTaseII.containing.BREX | TrbK->VirB6->VirB8->VirB9->VirB10->RHH-><-MODE-HTH+PBPII\|\|XIS-HTH->BrxB-BrxF-iSTAND->DUF6079*->Schlafen+HATPase+wHTH->MORC->N6-N4-METHYLASE->TUDOR+SF1-SF2-Helicase+iREase+CTD->PglZ->BrxA-DNA-binding->Phage_AlpA-><-Arm-DNA-bind_3+Phage_integrase | 1246 | Bacteria>Pseudomonadota>Alphaproteobacteria>Hyphomicrobiales>Parvibaculaceae>Parvibaculum | GCF_000017565.1 | Parvibaculum_lavamentivorans | PLAV_RS18035 | 3797139..3800879 | 256618 | DUF6079-family-protein-[Parvibaculum-lavamentivorans] |
| RWD51648.1 | Type3.MTaseII.containing.BREX | HTH_3->?->HTH_3->?-><-?\|\|?->BrxB-BrxF-iSTAND->DUF6079*->N6-N4-METHYLASE->TUDOR+SF1-SF2-Helicase+iREase+CTD->PglZ->BrxA-DNA-binding->?->?-><-HEPN<-?\|\|HTH_Tnp_1+HTH_21+rve+rve_3-> | 1241 | Bacteria>Pseudomonadota>Alphaproteobacteria>Hyphomicrobiales>Phyllobacteriaceae>Mesorhizobium | GCA_004017085.1 | Mesorhizobium_sp. | EOS75_31250 | 14991..18716 | 1871066 | ATP-binding-protein-[Mesorhizobium-sp.] |
| TIW29361.1 | Type3.MTaseII.containing.BREX | <-XIS-HTH\|\|?->Schlafen+HATPase+wHTH->PIWI-><-BrxA-DNA-binding<-TUDOR+SF1-SF2-Helicase+iREase+CTD<-?<-SbcC<-DUF6079*<-BrxB-BrxF-iSTAND<-XIS-HTH\|\|<-MODE-HTH+PBPII\|\|HTH_3-><-?<-Trypsin<-?\|\|UDG-> | 1258 | Bacteria>Pseudomonadota>Alphaproteobacteria>Hyphomicrobiales>Phyllobacteriaceae>Mesorhizobium | GCA_004964535.1 | Mesorhizobium_sp. | E5V63_01165 | 22561..26337 | 1871066 | ATP-binding-protein-[Mesorhizobium-sp.] |
| WP_023825443.1 | Type3.MTaseII.containing.BREX | <-HHH_4+HTH+wHTH+SF1-SF2-Helicase+REase<-?<-TUDOR+SF1-SF2-Helicase+iREase+CTD<-BrxA-DNA-binding<-PglZ<-TUDOR+SF1-SF2-Helicase+iREase+CTD<-N6-N4-METHYLASE<-HEPN<-DUF6079*<-BrxB-BrxF-iSTAND<-XIS-HTH<-?<-?<-?<-tRNA<-Peptidase-M15<-REC+NtrC-AAA+FIS-HTH\|\|Peptidase_M3_N+Peptidase_M3-> | 1229 | Bacteria>Pseudomonadota>Alphaproteobacteria>Hyphomicrobiales>Phyllobacteriaceae>Mesorhizobium | GCF_000502275.1 | Mesorhizobium_sp._L103C131B0 | X730_RS162110 | 447511..451200 | 1287089 | DUF6079-family-protein-[Mesorhizobium-sp.-L103C131B0] |
| WP_224642997.1 | Type3.MTaseII.containing.BREX | ABC-ATPase+SbcC+ABC-ATPase+SbcC->?->?->?->?->HNH->?->BrxB-BrxF-iSTAND->DUF6079*->N6-N4-METHYLASE->TUDOR+SF1-SF2-Helicase+iREase+CTD->PglZ->BrxA-DNA-binding->?->HTH+MPTase-><-NACHT\|\|ACET-> | 1241 | Bacteria>Pseudomonadota>Alphaproteobacteria>Hyphomicrobiales>Phyllobacteriaceae>Mesorhizobium | GCF_020165315.1 | Mesorhizobium_sp._BR115XR7A | LB557_RS18860 | 239767..243492 | 2876645 | DUF6079-family-protein-[Mesorhizobium-sp.-BR115XR7A] |
| WP_246365011.1 | Type3.MTaseII.containing.BREX | <-DEDD_Tnp_IS110<-BrxA-DNA-binding<-PglZ<-TUDOR+SF1-SF2-Helicase+iREase+CTD<-N6-N4-METHYLASE<-?<-DUF6079*<-BrxB-BrxF-iSTAND\|\|ParB->?->REase+ZnR->Classical-AAA->Classical-AAA->REase->?-><-Phage_integrase | 1243 | Bacteria>Pseudomonadota>Alphaproteobacteria>Hyphomicrobiales>Rhizobiaceae>Gellertiella | GCF_014197145.1 | Gellertiella_hungarica | GGR23_001421 | 542049..545780 | 1572859 | DUF6079-family-protein-[Gellertiella-hungarica] |
| WP_174047063.1 | Type3.MTaseII.containing.BREX | ABC-3C->SbcC-><-GDResolvase+HIN-HTH\|\|wHTH+WYL+WCX->DUF6079*->N6-N4-METHYLASE->TUDOR+SF1-SF2-Helicase+iREase+CTD->PglZ->BrxA-DNA-binding-><-PIN+OST-HTH\|\|?-><-?<-LexA-protease | 1241 | Bacteria>Pseudomonadota>Alphaproteobacteria>Hyphomicrobiales>Rhizobiaceae>Rhizobium/Agrobacterium_group | - | Agrobacterium_tumefaciens | G6M12_RS25340 | 82758..86483 | 358 | DUF6079-family-protein-[Agrobacterium-tumefaciens] |
| KQY13422.1 | Type3.MTaseII.containing.BREX | <-ABhydrolase\|\|?->?-><-CHTH\|\|GDResolvase+PPP1-HTH->ParB->ParB->wHTH+WYL+WCX->BrxB-BrxF-iSTAND->DUF6079*->N6-N4-METHYLASE->TUDOR+SF1-SF2-Helicase+iREase+CTD->PglZ->BrxA-DNA-binding-><-?<-?<-?<-?<-Glyco_hydro_53+Phage-tail_3 | 1241 | Bacteria>Pseudomonadota>Alphaproteobacteria>Hyphomicrobiales>Rhizobiaceae>Rhizobium/Agrobacterium_group | GCA_001426565.1 | Rhizobium_sp._Root482 | ASD31_13310 | 44054..47779 | 1736543 | ATPase-[Rhizobium-sp.-Root482] |
| PDS70271.1 | Type3.MTaseII.containing.BREX | BrxB-BrxF-iSTAND->DUF6079*->TUDOR+SF1-SF2-Helicase+iREase+CTD->PglZ->BrxA-DNA-binding-><-tRNA\|\|?->Thioredoxin-> | 1241 | Bacteria>Pseudomonadota>Alphaproteobacteria>Hyphomicrobiales>Rhizobiaceae>Rhizobium/Agrobacterium_group | GCA_002531715.1 | Rhizobium_phaseoli | CO651_19365 | 2097..5822 | 396 | ATPase-[Rhizobium-phaseoli] |
| TBE90170.1 | Type3.MTaseII.containing.BREX | <-Thioredoxin<-?\|\|tRNA-><-BrxA-DNA-binding<-PglZ<-TUDOR+SF1-SF2-Helicase+iREase+CTD<-N6-N4-METHYLASE<-DUF6079*<-BrxB-BrxF-iSTAND<-?<-TetR-HTH | 1241 | Bacteria>Pseudomonadota>Alphaproteobacteria>Hyphomicrobiales>Rhizobiaceae>Rhizobium/Agrobacterium_group | GCA_004305895.1 | Rhizobium_leguminosarum | ELG99_03805 | 766233..769958 | 384 | ATP-binding-protein-[Rhizobium-leguminosarum] |
| TBE09361.1 | Type3.MTaseII.containing.BREX | <-Thioredoxin<-?\|\|tRNA-><-BrxA-DNA-binding<-PglZ<-TUDOR+SF1-SF2-Helicase+iREase+CTD<-N6-N4-METHYLASE<-DUF6079*<-BrxB-BrxF-iSTAND<-?<-TetR-HTH | 1241 | Bacteria>Pseudomonadota>Alphaproteobacteria>Hyphomicrobiales>Rhizobiaceae>Rhizobium/Agrobacterium_group | GCA_004306185.1 | Rhizobium_leguminosarum | ELH12_03805 | 765413..769138 | 384 | ATP-binding-protein-[Rhizobium-leguminosarum] |
| TGE86678.1 | Type3.MTaseII.containing.BREX | <-Polbeta+HEPN\|\|Relaxase+SF1-SF2-Helicase-><-BrxA-DNA-binding<-PglZ<-TUDOR+SF1-SF2-Helicase+iREase+CTD<-N6-N4-METHYLASE<-MORC<-Schlafen+HATPase+wHTH<-DUF6079*<-BrxB-BrxF-iSTAND<-XIS-HTH\|\|McrB-AAA->McrC-NTD+REase->\|\|OLD-ABC+TOPRIM->SF1-SF2-Helicase-><-?<-ArsR-HTH\|\|HTH_3-> | 1246 | Bacteria>Pseudomonadota>Alphaproteobacteria>Hyphomicrobiales>Rhizobiaceae>Rhizobium/Agrobacterium_group | GCA_004766515.1 | Rhizobium_sp._SEMIA_4032 | C9418_21785 | 41425..45165 | 2137762 | ATPase-[Rhizobium-sp.-SEMIA-4032] |
| MBY5775964.1 | Type3.MTaseII.containing.BREX | <-BrxA-DNA-binding<-PglZ<-TUDOR+SF1-SF2-Helicase+iREase+CTD<-N6-N4-METHYLASE<-HEPN<-DUF6079*<-BrxB-BrxF-iSTAND<-?\|\|?-><-AbiEi<-URI-DNase<-REase+DUF5655 | 1241 | Bacteria>Pseudomonadota>Alphaproteobacteria>Hyphomicrobiales>Rhizobiaceae>Rhizobium/Agrobacterium_group | GCA_019799665.1 | Rhizobium_leguminosarum | HFN59_02345 | 481817..485542 | 384 | ATP-binding-protein-[Rhizobium-leguminosarum] |
| SCX35218.1 | Type3.MTaseII.containing.BREX | <-PIN+OST-HTH\|\|Relaxase+SF1-SF2-Helicase->Glyoxalase-><-BrxA-DNA-binding<-PglZ<-?<-SbcC<-N6-N4-METHYLASE<-DUF6079*<-BrxB-BrxF-iSTAND\|\|?-><-?<-?<-?<-?<-TIR<-SLATT<-TPR | 1168 | Bacteria>Pseudomonadota>Alphaproteobacteria>Hyphomicrobiales>Rhizobiaceae>Rhizobium/Agrobacterium_group | GCA_900100645.1 | Agrobacterium_rosae | DSM25559_4858 | 59277..62783 | 1972867 | hypothetical-protein-DSM25559_4858-[Agrobacterium-rosae] |
| WP_004111595.1 | Type3.MTaseII.containing.BREX | RNase->ACET+ACET-><-BrxA-DNA-binding<-PglZ<-TUDOR+SF1-SF2-Helicase+iREase+CTD<-N6-N4-METHYLASE<-MORC<-Schlafen+HATPase+wHTH<-DUF6079*<-BrxB-BrxF-iSTAND<-XIS-HTH<-DOC+HTH\|\|PhdYeFM_antitox->PIN->?->?->Phage_integrase-> | 1246 | Bacteria>Pseudomonadota>Alphaproteobacteria>Hyphomicrobiales>Rhizobiaceae>Rhizobium/Agrobacterium_group | GCF_000359745.1 | Rhizobium_freirei | RHSP_RS07185 | 17898..21638 | 1353277 | DUF6079-family-protein-[Rhizobium-freirei] |
| WP_244483902.1 | Type3.MTaseII.containing.BREX | <-ABhydrolase\|\|?->?-><-CHTH\|\|GDResolvase+PPP1-HTH->ParB->ParB->wHTH+WYL+WCX->BrxB-BrxF-iSTAND->DUF6079*->N6-N4-METHYLASE->TUDOR+SF1-SF2-Helicase+iREase+CTD->PglZ->BrxA-DNA-binding-><-?<-?<-?<-?<-Glyco_hydro_53+Phage-tail_3 | 1243 | Bacteria>Pseudomonadota>Alphaproteobacteria>Hyphomicrobiales>Rhizobiaceae>Rhizobium/Agrobacterium_group | GCF_001426565.1 | Rhizobium_sp._Root482 | ASD31_RS18785 | 44048..47779 | 1736543 | DUF6079-family-protein-[Rhizobium-sp.-Root482] |
| WP_245428179.1 | Type3.MTaseII.containing.BREX | BrxB-BrxF-iSTAND->DUF6079*->N6-N4-METHYLASE->N6-N4-METHYLASE->TUDOR+SF1-SF2-Helicase+iREase+CTD->PglZ->BrxA-DNA-binding-><-tRNA\|\|?->Thioredoxin-> | 1243 | Bacteria>Pseudomonadota>Alphaproteobacteria>Hyphomicrobiales>Rhizobiaceae>Rhizobium/Agrobacterium_group | GCF_002531715.1 | Rhizobium_phaseoli | CO651_RS19355 | 2091..5822 | 396 | DUF6079-family-protein-[Rhizobium-phaseoli] |
| WP_115156544.1 | Type3.MTaseII.containing.BREX | XIS-HTH->BrxB-BrxF-iSTAND->DUF6079*->OLD-ABC->TOPRIM->Schlafen+HATPase+wHTH->MORC->N6-N4-METHYLASE->TUDOR+SF1-SF2-Helicase+iREase+CTD->PglZ->BrxA-DNA-binding-><-NACHT+HEAT+REase | 1246 | Bacteria>Pseudomonadota>Alphaproteobacteria>Hyphomicrobiales>Rhizobiaceae>Rhizobium/Agrobacterium_group | GCF_004329915.1 | Rhizobium_leguminosarum | EOH49_RS18195 | 767..4507 | 384 | DUF6079-family-protein-[Rhizobium-leguminosarum] |
| WP_131702427.1 | Type3.MTaseII.containing.BREX | <-Polbeta+HEPN\|\|?->DOC+HTH->XIS-HTH->BrxB-BrxF-iSTAND->DUF6079*->OLD-ABC->TOPRIM->Schlafen+HATPase+wHTH->MORC->N6-N4-METHYLASE->TUDOR+SF1-SF2-Helicase+iREase+CTD->PglZ->BrxA-DNA-binding-><-NACHT+HEAT+REase | 1246 | Bacteria>Pseudomonadota>Alphaproteobacteria>Hyphomicrobiales>Rhizobiaceae>Rhizobium/Agrobacterium_group | GCF_004330525.1 | Rhizobium_leguminosarum | EOH64_RS12770 | 327665..331405 | 384 | DUF6079-family-protein-[Rhizobium-leguminosarum] |
| WP_236776373.1 | Type3.MTaseII.containing.BREX | <-HTH_3\|\|ArsR-HTH->?-><-SF1-SF2-Helicase<-OLD-ABC+TOPRIM<-McrC-NTD+REase<-McrB-AAA\|\|XIS-HTH->BrxB-BrxF-iSTAND->DUF6079*->Schlafen+HATPase+wHTH->MORC->N6-N4-METHYLASE->TUDOR+SF1-SF2-Helicase+iREase+CTD->PglZ->BrxA-DNA-binding-><-Relaxase+SF1-SF2-Helicase\|\|Polbeta+HEPN-> | 1248 | Bacteria>Pseudomonadota>Alphaproteobacteria>Hyphomicrobiales>Rhizobiaceae>Rhizobium/Agrobacterium_group | GCF_014200255.1 | Agrobacterium_tumefaciens | GGD52_RS24610 | 156089..159835 | 358 | DUF6079-family-protein-[Agrobacterium-tumefaciens] |
| WP_216330252.1 | Type3.MTaseII.containing.BREX | <-BrxA-DNA-binding<-PglZ<-TUDOR+SF1-SF2-Helicase+iREase+CTD<-N6-N4-METHYLASE<-MORC<-Schlafen+HATPase+wHTH<-DUF6079*<-BrxB-BrxF-iSTAND<-XIS-HTH<-RHH<-VirB10\|\|PhdYeFM_antitox->AAA-ATPase-><-DEDD_Tnp_IS110+Transposase_20\|\|?-><-LACI-HTH+PBPI | 1246 | Bacteria>Pseudomonadota>Alphaproteobacteria>Hyphomicrobiales>Rhizobiaceae>Rhizobium/Agrobacterium_group | GCF_018873185.1 | Rhizobium_sp._X9 | JZY08_RS11880 | 2438518..2442258 | 2815360 | DUF6079-family-protein-[Rhizobium-sp.-X9] |
| WP_164569833.1 | Type3.MTaseII.containing.BREX | <-MODE-HTH+PBPII\|\|CHTH-><-?<-HEPN<-?<-BrxA-DNA-binding<-PglZ<-TUDOR+SF1-SF2-Helicase+iREase+CTD<-N6-N4-METHYLASE<-DUF6079*<-BrxB-BrxF-iSTAND<-?<-SF1-SF2-Helicase<-MORC+DUF3684+REase<-PhdYeFM_antitox<-McrC-NTD+REase<-McrB-AAA<-REase+HTH+35exo+Small-B-barrel+SF1-SF2-Helicase<-?\|\|wHTH+WYL+WCX-> | 1249 | Bacteria>Pseudomonadota>Alphaproteobacteria>Hyphomicrobiales>Rhizobiaceae>Rhizobium/Agrobacterium_group | GCF_019798325.1 | Rhizobium_leguminosarum | HFN04_RS25070 | 142512..146261 | 384 | DUF6079-family-protein-[Rhizobium-leguminosarum] |
| WP_222285484.1 | Type3.MTaseII.containing.BREX | <-ADH_N+ADH_zinc_N\|\|AraC-HTH->GreA_GreB+SF1-SF2-Helicase->?->BrxB-BrxF-iSTAND->DUF6079*->N6-N4-METHYLASE->TUDOR+SF1-SF2-Helicase+iREase+CTD->PglZ->BrxA-DNA-binding->BirA-HTH->HIRAN->?->HisKin-><-REase | 1249 | Bacteria>Pseudomonadota>Alphaproteobacteria>Hyphomicrobiales>Rhizobiaceae>Rhizobium/Agrobacterium_group | GCF_019798555.1 | Rhizobium_leguminosarum | HFO88_RS26685 | 111734..115483 | 384 | DUF6079-family-protein-[Rhizobium-leguminosarum] |
| WP_261338694.1 | Type3.MTaseII.containing.BREX | <-BrxA-DNA-binding<-PglZ<-TUDOR+SF1-SF2-Helicase+iREase+CTD<-N6-N4-METHYLASE<-HEPN<-DUF6079*<-BrxB-BrxF-iSTAND<-?\|\|?-><-AbiEi<-URI-DNase<-REase+DUF5655 | 1243 | Bacteria>Pseudomonadota>Alphaproteobacteria>Hyphomicrobiales>Rhizobiaceae>Rhizobium/Agrobacterium_group | GCF_019799665.1 | Rhizobium_leguminosarum | HFN59_RS02305 | 481817..485548 | 384 | DUF6079-family-protein-[Rhizobium-leguminosarum] |
| WP_245581774.1 | Type3.MTaseII.containing.BREX | <-Thioredoxin<-?\|\|tRNA-><-BrxA-DNA-binding<-PglZ<-TUDOR+SF1-SF2-Helicase+iREase+CTD<-N6-N4-METHYLASE<-DUF6079*<-BrxB-BrxF-iSTAND<-?<-TetR-HTH | 1243 | Bacteria>Pseudomonadota>Alphaproteobacteria>Hyphomicrobiales>Rhizobiaceae>Rhizobium/Agrobacterium_group | - | Rhizobium_leguminosarum | ELG99_RS03805 | 766233..769964 | 384 | DUF6079-family-protein-[Rhizobium-leguminosarum] |
| WP_245497654.1 | Type3.MTaseII.containing.BREX | <-Thioredoxin<-?\|\|tRNA-><-BrxA-DNA-binding<-PglZ<-TUDOR+SF1-SF2-Helicase+iREase+CTD<-N6-N4-METHYLASE<-DUF6079*<-BrxB-BrxF-iSTAND<-?<-TetR-HTH | 1243 | Bacteria>Pseudomonadota>Alphaproteobacteria>Hyphomicrobiales>Rhizobiaceae>Rhizobium/Agrobacterium_group | - | Rhizobium_leguminosarum | ELH12_RS03800 | 765413..769144 | 384 | DUF6079-family-protein-[Rhizobium-leguminosarum] |
| WP_222282709.1 | Type3.MTaseII.containing.BREX | <-BrxA-DNA-binding<-PglZ<-TUDOR+SF1-SF2-Helicase+iREase+CTD<-N6-N4-METHYLASE<-HEPN<-DUF6079*<-BrxB-BrxF-iSTAND<-?\|\|?-><-AbiEi<-URI-DNase<-REase+DUF5655 | 1241 | Bacteria>Pseudomonadota>Alphaproteobacteria>Hyphomicrobiales>Rhizobiaceae>Rhizobium/Agrobacterium_group | - | Rhizobium_leguminosarum | HFN59_02345 | 481817..485542 | 384 | DUF6079-family-protein-[Rhizobium-leguminosarum] |
| WP_107026851.1 | Type3.MTaseII.containing.BREX | ParB->?->?->?->?-><-GDResolvase+HIN-HTH\|\|XIS-HTH->BrxB-BrxF-iSTAND->DUF6079*->N6-N4-METHYLASE->SbcC->?->TUDOR+SF1-SF2-Helicase+iREase+CTD->PglZ->BrxA-DNA-binding-> | 1256 | Bacteria>Pseudomonadota>Alphaproteobacteria>Hyphomicrobiales>Rhizobiaceae>Sinorhizobium/Ensifer_group | GCF_003024455.1 | Ensifer_sp._NM-2 | C6558_RS03465 | 697203..700973 | 2109730 | DUF6079-family-protein-[Ensifer-sp.-NM-2] |
| MBX3567502.1 | Type3.MTaseII.containing.BREX | <-ABhydrolase<-?<-GNTR-HTH\|\|?->JetB->SbcCD_C->DUF3323+TOPRIM->XIS-HTH->BrxB-BrxF-iSTAND->DUF6079*->Schlafen+HATPase+wHTH->MORC->N6-N4-METHYLASE->TUDOR+SF1-SF2-Helicase+iREase+CTD-p-> | 1246 | Bacteria>Pseudomonadota>Alphaproteobacteria>Hyphomicrobiales>Rhizobiaceae>unclassified_Rhizobiaceae | GCA_019635715.1 | Rhizobiaceae_bacterium | KF914_05550 | 63170..66910 | 1913961 | ATP-binding-protein-[Rhizobiaceae-bacterium] |
| MCC0042685.1 | Type3.MTaseII.containing.BREX | <-Schlafen+HATPase+wHTH<-DUF6079*<-BrxB-BrxF-iSTAND<-XIS-HTH\|\|MODE-HTH+PBPII-><-RHH<-VirB10<-VirB9<-VirB8<-VirB6<-TrbK | 1246 | Bacteria>Pseudomonadota>Alphaproteobacteria>Hyphomicrobiales>Rhodobiaceae>unclassified_Rhodobiaceae | GCA_020637975.1 | Rhodobiaceae_bacterium | H6880_10970 | 1838..5578 | 2026785 | ATP-binding-protein-[Rhodobiaceae-bacterium] |
| OSM08492.1 | Type3.MTaseII.containing.BREX | <-Y1_Tnp<-TPR<-SWIM\|\|wHTH+WYL+WCX->BrxB-BrxF-iSTAND->DUF6079*->Schlafen+HATPase+wHTH-><-?\|\|N6-N4-METHYLASE->TUDOR+SF1-SF2-Helicase+iREase+CTD->PglZ-><-Peptidase_G2 | 1249 | Bacteria>Pseudomonadota>Alphaproteobacteria>Magnetococcales>Magnetococcaceae>Magnetofaba | GCA_002109495.1 | Magnetofaba_australis_IT-1 | MAIT1_04664 | 7957..11706 | 1434232 | putative-ATPase,-VrlK-family-[Magnetofaba-australis-IT-1] |

| Protein-ID | System | Gene-Neighbourhoods | Length | Taxonomy | Assembly-ID | Strain | Locus-Tag | Genomic-coordinates | TaxID | Fasta-define |
|---|---|---|---|---|---|---|---|---|---|---|
| WP_085440117.1 | Type3.MTaseII.containing.BREX | <-Thioredoxin<-?<-Y1_Tnp<-TPR<-SWIM\|\|wHTH+WYL+WCX->BrxB-BrxF-iSTAND->DUF6079*->Schlafen+HATPase+wHTH->N6-N4-METHYLASE->TUDOR+SF1-SF2-Helicase+iREase+CTD->PglZ->BrxA-DNA-binding-><-Peptidase_G2 | 1243 | Bacteria>Pseudomonadota>Alphaproteobacteria>Magnetococcales>Magnetococcaceae>Magnetofaba | GCF_002109495.1 | Magnetofaba_australis | MAIT1_RS00700 | 7975..11706 | 1472297 | DUF6079-family-protein-[Magnetofaba-australis] |
| MBF0096963.1 | Type3.MTaseII.containing.BREX | <-TUDOR+SF1-SF2-Helicase+iREase+CTD<-?<-PIN<-SbcC<-Schlafen+HATPase+wHTH<-PHP+SbcC<-N6-N4-METHYLASE<-?<-?<-DUF6079*<-BrxB-BrxF-iSTAND\|\|SF1-SF2-Helicase->?->REase+HTH-> | 1243 | Bacteria>Pseudomonadota>Alphaproteobacteria>Magnetococcales>unclassified_Magnetococcales | GCA_015228825.1 | Magnetococcales_bacterium | HQM05_06450 | 17527..21258 | 2026759 | ATP-binding-protein-[Magnetococcales-bacterium] |
| MBF0133380.1 | Type3.MTaseII.containing.BREX | <-ROK-HTH\|\|REC+OmpR-HTH-><-?<-?\|\|BrxB-BrxF-iSTAND->DUF6079*->Schlafen+HATPase+wHTH->N6-N4-METHYLASE-> | 1250 | Bacteria>Pseudomonadota>Alphaproteobacteria>Magnetococcales>unclassified_Magnetococcales | GCA_015228925.1 | Magnetococcales_bacterium | HQL75_12430 | 13992..17744 | 2026759 | ATP-binding-protein-[Magnetococcales-bacterium] |
| MBF0181804.1 | Type3.MTaseII.containing.BREX | BrxB-BrxF-iSTAND->DUF6079*->Schlafen+HATPase+wHTH+HTH->N6-N4-METHYLASE->OLD-ABC->TOPRIM->TUDOR+SF1-SF2-Helicase+iREase+CTD->PglZ->BrxA-DNA-binding-><-SbcC | 1243 | Bacteria>Pseudomonadota>Alphaproteobacteria>Magnetococcales>unclassified_Magnetococcales | GCA_015231235.1 | Magnetococcales_bacterium | HQM03_17440 | 897..4628 | 2026759 | ATP-binding-protein-[Magnetococcales-bacterium] |
| MBF0213222.1 | Type3.MTaseII.containing.BREX | <-N6-N4-METHYLASE-p<-PIN<-SbcC<-N6-N4-METHYLASE<-PIN<-OLD-ABC<-DUF6079*<-BrxB-BrxF-iSTAND<-wHTH+WYL+WCX\|\|?->?-><-?<-?<-MoxR-AAA | 1246 | Bacteria>Pseudomonadota>Alphaproteobacteria>Magnetococcales>unclassified_Magnetococcales | GCA_015231295.1 | Magnetococcales_bacterium | HQM00_06620 | 7843..11583 | 2026759 | ATP-binding-protein-[Magnetococcales-bacterium] |
| MBF0308899.1 | Type3.MTaseII.containing.BREX | <-PglZ<-TUDOR+SF1-SF2-Helicase+iREase+CTD<-?<-OLD-ABC<-ParB<-N6-N4-METHYLASE<-Schlafen+HATPase+wHTH+HTH<-REase<-DUF6079*<-N6-N4-METHYLASE<-BrxB-BrxF-iSTAND<-wHTH+WYL+WCX\|\|glycosylase-><-rve_3\|\|?->?->SbcC-> | 1244 | Bacteria>Pseudomonadota>Alphaproteobacteria>Magnetococcales>unclassified_Magnetococcales | GCA_015231925.1 | Magnetococcales_bacterium | HQL56_05160 | 13287..17021 | 2026759 | ATP-binding-protein-[Magnetococcales-bacterium] |
| WP_028031177.1 | Type3.MTaseII.containing.BREX | GDResolvase+Recombinase+ZnR-><-?<-RADICAL-SAM<-?<-SWIM\|\|Arm-DNA-bind_3+Phage_int_SAM_3+Phage_integrase->wHTH+WYL+WCX->BrxB-BrxF-iSTAND->DUF6079-p*->TUDOR+SF1-SF2-Helicase+iREase+CTD->PglZ->BrxA-DNA-binding-> | 164 | Bacteria>Pseudomonadota>Alphaproteobacteria>Rhodobacterales>Paracoccaceae>Gemmobacter | GCF_000429765.1 | Gemmobacter_nectariphilus | G481_RS0117625 | 9893..10387 | 220343 | DUF6079-family-protein-[Gemmobacter-nectariphilus] |
| WP_211235232.1 | Type3.MTaseII.containing.BREX | <-HTH_Tnp_1<-PglZ-p<-TUDOR+SF1-SF2-Helicase+iREase+CTD<-N6-N4-METHYLASE<-?<-DUF6079*<-BrxB-BrxF-iSTAND<-wHTH+WYL+WCX | 1243 | Bacteria>Pseudomonadota>Alphaproteobacteria>Rhodobacterales>Paracoccaceae>Gemmobacter | GCF_000429765.1 | Gemmobacter_nectariphilus | G481_RS0117865 | 7055..10786 | 220343 | DUF6079-family-protein-[Gemmobacter-nectariphilus] |
| AUH66232.1 | Type3.MTaseII.containing.BREX | <-AbiGi<-GDResolvase+HIN-HTH\|\|?-><-BrxA-DNA-binding<-PglZ<-TUDOR+SF1-SF2-Helicase+iREase+CTD<-N6-N4-METHYLASE<-HEPN<-DUF6079*<-BrxB-BrxF-iSTAND<-XIS-HTH | 1241 | Bacteria>Pseudomonadota>Alphaproteobacteria>Rhodobacterales>Paracoccaceae>Paracoccus | GCA_002847445.1 | Paracoccus_zhejiangensis | CX676_07795 | 1579971..1583696 | 1077935 | ATPase-[Paracoccus-zhejiangensis] |
| TBN37292.1 | Type3.MTaseII.containing.BREX | Glyco_hydro_53+Phage-tail_3->?-><-BrxA-DNA-binding<-PglZ<-TUDOR+SF1-SF2-Helicase+iREase+CTD<-N6-N4-METHYLASE<-DUF6079*<-BrxB-BrxF-iSTAND<-wHTH+WYL+WCX<-Arm-DNA-bind_3+Phage_int_SAM_3+Phage_integrase\|\|SWIM-> | 1241 | Bacteria>Pseudomonadota>Alphaproteobacteria>Rhodobacterales>Paracoccaceae>Paracoccus | GCA_004310345.1 | Paracoccus_subflavus | EYE42_14755 | 52723..56448 | 2528244 | ATP-binding-protein-[Paracoccus-subflavus] |
| WP_198590304.1 | Type3.MTaseII.containing.BREX | HIN-HTH-><-AbiGi<-GDResolvase+HIN-HTH\|\|?-><-BrxA-DNA-binding<-PglZ<-TUDOR+SF1-SF2-Helicase+iREase+CTD<-N6-N4-METHYLASE<-HEPN<-DUF6079*<-BrxB-BrxF-iSTAND<-wHTH+WYL+WCX<-XIS-HTH | 1243 | Bacteria>Pseudomonadota>Alphaproteobacteria>Rhodobacterales>Paracoccaceae>Paracoccus | GCF_002847445.1 | Paracoccus_zhejiangensis | CX676_RS07800 | 1579971..1583702 | 1077935 | DUF6079-family-protein-[Paracoccus-zhejiangensis] |
| WP_198838502.1 | Type3.MTaseII.containing.BREX | TrbM->?-><-BrxA-DNA-binding<-PglZ<-TUDOR+SF1-SF2-Helicase+iREase+CTD<-N6-N4-METHYLASE<-DUF6079*<-BrxB-BrxF-iSTAND<-wHTH+WYL+WCX\|\|<-DOC+HTH\|\|<-?<-?<-Classical-AAA<-YTCD-HTH\|\|ADH_N+ADH_zinc_N_2-> | 1243 | Bacteria>Pseudomonadota>Alphaproteobacteria>Rhodobacterales>Paracoccaceae>Paracoccus | GCF_016405985.1 | Paracoccus_sp._IB05 | JC607_RS19250 | 38659..42390 | 2779367 | DUF6079-family-protein-[Paracoccus-sp.-IB05] |
| WP_211657955.1 | Type3.MTaseII.containing.BREX | Glyco_hydro_53+Phage-tail_3->?-><-BrxA-DNA-binding<-PglZ<-TUDOR+SF1-SF2-Helicase+iREase+CTD<-N6-N4-METHYLASE<-DUF6079*<-BrxB-BrxF-iSTAND<-wHTH+WYL+WCX<-Arm-DNA-bind_3+Phage_int_SAM_3+Phage_integrase\|\|SWIM-> | 1243 | Bacteria>Pseudomonadota>Alphaproteobacteria>Rhodobacterales>Paracoccaceae>Paracoccus | - | Paracoccus_subflavus | EYE42_RS14755 | 52723..56454 | 2528244 | DUF6079-family-protein-[Paracoccus-subflavus] |
| WP_217806501.1 | Type3.MTaseII.containing.BREX | HTH_3->Couple_hipA+HipA_C->?-><-SWIM<-BrxA-DNA-binding<-PglZ<-TUDOR+SF1-SF2-Helicase+iREase+CTD<-N6-N4-METHYLASE<-HEPN<-DUF6079*<-BrxB-BrxF-iSTAND<-wHTH+WYL+WCX\|\|ArdCN+MPTase->?->?->FurR-HTH->?->HIN-HTH+TraJ-RHH->TraI-VirD2-RCRE-> | 1243 | Bacteria>Pseudomonadota>Alphaproteobacteria>Rhodobacterales>Paracoccaceae>Planktotalea | GCF_002105285.1 | Planktotalea_arctica | B9P63_RS08325 | 222169..225900 | 1481893 | DUF6079-family-protein-[Planktotalea-arctica] |
| MCA3449867.1 | Type3.MTaseII.containing.BREX | XIS-HTH->wHTH+WYL+WCX->BrxB-BrxF-iSTAND->DUF6079*->Schlafen+HATPase+wHTH->N6-N4-METHYLASE->TUDOR+SF1-SF2-Helicase+iREase+CTD->PglZ->BrxA-DNA-binding-> | 1241 | Bacteria>Pseudomonadota>Alphaproteobacteria>Rhodobacterales>Paracoccaceae>Rhodobacter | GCA_020250135.1 | Rhodobacter_sp. | INF92_05750 | 5869..9594 | 1062 | ATP-binding-protein-[Rhodobacter-sp.] |
| WP_072244773.1 | Type3.MTaseII.containing.BREX | <-HIN-HTH+TraJ-RHH\|\|ParA-Soj-PloopNTPase->RHH->?->DOC+HTH-><-BrxA-DNA-binding<-PglZ<-TUDOR+SF1-SF2-Helicase+iREase+CTD<-N6-N4-METHYLASE<-DUF6079*<-BrxB-BrxF-iSTAND<-wHTH+WYL+WCX<-?<-?<-TOPRIM<-SAM-METHYLASE+SF1-SF2-Helicase | 1241 | Bacteria>Pseudomonadota>Alphaproteobacteria>Rhodobacterales>Paracoccaceae>Roseibaca | GCF_001517585.1 | Roseibaca_calidilacus | AWT76_RS02575 | 24099..27824 | 1666912 | DUF6079-family-protein-[Roseibaca-calidilacus] |
| WP_092891896.1 | Type3.MTaseII.containing.BREX | <-RNasePH<-?<-HIN-HTH\|\|<-BrxA-DNA-binding<-PglZ<-TUDOR+SF1-SF2-Helicase+iREase+CTD<-N6-N4-METHYLASE<-Schlafen+HATPase+wHTH<-DUF6079*<-BrxB-BrxF-iSTAND<-wHTH+WYL+WCX<-?<-?<-TOPRIM<-SAM-METHYLASE+SF1-SF2-Helicase | 1241 | Bacteria>Pseudomonadota>Alphaproteobacteria>Rhodobacterales>Paracoccaceae>Roseicitreum | GCF_014681765.1 | Roseicitreum_antarcticum | H9529_RS17210 | 41395..45120 | 564137 | DUF6079-family-protein-[Roseicitreum-antarcticum] |
| TGD60867.1 | Type3.MTaseII.containing.BREX | DnaK-><-BrxA-DNA-binding<-PglZ<-TUDOR+SF1-SF2-Helicase+iREase+CTD<-N6-N4-METHYLASE<-DUF6079*<-BrxB-BrxF-iSTAND | 1241 | Bacteria>Pseudomonadota>Alphaproteobacteria>Rhodobacterales>Paracoccaceae>Tabrizicola | GCA_004765575.1 | Tabrizicola_sp._WMC-M-20 | EYC08_19930 | 9961..13686 | 2528693 | ATP-binding-protein-[Tabrizicola-sp.-WMC-M-20] |
| WP_210114958.1 | Type3.MTaseII.containing.BREX | DnaK->?-><-BrxA-DNA-binding<-PglZ<-TUDOR+SF1-SF2-Helicase+iREase+CTD<-N6-N4-METHYLASE<-DUF6079*<-BrxB-BrxF-iSTAND<-wHTH+WYL+WCX | 1243 | Bacteria>Pseudomonadota>Alphaproteobacteria>Rhodobacterales>Paracoccaceae>Tabrizicola | - | Tabrizicola_sp._WMC-M-20 | EYC08_RS19940 | 9961..13692 | 2528693 | DUF6079-family-protein-[Tabrizicola-sp.-WMC-M-20] |
| KEP68194.1 | Type3.MTaseII.containing.BREX | <-HIN-HTH+TraJ-RHH\|\|ParA-Soj-PloopNTPase->RHH->?->DOC+HTH-><-BrxA-DNA-binding<-PglZ<-TUDOR+SF1-SF2-Helicase+iREase+CTD<-N6-N4-METHYLASE<-DUF6079*<-BrxB-BrxF-iSTAND<-wHTH+WYL+WCX<-?<-?<-TOPRIM | 1241 | Bacteria>Pseudomonadota>Alphaproteobacteria>Rhodobacterales>Paracoccaceae>Thioclava | GCA_000715505.1 | Thioclava_dalianensis | DL1_14480 | 15618..19343 | 1185766 | ATPase-[Thioclava-dalianensis] |
| OOY26688.1 | Type3.MTaseII.containing.BREX | CHTH->?-><-?\|\|SAM-METHYLASE+SF1-SF2-Helicase->TOPRIM->?->?->wHTH+WYL+WCX->BrxB-BrxF-iSTAND->DUF6079*->OLD-ABC->TOPRIM->N6-N4-METHYLASE->TUDOR+SF1-SF2-Helicase+iREase+CTD->PglZ->BrxA-DNA-binding-><-DOC+HTH<-RHH<-ParA-Soj-PloopNTPase | 1241 | Bacteria>Pseudomonadota>Alphaproteobacteria>Rhodobacterales>Paracoccaceae>Thioclava | GCA_002020275.1 | Thioclava_sp._L04-15 | BMI90_16680 | 37339..41064 | 1915318 | ATPase-[Thioclava-sp.-L04-15] |
| WP_210401976.1 | Type3.MTaseII.containing.BREX | CHTH->?-><-?\|\|SAM-METHYLASE+SF1-SF2-Helicase->TOPRIM->?->?->wHTH+WYL+WCX->BrxB-BrxF-iSTAND->DUF6079*->OLD-ABC->TOPRIM->N6-N4-METHYLASE->TUDOR+SF1-SF2-Helicase+iREase+CTD->PglZ->BrxA-DNA-binding-><-DOC+HTH<-?<-RHH | 1243 | Bacteria>Pseudomonadota>Alphaproteobacteria>Rhodobacterales>Paracoccaceae>Thioclava | GCF_002020275.1 | Thioclava_sp._L04-15 | BMI90_RS16610 | 37333..41064 | 1915318 | DUF6079-family-protein-[Thioclava-sp.-L04-15] |
| WP_211250644.1 | Type3.MTaseII.containing.BREX | <-HIN-HTH+TraJ-RHH\|\|ParA-Soj-PloopNTPase->RHH->?->DOC+HTH-><-BrxA-DNA-binding<-PglZ<-TUDOR+SF1-SF2-Helicase+iREase+CTD<-N6-N4-METHYLASE<-DUF6079*<-BrxB-BrxF-iSTAND<-wHTH+WYL+WCX<-?<-?<-TOPRIM | 1243 | Bacteria>Pseudomonadota>Alphaproteobacteria>Rhodobacterales>Paracoccaceae>Thioclava | GCF_900115035.1 | Thioclava_dalianensis | DL1_RS18500 | 15618..19349 | 1185766 | DUF6079-family-protein-[Thioclava-dalianensis] |
| WP_205321650.1 | Type3.MTaseII.containing.BREX | <-HIN-HTH+TraJ-RHH\|\|ParA-Soj-PloopNTPase->RHH->?->DOC+HTH-><-BrxA-DNA-binding<-PglZ<-TUDOR+SF1-SF2-Helicase+iREase+CTD<-N6-N4-METHYLASE<-DUF6079*<-BrxB-BrxF-iSTAND<-wHTH+WYL+WCX | 1243 | Bacteria>Pseudomonadota>Alphaproteobacteria>Rhodobacterales>Paracoccaceae>Tritonibacter | GCF_003347575.1 | Tritonibacter_mobilis | AOQ74_RS22550 | 81880..85611 | 379347 | DUF6079-family-protein-[Tritonibacter-mobilis] |
| MCC5993912.1 | Type3.MTaseII.containing.BREX | Peptidase-M15->?->?-><-DUF4236+TPR<-BrxA-DNA-binding<-PglZ<-TUDOR+SF1-SF2-Helicase+iREase+CTD<-N6-N4-METHYLASE<-?<-DUF6079*<-BrxB-BrxF-iSTAND<-wHTH+WYL+WCX<-RNase_T+35exo | 1243 | Bacteria>Pseudomonadota>Alphaproteobacteria>Rhodobacterales>Paracoccaceae>unclassified_Paracoccaceae | GCA_020832655.1 | Paracoccaceae_bacterium | JJT99_15470 | 35537..39268 | 1904441 | hypothetical-protein-[Paracoccaceae-bacterium] |
| MCR9110735.1 | Type3.MTaseII.containing.BREX | <-rve<-REase+wHTH<-?<-?\|\|?-><-BrxA-DNA-binding<-PglZ<-TUDOR+SF1-SF2-Helicase+iREase+CTD<-N6-N4-METHYLASE<-DUF6079-p* | 916 | Bacteria>Pseudomonadota>Alphaproteobacteria>Rhodobacterales>Paracoccaceae>unclassified_Paracoccaceae | GCA_024742895.1 | Paracoccaceae_bacterium | NXH94_17790 | 36972..39724 | 1904441 | DUF6079-family-protein-[Paracoccaceae-bacterium] |
| MCI5041378.1 | Type3.MTaseII.containing.BREX | RHH->?->DOC+HTH-><-?<-BrxA-DNA-binding<-PglZ<-TUDOR+SF1-SF2-Helicase+iREase+CTD<-N6-N4-METHYLASE<-DUF6079-p* | 737 | Bacteria>Pseudomonadota>Alphaproteobacteria>Rhodobacterales>Roseobacteraceae>Donghicola | GCA_022763195.1 | Donghicola_eburneus | MRY81_17050 | 11813..14026 | 393278 | DUF6079-family-protein-[Donghicola-eburneus] |
| AML53617.1 | Type3.MTaseII.containing.BREX | Couple_hipA+HipA_C->REase-><-?<-BrxA-DNA-binding<-PglZ<-TUDOR+SF1-SF2-Helicase+iREase+CTD<-?<-N6-N4-METHYLASE<-?<-DUF6079*<-BrxB-BrxF-iSTAND\|\|ArdCN+MPTase->?-><-?\|\|?->FurR-HTH->?->?->HIN-HTH+TraJ-RHH-> | 1241 | Bacteria>Pseudomonadota>Alphaproteobacteria>Rhodobacterales>Roseobacteraceae>Falsihalocynthiibacter | GCA_000812665.2 | Falsihalocynthiibacter_arcticus | RC74_09960 | 2004809..2008534 | 1579316 | ATPase-[Falsihalocynthiibacter-arcticus] |
| WP_218918124.1 | Type3.MTaseII.containing.BREX | Couple_hipA+HipA_C->REase-><-?<-BrxA-DNA-binding<-PglZ<-TUDOR+SF1-SF2-Helicase+iREase+CTD<-?<-N6-N4-METHYLASE<-?<-DUF6079*<-BrxB-BrxF-iSTAND<-wHTH+WYL+WCX\|\|ArdCN+MPTase->?-><-?\|\|?->FurR-HTH-> | 1243 | Bacteria>Pseudomonadota>Alphaproteobacteria>Rhodobacterales>Roseobacteraceae>Falsihalocynthiibacter | GCF_000812665.2 | Falsihalocynthiibacter_arcticus | RC74_RS09960 | 2004809..2008540 | 1579316 | DUF6079-family-protein-[Falsihalocynthiibacter-arcticus] |
| WP_264214692.1 | Type3.MTaseII.containing.BREX | SSB->?->REase->?->?->wHTH+WYL+WCX->BrxB-BrxF-iSTAND->DUF6079*->N6-N4-METHYLASE->TUDOR+SF1-SF2-Helicase+iREase+CTD->PglZ->BrxA-DNA-binding-><-RNasePH<-LZ_Tnp_IS66+zf-IS66+DDE_Tnp_IS66+DDE_Tnp_IS66_C<-?<-HIN-HTH<-DOC+HTH | 1241 | Bacteria>Pseudomonadota>Alphaproteobacteria>Rhodobacterales>Roseobacteraceae>Leisingera | GCF_025857195.1 | Leisingera_sp._BMJM1 | OKQ63_RS24995 | 24483..28208 | 2930199 | DUF6079-family-protein-[Leisingera-sp.-BMJM1] |
| TYB76568.1 | Type3.MTaseII.containing.BREX | CHTH->?-><-?\|\|SAM-METHYLASE+SF1-SF2-Helicase->TOPRIM->?->?->wHTH+WYL+WCX->BrxB-BrxF-iSTAND->DUF6079*->N6-N4-METHYLASE->TUDOR+SF1-SF2-Helicase+iREase+CTD->PglZ->BrxA-DNA-binding-><-DOC+HTH<-?<-RHH<-ParA-Soj-PloopNTPase\|\|HIN-HTH+TraJ-RHH-> | 1241 | Bacteria>Pseudomonadota>Alphaproteobacteria>Rhodobacterales>Roseobacteraceae>Muriiphilus | GCA_008086115.1 | Muriiphilus_fusiformis | FVF75_17215 | 37797..41522 | 2603819 | ATP-binding-protein-[Muriiphilus-fusiformis] |
| WP_222861238.1 | Type3.MTaseII.containing.BREX | CHTH->?-><-?\|\|SAM-METHYLASE+SF1-SF2-Helicase->TOPRIM->?->?->wHTH+WYL+WCX->BrxB-BrxF-iSTAND->DUF6079*->N6-N4-METHYLASE->TUDOR+SF1-SF2-Helicase+iREase+CTD->PglZ->BrxA-DNA-binding-><-DOC+HTH<-?<-RHH<-ParA-Soj-PloopNTPase\|\|HIN-HTH+TraJ-RHH-> | 1243 | Bacteria>Pseudomonadota>Alphaproteobacteria>Rhodobacterales>Roseobacteraceae>Muriiphilus | GCF_008086115.1 | Muriiphilus_fusiformis | FVF75_RS17210 | 37791..41522 | 2603819 | DUF6079-family-protein-[Muriiphilus-fusiformis] |
| WP_244691434.1 | Type3.MTaseII.containing.BREX | <-DOC+HTH<-HTH_3\|\|?-><-?\|\|TOPRIM->?->?->wHTH+WYL+WCX->BrxB-BrxF-iSTAND->DUF6079*->N6-N4-METHYLASE->TUDOR+SF1-SF2-Helicase+iREase+CTD-p->TUDOR+SF1-SF2-Helicase+iREase+CTD->PglZ-p->BrxA-DNA-binding-><-?<-RHH<-ParA-Soj-PloopNTPase\|\|HIN-HTH+TraJ-RHH-> | 1243 | Bacteria>Pseudomonadota>Alphaproteobacteria>Rhodobacterales>Roseobacteraceae>Phaeobacter | GCF_022867865.1 | Phaeobacter_sp._J2-8 | MUB49_RS21315 | 57708..61439 | 2931394 | DUF6079-family-protein-[Phaeobacter-sp.-J2-8] |
| GGE39036.1 | Type3.MTaseII.containing.BREX | <-DNA_methylase\|\|XIS-HTH->wHTH+WYL+WCX->BrxB-BrxF-iSTAND->DUF6079*->?->N6-N4-METHYLASE->TUDOR+SF1-SF2-Helicase+iREase+CTD->PglZ->BrxA-DNA-binding-><-?<-Glyco_hydro_53+Phage-tail_3 | 1241 | Bacteria>Pseudomonadota>Alphaproteobacteria>Rhodobacterales>Roseobacteraceae>Primorskyibacter | GCA_014638275.1 | Primorskyibacter_flagellatus | GCM10011360_28400 | 2984..6709 | 1387277 | hypothetical-protein-GCM10011360_28400-[Primorskyibacter-flagellatus] |
| WP_229737642.1 | Type3.MTaseII.containing.BREX | <-DNA_methylase\|\|XIS-HTH->wHTH+WYL+WCX->BrxB-BrxF-iSTAND->DUF6079*->?->N6-N4-METHYLASE->TUDOR+SF1-SF2-Helicase+iREase+CTD->PglZ->BrxA-DNA-binding-><-?<-Glyco_hydro_53+Phage-tail_3 | 1243 | Bacteria>Pseudomonadota>Alphaproteobacteria>Rhodobacterales>Roseobacteraceae>Primorskyibacter | GCF_014638275.1 | Primorskyibacter_flagellatus | IEV87_RS14250 | 2978..6709 | 1387277 | DUF6079-family-protein-[Primorskyibacter-flagellatus] |
| WP_109670728.1 | Type3.MTaseII.containing.BREX | <-HTH_3\|\|?-><-?\|\|SAM-METHYLASE+SF1-SF2-Helicase->TOPRIM->?->?->wHTH+WYL+WCX->BrxB-BrxF-iSTAND->DUF6079*->N6-N4-METHYLASE->TUDOR+SF1-SF2-Helicase+iREase+CTD->PglZ-p-> | 1241 | Bacteria>Pseudomonadota>Alphaproteobacteria>Rhodobacterales>Roseobacteraceae>Roseicyclus | GCF_003148775.1 | Roseicyclus_mahoneyensis | C7455_RS18355 | 66263..69988 | 164332 | DUF6079-family-protein-[Roseicyclus-mahoneyensis] |
| WP_210879912.1 | Type3.MTaseII.containing.BREX | ANK+ANK-><-?<-HHH_5+SF1-SF2-Helicase<-?\|\|RNase_T+35exo->wHTH+WYL+WCX->BrxB-BrxF-iSTAND->DUF6079*->HEPN->N6-N4-METHYLASE->TUDOR+SF1-SF2-Helicase+iREase+CTD->PglZ->BrxA-DNA-binding->?->DUF4236+TPR->?-><-MuB-ATPase | 1243 | Bacteria>Pseudomonadota>Alphaproteobacteria>Rhodobacterales>Roseobacteraceae>Roseovarius | GCF_018069895.1 | Roseovarius_autotrophicus | KAK09_RS15835 | 10081..13812 | 2824121 | DUF6079-family-protein-[Roseovarius-autotrophicus] |
| KIC47552.1 | Type3.MTaseII.containing.BREX | MODE-HTH+PBPII-><-MODE-HTH+PBPII\|\|?->?->?->?-><-ABhydrolase\|\|BrxB-BrxF-iSTAND->DUF6079*->N6-N4-METHYLASE->TUDOR+SF1-SF2-Helicase+iREase+CTD->PglZ->BrxA-DNA-binding-><-Phage_lambda<-XIS-HTH\|\|?-><-tRNA<-RNASE-EG | 1241 | Bacteria>Pseudomonadota>Alphaproteobacteria>Rhodobacterales>Roseobacteraceae>Ruegeria | GCA_000814005.1 | Ruegeria_sp._ANG-S4 | RA28_03065 | 590455..594180 | 1577904 | ATPase-[Ruegeria-sp.-ANG-S4] |
| NOD75957.1 | Type3.MTaseII.containing.BREX | DUF4096+DDE_Tnp_1+DDE_Tnp_1_2->XIS-HTH->HTH_36->Phage_lambda->REase-><-BrxA-DNA-binding<-PglZ<-TUDOR+SF1-SF2-Helicase+iREase+CTD<-N6-N4-METHYLASE<-DUF6079*<-BrxB-BrxF-iSTAND\|\|ABhydrolase-><-?\|\|?->TIR-><-?<-?<-?<-Aminotran_1_2 | 1241 | Bacteria>Pseudomonadota>Alphaproteobacteria>Rhodobacterales>Roseobacteraceae>Ruegeria | GCA_013110875.1 | Ruegeria_sp._HKCCD4332 | GS639_07235 | 417878..421603 | 2683021 | ATP-binding-protein-[Ruegeria-sp.-HKCCD4332] |
| WP_206539610.1 | Type3.MTaseII.containing.BREX | MODE-HTH+PBPII-><-MODE-HTH+PBPII\|\|?->?->?->?-><-ABhydrolase\|\|BrxB-BrxF-iSTAND->DUF6079*->N6-N4-METHYLASE->TUDOR+SF1-SF2-Helicase+iREase+CTD->PglZ->BrxA-DNA-binding-><-?<-Phage_lambda<-XIS-HTH\|\|?-><-Arm-DNA-bind_3+Phage_integrase | 1243 | Bacteria>Pseudomonadota>Alphaproteobacteria>Rhodobacterales>Roseobacteraceae>Ruegeria | GCF_000814005.1 | Ruegeria_sp._ANG-S4 | RA28_RS03060 | 590449..594180 | 1577904 | DUF6079-family-protein-[Ruegeria-sp.-ANG-S4] |
| WP_216601641.1 | Type3.MTaseII.containing.BREX | DUF4096+DDE_Tnp_1+DDE_Tnp_1_2->XIS-HTH->Phage_lambda->REase-><-BrxA-DNA-binding<-PglZ<-TUDOR+SF1-SF2-Helicase+iREase+CTD<-N6-N4-METHYLASE<-DUF6079*<-BrxB-BrxF-iSTAND\|\|ABhydrolase-><-?\|\|?->TIR-><-?<-?<-?<-Aminotran_1_2 | 1243 | Bacteria>Pseudomonadota>Alphaproteobacteria>Rhodobacterales>Roseobacteraceae>Ruegeria | GCF_013110875.1 | Ruegeria_sp._HKCCD4332 | GS639_RS07255 | 417878..421609 | 2683021 | DUF6079-family-protein-[Ruegeria-sp.-HKCCD4332] |
| WP_222458005.1 | Type3.MTaseII.containing.BREX | <-HIN-HTH<-?<-?<-MerR-HTH<-BrxA-DNA-binding<-PglZ<-TUDOR+SF1-SF2-Helicase+iREase+CTD<-N6-N4-METHYLASE<-HEPN<-DUF6079*<-BrxB-BrxF-iSTAND<-N6-METHYLASE<-DnaB<-?<-XIS-HTH<-?<-Arm-DNA-bind_3+Phage_integrase | 1243 | Bacteria>Pseudomonadota>Alphaproteobacteria>Rhodobacterales>Roseobacteraceae>Ruegeria | GCF_019801065.1 | Ruegeria_arenilitoris | KUW05_RS13435 | 236068..239799 | 1173585 | DUF6079-family-protein-[Ruegeria-arenilitoris] |
| WP_217650100.1 | Type3.MTaseII.containing.BREX | <-HTH_3\|\|?-><-?\|\|SAM-METHYLASE+SF1-SF2-Helicase->TOPRIM->?->?->wHTH+WYL+WCX->BrxB-BrxF-iSTAND->DUF6079*->N6-N4-METHYLASE->TUDOR+SF1-SF2-Helicase+iREase+CTD->PglZ->BrxA-DNA-binding-><-DOC+HTH<-?<-RHH<-ParA-Soj-PloopNTPase\|\|HIN-HTH+TraJ-RHH-> | 1243 | Bacteria>Pseudomonadota>Alphaproteobacteria>Rhodobacterales>Roseobacteraceae>Thalassobius | GCF_900110775.1 | Thalassobius_taeanensis | BSF79_RS17675 | 31913..35644 | 657014 | DUF6079-family-protein-[Thalassobius-taeanensis] |
| OYV51921.1 | Type3.MTaseII.containing.BREX | BrxB-BrxF-iSTAND->DUF6079*->Schlafen+HATPase+wHTH->N6-N4-METHYLASE->TUDOR+SF1-SF2-Helicase+iREase+CTD->PglZ-> | 1241 | Bacteria>Pseudomonadota>Alphaproteobacteria>Rhodospirillales>Acetobacteraceae>Acidocella | GCA_002255785.1 | Acidocella_sp._20-58-15 | B7Z77_03055 | 934..4659 | 1970302 | MAG:-ATPase-[Acidocella-sp.-20-58-15] |
| WP_183267040.1 | Type3.MTaseII.containing.BREX | TrbK->VirB6->VirB8->VirB9->VirB10->RHH-><-MODE-HTH+PBPII\|\|XIS-HTH->BrxB-BrxF-iSTAND->DUF6079*->Schlafen+HATPase+wHTH->MORC->N6-N4-METHYLASE->TUDOR+SF1-SF2-Helicase+iREase+CTD->PglZ->BrxA-DNA-binding-><-Arm-DNA-bind_3+Phage_int_SAM_3+Phage_integrase | 1246 | Bacteria>Pseudomonadota>Alphaproteobacteria>Rhodospirillales>Acetobacteraceae>Acidocella | GCF_014201825.1 | Acidocella_aromatica | HNP71_RS11360 | 77806..81546 | 1303579 | DUF6079-family-protein-[Acidocella-aromatica] |
| WP_227076189.1 | Type3.MTaseII.containing.BREX | DUF6079-p*->Schlafen+HATPase+wHTH->MORC->N6-N4-METHYLASE->TUDOR+SF1-SF2-Helicase+iREase+CTD->PglZ->BrxA-DNA-binding->Phage_AlpA-><-Arm-DNA-bind_3+Phage_integrase | 419 | Bacteria>Pseudomonadota>Alphaproteobacteria>Rhodospirillales>Acetobacteraceae>Acidocella | GCF_020567705.1 | Acidocella_sp._KAb_2-4 | LI953_RS11300 | 1..1261 | 2885158 | DUF6079-family-protein,-partial-[Acidocella-sp.-KAb-2-4] |
| WP_201727595.1 | Type3.MTaseII.containing.BREX | HTH+rve->AAA-ATPase->\|\|HIN-HTH->AAA-ATPase->\|\|<-BrxA-DNA-binding<-PglZ<-TUDOR+SF1-SF2-Helicase+iREase+CTD<-N6-N4-METHYLASE<-Schlafen+HATPase+wHTH<-DUF6079*<-BrxB-BrxF-iSTAND<-XIS-HTH\|\|Phage_AlpA-><-Arm-DNA-bind_3+Phage_integrase<-?<-ABhydrolase<-NUDIX<-PDZ+Peptidase | 1229 | Bacteria>Pseudomonadota>Alphaproteobacteria>Rhodospirillales>Acetobacteraceae>Acidocella | GCF_907164655.1 | Acidocella_sp._C78 | ACIDOCELLA78_RS32375 | 302326..306015 | 1671486 | DUF6079-family-protein-[Acidocella-sp.-C78] |
| SDB48716.1 | Type3.MTaseII.containing.BREX | <-Peptidase-M15<-?<-?<-?\|\|?->RAMA->GDResolvase+ZnR->?->DUF6079*->N6-N4-METHYLASE->TUDOR+SF1-SF2-Helicase+iREase+CTD->PglZ->BrxA-DNA-binding-p-><-tRNA\|\|?-><-REC<-REC | 1292 | Bacteria>Pseudomonadota>Alphaproteobacteria>Rhodospirillales>Acetobacteraceae>Belnapia | GCA_900104205.1 | Belnapia_rosea | SAMN02927895_01852 | 392011..395889 | 938405 | hypothetical-protein-SAMN02927895_01852-[Belnapia-rosea] |
| WP_218127383.1 | Type3.MTaseII.containing.BREX | <-Peptidase-M15<-?<-?<-?\|\|RAMA->GDResolvase+ZnR->?->PhdYeFM_antitox->BrxB-BrxF-iSTAND->DUF6079*->?->N6-N4-METHYLASE->TUDOR+SF1-SF2-Helicase+iREase+CTD->PglZ->BrxA-DNA-binding-><-tRNA\|\|?-><-REC<-REC | 1243 | Bacteria>Pseudomonadota>Alphaproteobacteria>Rhodospirillales>Acetobacteraceae>Belnapia | GCF_900104205.1 | Belnapia_rosea | BLR87_RS09110 | 392158..395889 | 938405 | DUF6079-family-protein-[Belnapia-rosea] |
| QDD97346.1 | Type3.MTaseII.containing.BREX | <-DDE_Tnp_ISL3<-Phage_integrase+Phage_integrase\|\|ParB+HNH-><-BrxA-DNA-binding<-PglZ<-TUDOR+SF1-SF2-Helicase+iREase+CTD<-N6-N4-METHYLASE<-Schlafen+HATPase+wHTH<-DUF6079*<-BrxB-BrxF-iSTAND<-PhdYeFM_antitox\|\|ParA-Soj-PloopNTPase->?-><-?\|\|?->?-><-REase | 1243 | Bacteria>Pseudomonadota>Alphaproteobacteria>Rhodospirillales>Acetobacteraceae>Roseomonas | GCA_006384895.1 | Roseomonas_mucosa | ADP8_05163 | 120547..124278 | 207340 | vrlK-[Roseomonas-mucosa] |
| KAA2211340.1 | Type3.MTaseII.containing.BREX | PhdYeFM_antitox->BrxB-BrxF-iSTAND->DUF6079*->Schlafen+HATPase+wHTH->N6-N4-METHYLASE->TUDOR+SF1-SF2-Helicase+iREase+CTD->PglZ->BrxA-DNA-binding-> | 1241 | Bacteria>Pseudomonadota>Alphaproteobacteria>Rhodospirillales>Acetobacteraceae>Roseomonas | GCA_008386565.1 | Roseomonas_oryzae | FOQ34_20610 | 1073..4798 | 1608942 | ATP-binding-protein-[Roseomonas-oryzae] |
| MCA3394955.1 | Type3.MTaseII.containing.BREX | <-BrxA-DNA-binding<-PglZ<-TUDOR+SF1-SF2-Helicase+iREase+CTD<-N6-N4-METHYLASE<-Schlafen+HATPase+wHTH<-DUF6079*<-BrxB-BrxF-iSTAND<-wHTH+WYL+WCX\|\|ArdCN+MPTase->ParB-> | 1241 | Bacteria>Pseudomonadota>Alphaproteobacteria>Rhodospirillales>Acetobacteraceae>Roseomonas | GCA_020249805.1 | Roseomonas_sp. | INF85_04840 | 23348..27073 | 1869190 | ATP-binding-protein-[Roseomonas-sp.] |
| WP_206668113.1 | Type3.MTaseII.containing.BREX | PhdYeFM_antitox->BrxB-BrxF-iSTAND->DUF6079*->Schlafen+HATPase+wHTH->N6-N4-METHYLASE->TUDOR+SF1-SF2-Helicase+iREase+CTD->PglZ->BrxA-DNA-binding-> | 1243 | Bacteria>Pseudomonadota>Alphaproteobacteria>Rhodospirillales>Acetobacteraceae>Roseomonas | GCF_008386565.1 | Roseomonas_oryzae | FOQ34_RS20615 | 1067..4798 | 1608942 | DUF6079-family-protein-[Roseomonas-oryzae] |
| AIB12890.1 | Type3.MTaseII.containing.BREX | ABC-ATPase+SbcC->?->Phage_integrase-><-PglZ<-TUDOR+SF1-SF2-Helicase+iREase+CTD<-N6-N4-METHYLASE<-HEPN<-DUF6079*<-wHTH+WYL+WCX<-?\|\|?->?->?-><-ZnR+TOPRIM+AAA-helicase+wHTH<-XIS-HTH<-MORC+DUF3684+DUF3883<-MAD-NTD+REase | 1241 | Bacteria>Pseudomonadota>Alphaproteobacteria>Rhodospirillales>Azospirillaceae>Azospirillum | GCA_000632475.2 | Azospirillum_argentinense | ABAZ39_13005 | 2719058..2722783 | 2970906 | ATPase-[Azospirillum-argentinense] |
| WP_200885622.1 | Type3.MTaseII.containing.BREX | ABC-ATPase+SbcC-><-?\|\|?->Phage_integrase-><-BrxA-DNA-binding<-PglZ<-TUDOR+SF1-SF2-Helicase+iREase+CTD<-N6-N4-METHYLASE<-HEPN<-DUF6079*<-BrxB-BrxF-iSTAND<-wHTH+WYL+WCX\|\|<-?\|\|?->?->?-><-ZnR+TOPRIM+AAA-helicase+wHTH<-XIS-HTH | 1243 | Bacteria>Pseudomonadota>Alphaproteobacteria>Rhodospirillales>Azospirillaceae>Azospirillum | GCF_000632475.1 | Azospirillum_argentinense | ABAZ39_RS12735 | 2719058..2722789 | 2970906 | DUF6079-family-protein-[Azospirillum-argentinense] |
| WP_264656683.1 | Type3.MTaseII.containing.BREX | Phage_int_SAM_5+Phage_integrase->?-><-?<-?<-Phage_integrase\|\|BrxB-BrxF-iSTAND->DUF6079*->N6-N4-METHYLASE->TUDOR+SF1-SF2-Helicase+iREase+CTD->PglZ->BrxA-DNA-binding->?->Phage_integrase-> | 1251 | Bacteria>Pseudomonadota>Alphaproteobacteria>Rhodospirillales>Azospirillaceae>Azospirillum | GCF_025961225.1 | Azospirillum_canadense | M2351_RS14165 | 2976535..2980290 | 403962 | DUF6079-family-protein-[Azospirillum-canadense] |
| MCB1832525.1 | Type3.MTaseII.containing.BREX | <-BrxA-DNA-binding<-PglZ<-TUDOR+SF1-SF2-Helicase+iREase+CTD<-N6-N4-METHYLASE<-MORC<-Schlafen+HATPase+wHTH<-TOPRIM<-OLD-ABC<-DUF6079-p* | 671 | Bacteria>Pseudomonadota>Alphaproteobacteria>Rhodospirillales>Geminicoccaceae>unclassified_Geminicoccaceae | GCA_020445435.1 | Geminicoccaceae_bacterium | KDH19_03625 | 19424..21441 | 2448052 | hypothetical-protein-[Geminicoccaceae-bacterium] |
| WP_173978271.1 | Type3.MTaseII.containing.BREX | <-XIS-HTH<-?<-?\|\|HTH->MPTase->BrxB-BrxF-iSTAND->DUF6079*->N6-N4-METHYLASE->?->TUDOR+SF1-SF2-Helicase+iREase+CTD->wHTH+AbiEi_2->Polbeta->PglZ->BrxA-DNA-binding-> | 1243 | Bacteria>Pseudomonadota>Alphaproteobacteria>Rhodospirillales>Rhodospirillaceae>Magnetospirillum | GCF_902729435.1 | Magnetospirillum_sp._UT-4 | MTBUT4_RS03465 | 59855..63586 | 2681467 | DUF6079-family-protein-[Magnetospirillum-sp.-UT-4] |
| WP_221238514.1 | Type3.MTaseII.containing.BREX | Recombinase+ZnR->Arm-DNA-bind_3+Phage_int_SAM_3+Phage_integrase->wHTH+WYL+WCX->BrxB-BrxF-iSTAND->DUF6079*->HEPN->N6-N4-METHYLASE->TUDOR+SF1-SF2-Helicase+iREase+CTD->PglZ->BrxA-DNA-binding-><-?<-Peptidase-M15 | 1257 | Bacteria>Pseudomonadota>Alphaproteobacteria>Rhodospirillales>Rhodospirillaceae>Roseospira | GCF_014197915.1 | Roseospira_visakhapatnamensis | GGD89_RS15995 | 4314..8087 | 390880 | DUF6079-family-protein-[Roseospira-visakhapatnamensis] |
| MXY42112.1 | Type3.MTaseII.containing.BREX | SIR2->?->?->BrxB-BrxF-iSTAND->DUF6079*->Schlafen+HATPase+wHTH->N6-N4-METHYLASE->PglZ->BrxA-DNA-binding-><-?<-RADICAL-SAM | 1242 | Bacteria>Pseudomonadota>Alphaproteobacteria>Rhodospirillales>Rhodospirillaceae>unclassified_Rhodospirillaceae | GCA_009838215.1 | Rhodospirillaceae_bacterium | F4Y62_20200 | 46242..49970 | 1898112 | ATP-binding-protein-[Rhodospirillaceae-bacterium] |
| MYK58411.1 | Type3.MTaseII.containing.BREX | DUF6079-p*->DUF6079-p->Schlafen+HATPase+wHTH->N6-N4-METHYLASE->PglZ->BrxA-DNA-binding-><-?<-RADICAL-SAM | 699 | Bacteria>Pseudomonadota>Alphaproteobacteria>Rhodospirillales>Rhodospirillaceae>unclassified_Rhodospirillaceae | GCA_009846065.1 | Rhodospirillaceae_bacterium | F4027_07305 | 74..2173 | 1898112 | hypothetical-protein-[Rhodospirillaceae-bacterium] |
| MYK58412.1 | Type3.MTaseII.containing.BREX | DUF6079-p->DUF6079-p*->Schlafen+HATPase+wHTH->N6-N4-METHYLASE->PglZ->BrxA-DNA-binding-><-?<-RADICAL-SAM | 461 | Bacteria>Pseudomonadota>Alphaproteobacteria>Rhodospirillales>Rhodospirillaceae>unclassified_Rhodospirillaceae | GCA_009846065.1 | Rhodospirillaceae_bacterium | F4027_07310 | 2115..3500 | 1898112 | hypothetical-protein-[Rhodospirillaceae-bacterium] |
| QJE74326.1 | Type3.MTaseII.containing.BREX | <-Thioredoxin\|\|tRNA->?->DUF6079-p->DUF6079-p*-><-?\|\|N6-N4-METHYLASE-p-><-HNH\|\|?-><-SF1-SF2-Helicase<-SF1-SF2-Helicase<-?<-Thioredoxin<-REC+NtrC-AAA+FIS-HTH | 196 | Bacteria>Pseudomonadota>Alphaproteobacteria>Rhodospirillales>Rhodospirillaceae>unclassified_Rhodospirillaceae | GCA_012913305.1 | Rhodospirillaceae_bacterium_B3 | HHL28_15700 | 3205701..3206291 | 2728875 | hypothetical-protein-HHL28_15700-[Rhodospirillaceae-bacterium-B3] |
| MBK8211737.1 | Type3.MTaseII.containing.BREX | <-BrxA-DNA-binding<-PglZ-p<-PglZ-p<-PglZ-p<-HATPase+wHTH<-Schlafen+HATPase+wHTH-p<-DUF6079-p<-?<-DUF6079-p*<-BrxB-BrxF-iSTAND<-XIS-HTH<-Arm-DNA-bind_3+Phage_int_SAM_3+Phage_integrase | 561 | Bacteria>Pseudomonadota>Alphaproteobacteria>Rhodospirillales>unclassified_Rhodospirillales | GCA_016712375.1 | Rhodospirillales_bacterium | IPK78_19080 | 31822..33507 | 2026786 | hypothetical-protein-[Rhodospirillales-bacterium] |
| MBK8211738.1 | Type3.MTaseII.containing.BREX | <-BrxA-DNA-binding<-PglZ-p<-PglZ-p<-PglZ-p<-HATPase+wHTH<-Schlafen+HATPase+wHTH-p<-DUF6079-p<-?<-DUF6079-p*<-BrxB-BrxF-iSTAND<-XIS-HTH<-Arm-DNA-bind_3+Phage_int_SAM_3+Phage_integrase | 259 | Bacteria>Pseudomonadota>Alphaproteobacteria>Rhodospirillales>unclassified_Rhodospirillales | GCA_016712375.1 | Rhodospirillales_bacterium | IPK78_19085 | 33576..34355 | 2026786 | hypothetical-protein-[Rhodospirillales-bacterium] |
| MBO6522093.1 | Type3.MTaseII.containing.BREX | ABC-ATPase+SbcC->?->Arm-DNA-bind_3+Phage_int_SAM_3+Phage_integrase->HTH_3->GDResolvase+PPP1-HTH->ParB->ParB->wHTH+WYL+WCX->BrxB-BrxF-iSTAND->DUF6079*->N6-N4-METHYLASE->TUDOR+SF1-SF2-Helicase+iREase+CTD->PglZ->BrxA-DNA-binding-><-?<-Peptidase | 1243 | Bacteria>Pseudomonadota>Alphaproteobacteria>Rhodospirillales>unclassified_Rhodospirillales | GCA_017640505.1 | Rhodospirillales_bacterium | JJ973_18765 | 226898..230629 | 2026786 | hypothetical-protein-[Rhodospirillales-bacterium] |
| WP_109907991.1 | Type3.MTaseII.containing.BREX | TPR->?->?->?-><-HIN-HTH<-DUF772+DDE_Tnp_1+DDE_Tnp_1_6\|\|XIS-HTH->BrxB-BrxF-iSTAND->DUF6079*->Schlafen+HATPase+wHTH->MORC->N6-N4-METHYLASE->OLD-ABC->REase->TUDOR+SF1-SF2-Helicase+iREase+CTD->PglZ->BrxA-DNA-binding-> | 1246 | Bacteria>Pseudomonadota>Alphaproteobacteria>Rhodospirillales>Zavarziniaceae>Zavarzinia | GCF_003173035.1 | Zavarzinia_aquatilis | DKG74_RS20205 | 21531..25271 | 2211142 | DUF6079-family-protein-[Zavarzinia-aquatilis] |
| USA60134.1 | Type3.MTaseII.containing.BREX | Thioredoxin->TraH->?-><-?<-SF1-SF2-Helicase<-BrxA-DNA-binding<-PglZ<-TUDOR+SF1-SF2-Helicase+iREase+CTD<-N6-N4-METHYLASE<-DUF6079*<-BrxB-BrxF-iSTAND<-wHTH+WYL+WCX\|\|LexA-protease->OBFOLD-NUCLEASE->?->?->?-><-MobF<-VirD4-TraG+VirD4-FtsK | 1243 | Bacteria>Pseudomonadota>Alphaproteobacteria>Sphingomonadales>Erythrobacteraceae>Qipengyuania | GCA_023702755.1 | Qipengyuania_citrea | NCF85_08350 | 1752461..1756192 | 225971 | DUF6079-family-protein-[Qipengyuania-citrea] |
| AOR78692.1 | Type3.MTaseII.containing.BREX | LZ_Tnp_IS66+zf-IS66+DDE_Tnp_IS66+DDE_Tnp_IS66_C-><-BrxA-DNA-binding<-PglZ<-TUDOR+SF1-SF2-Helicase+iREase+CTD<-N6-N4-METHYLASE<-DUF6079*<-BrxB-BrxF-iSTAND<-wHTH+WYL+WCX<-MODE-HTH+PBPII\|\|TetR-HTH-><-RHH<-VirB10<-VirB9<-VirB8<-VirB6 | 1241 | Bacteria>Pseudomonadota>Alphaproteobacteria>Sphingomonadales>Sphingomonadaceae>Novosphingobium | GCA_001742225.1 | Novosphingobium_resinovorum | BES08_14240 | 3028122..3031847 | 158500 | ATPase-[Novosphingobium-resinovorum] |
| WP_197524389.1 | Type3.MTaseII.containing.BREX | LZ_Tnp_IS66+zf-IS66+DDE_Tnp_IS66+DDE_Tnp_IS66_C-><-BrxA-DNA-binding<-PglZ<-TUDOR+SF1-SF2-Helicase+iREase+CTD<-N6-N4-METHYLASE<-DUF6079*<-BrxB-BrxF-iSTAND<-wHTH+WYL+WCX<-MODE-HTH+PBPII\|\|TetR-HTH-><-RHH<-VirB10<-VirB9<-VirB8<-VirB6 | 1243 | Bacteria>Pseudomonadota>Alphaproteobacteria>Sphingomonadales>Sphingomonadaceae>Novosphingobium | GCF_001742225.1 | Novosphingobium_resinovorum | BES08_RS14240 | 3028122..3031853 | 158500 | DUF6079-family-protein-[Novosphingobium-resinovorum] |
| WP_218033304.1 | Type3.MTaseII.containing.BREX | VirD4-TraG+VirD4-FtsK->MobF->?-><-?<-?<-OBFOLD-NUCLEASE<-LexA-protease\|\|wHTH+WYL+WCX->BrxB-BrxF-iSTAND->DUF6079*->N6-N4-METHYLASE->HTH_Tnp_1+HTH_28+HTH_32+rve+rve_3->SF1-SF2-Helicase->?-><-?<-TraH<-Thioredoxin<-TraN | 1243 | Bacteria>Pseudomonadota>Alphaproteobacteria>Sphingomonadales>Sphingomonadaceae>Novosphingobium | GCF_007991615.1 | Novosphingobium_sediminis | NSE01_RS18405 | 28027..31758 | 707214 | DUF6079-family-protein-[Novosphingobium-sediminis] |
| WP_201739862.1 | Type3.MTaseII.containing.BREX | VirD4-TraG+VirD4-FtsK->MobF->?-><-?<-?<-OBFOLD-NUCLEASE<-LexA-protease\|\|wHTH+WYL+WCX->BrxB-BrxF-iSTAND->DUF6079*->N6-N4-METHYLASE->TUDOR+SF1-SF2-Helicase+iREase+CTD->PglZ->BrxA-DNA-binding->SF1-SF2-Helicase->HIN-HTH-> | 1243 | Bacteria>Pseudomonadota>Alphaproteobacteria>Sphingomonadales>Sphingomonadaceae>Novosphingobium | GCF_027340905.1 | Novosphingobium_meiothermophilum | DKM43_RS12350 | 140057..143788 | 2202251 | DUF6079-family-protein-[Novosphingobium-meiothermophilum] |
| WP_021225977.1 | Type3.MTaseII.containing.BREX | <-BrxA-DNA-binding<-PglZ<-TUDOR+SF1-SF2-Helicase+iREase+CTD<-?<-SbcC<-N6-N4-METHYLASE<-DUF6079*<-BrxB-BrxF-iSTAND<-XIS-HTH<-?<-AraC-HTH+AraC-HTH\|\|?->Thermonuclease->PAPS-reduct->?->ParB-> | 1256 | Bacteria>Pseudomonadota>Alphaproteobacteria>Sphingomonadales>Sphingomonadaceae>Sphingobium | GCF_000445105.1 | Sphingobium_lactosutens | RLDS_RS39780 | 16923..20693 | 522773 | DUF6079-family-protein-[Sphingobium-lactosutens] |
| WP_066554193.1 | Type3.MTaseII.containing.BREX | VirD4-TraG+VirD4-FtsK->MobF->?-><-?<-?<-OBFOLD-NUCLEASE<-LexA-protease\|\|wHTH+WYL+WCX->BrxB-BrxF-iSTAND->DUF6079-p*->HIN-HTH->AAA-ATPase->?->PglZ->BrxA-DNA-binding->SF1-SF2-Helicase->?-><-?<-TraH | 418 | Bacteria>Pseudomonadota>Alphaproteobacteria>Sphingomonadales>Sphingomonadaceae>Sphingobium | GCF_001563265.1 | Sphingobium_sp._TKS | K426_RS04270 | 835830..837086 | 1315974 | DUF6079-family-protein-[Sphingobium-sp.-TKS] |
| MBZ6384174.1 | Type3.MTaseII.containing.BREX | PIN->DNA_METHYLASE->Phage_integrase+Phage_integrase->?-><-?\|\|?->URI-DNase->?->BrxB-BrxF-iSTAND->DUF6079*-> | 1032 | Bacteria>Pseudomonadota>Alphaproteobacteria>Sphingomonadales>Sphingomonadaceae>Sphingomonas | GCA_020093105.1 | Sphingomonas_sanguinis | LBE63_20795 | 6901..9999 | 33051 | DUF6079-family-protein-[Sphingomonas-sanguinis] |
| MCH4892413.1 | Type3.MTaseII.containing.BREX | VirD4-TraG+VirD4-FtsK->MobF->?-><-?<-?<-OBFOLD-NUCLEASE<-LexA-protease\|\|wHTH+WYL+WCX->BrxB-BrxF-iSTAND->DUF6079*->N6-N4-METHYLASE->TUDOR+SF1-SF2-Helicase+iREase+CTD->PglZ->BrxA-DNA-binding->SF1-SF2-Helicase->?-><-?<-TraH<-Thioredoxin | 1241 | Bacteria>Pseudomonadota>Alphaproteobacteria>Sphingomonadales>Sphingomonadaceae>Sphingomonas | GCA_022496715.1 | Sphingomonas_sp._SFZ2018-12 | GO308_04725 | 1025911..1029636 | 2683197 | ATP-binding-protein-[Sphingomonas-sp.-SFZ2018-12] |
| WP_168134696.1 | Type3.MTaseII.containing.BREX | SbcC->?->?->?->MORC+DNA_gyrase+TOPRIM-><-BrxA-DNA-binding<-PglZ<-TUDOR+SF1-SF2-Helicase+iREase+CTD<-N6-N4-METHYLASE<-DUF6079*<-BrxB-BrxF-iSTAND<-?<-HEPN | 1248 | Bacteria>Pseudomonadota>Alphaproteobacteria>Sphingomonadales>Sphingomonadaceae>Sphingomonas | GCF_012035195.1 | Sphingomonas_corticis | HBH26_RS11225 | 79093..82839 | 2722791 | DUF6079-family-protein-[Sphingomonas-corticis] |
| WP_170172272.1 | Type3.MTaseII.containing.BREX | <-DUF6079-p*<-BrxB-BrxF-iSTAND<-?<-URI-DNase<-?\|\|?-><-?<-Phage_integrase+Phage_integrase<-DNA_METHYLASE<-PIN | 980 | Bacteria>Pseudomonadota>Alphaproteobacteria>Sphingomonadales>Sphingomonadaceae>Sphingomonas | GCF_013002765.1 | Sphingomonas_sanguinis | HKX05_RS20025 | 1..2942 | 33051 | DUF6079-family-protein,-partial-[Sphingomonas-sanguinis] |
| WP_240884137.1 | Type3.MTaseII.containing.BREX | VirD4-TraG+VirD4-FtsK->MobF->?-><-?<-?<-OBFOLD-NUCLEASE<-LexA-protease\|\|wHTH+WYL+WCX->BrxB-BrxF-iSTAND->DUF6079*->N6-N4-METHYLASE->TUDOR+SF1-SF2-Helicase+iREase+CTD->PglZ->BrxA-DNA-binding->SF1-SF2-Helicase-><-?<-TraH<-Thioredoxin<-TraN | 1243 | Bacteria>Pseudomonadota>Alphaproteobacteria>Sphingomonadales>Sphingomonadaceae>Sphingomonas | GCF_022496715.1 | Sphingomonas_sp._SFZ2018-12 | GO308_RS04755 | 1025905..1029636 | 2683197 | DUF6079-family-protein-[Sphingomonas-sp.-SFZ2018-12] |
| AMG75218.1 | Type3.MTaseII.containing.BREX | LZ_Tnp_IS66+zf-IS66+DDE_Tnp_IS66+DDE_Tnp_IS66_C->rve->AAA-ATPase->?-><-N6-N4-METHYLASE<-N6-N4-METHYLASE\|\|HIN-HTH->HTH_Tnp_1+HTH_21+rve+rve_3-><-N6-N4-METHYLASE<-DUF6079*<-BrxB-BrxF-iSTAND<-wHTH+WYL+WCX\|\|LexA-protease->?->?-><-MobF<-VirD4-TraG+VirD4-FtsK<-TraJ-RHH | 1241 | Bacteria>Pseudomonadota>Alphaproteobacteria>Sphingomonadales>Sphingomonadaceae>Sphingopyxis | GCA_001559015.1 | Sphingopyxis_granuli | SGRAN_2870 | 3113278..3117003 | 267128 | ATPase-[Sphingopyxis-granuli] |
| AMU96668.1 | Type3.MTaseII.containing.BREX | TraN->Thioredoxin->TraH->?-><-SF1-SF2-Helicase<-BrxA-DNA-binding<-PglZ<-TUDOR+SF1-SF2-Helicase+iREase+CTD<-N6-N4-METHYLASE<-DUF6079*<-BrxB-BrxF-iSTAND<-wHTH+WYL+WCX<-HTH\|\|LexA-protease-><-?\|\|?-><-?<-MobF<-VirD4-TraG+VirD4-FtsK | 1241 | Bacteria>Pseudomonadota>Alphaproteobacteria>Sphingomonadales>Sphingomonadaceae>Sphingopyxis | GCA_001610975.1 | Sphingopyxis_terrae_subsp._terrae_NBRC_15098 | AOA14_18880 | 3958090..3961815 | 1219058 | ATPase-[Sphingopyxis-terrae-subsp.-terrae-NBRC-15098] |
| AMU88457.1 | Type3.MTaseII.containing.BREX | TraJ-RHH->VirD4-TraG+VirD4-FtsK->MobF->?-><-?<-?<-LexA-protease\|\|wHTH+WYL+WCX->DUF6079*->HEPN->N6-N4-METHYLASE->TUDOR+SF1-SF2-Helicase+iREase+CTD->PglZ->BrxA-DNA-binding->SF1-SF2-Helicase-><-DDE_Tnp_IS66+DDE_Tnp_IS66_C<-LZ_Tnp_IS66+zf-IS66+DDE_Tnp_IS66+DDE_Tnp_IS66_C | 1241 | Bacteria>Pseudomonadota>Alphaproteobacteria>Sphingomonadales>Sphingomonadaceae>Sphingopyxis | GCA_001610995.1 | Sphingopyxis_macrogoltabida | ATM17_05290 | 1078330..1082055 | 33050 | ATPase-[Sphingopyxis-macrogoltabida] |
| QCB56914.1 | Type3.MTaseII.containing.BREX | MobF->?-><-?<-?<-OBFOLD-NUCLEASE<-LexA-protease\|\|HTH->wHTH+WYL+WCX->BrxB-BrxF-iSTAND->DUF6079*->N6-N4-METHYLASE->TUDOR+SF1-SF2-Helicase+iREase+CTD->PglZ->BrxA-DNA-binding-><-?<-LZ_Tnp_IS66+zf-IS66+DDE_Tnp_IS66+DDE_Tnp_IS66_C<-?<-HIN-HTH\|\|SF1-SF2-Helicase-> | 1241 | Bacteria>Pseudomonadota>Alphaproteobacteria>Sphingomonadales>Sphingomonadaceae>Sphingopyxis | GCA_004795895.1 | Sphingopyxis_sp._PAMC25046 | E5675_09500 | 2019629..2023354 | 2565556 | ATP-binding-protein-[Sphingopyxis-sp.-PAMC25046] |
| WP_201258420.1 | Type3.MTaseII.containing.BREX | TraJ-RHH->VirD4-TraG+VirD4-FtsK->MobF->?-><-?<-?<-OBFOLD-NUCLEASE<-LexA-protease\|\|wHTH+WYL+WCX->BrxB-BrxF-iSTAND->DUF6079*->HEPN->N6-N4-METHYLASE-p-><-?\|\|DUF4096+DDE_Tnp_1+DDE_Tnp_1_2->N6-N4-METHYLASE->TUDOR+SF1-SF2-Helicase+iREase+CTD->PglZ->BrxA-DNA-binding->SF1-SF2-Helicase-> | 1243 | Bacteria>Pseudomonadota>Alphaproteobacteria>Sphingomonadales>Sphingomonadaceae>Sphingopyxis | GCF_001314325.1 | Sphingopyxis_macrogoltabida | LH19_RS05245 | 1076893..1080624 | 33050 | DUF6079-family-protein-[Sphingopyxis-macrogoltabida] |
| WP_202988337.1 | Type3.MTaseII.containing.BREX | Thioredoxin->TraH->?-><-?<-SF1-SF2-Helicase<-BrxA-DNA-binding<-PglZ<-TUDOR+SF1-SF2-Helicase+iREase+CTD<-N6-N4-METHYLASE<-DUF6079*<-BrxB-BrxF-iSTAND<-wHTH+WYL+WCX\|\|LexA-protease->OBFOLD-NUCLEASE->?->?->?-><-MobF<-VirD4-TraG+VirD4-FtsK | 1243 | Bacteria>Pseudomonadota>Alphaproteobacteria>Sphingomonadales>Sphingomonadaceae>Sphingopyxis | GCF_001610975.1 | Sphingopyxis_terrae | AOA14_RS18885 | 3958090..3961821 | 33052 | DUF6079-family-protein-[Sphingopyxis-terrae] |
| WP_210727634.1 | Type3.MTaseII.containing.BREX | VirD4-TraG+VirD4-FtsK->MobF->?-><-?<-?<-OBFOLD-NUCLEASE<-LexA-protease\|\|wHTH+WYL+WCX->BrxB-BrxF-iSTAND->DUF6079*->N6-N4-METHYLASE->TUDOR+SF1-SF2-Helicase+iREase+CTD->PglZ->BrxA-DNA-binding-><-?<-LZ_Tnp_IS66+zf-IS66+DDE_Tnp_IS66+DDE_Tnp_IS66_C<-?<-HIN-HTH | 1243 | Bacteria>Pseudomonadota>Alphaproteobacteria>Sphingomonadales>Sphingomonadaceae>Sphingopyxis | GCF_004795895.1 | Sphingopyxis_sp._PAMC25046 | E5675_RS09500 | 2019623..2023354 | 2565556 | DUF6079-family-protein-[Sphingopyxis-sp.-PAMC25046] |
| WP_251644434.1 | Type3.MTaseII.containing.BREX | <-SF1-SF2-Helicase<-DEDD_Tnp_IS110+Transposase_20<-DUF4096+DDE_Tnp_1+DDE_Tnp_1_2<-SF1-SF2-Helicase<-BrxA-DNA-binding<-PglZ<-TUDOR+SF1-SF2-Helicase+iREase+CTD<-N6-N4-METHYLASE<-DUF6079*<-BrxB-BrxF-iSTAND<-wHTH+WYL+WCX\|\|LexA-protease->OBFOLD-NUCLEASE->?->?->?-><-MobF<-VirD4-TraG+VirD4-FtsK | 1243 | Bacteria>Pseudomonadota>Alphaproteobacteria>Sphingomonadales>Sphingomonadaceae>Sphingopyxis | GCF_023712965.1 | Sphingopyxis_alaskensis | M3632_RS15935 | 35461..39192 | 117207 | DUF6079-family-protein-[Sphingopyxis-alaskensis] |
| WP_008603685.1 | Type3.MTaseII.containing.BREX | <-SF1-SF2-Helicase+iREase\|\|wHTH-><-?\|\|?->?->?-><-BrxA-DNA-binding<-PglZ<-N6-N4-METHYLASE<-DUF6079*<-BrxB-BrxF-iSTAND<-HTH_37\|\|DDE_Tnp_IS1595->?-><-?<-?<-HEPN+PAS+DUF4011+SF1-SF2-Helicase+REase<-?<-Zn_Tnp_IS91+Y2_Tnp<-Phage_int_SAM_4+Phage_integrase\|\|CHTH-> | 1246 | Bacteria>Pseudomonadota>Alphaproteobacteria>Sphingomonadales>Sphingosinicellaceae>Pacificimonas | GCF_000342165.1 | Pacificimonas_flava | C725_RS13250 | 48193..51933 | 1234595 | DUF6079-family-protein-[Pacificimonas-flava] |
| NQY81839.1 | Type3.MTaseII.containing.BREX | DDE_Tnp_IS1595-><-?\|\|?-><-?\|\|HIN-HTH->?->BrxB-BrxF-iSTAND->DUF6079*->N6-N4-METHYLASE->?->PglZ-><-?\|\|TraI-VirD2-RCRE-> | 1243 | Bacteria>Pseudomonadota>Alphaproteobacteria>unclassified_Alphaproteobacteria | GCA_013216065.1 | Alphaproteobacteria_bacterium | HRT36_02060 | 4558..8289 | 1913988 | ATP-binding-protein-[Alphaproteobacteria-bacterium] |
| MBI1186276.1 | Type3.MTaseII.containing.BREX | RAMA->GDResolvase+ZnR->?->PhdYeFM_antitox->BrxB-BrxF-iSTAND->DUF6079*->?->N6-N4-METHYLASE->TUDOR+SF1-SF2-Helicase+iREase+CTD->PglZ->BrxA-DNA-binding-> | 1241 | Bacteria>Pseudomonadota>Alphaproteobacteria>unclassified_Alphaproteobacteria | GCA_016124255.1 | Alphaproteobacteria_bacterium | GC206_02920 | 2831..6556 | 1913988 | ATP-binding-protein-[Alphaproteobacteria-bacterium] |
| KFB72158.1 | Type3.MTaseII.containing.BREX | DUF6079-p*->Schlafen+HATPase+wHTH->Schlafen+HATPase+wHTH->N6-N4-METHYLASE->OLD-ABC->TOPRIM->OLD-ABC->TOPRIM->TUDOR+SF1-SF2-Helicase+iREase+CTD-> | 388 | Bacteria>Pseudomonadota>Betaproteobacteria>Betaproteobacteria_incertae_sedis>Candidatus_Accumulibacter | GCA_000585035.2 | Candidatus_Accumulibacter_phosphatis | AW09_002650 | 30..1196 | 327160 | MAG:-hypothetical-protein-AW09_002650-[Candidatus-Accumulibacter-phosphatis] |
| EXI82568.1 | Type3.MTaseII.containing.BREX | <-McrC-NTD+REase<-McrB-AAA<-BrxA-DNA-binding<-PglZ<-TUDOR+SF1-SF2-Helicase+iREase+CTD<-N6-N4-METHYLASE<-?<-HerA/FtsK<-SIR2<-DUF6079*<-BrxB-BrxF-iSTAND<-wHTH+WYL+WCX\|\|HKD-DNase->?->?->?->?->?-><-ParB | 1246 | Bacteria>Pseudomonadota>Betaproteobacteria>Betaproteobacteria_incertae_sedis>Candidatus_Accumulibacter | GCA_000585055.1 | Candidatus_Accumulibacter_appositus | AW10_00354 | 44987..48727 | 1454003 | MAG:-hypothetical-protein-AW10_00354-[Candidatus-Accumulibacter-appositus] |
| RDE49561.1 | Type3.MTaseII.containing.BREX | <-HTH_3\|\|tRNA-><-?<-?<-?<-?<-?\|\|XIS-HTH->BrxB-BrxF-iSTAND->DUF6079*->HEPN->N6-N4-METHYLASE->TUDOR+SF1-SF2-Helicase+iREase+CTD->PglZ->BrxA-DNA-binding->?-><-?<-?<-TOPRIM | 1232 | Bacteria>Pseudomonadota>Betaproteobacteria>Betaproteobacteria_incertae_sedis>Candidatus_Accumulibacter | GCA_003332265.1 | Candidatus_Accumulibacter_meliphilus | DVS81_15885 | 8088..11786 | 2211374 | MAG:-ATP-binding-protein-[Candidatus-Accumulibacter-meliphilus] |
| QKS31658.1 | Type3.MTaseII.containing.BREX | Arm-DNA-bind_3+Phage_integrase-><-LysR-HTH\|\|HTH_3-><-MODE-HTH+PBPII\|\|wHTH+WYL+WCX->BrxB-BrxF-iSTAND->DUF6079*->SIR2->HerA/FtsK->?->N6-N4-METHYLASE->TUDOR+SF1-SF2-Helicase+iREase+CTD->PglZ->BrxA-DNA-binding-> | 1242 | Bacteria>Pseudomonadota>Betaproteobacteria>Betaproteobacteria_incertae_sedis>Candidatus_Accumulibacter | GCA_013347225.1 | Candidatus_Accumulibacter_similis | HT579_15245 | 3489535..3493263 | 2954388 | MAG:-ATP-binding-protein-[Candidatus-Accumulibacter-similis] |
| TLD44378.1 | Type3.MTaseII.containing.BREX | <-BrxA-DNA-binding<-PglZ<-TUDOR+SF1-SF2-Helicase+iREase+CTD<-N6-N4-METHYLASE<-?<-DUF6079*<-BrxB-BrxF-iSTAND<-HKD-DNase+DUF3150<-TOPRIM | 1241 | Bacteria>Pseudomonadota>Betaproteobacteria>Betaproteobacteria_incertae_sedis>Candidatus_Accumulibacter>unclassified_Candidatus_Accumulibacter | GCA_005524045.1 | Candidatus_Accumulibacter_sp. | FAZ92_03366 | 10235..13960 | 2053492 | MAG:-hypothetical-protein-FAZ92_03366-[Candidatus-Accumulibacter-sp.] |
| MCM8595100.1 | Type3.MTaseII.containing.BREX | <-BrxA-DNA-binding<-PglZ<-TUDOR+SF1-SF2-Helicase+iREase+CTD<-N6-N4-METHYLASE<-DUF6079*<-BrxB-BrxF-iSTAND<-?\|\|NtrC-AAA+FIS-HTH->RNase->?->STYKIN+CR-Kinase->SF1-SF2-Helicase->HTH+HEPN-> | 1241 | Bacteria>Pseudomonadota>Betaproteobacteria>Betaproteobacteria_incertae_sedis>Candidatus_Accumulibacter>unclassified_Candidatus_Accumulibacter | GCA_023806595.1 | Candidatus_Accumulibacter_sp. | NFV71_07035 | 98144..101869 | 2053492 | DUF6079-family-protein-[Candidatus-Accumulibacter-sp.] |
| MCM8625486.1 | Type3.MTaseII.containing.BREX | <-BrxA-DNA-binding<-PglZ<-TUDOR+SF1-SF2-Helicase+iREase+CTD<-REase<-OLD-ABC<-N6-N4-METHYLASE<-?<-DUF6079*<-BrxB-BrxF-iSTAND<-?\|\|NtrC-AAA+FIS-HTH->RNase->?->?->STYKIN+CR-Kinase->SF1-SF2-Helicase->HTH+HEPN-> | 1241 | Bacteria>Pseudomonadota>Betaproteobacteria>Betaproteobacteria_incertae_sedis>Candidatus_Accumulibacter>unclassified_Candidatus_Accumulibacter | GCA_023806625.1 | Candidatus_Accumulibacter_sp. | NFW05_05900 | 113658..117383 | 2053492 | DUF6079-family-protein-[Candidatus-Accumulibacter-sp.] |
| MCM8626700.1 | Type3.MTaseII.containing.BREX | <-BrxA-DNA-binding<-PglZ<-TUDOR+SF1-SF2-Helicase+iREase+CTD<-N6-N4-METHYLASE<-Schlafen+HATPase+wHTH<-DUF6079*<-BrxB-BrxF-iSTAND<-XIS-HTH<-?<-GDResolvase+ZnR | 1229 | Bacteria>Pseudomonadota>Betaproteobacteria>Betaproteobacteria_incertae_sedis>Candidatus_Accumulibacter>unclassified_Candidatus_Accumulibacter | GCA_023806625.1 | Candidatus_Accumulibacter_sp. | NFW05_12175 | 10591..14280 | 2053492 | DUF6079-family-protein-[Candidatus-Accumulibacter-sp.] |
| WP_156860858.1 | Type3.MTaseII.containing.BREX | <-BrxA-DNA-binding<-PglZ<-TUDOR+SF1-SF2-Helicase+iREase+CTD<-Schlafen+HATPase+wHTH<-N6-N4-METHYLASE<-?<-?<-HerA/FtsK<-SIR2<-DUF6079*<-BrxB-BrxF-iSTAND<-wHTH+WYL+WCX\|\|?->LysR-HTH-><-GNTR-HTH+Aminotran_1_2\|\|?->?->ACET->Phos_pyr_kin-> | 1246 | Bacteria>Pseudomonadota>Betaproteobacteria>Betaproteobacteria_incertae_sedis>Casimicrobiaceae>Casimicrobium | GCF_009746125.1 | Casimicrobium_huifangae | FKL89_RS00905 | 197066..200806 | 2591109 | DUF6079-family-protein-[Casimicrobium-huifangae] |
| WP_102774963.1 | Type3.MTaseII.containing.BREX | <-DNA_METHYLASE<-tRNA<-LysR-HTH\|\|HTH_3-><-Peptidase_S8<-Classical-AAA<-HTH_3<-MODE-HTH+PBPII\|\|wHTH+WYL+WCX->BrxB-BrxF-iSTAND->DUF6079*->SIR2->HerA/FtsK->?->N6-N4-METHYLASE-><-DUF772+DDE_Tnp_1\|\|HIN-HTH->N6-N4-METHYLASE->HTH+REase->HTH+REase-> | 1246 | Bacteria>Pseudomonadota>Betaproteobacteria>Burkholderiales>Alcaligenaceae>Achromobacter | - | Achromobacter_pulmonis | C1I89_23950 | 467033..470773 | 1389932 | DUF6079-family-protein-[Achromobacter-pulmonis] |
| PND31912.1 | Type3.MTaseII.containing.BREX | <-DNA_METHYLASE<-tRNA<-LysR-HTH\|\|HTH_3-><-Peptidase_S8<-Classical-AAA<-HTH_3<-MODE-HTH+PBPII\|\|wHTH+WYL+WCX->BrxB-BrxF-iSTAND->DUF6079*->SIR2->HerA/FtsK->?->N6-N4-METHYLASE-><-DUF772+DDE_Tnp_1\|\|HIN-HTH->N6-N4-METHYLASE->HTH+REase->HTH+REase-> | 1246 | Bacteria>Pseudomonadota>Betaproteobacteria>Burkholderiales>Alcaligenaceae>Achromobacter | GCA_002885955.2 | Achromobacter_pulmonis | C1I89_23950 | 467033..470773 | 1389932 | ATPase-[Achromobacter-pulmonis] |
| MCH1991581.1 | Type3.MTaseII.containing.BREX | <-TraE<-TraM<-TraM<-DUF3262<-?<-SMODS<-?\|\|wHTH+WYL+WCX->BrxB-BrxF-iSTAND->DUF6079*->?->N6-N4-METHYLASE->OLD-ABC->TOPRIM->TUDOR+SF1-SF2-Helicase+iREase+CTD->PglZ->BrxA-DNA-binding->?-><-DUF4400 | 1246 | Bacteria>Pseudomonadota>Betaproteobacteria>Burkholderiales>Alcaligenaceae>Achromobacter | GCA_022444305.1 | Achromobacter_xylosoxidans | L7Q78_02295 | 12685..16425 | 85698 | DUF6079-family-protein-[Achromobacter-xylosoxidans] |
| WP_258231514.1 | Type3.MTaseII.containing.BREX | <-DNA_METHYLASE<-tRNA<-LysR-HTH\|\|HTH_3-><-Peptidase_S8<-Classical-AAA<-HTH_3<-MODE-HTH+PBPII\|\|wHTH+WYL+WCX->BrxB-BrxF-iSTAND->DUF6079*->SIR2->HerA/FtsK->?->?->N6-N4-METHYLASE-><-DUF772+DDE_Tnp_1\|\|HIN-HTH->N6-N4-METHYLASE->HTH+REase-> | 1250 | Bacteria>Pseudomonadota>Betaproteobacteria>Burkholderiales>Alcaligenaceae>Achromobacter | GCF_002885955.2 | Achromobacter_pulmonis | C1I89_RS23930 | 467021..470773 | 1389932 | DUF6079-family-protein-[Achromobacter-pulmonis] |
| WP_125282998.1 | Type3.MTaseII.containing.BREX | <-Phos_pyr_kin<-ACET<-?<-?\|\|GNTR-HTH+Aminotran_1_2-><-LysR-HTH<-?\|\|wHTH+WYL+WCX->BrxB-BrxF-iSTAND->DUF6079*->SIR2->HerA/FtsK->?->?->N6-N4-METHYLASE->Schlafen+HATPase+wHTH->TUDOR+SF1-SF2-Helicase+iREase+CTD->PglZ->BrxA-DNA-binding-> | 1246 | Bacteria>Pseudomonadota>Betaproteobacteria>Burkholderiales>Alcaligenaceae>Achromobacter | GCF_003939895.1 | Achromobacter_denitrificans | EGU64_RS02450 | 220472..224212 | 32002 | DUF6079-family-protein-[Achromobacter-denitrificans] |
| WP_240294122.1 | Type3.MTaseII.containing.BREX | <-N6-N4-METHYLASE<-MORC<-Schlafen+HATPase+wHTH<-DUF6079*<-BrxB-BrxF-iSTAND<-XIS-HTH<-Phage_int_SAM_2+Integrase_1<-HTH_36<-?<-ABC-ATPase+SbcC+ABC-ATPase+SbcC<-?<-MarR-HTH | 1248 | Bacteria>Pseudomonadota>Betaproteobacteria>Burkholderiales>Alcaligenaceae>Achromobacter | GCF_022488505.1 | Achromobacter_xylosoxidans | L7A49_RS17320 | 6421..10167 | 85698 | DUF6079-family-protein-[Achromobacter-xylosoxidans] |
| WP_238920069.1 | Type3.MTaseII.containing.BREX | MarR-HTH->?->ABC-ATPase+SbcC+ABC-ATPase+SbcC->?->HTH_36->Phage_int_SAM_2+Integrase_1->XIS-HTH->BrxB-BrxF-iSTAND->DUF6079*->Schlafen+HATPase+wHTH->MORC->N6-N4-METHYLASE->TUDOR+SF1-SF2-Helicase+iREase+CTD->PglZ->BrxA-DNA-binding-><-Arm-DNA-bind_3+Phage_integrase<-sGTP | 1248 | Bacteria>Pseudomonadota>Betaproteobacteria>Burkholderiales>Alcaligenaceae>Achromobacter | GCF_022525265.1 | Achromobacter_xylosoxidans | MN082_RS16450 | 50800..54546 | 85698 | DUF6079-family-protein-[Achromobacter-xylosoxidans] |
| WP_175180616.1 | Type3.MTaseII.containing.BREX | sGTP->Arm-DNA-bind_3+Phage_integrase->Phage_integrase-><-BrxA-DNA-binding<-PglZ<-TUDOR+SF1-SF2-Helicase+iREase+CTD<-N6-N4-METHYLASE<-MORC<-Schlafen+HATPase+wHTH<-DUF6079*<-BrxB-BrxF-iSTAND<-XIS-HTH<-Phage_int_SAM_2+Integrase_1<-HTH_36<-?<-?<-MarR-HTH | 1248 | Bacteria>Pseudomonadota>Betaproteobacteria>Burkholderiales>Alcaligenaceae>Achromobacter | GCF_902859735.1 | Achromobacter_aegrifaciens | HV766_RS04070 | 908044..911790 | 1287736 | DUF6079-family-protein-[Achromobacter-aegrifaciens] |
| MCB4322560.1 | Type3.MTaseII.containing.BREX | DOC+HTH->SF1-SF2-Helicase->Phage_rep->HTH_3->Phage_int_SAM_2+Integrase_1->XIS-HTH->BrxB-BrxF-iSTAND->DUF6079*->MORC->?->?->N6-N4-METHYLASE->Schlafen+HATPase+wHTH->TUDOR+SF1-SF2-Helicase+iREase+CTD->PglZ->BrxA-DNA-binding->PspB-> | 1242 | Bacteria>Pseudomonadota>Betaproteobacteria>Burkholderiales>Alcaligenaceae>Alcaligenes | GCA_020496585.1 | Alcaligenes_sp._13f | KOE80_10155 | 22970..26698 | 2841924 | ATP-binding-protein-[Alcaligenes-sp.-13f] |
| WP_125277465.1 | Type3.MTaseII.containing.BREX | <-BrxA-DNA-binding<-PglZ<-TUDOR+SF1-SF2-Helicase+iREase+CTD<-HTH+REase<-N6-N4-METHYLASE<-HerA/FtsK<-SIR2<-PadR-HTH<-OLD-ABC+TOPRIM<-DUF6079*<-BrxB-BrxF-iSTAND<-wHTH+WYL+WCX\|\|MODE-HTH+PBPII-><-HTH_3\|\|LysR-HTH-><-DDE_Tnp_IS240<-ACET<-?\|\|<-MODE-HTH+PBPII | 1262 | Bacteria>Pseudomonadota>Betaproteobacteria>Burkholderiales>Alcaligenaceae>Alcaligenes | GCF_003938225.2 | Alcaligenes_aquatilis | EGK70_RS02760 | 606963..610751 | 323284 | DUF6079-family-protein-[Alcaligenes-aquatilis] |
| WP_226349060.1 | Type3.MTaseII.containing.BREX | DOC+HTH->SF1-SF2-Helicase->Phage_rep->HTH_3-><-?\|\|Phage_int_SAM_2+Integrase_1->XIS-HTH->BrxB-BrxF-iSTAND->DUF6079*->MORC->?->?->N6-N4-METHYLASE->Schlafen+HATPase+wHTH->TUDOR+SF1-SF2-Helicase+iREase+CTD->PglZ->BrxA-DNA-binding->PspB-> | 1246 | Bacteria>Pseudomonadota>Betaproteobacteria>Burkholderiales>Alcaligenaceae>Alcaligenes | GCF_020496585.1 | Alcaligenes_sp._13f | KOE80_RS10185 | 22958..26698 | 2841924 | DUF6079-family-protein-[Alcaligenes-sp.-13f] |
| OZI68382.1 | Type3.MTaseII.containing.BREX | <-HAMP+HISKIN<-REC+OmpR-HTH<-?\|\|MODE-HTH+PBPII->?-><-?\|\|ABhydrolase->XIS-HTH->BrxB-BrxF-iSTAND->DUF6079*->MORC->?->N6-N4-METHYLASE->Schlafen+HATPase+wHTH->TUDOR+SF1-SF2-Helicase+iREase+CTD->PglZ->BrxA-DNA-binding->SIR2-><-Arm-DNA-bind_3+Phage_integrase | 1242 | Bacteria>Pseudomonadota>Betaproteobacteria>Burkholderiales>Alcaligenaceae>Bordetella | GCA_002261435.1 | Bordetella_genomosp._1 | CAL27_02650 | 565236..568964 | 1395607 | ATPase-[Bordetella-genomosp.-1] |
| MCJ9707520.1 | Type3.MTaseII.containing.BREX | DOC+HTH->XIS-HTH-><-N6-N4-METHYLASE<-HerA/FtsK<-?\|\|XIS-HTH->BrxB-BrxF-iSTAND->DUF6079*->?->N6-N4-METHYLASE->TUDOR+SF1-SF2-Helicase+iREase+CTD->PglZ->BrxA-DNA-binding->?-><-Phage_lysozyme<-Phage_holin | 1241 | Bacteria>Pseudomonadota>Betaproteobacteria>Burkholderiales>Alcaligenaceae>Bordetella | GCA_023006425.1 | Bordetella_hinzii | LWS69_00545 | 7388..11113 | 103855 | DUF6079-family-protein-[Bordetella-hinzii] |
| WP_255032058.1 | Type3.MTaseII.containing.BREX | MODE-HTH+PBPII->?-><-?\|\|ABhydrolase->Phage_rep->?->Phage_int_SAM_2+Integrase_1->XIS-HTH->BrxB-BrxF-iSTAND->DUF6079*->MORC->?->?->N6-N4-METHYLASE->Schlafen+HATPase+wHTH->TUDOR+SF1-SF2-Helicase+iREase+CTD->PglZ->BrxA-DNA-binding->SIR2-> | 1246 | Bacteria>Pseudomonadota>Betaproteobacteria>Burkholderiales>Alcaligenaceae>Bordetella | GCF_002261435.1 | Bordetella_genomosp._1 | CAL27_RS02645 | 565224..568964 | 1395607 | DUF6079-family-protein-[Bordetella-genomosp.-1] |
| HBT33990.1 | Type3.MTaseII.containing.BREX | <-RelB<-BrxA-DNA-binding<-PglZ<-TUDOR+SF1-SF2-Helicase+iREase+CTD<-Schlafen+HATPase+wHTH<-N6-N4-METHYLASE<-PadR-HTH<-OLD-ABC+TOPRIM<-DUF6079-p<-DUF6079-p* | 161 | Bacteria>Pseudomonadota>Betaproteobacteria>Burkholderiales>Alcaligenaceae>Pusillimonas | GCA_003509985.1 | Pusillimonas_sp._(ex_Stolz_et_al._2005) | DEB15_14785 | 31746..32228 | 1979962 | ATPase-[Pusillimonas-sp.-(ex-Stolz-et-al.-2005)] |
| WP_105831338.1 | Type3.MTaseII.containing.BREX | XIS-HTH->?-><-?<-DUF932<-?\|\|?->HTH+MPTase->XIS-HTH->BrxB-BrxF-iSTAND->DUF6079*->Schlafen+HATPase+wHTH->N6-N4-METHYLASE->TUDOR+SF1-SF2-Helicase+iREase+CTD->PglZ->BrxA-DNA-binding-><-Phage_integrase | 1246 | Bacteria>Pseudomonadota>Betaproteobacteria>Burkholderiales>Burkholderiaceae>Burkholderia | - | Burkholderia_ambifaria | C6Q14_09295 | 40669..44409 | 152480 | DUF6079-family-protein-[Burkholderia-ambifaria] |
| WP_124920786.1 | Type3.MTaseII.containing.BREX | <-DUF932<-?\|\|?->HTH+MPTase->XIS-HTH->BrxB-BrxF-iSTAND->DUF6079*->Schlafen+HATPase+wHTH->N6-N4-METHYLASE->TUDOR+SF1-SF2-Helicase+iREase+CTD->PglZ->BrxA-DNA-binding-><-Phage_integrase | 1246 | Bacteria>Pseudomonadota>Betaproteobacteria>Burkholderiales>Burkholderiaceae>Burkholderia | - | Burkholderia_sp._Bp9099 | DIE17_07275 | 173126..176866 | 2184568 | DUF6079-family-protein-[Burkholderia-sp.-Bp9099] |
| ERJ33260.1 | Type3.MTaseII.containing.BREX | RFC-AAA+DNAA-HTH->?-><-?\|\|MORC+DNA_gyrase+TOPRIM->wHTH+WYL+WCX-p->DUF6079-p*-> | 790 | Bacteria>Pseudomonadota>Betaproteobacteria>Burkholderiales>Burkholderiaceae>Burkholderia | GCA_000466765.1 | Burkholderia_sp._AU4i | L810_4518 | 6539..8911 | 1335308 | Exonuclease-SbcC-[Burkholderia-sp.-AU4i] |
| KIX33653.1 | Type3.MTaseII.containing.BREX | ZnR+HTH_Tnp_ISL3+DDE_Tnp_ISL3-><-BrxA-DNA-binding<-PglZ<-TUDOR+SF1-SF2-Helicase+iREase+CTD<-N6-N4-METHYLASE<-TOPRIM<-OLD-ABC<-DUF6079*<-BrxB-BrxF-iSTAND\|\|DUF932-> | 1246 | Bacteria>Pseudomonadota>Betaproteobacteria>Burkholderiales>Burkholderiaceae>Burkholderia | GCA_000932075.1 | Burkholderia_pseudomallei | SY87_33360 | 11745..15485 | 28450 | ATPase-[Burkholderia-pseudomallei] |
| PRG07836.1 | Type3.MTaseII.containing.BREX | XIS-HTH->?-><-?<-DUF932<-?\|\|?->HTH+MPTase->XIS-HTH->BrxB-BrxF-iSTAND->DUF6079*->Schlafen+HATPase+wHTH->N6-N4-METHYLASE->TUDOR+SF1-SF2-Helicase+iREase+CTD->PglZ->BrxA-DNA-binding-><-Phage_integrase | 1246 | Bacteria>Pseudomonadota>Betaproteobacteria>Burkholderiales>Burkholderiaceae>Burkholderia | GCA_002981655.1 | Burkholderia_ambifaria | C6Q14_09295 | 40669..44409 | 152480 | ATPase-[Burkholderia-ambifaria] |
| RQZ50099.1 | Type3.MTaseII.containing.BREX | <-DUF932<-?\|\|?->HTH+MPTase->XIS-HTH->BrxB-BrxF-iSTAND->DUF6079*->Schlafen+HATPase+wHTH->N6-N4-METHYLASE->TUDOR+SF1-SF2-Helicase+iREase+CTD->PglZ->BrxA-DNA-binding-><-Phage_integrase | 1246 | Bacteria>Pseudomonadota>Betaproteobacteria>Burkholderiales>Burkholderiaceae>Burkholderia | GCA_003857805.1 | Burkholderia_sp._Bp9099 | DIE17_07275 | 173126..176866 | 2184568 | ATP-binding-protein-[Burkholderia-sp.-Bp9099] |
| MBH9659573.1 | Type3.MTaseII.containing.BREX | Phage_holin->Phage_lysozyme-><-?<-BrxA-DNA-binding<-PglZ<-TUDOR+SF1-SF2-Helicase+iREase+CTD<-N6-N4-METHYLASE<-DUF6079*<-BrxB-BrxF-iSTAND<-wHTH+WYL+WCX\|\|?->ARG->?->UDG-><-?<-?<-HTH_1+PBP_like | 1242 | Bacteria>Pseudomonadota>Betaproteobacteria>Burkholderiales>Burkholderiaceae>Burkholderia | GCA_016092255.1 | Burkholderia_multivorans | JAO06_00055 | 13327..17055 | 87883 | ATP-binding-protein-[Burkholderia-multivorans] |
| QQJ98611.1 | Type3.MTaseII.containing.BREX | XIS-HTH->?->?-><-?\|\|?->HTH+MPTase->XIS-HTH->BrxB-BrxF-iSTAND->DUF6079*->Schlafen+HATPase+wHTH->N6-N4-METHYLASE->TUDOR+SF1-SF2-Helicase+iREase+CTD->PglZ->BrxA-DNA-binding-><-Phage_integrase | 1246 | Bacteria>Pseudomonadota>Betaproteobacteria>Burkholderiales>Burkholderiaceae>Burkholderia | GCA_016458785.1 | Burkholderia_ambifaria | JG536_00765 | 175117..178857 | 152480 | ATP-binding-protein-[Burkholderia-ambifaria] |
| MBY4833604.1 | Type3.MTaseII.containing.BREX | XIS-HTH-><-DOC+HTH<-RNase<-?<-?\|\|DUF6079-p*->HEPN->N6-N4-METHYLASE->TUDOR+SF1-SF2-Helicase+iREase+CTD->PglZ->BrxA-DNA-binding->McrB-AAA->McrC-NTD+REase-> | 405 | Bacteria>Pseudomonadota>Betaproteobacteria>Burkholderiales>Burkholderiaceae>Burkholderia | GCA_019795915.1 | Burkholderia_dolosa | K6W55_25635 | 14267..15484 | 152500 | DUF6079-family-protein-[Burkholderia-dolosa] |
| WP_162175314.1 | Type3.MTaseII.containing.BREX | RFC-AAA+DNAA-HTH->?->MORC+DNA_gyrase+TOPRIM->wHTH+WYL+WCX->DUF6079-p*-> | 802 | Bacteria>Pseudomonadota>Betaproteobacteria>Burkholderiales>Burkholderiaceae>Burkholderia | GCF_000466765.1 | Burkholderia_sp._AU4i | L810_RS40980 | 6503..8911 | 1335308 | DUF6079-family-protein-[Burkholderia-sp.-AU4i] |
| WP_235357646.1 | Type3.MTaseII.containing.BREX | ZnR+HTH_Tnp_ISL3+DDE_Tnp_ISL3-><-BrxA-DNA-binding<-PglZ<-TUDOR+SF1-SF2-Helicase+iREase+CTD<-N6-N4-METHYLASE<-TOPRIM<-OLD-ABC<-DUF6079*<-BrxB-BrxF-iSTAND<-XIS-HTH\|\|?->?->DUF932-> | 1250 | Bacteria>Pseudomonadota>Betaproteobacteria>Burkholderiales>Burkholderiaceae>Burkholderia | GCF_000932075.1 | Burkholderia_pseudomallei | SY87_RS33410 | 11745..15497 | 28450 | DUF6079-family-protein-[Burkholderia-pseudomallei] |
| WP_059901856.1 | Type3.MTaseII.containing.BREX | <-BrxA-DNA-binding<-PglZ<-TUDOR+SF1-SF2-Helicase+iREase+CTD<-N6-N4-METHYLASE<-?<-DUF6079*<-BrxB-BrxF-iSTAND<-HEPN<-?<-MORC+DNA_gyrase+TOPRIM<-?<-RFC-AAA+DNAA-HTH\|\|?->RNaseP-> | 1241 | Bacteria>Pseudomonadota>Betaproteobacteria>Burkholderiales>Burkholderiaceae>Burkholderia | GCF_001526915.1 | Burkholderia_ubonensis | WJ99_RS10200 | 40174..43899 | 101571 | DUF6079-family-protein-[Burkholderia-ubonensis] |
| WP_059926577.1 | Type3.MTaseII.containing.BREX | <-RNaseP<-?\|\|RFC-AAA+DNAA-HTH->?->MORC+DNA_gyrase+TOPRIM->?->HEPN->BrxB-BrxF-iSTAND->DUF6079*->?->N6-N4-METHYLASE->TUDOR+SF1-SF2-Helicase+iREase+CTD->PglZ->BrxA-DNA-binding-> | 1241 | Bacteria>Pseudomonadota>Betaproteobacteria>Burkholderiales>Burkholderiaceae>Burkholderia | GCF_002880865.1 | Burkholderia_ubonensis | AHF08_RS12200 | 10606..14331 | 101571 | DUF6079-family-protein-[Burkholderia-ubonensis] |
| WP_225740305.1 | Type3.MTaseII.containing.BREX | HTH_3-><-?<-?<-RNaseP<-?\|\|RFC-AAA+DNAA-HTH->?->MORC+DNA_gyrase+TOPRIM->?->DUF6079-p*-> | 596 | Bacteria>Pseudomonadota>Betaproteobacteria>Burkholderiales>Burkholderiaceae>Burkholderia | GCF_002942785.2 | Burkholderia_cepacia | C5075_RS13600 | 38295..40085 | 292 | DUF6079-family-protein-[Burkholderia-cepacia] |
| WP_260439215.1 | Type3.MTaseII.containing.BREX | <-DUF932<-?<-?\|\|?->HTH+MPTase->XIS-HTH->BrxB-BrxF-iSTAND->DUF6079*->Schlafen+HATPase+wHTH->N6-N4-METHYLASE->TUDOR+SF1-SF2-Helicase+iREase+CTD->PglZ->BrxA-DNA-binding-><-Phage_integrase | 1250 | Bacteria>Pseudomonadota>Betaproteobacteria>Burkholderiales>Burkholderiaceae>Burkholderia | GCF_003857805.1 | Burkholderia_sp._Bp9099 | DIE17_RS07325 | 173114..176866 | 2184568 | DUF6079-family-protein-[Burkholderia-sp.-Bp9099] |
| WP_217105386.1 | Type3.MTaseII.containing.BREX | Phage_holin->Phage_lysozyme-><-BrxA-DNA-binding<-PglZ<-TUDOR+SF1-SF2-Helicase+iREase+CTD<-N6-N4-METHYLASE<-TOPRIM<-OLD-ABC<-DUF6079*<-BrxB-BrxF-iSTAND<-XIS-HTH\|\|<-?<-ParA-Soj-PloopNTPase+ParA-Soj-PloopNTPase | 1241 | Bacteria>Pseudomonadota>Betaproteobacteria>Burkholderiales>Burkholderiaceae>Burkholderia | GCF_019041275.1 | Burkholderia_multivorans | KTD09_RS06785 | 75280..79005 | 87883 | DUF6079-family-protein-[Burkholderia-multivorans] |
| WP_261316357.1 | Type3.MTaseII.containing.BREX | XIS-HTH->XIS-HTH+XIS-HTH-><-DOC+HTH<-RNase<-?<-?\|\|DUF6079-p*->HEPN->N6-N4-METHYLASE->TUDOR+SF1-SF2-Helicase+iREase+CTD->PglZ->BrxA-DNA-binding->McrB-AAA-> | 368 | Bacteria>Pseudomonadota>Betaproteobacteria>Burkholderiales>Burkholderiaceae>Burkholderia | GCF_019795915.1 | Burkholderia_dolosa | K6W55_RS25625 | 14378..15484 | 152500 | DUF6079-family-protein-[Burkholderia-dolosa] |
| WP_232356480.1 | Type3.MTaseII.containing.BREX | Phage_holin->Phage_lysozyme-><-?<-BrxA-DNA-binding<-PglZ<-TUDOR+SF1-SF2-Helicase+iREase+CTD<-N6-N4-METHYLASE<-DUF6079*<-BrxB-BrxF-iSTAND<-wHTH+WYL+WCX\|\|?->ARG->?->UDG-><-?<-?<-HTH_1+PBP_like | 1246 | Bacteria>Pseudomonadota>Betaproteobacteria>Burkholderiales>Burkholderiaceae>Burkholderia | GCF_023373625.1 | Burkholderia_multivorans | LOZ11_RS12580 | 2635149..2638889 | 87883 | DUF6079-family-protein-[Burkholderia-multivorans] |
| WP_249462784.1 | Type3.MTaseII.containing.BREX | RADICAL-SAM->?->Phage_holin->Phage_lysozyme-><-?<-BrxA-DNA-binding<-PglZ<-TUDOR+SF1-SF2-Helicase+iREase+CTD<-N6-N4-METHYLASE<-DUF6079*<-BrxB-BrxF-iSTAND<-wHTH+WYL+WCX\|\|ARG->?->UDG-><-?<-?<-HTH_1+PBP_like | 1246 | Bacteria>Pseudomonadota>Betaproteobacteria>Burkholderiales>Burkholderiaceae>Burkholderia | GCF_023374455.1 | Burkholderia_multivorans | LOZ09_RS11430 | 2414572..2418312 | 87883 | DUF6079-family-protein-[Burkholderia-multivorans] |
| WP_264838735.1 | Type3.MTaseII.containing.BREX | HDRnase-><-SIR2<-BrxA-DNA-binding<-PglZ<-TUDOR+SF1-SF2-Helicase+iREase+CTD<-Schlafen+HATPase+wHTH<-N6-N4-METHYLASE<-HEPN<-MORC<-DUF6079*<-BrxB-BrxF-iSTAND<-wHTH+WYL+WCX\|\|MODE-HTH+PBPII-><-HTH_3\|\|LysR-HTH->?-><-?\|\|HEPN-> | 1246 | Bacteria>Pseudomonadota>Betaproteobacteria>Burkholderiales>Burkholderiaceae>Burkholderia | GCF_025995855.1 | Burkholderia_cenocepacia | K6Y76_RS22795 | 78194..81934 | 95486 | DUF6079-family-protein-[Burkholderia-cenocepacia] |
| WP_122634120.1 | Type3.MTaseII.containing.BREX | <-BrxA-DNA-binding<-PglZ<-TUDOR+SF1-SF2-Helicase+iREase+CTD<-N6-N4-METHYLASE<-Schlafen+HATPase+wHTH<-DUF6079*<-BrxB-BrxF-iSTAND<-wHTH+WYL+WCX\|\|REase->?->DUF932->?->?->?-><-HTH_3 | 1241 | Bacteria>Pseudomonadota>Betaproteobacteria>Burkholderiales>Burkholderiaceae>Burkholderia | GCF_900596605.1 | Burkholderia_pseudomallei | EFR82_RS35455 | 10862..14587 | 28450 | DUF6079-family-protein-[Burkholderia-pseudomallei] |
| WP_233410152.1 | Type3.MTaseII.containing.BREX | XIS-HTH->?->?-><-?\|\|?->HTH+MPTase->XIS-HTH->BrxB-BrxF-iSTAND->DUF6079*->Schlafen+HATPase+wHTH->N6-N4-METHYLASE->TUDOR+SF1-SF2-Helicase+iREase+CTD->PglZ->BrxA-DNA-binding-><-Phage_integrase | 1250 | Bacteria>Pseudomonadota>Betaproteobacteria>Burkholderiales>Burkholderiaceae>Burkholderia | GCF_902829415.1 | Burkholderia_ambifaria | HWV76_RS23890 | 153198..156950 | 152480 | DUF6079-family-protein-[Burkholderia-ambifaria] |
| WP_244101683.1 | Type3.MTaseII.containing.BREX | SF1-SF2-Helicase-><-?\|\|tRNA-><-?\|\|DUF6079-p*->TUDOR+SF1-SF2-Helicase+iREase+CTD->PglZ->BrxA-DNA-binding-><-Phage_integrase | 663 | Bacteria>Pseudomonadota>Betaproteobacteria>Burkholderiales>Burkholderiaceae>Burkholderia | GCF_902829965.1 | Burkholderia_ambifaria | HWW18_RS26330 | 152521..154512 | 152480 | DUF6079-family-protein-[Burkholderia-ambifaria] |
| WP_244105277.1 | Type3.MTaseII.containing.BREX | Phage_integrase-><-BrxA-DNA-binding<-PglZ<-TUDOR+SF1-SF2-Helicase+iREase+CTD<-N6-N4-METHYLASE<-Schlafen+HATPase+wHTH<-DUF6079*<-BrxB-BrxF-iSTAND<-XIS-HTH<-HTH+MPTase<-?\|\|?->DUF932-><-?<-?<-XIS-HTH | 1250 | Bacteria>Pseudomonadota>Betaproteobacteria>Burkholderiales>Burkholderiaceae>Burkholderia | GCF_902830085.1 | Burkholderia_ambifaria | HWW14_RS24145 | 141364..145116 | 152480 | DUF6079-family-protein-[Burkholderia-ambifaria] |
| AOY94227.1 | Type3.MTaseII.containing.BREX | <-METHYLASE<-LysR-HTH\|\|HTH_3-><-MODE-HTH+PBPII\|\|wHTH+WYL+WCX->BrxB-BrxF-iSTAND->DUF6079*->OLD-ABC->TOPRIM->MORC->HEPN->N6-N4-METHYLASE-p->HIN-HTH->AAA-ATPase->HIN-HTH->DUF4158+DDE_Tnp_Tn3-> | 1242 | Bacteria>Pseudomonadota>Betaproteobacteria>Burkholderiales>Burkholderiaceae>Cupriavidus | GCA_001854305.1 | Cupriavidus_sp._USMAA2-4 | BKK79_15560 | 3504410..3508138 | 876364 | ATPase-[Cupriavidus-sp.-USMAA2-4] |
| WP_232346221.1 | Type3.MTaseII.containing.BREX | <-METHYLASE<-LysR-HTH\|\|HTH_3-><-MODE-HTH+PBPII\|\|wHTH+WYL+WCX->BrxB-BrxF-iSTAND->DUF6079*->OLD-ABC->TOPRIM->MORC->HEPN->N6-N4-METHYLASE-p->HIN-HTH->AAA-ATPase->DUF4158+DDE_Tnp_Tn3-><-SbcC+SbcC+ABC-ATPase+SbcC | 1246 | Bacteria>Pseudomonadota>Betaproteobacteria>Burkholderiales>Burkholderiaceae | GCF_001854305.1 | Cupriavidus | BKK79_RS15555 | 3504398..3508138 | 106589 | MULTISPECIES:-DUF6079-family-protein-[Cupriavidus] |
| WP_017772487.1 | Type3.MTaseII.containing.BREX | <-DUF932<-?<-?\|\|XIS-HTH->BrxB-BrxF-iSTAND->DUF6079*->OLD-ABC->TOPRIM->N6-N4-METHYLASE->TUDOR+SF1-SF2-Helicase+iREase+CTD->PglZ->BrxA-DNA-binding-><-LACI-HTH+PBPI | 1246 | Bacteria>Pseudomonadota>Betaproteobacteria>Burkholderiales>Burkholderiaceae>Paraburkholderia | GCF_000341045.1 | Paraburkholderia_kururiensis | G118_RS0102325 | 503731..507471 | 984307 | DUF6079-family-protein-[Paraburkholderia-kururiensis] |
| WP_236062649.1 | Type3.MTaseII.containing.BREX | TarH+HAMP+MA-><-?<-?\|\|AraC-HTH->Phage_integrase-><-BrxA-DNA-binding<-PglZ<-TUDOR+SF1-SF2-Helicase+iREase+CTD<-N6-N4-METHYLASE<-DUF6079*<-BrxB-BrxF-iSTAND<-wHTH+WYL+WCX<-?<-Phage_lambda_P<-HTH_36\|\|?-><-Y1_Tnp | 1246 | Bacteria>Pseudomonadota>Betaproteobacteria>Burkholderiales>Burkholderiaceae>Paraburkholderia | GCF_905220985.1 | Paraburkholderia_nemoris | KHO43_RS19855 | 63873..67613 | 2793076 | DUF6079-family-protein-[Paraburkholderia-nemoris] |
| WP_102080326.1 | Type3.MTaseII.containing.BREX | XIS-HTH->XIS-HTH+XIS-HTH-><-DOC+HTH<-RNase<-?<-?<-?\|\|wHTH+WYL+WCX->BrxB-BrxF-iSTAND->DUF6079*->HEPN->N6-N4-METHYLASE->TUDOR+SF1-SF2-Helicase+iREase+CTD->PglZ->BrxA-DNA-binding->McrB-AAA->McrC-NTD+REase-><-Phage_lysozyme<-Phage_holin | 1241 | Bacteria>Pseudomonadota>Betaproteobacteria>Burkholderiales>Burkholderiaceae>Ralstonia | GCF_002863525.1 | Ralstonia_mannitolilytica | CXP34_RS17665 | 53242..56967 | 105219 | DUF6079-family-protein-[Ralstonia-mannitolilytica] |

| Protein-ID | System | Gene-Neighbourhoods | Length | Taxonomy | Assembly-ID | Strain | Locus-Tag | Genomic-coordinates | TaxID | Fasta-defline |
|---|---|---|---|---|---|---|---|---|---|---|
| QLQ00182.1 | Type3.MTaseII.containing.BREX | Phage_lysozyme-><-BrxA-DNA-binding<-TUDOR+SF1-SF2-Helicase+iREase+CTD<-REase<-OLD-ABC<-N6-N4-METHYLASE<-OLD-ABC<-DUF6079*<-BrxB-BrxF-iST[illegible] | 1241 | Bacteria>Pseudomonadota>Betaproteobacteria>Burkholderiales>Burkholderiaceae>unclassified_Burkholderiaceae | GCA_013693755.1 | Burkholderiaceae_bacterium | HZY78_08275 | 1798535..1802260 | 2030806 | ATP-binding-protein-[Burkholderiaceae-bacterium] |
| MBX3589942.1 | Type3.MTaseII.containing.BREX | <-BrxA-DNA-binding<-PglZ<-ParB+HNH<-TUDOR+SF1-SF2-Helicase+iREase+CTD<-N6-N4-METHYLASE<-Schlafen+HATPase+wHTH<-DUF6079*<-BrxB-BrxF-iSTAN[illegible] | 1241 | Bacteria>Pseudomonadota>Betaproteobacteria>Burkholderiales>Burkholderiaceae>unclassified_Burkholderiaceae | GCA_019635275.1 | Burkholderiaceae_bacterium | KF755_03445 | 56011..59736 | 2030806 | ATP-binding-protein-[Burkholderiaceae-bacterium] |
| MBX7228949.1 | Type3.MTaseII.containing.BREX | Phage_int_SAM_1+Phage_integrase-><-BrxA-DNA-binding<-PglZ<-TUDOR+SF1-SF2-Helicase+iREase+CTD<-N6-N4-METHYLASE<-Schlafen+HATPase+wHTH<-DU[illegible] | 204 | Bacteria>Pseudomonadota>Betaproteobacteria>Burkholderiales>Burkholderiaceae>unclassified_Burkholderiaceae | GCA_019695155.1 | Burkholderiaceae_bacterium | K1X48_05035 | 19336..19952 | 2030806 | ATP-binding-protein-[Burkholderiaceae-bacterium] |
| MCD6733430.1 | Type3.MTaseII.containing.BREX | DUF932-><-Phage_int_SAM_2+Integrase_1<-?<-?<-?<-MerR-HTH\|\|XIS-HTH->BrxB-BrxF-iSTAND->DUF6079*->N6-N4-METHYLASE->TUDOR+SF1-SF2-Helicase+i[illegible] | 1241 | Bacteria>Pseudomonadota>Betaproteobacteria>Burkholderiales>Burkholderiaceae>unclassified_Burkholderiaceae | GCA_021166795.1 | Burkholderiaceae_bacterium | LT103_07535 | 131364..135089 | 2030806 | DUF6079-family-protein-[Burkholderiaceae-bacterium] |
| MCO5109351.1 | Type3.MTaseII.containing.BREX | Phage_integrase+Phage_integrase-><-BrxA-DNA-binding<-PglZ<-TUDOR+SF1-SF2-Helicase+iREase+CTD<-N6-N4-METHYLASE<-MORC<-Schlafen+HATPase+wH[illegible] | 1246 | Bacteria>Pseudomonadota>Betaproteobacteria>Burkholderiales>Burkholderiaceae>unclassified_Burkholderiaceae | GCA_023954095.1 | Burkholderiaceae_bacterium | M9929_00725 | 31681..35421 | 2030806 | DUF6079-family-protein-[Burkholderiaceae-bacterium] |
| MCP5271841.1 | Type3.MTaseII.containing.BREX | <-DNA_METHYLASE<-SbcC<-BrxA-DNA-binding<-PglZ<-TUDOR+SF1-SF2-Helicase+iREase+CTD<-?<-CHTH<-?<-N6-N4-METHYLASE<-DUF6079*<-BrxB-BrxF-iSTAN[illegible] | 1241 | Bacteria>Pseudomonadota>Betaproteobacteria>Burkholderiales>Burkholderiaceae>unclassified_Burkholderiaceae | GCA_024235855.1 | Burkholderiaceae_bacterium | H6932_11485 | 147063..150788 | 2030806 | ATP-binding-protein-[Burkholderiaceae-bacterium] |
| MCP5219704.1 | Type3.MTaseII.containing.BREX | HAD-><-35exo<-MODE-HTH+PBPII\|\|?-><-TetR-HTH+AbiJ-NTD3+REase<-?\|\|wHTH+WYL+WCX->BrxB-BrxF-iSTAND->DUF6079*->HIN-HTH+PAIREDC-HTH->HKD-DNase-[illegible] | 1242 | Bacteria>Pseudomonadota>Betaproteobacteria>Burkholderiales>Burkholderiaceae>unclassified_Burkholderiaceae | GCA_024235995.1 | Burkholderiaceae_bacterium | H6927_16565 | 3462616..3466344 | 2030806 | ATP-binding-protein-[Burkholderiaceae-bacterium] |
| ALT79667.1 | Type3.MTaseII.containing.BREX | Phage_lysozyme-><-?<-PspB\|\|LZ_Tnp_IS481+HTH_32+rve+rve_3-><-PspB<-BrxA-DNA-binding<-PglZ<-TUDOR+SF1-SF2-Helicase+iREase+CTD<-N6-N4-METHY[illegible] | 1246 | Bacteria>Pseudomonadota>Betaproteobacteria>Burkholderiales>Burkholderiales_genera_incertae_sedis>Paucibacter | GCA_001477625.1 | Paucibacter_sp._KCTC_42545 | AT984_05990 | 1349624..1353364 | 1768242 | ATPase-[Paucibacter-sp.-KCTC-42545] |
| WP_231741542.1 | Type3.MTaseII.containing.BREX | <-HNH<-ParB+HNH<-PspB\|\|LZ_Tnp_IS481+HTH_32+rve+rve_3-><-PspB<-BrxA-DNA-binding<-PglZ<-TUDOR+SF1-SF2-Helicase+iREase+CTD<-N6-N4-METHYLASE[illegible] | 1250 | Bacteria>Pseudomonadota>Betaproteobacteria>Burkholderiales>Burkholderiales_genera_incertae_sedis>Paucibacter | GCF_001477625.1 | Paucibacter_sp._KCTC_42545 | AT984_RS05990 | 1349624..1353376 | 1768242 | DUF6079-family-protein-[Paucibacter-sp.-KCTC-42545] |
| RZJ63032.1 | Type3.MTaseII.containing.BREX | Phage_lysozyme-><-SIR2<-McrC-NTD+REase<-McrB-AAA<-BrxA-DNA-binding<-PglZ<-TUDOR+SF1-SF2-Helicase+iREase+CTD<-N6-N4-METHYLASE<-HicA<-DUF6[illegible] | 1240 | Bacteria>Pseudomonadota>Betaproteobacteria>Burkholderiales>Comamonadaceae>Acidovorax | GCA_004211185.1 | Acidovorax_sp. | EON49_00335 | 64523..68245 | 1872122 | MAG:-ATP-binding-protein-[Acidovorax-sp.] |
| WP_235491831.1 | Type3.MTaseII.containing.BREX | Phage_holin->Phage_lysozyme-><-SIR2<-ParB+HNH<-BrxA-DNA-binding<-PglZ<-TUDOR+SF1-SF2-Helicase+iREase+CTD<-N6-N4-METHYLASE<-DUF6079*<-Brx[illegible] | 1246 | Bacteria>Pseudomonadota>Betaproteobacteria>Burkholderiales>Comamonadaceae>Acidovorax | GCF_001282395.1 | Acidovorax_avenae | AOG77_RS00390 | 81317..85057 | 80867 | DUF6079-family-protein-[Acidovorax-avenae] |
| SEN73398.1 | Type3.MTaseII.containing.BREX | VirB11->?->BrxB-BrxF-iSTAND->DUF6079*->N6-N4-METHYLASE->?->OLD-ABC->TOPRIM->TUDOR+SF1-SF2-Helicase+iREase+CTD->PglZ->BrxA-DNA-binding-><[illegible] | 1246 | Bacteria>Pseudomonadota>Betaproteobacteria>Burkholderiales>Comamonadaceae>Brachymonas | GCA_900110225.1 | Brachymonas_denitrificans_DSM_15123 | SAMN02745977_01903 | 66293..70033 | 1121117 | hypothetical-protein-SAMN02745977_01903-[Brachymonas-denitrificans-DSM-15123] |
| WP_091817240.1 | Type3.MTaseII.containing.BREX | VirB11->?->BrxB-BrxF-iSTAND->DUF6079*->N6-N4-METHYLASE->?->OLD-ABC->TOPRIM->TUDOR+SF1-SF2-Helicase+iREase+CTD->PglZ->BrxA-DNA-binding-><[illegible] | 1242 | Bacteria>Pseudomonadota>Betaproteobacteria>Burkholderiales>Comamonadaceae>Brachymonas | GCF_900110225.1 | Brachymonas_denitrificans | BMW28_RS09500 | 66305..70033 | 28220 | DUF6079-family-protein-[Brachymonas-denitrificans] |
| POR10230.1 | Type3.MTaseII.containing.BREX | DUF4400-><-?<-BrxA-DNA-binding<-PglZ<-TUDOR+SF1-SF2-Helicase+iREase+CTD<-N6-N4-METHYLASE<-?<-?<-MORC<-DUF6079*<-BrxB-BrxF-iSTAND<-wHTH+W[illegible] | 1242 | Bacteria>Pseudomonadota>Betaproteobacteria>Burkholderiales>Comamonadaceae>Diaphorobacter | GCA_002916785.1 | Diaphorobacter_sp._LR2014-1 | BV908_11590 | 35608..39336 | 1933219 | ATPase-[Diaphorobacter-sp.-LR2014-1] |
| MBF5004210.1 | Type3.MTaseII.containing.BREX | <-BrxA-DNA-binding<-PglZ<-TUDOR+SF1-SF2-Helicase+iREase+CTD<-Schlafen+HATPase+wHTH<-N6-N4-METHYLASE<-HEPN<-MORC<-TOPRIM<-OLD-ABC<-DUF607[illegible] | 1242 | Bacteria>Pseudomonadota>Betaproteobacteria>Burkholderiales>Comamonadaceae>Diaphorobacter | GCA_015354245.1 | Diaphorobacter_caeni | ISG32_08055 | 72029..75757 | 2784387 | ATP-binding-protein-[Diaphorobacter-caeni] |
| WP_246867495.1 | Type3.MTaseII.containing.BREX | DUF4400-><-?<-BrxA-DNA-binding<-PglZ<-TUDOR+SF1-SF2-Helicase+iREase+CTD<-N6-N4-METHYLASE<-?<-?<-MORC<-DUF6079*<-BrxB-BrxF-iSTAND<-wHTH+W[illegible] | 1246 | Bacteria>Pseudomonadota>Betaproteobacteria>Burkholderiales>Comamonadaceae>Diaphorobacter | GCF_002916785.1 | Diaphorobacter_sp._LR2014-1 | BV908_RS11585 | 35608..39348 | 1933219 | DUF6079-family-protein-[Diaphorobacter-sp.-LR2014-1] |
| WP_228529679.1 | Type3.MTaseII.containing.BREX | <-BrxA-DNA-binding<-PglZ<-TUDOR+SF1-SF2-Helicase+iREase+CTD<-Schlafen+HATPase+wHTH<-N6-N4-METHYLASE<-HEPN<-MORC<-TOPRIM<-OLD-ABC<-DUF607[illegible] | 1246 | Bacteria>Pseudomonadota>Betaproteobacteria>Burkholderiales>Comamonadaceae>Diaphorobacter | GCF_015354245.1 | Diaphorobacter_caeni | ISG32_RS08060 | 72029..75769 | 2784387 | DUF6079-family-protein-[Diaphorobacter-caeni] |
| QRR36380.1 | Type3.MTaseII.containing.BREX | Phage_holin->Phage_lysozyme-><-?<-ParB+HNH<-PspB<-BrxA-DNA-binding<-PglZ<-TUDOR+SF1-SF2-Helicase+iREase+CTD<-N6-N4-METHYLASE<-DUF6079*<-[illegible] | 1246 | Bacteria>Pseudomonadota>Betaproteobacteria>Burkholderiales>Comamonadaceae>Hydrogenophaga | GCA_016903055.1 | Hydrogenophaga_sp._YM1 | JNX00_02400 | 479305..483045 | 2806262 | ATP-binding-protein-[Hydrogenophaga-sp.-YM1] |
| MBS3911804.1 | Type3.MTaseII.containing.BREX | Phage_lysozyme-[illegible] | 1246 | Bacteria>Pseudomonadota>Betaproteobacteria>Burkholderiales>Comamonadaceae>Hydrogenophaga | GCA_018333605.1 | Hydrogenophaga_sp. | KGZ70_08240 | 56596..60336 | 1904254 | ATP-binding-protein-[Hydrogenophaga-sp.] |
| WP_239478241.1 | Type3.MTaseII.containing.BREX | Phage_holin->Ph[illegible] | 1250 | Bacteria>Pseudomonadota>Betaproteobacteria>Burkholderiales>Comamonadaceae>Hydrogenophaga | GCF_016903055.1 | Hydrogenophaga_sp._YM1 | JNX00_RS02400 | 479305..483057 | 2806262 | DUF6079-family-protein-[Hydrogenophaga-sp.-YM1] |
| WP_108412481.1 | Type3.MTaseII.containing.BREX | <-HORMA<-BrxA-[illegible] | 1241 | Bacteria>Pseudomonadota>Betaproteobacteria>Burkholderiales>Comamonadaceae>Limnohabitans | GCF_003063575.1 | Limnohabitans_sp._Jir72 | B9Z52_RS03490 | 121972..125697 | 1977909 | DUF6079-family-protein-[Limnohabitans-sp.-Jir72] |
| WP_231851188.1 | Type3.MTaseII.containing.BREX | Peptidase_M50->[illegible] | 1254 | Bacteria>Pseudomonadota>Betaproteobacteria>Burkholderiales>Comamonadaceae>Serpentinimonas | GCF_000828915.1 | Serpentinimonas_maccroryi | SMCB_RS08485 | 1836858..1840622 | 1458426 | DUF6079-family-protein-[Serpentinimonas-maccroryi] |
| PIQ51532.1 | Type3.MTaseII.containing.BREX | Phage_AlpA->Arm[illegible] | 1241 | Bacteria>Pseudomonadota>Betaproteobacteria>Burkholderiales>Comamonadaceae>unclassified_Comamonadaceae | GCA_002786915.1 | Comamonadaceae_bacterium_CG12_big_fil_rev_8_21_14_0_65_59_15 | COW02_14300 | 22759..26484 | 1973952 | ATPase-[Comamonadaceae-bacterium-CG12_big_fil_rev_8_21_14_0_65_59_15] |
| MBH2043716.1 | Type3.MTaseII.containing.BREX | <-TOPRIM<-OLD-A[illegible] | 1241 | Bacteria>Pseudomonadota>Betaproteobacteria>Burkholderiales>Comamonadaceae>unclassified_Comamonadaceae | GCA_016000285.1 | Comamonadaceae_bacterium | I8H87_09350 | 9585..13310 | 1871071 | ATP-binding-protein-[Comamonadaceae-bacterium] |
| WP_050467411.1 | Type3.MTaseII.containing.BREX | SF1-SF2-Helicas[illegible] | 1246 | Bacteria>Pseudomonadota>Betaproteobacteria>Burkholderiales>Oxalobacteraceae>Herbaspirillum | GCF_001571165.1 | Herbaspirillum_chlorophenolicum | HCH01S_RS12885 | 63057..66797 | 211589 | DUF6079-family-protein-[Herbaspirillum-chlorophenolicum] |
| WP_020656698.1 | Type3.MTaseII.containing.BREX | DDE_3-><-?\|\|wHT[illegible] | 1241 | Bacteria>Pseudomonadota>Betaproteobacteria>Burkholderiales>Oxalobacteraceae>Massilia_group | GCF_000382345.1 | Massilia_niastensis | H162_RS0127490 | 2592..6317 | 544911 | DUF6079-family-protein-[Massilia-niastensis] |
| WP_062194134.1 | Type3.MTaseII.containing.BREX | Phage_lysozyme-[illegible] | 1241 | Bacteria>Pseudomonadota>Betaproteobacteria>Burkholderiales>Sphaerotilaceae>Caldimonas | GCF_001592165.1 | Caldimonas_taiwanensis | CTA01S_RS07725 | 11112..14837 | 307483 | DUF6079-family-protein-[Caldimonas-taiwanensis] |
| MBN9203000.1 | Type3.MTaseII.containing.BREX | VirB9->VirB10->[illegible] | 1241 | Bacteria>Pseudomonadota>Betaproteobacteria>Burkholderiales>Sphaerotilaceae>Methylibium | GCA_017306535.1 | Methylibium_petroleiphilum | J0H28_00720 | 141559..145284 | 105560 | ATP-binding-protein-[Methylibium-petroleiphilum] |
| WP_273281156.1 | Type3.MTaseII.containing.BREX | VirB9->VirB10->[illegible] | 1241 | Bacteria>Pseudomonadota>Betaproteobacteria>Burkholderiales>Sphaerotilaceae>Methylibium | GCF_017306535.1 | Methylibium_petroleiphilum | J0H28_RS00725 | 141559..145284 | 105560 | DUF6079-family-protein-[Methylibium-petroleiphilum] |
| MBK7531541.1 | Type3.MTaseII.containing.BREX | VirD4-TraG+VirD[illegible] | 1242 | Bacteria>Pseudomonadota>Betaproteobacteria>Burkholderiales>Sphaerotilaceae>Piscinibacter | GCA_016706565.1 | Piscinibacter_sp. | IPI47_13325 | 105261..108989 | 1903157 | ATP-binding-protein-[Piscinibacter-sp.] |
| ODU09013.1 | Type3.MTaseII.containing.BREX | <-MetJarc-HTH\|\|[illegible] | 1246 | Bacteria>Pseudomonadota>Betaproteobacteria>Burkholderiales>Sphaerotilaceae>Rubrivivax | GCA_001724995.1 | Rubrivivax_sp._SCN_71-131 | ABS84_09895 | 9308..13048 | 1660138 | MAG:-ATPase-[Rubrivivax-sp.-SCN-71-131] |
| OJV49486.1 | Type3.MTaseII.containing.BREX | <-PglZ<-TUDOR+S[illegible] | 1246 | Bacteria>Pseudomonadota>Betaproteobacteria>Burkholderiales>unclassified_Burkholderiales | GCA_001897615.1 | Burkholderiales_bacterium_68-10 | BGO36_11460 | 29575..33315 | 1895730 | ATPase-[Burkholderiales-bacterium-68-10] |
| MCA3175340.1 | Type3.MTaseII.containing.BREX | wHTH+WYL+WCX-p-[illegible] | 1242 | Bacteria>Pseudomonadota>Betaproteobacteria>Burkholderiales>unclassified_Burkholderiales | GCA_020248585.1 | Burkholderiales_bacterium | ING36_07315 | 902..4630 | 1891238 | ATP-binding-protein-[Burkholderiales-bacterium] |
| WP_132764525.1 | Type3.MTaseII.containing.BREX | Phage_lysozyme-[illegible] | 1241 | Bacteria>Pseudomonadota>Betaproteobacteria | GCF_026057715.1 | Burkholderiales | ONS87_RS02500 | 493494..497219 | 80840 | MULTISPECIES:-DUF6079-family-protein-[Burkholderiales] |
| WP_022652891.1 | Type3.MTaseII.containing.BREX | <-BrxA-DNA-bind[illegible] | 1246 | Bacteria>Pseudomonadota>Betaproteobacteria>Neisseriales>Chromobacteriaceae>Aquaspirillum | GCF_000420525.1 | Aquaspirillum_serpens | G465_RS0102410 | 25502..29242 | 190 | DUF6079-family-protein-[Aquaspirillum-serpens] |
| MCG9054660.1 | Type3.MTaseII.containing.BREX | Phage_lysozyme-[illegible] | 1246 | Bacteria>Pseudomonadota>Betaproteobacteria>Neisseriales>Chromobacteriaceae>Laribacter | GCA_022394275.1 | Laribacter_hongkongensis | LH442_01420 | 284780..288520 | 168471 | DUF6079-family-protein-[Laribacter-hongkongensis] |
| WP_239882941.1 | Type3.MTaseII.containing.BREX | <-HEPN<-SF1-SF2[illegible] | 1243 | Bacteria>Pseudomonadota>Betaproteobacteria>Neisseriales>Chromobacteriaceae>Laribacter | GCF_022394215.1 | Laribacter_hongkongensis | LH420_RS00160 | 42696..46427 | 168471 | DUF6079-family-protein-[Laribacter-hongkongensis] |
| WP_239901909.1 | Type3.MTaseII.containing.BREX | <-HNH<-SMODS<-?[illegible] | 1242 | Bacteria>Pseudomonadota>Betaproteobacteria>Neisseriales>Chromobacteriaceae>Laribacter | GCF_022394235.1 | Laribacter_hongkongensis | LH447_RS07105 | 4517..8245 | 168471 | DUF6079-family-protein-[Laribacter-hongkongensis] |
| WP_003782592.1 | Type3.MTaseII.containing.BREX | <-HTH_6+SIS<-NU[illegible] | 1246 | Bacteria>Pseudomonadota>Betaproteobacteria>Neisseriales>Neisseriaceae>Kingella | GCF_016127355.1 | Kingella_denitrificans | I6I17_RS09765 | 1919649..1923389 | 502 | DUF6079-family-protein-[Kingella-denitrificans] |
| WP_205277212.1 | Type3.MTaseII.containing.BREX | <-DDE_Tnp_IS159[illegible] | 1246 | Bacteria>Pseudomonadota>Betaproteobacteria>Neisseriales>Neisseriaceae>Neisseria | GCF_003351545.1 | Neisseria_elongata | DV446_RS10120 | 1986248..1989988 | 495 | DUF6079-family-protein-[Neisseria-elongata] |
| HCJ50308.1 | Type3.MTaseII.containing.BREX | <-BrxA-DNA-bind[illegible] | 1245 | Bacteria>Pseudomonadota>Betaproteobacteria>Nitrosomonadales>Gallionellaceae>Gallionella | GCA_003530225.1 | Gallionella_sp. | DHV67_00400 | 12362..16099 | 2052161 | ATPase-[Gallionella-sp.] |
| QOJ10442.1 | Type3.MTaseII.containing.BREX | <-BrxA-DNA-bind[illegible] | 1242 | Bacteria>Pseudomonadota>Betaproteobacteria>Nitrosomonadales>Nitrosomonadaceae>Nitrosomonas | GCA_015709435.1 | Nitrosomonas_sp._H1_AOB3 | HRU73_09585 | 2063594..2067322 | 2741553 | ATP-binding-protein-[Nitrosomonas-sp.-H1_AOB3] |
| MCO6428551.1 | Type3.MTaseII.containing.BREX | DUF6079-p*->Sch[illegible] | 288 | Bacteria>Pseudomonadota>Betaproteobacteria>Nitrosomonadales>Nitrosomonadaceae>Nitrosomonas | GCA_024102715.1 | Nitrosomonas_communis | J5I83_10905 | 1..867 | 44574 | ATP-binding-protein-[Nitrosomonas-communis] |
| MCO6428679.1 | Type3.MTaseII.containing.BREX | <-Thr_synth_N+P[illegible] | 867 | Bacteria>Pseudomonadota>Betaproteobacteria>Nitrosomonadales>Nitrosomonadaceae>Nitrosomonas | GCA_024102715.1 | Nitrosomonas_communis | J5I83_11595 | 23923..26525 | 44574 | ATP-binding-protein-[Nitrosomonas-communis] |
| WP_172630389.1 | Type3.MTaseII.containing.BREX | <-METHYLASE<-?<[illegible] | 1287 | Bacteria>Pseudomonadota>Betaproteobacteria>Nitrosomonadales>Nitrosomonadaceae>Nitrosomonas | GCF_006742785.1 | Nitrosomonas_stercoris | Nstercoris_RS04280 | 893796..897659 | 1444684 | DUF6079-family-protein-[Nitrosomonas-stercoris] |
| WP_074721729.1 | Type3.MTaseII.containing.BREX | HNH-><-?\|\|HEPN-[illegible] | 1243 | Bacteria>Pseudomonadota>Betaproteobacteria>Nitrosomonadales>Nitrosomonadaceae>Nitrosomonas | GCF_900111055.1 | Nitrosomonas_ureae | BMW48_RS11200 | 12610..16341 | 44577 | DUF6079-family-protein-[Nitrosomonas-ureae] |
| WP_096293996.1 | Type3.MTaseII.containing.BREX | TraI-VirD2-RCRE[illegible] | 1243 | Bacteria>Pseudomonadota>Betaproteobacteria>Nitrosomonadales>Nitrosomonadaceae>Nitrosomonas | GCF_900206265.1 | Nitrosomonas_ureae | CPG39_RS13025 | 2801231..2804962 | 44577 | DUF6079-family-protein-[Nitrosomonas-ureae] |
| WP_218184832.1 | Type3.MTaseII.containing.BREX | <-ENDO3+NUDIX<-[illegible] | 1246 | Bacteria>Pseudomonadota>Betaproteobacteria>Nitrosomonadales>Nitrosomonadaceae>Nitrosovibrio | GCF_900119085.1 | Nitrosovibrio_sp._Nv17 | BT108_RS06120 | 109182..112922 | 1855339 | DUF6079-family-protein-[Nitrosovibrio-sp.-Nv17] |
| WP_008060486.1 | Type3.MTaseII.containing.BREX | ARG->NERD+SF1-S[illegible] | 1246 | Bacteria>Pseudomonadota>Betaproteobacteria>Nitrosomonadales>Sterolibacteriaceae>Methyloversatilis | GCF_000214035.2 | Methyloversatilis_universalis | METUNV1_RS07625 | 21792..25532 | 378211 | DUF6079-family-protein-[Methyloversatilis-universalis] |
| MBP9798940.1 | Type3.MTaseII.containing.BREX | PALP+Thr_dehydr[illegible] | 1246 | Bacteria>Pseudomonadota>Betaproteobacteria>Nitrosomonadales>Sterolibacteriaceae>Sterolibacterium | GCA_018062105.1 | Sterolibacterium_sp. | KBD39_02325 | 22013..25753 | 2819261 | ATP-binding-protein-[Sterolibacterium-sp.] |
| WP_198415290.1 | Type3.MTaseII.containing.BREX | Phage_holin->Ph[illegible] | 1250 | Bacteria>Pseudomonadota>Betaproteobacteria>Nitrosomonadales>Sulfuricellaceae>Sulfuriferula | GCF_009938015.1 | Sulfuriferula_plumbiphila | GZH91_RS10555 | 2028088..2031840 | 171865 | DUF6079-family-protein-[Sulfuriferula-plumbiphila] |
| WP_018076239.1 | Type3.MTaseII.containing.BREX | VirD4-TraG->DUF[illegible] | 1243 | Bacteria>Pseudomonadota>Betaproteobacteria>Nitrosomonadales>Thiobacillaceae>Thiobacillus | GCF_000376425.1 | Thiobacillus_denitrificans | B059_RS0100235 | 47866..51597 | 36861 | DUF6079-family-protein-[Thiobacillus-denitrificans] |
| ODT73869.1 | Type3.MTaseII.containing.BREX | DUF6079-p*->N6-[illegible] | 371 | Bacteria>Pseudomonadota>Betaproteobacteria>Nitrosomonadales>unclassified_Nitrosomonadales | GCA_001724915.1 | Nitrosomonadales_bacterium_SCN_54-20 | ABS69_13030 | 166..1281 | 1660164 | hypothetical-protein-ABS69_13030-[Nitrosomonadales-bacterium-SCN-54-20] |
| MBN8461437.1 | Type3.MTaseII.containing.BREX | AbiEi_4->AbiEii[illegible] | 1246 | Bacteria>Pseudomonadota>Betaproteobacteria>Rhodocyclales>Azonexaceae>Dechloromonas | GCA_017302355.1 | Dechloromonas_sp. | J0M01_01275 | 8315..12055 | 1917218 | hypothetical-protein-[Dechloromonas-sp.] |
| WP_153146309.1 | Type3.MTaseII.containing.BREX | REase-><-McrC-N[illegible] | 1242 | Bacteria>Pseudomonadota>Betaproteobacteria>Rhodocyclales>Azonexaceae>Dechloromonas | GCF_009469615.1 | Dechloromonas_sp._H13 | GH660_RS08865 | 103702..107430 | 2570193 | DUF6079-family-protein-[Dechloromonas-sp.-H13] |
| WP_211364234.1 | Type3.MTaseII.containing.BREX | <-LON-AAA<-TUDO[illegible] | 558 | Bacteria>Pseudomonadota>Betaproteobacteria>Rhodocyclales>Rhodocyclaceae>Propionivibrio | GCF_009469695.1 | Propionivibrio_limicola | GH659_RS12245 | 5858..7534 | 167645 | DUF6079-family-protein-[Propionivibrio-limicola] |
| WP_202636297.1 | Type3.MTaseII.containing.BREX | LPD32+LPD36+N6_[illegible] | 1241 | Bacteria>Pseudomonadota>Betaproteobacteria>Rhodocyclales>Rhodocyclaceae>Rugosibacter | GCF_000934545.1 | Rugosibacter_aromaticivorans | PG1C_RS04985 | 974601..978326 | 1565605 | DUF6079-family-protein-[Rugosibacter-aromaticivorans] |
| AUM01342.1 | Type3.MTaseII.containing.BREX | Integrase_1->Ph[illegible] | 1241 | Bacteria>Pseudomonadota>Betaproteobacteria>Rhodocyclales>Rhodocyclaceae>unclassified_Rhodocyclaceae | GCA_002863805.1 | Rhodocyclaceae_bacterium | B4966_15140 | 3176069..3179794 | 1898103 | ATPase-[Rhodocyclaceae-bacterium] |
| TXT28577.1 | Type3.MTaseII.containing.BREX | <-HD+PspB<-HKD-[illegible] | 1250 | Bacteria>Pseudomonadota>Betaproteobacteria>Rhodocyclales>Rhodocyclaceae>unclassified_Rhodocyclaceae | GCA_008080565.1 | Rhodocyclaceae_bacterium | FD131_3043 | 29190..32942 | 1898103 | hypothetical-protein-FD131_3043-[Rhodocyclaceae-bacterium] |
| MCA3121329.1 | Type3.MTaseII.containing.BREX | <-Methyltransf_[illegible] | 1240 | Bacteria>Pseudomonadota>Betaproteobacteria>Rhodocyclales>Rhodocyclaceae>unclassified_Rhodocyclaceae | GCA_020248305.1 | Rhodocyclaceae_bacterium | ING90_01685 | 255433..259155 | 1898103 | ATP-binding-protein-[Rhodocyclaceae-bacterium] |
| AKU11248.1 | Type3.MTaseII.containing.BREX | <-RNase<-DUF358[illegible] | 1150 | Bacteria>Pseudomonadota>Betaproteobacteria>Rhodocyclales>Zoogloeaceae>Azoarcus | GCA_001190925.1 | Azoarcus_sp._CIB | AzCIB_1343 | 1531839..1535291 | 198107 | hypothetical-protein-AzCIB_1343-[Azoarcus-sp.-CIB] |
| WP_114650027.1 | Type3.MTaseII.containing.BREX | HTH_36-><-DDE_T[illegible] | 1241 | Bacteria>Pseudomonadota>Betaproteobacteria>Rhodocyclales>Zoogloeaceae>Pseudothauera | GCF_003345255.1 | Pseudothauera_hydrothermalis | DIE29_RS12220 | 2549339..2553064 | 2184083 | DUF6079-family-protein-[Pseudothauera-hydrothermalis] |
| APR03213.1 | Type3.MTaseII.containing.BREX | wHTH+WYL+WCX->?[illegible] | 1246 | Bacteria>Pseudomonadota>Betaproteobacteria>Rhodocyclales>Zoogloeaceae>Thauera | GCA_001922305.1 | Thauera_chlorobenzoica | Tchl_0340 | 359729..363469 | 96773 | Exonuclease-SbcC-[Thauera-chlorobenzoica] |
| WP_043741317.1 | Type3.MTaseII.containing.BREX | <-BrxA-DNA-bind[illegible] | 1242 | Bacteria>Pseudomonadota>Betaproteobacteria>Rhodocyclales>Zoogloeaceae>Thauera | GCF_000831325.1 | Thauera_sp._SWB20 | PO78_4451 | 18675..22403 | 1572758 | DUF6079-family-protein-[Thauera-sp.-SWB20] |
| WP_068635434.1 | Type3.MTaseII.containing.BREX | VirD4-TraG+VirB[illegible] | 1241 | Bacteria>Pseudomonadota>Betaproteobacteria>Rhodocyclales>Zoogloeaceae>Thauera | GCF_001591165.1 | Thauera_butanivorans | TBU01S_RS04460 | 90960..94685 | 86174 | DUF6079-family-protein-[Thauera-butanivorans] |
| WP_075149527.1 | Type3.MTaseII.containing.BREX | wHTH+WYL+WCX->?[illegible] | 1242 | Bacteria>Pseudomonadota>Betaproteobacteria>Rhodocyclales>Zoogloeaceae>Thauera | GCF_001922305.1 | Thauera_chlorobenzoica | Tchl_RS01685 | 359729..363457 | 96773 | DUF6079-family-protein-[Thauera-chlorobenzoica] |
| WP_211309646.1 | Type3.MTaseII.containing.BREX | <-DUF3262<-?<-S[illegible] | 686 | Bacteria>Pseudomonadota>Betaproteobacteria>Rhodocyclales>Zoogloeaceae>Thauera | GCF_003030465.1 | Thauera_aromatica | Tharo_RS01595 | 342566..344626 | 59405 | DUF6079-family-protein-[Thauera-aromatica] |
| AVZ79926.1 | Type3.MTaseII.containing.BREX | <-BrxA-DNA-bind[illegible] | 1241 | Bacteria>Pseudomonadota>Betaproteobacteria>Rhodocyclales>Zoogloeaceae>unclassified_Zoogloeaceae | GCA_003054515.1 | Zoogloeaceae_bacteirum_Par-f-2 | C3497_11130 | 2343564..2347289 | 2080469 | ATPase-[Zoogloeaceae-bacteirum-Par-f-2] |
| MBK6654763.1 | Type3.MTaseII.containing.BREX | <-N6-N4-METHYLA[illegible] | 1245 | Bacteria>Pseudomonadota>Betaproteobacteria>Rhodocyclales>Zoogloeaceae>Zoogloea | GCA_016703645.1 | Zoogloea_sp. | IPG40_13145 | 3089..6826 | 49181 | ATP-binding-protein-[Zoogloea-sp.] |
| WP_153163450.1 | Type3.MTaseII.containing.BREX | <-Glycos_transf[illegible] | 1242 | Bacteria>Pseudomonadota>Betaproteobacteria>Rhodocyclales>Zoogloeaceae>Zoogloea | GCF_009469605.1 | Zoogloea_sp._1C4 | GH681_RS20745 | 16460..20188 | 2570190 | DUF6079-family-protein-[Zoogloea-sp.-1C4] |
| MBK7457970.1 | Type3.MTaseII.containing.BREX | <-Peptidase\|\|?-[illegible] | 1241 | Bacteria>Pseudomonadota>Betaproteobacteria>unclassified_Betaproteobacteria | GCA_016709385.1 | Betaproteobacteria_bacterium | IPJ42_06055 | 27399..31124 | 1891241 | ATP-binding-protein-[Betaproteobacteria-bacterium] |
| WP_118881598.1 | Type3.MTaseII.containing.BREX | DUF2787->?-><-F[illegible] | 258 | Bacteria>Pseudomonadota>Gammaproteobacteria>Aeromonadales>Aeromonadaceae>Aeromonas | - | Aeromonas_hydrophila | BFW41_14965 | 3349333..3350109 | 644 | DUF6079-family-protein-[Aeromonas-hydrophila] |
| WP_156852401.1 | Type3.MTaseII.containing.BREX | DUF2787->?->?->[illegible] | 1241 | Bacteria>Pseudomonadota>Gammaproteobacteria>Aeromonadales>Aeromonadaceae>Aeromonas | - | Aeromonas_jandaei | EF849_03870 | 853126..856851 | 650 | DUF6079-family-protein-[Aeromonas-jandaei] |
| AUV14856.1 | Type3.MTaseII.containing.BREX | wHTH+WYL+WCX->?[illegible] | 1242 | Bacteria>Pseudomonadota>Gammaproteobacteria>Aeromonadales>Aeromonadaceae>Aeromonas | GCA_002903405.1 | Aeromonas_sp._ASNIH3 | C2U39_15665 | 3331707..3335435 | 1636608 | ATPase-[Aeromonas-sp.-ASNIH3] |
| UXB14164.1 | Type3.MTaseII.containing.BREX | <-BrxA-DNA-bind[illegible] | 1242 | Bacteria>Pseudomonadota>Gammaproteobacteria>Aeromonadales>Aeromonadaceae>Aeromonas | GCA_025266835.1 | Aeromonas_dhakensis | GP476_17605 | 3854303..3858031 | 196024 | ATP-binding-protein-[Aeromonas-dhakensis] |
| WP_257119773.1 | Type3.MTaseII.containing.BREX | CbeA_antitoxin-[illegible] | 1241 | Bacteria>Pseudomonadota>Gammaproteobacteria>Aeromonadales>Aeromonadaceae>Aeromonas | GCA_030820635.1 | Aeromonas_hydrophila | NQU91_RS15385 | 3384091..3387816 | 644 | DUF6079-family-protein-[Aeromonas-hydrophila] |
| WP_042863015.1 | Type3.MTaseII.containing.BREX | DUF6079-p*->N6-[illegible] | 158 | Bacteria>Pseudomonadota>Gammaproteobacteria>Aeromonadales>Aeromonadaceae>Aeromonas | GCF_000586055.1 | Aeromonas_hydrophila | F881_RS24825 | 1..477 | 644 | DUF6079-family-protein,-partial-[Aeromonas-hydrophila] |
| WP_052814531.1 | Type3.MTaseII.containing.BREX | <-LexA-protease[illegible] | 1241 | Bacteria>Pseudomonadota>Gammaproteobacteria>Aeromonadales>Aeromonadaceae>Aeromonas | GCF_000820265.1 | Aeromonas_caviae | BN1141_RS02365 | 1219..4944 | 648 | DUF6079-family-protein-[Aeromonas-caviae] |
| WP_049048519.1 | Type3.MTaseII.containing.BREX | <-TRD+TRD<-N6-N[illegible] | 1241 | Bacteria>Pseudomonadota>Gammaproteobacteria>Aeromonadales>Aeromonadaceae>Aeromonas | GCF_001057275.1 | Aeromonas_hydrophila | ADB79_RS01365 | 81689..85414 | 644 | DUF6079-family-protein-[Aeromonas-hydrophila] |
| WP_199770375.1 | Type3.MTaseII.containing.BREX | <-BrxA-DNA-bind[illegible] | 1246 | Bacteria>Pseudomonadota>Gammaproteobacteria>Aeromonadales>Aeromonadaceae>Aeromonas | GCF_002903405.1 | Aeromonas_sp._ASNIH3 | C2U39_RS15665 | 3331707..3335447 | 1636608 | DUF6079-family-protein-[Aeromonas-sp.-ASNIH3] |
| WP_205421749.1 | Type3.MTaseII.containing.BREX | DUF932->?->?->?[illegible] | 986 | Bacteria>Pseudomonadota>Gammaproteobacteria>Aeromonadales>Aeromonadaceae>Aeromonas | GCF_003491245.1 | Aeromonas_hydrophila | BFW41_RS15345 | 3345046..3348006 | 644 | DUF6079-family-protein-[Aeromonas-hydrophila] |
| WP_160881875.1 | Type3.MTaseII.containing.BREX | <-TUDOR+SF1-SF2[illegible] | 1241 | Bacteria>Pseudomonadota>Gammaproteobacteria>Aeromonadales>Aeromonadaceae>Aeromonas | GCF_009834905.1 | Aeromonas_veronii | GS629_RS15370 | 5809..9534 | 654 | DUF6079-family-protein-[Aeromonas-veronii] |
| WP_167824812.1 | Type3.MTaseII.containing.BREX | GDResolvase+HIN[illegible] | 1241 | Bacteria>Pseudomonadota>Gammaproteobacteria>Aeromonadales>Aeromonadaceae>Aeromonas | GCF_012102435.1 | Aeromonas_caviae | HED45_RS02655 | 529489..533214 | 648 | DUF6079-family-protein-[Aeromonas-caviae] |
| WP_197970307.1 | Type3.MTaseII.containing.BREX | <-SIR2<-BrxA-DN[illegible] | 1246 | Bacteria>Pseudomonadota>Gammaproteobacteria>Aeromonadales>Aeromonadaceae>Aeromonas | GCF_014158455.1 | Aeromonas_caviae | WP3S18E02_14300 | 1546032..1549772 | 648 | DUF6079-family-protein-[Aeromonas-caviae] |
| WP_194759723.1 | Type3.MTaseII.containing.BREX | HNH-><-BrxA-DNA[illegible] | 1241 | Bacteria>Pseudomonadota>Gammaproteobacteria>Aeromonadales>Aeromonadaceae>Aeromonas | GCF_015353175.1 | Aeromonas_hydrophila | F3H83_RS12425 | 28743..32468 | 644 | DUF6079-family-protein-[Aeromonas-hydrophila] |
| WP_201940947.1 | Type3.MTaseII.containing.BREX | DUF2787->?->?->[illegible] | 1241 | Bacteria>Pseudomonadota>Gammaproteobacteria>Aeromonadales>Aeromonadaceae>Aeromonas | GCF_016730045.1 | Aeromonas_enteropelogenes | JD523_09010 | 534099..537824 | 29489 | DUF6079-family-protein-[Aeromonas-enteropelogenes] |
| WP_219264217.1 | Type3.MTaseII.containing.BREX | CbeA_antitoxin-[illegible] | 1241 | Bacteria>Pseudomonadota>Gammaproteobacteria>Aeromonadales>Aeromonadaceae>Aeromonas | GCF_019348735.1 | Aeromonas_veronii | GL272_RS08150 | 398371..402096 | 654 | DUF6079-family-protein-[Aeromonas-veronii] |
| WP_234657512.1 | Type3.MTaseII.containing.BREX | Phage_integrase[illegible] | 1241 | Bacteria>Pseudomonadota>Gammaproteobacteria>Aeromonadales>Aeromonadaceae>Aeromonas | GCF_021440765.1 | Aeromonas_caviae | LZ630_RS10960 | 7652..11377 | 648 | DUF6079-family-protein-[Aeromonas-caviae] |
| WP_244365015.1 | Type3.MTaseII.containing.BREX | CbeA_antitoxin-[illegible] | 1241 | Bacteria>Pseudomonadota>Gammaproteobacteria>Aeromonadales>Aeromonadaceae>Aeromonas | GCF_022836795.1 | Aeromonas_dhakensis | MRR68_RS06385 | 28219..31944 | 196024 | DUF6079-family-protein-[Aeromonas-dhakensis] |
| WP_260750840.1 | Type3.MTaseII.containing.BREX | <-BrxA-DNA-bind[illegible] | 1246 | Bacteria>Pseudomonadota>Gammaproteobacteria>Aeromonadales>Aeromonadaceae>Aeromonas | GCF_025266835.1 | Aeromonas_dhakensis | GP476_RS17630 | 3854303..3858043 | 196024 | DUF6079-family-protein-[Aeromonas-dhakensis] |
| WP_264787459.1 | Type3.MTaseII.containing.BREX | TetR-HTH->SbcC-[illegible] | 1175 | Bacteria>Pseudomonadota>Gammaproteobacteria>Aeromonadales>Aeromonadaceae>Aeromonas | GCF_026000055.1 | Aeromonas_caviae | OPI30_RS05975 | 1140878..1144405 | 648 | DUF6079-family-protein-[Aeromonas-caviae] |
| WP_268433388.1 | Type3.MTaseII.containing.BREX | HNH-><-BrxA-DNA[illegible] | 1241 | Bacteria>Pseudomonadota>Gammaproteobacteria>Aeromonadales>Aeromonadaceae>Aeromonas | GCF_026797875.1 | Aeromonas_media | OTH18_RS02155 | 466986..470711 | 651 | DUF6079-family-protein-[Aeromonas-media] |
| WP_274161900.1 | Type3.MTaseII.containing.BREX | <-LexA-protease[illegible] | 1241 | Bacteria>Pseudomonadota>Gammaproteobacteria>Aeromonadales>Aeromonadaceae>Aeromonas | GCF_028683275.1 | Aeromonas_veronii | LZT27_RS21700 | 1042..4767 | 654 | DUF6079-family-protein-[Aeromonas-veronii] |
| WP_275056624.1 | Type3.MTaseII.containing.BREX | HNH-><-BrxA-DNA[illegible] | 1241 | Bacteria>Pseudomonadota>Gammaproteobacteria>Aeromonadales>Aeromonadaceae>Aeromonas | GCF_029024985.1 | Aeromonas_allosaccharophila | PYU98_RS15735 | 3402895..3406620 | 656 | DUF6079-family-protein-[Aeromonas-allosaccharophila] |
| WP_275115563.1 | Type3.MTaseII.containing.BREX | Phage_int_SAM_4[illegible] | 1241 | Bacteria>Pseudomonadota>Gammaproteobacteria>Aeromonadales>Aeromonadaceae>Aeromonas | GCF_029025785.1 | Aeromonas_hydrophila | PY771_14825 | 3211006..3214731 | 644 | DUF6079-family-protein-[Aeromonas-hydrophila] |
| WP_139730802.1 | Type3.MTaseII.containing.BREX | <-TetR-HTH\|\|HH[illegible] | 1241 | Bacteria>Pseudomonadota>Gammaproteobacteria>Aeromonadales>Aeromonadaceae>Aeromonas | GCF_029975015.1 | Aeromonas_veronii | CF128_RS08315 | 222050..225775 | 654 | DUF6079-family-protein-[Aeromonas-veronii] |
| WP_108651333.1 | Type3.MTaseII.containing.BREX | <-HTH\|\|HNH-><-[illegible] | 1241 | Bacteria>Pseudomonadota>Gammaproteobacteria>Aeromonadales>Aeromonadaceae>Dongshaea | GCF_003072645.1 | Dongshaea_marina | DB847_RS14455 | 3054078..3057803 | 2047966 | DUF6079-family-protein-[Dongshaea-marina] |
| WP_163156127.1 | Type3.MTaseII.containing.BREX | HNH-><-BrxA-DNA[illegible] | 1241 | Bacteria>Pseudomonadota>Gammaproteobacteria>Aeromonadales>Aeromonadaceae | GCF_010974915.1 | Aeromonas | G4931_RS12740 | 204882..208607 | 642 | MULTISPECIES:-DUF6079-family-protein-[Aeromonas] |
| WP_180911160.1 | Type3.MTaseII.containing.BREX | TetR-HTH->SbcC-[illegible] | 1241 | Bacteria>Pseudomonadota>Gammaproteobacteria>Aeromonadales>Aeromonadaceae | GCF_013488005.1 | Aeromonas | OKC40_RS18425 | 21308..25033 | 642 | MULTISPECIES:-DUF6079-family-protein-[Aeromonas] |
| WP_106453122.1 | Type3.MTaseII.containing.BREX | DOC+HTH->HTH_36[illegible] | 1241 | Bacteria>Pseudomonadota>Gammaproteobacteria>Aeromonadales>Aeromonadaceae>Zobellella | GCF_003012795.1 | Zobellella_taiwanensis | C7I36_RS07620 | 82671..86396 | 347535 | DUF6079-family-protein-[Zobellella-taiwanensis] |
| WP_039222424.1 | Type3.MTaseII.containing.BREX | <-BrxA-DNA-bind[illegible] | 1240 | Bacteria>Pseudomonadota>Gammaproteobacteria>Alteromonadales>Alteromonadaceae>Alteromonas/Salinimonas_group | GCF_000808575.1 | Alteromonas_marina | RJ41_RS14760 | 12744..16466 | 203795 | DUF6079-family-protein-[Alteromonas-marina] |
| WP_073324035.1 | Type3.MTaseII.containing.BREX | REase+SF1-SF2-[illegible] | 1240 | Bacteria>Pseudomonadota>Gammaproteobacteria>Alteromonadales>Alteromonadaceae>Marisediminitalea | GCF_900129565.1 | Marisediminitalea_aggregata | BUB81_RS14520 | 60080..63802 | 634436 | DUF6079-family-protein-[Marisediminitalea-aggregata] |
| WP_274620656.1 | Type3.MTaseII.containing.BREX | HEPN->HEPN-p->[illegible] | 205 | Bacteria>Pseudomonadota>Gammaproteobacteria>Alteromonadales>Colwelliaceae>Colwellia | GCF_022669015.1 | Colwellia_maritima | L3081_RS27325 | 152696..153313 | 2912588 | DUF6079-family-protein-[Colwellia-maritima] |
| WP_274620657.1 | Type3.MTaseII.containing.BREX | HEPN->HEPN-p->[illegible] | 687 | Bacteria>Pseudomonadota>Gammaproteobacteria>Alteromonadales>Colwelliaceae>Colwellia | GCF_022669015.1 | Colwellia_maritima | L3081_RS27330 | 153243..155306 | 2912588 | DUF6079-family-protein-[Colwellia-maritima] |
| WP_274620658.1 | Type3.MTaseII.containing.BREX | HEPN->HEPN-p->[illegible] | 366 | Bacteria>Pseudomonadota>Gammaproteobacteria>Alteromonadales>Colwelliaceae>Colwellia | GCF_022669015.1 | Colwellia_maritima | L3081_RS27335 | 155326..156426 | 2912588 | DUF6079-family-protein-[Colwellia-maritima] |
| MCC5856369.1 | Type3.MTaseII.containing.BREX | <-MODE-HTH+PBP[illegible] | 1240 | Bacteria>Pseudomonadota>Gammaproteobacteria>Alteromonadales>Idiomarinaceae>Idiomarina | GCA_020831965.1 | Idiomarina_sp. | JJU10_11930 | 185517..189239 | 1874361 | hypothetical-protein-[Idiomarina-sp.] |
| KAF7780889.1 | Type3.MTaseII.containing.BREX | <-Phage_integra[illegible] | 1196 | Bacteria>Pseudomonadota>Gammaproteobacteria>Alteromonadales>Pseudoalteromonadaceae>Pseudoalteromonas | GCA_000238335.4 | Pseudoalteromonas_marina | PMAN_a1997 | 1867305..1870895 | 267375 | hypothetical-protein-PMAN_a1997-[Pseudoalteromonas-marina] |
| WP_010557259.1 | Type3.MTaseII.containing.BREX | <-Phage_integra[illegible] | 1240 | Bacteria>Pseudomonadota>Gammaproteobacteria>Alteromonadales>Pseudoalteromonadaceae>Pseudoalteromonas | GCF_000238335.3 | Pseudoalteromonas_marina | PMAN_RS08580 | 1867173..1870895 | 267375 | DUF6079-family-protein-[Pseudoalteromonas-marina] |
| WP_138595226.1 | Type3.MTaseII.containing.BREX | HNH-><-?<-BrxA-[illegible] | 1240 | Bacteria>Pseudomonadota>Gammaproteobacteria>Alteromonadales>Pseudoalteromonadaceae>Pseudoalteromonas | GCF_005887175.1 | Pseudoalteromonas_sp._S1612 | CWB78_RS09515 | 18150..21872 | 579507 | DUF6079-family-protein-[Pseudoalteromonas-sp.-S1612] |
| WP_256828522.1 | Type3.MTaseII.containing.BREX | HTH_3->HNH-><-[illegible] | 1240 | Bacteria>Pseudomonadota>Gammaproteobacteria>Alteromonadales>Pseudoalteromonadaceae>Pseudoalteromonas | GCF_024518675.1 | Pseudoalteromonas_shioyasakiensis | NQT69_RS15130 | 249668..253390 | 1190813 | DUF6079-family-protein-[Pseudoalteromonas-shioyasakiensis] |
| WP_256848669.1 | Type3.MTaseII.containing.BREX | HTH_3->HNH-><-[illegible] | 1240 | Bacteria>Pseudomonadota>Gammaproteobacteria>Alteromonadales>Pseudoalteromonadaceae>Pseudoalteromonas | GCF_024518825.1 | Pseudoalteromonas_carrageenovora | NQT72_RS15575 | 144009..147731 | 227 | DUF6079-family-protein-[Pseudoalteromonas-carrageenovora] |
| MBU76311.1 | Type3.MTaseII.containing.BREX | <-Phage_integra[illegible] | 1240 | Bacteria>Pseudomonadota>Gammaproteobacteria>Alteromonadales>Pseudoalteromonadaceae>unclassified_Pseudoalteromonadaceae | GCA_002731235.1 | Pseudoalteromonadaceae_bacterium | CMK63_04865 | 42457..46179 | 2026783 | hypothetical-protein-[Pseudoalteromonadaceae-bacterium] |
| MCF1431078.1 | Type3.MTaseII.containing.BREX | <-NERD+SF1-SF2-[illegible] | 236 | Bacteria>Pseudomonadota>Gammaproteobacteria>Alteromonadales>Shewanellaceae>Shewanella | GCA_021462555.1 | Shewanella_sp. | LPH20_09470 | 12421..13131 | 50422 | DUF6079-family-protein-[Shewanella-sp.] |
| WP_069455077.1 | Type3.MTaseII.containing.BREX | <-LexA-protease[illegible] | 1240 | Bacteria>Pseudomonadota>Gammaproteobacteria>Alteromonadales>Shewanellaceae>Shewanella | GCF_002074855.1 | Shewanella_xiamenensis | VO00_RS07310 | 30571..34293 | 332186 | DUF6079-family-protein-[Shewanella-xiamenensis] |
| WP_259569843.1 | Type3.MTaseII.containing.BREX | <-ANK+ANK+ANK<-[illegible] | 1240 | Bacteria>Pseudomonadota>Gammaproteobacteria>Alteromonadales>Shewanellaceae>Shewanella | GCF_024999565.1 | Shewanella_baltica | G3445_RS08200 | 194360..198082 | 62322 | DUF6079-family-protein-[Shewanella-baltica] |
| WP_263191773.1 | Type3.MTaseII.containing.BREX | <-ANK+ANK+ANK<-[illegible] | 1240 | Bacteria>Pseudomonadota>Gammaproteobacteria>Alteromonadales>Shewanellaceae>Shewanella | GCF_025653705.1 | Shewanella_sp._SM87 | L5M11_RS07375 | 37931..41653 | 2912808 | DUF6079-family-protein-[Shewanella-sp.-SM87] |
| KNZ40012.1 | Type3.MTaseII.containing.BREX | <-TOPRIM+Topois[illegible] | 123 | Bacteria>Pseudomonadota>Gammaproteobacteria>Cardiobacteriales>Cardiobacteriaceae>Dichelobacter | GCA_001263335.1 | Dichelobacter_nodosus | AKG33_00780 | 6863..7234 | 870 | hypothetical-protein-AKG33_00780-[Dichelobacter-nodosus] |
| WP_011927931.1 | Type3.MTaseII.containing.BREX | <-SF1-SF2-Helicase<-?<-BrxA-DNA-binding<-PglZ<-TUDOR+SF1-SF2-Helicase+iREase+CTD<-HEPN<-N6-N4-METHYLASE<-DUF6079*<-BrxB-BrxF-iSTAND<-XIS-HTH<-?<-?<-?<-?<-wHTH | 1243 | Bacteria>Pseudomonadota>Gammaproteobacteria>Cardiobacteriales>Cardiobacteriaceae>Dichelobacter | GCF_000015345.1 | Dichelobacter_nodosus | DNO_RS00875 | 195920..199651 | 870 | DUF6079-family-protein-[Dichelobacter-nodosus] |
| WP_235503142.1 | Type3.MTaseII.containing.BREX | <-TOPRIM+Topoisom_bac\|\|XIS-HTH->?->?->?->AAA+HTH-><-Arm-DNA-bind_3+Phage_integrase\|\|DUF6079-p*->N6-N4-METHYLASE->HEPN->TUDOR+SF1-SF2-Helicase+iREase+CTD->PglZ->BrxA[illegible] | 133 | Bacteria>Pseudomonadota>Gammaproteobacteria>Cardiobacteriales>Cardiobacteriaceae>Dichelobacter | GCF_001263335.1 | Dichelobacter_nodosus | AKG33_RS00780 | 6833..7234 | 870 | DUF6079-family-protein-[Dichelobacter-nodosus] |
| WP_235503143.1 | Type3.MTaseII.containing.BREX | <-HTH_OrfB_IS605+OrfB_IS605+OrfB_Zn_ribbon\|\|<-PhdYeFM_antitox<-?<-DUF6079*<-BrxB-BrxF-iSTAND<-XIS-HTH<-?<-?<-?<-?<-wHTH | 1113 | Bacteria>Pseudomonadota>Gammaproteobacteria>Cardiobacteriales>Cardiobacteriaceae>Dichelobacter | GCF_001263335.1 | Dichelobacter_nodosus | AKG33_RS00860 | 5171..8512 | 870 | DUF6079-family-protein-[Dichelobacter-nodosus] |
| MBU2886739.1 | Type3.MTaseII.containing.BREX | Arm-DNA-bind_3+Phage_integrase->?->XIS-HTH->?->?->BrxB-BrxF-iSTAND->DUF6079*->Schlafen+HATPase+wHTH->N6-N4-METHYLASE->TUDOR+SF1-SF2-Helicase+iREase+CTD->PglZ->BrxA-DNA[illegible] | 1248 | Bacteria>Pseudomonadota>Gammaproteobacteria>Cellvibrionales>Cellvibrionaceae>Gilvimarinus | GCA_018860575.1 | Gilvimarinus_agarilyticus | KO507_13275 | 315982..319728 | 679259 | hypothetical-protein-[Gilvimarinus-agarilyticus] |
| WP_266072734.1 | Type3.MTaseII.containing.BREX | Phage_integrase-><-?\|\|DOC+HTH->TraJ-RHH->VirB5->?->TrbL->wHTH+WYL+WCX->BrxB-BrxF-iSTAND->DUF6079*->N6-N4-METHYLASE->HKD-DNase->Schlafen+HATPase+wHTH->TUDOR+SF1-SF2-Hel[illegible] | 1241 | Bacteria>Pseudomonadota>Gammaproteobacteria>Cellvibrionales>Microbulbiferaceae>Microbulbifer | GCF_026241615.1 | Microbulbifer_thermotolerans | OQJ46_RS08120 | 146692..150417 | 252514 | DUF6079-family-protein-[Microbulbifer-thermotolerans] |
| MAT50696.1 | Type3.MTaseII.containing.BREX | VirB4-FtsK+A32->TrwJ->VirB6->VirB8->VirB9->VirB10->RHH->XIS-HTH->BrxB-BrxF-iSTAND->DUF6079*->?->N6-N4-METHYLASE->Classical-AAA->Peptidase_S8->TUDOR+SF1-SF2-Helicase+i[illegible] | 1246 | Bacteria>Pseudomonadota>Gammaproteobacteria>Cellvibrionales>Porticoccaceae>unclassified_Porticoccaceae | GCA_002702725.1 | Porticoccaceae_bacterium | CMK32_05880 | 13568..17308 | 2026782 | ATPase-[Porticoccaceae-bacterium] |
| BCU06243.1 | Type3.MTaseII.containing.BREX | <-OrfB_Zn_ribbon<-DDE_Tnp<-BrxA-DNA-binding<-PglZ<-TUDOR+SF1-SF2-Helicase+iREase+CTD-p<-TOPRIM<-OLD-ABC<-N6-N4-METHYLASE<-DUF6079*<-BrxB-BrxF-iSTAND<-ABhydrolase | 1246 | Bacteria>Pseudomonadota>Gammaproteobacteria>Chromatiales>Chromatiaceae>Allochromatium | GCA_018409545.1 | Allochromatium_tepidum | Atep_09200 | 1001877..1005617 | 553982 | hypothetical-protein-Atep_09200-[Allochromatium-tepidum] |
| WP_213380490.1 | Type3.MTaseII.containing.BREX | <-Amidohydro_1<-OrfB_Zn_ribbon<-BrxA-DNA-binding<-PglZ<-TUDOR+SF1-SF2-Helicase+iREase+CTD-p<-TOPRIM<-OLD-ABC<-N6-N4-METHYLASE<-DUF6079*<-BrxB-BrxF-iSTAND | 1242 | Bacteria>Pseudomonadota>Gammaproteobacteria>Chromatiales>Chromatiaceae>Allochromatium | GCF_018409545.1 | Allochromatium_tepidum | Atep_RS04630 | 1001877..1005605 | 553982 | DUF6079-family-protein-[Allochromatium-tepidum] |
| WP_201246603.1 | Type3.MTaseII.containing.BREX | HslUClpX-AAA+Classical-AAA->?-><-Pkinase+STYKIN<-?<-METHYLASE<-?<-?<-?\|\|BrxB-BrxF-iSTAND->DUF6079*->?->PemK_toxin->N6-N4-METHYLASE->OLD-ABC-> | 1266 | Bacteria>Pseudomonadota>Gammaproteobacteria>Chromatiales>Chromatiaceae>Halochromatium | GCF_016653295.1 | Halochromatium_salexigens | CCR82_RS14940 | 56680..60480 | 49447 | DUF6079-family-protein-[Halochromatium-salexigens] |
| WP_242519126.1 | Type3.MTaseII.containing.BREX | <-Pkinase+STYKIN<-?<-METHYLASE<-?<-?<-?<-REase\|\|BrxB-BrxF-iSTAND->DUF6079-p*-> | 284 | Bacteria>Pseudomonadota>Gammaproteobacteria>Chromatiales>Chromatiaceae>Halochromatium | GCF_016653315.1 | Halochromatium_roseum | CCR96_RS15560 | 61662..62514 | 391920 | DUF6079-family-protein,-partial-[Halochromatium-roseum] |
| WP_242519268.1 | Type3.MTaseII.containing.BREX | <-BrxA-DNA-binding<-PglZ\|\|?-><-TUDOR+SF1-SF2-Helicase+iREase+CTD<-TOPRIM<-OLD-ABC<-N6-N4-METHYLASE<-PemK_toxin<-?<-DUF6079-p* | 784 | Bacteria>Pseudomonadota>Gammaproteobacteria>Chromatiales>Chromatiaceae>Halochromatium | GCF_016653315.1 | Halochromatium_roseum | CCR96_RS21110 | 15785..18140 | 391920 | DUF6079-family-protein,-partial-[Halochromatium-roseum] |
| WP_020505853.1 | Type3.MTaseII.containing.BREX | <-Phage_integrase\|\|?-><-ACET<-BrxA-DNA-binding<-PglZ<-TOPRIM<-OLD-ABC<-N6-N4-METHYLASE<-Schlafen+HATPase+wHTH<-DUF6079*<-BrxB-BrxF-iSTAND | 1253 | Bacteria>Pseudomonadota>Gammaproteobacteria>Chromatiales>Chromatiaceae>Lamprocystis | GCF_000379525.1 | Lamprocystis_purpurea | A390_RS0116920 | 27902..31663 | 61598 | DUF6079-family-protein-[Lamprocystis-purpurea] |
| ADJ27036.1 | Type3.MTaseII.containing.BREX | <-Peptidase_M22\|\|?->?->ZnR+TOPRIM+TOPRIM->SIGMA-HTH+SIGMA-HTH+SIGMA-HTH->tRNA->BrxB-BrxF-iSTAND->DUF6079*->?->N6-N4-METHYLASE->STAS+HisKin->STAS-like->PIN->TUDOR+SF1-S[illegible] | 1326 | Bacteria>Pseudomonadota>Gammaproteobacteria>Chromatiales>Chromatiaceae>Nitrosococcus | GCA_000143085.1 | Nitrosococcus_watsonii_C-113 | Nwat_0045 | 44215..48195 | 105559 | conserved-hypothetical-protein-[Nitrosococcus-watsonii-C-113] |
| EDZ66011.1 | Type3.MTaseII.containing.BREX | HTH_36->?->TOPRIM+DnaB-><-?\|\|?->BrxB-BrxF-iSTAND->DUF6079*->Schlafen+HATPase+wHTH+HTH->REase->N6-N4-METHYLASE->STAS-like-><-?\|\|?->TUDOR+SF1-SF2-Helicase+iREase+CTD->Pg[illegible] | 1218 | Bacteria>Pseudomonadota>Gammaproteobacteria>Chromatiales>Chromatiaceae>Nitrosococcus | GCA_000155655.1 | Nitrosococcus_oceani_AFC27 | NOC27_2691 | 571553..575209 | 473788 | hypothetical-protein-NOC27_2691-[Nitrosococcus-oceani-AFC27] |
| KFI24074.1 | Type3.MTaseII.containing.BREX | <-Peptidase_M22\|\|?->?->ZnR+TOPRIM+TOPRIM->SIGMA-HTH+SIGMA-HTH+SIGMA-HTH->tRNA->BrxB-BrxF-iSTAND->DUF6079-p*-> | 486 | Bacteria>Pseudomonadota>Gammaproteobacteria>Chromatiales>Chromatiaceae>Nitrosococcus | GCA_000740735.1 | Nitrosococcus_oceani | HW44_00260 | 12938..14395 | 1229 | ATPase,-partial-[Nitrosococcus-oceani] |
| WP_011330215.1 | Type3.MTaseII.containing.BREX | HIN-HTH->DDE_3->?->HTH_36->?->TOPRIM+DnaB-><-?\|\|?->BrxB-BrxF-iSTAND->DUF6079*->Schlafen+HATPase+wHTH+HTH->REase->N6-N4-METHYLASE->STAS+HisKin->STAS-like->PIN->TUDOR+SF[illegible] | 1273 | Bacteria>Pseudomonadota>Gammaproteobacteria>Chromatiales>Chromatiaceae>Nitrosococcus | GCF_000012805.1 | Nitrosococcus_oceani | NOC_RS00500 | 54071..57892 | 1229 | DUF6079-family-protein-[Nitrosococcus-oceani] |
| WP_013219148.1 | Type3.MTaseII.containing.BREX | <-Peptidase_M22\|\|?->?->ZnR+TOPRIM+TOPRIM->SIGMA-HTH+SIGMA-HTH+SIGMA-HTH->tRNA->BrxB-BrxF-iSTAND->DUF6079*->?->N6-N4-METHYLASE->STAS+HisKin->STAS-like->PIN->TUDOR+SF1-S[illegible] | 1326 | Bacteria>Pseudomonadota>Gammaproteobacteria>Chromatiales>Chromatiaceae>Nitrosococcus | GCF_000143085.1 | Nitrosococcus_watsonii | NWAT_RS00235 | 44215..48195 | 473531 | DUF6079-family-protein-[Nitrosococcus-watsonii] |
| WP_269319872.1 | Type3.MTaseII.containing.BREX | <-Peptidase_M22<-?\|\|?->?->ZnR+TOPRIM+TOPRIM->SIGMA-HTH+SIGMA-HTH+SIGMA-HTH->tRNA->BrxB-BrxF-iSTAND->DUF6079-p*-> | 500 | Bacteria>Pseudomonadota>Gammaproteobacteria>Chromatiales>Chromatiaceae>Nitrosococcus | GCF_000740735.1 | Nitrosococcus_oceani | HW44_RS00240 | 12896..14395 | 1229 | DUF6079-family-protein,-partial-[Nitrosococcus-oceani] |
| WP_231561312.1 | Type3.MTaseII.containing.BREX | DUF6079-p*->N6-N4-METHYLASE->PHP+SbcC->TUDOR+SF1-SF2-Helicase+iREase+CTD->PglZ->BrxA-DNA-binding-><-?<-REase+SF1-SF2-Helicase\|\|?-><-ABC-ATPase+SbcC+SWACOS+SbcC | 796 | Bacteria>Pseudomonadota>Gammaproteobacteria>Chromatiales>Chromatiaceae>Nitrosococcus | GCF_000740735.1 | Nitrosococcus_oceani | HW44_RS00245 | 70..2460 | 1229 | DUF6079-family-protein-[Nitrosococcus-oceani] |
| WP_036498006.1 | Type3.MTaseII.containing.BREX | HTH_36->?->TOPRIM+DnaB-><-?\|\|?->BrxB-BrxF-iSTAND->DUF6079*->Schlafen+HATPase+wHTH+HTH->REase->N6-N4-METHYLASE->STAS+HisKin->STAS-like->PIN->TUDOR+SF1-SF2-Helicase+iREa[illegible] | 1248 | Bacteria>Pseudomonadota>Gammaproteobacteria>Chromatiales>Chromatiaceae>Nitrosococcus | GCF_000740745.1 | Nitrosococcus_oceani | IB75_RS00205 | 5920..9666 | 1229 | DUF6079-family-protein-[Nitrosococcus-oceani] |
| WP_244859996.1 | Type3.MTaseII.containing.BREX | HTH_36->?->TOPRIM+DnaB-><-?\|\|BrxB-BrxF-iSTAND->DUF6079*->Schlafen+HATPase+wHTH+HTH->REase->N6-N4-METHYLASE->STAS+HisKin->STAS-like->TUDOR+SF1-SF2-Helicase+iREase+CTD->[illegible] | 1231 | Bacteria>Pseudomonadota>Gammaproteobacteria>Chromatiales>Chromatiaceae>Nitrosococcus | - | Nitrosococcus_oceani | NONS58_31380 | 6077..9772 | 1229 | DUF6079-family-protein-[Nitrosococcus-oceani] |
| WP_246172808.1 | Type3.MTaseII.containing.BREX | SF1-SF2-Helicase->tRNA-><-TatD_DNase<-?\|\|?->?->?->?-><-?\|\|DUF6079-p*-><-RADICAL-SAM<-?<-?<-?<-NUDIX<-?\|\|NTP_transferase-> | 208 | Bacteria>Pseudomonadota>Gammaproteobacteria>Chromatiales>Chromatiaceae>Thermochromatium | GCF_009664085.1 | Thermochromatium_tepidum | E6P07_RS08910 | 1951030..1951656 | 1050 | DUF6079-family-protein-[Thermochromatium-tepidum] |
| QVL51256.1 | Type3.MTaseII.containing.BREX | <-TUDOR+SF1-SF2-Helicase+iREase+CTD<-TOPRIM<-OLD-ABC<-N6-N4-METHYLASE<-HEPN<-PIN<-OLD-ABC<-PIN<-PhdYeFM_antitox<-DUF6079*<-BrxB-BrxF-iSTAND\|\|?-><-?<-TOPRIM<-?<-?<-?[illegible] | [illegible] | [illegible] | [illegible] | [illegible] | [illegible] | [illegible] | [illegible] | [illegible] |
| WP_200249822.1 | Type3.MTaseII.containing.BREX | TOPRIM->?->TraJ-RHH->ACET+ACET->?->BrxB-BrxF-iSTAND->DUF6079*->Schlafen+HATPase+wHTH->?->HEPN->N6-N4-METHYLASE->HEPN->ParA-Soj-PloopNTPase+ORC-CDC6-like->TUDOR+SF1-SF2[illegible] | [illegible] | [illegible] | [illegible] | [illegible] | [illegible] | [illegible] | [illegible] | [illegible] |
| WP_201098798.1 | Type3.MTaseII.containing.BREX | [illegible] | [illegible] | [illegible] | [illegible] | [illegible] | [illegible] | [illegible] | [illegible] | [illegible] |
| WP_058557220.1 | Type3.MTaseII.containing.BREX | [illegible] | [illegible] | [illegible] | [illegible] | [illegible] | [illegible] | [illegible] | [illegible] | [illegible] |

| Protein-ID | System | Gene-Neighbourhoods | Length | Taxonomy | Assembly-ID | Strain | Locus-Tag | Genomic-coordinates | TaxID | Fasta-define |
|---|---|---|---|---|---|---|---|---|---|---|
| WP_150092059.1 | Type3.MTaseII.containing.BREX | <-GDResolvase+ZnR<-RelE-ParE<-PhdYeFM_antitox\|\|?->?->?->?->XIS-HTH->BrxB-BrxF-iSTAND->DUF6079*->?->N6-N4-METHYLASE->?->?->OLD-ABC->TOPRIM->TUDOR+SF1-SF2-Helicase+iREase+CTD->PglZ->BrxA-DNA-binding-> | 1261 | Bacteria>Pseudomonadota>Gammaproteobacteria>Chromatiales>Chromatiaceae>Thiohalocapsa | GCF_008632335.1 | Thiohalocapsa_marina | F2Q65_RS07725 | 154263..158048 | 424902 | DUF6079-family-protein-[Thiohalocapsa-marina] |
| WP_009146904.1 | Type3.MTaseII.containing.BREX | <-NERD+SF1-SF2-Helicase<-BrxA-DNA-binding<-PglZ<-TUDOR+SF1-SF2-Helicase+iREase+CTD<-?<-PIN<-SbcC<-N6-N4-METHYLASE<-HTH+REase<-DUF6079*<-BrxB-BrxF-iSTAND<-SF1-SF2-Helicase<-MORC+DUF3684+REase<-?<-MerR-HTH<-?<-?<-Radical_SAM<-MerR-HTH | 1250 | Bacteria>Pseudomonadota>Gammaproteobacteria>Chromatiales>Chromatiaceae>Thiorhodovibrio | GCF_000228725.2 | Thiorhodovibrio_sp._970 | THI970DRAFT_RS02035 | 44705..48457 | 631362 | DUF6079-family-protein-[Thiorhodovibrio-sp.-970] |
| MBK8637141.1 | Type3.MTaseII.containing.BREX | <-BrxA-DNA-binding<-PglZ<-TUDOR+SF1-SF2-Helicase+iREase+CTD<-TOPRIM<-OLD-ABC<-Schlafen+HATPase+wHTH<-DUF6079*<-BrxB-BrxF-iSTAND<-XIS-HTH\|\|?-><-HTH_Tnp_1 | 1245 | Bacteria>Pseudomonadota>Gammaproteobacteria>Chromatiales>Chromatiaceae>unclassified_Chromatiaceae | GCA_016714645.1 | Chromatiaceae_bacterium | IPN92_02265 | 482331..486068 | 2049432 | ATP-binding-protein-[Chromatiaceae-bacterium] |
| MBP6582546.1 | Type3.MTaseII.containing.BREX | <-N6-N4-METHYLASE<-Schlafen+HATPase+wHTH<-TOPRIM<-TOPRIM<-OLD-ABC<-DUF6079*<-BrxB-BrxF-iSTAND<-XIS-HTH | 1245 | Bacteria>Pseudomonadota>Gammaproteobacteria>Chromatiales>Chromatiaceae>unclassified_Chromatiaceae | GCA_017989555.1 | Chromatiaceae_bacterium | KA204_03605 | 5221..8958 | 2049432 | ATP-binding-protein-[Chromatiaceae-bacterium] |
| MBP6733371.1 | Type3.MTaseII.containing.BREX | <-OLD-ABC<-N6-N4-METHYLASE<-Schlafen+HATPase+wHTH<-TOPRIM<-TOPRIM<-OLD-ABC<-DUF6079*<-BrxB-BrxF-iSTAND<-XIS-HTH | 1245 | Bacteria>Pseudomonadota>Gammaproteobacteria>Chromatiales>Chromatiaceae>unclassified_Chromatiaceae | GCA_017990755.1 | Chromatiaceae_bacterium | KA142_01525 | 7507..11244 | 2049432 | ATP-binding-protein-[Chromatiaceae-bacterium] |
| MBP6806939.1 | Type3.MTaseII.containing.BREX | XIS-HTH->BrxB-BrxF-iSTAND->DUF6079*->OLD-ABC->TOPRIM->Schlafen+HATPase+wHTH->N6-N4-METHYLASE-> | 1245 | Bacteria>Pseudomonadota>Gammaproteobacteria>Chromatiales>Chromatiaceae>unclassified_Chromatiaceae | GCA_017991215.1 | Chromatiaceae_bacterium | KA125_01440 | 1631..5368 | 2049432 | ATP-binding-protein-[Chromatiaceae-bacterium] |
| MBP8282514.1 | Type3.MTaseII.containing.BREX | <-N6-N4-METHYLASE-p<-Y1_Tnp<-TOPRIM<-OLD-ABC<-N6-N4-METHYLASE<-Schlafen+HATPase+wHTH<-TOPRIM<-TOPRIM<-OLD-ABC<-DUF6079-p* | 223 | Bacteria>Pseudomonadota>Gammaproteobacteria>Chromatiales>Chromatiaceae>unclassified_Chromatiaceae | GCA_018007135.1 | Chromatiaceae_bacterium | KAX46_01135 | 9248..9920 | 2049432 | ATP-binding-protein-[Chromatiaceae-bacterium] |
| MCF8003110.1 | Type3.MTaseII.containing.BREX | <-BrxA-DNA-binding<-PglZ<-TUDOR+SF1-SF2-Helicase+iREase+CTD<-REase+AIPR<-N6-N4-METHYLASE<-HTH+REase<-DUF6079*<-BrxB-BrxF-iSTAND\|\|REase-> | 1295 | Bacteria>Pseudomonadota>Gammaproteobacteria>Chromatiales>Chromatiaceae>unclassified_Chromatiaceae | GCA_021736845.1 | Chromatiaceae_bacterium | K9L32_02700 | 11872..15759 | 2049432 | DUF6079-family-protein-[Chromatiaceae-bacterium] |
| MCP5408977.1 | Type3.MTaseII.containing.BREX | TraG_N-><-?<-HTH<-BrxA-DNA-binding<-PglZ<-N6-N4-METHYLASE<-Schlafen+HATPase+wHTH<-N6-N4-METHYLASE<-?<-DUF6079*<-BrxB-BrxF-iSTAND<-XIS-HTH\|\|SF1-SF2-Helicase->DrmB->SF1-SF2-Helicase->?->NERD+SF1-SF2-Helicase->\|\|?->DOC+HTH-> | 1241 | Bacteria>Pseudomonadota>Gammaproteobacteria>Chromatiales>Chromatiaceae>unclassified_Chromatiaceae | GCA_024235335.1 | Chromatiaceae_bacterium | H6963_06755 | 95301..99026 | 2049432 | ATP-binding-protein-[Chromatiaceae-bacterium] |
| MCP5307390.1 | Type3.MTaseII.containing.BREX | <-Peptidase\|\|Peptidase->?-><-?\|\|?->RHH_5->BrxB-BrxF-iSTAND->DUF6079*->HHH_4+HTH+wHTH+SF1-SF2-Helicase+REase->N6-N4-METHYLASE->TUDOR+SF1-SF2-Helicase+iREase+CTD->PglZ->BrxA-DNA-binding->HIN-HTH->?->LZ_Tnp_IS66+zf-IS66+DDE_Tnp_IS66+DDE_Tnp_IS66_C-> | 1236 | Bacteria>Pseudomonadota>Gammaproteobacteria>Chromatiales>Chromatiaceae>unclassified_Chromatiaceae | GCA_024235495.1 | Chromatiaceae_bacterium | H6953_18410 | 5242..8952 | 2049432 | ATP-binding-protein-[Chromatiaceae-bacterium] |
| WP_238621060.1 | Type3.MTaseII.containing.BREX | <-BrxA-DNA-binding<-PglZ<-TUDOR+SF1-SF2-Helicase+iREase+CTD<-Schlafen+HATPase+wHTH<-TOPRIM<-OLD-ABC<-N6-N4-METHYLASE<-DUF6079*<-BrxB-BrxF-iSTAND<-wHTH+WYL+WCX<-TrbL<-?<-VirB5<-TraJ-RHH<-DOC+HTH\|\|?-><-Phage_integrase | 1241 | Bacteria>Pseudomonadota>Gammaproteobacteria>Chromatiales>Ectothiorhodospiraceae>Ectothiorhodospira | - | Ectothiorhodospira_sp._B14B | LQ952_RS02380 | 48489..52214 | 2899127 | DUF6079-family-protein-[Ectothiorhodospira-sp.-B14B] |
| WP_008932926.1 | Type3.MTaseII.containing.BREX | <-PglZ<-TUDOR+SF1-SF2-Helicase+iREase+CTD<-Schlafen+HATPase+wHTH<-N6-N4-METHYLASE<-DUF6079*<-BrxB-BrxF-iSTAND<-wHTH+WYL+WCX<-TrbL<-tRNA<-SIGMA-HTH+SIGMA-HTH+SIGMA-HTH<-ZnR+TOPRIM+TOPRIM<-GspA-ATPase<-HD<-?<-SKI | 1241 | Bacteria>Pseudomonadota>Gammaproteobacteria>Chromatiales>Ectothiorhodospiraceae>Ectothiorhodospira | GCF_000225005.1 | Ectothiorhodospira_sp._PHS-1 | ECTPHS_RS11505 | 9601..13326 | 519989 | DUF6079-family-protein-[Ectothiorhodospira-sp.-PHS-1] |
| WP_123401498.1 | Type3.MTaseII.containing.BREX | <-SF1-SF2-Helicase\|\|Aminotran_1_2->?->?->tRNA->BrxB-BrxF-iSTAND->SF1-SF2-Helicase+PhospholipaseD->DUF6079*->Schlafen+HATPase+wHTH->N6-N4-METHYLASE->HEPN->TUDOR+SF1-SF2-Helicase+iREase+CTD->PglZ->BrxA-DNA-binding->ParB+HNH->DUF4236+TPR->DUF488-> | 1243 | Bacteria>Pseudomonadota>Gammaproteobacteria>Chromatiales>Ectothiorhodospiraceae>Inmirania | GCF_003751635.1 | Inmirania_thermothiophila | EDC57_RS08840 | 665304..669035 | 1750597 | DUF6079-family-protein-[Inmirania-thermothiophila] |
| WP_005002347.1 | Type3.MTaseII.containing.BREX | <-BrxA-DNA-binding<-PglZ<-TUDOR+SF1-SF2-Helicase+iREase+CTD<-?<-TOPRIM<-OLD-ABC<-N6-N4-METHYLASE<-HerA/FtsK<-SIR2<-DUF6079*<-BrxB-BrxF-iSTAND<-XIS-HTH | 1241 | Bacteria>Pseudomonadota>Gammaproteobacteria>Chromatiales>Ectothiorhodospiraceae>Nitrococcus | GCF_000153205.1 | Nitrococcus_mobilis | NB231_RS09995 | 2195577..2199302 | 35797 | DUF6079-family-protein-[Nitrococcus-mobilis] |
| QEP44630.1 | Type3.MTaseII.containing.BREX | Aminotran_1_2->?->Peptidase_M20->?->REC+SHELIX+HISKIN->XIS-HTH->BrxB-BrxF-iSTAND->DUF6079*->?->N6-N4-METHYLASE->PHP+SbcC->Schlafen+HATPase+wHTH->TUDOR+SF1-SF2-Helicase+iREase+CTD->PglZ->BrxA-DNA-binding->?-><-ABhydrolase | 1269 | Bacteria>Pseudomonadota>Gammaproteobacteria>Chromatiales>Ectothiorhodospiraceae>unclassified_Ectothiorhodospiraceae | GCA_008375315.1 | Ectothiorhodospiraceae_bacterium_BW-2 | D5085_16715 | 3562570..3566379 | 947515 | ATP-binding-protein-[Ectothiorhodospiraceae-bacterium-BW-2] |
| HH081872.1 | Type3.MTaseII.containing.BREX | XIS-HTH->SF1-SF2-Helicase+PRTase->SLOG+BirA-HTH->BrxB-BrxF-iSTAND->DUF6079-p*-> | 275 | Bacteria>Pseudomonadota>Gammaproteobacteria>Chromatiales>Halothiobacillaceae>Halothiobacillus | GCA_011372975.1 | Halothiobacillus_sp. | ENN01_01315 | 4732..5555 | 1891311 | ATP-binding-protein-[Halothiobacillus-sp.] |
| MBL8226335.1 | Type3.MTaseII.containing.BREX | <-REase\|\|ArdB->DUF932->?->TraJ-RHH->TraI-VirD2-RCRE->XIS-HTH->BrxB-BrxF-iSTAND->DUF6079*->N6-N4-METHYLASE->TUDOR+SF1-SF2-Helicase+iREase+CTD->PglZ->BrxA-DNA-binding-> | 1241 | Bacteria>Pseudomonadota>Gammaproteobacteria>Chromatiales>unclassified_Chromatiales | GCA_016789885.1 | Chromatiales_bacterium | JNM50_13505 | 141114..144839 | 2026725 | ATP-binding-protein-[Chromatiales-bacterium] |
| WP_130590275.1 | Type3.MTaseII.containing.BREX | <-Phage_GPD+Phage_base_V\|\|BrxB-BrxF-iSTAND->DUF6079*->N6-N4-METHYLASE->TUDOR+SF1-SF2-Helicase+iREase+CTD->PglZ->BrxA-DNA-binding->nSTAND1+GTPase-AIG+sGTP-> | 1241 | Bacteria>Pseudomonadota>Gammaproteobacteria>Enterobacterales>Budviciaceae>Jinshanibacter | GCF_004295645.1 | Jinshanibacter_zhutongyuii | EKN56_RS02005 | 530868..534593 | 2498113 | DUF6079-family-protein-[Jinshanibacter-zhutongyuii] |
| WP_086550271.1 | Type3.MTaseII.containing.BREX | XIS-HTH-><-REase<-Arm-DNA-bind_3+Phage_integrase\|\|tRNA->Arm-DNA-bind_3+Phage_integrase->Arm-DNA-bind_3+Phage_integrase->HNH->HIN-HTH-><-DUF6079*<-BrxB-BrxF-iSTAND<-HTH_23+HTH_32+HTH_33+DDE_3<-?<-REase<-?\|\|LACI-HTH+PBPI->?->?-><-Zn_Tnp_IS1+HTH_Tnp_IS1+DDE_Tnp_IS1 | 1224 | Bacteria>Pseudomonadota>Gammaproteobacteria>Enterobacterales>Enterobacteriaceae>Citrobacter | - | Citrobacter_freundii | AZ006_RS03430 | 660140..663814 | 546 | DUF6079-family-protein-[Citrobacter-freundii] |
| WP_109017752.1 | Type3.MTaseII.containing.BREX | Arm-DNA-bind_3+Phage_integrase->HNH-><-BrxA-DNA-binding<-BrxA-DNA-binding<-PglZ<-TUDOR+SF1-SF2-Helicase+iREase+CTD<-N6-N4-METHYLASE<-DUF6079-p<-HIN-HTH<-DUF6079*<-BrxB-BrxF-iSTAND<-?<-?<-?<-?\|\|PAS+AraC-HTH+AraC-HTH-> | 1127 | Bacteria>Pseudomonadota>Gammaproteobacteria>Enterobacterales>Enterobacteriaceae>Citrobacter | - | Citrobacter_freundii | DB684_RS11735 | 36435..39818 | 546 | DUF6079-family-protein-[Citrobacter-freundii] |
| WP_247751613.1 | Type3.MTaseII.containing.BREX | DsbC+DsbD+Thioredoxin_7->TetR-HTH->tRNA->Arm-DNA-bind_3+Phage_integrase->HNH-><-BrxA-DNA-binding<-BrxA-DNA-binding<-PglZ<-TUDOR+SF1-SF2-Helicase+iREase+CTD<-N6-N4-METHYLASE<-DUF6079-p* | 218 | Bacteria>Pseudomonadota>Gammaproteobacteria>Enterobacterales>Enterobacteriaceae>Citrobacter | - | Citrobacter_freundii | KKT92_RS23430 | 43282..43938 | 546 | DUF6079-family-protein-[Citrobacter-freundii] |
| EJW5372909.1 | Type3.MTaseII.containing.BREX | <-N6-N4-METHYLASE-p<-DUF6079*<-BrxB-BrxF-iSTAND<-?<-REase<-?<-?\|\|XIS-HTH->AAA+wHTH-> | 1241 | Bacteria>Pseudomonadota>Gammaproteobacteria>Enterobacterales>Enterobacteriaceae>Citrobacter | - | Citrobacter_freundii | OCH31_004483 | 448..4173 | 546 | hypothetical-protein-[Citrobacter-freundii] |
| OUE72492.1 | Type3.MTaseII.containing.BREX | XIS-HTH-><-REase<-Arm-DNA-bind_3+Phage_integrase\|\|tRNA->Arm-DNA-bind_3+Phage_integrase->Arm-DNA-bind_3+Phage_integrase->HNH->HIN-HTH->HTH_21+rve+rve_3-><-DUF6079*<-BrxB-BrxF-iSTAND<-HTH_23+HTH_32+HTH_33+DDE_3<-?<-REase<-?\|\|LACI-HTH+PBPI->?->?-><-DDE_Tnp_IS1 | 1224 | Bacteria>Pseudomonadota>Gammaproteobacteria>Enterobacterales>Enterobacteriaceae>Citrobacter | GCA_002151755.1 | Citrobacter_freundii | AZ006_000652 | 660140..663814 | 546 | hypothetical-protein-AZ006_000652-[Citrobacter-freundii] |
| HAT3908785.1 | Type3.MTaseII.containing.BREX | <-HerA/FtsK+A32<-?\|\|tRNA->Arm-DNA-bind_3+Phage_integrase->HNH-><-BrxA-DNA-binding<-BrxA-DNA-binding<-PglZ<-TUDOR+SF1-SF2-Helicase+iREase+CTD<-N6-N4-METHYLASE<-DUF6079*<-BrxB-BrxF-iSTAND\|\|?->ACET+ACET->\|\|AbiEi_4->AbiEii-><-wHTH(SBD)+HNH<-SymE_toxin | 1241 | Bacteria>Pseudomonadota>Gammaproteobacteria>Enterobacterales>Enterobacteriaceae>Citrobacter | GCA_015929465.1 | Citrobacter_freundii | I9Y28_003037 | 175326..179051 | 546 | hypothetical-protein-[Citrobacter-freundii] |
| HBI3680282.1 | Type3.MTaseII.containing.BREX | <-TUDOR+SF1-SF2-Helicase+iREase+CTD<-N6-N4-METHYLASE<-DUF6079*<-BrxB-BrxF-iSTAND<-SF1-SF2-Helicase<-HKD-DNase<-DUF6361<-sGTP<-?<-sGTP<-ZnR | 1241 | Bacteria>Pseudomonadota>Gammaproteobacteria>Enterobacterales>Enterobacteriaceae>Citrobacter | GCA_019570155.1 | Citrobacter_freundii | K3939_000893 | 6310..10035 | 546 | hypothetical-protein-[Citrobacter-freundii] |
| EIX7373173.1 | Type3.MTaseII.containing.BREX | HNH-><-BrxA-DNA-binding<-BrxA-DNA-binding<-PglZ<-TUDOR+SF1-SF2-Helicase+iREase+CTD<-N6-N4-METHYLASE<-DUF6079*<-BrxB-BrxF-iSTAND\|\|?->ACET+ACET->\|\|AbiEi_4->AbiEii-><-wHTH(SBD)+HNH<-SymE_toxin | 1241 | Bacteria>Pseudomonadota>Gammaproteobacteria>Enterobacterales>Enterobacteriaceae>Citrobacter | GCA_022400985.1 | Citrobacter_freundii | MK620_002547 | 178053..181778 | 546 | hypothetical-protein-[Citrobacter-freundii] |
| WP_137376011.1 | Type3.MTaseII.containing.BREX | <-nSTAND1+GTPase-AIG+sGTP\|\|?-><-AAA+wHTH<-XIS-HTH\|\|?->?->REase->?->BrxB-BrxF-iSTAND->DUF6079*->N6-N4-METHYLASE->TUDOR+SF1-SF2-Helicase+iREase+CTD->PglZ->BrxA-DNA-binding->BrxA-DNA-binding-><-HNH<-Arm-DNA-bind_3+Phage_integrase<-tRNA\|\|?-><-SKI | 1241 | Bacteria>Pseudomonadota>Gammaproteobacteria>Enterobacterales>Enterobacteriaceae>Citrobacter | GCA_030820635.1 | Citrobacter_sp._wls828 | FDW86_RS03800 | 41435..45160 | 2576413 | DUF6079-family-protein-[Citrobacter-sp.-wls828] |
| WP_135323298.1 | Type3.MTaseII.containing.BREX | SymE_toxin-><-MAD-NTD+HNH\|\|wHTH+SAD+HNH-><-ACET+ACET\|\|BrxB-BrxF-iSTAND->DUF6079*->N6-N4-METHYLASE->TUDOR+SF1-SF2-Helicase+iREase+CTD->PglZ->BrxA-DNA-binding->BrxA-DNA-binding-><-HNH<-NACHT+SF1-SF2-Helicase+HKD-DNase<-nSTAND1+HNH\|\|wHTH-> | 1241 | Bacteria>Pseudomonadota>Gammaproteobacteria>Enterobacterales>Enterobacteriaceae>Citrobacter | GCF_004684345.1 | Citrobacter_tructae | E4Z61_13985 | 2763162..2766887 | 2562449 | DUF6079-family-protein-[Citrobacter-tructae] |
| WP_182936542.1 | Type3.MTaseII.containing.BREX | <-nSTAND1+sGTP\|\|?-><-AAA+wHTH<-XIS-HTH\|\|?->?->REase->?->BrxB-BrxF-iSTAND->DUF6079*-><-HIN-HTH\|\|N6-N4-METHYLASE->TUDOR+SF1-SF2-Helicase+iREase+CTD->PglZ->BrxA-DNA-binding->BrxA-DNA-binding-><-HNH<-Arm-DNA-bind_3+Phage_integrase | 1232 | Bacteria>Pseudomonadota>Gammaproteobacteria>Enterobacterales>Enterobacteriaceae>Citrobacter | GCF_014169975.1 | Citrobacter_freundii | HS092_RS16800 | 3470050..3473748 | 546 | DUF6079-family-protein-[Citrobacter-freundii] |
| WP_199954350.1 | Type3.MTaseII.containing.BREX | ZnR->XIS-HTH->?->?->KAP_NTPase->?-><-HIN-HTH\|\|BrxB-BrxF-iSTAND->DUF6079*->N6-N4-METHYLASE->TUDOR+SF1-SF2-Helicase+iREase+CTD->PglZ->BrxA-DNA-binding->BrxA-DNA-binding-><-HNH<-Arm-DNA-bind_3+Phage_integrase<-tRNA\|\|?-><-SKI | 1241 | Bacteria>Pseudomonadota>Gammaproteobacteria>Enterobacterales>Enterobacteriaceae>Citrobacter | GCF_016505335.1 | Citrobacter_sp._FDAARGOS_156 | I5510_RS10650 | 334331..338056 | 1702170 | DUF6079-family-protein-[Citrobacter-sp.-FDAARGOS_156] |
| WP_204527022.1 | Type3.MTaseII.containing.BREX | <-METHYLASE\|\|SKI-><-?\|\|tRNA->HNH-><-BrxA-DNA-binding<-BrxA-DNA-binding<-PglZ<-TUDOR+SF1-SF2-Helicase+iREase+CTD<-DUF6079-p*<-BrxB-BrxF-iSTAND<-?<-URI-DNase\|\|<-?\|\|?->?->Phos_pyr_kin-> | 879 | Bacteria>Pseudomonadota>Gammaproteobacteria>Enterobacterales>Enterobacteriaceae>Citrobacter | GCF_016893745.1 | Citrobacter_cronae | JSN23_RS08120 | 1900683..1903319 | 1748967 | DUF6079-family-protein,-partial-[Citrobacter-cronae] |
| WP_044713057.1 | Type3.MTaseII.containing.BREX | <-ACET+ACET<-?\|\|BrxB-BrxF-iSTAND->DUF6079*->N6-N4-METHYLASE->TUDOR+SF1-SF2-Helicase+iREase+CTD->PglZ->BrxA-DNA-binding->BrxA-DNA-binding-> | 1241 | Bacteria>Pseudomonadota>Gammaproteobacteria>Enterobacterales>Enterobacteriaceae>Citrobacter | GCF_016905765.1 | Citrobacter_freundii | JTY75_RS22660 | 2431..6156 | 546 | DUF6079-family-protein-[Citrobacter-freundii] |
| WP_264363600.1 | Type3.MTaseII.containing.BREX | <-HerA/FtsK+A32<-?\|\|tRNA->Arm-DNA-bind_3+Phage_integrase->HNH-><-BrxA-DNA-binding<-BrxA-DNA-binding<-PglZ<-TUDOR+SF1-SF2-Helicase+iREase+CTD<-N6-N4-METHYLASE<-DUF6079*<-BrxB-BrxF-iSTAND<-?<-?<-sGTP\|\|<-?<-XIS-HTH | 1241 | Bacteria>Pseudomonadota>Gammaproteobacteria>Enterobacterales>Enterobacteriaceae>Citrobacter | GCF_025908555.1 | Citrobacter_freundii | OIU54_RS00800 | 166179..169904 | 546 | DUF6079-family-protein-[Citrobacter-freundii] |
| WP_275380015.1 | Type3.MTaseII.containing.BREX | <-HerA/FtsK+A32<-?\|\|tRNA->Arm-DNA-bind_3+Phage_integrase->HNH-><-BrxA-DNA-binding<-BrxA-DNA-binding<-PglZ<-TUDOR+SF1-SF2-Helicase+iREase+CTD<-N6-N4-METHYLASE<-DUF6079*<-BrxB-BrxF-iSTAND<-SF1-SF2-Helicase<-HKD-DNase<-DUF6361<-sGTP<-?<-sGTP\|\|OLD-ABC+TOPRIM->SF1-SF2-Helicase-> | 1241 | Bacteria>Pseudomonadota>Gammaproteobacteria>Enterobacterales>Enterobacteriaceae>Citrobacter | GCF_029104345.1 | Citrobacter_portucalensis | L2101_RS11645 | 154726..158451 | 1639133 | DUF6079-family-protein-[Citrobacter-portucalensis] |
| EGT5208817.1 | Type3.MTaseII.containing.BREX | <-BrxA-DNA-binding<-BrxA-DNA-binding<-PglZ<-TUDOR+SF1-SF2-Helicase+iREase+CTD<-N6-N4-METHYLASE<-DUF6079*<-BrxB-BrxF-iSTAND<-HTH_23+HTH_32+HTH_33+DDE_3<-?<-REase<-?<-?\|\|XIS-HTH->AAA+wHTH-> | 1241 | Bacteria>Pseudomonadota>Gammaproteobacteria>Enterobacterales>Enterobacteriaceae>Cronobacter | GCA_016081615.1 | Cronobacter_sakazakii | AGJ24_21620 | 9168..12893 | 28141 | hypothetical-protein-[Cronobacter-sakazakii] |
| WP_046886445.1 | Type3.MTaseII.containing.BREX | Arm-DNA-bind_3+Phage_integrase->HNH-><-BrxA-DNA-binding<-BrxA-DNA-binding<-PglZ<-TUDOR+SF1-SF2-Helicase+iREase+CTD<-N6-N4-METHYLASE<-DUF6079*<-BrxB-BrxF-iSTAND\|\|<-?<-DUF2754<-?<-SAM-METHYLASE<-wHTH(SBD)+HNH<-SymE_toxin | 1241 | Bacteria>Pseudomonadota>Gammaproteobacteria>Enterobacterales>Enterobacteriaceae>Enterobacter | - | Enterobacter_cloacae | OA46_RS16175 | 115286..119011 | 550 | DUF6079-family-protein-[Enterobacter-cloacae] |
| HAV1616224.1 | Type3.MTaseII.containing.BREX | <-BrxA-DNA-binding<-BrxA-DNA-binding<-PglZ<-TUDOR+SF1-SF2-Helicase+iREase+CTD<-N6-N4-METHYLASE<-DUF6079*<-BrxB-BrxF-iSTAND<-wHTH+SAD+HNH\|\|SymE_toxin-><-?<-?\|\|ASCH+Glyoxalase-><-LACI-HTH+PBPI | 1241 | Bacteria>Pseudomonadota>Gammaproteobacteria>Enterobacterales>Enterobacteriaceae>Enterobacter | GCA_016430465.1 | Enterobacter_hormaechei_subsp._xiangfangensis | JG867_004513 | 9296..13021 | 1296536 | hypothetical-protein-[Enterobacter-hormaechei-subsp.-xiangfangensis] |
| EHF4930923.1 | Type3.MTaseII.containing.BREX | <-wHTH\|\|nSTAND1+HNH->NACHT+SF1-SF2-Helicase+HKD-DNase->HNH-><-BrxA-DNA-binding<-BrxA-DNA-binding<-PglZ<-TUDOR+SF1-SF2-Helicase+iREase+CTD<-N6-N4-METHYLASE<-DUF6079*<-BrxB-BrxF-iSTAND\|\|ACET+ACET-><-wHTH+SAD+HNH\|\|SymE_toxin-><-?<-?\|\|ASCH+Glyoxalase-><-LACI-HTH+PBPI | 1241 | Bacteria>Pseudomonadota>Gammaproteobacteria>Enterobacterales>Enterobacteriaceae>Enterobacter | GCA_017328045.1 | Enterobacter_hormaechei | CLJ31_001997 | 206029..209754 | 158836 | hypothetical-protein-[Enterobacter-hormaechei] |
| HCA7855114.1 | Type3.MTaseII.containing.BREX | Arm-DNA-bind_3+Phage_integrase->HNH-><-BrxA-DNA-binding<-BrxA-DNA-binding<-PglZ<-TUDOR+SF1-SF2-Helicase+iREase+CTD<-N6-N4-METHYLASE<-DUF6079*<-BrxB-BrxF-iSTAND<-?<-XIS-HTH | 1241 | Bacteria>Pseudomonadota>Gammaproteobacteria>Enterobacterales>Enterobacteriaceae>Enterobacter | GCA_023000845.1 | Enterobacter_hormaechei | MX526_001465 | 158507..162232 | 158836 | hypothetical-protein-[Enterobacter-hormaechei] |
| HCM9194815.1 | Type3.MTaseII.containing.BREX | HNH-><-BrxA-DNA-binding<-BrxA-DNA-binding<-PglZ<-TUDOR+SF1-SF2-Helicase+iREase+CTD<-N6-N4-METHYLASE<-DUF6079*<-BrxB-BrxF-iSTAND<-wHTH(SBD)+HNH\|\|SymE_toxin->?->?->?-><-DUF2754<-?\|\|ASCH+Glyoxalase-> | 1241 | Bacteria>Pseudomonadota>Gammaproteobacteria>Enterobacterales>Enterobacteriaceae>Enterobacter | GCA_025209675.1 | Enterobacter_cloacae_subsp._dissolvens | N5K92_004842 | 14679..18404 | 69219 | hypothetical-protein-[Enterobacter-cloacae-subsp.-dissolvens] |
| WP_121528275.1 | Type3.MTaseII.containing.BREX | Arm-DNA-bind_3+Phage_integrase->HNH-><-BrxA-DNA-binding<-BrxA-DNA-binding<-PglZ<-TUDOR+SF1-SF2-Helicase+iREase+CTD<-N6-N4-METHYLASE<-DUF6079*<-BrxB-BrxF-iSTAND<-HIN-HTH<-?<-REase\|\|<-NTP_transferase<-?\|\|<-?<-ABC-ATPase+SbcC | 1241 | Bacteria>Pseudomonadota>Gammaproteobacteria>Enterobacterales>Enterobacteriaceae>Enterobacter | GCF_003665375.1 | Enterobacter_kobei | D9T11_RS25130 | 4904921..4908646 | 208224 | DUF6079-family-protein-[Enterobacter-kobei] |
| WP_126497372.1 | Type3.MTaseII.containing.BREX | XIS-HTH->?-><-?<-?<-SbcC\|\|BrxB-BrxF-iSTAND->DUF6079*->N6-N4-METHYLASE->TUDOR+SF1-SF2-Helicase+iREase+CTD->PglZ->BrxA-DNA-binding->BrxA-DNA-binding-><-HNH<-Arm-DNA-bind_3+Phage_integrase | 1241 | Bacteria>Pseudomonadota>Gammaproteobacteria>Enterobacterales>Enterobacteriaceae>Enterobacter | GCF_003965685.1 | Enterobacter_hormaechei | EKN63_RS10075 | 338811..342536 | 158836 | DUF6079-family-protein-[Enterobacter-hormaechei] |
| WP_200924817.1 | Type3.MTaseII.containing.BREX | Arm-DNA-bind_3+Phage_integrase->HNH-><-BrxA-DNA-binding<-BrxA-DNA-binding<-PglZ<-TUDOR+SF1-SF2-Helicase+iREase+CTD<-N6-N4-METHYLASE<-DUF6079*<-ZnR<-?<-XIS-HTH\|\|?->?->?->?->DUF932-> | 1241 | Bacteria>Pseudomonadota>Gammaproteobacteria>Enterobacterales>Enterobacteriaceae>Enterobacter | GCF_015136055.1 | Enterobacter_kobei | JLF39_RS03160 | 645294..649019 | 208224 | DUF6079-family-protein-[Enterobacter-kobei] |
| WP_248192361.1 | Type3.MTaseII.containing.BREX | XIS-HTH->ZnR->DUF6361->HKD-DNase->SF1-SF2-Helicase->BrxB-BrxF-iSTAND->DUF6079-p*-> | 390 | Bacteria>Pseudomonadota>Gammaproteobacteria>Enterobacterales>Enterobacteriaceae>Enterobacter | GCF_023153355.1 | Enterobacter_kobei | L8P93_RS24510 | 28368..29540 | 208224 | DUF6079-family-protein-[Enterobacter-kobei] |
| WP_262778588.1 | Type3.MTaseII.containing.BREX | LACI-HTH+PBPI-><-ASCH+Glyoxalase\|\|?->?-><-SymE_toxin\|\|wHTH(SBD)+HNH->BrxB-BrxF-iSTAND->DUF6079*->N6-N4-METHYLASE->TUDOR+SF1-SF2-Helicase+iREase+CTD->PglZ->BrxA-DNA-binding->BrxA-DNA-binding-><-HNH | 1241 | Bacteria>Pseudomonadota>Gammaproteobacteria>Enterobacterales>Enterobacteriaceae>Enterobacter | GCF_025535845.1 | Enterobacter_quasiroggenkampii | KW815_RS12600 | 303866..307591 | 2497436 | DUF6079-family-protein-[Enterobacter-quasiroggenkampii] |
| WP_262805832.1 | Type3.MTaseII.containing.BREX | HNH-><-BrxA-DNA-binding<-BrxA-DNA-binding<-PglZ<-TUDOR+SF1-SF2-Helicase+iREase+CTD<-N6-N4-METHYLASE<-DUF6079*<-BrxB-BrxF-iSTAND<-wHTH(SBD)+HNH\|\|SymE_toxin-><-?<-?\|\|ASCH+Glyoxalase-><-PBPI | 1241 | Bacteria>Pseudomonadota>Gammaproteobacteria>Enterobacterales>Enterobacteriaceae>Enterobacter | GCF_025536155.1 | Enterobacter_quasiroggenkampii | KSU18_RS16665 | 126264..129989 | 2497436 | DUF6079-family-protein-[Enterobacter-quasiroggenkampii] |
| WP_028016404.1 | Type3.MTaseII.containing.BREX | <-AAA+wHTH<-XIS-HTH\|\|?->?->DDE_Tnp_1_5+DDE_Tnp_1->REase->?->BrxB-BrxF-iSTAND->DUF6079*->N6-N4-METHYLASE->TUDOR+SF1-SF2-Helicase+iREase+CTD->PglZ->BrxA-DNA-binding->BrxA-DNA-binding-><-HNH<-Arm-DNA-bind_3+Phage_integrase<-tRNA<-TetR-HTH<-DsbC+DsbD+Thioredoxin_7 | 1241 | Bacteria>Pseudomonadota>Gammaproteobacteria>Enterobacterales>Enterobacteriaceae>Enterobacter | GCF_027688635.1 | Enterobacter_cloacae | P699_RS0102270 | 37920..41645 | 550 | DUF6079-family-protein-[Enterobacter-cloacae] |
| WP_274417362.1 | Type3.MTaseII.containing.BREX | Arm-DNA-bind_3+Phage_integrase->HNH-><-BrxA-DNA-binding<-BrxA-DNA-binding<-PglZ<-TUDOR+SF1-SF2-Helicase+iREase+CTD<-N6-N4-METHYLASE<-DUF6079*<-BrxB-BrxF-iSTAND<-?<-REase | 1241 | Bacteria>Pseudomonadota>Gammaproteobacteria>Enterobacterales>Enterobacteriaceae>Enterobacter | GCF_028765355.1 | Enterobacter_kobei | PVM09_RS09335 | 364975..368700 | 208224 | DUF6079-family-protein-[Enterobacter-kobei] |
| EEZ5786496.1 | Type3.MTaseII.containing.BREX | HNH-><-BrxA-DNA-binding<-BrxA-DNA-binding<-PglZ<-TUDOR+SF1-SF2-Helicase+iREase+CTD<-N6-N4-METHYLASE<-DUF6079*<-BrxB-BrxF-iSTAND | 1241 | Bacteria>Pseudomonadota>Gammaproteobacteria>Enterobacterales>Enterobacteriaceae>Escherichia | GCA_012214065.1 | Escherichia_coli_O107 | DBU15_004385 | 10406..14131 | 2162915 | hypothetical-protein-[Escherichia-coli-O107] |
| MBB9297699.1 | Type3.MTaseII.containing.BREX | Arm-DNA-bind_3+Phage_integrase->HNH-><-BrxA-DNA-binding<-BrxA-DNA-binding<-PglZ<-TUDOR+SF1-SF2-Helicase+iREase+CTD<-N6-N4-METHYLASE<-DUF6079-p* | 995 | Bacteria>Pseudomonadota>Gammaproteobacteria>Enterobacterales>Enterobacteriaceae>Escherichia | GCA_012376355.2 | Escherichia_coli | FSI29_018575 | 22589..25576 | 562 | hypothetical-protein-[Escherichia-coli] |
| HAX9835785.1 | Type3.MTaseII.containing.BREX | <-SF1-SF2-Helicase<-OLD-ABC+TOPRIM\|\|sGTP->?->sGTP->DUF6361->HKD-DNase->SF1-SF2-Helicase->BrxB-BrxF-iSTAND->DUF6079*->N6-N4-METHYLASE->TUDOR+SF1-SF2-Helicase+iREase+CTD->PglZ->BrxA-DNA-binding->BrxA-DNA-binding-><-HNH<-Arm-DNA-bind_3+Phage_integrase<-tRNA\|\|?-><-SKI | 1241 | Bacteria>Pseudomonadota>Gammaproteobacteria>Enterobacterales>Enterobacteriaceae>Escherichia | GCA_017073735.1 | Escherichia_coli | JS210_001190 | 46996..50721 | 562 | hypothetical-protein-[Escherichia-coli] |
| MCN9861063.1 | Type3.MTaseII.containing.BREX | <-DUF932<-?<-?<-?<-?\|\|XIS-HTH->?->\|\|BrxB-BrxF-iSTAND->DUF6079*->N6-N4-METHYLASE->TUDOR+SF1-SF2-Helicase+iREase+CTD->PglZ->BrxA-DNA-binding->BrxA-DNA-binding-><-HNH<-Arm-DNA-bind_3+Phage_integrase<-tRNA\|\|?->HerA/FtsK+A32-> | 1241 | Bacteria>Pseudomonadota>Gammaproteobacteria>Enterobacterales>Enterobacteriaceae>Escherichia | GCA_023889565.1 | Escherichia_coli | MLQ95_05995 | 11690..15415 | 562 | DUF6079-family-protein-[Escherichia-coli] |
| WP_097755484.1 | Type3.MTaseII.containing.BREX | DUF905->ArdB->?->CbeA_antitoxin->?->BrxB-BrxF-iSTAND->DUF6079*->N6-N4-METHYLASE->TUDOR+SF1-SF2-Helicase+iREase+CTD->PglZ->BrxA-DNA-binding->BrxA-DNA-binding-><-HNH<-HTH_Tnp_1+HTH_21+rve+rve_3 | 1241 | Bacteria>Pseudomonadota>Gammaproteobacteria>Enterobacterales>Enterobacteriaceae>Escherichia | GCF_005388805.1 | Escherichia_coli | FFF01_RS10370 | 151657..155382 | 562 | DUF6079-family-protein-[Escherichia-coli] |
| WP_038636828.1 | Type3.MTaseII.containing.BREX | SKI-><-?\|\|tRNA->Arm-DNA-bind_3+Phage_integrase->HNH-><-BrxA-DNA-binding<-BrxA-DNA-binding<-PglZ<-TUDOR+SF1-SF2-Helicase+iREase+CTD<-N6-N4-METHYLASE<-DUF6079*<-ZnR<-?<-XIS-HTH\|\|?->?->?->?->DUF932-> | 1241 | Bacteria>Pseudomonadota>Gammaproteobacteria>Enterobacterales>Enterobacteriaceae | GCF_001558935.2 | Citrobacter | CFNIH1_RS09050 | 1899594..1903319 | 544 | MULTISPECIES:-DUF6079-family-protein-[Citrobacter] |
| MCF0347731.1 | Type3.MTaseII.containing.BREX | <-wHTH+WYL<-?<-?\|\|?->XIS-HTH->ZnR->DUF6361->HKD-DNase->SF1-SF2-Helicase->DUF6079*->N6-N4-METHYLASE->TUDOR+SF1-SF2-Helicase+iREase+CTD->PglZ->BrxA-DNA-binding->BrxA-DNA-binding-><-HNH<-Arm-DNA-bind_3+Phage_integrase | 1241 | Bacteria>Pseudomonadota>Gammaproteobacteria>Enterobacterales>Enterobacteriaceae>Klebsiella/Raoultella_group | GCA_021455545.1 | Klebsiella_pneumoniae | LOF49_12125 | 63457..67182 | 573 | DUF6079-family-protein-[Klebsiella-pneumoniae] |
| HBS7446105.1 | Type3.MTaseII.containing.BREX | Arm-DNA-bind_3+Phage_integrase->HNH-><-BrxA-DNA-binding<-BrxA-DNA-binding<-PglZ<-TUDOR+SF1-SF2-Helicase+iREase+CTD<-N6-N4-METHYLASE<-DUF6079*<-BrxB-BrxF-iSTAND<-SF1-SF2-Helicase<-HKD-DNase<-DUF6361<-XIS-HTH<-?\|\|\|\|?->?->DUF905-> | 1241 | Bacteria>Pseudomonadota>Gammaproteobacteria>Enterobacterales>Enterobacteriaceae>Klebsiella/Raoultella_group | GCA_021978815.1 | Klebsiella_pneumoniae | MAQ81_001502 | 320914..324639 | 573 | hypothetical-protein-[Klebsiella-pneumoniae] |
| HBT3663674.1 | Type3.MTaseII.containing.BREX | Arm-DNA-bind_3+Phage_integrase->HNH-><-BrxA-DNA-binding<-BrxA-DNA-binding<-PglZ<-TUDOR+SF1-SF2-Helicase+iREase+CTD<-N6-N4-METHYLASE<-DUF6079*<-BrxB-BrxF-iSTAND | 1241 | Bacteria>Pseudomonadota>Gammaproteobacteria>Enterobacterales>Enterobacteriaceae>Klebsiella/Raoultella_group | GCA_022030495.1 | Klebsiella_pneumoniae | MBH96_005330 | 10616..14341 | 573 | hypothetical-protein-[Klebsiella-pneumoniae] |
| HBT9662835.1 | Type3.MTaseII.containing.BREX | <-nSTAND1+sGTP\|\|?-><-AAA+wHTH<-XIS-HTH\|\|?->?->REase->?->BrxB-BrxF-iSTAND->DUF6079*->N6-N4-METHYLASE->TUDOR+SF1-SF2-Helicase+iREase+CTD->PglZ->BrxA-DNA-binding->BrxA-DNA-binding-><-HNH<-Arm-DNA-bind_3+Phage_integrase | 1241 | Bacteria>Pseudomonadota>Gammaproteobacteria>Enterobacterales>Enterobacteriaceae>Klebsiella/Raoultella_group | GCA_022069925.1 | Klebsiella_pneumoniae | MCN56_004853 | 32761..36486 | 573 | hypothetical-protein-[Klebsiella-pneumoniae] |
| HBV9253299.1 | Type3.MTaseII.containing.BREX | XIS-HTH->BrxB-BrxF-iSTAND->DUF6079*->N6-N4-METHYLASE->TUDOR+SF1-SF2-Helicase+iREase+CTD->PglZ->BrxA-DNA-binding->BrxA-DNA-binding-> | 1241 | Bacteria>Pseudomonadota>Gammaproteobacteria>Enterobacterales>Enterobacteriaceae>Klebsiella/Raoultella_group | GCA_022163685.1 | Klebsiella_pneumoniae | MD920_005122 | 4004..7729 | 573 | hypothetical-protein-[Klebsiella-pneumoniae] |
| HBX6133595.1 | Type3.MTaseII.containing.BREX | Arm-DNA-bind_3+Phage_integrase->HNH-><-BrxA-DNA-binding<-PglZ<-TUDOR+SF1-SF2-Helicase+iREase+CTD<-N6-N4-METHYLASE<-DUF6079* | 1201 | Bacteria>Pseudomonadota>Gammaproteobacteria>Enterobacterales>Enterobacteriaceae>Klebsiella/Raoultella_group | GCA_022250115.1 | Klebsiella_pneumoniae | MH627_25405 | 28865..32470 | 573 | hypothetical-protein-[Klebsiella-pneumoniae] |
| HBX9795176.1 | Type3.MTaseII.containing.BREX | XIS-HTH->DUF6361->HKD-DNase->SF1-SF2-Helicase->BrxB-BrxF-iSTAND->DUF6079*-> | 1181 | Bacteria>Pseudomonadota>Gammaproteobacteria>Enterobacterales>Enterobacteriaceae>Klebsiella/Raoultella_group | GCA_022265715.1 | Klebsiella_pneumoniae | MIK97_005082 | 21145..24686 | 573 | hypothetical-protein-[Klebsiella-pneumoniae] |
| MCP6235010.1 | Type3.MTaseII.containing.BREX | Arm-DNA-bind_3+Phage_integrase->HNH-><-BrxA-DNA-binding<-BrxA-DNA-binding<-PglZ<-TUDOR+SF1-SF2-Helicase+iREase+CTD<-N6-N4-METHYLASE<-DUF6079*<-BrxB-BrxF-iSTAND<-SF1-SF2-Helicase<-HKD-DNase<-DUF6361<-ZnR<-XIS-HTH | 1241 | Bacteria>Pseudomonadota>Gammaproteobacteria>Enterobacterales>Enterobacteriaceae>Klebsiella/Raoultella_group | GCA_024239175.1 | Klebsiella_pneumoniae | NL436_15880 | 169849..173574 | 573 | DUF6079-family-protein-[Klebsiella-pneumoniae] |
| HCM4054959.1 | Type3.MTaseII.containing.BREX | DUF6079-p*->N6-N4-METHYLASE->TUDOR+SF1-SF2-Helicase+iREase+CTD->PglZ->BrxA-DNA-binding->BrxA-DNA-binding-><-HNH<-Arm-DNA-bind_3+Phage_integrase | 678 | Bacteria>Pseudomonadota>Gammaproteobacteria>Enterobacterales>Enterobacteriaceae>Klebsiella/Raoultella_group | GCA_025081835.1 | Klebsiella_quasipneumoniae | N3I99_003985 | 1..2037 | 1463165 | hypothetical-protein-[Klebsiella-quasipneumoniae] |
| HCM4056380.1 | Type3.MTaseII.containing.BREX | XIS-HTH->DUF6361->HKD-DNase->SF1-SF2-Helicase->BrxB-BrxF-iSTAND->DUF6079-p*-> | 563 | Bacteria>Pseudomonadota>Gammaproteobacteria>Enterobacterales>Enterobacteriaceae>Klebsiella/Raoultella_group | GCA_025081835.1 | Klebsiella_quasipneumoniae | N3N82_005459 | 32499..34188 | 1463165 | hypothetical-protein-[Klebsiella-quasipneumoniae] |
| HCM5386358.1 | Type3.MTaseII.containing.BREX | DUF6079-p*->N6-N4-METHYLASE->TUDOR+SF1-SF2-Helicase+iREase+CTD->PglZ->BrxA-DNA-binding->BrxA-DNA-binding-><-HNH<-Arm-DNA-bind_3+Phage_integrase | 676 | Bacteria>Pseudomonadota>Gammaproteobacteria>Enterobacterales>Enterobacteriaceae>Klebsiella/Raoultella_group | GCA_025094345.1 | Klebsiella_quasipneumoniae | N3N82_003980 | 1..2031 | 1463165 | hypothetical-protein-[Klebsiella-quasipneumoniae] |
| GKJ38632.1 | Type3.MTaseII.containing.BREX | Arm-DNA-bind_3+Phage_integrase->HNH-><-BrxA-DNA-binding<-BrxA-DNA-binding<-PglZ<-TUDOR+SF1-SF2-Helicase+iREase+CTD<-N6-N4-METHYLASE<-DUF6079*<-BrxB-BrxF-iSTAND<-?<-?<-XIS-HTH<-?\|\|?->?->wHTH+WYL-> | 1241 | Bacteria>Pseudomonadota>Gammaproteobacteria>Enterobacterales>Enterobacteriaceae>Klebsiella/Raoultella_group | GCA_026007675.1 | Klebsiella_variicola | NUKP24_47670 | 11433..15158 | 244366 | hypothetical-protein-NUKP24_47670-[Klebsiella-variicola] |
| WP_038435131.1 | Type3.MTaseII.containing.BREX | Arm-DNA-bind_3+Phage_integrase->HNH-><-BrxA-DNA-binding<-BrxA-DNA-binding<-PglZ<-TUDOR+SF1-SF2-Helicase+iREase+CTD<-N6-N4-METHYLASE<-DUF6079*<-BrxB-BrxF-iSTAND<-SF1-SF2-Helicase<-HKD-DNase<-DUF6361<-XIS-HTH | 1241 | Bacteria>Pseudomonadota>Gammaproteobacteria>Enterobacterales>Enterobacteriaceae>Klebsiella/Raoultella_group | GCF_000784945.1 | Klebsiella_pneumoniae | KPNIH29_RS02900 | 595912..599637 | 573 | DUF6079-family-protein-[Klebsiella-pneumoniae] |
| WP_055323323.1 | Type3.MTaseII.containing.BREX | <-ParB+BAM<-XIS-HTH<-wHTH\|\|BrxB-BrxF-iSTAND->DUF6079*->N6-N4-METHYLASE-> | 1241 | Bacteria>Pseudomonadota>Gammaproteobacteria>Enterobacterales>Enterobacteriaceae>Klebsiella/Raoultella_group | GCF_001317095.1 | Klebsiella_variicola | AN687_RS19930 | 3855..7580 | 244366 | DUF6079-family-protein-[Klebsiella-variicola] |
| WP_032738388.1 | Type3.MTaseII.containing.BREX | Arm-DNA-bind_3+Phage_integrase->HNH-><-BrxA-DNA-binding<-BrxA-DNA-binding<-PglZ<-TUDOR+SF1-SF2-Helicase+iREase+CTD<-N6-N4-METHYLASE<-DUF6079*<-BrxB-BrxF-iSTAND<-SF1-SF2-Helicase<-HKD-DNase<-DUF6361<-XIS-HTH | 1241 | Bacteria>Pseudomonadota>Gammaproteobacteria>Enterobacterales>Enterobacteriaceae>Klebsiella/Raoultella_group | GCF_002852945.3 | Klebsiella | C3378_RS10060 | 164999..168724 | 570 | MULTISPECIES:-DUF6079-family-protein-[Klebsiella] |
| WP_101825914.1 | Type3.MTaseII.containing.BREX | BrxB-BrxF-iSTAND->DUF6079*->N6-N4-METHYLASE->TUDOR+SF1-SF2-Helicase+iREase+CTD->PglZ->BrxA-DNA-binding->BrxA-DNA-binding-><-HNH<-Arm-DNA-bind_3+Phage_integrase | 1241 | Bacteria>Pseudomonadota>Gammaproteobacteria>Enterobacterales>Enterobacteriaceae>Klebsiella/Raoultella_group | GCF_002853035.1 | Klebsiella_variicola | CLQ66_RS24340 | 1116..4841 | 244366 | DUF6079-family-protein-[Klebsiella-variicola] |
| WP_103433591.1 | Type3.MTaseII.containing.BREX | XIS-HTH->DUF6361->HKD-DNase->SF1-SF2-Helicase->BrxB-BrxF-iSTAND->DUF6079*->N6-N4-METHYLASE->TUDOR+SF1-SF2-Helicase+iREase+CTD->PglZ->BrxA-DNA-binding->BrxA-DNA-binding-><-HNH<-Arm-DNA-bind_3+Phage_integrase<-tRNA\|\|wHTH(SBD)+HNH->SAM-METHYLASE-> | 1241 | Bacteria>Pseudomonadota>Gammaproteobacteria>Enterobacterales>Enterobacteriaceae>Klebsiella/Raoultella_group | GCF_002906395.1 | Klebsiella_oxytoca | C2U44_RS27535 | 5576240..5579965 | 571 | DUF6079-family-protein-[Klebsiella-oxytoca] |
| WP_117095259.1 | Type3.MTaseII.containing.BREX | DDE_Tnp_IS1->HNH-><-BrxA-DNA-binding<-BrxA-DNA-binding<-PglZ<-TUDOR+SF1-SF2-Helicase+iREase+CTD<-N6-N4-METHYLASE<-DUF6079*<-BrxB-BrxF-iSTAND\|\|ACET+ACET-><-wHTH+SAD+HNH\|\|MAD-NTD+HNH-><-Glyco_hydro_43+GH43_C2<-?\|\|AP_endonuc_2-> | 1241 | Bacteria>Pseudomonadota>Gammaproteobacteria>Enterobacterales>Enterobacteriaceae>Klebsiella/Raoultella_group | GCF_004312605.1 | Klebsiella_pneumoniae | EON34_RS25660 | 12944..16669 | 573 | DUF6079-family-protein-[Klebsiella-pneumoniae] |
| WP_131026262.1 | Type3.MTaseII.containing.BREX | <-wHTH+WYL<-?<-?\|\|?->XIS-HTH->ZnR->DUF6361->HKD-DNase->SF1-SF2-Helicase->DUF6079*->N6-N4-METHYLASE->TUDOR+SF1-SF2-Helicase+iREase+CTD->PglZ->BrxA-DNA-binding->BrxA-DNA-binding-><-HNH<-Arm-DNA-bind_3+Phage_integrase | 1241 | Bacteria>Pseudomonadota>Gammaproteobacteria>Enterobacterales>Enterobacteriaceae>Klebsiella/Raoultella_group | GCF_004312745.1 | Klebsiella_variicola | EON55_RS11865 | 53136..56861 | 244366 | DUF6079-family-protein-[Klebsiella-variicola] |
| WP_198072856.1 | Type3.MTaseII.containing.BREX | <-CbeA_antitoxin<-?<-ArdB\|\|Zn_Tnp_IS1+HTH_Tnp_IS1+DDE_Tnp_IS1->REase+SPF-cons->Herpes_LMP1+REase-><-?<-?<-HIN-HTH\|\|DUF6079-p*->DEDD_Tnp_IS110+Transposase_20->DDE_Tnp_1_5+DDE_Tnp_1->N6-N4-METHYLASE->TUDOR+SF1-SF2-Helicase+iREase+CTD->PglZ->BrxA-DNA-binding->BrxA-DNA-binding-><-HNH<-Arm-DNA-bind_3+Phage_integrase | 610 | Bacteria>Pseudomonadota>Gammaproteobacteria>Enterobacterales>Enterobacteriaceae>Klebsiella/Raoultella_group | GCF_016066695.1 | Klebsiella_quasipneumoniae | I8N79_RS22795 | 4658529..4660361 | 1463165 | DUF6079-family-protein-[Klebsiella-quasipneumoniae] |
| WP_087867049.1 | Type3.MTaseII.containing.BREX | <-wHTH+WYL<-?<-?\|\|?->XIS-HTH->ZnR->DUF6361->HKD-DNase->SF1-SF2-Helicase->DUF6079*->N6-N4-METHYLASE->TUDOR+SF1-SF2-Helicase+iREase+CTD->PglZ->BrxA-DNA-binding->BrxA-DNA-binding-><-HNH<-Arm-DNA-bind_3+Phage_integrase | 1241 | Bacteria>Pseudomonadota>Gammaproteobacteria>Enterobacterales>Enterobacteriaceae>Klebsiella/Raoultella_group | GCF_016762175.1 | Klebsiella_pneumoniae | H9R52_RS24840 | 41277..45002 | 573 | DUF6079-family-protein-[Klebsiella-pneumoniae] |
| WP_224257949.1 | Type3.MTaseII.containing.BREX | <-N6-N4-METHYLASE<-DUF6079*<-BrxB-BrxF-iSTAND\|\|ACET+ACET-><-?<-Glyoxalase<-SymE_toxin<-HNH\|\|?->LACI-HTH+PBPI-> | 1241 | Bacteria>Pseudomonadota>Gammaproteobacteria>Enterobacterales>Enterobacteriaceae>Klebsiella/Raoultella_group | GCF_020120515.1 | Klebsiella_oxytoca | FMJ84_RS26525 | 1240..4965 | 571 | DUF6079-family-protein-[Klebsiella-oxytoca] |
| WP_250099091.1 | Type3.MTaseII.containing.BREX | XIS-HTH->ZnR->DUF6361->HKD-DNase->SF1-SF2-Helicase->BrxB-BrxF-iSTAND->DUF6079*-> | 1199 | Bacteria>Pseudomonadota>Gammaproteobacteria>Enterobacterales>Enterobacteriaceae>Klebsiella/Raoultella_group | GCF_023544415.1 | Klebsiella_quasivariicola | M8363_RS25095 | 9201..12796 | 2026240 | DUF6079-family-protein,-partial-[Klebsiella-quasivariicola] |
| WP_255089518.1 | Type3.MTaseII.containing.BREX | DUF6361->HKD-DNase->SF1-SF2-Helicase->BrxB-BrxF-iSTAND->DUF6079*->N6-N4-METHYLASE->TUDOR+SF1-SF2-Helicase+iREase+CTD->PglZ->BrxA-DNA-binding->BrxA-DNA-binding-><-HNH<-Arm-DNA-bind_3+Phage_integrase | 1241 | Bacteria>Pseudomonadota>Gammaproteobacteria>Enterobacterales>Enterobacteriaceae>Klebsiella/Raoultella_group | GCF_024355505.1 | Klebsiella_pneumoniae | L1C10_RS16360 | 7315..11040 | 573 | DUF6079-family-protein-[Klebsiella-pneumoniae] |
| WP_086076407.1 | Type3.MTaseII.containing.BREX | XIS-HTH->DUF6361->HKD-DNase->SF1-SF2-Helicase->DUF6079*->N6-N4-METHYLASE->TUDOR+SF1-SF2-Helicase+iREase+CTD->PglZ->BrxA-DNA-binding->BrxA-DNA-binding-><-HNH<-Arm-DNA-bind_3+Phage_integrase<-tRNA\|\|Phage_integrase-><-PIWI | 1241 | Bacteria>Pseudomonadota>Gammaproteobacteria>Enterobacterales>Enterobacteriaceae>Klebsiella/Raoultella_group | GCF_024516485.1 | Klebsiella | JEK23_RS20160 | 34987..38712 | 570 | MULTISPECIES:-DUF6079-family-protein-[Klebsiella] |
| WP_265011148.1 | Type3.MTaseII.containing.BREX | Arm-DNA-bind_3+Phage_integrase->HNH-><-BrxA-DNA-binding<-BrxA-DNA-binding<-PglZ<-TUDOR+SF1-SF2-Helicase+iREase+CTD<-N6-N4-METHYLASE<-DUF6079*<-?<-XIS-HTH<-?\|\|?->?->wHTH+WYL-> | 1241 | Bacteria>Pseudomonadota>Gammaproteobacteria>Enterobacterales>Enterobacteriaceae>Klebsiella/Raoultella_group | GCF_026007675.1 | Klebsiella_variicola | OPB36_RS24690 | 11433..15158 | 244366 | DUF6079-family-protein-[Klebsiella-variicola] |
| WP_114266190.1 | Type3.MTaseII.containing.BREX | XIS-HTH->DUF6361->HKD-DNase->SF1-SF2-Helicase->BrxB-BrxF-iSTAND->DUF6079*->N6-N4-METHYLASE->TUDOR+SF1-SF2-Helicase+iREase+CTD->PglZ->BrxA-DNA-binding->BrxA-DNA-binding-><-HNH<-Arm-DNA-bind_3+Phage_integrase | 1241 | Bacteria>Pseudomonadota>Gammaproteobacteria>Enterobacterales>Enterobacteriaceae>Klebsiella/Raoultella_group | GCF_026010935.1 | Klebsiella_pneumoniae | OPE25_RS21535 | 23314..27039 | 573 | DUF6079-family-protein-[Klebsiella-pneumoniae] |
| WP_222859358.1 | Type3.MTaseII.containing.BREX | XIS-HTH->ZnR->DUF6361->HKD-DNase->SF1-SF2-Helicase->BrxB-BrxF-iSTAND->DUF6079*-> | 506 | Bacteria>Pseudomonadota>Gammaproteobacteria>Enterobacterales>Enterobacteriaceae>Klebsiella/Raoultella_group | GCF_900083795.1 | Klebsiella_oxytoca | A8T98_RS25255 | 76704..78224 | 571 | DUF6079-family-protein-[Klebsiella-oxytoca] |
| WP_064279013.1 | Type3.MTaseII.containing.BREX | Arm-DNA-bind_3+Phage_integrase->Arm-DNA-bind_3->HNH-><-BrxA-DNA-binding<-BrxA-DNA-binding<-PglZ<-N6-N4-METHYLASE-p<-N6-N4-METHYLASE<-DUF6079*<-BrxB-BrxF-iSTAND\|\|HNH->?->?-><-wHTH(SBD)+HNH<-Glyco_hydro_43+GH43_C2<-?\|\|AP_endonuc_2-> | 1241 | Bacteria>Pseudomonadota>Gammaproteobacteria>Enterobacterales>Enterobacteriaceae>Klebsiella/Raoultella_group | GCF_900084005.1 | Klebsiella_pneumoniae | SAMEA2273678_04067 | 185293..189018 | 573 | DUF6079-family-protein-[Klebsiella-pneumoniae] |
| WP_115385247.1 | Type3.MTaseII.containing.BREX | XIS-HTH->DUF6361->HKD-DNase->SF1-SF2-Helicase->BrxB-BrxF-iSTAND->DUF6079*->N6-N4-METHYLASE->TUDOR+SF1-SF2-Helicase+iREase+CTD->PglZ->BrxA-DNA-binding->BrxA-DNA-binding-><-HNH<-Arm-DNA-bind_3+Phage_integrase | 1241 | Bacteria>Pseudomonadota>Gammaproteobacteria>Enterobacterales>Enterobacteriaceae>Klebsiella/Raoultella_group | GCF_900451245.1 | Klebsiella_quasipneumoniae | DYI10_RS15560 | 3036028..3039753 | 1463165 | DUF6079-family-protein-[Klebsiella-quasipneumoniae] |
| WP_155011876.1 | Type3.MTaseII.containing.BREX | Glyco_hydro_43+GH43_C2->wHTH(SBD)+HNH-><-?<-?<-?<-HIN-HTH<-HNH<-?\|\|BrxB-BrxF-iSTAND->DUF6079*->N6-N4-METHYLASE->N6-N4-METHYLASE-p->PglZ->BrxA-DNA-binding->BrxA-DNA-binding-><-HNH\|\|TIR-> | 1241 | Bacteria>Pseudomonadota>Gammaproteobacteria>Enterobacterales>Enterobacteriaceae>Klebsiella/Raoultella_group | GCF_902165215.1 | Klebsiella_variicola | FN701_RS21035 | 12814..16539 | 244366 | DUF6079-family-protein-[Klebsiella-variicola] |
| WP_159174441.1 | Type3.MTaseII.containing.BREX | XIS-HTH->DUF6361->HKD-DNase->SF1-SF2-Helicase->DUF6079*->N6-N4-METHYLASE->TUDOR+SF1-SF2-Helicase+iREase+CTD->PglZ->BrxA-DNA-binding->BrxA-DNA-binding-><-HNH<-Arm-DNA-bind_3+Phage_integrase | 1241 | Bacteria>Pseudomonadota>Gammaproteobacteria>Enterobacterales>Enterobacteriaceae>Klebsiella/Raoultella_group | GCF_902704295.1 | Klebsiella_pneumoniae | FL632_RS17825 | 31829..35554 | 573 | DUF6079-family-protein-[Klebsiella-pneumoniae] |
| WP_181890665.1 | Type3.MTaseII.containing.BREX | XIS-HTH->DUF6361->HKD-DNase->SF1-SF2-Helicase->DUF6079*->N6-N4-METHYLASE->TUDOR+SF1-SF2-Helicase+iREase+CTD->PglZ->BrxA-DNA-binding->BrxA-DNA-binding-><-HNH<-Arm-DNA-bind_3+Phage_integrase | 1241 | Bacteria>Pseudomonadota>Gammaproteobacteria>Enterobacterales>Enterobacteriaceae>Klebsiella/Raoultella_group | GCF_902807195.1 | Raoultella_planticola | HUW11_RS11150 | 61321..65046 | 575 | DUF6079-family-protein-[Raoultella-planticola] |
| HCB9303670.1 | Type3.MTaseII.containing.BREX | <-AP_endonuc_2\|\|?->Glyco_hydro_43+GH43_C2-><-MAD-NTD+HNH\|\|wHTH+SAD+HNH-><-ACET+ACET\|\|BrxB-BrxF-iSTAND->DUF6079*-> | 1076 | Bacteria>Pseudomonadota>Gammaproteobacteria>Enterobacterales>Enterobacteriaceae>Klebsiella/Raoultella_group | - | Klebsiella_pneumoniae | M2P40_004872 | 30608..33837 | 573 | hypothetical-protein-[Klebsiella-pneumoniae] |
| WP_107147161.1 | Type3.MTaseII.containing.BREX | <-ArdB<-DUF932<-?<-?<-?<-?\|\|?->XIS-HTH->ZnR->DUF6079*->N6-N4-METHYLASE->TUDOR+SF1-SF2-Helicase+iREase+CTD->PglZ->BrxA-DNA-binding->BrxA-DNA-binding-><-HNH<-Arm-DNA-bind_3+Phage_integrase<-tRNA\|\|REC+LuxR-HTH-> | 1241 | Bacteria>Pseudomonadota>Gammaproteobacteria>Enterobacterales>Enterobacteriaceae>Kosakonia | GCF_003028445.1 | Kosakonia_sp._H7A | CWM66_RS20250 | 17461..21186 | 2054598 | DUF6079-family-protein-[Kosakonia-sp.-H7A] |
| MBD8580181.1 | Type3.MTaseII.containing.BREX | DUF6079*->N6-N4-METHYLASE->TUDOR+SF1-SF2-Helicase+iREase+CTD->PglZ->BrxA-DNA-binding->BrxA-DNA-binding-><-HNH<-Arm-DNA-bind_3+Phage_integrase | 1103 | Bacteria>Pseudomonadota>Gammaproteobacteria>Enterobacterales>Erwiniaceae>Pantoea | GCA_014841465.1 | Pantoea_agglomerans | IFU02_11635 | 198..3509 | 549 | hypothetical-protein-[Pantoea-agglomerans] |
| WP_033769541.1 | Type3.MTaseII.containing.BREX | Arm-DNA-bind_3+Phage_integrase->HNH-><-BrxA-DNA-binding<-BrxA-DNA-binding<-PglZ<-TUDOR+SF1-SF2-Helicase+iREase+CTD<-N6-N4-METHYLASE<-DUF6079*<-BrxB-BrxF-iSTAND<-SF1-SF2-Helicase<-HKD-DNase<-DUF6361<-XIS-HTH | 1241 | Bacteria>Pseudomonadota>Gammaproteobacteria>Enterobacterales>Erwiniaceae>Pantoea | GCF_000687245.1 | Pantoea_agglomerans | T296_RS18260 | 303869..307594 | 549 | DUF6079-family-protein-[Pantoea-agglomerans] |
| WP_065650234.1 | Type3.MTaseII.containing.BREX | XIS-HTH->?->sGTP->?->?->?->BrxB-BrxF-iSTAND->DUF6079*->N6-N4-METHYLASE->TUDOR+SF1-SF2-Helicase+iREase+CTD->PglZ->BrxA-DNA-binding->BrxA-DNA-binding-><-HNH<-Arm-DNA-bind_3+Phage_integrase<-tRNA\|\|?->DEDD_Tnp_IS110-> | 1241 | Bacteria>Pseudomonadota>Gammaproteobacteria>Enterobacterales>Erwiniaceae>Pantoea | GCF_001691555.1 | Pantoea_eucrina | BBA90_RS17380 | 2387197..2390922 | 472693 | DUF6079-family-protein-[Pantoea-eucrina] |
| WP_084884537.1 | Type3.MTaseII.containing.BREX | GNTR-HTH+Aminotran_1_2-><-?\|\|tRNA->Arm-DNA-bind_3+Phage_integrase->HNH-><-BrxA-DNA-binding<-BrxA-DNA-binding<-PglZ<-TUDOR+SF1-SF2-Helicase+iREase+CTD<-N6-N4-METHYLASE<-DUF6079* | 1241 | Bacteria>Pseudomonadota>Gammaproteobacteria>Enterobacterales>Erwiniaceae>Pantoea | GCF_002095575.1 | Pantoea_septica | HA46_RS12285 | 90403..94128 | 472695 | DUF6079-family-protein-[Pantoea-septica] |
| WP_187500220.1 | Type3.MTaseII.containing.BREX | <-SymE_toxin<-CHTH\|\|ColE7+HNH->MarR-HTH->MarR-HTH->\|\|BrxB-BrxF-iSTAND->DUF6079*->N6-N4-METHYLASE->TUDOR+SF1-SF2-Helicase+iREase+CTD->PglZ->BrxA-DNA-binding->BrxA-DNA-binding-><-HNH | 1241 | Bacteria>Pseudomonadota>Gammaproteobacteria>Enterobacterales>Erwiniaceae>Pantoea | GCF_014354155.1 | Pantoea_agglomerans | IBU41_RS09190 | 803357..807082 | 549 | DUF6079-family-protein-[Pantoea-agglomerans] |
| WP_192034226.1 | Type3.MTaseII.containing.BREX | sGTP->?->sGTP->DUF6361->HKD-DNase->SF1-SF2-Helicase->BrxB-BrxF-iSTAND->DUF6079-p*-> | 118 | Bacteria>Pseudomonadota>Gammaproteobacteria>Enterobacterales>Erwiniaceae>Pantoea | GCF_014841465.1 | Pantoea_agglomerans | IFU02_RS20350 | 110041..110395 | 549 | DUF6079-family-protein,-partial-[Pantoea-agglomerans] |
| WP_047715704.1 | Type3.MTaseII.containing.BREX | <-AAA+wHTH<-XIS-HTH\|\|?->?->REase->DDE_Tnp_1_5+DDE_Tnp_1->?->BrxB-BrxF-iSTAND->DUF6079*->N6-N4-METHYLASE->TUDOR+SF1-SF2-Helicase+iREase+CTD->PglZ->BrxA-DNA-binding->BrxA-DNA-binding-><-HNH<-Arm-DNA-bind_3+Phage_integrase | 1241 | Bacteria>Pseudomonadota>Gammaproteobacteria>Enterobacterales | GCF_001022015.1 | Enterobacteriaceae | ABY62_RS09035 | 1748148..1751873 | 543 | MULTISPECIES:-DUF6079-family-protein-[Enterobacteriaceae] |
| WP_088569507.1 | Type3.MTaseII.containing.BREX | BrxB-BrxF-iSTAND->DUF6079*->N6-N4-METHYLASE->TUDOR+SF1-SF2-Helicase+iREase+CTD->PglZ->BrxA-DNA-binding->BrxA-DNA-binding-><-HNH<-ORC-CDC6<-GDResolvase+HIN-HTH | 1241 | Bacteria>Pseudomonadota>Gammaproteobacteria>Enterobacterales | GCF_014103065.1 | Enterobacteriaceae | CEQ52_RS25735 | 1773..5498 | 543 | MULTISPECIES:-DUF6079-family-protein-[Enterobacteriaceae] |
| WP_064400984.1 | Type3.MTaseII.containing.BREX | Arm-DNA-bind_3+Phage_integrase->HNH-><-BrxA-DNA-binding<-BrxA-DNA-binding<-PglZ<-TUDOR+SF1-SF2-Helicase+iREase+CTD<-N6-N4-METHYLASE<-DUF6079*<-BrxB-BrxF-iSTAND<-SF1-SF2-Helicase<-HKD-DNase<-DUF6361<-ZnR<-XIS-HTH | 1241 | Bacteria>Pseudomonadota>Gammaproteobacteria>Enterobacterales | GCF_018255815.1 | Enterobacteriaceae | KFO54_RS02665 | 559330..563055 | 543 | MULTISPECIES:-DUF6079-family-protein-[Enterobacteriaceae] |
| WP_234625897.1 | Type3.MTaseII.containing.BREX | Arm-DNA-bind_3+Phage_integrase->HNH-><-BrxA-DNA-binding<-BrxA-DNA-binding<-PglZ<-TUDOR+SF1-SF2-Helicase+iREase+CTD<-N6-N4-METHYLASE<-DUF6079*<-BrxB-BrxF-iSTAND<-?<-REase | 1241 | Bacteria>Pseudomonadota>Gammaproteobacteria>Enterobacterales>Hafniaceae>Hafnia | GCF_021440185.1 | Hafnia_paralvei | LZ651_RS15655 | 36583..40308 | 546367 | DUF6079-family-protein-[Hafnia-paralvei] |
| WP_198860972.1 | Type3.MTaseII.containing.BREX | <-Peptidase_C80<-ACET<-?\|\|tRNA->Arm-DNA-bind_3+Phage_integrase->?->wHTH+WYL+WCX->BrxB-BrxF-iSTAND->DUF6079*->N6-N4-METHYLASE->TUDOR+SF1-SF2-Helicase+iREase+CTD-p->PglZ->BrxA-DNA-binding->BrxA-DNA-binding->?->?->?->REase-> | 1241 | Bacteria>Pseudomonadota>Gammaproteobacteria>Enterobacterales>Morganellaceae | GCF_022349885.1 | Providencia | MJ635_RS05620 | 34035..37760 | 586 | MULTISPECIES:-DUF6079-family-protein-[Providencia] |
| WP_186758892.1 | Type3.MTaseII.containing.BREX | <-SF1-SF2-Helicase<-?<-?<-MarR-HTH\|\|<-BrxA-DNA-binding<-BrxA-DNA-binding<-PglZ<-TUDOR+SF1-SF2-Helicase+iREase+CTD<-N6-N4-METHYLASE<-DUF6079*<-BrxB-BrxF-iSTAND<-XIS-HTH | 1241 | Bacteria>Pseudomonadota>Gammaproteobacteria>Enterobacterales>Morganellaceae>Morganella | GCF_014283995.1 | Morganella_morganii | H8R19_RS13740 | 175398..179123 | 582 | DUF6079-family-protein-[Morganella-morganii] |
| WP_079549421.1 | Type3.MTaseII.containing.BREX | XIS-HTH->BrxB-BrxF-iSTAND->DUF6079*->N6-N4-METHYLASE->TUDOR+SF1-SF2-Helicase+iREase+CTD-p->?->PglZ->BrxA-DNA-binding->BrxA-DNA-binding-><-Arm-DNA-bind_3+Phage_integrase<-tRNA\|\|Peptidase_M3-> | 1241 | Bacteria>Pseudomonadota>Gammaproteobacteria>Enterobacterales>Morganellaceae>Morganella | GCF_018475385.1 | Morganella_morganii | IS067_RS12990 | 4918..8643 | 582 | DUF6079-family-protein-[Morganella-morganii] |
| WP_111759512.1 | Type3.MTaseII.containing.BREX | Arm-DNA-bind_3+Phage_integrase->HNH-><-BrxA-DNA-binding<-BrxA-DNA-binding<-PglZ<-TUDOR+SF1-SF2-Helicase+iREase+CTD<-N6-N4-METHYLASE<-DUF6079*<-BrxB-BrxF-iSTAND<-?<-REase<-REase<-ZnR<-?<-XIS-HTH | 1241 | Bacteria>Pseudomonadota>Gammaproteobacteria>Enterobacterales>Morganellaceae>Morganella | GCF_902387845.1 | Morganella_morganii | FX120_RS01945 | 401947..405672 | 582 | DUF6079-family-protein-[Morganella-morganii] |
| GKW26535.1 | Type3.MTaseII.containing.BREX | <-nSTAND1+GTPase-AIG+sGTP<-CopG/RHH\|\|?->XIS-HTH->ZnR->?->?->LuxR-HTH->BrxB-BrxF-iSTAND->DUF6079*->N6-N4-METHYLASE->TUDOR+SF1-SF2-Helicase+iREase+CTD->PglZ->BrxA-DNA-binding->BrxA-DNA-binding-><-Arm-DNA-bind_3+Phage_integrase<-tRNA<-tRNA<-tRNA<-?<-URI-DNase+UVR+RNase+HHH<-REC+OmpR-HTH | 1241 | Bacteria>Pseudomonadota>Gammaproteobacteria>Enterobacterales>Pectobacteriaceae>Pectobacterium | GCA_024343625.1 | Pectobacterium_carotovorum_subsp._carotovorum | PEC311524_41290 | 13237..16962 | 555 | hypothetical-protein-PEC311524_41290-[Pectobacterium-carotovorum-subsp.-carotovorum] |
| WP_120162111.1 | Type3.MTaseII.containing.BREX | HNH-><-BrxA-DNA-binding<-BrxA-DNA-binding<-PglZ<-TUDOR+SF1-SF2-Helicase+iREase+CTD<-N6-N4-METHYLASE<-DUF6079*<-BrxB-BrxF-iSTAND<-SF1-SF2-Helicase<-SF1-SF2-Helicase<-?\|\|?-><-LuxR-HTH<-LuxR-HTH | 1241 | Bacteria>Pseudomonadota>Gammaproteobacteria>Enterobacterales>Yersiniaceae>Rahnella | GCF_003610915.1 | Rahnella_variigena | CKQ54_RS06125 | 1316224..1319949 | 574964 | DUF6079-family-protein-[Rahnella-variigena] |
| WP_129942186.1 | Type3.MTaseII.containing.BREX | Arm-DNA-bind_3+Phage_integrase->HNH-><-BrxA-DNA-binding<-BrxA-DNA-binding<-PglZ<-TUDOR+SF1-SF2-Helicase+iREase+CTD<-N6-N4-METHYLASE<-DUF6079*<-BrxB-BrxF-iSTAND<-SF1-SF2-Helicase<-HKD-DNase<-DUF6361<-ZnR<-XIS-HTH | 1241 | Bacteria>Pseudomonadota>Gammaproteobacteria>Enterobacterales>Yersiniaceae>Serratia | GCF_004153885.1 | Serratia_liquefaciens | BSR02_RS08575 | 211251..214976 | 614 | DUF6079-family-protein-[Serratia-liquefaciens] |
| TVR57519.1 | Type3.MTaseII.containing.BREX | <-DUF6079-p*<-REase<-BrxB-BrxF-iSTAND<-?\|\|?->Amidohydro_1->SAM-METHYLASE->HAD+HAD-> | 744 | Bacteria>Pseudomonadota>Gammaproteobacteria>Gammaproteobacteria_incertae_sedis>Candidatus_Competibacteraceae>unclassified_Candidatus_Competibacteraceae | GCA_007695245.1 | Candidatus_Competibacteraceae_bacterium | EA420_18880 | 1..2231 | 2053538 | ATP-binding-protein,-partial-[Candidatus-Competibacteraceae-bacterium] |
| MBK8753028.1 | Type3.MTaseII.containing.BREX | ParA-Soj-PloopNTPase->Peptidase->HNH-><-?\|\|?->BrxB-BrxF-iSTAND->DUF6079*-><-Peptidase_S8<-Classical-AAA\|\|N6-N4-METHYLASE->TUDOR+SF1-SF2-Helicase+iREase+CTD-> | 1229 | Bacteria>Pseudomonadota>Gammaproteobacteria>Gammaproteobacteria_incertae_sedis>Candidatus_Competibacteraceae>unclassified_Candidatus_Competibacteraceae | GCA_016710535.1 | Candidatus_Competibacteraceae_bacterium | IPL99_16005 | 6574..10263 | 2053538 | ATP-binding-protein-[Candidatus-Competibacteraceae-bacterium] |
| MCB1782191.1 | Type3.MTaseII.containing.BREX | DUF6079*->HHH_4+HTH+wHTH+SF1-SF2-Helicase+REase->N6-N4-METHYLASE->TUDOR+SF1-SF2-Helicase+iREase+CTD->PglZ->BrxA-DNA-binding->HIN-HTH-> | 1035 | Bacteria>Pseudomonadota>Gammaproteobacteria>Gammaproteobacteria_incertae_sedis>Candidatus_Competibacteraceae>unclassified_Candidatus_Competibacteraceae | GCA_020445025.1 | Candidatus_Competibacteraceae_bacterium | KDJ34_19295 | 1..3110 | 2053538 | ATP-binding-protein-[Candidatus-Competibacteraceae-bacterium] |
| PLY12604.1 | Type3.MTaseII.containing.BREX | XIS-HTH->BrxB-BrxF-iSTAND->DUF6079*->Schlafen+HATPase+wHTH->HEPN->N6-N4-METHYLASE->HEPN->?->OLD-ABC->TOPRIM->TUDOR+SF1-SF2-Helicase+iREase+CTD->PglZ-> | 1279 | Bacteria>Pseudomonadota>Gammaproteobacteria>Gammaproteobacteria_incertae_sedis>Sedimenticola>unclassified_Sedimenticola | GCA_002869585.1 | Sedimenticola_sp. | C0631_16515 | 1414..5253 | 1940285 | MAG:-ATPase-[Sedimenticola-sp.] |
| MBT4668298.1 | Type3.MTaseII.containing.BREX | XIS-HTH->BrxB-BrxF-iSTAND-><-PAIREDC+HTH+HIN-HTH\|\|DUF6079*->Schlafen+HATPase+wHTH->N6-N4-METHYLASE->TUDOR+SF1-SF2-Helicase+iREase+CTD->PglZ->BrxA-DNA-binding-> | 1247 | Bacteria>Pseudomonadota>Gammaproteobacteria>Gammaproteobacteria_incertae_sedis>sulfur-oxidizing_symbionts>Candidatus_Ruthia | GCA_018657745.1 | Candidatus_Ruthia_sp. | HOC17_02010 | 1923..5666 | 2735566 | ATP-binding-protein-[Candidatus-Ruthturnera-sp.] |
| ODJ87772.1 | Type3.MTaseII.containing.BREX | <-SbcC+CCDC158<-Calcineurin-SbcD<-SbcC<-?<-OBFOLD-NUCLEASE<-HNH\|\|wHTH+WYL+WCX->BrxB-BrxF-iSTAND->DUF6079*->DUF6079-p->N6-N4-METHYLASE->TUDOR+SF1-SF2-Helicase+iREase+CTD->PglZ->BrxA-DNA-binding->?->?-><-TraG_N | 1004 | Bacteria>Pseudomonadota>Gammaproteobacteria>Gammaproteobacteria_incertae_sedis>sulfur-oxidizing_symbionts>Candidatus_Thiodiazotropha | GCA_001715975.1 | Candidatus_Thiodiazotropha_endolucinida | CODIS_18800 | 29097..32111 | 1655433 | hypothetical-protein-CODIS_18800-[Candidatus-Thiodiazotropha-endolucinida] |
| ODJ87773.1 | Type3.MTaseII.containing.BREX | <-SbcC+CCDC158<-Calcineurin-SbcD<-SbcC<-?<-OBFOLD-NUCLEASE<-HNH\|\|wHTH+WYL+WCX->BrxB-BrxF-iSTAND->DUF6079-p*->N6-N4-METHYLASE->TUDOR+SF1-SF2-Helicase+iREase+CTD->PglZ->BrxA-DNA-binding->?->?-><-TraG_N<-?<-TraH | 212 | Bacteria>Pseudomonadota>Gammaproteobacteria>Gammaproteobacteria_incertae_sedis>sulfur-oxidizing_symbionts>Candidatus_Thiodiazotropha | GCA_001715975.1 | Candidatus_Thiodiazotropha_endolucinida | CODIS_18810 | 32185..32823 | 1655433 | hypothetical-protein-CODIS_18810-[Candidatus-Thiodiazotropha-endolucinida] |
| MCG7977135.1 | Type3.MTaseII.containing.BREX | <-SbcC+CCDC158<-Calcineurin-SbcD<-SbcC<-?<-OBFOLD-NUCLEASE<-HNH\|\|wHTH+WYL+WCX->BrxB-BrxF-iSTAND->DUF6079*->HEPN->N6-N4-METHYLASE->TUDOR+SF1-SF2-Helicase+iREase+CTD->PglZ->BrxA-DNA-binding->?->?-><-TraG_N | 1241 | Bacteria>Pseudomonadota>Gammaproteobacteria>Gammaproteobacteria_incertae_sedis>sulfur-oxidizing_symbionts>Candidatus_Thiodiazotropha | GCA_016842545.1 | Candidatus_Thiodiazotropha_taylori | JAY77_03165 | 13858..17583 | 2792791 | DUF6079-family-protein-[Candidatus-Thiodiazotropha-taylori] |
| MCG7969984.1 | Type3.MTaseII.containing.BREX | <-BrxA-DNA-binding<-PglZ<-TUDOR+SF1-SF2-Helicase+iREase+CTD<-?<-SbcC<-N6-N4-METHYLASE<-MORC<-DUF6079*<-BrxB-BrxF-iSTAND<-XIS-HTH<-RHH_5\|\|?->?->ZnR+TOPRIM+AAA-helicase+wHTH-> | 1243 | Bacteria>Pseudomonadota>Gammaproteobacteria>Gammaproteobacteria_incertae_sedis>sulfur-oxidizing_symbionts>Candidatus_Thiodiazotropha | GCA_016843285.1 | Candidatus_Thiodiazotropha_taylori | JBO36_04935 | 30051..33782 | 2792791 | DUF6079-family-protein-[Candidatus-Thiodiazotropha-taylori] |
| WP_061469286.1 | Type3.MTaseII.containing.BREX | PIN-><-XIS-HTH<-?<-?\|\|?->XIS-HTH->BrxB-BrxF-iSTAND->DUF6079*->N6-N4-METHYLASE->TUDOR+SF1-SF2-Helicase+iREase+CTD->PglZ->BrxA-DNA-binding-><-?<-TrwC+SF1-SF2-Helicase<-HerA/FtsK | 1244 | Bacteria>Pseudomonadota>Gammaproteobacteria>Legionellales>Legionellaceae>Legionella | GCF_001582325.1 | Legionella_pneumophila | AOU53_RS05715 | 45923..49657 | 446 | DUF6079-family-protein-[Legionella-pneumophila] |
| SJM92359.1 | Type3.MTaseII.containing.BREX | <-DUF6079-p<-DUF6079-p*<-HTH_21+rve+rve_3<-HTH_21+rve+rve_3<-HTH_Tnp_1 | 754 | Bacteria>Pseudomonadota>Gammaproteobacteria>Methylococcales>Crenotrichaceae>Crenothrix | GCA_900163745.1 | Crenothrix_polyspora | CRENPOLYSF2_2680003 | 1003..3267 | 360316 | conserved-hypothetical-protein-[Crenothrix-polyspora] |
| WP_218780277.1 | Type3.MTaseII.containing.BREX | <-Schlafen+HATPase+wHTH<-DUF6079-p*<-DUF6079-p<-DUF6079-p<-HTH_21+rve+rve_3<-HTH_21+rve+rve_3<-HTH_Tnp_1 | 191 | Bacteria>Pseudomonadota>Gammaproteobacteria>Methylococcales>Crenotrichaceae>Crenothrix | GCF_900163745.1 | Crenothrix_polyspora | CCY91_RS19335 | 438..1013 | 360316 | DUF6079-family-protein-[Crenothrix-polyspora] |
| WP_218780278.1 | Type3.MTaseII.containing.BREX | <-Schlafen+HATPase+wHTH<-DUF6079-p<-DUF6079-p*<-DUF6079-p<-HTH_21+rve+rve_3<-HTH_21+rve+rve_3<-HTH_Tnp_1 | 732 | Bacteria>Pseudomonadota>Gammaproteobacteria>Methylococcales>Crenotrichaceae>Crenothrix | GCF_900163745.1 | Crenothrix_polyspora | CCY91_RS19340 | 1003..3201 | 360316 | DUF6079-family-protein-[Crenothrix-polyspora] |
| WP_218780279.1 | Type3.MTaseII.containing.BREX | <-Schlafen+HATPase+wHTH<-DUF6079-p<-DUF6079-p<-DUF6079-p*<-HTH_21+rve+rve_3<-HTH_21+rve+rve_3<-HTH_Tnp_1 | 313 | Bacteria>Pseudomonadota>Gammaproteobacteria>Methylococcales>Crenotrichaceae>Crenothrix | GCF_900163745.1 | Crenothrix_polyspora | CCY91_RS19345 | 3245..4186 | 360316 | DUF6079-family-protein-[Crenothrix-polyspora] |
| MCF7967519.1 | Type3.MTaseII.containing.BREX | HigB_toxin->HTH_3-><-?\|\|XIS-HTH->BrxB-BrxF-iSTAND->DUF6079*->Schlafen+HATPase+wHTH->N6-N4-METHYLASE-> | 1246 | Bacteria>Pseudomonadota>Gammaproteobacteria>Methylococcales>Methylococcaceae>Methylobacter | GCA_021736725.1 | Methylobacter_tundripaludum | K9L79_18560 | 2994..6734 | 173365 | DUF6079-family-protein-[Methylobacter-tundripaludum] |
| WP_174627171.1 | Type3.MTaseII.containing.BREX | <-BrxA-DNA-binding<-PglZ<-TUDOR+SF1-SF2-Helicase+iREase+CTD<-HKD-DNase<-SF1-SF2-Helicase<-N6-N4-METHYLASE<-Schlafen+HATPase+wHTH<-DUF6079*<-BrxB-BrxF-iSTAND | 1240 | Bacteria>Pseudomonadota>Gammaproteobacteria>Methylococcales>Methylococcaceae>Methylobacter | GCF_902806695.1 | Candidatus_Methylobacter_favarea | HUX64_RS15760 | 14501..18223 | 2707345 | DUF6079-family-protein-[Candidatus-Methylobacter-favarea] |
| WP_017365797.1 | Type3.MTaseII.containing.BREX | <-MPTase<-HTH\|\|?->?->?-><-N6-N4-METHYLASE<-HerA/FtsK<-?\|\|BrxB-BrxF-iSTAND->DUF6079*->N6-N4-METHYLASE->TUDOR+SF1-SF2-Helicase+iREase+CTD->PglZ->BrxA-DNA-binding->?-><-Phage_lysozyme<-Phage_holin | 1241 | Bacteria>Pseudomonadota>Gammaproteobacteria>Methylococcales>Methylococcaceae>Methylococcus | GCF_000297615.1 | Methylococcus_capsulatus | B858_RS0111535 | 120844..124569 | 414 | DUF6079-family-protein-[Methylococcus-capsulatus] |
| WP_198322148.1 | Type3.MTaseII.containing.BREX | <-BrxA-DNA-binding<-PglZ<-TUDOR+SF1-SF2-Helicase+iREase+CTD<-REase<-OLD-ABC<-N6-N4-METHYLASE<-N6-N4-METHYLASE-p<-PIN<-PhdYeFM_antitox<-DUF6079*<-BrxB-BrxF-iSTAND<-RNA_METHYLASE<-Aminotran_3 | 1242 | Bacteria>Pseudomonadota>Gammaproteobacteria>Methylococcales>Methylococcaceae>Methylococcus | GCF_016106025.1 | Methylococcus_sp._BF19-07 | HUK90_RS04065 | 110959..114687 | 2743472 | DUF6079-family-protein-[Methylococcus-sp.-BF19-07] |
| WP_218808314.1 | Type3.MTaseII.containing.BREX | <-BrxA-DNA-binding<-PglZ<-TUDOR+SF1-SF2-Helicase+iREase+CTD<-REase<-OLD-ABC<-N6-N4-METHYLASE<-?<-PIN<-PhdYeFM_antitox<-DUF6079*<-BrxB-BrxF-iSTAND<-RNA_METHYLASE<-Aminotran_3<-?<-?<-NTP_transf_3+Hexapep+Hexapep | 1265 | Bacteria>Pseudomonadota>Gammaproteobacteria>Methylococcales>Methylococcaceae>Methylococcus | GCF_019285515.1 | Methylococcus_sp._Mc7 | KW115_RS06340 | 1279422..1283219 | 2860258 | DUF6079-family-protein-[Methylococcus-sp.-Mc7] |
| WP_218812148.1 | Type3.MTaseII.containing.BREX | Integrase_1->Phage_integrase-><-BrxA-DNA-binding<-PglZ<-TUDOR+SF1-SF2-Helicase+iREase+CTD<-Schlafen+HATPase+wHTH<-N6-N4-METHYLASE<-DUF6079*<-BrxB-BrxF-iSTAND\|\|?->HerA/FtsK->?->N6-N4-METHYLASE-><-Arm-DNA-bind_3+Phage_integrase<-sGTP | 1241 | Bacteria>Pseudomonadota>Gammaproteobacteria>Methylococcales>Methylococcaceae>Methylococcus | GCF_019285535.1 | Methylococcus_capsulatus | KW112_RS09060 | 1958527..1962252 | 414 | DUF6079-family-protein-[Methylococcus-capsulatus] |
| WP_218800686.1 | Type3.MTaseII.containing.BREX | RNaseP->?->?->sGTP->Arm-DNA-bind_3+Phage_integrase-><-N6-N4-METHYLASE<-HerA/FtsK<-?\|\|BrxB-BrxF-iSTAND->DUF6079*->OLD-ABC->TOPRIM->N6-N4-METHYLASE->Schlafen+HATPase+wHTH->TUDOR+SF1-SF2-Helicase+iREase+CTD->PglZ->BrxA-DNA-binding-><-Phage_integrase<-Integrase_1 | 1241 | Bacteria>Pseudomonadota>Gammaproteobacteria>Methylococcales>Methylococcaceae>Methylococcus | GCF_019285575.1 | Methylococcus_capsulatus | KW113_RS00555 | 116161..119886 | 414 | DUF6079-family-protein-[Methylococcus-capsulatus] |
| WP_249385185.1 | Type3.MTaseII.containing.BREX | Integrase_1->Phage_integrase-><-BrxA-DNA-binding<-PglZ<-TUDOR+SF1-SF2-Helicase+iREase+CTD<-Schlafen+HATPase+wHTH<-N6-N4-METHYLASE<-DUF6079*<-BrxB-BrxF-iSTAND\|\|?->HerA/FtsK->N6-N4-METHYLASE-><-Arm-DNA-bind_3+Phage_integrase<-sGTP<-?<-?<-RNaseP | 1241 | Bacteria>Pseudomonadota>Gammaproteobacteria>Methylococcales>Methylococcaceae>Methylococcus | GCF_023370095.1 | Methylococcus_capsulatus | M3M30_RS10805 | 2341456..2345181 | 414 | DUF6079-family-protein-[Methylococcus-capsulatus] |
| WP_266024638.1 | Type3.MTaseII.containing.BREX | DUF488-><-?\|\|HTH_29+HTH_32+HTH_33+DDE_3-><-ParB+HNH<-BrxA-DNA-binding<-PglZ<-TUDOR+SF1-SF2-Helicase+iREase+CTD<-N6-N4-METHYLASE<-DUF6079*<-BrxB-BrxF-iSTAND<-wHTH+WYL+WCX\|\|?->?->SMODS->HNH-><-?<-Phage_integrase | 1242 | Bacteria>Pseudomonadota>Gammaproteobacteria>Methylococcales>Methylococcaceae>Methylococcus | GCF_026247885.1 | Methylococcus_sp._16-5 | OOT43_RS09335 | 1977791..1981519 | 2993564 | DUF6079-family-protein-[Methylococcus-sp.-16-5] |
| MCF8007745.1 | Type3.MTaseII.containing.BREX | <-DUF6079-p*<-DUF6079-p<-BrxB-BrxF-iSTAND<-?\|\|?->ACET-><-?<-?<-?<-Aminotran_3 | 331 | Bacteria>Pseudomonadota>Gammaproteobacteria>Methylococcales>Methylococcaceae>Methylovulum | GCA_021736885.1 | Methylovulum_sp. | K9K84_10085 | 1..992 | 1916980 | DUF6079-family-protein-[Methylovulum-sp.] |
| MCF8007746.1 | Type3.MTaseII.containing.BREX | <-DUF6079-p<-DUF6079-p*<-BrxB-BrxF-iSTAND<-?\|\|?->ACET-><-?<-?<-?<-Aminotran_3 | 154 | Bacteria>Pseudomonadota>Gammaproteobacteria>Methylococcales>Methylococcaceae>Methylovulum | GCA_021736885.1 | Methylovulum_sp. | K9K84_10090 | 1005..1469 | 1916980 | DUF6079-family-protein-[Methylovulum-sp.] |
| PZN70748.1 | Type3.MTaseII.containing.BREX | <-BrxA-DNA-binding<-PglZ<-?<-TUDOR+SF1-SF2-Helicase+iREase+CTD<-?<-Schlafen+HATPase+wHTH<-PHP+SbcC<-N6-N4-METHYLASE<-Schlafen+HATPase+wHTH<-DUF6079*<-XIS-HTH | 1249 | Bacteria>Pseudomonadota>Gammaproteobacteria>Methylococcales>Methylococcales_incertae_sedis>Candidatus_Methylumidiphilus | GCA_003242955.1 | Candidatus_Methylumidiphilus_alinenensis | DM484_28025 | 24021..27770 | 2202197 | ATPase-[Candidatus-Methylumidiphilus-alinenensis] |
| EGY4091323.1 | Type3.MTaseII.containing.BREX | ZnR+ZnR->DUF932->?->?->XIS-HTH->SAM-METHYLASE-><-wHTH+HNH\|\|BrxB-BrxF-iSTAND->DUF6079*->N6-N4-METHYLASE->TUDOR+SF1-SF2-Helicase+iREase+CTD->PglZ->BrxA-DNA-binding-><-Arm-DNA-bind_3+Phage_integrase<-tRNA\|\|?-><-?\|\|REase-> | 1242 | Bacteria>Pseudomonadota>Gammaproteobacteria>Moraxellales>Moraxellaceae>Acinetobacter | - | Acinetobacter_baumannii | JIM75_000843 | 234969..238697 | 470 | ATP-binding-protein-[Acinetobacter-baumannii] |
| MBP8005395.1 | Type3.MTaseII.containing.BREX | Arm-DNA-bind_2+Phage_int_SAM_3+Phage_integrase->?->Phage_rep->AAA-ATPase-><-HTH_Tnp_1+HTH_21+rve+rve_3<-BrxA-DNA-binding<-PglZ<-TUDOR+SF1-SF2-Helicase+iREase+CTD<-N6-N4-METHYLASE<-DUF6079*<-BrxB-BrxF-iSTAND<-wHTH+HNH<-SAM-METHYLASE\|\|DUF4236->?-><-?\|\|<-SWIM+DUF3670+SF1-SF2-Helicase<-HTH_3<-HipA_C | 1242 | Bacteria>Pseudomonadota>Gammaproteobacteria>Moraxellales>Moraxellaceae>Acinetobacter | GCA_018004035.1 | Acinetobacter_sp. | KAZ18_00615 | 14503..18231 | 472 | ATP-binding-protein-[Acinetobacter-sp.] |
| WP_180049757.1 | Type3.MTaseII.containing.BREX | ICLR-HTH-><-BrxA-DNA-binding<-PglZ<-TUDOR+SF1-SF2-Helicase+iREase+CTD<-N6-N4-METHYLASE<-DUF6079*<-BrxB-BrxF-iSTAND<-wHTH(SBD)+HNH<-SAM-METHYLASE<-XIS-HTH<-?<-?<-DUF932 | 1242 | Bacteria>Pseudomonadota>Gammaproteobacteria>Moraxellales>Moraxellaceae>Acinetobacter | GCF_013419095.1 | Acinetobacter_sp._YH12105 | FTU44_RS05460 | 205234..208962 | 2601093 | DUF6079-family-protein-[Acinetobacter-sp.-YH12105] |
| MBQ0714975.1 | Type3.MTaseII.containing.BREX | <-HTH_Tnp<-N6-N4-METHYLASE<-DUF6079*<-BrxB-BrxF-iSTAND<-wHTH+WYL+WCX\|\|?->?->?->HerA/FtsK->TrwC+SF1-SF2-Helicase-> | 1242 | Bacteria>Pseudomonadota>Gammaproteobacteria>Moraxellales>Moraxellaceae>Paraperlucidibaca | GCA_018066805.1 | Paraperlucidibaca_sp. | KBT56_07365 | 1307..5035 | 2708021 | ATP-binding-protein-[Paraperlucidibaca-sp.] |
| WP_075107047.1 | Type3.MTaseII.containing.BREX | HTH_Tnp_IS630+DDE_3-><-?<-REase<-REase\|\|ZnR->McrB-AAA->McrC-NTD+REase->BrxB-BrxF-iSTAND->DUF6079*->N6-N4-METHYLASE->N6-N4-METHYLASE-><-HIN-HTH<-REase\|\|PglZ->BrxA-DNA-binding-><-Arm-DNA-bind_3+Phage_integrase | 1241 | Bacteria>Pseudomonadota>Gammaproteobacteria>Moraxellales>Moraxellaceae>Psychrobacter | GCF_001921765.1 | Psychrobacter_sp._C_20.9 | BTW00_RS08530 | 118940..122665 | 1926477 | DUF6079-family-protein-[Psychrobacter-sp.-C-20.9] |
| MCH2557387.1 | Type3.MTaseII.containing.BREX | <-BrxA-DNA-binding<-PglZ<-TUDOR+SF1-SF2-Helicase+iREase+CTD<-Schlafen+HATPase+wHTH<-N6-N4-METHYLASE<-HerA/FtsK<-SIR2<-OLD-ABC+TOPRIM<-DUF6079*<-BrxB-BrxF-iSTAND<-wHTH+WYL+WCX<-TraG_N<-?<-TraH<-TraU<-?<-?<-Thioredoxin_4 | 1241 | Bacteria>Pseudomonadota>Gammaproteobacteria>Oceanospirillales>Alcanivoracaceae>Alcanivorax | GCA_022449455.1 | Alcanivorax_sp. | MK005_08810 | 144603..148328 | 1872427 | DUF6079-family-protein-[Alcanivorax-sp.] |
| WP_008928712.1 | Type3.MTaseII.containing.BREX | <-BrxA-DNA-binding<-PglZ<-TUDOR+SF1-SF2-Helicase+iREase+CTD<-Schlafen+HATPase+wHTH<-N6-N4-METHYLASE<-HerA/FtsK<-SIR2<-OLD-ABC+TOPRIM<-DUF6079*<-BrxB-BrxF-iSTAND<-wHTH+WYL+WCX\|\|ARG->?->?-><-TraG_N<-?<-TraH<-TraU | 1241 | Bacteria>Pseudomonadota>Gammaproteobacteria>Oceanospirillales>Alcanivoracaceae>Alcanivorax | GCF_000300995.1 | Alcanivorax_hongdengensis | A11A3_RS07680 | 81332..85057 | 519051 | DUF6079-family-protein-[Alcanivorax-hongdengensis] |
| MCP5207854.1 | Type3.MTaseII.containing.BREX | NTP_transf_3+Hexapep+Hexapep->?->REase+wHTH->?->TnsC-ATPase->?->?->XIS-HTH->BrxB-BrxF-iSTAND->DUF6079*->Schlafen+HATPase+wHTH->N6-N4-METHYLASE->TUDOR+SF1-SF2-Helicase+iREase+CTD->PglZ->BrxA-DNA-binding-><-REase+SF1-SF2-Helicase<-?<-SF1-SF2-Helicase<-DrmB | 1243 | Bacteria>Pseudomonadota>Gammaproteobacteria>Oceanospirillales>Hahellaceae>unclassified_Hahellaceae | GCA_024234475.1 | Hahellaceae_bacterium | H7A01_11660 | 134013..137744 | 2048971 | ATP-binding-protein-[Hahellaceae-bacterium] |
| WP_136248423.1 | Type3.MTaseII.containing.BREX | <-DsbC+DsbD+Thioredoxin_7<-?\|\|XIS-HTH->?->DUF4236->Mrr-N->wHTH+WYL+WCX->BrxB-BrxF-iSTAND->DUF6079*->N6-N4-METHYLASE->Schlafen+HATPase+wHTH->TUDOR+SF1-SF2-Helicase+iREase+CTD->PglZ->BrxA-DNA-binding-> | 1241 | Bacteria>Pseudomonadota>Gammaproteobacteria>Oceanospirillales>Halomonadaceae>Halomonas | GCF_004798905.1 | Halomonas_borealis | ETC07_RS10250 | 48079..51804 | 2508710 | DUF6079-family-protein-[Halomonas-borealis] |
| WP_197566989.1 | Type3.MTaseII.containing.BREX | UDG->?->DUF932->?->?->?->?->wHTH+WYL+WCX->BrxB-BrxF-iSTAND->DUF6079*->N6-N4-METHYLASE->Schlafen+HATPase+wHTH->TUDOR+SF1-SF2-Helicase+iREase+CTD->PglZ->BrxA-DNA-binding-><-?<-?<-XIS-HTH<-CITB-HTH | 1241 | Bacteria>Pseudomonadota>Gammaproteobacteria>Oceanospirillales>Halomonadaceae>Halomonas | GCF_014931605.1 | Halomonas_sp._MCCC_1A13316 | HNO52_RS20455 | 4389739..4393464 | 2733487 | DUF6079-family-protein-[Halomonas-sp.-MCCC-1A13316] |
| WP_043221499.1 | Type3.MTaseII.containing.BREX | <-TraK<-TraE<-TraM<-TraM<-DUF3262<-?\|\|HipA_C->XIS-HTH->BrxB-BrxF-iSTAND->DUF6079*->N6-N4-METHYLASE->SbcC->?->TUDOR+SF1-SF2-Helicase+iREase+CTD->PglZ->BrxA-DNA-binding->Phage-HTH-><-DUF4400<-VirD4-TraG | 1243 | Bacteria>Pseudomonadota>Gammaproteobacteria>Pseudomonadales | GCF_929615065.1 | Pseudomonadaceae | MJP47_RS09275 | 951559..955290 | 135621 | MULTISPECIES:-DUF6079-family-protein-[Pseudomonadaceae] |

| Protein-ID | System | Gene-Neighbourhoods | Length | Taxonomy | Assembly-ID | Strain | Locus-Tag | Genomic-coordinates | TaxID | Fasta-defline |
|---|---|---|---|---|---|---|---|---|---|---|
| MAK49098.1 | Type3.MTaseII.containing.BREX | [illegible] | 1241 | Bacteria>Pseudomonadota>Gammaproteobacteria>Pseudomonadales>Marinobacteraceae>Marinobacter | GCA_002693995.1 | Marinobacter_sp. | CMG85_07410 | 15387..19112 | 50741 | ATPase-[Marinobacter-sp.] |
| WP_273070002.1 | Type3.MTaseII.containing.BREX | [illegible] | 1241 | Bacteria>Pseudomonadota>Gammaproteobacteria>Pseudomonadales>Marinobacteraceae>Marinobacter | GCF_002341575.1 | Marinobacter_sp. | CMG85_07410 | 15387..19112 | 50741 | DUF6079-family-protein-[Marinobacter-sp.] |
| WP_247116180.1 | Type3.MTaseII.containing.BREX | [illegible] | 1243 | Bacteria>Pseudomonadota>Gammaproteobacteria>Pseudomonadales>Marinobacteraceae>Marinobacter | - | Marinobacter_sp._S6332 | MWU62_RS18180 | 6270..10001 | 2926403 | DUF6079-family-protein-[Marinobacter-sp.-S6332] |
| WP_095610363.1 | Type3.MTaseII.containing.BREX | [illegible] | 1241 | Bacteria>Pseudomonadota>Gammaproteobacteria>Pseudomonadales>Marinobacteraceae>Tamilnaduibacter | GCF_002287355.1 | Tamilnaduibacter_salinus | CF392_RS04965 | 53834..57559 | 1484056 | DUF6079-family-protein-[Tamilnaduibacter-salinus] |
| WP_217641968.1 | Type3.MTaseII.containing.BREX | [illegible] | 1246 | Bacteria>Pseudomonadota>Gammaproteobacteria>Pseudomonadales>Pseudomonadaceae>Atopomonas | GCF_900109735.1 | Atopomonas_hussainii | BMX80_RS09305 | 101976..105716 | 1429083 | DUF6079-family-protein-[Atopomonas-hussainii] |
| WP_023116941.1 | Type3.MTaseII.containing.BREX | [illegible] | 1243 | Bacteria>Pseudomonadota>Gammaproteobacteria>Pseudomonadales>Pseudomonadaceae | GCF_000633495.1 | Pseudomonas | U864_RS0106985 | 410399..414130 | 286 | MULTISPECIES:-DUF6079-family-protein-[Pseudomonas] |
| KHO64388.1 | Type3.MTaseII.containing.BREX | [illegible] | 1242 | Bacteria>Pseudomonadota>Gammaproteobacteria>Pseudomonadales>Pseudomonadaceae>Pseudomonas | GCA_000806415.1 | Pseudomonas_flexibilis | PT85_13895 | 375883..379611 | 706570 | ATPase-[Pseudomonas-flexibilis] |
| KSL62728.1 | Type3.MTaseII.containing.BREX | [illegible] | 1242 | Bacteria>Pseudomonadota>Gammaproteobacteria>Pseudomonadales>Pseudomonadaceae>Pseudomonas | GCA_001452025.1 | Pseudomonas_aeruginosa | L5P76_005848 | 1168..4896 | 287 | ATPase-[Pseudomonas-aeruginosa] |
| PAA10532.1 | Type3.MTaseII.containing.BREX | [illegible] | 1248 | Bacteria>Pseudomonadota>Gammaproteobacteria>Pseudomonadales>Pseudomonadaceae>Pseudomonas | GCA_002269625.1 | Pseudomonas_fragi | CJU78_04000 | 74234..77980 | 296 | ATPase-[Pseudomonas-fragi] |
| RRI70507.1 | Type3.MTaseII.containing.BREX | [illegible] | 1246 | Bacteria>Pseudomonadota>Gammaproteobacteria>Pseudomonadales>Pseudomonadaceae>Pseudomonas | GCA_003863585.1 | Pseudomonas_aeruginosa | EIM10_18005 | 61537..65277 | 287 | ATP-binding-protein-[Pseudomonas-aeruginosa] |
| RTS56884.1 | Type3.MTaseII.containing.BREX | [illegible] | 1243 | Bacteria>Pseudomonadota>Gammaproteobacteria>Pseudomonadales>Pseudomonadaceae>Pseudomonas | GCA_003968225.1 | Pseudomonas_aeruginosa | DY943_06675 | 155490..159221 | 287 | ATP-binding-protein-[Pseudomonas-aeruginosa] |
| TRO09271.1 | Type3.MTaseII.containing.BREX | [illegible] | 1242 | Bacteria>Pseudomonadota>Gammaproteobacteria>Pseudomonadales>Pseudomonadaceae>Pseudomonas | GCA_007049815.1 | Pseudomonas_mendocina | EQ829_23215 | 20762..24490 | 300 | ATP-binding-protein-[Pseudomonas-mendocina] |
| TVT71190.1 | Type3.MTaseII.containing.BREX | [illegible] | 1246 | Bacteria>Pseudomonadota>Gammaproteobacteria>Pseudomonadales>Pseudomonadaceae>Pseudomonas | GCA_007713455.1 | Pseudomonas_sp. | FHK79_07520 | 336855..340595 | 306 | ATP-binding-protein-[Pseudomonas-sp.] |
| TXI26904.1 | Type3.MTaseII.containing.BREX | [illegible] | 1061 | Bacteria>Pseudomonadota>Gammaproteobacteria>Pseudomonadales>Pseudomonadaceae>Pseudomonas | GCA_008015435.1 | Pseudomonas_alcaligenes | E6Q69_18785 | 1..3184 | 43263 | ATP-binding-protein,-partial-[Pseudomonas-alcaligenes] |
| MUI85882.1 | Type3.MTaseII.containing.BREX | [illegible] | 1242 | Bacteria>Pseudomonadota>Gammaproteobacteria>Pseudomonadales>Pseudomonadaceae>Pseudomonas | GCA_009727325.1 | Pseudomonas_aeruginosa | GNQ43_32470 | 37788..41516 | 287 | ATP-binding-protein-[Pseudomonas-aeruginosa] |
| NNB48785.1 | Type3.MTaseII.containing.BREX | [illegible] | 1246 | Bacteria>Pseudomonadota>Gammaproteobacteria>Pseudomonadales>Pseudomonadaceae>Pseudomonas | GCA_012986825.1 | Pseudomonas_fragi | HBN89_05770 | 79154..82894 | 296 | ATP-binding-protein-[Pseudomonas-fragi] |
| MBA1269669.1 | Type3.MTaseII.containing.BREX | [illegible] | 1242 | Bacteria>Pseudomonadota>Gammaproteobacteria>Pseudomonadales>Pseudomonadaceae>Pseudomonas | GCA_013522615.1 | Pseudomonas_carnis | G7029_19575 | 130618..134346 | 2487355 | ATP-binding-protein-[Pseudomonas-carnis] |
| MBH9078684.1 | Type3.MTaseII.containing.BREX | [illegible] | 1243 | Bacteria>Pseudomonadota>Gammaproteobacteria>Pseudomonadales>Pseudomonadaceae>Pseudomonas | GCA_016090935.1 | Pseudomonas_aeruginosa | I5K41_09300 | 298308..302039 | 287 | ATP-binding-protein-[Pseudomonas-aeruginosa] |
| MBM2629143.1 | Type3.MTaseII.containing.BREX | [illegible] | 1250 | Bacteria>Pseudomonadota>Gammaproteobacteria>Pseudomonadales>Pseudomonadaceae>Pseudomonas | GCA_016836615.1 | Pseudomonas_aeruginosa | JQ488_19665 | 4235277..4239029 | 287 | ATP-binding-protein-[Pseudomonas-aeruginosa] |
| MBO3071654.1 | Type3.MTaseII.containing.BREX | [illegible] | 1246 | Bacteria>Pseudomonadota>Gammaproteobacteria>Pseudomonadales>Pseudomonadaceae>Pseudomonas | GCA_017582945.1 | Pseudomonas_aeruginosa | J4H77_25770 | 188215..191955 | 287 | ATP-binding-protein-[Pseudomonas-aeruginosa] |
| MBV6142874.1 | Type3.MTaseII.containing.BREX | [illegible] | 1246 | Bacteria>Pseudomonadota>Gammaproteobacteria>Pseudomonadales>Pseudomonadaceae>Pseudomonas | GCA_019167125.1 | Pseudomonas_aeruginosa | KUT38_04275 | 391807..395547 | 287 | ATP-binding-protein-[Pseudomonas-aeruginosa] |
| MBX6244464.1 | Type3.MTaseII.containing.BREX | [illegible] | 1262 | Bacteria>Pseudomonadota>Gammaproteobacteria>Pseudomonadales>Pseudomonadaceae>Pseudomonas | GCA_019686455.1 | Pseudomonas_aeruginosa | ISD49_17785 | 193059..196847 | 287 | ATP-binding-protein-[Pseudomonas-aeruginosa] |
| MBX6030772.1 | Type3.MTaseII.containing.BREX | [illegible] | 1241 | Bacteria>Pseudomonadota>Gammaproteobacteria>Pseudomonadales>Pseudomonadaceae>Pseudomonas | GCA_019688625.1 | Pseudomonas_aeruginosa | ISE67_08735 | 287568..291293 | 287 | ATP-binding-protein-[Pseudomonas-aeruginosa] |
| MCD2801270.1 | Type3.MTaseII.containing.BREX | [illegible] | 1246 | Bacteria>Pseudomonadota>Gammaproteobacteria>Pseudomonadales>Pseudomonadaceae>Pseudomonas | GCA_021060565.1 | Pseudomonas_aeruginosa | K4W73_30415 | 398718..402458 | 287 | ATP-binding-protein-[Pseudomonas-aeruginosa] |
| HBO1294566.1 | Type3.MTaseII.containing.BREX | [illegible] | 1250 | Bacteria>Pseudomonadota>Gammaproteobacteria>Pseudomonadales>Pseudomonadaceae>Pseudomonas | GCA_021597005.1 | Pseudomonas_aeruginosa | L4E86_006702 | 5308..9060 | 287 | ATP-binding-protein-[Pseudomonas-aeruginosa] |
| EIU3491391.1 | Type3.MTaseII.containing.BREX | [illegible] | 1246 | Bacteria>Pseudomonadota>Gammaproteobacteria>Pseudomonadales>Pseudomonadaceae>Pseudomonas | GCA_021644205.1 | Pseudomonas_aeruginosa | L4P75_002394 | 62876..66616 | 287 | ATP-binding-protein-[Pseudomonas-aeruginosa] |
| HBO5819655.1 | Type3.MTaseII.containing.BREX | [illegible] | 1242 | Bacteria>Pseudomonadota>Gammaproteobacteria>Pseudomonadales>Pseudomonadaceae>Pseudomonas | GCA_021668435.1 | Pseudomonas_aeruginosa | L4617_003481 | 113097..116825 | 287 | ATP-binding-protein-[Pseudomonas-aeruginosa] |
| HBO7582681.1 | Type3.MTaseII.containing.BREX | [illegible] | 1242 | Bacteria>Pseudomonadota>Gammaproteobacteria>Pseudomonadales>Pseudomonadaceae>Pseudomonas | GCA_021673555.1 | Pseudomonas_aeruginosa | L4920_006245 | 4928..8656 | 287 | ATP-binding-protein-[Pseudomonas-aeruginosa] |
| HBP0494148.1 | Type3.MTaseII.containing.BREX | [illegible] | 1246 | Bacteria>Pseudomonadota>Gammaproteobacteria>Pseudomonadales>Pseudomonadaceae>Pseudomonas | GCA_021685415.1 | Pseudomonas_aeruginosa | L5Q31_004998 | 30253..33993 | 287 | ATP-binding-protein-[Pseudomonas-aeruginosa] |
| EIU6855274.1 | Type3.MTaseII.containing.BREX | [illegible] | 1242 | Bacteria>Pseudomonadota>Gammaproteobacteria>Pseudomonadales>Pseudomonadaceae>Pseudomonas | GCA_021705865.1 | Pseudomonas_aeruginosa | L5557_000458 | 5822..9550 | 287 | ATP-binding-protein-[Pseudomonas-aeruginosa] |
| HBP6573981.1 | Type3.MTaseII.containing.BREX | [illegible] | 1242 | Bacteria>Pseudomonadota>Gammaproteobacteria>Pseudomonadales>Pseudomonadaceae>Pseudomonas | GCA_021710775.1 | Pseudomonas_aeruginosa | L6A42_23850 | 58047..61775 | 287 | ATP-binding-protein-[Pseudomonas-aeruginosa] |
| MCO1977065.1 | Type3.MTaseII.containing.BREX | [illegible] | 1242 | Bacteria>Pseudomonadota>Gammaproteobacteria>Pseudomonadales>Pseudomonadaceae>Pseudomonas | GCA_023905865.1 | Pseudomonas_aeruginosa | FA422_11970 | 22597..26325 | 287 | ATP-binding-protein-[Pseudomonas-aeruginosa] |
| MCO3958051.1 | Type3.MTaseII.containing.BREX | [illegible] | 1242 | Bacteria>Pseudomonadota>Gammaproteobacteria>Pseudomonadales>Pseudomonadaceae>Pseudomonas | GCA_023912735.1 | Pseudomonas_aeruginosa | FA108_20890 | 55174..58902 | 287 | ATP-binding-protein-[Pseudomonas-aeruginosa] |
| HCF1786781.1 | Type3.MTaseII.containing.BREX | [illegible] | 1250 | Bacteria>Pseudomonadota>Gammaproteobacteria>Pseudomonadales>Pseudomonadaceae>Pseudomonas | GCA_023942115.1 | Pseudomonas_aeruginosa | NH908_000689 | 60743..64495 | 287 | ATP-binding-protein-[Pseudomonas-aeruginosa] |
| HCF1890561.1 | Type3.MTaseII.containing.BREX | [illegible] | 1241 | Bacteria>Pseudomonadota>Gammaproteobacteria>Pseudomonadales>Pseudomonadaceae>Pseudomonas | GCA_023942415.1 | Pseudomonas_aeruginosa | NH920_001195 | 99271..102996 | 287 | ATP-binding-protein-[Pseudomonas-aeruginosa] |
| HCF2077872.1 | Type3.MTaseII.containing.BREX | [illegible] | 1241 | Bacteria>Pseudomonadota>Gammaproteobacteria>Pseudomonadales>Pseudomonadaceae>Pseudomonas | GCA_023942995.1 | Pseudomonas_aeruginosa | NH997_004587 | 20722..24447 | 287 | ATP-binding-protein-[Pseudomonas-aeruginosa] |
| HCF3378232.1 | Type3.MTaseII.containing.BREX | [illegible] | 1266 | Bacteria>Pseudomonadota>Gammaproteobacteria>Pseudomonadales>Pseudomonadaceae>Pseudomonas | GCA_023946955.1 | Pseudomonas_aeruginosa | NIC05_003401 | 65430..69230 | 287 | ATP-binding-protein-[Pseudomonas-aeruginosa] |
| HCF5128584.1 | Type3.MTaseII.containing.BREX | [illegible] | 1250 | Bacteria>Pseudomonadota>Gammaproteobacteria>Pseudomonadales>Pseudomonadaceae>Pseudomonas | GCA_023952375.1 | Pseudomonas_aeruginosa | NIG22_002413 | 2723..6475 | 287 | ATP-binding-protein-[Pseudomonas-aeruginosa] |
| HCG0728340.1 | Type3.MTaseII.containing.BREX | [illegible] | 1246 | Bacteria>Pseudomonadota>Gammaproteobacteria>Pseudomonadales>Pseudomonadaceae>Pseudomonas | GCA_024002515.1 | Pseudomonas_aeruginosa | NJJ48_005263 | 12717..16457 | 287 | ATP-binding-protein-[Pseudomonas-aeruginosa] |
| MCO7639785.1 | Type3.MTaseII.containing.BREX | [illegible] | 1242 | Bacteria>Pseudomonadota>Gammaproteobacteria>Pseudomonadales>Pseudomonadaceae>Pseudomonas | GCA_024128275.1 | Pseudomonas_sp._S_311-6 | NJI34_23730 | 51652..55380 | 2956727 | DUF6079-family-protein-[Pseudomonas-sp.-S-311-6] |
| GLO43967.1 | Type3.MTaseII.containing.BREX | [illegible] | 1246 | Bacteria>Pseudomonadota>Gammaproteobacteria>Pseudomonadales>Pseudomonadaceae>Pseudomonas | GCA_030159635.1 | Pseudomonas_putida | PPUN109347_05290 | 589827..593567 | 303 | hypothetical-protein-PPUN109347_05290-[Pseudomonas-putida] |
| WP_243736488.1 | Type3.MTaseII.containing.BREX | [illegible] | 861 | Bacteria>Pseudomonadota>Gammaproteobacteria>Pseudomonadales>Pseudomonadaceae>Pseudomonas | GCA_030820635.1 | Pseudomonas_aeruginosa | EQ618_RS03870 | 14161..16746 | 287 | DUF6079-family-protein,-partial-[Pseudomonas-aeruginosa] |
| WP_243736489.1 | Type3.MTaseII.containing.BREX | [illegible] | 384 | Bacteria>Pseudomonadota>Gammaproteobacteria>Pseudomonadales>Pseudomonadaceae>Pseudomonas | GCA_030820635.1 | Pseudomonas_aeruginosa | EQ618_RS37760 | 16774..17926 | 287 | DUF6079-family-protein,-partial-[Pseudomonas-aeruginosa] |
| WP_019751243.1 | Type3.MTaseII.containing.BREX | [illegible] | 1243 | Bacteria>Pseudomonadota>Gammaproteobacteria>Pseudomonadales>Pseudomonadaceae>Pseudomonas | GCF_000226035.2 | Pseudomonas_putida | KKK_RS20925 | 4609815..4613546 | 303 | DUF6079-family-protein-[Pseudomonas-putida] |
| WP_008016381.1 | Type3.MTaseII.containing.BREX | [illegible] | 1244 | Bacteria>Pseudomonadota>Gammaproteobacteria>Pseudomonadales>Pseudomonadaceae>Pseudomonas | GCF_000282395.1 | Pseudomonas_sp._GM55 | PMI31_RS03910 | 80959..84693 | 1144333 | DUF6079-family-protein-[Pseudomonas-sp.-GM55] |
| WP_021701537.1 | Type3.MTaseII.containing.BREX | [illegible] | 1246 | Bacteria>Pseudomonadota>Gammaproteobacteria>Pseudomonadales>Pseudomonadaceae>Pseudomonas | GCF_000467105.1 | Pseudomonas_alcaligenes | DYC96_RS14130 | 2987517..2991257 | 43263 | DUF6079-family-protein-[Pseudomonas-alcaligenes] |
| WP_023112387.1 | Type3.MTaseII.containing.BREX | [illegible] | 1246 | Bacteria>Pseudomonadota>Gammaproteobacteria>Pseudomonadales>Pseudomonadaceae>Pseudomonas | GCF_000481185.1 | Pseudomonas_aeruginosa | Q039_RS30195 | 963576..967316 | 287 | DUF6079-family-protein-[Pseudomonas-aeruginosa] |
| WP_033991628.1 | Type3.MTaseII.containing.BREX | [illegible] | 1246 | Bacteria>Pseudomonadota>Gammaproteobacteria>Pseudomonadales>Pseudomonadaceae>Pseudomonas | GCF_000795045.1 | Pseudomonas_aeruginosa | NR30_RS22645 | 72439..76179 | 287 | DUF6079-family-protein-[Pseudomonas-aeruginosa] |
| WP_202965933.1 | Type3.MTaseII.containing.BREX | [illegible] | 1250 | Bacteria>Pseudomonadota>Gammaproteobacteria>Pseudomonadales>Pseudomonadaceae>Pseudomonas | GCF_000796295.1 | Pseudomonas_aeruginosa | NQ68_RS16700 | 65424..69176 | 287 | DUF6079-family-protein-[Pseudomonas-aeruginosa] |
| WP_050481652.1 | Type3.MTaseII.containing.BREX | [illegible] | 1242 | Bacteria>Pseudomonadota>Gammaproteobacteria>Pseudomonadales>Pseudomonadaceae>Pseudomonas | GCF_001181125.1 | Pseudomonas_aeruginosa | J4H77_RS25765 | 188215..191943 | 287 | DUF6079-family-protein-[Pseudomonas-aeruginosa] |
| WP_058184960.1 | Type3.MTaseII.containing.BREX | [illegible] | 1242 | Bacteria>Pseudomonadota>Gammaproteobacteria>Pseudomonadales>Pseudomonadaceae>Pseudomonas | GCF_001453775.1 | Pseudomonas_aeruginosa | APB11_RS24340 | 63235..66963 | 287 | DUF6079-family-protein-[Pseudomonas-aeruginosa] |
| WP_197671916.1 | Type3.MTaseII.containing.BREX | [illegible] | 1246 | Bacteria>Pseudomonadota>Gammaproteobacteria>Pseudomonadales>Pseudomonadaceae>Pseudomonas | GCF_001548335.1 | Pseudomonas_aeruginosa | IOMTU133_RS11685 | 2532129..2535869 | 287 | DUF6079-family-protein-[Pseudomonas-aeruginosa] |
| WP_217363210.1 | Type3.MTaseII.containing.BREX | [illegible] | 1246 | Bacteria>Pseudomonadota>Gammaproteobacteria>Pseudomonadales>Pseudomonadaceae>Pseudomonas | GCF_001721765.1 | Pseudomonas_aeruginosa | A6681_RS28315 | 6159374..6163114 | 287 | DUF6079-family-protein-[Pseudomonas-aeruginosa] |
| WP_069377033.1 | Type3.MTaseII.containing.BREX | [illegible] | 1242 | Bacteria>Pseudomonadota>Gammaproteobacteria>Pseudomonadales>Pseudomonadaceae>Pseudomonas | GCF_001721825.1 | Pseudomonas_aeruginosa | A6701_RS28485 | 6198359..6202087 | 287 | DUF6079-family-protein-[Pseudomonas-aeruginosa] |
| WP_249920774.1 | Type3.MTaseII.containing.BREX | [illegible] | 1243 | Bacteria>Pseudomonadota>Gammaproteobacteria>Pseudomonadales>Pseudomonadaceae>Pseudomonas | GCF_001765455.1 | Pseudomonas_aeruginosa | BAW40_RS11025 | 50364..54095 | 287 | DUF6079-family-protein-[Pseudomonas-aeruginosa] |
| WP_070341174.1 | Type3.MTaseII.containing.BREX | [illegible] | 1241 | Bacteria>Pseudomonadota>Gammaproteobacteria>Pseudomonadales>Pseudomonadaceae>Pseudomonas | GCF_001765485.1 | Pseudomonas_aeruginosa | BAW37_RS11780 | 29962..33687 | 287 | DUF6079-family-protein-[Pseudomonas-aeruginosa] |
| WP_070331909.1 | Type3.MTaseII.containing.BREX | [illegible] | 1241 | Bacteria>Pseudomonadota>Gammaproteobacteria>Pseudomonadales>Pseudomonadaceae>Pseudomonas | GCF_001766035.1 | Pseudomonas_aeruginosa | BAW72_RS11835 | 79269..82994 | 287 | DUF6079-family-protein-[Pseudomonas-aeruginosa] |
| WP_196496122.1 | Type3.MTaseII.containing.BREX | [illegible] | 1246 | Bacteria>Pseudomonadota>Gammaproteobacteria>Pseudomonadales>Pseudomonadaceae>Pseudomonas | GCF_002242855.1 | Pseudomonas_aeruginosa | ACG91_RS23775 | 44073..47813 | 287 | DUF6079-family-protein-[Pseudomonas-aeruginosa] |
| WP_217902373.1 | Type3.MTaseII.containing.BREX | [illegible] | 1252 | Bacteria>Pseudomonadota>Gammaproteobacteria>Pseudomonadales>Pseudomonadaceae>Pseudomonas | GCF_002269625.1 | Pseudomonas_fragi | CJU78_RS04000 | 74222..77980 | 296 | DUF6079-family-protein-[Pseudomonas-fragi] |
| WP_255032603.1 | Type3.MTaseII.containing.BREX | [illegible] | 810 | Bacteria>Pseudomonadota>Gammaproteobacteria>Pseudomonadales>Pseudomonadaceae>Pseudomonas | GCF_002269625.1 | Pseudomonas_fragi | CJU78_RS22515 | 2983..5415 | 296 | DUF6079-family-protein-[Pseudomonas-fragi] |
| WP_140786438.1 | Type3.MTaseII.containing.BREX | [illegible] | 1241 | Bacteria>Pseudomonadota>Gammaproteobacteria>Pseudomonadales>Pseudomonadaceae>Pseudomonas | GCF_003626935.1 | Pseudomonas_aeruginosa | D7H65_RS11285 | 2305878..2309603 | 287 | DUF6079-family-protein-[Pseudomonas-aeruginosa] |
| WP_121372202.1 | Type3.MTaseII.containing.BREX | [illegible] | 1243 | Bacteria>Pseudomonadota>Gammaproteobacteria>Pseudomonadales>Pseudomonadaceae>Pseudomonas | GCF_003629765.1 | Pseudomonas_aeruginosa | DQI36_RS24190 | 17435..21166 | 287 | DUF6079-family-protein-[Pseudomonas-aeruginosa] |
| WP_121335902.1 | Type3.MTaseII.containing.BREX | [illegible] | 1242 | Bacteria>Pseudomonadota>Gammaproteobacteria>Pseudomonadales>Pseudomonadaceae>Pseudomonas | GCF_003633115.1 | Pseudomonas_aeruginosa | DQH02_RS27180 | 4878..8606 | 287 | DUF6079-family-protein-[Pseudomonas-aeruginosa] |
| WP_124182728.1 | Type3.MTaseII.containing.BREX | [illegible] | 1241 | Bacteria>Pseudomonadota>Gammaproteobacteria>Pseudomonadales>Pseudomonadaceae>Pseudomonas | GCF_003839865.1 | Pseudomonas_aeruginosa | IPC226_RS26580 | 200303..204028 | 287 | DUF6079-family-protein-[Pseudomonas-aeruginosa] |
| WP_124984784.1 | Type3.MTaseII.containing.BREX | [illegible] | 1241 | Bacteria>Pseudomonadota>Gammaproteobacteria>Pseudomonadales>Pseudomonadaceae>Pseudomonas | GCF_003863685.1 | Pseudomonas_aeruginosa | EIM22_RS03570 | 276481..280206 | 287 | DUF6079-family-protein-[Pseudomonas-aeruginosa] |
| WP_124988190.1 | Type3.MTaseII.containing.BREX | [illegible] | 1246 | Bacteria>Pseudomonadota>Gammaproteobacteria>Pseudomonadales>Pseudomonadaceae>Pseudomonas | GCF_003863815.1 | Pseudomonas_aeruginosa | EIM11_RS16920 | 77381..81121 | 287 | DUF6079-family-protein-[Pseudomonas-aeruginosa] |
| WP_221767937.1 | Type3.MTaseII.containing.BREX | [illegible] | 1247 | Bacteria>Pseudomonadota>Gammaproteobacteria>Pseudomonadales>Pseudomonadaceae>Pseudomonas | GCF_003968225.1 | Pseudomonas_aeruginosa | DY943_RS06675 | 155478..159221 | 287 | DUF6079-family-protein-[Pseudomonas-aeruginosa] |
| WP_221932157.1 | Type3.MTaseII.containing.BREX | [illegible] | 1246 | Bacteria>Pseudomonadota>Gammaproteobacteria>Pseudomonadales>Pseudomonadaceae>Pseudomonas | GCF_007049785.1 | Pseudomonas_mendocina | EQ836_RS24270 | 66157..69897 | 300 | DUF6079-family-protein-[Pseudomonas-mendocina] |
| WP_143503165.1 | Type3.MTaseII.containing.BREX | [illegible] | 1242 | Bacteria>Pseudomonadota>Gammaproteobacteria>Pseudomonadales>Pseudomonadaceae>Pseudomonas | GCF_007049885.1 | Pseudomonas_mendocina | EQ828_RS03320 | 274642..278370 | 300 | DUF6079-family-protein-[Pseudomonas-mendocina] |
| WP_223184022.1 | Type3.MTaseII.containing.BREX | [illegible] | 1250 | Bacteria>Pseudomonadota>Gammaproteobacteria>Pseudomonadales>Pseudomonadaceae>Pseudomonas | GCF_012986825.1 | Pseudomonas_fragi | HBN89_RS05745 | 79142..82894 | 296 | DUF6079-family-protein-[Pseudomonas-fragi] |
| WP_209007133.1 | Type3.MTaseII.containing.BREX | [illegible] | 1246 | Bacteria>Pseudomonadota>Gammaproteobacteria>Pseudomonadales>Pseudomonadaceae>Pseudomonas | GCF_013276295.1 | Pseudomonas_aeruginosa | GWK58_RS24945 | 23431..27171 | 287 | DUF6079-family-protein-[Pseudomonas-aeruginosa] |
| WP_181127557.1 | Type3.MTaseII.containing.BREX | [illegible] | 1242 | Bacteria>Pseudomonadota>Gammaproteobacteria>Pseudomonadales>Pseudomonadaceae>Pseudomonas | GCF_013522615.1 | Pseudomonas_carnis | G7029_RS19485 | 130618..134346 | 2487355 | DUF6079-family-protein-[Pseudomonas-carnis] |
| WP_200973278.1 | Type3.MTaseII.containing.BREX | [illegible] | 1246 | Bacteria>Pseudomonadota>Gammaproteobacteria>Pseudomonadales>Pseudomonadaceae>Pseudomonas | GCF_013912775.1 | Pseudomonas_aeruginosa | H2N73_RS00345 | 77595..81335 | 287 | DUF6079-family-protein-[Pseudomonas-aeruginosa] |
| WP_181859222.1 | Type3.MTaseII.containing.BREX | [illegible] | 1243 | Bacteria>Pseudomonadota>Gammaproteobacteria>Pseudomonadales>Pseudomonadaceae>Pseudomonas | GCF_013913095.1 | Pseudomonas_aeruginosa | H2063_RS20075 | 129101..132832 | 287 | DUF6079-family-protein-[Pseudomonas-aeruginosa] |
| WP_188867720.1 | Type3.MTaseII.containing.BREX | [illegible] | 1241 | Bacteria>Pseudomonadota>Gammaproteobacteria>Pseudomonadales>Pseudomonadaceae>Pseudomonas | GCF_014646755.1 | Pseudomonas_asuensis | IEX86_RS19080 | 192626..196351 | 1825787 | DUF6079-family-protein-[Pseudomonas-asuensis] |
| WP_202096817.1 | Type3.MTaseII.containing.BREX | [illegible] | 1266 | Bacteria>Pseudomonadota>Gammaproteobacteria>Pseudomonadales>Pseudomonadaceae>Pseudomonas | GCF_016745175.1 | Pseudomonas_aeruginosa | HUS13_RS26865 | 5697414..5701214 | 287 | DUF6079-family-protein-[Pseudomonas-aeruginosa] |
| WP_238999168.1 | Type3.MTaseII.containing.BREX | [illegible] | 1246 | Bacteria>Pseudomonadota>Gammaproteobacteria>Pseudomonadales>Pseudomonadaceae>Pseudomonas | GCF_016836685.1 | Pseudomonas_aeruginosa | JQ477_RS13000 | 2657563..2661303 | 287 | DUF6079-family-protein-[Pseudomonas-aeruginosa] |
| WP_228292836.1 | Type3.MTaseII.containing.BREX | [illegible] | 1243 | Bacteria>Pseudomonadota>Gammaproteobacteria>Pseudomonadales>Pseudomonadaceae>Pseudomonas | GCF_017741985.1 | Pseudomonas_aeruginosa | BAW68_RS10935 | 136915..140646 | 287 | DUF6079-family-protein-[Pseudomonas-aeruginosa] |
| WP_218424333.1 | Type3.MTaseII.containing.BREX | [illegible] | 1244 | Bacteria>Pseudomonadota>Gammaproteobacteria>Pseudomonadales>Pseudomonadaceae>Pseudomonas | GCF_019218965.1 | Pseudomonas_sp._PDM31 | KW839_RS11070 | 28010..31744 | 2854778 | DUF6079-family-protein-[Pseudomonas-sp.-PDM31] |
| WP_220803981.1 | Type3.MTaseII.containing.BREX | [illegible] | 1241 | Bacteria>Pseudomonadota>Gammaproteobacteria>Pseudomonadales>Pseudomonadaceae>Pseudomonas | GCF_019656335.1 | Pseudomonas_sp._NCCP-436 | NCCP436_RS10650 | 99901..103626 | 2842481 | DUF6079-family-protein-[Pseudomonas-sp.-NCCP-436] |
| WP_033990875.1 | Type3.MTaseII.containing.BREX | [illegible] | 1246 | Bacteria>Pseudomonadota>Gammaproteobacteria>Pseudomonadales>Pseudomonadaceae>Pseudomonas | GCF_020990445.1 | Pseudomonas_aeruginosa | LD755_RS25575 | 5366903..5370643 | 287 | DUF6079-family-protein-[Pseudomonas-aeruginosa] |
| WP_231281517.1 | Type3.MTaseII.containing.BREX | [illegible] | 1242 | Bacteria>Pseudomonadota>Gammaproteobacteria>Pseudomonadales>Pseudomonadaceae>Pseudomonas | GCF_021060565.1 | Pseudomonas_aeruginosa | K4W73_RS30470 | 398718..402446 | 287 | DUF6079-family-protein-[Pseudomonas-aeruginosa] |
| WP_239610961.1 | Type3.MTaseII.containing.BREX | [illegible] | 1250 | Bacteria>Pseudomonadota>Gammaproteobacteria>Pseudomonadales>Pseudomonadaceae>Pseudomonas | GCF_022355455.1 | Pseudomonas_sp._KSR10 | MCB86_RS11370 | 524669..528421 | 2916654 | DUF6079-family-protein-[Pseudomonas-sp.-KSR10] |
| WP_247258974.1 | Type3.MTaseII.containing.BREX | [illegible] | 1246 | Bacteria>Pseudomonadota>Gammaproteobacteria>Pseudomonadales>Pseudomonadaceae>Pseudomonas | GCF_023060325.1 | Pseudomonas_aeruginosa | MX104_RS19120 | 73167..76907 | 287 | DUF6079-family-protein-[Pseudomonas-aeruginosa] |
| WP_212646892.1 | Type3.MTaseII.containing.BREX | [illegible] | 1243 | Bacteria>Pseudomonadota>Gammaproteobacteria>Pseudomonadales>Pseudomonadaceae>Pseudomonas | GCF_023805305.1 | Pseudomonas_aeruginosa | KB219_RS25160 | 1317..5048 | 287 | DUF6079-family-protein-[Pseudomonas-aeruginosa] |
| WP_263477044.1 | Type3.MTaseII.containing.BREX | [illegible] | 1242 | Bacteria>Pseudomonadota>Gammaproteobacteria>Pseudomonadales>Pseudomonadaceae>Pseudomonas | GCF_025697865.1 | Pseudomonas_aeruginosa | KUC78_RS09980 | 112485..116213 | 287 | DUF6079-family-protein-[Pseudomonas-aeruginosa] |
| WP_058137613.1 | Type3.MTaseII.containing.BREX | [illegible] | 1242 | Bacteria>Pseudomonadota>Gammaproteobacteria>Pseudomonadales>Pseudomonadaceae>Pseudomonas | GCF_027856055.1 | Pseudomonas_aeruginosa | KKQ48_RS06875 | 1489509..1493237 | 287 | DUF6079-family-protein-[Pseudomonas-aeruginosa] |
| WP_270932592.1 | Type3.MTaseII.containing.BREX | [illegible] | 1246 | Bacteria>Pseudomonadota>Gammaproteobacteria>Pseudomonadales>Pseudomonadaceae>Pseudomonas | GCF_027856215.1 | Pseudomonas_aeruginosa | JR315_RS22265 | 5695..9435 | 287 | DUF6079-family-protein-[Pseudomonas-aeruginosa] |
| WP_058195288.1 | Type3.MTaseII.containing.BREX | [illegible] | 1246 | Bacteria>Pseudomonadota>Gammaproteobacteria>Pseudomonadales>Pseudomonadaceae>Pseudomonas | GCF_030034615.1 | Pseudomonas_aeruginosa | AO969_RS25430 | 34320..38060 | 287 | DUF6079-family-protein-[Pseudomonas-aeruginosa] |
| WP_197058174.1 | Type3.MTaseII.containing.BREX | [illegible] | 1246 | Bacteria>Pseudomonadota>Gammaproteobacteria>Pseudomonadales>Pseudomonadaceae>Pseudomonas | GCF_900101515.1 | Pseudomonas_flexibilis | BLP69_RS07640 | 71711..75451 | 706570 | DUF6079-family-protein-[Pseudomonas-flexibilis] |
| WP_031631068.1 | Type3.MTaseII.containing.BREX | [illegible] | 1241 | Bacteria>Pseudomonadota>Gammaproteobacteria>Pseudomonadales>Pseudomonadaceae>Pseudomonas | GCF_900707525.1 | Pseudomonas_aeruginosa | E4Z30_RS22025 | 15967..19692 | 287 | DUF6079-family-protein-[Pseudomonas-aeruginosa] |
| WP_244247695.1 | Type3.MTaseII.containing.BREX | [illegible] | 791 | Bacteria>Pseudomonadota>Gammaproteobacteria>Pseudomonadales>Pseudomonadaceae>Pseudomonas | GCF_900708635.1 | Pseudomonas_aeruginosa | E4068_RS14300 | 5262..7634 | 287 | DUF6079-family-protein,-partial-[Pseudomonas-aeruginosa] |
| WP_077565984.1 | Type3.MTaseII.containing.BREX | [illegible] | 1248 | Bacteria>Pseudomonadota>Gammaproteobacteria>Pseudomonadales>Pseudomonadaceae>Pseudomonas | - | Pseudomonas_aeruginosa | BM548_RS18915 | 99013..102759 | 287 | DUF6079-family-protein-[Pseudomonas-aeruginosa] |
| EIU1316754.1 | Type3.MTaseII.containing.BREX | [illegible] | 1250 | Bacteria>Pseudomonadota>Gammaproteobacteria>Pseudomonadales>Pseudomonadaceae>Pseudomonas | - | Pseudomonas_aeruginosa | L3560_003636 | 71973..75725 | 287 | ATP-binding-protein-[Pseudomonas-aeruginosa] |
| EIU1319524.1 | Type3.MTaseII.containing.BREX | [illegible] | 437 | Bacteria>Pseudomonadota>Gammaproteobacteria>Pseudomonadales>Pseudomonadaceae>Pseudomonas | - | Pseudomonas_aeruginosa | L3560_006461 | 1..1312 | 287 | ATP-binding-protein-[Pseudomonas-aeruginosa] |
| EJA2648340.1 | Type3.MTaseII.containing.BREX | [illegible] | 1246 | Bacteria>Pseudomonadota>Gammaproteobacteria>Pseudomonadales>Pseudomonadaceae>Pseudomonas | - | Pseudomonas_aeruginosa | MWT08_004828 | 9447..13187 | 287 | ATP-binding-protein-[Pseudomonas-aeruginosa] |
| EJO5623990.1 | Type3.MTaseII.containing.BREX | [illegible] | 1266 | Bacteria>Pseudomonadota>Gammaproteobacteria>Pseudomonadales>Pseudomonadaceae>Pseudomonas | - | Pseudomonas_aeruginosa | NRQ67_004692 | 65430..69230 | 287 | ATP-binding-protein-[Pseudomonas-aeruginosa] |
| WP_208235532.1 | Type3.MTaseII.containing.BREX | [illegible] | 1246 | Bacteria>Pseudomonadota>Gammaproteobacteria>Pseudomonadales>Pseudomonadaceae>Pseudomonas | - | Pseudomonas_aeruginosa | PAERUG_P36 | 25167..28907 | 287 | DUF6079-family-protein-[Pseudomonas-aeruginosa] |
| MCQ2032414.1 | Type3.MTaseII.containing.BREX | [illegible] | 1266 | Bacteria>Pseudomonadota>Gammaproteobacteria>Pseudomonadales>Pseudomonadaceae>Stutzerimonas | GCA_024397475.1 | Stutzerimonas_kunmingensis | NAU09_00540 | 118416..122216 | 1211807 | DUF6079-family-protein-[Stutzerimonas-kunmingensis] |
| WP_255847834.1 | Type3.MTaseII.containing.BREX | [illegible] | 1266 | Bacteria>Pseudomonadota>Gammaproteobacteria>Pseudomonadales>Pseudomonadaceae>Stutzerimonas | GCF_024397475.1 | Stutzerimonas_kunmingensis | NAU09_RS00540 | 118416..122216 | 1211807 | DUF6079-family-protein-[Stutzerimonas-kunmingensis] |
| WP_269263620.1 | Type3.MTaseII.containing.BREX | [illegible] | 1243 | Bacteria>Pseudomonadota>Gammaproteobacteria>Pseudomonadales>Pseudomonadaceae>Stutzerimonas | GCF_027158265.1 | Stutzerimonas_balearica | O3V18_RS20965 | 3018..6749 | 74829 | DUF6079-family-protein-[Stutzerimonas-balearica] |
| WP_019341816.1 | Type3.MTaseII.containing.BREX | [illegible] | 1262 | Bacteria>Pseudomonadota>Gammaproteobacteria>Pseudomonadales>Pseudomonadaceae>Stutzerimonas | - | Stutzerimonas_stutzeri | PSTU_RS0114850 | 50252..54040 | 316 | DUF6079-family-protein-[Stutzerimonas-stutzeri] |
| HEC58787.1 | Type3.MTaseII.containing.BREX | [illegible] | 592 | Bacteria>Pseudomonadota>Gammaproteobacteria>Thiotrichales>Piscirickettsiaceae>Methylophaga | GCA_011053005.1 | Methylophaga_sp. | ENI24_04380 | 43599..45377 | 2024840 | hypothetical-protein-[Methylophaga-sp.] |
| OQY55620.1 | Type3.MTaseII.containing.BREX | [illegible] | 1245 | Bacteria>Pseudomonadota>Gammaproteobacteria>Thiotrichales>Thiotrichaceae>Beggiatoa | GCA_002085445.1 | Beggiatoa_sp._4572_84 | B6247_07200 | 42863..46600 | 1972449 | MAG:-ATPase-[Beggiatoa-sp.-4572_84] |
| WP_171900971.1 | Type3.MTaseII.containing.BREX | [illegible] | 1003 | Bacteria>Pseudomonadota>Gammaproteobacteria>Thiotrichales>Thiotrichaceae>Candidatus_Marithrix | GCF_001723665.1 | Candidatus_Marithrix_sp._Canyon_246 | BHU67_RS17090 | 1..3012 | 1827136 | DUF6079-family-protein,-partial-[Candidatus-Marithrix-sp.-Canyon-246] |
| WP_069472954.1 | Type3.MTaseII.containing.BREX | [illegible] | 1001 | Bacteria>Pseudomonadota>Gammaproteobacteria>Thiotrichales>Thiotrichaceae>Candidatus_Marithrix | GCF_001723675.1 | Candidatus_Marithrix_sp._Canyon_246 | BHU66_RS17180 | 1..3008 | 1827136 | DUF6079-family-protein,-partial-[Candidatus-Marithrix-sp.-Canyon-246] |
| WP_069473694.1 | Type3.MTaseII.containing.BREX | [illegible] | 1157 | Bacteria>Pseudomonadota>Gammaproteobacteria>Thiotrichales>Thiotrichaceae>Candidatus_Marithrix | GCF_001723685.1 | Candidatus_Marithrix_sp._Canyon_246 | BHU68_RS03985 | 65893..69366 | 1827136 | DUF6079-family-protein-[Candidatus-Marithrix-sp.-Canyon-246] |
| MCK5717247.1 | Type3.MTaseII.containing.BREX | [illegible] | 529 | Bacteria>Pseudomonadota>Gammaproteobacteria>Thiotrichales>Thiotrichaceae>Thiomargarita | GCA_023144825.1 | Thiomargarita_sp. | KAH77_07135 | 9877..11463 | 2822435 | hypothetical-protein-[Thiomargarita-sp.] |
| BAP57136.1 | Type3.MTaseII.containing.BREX | [illegible] | 1248 | Bacteria>Pseudomonadota>Gammaproteobacteria>Thiotrichales>Thiotrichaceae>Thioploca | GCA_000828835.1 | Thioploca_ingrica | THII_2839 | 3401722..3405468 | 40754 | hypothetical-protein-THII_2839-[Thioploca-ingrica] |
| NJO17671.1 | Type3.MTaseII.containing.BREX | [illegible] | 293 | Bacteria>Pseudomonadota>Gammaproteobacteria>Thiotrichales>Thiotrichaceae>Thioploca | GCA_012032555.1 | Thioploca_sp. | HC877_18625 | 6169..7050 | 2268140 | hypothetical-protein-[Thioploca-sp.] |
| NJO17672.1 | Type3.MTaseII.containing.BREX | [illegible] | 957 | Bacteria>Pseudomonadota>Gammaproteobacteria>Thiotrichales>Thiotrichaceae>Thioploca | GCA_012032555.1 | Thioploca_sp. | HC877_18630 | 7071..9944 | 2268140 | ATP-binding-protein-[Thioploca-sp.] |
| MBL1294161.1 | Type3.MTaseII.containing.BREX | [illegible] | 1247 | Bacteria>Pseudomonadota>Gammaproteobacteria>Thiotrichales>unclassified_Thiotrichales | GCA_002401375.2 | Thiotrichales_bacterium | COB61_009840 | 15508..19251 | 2026796 | hypothetical-protein-[Thiotrichales-bacterium] |
| PKM04365.1 | Type3.MTaseII.containing.BREX | [illegible] | 1241 | Bacteria>Pseudomonadota>Gammaproteobacteria>unclassified_Gammaproteobacteria | GCA_002839965.1 | Gammaproteobacteria_bacterium_HGW-Gammaproteobacteria-6 | CVV16_05345 | 31508..35233 | 2013802 | ATPase-[Gammaproteobacteria-bacterium-HGW-Gammaproteobacteria-6] |
| RKZ51977.1 | Type3.MTaseII.containing.BREX | [illegible] | 1250 | Bacteria>Pseudomonadota>Gammaproteobacteria>unclassified_Gammaproteobacteria | GCA_003645185.1 | Gammaproteobacteria_bacterium | DRR00_09870 | 7231..10983 | 1913989 | ATP-binding-protein-[Gammaproteobacteria-bacterium] |
| MYA37130.1 | Type3.MTaseII.containing.BREX | [illegible] | 1243 | Bacteria>Pseudomonadota>Gammaproteobacteria>unclassified_Gammaproteobacteria | GCA_009838725.1 | Gammaproteobacteria_bacterium | F4Y34_10780 | 18894..22625 | 1913989 | ATP-binding-protein-[Gammaproteobacteria-bacterium] |
| HDH16018.1 | Type3.MTaseII.containing.BREX | [illegible] | 232 | Bacteria>Pseudomonadota>Gammaproteobacteria>unclassified_Gammaproteobacteria | GCA_011040115.1 | Gammaproteobacteria_bacterium | ENG90_06030 | 7201..7901 | 1913989 | ATP-binding-protein-[Gammaproteobacteria-bacterium] |
| NIP47373.1 | Type3.MTaseII.containing.BREX | [illegible] | 348 | Bacteria>Pseudomonadota>Gammaproteobacteria>unclassified_Gammaproteobacteria | GCA_011774445.1 | Gammaproteobacteria_bacterium | GWD29_13500 | 143386..144432 | 1913989 | hypothetical-protein-[Gammaproteobacteria-bacterium] |
| NKB61264.1 | Type3.MTaseII.containing.BREX | [illegible] | 1247 | Bacteria>Pseudomonadota>Gammaproteobacteria>unclassified_Gammaproteobacteria | GCA_012103415.1 | Gammaproteobacteria_bacterium | GKR95_03675 | 840288..844031 | 1913989 | ATP-binding-protein-[Gammaproteobacteria-bacterium] |
| NKC13928.1 | Type3.MTaseII.containing.BREX | [illegible] | 319 | Bacteria>Pseudomonadota>Gammaproteobacteria>unclassified_Gammaproteobacteria | GCA_012103455.1 | Gammaproteobacteria_bacterium | GKR94_17600 | 3530439..3531398 | 1913989 | hypothetical-protein-[Gammaproteobacteria-bacterium] |
| MBU0782760.1 | Type3.MTaseII.containing.BREX | [illegible] | 1246 | Bacteria>Pseudomonadota>Gammaproteobacteria>unclassified_Gammaproteobacteria | GCA_018815355.1 | Gammaproteobacteria_bacterium | KJ979_02525 | 62574..66314 | 1913989 | ATP-binding-protein-[Gammaproteobacteria-bacterium] |
| MBU1223667.1 | Type3.MTaseII.containing.BREX | [illegible] | 1241 | Bacteria>Pseudomonadota>Gammaproteobacteria>unclassified_Gammaproteobacteria | GCA_018819475.1 | Gammaproteobacteria_bacterium | KKI05_07425 | 77391..81116 | 1913989 | ATP-binding-protein-[Gammaproteobacteria-bacterium] |
| WP_102530608.1 | Type3.MTaseII.containing.BREX | [illegible] | 1241 | Bacteria>Pseudomonadota>Gammaproteobacteria>Vibrionales>Vibrionaceae | GCF_002878515.1 | Vibrio | BCU87_RS25110 | 12426..16151 | 662 | MULTISPECIES:-DUF6079-family-protein-[Vibrio] |
| WP_077679391.1 | Type3.MTaseII.containing.BREX | [illegible] | 1241 | Bacteria>Pseudomonadota>Gammaproteobacteria>Vibrionales>Vibrionaceae>Salinivibrio | GCF_001996265.1 | Salinivibrio_sp._MA440 | BZG80_11580 | 16826..20551 | 1909456 | DUF6079-family-protein-[Salinivibrio-sp.-MA440] |
| HAS6049575.1 | Type3.MTaseII.containing.BREX | [illegible] | 1241 | Bacteria>Pseudomonadota>Gammaproteobacteria>Vibrionales>Vibrionaceae>Vibrio | GCA_015776375.1 | Vibrio_vulnificus | I7138_16200 | 19766..23491 | 672 | hypothetical-protein-[Vibrio-vulnificus] |
| EGR2106512.1 | Type3.MTaseII.containing.BREX | [illegible] | 1241 | Bacteria>Pseudomonadota>Gammaproteobacteria>Vibrionales>Vibrionaceae>Vibrio | GCA_015802515.1 | Vibrio_cholerae | DYF88_13630 | 10389..14114 | 666 | hypothetical-protein-[Vibrio-cholerae] |
| EJG0684220.1 | Type3.MTaseII.containing.BREX | [illegible] | 1241 | Bacteria>Pseudomonadota>Gammaproteobacteria>Vibrionales>Vibrionaceae>Vibrio | GCA_015814245.2 | Vibrio_parahaemolyticus | C4G75_RS05935 | 47412..51137 | 670 | hypothetical-protein-[Vibrio-parahaemolyticus] |
| EGU0166216.1 | Type3.MTaseII.containing.BREX | [illegible] | 1241 | Bacteria>Pseudomonadota>Gammaproteobacteria>Vibrionales>Vibrionaceae>Vibrio | GCA_016147415.1 | Vibrio_parahaemolyticus | JFQ74_002030 | 212242..215967 | 670 | hypothetical-protein-[Vibrio-parahaemolyticus] |
| EHC7288062.1 | Type3.MTaseII.containing.BREX | [illegible] | 1241 | Bacteria>Pseudomonadota>Gammaproteobacteria>Vibrionales>Vibrionaceae>Vibrio | GCA_017078635.1 | Vibrio_parahaemolyticus | JS475_001525 | 801180..804905 | 670 | hypothetical-protein-[Vibrio-parahaemolyticus] |
| EHD1694795.1 | Type3.MTaseII.containing.BREX | [illegible] | 1241 | Bacteria>Pseudomonadota>Gammaproteobacteria>Vibrionales>Vibrionaceae>Vibrio | GCA_017123195.1 | Vibrio_vulnificus | JR728_000349 | 364709..368434 | 672 | hypothetical-protein-[Vibrio-vulnificus] |
| EHK2855481.1 | Type3.MTaseII.containing.BREX | [illegible] | 1241 | Bacteria>Pseudomonadota>Gammaproteobacteria>Vibrionales>Vibrionaceae>Vibrio | GCA_018017035.1 | Vibrio_parahaemolyticus | J7X11_003955 | 8440..12165 | 670 | hypothetical-protein-[Vibrio-parahaemolyticus] |
| HBC3410538.1 | Type3.MTaseII.containing.BREX | [illegible] | 1240 | Bacteria>Pseudomonadota>Gammaproteobacteria>Vibrionales>Vibrionaceae>Vibrio | GCA_018118845.1 | Vibrio_parahaemolyticus | KDY13_002790 | 91232..94954 | 670 | hypothetical-protein-[Vibrio-parahaemolyticus] |
| EHR6926057.1 | Type3.MTaseII.containing.BREX | [illegible] | 1241 | Bacteria>Pseudomonadota>Gammaproteobacteria>Vibrionales>Vibrionaceae>Vibrio | GCA_019067225.1 | Vibrio_parahaemolyticus | KUK83_003900 | 12534..16259 | 670 | hypothetical-protein-[Vibrio-parahaemolyticus] |
| EIA1331990.1 | Type3.MTaseII.containing.BREX | [illegible] | 1241 | Bacteria>Pseudomonadota>Gammaproteobacteria>Vibrionales>Vibrionaceae>Vibrio | GCA_019764355.1 | Vibrio_parahaemolyticus | K6675_002577 | 8440..12165 | 670 | hypothetical-protein-[Vibrio-parahaemolyticus] |
| EIF5159839.1 | Type3.MTaseII.containing.BREX | [illegible] | 1241 | Bacteria>Pseudomonadota>Gammaproteobacteria>Vibrionales>Vibrionaceae>Vibrio | GCA_020260965.1 | Vibrio_cholerae | LFW18_002408 | 87962..91687 | 666 | hypothetical-protein-[Vibrio-cholerae] |
| EII3096499.1 | Type3.MTaseII.containing.BREX | [illegible] | 1009 | Bacteria>Pseudomonadota>Gammaproteobacteria>Vibrionales>Vibrionaceae>Vibrio | GCA_020410925.1 | Vibrio_parahaemolyticus | CWL41_002108 | 340153..343184 | 670 | hypothetical-protein-[Vibrio-parahaemolyticus] |
| HBK7260599.1 | Type3.MTaseII.containing.BREX | [illegible] | 1241 | Bacteria>Pseudomonadota>Gammaproteobacteria>Vibrionales>Vibrionaceae>Vibrio | GCA_020795255.1 | Vibrio_cholerae | LNF50_003076 | 2675..6400 | 666 | hypothetical-protein-[Vibrio-cholerae] |
| EIN6339961.1 | Type3.MTaseII.containing.BREX | [illegible] | 1241 | Bacteria>Pseudomonadota>Gammaproteobacteria>Vibrionales>Vibrionaceae>Vibrio | GCA_020908185.1 | Vibrio_parahaemolyticus | LPQ31_001314 | 349939..353664 | 670 | hypothetical-protein-[Vibrio-parahaemolyticus] |
| EJB8504845.1 | Type3.MTaseII.containing.BREX | [illegible] | 1241 | Bacteria>Pseudomonadota>Gammaproteobacteria>Vibrionales>Vibrionaceae>Vibrio | GCA_022967155.1 | Vibrio_parahaemolyticus | MW364_003023 | 7653..11378 | 670 | hypothetical-protein-[Vibrio-parahaemolyticus] |
| HCE2000767.1 | Type3.MTaseII.containing.BREX | [illegible] | 1241 | Bacteria>Pseudomonadota>Gammaproteobacteria>Vibrionales>Vibrionaceae>Vibrio | GCA_023810945.1 | Vibrio_parahaemolyticus | NGT00_003030 | 162203..165928 | 670 | hypothetical-protein-[Vibrio-parahaemolyticus] |
| HCE5247588.1 | Type3.MTaseII.containing.BREX | [illegible] | 1242 | Bacteria>Pseudomonadota>Gammaproteobacteria>Vibrionales>Vibrionaceae>Vibrio | GCA_023836695.1 | Vibrio_parahaemolyticus | NG505_004023 | 93782..97510 | 670 | hypothetical-protein-[Vibrio-parahaemolyticus] |
| HCG6379153.1 | Type3.MTaseII.containing.BREX | [illegible] | 1241 | Bacteria>Pseudomonadota>Gammaproteobacteria>Vibrionales>Vibrionaceae>Vibrio | GCA_024039965.1 | Vibrio_parahaemolyticus | NJ203_004422 | 3538..7263 | 670 | hypothetical-protein-[Vibrio-parahaemolyticus] |
| HCH5614387.1 | Type3.MTaseII.containing.BREX | [illegible] | 1241 | Bacteria>Pseudomonadota>Gammaproteobacteria>Vibrionales>Vibrionaceae>Vibrio | GCA_024095595.1 | Vibrio_parahaemolyticus | NKZ26_003757 | 148723..152448 | 670 | hypothetical-protein-[Vibrio-parahaemolyticus] |
| EJS1625578.1 | Type3.MTaseII.containing.BREX | [illegible] | 1241 | Bacteria>Pseudomonadota>Gammaproteobacteria>Vibrionales>Vibrionaceae>Vibrio | GCA_024944445.1 | Vibrio_cholerae | NXA67_002434 | 122275..126000 | 666 | hypothetical-protein-[Vibrio-cholerae] |

| Protein-ID | System | Gene-Neighbourhoods | Length | Taxonomy | Assembly-ID | Strain | Locus-Tag | Genomic-coordinates | TaxID | Fasta-defline |
|---|---|---|---|---|---|---|---|---|---|---|
| WP_005593745.1 | Type3.MTaseII.containing.BREX | HNH-><-BrxA-DNA-binding\|\|?-><-BrxA-DNA-binding<-PglZ<-TUDOR+SF1-SF2-Helicase+iREase+CTD<-N6-N4-METHYLASE<-DUF6079*<-BrxB-BrxF-iSTAND<-wHTH+WYL+WCX\|\|?-><-HNH<-?<-DEOR-H [illegible] | 1241 | Bacteria>Pseudomonadota>Gammaproteobacteria>Vibrionales>Vibrionaceae>Vibrio | GCF_000222585.1 | Vibrio_scophthalmi | RH88_RS04470 | 9667..13392 | 45658 | DUF6079-family-protein-[Vibrio-scophthalmi] |
| WP_065206911.1 | Type3.MTaseII.containing.BREX | Arm-DNA-bind_3+Phage_integrase-><-XIS-HTH\|\|HNH-><-BrxA-DNA-binding<-BrxA-DNA-binding<-PglZ<-TUDOR+SF1-SF2-Helicase+iREase+CTD<-N6-N4-METHYLASE\|\|tRNA->tRNA->tRNA->tRNA- [illegible] | 1241 | Bacteria>Pseudomonadota>Gammaproteobacteria>Vibrionales>Vibrionaceae>Vibrio | GCF_001676015.1 | Vibrio_splendidus | A9262_RS18605 | 305898..309623 | 29497 | DUF6079-family-protein-[Vibrio-splendidus] |
| WP_095469002.1 | Type3.MTaseII.containing.BREX | <-TPR<-?<-?<-?<-?<-wHTH+WYL+WCX\|\|REase+DUF5401->XIS-HTH->BrxB-BrxF-iSTAND->DUF6079*->N6-N4-METHYLASE->TUDOR+SF1-SF2-Helicase+iREase+CTD->PglZ->BrxA-DNA-binding->BrxA-D [illegible] | 1241 | Bacteria>Pseudomonadota>Gammaproteobacteria>Vibrionales>Vibrionaceae>Vibrio | GCF_002283905.1 | Vibrio_metoecus | CGT97_RS16085 | 12636..16361 | 1481663 | DUF6079-family-protein-[Vibrio-metoecus] |
| WP_102397786.1 | Type3.MTaseII.containing.BREX | Arm-DNA-bind_3+Phage_integrase->HNH-><-BrxA-DNA-binding<-BrxA-DNA-binding<-PglZ<-TUDOR+SF1-SF2-Helicase+iREase+CTD<-RecR<-N6-N4-METHYLASE<-DUF6079*<-BrxB-BrxF-iSTAND<- [illegible] | 1241 | Bacteria>Pseudomonadota>Gammaproteobacteria>Vibrionales>Vibrionaceae>Vibrio | GCF_002873235.1 | Vibrio_cyclitrophicus | BCU59_RS21850 | 110758..114483 | 47951 | DUF6079-family-protein-[Vibrio-cyclitrophicus] |
| WP_069513733.1 | Type3.MTaseII.containing.BREX | <-HAMP+MA+MA<-ParB+HNH\|\|HNH-><-BrxA-DNA-binding<-BrxA-DNA-binding<-PglZ<-TUDOR+SF1-SF2-Helicase+iREase+CTD<-N6-N4-METHYLASE<-DUF6079*<-BrxB-BrxF-iSTAND<-wHTH+WYL+WCX\|\| [illegible] | 1241 | Bacteria>Pseudomonadota>Gammaproteobacteria>Vibrionales>Vibrionaceae>Vibrio | GCF_003691525.1 | Vibrio_parahaemolyticus | D0871_RS11680 | 2311889..2315614 | 670 | DUF6079-family-protein-[Vibrio-parahaemolyticus] |
| WP_123322066.1 | Type3.MTaseII.containing.BREX | MODE-HTH+PBPII-><-ParB+HNH\|\|HNH-><-BrxA-DNA-binding<-BrxA-DNA-binding<-PglZ<-TUDOR+SF1-SF2-Helicase+iREase+CTD<-?<-N6-N4-METHYLASE<-DUF6079*<-BrxB-BrxF-iSTAND<-XIS-HTH [illegible] | 1241 | Bacteria>Pseudomonadota>Gammaproteobacteria>Vibrionales>Vibrionaceae>Vibrio | GCF_003751765.1 | Vibrio_crassostreae | EDB73_RS01705 | 379275..383000 | 246167 | DUF6079-family-protein-[Vibrio-crassostreae] |
| WP_129147714.1 | Type3.MTaseII.containing.BREX | Arm-DNA-bind_3+Phage_integrase->HNH-><-BrxA-DNA-binding<-BrxA-DNA-binding<-PglZ<-TUDOR+SF1-SF2-Helicase+iREase+CTD<-RecR<-N6-N4-METHYLASE<-DUF6079*<-BrxB-BrxF-iSTAND<- [illegible] | 1241 | Bacteria>Pseudomonadota>Gammaproteobacteria>Vibrionales>Vibrionaceae>Vibrio | GCF_004121285.1 | Vibrio_parahaemolyticus | EGL68_RS05395 | 1123769..1127494 | 670 | DUF6079-family-protein-[Vibrio-parahaemolyticus] |
| WP_132941357.1 | Type3.MTaseII.containing.BREX | <-ParB+HNH\|\|HNH-><-BrxA-DNA-binding<-BrxA-DNA-binding<-PglZ<-TUDOR+SF1-SF2-Helicase+iREase+CTD\|\|REase-><-N6-N4-METHYLASE<-N6-N4-METHYLASE<-DUF6079*<-BrxB-BrxF-iSTAND<- [illegible] | 1241 | Bacteria>Pseudomonadota>Gammaproteobacteria>Vibrionales>Vibrionaceae>Vibrio | GCF_004345555.1 | Vibrio_crassostreae | EDB60_RS00610 | 124768..128493 | 246167 | DUF6079-family-protein-[Vibrio-crassostreae] |
| WP_194592798.1 | Type3.MTaseII.containing.BREX | RNase_H-><-DUF2787<-?<-?\|\|XIS-HTH-><-wHTH+WYL+WCX\|\|REase+DUF5401->XIS-HTH->BrxB-BrxF-iSTAND->DUF6079*->N6-N4-METHYLASE->TUDOR+SF1-SF2-Helicase+iREase+CTD-p->TUDOR+SF1- [illegible] | 1241 | Bacteria>Pseudomonadota>Gammaproteobacteria>Vibrionales>Vibrionaceae>Vibrio | GCF_015351545.2 | Vibrio_vulnificus | IRL01_RS02825 | 610034..613759 | 672 | DUF6079-family-protein-[Vibrio-vulnificus] |
| WP_206445947.1 | Type3.MTaseII.containing.BREX | Arm-DNA-bind_3+Phage_integrase->HNH-><-BrxA-DNA-binding<-BrxA-DNA-binding<-PglZ<-TUDOR+SF1-SF2-Helicase+iREase+CTD<-RecR<-N6-N4-METHYLASE<-DUF6079*<-BrxB-BrxF-iSTAND<- [illegible] | 1241 | Bacteria>Pseudomonadota>Gammaproteobacteria>Vibrionales>Vibrionaceae>Vibrio | GCF_017169515.1 | Vibrio_paracholerae | GU254_RS11830 | 87056..90781 | 650003 | DUF6079-family-protein-[Vibrio-paracholerae] |
| WP_206959706.1 | Type3.MTaseII.containing.BREX | Arm-DNA-bind_3+Phage_integrase-><-XIS-HTH\|\|HNH-><-BrxA-DNA-binding<-BrxA-DNA-binding<-PglZ<-TUDOR+SF1-SF2-Helicase+iREase+CTD<-N6-N4-METHYLASE<-DUF6079*<-BrxB-BrxF-iST [illegible] | 1241 | Bacteria>Pseudomonadota>Gammaproteobacteria>Vibrionales>Vibrionaceae>Vibrio | GCF_017313725.1 | Vibrio_sp._Vb0592 | J0671_RS16335 | 111003..114728 | 2816072 | DUF6079-family-protein-[Vibrio-sp.-Vb0592] |
| WP_132761830.1 | Type3.MTaseII.containing.BREX | XIS-HTH->RNase_H->SAM-METHYLASE->wHTH(SBD)+HNH-><-wHTH+WYL+WCX\|\|REase+DUF5401->XIS-HTH->BrxB-BrxF-iSTAND->DUF6079*->N6-N4-METHYLASE->?->TUDOR+SF1-SF2-Helicase+iREase+C [illegible] | 1241 | Bacteria>Pseudomonadota>Gammaproteobacteria>Vibrionales>Vibrionaceae>Vibrio | GCF_017918175.1 | Vibrio_crassostreae | EDB51_RS02165 | 53539..57264 | 246167 | DUF6079-family-protein-[Vibrio-crassostreae] |
| WP_124009095.1 | Type3.MTaseII.containing.BREX | RNA_METHYLASE->HNH-><-BrxA-DNA-binding<-BrxA-DNA-binding<-PglZ<-TUDOR+SF1-SF2-Helicase+iREase+CTD<-?<-N6-N4-METHYLASE<-DUF6079*<-BrxB-BrxF-iSTAND<-wHTH+WYL+WCX<-XIS-HT [illegible] | 1241 | Bacteria>Pseudomonadota>Gammaproteobacteria>Vibrionales>Vibrionaceae>Vibrio | GCF_018458505.1 | Vibrio_harveyi_group | J4H11_RS21330 | 16878..20603 | 717610 | MULTISPECIES:-DUF6079-family-protein-[Vibrio-harveyi-group] |
| WP_230666600.1 | Type3.MTaseII.containing.BREX | <-TPR<-?<-?<-?<-?<-wHTH+WYL+WCX\|\|REase+DUF5401->XIS-HTH->BrxB-BrxF-iSTAND->DUF6079*->N6-N4-METHYLASE->TUDOR+SF1-SF2-Helicase+iREase+CTD->PglZ->BrxA-DNA-binding->BrxA-D [illegible] | 1241 | Bacteria>Pseudomonadota>Gammaproteobacteria>Vibrionales>Vibrionaceae>Vibrio | GCF_021014645.1 | Vibrio_cholerae | B4944_RS12115 | 17627..21352 | 666 | DUF6079-family-protein-[Vibrio-cholerae] |
| WP_233934109.1 | Type3.MTaseII.containing.BREX | Arm-DNA-bind_3+Phage_integrase->HNH-><-BrxA-DNA-binding<-BrxA-DNA-binding<-PglZ<-TUDOR+SF1-SF2-Helicase+iREase+CTD<-?<-N6-N4-METHYLASE<-DUF6079*<-BrxB-BrxF-iSTAND<-wHT [illegible] | 1241 | Bacteria>Pseudomonadota>Gammaproteobacteria>Vibrionales>Vibrionaceae>Vibrio | GCF_021394995.1 | Vibrio_fluvialis | LZT09_RS15360 | 26135..29860 | 676 | DUF6079-family-protein-[Vibrio-fluvialis] |
| WP_243191699.1 | Type3.MTaseII.containing.BREX | <-DUF2787<-?<-?\|\|XIS-HTH->wHTH+WYL+WCX->BrxB-BrxF-iSTAND->DUF6079*->N6-N4-METHYLASE->TUDOR+SF1-SF2-Helicase+iREase+CTD->PglZ->BrxA-DNA-binding->BrxA-DNA-binding-><-HNH [illegible] | 1240 | Bacteria>Pseudomonadota>Gammaproteobacteria>Vibrionales>Vibrionaceae>Vibrio | GCF_022802175.1 | Vibrio_parahaemolyticus | JL838_RS10850 | 15265..18987 | 670 | DUF6079-family-protein-[Vibrio-parahaemolyticus] |
| WP_258672263.1 | Type3.MTaseII.containing.BREX | Arm-DNA-bind_3+Phage_integrase->HNH-><-BrxA-DNA-binding<-BrxA-DNA-binding<-PglZ<-TUDOR+SF1-SF2-Helicase+iREase+CTD<-RecR<-N6-N4-METHYLASE<-DUF6079*<-BrxB-BrxF-iSTAND<- [illegible] | 1241 | Bacteria>Pseudomonadota>Gammaproteobacteria>Vibrionales>Vibrionaceae>Vibrio | GCF_024747115.1 | Vibrio_parahaemolyticus | NDJ77_RS09340 | 109446..113171 | 670 | DUF6079-family-protein-[Vibrio-parahaemolyticus] |
| WP_148514678.1 | Type3.MTaseII.containing.BREX | PTSEIIA1+BLBD->?->tRNA->Arm-DNA-bind_3+Phage_integrase->HNH-><-BrxA-DNA-binding<-BrxA-DNA-binding<-PglZ<-TUDOR+SF1-SF2-Helicase+iREase+CTD<-N6-N4-METHYLASE<-DUF6079*<- [illegible] | 1241 | Bacteria>Pseudomonadota>Gammaproteobacteria>Vibrionales>Vibrionaceae>Vibrio | GCF_026613655.1 | Vibrio_cholerae | MRS66_RS05695 | 543936..547661 | 666 | DUF6079-family-protein-[Vibrio-cholerae] |
| WP_274676098.1 | Type3.MTaseII.containing.BREX | ZnR+TOPRIM+TOPRIM->SIGMA-HTH+SIGMA-HTH+SIGMA-HTH->tRNA->Arm-DNA-bind_3+Phage_integrase->HNH-><-BrxA-DNA-binding<-BrxA-DNA-binding<-PglZ<-TUDOR+SF1-SF2-Helicase+iREase+ [illegible] | 1241 | Bacteria>Pseudomonadota>Gammaproteobacteria>Vibrionales>Vibrionaceae>Vibrio | GCF_028858035.1 | Vibrio_aestuarianus | L9X51_RS11535 | 32363..36088 | 28171 | DUF6079-family-protein-[Vibrio-aestuarianus] |
| WP_182037014.1 | Type3.MTaseII.containing.BREX | <-wHTH+WYL+WCX<-DUF2787<-?<-?\|\|XIS-HTH->HNH-><-HEPN\|\|wHTH+WYL+WCX->BrxB-BrxF-iSTAND->DUF6079*->N6-N4-METHYLASE-><-REase\|\|TUDOR+SF1-SF2-Helicase+iREase+CTD->PglZ->BrxA- [illegible] | 1241 | Bacteria>Pseudomonadota>Gammaproteobacteria>Vibrionales>Vibrionaceae>Vibrio | GCF_903986725.1 | Vibrio_diabolicus | H7A60_RS18185 | 117894..121619 | 50719 | DUF6079-family-protein-[Vibrio-diabolicus] |
| WP_229626017.1 | Type3.MTaseII.containing.BREX | Arm-DNA-bind_3+Phage_integrase-><-XIS-HTH\|\|HNH-><-BrxA-DNA-binding<-BrxA-DNA-binding<-PglZ<-TUDOR+SF1-SF2-Helicase+iREase+CTD<-N6-N4-METHYLASE<-DUF6079*<-BrxB-BrxF-iS [illegible] | 1240 | Bacteria>Pseudomonadota>Gammaproteobacteria>Vibrionales>Vibrionaceae>Vibrio | GCF_905331745.1 | Vibrio_parahaemolyticus | LPS00_RS14565 | 121441..125163 | 670 | DUF6079-family-protein-[Vibrio-parahaemolyticus] |
| MBX3699888.1 | Type3.MTaseII.containing.BREX | [illegible] | 1246 | Bacteria>Pseudomonadota>Gammaproteobacteria>Xanthomonadales>Rhodanobacteraceae>Dokdonella | GCA_019634845.1 | Dokdonella_sp. | KF903_02665 | 9380..13120 | 2291710 | hypothetical-protein-[Dokdonella-sp.] |
| MBL8299346.1 | Type3.MTaseII.containing.BREX | [illegible] | 1246 | Bacteria>Pseudomonadota>Gammaproteobacteria>Xanthomonadales>Rhodanobacteraceae>unclassified_Rhodanobacteraceae | GCA_016790075.1 | Rhodanobacteraceae_bacterium | JNN30_13490 | 4880..8620 | 2052182 | hypothetical-protein-[Rhodanobacteraceae-bacterium] |
| MBK9496395.1 | Type3.MTaseII.containing.BREX | [illegible] | 1242 | Bacteria>Pseudomonadota>Gammaproteobacteria>Xanthomonadales>unclassified_Xanthomonadales | GCA_016718605.1 | Xanthomonadales_bacterium | IPO08_18210 | 258206..261934 | 2006849 | ATP-binding-protein-[Xanthomonadales-bacterium] |
| WP_075676612.1 | Type3.MTaseII.containing.BREX | [illegible] | 1246 | Bacteria>Pseudomonadota>Gammaproteobacteria>Xanthomonadales>Xanthomonadaceae | GCF_001939825.1 | Stenotrophomonas | BKE09_RS06610 | 1401977..1405717 | 40323 | MULTISPECIES:-DUF6079-family-protein-[Stenotrophomonas] |
| WP_144815719.1 | Type3.MTaseII.containing.BREX | [illegible] | 1243 | Bacteria>Pseudomonadota>Gammaproteobacteria>Xanthomonadales>Xanthomonadaceae>Lysobacter | GCF_007830115.1 | Lysobacter_ruishenii | IP93_RS11165 | 70444..74175 | 686800 | DUF6079-family-protein-[Lysobacter-ruishenii] |
| WP_228464986.1 | Type3.MTaseII.containing.BREX | [illegible] | 1246 | Bacteria>Pseudomonadota>Gammaproteobacteria>Xanthomonadales>Xanthomonadaceae>Lysobacter | GCF_014779555.2 | Lysobacter_capsici | IEQ11_RS15915 | 3926369..3930109 | 435897 | DUF6079-family-protein-[Lysobacter-capsici] |
| ALJ26653.1 | Type3.MTaseII.containing.BREX | [illegible] | 1246 | Bacteria>Pseudomonadota>Gammaproteobacteria>Xanthomonadales>Xanthomonadaceae>Stenotrophomonas | GCA_001314305.1 | Stenotrophomonas_acidaminiphila | AOT14_01920 | 231274..235014 | 128780 | ATPase-[Stenotrophomonas-acidaminiphila] |
| QNG75234.1 | Type3.MTaseII.containing.BREX | [illegible] | 1246 | Bacteria>Pseudomonadota>Gammaproteobacteria>Xanthomonadales>Xanthomonadaceae>Stenotrophomonas | GCA_014236105.1 | Stenotrophomonas_maltophilia | EIELFIGP_04091 | 4367916..4371656 | 40324 | hypothetical-protein-EIELFIGP_04091-[Stenotrophomonas-maltophilia] |
| MBH1645065.1 | Type3.MTaseII.containing.BREX | [illegible] | 1242 | Bacteria>Pseudomonadota>Gammaproteobacteria>Xanthomonadales>Xanthomonadaceae>Stenotrophomonas | GCA_015997905.1 | Stenotrophomonas_maltophilia | I5U75_16585 | 22909..26637 | 40324 | hypothetical-protein-[Stenotrophomonas-maltophilia] |
| MBH1712514.1 | Type3.MTaseII.containing.BREX | [illegible] | 1242 | Bacteria>Pseudomonadota>Gammaproteobacteria>Xanthomonadales>Xanthomonadaceae>Stenotrophomonas | GCA_015998125.1 | Stenotrophomonas_maltophilia | I5U96_12260 | 1756..5484 | 40324 | hypothetical-protein-[Stenotrophomonas-maltophilia] |
| MBN5005304.1 | Type3.MTaseII.containing.BREX | [illegible] | 1258 | Bacteria>Pseudomonadota>Gammaproteobacteria>Xanthomonadales>Xanthomonadaceae>Stenotrophomonas | GCA_017149675.1 | Stenotrophomonas_maltophilia | JY447_14725 | 10675..14451 | 40324 | hypothetical-protein-[Stenotrophomonas-maltophilia] |
| WP_054668533.1 | Type3.MTaseII.containing.BREX | [illegible] | 1242 | Bacteria>Pseudomonadota>Gammaproteobacteria>Xanthomonadales>Xanthomonadaceae>Stenotrophomonas | GCF_001314305.1 | Stenotrophomonas_acidaminiphila | AOT14_RS00935 | 231286..235014 | 128780 | DUF6079-family-protein-[Stenotrophomonas-acidaminiphila] |
| WP_070427208.1 | Type3.MTaseII.containing.BREX | [illegible] | 1242 | Bacteria>Pseudomonadota>Gammaproteobacteria>Xanthomonadales>Xanthomonadaceae>Stenotrophomonas | GCF_001806305.1 | Stenotrophomonas_sp._LM091 | BIZ42_RS17705 | 3964619..3968347 | 1904944 | DUF6079-family-protein-[Stenotrophomonas-sp.-LM091] |
| WP_199122707.1 | Type3.MTaseII.containing.BREX | [illegible] | 1250 | Bacteria>Pseudomonadota>Gammaproteobacteria>Xanthomonadales>Xanthomonadaceae>Stenotrophomonas | GCF_002739155.1 | Stenotrophomonas_maltophilia | BRB81_RS09920 | 103620..107372 | 40324 | DUF6079-family-protein-[Stenotrophomonas-maltophilia] |
| WP_111188909.1 | Type3.MTaseII.containing.BREX | [illegible] | 1244 | Bacteria>Pseudomonadota>Gammaproteobacteria>Xanthomonadales>Xanthomonadaceae>Stenotrophomonas | GCF_003244415.1 | Stenotrophomonas_maltophilia | A7X66_04325 | 19797..23531 | 40324 | DUF6079-family-protein-[Stenotrophomonas-maltophilia] |
| WP_049407028.1 | Type3.MTaseII.containing.BREX | [illegible] | 1242 | Bacteria>Pseudomonadota>Gammaproteobacteria>Xanthomonadales>Xanthomonadaceae>Stenotrophomonas | GCF_009676585.1 | Stenotrophomonas_maltophilia | FE088_RS04735 | 1035620..1039348 | 40324 | DUF6079-family-protein-[Stenotrophomonas-maltophilia] |
| WP_253077533.1 | Type3.MTaseII.containing.BREX | [illegible] | 1246 | Bacteria>Pseudomonadota>Gammaproteobacteria>Xanthomonadales>Xanthomonadaceae>Stenotrophomonas | GCF_024126795.1 | Stenotrophomonas_maltophilia | NJG15_RS04315 | 85252..88992 | 40324 | DUF6079-family-protein-[Stenotrophomonas-maltophilia] |
| WP_135965894.1 | Type3.MTaseII.containing.BREX | [illegible] | 1258 | Bacteria>Pseudomonadota>Gammaproteobacteria>Xanthomonadales>Xanthomonadaceae>Stenotrophomonas | GCF_029839415.1 | Stenotrophomonas_maltophilia | N5C11_RS09555 | 28442..32218 | 40324 | MULTISPECIES:-DUF6079-family-protein-[Stenotrophomonas] |
| WP_100449517.1 | Type3.MTaseII.containing.BREX | [illegible] | 1258 | Bacteria>Pseudomonadota>Gammaproteobacteria>Xanthomonadales>Xanthomonadaceae>Stenotrophomonas | - | Stenotrophomonas_maltophilia | B9Y88_RS03200 | 89921..93697 | 40324 | DUF6079-family-protein-[Stenotrophomonas-maltophilia] |
| WP_046429279.1 | Type3.MTaseII.containing.BREX | [illegible] | 1244 | Bacteria>Pseudomonadota>Gammaproteobacteria>Xanthomonadales>Xanthomonadaceae>Stenotrophomonas | - | Stenotrophomonas_maltophilia | XY58_RS04945 | 196831..200565 | 40324 | DUF6079-family-protein-[Stenotrophomonas-maltophilia] |
| WP_271679097.1 | Type3.MTaseII.containing.BREX | [illegible] | 1241 | Bacteria>Pseudomonadota>Gammaproteobacteria>Xanthomonadales>Xanthomonadaceae>Thermomonas | GCF_028201335.1 | Thermomonas_sp._2C3345 | ON759_RS04090 | 81596..85321 | 2993316 | DUF6079-family-protein-[Thermomonas-sp.-2C3345] |
| MBH2009268.1 | Type3.MTaseII.containing.BREX | [illegible] | 1074 | Bacteria>Pseudomonadota>Gammaproteobacteria>Xanthomonadales>Xanthomonadaceae>unclassified_Xanthomonadaceae | GCA_015999665.1 | Xanthomonadaceae_bacterium | I8H71_06175 | 1..3221 | 1926873 | hypothetical-protein-[Xanthomonadaceae-bacterium] |
| WP_033836879.1 | Type3.MTaseII.containing.BREX | [illegible] | 1248 | Bacteria>Pseudomonadota>Gammaproteobacteria>Xanthomonadales>Xanthomonadaceae>Xanthomonas | GCF_000454525.1 | Xanthomonas_citri | XMLX20_RS19135 | 4482061..4485807 | 346 | DUF6079-family-protein-[Xanthomonas-citri] |
| WP_230434266.1 | Type3.MTaseII.containing.BREX | [illegible] | 1246 | Bacteria>Pseudomonadota>Gammaproteobacteria>Xanthomonadales>Xanthomonadaceae>Xanthomonas | GCF_020990895.1 | Xanthomonas_melonis | JWH05_RS03165 | 14918..18658 | 56456 | DUF6079-family-protein-[Xanthomonas-melonis] |
| WP_065846576.1 | Type3.MTaseII.containing.BREX | [illegible] | 1246 | Bacteria>Pseudomonadota | GCF_011762225.1 | Alphaproteobacteria | FHS82_RS15730 | 23668..27408 | 28211 | MULTISPECIES:-DUF6079-family-protein-[Alphaproteobacteria] |
| WP_119335620.1 | Type3.MTaseII.containing.BREX | [illegible] | 1246 | Bacteria>Pseudomonadota>Hydrogenophilia>Hydrogenophilales>Hydrogenophilaceae>Hydrogenophilus | GCF_003574215.1 | Hydrogenophilus_thermoluteolus | HPTL_1674 | 1775492..1779232 | 297 | DUF6079-family-protein-[Hydrogenophilus-thermoluteolus] |
| WP_217125923.1 | Type3.MTaseII.containing.BREX | [illegible] | 1243 | Bacteria>Pseudomonadota>Hydrogenophilia>Hydrogenophilales>Hydrogenophilaceae>Hydrogenophilus | GCF_019049855.1 | Hydrogenophilus_thiooxidans | KUL93_RS04625 | 87236..90967 | 2820326 | DUF6079-family-protein-[Hydrogenophilus-thiooxidans] |
| WP_142810711.1 | Type3.MTaseII.containing.BREX | [illegible] | 1241 | Bacteria>Pseudomonadota>Hydrogenophilia>Hydrogenophilales>Hydrogenophilaceae>Tepidiphilus | GCF_006980705.1 | Tepidiphilus_olei | EPD80_RS07330 | 156841..160566 | 2502184 | DUF6079-family-protein-[Tepidiphilus-olei] |
| MBK6849133.1 | Type3.MTaseII.containing.BREX | [illegible] | 1232 | Bacteria>Pseudomonadota>unclassified_Pseudomonadota | GCA_016702545.1 | Pseudomonadota_bacterium | IPG96_16905 | 43585..47283 | 1977087 | ATP-binding-protein-[Pseudomonadota-bacterium] |
| MBS0577369.1 | Type3.MTaseII.containing.BREX | [illegible] | 1246 | Bacteria>Pseudomonadota>unclassified_Pseudomonadota | GCA_018240435.1 | Pseudomonadota_bacterium | JSS45_13265 | 2298..6038 | 1977087 | ATP-binding-protein-[Pseudomonadota-bacterium] |
| MBS0352305.1 | Type3.MTaseII.containing.BREX | [illegible] | 710 | Bacteria>Pseudomonadota>unclassified_Pseudomonadota | GCA_018241725.1 | Pseudomonadota_bacterium | JSR83_00145 | 1..2133 | 1977087 | ATP-binding-protein-[Pseudomonadota-bacterium] |
| MBU0995361.1 | Type3.MTaseII.containing.BREX | [illegible] | 1445 | Bacteria>Pseudomonadota>unclassified_Pseudomonadota | GCA_018816735.1 | Pseudomonadota_bacterium | KJ737_22930 | 3653..7990 | 1977087 | hypothetical-protein-[Pseudomonadota-bacterium] |
| MBU4427040.1 | Type3.MTaseII.containing.BREX | [illegible] | 456 | Bacteria>Pseudomonadota>unclassified_Pseudomonadota | GCA_018895815.1 | Pseudomonadota_bacterium | KKB35_09910 | 1..1370 | 1977087 | ATP-binding-protein-[Pseudomonadota-bacterium] |
| MCA0325062.1 | Type3.MTaseII.containing.BREX | [illegible] | 1246 | Bacteria>Pseudomonadota>unclassified_Pseudomonadota | GCA_020161215.1 | Pseudomonadota_bacterium | LCH89_05600 | 7182..10922 | 1977087 | DUF6079-family-protein-[Pseudomonadota-bacterium] |
| MCA0212097.1 | Type3.MTaseII.containing.BREX | [illegible] | 1242 | Bacteria>Pseudomonadota>unclassified_Pseudomonadota | GCA_020161775.1 | Pseudomonadota_bacterium | LCH79_02850 | 268802..272530 | 1977087 | DUF6079-family-protein-[Pseudomonadota-bacterium] |
| RJP35075.1 | Type3.MTaseII.containing.BREX | [illegible] | 1445 | Bacteria>PVC_group>Candidatus_Omnitrophica>unclassified_Candidatus_Omnitrophica | GCA_003598175.1 | Candidatus_Omnitrophica_bacterium | C4527_01180 | 9487..13824 | 2035772 | MAG:-hypothetical-protein-C4527_01180-[Candidatus-Omnitrophota-bacterium] |
| MBK7497323.1 | Type3.MTaseII.containing.BREX | [illegible] | 1241 | Bacteria>PVC_group>Candidatus_Omnitrophica>unclassified_Candidatus_Omnitrophica | GCA_016706995.1 | Candidatus_Omnitrophica_bacterium | IPI28_18700 | 38924..42649 | 2035772 | ATP-binding-protein-[Candidatus-Omnitrophota-bacterium] |
| NLF62180.1 | Type3.MTaseII.containing.BREX | [illegible] | 1251 | Bacteria>PVC_group>Lentisphaerota>unclassified_Lentisphaerota | GCA_012515215.1 | Lentisphaerota_bacterium | GX574_13635 | 31199..34954 | 1932692 | ATP-binding-protein-[Lentisphaerota-bacterium] |
| MBT3379341.1 | Type3.MTaseII.containing.BREX | [illegible] | 1252 | Bacteria>PVC_group>Lentisphaerota>unclassified_Lentisphaerota | GCA_018648355.1 | Lentisphaerota_bacterium | HN406_27320 | 7239..10997 | 1932692 | ATP-binding-protein-[Lentisphaerota-bacterium] |
| MBT4818762.1 | Type3.MTaseII.containing.BREX | [illegible] | 298 | Bacteria>PVC_group>Lentisphaerota>unclassified_Lentisphaerota | GCA_018664235.1 | Lentisphaerota_bacterium | HON70_23840 | 8568..9465 | 1932692 | ATP-binding-protein-[Lentisphaerota-bacterium] |
| MBT5606091.1 | Type3.MTaseII.containing.BREX | [illegible] | 722 | Bacteria>PVC_group>Lentisphaerota>unclassified_Lentisphaerota | GCA_018666605.1 | Lentisphaerota_bacterium | HOJ57_09130 | 7357..9522 | 1932692 | ATP-binding-protein-[Lentisphaerota-bacterium] |
| MBT5608619.1 | Type3.MTaseII.containing.BREX | [illegible] | 718 | Bacteria>PVC_group>Lentisphaerota>unclassified_Lentisphaerota | GCA_018666605.1 | Lentisphaerota_bacterium | HOJ57_21965 | 15839..17992 | 1932692 | ATP-binding-protein-[Lentisphaerota-bacterium] |
| MBT5612037.1 | Type3.MTaseII.containing.BREX | [illegible] | 312 | Bacteria>PVC_group>Lentisphaerota>unclassified_Lentisphaerota | GCA_018666605.1 | Lentisphaerota_bacterium | HOJ57_39220 | 1..940 | 1932692 | ATP-binding-protein-[Lentisphaerota-bacterium] |
| KAA0248059.1 | Type3.MTaseII.containing.BREX | [illegible] | 1244 | Bacteria>PVC_group>Planctomycetota>Candidatus_Brocadiia>Candidatus_Brocadiales>Candidatus_Brocadiaceae | GCA_008363445.1 | Candidatus_Jettenia_sp._AMX1 | EDM77_13475 | 2414..6148 | 2293637 | MAG:-ATP-binding-protein-[Candidatus-Jettenia-sp.-AMX1] |
| NUN22618.1 | Type3.MTaseII.containing.BREX | [illegible] | 1244 | Bacteria>PVC_group>Planctomycetota>Candidatus_Brocadiia>Candidatus_Brocadiales>Candidatus_Brocadiaceae | GCA_013360885.1 | Candidatus_Jettenia_caeni | HUU09_04020 | 13155..16889 | 247490 | ATP-binding-protein-[Candidatus-Jettenia-caeni] |
| MBT6049514.1 | Type3.MTaseII.containing.BREX | [illegible] | 1248 | Bacteria>PVC_group>Planctomycetota>Candidatus_Brocadiia>Candidatus_Brocadiales>Candidatus_Scalinduaceae | GCA_018697755.1 | Candidatus_Scalindua_sp. | HOG49_22170 | 17519..21265 | 2268189 | ATP-binding-protein-[Candidatus-Scalindua-sp.] |
| WP_096895081.1 | Type3.MTaseII.containing.BREX | [illegible] | 1248 | Bacteria>PVC_group>Planctomycetota>Candidatus_Brocadiia>Candidatus_Brocadiales>Candidatus_Scalinduaceae | GCF_002443295.1 | Candidatus_Scalindua_japonica | SCALIN_RS13975 | 114317..118063 | 1284222 | DUF6079-family-protein-[Candidatus-Scalindua-japonica] |
| NQV34261.1 | Type3.MTaseII.containing.BREX | [illegible] | 1244 | Bacteria>PVC_group>Planctomycetota>Phycisphaerae>Phycisphaerales>Phycisphaeraceae | GCA_013203945.1 | Phycisphaeraceae_bacterium | HQ515_16320 | 6009..9743 | 2026777 | ATP-binding-protein-[Phycisphaeraceae-bacterium] |
| MCA9303594.1 | Type3.MTaseII.containing.BREX | [illegible] | 516 | Bacteria>PVC_group>Planctomycetota>Phycisphaerae>Phycisphaerales>unclassified_Phycisphaerales | GCA_020427745.1 | Phycisphaerales_bacterium | KC996_05690 | 22249..23798 | 2052180 | hypothetical-protein-[Phycisphaerales-bacterium] |
| MBN2137808.1 | Type3.MTaseII.containing.BREX | [illegible] | 286 | Bacteria>PVC_group>Planctomycetota>Phycisphaerae>Sedimentisphaerales>unclassified_Sedimentisphaerales | GCA_016932835.1 | Sedimentisphaerales_bacterium | JW720_08375 | 10142..11004 | 2813034 | hypothetical-protein-[Sedimentisphaerales-bacterium] |
| MBC04158.1 | Type3.MTaseII.containing.BREX | [illegible] | 1243 | Bacteria>PVC_group>Planctomycetota>Phycisphaerae>unclassified_Phycisphaerae | GCA_002709295.1 | Phycisphaerae_bacterium | CMJ34_12780 | 71480..75211 | 2026778 | ATPase-[Phycisphaerae-bacterium] |
| MBL9119762.1 | Type3.MTaseII.containing.BREX | [illegible] | 526 | Bacteria>PVC_group>Planctomycetota>Phycisphaerae>unclassified_Phycisphaerae | GCA_016795065.1 | Phycisphaerae_bacterium | JNL80_07605 | 27199..28779 | 2026778 | hypothetical-protein-[Phycisphaerae-bacterium] |
| WP_088253237.1 | Type3.MTaseII.containing.BREX | [illegible] | 1420 | Bacteria>PVC_group>Planctomycetota>Planctomycetia>Gemmatales>Gemmataceae | GCF_002197845.1 | Fimbriiglobus_ruber | FRUB_RS08795 | 650711..654973 | 1908690 | hypothetical-protein-[Fimbriiglobus-ruber] |
| MCO6458543.1 | Type3.MTaseII.containing.BREX | [illegible] | 1446 | Bacteria>PVC_group>Planctomycetota>Planctomycetia>Pirellulales>Pirellulaceae | GCA_024102795.1 | Pirellulaceae_bacterium | J5I93_24825 | 24105..28445 | 2699754 | hypothetical-protein-[Pirellulaceae-bacterium] |
| WP_146535598.1 | Type3.MTaseII.containing.BREX | [illegible] | 1234 | Bacteria>PVC_group>Planctomycetota>Planctomycetia>Pirellulales>Pirellulaceae | GCF_007860175.1 | Rubripirellula_reticaptiva | Poly59_RS18475 | 382484..386188 | 2528013 | DUF6079-family-protein-[Rubripirellula-reticaptiva] |
| WP_150079789.1 | Type3.MTaseII.containing.BREX | [illegible] | 1243 | Bacteria>PVC_group>Planctomycetota>Planctomycetia>Pirellulales>Pirellulaceae | GCF_008629675.1 | Roseiconus_nitratireducens | FYK55_RS27325 | 17824..21555 | 2605748 | DUF6079-family-protein-[Roseiconus-nitratireducens] |
| ADY58758.1 | Type3.MTaseII.containing.BREX | [illegible] | 1251 | Bacteria>PVC_group>Planctomycetota>Planctomycetia>Planctomycetales>Planctomycetaceae | GCA_000165715.3 | Rubinisphaera_brasiliensis_DSM_5305 | Plabr_1142 | 1331346..1335101 | 756272 | hypothetical-protein-Plabr_1142-[Rubinisphaera-brasiliensis-DSM-5305] |
| WP_041397812.1 | Type3.MTaseII.containing.BREX | [illegible] | 1232 | Bacteria>PVC_group>Planctomycetota>Planctomycetia>Planctomycetales>Planctomycetaceae | GCF_000165715.2 | Rubinisphaera_brasiliensis | PLABR_RS05750 | 1331346..1335044 | 119 | DUF6079-family-protein-[Rubinisphaera-brasiliensis] |
| WP_145380206.1 | Type3.MTaseII.containing.BREX | [illegible] | 1234 | Bacteria>PVC_group>Planctomycetota>Planctomycetia>Planctomycetales>Planctomycetaceae | GCF_007747995.1 | Symmachiella_dynata | Mal52_RS28010 | 7362136..7365840 | 2527995 | DUF6079-family-protein-[Symmachiella-dynata] |
| MBE7447013.1 | Type3.MTaseII.containing.BREX | [illegible] | 1244 | Bacteria>PVC_group>Planctomycetota>Planctomycetia>unclassified_Planctomycetia | GCA_015075005.1 | Planctomycetia_bacterium | HS132_18070 | 382904..386638 | 2052181 | ATP-binding-protein-[Planctomycetia-bacterium] |
| OQZ06364.1 | Type3.MTaseII.containing.BREX | [illegible] | 1231 | Bacteria>PVC_group>Planctomycetota>unclassified_Planctomycetota | GCA_002050205.1 | Planctomycetes_bacterium_UTPLA1 | B6D36_05375 | 2828..6523 | 1961112 | ATPase-[Planctomycetes-bacterium-UTPLA1] |
| RLT19656.1 | Type3.MTaseII.containing.BREX | [illegible] | 987 | Bacteria>PVC_group>Planctomycetota>unclassified_Planctomycetota | GCA_003672035.1 | Planctomycetota_bacterium | DWI28_02425 | 16452..19415 | 2026780 | MAG:-ATP-binding-protein-[Planctomycetota-bacterium] |
| MBI4882201.1 | Type3.MTaseII.containing.BREX | [illegible] | 1237 | Bacteria>PVC_group>Planctomycetota>unclassified_Planctomycetota | GCA_016213895.1 | Planctomycetota_bacterium | HY812_21445 | 2712..6425 | 2026780 | ATP-binding-protein-[Planctomycetota-bacterium] |
| MBN8597958.1 | Type3.MTaseII.containing.BREX | [illegible] | 1232 | Bacteria>PVC_group>Planctomycetota>unclassified_Planctomycetota | GCA_017303555.1 | Planctomycetota_bacterium | JOL78_09815 | 34323..38021 | 2026780 | ATP-binding-protein-[Planctomycetota-bacterium] |
| ACD82384.1 | Type3.MTaseII.containing.BREX | [illegible] | 1243 | Bacteria>PVC_group>Verrucomicrobiota>Methylacidiphilae>Methylacidiphilales>Methylacidiphilaceae | GCA_000019665.1 | Methylacidiphilum_infernorum_V4 | Minf_0326 | 329568..333299 | 481448 | Predicted-ATPase,-VrlK-family-[Methylacidiphilum-infernorum-V4] |
| WP_012462666.1 | Type3.MTaseII.containing.BREX | [illegible] | 1243 | Bacteria>PVC_group>Verrucomicrobiota>Methylacidiphilae>Methylacidiphilales>Methylacidiphilaceae | GCF_000019665.1 | Methylacidiphilum_infernorum | MINF_RS01440 | 329568..333299 | 511746 | DUF6079-family-protein-[Methylacidiphilum-infernorum] |
| MBL9185851.1 | Type3.MTaseII.containing.BREX | [illegible] | 1236 | Bacteria>PVC_group>Verrucomicrobiota>Opitutae>Opitutales>Opitutaceae | GCA_016794785.1 | Opitutaceae_bacterium | JNK23_00090 | 29545..33255 | 2006848 | hypothetical-protein-[Opitutaceae-bacterium] |
| HCF94153.1 | Type3.MTaseII.containing.BREX | [illegible] | 1288 | Bacteria>PVC_group>Verrucomicrobiota>unclassified_Verrucomicrobiota | GCA_003534995.1 | Verrucomicrobiota_bacterium | DEW46_03760 | 1029..4895 | 2026799 | ATPase-[Verrucomicrobiota-bacterium] |
| NMD20861.1 | Type3.MTaseII.containing.BREX | [illegible] | 1236 | Bacteria>PVC_group>Verrucomicrobiota>unclassified_Verrucomicrobiota | GCA_012797305.1 | Verrucomicrobiota_bacterium | GYA76_11470 | 14764..18474 | 2026799 | ATP-binding-protein-[Verrucomicrobiota-bacterium] |
| MBI4023511.1 | Type3.MTaseII.containing.BREX | [illegible] | 1371 | Bacteria>PVC_group>Verrucomicrobiota>unclassified_Verrucomicrobiota | GCA_016198745.1 | Verrucomicrobiota_bacterium | HY360_00925 | 8259..12374 | 2026799 | hypothetical-protein-[Verrucomicrobiota-bacterium] |
| MBP7946969.1 | Type3.MTaseII.containing.BREX | [illegible] | 1351 | Bacteria>PVC_group>Verrucomicrobiota>unclassified_Verrucomicrobiota | GCA_018003215.1 | Verrucomicrobiota_bacterium | KA191_12990 | 22322..26377 | 2026799 | hypothetical-protein-[Verrucomicrobiota-bacterium] |
| MCF7763181.1 | Type3.MTaseII.containing.BREX | [illegible] | 1257 | Bacteria>PVC_group>Verrucomicrobiota>unclassified_Verrucomicrobiota | GCA_021733945.1 | Verrucomicrobiota_bacterium | K9N62_05860 | 12732..16505 | 2026799 | DUF6079-family-protein-[Verrucomicrobiota-bacterium] |
| MCU0771894.1 | Type3.MTaseII.containing.BREX | [illegible] | 1236 | Bacteria>PVC_group>Verrucomicrobiota>unclassified_Verrucomicrobiota | GCA_025457615.1 | Verrucomicrobiota_bacterium | MUE94_09045 | 7531..11241 | 2026799 | DUF6079-family-protein-[Verrucomicrobiota-bacterium] |
| SPE50997.1 | Type3.MTaseII.containing.BREX | [illegible] | 1442 | Bacteria>PVC_group>Verrucomicrobiota>unclassified_Verrucomicrobiota | GCA_900290355.1 | Verrucomicrobiota_bacterium | SBV1_1250002 | 992..5320 | 2026799 | conserved-hypothetical-protein-[Verrucomicrobiota-bacterium] |
| MBS5507937.1 | Type3.MTaseII.containing.BREX | [illegible] | 1242 | Bacteria>PVC_group>Verrucomicrobiota>Verrucomicrobiae>Verrucomicrobiales>Akkermansiaceae | GCA_018368805.1 | Akkermansia_sp. | KHX31_04810 | 60826..64554 | 1872421 | hypothetical-protein-[Akkermansia-sp.] |
| MBX3742594.1 | Type3.MTaseII.containing.BREX | [illegible] | 1243 | Bacteria>PVC_group>Verrucomicrobiota>Verrucomicrobiae>Verrucomicrobiales>Akkermansiaceae | GCA_019634595.1 | Akkermansiaceae_bacterium | KF712_16525 | 314704..318435 | 2562705 | hypothetical-protein-[Akkermansiaceae-bacterium] |
| MCD8271875.1 | Type3.MTaseII.containing.BREX | [illegible] | 1242 | Bacteria>PVC_group>Verrucomicrobiota>Verrucomicrobiae>Verrucomicrobiales>Akkermansiaceae | GCA_021204505.1 | Akkermansia_sp. | LUC46_04005 | 29201..32929 | 1872421 | DUF6079-family-protein-[Akkermansia-sp.] |
| WP_067776555.1 | Type3.MTaseII.containing.BREX | [illegible] | 1243 | Bacteria>PVC_group>Verrucomicrobiota>Verrucomicrobiae>Verrucomicrobiales>Akkermansiaceae | GCF_001683795.1 | Akkermansia_glycaniphila | AC781_RS08510 | 117484..121215 | 1679444 | DUF6079-family-protein-[Akkermansia-glycaniphila] |
| WP_102758537.1 | Type3.MTaseII.containing.BREX | [illegible] | 1242 | Bacteria>PVC_group>Verrucomicrobiota>Verrucomicrobiae>Verrucomicrobiales>Akkermansiaceae | GCF_001917295.1 | Akkermansia_muciniphila | CXU07_RS02965 | 669523..673251 | 239935 | DUF6079-family-protein-[Akkermansia-muciniphila] |
| WP_123052213.1 | Type3.MTaseII.containing.BREX | [illegible] | 1242 | Bacteria>PVC_group>Verrucomicrobiota>Verrucomicrobiae>Verrucomicrobiales>Akkermansiaceae | GCF_003716915.1 | Akkermansia_muciniphila | CUB89_RS06295 | 1401029..1404757 | 239935 | DUF6079-family-protein-[Akkermansia-muciniphila] |
| WP_215841664.1 | Type3.MTaseII.containing.BREX | [illegible] | 1242 | Bacteria>PVC_group>Verrucomicrobiota>Verrucomicrobiae>Verrucomicrobiales>Akkermansiaceae | GCF_018847055.1 | Akkermansia_muciniphila | J5W61_RS00450 | 112413..116141 | 239935 | DUF6079-family-protein-[Akkermansia-muciniphila] |
| WP_215833375.1 | Type3.MTaseII.containing.BREX | [illegible] | 1242 | Bacteria>PVC_group>Verrucomicrobiota>Verrucomicrobiae>Verrucomicrobiales>Akkermansiaceae | GCF_018847255.1 | Akkermansia_muciniphila | J5W48_RS06490 | 1515271..1518999 | 239935 | DUF6079-family-protein-[Akkermansia-muciniphila] |
| RFU95875.1 | Type3.MTaseII.containing.BREX | [illegible] | 1246 | Bacteria>Spirochaetota>Spirochaetia>Spirochaetales>Sphaerochaetaceae>Sphaerochaeta | GCA_003429665.1 | Sphaerochaeta_halotolerans | DYP60_02410 | 179528..183268 | 2293840 | ATP-binding-protein-[Sphaerochaeta-halotolerans] |
| WP_206363090.1 | Type3.MTaseII.containing.BREX | [illegible] | 1255 | Bacteria>Spirochaetota>Spirochaetia>Spirochaetales>Sphaerochaetaceae>Sphaerochaeta | GCF_003429665.1 | Sphaerochaeta_halotolerans | DYP60_RS02405 | 179501..183268 | 2293840 | DUF6079-family-protein-[Sphaerochaeta-halotolerans] |
| WP_020614466.1 | Type3.MTaseII.containing.BREX | [illegible] | 1241 | Bacteria>Spirochaetota>Spirochaetia>Spirochaetales>Spirochaetaceae>Sediminispirochaeta | GCF_000378205.1 | Sediminispirochaeta_bajacaliforniensis | F459_RS0120160 | 20907..24632 | 148 | DUF6079-family-protein-[Sediminispirochaeta-bajacaliforniensis] |
| OQY33159.1 | Type3.MTaseII.containing.BREX | [illegible] | 1241 | Bacteria>Spirochaetota>Spirochaetia>Spirochaetales>Spirochaetaceae>unclassified_Spirochaetaceae | GCA_002084805.1 | Spirochaetaceae_bacterium_4572_59 | B6241_08630 | 14589..18314 | 1971727 | hypothetical-protein-B6241_08630-[Spirochaetaceae-bacterium-4572_59] |
| MBO7485543.1 | Type3.MTaseII.containing.BREX | [illegible] | 1226 | Bacteria>Spirochaetota>Spirochaetia>Spirochaetales>Spirochaetaceae>unclassified_Spirochaetaceae | GCA_017649985.1 | Spirochaetaceae_bacterium | J6T84_05730 | 14536..18216 | 1898206 | hypothetical-protein-[Spirochaetaceae-bacterium] |
| WP_010252922.1 | Type3.MTaseII.containing.BREX | [illegible] | 1224 | Bacteria>Spirochaetota>Spirochaetia>Spirochaetales>Treponemataceae>Treponema | GCF_000297095.1 | Treponema_primitia | TPRIMZ1_RS18025 | 17306..20980 | 88058 | DUF6079-family-protein-[Treponema-primitia] |
| WP_132780483.1 | Type3.MTaseII.containing.BREX | [illegible] | 1243 | Bacteria>Spirochaetota>Spirochaetia>Spirochaetales>Treponemataceae>Treponema | GCF_004343725.1 | Treponema_sp._J25 | C5022_RS06925 | 72481..76212 | 2094121 | DUF6079-family-protein-[Treponema-sp.-J25] |
| WP_198443345.1 | Type3.MTaseII.containing.BREX | [illegible] | 1225 | Bacteria>Spirochaetota>Spirochaetia>Spirochaetales>Treponemataceae>Treponema | GCF_016117655.1 | Treponema_peruense | IWA51_RS04150 | 893200..896877 | 2787628 | DUF6079-family-protein-[Treponema-peruense] |
| MBN2531353.1 | Type3.MTaseII.containing.BREX | [illegible] | 1396 | Bacteria>Spirochaetota>Spirochaetia>Spirochaetales>unclassified_Spirochaetales | GCA_016935735.1 | Spirochaetales_bacterium | JXB88_00605 | 786..4976 | 2026792 | hypothetical-protein-[Spirochaetales-bacterium] |
| MBN2531851.1 | Type3.MTaseII.containing.BREX | [illegible] | 1446 | Bacteria>Spirochaetota>Spirochaetia>Spirochaetales>unclassified_Spirochaetales | GCA_016935735.1 | Spirochaetales_bacterium | JXB88_03110 | 8268..12608 | 2026792 | hypothetical-protein-[Spirochaetales-bacterium] |
| MBQ3728152.1 | Type3.MTaseII.containing.BREX | [illegible] | 1237 | Bacteria>Spirochaetota>Spirochaetia>Spirochaetales>unclassified_Spirochaetales | GCA_017412555.1 | Spirochaetales_bacterium | II903_01460 | 5771..9484 | 2026792 | hypothetical-protein-[Spirochaetales-bacterium] |
| GHV83239.1 | Type3.MTaseII.containing.BREX | [illegible] | 1246 | Bacteria>Spirochaetota>Spirochaetia>unclassified_Spirochaetia | GCA_025997755.1 | Spirochaetia_bacterium | AGMMS50212_05790 | 691..4431 | 2053615 | hypothetical-protein-AGMMS50212_05790-[Spirochaetia-bacterium] |
| GHT84073.1 | Type3.MTaseII.containing.BREX | [illegible] | 177 | Bacteria>Spirochaetota>Spirochaetia>unclassified_Spirochaetia | GCA_026002105.1 | Spirochaetia_bacterium | FACS1894137_06670 | 4966..5499 | 2053615 | hypothetical-protein-FACS1894137_06670-[Spirochaetia-bacterium] |
| GHT84077.1 | Type3.MTaseII.containing.BREX | [illegible] | 180 | Bacteria>Spirochaetota>Spirochaetia>unclassified_Spirochaetia | GCA_026002105.1 | Spirochaetia_bacterium | FACS1894137_06680 | 5766..6308 | 2053615 | hypothetical-protein-FACS1894137_06680-[Spirochaetia-bacterium] |
| GHU24474.1 | Type3.MTaseII.containing.BREX | [illegible] | 1244 | Bacteria>Spirochaetota>Spirochaetia>unclassified_Spirochaetia | GCA_026002265.1 | Spirochaetia_bacterium | FACS1894164_11200 | 5785..9519 | 2053615 | hypothetical-protein-FACS1894164_11200-[Spirochaetia-bacterium] |
| OQB94794.1 | Type3.MTaseII.containing.BREX | [illegible] | 1248 | Bacteria>Spirochaetota>unclassified_Spirochaetota | GCA_002070755.1 | Spirochaetes_bacterium_ADurb.Bin110 | BWX81_02285 | 880..4626 | 1852908 | hypothetical-protein-BWX81_02285-[Spirochaetes-bacterium-ADurb.Bin110] |
| OQB94579.1 | Type3.MTaseII.containing.BREX | [illegible] | 1246 | Bacteria>Spirochaetota>unclassified_Spirochaetota | GCA_002070755.1 | Spirochaetes_bacterium_ADurb.Bin110 | BWX81_02438 | 21902..25642 | 1852908 | hypothetical-protein-BWX81_02438-[Spirochaetes-bacterium-ADurb.Bin110] |
| TXT44742.1 | Type3.MTaseII.containing.BREX | [illegible] | 488 | Bacteria>Spirochaetota>unclassified_Spirochaetota | GCA_008080685.1 | Spirochaetota_bacterium | FD137_1517 | 14949..16415 | 2202144 | hypothetical-protein-FD137_1517-[Spirochaetota-bacterium] |
| TXT50777.1 | Type3.MTaseII.containing.BREX | [illegible] | 1244 | Bacteria>Spirochaetota>unclassified_Spirochaetota | GCA_008080685.1 | Spirochaetota_bacterium | FD137_232 | 19223..22957 | 2202144 | hypothetical-protein-FD137_232-[Spirochaetota-bacterium] |
| SIN70165.1 | Type3.MTaseII.containing.BREX | [illegible] | 1265 | Bacteria>Synergistota>Synergistia>Synergistales>Acetomicrobiaceae>Acetomicrobium | GCA_900129645.1 | Acetomicrobium_flavidum | SAMN05444368_1310 | 1282728..1286525 | 49896 | hypothetical-protein-SAMN05444368_1310-[Acetomicrobium-flavidum] |
| WP_014807685.1 | Type3.MTaseII.containing.BREX | [illegible] | 1263 | Bacteria>Synergistota>Synergistia>Synergistales>Acetomicrobiaceae>Acetomicrobium | GCF_000266925.1 | Acetomicrobium_mobile | Anamo_1876 | 1904829..1908620 | 97477 | DUF6079-family-protein-[Acetomicrobium-mobile] |
| MBL3538350.1 | Type3.MTaseII.containing.BREX | [illegible] | 1442 | Bacteria>Synergistota>Synergistia>Synergistales>Aminobacteriaceae>Aminivibrio | GCA_016756745.1 | Aminivibrio_sp. | JMJ95_01550 | 21005..25333 | 1872489 | hypothetical-protein-[Aminivibrio-sp.] |
| MBL3538483.1 | Type3.MTaseII.containing.BREX | [illegible] | 1252 | Bacteria>Synergistota>Synergistia>Synergistales>Aminobacteriaceae>Aminivibrio | GCA_016756745.1 | Aminivibrio_sp. | JMJ95_02220 | 62169..65927 | 1872489 | ATP-binding-protein-[Aminivibrio-sp.] |
| MBL3538842.1 | Type3.MTaseII.containing.BREX | [illegible] | 279 | Bacteria>Synergistota>Synergistia>Synergistales>Aminobacteriaceae>Aminivibrio | GCA_016756745.1 | Aminivibrio_sp. | JMJ95_04075 | 1..839 | 1872489 | hypothetical-protein-[Aminivibrio-sp.] |
| WP_005662440.1 | Type3.MTaseII.containing.BREX | [illegible] | 1247 | Bacteria>Synergistota>Synergistia>Synergistales>Dethiosulfovibrionaceae>Dethiosulfovibrio | GCF_000172975.1 | Dethiosulfovibrio_peptidovorans | DPEP_RS12015 | 2485235..2488978 | 47055 | DUF6079-family-protein-[Dethiosulfovibrio-peptidovorans] |
| WP_236100997.1 | Type3.MTaseII.containing.BREX | [illegible] | 1246 | Bacteria>Synergistota>Synergistia>Synergistales>Dethiosulfovibrionaceae>Dethiosulfovibrio | GCF_021568795.1 | Dethiosulfovibrio_faecalis | L2W58_RS00505 | 84364..88104 | 2720018 | DUF6079-family-protein-[Dethiosulfovibrio-faecalis] |
| WP_006299685.1 | Type3.MTaseII.containing.BREX | [illegible] | 1243 | Bacteria>Synergistota>Synergistia>Synergistales>Synergistaceae>Aminomonas | GCF_000165795.1 | Aminomonas_paucivorans | APAU_RS00530 | 111006..114737 | 81412 | DUF6079-family-protein-[Aminomonas-paucivorans] |
| WP_012870232.1 | Type3.MTaseII.containing.BREX | [illegible] | 1272 | Bacteria>Synergistota>Synergistia>Synergistales>Synergistaceae>Thermanaerovibrio | GCF_000024905.1 | Thermanaerovibrio_acidaminovorans | TACI_RS07710 | 1563888..1567706 | 81462 | DUF6079-family-protein-[Thermanaerovibrio-acidaminovorans] |
| NLD05176.1 | Type3.MTaseII.containing.BREX | [illegible] | 1242 | Bacteria>Synergistota>Synergistia>Synergistales>Synergistaceae>unclassified_Synergistaceae | GCA_012513245.1 | Synergistaceae_bacterium | GX672_04650 | 4463..8191 | 2053624 | ATP-binding-protein-[Synergistaceae-bacterium] |
| MBL3593375.1 | Type3.MTaseII.containing.BREX | [illegible] | 1249 | Bacteria>Synergistota>Synergistia>Synergistales>Synergistaceae>unclassified_Synergistaceae | GCA_016756725.1 | Synergistaceae_bacterium | JMJ93_07675 | 80117..83866 | 2053624 | ATP-binding-protein-[Synergistaceae-bacterium] |
| MCK9435717.1 | Type3.MTaseII.containing.BREX | [illegible] | 1219 | Bacteria>Synergistota>Synergistia>Synergistales>Synergistaceae>unclassified_Synergistaceae | GCA_023229345.1 | Synergistaceae_bacterium | MOQ12_00750 | 4305..7964 | 2053624 | DUF6079-family-protein-[Synergistaceae-bacterium] |
| GHV29554.1 | Type3.MTaseII.containing.BREX | [illegible] | 1230 | Bacteria>Synergistota>Synergistia>Synergistales>unclassified_Synergistales | GCA_026002695.1 | Synergistales_bacterium | FACS1894167_09040 | 12998..16690 | 2268201 | hypothetical-protein-FACS1894167_09040-[Synergistales-bacterium] |
| MXW95274.1 | Type3.MTaseII.containing.BREX | [illegible] | 1228 | Bacteria>Terrabacteria_group>Actinomycetota>Acidimicrobiia>Acidimicrobiales>Acidimicrobiaceae | GCA_009836165.1 | Acidimicrobiaceae_bacterium | F4Z64_07180 | 9801..13487 | 2024894 | ATP-binding-protein-[Acidimicrobiaceae-bacterium] |
| MYH43036.1 | Type3.MTaseII.containing.BREX | [illegible] | 1228 | Bacteria>Terrabacteria_group>Actinomycetota>Acidimicrobiia>Acidimicrobiales>Acidimicrobiaceae | GCA_009843995.1 | Acidimicrobiaceae_bacterium | F4131_04965 | 9076..12762 | 2024894 | ATP-binding-protein-[Acidimicrobiaceae-bacterium] |
| GIU85384.1 | Type3.MTaseII.containing.BREX | [illegible] | 371 | Bacteria>Terrabacteria_group>Actinomycetota>Acidimicrobiia>Acidimicrobiales>unclassified_Acidimicrobiales | GCA_026002915.1 | Acidimicrobiales_bacterium | KatS3mg008_2159 | 241814..242929 | 2201156 | MAG:-hypothetical-protein-KatS3mg008_2159-[Acidimicrobiales-bacterium] |
| MYB24177.1 | Type3.MTaseII.containing.BREX | [illegible] | 1228 | Bacteria>Terrabacteria_group>Actinomycetota>Acidimicrobiia>unclassified_Acidimicrobiia | GCA_009840735.1 | Acidimicrobiia_bacterium | F4X37_03560 | 269..3955 | 2080302 | ATP-binding-protein-[Acidimicrobiia-bacterium] |
| WP_011719648.1 | Type3.MTaseII.containing.BREX | [illegible] | 1243 | Bacteria>Terrabacteria_group>Actinomycetota>Actinomycetes>Acidothermales>Acidothermaceae | GCF_000015025.1 | Acidothermus_cellulolyticus | ACEL_RS04195 | 896448..900179 | 28049 | DUF6079-family-protein-[Acidothermus-cellulolyticus] |
| WP_030992070.1 | Type3.MTaseII.containing.BREX | [illegible] | 1228 | Bacteria>Terrabacteria_group>Actinomycetota>Actinomycetes>Kitasatosporales>Streptomycetaceae | GCF_000717895.1 | Streptomyces_sp._NRRL_WC-3744 | IF36_RS0138665 | 24616..28302 | 1463935 | DUF6079-family-protein-[Streptomyces-sp.-NRRL-WC-3744] |
| WP_227569851.1 | Type3.MTaseII.containing.BREX | [illegible] | 1229 | Bacteria>Terrabacteria_group>Actinomycetota>Actinomycetes>Micrococcales>Cellulomonadaceae | GCF_024508335.1 | unclassified_Cellulomonas | NP064_RS16070 | 3464938..3468627 | 2620175 | MULTISPECIES:-DUF6079-family-protein-[unclassified-Cellulomonas] |
| SLK50155.1 | Type3.MTaseII.containing.BREX | [illegible] | 1233 | Bacteria>Terrabacteria_group>Actinomycetota>Actinomycetes>Mycobacteriales>Mycobacteriaceae | GCA_900132645.1 | Mycobacteroides_abscessus_subsp._abscessus | SAMEA2071448_00696 | 756207..759908 | 1185650 | Uncharacterised-protein-[Mycobacteroides-abscessus-subsp.-abscessus] |

| Protein-ID | System | Gene-Neighbourhoods | Length | Taxonomy | Assembly-ID | Strain | Locus-Tag | Genomic-coordinates | TaxID | Fasta-define |
|---|---|---|---|---|---|---|---|---|---|---|
| WP_067936768.1 | Type3.MTaseII.containing.BREX | <-TUDOR+SF1-SF2-Helicase+iREase+CTD<-N6-N4-METHYLASE<-DUF6079*<-BrxB-BrxF-iSTAND<-?<-?\|\|BirA-HTH->?->Cox-HTH->Primpol-> | 1230 | Bacteria>Terrabacteria_group>Actinomycetota>Actinomycetes>Mycobacteriales>Mycobacteriaceae | GCF_001667925.1 | Mycobacterium_sp._E2479 | A5686_RS17380 | 5462..9154 | 1834134 | DUF6079-family-protein-[Mycobacterium-sp.-E2479] |
| WP_056925870.1 | Type3.MTaseII.containing.BREX | <-RnaseH<-?\|\|SIGMA-HTH->?->?-><-BrxA-DNA-binding<-PglZ<-TUDOR+SF1-SF2-Helicase+iREase+CTD<-N6-N4-METHYLASE<-DUF6079*<-BrxB-BrxF-iSTAND\|\|SF1-SF2-Helicase->NERD+Pkinase+Pkinase+DndF->DndG->DndH->SF1-SF2-Helicase+HTH->?-><-DNA-glycosylase | 1229 | Bacteria>Terrabacteria_group>Actinomycetota>Actinomycetes>Propionibacteriales>Nocardioidaceae | GCF_001429625.1 | Nocardioides_sp._Soil805 | ASG94_RS08685 | 1773281..1776970 | 1736416 | DUF6079-family-protein-[Nocardioides-sp.-Soil805] |
| WP_182526621.1 | Type3.MTaseII.containing.BREX | <-STAS<-?<-?<-?<-?\|\|?->Acetate_kinase-><-tRNA<-tRNA<-tRNA\|\|Aminotran_4->tRNA->BrxB-BrxF-iSTAND->DUF6079*->N6-N4-METHYLASE->TUDOR+SF1-SF2-Helicase+iREase+CTD->PglZ->BrxA-DNA-binding-><-?\|\|TrwC+SF1-SF2-Helicase->XIS-HTH->XIS-HTH-> | 1229 | Bacteria>Terrabacteria_group>Actinomycetota>Actinomycetes>Propionibacteriales>Nocardioidaceae | GCF_014127485.1 | Nocardioides_dongkuii | H4O22_RS08820 | 1819921..1823610 | 2760089 | DUF6079-family-protein-[Nocardioides-dongkuii] |
| WP_262852283.1 | Type3.MTaseII.containing.BREX | <-Peptidase_M23<-ParB<-?<-?<-Primpol+PriCT1\|\|VirB11+MuB-ATPase->?-><-?\|\|?->DUF6079*->N6-N4-METHYLASE->TUDOR+SF1-SF2-Helicase+iREase+CTD->PglZ->BrxA-DNA-binding-><-?<-?<-SbcC+CCDC158\|\|JetB->SbcC-> | 1228 | Bacteria>Terrabacteria_group>Actinomycetota>Actinomycetes>Propionibacteriales>Nocardioidaceae | GCF_025506335.1 | Mumia_sp._NEAU-365 | NYZ95_RS15800 | 255507..259193 | 2976125 | DUF6079-family-protein-[Mumia-quercus] |
| KUK47744.1 | Type3.MTaseII.containing.BREX | IG+IG->?->BrxB-BrxF-iSTAND->DUF6079*->N6-N4-METHYLASE->TUDOR+SF1-SF2-Helicase+iREase+CTD->PglZ->BrxA-DNA-binding-> | 1226 | Bacteria>Terrabacteria_group>Actinomycetota>Actinomycetes>unclassified_Actinomycetes | GCA_001508275.1 | Actinobacteria_bacterium_66_15 | XD74_1657 | 1866..5546 | 1635289 | Uncharacterized-protein-XD74_1657-[Actinobacteria-bacterium-66_15] |
| MBS3909265.1 | Type3.MTaseII.containing.BREX | DNA_pol3_beta+DNA_pol3_beta_2+DNA_pol3_beta_3-><-REase\|\|?->METHYLASE+SF1-SF2-Helicase-><-N6-N4-METHYLASE<-?<-SIGMA+OB-Fold+AAA-ATPase-BrxL<-PglZ<-DUF6079*<-LON-AAA<-DUF4007<-N6-METHYLASE<-PHP+SbcC+PspA<-XIS-HTH<-CHTH<-LuxR-HTH\|\|?->XIS-HTH-> | 1374 | Bacteria>Terrabacteria_group>Actinomycetota>Actinomycetes>unclassified_Actinomycetes | GCA_018333515.1 | Actinomycetia_bacterium | KGZ93_06520 | 24417..28541 | 1883427 | hypothetical-protein-[Actinomycetia-bacterium] |
| MBS3974431.1 | Type3.MTaseII.containing.BREX | <-N6-N4-METHYLASE<-Schlafen+HATPase+wHTH<-DUF6079*<-BrxB-BrxF-iSTAND<-?<-?<-CHTH<-?<-RhoN+RhoN+RNase+S1\|\|?-><-PDZ+Peptidase | 1244 | Bacteria>Terrabacteria_group>Actinomycetota>Actinomycetes>unclassified_Actinomycetes | GCA_018334075.1 | Actinomycetia_bacterium | KGZ89_06140 | 2651..6385 | 1883427 | ATP-binding-protein-[Actinomycetia-bacterium] |
| MBU4174457.1 | Type3.MTaseII.containing.BREX | <-Glyco_hydro_35\|\|XIS-HTH->BrxB-BrxF-iSTAND->DUF6079*->Schlafen+HATPase+wHTH->SbcC->N6-N4-METHYLASE->SbcC->?->?->PglZ->BrxA-DNA-binding->GDResolvase-> | 1242 | Bacteria>Terrabacteria_group>Actinomycetota>Actinomycetes>unclassified_Actinomycetes | GCA_018898375.1 | Actinomycetia_bacterium | KJ686_06080 | 21233..24961 | 1883427 | ATP-binding-protein-[Actinomycetia-bacterium] |
| GFP32117.1 | Type3.MTaseII.containing.BREX | <-PglZ<-HARE-HTH<-N6-N4-METHYLASE<-DUF6079* | 1276 | Bacteria>Terrabacteria_group>Actinomycetota>Actinomycetota_incertae_sedis>Candidatus_Hakubanella | GCA_013281995.1 | Candidatus_Hakubanella_thermoalkaliphilus | HKBW3S42_00422 | 4200..8030 | 2754717 | hypothetical-protein-HKBW3S42_00422-[Candidatus-Hakubanella-thermoalkaliphilus] |
| MBR3317259.1 | Type3.MTaseII.containing.BREX | HD-><-?<-?\|\|HTH_3->ACET-><-HTH_3\|\|BrxB-BrxF-iSTAND->DUF6079*->N6-N4-METHYLASE->TUDOR+SF1-SF2-Helicase+iREase+CTD->PglZ->BrxA-DNA-binding-><-PIN<-MPTase\|\|TraJ-RHH->RNase->DEDD_Tnp_IS110+Transposase_20-> | 1225 | Bacteria>Terrabacteria_group>Actinomycetota>Coriobacteriia>Coriobacteriales>Atopobiaceae | GCA_017510785.1 | Atopobiaceae_bacterium | IKG21_05515 | 88091..91768 | 2485926 | hypothetical-protein-[Atopobiaceae-bacterium] |
| RRF92484.1 | Type3.MTaseII.containing.BREX | <-Acyl_transf_3\|\|?->?->?-><-?<-Peptidase_C26\|\|zf-ISL3->BrxB-BrxF-iSTAND->DUF6079*->N6-N4-METHYLASE-> | 1226 | Bacteria>Terrabacteria_group>Actinomycetota>Coriobacteriia>Coriobacteriales>Coriobacteriaceae | GCA_003862165.1 | Coriobacteriaceae_bacterium | DUD31_07560 | 9244..12924 | 2011094 | hypothetical-protein-DUD31_07560-[Coriobacteriaceae-bacterium] |
| WP_087187058.1 | Type3.MTaseII.containing.BREX | <-ACET\|\|Phage-portal->BrxB-BrxF-iSTAND->DUF6079*->N6-N4-METHYLASE->TUDOR+SF1-SF2-Helicase+iREase+CTD-> | 1225 | Bacteria>Terrabacteria_group>Actinomycetota>Coriobacteriia>Coriobacteriales>Coriobacteriaceae | GCF_002159295.1 | Enorma_massiliensis | B5G21_RS10120 | 3408..7085 | 1472761 | DUF6079-family-protein-[Enorma-massiliensis] |
| WP_087190343.1 | Type3.MTaseII.containing.BREX | <-ABhydrolase<-?\|\|?->?->?->REase+SPF-cons-><-wHTH+WYL\|\|?->BrxB-BrxF-iSTAND-p->DUF6079*->N6-N4-METHYLASE->TUDOR+SF1-SF2-Helicase+iREase+CTD->PglZ->BrxA-DNA-binding->ParB+HNH->?->?-><-HTH_Tnp_1+HTH_21+rve+rve_3 | 1225 | Bacteria>Terrabacteria_group>Actinomycetota>Coriobacteriia>Eggerthellales>Eggerthellaceae | GCF_002159895.1 | Gordonibacter | B5F44_RS03995 | 62178..65855 | 644652 | MULTISPECIES:-DUF6079-family-protein-[Gordonibacter] |
| WP_273124462.1 | Type3.MTaseII.containing.BREX | <-ACET<-ARG\|\|STYKinase-like->ACET-><-?\|\|Phage-portal->BrxB-BrxF-iSTAND-p->DUF6079-p*-> | 332 | Bacteria>Terrabacteria_group>Actinomycetota>Coriobacteriia>Eggerthellales>Eggerthellaceae | - | Slackia_equolifaciens | K8U77_09670 | 12740..13735 | 498718 | DUF6079-family-protein,-partial-[Slackia-equolifaciens] |
| MBX6763713.1 | Type3.MTaseII.containing.BREX | SIGMA-HTH->?->Primpol->?->?->?->Thermonuclease->XIS-HTH->BrxB-BrxF-iSTAND->DUF6079-p*-> | 178 | Bacteria>Terrabacteria_group>Actinomycetota>Rubrobacteria>Rubrobacterales>Rubrobacteraceae | GCA_019690035.1 | Rubrobacteraceae_bacterium | IRY88_08530 | 17576..18109 | 2740537 | ATP-binding-protein-[Rubrobacteraceae-bacterium] |
| WP_273888494.1 | Type3.MTaseII.containing.BREX | SIGMA-HTH->?->Primpol->?->?->?->Thermonuclease->XIS-HTH->BrxB-BrxF-iSTAND->DUF6079*->Schlafen+HATPase+wHTH->N6-N4-METHYLASE->nSTAND->TUDOR+SF1-SF2-Helicase+iREase+CTD->PglZ->BrxA-DNA-binding-><-?\|\|?->HerA/FtsK-> | 1245 | Bacteria>Terrabacteria_group>Actinomycetota>Rubrobacteria>Rubrobacterales>Rubrobacteraceae | GCF_028617085.1 | Rubrobacter_naiadicus | PJB25_RS10020 | 42125..45862 | 1392641 | DUF6079-family-protein-[Rubrobacter-naiadicus] |
| MBV6489844.1 | Type3.MTaseII.containing.BREX | <-Schlafen+REase<-?<-BrxA-DNA-binding<-PglZ<-TUDOR+SF1-SF2-Helicase+iREase+CTD<-URI-DNase<-N6-N4-METHYLASE<-?<-?<-DUF6079*\|\|HTH->?->?->ArdCN+MPTase-> | 1250 | Bacteria>Terrabacteria_group>Armatimonadota>Fimbriimonadia>Fimbriimonadales>Fimbriimonadaceae | GCA_019187035.1 | Fimbriimonadaceae_bacterium | CNCCGFBP_00218 | 43734..47486 | 2682144 | hypothetical-protein-[Fimbriimonadaceae-bacterium] |
| MCK6631903.1 | Type3.MTaseII.containing.BREX | <-Schlafen+REase<-?<-BrxA-DNA-binding<-PglZ<-TUDOR+SF1-SF2-Helicase+iREase+CTD<-URI-DNase<-N6-N4-METHYLASE<-HEPN<-DUF6079*\|\|HTH->?->?->MPTase->?->REase+35exo-> | 1250 | Bacteria>Terrabacteria_group>Armatimonadota>Fimbriimonadia>Fimbriimonadales>Fimbriimonadaceae | GCA_023151345.1 | Fimbriimonadaceae_bacterium | L6Q31_05960 | 30666..34418 | 2682144 | DUF6079-family-protein-[Fimbriimonadaceae-bacterium] |
| GIV04203.1 | Type3.MTaseII.containing.BREX | DUF6079*->N6-N4-METHYLASE-><-?\|\|nSTAND->TUDOR+SF1-SF2-Helicase+iREase+CTD->PglZ->BrxA-DNA-binding-> | 1243 | Bacteria>Terrabacteria_group>Armatimonadota>Fimbriimonadia>Fimbriimonadales>unclassified_Fimbriimonadales | GCA_026003055.1 | Fimbriimonadales_bacterium | KatS3mg015_3033 | 865..4596 | 2682145 | MAG:-hypothetical-protein-KatS3mg015_3033-[Fimbriimonadales-bacterium] |
| NUM38593.1 | Type3.MTaseII.containing.BREX | <-REase+35exo<-?<-ArdCN+MPTase<-?<-?<-HTH\|\|DUF6079*->HEPN->N6-N4-METHYLASE->URI-DNase->TUDOR+SF1-SF2-Helicase+iREase+CTD->PglZ->BrxA-DNA-binding->?->Schlafen+REase-> | 1250 | Bacteria>Terrabacteria_group>Armatimonadota>unclassified_Armatimonadota | GCA_013359165.1 | Armatimonadota_bacterium | HUU52_05595 | 233692..237444 | 2033014 | hypothetical-protein-[Armatimonadota-bacterium] |
| MBN1459900.1 | Type3.MTaseII.containing.BREX | BrxB-BrxF-iSTAND->DUF6079*->N6-N4-METHYLASE->TUDOR+SF1-SF2-Helicase+iREase+CTD->PglZ->BrxA-DNA-binding->Classical-AAA->Peptidase_S8-> | 1243 | Bacteria>Terrabacteria_group>Armatimonadota>unclassified_Armatimonadota | GCA_016928475.1 | Armatimonadota_bacterium | JXA57_10195 | 1557..5288 | 2033014 | ATP-binding-protein-[Armatimonadota-bacterium] |
| MBN1461881.1 | Type3.MTaseII.containing.BREX | <-SF1-SF2-Helicase<-REase+SF1-SF2-Helicase\|\|XIS-HTH->BrxB-BrxF-iSTAND->DUF6079*->N6-N4-METHYLASE->?->?->?->TUDOR+SF1-SF2-Helicase+iREase+CTD->PglZ->BrxA-DNA-binding->SF1-SF2-Helicase-> | 1249 | Bacteria>Terrabacteria_group>Armatimonadota>unclassified_Armatimonadota | GCA_016928475.1 | Armatimonadota_bacterium | JXA57_20310 | 5275..9024 | 2033014 | ATP-binding-protein-[Armatimonadota-bacterium] |
| BCW97487.1 | Type3.MTaseII.containing.BREX | SMS-KaiC->NUDIX->?->BirA-HTH->His_Phos_1->tRNA->BrxB-BrxF-iSTAND->DUF6079*->N6-N4-METHYLASE->N6-N4-METHYLASE->HEPN->TUDOR+SF1-SF2-Helicase+iREase+CTD->?->PglZ->BrxA-DNA-binding->DUF4236+TPR-> | 1252 | Bacteria>Terrabacteria_group>Armatimonadota>unclassified_Armatimonadota | GCA_025998755.1 | Armatimonadota_bacterium | KatS3mg024_0314 | 379484..383242 | 2033014 | MAG:-hypothetical-protein-KatS3mg024_0314-[Armatimonadota-bacterium] |
| WP_236819147.1 | Type3.MTaseII.containing.BREX | <-BrxA-DNA-binding<-PglZ<-N6-N4-METHYLASE<-DUF6079*<-?<-?<-?<-?<-?<-?<-SF1-SF2-Helicase+REase-p | 1245 | Bacteria>Terrabacteria_group>Bacillota>Bacilli>Bacillales>Alicyclobacillaceae | - | Alicyclobacillus_tolerans | LLE49_24515 | 6515..10252 | 90970 | DUF6079-family-protein-[Alicyclobacillus-tolerans] |
| MCF8567890.1 | Type3.MTaseII.containing.BREX | <-BrxA-DNA-binding<-PglZ<-N6-N4-METHYLASE<-DUF6079*<-?<-?<-?<-?<-?<-?<-SF1-SF2-Helicase+REase-p | 1245 | Bacteria>Terrabacteria_group>Bacillota>Bacilli>Bacillales>Alicyclobacillaceae | GCA_021739485.1 | Alicyclobacillus_tolerans | LLE49_24515 | 6515..10252 | 90970 | DUF6079-family-protein-[Alicyclobacillus-tolerans] |
| WP_052300518.1 | Type3.MTaseII.containing.BREX | Pkinase+STYKIN->?-><-PIN<-PhdYeFM_antitox\|\|?->?->BrxB-BrxF-iSTAND->DUF6079-p*-><-HTH_Tnp_1+HTH_21+rve+rve_3\|\|DUF6079->N6-N4-METHYLASE->PglZ->BrxA-DNA-binding-><-DDE_Tnp | 495 | Bacteria>Terrabacteria_group>Bacillota>Bacilli>Bacillales>Alicyclobacillaceae | GCF_000092905.1 | Kyrpidia_tusciae | BTUS_RS01970 | 375338..376825 | 33943 | DUF6079-family-protein-[Kyrpidia-tusciae] |
| WP_026975941.1 | Type3.MTaseII.containing.BREX | DDE_Tnp_IS66+DDE_Tnp_IS66_C-><-DUF6079*<-BrxB-BrxF-iSTAND<-ZnR+TOPRIM+AAA-helicase+wHTH<-XIS-HTH<-NUDIX | 1243 | Bacteria>Terrabacteria_group>Bacillota>Bacilli>Bacillales>Alicyclobacillaceae | GCF_000429525.1 | Alicyclobacillus_contaminans | G463_RS0115865 | 2132..5863 | 392016 | DUF6079-family-protein-[Alicyclobacillus-contaminans] |
| WP_272908837.1 | Type3.MTaseII.containing.BREX | <-BrxA-DNA-binding<-PglZ<-N6-N4-METHYLASE<-DUF6079*<-?<-?<-?<-?<-?<-?<-SF1-SF2-Helicase+REase-p | 1246 | Bacteria>Terrabacteria_group>Bacillota>Bacilli>Bacillales>Alicyclobacillaceae | GCF_021739485.1 | Alicyclobacillus_tolerans | LLE49_RS24360 | 6515..10255 | 90970 | DUF6079-family-protein-[Alicyclobacillus-tolerans] |
| WP_077114048.1 | Type3.MTaseII.containing.BREX | <-ABC-ATPase+SbcC\|\|DNA_METHYLASE-><-REase<-AAA-ATPase-p<-TUDOR+SF1-SF2-Helicase+iREase+CTD+REase<-BrxA-DNA-binding<-PglZ<-N6-N4-METHYLASE<-DUF6079*<-BrxB-BrxF-iSTAND<-DDE_5+DDE_Tnp_1<-DrmB<-DrmA<-HTH+HTH+SF1-SF2-Helicase\|\|?-><-?<-Phage_CRI+Phage_X | 1233 | Bacteria>Terrabacteria_group>Bacillota>Bacilli>Bacillales>Bacillaceae | - | Bacillus_sp._VT-16-64 | BLX87_RS22765 | 22398..26099 | 1917019 | DUF6079-family-protein-[Bacillus-sp.-VT-16-64] |
| EDX56188.1 | Type3.MTaseII.containing.BREX | <-MerR-HTH<-?<-?\|\|REase-><-TUDOR+SF1-SF2-Helicase+iREase+CTD+REase<-BrxA-DNA-binding<-PglZ<-N6-N4-METHYLASE<-DUF6079*<-BrxB-BrxF-iSTAND<-DrmB<-DrmA<-wHTH+WYL+WCX<-?<-?<-REC+CITB-HTH+ZDNA-HTH<-IONCHANNEL+PAS+HISKIN+MORC | 1232 | Bacteria>Terrabacteria_group>Bacillota>Bacilli>Bacillales>Bacillaceae | GCA_000181595.1 | Bacillus_cereus_W | BCW_0922 | 72761..76459 | 405917 | virulence-associated-protein-[Bacillus-cereus-W] |
| AKM18684.1 | Type3.MTaseII.containing.BREX | Y1_Tnp->HTH_OrfB_IS605+OrfB_IS605+OrfB_Zn_ribbon-><-?\|\|BrxB-BrxF-iSTAND->DUF6079*->N6-N4-METHYLASE->PglZ->BrxA-DNA-binding->TUDOR+SF1-SF2-Helicase+iREase+CTD+REase->REase-><-DDE_Tnp_1 | 1232 | Bacteria>Terrabacteria_group>Bacillota>Bacilli>Bacillales>Bacillaceae | GCA_001028085.1 | Geobacillus_sp._12AMOR1 | GARCT_01391 | 1367234..1370932 | 1629723 | hypothetical-protein-GARCT_01391-[Geobacillus-sp.-12AMOR1] |
| KZE90469.1 | Type3.MTaseII.containing.BREX | HTH+MPTase->PIN-><-?<-?\|\|BrxB-BrxF-iSTAND->DUF6079*->N6-N4-METHYLASE->PglZ->BrxA-DNA-binding->TUDOR+SF1-SF2-Helicase+iREase+CTD+REase->HTH_Tnp_ISL3+DDE_Tnp_ISL3->?->?-><-OrfB_IS605+OrfB_Zn_ribbon\|\|HTH_3-> | 1233 | Bacteria>Terrabacteria_group>Bacillota>Bacilli>Bacillales>Bacillaceae | GCA_001620045.1 | Geobacillus_stearothermophilus | AVP43_03286 | 55510..59211 | 1422 | hypothetical-protein-AVP43_03286-[Geobacillus-stearothermophilus] |
| ASN07261.1 | Type3.MTaseII.containing.BREX | SbcC-><-?<-PADR-HTH\|\|SPOUT-><-?\|\|BrxB-BrxF-iSTAND->DUF6079*->N6-N4-METHYLASE->PglZ->BrxA-DNA-binding->?->HEPN->MarR-HTH->?->Glyoxalase-> | 1233 | Bacteria>Terrabacteria_group>Bacillota>Bacilli>Bacillales>Bacillaceae | GCA_002224365.1 | Virgibacillus_necropolis | CFK40_01940 | 358932..362633 | 163877 | ATPase-[Virgibacillus-necropolis] |
| PAD23515.1 | Type3.MTaseII.containing.BREX | <-Arm-DNA-bind_4+Phage_int_SAM_5+Phage_integrase<-?\|\|GDResolvase+HIN-HTH->BrxB-BrxF-iSTAND->DUF6079*->N6-N4-METHYLASE->PglZ->BrxA-DNA-binding->TUDOR+SF1-SF2-Helicase+iREase+CTD+REase->SIR2->HerA/FtsK->HKD-DNase->SbcC->HNH-> | 1234 | Bacteria>Terrabacteria_group>Bacillota>Bacilli>Bacillales>Bacillaceae | GCA_002271635.1 | Niallia_circulans | CHH62_21825 | 4343..8047 | 1397 | ATPase-[Niallia-circulans] |
| PED47023.1 | Type3.MTaseII.containing.BREX | <-DnaB+TnsC-ATPase<-DnaB_2\|\|?-><-?<-TUDOR+SF1-SF2-Helicase+iREase+CTD+REase<-BrxA-DNA-binding<-PglZ<-N6-N4-METHYLASE<-DUF6079*<-BrxB-BrxF-iSTAND<-DrmB<-DrmA<-wHTH+WYL+WCX<-?<-?<-REC+CITB-HTH+ZDNA-HTH<-IONCHANNEL+PAS+HISKIN+MORC | 1232 | Bacteria>Terrabacteria_group>Bacillota>Bacilli>Bacillales>Bacillaceae | GCA_002552505.1 | Bacillus_cereus | CON49_25665 | 28116..31814 | 1396 | ATPase-[Bacillus-cereus] |
| PEY26875.1 | Type3.MTaseII.containing.BREX | IONCHANNEL+PAS+HISKIN+MORC->REC+CITB-HTH+ZDNA-HTH->?->?->wHTH+WYL+WCX->DrmA->DrmB->BrxB-BrxF-iSTAND->DUF6079*->N6-N4-METHYLASE->PglZ->BrxA-DNA-binding->TUDOR+SF1-SF2-Helicase+iREase+CTD+REase-><-REase\|\|?->?->MerR-HTH-> | 1232 | Bacteria>Terrabacteria_group>Bacillota>Bacilli>Bacillales>Bacillaceae | GCA_002559605.1 | Bacillus_anthracis | CN340_12720 | 51340..55038 | 1392 | ATPase-[Bacillus-anthracis] |
| PFA00385.1 | Type3.MTaseII.containing.BREX | IONCHANNEL+PAS+HISKIN+MORC->REC+CITB-HTH+ZDNA-HTH->?->?->wHTH+WYL+WCX->DrmA->DrmB->BrxB-BrxF-iSTAND->DUF6079*->N6-N4-METHYLASE->PglZ->BrxA-DNA-binding->TUDOR+SF1-SF2-Helicase+iREase+CTD+REase->?-><-?\|\|DnaB_2->DnaB+TnsC-ATPase-> | 1232 | Bacteria>Terrabacteria_group>Bacillota>Bacilli>Bacillales>Bacillaceae | GCA_002561255.1 | Bacillus_cereus | CN375_05445 | 15872..19570 | 1396 | ATPase-[Bacillus-cereus] |
| PFT90557.1 | Type3.MTaseII.containing.BREX | <-DnaB+TnsC-ATPase<-DnaB_2\|\|?-><-?<-TUDOR+SF1-SF2-Helicase+iREase+CTD+REase<-BrxA-DNA-binding<-PglZ<-N6-N4-METHYLASE<-DUF6079*<-BrxB-BrxF-iSTAND<-DrmB<-DrmA<-wHTH+WYL+WCX<-?<-?<-REC+CITB-HTH+ZDNA-HTH<-IONCHANNEL+PAS+HISKIN+MORC | 1232 | Bacteria>Terrabacteria_group>Bacillota>Bacilli>Bacillales>Bacillaceae | GCA_002568375.1 | Bacillus_cereus | COK66_16930 | 34253..37951 | 1396 | ATPase-[Bacillus-cereus] |
| PGQ77680.1 | Type3.MTaseII.containing.BREX | <-RNase<-wHTH+WYL+WCX-p<-?<-TUDOR+SF1-SF2-Helicase+iREase+CTD+REase<-BrxA-DNA-binding<-PglZ<-N6-N4-METHYLASE<-DUF6079*<-BrxB-BrxF-iSTAND<-DrmB<-DrmA<-wHTH+WYL+WCX<-?<-REase<-?<-REC+CITB-HTH+ZDNA-HTH<-IONCHANNEL+PAS+HISKIN+MORC | 1232 | Bacteria>Terrabacteria_group>Bacillota>Bacilli>Bacillales>Bacillaceae | GCA_002576945.1 | Bacillus_anthracis | COA15_22160 | 380582..384280 | 1392 | ATPase-[Bacillus-anthracis] |
| MBG0973007.1 | Type3.MTaseII.containing.BREX | REase->?->wHTH+WYL+WCX->DrmA->DrmB->BrxB-BrxF-iSTAND->DUF6079*->N6-N4-METHYLASE->PglZ->BrxA-DNA-binding->TUDOR+SF1-SF2-Helicase+iREase+CTD+REase->?->SbcC->?->DnaB_2->DnaB+TnsC-ATPase-> | 1232 | Bacteria>Terrabacteria_group>Bacillota>Bacilli>Bacillales>Bacillaceae | GCA_015712745.1 | Bacillus_sp._SRB3LM | CN504_21520 | 9373..13071 | 2608689 | ATP-binding-protein-[Bacillus-sp.-SRB3LM] |
| MBL4952760.1 | Type3.MTaseII.containing.BREX | ABC-ATPase+SbcC->ABC-ATPase+SbcC-><-SpoOE-like\|\|DsbD->?-><-?\|\|tRNA->BrxB-BrxF-iSTAND->DUF6079*->N6-N4-METHYLASE->PglZ->BrxA-DNA-binding->?->?->?->?->SbcC-> | 1233 | Bacteria>Terrabacteria_group>Bacillota>Bacilli>Bacillales>Bacillaceae | GCA_016765675.1 | Neobacillus_paridis | JK635_11110 | 423062..426763 | 2803862 | ATP-binding-protein-[Neobacillus-paridis] |
| MBO0997207.1 | Type3.MTaseII.containing.BREX | <-RVT<-?\|\|Phage_GPA->XIS-HTH->?-><-BrxA-DNA-binding<-PglZ<-N6-N4-METHYLASE<-DUF6079*<-BrxB-BrxF-iSTAND\|\|XIS-HTH->Arm-DNA-bind_4+Phage_int_SAM_3+Phage_integrase-><-tRNA<-?\|\|wHTH+WYL+WCX-p-><-?\|\|ABhydrolase->?->ACET-> | 1233 | Bacteria>Terrabacteria_group>Bacillota>Bacilli>Bacillales>Bacillaceae | GCA_017355165.1 | Bacillus_sp._SD075 | IOC57_05435 | 198059..201760 | 2781732 | ATP-binding-protein-[Bacillus-sp.-SD075] |
| MBO8173526.1 | Type3.MTaseII.containing.BREX | <-HIN-HTH<-TUDOR+SF1-SF2-Helicase+iREase+CTD+REase<-BrxA-DNA-binding<-PglZ<-N6-N4-METHYLASE<-DUF6079*<-BrxB-BrxF-iSTAND<-35exo | 1232 | Bacteria>Terrabacteria_group>Bacillota>Bacilli>Bacillales>Bacillaceae | GCA_017656255.1 | Bacillaceae_bacterium | HOZ33_16790 | 11282..14980 | 1889774 | ATP-binding-protein-[Bacillaceae-bacterium] |
| MCD8511941.1 | Type3.MTaseII.containing.BREX | <-ABC-ATPase+SbcC<-?<-?<-BrxA-DNA-binding<-PglZ<-N6-N4-METHYLASE<-DUF6079-p<-?<-DUF6079-p*<-?<-ABC-ATPase+SbcC<-ACET<-HD<-HKD-DNase+HKD-DNase<-DUF4007\|\|SF1-SF2-Helicase-><-?<-HISKIN | 750 | Bacteria>Terrabacteria_group>Bacillota>Bacilli>Bacillales>Bacillaceae | GCA_021208645.1 | Bacillus_sp._(in:_firmicutes) | LRY73_20215 | 22163..24415 | 1409 | DUF6079-family-protein-[Bacillus-sp.-(in:-firmicutes)] |
| WP_007504072.1 | Type3.MTaseII.containing.BREX | RNase->?->?->?-><-?\|\|?-><-?<-?\|\|BrxB-BrxF-iSTAND->DUF6079-p*->DUF6079->N6-N4-METHYLASE-> | 608 | Bacteria>Terrabacteria_group>Bacillota>Bacilli>Bacillales>Bacillaceae | GCF_000218765.1 | Caldalkalibacillus_thermarum | CATHTA2_RS16890 | 15578..17404 | 296745 | DUF6079-family-protein-[Caldalkalibacillus-thermarum] |
| WP_003349666.1 | Type3.MTaseII.containing.BREX | BrxB-BrxF-iSTAND->DUF6079*->N6-N4-METHYLASE->PglZ->BrxA-DNA-binding->TUDOR+SF1-SF2-Helicase+iREase+CTD+REase->?-><-DDE_Tnp<-DDE_Tnp_ISL3<-?\|\|RVT-> | 1235 | Bacteria>Terrabacteria_group>Bacillota>Bacilli>Bacillales>Bacillaceae | GCF_000262735.1 | Bacillus_methanolicus | MGA3_RS15495 | 40396..44103 | 1471 | DUF6079-family-protein-[Bacillus-methanolicus] |
| WP_000966765.1 | Type3.MTaseII.containing.BREX | <-Resolvase+Recombinase+ZnR<-?\|\|?-><-?\|\|GDResolvase+Recombinase-><-TUDOR+SF1-SF2-Helicase+iREase+CTD+REase<-BrxA-DNA-binding<-PglZ<-N6-N4-METHYLASE<-DUF6079*<-BrxB-BrxF-iSTAND<-?\|\|<-?<-SWACOS<-PAS+PAS+GGDEF<-SPOUT | 1234 | Bacteria>Terrabacteria_group>Bacillota>Bacilli>Bacillales>Bacillaceae | GCF_000291665.1 | Bacillus_cereus | ICU_RS21145 | 4139883..4143587 | 1396 | DUF6079-family-protein-[Bacillus-cereus] |
| WP_019382828.1 | Type3.MTaseII.containing.BREX | <-HNH\|\|HTH_Tnp+HTH_Tnp-><-HIN-HTH\|\|PadR-HTH-><-TUDOR+SF1-SF2-Helicase+iREase+CTD+REase<-BrxA-DNA-binding<-PglZ<-N6-N4-METHYLASE<-DUF6079*<-BrxB-BrxF-iSTAND<-GDResolvase+HIN-HTH<-NUDIX\|\|?->GDResolvase+Recombinase+ZnR-><-SPOUT<-?<-Trypsin+PDZ | 1234 | Bacteria>Terrabacteria_group>Bacillota>Bacilli>Bacillales>Bacillaceae | GCF_000294775.2 | Cytobacillus_oceanisediminis | A361_RS26950 | 5414558..5418262 | 665099 | DUF6079-family-protein-[Cytobacillus-oceanisediminis] |
| WP_034748802.1 | Type3.MTaseII.containing.BREX | TIR->?->?-><-?<-HTH_3\|\|HEPN->REase+SF1-SF2-Helicase->REase->BrxB-BrxF-iSTAND->DUF6079*->N6-N4-METHYLASE->PglZ-p->PglZ-p->BrxA-DNA-binding->TUDOR+SF1-SF2-Helicase+iREase+CTD+REase-p->TUDOR+SF1-SF2-Helicase+iREase+CTD+REase-p->TUDOR+SF1-SF2-Helicase+iREase+CTD+REase-p->REase->ParB+HNH-> | 1233 | Bacteria>Terrabacteria_group>Bacillota>Bacilli>Bacillales>Bacillaceae | GCF_000513095.1 | Halalkalibacter_wakoensis | JCM9140_RS16205 | 9393..13094 | 127891 | DUF6079-family-protein-[Halalkalibacter-wakoensis] |
| WP_036836008.1 | Type3.MTaseII.containing.BREX | SPOUT-><-?<-wHTH+HNH<-?<-wHTH\|\|?->BrxB-BrxF-iSTAND->DUF6079*->N6-N4-METHYLASE->PglZ->BrxA-DNA-binding->TUDOR+SF1-SF2-Helicase+iREase+CTD+REase-p->TUDOR+SF1-SF2-Helicase+iREase+CTD+REase-p-> | 1226 | Bacteria>Terrabacteria_group>Bacillota>Bacilli>Bacillales>Bacillaceae | GCF_000775615.1 | Pontibacillus_litoralis | N784_RS14330 | 18020..21700 | 516703 | DUF6079-family-protein-[Pontibacillus-litoralis] |
| WP_001994602.1 | Type3.MTaseII.containing.BREX | IONCHANNEL+PAS+HISKIN+MORC->REC+CITB-HTH+ZDNA-HTH->?->REase->?->wHTH+WYL+WCX->DrmA->DrmB->BrxB-BrxF-iSTAND->DUF6079*->N6-N4-METHYLASE->PglZ->BrxA-DNA-binding->TUDOR+SF1-SF2-Helicase+iREase+CTD+REase->?->?->RNase-> | 1232 | Bacteria>Terrabacteria_group>Bacillota>Bacilli>Bacillales>Bacillaceae | GCF_000832865.1 | Bacillus_cereus | AK40_RS29415 | 4972014..4975712 | 1396 | DUF6079-family-protein-[Bacillus-cereus] |
| WP_044893683.1 | Type3.MTaseII.containing.BREX | <-MazG<-HNH<-?\|\|?-><-?<-TUDOR+SF1-SF2-Helicase+iREase+CTD+REase<-BrxA-DNA-binding<-PglZ<-N6-N4-METHYLASE<-DUF6079*<-BrxB-BrxF-iSTAND | 1233 | Bacteria>Terrabacteria_group>Bacillota>Bacilli>Bacillales>Bacillaceae | GCF_000948175.1 | Bacillus_alveayuensis | LG50_RS18070 | 22446..26147 | 279215 | DUF6079-family-protein-[Bacillus-alveayuensis] |
| WP_082798212.1 | Type3.MTaseII.containing.BREX | OrfB_Zn_ribbon->HIRAN->?->BrxB-BrxF-iSTAND->DUF6079*->N6-N4-METHYLASE->PglZ->BrxA-DNA-binding->TUDOR+SF1-SF2-Helicase+iREase+CTD+REase->MazG->REase+SF1-SF2-Helicase->zf-ISL3+HTH_Tnp_ISL3-> | 1233 | Bacteria>Terrabacteria_group>Bacillota>Bacilli>Bacillales>Bacillaceae | GCF_001587455.1 | Parageobacillus_toebii | B4110_RS17500 | 3743..7444 | 153151 | DUF6079-family-protein-[Parageobacillus-toebii] |
| WP_061808755.1 | Type3.MTaseII.containing.BREX | HTH_3->?->SF1-SF2-Helicase+REase->?->?->HKD-DNase-><-CHTH\|\|REase+SF1-SF2-Helicase->BrxB-BrxF-iSTAND->DUF6079*->N6-N4-METHYLASE->PglZ->BrxA-DNA-binding->TUDOR+SF1-SF2-Helicase+iREase+CTD+REase->ParB+HNH->?->OLD-ABC->HNH-> | 1226 | Bacteria>Terrabacteria_group>Bacillota>Bacilli>Bacillales>Bacillaceae | GCF_001591825.1 | Rossellomorea_vietnamensis | BVI01S_RS00200 | 36378..40058 | 218284 | DUF6079-family-protein-[Rossellomorea-vietnamensis] |
| WP_078596332.1 | Type3.MTaseII.containing.BREX | <-HTH_3\|\|NUDIX-><-SAD+HNH<-NERD+SF1-SF2-Helicase\|\|?-><-TUDOR+SF1-SF2-Helicase+iREase+CTD+REase<-BrxA-DNA-binding<-PglZ<-N6-N4-METHYLASE<-DUF6079*<-BrxB-BrxF-iSTAND<-DrmB<-DrmA<-HTH+HTH+SF1-SF2-Helicase+HTH+HTH+SRA\|\|?-><-TIR<-?<-REase | 1233 | Bacteria>Terrabacteria_group>Bacillota>Bacilli>Bacillales>Bacillaceae | GCF_002019695.1 | Evansella_clarkii | BK582_RS17790 | 3802313..3806014 | 79879 | DUF6079-family-protein-[Evansella-clarkii] |
| WP_227001846.1 | Type3.MTaseII.containing.BREX | SbcC-><-?<-PADR-HTH\|\|SPOUT-><-?\|\|?->BrxB-BrxF-iSTAND->DUF6079*->N6-N4-METHYLASE->PglZ->BrxA-DNA-binding->?->HEPN->MarR-HTH->?->Glyoxalase-> | 1234 | Bacteria>Terrabacteria_group>Bacillota>Bacilli>Bacillales>Bacillaceae | GCF_002224365.1 | Virgibacillus_necropolis | CFK40_RS01940 | 358929..362633 | 163877 | DUF6079-family-protein-[Virgibacillus-necropolis] |
| WP_235850150.1 | Type3.MTaseII.containing.BREX | <-Arm-DNA-bind_4+Phage_int_SAM_5+Phage_integrase<-?\|\|GDResolvase+HIN-HTH->BrxB-BrxF-iSTAND->DUF6079*->N6-N4-METHYLASE->PglZ->BrxA-DNA-binding->TUDOR+SF1-SF2-Helicase+iREase+CTD+REase->?->SIR2->HerA/FtsK->HKD-DNase->SbcC-> | 1260 | Bacteria>Terrabacteria_group>Bacillota>Bacilli>Bacillales>Bacillaceae | GCF_002271635.1 | Niallia_circulans | CHH62_RS21835 | 4265..8047 | 1397 | DUF6079-family-protein-[Niallia-circulans] |
| WP_098172532.1 | Type3.MTaseII.containing.BREX | XIS-HTH->ZnR+TOPRIM->ZnR+TOPRIM+AAA-helicase+wHTH->?->?-><-?\|\|DrmA->DrmB->BrxB-BrxF-iSTAND->DUF6079*->N6-N4-METHYLASE->PglZ->BrxA-DNA-binding->TUDOR+SF1-SF2-Helicase+iREase+CTD+REase->NTP_transf->HTH->\|\|?-><-MerR-HTH | 1231 | Bacteria>Terrabacteria_group>Bacillota>Bacilli>Bacillales>Bacillaceae | GCF_002555445.1 | Bacillus_sp._AFS017336 | CN601_RS23830 | 64001..67696 | 2033489 | DUF6079-family-protein-[Bacillus-sp.-AFS017336] |
| WP_098349130.1 | Type3.MTaseII.containing.BREX | IONCHANNEL+PAS+HISKIN+MORC->REC+CITB-HTH+ZDNA-HTH->?->?->wHTH+WYL+WCX->DrmA->DrmB->BrxB-BrxF-iSTAND->DUF6079*->N6-N4-METHYLASE->PglZ->BrxA-DNA-binding->TUDOR+SF1-SF2-Helicase+iREase+CTD+REase->?->?-><-?\|\|DnaB_2->DnaB+TnsC-ATPase-> | 1232 | Bacteria>Terrabacteria_group>Bacillota>Bacilli>Bacillales>Bacillaceae | GCF_002561255.1 | Bacillus_cereus | CN375_RS05435 | 15872..19570 | 1396 | DUF6079-family-protein-[Bacillus-cereus] |
| WP_098658108.1 | Type3.MTaseII.containing.BREX | <-PALP+ACT_4<-?<-?\|\|<-?<-PADR-HTH<-BrxA-DNA-binding<-PglZ<-?<-N6-N4-METHYLASE<-DUF6079*<-BrxB-BrxF-iSTAND<-TOPRIM+Topoisom_bac<-HIN-HTH\|\|HTH_3-><-?<-Phage_integrase | 1228 | Bacteria>Terrabacteria_group>Bacillota>Bacilli>Bacillales>Bacillaceae | GCF_002570105.1 | Bacillus_toyonensis | COM09_RS29270 | 36276..39962 | 155322 | DUF6079-family-protein-[Bacillus-toyonensis] |
| WP_098582494.1 | Type3.MTaseII.containing.BREX | IONCHANNEL+PAS+HISKIN+MORC->REC+CITB-HTH+ZDNA-HTH->?->?->wHTH+WYL+WCX->DrmA->DrmB->BrxB-BrxF-iSTAND->DUF6079*->N6-N4-METHYLASE->PglZ->BrxA-DNA-binding->TUDOR+SF1-SF2-Helicase+iREase+CTD+REase->?->?->?-><-?\|\|DnaB_2->DnaB+TnsC-ATPase-> | 1232 | Bacteria>Terrabacteria_group>Bacillota>Bacilli>Bacillales>Bacillaceae | GCF_002574415.1 | Bacillus | CN915_RS11630 | 19729..23427 | 1386 | MULTISPECIES:-DUF6079-family-protein-[Bacillus] |
| WP_098606674.1 | Type3.MTaseII.containing.BREX | IONCHANNEL+PAS+HISKIN+MORC->REC+CITB-HTH+ZDNA-HTH->?->?->wHTH+WYL+WCX->DrmA->DrmB->BrxB-BrxF-iSTAND->DUF6079*->N6-N4-METHYLASE->PglZ->BrxA-DNA-binding->TUDOR+SF1-SF2-Helicase+iREase+CTD+REase->?->?->?-><-?\|\|DnaB_2->DnaB+TnsC-ATPase-> | 1232 | Bacteria>Terrabacteria_group>Bacillota>Bacilli>Bacillales>Bacillaceae | GCF_002575835.1 | Bacillus_cereus | COA03_RS10365 | 67720..71418 | 1396 | DUF6079-family-protein-[Bacillus-cereus] |
| WP_116555899.1 | Type3.MTaseII.containing.BREX | BrxB-BrxF-iSTAND->DUF6079*->N6-N4-METHYLASE->PglZ->BrxA-DNA-binding->McrB-AAA->McrC-NTD+REase->PadR-HTH->?->TPR-><-DSBH | 1233 | Bacteria>Terrabacteria_group>Bacillota>Bacilli>Bacillales>Bacillaceae | GCF_003097615.1 | Pueribacillus_theae | DCC39_RS15950 | 661..4362 | 2171751 | DUF6079-family-protein-[Pueribacillus-theae] |
| WP_208731622.1 | Type3.MTaseII.containing.BREX | IONCHANNEL+PAS+HISKIN+MORC->REC+CITB-HTH+ZDNA-HTH->?->?->wHTH+WYL+WCX->DrmA->DrmB->BrxB-BrxF-iSTAND->DUF6079*->N6-N4-METHYLASE->PglZ->BrxA-DNA-binding->TUDOR+SF1-SF2-Helicase+iREase+CTD+REase->?->wHTH+WYL+WCX-p->?-><-?\|\|DnaB_2-> | 1232 | Bacteria>Terrabacteria_group>Bacillota>Bacilli>Bacillales>Bacillaceae | GCF_003355115.1 | Bacillus_sp._COPE52 | DPQ31_RS04895 | 984162..987860 | 2233998 | DUF6079-family-protein-[Bacillus-sp.-COPE52] |
| WP_116277929.1 | Type3.MTaseII.containing.BREX | BrxB-BrxF-iSTAND->DUF6079*->N6-N4-METHYLASE->PglZ->BrxA-DNA-binding->?->?->?->?->ABC-ATPase+SbcC-> | 1233 | Bacteria>Terrabacteria_group>Bacillota>Bacilli>Bacillales>Bacillaceae | GCF_003399705.1 | Virgibacillus_dokdonensis | CAI16_RS07730 | 2030..5731 | 302167 | DUF6079-family-protein-[Virgibacillus-dokdonensis] |
| WP_119115140.1 | Type3.MTaseII.containing.BREX | <-D5ATPase<-?<-CHTH\|\|?-><-TUDOR+SF1-SF2-Helicase+iREase+CTD+REase<-BrxA-DNA-binding<-PglZ<-N6-N4-METHYLASE<-DUF6079*<-BrxB-BrxF-iSTAND<-?<-ParB+HNH<-?<-?<-?<-GDResolvase+Recombinase+ZnR | 1233 | Bacteria>Terrabacteria_group>Bacillota>Bacilli>Bacillales>Bacillaceae | GCF_003570725.1 | Peribacillus_asahii | D1953_RS00175 | 14623..18324 | 228899 | DUF6079-family-protein-[Peribacillus-asahii] |
| WP_126294755.1 | Type3.MTaseII.containing.BREX | <-HEPN+HTH<-TUDOR+SF1-SF2-Helicase+iREase+CTD+REase<-BrxA-DNA-binding<-PglZ<-N6-N4-METHYLASE<-N6-N4-METHYLASE<-DUF6079*<-BrxB-BrxF-iSTAND<-HHH_4+HTH+wHTH+SF1-SF2-Helicase+REase<-?<-?<-?\|\|HTH_3-> | 1233 | Bacteria>Terrabacteria_group>Bacillota>Bacilli>Bacillales>Bacillaceae | GCF_003966145.1 | Lysinibacillus_telephonicus | EKG35_RS12280 | 21521..25222 | 1714840 | DUF6079-family-protein-[Lysinibacillus-telephonicus] |
| WP_131246810.1 | Type3.MTaseII.containing.BREX | IONCHANNEL+PAS+HISKIN+MORC->REC+CITB-HTH+ZDNA-HTH->?->REase->wHTH+WYL+WCX->DrmA->DrmB->BrxB-BrxF-iSTAND->DUF6079*->?->N6-N4-METHYLASE->PglZ->BrxA-DNA-binding->TUDOR+SF1-SF2-Helicase+iREase+CTD+REase->?->?-><-?\|\|DnaB_2-> | 1232 | Bacteria>Terrabacteria_group>Bacillota>Bacilli>Bacillales>Bacillaceae | GCF_004330345.1 | Bacillus_mycoides | EOM27_RS18685 | 84585..88283 | 1405 | DUF6079-family-protein-[Bacillus-mycoides] |
| WP_146817812.1 | Type3.MTaseII.containing.BREX | <-GDResolvase+HIN-HTH<-?<-TUDOR+SF1-SF2-Helicase+iREase+CTD+REase<-BrxA-DNA-binding<-PglZ<-N6-N4-METHYLASE<-DUF6079*<-BrxB-BrxF-iSTAND<-?<-HTH_3<-?<-ZnR+TOPRIM+AAA-helicase+wHTH<-?\|\|HTH_3->Arm-DNA-bind_4+Phage_int_SAM_5+Phage_integrase-> | 1227 | Bacteria>Terrabacteria_group>Bacillota>Bacilli>Bacillales>Bacillaceae | GCF_007991275.1 | Alkalibacillus_haloalkaliphilus | AHA02nite_RS12530 | 64321..68004 | 94136 | DUF6079-family-protein-[Alkalibacillus-haloalkaliphilus] |
| WP_152155470.1 | Type3.MTaseII.containing.BREX | <-MerR-HTH<-?<-?<-?<-TUDOR+SF1-SF2-Helicase+iREase+CTD+REase<-BrxA-DNA-binding<-PglZ<-N6-N4-METHYLASE<-DUF6079*<-BrxB-BrxF-iSTAND<-DrmB<-DrmA<-wHTH+WYL+WCX<-?<-REase<-?<-REC+CITB-HTH+ZDNA-HTH<-IONCHANNEL+PAS+HISKIN+MORC | 1232 | Bacteria>Terrabacteria_group>Bacillota>Bacilli>Bacillales>Bacillaceae | GCF_009183765.1 | Bacillus_sp._B3-WWTP-C-10-D-3 | GBN83_RS14870 | 21828..25526 | 2653217 | DUF6079-family-protein-[Bacillus-sp.-B3-WWTP-C-10-D-3] |
| WP_153253563.1 | Type3.MTaseII.containing.BREX | <-VirD4-TraG+VirD4-FtsK<-?\|\|MerR-HTH-><-?\|\|HTH_3-><-?\|\|?->BrxB-BrxF-iSTAND->DUF6079*->N6-N4-METHYLASE->PglZ->BrxA-DNA-binding->TUDOR+SF1-SF2-Helicase+iREase+CTD+REase-><-SpoOE-like<-?<-HAD | 1234 | Bacteria>Terrabacteria_group>Bacillota>Bacilli>Bacillales>Bacillaceae | GCF_009497105.1 | Priestia_megaterium | GFV16_13160 | 12206..15910 | 1404 | DUF6079-family-protein-[Priestia-megaterium] |
| WP_160269222.1 | Type3.MTaseII.containing.BREX | BrxB-BrxF-iSTAND->DUF6079*->N6-N4-METHYLASE->PglZ->BrxA-DNA-binding->TUDOR+SF1-SF2-Helicase+iREase+CTD+REase->MazG-><-DDE_5+DDE_Tnp_2 | 1233 | Bacteria>Terrabacteria_group>Bacillota>Bacilli>Bacillales>Bacillaceae | GCF_009914195.1 | Geobacillus_stearothermophilus | EPB69_RS06760 | 1331297..1334998 | 1422 | DUF6079-family-protein-[Geobacillus-stearothermophilus] |
| WP_161986542.1 | Type3.MTaseII.containing.BREX | ADH_N+ADH_zinc_N-><-?\|\|?->SPOUT->BrxB-BrxF-iSTAND->DUF6079*->N6-N4-METHYLASE->PglZ->BrxA-DNA-binding->TUDOR+SF1-SF2-Helicase+iREase+CTD+REase-><-?\|\|ABC-ATPase-> | 1234 | Bacteria>Terrabacteria_group>Bacillota>Bacilli>Bacillales>Bacillaceae | GCF_009937745.1 | Bacillus_subtilis | GPJ55_RS17625 | 3379797..3383501 | 1423 | DUF6079-family-protein-[Bacillus-subtilis] |
| WP_168409848.1 | Type3.MTaseII.containing.BREX | Phage_integrase->?->Phage_integrase->?-><-?<-HTH_3\|\|HIN-HTH->TOPRIM+Topoisom_bac->BrxB-BrxF-iSTAND->DUF6079*->N6-N4-METHYLASE->?->PglZ->BrxA-DNA-binding->\|\|?->?-><-?\|\|HAMP+MA-> | 1228 | Bacteria>Terrabacteria_group>Bacillota>Bacilli>Bacillales>Bacillaceae | GCF_012395385.1 | Bacillus_cereus | HB669_RS25765 | 125607..129293 | 1396 | DUF6079-family-protein-[Bacillus-cereus] |
| WP_169099403.1 | Type3.MTaseII.containing.BREX | <-N6-METHYLASE\|\|?-><-TUDOR+SF1-SF2-Helicase+iREase+CTD+REase<-BrxA-DNA-binding<-PglZ<-N6-N4-METHYLASE<-DUF6079*<-BrxB-BrxF-iSTAND<-McrC-NTD+REase<-?<-SbcC+ABC-ATPase<-RADICAL-SAM | 1233 | Bacteria>Terrabacteria_group>Bacillota>Bacilli>Bacillales>Bacillaceae | GCF_012843265.1 | Bacillus_sp._DNRA2 | HHD41_RS13370 | 109180..112881 | 2723053 | DUF6079-family-protein-[Bacillus-sp.-DNRA2] |
| WP_183186397.1 | Type3.MTaseII.containing.BREX | <-AAA-ATPase<-HTH_7+rve<-SF1-SF2-Helicase<-OLD-ABC+TOPRIM<-?<-TUDOR+SF1-SF2-Helicase+iREase+CTD+REase<-BrxA-DNA-binding<-PglZ<-N6-N4-METHYLASE<-DUF6079*<-BrxB-BrxF-iSTAND\|\|?->?-><-MODE-HTH+PBPII | 1233 | Bacteria>Terrabacteria_group>Bacillota>Bacilli>Bacillales>Bacillaceae | GCF_014196195.1 | Anoxybacillus_rupiensis | GGQ95_000812 | 79054..82755 | 311460 | DUF6079-family-protein-[Anoxybacillus-rupiensis] |
| WP_188624106.1 | Type3.MTaseII.containing.BREX | RNase->?->?->?-><-?\|\|?-><-?<-?\|\|BrxB-BrxF-iSTAND->DUF6079*->N6-N4-METHYLASE->PglZ->BrxA-DNA-binding->ParB+HNH->?->?->SWIM+DUF3670+SF1-SF2-Helicase->McrC-NTD+REase-> | 1236 | Bacteria>Terrabacteria_group>Bacillota>Bacilli>Bacillales>Bacillaceae | GCF_014644735.1 | Caldalkalibacillus_thermarum | IEW48_RS12860 | 15714..19424 | 296745 | DUF6079-family-protein-[Caldalkalibacillus-thermarum] |
| WP_191274572.1 | Type3.MTaseII.containing.BREX | <-SbcC<-?<-?<-?<-?<-TUDOR+SF1-SF2-Helicase+iREase+CTD+REase<-BrxA-DNA-binding<-PglZ<-N6-N4-METHYLASE<-DUF6079*<-BrxB-BrxF-iSTAND\|\|?->?->Resolvase+Recombinase+ZnR->GDResolvase+Recombinase+ZnR->XIS-HTH->Arm-DNA-bind_3+Phage_integrase->Phage_int_SAM_3+Phage_integrase->Phage_integrase-> | 1234 | Bacteria>Terrabacteria_group>Bacillota>Bacilli>Bacillales>Bacillaceae | GCF_014656545.1 | Neobacillus_kokaensis | IE222_RS16245 | 114204..117908 | 2759023 | DUF6079-family-protein-[Neobacillus-kokaensis] |
| WP_190997469.1 | Type3.MTaseII.containing.BREX | <-HEPN+PAS+DUF4011+SF1-SF2-Helicase+REase<-?<-?<-?<-REase+SF1-SF2-Helicase<-TUDOR+SF1-SF2-Helicase+iREase+CTD+REase<-BrxA-DNA-binding<-PglZ<-N6-N4-METHYLASE<-DUF6079*<-BrxB-BrxF-iSTAND<-DrmB<-DrmA<-HKD-DNase+SF1-SF2-Helicase+HTH+HTH+SRA<-?<-?\|\|?->HAD-> | 1232 | Bacteria>Terrabacteria_group>Bacillota>Bacilli>Bacillales>Bacillaceae | GCF_014712675.1 | Peribacillus_faecalis | IEO70_RS06060 | 58826..62524 | 2772559 | DUF6079-family-protein-[Peribacillus-faecalis] |
| WP_134783939.1 | Type3.MTaseII.containing.BREX | <-ABC-ATPase+SbcC<-?\|\|?-><-?<-DndH<-DndG<-NERD+Pkinase+Pkinase+DndF<-DUF6079*<-DUF4007<-REase<-SF1-SF2-Helicase<-?<-N6-METHYLASE\|\|HTH_3-> | 1309 | Bacteria>Terrabacteria_group>Bacillota>Bacilli>Bacillales>Bacillaceae | GCF_014876495.1 | Peribacillus_frigoritolerans | IMY49_RS24145 | 4156939..4160868 | 450367 | hypothetical-protein-[Peribacillus-frigoritolerans] |
| WP_192949168.1 | Type3.MTaseII.containing.BREX | <-RADICAL-SAM\|\|?->?->?->?->TRAM+METHYLASE->\|\|BrxB-BrxF-iSTAND->DUF6079*->N6-N4-METHYLASE-><-DDE_Tnp | 1234 | Bacteria>Terrabacteria_group>Bacillota>Bacilli>Bacillales>Bacillaceae | GCF_014890275.1 | Anoxybacillus_flavithermus | HPK06_RS06765 | 42738..46442 | 33934 | DUF6079-family-protein-[Anoxybacillus-flavithermus] |
| WP_192952464.1 | Type3.MTaseII.containing.BREX | <-NUDIX<-?<-?<-TUDOR+SF1-SF2-Helicase+iREase+CTD+REase<-BrxA-DNA-binding<-PglZ<-N6-N4-METHYLASE<-DUF6079*<-BrxB-BrxF-iSTAND\|\|<-TRAM+METHYLASE<-?<-?<-?<-?\|\|RADICAL-SAM-> | 1234 | Bacteria>Terrabacteria_group>Bacillota>Bacilli>Bacillales>Bacillaceae | GCF_014890335.1 | Anoxybacillus_flavithermus | HPK03_RS06785 | 13196..16900 | 33934 | DUF6079-family-protein-[Anoxybacillus-flavithermus] |
| WP_098215393.1 | Type3.MTaseII.containing.BREX | <-DnaB+TnsC-ATPase<-DnaB_2\|\|?-><-?<-TUDOR+SF1-SF2-Helicase+iREase+CTD+REase<-BrxA-DNA-binding<-PglZ<-N6-N4-METHYLASE<-DUF6079*<-BrxB-BrxF-iSTAND<-DrmB<-DrmA<-wHTH+WYL+WCX<-?<-?<-REC+CITB-HTH+ZDNA-HTH<-IONCHANNEL+PAS+HISKIN+MORC | 1232 | Bacteria>Terrabacteria_group>Bacillota>Bacilli>Bacillales>Bacillaceae | GCF_015712745.1 | Bacillus_sp._SRB3LM | F3I02_RS30530 | 659881..663579 | 2608689 | DUF6079-family-protein-[Bacillus-sp.-SRB3LM] |
| WP_098109212.1 | Type3.MTaseII.containing.BREX | <-TOPRIM+Topoisom_bac\|\|Phage_integrase->?->Phage_integrase->?-><-HTH_3\|\|HIN-HTH->TOPRIM+Topoisom_bac->BrxB-BrxF-iSTAND->DUF6079*->N6-N4-METHYLASE->?->PglZ->BrxA-DNA-binding->\|\|?->?-><-?\|\|HAMP+MA-> | 1228 | Bacteria>Terrabacteria_group>Bacillota>Bacilli>Bacillales>Bacillaceae | GCF_015880975.1 | Bacillus_toyonensis | ABE55_RS07715 | 118904..122590 | 155322 | MULTISPECIES:-DUF6079-family-protein-[Bacillus-cereus-group] |
| WP_098016179.1 | Type3.MTaseII.containing.BREX | <-Resolvase+Recombinase+ZnR<-?\|\|?-><-?\|\|GDResolvase+Recombinase-><-TUDOR+SF1-SF2-Helicase+iREase+CTD+REase<-BrxA-DNA-binding<-PglZ<-N6-N4-METHYLASE<-DUF6079*<-BrxB-BrxF-iSTAND\|\|<-?<-SWACOS<-PAS+PAS+GGDEF<-?<-?<-tRNA<-?<-?<-NUDIX | 1234 | Bacteria>Terrabacteria_group>Bacillota>Bacilli>Bacillales>Bacillaceae | GCF_016483725.1 | Bacillus_cereus_group | JDS69_RS23095 | 101937..105641 | 86661 | MULTISPECIES:-DUF6079-family-protein-[Bacillus-cereus-group] |
| WP_202283221.1 | Type3.MTaseII.containing.BREX | <-DnaB_2<-?<-MerR-HTH<-?<-?<-TUDOR+SF1-SF2-Helicase+iREase+CTD+REase<-BrxA-DNA-binding<-PglZ<-N6-N4-METHYLASE<-DUF6079*<-BrxB-BrxF-iSTAND<-DrmB<-DrmA<-wHTH+WYL+WCX<-?<-REase<-?<-REC+CITB-HTH+ZDNA-HTH<-IONCHANNEL+PAS+HISKIN+MORC | 1232 | Bacteria>Terrabacteria_group>Bacillota>Bacilli>Bacillales>Bacillaceae | GCF_016757895.1 | Bacillus_cereus | IRV09_RS02780 | 86749..90447 | 1396 | DUF6079-family-protein-[Bacillus-cereus] |
| WP_207452355.1 | Type3.MTaseII.containing.BREX | <-AAA-ATPase<-BrxA-DNA-binding<-PglZ<-N6-N4-METHYLASE<-DUF6079*<-BrxB-BrxF-iSTAND<-?<-DrmA<-DDE_Tnp_ISL3<-DDE_Tnp_ISL3<-NAD_binding_10<-Glyoxalase\|\|SIGMA-HTH-> | 1233 | Bacteria>Terrabacteria_group>Bacillota>Bacilli>Bacillales>Bacillaceae | GCF_017355205.1 | Bacillus_sp._SD088 | IPU53_RS17755 | 8287..11988 | 2782012 | DUF6079-family-protein-[Bacillus-sp.-SD088] |
| WP_066398773.1 | Type3.MTaseII.containing.BREX | <-ParB+HNH<-TUDOR+SF1-SF2-Helicase+iREase+CTD+REase<-BrxA-DNA-binding<-PglZ<-N6-N4-METHYLASE<-DUF6079*\|\|wHTH->?->wHTH(SBD)+HNH->?-><-SPOUT | 1227 | Bacteria>Terrabacteria_group>Bacillota>Bacilli>Bacillales>Bacillaceae | GCF_017874625.1 | Cytobacillus_eiseniae | J2Z40_RS15720 | 24192..27875 | 762947 | DUF6079-family-protein-[Cytobacillus-eiseniae] |
| WP_213145782.1 | Type3.MTaseII.containing.BREX | <-SF1-SF2-Helicase<-?<-SF1-SF2-Helicase<-REase+SF1-SF2-Helicase<-REase<-TUDOR+SF1-SF2-Helicase+iREase+CTD+REase<-BrxA-DNA-binding<-PglZ<-N6-N4-METHYLASE<-DUF6079*<-BrxB-BrxF-iSTAND<-pPIWI+MID+RNaseH<-pPIWI<-pPIWI+pPIWI<-wHTH+WYL+WCX<-Thermonuclease<-?<-DnaB+TnsC-ATPase | 1233 | Bacteria>Terrabacteria_group>Bacillota>Bacilli>Bacillales>Bacillaceae | GCF_018343545.2 | Neobacillus_citreus | KHB02_RS07300 | 40729..44430 | 2833578 | DUF6079-family-protein-[Neobacillus-citreus] |
| WP_225002267.1 | Type3.MTaseII.containing.BREX | Trypsin+PDZ->?->SPOUT->?-><-?<-wHTH\|\|DUF6079*->N6-N4-METHYLASE->PglZ->BrxA-DNA-binding->TUDOR+SF1-SF2-Helicase+iREase+CTD+REase->ParB+HNH-> | 1226 | Bacteria>Terrabacteria_group>Bacillota>Bacilli>Bacillales>Bacillaceae | GCF_020176725.1 | Priestia_flexa | LC052_RS15430 | 42054..45734 | 86664 | DUF6079-family-protein-[Priestia-flexa] |
| WP_248893726.1 | Type3.MTaseII.containing.BREX | BrxB-BrxF-iSTAND->DUF6079*->N6-N4-METHYLASE->PglZ->BrxA-DNA-binding->TUDOR+SF1-SF2-Helicase+iREase+CTD+REase->?->\|\|zf-ISL3+HTH_Tnp_ISL3+DDE_Tnp_ISL3->?->ABC-ATPase+SbcC-> | 1235 | Bacteria>Terrabacteria_group>Bacillota>Bacilli>Bacillales>Bacillaceae | GCF_023278615.1 | Bacillus_methanolicus | C0971_RS07440 | 1482355..1486062 | 1471 | DUF6079-family-protein-[Bacillus-methanolicus] |
| WP_251555692.1 | Type3.MTaseII.containing.BREX | <-GTPase-AIG+IIGP-C<-HIN-HTH\|\|<-HNH<-OLD-ABC<-GDResolvase<-TUDOR+SF1-SF2-Helicase+iREase+CTD+REase<-BrxA-DNA-binding<-PglZ<-N6-N4-METHYLASE<-DUF6079*<-BrxB-BrxF-iSTAND<-HicB<-?<-?<-?<-CR-Kinase<-Phage_integrase | 1233 | Bacteria>Terrabacteria_group>Bacillota>Bacilli>Bacillales>Bacillaceae | GCF_023713305.1 | Psychrobacillus_sp._MER_TA_171 | M3627_11610 | 24549..28250 | 2939577 | DUF6079-family-protein-[Psychrobacillus-sp.-MER-TA-171] |
| WP_251162457.1 | Type3.MTaseII.containing.BREX | SPOUT->?->?->?->RVT->SF1-SF2-Helicase+HKD-DNase-><-NUDIX\|\|BrxB-BrxF-iSTAND->DUF6079*->N6-N4-METHYLASE->PglZ->BrxA-DNA-binding->TUDOR+SF1-SF2-Helicase+iREase+CTD+REase->ParB+HNH->?->?->ZnR-> | 1226 | Bacteria>Terrabacteria_group>Bacillota>Bacilli>Bacillales>Bacillaceae | GCF_023713445.1 | Cytobacillus_kochii | M3670_RS08680 | 58459..62139 | 859143 | DUF6079-family-protein-[Cytobacillus-kochii] |
| WP_257961947.1 | Type3.MTaseII.containing.BREX | ABhydrolase->RhoN+RhoN+RNase+S1->?-><-Arm-DNA-bind_4+Phage_int_SAM_3+Phage_integrase<-Phage_int_SAM_3+Phage_integrase<-XIS-HTH<-ZnR+TOPRIM+AAA-helicase+wHTH<-?\|\|BrxB-BrxF-iSTAND->DUF6079*->N6-N4-METHYLASE->PglZ->BrxA-DNA-binding->TUDOR+SF1-SF2-Helicase+iREase+CTD+REase->JAB->HEPN+HTH-><-ParB<-ParB | 1231 | Bacteria>Terrabacteria_group>Bacillota>Bacilli>Bacillales>Bacillaceae | GCF_024705655.1 | Lysinibacillus_capsici | NUA17_RS16970 | 35936..39631 | 2115968 | DUF6079-family-protein-[Lysinibacillus-capsici] |
| WP_242290722.1 | Type3.MTaseII.containing.BREX | IONCHANNEL+PAS+HISKIN+MORC->REC+CITB-HTH+ZDNA-HTH->REase->?->wHTH+WYL+WCX->DrmA->DrmB->BrxB-BrxF-iSTAND->DUF6079*->N6-N4-METHYLASE->PglZ->BrxA-DNA-binding->TUDOR+SF1-SF2-Helicase+iREase+CTD+REase->?->?->?->MerR-HTH-> | 1232 | Bacteria>Terrabacteria_group>Bacillota>Bacilli>Bacillales>Bacillaceae | GCF_026684175.1 | Bacillus_cereus_group | OU819_RS04775 | 931389..935087 | 86661 | MULTISPECIES:-DUF6079-family-protein-[Bacillus-cereus-group] |
| WP_268328395.1 | Type3.MTaseII.containing.BREX | ADH_N+ADH_zinc_N-><-?\|\|?->SPOUT->BrxB-BrxF-iSTAND->DUF6079*->N6-N4-METHYLASE->PglZ->BrxA-DNA-binding->TUDOR+SF1-SF2-Helicase+iREase+CTD+REase-><-GNTR-HTH+UTRA\|\|?->Glyco_hydro_1-> | 1234 | Bacteria>Terrabacteria_group>Bacillota>Bacilli>Bacillales>Bacillaceae | GCF_026793125.1 | Bacillus_inaquosorum | MOD47_RS12330 | 91172..94876 | 483913 | DUF6079-family-protein-[Bacillus-inaquosorum] |
| WP_270691671.1 | Type3.MTaseII.containing.BREX | <-RNase<-?<-TUDOR+SF1-SF2-Helicase+iREase+CTD+REase<-BrxA-DNA-binding<-PglZ<-N6-N4-METHYLASE<-DUF6079*<-BrxB-BrxF-iSTAND<-DrmB<-DrmA<-wHTH+WYL+WCX<-?<-REase<-?<-REC+CITB-HTH+ZDNA-HTH<-IONCHANNEL+PAS+HISKIN+MORC | 1232 | Bacteria>Terrabacteria_group>Bacillota>Bacilli>Bacillales>Bacillaceae | GCF_027715825.1 | Bacillus_cereus_group_sp._Bc061 | PDQ40_RS23465 | 111028..114726 | 3018117 | DUF6079-family-protein-[Bacillus-cereus-group-sp.-Bc061] |
| WP_272767828.1 | Type3.MTaseII.containing.BREX | <-SF1-SF2-Helicase<-OLD-ABC+TOPRIM<-REase<-TUDOR+SF1-SF2-Helicase+iREase+CTD+REase<-BrxA-DNA-binding<-PglZ<-N6-N4-METHYLASE<-DUF6079*<-BrxB-BrxF-iSTAND<-DrmB<-DrmA | 1232 | Bacteria>Terrabacteria_group>Bacillota>Bacilli>Bacillales>Bacillaceae | GCF_028548275.1 | Priestia_megaterium | PNF31_RS27520 | 33543..37241 | 1404 | DUF6079-family-protein-[Priestia-megaterium] |
| WP_275191614.1 | Type3.MTaseII.containing.BREX | <-SF1-SF2-Helicase<-OLD-ABC+TOPRIM<-TUDOR+SF1-SF2-Helicase+iREase+CTD+REase<-BrxA-DNA-binding<-PglZ<-N6-N4-METHYLASE<-DUF6079*<-BrxB-BrxF-iSTAND\|\|ZnR-><-MODE-HTH+PBPII | 1233 | Bacteria>Terrabacteria_group>Bacillota>Bacilli>Bacillales>Bacillaceae | GCF_029077325.1 | Anoxybacillus_rupiensis | PNH38_RS00045 | 14132..17833 | 311460 | DUF6079-family-protein-[Anoxybacillus-rupiensis] |
| WP_090233733.1 | Type3.MTaseII.containing.BREX | <-HTH_3\|\|HTH_36->?-><-?<-wHTH+HNH<-?<-wHTH\|\|?->BrxB-BrxF-iSTAND->DUF6079*->N6-N4-METHYLASE->PglZ->BrxA-DNA-binding->Classical-AAA->Peptidase_S8->?->?->?->ADH_N+ADH_zinc_N-> | 1227 | Bacteria>Terrabacteria_group>Bacillota>Bacilli>Bacillales>Bacillaceae | GCF_900112045.1 | Lentibacillus_halodurans | BM195_RS03570 | 169567..173250 | 237679 | DUF6079-family-protein-[Lentibacillus-halodurans] |
| WP_254843978.1 | Type3.MTaseII.containing.BREX | <-ABC-ATPase+SbcC<-?<-?<-?<-?<-?<-BrxA-DNA-binding<-PglZ<-N6-N4-METHYLASE<-DUF6079*<-BrxB-BrxF-iSTAND<-?<-?<-?<-?<-?<-?<-?<-PGK | 1238 | Bacteria>Terrabacteria_group>Bacillota>Bacilli>Bacillales>Bacillaceae | GCF_900166595.1 | Virgibacillus_dokdonensis | B2C77_RS14515 | 2746432..2750148 | 302167 | DUF6079-family-protein-[Virgibacillus-dokdonensis] |
| WP_086717959.1 | Type3.MTaseII.containing.BREX | <-MerR-HTH<-?<-?<-TUDOR+SF1-SF2-Helicase+iREase+CTD+REase<-BrxA-DNA-binding<-PglZ<-N6-N4-METHYLASE<-DUF6079*<-BrxB-BrxF-iSTAND\|\|?-><-wHTH+WYL+WCX<-?<-REase<-?<-REC+CITB-HTH+ZDNA-HTH<-IONCHANNEL+PAS+HISKIN+MORC | 1232 | Bacteria>Terrabacteria_group>Bacillota>Bacilli>Bacillales>Bacillaceae | GCF_900176835.1 | Bacillus_cereus_group | BACERE00188_RS21780 | 68079..71777 | 86661 | MULTISPECIES:-DUF6079-family-protein-[Bacillus-cereus-group] |
| WP_102026198.1 | Type3.MTaseII.containing.BREX | <-HTH_3\|\|?->SPOUT-><-?<-wHTH(SBD)+HNH<-?<-wHTH\|\|BrxB-BrxF-iSTAND->DUF6079*->N6-N4-METHYLASE->PglZ->BrxA-DNA-binding->TUDOR+SF1-SF2-Helicase+iREase+CTD+REase->ParB+HNH-> | 1227 | Bacteria>Terrabacteria_group>Bacillota>Bacilli>Bacillales>Bacillaceae | GCF_900240405.1 | Salirhabdus_sp._Marseille-P4669 | C1A05_RS00395 | 77615..81298 | 2042310 | DUF6079-family-protein-[Salirhabdus-sp.-Marseille-P4669] |
| WP_084780292.1 | Type3.MTaseII.containing.BREX | ADH_N+ADH_zinc_N-><-HHH+DNA_pol_B+RNase<-ACET<-?\|\|?-><-BrxA-DNA-binding<-PglZ<-N6-N4-METHYLASE<-N6-N4-METHYLASE<-DUF6079*<-BrxB-BrxF-iSTAND<-?<-?<-?<-?<-REC+OmpR-HTH<-?<-sGTP | 1235 | Bacteria>Terrabacteria_group>Bacillota>Bacilli>Bacillales>Bacillaceae | GCF_902386855.1 | Bacillus_massilionigeriensis | BQ6100_RS08540 | 1710832..1714539 | 1805475 | DUF6079-family-protein-[Bacillus-massilionigeriensis] |
| WP_075981302.1 | Type3.MTaseII.containing.BREX | <-METHYLASE<-?\|\|?-><-Peptidase_C14<-TUDOR+SF1-SF2-Helicase+iREase+CTD+REase<-BrxA-DNA-binding<-PglZ<-N6-N4-METHYLASE<-N6-N4-METHYLASE<-DUF6079*<-BrxB-BrxF-iSTAND<-DrmB<-DrmA<-HTH+HTH+SF1-SF2-Helicase<-?\|\|?-><-?<-?<-MerR-HTH | 1233 | Bacteria>Terrabacteria_group>Bacillota>Bacilli>Bacillales>Bacillaceae | GCF_902386855.1 | Bacillus_massilionigeriensis | BQ6100_RS08730 | 1758290..1761991 | 1805475 | DUF6079-family-protein-[Bacillus-massilionigeriensis] |
| WP_075982597.1 | Type3.MTaseII.containing.BREX | <-PolB<-?<-TUDOR+SF1-SF2-Helicase+iREase+CTD+REase<-BrxA-DNA-binding<-PglZ<-N6-N4-METHYLASE<-DUF6079*<-BrxB-BrxF-iSTAND<-GDResolvase+HIN-HTH | 1234 | Bacteria>Terrabacteria_group>Bacillota>Bacilli>Bacillales>Bacillaceae | GCF_902386855.1 | Bacillus_massilionigeriensis | FX071_RS15940 | 3241888..3245592 | 1805475 | DUF6079-family-protein-[Bacillus-massilionigeriensis] |
| WP_174496250.1 | Type3.MTaseII.containing.BREX | McrB-AAA->REase->SF1-SF2-Helicase-><-SF1-SF2-Helicase<-SF1-SF2-Helicase\|\|HKD-DNase->DNA_METHYLASE->BrxB-BrxF-iSTAND->DUF6079*->N6-N4-METHYLASE->PglZ->BrxA-DNA-binding->TUDOR+SF1-SF2-Helicase+iREase+CTD+REase->?->?->?->N6-N4-METHYLASE-> | 1226 | Bacteria>Terrabacteria_group>Bacillota>Bacilli>Bacillales>Bacillaceae | GCF_902809875.1 | Salirhabdus_euzebyi | HNQ94_RS00270 | 50426..54106 | 394506 | DUF6079-family-protein-[Salirhabdus-euzebyi] |
| WP_191566563.1 | Type3.MTaseII.containing.BREX | <-GDResolvase+Recombinase+ZnR<-?<-HTH_3\|\|HTH_36->?-><-?<-wHTH(SBD)+HNH<-?\|\|BrxB-BrxF-iSTAND->DUF6079*->N6-N4-METHYLASE->PglZ->BrxA-DNA-binding->TUDOR+SF1-SF2-Helicase+iREase+CTD+REase->ParB+HNH->?->?->?->ZnR-> | 1226 | Bacteria>Terrabacteria_group>Bacillota>Bacilli>Bacillales>Bacillaceae | GCF_904423835.1 | Metabacillus_idriensis | IHQ32_RS17375 | 3417023..3420703 | 324768 | DUF6079-family-protein-[Metabacillus-idriensis] |
| WP_017552551.1 | Type3.MTaseII.containing.BREX | transposase->HTH+HTH+SF1-SF2-Helicase-p->DrmA->DrmB->BrxB-BrxF-iSTAND->DUF6079-p*-> | 496 | Bacteria>Terrabacteria_group>Bacillota>Bacilli>Bacillales>Bacillaceae | - | Weizmannia_coagulans | C893_RS0101435 | 7139..8629 | 1398 | DUF6079-family-protein-[Weizmannia-coagulans] |
| STO11899.1 | Type3.MTaseII.containing.BREX | Y1_Tnp->HTH_OrfB_IS605+OrfB_IS605+OrfB_Zn_ribbon-><-?\|\|BrxB-BrxF-iSTAND->DUF6079*->N6-N4-METHYLASE->PglZ->BrxA-DNA-binding->TUDOR+SF1-SF2-Helicase+iREase+CTD+REase-p->TUDOR+SF1-SF2-Helicase+iREase+CTD+REase->REase-> | 1232 | Bacteria>Terrabacteria_group>Bacillota>Bacilli>Bacillales>Bacillales_incertae_sedis | GCA_900450595.1 | [Flavobacterium]_thermophilum | NCTC11526_00565 | 545789..549487 | 1414643 | Uncharacterised-protein-[[Flavobacterium] |
| WP_047391744.1 | Type3.MTaseII.containing.BREX | Phage_holin-><-RNase\|\|?->HKD-DNase+SF1-SF2-Helicase+HTH+HTH+SRA->?-><-TUDOR+SF1-SF2-Helicase+iREase+CTD+REase<-BrxA-DNA-binding<-PglZ<-N6-N4-METHYLASE<-DUF6079*<-BrxB-BrxF-iSTAND<-DrmB<-DrmA\|\|SIGMA-HTH+SIGMA-HTH-><-MazG<-NERD+SF1-SF2-Helicase\|\|?-><-Glyoxalase\|\|LACI-HTH+PBPI-> | 1234 | Bacteria>Terrabacteria_group>Bacillota>Bacilli>Bacillales>Bacillales_incertae_sedis | GCF_000798935.2 | Exiguobacterium_sp._ZWU0009 | L985_RS01925 | 348423..352127 | 1224749 | DUF6079-family-protein-[Exiguobacterium-sp.-ZWU0009] |
| WP_214689527.1 | Type3.MTaseII.containing.BREX | <-PIWI<-REase+DUF5655<-SF1-SF2-Helicase<-?<-?<-SIGMA-HTH+SIGMA-HTH\|\|DrmA->DrmB->BrxB-BrxF-iSTAND->DUF6079*->N6-N4-METHYLASE->PglZ->BrxA-DNA-binding->TUDOR+SF1-SF2-Helicase+iREase+CTD+REase-><-REase+SF1-SF2-Helicase<-wHTH+HNH | 1234 | Bacteria>Terrabacteria_group>Bacillota>Bacilli>Bacillales>Bacillales_incertae_sedis | GCF_018617135.1 | Exiguobacterium_sp._s163 | KNO71_RS16205 | 79478..83182 | 2751287 | DUF6079-family-protein-[Exiguobacterium-sp.-s163] |
| WP_214788925.1 | Type3.MTaseII.containing.BREX | JetA->?->?->?->REase+SF1-SF2-Helicase-><-TUDOR+SF1-SF2-Helicase+iREase+CTD+REase<-BrxA-DNA-binding<-PglZ<-N6-N4-METHYLASE<-DUF6079*<-BrxB-BrxF-iSTAND<-DrmB<-DrmA\|\|SIGMA-HTH+SIGMA-HTH->?->?->SF1-SF2-Helicase->REase+DUF5655->PIWI-> | 1234 | Bacteria>Terrabacteria_group>Bacillota>Bacilli>Bacillales>Bacillales_incertae_sedis | GCF_018617975.1 | Exiguobacterium_sp._s21 | KNP06_RS10885 | 43669..47373 | 2751244 | DUF6079-family-protein-[Exiguobacterium-sp.-s21] |
| WP_214805994.1 | Type3.MTaseII.containing.BREX | HARE-HTH-><-RNase\|\|?->HKD-DNase+SF1-SF2-Helicase+HTH+HTH+SRA->?-><-TUDOR+SF1-SF2-Helicase+iREase+CTD+REase<-BrxA-DNA-binding<-PglZ<-N6-N4-METHYLASE<-DUF6079*<-BrxB-BrxF-iSTAND<-DrmB<-DrmA\|\|SIGMA-HTH+SIGMA-HTH-><-MazG\|\|?-><-?<-Glyoxalase\|\|LACI-HTH+PBPI-> | 1234 | Bacteria>Terrabacteria_group>Bacillota>Bacilli>Bacillales>Bacillales_incertae_sedis | GCF_018618875.1 | Exiguobacterium_sp._s46 | KNQ08_RS12560 | 192261..195965 | 2751200 | DUF6079-family-protein-[Exiguobacterium-sp.-s46] |
| WP_271306165.1 | Type3.MTaseII.containing.BREX | <-MazG<-SF1-SF2-Helicase<-?<-?<-SIGMA-HTH+SIGMA-HTH\|\|DrmA->DrmB->BrxB-BrxF-iSTAND->DUF6079*->N6-N4-METHYLASE->PglZ->BrxA-DNA-binding->TUDOR+SF1-SF2-Helicase+iREase+CTD+REase->?-><-?<-?\|\|Abi-> | 1234 | Bacteria>Terrabacteria_group>Bacillota>Bacilli>Bacillales>Bacillales_incertae_sedis | GCF_027947425.1 | Exiguobacterium_sp._MMG028 | PJK55_RS13660 | 63875..67579 | 3021979 | DUF6079-family-protein-[Exiguobacterium-sp.-MMG028] |
| WP_275060104.1 | Type3.MTaseII.containing.BREX | <-PIWI<-REase+DUF5655<-SF1-SF2-Helicase<-?<-?<-SIGMA-HTH+SIGMA-HTH\|\|DrmA->DrmB->BrxB-BrxF-iSTAND->DUF6079*->N6-N4-METHYLASE->PglZ->BrxA-DNA-binding->TUDOR+SF1-SF2-Helicase+iREase+CTD+REase-><-REase+SF1-SF2-Helicase<-wHTH+HNH | 1234 | Bacteria>Terrabacteria_group>Bacillota>Bacilli>Bacillales>Bacillales_incertae_sedis | GCF_029026745.1 | Exiguobacterium_profundum | OE059_RS13135 | 2522786..2526490 | 307643 | DUF6079-family-protein-[Exiguobacterium-profundum] |
| WP_227092780.1 | Type3.MTaseII.containing.BREX | <-BrxA-DNA-binding<-PglZ<-N6-N4-METHYLASE<-DUF6079*<-BrxB-BrxF-iSTAND\|\|<-Rrf2-HTH | 1233 | Bacteria>Terrabacteria_group>Bacillota>Bacilli>Bacillales | GCF_020554445.1 | Bacillaceae | LIO12_RS14010 | 6021..9722 | 186817 | MULTISPECIES:-DUF6079-family-protein-[Bacillaceae] |
| EUJ41574.1 | Type3.MTaseII.containing.BREX | <-HIN-HTH\|\|?->BrxB-BrxF-iSTAND->DUF6079-p*->DUF6079-p->N6-N4-METHYLASE->PglZ->BrxA-DNA-binding-><-?\|\|DSBH+AraC-HTH-> | 954 | Bacteria>Terrabacteria_group>Bacillota>Bacilli>Bacillales>Listeriaceae | GCA_000525915.1 | Brochothrix_campestris_FSL_F6-1037 | BCAMP_03295 | 1378..4242 | 1265861 | hypothetical-protein-BCAMP_03295-[Brochothrix-campestris-FSL-F6-1037] |
| EAG7745520.1 | Type3.MTaseII.containing.BREX | <-MerR-HTH<-?\|\|?-><-RVT<-BrxA-DNA-binding<-PglZ<-N6-N4-METHYLASE<-DUF6079*<-BrxB-BrxF-iSTAND<-HTH_3\|\|?->Phage_GPA->XIS-HTH->Phage_int_SAM_3+Phage_integrase-> | 1239 | Bacteria>Terrabacteria_group>Bacillota>Bacilli>Bacillales>Listeriaceae | GCA_004719905.1 | Listeria_monocytogenes | BLI34_06105 | 13482..17201 | 1639 | ATP-binding-protein-[Listeria-monocytogenes] |
| WP_198015686.1 | Type3.MTaseII.containing.BREX | <-HIN-HTH\|\|?->BrxB-BrxF-iSTAND->DUF6079-p*->DUF6079-p->N6-N4-METHYLASE->PglZ->BrxA-DNA-binding-><-?\|\|DSBH+AraC-HTH->?->?->Glyco_hydro_3_C+Fn3-like+Glyco_hydro_3-> | 954 | Bacteria>Terrabacteria_group>Bacillota>Bacilli>Bacillales>Listeriaceae | GCF_000525915.1 | Brochothrix_campestris | BCAMP_RS02945 | 1378..4242 | 2757 | DUF6079-family-protein-[Brochothrix-campestris] |
| GIO15806.1 | Type3.MTaseII.containing.BREX | <-ParA-Soj-PloopNTPase<-HTH_Tnp_1+HTH_21+rve+rve_3<-?<-?<-OLD-ABC+TOPRIM<-TUDOR+SF1-SF2-Helicase+iREase+CTD+REase<-BrxA-DNA-binding<-PglZ<-N6-N4-METHYLASE<-DUF6079-p* | 821 | Bacteria>Terrabacteria_group>Bacillota>Bacilli>Bacillales>Paenibacillaceae | GCA_018333175.1 | Cohnella_xylanilytica | J19TS2_53610 | 56514..58979 | 557555 | hypothetical-protein-J19TS2_53610-[Cohnella-xylanilytica] |
| GIO16852.1 | Type3.MTaseII.containing.BREX | <-DUF6079-p*<-BrxB-BrxF-iSTAND<-?<-DUF772+DDE_Tnp_1_2<-?<-?<-XIS-HTH\|\|HTH_3-> | 394 | Bacteria>Terrabacteria_group>Bacillota>Bacilli>Bacillales>Paenibacillaceae | GCA_018333175.1 | Cohnella_xylanilytica | J19TS2_64070 | 139..1323 | 557555 | hypothetical-protein-J19TS2_64070-[Cohnella-xylanilytica] |
| UUZ84567.1 | Type3.MTaseII.containing.BREX | N-OB+HDRnase->?->wHTH+WYL+WCX->?->SF1-SF2-Helicase+REase-p->SF1-SF2-Helicase+REase->REase->BrxB-BrxF-iSTAND->DUF6079*->N6-N4-METHYLASE-p->N6-N4-METHYLASE->PglZ->BrxA-DNA-binding->TUDOR+SF1-SF2-Helicase+iREase+CTD+REase->OLD-ABC+TOPRIM->SF1-SF2-Helicase->?-><-DUF772+DDE_Tnp_1 | 1226 | Bacteria>Terrabacteria_group>Bacillota>Bacilli>Bacillales>Paenibacillaceae | GCA_024703795.1 | Paenibacillus_sp._P26 | LJK88_13395 | 2157947..2161627 | 2885204 | ATP-binding-protein-[Paenibacillus-sp.-P26] |
| WP_023554877.1 | Type3.MTaseII.containing.BREX | <-HTH+McrB-AAA<-HD+PspB<-SF1-SF2-Helicase<-SF1-SF2-Helicase<-OLD-ABC+TOPRIM<-TUDOR+SF1-SF2-Helicase+iREase+CTD+REase<-BrxA-DNA-binding<-PglZ<-N6-N4-METHYLASE<-DUF6079*<-BrxB-BrxF-iSTAND<-SF1-SF2-Helicase+REase<-Exonuc_V_gamma+REase<-wHTH+WYL+WCX<-?<-DOC+HTH<-HDRnase\|\|?->LZ_Tnp_IS66+zf-IS66+DDE_Tnp_IS66+DDE_Tnp_IS66_C-> | 1226 | Bacteria>Terrabacteria_group>Bacillota>Bacilli>Bacillales>Paenibacillaceae | GCF_000503775.1 | Brevibacillus_panacihumi | T458_RS33500 | 831264..834944 | 497735 | DUF6079-family-protein-[Brevibacillus-panacihumi] |
| WP_042184800.1 | Type3.MTaseII.containing.BREX | TRAM+METHYLASE-><-Phage_int_SAM_3+Phage_integrase<-XIS-HTH<-D5N+viral-AAA+CDC6-HTH<-?<-?<-XIS-HTH<-?\|\|BrxB-BrxF-iSTAND->DUF6079*->N6-N4-METHYLASE->PglZ->BrxA-DNA-binding->TUDOR+SF1-SF2-Helicase+iREase+CTD+REase->REase->SIR2->?-><-?\|\|dCache_1+HAMP+HisKin+HATPase_c->REC+AraC-HTH+AraC-HTH-> | 1228 | Bacteria>Terrabacteria_group>Bacillota>Bacilli>Bacillales>Paenibacillaceae | GCF_000758545.1 | Paenibacillus_sp._FSL_H7-0737 | H70737_RS03460 | 769371..773057 | 1536775 | DUF6079-family-protein-[Paenibacillus-sp.-FSL-H7-0737] |
| WP_076118191.1 | Type3.MTaseII.containing.BREX | <-SF1-SF2-Helicase+REase+HTH<-SbcC+SbcC<-Calcineurin<-TUDOR+SF1-SF2-Helicase+iREase+CTD+REase<-BrxA-DNA-binding<-PglZ<-N6-N4-METHYLASE<-DUF6079*<-BrxB-BrxF-iSTAND<-SF1-SF2-Helicase+REase<-SF1-SF2-Helicase+REase<-wHTH+WYL+WCX\|\|?-><-tRNA<-HTH_3\|\|?-><-tRNA<-tRNA<-?<-GNTR-HTH | 1226 | Bacteria>Terrabacteria_group>Bacillota>Bacilli>Bacillales>Paenibacillaceae | GCF_001954425.1 | Paenibacillus_odorifer | BSK52_RS07875 | 302005..305685 | 189426 | DUF6079-family-protein-[Paenibacillus-odorifer] |
| WP_113032646.1 | Type3.MTaseII.containing.BREX | <-Amidohydro_2<-?<-?<-?<-REC+AraC-HTH+AraC-HTH<-dCache_1+HAMP+HisKin+HATPase_c<-BrxA-DNA-binding<-PglZ<-N6-N4-METHYLASE<-DUF6079*<-BrxB-BrxF-iSTAND\|\|?->?->?->?->?-><-PemK_toxin<-MazE_antitoxin<-IG+IG | 1226 | Bacteria>Terrabacteria_group>Bacillota>Bacilli>Bacillales>Paenibacillaceae | GCF_003285015.1 | Paenibacillus_contaminans | DQG23_RS19640 | 188571..192251 | 450362 | DUF6079-family-protein-[Paenibacillus-contaminans] |
| WP_114378272.1 | Type3.MTaseII.containing.BREX | HAMP+HisKin+HATPase_c->REC+AraC-HTH+AraC-HTH->?-><-?<-?\|\|REase+SF1-SF2-Helicase->?->?->BrxB-BrxF-iSTAND->DUF6079*->N6-N4-METHYLASE->PglZ->BrxA-DNA-binding->TUDOR+SF1-SF2-Helicase+iREase+CTD+REase-><-TnpB_IS66\|\|?-><-?\|\|ABhydrolase-> | 1228 | Bacteria>Terrabacteria_group>Bacillota>Bacilli>Bacillales>Paenibacillaceae | GCF_003337375.1 | Paenibacillus_prosopidis | DFP97_RS01955 | 467287..470973 | 630520 | DUF6079-family-protein-[Paenibacillus-prosopidis] |
| WP_138789730.1 | Type3.MTaseII.containing.BREX | TRPR-HTH->?->?->TRAM+METHYLASE->\|\|BrxB-BrxF-iSTAND->DUF6079*->N6-N4-METHYLASE->PglZ->BrxA-DNA-binding->TUDOR+SF1-SF2-Helicase+iREase+CTD+REase->REase-> | 1226 | Bacteria>Terrabacteria_group>Bacillota>Bacilli>Bacillales>Paenibacillaceae | GCF_005938195.1 | Paenibacillus_thermoaerophilus | FE781_RS11300 | 101668..105348 | 1215385 | DUF6079-family-protein-[Paenibacillus-thermoaerophilus] |
| WP_122960163.1 | Type3.MTaseII.containing.BREX | <-sGTP<-?<-sGTP<-SbcC+SbcC<-Calcineurin<-TUDOR+SF1-SF2-Helicase+iREase+CTD+REase<-BrxA-DNA-binding<-PglZ<-N6-N4-METHYLASE<-DUF6079*<-BrxB-BrxF-iSTAND<-SF1-SF2-Helicase+REase<-SF1-SF2-Helicase+REase<-wHTH+WYL+WCX<-pPIWI+MID+RNaseH<-pPIWI<-pPIWI+pPIWI<-TRAM+METHYLASE | 1226 | Bacteria>Terrabacteria_group>Bacillota>Bacilli>Bacillales>Paenibacillaceae | GCF_006540045.1 | Brevibacillus_centrosporus | BCE02nite_RS06965 | 43233..46913 | 54910 | DUF6079-family-protein-[Brevibacillus-centrosporus] |
| WP_173598487.1 | Type3.MTaseII.containing.BREX | N6-METHYLASE->?-><-SF1-SF2-Helicase<-?<-?\|\|?->?->HTH_Tnp_1+HTH_21+rve+rve_3->BrxB-BrxF-iSTAND->DUF6079*->N6-N4-METHYLASE->PglZ->BrxA-DNA-binding->TUDOR+SF1-SF2-Helicase+iREase+CTD+REase-><-AAA-ATPase<-HIN-HTH | 1228 | Bacteria>Terrabacteria_group>Bacillota>Bacilli>Bacillales>Paenibacillaceae | GCF_013284375.2 | Brevibacillus | HP435_RS06005 | 1182044..1185730 | 55080 | MULTISPECIES:-DUF6079-family-protein-[Brevibacillus] |
| WP_189000169.1 | Type3.MTaseII.containing.BREX | wHTH+WYL+WCX->?->?->?->SF1-SF2-Helicase+REase->SF1-SF2-Helicase+REase->BrxB-BrxF-iSTAND->DUF6079*->N6-N4-METHYLASE->PglZ->BrxA-DNA-binding->TUDOR+SF1-SF2-Helicase+iREase+CTD+REase->SF1-SF2-Helicase+REase+HTH->SF1-SF2-Helicase->SF1-SF2-Helicase->HTH+McrB-AAA->McrC-NTD+REase-> | 1227 | Bacteria>Terrabacteria_group>Bacillota>Bacilli>Bacillales>Paenibacillaceae | GCF_014641075.1 | Paenibacillus_nasutitermitis | IEY46_RS33895 | 11702..15385 | 1652958 | DUF6079-family-protein-[Paenibacillus-nasutitermitis] |
| WP_198826837.1 | Type3.MTaseII.containing.BREX | <-rve<-HTH_Tnp_1+HTH_21+rve+rve_3<-HTH_Tnp_1<-SF1-SF2-Helicase<-OLD-ABC+TOPRIM<-TUDOR+SF1-SF2-Helicase+iREase+CTD+REase<-BrxA-DNA-binding<-PglZ<-N6-N4-METHYLASE<-DUF6079*<-BrxB-BrxF-iSTAND\|\|DUF4372+DDE_Tnp_1-><-pPIWI+pPIWI<-?<-SWIM<-ParB\|\|<-PIN | 1226 | Bacteria>Terrabacteria_group>Bacillota>Bacilli>Bacillales>Paenibacillaceae | GCF_016406105.1 | Brevibacillus_composti | KDJ56_RS14960 | 2939715..2943395 | 2796470 | DUF6079-family-protein-[Brevibacillus-composti] |
| WP_209876427.1 | Type3.MTaseII.containing.BREX | ABC-ATPase+ABC-ATPase+ABC-ATPase+SbcC->ABC-ATPase+SbcC->?->?->tRNA->tRNA->wHTH+WYL+WCX->?->SF1-SF2-Helicase+REase->SF1-SF2-Helicase+REase->BrxB-BrxF-iSTAND->DUF6079*->N6-N4-METHYLASE->PglZ->BrxA-DNA-binding->TUDOR+SF1-SF2-Helicase+iREase+CTD+REase->Calcineurin->SbcC+SbcC->?->pPIWI+pPIWI->pPIWI-> | 1226 | Bacteria>Terrabacteria_group>Bacillota>Bacilli>Bacillales>Paenibacillaceae | GCF_017874735.1 | Paenibacillus_silagei | J2Z70_RS20895 | 40400..44080 | 1670801 | DUF6079-family-protein-[Paenibacillus-silagei] |
| WP_246578858.1 | Type3.MTaseII.containing.BREX | <-ParB<-ParA-Soj-PloopNTPase<-?<-?<-OLD-ABC+TOPRIM<-TUDOR+SF1-SF2-Helicase+iREase+CTD+REase<-BrxA-DNA-binding<-PglZ<-N6-N4-METHYLASE<-DUF6079-p* | 845 | Bacteria>Terrabacteria_group>Bacillota>Bacilli>Bacillales>Paenibacillaceae | GCF_018333175.1 | Cohnella_xylanilytica | KI376_RS26605 | 56514..59051 | 557555 | DUF6079-family-protein-[Cohnella-xylanilytica] |
| WP_212963288.1 | Type3.MTaseII.containing.BREX | <-DUF6079-p*<-BrxB-BrxF-iSTAND<-?<-DUF772+DDE_Tnp_1_2<-?<-?<-XIS-HTH\|\|HTH_3-> | 391 | Bacteria>Terrabacteria_group>Bacillota>Bacilli>Bacillales>Paenibacillaceae | GCF_018333175.1 | Cohnella_xylanilytica | KI376_RS31905 | 139..1314 | 557555 | DUF6079-family-protein-[Cohnella-xylanilytica] |
| WP_221842513.1 | Type3.MTaseII.containing.BREX | wHTH+WYL+WCX->Exonuc_V_gamma+REase->SF1-SF2-Helicase+REase->BrxB-BrxF-iSTAND->DUF6079*->N6-N4-METHYLASE->PglZ->BrxA-DNA-binding->TUDOR+SF1-SF2-Helicase+iREase+CTD+REase->OLD-ABC+TOPRIM->SF1-SF2-Helicase-><-DUF772+DDE_Tnp_1\|\|HIN-HTH-> | 1226 | Bacteria>Terrabacteria_group>Bacillota>Bacilli>Bacillales>Paenibacillaceae | GCF_019749055.1 | Brevibacillus_agri | H7K32_RS04485 | 8422..12102 | 51101 | DUF6079-family-protein-[Brevibacillus-agri] |
| WP_054819688.1 | Type3.MTaseII.containing.BREX | HDRnase->?->URI-DNase->DOC+HTH->?->wHTH+WYL+WCX->Exonuc_V_gamma+REase->SF1-SF2-Helicase+REase->BrxB-BrxF-iSTAND->DUF6079*->N6-N4-METHYLASE->PglZ->BrxA-DNA-binding->TUDOR+SF1-SF2-Helicase+iREase+CTD+REase->OLD-ABC+TOPRIM->SF1-SF2-Helicase-> | 1226 | Bacteria>Terrabacteria_group>Bacillota>Bacilli>Bacillales>Paenibacillaceae | GCF_024519685.1 | unclassified_Paenibacillus | NR121_RS23310 | 15408..19088 | 185978 | MULTISPECIES:-DUF6079-family-protein-[unclassified-Paenibacillus] |
| WP_274359443.1 | Type3.MTaseII.containing.BREX | wHTH+WYL+WCX->?->?->?->SF1-SF2-Helicase+REase->SF1-SF2-Helicase+REase->BrxB-BrxF-iSTAND->DUF6079*->N6-N4-METHYLASE->PglZ->BrxA-DNA-binding->TUDOR+SF1-SF2-Helicase+iREase+CTD+REase->?->McrB-AAA->McrC-NTD+REase->HDRnase->SIR2-> | 1227 | Bacteria>Terrabacteria_group>Bacillota>Bacilli>Bacillales>Paenibacillaceae | GCF_028747665.1 | Paenibacillus_urinalis | PUW23_03440 | 763405..767088 | 521520 | DUF6079-family-protein-[Paenibacillus-urinalis] |
| WP_090826254.1 | Type3.MTaseII.containing.BREX | SF1-SF2-Helicase+REase->?->?->?->35exo\|\|\|\|?-><-?<-?<-?\|\|BrxB-BrxF-iSTAND->DUF6079*->N6-N4-METHYLASE->PglZ->BrxA-DNA-binding->TUDOR+SF1-SF2-Helicase+iREase+CTD+REase->Calcineurin->SbcC+SbcC->SIR2-> | 1226 | Bacteria>Terrabacteria_group>Bacillota>Bacilli>Bacillales>Paenibacillaceae | GCF_900103825.1 | Paenibacillus_sp._yr247 | BLT91_RS25150 | 19136..22816 | 1761880 | DUF6079-family-protein-[Paenibacillus-sp.-yr247] |
| WP_085168718.1 | Type3.MTaseII.containing.BREX | HDRnase->DOC+HTH->?->wHTH+WYL+WCX->Exonuc_V_gamma+REase->SF1-SF2-Helicase+REase->BrxB-BrxF-iSTAND->DUF6079*->N6-N4-METHYLASE->PglZ->BrxA-DNA-binding->TUDOR+SF1-SF2-Helicase+iREase+CTD+REase->OLD-ABC+TOPRIM->SF1-SF2-Helicase->SF1-SF2-Helicase->HD+PspB->HTH+McrB-AAA-> | 1226 | Bacteria>Terrabacteria_group>Bacillota>Bacilli>Bacillales>Paenibacillaceae | GCF_900177285.1 | Paenibacillus_barengoltzii | B9N82_RS00040 | 11271..14951 | 343517 | DUF6079-family-protein-[Paenibacillus-barengoltzii] |
| WP_058302024.1 | Type3.MTaseII.containing.BREX | <-pPIWI<-pPIWI+pPIWI<-McrC-NTD+REase<-McrB-AAA<-?<-TUDOR+SF1-SF2-Helicase+iREase+CTD+REase<-BrxA-DNA-binding<-PglZ<-N6-N4-METHYLASE<-DUF6079*<-BrxB-BrxF-iSTAND<-SF1-SF2-Helicase+REase<-SF1-SF2-Helicase+REase<-?<-wHTH+WYL+WCX<-?<-DOC+HTH<-DEDD_Tnp_IS110+Transposase_20<-RVT | 1227 | Bacteria>Terrabacteria_group>Bacillota>Bacilli>Bacillales>Paenibacillaceae | GCF_902375625.1 | Gorillibacterium_timonense | BN2684_RS09980 | 487147..490830 | 1689269 | DUF6079-family-protein-[Gorillibacterium-timonense] |
| WP_019415999.1 | Type3.MTaseII.containing.BREX | TPR-><-?<-HKD-DNase+SF1-SF2-Helicase+HTH+HTH+SRA<-?\|\|Phage_integrase->HicB->BrxB-BrxF-iSTAND->DUF6079*->N6-N4-METHYLASE->PglZ->BrxA-DNA-binding->TUDOR+SF1-SF2-Helicase+iREase+CTD+REase->SF1-SF2-Helicase+Z1-> | 1233 | Bacteria>Terrabacteria_group>Bacillota>Bacilli>Bacillales>Planococcaceae | GCF_000286315.1 | Paenisporosarcina_sp._TG20 | B371_RS0119450 | 28228..31929 | 1211706 | DUF6079-family-protein-[Paenisporosarcina-sp.-TG20] |

| Protein-ID | System | Gene-Neighbourhoods | Length | Taxonomy | Assembly-ID | Strain | Locus-Tag | Genomic-coordinates | TaxID | Fasta-define |
|---|---|---|---|---|---|---|---|---|---|---|
| WP_065525137.1 | Type3.MTaseII.containing.BREX | <-HKD-DNase+SF1-SF2-Helicase+HTH+HTH+SRA<-?<-?<-SF1-SF2-Helicase+Z1<-?<-TUDOR+SF1-SF2-Helicase+iREase+CTD+REase<-BrxA-DNA-binding<-PglZ<-N6-N4-METHYLASE<-DUF6079*<-BrxB-BrxF-iSTAND<-HicB<-?<-REase+ZnR<-McrB-AAA | 1233 | Bacteria>Terrabacteria_group>Bacillota>Bacilli>Bacillales>Planococcaceae | GCF_001687665.2 | Planococcus_donghaensis | BCM40_RS15180 | 3100969..3104670 | 414778 | DUF6079-family-protein-[Planococcus-donghaensis] |
| WP_149457005.1 | Type3.MTaseII.containing.BREX | Phage_integrase-><-?\|\|?->?-><-RelE-ParE<-TraJ-RHH\|\|?->HicB->BrxB-BrxF-iSTAND->DUF6079*->N6-N4-METHYLASE->PglZ->BrxA-DNA-binding->TUDOR+SF1-SF2-Helicase+iREase+CTD+REase->?-><-?<-Trypsin<-Schlafen+HATPase+wHTH\|\|SAVED-> | 1233 | Bacteria>Terrabacteria_group>Bacillota>Bacilli>Bacillales>Planococcaceae | GCF_008369335.1 | Planococcus_sp._ANT_H30 | FQ085_RS14925 | 107901..111602 | 2597347 | DUF6079-family-protein-[Planococcus-sp.-ANT_H30] |
| WP_192599219.1 | Type3.MTaseII.containing.BREX | DDE_Tnp_1-><-Small-B-barrel+SF1-SF2-Helicase<-REase+SF1-SF2-Helicase<-TUDOR+SF1-SF2-Helicase+iREase+CTD+REase<-BrxA-DNA-binding<-PglZ<-N6-N4-METHYLASE<-DUF6079*<-BrxB-BrxF-iSTAND\|\|HTH_3->?->?->RelE-ParE->HTH_3-><-?<-SF1-SF2-Helicase<-PIN | 1233 | Bacteria>Terrabacteria_group>Bacillota>Bacilli>Bacillales>Planococcaceae | GCF_014873855.1 | Sporosarcina_limicola | H4683_RS12805 | 42825..46526 | 34101 | DUF6079-family-protein-[Sporosarcina-limicola] |
| WP_215106202.1 | Type3.MTaseII.containing.BREX | <-Phage_integrase<-TUDOR+SF1-SF2-Helicase+iREase+CTD+REase<-BrxA-DNA-binding<-PglZ<-N6-N4-METHYLASE<-DUF6079*<-BrxB-BrxF-iSTAND<-HicB<-?<-?<-HKD-DNase+SF1-SF2-Helicase+HTH+HTH+SRA<-NUDIX\|\|?-><-McrC-NTD+REase<-SF1-SF2-Helicase+McrB-AAA | 1233 | Bacteria>Terrabacteria_group>Bacillota>Bacilli>Bacillales>Planococcaceae | GCF_018613655.1 | Planococcus_sp._ISL-109 | J7E21_RS07380 | 9923..13624 | 2819166 | DUF6079-family-protein-[Planococcus-sp.-ISL-109] |
| WP_239738037.1 | Type3.MTaseII.containing.BREX | <-Glyoxalase<-HTH_26<-?<-?<-?\|\|ABC-ATPase+SbcC-><-BrxA-DNA-binding<-PglZ<-N6-N4-METHYLASE<-DUF6079*<-BrxB-BrxF-iSTAND<-CHTH+CHTH\|\|SIGMA-HTH->?->HerA/FtsK->?->?->XIS-HTH->Phage_integrase-> | 1240 | Bacteria>Terrabacteria_group>Bacillota>Bacilli>Bacillales>Staphylococcaceae | GCF_022386515.1 | Mammaliicoccus_sp._H-M32 | LQ616_RS02785 | 121216..124938 | 2898691 | DUF6079-family-protein-[Mammaliicoccus-sp.-H-M32] |
| WP_106343732.1 | Type3.MTaseII.containing.BREX | <-ABhydrolase+ABhydrolase<-?<-?<-BrxA-DNA-binding<-PglZ<-PIWI<-?<-N6-N4-METHYLASE<-DUF6079*<-BrxB-BrxF-iSTAND<-?<-Peptidase_M28 | 1237 | Bacteria>Terrabacteria_group>Bacillota>Bacilli>Bacillales>Thermoactinomycetaceae | GCF_003001905.1 | Planifilum_fimeticola | CLV97_RS00260 | 62544..66257 | 201975 | DUF6079-family-protein-[Planifilum-fimeticola] |
| WP_181742021.1 | Type3.MTaseII.containing.BREX | <-BrxA-DNA-binding<-PglZ<-N6-N4-METHYLASE<-DUF6079*<-BrxB-BrxF-iSTAND<-REase<-SF1-SF2-Helicase+REase-p | 1225 | Bacteria>Terrabacteria_group>Bacillota>Bacilli>Bacillales>Thermoactinomycetaceae | GCF_013868055.1 | Thermoactinomyces_mirandus | H2C83_RS15065 | 6174..9851 | 2756294 | DUF6079-family-protein-[Thermoactinomyces-mirandus] |
| WP_229736080.1 | Type3.MTaseII.containing.BREX | TRAM+METHYLASE-><-?\|\|REase->?->REase->?-><-?\|\|BrxB-BrxF-iSTAND->DUF6079*->N6-N4-METHYLASE->PglZ->BrxA-DNA-binding->?-><-TPR+TPR+TPR+TPR | 1245 | Bacteria>Terrabacteria_group>Bacillota>Bacilli>Bacillales>Thermoactinomycetaceae | GCF_014637345.1 | Kroppenstedtia_guangzhouensis | IEV46_RS12090 | 12192..15929 | 1274356 | DUF6079-family-protein-[Kroppenstedtia-guangzhouensis] |
| WP_044640701.1 | Type3.MTaseII.containing.BREX | <-BrxA-DNA-binding<-PglZ<-N6-N4-METHYLASE<-DUF6079*<-BrxB-BrxF-iSTAND<-HEPN<-REase<-SF1-SF2-Helicase<-?<-HAD+PRTase+PRTase+REase+SF1-SF2-Helicase | 1225 | Bacteria>Terrabacteria_group>Bacillota>Bacilli>Bacillales>Thermoactinomycetaceae | GCF_902375465.1 | Risungbinella_massiliensis | VJ09_RS06145 | 1166148..1169825 | 1329796 | DUF6079-family-protein-[Risungbinella-massiliensis] |
| WP_016896589.1 | Type3.MTaseII.containing.BREX | Peptidase_M3_N+Peptidase_M3-><-?\|\|MODE-HTH+PBPII->?->?-><-Phage_integrase\|\|BrxB-BrxF-iSTAND->DUF6079*->N6-N4-METHYLASE->PglZ->BrxA-DNA-binding->?->ZnR-> | 1240 | Bacteria>Terrabacteria_group>Bacillota>Bacilli>Lactobacillales>Aerococcaceae | GCF_000262085.1 | Aerococcus_viridans | Y15_RS0101355 | 83024..86746 | 1377 | DUF6079-family-protein-[Aerococcus-viridans] |
| MBP8682477.1 | Type3.MTaseII.containing.BREX | Trypsin+PDZ-><-?<-TrkH\|\|SPOUT->XIS-HTH->BrxB-BrxF-iSTAND->DUF6079*->N6-N4-METHYLASE->TUDOR+SF1-SF2-Helicase+iREase+CTD->PglZ->BrxA-DNA-binding->N6-METHYLASE-p->?->?->?->UDG-conflict-> | 1236 | Bacteria>Terrabacteria_group>Bacillota>Bacilli>Lactobacillales>Carnobacteriaceae | GCA_018053525.1 | Trichococcus_sp. | KBH68_01095 | 40360..44070 | 1985464 | hypothetical-protein-[Trichococcus-sp.] |
| MCA9765537.1 | Type3.MTaseII.containing.BREX | REase+N6_METHYLASE->TRD->REase+SF1-SF2-Helicase-><-?<-MarR-HTH\|\|GDResolvase+HIN-HTH-><-BrxA-DNA-binding<-PglZ<-N6-N4-METHYLASE<-DUF6079*<-BrxB-BrxF-iSTAND<-AIPR | 1239 | Bacteria>Terrabacteria_group>Bacillota>Bacilli>Lactobacillales>Carnobacteriaceae | GCA_020431645.1 | Carnobacterium_sp. | KC455_03870 | 18945..22664 | 48221 | ATP-binding-protein-[Carnobacterium-sp.] |
| WP_028274683.1 | Type3.MTaseII.containing.BREX | BrxB-BrxF-iSTAND->DUF6079*->N6-N4-METHYLASE->PglZ->BrxA-DNA-binding-> | 1240 | Bacteria>Terrabacteria_group>Bacillota>Bacilli>Lactobacillales>Carnobacteriaceae | GCF_000429585.1 | Atopococcus_tabaci | G470_RS0109895 | 1044..4766 | 269774 | DUF6079-family-protein-[Atopococcus-tabaci] |
| WP_034538267.1 | Type3.MTaseII.containing.BREX | ABC-ATPase+SbcC->?->Cysteine-Desulfurase->?->?->?->?->?->BrxB-BrxF-iSTAND->DUF6079*->N6-N4-METHYLASE->PglZ->BrxA-DNA-binding->HTH_3-><-SF1-SF2-Helicase\|\|?->?->ABC-ATPase+SbcC-> | 1239 | Bacteria>Terrabacteria_group>Bacillota>Bacilli>Lactobacillales>Carnobacteriaceae | GCF_000746825.1 | Carnobacterium_inhibens | BR65_RS11415 | 2140041..2143760 | 147709 | DUF6079-family-protein-[Carnobacterium-inhibens] |
| WP_106450120.1 | Type3.MTaseII.containing.BREX | Glyco_hydro_4+Glyco_hydro_4C->?-><-?\|\|?->HAD+HAD->?->LACI-HTH+PBPI->BrxB-BrxF-iSTAND->DUF6079*->N6-N4-METHYLASE->PglZ->BrxA-DNA-binding->TUDOR+SF1-SF2-Helicase+iREase+CTD+REase->?->TPR->Peptidase_M42->SIGMA-HTH+SIGMA-HTH+SIGMA-HTH->35exo+wHTH+SF1-SF2-Helicase+HTH-> | 1234 | Bacteria>Terrabacteria_group>Bacillota>Bacilli>Lactobacillales>Carnobacteriaceae | GCF_003008695.1 | Trichococcus_alkaliphilus | C7R73_RS08650 | 50442..54146 | 2052943 | DUF6079-family-protein-[Trichococcus-alkaliphilus] |
| WP_091481844.1 | Type3.MTaseII.containing.BREX | ACET+ACET->?->?->ABC-ATPase+SbcC+ABC-ATPase+SbcC->?->?->?->\|\|BrxB-BrxF-iSTAND->DUF6079*->N6-N4-METHYLASE->PglZ->BrxA-DNA-binding-> | 1240 | Bacteria>Terrabacteria_group>Bacillota>Bacilli>Lactobacillales>Carnobacteriaceae | GCF_007991315.1 | Alkalibacterium_pelagium | APE02nite_RS09095 | 86351..90073 | 426702 | DUF6079-family-protein-[Alkalibacterium-pelagium] |
| WP_160806092.1 | Type3.MTaseII.containing.BREX | <-Arm-DNA-bind_4+Phage_int_SAM_3+Phage_integrase<-XIS-HTH<-Phage_GPA<-?\|\|HTH_3->BrxB-BrxF-iSTAND->DUF6079*->N6-N4-METHYLASE->PglZ->BrxA-DNA-binding->RVT->?->?->?->?->Glyco_hydro_4+Glyco_hydro_4C-> | 1239 | Bacteria>Terrabacteria_group>Bacillota>Bacilli>Lactobacillales>Enterococcaceae | - | Enterococcus | PG785_RS12680 | 21021..24740 | 1350 | MULTISPECIES:-DUF6079-family-protein-[Enterococcus] |
| EGP4935252.1 | Type3.MTaseII.containing.BREX | <-BrxA-DNA-binding<-PglZ<-N6-N4-METHYLASE<-DUF6079*<-BrxB-BrxF-iSTAND | 1240 | Bacteria>Terrabacteria_group>Bacillota>Bacilli>Lactobacillales>Enterococcaceae | GCA_015504425.1 | Enterococcus_faecium | FEW45_002553 | 4787..8509 | 1352 | ATP-binding-protein-[Enterococcus-faecium] |
| EGP5109070.1 | Type3.MTaseII.containing.BREX | HTH_3->BrxB-BrxF-iSTAND->DUF6079*->N6-N4-METHYLASE->PglZ->BrxA-DNA-binding->RVT-> | 1239 | Bacteria>Terrabacteria_group>Bacillota>Bacilli>Lactobacillales>Enterococcaceae | GCA_015505085.1 | Enterococcus_faecium | EWX82_13105 | 1939..5658 | 1352 | ATP-binding-protein-[Enterococcus-faecium] |
| MBG7676594.1 | Type3.MTaseII.containing.BREX | <-ZnR<-?<-BrxA-DNA-binding<-PglZ<-N6-N4-METHYLASE<-DUF6079-p* | 727 | Bacteria>Terrabacteria_group>Bacillota>Bacilli>Lactobacillales>Enterococcaceae | GCA_015768365.1 | Enterococcus_faecium | I5V21_12705 | 6623..8806 | 1352 | hypothetical-protein-[Enterococcus-faecium] |
| MBG8188679.1 | Type3.MTaseII.containing.BREX | <-ZnR<-?<-BrxA-DNA-binding<-PglZ<-N6-N4-METHYLASE<-DUF6079-p* | 748 | Bacteria>Terrabacteria_group>Bacillota>Bacilli>Lactobacillales>Enterococcaceae | GCA_015771775.1 | Enterococcus_faecium | I5079_12220 | 6496..8744 | 1352 | ATP-binding-protein-[Enterococcus-faecium] |
| MBH0871427.1 | Type3.MTaseII.containing.BREX | DDE_Tnp_1_3->?->BrxB-BrxF-iSTAND->DUF6079*->N6-N4-METHYLASE->PglZ->BrxA-DNA-binding->?->ZnR-> | 1240 | Bacteria>Terrabacteria_group>Bacillota>Bacilli>Lactobacillales>Enterococcaceae | GCA_015992865.1 | Enterococcus_faecium | I5R84_11420 | 2044..5766 | 1352 | ATP-binding-protein-[Enterococcus-faecium] |
| EIB3113460.1 | Type3.MTaseII.containing.BREX | <-DUF6079-p*<-BrxB-BrxF-iSTAND<-CHTH+CHTH\|\|SIGMA-HTH->?->HerA/FtsK->?->?->XIS-HTH->Phage_integrase-> | 628 | Bacteria>Terrabacteria_group>Bacillota>Bacilli>Lactobacillales>Enterococcaceae | GCA_019875925.1 | Enterococcus_faecalis | K8S68_002277 | 1..1885 | 1351 | ATP-binding-protein-[Enterococcus-faecalis] |
| EJI7171896.1 | Type3.MTaseII.containing.BREX | <-DUF6079-p*<-BrxB-BrxF-iSTAND<-CHTH+CHTH<-CHTH\|\|SIGMA-HTH->?->?->?->XIS-HTH->Phage_integrase-> | 954 | Bacteria>Terrabacteria_group>Bacillota>Bacilli>Lactobacillales>Enterococcaceae | GCA_023804595.1 | Enterococcus_faecalis | NFW91_002051 | 1..2863 | 1351 | ATP-binding-protein-[Enterococcus-faecalis] |
| WP_010718371.1 | Type3.MTaseII.containing.BREX | SF1-SF2-Helicase+REase->CR-Kinase->?-><-Phage_GPA<-?\|\|HTH_3->BrxB-BrxF-iSTAND->DUF6079*->N6-N4-METHYLASE->PglZ->BrxA-DNA-binding->RVT-><-DDE_Tnp_IS240\|\|<-ROK-HTH<-?\|\|?->LACI-HTH+PBPI-> | 1239 | Bacteria>Terrabacteria_group>Bacillota>Bacilli>Lactobacillales>Enterococcaceae | GCF_000391745.1 | Enterococcus_faecium | SCW_RS15875 | 200040..203759 | 1352 | DUF6079-family-protein-[Enterococcus-faecium] |
| WP_095803873.1 | Type3.MTaseII.containing.BREX | <-ABC-ATPase+SbcC<-?<-?<-RVT<-BrxA-DNA-binding<-PglZ<-N6-N4-METHYLASE<-DUF6079*<-BrxB-BrxF-iSTAND<-HTH_3\|\|?->Phage_GPA->XIS-HTH->Phage_int_SAM_3+Phage_integrase-><-DOC+HTH<-?<-ABC-ATPase+SbcC | 1239 | Bacteria>Terrabacteria_group>Bacillota>Bacilli>Lactobacillales>Enterococcaceae | GCF_002290025.1 | Enterococcus_thailandicus | CK496_RS12070 | 2555880..2559599 | 417368 | DUF6079-family-protein-[Enterococcus-thailandicus] |
| WP_257957656.1 | Type3.MTaseII.containing.BREX | <-ZnR<-?<-BrxA-DNA-binding<-PglZ<-N6-N4-METHYLASE<-DUF6079-p*<-BrxB-BrxF-iSTAND | 969 | Bacteria>Terrabacteria_group>Bacillota>Bacilli>Lactobacillales>Enterococcaceae | GCF_002859045.1 | Enterococcus_faecium | CYQ99_RS14345 | 7365..10274 | 1352 | DUF6079-family-protein-[Enterococcus-faecium] |
| WP_104889221.1 | Type3.MTaseII.containing.BREX | <-Phage_int_SAM_3+Phage_integrase<-XIS-HTH<-Phage_GPA<-?\|\|HTH_3->?->BrxB-BrxF-iSTAND->DUF6079*->N6-N4-METHYLASE->PglZ->BrxA-DNA-binding->?->ZnR->\|\|SbcC->?->ABC-ATPase+SbcC-> | 1240 | Bacteria>Terrabacteria_group>Bacillota>Bacilli>Lactobacillales>Enterococcaceae | GCF_002949075.1 | Enterococcus | CUS48_RS02210 | 37761..41483 | 1350 | MULTISPECIES:-DUF6079-family-protein-[Enterococcus] |
| WP_128704481.1 | Type3.MTaseII.containing.BREX | <-RVT<-BrxA-DNA-binding<-PglZ<-N6-N4-METHYLASE<-DUF6079*<-BrxB-BrxF-iSTAND<-HTH_3 | 1239 | Bacteria>Terrabacteria_group>Bacillota>Bacilli>Lactobacillales>Enterococcaceae | GCF_004103165.1 | Enterococcus_faecium | EG862_RS12265 | 7859..11578 | 1352 | DUF6079-family-protein-[Enterococcus-faecium] |
| WP_159162355.1 | Type3.MTaseII.containing.BREX | <-Phage_integrase<-XIS-HTH<-?<-?<-?<-SIGMA-HTH\|\|CHTH->CHTH+CHTH->BrxB-BrxF-iSTAND->DUF6079*->N6-N4-METHYLASE->PglZ->BrxA-DNA-binding->NUDIX->HKD-DNase+SF1-SF2-Helicase+HTH+HTH+SRA-><-NtrC-AAA+FIS-HTH | 1240 | Bacteria>Terrabacteria_group>Bacillota>Bacilli>Lactobacillales>Enterococcaceae | GCF_009830655.1 | Enterococcus_faecalis | F1765_RS09375 | 70659..74381 | 1351 | DUF6079-family-protein-[Enterococcus-faecalis] |
| WP_104802017.1 | Type3.MTaseII.containing.BREX | <-Phage_integrase<-XIS-HTH<-?<-?<-HerA/FtsK<-?<-SIGMA-HTH\|\|CHTH+CHTH->BrxB-BrxF-iSTAND->DUF6079*->N6-N4-METHYLASE->PglZ->BrxA-DNA-binding->NUDIX->HKD-DNase+SF1-SF2-Helicase+HTH+HTH+SRA-><-NtrC-AAA+FIS-HTH | 1240 | Bacteria>Terrabacteria_group>Bacillota>Bacilli>Lactobacillales>Enterococcaceae | GCF_015547115.1 | Enterococcus_faecalis | PMV45_RS09150 | 27614..31336 | 1351 | DUF6079-family-protein-[Enterococcus-faecalis] |
| WP_202065241.1 | Type3.MTaseII.containing.BREX | <-REase\|\|?-><-RVT<-BrxA-DNA-binding<-PglZ<-N6-N4-METHYLASE<-DUF6079*<-BrxB-BrxF-iSTAND<-HTH_3\|\|?->Phage_GPA->XIS-HTH->Phage_int_SAM_3+Phage_integrase-><-?<-?<-ACET | 1239 | Bacteria>Terrabacteria_group>Bacillota>Bacilli>Lactobacillales>Enterococcaceae | GCF_016745275.1 | Enterococcus | IW492_RS03495 | 90710..94429 | 1350 | MULTISPECIES:-DUF6079-family-protein-[Enterococcus] |
| WP_227259064.1 | Type3.MTaseII.containing.BREX | <-Glyco_hydro_1<-?<-?<-?<-ZnR<-?<-BrxA-DNA-binding<-PglZ<-N6-N4-METHYLASE<-DUF6079*<-BrxB-BrxF-iSTAND<-?\|\|SIGMA-HTH->?->HerA/FtsK->?->?->XIS-HTH->Phage_integrase-> | 1240 | Bacteria>Terrabacteria_group>Bacillota>Bacilli>Lactobacillales>Enterococcaceae | GCF_020628455.1 | Vagococcus_fluvialis | K5K99_RS01985 | 381627..385349 | 2738 | DUF6079-family-protein-[Vagococcus-fluvialis] |
| WP_221685687.1 | Type3.MTaseII.containing.BREX | <-BrxA-DNA-binding<-PglZ<-N6-N4-METHYLASE<-DUF6079*<-BrxB-BrxF-iSTAND | 1240 | Bacteria>Terrabacteria_group>Bacillota>Bacilli>Lactobacillales>Enterococcaceae | GCF_021560195.1 | Enterococcus_faecium | HCH26_11475 | 6041..9763 | 1352 | DUF6079-family-protein-[Enterococcus-faecium] |
| WP_250864578.1 | Type3.MTaseII.containing.BREX | <-ZnR<-?<-BrxA-DNA-binding<-PglZ<-N6-N4-METHYLASE<-DUF6079*<-BrxB-BrxF-iSTAND<-CHTH+CHTH\|\|SIGMA-HTH->?->HerA/FtsK->?->?->XIS-HTH->Phage_integrase-> | 1240 | Bacteria>Terrabacteria_group>Bacillota>Bacilli>Lactobacillales>Enterococcaceae | GCF_023698675.1 | Enterococcus_faecalis | NEI57_RS08695 | 6712..10434 | 1351 | DUF6079-family-protein-[Enterococcus-faecalis] |
| WP_251845434.1 | Type3.MTaseII.containing.BREX | REC+OmpR-HTH->HAMP+HISKIN->?-><-zf-ISL3+HTH_Tnp_ISL3+DDE_Tnp_ISL3\|\|?->HTH_3-><-MULE-TRANSPOSASE\|\|BrxB-BrxF-iSTAND->DUF6079*->N6-N4-METHYLASE->PglZ->BrxA-DNA-binding->RVT->?->ClpABC-AAA-><-Phage_integrase<-MULE-TRANSPOSASE<-HIN-HTH | 1239 | Bacteria>Terrabacteria_group>Bacillota>Bacilli>Lactobacillales>Enterococcaceae | GCF_023743525.1 | Enterococcus_faecium | NE259_RS00795 | 135954..139673 | 1352 | DUF6079-family-protein-[Enterococcus-faecium] |
| WP_085837461.1 | Type3.MTaseII.containing.BREX | <-LACI-HTH+PBPI<-LZ_Tnp_IS66+zf-IS66+DDE_Tnp_IS66+DDE_Tnp_IS66_C<-?<-?<-ZnR<-?<-BrxA-DNA-binding<-PglZ<-N6-N4-METHYLASE<-DUF6079*<-BrxB-BrxF-iSTAND | 1240 | Bacteria>Terrabacteria_group>Bacillota>Bacilli>Lactobacillales>Enterococcaceae | GCF_900178965.1 | Enterococcus_faecium | CAP04_RS11535 | 13715..17437 | 1352 | DUF6079-family-protein-[Enterococcus-faecium] |
| WP_142971906.1 | Type3.MTaseII.containing.BREX | <-HTH_26\|\|Classical-AAA->Peptidase_S8->?-><-BrxA-DNA-binding<-PglZ<-N6-N4-METHYLASE<-DUF6079*<-BrxB-BrxF-iSTAND<-CHTH+CHTH\|\|SIGMA-HTH->?->HerA/FtsK->?->?->XIS-HTH->Phage_integrase-> | 1240 | Bacteria>Terrabacteria_group>Bacillota>Bacilli>Lactobacillales>Enterococcaceae | GCF_902159265.1 | Enterococcus_gallinarum | FN047_RS13415 | 79668..83390 | 1353 | DUF6079-family-protein-[Enterococcus-gallinarum] |
| WP_057086594.1 | Type3.MTaseII.containing.BREX | <-Phage_integrase<-XIS-HTH<-?<-?<-?<-SIGMA-HTH\|\|CHTH->CHTH+CHTH->BrxB-BrxF-iSTAND->DUF6079*->N6-N4-METHYLASE->PglZ->BrxA-DNA-binding->NUDIX->HKD-DNase+SF1-SF2-Helicase+HTH+HTH+SRA-><-NtrC-AAA+FIS-HTH | 1240 | Bacteria>Terrabacteria_group>Bacillota>Bacilli>Lactobacillales>Enterococcaceae | GCF_902160495.1 | Enterococcus_faecalis | FQS91_RS11170 | 89842..93564 | 1351 | DUF6079-family-protein-[Enterococcus-faecalis] |
| WP_142425732.1 | Type3.MTaseII.containing.BREX | <-Phage_integrase<-XIS-HTH<-?<-?<-HerA/FtsK<-?<-SIGMA-HTH\|\|CHTH+CHTH->BrxB-BrxF-iSTAND->DUF6079*->N6-N4-METHYLASE->PglZ->BrxA-DNA-binding->?->ZnR-> | 1240 | Bacteria>Terrabacteria_group>Bacillota>Bacilli>Lactobacillales>Enterococcaceae | GCF_902161145.1 | Enterococcus_faecalis | FN270_RS12040 | 67344..71066 | 1351 | DUF6079-family-protein-[Enterococcus-faecalis] |
| WP_002307575.1 | Type3.MTaseII.containing.BREX | <-RVT<-BrxA-DNA-binding<-PglZ<-N6-N4-METHYLASE<-DUF6079*<-BrxB-BrxF-iSTAND | 1239 | Bacteria>Terrabacteria_group>Bacillota>Bacilli>Lactobacillales>Enterococcaceae | GCF_947051815.1 | Enterococcus_faecium | PG973_RS13380 | 6760..10479 | 1352 | DUF6079-family-protein-[Enterococcus-faecium] |
| KRN24181.1 | Type3.MTaseII.containing.BREX | <-GDResolvase+HIN-HTH<-?<-?<-BrxA-DNA-binding-p<-PglZ<-TUDOR+SF1-SF2-Helicase+iREase+CTD<-N6-N4-METHYLASE<-DUF6079*<-BrxB-BrxF-iSTAND-p | 1198 | Bacteria>Terrabacteria_group>Bacillota>Bacilli>Lactobacillales>Lactobacillaceae | GCA_001436615.1 | Lacticaseibacillus_camelliae_DSM_22697_=_JCM_13995 | FC75_GL001360 | 56871..60467 | 1423730 | hypothetical-protein-FC75_GL001360-[Lacticaseibacillus-camelliae-DSM-22697-=-JCM-13995] |
| HAM87243.1 | Type3.MTaseII.containing.BREX | <-SPOUT\|\|?-><-BrxA-DNA-binding<-PglZ<-N6-N4-METHYLASE<-DUF6079*<-BrxB-BrxF-iSTAND<-CHTH<-?\|\|Tox-RelA->REC+OmpR-HTH->HISKIN-><-?<-Sulfatase+Phosphodiest<-SF1-SF2-Helicase | 1226 | Bacteria>Terrabacteria_group>Bacillota>Bacilli>Lactobacillales>Lactobacillaceae | GCA_003452595.1 | Lactobacillus_sp. | DCP87_06820 | 91663..95343 | 1591 | ATPase-[Lactobacillus-sp.] |
| MBT8908682.1 | Type3.MTaseII.containing.BREX | NUDIX->?-><-AraC-HTH+O6MGT-HTH<-NERD+SF1-SF2-Helicase<-?<-BrxA-DNA-binding-p<-PglZ<-TUDOR+SF1-SF2-Helicase+iREase+CTD<-N6-N4-METHYLASE<-DUF6079-p*<-BrxB-BrxF-iSTAND<-?<-SF1-SF2-Helicase+REase+HTH<-?<-?\|\|?-><-HAD+HAD<-SIGMA-HTH+SIGMA-HTH<-SIGMA-HTH+SIGMA-HTH+SIGMA-HTH | 941 | Bacteria>Terrabacteria_group>Bacillota>Bacilli>Lactobacillales>Lactobacillaceae | GCA_018720765.1 | Lactobacillus_delbrueckii_subsp._bulgaricus | BTI62_02310 | 20564..23389 | 1585 | hypothetical-protein-[Lactobacillus-delbrueckii-subsp.-bulgaricus] |
| WP_263849636.1 | Type3.MTaseII.containing.BREX | <-GDResolvase+HIN-HTH<-?<-BrxA-DNA-binding<-PglZ-p<-PglZ-p<-TUDOR+SF1-SF2-Helicase+iREase+CTD<-N6-N4-METHYLASE<-DUF6079-p*<-DUF6079-p<-DUF6079-p<-BrxB-BrxF-iSTAND | 193 | Bacteria>Terrabacteria_group>Bacillota>Bacilli>Lactobacillales>Lactobacillaceae | GCF_001311195.1 | Lacticaseibacillus_camelliae | JCM13995_RS17045 | 12766..13347 | 381742 | DUF6079-family-protein-[Lacticaseibacillus-camelliae] |
| WP_054664892.1 | Type3.MTaseII.containing.BREX | <-BrxA-DNA-binding<-PglZ-p<-PglZ-p<-TUDOR+SF1-SF2-Helicase+iREase+CTD<-N6-N4-METHYLASE<-DUF6079-p<-DUF6079-p<-DUF6079-p*<-BrxB-BrxF-iSTAND | 317 | Bacteria>Terrabacteria_group>Bacillota>Bacilli>Lactobacillales>Lactobacillaceae | GCF_001311195.1 | Lacticaseibacillus_camelliae | JCM13995_RS17060 | 15502..16455 | 381742 | DUF6079-family-protein-[Lacticaseibacillus-camelliae] |
| WP_056989286.1 | Type3.MTaseII.containing.BREX | <-GDResolvase+HIN-HTH<-?<-BrxA-DNA-binding<-PglZ<-TUDOR+SF1-SF2-Helicase+iREase+CTD<-N6-N4-METHYLASE<-DUF6079*<-BrxB-BrxF-iSTAND<-HTH_26 | 1229 | Bacteria>Terrabacteria_group>Bacillota>Bacilli>Lactobacillales>Lactobacillaceae | GCF_001436615.1 | Lacticaseibacillus_camelliae | FC75_RS04945 | 56871..60560 | 381742 | DUF6079-family-protein-[Lacticaseibacillus-camelliae] |
| WP_199799129.1 | Type3.MTaseII.containing.BREX | MobA_MobL-><-?<-BrxA-DNA-binding<-PglZ<-TUDOR+SF1-SF2-Helicase+iREase+CTD<-N6-N4-METHYLASE<-DUF6079*<-BrxB-BrxF-iSTAND<-DUF4096+DDE_Tnp_1+DDE_Tnp_1_2<-?\|\|TetR-HTH->?->GDResolvase+HIN-HTH-><-?<-HTH_Tnp_1+HTH_21+rve+rve_3<-HTH_Tnp_1 | 1229 | Bacteria>Terrabacteria_group>Bacillota>Bacilli>Lactobacillales>Lactobacillaceae | GCF_002849955.1 | Lactobacillus_helveticus | Lh22155_RS09850 | 1898512..1902201 | 1587 | DUF6079-family-protein-[Lactobacillus-helveticus] |
| WP_146342557.1 | Type3.MTaseII.containing.BREX | <-ABC-ATPase+SbcC+ABC-ATPase+SbcC<-?<-BrxA-DNA-binding<-PglZ<-N6-N4-METHYLASE<-DUF6079*<-BrxB-BrxF-iSTAND<-?<-?<-?<-?<-ABC-ATPase+SbcC<-ABC-ATPase+SbcC<-ACET | 1241 | Bacteria>Terrabacteria_group>Bacillota>Bacilli>Lactobacillales>Lactobacillaceae | GCF_007922695.1 | Dellaglioa_algida | LABALGNA3A7_RS03365 | 274772..278497 | 105612 | DUF6079-family-protein-[Dellaglioa-algida] |
| WP_254672326.1 | Type3.MTaseII.containing.BREX | <-BrxA-DNA-binding<-PglZ<-N6-N4-METHYLASE<-DUF6079*<-BrxB-BrxF-iSTAND | 1240 | Bacteria>Terrabacteria_group>Bacillota>Bacilli>Lactobacillales>Lactobacillaceae | GCF_024272965.1 | Latilactobacillus_curvatus | LOX55_RS08145 | 6545..10267 | 28038 | DUF6079-family-protein-[Latilactobacillus-curvatus] |
| WP_270641084.1 | Type3.MTaseII.containing.BREX | HerA/FtsK->?->AAA-ATPase->?->Phage_int_SAM_3+Phage_integrase->CHTH->BrxB-BrxF-iSTAND->DUF6079*->N6-N4-METHYLASE->PglZ->BrxA-DNA-binding-><-?\|\|SPOUT-> | 1226 | Bacteria>Terrabacteria_group>Bacillota>Bacilli>Lactobacillales>Lactobacillaceae | GCF_027693785.1 | Limosilactobacillus_mucosae | PG719_RS03100 | 49653..53333 | 97478 | DUF6079-family-protein-[Limosilactobacillus-mucosae] |
| WP_106483446.1 | Type3.MTaseII.containing.BREX | <-TetR-HTH\|\|Peptidase_M29->?->ZnR->ZnR->?->?->BrxB-BrxF-iSTAND->DUF6079*->N6-N4-METHYLASE-><-?<-?<-?<-?<-Amidohydro_1 | 1239 | Bacteria>Terrabacteria_group>Bacillota>Bacilli>Lactobacillales>Lactobacillaceae | GCF_900258455.1 | Latilactobacillus_fuchuensis | LFUMFP_RS08315 | 1739454..1743173 | 164393 | DUF6079-family-protein-[Latilactobacillus-fuchuensis] |
| WP_270631004.1 | Type3.MTaseII.containing.BREX | HerA/FtsK->?->AAA-ATPase->?->Phage_int_SAM_3+Phage_integrase->CHTH->BrxB-BrxF-iSTAND->DUF6079*->N6-N4-METHYLASE->PglZ->BrxA-DNA-binding-><-?\|\|SPOUT-> | 1226 | Bacteria>Terrabacteria_group>Bacillota>Bacilli>Lactobacillales>Lactobacillaceae | GCF_905194075.1 | Limosilactobacillus_mucosae | QDG36_RS02675 | 49996..53676 | 97478 | DUF6079-family-protein-[Limosilactobacillus-mucosae] |
| MBR2506166.1 | Type3.MTaseII.containing.BREX | <-HTH+HTH\|\|?->HTH_36->?->?->TraJ-RHH->TraI-VirD2-RCRE->?->BrxB-BrxF-iSTAND->DUF6079*->OLD-ABC->HNH->N6-N4-METHYLASE->AIPR->Schlafen+HATPase+wHTH->TUDOR+SF1-SF2-Helicase+iREase+CTD->PglZ->BrxA-DNA-binding-> | 1225 | Bacteria>Terrabacteria_group>Bacillota>Bacilli>unclassified_Bacilli | GCA_017502465.1 | Bacilli_bacterium | IKB70_04475 | 8185..11862 | 1903720 | hypothetical-protein-[Bacilli-bacterium] |
| NPV81613.1 | Type3.MTaseII.containing.BREX | BrxB-BrxF-iSTAND->DUF6079*->ParB+HNH->N6-N4-METHYLASE->SIR2->HerA/FtsK->TUDOR+SF1-SF2-Helicase+iREase+CTD->PglZ->BrxA-DNA-binding-> | 1243 | Bacteria>Terrabacteria_group>Bacillota | - | Bacillota_bacterium | HPY52_15350 | 1366..5097 | 1879010 | ATP-binding-protein-[Bacillota-bacterium] |
| WP_252226012.1 | Type3.MTaseII.containing.BREX | <-BrxA-DNA-binding<-PglZ<-TUDOR+SF1-SF2-Helicase+iREase+CTD<-N6-N4-METHYLASE<-Ntox45<-HEPN<-DUF6079*<-BrxB-BrxF-iSTAND\|\|?-><-MACRODOMAIN<-?\|\|?-><-?<-HMA+HAD+HAD<-?<-MarR_2+ROK-HTH | 1228 | Bacteria>Terrabacteria_group>Bacillota>Clostridia>Eubacteriales>Caldicoprobacteraceae | GCF_023838525.1 | Caldicoprobacter_algeriensis | KVG29_RS01735 | 103760..107446 | 699281 | DUF6079-family-protein-[Caldicoprobacter-algeriensis] |
| HBM74930.1 | Type3.MTaseII.containing.BREX | <-N6-N4-METHYLASE-p<-DUF6079-p<-DUF6079-p<-?<-DUF6079-p*<-BrxB-BrxF-iSTAND | 538 | Bacteria>Terrabacteria_group>Bacillota>Clostridia>Eubacteriales>Clostridiaceae | GCA_003503385.1 | Clostridiaceae_bacterium | DD429_05165 | 2962..4576 | 1898204 | exonuclease-SbcC-[Clostridiaceae-bacterium] |
| HHW00112.1 | Type3.MTaseII.containing.BREX | <-XIS-HTH<-BrxA-DNA-binding-p<-PglZ<-TUDOR+SF1-SF2-Helicase+iREase+CTD<-N6-N4-METHYLASE<-DUF6079-p*<-BrxB-BrxF-iSTAND<-HTH_26<-?<-Radical_SAM<-ABC-ATPase+SbcC | 942 | Bacteria>Terrabacteria_group>Bacillota>Clostridia>Eubacteriales>Clostridiaceae | GCA_012840085.1 | Clostridiaceae_bacterium | GXX36_11210 | 108319..111147 | 1898204 | hypothetical-protein-[Clostridiaceae-bacterium] |
| NMM63086.1 | Type3.MTaseII.containing.BREX | wHTH+WYL+WCX->?->HKD-DNase+SF1-SF2-Helicase+HTH+HTH+SRA->?->XIS-HTH->BrxB-BrxF-iSTAND->DUF6079-p*->N6-N4-METHYLASE->TUDOR+SF1-SF2-Helicase+iREase+CTD->PglZ->BrxA-DNA-binding-p-><-?<-?\|\|?->?->SF1-SF2-Helicase+REase-> | 942 | Bacteria>Terrabacteria_group>Bacillota>Clostridia>Eubacteriales>Clostridiaceae | GCA_012926525.1 | Clostridium_muellerianum | HBE96_10315 | 79594..82422 | 2716538 | hypothetical-protein-[Clostridium-muellerianum] |
| MBS3994302.1 | Type3.MTaseII.containing.BREX | <-PglZ<-TUDOR+SF1-SF2-Helicase+iREase+CTD<-N6-N4-METHYLASE<-DUF6079* | 1123 | Bacteria>Terrabacteria_group>Bacillota>Clostridia>Eubacteriales>Clostridiaceae | GCA_018335765.1 | Alkaliphilus_sp. | KGZ33_00785 | 6373..9744 | 1872455 | hypothetical-protein-[Alkaliphilus-sp.] |
| MCI7012744.1 | Type3.MTaseII.containing.BREX | ArdA+LPD40+LPD18+LPD17+LPD2->?-><-BrxA-DNA-binding<-PglZ<-TUDOR+SF1-SF2-Helicase+iREase+CTD<-TIR<-Peptidase_C14<-N6-N4-METHYLASE<-DUF6079*<-BrxB-BrxF-iSTAND<-HTH_26<-TraI-VirD2-RCRE<-TraJ-RHH | 1225 | Bacteria>Terrabacteria_group>Bacillota>Clostridia>Eubacteriales>Clostridiaceae | GCA_022775205.1 | Clostridium_sp. | MR899_03245 | 10846..14523 | 1506 | DUF6079-family-protein-[Clostridium-sp.] |
| MCI9500566.1 | Type3.MTaseII.containing.BREX | <-REase<-BrxA-DNA-binding<-PglZ<-TUDOR+SF1-SF2-Helicase+iREase+CTD<-N6-N4-METHYLASE<-?<-DUF6079-p*<-BrxB-BrxF-iSTAND<-RADICAL-SAM<-METHYLASE<-HTH_26\|\|MPTase->?-><-?<-SymE_toxin<-TRAM+METHYLASE | 941 | Bacteria>Terrabacteria_group>Bacillota>Clostridia>Eubacteriales>Clostridiaceae | GCA_022800895.1 | Hungatella_sp. | HFG76_00915 | 32675..35500 | 2613924 | hypothetical-protein-[Hungatella-sp.] |
| MCM1450901.1 | Type3.MTaseII.containing.BREX | Glycos_transf_4->Glycos_transf->RADICAL-SAM->Glycos_transf->tRNA-><-AbiGi<-BrxA-DNA-binding<-PglZ<-N6-N4-METHYLASE<-DUF6079*<-BrxB-BrxF-iSTAND\|\|?->DADA-Ligase-><-?\|\|tRNA->?->Tox-RelE-><-?\|\|METHYLASE->SF1-SF2-Helicase-> | 1231 | Bacteria>Terrabacteria_group>Bacillota>Clostridia>Eubacteriales>Clostridiaceae | GCA_023666615.1 | Clostridium_sp. | NC102_01400 | 304803..308498 | 1506 | DUF6079-family-protein-[Clostridium-sp.] |
| SCI20961.1 | Type3.MTaseII.containing.BREX | ArdCN+LPD32+LPD36+N6_Mtase+SF1-SF2-Helicase->TraJ-RHH->TraM->?->?->?->HTH_26->BrxB-BrxF-iSTAND->DUF6079*->N6-N4-METHYLASE->Polbeta->?->TUDOR+SF1-SF2-Helicase+iREase+CTD->PglZ->BrxA-DNA-binding->?->TERNS3+HEPN-> | 1224 | Bacteria>Terrabacteria_group>Bacillota>Clostridia>Eubacteriales>Clostridiaceae | GCA_900066055.1 | uncultured_Clostridium_sp. | SAMEA2393089_01916 | 116920..120594 | 59620 | Uncharacterised-protein-[uncultured-Clostridium-sp.] |
| WP_008909101.1 | Type3.MTaseII.containing.BREX | <-RelE-ParE<-PhdYeFM_antitox\|\|TPR+TPR->?->?->?->BrxB-BrxF-iSTAND->DUF6079*->N6-N4-METHYLASE->N6-N4-METHYLASE-p->N6-N4-METHYLASE->TUDOR+SF1-SF2-Helicase+iREase+CTD->PglZ->BrxA-DNA-binding->RelE->HTH_3-> | 1228 | Bacteria>Terrabacteria_group>Bacillota>Clostridia>Eubacteriales>Clostridiaceae | GCF_000297115.1 | Caloramator_australicus | CAAU_RS08850 | 18042..21728 | 515264 | DUF6079-family-protein-[Caloramator-australicus] |
| WP_045091538.1 | Type3.MTaseII.containing.BREX | <-TERNS3+HEPN<-?<-BrxA-DNA-binding<-PglZ<-TUDOR+SF1-SF2-Helicase+iREase+CTD<-?<-SMODS<-HNH+SAVED<-N6-N4-METHYLASE<-DUF6079*<-BrxB-BrxF-iSTAND<-HTH_26<-?<-MACRODOMAIN<-?<-?<-TraM<-TraJ-RHH<-LPD32+LPD36+N6_Mtase+SF1-SF2-Helicase | 1224 | Bacteria>Terrabacteria_group>Bacillota>Clostridia>Eubacteriales>Clostridiaceae | GCF_000954015.1 | Clostridium_sp._FS41 | CLFS41_RS04045 | 44721..48395 | 1609975 | DUF6079-family-protein-[Clostridium-sp.-FS41] |
| WP_119865507.1 | Type3.MTaseII.containing.BREX | LZ_Tnp_IS66+zf-IS66+DDE_Tnp_IS66+DDE_Tnp_IS66_C-><-MORC+DUF3684+SF1-SF2-Helicase<-HHH_4+HTH+wHTH+SF1-SF2-Helicase+REase<-REase<-REase<-TUDOR+SF1-SF2-Helicase+iREase+CTD+REase<-BrxA-DNA-binding<-PglZ<-N6-N4-METHYLASE<-DUF6079*<-BrxB-BrxF-iSTAND<-?\|\|RNase->?-><-LZ_Tnp_IS66+zf-IS66+DDE_Tnp_IS66+DDE_Tnp_IS66_C<-?<-HTH_Tnp | 1229 | Bacteria>Terrabacteria_group>Bacillota>Clostridia>Eubacteriales>Clostridiaceae | GCF_002285495.1 | Clostridium_isatidis | BEN51_RS07785 | 1650167..1653856 | 182773 | DUF6079-family-protein-[Clostridium-isatidis] |
| WP_243186892.1 | Type3.MTaseII.containing.BREX | wHTH+WYL+WCX->?->HKD-DNase+SF1-SF2-Helicase+HTH+HTH+SRA->?->BrxB-BrxF-iSTAND->DUF6079*->N6-N4-METHYLASE->TUDOR+SF1-SF2-Helicase+iREase+CTD->PglZ->BrxA-DNA-binding-><-?<-?\|\|?->?->SF1-SF2-Helicase+REase-> | 1225 | Bacteria>Terrabacteria_group>Bacillota>Clostridia>Eubacteriales>Clostridiaceae | GCF_012926525.1 | Clostridium_muellerianum | HBE96_RS10315 | 78745..82422 | 2716538 | DUF6079-family-protein-[Clostridium-muellerianum] |
| WP_191750181.1 | Type3.MTaseII.containing.BREX | <-XIS-HTH<-?<-Herpes_LMP1+REase<-?<-?<-BrxA-DNA-binding<-PglZ<-TUDOR+SF1-SF2-Helicase+iREase+CTD<-N6-N4-METHYLASE<-DUF6079*<-BrxB-BrxF-iSTAND<-?<-?<-XIS-HTH\|\|REase->?-><-?<-?\|\|LexA-protease-> | 1244 | Bacteria>Terrabacteria_group>Bacillota>Clostridia>Eubacteriales>Clostridiaceae | GCF_014836325.1 | Clostridium_gallinarum | H9660_RS09695 | 17179..20913 | 2762246 | DUF6079-family-protein-[Clostridium-gallinarum] |
| WP_195245441.1 | Type3.MTaseII.containing.BREX | <-HIRAN<-BrxA-DNA-binding<-PglZ<-TUDOR+SF1-SF2-Helicase+iREase+CTD<-N6-N4-METHYLASE<-DUF6079*<-BrxB-BrxF-iSTAND<-Cas4+SF1-SF2-Helicase<-?<-GDResolvase+Recombinase+ZnR<-MPTase\|\|Zn_Tnp_IS1+DDE_Tnp_IS1595-> | 1241 | Bacteria>Terrabacteria_group>Bacillota>Clostridia>Eubacteriales>Clostridiaceae | GCF_015547475.1 | Clostridium_celatum | I2A54_RS14225 | 9217..12942 | 36834 | DUF6079-family-protein-[Clostridium-celatum] |
| WP_195923517.1 | Type3.MTaseII.containing.BREX | VWA+SIGMA-HTH->TOPRIM+Topoisom_bac->DNA_METHYLASE->?->JAB->?->BrxB-BrxF-iSTAND->DUF6079*->N6-N4-METHYLASE->TUDOR+SF1-SF2-Helicase+iREase+CTD->PglZ->BrxA-DNA-binding->SF1-SF2-Helicase->DrmB->HKD-DNase->DNA_METHYLASE-><-TraI-VirD2-RCRE | 1245 | Bacteria>Terrabacteria_group>Bacillota>Clostridia>Eubacteriales>Clostridiaceae | GCF_015667975.1 | Clostridium_tepidum | I4U60_RS07000 | 31067..34804 | 1962263 | DUF6079-family-protein-[Clostridium-tepidum] |
| WP_226104522.1 | Type3.MTaseII.containing.BREX | <-ACET<-?<-GDResolvase+Recombinase+Zn_ribbon_recom<-GDResolvase<-?<-BrxA-DNA-binding<-PglZ<-TUDOR+SF1-SF2-Helicase+iREase+CTD<-N6-N4-METHYLASE<-DUF6079*<-BrxB-BrxF-iSTAND<-XIS-HTH<-?<-ABC-ATPase+SbcC+ABC-ATPase+SbcC<-?\|\|METHYLASE-> | 1244 | Bacteria>Terrabacteria_group>Bacillota>Clostridia>Eubacteriales>Clostridiaceae | GCF_020443655.1 | Clostridium_algidicarnis | LGK99_RS07250 | 74205..77939 | 37659 | DUF6079-family-protein-[Clostridium-algidicarnis] |
| WP_226104829.1 | Type3.MTaseII.containing.BREX | ParB+HNH->ZnR+TOPRIM->VirD4-TraG+VirD4-FtsK->HTH_36->HTH_26->BrxB-BrxF-iSTAND->DUF6079*->N6-N4-METHYLASE->?->?->TUDOR+SF1-SF2-Helicase+iREase+CTD->PglZ->BrxA-DNA-binding->HKD-DNase-> | 1230 | Bacteria>Terrabacteria_group>Bacillota>Clostridia>Eubacteriales>Clostridiaceae | GCF_020443655.1 | Clostridium_algidicarnis | LGK99_RS12760 | 10487..14179 | 37659 | DUF6079-family-protein-[Clostridium-algidicarnis] |
| WP_236916068.1 | Type3.MTaseII.containing.BREX | SF1-SF2-Helicase->?->JAB->?->BrxB-BrxF-iSTAND->DUF6079*->N6-N4-METHYLASE->TUDOR+SF1-SF2-Helicase+iREase+CTD->PglZ->BrxA-DNA-binding->DrmB->HKD-DNase->DNA_METHYLASE-><-TraI-VirD2-RCRE<-TraJ-RHH | 1245 | Bacteria>Terrabacteria_group>Bacillota>Clostridia>Eubacteriales>Clostridiaceae | GCF_021655075.1 | Clostridium_sp._Cult2 | C3E90_RS12825 | 4884..8621 | 2079003 | DUF6079-family-protein-[Clostridium-sp.-Cult2] |
| WP_261669784.1 | Type3.MTaseII.containing.BREX | D5ATPase-><-PADR-HTH+MarR-HTH\|\|MULE-TRANSPOSASE->HTH_Tnp_1+HTH_21+rve+rve_3->HTH_Tnp_1+HTH_21+rve+rve_3-><-?<-RNase\|\|BrxB-BrxF-iSTAND->DUF6079*->N6-N4-METHYLASE->PglZ->BrxA-DNA-binding->TUDOR+SF1-SF2-Helicase+iREase+CTD+REase->REase->HHH_4+HTH+wHTH+SF1-SF2-Helicase+REase->?->XIS-HTH->MPTase-> | 1229 | Bacteria>Terrabacteria_group>Bacillota>Clostridia>Eubacteriales>Clostridiaceae | GCF_025398335.1 | Clostridium_sp._CX1 | N4T77_RS00670 | 140475..144164 | 2978346 | DUF6079-family-protein-[Clostridium-sp.-CX1] |
| WP_270566901.1 | Type3.MTaseII.containing.BREX | wHTH+HNH->?->REase->wHTH+WYL+WCX->SF1-SF2-Helicase+REase->?->XIS-HTH->BrxB-BrxF-iSTAND->DUF6079*->N6-N4-METHYLASE-> | 1243 | Bacteria>Terrabacteria_group>Bacillota>Clostridia>Eubacteriales>Clostridiaceae | GCF_027687085.1 | Clostridium_beijerinckii | PG029_RS16420 | 12525..16256 | 1520 | DUF6079-family-protein-[Clostridium-beijerinckii] |
| WP_082812355.1 | Type3.MTaseII.containing.BREX | <-Phage_int_SAM_4+Phage_integrase<-?<-TRD+TRD<-?<-URI-DNase+RAMA<-BrxA-DNA-binding<-PglZ<-TUDOR+SF1-SF2-Helicase+iREase+CTD<-N6-N4-METHYLASE<-DUF6079*<-BrxB-BrxF-iSTAND<-XIS-HTH<-NACHT+SNaCT\|\|SIGMA-HTH->?->REase-> | 1230 | Bacteria>Terrabacteria_group>Bacillota>Clostridia>Eubacteriales>Clostridiaceae | GCF_900078195.1 | Clostridium_sp._Marseille-P299 | BN4220_RS16460 | 1970098..1973790 | 1805477 | DUF6079-family-protein-[Clostridium-sp.-Marseille-P299] |
| WP_114648808.1 | Type3.MTaseII.containing.BREX | <-DNA_METHYLASE<-HKD-DNase<-DrmB<-SF1-SF2-Helicase<-SF1-SF2-Helicase<-BrxA-DNA-binding<-PglZ<-TUDOR+SF1-SF2-Helicase+iREase+CTD<-N6-N4-METHYLASE<-DUF6079*<-BrxB-BrxF-iSTAND<-?<-JAB<-?<-ArdCN+LPD32+LPD36+N6_Mtase+SF1-SF2-Helicase<-DNA_METHYLASE<-TOPRIM+Topoisom_bac<-VWA+SIGMA-HTH | 1245 | Bacteria>Terrabacteria_group>Bacillota>Clostridia>Eubacteriales>Clostridiaceae | GCF_900446985.1 | Clostridium_cochlearium | DXE95_RS10410 | 2107504..2111241 | 1494 | DUF6079-family-protein-[Clostridium-cochlearium] |
| WP_195713928.1 | Type3.MTaseII.containing.BREX | <-DsbD<-?<-?<-SIR2<-?<-DUF6079-p*<-BrxB-BrxF-iSTAND-p\|\|?-><-RelE-ParE<-PhdYeFM_antitox<-?<-?<-NtrC-AAA+FIS-HTH | 154 | Bacteria>Terrabacteria_group>Bacillota>Clostridia>Eubacteriales>Desulfallaceae | GCF_015594625.1 | Desulfallas_sp._Bu1-1 | IT084_RS16190 | 15735..16199 | 2787620 | DUF6079-family-protein-[Desulfallas-sp.-Bu1-1] |
| MCM1566961.1 | Type3.MTaseII.containing.BREX | REase+wHTH->rve+Mu-transpos_C->TnsC-ATPase->?->DUF6079*->N6-N4-METHYLASE->TUDOR+SF1-SF2-Helicase+iREase+CTD-> | 1249 | Bacteria>Terrabacteria_group>Bacillota>Clostridia>Eubacteriales>Desulfitobacteriaceae | GCA_023667845.1 | Dehalobacter_sp. | NC238_13700 | 8149..11898 | 1962289 | DUF6079-family-protein-[Dehalobacter-sp.] |
| WP_068882975.1 | Type3.MTaseII.containing.BREX | <-AAA-ATPase<-HTH+rve<-?<-REase<-Preader17<-BrxA-DNA-binding<-PglZ<-TUDOR+SF1-SF2-Helicase+iREase+CTD<-N6-N4-METHYLASE<-DUF6079*<-BrxB-BrxF-iSTAND<-SIGMA-HTH<-Radical_SAM<-?<-Peptidase_U32<-RADICAL-SAM | 1202 | Bacteria>Terrabacteria_group>Bacillota>Clostridia>Eubacteriales>Desulfitobacteriaceae | GCF_001707455.1 | Dehalobacter_sp._TeCB1 | A7D23_05610 | 38739..42347 | 1843715 | DUF6079-family-protein-[Dehalobacter-sp.-TeCB1] |
| BAF58887.1 | Type3.MTaseII.containing.BREX | REase-><-?<-SIR2<-DEOR-HTH\|\|?->MazE_antitoxin->PIN->?-><-wHTH\|\|DUF6079*->RAMPs->RAMPs->HD-><-DEDD_Tnp_IS110+Transposase_20<-DEDD_Tnp_IS110\|\|RAMPs->?->RAMPs->N6-N4-METHYLASE-> | 1226 | Bacteria>Terrabacteria_group>Bacillota>Clostridia>Eubacteriales>Desulfotomaculaceae | GCA_000010565.1 | Pelotomaculum_thermopropionicum_SI | PTH_0706 | 698519..702199 | 370438 | hypothetical-protein-PTH_0706-[Pelotomaculum-thermopropionicum-SI] |
| KUK64703.1 | Type3.MTaseII.containing.BREX | BrxA-DNA-binding->BrxB-BrxF-iSTAND->DUF6079*->PglZ->N6-N4-METHYLASE->SIGMA+OB-Fold+AAA-ATPase+Lon-Protease-BrxL->HKD-DNase+DUF3150->?->RADICAL-SAM->?->Aminotran_1_2-> | 1437 | Bacteria>Terrabacteria_group>Bacillota>Clostridia>Eubacteriales>Desulfotomaculaceae | GCA_001508995.1 | Desulfotomaculum_sp._46_80 | XD84_1194 | 2070..6383 | 1641375 | MAG:-Uncharacterized-protein-XD84_1194-[Desulfotomaculum-sp.-46_80] |
| OPX93595.1 | Type3.MTaseII.containing.BREX | <-ZnR+Fer2_BFD<-?<-ZnR+HARE-HTH<-REase+iREase<-BrxA-DNA-binding<-PglZ<-TUDOR+SF1-SF2-Helicase+iREase+CTD<-SbcC<-N6-N4-METHYLASE<-DUF6079*<-BrxB-BrxF-iSTAND<-HTH_26<-?<-?<-HTH_36<-VirD4-TraG+VirD4-FtsK<-ZnR+TOPRIM | 1229 | Bacteria>Terrabacteria_group>Bacillota>Clostridia>Eubacteriales>Desulfotomaculaceae | GCA_002067205.1 | Pelotomaculum_sp._PtaB.Bin104 | A4E53_00238 | 25374..29063 | 1811693 | MAG:-hypothetical-protein-A4E53_00238-[Pelotomaculum-sp.-PtaB.Bin104] |
| WP_003539947.1 | Type3.MTaseII.containing.BREX | <-AA_permease<-?\|\|<-BrxA-DNA-binding<-PglZ<-?<-?<-N6-N4-METHYLASE<-DUF6079*<-BrxB-BrxF-iSTAND<-?<-?\|\|?-><-?<-HISKIN<-?<-DsbD | 1224 | Bacteria>Terrabacteria_group>Bacillota>Clostridia>Eubacteriales>Desulfotomaculaceae | GCF_000189755.2 | Desulfotomaculum_nigrificans | DESNIDRAFT_RS0209520 | 1819828..1823502 | 1565 | DUF6079-family-protein-[Desulfotomaculum-nigrificans] |
| WP_134213462.1 | Type3.MTaseII.containing.BREX | SMODS->METHYLASE->?->?->ncRNA->BrxB-BrxF-iSTAND->DUF6079*->N6-N4-METHYLASE->HEPN->Polbeta->PglZ->BrxA-DNA-binding->SF1-SF2-Helicase+REase-p->?-><-wHTH+WYL+WCX | 1225 | Bacteria>Terrabacteria_group>Bacillota>Clostridia>Eubacteriales>Desulfotomaculaceae | GCF_004369225.1 | Pelotomaculum_propionicicum | Pmgp_RS07945 | 51068..54745 | 258475 | DUF6079-family-protein-[Pelotomaculum-propionicicum] |
| MBR2547797.1 | Type3.MTaseII.containing.BREX | <-HISKIN+REC+REC<-ACET<-Abi<-ASCH<-BrxA-DNA-binding<-PglZ<-TUDOR+SF1-SF2-Helicase+iREase+CTD<-N6-N4-METHYLASE<-DUF6079*<-BrxB-BrxF-iSTAND<-HTH_26\|\|nSTAND3->ACET+EVE->ASCH->HTH_31->MPTase->SIGMA-HTH->MazG-> | 1225 | Bacteria>Terrabacteria_group>Bacillota>Clostridia>Eubacteriales>Eubacteriaceae | GCA_017533915.1 | Eubacterium_sp. | IKF07_06380 | 191152..194829 | 142586 | hypothetical-protein-[Eubacterium-sp.] |
| PSR33881.1 | Type3.MTaseII.containing.BREX | HISKIN->?-><-?<-Calcineurin\|\|?->REase->HEPN+PAS+DUF4011+SF1-SF2-Helicase+REase->BrxB-BrxF-iSTAND->DUF6079*->N6-N4-METHYLASE->TUDOR+SF1-SF2-Helicase+iREase+CTD->PglZ->BrxA-DNA-binding->?->?->?->SF1-SF2-Helicase->SF1-SF2-Helicase-> | 1224 | Bacteria>Terrabacteria_group>Bacillota>Clostridia>Eubacteriales>Eubacteriales_incertae_sedis | GCA_003023695.1 | Sulfobacillus_benefaciens | C7B46_07770 | 13431..17105 | 453960 | hypothetical-protein-C7B46_07770-[Sulfobacillus-benefaciens] |
| NPE22818.1 | Type3.MTaseII.containing.BREX | LuxR-HTH-><-wHTH+WYL+WCX<-BrxA-DNA-binding-p<-PglZ<-TUDOR+SF1-SF2-Helicase+iREase+CTD<-N6-N4-METHYLASE<-DUF6079-p*<-BrxB-BrxF-iSTAND<-HTH_36<-SMS-KaiC<-?<-TOPRIM<-HKD-DNase+SF1-SF2-Helicase+HTH+HTH+SRA<-HKD-DNase | 942 | Bacteria>Terrabacteria_group>Bacillota>Clostridia>Eubacteriales>Eubacteriales_incertae_sedis | GCA_013166675.1 | Fusibacter_sp._A1 | HNV05_13335 | 230680..233508 | 2283630 | hypothetical-protein-[Fusibacter-sp.-A1] |
| MBT9676182.1 | Type3.MTaseII.containing.BREX | <-TERNS3+HEPN<-?<-BrxA-DNA-binding<-PglZ<-TUDOR+SF1-SF2-Helicase+iREase+CTD<-?<-Polbeta<-N6-N4-METHYLASE<-DUF6079*<-BrxB-BrxF-iSTAND<-HTH_26<-?<-?<-TraM<-TraJ-RHH<-RVT+HNH | 1224 | Bacteria>Terrabacteria_group>Bacillota>Clostridia>Eubacteriales>Eubacteriales_incertae_sedis | GCA_018784445.1 | Gemmiger_formicilis | GPK89_15935 | 141062..144736 | 745368 | hypothetical-protein-[Gemmiger-formicilis] |
| MCG4528059.1 | Type3.MTaseII.containing.BREX | LPD32+LPD36+N6_Mtase+SF1-SF2-Helicase->TraJ-RHH->TraM->?->?->HTH_26->BrxB-BrxF-iSTAND->DUF6079*->N6-N4-METHYLASE->Polbeta->?->TUDOR+SF1-SF2-Helicase+iREase+CTD->PglZ->BrxA-DNA-binding->?->TERNS3+HEPN-> | 1224 | Bacteria>Terrabacteria_group>Bacillota>Clostridia>Eubacteriales>Eubacteriales_incertae_sedis | GCA_022132365.1 | Intestinimonas_massiliensis | LOP79_13445 | 32491..36165 | 1673721 | DUF6079-family-protein-[Intestinimonas-massiliensis] |
| WP_020373658.1 | Type3.MTaseII.containing.BREX | SWIM+DUF3670+SF1-SF2-Helicase->?->?->BrxB-BrxF-iSTAND->DUF6079*->N6-N4-METHYLASE->DEDD_Tnp_IS110->Transposase_20->N6-N4-METHYLASE->TUDOR+SF1-SF2-Helicase+iREase+CTD->PglZ->BrxA-DNA-binding->?->SIGMA-HTH-> | 1236 | Bacteria>Terrabacteria_group>Bacillota>Clostridia>Eubacteriales>Eubacteriales_incertae_sedis | GCF_000294425.1 | Sulfobacillus_thermosulfidooxidans | B471_RS0103800 | 142039..145749 | 28034 | DUF6079-family-protein-[Sulfobacillus-thermosulfidooxidans] |
| WP_053959405.1 | Type3.MTaseII.containing.BREX | <-SIGMA-HTH<-?<-BrxA-DNA-binding<-PglZ<-TUDOR+SF1-SF2-Helicase+iREase+CTD<-N6-N4-METHYLASE<-DDE_Tnp<-DUF6079*<-BrxB-BrxF-iSTAND<-SWIM+DUF3670+SF1-SF2-Helicase | 1236 | Bacteria>Terrabacteria_group>Bacillota>Clostridia>Eubacteriales>Eubacteriales_incertae_sedis | GCF_001280565.1 | Sulfobacillus_thermosulfidooxidans | A0A63_RS09120 | 1818077..1821787 | 28034 | DUF6079-family-protein-[Sulfobacillus-thermosulfidooxidans] |
| WP_076005773.1 | Type3.MTaseII.containing.BREX | SWIM+DUF3670+SF1-SF2-Helicase->?->BrxB-BrxF-iSTAND->DUF6079*->N6-N4-METHYLASE->TUDOR+SF1-SF2-Helicase+iREase+CTD->PglZ->BrxA-DNA-binding-> | 1236 | Bacteria>Terrabacteria_group>Bacillota>Clostridia>Eubacteriales>Eubacteriales_incertae_sedis | GCF_001953285.1 | Sulfobacillus_thermosulfidooxidans | BFX05_RS03700 | 56347..60057 | 28034 | DUF6079-family-protein-[Sulfobacillus-thermosulfidooxidans] |
| WP_243118983.1 | Type3.MTaseII.containing.BREX | LuxR-HTH-><-wHTH+WYL+WCX<-BrxA-DNA-binding<-PglZ<-TUDOR+SF1-SF2-Helicase+iREase+CTD<-N6-N4-METHYLASE<-DUF6079*<-BrxB-BrxF-iSTAND<-HTH_36<-SMS-KaiC<-?<-HKD-DNase+SF1-SF2-Helicase+HTH+HTH+SRA | 1225 | Bacteria>Terrabacteria_group>Bacillota>Clostridia>Eubacteriales>Eubacteriales_incertae_sedis | GCF_013166675.1 | unclassified_Fusibacter | HNV05_RS13330 | 230680..234357 | 2624464 | MULTISPECIES:-DUF6079-family-protein-[unclassified-Fusibacter] |
| WP_252261727.1 | Type3.MTaseII.containing.BREX | <-TERNS3+HEPN<-?<-BrxA-DNA-binding<-PglZ<-TUDOR+SF1-SF2-Helicase+iREase+CTD<-?<-Polbeta<-N6-N4-METHYLASE<-DUF6079*<-BrxB-BrxF-iSTAND<-HTH_26<-?<-TraM<-TraJ-RHH<-SF1-SF2-Helicase<-RVT+HNH | 1219 | Bacteria>Terrabacteria_group>Bacillota>Clostridia>Eubacteriales>Eubacteriales_incertae_sedis | GCF_018784445.1 | Gemmiger_formicilis | GPK89_RS15985 | 141062..144721 | 745368 | DUF6079-family-protein-[Gemmiger-formicilis] |
| WP_269190175.1 | Type3.MTaseII.containing.BREX | LPD32+LPD36+N6_Mtase+SF1-SF2-Helicase->TraJ-RHH->TraM->?->?->HTH_26->BrxB-BrxF-iSTAND->DUF6079*->N6-N4-METHYLASE->Polbeta->?->TUDOR+SF1-SF2-Helicase+iREase+CTD->PglZ->BrxA-DNA-binding->?->TERNS3+HEPN-> | 1219 | Bacteria>Terrabacteria_group>Bacillota>Clostridia>Eubacteriales>Eubacteriales_incertae_sedis | GCF_022132365.1 | Intestinimonas_massiliensis | LOP79_RS13450 | 32506..36165 | 1673721 | DUF6079-family-protein-[Intestinimonas-massiliensis] |
| WP_089285463.1 | Type3.MTaseII.containing.BREX | APATPase+FtsK+VirB4->?->SMS-KaiC->HTH_36->?->BrxB-BrxF-iSTAND->DUF6079*->N6-N4-METHYLASE->TUDOR+SF1-SF2-Helicase+iREase+CTD->PglZ->BrxA-DNA-binding->SF1-SF2-Helicase-> | 1245 | Bacteria>Terrabacteria_group>Bacillota>Clostridia>Eubacteriales>Eubacteriales_incertae_sedis | GCF_900188145.1 | Anaerovirgula_multivorans | CHB72_RS24325 | 4574..8311 | 312168 | DUF6079-family-protein-[Anaerovirgula-multivorans] |
| WP_238074554.1 | Type3.MTaseII.containing.BREX | LPD32+LPD36+N6_Mtase+SF1-SF2-Helicase->TraJ-RHH->TraM->?->?->HTH_26->BrxB-BrxF-iSTAND->DUF6079*->N6-N4-METHYLASE->Polbeta->?->TUDOR+SF1-SF2-Helicase+iREase+CTD->PglZ->BrxA-DNA-binding->?->TERNS3+HEPN-> | 1224 | Bacteria>Terrabacteria_group>Bacillota>Clostridia>Eubacteriales>Eubacteriales_incertae_sedis | - | Intestinimonas_massiliensis | LOP79_13445 | 32491..36165 | 1673721 | DUF6079-family-protein-[Intestinimonas-massiliensis] |
| WP_256170312.1 | Type3.MTaseII.containing.BREX | SF1-SF2-Helicase->TraJ-RHH->TraM->?->?->HTH_26->BrxB-BrxF-iSTAND->DUF6079*->HTH+REase->N6-N4-METHYLASE->Polbeta->?->TUDOR+SF1-SF2-Helicase+iREase+CTD->PglZ->BrxA-DNA-binding->?->TERNS3+HEPN-> | 1224 | Bacteria>Terrabacteria_group>Bacillota>Clostridia>Eubacteriales>Lachnospiraceae | - | Enterocloster_bolteae | NE538_26270 | 5581..9255 | 208479 | DUF6079-family-protein-[Enterocloster-bolteae] |
| MQN00460.1 | Type3.MTaseII.containing.BREX | <-Tox-RelA<-REase+SF1-SF2-Helicase<-TRD+TRD<-N6-METHYLASE-p<-TERNS3+HEPN<-BrxA-DNA-binding<-PglZ<-TUDOR+SF1-SF2-Helicase+iREase+CTD<-N6-N4-METHYLASE<-DUF6079-p*<-BrxB-BrxF-iSTAND<-SbcC<-XIS-HTH<-?<-?\|\|SIGMA-HTH->MazG->?->REase-> | 944 | Bacteria>Terrabacteria_group>Bacillota>Clostridia>Eubacteriales>Lachnospiraceae | GCA_009493715.1 | Candidatus_Weimeria_bifida | FRC54_00430 | 18825..21659 | 2599074 | hypothetical-protein-[Candidatus-Weimeria-bifida] |
| MST58790.1 | Type3.MTaseII.containing.BREX | <-LexA-protease<-SbcC<-wHTH+WYL+WCX<-HNH<-REase<-BrxA-DNA-binding<-PglZ<-TUDOR+SF1-SF2-Helicase+iREase+CTD<-N6-N4-METHYLASE<-DUF6079-p*<-BrxB-BrxF-iSTAND<-XIS-HTH<-RNase<-SF1-SF2-Helicase+REase<-HTH_3<-?<-?<-TraJ-RHH<-LPD2 | 943 | Bacteria>Terrabacteria_group>Bacillota>Clostridia>Eubacteriales>Lachnospiraceae | GCA_009695995.1 | Waltera_intestinalis | FYJ59_11185 | 33716..36547 | 2606635 | hypothetical-protein-[Waltera-intestinalis] |
| MBD9124682.1 | Type3.MTaseII.containing.BREX | <-MazG<-BrxA-DNA-binding<-PglZ<-TUDOR+SF1-SF2-Helicase+iREase+CTD<-?<-SMODS<-HNH+SAVED<-N6-N4-METHYLASE<-HTH+REase<-DUF6079*<-BrxB-BrxF-iSTAND<-HTH_26<-Rrf2-HTH\|\|?-><-Schlafen+HATPase+wHTH | 1224 | Bacteria>Terrabacteria_group>Bacillota>Clostridia>Eubacteriales>Lachnospiraceae | GCA_014845505.1 | Lachnospiraceae_bacterium | EGP68_08015 | 66506..70180 | 1898203 | hypothetical-protein-[Lachnospiraceae-bacterium] |
| MBP3664797.1 | Type3.MTaseII.containing.BREX | <-TUDOR+SF1-SF2-Helicase+iREase+CTD<-Schlafen+HATPase+wHTH<-N6-N4-METHYLASE<-DUF6079*<-BrxB-BrxF-iSTAND<-HTH_3 | 1221 | Bacteria>Terrabacteria_group>Bacillota>Clostridia>Eubacteriales>Lachnospiraceae | GCA_017939005.1 | Tyzzerella_sp. | J6J03_06510 | 6975..10640 | 2053632 | hypothetical-protein-[Tyzzerella-sp.] |
| MCC8150845.1 | Type3.MTaseII.containing.BREX | <-OLD-ABC<-N6-N4-METHYLASE<-?<-HTH+REase<-DUF6079*<-BrxB-BrxF-iSTAND<-HIRAN<-DnaJ | 1233 | Bacteria>Terrabacteria_group>Bacillota>Clostridia>Eubacteriales>Lachnospiraceae | GCA_020860745.1 | Lachnospiraceae_bacterium | LI096_05190 | 5982..9683 | 1898203 | DUF6079-family-protein-[Lachnospiraceae-bacterium] |
| MCD7957103.1 | Type3.MTaseII.containing.BREX | <-ABC-ATPase+SbcC<-?<-?<-?<-?<-PglZ-p<-N6-METHYLASE-p<-HTH+REase<-DUF6079-p*<-McrB-AAA<-?<-?<-?<-?<-?<-IG+IG<-?<-LuxR-HTH | 221 | Bacteria>Terrabacteria_group>Bacillota>Clostridia>Eubacteriales>Lachnospiraceae | GCA_021201665.1 | Lachnospiraceae_bacterium | LUG93_15410 | 11504..12169 | 1898203 | DUF6079-family-protein-[Lachnospiraceae-bacterium] |
| MCD8067718.1 | Type3.MTaseII.containing.BREX | DUF6079-p*->HTH+REase->TUDOR+SF1-SF2-Helicase+iREase+CTD-p->ACET->?->HTH_42-><-PARG+MACRODOMAIN<-SIR2 | 945 | Bacteria>Terrabacteria_group>Bacillota>Clostridia>Eubacteriales>Lachnospiraceae | GCA_021202575.1 | Lachnospiraceae_bacterium | LUE87_02260 | 7..2844 | 1898203 | DUF6079-family-protein-[Lachnospiraceae-bacterium] |
| MCI7595222.1 | Type3.MTaseII.containing.BREX | TRAM+METHYLASE->SymE_toxin-><-XIS-HTH<-?\|\|HNH->?->HTH_26->BrxB-BrxF-iSTAND->DUF6079*->N6-N4-METHYLASE->Schlafen+BirA-HTH->TUDOR+SF1-SF2-Helicase+iREase+CTD->PglZ->BrxA-DNA-binding-> | 1230 | Bacteria>Terrabacteria_group>Bacillota>Clostridia>Eubacteriales>Lachnospiraceae | GCA_022786115.1 | Lachnospiraceae_bacterium | MSS92_03230 | 13246..16938 | 1898203 | DUF6079-family-protein-[Lachnospiraceae-bacterium] |
| MCI8530494.1 | Type3.MTaseII.containing.BREX | <-BrxA-DNA-binding<-PglZ<-N6-N4-METHYLASE<-DUF6079-p*<-BrxB-BrxF-iSTAND<-XIS-HTH<-NACHT+SNaCT | 947 | Bacteria>Terrabacteria_group>Bacillota>Clostridia>Eubacteriales>Lachnospiraceae | GCA_022793065.1 | Lachnospiraceae_bacterium | HFH82_15375 | 7218..10061 | 1898203 | hypothetical-protein-[Lachnospiraceae-bacterium] |
| MCM1551089.1 | Type3.MTaseII.containing.BREX | <-ACET<-Peptidase_M22<-?<-?<-?<-DUF6079-p*<-BrxB-BrxF-iSTAND\|\|Calcineurin->SbcC+SbcC->?->?->HNH+ADA-DBD-><-?<-tRNA\|\|METHYLASE-> | 181 | Bacteria>Terrabacteria_group>Bacillota>Clostridia>Eubacteriales>Lachnospiraceae | GCA_023667795.1 | Butyrivibrio_sp. | NC092_00170 | 31699..32242 | 28121 | DUF6079-family-protein-[Butyrivibrio-sp.] |
| MCQ4758732.1 | Type3.MTaseII.containing.BREX | SF1-SF2-Helicase->TraJ-RHH->TraM->?->HTH_26->BrxB-BrxF-iSTAND->DUF6079*->HTH+REase->N6-N4-METHYLASE->Polbeta->?->TUDOR+SF1-SF2-Helicase+iREase+CTD->PglZ->BrxA-DNA-binding->?->TERNS3+HEPN-> | 1224 | Bacteria>Terrabacteria_group>Bacillota>Clostridia>Eubacteriales>Lachnospiraceae | GCA_024460255.1 | Enterocloster_bolteae | NE538_26270 | 5581..9255 | 208479 | DUF6079-family-protein-[Enterocloster-bolteae] |
| WP_028243926.1 | Type3.MTaseII.containing.BREX | <-SF1-SF2-Helicase+REase<-SF1-SF2-Helicase<-?<-BrxA-DNA-binding<-PglZ<-TUDOR+SF1-SF2-Helicase+iREase+CTD<-N6-N4-METHYLASE<-N6-N4-METHYLASE<-DUF6079*<-BrxB-BrxF-iSTAND<-HTH_26<-?<-wHTH+WYL+WCX<-?<-?<-?<-?\|\|N6-METHYLASE-> | 1227 | Bacteria>Terrabacteria_group>Bacillota>Clostridia>Eubacteriales>Lachnospiraceae | GCF_000424305.1 | Pseudobutyrivibrio_ruminis | G611_RS0112540 | 501700..505383 | 46206 | DUF6079-family-protein-[Pseudobutyrivibrio-ruminis] |
| WP_031552730.1 | Type3.MTaseII.containing.BREX | <-SIGMA-HTH+SIGMA-HTH<-?<-ZnR+ZnR<-?<-BrxA-DNA-binding<-PglZ<-TUDOR+SF1-SF2-Helicase+iREase+CTD<-N6-N4-METHYLASE<-DUF6079*<-BrxB-BrxF-iSTAND<-HTH_26 | 1234 | Bacteria>Terrabacteria_group>Bacillota>Clostridia>Eubacteriales>Lachnospiraceae | GCF_000701645.1 | Oribacterium_sp._FC2011 | T357_RS19305 | 67923..71627 | 1408311 | DUF6079-family-protein-[Oribacterium-sp.-FC2011] |
| WP_205839527.1 | Type3.MTaseII.containing.BREX | <-LexA-protease<-SbcC<-wHTH+WYL+WCX<-HNH<-REase<-BrxA-DNA-binding<-PglZ<-TUDOR+SF1-SF2-Helicase+iREase+CTD<-N6-N4-METHYLASE<-DUF6079*<-BrxB-BrxF-iSTAND<-XIS-HTH<-RNase<-SF1-SF2-Helicase+REase<-HTH_3<-?<-?<-TraJ-RHH<-LPD2 | 1230 | Bacteria>Terrabacteria_group>Bacillota>Clostridia>Eubacteriales>Lachnospiraceae | GCF_009695995.1 | Waltera_intestinalis | FYJ59_RS11205 | 33716..37408 | 2606635 | DUF6079-family-protein-[Waltera-intestinalis] |
| WP_243273543.1 | Type3.MTaseII.containing.BREX | MPTase-><-?<-TERNS3+HEPN<-?<-BrxA-DNA-binding<-PglZ<-N6-N4-METHYLASE<-HTH+REase<-DUF6079*<-BrxB-BrxF-iSTAND<-HTH_26<-?<-?<-TraM<-TraJ-RHH<-LPD32+LPD36+N6_Mtase+SF1-SF2-Helicase | 1219 | Bacteria>Terrabacteria_group>Bacillota>Clostridia>Eubacteriales>Lachnospiraceae | GCF_015556105.1 | Coprococcus_comes | I2F75_RS13170 | 32473..36132 | 410072 | DUF6079-family-protein-[Coprococcus-comes] |
| WP_243285215.1 | Type3.MTaseII.containing.BREX | LPD32+LPD36+N6_Mtase+SF1-SF2-Helicase->RVT+HNH->TraJ-RHH->TraM->?->?->HTH_26->BrxB-BrxF-iSTAND->DUF6079-p*-> | 489 | Bacteria>Terrabacteria_group>Bacillota>Clostridia>Eubacteriales>Lachnospiraceae | GCF_015558345.1 | Enterocloster_bolteae | I2I03_RS19155 | 145904..147373 | 208479 | DUF6079-family-protein-[Enterocloster-bolteae] |
| WP_226978790.1 | Type3.MTaseII.containing.BREX | <-MPTase<-HTH_31<-ASCH<-ACET+EVE<-nSTAND3\|\|HTH_26->BrxB-BrxF-iSTAND->DUF6079*->?->N6-N4-METHYLASE->TUDOR+SF1-SF2-Helicase+iREase+CTD->PglZ->BrxA-DNA-binding->ZnR+NACHT-><-?<-?<-REase | 1233 | Bacteria>Terrabacteria_group>Bacillota>Clostridia>Eubacteriales>Lachnospiraceae | GCF_020537245.1 | [Ruminococcus]_lactaris | LIP51_RS01680 | 26782..30483 | 46228 | DUF6079-family-protein-[[Ruminococcus] |
| WP_262586687.1 | Type3.MTaseII.containing.BREX | Glycoside_Hydrolase-><-?\|\|ACET->?->Schlafen+HATPase+wHTH-><-?\|\|Rrf2-HTH->HTH_26->BrxB-BrxF-iSTAND->DUF6079*->HTH+REase->N6-N4-METHYLASE->HNH+SAVED->SMODS->?->TUDOR+SF1-SF2-Helicase+iREase+CTD->PglZ->BrxA-DNA-binding-> | 1224 | Bacteria>Terrabacteria_group>Bacillota>Clostridia>Eubacteriales>Lachnospiraceae | GCF_025567525.1 | Gallintestinimicrobium_propionicum | OCV62_RS11070 | 8721..12395 | 2981770 | DUF6079-family-protein-[Gallintestinimicrobium-propionicum] |
| WP_073587001.1 | Type3.MTaseII.containing.BREX | ACET-><-Glyco_hydro_3+Glyco_hydro_3_C+Fn3-like\|\|TRAM+METHYLASE-><-XIS-HTH\|\|NACHT+SNaCT->XIS-HTH->BrxB-BrxF-iSTAND->DUF6079*->N6-N4-METHYLASE->TUDOR+SF1-SF2-Helicase+iREase+CTD->PglZ->BrxA-DNA-binding-><-?<-?<-?<-REase | 1236 | Bacteria>Terrabacteria_group>Bacillota>Clostridia>Eubacteriales>Lachnospiraceae | GCF_900143645.1 | Anaerocolumna_xylanovorans | BUF11_RS01300 | 284575..288285 | 100134 | DUF6079-family-protein-[Anaerocolumna-xylanovorans] |
| WP_009252807.1 | Type3.MTaseII.containing.BREX | <-HTH_3\|\|D5N+D5ATPase->?->?->TraJ-RHH->?->?->XIS-HTH->BrxB-BrxF-iSTAND->DUF6079*->N6-N4-METHYLASE->N6-N4-METHYLASE->TUDOR+SF1-SF2-Helicase+iREase+CTD->PglZ->BrxA-DNA-binding-><-?\|\|?->?->PhoH-> | 1231 | Bacteria>Terrabacteria_group>Bacillota>Clostridia>Eubacteriales>Lachnospiraceae | GCF_902385915.1 | Eisenbergiella_tayi | FYC33_RS15820 | 47046..50741 | 1432052 | DUF6079-family-protein-[Eisenbergiella-tayi] |
| ALX09382.1 | Type3.MTaseII.containing.BREX | <-ParB\|\|?->?-><-BrxA-DNA-binding-p<-BrxA-DNA-binding<-PglZ<-TUDOR+SF1-SF2-Helicase+iREase+CTD<-N6-N4-METHYLASE<-DUF6079-p*<-HTH_Tnp_1+HTH_21+rve+rve_3<-HTH_Tnp_1<-?<-?<-HARE-HTH<-SAD+HNH<-NUDIX<-HIN-HTH+PAIREDC-HTH<-Resolvase+Recombinase+ZnR | 946 | Bacteria>Terrabacteria_group>Bacillota>Clostridia>Eubacteriales>Oscillospiraceae | GCA_000255615.3 | Acetivibrio_thermocellus_AD2 | LQRI_2395 | 2785039..2787879 | 1138384 | hypothetical-protein-AD2_02396-[Acetivibrio-thermocellus-AD2] |
| HBR02120.1 | Type3.MTaseII.containing.BREX | DDE_Tnp_1->?->?-><-?<-BrxA-DNA-binding-p<-PglZ<-TUDOR+SF1-SF2-Helicase+iREase+CTD<-REase<-N6-N4-METHYLASE<-DUF6079-p*<-DUF6079-p*<-BrxB-BrxF-iSTAND\|\|Trypsin+PDZ-> | 942 | Bacteria>Terrabacteria_group>Bacillota>Clostridia>Eubacteriales>Oscillospiraceae | GCA_003535195.1 | Ruminiclostridium_sp. | DD738_05890 | 21181..24009 | 2053608 | hypothetical-protein-[Ruminiclostridium-sp.] |
| HIT07770.1 | Type3.MTaseII.containing.BREX | <-GDResolvase<-DUF6079-p*<-BrxB-BrxF-iSTAND<-XIS-HTH<-?<-NACHT+SNaCT\|\|SIGMA-HTH->Vrr-REase->SF1-SF2-Helicase-p->?->AraC-HTH+AraC-HTH-> | 801 | Bacteria>Terrabacteria_group>Bacillota>Clostridia>Eubacteriales>Oscillospiraceae | GCA_018716285.1 | Candidatus_Merdivicinus_faecavium | IAB55_01615 | 296..2701 | 2840868 | hypothetical-protein-[Candidatus-Merdivicinus-faecavium] |
| MCG4604364.1 | Type3.MTaseII.containing.BREX | <-HAMP+HISKIN<-REC+OmpR-HTH<-HKD-DNase+HKD-DNase<-HKD-DNase+HKD-DNase<-Tox-RelA<-?<-?<-TERNS3+HEPN<-DUF6079-p*<-BrxB-BrxF-iSTAND<-HTH_26<-?<-?<-TraM<-TraJ-RHH | 402 | Bacteria>Terrabacteria_group>Bacillota>Clostridia>Eubacteriales>Oscillospiraceae | GCA_022133545.1 | Faecalibacterium_prausnitzii | LON24_12700 | 6832..8040 | 853 | DUF6079-family-protein-[Faecalibacterium-prausnitzii] |
| WP_004621790.1 | Type3.MTaseII.containing.BREX | SIGMA-HTH+SIGMA-HTH+SIGMA-HTH->tRNA->tRNA-><-?\|\|METHYLASE->?->ncRNA->Phage_integrase->?->SbcC-><-SbcC\|\|BrxB-BrxF-iSTAND->DUF6079*->N6-N4-METHYLASE->TUDOR+SF1-SF2-Helicase+iREase+CTD->PglZ->BrxA-DNA-binding-><-REase\|\|HHH_4+SF1-SF2-Helicase+REase->URI-DNase-> | 1222 | Bacteria>Terrabacteria_group>Bacillota>Clostridia>Eubacteriales>Oscillospiraceae | GCF_000175795.2 | Ruminiclostridium_papyrosolvens | P0092_RS08120 | 1816637..1820305 | 29362 | DUF6079-family-protein-[Ruminiclostridium-papyrosolvens] |
| WP_087378434.1 | Type3.MTaseII.containing.BREX | TraJ-RHH->TraI-VirD2-RCRE->HTH_26->BrxB-BrxF-iSTAND->DUF6079*->N6-N4-METHYLASE->SbcC->SF1-SF2-Helicase->TUDOR+SF1-SF2-Helicase+iREase+CTD->PglZ->BrxA-DNA-binding->TERNS3+HEPN-><-?<-HTH_3 | 1230 | Bacteria>Terrabacteria_group>Bacillota>Clostridia>Eubacteriales>Oscillospiraceae | GCF_002160795.1 | Flavonifractor_sp._An135 | B5E80_RS08695 | 45549..49241 | 1965558 | DUF6079-family-protein-[Flavonifractor-sp.-An135] |
| WP_223382969.1 | Type3.MTaseII.containing.BREX | <-CHTH+Tox-ART-HYD2<-BrxA-DNA-binding<-PglZ<-TUDOR+SF1-SF2-Helicase+iREase+CTD<-?<-SMODS<-HNH+SAVED<-N6-N4-METHYLASE<-DUF6079*<-BrxB-BrxF-iSTAND<-HTH_26\|\|DUF932->?->?->CR-ATPase5+REase->?->CR-ATPase8+REase-> | 1224 | Bacteria>Terrabacteria_group>Bacillota>Clostridia>Eubacteriales>Oscillospiraceae | GCF_019967995.1 | Faecalibacterium_prausnitzii | I5P97_RS02615 | 537900..541574 | 853 | DUF6079-family-protein-[Faecalibacterium-prausnitzii] |
| WP_263291086.1 | Type3.MTaseII.containing.BREX | <-CHTH+Tox-ART-HYD2<-BrxA-DNA-binding<-PglZ<-TUDOR+SF1-SF2-Helicase+iREase+CTD<-?<-SMODS<-HNH+SAVED<-N6-N4-METHYLASE<-DUF6079*<-BrxB-BrxF-iSTAND<-HTH_26\|\|DUF932->?->?->CR-ATPase5+REase->?->CR-ATPase8+REase-> | 1230 | Bacteria>Terrabacteria_group>Bacillota>Clostridia>Eubacteriales>Oscillospiraceae | GCF_019969425.1 | Faecalibacterium_prausnitzii | LC356_RS02735 | 537900..541592 | 853 | DUF6079-family-protein-[Faecalibacterium-prausnitzii] |
| WP_273700773.1 | Type3.MTaseII.containing.BREX | <-HAMP+HISKIN<-REC+OmpR-HTH<-HKD-DNase+HKD-DNase<-HKD-DNase+HKD-DNase<-Tox-RelA<-?<-?<-TERNS3+HEPN<-DUF6079-p*<-BrxB-BrxF-iSTAND<-HTH_26<-?<-?<-TraM<-TraJ-RHH | 397 | Bacteria>Terrabacteria_group>Bacillota>Clostridia>Eubacteriales>Oscillospiraceae | GCF_022133545.1 | Faecalibacterium_prausnitzii | LON24_RS12700 | 6832..8025 | 853 | DUF6079-family-protein-[Faecalibacterium-prausnitzii] |
| WP_260489645.1 | Type3.MTaseII.containing.BREX | <-ParB\|\|?->?-><-BrxA-DNA-binding<-PglZ<-DUF6079-p<-TUDOR+SF1-SF2-Helicase+iREase+CTD<-N6-N4-METHYLASE<-DUF6079*<-BrxB-BrxF-iSTAND<-ABC-ATPase+SbcC<-?<-REC+OmpR-HTH<-?\|\|?-><-?<-?<-ZnR+Phage_GP20 | 1229 | Bacteria>Terrabacteria_group>Bacillota>Clostridia>Eubacteriales>Oscillospiraceae | GCF_025213355.1 | Acetivibrio_thermocellus | N1236_RS12350 | 2845166..2848855 | 1515 | DUF6079-family-protein-[Acetivibrio-thermocellus] |
| WP_271938551.1 | Type3.MTaseII.containing.BREX | LPD32+LPD36+N6_Mtase+SF1-SF2-Helicase->TraJ-RHH->TraM->?->?->HTH_26->BrxB-BrxF-iSTAND->DUF6079*->HTH+REase->N6-N4-METHYLASE-><-TraJ-RHH<-HTH_3\|\|?->RADICAL-SAM->SIGMA-HTH->HTH_16-> | 1224 | Bacteria>Terrabacteria_group>Bacillota>Clostridia>Eubacteriales>Oscillospiraceae | GCF_028321805.1 | Flavonifractor_plautii | PND94_RS10175 | 43583..47257 | 292800 | DUF6079-family-protein-[Flavonifractor-plautii] |
| WP_274235722.1 | Type3.MTaseII.containing.BREX | <-CR-ATPase8+REase<-?<-CR-ATPase5+REase<-?<-?<-DUF932\|\|HTH_26->BrxB-BrxF-iSTAND->DUF6079*->N6-N4-METHYLASE->?->?->TUDOR+SF1-SF2-Helicase+iREase+CTD->PglZ->BrxA-DNA-binding->CHTH+Tox-ART-HYD2->?->Peptidase-> | 1224 | Bacteria>Terrabacteria_group>Bacillota>Clostridia>Eubacteriales>Oscillospiraceae | GCF_028743395.1 | Faecalibacterium_prausnitzii | PT301_RS03130 | 649364..653038 | 853 | DUF6079-family-protein-[Faecalibacterium-prausnitzii] |
| WP_253098837.1 | Type3.MTaseII.containing.BREX | LPD32+LPD36+N6_Mtase+SF1-SF2-Helicase->TraJ-RHH->TraM->?->?->HTH_26->BrxB-BrxF-iSTAND->DUF6079*->N6-N4-METHYLASE->Polbeta->?->TUDOR+SF1-SF2-Helicase+iREase+CTD->PglZ->BrxA-DNA-binding->?->TERNS3+HEPN-> | 1224 | Bacteria>Terrabacteria_group>Bacillota>Clostridia>Eubacteriales>Oscillospiraceae | - | Oscillibacter_valericigenes | NIA13_RS09300 | 67071..70745 | 351091 | DUF6079-family-protein-[Oscillibacter-valericigenes] |
| KFD41618.1 | Type3.MTaseII.containing.BREX | CASPASE->?->?-><-BrxA-DNA-binding<-PglZ<-?<-SIR2<-?<-DUF6079*\|\|?->?->Peptidase_M50->?->?-><-HEPN<-?<-HicB_lk_antitox | 1215 | Bacteria>Terrabacteria_group>Bacillota>Clostridia>Eubacteriales>Peptococcaceae | GCA_000681355.2 | Peptococcaceae_bacterium_SCADC1_2_3 | DK28_0207245 | 14116..17763 | 1487582 | hypothetical-protein-DK28_0207245-[Peptococcaceae-bacterium-SCADC1_2_3] |
| KFD41493.1 | Type3.MTaseII.containing.BREX | <-RAMPs<-HD<-SF1-SF2-Helicase<-N6-N4-METHYLASE<-REase<-N6-METHYLASE<-DUF6079-p* | 112 | Bacteria>Terrabacteria_group>Bacillota>Clostridia>Eubacteriales>Peptococcaceae | GCA_000681355.2 | Peptococcaceae_bacterium_SCADC1_2_3 | DK28_0208715 | 20340..20678 | 1487582 | hypothetical-protein-DK28_0208715-[Peptococcaceae-bacterium-SCADC1_2_3] |
| KFI36375.1 | Type3.MTaseII.containing.BREX | wHTH->?->HicB_lk_antitox->?->HEPN-><-?<-?<-Peptidase_M50<-?\|\|DUF6079*->?->?->?->SIR2->PglZ->BrxA-DNA-binding->TPR+TPR+TPR+TPR+TPR->HEPN-> | 1215 | Bacteria>Terrabacteria_group>Bacillota>Clostridia>Eubacteriales>Peptococcaceae | GCA_000740805.1 | Peptococcaceae_bacterium_SCADC1_2_3 | HY00_00360 | 42493..46140 | 1487582 | hypothetical-protein-HY00_00360-[Peptococcaceae-bacterium-SCADC1_2_3] |
| KFI36120.1 | Type3.MTaseII.containing.BREX | <-RAMPs<-HD<-SF1-SF2-Helicase<-N6-N4-METHYLASE<-REase<-N6-METHYLASE<-DUF6079-p*<-BrxB-BrxF-iSTAND\|\|GNTR-HTH+UTRA->?->HNH->?->ABC-ATPase+SbcC-><-?<-?<-DSBH+DSBH | 808 | Bacteria>Terrabacteria_group>Bacillota>Clostridia>Eubacteriales>Peptococcaceae | GCA_000740805.1 | Peptococcaceae_bacterium_SCADC1_2_3 | HY00_07510 | 43328..45754 | 1487582 | hypothetical-protein-HY00_07510-[Peptococcaceae-bacterium-SCADC1_2_3] |
| HBT46725.1 | Type3.MTaseII.containing.BREX | ACET->Aminotran_3->?->?->?->?->BrxB-BrxF-iSTAND->DUF6079*->HD->ABC-3C->N6-N4-METHYLASE->TUDOR+SF1-SF2-Helicase+iREase+CTD->PglZ->BrxA-DNA-binding-><-?<-HIN-HTH | 1243 | Bacteria>Terrabacteria_group>Bacillota>Clostridia>Eubacteriales>Peptococcaceae | GCA_003509545.1 | Peptococcaceae_bacterium | DEA73_02415 | 25203..28934 | 2052179 | ATPase-[Peptococcaceae-bacterium] |
| RJX22765.1 | Type3.MTaseII.containing.BREX | <-BrxA-DNA-binding-p<-PglZ<-TUDOR+SF1-SF2-Helicase+iREase+CTD<-RNaseH<-N6-N4-METHYLASE<-DUF6079*<-BrxB-BrxF-iSTAND<-ArdCN+MPTase-inact<-?<-?<-?<-OrfB_Zn_ribbon | 1242 | Bacteria>Terrabacteria_group>Bacillota>Clostridia>Eubacteriales>Peptococcaceae | GCA_003604975.1 | Candidatus_Desulforudis_sp. | C4575_00525 | 8787..12515 | 2093368 | MAG:-ATP-binding-protein-[Candidatus-Desulforudis-sp.] |
| NHM28431.1 | Type3.MTaseII.containing.BREX | <-OLD-ABC<-ParB<-BrxA-DNA-binding<-PglZ<-?<-REase<-N6-N4-METHYLASE<-DUF6079*<-BrxB-BrxF-iSTAND | 1226 | Bacteria>Terrabacteria_group>Bacillota>Clostridia>Eubacteriales>Peptococcaceae | GCA_011393055.1 | Desulfofundulus_sp._TPOSR | G7K71_15925 | 30486..34166 | 2714340 | ATP-binding-protein-[Desulfofundulus-sp.-TPOSR] |
| NLI12136.1 | Type3.MTaseII.containing.BREX | SMODS->METHYLASE->?->?->ncRNA->BrxB-BrxF-iSTAND->DUF6079*->N6-N4-METHYLASE-> | 1225 | Bacteria>Terrabacteria_group>Bacillota>Clostridia>Eubacteriales>Peptococcaceae | GCA_012517685.1 | Peptococcaceae_bacterium | GX425_05840 | 8823..12500 | 2052179 | ATP-binding-protein-[Peptococcaceae-bacterium] |
| HHU86035.1 | Type3.MTaseII.containing.BREX | <-N6-N4-METHYLASE<-PglZ<-DUF6079*<-BrxB-BrxF-iSTAND<-BrxA-DNA-binding\|\|HNH+NERD+ZnR->NERD-> | 1437 | Bacteria>Terrabacteria_group>Bacillota>Clostridia>Eubacteriales>Peptococcaceae | GCA_012839155.1 | Peptococcaceae_bacterium | GXZ25_04390 | 4519..8832 | 2052179 | hypothetical-protein-[Peptococcaceae-bacterium] |
| MCR4442915.1 | Type3.MTaseII.containing.BREX | <-HAMP+PAS+HISKIN<-REC+OmpR-HTH\|\|?->?->HTH_3->MPTase-><-DUF6079-p<-DUF6079-p*<-BrxB-BrxF-iSTAND<-SF1-SF2-Helicase+DUF1998<-?<-wHTH+WYL+WCX | 429 | Bacteria>Terrabacteria_group>Bacillota>Clostridia>Eubacteriales>Peptococcaceae | GCA_024655825.1 | Peptococcaceae_bacterium | NUV48_12275 | 5866..7155 | 2052179 | DUF6079-family-protein-[Peptococcaceae-bacterium] |

| Protein-ID | System | Gene-Neighbourhoods | Length | Taxonomy | Assembly-ID | Strain | Locus-Tag | Genomic-coordinates | TaxID | Fasta-define |
|---|---|---|---|---|---|---|---|---|---|---|
| WP_028308210.1 | Type3.MTaseII.containing.BREX | CopG/RHH->RHH_8->?-><-DEDD_Tnp_IS110+Transposase_20\|\|ZnR+TOPRIM->VirD4-TraG+VirD4-FtsK->HTH_36->HTH_26->BrxB-BrxF-iSTAND->DUF6079*->N6-N4-METHYLASE->?->TUDOR+SF1-SF2-Helicase+iREase+CTD->PglZ->BrxA-DNA-binding->REase+iREase->Preader+URI-DNase->ZnR+HARE-HTH-> | 1229 | Bacteria>Terrabacteria_group>Bacillota>Clostridia>Eubacteriales>Peptococcaceae | GCF_000620305.1 | Desulfitibacter_alkalitolerans | K364_RS0111915 | 37253..40942 | 264641 | DUF6079-family-protein-[Desulfitibacter-alkalitolerans] |
| WP_028309574.1 | Type3.MTaseII.containing.BREX | wHTH+WYL+WCX->?->?->?->SF1-SF2-Helicase-><-HTH_Tnp_1+HTH_21+rve+rve_3\|\|SF1-SF2-Helicase+DUF1998->BrxB-BrxF-iSTAND->DUF6079*->N6-N4-METHYLASE->TUDOR+SF1-SF2-Helicase+iREase+CTD->PglZ->BrxA-DNA-binding->JetA-> | 1222 | Bacteria>Terrabacteria_group>Bacillota>Clostridia>Eubacteriales>Peptococcaceae | GCF_000620305.1 | Desulfitibacter_alkalitolerans | K364_RS0120625 | 201380..205048 | 264641 | DUF6079-family-protein-[Desulfitibacter-alkalitolerans] |
| WP_152946296.1 | Type3.MTaseII.containing.BREX | <-RAMPs<-?<-RAMPs<-HD<-RAMPs<-TOPRIM<-?<-RAMPs<-N6-N4-METHYLASE-p<-DUF6079*<-BrxB-BrxF-iSTAND<-?<-DOC+zhth<-REase+HTH<-ABC-ATPase+SbcC | 1243 | Bacteria>Terrabacteria_group>Bacillota>Clostridia>Eubacteriales>Peptococcaceae | GCF_009428905.1 | Desulfofundulus_thermobenzoicus | GFC01_RS08615 | 53615..57346 | 29376 | DUF6079-family-protein-[Desulfofundulus-thermobenzoicus] |
| WP_259498357.1 | Type3.MTaseII.containing.BREX | PIN-><-DUF5086<-BrxA-DNA-binding<-PglZ<-TUDOR+SF1-SF2-Helicase+iREase+CTD-p<-SF1-SF2-Helicase<-OLD-ABC+TOPRIM<-N6-N4-METHYLASE<-DUF6079*<-BrxB-BrxF-iSTAND | 1226 | Bacteria>Terrabacteria_group>Bacillota>Clostridia>Eubacteriales>Peptococcaceae | GCF_024958695.1 | Desulfofundulus_thermocisternus | NZJ93_RS02610 | 113330..117010 | 42471 | DUF6079-family-protein-[Desulfofundulus-thermocisternus] |
| WP_242945693.1 | Type3.MTaseII.containing.BREX | BrxB-BrxF-iSTAND->DUF6079*->N6-N4-METHYLASE->SbcC->TUDOR+SF1-SF2-Helicase+iREase+CTD->PglZ->BrxA-DNA-binding->BirA-HTH->?->Peptidase->Peptidase_M16+Peptidase_M16_C-> | 1228 | Bacteria>Terrabacteria_group>Bacillota>Clostridia>Eubacteriales>Proteinivoraceae | GCF_900111575.1 | Anaerobranca_gottschalkii | BMX60_RS03100 | 20741..24427 | 108328 | DUF6079-family-protein-[Anaerobranca-gottschalkii] |
| HHW29753.1 | Type3.MTaseII.containing.BREX | <-SbcC<-?<-?<-RNase<-BrxA-DNA-binding<-PglZ<-N6-N4-METHYLASE<-DUF6079*<-BrxB-BrxF-iSTAND | 1224 | Bacteria>Terrabacteria_group>Bacillota>Clostridia>Eubacteriales>Syntrophomonadaceae | GCA_012840405.1 | Syntrophomonadaceae_bacterium | GXX21_09420 | 13995..17669 | 2093811 | ATP-binding-protein-[Syntrophomonadaceae-bacterium] |
| NSW83904.1 | Type3.MTaseII.containing.BREX | HTH+REase->\|\|AIPR->?->?->HEPN->BrxB-BrxF-iSTAND->DUF6079*->N6-N4-METHYLASE->TUDOR+SF1-SF2-Helicase+iREase+CTD->PglZ->BrxA-DNA-binding-> | 1203 | Bacteria>Terrabacteria_group>Bacillota>Clostridia>Eubacteriales>Syntrophomonadaceae | GCA_013314835.1 | Syntrophothermus_sp. | HPY90_11660 | 7329..10940 | 2736299 | hypothetical-protein-[Syntrophothermus-sp.] |
| MBS3887625.1 | Type3.MTaseII.containing.BREX | <-N6-N4-METHYLASE-p<-DUF6079*<-BrxB-BrxF-iSTAND<-HTH_26<-?<-HNH\|\|SIGMA-HTH->MazG->?->REase-> | 1231 | Bacteria>Terrabacteria_group>Bacillota>Clostridia>Eubacteriales>Syntrophomonadaceae | GCA_018333455.1 | Dethiobacter_sp. | KGZ56_11280 | 619..4314 | 2093370 | hypothetical-protein-[Dethiobacter-sp.] |
| MBT9173637.1 | Type3.MTaseII.containing.BREX | PIN->?->?-><-CN_hydrolase+NAD_synthase\|\|ABhydrolase+ABhydrolase->?-><-RelE-ParE<-PhdYeFM_antitox\|\|BrxB-BrxF-iSTAND->DUF6079*->HD->ABC-3C->N6-N4-METHYLASE->PhdYeFM_antitox->RelE-ParE->DUF488-1->TUDOR+SF1-SF2-Helicase+iREase+CTD->?->HerA/FtsK-> | 1248 | Bacteria>Terrabacteria_group>Bacillota>Clostridia>Eubacteriales>Syntrophomonadaceae | GCA_018725565.1 | Syntrophomonadaceae_bacterium | DDT21_02038 | 9261..13007 | 2093811 | hypothetical-protein-[Syntrophomonadaceae-bacterium] |
| WP_013175448.1 | Type3.MTaseII.containing.BREX | PIN->?-><-wHTH<-?<-OLD-ABC+TOPRIM<-HEPN<-BrxA-DNA-binding<-PglZ<-N6-N4-METHYLASE<-DUF6079*<-BrxB-BrxF-iSTAND<-?\|\|?->PIN->?-><-Amidohydro_1 | 1226 | Bacteria>Terrabacteria_group>Bacillota>Clostridia>Eubacteriales>Syntrophomonadaceae | GCF_000092405.1 | Syntrophothermus_lipocalidus | SLIP_RS06315 | 1312797..1316477 | 86170 | DUF6079-family-protein-[Syntrophothermus-lipocalidus] |
| WP_046495069.1 | Type3.MTaseII.containing.BREX | <-SF1-SF2-Helicase<-BrxA-DNA-binding<-PglZ<-TUDOR+SF1-SF2-Helicase+iREase+CTD<-N6-N4-METHYLASE<-DUF6079*<-BrxB-BrxF-iSTAND<-?\|\|HD->ABC-3C-><-XIS-HTH<-tRNA<-?<-?\|\|GntR-HTH->ABC-ATPase+SbcC-> | 1203 | Bacteria>Terrabacteria_group>Bacillota>Clostridia>Eubacteriales>Syntrophomonadaceae | GCF_000983115.1 | Syntrophomonas_zehnderi | AAV90_RS05050 | 66936..70547 | 404335 | DUF6079-family-protein-[Syntrophomonas-zehnderi] |
| WP_061214834.1 | Type3.MTaseII.containing.BREX | <-SIGMA-HTH+SIGMA-HTH<-HEPN+REase+SF1-SF2-Helicase<-?<-TRD+TRD<-BrxA-DNA-binding<-PglZ<-TUDOR+SF1-SF2-Helicase+iREase+CTD<-N6-N4-METHYLASE<-DUF6079*<-BrxB-BrxF-iSTAND<-XIS-HTH<-?<-GspA-ATPase | 1236 | Bacteria>Terrabacteria_group>Bacillota>Clostridia>Eubacteriales>Syntrophomonadaceae | GCF_001570625.1 | Syntrophomonas_wolfei | AWB54_RS11995 | 64212..67922 | 863 | DUF6079-family-protein-[Syntrophomonas-wolfei] |
| KKM12037.1 | Type3.MTaseII.containing.BREX | <-BrxA-DNA-binding<-PglZ<-N6-N4-METHYLASE<-DUF6079*<-BrxB-BrxF-iSTAND\|\|?->?->?-><-NAD_binding_10<-MarR-HTH<-?<-?<-ACET | 1234 | Bacteria>Terrabacteria_group>Bacillota>Clostridia>Eubacteriales>unclassified_Eubacteriales | GCA_000987045.1 | Clostridiales_bacterium_PH28_bin88 | SY88_05475 | 7325..11029 | 1605376 | ATPase-[Clostridiales-bacterium-PH28_bin88] |
| KKM08525.1 | Type3.MTaseII.containing.BREX | <-DDE_Tnp\|\|?->BrxB-BrxF-iSTAND->DUF6079*->?->N6-N4-METHYLASE->SbcC->?->TUDOR+SF1-SF2-Helicase+iREase+CTD->PglZ->BrxA-DNA-binding->DOC+HTH->PadR-HTH-> | 1243 | Bacteria>Terrabacteria_group>Bacillota>Clostridia>Eubacteriales>unclassified_Eubacteriales | GCA_000987045.1 | Clostridiales_bacterium_PH28_bin88 | SY88_23360 | 4250..7981 | 1605376 | ATPase-[Clostridiales-bacterium-PH28_bin88] |
| HCV69871.1 | Type3.MTaseII.containing.BREX | <-HTH_Tnp<-?<-BrxA-DNA-binding<-PglZ<-TUDOR+SF1-SF2-Helicase+iREase+CTD<-?<-SMODS<-SAVED<-N6-N4-METHYLASE<-DUF6079-p*<-BrxB-BrxF-iSTAND<-HTH_26 | 943 | Bacteria>Terrabacteria_group>Bacillota>Clostridia>Eubacteriales>unclassified_Eubacteriales | GCA_003538415.1 | Clostridiales_bacterium | DGW67_10740 | 113882..116713 | 1898207 | hypothetical-protein-[Clostridiales-bacterium] |
| NMA94940.1 | Type3.MTaseII.containing.BREX | <-N6-N4-METHYLASE<-DUF6079*<-BrxB-BrxF-iSTAND<-HTH_Tnp_1+HTH_21+rve+rve_3 | 1250 | Bacteria>Terrabacteria_group>Bacillota>Clostridia>Eubacteriales>unclassified_Eubacteriales | GCA_012799205.1 | Clostridiales_bacterium | GX974_02750 | 1461..5213 | 1898207 | hypothetical-protein-[Clostridiales-bacterium] |
| MBE5774055.1 | Type3.MTaseII.containing.BREX | <-Phage_int_SAM_3+Phage_integrase<-HTH_3\|\|XIS-HTH->?-><-?\|\|BrxB-BrxF-iSTAND->DUF6079*->?->?->?->N6-N4-METHYLASE-> | 1231 | Bacteria>Terrabacteria_group>Bacillota>Clostridia>Eubacteriales>unclassified_Eubacteriales | GCA_015055915.1 | Clostridiales_bacterium | E7337_08995 | 5564..9259 | 1898207 | exonuclease-SbcC-[Clostridiales-bacterium] |
| MBS4886749.1 | Type3.MTaseII.containing.BREX | ABC-ATPase+SbcC->?->?->?->?->MACRODOMAIN->?->HTH_26->BrxB-BrxF-iSTAND->DUF6079*->HTH+REase->N6-N4-METHYLASE->?->?->TUDOR+SF1-SF2-Helicase+iREase+CTD->PglZ->BrxA-DNA-binding-> | 1224 | Bacteria>Terrabacteria_group>Bacillota>Clostridia>Eubacteriales>unclassified_Eubacteriales | GCA_018363765.1 | Clostridiales_bacterium | KH027_08810 | 19109..22783 | 1898207 | hypothetical-protein-[Clostridiales-bacterium] |
| MBZ4688268.1 | Type3.MTaseII.containing.BREX | RNase->?->SPOUT->PIN->SIGMA-HTH+LuxR-HTH->HTH+MPTase->PIN->?->BrxB-BrxF-iSTAND->DUF6079*->N6-N4-METHYLASE-> | 1229 | Bacteria>Terrabacteria_group>Bacillota>Clostridia>Eubacteriales>unclassified_Eubacteriales | GCA_020063565.1 | Clostridiales_bacterium | JG764_1901 | 61044..64733 | 1898207 | ATPase-[Clostridiales-bacterium] |
| MCI6377637.1 | Type3.MTaseII.containing.BREX | <-SF1-SF2-Helicase-p\|\|?-><-?<-DEOR-HTH+ISOCOT<-?<-?<-?<-N6-N4-METHYLASE-p<-DUF6079*<-BrxB-BrxF-iSTAND<-HTH_26<-tRNA\|\|?->MORC+DNA_gyrase+TOPRIM-> | 1231 | Bacteria>Terrabacteria_group>Bacillota>Clostridia>Eubacteriales>unclassified_Eubacteriales | GCA_022776425.1 | Clostridiales_bacterium | MR842_07770 | 47561..51256 | 1898207 | DUF6079-family-protein-[Clostridiales-bacterium] |
| WP_114474457.1 | Type3.MTaseII.containing.BREX | Peptidase_M42+Peptidase_M28->?->?->?->HEPN+REase+SF1-SF2-Helicase->SF1-SF2-Helicase+REase+HTH->?->ParB+HNH->BrxB-BrxF-iSTAND->DUF6079*-><-SIGMA-HTH+LuxR-HTH\|\|N6-N4-METHYLASE->TUDOR+SF1-SF2-Helicase+iREase+CTD->PglZ->BrxA-DNA-binding->MazG->EVE+McrB-AAA-> | 1218 | Bacteria>Terrabacteria_group>Bacillota>Clostridia>Halanaerobiales>Halanaerobiaceae | GCF_003338615.1 | Halanaerobium_sp._DL-01 | DER71_RS07030 | 29697..33353 | 1653064 | DUF6079-family-protein-[Halanaerobium-sp.-DL-01] |
| WP_133505387.1 | Type3.MTaseII.containing.BREX | <-Phage_int_SAM_1+Phage_integrase<-BrxA-DNA-binding<-PglZ<-TUDOR+SF1-SF2-Helicase+iREase+CTD<-N6-N4-METHYLASE<-DUF6079*<-BrxB-BrxF-iSTAND<-?<-?<-?<-ParA-Soj-PloopNTPase<-?<-?<-HNH | 1218 | Bacteria>Terrabacteria_group>Bacillota>Clostridia>Halanaerobiales>Halanaerobiaceae | GCF_004362065.1 | Halanaerobium_congolense | C8C79_10233 | 27115..30771 | 54121 | DUF6079-family-protein-[Halanaerobium-congolense] |
| WP_134059933.1 | Type3.MTaseII.containing.BREX | SF1-SF2-Helicase+REase-p->?->ParB+HNH->BrxB-BrxF-iSTAND->DUF6079*->N6-N4-METHYLASE->TUDOR+SF1-SF2-Helicase+iREase+CTD->PglZ->BrxA-DNA-binding->MazG-><-HIN-HTH\|\|?->?->RAMA-> | 1218 | Bacteria>Terrabacteria_group>Bacillota>Clostridia>Halanaerobiales>Halanaerobiaceae | GCF_004366375.1 | Halanaerobium_congolense | C7954_RS10330 | 6715..10371 | 54121 | DUF6079-family-protein-[Halanaerobium-congolense] |
| WP_073159266.1 | Type3.MTaseII.containing.BREX | HIN-HTH-><-MazG<-BrxA-DNA-binding<-PglZ<-TUDOR+SF1-SF2-Helicase+iREase+CTD<-N6-N4-METHYLASE\|\|SIGMA-HTH+LuxR-HTH-><-DUF6079*<-BrxB-BrxF-iSTAND<-ParB+HNH<-?<-?<-SF1-SF2-Helicase+REase<-HEPN+REase+SF1-SF2-Helicase<-?<-?<-MULE-TRANSPOSASE | 1218 | Bacteria>Terrabacteria_group>Bacillota>Clostridia>Halanaerobiales>Halanaerobiaceae | GCF_900102605.1 | Halanaerobium_congolense | F3G65_RS12720 | 15263..18919 | 54121 | DUF6079-family-protein-[Halanaerobium-congolense] |
| WP_013278344.1 | Type3.MTaseII.containing.BREX | <-HEPN<-Polbeta<-DSBH<-SIR2<-BrxA-DNA-binding<-PglZ<-N6-N4-METHYLASE<-DUF6079*<-BrxB-BrxF-iSTAND<-Peptidase_S8<-Classical-AAA\|\|?-><-?\|\|Peptidase_M20->?-><-SF1-SF2-Helicase+REase<-SF1-SF2-Helicase+REase | 1215 | Bacteria>Terrabacteria_group>Bacillota>Clostridia>Halanaerobiales>Halobacteroidaceae | GCF_000144695.1 | Acetohalobium_arabaticum | acear_RS07195 | 1477107..1480754 | 28187 | DUF6079-family-protein-[Acetohalobium-arabaticum] |
| EGD52432.1 | Type3.MTaseII.containing.BREX | <-McrB-AAA<-McrB-AAA<-?<-?<-BrxA-DNA-binding<-PglZ<-TUDOR+SF1-SF2-Helicase+iREase+CTD<-N6-N4-METHYLASE\|\|DUF772+DDE_Tnp_1-><-DUF6079*<-BrxB-BrxF-iSTAND<-RAMPs<-?<-RAMPs<-Cas10/Csm1<-Cas10/Csm1<-RAMPs<-RAMPs\|\|DEDD_Tnp_IS110+Transposase_20-> | 1253 | Bacteria>Terrabacteria_group>Bacillota>Clostridia>Thermoanaerobacterales>Thermoanaerobacteraceae | GCA_000192295.2 | Thermoanaerobacter_ethanolicus_JW_200 | TheetDRAFT_0743 | 44877..48638 | 509192 | hypothetical-protein-TheetDRAFT_0743-[Thermoanaerobacter-ethanolicus-JW-200] |
| KUJ90862.1 | Type3.MTaseII.containing.BREX | CARF+HEPN->?->Cas10->?->?->?->?->35exo+wHTH+SF1-SF2-Helicase+HTH->BrxB-BrxF-iSTAND->DUF6079*->N6-N4-METHYLASE->PglZ->BrxA-DNA-binding->DUF4236-><-REase<-REase\|\|Peptidase_A8->S4+PSYN-> | 1227 | Bacteria>Terrabacteria_group>Bacillota>Clostridia>Thermoanaerobacterales>Thermoanaerobacteraceae | GCA_001508735.1 | Thermoanaerobacter_thermocopriae | XD37_0930 | 27167..30850 | 29350 | hypothetical-protein-XD37_0930-[Thermoanaerobacter-thermocopriae] |
| RKL62727.1 | Type3.MTaseII.containing.BREX | <-HEPN<-PolB<-?<-DUF4236<-HEPN<-ParB+HNH<-BrxA-DNA-binding<-PglZ<-N6-N4-METHYLASE<-DUF6079*<-BrxB-BrxF-iSTAND<-REase+HTH | 1223 | Bacteria>Terrabacteria_group>Bacillota>Clostridia>Thermoanaerobacterales>Thermoanaerobacteraceae | GCA_003615265.1 | Thermoanaerobacteraceae_bacterium_SP2 | DXT63_09740 | 11701..15372 | 2292448 | exonuclease-SbcC-[Thermoanaerobacteraceae-bacterium-SP2] |
| SFE40605.1 | Type3.MTaseII.containing.BREX | <-S4+PSYN<-Peptidase_A8\|\|REase-><-DUF4236+TPR<-BrxA-DNA-binding<-PglZ<-?<-N6-N4-METHYLASE<-DUF6079*<-BrxB-BrxF-iSTAND<-RAMPs<-?<-RAMPs<-Cas10/Csm1<-Cas10/Csm1<-RAMPs<-RAMPs | 1227 | Bacteria>Terrabacteria_group>Bacillota>Clostridia>Thermoanaerobacterales>Thermoanaerobacteraceae | GCA_900112905.1 | Thermoanaerobacter_thermohydrosulfuricus | SAMN04324257_01605 | 19229..22912 | 1516 | hypothetical-protein-SAMN04324257_01605-[Thermoanaerobacter-thermohydrosulfuricus] |
| WP_012269233.1 | Type3.MTaseII.containing.BREX | CARF+HEPN->?->Cas10->?->?->?->?->35exo+wHTH+SF1-SF2-Helicase+HTH->BrxB-BrxF-iSTAND->DUF6079*->N6-N4-METHYLASE->TUDOR+SF1-SF2-Helicase+iREase+CTD->PglZ->BrxA-DNA-binding->wHTH+McrB-AAA->HIN-HTH+PAIREDC-HTH->McrB-AAA->wHTH+REase-p->DDE_Tnp_1_4-> | 1227 | Bacteria>Terrabacteria_group>Bacillota>Clostridia>Thermoanaerobacterales>Thermoanaerobacteraceae | GCF_000019085.1 | Thermoanaerobacter | THEBR_RS04735 | 974574..978257 | 1754 | MULTISPECIES:-DUF6079-family-protein-[Thermoanaerobacter] |
| WP_012995396.1 | Type3.MTaseII.containing.BREX | <-PRTase\|\|?-><-S4+PSYN<-Peptidase_A8\|\|<-DUF4236+TPR<-BrxA-DNA-binding<-PglZ<-?<-N6-N4-METHYLASE<-DUF6079*<-BrxB-BrxF-iSTAND<-RAMPs<-?<-RAMPs<-Cas10/Csm1<-RAMPs<-RAMPs\|\|DEDD_Tnp_IS110+Transposase_20-> | 1227 | Bacteria>Terrabacteria_group>Bacillota>Clostridia>Thermoanaerobacterales>Thermoanaerobacteraceae | GCF_000025645.1 | Thermoanaerobacter_italicus | THIT_RS06730 | 1375258..1378941 | 108150 | DUF6079-family-protein-[Thermoanaerobacter-italicus] |
| WP_043883710.1 | Type3.MTaseII.containing.BREX | DsbD->?-><-?\|\|?->tRNA->Phage_integrase->PhdYeFM_antitox->PIN->XIS-HTH->\|\|BrxB-BrxF-iSTAND->DUF6079*->N6-N4-METHYLASE->TUDOR+SF1-SF2-Helicase+iREase+CTD->PglZ->BrxA-DNA-binding->SF1-SF2-Helicase+REase+HTH->ParB+HNH-> | 1235 | Bacteria>Terrabacteria_group>Bacillota>Clostridia>Thermoanaerobacterales>Thermoanaerobacteraceae | GCF_000156275.2 | Caldanaerobacter_subterraneus | CDSM653_RS04915 | 16153..19860 | 911092 | DUF6079-family-protein-[Caldanaerobacter-subterraneus] |
| WP_269207978.1 | Type3.MTaseII.containing.BREX | <-S4+PSYN<-Peptidase_A8\|\|REase-><-DUF4236+TPR<-BrxA-DNA-binding<-PglZ<-TUDOR+SF1-SF2-Helicase+iREase+CTD<-DUF6079-p<-DUF6079-p*<-BrxB-BrxF-iSTAND<-RAMPs<-?<-RAMPs<-Cas10/Csm1<-RAMPs<-RAMPs<-?\|\|DEDD_Tnp_IS110+Transposase_20-> | 945 | Bacteria>Terrabacteria_group>Bacillota>Clostridia>Thermoanaerobacterales>Thermoanaerobacteraceae | GCF_000373165.1 | Thermoanaerobacter_indiensis | B044_RS14345 | 1544650..1547487 | 1125974 | DUF6079-family-protein-[Thermoanaerobacter-indiensis] |
| WP_269207977.1 | Type3.MTaseII.containing.BREX | <-PRTase\|\|?-><-S4+PSYN<-Peptidase_A8\|\|REase-><-DUF4236+TPR<-BrxA-DNA-binding<-PglZ<-?<-DUF6079-p<-DUF6079-p*<-BrxB-BrxF-iSTAND<-RAMPs<-?<-RAMPs<-Cas10/Csm1<-RAMPs<-RAMPs | 266 | Bacteria>Terrabacteria_group>Bacillota>Clostridia>Thermoanaerobacterales>Thermoanaerobacteraceae | GCF_000373165.1 | Thermoanaerobacter_indiensis | B044_RS15690 | 1543815..1544615 | 1125974 | DUF6079-family-protein-[Thermoanaerobacter-indiensis] |
| WP_026486463.1 | Type3.MTaseII.containing.BREX | <-HEPN<-PolB<-?<-BrxA-DNA-binding<-PglZ<-TUDOR+SF1-SF2-Helicase+iREase+CTD<-N6-N4-METHYLASE<-HicB_lk_antitox<-RRXRR+HNH<-DUF6079*<-BrxB-BrxF-iSTAND<-Trypsin+PDZ\|\|?-><-?<-?<-?<-?<-ABC-ATPase+SbcC+ABC-ATPase+SbcC | 1236 | Bacteria>Terrabacteria_group>Bacillota>Clostridia>Thermoanaerobacterales>Thermoanaerobacteraceae | GCA_000427425.1 | Caldanaerobius_polysaccharolyticus | CALPO_RS0105665 | 974700..978410 | 44256 | DUF6079-family-protein-[Caldanaerobius-polysaccharolyticus] |
| WP_028992367.1 | Type3.MTaseII.containing.BREX | RAMPs->RAMPs->Cas10/Csm1->RAMPs->?->RAMPs->BrxB-BrxF-iSTAND->DUF6079*->N6-N4-METHYLASE->?->PglZ->BrxA-DNA-binding->DUF4236->DUF4236->\|\|Peptidase_A8->S4+PSYN-> | 1227 | Bacteria>Terrabacteria_group>Bacillota>Clostridia>Thermoanaerobacterales>Thermoanaerobacteraceae | GCF_000518565.1 | Thermoanaerobacter_thermocopriae | L887_RS0109270 | 9659..13342 | 29350 | DUF6079-family-protein-[Thermoanaerobacter-thermocopriae] |
| WP_042833177.1 | Type3.MTaseII.containing.BREX | <-PRTase\|\|?-><-S4+PSYN<-Peptidase_A8\|\|REase-><-DUF4236+TPR<-PglZ-p<-TUDOR+SF1-SF2-Helicase+iREase+CTD<-N6-N4-METHYLASE<-DUF6079*<-BrxB-BrxF-iSTAND\|\|<-AAA-ATPase<-HTH_7+rve+rve_3\|\|<-RAMPs<-?<-RAMPs<-Cas10/Csm1<-RAMPs<-RAMPs | 1227 | Bacteria>Terrabacteria_group>Bacillota>Clostridia>Thermoanaerobacterales>Thermoanaerobacteraceae | GCF_000806225.2 | Thermoanaerobacter_sp._YS13 | THYS13_RS03510 | 675539..679222 | 1511746 | DUF6079-family-protein-[Thermoanaerobacter-sp.-YS13] |
| WP_256201197.1 | Type3.MTaseII.containing.BREX | RAMPs->Cas10/Csm1->Cas10/Csm1->Cas10/Csm1->RAMPs->?->RAMPs->RAMPs->BrxB-BrxF-iSTAND->DUF6079-p*->?->PglZ-p->PglZ-p->?->PglZ-p->BrxA-DNA-binding->DUF4236->DUF4236->\|\|S4+PSYN-> | 217 | Bacteria>Terrabacteria_group>Bacillota>Clostridia>Thermoanaerobacterales>Thermoanaerobacteraceae | GCF_001310975.1 | Thermoanaerobacter_thermocopriae | JCM7501_RS14380 | 44306..44959 | 29350 | DUF6079-family-protein-[Thermoanaerobacter-thermocopriae] |
| WP_071520604.1 | Type3.MTaseII.containing.BREX | HHH+DNA_pol_B+RNase->?-><-TOPRIM<-OLD-ABC\|\|BrxB-BrxF-iSTAND->DUF6079*->N6-N4-METHYLASE->TUDOR+SF1-SF2-Helicase+iREase+CTD->?->PglZ->BrxA-DNA-binding-><-PhdYeFM_antitox\|\|MNS-> | 1226 | Bacteria>Terrabacteria_group>Bacillota>Clostridia>Thermoanaerobacterales>Thermoanaerobacteraceae | GCF_001874065.1 | Moorella_thermoacetica | MOOR_RS04485 | 140979..144659 | 1525 | DUF6079-family-protein-[Moorella-thermoacetica] |
| WP_106004662.1 | Type3.MTaseII.containing.BREX | <-REase<-REase+EVE<-PIN<-SIR2<-BrxA-DNA-binding<-PglZ<-TUDOR+SF1-SF2-Helicase+iREase+CTD<-N6-N4-METHYLASE<-DUF6079*<-BrxB-BrxF-iSTAND\|\|?->?-><-?\|\|?->?->?-><-LysM+LysM<-AA_kinase+ACT+ACT_7 | 1226 | Bacteria>Terrabacteria_group>Bacillota>Clostridia>Thermoanaerobacterales>Thermoanaerobacteraceae | GCF_002995755.1 | Moorella_humiferrea | MOHU_RS03240 | 30215..33895 | 676965 | DUF6079-family-protein-[Moorella-humiferrea] |
| WP_115793167.1 | Type3.MTaseII.containing.BREX | BrxB-BrxF-iSTAND->DUF6079*->N6-N4-METHYLASE->OLD-ABC+TOPRIM->TUDOR+SF1-SF2-Helicase+iREase+CTD->PglZ->BrxA-DNA-binding->ARG-><-?\|\|?->PRTase-> | 1239 | Bacteria>Terrabacteria_group>Bacillota>Clostridia>Thermoanaerobacterales>Thermoanaerobacteraceae | GCF_003368535.1 | Ammonifex_thiophilus | DXX99_RS09060 | 2331..6050 | 444093 | DUF6079-family-protein-[Ammonifex-thiophilus] |
| WP_148854792.1 | Type3.MTaseII.containing.BREX | LACI-HTH+PBPI->HIN-HTH->?->?->SMODS->BrxB-BrxF-iSTAND->DUF6079*->N6-N4-METHYLASE->TUDOR+SF1-SF2-Helicase+iREase+CTD->PglZ->BrxA-DNA-binding->SIR2->?-><-IstB_IS21 | 1203 | Bacteria>Terrabacteria_group>Bacillota>Clostridia>Thermoanaerobacterales>Thermoanaerobacteraceae | GCF_008121875.1 | Moorella_thermoacetica | MOLA_RS10055 | 195437..199048 | 1525 | DUF6079-family-protein-[Moorella-thermoacetica] |
| WP_149544903.1 | Type3.MTaseII.containing.BREX | LACI-HTH+PBPI->?-><-DNA_METHYLASE\|\|RAMPs->RAMPs->?->?-><-?\|\|BrxB-BrxF-iSTAND->DUF6079*->N6-N4-METHYLASE->DUF6079->TUDOR+SF1-SF2-Helicase+iREase+CTD->PglZ->BrxA-DNA-binding->N6-N4-METHYLASE->?->Radical_SAM-> | 1219 | Bacteria>Terrabacteria_group>Bacillota>Clostridia>Thermoanaerobacterales>Thermoanaerobacteraceae | GCF_008274215.1 | Calorimonas_adulescens | FWJ32_RS05120 | 20661..24320 | 2606906 | DUF6079-family-protein-[Calorimonas-adulescens] |
| WP_149544905.1 | Type3.MTaseII.containing.BREX | <-DNA_METHYLASE\|\|RAMPs->RAMPs->?->?-><-?\|\|BrxB-BrxF-iSTAND->DUF6079->N6-N4-METHYLASE->DUF6079*->TUDOR+SF1-SF2-Helicase+iREase+CTD->PglZ->BrxA-DNA-binding->N6-N4-METHYLASE->?->Radical_SAM-> | 1202 | Bacteria>Terrabacteria_group>Bacillota>Clostridia>Thermoanaerobacterales>Thermoanaerobacteraceae | GCF_008274215.1 | Calorimonas_adulescens | FWJ32_RS05130 | 26921..30529 | 2606906 | DUF6079-family-protein-[Calorimonas-adulescens] |
| WP_214495594.1 | Type3.MTaseII.containing.BREX | <-DEDD_Tnp_IS110+Transposase_20\|\|RAMPs->RAMPs->Cas10/Csm1->RAMPs->?->RAMPs->BrxB-BrxF-iSTAND->DUF6079*->N6-N4-METHYLASE->?->PglZ->BrxA-DNA-binding->DUF4236+TPR->\|\|Peptidase_A8->S4+PSYN-><-?\|\|PRTase-> | 1227 | Bacteria>Terrabacteria_group>Bacillota>Clostridia>Thermoanaerobacterales>Thermoanaerobacteraceae | GCF_018588165.1 | Thermoanaerobacter_sp._CM-CNRG_TB177 | I4N53_RS03585 | 103174..106857 | 2800659 | DUF6079-family-protein-[Thermoanaerobacter-sp.-CM-CNRG-TB177] |
| WP_109208176.1 | Type3.MTaseII.containing.BREX | <-AbiEii<-?<-?<-BrxA-DNA-binding<-PglZ<-TUDOR+SF1-SF2-Helicase+iREase+CTD<-NACHT<-HEPN<-N6-N4-METHYLASE<-DUF6079*<-BrxB-BrxF-iSTAND<-?<-?<-Polbeta\|\|DDE_Tnp_1-><-?<-ArsR-HTH | 1239 | Bacteria>Terrabacteria_group>Bacillota>Clostridia>Thermoanaerobacterales>Thermoanaerobacteraceae | GCA_023734095.1 | Moorella_sp._Hama-1 | NGH78_RS14130 | 2894050..2897769 | 2138101 | DUF6079-family-protein-[Moorella-sp.-Hama-1] |
| WP_266037231.1 | Type3.MTaseII.containing.BREX | <-S4+PSYN\|\|REase-><-DUF4236+TPR<-DUF4236<-BrxA-DNA-binding<-BrxA-DNA-binding-p<-PglZ<-TUDOR+SF1-SF2-Helicase+iREase+CTD<-N6-N4-METHYLASE<-DUF6079*<-BrxB-BrxF-iSTAND<-RAMPs<-?<-RAMPs<-Cas10/Csm1<-RAMPs<-RAMPs<-RAMPs | 1227 | Bacteria>Terrabacteria_group>Bacillota>Clostridia>Thermoanaerobacterales>Thermoanaerobacteraceae | GCF_026240795.1 | Thermoanaerobacter_sp._RKWS2 | OEI98_RS08775 | 1653038..1656721 | 2983842 | DUF6079-family-protein-[Thermoanaerobacter-sp.-RKWS2] |
| WP_011392173.1 | Type3.MTaseII.containing.BREX | IstB_IS21<-?<-SIR2<-BrxA-DNA-binding<-PglZ<-TUDOR+SF1-SF2-Helicase+iREase+CTD<-N6-N4-METHYLASE<-DUF6079*<-BrxB-BrxF-iSTAND<-SMODS<-?<-?<-?<-HIN-HTH<-LACI-HTH+PBPI | 1203 | Bacteria>Terrabacteria_group>Bacillota>Clostridia>Thermoanaerobacterales>Thermoanaerobacteraceae | GCF_027925125.1 | Moorella_thermoacetica | QJ070_RS08455 | 11985..15596 | 1525 | DUF6079-family-protein-[Moorella-thermoacetica] |
| WP_074592540.1 | Type3.MTaseII.containing.BREX | RAMPs->RAMPs->Cas10/Csm1->RAMPs->?->RAMPs->BrxB-BrxF-iSTAND->DUF6079*->N6-N4-METHYLASE->?->PglZ->BrxA-DNA-binding->DUF4236+TPR-><-REase\|\|Peptidase_A8->S4+PSYN-> | 1227 | Bacteria>Terrabacteria_group>Bacillota>Clostridia>Thermoanaerobacterales>Thermoanaerobacteraceae | GCF_900102395.1 | Thermoanaerobacter_thermohydrosulfuricus | BLS86_RS06330 | 10439..14122 | 1516 | DUF6079-family-protein-[Thermoanaerobacter-thermohydrosulfuricus] |
| CEO89246.1 | Type3.MTaseII.containing.BREX | <-RNase<-HEPN<-PolB<-BrxA-DNA-binding<-PglZ-p<-PglZ-p<-N6-N4-METHYLASE<-N6-N4-METHYLASE-p<-DUF6079-p*<-BrxB-BrxF-iSTAND | 819 | Bacteria>Terrabacteria_group>Bacillota>Clostridia>Thermoanaerobacterales>Thermoanaerobacterales_Family_III._Incertae_Sedis | GCA_000946815.1 | Syntrophaceticus_schinkii | SSCH_420039 | 38369..40828 | 499207 | hypothetical-protein-SSCH_420039-[Syntrophaceticus-schinkii] |
| WP_013432000.1 | Type3.MTaseII.containing.BREX | Thr_synth_N+PALP->?->?->\|\|?->?->?->ABC-ATPase+SbcC->?->BrxB-BrxF-iSTAND->DUF6079*->N6-N4-METHYLASE->TUDOR+SF1-SF2-Helicase+iREase+CTD->?->PglZ->BrxA-DNA-binding->?->HD->?->Schlafen+HATPase+wHTH-> | 1230 | Bacteria>Terrabacteria_group>Bacillota>Clostridia>Thermoanaerobacterales>Thermoanaerobacterales_Family_III._Incertae_Sedis | GCF_000166695.1 | Caldicellulosiruptor_acetigenus | CALKR_RS03100 | 672126..675818 | 301953 | DUF6079-family-protein-[Caldicellulosiruptor-acetigenus] |
| WP_013787753.1 | Type3.MTaseII.containing.BREX | ACET->HTH_36+DnaB_2+DnaB_2->SIGMA-HTH+SIGMA-HTH+SIGMA-HTH->35exo+wHTH+SF1-SF2-Helicase+HTH->BrxB-BrxF-iSTAND->DUF6079*->N6-N4-METHYLASE->TUDOR+SF1-SF2-Helicase+iREase+CTD->PglZ->BrxA-DNA-binding->ParA-Soj-PloopNTPase+McrB-AAA->McrC-NTD+REase->5TMR-LYT+GAF+MORC->REC+LYTTR-> | 1227 | Bacteria>Terrabacteria_group>Bacillota>Clostridia>Thermoanaerobacterales>Thermoanaerobacterales_Family_III._Incertae_Sedis | GCF_000189775.2 | Thermoanaerobacterium_xylanolyticum | THEXY_RS05040 | 1037327..1041010 | 29329 | DUF6079-family-protein-[Thermoanaerobacterium-xylanolyticum] |
| WP_014759120.1 | Type3.MTaseII.containing.BREX | <-REC+LYTTR<-5TMR-LYT+GAF+MORC<-REase<-McrC-NTD+REase<-ParA-Soj-PloopNTPase+McrB-AAA<-BrxA-DNA-binding<-PglZ<-TUDOR+SF1-SF2-Helicase+iREase+CTD<-N6-N4-METHYLASE<-DUF6079*<-BrxB-BrxF-iSTAND<-?<-35exo+wHTH+SF1-SF2-Helicase+HTH<-SIGMA-HTH+SIGMA-HTH<-HTH_36+DnaB_2+DnaB_2<-ACET | 1227 | Bacteria>Terrabacteria_group>Bacillota>Clostridia>Thermoanaerobacterales>Thermoanaerobacterales_Family_III._Incertae_Sedis | GCF_000307585.2 | Thermoanaerobacterium | TSAC_RS11555 | 2333389..2337072 | 28895 | MULTISPECIES:-DUF6079-family-protein-[Thermoanaerobacterium] |
| WP_015312001.1 | Type3.MTaseII.containing.BREX | <-REC+LYTTR<-5TMR-LYT+GAF+MORC<-REase<-McrC-NTD+REase<-?<-BrxA-DNA-binding<-PglZ<-TUDOR+SF1-SF2-Helicase+iREase+CTD<-N6-N4-METHYLASE<-DUF6079*<-BrxB-BrxF-iSTAND<-?<-HNH+NERD+ZnR<-HTH_36+DnaB_2+DnaB_2<-ACET\|\|?->?->HEPN-> | 1227 | Bacteria>Terrabacteria_group>Bacillota>Clostridia>Thermoanaerobacterales>Thermoanaerobacterales_Family_III._Incertae_Sedis | GCF_000328545.1 | Thermoanaerobacterium_thermosaccharolyticum | THETHE_RS09155 | 1853967..1857650 | 1517 | DUF6079-family-protein-[Thermoanaerobacterium-thermosaccharolyticum] |
| WP_231560571.1 | Type3.MTaseII.containing.BREX | <-REC+LYTTR<-5TMR-LYT+GAF+MORC<-REase<-McrC-NTD+REase<-ParA-Soj-PloopNTPase+McrB-AAA<-BrxA-DNA-binding<-PglZ<-TUDOR+SF1-SF2-Helicase+iREase+CTD<-N6-N4-METHYLASE<-DUF6079-p*<-DUF6079-p<-BrxB-BrxF-iSTAND<-?<-SF1-SF2-Helicase+HTH<-35exo+wHTH+SF1-SF2-Helicase<-SIGMA-HTH<-SIGMA-HTH+SIGMA-HTH<-HTH_36+DnaB_2+DnaB_2<-ACET | 958 | Bacteria>Terrabacteria_group>Bacillota>Clostridia>Thermoanaerobacterales>Thermoanaerobacterales_Family_III._Incertae_Sedis | GCF_000747665.1 | Thermoanaerobacterium_saccharolyticum | NTOU1_RS07135 | 43730..46606 | 28896 | DUF6079-family-protein-[Thermoanaerobacterium-saccharolyticum] |
| WP_231560568.1 | Type3.MTaseII.containing.BREX | <-5TMR-LYT+GAF+MORC<-REase<-McrC-NTD+REase<-ParA-Soj-PloopNTPase+McrB-AAA<-BrxA-DNA-binding<-PglZ<-TUDOR+SF1-SF2-Helicase+iREase+CTD<-N6-N4-METHYLASE<-DUF6079-p<-DUF6079-p*<-BrxB-BrxF-iSTAND<-?<-SF1-SF2-Helicase+HTH<-35exo+wHTH+SF1-SF2-Helicase<-SIGMA-HTH<-SIGMA-HTH+SIGMA-HTH<-HTH_36+DnaB_2+DnaB_2<-ACET | 263 | Bacteria>Terrabacteria_group>Bacillota>Clostridia>Thermoanaerobacterales>Thermoanaerobacterales_Family_III._Incertae_Sedis | GCF_000747665.1 | Thermoanaerobacterium_saccharolyticum | NTOU1_RS15720 | 46623..47414 | 28896 | DUF6079-family-protein-[Thermoanaerobacterium-saccharolyticum] |
| WP_156972215.1 | Type3.MTaseII.containing.BREX | <-RNase<-HEPN<-PolB<-BrxA-DNA-binding<-PglZ-p<-PglZ-p<-DUF6079-p*<-DUF6079-p<-BrxB-BrxF-iSTAND | 789 | Bacteria>Terrabacteria_group>Bacillota>Clostridia>Thermoanaerobacterales>Thermoanaerobacterales_Family_III._Incertae_Sedis | GCF_000946815.1 | Syntrophaceticus_schinkii | SSCH_RS16730 | 38369..40738 | 499207 | DUF6079-family-protein-[Syntrophaceticus-schinkii] |
| WP_052835524.1 | Type3.MTaseII.containing.BREX | <-RNase<-HEPN<-PolB<-BrxA-DNA-binding<-PglZ-p<-PglZ-p<-DUF6079-p<-DUF6079-p*<-BrxB-BrxF-iSTAND | 466 | Bacteria>Terrabacteria_group>Bacillota>Clostridia>Thermoanaerobacterales>Thermoanaerobacterales_Family_III._Incertae_Sedis | GCF_000946815.1 | Syntrophaceticus_schinkii | SSCH_RS20240 | 40644..42044 | 499207 | DUF6079-family-protein-[Syntrophaceticus-schinkii] |
| WP_045174402.1 | Type3.MTaseII.containing.BREX | ABC-ATPase+SbcC->?->BrxB-BrxF-iSTAND->DUF6079*->TUDOR+SF1-SF2-Helicase+iREase+CTD->PglZ->BrxA-DNA-binding->HATPase+wHTH->PolB->HEPN->?-><-SH3 | 1230 | Bacteria>Terrabacteria_group>Bacillota>Clostridia>Thermoanaerobacterales>Thermoanaerobacterales_Family_III._Incertae_Sedis | GCF_000955725.1 | Caldicellulosiruptor_danielii | N913_RS05165 | 1164388..1168080 | 1387557 | DUF6079-family-protein-[Caldicellulosiruptor-danielii] |
| WP_094045777.1 | Type3.MTaseII.containing.BREX | <-HEPN<-?<-?\|\|ACET->HTH_36+DnaB_2+DnaB_2->SIGMA-HTH+SIGMA-HTH+SIGMA-HTH->35exo+wHTH+SF1-SF2-Helicase+HTH->?->BrxB-BrxF-iSTAND->DUF6079*->N6-N4-METHYLASE->TUDOR+SF1-SF2-Helicase+iREase+CTD->PglZ->BrxA-DNA-binding->?->McrC-NTD+REase->?->5TMR-LYT+GAF+MORC->REC+LYTTR-> | 1227 | Bacteria>Terrabacteria_group>Bacillota>Clostridia>Thermoanaerobacterales>Thermoanaerobacterales_Family_III._Incertae_Sedis | GCF_002238085.1 | Thermoanaerobacterium_thermosaccharolyticum | CE561_RS09515 | 11232..14915 | 1517 | DUF6079-family-protein-[Thermoanaerobacterium-thermosaccharolyticum] |
| WP_150200391.1 | Type3.MTaseII.containing.BREX | <-REC+LYTTR<-5TMR-LYT+GAF+MORC<-REase<-McrC-NTD+REase<-?<-BrxA-DNA-binding<-PglZ<-TUDOR+SF1-SF2-Helicase+iREase+CTD<-N6-N4-METHYLASE<-DUF6079*<-BrxB-BrxF-iSTAND<-?<-HNH+NERD+ZnR<-N6-N4-METHYLASE<-?\|\|HTH_7+rve+rve_3->AAA-ATPase-><-REase<-DNA_METHYLASE | 1229 | Bacteria>Terrabacteria_group>Bacillota>Clostridia>Thermoanaerobacterales>Thermoanaerobacterales_Family_III._Incertae_Sedis | GCF_008632855.1 | Thermoanaerobacterium_thermosaccharolyticum | F1655_RS01835 | 39469..43158 | 1517 | DUF6079-family-protein-[Thermoanaerobacterium-thermosaccharolyticum] |
| WP_045165709.1 | Type3.MTaseII.containing.BREX | <-HATPase+wHTH<-Schlafen+HATPase+wHTH<-HD<-HD<-?<-BrxA-DNA-binding<-PglZ<-TUDOR+SF1-SF2-Helicase+iREase+CTD<-N6-N4-METHYLASE<-DUF6079*<-BrxB-BrxF-iSTAND<-?<-ABC-ATPase+SbcC<-?<-?<-?\|\|Aminotran_1_2-> | 1230 | Bacteria>Terrabacteria_group>Bacillota>Clostridia>Thermoanaerobacterales>Thermoanaerobacterales_Family_III._Incertae_Sedis | GCF_026914285.1 | Caldicellulosiruptor_naganoensis | OTJ99_RS09170 | 1832428..1836120 | 29324 | DUF6079-family-protein-[Caldicellulosiruptor-naganoensis] |
| MCR4431939.1 | Type3.MTaseII.containing.BREX | BrxB-BrxF-iSTAND->DUF6079*->N6-N4-METHYLASE->PglZ->BrxA-DNA-binding->?->?->HEPN-> | 1223 | Bacteria>Terrabacteria_group>Bacillota>Clostridia>Thermosediminibacterales>Tepidanaerobacteraceae | GCA_024655905.1 | Tepidanaerobacteraceae_bacterium | NUV45_13095 | 103146..106817 | 2963290 | DUF6079-family-protein-[Tepidanaerobacteraceae-bacterium] |
| WP_013778923.1 | Type3.MTaseII.containing.BREX | PTSEIIC->?->?->Tox-ABhydrolase-><-NtrC-AAA+FIS-HTH<-?<-BrxA-DNA-binding<-PglZ<-N6-N4-METHYLASE<-DUF6079*<-BrxB-BrxF-iSTAND\|\|?-><-?<-?<-PAS+NtrC-AAA+FIS-HTH | 1221 | Bacteria>Terrabacteria_group>Bacillota>Clostridia>Thermosediminibacterales>Tepidanaerobacteraceae | GCF_000213235.1 | Tepidanaerobacter_acetatoxydans | TEPRE1_RS09335 | 1938923..1942588 | 499229 | DUF6079-family-protein-[Tepidanaerobacter-acetatoxydans] |
| WP_059033512.1 | Type3.MTaseII.containing.BREX | BrxB-BrxF-iSTAND->DUF6079*->N6-N4-METHYLASE->PglZ->BrxA-DNA-binding->?->HAMP+MA+MA->?->?->SIGMA-HTH-> | 1221 | Bacteria>Terrabacteria_group>Bacillota>Clostridia>Thermosediminibacterales>Tepidanaerobacteraceae | GCF_001485475.2 | Tepidanaerobacter_syntrophicus | TSYNT_RS09600 | 148625..152290 | 224999 | DUF6079-family-protein-[Tepidanaerobacter-syntrophicus] |
| WP_122014530.1 | Type3.MTaseII.containing.BREX | REase+HTH->BrxB-BrxF-iSTAND->DUF6079*->N6-N4-METHYLASE->N6-N4-METHYLASE->PglZ->BrxA-DNA-binding->HEPN->HTH+REase->?->PolB-> | 1223 | Bacteria>Terrabacteria_group>Bacillota>Clostridia>Thermosediminibacterales>Tepidanaerobacteraceae | GCF_003691585.1 | Biomaibacter_acetigenes | D2962_RS06505 | 1331305..1334976 | 2316383 | DUF6079-family-protein-[Biomaibacter-acetigenes] |
| WP_273702429.1 | Type3.MTaseII.containing.BREX | Cysteine-Desulfurase->?->?->?->?-><-?\|\|BrxB-BrxF-iSTAND->DUF6079*->N6-N4-METHYLASE->BrxA-DNA-binding->?->HAMP+MA+MA->?->?->SIGMA-HTH->?->NtrC-AAA+FIS-HTH-> | 1231 | Bacteria>Terrabacteria_group>Bacillota>Clostridia>Thermosediminibacterales>Tepidanaerobacteraceae | GCF_018436545.1 | Tepidanaerobacter_syntrophicus | KKR28_RS04975 | 15594..19289 | 224999 | DUF6079-family-protein-[Tepidanaerobacter-syntrophicus] |
| MCF6096371.1 | Type3.MTaseII.containing.BREX | <-HTH_3<-RelE<-ParB+HNH<-DUF4011+SF1-SF2-Helicase+REase<-BrxA-DNA-binding<-PglZ<-TUDOR+SF1-SF2-Helicase+iREase+CTD<-N6-N4-METHYLASE<-DUF6079*<-BrxB-BrxF-iSTAND\|\|DEOR-HTH+ISOCOT->?->?->?-><-?<-?<-?<-Glycos_transf | 1232 | Bacteria>Terrabacteria_group>Bacillota>Clostridia>Thermosediminibacterales>Thermosediminibacteraceae | GCA_021608145.1 | Thermovorax_subterraneus | L1766_05030 | 56890..60588 | 496831 | DUF6079-family-protein-[Thermovorax-subterraneus] |
| MCG0276139.1 | Type3.MTaseII.containing.BREX | <-N6-N4-METHYLASE<-DUF6079*<-BrxB-BrxF-iSTAND<-Cysteine-Desulfurase<-DUF6079<-DUF4007<-PAPS-reduct+Fer4_7<-DndE<-DndD<-viral-protein<-HKD-DNase+SF1-SF2-Helicase | 1234 | Bacteria>Terrabacteria_group>Bacillota>Clostridia>Thermosediminibacterales>Thermosediminibacteraceae | GCA_021890755.1 | Thermosediminibacteraceae_bacterium | L5655_08270 | 2182..5886 | 2885163 | DUF6079-family-protein-[Thermosediminibacteraceae-bacterium] |
| NCB91957.1 | Type3.MTaseII.containing.BREX | <-REase<-?<-SIGMA-HTH\|\|NACHT+SNaCT->XIS-HTH->BrxB-BrxF-iSTAND->DUF6079*->N6-N4-METHYLASE->TUDOR+SF1-SF2-Helicase+iREase+CTD->PglZ->N6-N4-METHYLASE-p-> | 1230 | Bacteria>Terrabacteria_group>Bacillota>Clostridia>unclassified_Clostridia | GCA_009929525.1 | Clostridia_bacterium | EDM40_05205 | 33450..37142 | 2044939 | hypothetical-protein-[Clostridia-bacterium] |
| MBQ4455501.1 | Type3.MTaseII.containing.BREX | Classical-AAA->?->?->NERD+SF1-SF2-Helicase->?->BrxB-BrxF-iSTAND->DUF6079-p*-> | 356 | Bacteria>Terrabacteria_group>Bacillota>Clostridia>unclassified_Clostridia | GCA_017394825.1 | Clostridia_bacterium | II895_12170 | 7685..8754 | 2044939 | hypothetical-protein-[Clostridia-bacterium] |
| MBQ3268373.1 | Type3.MTaseII.containing.BREX | <-SF1-SF2-Helicase<-Vrr-REase<-PriCT1+D5N+D5ATPase\|\|?->?->REase->?-><-?<-DUF6079-p*<-BrxB-BrxF-iSTAND\|\|HTH_3->?->?->XIS-HTH->Phage_int_SAM_3+Phage_integrase-><-tRNA<-tRNA\|\|GGDEF-> | 308 | Bacteria>Terrabacteria_group>Bacillota>Clostridia>unclassified_Clostridia | GCA_017404615.1 | Clostridia_bacterium | IJH09_00480 | 39487..40413 | 2044939 | hypothetical-protein-[Clostridia-bacterium] |
| MBQ5956552.1 | Type3.MTaseII.containing.BREX | Classical-AAA->?->?->NERD+SF1-SF2-Helicase->?->BrxB-BrxF-iSTAND->DUF6079-p*-> | 196 | Bacteria>Terrabacteria_group>Bacillota>Clostridia>unclassified_Clostridia | GCA_017439085.1 | Clostridia_bacterium | IJL46_03155 | 63510..64097 | 2044939 | hypothetical-protein-[Clostridia-bacterium] |
| MBR6410257.1 | Type3.MTaseII.containing.BREX | <-DUF6079-p*<-BrxB-BrxF-iSTAND<-HTH_26\|\|?-><-ABhydrolase+ABhydrolase<-?<-Glyco_hydro_43<-?<-HATPase_c | 480 | Bacteria>Terrabacteria_group>Bacillota>Clostridia>unclassified_Clostridia | GCA_017539425.1 | Clostridia_bacterium | IKS35_02600 | 1..1440 | 2044939 | hypothetical-protein-[Clostridia-bacterium] |
| MBO2493599.1 | Type3.MTaseII.containing.BREX | HMA+HAD+HAD->?-><-?\|\|?->MACRODOMAIN->BrxB-BrxF-iSTAND->DUF6079*->HEPN->Ntox45->N6-N4-METHYLASE->TUDOR+SF1-SF2-Helicase+iREase+CTD->PglZ->BrxA-DNA-binding-> | 1228 | Bacteria>Terrabacteria_group>Bacillota>Clostridia>unclassified_Clostridia | GCA_017577715.1 | Clostridia_bacterium | COP72_02085 | 143333..147019 | 2044939 | exonuclease-SbcC-[Clostridia-bacterium] |
| MBP3452735.1 | Type3.MTaseII.containing.BREX | Vrr-REase->SF1-SF2-Helicase->?->?-><-MACRODOMAIN<-BrxA-DNA-binding<-PglZ<-TUDOR+SF1-SF2-Helicase+iREase+CTD<-N6-N4-METHYLASE<-DUF6079*<-BrxB-BrxF-iSTAND<-CHTH<-Calcineurin\|\|ParB+HNH->ACET->TERNS3+HEPN-> | 1229 | Bacteria>Terrabacteria_group>Bacillota>Clostridia>unclassified_Clostridia | GCA_017936765.1 | Clostridia_bacterium | J6M20_03340 | 17195..20884 | 2044939 | exonuclease-SbcC-[Clostridia-bacterium] |
| MBS4024829.1 | Type3.MTaseII.containing.BREX | XIS-HTH->BrxB-BrxF-iSTAND->DUF6079-p*->N6-N4-METHYLASE->TUDOR+SF1-SF2-Helicase+iREase+CTD->PglZ->BrxA-DNA-binding-p->?->DUF4011+SF1-SF2-Helicase+REase->ParB+HNH-> | 942 | Bacteria>Terrabacteria_group>Bacillota>Clostridia>unclassified_Clostridia | GCA_018335815.1 | Clostridia_bacterium | KGZ96_04035 | 1795..4623 | 2044939 | hypothetical-protein-[Clostridia-bacterium] |
| MCD8293625.1 | Type3.MTaseII.containing.BREX | ArdCN+MPTase->BrxB-BrxF-iSTAND->DUF6079*->N6-N4-METHYLASE->TUDOR+SF1-SF2-Helicase+iREase+CTD->PglZ->BrxA-DNA-binding-> | 1225 | Bacteria>Terrabacteria_group>Bacillota>Clostridia>unclassified_Clostridia | GCA_021204625.1 | Clostridia_bacterium | LUE27_00050 | 2544..6221 | 2044939 | DUF6079-family-protein-[Clostridia-bacterium] |
| CDC76539.1 | Type3.MTaseII.containing.BREX | ACET->?->?->?->TraJ-RHH->TraI-VirD2-RCRE->HTH_26->BrxB-BrxF-iSTAND->DUF6079*->N6-N4-METHYLASE->SIR2->HerA/FtsK->TUDOR+SF1-SF2-Helicase+iREase+CTD->PglZ->BrxA-DNA-binding->?-><-?<-ArdA+LPD40+LPD18+LPD17+LPD2 | 1230 | Bacteria>Terrabacteria_group>Bacillota>environmental_samples | GCA_000433515.1 | Firmicutes_bacterium_CAG:272 | BN580_00173 | 10293..13985 | 1263015 | putative-uncharacterized-protein-[Firmicutes-bacterium-CAG:272] |
| WP_209451817.1 | Type3.MTaseII.containing.BREX | ArdA->?->?->TraJ-RHH->TraM->?-><-HEPN\|\|HTH_26->BrxB-BrxF-iSTAND->DUF6079*->N6-N4-METHYLASE->?->?->TUDOR+SF1-SF2-Helicase+iREase+CTD->PglZ->BrxA-DNA-binding->?->wHTH+TPR->HTH_3-> | 1241 | Bacteria>Terrabacteria_group>Bacillota>Erysipelotrichia>Erysipelotrichales>Erysipelotrichaceae | GCF_900343155.1 | Lactimicrobium_massiliense | DHU66_RS10865 | 52399..56124 | 2161814 | DUF6079-family-protein-[Lactimicrobium-massiliense] |
| KAF0196954.1 | Type3.MTaseII.containing.BREX | <-HEPN+PAS+DUF4011+SF1-SF2-Helicase+REase\|\|PBPI->DUF6079-p*->DUF6079*->N6-N4-METHYLASE->TUDOR+SF1-SF2-Helicase+iREase+CTD->PglZ->BrxA-DNA-binding-> | 1159 | Bacteria>Terrabacteria_group>Bacillota | GCA_009773495.1 | Bacillota_bacterium | FD169_626 | 6548..10027 | 1879010 | MAG:-hypothetical-protein-FD169_626-[Bacillota-bacterium] |
| NMB01453.1 | Type3.MTaseII.containing.BREX | <-RNase<-?<-BrxA-DNA-binding<-PglZ<-TUDOR+SF1-SF2-Helicase+iREase+CTD<-N6-N4-METHYLASE<-DUF6079*<-BrxB-BrxF-iSTAND<-?<-?<-?<-?<-ABC-ATPase+SbcC | 1239 | Bacteria>Terrabacteria_group>Bacillota | GCA_012799255.1 | Bacillota_bacterium | GX971_08065 | 11511..15230 | 1879010 | hypothetical-protein-[Bacillota-bacterium] |
| NMB02483.1 | Type3.MTaseII.containing.BREX | DUF6079-p*->N6-N4-METHYLASE->TUDOR+SF1-SF2-Helicase+iREase+CTD->PglZ->BrxA-DNA-binding-> | 335 | Bacteria>Terrabacteria_group>Bacillota | GCA_012799255.1 | Bacillota_bacterium | GX971_13360 | 1..1010 | 1879010 | hypothetical-protein-[Bacillota-bacterium] |
| HHX26369.1 | Type3.MTaseII.containing.BREX | <-DUF6079-p*<-BrxB-BrxF-iSTAND\|\|?->?-><-?<-Polbeta\|\|MULE-TRANSPOSASE->MULE-TRANSPOSASE->?->HTH_Tnp_1+HTH_21+rve+rve_3->MazE_antitoxin-> | 186 | Bacteria>Terrabacteria_group>Bacillota | GCA_012841305.1 | Bacillota_bacterium | GX721_06840 | 1..559 | 1879010 | ATP-binding-protein-[Bacillota-bacterium] |
| HHY32854.1 | Type3.MTaseII.containing.BREX | 35exo->Ntox45->?->ABhydrolase+ABhydrolase-><-BrxA-DNA-binding<-PglZ<-TUDOR+SF1-SF2-Helicase+iREase+CTD<-N6-N4-METHYLASE<-DUF6079*<-BrxB-BrxF-iSTAND<-?<-?<-?<-?<-?\|\|Calcineurin-> | 1203 | Bacteria>Terrabacteria_group>Bacillota | GCA_012842395.1 | Bacillota_bacterium | GX515_07525 | 73870..77481 | 1879010 | hypothetical-protein-[Bacillota-bacterium] |
| HHY35079.1 | Type3.MTaseII.containing.BREX | RNase->MPTase->PIN-><-BrxA-DNA-binding<-PglZ<-TUDOR+SF1-SF2-Helicase+iREase+CTD<-N6-N4-METHYLASE<-DUF6079*<-BrxB-BrxF-iSTAND\|\|?-><-ABC-ATPase+SbcC<-ABC-ATPase+SbcC<-?<-?<-?<-?<-PALP | 1226 | Bacteria>Terrabacteria_group>Bacillota | GCA_012842495.1 | Bacillota_bacterium | GX510_05530 | 23147..26827 | 1879010 | ATP-binding-protein-[Bacillota-bacterium] |
| NPV30272.1 | Type3.MTaseII.containing.BREX | <-TUDOR+SF1-SF2-Helicase+iREase+CTD<-HTH+REase<-N6-N4-METHYLASE<-RAMPs<-?<-RAMPs<-HD<-RAMPs<-RAMPs<-DUF6079*<-BrxB-BrxF-iSTAND\|\|?->HTH_3-> | 1226 | Bacteria>Terrabacteria_group>Bacillota | GCA_013177945.1 | Bacillota_bacterium | HPY58_11625 | 17614..21294 | 1879010 | ATP-binding-protein-[Bacillota-bacterium] |
| NPV81513.1 | Type3.MTaseII.containing.BREX | <-BrxA-DNA-binding<-PglZ<-TUDOR+SF1-SF2-Helicase+iREase+CTD<-N6-N4-METHYLASE<-DUF6079*<-BrxB-BrxF-iSTAND<-?<-SbcC<-TRD+TRD<-?\|\|?->?->PIN-><-HTH_26 | 1203 | Bacteria>Terrabacteria_group>Bacillota | GCA_013178415.1 | Bacillota_bacterium | HPY52_14840 | 8262..11873 | 1879010 | hypothetical-protein-[Bacillota-bacterium] |
| NSW90670.1 | Type3.MTaseII.containing.BREX | SIGMA-HTH+LuxR-HTH->tRNA->tRNA->?->BrxB-BrxF-iSTAND->DUF6079*->N6-N4-METHYLASE->TUDOR+SF1-SF2-Helicase+iREase+CTD->PglZ->BrxA-DNA-binding->?->?-><-RelE-ParE<-MazE_antitoxin | 1229 | Bacteria>Terrabacteria_group>Bacillota | GCA_013314855.1 | Bacillota_bacterium | HPY74_08355 | 2908..6597 | 1879010 | exonuclease-SbcC-[Bacillota-bacterium] |
| MBA1336325.1 | Type3.MTaseII.containing.BREX | <-BrxA-DNA-binding<-PglZ<-TUDOR+SF1-SF2-Helicase+iREase+CTD<-Schlafen+HATPase+wHTH-p<-Schlafen+HATPase+wHTH-p<-N6-N4-METHYLASE<-DUF6079*<-BrxB-BrxF-iSTAND\|\|DDE_Tnp_1+DDE_Tnp_1-> | 1223 | Bacteria>Terrabacteria_group>Bacillota | GCA_013540605.1 | Bacillota_bacterium | HPY66_2760 | 63534..67205 | 1879010 | hypothetical-protein-[Bacillota-bacterium] |
| MBQ5959619.1 | Type3.MTaseII.containing.BREX | VirD4-TraG+VirD4-FtsK->TraM->TrwI->?->?->VirB4-FtsK->Lysozyme+NLPC->HTH_26->BrxB-BrxF-iSTAND->DUF6079*->N6-N4-METHYLASE->Polbeta->?->TUDOR+SF1-SF2-Helicase+iREase+CTD->PglZ-> | 1230 | Bacteria>Terrabacteria_group>Bacillota | GCA_017441925.1 | Bacillota_bacterium | IJL53_03020 | 301803..305495 | 1879010 | hypothetical-protein-[Bacillota-bacterium] |
| MBP8620504.1 | Type3.MTaseII.containing.BREX | TRAM+METHYLASE->RVT->HTH_26->BrxB-BrxF-iSTAND->DUF6079-p*->N6-N4-METHYLASE->Polbeta->?->TUDOR+SF1-SF2-Helicase+iREase+CTD-p-> | 942 | Bacteria>Terrabacteria_group>Bacillota | GCA_018052475.1 | Bacillota_bacterium | KBI39_03445 | 79014..81842 | 1879010 | hypothetical-protein-[Bacillota-bacterium] |
| MBS6916850.1 | Type3.MTaseII.containing.BREX | <-BrxA-DNA-binding<-PglZ<-TUDOR+SF1-SF2-Helicase+iREase+CTD<-?<-SMODS<-HNH+SAVED<-N6-N4-METHYLASE<-HTH+REase<-DUF6079*<-BrxB-BrxF-iSTAND<-HTH_26<-Rrf2-HTH\|\|?-><-Schlafen+HATPase+wHTH<-?<-ACET<-GDResolvase+Recombinase+ZnR | 1224 | Bacteria>Terrabacteria_group>Bacillota | GCA_018380515.1 | Bacillota_bacterium | KH272_06725 | 41402..45076 | 1879010 | hypothetical-protein-[Bacillota-bacterium] |
| MBT9153586.1 | Type3.MTaseII.containing.BREX | <-DUF6079*<-BrxB-BrxF-iSTAND<-XIS-HTH\|\|?-><-?<-ABC-ATPase+SbcC<-?<-?<-?<-CDP-OH_P_transf | 1237 | Bacteria>Terrabacteria_group>Bacillota | GCA_018725105.1 | Bacillota_bacterium | DDT39_00243 | 1298..5011 | 1879010 | hypothetical-protein-[Bacillota-bacterium] |
| MCI6854478.1 | Type3.MTaseII.containing.BREX | HTH_26->BrxB-BrxF-iSTAND->DUF6079*->HTH+REase->N6-N4-METHYLASE->Polbeta->?->TUDOR+SF1-SF2-Helicase+iREase+CTD->PglZ->BrxA-DNA-binding-> | 1224 | Bacteria>Terrabacteria_group>Bacillota | GCA_022776865.1 | Bacillota_bacterium | MR908_04035 | 2221..5895 | 1879010 | DUF6079-family-protein-[Bacillota-bacterium] |
| MCI6854911.1 | Type3.MTaseII.containing.BREX | DUF6079-p*->N6-N4-METHYLASE->TUDOR+SF1-SF2-Helicase+iREase+CTD->PglZ->BrxA-DNA-binding->TRD+TRD->HEPN+SF1-SF2-Helicase->?->GTPase-AIG-> | 570 | Bacteria>Terrabacteria_group>Bacillota | GCA_022776865.1 | Bacillota_bacterium | MR908_06285 | 1..1714 | 1879010 | DUF6079-family-protein-[Bacillota-bacterium] |
| MCI6593671.1 | Type3.MTaseII.containing.BREX | HNH->?->?->HTH_26->BrxB-BrxF-iSTAND->DUF6079*->N6-N4-METHYLASE->TUDOR+SF1-SF2-Helicase+iREase+CTD->PglZ-> | 1230 | Bacteria>Terrabacteria_group>Bacillota | GCA_022778915.1 | Bacillota_bacterium | MSJ87_00260 | 3057..6749 | 1879010 | DUF6079-family-protein-[Bacillota-bacterium] |
| MCL4514314.1 | Type3.MTaseII.containing.BREX | <-PglZ<-TUDOR+SF1-SF2-Helicase+iREase+CTD<-RAMPs<-?<-RAMPs<-HD<-RAMPs<-RAMPs<-N6-N4-METHYLASE<-DUF6079*<-BrxB-BrxF-iSTAND | 1235 | Bacteria>Terrabacteria_group>Bacillota | GCA_023381695.1 | Bacillota_bacterium | M1379_01765 | 35011..38718 | 1879010 | DUF6079-family-protein-[Bacillota-bacterium] |
| MCL5779814.1 | Type3.MTaseII.containing.BREX | <-Acetate_kinase<-?\|\|?->HTH_26->BrxB-BrxF-iSTAND->DUF6079*->N6-N4-METHYLASE->?->TUDOR+SF1-SF2-Helicase+iREase+CTD->PglZ->BrxA-DNA-binding-> | 1226 | Bacteria>Terrabacteria_group>Bacillota | GCA_023487165.1 | Bacillota_bacterium | M1119_02500 | 39189..42869 | 1879010 | DUF6079-family-protein-[Bacillota-bacterium] |
| MCL6590794.1 | Type3.MTaseII.containing.BREX | REC->?->?->?-><-?\|\|ACET+ACET->?->?->BrxB-BrxF-iSTAND->DUF6079*->N6-N4-METHYLASE->SbcC->?->PglZ->BrxA-DNA-binding->Thioredoxin-> | 1244 | Bacteria>Terrabacteria_group>Bacillota | GCA_023512195.1 | Bacillota_bacterium | K6U80_12665 | 24320..28054 | 1879010 | DUF6079-family-protein-[Bacillota-bacterium] |
| MCK9221196.1 | Type3.MTaseII.containing.BREX | <-ABC-ATPase+SbcC<-HKD-DNase-p\|\|BrxB-BrxF-iSTAND->DUF6079*->N6-N4-METHYLASE->TUDOR+SF1-SF2-Helicase+iREase+CTD->PglZ->BrxA-DNA-binding->Calcineurin->SbcC-> | 1230 | Bacteria>Terrabacteria_group>Bacillota>Limnochordia>unclassified_Limnochordia | GCA_023230925.1 | Limnochordia_bacterium | MOQ40_00990 | 25393..29085 | 2497621 | DUF6079-family-protein-[Limnochordia-bacterium] |
| MCD8175748.1 | Type3.MTaseII.containing.BREX | <-wHTH+TPR<-HTH_3\|\|HTH_26->BrxB-BrxF-iSTAND->DUF6079*->N6-N4-METHYLASE->RNase+PHP-> | 1228 | Bacteria>Terrabacteria_group>Bacillota>Negativicutes>Acidaminococcales>Acidaminococcaceae | GCA_021203645.1 | Phascolarctobacterium_sp. | LUD41_07560 | 2922..6608 | 2049039 | DUF6079-family-protein-[Phascolarctobacterium-sp.] |
| MCH4095444.1 | Type3.MTaseII.containing.BREX | <-REase<-?<-MazG<-SIGMA-HTH\|\|HNH->?->SbcC->HTH_26->BrxB-BrxF-iSTAND->DUF6079*->N6-N4-METHYLASE->TUDOR+SF1-SF2-Helicase+iREase+CTD->PglZ->BrxA-DNA-binding->?-><-SbcC<-?<-HNH\|\|HTH+rve-> | 1230 | Bacteria>Terrabacteria_group>Bacillota>Negativicutes>Acidaminococcales>Acidaminococcaceae | GCA_022484025.1 | Acidaminococcus_provencensis | LKE88_02180 | 456929..460621 | 2058289 | DUF6079-family-protein-[Acidaminococcus-provencensis] |
| WP_196601800.1 | Type3.MTaseII.containing.BREX | <-Resolvase+Recombinase<-?<-?<-BrxA-DNA-binding<-PglZ<-TUDOR+SF1-SF2-Helicase+iREase+CTD<-N6-N4-METHYLASE<-?<-DUF6079*<-BrxB-BrxF-iSTAND<-HTH_26<-?<-SbcC<-?<-HNH-p<-AbiEii<-AbiEi_4<-TRAM+METHYLASE | 1231 | Bacteria>Terrabacteria_group>Bacillota>Negativicutes>Selenomonadales>Selenomonadaceae | GCF_015732325.1 | Pectinatus_frisingensis | I6752_RS08745 | 13360..17055 | 865 | DUF6079-family-protein-[Pectinatus-frisingensis] |
| WP_270837224.1 | Type3.MTaseII.containing.BREX | ChlI+Lon_C+AAA->SF1-SF2-Helicase->?->?->?->HAD+HAD->TRAM+METHYLASE-><-?\|\|BrxB-BrxF-iSTAND->DUF6079-p*-> | 375 | Bacteria>Terrabacteria_group>Bacillota>Negativicutes>Selenomonadales>Selenomonadaceae | GCF_027670005.1 | Mitsuokella_multacida | PHA28_RS01560 | 158316..159443 | 52226 | DUF6079-family-protein-[Mitsuokella-multacida] |
| WP_270838224.1 | Type3.MTaseII.containing.BREX | DUF6079-p*->N6-N4-METHYLASE->TUDOR+SF1-SF2-Helicase+iREase+CTD->PglZ->BrxA-DNA-binding->ZnR->PNPase-> | 874 | Bacteria>Terrabacteria_group>Bacillota>Negativicutes>Selenomonadales>Selenomonadaceae | GCF_027670005.1 | Mitsuokella_multacida | PHA28_RS10410 | 19..2643 | 52226 | DUF6079-family-protein-[Mitsuokella-multacida] |
| WP_270400695.1 | Type3.MTaseII.containing.BREX | <-OrfB_IS605+OrfB_Zn_ribbon<-XIS-HTH+GDResolvase<-?<-PNPase<-ZnR<-BrxA-DNA-binding<-PglZ<-TUDOR+SF1-SF2-Helicase+iREase+CTD<-N6-N4-METHYLASE<-DUF6079* | 1150 | Bacteria>Terrabacteria_group>Bacillota>Negativicutes>Selenomonadales>Selenomonadaceae | GCF_027670365.1 | Mitsuokella_multacida | PGS78_RS10195 | 12493..15945 | 52226 | DUF6079-family-protein-[Mitsuokella-multacida] |
| WP_021167322.1 | Type3.MTaseII.containing.BREX | <-DOC+HTH<-?\|\|ArsR-HTH->?->?->DOC+HTH->HD-><-CR-ATPase5+REase\|\|BrxB-BrxF-iSTAND->DUF6079*->N6-N4-METHYLASE->SbcC->?->PglZ->BrxA-DNA-binding->REase->\|\|GDResolvase+Recombinase+ZnR->SIGMA-HTH+SIGMA-HTH-> | 1242 | Bacteria>Terrabacteria_group>Bacillota>Negativicutes>Selenomonadales>Sporomusaceae | GCF_000445445.1 | Sporomusa_ovata | SOV_RS04400 | 941244..944972 | 2378 | DUF6079-family-protein-[Sporomusa-ovata] |
| NCD09233.1 | Type3.MTaseII.containing.BREX | BrxB-BrxF-iSTAND->DUF6079*->N6-N4-METHYLASE->Schlafen+HATPase+wHTH->TUDOR+SF1-SF2-Helicase+iREase+CTD->PglZ->BrxA-DNA-binding-> | 1222 | Bacteria>Terrabacteria_group>Bacillota>Negativicutes>unclassified_Negativicutes | GCA_009930275.1 | Negativicutes_bacterium | EOL98_07380 | 621..4289 | 2499140 | hypothetical-protein-[Negativicutes-bacterium] |
| NMC34869.1 | Type3.MTaseII.containing.BREX | 35exo+TUDOR+SF1-SF2-Helicase+iREase+CTD->SF1-SF2-Helicase->BrxB-BrxF-iSTAND->DUF6079*->N6-N4-METHYLASE->TUDOR+SF1-SF2-Helicase+iREase+CTD->PglZ->BrxA-DNA-binding-><-DDE_Tnp_1<-?<-VirB11+MuB-ATPase | 1228 | Bacteria>Terrabacteria_group>Bacillota>Negativicutes>Veillonellales>Veillonellaceae | GCA_012798135.1 | Veillonellaceae_bacterium | GYA36_20810 | 6163..9849 | 2049049 | ATP-binding-protein-[Veillonellaceae-bacterium] |
| WP_273012836.1 | Type3.MTaseII.containing.BREX | DUF6079*->HTH+REase->N6-N4-METHYLASE->SbcC->SbcC->TUDOR+SF1-SF2-Helicase+iREase+CTD->PglZ->BrxA-DNA-binding->?->Recombinase+ZnR-> | 1082 | Bacteria>Terrabacteria_group>Bacillota>Negativicutes>Veillonellales>Veillonellaceae | GCF_014872435.1 | Dialister_invisus | M3287_RS05160 | 60..3308 | 218538 | DUF6079-family-protein-[Dialister-invisus] |
| WP_071060853.1 | Type3.MTaseII.containing.BREX | <-HEPN\|\|HTH_Tnp_1->rve+rve_3-><-?<-BrxA-DNA-binding<-PglZ<-TUDOR+SF1-SF2-Helicase+iREase+CTD<-N6-N4-METHYLASE<-DUF6079*<-BrxB-BrxF-iSTAND<-TRAM+METHYLASE\|\|RADICAL-SAM-><-Peptidase | 1225 | Bacteria>Terrabacteria_group>Bacillota>Tissierellia>Tissierellales>Gottschalkiaceae | GCF_001855385.1 | Andreesenia_angusta | EUAN_RS01300 | 271927..275604 | 39480 | DUF6079-family-protein-[Andreesenia-angusta] |
| WP_071138485.1 | Type3.MTaseII.containing.BREX | VWA+SIGMA-HTH->TOPRIM+Topoisom_bac->DNA_METHYLASE->ArdCN+LPD32+LPD36+N6_Mtase+SF1-SF2-Helicase->?->JAB->?->BrxB-BrxF-iSTAND->DUF6079*->N6-N4-METHYLASE->TUDOR+SF1-SF2-Helicase+iREase+CTD->PglZ->BrxA-DNA-binding->SF1-SF2-Helicase->DrmB->HKD-DNase->DNA_METHYLASE-><-TraI-VirD2-RCRE | 1245 | Bacteria>Terrabacteria_group>Bacillota>Tissierellia>Tissierellales>Sporanaerobacteraceae | GCF_900095615.1 | Sporanaerobacter_sp._PP17-6a | PP176A_RS00505 | 115965..119702 | 1891289 | DUF6079-family-protein-[Sporanaerobacter-sp.-PP17-6a] |
| HHV38843.1 | Type3.MTaseII.containing.BREX | ParBC-><-?\|\|TRAM+METHYLASE-><-BrxA-DNA-binding<-PglZ<-TUDOR+SF1-SF2-Helicase+iREase+CTD<-N6-N4-METHYLASE<-DUF6079*<-BrxB-BrxF-iSTAND | 1246 | Bacteria>Terrabacteria_group>Bacillota>Tissierellia>Tissierellales>Tepidimicrobiaceae | GCA_012839395.1 | Tepidimicrobium_sp. | GXX70_05035 | 11312..15052 | 2184419 | hypothetical-protein-[Tepidimicrobium-sp.] |
| WP_132029685.1 | Type3.MTaseII.containing.BREX | ParA-Soj-PloopNTPase->TRPR-HTH->?->?->BrxB-BrxF-iSTAND->McrB-NTD+McrB-AAA->BrxB-BrxF-iSTAND->DUF6079*->N6-N4-METHYLASE->TUDOR+SF1-SF2-Helicase+iREase+CTD->PglZ->BrxA-DNA-binding->SAM-METHYLASE->?->?->?->ABC-ATPase+SbcC+ABC-ATPase+SbcC-> | 1246 | Bacteria>Terrabacteria_group>Bacillota>Tissierellia>Tissierellales>Tepidimicrobiaceae | GCF_016767175.1 | Keratinibaculum_paraultunense | JL105_RS02220 | 481272..485012 | 1278232 | DUF6079-family-protein-[Keratinibaculum-paraultunense] |
| WP_075724820.1 | Type3.MTaseII.containing.BREX | TraI-VirD2-RCRE-><-DNA_METHYLASE<-HKD-DNase<-DrmB<-SF1-SF2-Helicase<-BrxA-DNA-binding<-PglZ<-TUDOR+SF1-SF2-Helicase+iREase+CTD<-N6-N4-METHYLASE<-DUF6079*<-BrxB-BrxF-iSTAND | 1245 | Bacteria>Terrabacteria_group>Bacillota>Tissierellia>Tissierellales>Tissierellaceae | GCF_001940565.1 | Tissierella_creatinophila | TICRE_RS02550 | 58874..62611 | 79681 | DUF6079-family-protein-[Tissierella-creatinophila] |
| KUK98165.1 | Type3.MTaseII.containing.BREX | XIS-HTH->BrxA-DNA-binding->BrxB-BrxF-iSTAND->DUF6079*->PglZ->SIGMA+OB-Fold+AAA-ATPase+Lon-Protease-BrxL->N6-N4-METHYLASE->SF1-SF2-Helicase-> | 1445 | Bacteria>Terrabacteria_group>Chloroflexota>Anaerolineae>Anaerolineales>Anaerolineaceae | GCA_001508595.1 | Anaerolineaceae_bacterium_46_22 | XE06_0076 | 6025..10362 | 1635294 | Uncharacterized-protein-XE06_0076-[Anaerolineaceae-bacterium-46_22] |
| MAT45421.1 | Type3.MTaseII.containing.BREX | <-XIS-HTH<-?\|\|LuxR-HTH->CHTH->REase->HEPN->XIS-HTH->BrxA-DNA-binding->BrxB-BrxF-iSTAND->DUF6079*->N6-N4-METHYLASE->N6-N4-METHYLASE->PglZ->SF1-SF2-Helicase->SIGMA+OB-Fold+AAA-ATPase+Lon-Protease-BrxL->?->?->?->TPR+TPR-> | 1433 | Bacteria>Terrabacteria_group>Chloroflexota>Anaerolineae>Anaerolineales>Anaerolineaceae | GCA_002702705.1 | Anaerolineaceae_bacterium | CL609_24095 | 114473..118774 | 2024896 | hypothetical-protein-[Anaerolineaceae-bacterium] |
| MBI9046675.1 | Type3.MTaseII.containing.BREX | <-DUF6079*<-BrxB-BrxF-iSTAND<-BrxA-DNA-binding<-XIS-HTH<-HTH_3\|\|GDResolvase+Recombinase-><-tRNA<-?<-TPR+TPR+TPR+TPR<-SIGMA-HTH+SIGMA-HTH | 1107 | Bacteria>Terrabacteria_group>Chloroflexota>Anaerolineae>Anaerolineales>Anaerolineaceae | GCA_016342475.1 | Anaerolineaceae_bacterium | JEZ06_19445 | 1..3323 | 2024896 | hypothetical-protein-[Anaerolineaceae-bacterium] |
| GER78906.1 | Type3.MTaseII.containing.BREX | METHYLASE+SF1-SF2-Helicase->?-><-BrxA-DNA-binding<-SF1-SF2-Helicase+DUF1998<-viral-protein<-N6-N4-METHYLASE<-DUF6079*<-NERD+RNA-helicase<-RVT\|\|MULE-TRANSPOSASE-><-?<-?\|\|?->?->MPTase-><-MarR-HTH | 1235 | Bacteria>Terrabacteria_group>Chloroflexota>Anaerolineae>Anaerolineales>Anaerolineaceae | GCA_017312345.1 | Candidatus_Denitrolinea_symbiosum | DIM_09870 | 1052900..1056607 | 2608986 | conserved-hypothetical-protein-[Candidatus-Denitrolinea-symbiosum] |
| MBQ6482912.1 | Type3.MTaseII.containing.BREX | <-TUDOR+SF1-SF2-Helicase+iREase+CTD<-?<-?<-SMODS<-HNH+SAVED<-N6-N4-METHYLASE<-DUF6079-p* | 675 | Bacteria>Terrabacteria_group>Chloroflexota>Anaerolineae>Anaerolineales>Anaerolineaceae | GCA_017432945.1 | Anaerolineaceae_bacterium | IJI45_17550 | 8823..10851 | 2024896 | hypothetical-protein-[Anaerolineaceae-bacterium] |
| MBP9040766.1 | Type3.MTaseII.containing.BREX | <-N6-N4-METHYLASE<-HEPN<-ParB+HNH<-DUF6079*<-BrxB-BrxF-iSTAND<-BrxA-DNA-binding<-XIS-HTH\|\|XIS-HTH-> | 1434 | Bacteria>Terrabacteria_group>Chloroflexota>Anaerolineae>Anaerolineales>Anaerolineaceae | GCA_018057145.1 | Anaerolineaceae_bacterium | KBF64_03225 | 4239..8543 | 2024896 | hypothetical-protein-[Anaerolineaceae-bacterium] |
| WP_061919060.1 | Type3.MTaseII.containing.BREX | <-BrxA-DNA-binding<-SF1-SF2-Helicase+DUF1998<-viral-protein<-N6-N4-METHYLASE<-DUF6079*<-NERD+RNA-helicase\|\|Phage_int_SAM_1+Phage_integrase->NtrC-AAA+FIS-HTH->?->?->?->PAS+HISKIN+REC->?->PADR-HTH-> | 1235 | Bacteria>Terrabacteria_group>Chloroflexota>Anaerolineae>Anaerolineales>Anaerolineaceae | GCF_001050215.2 | Bellilinea_caldifistulae | BECAL_RS15120 | 3268182..3271889 | 360411 | DUF6079-family-protein-[Bellilinea-caldifistulae] |
| WP_075072883.1 | Type3.MTaseII.containing.BREX | <-XIS-HTH<-AbiGi+seqA+RAMA<-BrxA-DNA-binding<-PglZ<-?<-?<-?<-N6-N4-METHYLASE<-?<-RNaseH+RNaseH<-DUF6079*<-BrxB-BrxF-iSTAND<-MPTase<-HTH_3\|\|?-><-?<-TetR-HTH<-?\|\|LACI-HTH+PBPI-> | 1258 | Bacteria>Terrabacteria_group>Chloroflexota>Anaerolineae>Anaerolineales>Anaerolineaceae | GCF_001050235.1 | Longilinea_arvoryzae | LARV_RS06550 | 1459635..1463411 | 360412 | DUF6079-family-protein-[Longilinea-arvoryzae] |
| WP_062416901.1 | Type3.MTaseII.containing.BREX | <-LuxR-HTH<-SIGMA+OB-Fold+AAA-ATPase+Lon-Protease-BrxL<-?<-TIR<-PglZ<-TOPRIM<-OLD-ABC<-N6-N4-METHYLASE<-?<-DUF6079*<-BrxB-BrxF-iSTAND<-BrxA-DNA-binding<-XIS-HTH<-OLD-ABC+TOPRIM<-?\|\|HicA->HicB-> | 1444 | Bacteria>Terrabacteria_group>Chloroflexota>Anaerolineae>Anaerolineales>Anaerolineaceae | GCF_001050255.2 | Levilinea_saccharolytica | LSAC_RS05170 | 242785..247119 | 229921 | DUF6079-family-protein-[Levilinea-saccharolytica] |
| WP_062280357.1 | Type3.MTaseII.containing.BREX | ACET->?->?->?->BrxB-BrxF-iSTAND->DUF6079*->N6-N4-METHYLASE->PIN->PIN->PglZ->BrxA-DNA-binding-> | 1258 | Bacteria>Terrabacteria_group>Chloroflexota>Anaerolineae>Anaerolineales>Anaerolineaceae | GCF_001192795.1 | Flexilinea_flocculi | ATC1_RS08435 | 808152..811928 | 1678840 | DUF6079-family-protein-[Flexilinea-flocculi] |
| WP_075063463.1 | Type3.MTaseII.containing.BREX | RAMPs->?->?->CRISPR_Cas6->RVT->Cas_Cas1->?->HTH_3->BrxB-BrxF-iSTAND->DUF6079*->HKD-DNase->N6-N4-METHYLASE->REase->PglZ->BrxA-DNA-binding-> | 1258 | Bacteria>Terrabacteria_group>Chloroflexota>Anaerolineae>Anaerolineales>Anaerolineaceae | GCF_001306115.1 | Ornatilinea_apprima | ADN00_RS13180 | 17619..21395 | 1134406 | DUF6079-family-protein-[Ornatilinea-apprima] |
| NLF03150.1 | Type3.MTaseII.containing.BREX | BrxA-DNA-binding->?->DUF6079*->N6-N4-METHYLASE->AbiEi_4->AbiEii->REase-> | 1317 | Bacteria>Terrabacteria_group>Chloroflexota>Anaerolineae>Anaerolineales>unclassified_Anaerolineales | - | Anaerolineales_bacterium | GX601_19475 | 1931..5884 | 2073117 | hypothetical-protein-[Anaerolineales-bacterium] |
| TEU16812.1 | Type3.MTaseII.containing.BREX | DUF6079*->?->N6-N4-METHYLASE->HARE-HTH->PglZ->SIGMA+OB-Fold+AAA-ATPase+Lon-Protease-BrxL->?->?->?->PGK-> | 1313 | Bacteria>Terrabacteria_group>Chloroflexota>Anaerolineae>Anaerolineales>unclassified_Anaerolineales | GCA_004377365.1 | Anaerolineales_bacterium | E3J21_09750 | 731..4672 | 2073117 | hypothetical-protein-E3J21_09750-[Anaerolineales-bacterium] |
| TEU11065.1 | Type3.MTaseII.containing.BREX | Acyltransferase->?->URI-DNase->?->BrxB-BrxF-iSTAND->DUF6079*->N6-N4-METHYLASE->PglZ->BrxA-DNA-binding-> | 1256 | Bacteria>Terrabacteria_group>Chloroflexota>Anaerolineae>Anaerolineales>unclassified_Anaerolineales | GCA_004377365.1 | Anaerolineales_bacterium | E3J21_24555 | 6878..10648 | 2073117 | ATP-binding-protein-[Anaerolineales-bacterium] |
| MBN1316405.1 | Type3.MTaseII.containing.BREX | DUF6079-p*->TUDOR+SF1-SF2-Helicase+iREase+CTD->N6-N4-METHYLASE->MORC+HISKIN->?->REase->REase+NACHT->N6-N4-METHYLASE-p->HTH_Tnp_1->HTH_21+rve+rve_3-> | 703 | Bacteria>Terrabacteria_group>Chloroflexota>Anaerolineae>Anaerolineales>unclassified_Anaerolineales | GCA_016927355.1 | Anaerolineales_bacterium | JXA42_13100 | 1..2113 | 2073117 | hypothetical-protein-[Anaerolineales-bacterium] |
| OIO98951.1 | Type3.MTaseII.containing.BREX | BrxA-DNA-binding->?->DUF6079*->N6-N4-METHYLASE->Y1_Tnp->HARE-HTH->PglZ-> | 1309 | Bacteria>Terrabacteria_group>Chloroflexota>Anaerolineae>unclassified_Anaerolineae | GCA_001871755.1 | Anaerolineae_bacterium_CG2_30_64_16 | AUK03_00385 | 2801..6730 | 1805005 | MAG:-hypothetical-protein-AUK03_00385-[Anaerolineae-bacterium-CG2_30_64_16] |
| MBL7200858.1 | Type3.MTaseII.containing.BREX | <-N6-N4-METHYLASE<-PIN<-RHH<-PglZ<-DUF6079*<-BrxB-BrxF-iSTAND<-BrxA-DNA-binding<-tRNA<-AAA-ATPase\|\|PHP->RADICAL-SAM-><-TP_METHYLASE\|\|MazG->TPR->Calcineurin-> | 1427 | Bacteria>Terrabacteria_group>Chloroflexota>Anaerolineae>unclassified_Anaerolineae | GCA_016783145.1 | Anaerolineae_bacterium | ISS56_11960 | 5308..9591 | 2052143 | hypothetical-protein-[Anaerolineae-bacterium] |
| MBL7065771.1 | Type3.MTaseII.containing.BREX | <-HTH\|\|BrxB-BrxF-iSTAND->DUF6079*->?->N6-N4-METHYLASE->Schlafen+HATPase+wHTH->N6-N4-METHYLASE-p->AbiEi_4+AbiEi_1->AbiEii+ZnR-> | 1237 | Bacteria>Terrabacteria_group>Chloroflexota>Anaerolineae>unclassified_Anaerolineae | GCA_016783285.1 | Anaerolineae_bacterium | ISS49_16465 | 1306..5019 | 2052143 | ATP-binding-protein-[Anaerolineae-bacterium] |
| MBN1136469.1 | Type3.MTaseII.containing.BREX | <-HNH<-BrxA-DNA-binding<-PglZ<-N6-N4-METHYLASE<-Schlafen+HATPase+wHTH<-DUF6079*<-BrxB-BrxF-iSTAND\|\|?->HNH->REase->PolB->?->REase-> | 1242 | Bacteria>Terrabacteria_group>Chloroflexota>Anaerolineae>unclassified_Anaerolineae | GCA_016926595.1 | Anaerolineae_bacterium | JXM73_07780 | 8570..12298 | 2052143 | ATP-binding-protein-[Anaerolineae-bacterium] |
| MBN1247678.1 | Type3.MTaseII.containing.BREX | <-BrxA-DNA-binding<-PglZ<-NERD+RNA-helicase<-SF1-SF2-Helicase+DUF1998<-viral-protein<-N6-N4-METHYLASE<-MORC+DUF3684+REase<-DUF6079*<-BrxB-BrxF-iSTAND | 1229 | Bacteria>Terrabacteria_group>Chloroflexota>Anaerolineae>unclassified_Anaerolineae | GCA_016927095.1 | Anaerolineae_bacterium | JXC32_08475 | 24187..27876 | 2052143 | ATP-binding-protein-[Anaerolineae-bacterium] |
| MBN1874773.1 | Type3.MTaseII.containing.BREX | ACET->STYKIN->tRNA-><-BrxA-DNA-binding<-PglZ<-N6-N4-METHYLASE<-Y1_Tnp<-DUF6079*<-BrxB-BrxF-iSTAND<-?<-Peptidase_M20 | 1263 | Bacteria>Terrabacteria_group>Chloroflexota>Anaerolineae>unclassified_Anaerolineae | GCA_016931185.1 | Anaerolineae_bacterium | JXA33_11135 | 8271..12062 | 2052143 | ATP-binding-protein-[Anaerolineae-bacterium] |
| MBN2003264.1 | Type3.MTaseII.containing.BREX | <-AbiEi_4+AbiEi_1<-N6-N4-METHYLASE<-NERD+RNA-helicase<-SF1-SF2-Helicase+DUF1998<-viral-protein<-DUF6079*<-BrxB-BrxF-iSTAND<-?<-HTH_3\|\|?->Phage_int_SAM_4+Phage_integrase-> | 1234 | Bacteria>Terrabacteria_group>Chloroflexota>Anaerolineae>unclassified_Anaerolineae | GCA_016931835.1 | Anaerolineae_bacterium | JXA21_07905 | 14338..18042 | 2052143 | ATP-binding-protein-[Anaerolineae-bacterium] |
| MCB0191463.1 | Type3.MTaseII.containing.BREX | <-GDResolvase+HIN-HTH\|\|BrxB-BrxF-iSTAND->DUF6079*->?->CR-ATPase9->N6-N4-METHYLASE->TUDOR+SF1-SF2-Helicase+iREase+CTD->REase->?->?->MORC+DUF3684+REase->SF1-SF2-Helicase-> | 1225 | Bacteria>Terrabacteria_group>Chloroflexota>Anaerolineae>unclassified_Anaerolineae | GCA_020433805.1 | Anaerolineae_bacterium | KDJ65_05920 | 1466..5143 | 2052143 | hypothetical-protein-[Anaerolineae-bacterium] |
| MCK6629895.1 | Type3.MTaseII.containing.BREX | Glyoxalase->?->AraC-HTH->?->?->BrxB-BrxF-iSTAND->DUF6079*->N6-N4-METHYLASE->REase->PglZ->BrxA-DNA-binding-> | 1241 | Bacteria>Terrabacteria_group>Chloroflexota>Anaerolineae>unclassified_Anaerolineae | GCA_023151115.1 | Anaerolineae_bacterium | L6R45_32560 | 7534..11259 | 2052143 | DUF6079-family-protein-[Anaerolineae-bacterium] |
| MCL4298469.1 | Type3.MTaseII.containing.BREX | <-TUDOR+SF1-SF2-Helicase+iREase+CTD<-?<-N6-N4-METHYLASE<-?<-DnaJ<-?<-?<-?<-Trypsin+PDZ<-DUF6079*<-BrxB-BrxF-iSTAND<-?<-HTH_Tnp_1\|\|?->?->?-><-?\|\|LuxR-HTH-> | 1239 | Bacteria>Terrabacteria_group>Chloroflexota>Anaerolineae>unclassified_Anaerolineae | GCA_023382535.1 | Anaerolineae_bacterium | KJ077_22220 | 132971..136690 | 2052143 | hypothetical-protein-[Anaerolineae-bacterium] |
| MCQ3973388.1 | Type3.MTaseII.containing.BREX | ABhydrolase->SF1-SF2-Helicase-><-REase<-PhdYeFM_antitox<-BrxA-DNA-binding<-PglZ<-REase<-N6-N4-METHYLASE<-?<-DUF6079*<-BrxB-BrxF-iSTAND<-XIS-HTH<-?<-?<-wHTH<-?<-?<-AraC-HTH | 1241 | Bacteria>Terrabacteria_group>Chloroflexota>Anaerolineae>unclassified_Anaerolineae | GCA_024434315.1 | Anaerolineae_bacterium | DPW09_08095 | 27820..31545 | 2052143 | ATP-binding-protein-[Anaerolineae-bacterium] |
| MCR4407612.1 | Type3.MTaseII.containing.BREX | <-PhdYeFM_antitox<-?<-?<-?<-?<-?<-N6-N4-METHYLASE-p<-NTP_transf_5<-?<-DUF6079-p*<-BrxB-BrxF-iSTAND<-REase<-?\|\|?-><-RelE-ParE<-Phd-YefM<-Phage_int_SAM_4+Phage_integrase\|\|MazE_antitoxin->PIN-> | 600 | Bacteria>Terrabacteria_group>Chloroflexota>Anaerolineae>unclassified_Anaerolineae | GCA_024653325.1 | Anaerolineae_bacterium | NUW24_11945 | 13415..15217 | 2052143 | DUF6079-family-protein-[Anaerolineae-bacterium] |

| Protein-ID | System | Gene-Neighbourhoods | Length | Taxonomy | Assembly-ID | Strain | Locus-Tag | Genomic-coordinates | TaxID | Fasta-defline |
|---|---|---|---|---|---|---|---|---|---|---|
| MCE7981875.1 | Type3.MTaseII.containing.BREX | [illegible] | 1044 | Bacteria>Terrabacteria_group>Chloroflexota>Caldilineae>Caldilineales>Caldilineaceae | GCA_021405325.1 | Caldilinea_sp._CFX5 | DYG89_11820 | 173689..176823 | 2293554 | ATP-binding-protein-[Caldilinea-sp.-CFX5] |
| MCE7981876.1 | Type3.MTaseII.containing.BREX | [illegible] | 195 | Bacteria>Terrabacteria_group>Chloroflexota>Caldilineae>Caldilineales>Caldilineaceae | GCA_021405325.1 | Caldilinea_sp._CFX5 | DYG89_11825 | 176808..177395 | 2293554 | hypothetical-protein-[Caldilinea-sp.-CFX5] |
| MCE7988231.1 | Type3.MTaseII.containing.BREX | [illegible] | 718 | Bacteria>Terrabacteria_group>Chloroflexota>Caldilineae>Caldilineales>Caldilineaceae | GCA_021405325.1 | Caldilinea_sp._CFX5 | DYG89_44295 | 35554..37709 | 2293554 | hypothetical-protein-[Caldilinea-sp.-CFX5] |
| MCL4834878.1 | Type3.MTaseII.containing.BREX | [illegible] | 1260 | Bacteria>Terrabacteria_group>Chloroflexota>Caldilineae>Caldilineales>Caldilineaceae | GCA_023384455.1 | Caldilineaceae_bacterium | KJZ86_20725 | 10548..14330 | 2614250 | hypothetical-protein-[Caldilineaceae-bacterium] |
| GIV90317.1 | Type3.MTaseII.containing.BREX | [illegible] | 1153 | Bacteria>Terrabacteria_group>Chloroflexota>Chloroflexia>Chloroflexales>Chloroflexineae | GCA_026003795.1 | Chloroflexus_sp. | KatS3mg055_2835 | 363038..366499 | 1904827 | MAG:-hypothetical-protein-KatS3mg055_2835-[Chloroflexus-sp.] |
| GIV90321.1 | Type3.MTaseII.containing.BREX | [illegible] | 1250 | Bacteria>Terrabacteria_group>Chloroflexota>Chloroflexia>Chloroflexales>Chloroflexineae | GCA_026003795.1 | Chloroflexus_sp. | KatS3mg055_2839 | 369317..373069 | 1904827 | MAG:-hypothetical-protein-KatS3mg055_2839-[Chloroflexus-sp.] |
| WP_012616394.1 | Type3.MTaseII.containing.BREX | [illegible] | 1242 | Bacteria>Terrabacteria_group>Chloroflexota>Chloroflexia>Chloroflexales>Chloroflexineae | GCF_000021945.1 | Chloroflexus_aggregans | CAGG_RS05545 | 1379812..1383540 | 152260 | DUF6079-family-protein-[Chloroflexus-aggregans] |
| WP_217352396.1 | Type3.MTaseII.containing.BREX | [illegible] | 741 | Bacteria>Terrabacteria_group>Chloroflexota>Chloroflexia>Chloroflexales>Chloroflexineae | GCF_002532535.1 | Candidatus_Viridilinea_mediisalina | CJ255_RS03915 | 76155..78380 | 2024553 | hypothetical-protein-[Candidatus-Viridilinea-mediisalina] |
| WP_135481574.1 | Type3.MTaseII.containing.BREX | [illegible] | 1262 | Bacteria>Terrabacteria_group>Chloroflexota>Chloroflexia>Chloroflexales>Chloroflexineae | GCF_004762035.2 | Candidatus_Chloroploca_sp._M50-1 | EYB53_RS23165 | 7974..11762 | 2528176 | DUF6079-family-protein-[Candidatus-Chloroploca-sp.-M50-1] |
| OBW63497.1 | Type3.MTaseII.containing.BREX | [illegible] | 1237 | Bacteria>Terrabacteria_group>Chloroflexota>Dehalococcoidia>Dehalococcoidales>Dehalococcoidaceae | GCA_001678035.1 | Dehalococcoides_mccartyi | A9183_07180 | 25380..29093 | 61435 | ATPase-[Dehalococcoides-mccartyi] |
| MCF7634650.1 | Type3.MTaseII.containing.BREX | [illegible] | 1446 | Bacteria>Terrabacteria_group>Chloroflexota>Dehalococcoidia>Dehalococcoidales>Dehalococcoidaceae | GCA_021732865.1 | Dehalococcoides_mccartyi | LLF82_000114 | 104665..109005 | 61435 | hypothetical-protein-[Dehalococcoides-mccartyi] |
| WP_015406900.1 | Type3.MTaseII.containing.BREX | [illegible] | 1447 | Bacteria>Terrabacteria_group>Chloroflexota>Dehalococcoidia>Dehalococcoidales>Dehalococcoidaceae | GCF_000341655.1 | Dehalococcoides_mccartyi | DCMB_RS01625 | 325612..329955 | 61435 | hypothetical-protein-[Dehalococcoides-mccartyi] |
| MBT9148683.1 | Type3.MTaseII.containing.BREX | [illegible] | 1317 | Bacteria>Terrabacteria_group>Chloroflexota>Dehalococcoidia>unclassified_Dehalococcoidia | GCA_018725325.1 | Dehalococcoidia_bacterium | DDT28_00095 | 1544..5497 | 2026734 | hypothetical-protein-[Dehalococcoidia-bacterium] |
| MBT9159326.1 | Type3.MTaseII.containing.BREX | [illegible] | 1317 | Bacteria>Terrabacteria_group>Chloroflexota>Dehalococcoidia>unclassified_Dehalococcoidia | GCA_018725405.1 | Dehalococcoidia_bacterium | DDT26_00584 | 4420..8373 | 2026734 | hypothetical-protein-[Dehalococcoidia-bacterium] |
| MBN1887191.1 | Type3.MTaseII.containing.BREX | [illegible] | 1235 | Bacteria>Terrabacteria_group>Chloroflexota>Thermoflexia>Thermoflexales>unclassified_Thermoflexales | GCA_016931085.1 | Thermoflexales_bacterium | JW850_04355 | 27356..31063 | 2497639 | hypothetical-protein-[Thermoflexales-bacterium] |
| NMB57621.1 | Type3.MTaseII.containing.BREX | [illegible] | 1428 | Bacteria>Terrabacteria_group>Chloroflexota>unclassified_Chloroflexota | - | Chloroflexota_bacterium | GYA12_00495 | 2698..6984 | 2026724 | hypothetical-protein-[Chloroflexota-bacterium] |
| OGN72032.1 | Type3.MTaseII.containing.BREX | [illegible] | 1254 | Bacteria>Terrabacteria_group>Chloroflexota>unclassified_Chloroflexota | GCA_001796335.1 | Chloroflexi_bacterium_GWB2_49_20 | A2X25_12855 | 139097..142861 | 1797612 | ATPase-[Chloroflexi-bacterium-GWB2_49_20] |
| OJX40546.1 | Type3.MTaseII.containing.BREX | [illegible] | 1258 | Bacteria>Terrabacteria_group>Chloroflexota>unclassified_Chloroflexota | GCA_001899005.1 | Chloroflexi_bacterium_44-23 | BGO78_06045 | 155438..159214 | 1895927 | hypothetical-protein-BGO78_06045-[Chloroflexi-bacterium-44-23] |
| PKO03865.1 | Type3.MTaseII.containing.BREX | [illegible] | 1236 | Bacteria>Terrabacteria_group>Chloroflexota>unclassified_Chloroflexota | GCA_002842015.1 | Chloroflexi_bacterium_HGW-Chloroflexi-5 | CVU43_00685 | 60388..64098 | 2013728 | hypothetical-protein-CVU43_00685-[Chloroflexi-bacterium-HGW-Chloroflexi-5] |
| PKO02179.1 | Type3.MTaseII.containing.BREX | [illegible] | 1222 | Bacteria>Terrabacteria_group>Chloroflexota>unclassified_Chloroflexota | GCA_002842015.1 | Chloroflexi_bacterium_HGW-Chloroflexi-5 | CVU43_09165 | 5258..8926 | 2013728 | hypothetical-protein-CVU43_09165-[Chloroflexi-bacterium-HGW-Chloroflexi-5] |
| PKO06212.1 | Type3.MTaseII.containing.BREX | [illegible] | 1256 | Bacteria>Terrabacteria_group>Chloroflexota>unclassified_Chloroflexota | GCA_002842045.1 | Chloroflexi_bacterium_HGW-Chloroflexi-3 | CVU41_08900 | 234347..238117 | 2013726 | ATPase-[Chloroflexi-bacterium-HGW-Chloroflexi-3] |
| HAJ37908.1 | Type3.MTaseII.containing.BREX | [illegible] | 1436 | Bacteria>Terrabacteria_group>Chloroflexota>unclassified_Chloroflexota | GCA_003451595.1 | Chloroflexota_bacterium | DCL15_19735 | 113168..117478 | 2026724 | hypothetical-protein-[Chloroflexota-bacterium] |
| MSP14286.1 | Type3.MTaseII.containing.BREX | [illegible] | 1257 | Bacteria>Terrabacteria_group>Chloroflexota>unclassified_Chloroflexota | GCA_009692745.1 | Chloroflexota_bacterium | EXR62_15195 | 8774..12547 | 2026724 | ATP-binding-protein-[Chloroflexota-bacterium] |
| HDQ73860.1 | Type3.MTaseII.containing.BREX | [illegible] | 1280 | Bacteria>Terrabacteria_group>Chloroflexota>unclassified_Chloroflexota | GCA_011046405.1 | Chloroflexota_bacterium | ENN19_17455 | 10020..13862 | 2026724 | ATP-binding-protein-[Chloroflexota-bacterium] |
| HEC34266.1 | Type3.MTaseII.containing.BREX | [illegible] | 1308 | Bacteria>Terrabacteria_group>Chloroflexota>unclassified_Chloroflexota | GCA_011052755.1 | Chloroflexota_bacterium | ENI37_06065 | 1523..5449 | 2026724 | hypothetical-protein-[Chloroflexota-bacterium] |
| NMA12761.1 | Type3.MTaseII.containing.BREX | [illegible] | 1233 | Bacteria>Terrabacteria_group>Chloroflexota>unclassified_Chloroflexota | GCA_012799995.1 | Chloroflexota_bacterium | GX933_06230 | 36696..40397 | 2026724 | ATP-binding-protein-[Chloroflexota-bacterium] |
| HHY54153.1 | Type3.MTaseII.containing.BREX | [illegible] | 767 | Bacteria>Terrabacteria_group>Chloroflexota>unclassified_Chloroflexota | GCA_012842785.1 | Chloroflexota_bacterium | GYA08_01840 | 7589..9892 | 2026724 | hypothetical-protein-[Chloroflexota-bacterium] |
| MBI2332378.1 | Type3.MTaseII.containing.BREX | [illegible] | 865 | Bacteria>Terrabacteria_group>Chloroflexota>unclassified_Chloroflexota | GCA_016183455.1 | Chloroflexota_bacterium | HYU84_09520 | 13254..15851 | 2026724 | hypothetical-protein-[Chloroflexota-bacterium] |
| MBI2332379.1 | Type3.MTaseII.containing.BREX | [illegible] | 158 | Bacteria>Terrabacteria_group>Chloroflexota>unclassified_Chloroflexota | GCA_016183455.1 | Chloroflexota_bacterium | HYU84_09525 | 15859..16335 | 2026724 | ATP-binding-protein-[Chloroflexota-bacterium] |
| MBI2332380.1 | Type3.MTaseII.containing.BREX | [illegible] | 153 | Bacteria>Terrabacteria_group>Chloroflexota>unclassified_Chloroflexota | GCA_016183455.1 | Chloroflexota_bacterium | HYU84_09530 | 16364..16822 | 2026724 | ATP-binding-protein-[Chloroflexota-bacterium] |
| MBI2939062.1 | Type3.MTaseII.containing.BREX | [illegible] | 1468 | Bacteria>Terrabacteria_group>Chloroflexota>unclassified_Chloroflexota | GCA_016190465.1 | Chloroflexota_bacterium | HYY04_01385 | 8416..12822 | 2026724 | hypothetical-protein-[Chloroflexota-bacterium] |
| MBI2939757.1 | Type3.MTaseII.containing.BREX | [illegible] | 1237 | Bacteria>Terrabacteria_group>Chloroflexota>unclassified_Chloroflexota | GCA_016190465.1 | Chloroflexota_bacterium | HYY04_04900 | 582..4295 | 2026724 | ATP-binding-protein-[Chloroflexota-bacterium] |
| MBI3175730.1 | Type3.MTaseII.containing.BREX | [illegible] | 1234 | Bacteria>Terrabacteria_group>Chloroflexota>unclassified_Chloroflexota | GCA_016191845.1 | Chloroflexota_bacterium | HYZ25_18570 | 129078..132782 | 2026724 | ATP-binding-protein-[Chloroflexota-bacterium] |
| MBI3973657.1 | Type3.MTaseII.containing.BREX | [illegible] | 770 | Bacteria>Terrabacteria_group>Chloroflexota>unclassified_Chloroflexota | GCA_016199265.1 | Chloroflexota_bacterium | HY332_20470 | 1..2310 | 2026724 | ATP-binding-protein-[Chloroflexota-bacterium] |
| MBI4318416.1 | Type3.MTaseII.containing.BREX | [illegible] | 1447 | Bacteria>Terrabacteria_group>Chloroflexota>unclassified_Chloroflexota | GCA_016210495.1 | Chloroflexota_bacterium | HY675_07990 | 50665..55008 | 2026724 | hypothetical-protein-[Chloroflexota-bacterium] |
| MBI4321182.1 | Type3.MTaseII.containing.BREX | [illegible] | 1255 | Bacteria>Terrabacteria_group>Chloroflexota>unclassified_Chloroflexota | GCA_016210495.1 | Chloroflexota_bacterium | HY675_22045 | 10530..14297 | 2026724 | ATP-binding-protein-[Chloroflexota-bacterium] |
| MBI5824577.1 | Type3.MTaseII.containing.BREX | [illegible] | 1234 | Bacteria>Terrabacteria_group>Chloroflexota>unclassified_Chloroflexota | GCA_016235445.1 | Chloroflexota_bacterium | HZB18_11155 | 1964..5668 | 2026724 | ATP-binding-protein-[Chloroflexota-bacterium] |
| MBK8034669.1 | Type3.MTaseII.containing.BREX | [illegible] | 1236 | Bacteria>Terrabacteria_group>Chloroflexota>unclassified_Chloroflexota | GCA_016713355.1 | Chloroflexota_bacterium | IPK17_35240 | 1112919..1116629 | 2026724 | ATP-binding-protein-[Chloroflexota-bacterium] |
| MCL5960615.1 | Type3.MTaseII.containing.BREX | [illegible] | 1447 | Bacteria>Terrabacteria_group>Chloroflexota>unclassified_Chloroflexota | GCA_023489135.1 | Chloroflexota_bacterium | M1358_15150 | 7286..11629 | 2026724 | hypothetical-protein-[Chloroflexota-bacterium] |
| PKL79310.1 | Type3.MTaseII.containing.BREX | [illegible] | 1236 | Bacteria>Terrabacteria_group>Cyanobacteriota/Melainabacteria_group>Candidatus_Melainabacteria>unclassified_Candidatus_Melainabacteria | GCA_002839735.1 | Candidatus_Melainabacteria_bacterium_HGW-Melainabacteria-1 | CVV27_02010 | 5841..9551 | 2013813 | MAG:-ATPase-[Candidatus-Melainabacteria-bacterium-HGW-Melainabacteria-1] |
| MBR8836037.1 | Type3.MTaseII.containing.BREX | [illegible] | 493 | Bacteria>Terrabacteria_group>Cyanobacteriota/Melainabacteria_group>Cyanobacteriota>Cyanophyceae>Nostocales | GCA_016903675.1 | Stigonema_ocellatum_SAG_48.90_=_DSM_106950 | DSM106950_18940 | 32709..34190 | 221294 | hypothetical-protein-[Stigonema-ocellatum-SAG-48.90-=-DSM-106950] |
| MBW4477582.1 | Type3.MTaseII.containing.BREX | [illegible] | 264 | Bacteria>Terrabacteria_group>Cyanobacteriota/Melainabacteria_group>Cyanobacteriota>Cyanophyceae>Nostocales | GCA_019358985.1 | Tolypothrix_brevis_GSE-NOS-MK-07-07A | KME54_12080 | 53329..54123 | 2835095 | hypothetical-protein-[Tolypothrix-brevis-GSE-NOS-MK-07-07A] |
| WP_026134770.1 | Type3.MTaseII.containing.BREX | [illegible] | 76 | Bacteria>Terrabacteria_group>Cyanobacteriota/Melainabacteria_group>Cyanobacteriota>Cyanophyceae>Nostocales | GCF_000346485.2 | Scytonema_hofmannii | WA1_RS22390 | 5399151..5399381 | 34078 | DUF6079-family-protein-[Scytonema-hofmannii] |
| WP_096597413.1 | Type3.MTaseII.containing.BREX | [illegible] | 1236 | Bacteria>Terrabacteria_group>Cyanobacteriota/Melainabacteria_group>Cyanobacteriota>Cyanophyceae>Nostocales | GCF_002368195.1 | Calothrix_sp._NIES-2100 | CA734_RS02095 | 551989..555699 | 1954172 | DUF6079-family-protein-[Calothrix-sp.-NIES-2100] |
| WP_096683471.1 | Type3.MTaseII.containing.BREX | [illegible] | 1236 | Bacteria>Terrabacteria_group>Cyanobacteriota/Melainabacteria_group>Cyanobacteriota>Cyanophyceae>Nostocales | GCF_002368215.1 | Nostoc_sp._NIES-2111 | CA737_RS32345 | 59866..63576 | 1973475 | DUF6079-family-protein-[Nostoc-sp.-NIES-2111] |
| WP_168540454.1 | Type3.MTaseII.containing.BREX | [illegible] | 1238 | Bacteria>Terrabacteria_group>Cyanobacteriota/Melainabacteria_group>Cyanobacteriota>Cyanophyceae>Nostocales | GCF_009712085.1 | Anabaena_sp._UHCC_0253 | FJR38_26290 | 50119..53835 | 2590019 | DUF6079-family-protein-[Anabaena-sp.-UHCC-0253] |
| WP_155743287.1 | Type3.MTaseII.containing.BREX | [illegible] | 1237 | Bacteria>Terrabacteria_group>Cyanobacteriota/Melainabacteria_group>Cyanobacteriota>Cyanophyceae>Nostocales | GCF_009725235.1 | Scytonema_sp._UIC_10036 | F7734_RS02805 | 108764..112477 | 2304196 | DUF6079-family-protein-[Scytonema-sp.-UIC-10036] |
| WP_190937388.1 | Type3.MTaseII.containing.BREX | [illegible] | 1237 | Bacteria>Terrabacteria_group>Cyanobacteriota/Melainabacteria_group>Cyanobacteriota>Cyanophyceae>Nostocales | GCF_014698055.1 | Nostoc_sp._FACHB-133 | H6G86_RS30115 | 33879..37592 | 2692835 | DUF6079-family-protein-[Nostoc-sp.-FACHB-133] |
| WP_190509459.1 | Type3.MTaseII.containing.BREX | [illegible] | 1244 | Bacteria>Terrabacteria_group>Cyanobacteriota/Melainabacteria_group>Cyanobacteriota>Cyanophyceae>Oscillatoriophycideae | GCF_014697545.1 | Oscillatoria_sp._FACHB-1407 | H6G89_RS19405 | 686..4420 | 2692847 | DUF6079-family-protein-[Oscillatoria-sp.-FACHB-1407] |
| PZV08803.1 | Type3.MTaseII.containing.BREX | [illegible] | 1237 | Bacteria>Terrabacteria_group>Cyanobacteriota/Melainabacteria_group>Cyanobacteriota>Cyanophyceae>Pseudanabaenales | GCA_003249105.1 | Leptolyngbya_sp. | DCF32_03945 | 21297..25010 | 47254 | ATPase-[Leptolyngbya-sp.] |
| WP_255112871.1 | Type3.MTaseII.containing.BREX | [illegible] | 1232 | Bacteria>Terrabacteria_group>Cyanobacteriota/Melainabacteria_group>Cyanobacteriota>Cyanophyceae>Synechococcales | GCF_024345735.1 | Synechococcus_lacustris | KBY64_RS06860 | 13686..17384 | 2116544 | DUF6079-family-protein-[Synechococcus-lacustris] |
| WP_255117825.1 | Type3.MTaseII.containing.BREX | [illegible] | 1242 | Bacteria>Terrabacteria_group>Cyanobacteriota/Melainabacteria_group>Cyanobacteriota>Cyanophyceae>Synechococcales | GCF_024345835.1 | Synechococcus_sp._HJ21-Hayes | KBZ14_RS08140 | 86825..90553 | 2823736 | DUF6079-family-protein-[Synechococcus-sp.-HJ21-Hayes] |
| WP_254950517.1 | Type3.MTaseII.containing.BREX | [illegible] | 1233 | Bacteria>Terrabacteria_group>Cyanobacteriota/Melainabacteria_group>Cyanobacteriota>Cyanophyceae>Synechococcales | GCF_024345955.1 | unclassified_Cyanobium | KBZ34_RS03540 | 162386..166087 | 2627006 | MULTISPECIES:-DUF6079-family-protein-[unclassified-Cyanobium] |
| KMY65633.1 | Type3.MTaseII.containing.BREX | [illegible] | 1240 | Bacteria>Thermodesulfobacteriota>Desulfarculia>Desulfarculales>Desulfarculaceae>Desulfocarbo | GCA_001184205.1 | Desulfocarbo_indianensis | AAU61_21215 | 11918..15640 | 1348163 | ATPase-[Desulfocarbo-indianensis] |
| WP_073478300.1 | Type3.MTaseII.containing.BREX | [illegible] | 1242 | Bacteria>Thermodesulfobacteriota>Desulfobacteria>Desulfobacterales>Desulfatibacillaceae>Desulfatibacillum | GCF_900142135.1 | Desulfatibacillum_alkenivorans | BUE18_RS21830 | 56360..60088 | 259354 | DUF6079-family-protein-[Desulfatibacillum-alkenivorans] |
| WP_073478764.1 | Type3.MTaseII.containing.BREX | [illegible] | 1242 | Bacteria>Thermodesulfobacteriota>Desulfobacteria>Desulfobacterales>Desulfatibacillaceae>Desulfatibacillum | GCF_900142135.1 | Desulfatibacillum_alkenivorans | BUE18_RS24405 | 18194..21922 | 259354 | DUF6079-family-protein-[Desulfatibacillum-alkenivorans] |
| SLM29542.1 | Type3.MTaseII.containing.BREX | [illegible] | 1248 | Bacteria>Thermodesulfobacteriota>Desulfobacteria>Desulfobacterales>Desulfobacteraceae>Desulfamplus | GCA_900170035.1 | Desulfamplus_magnetovallimortis | MTBBW1_1860032 | 36511..40257 | 1246637 | conserved-hypothetical-protein-[Desulfamplus-magnetovallimortis] |
| WP_080806571.1 | Type3.MTaseII.containing.BREX | [illegible] | 1246 | Bacteria>Thermodesulfobacteriota>Desulfobacteria>Desulfobacterales>Desulfobacteraceae>Desulfamplus | GCF_900170035.1 | Desulfamplus_magnetovallimortis | MTBBW1_RS20320 | 62532..66272 | 1246637 | DUF6079-family-protein-[Desulfamplus-magnetovallimortis] |
| VBB41622.1 | Type3.MTaseII.containing.BREX | [illegible] | 1447 | Bacteria>Thermodesulfobacteriota>Desulfobacteria>Desulfobacterales>Desulfobacteraceae>Desulfatiglans | GCA_900498135.1 | uncultured_Desulfatiglans_sp. | TRIP_B120057 | 39977..44320 | 1748965 | conserved-hypothetical-protein-[uncultured-Desulfatiglans-sp.] |
| VBB44945.1 | Type3.MTaseII.containing.BREX | [illegible] | 1445 | Bacteria>Thermodesulfobacteriota>Desulfobacteria>Desulfobacterales>Desulfobacteraceae>Desulfatiglans | GCA_900498135.1 | uncultured_Desulfatiglans_sp. | TRIP_B350116 | 4274729..4279066 | 1748965 | conserved-hypothetical-protein-[uncultured-Desulfatiglans-sp.] |
| WP_028321661.1 | Type3.MTaseII.containing.BREX | [illegible] | 1242 | Bacteria>Thermodesulfobacteriota>Desulfobacteria>Desulfobacterales>Desulfobacteraceae>Desulfatiglans | GCF_000422285.1 | Desulfatiglans_anilini | H567_RS0112520 | 1880..5608 | 90728 | DUF6079-family-protein-[Desulfatiglans-anilini] |
| HBT88457.1 | Type3.MTaseII.containing.BREX | [illegible] | 1246 | Bacteria>Thermodesulfobacteriota>Desulfobacteria>Desulfobacterales>Desulfobacteraceae>Desulfobacter | GCA_003510365.1 | Desulfobacter_sp. | DEB50_08330 | 33339..37079 | 2294 | ATPase-[Desulfobacter-sp.] |
| WP_246293874.1 | Type3.MTaseII.containing.BREX | [illegible] | 1068 | Bacteria>Thermodesulfobacteriota>Desulfobacteria>Desulfobacterales>Desulfobacteraceae>Desulfobacter | GCF_013389365.1 | Desulfobacter_latus | HXW94_RS11125 | 9686..12892 | 2292 | DUF6079-family-protein-[Desulfobacter-latus] |
| WP_144683374.1 | Type3.MTaseII.containing.BREX | [illegible] | 1243 | Bacteria>Thermodesulfobacteriota>Desulfobacteria>Desulfobacterales>Desulfobacteraceae>Desulfobotulus | GCF_007830435.1 | Desulfobotulus_alkaliphilus | LZ24_RS05840 | 46533..50264 | 622671 | DUF6079-family-protein-[Desulfobotulus-alkaliphilus] |
| RJQ83728.1 | Type3.MTaseII.containing.BREX | [illegible] | 1285 | Bacteria>Thermodesulfobacteriota>Desulfobacteria>Desulfobacterales>Desulfobacteraceae>unclassified_Desulfobacteraceae | GCA_003599575.1 | Desulfobacteraceae_bacterium | C4519_06260 | 1729..5586 | 2049433 | ATP-binding-protein-[Desulfobacteraceae-bacterium] |
| RJR29735.1 | Type3.MTaseII.containing.BREX | [illegible] | 1244 | Bacteria>Thermodesulfobacteriota>Desulfobacteria>Desulfobacterales>Desulfobacteraceae>unclassified_Desulfobacteraceae | GCA_003599615.1 | Desulfobacteraceae_bacterium | C4576_34875 | 7434..11168 | 2049433 | ATP-binding-protein-[Desulfobacteraceae-bacterium] |
| MBC2732793.1 | Type3.MTaseII.containing.BREX | [illegible] | 1242 | Bacteria>Thermodesulfobacteriota>Desulfobacteria>Desulfobacterales>Desulfobacteraceae>unclassified_Desulfobacteraceae | GCA_014237205.1 | Desulfobacteraceae_bacterium | HF981_00360 | 90454..94182 | 2049433 | ATP-binding-protein-[Desulfobacteraceae-bacterium] |
| MBL7172016.1 | Type3.MTaseII.containing.BREX | [illegible] | 1242 | Bacteria>Thermodesulfobacteriota>Desulfobacteria>Desulfobacterales>Desulfobacteraceae>unclassified_Desulfobacteraceae | GCA_016782905.1 | Desulfobacteraceae_bacterium | ISS68_04195 | 1503..5231 | 2049433 | ATP-binding-protein-[Desulfobacteraceae-bacterium] |
| MCP4110753.1 | Type3.MTaseII.containing.BREX | [illegible] | 1399 | Bacteria>Thermodesulfobacteriota>Desulfobacteria>Desulfobacterales>Desulfobacteraceae>unclassified_Desulfobacteraceae | GCA_024225235.1 | Desulfobacteraceae_bacterium | GY749_35375 | 575..4774 | 2049433 | hypothetical-protein-[Desulfobacteraceae-bacterium] |
| MCP3942933.1 | Type3.MTaseII.containing.BREX | [illegible] | 1204 | Bacteria>Thermodesulfobacteriota>Desulfobacteria>Desulfobacterales>Desulfobacteraceae>unclassified_Desulfobacteraceae | GCA_024226935.1 | Desulfobacteraceae_bacterium | GY710_15805 | 766..4380 | 2049433 | BREX-system-P-loop-protein-BrxC-[Desulfobacteraceae-bacterium] |
| WP_020876159.1 | Type3.MTaseII.containing.BREX | [illegible] | 1243 | Bacteria>Thermodesulfobacteriota>Desulfobacteria>Desulfobacterales>Desulfococcaceae>Desulfococcus | GCF_000422185.1 | Desulfococcus_multivorans | B2D07_RS03090 | 705815..709546 | 897 | DUF6079-family-protein-[Desulfococcus-multivorans] |
| WP_207691048.1 | Type3.MTaseII.containing.BREX | [illegible] | 1241 | Bacteria>Thermodesulfobacteriota>Desulfobacteria>Desulfobacterales>Desulfococcaceae>Desulfonema | GCF_017377355.1 | Desulfonema_limicola | dnl_RS07085 | 1641549..1645274 | 45656 | DUF6079-family-protein-[Desulfonema-limicola] |
| MBC8362871.1 | Type3.MTaseII.containing.BREX | [illegible] | 1209 | Bacteria>Thermodesulfobacteriota>Desulfobacteria>Desulfobacterales>unclassified_Desulfobacterales | GCA_014382665.1 | Desulfobacterales_bacterium | H8E23_15910 | 40..3669 | 2044940 | BREX-system-P-loop-protein-BrxC-[Desulfobacterales-bacterium] |
| MBF0121090.1 | Type3.MTaseII.containing.BREX | [illegible] | 1399 | Bacteria>Thermodesulfobacteriota>Desulfobacteria>Desulfobacterales>unclassified_Desulfobacterales | GCA_015228685.1 | Desulfobacterales_bacterium | HQK79_19845 | 16803..21002 | 2044940 | hypothetical-protein-[Desulfobacterales-bacterium] |
| MBF0225032.1 | Type3.MTaseII.containing.BREX | [illegible] | 1398 | Bacteria>Thermodesulfobacteriota>Desulfobacteria>Desulfobacterales>unclassified_Desulfobacterales | GCA_015231595.1 | Desulfobacterales_bacterium | HQK76_06210 | 65605..69801 | 2044940 | hypothetical-protein-[Desulfobacterales-bacterium] |
| MBL7208341.1 | Type3.MTaseII.containing.BREX | [illegible] | 538 | Bacteria>Thermodesulfobacteriota>Desulfobacteria>Desulfobacterales>unclassified_Desulfobacterales | GCA_016782935.1 | Desulfobacterales_bacterium | ISS67_07480 | 3997..5611 | 2044940 | BREX-system-P-loop-protein-BrxC-[Desulfobacterales-bacterium] |
| MBL7181321.1 | Type3.MTaseII.containing.BREX | [illegible] | 918 | Bacteria>Thermodesulfobacteriota>Desulfobacteria>Desulfobacterales>unclassified_Desulfobacterales | GCA_016782965.1 | Desulfobacterales_bacterium | ISS65_14115 | 3135..5888 | 2044940 | ATP-binding-protein-[Desulfobacterales-bacterium] |
| KGO32881.1 | Type3.MTaseII.containing.BREX | [illegible] | 754 | Bacteria>Thermodesulfobacteriota>Desulfobulbia>Desulfobulbales>Desulfobulbaceae>Desulfobulbus | GCA_000769715.1 | Desulfobulbus_sp._Tol-SR | JT06_17820 | 7334..9599 | 1536652 | ATPase,-partial-[Desulfobulbus-sp.-Tol-SR] |
| KGO32875.1 | Type3.MTaseII.containing.BREX | [illegible] | 789 | Bacteria>Thermodesulfobacteriota>Desulfobulbia>Desulfobulbales>Desulfobulbaceae>Desulfobulbus | GCA_000769715.1 | Desulfobulbus_sp._Tol-SR | JT06_17890 | 17101..19470 | 1536652 | ATPase,-partial-[Desulfobulbus-sp.-Tol-SR] |
| HHD64075.1 | Type3.MTaseII.containing.BREX | [illegible] | 462 | Bacteria>Thermodesulfobacteriota>Desulfobulbia>Desulfobulbales>Desulfobulbaceae>unclassified_Desulfobulbaceae | GCA_011372665.1 | Desulfobulbaceae_bacterium | ENK96_06915 | 1..1391 | 2053307 | ATP-binding-protein-[Desulfobulbaceae-bacterium] |
| HBH32876.1 | Type3.MTaseII.containing.BREX | [illegible] | 155 | Bacteria>Thermodesulfobacteriota>Desulfobulbia>Desulfobulbales>Desulfocapsaceae>Desulfofustis | GCA_003499505.1 | Desulfofustis_sp. | DDX81_13445 | 1..470 | 2053546 | ATPase-[Desulfofustis-sp.] |
| MBL0380281.1 | Type3.MTaseII.containing.BREX | [illegible] | 1243 | Bacteria>Thermodesulfobacteriota>Desulfobulbia>Desulfobulbales>Desulfocapsaceae>Desulfofustis | GCA_016722945.1 | Desulfofustis_sp._PB-SRB1 | JKP90_11585 | 146764..150495 | 1385624 | ATP-binding-protein-[Desulfofustis-sp.-PB-SRB1] |
| MBL0380707.1 | Type3.MTaseII.containing.BREX | [illegible] | 1242 | Bacteria>Thermodesulfobacteriota>Desulfobulbia>Desulfobulbales>Desulfocapsaceae>Desulfofustis | GCA_016722945.1 | Desulfofustis_sp._PB-SRB1 | JKP90_13820 | 384202..387930 | 1385624 | ATP-binding-protein-[Desulfofustis-sp.-PB-SRB1] |
| WP_205217413.1 | Type3.MTaseII.containing.BREX | [illegible] | 1249 | Bacteria>Thermodesulfobacteriota>Desulfobulbia>Desulfobulbales>Desulfocapsaceae>Desulforhopalus | GCF_016918505.1 | Desulforhopalus_vacuolatus | JWG39_RS00650 | 155250..158999 | 40414 | DUF6079-family-protein-[Desulforhopalus-vacuolatus] |
| WP_107736576.1 | Type3.MTaseII.containing.BREX | [illegible] | 1244 | Bacteria>Thermodesulfobacteriota>Desulfovibrionia>Desulfovibrionales>Desulfonatronaceae>Desulfonatronum | GCF_003046795.1 | Desulfonatronum_sp._SC1 | C6366_RS06755 | 68934..72668 | 2109626 | DUF6079-family-protein-[Desulfonatronum-sp.-SC1] |
| WP_045217268.1 | Type3.MTaseII.containing.BREX | [illegible] | 1254 | Bacteria>Thermodesulfobacteriota>Desulfovibrionia>Desulfovibrionales>Desulfonatronovibrionaceae>Desulfonatronovibrio | GCF_000934755.1 | Desulfonatronovibrio_magnus | LZ23_RS20570 | 26027..29791 | 698827 | DUF6079-family-protein-[Desulfonatronovibrio-magnus] |
| WP_270570163.1 | Type3.MTaseII.containing.BREX | [illegible] | 346 | Bacteria>Thermodesulfobacteriota>Desulfovibrionia>Desulfovibrionales>Desulfovibrionaceae>Bilophila | GCA_030820635.1 | Bilophila_wadsworthia | PG042_RS02170 | 68996..70036 | 35833 | DUF6079-family-protein-[Bilophila-wadsworthia] |
| WP_270570164.1 | Type3.MTaseII.containing.BREX | [illegible] | 445 | Bacteria>Thermodesulfobacteriota>Desulfovibrionia>Desulfovibrionales>Desulfovibrionaceae>Bilophila | GCA_030820635.1 | Bilophila_wadsworthia | PG042_RS02180 | 70866..72203 | 35833 | DUF6079-family-protein-[Bilophila-wadsworthia] |
| WP_270570100.1 | Type3.MTaseII.containing.BREX | [illegible] | 191 | Bacteria>Thermodesulfobacteriota>Desulfovibrionia>Desulfovibrionales>Desulfovibrionaceae>Bilophila | GCA_030820635.1 | Bilophila_wadsworthia | PG042_RS02185 | 72149..72724 | 35833 | DUF6079-family-protein-[Bilophila-wadsworthia] |
| WP_028587811.1 | Type3.MTaseII.containing.BREX | [illegible] | 1243 | Bacteria>Thermodesulfobacteriota>Desulfovibrionia>Desulfovibrionales>Desulfovibrionaceae>Desulfocurvus | GCF_000519125.1 | Desulfocurvus_vexinensis | G495_RS0110700 | 70932..74663 | 399548 | DUF6079-family-protein-[Desulfocurvus-vexinensis] |
| WP_029894986.1 | Type3.MTaseII.containing.BREX | [illegible] | 1243 | Bacteria>Thermodesulfobacteriota>Desulfovibrionia>Desulfovibrionales>Desulfovibrionaceae>Desulfohalovibrio | GCF_000711295.1 | Desulfohalovibrio_reitneri | N911_RS0105065 | 956113..959844 | 1307759 | DUF6079-family-protein-[Desulfohalovibrio-reitneri] |
| WP_256267769.1 | Type3.MTaseII.containing.BREX | [illegible] | 1243 | Bacteria>Thermodesulfobacteriota>Desulfovibrionia>Desulfovibrionales>Desulfovibrionaceae>Desulfovibrio | - | Desulfovibrio_desulfuricans | NE637_14445 | 12990..16721 | 876 | DUF6079-family-protein-[Desulfovibrio-desulfuricans] |
| RHH19729.1 | Type3.MTaseII.containing.BREX | [illegible] | 1243 | Bacteria>Thermodesulfobacteriota>Desulfovibrionia>Desulfovibrionales>Desulfovibrionaceae>Desulfovibrio | GCA_003470605.1 | Desulfovibrio_sp._AM18-2 | DW219_10435 | 17932..21663 | 2292040 | ATP-binding-protein-[Desulfovibrio-sp.-AM18-2] |
| MBQ9405388.1 | Type3.MTaseII.containing.BREX | [illegible] | 1241 | Bacteria>Thermodesulfobacteriota>Desulfovibrionia>Desulfovibrionales>Desulfovibrionaceae>Desulfovibrio | GCA_017454855.1 | Desulfovibrio_sp. | IJU37_01455 | 3099..6824 | 885 | ATP-binding-protein-[Desulfovibrio-sp.] |
| MCQ4862360.1 | Type3.MTaseII.containing.BREX | [illegible] | 1243 | Bacteria>Thermodesulfobacteriota>Desulfovibrionia>Desulfovibrionales>Desulfovibrionaceae>Desulfovibrio | GCA_024460775.1 | Desulfovibrio_desulfuricans | NE637_14445 | 12990..16721 | 876 | DUF6079-family-protein-[Desulfovibrio-desulfuricans] |
| MCQ5218961.1 | Type3.MTaseII.containing.BREX | [illegible] | 1014 | Bacteria>Thermodesulfobacteriota>Desulfovibrionia>Desulfovibrionales>Desulfovibrionaceae>Desulfovibrio | GCA_024463415.1 | Desulfovibrio_desulfuricans | NE596_12380 | 115185..118231 | 876 | DUF6079-family-protein-[Desulfovibrio-desulfuricans] |
| MCQ5219595.1 | Type3.MTaseII.containing.BREX | [illegible] | 852 | Bacteria>Thermodesulfobacteriota>Desulfovibrionia>Desulfovibrionales>Desulfovibrionaceae>Desulfovibrio | GCA_024463415.1 | Desulfovibrio_desulfuricans | NE596_15655 | 1..2561 | 876 | DUF6079-family-protein-[Desulfovibrio-desulfuricans] |
| CAI3239427.1 | Type3.MTaseII.containing.BREX | [illegible] | 1045 | Bacteria>Thermodesulfobacteriota>Desulfovibrionia>Desulfovibrionales>Desulfovibrionaceae>Desulfovibrio | GCA_902498985.2 | Desulfovibrio_diazotrophicus | DWUX_2110 | 2221222..2224359 | 2609801 | hypothetical-protein-BVU_3741-[Desulfovibrio-diazotrophicus] |
| WP_250783862.1 | Type3.MTaseII.containing.BREX | [illegible] | 1243 | Bacteria>Thermodesulfobacteriota>Desulfovibrionia>Desulfovibrionales>Desulfovibrionaceae>Desulfovibrio | GCF_023660105.1 | Desulfovibrio_aminophilus | M7784_08630 | 72453..76184 | 81425 | DUF6079-family-protein-[Desulfovibrio-aminophilus] |
| WP_256186046.1 | Type3.MTaseII.containing.BREX | [illegible] | 1014 | Bacteria>Thermodesulfobacteriota>Desulfovibrionia>Desulfovibrionales>Desulfovibrionaceae>Desulfovibrio | GCF_024463415.1 | Desulfovibrio_desulfuricans | NE596_RS12370 | 115185..118231 | 876 | DUF6079-family-protein,-partial-[Desulfovibrio-desulfuricans] |
| WP_256186110.1 | Type3.MTaseII.containing.BREX | [illegible] | 852 | Bacteria>Thermodesulfobacteriota>Desulfovibrionia>Desulfovibrionales>Desulfovibrionaceae>Desulfovibrio | GCF_024463415.1 | Desulfovibrio_desulfuricans | NE596_RS15640 | 1..2561 | 876 | DUF6079-family-protein,-partial-[Desulfovibrio-desulfuricans] |
| WP_118230669.1 | Type3.MTaseII.containing.BREX | [illegible] | 1243 | Bacteria>Thermodesulfobacteriota>Desulfovibrionia>Desulfovibrionales>Desulfovibrionaceae>Desulfovibrio | GCF_027671995.1 | Desulfovibrio_legallii | PGT02_RS09615 | 17960..21691 | 571438 | DUF6079-family-protein-[Desulfovibrio-legallii] |
| WP_215646936.1 | Type3.MTaseII.containing.BREX | [illegible] | 1243 | Bacteria>Thermodesulfobacteriota>Desulfovibrionia>Desulfovibrionales>Desulfovibrionaceae>Desulfovibrio | GCF_934502845.1 | Desulfovibrio_desulfuricans | GPK78_RS01810 | 338638..342369 | 876 | DUF6079-family-protein-[Desulfovibrio-desulfuricans] |
| WP_009368945.1 | Type3.MTaseII.containing.BREX | [illegible] | 1243 | Bacteria>Thermodesulfobacteriota>Desulfovibrionia>Desulfovibrionales>Desulfovibrionaceae | GCF_020592975.1 | Bilophila | LJE06_RS05510 | 48022..51753 | 35832 | MULTISPECIES:-DUF6079-family-protein-[Bilophila] |
| HJA07900.1 | Type3.MTaseII.containing.BREX | [illegible] | 1243 | Bacteria>Thermodesulfobacteriota>Desulfovibrionia>Desulfovibrionales>Desulfovibrionaceae>Mailhella | GCA_019117045.1 | Candidatus_Mailhella_merdigallinarum | H9962_01740 | 210661..214392 | 2838658 | ATP-binding-protein-[Candidatus-Mailhella-merdigallinarum] |
| MCG2733856.1 | Type3.MTaseII.containing.BREX | [illegible] | 1243 | Bacteria>Thermodesulfobacteriota>Desulfovibrionia>Desulfovibrionales>Desulfovibrionaceae>Pseudodesulfovibrio | GCA_022013435.1 | Pseudodesulfovibrio_aespoeensis | L6357_12305 | 5896..9627 | 182210 | DUF6079-family-protein-[Pseudodesulfovibrio-aespoeensis] |
| WP_013514583.1 | Type3.MTaseII.containing.BREX | [illegible] | 1243 | Bacteria>Thermodesulfobacteriota>Desulfovibrionia>Desulfovibrionales>Desulfovibrionaceae>Pseudodesulfovibrio | GCF_000176915.2 | Pseudodesulfovibrio_aespoeensis | DAES_RS08285 | 1822612..1826343 | 182210 | DUF6079-family-protein-[Pseudodesulfovibrio-aespoeensis] |
| WP_273435850.1 | Type3.MTaseII.containing.BREX | [illegible] | 1243 | Bacteria>Thermodesulfobacteriota>Desulfovibrionia>Desulfovibrionales>Desulfovibrionaceae>Pseudodesulfovibrio | GCF_022013435.1 | Pseudodesulfovibrio_aespoeensis | L6357_12305 | 5896..9627 | 182210 | DUF6079-family-protein-[Pseudodesulfovibrio-aespoeensis] |
| WP_097010673.1 | Type3.MTaseII.containing.BREX | [illegible] | 1243 | Bacteria>Thermodesulfobacteriota>Desulfovibrionia>Desulfovibrionales>Desulfovibrionaceae>Pseudodesulfovibrio | GCF_900217235.1 | Pseudodesulfovibrio_profundus | DPRO_RS02630 | 528076..531807 | 57320 | DUF6079-family-protein-[Pseudodesulfovibrio-profundus] |
| WP_129350664.1 | Type3.MTaseI.containing.BREX | [illegible] | 1243 | Bacteria>Thermodesulfobacteriota>Desulfovibrionia>Desulfovibrionales>Desulfovibrionaceae>Solidesulfovibrio | GCF_004135975.1 | Solidesulfovibrio_carbinolicus | C3Y92_RS06065 | 1371695..1375426 | 296842 | DUF6079-family-protein-[Solidesulfovibrio-carbinolicus] |
| WP_092055244.1 | Type3.MTaseII.containing.BREX | [illegible] | 1267 | Bacteria>Thermodesulfobacteriota>Desulfuromonadia>Desulfuromonadales>Desulfuromonadaceae>Desulfuromonas | GCF_900111775.1 | Desulfuromonas_acetexigens | BQ4888_RS06420 | 13598..17401 | 38815 | DUF6079-family-protein-[Desulfuromonas-acetexigens] |
| MBW6511064.1 | Type3.MTaseII.containing.BREX | [illegible] | 69 | Bacteria>Thermodesulfobacteriota>Desulfuromonadia>Desulfuromonadales>Desulfuromonadaceae>unclassified_Desulfuromonadaceae | GCA_019429445.1 | Desulfuromonadaceae_bacterium | K0A93_02955 | 88186..88395 | 2099677 | hypothetical-protein-[Desulfuromonadaceae-bacterium] |
| MBW6511065.1 | Type3.MTaseII.containing.BREX | [illegible] | 1290 | Bacteria>Thermodesulfobacteriota>Desulfuromonadia>Desulfuromonadales>Desulfuromonadaceae>unclassified_Desulfuromonadaceae | GCA_019429445.1 | Desulfuromonadaceae_bacterium | K0A93_02960 | 88337..92209 | 2099677 | hypothetical-protein-[Desulfuromonadaceae-bacterium] |
| WP_040199369.1 | Type3.MTaseII.containing.BREX | [illegible] | 1287 | Bacteria>Thermodesulfobacteriota>Desulfuromonadia>Desulfuromonadales>Geoalkalibacteraceae>Geoalkalibacter | GCF_000827125.1 | Geoalkalibacter_subterraneus | GSUB_04365 | 972871..976734 | 483547 | DUF6079-family-protein-[Geoalkalibacter-subterraneus] |
| OPY17362.1 | Type3.MTaseII.containing.BREX | [illegible] | 1378 | Bacteria>Thermodesulfobacteriota>Syntrophia>Syntrophales>Syntrophaceae>Syntrophus | GCA_002067245.1 | Syntrophus_sp._PtaB.Bin138 | A4E69_00025 | 12131..16267 | 1811711 | MAG:-hypothetical-protein-A4E69_00025-[Syntrophus-sp.-PtaB.Bin138] |
| SMC23767.1 | Type3.MTaseII.containing.BREX | [illegible] | 242 | Bacteria>Thermodesulfobacteriota>Syntrophobacteria>Syntrophobacterales>Syntrophobacteraceae>Desulfacinum | GCA_900176285.1 | Desulfacinum_hydrothermale_DSM_13146 | SAMN02746041_01839 | 170359..171087 | 1121390 | hypothetical-protein-SAMN02746041_01839-[Desulfacinum-hydrothermale-DSM-13146] |
| WP_212636882.1 | Type3.MTaseII.containing.BREX | [illegible] | 220 | Bacteria>Thermodesulfobacteriota>Syntrophobacteria>Syntrophobacterales>Syntrophobacteraceae>Desulfacinum | GCF_900176285.1 | Desulfacinum_hydrothermale | B9A12_RS17330 | 170359..171021 | 109258 | DUF6079-family-protein-[Desulfacinum-hydrothermale] |
| WP_212636883.1 | Type3.MTaseII.containing.BREX | [illegible] | 353 | Bacteria>Thermodesulfobacteriota>Syntrophobacteria>Syntrophobacterales>Syntrophobacteraceae>Desulfacinum | GCF_900176285.1 | Desulfacinum_hydrothermale | B9A12_RS17335 | 171027..172088 | 109258 | DUF6079-family-protein-[Desulfacinum-hydrothermale] |
| MCK8604085.1 | Type3.MTaseII.containing.BREX | [illegible] | 480 | Bacteria>Thermodesulfobacteriota>Syntrophobacteria>Syntrophobacterales>Syntrophobacteraceae>Desulfoferrobacter | GCA_022603395.1 | Desulfoferrobacter_suflitae | K1V27_20485 | 64658..66100 | 2865782 | DUF6079-family-protein-[Desulfoferrobacter-suflitae] |
| WP_258932099.1 | Type3.MTaseII.containing.BREX | [illegible] | 174 | Bacteria>Thermodesulfobacteriota>Syntrophobacteria>Syntrophobacterales>Syntrophobacteraceae>Desulfoferrobacter | GCF_022603395.1 | Desulfoferrobacter_suflitae | K1V27_RS20480 | 64243..64767 | 2865782 | DUF6079-family-protein-[Desulfoferrobacter-suflitae] |
| WP_258932100.1 | Type3.MTaseII.containing.BREX | [illegible] | 503 | Bacteria>Thermodesulfobacteriota>Syntrophobacteria>Syntrophobacterales>Syntrophobacteraceae>Desulfoferrobacter | GCF_022603395.1 | Desulfoferrobacter_suflitae | K1V27_RS20485 | 64658..66169 | 2865782 | DUF6079-family-protein-[Desulfoferrobacter-suflitae] |
| WP_258932103.1 | Type3.MTaseII.containing.BREX | [illegible] | 660 | Bacteria>Thermodesulfobacteriota>Syntrophobacteria>Syntrophobacterales>Syntrophobacteraceae>Desulfoferrobacter | GCF_022603395.1 | Desulfoferrobacter_suflitae | K1V27_RS20505 | 67471..69453 | 2865782 | DUF6079-family-protein-[Desulfoferrobacter-suflitae] |
| WP_137424076.1 | Type3.MTaseII.containing.BREX | [illegible] | 1276 | Bacteria>Thermodesulfobacteriota>Syntrophobacteria>Syntrophobacterales>Syntrophobacteraceae>Desulfoglaeba | GCF_005377625.1 | Desulfoglaeba_alkanexedens | FDQ92_RS07965 | 1756527..1760357 | 361111 | DUF6079-family-protein-[Desulfoglaeba-alkanexedens] |
| WP_211341213.1 | Type3.MTaseII.containing.BREX | [illegible] | 242 | Bacteria>Thermodesulfobacteriota>Syntrophobacteria>Syntrophobacterales>Syntrophobacteraceae>Desulfoglaeba | GCF_005377625.1 | Desulfoglaeba_alkanexedens | FDQ92_RS08055 | 1782265..1782993 | 361111 | DUF6079-family-protein-[Desulfoglaeba-alkanexedens] |
| GKT09455.1 | Type3.MTaseII.containing.BREX | [illegible] | 1248 | Bacteria>Thermodesulfobacteriota>Syntrophobacteria>Syntrophobacterales>Syntrophobacteraceae>Desulforhabdus | GCA_022836495.1 | Desulforhabdus_sp._TSK | DSTSK_27600 | 41946..45692 | 2925014 | hypothetical-protein-DSTSK_27600-[Desulforhabdus-sp.-TSK] |
| WP_244082336.1 | Type3.MTaseII.containing.BREX | [illegible] | 1245 | Bacteria>Thermodesulfobacteriota>Syntrophobacteria>Syntrophobacterales>Syntrophobacteraceae>Desulforhabdus | GCF_022836495.1 | Desulforhabdus_sp._TSK | MWC30_RS01305 | 286103..289840 | 2925014 | DUF6079-family-protein-[Desulforhabdus-sp.-TSK] |
| WP_244084753.1 | Type3.MTaseII.containing.BREX | [illegible] | 1235 | Bacteria>Thermodesulfobacteriota>Syntrophobacteria>Syntrophobacterales>Syntrophobacteraceae>Desulforhabdus | GCF_022836495.1 | Desulforhabdus_sp._TSK | MWC30_RS13665 | 41946..45653 | 2925014 | DUF6079-family-protein-[Desulforhabdus-sp.-TSK] |
| WP_123290345.1 | Type3.MTaseII.containing.BREX | [illegible] | 1266 | Bacteria>Thermodesulfobacteriota>Syntrophobacteria>Syntrophobacterales>Syntrophobacteraceae>Desulfosoma | GCF_003751385.1 | Desulfosoma_caldarium | EDC27_RS09315 | 252284..256084 | 610254 | DUF6079-family-protein-[Desulfosoma-caldarium] |
| NSW86901.1 | Type3.MTaseII.containing.BREX | [illegible] | 385 | Bacteria>Thermodesulfobacteriota>Syntrophobacteria>Syntrophobacterales>Syntrophobacteraceae>unclassified_Syntrophobacteraceae | GCA_013314815.1 | Syntrophobacteraceae_bacterium | HPY84_11335 | 4225..5382 | 2052185 | hypothetical-protein-[Syntrophobacteraceae-bacterium] |
| MBP8646529.1 | Type3.MTaseII.containing.BREX | [illegible] | 591 | Bacteria>Thermodesulfobacteriota>Syntrophobacteria>Syntrophobacterales>Syntrophobacteraceae>unclassified_Syntrophobacteraceae | GCA_018053185.1 | Syntrophobacteraceae_bacterium | KBH99_10480 | 8770..10544 | 2052185 | hypothetical-protein-[Syntrophobacteraceae-bacterium] |
| OHE16646.1 | Type3.MTaseII.containing.BREX | [illegible] | 1186 | Bacteria>Thermodesulfobacteriota>Syntrophobacteria>Syntrophobacterales>unclassified_Syntrophobacterales | GCA_001830835.1 | Syntrophobacterales_bacterium_GWC2_56_13 | A2X96_09765 | 10734..14294 | 1802287 | ATPase-[Syntrophobacterales-bacterium-GWC2_56_13] |
| MBP6974014.1 | Type3.MTaseII.containing.BREX | [illegible] | 1236 | Bacteria>Thermodesulfobacteriota>Syntrophorhabdia>Syntrophorhabdales>Syntrophorhabdaceae>Syntrophorhabdus | GCA_017992995.1 | Syntrophorhabdus_sp. | KBB67_13225 | 35..3745 | 2699756 | hypothetical-protein-[Syntrophorhabdus-sp.] |
| MBP7089501.1 | Type3.MTaseII.containing.BREX | [illegible] | 1262 | Bacteria>Thermodesulfobacteriota>Syntrophorhabdia>Syntrophorhabdales>Syntrophorhabdaceae>unclassified_Syntrophorhabdaceae | - | Syntrophorhabdaceae_bacterium | KBA98_01535 | 4905..8693 | 2660714 | hypothetical-protein-[Syntrophorhabdaceae-bacterium] |
| WP_068347596.1 | Type3.MTaseII.containing.BREX | [illegible] | 1239 | Bacteria>Thermotogota>Thermotogae>Kosmotogales>Kosmotogaceae>Kosmotoga | GCF_001636545.1 | Kosmotoga_arenicorallina | AT15_RS06370 | 8494..12213 | 688066 | DUF6079-family-protein-[Kosmotoga-arenicorallina] |
| WP_047754172.1 | Type3.MTaseII.containing.BREX | [illegible] | 1241 | Bacteria>Thermotogota>Thermotogae>Kosmotogales>Kosmotogaceae>Kosmotoga | GCF_910586115.1 | Kosmotoga_pacifica | IX53_RS03475 | 736249..739974 | 1330330 | DUF6079-family-protein-[Kosmotoga-pacifica] |
| CCU85049.1 | Type3.MTaseII.containing.BREX | [illegible] | 1239 | Bacteria>Thermotogota>Thermotogae>Kosmotogales>Kosmotogaceae>Mesotoga | GCA_000367705.1 | Mesotoga_infera | PHOSAC3_150049 | 70008..73727 | 1236046 | Predicted-ATPase,-VrlK-family-[Mesotoga-infera] |
| WP_103135224.1 | Type3.MTaseII.containing.BREX | [illegible] | 1239 | Bacteria>Thermotogota>Thermotogae>Kosmotogales>Kosmotogaceae>Mesotoga | GCF_002896535.1 | Mesotoga_sp._B105.6.4 | RJ60_RS14495 | 2748..6467 | 1582224 | DUF6079-family-protein-[Mesotoga-sp.-B105.6.4] |
| WP_121510563.1 | Type3.MTaseII.containing.BREX | [illegible] | 1239 | Bacteria>Thermotogota>Thermotogae>Kosmotogales>Kosmotogaceae>Mesotoga | GCF_003664935.1 | Mesotoga_sp._H07pep.5.4 | Y696_RS11485 | 59441..63160 | 1463664 | DUF6079-family-protein-[Mesotoga-sp.-H07pep.5.4] |
| WP_121526195.1 | Type3.MTaseII.containing.BREX | [illegible] | 1239 | Bacteria>Thermotogota>Thermotogae>Kosmotogales>Kosmotogaceae>Mesotoga | [illegible] | [illegible] | [illegible] | 22219..25938 | [illegible] | [illegible] |
| NLI08750.1 | Type3.MTaseII.containing.BREX | [illegible] | [illegible] | [illegible] | [illegible] | [illegible] | [illegible] | [illegible] | [illegible] | [illegible] |
| QQA16944.1 | Type3.MTaseII.containing.BREX | [illegible] | 835 | Bacteria>unclassified_Bacteria | GCA_002069435.1 | bacterium_ADurb.Bin363 | BWY64_02361 | 5209..7716 | 1866927 | hypothetical-protein-BWY64_02361-[bacterium-ADurb.Bin363] |
| [illegible] | Type3.MTaseII.containing.BREX | [illegible] | 1232 | Bacteria>unclassified_Bacteria | GCA_002069295.1 | bacterium_ADurb.Bin236 | BWY28_02297 | 29686..33414 | 1866925 | hypothetical-protein-BWY28_02297-[bacterium-ADurb.Bin236] |
| [illegible] | Type3.MTaseII.containing.BREX | [illegible] | 1238 | Bacteria>unclassified_Bacteria | GCA_002897895.1 | bacterium_BMS3Abin06 | BMS3Abin06_01802 | 10683..14699 | 2005714 | hypothetical-protein-BMS3Abin06_01802-[bacterium-BMS3Abin06] |

| Protein-ID | System | Gene-Neighbourhoods | Length | Taxonomy | Assembly-ID | Strain | Locus-Tag | Genomic-coordinates | TaxID | Fasta-defline |
|---|---|---|---|---|---|---|---|---|---|---|
| MBC7326963.1 | Type3.MTaseII.containing.BREX | RNase->?->?->?-><-ORC-AAA<-N6-N4-METHYLASE\|\|?->DUF6079*->PglZ->TPR+TPR+TPR+TPR->?->?->ArsR-HTH-> | 1239 | Bacteria>unclassified_Bacteria | GCA_014360465.1 | bacterium | H5T87_02470 | 27042..30761 | 1869227 | hypothetical-protein-[bacterium] |
| MBC7329234.1 | Type3.MTaseII.containing.BREX | <-Glyco_hydro_4+Glyco_hydro_4C\|\|?->?->?->?->ROK-HTH->?->DUF6079*->PglZ->?-><-?\|\|?-><-ORC-AAA<-N6-N4-METHYLASE<-Mrr_N | 1240 | Bacteria>unclassified_Bacteria | GCA_014360495.1 | bacterium | H5T88_02635 | 233811..237533 | 1869227 | hypothetical-protein-[bacterium] |
| MBI9032245.1 | Type3.MTaseII.containing.BREX | ParA-Soj-PloopNTPase->BrxB-BrxF-iSTAND->DUF6079*->N6-N4-METHYLASE->TUDOR+SF1-SF2-Helicase+iREase+CTD->PglZ->BrxA-DNA-binding->?->URI-DNase-><-HNH<-Phage_integrase\|\|AbiEi_4-> | 1216 | Bacteria>unclassified_Bacteria | GCA_016342615.1 | bacterium | JEZ13_09640 | 6548..10198 | 1869227 | hypothetical-protein-[bacterium] |
| MBN2056796.1 | Type3.MTaseII.containing.BREX | Arm-DNA-bind_3->DOC+HTH->?->BrxB-BrxF-iSTAND->DUF6079*->Schlafen+HATPase+wHTH->N6-N4-METHYLASE-p-> | 1261 | Bacteria>unclassified_Bacteria | GCA_016932295.1 | bacterium | JW905_17845 | 3771..7556 | 1869227 | ATP-binding-protein-[bacterium] |
| MCG3155095.1 | Type3.MTaseII.containing.BREX | <-Phage_integrase<-XIS-HTH<-?<-?<-DNA_METHYLASE<-ParB<-DnaB+TnsC-ATPase\|\|BrxB-BrxF-iSTAND->DUF6079*->N6-N4-METHYLASE->N6-N4-METHYLASE->DOC+HTH->TUDOR+SF1-SF2-Helicase+iREase+CTD->PglZ->BrxA-DNA-binding->REC-> | 1245 | Bacteria>unclassified_Bacteria | GCA_022072305.1 | bacterium | DKINENOH_01697 | 42467..46204 | 1869227 | hypothetical-protein-[bacterium] |
| MCP4405112.1 | Type3.MTaseII.containing.BREX | <-DUF6079-p*<-BrxB-BrxF-iSTAND<-PadR-HTH<-Schlafen+HATPase+wHTH<-?<-?<-RelE-ParE<-HEPN<-MoxR-AAA | 120 | Bacteria>unclassified_Bacteria | GCA_024222835.1 | bacterium | GY801_48400 | 1..359 | 1869227 | hypothetical-protein-[bacterium] |
| MCP4662276.1 | Type3.MTaseII.containing.BREX | BrxB-BrxF-iSTAND->DUF6079*->OLD-ABC->TOPRIM->N6-N4-METHYLASE->TUDOR+SF1-SF2-Helicase+iREase+CTD->PglZ->BrxA-DNA-binding-> | 1255 | Bacteria>unclassified_Bacteria | GCA_024231315.1 | bacterium | GY856_43295 | 2092..5859 | 1869227 | ATP-binding-protein-[bacterium] |
| WP_142434925.1 | Type3.MTaseII.containing.BREX | <-RVT<-BrxA-DNA-binding<-PglZ<-N6-N4-METHYLASE<-DUF6079*<-BrxB-BrxF-iSTAND<-HTH_3\|\|?->Phage_GPA->XIS-HTH->Arm-DNA-bind_4+Phage_int_SAM_3+Phage_integrase-> | 1239 | - | GCF_902159105.1 | Bacteria | FN024_RS14190 | 7834..11553 | 2 | DUF6079-family-protein-[Enterococcus-gallinarum] |

| Protein-ID | System | Gene-Neighbourhoods | Length | Taxonomy | Assembly-ID | Strain | Locus-Tag | Genomic-coordinates | TaxID | Fasta-define |
|---|---|---|---|---|---|---|---|---|---|---|
| -- | | | | | | | | | | |
| --Type4.BREX.Systems | | | | | | | | | | |
| -- | | | | | | | | | | |
| MBD3337553.1 | Type4.PglZ.containing.BREX | <-MazE_antitoxin<-N6-METHYLASE\|\|PAPS-reduct+DUF4007->DUF6079*->iSTAND+PglZ-> | 1203 | Archaea>Asgard_group>Candidatus_Lokiarchaeota>unclassified_Lokiarchaeota | GCA_014729315.1 | Candidatus_Lokiarchaeota_archaeon | GF353_00500 | 4154..7765 | 2053489 | PKD-domain-containing-protein-[Candidatus-Lokiarchaeota-archaeon] |
| WP_015504574.1 | Type4.PglZ.containing.BREX | <-DDE_Tnp_1\|\|?-><-Cysteine-Desulfurase<-SIGMA+OB-Fold+AAA-ATPase-BrxL<-PglZ<-?<-?<-?<-DUF6079*<-PAPS-reduct+a+b-domain+Fer4_9+DUF4007<-RAMA<-?\|\|GDResolvase->SF1-SF2-Helicase-><-?<-RADICAL-SAM\|\|?-><-REase | 1046 | Archaea>Candidatus_Thermoplasmatota>Thermoplasmata>Methanomassiliicoccales>Candidatus_Methanomethylophilaceae>Candidatus_Methanomethylophilus | GCF_000300255.2 | Candidatus_Methanomethylophilus_alvus | FX051_RS03360 | 653642..656782 | 1291540 | hypothetical-protein-[Candidatus-Methanomethylophilus-alvus] |
| KUK07754.1 | Type4.PglZ.containing.BREX | HEPN->?->?->?->?->?-><-SIGMA+OB-Fold+AAA-ATPase-BrxL<-iSTAND+PglZ<-DUF6079*<-?<-PAPS-reduct+a+b-domain+Fer4_9+DUF4007\|\|?->MazE_antitoxin->PIN->?-><-LexA-protease<-ORC-CDC6+RuvB-AAA<-Phage_integrase | 1419 | Archaea>Euryarchaeota>Archaeoglobi>Archaeoglobales>Archaeoglobaceae>Archaeoglobus | GCA_001508815.1 | Archaeoglobus_fulgidus | XD48_0089 | 68594..72853 | 2234 | Uncharacterized-protein-XD48_0089-[Archaeoglobus-fulgidus] |
| WP_048095117.1 | Type4.PglZ.containing.BREX | MazE_antitoxin->Thioredoxin->?-><-?<-?<-SIGMA+OB-Fold+AAA-ATPase-BrxL<-iSTAND+PglZ<-DUF6079*<-PAPS-reduct+a+b-domain+Fer4_9+DUF4007<-?\|\|?->?->HEPN->?->?->SLOG-> | 1419 | Archaea>Euryarchaeota>Archaeoglobi>Archaeoglobales>Archaeoglobaceae>Archaeoglobus | GCF_000734035.1 | Archaeoglobus_fulgidus | AFULGI_RS02805 | 507151..511410 | 2234 | hypothetical-protein-[Archaeoglobus-fulgidus] |
| WP_048175326.1 | Type4.PglZ.containing.BREX | <-HEPN+PAS+DUF4011+SF1-SF2-Helicase+REase<-?<-?<-35exo<-SF1-SF2-Helicase+wHTH+Small-B-barrel+DUF1998\|\|?->Thermonuclease-><-SIGMA+OB-Fold+AAA-ATPase-BrxL<-iSTAND+PglZ<-DUF6079*<-PAPS-reduct+a+b-domain+Fer4_9+DUF4007\|\|McrB-AAA->McrC-NTD+REase-><-N6-METHYLASE<-REase | 1122 | Archaea>Euryarchaeota>Methanomada_group>Methanobacteria>Methanobacteriales>Methanobacteriaceae | GCF_000828575.1 | Methanothermobacter_sp._CaT2 | MTCT_RS02130 | 407939..411307 | 866790 | hypothetical-protein-[Methanothermobacter-sp.-CaT2] |
| WP_147671926.1 | Type4.PglZ.containing.BREX | <-PSBP\|\|?->?-><-?\|\|?->?-><-McrC-NTD+REase<-McrB-NTD+McrB-AAA\|\|PAPS-reduct+DUF4007->DUF6079*->iSTAND+PglZ->SIGMA+OB-Fold+AAA-ATPase-BrxL-><-?<-Thermonuclease\|\|SF1-SF2-Helicase+wHTH+Small-B-barrel+DUF1998->35exo->?->?->HEPN+PAS+DUF4011+SF1-SF2-Helicase+REase-> | 1116 | Archaea>Euryarchaeota>Methanomada_group>Methanobacteria>Methanobacteriales>Methanobacteriaceae | GCF_008033705.1 | Methanothermobacter_sp._KEPCO-1 | FVF72_RS08995 | 1680163..1683513 | 2603820 | hypothetical-protein-[Methanothermobacter-sp.-KEPCO-1] |
| WP_191216538.1 | Type4.PglZ.containing.BREX | <-SF1-SF2-Helicase+SMS-KaiC<-?<-?<-35exo<-TPR<-SF1-SF2-Helicase+wHTH+Small-B-barrel+DUF1998\|\|Thermonuclease-><-SIGMA+OB-Fold+AAA-ATPase-BrxL<-iSTAND+PglZ<-DUF6079*<-PAPS-reduct+a+b-domain+Fer4_9+DUF4007<-DUF4007\|\|McrB-NTD->McrB-NTD+McrB-AAA->McrC-NTD+REase-><-?<-?\|\|?-><-SF1-SF2-Helicase+REase | 1118 | Archaea>Euryarchaeota>Methanomada_group>Methanobacteria>Methanobacteriales>Methanobacteriaceae | GCF_009914355.1 | Methanothermobacter_sp._THM-1 | FZP57_RS04205 | 797900..801256 | 2606911 | hypothetical-protein-[Methanothermobacter-sp.-THM-1] |
| MBB6067915.1 | Type4.PglZ.containing.BREX | TPR+TPR+TPR+TPR->GDResolvase+HIN-HTH-><-SF1-SF2-Helicase<-SIGMA+OB-Fold+AAA-ATPase-BrxL<-iSTAND+PglZ<-DUF6079*<-PAPS-reduct+a+b-domain+Fer4_9+DUF4007<-?\|\|?->N6-N4-METHYLASE-><-TPR+TPR+TPR\|\|Schlafen+HATPase+wHTH-><-PadR-HTH | 1099 | Archaea>Euryarchaeota>Methanomada_group>Methanococci>Methanococcales>Methanococcaceae | GCA_014203295.1 | Methanococcus_maripaludis | HNP97_001425 | 406681..409980 | 39152 | putative-house-cleaning-noncanonical-NTP-pyrophosphatase-(MazG-superfamily)-[Methanococcus-maripaludis] |
| HIQ39047.1 | Type4.PglZ.containing.BREX | <-SIGMA+OB-Fold+AAA-ATPase-BrxL<-iSTAND+PglZ<-DUF6079*<-PAPS-reduct+a+b-domain+Fer4_9+DUF4007 | 1140 | Archaea>Euryarchaeota>Methanomada_group>Methanococci>Methanococcales>Methanococcaceae | GCA_015661585.1 | Methanothermococcus_okinawensis | EYH53_03460 | 3935..7357 | 155863 | hypothetical-protein-[Methanothermococcus-okinawensis] |
| WP_183547001.1 | Type4.PglZ.containing.BREX | TPR+TPR+TPR+TPR->GDResolvase+HIN-HTH-><-SF1-SF2-Helicase<-SIGMA+OB-Fold+AAA-ATPase-BrxL<-iSTAND+PglZ<-?<-DUF6079*<-PAPS-reduct+a+b-domain+Fer4_9+DUF4007<-?\|\|?->N6-N4-METHYLASE-><-TPR+TPR+TPR\|\|Schlafen+HATPase+wHTH-><-PadR-HTH | 1064 | Archaea>Euryarchaeota>Methanomada_group>Methanococci>Methanococcales>Methanococcaceae | GCF_014203295.1 | Methanococcus_maripaludis | HNP97_RS07065 | 406788..409980 | 39152 | hypothetical-protein,-partial-[Methanococcus-maripaludis] |
| WP_229125940.1 | Type4.PglZ.containing.BREX | <-Phage_integrase\|\|?->?->?->DUF4007->DnaA->DUF6079*->PglZ-p->HerA/FtsK->ACET-><-?\|\|?-><-Phage_integrase+Phage_integr_3<-DDE_3<-HTH_23+HTH_32+HTH_33<-tRNA\|\|HTH_3-> | 1325 | Archaea>Euryarchaeota>Stenosarchaea_group>Halobacteria>Halobacteriales>Haloarculaceae | GCF_017094445.1 | Halapricum_desulfuricans | HSBGL_RS04755 | 937014..940991 | 2841257 | hypothetical-protein-[Halapricum-desulfuricans] |
| WP_092695620.1 | Type4.PglZ.containing.BREX | ORC-CDC6+HTH->?-><-?\|\|?->?->DUF4007->DnaA->DUF6079*->PglZ-p->HerA/FtsK->ACET-> | 1325 | Archaea>Euryarchaeota>Stenosarchaea_group>Halobacteria>Halobacteriales>Haloarculaceae | GCF_900102305.1 | Halorientalis_regularis | BLQ72_RS19855 | 7350..11327 | 660518 | hypothetical-protein-[Halorientalis-regularis] |
| WP_232571134.1 | Type4.PglZ.containing.BREX | HAD+HAD->?-><-?<-?\|\|?->?->DUF4007->DnaA->DUF6079*->PglZ-p->HerA/FtsK->ACET-> | 1334 | Archaea>Euryarchaeota>Stenosarchaea_group>Halobacteria>Halobacteriales>Halobacteriaceae | GCF_021233415.1 | Halobacterium_litoreum | LT972_RS14690 | 2690581..2694585 | 2039234 | hypothetical-protein-[Halobacterium-litoreum] |
| WP_147270938.1 | Type4.PglZ.containing.BREX | <-RADICAL-SAM<-?<-?\|\|?->?->DUF4007->DnaA->DUF6079*->PglZ-p->HerA/FtsK->ACET->?-><-REase\|\|?->TUDOR+SF1-SF2-Helicase+iREase+CTD+REase-p->?->TUDOR+SF1-SF2-Helicase+iREase+CTD-> | 1322 | Archaea>Euryarchaeota>Stenosarchaea_group>Halobacteria>Haloferacales>Haloferacaceae | GCF_003336245.1 | Haloplanus_salinus | DU504_RS14980 | 2902089..2906057 | 1126245 | hypothetical-protein-[Haloplanus-salinus] |
| WP_232819696.1 | Type4.PglZ.containing.BREX | MACRODOMAIN->?->?-><-McrC-NTD+REase<-McrB-AAA\|\|?->DUF4007->DUF6079*->PglZ-p->HerA/FtsK->ACET-><-?\|\|?-><-?<-SF1-SF2-Helicase+REase<-SF1-SF2-Helicase+REase<-PIN | 1317 | Archaea>Euryarchaeota>Stenosarchaea_group>Halobacteria>Haloferacales>Haloferacaceae | GCF_003439925.1 | Haloprofundus_halophilus | DV709_RS05700 | 1124449..1128402 | 2283527 | hypothetical-protein-[Haloprofundus-halophilus] |
| WP_089673072.1 | Type4.PglZ.containing.BREX | <-TBP+TBP<-ACET<-?<-?\|\|?->?-><-ACET<-HerA/FtsK<-PglZ-p<-DUF6079*<-DnaA<-DUF4007 | 1322 | Archaea>Euryarchaeota>Stenosarchaea_group>Halobacteria>Haloferacales>Halorubraceae | GCF_002788215.1 | Halohasta_litchfieldiae | BMX08_RS11270 | 26292..30260 | 1073996 | hypothetical-protein-[Halohasta-litchfieldiae] |
| MCP1662128.1 | Type4.PglZ.containing.BREX | <-HTH_OrfB_IS605+OrfB_IS605+OrfB_Zn_ribbon<-Y1_Tnp\|\|?->HTH+REase-><-REase\|\|?-><-Cysteine-Desulfurase<-SIGMA+OB-Fold+AAA-ATPase-BrxL<-iSTAND+PglZ<-DUF6079*<-PAPS-reduct+a+b-domain+Fer4_9+DUF4007<-HARE-HTH+E3_UFM1_ligase\|\|?-><-?<-?<-PIN<-AbrB-HTH\|\|PolB-> | 1384 | Archaea>Euryarchaeota>Stenosarchaea_group>Methanomicrobia>Methanomicrobiales>Methanocalculaceae | GCA_024170265.1 | Methanocalculus_sp._AMF5 | J2T61_000799 | 111729..115883 | 1198257 | hypothetical-protein-[Methanocalculus-sp.-AMF5] |
| WP_253457733.1 | Type4.PglZ.containing.BREX | <-ASCH<-HTH_OrfB_IS605+OrfB_IS605+OrfB_Zn_ribbon<-Y1_Tnp\|\|HTH+REase-><-REase\|\|?-><-Cysteine-Desulfurase<-SIGMA+OB-Fold+AAA-ATPase-BrxL<-iSTAND+PglZ<-DUF6079*<-PAPS-reduct+a+b-domain+Fer4_9+DUF4007<-HARE-HTH+E3_UFM1_ligase\|\|?-><-?<-?<-PIN<-AbrB-HTH\|\|PolB-> | 1387 | Archaea>Euryarchaeota>Stenosarchaea_group>Methanomicrobia>Methanomicrobiales>Methanocalculaceae | GCF_024170265.1 | Methanocalculus_sp._AMF5 | J2T61_RS03890 | 111729..115892 | 1198257 | hypothetical-protein-[Methanocalculus-sp.-AMF5] |
| MCQ2357400.1 | Type4.PglZ.containing.BREX | <-HTH+REase<-?<-?\|\|?->REase->?->PAPS-reduct+DUF4007->GspA-ATPase->DUF6079-p*->iSTAND+PglZ->SIGMA+OB-Fold+AAA-ATPase-BrxL->ZnR->Cysteine-Desulfurase->?-><-wHTH(SBD)+HNH\|\|?->?->MORC-> | 716 | Archaea>Euryarchaeota>Stenosarchaea_group>Methanomicrobia>Methanomicrobiales>Methanocorpusculaceae | GCA_024405795.1 | Methanocorpusculum_sp. | MJ005_01325 | 29309..31459 | 2058474 | hypothetical-protein-[Methanocorpusculum-sp.] |
| KQC03544.1 | Type4.PglZ.containing.BREX | <-ASCH\|\|REase->?->?->HARE-HTH+E3_UFM1_ligase->PAPS-reduct+a+b-domain+Fer4_9+DUF4007->DUF6079*->iSTAND+PglZ->SIGMA+OB-Fold+AAA-ATPase-BrxL->ZnR->Cysteine-Desulfurase-><-DDE_Tnp_1<-ORC-CDC6+RuvB-AAA | 1386 | Archaea>Euryarchaeota>Stenosarchaea_group>Methanomicrobia>Methanomicrobiales>Methanomicrobiaceae | GCA_001412355.1 | Methanoculleus_sp._SDB | APR53_04960 | 13163..17323 | 1735326 | MAG:-hypothetical-protein-APR53_04960-[Methanoculleus-sp.-SDB] |
| WP_176788577.1 | Type4.PglZ.containing.BREX | <-DDE_Tnp_1\|\|MULE-TRANSPOSASE->\|\|HARE-HTH+E3_UFM1_ligase->?->?-><-MULE-TRANSPOSASE\|\|PAPS-reduct+DUF4007->DUF6079*->iSTAND+PglZ->SIGMA+OB-Fold+AAA-ATPase-BrxL-><-PadR-HTH<-?<-?<-?<-?<-?<-DNA_METHYLASE | 1384 | Archaea>Euryarchaeota>Stenosarchaea_group>Methanomicrobia>Methanomicrobiales>Methanomicrobiaceae | GCF_013377755.1 | Methanofollis_tationis | HWN36_RS06325 | 1207344..1211498 | 81417 | hypothetical-protein-[Methanofollis-tationis] |
| MBU7048060.1 | Type4.PglZ.containing.BREX | <-ZnR\|\|?->PAPS-reduct+a+b-domain+Fer4_9+DUF4007->DUF6079*->iSTAND+PglZ->SIGMA+OB-Fold+AAA-ATPase-BrxL->TPR-><-?\|\|?->Thermonuclease-> | 1150 | Archaea>Euryarchaeota>Theionarchaea>unclassified_Theionarchaea | GCA_019008725.1 | Theionarchaea_archaeon | HXS54_16620 | 6350..9802 | 2747605 | hypothetical-protein-[Theionarchaea-archaeon] |
| NLB77117.1 | Type4.PglZ.containing.BREX | <-Cysteine-Desulfurase-p<-SIGMA+OB-Fold+AAA-ATPase-BrxL<-ZnR<-iSTAND+PglZ<-DUF6079*<-PAPS-reduct+a+b-domain+Fer4_9+DUF4007<-HARE-HTH+E3_UFM1_ligase | 1398 | Archaea>TACK_group>Thermoproteota>unclassified_Thermoproteota | GCA_012511995.1 | Thermoproteota_archaeon | GX799_11760 | 4150..8346 | 2056631 | hypothetical-protein-[Thermoproteota-archaeon] |
| WP_149796608.1 | Type4.PglZ.containing.BREX | <-HHH_4+SF1-SF2-Helicase\|\|?->PAS+PAS+GGDEF->REase+DUF4268->?->N6-METHYLASE->DUF4007->BrxB-BrxF-iSTAND->DUF6079*->PglZ->SIGMA+OB-Fold+AAA-ATPase-BrxL->PHP+SbcC->Phage_integrase->TRPR-HTH->tRNA->?->tRNA->tRNA->ncRNA->RFC-AAA+RuvB-AAA->?->HHH_5+RecR+TOPRIM-> | 1357 | Bacteria>Terrabacteria_group>Bacillota>Clostridia>Halanaerobiales>Halanaerobiaceae | GCF_900102605.1 | Halanaerobium_congolense | F3G65_RS02025 | 91352..95425 | 54121 | hypothetical-protein-[Halanaerobium-congolense] |
| PXF58772.1 | Type4.PglZ.containing.BREX | <-GAF+GAF+HISKIN\|\|?-><-?\|\|Aminotran_1_2-><-RHH<-SIGMA+OB-Fold+AAA-ATPase-BrxL<-PglZ<-DDE_Tnp_IS66<-PglZ<-DUF6079*<-?\|\|?-><-TOPRIM<-OLD-ABC<-N6-METHYLASE<-LON-AAA<-DUF4007<-DNA_METHYLASE | 1354 | Bacteria>delta/epsilon_subdivisions>Deltaproteobacteria>unclassified_Deltaproteobacteria | GCA_003194495.1 | Deltaproteobacteria_bacterium | C4B58_05630 | 57211..61275 | 2026735 | hypothetical-protein-C4B58_05630-[Deltaproteobacteria-bacterium] |
| KUK66683.1 | Type4.PglZ.containing.BREX | <-SIGMA+OB-Fold+AAA-ATPase-BrxL<-iSTAND+PglZ<-DUF6079*<-DUF4007<-PAPS-reduct+a+b-domain+Fer4_9+DUF4007 | 1400 | Bacteria>Bacteria_incertae_sedis>Bacteria_candidate_phyla>Patescibacteria_group>Parcubacteria_group>Candidatus_Parcubacteria | GCA_001508405.1 | Parcubacteria_bacterium_34_609 | XD85_0045 | 3984..8186 | 1641386 | hypothetical-protein-XD85_0045-[Parcubacteria-bacterium-34_609] |
| MBU4000565.1 | Type4.PglZ.containing.BREX | DUF6079-p*->PAPS-reduct->DUF4007->DnaA->DUF6079*->PglZ-p->HerA/FtsK->?-><-?\|\|MPTase-> | 151 | Bacteria>Bacteria_incertae_sedis>Bacteria_candidate_phyla>Patescibacteria_group>unclassified_Patescibacteria_group | GCA_018900475.1 | Patescibacteria_group_bacterium | KKG29_05365 | 1..457 | 2052139 | hypothetical-protein-[Patescibacteria-group-bacterium] |
| MBU4000569.1 | Type4.PglZ.containing.BREX | DUF6079-p->PAPS-reduct->DUF4007->DnaA->DUF6079*->PglZ-p->HerA/FtsK->?-><-?\|\|MPTase-><-HTH_3<-SIGMA-HTH\|\|HerA+REase-> | 1313 | Bacteria>Bacteria_incertae_sedis>Bacteria_candidate_phyla>Patescibacteria_group>unclassified_Patescibacteria_group | GCA_018900475.1 | Patescibacteria_group_bacterium | KKG29_05385 | 2720..6661 | 2052139 | hypothetical-protein-[Patescibacteria-group-bacterium] |
| HIS35237.1 | Type4.PglZ.containing.BREX | PAPS-reduct+a+b-domain+Fer4_9+DUF4007->DUF6079*->iSTAND+PglZ->SIGMA+OB-Fold+AAA-ATPase-BrxL->ZnR->Cysteine-Desulfurase-><-?\|\|AbiEi_4+AbiEi_1-><-GspA-ATPase+REase | 1418 | Bacteria>Bacteria_incertae_sedis>Candidatus_Scatousia | GCA_018711035.1 | Candidatus_Scatousia_excrementigallinarum | IAC10_01210 | 6584..10840 | 2840935 | hypothetical-protein-[Candidatus-Scatousia-excrementigallinarum] |
| WP_165846692.1 | Type4.PglZ.containing.BREX | HNH-p->?->?->?-><-SIGMA+OB-Fold+AAA-ATPase-BrxL<-iSTAND+PglZ<-DUF6079*<-?\|\|?-><-PAPS-reduct+a+b-domain+Fer4_9+DUF4007<-REase+SF1-SF2-Helicase-><-?<-TPR | 1272 | Bacteria>Caldisericota/Cryosericota_group>Candidatus_Cryosericota>Candidatus_Cryosericia>Candidatus_Cryosericales>Candidatus_Cryosericaceae | GCF_003570985.1 | Candidatus_Cryosericum_hinesii | SMC3_RS09445 | 19815..23633 | 2290915 | DUF6079-family-protein-[Candidatus-Cryosericum-hinesii] |
| NLT83665.1 | Type4.PglZ.containing.BREX | <-SIGMA+OB-Fold+AAA-ATPase-BrxL<-iSTAND+PglZ<-DUF6079*<-DUF4007<-PAPS-reduct+a+b-domain+Fer4_9+DUF4007 | 1419 | Bacteria>Coprothermobacterota>Coprothermobacteria>Coprothermobacterales>Coprothermobacteraceae>Coprothermobacter | GCA_012719895.1 | Coprothermobacter_proteolyticus | GXW92_02630 | 4312..8571 | 35786 | hypothetical-protein-[Coprothermobacter-proteolyticus] |
| WP_012544415.1 | Type4.PglZ.containing.BREX | <-ABC-ATPase+ABC-ATPase<-?<-?<-?<-SIGMA+OB-Fold+AAA-ATPase-BrxL<-iSTAND+PglZ<-DUF6079*<-DUF4007<-PAPS-reduct+a+b-domain+Fer4_9+DUF4007\|\|?->Ferredoxin-RRM+Radical_SAM-><-?\|\|Peripla_BP_2-><-?\|\|ABC-ATPase+SbcC-> | 1419 | Bacteria>Coprothermobacterota>Coprothermobacteria>Coprothermobacterales>Coprothermobacteraceae>Coprothermobacter | GCF_000020945.1 | Coprothermobacter_proteolyticus | COPRO5265_RS07125 | 1377545..1381804 | 35786 | hypothetical-protein-[Coprothermobacter-proteolyticus] |
| WP_018963724.1 | Type4.PglZ.containing.BREX | <-MarR-HTH<-GNTR-HTH\|\|?->?-><-?<-?<-?<-SIGMA+OB-Fold+AAA-ATPase-BrxL<-iSTAND+PglZ<-DUF6079*<-DUF4007<-PAPS-reduct+a+b-domain+Fer4_9+DUF4007\|\|Peripla_BP_2-><-?\|\|?->?-><-?<-?<-Peptidase_S26 | 1415 | Bacteria>Coprothermobacterota>Coprothermobacteria>Coprothermobacterales>Coprothermobacteraceae>Coprothermobacter | GCF_000378005.1 | Coprothermobacter_platensis | D890_RS0106570 | 1225420..1229667 | 108819 | hypothetical-protein-[Coprothermobacter-platensis] |
| WP_107689600.1 | Type4.PglZ.containing.BREX | MazE_antitoxin->RelE-ParE-><-?<-?<-?<-?<-?<-SIGMA+OB-Fold+AAA-ATPase-BrxL<-iSTAND+PglZ<-DUF6079*<-DUF4007<-PAPS-reduct+a+b-domain+Fer4_9+DUF4007\|\|?->Ferredoxin-RRM+Radical_SAM-><-?\|\|Peripla_BP_2-> | 1419 | Bacteria>Coprothermobacterota>Coprothermobacteria>Coprothermobacterales>Coprothermobacteraceae>Coprothermobacter | GCF_003052365.1 | Coprothermobacter_proteolyticus | DCP27_RS03020 | 210727..214986 | 35786 | hypothetical-protein-[Coprothermobacter-proteolyticus] |
| WP_107764156.1 | Type4.PglZ.containing.BREX | MazE_antitoxin->RelE-ParE-><-?<-?<-?<-?<-?<-SIGMA+OB-Fold+AAA-ATPase-BrxL<-iSTAND+PglZ<-DUF6079*<-DUF4007<-PAPS-reduct+a+b-domain+Fer4_9+DUF4007\|\|?->Ferredoxin-RRM+Radical_SAM-><-?<-?<-Peptidase_S26<-Peptidase\|\|Peripla_BP_2-> | 1419 | Bacteria>Coprothermobacterota>Coprothermobacteria>Coprothermobacterales>Coprothermobacteraceae>Coprothermobacter | GCF_003052375.1 | Coprothermobacter_proteolyticus | DCP34_RS06365 | 31435..35694 | 35786 | hypothetical-protein-[Coprothermobacter-proteolyticus] |
| WP_013009914.1 | Type4.PglZ.containing.BREX | <-TraG_N<-TraH\|\|?->?->?-><-35exo<-wHTH+WYL+WCX\|\|HARE-HTH+E3_UFM1_ligase->PAPS-reduct+a+b-domain+Fer4_9+DUF4007->DUF6079*->iSTAND+PglZ->SIGMA+OB-Fold+AAA-ATPase-BrxL->ZnR-><-?<-?<-?<-ArdCN+MPTase | 1376 | Bacteria>Deferribacterota>Deferribacteres>Deferribacterales>Geovibrionaceae>Denitrovibrio | GCF_000025725.1 | Denitrovibrio_acetiphilus | DACET_RS02880 | 585751..589881 | 118000 | hypothetical-protein-[Denitrovibrio-acetiphilus] |
| MBS7379910.1 | Type4.PglZ.containing.BREX | PAPS-reduct+a+b-domain+Fer4_9+DUF4007->DUF6079*->iSTAND+PglZ->SIGMA+OB-Fold+AAA-ATPase-BrxL->ZnR-> | 1402 | Bacteria>delta/epsilon_subdivisions>Deltaproteobacteria>Bradymonadales>unclassified_Bradymonadales | GCA_018384915.1 | Bradymonadales_bacterium | KIG72_01410 | 1812..6020 | 2099667 | hypothetical-protein-[Bradymonadales-bacterium] |
| EFK09464.1 | Type4.PglZ.containing.BREX | METHYLASE->?-><-?\|\|?->?-><-SIGMA+OB-Fold+AAA-ATPase-BrxL<-PglZ<-DUF6079*<-?<-HTH+REase<-DUF4007<-PAPS-reduct+a+b-domain+Fer4_9<-SIGMA-HTH+SIGMA-HTH+HARE-HTH\|\|?->MetJarc-HTH-><-?<-DDE_5+DDE_Tnp_1 | 1257 | Bacteria>delta/epsilon_subdivisions>Deltaproteobacteria>unclassified_Deltaproteobacteria | GCA_000179315.1 | delta_proteobacterium_NaphS2 | NPH_3107 | 44675..48448 | 88274 | conserved-hypothetical-protein-[delta-proteobacterium-NaphS2] |
| OGQ95915.1 | Type4.PglZ.containing.BREX | XIS-HTH->?->ParA-Soj-PloopNTPase->?->AbiEi_4+AbiEi_1->AbiEii+ZnR->?->HARE-HTH+E3_UFM1_ligase->PAPS-reduct+a+b-domain+Fer4_9+DUF4007->DUF6079*->iSTAND+PglZ->SIGMA+OB-Fold+AAA-ATPase-BrxL->ZnR->Cysteine-Desulfurase->HTH+HTH->?->ZnR-> | 1204 | Bacteria>delta/epsilon_subdivisions>Deltaproteobacteria>unclassified_Deltaproteobacteria | GCA_001799275.1 | Deltaproteobacteria_bacterium_RIFOXYD12_FULL_57_12 | A2521_02980 | 25708..29322 | 1797906 | hypothetical-protein-A2521_02980-[Deltaproteobacteria-bacterium-RIFOXYD12_FULL_57_12] |
| PKN32801.1 | Type4.PglZ.containing.BREX | HISKIN+MORC+REC-><-Cysteine-Desulfurase<-ZnR<-SIGMA+OB-Fold+AAA-ATPase-BrxL<-iSTAND+PglZ<-DUF6079*<-PAPS-reduct+a+b-domain+Fer4_9+DUF4007<-RAMA<-NtrC-AAA+FIS-HTH | 1384 | Bacteria>delta/epsilon_subdivisions>Deltaproteobacteria>unclassified_Deltaproteobacteria | GCA_002841865.1 | Deltaproteobacteria_bacterium_HGW-Deltaproteobacteria-19 | CVU61_16495 | 6054..10208 | 2013746 | transposase-[Deltaproteobacteria-bacterium-HGW-Deltaproteobacteria-19] |
| RJP56333.1 | Type4.PglZ.containing.BREX | PAPS-reduct->DUF4007->DnaA->DUF6079*->PglZ-p->HerA/FtsK-> | 1347 | Bacteria>delta/epsilon_subdivisions>Deltaproteobacteria>unclassified_Deltaproteobacteria | GCA_003599365.1 | Deltaproteobacteria_bacterium | C4549_06710 | 1956..5999 | 2026735 | hypothetical-protein-C4549_06710-[Deltaproteobacteria-bacterium] |
| MRR05466.1 | Type4.PglZ.containing.BREX | <-Cysteine-Desulfurase<-ZnR<-SIGMA+OB-Fold+AAA-ATPase-BrxL<-iSTAND+PglZ<-DUF6079*<-PAPS-reduct+a+b-domain+Fer4_9+DUF4007<-HARE-HTH+E3_UFM1_ligase<-XIS-HTH<-MarR-HTH<-HATPase+wHTH | 1385 | Bacteria>delta/epsilon_subdivisions>Deltaproteobacteria>unclassified_Deltaproteobacteria | GCA_009668845.1 | Deltaproteobacteria_bacterium | EG828_00750 | 16051..20208 | 2026735 | hypothetical-protein-[Deltaproteobacteria-bacterium] |
| MBI2377782.1 | Type4.PglZ.containing.BREX | DUF4007->DUF6079->PglZ->SIGMA+OB-Fold+AAA-ATPase-BrxL-><-SAM-METHYLASE\|\|SAM-METHYLASE-><-DndB<-URI-DNase<-PAPS-reduct<-DUF6079*<-DUF4007<-?\|\|?->AAA-ATPase-p-> | 1190 | Bacteria>delta/epsilon_subdivisions>Deltaproteobacteria>unclassified_Deltaproteobacteria | GCA_016183035.1 | Deltaproteobacteria_bacterium | HYV07_27515 | 41194..44766 | 2026735 | hypothetical-protein-[Deltaproteobacteria-bacterium] |
| MBM4284316.1 | Type4.PglZ.containing.BREX | Nudix_N_2+NUDIX-><-Thioredoxin\|\|GGDEF->ParA-Soj-PloopNTPase->?->?->tRNA->DDE_Tnp->DUF4007->DUF6079*->PAPS-reduct->DUF4007->DnaA->DUF6079->PglZ-p->HerA/FtsK-><-HTH_3<-?<-TUDOR+SF1-SF2-Helicase+iREase+CTD+REase-p | 1196 | Bacteria>delta/epsilon_subdivisions>Deltaproteobacteria>unclassified_Deltaproteobacteria | GCA_016874975.1 | Deltaproteobacteria_bacterium | FJ128_03560 | 20498..24088 | 2026735 | hypothetical-protein-[Deltaproteobacteria-bacterium] |
| MBM4284320.1 | Type4.PglZ.containing.BREX | ParA-Soj-PloopNTPase->?->?->tRNA->DDE_Tnp->DUF4007->DUF6079->PAPS-reduct->DUF4007->DnaA->DUF6079*->PglZ-p->HerA/FtsK-><-HTH_3<-?<-TUDOR+SF1-SF2-Helicase+iREase+CTD+REase-p\|\|<-?<-?<-TOPRIM+Primase | 1348 | Bacteria>delta/epsilon_subdivisions>Deltaproteobacteria>unclassified_Deltaproteobacteria | GCA_016874975.1 | Deltaproteobacteria_bacterium | FJ128_03580 | 26257..30303 | 2026735 | hypothetical-protein-[Deltaproteobacteria-bacterium] |
| MBW2003213.1 | Type4.PglZ.containing.BREX | <-RelE-ParE\|\|HKD-DNase->PAPS-reduct+a+b-domain+Fer4_9+DUF4007->DUF4007->DUF6079*->iSTAND+PglZ->SIGMA+OB-Fold+AAA-ATPase-BrxL-><-His_biosynth<-?<-CN_hydrolase+NAD_synthase<-PRTase | 1139 | Bacteria>delta/epsilon_subdivisions>Deltaproteobacteria>unclassified_Deltaproteobacteria | GCA_019308645.1 | Deltaproteobacteria_bacterium | JRI72_01160 | 9596..13015 | 2026735 | hypothetical-protein-[Deltaproteobacteria-bacterium] |
| PMQ01344.1 | Type4.PglZ.containing.BREX | TPR-><-SIGMA-HTH+SIGMA-HTH<-SIGMA+OB-Fold+AAA-ATPase-BrxL<-iSTAND+PglZ<-DUF6079*<-PAPS-reduct+a+b-domain+Fer4_9+DUF4007\|\|?-><-?<-?<-TetR-HTH | 1414 | Bacteria>Dictyoglomota>Dictyoglomia>Dictyoglomales>Dictyoglomaceae>Dictyoglomus | GCF_002878375.1 | Dictyoglomus_sp._NZ13-RE01 | CBR30_06730 | 77755..81999 | 2010949 | MAG:-hypothetical-protein-CBR30_06730-[Dictyoglomus-sp.-NZ13-RE01] |
| WP_012547648.1 | Type4.PglZ.containing.BREX | PAPS-reduct+Fer4_9-p->DUF4007->DUF4007->DUF6079*->iSTAND+PglZ->SIGMA+OB-Fold+AAA-ATPase-BrxL->Mrr_N->SIGMA-HTH-><-?<-?<-?<-DEOR-HTH+ISOCOT | 1393 | Bacteria>Dictyoglomota>Dictyoglomia>Dictyoglomales>Dictyoglomaceae>Dictyoglomus | GCF_000020965.1 | Dictyoglomus_thermophilum | DICTH_RS00520 | 116037..120218 | 14 | hypothetical-protein-[Dictyoglomus-thermophilum] |
| WP_149123247.1 | Type4.PglZ.containing.BREX | DEOR-HTH+ISOCOT->?->?->?-><-SIGMA-HTH<-Mrr_N<-SIGMA+OB-Fold+AAA-ATPase-BrxL<-iSTAND+PglZ<-DUF6079*<-DUF4007<-DUF4007<-PAPS-reduct+Fer4_9-p | 1393 | Bacteria>Dictyoglomota>Dictyoglomia>Dictyoglomales>Dictyoglomaceae>Dictyoglomus | GCF_008244535.1 | Dictyoglomus_thermophilum | FY122_08690 | 45431..49612 | 14 | hypothetical-protein-[Dictyoglomus-thermophilum] |
| WP_149122444.1 | Type4.PglZ.containing.BREX | <-SIGMA-HTH+SIGMA-HTH+SIGMA-HTH<-SIGMA+OB-Fold+AAA-ATPase-BrxL<-iSTAND+PglZ<-DUF6079*<-DUF4007<-DUF4007<-PAPS-reduct+Fer4_9-p | 1392 | Bacteria>Dictyoglomota>Dictyoglomia>Dictyoglomales>Dictyoglomaceae>Dictyoglomus | GCF_008244535.1 | Dictyoglomus_thermophilum | FY122_RS02095 | 466758..470936 | 14 | hypothetical-protein-[Dictyoglomus-thermophilum] |
| CDD83746.1 | Type4.PglZ.containing.BREX | HTH_3->PAPS-reduct+a+b-domain+Fer4_9+DUF4007->DUF6079*->iSTAND+PglZ-><-?\|\|SIGMA+OB-Fold+AAA-ATPase-BrxL->Schlafen+HATPase+wHTH-><-?<-Relaxase<-TraJ-RHH | 1160 | Bacteria>FCB_group>Bacteroidetes/Chlorobi_group>Bacteroidota>Bacteroidia>Bacteroidales | GCA_000435095.1 | Bacteroides_sp._CAG:462 | BN666_00387 | 3677..7159 | 1262740 | putative-uncharacterized-protein-[Bacteroides-sp.-CAG:462] |
| CDE58178.1 | Type4.PglZ.containing.BREX | DUF6079*->iSTAND+PglZ->SIGMA+OB-Fold+AAA-ATPase-BrxL-><-?\|\|?->?->?->Peptidase_M16+Peptidase_M16_C->JAB+sGTP-> | 1096 | Bacteria>FCB_group>Bacteroidetes/Chlorobi_group>Bacteroidota>Bacteroidia>Bacteroidales | GCA_000436235.1 | Prevotella_sp._CAG:873 | BN799_00258 | 42..3332 | 1262936 | putative-uncharacterized-protein-[Prevotella-sp.-CAG:873] |
| CDE63279.1 | Type4.PglZ.containing.BREX | Mob_Pre->HTH_3->PAPS-reduct+a+b-domain+Fer4_9+DUF4007->DUF6079*->iSTAND+PglZ->SIGMA+OB-Fold+AAA-ATPase-BrxL->SeqA->SeqA->Schlafen+HATPase+wHTH->?->?->Calcineurin-SbcD->ClpABN-AAA+ClpABC-AAA-> | 1145 | Bacteria>FCB_group>Bacteroidetes/Chlorobi_group>Bacteroidota>Bacteroidia>Bacteroidales | GCA_000436355.1 | Alistipes_putredinis_CAG:67 | BN752_00056 | 4952..8389 | 1263036 | putative-uncharacterized-protein-[Alistipes-putredinis-CAG:67] |
| ERK01021.1 | Type4.PglZ.containing.BREX | Phage_int_SAM_5+Phage_integrase->PAPS-reduct+a+b-domain+Fer4_9+DUF4007->wHTH->DUF6079*->iSTAND+PglZ->SIGMA+OB-Fold+AAA-ATPase-BrxL->?->Schlafen+BirA-HTH-><-FurR-HTH<-?<-TraJ-RHH<-ZnR+TOPRIM | 1122 | Bacteria>FCB_group>Bacteroidetes/Chlorobi_group>Bacteroidota>Bacteroidia>Bacteroidales | GCA_000468135.1 | Prevotella_pleuritidis_F0068 | HMPREF1218_0904 | 14576..17944 | 1081904 | hypothetical-protein-HMPREF1218_0904-[Prevotella-pleuritidis-F0068] |
| OYP42213.1 | Type4.PglZ.containing.BREX | <-REase<-SIGMA+OB-Fold+AAA-ATPase-BrxL<-iSTAND+PglZ<-DUF6079*<-PAPS-reduct+a+b-domain+Fer4_9+DUF4007 | 1159 | Bacteria>FCB_group>Bacteroidetes/Chlorobi_group>Bacteroidota>Bacteroidia>Bacteroidales | GCA_002251295.1 | Prevotella_sp._P5-50 | CIK88_02545 | 5600..9079 | 2024217 | hypothetical-protein-CIK88_02545-[Prevotella-sp.-P5-50] |
| RGS50057.1 | Type4.PglZ.containing.BREX | <-TIR<-SIGMA+OB-Fold+AAA-ATPase-BrxL<-PglZ<-DUF6079*<-wHTH<-PAPS-reduct+a+b-domain+Fer4_9+DUF4007<-HTH_3<-STYKinase-like<-TRPR-HTH\|\|SPOUT-><-?<-ADH_N+ADH_zinc_N+METHYLASE | 1145 | Bacteria>FCB_group>Bacteroidetes/Chlorobi_group>Bacteroidota>Bacteroidia>Bacteroidales | GCA_003458755.1 | Bacteroides_xylanisolvens | DWX88_20865 | 4657..8094 | 371601 | hypothetical-protein-DWX88_20865-[Bacteroides-xylanisolvens] |
| RGX95621.1 | Type4.PglZ.containing.BREX | <-HHH+DNA_pol_B+RNase\|\|<-HU-HIG+DEOR-HTH<-HTH_3<-SIGMA+OB-Fold+AAA-ATPase-BrxL<-iSTAND+PglZ<-DUF6079*<-wHTH<-PAPS-reduct+a+b-domain+Fer4_9+DUF4007<-?\|\|?-><-?<-?<-?\|\|?->S1+RNase-> | 1152 | Bacteria>FCB_group>Bacteroidetes/Chlorobi_group>Bacteroidota>Bacteroidia>Bacteroidales | GCA_003463005.1 | Prevotella_copri | DXA63_07170 | 10328..13786 | 165179 | hypothetical-protein-DXA63_07170-[Prevotella-copri] |
| HCE48067.1 | Type4.PglZ.containing.BREX | <-iSTAND+PglZ<-DUF6079*<-PAPS-reduct+a+b-domain+Fer4_9+DUF4007<-?<-?\|\|GDResolvase-> | 1144 | Bacteria>FCB_group>Bacteroidetes/Chlorobi_group>Bacteroidota>Bacteroidia>Bacteroidales | GCA_003516685.1 | Prevotellaceae_bacterium | DEQ84_05375 | 621..4055 | 2049047 | hypothetical-protein-[Prevotellaceae-bacterium] |
| HBZ34375.1 | Type4.PglZ.containing.BREX | <-SIGMA+OB-Fold+AAA-ATPase-BrxL<-iSTAND+PglZ<-DUF6079*<-PAPS-reduct+a+b-domain+Fer4_9+DUF4007<-HNH+SeqA<-NERD<-OLD-ABC<-?<-?\|\|PRTase->?->sGTP-> | 1189 | Bacteria>FCB_group>Bacteroidetes/Chlorobi_group>Bacteroidota>Bacteroidia>Bacteroidales | GCA_003520485.1 | Bacteroidales_bacterium | DEO38_04655 | 4611..8180 | 2030927 | hypothetical-protein-[Bacteroidales-bacterium] |
| ROT09369.1 | Type4.PglZ.containing.BREX | SIGMA-HTH->?->O6MGT-HTH-><-?<-DndE<-DndD<-RNase<-SIGMA+OB-Fold+AAA-ATPase-BrxL<-iSTAND+PglZ<-DUF6079*<-wHTH<-PAPS-reduct+a+b-domain+Fer4_9+DUF4007<-REase\|\|?->Aminotran_1_2->?->HigB->HTH_3-><-HD | 1160 | Bacteria>FCB_group>Bacteroidetes/Chlorobi_group>Bacteroidota>Bacteroidia>Bacteroidales | GCA_003762615.1 | Muribaculaceae_bacterium_Isolate-037_(Harlan) | EEL33_02915 | 250198..253680 | 2486461 | hypothetical-protein-EEL33_02915-[Muribaculaceae-bacterium-Isolate-037-(Harlan)] |
| ROT09402.1 | Type4.PglZ.containing.BREX | Tox-RelE-><-?\|\|ABhydrolase->AA_kinase->?-><-Cysteine-Desulfurase<-ZnR<-SIGMA+OB-Fold+AAA-ATPase-BrxL<-iSTAND+PglZ<-DUF6079*<-PAPS-reduct+a+b-domain+Fer4_9+DUF4007 | 1387 | Bacteria>FCB_group>Bacteroidetes/Chlorobi_group>Bacteroidota>Bacteroidia>Bacteroidales | GCA_003762615.1 | Muribaculaceae_bacterium_Isolate-037_(Harlan) | EEL33_03100 | 299295..303458 | 2486461 | hypothetical-protein-EEL33_03100-[Muribaculaceae-bacterium-Isolate-037-(Harlan)] |
| KAA4699236.1 | Type4.PglZ.containing.BREX | Arm-DNA-bind_5+Phage_int_SAM_5+Phage_integrase->?->XIS-HTH->SMS-KaiC-><-?<-LRR+LRR<-ACYC<-SIGMA+OB-Fold+AAA-ATPase-BrxL<-iSTAND+PglZ<-DUF6079*<-wHTH<-PAPS-reduct+a+b-domain+Fer4_9+DUF4007<-HTH_3<-?<-AbiEii<-AbiEi_4+AbiEi_1<-STYKinase-like<-TRPR-HTH | 1106 | Bacteria>FCB_group>Bacteroidetes/Chlorobi_group>Bacteroidota>Bacteroidia>Bacteroidales | GCA_008569055.1 | Bacteroides_fragilis | F3B26_17335 | 35427..38747 | 817 | hypothetical-protein-F3B26_17335-[Bacteroides-fragilis] |
| GFI35177.1 | Type4.PglZ.containing.BREX | <-Arm-DNA-bind_5+Phage_int_SAM_5+Phage_integrase<-XIS-HTH+XIS-HTH+XIS-HTH+XIS-HTH<-?<-?\|\|TRPR-HTH->MNS+HTH->?->PAPS-reduct+a+b-domain+Fer4_9+DUF4007->DUF6079-p->DUF6079-p*->iSTAND+PglZ->SIGMA+OB-Fold+AAA-ATPase-BrxL->SeqA->?-><-?<-Relaxase<-TraJ-RHH<-SMS-KaiC<-XIS-HTH | 962 | Bacteria>FCB_group>Bacteroidetes/Chlorobi_group>Bacteroidota>Bacteroidia>Bacteroidales | GCA_011960185.1 | Bacteroidaceae_bacterium | IMSAGC014_01691 | 8937..11825 | 2212467 | hypothetical-protein-IMSAGC014_01691-[Bacteroidaceae-bacterium] |
| NLE35797.1 | Type4.PglZ.containing.BREX | <-LexA-protease<-Classical-AAA<-?<-?<-wHTH+WYL+WCX<-Cysteine-Desulfurase<-ZnR<-SIGMA+OB-Fold+AAA-ATPase-BrxL<-iSTAND+PglZ<-DUF6079*<-PAPS-reduct+a+b-domain+Fer4_9+DUF4007<-SeqA+Phage_int+SeqA+E3_UFM1_ligase<-HTH_26<-?<-TraJ-RHH<-LPD2 | 1391 | Bacteria>FCB_group>Bacteroidetes/Chlorobi_group>Bacteroidota>Bacteroidia>Bacteroidales | GCA_012514255.1 | Bacteroidales_bacterium | GX622_11890 | 11999..16174 | 2030927 | hypothetical-protein-[Bacteroidales-bacterium] |
| NLE36132.1 | Type4.PglZ.containing.BREX | <-iSTAND+PglZ<-DUF6079*<-PAPS-reduct+a+b-domain+Fer4_9+DUF4007<-REase<-DNA_METHYLASE<-?<-?\|\|?->RADICAL-SAM-><-His_Phos_1 | 1127 | Bacteria>FCB_group>Bacteroidetes/Chlorobi_group>Bacteroidota>Bacteroidia>Bacteroidales | GCA_012514255.1 | Bacteroidales_bacterium | GX622_13610 | 555..3938 | 2030927 | hypothetical-protein-[Bacteroidales-bacterium] |
| NMA75533.1 | Type4.PglZ.containing.BREX | REC+AraC-HTH-><-?\|\|?-><-SIGMA+OB-Fold+AAA-ATPase-BrxL<-iSTAND+PglZ<-DUF6079*<-PAPS-reduct+a+b-domain+Fer4_9+DUF4007 | 1376 | Bacteria>FCB_group>Bacteroidetes/Chlorobi_group>Bacteroidota>Bacteroidia>Bacteroidales | GCA_012799395.1 | Bacteroidales_bacterium | GX963_15590 | 16861..20991 | 2030927 | hypothetical-protein-[Bacteroidales-bacterium] |
| MBA7488708.1 | Type4.PglZ.containing.BREX | Phage_int_SAM_5+Phage_integrase->HTH_3->PAPS-reduct+a+b-domain+Fer4_9+DUF4007->wHTH->DUF6079*->iSTAND+PglZ->SIGMA+OB-Fold+AAA-ATPase-BrxL->?->Schlafen+BirA-HTH-> | 1134 | Bacteria>FCB_group>Bacteroidetes/Chlorobi_group>Bacteroidota>Bacteroidia>Bacteroidales | GCA_013333935.1 | Prevotella_sp. | HG422_06745 | 4580..7984 | 59823 | hypothetical-protein-[Prevotella-sp.] |
| MBD5262116.1 | Type4.PglZ.containing.BREX | Tox-ART-HYD2->?->?-><-?\|\|?-><-Schlafen+HATPase+wHTH\|\|PAPS-reduct+DUF4007->DUF6079*->iSTAND+PglZ->SIGMA+OB-Fold+AAA-ATPase-BrxL-><-?<-SIGMA-HTH | 1182 | Bacteria>FCB_group>Bacteroidetes/Chlorobi_group>Bacteroidota>Bacteroidia>Bacteroidales | GCA_014801315.1 | Bacteroides_sp. | HDS38_08315 | 60698..64246 | 29523 | hypothetical-protein-[Bacteroides-sp.] |
| MBD5285709.1 | Type4.PglZ.containing.BREX | TraJ-RHH->TraI-VirD2-RCRE->HTH_3->PAPS-reduct+a+b-domain+Fer4_9+DUF4007->DUF6079*->iSTAND+PglZ->SIGMA+OB-Fold+AAA-ATPase-BrxL->REase->Phage_holin->DOC+HTH->HTH_31->?->RNase->Calcineurin-> | 1179 | Bacteria>FCB_group>Bacteroidetes/Chlorobi_group>Bacteroidota>Bacteroidia>Bacteroidales | GCA_014801525.1 | Bacteroides_sp. | HDS29_05470 | 5866..9405 | 29523 | hypothetical-protein-[Bacteroides-sp.] |
| MBD5332113.1 | Type4.PglZ.containing.BREX | <-KAP-NTPase<-HerA/FtsK<-SIR2<-SIGMA+OB-Fold+AAA-ATPase-BrxL<-iSTAND+PglZ<-DUF6079*<-PAPS-reduct+a+b-domain+Fer4_9+DUF4007<-HTH_3 | 1182 | Bacteria>FCB_group>Bacteroidetes/Chlorobi_group>Bacteroidota>Bacteroidia>Bacteroidales | GCA_014802045.1 | Bacteroides_sp. | HDS00_09520 | 10265..13813 | 29523 | hypothetical-protein-[Bacteroides-sp.] |
| MBD5420816.1 | Type4.PglZ.containing.BREX | sGTP->?-><-?<-DndE<-DndD<-RNase<-SIGMA+OB-Fold+AAA-ATPase-BrxL<-iSTAND+PglZ<-DUF6079*<-wHTH<-PAPS-reduct+a+b-domain+Fer4_9+DUF4007<-?\|\|?->?-><-?<-?<-CNMP+CRP-HTH<-REase+HTH | 1143 | Bacteria>FCB_group>Bacteroidetes/Chlorobi_group>Bacteroidota>Bacteroidia>Bacteroidales | GCA_014803785.1 | Bacteroides_sp. | HDR47_06255 | 15095..18526 | 29523 | hypothetical-protein-[Bacteroides-sp.] |
| MBD9229349.1 | Type4.PglZ.containing.BREX | <-SF1-SF2-Helicase+RNase+RNase<-SIGMA+OB-Fold+AAA-ATPase-BrxL<-iSTAND+PglZ<-DUF6079*<-wHTH<-PAPS-reduct+a+b-domain+Fer4_9+DUF4007\|\|?->Peptidase_M3-> | 1166 | Bacteria>FCB_group>Bacteroidetes/Chlorobi_group>Bacteroidota>Bacteroidia>Bacteroidales | GCA_014846805.1 | Prevotella_sp. | EGQ31_01705 | 81403..84903 | 59823 | hypothetical-protein-[Prevotella-sp.] |
| MBD8978002.1 | Type4.PglZ.containing.BREX | <-SIGMA+OB-Fold+AAA-ATPase-BrxL<-iSTAND+PglZ<-DUF6079*<-PAPS-reduct+a+b-domain+Fer4_9+DUF4007<-?\|\|PhoH-> | 1160 | Bacteria>FCB_group>Bacteroidetes/Chlorobi_group>Bacteroidota>Bacteroidia>Bacteroidales | GCA_014847385.1 | Prevotellamassilia_timonensis | EGQ88_05395 | 14012..17494 | 1852370 | hypothetical-protein-[Prevotellamassilia-timonensis] |
| MBD9293002.1 | Type4.PglZ.containing.BREX | PAPS-reduct+DUF4007->DUF6079-p*->?->iSTAND+PglZ->SIGMA+OB-Fold+AAA-ATPase-BrxL->ZnR->Cysteine-Desulfurase->wHTH+WYL+WCX-> | 898 | Bacteria>FCB_group>Bacteroidetes/Chlorobi_group>Bacteroidota>Bacteroidia>Bacteroidales | GCA_014848995.1 | Bacteroidales_bacterium | EGS78_00050 | 393..3089 | 2030927 | hypothetical-protein-[Bacteroidales-bacterium] |
| MBE6296482.1 | Type4.PglZ.containing.BREX | <-iSTAND+PglZ<-DUF6079*<-wHTH<-PAPS-reduct+a+b-domain+Fer4_9+DUF4007<-HTH_3\|\|MNS+CR-ATPase5->?-><-?<-?<-?<-ADH_N+ADH_zinc_N+METHYLASE | 1143 | Bacteria>FCB_group>Bacteroidetes/Chlorobi_group>Bacteroidota>Bacteroidia>Bacteroidales | GCA_015060965.1 | Bacteroidales_bacterium | E7086_08280 | 1346..4777 | 2030927 | hypothetical-protein-[Bacteroidales-bacterium] |
| MBE6328327.1 | Type4.PglZ.containing.BREX | CR-ATPase5+REase-><-?\|\|?-><-?<-Glycos_transf<-?<-?<-SIGMA+OB-Fold+AAA-ATPase-BrxL<-iSTAND+PglZ<-DUF6079*<-PAPS-reduct+a+b-domain+Fer4_9+DUF4007 | 1184 | Bacteria>FCB_group>Bacteroidetes/Chlorobi_group>Bacteroidota>Bacteroidia>Bacteroidales | GCA_015061115.1 | Bacteroidales_bacterium | E7077_14990 | 31086..34640 | 2030927 | hypothetical-protein-[Bacteroidales-bacterium] |
| MBF1451715.1 | Type4.PglZ.containing.BREX | URI-DNase+RAMA->?->?->Phage_int_SAM_5+Phage_integrase->PAPS-reduct+a+b-domain+Fer4_9+DUF4007->wHTH->DUF6079*->iSTAND+PglZ->SIGMA+OB-Fold+AAA-ATPase-BrxL->?->Schlafen+BirA-HTH-><-FurR-HTH | 1133 | Bacteria>FCB_group>Bacteroidetes/Chlorobi_group>Bacteroidota>Bacteroidia>Bacteroidales | GCA_015261915.1 | Prevotella_pallens | HXN61_RS08010 | 23695..27096 | 60133 | hypothetical-protein-[Prevotella-pallens] |
| MBQ3618950.1 | Type4.PglZ.containing.BREX | PAPS-reduct+a+b-domain+Fer4_9+DUF4007->DUF6079*->iSTAND+PglZ->SIGMA+OB-Fold+AAA-ATPase-BrxL-> | 1184 | Bacteria>FCB_group>Bacteroidetes/Chlorobi_group>Bacteroidota>Bacteroidia>Bacteroidales | GCA_017394445.1 | Bacteroidales_bacterium | II939_12420 | 3317..6871 | 2030927 | hypothetical-protein-[Bacteroidales-bacterium] |
| MBQ4420211.1 | Type4.PglZ.containing.BREX | <-Aminotran_1_2<-?<-?<-?\|\|?->?-><-Peptidase_M20\|\|PAPS-reduct+DUF4007->DUF6079*->iSTAND+PglZ->SIGMA+OB-Fold+AAA-ATPase-BrxL->?-><-?\|\|?-><-?<-HNH | 1149 | Bacteria>FCB_group>Bacteroidetes/Chlorobi_group>Bacteroidota>Bacteroidia>Bacteroidales | GCA_017394975.1 | Bacteroidales_bacterium | II878_01540 | 63178..66627 | 2030927 | hypothetical-protein-[Bacteroidales-bacterium] |
| MBQ3604732.1 | Type4.PglZ.containing.BREX | <-OLD-ABC+TOPRIM<-SIGMA+OB-Fold+AAA-ATPase-BrxL<-iSTAND+PglZ<-DUF6079*<-wHTH<-PAPS-reduct+a+b-domain+Fer4_9+DUF4007<-HTH_3<-BrxD<-RVT<-HTH_3<-?\|\|ACET-> | 1143 | Bacteria>FCB_group>Bacteroidetes/Chlorobi_group>Bacteroidota>Bacteroidia>Bacteroidales | GCA_017399915.1 | Muribaculaceae_bacterium | II990_00560 | 6286..9717 | 2498093 | hypothetical-protein-[Muribaculaceae-bacterium] |
| MBQ6275504.1 | Type4.PglZ.containing.BREX | REC+PglZ->lipoprotein-><-BirA-HTH<-Schlafen+HATPase+wHTH<-HTH+REase<-SeqA<-SIGMA+OB-Fold+AAA-ATPase-BrxL<-iSTAND+PglZ<-DUF6079*<-PAPS-reduct+a+b-domain+Fer4_9+DUF4007<-?<-TOPRIM<-SbcC<-DUF4007<-?<-Couple_hipA+HipA_C<-HTH_3 | 1144 | Bacteria>FCB_group>Bacteroidetes/Chlorobi_group>Bacteroidota>Bacteroidia>Bacteroidales | GCA_017428605.1 | Bacteroidales_bacterium | IJK62_02205 | 79849..83283 | 2030927 | hypothetical-protein-[Bacteroidales-bacterium] |
| MBQ6210586.1 | Type4.PglZ.containing.BREX | Glycos_transf->?->Glycos_transf-><-?<-?\|\|?-><-ABC-ATPase+SbcC<-SIGMA+OB-Fold+AAA-ATPase-BrxL<-iSTAND+PglZ<-DUF6079*<-PAPS-reduct+a+b-domain+Fer4_9+DUF4007\|\|?-><-35exo<-?<-RhoN+RNase+S1\|\|?->?->RADICAL-SAM+TRAM-><-Glyco_hydro_88 | 1145 | Bacteria>FCB_group>Bacteroidetes/Chlorobi_group>Bacteroidota>Bacteroidia>Bacteroidales | GCA_017428995.1 | Prevotella_sp. | IJK42_12575 | 65363..68800 | 59823 | hypothetical-protein-[Prevotella-sp.] |
| MBQ7062982.1 | Type4.PglZ.containing.BREX | <-NTP_transferase+MannoseP_isomer<-?<-?<-?\|\|?->PAPS-reduct+a+b-domain+Fer4_9+DUF4007->wHTH->CR-ATPase5+REase->DUF6079*->iSTAND+PglZ->SIGMA+OB-Fold+AAA-ATPase-BrxL-><-?\|\|?->RelE-ParE-> | 1153 | Bacteria>FCB_group>Bacteroidetes/Chlorobi_group>Bacteroidota>Bacteroidia>Bacteroidales | GCA_017442615.1 | Bacteroidales_bacterium | IJM88_06815 | 35695..39156 | 2030927 | hypothetical-protein-[Bacteroidales-bacterium] |
| MBQ6741271.1 | Type4.PglZ.containing.BREX | HTH_3->Couple_hipA+HipA_C->?->HTH_3->PAPS-reduct+a+b-domain+Fer4_9+DUF4007->DUF6079*->iSTAND+PglZ->SIGMA+OB-Fold+AAA-ATPase-BrxL->NACHT+SNaCT->HTH+REase->CR-ATPase5+REase-> | 1157 | Bacteria>FCB_group>Bacteroidetes/Chlorobi_group>Bacteroidota>Bacteroidia>Bacteroidales | GCA_017442975.1 | Bacteroidales_bacterium | IJR04_01610 | 122369..125842 | 2030927 | hypothetical-protein-[Bacteroidales-bacterium] |
| MBQ7238833.1 | Type4.PglZ.containing.BREX | SIGMA-HTH->?->?-><-CR-ATPase5+REase\|\|PAPS-reduct+DUF4007->DUF6079*->iSTAND+PglZ->SIGMA+OB-Fold+AAA-ATPase-BrxL->SeqA->SeqA->wHTH->TRPR-HTH-> | 1143 | Bacteria>FCB_group>Bacteroidetes/Chlorobi_group>Bacteroidota>Bacteroidia>Bacteroidales | GCA_017445605.1 | Bacteroidales_bacterium | IJS20_08580 | 20484..23915 | 2030927 | hypothetical-protein-[Bacteroidales-bacterium] |
| MBR0038055.1 | Type4.PglZ.containing.BREX | <-Schlafen+HATPase+wHTH<-SeqA<-SIGMA+OB-Fold+AAA-ATPase-BrxL<-iSTAND+PglZ<-DUF6079*<-PAPS-reduct+a+b-domain+Fer4_9+DUF4007<-DOC+HTH | 1139 | Bacteria>FCB_group>Bacteroidetes/Chlorobi_group>Bacteroidota>Bacteroidia>Bacteroidales | GCA_017449065.1 | Bacteroidales_bacterium | IJP70_10530 | 41390..44809 | 2030927 | hypothetical-protein-[Bacteroidales-bacterium] |
| MBQ7423401.1 | Type4.PglZ.containing.BREX | Peptidase->?->?-><-?<-?\|\|?->?->PAPS-reduct+a+b-domain+Fer4_9+DUF4007->DUF6079*->iSTAND+PglZ->SIGMA+OB-Fold+AAA-ATPase-BrxL->wHTH->SeqA-><-CR-ATPase5+REase<-CR-ATPase5+REase\|\|CR-ATPase5+REase->REase->CR-ATPase5-> | 1143 | Bacteria>FCB_group>Bacteroidetes/Chlorobi_group>Bacteroidota>Bacteroidia>Bacteroidales | GCA_017449665.1 | Prevotella_sp. | IJV19_01510 | 35528..38959 | 59823 | hypothetical-protein-[Prevotella-sp.] |
| MBQ7712017.1 | Type4.PglZ.containing.BREX | <-Schlafen+HATPase+wHTH<-Schlafen+HATPase+wHTH<-SF1-SF2-Helicase<-OLD-ABC+TOPRIM<-?<-Cysteine-Desulfurase<-ZnR<-SIGMA+OB-Fold+AAA-ATPase-BrxL<-iSTAND+PglZ<-DUF6079*<-PAPS-reduct+a+b-domain+Fer4_9+DUF4007<-?<-HTH_3\|\|MNS+REase->CR-ATPase5+REase-> | 1392 | Bacteria>FCB_group>Bacteroidetes/Chlorobi_group>Bacteroidota>Bacteroidia>Bacteroidales | GCA_017449975.1 | Bacteroidales_bacterium | IJT39_09370 | 54337..58515 | 2030927 | hypothetical-protein-[Bacteroidales-bacterium] |
| MBQ7361758.1 | Type4.PglZ.containing.BREX | <-DndE<-DndD<-RNase<-HKD-DNase+SF1-SF2-Helicase-p<-SIGMA+OB-Fold+AAA-ATPase-BrxL<-iSTAND+PglZ<-DUF6079*<-wHTH<-PAPS-reduct+a+b-domain+Fer4_9+DUF4007<-HTH_3<-?<-?<-AbiEii<-wHTH+AbiEi_2<-?\|\|HTH_3-> | 1137 | Bacteria>FCB_group>Bacteroidetes/Chlorobi_group>Bacteroidota>Bacteroidia>Bacteroidales | GCA_017452965.1 | Bacteroidaceae_bacterium | IJW68_04630 | 10917..14330 | 2212467 | hypothetical-protein-[Bacteroidaceae-bacterium] |
| MBQ9191708.1 | Type4.PglZ.containing.BREX | <-SeqA<-SIGMA+OB-Fold+AAA-ATPase-BrxL<-iSTAND+PglZ<-DUF6079*<-PAPS-reduct+a+b-domain+Fer4_9+DUF4007<-?\|\|?->RADICAL-SAM->?->?->ADH_N+ADH_zinc_N->YTCD-HTH-> | 1144 | Bacteria>FCB_group>Bacteroidetes/Chlorobi_group>Bacteroidota>Bacteroidia>Bacteroidales | GCA_017459245.1 | Bacteroidaceae_bacterium | IJ142_08980 | 7614..11048 | 2212467 | hypothetical-protein-[Bacteroidaceae-bacterium] |
| MBQ8968316.1 | Type4.PglZ.containing.BREX | TraJ-RHH->?->?->ParA-Soj-PloopNTPase->CopG/RHH->Classical-AAA->Peptidase_S8->PAPS-reduct+a+b-domain+Fer4_9+DUF4007->DUF6079*->iSTAND+PglZ->SIGMA+OB-Fold+AAA-ATPase-BrxL->wHTH->SeqA->?-><-REase | 1140 | Bacteria>FCB_group>Bacteroidetes/Chlorobi_group>Bacteroidota>Bacteroidia>Bacteroidales | GCA_017464345.1 | Bacteroidaceae_bacterium | IJ064_01080 | 10715..14137 | 2212467 | hypothetical-protein-[Bacteroidaceae-bacterium] |
| MBQ8722027.1 | Type4.PglZ.containing.BREX | <-REC+OmpR-HTH<-HISKIN<-?<-?\|\|METHYLASE->?->?->PAPS-reduct+DUF4007->wHTH->DUF6079*->iSTAND+PglZ->SIGMA+OB-Fold+AAA-ATPase-BrxL->?-><-?<-SIGMA-HTH | 1151 | Bacteria>FCB_group>Bacteroidetes/Chlorobi_group>Bacteroidota>Bacteroidia>Bacteroidales | GCA_017467185.1 | Paludibacteraceae_bacterium | IJY67_07785 | 35980..39435 | 2748031 | hypothetical-protein-[Paludibacteraceae-bacterium] |
| MBQ9309284.1 | Type4.PglZ.containing.BREX | <-AT-HTH-HYD2<-Tox-ART-HYD2<-AT-HTH-HYD2<-DndB<-URI-DNase<-SIGMA+OB-Fold+AAA-ATPase-BrxL<-iSTAND+PglZ<-DUF6079*<-wHTH<-PAPS-reduct+a+b-domain+Fer4_9+DUF4007<-SIGMA-HTH<-?<-?<-Schlafen+HATPase+wHTH<-Phage_integrase\|\|?->SF1-SF2-Helicase-> | 1135 | Bacteria>FCB_group>Bacteroidetes/Chlorobi_group>Bacteroidota>Bacteroidia>Bacteroidales | GCA_017469425.1 | Bacteroidales_bacterium | IJ222_00280 | 59502..62909 | 2030927 | hypothetical-protein-[Bacteroidales-bacterium] |
| MBR1683401.1 | Type4.PglZ.containing.BREX | NTP_transf+HDRnase-><-?<-?\|\|?->?->PAPS-reduct+a+b-domain+Fer4_9+DUF4007->wHTH->DUF6079*->iSTAND+PglZ->SIGMA+OB-Fold+AAA-ATPase-BrxL-> | 1140 | Bacteria>FCB_group>Bacteroidetes/Chlorobi_group>Bacteroidota>Bacteroidia>Bacteroidales | GCA_017470425.1 | Bacteroidaceae_bacterium | IJ700_08660 | 141228..144650 | 2212467 | hypothetical-protein-[Bacteroidaceae-bacterium] |
| MBR1595281.1 | Type4.PglZ.containing.BREX | DUF6079-p*->iSTAND+PglZ->SIGMA+OB-Fold+AAA-ATPase-BrxL->URI-DNase->DndB->KAP-NTPase->AbiGi+seqA+RAMA-><-tRNA\|\|?->HAD+HAD-> | 590 | Bacteria>FCB_group>Bacteroidetes/Chlorobi_group>Bacteroidota>Bacteroidia>Bacteroidales | GCA_017470885.1 | Phocaeicola_sp. | IJ665_01115 | 1..1774 | 2773926 | hypothetical-protein-[Phocaeicola-sp.] |
| MBR2184414.1 | Type4.PglZ.containing.BREX | <-Acyl_transf_3<-Glyco_hydro_43<-HTH_3\|\|ABhydrolase-><-?<-SIGMA+OB-Fold+AAA-ATPase-BrxL<-iSTAND+PglZ<-DUF6079*<-wHTH<-PAPS-reduct+a+b-domain+Fer4_9+DUF4007\|\|?->?->REase+RNase-> | 1134 | Bacteria>FCB_group>Bacteroidetes/Chlorobi_group>Bacteroidota>Bacteroidia>Bacteroidales | GCA_017475005.1 | Prevotella_sp. | IJ897_03200 | 10144..13548 | 59823 | hypothetical-protein-[Prevotella-sp.] |
| MBR8703686.1 | Type4.PglZ.containing.BREX | <-ACET<-?<-?<-ACET<-DUF1016+REase<-CR-ATPase5+REase<-SIGMA+OB-Fold+AAA-ATPase-BrxL<-iSTAND+PglZ<-DUF6079*<-CR-ATPase5+REase<-wHTH<-PAPS-reduct+a+b-domain+Fer4_9+DUF4007<-CR-ATPase5+REase\|\|?-><-SIR2<-?<-?<-RNase | 1154 | Bacteria>FCB_group>Bacteroidetes/Chlorobi_group>Bacteroidota>Bacteroidia>Bacteroidales | GCA_017480785.1 | Bacteroidales_bacterium | IJ524_04875 | 122008..125472 | 2030927 | hypothetical-protein-[Bacteroidales-bacterium] |
| MBQ7786644.1 | Type4.PglZ.containing.BREX | <-ABhydrolase<-ABhydrolase\|\|?->ABhydrolase->?->?->DUF6079-p*->iSTAND+PglZ->SIGMA+OB-Fold+AAA-ATPase-BrxL-><-RADICAL-SAM<-Peptidase | 473 | Bacteria>FCB_group>Bacteroidetes/Chlorobi_group>Bacteroidota>Bacteroidia>Bacteroidales | GCA_017482855.1 | Alistipes_sp. | IJ378_00570 | 111105..112526 | 1872444 | hypothetical-protein-[Alistipes-sp.] |
| MBR3434196.1 | Type4.PglZ.containing.BREX | <-Glyco_hydro_92N+Glyco_hydro_92<-?<-?<-METHYLASE<-?\|\|tRNA->HTH_3->PAPS-reduct+a+b-domain+Fer4_9+DUF4007->DUF6079*->iSTAND+PglZ->SIGMA+OB-Fold+AAA-ATPase-BrxL->SeqA->SeqA->SeqA->wHTH->?->Peptidase->Schlafen+HATPase+wHTH-> | 1141 | Bacteria>FCB_group>Bacteroidetes/Chlorobi_group>Bacteroidota>Bacteroidia>Bacteroidales | GCA_017509465.1 | Bacteroidaceae_bacterium | IKH05_11770 | 13465..16890 | 2212467 | hypothetical-protein-[Bacteroidaceae-bacterium] |
| MBR3078191.1 | Type4.PglZ.containing.BREX | REase->LRR_5+LRR_5+LRR_5+LRR_5->HTH_3->PAPS-reduct+a+b-domain+Fer4_9+DUF4007->wHTH->DUF6079*->iSTAND+PglZ->SIGMA+OB-Fold+AAA-ATPase-BrxL->RNase->DndD->DndE->SF1-SF2-Helicase+PRTase->?->?->NUDIX-> | 1146 | Bacteria>FCB_group>Bacteroidetes/Chlorobi_group>Bacteroidota>Bacteroidia>Bacteroidales | GCA_017509495.1 | Prevotella_sp. | IKH01_00060 | 8883..12323 | 59823 | hypothetical-protein-[Prevotella-sp.] |
| MBR6930698.1 | Type4.PglZ.containing.BREX | <-TOPRIM<-SbcC<-SIGMA+OB-Fold+AAA-ATPase-BrxL<-iSTAND+PglZ<-DUF6079*<-PAPS-reduct+a+b-domain+Fer4_9+DUF4007<-SIR2 | 1188 | Bacteria>FCB_group>Bacteroidetes/Chlorobi_group>Bacteroidota>Bacteroidia>Bacteroidales | GCA_017519875.1 | Bacteroidales_bacterium | IKH61_10820 | 63853..67419 | 2030927 | hypothetical-protein-[Bacteroidales-bacterium] |
| MBR3710350.1 | Type4.PglZ.containing.BREX | SIR2-><-RelE-ParE<-?\|\|?->?->?-><-?\|\|PAPS-reduct+DUF4007->DUF6079*->iSTAND+PglZ->SIGMA+OB-Fold+AAA-ATPase-BrxL->SIGMA-HTH->?->wHTH-> | 1160 | Bacteria>FCB_group>Bacteroidetes/Chlorobi_group>Bacteroidota>Bacteroidia>Bacteroidales | GCA_017530365.1 | Bacteroidales_bacterium | IKM99_05240 | 9571..13053 | 2030927 | hypothetical-protein-[Bacteroidales-bacterium] |
| MBR6041787.1 | Type4.PglZ.containing.BREX | <-SF1-SF2-Helicase+TUDOR<-?<-?<-?\|\|CR-ATPase8+REase->PAPS-reduct+a+b-domain+Fer4_9+DUF4007->wHTH->DUF6079*->iSTAND+PglZ->SIGMA+OB-Fold+AAA-ATPase-BrxL->?->REase-><-PRTase\|\|Aminotran_1_2-> | 1147 | Bacteria>FCB_group>Bacteroidetes/Chlorobi_group>Bacteroidota>Bacteroidia>Bacteroidales | GCA_017534895.1 | Paludibacteraceae_bacterium | IKP37_04040 | 49742..53185 | 2748031 | hypothetical-protein-[Paludibacteraceae-bacterium] |
| MBR6489767.1 | Type4.PglZ.containing.BREX | PAPS-reduct+DUF4007->wHTH->DUF6079*->iSTAND+PglZ->SIGMA+OB-Fold+AAA-ATPase-BrxL->RNase->DndD->DndD->DndE-><-HNH<-?<-SbcC | 1138 | Bacteria>FCB_group>Bacteroidetes/Chlorobi_group>Bacteroidota>Bacteroidia>Bacteroidales | GCA_017538045.1 | Muribaculaceae_bacterium | IKT03_04450 | 4842..8258 | 2498093 | hypothetical-protein-[Muribaculaceae-bacterium] |
| MBR5214693.1 | Type4.PglZ.containing.BREX | <-Schlafen+HATPase+wHTH<-NACHT+SNaCT<-Phage_int_SAM_5+Phage_integrase<-OLD-ABC+TOPRIM<-SeqA<-SIGMA+OB-Fold+AAA-ATPase-BrxL<-iSTAND+PglZ<-DUF6079*<-PAPS-reduct+a+b-domain+Fer4_9+DUF4007 | 1143 | Bacteria>FCB_group>Bacteroidetes/Chlorobi_group>Bacteroidota>Bacteroidia>Bacteroidales | GCA_017549605.1 | Bacteroidales_bacterium | IKV80_00540 | 11509..14940 | 2030927 | hypothetical-protein-[Bacteroidales-bacterium] |
| MBR5216328.1 | Type4.PglZ.containing.BREX | <-Glyco_hydro_3\|\|?->Peptidase_M13_N+Peptidase_M13-><-?\|\|Calcineurin->?->HTH_3->PAPS-reduct+a+b-domain+Fer4_9+DUF4007->wHTH->DUF6079*->iSTAND+PglZ->SIGMA+OB-Fold+AAA-ATPase-BrxL->URI-DNase->DndB->?->?->OLD-ABC->TOPRIM-> | 1146 | Bacteria>FCB_group>Bacteroidetes/Chlorobi_group>Bacteroidota>Bacteroidia>Bacteroidales | GCA_017549605.1 | Bacteroidales_bacterium | IKV80_08945 | 9711..13151 | 2030927 | hypothetical-protein-[Bacteroidales-bacterium] |
| MBR5399198.1 | Type4.PglZ.containing.BREX | <-HTH+REase\|\|PAPS-reduct+DUF4007->DUF6079*->iSTAND+PglZ->SIGMA+OB-Fold+AAA-ATPase-BrxL->?->?->?-><-?<-?<-?<-SIGMA-HTH | 1143 | Bacteria>FCB_group>Bacteroidetes/Chlorobi_group>Bacteroidota>Bacteroidia>Bacteroidales | GCA_017550655.1 | Bacteroidales_bacterium | IK103_05335 | 51836..55267 | 2030927 | hypothetical-protein-[Bacteroidales-bacterium] |
| MBR5972234.1 | Type4.PglZ.containing.BREX | PAPS-reduct+a+b-domain+Fer4_9+DUF4007->DUF6079*->iSTAND+PglZ->SIGMA+OB-Fold+AAA-ATPase-BrxL->SIGMA-HTH-> | 1155 | Bacteria>FCB_group>Bacteroidetes/Chlorobi_group>Bacteroidota>Bacteroidia>Bacteroidales | GCA_017551675.1 | Paludibacteraceae_bacterium | IK017_06235 | 4335..7802 | 2748031 | hypothetical-protein-[Paludibacteraceae-bacterium] |
| MBR4838438.1 | Type4.PglZ.containing.BREX | NTP_transf+HDRnase->?->HAD+HAD-><-?<-ARG<-?<-SIGMA+OB-Fold+AAA-ATPase-BrxL<-iSTAND+PglZ<-DUF6079*<-wHTH<-PAPS-reduct+a+b-domain+Fer4_9+DUF4007<-?<-?<-HNH | 1194 | Bacteria>FCB_group>Bacteroidetes/Chlorobi_group>Bacteroidota>Bacteroidia>Bacteroidales | GCA_017553625.1 | Bacteroidales_bacterium | IK004_08435 | 36887..40471 | 2030927 | hypothetical-protein-[Bacteroidales-bacterium] |
| MBR5695142.1 | Type4.PglZ.containing.BREX | METHYLASE->Peptidase_M48_N+Peptidase_M48->RADICAL-SAM+TRAM->Thermonuclease->?->?->PAPS-reduct+a+b-domain+Fer4_9+DUF4007->wHTH->DUF6079*->iSTAND+PglZ->SIGMA+OB-Fold+AAA-ATPase-BrxL-><-?\|\|?->?->?->TetR-HTH-> | 1156 | Bacteria>FCB_group>Bacteroidetes/Chlorobi_group>Bacteroidota>Bacteroidia>Bacteroidales | GCA_017557245.1 | Paludibacteraceae_bacterium | IKX43_02780 | 66059..69529 | 2748031 | hypothetical-protein-[Paludibacteraceae-bacterium] |
| MBR5673318.1 | Type4.PglZ.containing.BREX | RADICAL-SAM->?->?->HEPN->DUF772+DDE_Tnp_1-><-OLD-ABC<-SIGMA+OB-Fold+AAA-ATPase-BrxL<-iSTAND+PglZ<-DUF6079*<-PAPS-reduct+a+b-domain+Fer4_9+DUF4007<-HTH_3<-ZnR+TOPRIM<-ZnR+TOPRIM+AAA-helicase+wHTH<-XIS-HTH<-?\|\|DOC+HTH->?-><-Phage_integrase | 1148 | Bacteria>FCB_group>Bacteroidetes/Chlorobi_group>Bacteroidota>Bacteroidia>Bacteroidales | GCA_017557425.1 | Muribaculaceae_bacterium | IKW97_02875 | 38451..41897 | 2498093 | hypothetical-protein-[Muribaculaceae-bacterium] |
| MBO4331558.1 | Type4.PglZ.containing.BREX | <-ClpABN-AAA+ClpABC-AAA<-?<-?<-Aminotran_1_2\|\|PRTase-><-REase<-?<-SIGMA+OB-Fold+AAA-ATPase-BrxL<-iSTAND+PglZ<-DUF6079*<-wHTH<-PAPS-reduct+a+b-domain+Fer4_9+DUF4007<-CR-ATPase8+REase\|\|?->?->?->SF1-SF2-Helicase+TUDOR-> | 1143 | Bacteria>FCB_group>Bacteroidetes/Chlorobi_group>Bacteroidota>Bacteroidia>Bacteroidales | GCA_017613615.1 | Paludibacteraceae_bacterium | J5875_00130 | 33324..36755 | 2748031 | hypothetical-protein-[Paludibacteraceae-bacterium] |
| MBO4529016.1 | Type4.PglZ.containing.BREX | SH3->?->?->?->?->?->?->?->PAPS-reduct+a+b-domain+Fer4_9+DUF4007->DUF6079*->iSTAND+PglZ->SIGMA+OB-Fold+AAA-ATPase-BrxL->DndE->?->SH3->DOC+HTH-><-NAD_synthase<-?<-Aminotran_1_2 | 1185 | Bacteria>FCB_group>Bacteroidetes/Chlorobi_group>Bacteroidota>Bacteroidia>Bacteroidales | GCA_017616155.1 | Paludibacteraceae_bacterium | J5767_00080 | 11689..15246 | 2748031 | hypothetical-protein-[Paludibacteraceae-bacterium] |
| MBO4818256.1 | Type4.PglZ.containing.BREX | <-HARE-HTH<-?<-?<-wHTH+WYL+WCX<-SIGMA+OB-Fold+AAA-ATPase-BrxL<-iSTAND+PglZ<-DUF6079*<-wHTH<-PAPS-reduct+a+b-domain+Fer4_9+DUF4007<-SIGMA-HTH<-?<-?<-?<-?<-?<-Arm-DNA-bind_5+Phage_int_SAM_5 | 1135 | Bacteria>FCB_group>Bacteroidetes/Chlorobi_group>Bacteroidota>Bacteroidia>Bacteroidales | GCA_017619415.1 | Bacteroidales_bacterium | J5498_08280 | 8666..12073 | 2030927 | hypothetical-protein-[Bacteroidales-bacterium] |
| MBO5133522.1 | Type4.PglZ.containing.BREX | AbiEii->?->?-><-MNS+REase<-?\|\|HTH_3->PAPS-reduct+a+b-domain+Fer4_9+DUF4007->wHTH->DUF6079*->iSTAND+PglZ->SIGMA+OB-Fold+AAA-ATPase-BrxL->DndB->HEPN-><-GNAT<-?<-TraJ-RHH<-ZnR+TOPRIM<-CR-ATPase4 | 1143 | Bacteria>FCB_group>Bacteroidetes/Chlorobi_group>Bacteroidota>Bacteroidia>Bacteroidales | GCA_017622315.1 | Bacteroidaceae_bacterium | J6C15_00045 | 7922..11353 | 2212467 | hypothetical-protein-[Bacteroidaceae-bacterium] |
| MBO5614530.1 | Type4.PglZ.containing.BREX | <-Schlafen+HATPase+wHTH<-TIR<-KAP-NTPase<-DndB<-URI-DNase<-SIGMA+OB-Fold+AAA-ATPase-BrxL<-iSTAND+PglZ<-DUF6079*<-wHTH<-PAPS-reduct+a+b-domain+Fer4_9+DUF4007<-HTH_3\|\|XIS-HTH->?->?-><-Phage_int_SAM_5+Phage_integrase\|\|REase->XIS-HTH-> | 1144 | Bacteria>FCB_group>Bacteroidetes/Chlorobi_group>Bacteroidota>Bacteroidia>Bacteroidales | GCA_017627755.1 | Prevotella_sp. | J5932_00210 | 16056..19490 | 59823 | hypothetical-protein-[Prevotella-sp.] |
| MBO5957873.1 | Type4.PglZ.containing.BREX | ZnR+TOPRIM->TraJ-RHH->?-><-?<-HEPN<-DndB<-SIGMA+OB-Fold+AAA-ATPase-BrxL<-iSTAND+PglZ<-DUF6079*<-wHTH<-PAPS-reduct+a+b-domain+Fer4_9+DUF4007<-HTH_3\|\|?->?-><-Calcineurin<-?<-?<-STAS+HisKin | 1143 | Bacteria>FCB_group>Bacteroidetes/Chlorobi_group>Bacteroidota>Bacteroidia>Bacteroidales | GCA_017631995.1 | Bacteroidales_bacterium | J6Q39_10010 | 9257..12688 | 2030927 | hypothetical-protein-[Bacteroidales-bacterium] |
| MBO6251939.1 | Type4.PglZ.containing.BREX | <-DndE<-DndD<-RNase<-SIGMA+OB-Fold+AAA-ATPase-BrxL<-iSTAND+PglZ<-DUF6079*<-wHTH<-PAPS-reduct+a+b-domain+Fer4_9+DUF4007 | 1136 | Bacteria>FCB_group>Bacteroidetes/Chlorobi_group>Bacteroidota>Bacteroidia>Bacteroidales | GCA_017635355.1 | Muribaculaceae_bacterium | J6N71_11410 | 11513..14923 | 2498093 | hypothetical-protein-[Muribaculaceae-bacterium] |
| MBO7067674.1 | Type4.PglZ.containing.BREX | NUDIX->MNS+REase->CR-ATPase5+REase->?-><-XIS-HTH\|\|HTH_3->PAPS-reduct+a+b-domain+Fer4_9+DUF4007->wHTH->DUF6079*->iSTAND+PglZ->SIGMA+OB-Fold+AAA-ATPase-BrxL->URI-DNase->DndB->Peptidase->Tox-RES->Schlafen+HATPase+wHTH-><-tRNA\|\|?->METHYLASE-> | 1144 | Bacteria>FCB_group>Bacteroidetes/Chlorobi_group>Bacteroidota>Bacteroidia>Bacteroidales | GCA_017645305.1 | Bacteroidaceae_bacterium | J6W52_03195 | 64285..67719 | 2212467 | hypothetical-protein-[Bacteroidaceae-bacterium] |
| MBO7636113.1 | Type4.PglZ.containing.BREX | OLD-ABC->?->PAPS-reduct+a+b-domain+Fer4_9+DUF4007->wHTH->DUF6079*->iSTAND+PglZ->SIGMA+OB-Fold+AAA-ATPase-BrxL->SeqA-><-KAP-NTPase\|\|SIGMA-HTH+MA-> | 1197 | Bacteria>FCB_group>Bacteroidetes/Chlorobi_group>Bacteroidota>Bacteroidia>Bacteroidales | GCA_017651825.1 | Paludibacteraceae_bacterium | J6S89_06000 | 5511..9104 | 2748031 | hypothetical-protein-[Paludibacteraceae-bacterium] |
| MBP3245577.1 | Type4.PglZ.containing.BREX | Arm-DNA-bind_5+Phage_int_SAM_5+Phage_integrase->PAPS-reduct+a+b-domain+Fer4_9+DUF4007->DUF6079*->iSTAND+PglZ->SIGMA+OB-Fold+AAA-ATPase-BrxL->SeqA->SeqA-> | 1149 | Bacteria>FCB_group>Bacteroidetes/Chlorobi_group>Bacteroidota>Bacteroidia>Bacteroidales | GCA_017934435.1 | Bacteroidaceae_bacterium | J6M59_10820 | 4488..7937 | 2212467 | hypothetical-protein-[Bacteroidaceae-bacterium] |
| MBP3256967.1 | Type4.PglZ.containing.BREX | sGTP->XIS-HTH->Arm-DNA-bind_5+Phage_int_SAM_5+Phage_integrase->?->XIS-HTH->?->?->?->PAPS-reduct+a+b-domain+Fer4_9+DUF4007->DUF6079*->iSTAND+PglZ->SIGMA+OB-Fold+AAA-ATPase-BrxL->SIGMA-HTH->PIWI->HARE-HTH-> | 1155 | Bacteria>FCB_group>Bacteroidetes/Chlorobi_group>Bacteroidota>Bacteroidia>Bacteroidales | GCA_017934575.1 | Bacteroidales_bacterium | J6M23_03055 | 94852..98319 | 2030927 | hypothetical-protein-[Bacteroidales-bacterium] |
| MBP3716608.1 | Type4.PglZ.containing.BREX | <-ClpABN-AAA+ClpABC-AAA<-?<-?<-Aminotran_1_2\|\|PRTase-><-REase<-?<-SIGMA+OB-Fold+AAA-ATPase-BrxL<-iSTAND+PglZ<-DUF6079*<-wHTH<-PAPS-reduct+a+b-domain+Fer4_9+DUF4007\|\|?->?->SF1-SF2-Helicase+TUDOR->?->?-><-HAD+HAD | 1147 | Bacteria>FCB_group>Bacteroidetes/Chlorobi_group>Bacteroidota>Bacteroidia>Bacteroidales | GCA_017940035.1 | Paludibacteraceae_bacterium | J6I79_03805 | 359807..363250 | 2748031 | hypothetical-protein-[Paludibacteraceae-bacterium] |
| MBP3741081.1 | Type4.PglZ.containing.BREX | MarR-HTH-><-?<-?\|\|tRNA->Phage_int_SAM_5+Phage_integrase->XIS-HTH->?->?-><-SIGMA+OB-Fold+AAA-ATPase-BrxL<-iSTAND+PglZ<-DUF6079*<-wHTH<-PAPS-reduct+a+b-domain+Fer4_9+DUF4007<-?<-CR-ATPase5+REase<-?<-AraC-HTH+AraC-HTH | 1144 | Bacteria>FCB_group>Bacteroidetes/Chlorobi_group>Bacteroidota>Bacteroidia>Bacteroidales | GCA_017940305.1 | Bacteroidales_bacterium | J6I41_08200 | 65369..68803 | 2030927 | hypothetical-protein-[Bacteroidales-bacterium] |
| MBP8994079.1 | Type4.PglZ.containing.BREX | <-SIGMA+OB-Fold+AAA-ATPase-BrxL<-iSTAND+PglZ<-DUF6079*<-PAPS-reduct+DUF4007 | 1133 | Bacteria>FCB_group>Bacteroidetes/Chlorobi_group>Bacteroidota>Bacteroidia>Bacteroidales | GCA_018056525.1 | Bacteroidales_bacterium | KBG30_09750 | 2779..6180 | 2030927 | hypothetical-protein-[Bacteroidales-bacterium] |
| MBQ0053428.1 | Type4.PglZ.containing.BREX | <-DndE<-DndD<-RNase<-HKD-DNase+SF1-SF2-Helicase-p<-SIGMA+OB-Fold+AAA-ATPase-BrxL<-iSTAND+PglZ<-DUF6079*<-wHTH<-PAPS-reduct+a+b-domain+Fer4_9+DUF4007<-ACET | 1138 | Bacteria>FCB_group>Bacteroidetes/Chlorobi_group>Bacteroidota>Bacteroidia>Bacteroidales | GCA_018065035.1 | Bacteroidales_bacterium | KBS89_03105 | 8485..11901 | 2030927 | hypothetical-protein-[Bacteroidales-bacterium] |
| MBQ0062830.1 | Type4.PglZ.containing.BREX | <-TRPR-HTH<-?<-?<-?<-?\|\|?->PAPS-reduct+a+b-domain+Fer4_9+DUF4007->wHTH->DUF6079*->iSTAND+PglZ->SIGMA+OB-Fold+AAA-ATPase-BrxL->RNase->DndD-><-DDE_Tnp_1_3<-HTH_29+HTH_32+DDE_3 | 1154 | Bacteria>FCB_group>Bacteroidetes/Chlorobi_group>Bacteroidota>Bacteroidia>Bacteroidales | GCA_018065185.1 | Prevotella_sp. | KBS94_02235 | 20608..24072 | 59823 | hypothetical-protein-[Prevotella-sp.] |
| MBQ0159481.1 | Type4.PglZ.containing.BREX | HisKin->REC+LYTTR-><-?\|\|?->REC+OmpR-HTH->?-><-?<-SIGMA+OB-Fold+AAA-ATPase-BrxL<-iSTAND+PglZ<-DUF6079*<-PAPS-reduct+a+b-domain+Fer4_9+DUF4007 | 1138 | Bacteria>FCB_group>Bacteroidetes/Chlorobi_group>Bacteroidota>Bacteroidia>Bacteroidales | GCA_018066145.1 | Bacteroidales_bacterium | KBT28_02495 | 19278..22694 | 2030927 | hypothetical-protein-[Bacteroidales-bacterium] |
| MBS5552423.1 | Type4.PglZ.containing.BREX | AraC-HTH->HMA+HAD+HAD-><-?<-SIGMA+OB-Fold+AAA-ATPase-BrxL<-iSTAND+PglZ<-DUF6079*<-PAPS-reduct+a+b-domain+Fer4_9+DUF4007 | 1151 | Bacteria>FCB_group>Bacteroidetes/Chlorobi_group>Bacteroidota>Bacteroidia>Bacteroidales | GCA_018369195.1 | Bacteroides_sp. | KHX53_10345 | 8275..11730 | 29523 | hypothetical-protein-[Bacteroides-sp.] |
| MBS6966832.1 | Type4.PglZ.containing.BREX | TRPR-HTH->MNS+HTH->?->TRPR-HTH->STYKinase-like->?->PAPS-reduct+a+b-domain+Fer4_9+DUF4007->DUF6079*->iSTAND+PglZ->SIGMA+OB-Fold+AAA-ATPase-BrxL->ZnR->Cysteine-Desulfurase->?-><-?<-Relaxase<-TraJ-RHH<-SMS-KaiC | 1395 | Bacteria>FCB_group>Bacteroidetes/Chlorobi_group>Bacteroidota>Bacteroidia>Bacteroidales | GCA_018381015.1 | Bacteroides_sp. | KH193_15175 | 27111..31298 | 29523 | hypothetical-protein-[Bacteroides-sp.] |
| MBS7314725.1 | Type4.PglZ.containing.BREX | <-DOC+HTH<-?<-SIGMA+OB-Fold+AAA-ATPase-BrxL<-iSTAND+PglZ<-DUF6079*<-wHTH<-PAPS-reduct+a+b-domain+Fer4_9+DUF4007<-?<-SbcC<-?<-?<-?<-S4+PSYN | 1179 | Bacteria>FCB_group>Bacteroidetes/Chlorobi_group>Bacteroidota>Bacteroidia>Bacteroidales | GCA_018384335.1 | Bacteroidales_bacterium | KIG54_08120 | 6238..9777 | 2030927 | hypothetical-protein-[Bacteroidales-bacterium] |
| MBZ4674435.1 | Type4.PglZ.containing.BREX | DDE_Tnp_1->HTH->HNH+SeqA->PAPS-reduct+a+b-domain+Fer4_9+DUF4007->DUF6079*->iSTAND+PglZ->SIGMA+OB-Fold+AAA-ATPase-BrxL->DEOR-HTH-> | 1146 | Bacteria>FCB_group>Bacteroidetes/Chlorobi_group>Bacteroidota>Bacteroidia>Bacteroidales | GCA_020055075.1 | Dysgonamonadaceae_bacterium | JG772_123 | 6963..10403 | 2065189 | hypothetical-protein-[Dysgonamonadaceae-bacterium] |
| HJH39685.1 | Type4.PglZ.containing.BREX | HTH_26->?->PAPS-reduct+a+b-domain+Fer4_9+DUF4007->DUF6079*->iSTAND+PglZ->SIGMA+OB-Fold+AAA-ATPase-BrxL->ZnR->Cysteine-Desulfurase-><-HTH_26 | 1379 | Bacteria>FCB_group>Bacteroidetes/Chlorobi_group>Bacteroidota>Bacteroidia>Bacteroidales | GCA_020809995.1 | Bacteroides_ovatus | K8V76_10925 | 6615..10754 | 28116 | hypothetical-protein-[Bacteroides-ovatus] |
| MCD8290137.1 | Type4.PglZ.containing.BREX | HTH_3->PAPS-reduct+a+b-domain+Fer4_9+DUF4007->DUF6079*->iSTAND+PglZ->SIGMA+OB-Fold+AAA-ATPase-BrxL->SeqA-> | 1144 | Bacteria>FCB_group>Bacteroidetes/Chlorobi_group>Bacteroidota>Bacteroidia>Bacteroidales | GCA_021204735.1 | Prevotella_sp. | LUC91_01385 | 3195..6629 | 59823 | hypothetical-protein-[Prevotella-sp.] |
| MCD8306336.1 | Type4.PglZ.containing.BREX | HKD-DNase+HKD-DNase->HISKIN->?->?->?-><-Peptidase_M6\|\|tRNA->PAPS-reduct+DUF4007->wHTH->DUF6079*->iSTAND+PglZ->SIGMA+OB-Fold+AAA-ATPase-BrxL-><-?<-Relaxase<-TraJ-RHH<-?<-XIS-HTH | 1143 | Bacteria>FCB_group>Bacteroidetes/Chlorobi_group>Bacteroidota>Bacteroidia>Bacteroidales | GCA_021204905.1 | Prevotella_sp. | LUC49_06760 | 87929..91360 | 59823 | hypothetical-protein-[Prevotella-sp.] |
| MCH4147877.1 | Type4.PglZ.containing.BREX | <-TRPR-HTH<-?<-ParA-Soj-PloopNTPase\|\|HTH_3->PAPS-reduct+a+b-domain+Fer4_9+DUF4007->wHTH->DUF6079*->iSTAND+PglZ->SIGMA+OB-Fold+AAA-ATPase-BrxL->OLD-ABC+TOPRIM->SF1-SF2-Helicase->DDE_Tnp_1-> | 1131 | Bacteria>FCB_group>Bacteroidetes/Chlorobi_group>Bacteroidota>Bacteroidia>Bacteroidales | GCA_022484605.1 | Prevotella_sp. | LKF33_05565 | 19991..23386 | 59823 | hypothetical-protein-[Prevotella-sp.] |
| MCH4183502.1 | Type4.PglZ.containing.BREX | <-DndE<-DndD<-RNase<-SIGMA+OB-Fold+AAA-ATPase-BrxL<-iSTAND+PglZ<-DUF6079*<-wHTH<-PAPS-reduct+a+b-domain+Fer4_9+DUF4007 | 1144 | Bacteria>FCB_group>Bacteroidetes/Chlorobi_group>Bacteroidota>Bacteroidia>Bacteroidales | GCA_022486515.1 | Prevotella_sp. | LKF48_10145 | 7616..11050 | 59823 | hypothetical-protein-[Prevotella-sp.] |
| MCI6048552.1 | Type4.PglZ.containing.BREX | CR-ATPase8+REase-><-?<-Peptidase_C69\|\|?-><-SIGMA+OB-Fold+AAA-ATPase-BrxL<-iSTAND+PglZ<-DUF6079*<-wHTH<-PAPS-reduct+a+b-domain+Fer4_9+DUF4007\|\|wHTH->REase-> | 1151 | Bacteria>FCB_group>Bacteroidetes/Chlorobi_group>Bacteroidota>Bacteroidia>Bacteroidales | GCA_022770845.1 | Prevotella_pectinovora | MR736_08345 | 11854..15309 | 1602169 | hypothetical-protein-[Prevotella-pectinovora] |
| MCI6106023.1 | Type4.PglZ.containing.BREX | PAPS-reduct+a+b-domain+Fer4_9+DUF4007->wHTH->DUF6079*->iSTAND+PglZ->SIGMA+OB-Fold+AAA-ATPase-BrxL->SeqA-><-?<-?<-?<-Thioredoxin | 1147 | Bacteria>FCB_group>Bacteroidetes/Chlorobi_group>Bacteroidota>Bacteroidia>Bacteroidales | GCA_022771345.1 | Prevotella_sp. | MR703_04050 | 3960..7403 | 59823 | hypothetical-protein-[Prevotella-sp.] |
| MCI6428296.1 | Type4.PglZ.containing.BREX | <-SeqA<-SIGMA+OB-Fold+AAA-ATPase-BrxL<-iSTAND+PglZ<-DUF6079*<-PAPS-reduct+a+b-domain+Fer4_9+DUF4007 | 1141 | Bacteria>FCB_group>Bacteroidetes/Chlorobi_group>Bacteroidota>Bacteroidia>Bacteroidales | GCA_022777205.1 | Rikenellaceae_bacterium | MR809_05345 | 6027..9452 | 2049048 | hypothetical-protein-[Rikenellaceae-bacterium] |

| Protein-ID | System | Gene-Neighbourhoods | Length | Taxonomy | Assembly-ID | Strain | Locus-Tag | Genomic-coordinates | TaxID | Fasta-define |
|---|---|---|---|---|---|---|---|---|---|---|
| MCI6342615.1 | Type4.PglZ.containing.BREX | TOPRIM+Topoisom_bac-><-?<-REC+PglZ<-?<-?<-?<-?<-wHTH\|\|DUF4007->DUF6079*->PAPS-reduct->HKD-DNase+SF1-SF2-Helicase->?->DndD->DndE-><-?<-?<-?<-Thioredoxin | 1073 | Bacteria>FCB_group>Bacteroidetes/Chlorobi_group>Bacteroidota>Bacteroidia>Bacteroidales | GCA_022777695.1 | Prevotella_sp. | MR788_08715 | 48810..52031 | 59823 | ATP-binding-protein-[Prevotella-sp.] |
| MCI7182580.1 | Type4.PglZ.containing.BREX | <-Thioredoxin\|\|LexA-protease->?->ACET-><-SeqA<-SIGMA+OB-Fold+AAA-ATPase-BrxL<-iSTAND+PglZ<-DUF6079*\|\|?->?-><-TRD+TRD<-?<-Four-helical-protein<-REase+SF1-SF2-Helicase<-Thr_synth_N+PALP<-?<-AA_kinase+ACT_7 | 1147 | Bacteria>FCB_group>Bacteroidetes/Chlorobi_group>Bacteroidota>Bacteroidia>Bacteroidales | GCA_022782505.1 | Prevotella_sp. | MSA05_00150 | 35510..38953 | 59823 | hypothetical-protein-[Prevotella-sp.] |
| MCI7370543.1 | Type4.PglZ.containing.BREX | <-transposase<-SIGMA+OB-Fold+AAA-ATPase-BrxL<-SIGMA+OB-Fold+AAA-ATPase-BrxL<-iSTAND+PglZ<-DUF6079*<-wHTH<-PAPS-reduct+a+b-domain+Fer4_9+DUF4007<-?<-CR-ATPase5+REase<-RNase<-SbcC<-JetB<-?\|\|RNase-> | 1161 | Bacteria>FCB_group>Bacteroidetes/Chlorobi_group>Bacteroidota>Bacteroidia>Bacteroidales | GCA_022784205.1 | Prevotella_sp. | MSH42_01135 | 4304..7789 | 59823 | hypothetical-protein-[Prevotella-sp.] |
| MCI7376845.1 | Type4.PglZ.containing.BREX | <-FURR-HTH\|\|PAPS-reduct+DUF4007->DUF6079*->iSTAND+PglZ->SIGMA+OB-Fold+AAA-ATPase-BrxL-> | 1137 | Bacteria>FCB_group>Bacteroidetes/Chlorobi_group>Bacteroidota>Bacteroidia>Bacteroidales | GCA_022784405.1 | Bacteroidales_bacterium | MSH43_02240 | 22233..25646 | 2030927 | hypothetical-protein-[Bacteroidales-bacterium] |
| MCI7482635.1 | Type4.PglZ.containing.BREX | <-SeqA<-SIGMA+OB-Fold+AAA-ATPase-BrxL<-iSTAND+PglZ<-DUF6079*<-wHTH<-PAPS-reduct+a+b-domain+Fer4_9+DUF4007\|\|?->?->?->REC+PglZ->?-><-TOPRIM+Topoisom_bac | 1147 | Bacteria>FCB_group>Bacteroidetes/Chlorobi_group>Bacteroidota>Bacteroidia>Bacteroidales | GCA_022785245.1 | Prevotella_sp. | MSA97_01395 | 5486..8929 | 59823 | hypothetical-protein-[Prevotella-sp.] |
| MCI7465921.1 | Type4.PglZ.containing.BREX | PAPS-reduct+a+b-domain+Fer4_9+DUF4007->wHTH->DUF6079*->iSTAND+PglZ->SIGMA+OB-Fold+AAA-ATPase-BrxL->?->DOC+HTH-><-?<-RADICAL-SAM<-?<-?<-MMR_HSR1+sGTP+sGTP | 1179 | Bacteria>FCB_group>Bacteroidetes/Chlorobi_group>Bacteroidota>Bacteroidia>Bacteroidales | GCA_022785405.1 | Bacteroidales_bacterium | MSB01_06485 | 2594..6133 | 2030927 | hypothetical-protein-[Bacteroidales-bacterium] |
| MCI7634857.1 | Type4.PglZ.containing.BREX | SIGMA-HTH+LuxR-HTH->?->HATPase+wHTH->?-><-?\|\|SIGMA-HTH->PAPS-reduct+a+b-domain+Fer4_9+DUF4007->wHTH->DUF6079*->iSTAND+PglZ->SIGMA+OB-Fold+AAA-ATPase-BrxL-><-tRNA\|\|?->?-><-Glyco_hydro_3\|\|?-><-Peptidase_S46+Trypsin_2 | 1135 | Bacteria>FCB_group>Bacteroidetes/Chlorobi_group>Bacteroidota>Bacteroidia>Bacteroidales | GCA_022787815.1 | Bacteroidales_bacterium | MSS78_04230 | 20004..23411 | 2030927 | hypothetical-protein-[Bacteroidales-bacterium] |
| MCM1139530.1 | Type4.PglZ.containing.BREX | <-ADH_N+ADH_zinc_N+METHYLASE<-?<-TIR<-Cysteine-Desulfurase<-ZnR<-SIGMA+OB-Fold+AAA-ATPase-BrxL<-iSTAND+PglZ<-DUF6079*<-PAPS-reduct+a+b-domain+Fer4_9+DUF4007<-?<-Phage_lysozyme | 1386 | Bacteria>FCB_group>Bacteroidetes/Chlorobi_group>Bacteroidota>Bacteroidia>Bacteroidales | GCA_023663885.1 | Muribaculum_sp. | NC453_13245 | 8928..13088 | 1918611 | hypothetical-protein-[Muribaculum-sp.] |
| MCM1373150.1 | Type4.PglZ.containing.BREX | HTH_3->PAPS-reduct+a+b-domain+Fer4_9+DUF4007->DUF6079*->iSTAND+PglZ->SIGMA+OB-Fold+AAA-ATPase-BrxL->?->Schlafen+HATPase+wHTH->?-><-?<-?<-ParBC | 1137 | Bacteria>FCB_group>Bacteroidetes/Chlorobi_group>Bacteroidota>Bacteroidia>Bacteroidales | GCA_023665805.1 | Bacteroides_sp. | NC208_09330 | 3566..6979 | 29523 | hypothetical-protein-[Bacteroides-sp.] |
| MCM1529336.1 | Type4.PglZ.containing.BREX | BrxA-DNA-binding->Phage_integrase+wHTH->PAPS-reduct+a+b-domain+Fer4_9+DUF4007->DUF6079*->iSTAND+PglZ->SIGMA+OB-Fold+AAA-ATPase-BrxL->ZnR->Cysteine-Desulfurase-><-?<-TRAM+METHYLASE<-RADICAL-SAM<-?<-LRR_5+LRR_5+LRR_5+LRR_5 | 1400 | Bacteria>FCB_group>Bacteroidetes/Chlorobi_group>Bacteroidota>Bacteroidia>Bacteroidales | GCA_023667265.1 | Alistipes_sp. | NC093_04995 | 1112262..1116464 | 1872444 | hypothetical-protein-[Alistipes-sp.] |
| MCQ2337979.1 | Type4.PglZ.containing.BREX | <-iSTAND+PglZ<-DUF6079*<-wHTH<-PAPS-reduct+a+b-domain+Fer4_9+DUF4007<-?<-PolB<-sGTP<-?<-DUF4369+Thioredoxin_8<-?\|\|TPR+TPR-> | 1139 | Bacteria>FCB_group>Bacteroidetes/Chlorobi_group>Bacteroidota>Bacteroidia>Bacteroidales | GCA_024399515.1 | Paludibacteraceae_bacterium | MJ001_03510 | 1053..4472 | 2748031 | hypothetical-protein-[Paludibacteraceae-bacterium] |
| MCQ2340465.1 | Type4.PglZ.containing.BREX | <-D5ATPase+DUF3874<-sGTP<-?<-SIGMA+OB-Fold+AAA-ATPase-BrxL<-iSTAND+PglZ<-DUF6079*<-PAPS-reduct+a+b-domain+Fer4_9+DUF4007<-tRNA<-?<-Schlafen+HATPase+wHTH<-?<-?<-?<-Peptidase_M20<-?\|\|REC+OmpR-HTH-> | 1153 | Bacteria>FCB_group>Bacteroidetes/Chlorobi_group>Bacteroidota>Bacteroidia>Bacteroidales | GCA_024399955.1 | Paludibacteraceae_bacterium | MJZ79_06790 | 9684..13145 | 2748031 | hypothetical-protein-[Paludibacteraceae-bacterium] |
| MCQ2306876.1 | Type4.PglZ.containing.BREX | REase->?->Peptidase_M20->HTH+REase->HNH+SeqA->PAPS-reduct+a+b-domain+Fer4_9+DUF4007->Schlafen+HATPase+wHTH->DUF6079*->iSTAND+PglZ->SIGMA+OB-Fold+AAA-ATPase-BrxL-><-?\|\|CR-ATPase5+REase-><-?<-?\|\|TPR+TPR+TPR+TPR-><-RADICAL-SAM+TRAM | 1184 | Bacteria>FCB_group>Bacteroidetes/Chlorobi_group>Bacteroidota>Bacteroidia>Bacteroidales | GCA_024400305.1 | Bacteroidales_bacterium | MJ000_04835 | 42478..46032 | 2030927 | hypothetical-protein-[Bacteroidales-bacterium] |
| MCQ2302016.1 | Type4.PglZ.containing.BREX | <-Schlafen+HATPase+wHTH<-SeqA<-wHTH(SBD)+HNH<-SIGMA+OB-Fold+AAA-ATPase-BrxL<-iSTAND+PglZ<-DUF6079*<-PAPS-reduct+a+b-domain+Fer4_9+DUF4007<-?<-TrwF<-wHTH+WYL+WCX\|\|?->?->?-><-LRR_5+LRR_5+LRR_5+LRR_5 | 1121 | Bacteria>FCB_group>Bacteroidetes/Chlorobi_group>Bacteroidota>Bacteroidia>Bacteroidales | GCA_024400425.1 | Bacteroidales_bacterium | MJZ94_05255 | 16013..19378 | 2030927 | hypothetical-protein-[Bacteroidales-bacterium] |
| MCQ2228644.1 | Type4.PglZ.containing.BREX | <-ABhydrolase<-?<-?<-?\|\|?-><-?<-SIGMA+OB-Fold+AAA-ATPase-BrxL<-iSTAND+PglZ<-DUF6079*<-PAPS-reduct+a+b-domain+Fer4_9+DUF4007<-TOPRIM<-OLD-ABC<-Calcineurin<-Polbeta<-MarR-HTH\|\|?->GH3->HAD-> | 1149 | Bacteria>FCB_group>Bacteroidetes/Chlorobi_group>Bacteroidota>Bacteroidia>Bacteroidales | GCA_024401335.1 | Bacteroidales_bacterium | MJZ13_02730 | 163288..166737 | 2030927 | hypothetical-protein-[Bacteroidales-bacterium] |
| MCR4857685.1 | Type4.PglZ.containing.BREX | <-MNS+REase<-?<-SeqA<-SeqA<-SIGMA+OB-Fold+AAA-ATPase-BrxL<-iSTAND+PglZ<-DUF6079*<-PAPS-reduct+a+b-domain+Fer4_9+DUF4007<-?<-RhoN+RNase+S1 | 1167 | Bacteria>FCB_group>Bacteroidetes/Chlorobi_group>Bacteroidota>Bacteroidia>Bacteroidales | GCA_024681675.1 | Bacteroidales_bacterium | K5890_05755 | 7652..11155 | 2030927 | hypothetical-protein-[Bacteroidales-bacterium] |
| MCR4768684.1 | Type4.PglZ.containing.BREX | <-TOPRIM<-OLD-ABC\|\|HTH_3->DUF1016+REase->PAPS-reduct+a+b-domain+Fer4_9+DUF4007->wHTH->DUF6079*->iSTAND+PglZ->SIGMA+OB-Fold+AAA-ATPase-BrxL->URI-DNase->DndB-> | 1135 | Bacteria>FCB_group>Bacteroidetes/Chlorobi_group>Bacteroidota>Bacteroidia>Bacteroidales | GCA_024682395.1 | Bacteroidaceae_bacterium | K5874_00480 | 7551..10958 | 2212467 | hypothetical-protein-[Bacteroidaceae-bacterium] |
| MCR4564664.1 | Type4.PglZ.containing.BREX | ZnR+TOPRIM->?->SIGMA-HTH->PAPS-reduct+a+b-domain+Fer4_9+DUF4007->wHTH->DUF6079*->iSTAND+PglZ->SIGMA+OB-Fold+AAA-ATPase-BrxL-> | 1135 | Bacteria>FCB_group>Bacteroidetes/Chlorobi_group>Bacteroidota>Bacteroidia>Bacteroidales | GCA_024683825.1 | Bacteroidales_bacterium | K5651_01085 | 6033..9440 | 2030927 | hypothetical-protein-[Bacteroidales-bacterium] |
| MCR5064645.1 | Type4.PglZ.containing.BREX | PAPS-reduct+a+b-domain+Fer4_9+DUF4007->DUF6079*->iSTAND+PglZ->SIGMA+OB-Fold+AAA-ATPase-BrxL->CR-ATPase5+REase->?->wHTH+AbiEi_2->AbiEii->Calcineurin-> | 1152 | Bacteria>FCB_group>Bacteroidetes/Chlorobi_group>Bacteroidota>Bacteroidia>Bacteroidales | GCA_024688465.1 | Bacteroidales_bacterium | K6A67_02640 | 2312..5770 | 2030927 | hypothetical-protein-[Bacteroidales-bacterium] |
| MCR5643396.1 | Type4.PglZ.containing.BREX | <-XIS-HTH\|\|TraJ-RHH->HTH_3->PAPS-reduct+a+b-domain+Fer4_9+DUF4007->DUF6079*->iSTAND+PglZ->SIGMA+OB-Fold+AAA-ATPase-BrxL->URI-DNase->DndB->?->Schlafen+HATPase+wHTH->?->?-><-GTPase-AIG | 1147 | Bacteria>FCB_group>Bacteroidetes/Chlorobi_group>Bacteroidota>Bacteroidia>Bacteroidales | GCA_024690905.1 | Prevotella_sp. | K6G32_08685 | 4225..7668 | 59823 | hypothetical-protein-[Prevotella-sp.] |
| MCS2440786.1 | Type4.PglZ.containing.BREX | <-Phage_integrase\|\|Arm-DNA-bind_5+Phage_int_SAM_5+Phage_integrase->Virulence_RhuM->?-><-XIS-HTH\|\|TraJ-RHH-><-SIGMA+OB-Fold+AAA-ATPase-BrxL<-PglZ<-PglZ-p<-DUF6079-p*<-DUF6079-p*<-PAPS-reduct+a+b-domain+Fer4_9+DUF4007<-PAPS-reduct<-PAPS-reduct+Fer4_9-p<-Arm-DNA-bind_5+Phage_int_SAM_5+Phage_integrase<-tRNA\|\|TP_METHYLASE->?->HAD+HAD-> | 639 | Bacteria>FCB_group>Bacteroidetes/Chlorobi_group>Bacteroidota>Bacteroidia>Bacteroidales | GCA_024788655.1 | Bacteroides_uniformis | NXX09_06240 | 1386576..1388495 | 820 | hypothetical-protein-[Bacteroides-uniformis] |
| MCS2440787.1 | Type4.PglZ.containing.BREX | Arm-DNA-bind_5+Phage_int_SAM_5+Phage_integrase->Virulence_RhuM->?-><-XIS-HTH\|\|TraJ-RHH-><-SIGMA+OB-Fold+AAA-ATPase-BrxL<-PglZ<-PglZ-p<-DUF6079-p*<-DUF6079-p*<-PAPS-reduct+a+b-domain+Fer4_9+DUF4007<-PAPS-reduct<-PAPS-reduct+Fer4_9-p<-Arm-DNA-bind_5+Phage_int_SAM_5+Phage_integrase<-tRNA\|\|TP_METHYLASE->?->HAD+HAD-> | 514 | Bacteria>FCB_group>Bacteroidetes/Chlorobi_group>Bacteroidota>Bacteroidia>Bacteroidales | GCA_024788655.1 | Bacteroides_uniformis | NXX09_06245 | 1388488..1390032 | 820 | hypothetical-protein-[Bacteroides-uniformis] |
| WP_004344914.1 | Type4.PglZ.containing.BREX | Phage_int_SAM_5+Phage_integrase->PAPS-reduct+a+b-domain+Fer4_9+DUF4007->wHTH->DUF6079*->iSTAND+PglZ->SIGMA+OB-Fold+AAA-ATPase-BrxL->?->Schlafen+BirA-HTH-><-FurR-HTH<-?<-TraJ-RHH<-ZnR+TOPRIM<-D5ATPase | 1133 | Bacteria>FCB_group>Bacteroidetes/Chlorobi_group>Bacteroidota>Bacteroidia>Bacteroidales | GCF_000184945.1 | Prevotella_buccae | HMPREF6485_RS04250 | 1122248..1125649 | 28126 | hypothetical-protein-[Prevotella-buccae] |
| WP_013670603.1 | Type4.PglZ.containing.BREX | SF1-SF2-Helicase-><-?<-DDE_Tnp<-DDE_Tnp\|\|Phage_int_SAM_5+Phage_integrase->HTH_3->PAPS-reduct+a+b-domain+Fer4_9+DUF4007->wHTH->DUF6079*->iSTAND+PglZ->SIGMA+OB-Fold+AAA-ATPase-BrxL->?->Schlafen+BirA-HTH->HEPN+REase->DDE_Tnp_1_3-><-FurR-HTH<-?<-TraJ-RHH | 1133 | Bacteria>FCB_group>Bacteroidetes/Chlorobi_group>Bacteroidota>Bacteroidia>Bacteroidales | GCF_000193395.1 | Prevotella_denticola | HMPREF9137_RS01730 | 417083..420484 | 28129 | hypothetical-protein-[Prevotella-denticola] |
| WP_016268715.1 | Type4.PglZ.containing.BREX | CopG/RHH-><-AbrB-HTH<-?\|\|XIS-HTH->XIS-HTH-><-?<-Schlafen+HATPase+wHTH<-SIGMA+OB-Fold+AAA-ATPase-BrxL<-iSTAND+PglZ<-DUF6079*<-PAPS-reduct+a+b-domain+Fer4_9+DUF4007<-HTH_3\|\|BirA-HTH->?-><-Arm-DNA-bind_5+Phage_int_SAM_5+Phage_integrase<-Arm-DNA-bind_5+Phage_int_SAM_5+Phage_integrase\|\|?->?->ArdA-> | 1140 | Bacteria>FCB_group>Bacteroidetes/Chlorobi_group>Bacteroidota>Bacteroidia>Bacteroidales | GCF_000403155.2 | Bacteroides_thetaiotaomicron | C799_RS09485 | 3549049..3552471 | 818 | hypothetical-protein-[Bacteroides-thetaiotaomicron] |
| WP_039876352.1 | Type4.PglZ.containing.BREX | Phage_int_SAM_5+Phage_integrase->PAPS-reduct+a+b-domain+Fer4_9+DUF4007->wHTH->DUF6079*->iSTAND+PglZ->SIGMA+OB-Fold+AAA-ATPase-BrxL->?->Schlafen+BirA-HTH-><-FurR-HTH<-?<-TraJ-RHH<-ZnR+TOPRIM<-D5ATPase | 1133 | Bacteria>FCB_group>Bacteroidetes/Chlorobi_group>Bacteroidota>Bacteroidia>Bacteroidales | GCF_000468135.1 | Prevotellaceae | HMPREF1218_RS06320 | 14543..17944 | 171552 | MULTISPECIES:-hypothetical-protein-[Prevotellaceae] |
| WP_028899576.1 | Type4.PglZ.containing.BREX | XIS-HTH-><-Virulence_RhuM<-Arm-DNA-bind_5+Phage_int_SAM_5+Phage_integrase\|\|?-><-Peptidase_S8<-?\|\|Phage_int_SAM_5+Phage_integrase->PAPS-reduct+a+b-domain+Fer4_9+DUF4007->wHTH->DUF6079*->iSTAND+PglZ->SIGMA+OB-Fold+AAA-ATPase-BrxL->?->Schlafen+BirA-HTH-><-FurR-HTH\|\|?->?->ZnR-><-XIS-HTH | 1133 | Bacteria>FCB_group>Bacteroidetes/Chlorobi_group>Bacteroidota>Bacteroidia>Bacteroidales | GCF_000599605.1 | Prevotella_sp._HJM029 | X919_RS0107460 | 47159..50560 | 1433844 | hypothetical-protein-[Prevotella-sp.-HJM029] |
| WP_021669284.1 | Type4.PglZ.containing.BREX | HTH_3->PAPS-reduct+Fer4_9-p->PAPS-reduct+a+b-domain+Fer4_9->PAPS-reduct+DUF4007->wHTH->DUF6079*->iSTAND+PglZ->SIGMA+OB-Fold+AAA-ATPase-BrxL->?-><-FurR-HTH | 1133 | Bacteria>FCB_group>Bacteroidetes/Chlorobi_group>Bacteroidota>Bacteroidia>Bacteroidales | GCF_000613345.1 | Prevotella_disiens | JCM6334_RS10425 | 3157..6558 | 28130 | hypothetical-protein-[Prevotella-disiens] |
| WP_232220146.1 | Type4.PglZ.containing.BREX | D5ATPase->ZnR+TOPRIM->TraJ-RHH->?->FurR-HTH-><-Schlafen+BirA-HTH<-?<-SIGMA+OB-Fold+AAA-ATPase-BrxL<-iSTAND+PglZ<-DUF6079-p*<-wHTH<-PAPS-reduct+a+b-domain+Fer4_9+DUF4007<-PAPS-reduct+Fer4_9-p<-HTH_3<-?<-?<-N6-METHYLASE<-SF1-SF2-Helicase | 900 | Bacteria>FCB_group>Bacteroidetes/Chlorobi_group>Bacteroidota>Bacteroidia>Bacteroidales | GCF_000613945.1 | Prevotella_falsenii | JCM15124_RS13945 | 13149..15851 | 515414 | hypothetical-protein-[Prevotella-falsenii] |
| WP_232220147.1 | Type4.PglZ.containing.BREX | ZnR+TOPRIM->TraJ-RHH->?->FurR-HTH-><-Schlafen+BirA-HTH<-?<-SIGMA+OB-Fold+AAA-ATPase-BrxL<-iSTAND+PglZ<-DUF6079-p*<-wHTH<-PAPS-reduct+a+b-domain+Fer4_9+DUF4007<-PAPS-reduct+Fer4_9-p<-HTH_3<-?<-?<-N6-METHYLASE<-SF1-SF2-Helicase | 231 | Bacteria>FCB_group>Bacteroidetes/Chlorobi_group>Bacteroidota>Bacteroidia>Bacteroidales | GCF_000613945.1 | Prevotella_falsenii | JCM15124_RS16690 | 15832..16527 | 515414 | DUF6079-family-protein-[Prevotella-falsenii] |
| WP_234397399.1 | Type4.PglZ.containing.BREX | ZnR+TOPRIM->TraJ-RHH->?-><-?<-SIGMA+OB-Fold+AAA-ATPase-BrxL<-PglZ-p<-iSTAND+PglZ<-DUF6079*<-wHTH<-PAPS-reduct+a+b-domain+Fer4_9+DUF4007<-?<-?<-?\|\|?-><-METHYLASE<-?\|\|SprT-like-> | 1077 | Bacteria>FCB_group>Bacteroidetes/Chlorobi_group>Bacteroidota>Bacteroidia>Bacteroidales | GCF_000614025.1 | Prevotella_aurantiaca | JCM15754_RS13660 | 51590..54823 | 596085 | hypothetical-protein-[Prevotella-aurantiaca] |
| WP_028902856.1 | Type4.PglZ.containing.BREX | Relaxase->?-><-?<-REase<-DndE<-DndD<-RNase<-SIGMA+OB-Fold+AAA-ATPase-BrxL<-iSTAND+PglZ<-DUF6079*<-wHTH<-PAPS-reduct+a+b-domain+Fer4_9+DUF4007<-HTH_3<-LRR_5<-REase\|\|?->?->?->REase-> | 1145 | Bacteria>FCB_group>Bacteroidetes/Chlorobi_group>Bacteroidota>Bacteroidia>Bacteroidales | GCF_000619985.1 | Prevotella_sp._P6B4 | T491_RS0104670 | 53738..57175 | 1410614 | hypothetical-protein-[Prevotella-sp.-P6B4] |
| WP_044080181.1 | Type4.PglZ.containing.BREX | REase-><-Peptidase_C69\|\|?-><-Cysteine-Desulfurase<-ZnR<-SIGMA+OB-Fold+AAA-ATPase-BrxL<-iSTAND+PglZ<-DUF6079*<-PAPS-reduct+a+b-domain+Fer4_9+DUF4007<-?\|\|wHTH->REase->?-><-?\|\|?->RADICAL-SAM-> | 1390 | Bacteria>FCB_group>Bacteroidetes/Chlorobi_group>Bacteroidota>Bacteroidia>Bacteroidales | GCF_000833955.1 | Prevotella_pectinovora | ST41_RS06630 | 10771..14943 | 1602169 | hypothetical-protein-[Prevotella-pectinovora] |
| WP_044072855.1 | Type4.PglZ.containing.BREX | <-RADICAL-SAM<-?\|\|?-><-?<-?<-wHTH\|\|PAPS-reduct+DUF4007->wHTH->DUF6079*->iSTAND+PglZ-> | 1151 | Bacteria>FCB_group>Bacteroidetes/Chlorobi_group>Bacteroidota>Bacteroidia>Bacteroidales | GCF_000834015.1 | Prevotella_pectinovora | ST42_RS06090 | 46336..49791 | 1602169 | hypothetical-protein-[Prevotella-pectinovora] |
| WP_087402065.1 | Type4.PglZ.containing.BREX | <-SIGMA+OB-Fold+AAA-ATPase-BrxL<-iSTAND+PglZ<-DUF6079*<-wHTH<-PAPS-reduct+a+b-domain+Fer4_9+DUF4007<-HTH_3<-STYKinase-like<-TRPR-HTH<-?<-?<-?<-DOC+HTH | 1151 | Bacteria>FCB_group>Bacteroidetes/Chlorobi_group>Bacteroidota>Bacteroidia>Bacteroidales | GCF_002159215.1 | Alistipes_onderdonkii | B5G41_RS07005 | 125877..129332 | 328813 | hypothetical-protein-[Alistipes-onderdonkii] |
| WP_087394606.1 | Type4.PglZ.containing.BREX | <-SAVED<-TOPRIM<-SbcC\|\|Calcineurin->?->ACET->HTH_3->?->PAPS-reduct+a+b-domain+Fer4_9+DUF4007->DUF6079*->iSTAND+PglZ->SIGMA+OB-Fold+AAA-ATPase-BrxL->ZnR->Cysteine-Desulfurase->?->?->?->ART-DarT-> | 1396 | Bacteria>FCB_group>Bacteroidetes/Chlorobi_group>Bacteroidota>Bacteroidia>Bacteroidales | GCF_002161635.1 | Odoribacter_splanchnicus | B5F99_RS09425 | 101934..106124 | 28118 | hypothetical-protein-[Odoribacter-splanchnicus] |
| WP_087426296.1 | Type4.PglZ.containing.BREX | XIS-HTH+XIS-HTH+XIS-HTH+XIS-HTH+XIS-HTH+XIS-HTH+XIS-HTH->Arm-DNA-bind_5+Phage_int_SAM_5+Phage_integrase->DUF4007->XIS-HTH->SMS-KaiC-><-?<-SeqA<-SIGMA+OB-Fold+AAA-ATPase-BrxL<-iSTAND+PglZ<-DUF6079*<-PAPS-reduct+a+b-domain+Fer4_9+DUF4007<-HTH_3<-?<-MNS+HTH\|\|?->SPOUT-><-ADH_N+ADH_zinc_N+METHYLASE | 1147 | Bacteria>FCB_group>Bacteroidetes/Chlorobi_group>Bacteroidota>Bacteroidia>Bacteroidales | GCF_002161685.1 | Bacteroides_clarus | B5F97_RS11100 | 142848..146291 | 626929 | hypothetical-protein-[Bacteroides-clarus] |
| WP_094434346.1 | Type4.PglZ.containing.BREX | TetR-HTH->?->?->?-><-REase<-SIGMA+OB-Fold+AAA-ATPase-BrxL<-iSTAND+PglZ<-DUF6079*<-PAPS-reduct+a+b-domain+Fer4_9+DUF4007 | 1157 | Bacteria>FCB_group>Bacteroidetes/Chlorobi_group>Bacteroidota>Bacteroidia>Bacteroidales | GCF_002251185.1 | unclassified_Prevotella | CIK90_RS10290 | 69855..73328 | 2638335 | MULTISPECIES:-hypothetical-protein-[unclassified-Prevotella] |
| WP_022309959.1 | Type4.PglZ.containing.BREX | PAPS-reduct+DUF4007->DUF6079*->iSTAND+PglZ->SIGMA+OB-Fold+AAA-ATPase-BrxL-> | 1161 | Bacteria>FCB_group>Bacteroidetes/Chlorobi_group>Bacteroidota>Bacteroidia>Bacteroidales | GCF_002251325.1 | Prevotella_sp._P4-67 | CIK92_RS12670 | 19690..23175 | 2024227 | hypothetical-protein-[Prevotella-sp.-P4-67] |
| WP_104404274.1 | Type4.PglZ.containing.BREX | <-Arm-DNA-bind_5+Phage_int_SAM_5+Phage_integrase<-XIS-HTH+XIS-HTH+XIS-HTH+XIS-HTH\|\|TRPR-HTH->MNS+HTH->?->HTH_3->PAPS-reduct+a+b-domain+Fer4_9+DUF4007->DUF6079-p*->DUF6079-p->iSTAND+PglZ->SIGMA+OB-Fold+AAA-ATPase-BrxL->SeqA->?-><-?<-Relaxase<-TraJ-RHH<-SMS-KaiC | 168 | Bacteria>FCB_group>Bacteroidetes/Chlorobi_group>Bacteroidota>Bacteroidia>Bacteroidales | GCF_002933955.1 | Prevotella_sp._MGM2 | C5S62_RS04720 | 8456..8962 | 2033406 | hypothetical-protein-[Prevotella-sp.-MGM2] |
| WP_104404275.1 | Type4.PglZ.containing.BREX | <-Arm-DNA-bind_5+Phage_int_SAM_5+Phage_integrase<-XIS-HTH+XIS-HTH+XIS-HTH+XIS-HTH\|\|TRPR-HTH->MNS+HTH->?->HTH_3->PAPS-reduct+a+b-domain+Fer4_9+DUF4007->DUF6079-p->DUF6079-p*->iSTAND+PglZ->SIGMA+OB-Fold+AAA-ATPase-BrxL->SeqA->?-><-?<-Relaxase<-TraJ-RHH<-SMS-KaiC<-XIS-HTH | 948 | Bacteria>FCB_group>Bacteroidetes/Chlorobi_group>Bacteroidota>Bacteroidia>Bacteroidales | GCF_002933955.1 | Prevotella_sp._MGM2 | C5S62_RS04725 | 8979..11825 | 2033406 | hypothetical-protein-[Prevotella-sp.-MGM2] |
| WP_117856025.1 | Type4.PglZ.containing.BREX | TRPR-HTH->MNS+HTH->?->TRPR-HTH->STYKinase-like->?->PAPS-reduct+a+b-domain+Fer4_9+DUF4007->DUF6079-p*->DUF6079-p->iSTAND+PglZ->SIGMA+OB-Fold+AAA-ATPase-BrxL->ZnR->Cysteine-Desulfurase-><-?<-Relaxase<-TraJ-RHH | 745 | Bacteria>FCB_group>Bacteroidetes/Chlorobi_group>Bacteroidota>Bacteroidia>Bacteroidales | GCF_003468295.1 | Phocaeicola_vulgatus | DW874_RS12310 | 10805..13042 | 821 | hypothetical-protein-[Phocaeicola-vulgatus] |
| WP_118866086.1 | Type4.PglZ.containing.BREX | SF1-SF2-Helicase->?->SF1-SF2-Helicase-><-?<-DDE_Tnp\|\|Phage_int_SAM_5+Phage_integrase->HTH_3->PAPS-reduct+a+b-domain+Fer4_9+DUF4007->wHTH->DUF6079*->iSTAND+PglZ->SIGMA+OB-Fold+AAA-ATPase-BrxL->?->Schlafen+BirA-HTH-><-FurR-HTH<-?<-TraJ-RHH<-ZnR+TOPRIM<-D5ATPase | 1134 | Bacteria>FCB_group>Bacteroidetes/Chlorobi_group>Bacteroidota>Bacteroidia>Bacteroidales | GCF_003515045.1 | Prevotella_denticola | DYJ25_RS04070 | 960935..964339 | 28129 | hypothetical-protein-[Prevotella-denticola] |
| WP_119962087.1 | Type4.PglZ.containing.BREX | XIS-HTH->?->XIS-HTH-><-?\|\|Arm-DNA-bind_5+Phage_int_SAM_5+Phage_integrase->Arm-DNA-bind_5+Phage_int_SAM_5+Phage_integrase-><-BirA-HTH\|\|HTH_3->PAPS-reduct+a+b-domain+Fer4_9+DUF4007->DUF6079*->iSTAND+PglZ->SIGMA+OB-Fold+AAA-ATPase-BrxL->Schlafen+HATPase+wHTH->?-><-XIS-HTH<-XIS-HTH<-?\|\|?->AbrB-HTH-> | 1140 | Bacteria>FCB_group>Bacteroidetes/Chlorobi_group>Bacteroidota>Bacteroidia>Bacteroidales | GCF_003602965.1 | unclassified_Bacteroides | DWY57_RS16330 | 92363..95785 | 2646097 | MULTISPECIES:-hypothetical-protein-[unclassified-Bacteroides] |
| WP_121766191.1 | Type4.PglZ.containing.BREX | SMS-KaiC->TraJ-RHH->Relaxase->?-><-?<-Cysteine-Desulfurase<-ZnR<-SIGMA+OB-Fold+AAA-ATPase-BrxL<-iSTAND+PglZ<-DUF6079*<-PAPS-reduct+a+b-domain+Fer4_9+DUF4007<-?<-?<-STYKinase-like<-TRPR-HTH<-?<-MNS+HTH<-TRPR-HTH | 1395 | Bacteria>FCB_group>Bacteroidetes/Chlorobi_group>Bacteroidota>Bacteroidia>Bacteroidales | GCF_003671855.1 | Bacteroides_acidifaciens | D7Y07_RS10600 | 22072..26259 | 85831 | hypothetical-protein-[Bacteroides-acidifaciens] |
| WP_124140263.1 | Type4.PglZ.containing.BREX | <-TOPRIM<-ArdCN+MPTase<-?<-PRTase+MutS<-?\|\|PTSEIIA1->HTH_3->?->PAPS-reduct+a+b-domain+Fer4_9+DUF4007->DUF6079*->iSTAND+PglZ->SIGMA+OB-Fold+AAA-ATPase-BrxL->?\|\|LRR_5-> | 1136 | Bacteria>FCB_group>Bacteroidetes/Chlorobi_group>Bacteroidota>Bacteroidia>Bacteroidales | GCF_003838285.1 | Prevotella_intermedia | D2S45_RS11175 | 14406..17816 | 28131 | hypothetical-protein-[Prevotella-intermedia] |
| WP_126679193.1 | Type4.PglZ.containing.BREX | <-DOC+HTH<-GspA-ATPase+REase<-?<-HTH<-Schlafen+HATPase+wHTH<-?<-SIGMA+OB-Fold+AAA-ATPase-BrxL<-iSTAND+PglZ<-DUF6079*<-wHTH<-PAPS-reduct+a+b-domain+Fer4_9+DUF4007<-?<-SF1-SF2-Helicase\|\|CR-ATPase5+REase-> | 1144 | Bacteria>FCB_group>Bacteroidetes/Chlorobi_group>Bacteroidota>Bacteroidia>Bacteroidales | GCF_003977605.1 | Prevotella_koreensis | EHV08_RS10340 | 2429244..2432678 | 2490854 | hypothetical-protein-[Prevotella-koreensis] |
| WP_131927120.1 | Type4.PglZ.containing.BREX | XIS-HTH->D5ATPase->ZnR+TOPRIM->TraJ-RHH->?-><-Schlafen+BirA-HTH<-SIGMA+OB-Fold+AAA-ATPase-BrxL<-iSTAND+PglZ<-DUF6079*<-wHTH<-PAPS-reduct+a+b-domain+Fer4_9+DUF4007<-HTH_3<-Phage_int_SAM_5+Phage_integrase | 1132 | Bacteria>FCB_group>Bacteroidetes/Chlorobi_group>Bacteroidota>Bacteroidia>Bacteroidales | GCF_004342845.1 | Bacteroides_heparinolyticus | EV202_RS11185 | 61051..64449 | 28113 | hypothetical-protein-[Bacteroides-heparinolyticus] |
| WP_134849806.1 | Type4.PglZ.containing.BREX | <-SIGMA+OB-Fold+AAA-ATPase-BrxL<-iSTAND+PglZ<-DUF6079*<-PAPS-reduct+a+b-domain+Fer4_9+DUF4007\|\|DUF4099+DUF3945+TOPRIM+TOPRIM->TOPRIM+Topoisom_bac->N6-N4-METHYLASE-> | 1152 | Bacteria>FCB_group>Bacteroidetes/Chlorobi_group>Bacteroidota>Bacteroidia>Bacteroidales | GCF_004551515.1 | Porphyromonas_levii | E4P48_RS08045 | 4254..7712 | 28114 | hypothetical-protein-[Porphyromonas-levii] |
| WP_135993957.1 | Type4.PglZ.containing.BREX | <-SIGMA+OB-Fold+AAA-ATPase-BrxL<-iSTAND+PglZ<-DUF6079*<-PAPS-reduct+DUF4007 | 1125 | Bacteria>FCB_group>Bacteroidetes/Chlorobi_group>Bacteroidota>Bacteroidia>Bacteroidales | GCF_004793655.1 | Muribaculum_intestinale | E5333_RS14435 | 6333..9710 | 1796646 | hypothetical-protein-[Muribaculum-intestinale] |
| WP_136000316.1 | Type4.PglZ.containing.BREX | Calcineurin->Calcineurin->?->TRPR-HTH->STYKinase-like->?->HTH_3->PAPS-reduct+a+b-domain+Fer4_9+DUF4007->DUF6079*->iSTAND+PglZ->SIGMA+OB-Fold+AAA-ATPase-BrxL->SeqA-> | 1147 | Bacteria>FCB_group>Bacteroidetes/Chlorobi_group>Bacteroidota>Bacteroidia>Bacteroidales | GCF_004793675.1 | Bacteroides_caecimuris | E5353_RS15755 | 8657..12100 | 1796613 | hypothetical-protein-[Bacteroides-caecimuris] |
| WP_123481399.1 | Type4.PglZ.containing.BREX | SIGMA-HTH->?->O6MGT-HTH-><-?<-DndE<-DndD<-RNase<-SIGMA+OB-Fold+AAA-ATPase-BrxL<-iSTAND+PglZ<-DUF6079*<-wHTH<-PAPS-reduct+a+b-domain+Fer4_9+DUF4007<-REase\|\|?->Aminotran_1_2->?->?-><-HTH+REase<-HD | 1160 | Bacteria>FCB_group>Bacteroidetes/Chlorobi_group>Bacteroidota>Bacteroidia>Bacteroidales | GCF_004793975.1 | Lepagella_muris | E5331_RS10660 | 44917..48399 | 3032870 | hypothetical-protein-[Lepagella-muris] |
| WP_123481433.1 | Type4.PglZ.containing.BREX | Tox-RelE-><-?\|\|ABhydrolase->AA_kinase->?-><-Cysteine-Desulfurase<-ZnR<-SIGMA+OB-Fold+AAA-ATPase-BrxL<-iSTAND+PglZ<-DUF6079*<-PAPS-reduct+DUF4007<-PBPI | 1387 | Bacteria>FCB_group>Bacteroidetes/Chlorobi_group>Bacteroidota>Bacteroidia>Bacteroidales | GCF_004793975.1 | Lepagella_muris | E5331_RS10855 | 95524..99687 | 3032870 | hypothetical-protein-[Lepagella-muris] |
| WP_154480799.1 | Type4.PglZ.containing.BREX | <-AT-HTH-HYD2<-?\|\|?->RelE-ParE-><-?<-?<-Phage_integrase<-SIGMA+OB-Fold+AAA-ATPase-BrxL<-iSTAND+PglZ<-DUF6079*<-PAPS-reduct+a+b-domain+Fer4_9+DUF4007\|\|?->D5ATPase+DUF3874-><-?\|\|CR-ATPase5+REase->MNS-><-CR-ATPase8+REase | 1144 | Bacteria>FCB_group>Bacteroidetes/Chlorobi_group>Bacteroidota>Bacteroidia>Bacteroidales | GCF_009695805.1 | Prevotella_copri | FYJ72_RS06390 | 22860..26294 | 165179 | hypothetical-protein-[Prevotella-copri] |
| WP_155716074.1 | Type4.PglZ.containing.BREX | REase+RNase-><-?<-ANK<-?\|\|Phage_int_SAM_5+Phage_integrase->PAPS-reduct+a+b-domain+Fer4_9+DUF4007->wHTH->DUF6079*->iSTAND+PglZ->SIGMA+OB-Fold+AAA-ATPase-BrxL->?->Schlafen+BirA-HTH-><-FurR-HTH<-?<-TraJ-RHH<-ZnR+TOPRIM<-D5ATPase | 1133 | Bacteria>FCB_group>Bacteroidetes/Chlorobi_group>Bacteroidota>Bacteroidia>Bacteroidales | GCF_009728485.1 | Prevotella_vespertina | F0475_RS07095 | 124515..127916 | 2608404 | hypothetical-protein-[Prevotella-vespertina] |
| WP_165044724.1 | Type4.PglZ.containing.BREX | TraI-VirD2-RCRE-><-MerR-HTH\|\|?->?->?->?->\|\|VirD4-TraG+VirD4-FtsK-><-SIGMA+OB-Fold+AAA-ATPase-BrxL<-iSTAND+PglZ<-DUF6079*<-PAPS-reduct+a+b-domain+Fer4_9+DUF4007\|\|DUF4099+DUF3945+TOPRIM+TOPRIM-> | 1135 | Bacteria>FCB_group>Bacteroidetes/Chlorobi_group>Bacteroidota>Bacteroidia>Bacteroidales | GCF_011039195.1 | Dysgonomonas_sp._ZJ709 | G7Y56_RS15020 | 33443..36850 | 2709797 | hypothetical-protein-[Dysgonomonas-sp.-ZJ709] |
| WP_167963304.1 | Type4.PglZ.containing.BREX | <-VbhA+DOC+HTH\|\|?-><-RNase\|\|HTH_3->Couple_hipA->HipA_C->CR-ATPase5+REase->?->PAPS-reduct+a+b-domain+Fer4_9+DUF4007->DUF6079*->iSTAND+PglZ->SIGMA+OB-Fold+AAA-ATPase-BrxL->Cysteine-Desulfurase->TetR-HTH-> | 1392 | Bacteria>FCB_group>Bacteroidetes/Chlorobi_group>Bacteroidota>Bacteroidia>Bacteroidales | GCF_012113595.1 | Bacteroides_faecium | BacF7301_RS12655 | 3544858..3549036 | 2715212 | hypothetical-protein-[Bacteroides-faecium] |
| WP_176468401.1 | Type4.PglZ.containing.BREX | Arm-DNA-bind_5+Phage_int_SAM_5+Phage_integrase-><-?\|\|PAPS-reduct+DUF4007->DUF6079*->iSTAND+PglZ->SIGMA+OB-Fold+AAA-ATPase-BrxL-><-?<-TraJ-RHH<-CopG/RHH\|\|XIS-HTH->XIS-HTH->?-><-Virulence_RhuM | 1151 | Bacteria>FCB_group>Bacteroidetes/Chlorobi_group>Bacteroidota>Bacteroidia>Bacteroidales | GCF_013373675.1 | Bacteroides_sp._L10-4 | HU824_RS06390 | 68246..71701 | 2746063 | hypothetical-protein-[Bacteroides-sp.-L10-4] |
| WP_195207461.1 | Type4.PglZ.containing.BREX | D5ATPase->?->?->TraJ-RHH->Relaxase->?-><-Schlafen+HATPase+wHTH<-SIGMA+OB-Fold+AAA-ATPase-BrxL<-iSTAND+PglZ<-DUF6079*<-wHTH<-PAPS-reduct+a+b-domain+Fer4_9+DUF4007<-HTH_3\|\|?->LYTTR->Peptidase_M15->METHYLASE->?->Calcineurin+Calcineurin-conf-> | 1161 | Bacteria>FCB_group>Bacteroidetes/Chlorobi_group>Bacteroidota>Bacteroidia>Bacteroidales | GCF_015547015.1 | Bacteroides_clarus | I1P93_RS11595 | 64548..68033 | 626929 | hypothetical-protein-[Bacteroides-clarus] |
| WP_195342613.1 | Type4.PglZ.containing.BREX | Phage_integrase->Phage_integrase->?->XIS-HTH->ZnR+TOPRIM+AAA-helicase+wHTH->ZnR+TOPRIM->Mob_Pre->HTH_3->PAPS-reduct+a+b-domain+Fer4_9+DUF4007->DUF6079*->iSTAND+PglZ->SIGMA+OB-Fold+AAA-ATPase-BrxL->SeqA->Schlafen+HATPase+wHTH->SF1-SF2-Helicase->DOC+zhth->?->Cysteine-Desulfurase->TPR+TPR+TPR+TPR-> | 1145 | Bacteria>FCB_group>Bacteroidetes/Chlorobi_group>Bacteroidota>Bacteroidia>Bacteroidales | GCF_015551465.1 | Barnesiella_intestinihominis | I2D16_RS03535 | 160234..163671 | 487174 | hypothetical-protein-[Barnesiella-intestinihominis] |
| WP_195420883.1 | Type4.PglZ.containing.BREX | XIS-HTH->SMS-KaiC->TraJ-RHH->Relaxase->?-><-?<-SIGMA+OB-Fold+AAA-ATPase-BrxL<-iSTAND+PglZ<-DUF6079*<-PAPS-reduct+a+b-domain+Fer4_9+DUF4007<-HTH_3<-STYKinase-like<-TRPR-HTH<-?<-Calcineurin<-Calcineurin | 1144 | Bacteria>FCB_group>Bacteroidetes/Chlorobi_group>Bacteroidota>Bacteroidia>Bacteroidales | GCF_015551975.1 | Phocaeicola_vulgatus | I2E38_RS16540 | 31417..34851 | 821 | hypothetical-protein-[Phocaeicola-vulgatus] |
| WP_195554878.1 | Type4.PglZ.containing.BREX | XIS-HTH+XIS-HTH+XIS-HTH+XIS-HTH->Phage_int_SAM_5+Phage_integrase->?->XIS-HTH->SMS-KaiC-><-?<-TIR<-SIGMA+OB-Fold+AAA-ATPase-BrxL<-iSTAND+PglZ<-DUF6079*<-wHTH<-PAPS-reduct+a+b-domain+Fer4_9+DUF4007<-HTH_3<-STYKinase-like<-TRPR-HTH<-?<-ParB+HNH<-CR-ATPase5+REase\|\|MULE-TRANSPOSASE-><-URI-DNase+RAMA | 1152 | Bacteria>FCB_group>Bacteroidetes/Chlorobi_group>Bacteroidota>Bacteroidia>Bacteroidales | GCF_015556345.1 | Barnesiella_intestinihominis | I2F72_RS04925 | 148617..152075 | 487174 | hypothetical-protein-[Barnesiella-intestinihominis] |
| WP_138286707.1 | Type4.PglZ.containing.BREX | TRPR-HTH->MNS+HTH->?->TRPR-HTH->STYKinase-like->?->PAPS-reduct+a+b-domain+Fer4_9+DUF4007->GspA-ATPase->DUF6079*->iSTAND+PglZ->SIGMA+OB-Fold+AAA-ATPase-BrxL->ZnR->Cysteine-Desulfurase-><-?<-Relaxase<-TraJ-RHH<-SMS-KaiC<-XIS-HTH | 1188 | Bacteria>FCB_group>Bacteroidetes/Chlorobi_group>Bacteroidota>Bacteroidia>Bacteroidales | GCF_015667255.1 | Phocaeicola_vulgatus | FGQ91_RS06545 | 106226..109792 | 821 | hypothetical-protein-[Phocaeicola-vulgatus] |
| WP_196042835.1 | Type4.PglZ.containing.BREX | <-HTH_Tnp<-?<-?<-?\|\|HTH_3-><-?\|\|DUF6079-p*->iSTAND+PglZ->SIGMA+OB-Fold+AAA-ATPase-BrxL-><-HTH<-Schlafen+HATPase+wHTH<-rRNA<-Classical-AAA<-?<-SF1-SF2-Helicase<-TPR+TPR | 791 | Bacteria>FCB_group>Bacteroidetes/Chlorobi_group>Bacteroidota>Bacteroidia>Bacteroidales | GCF_015670365.1 | Parabacteroides_merdae | PN609_RS10245 | 28915..31290 | 46503 | hypothetical-protein-[Parabacteroides-merdae] |
| WP_205109180.1 | Type4.PglZ.containing.BREX | <-AraC-HTH+AraC-HTH<-REC<-HISKIN\|\|?->?->?-><-Schlafen+HATPase+wHTH<-SIGMA+OB-Fold+AAA-ATPase-BrxL<-iSTAND+PglZ<-DUF6079*<-PAPS-reduct+a+b-domain+Fer4_9+DUF4007<-HTH_3<-FurR-HTH<-?<-TraJ-RHH<-CopG/RHH\|\|XIS-HTH->XIS-HTH->\|\|Arm-DNA-bind_5+Phage_int_SAM_5+Phage_integrase-> | 1183 | Bacteria>FCB_group>Bacteroidetes/Chlorobi_group>Bacteroidota>Bacteroidia>Bacteroidales | GCF_016899955.1 | Marseilla_massiliensis | H6B30_RS07350 | 50180..53731 | 1841864 | hypothetical-protein-[Marseilla-massiliensis] |
| WP_008824212.1 | Type4.PglZ.containing.BREX | REase+RNase-><-?<-?\|\|?->PAPS-reduct+DUF4007->wHTH->DUF6079*->iSTAND+PglZ->SIGMA+OB-Fold+AAA-ATPase-BrxL->?-><-tRNA\|\|NTP_transferase+MannoseP_isomer-><-HIT | 1133 | Bacteria>FCB_group>Bacteroidetes/Chlorobi_group>Bacteroidota>Bacteroidia>Bacteroidales | GCF_018128125.1 | Prevotella_histicola | J5A62_RS00890 | 248148..251549 | 470565 | hypothetical-protein-[Prevotella-histicola] |
| WP_272194288.1 | Type4.PglZ.containing.BREX | SMS-KaiC->TraJ-RHH->Relaxase->?-><-?<-Cysteine-Desulfurase<-ZnR<-SIGMA+OB-Fold+AAA-ATPase-BrxL<-iSTAND+PglZ<-DUF6079*<-PAPS-reduct+DUF4007<-PAPS-reduct+a+b-domain+Fer4_9<-?<-STYKinase-like<-TRPR-HTH<-?<-MNS+HTH<-TRPR-HTH | 1395 | Bacteria>FCB_group>Bacteroidetes/Chlorobi_group>Bacteroidota>Bacteroidia>Bacteroidales | GCF_018381015.1 | Bacteroides_uniformis | POY93_RS01295 | 283800..287987 | 820 | hypothetical-protein-[Bacteroides-uniformis] |
| WP_214462063.1 | Type4.PglZ.containing.BREX | DUF4372+DDE_Tnp_1->\|\|?->DDE_Tnp_1_4-><-Arm-DNA-bind_5+Phage_int_SAM_5+Phage_integrase<-Tox-ART-HYD3<-Schlafen+HATPase+wHTH<-SIGMA+OB-Fold+AAA-ATPase-BrxL<-iSTAND+PglZ<-DUF6079*<-PAPS-reduct+a+b-domain+Fer4_9+DUF4007<-HTH_3<-FurR-HTH<-?<-TraJ-RHH<-CopG/RHH<-?\|\|XIS-HTH->XIS-HTH-> | 1183 | Bacteria>FCB_group>Bacteroidetes/Chlorobi_group>Bacteroidota>Bacteroidia>Bacteroidales | GCF_018588025.1 | Bacteroidaceae | JQN06_RS10465 | 618028..621579 | 815 | MULTISPECIES:-hypothetical-protein-[Bacteroidaceae] |
| WP_214455629.1 | Type4.PglZ.containing.BREX | Arm-DNA-bind_5+Phage_int_SAM_5+Phage_integrase->Arm-DNA-bind_5+Phage_int_SAM_5+Phage_integrase->Virulence_RhuM->?-><-XIS-HTH<-XIS-HTH\|\|TraJ-RHH-><-SIGMA+OB-Fold+AAA-ATPase-BrxL<-iSTAND+PglZ<-DUF6079*<-PAPS-reduct+a+b-domain+Fer4_9+DUF4007<-Arm-DNA-bind_5+Phage_int_SAM_5+Phage_integrase | 1151 | Bacteria>FCB_group>Bacteroidetes/Chlorobi_group>Bacteroidota>Bacteroidia>Bacteroidales | GCF_018588075.1 | Phocaeicola_dorei | JQN03_RS15165 | 93798..97253 | 357276 | hypothetical-protein-[Phocaeicola-dorei] |
| WP_215683073.1 | Type4.PglZ.containing.BREX | <-XIS-HTH<-XIS-HTH\|\|CopG/RHH-><-?\|\|TraJ-RHH->?->?->HTH_3->PAPS-reduct+a+b-domain+Fer4_9+DUF4007->DUF6079*->iSTAND+PglZ->SIGMA+OB-Fold+AAA-ATPase-BrxL->Schlafen+HATPase+wHTH-> | 1140 | Bacteria>FCB_group>Bacteroidetes/Chlorobi_group>Bacteroidota>Bacteroidia>Bacteroidales | GCF_018785945.1 | Bacteroides_salyersiae | GPL06_RS15285 | 15394..18816 | 291644 | hypothetical-protein-[Bacteroides-salyersiae] |
| WP_218484873.1 | Type4.PglZ.containing.BREX | HTH_31->?-><-MNS<-CR-ATPase5+REase\|\|?-><-D5ATPase+DUF3874<-?\|\|PAPS-reduct+DUF4007->DUF6079*->iSTAND+PglZ->SIGMA+OB-Fold+AAA-ATPase-BrxL->Phage_integrase->?-><-CR-ATPase5+REase<-35exo<-CR-ATPase5+REase | 1143 | Bacteria>FCB_group>Bacteroidetes/Chlorobi_group>Bacteroidota>Bacteroidia>Bacteroidales | GCF_019249635.1 | Prevotella_copri | KUA42_12150 | 24442..27873 | 165179 | hypothetical-protein-[Prevotella-copri] |
| WP_225722473.1 | Type4.PglZ.containing.BREX | AbiGi->?->HTH_3->Couple_hipA->HipA_C->?->AraC-HTH+AraC-HTH->HTH_3->PAPS-reduct+a+b-domain+Fer4_9+DUF4007->DUF6079*->iSTAND+PglZ->SIGMA+OB-Fold+AAA-ATPase-BrxL->SeqA->SeqA-><-zf-ISL3+HTH_Tnp_ISL3\|\|TetR-HTH->?->AraC-HTH+AraC-HTH-> | 1155 | Bacteria>FCB_group>Bacteroidetes/Chlorobi_group>Bacteroidota>Bacteroidia>Bacteroidales | GCF_020313435.1 | Bacteroides_thetaiotaomicron | KFX44_RS14795 | 88138..91605 | 818 | hypothetical-protein-[Bacteroides-thetaiotaomicron] |
| WP_234129192.1 | Type4.PglZ.containing.BREX | <-XIS-HTH<-XIS-HTH\|\|CopG/RHH->TraJ-RHH->?->FurR-HTH-><-SIGMA+OB-Fold+AAA-ATPase-BrxL<-iSTAND+PglZ<-DUF6079*<-PAPS-reduct+a+b-domain+Fer4_9+DUF4007\|\|?-><-Arm-DNA-bind_5+Phage_int_SAM_5+Phage_integrase<-XIS-HTH<-?<-SIGMA-HTH<-?<-Glyco_hydro_88 | 1151 | Bacteria>FCB_group>Bacteroidetes/Chlorobi_group>Bacteroidota>Bacteroidia>Bacteroidales | GCF_021411885.1 | Bacteroides_thetaiotaomicron | KOH07_RS22195 | 31143..34598 | 818 | hypothetical-protein-[Bacteroides-thetaiotaomicron] |
| WP_235392644.1 | Type4.PglZ.containing.BREX | <-MODE-HTH+PBPII<-?\|\|CR-ATPase8+REase<-?<-wHTH\|\|?->PAPS-reduct+a+b-domain+Fer4_9+DUF4007->DUF6079*->iSTAND+PglZ->SIGMA+OB-Fold+AAA-ATPase-BrxL->ZnR->Cysteine-Desulfurase-><-CR-ATPase8+REase<-REase\|\|SIGMA-HTH-> | 1392 | Bacteria>FCB_group>Bacteroidetes/Chlorobi_group>Bacteroidota>Bacteroidia>Bacteroidales | GCF_021531355.1 | Prevotella_stercorea | I6E38_RS03195 | 44898..49076 | 363265 | hypothetical-protein-[Prevotella-stercorea] |
| WP_235402044.1 | Type4.PglZ.containing.BREX | <-RNase\|\|?->JetB->SbcC->RNase-><-CR-ATPase5+REase->?->PAPS-reduct+a+b-domain+Fer4_9+DUF4007->wHTH->DUF6079*->iSTAND+PglZ->SIGMA+OB-Fold+AAA-ATPase-BrxL-> | 1161 | Bacteria>FCB_group>Bacteroidetes/Chlorobi_group>Bacteroidota>Bacteroidia>Bacteroidales | GCF_021531655.1 | Prevotella_stercorea | I6E49_RS09825 | 36590..40075 | 363265 | hypothetical-protein-[Prevotella-stercorea] |
| WP_151259905.1 | Type4.PglZ.containing.BREX | SMS-KaiC->TraJ-RHH->Relaxase->?-><-?<-Cysteine-Desulfurase<-ZnR<-SIGMA+OB-Fold+AAA-ATPase-BrxL<-iSTAND+PglZ<-DUF6079*<-PAPS-reduct+a+b-domain+Fer4_9+DUF4007<-?<-STYKinase-like<-TRPR-HTH<-?<-MNS+HTH<-TRPR-HTH | 1395 | Bacteria>FCB_group>Bacteroidetes/Chlorobi_group>Bacteroidota>Bacteroidia>Bacteroidales | GCF_021890855.1 | Phocaeicola_vulgatus | L4X28_RS13310 | 49264..53451 | 821 | hypothetical-protein-[Phocaeicola-vulgatus] |
| WP_060940051.1 | Type4.PglZ.containing.BREX | DDE_Tnp_1_3->CR-ATPase5+REase->?->?->PAPS-reduct+Fer4_9-p->DUF6079*->iSTAND+PglZ->SIGMA+OB-Fold+AAA-ATPase-BrxL-><-?<-?\|\|?-><-Peptidase_M13_N+Peptidase_M13<-GNTR-HTH | 1160 | Bacteria>FCB_group>Bacteroidetes/Chlorobi_group>Bacteroidota>Bacteroidia>Bacteroidales | GCF_022713225.1 | Prevotella_corporis | P6744_RS07155 | 6822..10304 | 28128 | hypothetical-protein-[Prevotella-corporis] |
| WP_257421405.1 | Type4.PglZ.containing.BREX | DSBH+AraC-HTH-><-?\|\|XIS-HTH+XIS-HTH+XIS-HTH+XIS-HTH->Arm-DNA-bind_5+Phage_int_SAM_5+Phage_integrase-><-?\|\|PAPS-reduct+a+b-domain+Fer4_9+DUF4007->DUF6079*->iSTAND+PglZ->SIGMA+OB-Fold+AAA-ATPase-BrxL-><-FurR-HTH<-?<-TraJ-RHH<-CopG/RHH\|\|XIS-HTH->XIS-HTH-> | 1151 | Bacteria>FCB_group>Bacteroidetes/Chlorobi_group>Bacteroidota>Bacteroidia>Bacteroidales | GCF_024623115.1 | Bacteroides_acidifaciens | NSB20_RS19010 | 12273..15728 | 85831 | hypothetical-protein-[Bacteroides-acidifaciens] |
| WP_258901871.1 | Type4.PglZ.containing.BREX | <-NERD+ZnR<-?\|\|Arm-DNA-bind_5+Phage_int_SAM_5+Phage_integrase->Virulence_RhuM->?->PAPS-reduct+a+b-domain+Fer4_9+DUF4007->DUF6079*->iSTAND+PglZ->SIGMA+OB-Fold+AAA-ATPase-BrxL-><-?<-?<-GDResolvase<-?<-?<-Peptidase_M42 | 1157 | Bacteria>FCB_group>Bacteroidetes/Chlorobi_group>Bacteroidota>Bacteroidia>Bacteroidales | GCF_024760245.1 | Bacteroides_thetaiotaomicron | NXY23_RS10185 | 2745346..2748819 | 818 | hypothetical-protein-[Bacteroides-thetaiotaomicron] |
| WP_258983772.1 | Type4.PglZ.containing.BREX | GDResolvase+HIN-HTH-><-TetR-HTH<-?<-?\|\|?->SIGMA-HTH+MA->REase->REase->PAPS-reduct+a+b-domain+Fer4_9+DUF4007->DUF6079*->iSTAND+PglZ->SIGMA+OB-Fold+AAA-ATPase-BrxL->?-><-?<-?<-?<-SF1-SF2-Helicase+35exo | 1256 | Bacteria>FCB_group>Bacteroidetes/Chlorobi_group>Bacteroidota>Bacteroidia>Bacteroidales | GCF_024788465.1 | Bacteroides_thetaiotaomicron | NXV24_RS26455 | 35978..39748 | 818 | hypothetical-protein-[Bacteroides-thetaiotaomicron] |
| WP_259029872.1 | Type4.PglZ.containing.BREX | <-Glyco_hydro_43+GH43_C2<-Schlafen+HATPase+wHTH<-SIGMA+OB-Fold+AAA-ATPase-BrxL<-iSTAND+PglZ<-DUF6079*<-wHTH<-PAPS-reduct+a+b-domain+Fer4_9+DUF4007<-HTH_3<-?<-?<-CopG/RHH\|\|XIS-HTH->XIS-HTH-><-Arm-DNA-bind_5+Phage_int_SAM_5+Phage_integrase | 1181 | Bacteria>FCB_group>Bacteroidetes/Chlorobi_group>Bacteroidota>Bacteroidia>Bacteroidales | GCF_024792385.1 | Bacteroides_salyersiae | NXY20_RS18880 | 4719748..4723293 | 291644 | hypothetical-protein-[Bacteroides-salyersiae] |
| WP_260776559.1 | Type4.PglZ.containing.BREX | ADH_N+ADH_zinc_N+METHYLASE->SPOUT-><-?\|\|TRPR-HTH->MNS+HTH->HTH_3->?->PAPS-reduct+a+b-domain+Fer4_9+DUF4007->DUF6079*->?->iSTAND+PglZ->SIGMA+OB-Fold+AAA-ATPase-BrxL->ZnR->Cysteine-Desulfurase->?-><-?<-Relaxase<-TraJ-RHH | 1336 | Bacteria>FCB_group>Bacteroidetes/Chlorobi_group>Bacteroidota>Bacteroidia>Bacteroidales | GCF_025263845.1 | Bacteroides_cellulosilyticus | NWT25_RS14255 | 3996268..4000278 | 246787 | hypothetical-protein-[Bacteroides-cellulosilyticus] |
| WP_264966907.1 | Type4.PglZ.containing.BREX | <-SIGMA+OB-Fold+AAA-ATPase-BrxL<-DDE_5+DDE_Tnp_1<-SIGMA+OB-Fold+AAA-ATPase+Lon-Protease-BrxL-p<-iSTAND+PglZ<-DUF6079-p*\|\|DDE_Tnp_1_4-><-DUF6079-p<-PAPS-reduct+a+b-domain+Fer4_9+DUF4007<-PAPS-reduct+Fer4_9-p<-HU-HIG+DEOR-HTH<-CR-ATPase5+REase<-?<-wHTH+WYL+WCX<-URI-DNase | 922 | Bacteria>FCB_group>Bacteroidetes/Chlorobi_group>Bacteroidota>Bacteroidia>Bacteroidales | GCF_026015685.1 | Prevotella_copri | ONT16_RS14450 | 326301..329069 | 165179 | hypothetical-protein-[Prevotella-copri] |
| WP_264966908.1 | Type4.PglZ.containing.BREX | <-SIGMA+OB-Fold+AAA-ATPase-BrxL<-DDE_5+DDE_Tnp_1<-SIGMA+OB-Fold+AAA-ATPase+Lon-Protease-BrxL-p<-iSTAND+PglZ<-DUF6079-p\|\|DDE_Tnp_1_4-><-DUF6079-p*<-PAPS-reduct+a+b-domain+Fer4_9+DUF4007<-PAPS-reduct+Fer4_9-p<-HU-HIG+DEOR-HTH<-CR-ATPase5+REase<-?<-wHTH+WYL+WCX<-URI-DNase<-HU-HIG+DEOR-HTH<-DUF1016+REase | 304 | Bacteria>FCB_group>Bacteroidetes/Chlorobi_group>Bacteroidota>Bacteroidia>Bacteroidales | GCF_026015685.1 | Prevotella_copri | ONT16_RS14460 | 330484..331398 | 165179 | hypothetical-protein-[Prevotella-copri] |
| WP_270833287.1 | Type4.PglZ.containing.BREX | ADH_N+ADH_zinc_N+METHYLASE->SPOUT-><-?\|\|TRPR-HTH->MNS+HTH->HTH_3->?->PAPS-reduct+a+b-domain+Fer4_9+DUF4007->DUF6079*->iSTAND+PglZ->SIGMA+OB-Fold+AAA-ATPase-BrxL->ZnR->Cysteine-Desulfurase->?-><-?<-Relaxase<-TraJ-RHH<-SMS-KaiC | 1395 | Bacteria>FCB_group>Bacteroidetes/Chlorobi_group>Bacteroidota>Bacteroidia>Bacteroidales | GCF_027663765.1 | Bacteroides_cellulosilyticus | PHA19_RS02430 | 16212..20399 | 246787 | hypothetical-protein-[Bacteroides-cellulosilyticus] |
| WP_068855222.1 | Type4.PglZ.containing.BREX | <-SIGMA+OB-Fold+AAA-ATPase-BrxL<-iSTAND+PglZ<-DUF6079*<-wHTH<-PAPS-reduct+Fer4_9-p<-PAPS-reduct+a+b-domain+Fer4_9<-HTH_3<-GNAT<-?<-TraJ-RHH<-ZnR+TOPRIM<-CR-ATPase4 | 1135 | Bacteria>FCB_group>Bacteroidetes/Chlorobi_group>Bacteroidota>Bacteroidia>Bacteroidales | GCF_900079775.1 | Prevotella_lascolaii | BN4609_RS03055 | 424682..428089 | 1776379 | hypothetical-protein-[Prevotella-lascolaii] |
| WP_071148139.1 | Type4.PglZ.containing.BREX | ParB+HNH->OLD-ABC->?->?->?->Phage_holin->tRNA->HTH_3->PAPS-reduct+a+b-domain+Fer4_9+DUF4007->DUF6079*->iSTAND+PglZ->SIGMA+OB-Fold+AAA-ATPase-BrxL->Schlafen+HATPase+wHTH-><-?<-Relaxase<-TraJ-RHH<-?<-?<-D5ATPase | 1182 | Bacteria>FCB_group>Bacteroidetes/Chlorobi_group>Bacteroidota>Bacteroidia>Bacteroidales | GCF_900108345.1 | Bacteroides_ndongoniae | BQ5591_RS01340 | 248814..252362 | 1903262 | hypothetical-protein-[Bacteroides-ndongoniae] |
| WP_073047778.1 | Type4.PglZ.containing.BREX | <-DndE<-DndD<-RNase<-SIGMA+OB-Fold+AAA-ATPase-BrxL<-iSTAND+PglZ<-DUF6079*<-wHTH<-PAPS-reduct+a+b-domain+Fer4_9+DUF4007\|\|?-><-HicB<-?\|\|?->?-><-?<-HAD+HAD | 1145 | Bacteria>FCB_group>Bacteroidetes/Chlorobi_group>Bacteroidota>Bacteroidia>Bacteroidales | GCF_900112275.1 | Prevotella_ruminicola | BM023_RS13275 | 8117..11554 | 839 | hypothetical-protein-[Prevotella-ruminicola] |
| WP_073046231.1 | Type4.PglZ.containing.BREX | <-REase<-?<-?<-?\|\|REase->LRR_5->HTH_3->PAPS-reduct+a+b-domain+Fer4_9+DUF4007->wHTH->DUF6079*->iSTAND+PglZ-> | 1145 | Bacteria>FCB_group>Bacteroidetes/Chlorobi_group>Bacteroidota>Bacteroidia>Bacteroidales | GCF_900142855.1 | Prevotella_ruminicola | BUB50_RS10445 | 275764..279201 | 839 | hypothetical-protein-[Prevotella-ruminicola] |
| WP_075965318.1 | Type4.PglZ.containing.BREX | DEDD_Tnp_IS110+Transposase_20->HATPase+wHTH->HTH-><-?<-?<-TIR<-SIGMA+OB-Fold+AAA-ATPase-BrxL<-iSTAND+PglZ<-DUF6079*<-wHTH<-PAPS-reduct+a+b-domain+Fer4_9+DUF4007<-HTH_3<-STYKinase-like<-TRPR-HTH<-DEDD_Tnp_IS110+Transposase_20<-SbcC<-JetB | 1145 | Bacteria>FCB_group>Bacteroidetes/Chlorobi_group>Bacteroidota>Bacteroidia>Bacteroidales | GCF_900155425.1 | Parabacteroides_massiliensis | BQ9893_RS06875 | 1631215..1634652 | 1750560 | hypothetical-protein-[Parabacteroides-massiliensis] |
| WP_005930488.1 | Type4.PglZ.containing.BREX | Arm-DNA-bind_5+Phage_int_SAM_5+Phage_integrase-><-XIS-HTH<-XIS-HTH\|\|CopG/RHH->?->?->HTH_3->PAPS-reduct+a+b-domain+Fer4_9+DUF4007->wHTH->DUF6079*->iSTAND+PglZ->SIGMA+OB-Fold+AAA-ATPase-BrxL->Schlafen+HATPase+wHTH->Glyco_hydro_43+GH43_C2-> | 1181 | Bacteria>FCB_group>Bacteroidetes/Chlorobi_group>Bacteroidota>Bacteroidia>Bacteroidales | GCF_902374995.1 | Bacteroides_salyersiae | HMPREF1532_RS13495 | 521965..525510 | 291644 | hypothetical-protein-[Bacteroides-salyersiae] |
| WP_091533684.1 | Type4.PglZ.containing.BREX | DSBH+AraC-HTH-><-N6-N4-METHYLASE<-SIGMA-HTH<-SIGMA+OB-Fold+AAA-ATPase-BrxL<-iSTAND+PglZ<-DUF6079*<-PAPS-reduct+a+b-domain+Fer4_9+DUF4007\|\|?->XIS-HTH->MarR-HTH->AAA+->?->Phage_int_SAM_1+Phage_integrase-> | 1160 | Bacteria>FCB_group>Bacteroidetes/Chlorobi_group>Bacteroidota>Bacteroidia>Bacteroidales | GCF_904381355.1 | Alistipes_sp._CHKCI003 | BN3659_00095 | 18669..22151 | 1780376 | hypothetical-protein-[Alistipes-sp.-CHKCI003] |
| WP_195558544.1 | Type4.PglZ.containing.BREX | <-TetR-HTH<-?\|\|?->SIGMA-HTH+MA->REase->REase->HTH_3->PAPS-reduct+a+b-domain+Fer4_9+DUF4007->DUF6079*->iSTAND+PglZ->SIGMA+OB-Fold+AAA-ATPase-BrxL->?-><-?<-?<-?<-SF1-SF2-Helicase+35exo | 1273 | Bacteria>FCB_group>Bacteroidetes/Chlorobi_group>Bacteroidota>Bacteroidia>Bacteroidales | GCF_918279245.1 | Bacteroides | NMG78_RS00315 | 41758..45579 | 816 | MULTISPECIES:-hypothetical-protein-[Bacteroides] |
| WP_057099388.1 | Type4.PglZ.containing.BREX | <-SIR2<-?<-ACET<-Schlafen+HATPase+wHTH<-SIGMA+OB-Fold+AAA-ATPase-BrxL<-iSTAND+PglZ<-DUF6079*<-PAPS-reduct+a+b-domain+Fer4_9+DUF4007<-HTH_3<-?<-TraJ-RHH\|\|?->?->CopG/RHH\|\|XIS-HTH->XIS-HTH-> | 1140 | Bacteria>FCB_group>Bacteroidetes/Chlorobi_group>Bacteroidota>Bacteroidia>Bacteroidales | GCF_932186025.1 | Bacteroides_acidifaciens | QDP01_RS07415 | 40613..44035 | 85831 | hypothetical-protein-[Bacteroides-acidifaciens] |
| WP_136416193.1 | Type4.PglZ.containing.BREX | <-SeqA<-SIGMA+OB-Fold+AAA-ATPase-BrxL<-iSTAND+PglZ<-DUF4372+DDE_Tnp_1<-?<-DUF6079-p<-DUF6079-p*<-PAPS-reduct+a+b-domain+Fer4_9+DUF4007<-HTH_3<-?<-STYKinase-like<-TRPR-HTH<-?<-Calcineurin<-Calcineurin | 159 | Bacteria>FCB_group>Bacteroidetes/Chlorobi_group>Bacteroidota>Bacteroidia>Bacteroidales | GCF_949104925.1 | Duncaniella_dubosii | E7747_RS12100 | 2715438..2715917 | 2518971 | hypothetical-protein-[Duncaniella-dubosii] |
| WP_216651884.1 | Type4.PglZ.containing.BREX | XIS-HTH+XIS-HTH->?-><-?\|\|PAPS-reduct+a+b-domain+Fer4_9+DUF4007->DUF6079*->iSTAND+PglZ->?->?->?->?-><-?<-Zn_Tnp_IS1595+DDE_Tnp_IS1595\|\|XIS-HTH-> | 1073 | Bacteria>FCB_group>Bacteroidetes/Chlorobi_group>Bacteroidota>Bacteroidia>Bacteroidales | GCF_949526175.1 | Prevotella_sp._PTAC | HPS53_RS04320 | 189281..192502 | 2736295 | hypothetical-protein-[Prevotella-sp.-PTAC] |
| WP_195241298.1 | Type4.PglZ.containing.BREX | <-AbrB-HTH<-?\|\|?->XIS-HTH->XIS-HTH-><-?<-Schlafen+HATPase+wHTH<-SIGMA+OB-Fold+AAA-ATPase-BrxL<-iSTAND+PglZ<-DUF6079*<-PAPS-reduct+a+b-domain+Fer4_9+DUF4007<-HTH_3\|\|BirA-HTH-><-Arm-DNA-bind_5+Phage_int_SAM_5+Phage_integrase<-Arm-DNA-bind_5+Phage_int_SAM_5+Phage_integrase\|\|Phage_integrase-><-TIR<-Peptidase_C14<-TIR | 1140 | Bacteria>FCB_group>Bacteroidetes/Chlorobi_group>Bacteroidota>Bacteroidia>Bacteroidales | GCF_958425225.1 | Odoribacter_laneus | I2A63_RS11780 | 58644..62066 | 626933 | hypothetical-protein-[Odoribacter-laneus] |
| WP_160246700.1 | Type4.PglZ.containing.BREX | CopG/RHH-><-AbrB-HTH<-?\|\|XIS-HTH->XIS-HTH-><-?<-Schlafen+HATPase+wHTH<-SIGMA+OB-Fold+AAA-ATPase-BrxL<-iSTAND+PglZ<-DUF6079*<-PAPS-reduct+a+b-domain+Fer4_9+DUF4007<-HTH_3\|\|<-Arm-DNA-bind_5+Phage_int_SAM_5+Phage_integrase<-Arm-DNA-bind_5+Phage_int_SAM_5+Phage_integrase\|\|Phage_integrase->XIS-HTH-> | 1140 | Bacteria>FCB_group>Bacteroidetes/Chlorobi_group>Bacteroidota>Bacteroidia>Bacteroidales | - | Phocaeicola_sartorii | D3Z46_05570 | 68877..72299 | 671267 | hypothetical-protein-[Phocaeicola-sartorii] |
| WP_153112824.1 | Type4.PglZ.containing.BREX | ABC-ATPase+SbcC+SWACOS+SbcC->REase-><-Cysteine-Desulfurase<-ZnR<-SIGMA+OB-Fold+AAA-ATPase-BrxL<-iSTAND+PglZ<-DUF6079*<-PAPS-reduct+a+b-domain+Fer4_9+DUF4007<-?\|\|wHTH-> | 1390 | Bacteria>FCB_group>Bacteroidetes/Chlorobi_group>Bacteroidota>Bacteroidia>Bacteroidales | - | Prevotella_copri | F7D34_RS04375 | 44361..48533 | 165179 | hypothetical-protein-[Prevotella-copri] |
| WP_153133795.1 | Type4.PglZ.containing.BREX | ABC-ATPase+SbcC+SWACOS+SbcC->REase-><-Cysteine-Desulfurase<-SIGMA+OB-Fold+AAA-ATPase-BrxL<-iSTAND+PglZ<-DUF6079*<-PAPS-reduct+a+b-domain+Fer4_9+DUF4007<-?\|\|wHTH-> | 758 | Bacteria>FCB_group>Bacteroidetes/Chlorobi_group>Bacteroidota>Bacteroidia>Bacteroidales | - | Prevotella_copri | F7D59_RS16355 | 52723..54999 | 165179 | hypothetical-protein-[Prevotella-copri] |
| CDC25991.1 | Type4.PglZ.containing.BREX | PAPS-reduct+DUF4007->DUF6079*->iSTAND+PglZ->SIGMA+OB-Fold+AAA-ATPase-BrxL->SIGMA+OB-Fold+AAA-ATPase-BrxL->HTH_3-><-?<-?\|\|LuxR-HTH-> | 1166 | Bacteria>FCB_group>Bacteroidetes/Chlorobi_group>Bacteroidota>Bacteroidia>Bacteroidales | - | Prevotella_sp._CAG:386 | BN637_01806 | 657..4157 | 1262925 | putative-uncharacterized-protein-[Prevotella-sp.-CAG:386] |
| WP_195485920.1 | Type4.PglZ.containing.BREX | SIR2->HerA/FtsK->?->REase->ACET-><-?<-SeqA<-SIGMA+OB-Fold+AAA-ATPase-BrxL<-iSTAND+PglZ<-DUF6079*<-PAPS-reduct+a+b-domain+Fer4_9+DUF4007<-HTH_3<-?<-STYKinase-like<-TRPR-HTH\|\|SPOUT-><-?<-ADH_N+ADH_zinc_N+METHYLASE\|\|HTH+REase-> | 1134 | Bacteria>FCB_group>Bacteroidetes/Chlorobi_group>Bacteroidota>Bacteroidia | GCF_015559535.1 | Bacteroidales | I2I08_RS05855 | 97983..101387 | 171549 | MULTISPECIES:-hypothetical-protein-[Bacteroidales] |
| WP_016212911.1 | Type4.PglZ.containing.BREX | ABC-ATPase+SbcC->?->?->?-><-?<-?\|\|Arm-DNA-bind_5+Phage_int_SAM_5+Phage_integrase-><-?\|\|PAPS-reduct+a+b-domain+Fer4_9+DUF4007->DUF6079*->iSTAND+PglZ->SIGMA+OB-Fold+AAA-ATPase-BrxL-><-FurR-HTH<-?<-TraJ-RHH<-CopG/RHH\|\|XIS-HTH->XIS-HTH-> | 1151 | Bacteria>FCB_group>Bacteroidetes/Chlorobi_group>Bacteroidota>Bacteroidia | GCF_947368045.1 | Bacteroidales | QDP13_RS02715 | 27965..31420 | 171549 | MULTISPECIES:-hypothetical-protein-[Bacteroidales] |
| WP_178291773.1 | Type4.PglZ.containing.BREX | <-CNMP+CRP-HTH\|\|?->?->XIS-HTH->TOPRIM->XIS-HTH->SbcC-><-?\|\|PAPS-reduct+a+b-domain+Fer4_9+DUF4007->DUF6079*->iSTAND+PglZ->SIGMA+OB-Fold+AAA-ATPase-BrxL-><-FurR-HTH<-?<-TraJ-RHH<-CopG/RHH\|\|XIS-HTH->XIS-HTH-> | 1151 | Bacteria>FCB_group>Bacteroidetes/Chlorobi_group>Bacteroidota>Bacteroidia | GCF_948480705.1 | Bacteroidales | K6V28_RS12855 | 2860583..2864038 | 171549 | MULTISPECIES:-hypothetical-protein-[Bacteroidales] |
| NCB24126.1 | Type4.PglZ.containing.BREX | HTH_7+rve+rve_3->AAA-ATPase-><-?<-OLD-ABC+TOPRIM-p<-?<-wHTH+WYL+WCX<-Cysteine-Desulfurase<-ZnR<-SIGMA+OB-Fold+AAA-ATPase-BrxL<-iSTAND+PglZ<-DUF6079*<-PAPS-reduct+a+b-domain+Fer4_9+DUF4007<-SeqA+Phage_int+SeqA+E3_UFM1_ligase<-HTH_26<-?<-?<-TraJ-RHH<-?<-ArdCN+LPD32+LPD36+N6_Mtase+SF1-SF2-Helicase<-URI-DNase | 1381 | Bacteria>FCB_group>Bacteroidetes/Chlorobi_group>Bacteroidota>Bacteroidia>unclassified_Bacteroidia | GCA_009929165.1 | Bacteroidia_bacterium | EOM62_01415 | 22073..26218 | 2044936 | hypothetical-protein-[Bacteroidia-bacterium] |
| NCB25245.1 | Type4.PglZ.containing.BREX | HARE-HTH+E3_UFM1_ligase->HNH->PAPS-reduct+a+b-domain+Fer4_9+DUF4007->DUF6079*->iSTAND+PglZ->SIGMA+OB-Fold+AAA-ATPase-BrxL->Cysteine-Desulfurase-> | 1397 | Bacteria>FCB_group>Bacteroidetes/Chlorobi_group>Bacteroidota>Bacteroidia>unclassified_Bacteroidia | GCA_009929165.1 | Bacteroidia_bacterium | EOM62_07190 | 5577..9770 | 2044936 | hypothetical-protein-[Bacteroidia-bacterium] |
| OGX83058.1 | Type4.PglZ.containing.BREX | HTH_36->DnaB+TnsC-ATPase->STYKinase-like->?->DUF4007->DUF6079*->PAPS-reduct->?->DUF6079->PglZ->SIGMA+OB-Fold+AAA-ATPase-BrxL->Cysteine-Desulfurase->SF1-SF2-Helicase-><-?<-HTH_Tnp_1+HTH_21+rve+rve_3 | 1109 | Bacteria>FCB_group>Bacteroidetes/Chlorobi_group>Bacteroidota>Cytophagia>Cytophagales | GCA_001816165.1 | Hymenobacter_glacialis | BEN48_04715 | 162359..165688 | 1908236 | hypothetical-protein-BEN48_04715-[Hymenobacter-glacialis] |
| TPG64627.1 | Type4.PglZ.containing.BREX | <-DndB<-SIGMA+OB-Fold+AAA-ATPase-BrxL<-PglZ<-DUF6079*<-DUF4007<-PAPS-reduct<-DUF6079*<-DUF4007<-Phage_integrase+Phage_integrase\|\|?->?->?-><-tRNA\|\|?->ACET+ACET->?-><-N6-METHYLASE | 1041 | Bacteria>FCB_group>Bacteroidetes/Chlorobi_group>Bacteroidota>Cytophagia>Cytophagales | GCA_006439025.1 | Hymenobacter_nivis | EAH73_15805 | 346011..349136 | 1850093 | hypothetical-protein-EAH73_15805-[Hymenobacter-nivis] |
| MBC7570527.1 | Type4.PglZ.containing.BREX | XIS-HTH->?->?->?-><-Peptidase_M28\|\|?-><-SIGMA+OB-Fold+AAA-ATPase-BrxL<-PglZ<-DUF6079*<-?<-DUF4007<-PAPS-reduct | 1552 | Bacteria>FCB_group>Bacteroidetes/Chlorobi_group>Bacteroidota>Cytophagia>Cytophagales | GCA_014377625.1 | Spirosoma_sp. | H7319_12430 | 32428..37086 | 1899569 | hypothetical-protein-[Spirosoma-sp.] |
| MBH8560397.1 | Type4.PglZ.containing.BREX | <-REC<-?<-?\|\|Phage_integrase+Phage_integrase->?->?-><-?\|\|DUF4007->DUF6079*->PAPS-reduct->wHTH+URI-DNase->DUF4007->?->DUF6079->PglZ->SIGMA+OB-Fold+AAA-ATPase-BrxL-><-Sp<-TOPRIM | 1075 | Bacteria>FCB_group>Bacteroidetes/Chlorobi_group>Bacteroidota>Cytophagia>Cytophagales | GCA_016056375.1 | Hymenobacter_negativus | I7X13_20215 | 50672..53899 | 2795026 | hypothetical-protein-[Hymenobacter-negativus] |
| WP_035559952.1 | Type4.PglZ.containing.BREX | Peptidase_M15_4->LexA-protease->?->?-><-Cysteine-Desulfurase<-SIGMA+OB-Fold+AAA-ATPase-BrxL<-PglZ<-DUF6079*<-?<-DUF4007<-PAPS-reduct<-REase<-?<-DUF6079*<-DUF4007\|\|DOC+HTH-> | 1453 | Bacteria>FCB_group>Bacteroidetes/Chlorobi_group>Bacteroidota>Cytophagia>Cytophagales | GCF_000715495.1 | Hymenobacter_sp._IS2118 | GM79_RS04550 | 15393..19754 | 1505605 | hypothetical-protein-[Hymenobacter-sp.-IS2118] |
| WP_035559967.1 | Type4.PglZ.containing.BREX | <-Cysteine-Desulfurase<-SIGMA+OB-Fold+AAA-ATPase-BrxL<-PglZ<-DUF6079*<-DUF4007<-PAPS-reduct<-REase<-?<-DUF6079*<-DUF4007\|\|DOC+HTH-><-?<-DnaB+TnsC-ATPase<-HTH_36<-?<-XIS-HTH<-?\|\|CHTH+LexA-protease-> | 1127 | Bacteria>FCB_group>Bacteroidetes/Chlorobi_group>Bacteroidota>Cytophagia>Cytophagales | GCF_000715495.1 | Hymenobacter_sp._IS2118 | GM79_RS04580 | 23577..26960 | 1505605 | hypothetical-protein-[Hymenobacter-sp.-IS2118] |
| WP_125932665.1 | Type4.PglZ.containing.BREX | HTH_36->DnaB+TnsC-ATPase->STYKinase-like->?->DUF4007->DUF6079*->PAPS-reduct->DUF4007->?->DUF6079->PglZ->SIGMA+OB-Fold+AAA-ATPase-BrxL->Cysteine-Desulfurase->SF1-SF2-Helicase-> | 1123 | Bacteria>FCB_group>Bacteroidetes/Chlorobi_group>Bacteroidota>Cytophagia>Cytophagales | GCF_001816165.1 | Hymenobacter_glacialis | BEN48_RS14520 | 162317..165688 | 1908236 | hypothetical-protein-[Hymenobacter-glacialis] |
| WP_135462782.1 | Type4.PglZ.containing.BREX | DrHyd->Schlafen+HATPase+wHTH-><-?<-?<-DEDD_Tnp_IS110<-XIS-HTH<-Cysteine-Desulfurase<-SIGMA+OB-Fold+AAA-ATPase-BrxL<-PglZ<-DUF6079*<-DUF4007<-PAPS-reduct<-DUF6079-p<-DUF6079-p<-DUF4007\|\|REase+NACHT-><-MORC+DUF3684+REase<-Tox-RES<-HTH_3 | 1444 | Bacteria>FCB_group>Bacteroidetes/Chlorobi_group>Bacteroidota>Cytophagia>Cytophagales | GCF_004765605.1 | Hymenobacter_aquaticus | E5K00_RS08395 | 2047536..2051870 | 1867101 | hypothetical-protein-[Hymenobacter-aquaticus] |
| WP_135462784.1 | Type4.PglZ.containing.BREX | <-DEDD_Tnp_IS110<-XIS-HTH<-Cysteine-Desulfurase<-SIGMA+OB-Fold+AAA-ATPase-BrxL<-PglZ<-DUF6079*<-DUF4007<-PAPS-reduct<-DUF6079-p*<-DUF6079-p<-DUF4007\|\|REase+NACHT-><-MORC+DUF3684+REase<-Tox-RES<-HTH_3\|\|GDResolvase+Recombinase+ZnR->?-><-tRNA\|\|HAD+HAD-> | 860 | Bacteria>FCB_group>Bacteroidetes/Chlorobi_group>Bacteroidota>Cytophagia>Cytophagales | GCF_004765605.1 | Hymenobacter_aquaticus | E5K00_RS08410 | 2054589..2057171 | 1867101 | hypothetical-protein-[Hymenobacter-aquaticus] |
| WP_135462785.1 | Type4.PglZ.containing.BREX | <-DEDD_Tnp_IS110<-XIS-HTH<-Cysteine-Desulfurase<-SIGMA+OB-Fold+AAA-ATPase-BrxL<-PglZ<-DUF6079*<-DUF4007<-PAPS-reduct<-DUF6079-p*<-DUF6079-p<-DUF4007\|\|REase+NACHT-><-MORC+DUF3684+REase<-Tox-RES<-HTH_3\|\|GDResolvase+Recombinase+ZnR->?-><-tRNA\|\|HAD+HAD->Peptidase_M50-> | 246 | Bacteria>FCB_group>Bacteroidetes/Chlorobi_group>Bacteroidota>Cytophagia>Cytophagales | GCF_004765605.1 | Hymenobacter_aquaticus | E5K00_RS08415 | 2057214..2057954 | 1867101 | hypothetical-protein-[Hymenobacter-aquaticus] |
| WP_139515175.1 | Type4.PglZ.containing.BREX | RVT-><-Cysteine-Desulfurase<-SIGMA+OB-Fold+AAA-ATPase-BrxL<-PglZ<-DUF6079*<-?<-DUF4007<-PAPS-reduct<-DUF6079<-DUF4007<-?<-?<-D5N+viral-AAA+CDC6-HTH<-XIS-HTH | 1453 | Bacteria>FCB_group>Bacteroidetes/Chlorobi_group>Bacteroidota>Cytophagia>Cytophagales | GCF_006337165.1 | Hymenobacter_jejuensis | FHG12_RS07675 | 1893030..1897391 | 2502781 | hypothetical-protein-[Hymenobacter-jejuensis] |
| WP_139515178.1 | Type4.PglZ.containing.BREX | RVT-><-Cysteine-Desulfurase<-SIGMA+OB-Fold+AAA-ATPase-BrxL<-PglZ<-DUF6079<-?<-DUF4007<-PAPS-reduct<-DUF6079*<-DUF4007<-?<-?<-D5N+viral-AAA+CDC6-HTH<-XIS-HTH<-?<-Arm-DNA-bind_5+Phage_int_SAM_5+Phage_integrase | 1113 | Bacteria>FCB_group>Bacteroidetes/Chlorobi_group>Bacteroidota>Cytophagia>Cytophagales | GCF_006337165.1 | Hymenobacter_jejuensis | FHG12_RS07695 | 1899664..1903005 | 2502781 | hypothetical-protein-[Hymenobacter-jejuensis] |
| WP_140468254.1 | Type4.PglZ.containing.BREX | TraJ-RHH->TraI-VirD2-RCRE->?-><-?<-?<-?<-DndB<-SIGMA+OB-Fold+AAA-ATPase-BrxL<-PglZ<-DUF6079*<-DUF4007<-PAPS-reduct<-DUF6079<-DUF4007<-Phage_integrase+Phage_integrase | 1563 | Bacteria>FCB_group>Bacteroidetes/Chlorobi_group>Bacteroidota>Cytophagia>Cytophagales | GCF_006439025.1 | Hymenobacter_nivis | EAH73_RS15850 | 338987..343678 | 1850093 | hypothetical-protein-[Hymenobacter-nivis] |
| WP_140468259.1 | Type4.PglZ.containing.BREX | <-DndB<-SIGMA+OB-Fold+AAA-ATPase-BrxL<-PglZ<-DUF6079<-DUF4007<-PAPS-reduct<-DUF6079*<-DUF4007<-Phage_integrase+Phage_integrase\|\|?->?->?-><-tRNA\|\|?->ACET+ACET-> | 1075 | Bacteria>FCB_group>Bacteroidetes/Chlorobi_group>Bacteroidota>Cytophagia>Cytophagales | GCF_006439025.1 | Hymenobacter_nivis | EAH73_RS15865 | 346011..349238 | 1850093 | hypothetical-protein-[Hymenobacter-nivis] |
| WP_150087478.1 | Type4.PglZ.containing.BREX | <-Cysteine-Desulfurase<-SIGMA+OB-Fold+AAA-ATPase-BrxL<-PglZ<-DUF6079*<-DUF4007<-PAPS-reduct<-DUF6079<-DUF4007<-?<-SF1-SF2-Helicase<-?\|\|ABhydrolase-> | 1456 | Bacteria>FCB_group>Bacteroidetes/Chlorobi_group>Bacteroidota>Cytophagia>Cytophagales | GCF_008629685.1 | Adhaeribacter_rhizoryzae | F0145_RS06375 | 322504..326874 | 2607907 | hypothetical-protein-[Adhaeribacter-rhizoryzae] |
| WP_150087481.1 | Type4.PglZ.containing.BREX | <-Cysteine-Desulfurase<-SIGMA+OB-Fold+AAA-ATPase-BrxL<-PglZ<-DUF6079<-DUF4007<-PAPS-reduct<-DUF6079*<-DUF4007<-?<-SF1-SF2-Helicase<-?\|\|ABhydrolase-><-?<-?\|\|?-><-HTH_3 | 1125 | Bacteria>FCB_group>Bacteroidetes/Chlorobi_group>Bacteroidota>Cytophagia>Cytophagales | GCF_008629685.1 | Adhaeribacter_rhizoryzae | F0145_RS06390 | 329614..332991 | 2607907 | hypothetical-protein-[Adhaeribacter-rhizoryzae] |
| WP_171591910.1 | Type4.PglZ.containing.BREX | DnaB+TnsC-ATPase->HEPN-><-Classical-AAA<-?<-?<-?<-?<-?\|\|DUF4007->DUF6079*->PAPS-reduct->DUF4007->?->DUF6079->PglZ->SIGMA+OB-Fold+AAA-ATPase-BrxL->Cysteine-Desulfurase-> | 1116 | Bacteria>FCB_group>Bacteroidetes/Chlorobi_group>Bacteroidota>Cytophagia>Cytophagales | GCF_013137895.1 | Hymenobacter_taeanensis | HMJ29_RS13045 | 3059346..3062696 | 2735321 | hypothetical-protein-[Hymenobacter-taeanensis] |
| WP_171591913.1 | Type4.PglZ.containing.BREX | DUF4007->DUF6079->PAPS-reduct->DUF4007->?->DUF6079*->PglZ->SIGMA+OB-Fold+AAA-ATPase-BrxL->Cysteine-Desulfurase-> | 1445 | Bacteria>FCB_group>Bacteroidetes/Chlorobi_group>Bacteroidota>Cytophagia>Cytophagales | GCF_013137895.1 | Hymenobacter_taeanensis | HMJ29_RS13065 | 3065021..3069358 | 2735321 | hypothetical-protein-[Hymenobacter-taeanensis] |
| WP_192822483.1 | Type4.PglZ.containing.BREX | <-SIGMA+OB-Fold+AAA-ATPase-BrxL<-PglZ<-DUF6079*<-?<-DUF4007<-PAPS-reduct<-DUF6079<-DUF4007<-?\|\|HNH-> | 1456 | Bacteria>FCB_group>Bacteroidetes/Chlorobi_group>Bacteroidota>Cytophagia>Cytophagales | GCF_014876175.1 | Rufibacter_sp._LB8 | IMY37_RS12980 | 3071589..3075959 | 2777781 | hypothetical-protein-[Rufibacter-sp.-LB8] |
| WP_192822487.1 | Type4.PglZ.containing.BREX | <-SIGMA+OB-Fold+AAA-ATPase-BrxL<-PglZ<-DUF6079<-?<-DUF4007<-PAPS-reduct<-DUF6079*<-DUF4007<-?\|\|HNH->?-><-?\|\|?-><-?<-Calcineurin\|\|Phage_holin_3_6-> | 1073 | Bacteria>FCB_group>Bacteroidetes/Chlorobi_group>Bacteroidota>Cytophagia>Cytophagales | GCF_014876175.1 | Rufibacter_sp._LB8 | IMY37_RS13000 | 3078586..3081807 | 2777781 | hypothetical-protein-[Rufibacter-sp.-LB8] |
| WP_198076871.1 | Type4.PglZ.containing.BREX | <-REC<-?<-?\|\|Phage_integrase+Phage_integrase->?->?-><-?\|\|DUF4007->DUF6079*->PAPS-reduct->wHTH+URI-DNase->DUF4007->?->DUF6079->PglZ->SIGMA+OB-Fold+AAA-ATPase-BrxL-><-Sp<-TOPRIM | 1071 | Bacteria>FCB_group>Bacteroidetes/Chlorobi_group>Bacteroidota>Cytophagia>Cytophagales | GCF_016056375.1 | Hymenobacter_negativus | I7X13_RS20240 | 50684..53899 | 2795026 | hypothetical-protein-[Hymenobacter-negativus] |
| WP_198076876.1 | Type4.PglZ.containing.BREX | DUF4007->DUF6079->PAPS-reduct->wHTH+URI-DNase->DUF4007->?->DUF6079*->PglZ->SIGMA+OB-Fold+AAA-ATPase-BrxL-><-Sp<-TOPRIM<-CHTH<-HEPN | 1553 | Bacteria>FCB_group>Bacteroidetes/Chlorobi_group>Bacteroidota>Cytophagia>Cytophagales | GCF_016056375.1 | Hymenobacter_negativus | I7X13_RS20265 | 57107..61768 | 2795026 | hypothetical-protein-[Hymenobacter-negativus] |
| WP_215594963.1 | Type4.PglZ.containing.BREX | STYKinase-like->?->DUF4007->DUF6079*->PAPS-reduct->DUF4007->?->DUF6079->PglZ->SIGMA+OB-Fold+AAA-ATPase-BrxL->Cysteine-Desulfurase-> | 1119 | Bacteria>FCB_group>Bacteroidetes/Chlorobi_group>Bacteroidota>Cytophagia>Cytophagales | GCF_018760735.1 | Hymenobacter_piscis | KLP40_RS16680 | 3218..6577 | 2839984 | hypothetical-protein-[Hymenobacter-piscis] |
| WP_219156489.1 | Type4.PglZ.containing.BREX | <-LexA-protease\|\|?-><-APATPase<-AbiEii<-wHTH+AbiEi_2<-SLATT<-Cysteine-Desulfurase<-SIGMA+OB-Fold+AAA-ATPase-BrxL<-PglZ<-DUF6079*<-?<-DUF4007<-PAPS-reduct<-DUF6079<-DUF4007<-?<-STYKinase-like<-?<-DnaB+TnsC-ATPase | 1456 | Bacteria>FCB_group>Bacteroidetes/Chlorobi_group>Bacteroidota>Cytophagia>Cytophagales | GCF_019334315.1 | Hymenobacter_profundi | KYK14_RS01920 | 88615..92985 | 1982110 | hypothetical-protein-[Hymenobacter-profundi] |
| WP_219156495.1 | Type4.PglZ.containing.BREX | <-wHTH+AbiEi_2<-SLATT<-Cysteine-Desulfurase<-SIGMA+OB-Fold+AAA-ATPase-BrxL<-PglZ<-DUF6079<-?<-DUF4007<-PAPS-reduct<-DUF6079*<-DUF4007<-?<-STYKinase-like<-?<-DnaB+TnsC-ATPase<-HTH_36<-?<-XIS-HTH | 1119 | Bacteria>FCB_group>Bacteroidetes/Chlorobi_group>Bacteroidota>Cytophagia>Cytophagales | GCF_019334315.1 | Hymenobacter_profundi | KYK14_RS01940 | 95233..98592 | 1982110 | hypothetical-protein-[Hymenobacter-profundi] |
| WP_235137403.1 | Type4.PglZ.containing.BREX | ABC-ATPase+SbcC->Glyco_hydro_3+Glyco_hydro_3_C+Fn3-like->?->?->?->tRNA->Arm-DNA-bind_5+Phage_int_SAM_5+Phage_integrase->DUF4007->DUF6079*->PAPS-reduct->DUF4007->?->DUF6079->PglZ->SIGMA+OB-Fold+AAA-ATPase-BrxL-><-Cysteine-Desulfurase<-?<-MORC+DUF3684+REase | 1079 | Bacteria>FCB_group>Bacteroidetes/Chlorobi_group>Bacteroidota>Cytophagia>Cytophagales | GCF_021502735.1 | Dyadobacter_fanqingshengii | L0663_RS10270 | 2404601..2407840 | 2906443 | hypothetical-protein-[Dyadobacter-fanqingshengii] |
| WP_235137407.1 | Type4.PglZ.containing.BREX | Arm-DNA-bind_5+Phage_int_SAM_5+Phage_integrase->DUF4007->DUF6079->PAPS-reduct->DUF4007->?->DUF6079*->PglZ->SIGMA+OB-Fold+AAA-ATPase-BrxL-><-Cysteine-Desulfurase<-?<-MORC+DUF3684+REase<-VirD4-TraG+VirD4-FtsK<-?\|\|DOC+HTH->ANK+ANK-> | 1552 | Bacteria>FCB_group>Bacteroidetes/Chlorobi_group>Bacteroidota>Cytophagia>Cytophagales | GCF_021502735.1 | Dyadobacter_fanqingshengii | L0663_RS10290 | 2410166..2414824 | 2906443 | hypothetical-protein-[Dyadobacter-fanqingshengii] |
| WP_243797657.1 | Type4.PglZ.containing.BREX | HTH+REase->?->?-><-DDE_Tnp<-TIR<-HTH_3\|\|?->?->DUF4007->DUF6079*->PAPS-reduct->DUF4007->?->DUF6079->PglZ->SIGMA+OB-Fold+AAA-ATPase-BrxL->Cysteine-Desulfurase->SF1-SF2-Helicase-> | 1123 | Bacteria>FCB_group>Bacteroidetes/Chlorobi_group>Bacteroidota>Cytophagia>Cytophagales | GCF_022827545.1 | Hymenobacter_tibetensis | MTX78_RS19760 | 4898444..4901815 | 497967 | hypothetical-protein-[Hymenobacter-tibetensis] |
| WP_243797663.1 | Type4.PglZ.containing.BREX | <-TIR<-HTH_3\|\|?->?->DUF4007->DUF6079->PAPS-reduct->DUF4007->?->DUF6079*->PglZ->SIGMA+OB-Fold+AAA-ATPase-BrxL->Cysteine-Desulfurase->SF1-SF2-Helicase->?->N6-METHYLASE->?-><-Arm-DNA-bind_5+Phage_int_SAM_5+Phage_integrase | 1449 | Bacteria>FCB_group>Bacteroidetes/Chlorobi_group>Bacteroidota>Cytophagia>Cytophagales | GCF_022827545.1 | Hymenobacter_tibetensis | MTX78_RS19780 | 4904124..4908473 | 497967 | hypothetical-protein-[Hymenobacter-tibetensis] |
| WP_092740295.1 | Type4.PglZ.containing.BREX | HTH_36->DnaB+TnsC-ATPase->DUF4007->DUF6079-p->wHTH(SBD)+HNH->SAM-METHYLASE+HIT-><-?<-REase\|\|DUF6079*->PglZ->SIGMA+OB-Fold+AAA-ATPase-BrxL->Cysteine-Desulfurase->MORC+DUF3684+REase->wHTH+AbiEi_2->AbiEii-><-ParB<-?<-Peptidase_M15_4 | 1353 | Bacteria>FCB_group>Bacteroidetes/Chlorobi_group>Bacteroidota>Cytophagia>Cytophagales | GCF_900107135.1 | Hymenobacter_psychrophilus | BLW21_RS11955 | 114723..118784 | 651662 | hypothetical-protein-[Hymenobacter-psychrophilus] |
| WP_194112249.1 | Type4.PglZ.containing.BREX | <-VirD4-TraG+VirD4-FtsK<-TraI-VirD2-RCRE<-TraJ-RHH<-?<-XIS-HTH\|\|DUF4007->DUF6079*->PAPS-reduct->DUF4007->?->DUF6079->PglZ->SIGMA+OB-Fold+AAA-ATPase-BrxL-><-Cysteine-Desulfurase<-HARE-HTH | 1079 | Bacteria>FCB_group>Bacteroidetes/Chlorobi_group>Bacteroidota>Sphingobacteriia>Sphingobacteriales | GCF_015222005.1 | Mucilaginibacter_myungsuensis | IRJ16_RS14095 | 129346..132585 | 649104 | hypothetical-protein-[Mucilaginibacter-myungsuensis] |
| WP_194112253.1 | Type4.PglZ.containing.BREX | <-TraI-VirD2-RCRE<-TraJ-RHH<-?<-XIS-HTH\|\|DUF4007->DUF6079->PAPS-reduct->DUF4007->?->DUF6079*->PglZ->SIGMA+OB-Fold+AAA-ATPase-BrxL-><-Cysteine-Desulfurase<-HARE-HTH\|\|?->?-><-Arm-DNA-bind_5+Phage_int_SAM_5+Phage_integrase | 1550 | Bacteria>FCB_group>Bacteroidetes/Chlorobi_group>Bacteroidota>Sphingobacteriia>Sphingobacteriales | GCF_015222005.1 | Mucilaginibacter_myungsuensis | IRJ16_RS14115 | 134911..139563 | 649104 | hypothetical-protein-[Mucilaginibacter-myungsuensis] |
| OQC38485.1 | Type4.PglZ.containing.BREX | <-SIGMA+OB-Fold+AAA-ATPase-BrxL<-iSTAND+PglZ<-DUF6079*<-PAPS-reduct+a+b-domain+Fer4_9+DUF4007<-Schlafen+HATPase+wHTH\|\|?->?->?->HNH-> | 1201 | Bacteria>FCB_group>Bacteroidetes/Chlorobi_group>Bacteroidota>unclassified_Bacteroidota | GCA_002071145.1 | Bacteroidetes_bacterium_ADurb.Bin041 | BWX63_00120 | 3698..7303 | 1852801 | hypothetical-protein-BWX63_00120-[Bacteroidetes-bacterium-ADurb.Bin041] |
| OQC39308.1 | Type4.PglZ.containing.BREX | ABhydrolase-><-ABhydrolase<-?<-?<-DOC+HTH\|\|?->?-><-SIGMA+OB-Fold+AAA-ATPase-BrxL<-iSTAND+PglZ<-DUF6079*<-PAPS-reduct+a+b-domain+Fer4_9+DUF4007<-?<-REase<-?\|\|tRNA->?->LuxR-HTH->LuxR-HTH-> | 1129 | Bacteria>FCB_group>Bacteroidetes/Chlorobi_group>Bacteroidota>unclassified_Bacteroidota | GCA_002071175.1 | Bacteroidetes_bacterium_ADurb.Bin037 | BWX62_00954 | 67069..70458 | 1852800 | hypothetical-protein-BWX62_00954-[Bacteroidetes-bacterium-ADurb.Bin037] |
| MBV5327599.1 | Type4.PglZ.containing.BREX | <-Calcineurin<-?<-?<-?\|\|Phage_integrase-><-Cysteine-Desulfurase\|\|DUF4007->DUF6079*->PAPS-reduct-><-SIGMA+OB-Fold+AAA-ATPase-BrxL<-PglZ<-DUF6079<-?<-DUF4007<-GreA_GreB+SF1-SF2-Helicase<-PIN<-wHTH+WYL+WCX-p | 1116 | Bacteria>FCB_group>Bacteroidetes/Chlorobi_group>Chlorobiota>Chlorobiia>Chlorobiales | GCA_019162945.1 | Chlorobium_sp. | JZU65_08155 | 102097..105447 | 1095 | hypothetical-protein-[Chlorobium-sp.] |
| NLN94368.1 | Type4.PglZ.containing.BREX | <-Cysteine-Desulfurase<-ZnR<-SIGMA+OB-Fold+AAA-ATPase-BrxL<-iSTAND+PglZ<-DUF6079*<-PAPS-reduct+a+b-domain+Fer4_9+DUF4007 | 1388 | Bacteria>FCB_group>Candidatus_Hydrogenedentes>Candidatus_Hydrogenedens>unclassified_Candidatus_Hydrogenedens | GCA_012522755.1 | Candidatus_Hydrogenedens_sp. | GX130_13835 | 4950..9116 | 2654184 | hypothetical-protein-[Candidatus-Hydrogenedens-sp.] |
| MBO8152008.1 | Type4.PglZ.containing.BREX | <-KAP-NTPase<-SIGMA+OB-Fold+AAA-ATPase-BrxL<-iSTAND+PglZ<-DUF6079*<-PAPS-reduct+a+b-domain+Fer4_9+DUF4007\|\|?->?->?->?->HAD+HAD->CBS+CBS->?->ABC-ATPase+SbcC-> | 1124 | Bacteria>FCB_group>Candidatus_Marinimicrobia | GCA_017656455.1 | Candidatus_Marinimicrobia_bacterium | H0Z30_05895 | 8091..11465 | 2026760 | hypothetical-protein-[Candidatus-Marinimicrobia-bacterium] |

| Protein-ID | System | Gene-Neighbourhoods | Length | Taxonomy | Assembly-ID | Strain | Locus-Tag | Genomic-coordinates | TaxID | Fasta-defline |
|---|---|---|---|---|---|---|---|---|---|---|
| MCF0215815.1 | Type4.PglZ.containing.BREX | <-RADICAL-SAM<-?<-?<-?<-?\|\|HNH->JetC->\|\|?->PAPS-reduct+a+b-domain+Fer4_9+DUF4007->DUF6079*->iSTAND+PglZ->SIGMA+OB-Fold+AAA-ATPase-BrxL->ZnR->Cysteine-Desulfurase->SAM-METHYLASE-> | 1393 | Bacteria>FCB_group>Fibrobacterota>Fibrobacteria>Fibrobacterales>Fibrobacteraceae | GCA_021443045.1 | Fibrobacteraceae_bacterium | HUK21_05025 | 47335..51516 | 2652288 | hypothetical-protein-[Fibrobacteraceae-bacterium] |
| WP_073324032.1 | Type4.PglZ.containing.BREX | <-Cysteine-Desulfurase<-ZnR<-SIGMA+OB-Fold+AAA-ATPase-BrxL<-iSTAND+PglZ<-DUF6079*<-PAPS-reduct+a+b-domain+Fer4_9+DUF4007<-?\|\|MNS+REase-> | 1393 | Bacteria>FCB_group>Fibrobacterota>Fibrobacteria>Fibrobacterales>Fibrobacteraceae | GCF_900142495.1 | Fibrobacter_sp._UWOV1 | BUC02_RS15315 | 5401..9582 | 1896215 | hypothetical-protein-[Fibrobacter-sp.-UWOV1] |
| WP_185891394.1 | Type4.PglZ.containing.BREX | Cysteine-Desulfurase->?->?->PAPS-reduct+a+b-domain+Fer4_9->DUF4007->DUF6079*->wHTH+HNH-p->wHTH+HNH-p->DUF4007->?->DUF6079->PglZ+Phosphodiest->SIGMA+OB-Fold+AAA-ATPase-BrxL->TPR+S1->SF1-SF2-Helicase-> | 1100 | Bacteria>Fusobacteriota>Fusobacteriia>Fusobacteriales>Fusobacteriaceae>Cetobacterium | GCF_014250655.1 | Cetobacterium_sp._2G_large | H5J23_RS06535 | 772427..775729 | 2759680 | hypothetical-protein-[Cetobacterium-sp.-2G-large] |
| WP_185891404.1 | Type4.PglZ.containing.BREX | PAPS-reduct+a+b-domain+Fer4_9->DUF4007->DUF6079->wHTH+HNH-p->wHTH+HNH-p->DUF4007->?->DUF6079*->PglZ+Phosphodiest->SIGMA+OB-Fold+AAA-ATPase-BrxL->TPR+S1->SF1-SF2-Helicase->NADAR->GrpE->?->TPR+TPR+TPR->DnaJ-> | 1321 | Bacteria>Fusobacteriota>Fusobacteriia>Fusobacteriales>Fusobacteriaceae>Cetobacterium | GCF_014250655.1 | Cetobacterium_sp._2G_large | H5J23_RS06560 | 777659..781624 | 2759680 | hypothetical-protein-[Cetobacterium-sp.-2G-large] |
| WP_118127458.1 | Type4.PglZ.containing.BREX | <-HTH_3\|\|?->HNH-><-CR-ATPase8+REase\|\|Cysteine-Desulfurase->?->?->PAPS-reduct+a+b-domain+Fer4_9->DUF4007->DUF6079*->wHTH+HNH-p->DUF4007->?->DUF6079->PglZ+Phosphodiest->SIGMA+OB-Fold+AAA-ATPase-BrxL-> | 1117 | Bacteria>Fusobacteriota>Fusobacteriia>Fusobacteriales>Fusobacteriaceae>Fusobacterium | GCF_003470285.1 | Fusobacterium_mortiferum | DW670_RS09740 | 32239..35592 | 850 | AAA-family-ATPase-[Fusobacterium-mortiferum] |
| WP_118127466.1 | Type4.PglZ.containing.BREX | Cysteine-Desulfurase->?->?->PAPS-reduct+a+b-domain+Fer4_9->DUF4007->DUF6079->wHTH+HNH-p->DUF4007->?->DUF6079*->PglZ+Phosphodiest->SIGMA+OB-Fold+AAA-ATPase-BrxL-><-?<-?<-?\|\|Aminotran_4->?->?-><-GNTR-HTH | 1321 | Bacteria>Fusobacteriota>Fusobacteriia>Fusobacteriales>Fusobacteriaceae>Fusobacterium | GCF_003470285.1 | Fusobacterium_mortiferum | DW670_RS09760 | 37530..41495 | 850 | DUF2791-family-P-loop-domain-containing-protein-[Fusobacterium-mortiferum] |
| WP_118234045.1 | Type4.PglZ.containing.BREX | <-HTH_3\|\|?->HNH-><-CR-ATPase8+REase\|\|Cysteine-Desulfurase->?->?->PAPS-reduct+a+b-domain+Fer4_9->DUF4007->DUF6079*->wHTH+HNH-p->DUF4007->?->DUF6079->PglZ+Phosphodiest->SIGMA+OB-Fold+AAA-ATPase-BrxL-> | 1117 | Bacteria>Fusobacteriota>Fusobacteriia>Fusobacteriales>Fusobacteriaceae>Fusobacterium | GCF_003470805.1 | Fusobacterium_mortiferum | DW663_RS02810 | 250706..254059 | 850 | AAA-family-ATPase-[Fusobacterium-mortiferum] |
| WP_118234047.1 | Type4.PglZ.containing.BREX | Cysteine-Desulfurase->?->?->PAPS-reduct+a+b-domain+Fer4_9->DUF4007->DUF6079->wHTH+HNH-p->DUF4007->?->DUF6079*->PglZ+Phosphodiest->SIGMA+OB-Fold+AAA-ATPase-BrxL-><-?<-?<-?\|\|Aminotran_4->?->?-><-GNTR-HTH | 1321 | Bacteria>Fusobacteriota>Fusobacteriia>Fusobacteriales>Fusobacteriaceae>Fusobacterium | GCF_003470805.1 | Fusobacterium_mortiferum | DW663_RS02830 | 255997..259962 | 850 | DUF2791-family-P-loop-domain-containing-protein-[Fusobacterium-mortiferum] |
| WP_211494048.1 | Type4.PglZ.containing.BREX | SF1-SF2-Helicase->?->PAPS-reduct+a+b-domain+Fer4_9+DUF4007->DUF6079*->iSTAND+PglZ->ZnR->Cysteine-Desulfurase-> | 1399 | Bacteria>Fusobacteriota>Fusobacteriia>Fusobacteriales>Fusobacteriaceae>Fusobacterium | GCF_018206885.1 | unclassified_Fusobacterium | IX316_RS06940 | 10243..14442 | 2648384 | MULTISPECIES:-hypothetical-protein-[unclassified-Fusobacterium] |
| MBP6281401.1 | Type4.PglZ.containing.BREX | Cysteine-Desulfurase->SIGMA-HTH+HARE-HTH->PAPS-reduct+a+b-domain+Fer4_9->DUF4007->DUF6079*->wHTH+HNH-p->DUF4007->?->DUF6079->PglZ+Phosphodiest-> | 1125 | Bacteria>Fusobacteriota>Fusobacteriia>Fusobacteriales>Leptotrichiaceae>unclassified_Leptotrichiaceae | GCA_017986595.1 | Leptotrichiaceae_bacterium | KA384_05040 | 6410..9787 | 2782225 | hypothetical-protein-[Leptotrichiaceae-bacterium] |
| MBP6281405.1 | Type4.PglZ.containing.BREX | Cysteine-Desulfurase->SIGMA-HTH+HARE-HTH->PAPS-reduct+a+b-domain+Fer4_9->DUF4007->DUF6079->wHTH+HNH-p->DUF4007->?->DUF6079*->PglZ+Phosphodiest-> | 1334 | Bacteria>Fusobacteriota>Fusobacteriia>Fusobacteriales>Leptotrichiaceae>unclassified_Leptotrichiaceae | GCA_017986595.1 | Leptotrichiaceae_bacterium | KA384_05060 | 11776..15780 | 2782225 | hypothetical-protein-[Leptotrichiaceae-bacterium] |
| KJU83051.1 | Type4.PglZ.containing.BREX | PAPS-reduct+DUF4007->DUF6079*->iSTAND+PglZ-><-?\|\|?-><-RADICAL-SAM | 1078 | Bacteria>Nitrospirota>Nitrospiria>Nitrospirales>Nitrospiraceae>Candidatus_Magnetobacterium | GCA_000963235.1 | Candidatus_Magnetobacterium_bavaricum | MBAV_004751 | 2988..6224 | 29290 | hypothetical-protein-MBAV_004751-[Candidatus-Magnetobacterium-bavaricum] |
| WP_085053835.1 | Type4.PglZ.containing.BREX | TPR-><-?<-?\|\|?->PAPS-reduct+a+b-domain+Fer4_9+DUF4007->DUF6079*->iSTAND+PglZ->SIGMA+OB-Fold+AAA-ATPase-BrxL-><-?<-?<-?<-?<-SAM-METHYLASE | 1385 | Bacteria>Nitrospirota>Nitrospiria>Nitrospirales>Nitrospiraceae>Candidatus_Magnetominusculus | - | Candidatus_Magnetominusculus_xianensis | HQL00_07110 | 53290..57447 | 1748249 | hypothetical-protein-[Nitrospirae-bacterium-HCH-1] |
| HHW21153.1 | Type4.PglZ.containing.BREX | HTH_Tnp_1+HTH_21+rve+rve_3-><-SIGMA+OB-Fold+AAA-ATPase-BrxL<-iSTAND+PglZ<-DUF6079*<-PAPS-reduct+a+b-domain+Fer4_9+DUF4007\|\|?->?-><-?<-?<-?<-?\|\|tRNA->?->Amidohydro_2-> | 1123 | Bacteria>Nitrospirota>Thermodesulfovibrionia>Thermodesulfovibrionales>Thermodesulfovibrionaceae>Thermodesulfovibrio | GCA_012840265.1 | Thermodesulfovibrio_thiophilus | GXX27_08530 | 4199..7570 | 340095 | hypothetical-protein-[Thermodesulfovibrio-thiophilus] |
| MBF0318625.1 | Type4.PglZ.containing.BREX | <-Peptidase<-?<-?<-?<-Peptidase_M23<-?\|\|PAPS-reduct+DUF4007->DUF6079*->iSTAND+PglZ->SIGMA+OB-Fold+AAA-ATPase-BrxL->NTP_transf->?-><-SAM-METHYLASE\|\|HAD->?->XIS-HTH-> | 1397 | Bacteria>Nitrospirota>unclassified_Nitrospirota | GCA_015232115.1 | Nitrospirota_bacterium | HQL01_02300 | 22499..26692 | 2026887 | hypothetical-protein-[Nitrospirota-bacterium] |
| MBF0554175.1 | Type4.PglZ.containing.BREX | SIGMA-HTH+E3_UFM1_ligase->PAPS-reduct+a+b-domain+Fer4_9+DUF4007->DUF6079*->iSTAND+PglZ->SIGMA+OB-Fold+AAA-ATPase-BrxL-> | 1394 | Bacteria>Nitrospirota>unclassified_Nitrospirota | GCA_015233485.1 | Nitrospirota_bacterium | HQK96_06390 | 4648..8832 | 2026887 | hypothetical-protein-[Nitrospirota-bacterium] |
| NJO81811.1 | Type4.PglZ.containing.BREX | <-SIGMA+OB-Fold+AAA-ATPase-BrxL<-PglZ<-DUF6079-p*<-DUF6079-p<-CR-ATPase5<-DUF4007<-PAPS-reduct<-?<-DUF6079-p | 925 | Bacteria>Pseudomonadota>Alphaproteobacteria>Hyphomicrobiales>Blastochloridaceae>Blastochloris | GCA_012033475.1 | Blastochloris_sp. | HC828_02890 | 6893..9670 | 2066513 | hypothetical-protein-[Blastochloris-sp.] |
| NJO81812.1 | Type4.PglZ.containing.BREX | <-SIGMA+OB-Fold+AAA-ATPase-BrxL<-PglZ<-DUF6079-p<-DUF6079-p*<-CR-ATPase5<-DUF4007<-PAPS-reduct<-?<-DUF6079-p | 759 | Bacteria>Pseudomonadota>Alphaproteobacteria>Hyphomicrobiales>Blastochloridaceae>Blastochloris | GCA_012033475.1 | Blastochloris_sp. | HC828_02895 | 9670..11949 | 2066513 | hypothetical-protein-[Blastochloris-sp.] |
| MBQ2883389.1 | Type4.PglZ.containing.BREX | ZnR+TOPRIM->TraJ-RHH->?->GNAT-><-?<-DndB<-URI-DNase<-SIGMA+OB-Fold+AAA-ATPase-BrxL<-iSTAND+PglZ<-DUF6079*<-wHTH<-PAPS-reduct+a+b-domain+Fer4_9+DUF4007<-HTH_3\|\|?->MNS+REase-> | 1145 | Bacteria>Pseudomonadota>Alphaproteobacteria>unclassified_Alphaproteobacteria | GCA_017409945.1 | Alphaproteobacteria_bacterium | IJE43_06425 | 9636..13073 | 1913988 | hypothetical-protein-[Alphaproteobacteria-bacterium] |
| WP_194715672.1 | Type4.PglZ.containing.BREX | <-SAM-METHYLASE\|\|<-PAPS-reduct<-DUF6079*<-DUF4007<-SIGMA+OB-Fold+AAA-ATPase-BrxL<-PglZ<-DUF6079-p | 1101 | Bacteria>Pseudomonadota>Betaproteobacteria>Burkholderiales>Oxalobacteraceae>Noviherbaspirillum | GCF_015352955.1 | Noviherbaspirillum_soli | IGW04_RS24240 | 5612..8917 | 1064518 | ATP-binding-protein-[Noviherbaspirillum-soli] |
| WP_031492453.1 | Type4.PglZ.containing.BREX | <-Cysteine-Desulfurase<-ZnR<-SIGMA+OB-Fold+AAA-ATPase-BrxL<-iSTAND+PglZ<-DUF6079*<-PAPS-reduct+a+b-domain+Fer4_9+DUF4007<-Phage_integrase+E3_UFM1_ligase<-?<-OrfB_Zn_ribbon | 1398 | Bacteria>Pseudomonadota>Gammaproteobacteria>Aeromonadales>Succinivibrionaceae>Succinivibrio | GCF_000702045.1 | Succinivibrio_dextrinosolvens | T508_RS0109495 | 7116..11312 | 83771 | hypothetical-protein-[Succinivibrio-dextrinosolvens] |
| MBQ8976517.1 | Type4.PglZ.containing.BREX | <-Cysteine-Desulfurase<-ZnR<-SIGMA+OB-Fold+AAA-ATPase-BrxL<-iSTAND+PglZ<-DUF6079*<-PAPS-reduct+a+b-domain+Fer4_9+DUF4007<-SeqA+Phage_int+SeqA+E3_UFM1_ligase | 1401 | Bacteria>Pseudomonadota>Gammaproteobacteria>Aeromonadales>Succinivibrionaceae>unclassified_Succinivibrionaceae | GCA_017460495.1 | Succinivibrionaceae_bacterium | IJ078_03745 | 9519..13724 | 2053620 | hypothetical-protein-[Succinivibrionaceae-bacterium] |
| WP_182216823.1 | Type4.PglZ.containing.BREX | Arm-DNA-bind_3+Phage_integrase-><-?<-?\|\|Integrase_1->?->?-><-?\|\|Cysteine-Desulfurase-><-PAPS-reduct<-DUF6079*<-DUF4007\|\|DUF4007->?->DUF6079-p->DUF6079-p->PglZ->SIGMA+OB-Fold+AAA-ATPase-BrxL-> | 1076 | Bacteria>Pseudomonadota>Gammaproteobacteria>Alteromonadales>Colwelliaceae>Colwellia | GCF_014077305.1 | unclassified_Colwellia | H4J49_RS11080 | 49749..52979 | 196834 | MULTISPECIES:-AAA-family-ATPase-[unclassified-Colwellia] |
| WP_182231328.1 | Type4.PglZ.containing.BREX | Cysteine-Desulfurase-><-PAPS-reduct<-DUF6079<-DUF4007\|\|DUF4007->?->DUF6079*->PglZ->SIGMA+OB-Fold+AAA-ATPase-BrxL-><-SAM-METHYLASE\|\|KAP_NTPase->?-><-SAM-METHYLASE | 1229 | Bacteria>Pseudomonadota>Gammaproteobacteria>Alteromonadales>Colwelliaceae>Colwellia | GCF_014077325.1 | unclassified_Colwellia | H4J62_RS08490 | 55638..59327 | 196834 | MULTISPECIES:-hypothetical-protein-[unclassified-Colwellia] |
| PIQ01000.1 | Type4.PglZ.containing.BREX | <-GAF+PAS+PAS+PAS+PAS+HISKIN<-REC\|\|?-><-?\|\|SAM-METHYLASE->SAM-METHYLASE-><-SIGMA+OB-Fold+AAA-ATPase-BrxL<-PglZ<-DUF6079*<-?<-DUF4007<-?\|\|DUF4007->DUF6079->PAPS-reduct-><-Cysteine-Desulfurase | 1234 | Bacteria>Pseudomonadota>Gammaproteobacteria>Alteromonadales>Shewanellaceae>Shewanella | GCA_002770875.1 | Shewanella_sp._CG18_big_fil_WC_8_21_14_2_50_42_11 | COW76_07770 | 157907..161611 | 1975538 | hypothetical-protein-COW76_07770-[Shewanella-sp.-CG18_big_fil_WC_8_21_14_2_50_42_11] |
| NCO70514.1 | Type4.PglZ.containing.BREX | <-EAL<-?<-?<-?<-?\|\|?-><-?\|\|Cysteine-Desulfurase-><-PAPS-reduct<-DUF6079*<-DUF4007\|\|?->DUF4007->?->PglZ->SIGMA+OB-Fold+AAA-ATPase-BrxL-><-SAM-METHYLASE<-SAM-METHYLASE | 1134 | Bacteria>Pseudomonadota>Gammaproteobacteria>Alteromonadales>Shewanellaceae>Shewanella | GCA_009993165.1 | Shewanella_vesiculosa | GW877_03695 | 118842..122246 | 518738 | hypothetical-protein-[Shewanella-vesiculosa] |
| WP_273058118.1 | Type4.PglZ.containing.BREX | <-EAL<-?<-?<-?<-?\|\|?-><-?\|\|Cysteine-Desulfurase-><-PAPS-reduct<-DUF6079*<-DUF4007\|\|?->DUF4007->?->DUF6079->PglZ->SIGMA+OB-Fold+AAA-ATPase-BrxL-><-SAM-METHYLASE<-XIS-HTH | 1134 | Bacteria>Pseudomonadota>Gammaproteobacteria>Alteromonadales>Shewanellaceae>Shewanella | GCA_009994445.1 | Shewanella_vesiculosa | GW807_RS03095 | 234894..238298 | 518738 | hypothetical-protein-[Shewanella-vesiculosa] |
| WP_020504539.1 | Type4.PglZ.containing.BREX | DUF4007->DUF6079*->PAPS-reduct->DUF4007->?->DUF6079->PglZ->SIGMA+OB-Fold+AAA-ATPase-BrxL->?->RHH_5->N6-METHYLASE-p-> | 1135 | Bacteria>Pseudomonadota>Gammaproteobacteria>Chromatiales>Chromatiaceae>Lamprocystis | GCF_000379525.1 | Lamprocystis_purpurea | A390_RS0110110 | 143015..146422 | 61598 | hypothetical-protein-[Lamprocystis-purpurea] |
| WP_020504542.1 | Type4.PglZ.containing.BREX | DUF4007->DUF6079->PAPS-reduct->DUF4007->?->DUF6079*->PglZ->SIGMA+OB-Fold+AAA-ATPase-BrxL->?->RHH_5->N6-METHYLASE-p->Schlafen+HATPase+wHTH->?-><-?<-SF1-SF2-Helicase | 1233 | Bacteria>Pseudomonadota>Gammaproteobacteria>Chromatiales>Chromatiaceae>Lamprocystis | GCF_000379525.1 | Lamprocystis_purpurea | A390_RS0110125 | 148614..152315 | 61598 | hypothetical-protein-[Lamprocystis-purpurea] |
| EGV17033.1 | Type4.PglZ.containing.BREX | <-SF1-SF2-Helicase+wHTH+Small-B-barrel+DUF1998<-BrxD<-BrxD-AAA<-BrxD-AAA<-?<-?<-DndH<-SIGMA+OB-Fold+AAA-ATPase-BrxL<-PglZ<-DUF6079-p* | 844 | Bacteria>Pseudomonadota>Gammaproteobacteria>Chromatiales>Chromatiaceae>Thiocapsa | GCA_000223985.2 | Thiocapsa_marina_5811 | ThimaDRAFT_3862 | 209877..212411 | 768671 | hypothetical-protein-ThimaDRAFT_3862-[Thiocapsa-marina-5811] |
| WP_139052913.1 | Type4.PglZ.containing.BREX | <-SF1-SF2-Helicase+wHTH+Small-B-barrel+DUF1998<-BrxD<-BrxD-AAA<-BrxD-AAA<-?<-?<-DndH<-SIGMA+OB-Fold+AAA-ATPase-BrxL<-PglZ<-DUF6079-p* | 961 | Bacteria>Pseudomonadota>Gammaproteobacteria>Chromatiales>Chromatiaceae>Thiocapsa | GCF_000223985.1 | Thiocapsa_marina | THIMADRAFT_RS19070 | 209877..212762 | 244573 | hypothetical-protein-[Thiocapsa-marina] |
| QEP42413.1 | Type4.PglZ.containing.BREX | REC+OmpR-HTH->HISKIN->SAM-METHYLASE-><-Transposase_31<-Transposase_31<-SIGMA+OB-Fold+AAA-ATPase-BrxL<-PglZ<-DUF6079*<-?<-DUF4007<-PAPS-reduct<-HEPN<-Polbeta<-DUF6079<-DUF4007<-Cysteine-Desulfurase<-SAM-METHYLASE | 1312 | Bacteria>Pseudomonadota>Gammaproteobacteria>Chromatiales>Ectothiorhodospiraceae>unclassified_Ectothiorhodospiraceae | GCA_008375315.1 | Ectothiorhodospiraceae_bacterium_BW-2 | D5085_04220 | 905628..909566 | 947515 | hypothetical-protein-D5085_04220-[Ectothiorhodospiraceae-bacterium-BW-2] |
| QEP42419.1 | Type4.PglZ.containing.BREX | <-Transposase_31<-SIGMA+OB-Fold+AAA-ATPase-BrxL<-PglZ<-DUF6079<-?<-DUF4007<-PAPS-reduct<-HEPN<-Polbeta<-DUF6079*<-DUF4007<-Cysteine-Desulfurase<-SAM-METHYLASE\|\|?->?-><-ABhydrolase<-?<-?<-RADICAL-SAM | 1117 | Bacteria>Pseudomonadota>Gammaproteobacteria>Chromatiales>Ectothiorhodospiraceae>unclassified_Ectothiorhodospiraceae | GCA_008375315.1 | Ectothiorhodospiraceae_bacterium_BW-2 | D5085_04250 | 913304..916657 | 947515 | hypothetical-protein-D5085_04250-[Ectothiorhodospiraceae-bacterium-BW-2] |
| MBL1007423.1 | Type4.PglZ.containing.BREX | RVT-><-HTH_3<-?\|\|HU-HIG+DEOR-HTH->?->PAPS-reduct+a+b-domain+Fer4_9+DUF4007->DUF6079*->iSTAND+PglZ->SIGMA+OB-Fold+AAA-ATPase-BrxL->HTH_3->?->NUDIX-> | 1165 | Bacteria>Pseudomonadota>Gammaproteobacteria>Enterobacterales>Enterobacteriaceae>Escherichia | GCA_016742525.1 | Escherichia_coli | JJO71_27710 | 7246..10743 | 562 | hypothetical-protein-[Escherichia-coli] |
| EBT6907986.1 | Type4.PglZ.containing.BREX | <-TraI-VirD2-RCRE<-TraJ-RHH<-ANK+ANK\|\|ParB+HNH-><-PAPS-reduct<-DUF6079*<-DUF4007\|\|MORC+HISKIN->Response_reg->NACHT+SF1-SF2-Helicase+HKD-DNase->REase-><-?<-SIGMA+OB-Fold+AAA-ATPase+Lon-Protease-BrxL<-PglZ<-TOPRIM | 1099 | Bacteria>Pseudomonadota>Gammaproteobacteria>Enterobacterales>Enterobacteriaceae>Salmonella | GCA_007067565.1 | Salmonella_enterica | CLA26_04485 | 182280..185579 | 28901 | ATP-binding-protein-[Salmonella-enterica] |
| EDX0369816.1 | Type4.PglZ.containing.BREX | PglZ->SIGMA+OB-Fold+AAA-ATPase+Lon-Protease-BrxL->HKD-DNase+OGG-like-><-REase<-NACHT+SF1-SF2-Helicase+HKD-DNase<-Arm-DNA-bind_3<-Response_reg<-MORC+HISKIN\|\|DUF4007->DUF6079*->PAPS-reduct-><-ParB+HNH\|\|ANK+ANK->TraJ-RHH->TraI-VirD2-RCRE-> | 1099 | Bacteria>Pseudomonadota>Gammaproteobacteria>Enterobacterales>Enterobacteriaceae>Salmonella | GCA_010742735.1 | Salmonella_enterica | GTL21_000374 | 413346..416645 | 28901 | ATP-binding-protein-[Salmonella-enterica] |
| EEC5291039.1 | Type4.PglZ.containing.BREX | <-TraI-VirD2-RCRE<-TraJ-RHH<-ANK+ANK\|\|ParB+HNH-><-PAPS-reduct<-DUF6079*<-DUF4007\|\|MORC+HISKIN->Response_reg->NACHT+SF1-SF2-Helicase+HKD-DNase->REase-><-?<-SIGMA+OB-Fold+AAA-ATPase+Lon-Protease-BrxL<-PglZ<-TOPRIM | 1099 | Bacteria>Pseudomonadota>Gammaproteobacteria>Enterobacterales>Enterobacteriaceae>Salmonella | GCA_011001995.1 | Salmonella_enterica_subsp._enterica | ZD54_001383 | 9715..13014 | 59201 | ATP-binding-protein-[Salmonella-enterica-subsp.-enterica] |
| WP_045341604.1 | Type4.PglZ.containing.BREX | <-TraI-VirD2-RCRE<-TraJ-RHH<-ANK+ANK\|\|ParB+HNH-><-PAPS-reduct<-DUF6079*<-DUF4007\|\|MORC+HISKIN->Response_reg->NACHT+SF1-SF2-Helicase+HKD-DNase->REase-><-?<-SIGMA+OB-Fold+AAA-ATPase+Lon-Protease-BrxL<-PglZ<-TOPRIM | 1099 | Bacteria>Pseudomonadota>Gammaproteobacteria>Enterobacterales | GCF_014518275.1 | Enterobacteriaceae | IBX74_RS21060 | 336197..339496 | 543 | MULTISPECIES:-ATP-binding-protein-[Enterobacteriaceae] |
| WP_087869662.1 | Type4.PglZ.containing.BREX | DDE_Tnp_IS240->\|\|?-><-?\|\|?->\|\|PAPS-reduct->Zn_Tnp_IS1+HTH_Tnp_IS1+DDE_Tnp_IS1->?->DUF4007->?->DUF6079*->PglZ->SIGMA+OB-Fold+AAA-ATPase-BrxL-><-Cysteine-Desulfurase\|\|ParA-Soj-PloopNTPase-> | 1220 | Bacteria>Pseudomonadota>Gammaproteobacteria | GCF_002189065.1 | unclassified_Gammaproteobacteria | CDZ78_RS08660 | 8369..12031 | 33811 | MULTISPECIES:-hypothetical-protein-[unclassified-Gammaproteobacteria] |
| MCK9608995.1 | Type4.PglZ.containing.BREX | <-SSB\|\|?->?->?-><-PIN+HTH<-wHTH+WYL+WCX\|\|HNH->DUF4007->DUF6079*->PAPS-reduct->DUF4007->?->DUF6079->PglZ->SIGMA+OB-Fold+AAA-ATPase-BrxL-><-SAM-METHYLASE<-Cysteine-Desulfurase | 1135 | Bacteria>Pseudomonadota>Gammaproteobacteria>Methylococcales>Methylococcaceae>Methylomonas | GCA_023228165.1 | Methylomonas_sp. | MOR33_21370 | 16404..19811 | 418 | DUF6079-family-protein-[Methylomonas-sp.] |
| MCK9608999.1 | Type4.PglZ.containing.BREX | <-PIN+HTH<-wHTH+WYL+WCX\|\|HNH->DUF4007->DUF6079->PAPS-reduct->DUF4007->?->DUF6079*->PglZ->SIGMA+OB-Fold+AAA-ATPase-BrxL-><-SAM-METHYLASE<-Cysteine-Desulfurase\|\|?-><-TraN | 1237 | Bacteria>Pseudomonadota>Gammaproteobacteria>Methylococcales>Methylococcaceae>Methylomonas | GCA_023228165.1 | Methylomonas_sp. | MOR33_21390 | 22122..25835 | 418 | hypothetical-protein-[Methylomonas-sp.] |
| WP_017840084.1 | Type4.PglZ.containing.BREX | Polbeta->HEPN->PAPS-reduct-><-DDE_Tnp_IS66\|\|?->DUF4007->?->PemK_toxin->?->DUF6079*->PglZ->SIGMA+OB-Fold+AAA-ATPase-BrxL->?->?->CopG/RHH->RelE-ParE->BrnT_toxin->BrnA_antitoxin->REase-> | 1205 | Bacteria>Pseudomonadota>Gammaproteobacteria>Methylococcales>Methylococcaceae>Methylotuvimicrobium | GCF_005931095.1 | Methylotuvimicrobium_buryatense | EQU24_RS07510 | 1654570..1658187 | 95641 | hypothetical-protein-[Methylotuvimicrobium-buryatense] |
| TSA39893.1 | Type4.PglZ.containing.BREX | PAPS-reduct->DUF4007->?->RelE-ParE->?->DUF6079*->PglZ-> | 1217 | Bacteria>Pseudomonadota>Gammaproteobacteria>Methylococcales>Methylococcaceae>unclassified_Methylococcaceae | GCA_007280895.1 | Methylococcaceae_bacterium | D4R63_07735 | 3187..6840 | 1933926 | hypothetical-protein-D4R63_07735-[Methylococcaceae-bacterium] |
| TXK94054.1 | Type4.PglZ.containing.BREX | DDE_Tnp-><-SIGMA+OB-Fold+AAA-ATPase-BrxL<-PglZ<-DUF6079*<-?<-DUF4007<-?<-PAPS-reduct | 1220 | Bacteria>Pseudomonadota>Gammaproteobacteria>Methylococcales>Methylococcaceae>unclassified_Methylococcaceae | GCA_008030135.1 | Methylococcaceae_bacterium_CS4 | BMR10_14205 | 4315..7977 | 1916069 | hypothetical-protein-BMR10_14205-[Methylococcaceae-bacterium-CS4] |
| MCK4843012.1 | Type4.PglZ.containing.BREX | <-HerA/FtsK\|\|SAM-METHYLASE->SAM-METHYLASE->SKI->?->Cysteine-Desulfurase-><-ParB+HNH<-SIGMA+OB-Fold+AAA-ATPase-BrxL<-PglZ<-DUF6079*<-?<-DUF4007<-PAPS-reduct | 1241 | Bacteria>Pseudomonadota>Gammaproteobacteria>Methylococcales>unclassified_Methylococcales | GCA_023134955.1 | Methylococcales_bacterium | KAT04_14215 | 26352..30077 | 2304002 | hypothetical-protein-[Methylococcales-bacterium] |
| MCK5872692.1 | Type4.PglZ.containing.BREX | <-SIGMA+OB-Fold+AAA-ATPase-BrxL<-PglZ<-DUF6079*<-?<-DUF4007<-?<-PAPS-reduct-p | 1220 | Bacteria>Pseudomonadota>Gammaproteobacteria>Methylococcales>unclassified_Methylococcales | GCA_023146315.1 | Methylococcales_bacterium | KAG26_07680 | 3979..7641 | 2304002 | hypothetical-protein-[Methylococcales-bacterium] |
| MCK5897799.1 | Type4.PglZ.containing.BREX | DUF4007->DUF6079*->PAPS-reduct->DUF4007->?->DUF6079->PglZ->SIGMA+OB-Fold+AAA-ATPase-BrxL-><-SAM-METHYLASE<-Cysteine-Desulfurase<-ACET | 1127 | Bacteria>Pseudomonadota>Gammaproteobacteria>Methylococcales>unclassified_Methylococcales | GCA_023146575.1 | Methylococcales_bacterium | KAG06_01855 | 1447..4830 | 2304002 | hypothetical-protein-[Methylococcales-bacterium] |
| MCK5897803.1 | Type4.PglZ.containing.BREX | DUF4007->DUF6079->PAPS-reduct->DUF4007->?->DUF6079*->PglZ->SIGMA+OB-Fold+AAA-ATPase-BrxL-><-SAM-METHYLASE<-Cysteine-Desulfurase<-ACET | 1208 | Bacteria>Pseudomonadota>Gammaproteobacteria>Methylococcales>unclassified_Methylococcales | GCA_023146575.1 | Methylococcales_bacterium | KAG06_01875 | 7066..10692 | 2304002 | hypothetical-protein-[Methylococcales-bacterium] |
| SEH06867.1 | Type4.PglZ.containing.BREX | <-SIGMA+OB-Fold+AAA-ATPase-BrxL<-PglZ<-DUF6079*<-?<-DUF4007<-PAPS-reduct<-wHTH(SBD)+HNH<-DUF6079<-DUF4007 | 1241 | Bacteria>Pseudomonadota>Gammaproteobacteria>Thiotrichales>unclassified_Thiotrichales | GCA_900099115.2 | Thiotrichales_bacterium_HS_08 | MBHS_02733 | 4364..8089 | 1899563 | Uncharacterised-protein-[Thiotrichales-bacterium-HS_08] |
| SEH06872.1 | Type4.PglZ.containing.BREX | <-SIGMA+OB-Fold+AAA-ATPase-BrxL<-PglZ<-DUF6079<-?<-DUF4007<-PAPS-reduct<-wHTH(SBD)+HNH<-DUF6079*<-DUF4007 | 1141 | Bacteria>Pseudomonadota>Gammaproteobacteria>Thiotrichales>unclassified_Thiotrichales | GCA_900099115.2 | Thiotrichales_bacterium_HS_08 | MBHS_02738 | 11220..14645 | 1899563 | Uncharacterised-protein-[Thiotrichales-bacterium-HS_08] |
| WP_174482024.1 | Type4.PglZ.containing.BREX | BrxB-BrxF-iSTAND->ACET->Cysteine-Desulfurase-><-DndE<-DndD<-?<-HKD-DNase+SF1-SF2-Helicase<-SIGMA+OB-Fold+AAA-ATPase-BrxL<-PglZ<-DUF6079*\|\|SAM-METHYLASE-><-wHTH+HNH-p | 1162 | Bacteria>Pseudomonadota>Gammaproteobacteria>unclassified_Gammaproteobacteria | GCF_900128525.1 | methanotrophic_endosymbiont_of_Bathymodiolus_puteoserpentis_(Logatchev) | BQ7314_RS01950 | 13427..16915 | 343235 | hypothetical-protein-[methanotrophic-endosymbiont-of-Bathymodiolus-puteoserpentis-(Logatchev)] |
| WP_102456724.1 | Type4.PglZ.containing.BREX | <-Peptidase_M15_4<-NUDIX+DUF429<-?\|\|SAM-METHYLASE->SAM-METHYLASE->ParB+HNH->Cysteine-Desulfurase-><-SIGMA+OB-Fold+AAA-ATPase-BrxL<-PglZ<-DUF6079*<-?<-DUF4007<-PAPS-reduct<-DUF6079<-DUF4007\|\|HNH->Acyltransferase-> | 1254 | Bacteria>Pseudomonadota>Gammaproteobacteria>Vibrionales>Vibrionaceae | GCF_002877415.1 | Vibrio | BCU08_RS19910 | 81955..85719 | 662 | MULTISPECIES:-DUF6079-family-protein-[Vibrio] |
| WP_099078661.1 | Type4.PglZ.containing.BREX | <-wHTH+WYL+WCX\|\|HEPN->URI-DNase->ARG->?->PAPS-reduct->ParB+HNH->DUF4007->DUF6079*->PAPS-reduct->DUF4007->?->DUF6079->PglZ->SIGMA+OB-Fold+AAA-ATPase-BrxL-><-Cysteine-Desulfurase<-SAM-METHYLASE<-SAM-METHYLASE | 1079 | Bacteria>Pseudomonadota>Gammaproteobacteria>Vibrionales>Vibrionaceae>Vibrio | GCF_002608565.1 | Vibrio_sp._PID17_43 | AK965_RS05465 | 175401..178640 | 1583451 | hypothetical-protein-[Vibrio-sp.-PID17_43] |
| WP_099078665.1 | Type4.PglZ.containing.BREX | ARG->?->PAPS-reduct->ParB+HNH->DUF4007->DUF6079->PAPS-reduct->DUF4007->?->DUF6079*->PglZ->SIGMA+OB-Fold+AAA-ATPase-BrxL-><-Cysteine-Desulfurase<-SAM-METHYLASE<-SAM-METHYLASE | 1258 | Bacteria>Pseudomonadota>Gammaproteobacteria>Vibrionales>Vibrionaceae>Vibrio | GCF_002608565.1 | Vibrio_sp._PID17_43 | AK965_RS05485 | 180971..184747 | 1583451 | hypothetical-protein-[Vibrio-sp.-PID17_43] |
| WP_102456728.1 | Type4.PglZ.containing.BREX | SAM-METHYLASE->ParB+HNH->Cysteine-Desulfurase-><-SIGMA+OB-Fold+AAA-ATPase-BrxL<-PglZ<-DUF6079<-?<-DUF4007<-PAPS-reduct<-DUF6079*<-DUF4007\|\|HNH->Acyltransferase->?->?-><-?<-?<-Glyco_hydro_19+CBM_5_12 | 1085 | Bacteria>Pseudomonadota>Gammaproteobacteria>Vibrionales>Vibrionaceae>Vibrio | GCF_002877315.1 | Vibrio_splendidus | BCU08_RS19930 | 88041..91298 | 29497 | hypothetical-protein-[Vibrio-splendidus] |
| WP_102514132.1 | Type4.PglZ.containing.BREX | SAM-METHYLASE->ParB+HNH->Cysteine-Desulfurase-><-SIGMA+OB-Fold+AAA-ATPase-BrxL<-PglZ<-DUF6079<-?<-DUF4007<-PAPS-reduct<-DUF6079*<-DUF4007\|\|HNH->Acyltransferase-> | 1085 | Bacteria>Pseudomonadota>Gammaproteobacteria>Vibrionales>Vibrionaceae>Vibrio | GCF_002877415.1 | Vibrio_lentus | BCU13_RS17470 | 198396..201653 | 136468 | hypothetical-protein-[Vibrio-lentus] |
| NCC04461.1 | Type4.PglZ.containing.BREX | Calcineurin->AbiEi_4->?->Calcineurin-SbcD->Calcineurin-SbcD->?->RAMA->PAPS-reduct+a+b-domain+Fer4_9+DUF4007->DUF6079*->PglZ->SIGMA+OB-Fold+AAA-ATPase-BrxL->ZnR->?->STYKIN->?->SIR2->HerA/FtsK-> | 1383 | Bacteria>Pseudomonadota>unclassified_Pseudomonadota | GCA_009929575.1 | Pseudomonadota_bacterium | EDM37_10550 | 10272..14423 | 1977087 | transposase-[Pseudomonadota-bacterium] |
| HED00229.1 | Type4.PglZ.containing.BREX | <-DSBH<-NTP_transferase\|\|?->Peptidase_M23->PAPS-reduct+a+b-domain+Fer4_9+DUF4007->DUF4007->DUF6079*->iSTAND+PglZ->SIGMA+OB-Fold+AAA-ATPase-BrxL-> | 1386 | Bacteria>Pseudomonadota>unclassified_Pseudomonadota | GCA_011046395.1 | Pseudomonadota_bacterium | ENN18_07590 | 18713..22873 | 1977087 | hypothetical-protein-[Pseudomonadota-bacterium] |
| MBU0967620.1 | Type4.PglZ.containing.BREX | TraH->TraG_N-><-ZnR<-Schlafen+HATPase+wHTH<-SIGMA+OB-Fold+AAA-ATPase-BrxL<-PglZ<-DUF6079*<-?<-DUF4007<-PAPS-reduct+a+b-domain+Fer4_9<-SIGMA-HTH+SIGMA-HTH+HARE-HTH<-?<-PadR-HTH | 1260 | Bacteria>Pseudomonadota>unclassified_Pseudomonadota | GCA_018816555.1 | Pseudomonadota_bacterium | KKA54_14705 | 31546..35328 | 1977087 | hypothetical-protein-[Pseudomonadota-bacterium] |
| MBU1057840.1 | Type4.PglZ.containing.BREX | SSB->?->DUF932->AbiEi_4+AbiEi_1->AbiEii+ZnR->XIS-HTH->SF1-SF2-Helicase->RAMA->PAPS-reduct+a+b-domain+Fer4_9+DUF4007->DUF6079*->PglZ->SIGMA+OB-Fold+AAA-ATPase-BrxL-><-SAM-METHYLASE<-BrxB-BrxF-iSTAND<-?\|\|DEOR-HTH->?-><-TraG_N<-TraH | 1383 | Bacteria>Pseudomonadota>unclassified_Pseudomonadota | GCA_018817385.1 | Pseudomonadota_bacterium | KJ804_05920 | 8730..12881 | 1977087 | transposase-[Pseudomonadota-bacterium] |
| MBU4261962.1 | Type4.PglZ.containing.BREX | <-PAS+SHELIX+PAS+PAS+PAS<-tRNA<-MPTase<-HTH_19\|\|BrnT_toxin->CopG_antitoxin->SIGMA-HTH+SIGMA-HTH+HARE-HTH->PAPS-reduct+a+b-domain+Fer4_9->DUF4007->?->DUF6079*->PglZ->SIGMA+OB-Fold+AAA-ATPase-BrxL->REase->HEPN->DnaJ->?-><-?<-HTH_3\|\|HTH_3-> | 1267 | Bacteria>Pseudomonadota>unclassified_Pseudomonadota | GCA_018897275.1 | Pseudomonadota_bacterium | KKC76_08830 | 21089..24892 | 1977087 | hypothetical-protein-[Pseudomonadota-bacterium] |
| HCG26922.1 | Type4.PglZ.containing.BREX | SIGMA-HTH-><-?<-?\|\|wHTH+WYL+WCX-><-Couple_hipA+HipA_C<-HTH_3\|\|?->HARE-HTH+E3_UFM1_ligase->PAPS-reduct+a+b-domain+Fer4_9+DUF4007->DUF6079*->iSTAND+PglZ->ZnR->SIGMA+OB-Fold+AAA-ATPase-BrxL->Cysteine-Desulfurase-> | 1398 | Bacteria>PVC_group>Lentisphaerota>Lentisphaeria>unclassified_Lentisphaeria | GCA_003526375.1 | Lentisphaeria_bacterium | DER70_15205 | 54616..58812 | 2053569 | hypothetical-protein-[Lentisphaeria-bacterium] |
| MBE6363378.1 | Type4.PglZ.containing.BREX | HTH_26->Preader17->PAPS-reduct+a+b-domain+Fer4_9+DUF4007->DUF6079*->PglZ->SIGMA+OB-Fold+AAA-ATPase-BrxL->ZnR->Cysteine-Desulfurase->STYKinase-like->O6MGT-HTH-><-?<-MqsA_antitoxin+DUF4065 | 1393 | Bacteria>PVC_group>Lentisphaerota>unclassified_Lentisphaerota | GCA_015061605.1 | Lentisphaerota_bacterium | E7054_06955 | 8438..12619 | 1932692 | transposase-[Lentisphaerota-bacterium] |
| MBX3399558.1 | Type4.PglZ.containing.BREX | Glycos_transf->?->Glycos_transf-><-?\|\|REase-><-SAM-METHYLASE\|\|DUF4007->DUF6079*->PAPS-reduct->DUF4007->?->DUF6079->PglZ->SIGMA+OB-Fold+AAA-ATPase-BrxL->?->REase->REase+SF1-SF2-Helicase-> | 1169 | Bacteria>PVC_group>Planctomycetota>Planctomycetia>Gemmatales>Gemmataceae | GCA_019636445.1 | Gemmataceae_bacterium | KF873_12515 | 94876..98385 | 2052164 | hypothetical-protein-[Gemmataceae-bacterium] |
| MBX3399562.1 | Type4.PglZ.containing.BREX | Glycos_transf-><-?\|\|REase-><-SAM-METHYLASE\|\|DUF4007->DUF6079->PAPS-reduct->DUF4007->?->DUF6079*->PglZ->SIGMA+OB-Fold+AAA-ATPase-BrxL->?->REase->REase+SF1-SF2-Helicase-> | 1264 | Bacteria>PVC_group>Planctomycetota>Planctomycetia>Gemmatales>Gemmataceae | GCA_019636445.1 | Gemmataceae_bacterium | KF873_12535 | 100604..104398 | 2052164 | hypothetical-protein-[Gemmataceae-bacterium] |
| WP_210653444.1 | Type4.PglZ.containing.BREX | <-URI-DNase<-MORC<-SIGMA+OB-Fold+AAA-ATPase-BrxL<-PglZ<-DUF6079*<-?<-DUF4007<-PAPS-reduct<-DUF6079<-DUF4007\|\|SAM-METHYLASE-> | 1262 | Bacteria>PVC_group>Planctomycetota>Planctomycetia>Gemmatales>Gemmataceae | GCF_017939745.1 | Gemmata_palustris | J8F10_RS08725 | 2215604..2219392 | 2822762 | hypothetical-protein-[Gemmata-palustris] |
| WP_210653448.1 | Type4.PglZ.containing.BREX | <-URI-DNase<-MORC<-SIGMA+OB-Fold+AAA-ATPase-BrxL<-PglZ<-DUF6079<-?<-DUF4007<-PAPS-reduct<-DUF6079*<-DUF4007\|\|SAM-METHYLASE->?->?->?->?->?-><-?<-LPD32+LPD36+N6_Mtase+SF1-SF2-Helicase | 1168 | Bacteria>PVC_group>Planctomycetota>Planctomycetia>Gemmatales>Gemmataceae | GCF_017939745.1 | Gemmata_palustris | J8F10_RS08745 | 2221600..2225106 | 2822762 | hypothetical-protein-[Gemmata-palustris] |
| NLY01889.1 | Type4.PglZ.containing.BREX | <-ZnR<-SIGMA+OB-Fold+AAA-ATPase-BrxL<-iSTAND+PglZ<-DUF6079*<-PAPS-reduct+a+b-domain+Fer4_9+DUF4007<-RAMA<-DUF4007<-XIS-HTH<-DUF932 | 1389 | Bacteria>PVC_group>Planctomycetota>Planctomycetia>Pirellulales>Pirellulaceae | GCA_012728855.1 | Rhodopirellula_sp. | GXY83_37875 | 5317..9486 | 2024855 | transposase-[Rhodopirellula-sp.] |
| MBQ4143519.1 | Type4.PglZ.containing.BREX | PAPS-reduct+a+b-domain+Fer4_9+DUF4007->DUF6079*->iSTAND+PglZ->SIGMA+OB-Fold+AAA-ATPase-BrxL->ZnR->Cysteine-Desulfurase-><-?<-SIGMA-HTH+SIGMA-HTH+SIGMA-HTH\|\|?->DnaB+SMS-KaiC-> | 1388 | Bacteria>PVC_group>Planctomycetota>Planctomycetia>Pirellulales>Thermoguttaceae | GCA_017403345.1 | Thermoguttaceae_bacterium | IJD43_08600 | 3436..7602 | 2691551 | hypothetical-protein-[Thermoguttaceae-bacterium] |
| MBQ7110120.1 | Type4.PglZ.containing.BREX | <-ABhydrolase<-Classical-AAA\|\|?->?->?-><-Cysteine-Desulfurase<-ZnR<-SIGMA+OB-Fold+AAA-ATPase-BrxL<-iSTAND+PglZ<-DUF6079*<-PAPS-reduct+a+b-domain+Fer4_9+DUF4007 | 1385 | Bacteria>PVC_group>Planctomycetota>Planctomycetia>Pirellulales>Thermoguttaceae | GCA_017436025.1 | Thermoguttaceae_bacterium | IJD06_02770 | 19623..23780 | 2691551 | hypothetical-protein-[Thermoguttaceae-bacterium] |
| MBQ7111119.1 | Type4.PglZ.containing.BREX | SbcC-><-Cysteine-Desulfurase<-ZnR<-SIGMA+OB-Fold+AAA-ATPase-BrxL<-iSTAND+PglZ<-DUF6079*<-PAPS-reduct+a+b-domain+Fer4_9+DUF4007<-?<-?\|\|?->?->SF1-SF2-Helicase-><-SF1-SF2-Helicase<-OLD-ABC+TOPRIM<-HEPN | 1386 | Bacteria>PVC_group>Planctomycetota>Planctomycetia>Pirellulales>Thermoguttaceae | GCA_017436025.1 | Thermoguttaceae_bacterium | IJD06_07855 | 9692..13852 | 2691551 | hypothetical-protein-[Thermoguttaceae-bacterium] |
| MBR0237621.1 | Type4.PglZ.containing.BREX | PAPS-reduct+a+b-domain+Fer4_9+DUF4007->DUF6079*->PglZ->SIGMA+OB-Fold+AAA-ATPase-BrxL->ZnR->Cysteine-Desulfurase-><-tRNA<-Phosphodiest | 1411 | Bacteria>PVC_group>Planctomycetota>Planctomycetia>Pirellulales>Thermoguttaceae | GCA_017447745.1 | Thermoguttaceae_bacterium | IJQ39_06000 | 10475..14710 | 2691551 | hypothetical-protein-[Thermoguttaceae-bacterium] |
| MBQ9371615.1 | Type4.PglZ.containing.BREX | PAPS-reduct+a+b-domain+Fer4_9+DUF4007->DUF6079*->iSTAND+PglZ->SIGMA+OB-Fold+AAA-ATPase-BrxL->ZnR-> | 1426 | Bacteria>PVC_group>Planctomycetota>Planctomycetia>Pirellulales>Thermoguttaceae | GCA_017458405.1 | Thermoguttaceae_bacterium | IJU03_05440 | 17929..22209 | 2691551 | hypothetical-protein-[Thermoguttaceae-bacterium] |
| MBR2693833.1 | Type4.PglZ.containing.BREX | ZnR->Acyltransferase->?->?->PAPS-reduct+a+b-domain+Fer4_9+DUF4007->DUF6079*->iSTAND+PglZ->SIGMA+OB-Fold+AAA-ATPase-BrxL->ZnR->Cysteine-Desulfurase-> | 1392 | Bacteria>PVC_group>Planctomycetota>Planctomycetia>Pirellulales>Thermoguttaceae | GCA_017515305.1 | Thermoguttaceae_bacterium | IKE69_06430 | 40642..44820 | 2691551 | hypothetical-protein-[Thermoguttaceae-bacterium] |
| MBR4835363.1 | Type4.PglZ.containing.BREX | SF1-SF2-Helicase+DUF1998-><-wHTH(SBD)+HNH<-Cysteine-Desulfurase<-ZnR<-SIGMA+OB-Fold+AAA-ATPase-BrxL<-PglZ<-DUF6079*<-PAPS-reduct+a+b-domain+Fer4_9+DUF4007<-RAMA | 1411 | Bacteria>PVC_group>Planctomycetota>Planctomycetia>Pirellulales>Thermoguttaceae | GCA_017559825.1 | Thermoguttaceae_bacterium | IKU86_13690 | 23842..28077 | 2691551 | hypothetical-protein-[Thermoguttaceae-bacterium] |
| MBP3531683.1 | Type4.PglZ.containing.BREX | SF1-SF2-Helicase+REase->HEPN->OLD-ABC+TOPRIM->SF1-SF2-Helicase-><-Cysteine-Desulfurase<-ZnR<-SIGMA+OB-Fold+AAA-ATPase-BrxL<-PglZ<-DUF6079*<-PAPS-reduct+a+b-domain+Fer4_9+DUF4007<-RAMA | 1411 | Bacteria>PVC_group>Planctomycetota>Planctomycetia>Pirellulales>Thermoguttaceae | GCA_017937485.1 | Thermoguttaceae_bacterium | J6K25_10960 | 13859..18094 | 2691551 | hypothetical-protein-[Thermoguttaceae-bacterium] |
| MBP3693065.1 | Type4.PglZ.containing.BREX | PAPS-reduct+a+b-domain+Fer4_9+DUF4007->DUF6079*->iSTAND+PglZ->SIGMA+OB-Fold+AAA-ATPase-BrxL->ZnR->Cysteine-Desulfurase-> | 1387 | Bacteria>PVC_group>Planctomycetota>Planctomycetia>Pirellulales>Thermoguttaceae | GCA_017939795.1 | Thermoguttaceae_bacterium | J6J31_01340 | 3330..7493 | 2691551 | hypothetical-protein-[Thermoguttaceae-bacterium] |
| HBJ34335.1 | Type4.PglZ.containing.BREX | Cysteine-Desulfurase->DUF4007->DUF6079-p*->?->PAPS-reduct->DUF4007->?->DUF6079->PglZ->SIGMA+OB-Fold+AAA-ATPase-BrxL-><-SAM-METHYLASE\|\|XIS-HTH-> | 917 | Bacteria>PVC_group>Planctomycetota>Planctomycetia>Planctomycetales>Planctomycetaceae | GCA_003506095.1 | Planctomycetaceae_bacterium | DDZ51_06145 | 3301..6054 | 2026779 | hypothetical-protein-[Planctomycetaceae-bacterium] |
| HBJ34340.1 | Type4.PglZ.containing.BREX | Cysteine-Desulfurase->DUF4007->DUF6079-p->?->PAPS-reduct->DUF4007->?->DUF6079*->PglZ->SIGMA+OB-Fold+AAA-ATPase-BrxL-><-SAM-METHYLASE\|\|XIS-HTH-><-wHTH(SBD)+HNH | 1291 | Bacteria>PVC_group>Planctomycetota>Planctomycetia>Planctomycetales>Planctomycetaceae | GCA_003506095.1 | Planctomycetaceae_bacterium | DDZ51_06170 | 8781..12656 | 2026779 | hypothetical-protein-[Planctomycetaceae-bacterium] |
| MBE6424426.1 | Type4.PglZ.containing.BREX | <-DnaB+SMS-KaiC<-?\|\|SIGMA-HTH+SIGMA-HTH+SIGMA-HTH->?-><-Cysteine-Desulfurase<-ZnR<-SIGMA+OB-Fold+AAA-ATPase-BrxL<-iSTAND+PglZ<-DUF6079*<-PAPS-reduct+a+b-domain+Fer4_9+DUF4007<-?\|\|BrxA-DNA-binding->BrxB-BrxF-iSTAND->DUF6079-p-> | 1388 | Bacteria>PVC_group>Planctomycetota>Planctomycetia>Planctomycetales>Planctomycetaceae | GCA_015062085.1 | Planctomycetaceae_bacterium | E7029_00260 | 77820..81986 | 2026779 | hypothetical-protein-[Planctomycetaceae-bacterium] |
| MBE6424431.1 | Type4.PglZ.containing.BREX | <-Cysteine-Desulfurase<-ZnR<-SIGMA+OB-Fold+AAA-ATPase-BrxL<-iSTAND+PglZ<-DUF6079<-PAPS-reduct+a+b-domain+Fer4_9+DUF4007<-?\|\|BrxA-DNA-binding->BrxB-BrxF-iSTAND->DUF6079-p*-> | 716 | Bacteria>PVC_group>Planctomycetota>Planctomycetia>Planctomycetales>Planctomycetaceae | GCA_015062085.1 | Planctomycetaceae_bacterium | E7029_00285 | 88488..90634 | 2026779 | BREX-system-P-loop-protein-BrxC-[Planctomycetaceae-bacterium] |
| MCE5327910.1 | Type4.PglZ.containing.BREX | DUF4007->DUF6079*->PAPS-reduct->DUF4007->?->DUF6079->PglZ->SIGMA+OB-Fold+AAA-ATPase-BrxL->Cysteine-Desulfurase-><-HEPN<-wHTH+WYL+WCX | 1136 | Bacteria>PVC_group>Planctomycetota>Planctomycetia>Planctomycetales>Planctomycetaceae | GCA_021371965.1 | Planctomycetaceae_bacterium | LLG01_16000 | 69782..73192 | 2026779 | hypothetical-protein-[Planctomycetaceae-bacterium] |
| MBT3194523.1 | Type4.PglZ.containing.BREX | PAPS-reduct+Fer4_9-p->DUF6079*->PglZ->SIGMA+OB-Fold+AAA-ATPase-BrxL->ZnR->Cysteine-Desulfurase-><-SAM-METHYLASE | 1384 | Bacteria>PVC_group>Verrucomicrobiota>unclassified_Verrucomicrobiota | GCA_018645275.1 | Verrucomicrobiota_bacterium | HN341_18405 | 406..4560 | 2026799 | transposase-[Verrucomicrobiota-bacterium] |
| MCC8021078.1 | Type4.PglZ.containing.BREX | <-SIGMA+OB-Fold+AAA-ATPase-BrxL<-iSTAND+PglZ<-DUF6079*<-PAPS-reduct+a+b-domain+Fer4_9+DUF4007\|\|MACRODOMAIN-> | 1160 | Bacteria>PVC_group>Verrucomicrobiota>Verrucomicrobiae>Verrucomicrobiales>Akkermansiaceae | GCA_020859585.1 | Akkermansia_sp. | LIO63_03745 | 4074..7556 | 1872421 | DUF6079-family-protein-[Akkermansia-sp.] |
| MCD7797871.1 | Type4.PglZ.containing.BREX | <-SIGMA+OB-Fold+AAA-ATPase-BrxL<-iSTAND+PglZ<-DUF6079*<-PAPS-reduct+a+b-domain+Fer4_9+DUF4007\|\|MACRODOMAIN-> | 1161 | Bacteria>PVC_group>Verrucomicrobiota>Verrucomicrobiae>Verrucomicrobiales>Akkermansiaceae | GCA_021200315.1 | Akkermansiaceae_bacterium | LUG84_00465 | 4044..7529 | 2562705 | DUF6079-family-protein-[Akkermansiaceae-bacterium] |
| MCD7799233.1 | Type4.PglZ.containing.BREX | SeqA+Phage_int+SeqA+E3_UFM1_ligase->PAPS-reduct+a+b-domain+Fer4_9+DUF4007->DUF6079*->iSTAND+PglZ->SIGMA+OB-Fold+AAA-ATPase-BrxL->ZnR->Cysteine-Desulfurase->?->Recombinase+ZnR-><-?<-PAS+HISKIN<-PDZ+Peptidase | 1388 | Bacteria>PVC_group>Verrucomicrobiota>Verrucomicrobiae>Verrucomicrobiales>Akkermansiaceae | GCA_021200315.1 | Akkermansiaceae_bacterium | LUG84_07525 | 6371..10537 | 2562705 | hypothetical-protein-[Akkermansiaceae-bacterium] |
| MCI6981217.1 | Type4.PglZ.containing.BREX | PAPS-reduct+a+b-domain+Fer4_9+DUF4007->DUF6079*->iSTAND+PglZ->SIGMA+OB-Fold+AAA-ATPase-BrxL-><-tRNA\|\|?->ACET-> | 1153 | Bacteria>PVC_group>Verrucomicrobiota>Verrucomicrobiae>Verrucomicrobiales>Akkermansiaceae | GCA_022775565.1 | Akkermansia_muciniphila | MR894_03270 | 5583..9044 | 239935 | hypothetical-protein-[Akkermansia-muciniphila] |
| TAL39060.1 | Type4.PglZ.containing.BREX | <-Cysteine-Desulfurase<-ZnR<-SIGMA+OB-Fold+AAA-ATPase-BrxL<-iSTAND+PglZ<-DUF6079*<-PAPS-reduct+a+b-domain+Fer4_9+DUF4007<-HARE-HTH+E3_UFM1_ligase\|\|?->?-><-?<-?<-?<-METHYLASE | 1386 | Bacteria>Spirochaetota>unclassified_Spirochaetota | GCA_004297825.1 | Spirochaetota_bacterium | EPN93_03015 | 109350..113510 | 2202144 | hypothetical-protein-EPN93_03015-[Spirochaetota-bacterium] |
| AFM22178.1 | Type4.PglZ.containing.BREX | <-PAS+PAS+GGDEF<-?<-?<-?<-SIGMA+OB-Fold+AAA-ATPase-BrxL<-iSTAND+PglZ<-DUF6079*<-DUF4007<-PAPS-reduct+a+b-domain+Fer4_9+DUF4007<-?<-?<-?\|\|Radical_SAM-> | 1387 | Bacteria>Synergistota>Synergistia>Synergistales>Acetomicrobiaceae>Acetomicrobium | GCA_000266925.1 | Acetomicrobium_mobile_DSM_13181 | Anamo_1580 | 1578434..1582597 | 891968 | hypothetical-protein-Anamo_1580-[Acetomicrobium-mobile-DSM-13181] |
| WP_169312880.1 | Type4.PglZ.containing.BREX | <-PAS+PAS+GGDEF\|\|?->HAMP+MA+MA-><-?<-?<-?<-SIGMA+OB-Fold+AAA-ATPase-BrxL<-iSTAND+PglZ<-DUF6079*<-DUF4007<-PAPS-reduct+a+b-domain+Fer4_9+DUF4007<-SIGMA-HTH<-?<-?<-?\|\|Radical_SAM-> | 1380 | Bacteria>Synergistota>Synergistia>Synergistales>Acetomicrobiaceae>Acetomicrobium | GCF_000266925.1 | Acetomicrobium_mobile | ANAMO_RS07785 | 1578434..1582576 | 97477 | ATP-binding-protein-[Acetomicrobium-mobile] |
| WP_198469750.1 | Type4.PglZ.containing.BREX | MULE-TRANSPOSASE->MULE-TRANSPOSASE->TPR-><-?<-?<-SIGMA+OB-Fold+AAA-ATPase-BrxL<-iSTAND+PglZ<-DUF6079*<-DUF4007<-PAPS-reduct+a+b-domain+Fer4_9+DUF4007<-SIGMA-HTH+SIGMA-HTH\|\|?->?-><-Peptidase_M20<-?<-?<-Peptidase_M42+Peptidase_M28 | 1397 | Bacteria>Synergistota>Synergistia>Synergistales>Acetomicrobiaceae>Acetomicrobium | GCF_016125955.1 | Acetomicrobium_sp._S15_=_DSM_107314 | EZM41_RS04010 | 10315..14508 | 2529858 | hypothetical-protein-[Acetomicrobium-sp.-S15-=-DSM-107314] |
| WP_133956945.1 | Type4.PglZ.containing.BREX | <-MetJarc-HTH\|\|?->DUF772+DDE_Tnp_1->?->?-><-Cysteine-Desulfurase<-SIGMA+OB-Fold+AAA-ATPase-BrxL<-iSTAND+PglZ<-DUF6079*<-PAPS-reduct+a+b-domain+Fer4_9+DUF4007<-HNH<-HARE-HTH+E3_UFM1_ligase | 1396 | Bacteria>Synergistota>Synergistia>Synergistales>Aminobacteriaceae>Aminivibrio | GCF_004366815.1 | Aminivibrio_pyruvatiphilus | C8D99_RS04780 | 128248..132438 | 1005740 | hypothetical-protein-[Aminivibrio-pyruvatiphilus] |
| MCI5864797.1 | Type4.PglZ.containing.BREX | HTH_26->SeqA+Phage_int+SeqA+E3_UFM1_ligase->PAPS-reduct+a+b-domain+Fer4_9+DUF4007->DUF6079*->iSTAND+PglZ->SIGMA+OB-Fold+AAA-ATPase-BrxL->ZnR->Cysteine-Desulfurase->?->TERNS3+HEPN->?-><-MobA_MobL | 1388 | Bacteria>Synergistota>Synergistia>Synergistales>Synergistaceae>Cloacibacillus | GCF_022771785.1 | Cloacibacillus_porcorum | MR000_06180 | 158443..162609 | 1197717 | hypothetical-protein-[Cloacibacillus-porcorum] |
| WP_273309660.1 | Type4.PglZ.containing.BREX | HTH_26->SeqA+Phage_int+SeqA+E3_UFM1_ligase->PAPS-reduct+a+b-domain+Fer4_9+DUF4007->DUF6079*->iSTAND+PglZ->SIGMA+OB-Fold+AAA-ATPase-BrxL->ZnR->Cysteine-Desulfurase->?->TERNS3+HEPN->?-><-MobA_MobL | 1388 | Bacteria>Synergistota>Synergistia>Synergistales>Synergistaceae>Cloacibacillus | GCF_022771785.1 | Cloacibacillus_porcorum | MR000_RS06245 | 158443..162609 | 1197717 | hypothetical-protein-[Cloacibacillus-porcorum] |
| GHS97043.1 | Type4.PglZ.containing.BREX | CRISPR_Cas6->Cas_Cas1->?-><-?<-ZnR<-SIGMA+OB-Fold+AAA-ATPase-BrxL<-iSTAND+PglZ<-DUF6079*<-PAPS-reduct+a+b-domain+Fer4_9+DUF4007<-HARE-HTH+E3_UFM1_ligase | 1389 | Bacteria>Synergistota>Synergistia>Synergistales>unclassified_Synergistales | GCA_025997955.1 | Synergistales_bacterium | AGMMS50276_16420 | 14134..18303 | 2268201 | hypothetical-protein-AGMMS50276_16420-[Synergistales-bacterium] |
| GHV54672.1 | Type4.PglZ.containing.BREX | <-35exo<-?\|\|?->PAPS-reduct+a+b-domain+Fer4_9+DUF4007->DUF6079*->iSTAND+PglZ->SIGMA+OB-Fold+AAA-ATPase-BrxL->ZnR-> | 1394 | Bacteria>Synergistota>Synergistia>Synergistales>unclassified_Synergistales | GCA_026002755.1 | Synergistales_bacterium | FACS1894216_15170 | 7753..11937 | 2268201 | hypothetical-protein-FACS1894216_15170-[Synergistales-bacterium] |
| WP_105482022.1 | Type4.PglZ.containing.BREX | <-NACHT<-DUF6079*<-DUF4007\|\|wHTH(SBD)+HNH->?-><-?\|\|DUF6079->PglZ->SIGMA+OB-Fold+AAA-ATPase-BrxL->Cysteine-Desulfurase-> | 1152 | Bacteria>Terrabacteria_group>Abditibacteriota>Abditibacteria>Abditibacteriales>Abitibacteriaceae | GCF_002973605.1 | Abditibacterium_utsteinense | B1R32_RS00015 | 3258..6716 | 1960156 | hypothetical-protein-[Abditibacterium-utsteinense] |
| MBQ7256248.1 | Type4.PglZ.containing.BREX | <-LACI-HTH+PBPI<-?<-Phage_integrase<-?<-?\|\|?->PAPS-reduct+a+b-domain+Fer4_9+DUF4007->DUF6079*->iSTAND+PglZ->SIGMA+OB-Fold+AAA-ATPase-BrxL->ZnR->OLD-ABC+TOPRIM->SF1-SF2-Helicase-> | 1403 | Bacteria>Terrabacteria_group>Abditibacteriota>unclassified_Abditibacteriota | GCA_017445025.1 | Abditibacteriota_bacterium | IJS60_00965 | 22785..26996 | 2783868 | hypothetical-protein-[Abditibacteriota-bacterium] |
| WP_050340978.1 | Type4.PglZ.containing.BREX | <-REase<-DDE_Tnp_1\|\|rve->AAA+-><-HEPN<-Cysteine-Desulfurase<-ZnR<-SIGMA+OB-Fold+AAA-ATPase-BrxL<-iSTAND+PglZ<-DUF6079*<-PAPS-reduct+a+b-domain+Fer4_9+DUF4007<-?\|\|HTH_3->Couple_hipA+HipA_C->CR-ATPase5-> | 1390 | Bacteria>Terrabacteria_group>Actinomycetota>Coriobacteriia>Coriobacteriales>Atopobiaceae | GCF_001189515.2 | Olsenella_sp._oral_taxon_807 | ADJ70_RS09690 | 2263054..2267226 | 712411 | hypothetical-protein-[Olsenella-sp.-oral-taxon-807] |
| WP_114551671.1 | Type4.PglZ.containing.BREX | PAPS-reduct+DUF4007->DUF6079*->PglZ->SIGMA+OB-Fold+AAA-ATPase-BrxL->Cysteine-Desulfurase-><-?<-Phage_integrase<-?\|\|HTH_3-><-Phage_integrase | 1393 | Bacteria>Terrabacteria_group>Actinomycetota>Coriobacteriia>Eggerthellales>Eggerthellaceae | GCF_015561125.1 | Eggerthella_lenta | I2I24_RS08680 | 714..4895 | 84112 | hypothetical-protein-[Eggerthella-lenta] |
| NLN77497.1 | Type4.PglZ.containing.BREX | SF1-SF2-Helicase+REase->Calcineurin->SbcC-><-rve<-HTH_Tnp_1\|\|HARE-HTH+E3_UFM1_ligase->PAPS-reduct+a+b-domain+Fer4_9+DUF4007->DUF6079*->iSTAND+PglZ->SIGMA+OB-Fold+AAA-ATPase-BrxL->Cysteine-Desulfurase-> | 1380 | Bacteria>Terrabacteria_group>Armatimonadota>unclassified_Armatimonadota | GCA_012522555.1 | Armatimonadota_bacterium | GX141_01075 | 12642..16784 | 2033014 | hypothetical-protein-[Armatimonadota-bacterium] |
| MBI2842129.1 | Type4.PglZ.containing.BREX | <-BrxB-BrxF-iSTAND<-MPTase<-HTH<-?<-MORC+REase<-HerA/FtsK<-PglZ-p<-DUF6079*<-DnaA<-DUF4007<-wHTH<-PAPS-reduct<-DUF6079<-DUF4007<-XIS-HTH<-GDResolvase+Recombinase+ZnR | 1344 | Bacteria>Terrabacteria_group>Armatimonadota>unclassified_Armatimonadota | GCA_016190935.1 | Armatimonadota_bacterium | HYX78_01885 | 41359..45393 | 2033014 | hypothetical-protein-[Armatimonadota-bacterium] |
| MBI2842134.1 | Type4.PglZ.containing.BREX | <-MORC+REase<-HerA/FtsK<-PglZ-p<-DUF6079<-DnaA<-DUF4007<-wHTH<-PAPS-reduct<-DUF6079*<-DUF4007<-XIS-HTH<-GDResolvase+Recombinase+ZnR<-?\|\|PHP->?-><-?<-?<-MoxR-AAA | 1147 | Bacteria>Terrabacteria_group>Armatimonadota>unclassified_Armatimonadota | GCA_016190935.1 | Armatimonadota_bacterium | HYX78_01910 | 48099..51542 | 2033014 | hypothetical-protein-[Armatimonadota-bacterium] |
| MCL6576916.1 | Type4.PglZ.containing.BREX | <-DDE_Tnp_1\|\|HIN-HTH->TnpB_IS66-><-AAA-ATPase<-HIN-HTH<-DEDD_Tnp_IS110\|\|DUF4007->DUF6079*->PAPS-reduct->DUF4007->?->DUF6079->PglZ->SIGMA+OB-Fold+AAA-ATPase-BrxL->Cysteine-Desulfurase-><-?<-DDE_Tnp_IS66+DDE_Tnp_IS66_C | 1045 | Bacteria>Terrabacteria_group>Bacillota>Bacilli>Bacillales>Alicyclobacillaceae | GCA_023512215.1 | Kyrpidia_sp. | K6T64_12345 | 15300..18437 | 2073077 | hypothetical-protein-[Kyrpidia-sp.] |
| MCL6576920.1 | Type4.PglZ.containing.BREX | TnpB_IS66-><-AAA-ATPase<-HIN-HTH<-DEDD_Tnp_IS110\|\|DUF4007->DUF6079->PAPS-reduct->DUF4007->?->DUF6079*->PglZ->SIGMA+OB-Fold+AAA-ATPase-BrxL->Cysteine-Desulfurase-><-?<-DDE_Tnp_IS66+DDE_Tnp_IS66_C | 1516 | Bacteria>Terrabacteria_group>Bacillota>Bacilli>Bacillales>Alicyclobacillaceae | GCA_023512215.1 | Kyrpidia_sp. | K6T64_12365 | 20710..25260 | 2073077 | hypothetical-protein-[Kyrpidia-sp.] |
| MBR3117631.1 | Type4.PglZ.containing.BREX | <-SMODS<-NERD+SF1-SF2-Helicase\|\|?->BirA-HTH->?->REase+SF1-SF2-Helicase->HTH_26->PAPS-reduct+a+b-domain+Fer4_9+DUF4007->ZnR->DUF6079*->iSTAND+PglZ->SIGMA+OB-Fold+AAA-ATPase-BrxL->?->Cysteine-Desulfurase->ParB+HTH->?->MerR-HTH-> | 1173 | Bacteria>Terrabacteria_group>Bacillota>Bacilli>Bacillales>Bacillaceae | GCA_017514815.1 | Oceanobacillus_sp. | IKF29_00175 | 12273..15794 | 1871622 | hypothetical-protein-[Oceanobacillus-sp.] |
| HIT17410.1 | Type4.PglZ.containing.BREX | <-HEPN<-SMODS<-Cysteine-Desulfurase<-ZnR<-SIGMA+OB-Fold+AAA-ATPase-BrxL<-iSTAND+PglZ<-DUF6079*<-PAPS-reduct+a+b-domain+Fer4_9+DUF4007<-HNH<-SIGMA-HTH | 1394 | Bacteria>Terrabacteria_group>Bacillota>Bacillota_incertae_sedis>Candidatus_Caccosoma | GCA_018716205.1 | Candidatus_Caccosoma_faecigallinarum | IAD04_03385 | 6723..10907 | 2840720 | hypothetical-protein-[Candidatus-Caccosoma-faecigallinarum] |
| CDC05527.1 | Type4.PglZ.containing.BREX | HEPN->MNS+REase-><-Cysteine-Desulfurase<-ZnR<-SIGMA+OB-Fold+AAA-ATPase-BrxL<-iSTAND+PglZ<-DUF6079*<-PAPS-reduct+a+b-domain+Fer4_9+DUF4007<-SeqA+Phage_int+SeqA+E3_UFM1_ligase<-?<-XIS-HTH\|\|XIS-HTH->?->?->DnaB->TraJ-RHH-> | 1391 | Bacteria>Terrabacteria_group>Bacillota>Clostridia>Eubacteriales>Clostridiaceae | GCA_000431955.1 | Clostridium_leptum_CAG:27 | BN578_00994 | 8520..12695 | 1263068 | putative-uncharacterized-protein-[Clostridium-leptum-CAG:27] |
| CDC79176.1 | Type4.PglZ.containing.BREX | <-HIN-HTH+PAIREDC-HTH\|\|Schlafen+HATPase+wHTH->SF1-SF2-Helicase+PRTase->SeqA+Phage_int+SeqA+E3_UFM1_ligase->PAPS-reduct+a+b-domain+Fer4_9+DUF4007->DUF6079*->iSTAND+PglZ->SIGMA+OB-Fold+AAA-ATPase-BrxL->Cysteine-Desulfurase-><-?<-?<-?<-?\|\|DEDD_Tnp_IS110-> | 1390 | Bacteria>Terrabacteria_group>Bacillota>Clostridia>Eubacteriales>Clostridiaceae | GCA_000435335.1 | Clostridium_sp._CAG:964 | BN818_00604 | 13029..17201 | 1262848 | putative-uncharacterized-protein-[Clostridium-sp.-CAG:964] |
| NBI83182.1 | Type4.PglZ.containing.BREX | HTH_32+DDE_3-><-?\|\|RAMA->PAPS-reduct+a+b-domain+Fer4_9+DUF4007->DUF6079*->iSTAND+PglZ->SIGMA+OB-Fold+AAA-ATPase-BrxL->ZnR->Cysteine-Desulfurase-> | 1383 | Bacteria>Terrabacteria_group>Bacillota>Clostridia>Eubacteriales>Clostridiaceae | GCA_009911335.1 | Clostridiaceae_bacterium | D3Z48_14050 | 4732..8883 | 1898204 | hypothetical-protein-[Clostridiaceae-bacterium] |
| NLE25893.1 | Type4.PglZ.containing.BREX | HARE-HTH+E3_UFM1_ligase->HNH->PAPS-reduct+a+b-domain+Fer4_9+DUF4007->DUF6079*->iSTAND+PglZ->ZnR->SIGMA+OB-Fold+AAA-ATPase-BrxL->Cysteine-Desulfurase-> | 1396 | Bacteria>Terrabacteria_group>Bacillota>Clostridia>Eubacteriales>Clostridiaceae | GCA_012514215.1 | Clostridiaceae_bacterium | GX625_11245 | 6111..10301 | 1898204 | hypothetical-protein-[Clostridiaceae-bacterium] |
| NLM18925.1 | Type4.PglZ.containing.BREX | MPTase-><-wHTH+WYL+WCX<-Cysteine-Desulfurase<-ZnR<-SIGMA+OB-Fold+AAA-ATPase-BrxL<-iSTAND+PglZ<-DUF6079*<-PAPS-reduct+a+b-domain+Fer4_9+DUF4007<-SeqA+Phage_int+SeqA+Phage_int+E3_UFM1_ligase | 1398 | Bacteria>Terrabacteria_group>Bacillota>Clostridia>Eubacteriales>Clostridiaceae | GCA_012521015.1 | Clostridiaceae_bacterium | GX217_02745 | 6813..11009 | 1898204 | hypothetical-protein-[Clostridiaceae-bacterium] |
| QUO34989.1 | Type4.PglZ.containing.BREX | GDResolvase+Recombinase+ZnR-><-?\|\|HTH_28+rve+rve_3->?-><-DEDD_Tnp_IS110+Transposase_20<-?\|\|HTH_26->SeqA+Phage_int+SeqA+Phage_int+E3_UFM1_ligase->PAPS-reduct+a+b-domain+Fer4_9+DUF4007->DUF6079*->iSTAND+PglZ->SIGMA+OB-Fold+AAA-ATPase-BrxL->ZnR->Cysteine-Desulfurase-> | 1402 | Bacteria>Terrabacteria_group>Bacillota>Clostridia>Eubacteriales>Clostridiaceae | GCA_018228685.1 | Clostridiaceae_bacterium_Marseille-Q4149 | KFE16_14110 | 2776119..2780327 | 2830659 | hypothetical-protein-KFE16_14110-[Clostridiaceae-bacterium-Marseille-Q4149] |
| MBS6228511.1 | Type4.PglZ.containing.BREX | HTH_3->MPTase-><-?<-?<-Cysteine-Desulfurase<-SIGMA+OB-Fold+AAA-ATPase-BrxL<-ZnR<-iSTAND+PglZ<-DUF6079*<-PAPS-reduct+a+b-domain+Fer4_9+DUF4007<-HNH<-?<-HTH_26<-TraM<-TraJ-RHH<-ArdCN+LPD32+LPD36+N6_Mtase+SF1-SF2-Helicase | 1399 | Bacteria>Terrabacteria_group>Bacillota>Clostridia>Eubacteriales>Clostridiaceae | GCA_018374725.1 | Clostridium_sp. | KH509_01900 | 55769..59968 | 1506 | hypothetical-protein-[Clostridium-sp.] |
| MBS6266252.1 | Type4.PglZ.containing.BREX | <-LZ_Tnp_IS66+DDE_Tnp_IS66+DDE_Tnp_IS66_C<-?<-HIN-HTH<-Cysteine-Desulfurase<-SIGMA+OB-Fold+AAA-ATPase-BrxL<-ZnR<-PglZ<-iSTAND+PglZ<-DUF6079*<-PAPS-reduct+a+b-domain+Fer4_9+DUF4007\|\|?-><-?<-HTH_26 | 1399 | Bacteria>Terrabacteria_group>Bacillota>Clostridia>Eubacteriales>Clostridiaceae | GCA_018374955.1 | Clostridium_sp. | KH611_10765 | 10729..14928 | 1506 | hypothetical-protein-[Clostridium-sp.] |
| MCI8605659.1 | Type4.PglZ.containing.BREX | GGDEF-><-?<-?<-?<-tRNA\|\|OLD-ABC->HNH->SeqA+E3_UFM1_ligase->PAPS-reduct+a+b-domain+Fer4_9+DUF4007-><-?\|\|DUF6079*->iSTAND+PglZ->SIGMA+OB-Fold+AAA-ATPase-BrxL->ZnR->Cysteine-Desulfurase->ParB+HNH->VirB11+SbcC->HNH-> | 1121 | Bacteria>Terrabacteria_group>Bacillota>Clostridia>Eubacteriales>Clostridiaceae | GCA_022793615.1 | Hungatella_sp. | HFG72_02125 | 13672..17037 | 2613924 | hypothetical-protein-[Hungatella-sp.] |
| MCI8664738.1 | Type4.PglZ.containing.BREX | <-MarR-HTH\|\|?->?->?->?->NUDIX->?->PAPS-reduct+a+b-domain+Fer4_9+DUF4007->DUF6079*->iSTAND+PglZ->SIGMA+OB-Fold+AAA-ATPase-BrxL->Preader8->Cysteine-Desulfurase->ParB+HNH->VirB11+SbcC->HNH-> | 1165 | Bacteria>Terrabacteria_group>Bacillota>Clostridia>Eubacteriales>Clostridiaceae | GCA_022794315.1 | Hungatella_sp. | HFG69_16105 | 97871..101368 | 2613924 | hypothetical-protein-[Hungatella-sp.] |
| MCM1227008.1 | Type4.PglZ.containing.BREX | XIS-HTH->?->?->TraJ-RHH->XIS-HTH-><-Cysteine-Desulfurase<-SIGMA+OB-Fold+AAA-ATPase-BrxL<-ZnR<-iSTAND+PglZ<-DUF6079*<-PAPS-reduct+a+b-domain+Fer4_9+DUF4007<-?<-?<-CHTH<-?\|\|HTH->?-><-Tox-RES<-Tox-RES | 1399 | Bacteria>Terrabacteria_group>Bacillota>Clostridia>Eubacteriales>Clostridiaceae | GCA_023664445.1 | Clostridium_sp. | NC320_06220 | 22495..26694 | 1506 | hypothetical-protein-[Clostridium-sp.] |
| SCH09324.1 | Type4.PglZ.containing.BREX | Abi->DDE_Tnp_1->?-><-?<-?<-Cysteine-Desulfurase<-SIGMA+OB-Fold+AAA-ATPase-BrxL<-ZnR<-iSTAND+PglZ<-DUF6079*<-?<-PAPS-reduct+a+b-domain+Fer4_9+DUF4007<-HNH | 1355 | Bacteria>Terrabacteria_group>Bacillota>Clostridia>Eubacteriales>Clostridiaceae | GCA_900066525.1 | uncultured_Clostridium_sp. | SAMEA3545335_00161 | 175840..179907 | 59620 | Uncharacterised-protein-[uncultured-Clostridium-sp.] |
| WP_045091744.1 | Type4.PglZ.containing.BREX | <-ABC-ATPase+SbcC+ABC-ATPase+SbcC<-PBPI\|\|LACI-HTH+PBPI-><-tRNA\|\|HAD+HAD->?->?->\|\|TRAM+METHYLASE->?->PAPS-reduct+a+b-domain+Fer4_9+DUF4007->DUF6079*->iSTAND+PglZ->SIGMA+OB-Fold+AAA-ATPase-BrxL->ZnR->Cysteine-Desulfurase->?->SF1-SF2-Helicase->?->HD+SF1-SF2-Helicase-> | 1378 | Bacteria>Terrabacteria_group>Bacillota>Clostridia>Eubacteriales>Clostridiaceae | GCF_000954015.1 | Clostridium_sp._FS41 | CLFS41_RS05995 | 67116..71252 | 1609975 | hypothetical-protein-[Clostridium-sp.-FS41] |
| WP_103226893.1 | Type4.PglZ.containing.BREX | <-RVT<-?<-?<-?<-Cysteine-Desulfurase<-SIGMA+OB-Fold+AAA-ATPase-BrxL<-iSTAND+PglZ<-DUF6079*<-PAPS-reduct+a+b-domain+Fer4_9+DUF4007<-?<-HEPN<-HTH_3<-?<-Peptidase_S8<-Classical-AAA<-?<-ACET | 1415 | Bacteria>Terrabacteria_group>Bacillota>Clostridia>Eubacteriales>Clostridiaceae | GCF_002899675.1 | Clostridium_sp._chh4-2 | C0033_RS05605 | 146243..150490 | 2067550 | hypothetical-protein-[Clostridium-sp.-chh4-2] |
| WP_117621173.1 | Type4.PglZ.containing.BREX | REC+AraC-HTH+AraC-HTH->?-><-Cysteine-Desulfurase<-Preader8<-SIGMA+OB-Fold+AAA-ATPase-BrxL<-iSTAND+PglZ<-DUF6079*<-ZnR<-PAPS-reduct+a+b-domain+Fer4_9+DUF4007\|\|DDE_Tnp_1->?->?-><-?<-McrB-NTD+HNH<-?<-Calcineurin | 1165 | Bacteria>Terrabacteria_group>Bacillota>Clostridia>Eubacteriales>Clostridiaceae | GCF_003439535.1 | Hungatella_hathewayi | DXB08_RS05735 | 287141..290638 | 154046 | hypothetical-protein-[Hungatella-hathewayi] |
| WP_125003657.1 | Type4.PglZ.containing.BREX | <-SIR2<-SF1-SF2-Helicase<-?\|\|<-HKD-DNase+SF1-SF2-Helicase+HTH+HTH+SRA-p<-?<-REase<-Preader4<-SIGMA+OB-Fold+AAA-ATPase-BrxL<-PglZ+Phosphodiest<-DUF6079*<-?<-DUF4007<-DUF6079<-DUF4007<-PAPS-reduct+a+b-domain+Fer4_9<-?<-Cysteine-Desulfurase<-wHTH(SBD)+HNH | 1323 | Bacteria>Terrabacteria_group>Bacillota>Clostridia>Eubacteriales>Clostridiaceae | GCF_003865095.1 | Clostridium_tagluense | Ctaglu_RS16485 | 75738..79709 | 360422 | DUF6079-family-protein-[Clostridium-tagluense] |
| WP_125003663.1 | Type4.PglZ.containing.BREX | <-HKD-DNase+SF1-SF2-Helicase+HTH+HTH+SRA-p<-?<-REase<-Preader4<-SIGMA+OB-Fold+AAA-ATPase-BrxL<-PglZ+Phosphodiest<-DUF6079<-?<-DUF4007<-DUF6079*<-DUF4007<-PAPS-reduct+a+b-domain+Fer4_9<-?<-Cysteine-Desulfurase<-wHTH(SBD)+HNH | 1118 | Bacteria>Terrabacteria_group>Bacillota>Clostridia>Eubacteriales>Clostridiaceae | GCF_003865095.1 | Clostridium_tagluense | Ctaglu_RS16500 | 81116..84472 | 360422 | hypothetical-protein-[Clostridium-tagluense] |
| WP_187018530.1 | Type4.PglZ.containing.BREX | <-HTH\|\|PAS+GGDEF-><-?\|\|TRAM+METHYLASE->SIGMA-HTH->HNH->PAPS-reduct+a+b-domain+Fer4_9+DUF4007->DUF6079*->iSTAND+PglZ->SIGMA+OB-Fold+AAA-ATPase-BrxL->Cysteine-Desulfurase->wHTH+WYL+WCX->?->VbhA+DOC+HTH-><-?<-SIGMA-HTH<-ACET | 1402 | Bacteria>Terrabacteria_group>Bacillota>Clostridia>Eubacteriales>Clostridiaceae | GCF_014288005.1 | Hungatella_sp._L12 | H8S75_RS00675 | 149912..154120 | 2763050 | hypothetical-protein-[Hungatella-sp.-L12] |
| WP_194611685.1 | Type4.PglZ.containing.BREX | <-TraI-VirD2-RCRE<-TraJ-RHH<-LexA-protease\|\|wHTH+WYL+WCX->SeqA+Phage_int+SeqA+Phage_int+E3_UFM1_ligase->PAPS-reduct+a+b-domain+Fer4_9+DUF4007->DUF6079*->iSTAND+PglZ->SIGMA+OB-Fold+AAA-ATPase-BrxL->ZnR->Cysteine-Desulfurase->Schlafen+BirA-HTH->SLATT->SMODS-><-MPTase<-HTH_3 | 1405 | Bacteria>Terrabacteria_group>Bacillota>Clostridia>Eubacteriales>Clostridiaceae | GCF_015351765.1 | Clostridium_vitabionis | ISF27_RS11770 | 47072..51289 | 2784388 | hypothetical-protein-[Clostridium-vitabionis] |
| WP_195464807.1 | Type4.PglZ.containing.BREX | <-Cysteine-Desulfurase<-ZnR<-SIGMA+OB-Fold+AAA-ATPase-BrxL<-iSTAND+PglZ<-DUF6079*<-PAPS-reduct+a+b-domain+Fer4_9+DUF4007<-HNH<-SIGMA-HTH\|\|NtrC-AAA+FIS-HTH-><-?<-PADR-HTH | 1408 | Bacteria>Terrabacteria_group>Bacillota>Clostridia>Eubacteriales>Clostridiaceae | GCF_015554605.1 | Clostridium_jeddahense | I2E80_RS10445 | 132208..136434 | 1414721 | hypothetical-protein-[Clostridium-jeddahense] |
| WP_204907614.1 | Type4.PglZ.containing.BREX | <-PemK_toxin<-HTH_3<-SIGMA-HTH\|\|HTH_3->MPTase->AT-HTH-HYD2-><-Cysteine-Desulfurase<-SIGMA+OB-Fold+AAA-ATPase-BrxL<-iSTAND+PglZ<-DUF6079*<-PAPS-reduct+a+b-domain+Fer4_9+DUF4007<-SeqA+Phage_int+SeqA+E3_UFM1_ligase<-XIS-HTH<-HTH\|\|?-><-?<-?<-TraJ-RHH<-RVT | 1401 | Bacteria>Terrabacteria_group>Bacillota>Clostridia>Eubacteriales>Clostridiaceae | GCF_016902415.1 | Mordavella_massiliensis | H6A13_RS00265 | 58679..62884 | 1871024 | hypothetical-protein-[Mordavella-massiliensis] |
| WP_215704540.1 | Type4.PglZ.containing.BREX | <-METHYLASE<-SAM-METHYLASE\|\|MPTase-><-?<-?<-Cysteine-Desulfurase<-SIGMA+OB-Fold+AAA-ATPase-BrxL<-ZnR<-iSTAND+PglZ<-DUF6079*<-PAPS-reduct+DUF4007 | 1399 | Bacteria>Terrabacteria_group>Bacillota>Clostridia>Eubacteriales>Clostridiaceae | GCF_018785465.1 | Clostridium_sp._MCC328 | GPK83_RS08485 | 28075..32274 | 2592642 | hypothetical-protein-[Clostridium-sp.-MCC328] |
| WP_216157725.1 | Type4.PglZ.containing.BREX | AbiEii->Tox-ART-HYD3->?->?-><-?\|\|?-><-?\|\|?->PAPS-reduct+a+b-domain+Fer4_9+DUF4007->DUF6079*->iSTAND+PglZ->ZnR->GDResolvase->Tox-RES->?->PIN->\|\|?-><-?<-SH3 | 1385 | Bacteria>Terrabacteria_group>Bacillota>Clostridia>Eubacteriales>Clostridiaceae | GCF_018861315.1 | Clostridium_bowmanii | KPL51_RS04035 | 114654..118811 | 132925 | hypothetical-protein-[Clostridium-bowmanii] |
| WP_216195025.1 | Type4.PglZ.containing.BREX | HEPN->Phage_integrase-><-?<-HicB_lk_antitox\|\|?->PAPS-reduct+a+b-domain+Fer4_9+DUF4007->DUF6079*->iSTAND+PglZ->SIGMA+OB-Fold+AAA-ATPase-BrxL->ZnR-><-?<-?<-?<-Phage_integrase\|\|?->REase-> | 1386 | Bacteria>Terrabacteria_group>Bacillota>Clostridia>Eubacteriales>Clostridiaceae | GCF_018861375.1 | Clostridium_estertheticum | KPL45_20640 | 48461..52621 | 238834 | hypothetical-protein-[Clostridium-estertheticum] |
| WP_216471319.1 | Type4.PglZ.containing.BREX | <-Phage_int_SAM_3+Phage_integrase<-XIS-HTH\|\|?->?->?->PAPS-reduct+a+b-domain+Fer4_9+DUF4007->DUF6079*->iSTAND+PglZ->SIGMA+OB-Fold+AAA-ATPase-BrxL->ZnR-> | 1400 | Bacteria>Terrabacteria_group>Bacillota>Clostridia>Eubacteriales>Clostridiaceae | GCF_018918345.1 | Butyricicoccus_intestinisimiae | KQI75_RS13305 | 11265..15467 | 2841509 | hypothetical-protein-[Butyricicoccus-intestinisimiae] |
| WP_118522381.1 | Type4.PglZ.containing.BREX | Glyoxalase_2->PIN->?->MACRODOMAIN->XIS-HTH->SeqA+Phage_int+SeqA+E3_UFM1_ligase->?->PAPS-reduct+a+b-domain+Fer4_9+DUF4007->DUF6079*->iSTAND+PglZ->SIGMA+OB-Fold+AAA-ATPase-BrxL->Cysteine-Desulfurase->IstB_IS21-> | 1401 | Bacteria>Terrabacteria_group>Bacillota>Clostridia>Eubacteriales>Clostridiaceae | GCF_027660525.1 | Clostridium_fessum | PGP72_RS10725 | 129729..133934 | 2126740 | hypothetical-protein-[Clostridium-fessum] |
| WP_117633843.1 | Type4.PglZ.containing.BREX | REC+AraC-HTH+AraC-HTH->?-><-Cysteine-Desulfurase<-Preader8<-SIGMA+OB-Fold+AAA-ATPase-BrxL<-iSTAND+PglZ<-DUF6079*<-?<-?<-?<-?<-HTH_3\|\|?->Mob_Pre->?->GDResolvase+Recombinase+ZnR-> | 1079 | Bacteria>Terrabacteria_group>Bacillota>Clostridia>Eubacteriales>Clostridiaceae | GCF_027662785.1 | Hungatella | PGR72_RS05990 | 317870..321109 | 1649459 | MULTISPECIES:-hypothetical-protein-[Hungatella] |
| WP_073340811.1 | Type4.PglZ.containing.BREX | <-Tox-ART-HYD2<-SIGMA+OB-Fold+AAA-ATPase-BrxL<-PglZ+Phosphodiest<-DUF6079*<-?<-DUF4007<-wHTH+HNH-p<-DUF6079<-DUF4007<-PAPS-reduct+a+b-domain+Fer4_9<-?<-Cysteine-Desulfurase | 1323 | Bacteria>Terrabacteria_group>Bacillota>Clostridia>Eubacteriales>Clostridiaceae | GCF_900129965.1 | Clostridium_grantii | BUB83_RS19620 | 4604..8575 | 40575 | hypothetical-protein-[Clostridium-grantii] |
| WP_073340815.1 | Type4.PglZ.containing.BREX | <-Tox-ART-HYD2<-SIGMA+OB-Fold+AAA-ATPase-BrxL<-PglZ+Phosphodiest<-DUF6079<-?<-DUF4007<-wHTH+HNH-p<-DUF6079*<-DUF4007<-PAPS-reduct+a+b-domain+Fer4_9<-?<-Cysteine-Desulfurase | 1118 | Bacteria>Terrabacteria_group>Bacillota>Clostridia>Eubacteriales>Clostridiaceae | GCF_900129965.1 | Clostridium_grantii | BUB83_RS19640 | 10582..13938 | 40575 | hypothetical-protein-[Clostridium-grantii] |
| MCL2499300.1 | Type4.PglZ.containing.BREX | <-RADICAL-SAM<-?<-HTH_26<-ZnR<-SIGMA+OB-Fold+AAA-ATPase-BrxL<-iSTAND+PglZ<-DUF6079*<-PAPS-reduct+a+b-domain+Fer4_9+DUF4007<-HARE-HTH+E3_UFM1_ligase<-HTH_26<-?<-?<-primpol+D5N+D5ATPase | 1383 | Bacteria>Terrabacteria_group>Bacillota>Clostridia>Eubacteriales>Defluviitaleaceae | GCA_009779755.1 | Defluviitaleaceae_bacterium | FWE90_03070 | 105821..109972 | 2660712 | hypothetical-protein-[Defluviitaleaceae-bacterium] |
| MCL2016625.1 | Type4.PglZ.containing.BREX | ZnR->HARE-HTH+E3_UFM1_ligase->PAPS-reduct+a+b-domain+Fer4_9+DUF4007->DUF6079*->iSTAND+PglZ->SIGMA+OB-Fold+AAA-ATPase-BrxL-> | 1404 | Bacteria>Terrabacteria_group>Bacillota>Clostridia>Eubacteriales>Defluviitaleaceae | GCA_009782245.1 | Defluviitaleaceae_bacterium | FWG68_10315 | 5487..9701 | 2660712 | hypothetical-protein-[Defluviitaleaceae-bacterium] |
| WP_092467481.1 | Type4.PglZ.containing.BREX | Phage_int_SAM_1+Phage_integrase->35exo+wHTH+SF1-SF2-Helicase+HTH->HEPN+ZNR-><-AAA-ATPase<-HTH_7+rve\|\|SIR2-><-HNH+CXC3\|\|HARE-HTH+E3_UFM1_ligase->PAPS-reduct+a+b-domain+Fer4_9+DUF4007->DUF6079*->iSTAND+PglZ->SIGMA+OB-Fold+AAA-ATPase-BrxL->ZnR->Cysteine-Desulfurase->AAA-ATPase-p->?-><-AAA-ATPase<-HTH+rve | 1383 | Bacteria>Terrabacteria_group>Bacillota>Clostridia>Eubacteriales>Desulfallaceae | GCF_900113335.1 | Desulfotruncus_arcticus | SAMN05660649_00012 | 18170..22321 | 341036 | hypothetical-protein-[Desulfotruncus-arcticus] |
| NBJ16968.1 | Type4.PglZ.containing.BREX | <-GDResolvase+Recombinase+Zn_ribbon_recom<-?<-MACRODOMAIN<-Cysteine-Desulfurase<-ZnR<-SIGMA+OB-Fold+AAA-ATPase-BrxL<-iSTAND+PglZ<-DUF6079*<-PAPS-reduct+a+b-domain+Fer4_9+DUF4007<-HARE-HTH+E3_UFM1_ligase<-SIGMA-HTH\|\|HTH_3-><-Radical_SAM<-?<-Peptidase_U32<-RADICAL-SAM | 1398 | Bacteria>Terrabacteria_group>Bacillota>Clostridia>Eubacteriales>Desulfitobacteriaceae | GCA_009910795.1 | Dehalobacter_sp._4CP | FNP40_15715 | 33739..37935 | 2594474 | hypothetical-protein-[Dehalobacter-sp.-4CP] |
| MCO5388571.1 | Type4.PglZ.containing.BREX | <-ParB<-?<-?<-HKD-DNase+SF1-SF2-Helicase+HTH+HTH+SRA-p<-Cysteine-Desulfurase<-ZnR<-SIGMA+OB-Fold+AAA-ATPase-BrxL<-iSTAND+PglZ<-DUF6079-p<-DUF6079*<-DUF4007<-PAPS-reduct+a+b-domain+Fer4_9<-?<-HARE-HTH+E3_UFM1_ligase<-HTH_26<-HTH_38<-REase+SF1-SF2-Helicase | 1244 | Bacteria>Terrabacteria_group>Bacillota>Clostridia>Eubacteriales>Desulfitobacteriaceae | GCA_023955555.1 | Desulfosporosinus_sp. | NHB14_25450 | 97412..101146 | 157907 | hypothetical-protein-[Desulfosporosinus-sp.] |
| WP_015261292.1 | Type4.PglZ.containing.BREX | HAD+HAD->?->RAMA->PAPS-reduct+a+b-domain+Fer4_9+DUF4007->DUF6079*->PglZ->SIGMA+OB-Fold+AAA-ATPase-BrxL->ZnR->HNH+NERD+ZnR-><-HTH_3 | 1402 | Bacteria>Terrabacteria_group>Bacillota>Clostridia>Eubacteriales>Desulfitobacteriaceae | GCF_000243135.2 | Desulfitobacterium_dichloroeliminans | DESDI_RS03685 | 796259..800467 | 233055 | hypothetical-protein-[Desulfitobacterium-dichloroeliminans] |
| WP_229249285.1 | Type4.PglZ.containing.BREX | <-Cysteine-Desulfurase<-ZnR<-SIGMA+OB-Fold+AAA-ATPase-BrxL<-iSTAND+PglZ<-DUF6079*<-PAPS-reduct+a+b-domain+Fer4_9+DUF4007<-HARE-HTH+E3_UFM1_ligase<-HTH_3\|\|DUF772+DDE_Tnp_1+DDE_Tnp_1_6-><-TRAM+METHYLASE | 1379 | Bacteria>Terrabacteria_group>Bacillota>Clostridia>Eubacteriales>Desulfitobacteriaceae | GCF_020595055.1 | Desulfosporosinus_sp._SRJS8 | LJE72_RS17880 | 5352..9491 | 2885154 | hypothetical-protein-[Desulfosporosinus-sp.-SRJS8] |

| Protein-ID | System | Gene-Neighbourhoods | Length | Taxonomy | Assembly-ID | Strain | Locus-Tag | Genomic-coordinates | TaxID | Fasta-define |
|---|---|---|---|---|---|---|---|---|---|---|
| WP_073031289.1 | Type4.PglZ.containing.BREX | <-Cysteine-Desulfurase<-ZnR<-SIGMA+OB-Fold+AAA-ATPase-BrxL<-iSTAND+PglZ<-DUF6079*<-PAPS-reduct+a+b-domain+Fer4_9+DUF4007<-SeqA+Phage_int+SeqA+Phage_int+E3_UFM1_ligase<-DDE_Tnp_1<-?\|\|Phage_integrase->Zn_Tnp_IS91+Y2_Tnp-><-HIN-HTH | 1396 | Bacteria>Terrabacteria_group>Bacillota>Clostridia>Eubacteriales>Desulfitobacteriaceae | GCF_900129935.1 | Desulfosporosinus_lacus | BUB11_RS18755 | 6357..10547 | 329936 | hypothetical-protein-[Desulfosporosinus-lacus] |
| OPY55979.1 | Type4.PglZ.containing.BREX | <-TPR+TPR+TPR+TPR<-HAD<-LysM+PTSEIIA1<-?<-ZnR<-RNaseH\|\|tRNA->XIS-HTH->RAMA->PAPS-reduct+a+b-domain+Fer4_9+DUF4007->DUF6079*->PglZ->SIGMA+OB-Fold+AAA-ATPase-BrxL->ZnR->?-><-?\|\|?->RHH_8->?->GDResolvase+Recombinase+ZnR-> | 1404 | Bacteria>Terrabacteria_group>Bacillota>Clostridia>Eubacteriales>Desulfotomaculaceae | GCA_002067965.1 | Pelotomaculum_sp._PtaU1.Bin035 | A4E55_02300 | 29979..34193 | 1811695 | MAG:-hypothetical-protein-A4E55_02300-[Pelotomaculum-sp.-PtaU1.Bin035] |
| HBQ28532.1 | Type4.PglZ.containing.BREX | PAPS-reduct+DUF4007->DUF6079*->iSTAND+PglZ->SIGMA+OB-Fold+AAA-ATPase-BrxL->ZnR->HicB_lk_antitox->HicA_toxin->?->?-><-URI-DNase | 1394 | Bacteria>Terrabacteria_group>Bacillota>Clostridia>Eubacteriales>Desulfotomaculaceae | GCA_003535365.1 | Desulfotomaculum_sp. | DD719_03850 | 6237..10421 | 41211 | hypothetical-protein-[Desulfotomaculum-sp.] |
| WP_134212086.1 | Type4.PglZ.containing.BREX | <-RelE-ParE<-PhdYeFM_antitox<-?\|\|OLD-ABC->PIN-><-?\|\|?->PAPS-reduct+a+b-domain+Fer4_9+DUF4007->DUF4007->DUF6079*->iSTAND+PglZ->SIGMA+OB-Fold+AAA-ATPase-BrxL-> | 1375 | Bacteria>Terrabacteria_group>Bacillota>Clostridia>Eubacteriales>Desulfotomaculaceae | GCF_004369225.1 | Pelotomaculum_propionicicum | Pmgp_RS00985 | 191364..195491 | 258475 | ATP-binding-protein-[Pelotomaculum-propionicicum] |
| MBQ8951403.1 | Type4.PglZ.containing.BREX | <-CarD+HTH<-HTH_3<-?<-DndB<-wHTH(SBD)+HNH<-Cysteine-Desulfurase<-SIGMA+OB-Fold+AAA-ATPase-BrxL<-iSTAND+PglZ<-DUF6079*<-ZnR<-PAPS-reduct+a+b-domain+Fer4_9+DUF4007<-XIS-HTH | 1195 | Bacteria>Terrabacteria_group>Bacillota>Clostridia>Eubacteriales>Eubacteriaceae | GCA_017460785.1 | Eubacterium_sp. | IJ065_09690 | 25576..29163 | 142586 | hypothetical-protein-[Eubacterium-sp.] |
| MBO5175449.1 | Type4.PglZ.containing.BREX | DnaJ->AbiEi_4->AbiEii->NUDIX->?->XIS-HTH->PAPS-reduct+a+b-domain+Fer4_9+DUF4007->ZnR->DUF6079*->iSTAND+PglZ->SIGMA+OB-Fold+AAA-ATPase-BrxL->Cysteine-Desulfurase->?->?->Peptidase_M18+Peptidase_M42->?->?->TatD_DNase-> | 1196 | Bacteria>Terrabacteria_group>Bacillota>Clostridia>Eubacteriales>Eubacteriaceae | GCA_017623035.1 | Eubacterium_sp. | J6B84_09950 | 12194..15784 | 142586 | hypothetical-protein-[Eubacterium-sp.] |
| MBS6586268.1 | Type4.PglZ.containing.BREX | Phage-portal->?->?-><-HTH_3\|\|SeqA+Phage_int+SeqA+E3_UFM1_ligase->PAPS-reduct+a+b-domain+Fer4_9->DUF4007->DUF6079*->iSTAND+PglZ->SIGMA+OB-Fold+AAA-ATPase-BrxL->ZnR->Cysteine-Desulfurase-> | 1408 | Bacteria>Terrabacteria_group>Bacillota>Clostridia>Eubacteriales>Eubacteriaceae | GCA_018377805.1 | Eubacterium_sp. | KH345_08940 | 91074..95300 | 142586 | hypothetical-protein-[Eubacterium-sp.] |
| MCC8015067.1 | Type4.PglZ.containing.BREX | RAMA->PAPS-reduct+a+b-domain+Fer4_9+DUF4007->DUF6079*->PglZ->SIGMA+OB-Fold+AAA-ATPase+Lon-Protease-BrxL-p-> | 1397 | Bacteria>Terrabacteria_group>Bacillota>Clostridia>Eubacteriales>Eubacteriaceae | GCA_020859645.1 | Eubacterium_sp. | LI087_07710 | 3437..7630 | 142586 | hypothetical-protein-[Eubacterium-sp.] |
| MCI9127450.1 | Type4.PglZ.containing.BREX | REase->?->?->XIS-HTH->PAPS-reduct+a+b-domain+Fer4_9+DUF4007->ZnR->DUF6079*->iSTAND+PglZ->SIGMA+OB-Fold+AAA-ATPase-BrxL->Cysteine-Desulfurase->wHTH(SBD)+HNH->DndB->?->?->?->SIGMA-HTH-> | 1199 | Bacteria>Terrabacteria_group>Bacillota>Clostridia>Eubacteriales>Eubacteriaceae | GCA_022797855.1 | Eubacterium_sp. | HFG28_09665 | 5456..9055 | 142586 | hypothetical-protein-[Eubacterium-sp.] |
| MCM1063037.1 | Type4.PglZ.containing.BREX | <-GrpE<-?<-DndB<-URI-DNase<-Cysteine-Desulfurase<-Preader8<-SIGMA+OB-Fold+AAA-ATPase-BrxL<-iSTAND+PglZ<-DUF6079*<-PAPS-reduct+a+b-domain+Fer4_9+DUF4007<-XIS-HTH<-?<-?<-?<-Aminotran_1_2 | 1171 | Bacteria>Terrabacteria_group>Bacillota>Clostridia>Eubacteriales>Eubacteriaceae | GCA_023663325.1 | Eubacterium_sp. | NC420_00970 | 232065..235580 | 142586 | hypothetical-protein-[Eubacterium-sp.] |
| SEG30141.1 | Type4.PglZ.containing.BREX | <-HEPN<-?<-?<-McrC-NTD+REase<-McrB-AAA<-Cysteine-Desulfurase<-ZnR<-SIGMA+OB-Fold+AAA-ATPase-BrxL<-iSTAND+PglZ<-DUF6079*<-PAPS-reduct+a+b-domain+Fer4_9+DUF4007<-SeqA+Phage_int+SeqA+E3_UFM1_ligase<-wHTH+WYL+WCX<-?<-?<-TIR | 1399 | Bacteria>Terrabacteria_group>Bacillota>Clostridia>Eubacteriales>Eubacteriaceae | GCA_900108205.1 | Eubacterium_ruminantium | SAMN04487934_11447 | 47901..52100 | 42322 | hypothetical-protein-SAMN04487934_11447-[Eubacterium-ruminantium] |
| WP_006599851.1 | Type4.PglZ.containing.BREX | <-ACET<-ACET<-?<-N6-METHYLASE-p<-?<-Cysteine-Desulfurase<-ZnR<-SIGMA+OB-Fold+AAA-ATPase-BrxL<-iSTAND+PglZ<-DUF6079*<-PAPS-reduct+DUF4007<-PAPS-reduct+a+b-domain+Fer4_9<-SeqA+Phage_int+SeqA+Phage_int+E3_UFM1_ligase<-HTH_26<-SbcC<-?<-HNH\|\|?->SF1-SF2-Helicase-> | 1409 | Bacteria>Terrabacteria_group>Bacillota>Clostridia>Eubacteriales>Eubacteriaceae | GCF_000185505.1 | Pseudoramibacter_alactolyticus | HMPREF0721_RS10900 | 2289610..2293839 | 113287 | hypothetical-protein-[Pseudoramibacter-alactolyticus] |
| WP_228883703.1 | Type4.PglZ.containing.BREX | <-REC+HDGYP\|\|?->?->?->?->wHTH+WYL+WCX->?->SeqA+Phage_int+SeqA+Phage_int+E3_UFM1_ligase->PAPS-reduct+a+b-domain+Fer4_9+DUF4007->DUF6079*->iSTAND+PglZ-> | 1399 | Bacteria>Terrabacteria_group>Bacillota>Clostridia>Eubacteriales>Eubacteriaceae | GCF_902175055.1 | Acetobacterium_wieringae | LPA28_RS18135 | 161486..165685 | 52694 | hypothetical-protein-[Acetobacterium-wieringae] |
| MCC2258802.1 | Type4.PglZ.containing.BREX | <-TERNS3+HEPN<-Preader+URI-DNase<-?<-GDSL<-NUDIX<-?\|\|?->HNH->PAPS-reduct+a+b-domain+Fer4_9+DUF4007->DUF6079*->iSTAND+PglZ->ZnR->SIGMA+OB-Fold+AAA-ATPase-BrxL->Cysteine-Desulfurase->wHTH+WYL+WCX->HEPN-> | 1403 | Bacteria>Terrabacteria_group>Bacillota>Clostridia>Eubacteriales>Eubacteriales_incertae_sedis | GCA_020687605.1 | Intestinimonas_sp._CLA-AA-H199 | LKD74_12670 | 10236..14447 | 2881257 | DUF6079-family-protein-[Intestinimonas-sp.-CLA-AA-H199] |
| WP_206458293.1 | Type4.PglZ.containing.BREX | <-SIGMA-HTH<-SIGMA-HTH\|\|HTH_3->MPTase-><-Cysteine-Desulfurase<-ZnR<-SIGMA+OB-Fold+AAA-ATPase-BrxL<-PglZ<-iSTAND+PglZ<-DUF6079*<-PAPS-reduct+a+b-domain+Fer4_9+DUF4007<-SeqA+E3_UFM1_ligase<-HTH_26<-HTH_36<-RVT+HNH\|\|?-><-ZnR+TOPRIM | 1393 | Bacteria>Terrabacteria_group>Bacillota>Clostridia>Eubacteriales>Eubacteriales_incertae_sedis | GCF_017168145.1 | Anaerovorax_sp._IOR16 | I2490_RS02970 | 200700..204881 | 2773458 | hypothetical-protein-[Anaerovorax-sp.-IOR16] |
| WP_215692602.1 | Type4.PglZ.containing.BREX | XIS-HTH+XIS-HTH->\|\|?->?->HTH_26->?->PAPS-reduct+Fer4_9-p->PAPS-reduct+a+b-domain+Fer4_9+DUF4007->DUF6079*->iSTAND+PglZ->SIGMA+OB-Fold+AAA-ATPase-BrxL->ZnR->Cysteine-Desulfurase->wHTH+WYL+WCX-><-HTH_3\|\|XIS-HTH->?->TraJ-RHH-> | 1399 | Bacteria>Terrabacteria_group>Bacillota>Clostridia>Eubacteriales>Eubacteriales_incertae_sedis | GCF_017884445.1 | Gemmiger_formicilis | GPK89_RS09430 | 79487..83686 | 745368 | hypothetical-protein-[Gemmiger-formicilis] |
| WP_071430727.1 | Type4.PglZ.containing.BREX | HTH+rve->AAA-ATPase-><-?<-TERNS3+HEPN<-?<-?<-ZnR<-iSTAND+PglZ<-DUF6079*<-PAPS-reduct+a+b-domain+Fer4_9+DUF4007<-SeqA+Phage_int+SeqA+Phage_int+E3_UFM1_ligase<-HTH_26<-HTH_36<-VirD4-TraG+VirD4-FtsK<-ZnR+TOPRIM<-?<-RHH_8 | 1401 | Bacteria>Terrabacteria_group>Bacillota>Clostridia>Eubacteriales | GCF_015554745.1 | Oscillospiraceae | I2E77_RS04130 | 59703..63908 | 216572 | MULTISPECIES:-hypothetical-protein-[Oscillospiraceae] |
| CCY54306.1 | Type4.PglZ.containing.BREX | <-AAA-ATPase<-HIN-HTH<-DndB<-URI-DNase<-Cysteine-Desulfurase<-ZnR<-SIGMA+OB-Fold+AAA-ATPase-BrxL<-iSTAND+PglZ<-DUF6079*<-PAPS-reduct+a+b-domain+Fer4_9+DUF4007<-?<-HTH_26<-?<-?<-?<-?<-SIGMA-HTH<-Pkinase+Pkinase | 1378 | Bacteria>Terrabacteria_group>Bacillota>Clostridia>Eubacteriales>Lachnospiraceae | GCA_000433075.1 | Coprococcus_sp._CAG:782 | BN781_00828 | 17633..21769 | 1262863 | putative-uncharacterized-protein-[Coprococcus-sp.-CAG:782] |
| CRL42117.1 | Type4.PglZ.containing.BREX | SMODS->VirB4-FtsK->?->?->Lysozyme+NLPC->HTH_26->Phage_integrase+E3_UFM1_ligase->PAPS-reduct+a+b-domain+Fer4_9+DUF4007->DUF6079*->iSTAND+PglZ->SIGMA+OB-Fold+AAA-ATPase-BrxL->ZnR->Cysteine-Desulfurase->?->wHTH+WYL+WCX->HTH+REase->LexA-protease-> | 1394 | Bacteria>Terrabacteria_group>Bacillota>Clostridia>Eubacteriales>Lachnospiraceae | GCA_001406835.1 | [Eubacterium]_rectale | T1815_00811 | 85812..89996 | 39491 | hypothetical-protein-T1815_00811-[[Eubacterium] |
| OKZ93162.1 | Type4.PglZ.containing.BREX | Aminotran_3->?->HTH_3->?->?->HTH_26->?->PAPS-reduct+a+b-domain+Fer4_9+DUF4007->DUF6079*->iSTAND+PglZ->SIGMA+OB-Fold+AAA-ATPase-BrxL->ZnR->Cysteine-Desulfurase->URI-DNase->DndB->ZnR->HTH_3-> | 1378 | Bacteria>Terrabacteria_group>Bacillota>Clostridia>Eubacteriales>Lachnospiraceae | GCA_001916305.1 | Coprococcus_sp._CAG:131_42_139 | BHW15_01205 | 80914..85050 | 1896997 | MAG:-hypothetical-protein-BHW15_01205-[Coprococcus-sp.-CAG:131_42_139] |
| ONI45222.1 | Type4.PglZ.containing.BREX | GGDEF->?-><-?<-?<-?<-?<-ZnR<-SIGMA+OB-Fold+AAA-ATPase-BrxL<-iSTAND+PglZ<-DUF6079*<-PAPS-reduct+a+b-domain+Fer4_9+DUF4007 | 1388 | Bacteria>Terrabacteria_group>Bacillota>Clostridia>Eubacteriales>Lachnospiraceae | GCA_001983555.1 | Epulopiscium_sp._SCG-C07WGA-EpuloA2 | AN641_04455 | 13048..17214 | 1712378 | hypothetical-protein-AN641_04455-[Epulopiscium-sp.-SCG-C07WGA-EpuloA2] |
| RGE15632.1 | Type4.PglZ.containing.BREX | Methyltransf_25->?->?-><-?\|\|MACRODOMAIN->HTH_26->?->HNH->PAPS-reduct+a+b-domain+Fer4_9+DUF4007->DUF6079*->iSTAND+PglZ->ZnR->SIGMA+OB-Fold+AAA-ATPase-BrxL->Cysteine-Desulfurase->?->?-><-MPTase\|\|SAM-METHYLASE->METHYLASE-> | 1399 | Bacteria>Terrabacteria_group>Bacillota>Clostridia>Eubacteriales>Lachnospiraceae | GCA_003435545.1 | Lachnospiraceae_bacterium_OF11-28 | DXB03_06330 | 29137..33336 | 2305244 | hypothetical-protein-DXB03_06330-[Lachnospiraceae-bacterium-OF11-28] |
| RGF12868.1 | Type4.PglZ.containing.BREX | Lysozyme+NLPC->HTH_26->Phage_integrase+E3_UFM1_ligase->PAPS-reduct+a+b-domain+Fer4_9+DUF4007->DUF6079*->iSTAND+PglZ->SIGMA+OB-Fold+AAA-ATPase-BrxL->ZnR->Cysteine-Desulfurase->wHTH+WYL+WCX->HTH+REase->LexA-protease->MPTase->ZnR-> | 1394 | Bacteria>Terrabacteria_group>Bacillota>Clostridia>Eubacteriales>Lachnospiraceae | GCA_003435775.1 | Blautia_sp._AM16-16B | DW177_12610 | 7334..11518 | 2292969 | hypothetical-protein-DW177_12610-[Blautia-sp.-AM16-16B] |
| RGO24546.1 | Type4.PglZ.containing.BREX | <-Phage_int_SAM_3+Phage_integrase\|\|Lysozyme+NLPC->HTH_26->Phage_integrase+E3_UFM1_ligase->PAPS-reduct+a+b-domain+Fer4_9+DUF4007->GspA-ATPase->DUF6079-p*->iSTAND+PglZ->SIGMA+OB-Fold+AAA-ATPase-BrxL->ZnR->Cysteine-Desulfurase->wHTH+WYL+WCX->HTH+REase-> | 858 | Bacteria>Terrabacteria_group>Bacillota>Clostridia>Eubacteriales>Lachnospiraceae | GCA_003439775.1 | Dorea_sp._OM02-2LB | DXB23_09190 | 29363..31939 | 2292347 | hypothetical-protein-DXB23_09190-[Dorea-sp.-OM02-2LB] |
| RGS73808.1 | Type4.PglZ.containing.BREX | SMODS->?->?->Lysozyme+NLPC->HTH_26->Phage_integrase+E3_UFM1_ligase->PAPS-reduct+Fer4_9-p->PAPS-reduct+a+b-domain+Fer4_9+DUF4007->DUF6079*->iSTAND+PglZ-> | 1394 | Bacteria>Terrabacteria_group>Bacillota>Clostridia>Eubacteriales>Lachnospiraceae | GCA_003458845.1 | Dorea_formicigenerans | DWX78_00790 | 74696..78880 | 39486 | hypothetical-protein-DWX78_00790-[Dorea-formicigenerans] |
| HAX53679.1 | Type4.PglZ.containing.BREX | <-ROK-HTH<-Glyco_hydro_1<-?\|\|?-><-?<-Cysteine-Desulfurase<-ZnR<-SIGMA+OB-Fold+AAA-ATPase-BrxL<-iSTAND+PglZ<-DUF6079*<-PAPS-reduct+a+b-domain+Fer4_9+DUF4007<-?<-HTH_26<-?<-?<-?<-?<-HNH+3C-ABC\|\|SIGMA-HTH-> | 1379 | Bacteria>Terrabacteria_group>Bacillota>Clostridia>Eubacteriales>Lachnospiraceae | GCA_003487345.1 | Lachnospiraceae_bacterium | DCX82_17395 | 15220..19359 | 1898203 | hypothetical-protein-[Lachnospiraceae-bacterium] |
| HCH97636.1 | Type4.PglZ.containing.BREX | HTH+LexA-protease->HTH_3->?-><-?<-Cysteine-Desulfurase<-SIGMA+OB-Fold+AAA-ATPase-BrxL<-ZnR<-iSTAND+PglZ<-DUF6079*<-PAPS-reduct+a+b-domain+Fer4_9+DUF4007<-HARE-HTH+E3_UFM1_ligase<-?<-HTH_26<-PolB<-?<-TraJ-RHH<-LPD2 | 1404 | Bacteria>Terrabacteria_group>Bacillota>Clostridia>Eubacteriales>Lachnospiraceae | GCA_003521595.1 | Lachnospiraceae_bacterium | DFI63_06420 | 14994..19208 | 1898203 | hypothetical-protein-[Lachnospiraceae-bacterium] |
| GFI28164.1 | Type4.PglZ.containing.BREX | TraJ-RHH->TraI-VirD2-RCRE->XIS-HTH->Phage_integrase+E3_UFM1_ligase->PAPS-reduct+a+b-domain+Fer4_9+DUF4007->DUF6079*->DUF6079-p->iSTAND+PglZ->SIGMA+OB-Fold+AAA-ATPase-BrxL->ZnR->Cysteine-Desulfurase->Schlafen+HATPase+wHTH->?->?->SF1-SF2-Helicase-> | 1001 | Bacteria>Terrabacteria_group>Bacillota>Clostridia>Eubacteriales>Lachnospiraceae | GCA_011959925.1 | Lachnospiraceae_bacterium | IMSAGC012_03295 | 7920..10925 | 1898203 | hypothetical-protein-IMSAGC012_03295-[Lachnospiraceae-bacterium] |
| MBD5156842.1 | Type4.PglZ.containing.BREX | Thymidylate_kin-><-CHTH<-SMODS\|\|HTH_3->SeqA+Phage_int+SeqA+E3_UFM1_ligase->PAPS-reduct+a+b-domain+Fer4_9+DUF4007->DUF6079*->iSTAND+PglZ->SIGMA+OB-Fold+AAA-ATPase-BrxL->ZnR->Cysteine-Desulfurase->URI-DNase->DndB-><-SMODS<-?\|\|PadR-HTH-> | 1381 | Bacteria>Terrabacteria_group>Bacillota>Clostridia>Eubacteriales>Lachnospiraceae | GCA_014799955.1 | Butyrivibrio_sp. | HDT13_04330 | 35388..39533 | 28121 | hypothetical-protein-[Butyrivibrio-sp.] |
| MBD5471182.1 | Type4.PglZ.containing.BREX | TraI-VirD2-RCRE->?->HTH_26->SeqA+Phage_int+SeqA+E3_UFM1_ligase->PAPS-reduct+a+b-domain+Fer4_9+DUF4007->DUF6079*->iSTAND+PglZ->SIGMA+OB-Fold+AAA-ATPase-BrxL-> | 1405 | Bacteria>Terrabacteria_group>Bacillota>Clostridia>Eubacteriales>Lachnospiraceae | GCA_014804345.1 | Lachnospiraceae_bacterium | HDR19_08755 | 5661..9878 | 1898203 | hypothetical-protein-[Lachnospiraceae-bacterium] |
| MBD5489945.1 | Type4.PglZ.containing.BREX | <-Peptidase_M18+Peptidase_M42<-CR-ATPase8+REase<-?<-Cysteine-Desulfurase<-Preader8<-SIGMA+OB-Fold+AAA-ATPase-BrxL<-iSTAND+PglZ<-DUF6079*<-ZnR<-PAPS-reduct+a+b-domain+Fer4_9+DUF4007<-PIN<-HTH+MPTase<-MNS+REase | 1160 | Bacteria>Terrabacteria_group>Bacillota>Clostridia>Eubacteriales>Lachnospiraceae | GCA_014804485.1 | Lachnospiraceae_bacterium | HDR13_14335 | 23664..27146 | 1898203 | hypothetical-protein-[Lachnospiraceae-bacterium] |
| MBD5550155.1 | Type4.PglZ.containing.BREX | XIS-HTH->PAPS-reduct+a+b-domain+Fer4_9+DUF4007->DUF6079*->iSTAND+PglZ->SIGMA+OB-Fold+AAA-ATPase-BrxL->Preader8->Cysteine-Desulfurase->ParB+HNH->VirB11+SbcC->HNH->SF1-SF2-Helicase->Schlafen+HATPase+wHTH-> | 1166 | Bacteria>Terrabacteria_group>Bacillota>Clostridia>Eubacteriales>Lachnospiraceae | GCA_014804795.1 | Lachnospiraceae_bacterium | HDQ96_03115 | 108470..111970 | 1898203 | hypothetical-protein-[Lachnospiraceae-bacterium] |
| MBQ8803650.1 | Type4.PglZ.containing.BREX | <-Integrase_DNA+Phage_integrase<-XIS-HTH<-?<-?<-Cysteine-Desulfurase<-ZnR<-SIGMA+OB-Fold+AAA-ATPase-BrxL<-iSTAND+PglZ<-DUF6079*<-PAPS-reduct+a+b-domain+Fer4_9+DUF4007<-?<-MazG<-NERD+SF1-SF2-Helicase<-OLD-ABC<-?<-CHTH | 1413 | Bacteria>Terrabacteria_group>Bacillota>Clostridia>Eubacteriales>Lachnospiraceae | GCA_017465565.1 | Tyzzerella_sp. | IJZ53_08465 | 137120..141361 | 2053632 | hypothetical-protein-[Tyzzerella-sp.] |
| MBR3103810.1 | Type4.PglZ.containing.BREX | PAPS-reduct+a+b-domain+Fer4_9+DUF4007->DUF6079*->iSTAND+PglZ->SIGMA+OB-Fold+AAA-ATPase-BrxL->ZnR->Cysteine-Desulfurase-> | 1320 | Bacteria>Terrabacteria_group>Bacillota>Clostridia>Eubacteriales>Lachnospiraceae | GCA_017508825.1 | Lachnospiraceae_bacterium | IKQ27_11075 | 1374..5336 | 1898203 | hypothetical-protein-[Lachnospiraceae-bacterium] |
| MBR3507814.1 | Type4.PglZ.containing.BREX | PRTase->Classical-AAA->?->tRNA->tRNA-><-Phage_integrase\|\|HTH_3->XIS-HTH->HARE-HTH+E3_UFM1_ligase->PAPS-reduct+a+b-domain+Fer4_9+DUF4007->DUF6079*->iSTAND+PglZ->ZnR->SIGMA+OB-Fold+AAA-ATPase-BrxL->Cysteine-Desulfurase->Schlafen+HATPase+wHTH-><-Phage_integrase\|\|Classical-AAA->Peptidase_S8->SF1-SF2-Helicase+REase-> | 1398 | Bacteria>Terrabacteria_group>Bacillota>Clostridia>Eubacteriales>Lachnospiraceae | GCA_017527205.1 | Lachnospiraceae_bacterium | IKO03_03350 | 38351..42547 | 1898203 | hypothetical-protein-[Lachnospiraceae-bacterium] |
| MBR4981305.1 | Type4.PglZ.containing.BREX | <-HNH<-VirB11+SbcC<-ParB+HNH<-Cysteine-Desulfurase<-Preader8<-SIGMA+OB-Fold+AAA-ATPase-BrxL<-iSTAND+PglZ<-DUF6079*<-ZnR<-PAPS-reduct+a+b-domain+Fer4_9+DUF4007<-XIS-HTH<-?<-?<-HEPN<-ZetaToxin | 1166 | Bacteria>Terrabacteria_group>Bacillota>Clostridia>Eubacteriales>Lachnospiraceae | GCA_017555455.1 | Lachnospiraceae_bacterium | IKY94_01945 | 9438..12938 | 1898203 | hypothetical-protein-[Lachnospiraceae-bacterium] |
| MBP3458816.1 | Type4.PglZ.containing.BREX | <-TrkH<-?\|\|?->?->ParB+HNH->HEPN->Phage_integrase+E3_UFM1_ligase->PAPS-reduct+a+b-domain+Fer4_9+DUF4007->DUF6079*->iSTAND+PglZ-> | 1394 | Bacteria>Terrabacteria_group>Bacillota>Clostridia>Eubacteriales>Lachnospiraceae | GCA_017936785.1 | Lachnospiraceae_bacterium | J6K58_06375 | 32294..36478 | 1898203 | hypothetical-protein-[Lachnospiraceae-bacterium] |
| MBS5262590.1 | Type4.PglZ.containing.BREX | <-DndB<-URI-DNase<-Cysteine-Desulfurase<-ZnR<-SIGMA+OB-Fold+AAA-ATPase-BrxL<-iSTAND+PglZ<-DUF6079*<-PAPS-reduct+a+b-domain+Fer4_9+DUF4007<-?<-HTH_26<-?<-?<-?<-?<-SbcC | 1378 | Bacteria>Terrabacteria_group>Bacillota>Clostridia>Eubacteriales>Lachnospiraceae | GCA_018366895.1 | Roseburia_sp. | KHY54_12575 | 7092..11228 | 2049040 | hypothetical-protein-[Roseburia-sp.] |
| HIV92079.1 | Type4.PglZ.containing.BREX | CHTH-><-?<-HISKIN<-Cysteine-Desulfurase<-ZnR<-SIGMA+OB-Fold+AAA-ATPase-BrxL<-iSTAND+PglZ<-DUF6079*<-PAPS-reduct+a+b-domain+Fer4_9+DUF4007<-SeqA+Phage_int+SeqA+E3_UFM1_ligase | 1407 | Bacteria>Terrabacteria_group>Bacillota>Clostridia>Eubacteriales>Lachnospiraceae | GCA_019113175.1 | Candidatus_Eisenbergiella_stercoravium | H9744_10440 | 81665..85888 | 2838556 | hypothetical-protein-[Candidatus-Eisenbergiella-stercoravium] |
| HJC24805.1 | Type4.PglZ.containing.BREX | ACET-><-Transposase_31<-?<-Peptidase<-Peptidase_S66<-MarR-HTH\|\|RAMA->PAPS-reduct+a+b-domain+Fer4_9+DUF4007->DUF6079*->PglZ->SIGMA+OB-Fold+AAA-ATPase+Lon-Protease-BrxL-p-><-Phage_int_SAM_3+Phage_integrase<-XIS-HTH<-HTH_16<-SIGMA-HTH<-?<-Thioredoxin_3<-DsbD | 1385 | Bacteria>Terrabacteria_group>Bacillota>Clostridia>Eubacteriales>Lachnospiraceae | GCA_019119595.1 | Candidatus_Eisenbergiella_merdavium | H9761_14060 | 62475..66632 | 2838551 | hypothetical-protein-[Candidatus-Eisenbergiella-merdavium] |
| MCD7884237.1 | Type4.PglZ.containing.BREX | <-Cysteine-Desulfurase<-ZnR<-SIGMA+OB-Fold+AAA-ATPase-BrxL<-iSTAND+PglZ<-DUF6079*<-PAPS-reduct+a+b-domain+Fer4_9+DUF4007<-SeqA+Phage_int+SeqA+Phage_int+E3_UFM1_ligase<-?<-wHTH+WYL+WCX | 1400 | Bacteria>Terrabacteria_group>Bacillota>Clostridia>Eubacteriales>Lachnospiraceae | GCA_021201095.1 | Lachnospiraceae_bacterium | LUI87_11135 | 15344..19546 | 1898203 | hypothetical-protein-[Lachnospiraceae-bacterium] |
| MCD8054955.1 | Type4.PglZ.containing.BREX | <-Cysteine-Desulfurase<-ZnR<-SIGMA+OB-Fold+AAA-ATPase-BrxL<-iSTAND+PglZ<-DUF6079*<-PAPS-reduct+a+b-domain+Fer4_9+DUF4007<-SeqA+Phage_int+SeqA+Phage_int+E3_UFM1_ligase<-wHTH+WYL+WCX<-Lysozyme+NLPC<-VirB4-FtsK<-?<-TrwI<-TraM | 1400 | Bacteria>Terrabacteria_group>Bacillota>Clostridia>Eubacteriales>Lachnospiraceae | GCA_021202465.1 | Lachnospiraceae_bacterium | LUF00_13175 | 17269..21471 | 1898203 | hypothetical-protein-[Lachnospiraceae-bacterium] |
| MCI6886009.1 | Type4.PglZ.containing.BREX | <-SIGMA+OB-Fold+AAA-ATPase-BrxL<-iSTAND+PglZ<-DUF6079*<-GspA-ATPase<-PAPS-reduct+a+b-domain+Fer4_9+DUF4007<-SeqA+Phage_int+SeqA+E3_UFM1_ligase<-XIS-HTH | 1084 | Bacteria>Terrabacteria_group>Bacillota>Clostridia>Eubacteriales>Lachnospiraceae | GCA_022776445.1 | Lachnospiraceae_bacterium | MR868_01980 | 3882..7136 | 1898203 | hypothetical-protein-[Lachnospiraceae-bacterium] |
| MCI7180753.1 | Type4.PglZ.containing.BREX | MACRODOMAIN->?->PIN->ZnR+ZnR+ZnR->XIS-HTH->SeqA+Phage_int+SeqA+E3_UFM1_ligase->?->?->PAPS-reduct+a+b-domain+Fer4_9+DUF4007->DUF6079*->iSTAND+PglZ->SIGMA+OB-Fold+AAA-ATPase-BrxL->ZnR->Cysteine-Desulfurase->?->STYKIN-> | 1401 | Bacteria>Terrabacteria_group>Bacillota>Clostridia>Eubacteriales>Lachnospiraceae | GCA_022782435.1 | Lachnospiraceae_bacterium | MSA26_09330 | 14283..18488 | 1898203 | hypothetical-protein-[Lachnospiraceae-bacterium] |
| MCI8507239.1 | Type4.PglZ.containing.BREX | MerR-HTH->GspA-ATPase+REase->?->?->?->SeqA+Phage_int+SeqA+E3_UFM1_ligase->PAPS-reduct+a+b-domain+Fer4_9+DUF4007->DUF6079*->iSTAND+PglZ->SIGMA+OB-Fold+AAA-ATPase-BrxL->ZnR->Cysteine-Desulfurase->?->ACET+ACET-> | 1380 | Bacteria>Terrabacteria_group>Bacillota>Clostridia>Eubacteriales>Lachnospiraceae | GCA_022792915.1 | Lachnospiraceae_bacterium | HFJ06_01555 | 39671..43813 | 1898203 | hypothetical-protein-[Lachnospiraceae-bacterium] |
| MCI8565873.1 | Type4.PglZ.containing.BREX | <-RVT<-?<-?<-MORC<-Cysteine-Desulfurase<-ZnR<-SIGMA+OB-Fold+AAA-ATPase-BrxL<-iSTAND+PglZ<-DUF6079*<-PAPS-reduct+a+b-domain+Fer4_9+DUF4007<-?<-?<-HHH+DNA_pol_B+RNase\|\|?-><-?<-?<-MerR-HTH+SHS2+SHS2\|\|DSBH+AraC-HTH+AraC-HTH-> | 1394 | Bacteria>Terrabacteria_group>Bacillota>Clostridia>Eubacteriales>Lachnospiraceae | GCA_022793375.1 | Lachnospiraceae_bacterium | HFI39_06080 | 27700..31884 | 1898203 | hypothetical-protein-[Lachnospiraceae-bacterium] |
| MCI8950348.1 | Type4.PglZ.containing.BREX | <-Cysteine-Desulfurase<-ZnR<-SIGMA+OB-Fold+AAA-ATPase-BrxL<-iSTAND+PglZ<-DUF6079*<-PAPS-reduct+a+b-domain+Fer4_9+DUF4007<-SeqA+Phage_int+SeqA+E3_UFM1_ligase<-SeqA+Phage_int+SeqA+E3_UFM1_ligase<-HTH_26<-?<-HNH<-REC<-MORC+REase<-N6-METHYLASE | 1407 | Bacteria>Terrabacteria_group>Bacillota>Clostridia>Eubacteriales>Lachnospiraceae | GCA_022796475.1 | Lachnospiraceae_bacterium | HFG49_09935 | 6060..10283 | 1898203 | hypothetical-protein-[Lachnospiraceae-bacterium] |
| MCI9323607.1 | Type4.PglZ.containing.BREX | <-HNH<-VirB11+SbcC<-ParB+HNH<-Cysteine-Desulfurase<-Preader8<-SIGMA+OB-Fold+AAA-ATPase-BrxL<-iSTAND+PglZ<-DUF6079*<-PAPS-reduct+a+b-domain+Fer4_9+DUF4007\|\|ABhydrolase-><-?<-SbcC+SbcC<-?<-?<-?<-CR-ATPase8+REase | 1171 | Bacteria>Terrabacteria_group>Bacillota>Clostridia>Eubacteriales>Lachnospiraceae | GCA_022799595.1 | Lachnospiraceae_bacterium | HFH81_10180 | 61695..65210 | 1898203 | hypothetical-protein-[Lachnospiraceae-bacterium] |
| MCI9486770.1 | Type4.PglZ.containing.BREX | PAPS-reduct+a+b-domain+Fer4_9+DUF4007->DUF6079*->iSTAND+PglZ->SIGMA+OB-Fold+AAA-ATPase-BrxL->ZnR->Cysteine-Desulfurase->REase->SF1-SF2-Helicase->SF1-SF2-Helicase->DrmB->NERD+SF1-SF2-Helicase-> | 1400 | Bacteria>Terrabacteria_group>Bacillota>Clostridia>Eubacteriales>Lachnospiraceae | GCA_022800815.1 | Lachnospiraceae_bacterium | HFI64_07295 | 4690..8892 | 1898203 | hypothetical-protein-[Lachnospiraceae-bacterium] |
| MCI9535149.1 | Type4.PglZ.containing.BREX | <-HNH<-VirB11+SbcC<-ParB+HNH<-Cysteine-Desulfurase<-Preader8<-SIGMA+OB-Fold+AAA-ATPase-BrxL<-iSTAND+PglZ<-DUF6079*<-PAPS-reduct+a+b-domain+Fer4_9+DUF4007<-XIS-HTH\|\|?-><-?\|\|GNTR-HTH-> | 1171 | Bacteria>Terrabacteria_group>Bacillota>Clostridia>Eubacteriales>Lachnospiraceae | GCA_022801195.1 | Lachnospiraceae_bacterium | HFG53_10180 | 52808..56323 | 1898203 | hypothetical-protein-[Lachnospiraceae-bacterium] |
| MCI9559903.1 | Type4.PglZ.containing.BREX | <-DndB<-URI-DNase<-Cysteine-Desulfurase<-ZnR<-SIGMA+OB-Fold+AAA-ATPase-BrxL<-iSTAND+PglZ<-DUF6079*<-PAPS-reduct+a+b-domain+Fer4_9+DUF4007<-SeqA+Phage_int+SeqA+E3_UFM1_ligase\|\|?-><-Amidohydro_1<-BPD_transp_2<-?<-ABC-ATPase+SbcC+ABC-ATPase+SbcC | 1411 | Bacteria>Terrabacteria_group>Bacillota>Clostridia>Eubacteriales>Lachnospiraceae | GCA_022801375.1 | Lachnospiraceae_bacterium | HFG52_11805 | 7188..11423 | 1898203 | hypothetical-protein-[Lachnospiraceae-bacterium] |
| BDF48894.1 | Type4.PglZ.containing.BREX | LZ_Tnp_IS66+zf-IS66+DDE_Tnp_IS66+DDE_Tnp_IS66_C->DDE_Tnp_IS66+DDE_Tnp_IS66_C-><-?<-MULE-TRANSPOSASE<-wHTH+WYL+WCX<-Cysteine-Desulfurase<-ZnR<-SIGMA+OB-Fold+AAA-ATPase-BrxL<-iSTAND+PglZ<-DUF6079*<-PAPS-reduct+a+b-domain+Fer4_9+DUF4007<-HNH<-HNH<-SIGMA-HTH<-TRAM+METHYLASE<-N-OB+HDRnase<-Trypsin+PDZ<-ACET | 1402 | Bacteria>Terrabacteria_group>Bacillota>Clostridia>Eubacteriales>Lachnospiraceae | GCA_022845955.1 | Lachnospiraceae_bacterium | CE91St56_60170 | 6872214..6876422 | 1898203 | hypothetical-protein-CE91St56_60170-[Lachnospiraceae-bacterium] |
| MCR4802121.1 | Type4.PglZ.containing.BREX | HTH_26->?->PAPS-reduct+a+b-domain+Fer4_9+DUF4007->DUF6079*->iSTAND+PglZ->SIGMA+OB-Fold+AAA-ATPase-BrxL->ZnR->Cysteine-Desulfurase->URI-DNase->DndB->AbiGi+seqA+RAMA-> | 1378 | Bacteria>Terrabacteria_group>Bacillota>Clostridia>Eubacteriales>Lachnospiraceae | GCA_024681895.1 | Lachnospiraceae_bacterium | K5895_03850 | 4502..8638 | 1898203 | hypothetical-protein-[Lachnospiraceae-bacterium] |
| SDJ32367.1 | Type4.PglZ.containing.BREX | CHTH->?->?->?->?->HTH_26->Phage_integrase+E3_UFM1_ligase->PAPS-reduct+a+b-domain+Fer4_9+DUF4007->DUF6079*->iSTAND+PglZ->SIGMA+OB-Fold+AAA-ATPase-BrxL->ZnR->Cysteine-Desulfurase->HTH_3-><-TraI-VirD2-RCRE<-TraJ-RHH\|\|HTH_3-> | 1394 | Bacteria>Terrabacteria_group>Bacillota>Clostridia>Eubacteriales>Lachnospiraceae | GCA_900100245.1 | Lachnospiraceae_bacterium_G41 | SAMN04487760_10564 | 50138..54322 | 1200749 | hypothetical-protein-SAMN04487760_10564-[Lachnospiraceae-bacterium-G41] |
| WP_029201261.1 | Type4.PglZ.containing.BREX | DDE_Tnp_ISL3-><-iSTAND+PglZ<-DUF6079*<-ZnR<-PAPS-reduct+a+b-domain+Fer4_9+DUF4007<-XIS-HTH<-ParB<-ParA-Soj-PloopNTPase<-SymE_toxin | 1193 | Bacteria>Terrabacteria_group>Bacillota>Clostridia>Eubacteriales>Lachnospiraceae | GCF_000424445.1 | Oribacterium_sp._NK2B42 | G625_RS0111710 | 1739..5320 | 689781 | DUF6079-family-protein-[Oribacterium-sp.-NK2B42] |
| WP_027209045.1 | Type4.PglZ.containing.BREX | nSTAND->URI-DNase->HTH_26->?->PAPS-reduct+a+b-domain+Fer4_9+DUF4007->DUF6079*->iSTAND+PglZ->SIGMA+OB-Fold+AAA-ATPase-BrxL->ZnR->Cysteine-Desulfurase->?->DnaJ->HIRAN->BirA-HTH-> | 1381 | Bacteria>Terrabacteria_group>Bacillota>Clostridia>Eubacteriales>Lachnospiraceae | GCF_000424465.1 | Butyrivibrio_hungatei | G628_RS0109900 | 12562..16707 | 185008 | hypothetical-protein-[Butyrivibrio-hungatei] |
| WP_024292380.1 | Type4.PglZ.containing.BREX | LZ_Tnp_IS66+zf-IS66+DDE_Tnp_IS66+DDE_Tnp_IS66_C-><-HNH<-VirB11+SbcC<-ParB+HNH<-Cysteine-Desulfurase<-Preader8<-SIGMA+OB-Fold+AAA-ATPase-BrxL<-iSTAND+PglZ<-DUF6079*<-PAPS-reduct+a+b-domain+Fer4_9+DUF4007<-XIS-HTH<-?<-CR-ATPase5+REase<-?<-?<-?<-PhdYeFM_antitox | 1168 | Bacteria>Terrabacteria_group>Bacillota>Clostridia>Eubacteriales>Lachnospiraceae | GCF_000526995.1 | Lacrimispora_indolis | K401_RS0107520 | 1580339..1583845 | 69825 | hypothetical-protein-[Lacrimispora-indolis] |
| WP_026654575.1 | Type4.PglZ.containing.BREX | GNTR-HTH->Aminotran_1_2->?->XIS-HTH->PAPS-reduct+a+b-domain+Fer4_9+DUF4007->ZnR->DUF6079*->iSTAND+PglZ->SIGMA+OB-Fold+AAA-ATPase-BrxL->Cysteine-Desulfurase-p->Cysteine-Desulfurase->ParB+HNH->VirB11+SbcC->HNH-> | 1192 | Bacteria>Terrabacteria_group>Bacillota>Clostridia>Eubacteriales>Lachnospiraceae | GCF_000686065.1 | Butyrivibrio_sp._AE3003 | CD00_RS0103055 | 628547..632125 | 1496721 | hypothetical-protein-[Butyrivibrio-sp.-AE3003] |
| WP_038284935.1 | Type4.PglZ.containing.BREX | <-DndB<-URI-DNase<-Cysteine-Desulfurase<-ZnR<-SIGMA+OB-Fold+AAA-ATPase-BrxL<-iSTAND+PglZ<-DUF6079*<-PAPS-reduct+a+b-domain+Fer4_9+DUF4007<-SeqA+Phage_int+SeqA+E3_UFM1_ligase\|\|?->?-><-RelE-ParE<-PhdYeFM_antitox<-?<-?<-DUF932 | 1404 | Bacteria>Terrabacteria_group>Bacillota>Clostridia>Eubacteriales>Lachnospiraceae | GCF_000732605.1 | Lacrimispora_celerecrescens | IO98_RS22120 | 7610..11824 | 29354 | hypothetical-protein-[Lacrimispora-celerecrescens] |
| WP_057572019.1 | Type4.PglZ.containing.BREX | HAD+HAD->?->TRAM+METHYLASE->HTH_26->SeqA+Phage_int+SeqA+E3_UFM1_ligase->PAPS-reduct+a+b-domain+Fer4_9+DUF4007->DUF6079*->iSTAND+PglZ->SIGMA+OB-Fold+AAA-ATPase-BrxL->ZnR->Cysteine-Desulfurase->Phage_integrase->Phage_integrase->Phage_int_SAM_4+Phage_integrase-> | 1407 | Bacteria>Terrabacteria_group>Bacillota>Clostridia>Eubacteriales>Lachnospiraceae | GCF_001405335.1 | Enterocloster_clostridioformis | ARA87_RS13265 | 53989..58212 | 1531 | hypothetical-protein-[Enterocloster-clostridioformis] |
| WP_055301605.1 | Type4.PglZ.containing.BREX | <-DndB<-URI-DNase<-Cysteine-Desulfurase<-Preader8<-SIGMA+OB-Fold+AAA-ATPase-BrxL<-iSTAND+PglZ<-DUF6079*<-PAPS-reduct+a+b-domain+Fer4_9+DUF4007<-XIS-HTH<-?<-ABhydrolase<-MA | 1174 | Bacteria>Terrabacteria_group>Bacillota>Clostridia>Eubacteriales>Lachnospiraceae | GCF_001406675.1 | Roseburia_inulinivorans | ARA14_RS04785 | 53366..56890 | 360807 | hypothetical-protein-[Roseburia-inulinivorans] |
| WP_087307206.1 | Type4.PglZ.containing.BREX | LON-AAA-><-ABC-ATPase+SbcC<-TetR-HTH<-?\|\|?-><-?\|\|SAM-METHYLASE->?->PAPS-reduct+a+b-domain+Fer4_9+DUF4007->DUF6079*->iSTAND+PglZ->SIGMA+OB-Fold+AAA-ATPase-BrxL->ZnR->Cysteine-Desulfurase->?->METHYLASE-> | 1379 | Bacteria>Terrabacteria_group>Bacillota>Clostridia>Eubacteriales>Lachnospiraceae | GCF_002160765.1 | Lachnoclostridium_sp._An138 | B5E82_RS08875 | 137282..141421 | 1965560 | hypothetical-protein-[Lachnoclostridium-sp.-An138] |
| WP_087989664.1 | Type4.PglZ.containing.BREX | HTH_36->?-><-?<-Cysteine-Desulfurase<-ZnR<-SIGMA+OB-Fold+AAA-ATPase-BrxL<-iSTAND+PglZ<-DUF6079*<-PAPS-reduct+a+b-domain+Fer4_9+DUF4007<-SeqA+Phage_int+SeqA+Phage_int+E3_UFM1_ligase<-HTH_26<-?<-?<-?<-?\|\|DEDD_Tnp_IS110+Transposase_20-><-Preader17 | 1401 | Bacteria>Terrabacteria_group>Bacillota>Clostridia>Eubacteriales>Lachnospiraceae | GCF_002161375.1 | Anaerotignum_lactatifermentans | B5G26_RS11050 | 25727..29932 | 160404 | hypothetical-protein-[Anaerotignum-lactatifermentans] |
| WP_110290602.1 | Type4.PglZ.containing.BREX | sGTP->?->?->?->?->?->HTH_3->HARE-HTH+E3_UFM1_ligase->PAPS-reduct+a+b-domain+Fer4_9+DUF4007->DUF6079*->iSTAND+PglZ->SIGMA+OB-Fold+AAA-ATPase-BrxL->ZnR->Cysteine-Desulfurase->HTH_31->MPTase->?->SIGMA-HTH-> | 1393 | Bacteria>Terrabacteria_group>Bacillota>Clostridia>Eubacteriales>Lachnospiraceae | GCF_003201285.1 | Lachnotalea_glycerini | C8E03_RS06940 | 557759..561940 | 1763509 | hypothetical-protein-[Lachnotalea-glycerini] |
| WP_117841749.1 | Type4.PglZ.containing.BREX | <-DndB<-URI-DNase<-Cysteine-Desulfurase<-ZnR<-SIGMA+OB-Fold+AAA-ATPase-BrxL<-iSTAND+PglZ<-DUF6079*<-PAPS-reduct+a+b-domain+Fer4_9+DUF4007<-?<-HTH_26\|\|?-><-wHTH+WYL+WCX\|\|?-><-?<-ACET+ACET<-ACET+ACET | 1378 | Bacteria>Terrabacteria_group>Bacillota>Clostridia>Eubacteriales>Lachnospiraceae | GCF_003462625.1 | unclassified_Roseburia | DXB39_RS10385 | 8256..12392 | 2637578 | MULTISPECIES:-hypothetical-protein-[unclassified-Roseburia] |
| WP_118093056.1 | Type4.PglZ.containing.BREX | MA->ABhydrolase->?->HTH_38->?->HTH_26->Phage_integrase+E3_UFM1_ligase->PAPS-reduct+a+b-domain+Fer4_9+DUF4007->DUF6079*->iSTAND+PglZ->SIGMA+OB-Fold+AAA-ATPase-BrxL->ZnR->Cysteine-Desulfurase->HTH+REase-> | 1397 | Bacteria>Terrabacteria_group>Bacillota>Clostridia>Eubacteriales>Lachnospiraceae | GCF_003468625.1 | Roseburia_inulinivorans | DW813_RS09690 | 54941..59134 | 360807 | hypothetical-protein-[Roseburia-inulinivorans] |
| WP_173896440.1 | Type4.PglZ.containing.BREX | <-MarR-HTH\|\|?-><-Cysteine-Desulfurase<-ZnR<-SIGMA+OB-Fold+AAA-ATPase-BrxL<-iSTAND+PglZ<-DUF6079*<-PAPS-reduct+a+b-domain+Fer4_9+DUF4007<-SeqA+Phage_int+SeqA+Phage_int+E3_UFM1_ligase<-XIS-HTH<-?<-ArdCN+MPTase-inact+LPD19 | 1398 | Bacteria>Terrabacteria_group>Bacillota>Clostridia>Eubacteriales>Lachnospiraceae | GCF_013300195.1 | Faecalicatena_fissicatena | HFM84_RS09860 | 22649..26845 | 290055 | P-loop-NTPase-fold-protein-[Faecalicatena-fissicatena] |
| WP_186847752.1 | Type4.PglZ.containing.BREX | PBPI->?->PIN-><-Cysteine-Desulfurase<-ZnR<-SIGMA+OB-Fold+AAA-ATPase-BrxL<-iSTAND+PglZ<-DUF6079*<-GspA-ATPase<-PAPS-reduct+a+b-domain+Fer4_9+DUF4007<-Phage_integrase+E3_UFM1_ligase<-?<-?<-?<-VirD4-TraG+VirD4-FtsK | 1267 | Bacteria>Terrabacteria_group>Bacillota>Clostridia>Eubacteriales>Lachnospiraceae | GCF_014287735.1 | Coprococcus_sp._NSJ-10 | H8S09_RS09190 | 26541..30344 | 2763039 | hypothetical-protein-[Coprococcus-sp.-NSJ-10] |
| WP_138262164.1 | Type4.PglZ.containing.BREX | <-GDResolvase+Recombinase+ZnR<-UDG-conflict<-TOPRIM<-OLD-ABC<-?<-Cysteine-Desulfurase<-ZnR<-SIGMA+OB-Fold+AAA-ATPase-BrxL<-iSTAND+PglZ<-DUF6079*<-PAPS-reduct+a+b-domain+Fer4_9+DUF4007<-SeqA+Phage_int+SeqA+E3_UFM1_ligase\|\|REC+AraC-HTH->HisKin+HATPase_c->?-><-?<-PBPI<-ABhydrolase | 1381 | Bacteria>Terrabacteria_group>Bacillota>Clostridia>Eubacteriales>Lachnospiraceae | GCF_015550395.1 | [Clostridium]_hylemonae | I2C33_RS01925 | 56979..61124 | 89153 | hypothetical-protein-[[Clostridium] |
| WP_195565920.1 | Type4.PglZ.containing.BREX | PIN->OLD-ABC->NERD->URI-DNase->?->HDRnase->HTH_26->Phage_integrase+E3_UFM1_ligase->PAPS-reduct+a+b-domain+Fer4_9+DUF4007->DUF6079*->iSTAND+PglZ->SIGMA+OB-Fold+AAA-ATPase-BrxL->ZnR->Cysteine-Desulfurase->AAA-ATPase-p->?->?->?->CR-ATPase8+REase-> | 1394 | Bacteria>Terrabacteria_group>Bacillota>Clostridia>Eubacteriales>Lachnospiraceae | GCF_015556425.1 | Blautia_wexlerae | I2F87_RS07370 | 148804..152988 | 418240 | hypothetical-protein-[Blautia-wexlerae] |
| WP_195942878.1 | Type4.PglZ.containing.BREX | <-ABC-ATPase+SbcC<-?<-LexA-protease<-HTH+REase<-wHTH+WYL+WCX<-Cysteine-Desulfurase<-ZnR<-SIGMA+OB-Fold+AAA-ATPase-BrxL<-iSTAND+PglZ<-DUF6079*<-PAPS-reduct+a+b-domain+Fer4_9+DUF4007<-Phage_integrase+E3_UFM1_ligase<-HTH_26<-Lysozyme+NLPC<-?<-?<-Integrase_DNA+Phage_integrase<-TraJ-RHH | 1394 | Bacteria>Terrabacteria_group>Bacillota>Clostridia>Eubacteriales>Lachnospiraceae | GCF_015668255.1 | Mediterraneibacter_faecis | I4U97_RS14630 | 14129..18313 | 592978 | hypothetical-protein-[Mediterraneibacter-faecis] |
| WP_238014881.1 | Type4.PglZ.containing.BREX | HTH_26->Phage_integrase+E3_UFM1_ligase->PAPS-reduct+a+b-domain+Fer4_9->PAPS-reduct+DUF4007->DUF6079*->iSTAND+PglZ->SIGMA+OB-Fold+AAA-ATPase-BrxL->ZnR->Cysteine-Desulfurase->wHTH+WYL+WCX->HTH+REase-> | 1394 | Bacteria>Terrabacteria_group>Bacillota>Clostridia>Eubacteriales>Lachnospiraceae | GCF_022132665.1 | Mediterraneibacter_faecis | LOP77_RS14920 | 5513..9697 | 592978 | hypothetical-protein-[Mediterraneibacter-faecis] |
| WP_073992527.1 | Type4.PglZ.containing.BREX | <-UDG-conflict<-?<-Cysteine-Desulfurase<-ZnR<-SIGMA+OB-Fold+AAA-ATPase-BrxL<-iSTAND+PglZ<-DUF6079*<-PAPS-reduct+a+b-domain+Fer4_9+DUF4007<-HARE-HTH+E3_UFM1_ligase | 1405 | Bacteria>Terrabacteria_group>Bacillota>Clostridia>Eubacteriales>Lachnospiraceae | GCF_900141895.1 | Parasporobacterium_paucivorans | BUA79_RS01130 | 11911..16128 | 115544 | hypothetical-protein-[Parasporobacterium-paucivorans] |
| RGF94077.1 | Type4.PglZ.containing.BREX | ParB->ParB->?->?->?->?->Phage_integrase+wHTH->PAPS-reduct+a+b-domain+Fer4_9+DUF4007->DUF6079*->iSTAND+PglZ->SIGMA+OB-Fold+AAA-ATPase-BrxL->ZnR->Cysteine-Desulfurase->ZnR-><-RADICAL-SAM<-SF1-SF2-Helicase<-LPD32+LPD36+N6_Mtase+SF1-SF2-Helicase | 1405 | Bacteria>Terrabacteria_group>Bacillota>Clostridia>Eubacteriales>Oscillospiraceae | GCA_003460765.1 | Ruminococcus_sp._AM54-1NS | DXA02_04830 | 56881..61098 | 2293226 | hypothetical-protein-DXA02_04830-[Ruminococcus-sp.-AM54-1NS] |
| RGH34271.1 | Type4.PglZ.containing.BREX | <-SF1-SF2-Helicase+REase<-N6-N4-METHYLASE\|\|?->SF1-SF2-Helicase->RADICAL-SAM-><-ZnR<-Cysteine-Desulfurase<-ZnR<-iSTAND+PglZ<-DUF6079*<-PAPS-reduct+a+b-domain+Fer4_9+DUF4007<-Phage_integrase+wHTH<-?<-rve+rve_3<-HTH_Tnp_1 | 1405 | Bacteria>Terrabacteria_group>Bacillota>Clostridia>Eubacteriales>Oscillospiraceae | GCA_003461775.1 | Ruminococcus_sp._AM47-2BH | DW964_01575 | 149937..154154 | 2293221 | hypothetical-protein-DW964_01575-[Ruminococcus-sp.-AM47-2BH] |
| TCL45228.1 | Type4.PglZ.containing.BREX | <-RHH_8<-?<-HTH_36<-SIGMA-HTH<-?<-Cysteine-Desulfurase<-ZnR<-SIGMA+OB-Fold+AAA-ATPase-BrxL<-iSTAND+PglZ<-DUF6079*<-PAPS-reduct+a+b-domain+Fer4_9+DUF4007<-SeqA+Phage_int+SeqA+Phage_int+E3_UFM1_ligase<-Phage_int_SAM_3+Phage_integrase<-Phage_int_SAM_3\|\|?->?->PIN->?->SIGMA-HTH-> | 1401 | Bacteria>Terrabacteria_group>Bacillota>Clostridia>Eubacteriales>Oscillospiraceae | GCA_004340125.1 | Harryflintia_acetispora | EDD78_101210 | 218405..222610 | 1849041 | hypothetical-protein-EDD78_101210-[Harryflintia-acetispora] |
| MBD5142973.1 | Type4.PglZ.containing.BREX | DDE_Tnp-><-DDE_Tnp_4<-HTH_Tnp<-DDE_Tnp_1_3<-?\|\|Phage_integrase+wHTH->PAPS-reduct+a+b-domain+Fer4_9+DUF4007->DUF6079*->iSTAND+PglZ->SIGMA+OB-Fold+AAA-ATPase-BrxL->ZnR->Cysteine-Desulfurase-><-?\|\|?->RHH_8->?->GDResolvase+Recombinase+ZnR-> | 1399 | Bacteria>Terrabacteria_group>Bacillota>Clostridia>Eubacteriales>Oscillospiraceae | GCA_014799805.1 | Ruminococcus_sp. | HDT22_05090 | 13208..17407 | 41978 | hypothetical-protein-[Ruminococcus-sp.] |
| MBE6838776.1 | Type4.PglZ.containing.BREX | <-HIN-HTH\|\|ArdCN+MPTase->?->?->?->VirD4-TraG+VirD4-FtsK->HTH_26->PAPS-reduct+a+b-domain+Fer4_9+DUF4007->ZnR->DUF6079*->iSTAND+PglZ->SIGMA+OB-Fold+AAA-ATPase-BrxL->Cysteine-Desulfurase->?->ArsR-HTH->SAM-METHYLASE-> | 1188 | Bacteria>Terrabacteria_group>Bacillota>Clostridia>Eubacteriales>Oscillospiraceae | GCA_015066785.1 | Ruminococcus_sp. | E7507_04455 | 136451..140017 | 41978 | hypothetical-protein-[Ruminococcus-sp.] |
| MBQ3756408.1 | Type4.PglZ.containing.BREX | <-ZnR\|\|?-><-?\|\|wHTH+WYL+WCX->?->HNH->PAPS-reduct+a+b-domain+Fer4_9+DUF4007->DUF6079*->iSTAND+PglZ->ZnR->SIGMA+OB-Fold+AAA-ATPase-BrxL->Cysteine-Desulfurase->RFC-AAA->DnaJ->HIRAN->wHTH+WYL+WCX->LexA-protease-> | 1401 | Bacteria>Terrabacteria_group>Bacillota>Clostridia>Eubacteriales>Oscillospiraceae | GCA_017395085.1 | Oscillospiraceae_bacterium | II873_04770 | 29507..33712 | 2485925 | hypothetical-protein-[Oscillospiraceae-bacterium] |
| MBQ3262726.1 | Type4.PglZ.containing.BREX | <-D5N+viral-AAA+CDC6-HTH<-?<-?<-?\|\|CR-ATPase5+REase-><-?<-DndB\|\|HARE-HTH+E3_UFM1_ligase->PAPS-reduct+a+b-domain+Fer4_9+DUF4007->DUF6079*->iSTAND+PglZ-> | 1397 | Bacteria>Terrabacteria_group>Bacillota>Clostridia>Eubacteriales>Oscillospiraceae | GCA_017411225.1 | Oscillospiraceae_bacterium | IJH52_06090 | 85829..90022 | 2485925 | hypothetical-protein-[Oscillospiraceae-bacterium] |
| MBQ6207831.1 | Type4.PglZ.containing.BREX | DUF4007->GspA-ATPase->DUF6079*->iSTAND+PglZ->SIGMA+OB-Fold+AAA-ATPase-BrxL->ZnR->Cysteine-Desulfurase-p-> | 1311 | Bacteria>Terrabacteria_group>Bacillota>Clostridia>Eubacteriales>Oscillospiraceae | GCA_017428815.1 | Oscillospiraceae_bacterium | IJK52_12220 | 728..4663 | 2485925 | hypothetical-protein-[Oscillospiraceae-bacterium] |
| MBQ6632813.1 | Type4.PglZ.containing.BREX | <-XIS-HTH<-TraI-VirD2-RCRE<-TraJ-RHH<-?\|\|HTH->?->?->SeqA+Phage_int+SeqA+Phage_int+E3_UFM1_ligase->PAPS-reduct+a+b-domain+Fer4_9+DUF4007->DUF6079*->iSTAND+PglZ->SIGMA+OB-Fold+AAA-ATPase-BrxL->ZnR->Cysteine-Desulfurase->ARG->Schlafen+HATPase+wHTH-><-?<-UDG | 1396 | Bacteria>Terrabacteria_group>Bacillota>Clostridia>Eubacteriales>Oscillospiraceae | GCA_017434205.1 | Ruminococcus_sp. | IJH80_01280 | 17248..21438 | 41978 | hypothetical-protein-[Ruminococcus-sp.] |
| MBQ7681483.1 | Type4.PglZ.containing.BREX | DUF6079*->iSTAND+PglZ->SIGMA+OB-Fold+AAA-ATPase-BrxL->Cysteine-Desulfurase-><-?<-MODE-HTH+PBPII<-?<-?<-?<-SPOUT | 1073 | Bacteria>Terrabacteria_group>Bacillota>Clostridia>Eubacteriales>Oscillospiraceae | GCA_017443745.1 | Oscillibacter_sp. | IJT31_04655 | 1..3222 | 1945593 | hypothetical-protein-[Oscillibacter-sp.] |
| MBR6984229.1 | Type4.PglZ.containing.BREX | <-HEPN<-MPTase<-HTH_3\|\|SIGMA-HTH->TraJ-RHH->TraI-VirD2-RCRE-><-Cysteine-Desulfurase<-SIGMA+OB-Fold+AAA-ATPase-BrxL<-PglZ<-DUF6079*<-PAPS-reduct+a+b-domain+Fer4_9+DUF4007<-RAMA\|\|RADICAL-SAM->DUF3670+SF1-SF2-Helicase-><-SMODS<-ZnR+TOPRIM<-LPD32+LPD36+N6_Mtase+SF1-SF2-Helicase | 1404 | Bacteria>Terrabacteria_group>Bacillota>Clostridia>Eubacteriales>Oscillospiraceae | GCA_017519475.1 | Ruminococcus_sp. | IKH75_12020 | 30268..34482 | 41978 | hypothetical-protein-[Ruminococcus-sp.] |
| MBR4291498.1 | Type4.PglZ.containing.BREX | <-HTH_36<-XIS-HTH\|\|?-><-N6-METHYLASE-p<-Cysteine-Desulfurase<-SIGMA+OB-Fold+AAA-ATPase-BrxL<-iSTAND+PglZ<-DUF6079*<-PAPS-reduct+a+b-domain+Fer4_9+DUF4007 | 1400 | Bacteria>Terrabacteria_group>Bacillota>Clostridia>Eubacteriales>Oscillospiraceae | GCA_017537045.1 | Oscillospiraceae_bacterium | IKT52_12805 | 6995..11197 | 2485925 | hypothetical-protein-[Oscillospiraceae-bacterium] |
| MBP3335378.1 | Type4.PglZ.containing.BREX | HARE-HTH->PAPS-reduct+a+b-domain+Fer4_9+DUF4007->DUF6079*->iSTAND+PglZ->SIGMA+OB-Fold+AAA-ATPase-BrxL->ZnR->Cysteine-Desulfurase-><-?<-?<-ORC-CDC6<-tRNA<-?<-HTH_3 | 1382 | Bacteria>Terrabacteria_group>Bacillota>Clostridia>Eubacteriales>Oscillospiraceae | GCA_017935365.1 | Ruminiclostridium_sp. | J6L61_04475 | 2538..6686 | 2053608 | hypothetical-protein-[Ruminiclostridium-sp.] |
| MBS7204849.1 | Type4.PglZ.containing.BREX | REase->Tox-RES->?->?->?->MACRODOMAIN->?->HNH->PAPS-reduct+a+b-domain+Fer4_9+DUF4007->DUF6079*->iSTAND+PglZ->ZnR->SIGMA+OB-Fold+AAA-ATPase-BrxL->Cysteine-Desulfurase->Schlafen+HATPase+wHTH-p->?->BirA-HTH->Phage_integrase-><-CHTH | 1402 | Bacteria>Terrabacteria_group>Bacillota>Clostridia>Eubacteriales>Oscillospiraceae | GCA_018383015.1 | Ruminococcus_bicirculans | KH048_01075 | 243513..247721 | 1160721 | hypothetical-protein-[Ruminococcus-bicirculans] |
| HJA48126.1 | Type4.PglZ.containing.BREX | SIGMA-HTH->?->TRAM+METHYLASE->HTH_26->SeqA+Phage_int+SeqA+E3_UFM1_ligase->PAPS-reduct+a+b-domain+Fer4_9+DUF4007->DUF6079*->iSTAND+PglZ->SIGMA+OB-Fold+AAA-ATPase-BrxL-> | 1407 | Bacteria>Terrabacteria_group>Bacillota>Clostridia>Eubacteriales>Oscillospiraceae | GCA_019117485.1 | Candidatus_Agathobaculum_intestinipullorum | H9790_02170 | 36292..40515 | 2838422 | hypothetical-protein-[Candidatus-Agathobaculum-intestinipullorum] |
| MCD7785561.1 | Type4.PglZ.containing.BREX | <-TIR+TPR<-TIR+TPR\|\|?-><-SIGMA-HTH+SIGMA-HTH<-GDResolvase+Recombinase+ZnR<-KAP-NTPase<-REase<-SIGMA+OB-Fold+AAA-ATPase-BrxL<-iSTAND+PglZ<-DUF6079*<-PAPS-reduct+a+b-domain+Fer4_9+DUF4007<-SeqA+Phage_int+SeqA+E3_UFM1_ligase\|\|TraJ-RHH->RepA_N+HTH->?->MACRODOMAIN-><-GDResolvase<-HipA_C+STYKinase\|\|HTH_3-> | 1382 | Bacteria>Terrabacteria_group>Bacillota>Clostridia>Eubacteriales>Oscillospiraceae | GCA_021199855.1 | Oscillospiraceae_bacterium | LUH18_08335 | 20236..24384 | 2485925 | hypothetical-protein-[Oscillospiraceae-bacterium] |
| MCD7847862.1 | Type4.PglZ.containing.BREX | <-AAA+<-AAA+<-RepA_N+HTH<-?<-TraJ-RHH\|\|?->HNH->PAPS-reduct+a+b-domain+Fer4_9+DUF4007->DUF6079*->iSTAND+PglZ->SIGMA+OB-Fold+AAA-ATPase-BrxL-> | 1405 | Bacteria>Terrabacteria_group>Bacillota>Clostridia>Eubacteriales>Oscillospiraceae | GCA_021200775.1 | Oscillospiraceae_bacterium | LUG49_07570 | 34177..38394 | 2485925 | DUF6079-family-protein-[Oscillospiraceae-bacterium] |
| MCD8005837.1 | Type4.PglZ.containing.BREX | <-SIGMA+OB-Fold+AAA-ATPase-BrxL<-iSTAND+PglZ<-DUF6079*<-PAPS-reduct+a+b-domain+Fer4_9+DUF4007<-SeqA+Phage_int+SeqA+Phage_int+E3_UFM1_ligase<-ACET<-?\|\|?-><-?<-?\|\|AAA-ATPase->AbiEi_4-> | 1405 | Bacteria>Terrabacteria_group>Bacillota>Clostridia>Eubacteriales>Oscillospiraceae | GCA_021202155.1 | Oscillospiraceae_bacterium | LUF29_02525 | 16962..21179 | 2485925 | hypothetical-protein-[Oscillospiraceae-bacterium] |
| MCI6195361.1 | Type4.PglZ.containing.BREX | <-OLD-ABC<-?<-HIN-HTH<-Cysteine-Desulfurase<-ZnR<-SIGMA+OB-Fold+AAA-ATPase-BrxL<-iSTAND+PglZ<-DUF6079*<-PAPS-reduct+a+b-domain+Fer4_9+DUF4007<-SeqA+Phage_int+SeqA+E3_UFM1_ligase<-?\|\|MPTase-> | 1411 | Bacteria>Terrabacteria_group>Bacillota>Clostridia>Eubacteriales>Oscillospiraceae | GCA_022772255.1 | Faecalibacterium_sp. | MR651_10350 | 8163..12398 | 1971605 | hypothetical-protein-[Faecalibacterium-sp.] |
| MCI6401851.1 | Type4.PglZ.containing.BREX | LPD32+LPD36+N6_Mtase+SF1-SF2-Helicase->?->TraJ-RHH->?->?->SeqA+Phage_int+SeqA+Phage_int+E3_UFM1_ligase->PAPS-reduct+a+b-domain+Fer4_9+DUF4007->DUF6079*->iSTAND+PglZ->SIGMA+OB-Fold+AAA-ATPase-BrxL->Cysteine-Desulfurase->wHTH+WYL+WCX->?->?->?->Calcineurin-SbcD->HTH-> | 1399 | Bacteria>Terrabacteria_group>Bacillota>Clostridia>Eubacteriales>Oscillospiraceae | GCA_022777155.1 | Oscillospiraceae_bacterium | MR815_08540 | 38381..42580 | 2485925 | YqaJ-viral-recombinase-family-protein-[Oscillospiraceae-bacterium] |
| MCI8423043.1 | Type4.PglZ.containing.BREX | <-SIGMA-HTH<-ABC-ATPase+ABC-ATPase<-?<-?\|\|MORC-><-Cysteine-Desulfurase<-ZnR<-SIGMA+OB-Fold+AAA-ATPase-BrxL<-iSTAND+PglZ<-DUF6079*<-PAPS-reduct+a+b-domain+Fer4_9+DUF4007<-HARE-HTH+E3_UFM1_ligase<-?<-?<-?<-?<-?<-HD+SF1-SF2-Helicase | 1384 | Bacteria>Terrabacteria_group>Bacillota>Clostridia>Eubacteriales>Oscillospiraceae | GCA_022791995.1 | Lawsonibacter_sp. | HFF50_05885 | 23728..27882 | 2185275 | hypothetical-protein-[Lawsonibacter-sp.] |
| MCI8620198.1 | Type4.PglZ.containing.BREX | <-HTH_36<-VirD4-TraG+VirD4-FtsK<-ZnR+TOPRIM<-RVT<-Cysteine-Desulfurase<-ZnR<-SIGMA+OB-Fold+AAA-ATPase-BrxL<-iSTAND+PglZ<-DUF6079*<-PAPS-reduct+a+b-domain+Fer4_9+DUF4007<-HARE-HTH+E3_UFM1_ligase<-XIS-HTH\|\|DndB-><-tRNA<-RHS+RHS+HNH<-?<-RHS+RHS+HNH<-?<-RHS+RHS+HNH | 1397 | Bacteria>Terrabacteria_group>Bacillota>Clostridia>Eubacteriales>Oscillospiraceae | GCA_022793835.1 | Oscillospiraceae_bacterium | HFG44_09070 | 10421..14614 | 2485925 | hypothetical-protein-[Oscillospiraceae-bacterium] |
| MCI9347611.1 | Type4.PglZ.containing.BREX | <-LACI-HTH+PBPI<-?\|\|ADH_N+ADH_zinc_N->ABC-ATPase+SbcC+ABC-ATPase+SbcC->?->?->?->HNH->PAPS-reduct+a+b-domain+Fer4_9+DUF4007->DUF6079*->iSTAND+PglZ->ZnR->SIGMA+OB-Fold+AAA-ATPase-BrxL->Cysteine-Desulfurase->?->?->GDResolvase+Recombinase+ZnR-> | 1399 | Bacteria>Terrabacteria_group>Bacillota>Clostridia>Eubacteriales>Oscillospiraceae | GCA_022799735.1 | Oscillibacter_sp. | HFF83_00080 | 19216..23415 | 1945593 | hypothetical-protein-[Oscillibacter-sp.] |
| MCI9677950.1 | Type4.PglZ.containing.BREX | <-DDE_Tnp<-?<-wHTH\|\|tRNA-><-DDE_Tnp<-?<-Cysteine-Desulfurase<-ZnR<-SIGMA+OB-Fold+AAA-ATPase-BrxL<-iSTAND+PglZ<-DUF6079*<-PAPS-reduct+a+b-domain+Fer4_9+DUF4007<-?\|\|DndB->?->?-><-?<-?\|\|FURR-HTH-> | 1393 | Bacteria>Terrabacteria_group>Bacillota>Clostridia>Eubacteriales>Oscillospiraceae | GCA_022804995.1 | Oscillospiraceae_bacterium | HFF77_03105 | 16778..20959 | 2485925 | hypothetical-protein-[Oscillospiraceae-bacterium] |
| MCM1232922.1 | Type4.PglZ.containing.BREX | <-HIN-HTH<-Cysteine-Desulfurase<-ZnR<-SIGMA+OB-Fold+AAA-ATPase-BrxL<-iSTAND+PglZ<-DUF6079*<-PAPS-reduct+a+b-domain+Fer4_9+DUF4007<-SeqA+Phage_int+SeqA+Phage_int+E3_UFM1_ligase<-Mrr_N+ACET | 1400 | Bacteria>Terrabacteria_group>Bacillota>Clostridia>Eubacteriales>Oscillospiraceae | GCA_023664505.1 | Ruminococcus_flavefaciens | NC489_22580 | 5740..9942 | 1265 | hypothetical-protein-[Ruminococcus-flavefaciens] |
| MCM1531711.1 | Type4.PglZ.containing.BREX | Phage_int_SAM_1+Phage_integrase->?->sGTP->?-><-ADH_N+ADH_zinc_N+METHYLASE<-?\|\|PAPS-reduct+DUF4007->DUF6079*->iSTAND+PglZ->SIGMA+OB-Fold+AAA-ATPase-BrxL->\|\|?-><-URI-DNase+RAMA<-?\|\|TonBC->?->?-><-?<-CR-ATPase5+REase | 1174 | Bacteria>Terrabacteria_group>Bacillota>Clostridia>Eubacteriales>Oscillospiraceae | GCA_023667315.1 | Ruminococcus_flavefaciens | NC114_05510 | 186850..190374 | 1265 | hypothetical-protein-[Ruminococcus-flavefaciens] |
| MCQ4784087.1 | Type4.PglZ.containing.BREX | <-HTH_36<-XIS-HTH\|\|DEDD_Tnp_IS110+Transposase_20->?-><-TERNS3+HEPN<-Preader+URI-DNase<-?<-GDSL\|\|DUF6079*->iSTAND+PglZ->ZnR->SIGMA+OB-Fold+AAA-ATPase-BrxL->Cysteine-Desulfurase->wHTH+WYL+WCX->HEPN-><-?<-GDResolvase+Recombinase+ZnR | 1325 | Bacteria>Terrabacteria_group>Bacillota>Clostridia>Eubacteriales>Oscillospiraceae | GCA_024460695.1 | Flavonifractor_plautii | NE670_02185 | 68841..72818 | 292800 | hypothetical-protein-[Flavonifractor-plautii] |
| WP_117523970.1 | Type4.PglZ.containing.BREX | <-Cysteine-Desulfurase<-ZnR<-SIGMA+OB-Fold+AAA-ATPase-BrxL<-?<-iSTAND+PglZ<-DUF6079*<-PAPS-reduct+a+b-domain+Fer4_9+DUF4007<-SeqA+Phage_int+SeqA+E3_UFM1_ligase<-SF1-SF2-Helicase+PRTase<-Schlafen+HATPase+wHTH<-?<-?<-TetR-HTH\|\|RelB-> | 1390 | Bacteria>Terrabacteria_group>Bacillota>Clostridia>Eubacteriales>Oscillospiraceae | GCA_030820635.1 | Ruminococcus_sp._AF42-10 | DW050_RS08760 | 15777..19949 | 2293180 | hypothetical-protein-[Ruminococcus-sp.-AF42-10] |
| WP_118161974.1 | Type4.PglZ.containing.BREX | <-SF1-SF2-Helicase+REase<-N6-N4-METHYLASE\|\|?->SF1-SF2-Helicase->RADICAL-SAM-><-ZnR<-Cysteine-Desulfurase<-ZnR<-iSTAND+PglZ<-DUF6079*<-PAPS-reduct+a+b-domain+Fer4_9+DUF4007<-Phage_integrase+wHTH<-?<-rve+rve_3<-HTH_Tnp_1 | 1405 | Bacteria>Terrabacteria_group>Bacillota>Clostridia>Eubacteriales>Oscillospiraceae | GCA_030820635.1 | Ruminococcus_sp._AM47-2BH | DW964_01575 | 149937..154154 | 2293221 | hypothetical-protein-[Ruminococcus-sp.-AM47-2BH] |
| SCJ21704.1 | Type4.PglZ.containing.BREX | Phage_integrase+wHTH->PAPS-reduct+a+b-domain+Fer4_9+DUF4007->DUF6079*->iSTAND+PglZ->SIGMA+OB-Fold+AAA-ATPase-BrxL->ZnR->Cysteine-Desulfurase->ZnR-><-LPD32+LPD36+N6_Mtase+SF1-SF2-Helicase<-TOPRIM | 1405 | Bacteria>Terrabacteria_group>Bacillota>Clostridia>Eubacteriales>Oscillospiraceae | GCA_900067025.1 | uncultured_Ruminococcus_sp. | SAMEA3545436_04964 | 5452..9669 | 165186 | Uncharacterised-protein-[uncultured-Ruminococcus-sp.] |
| WP_087233702.1 | Type4.PglZ.containing.BREX | RHH_8->?->ZnR+TOPRIM->VirD4-TraG+VirD4-FtsK->HTH_36->HTH_26->SeqA+Phage_int+SeqA+Phage_int+E3_UFM1_ligase->PAPS-reduct+a+b-domain+Fer4_9+DUF4007->DUF6079*->iSTAND+PglZ->SIGMA+OB-Fold+AAA-ATPase-BrxL->ZnR->Cysteine-Desulfurase->?->?->TERNS3+HEPN-> | 1401 | Bacteria>Terrabacteria_group>Bacillota>Clostridia>Eubacteriales>Oscillospiraceae | GCF_002159845.1 | Anaeromassilibacillus_sp._An250 | B5F54_RS09660 | 80462..84667 | 1965604 | hypothetical-protein-[Anaeromassilibacillus-sp.-An250] |
| WP_087292451.1 | Type4.PglZ.containing.BREX | <-RHH_8<-HTH_36<-SIGMA-HTH<-?<-?<-Cysteine-Desulfurase<-ZnR<-SIGMA+OB-Fold+AAA-ATPase-BrxL<-iSTAND+PglZ<-DUF6079*<-PAPS-reduct+a+b-domain+Fer4_9+DUF4007<-SeqA+Phage_int+SeqA+Phage_int+E3_UFM1_ligase<-HTH_26<-?<-Phage_int_SAM_3+Phage_integrase<-Phage_int_SAM_3<-CR-ATPase5+REase\|\|tRNA->?-><-Phage_int_SAM_3+Phage_integrase | 1401 | Bacteria>Terrabacteria_group>Bacillota>Clostridia>Eubacteriales>Oscillospiraceae | GCF_002160015.1 | Anaerofilum_sp._An201 | B5F36_RS06485 | 32311..36516 | 1965588 | hypothetical-protein-[Anaerofilum-sp.-An201] |
| WP_097779439.1 | Type4.PglZ.containing.BREX | <-Phage_int_SAM_5+Phage_integrase<-XIS-HTH<-PIN<-?<-Cysteine-Desulfurase<-SIGMA+OB-Fold+AAA-ATPase-BrxL<-iSTAND+PglZ<-DUF6079*<-PAPS-reduct+a+b-domain+Fer4_9+DUF4007<-?<-DOC+HTH<-?<-STYKinase-like<-OLD-ABC<-?<-PIN | 1398 | Bacteria>Terrabacteria_group>Bacillota>Clostridia>Eubacteriales>Oscillospiraceae | GCF_002550015.1 | Faecalibacterium_prausnitzii | CGS59_RS07225 | 164155..168351 | 853 | hypothetical-protein-[Faecalibacterium-prausnitzii] |
| WP_102365412.1 | Type4.PglZ.containing.BREX | MODE-HTH+PBPII->HTH_26->SeqA+Phage_int+SeqA+Phage_int+E3_UFM1_ligase->PAPS-reduct+a+b-domain+Fer4_9+DUF4007->GspA-ATPase->DUF6079-p*->iSTAND+PglZ->SIGMA+OB-Fold+AAA-ATPase-BrxL->ZnR->Cysteine-Desulfurase->wHTH+WYL+WCX-><-?<-TraI-VirD2-RCRE<-TraJ-RHH | 890 | Bacteria>Terrabacteria_group>Bacillota>Clostridia>Eubacteriales>Oscillospiraceae | GCF_002874775.1 | Monoglobus_pectinilyticus | B9O19_RS05180 | 1163342..1166014 | 1981510 | hypothetical-protein-[Monoglobus-pectinilyticus] |
| WP_112148251.1 | Type4.PglZ.containing.BREX | <-GDResolvase+Recombinase+ZnR\|\|?->DOC+HTH-><-?<-XIS-HTH<-?<-SF1-SF2-Helicase\|\|?->PAPS-reduct+a+b-domain+Fer4_9+DUF4007->DUF6079-p*->DUF6079-p->iSTAND+PglZ->SIGMA+OB-Fold+AAA-ATPase-BrxL->ZnR->Cysteine-Desulfurase->wHTH+WYL+WCX->SIGMA-HTH->?->HTH_3-> | 812 | Bacteria>Terrabacteria_group>Bacillota>Clostridia>Eubacteriales>Oscillospiraceae | GCF_003287495.1 | Faecalibacterium_prausnitzii | C4N22_RS05435 | 430909..433347 | 853 | hypothetical-protein-[Faecalibacterium-prausnitzii] |
| WP_118124222.1 | Type4.PglZ.containing.BREX | LPD32+LPD36+N6_Mtase+SF1-SF2-Helicase->?-><-Cysteine-Desulfurase<-ZnR<-SIGMA+OB-Fold+AAA-ATPase-BrxL<-iSTAND+PglZ<-DUF6079*<-PAPS-reduct+a+b-domain+Fer4_9+DUF4007<-Phage_integrase+wHTH<-?<-?<-?<-?<-ParB | 1405 | Bacteria>Terrabacteria_group>Bacillota>Clostridia>Eubacteriales>Oscillospiraceae | GCF_003461785.1 | Ruminococcus_sp._AM43-6 | DW938_RS06230 | 89478..93695 | 2293216 | hypothetical-protein-[Ruminococcus-sp.-AM43-6] |
| WP_174403076.1 | Type4.PglZ.containing.BREX | Resolvase+Recombinase+ZnR->Resolvase+Recombinase+ZnR-><-Cysteine-Desulfurase<-SIGMA+OB-Fold+AAA-ATPase-BrxL<-iSTAND+PglZ<-DUF6079*<-PAPS-reduct+a+b-domain+Fer4_9+DUF4007<-SeqA+Phage_int+SeqA+Phage_int+E3_UFM1_ligase<-wHTH+WYL+WCX<-TRAM+METHYLASE\|\|?->HAD+HAD-> | 1407 | Bacteria>Terrabacteria_group>Bacillota>Clostridia>Eubacteriales>Oscillospiraceae | GCF_013340725.1 | Caproicibacterium_lactatifermentans | GKP14_RS03905 | 815047..819270 | 2666138 | hypothetical-protein-[Caproicibacterium-lactatifermentans] |
| WP_204883670.1 | Type4.PglZ.containing.BREX | DUF4096+DDE_Tnp_1+DDE_Tnp_1_2-><-Cysteine-Desulfurase<-SIGMA+OB-Fold+AAA-ATPase-BrxL<-PglZ<-DUF6079*<-PAPS-reduct+a+b-domain+Fer4_9+DUF4007<-RAMA\|\|rve+rve_3-><-DOC+HTH<-?<-?<-?<-TraJ-RHH<-Arm-DNA-bind_4+Phage_int_SAM_3+Phage_integrase | 1390 | Bacteria>Terrabacteria_group>Bacillota>Clostridia>Eubacteriales>Oscillospiraceae | GCF_016901955.1 | Pseudoflavonifractor_phocaeensis | H6B51_RS10645 | 5549..9721 | 1870988 | hypothetical-protein-[Pseudoflavonifractor-phocaeensis] |
| WP_212792178.1 | Type4.PglZ.containing.BREX | <-Phage_int_SAM_5+Phage_integrase<-XIS-HTH<-PIN<-?<-Cysteine-Desulfurase<-SIGMA+OB-Fold+AAA-ATPase-BrxL<-iSTAND+PglZ<-DUF6079*<-PAPS-reduct+a+b-domain+Fer4_9+DUF4007<-?<-DOC+HTH<-?<-STYKinase-like<-?<-HTH_36<-?<-PIN | 1398 | Bacteria>Terrabacteria_group>Bacillota>Clostridia>Eubacteriales>Oscillospiraceae | GCF_018327745.1 | Faecalibacterium_prausnitzii | MCC2041_RS04450 | 18680..22876 | 853 | hypothetical-protein-[Faecalibacterium-prausnitzii] |
| WP_215649152.1 | Type4.PglZ.containing.BREX | HNH->PAPS-reduct+a+b-domain+Fer4_9+DUF4007->DUF6079*->iSTAND+PglZ->ZnR->SIGMA+OB-Fold+AAA-ATPase-BrxL->Cysteine-Desulfurase->Schlafen+HATPase+wHTH-p->?->BirA-HTH->Phage_integrase-><-CHTH | 1402 | Bacteria>Terrabacteria_group>Bacillota>Clostridia>Eubacteriales>Oscillospiraceae | GCF_018784005.1 | Ruminococcus_bicirculans | GPK54_04625 | 6087..10295 | 1160721 | hypothetical-protein-[Ruminococcus-bicirculans] |
| WP_216553765.1 | Type4.PglZ.containing.BREX | <-LexA-protease<-CHTH<-HTH+REase<-wHTH+WYL+WCX<-DEDD_Tnp_IS110+Transposase_20<-Cysteine-Desulfurase<-ZnR<-SIGMA+OB-Fold+AAA-ATPase-BrxL<-iSTAND+PglZ<-DUF6079*<-PAPS-reduct+a+b-domain+Fer4_9+DUF4007<-Phage_integrase+E3_UFM1_ligase<-HTH_26<-Lysozyme+NLPC<-ZnR<-CHTH | 1394 | Bacteria>Terrabacteria_group>Bacillota>Clostridia>Eubacteriales>Oscillospiraceae | GCF_018917825.1 | Ruminococcus_sp._MSJ-25 | KQH97_RS08315 | 272186..276370 | 2841536 | hypothetical-protein-[Ruminococcus-sp.-MSJ-25] |
| WP_216566930.1 | Type4.PglZ.containing.BREX | TraM->TrwI->?->VirB4-FtsK->?->Lysozyme+NLPC->HTH_26->Phage_integrase+E3_UFM1_ligase->PAPS-reduct+a+b-domain+Fer4_9+DUF4007->DUF6079*->iSTAND+PglZ->SIGMA+OB-Fold+AAA-ATPase-BrxL->ZnR->Cysteine-Desulfurase->wHTH+WYL+WCX->LexA-protease->?->?-><-HTH_3 | 1394 | Bacteria>Terrabacteria_group>Bacillota>Clostridia>Eubacteriales>Oscillospiraceae | GCF_018918045.1 | Pseudoflavonifractor_sp._MSJ-30 | KQI10_RS08270 | 155136..159320 | 2841525 | hypothetical-protein-[Pseudoflavonifractor-sp.-MSJ-30] |
| WP_227612027.1 | Type4.PglZ.containing.BREX | HTH_36->STYKinase-like->?->DOC+HTH->?->PAPS-reduct+a+b-domain+Fer4_9+DUF4007->DUF6079*->iSTAND+PglZ->SIGMA+OB-Fold+AAA-ATPase-BrxL->Cysteine-Desulfurase->?->PIN-> | 1398 | Bacteria>Terrabacteria_group>Bacillota>Clostridia>Eubacteriales>Oscillospiraceae | GCF_020686905.1 | Faecalibacterium_sp._CLA-AA-H283 | LKD27_RS12610 | 9890..14086 | 2881278 | hypothetical-protein-[Faecalibacterium-sp.-CLA-AA-H283] |
| WP_235396409.1 | Type4.PglZ.containing.BREX | HTH_3->?-><-?<-TERNS3+HEPN<-?<-?<-ZnR<-iSTAND+PglZ<-DUF6079*<-PAPS-reduct+a+b-domain+Fer4_9+DUF4007<-SeqA+Phage_int+SeqA+Phage_int+E3_UFM1_ligase<-HTH_26<-HTH_36<-VirD4-TraG+VirD4-FtsK<-ZnR+TOPRIM | 1401 | Bacteria>Terrabacteria_group>Bacillota>Clostridia>Eubacteriales>Oscillospiraceae | GCF_021531955.1 | Pseudoflavonifractor_phocaeensis | I6E44_RS02825 | 56517..60722 | 1870988 | hypothetical-protein-[Pseudoflavonifractor-phocaeensis] |
| WP_249377195.1 | Type4.PglZ.containing.BREX | <-SIGMA-HTH<-?<-?<-DndB\|\|XIS-HTH->HARE-HTH+E3_UFM1_ligase->PAPS-reduct+a+b-domain+Fer4_9+DUF4007->DUF6079*->iSTAND+PglZ->SIGMA+OB-Fold+AAA-ATPase-BrxL->Cysteine-Desulfurase->TRD+TRD->Phage_int_SAM_4+Phage_integrase->TRD+TRD->TRD->TIR->REase+SF1-SF2-Helicase-> | 1398 | Bacteria>Terrabacteria_group>Bacillota>Clostridia>Eubacteriales>Oscillospiraceae | GCF_023369675.1 | Ruminococcus_bromii | E2N93_RS09935 | 123130..127326 | 40518 | DUF6079-family-protein-[Ruminococcus-bromii] |
| WP_268058969.1 | Type4.PglZ.containing.BREX | TraJ-RHH->?-><-?<-TetR-HTH\|\|?-><-Cysteine-Desulfurase<-ZnR<-SIGMA+OB-Fold+AAA-ATPase-BrxL<-iSTAND+PglZ<-DUF6079*<-PAPS-reduct+a+b-domain+Fer4_9+DUF4007<-SeqA+Phage_int+SeqA+E3_UFM1_ligase<-XIS-HTH<-PadR-HTH\|\|?->TetR-HTH->?-><-ABhydrolase\|\|TetR-HTH-> | 1414 | Bacteria>Terrabacteria_group>Bacillota>Clostridia>Eubacteriales>Oscillospiraceae | GCF_026672255.1 | Caproiciproducens_galactitolivorans | OUY18_RS11685 | 63176..67420 | 642589 | hypothetical-protein-[Caproiciproducens-galactitolivorans] |
| WP_092639168.1 | Type4.PglZ.containing.BREX | Resolvase+Recombinase+ZnR->Recombinase->Resolvase+Recombinase+ZnR-><-?<-ZnR+Fer2_BFD<-Cysteine-Desulfurase<-SIGMA+OB-Fold+AAA-ATPase-BrxL<-iSTAND+PglZ<-DUF6079*<-PAPS-reduct+a+b-domain+Fer4_9+DUF4007<-SeqA+Phage_int+SeqA+Phage_int+E3_UFM1_ligase<-?\|\|HTH_31->MPTase->SIGMA-HTH->?->?->SF1-SF2-Helicase-p-> | 1403 | Bacteria>Terrabacteria_group>Bacillota>Clostridia>Eubacteriales>Oscillospiraceae | GCF_900103835.1 | Acetanaerobacterium_elongatum | BLR50_RS04950 | 69653..73864 | 258515 | hypothetical-protein-[Acetanaerobacterium-elongatum] |
| WP_072949693.1 | Type4.PglZ.containing.BREX | SeqA+Phage_int+SeqA+E3_UFM1_ligase->PAPS-reduct+a+b-domain+Fer4_9+DUF4007->DUF6079*->iSTAND+PglZ->SIGMA+OB-Fold+AAA-ATPase-BrxL->Cysteine-Desulfurase->UDG-conflict->SF1-SF2-Helicase+TOPC->?->HTH+HTH+SF1-SF2-Helicase-> | 1390 | Bacteria>Terrabacteria_group>Bacillota>Clostridia>Eubacteriales>Oscillospiraceae | GCF_900108555.1 | Ruminococcus_flavefaciens | BLW60_RS12070 | 5201..9373 | 1265 | hypothetical-protein-[Ruminococcus-flavefaciens] |
| WP_106764690.1 | Type4.PglZ.containing.BREX | Lysozyme+NLPC-><-HTH_3\|\|?->HTH_26->?->PAPS-reduct+a+b-domain+Fer4_9+DUF4007->DUF6079*->iSTAND+PglZ->SIGMA+OB-Fold+AAA-ATPase-BrxL->ZnR->Cysteine-Desulfurase->wHTH+WYL+WCX->LexA-protease-> | 1404 | Bacteria>Terrabacteria_group>Bacillota>Clostridia>Eubacteriales>Oscillospiraceae | GCF_900291955.1 | Pseudoruminococcus_massiliensis | C9914_RS08220 | 11258..15472 | 2086583 | hypothetical-protein-[Pseudoruminococcus-massiliensis] |
| WP_117927146.1 | Type4.PglZ.containing.BREX | ParB->ParB->?->?->?->?->Phage_integrase+wHTH->PAPS-reduct+a+b-domain+Fer4_9+DUF4007->DUF6079*->iSTAND+PglZ->SIGMA+OB-Fold+AAA-ATPase-BrxL->ZnR->Cysteine-Desulfurase->ZnR-><-RADICAL-SAM<-SF1-SF2-Helicase<-LPD32+LPD36+N6_Mtase+SF1-SF2-Helicase | 1405 | Bacteria>Terrabacteria_group>Bacillota>Clostridia>Eubacteriales>Oscillospiraceae | GCF_946062445.1 | unclassified_Ruminococcus | DXA02_04830 | 56881..61098 | 2608920 | MULTISPECIES:-hypothetical-protein-[Ruminococcus] |
| WP_162208121.1 | Type4.PglZ.containing.BREX | DUF6079-p*->?->?->iSTAND+PglZ->PglZ->SIGMA+OB-Fold+AAA-ATPase-BrxL->SIGMA+OB-Fold+AAA-ATPase-BrxL->ZnR->Cysteine-Desulfurase->wHTH+WYL+WCX-> | 785 | Bacteria>Terrabacteria_group>Bacillota>Clostridia>Eubacteriales>Oscillospiraceae | - | Ruminococcus_champanellensis | JCM17042_RS14580 | 1..2359 | 1161942 | hypothetical-protein,-partial-[Ruminococcus-champanellensis] |
| KLU60083.1 | Type4.PglZ.containing.BREX | ABC-ATPase+SbcC->ABC-ATPase+SbcC->GNTR-HTH->Thioredoxin-><-ZnR<-SIGMA+OB-Fold+AAA-ATPase-BrxL<-iSTAND+PglZ<-DUF6079*<-PAPS-reduct+a+b-domain+Fer4_9+DUF4007<-RAMA | 1381 | Bacteria>Terrabacteria_group>Bacillota>Clostridia>Eubacteriales>Peptococcaceae | GCA_001029295.1 | Peptococcaceae_bacterium_CEB3 | CEB3_c35140 | 140776..144921 | 999898 | hypothetical-protein-CEB3_c35140-[Peptococcaceae-bacterium-CEB3] |
| MCI9044303.1 | Type4.PglZ.containing.BREX | ACET->ABhydrolase->?->TPR+TPR->SeqA+Phage_int+SeqA+E3_UFM1_ligase->PAPS-reduct+a+b-domain+Fer4_9+DUF4007->GspA-ATPase->DUF6079*->iSTAND+PglZ->SIGMA+OB-Fold+AAA-ATPase-BrxL->ZnR->Cysteine-Desulfurase->XIS-HTH->Phage_int_SAM_5+Phage_integrase-> | 1114 | Bacteria>Terrabacteria_group>Bacillota>Clostridia>Eubacteriales>Peptococcaceae | GCA_022797295.1 | Peptococcaceae_bacterium | HFJ96_02660 | 10678..14022 | 2052179 | hypothetical-protein-[Peptococcaceae-bacterium] |
| SHE91973.1 | Type4.PglZ.containing.BREX | ABC-ATPase+SbcC-><-TOPRIM<-OLD-ABC<-SIGMA+OB-Fold+AAA-ATPase-BrxL<-iSTAND+PglZ<-DUF6079*<-DUF4007<-DUF4007<-PAPS-reduct+a+b-domain+Fer4_9+DUF4007<-LuxR-HTH | 1398 | Bacteria>Terrabacteria_group>Bacillota>Clostridia>Eubacteriales>Peptococcaceae | GCA_900129285.1 | Desulfofundulus_australicus_DSM_11792 | SAMN02745218_01069 | 50271..54467 | 1121425 | hypothetical-protein-SAMN02745218_01069-[Desulfofundulus-australicus-DSM-11792] |
| WP_157832982.1 | Type4.PglZ.containing.BREX | ABC-ATPase+SbcC-><-OLD-ABC<-SIGMA+OB-Fold+AAA-ATPase-BrxL<-iSTAND+PglZ<-DUF6079*\|\|?-><-DUF4007<-PAPS-reduct+a+b-domain+Fer4_9+DUF4007<-LuxR-HTH<-MORC | 1121 | Bacteria>Terrabacteria_group>Bacillota>Clostridia>Eubacteriales>Peptococcaceae | GCF_000686645.1 | Desulfofundulus_thermocisternus | BR83_RS0107745 | 50508..53873 | 42471 | hypothetical-protein-[Desulfofundulus-thermocisternus] |
| WP_240986375.1 | Type4.PglZ.containing.BREX | XIS-HTH->HARE-HTH+E3_UFM1_ligase->PAPS-reduct+a+b-domain+Fer4_9+DUF4007->DUF6079*->iSTAND+PglZ->SIGMA+OB-Fold+AAA-ATPase-BrxL->Cysteine-Desulfurase->?->RelE-ParE-><-?\|\|DOC+HTH->?->HEPN+PAS+DUF4011+SF1-SF2-Helicase+REase-> | 1400 | Bacteria>Terrabacteria_group>Bacillota>Clostridia>Eubacteriales>Peptococcaceae | GCF_900016645.1 | Acididesulfobacillus_acetoxydans | OCJ20_RS00105 | 17835..22037 | 1561005 | hypothetical-protein-[Acididesulfobacillus-acetoxydans] |
| WP_131821581.1 | Type4.PglZ.containing.BREX | ABC-ATPase+SbcC-><-TOPRIM<-OLD-ABC<-SIGMA+OB-Fold+AAA-ATPase-BrxL<-iSTAND+PglZ<-DUF6079*\|\|?-><-DUF4007<-DUF4007<-PAPS-reduct+a+b-domain+Fer4_9+DUF4007<-LuxR-HTH | 1132 | Bacteria>Terrabacteria_group>Bacillota>Clostridia>Eubacteriales>Peptococcaceae | GCF_900129285.1 | Desulfofundulus_australicus | BUB44_RS05275 | 50271..53669 | 1566 | hypothetical-protein-[Desulfofundulus-australicus] |
| MZQ76088.1 | Type4.PglZ.containing.BREX | <-ZnR\|\|?-><-?<-NUDIX+DUF429<-?<-ZnR<-SIGMA+OB-Fold+AAA-ATPase-BrxL<-iSTAND+PglZ<-DUF6079*<-PAPS-reduct+a+b-domain+Fer4_9+DUF4007<-?<-?<-GGDEF<-?<-?<-?<-Aminotran_4 | 1390 | Bacteria>Terrabacteria_group>Bacillota>Clostridia>Eubacteriales>Peptostreptococcaceae | GCA_009881055.1 | Peptoclostridium_sp. | GT589_08075 | 119308..123480 | 1904860 | hypothetical-protein-[Peptoclostridium-sp.] |
| MBK5260962.1 | Type4.PglZ.containing.BREX | <-SIGMA+OB-Fold+AAA-ATPase-BrxL<-iSTAND+PglZ<-DUF6079*<-PAPS-reduct+a+b-domain+Fer4_9+DUF4007<-SIGMA-HTH+E3_UFM1_ligase<-SbcC+SbcC | 1393 | Bacteria>Terrabacteria_group>Bacillota>Clostridia>Eubacteriales>Peptostreptococcaceae | GCA_016650135.1 | Peptostreptococcaceae_bacterium | JJE17_00125 | 25769..29950 | 1904861 | hypothetical-protein-[Peptostreptococcaceae-bacterium] |
| NLV22349.1 | Type4.PglZ.containing.BREX | <-Cysteine-Desulfurase<-ZnR<-SIGMA+OB-Fold+AAA-ATPase-BrxL<-PglZ<-DUF6079*<-PAPS-reduct+a+b-domain+Fer4_9+DUF4007<-RAMA | 1388 | Bacteria>Terrabacteria_group>Bacillota>Clostridia>Eubacteriales>Syntrophomonadaceae | GCA_012729495.1 | Syntrophomonadaceae_bacterium | GXY49_10235 | 6703..10869 | 2093811 | hypothetical-protein-[Syntrophomonadaceae-bacterium] |
| HHV76575.1 | Type4.PglZ.containing.BREX | Glycos_transf-><-?<-?<-DDE_Tnp<-SIGMA+OB-Fold+AAA-ATPase-BrxL<-iSTAND+PglZ<-DUF6079*<-DUF4007<-PAPS-reduct+a+b-domain+Fer4_9+DUF4007<-?<-?<-?<-wHTH+WYL+WCX<-PIN<-?<-ARG | 1389 | Bacteria>Terrabacteria_group>Bacillota>Clostridia>Eubacteriales>Syntrophomonadaceae | GCA_012840045.1 | Syntrophothermus_lipocalidus | GXX39_04275 | 25444..29613 | 86170 | hypothetical-protein-[Syntrophothermus-lipocalidus] |
| HCG68748.1 | Type4.PglZ.containing.BREX | <-HIN-HTH<-Calcineurin-SbcD<-?<-?<-wHTH+WYL+WCX<-Cysteine-Desulfurase<-ZnR<-SIGMA+OB-Fold+AAA-ATPase-BrxL<-iSTAND+PglZ<-DUF6079*<-PAPS-reduct+a+b-domain+Fer4_9+DUF4007<-SeqA+E3_UFM1_ligase<-?<-?<-PDZ+Peptidase<-PTSEIIA1<-?<-ABC-ATPase+SbcC<-DNA-glycosylase | 1399 | Bacteria>Terrabacteria_group>Bacillota>Clostridia>Eubacteriales>unclassified_Eubacteriales | GCA_003523225.1 | Clostridiales_bacterium | DEW22_08745 | 12492..16691 | 1898207 | hypothetical-protein-[Clostridiales-bacterium] |
| HCS73735.1 | Type4.PglZ.containing.BREX | DUF4007->DUF6079->SIGMA-HTH->HARE-HTH->?->PAPS-reduct+a+b-domain+Fer4_9->DUF4007->BrxB-BrxF-iSTAND->DUF6079*->?->PglZ->SIGMA+OB-Fold+AAA-ATPase-BrxL->Cysteine-Desulfurase->?->URI-DNase->DndB-> | 1032 | Bacteria>Terrabacteria_group>Bacillota>Clostridia>Eubacteriales>unclassified_Eubacteriales | GCA_003524825.1 | Clostridiales_bacterium | DIW17_07660 | 11456..14551 | 1898207 | hypothetical-protein-[Clostridiales-bacterium] |
| NLN48282.1 | Type4.PglZ.containing.BREX | HKD-DNase->?->HTH_Tnp_1-><-SIGMA+OB-Fold+AAA-ATPase-BrxL<-iSTAND+PglZ<-DUF6079*<-PAPS-reduct+a+b-domain+Fer4_9+DUF4007<-HNH+SeqA<-HTH | 1138 | Bacteria>Terrabacteria_group>Bacillota>Clostridia>Eubacteriales>unclassified_Eubacteriales | GCA_012522255.1 | Clostridiales_bacterium | GX154_04185 | 7445..10861 | 1898207 | hypothetical-protein-[Clostridiales-bacterium] |
| NLW00223.1 | Type4.PglZ.containing.BREX | <-HTH_3\|\|?->PAPS-reduct+a+b-domain+Fer4_9+DUF4007->DUF6079*->iSTAND+PglZ->SIGMA+OB-Fold+AAA-ATPase-BrxL-> | 1377 | Bacteria>Terrabacteria_group>Bacillota>Clostridia>Eubacteriales>unclassified_Eubacteriales | GCA_012727765.1 | Clostridiales_bacterium | GX041_03380 | 3853..7986 | 1898207 | hypothetical-protein-[Clostridiales-bacterium] |
| HHY53749.1 | Type4.PglZ.containing.BREX | PAPS-reduct+a+b-domain+Fer4_9+DUF4007->DUF4007->DUF6079*->iSTAND+PglZ-> | 1416 | Bacteria>Terrabacteria_group>Bacillota>Clostridia>Eubacteriales>unclassified_Eubacteriales | GCA_012842695.1 | Clostridiales_bacterium | GX499_11045 | 1777..6027 | 1898207 | hypothetical-protein-[Clostridiales-bacterium] |
| MBE5813905.1 | Type4.PglZ.containing.BREX | <-Cap4_nuclease<-Cysteine-Desulfurase<-ZnR<-SIGMA+OB-Fold+AAA-ATPase-BrxL<-iSTAND+PglZ<-DUF6079*<-PAPS-reduct+a+b-domain+Fer4_9+DUF4007<-HNH<-SIGMA-HTH<-HTH_26<-?<-?<-HTH_3\|\|LRR_5+LRR_5+LRR_5+LRR_5-> | 1394 | Bacteria>Terrabacteria_group>Bacillota>Clostridia>Eubacteriales>unclassified_Eubacteriales | GCA_015056315.1 | Clostridiales_bacterium | E7320_01680 | 119434..123618 | 1898207 | hypothetical-protein-[Clostridiales-bacterium] |
| MBS5594067.1 | Type4.PglZ.containing.BREX | <-VbhA+DOC+HTH<-TERNS3+HEPN<-?<-Cysteine-Desulfurase<-SIGMA+OB-Fold+AAA-ATPase-BrxL<-iSTAND+PglZ<-DUF6079*<-PAPS-reduct+a+b-domain+Fer4_9+DUF4007<-SeqA+Phage_int+SeqA+Phage_int+E3_UFM1_ligase<-HTH_26<-?<-HNH\|\|SIGMA-HTH->?->?->Recombinase-> | 1401 | Bacteria>Terrabacteria_group>Bacillota>Clostridia>Eubacteriales>unclassified_Eubacteriales | GCA_018369595.1 | Clostridiales_bacterium | KHX08_01355 | 31404..35609 | 1898207 | hypothetical-protein-[Clostridiales-bacterium] |
| MBS7306476.1 | Type4.PglZ.containing.BREX | <-TERNS3+HEPN<-?<-Cysteine-Desulfurase<-?<-ZnR<-SIGMA+OB-Fold+AAA-ATPase-BrxL<-iSTAND+PglZ<-DUF6079*<-PAPS-reduct+a+b-domain+Fer4_9+DUF4007<-SeqA+Phage_int+SeqA+Phage_int+E3_UFM1_ligase<-?<-XIS-HTH<-TraI-VirD2-RCRE<-TraJ-RHH | 1400 | Bacteria>Terrabacteria_group>Bacillota>Clostridia>Eubacteriales>unclassified_Eubacteriales | GCA_018384315.1 | Eubacteriales_bacterium | KIG43_00140 | 25153..29355 | 2831996 | hypothetical-protein-[Eubacteriales-bacterium] |
| MCD7774058.1 | Type4.PglZ.containing.BREX | <-Cysteine-Desulfurase<-SIGMA+OB-Fold+AAA-ATPase-BrxL<-ZnR<-iSTAND+PglZ<-DUF6079*<-PAPS-reduct+a+b-domain+Fer4_9+DUF4007<-HNH | 1402 | Bacteria>Terrabacteria_group>Bacillota>Clostridia>Eubacteriales>unclassified_Eubacteriales | GCA_021199775.1 | Clostridiales_bacterium | LUH40_00545 | 5553..9761 | 1898207 | DUF6079-family-protein-[Clostridiales-bacterium] |

| Protein-ID | System | Gene-Neighbourhoods | Length | Taxonomy | Assembly-ID | Strain | Locus-Tag | Genomic-coordinates | TaxID | Fasta-defline |
|---|---|---|---|---|---|---|---|---|---|---|
| MCD7827861.1 | Type4.PglZ.containing.BREX | <-Cysteine-Desulfurase<-SIGMA+OB-Fold+AAA-ATPase-BrxL<-ZnR<-iSTAND+PglZ<-DUF6079*<-PAPS-reduct+a+b-domain+Fer4_9+DUF4007 | 1402 | Bacteria>Terrabacteria_group>Bacillota>Clostridia>Eubacteriales>unclassified_Eubacteriales | GCA_021200555.1 | Clostridiales_bacterium | LUG85_04930 | 5416..9624 | 1898207 | DUF6079-family-protein-[Clostridiales-bacterium] |
| MCD7828248.1 | Type4.PglZ.containing.BREX | SeqA+Phage_int+SeqA+Phage_int+E3_UFM1_ligase->PAPS-reduct+a+b-domain+Fer4_9+DUF4007->DUF6079*->iSTAND+PglZ->SIGMA+OB-Fold+AAA-ATPase-BrxL->ZnR->Cysteine-Desulfurase->LRR_5->LRR_5+LRR_5+LRR_5+LRR_5-><-?\|\|?->REC+OmpR-HTH-> | 1402 | Bacteria>Terrabacteria_group>Bacillota>Clostridia>Eubacteriales>unclassified_Eubacteriales | GCA_021200555.1 | Clostridiales_bacterium | LUG85_06920 | 6745..10953 | 1898207 | hypothetical-protein-[Clostridiales-bacterium] |
| MCD8238627.1 | Type4.PglZ.containing.BREX | RNA_METHYLASE->?->REase-><-?<-ABC-ATPase<-GntR-HTH\|\|?->HNH->PAPS-reduct+a+b-domain+Fer4_9+DUF4007->DUF6079-p*->?->iSTAND+PglZ->ZnR->SIGMA+OB-Fold+AAA-ATPase-BrxL->Cysteine-Desulfurase-><-HTH+HTH+SF1-SF2-Helicase<-wHTH+WYL+WCX | 957 | Bacteria>Terrabacteria_group>Bacillota>Clostridia>Eubacteriales>unclassified_Eubacteriales | GCA_021204175.1 | Clostridiales_bacterium | LUC92_04725 | 25081..27954 | 1898207 | DUF6079-family-protein-[Clostridiales-bacterium] |
| MCI8471974.1 | Type4.PglZ.containing.BREX | ADH_N+ADH_zinc_N+METHYLASE->?->HTH_3->?->?->XIS-HTH->HARE-HTH+E3_UFM1_ligase->PAPS-reduct+a+b-domain+Fer4_9+DUF4007->DUF6079*->iSTAND+PglZ->SIGMA+OB-Fold+AAA-ATPase-BrxL->Cysteine-Desulfurase-><-?<-GDResolvase+Recombinase+ZnR\|\|?->tRNA-><-?<-HISKIN<-REC+OmpR-HTH | 1399 | Bacteria>Terrabacteria_group>Bacillota>Clostridia>Eubacteriales>unclassified_Eubacteriales | GCA_022792475.1 | Clostridiales_bacterium | HFE65_02590 | 168336..172535 | 1898207 | hypothetical-protein-[Clostridiales-bacterium] |
| WP_134114886.1 | Type4.PglZ.containing.BREX | <-SF1-SF2-Helicase+REase<-SF1-SF2-Helicase+REase<-?<-Cysteine-Desulfurase<-SIGMA+OB-Fold+AAA-ATPase-BrxL<-PglZ+Phosphodiest<-DUF6079<-BrxB-BrxF-iSTAND<-DUF4007<-DUF6079*<-DUF4007<-PAPS-reduct+a+b-domain+Fer4_9<-?<-SIGMA-HTH+HARE-HTH<-SIGMA-HTH+HARE-HTH<-?<-Peptidase_S78 | 1189 | Bacteria>Terrabacteria_group>Bacillota>Clostridia>Halanaerobiales>Halobacteroidaceae | GCF_004365685.1 | Orenia_marismortui | C7959_RS02420 | 119047..122616 | 46469 | hypothetical-protein-[Orenia-marismortui] |
| WP_089748436.1 | Type4.PglZ.containing.BREX | <-Cysteine-Desulfurase<-SIGMA+OB-Fold+AAA-ATPase-BrxL<-PglZ+Phosphodiest<-DUF6079<-?<-DUF4007<-DUF6079*<-DUF4007<-PAPS-reduct+a+b-domain+Fer4_9<-?<-?<-SIGMA-HTH+HARE-HTH<-SIGMA-HTH+HARE-HTH<-?<-SSB<-HEPN | 1192 | Bacteria>Terrabacteria_group>Bacillota>Clostridia>Halanaerobiales>Halobacteroidaceae | GCF_900101375.1 | unclassified_Candidatus_Frackibacter | BLT14_RS04865 | 91742..95320 | 2648818 | MULTISPECIES:-hypothetical-protein-[unclassified-Candidatus-Frackibacter] |
| RJX19215.1 | Type4.PglZ.containing.BREX | <-SIR2<-SIGMA+OB-Fold+AAA-ATPase-BrxL<-iSTAND+PglZ<-DUF6079*<-DUF4007<-PAPS-reduct+a+b-domain+Fer4_9+DUF4007<-?<-?\|\|HicA_toxin->HicB_lk_antitox-> | 1377 | Bacteria>Terrabacteria_group>Bacillota>Clostridia>Thermoanaerobacterales>Thermoanaerobacteraceae | GCA_003604985.1 | Ammonifex_sp. | C4570_05550 | 5055..9188 | 1872493 | MAG:-ATP-binding-protein-[Ammonifex-sp.] |
| WP_028051764.1 | Type4.PglZ.containing.BREX | PAPS-reduct+a+b-domain+Fer4_9+DUF4007->DUF4007->DUF6079*->iSTAND+PglZ->SIGMA+OB-Fold+AAA-ATPase-BrxL->DUF4236-><-?\|\|CHTH->PolB->?->GNTR-HTH+UTRA-> | 1372 | Bacteria>Terrabacteria_group>Bacillota>Clostridia>Thermoanaerobacterales>Thermoanaerobacteraceae | GCF_000427565.1 | Carboxydothermus_ferrireducens | CARFE_RS0103565 | 10424..14542 | 54265 | hypothetical-protein-[Carboxydothermus-ferrireducens] |
| WP_062284551.1 | Type4.PglZ.containing.BREX | <-DrsE<-HTH+HTH+CHTH<-rve\|\|?-><-SIGMA+OB-Fold+AAA-ATPase-BrxL<-iSTAND+PglZ<-DUF6079*<-DUF4007<-?<-PAPS-reduct+a+b-domain+Fer4_9+DUF4007<-SIGMA-HTH+SIGMA-HTH+SIGMA-HTH<-?\|\|REase+HTH-><-tRNA<-tRNA<-tRNA<-tRNA<-tRNA<-?<-RNasePH | 1373 | Bacteria>Terrabacteria_group>Bacillota>Clostridia>Thermoanaerobacterales>Thermoanaerobacteraceae | GCF_001594015.1 | Moorella_mulderi | MOMUL_RS09875 | 86448..90569 | 202604 | ATP-binding-protein-[Moorella-mulderi] |
| WP_071521068.1 | Type4.PglZ.containing.BREX | <-SIGMA+OB-Fold+AAA-ATPase-BrxL<-iSTAND+PglZ<-DUF6079*<-DUF4007<-?\|\|?-><-PAPS-reduct+a+b-domain+Fer4_9+DUF4007<-SIGMA-HTH+SIGMA-HTH\|\|?-><-tRNA<-tRNA<-tRNA<-tRNA<-tRNA<-tRNA<-?<-RNasePH | 1376 | Bacteria>Terrabacteria_group>Bacillota>Clostridia>Thermoanaerobacterales>Thermoanaerobacteraceae | GCF_001874065.1 | Moorella_thermoacetica | MOOR_RS08745 | 73773..77903 | 1525 | ATP-binding-protein-[Moorella-thermoacetica] |
| NCD03519.1 | Type4.PglZ.containing.BREX | <-SIGMA+OB-Fold+AAA-ATPase-BrxL<-iSTAND+PglZ<-DUF6079*<-PAPS-reduct+a+b-domain+Fer4_9+DUF4007<-SeqA+Phage_int+SeqA+Phage_int+E3_UFM1_ligase<-HTH_26\|\|?-><-?<-?<-TraJ-RHH<-ArdCN+LPD32+LPD36+N6_Mtase+SF1-SF2-Helicase | 1402 | Bacteria>Terrabacteria_group>Bacillota>Clostridia>unclassified_Clostridia | GCA_009930255.1 | Clostridia_bacterium | EOM00_11135 | 3236..7444 | 2044939 | hypothetical-protein-[Clostridia-bacterium] |
| MBD5583500.1 | Type4.PglZ.containing.BREX | <-HATPase_c<-?<-REase+35exo<-NUDIX<-RNase+DSRBD<-Cysteine-Desulfurase<-?<-SIGMA+OB-Fold+AAA-ATPase-BrxL<-iSTAND+PglZ<-DUF6079*<-ZnR<-PAPS-reduct+a+b-domain+Fer4_9+DUF4007\|\|Mob_Pre->Phage_integrase-><-tRNA<-?\|\|?-><-LRR_5+LRR_5+LRR_5+LRR_5<-SIGMA-HTH<-BirA-HTH+3H | 1169 | Bacteria>Terrabacteria_group>Bacillota>Clostridia>unclassified_Clostridia | GCA_014804995.1 | Clostridia_bacterium | HDQ88_00210 | 24831..28340 | 2044939 | hypothetical-protein-[Clostridia-bacterium] |
| MBQ4585186.1 | Type4.PglZ.containing.BREX | <-DndB<-URI-DNase<-Cysteine-Desulfurase<-?<-SIGMA+OB-Fold+AAA-ATPase-BrxL<-iSTAND+PglZ<-DUF6079*<-PAPS-reduct+a+b-domain+Fer4_9+DUF4007<-?\|\|GDResolvase+Recombinase+ZnR-><-?\|\|?->DSBH+AraC-HTH+AraC-HTH-> | 1170 | Bacteria>Terrabacteria_group>Bacillota>Clostridia>unclassified_Clostridia | GCA_017399145.1 | Clostridia_bacterium | IJA82_03155 | 24176..27688 | 2044939 | hypothetical-protein-[Clostridia-bacterium] |
| MBQ6067434.1 | Type4.PglZ.containing.BREX | CR-ATPase5+REase->?->?-><-CN_hydrolase<-XIS-HTH<-?\|\|XIS-HTH->HARE-HTH+E3_UFM1_ligase->PAPS-reduct+a+b-domain+Fer4_9+DUF4007->DUF6079*->iSTAND+PglZ->SIGMA+OB-Fold+AAA-ATPase-BrxL->Cysteine-Desulfurase->NERD+SF1-SF2-Helicase->SF1-SF2-Helicase+REase-p-> | 1397 | Bacteria>Terrabacteria_group>Bacillota>Clostridia>unclassified_Clostridia | GCA_017428085.1 | Clostridia_bacterium | IJK89_11515 | 53896..58089 | 2044939 | hypothetical-protein-[Clostridia-bacterium] |
| MBR1677452.1 | Type4.PglZ.containing.BREX | <-TraJ-RHH<-DEOR-HTH<-Cysteine-Desulfurase<-ZnR<-SIGMA+OB-Fold+AAA-ATPase-BrxL<-iSTAND+PglZ<-DUF6079*<-PAPS-reduct+a+b-domain+Fer4_9+DUF4007<-HNH<-HNH<-SIGMA-HTH<-?<-HipA_C<-Couple_hipA<-HTH_3<-PAIREDC-HTH | 1394 | Bacteria>Terrabacteria_group>Bacillota>Clostridia>unclassified_Clostridia | GCA_017476895.1 | Clostridia_bacterium | IJ706_09110 | 25756..29940 | 2044939 | hypothetical-protein-[Clostridia-bacterium] |
| MBR3817931.1 | Type4.PglZ.containing.BREX | <-MPTase<-HTH_19<-?\|\|?->REase+RNase->HEPN-><-Cysteine-Desulfurase<-SIGMA+OB-Fold+AAA-ATPase-BrxL<-iSTAND+PglZ<-DUF6079*<-PAPS-reduct+a+b-domain+Fer4_9+DUF4007<-?\|\|Classical-AAA->PIN->HEPN-> | 1389 | Bacteria>Terrabacteria_group>Bacillota>Clostridia>unclassified_Clostridia | GCA_017517665.1 | Clostridia_bacterium | IKJ41_02180 | 15710..19879 | 2044939 | hypothetical-protein-[Clostridia-bacterium] |
| MBR6632684.1 | Type4.PglZ.containing.BREX | <-PARG+MACRODOMAIN<-SLATT<-Cysteine-Desulfurase<-SIGMA+OB-Fold+AAA-ATPase-BrxL<-iSTAND+PglZ<-DUF6079*<-PAPS-reduct+a+b-domain+Fer4_9+DUF4007<-SeqA+Phage_int+SeqA+E3_UFM1_ligase<-HTH_26<-XIS-HTH+XIS-HTH<-XIS-HTH<-HNH | 1382 | Bacteria>Terrabacteria_group>Bacillota>Clostridia>unclassified_Clostridia | GCA_017520825.1 | Clostridia_bacterium | IKL05_00090 | 8302..12450 | 2044939 | hypothetical-protein-[Clostridia-bacterium] |
| MBP3656734.1 | Type4.PglZ.containing.BREX | <-XIS-HTH\|\|HNH->?->?->HTH_26->OLD-ABC->TOPRIM->?->PAPS-reduct+a+b-domain+Fer4_9+DUF4007->DUF6079*->iSTAND+PglZ->SIGMA+OB-Fold+AAA-ATPase-BrxL->Cysteine-Desulfurase-> | 1386 | Bacteria>Terrabacteria_group>Bacillota>Clostridia>unclassified_Clostridia | GCA_017938795.1 | Clostridia_bacterium | J6K32_08570 | 28511..32671 | 2044939 | hypothetical-protein-[Clostridia-bacterium] |
| MCD8003208.1 | Type4.PglZ.containing.BREX | <-HIN-HTH<-?<-?\|\|DUF772+DDE_Tnp_1->?-><-SIGMA+OB-Fold+AAA-ATPase-BrxL<-iSTAND+PglZ<-DUF6079*<-PAPS-reduct+a+b-domain+Fer4_9+DUF4007<-HNH<-?\|\|?->ACET-><-CMP-hydrolase<-CMP-hydrolase<-?\|\|PDZ+RADICAL-SAM-> | 1406 | Bacteria>Terrabacteria_group>Bacillota>Clostridia>unclassified_Clostridia | GCA_021202035.1 | Clostridia_bacterium | LUG88_04775 | 11357..15577 | 2044939 | hypothetical-protein-[Clostridia-bacterium] |
| MCD8049835.1 | Type4.PglZ.containing.BREX | <-Cysteine-Desulfurase<-ZnR<-SIGMA+OB-Fold+AAA-ATPase-BrxL<-iSTAND+PglZ<-DUF6079*<-PAPS-reduct+a+b-domain+Fer4_9+DUF4007<-HARE-HTH | 1244 | Bacteria>Terrabacteria_group>Bacillota>Clostridia>unclassified_Clostridia | GCA_021202495.1 | Clostridia_bacterium | LUG52_09665 | 6156..9890 | 2044939 | hypothetical-protein-[Clostridia-bacterium] |
| MCD8287174.1 | Type4.PglZ.containing.BREX | <-SIGMA+OB-Fold+AAA-ATPase-BrxL<-iSTAND+PglZ<-DUF6079-p<-DUF6079-p*<-PAPS-reduct+a+b-domain+Fer4_9+DUF4007 | 556 | Bacteria>Terrabacteria_group>Bacillota>Clostridia>unclassified_Clostridia | GCA_021204585.1 | Clostridia_bacterium | LUD50_08130 | 5791..7461 | 2044939 | hypothetical-protein-[Clostridia-bacterium] |
| MCI8555500.1 | Type4.PglZ.containing.BREX | <-NTP_transf_3+Hexapep+Hexapep\|\|tRNA->?-><-YTCD-HTH\|\|DSBH->?-><-?\|\|DOC+HTH->?->PAPS-reduct+a+b-domain+Fer4_9+DUF4007->DUF6079*->iSTAND+PglZ->SIGMA+OB-Fold+AAA-ATPase-BrxL->ZnR-><-Cysteine-Desulfurase\|\|rve->?->AAA-ATPase-> | 1396 | Bacteria>Terrabacteria_group>Bacillota>Clostridia>unclassified_Clostridia | GCA_022793335.1 | Clostridia_bacterium | HFI85_02890 | 53624..57814 | 2044939 | hypothetical-protein-[Clostridia-bacterium] |
| MCR5186519.1 | Type4.PglZ.containing.BREX | <-CHTH<-Cysteine-Desulfurase<-ZnR<-SIGMA+OB-Fold+AAA-ATPase-BrxL<-iSTAND+PglZ<-DUF6079*<-PAPS-reduct+a+b-domain+Fer4_9+DUF4007<-?<-XIS-HTH | 1390 | Bacteria>Terrabacteria_group>Bacillota>Clostridia>unclassified_Clostridia | GCA_024697005.1 | Clostridia_bacterium | K6D97_05340 | 5557..9729 | 2044939 | hypothetical-protein-[Clostridia-bacterium] |
| GHU63478.1 | Type4.PglZ.containing.BREX | <-Schlafen+HATPase+wHTH<-ZnR<-SIGMA+OB-Fold+AAA-ATPase-BrxL<-iSTAND+PglZ<-DUF6079*<-PAPS-reduct+a+b-domain+Fer4_9+DUF4007<-HARE-HTH+E3_UFM1_ligase<-?<-AAA-ATPase<-rve_3<-?<-STYKIN<-sGTP | 1385 | Bacteria>Terrabacteria_group>Bacillota>Clostridia>unclassified_Clostridia | GCA_025997195.1 | Clostridia_bacterium | AGMMS49983_07010 | 5945..10102 | 2044939 | hypothetical-protein-AGMMS49983_07010-[Clostridia-bacterium] |
| WP_216880784.1 | Type4.PglZ.containing.BREX | <-MPTase<-LexA-protease<-HTH+REase<-wHTH+WYL+WCX<-Cysteine-Desulfurase<-ZnR<-SIGMA+OB-Fold+AAA-ATPase-BrxL<-iSTAND+PglZ<-DUF6079-p<-DUF6079*<-PAPS-reduct+a+b-domain+Fer4_9+DUF4007<-Phage_integrase+E3_UFM1_ligase<-HTH_26<-Lysozyme+NLPC<-?<-?<-VirB4-FtsK<-?<-TrwI | 1065 | Bacteria>Terrabacteria_group>Bacillota>Erysipelotrichia>Erysipelotrichales>Coprobacillaceae | GCF_019041895.1 | [Clostridium]_innocuum | KTG08_RS07105 | 40912..44109 | 1522 | hypothetical-protein-[[Clostridium] |
| WP_273401279.1 | Type4.PglZ.containing.BREX | ABhydrolase->?->HTH_26->SeqA+Phage_int+SeqA+Phage_int+E3_UFM1_ligase->PAPS-reduct+a+b-domain+Fer4_9+DUF4007->DUF6079*->iSTAND+PglZ->SIGMA+OB-Fold+AAA-ATPase-BrxL->ZnR->Cysteine-Desulfurase-> | 1398 | Bacteria>Terrabacteria_group>Bacillota>Erysipelotrichia>Erysipelotrichales>Erysipelotrichaceae | GCF_944324785.1 | Traorella_massiliensis | PUQ26_RS07405 | 37177..41373 | 1903263 | hypothetical-protein-[Traorella-massiliensis] |
| PWL70024.1 | Type4.PglZ.containing.BREX | <-STYKIN<-?<-Cysteine-Desulfurase<-ZnR<-SIGMA+OB-Fold+AAA-ATPase-BrxL<-iSTAND+PglZ<-DUF6079*<-PAPS-reduct+a+b-domain+Fer4_9+DUF4007<-SeqA+Phage_int+SeqA+E3_UFM1_ligase<-HTH_26\|\|LexA-protease-> | 1401 | Bacteria>Terrabacteria_group>Bacillota | GCA_003149915.1 | Bacillota_bacterium | DBY23_05625 | 10514..14719 | 1879010 | MAG:-hypothetical-protein-DBY23_05625-[Bacillota-bacterium] |
| HCF89833.1 | Type4.PglZ.containing.BREX | LACI-HTH+PBPI->?->?->?->?->?->HARE-HTH+E3_UFM1_ligase->PAPS-reduct+a+b-domain+Fer4_9+DUF4007->DUF6079*->iSTAND+PglZ->ZnR->SIGMA+OB-Fold+AAA-ATPase-BrxL->SF1-SF2-Helicase-> | 1395 | Bacteria>Terrabacteria_group>Bacillota | GCA_003534975.1 | Bacillota_bacterium | DER41_07745 | 16768..20955 | 1879010 | hypothetical-protein-[Bacillota-bacterium] |
| NLI70096.1 | Type4.PglZ.containing.BREX | <-SIR2<-?\|\|SF1-SF2-Helicase-><-SIGMA+OB-Fold+AAA-ATPase-BrxL<-iSTAND+PglZ<-DUF6079*<-PAPS-reduct+a+b-domain+Fer4_9+DUF4007<-HARE-HTH+E3_UFM1_ligase | 1387 | Bacteria>Terrabacteria_group>Bacillota | GCA_012518215.1 | Bacillota_bacterium | GX364_04445 | 64498..68661 | 1879010 | hypothetical-protein-[Bacillota-bacterium] |
| NLJ56152.1 | Type4.PglZ.containing.BREX | PAPS-reduct+a+b-domain+Fer4_9+DUF4007->DUF6079*->iSTAND+PglZ->SIGMA+OB-Fold+AAA-ATPase-BrxL->?->tRNA->tRNA-><-?<-?<-tRNA\|\|Calcineurin->?->MazE_antitoxin+SpoVT_C-> | 1375 | Bacteria>Terrabacteria_group>Bacillota | GCA_012518675.1 | Bacillota_bacterium | GX334_03760 | 1385..5512 | 1879010 | hypothetical-protein-[Bacillota-bacterium] |
| HHW17414.1 | Type4.PglZ.containing.BREX | <-Phage_integrase<-?\|\|N6-N4-METHYLASE-><-?\|\|?->?->PAPS-reduct+a+b-domain+Fer4_9+DUF4007->DUF6079*->iSTAND+PglZ->SIGMA+OB-Fold+AAA-ATPase-BrxL->ZnR-> | 1381 | Bacteria>Terrabacteria_group>Bacillota | GCA_012840505.1 | Bacillota_bacterium | GXX30_00715 | 37190..41335 | 1879010 | hypothetical-protein-[Bacillota-bacterium] |
| HHY36028.1 | Type4.PglZ.containing.BREX | <-ACET<-SIGMA+OB-Fold+AAA-ATPase-BrxL<-iSTAND+PglZ<-DUF6079*<-DUF4007<-PAPS-reduct+a+b-domain+Fer4_9+DUF4007<-?<-?\|\|?->Glycos_transf-><-PIN<-MazE_antitoxin<-ABC-ATPase+SbcC | 1384 | Bacteria>Terrabacteria_group>Bacillota | GCA_012842495.1 | Bacillota_bacterium | GX510_10465 | 6005..10159 | 1879010 | hypothetical-protein-[Bacillota-bacterium] |
| NSW92344.1 | Type4.PglZ.containing.BREX | <-ZnR<-SIGMA+OB-Fold+AAA-ATPase-BrxL<-iSTAND+PglZ<-DUF6079*<-PAPS-reduct+a+b-domain+Fer4_9+DUF4007<-HNH | 1385 | Bacteria>Terrabacteria_group>Bacillota | GCA_013314855.1 | Bacillota_bacterium | HPY74_17040 | 4834..8991 | 1879010 | hypothetical-protein-[Bacillota-bacterium] |
| QUL97787.1 | Type4.PglZ.containing.BREX | <-Classical-AAA<-?<-?\|\|?->?-><-HTH_Tnp<-ZnR<-SIGMA+OB-Fold+AAA-ATPase-BrxL<-iSTAND+PglZ<-DUF6079*<-PAPS-reduct+a+b-domain+Fer4_9+DUF4007<-?\|\|?->AAA-ATPase-p-><-N6-N4-METHYLASE | 1381 | Bacteria>Terrabacteria_group>Bacillota | GCA_018208355.1 | Bacillota_bacterium | IMF26_06655 | 1371991..1376136 | 1879010 | MAG:-hypothetical-protein-IMF26_06655-[Bacillota-bacterium] |
| MBQ7883783.1 | Type4.PglZ.containing.BREX | <-DOC+HTH<-REase\|\|HTH_26->SeqA+Phage_int+SeqA+Phage_int+E3_UFM1_ligase->PAPS-reduct+a+b-domain+Fer4_9+DUF4007->DUF6079*->iSTAND+PglZ->PglZ->SIGMA+OB-Fold+AAA-ATPase-BrxL->ZnR->Cysteine-Desulfurase->?->?->Schlafen+HATPase+wHTH-> | 1401 | Bacteria>Terrabacteria_group>Bacillota>Negativicutes>Acidaminococcales>Acidaminococcaceae | GCA_017467905.1 | Phascolarctobacterium_sp. | IJ320_05450 | 9669..13874 | 2049039 | hypothetical-protein-[Phascolarctobacterium-sp.] |
| MBR6862437.1 | Type4.PglZ.containing.BREX | HTH_3->SIGMA-HTH-><-?<-?<-DOC+HTH<-?<-DndB\|\|HARE-HTH+E3_UFM1_ligase->PAPS-reduct+a+b-domain+Fer4_9+DUF4007->DUF6079*->iSTAND+PglZ->ZnR->SIGMA+OB-Fold+AAA-ATPase-BrxL->Cysteine-Desulfurase->Schlafen+HATPase+wHTH-> | 1397 | Bacteria>Terrabacteria_group>Bacillota>Negativicutes>Acidaminococcales>Acidaminococcaceae | GCA_017531245.1 | Acidaminococcaceae_bacterium | IKM73_14065 | 182681..186874 | 2049052 | hypothetical-protein-[Acidaminococcaceae-bacterium] |
| MBE8952290.1 | Type4.PglZ.containing.BREX | <-SIGMA-HTH<-ABC-ATPase+SbcC<-Peptidase-C39+ABC-ATPase+SbcC<-?<-?<-Cysteine-Desulfurase<-ZnR<-SIGMA+OB-Fold+AAA-ATPase-BrxL<-iSTAND+PglZ<-DUF6079*<-PAPS-reduct+a+b-domain+Fer4_9+DUF4007<-?\|\|?->?->?->?->?->?->ABC-ATPase+SbcC-> | 1395 | Bacteria>Terrabacteria_group>Bacillota>Negativicutes>Negativicutes_incertae_sedis>Quinella | GCA_015206805.1 | Quinella_sp._1Q7 | SR1Q7_03985 | 12719..16906 | 1958789 | hypothetical-protein-[Quinella-sp.-1Q7] |
| MBE8949589.1 | Type4.PglZ.containing.BREX | REase+N6_METHYLASE->TRD+TRD->REase+SF1-SF2-Helicase-><-Cysteine-Desulfurase<-ZnR<-SIGMA+OB-Fold+AAA-ATPase-BrxL<-iSTAND+PglZ<-DUF6079*<-PAPS-reduct+a+b-domain+Fer4_9+DUF4007<-?\|\|CARF-><-wHTH+WYL+WCX<-wHTH+WYL+WCX<-CARF<-?<-Cas_Cas1<-CRISPR_Cas6 | 1400 | Bacteria>Terrabacteria_group>Bacillota>Negativicutes>Negativicutes_incertae_sedis>Quinella | GCA_015206825.1 | Quinella_sp._3Q1 | SR3Q1_03195 | 54730..58932 | 1958791 | hypothetical-protein-[Quinella-sp.-3Q1] |
| MBE6101263.1 | Type4.PglZ.containing.BREX | RADICAL-SAM+TRAM->?->MORC->?->RelB->PIN->?->?->PAPS-reduct+a+b-domain+Fer4_9+DUF4007->DUF6079*->iSTAND+PglZ->SIGMA+OB-Fold+AAA-ATPase-BrxL->ZnR->?->Cysteine-Desulfurase->Phd-YefM->PIN->CR-ATPase8+REase-> | 1381 | Bacteria>Terrabacteria_group>Bacillota>Negativicutes>Selenomonadales>Selenomonadaceae | GCA_015058655.1 | Selenomonas_ruminantium | E7200_04070 | 39739..43884 | 971 | hypothetical-protein-[Selenomonas-ruminantium] |
| MBQ3442988.1 | Type4.PglZ.containing.BREX | <-Peptidase_M15_4\|\|?-><-PglZ<-iSTAND+PglZ<-DUF6079-p*<-PAPS-reduct+DUF4007<-SIGMA-HTH<-AAA+HTH | 962 | Bacteria>Terrabacteria_group>Bacillota>Negativicutes>Selenomonadales>Selenomonadaceae | GCA_017406175.1 | Selenomonadaceae_bacterium | IJG33_07070 | 8183..11071 | 2652289 | hypothetical-protein-[Selenomonadaceae-bacterium] |
| MBQ3725451.1 | Type4.PglZ.containing.BREX | DSBH+AraC-HTH+AraC-HTH-><-Cysteine-Desulfurase<-ZnR<-SIGMA+OB-Fold+AAA-ATPase-BrxL<-iSTAND+PglZ<-DUF6079*<-PAPS-reduct+a+b-domain+Fer4_9+DUF4007<-?<-ABC-ATPase+SbcC | 1397 | Bacteria>Terrabacteria_group>Bacillota>Negativicutes>Selenomonadales>Selenomonadaceae | GCA_017412545.1 | Selenomonadaceae_bacterium | II902_00100 | 22345..26538 | 2652289 | hypothetical-protein-[Selenomonadaceae-bacterium] |
| MBQ4403148.1 | Type4.PglZ.containing.BREX | <-DOC+HTH<-Cysteine-Desulfurase<-ZnR<-SIGMA+OB-Fold+AAA-ATPase-BrxL<-iSTAND+PglZ<-DUF6079*<-PAPS-reduct+a+b-domain+Fer4_9+DUF4007 | 1401 | Bacteria>Terrabacteria_group>Bacillota>Negativicutes>Selenomonadales>Selenomonadaceae | GCA_017412925.1 | Selenomonadaceae_bacterium | II857_01935 | 5510..9715 | 2652289 | hypothetical-protein-[Selenomonadaceae-bacterium] |
| MBQ6296992.1 | Type4.PglZ.containing.BREX | <-PRTase\|\|?->PAPS-reduct+a+b-domain+Fer4_9+DUF4007->DUF6079*->iSTAND+PglZ->HTH_OrfB_IS605-> | 1400 | Bacteria>Terrabacteria_group>Bacillota>Negativicutes>Selenomonadales>Selenomonadaceae | GCA_017428215.1 | Selenomonadaceae_bacterium | IJK81_04780 | 39302..43504 | 2652289 | hypothetical-protein-[Selenomonadaceae-bacterium] |
| MBQ7630271.1 | Type4.PglZ.containing.BREX | <-SIGMA+OB-Fold+AAA-ATPase-BrxL<-iSTAND+PglZ<-DUF6079*<-?<-PAPS-reduct+DUF4007 | 1228 | Bacteria>Terrabacteria_group>Bacillota>Negativicutes>Selenomonadales>Selenomonadaceae | GCA_017444605.1 | Selenomonadaceae_bacterium | IJS81_08690 | 4143..7829 | 2652289 | hypothetical-protein-[Selenomonadaceae-bacterium] |
| MBR0062286.1 | Type4.PglZ.containing.BREX | PAPS-reduct+a+b-domain+Fer4_9+DUF4007->DUF6079*->iSTAND+PglZ->SIGMA+OB-Fold+AAA-ATPase-BrxL->ZnR->Cysteine-Desulfurase-> | 1401 | Bacteria>Terrabacteria_group>Bacillota>Negativicutes>Selenomonadales>Selenomonadaceae | GCA_017449105.1 | Selenomonadaceae_bacterium | IJP68_12495 | 5499..9704 | 2652289 | hypothetical-protein-[Selenomonadaceae-bacterium] |
| MBQ9486954.1 | Type4.PglZ.containing.BREX | <-RADICAL-SAM<-?<-?<-Cysteine-Desulfurase<-ZnR<-SIGMA+OB-Fold+AAA-ATPase-BrxL<-iSTAND+PglZ<-DUF6079*<-GspA-ATPase<-PAPS-reduct+a+b-domain+Fer4_9+DUF4007<-?<-HAMP+MA | 1208 | Bacteria>Terrabacteria_group>Bacillota>Negativicutes>Selenomonadales>Selenomonadaceae | GCA_017453765.1 | Selenomonadaceae_bacterium | IJU91_04005 | 7962..11588 | 2652289 | hypothetical-protein-[Selenomonadaceae-bacterium] |
| MBQ7704941.1 | Type4.PglZ.containing.BREX | PALP->?->PAPS-reduct+a+b-domain+Fer4_9+DUF4007->DUF6079*->iSTAND+PglZ->SIGMA+OB-Fold+AAA-ATPase-BrxL->ZnR->Cysteine-Desulfurase-> | 1399 | Bacteria>Terrabacteria_group>Bacillota>Negativicutes>Selenomonadales>Selenomonadaceae | GCA_017455645.1 | Selenomonadaceae_bacterium | IJT73_05910 | 5245..9444 | 2652289 | hypothetical-protein-[Selenomonadaceae-bacterium] |
| MBQ7705973.1 | Type4.PglZ.containing.BREX | <-iSTAND+PglZ<-DUF6079*<-PAPS-reduct+a+b-domain+Fer4_9+DUF4007<-SIGMA-HTH<-?<-PRTase<-Phos_pyr_kin<-?<-Peptidase_M22<-ACET<-ACET | 1395 | Bacteria>Terrabacteria_group>Bacillota>Negativicutes>Selenomonadales>Selenomonadaceae | GCA_017455645.1 | Selenomonadaceae_bacterium | IJT73_11225 | 2093..6280 | 2652289 | hypothetical-protein-[Selenomonadaceae-bacterium] |
| MBR2215718.1 | Type4.PglZ.containing.BREX | PIN->?->Glyco_transf_9->E3_UFM1_ligase+RAMA->E3_UFM1_ligase+RAMA->Ntox45->HEPN->DOC+HTH->PAPS-reduct+a+b-domain+Fer4_9+DUF4007->DUF6079*->iSTAND+PglZ->SIGMA+OB-Fold+AAA-ATPase-BrxL->ZnR->Cysteine-Desulfurase->?-><-?<-?<-?<-PRTase | 1407 | Bacteria>Terrabacteria_group>Bacillota>Negativicutes>Selenomonadales>Selenomonadaceae | GCA_017476005.1 | Selenomonadaceae_bacterium | IJ849_08180 | 106784..111007 | 2652289 | hypothetical-protein-[Selenomonadaceae-bacterium] |
| MBR1421803.1 | Type4.PglZ.containing.BREX | <-Glycos_transf\|\|?->PAPS-reduct+a+b-domain+Fer4_9+DUF4007->DUF6079*->iSTAND+PglZ->SIGMA+OB-Fold+AAA-ATPase-BrxL->Cysteine-Desulfurase-><-?<-RADICAL-SAM<-?<-CheW+REC\|\|?->TPR+TPR-> | 1396 | Bacteria>Terrabacteria_group>Bacillota>Negativicutes>Selenomonadales>Selenomonadaceae | GCA_017484005.1 | Selenomonadaceae_bacterium | IJ575_12220 | 14581..18771 | 2652289 | hypothetical-protein-[Selenomonadaceae-bacterium] |
| MBR2520311.1 | Type4.PglZ.containing.BREX | PAPS-reduct+a+b-domain+Fer4_9+DUF4007->DUF6079*->iSTAND+PglZ->SIGMA+OB-Fold+AAA-ATPase-BrxL->ZnR->Cysteine-Desulfurase->?->?-><-SAM-METHYLASE<-?<-SPOUT | 1406 | Bacteria>Terrabacteria_group>Bacillota>Negativicutes>Selenomonadales>Selenomonadaceae | GCA_017515635.1 | Selenomonadaceae_bacterium | IKE46_11095 | 4107..8327 | 2652289 | hypothetical-protein-[Selenomonadaceae-bacterium] |
| MBR7025508.1 | Type4.PglZ.containing.BREX | <-STAS<-?<-Cysteine-Desulfurase<-ZnR<-SIGMA+OB-Fold+AAA-ATPase-BrxL<-iSTAND+PglZ<-DUF6079*<-PAPS-reduct+a+b-domain+Fer4_9+DUF4007<-SIGMA-HTH | 1390 | Bacteria>Terrabacteria_group>Bacillota>Negativicutes>Selenomonadales>Selenomonadaceae | GCA_017518905.1 | Selenomonadaceae_bacterium | IKI08_05815 | 6338..10510 | 2652289 | hypothetical-protein-[Selenomonadaceae-bacterium] |
| MBR7025837.1 | Type4.PglZ.containing.BREX | <-iSTAND+PglZ<-DUF6079*<-PAPS-reduct+DUF4007<-?\|\|RADICAL-SAM->?->?->?-><-?\|\|TUDOR+SF1-SF2-Helicase+iREase+CTD+REase->N6-METHYLASE-> | 1401 | Bacteria>Terrabacteria_group>Bacillota>Negativicutes>Selenomonadales>Selenomonadaceae | GCA_017518905.1 | Selenomonadaceae_bacterium | IKI08_07535 | 7948..12153 | 2652289 | hypothetical-protein-[Selenomonadaceae-bacterium] |
| MBR6887677.1 | Type4.PglZ.containing.BREX | CARF->wHTH+WYL+WCX->wHTH+WYL+WCX-><-CARF\|\|?->PAPS-reduct+a+b-domain+Fer4_9+DUF4007->DUF6079*->iSTAND+PglZ->SIGMA+OB-Fold+AAA-ATPase-BrxL->SIGMA+OB-Fold+AAA-ATPase-BrxL->ZnR->Cysteine-Desulfurase-><-REase+SF1-SF2-Helicase<-TRD+TRD<-REase+N6_METHYLASE | 1400 | Bacteria>Terrabacteria_group>Bacillota>Negativicutes>Selenomonadales>Selenomonadaceae | GCA_017529935.1 | Selenomonadaceae_bacterium | IKN16_04455 | 10643..14845 | 2652289 | hypothetical-protein-[Selenomonadaceae-bacterium] |
| MBR4643031.1 | Type4.PglZ.containing.BREX | <-ABC-ATPase+SbcC<-?<-?<-Glycos_transf<-?\|\|?-><-SIGMA+OB-Fold+AAA-ATPase-BrxL<-iSTAND+PglZ<-DUF6079*<-?<-PAPS-reduct+a+b-domain+Fer4_9<-SIGMA-HTH<-?<-?\|\|?-><-Calcineurin<-?<-DEOR-HTH | 1275 | Bacteria>Terrabacteria_group>Bacillota>Negativicutes>Selenomonadales>Selenomonadaceae | GCA_017535925.1 | Selenomonadaceae_bacterium | IKO74_10015 | 37982..41809 | 2652289 | hypothetical-protein-[Selenomonadaceae-bacterium] |
| MBR4643448.1 | Type4.PglZ.containing.BREX | <-GAF+GGDEF<-?<-CHASE2+ACYC<-?<-PAS+GGDEF<-DnaB\|\|?->?-><-iSTAND+PglZ<-DUF6079*<-?<-ABC-ATPase+SbcC<-ABC-ATPase+SbcC<-?<-?<-?<-Peptidase_M15_4 | 1401 | Bacteria>Terrabacteria_group>Bacillota>Negativicutes>Selenomonadales>Selenomonadaceae | GCA_017535925.1 | Selenomonadaceae_bacterium | IKO74_12145 | 80410..84615 | 2652289 | hypothetical-protein-[Selenomonadaceae-bacterium] |
| MBR2732785.1 | Type4.PglZ.containing.BREX | <-ABC-ATPase<-Peptidase-C39+ABC-ATPase+SbcC<-?<-?<-Cysteine-Desulfurase<-ZnR<-SIGMA+OB-Fold+AAA-ATPase-BrxL<-iSTAND+PglZ<-DUF6079*<-PAPS-reduct+a+b-domain+Fer4_9+DUF4007 | 1395 | Bacteria>Terrabacteria_group>Bacillota>Negativicutes>Selenomonadales>Selenomonadaceae | GCA_017536785.1 | Selenomonadaceae_bacterium | IKD80_00830 | 11293..15480 | 2652289 | hypothetical-protein-[Selenomonadaceae-bacterium] |
| MBR6012951.1 | Type4.PglZ.containing.BREX | <-Cysteine-Desulfurase<-ZnR<-SIGMA+OB-Fold+AAA-ATPase-BrxL\|\|?->SIGMA+OB-Fold+AAA-ATPase-BrxL-><-iSTAND+PglZ<-DUF6079*<-PAPS-reduct+a+b-domain+Fer4_9+DUF4007<-?<-?<-DnaB+SMS-KaiC\|\|HTH_3-> | 1399 | Bacteria>Terrabacteria_group>Bacillota>Negativicutes>Selenomonadales>Selenomonadaceae | GCA_017553205.1 | Selenomonadaceae_bacterium | IK062_04105 | 6931..11130 | 2652289 | hypothetical-protein-[Selenomonadaceae-bacterium] |
| MBR5913419.1 | Type4.PglZ.containing.BREX | METHYLASE-><-?\|\|?->?->S1+GTPase-AIG->?->SIGMA-HTH->PAPS-reduct+a+b-domain+Fer4_9+DUF4007->DUF6079*->iSTAND+PglZ->SIGMA+OB-Fold+AAA-ATPase-BrxL->ZnR->Cysteine-Desulfurase-> | 1395 | Bacteria>Terrabacteria_group>Bacillota>Negativicutes>Selenomonadales>Selenomonadaceae | GCA_017554235.1 | Selenomonadaceae_bacterium | IKZ58_03510 | 94120..98307 | 2652289 | hypothetical-protein-[Selenomonadaceae-bacterium] |
| MBO4400231.1 | Type4.PglZ.containing.BREX | PAPS-reduct+a+b-domain+Fer4_9+DUF4007->DUF6079*->iSTAND+PglZ->SIGMA+OB-Fold+AAA-ATPase-BrxL->ZnR->Cysteine-Desulfurase-> | 1396 | Bacteria>Terrabacteria_group>Bacillota>Negativicutes>Selenomonadales>Selenomonadaceae | GCA_017614505.1 | Selenomonadaceae_bacterium | J5809_00100 | 24568..28758 | 2652289 | hypothetical-protein-[Selenomonadaceae-bacterium] |
| MCI6086972.1 | Type4.PglZ.containing.BREX | <-Cysteine-Desulfurase<-SIGMA+OB-Fold+AAA-ATPase-BrxL<-ZnR<-iSTAND+PglZ<-DUF6079*<-PAPS-reduct+a+b-domain+Fer4_9+DUF4007 | 1406 | Bacteria>Terrabacteria_group>Bacillota>Negativicutes>Selenomonadales>Selenomonadaceae | GCA_022771645.1 | Selenomonas_sp. | MR711_12165 | 5740..9960 | 2053611 | hypothetical-protein-[Selenomonas-sp.] |
| MCR5834345.1 | Type4.PglZ.containing.BREX | TPR->?->?->LRR_5-><-?\|\|?->PAPS-reduct+a+b-domain+Fer4_9+DUF4007->DUF6079*->iSTAND+PglZ->SIGMA+OB-Fold+AAA-ATPase-BrxL->ZnR->Cysteine-Desulfurase-><-MORC+SF1-SF2-Helicase<-?<-?<-REase<-DNA_METHYLASE | 1397 | Bacteria>Terrabacteria_group>Bacillota>Negativicutes>Selenomonadales>Selenomonadaceae | GCA_024698045.1 | Selenomonadaceae_bacterium | K6G55_06840 | 68910..73103 | 2652289 | hypothetical-protein-[Selenomonadaceae-bacterium] |
| SEH28739.1 | Type4.PglZ.containing.BREX | MULE-TRANSPOSASE-><-HAD<-Cysteine-Desulfurase<-ZnR<-SIGMA+OB-Fold+AAA-ATPase-BrxL<-iSTAND+PglZ<-DUF6079*<-PAPS-reduct+a+b-domain+Fer4_9+DUF4007<-E3_UFM1_ligase+RAMA\|\|?-><-HTH_OrfB_IS605+OrfB_IS605+OrfB_Zn_ribbon<-Y1_Tnp | 1403 | Bacteria>Terrabacteria_group>Bacillota>Negativicutes>Selenomonadales>Selenomonadaceae | GCA_900108415.1 | Selenomonas_ruminantium | SAMN05216583_1127 | 7179..11390 | 971 | hypothetical-protein-SAMN05216583_1127-[Selenomonas-ruminantium] |
| WP_006692207.1 | Type4.PglZ.containing.BREX | <-PAS+GGDEF<-?\|\|?-><-?<-Cysteine-Desulfurase<-ZnR<-SIGMA+OB-Fold+AAA-ATPase-BrxL<-iSTAND+PglZ<-DUF6079*<-PAPS-reduct+a+b-domain+Fer4_9+DUF4007<-?<-MACRODOMAIN | 1381 | Bacteria>Terrabacteria_group>Bacillota>Negativicutes>Selenomonadales>Selenomonadaceae | GCF_000234095.1 | Selenomonas_infelix | HMPREF9334_RS03955 | 143269..147414 | 135082 | hypothetical-protein-[Selenomonas-infelix] |
| WP_021772229.1 | Type4.PglZ.containing.BREX | TOPRIM-><-Cysteine-Desulfurase<-SIGMA+OB-Fold+AAA-ATPase-BrxL<-ZnR<-iSTAND+PglZ<-DUF6079*<-PAPS-reduct+a+b-domain+Fer4_9+DUF4007<-?<-?<-MPTase<-HTH_3<-rve+rve_3<-wHTH+WYL | 1395 | Bacteria>Terrabacteria_group>Bacillota>Negativicutes>Selenomonadales>Selenomonadaceae | GCF_000469545.1 | Mitsuokella_sp._oral_taxon_131 | HMPREF1985_RS08730 | 190570..194757 | 1321780 | hypothetical-protein-[Mitsuokella-sp.-oral-taxon-131] |
| WP_037355507.1 | Type4.PglZ.containing.BREX | <-RuvB-AAA+RuvB-HTH<-?<-?<-HEPN<-Cysteine-Desulfurase<-ZnR<-SIGMA+OB-Fold+AAA-ATPase-BrxL<-iSTAND+PglZ<-DUF6079*<-PAPS-reduct+a+b-domain+Fer4_9+DUF4007<-?\|\|HTH_3-><-HKD-DNase+SF1-SF2-Helicase+HTH+HTH+SRA<-SAM-METHYLASE | 1410 | Bacteria>Terrabacteria_group>Bacillota>Negativicutes>Selenomonadales>Selenomonadaceae | GCF_000701565.1 | Selenomonas_sp._FC4001 | T320_RS10205 | 100764..104996 | 1408313 | hypothetical-protein-[Selenomonas-sp.-FC4001] |
| WP_118016040.1 | Type4.PglZ.containing.BREX | <-Cysteine-Desulfurase<-ZnR<-SIGMA+OB-Fold+AAA-ATPase-BrxL<-iSTAND+PglZ<-DUF6079*<-PAPS-reduct+a+b-domain+Fer4_9+DUF4007<-SeqA+E3_UFM1_ligase<-?<-MODE-HTH+PBPII<-35exo | 1409 | Bacteria>Terrabacteria_group>Bacillota>Negativicutes>Selenomonadales>Selenomonadaceae | GCF_003458855.1 | Mitsuokella_sp._AF21-1AC | DWX75_RS09240 | 5529..9758 | 2292235 | hypothetical-protein-[Mitsuokella-sp.-AF21-1AC] |
| WP_270379983.1 | Type4.PglZ.containing.BREX | <-SF1-SF2-Helicase+REase<-Exonuc_V_gamma+REase<-SbcC+SbcC<-Calcineurin<-tRNA<-HTH_32+DDE_3<-HTH_32+DDE_3<-Cysteine-Desulfurase<-SIGMA+OB-Fold+AAA-ATPase-BrxL<-iSTAND+PglZ<-DUF6079* | 1409 | Bacteria>Terrabacteria_group>Bacillota>Negativicutes>Selenomonadales>Selenomonadaceae | GCF_027670365.1 | Mitsuokella_multacida | PGS78_RS09130 | 23402..27631 | 52226 | hypothetical-protein-[Mitsuokella-multacida] |
| WP_074573569.1 | Type4.PglZ.containing.BREX | <-Cysteine-Desulfurase<-ZnR<-SIGMA+OB-Fold+AAA-ATPase-BrxL<-iSTAND+PglZ<-DUF6079*<-PAPS-reduct+a+b-domain+Fer4_9+DUF4007<-?\|\|HTH_3->HTH_28-> | 1409 | Bacteria>Terrabacteria_group>Bacillota>Negativicutes>Selenomonadales>Selenomonadaceae | GCF_900104055.1 | Selenomonas_ruminantium | BLQ59_RS14670 | 5312..9541 | 971 | hypothetical-protein-[Selenomonas-ruminantium] |
| WP_074670413.1 | Type4.PglZ.containing.BREX | <-RuvB-AAA+RuvB-HTH<-?<-?<-HEPN<-Cysteine-Desulfurase<-SIGMA+OB-Fold+AAA-ATPase-BrxL<-iSTAND+PglZ<-DUF6079*<-PAPS-reduct+a+b-domain+Fer4_9+DUF4007<-?\|\|HTH_3-><-HKD-DNase+SF1-SF2-Helicase+HTH+HTH+SRA<-SAM-METHYLASE | 1410 | Bacteria>Terrabacteria_group>Bacillota>Negativicutes>Selenomonadales>Selenomonadaceae | GCF_900107615.1 | Selenomonas_ruminantium | BLV69_RS01695 | 324442..328674 | 971 | hypothetical-protein-[Selenomonas-ruminantium] |
| WP_073090449.1 | Type4.PglZ.containing.BREX | <-RuvB-AAA+RuvB-HTH<-?<-HEPN<-Cysteine-Desulfurase<-ZnR<-SIGMA+OB-Fold+AAA-ATPase-BrxL<-iSTAND+PglZ<-DUF6079*<-PAPS-reduct+a+b-domain+Fer4_9+DUF4007<-?\|\|HTH_3-><-HKD-DNase+SF1-SF2-Helicase+HTH+HTH+SRA<-SAM-METHYLASE<-?<-VirB11 | 1410 | Bacteria>Terrabacteria_group>Bacillota>Negativicutes>Selenomonadales>Selenomonadaceae | GCF_900142515.1 | Selenomonas_ruminantium | SAMN05216582_11738 | 34242..38474 | 971 | hypothetical-protein-[Selenomonas-ruminantium] |
| WP_093795253.1 | Type4.PglZ.containing.BREX | <-ZnR<-SIGMA+OB-Fold+AAA-ATPase-BrxL<-iSTAND+PglZ<-DUF6079*<-PAPS-reduct+a+b-domain+Fer4_9+DUF4007<-SIGMA-HTH+E3_UFM1_ligase<-?<-?<-MPTase<-HTH_3 | 1393 | Bacteria>Terrabacteria_group>Bacillota>Negativicutes>Selenomonadales>Sporomusaceae | GCF_900101845.1 | Sporomusa_acidovorans | BLR65_RS16325 | 5121..9302 | 112900 | hypothetical-protein-[Sporomusa-acidovorans] |
| HAD77166.1 | Type4.PglZ.containing.BREX | REase->REase->REase->HKD-DNase+SF1-SF2-Helicase+HTH+HTH+SRA->?->?->PAPS-reduct+a+b-domain+Fer4_9+DUF4007->DUF6079*->iSTAND+PglZ->SIGMA+OB-Fold+AAA-ATPase-BrxL->ZnR->Cysteine-Desulfurase->Aminotran_1_2->sGTP->?->ABC-ATPase+SbcC-> | 1397 | Bacteria>Terrabacteria_group>Bacillota>Negativicutes>Veillonellales>Veillonellaceae | GCA_003447275.1 | Dialister_sp. | DCG08_04065 | 35312..39505 | 1955814 | hypothetical-protein-[Dialister-sp.] |
| MBD8953984.1 | Type4.PglZ.containing.BREX | <-ABC-ATPase+SbcC<-?\|\|?-><-sGTP<-Aminotran_1_2<-Cysteine-Desulfurase<-ZnR<-SIGMA+OB-Fold+AAA-ATPase-BrxL<-iSTAND+PglZ<-DUF6079*<-PAPS-reduct+a+b-domain+Fer4_9+DUF4007<-?<-?<-HKD-DNase+SF1-SF2-Helicase+HTH+HTH+SRA<-REase | 1397 | Bacteria>Terrabacteria_group>Bacillota>Negativicutes>Veillonellales>Veillonellaceae | GCA_014847205.1 | Dialister_sp. | EGQ71_02915 | 42379..46572 | 1955814 | hypothetical-protein-[Dialister-sp.] |
| MCQ4113132.1 | Type4.PglZ.containing.BREX | PRTase-><-?<-?\|\|REase->?->XIS-HTH->SeqA+E3_UFM1_ligase->PAPS-reduct+a+b-domain+Fer4_9+DUF4007->DUF6079*->iSTAND+PglZ->SIGMA+OB-Fold+AAA-ATPase-BrxL->ZnR->Cysteine-Desulfurase-> | 1405 | Bacteria>Terrabacteria_group>Bacillota>Negativicutes>Veillonellales>Veillonellaceae | GCA_024436295.1 | Megasphaera_sp._SC8-1 | NOS26_07925 | 50940..55157 | 2965102 | hypothetical-protein-[Megasphaera-sp.-SC8-1] |
| WP_054335796.1 | Type4.PglZ.containing.BREX | <-MODE-HTH+PBPII\|\|?-><-Cysteine-Desulfurase<-SIGMA+OB-Fold+AAA-ATPase-BrxL<-iSTAND+PglZ<-DUF6079*<-PAPS-reduct+a+b-domain+Fer4_9+DUF4007<-SeqA+E3_UFM1_ligase<-SIGMA-HTH | 1402 | Bacteria>Terrabacteria_group>Bacillota>Negativicutes>Veillonellales>Veillonellaceae | GCF_001304715.1 | Megasphaera_elsdenii | AZ49_RS00095 | 24983..29191 | 907 | hypothetical-protein-[Megasphaera-elsdenii] |
| WP_255917521.1 | Type4.PglZ.containing.BREX | <-Cysteine-Desulfurase<-ZnR<-SIGMA+OB-Fold+AAA-ATPase-BrxL<-iSTAND+PglZ<-DUF6079*<-PAPS-reduct+a+b-domain+Fer4_9+DUF4007<-SeqA+E3_UFM1_ligase<-XIS-HTH<-REase\|\|?->?->?->PRTase | 1405 | Bacteria>Terrabacteria_group>Bacillota>Negativicutes>Veillonellales>Veillonellaceae | GCF_902760425.1 | Megasphaera_sp._SC8-1 | QDP60_RS02680 | 15576..19793 | 2965102 | MULTISPECIES:-hypothetical-protein-[Megasphaera] |
| WP_132995095.1 | Type4.PglZ.containing.BREX | <-ABC-ATPase+SbcC\|\|DSBH-><-?<-?<-SIGMA+OB-Fold+AAA-ATPase-BrxL<-PglZ+Phosphodiest<-DUF6079*<-BrxB-BrxF-iSTAND<-DUF4007<-DUF6079<-DUF4007<-PAPS-reduct+a+b-domain+Fer4_9<-?<-SIGMA-HTH\|\|DSBH-> | 1332 | Bacteria>Terrabacteria_group>Bacillota>Tissierellia>Tissierellales>Sporanaerobacteraceae | GCF_004349435.1 | Sporanaerobacter_acetigenes | EIC75_RS01970 | 156657..160655 | 165813 | hypothetical-protein-[Sporanaerobacter-acetigenes] |
| WP_132995101.1 | Type4.PglZ.containing.BREX | <-ABC-ATPase+SbcC\|\|DSBH-><-?<-?<-SIGMA+OB-Fold+AAA-ATPase-BrxL<-PglZ+Phosphodiest<-DUF6079*<-BrxB-BrxF-iSTAND<-DUF4007<-DUF6079*<-DUF4007<-PAPS-reduct+a+b-domain+Fer4_9<-?<-SIGMA-HTH\|\|DSBH-><-?<-?<-?<-HKD-DNase+SF1-SF2-Helicase+HTH+HTH+SRA-p | 1176 | Bacteria>Terrabacteria_group>Bacillota>Tissierellia>Tissierellales>Sporanaerobacteraceae | GCF_004349435.1 | Sporanaerobacter_acetigenes | EIC75_RS01985 | 162032..165562 | 165813 | hypothetical-protein-[Sporanaerobacter-acetigenes] |
| WP_077369755.1 | Type4.PglZ.containing.BREX | Peptidase_M42->Aminotran_1_2->DUF4007->DUF6079->HARE-HTH->?->PAPS-reduct+a+b-domain+Fer4_9->DUF4007->AAA-ATPase->DUF6079*->PglZ->SIGMA+OB-Fold+AAA-ATPase-BrxL->Cysteine-Desulfurase->?->Y1_Tnp->?-><-HTH_6+SIS | 1367 | Bacteria>Terrabacteria_group>Bacillota>Tissierellia>Tissierellales>Sporanaerobacteraceae | GCF_900155565.1 | Anaerosalibacter_sp._Marseille-P3206 | BQ9840_RS10580 | 1903576..1907679 | 1871005 | hypothetical-protein-[Anaerosalibacter-sp.-Marseille-P3206] |
| NLD15978.1 | Type4.PglZ.containing.BREX | REase-><-Cysteine-Desulfurase<-ZnR<-SIGMA+OB-Fold+AAA-ATPase-BrxL<-iSTAND+PglZ<-DUF6079*<-PAPS-reduct+a+b-domain+Fer4_9+DUF4007<-SeqA+E3_UFM1_ligase<-HTH_26<-JAB | 1401 | Bacteria>Terrabacteria_group>Bacillota>Tissierellia>unclassified_Tissierellia | GCA_012513395.1 | Tissierellia_bacterium | GX666_00155 | 5529..9734 | 2049431 | hypothetical-protein-[Tissierellia-bacterium] |
| NMB09044.1 | Type4.PglZ.containing.BREX | SeqA+E3_UFM1_ligase->PAPS-reduct+a+b-domain+Fer4_9+DUF4007->DUF6079*->iSTAND+PglZ->SIGMA+OB-Fold+AAA-ATPase-BrxL->ZnR->Cysteine-Desulfurase-p-> | 1401 | Bacteria>Terrabacteria_group>Bacillota>Tissierellia>unclassified_Tissierellia | GCA_012799055.1 | Tissierellia_bacterium | GX981_11740 | 3591..7796 | 2049431 | hypothetical-protein-[Tissierellia-bacterium] |
| OQB25825.1 | Type4.PglZ.containing.BREX | TRAM+METHYLASE-><-Cysteine-Desulfurase<-SIGMA+OB-Fold+AAA-ATPase-BrxL<-ZnR<-iSTAND+PglZ<-DUF6079*<-PAPS-reduct+a+b-domain+Fer4_9+DUF4007<-HARE-HTH+E3_UFM1_ligase<-XIS-HTH<-MPTase<-HTH_3\|\|?->?->SIGMA-HTH-> | 1411 | Bacteria>Terrabacteria_group>Bacillota>unclassified_Bacillota | GCA_002070205.1 | Firmicutes_bacterium_ADurb.Bin182 | BWY11_00203 | 65417..69652 | 1852884 | hypothetical-protein-BWY11_00203-[Firmicutes-bacterium-ADurb.Bin182] |
| PKM78481.1 | Type4.PglZ.containing.BREX | <-HIN-HTH<-Cysteine-Desulfurase<-ZnR<-SIGMA+OB-Fold+AAA-ATPase-BrxL<-iSTAND+PglZ<-DUF6079*<-PAPS-reduct+a+b-domain+Fer4_9+DUF4007<-HARE-HTH+E3_UFM1_ligase<-HKD-DNase+SF1-SF2-Helicase+HTH+HTH+SRA<-NUDIX<-PIN<-?<-?\|\|REC+OmpR-HTH->HISKIN+MORC-> | 1388 | Bacteria>Terrabacteria_group>Bacillota>unclassified_Bacillota | GCA_002840245.1 | Firmicutes_bacterium_HGW-Firmicutes-15 | CVU90_01710 | 40231..44397 | 2013776 | hypothetical-protein-CVU90_01710-[Firmicutes-bacterium-HGW-Firmicutes-15] |
| RHS79257.1 | Type4.PglZ.containing.BREX | <-MPTase<-LexA-protease<-HTH+REase<-wHTH+WYL+WCX<-?<-Cysteine-Desulfurase<-ZnR<-SIGMA+OB-Fold+AAA-ATPase-BrxL<-iSTAND+PglZ<-DUF6079*<-PAPS-reduct+a+b-domain+Fer4_9+DUF4007<-Phage_integrase+E3_UFM1_ligase<-HTH_26<-Lysozyme+NLPC<-?<-?<-VirB4-FtsK | 1394 | Bacteria>Terrabacteria_group>Bacillota>unclassified_Bacillota | GCA_003480065.1 | Firmicutes_bacterium_AM43-11BH | DW928_10670 | 91212..95396 | 2292895 | hypothetical-protein-DW928_10670-[Firmicutes-bacterium-AM43-11BH] |
| MAU00653.1 | Type4.PglZ.containing.BREX | <-ParA-Soj-PloopNTPase<-?\|\|PAPS-reduct+Fer4_7->DUF4007->DUF6079*->Cysteine-Desulfurase->SIGMA-HTH+SIGMA-HTH->DUF4007->?->DUF6079->?->PglZ->SIGMA+OB-Fold+AAA-ATPase-BrxL-> | 1257 | Bacteria>Terrabacteria_group>Chloroflexota>Anaerolineae>Anaerolineales>Anaerolineaceae | GCA_002699125.1 | Anaerolineaceae_bacterium | CL608_26205 | 91832..95605 | 2024896 | hypothetical-protein-[Anaerolineaceae-bacterium] |
| MBI9047161.1 | Type4.PglZ.containing.BREX | DDE_Tnp_IS66+DDE_Tnp_IS66_C->DDE_Tnp_IS66+DDE_Tnp_IS66_C-><-SIGMA+OB-Fold+AAA-ATPase-BrxL<-PglZ<-DUF6079* | 1149 | Bacteria>Terrabacteria_group>Chloroflexota>Anaerolineae>Anaerolineales>Anaerolineaceae | GCA_016342475.1 | Anaerolineaceae_bacterium | JEZ06_21925 | 4627..8078 | 2024896 | hypothetical-protein-[Anaerolineaceae-bacterium] |
| MBN2258343.1 | Type4.PglZ.containing.BREX | <-ABC-ATPase+SbcC\|\|RNase-><-DndB<-URI-DNase<-Cysteine-Desulfurase<-SIGMA+OB-Fold+AAA-ATPase-BrxL<-PglZ<-DUF6079*<-?<-DUF4007 | 1422 | Bacteria>Terrabacteria_group>Chloroflexota>Anaerolineae>Anaerolineales>Anaerolineaceae | GCA_016933925.1 | Anaerolineaceae_bacterium | JW704_11050 | 24678..28946 | 2024896 | hypothetical-protein-[Anaerolineaceae-bacterium] |
| MBN2678363.1 | Type4.PglZ.containing.BREX | AbiEii-><-?<-?<-ABC-ATPase+SbcC<-DndB<-URI-DNase<-Cysteine-Desulfurase<-SIGMA+OB-Fold+AAA-ATPase-BrxL<-PglZ<-DUF6079*<-?<-DUF4007<-wHTH<-PAPS-reduct<-DUF6079<-DUF4007 | 1422 | Bacteria>Terrabacteria_group>Chloroflexota>Anaerolineae>Anaerolineales>Anaerolineaceae | GCA_016936755.1 | Anaerolineaceae_bacterium | JXR32_09930 | 14087..18355 | 2024896 | hypothetical-protein-[Anaerolineaceae-bacterium] |
| WP_162787802.1 | Type4.PglZ.containing.BREX | <-Glyco_hydro_36N+Melibiase+Glyco_hydro_36C<-?\|\|?-><-?<-?<-Phage_int_SAM_4+Phage_integrase\|\|DUF4007->DUF6079-p->DUF6079-p*->PAPS-reduct->wHTH->DUF4007->?->DUF6079->PglZ->SIGMA+OB-Fold+AAA-ATPase-BrxL->Cysteine-Desulfurase-> | 846 | Bacteria>Terrabacteria_group>Chloroflexota>Anaerolineae>Anaerolineales>Anaerolineaceae | GCF_001050195.2 | Anaerolinea_thermolimosa | ATHL_RS17765 | 101882..104422 | 229919 | hypothetical-protein,-partial-[Anaerolinea-thermolimosa] |
| WP_062196491.1 | Type4.PglZ.containing.BREX | <-Phage_int_SAM_4+Phage_integrase\|\|DUF4007->DUF6079-p->PAPS-reduct->wHTH->DUF4007->?->DUF6079*->PglZ->SIGMA+OB-Fold+AAA-ATPase-BrxL->Cysteine-Desulfurase-> | 1403 | Bacteria>Terrabacteria_group>Chloroflexota>Anaerolineae>Anaerolineales>Anaerolineaceae | GCF_001050195.2 | Anaerolinea_thermolimosa | ATHL_RS17790 | 107066..111277 | 229919 | hypothetical-protein-[Anaerolinea-thermolimosa] |
| WP_162787801.1 | Type4.PglZ.containing.BREX | <-Glyco_hydro_36N+Melibiase+Glyco_hydro_36C<-?\|\|?-><-?<-?<-Phage_int_SAM_4+Phage_integrase\|\|DUF4007->DUF6079-p*->DUF6079-p->PAPS-reduct->wHTH->DUF4007->?->DUF6079->PglZ->SIGMA+OB-Fold+AAA-ATPase-BrxL->Cysteine-Desulfurase-> | 285 | Bacteria>Terrabacteria_group>Chloroflexota>Anaerolineae>Anaerolineales>Anaerolineaceae | GCF_001050195.2 | Anaerolinea_thermolimosa | ATHL_RS19050 | 99708..100565 | 229919 | hypothetical-protein-[Anaerolinea-thermolimosa] |
| WP_062417761.1 | Type4.PglZ.containing.BREX | <-Phage_int_SAM_3+Phage_integrase\|\|HTH_3-><-XIS-HTH\|\|HTH_3-><-?<-Cysteine-Desulfurase<-SIGMA+OB-Fold+AAA-ATPase-BrxL<-PglZ<-DUF6079<-?<-DUF4007<-PAPS-reduct<-DUF6079*<-DUF4007\|\|?->DEDD_Tnp_IS110+Transposase_20->AbiEi_4+AbiEi_1->AbiEii-> | 1410 | Bacteria>Terrabacteria_group>Chloroflexota>Anaerolineae>Anaerolineales>Anaerolineaceae | GCF_001050255.2 | Levilinea_saccharolytica | LSAC_RS09855 | 1334908..1339140 | 229921 | hypothetical-protein-[Levilinea-saccharolytica] |
| WP_062417765.1 | Type4.PglZ.containing.BREX | <-XIS-HTH\|\|HTH_3-><-?<-Cysteine-Desulfurase<-SIGMA+OB-Fold+AAA-ATPase-BrxL<-PglZ<-DUF6079<-?<-DUF4007<-PAPS-reduct<-DUF6079*<-DUF4007\|\|?->DEDD_Tnp_IS110+Transposase_20->AbiEi_4+AbiEi_1->AbiEii->SF1-SF2-Helicase+REase->ParB+HNH+Schlafen->N6-N4-METHYLASE-> | 1141 | Bacteria>Terrabacteria_group>Chloroflexota>Anaerolineae>Anaerolineales>Anaerolineaceae | GCF_001050255.2 | Levilinea_saccharolytica | LSAC_RS09875 | 1341322..1344747 | 229921 | phage-tail-tape-measure-protein-[Levilinea-saccharolytica] |
| WP_062281142.1 | Type4.PglZ.containing.BREX | <-DnaB+SMS-KaiC<-?<-?<-?<-XIS-HTH\|\|?-><-Cysteine-Desulfurase<-SIGMA+OB-Fold+AAA-ATPase-BrxL<-iSTAND+PglZ<-DUF6079*<-PAPS-reduct+a+b-domain+Fer4_9+DUF4007<-?<-XIS-HTH\|\|HTH+MPTase->PIN->?->McrB-AAA->REase-> | 1401 | Bacteria>Terrabacteria_group>Chloroflexota>Anaerolineae>Anaerolineales>Anaerolineaceae | GCF_001192795.1 | Flexilinea_flocculi | ATC1_RS09745 | 1119507..1123712 | 1678840 | hypothetical-protein-[Flexilinea-flocculi] |
| WP_062282329.1 | Type4.PglZ.containing.BREX | FGS-><-?<-?<-?\|\|?-><-?<-Cysteine-Desulfurase<-SIGMA+OB-Fold+AAA-ATPase-BrxL<-iSTAND+PglZ<-DUF6079*<-PAPS-reduct+a+b-domain+Fer4_9+DUF4007<-HARE-HTH+E3_UFM1_ligase\|\|?->?->?-><-tRNA<-?<-?<-HARE-HTH<-LysM+PTSEIIA1 | 1382 | Bacteria>Terrabacteria_group>Chloroflexota>Anaerolineae>Anaerolineales>Anaerolineaceae | GCF_001192795.1 | Flexilinea_flocculi | ATC1_RS11785 | 1650028..1654176 | 1678840 | hypothetical-protein-[Flexilinea-flocculi] |
| WP_160318459.1 | Type4.PglZ.containing.BREX | <-MULE-TRANSPOSASE\|\|DUF4007->DUF6079->PAPS-reduct->DUF4007->?->DUF6079*->PglZ->SIGMA+OB-Fold+AAA-ATPase-BrxL->Cysteine-Desulfurase-> | 1402 | Bacteria>Terrabacteria_group>Chloroflexota>Anaerolineae>Anaerolineales>Anaerolineaceae | GCF_001306035.1 | Levilinea_saccharolytica | ADN01_RS16860 | 7210..11418 | 229921 | hypothetical-protein-[Levilinea-saccharolytica] |
| TEU15946.1 | Type4.PglZ.containing.BREX | <-HerA/FtsK<-PglZ-p<-DUF6079*<-DnaA<-DUF4007<-HTH_3<-DndB<-Cysteine-Desulfurase | 1350 | Bacteria>Terrabacteria_group>Chloroflexota>Anaerolineae>Anaerolineales>unclassified_Anaerolineales | GCA_004377365.1 | Anaerolineales_bacterium | E3J21_11775 | 2704..6756 | 2073117 | hypothetical-protein-E3J21_11775-[Anaerolineales-bacterium] |
| MCA9963061.1 | Type4.PglZ.containing.BREX | ACET->ABC-ATPase+ABC-ATPase+ABC-ATPase+SbcC->DUF4007->DUF6079*->PAPS-reduct->DUF4007->?->DUF6079->PglZ->SIGMA+OB-Fold+AAA-ATPase-BrxL->Cysteine-Desulfurase-> | 1131 | Bacteria>Terrabacteria_group>Chloroflexota>Anaerolineae>Anaerolineales>unclassified_Anaerolineales | GCA_020432595.1 | Anaerolineales_bacterium | KC423_02415 | 5096..8491 | 2073117 | hypothetical-protein-[Anaerolineales-bacterium] |
| MCC6188629.1 | Type4.PglZ.containing.BREX | <-SIGMA+OB-Fold+AAA-ATPase-BrxL<-PglZ<-DUF6079*<-iSTAND+PglZ<-DUF4007<-PAPS-reduct<-DUF6079<-DUF4007 | 1395 | Bacteria>Terrabacteria_group>Chloroflexota>Anaerolineae>Anaerolineales>unclassified_Anaerolineales | GCA_020845955.1 | Anaerolineales_bacterium | IT318_06315 | 4265..8452 | 2073117 | hypothetical-protein-[Anaerolineales-bacterium] |
| MCC6188633.1 | Type4.PglZ.containing.BREX | <-SIGMA+OB-Fold+AAA-ATPase-BrxL<-PglZ<-DUF6079<-iSTAND+PglZ<-DUF4007<-PAPS-reduct<-DUF6079*<-DUF4007 | 1120 | Bacteria>Terrabacteria_group>Chloroflexota>Anaerolineae>Anaerolineales>unclassified_Anaerolineales | GCA_020845955.1 | Anaerolineales_bacterium | IT318_06335 | 10698..14060 | 2073117 | hypothetical-protein-[Anaerolineales-bacterium] |
| MCL4273187.1 | Type4.PglZ.containing.BREX | <-HTH_3<-?<-Cysteine-Desulfurase<-SIGMA+OB-Fold+AAA-ATPase-BrxL<-PglZ<-DUF6079<-?<-DUF4007<-PAPS-reduct<-DUF6079*<-DUF4007<-?<-?<-NERD+ZnR<-?<-?<-ARG<-Thermonuclease | 1131 | Bacteria>Terrabacteria_group>Chloroflexota>Anaerolineae>Anaerolineales>unclassified_Anaerolineales | GCA_023382595.1 | Anaerolineales_bacterium | KJZ77_04890 | 1008497..1011892 | 2073117 | hypothetical-protein-[Anaerolineales-bacterium] |
| NPV08139.1 | Type4.PglZ.containing.BREX | XIS-HTH->?->?->?->?->MULE-TRANSPOSASE-><-SIGMA+OB-Fold+AAA-ATPase-BrxL<-iSTAND+PglZ<-DUF6079*<-DUF4007<-PAPS-reduct+a+b-domain+Fer4_9+DUF4007<-SIGMA-HTH+SIGMA-HTH | 1378 | Bacteria>Terrabacteria_group>Chloroflexota>Anaerolineae>unclassified_Anaerolineae | GCA_013178015.1 | Anaerolineae_bacterium | HPY83_09275 | 50503..54639 | 2052143 | hypothetical-protein-[Anaerolineae-bacterium] |
| NPV84908.1 | Type4.PglZ.containing.BREX | <-NAD_binding_3+Homoserine_dh<-STYKinase-like<-?<-Peptidase_C26<-?\|\|DUF4007->DUF6079*->PAPS-reduct->DUF4007->?-><-?\|\|DUF6079->PglZ->SIGMA+OB-Fold+AAA-ATPase-BrxL->Cysteine-Desulfurase-><-DUF772+DDE_Tnp_1 | 1116 | Bacteria>Terrabacteria_group>Chloroflexota>Anaerolineae>unclassified_Anaerolineae | GCA_013178295.1 | Anaerolineae_bacterium | HPY45_02710 | 245912..249262 | 2052143 | hypothetical-protein-[Anaerolineae-bacterium] |
| NPV84913.1 | Type4.PglZ.containing.BREX | <-Peptidase_C26<-?\|\|DUF4007->DUF6079->PAPS-reduct->DUF4007->?-><-?\|\|DUF6079*->PglZ->SIGMA+OB-Fold+AAA-ATPase-BrxL->Cysteine-Desulfurase-><-DUF772+DDE_Tnp_1 | 1134 | Bacteria>Terrabacteria_group>Chloroflexota>Anaerolineae>unclassified_Anaerolineae | GCA_013178295.1 | Anaerolineae_bacterium | HPY45_02735 | 252242..255646 | 2052143 | hypothetical-protein-[Anaerolineae-bacterium] |
| MCG3139195.1 | Type4.PglZ.containing.BREX | <-Cysteine-Desulfurase<-SIGMA+OB-Fold+AAA-ATPase-BrxL<-PglZ<-DUF6079<-?<-DUF4007<-wHTH<-PAPS-reduct<-DUF6079*<-DUF4007<-?<-?<-?<-MarR-HTH<-PTSEIIA1 | 1131 | Bacteria>Terrabacteria_group>Chloroflexota>Anaerolineae>unclassified_Anaerolineae | GCA_022072245.1 | Anaerolineae_bacterium | HDKAJFGB_00057 | 19305..22700 | 2052143 | hypothetical-protein-[Anaerolineae-bacterium] |
| GIV81897.1 | Type4.PglZ.containing.BREX | HEPN->REase+SF1-SF2-Helicase->DUF4007->DUF6079-p*->DUF6079-p->wHTH(SBD)+HNH->SAM-METHYLASE+HIT->?->DUF6079-p->PglZ->PglZ-> | 404 | Bacteria>Terrabacteria_group>Chloroflexota>Anaerolineae>unclassified_Anaerolineae | GCA_026003715.1 | Anaerolineae_bacterium | KatS3mg051_1251 | 6189..7403 | 2052143 | MAG:-hypothetical-protein-KatS3mg051_1251-[Anaerolineae-bacterium] |
| MBX7251231.1 | Type4.PglZ.containing.BREX | <-TPR+TPR+TPR+TPR+TPR+TPR<-?<-TPR+TPR+TPR<-Cysteine-Desulfurase<-SIGMA+OB-Fold+AAA-ATPase-BrxL<-PglZ<-DUF6079*<-?<-DUF4007<-PAPS-reduct | 1413 | Bacteria>Terrabacteria_group>Chloroflexota>Ardenticatenia>Ardenticatenia_incertae_sedis>Candidatus_Promineifilum | GCA_019695075.1 | Candidatus_Promineofilum_sp. | K1X50_04545 | 11533..15774 | 2664178 | hypothetical-protein-[Candidatus-Promineofilum-sp.] |
| NJL05929.1 | Type4.PglZ.containing.BREX | <-SIGMA+OB-Fold+AAA-ATPase-BrxL<-PglZ<-DUF6079*<-BrxB-BrxF-iSTAND<-DUF4007<-PAPS-reduct<-DUF6079<-DUF4007 | 1706 | Bacteria>Terrabacteria_group>Chloroflexota>Chloroflexia>Chloroflexales>Chloroflexineae | GCA_012031225.1 | Chloroflexaceae_bacterium | HC911_13690 | 3825..8945 | 2562117 | hypothetical-protein-[Chloroflexaceae-bacterium] |
| NJL05933.1 | Type4.PglZ.containing.BREX | <-SIGMA+OB-Fold+AAA-ATPase-BrxL<-PglZ<-DUF6079<-BrxB-BrxF-iSTAND<-DUF4007<-PAPS-reduct<-DUF6079*<-DUF4007 | 1150 | Bacteria>Terrabacteria_group>Chloroflexota>Chloroflexia>Chloroflexales>Chloroflexineae | GCA_012031225.1 | Chloroflexaceae_bacterium | HC911_13710 | 11059..14511 | 2562117 | hypothetical-protein-[Chloroflexaceae-bacterium] |
| NJN65900.1 | Type4.PglZ.containing.BREX | SAM-METHYLASE-><-MuB-ATPase<-?<-rve<-?<-?<-?\|\|DUF4007->BrxB-BrxF-iSTAND->DUF6079->DUF6079-p->PglZ->SIGMA+OB-Fold+AAA-ATPase-BrxL->CASPASE->CASPASE->?->?->RAMPs->RAMPs-> | 1164 | Bacteria>Terrabacteria_group>Chloroflexota>Chloroflexia>Chloroflexales>Chloroflexineae | GCA_012032475.1 | Chloroflexaceae_bacterium | HC884_03880 | 24938..28432 | 2562117 | hypothetical-protein-[Chloroflexaceae-bacterium] |
| WP_097643874.1 | Type4.PglZ.containing.BREX | <-DNA_METHYLASE+DCM<-DndG<-?<-DndF<-SIGMA+OB-Fold+AAA-ATPase-BrxL<-PglZ<-DUF6079*<-BrxB-BrxF-iSTAND<-DUF4007<-PAPS-reduct\|\|PALP-><-?\|\|PAS+STAS-> | 1693 | Bacteria>Terrabacteria_group>Chloroflexota>Chloroflexia>Chloroflexales>Chloroflexineae | GCF_002532535.1 | Candidatus_Viridilinea_mediisalina | CJ255_RS09535 | 7330..12411 | 2024553 | hypothetical-protein-[Candidatus-Viridilinea-mediisalina] |
| WP_129675132.1 | Type4.PglZ.containing.BREX | <-ACET<-PIN<-CopG/RHH<-ABhydrolase<-PADR-HTH\|\|?->DUF4007->DUF6079*->PAPS-reduct->DUF4007->CR-ATPase5->DUF6079->PglZ->SIGMA+OB-Fold+AAA-ATPase-BrxL->?-><-PIN<-HTH+MPTase | 1154 | Bacteria>Terrabacteria_group>Chloroflexota>Chloroflexia>Chloroflexales>Chloroflexineae | GCF_004138165.1 | Candidatus_Chloroploca_sp._Khr17 | EKD05_RS17615 | 27349..30813 | 2496869 | hypothetical-protein-[Candidatus-Chloroploca-sp.-Khr17] |
| WP_129675138.1 | Type4.PglZ.containing.BREX | <-CopG/RHH<-ABhydrolase<-PADR-HTH\|\|?->DUF4007->DUF6079->PAPS-reduct->DUF4007->CR-ATPase5->DUF6079*->PglZ->SIGMA+OB-Fold+AAA-ATPase-BrxL->?-><-PIN<-HTH+MPTase\|\|REase-> | 1697 | Bacteria>Terrabacteria_group>Chloroflexota>Chloroflexia>Chloroflexales>Chloroflexineae | GCF_004138165.1 | Candidatus_Chloroploca_sp._Khr17 | EKD05_RS17625 | 32935..38028 | 2496869 | hypothetical-protein-[Candidatus-Chloroploca-sp.-Khr17] |
| MBK9943584.1 | Type4.PglZ.containing.BREX | <-LACI-HTH+PBPI<-?<-?\|\|DUF4007->DUF6079*->PAPS-reduct->DUF4007->BrxB-BrxF-iSTAND->DUF6079->PglZ->SIGMA+OB-Fold+AAA-ATPase-BrxL-><-ABC-ATPase+SbcC | 1154 | Bacteria>Terrabacteria_group>Chloroflexota>Chloroflexia>Chloroflexales>Roseiflexineae | GCA_016722075.1 | Kouleothrix_sp. | IPP13_18415 | 4645779..4649243 | 2779161 | hypothetical-protein-[Kouleothrix-sp.] |
| MBK9943588.1 | Type4.PglZ.containing.BREX | <-LACI-HTH+PBPI<-?<-?\|\|DUF4007->DUF6079->PAPS-reduct->DUF4007->BrxB-BrxF-iSTAND->DUF6079*->PglZ->SIGMA+OB-Fold+AAA-ATPase-BrxL-><-ABC-ATPase+SbcC<-?<-?\|\|ACET-><-Glyco_hydro_130<-?<-ABC-ATPase+SbcC | 1685 | Bacteria>Terrabacteria_group>Chloroflexota>Chloroflexia>Chloroflexales>Roseiflexineae | GCA_016722075.1 | Kouleothrix_sp. | IPP13_18435 | 4651728..4656785 | 2779161 | hypothetical-protein-[Kouleothrix-sp.] |
| NTU79112.1 | Type4.PglZ.containing.BREX | PAPS-reduct->DUF4007->BrxB-BrxF-iSTAND->DUF6079*->PglZ->SIGMA+OB-Fold+AAA-ATPase-BrxL-><-?\|\|?-><-SF1-SF2-Helicase | 1690 | Bacteria>Terrabacteria_group>Chloroflexota>Chloroflexia>Chloroflexales>unclassified_Chloroflexales | GCA_013336035.1 | Chloroflexales_bacterium | HGA45_06880 | 2541..7613 | 2725990 | hypothetical-protein-[Chloroflexales-bacterium] |
| WP_078022411.1 | Type4.PglZ.containing.BREX | <-MORC\|\|SbcC->Recombinase+ZnR-><-?<-Resolvase+Recombinase<-Cysteine-Desulfurase<-ZnR<-SIGMA+OB-Fold+AAA-ATPase-BrxL<-iSTAND+PglZ<-DUF6079*<-PAPS-reduct+a+b-domain+Fer4_9+DUF4007<-HARE-HTH+E3_UFM1_ligase<-HTH_26<-SbcC<-?<-HNH\|\|XIS-HTH-> | 1387 | Bacteria>Terrabacteria_group>Chloroflexota>Dehalococcoidia>Dehalococcoidales>Dehalococcoidaceae | GCF_002007845.1 | Dehalococcoides_mccartyi | B1777_RS01280 | 239694..243857 | 61435 | hypothetical-protein-[Dehalococcoides-mccartyi] |
| WP_077997895.1 | Type4.PglZ.containing.BREX | <-MORC\|\|SbcC->Recombinase+ZnR-><-?<-Resolvase+Recombinase<-Cysteine-Desulfurase<-ZnR<-SIGMA+OB-Fold+AAA-ATPase-BrxL<-iSTAND+PglZ<-DUF6079*<-PAPS-reduct+a+b-domain+Fer4_9+DUF4007<-HARE-HTH+E3_UFM1_ligase<-HTH_26<-SbcC<-?<-HNH\|\|XIS-HTH-> | 1387 | Bacteria>Terrabacteria_group>Chloroflexota>Dehalococcoidia>Dehalococcoidales>Dehalococcoidaceae | GCF_002007865.1 | Dehalococcoides_mccartyi | B1778_RS01135 | 216794..220957 | 61435 | hypothetical-protein-[Dehalococcoides-mccartyi] |
| WP_076004840.1 | Type4.PglZ.containing.BREX | SIGMA-HTH->DNA_pol3_beta_2+DNA_pol3_beta_3->?->?->?->?-><-ZnR<-SIGMA+OB-Fold+AAA-ATPase-BrxL<-iSTAND+PglZ<-DUF6079-p*<-GspA-ATPase<-PAPS-reduct+a+b-domain+Fer4_9+DUF4007<-HARE-HTH+E3_UFM1_ligase\|\|?->?-><-HD | 767 | Bacteria>Terrabacteria_group>Chloroflexota>Dehalococcoidia>Dehalogenimonas | GCF_001953175.1 | Dehalogenimonas_formicexedens | Dform_RS10005 | 1909335..1911638 | 1839801 | hypothetical-protein-[Dehalogenimonas-formicexedens] |
| WP_126601676.1 | Type4.PglZ.containing.BREX | ABhydrolase+ABhydrolase->zf-IS66+DDE_Tnp_IS66-><-Cysteine-Desulfurase<-DndH<-DndG<-DndF<-PAPS-reduct<-DUF6079*<-DUF4007\|\|zf-ISL3+HTH_Tnp_ISL3-><-zf-ISL3+HTH_Tnp_ISL3<-SIGMA+OB-Fold+AAA-ATPase-BrxL<-PglZ<-DUF6079<-BrxB-BrxF-iSTAND<-DUF4007\|\|SF1-SF2-Helicase+wHTH+Small-B-barrel+DUF1998-p-> | 1165 | Bacteria>Terrabacteria_group>Chloroflexota>Ktedonobacteria>Ktedonobacterales>Dictyobacteraceae | GCF_003967515.1 | Dictyobacter_aurantiacus | KTT_RS31760 | 1724003..1727500 | 1936993 | hypothetical-protein-[Dictyobacter-aurantiacus] |
| WP_126601680.1 | Type4.PglZ.containing.BREX | <-DndG<-DndF<-PAPS-reduct<-DUF6079<-DUF4007\|\|zf-ISL3+HTH_Tnp_ISL3-><-zf-ISL3+HTH_Tnp_ISL3<-SIGMA+OB-Fold+AAA-ATPase-BrxL<-PglZ<-DUF6079*<-BrxB-BrxF-iSTAND<-DUF4007\|\|SF1-SF2-Helicase+wHTH+Small-B-barrel+DUF1998-p->?-><-URI-DNase+URI-DNase<-?<-?\|\|HTH_29+HTH_32+DDE_3-> | 1322 | Bacteria>Terrabacteria_group>Chloroflexota>Ktedonobacteria>Ktedonobacterales>Dictyobacteraceae | GCF_003967515.1 | Dictyobacter_aurantiacus | KTT_RS31795 | 1737480..1741448 | 1936993 | hypothetical-protein-[Dictyobacter-aurantiacus] |
| WP_151757600.1 | Type4.PglZ.containing.BREX | NACHT+GUN4-><-?<-?<-Cysteine-Desulfurase<-DndH<-DndG-p<-DndF<-PAPS-reduct<-DUF6079<-DUF4007<-SIGMA+OB-Fold+AAA-ATPase-BrxL\|\|DDE_Tnp-><-PglZ\|\|HTH_29+HTH_32+DDE_3-><-?<-DUF6079-p<-BrxB-BrxF-iSTAND | 1165 | Bacteria>Terrabacteria_group>Chloroflexota>Ktedonobacteria>Ktedonobacterales>Dictyobacteraceae | GCF_008974265.1 | Dictyobacter_vulcani | KDW_RS19240 | 284962..288459 | 2607529 | hypothetical-protein-[Dictyobacter-vulcani] |
| WP_007916553.1 | Type4.PglZ.containing.BREX | <-OLD-ABC+TOPRIM<-TRD+TRD<-N6-METHYLASE<-REase+SF1-SF2-Helicase<-Cysteine-Desulfurase<-DndH<-DndG<-DndF<-PAPS-reduct<-DUF6079*<-DUF4007<-SIGMA+OB-Fold+AAA-ATPase-BrxL<-PglZ<-DUF6079<-BrxB-BrxF-iSTAND<-DUF4007\|\|?->RelE-ParE+SF1-SF2-Helicase-> | 1164 | Bacteria>Terrabacteria_group>Chloroflexota>Ktedonobacteria>Ktedonobacterales>Ktedonobacteraceae | GCF_000178855.1 | Ktedonobacter_racemifer | KRAC_RS37925 | 2033471..2036965 | 363277 | hypothetical-protein-[Ktedonobacter-racemifer] |
| WP_007916557.1 | Type4.PglZ.containing.BREX | <-Cysteine-Desulfurase<-DndH<-DndG<-DndF<-PAPS-reduct<-DUF6079<-DUF4007<-SIGMA+OB-Fold+AAA-ATPase-BrxL<-PglZ<-DUF6079*<-BrxB-BrxF-iSTAND<-DUF4007\|\|?->RelE-ParE+SF1-SF2-Helicase->?->HHH_4+SF1-SF2-Helicase+REase-><-?<-NERD+RNA-helicase<-SF1-SF2-Helicase+DUF1998 | 1322 | Bacteria>Terrabacteria_group>Chloroflexota>Ktedonobacteria>Ktedonobacterales>Ktedonobacteraceae | GCF_000178855.1 | Ktedonobacter_racemifer | KRAC_RS37945 | 2043720..2047688 | 363277 | hypothetical-protein-[Ktedonobacter-racemifer] |
| WP_201382002.1 | Type4.PglZ.containing.BREX | <-HerA/FtsK<-HerA/FtsK\|\|ClpABN-AAA+ClpABC-AAA->REC+OmpR-HTH-><-?\|\|ZnR->MORC+DNA_gyrase+TOPRIM->DUF4007->BrxB-BrxF-iSTAND->DUF6079*->PglZ->SIGMA+OB-Fold+AAA-ATPase-BrxL-><-Cysteine-Desulfurase<-DUF6079<-DUF4007<-PAPS-reduct+a+b-domain+Fer4_9<-?<-TnsC-ATPase<-rve+Mu-transpos_C | 1322 | Bacteria>Terrabacteria_group>Chloroflexota>Ktedonobacteria>Ktedonobacterales>Ktedonobacteraceae | GCF_016587415.1 | Ktedonobacter_sp._SOSP1-85 | KTE_RS14510 | 3478226..3482194 | 2778367 | hypothetical-protein-[Ktedonobacter-sp.-SOSP1-85] |
| WP_201382007.1 | Type4.PglZ.containing.BREX | ZnR->MORC+DNA_gyrase+TOPRIM->DUF4007->BrxB-BrxF-iSTAND->DUF6079->PglZ->SIGMA+OB-Fold+AAA-ATPase-BrxL-><-Cysteine-Desulfurase<-DUF6079*<-DUF4007<-PAPS-reduct+a+b-domain+Fer4_9<-?<-TnsC-ATPase<-rve+Mu-transpos_C<-?\|\|XIS-HTH-><-?\|\|MPTase-> | 1239 | Bacteria>Terrabacteria_group>Chloroflexota>Ktedonobacteria>Ktedonobacterales>Ktedonobacteraceae | GCF_016587415.1 | Ktedonobacter_sp._SOSP1-85 | KTE_RS14530 | 3487172..3490891 | 2778367 | hypothetical-protein-[Ktedonobacter-sp.-SOSP1-85] |
| MBP7042334.1 | Type4.PglZ.containing.BREX | <-Cysteine-Desulfurase<-SIGMA+OB-Fold+AAA-ATPase-BrxL<-PglZ<-?<-SAM-METHYLASE+HIT<-wHTH(SBD)+HNH<-DUF6079-p*<-DUF4007\|\|?->?->SIGMA-HTH-><-HicB<-HicA_toxin | 119 | Bacteria>Terrabacteria_group>Chloroflexota>unclassified_Chloroflexota | GCA_017993775.1 | Chloroflexota_bacterium | KBE23_06295 | 11926..12285 | 2026724 | hypothetical-protein-[Chloroflexota-bacterium] |
| MBP8002222.1 | Type4.PglZ.containing.BREX | <-HTH_3<-RelE-ParE\|\|DUF4007->DUF6079-p*->wHTH(SBD)+HNH->SAM-METHYLASE+HIT->DUF6079->PglZ->SIGMA+OB-Fold+AAA-ATPase-BrxL->Cysteine-Desulfurase->N6-METHYLASE->?-><-tRNA<-SF1-SF2-Helicase | 757 | Bacteria>Terrabacteria_group>Chloroflexota>unclassified_Chloroflexota | GCA_018004095.1 | Chloroflexota_bacterium | KA338_23440 | 4412..6685 | 2026724 | hypothetical-protein-[Chloroflexota-bacterium] |
| MBP8002225.1 | Type4.PglZ.containing.BREX | <-HTH_3<-RelE-ParE\|\|DUF4007->DUF6079-p->wHTH(SBD)+HNH->SAM-METHYLASE+HIT->DUF6079*->PglZ->SIGMA+OB-Fold+AAA-ATPase-BrxL->Cysteine-Desulfurase->N6-METHYLASE->?-><-tRNA<-SF1-SF2-Helicase\|\|?->?->ZnR-> | 1304 | Bacteria>Terrabacteria_group>Chloroflexota>unclassified_Chloroflexota | GCA_018004095.1 | Chloroflexota_bacterium | KA338_23455 | 8870..12784 | 2026724 | hypothetical-protein-[Chloroflexota-bacterium] |
| MBU6335171.1 | Type4.PglZ.containing.BREX | <-SIGMA+OB-Fold+AAA-ATPase-BrxL<-PglZ<-DUF6079*<-BrxB-BrxF-iSTAND<-DUF4007<-PAPS-reduct<-DUF6079<-DUF4007 | 1131 | Bacteria>Terrabacteria_group>Chloroflexota>unclassified_Chloroflexota | GCA_018969485.1 | Chloroflexota_bacterium | KGS47_12335 | 13669..17064 | 2026724 | hypothetical-protein-[Chloroflexota-bacterium] |
| WP_190709582.1 | Type4.PglZ.containing.BREX | DnaB->\|\|wHTH(SBD)+HNH->DUF4007->viral-protein->DUF6079*->PglZ->SIGMA+OB-Fold+AAA-ATPase-BrxL-><-?<-?\|\|?-><-HNH\|\|RHH_9->Phage_integrase-> | 1810 | Bacteria>Terrabacteria_group>Cyanobacteriota/Melainabacteria_group>Cyanobacteriota>Cyanophyceae>Nostocales | - | Anabaena_minutissima | H6G36_27855 | 3815..9247 | 2303152 | hypothetical-protein-[Anabaena-minutissima] |
| WP_224177031.1 | Type4.PglZ.containing.BREX | HNH->?-><-?\|\|?-><-35exo\|\|SF1-SF2-Helicase->DUF4007->viral-protein->DUF6079*->NERD+Pkinase+Pkinase->PglZ->SIGMA+OB-Fold+AAA-ATPase-BrxL-><-REase+HTH<-ABhydrolase<-?<-?\|\|?-><-CASPASE | 1801 | Bacteria>Terrabacteria_group>Cyanobacteriota/Melainabacteria_group>Cyanobacteriota>Cyanophyceae>Nostocales | - | Calothrix_sp._PCC_7716 | CAL7716_003600 | 406323..411728 | 2853227 | hypothetical-protein-[Calothrix-sp.-PCC-7716] |
| OBQ15938.1 | Type4.PglZ.containing.BREX | ABC-ATPase+ABC-ATPase->?->tRNA->?-><-SIGMA+OB-Fold+AAA-ATPase-BrxL<-PglZ<-DUF6079*<-viral-protein<-DUF4007<-?<-Glyco_transf_22<-REase | 1545 | Bacteria>Terrabacteria_group>Cyanobacteriota/Melainabacteria_group>Cyanobacteriota>Cyanophyceae>Nostocales | GCA_001672085.1 | Anabaena_sp._AL93 | AN486_21110 | 9967..14604 | 1678133 | MAG:-hypothetical-protein-AN486_21110-[Anabaena-sp.-AL93] |
| BAY31213.1 | Type4.PglZ.containing.BREX | DDE_Tnp-><-?\|\|?-><-Methyltransf_33<-Glyco_hydro_63<-?\|\|DndD-><-SIGMA+OB-Fold+AAA-ATPase-BrxL<-PglZ<-DUF6079*<-viral-protein<-DUF4007\|\|DndE->SAM-METHYLASE->?->TPR->ACET->HEAT->DDE_Tnp-> | 1534 | Bacteria>Terrabacteria_group>Cyanobacteriota/Melainabacteria_group>Cyanobacteriota>Cyanophyceae>Nostocales | GCA_002368155.1 | Nostoc_carneum_NIES-2107 | NIES2107_30720 | 3836836..3841440 | 1973483 | hypothetical-protein-NIES2107_30720-[Nostoc-carneum-NIES-2107] |
| PMB40674.1 | Type4.PglZ.containing.BREX | <-NTP_transf_6<-?\|\|?->DndE-><-DndB\|\|PAPS-reduct+Sp->DndD-><-SIGMA+OB-Fold+AAA-ATPase-BrxL<-PglZ<-DUF6079*<-viral-protein<-DUF4007\|\|DndE->SAM-METHYLASE->?->TPR->?->ACET->HEAT-> | 1539 | Bacteria>Terrabacteria_group>Cyanobacteriota/Melainabacteria_group>Cyanobacteriota>Cyanophyceae>Nostocales | GCA_002870745.1 | Fischerella_thermalis_CCMEE_5205 | CEN40_21900 | 47142..51761 | 2019671 | hypothetical-protein-CEN40_21900-[Fischerella-thermalis-CCMEE-5205] |
| MBF2068020.1 | Type4.PglZ.containing.BREX | ACET->?-><-Aminotran_1_2<-ABhydrolase+ABhydrolase<-?<-?<-?<-SIGMA+OB-Fold+AAA-ATPase-BrxL<-NERD+iPkinase+Pkinase+PglZ<-DUF6079*<-viral-protein<-DUF4007<-SF1-SF2-Helicase\|\|35exo-><-?<-Transposase\|\|?-><-ABhydrolase | 1802 | Bacteria>Terrabacteria_group>Cyanobacteriota/Melainabacteria_group>Cyanobacteriota>Cyanophyceae>Nostocales | GCA_015272395.1 | Calothrix_sp._C42_A2020_038 | IGS39_26935 | 19946..25354 | 2774333 | hypothetical-protein-[Calothrix-sp.-C42_A2020_038] |
| MBI1240662.1 | Type4.PglZ.containing.BREX | <-DndE\|\|DUF4007->viral-protein->DUF6079*->PglZ->SIGMA+OB-Fold+AAA-ATPase-BrxL-><-DndD<-PAPS-reduct+Sp\|\|DndB-> | 1543 | Bacteria>Terrabacteria_group>Cyanobacteriota/Melainabacteria_group>Cyanobacteriota>Cyanophyceae>Nostocales | GCA_016124525.1 | Nostoc_sp._RI_552 | GC195_03465 | 2339..6970 | 2654185 | hypothetical-protein-[Nostoc-sp.-RI_552] |
| MBL1198056.1 | Type4.PglZ.containing.BREX | DndE->wHTH+SAD+HNH-><-DndB\|\|PAPS-reduct+Sp-><-HTH\|\|DUF3970->DndD-><-SIGMA+OB-Fold+AAA-ATPase-BrxL<-PglZ<-DUF6079*<-viral-protein<-DUF4007\|\|DndE->SAM-METHYLASE->?->TPR->?->ACET->HEAT-> | 1539 | Bacteria>Terrabacteria_group>Cyanobacteriota/Melainabacteria_group>Cyanobacteriota>Cyanophyceae>Nostocales | GCA_016743225.1 | Nostoc_sp._GBBB01 | FWK04_02950 | 279898..284517 | 2604138 | hypothetical-protein-[Nostoc-sp.-GBBB01] |
| MBN4003526.1 | Type4.PglZ.containing.BREX | DndE->URI-DNase-><-DndB\|\|PAPS-reduct+Sp-><-HTH\|\|DUF3970->DndD-><-SIGMA+OB-Fold+AAA-ATPase-BrxL<-PglZ<-DUF6079*<-viral-protein<-DUF4007\|\|DndE->SAM-METHYLASE->?->TPR->?->ACET->HEAT-> | 1331 | Bacteria>Terrabacteria_group>Cyanobacteriota/Melainabacteria_group>Cyanobacteriota>Cyanophyceae>Nostocales | GCA_017115165.1 | Nostoc_sp._LPT | HWQ58_17650 | 103712..107707 | 2815387 | hypothetical-protein-[Nostoc-sp.-LPT] |
| MBW4641838.1 | Type4.PglZ.containing.BREX | RHH_5->HTH_3-><-DOC+HTH<-?<-?\|\|Mob_Pre-><-SAM-METHYLASE\|\|DUF4007->DUF6079*->PAPS-reduct->DUF4007->viral-protein->DUF6079->NERD+iPkinase+Pkinase+PglZ->SIGMA+OB-Fold+AAA-ATPase-BrxL->Cysteine-Desulfurase-> | 1153 | Bacteria>Terrabacteria_group>Cyanobacteriota/Melainabacteria_group>Cyanobacteriota>Cyanophyceae>Nostocales | GCA_019359495.1 | Goleter_apudmare_HA4340-LM2 | KME23_02240 | 23265..26726 | 1382955 | hypothetical-protein-[Goleter-apudmare-HA4340-LM2] |
| MBW4641842.1 | Type4.PglZ.containing.BREX | Mob_Pre-><-SAM-METHYLASE\|\|DUF4007->DUF6079->PAPS-reduct->DUF4007->viral-protein->DUF6079*->NERD+iPkinase+Pkinase+PglZ->SIGMA+OB-Fold+AAA-ATPase-BrxL->Cysteine-Desulfurase-> | 1765 | Bacteria>Terrabacteria_group>Cyanobacteriota/Melainabacteria_group>Cyanobacteriota>Cyanophyceae>Nostocales | GCA_019359495.1 | Goleter_apudmare_HA4340-LM2 | KME23_02260 | 29054..34351 | 1382955 | hypothetical-protein-[Goleter-apudmare-HA4340-LM2] |
| WP_016868044.1 | Type4.PglZ.containing.BREX | <-TPR<-?<-?<-SAM-METHYLASE<-?<-DndE\|\|DUF4007->viral-protein->?->DUF6079*->PglZ->SIGMA+OB-Fold+AAA-ATPase-BrxL-><-?<-DndD<-DUF3970\|\|HTH-><-PAPS-reduct+Sp\|\|DndB-><-DndE | 1540 | Bacteria>Terrabacteria_group>Cyanobacteriota/Melainabacteria_group>Cyanobacteriota>Cyanophyceae>Nostocales | GCF_000317205.1 | Fischerella | UYI_RS0118020 | 44597..49219 | 1190 | MULTISPECIES:-hypothetical-protein-[Fischerella] |

<table>
<tr><th>Protein-ID</th><th>System</th><th>Gene-Neighbourhoods</th><th>Length</th><th>Taxonomy</th><th>Assembly-ID</th><th>Strain</th><th>Locus-Tag</th><th>Genomic-coordinates</th><th>TaxID</th><th>Fasta-define</th></tr>
<tr><td>WP_137986287.1</td><td>Type4.PglZ.containing.BREX</td><td><-Glycos_transf||?->?-><-MerR-HTH<-DSBH<-SIGMA+OB-Fold+AAA-ATPase-BrxL||?->?-><-NERD+iPkinase+Pkinase+PglZ<-DUF6079*<-viral-protein||?-><-DUF4007<-?||Peptidase_M48+TerB-><-DUF4007<-VapB_antitoxin<-Cysteine-Desulfurase<-ORC-CDC6</td><td>1796</td><td>Bacteria>Terrabacteria_group>Cyanobacteriota/Melainabacteria_group>Cyanobacteriota>Cyanophyceae>Nostocales</td><td>GCF_000760695.4</td><td>Tolypothrix_bouteillei</td><td>HC643_RS14195</td><td>3587892..3593282</td><td>1246981</td><td>hypothetical-protein-[Tolypothrix-bouteillei]</td></tr>
<tr><td>WP_053538156.1</td><td>Type4.PglZ.containing.BREX</td><td>ABC-ATPase+ABC-ATPase->?->tRNA->?-><-BrxL-p<-Lon_2<-PglZ<-DUF6079*<-viral-protein<-DUF4007<-?<-Glyco_transf_22<-REase||Glyco_transf-></td><td>1325</td><td>Bacteria>Terrabacteria_group>Cyanobacteriota/Melainabacteria_group>Cyanobacteriota>Cyanophyceae>Nostocales</td><td>GCF_001277295.1</td><td>Anabaena_sp._WA102</td><td>AA650_RS04645</td><td>1131316..1135293</td><td>1647413</td><td>hypothetical-protein-[Anabaena-sp.-WA102]</td></tr>
<tr><td>WP_062244415.1</td><td>Type4.PglZ.containing.BREX</td><td><-HEAT<-ACET<-?<-TPR<-?<-SAM-METHYLASE<-DndE||DUF4007->viral-protein->DUF6079*->PglZ->SIGMA+OB-Fold+AAA-ATPase-BrxL-><-DndD<-PAPS-reduct+Sp||DndB->?-><-DndE<-REase</td><td>1541</td><td>Bacteria>Terrabacteria_group>Cyanobacteriota/Melainabacteria_group>Cyanobacteriota>Cyanophyceae>Nostocales</td><td>GCF_001548455.1</td><td>Fischerella_sp._NIES-3754</td><td>FIS3754_RS04725</td><td>1081097..1085722</td><td>1752063</td><td>hypothetical-protein-[Fischerella-sp.-NIES-3754]</td></tr>
<tr><td>WP_073556530.1</td><td>Type4.PglZ.containing.BREX</td><td>DndE-><-DndB||PAPS-reduct+Sp->DndD-><-SIGMA+OB-Fold+AAA-ATPase-BrxL<-PglZ<-DUF6079*<-viral-protein<-DUF4007||DndE->SAM-METHYLASE->?->TPR->?->ACET->HEAT-></td><td>1542</td><td>Bacteria>Terrabacteria_group>Cyanobacteriota/Melainabacteria_group>Cyanobacteriota>Cyanophyceae>Nostocales</td><td>GCF_001904645.1</td><td>Fischerella_major</td><td>NIES592_RS18590</td><td>35296..39924</td><td>210993</td><td>hypothetical-protein-[Fischerella-major]</td></tr>
<tr><td>WP_086759833.1</td><td>Type4.PglZ.containing.BREX</td><td><-MerR-HTH<-Cysteine-Desulfurase<-SIGMA+OB-Fold+AAA-ATPase-BrxL<-PglZ<-DUF6079*<-viral-protein<-DUF4007<-ParB+HNH<-PAPS-reduct<-DUF6079<-DUF4007||SAM-METHYLASE->?-><-METHYLASE</td><td>1815</td><td>Bacteria>Terrabacteria_group>Cyanobacteriota/Melainabacteria_group>Cyanobacteriota>Cyanophyceae>Nostocales</td><td>GCF_002154725.1</td><td>Nostoc_sp._106C</td><td>BV375_RS09695</td><td>43659..49106</td><td>1932667</td><td>hypothetical-protein-[Nostoc-sp.-106C]</td></tr>
<tr><td>WP_157750035.1</td><td>Type4.PglZ.containing.BREX</td><td><-AA_kinase+ACT+ACT_7||?->?->?->ABC-ATPase+ABC-ATPase->?->tRNA->?-><-SIGMA+OB-Fold+AAA-ATPase-BrxL<-PglZ<-DUF6079*<-viral-protein<-DUF4007<-?<-Glyco_transf_22<-REase||Glyco_transf-></td><td>1566</td><td>Bacteria>Terrabacteria_group>Cyanobacteriota/Melainabacteria_group>Cyanobacteriota>Cyanophyceae>Nostocales</td><td>GCF_002368115.1</td><td>Dolichospermum_compactum</td><td>CA730_RS20995</td><td>4516895..4521595</td><td>136073</td><td>hypothetical-protein-[Dolichospermum-compactum]</td></tr>
<tr><td>WP_096587560.1</td><td>Type4.PglZ.containing.BREX</td><td>REase->?->?-><-HTH_32+DDE_3||DndD-><-SIGMA+OB-Fold+AAA-ATPase-BrxL<-PglZ<-DUF6079*<-viral-protein<-DUF4007||DndE->SAM-METHYLASE-><-HAMP+PAS+HISKIN<-?||?-><-REase</td><td>1535</td><td>Bacteria>Terrabacteria_group>Cyanobacteriota/Melainabacteria_group>Cyanobacteriota>Cyanophyceae>Nostocales</td><td>GCF_002368175.1</td><td>Calothrix_sp._NIES-2098</td><td>CA733_RS06225</td><td>1488383..1492990</td><td>1954171</td><td>hypothetical-protein-[Calothrix-sp.-NIES-2098]</td></tr>
<tr><td>WP_096691989.1</td><td>Type4.PglZ.containing.BREX</td><td>Peptidase_S51-><-?||RelE-ParE->?->||<-?<-HTH||HNH-><-SIGMA+OB-Fold+AAA-ATPase-BrxL<-NERD+iPkinase+Pkinase+PglZ<-DUF6079*<-viral-protein<-DUF4007<-SF1-SF2-Helicase||35exo-><-MerR-HTH||N6-METHYLASE-></td><td>1799</td><td>Bacteria>Terrabacteria_group>Cyanobacteriota/Melainabacteria_group>Cyanobacteriota>Cyanophyceae>Nostocales</td><td>GCF_002368455.1</td><td>unclassified_Calothrix</td><td>CDC41_RS40685</td><td>9230987..9236386</td><td>2619626</td><td>MULTISPECIES:-hypothetical-protein-[unclassified-Calothrix]</td></tr>
<tr><td>WP_102182916.1</td><td>Type4.PglZ.containing.BREX</td><td>DndE-><-DndB||PAPS-reduct+Sp->DndD-><-SIGMA+OB-Fold+AAA-ATPase-BrxL<-PglZ<-DUF6079*<-?<-viral-protein<-DUF4007||DndE->SAM-METHYLASE-></td><td>1539</td><td>Bacteria>Terrabacteria_group>Cyanobacteriota/Melainabacteria_group>Cyanobacteriota>Cyanophyceae>Nostocales</td><td>GCF_002870675.1</td><td>Fischerella_thermalis</td><td>CEN46_RS20455</td><td>50195..54814</td><td>372787</td><td>hypothetical-protein-[Fischerella-thermalis]</td></tr>
<tr><td>WP_102206594.1</td><td>Type4.PglZ.containing.BREX</td><td>REase->DndE-><-?<-DndB||PAPS-reduct+Sp->DndD-><-SIGMA+OB-Fold+AAA-ATPase-BrxL<-PglZ<-DUF6079*<-viral-protein<-DUF4007||DndE->SAM-METHYLASE-></td><td>1541</td><td>Bacteria>Terrabacteria_group>Cyanobacteriota/Melainabacteria_group>Cyanobacteriota>Cyanophyceae>Nostocales</td><td>GCF_002870725.1</td><td>Fischerella_thermalis</td><td>CEN41_RS06915</td><td>14951..19576</td><td>372787</td><td>hypothetical-protein-[Fischerella-thermalis]</td></tr>
<tr><td>WP_168492336.1</td><td>Type4.PglZ.containing.BREX</td><td><-URI-DNase<-HEPN<-TRD+TRD<-N6-METHYLASE<-Peptidase_M17_N+Peptidase_M17||?-><-?<-SIGMA+OB-Fold+AAA-ATPase-BrxL<-PglZ<-DUF6079*<-viral-protein<-DUF4007<-Glyco_transf_22||?->METHYLASE->?->?->CBS+CBS-></td><td>1565</td><td>Bacteria>Terrabacteria_group>Cyanobacteriota/Melainabacteria_group>Cyanobacteriota>Cyanophyceae>Nostocales</td><td>GCF_009711975.1</td><td>Anabaena_sp._UHCC_0204</td><td>FJR04_RS04930</td><td>130875..135572</td><td>2590009</td><td>hypothetical-protein-[Anabaena-sp.-UHCC-0204]</td></tr>
<tr><td>WP_168535727.1</td><td>Type4.PglZ.containing.BREX</td><td><-CBS+CBS<-?<-?<-METHYLASE<-?||Glyco_transf_22->DUF4007->viral-protein->DUF6079-p->DUF6079-p*->PglZ->SIGMA+OB-Fold+AAA-ATPase-BrxL->?-><-?||Peptidase_M17_N+Peptidase_M17->N6-METHYLASE->TRD+TRD->HEPN->URI-DNase-></td><td>701</td><td>Bacteria>Terrabacteria_group>Cyanobacteriota/Melainabacteria_group>Cyanobacteriota>Cyanophyceae>Nostocales</td><td>GCF_009712085.1</td><td>Anabaena_sp._UHCC_0253</td><td>FJR38_RS04700</td><td>127202..129307</td><td>2590019</td><td>hypothetical-protein-[Anabaena-sp.-UHCC-0253]</td></tr>
<tr><td>WP_155748747.1</td><td>Type4.PglZ.containing.BREX</td><td><-ParA-Soj-PloopNTPase||?->Glycos_transf->Glycos_transf->Glycos_transf-><-?||MetJarc-HTH->?->viral-protein->DUF6079*->NERD+iPkinase+Pkinase+PglZ-><-DOC+HTH||SIGMA+OB-Fold+AAA-ATPase-BrxL->?-><-?<-?||MerR-HTH-></td><td>1592</td><td>Bacteria>Terrabacteria_group>Cyanobacteriota/Melainabacteria_group>Cyanobacteriota>Cyanophyceae>Nostocales</td><td>GCF_009725235.1</td><td>Scytonema_sp._UIC_10036</td><td>F7734_RS32875</td><td>19370..24148</td><td>2304196</td><td>hypothetical-protein-[Scytonema-sp.-UIC-10036]</td></tr>
<tr><td>WP_190936010.1</td><td>Type4.PglZ.containing.BREX</td><td><-HEAT<-ACET<-?<-TPR<-?<-SAM-METHYLASE<-DndE||DUF4007->viral-protein->DUF6079*->PglZ->SIGMA+OB-Fold+AAA-ATPase-BrxL-><-DndD<-REase<-PAPS-reduct+Sp||DndB-><-wHTH+SAD+HNH<-DndE<-REase</td><td>1439</td><td>Bacteria>Terrabacteria_group>Cyanobacteriota/Melainabacteria_group>Cyanobacteriota>Cyanophyceae>Nostocales</td><td>GCF_014698055.1</td><td>Nostoc_sp._FACHB-133</td><td>H6G86_RS21500</td><td>24399..28718</td><td>2692835</td><td>hypothetical-protein-[Nostoc-sp.-FACHB-133]</td></tr>
<tr><td>WP_190950967.1</td><td>Type4.PglZ.containing.BREX</td><td><-HEAT<-ACET<-?<-TPR<-?<-SAM-METHYLASE<-DndE||DUF4007->viral-protein->DUF6079*->PglZ->SIGMA+OB-Fold+AAA-ATPase-BrxL-><-DndD<-DUF3970||HTH-><-PAPS-reduct+Sp||DndB-><-wHTH+SAD+HNH<-DndE</td><td>1539</td><td>Bacteria>Terrabacteria_group>Cyanobacteriota/Melainabacteria_group>Cyanobacteriota>Cyanophyceae>Nostocales</td><td>GCF_014698495.1</td><td>Nostoc_punctiforme</td><td>H6G94_RS21265</td><td>186521..191140</td><td>272131</td><td>hypothetical-protein-[Nostoc-punctiforme]</td></tr>
<tr><td>WP_190826790.1</td><td>Type4.PglZ.containing.BREX</td><td><-SAM-METHYLASE<-Glyoxalase||MqsA_antitoxin+DUF4065->wHTH->?->DUF4007->viral-protein->DUF6079*->PglZ->SIGMA+OB-Fold+AAA-ATPase-BrxL->?-><-?<-?<-?||<-HNH</td><td>1547</td><td>Bacteria>Terrabacteria_group>Cyanobacteriota/Melainabacteria_group>Cyanobacteriota>Cyanophyceae>Nostocales</td><td>GCF_014698965.1</td><td>Iningainema_tapete</td><td>ICL16_RS09725</td><td>10324..14967</td><td>2806730</td><td>hypothetical-protein-[Iningainema-tapete]</td></tr>
<tr><td>WP_260504429.1</td><td>Type4.PglZ.containing.BREX</td><td><-DNA_METHYLASE<-?<-?||?->?->METHYLASE-><-?<-SAM-METHYLASE||DUF4007->DUF6079-p*-><-?||wHTH(SBD)+HNH-><-?||DndB->MORC->URI-DNase-><-SIGMA+OB-Fold+AAA-ATPase-BrxL<-PglZ<-DUF6079</td><td>931</td><td>Bacteria>Terrabacteria_group>Cyanobacteriota/Melainabacteria_group>Cyanobacteriota>Cyanophyceae>Nostocales</td><td>GCF_019734875.1</td><td>Desmonostoc_muscorum</td><td>H1Q63_RS02880</td><td>25424..28219</td><td>1179</td><td>hypothetical-protein-[Desmonostoc-muscorum]</td></tr>
<tr><td>WP_221704725.1</td><td>Type4.PglZ.containing.BREX</td><td>DUF6079-p-><-?||wHTH(SBD)+HNH-><-?||DndB->MORC->URI-DNase-><-SIGMA+OB-Fold+AAA-ATPase-BrxL<-PglZ<-DUF6079*<-BrxB-BrxF-iSTAND<-DUF4007||Cysteine-Desulfurase->?->ACET->Beta-lactamase-><-?<-Glyoxalase<-Glyoxalase</td><td>1303</td><td>Bacteria>Terrabacteria_group>Cyanobacteriota/Melainabacteria_group>Cyanobacteriota>Cyanophyceae>Nostocales</td><td>GCF_019734875.1</td><td>Desmonostoc_muscorum</td><td>H1Q63_RS02925</td><td>38336..42247</td><td>1179</td><td>hypothetical-protein-[Desmonostoc-muscorum]</td></tr>
<tr><td>WP_224094424.1</td><td>Type4.PglZ.containing.BREX</td><td>DDE_Tnp-><-tRNA<-tRNA<-?<-REC+HISKIN+MORC<-SHELIX+HISKIN+REC<-?<-Methyltransf_33||DndD-><-SIGMA+OB-Fold+AAA-ATPase-BrxL<-PglZ<-DUF6079*<-viral-protein<-DUF4007||DndE->SAM-METHYLASE-></td><td>1542</td><td>Bacteria>Terrabacteria_group>Cyanobacteriota/Melainabacteria_group>Cyanobacteriota>Cyanophyceae>Nostocales</td><td>GCF_019976755.1</td><td>Nostoc_sp._MS1</td><td>NSMS1_RS14245</td><td>3290873..3295501</td><td>2764711</td><td>hypothetical-protein-[Nostoc-sp.-MS1]</td></tr>
<tr><td>WP_236141666.1</td><td>Type4.PglZ.containing.BREX</td><td><-Cysteine-Desulfurase-p<-Cysteine-Desulfurase-p<-SIGMA+OB-Fold+AAA-ATPase-BrxL<-NERD+iPkinase+Pkinase+PglZ<-DUF6079*<-viral-protein<-DUF4007<-PAPS-reduct<-DUF6079<-DUF4007<-?||HTH_36-><-Phage_integrase+Phage_integrase<-Phage_integrase+Phage_integrase</td><td>1804</td><td>Bacteria>Terrabacteria_group>Cyanobacteriota/Melainabacteria_group>Cyanobacteriota>Cyanophyceae>Nostocales</td><td>GCF_021596505.1</td><td>Nostoc_sp._CMAA1605</td><td>CV014_RS08620</td><td>8960..14374</td><td>2055159</td><td>hypothetical-protein-[Nostoc-sp.-CMAA1605]</td></tr>
<tr><td>WP_236141670.1</td><td>Type4.PglZ.containing.BREX</td><td><-Cysteine-Desulfurase-p<-Cysteine-Desulfurase-p<-SIGMA+OB-Fold+AAA-ATPase-BrxL<-NERD+iPkinase+Pkinase+PglZ<-DUF6079<-viral-protein<-DUF4007<-PAPS-reduct<-DUF6079*<-DUF4007<-?||HTH_36-><-Phage_integrase+Phage_integrase<-Phage_integrase+Phage_integrase||<-?<-?||<-AAA-ATPase</td><td>1168</td><td>Bacteria>Terrabacteria_group>Cyanobacteriota/Melainabacteria_group>Cyanobacteriota>Cyanophyceae>Nostocales</td><td>GCF_021596505.1</td><td>Nostoc_sp._CMAA1605</td><td>CV014_RS08640</td><td>16536..20042</td><td>2055159</td><td>hypothetical-protein-[Nostoc-sp.-CMAA1605]</td></tr>
<tr><td>WP_251958553.1</td><td>Type4.PglZ.containing.BREX</td><td><-HEAT<-ACET<-?<-TPR<-?<-SAM-METHYLASE<-DndE||DUF4007->viral-protein->DUF6079-p*->?->?->NERD+Pkinase+Pkinase->PglZ-p->PglZ->SIGMA+OB-Fold+AAA-ATPase-BrxL-><-REase<-PAPS-reduct+Sp||DndB-></td><td>936</td><td>Bacteria>Terrabacteria_group>Cyanobacteriota/Melainabacteria_group>Cyanobacteriota>Cyanophyceae>Nostocales</td><td>GCF_023734775.1</td><td>Nostoc_commune</td><td>ANS036C_RS04395</td><td>979875..982685</td><td>1178</td><td>hypothetical-protein-[Nostoc-commune]</td></tr>
<tr><td>WP_086759843.1</td><td>Type4.PglZ.containing.BREX</td><td><-MerR-HTH<-Cysteine-Desulfurase<-SIGMA+OB-Fold+AAA-ATPase-BrxL<-PglZ<-DUF6079<-viral-protein<-DUF4007<-ParB+HNH<-PAPS-reduct<-DUF6079*<-DUF4007||SAM-METHYLASE->?-><-METHYLASE<-?<-?||?->?->DNA_METHYLASE-></td><td>1170</td><td>Bacteria>Terrabacteria_group>Cyanobacteriota/Melainabacteria_group>Cyanobacteriota>Cyanophyceae>Nostocales</td><td>-</td><td>Nostoc_sp._106C</td><td>BV375_RS09720</td><td>53554..57066</td><td>1932667</td><td>hypothetical-protein-[Nostoc-sp.-106C]</td></tr>
<tr><td>WP_206265997.1</td><td>Type4.PglZ.containing.BREX</td><td><-HEAT<-ACET<-?<-TPR<-?<-SAM-METHYLASE<-DndE||DUF4007->viral-protein->DUF6079*->PglZ->SIGMA+OB-Fold+AAA-ATPase-BrxL-><-PemK_toxin<-?<-DndD<-Y1_Tnp<-DUF3970||HTH-></td><td>1529</td><td>Bacteria>Terrabacteria_group>Cyanobacteriota/Melainabacteria_group>Cyanobacteriota>Cyanophyceae>Nostocales</td><td>-</td><td>Nostoc_sp._UHCC_0702</td><td>JYQ62_02970</td><td>615044..619633</td><td>2814655</td><td>hypothetical-protein-[Nostoc-sp.-UHCC-0702]</td></tr>
<tr><td>WP_223980148.1</td><td>Type4.PglZ.containing.BREX</td><td>HNH->?-><-?||?-><-35exo||SF1-SF2-Helicase->DUF4007->viral-protein->DUF6079*->NERD+iPkinase+Pkinase+PglZ->SIGMA+OB-Fold+AAA-ATPase-BrxL-><-REase+HTH<-ABhydrolase<-?<-?||?-><-CASPASE</td><td>1805</td><td>Bacteria>Terrabacteria_group>Cyanobacteriota/Melainabacteria_group>Cyanobacteriota>Cyanophyceae>Nostocales</td><td>-</td><td>Rivularia_sp._IAM_M-261</td><td>RIVM261_050350</td><td>43225..48642</td><td>448371</td><td>hypothetical-protein-[Rivularia-sp.-IAM-M-261]</td></tr>
<tr><td>AFZ48539.1</td><td>Type4.PglZ.containing.BREX</td><td>ABhydrolase->?-><-?<-Cysteine-Desulfurase||VapB_antitoxin->PIN-><-SIGMA+OB-Fold+AAA-ATPase-BrxL<-PglZ<-DUF6079*<-viral-protein<-DUF4007<-PAPS-reduct<-wHTH(SBD)+HNH<-DUF6079<-DUF4007</td><td>1565</td><td>Bacteria>Terrabacteria_group>Cyanobacteriota/Melainabacteria_group>Cyanobacteriota>Cyanophyceae>Oscillatoriophycideae</td><td>GCA_000317655.1</td><td>Cyanobacterium_stanieri_PCC_7202</td><td>Cyast_2596</td><td>2851603..2856300</td><td>292563</td><td>hypothetical-protein-Cyast_2596-[Cyanobacterium-stanieri-PCC-7202]</td></tr>
<tr><td>AFZ48544.1</td><td>Type4.PglZ.containing.BREX</td><td>VapB_antitoxin->PIN-><-SIGMA+OB-Fold+AAA-ATPase-BrxL<-PglZ<-DUF6079<-viral-protein<-DUF4007<-PAPS-reduct<-wHTH(SBD)+HNH<-DUF6079*<-DUF4007<-?||?->?->?->?->?->ABC-ATPase+SbcC-><-TPR+TPR</td><td>1165</td><td>Bacteria>Terrabacteria_group>Cyanobacteriota/Melainabacteria_group>Cyanobacteriota>Cyanophyceae>Oscillatoriophycideae</td><td>GCA_000317655.1</td><td>Cyanobacterium_stanieri_PCC_7202</td><td>Cyast_2601</td><td>2859880..2863377</td><td>292563</td><td>hypothetical-protein-Cyast_2601-[Cyanobacterium-stanieri-PCC-7202]</td></tr>
<tr><td>KPQ40624.1</td><td>Type4.PglZ.containing.BREX</td><td><-ParB||?->?->ParD_antitoxin->REase->?->PIN->DUF4007->DUF6079*->PAPS-reduct->ParB+HNH->DUF4007->viral-protein->DUF6079->PglZ->SIGMA+OB-Fold+AAA-ATPase-BrxL->?-><-HEPN</td><td>1162</td><td>Bacteria>Terrabacteria_group>Cyanobacteriota/Melainabacteria_group>Cyanobacteriota>Cyanophyceae>Oscillatoriophycideae</td><td>GCA_001314905.1</td><td>Phormidium_sp._OSCR</td><td>HLUCC016_02250</td><td>106799..110287</td><td>1666905</td><td>hypothetical-protein-HLUCC016_02250-[Phormidium-sp.-OSCR]</td></tr>
<tr><td>KPQ40629.1</td><td>Type4.PglZ.containing.BREX</td><td>REase->?->PIN->DUF4007->DUF6079->PAPS-reduct->ParB+HNH->DUF4007->viral-protein->DUF6079*->PglZ->SIGMA+OB-Fold+AAA-ATPase-BrxL->?-><-HEPN<-PolB||Cysteine-Desulfurase-></td><td>1552</td><td>Bacteria>Terrabacteria_group>Cyanobacteriota/Melainabacteria_group>Cyanobacteriota>Cyanophyceae>Oscillatoriophycideae</td><td>GCA_001314905.1</td><td>Phormidium_sp._OSCR</td><td>HLUCC016_02275</td><td>114944..119602</td><td>1666905</td><td>Protein-of-unknown-function-(DUF3584)-[Phormidium-sp.-OSCR]</td></tr>
<tr><td>OAB59992.1</td><td>Type4.PglZ.containing.BREX</td><td>ParD_antitoxin->REase->?->PIN->DUF4007->DUF6079*->PAPS-reduct->ParB+HNH->DUF4007->viral-protein->DUF6079->PglZ->SIGMA+OB-Fold+AAA-ATPase-BrxL-><-PolB||Cysteine-Desulfurase-></td><td>1161</td><td>Bacteria>Terrabacteria_group>Cyanobacteriota/Melainabacteria_group>Cyanobacteriota>Cyanophyceae>Oscillatoriophycideae</td><td>GCA_001637315.1</td><td>Phormidium_willei_BDU_130791</td><td>AY600_15255</td><td>4542..8027</td><td>399101</td><td>hypothetical-protein-AY600_15255-[Phormidium-willei-BDU-130791]</td></tr>
<tr><td>OAB59997.1</td><td>Type4.PglZ.containing.BREX</td><td>REase->?->PIN->DUF4007->DUF6079->PAPS-reduct->ParB+HNH->DUF4007->viral-protein->DUF6079*->PglZ->SIGMA+OB-Fold+AAA-ATPase-BrxL-><-PolB||Cysteine-Desulfurase->?->?->?->PIN-><-SF1-SF2-Helicase</td><td>1555</td><td>Bacteria>Terrabacteria_group>Cyanobacteriota/Melainabacteria_group>Cyanobacteriota>Cyanophyceae>Oscillatoriophycideae</td><td>GCA_001637315.1</td><td>Phormidium_willei_BDU_130791</td><td>AY600_15280</td><td>12637..17304</td><td>399101</td><td>hypothetical-protein-AY600_15280-[Phormidium-willei-BDU-130791]</td></tr>
<tr><td>OIP72761.1</td><td>Type4.PglZ.containing.BREX</td><td>DNA_METHYLASE-><-DUF29<-?||?-><-?||SPOUT->?-><-SIGMA+OB-Fold+AAA-ATPase-BrxL<-PglZ<-DUF6079*<-viral-protein<-DUF4007<-?<-?<-AbrB-HTH<-?||?->SIR2->ChlI+Lon_C+AAA-></td><td>1442</td><td>Bacteria>Terrabacteria_group>Cyanobacteriota/Melainabacteria_group>Cyanobacteriota>Cyanophyceae>Oscillatoriophycideae</td><td>GCA_001873365.1</td><td>Oscillatoriales_cyanobacterium_CG2_30_40_61</td><td>AUK43_02005</td><td>14388..18716</td><td>1805291</td><td>hypothetical-protein-AUK43_02005-[Oscillatoriales-cyanobacterium-CG2_30_40_61]</td></tr>
<tr><td>MCC3434391.1</td><td>Type4.PglZ.containing.BREX</td><td><-TraI-VirD2-RCRE<-TraJ-RHH<-?||zf-ISL3+HTH_Tnp_ISL3+DDE_Tnp_ISL3-><-?<-Cysteine-Desulfurase||DUF4007->BrxB-BrxF-iSTAND->DUF6079*->PglZ->SIGMA+OB-Fold+AAA-ATPase-BrxL-><-ParB+HNH<-PAPS-reduct<-DOC+HTH<-DUF6079<-DUF4007<-?||NACHT+GUN4-></td><td>1316</td><td>Bacteria>Terrabacteria_group>Cyanobacteriota/Melainabacteria_group>Cyanobacteriota>Cyanophyceae>Oscillatoriophycideae</td><td>GCA_020738785.1</td><td>Microcoleus_sp._PH2017_05_CCC_O_A</td><td>JGK20_02525</td><td>12906..16856</td><td>2798816</td><td>hypothetical-protein-[Microcoleus-sp.-PH2017_05_CCC_O_A]</td></tr>
<tr><td>MCC3434397.1</td><td>Type4.PglZ.containing.BREX</td><td><-Cysteine-Desulfurase||DUF4007->BrxB-BrxF-iSTAND->DUF6079->PglZ->SIGMA+OB-Fold+AAA-ATPase-BrxL-><-ParB+HNH<-PAPS-reduct<-DOC+HTH<-DUF6079*<-DUF4007<-?||NACHT+GUN4-><-McrC-NTD+REase<-HTH+McrB-AAA<-ASCH<-ACET+PIN+ACET||Phage_integrase-><-CopG/RHH</td><td>1175</td><td>Bacteria>Terrabacteria_group>Cyanobacteriota/Melainabacteria_group>Cyanobacteriota>Cyanophyceae>Oscillatoriophycideae</td><td>GCA_020738785.1</td><td>Microcoleus_sp._PH2017_05_CCC_O_A</td><td>JGK20_02555</td><td>25251..28778</td><td>2798816</td><td>hypothetical-protein-[Microcoleus-sp.-PH2017_05_CCC_O_A]</td></tr>
<tr><td>MCP9293151.1</td><td>Type4.PglZ.containing.BREX</td><td><-ChlI+Lon_C+AAA<-?||?-><-?||AbrB-HTH->DUF4007->viral-protein->DUF6079*->PglZ->SIGMA+OB-Fold+AAA-ATPase-BrxL->REase-><-?<-SPOUT</td><td>1438</td><td>Bacteria>Terrabacteria_group>Cyanobacteriota/Melainabacteria_group>Cyanobacteriota>Cyanophyceae>Oscillatoriophycideae</td><td>GCA_024320855.1</td><td>Planktothrix_agardhii_LY1</td><td>NDM07_00465</td><td>8150..12466</td><td>2950590</td><td>hypothetical-protein-[Planktothrix-agardhii-LY1]</td></tr>
<tr><td>WP_235767360.1</td><td>Type4.PglZ.containing.BREX</td><td>AbrB-HTH->DUF4007->DUF6079-p->DUF6079*->PglZ->SIGMA+OB-Fold+AAA-ATPase-BrxL->REase-><-?<-SPOUT||?->?->||?->DUF29-></td><td>1177</td><td>Bacteria>Terrabacteria_group>Cyanobacteriota/Melainabacteria_group>Cyanobacteriota>Cyanophyceae>Oscillatoriophycideae</td><td>GCA_030820635.1</td><td>Planktothrix_agardhii</td><td>L2E80_RS19275</td><td>4094890..4098423</td><td>1160</td><td>hypothetical-protein-[Planktothrix-agardhii]</td></tr>
<tr><td>CAD5929387.1</td><td>Type4.PglZ.containing.BREX</td><td><-DUF29<-?||?-><-?||SPOUT->?-><-REase<-SIGMA+OB-Fold+AAA-ATPase-BrxL<-PglZ<-DUF6079-p*<-HTH_Tnp_4+DDE_Tnp_4<-DUF6079-p<-viral-protein<-DUF4007<-AbrB-HTH||?-><-?||?->ChlI+Lon_C+AAA-></td><td>704</td><td>Bacteria>Terrabacteria_group>Cyanobacteriota/Melainabacteria_group>Cyanobacteriota>Cyanophyceae>Oscillatoriophycideae</td><td>GCA_904830855.1</td><td>Planktothrix_agardhii</td><td>PANO66_01242</td><td>1443350..1445464</td><td>1160</td><td>putative-DNA-double-strand-break-repair-Rad50-ATPase-[Planktothrix-agardhii]</td></tr>
<tr><td>WP_006100228.1</td><td>Type4.PglZ.containing.BREX</td><td><-ABC-ATPase+SbcC<-?<-tRNA||?-><-?<-?<-ACET+ACET<-ACET||<-ACET+ACET||DUF4007->DUF6079*->PAPS-reduct->ParB+HNH->DUF4007->viral-protein->DUF6079->PglZ->SIGMA+OB-Fold+AAA-ATPase-BrxL->Cysteine-Desulfurase-></td><td>1187</td><td>Bacteria>Terrabacteria_group>Cyanobacteriota/Melainabacteria_group>Cyanobacteriota>Cyanophyceae>Oscillatoriophycideae</td><td>GCF_000155555.1</td><td>Coleofasciculus_chthonoplastes</td><td>MC7420_RS09345</td><td>54832..58395</td><td>64178</td><td>hypothetical-protein-[Coleofasciculus-chthonoplastes]</td></tr>
<tr><td>WP_006100246.1</td><td>Type4.PglZ.containing.BREX</td><td><-ACET+ACET<-ACET||<-ACET+ACET||DUF4007->DUF6079->PAPS-reduct->ParB+HNH->DUF4007->viral-protein->DUF6079*->PglZ->SIGMA+OB-Fold+AAA-ATPase-BrxL->Cysteine-Desulfurase-></td><td>1430</td><td>Bacteria>Terrabacteria_group>Cyanobacteriota/Melainabacteria_group>Cyanobacteriota>Cyanophyceae>Oscillatoriophycideae</td><td>GCF_000155555.1</td><td>Coleofasciculus_chthonoplastes</td><td>MC7420_RS09370</td><td>63187..67479</td><td>64178</td><td>hypothetical-protein-[Coleofasciculus-chthonoplastes]</td></tr>
<tr><td>WP_015181437.1</td><td>Type4.PglZ.containing.BREX</td><td>REase-><-Glycos_transf<-?||MORC+DNA_gyrase+TOPRIM->DUF4007->viral-protein->DUF6079*->PglZ->SIGMA+OB-Fold+AAA-ATPase-BrxL-></td><td>1797</td><td>Bacteria>Terrabacteria_group>Cyanobacteriota/Melainabacteria_group>Cyanobacteriota>Cyanophyceae>Oscillatoriophycideae</td><td>GCF_000317515.1</td><td>Allocoleopsis_franciscana</td><td>MIC7113_RS06770</td><td>1598027..1603420</td><td>2886352</td><td>hypothetical-protein-[Allocoleopsis-franciscana]</td></tr>
<tr><td>WP_027249366.1</td><td>Type4.PglZ.containing.BREX</td><td><-ChlI+Lon_C+AAA<-?||?->AbrB-HTH->DUF4007->viral-protein->DUF6079*->PglZ->SIGMA+OB-Fold+AAA-ATPase-BrxL->REase-><-?<-SPOUT||?-><-?||?->DUF29-></td><td>1447</td><td>Bacteria>Terrabacteria_group>Cyanobacteriota/Melainabacteria_group>Cyanobacteriota>Cyanophyceae>Oscillatoriophycideae</td><td>GCF_000464665.1</td><td>Planktothrix_agardhii</td><td>N318_RS0111070</td><td>2475843..2480186</td><td>1160</td><td>hypothetical-protein-[Planktothrix-agardhii]</td></tr>
<tr><td>WP_026793814.1</td><td>Type4.PglZ.containing.BREX</td><td>SPOUT->?-><-REase<-SIGMA+OB-Fold+AAA-ATPase-BrxL<-PglZ<-DUF6079-p<-DUF6079-p*<-viral-protein<-DUF4007<-AbrB-HTH<-?||?->ChlI+Lon_C+AAA-></td><td>920</td><td>Bacteria>Terrabacteria_group>Cyanobacteriota/Melainabacteria_group>Cyanobacteriota>Cyanophyceae>Oscillatoriophycideae</td><td>GCF_000464725.1</td><td>Planktothrix</td><td>N317_RS0104230</td><td>886317..889079</td><td>54304</td><td>MULTISPECIES:-hypothetical-protein-[Planktothrix]</td></tr>
<tr><td>WP_026786969.1</td><td>Type4.PglZ.containing.BREX</td><td><-ChlI+Lon_C+AAA<-?||?-><-?||AbrB-HTH->DUF4007->viral-protein->DUF6079*->PglZ->SIGMA+OB-Fold+AAA-ATPase-BrxL->REase-><-?<-SPOUT</td><td>1438</td><td>Bacteria>Terrabacteria_group>Cyanobacteriota/Melainabacteria_group>Cyanobacteriota>Cyanophyceae>Oscillatoriophycideae</td><td>GCF_000464785.1</td><td>Planktothrix_rubescens</td><td>N313_RS0110825</td><td>2367475..2371791</td><td>59512</td><td>hypothetical-protein-[Planktothrix-rubescens]</td></tr>
<tr><td>WP_042151520.1</td><td>Type4.PglZ.containing.BREX</td><td>SPOUT->?-><-REase<-SIGMA+OB-Fold+AAA-ATPase-BrxL<-PglZ<-DUF6079*<-viral-protein<-DUF4007<-AbrB-HTH||?-><-?||?->ChlI+Lon_C+AAA-></td><td>1322</td><td>Bacteria>Terrabacteria_group>Cyanobacteriota/Melainabacteria_group>Cyanobacteriota>Cyanophyceae>Oscillatoriophycideae</td><td>GCF_000710505.1</td><td>Planktothrix_agardhii</td><td>A19Y_RS01335</td><td>297140..301108</td><td>1160</td><td>hypothetical-protein-[Planktothrix-agardhii]</td></tr>
<tr><td>WP_141294168.1</td><td>Type4.PglZ.containing.BREX</td><td><-ChlI+Lon_C+AAA<-?||?-><-?||AbrB-HTH->DUF4007->viral-protein->DUF6079*->PglZ->SIGMA+OB-Fold+AAA-ATPase-BrxL->REase-></td><td>1438</td><td>Bacteria>Terrabacteria_group>Cyanobacteriota/Melainabacteria_group>Cyanobacteriota>Cyanophyceae>Oscillatoriophycideae</td><td>GCF_005402765.1</td><td>Planktothrix_agardhii</td><td>FLZ02_RS12440</td><td>118679..122995</td><td>1160</td><td>hypothetical-protein-[Planktothrix-agardhii]</td></tr>
<tr><td>WP_168570765.1</td><td>Type4.PglZ.containing.BREX</td><td><-HTH_23+HTH_32||HTH_OrfB_IS605+OrfB_Zn_ribbon->Aminotran_1_2->?-><-?<-?||?-><-?<-SIGMA+OB-Fold+AAA-ATPase-BrxL<-PglZ<-DUF6079*<-viral-protein<-DUF4007<-?<-?<-TPR+TPR+TPR+TPR+TPR+TPR<-OrfB_Zn_ribbon<-OrfB_IS605<-HTH_OrfB_IS605+OrfB_IS605+OrfB_Zn_ribbon</td><td>1557</td><td>Bacteria>Terrabacteria_group>Cyanobacteriota/Melainabacteria_group>Cyanobacteriota>Cyanophyceae>Oscillatoriophycideae</td><td>GCF_012295525.1</td><td>Oxynema_aestuarii</td><td>HCG48_RS20145</td><td>5031560..5036233</td><td>2874213</td><td>hypothetical-protein-[Oxynema-aestuarii]</td></tr>
<tr><td>WP_190414234.1</td><td>Type4.PglZ.containing.BREX</td><td>TPR+TPR+TPR->TPR+TPR+TPR+TPR+TPR+TPR-><-Phage_integrase||Phage_integrase-><-CopG/RHH<-ParA-Soj-PloopNTPase<-Cysteine-Desulfurase<-SIGMA+OB-Fold+AAA-ATPase-BrxL<-PglZ<-DUF6079*<-viral-protein<-DUF4007<-PAPS-reduct<-wHTH(SBD)+HNH||SAM-METHYLASE-><-PrimPol<-?||?->Phage_integrase-></td><td>1801</td><td>Bacteria>Terrabacteria_group>Cyanobacteriota/Melainabacteria_group>Cyanobacteriota>Cyanophyceae>Oscillatoriophycideae</td><td>GCF_014695485.1</td><td>Coleofasciculus_sp._FACHB-125</td><td>H6F83_RS19380</td><td>40084..45489</td><td>2692784</td><td>hypothetical-protein-[Coleofasciculus-sp.-FACHB-125]</td></tr>
<tr><td>WP_192219076.1</td><td>Type4.PglZ.containing.BREX</td><td>ParA-Soj-PloopNTPase-><-REase+RNase||?-><-CASPASE<-?||DUF4007->DUF6079*->PAPS-reduct->?->DUF4007->viral-protein->DUF6079->NERD+iPkinase+Pkinase+PglZ->SIGMA+OB-Fold+AAA-ATPase-BrxL-></td><td>1190</td><td>Bacteria>Terrabacteria_group>Cyanobacteriota/Melainabacteria_group>Cyanobacteriota>Cyanophyceae>Oscillatoriophycideae</td><td>GCF_014849525.1</td><td>Microcoleus_sp._FACHB-SPT15</td><td>H6F98_RS12785</td><td>37537..41109</td><td>2692830</td><td>hypothetical-protein-[Microcoleus-sp.-FACHB-SPT15]</td></tr>
<tr><td>WP_192219081.1</td><td>Type4.PglZ.containing.BREX</td><td><-CASPASE<-?||DUF4007->DUF6079->PAPS-reduct->?->DUF4007->viral-protein->DUF6079*->NERD+iPkinase+Pkinase+PglZ->SIGMA+OB-Fold+AAA-ATPase-BrxL-></td><td>1819</td><td>Bacteria>Terrabacteria_group>Cyanobacteriota/Melainabacteria_group>Cyanobacteriota>Cyanophyceae>Oscillatoriophycideae</td><td>GCF_014849525.1</td><td>Microcoleus_sp._FACHB-SPT15</td><td>H6F98_RS12810</td><td>43356..48815</td><td>2692830</td><td>hypothetical-protein-[Microcoleus-sp.-FACHB-SPT15]</td></tr>
<tr><td>WP_228059560.1</td><td>Type4.PglZ.containing.BREX</td><td><-SAM-METHYLASE||DUF4007->DUF6079*-><-?||wHTH(SBD)+HNH->DUF4007->viral-protein->DUF6079->PglZ->SIGMA+OB-Fold+AAA-ATPase-BrxL->Cysteine-Desulfurase->ParA-Soj-PloopNTPase-></td><td>1152</td><td>Bacteria>Terrabacteria_group>Cyanobacteriota/Melainabacteria_group>Cyanobacteriota>Cyanophyceae>Oscillatoriophycideae</td><td>GCF_015207665.1</td><td>Plectonema_radiosum</td><td>IQ247_RS07910</td><td>5741..9199</td><td>945768</td><td>hypothetical-protein-[Plectonema-radiosum]</td></tr>
<tr><td>WP_193918771.1</td><td>Type4.PglZ.containing.BREX</td><td><-SAM-METHYLASE||DUF4007->DUF6079-><-?||wHTH(SBD)+HNH->DUF4007->viral-protein->DUF6079*->PglZ->SIGMA+OB-Fold+AAA-ATPase-BrxL->Cysteine-Desulfurase->ParA-Soj-PloopNTPase->ParB+HTH->?->PIN->SF1-SF2-Helicase-><-REase</td><td>1809</td><td>Bacteria>Terrabacteria_group>Cyanobacteriota/Melainabacteria_group>Cyanobacteriota>Cyanophyceae>Oscillatoriophycideae</td><td>GCF_015207665.1</td><td>Plectonema_radiosum</td><td>IQ247_RS07935</td><td>13410..18839</td><td>945768</td><td>hypothetical-protein-[Plectonema-radiosum]</td></tr>
<tr><td>WP_227349999.1</td><td>Type4.PglZ.containing.BREX</td><td><-DUF29<-?||<-?<-?||SPOUT->?-><-REase<-SIGMA+OB-Fold+AAA-ATPase-BrxL<-PglZ<-DUF6079*<-viral-protein<-DUF4007<-AbrB-HTH||?-><-?||?->ChlI+Lon_C+AAA-></td><td>1435</td><td>Bacteria>Terrabacteria_group>Cyanobacteriota/Melainabacteria_group>Cyanobacteriota>Cyanophyceae>Oscillatoriophycideae</td><td>GCF_020594555.1</td><td>Planktothrix_agardhii</td><td>LJE14_RS01380</td><td>293264..297571</td><td>1160</td><td>hypothetical-protein-[Planktothrix-agardhii]</td></tr>
<tr><td>WP_227382110.1</td><td>Type4.PglZ.containing.BREX</td><td><-ChlI+Lon_C+AAA<-?||?-><-?||AbrB-HTH->DUF4007->viral-protein->DUF6079*->PglZ->SIGMA+OB-Fold+AAA-ATPase-BrxL->REase-><-?<-SPOUT||?->?->||?->DUF29-></td><td>1438</td><td>Bacteria>Terrabacteria_group>Cyanobacteriota/Melainabacteria_group>Cyanobacteriota>Cyanophyceae>Oscillatoriophycideae</td><td>GCF_020594575.1</td><td>Planktothrix_agardhii</td><td>LJE16_RS18160</td><td>3953873..3958189</td><td>1160</td><td>hypothetical-protein-[Planktothrix-agardhii]</td></tr>
<tr><td>WP_235765519.1</td><td>Type4.PglZ.containing.BREX</td><td><-DUF29<-?||<-?<-?||SPOUT->?-><-REase<-SIGMA+OB-Fold+AAA-ATPase-BrxL<-PglZ<-DUF6079*<-viral-protein<-DUF4007<-AbrB-HTH||?-><-?||?->ChlI+Lon_C+AAA-></td><td>1441</td><td>Bacteria>Terrabacteria_group>Cyanobacteriota/Melainabacteria_group>Cyanobacteriota>Cyanophyceae>Oscillatoriophycideae</td><td>GCF_021554705.1</td><td>Planktothrix_agardhii</td><td>L2E76_RS02370</td><td>508842..513167</td><td>1160</td><td>hypothetical-protein-[Planktothrix-agardhii]</td></tr>
<tr><td>WP_235750932.1</td><td>Type4.PglZ.containing.BREX</td><td><-DUF29<-?||<-?<-?||SPOUT->?-><-REase<-SIGMA+OB-Fold+AAA-ATPase-BrxL<-PglZ<-DUF6079*<-viral-protein<-DUF4007<-AbrB-HTH||?-><-?||?->ChlI+Lon_C+AAA-></td><td>1441</td><td>Bacteria>Terrabacteria_group>Cyanobacteriota/Melainabacteria_group>Cyanobacteriota>Cyanophyceae>Oscillatoriophycideae</td><td>GCF_021554875.1</td><td>Planktothrix_agardhii</td><td>L2E65_RS00980</td><td>208238..212563</td><td>1160</td><td>hypothetical-protein-[Planktothrix-agardhii]</td></tr>
<tr><td>WP_275570982.1</td><td>Type4.PglZ.containing.BREX</td><td>SF1-SF2-Helicase->Phage_integrase-><-?<-?<-?||ParA-Soj-PloopNTPase->?->DUF4007->DUF6079*->Tn7_Tnp_TnsA_N+Tn7_Tnp_TnsA_C+rve+Mu-transpos_C->TnsC-ATPase->?->DOC+HTH->PAPS-reduct->ParB+HNH-><-SIGMA+OB-Fold+AAA-ATPase-BrxL<-PglZ<-DUF6079</td><td>1175</td><td>Bacteria>Terrabacteria_group>Cyanobacteriota/Melainabacteria_group>Cyanobacteriota>Cyanophyceae>Oscillatoriophycideae</td><td>GCF_029167235.1</td><td>Kamptonema_sp._UHCC_0994</td><td>PY364_RS28020</td><td>9183..12710</td><td>3031329</td><td>hypothetical-protein-[Kamptonema-sp.-UHCC-0994]</td></tr>
<tr><td>WP_083623965.1</td><td>Type4.PglZ.containing.BREX</td><td>SPOUT->?-><-REase<-?<-?<-SIGMA+OB-Fold+AAA-ATPase-BrxL<-PglZ<-DUF6079*<-viral-protein<-DUF4007<-AbrB-HTH<-?||?->SIR2->ChlI+Lon_C+AAA-></td><td>1437</td><td>Bacteria>Terrabacteria_group>Cyanobacteriota/Melainabacteria_group>Cyanobacteriota>Cyanophyceae>Oscillatoriophycideae</td><td>GCF_900009265.2</td><td>Planktothrix_paucivesiculata</td><td>PL9631_RS22955</td><td>536639..540952</td><td>1678308</td><td>hypothetical-protein-[Planktothrix-paucivesiculata]</td></tr>
<tr><td>WP_214560754.1</td><td>Type4.PglZ.containing.BREX</td><td><-DUF29<-?||?-><-?||SPOUT->?-><-REase<-SIGMA+OB-Fold+AAA-ATPase-BrxL<-PglZ<-DUF6079*<-viral-protein<-DUF4007<-AbrB-HTH<-?||?->ChlI+Lon_C+AAA-></td><td>1447</td><td>Bacteria>Terrabacteria_group>Cyanobacteriota/Melainabacteria_group>Cyanobacteriota>Cyanophyceae>Oscillatoriophycideae</td><td>GCF_903969095.1</td><td>Planktothrix_agardhii</td><td>KM073_RS19550</td><td>15976..20319</td><td>1160</td><td>hypothetical-protein-[Planktothrix-agardhii]</td></tr>
<tr><td>WP_254033533.1</td><td>Type4.PglZ.containing.BREX</td><td><-ChlI+Lon_C+AAA<-?||?-><-?||DUF4007->viral-protein->DUF6079*->PglZ->BrxL-p->REase-><-?<-SPOUT</td><td>1437</td><td>Bacteria>Terrabacteria_group>Cyanobacteriota/Melainabacteria_group>Cyanobacteriota>Cyanophyceae>Oscillatoriophycideae</td><td>GCF_904830845.1</td><td>Planktothrix_agardhii</td><td>NMG87_RS10510</td><td>2379967..2384280</td><td>1160</td><td>hypothetical-protein-[Planktothrix-agardhii]</td></tr>
<tr><td>WP_254032199.1</td><td>Type4.PglZ.containing.BREX</td><td>SPOUT->?-><-REase<-SIGMA+OB-Fold+AAA-ATPase-BrxL<-PglZ<-DUF6079-p<-HTH_Tnp_4+DDE_Tnp_4<-DUF6079-p*<-viral-protein<-DUF4007<-AbrB-HTH||?-><-?||?->ChlI+Lon_C+AAA-></td><td>768</td><td>Bacteria>Terrabacteria_group>Cyanobacteriota/Melainabacteria_group>Cyanobacteriota>Cyanophyceae>Oscillatoriophycideae</td><td>GCF_904830855.1</td><td>Planktothrix_agardhii</td><td>NMG84_RS06245</td><td>1446496..1448802</td><td>1160</td><td>hypothetical-protein-[Planktothrix-agardhii]</td></tr>
<tr><td>WP_254035079.1</td><td>Type4.PglZ.containing.BREX</td><td>SPOUT->?-><-REase<-SIGMA+OB-Fold+AAA-ATPase-BrxL<-PglZ<-DUF6079*<-viral-protein<-DUF4007||?-><-?||?->ChlI+Lon_C+AAA->?-><-?<-Peptidase</td><td>1438</td><td>Bacteria>Terrabacteria_group>Cyanobacteriota/Melainabacteria_group>Cyanobacteriota>Cyanophyceae>Oscillatoriophycideae</td><td>GCF_904830935.1</td><td>Planktothrix_agardhii</td><td>NMG88_RS02425</td><td>528904..533220</td><td>1160</td><td>hypothetical-protein-[Planktothrix-agardhii]</td></tr>
<tr><td>WP_254031240.1</td><td>Type4.PglZ.containing.BREX</td><td>SPOUT->?-><-REase<-SIGMA+OB-Fold+AAA-ATPase-BrxL<-PglZ<-DUF6079-p<-DUF6079-p*<-viral-protein<-DUF4007<-AbrB-HTH<-?||?->ChlI+Lon_C+AAA-></td><td>919</td><td>Bacteria>Terrabacteria_group>Cyanobacteriota/Melainabacteria_group>Cyanobacteriota>Cyanophyceae>Oscillatoriophycideae</td><td>GCF_904830945.1</td><td>Planktothrix_agardhii</td><td>NMG75_RS11300</td><td>2470908..2473667</td><td>1160</td><td>hypothetical-protein-[Planktothrix-agardhii]</td></tr>
<tr><td>OUC12890.1</td><td>Type4.PglZ.containing.BREX</td><td><-SAM-METHYLASE||DUF4007->DUF6079*->REase->PAPS-reduct->DUF4007->viral-protein->DUF6079->PglZ->SIGMA+OB-Fold+AAA-ATPase-BrxL->Cysteine-Desulfurase-></td><td>1172</td><td>Bacteria>Terrabacteria_group>Cyanobacteriota/Melainabacteria_group>Cyanobacteriota>Cyanophyceae>Pseudanabaenales</td><td>GCA_002148405.1</td><td>Alkalinema_sp._CACIAM_70d</td><td>BOA82_20145</td><td>16127..19645</td><td>1934309</td><td>MAG:-hypothetical-protein-BOA82_20145-[Alkalinema-sp.-CACIAM-70d]</td></tr>
<tr><td>OUC12874.1</td><td>Type4.PglZ.containing.BREX</td><td><-SAM-METHYLASE||DUF4007->DUF6079->REase->PAPS-reduct->DUF4007->viral-protein->DUF6079*->PglZ->SIGMA+OB-Fold+AAA-ATPase-BrxL->Cysteine-Desulfurase->?->?->DNA_METHYLASE-></td><td>1814</td><td>Bacteria>Terrabacteria_group>Cyanobacteriota/Melainabacteria_group>Cyanobacteriota>Cyanophyceae>Pseudanabaenales</td><td>GCA_002148405.1</td><td>Alkalinema_sp._CACIAM_70d</td><td>BOA82_20170</td><td>23110..28554</td><td>1934309</td><td>MAG:-hypothetical-protein-BOA82_20170-[Alkalinema-sp.-CACIAM-70d]</td></tr>
<tr><td>PSB31008.1</td><td>Type4.PglZ.containing.BREX</td><td>TnsC-ATPase->?-><-Cysteine-Desulfurase<-SIGMA+OB-Fold+AAA-ATPase-BrxL<-NERD+iPkinase+Pkinase+PglZ<-DUF6079*<-viral-protein<-DUF4007<-PAPS-reduct<-DUF6079<-DUF4007||SAM-METHYLASE->REase+SF1-SF2-Helicase->||REase+RNase->||Tn7_Tnp_TnsA_N+Tn7_Tnp_TnsA_C+rve+Mu-transpos_C-></td><td>1811</td><td>Bacteria>Terrabacteria_group>Cyanobacteriota/Melainabacteria_group>Cyanobacteriota>Cyanophyceae>Pseudanabaenales</td><td>GCA_003003795.1</td><td>Stenomitos_frigidus_ULC18</td><td>C7B82_07770</td><td>10215..15650</td><td>2107698</td><td>hypothetical-protein-C7B82_07770-[Stenomitos-frigidus-ULC18]</td></tr>
<tr><td>HBC40336.1</td><td>Type4.PglZ.containing.BREX</td><td>PAPS-reduct-p->AbrB-HTH->PIN->REase->HicB_lk_antitox->HicA_toxin->DUF4007->viral-protein->DUF6079*->?->NERD+iPkinase+Pkinase+PglZ->SIGMA+OB-Fold+AAA-ATPase-BrxL->ANK->wHTH->PolB->HEPN->Cysteine-Desulfurase->Cysteine-Desulfurase-p-></td><td>1805</td><td>Bacteria>Terrabacteria_group>Cyanobacteriota/Melainabacteria_group>Cyanobacteriota>Cyanophyceae>Pseudanabaenales</td><td>GCA_003488605.1</td><td>Pseudanabaena_sp.</td><td>DCZ88_00125</td><td>3477..8894</td><td>1153</td><td>hypothetical-protein-[Pseudanabaena-sp.]</td></tr>
<tr><td>NEQ47451.1</td><td>Type4.PglZ.containing.BREX</td><td>Phage_integrase+Phage_integrase-><-?||SAM-METHYLASE-><-?<-?<-?<-SAM-METHYLASE||DUF4007->DUF6079*->REase->DUF29->Preader1+REase->PAPS-reduct->DUF4007->viral-protein->DUF6079->PglW->PglZ-></td><td>1181</td><td>Bacteria>Terrabacteria_group>Cyanobacteriota/Melainabacteria_group>Cyanobacteriota>Cyanophyceae>Pseudanabaenales</td><td>GCA_010671905.1</td><td>Leptolyngbya_sp._SIOISBB</td><td>F6K00_29485</td><td>6889470..6893015</td><td>2607771</td><td>hypothetical-protein-[Leptolyngbya-sp.-SIOISBB]</td></tr>
<tr><td>NEQ47458.1</td><td>Type4.PglZ.containing.BREX</td><td><-SAM-METHYLASE||DUF4007->DUF6079->REase->DUF29->Preader1+REase->PAPS-reduct->DUF4007->viral-protein->DUF6079*->PglW->PglZ->SIGMA+OB-Fold+AAA-ATPase-BrxL-><-Nuc_deoxyri_tr2||Cysteine-Desulfurase->?-><-DUF3970||HTH_26-></td><td>1826</td><td>Bacteria>Terrabacteria_group>Cyanobacteriota/Melainabacteria_group>Cyanobacteriota>Cyanophyceae>Pseudanabaenales</td><td>GCA_010671905.1</td><td>Leptolyngbya_sp._SIOISBB</td><td>F6K00_29525</td><td>6898266..6903746</td><td>2607771</td><td>hypothetical-protein-[Leptolyngbya-sp.-SIOISBB]</td></tr>
<tr><td>WP_121970835.1</td><td>Type4.PglZ.containing.BREX</td><td>NUDIX->CN_hydrolase+NAD_synthase-><-?||RHH_6-><-Cysteine-Desulfurase<-SIGMA+OB-Fold+AAA-ATPase-BrxL<-NERD+iPkinase+Pkinase+PglZ<-DUF6079*</td><td>1490</td><td>Bacteria>Terrabacteria_group>Cyanobacteriota/Melainabacteria_group>Cyanobacteriota>Cyanophyceae>Pseudanabaenales</td><td>GCF_002286735.1</td><td>Leptolyngbya_sp._BC1307</td><td>CKL79_RS17360</td><td>33040..37512</td><td>2029589</td><td>hypothetical-protein-[Leptolyngbya-sp.-BC1307]</td></tr>
<tr><td>WP_146141087.1</td><td>Type4.PglZ.containing.BREX</td><td>TnsC-ATPase->?-><-Cysteine-Desulfurase<-SIGMA+OB-Fold+AAA-ATPase-BrxL<-NERD+iPkinase+Pkinase+PglZ<-DUF6079*<-viral-protein<-DUF4007<-PAPS-reduct<-DUF6079<-DUF4007||SAM-METHYLASE->REase+SF1-SF2-Helicase-><-SAM-METHYLASE||REase+RNase-></td><td>1827</td><td>Bacteria>Terrabacteria_group>Cyanobacteriota/Melainabacteria_group>Cyanobacteriota>Cyanophyceae>Pseudanabaenales</td><td>GCF_003003795.1</td><td>Stenomitos_frigidus</td><td>C7B82_RS07800</td><td>10215..15698</td><td>1886765</td><td>hypothetical-protein-[Stenomitos-frigidus]</td></tr>
<tr><td>WP_106255742.1</td><td>Type4.PglZ.containing.BREX</td><td>TnsC-ATPase->?-><-Cysteine-Desulfurase<-SIGMA+OB-Fold+AAA-ATPase-BrxL<-NERD+iPkinase+Pkinase+PglZ<-DUF6079<-viral-protein<-DUF4007<-PAPS-reduct<-DUF6079*<-DUF4007||SAM-METHYLASE->REase+SF1-SF2-Helicase-><-SAM-METHYLASE||REase+RNase-><-?||Tn7_Tnp_TnsA_N+Tn7_Tnp_TnsA_C+rve+Mu-transpos_C->TnsC-ATPase-></td><td>1167</td><td>Bacteria>Terrabacteria_group>Cyanobacteriota/Melainabacteria_group>Cyanobacteriota>Cyanophyceae>Pseudanabaenales</td><td>GCF_003003795.1</td><td>Stenomitos_frigidus</td><td>C7B82_RS07820</td><td>17910..21413</td><td>1886765</td><td>hypothetical-protein-[Stenomitos-frigidus]</td></tr>
<tr><td>WP_169366114.1</td><td>Type4.PglZ.containing.BREX</td><td>DUF6079->REase->REase->REase->HARE-HTH->PAPS-reduct->REase->DUF4007->viral-protein->DUF6079*->NERD+iPkinase+Pkinase+PglZ->SIGMA+OB-Fold+AAA-ATPase-BrxL->Cysteine-Desulfurase->ParA-Soj-PloopNTPase->CopG/RHH-><-TOPRIM+SF1-SF2-Helicase</td><td>1805</td><td>Bacteria>Terrabacteria_group>Cyanobacteriota/Melainabacteria_group>Cyanobacteriota>Cyanophyceae>Pseudanabaenales</td><td>GCF_012863495.1</td><td>Pseudanabaena_yagii</td><td>HC246_RS24875</td><td>14182..19599</td><td>2661615</td><td>hypothetical-protein-[Pseudanabaena-yagii]</td></tr>
<tr><td>WP_190351065.1</td><td>Type4.PglZ.containing.BREX</td><td><-Cysteine-Desulfurase<-HEPN<-PolB<-SIGMA+OB-Fold+AAA-ATPase-BrxL<-NERD+iPkinase+Pkinase+PglZ<-?<-?<-REase<-DUF6079*||?->?-><-viral-protein<-DUF4007<-HicB_lk_antitox<-REase<-PIN</td><td>1594</td><td>Bacteria>Terrabacteria_group>Cyanobacteriota/Melainabacteria_group>Cyanobacteriota>Cyanophyceae>Pseudanabaenales</td><td>GCF_014696345.1</td><td>Pseudanabaena_sp._FACHB-1277</td><td>H6F44_RS11335</td><td>40802..45586</td><td>1540896</td><td>hypothetical-protein-[Pseudanabaena-sp.-FACHB-1277]</td></tr>
<tr><td>WP_204150070.1</td><td>Type4.PglZ.containing.BREX</td><td><-Cysteine-Desulfurase-p<-SIGMA+OB-Fold+AAA-ATPase-BrxL<-NERD+iPkinase+Pkinase+PglZ<-DUF6079*<-viral-protein<-DUF4007<-PAPS-reduct<-Preader1+REase||DUF4007-></td><td>1816</td><td>Bacteria>Terrabacteria_group>Cyanobacteriota/Melainabacteria_group>Cyanobacteriota>Cyanophyceae>Pseudanabaenales</td><td>GCF_016888135.1</td><td>Leptolyngbya_sp._CCY15150</td><td>JUJ53_RS00715</td><td>6487..11937</td><td>2767772</td><td>hypothetical-protein-[Leptolyngbya-sp.-CCY15150]</td></tr>
<tr><td>WP_204104298.1</td><td>Type4.PglZ.containing.BREX</td><td><-REase||DUF4007->DUF6079-p*->wHTH(SBD)+HNH->PAPS-reduct->REase->DUF4007->viral-protein->DUF6079->PglZ->SIGMA+OB-Fold+AAA-ATPase-BrxL->Cysteine-Desulfurase-></td><td>882</td><td>Bacteria>Terrabacteria_group>Cyanobacteriota/Melainabacteria_group>Cyanobacteriota>Cyanophyceae>Spirulinales</td><td>GCF_016888125.1</td><td>Spirulina_major</td><td>JUI58_RS13385</td><td>3450..6098</td><td>270636</td><td>hypothetical-protein-[Spirulina-major]</td></tr>
<tr><td>WP_204104304.1</td><td>Type4.PglZ.containing.BREX</td><td><-REase||DUF4007->DUF6079-p->wHTH(SBD)+HNH->PAPS-reduct->REase->DUF4007->viral-protein->DUF6079*->PglZ->SIGMA+OB-Fold+AAA-ATPase-BrxL->Cysteine-Desulfurase->PTSEIIA1-><-?<-?||METHYLASE-></td><td>1570</td><td>Bacteria>Terrabacteria_group>Cyanobacteriota/Melainabacteria_group>Cyanobacteriota>Cyanophyceae>Spirulinales</td><td>GCF_016888125.1</td><td>Spirulina_major</td><td>JUI58_RS13415</td><td>10116..14828</td><td>270636</td><td>hypothetical-protein-[Spirulina-major]</td></tr>
<tr><td>MBF2075132.1</td><td>Type4.PglZ.containing.BREX</td><td><-Cysteine-Desulfurase<-REase||REase->?->SWACOS+HerA/FtsK-><-REase<-SIGMA+OB-Fold+AAA-ATPase-BrxL<-NERD+iPkinase+Pkinase+PglZ<-DUF6079*<-viral-protein<-DUF4007<-ParB+HNH<-PAPS-reduct<-Preader1+REase<-METHYLASE<-Radical-SAM<-DUF6079<-DUF4007</td><td>1797</td><td>Bacteria>Terrabacteria_group>Cyanobacteriota/Melainabacteria_group>Cyanobacteriota>Cyanophyceae>Synechococcales</td><td>GCA_015272325.1</td><td>Synechococcales_cyanobacterium_C42_A2020_086</td><td>IGS50_15425</td><td>20530..25923</td><td>2774344</td><td>hypothetical-protein-[Synechococcales-cyanobacterium-C42_A2020_086]</td></tr>
<tr><td>MBF2075140.1</td><td>Type4.PglZ.containing.BREX</td><td><-NERD+iPkinase+Pkinase+PglZ<-DUF6079<-viral-protein<-DUF4007<-ParB+HNH<-PAPS-reduct<-Preader1+REase<-METHYLASE<-Radical-SAM<-DUF6079*<-DUF4007<-REase||REase->SAM-METHYLASE-><-DNA_pol3_beta+DNA_pol3_beta_2+DNA_pol3_beta_3||sGTP-></td><td>1165</td><td>Bacteria>Terrabacteria_group>Cyanobacteriota/Melainabacteria_group>Cyanobacteriota>Cyanophyceae>Synechococcales</td><td>GCA_015272325.1</td><td>Synechococcales_cyanobacterium_C42_A2020_086</td><td>IGS50_15465</td><td>33423..36920</td><td>2774344</td><td>hypothetical-protein-[Synechococcales-cyanobacterium-C42_A2020_086]</td></tr>
<tr><td>KEF34342.1</td><td>Type4.PglZ.containing.BREX</td><td><-NTP_transf+HDRnase||tRNA-><-Cysteine-Desulfurase<-SIGMA+OB-Fold+AAA-ATPase-BrxL<-PglZ<-DUF6079<-wHTH(SBD)+HNH<-DUF6079*<-DUF4007||?->ZnR+Pkinase+pi3KINASE->ZnR+FHA->?->?->?-><-NAD_binding_3+Homoserine_dh</td><td>1091</td><td>Bacteria>Terrabacteria_group>Deinococcota>Deinococci>Deinococcales>Deinococcaceae</td><td>GCA_000708185.1</td><td>Deinococcus_sp._RL</td><td>RDMS_08055</td><td>112880..116155</td><td>1489678</td><td>hypothetical-protein-RDMS_08055-[Deinococcus-sp.-RL]</td></tr>
<tr><td>MCD0174458.1</td><td>Type4.PglZ.containing.BREX</td><td><-PglZ<-DUF6079*<-?<-DUF4007<-DndH<-DndG<-DndF<-Cysteine-Desulfurase<-PAPS-reduct<-DUF6079*<-DUF4007<-ParB+HTH<-ParA-Soj-PloopNTPase||?-><-wHTH+WYL+WCX<-?<-SF1-SF2-Helicase+REase</td><td>1094</td><td>Bacteria>Terrabacteria_group>Deinococcota>Deinococci>Deinococcales>Deinococcaceae</td><td>GCA_020984885.1</td><td>Deinococcus_sp._14RED07</td><td>IHN32_00610</td><td>22148..25432</td><td>2745874</td><td>hypothetical-protein-[Deinococcus-sp.-14RED07]</td></tr>
<tr><td>WP_034404733.1</td><td>Type4.PglZ.containing.BREX</td><td><-NTP_transf+HDRnase||tRNA-><-Cysteine-Desulfurase<-SIGMA+OB-Fold+AAA-ATPase-BrxL<-PglZ<-DUF6079*<-HTH_OrfB_IS605+OrfB_IS605+OrfB_Zn_ribbon<-wHTH(SBD)+HNH<-DUF6079<-DUF4007||?->Pkinase+pi3KINASE->ZnR+FHA-></td><td>1726</td><td>Bacteria>Terrabacteria_group>Deinococcota>Deinococci>Deinococcales>Deinococcaceae</td><td>GCF_000708185.1</td><td>Deinococcus_sp._RL</td><td>RDMS_RS07910</td><td>105181..110361</td><td>1489678</td><td>hypothetical-protein-[Deinococcus-sp.-RL]</td></tr>
<tr><td>WP_152550631.1</td><td>Type4.PglZ.containing.BREX</td><td><-NTP_transf+HDRnase||tRNA-><-Cysteine-Desulfurase<-SIGMA+OB-Fold+AAA-ATPase-BrxL<-PglZ<-DUF6079<-HTH_OrfB_IS605+OrfB_IS605+OrfB_Zn_ribbon<-wHTH(SBD)+HNH<-DUF6079*<-DUF4007||?->Pkinase+pi3KINASE->ZnR+FHA->?->?->?-><-NAD_binding_3+Homoserine_dh</td><td>1111</td><td>Bacteria>Terrabacteria_group>Deinococcota>Deinococci>Deinococcales>Deinococcaceae</td><td>GCF_000708185.1</td><td>Deinococcus_sp._RL</td><td>RDMS_RS07920</td><td>112880..116215</td><td>1489678</td><td>hypothetical-protein-[Deinococcus-sp.-RL]</td></tr>
<tr><td>WP_102126051.1</td><td>Type4.PglZ.containing.BREX</td><td><-N6-METHYLASE<-?<-Calcineurin<-Peptidase_S66+Peptidase_S66C||DDE_Tnp_1-><-virion-envelope<-Cysteine-Desulfurase<-SIGMA+OB-Fold+AAA-ATPase-BrxL<-PglZ<-DUF6079*<-?<-DUF4007<-PAPS-reduct<-DUF6079<-DUF4007||DndB-></td><td>1747</td><td>Bacteria>Terrabacteria_group>Deinococcota>Deinococci>Deinococcales>Deinococcaceae</td><td>GCF_002869765.1</td><td>Deinococcus_planocerae</td><td>A7B18_RS07415</td><td>27365..32608</td><td>1737569</td><td>hypothetical-protein-[Deinococcus-planocerae]</td></tr>
<tr><td>WP_146009483.1</td><td>Type4.PglZ.containing.BREX</td><td>DDE_Tnp_1-><-virion-envelope<-Cysteine-Desulfurase<-SIGMA+OB-Fold+AAA-ATPase-BrxL<-PglZ<-DUF6079<-?<-DUF4007<-PAPS-reduct<-DUF6079*<-DUF4007||DndB-><-?<-?<-?<-?||HTH_3-></td><td>1094</td><td>Bacteria>Terrabacteria_group>Deinococcota>Deinococci>Deinococcales>Deinococcaceae</td><td>GCF_002869765.1</td><td>Deinococcus_planocerae</td><td>A7B18_RS07435</td><td>34864..38148</td><td>1737569</td><td>hypothetical-protein-[Deinococcus-planocerae]</td></tr>
<tr><td>WP_103311070.1</td><td>Type4.PglZ.containing.BREX</td><td>Peptidase_M20-><-LACI-HTH+PBP1||?-><-?<-Cysteine-Desulfurase<-SIGMA+OB-Fold+AAA-ATPase-BrxL<-PglZ<-DUF6079*<-?<-DUF4007<-PAPS-reduct<-DUF6079<-DUF4007<-ParB+HNH</td><td>1747</td><td>Bacteria>Terrabacteria_group>Deinococcota>Deinococci>Deinococcales>Deinococcaceae</td><td>GCF_002901445.1</td><td>Deinococcus_koreensis</td><td>CVO96_05115</td><td>1082261..1087504</td><td>2054903</td><td>hypothetical-protein-[Deinococcus-koreensis]</td></tr>
<tr><td>WP_103311079.1</td><td>Type4.PglZ.containing.BREX</td><td><-Cysteine-Desulfurase<-SIGMA+OB-Fold+AAA-ATPase-BrxL<-PglZ<-DUF6079<-?<-DUF4007<-wHTH<-PAPS-reduct<-DUF6079*<-DUF4007<-ParB+HNH<-?||?->?-><-tRNA<-rRNA<-rRNA<-?||Peptidase_A8-></td><td>1115</td><td>Bacteria>Terrabacteria_group>Deinococcota>Deinococci>Deinococcales>Deinococcaceae</td><td>GCF_002901445.1</td><td>Deinococcus_koreensis</td><td>CVO96_RS05135</td><td>1090249..1093596</td><td>2054903</td><td>hypothetical-protein-[Deinococcus-koreensis]</td></tr>
<tr><td>WP_104989993.1</td><td>Type4.PglZ.containing.BREX</td><td><-DDE_Tnp||OrfB_Zn_ribbon-><-GDResolvase+HIN-HTH||Primpol->?->?-><-Cysteine-Desulfurase<-SIGMA+OB-Fold+AAA-ATPase-BrxL<-PglZ<-DUF6079*<-?<-DUF4007<-PAPS-reduct<-DUF6079<-DUF4007||?->ZnR+Pkinase+pi3KINASE->ZnR+FHA-></td><td>1748</td><td>Bacteria>Terrabacteria_group>Deinococcota>Deinococci>Deinococcales>Deinococcaceae</td><td>GCF_002953415.1</td><td>Deinococcus_sp._NW-56</td><td>C3K08_RS03145</td><td>599930..605176</td><td>2080419</td><td>hypothetical-protein-[Deinococcus-sp.-NW-56]</td></tr>
<tr><td>WP_104989997.1</td><td>Type4.PglZ.containing.BREX</td><td><-Cysteine-Desulfurase<-SIGMA+OB-Fold+AAA-ATPase-BrxL<-PglZ<-DUF6079<-?<-DUF4007<-PAPS-reduct<-DUF6079*<-DUF4007||?->ZnR+Pkinase+pi3KINASE->ZnR+FHA-></td><td>1118</td><td>Bacteria>Terrabacteria_group>Deinococcota>Deinococci>Deinococcales>Deinococcaceae</td><td>GCF_002953415.1</td><td>Deinococcus_sp._NW-56</td><td>C3K08_RS03165</td><td>607914..611270</td><td>2080419</td><td>hypothetical-protein-[Deinococcus-sp.-NW-56]</td></tr>
<tr><td>WP_139401673.1</td><td>Type4.PglZ.containing.BREX</td><td><-HTH+HTH+SF1-SF2-Helicase||?-><-?<-?<-?<-ZnR+FHA<-Pkinase+pi3KINASE<-?||DUF4007->DUF6079*->PAPS-reduct->DUF4007->?->DUF6079->PglZ->SIGMA+OB-Fold+AAA-ATPase-BrxL->Cysteine-Desulfurase->?-><-DDE_Tnp_1</td><td>1117</td><td>Bacteria>Terrabacteria_group>Deinococcota>Deinococci>Deinococcales>Deinococcaceae</td><td>GCF_014201625.1</td><td>Deinococcus_radiopugnans</td><td>HNQ04_RS13215</td><td>20790..24143</td><td>57497</td><td>hypothetical-protein-[Deinococcus-radiopugnans]</td></tr>
<tr><td>WP_139401681.1</td><td>Type4.PglZ.containing.BREX</td><td><-ZnR+FHA<-Pkinase+pi3KINASE<-?||DUF4007->DUF6079->PAPS-reduct->DUF4007<-?||DUF6079*->PglZ->SIGMA+OB-Fold+AAA-ATPase-BrxL->Cysteine-Desulfurase->?-><-DDE_Tnp_1||?->DOC+HTH-></td><td>1749</td><td>Bacteria>Terrabacteria_group>Deinococcota>Deinococci>Deinococcales>Deinococcaceae</td><td>GCF_014201625.1</td><td>Deinococcus_radiopugnans</td><td>HNQ04_RS13235</td><td>27254..32503</td><td>57497</td><td>hypothetical-protein-[Deinococcus-radiopugnans]</td></tr>
<tr><td>WP_184137439.1</td><td>Type4.PglZ.containing.BREX</td><td>TPR+TPR+TPR->DUF6079*->PAPS-reduct->wHTH->DUF4007->?->DUF6079->PglZ->SIGMA+OB-Fold+AAA-ATPase-BrxL->Cysteine-Desulfurase-></td><td>1111</td><td>Bacteria>Terrabacteria_group>Deinococcota>Deinococci>Deinococcales>Deinococcaceae</td><td>GCF_014201875.1</td><td>Deinococcus_humi</td><td>HNQ08_RS23380</td><td>6394..9729</td><td>662880</td><td>hypothetical-protein-[Deinococcus-humi]</td></tr>
<tr><td>WP_184137449.1</td><td>Type4.PglZ.containing.BREX</td><td><-ABhydrolase<-?||?-><-Cysteine-Desulfurase<-SIGMA+OB-Fold+AAA-ATPase-BrxL<-PglZ<-DUF6079*<-?<-DUF4007<-wHTH<-PAPS-reduct<-DUF6079<-DUF4007<-TPR+TPR+TPR</td><td>1744</td><td>Bacteria>Terrabacteria_group>Deinococcota>Deinococci>Deinococcales>Deinococcaceae</td><td>GCF_014201875.1</td><td>Deinococcus_humi</td><td>HNQ08_RS23405</td><td>12449..17683</td><td>662880</td><td>hypothetical-protein-[Deinococcus-humi]</td></tr>
<tr><td>WP_189070975.1</td><td>Type4.PglZ.containing.BREX</td><td><-SIGMA+OB-Fold+AAA-ATPase-BrxL<-PglZ<-DUF6079*<-?<-DUF4007||RNase->Exonuc_V_gamma+REase->SF1-SF2-Helicase+REase->?->wHTH+WYL+WCX-></td><td>1753</td><td>Bacteria>Terrabacteria_group>Deinococcota>Deinococci>Deinococcales>Deinococcaceae</td><td>GCF_014647435.1</td><td>Deinococcus_radiotolerans</td><td>IEY63_RS21095</td><td>4460..9721</td><td>1309407</td><td>hypothetical-protein-[Deinococcus-radiotolerans]</td></tr>
<tr><td>WP_230285997.1</td><td>Type4.PglZ.containing.BREX</td><td><-PIN<-PhdYeFM_antitox<-?||ParA-Soj-PloopNTPase->ParB+HTH->?->DUF6079*->PAPS-reduct->Cysteine-Desulfurase->DndF->DndG->DndH->DUF4007->?->DUF6079->PglZ-></td><td>1110</td><td>Bacteria>Terrabacteria_group>Deinococcota>Deinococci>Deinococcales>Deinococcaceae</td><td>GCF_020984835.1</td><td>Deinococcus_sp._12RED42</td><td>IHN58_RS00685</td><td>7094..10426</td><td>2745872</td><td>hypothetical-protein-[Deinococcus-sp.-12RED42]</td></tr>
<tr><td>WP_230286005.1</td><td>Type4.PglZ.containing.BREX</td><td>DUF6079->PAPS-reduct->Cysteine-Desulfurase->DndF->DndG->DndH->DUF4007->?->DUF6079*->PglZ->SIGMA+OB-Fold+AAA-ATPase-BrxL-></td><td>1746</td><td>Bacteria>Terrabacteria_group>Deinococcota>Deinococci>Deinococcales>Deinococcaceae</td><td>GCF_020984835.1</td><td>Deinococcus_sp._12RED42</td><td>IHN58_RS00725</td><td>23805..29045</td><td>2745872</td><td>hypothetical-protein-[Deinococcus-sp.-12RED42]</td></tr>
<tr><td>WP_230288670.1</td><td>Type4.PglZ.containing.BREX</td><td><-SIGMA+OB-Fold+AAA-ATPase-BrxL<-PglZ<-DUF6079*<-?<-DUF4007<-DndH<-DndG<-DndF<-Cysteine-Desulfurase<-PAPS-reduct<-DUF6079<-ParB+HTH</td><td>1746</td><td>Bacteria>Terrabacteria_group>Deinococcota>Deinococci>Deinococcales>Deinococcaceae</td><td>GCF_020984885.1</td><td>Deinococcus_sp._14RED07</td><td>IHN32_RS00575</td><td>3862..9102</td><td>2745874</td><td>hypothetical-protein-[Deinococcus-sp.-14RED07]</td></tr>
<tr><td>WP_230288676.1</td><td>Type4.PglZ.containing.BREX</td><td><-PglZ<-DUF6079<-?<-DUF4007<-DndH<-DndG<-DndF<-Cysteine-Desulfurase<-PAPS-reduct<-DUF6079*<-ParB+HTH||?-><-wHTH+WYL+WCX<-?<-SF1-SF2-Helicase+REase</td><td>1110</td><td>Bacteria>Terrabacteria_group>Deinococcota>Deinococci>Deinococcales>Deinococcaceae</td><td>GCF_020984885.1</td><td>Deinococcus_sp._14RED07</td><td>IHN32_RS00615</td><td>22148..25480</td><td>2745874</td><td>hypothetical-protein-[Deinococcus-sp.-14RED07]</td></tr>
<tr><td>WP_092265764.1</td><td>Type4.PglZ.containing.BREX</td><td><-Cysteine-Desulfurase<-SIGMA+OB-Fold+AAA-ATPase-BrxL<-PglZ<-DUF6079*<-?<-DUF4007<-PAPS-reduct</td><td>1748</td><td>Bacteria>Terrabacteria_group>Deinococcota>Deinococci>Deinococcales>Deinococcaceae</td><td>GCF_900109185.1</td><td>Deinococcus_reticulitermitis</td><td>BMY43_RS15955</td><td>10500..15746</td><td>856736</td><td>hypothetical-protein-[Deinococcus-reticulitermitis]</td></tr>
<tr><td>WP_084784956.1</td><td>Type4.PglZ.containing.BREX</td><td><-TRD+TRD||?-><-ABhydrolase||OrfB_IS605+OrfB_Zn_ribbon-><-TPR+TPR+TPR+TPR+TPR<-?<-MarR-HTH<-?||DUF4007->DUF6079*->PAPS-reduct->DUF4007->?->DUF6079->PglZ->SIGMA+OB-Fold+AAA-ATPase-BrxL->Cysteine-Desulfurase-><-DsbD</td><td>1105</td><td>Bacteria>Terrabacteria_group>Deinococcota>Deinococci>Thermales>Thermaceae</td><td>GCF_000373205.1</td><td>Calidithermus_timidus</td><td>B047_RS0106605</td><td>16642..19959</td><td>307124</td><td>hypothetical-protein-[Calidithermus-timidus]</td></tr>
<tr><td>WP_018466167.1</td><td>Type4.PglZ.containing.BREX</td><td><-TPR+TPR+TPR+TPR+TPR<-?<-MarR-HTH<-?||DUF4007->DUF6079->PAPS-reduct->DUF4007->?->DUF6079*->PglZ->SIGMA+OB-Fold+AAA-ATPase-BrxL->Cysteine-Desulfurase-><-DsbD</td><td>1857</td><td>Bacteria>Terrabacteria_group>Deinococcota>Deinococci>Thermales>Thermaceae</td><td>GCF_000373205.1</td><td>Calidithermus_timidus</td><td>B047_RS0106625</td><td>22582..28155</td><td>307124</td><td>hypothetical-protein-[Calidithermus-timidus]</td></tr>
<tr><td>WP_027878018.1</td><td>Type4.PglZ.containing.BREX</td><td>35exo+SF1-SF2-Helicase->?->DDE_5+DDE_Tnp_1-><-Phage_int_SAM_1+Phage_integrase<-Cysteine-Desulfurase<-SIGMA+OB-Fold+AAA-ATPase-BrxL<-PglZ<-DUF6079*<-?<-DUF4007<-PAPS-reduct<-DUF6079-p<-DUF4007</td><td>1850</td><td>Bacteria>Terrabacteria_group>Deinococcota>Deinococci>Thermales>Thermaceae</td><td>GCF_000620065.1</td><td>Meiothermus_cerbereus</td><td>Q355_RS0111985</td><td>14698..20250</td><td>65552</td><td>hypothetical-protein-[Meiothermus-cerbereus]</td></tr>
<tr><td>WP_245597570.1</td><td>Type4.PglZ.containing.BREX</td><td>DDE_5+DDE_Tnp_1-><-Phage_int_SAM_1+Phage_integrase<-Cysteine-Desulfurase<-SIGMA+OB-Fold+AAA-ATPase-BrxL<-PglZ<-DUF6079<-?<-DUF4007<-PAPS-reduct<-DUF6079-p*<-DUF4007</td><td>986</td><td>Bacteria>Terrabacteria_group>Deinococcota>Deinococci>Thermales>Thermaceae</td><td>GCF_000620065.1</td><td>Meiothermus_cerbereus</td><td>Q355_RS0112000</td><td>22922..25882</td><td>65552</td><td>hypothetical-protein-[Meiothermus-cerbereus]</td></tr>
<tr><td>HAR58250.1</td><td>Type4.PglZ.containing.BREX</td><td><-SF1-SF2-Helicase<-SF1-SF2-Helicase+REase||HTH_7+rve+rve_3->SMODS-><-RNase+DSRBD<-Cysteine-Desulfurase<-?<-SIGMA+OB-Fold+AAA-ATPase-BrxL<-iSTAND+PglZ<-DUF6079*<-ZnR<-PAPS-reduct+a+b-domain+Fer4_9+DUF4007<-?<-?<-MarR-HTH</td><td>1175</td><td>Bacteria>Terrabacteria_group>Mycoplasmatota>Mollicutes>Acholeplasmatales>unclassified_Acholeplasmatales</td><td>GCA_003476205.1</td><td>Acholeplasmatales_bacterium</td><td>DCR62_05855</td><td>181934..185461</td><td>2053494</td><td>hypothetical-protein-[Acholeplasmatales-bacterium]</td></tr>
<tr><td>RIA77964.1</td><td>Type4.PglZ.containing.BREX</td><td><-HTH_3<-?<-?<-CR-Kinase<-Cysteine-Desulfurase<-ZnR<-SIGMA+OB-Fold+AAA-ATPase-BrxL<-iSTAND+PglZ<-DUF6079-p<-DUF6079-p*<-PAPS-reduct+a+b-domain+Fer4_9+DUF4007<-?<-NUDIX<-NERD+SF1-SF2-Helicase<-TRAM+METHYLASE<-SF1-SF2-Helicase<-RnaseH+HEPN<-HD</td><td>968</td><td>Bacteria>Terrabacteria_group>Mycoplasmatota>Mollicutes>Anaeroplasmatales>Anaeroplasmataceae</td><td>GCA_003550015.1</td><td>Anaeroplasma_bactoclasticum</td><td>EI71_00747</td><td>88215..91121</td><td>2088</td><td>hypothetical-protein-EI71_00747-[Anaeroplasma-bactoclasticum]</td></tr>
<tr><td>MCM1514005.1</td><td>Type4.PglZ.containing.BREX</td><td>RhoN+RhoN+RNase+S1->?->?-><-?||?->tRNA->CHTH->?->PAPS-reduct+a+b-domain+Fer4_9+DUF4007->DUF6079*->iSTAND+PglZ->SIGMA+OB-Fold+AAA-ATPase-BrxL->ZnR->Cysteine-Desulfurase->rve-></td><td>1399</td><td>Bacteria>Terrabacteria_group>Mycoplasmatota>Mollicutes>Anaeroplasmatales>Anaeroplasmataceae</td><td>GCA_023667245.1</td><td>Anaeroplasma_bactoclasticum</td><td>NC090_03365</td><td>35773..39972</td><td>2088</td><td>hypothetical-protein-[Anaeroplasma-bactoclasticum]</td></tr>
<tr><td>WP_162849737.1</td><td>Type4.PglZ.containing.BREX</td><td><-HTH_3<-?<-?<-CR-Kinase<-Cysteine-Desulfurase<-ZnR<-SIGMA+OB-Fold+AAA-ATPase-BrxL<-iSTAND+PglZ<-DUF6079-p<-DUF6079-p*<-PAPS-reduct+a+b-domain+Fer4_9+DUF4007<-?<-NUDIX<-NERD+SF1-SF2-Helicase<-TRAM+METHYLASE<-SF1-SF2-Helicase<-RnaseH+HEPN<-HD</td><td>959</td><td>Bacteria>Terrabacteria_group>Mycoplasmatota>Mollicutes>Anaeroplasmatales>Anaeroplasmataceae</td><td>GCF_003550015.1</td><td>Anaeroplasma_bactoclasticum</td><td>EI71_RS03700</td><td>88242..91121</td><td>2088</td><td>hypothetical-protein-[Anaeroplasma-bactoclasticum]</td></tr>
<tr><td>ACM07070.1</td><td>Type4.PglZ.containing.BREX</td><td>DUF4007->DUF6079->DUF4007->?->DUF6079*->PglZ->SIGMA+OB-Fold+AAA-ATPase-BrxL->Cysteine-Desulfurase-><-CARF+HEPN<-RAMPs<-?<-RAMPs</td><td>1382</td><td>Bacteria>Terrabacteria_group>Thermomicrobiota>Thermomicrobia>Thermomicrobiales>Thermomicrobiaceae</td><td>GCA_000021685.1</td><td>Thermomicrobium_roseum_DSM_5159</td><td>trd_A0106</td><td>96854..101002</td><td>309801</td><td>hypothetical-protein-trd_A0106-[Thermomicrobium-roseum-DSM-5159]</td></tr>
<tr><td>MBO9307693.1</td><td>Type4.PglZ.containing.BREX</td><td>DUF6079-p->PAPS-reduct->DUF4007->?->DUF6079*->PglZ-></td><td>1382</td><td>Bacteria>Terrabacteria_group>Thermomicrobiota>Thermomicrobia>Thermomicrobiales>Thermomicrobiaceae</td><td>GCA_017743575.1</td><td>Thermomicrobium_sp.</td><td>J7451_10715</td><td>3318..7466</td><td>1969469</td><td>hypothetical-protein-[Thermomicrobium-sp.]</td></tr>
<tr><td>WP_012643011.1</td><td>Type4.PglZ.containing.BREX</td><td>ABC-ATPase+SbcC->?-><-?<-?||?->?->?-><-?||DUF4007->DUF6079*->PAPS-reduct->DUF4007->?->DUF6079->PglZ->SIGMA+OB-Fold+AAA-ATPase-BrxL->Cysteine-Desulfurase-><-CARF+HEPN<-RAMPs</td><td>1108</td><td>Bacteria>Terrabacteria_group>Thermomicrobiota>Thermomicrobia>Thermomicrobiales>Thermomicrobiaceae</td><td>GCF_000021685.1</td><td>Thermomicrobium_roseum</td><td>TRD_RS09735</td><td>91162..94488</td><td>500</td><td>hypothetical-protein-[Thermomicrobium-roseum]</td></tr>
<tr><td>WP_143714756.1</td><td>Type4.PglZ.containing.BREX</td><td>DUF4007->DUF6079->PAPS-reduct->DUF4007->?->DUF6079*->PglZ->SIGMA+OB-Fold+AAA-ATPase-BrxL->Cysteine-Desulfurase-><-CARF+HEPN<-RAMPs<-?<-RAMPs</td><td>1401</td><td>Bacteria>Terrabacteria_group>Thermomicrobiota>Thermomicrobia>Thermomicrobiales>Thermomicrobiaceae</td><td>GCF_000021685.1</td><td>Thermomicrobium_roseum</td><td>TRD_RS09755</td><td>96797..101002</td><td>500</td><td>hypothetical-protein-[Thermomicrobium-roseum]</td></tr>
<tr><td>MBF0303406.1</td><td>Type4.PglZ.containing.BREX</td><td><-SIGMA+OB-Fold+AAA-ATPase-BrxL<-PglZ<-DUF6079*<-CHTH<-RelE<-?<-DUF4007<-PAPS-reduct+a+b-domain+Fer4_9</td><td>1285</td><td>Bacteria>Thermodesulfobacteriota>Desulfobacteria>Desulfobacterales>Desulfobacteraceae>Desulfamplus</td><td>GCA_015232035.1</td><td>Desulfamplus_sp.</td><td>HQK73_10215</td><td>4373..8230</td><td>2006184</td><td>hypothetical-protein-[Desulfamplus-sp.]</td></tr>
<tr><td>WP_211221097.1</td><td>Type4.PglZ.containing.BREX</td><td><-RelE||?->CHTH->Unstab_antitox->ParB->wHTH+WYL+WCX->DUF4007->RAMA->PAPS-reduct+a+b-domain+Fer4_9+DUF4007->DUF6079-p*->?->PglZ->SIGMA+OB-Fold+AAA-ATPase-BrxL->ZnR-><-SF1-SF2-Helicase<-SF1-SF2-Helicase<-?<-PIN<-wHTH</td><td>970</td><td>Bacteria>Thermodesulfobacteriota>Desulfobacteria>Desulfobacterales>Desulfobacteraceae>Desulfotignum</td><td>GCF_000421285.1</td><td>Desulfotignum_balticum</td><td>K365_RS25475</td><td>195447..198359</td><td>115781</td><td>hypothetical-protein-[Desulfotignum-balticum]</td></tr>
<tr><td>WP_020878637.1</td><td>Type4.PglZ.containing.BREX</td><td>XIS-HTH->RAMA->PAPS-reduct+a+b-domain+Fer4_9+DUF4007->DUF6079*->PglZ->SIGMA+OB-Fold+AAA-ATPase-BrxL->?->?-><-?<-?<-?<-D5ATPase</td><td>1384</td><td>Bacteria>Thermodesulfobacteriota>Desulfobacteria>Desulfobacterales>Desulfococcaceae>Desulfococcus</td><td>GCF_000422185.1</td><td>Desulfococcus_multivorans</td><td>DSMV_RS18975</td><td>4075..8229</td><td>897</td><td>hypothetical-protein-[Desulfococcus-multivorans]</td></tr>
<tr><td>WP_207678198.1</td><td>Type4.PglZ.containing.BREX</td><td>DDE_Tnp->?->?->OLD-ABC->TOPRIM-><-?||HKD-DNase+SF1-SF2-Helicase+HTH+HTH+SRA->PAPS-reduct+a+b-domain+Fer4_9+DUF4007->DUF6079*->iSTAND+PglZ->SIGMA+OB-Fold+AAA-ATPase-BrxL->wHTH(SBD)+HNH->BrxD-AAA->PIN->ParA-Soj-PloopNTPase-></td><td>1129</td><td>Bacteria>Thermodesulfobacteriota>Desulfobacteria>Desulfobacterales>Desulfococcaceae>Desulfonema</td><td>GCF_017377395.1</td><td>Desulfonema_magnum</td><td>dnm_RS17465</td><td>4564753..4568142</td><td>45655</td><td>hypothetical-protein-[Desulfonema-magnum]</td></tr>
<tr><td>WP_223921766.1</td><td>Type4.PglZ.containing.BREX</td><td><-HTH_3||?->?->RAMA->PAPS-reduct+a+b-domain+Fer4_9+DUF4007->DUF6079*->PglZ->SIGMA+OB-Fold+AAA-ATPase-BrxL->ZnR->Cysteine-Desulfurase->?->?-><-tRNA<-IG+IG<-TPR||METHYLASE-></td><td>1384</td><td>Bacteria>Thermodesulfobacteriota>Desulfuromonadia>Geobacterales>Geobacteraceae>Geobacter</td><td>GCF_019972295.1</td><td>Geobacter_sp._AOG2</td><td>LDN12_RS05980</td><td>1299043..1303197</td><td>1566347</td><td>transposase-[Geobacter-sp.-AOG2]</td></tr>
<tr><td>WP_136523554.1</td><td>Type4.PglZ.containing.BREX</td><td><-Phage_integrase||?-><-HTH_3||?->HNH->?->?-><-SIGMA+OB-Fold+AAA-ATPase-BrxL<-PglZ<-DUF6079*<-PAPS-reduct+a+b-domain+Fer4_9+DUF4007<-RAMA<-?<-TIR<-Peptidase_C14<-TIR</td><td>1384</td><td>Bacteria>Thermodesulfobacteriota>Desulfuromonadia>Geobacterales>Geobacteraceae>Geomonas</td><td>GCF_004917065.1</td><td>Geomonas_ferrireducens</td><td>E8L22_RS01670</td><td>362651..366805</td><td>2570227</td><td>transposase-[Geomonas-ferrireducens]</td></tr>
<tr><td>OGU05541.1</td><td>Type4.PglZ.containing.BREX</td><td><-McrC-NTD+REase<-McrB-AAA||?-><-DndB||?-><-SIGMA+OB-Fold+AAA-ATPase-BrxL<-iSTAND+PglZ<-DUF6079*<-PAPS-reduct+a+b-domain+Fer4_9+DUF4007<-?<-AbiEii+ZnR<-AbiEi_4+AbiEi_1||PVC-AAA->?->?-><-tRNA<-?<-HAD+HAD</td><td>1390</td><td>Bacteria>Thermodesulfobacteriota>Desulfuromonadia>Geobacterales>Geobacteraceae>unclassified_Geobacteraceae</td><td>GCA_001802135.1</td><td>Geobacteraceae_bacterium_GWC2_58_44</td><td>A2075_23910</td><td>25996..30168</td><td>1798318</td><td>hypothetical-protein-A2075_23910-[Geobacteraceae-bacterium-GWC2_58_44]</td></tr>
<tr><td>MCK9390122.1</td><td>Type4.PglZ.containing.BREX</td><td><-AbiEii<-?||HIN-HTH->?->?->SIR2->HerA/FtsK->HARE-HTH+E3_UFM1_ligase->PAPS-reduct+a+b-domain+Fer4_9+DUF4007->DUF6079*->iSTAND+PglZ->SIGMA+OB-Fold+AAA-ATPase-BrxL->ZnR->Cysteine-Desulfurase-></td><td>1388</td><td>Bacteria>Thermodesulfobacteriota>Syntrophia>Syntrophales>unclassified_Syntrophales</td><td>GCA_023229565.1</td><td>Syntrophales_bacterium</td><td>MOQ01_00975</td><td>17161..21327</td><td>2928017</td><td>hypothetical-protein-[Syntrophales-bacterium]</td></tr>
<tr><td>WP_011944159.1</td><td>Type4.PglZ.containing.BREX</td><td>AraC-HTH->?->ArsR-HTH->?->Thioredoxin_3->PAPS-reduct+a+b-domain+Fer4_9+DUF4007->PAPS-reduct+DUF4007->DUF6079*->iSTAND+PglZ->SIGMA+OB-Fold+AAA-ATPase-BrxL-><-?<-?<-?||?->?->?->DrsE-></td><td>1123</td><td>Bacteria>Thermotogota>Thermotogae>Thermotogales>Thermotogaceae>Thermotoga</td><td>GCF_000016785.1</td><td>Thermotoga_petrophila</td><td>TPET_RS08935</td><td>1749551..1752922</td><td>93929</td><td>hypothetical-protein-[Thermotoga-petrophila]</td></tr>
<tr><td>NLI07997.1</td><td>Type4.PglZ.containing.BREX</td><td><-SAD+HNH<-Cysteine-Desulfurase<-ZnR<-SIGMA+OB-Fold+AAA-ATPase-BrxL<-iSTAND+PglZ<-DUF6079*<-PAPS-reduct+a+b-domain+Fer4_9+DUF4007<-HARE-HTH+E3_UFM1_ligase<-ZetaToxin</td><td>1379</td><td>Bacteria>Thermotogota>Thermotogae>Thermotogales>Thermotogaceae>unclassified_Thermotogaceae</td><td>GCA_012517055.1</td><td>Thermotogaceae_bacterium</td><td>GX457_13045</td><td>6174..10313</td><td>2268206</td><td>hypothetical-protein-[Thermotogaceae-bacterium]</td></tr>
<tr><td>MBP6982276.1</td><td>Type4.PglZ.containing.BREX</td><td><-Cysteine-Desulfurase<-ZnR<-SIGMA+OB-Fold+AAA-ATPase-BrxL<-iSTAND+PglZ<-DUF6079*<-PAPS-reduct+a+b-domain+Fer4_9+DUF4007<-?||DTXR-HTH->?->?->TatD_DNase-></td><td>1379</td><td>Bacteria>Thermotogota>unclassified_Thermotogota</td><td>GCA_017993175.1</td><td>Thermotogota_bacterium</td><td>KBC37_01725</td><td>5376..9515</td><td>2053689</td><td>hypothetical-protein-[Thermotogota-bacterium]</td></tr>
<tr><td>RKJ03002.1</td><td>Type4.PglZ.containing.BREX</td><td><-DndB<-URI-DNase<-Cysteine-Desulfurase<-ZnR<-SIGMA+OB-Fold+AAA-ATPase-BrxL<-iSTAND+PglZ<-DUF6079*<-PAPS-reduct+a+b-domain+Fer4_9+DUF4007<-SeqA+Phage_int+SeqA+E3_UFM1_ligase<-?||GNTR-HTH-></td><td>1411</td><td>Bacteria>unclassified_Bacteria</td><td>GCA_003611905.1</td><td>bacterium_D16-54</td><td>D7X87_15375</td><td>10196..14431</td><td>2320104</td><td>hypothetical-protein-D7X87_15375-[bacterium-D16-54]</td></tr>
<tr><td>RKI68143.1</td><td>Type4.PglZ.containing.BREX</td><td><-SIGMA-HTH+SIGMA-HTH<-GDResolvase+Recombinase+ZnR<-ACET<-DOC+HTH||?->?->PAPS-reduct+a+b-domain+Fer4_9+DUF4007->DUF6079*->iSTAND+PglZ->SIGMA+OB-Fold+AAA-ATPase-BrxL->ZnR->Cysteine-Desulfurase-><-HTH_3||?->?->ZnR-></td><td>1394</td><td>Bacteria>unclassified_Bacteria</td><td>GCA_003612335.1</td><td>bacterium_1xD42-67</td><td>D7V91_08085</td><td>12343..16527</td><td>2320107</td><td>hypothetical-protein-D7V91_08085-[bacterium-1xD42-67]</td></tr>
<tr><td>NBK77201.1</td><td>Type4.PglZ.containing.BREX</td><td>SIGMA-HTH->Resolvase+Recombinase+ZnR->HEPN->?->PAPS-reduct+a+b-domain+Fer4_9+DUF4007->DUF6079*->iSTAND+PglZ->SIGMA+OB-Fold+AAA-ATPase-BrxL->ZnR->Cysteine-Desulfurase-><-XIS-HTH+XIS-HTH<-Phage_int_SAM_5+Phage_integrase<-XIS-HTH<-TraI-VirD2-RCRE<-TraJ-RHH</td><td>1435</td><td>Bacteria>unclassified_Bacteria</td><td>GCA_009917445.1</td><td>bacterium_D16-76</td><td>D5272_00940</td><td>181388..185695</td><td>2320886</td><td>hypothetical-protein-[bacterium-D16-76]</td></tr>
<tr><td>MBO6272668.1</td><td>Type4.PglZ.containing.BREX</td><td>PAPS-reduct+a+b-domain+Fer4_9+DUF4007->DUF6079*->iSTAND+PglZ->SIGMA+OB-Fold+AAA-ATPase-BrxL->ZnR->Cysteine-Desulfurase->?-><-?||GDResolvase+HIN-HTH-><-CR-ATPase5+REase</td><td>1400</td><td>Bacteria>unclassified_Bacteria</td><td>GCA_017635555.1</td><td>bacterium</td><td>J6048_07790</td><td>3341..7543</td><td>1869227</td><td>hypothetical-protein-[bacterium]</td></tr>
<tr><td>MBP9995129.1</td><td>Type4.PglZ.containing.BREX</td><td><-Glycos_transf<-?||?->DUF4007->DUF1016+REase->HNH+SeqA->PAPS-reduct+a+b-domain+Fer4_9+DUF4007->Schlafen+HATPase+wHTH->HATPase+wHTH->DUF6079*->iSTAND+PglZ->?->SIGMA+OB-Fold+AAA-ATPase-BrxL-></td><td>1184</td><td>Bacteria>unclassified_Bacteria</td><td>GCA_018064605.1</td><td>bacterium</td><td>KBT67_09380</td><td>71344..74898</td><td>1869227</td><td>hypothetical-protein-[bacterium]</td></tr>
</table>

| Protein-ID | DUF499-architecture | System | Gene-Neighbourhoods | Length | Taxonomy | Assembly-ID | Strain | Locus-Tag | Genomic-coordinates | TaxID | Fasta-defline |
|---|---|---|---|---|---|---|---|---|---|---|---|
| -- | | | | | | | | | | | |
| --BR-1.RM.like.systems | | | | | | | | | | | |
| -- | | | | | | | | | | | |
| UCG02108.1 | AAA+ATPase+wHTH+Alpha+Beta-domain+wHTH+RRM | Type-1_BR.RM.like.systems | <-ABhydrolase<-?\|\|DUF499*-><-URI-DNase+HhH-RADC<-ACET\|\|?-><-?<-GTPase-AIG\|\|SF1-SF2-Helicase-> | 889 | Archaea>Asgard_group>Candidatus_Heimdallarchaeota>unclassified_Candidatus_Heimdallarchaeota | GCA_020353515.1 | Candidatus_Heimdallarchaeota_archaeon | JSW11_21265 | 4620650..4623319 | 2026747 | MAG:-ATP-binding-protein-[Candidatus-Heimdallarchaeota-archaeon] |
| PWI48343.1 | AAA+ATPase+wHTH+Alpha+Beta-domain+wHTH+RRM | Type-1_BR.RM.like.systems | <-SF1-SF2-Helicase\|\|GTPase-AIG-><-?\|\|URI-DNase+HhH-RADC-><-DUF499*\|\|?->DSBH->ABhydrolase-> | 863 | Archaea>Asgard_group>Candidatus_Heimdallarchaeota>unclassified_Candidatus_Heimdallarchaeota | GCA_003144275.1 | Candidatus_Heimdallarchaeota_archaeon_B3_Heim | CEE45_07040 | 29302..31893 | 2012493 | hypothetical-protein-CEE45_07040-[Candidatus-Heimdallarchaeota-archaeon-B3_Heim] |
| NHJ00877.1 | AAA+ATPase+wHTH+Alpha+Beta-domain+wHTH+RRM | Type-1_BR.RM.like.systems | <-SF1-SF2-Helicase\|\|GTPase-AIG-><-?\|\|URI-DNase+HhH-RADC-><-DUF499*\|\|?->DSBH->ABhydrolase-> | 848 | Archaea>Asgard_group>Candidatus_Heimdallarchaeota>unclassified_Candidatus_Heimdallarchaeota | GCA_011364965.1 | Candidatus_Heimdallarchaeota_archaeon | EAX86_01990 | 139948..142494 | 2026747 | ATP-binding-protein-[Candidatus-Heimdallarchaeota-archaeon] |
| UCE13226.1 | AAA+ATPase+wHTH+Alpha+Beta-domain+wHTH+RRM | Type-1_BR.RM.like.systems | <-ABhydrolase<-?\|\|DUF499*->ZnR-><-URI-DNase+HhH-RADC\|\|?-><-GTPase-AIG\|\|SF1-SF2-Helicase-> | 882 | Archaea>Asgard_group>Candidatus_Heimdallarchaeota>unclassified_Candidatus_Heimdallarchaeota | GCA_020348965.1 | Candidatus_Heimdallarchaeota_archaeon | JSV04_13700 | 2755433..2758081 | 2026747 | MAG:-ATP-binding-protein-[Candidatus-Heimdallarchaeota-archaeon] |
| UCG90052.1 | AAA+ATPase+wHTH+Alpha+Beta-domain+wHTH+RRM | Type-1_BR.RM.like.systems | <-DSBH<-?\|\|DUF499*->?->?-><-URI-DNase+HhH-RADC<-ACET\|\|?-><-GTPase-AIG | 892 | Archaea>Asgard_group>Candidatus_Heimdallarchaeota>unclassified_Candidatus_Heimdallarchaeota | GCA_020351745.1 | Candidatus_Heimdallarchaeota_archaeon | JSU57_06205 | 1279746..1282424 | 2026747 | MAG:-ATP-binding-protein-[Candidatus-Heimdallarchaeota-archaeon] |
| OLS21118.1 | AAA+ATPase+wHTH+Alpha+Beta-domain+wHTH+RRM | Type-1_BR.RM.like.systems | <-DUF499*<-METHYLASE<-?<-?\|\|?-><-MPTase | 868 | Archaea>Asgard_group>Candidatus_Heimdallarchaeota>unclassified_Candidatus_Heimdallarchaeota | GCA_001940645.1 | Candidatus_Heimdallarchaeota_archaeon_LC_3 | HeimC3_37120 | 6101..8707 | 1841598 | hypothetical-protein-HeimC3_37120-[Candidatus-Heimdallarchaeota-archaeon-LC_3] |
| MBD3343284.1 | AAA+ATPase+wHTH+Alpha+Beta-domain+wHTH+RRM | Type-1_BR.RM.like.systems | TUDOR+SF1-SF2-Helicase+iREase+CTD+REase->METHYLASE->DUF499*->?->PAPS_reduct->ZnR-> | 839 | Archaea>Asgard_group>Candidatus_Lokiarchaeota>unclassified_Candidatus_Lokiarchaeota | GCA_014729315.1 | Candidatus_Lokiarchaeota_archaeon | GF353_29580 | 5892..8411 | 2053489 | DUF499-domain-containing-protein-[Candidatus-Lokiarchaeota-archaeon] |
| RLI54042.1 | AAA+ATPase+wHTH+Alpha+Beta-domain+wHTH+RRM | Type-1_BR.RM.like.systems | TUDOR+SF1-SF2-Helicase+iREase+CTD+REase->REase->METHYLASE->DUF499*-> | 866 | Archaea>Asgard_group>Candidatus_Thorarchaeota>unclassified_Candidatus_Thorarchaeota | GCA_003662775.1 | Candidatus_Thorarchaeota_archaeon | DRP09_14185 | 7004..9603 | 2053491 | AAA-family-ATPase,-partial-[Candidatus-Thorarchaeota-archaeon] |
| MBC7109130.1 | AAA+ATPase+wHTH+Alpha+Beta-domain+wHTH+RRM | Type-1_BR.RM.like.systems | <-SF1-SF2-Helicase<-DUF499-p<-DUF499*<-METHYLASE<-TUDOR+SF1-SF2-Helicase+iREase+CTD+REase | 615 | Archaea>Candidatus_Thermoplasmatota>Thermoplasmata>Methanomassiliicoccales>unclassified_Methanomassiliicoccales | GCA_014361295.1 | Methanomassiliicoccales_archaeon | H5T41_10180 | 1517..3364 | 1906667 | ATP-binding-protein-[Methanomassiliicoccales-archaeon] |
| MDA8055456.1 | AAA+ATPase+wHTH+Alpha+Beta-domain+wHTH+RRM | Type-1_BR.RM.like.systems | <-DUF499*<-METHYLASE<-TUDOR+SF1-SF2-Helicase+iREase+CTD+REase | 844 | Archaea>Candidatus_Thermoplasmatota>Thermoplasmata>Thermoplasmatales>unclassified_Thermoplasmatales | GCA_028063715.1 | Thermoplasmatales_archaeon | MOZ77_07395 | 61..2595 | 2268204 | MAG:-hypothetical-protein-MOZ77_07395-[Thermoplasmatales-archaeon] |
| MBE3122676.1 | AAA+ATPase+wHTH+Alpha+Beta-domain+wHTH+RRM | Type-1_BR.RM.like.systems | TPR+TPR->?->TUDOR+SF1-SF2-Helicase+iREase+CTD+REase->METHYLASE->DUF499*-><-?<-?<-SMODS | 921 | Archaea>Candidatus_Thermoplasmatota>Thermoplasmata>unclassified_Thermoplasmata | GCA_014894395.1 | Thermoplasmata_archaeon | IMZ58_10795 | 35310..38075 | 1906666 | ATP-binding-protein-[Thermoplasmata-archaeon] |
| RLF71961.1 | AAA+ATPase+wHTH+Alpha+Beta-domain+wHTH+RRM | Type-1_BR.RM.like.systems | <-DUF499*<-TOPRIM<-OLD-ABC<-METHYLASE<-?<-SbcC<-TUDOR+SF1-SF2-Helicase+iREase+CTD+REase | 691 | Archaea>Candidatus_Thermoplasmatota>Thermoplasmata>unclassified_Thermoplasmata | GCA_003663625.1 | Thermoplasmata_archaeon | DRN40_01330 | 1..2074 | 1906666 | MAG:-AAA-family-ATPase,-partial-[Thermoplasmata-archaeon] |
| RLF61780.1 | AAA+ATPase+wHTH+Alpha+Beta-domain+wHTH+RRM | Type-1_BR.RM.like.systems | METHYLASE-p->PolB->HEPN->DUF499*-> | 863 | Archaea>Candidatus_Thermoplasmatota>Thermoplasmata>unclassified_Thermoplasmata | GCA_003650005.1 | Thermoplasmata_archaeon | DRN16_03190 | 1440..4028 | 1906666 | MAG:-hypothetical-protein-DRN16_03190,-partial-[Thermoplasmata-archaeon] |
| RLI98027.1 | AAA+ATPase+wHTH+Alpha+Beta-domain+wHTH+RRM | Type-1_BR.RM.like.systems | TUDOR+SF1-SF2-Helicase+iREase+CTD+REase-p->METHYLASE->DUF499*-> | 402 | Archaea>DPANN_group>Candidatus_Aenigmarchaeota>unclassified_Candidatus_Aenigmarchaeota | GCA_003663235.1 | Candidatus_Aenigmarchaeota_archaeon | DRP00_02915 | 3390..4595 | 2093792 | hypothetical-protein-DRP00_02915,-partial-[Candidatus-Aenigmarchaeota-archaeon] |
| MCD6477944.1 | AAA+ATPase+wHTH+Alpha+Beta-domain+wHTH+RRM | Type-1_BR.RM.like.systems | <-DUF499*<-METHYLASE<-TUDOR+SF1-SF2-Helicase+iREase+CTD+REase | 886 | Archaea>DPANN_group>Candidatus_Aenigmarchaeota>unclassified_Candidatus_Aenigmarchaeota | GCA_021162085.1 | Candidatus_Aenigmarchaeota_archaeon | J7K87_02990 | 442..3102 | 2093792 | ATP-binding-protein-[Candidatus-Aenigmarchaeota-archaeon] |
| HDN83604.1 | AAA+ATPase+wHTH+Alpha+Beta-domain+wHTH+RRM | Type-1_BR.RM.like.systems | BirA-HTH->?-><-REase\|\|HTH->?->HKD-DNase-><-DUF499*<-METHYLASE-p | 871 | Archaea>DPANN_group>Candidatus_Altarchaeota>Candidatus_Altarchaeales>unclassified_Candidatus_Altiarchaeales | GCA_011041525.1 | Candidatus_Altiarchaeales_archaeon | ENG50_04490 | 4161..6776 | 2250256 | ATP-binding-protein-[Candidatus-Altiarchaeales-archaeon] |
| MCX6771634.1 | AAA+ATPase+wHTH+Alpha+Beta-domain+wHTH+RRM | Type-1_BR.RM.like.systems | TUDOR+SF1-SF2-Helicase+iREase+CTD+REase-p->METHYLASE->DUF499*->ORC-CDC6+HTH->?-><-VirB4-FtsK<-?<-?<-Methyltransf | 911 | Archaea>DPANN_group>Candidatus_Micrarchaeota>unclassified_Candidatus_Micrarchaeota | GCA_026396415.1 | Candidatus_Micrarchaeota_archaeon | NTX79_06270 | 3316..6051 | 2250274 | MAG:-DUF499-domain-containing-protein-[Candidatus-Micrarchaeota-archaeon] |
| MBM3202348.1 | AAA+ATPase+wHTH+Alpha+Beta-domain+wHTH+RRM | Type-1_BR.RM.like.systems | <-TP_methylase\|\|?->REase->?-><-?<-DUF499* | 944 | Archaea>DPANN_group>Candidatus_Woesearchaeota>unclassified_Candidatus_Woesearchaeota | GCA_016866615.1 | Candidatus_Woesearchaeota_archaeon | FJZ55_00360 | 7583..10417 | 2026803 | ATP-binding-protein-[Candidatus-Woesearchaeota-archaeon] |
| MBU4502274.1 | AAA+ATPase+wHTH+Alpha+Beta-domain+wHTH+RRM | Type-1_BR.RM.like.systems | MACRODOMAIN->?->TUDOR+SF1-SF2-Helicase+iREase+CTD+REase->?->METHYLASE->DUF499*-> | 835 | Archaea>DPANN_group>Nanoarchaeota>unclassified_Nanoarchaeota | GCA_018894775.1 | Nanoarchaeota_archaeon | KKA79_06765 | 20975..23482 | 2026764 | DUF499-domain-containing-protein-[Nanoarchaeota-archaeon] |
| MBU1103405.1 | AAA+ATPase+wHTH+Alpha+Beta-domain+wHTH+RRM | Type-1_BR.RM.like.systems | <-REase+35exo<-DUF499*<-?<-METHYLASE<-REase | 918 | Archaea>DPANN_group>Nanoarchaeota>unclassified_Nanoarchaeota | GCA_018817905.1 | Nanoarchaeota_archaeon | KJ600_02510 | 1498..4254 | 2026764 | hypothetical-protein-[Nanoarchaeota-archaeon] |
| CBH38392.1 | AAA+ATPase+wHTH+Alpha+Beta-domain+wHTH+RRM | Type-1_BR.RM.like.systems | <-HicB_lk_antitox\|\|?->METHYLASE-p->TUDOR+SF1-SF2-Helicase+iREase+CTD+REase->Schlafen->METHYLASE->DUF499*->?-><-?<-?<-?<-SPOUT | 900 | Archaea>environmental_samples | - | uncultured_archaeon | BSM_18690 | 1759679..1762381 | 115547 | conserved-hypothetical-protein-[uncultured-archaeon] |
| VVB90090.1 | AAA+ATPase+wHTH+Alpha+Beta-domain+wHTH+RRM | Type-1_BR.RM.like.systems | METHYLASE-p->PolB->HEPN->?->Polbeta->?->DUF499*-><-METHYLASE-p<-ParA-Soj-PloopNTPase | 845 | Archaea>environmental_samples | GCA_902386115.1 | uncultured_archaeon | LFW28022_00224 | 2135..4672 | 115547 | Uncharacterised-protein-[uncultured-archaeon] |
| WP_012966137.1 | AAA+ATPase+wHTH+Alpha+Beta-domain+wHTH+RRM | Type-1_BR.RM.like.systems | <-ABC-ATPase+SbcC<-?<-?\|\|DUF499*-><-METHYLASE<-?\|\|PADR-HTH->PIN->REase->SF1-SF2-Helicase-> | 977 | Archaea>Euryarchaeota>Archaeoglobi>Archaeoglobales>Archaeoglobaceae | GCF_000025505.1 | Ferroglobus_placidus | FERP_RS08250 | 1427325..1430258 | 54261 | ATP-binding-protein-[Ferroglobus-placidus] |
| MCS7144689.1 | AAA+ATPase+wHTH+Alpha+Beta-domain+wHTH+RRM | Type-1_BR.RM.like.systems | METHYLASE->DUF499*->?->TUDOR+SF1-SF2-Helicase+iREase+CTD+REase-> | 1110 | Archaea>Euryarchaeota>Archaeoglobi>Archaeoglobales>Archaeoglobaceae | GCA_025059545.1 | Archaeoglobaceae_archaeon | NZ879_06680 | 2440..5772 | 2511821 | ATP-binding-protein-[Archaeoglobaceae-archaeon] |
| MBO8180197.1 | AAA+ATPase+wHTH+Alpha+Beta-domain+wHTH+RRM | Type-1_BR.RM.like.systems | <-HEPN<-PolB<-?<-?<-EVE<-DUF499*<-HEPN<-Polbeta\|\|?-><-DEDD_Tnp_IS110+Transposase_20 | 875 | Archaea>Euryarchaeota>Archaeoglobi>Archaeoglobales>Archaeoglobaceae | GCA_017656475.1 | Archaeoglobus_sp. | HOZ19_06905 | 5134..7761 | 1872626 | ATP-binding-protein-[Archaeoglobus-sp.] |
| RLI82780.1 | AAA+ATPase+wHTH+Alpha+Beta-domain+wHTH+RRM | Type-1_BR.RM.like.systems | <-TUDOR+SF1-SF2-Helicase+iREase+CTD+REase<-?<-OLD-ABC<-?<-DUF499*<-METHYLASE-p | 1128 | Archaea>Euryarchaeota>Archaeoglobi>Archaeoglobales>unclassified_Archaeoglobales | GCA_003662925.1 | Archaeoglobales_archaeon | DRP04_03205 | 5696..9082 | 2250258 | hypothetical-protein-DRP04_03205-[Archaeoglobales-archaeon] |
| RLI77774.1 | AAA+ATPase+wHTH+Alpha+Beta-domain+wHTH+RRM | Type-1_BR.RM.like.systems | METHYLASE-p-><-TOPRIM<-OLD-ABC\|\|MazE_antitoxin->PIN->DUF499*-> | 227 | Archaea>Euryarchaeota>Archaeoglobi>Archaeoglobales>unclassified_Archaeoglobales | GCA_003662925.1 | Archaeoglobales_archaeon | DRP04_10895 | 3032..3711 | 2250258 | hypothetical-protein-DRP04_10895,-partial-[Archaeoglobales-archaeon] |
| RLI80545.1 | AAA+ATPase+wHTH+Alpha+Beta-domain+wHTH+RRM | Type-1_BR.RM.like.systems | METHYLASE-p->?->?->DUF499*->EVE-> | 877 | Archaea>Euryarchaeota>Archaeoglobi>Archaeoglobales>unclassified_Archaeoglobales | GCA_003662925.1 | Archaeoglobales_archaeon | DRP04_08060 | 1906..4539 | 2250258 | AAA-family-ATPase-[Archaeoglobales-archaeon] |
| WP_229235289.1 | AAA+ATPase+wHTH+Alpha+Beta-domain+wHTH+RRM | Type-1_BR.RM.like.systems | <-DUF499*<-METHYLASE<-METHYLASE-p<-TUDOR+SF1-SF2-Helicase+iREase+CTD+REase<-?<-tRNA<-?<-Endonuclease_5 | 856 | Archaea>Euryarchaeota>Candidatus_Methanoliparia>Candidatus_Methanoliparales>Candidatus_Methanoliparaceae | GCF_020885915.1 | Candidatus_Methanoliparum_sp._LAM-1 | LPC15_RS03270 | 585052..587622 | 2874846 | DUF499-domain-containing-protein-[Candidatus-Methanoliparum-sp.-LAM-1] |
| RZN64443.1 | AAA+ATPase+wHTH+Alpha+Beta-domain+wHTH+RRM | Type-1_BR.RM.like.systems | Endonuclease_5->?->tRNA->TUDOR+SF1-SF2-Helicase+iREase+CTD+REase->METHYLASE->DUF499*-> | 858 | Archaea>Euryarchaeota>Candidatus_Methanoliparia>Candidatus_Methanoliparales>Candidatus_Methanoliparaceae | GCA_004212075.1 | Candidatus_Methanoliparum_thermophilum | EF806_03620 | 30329..32905 | 2491083 | ATP-binding-protein-[Candidatus-Methanoliparum-thermophilum] |
| KXB06557.1 | AAA+ATPase+wHTH+Alpha+Beta-domain+wHTH+RRM | Type-1_BR.RM.like.systems | PIN-><-DUF499*<-METHYLASE<-TUDOR+SF1-SF2-Helicase+iREase+CTD+REase-p | 953 | Archaea>Euryarchaeota>Euryarchaeota_incertae_sedis>candidate_division_MSBL1 | GCA_001549275.1 | candidate_division_MSBL1_archaeon_SCGC-AAA382A20 | AKJ51_03505 | 2453..5314 | 1698280 | hypothetical-protein-AKJ51_03505-[candidate-division-MSBL1-archaeon-SCGC-AAA382A20] |
| EKQ50758.1 | AAA+ATPase+wHTH+Alpha+Beta-domain+wHTH+RRM | Type-1_BR.RM.like.systems | <-EVE<-?\|\|PIN-><-?<-DUF499*\|\|MORC->REase->ParB+HNH-><-SbcC+SbcC | 701 | Archaea>Euryarchaeota>Methanomada_group>Methanobacteria>Methanobacteriales | GCA_000309865.1 | Methanobacterium_sp._Maddingley_MBC34 | B655_2273 | 4021..6126 | 1220534 | MAG:-putative-ATPase-(AAA+-superfamily)-[Methanobacterium-sp.-Maddingley-MBC34] |
| WP_013295752.1 | AAA+ATPase+wHTH+Alpha+Beta-domain+wHTH+RRM | Type-1_BR.RM.like.systems | <-SbcC+SbcC<-Calcineurin<-FtsK+VirB4-FtsK+HerA<-NurA<-?\|\|Glyco_transf-><-DUF499*<-METHYLASE<-TUDOR+SF1-SF2-Helicase+iREase+CTD+REase<-TUDOR+SF1-SF2-Helicase+iREase+CTD+REase-p\|\|?-><-TPR+TPR | 904 | Archaea>Euryarchaeota>Methanomada_group>Methanobacteria>Methanobacteriales | GCF_000145295.1 | Methanothermobacter_marburgensis | MTBMA_RS04570 | 869962..872676 | 145263 | DUF499-domain-containing-protein-[Methanothermobacter-marburgensis] |
| WP_237779669.1 | AAA+ATPase+wHTH+Alpha+Beta-domain+wHTH+RRM | Type-1_BR.RM.like.systems | <-Peptidase<-SbcC+SbcC<-Calcineurin<-FtsK+VirB4-FtsK+HerA<-NurA\|\|Glyco_transf-><-DUF499*<-METHYLASE<-TUDOR+SF1-SF2-Helicase+iREase+CTD+REase\|\|?-><-?<-?\|\|METHYLASE-p-> | 896 | Archaea>Euryarchaeota>Methanomada_group>Methanobacteria>Methanobacteriales | GCF_022014235.1 | Methanothermobacter_sp._K4 | L5462_RS04755 | 204522..207212 | 2913262 | DUF499-domain-containing-protein-[Methanothermobacter-sp.-K4] |
| AXV39094.1 | AAA+ATPase+wHTH+Alpha+Beta-domain+wHTH+RRM | Type-1_BR.RM.like.systems | <-CR-ATPase9<-?<-?<-METHYLASE-p<-TUDOR+SF1-SF2-Helicase+iREase+CTD+REase<-DUF499*<-METHYLASE<-TUDOR+SF1-SF2-Helicase+iREase+CTD+REase\|\|HHH_4+SF1-SF2-Helicase+REase-><-?<-N6_MTase+TRD+HTL1 | 1017 | Archaea>Euryarchaeota>Methanomada_group>Methanobacteria>Methanobacteriales | GCA_003491305.1 | Methanobacterium_sp._BAmetb5 | CIT02_01575 | 337652..340705 | 2025351 | MAG:-hypothetical-protein-CIT02_01575-[Methanobacterium-sp.-BAmetb5] |
| SCM57147.1 | AAA+ATPase+wHTH+Alpha+Beta-domain+wHTH+RRM | Type-1_BR.RM.like.systems | <-SbcC+SbcC<-Calcineurin<-FtsK+VirB4-FtsK+HerA<-NurA<-?\|\|Glyco_transf-><-DUF499*<-METHYLASE<-TUDOR+SF1-SF2-Helicase+iREase+CTD+REase\|\|?-><-?<-Hpr-Kinase-N | 903 | Archaea>Euryarchaeota>Methanomada_group>Methanobacteria>Methanobacteriales | GCA_900095815.1 | Methanothermobacter_wolfeii | MWSIV6_0911 | 858412..861123 | 145261 | ATPase-{ECO:0000313\|EMBL:BAM69720,1}-[Methanothermobacter-wolfeii] |
| WP_261599343.1 | AAA+ATPase+wHTH+Alpha+Beta-domain+wHTH+RRM | Type-1_BR.RM.like.systems | <-Peptidase<-SbcC+SbcC<-Calcineurin<-FtsK+VirB4-FtsK+HerA<-NurA<-?<-DUF499*<-METHYLASE<-TUDOR+SF1-SF2-Helicase+iREase+CTD+REase\|\|?-><-?\|\|?-><-Hpr-Kinase-N | 855 | Archaea>Euryarchaeota>Methanomada_group>Methanobacteria>Methanobacteriales | GCF_025397995.1 | Methanothermobacter_wolfeii | N5910_RS04690 | 870325..872892 | 145261 | DUF499-domain-containing-protein-[Methanothermobacter-wolfeii] |
| WP_060537629.1 | AAA+ATPase+wHTH+Alpha+Beta-domain+wHTH+RRM | Type-1_BR.RM.like.systems | N6_MTase+TRD+HTL1->?-><-?<-?<-SF1-SF2-Helicase\|\|TUDOR+SF1-SF2-Helicase+iREase+CTD+REase->DUF499*->METHYLASE->METHYLASE-p->METHYLASE-p->HNH->McrC-NTD+REase-> | 1010 | Archaea>Euryarchaeota>Methanomada_group>Methanobacteria>Methanobacteriales | GCF_001458655.1 | Methanobacterium_formicicum | MB9_RS05635 | 1192394..1195426 | 2162 | DUF499-domain-containing-protein-[Methanobacterium-formicicum] |
| MBW4257496.1 | AAA+ATPase+wHTH+Alpha+Beta-domain+wHTH+RRM | Type-1_BR.RM.like.systems | <-ParB+HNH\|\|N6_MTase+TRD+HTL1-><-?\|\|?->TUDOR+SF1-SF2-Helicase+iREase+CTD+REase->METHYLASE->DUF499*->TUDOR+SF1-SF2-Helicase+iREase+CTD+REase->METHYLASE-p->?->?->CR-ATPase9-> | 1017 | Archaea>Euryarchaeota>Methanomada_group>Methanobacteria>Methanobacteriales | GCA_019351055.1 | Methanobacterium_sp._YSL | KTG15_07295 | 1502044..1505097 | 2841655 | hypothetical-protein-[Methanobacterium-sp.-YSL] |
| WP_048175355.1 | AAA+ATPase+wHTH+Alpha+Beta-domain+wHTH+RRM | Type-1_BR.RM.like.systems | <-SbcC+SbcC<-Calcineurin<-FtsK+VirB4-FtsK+HerA<-NurA<-?\|\|Glyco_transf-><-DUF499*<-METHYLASE<-TUDOR+SF1-SF2-Helicase+iREase+CTD+REase\|\|?-><-TPR+TPR | 904 | Archaea>Euryarchaeota>Methanomada_group>Methanobacteria>Methanobacteriales | GCF_000828575.1 | Methanothermobacter_sp._CaT2 | MTCT_RS02355 | 461521..464235 | 866790 | DUF499-domain-containing-protein-[Methanothermobacter-sp.-CaT2] |
| WP_191216553.1 | AAA+ATPase+wHTH+Alpha+Beta-domain+wHTH+RRM | Type-1_BR.RM.like.systems | <-Peptidase<-SbcC+SbcC<-Calcineurin<-FtsK+VirB4-FtsK+HerA<-NurA<-?<-DUF499*<-METHYLASE<-TUDOR+SF1-SF2-Helicase+iREase+CTD+REase | 855 | Archaea>Euryarchaeota>Methanomada_group>Methanobacteria>Methanobacteriales | GCF_009914355.1 | Methanothermobacter_sp._THM-1 | FZP57_RS04385 | 843523..846090 | 2606911 | DUF499-domain-containing-protein-[Methanothermobacter-sp.-THM-1] |
| WP_160323097.1 | AAA+ATPase+wHTH+Alpha+Beta-domain+wHTH+RRM | Type-1_BR.RM.like.systems | <-FtsK+VirB4-FtsK+HerA<-NurA<-?<-?\|\|Glyco_transf-><-DUF499-p<-DUF499*<-METHYLASE<-TUDOR+SF1-SF2-Helicase+iREase+CTD+REase\|\|?-><-TPR+TPR | 328 | Archaea>Euryarchaeota>Methanomada_group>Methanobacteria>Methanobacteriales | GCF_009917665.1 | Methanothermobacter_sp._THM-2 | FZP68_RS04845 | 935607..936593 | 2606912 | DUF499-domain-containing-protein-[Methanothermobacter-sp.-THM-2] |
| WP_115892292.1 | AAA+ATPase+wHTH+Alpha+Beta-domain+wHTH+RRM | Type-1_BR.RM.like.systems | TUDOR+SF1-SF2-Helicase+iREase+CTD+REase->METHYLASE->DUF499*->?->NurA->FtsK+VirB4-FtsK+HerA->Calcineurin->SbcC+SbcC->Peptidase-> | 855 | Archaea>Euryarchaeota>Methanomada_group>Methanobacteria>Methanobacteriales | GCF_003385755.1 | Methanothermobacter_defluvii | C7452_RS04080 | 750536..753103 | 49339 | DUF499-domain-containing-protein-[Methanothermobacter-defluvii] |
| TMS41412.1 | AAA+ATPase+wHTH+Alpha+Beta-domain+wHTH+RRM | Type-1_BR.RM.like.systems | Primpol+PriCT1->?->?->?->?->TUDOR+SF1-SF2-Helicase+iREase+CTD+REase->DUF499*->?->METHYLASE->?->METHYLASE-p->?->SF1-SF2-Helicase+HNH-> | 1011 | Archaea>Euryarchaeota>Methanomada_group>Methanobacteria>Methanobacteriales | GCA_005924185.1 | Methanobacterium_sp. | FGO69_08395 | 12026..15061 | 2164 | MAG:-ATP-binding-protein-[Methanobacterium-sp.] |
| WP_219304522.1 | AAA+ATPase+wHTH+Alpha+Beta-domain+wHTH+RRM | Type-1_BR.RM.like.systems | <-CR-ATPase9<-?<-?<-METHYLASE-p<-TUDOR+SF1-SF2-Helicase+iREase+CTD+REase<-DUF499*<-METHYLASE<-TUDOR+SF1-SF2-Helicase+iREase+CTD+REase-p | 1017 | Archaea>Euryarchaeota>Methanomada_group>Methanobacteria>Methanobacteriales | - | Methanobacterium_sp._YSL | DSECCO2_272550 | 16298..19351 | 2841655 | hypothetical-protein-[Methanobacterium-sp.-YSL] |
| NLM03057.1 | AAA+ATPase+wHTH+Alpha+Beta-domain+wHTH+RRM | Type-1_BR.RM.like.systems | DUF499*->?->NurA->FtsK+VirB4-FtsK+HerA->Calcineurin->SbcC+SbcC->Peptidase-> | 855 | Archaea>Euryarchaeota>Methanomada_group>Methanobacteria>Methanobacteriales | GCA_012521115.1 | Methanothermobacter_wolfeii | GX212_06825 | 89..2656 | 145261 | ATP-binding-protein-[Methanothermobacter-wolfeii] |
| MBC7110764.1 | AAA+ATPase+wHTH+Alpha+Beta-domain+wHTH+RRM | Type-1_BR.RM.like.systems | <-DUF499*<-METHYLASE<-TUDOR+SF1-SF2-Helicase+iREase+CTD+REase\|\|?-><-TPR+TPR | 641 | Archaea>Euryarchaeota>Methanomada_group>Methanobacteria>Methanobacteriales | GCA_014361435.1 | Methanothermobacter_sp. | H5T35_00885 | 1..1924 | 1884223 | ATP-binding-protein-[Methanothermobacter-sp.] |
| WP_007043961.1 | AAA+ATPase+wHTH+Alpha+Beta-domain+wHTH+RRM | Type-1_BR.RM.like.systems | METHYLASE-p->METHYLASE-p->MNS->DUF499*->Cluster35_66clades->TUDOR+SF1-SF2-Helicase+iREase+CTD+REase-p->TUDOR+SF1-SF2-Helicase+iREase+CTD+REase->REase->EVE-> | 1104 | Archaea>Euryarchaeota>Methanomada_group>Methanococci>Methanococcales | GCF_000243455.1 | Methanotorris_formicicus | METFODRAFT_RS02510 | 35682..38996 | 213185 | DUF499-domain-containing-protein-[Methanotorris-formicicus] |
| WP_015733362.1 | AAA+ATPase+wHTH+Alpha+Beta-domain+wHTH+RRM | Type-1_BR.RM.like.systems | METHYLASE->MNS->DUF499*->Cluster35_66clades->TUDOR+SF1-SF2-Helicase+iREase+CTD+REase->HEPN->RAMPs->HDRnase-> | 1104 | Archaea>Euryarchaeota>Methanomada_group>Methanococci>Methanococcales | GCF_000024625.1 | Methanocaldococcus_vulcanius | METVU_RS06315 | 1284462..1287776 | 73913 | DUF499-domain-containing-protein-[Methanocaldococcus-vulcanius] |
| WP_017981059.1 | AAA+ATPase+wHTH+Alpha+Beta-domain+wHTH+RRM | Type-1_BR.RM.like.systems | DUF499*->Cluster35_66clades->TUDOR+SF1-SF2-Helicase+iREase+CTD+REase->?->Hydrolase_3-><-SF1-SF2-Helicase | 1137 | Archaea>Euryarchaeota>Methanomada_group>Methanococci>Methanococcales | GCF_000371805.1 | Methanocaldococcus_villosus | METVI_RS0104265 | 721661..725074 | 667126 | DUF499-domain-containing-protein-[Methanocaldococcus-villosus] |
| WP_004592902.1 | AAA+ATPase+wHTH+Alpha+Beta-domain+wHTH+RRM | Type-1_BR.RM.like.systems | <-TUDOR+SF1-SF2-Helicase+iREase+CTD+REase<-Cluster35_66clades<-DUF499* | 1128 | Archaea>Euryarchaeota>Methanomada_group>Methanococci>Methanococcales | GCF_000363885.1 | Methanocaldococcus_villosus | J422_RS05570 | 4111..7497 | 667126 | DUF499-domain-containing-protein-[Methanocaldococcus-villosus] |
| WP_214400399.1 | AAA+ATPase+wHTH+Alpha+Beta-domain+wHTH+RRM | Type-1_BR.RM.like.systems | HEPN->METHYLASE->SbcC-><-TOPRIM<-TOPRIM<-OLD-ABC\|\|DUF499*-><-?<-?\|\|MarR-HTH->?-><-Peptidase_M42 | 878 | Archaea>Euryarchaeota>Methanomada_group>Methanococci>Methanococcales | GCF_902827225.1 | Methanocaldococcus_lauensis | KMP69_RS02590 | 459984..462620 | 2546128 | DUF499-domain-containing-protein-[Methanocaldococcus-sp.-SG7] |
| WP_048201423.1 | AAA+ATPase+wHTH+Alpha+Beta-domain+wHTH+RRM | Type-1_BR.RM.like.systems | PADR-HTH->PIN->?->METHYLASE->?->MNS->DUF499*->?-><-?\|\|?-><-ABC-ATPase+SbcC<-Peptidase | 887 | Archaea>Euryarchaeota>Methanomada_group>Methanococci>Methanococcales | GCF_000739065.1 | Methanocaldococcus_bathoardescens | JH146_RS01935 | 372793..375456 | 1301915 | DUF499-domain-containing-protein-[Methanocaldococcus-bathoardescens] |
| WP_148119962.1 | AAA+ATPase+wHTH+Alpha+Beta-domain+wHTH+RRM | Type-1_BR.RM.like.systems | TUDOR+SF1-SF2-Helicase+iREase+CTD+REase->METHYLASE->DUF499*->?->?-><-?<-?<-Peptidase_M22<-HD | 852 | Archaea>Euryarchaeota>Methanomada_group>Methanococci>Methanococcales | GCF_003351865.1 | Methanofervidicoccus_sp._A16 | CFE53_RS00575 | 131182..133740 | 2607662 | ATP-binding-protein-[Methanofervidicoccus-sp.-A16] |
| WP_214400221.1 | AAA+ATPase+wHTH+Alpha+Beta-domain+wHTH+RRM | Type-1_BR.RM.like.systems | DUF499*-><-?<-?\|\|?-><-?\|\|ParA-Soj-PloopNTPase-><-Radical-SAM | 250 | Archaea>Euryarchaeota>Methanomada_group>Methanococci>Methanococcales | GCF_902827225.1 | Methanocaldococcus_lauensis | KMP69_RS01560 | 269632..270384 | 2546128 | DUF499-domain-containing-protein-[Methanocaldococcus-lauensis] |
| MBW9223096.1 | AAA+ATPase+wHTH+Alpha+Beta-domain+wHTH+RRM | Type-1_BR.RM.like.systems | METHYLASE-p->DUF499*->?->?-><-?<-Peptidase_M22<-HD | 851 | Archaea>Euryarchaeota>Methanomada_group>Methanococci>Methanococcales | GCA_019561615.1 | Methanothermococcus_sp._SCGC_AD-155-E23 | KKP90_00650 | 638..3193 | 2838162 | hypothetical-protein-[Methanothermococcus-sp.-SCGC-AD-155-E23] |
| HIQ32859.1 | AAA+ATPase+wHTH+Alpha+Beta-domain+wHTH+RRM | Type-1_BR.RM.like.systems | <-DUF499*<-METHYLASE-p | 849 | Archaea>Euryarchaeota>Methanomada_group>Methanococci>Methanococcales | GCA_015661555.1 | Methanothermococcus_okinawensis | EYH55_05210 | 3856..6405 | 155863 | ATP-binding-protein-[Methanothermococcus-okinawensis] |
| WP_011019628.1 | AAA+ATPase+wHTH+Alpha+Beta-domain+wHTH+RRM | Type-1_BR.RM.like.systems | PIN-><-?<-?\|\|Mbetalac-><-METHYLASE\|\|DUF499*->?->?->?->ParA-Soj-PloopNTPase-> | 722 | Archaea>Euryarchaeota>Methanomada_group>Methanopyri>Methanopyrales | GCF_000007185.1 | Methanopyrus_kandleri | MK_RS06710 | 1248593..1250761 | 2320 | ATP-binding-protein-[Methanopyrus-kandleri] |
| WP_269070304.1 | AAA+ATPase+wHTH+Alpha+Beta-domain+wHTH+RRM | Type-1_BR.RM.like.systems | Zn_Tnp_IS1595-><-HEPN<-METHYLASE\|\|?->Zn_Tnp_IS1595+DDE_Tnp_IS1595-><-PADR-HTH<-DUF499*<-?\|\|SF1-SF2-Helicase->?-><-SF1-SF2-Helicase+REase<-REase | 1071 | Archaea>Euryarchaeota>Stenosarchaea_group>Halobacteria>Halobacteriales | GCF_021028995.1 | Halobacterium_sp._KA-4 | LPA44_RS15370 | 20251..23466 | 2896367 | DUF499-domain-containing-protein-[Halobacterium-sp.-KA-4] |
| WP_230890402.1 | AAA+ATPase+wHTH+Alpha+Beta-domain+wHTH+RRM | Type-1_BR.RM.like.systems | Zn_Tnp_IS1595-><-HEPN<-METHYLASE\|\|?->Zn_Tnp_IS1595+DDE_Tnp_IS1595-><-PADR-HTH<-DUF499*\|\|SF1-SF2-Helicase->?-><-SF1-SF2-Helicase+REase<-REase | 1057 | Archaea>Euryarchaeota>Stenosarchaea_group>Halobacteria>Halobacteriales | - | Halobacterium_sp._KA-4 | LPA44_15370 | 20251..23424 | 2896367 | DUF499-domain-containing-protein-[Halobacterium-sp.-KA-4] |
| MCD2201253.1 | AAA+ATPase+wHTH+Alpha+Beta-domain+wHTH+RRM | Type-1_BR.RM.like.systems | Zn_Tnp_IS1595-><-HEPN<-METHYLASE\|\|?->Zn_Tnp_IS1595+DDE_Tnp_IS1595-><-PADR-HTH<-DUF499*\|\|SF1-SF2-Helicase->?-><-SF1-SF2-Helicase+REase<-REase | 1057 | Archaea>Euryarchaeota>Stenosarchaea_group>Halobacteria>Halobacteriales | GCA_021028995.1 | Halobacterium_sp._KA-4 | LPA44_15370 | 20251..23424 | 2896367 | DUF499-domain-containing-protein-[Halobacterium-sp.-KA-4] |
| WP_181685013.1 | AAA+ATPase+wHTH+Alpha+Beta-domain+wHTH+RRM | Type-1_BR.RM.like.systems | <-MarR-HTH\|\|?->?-><-SF1-SF2-Helicase<-?<-SF1-SF2-Helicase\|\|DUF499*-><-?<-ParB+HNH\|\|?->METHYLASE-><-DNA-glycosylase\|\|ParB+HNH-> | 1068 | Archaea>Euryarchaeota>Stenosarchaea_group>Halobacteria>Halobacteriales | GCF_013839425.1 | Halorhabdus_salina | HZR20_RS02465 | 491897..495103 | 2750670 | DUF499-domain-containing-protein-[Halorhabdus-salina] |
| WP_237006757.1 | AAA+ATPase+wHTH+Alpha+Beta-domain+wHTH+RRM | Type-1_BR.RM.like.systems | <-HEPN\|\|REase->SF1-SF2-Helicase+REase-><-?<-SF1-SF2-Helicase\|\|?->DUF499*-><-DOC+HTH<-?<-?<-?<-?\|\|METHYLASE-> | 1068 | Archaea>Euryarchaeota>Stenosarchaea_group>Halobacteria>Halobacteriales | GCF_021729015.1 | Halobacterium | QFK75_RS08735 | 232135..235341 | 2239 | DUF499-domain-containing-protein-[Halobacterium-sp.-TGN-42-S1] |
| WP_220599369.1 | AAA+ATPase+wHTH+Alpha+Beta-domain+wHTH+RRM | Type-1_BR.RM.like.systems | <-PADR-HTH<-?\|\|ParB+HNH->MarR-HTH->?-><-DUF499*<-?\|\|PADR-HTH->SF1-SF2-Helicase->?-><-SF1-SF2-Helicase<-Resolvase | 1070 | Archaea>Euryarchaeota>Stenosarchaea_group>Halobacteria>Halobacteriales | GCF_019599485.1 | Halomicroarcula_salinisoli | EGH22_RS09775 | 143236..146448 | 2487746 | DUF499-domain-containing-protein-[Halomicroarcula-salinisoli] |
| WP_151104334.1 | AAA+ATPase+wHTH+Alpha+Beta-domain+wHTH+RRM | Type-1_BR.RM.like.systems | <-SF1-SF2-Helicase\|\|?->DUF499*-><-?<-MarR-HTH<-ParB+HNH<-PHP+SbcC+SidE+CR-ATPase9<-RHH+RAMA\|\|METHYLASE-> | 1070 | Archaea>Euryarchaeota>Stenosarchaea_group>Halobacteria>Halobacteriales | GCF_008729095.1 | Haloarcula_hispanica | EGO51_RS19160 | 76643..79855 | 51589 | DUF499-domain-containing-protein-[Haloarcula-hispanica] |
| WP_146418283.1 | AAA+ATPase+wHTH+Alpha+Beta-domain+wHTH+RRM | Type-1_BR.RM.like.systems | <-RNase<-METHYLASE\|\|ParB+HNH-><-DUF499*\|\|SF1-SF2-Helicase->?-><-ORC/CDC6+HTH | 1068 | Archaea>Euryarchaeota>Stenosarchaea_group>Halobacteria>Halobacteriales | GCF_902209435.1 | Haloarcula_hispanica | FUT35_RS18485 | 5617..8823 | 51589 | DUF499-domain-containing-protein-[Haloarcula-hispanica] |
| WP_240147535.1 | AAA+ATPase+wHTH+Alpha+Beta-domain+wHTH+RRM | Type-1_BR.RM.like.systems | <-METHYLASE<-?\|\|?->?-><-DUF499*\|\|SF1-SF2-Helicase->?-><-?\|\|?-><-TIR<-SLATT | 1062 | Archaea>Euryarchaeota>Stenosarchaea_group>Halobacteria>Halobacteriales | GCF_011440385.1 | Halorussus | G9466_RS13305 | 934040..937228 | 1070314 | MULTISPECIES:-DUF499-domain-containing-protein-[Halorussus] |
| NHN60046.1 | AAA+ATPase+wHTH+Alpha+Beta-domain+wHTH+RRM | Type-1_BR.RM.like.systems | <-METHYLASE<-?\|\|?->?-><-DUF499*\|\|SF1-SF2-Helicase->?-><-?\|\|?->?-><-TIR | 1068 | Archaea>Euryarchaeota>Stenosarchaea_group>Halobacteria>Halobacteriales | GCA_011440385.1 | Halorussus_sp._JP-T4 | G9466_13305 | 934040..937246 | 2716718 | DUF499-domain-containing-protein-[Halorussus-sp.-JP-T4] |
| WP_188854179.1 | AAA+ATPase+wHTH+Alpha+Beta-domain+wHTH+RRM | Type-1_BR.RM.like.systems | <-PIN<-?<-?<-METHYLASE\|\|?-><-?<-DUF499*\|\|SF1-SF2-Helicase-> | 1068 | Archaea>Euryarchaeota>Stenosarchaea_group>Halobacteria>Halobacteriales | GCF_014647095.1 | Haloarcula_argentinensis | IEY10_RS18745 | 18274..21480 | 43776 | DUF499-domain-containing-protein-[Haloarcula-salaria] |
| ERH10996.1 | AAA+ATPase+wHTH+Alpha+Beta-domain+wHTH+RRM | Type-1_BR.RM.like.systems | <-SF1-SF2-Helicase\|\|DUF499*-><-?<-?\|\|?->?-><-?\|\|ParB+HNH-p-> | 565 | Archaea>Euryarchaeota>Stenosarchaea_group>Halobacteria>Halobacteriales | GCA_000416085.1 | halophilic_archaeon_J07HX64 | J07HX64_02775 | 2657984..2659681 | 1085028 | hypothetical-protein-J07HX64_02775-[halophilic-archaeon-J07HX64] |
| WP_092695686.1 | AAA+ATPase+wHTH+Alpha+Beta-domain+wHTH+RRM | Type-1_BR.RM.like.systems | DDE_Tnp_1-><-?<-REase<-METHYLASE\|\|DUF499*-> | 805 | Archaea>Euryarchaeota>Stenosarchaea_group>Halobacteria>Halobacteriales | GCF_900102305.1 | Halorientalis_regularis | BLQ72_RS20055 | 9401..11817 | 660518 | DUF499-domain-containing-protein,-partial-[Halorientalis-regularis] |
| WP_049920397.1 | AAA+ATPase+wHTH+Alpha+Beta-domain+wHTH+RRM | Type-1_BR.RM.like.systems | <-METHYLASE<-MORC+DUF3684+REase\|\|ParB+HNH-><-?\|\|?-><-Zn_Tnp_IS1595+DDE_Tnp_IS1595<-DUF499*\|\|SF1-SF2-Helicase->?-><-RHH_6+RHH_6<-?<-Phage_integrase<-SF1-SF2-Helicase | 1071 | Archaea>Euryarchaeota>Stenosarchaea_group>Halobacteria>Haloferacales | GCF_000337835.1 | Haloferax_sulfurifontis | C441_RS03265 | 199646..202861 | 255616 | DUF499-domain-containing-protein-[Haloferax-sulfurifontis] |
| ELZ97832.1 | AAA+ATPase+wHTH+Alpha+Beta-domain+wHTH+RRM | Type-1_BR.RM.like.systems | <-METHYLASE<-MORC+DUF3684+REase\|\|?->ParB+HNH->?-><-Zn_Tnp_IS1595+DDE_Tnp_IS1595<-DUF499*\|\|?->SF1-SF2-Helicase-><-?<-RHH_6+RHH_6<-Phage_integrase<-SF1-SF2-Helicase | 940 | Archaea>Euryarchaeota>Stenosarchaea_group>Halobacteria>Haloferacales | GCA_000337835.1 | Haloferax_sulfurifontis_ATCC_BAA-897 | C441_03337 | 199646..202468 | 662480 | ATPase-(AAA+-superfamily)-like-protein-[Haloferax-sulfurifontis-ATCC-BAA-897] |
| WP_224338217.1 | AAA+ATPase+wHTH+Alpha+Beta-domain+wHTH+RRM | Type-1_BR.RM.like.systems | <-PIWI<-SF1-SF2-Helicase+REase<-Exonuc_V_gamma+REase<-?<-?<-SF1-SF2-Helicase\|\|DUF499*->?-><-ParB+HNH<-ParB+HNH-p<-?\|\|METHYLASE->RNase-> | 1069 | Archaea>Euryarchaeota>Stenosarchaea_group>Halobacteria>Haloferacales | GCF_020097835.1 | Haloprofundus_halobius | LAQ74_RS18925 | 18862..22071 | 2876194 | DUF499-domain-containing-protein-[Haloprofundus-halobius] |
| WP_049905502.1 | AAA+ATPase+wHTH+Alpha+Beta-domain+wHTH+RRM | Type-1_BR.RM.like.systems | <-SF1-SF2-Helicase<-HTH+ACET+HNH\|\|DUF499*->METHYLASE->DUF429-><-DUF772+DDE_Tnp_1\|\|<-BirA-HTH | 1068 | Archaea>Euryarchaeota>Stenosarchaea_group>Halobacteria>Haloferacales | GCF_000336875.1 | Halorubrum_californiense | C463_RS04080 | 15995..19201 | 416585 | DUF499-domain-containing-protein-[Halorubrum-californiense] |
| WP_241211153.1 | AAA+ATPase+wHTH+Alpha+Beta-domain+wHTH+RRM | Type-1_BR.RM.like.systems | Phage_integrase->?->RHH_6+RHH_6-><-?<-SF1-SF2-Helicase<-?\|\|DUF499*-><-?\|\|?->METHYLASE->?->CITB-HTH-> | 1033 | Archaea>Euryarchaeota>Stenosarchaea_group>Halobacteria>Haloferacales | - | Halorubrum_lacusprofundi | HLAC_RS13700 | 12931..16032 | 2247 | DUF499-domain-containing-protein-[Halorubrum-lacusprofundi] |
| WP_004062079.1 | AAA+ATPase+wHTH+Alpha+Beta-domain+wHTH+RRM | Type-1_BR.RM.like.systems | ORC/CDC6+HTH->?-><-?<-SF1-SF2-Helicase\|\|DUF499*-><-?\|\|?->?->METHYLASE->DUF429->ParB+HNH-> | 1039 | Archaea>Euryarchaeota>Stenosarchaea_group>Halobacteria>Haloferacales | GCF_000336795.1 | Haloferax_lucentense | C456_RS02980 | 68527..71646 | 2254 | DUF499-domain-containing-protein-[Haloferax-lucentense] |
| ACM58329.1 | AAA+ATPase+wHTH+Alpha+Beta-domain+wHTH+RRM | Type-1_BR.RM.like.systems | RHH_6+RHH_6-><-?<-SF1-SF2-Helicase<-?\|\|DUF499*-><-?\|\|?->METHYLASE->?->CITB-HTH-> | 1068 | Archaea>Euryarchaeota>Stenosarchaea_group>Halobacteria>Haloferacales | GCA_000022205.1 | Halorubrum_lacusprofundi_ATCC_49239 | Hlac_2758 | 12826..16032 | 416348 | ATPase-(AAA+-superfamily)-like-protein-[Halorubrum-lacusprofundi-ATCC-49239] |
| WP_159663625.1 | AAA+ATPase+wHTH+Alpha+Beta-domain+wHTH+RRM | Type-1_BR.RM.like.systems | <-PIN<-METHYLASE<-?\|\|ParB+HNH->?-><-DUF499*\|\|SF1-SF2-Helicase-> | 1071 | Archaea>Euryarchaeota>Stenosarchaea_group>Halobacteria>Haloferacales | - | Halobaculum_saliterrae | GRX01_RS03890 | 204032..207247 | 2073113 | DUF499-domain-containing-protein-[Halobaculum-saliterrae] |
| WP_092814613.1 | AAA+ATPase+wHTH+Alpha+Beta-domain+wHTH+RRM | Type-1_BR.RM.like.systems | <-SF1-SF2-Helicase\|\|DUF499*-><-?<-?<-?\|\|METHYLASE->PIN-> | 1069 | Archaea>Euryarchaeota>Stenosarchaea_group>Halobacteria>Haloferacales | GCF_900108505.1 | Halopenitus_malekzadehii | BLW38_RS04065 | 817111..820320 | 1267564 | DUF499-domain-containing-protein-[Halopenitus-malekzadehii] |
| WP_098723992.1 | AAA+ATPase+wHTH+Alpha+Beta-domain+wHTH+RRM | Type-1_BR.RM.like.systems | Phage_integrase->?->RHH_6+RHH_6-><-?<-SF1-SF2-Helicase\|\|DUF499*-><-?\|\|MTaX1+HNH-><-MethyltransfD12\|\|METHYLASE-> | 1068 | Archaea>Euryarchaeota>Stenosarchaea_group>Halobacteria>Natrialbales | GCF_002572525.1 | Natrinema_sp._CBA1119 | CP556_RS01420 | 289051..292257 | 1608465 | DUF499-domain-containing-protein-[Natrinema-sp.-CBA1119] |
| WP_226007074.1 | AAA+ATPase+wHTH+Alpha+Beta-domain+wHTH+RRM | Type-1_BR.RM.like.systems | <-REase\|\|Abi->?-><-DUF499* | 1071 | Archaea>Euryarchaeota>Stenosarchaea_group>Halobacteria>Natrialbales | GCF_020405205.1 | Natrinema_salinisoli | LDB05_RS06295 | 1275402..1278617 | 2878535 | DUF499-domain-containing-protein-[Natrinema-salinisoli] |
| HIH77424.1 | AAA+ATPase+wHTH+Alpha+Beta-domain+wHTH+RRM | Type-1_BR.RM.like.systems | DUF499*->Cluster35_66clades->TUDOR+SF1-SF2-Helicase+iREase+CTD+REase-> | 830 | Archaea>Euryarchaeota>Stenosarchaea_group>Halobacteria>unclassified_Halobacteria | GCA_013329575.1 | Halobacteria_archaeon | HA341_00655 | 1..2495 | 2670410 | DUF499-domain-containing-protein-[Halobacteria-archaeon] |
| MCE5337191.1 | AAA+ATPase+wHTH+Alpha+Beta-domain+wHTH+RRM | Type-1_BR.RM.like.systems | ORC/CDC6+HTH->DOC+HTH-><-?<-PolB<-?<-PolB<-DUF499*<-ParB+HNH<-METHYLASE<-?<-wHTH+REase<-METHYLASE-p<-TUDOR+SF1-SF2-Helicase+iREase+CTD+REase | 944 | Archaea>Euryarchaeota>Stenosarchaea_group>Methanomicrobia>Methanomicrobiales | GCF_021371835.1 | Methanomicrobiaceae_archaeon | LLF90_00675 | 34057..36891 | 2716262 | DUF499-domain-containing-protein-[Methanomicrobiaceae-archaeon] |
| WP_156147608.1 | AAA+ATPase+wHTH+Alpha+Beta-domain+wHTH+RRM | Type-1_BR.RM.like.systems | TUDOR+SF1-SF2-Helicase+iREase+CTD+REase->METHYLASE-p->wHTH+REase->?->METHYLASE->ParB+HNH->DUF499*->PolB->HEPN-><-?<-Ntox49 | 942 | Archaea>Euryarchaeota>Stenosarchaea_group>Methanomicrobia>Methanomicrobiales | GCF_000304355.2 | Methanoculleus_bourgensis | BN140_RS09865 | 2039585..2042413 | 83986 | DUF499-domain-containing-protein-[Methanoculleus-bourgensis] |
| WP_004078075.1 | AAA+ATPase+wHTH+Alpha+Beta-domain+wHTH+RRM | Type-1_BR.RM.like.systems | <-DUF499*<-METHYLASE<-HicB_lk_antitox<-TUDOR+SF1-SF2-Helicase+iREase+CTD+REase-p<-TUDOR+SF1-SF2-Helicase+iREase+CTD+REase | 948 | Archaea>Euryarchaeota>Stenosarchaea_group>Methanomicrobia>Methanomicrobiales | GCF_000243255.1 | Methanoplanus_limicola | METLIM_RS09500 | 1961623..1964469 | 2315 | DUF499-domain-containing-protein-[Methanoplanus-limicola] |
| CCJ36896.1 | AAA+ATPase+wHTH+Alpha+Beta-domain+wHTH+RRM | Type-1_BR.RM.like.systems | TUDOR+SF1-SF2-Helicase+iREase+CTD+REase->METHYLASE-p->wHTH+REase->METHYLASE->ParB+HNH->DUF499*->?->PolB->HEPN-> | 944 | Archaea>Euryarchaeota>Stenosarchaea_group>Methanomicrobia>Methanomicrobiales | GCA_000304355.2 | Methanoculleus_bourgensis_MS2 | BN140_1973 | 2039579..2042413 | 1201294 | ATPase-[Methanoculleus-bourgensis-MS2] |
| WP_235808115.1 | AAA+ATPase+wHTH+Alpha+Beta-domain+wHTH+RRM | Type-1_BR.RM.like.systems | <-SF1-SF2-Helicase<-?<-?\|\|TUDOR+SF1-SF2-Helicase+iREase+CTD+REase->METHYLASE->ParB+HNH->DUF499*-><-?<-?<-?<-?<-?<-XIS-HTH | 945 | Archaea>Euryarchaeota>Stenosarchaea_group>Methanomicrobia>Methanomicrobiales | GCF_001602375.1 | Methanoculleus_horonobensis | MCUHO_RS01275 | 28363..31200 | 528314 | DUF499-domain-containing-protein-[Methanoculleus-horonobensis] |
| WP_253456790.1 | AAA+ATPase+wHTH+Alpha+Beta-domain+wHTH+RRM | Type-1_BR.RM.like.systems | TUDOR+SF1-SF2-Helicase+iREase+CTD+REase->METHYLASE->ParB+HNH->DUF499*-><-?<-tRNA<-?\|\|Radical-SAM->?->tRNA-><-METHYLASE | 947 | Archaea>Euryarchaeota>Stenosarchaea_group>Methanomicrobia>Methanomicrobiales | GCF_024170265.1 | Methanocalculus_sp._AMF5 | J2T61_RS02245 | 78750..81593 | 1198257 | DUF499-domain-containing-protein-[Methanocalculus-sp.-AMF5] |
| WP_206633427.1 | AAA+ATPase+wHTH+Alpha+Beta-domain+wHTH+RRM | Type-1_BR.RM.like.systems | ORC/CDC6+HTH-><-DUF499*<-ParB+HNH<-METHYLASE<-TUDOR+SF1-SF2-Helicase+iREase+CTD+REase\|\|REase-> | 937 | Archaea>Euryarchaeota>Stenosarchaea_group>Methanomicrobia>Methanomicrobiales | GCF_004102725.1 | Methanoculleus_taiwanensis | ABH15_RS07615 | 552550..555363 | 1550565 | DUF499-domain-containing-protein-[Methanoculleus-taiwanensis] |
| RXE56548.1 | AAA+ATPase+wHTH+Alpha+Beta-domain+wHTH+RRM | Type-1_BR.RM.like.systems | ORC/CDC6+HTH-><-DUF499*<-ParB+HNH<-METHYLASE<-TUDOR+SF1-SF2-Helicase+iREase+CTD+REase\|\|REase-> | 935 | Archaea>Euryarchaeota>Stenosarchaea_group>Methanomicrobia>Methanomicrobiales | GCA_004102725.1 | Methanoculleus_taiwanensis | ABH15_07530 | 552550..555357 | 1550565 | hypothetical-protein-ABH15_07530-[Methanoculleus-taiwanensis] |
| NQS77681.1 | AAA+ATPase+wHTH+Alpha+Beta-domain+wHTH+RRM | Type-1_BR.RM.like.systems | <-ArdCN+MPTase<-?<-?<-?<-?\|\|?-><-DUF499*<-METHYLASE<-TUDOR+SF1-SF2-Helicase+iREase+CTD+REase | 943 | Archaea>Euryarchaeota>Stenosarchaea_group>Methanomicrobia>Methanomicrobiales | GCA_013201445.1 | Methanoculleus_bourgensis | HQQ74_03005 | 9556..12387 | 83986 | ATP-binding-protein-[Methanoculleus-bourgensis] |
| WP_167815556.1 | AAA+ATPase+wHTH+Alpha+Beta-domain+wHTH+RRM | Type-1_BR.RM.like.systems | TUDOR+SF1-SF2-Helicase+iREase+CTD+REase->METHYLASE->DUF2971->DUF499*-> | 935 | Archaea>Euryarchaeota>Stenosarchaea_group>Methanomicrobia>Methanomicrobiales | GCF_012062385.1 | Methanocorpusculum_sp._GPch4 | EYA77_RS00585 | 22517..25324 | 2527877 | DUF499-domain-containing-protein-[Methanocorpusculum-sp.-GPch4] |
| WP_042699664.1 | AAA+ATPase+wHTH+Alpha+Beta-domain+wHTH+RRM | Type-1_BR.RM.like.systems | TUDOR+SF1-SF2-Helicase+iREase+CTD+REase->METHYLASE->DUF2971->DUF499*-> | 935 | Archaea>Euryarchaeota>Stenosarchaea_group>Methanomicrobia>Methanomicrobiales | GCF_000430905.1 | Methanocorpusculum_bavaricum | H575_RS06345 | 9221..12028 | 71518 | DUF499-domain-containing-protein-[Methanocorpusculum-bavaricum] |
| MBP7411338.1 | AAA+ATPase+wHTH+Alpha+Beta-domain+wHTH+RRM | Type-1_BR.RM.like.systems | TUDOR+SF1-SF2-Helicase+iREase+CTD+REase-p->METHYLASE->DUF499*-> | 741 | Archaea>Euryarchaeota>Stenosarchaea_group>Methanomicrobia>Methanomicrobiales | GCA_017997385.1 | Methanoculleus_sp. | KA818_07355 | 4684..6906 | 90427 | ATP-binding-protein-[Methanoculleus-sp.] |
| KUL01263.1 | AAA+ATPase+wHTH+Alpha+Beta-domain+wHTH+RRM | Type-1_BR.RM.like.systems | TUDOR+SF1-SF2-Helicase+iREase+CTD+REase-p->METHYLASE->DUF499*-> | 490 | Archaea>Euryarchaeota>Stenosarchaea_group>Methanomicrobia>Methanomicrobiales | GCA_001508695.1 | Methanomicrobiales_archaeon_53_19 | XE11_2189 | 3171..4641 | 1641394 | Uncharacterized-protein-XE11_2189,-partial-[Methanomicrobiales-archaeon-53_19] |
| NLM31070.1 | AAA+ATPase+wHTH+Alpha+Beta-domain+wHTH+RRM | Type-1_BR.RM.like.systems | METHYLASE->ParB+HNH->DUF499*-> | 942 | Archaea>Euryarchaeota>Stenosarchaea_group>Methanomicrobia>Methanomicrobiales | GCA_012521035.1 | Methanomicrobiales_archaeon | GX216_11755 | 4804..7632 | 2184052 | ATP-binding-protein-[Methanomicrobiales-archaeon] |
| NLV27387.1 | AAA+ATPase+wHTH+Alpha+Beta-domain+wHTH+RRM | Type-1_BR.RM.like.systems | <-XIS-HTH<-?\|\|TUDOR+SF1-SF2-Helicase+iREase+CTD+REase->METHYLASE->?->DUF499*->?-><-?<-SF1-SF2-Helicase | 972 | Archaea>Euryarchaeota>Stenosarchaea_group>Methanomicrobia>Methanomicrobiales | GCA_012729535.1 | Methanomicrobiales_archaeon | GXY48_09515 | 11881..14799 | 2184052 | ATP-binding-protein-[Methanomicrobiales-archaeon] |
| MBP7298783.1 | AAA+ATPase+wHTH+Alpha+Beta-domain+wHTH+RRM | Type-1_BR.RM.like.systems | SF1-SF2-Helicase->?->XIS-HTH->TUDOR+SF1-SF2-Helicase+iREase+CTD+REase-><-?\|\|METHYLASE->DUF499*-> | 988 | Archaea>Euryarchaeota>Stenosarchaea_group>Methanomicrobia>Methanomicrobiales | GCA_017996475.1 | Methanoculleus_sp. | KA967_00055 | 14120..17086 | 90427 | DUF499-domain-containing-protein-[Methanoculleus-sp.] |
| WP_062262810.1 | AAA+ATPase+wHTH+Alpha+Beta-domain+wHTH+RRM | Type-1_BR.RM.like.systems | PolB->HEPN-><-?<-DUF499-p<-DUF499*<-?<-?<-?<-?<-SF1-SF2-Helicase<-METHYLASE | 567 | Archaea>Euryarchaeota>Stenosarchaea_group>Methanomicrobia>Methanomicrobiales | GCF_900036045.1 | Methanoculleus_bourgensis | MAB1_RS15240 | 1008706..1010409 | 83986 | DUF499-domain-containing-protein-[Methanoculleus-bourgensis] |
| NLL10045.1 | AAA+ATPase+wHTH+Alpha+Beta-domain+wHTH+RRM | Type-1_BR.RM.like.systems | METHYLASE-p->OLD-ABC->TOPRIM->DUF499*-><-?\|\|Transposase_31+DUF4351-> | 970 | Archaea>Euryarchaeota>Stenosarchaea_group>Methanomicrobia>Methanomicrobiales | GCA_012520015.1 | Methanomicrobiales_archaeon | GX268_03955 | 2465..5377 | 2184052 | ATP-binding-protein-[Methanomicrobiales-archaeon] |
| WP_233504506.1 | AAA+ATPase+wHTH+Alpha+Beta-domain+wHTH+RRM | Type-1_BR.RM.like.systems | TUDOR+SF1-SF2-Helicase+iREase+CTD+REase->METHYLASE->?->PADR-HTH->PIN->DUF499*-> | 867 | Archaea>Euryarchaeota>Stenosarchaea_group>Methanomicrobia>Methanomicrobiales | GCF_000327485.1 | Methanoregula_formicica | METFOR_RS11875 | 2388618..2391221 | 882104 | DUF499-domain-containing-protein-[Methanoregula-formicica] |
| AGB03431.1 | AAA+ATPase+wHTH+Alpha+Beta-domain+wHTH+RRM | Type-1_BR.RM.like.systems | TUDOR+SF1-SF2-Helicase+iREase+CTD+REase->METHYLASE->?->PADR-HTH->PIN->DUF499*-> | 981 | Archaea>Euryarchaeota>Stenosarchaea_group>Methanomicrobia>Methanomicrobiales | GCA_000327485.1 | Methanoregula_formicica_SMSP | Metfor_2431 | 2388276..2391221 | 593750 | putative-ATPase-(AAA+-superfamily)-[Methanoregula-formicica-SMSP] |
| MDD3135451.1 | AAA+ATPase+wHTH+Alpha+Beta-domain+wHTH+RRM | Type-1_BR.RM.like.systems | <-HicB_lk_antitox<-DUF499*<-?\|\|REase+ParB-><-?<-?<-Schlafen+HATPase+HTH | 312 | Archaea>Euryarchaeota>Stenosarchaea_group>Methanomicrobia>Methanomicrobiales | GCA_028690975.1 | Methanoregula_sp. | PHF64_03010 | 279..1217 | 2052170 | MAG:-DUF499-domain-containing-protein-[Methanoregula-sp.] |
| MDD5047888.1 | AAA+ATPase+wHTH+Alpha+Beta-domain+wHTH+RRM | Type-1_BR.RM.like.systems | XIS-HTH-><-?<-DUF499*<-?<-METHYLASE\|\|?->?-><-?<-TUDOR+SF1-SF2-Helicase+iREase+CTD+REase | 979 | Archaea>Euryarchaeota>Stenosarchaea_group>Methanomicrobia>Methanomicrobiales | GCA_028710825.1 | Methanoregulaceae_archaeon | PHH09_03080 | 1457..4396 | 2485498 | MAG:-DUF499-domain-containing-protein-[Methanoregulaceae-archaeon] |
| MDD5187496.1 | AAA+ATPase+wHTH+Alpha+Beta-domain+wHTH+RRM | Type-1_BR.RM.like.systems | <-REase+ParB\|\|?-><-?\|\|DUF499*->HicB_lk_antitox->HicA_toxin-> | 290 | Archaea>Euryarchaeota>Stenosarchaea_group>Methanomicrobia>Methanomicrobiales | GCA_028713305.1 | Methanoregula_sp. | PHF57_04740 | 3538..4410 | 2052170 | MAG:-DUF499-domain-containing-protein-[Methanoregula-sp.] |
| OPX61890.1 | AAA+ATPase+wHTH+Alpha+Beta-domain+wHTH+RRM | Type-1_BR.RM.like.systems | TUDOR+SF1-SF2-Helicase+iREase+CTD+REase->METHYLASE->DUF499*-><-?<-?<-SMODS | 845 | Archaea>Euryarchaeota>Stenosarchaea_group>Methanomicrobia>Methanomicrobiales | GCA_002067035.1 | Methanoregula_sp._PtaB.Bin085 | A4E33_02487 | 8643..11180 | 1811680 | MAG:-hypothetical-protein-A4E33_02487-[Methanoregula-sp.-PtaB.Bin085] |
| MDD5024653.1 | AAA+ATPase+wHTH+Alpha+Beta-domain+wHTH+RRM | Type-1_BR.RM.like.systems | <-HicB_lk_antitox<-DUF499*<-?<-?\|\|REase+ParB-> | 298 | Archaea>Euryarchaeota>Stenosarchaea_group>Methanomicrobia>Methanomicrobiales | GCA_028711405.1 | Methanoregula_sp. | PHN79_06075 | 2121..3017 | 2052170 | MAG:-DUF499-domain-containing-protein-[Methanoregula-sp.] |
| MDD4255954.1 | AAA+ATPase+wHTH+Alpha+Beta-domain+wHTH+RRM | Type-1_BR.RM.like.systems | <-DUF499*<-?<-METHYLASE-p | 988 | Archaea>Euryarchaeota>Stenosarchaea_group>Methanomicrobia>Methanomicrobiales | GCA_028702905.1 | Methanofollis_sp. | PHP59_11340 | 544..3510 | 2052835 | MAG:-DUF499-domain-containing-protein-[Methanofollis-sp.] |
| MCW3130203.1 | AAA+ATPase+wHTH+Alpha+Beta-domain+wHTH+RRM | Type-1_BR.RM.like.systems | HhH-RADC+JAB->REase+SF1-SF2-Helicase->HicB_lk_antitox->HicA_toxin->REase-><-?<-DUF499*<-?<-?<-?<-METHYLASE-p<-HEPN<-PolB | 934 | Archaea>Euryarchaeota>Stenosarchaea_group>Methanomicrobia>Methanophagales | GCA_025966515.1 | Methanophagales_archaeon | N2V75_08925 | 8266..11070 | 2056316 | MAG:-DUF499-domain-containing-protein-[Methanophagales-archaeon] |
| MCW3134161.1 | AAA+ATPase+wHTH+Alpha+Beta-domain+wHTH+RRM | Type-1_BR.RM.like.systems | <-DUF499*<-TOPRIM<-OLD-ABC<-METHYLASE-p<-METHYLASE<-SF1-SF2-Helicase | 434 | Archaea>Euryarchaeota>Stenosarchaea_group>Methanomicrobia>Methanophagales | GCA_025966595.1 | Methanophagales_archaeon | N2V78_07540 | 1..1304 | 2056316 | MAG:-DUF499-domain-containing-protein,-partial-[Methanophagales-archaeon] |
| MCW3129356.1 | AAA+ATPase+wHTH+Alpha+Beta-domain+wHTH+RRM | Type-1_BR.RM.like.systems | METHYLASE->OLD-ABC->TOPRIM->DUF499*-><-?<-?<-Aminotran_1_2<-?<-?<-sGTP | 843 | Archaea>Euryarchaeota>Stenosarchaea_group>Methanomicrobia>Methanophagales | GCA_025966515.1 | Methanophagales_archaeon | N2V75_04565 | 3652..6183 | 2056316 | MAG:-hypothetical-protein-N2V75_04565-[Methanophagales-archaeon] |
| MCD6456992.1 | AAA+ATPase+wHTH+Alpha+Beta-domain+wHTH+RRM | Type-1_BR.RM.like.systems | <-Cysteine_desulfurase<-?<-DUF499*<-METHYLASE-p | 843 | Archaea>Euryarchaeota>Stenosarchaea_group>Methanomicrobia>Methanophagales | GCA_021161765.1 | Methanophagales_archaeon | J7K81_09440 | 1136..3667 | 2056316 | ATP-binding-protein-[Methanophagales-archaeon] |
| NQE46165.1 | AAA+ATPase+wHTH+Alpha+Beta-domain+wHTH+RRM | Type-1_BR.RM.like.systems | DDE_Tnp_1->METHYLASE-p->DUF499*->?->HEPN-> | 844 | Archaea>Euryarchaeota>Stenosarchaea_group>Methanomicrobia>Methanophagales | GCA_013180585.1 | ANME-1_cluster_archaeon_GoMg2 | C5S31_09110 | 1445..3979 | 2081358 | hypothetical-protein-[ANME-1-cluster-archaeon-GoMg2] |
| RLG39156.1 | AAA+ATPase+wHTH+Alpha+Beta-domain+wHTH+RRM | Type-1_BR.RM.like.systems | <-TUDOR+SF1-SF2-Helicase+iREase+CTD+REase<-Cluster35_66clades<-DUF499*<-METHYLASE-p | 1129 | Archaea>Euryarchaeota>Stenosarchaea_group>Methanomicrobia>Methanosarcinales | GCA_003661195.1 | Methanosarcinales_archaeon | DRN79_00405 | 3086..6475 | 2250255 | hypothetical-protein-DRN79_00405-[Methanosarcinales-archaeon] |
| MCZ7362504.1 | AAA+ATPase+wHTH+Alpha+Beta-domain+wHTH+RRM | Type-1_BR.RM.like.systems | <-DUF499*\|\|?->TPR+TPR+TPR+TPR+TPR+TPR+TPR+TPR-> | 980 | Archaea>Euryarchaeota>Stenosarchaea_group>Methanomicrobia>Methanosarcinales | GCA_027460385.1 | Candidatus_Methanoperedens_sp. | O8C58_04085 | 67..3009 | 2032688 | MAG:-DUF499-domain-containing-protein-[Candidatus-Methanoperedens-sp.] |
| AKB47809.1 | AAA+ATPase+wHTH+Alpha+Beta-domain+wHTH+RRM | Type-1_BR.RM.like.systems | TUDOR+SF1-SF2-Helicase+iREase+CTD+REase->METHYLASE->REase->DUF499*-><-Radical-SAM<-?<-Radical-SAM<-Radical-SAM<-?\|\|DDE_Tnp_1_2-> | 939 | Archaea>Euryarchaeota>Stenosarchaea_group>Methanomicrobia>Methanosarcinales | GCA_000969945.1 | Methanosarcina_sp._Kolksee | MSKOL_2032 | 2634692..2637511 | 1434099 | putative-ATPase-(AAA+-superfamily)-[Methanosarcina-sp.-Kolksee] |
| PXF61726.1 | AAA+ATPase+wHTH+Alpha+Beta-domain+wHTH+RRM | Type-1_BR.RM.like.systems | TUDOR+SF1-SF2-Helicase+iREase+CTD+REase->HD->DUF4065->REase->PolB->HEPN->DUF499*-><-Mbetalac<-?<-?<-?<-CR-Trypsin1 | 968 | Archaea>Euryarchaeota>Stenosarchaea_group>Methanomicrobia>Methanosarcinales | GCA_003194445.1 | ANME-2_cluster_archaeon | C4B59_02400 | 71391..74297 | 2056317 | AAA-family-ATPase-[ANME-2-cluster-archaeon] |
| WP_197078759.1 | AAA+ATPase+wHTH+Alpha+Beta-domain+wHTH+RRM | Type-1_BR.RM.like.systems | TUDOR+SF1-SF2-Helicase+iREase+CTD+REase->?->METHYLASE->REase->DUF499*-><-Radical-SAM<-?<-Radical-SAM<-Radical-SAM | 943 | Archaea>Euryarchaeota>Stenosarchaea_group>Methanomicrobia>Methanosarcinales | GCF_000969945.1 | Methanosarcina_sp._Kolksee | MSKOL_RS10880 | 2634680..2637511 | 1434099 | DUF499-domain-containing-protein-[Methanosarcina-sp.-Kolksee] |
| RZB29403.1 | AAA+ATPase+wHTH+Alpha+Beta-domain+wHTH+RRM | Type-1_BR.RM.like.systems | <-DUF499*<-PolB<-METHYLASE-p<-METHYLASE-p<-METHYLASE-p<-TUDOR+SF1-SF2-Helicase+iREase+CTD+REase<-TUDOR+SF1-SF2-Helicase+iREase+CTD+REase-p | 967 | Archaea>Euryarchaeota>Stenosarchaea_group>Methanomicrobia>Methanosarcinales | GCA_004193545.1 | Candidatus_Argoarchaeum_ethanivorans | AEth_01220 | 45492..48395 | 2608793 | hypothetical-protein-AEth_01220-[Candidatus-Argoarchaeum-ethanivorans] |
| QNO47559.1 | AAA+ATPase+wHTH+Alpha+Beta-domain+wHTH+RRM | Type-1_BR.RM.like.systems | <-DDE_Tnp_1\|\|METHYLASE-p->METHYLASE-p->OLD-ABC->TOPRIM->DUF499*->?->?-><-?<-?<-?<-Phage_integrase | 854 | Archaea>Euryarchaeota>Stenosarchaea_group>Methanomicrobia>Methanosarcinales | - | Methanosarcinales_archaeon_ANME-2c_ERB4 | GGGHDLIA_00049 | 33141..35705 | 2759911 | hypothetical-protein-GGGHDLIA_00049-[Methanosarcinales-archaeon-ANME-2c-ERB4] |
| CAD7774787.1 | AAA+ATPase+wHTH+Alpha+Beta-domain+wHTH+RRM | Type-1_BR.RM.like.systems | DUF499*->DUF499-p-><-?<-?\|\|?->Glyco_hydro_57+DUF3536->SAM-methylase-> | 375 | Archaea>Euryarchaeota>Stenosarchaea_group>Methanomicrobia>Methanosarcinales | GCA_905171735.1 | Candidatus_Methanoperedenaceae_archaeon_GB50 | KIIPBIDF_00607 | 67..1194 | 2691038 | hypothetical-protein-KIIPBIDF_00607-[Candidatus-Methanoperedenaceae-archaeon-GB50] |

| Protein-ID | DUF499-architecture | System | Gene-Neighbourhoods | Length | Taxonomy | Assembly-ID | Strain | Locus-Tag | Genomic-coordinates | TaxID | Fasta-defline |
|---|---|---|---|---|---|---|---|---|---|---|---|
| OFV65717.1 | AAA+ATPase+wHTH+Alpha+Beta-domain+wHTH+RRM | Type-1_BR.RM.like.systems | <-DUF499*<-TOPRIM<-OLD-ABC<-METHYLASE<-?\|\|PADR-HTH->PIN-> | 860 | Archaea>Euryarchaeota>Stenosarchaea_group>Methanomicrobia>Methanosarcinales | GCA_001766825.1 | Candidatus_Syntrophoarchaeum_butanivorans | SBU_001300 | 796..3378 | 1839936 | ATPase-AAA-[Candidatus-Syntrophoarchaeum-butanivorans] |
| MQY54544.1 | AAA+ATPase+wHTH+Alpha+Beta-domain+wHTH+RRM | Type-1_BR.RM.like.systems | PolB->HEPN->DUF499*->Cysteine_desulfurase-> | 840 | Archaea>Euryarchaeota>Stenosarchaea_group>Methanomicrobia>Methanosarcinales | GCA_009618475.1 | Methanosarcinales_archaeon | GH149_04580 | 2436..4958 | 2250255 | AAA-family-ATPase-[Methanosarcinales-archaeon] |
| RLG35314.1 | AAA+ATPase+wHTH+Alpha+Beta-domain+wHTH+RRM | Type-1_BR.RM.like.systems | METHYLASE-p->HEPN->DUF499*-> | 886 | Archaea>Euryarchaeota>Stenosarchaea_group>Methanomicrobia>Methanosarcinales | GCA_003661135.1 | Methanosarcinales_archaeon | DRN98_01510 | 3349..6009 | 2250255 | hypothetical-protein-DRN98_01510-[Methanosarcinales-archaeon] |
| MCG7853928.1 | AAA+ATPase+wHTH+Alpha+Beta-domain+wHTH+RRM | Type-1_BR.RM.like.systems | <-DUF499*<-METHYLASE<-?<-Schlafen+HATPase+HTH<-TUDOR+SF1-SF2-Helicase+iREase+CTD+REase | 377 | Archaea>Euryarchaeota>Stenosarchaea_group>Methanomicrobia>Methanosarcinales | GCA_022353215.1 | Methanosarcinaceae_archaeon | MI092_15510 | 1..1132 | 1945595 | DUF499-domain-containing-protein-[Methanosarcinaceae-archaeon] |
| MDD3564060.1 | AAA+ATPase+wHTH+Alpha+Beta-domain+wHTH+RRM | Type-1_BR.RM.like.systems | DUF6361->HKD-DNase->SF1-SF2-Helicase-><-SF1-SF2-Helicase+REase<-REase<-?<-DUF499*<-REase<-METHYLASE<-DUF4065<-HD<-HD<-TUDOR+SF1-SF2-Helicase+iREase+CTD+REase | 1047 | Archaea>Euryarchaeota>Stenosarchaea_group>Methanomicrobia>Methanotrichales | GCA_028695755.1 | Methanothrix_sp. | PHN90_00125 | 45442..48585 | 90426 | MAG:-DUF499-domain-containing-protein-[Methanothrix-sp.] |
| WP_048144800.1 | AAA+ATPase+wHTH+Alpha+Beta-domain+wHTH+RRM | Type-1_BR.RM.like.systems | TUDOR+SF1-SF2-Helicase+iREase+CTD+REase->Schlafen+HATPase+HTH->METHYLASE->REase-><-HicB_lk_antitox\|\|DUF499*->RelB->HicB_lk_antitox-> | 987 | Archaea>Euryarchaeota>Stenosarchaea_group>Methanomicrobia>Methanotrichales | GCF_000235565.1 | Methanothrix_harundinacea | MHAR_RS03425 | 698773..701736 | 301375 | DUF499-domain-containing-protein-[Methanothrix-harundinacea] |
| AET64040.1 | AAA+ATPase+wHTH+Alpha+Beta-domain+wHTH+RRM | Type-1_BR.RM.like.systems | TUDOR+SF1-SF2-Helicase+iREase+CTD+REase->Schlafen+HATPase+HTH->METHYLASE->REase-><-HicB_lk_antitox\|\|DUF499*->HicB_lk_antitox-> | 1010 | Archaea>Euryarchaeota>Stenosarchaea_group>Methanomicrobia>Methanotrichales | GCA_000235565.1 | Methanothrix_harundinacea_6Ac | Mhar_0662 | 698704..701736 | 1110509 | ATPase-(AAA+-superfamily)-like-protein-[Methanosaeta-harundinacea-6Ac] |
| KUK43676.1 | AAA+ATPase+wHTH+Alpha+Beta-domain+wHTH+RRM | Type-1_BR.RM.like.systems | TUDOR+SF1-SF2-Helicase+iREase+CTD+REase->HD->DUF4065->METHYLASE->REase->DUF499*->RelB->?-><-HicB_lk_antitox | 987 | Archaea>Euryarchaeota>Stenosarchaea_group>Methanomicrobia>Methanotrichales | GCA_001509375.1 | Methanothrix_harundinacea | XD72_1943 | 10548..13511 | 301375 | ATPase-(AAA+-superfamily)-like-protein-[Methanothrix-harundinacea] |
| MBN1235613.1 | AAA+ATPase+wHTH+Alpha+Beta-domain+wHTH+RRM | Type-1_BR.RM.like.systems | <-DUF499*<-REase<-ASCH<-PIN+ACET+EVE<-METHYLASE<-Schlafen+HATPase+HTH<-TUDOR+SF1-SF2-Helicase+iREase+CTD+REase | 1051 | Archaea>Euryarchaeota>Stenosarchaea_group>Methanomicrobia>Methanotrichales | GCA_016927055.1 | Methanotrichaceae_archaeon | JW999_06125 | 575..3730 | 2595067 | DUF499-domain-containing-protein-[Methanotrichaceae-archaeon] |
| WP_013719923.1 | AAA+ATPase+wHTH+Alpha+Beta-domain+wHTH+RRM | Type-1_BR.RM.like.systems | PADR-HTH->?->?->METHYLASE->?->REase->DUF499*-><-?<-SbcC+SbcC\|\|RADICAL-SAM->Radical-SAM-> | 1047 | Archaea>Euryarchaeota>Stenosarchaea_group>Methanomicrobia>Methanotrichales | GCF_000204415.1 | Methanothrix | QVD88_RS00980 | 24822..27965 | 2222 | DUF499-domain-containing-protein-[Methanothrix-soehngenii] |
| NLX39269.1 | AAA+ATPase+wHTH+Alpha+Beta-domain+wHTH+RRM | Type-1_BR.RM.like.systems | Y1_Tnp-><-DUF499*<-REase<-METHYLASE<-TUDOR+SF1-SF2-Helicase+iREase+CTD+REase | 1044 | Archaea>Euryarchaeota>Stenosarchaea_group>Methanomicrobia>Methanotrichales | GCA_012729025.1 | Methanothrix_sp. | GXY73_07140 | 1114..4248 | 90426 | ATP-binding-protein-[Methanothrix-sp.] |
| MCX6669299.1 | AAA+ATPase+wHTH+Alpha+Beta-domain+wHTH+RRM | Type-1_BR.RM.like.systems | METHYLASE-p->PIN->PIN+ACET+EVE->ASCH->REase->DUF499*-> | 1023 | Archaea>Euryarchaeota>Stenosarchaea_group>Methanomicrobia>Methanotrichales | GCA_026394475.1 | Methanothrix_sp. | NTV25_05770 | 3501..6572 | 90426 | MAG:-DUF499-domain-containing-protein-[Methanothrix-sp.] |
| MCP1392941.1 | AAA+ATPase+wHTH+Alpha+Beta-domain+wHTH+RRM | Type-1_BR.RM.like.systems | <-RelB<-DUF499*\|\|HicB_lk_antitox-><-REase<-METHYLASE-p | 1008 | Archaea>Euryarchaeota>Stenosarchaea_group>Methanomicrobia>Methanotrichales | GCA_024166385.1 | Methanothrix_harundinacea | J5I35_09320 | 253..3279 | 301375 | ATP-binding-protein-[Methanothrix-harundinacea] |
| WP_011011684.1 | AAA+ATPase+wHTH+Alpha+Beta-domain+wHTH+RRM | Type-1_BR.RM.like.systems | DUF2791->DDE_Tnp_IS240->DUF2791-><-ncRNA<-TUDOR+SF1-SF2-Helicase+iREase+CTD+REase<-Cluster35_66clades<-DUF499*<-METHYLASE<-METHYLASE-p\|\|EVE-><-TPR<-TPR\|\|SF1-SF2-Helicase-> | 1105 | Archaea>Euryarchaeota>Thermococci>Thermococcales>Thermococcaceae | GCF_000007305.1 | Pyrococcus_furiosus | PFDSM3638_RS02850 | 566793..570110 | 2261 | ATP-binding-protein-[Pyrococcus-furiosus] |
| WP_013748669.1 | AAA+ATPase+wHTH+Alpha+Beta-domain+wHTH+RRM | Type-1_BR.RM.like.systems | REase->HTH+IclR-HTH->NTP_transf+wHTH-><-EVE<-EVE\|\|METHYLASE->DUF499*->Cluster35_66clades->TUDOR+SF1-SF2-Helicase+iREase+CTD+REase->tRNA->MNS+HTH+REase-><-?<-Peptidase | 1105 | Archaea>Euryarchaeota>Thermococci>Thermococcales>Thermococcaceae | GCF_000211475.1 | Pyrococcus_sp._NA2 | PNA2_RS06085 | 1101176..1104493 | 342949 | DUF499-domain-containing-protein-[Pyrococcus-sp.-NA2] |
| WP_010884994.1 | AAA+ATPase+wHTH+Alpha+Beta-domain+wHTH+RRM | Type-1_BR.RM.like.systems | PIN-><-MazE_antitoxin<-SF1-SF2-Helicase\|\|TPR-><-EVE\|\|METHYLASE->DUF499*->Cluster35_66clades->TUDOR+SF1-SF2-Helicase+iREase+CTD+REase->ncRNA-><-DUF2791 | 1105 | Archaea>Euryarchaeota>Thermococci>Thermococcales>Thermococcaceae | GCF_000011105.1 | Pyrococcus_horikoshii | HA331_RS05800 | 71976..75293 | 53953 | ATP-binding-protein-[Pyrococcus-horikoshii] |
| WP_058937731.1 | AAA+ATPase+wHTH+Alpha+Beta-domain+wHTH+RRM | Type-1_BR.RM.like.systems | ParB+HNH->BirA-HTH-><-Phage_integrase<-EVE<-TUDOR+SF1-SF2-Helicase+iREase+CTD+REase<-Cluster35_66clades<-DUF499*<-METHYLASE<-?\|\|EVE->PADR-HTH->PIN-> | 1128 | Archaea>Euryarchaeota>Thermococci>Thermococcales>Thermococcaceae | GCF_001484195.1 | Thermococcus_celericrescens | APY94_RS00205 | 42045..45431 | 227598 | ATP-binding-protein-[Thermococcus-celericrescens] |
| WP_013468121.1 | AAA+ATPase+wHTH+Alpha+Beta-domain+wHTH+RRM | Type-1_BR.RM.like.systems | Phage_integrase-><-BirA-HTH+CR-ATPase6<-?\|\|?-><-TUDOR+SF1-SF2-Helicase+iREase+CTD+REase<-Cluster35_66clades<-DUF499*<-?<-AbiEii<-CRP-HTH<-?<-METHYLASE | 1129 | Archaea>Euryarchaeota>Thermococci>Thermococcales>Thermococcaceae | GCF_000151105.2 | Thermococcus_barophilus | TERMP_RS09145 | 1635861..1639250 | 55802 | DUF499-domain-containing-protein-[Thermococcus-barophilus] |
| WP_010868001.1 | AAA+ATPase+wHTH+Alpha+Beta-domain+wHTH+RRM | Type-1_BR.RM.like.systems | PIN-><-Peptidase_M24<-EVE<-EVE\|\|METHYLASE->DUF499*->Cluster35_66clades->TUDOR+SF1-SF2-Helicase+iREase+CTD+REase->SF1-SF2-Helicase-> | 1109 | Archaea>Euryarchaeota>Thermococci>Thermococcales>Thermococcaceae | GCF_000195935.2 | Pyrococcus_abyssi | PAB_RS04690 | 840711..844040 | 29292 | ATP-binding-protein-[Pyrococcus-abyssi] |
| WP_004069029.1 | AAA+ATPase+wHTH+Alpha+Beta-domain+wHTH+RRM | Type-1_BR.RM.like.systems | <-TUDOR+SF1-SF2-Helicase+iREase+CTD+REase<-Cluster35_66clades<-DUF499*<-METHYLASE\|\|EVE-><-?\|\|BirA-HTH+CR-ATPase6-><-SLATT\|\|NACHT-> | 1131 | Archaea>Euryarchaeota>Thermococci>Thermococcales>Thermococcaceae | GCF_000246985.2 | Thermococcus_litoralis | OCC_RS01665 | 312029..315424 | 2265 | DUF499-domain-containing-protein-[Thermococcus-litoralis] |
| WP_167915555.1 | AAA+ATPase+wHTH+Alpha+Beta-domain+wHTH+RRM | Type-1_BR.RM.like.systems | <-BirA-HTH+CR-ATPase6<-?\|\|?-><-?<-TUDOR+SF1-SF2-Helicase+iREase+CTD+REase<-Cluster35_66clades<-DUF499*<-?<-AbiEii<-CRP-HTH<-?<-METHYLASE | 1129 | Archaea>Euryarchaeota>Thermococci>Thermococcales>Thermococcaceae | GCF_012027375.1 | Thermococcus_sp._LS2 | E3E24_RS01880 | 361390..364779 | 1638260 | DUF499-domain-containing-protein-[Thermococcus-sp.-LS2] |
| WP_068665282.1 | AAA+ATPase+wHTH+Alpha+Beta-domain+wHTH+RRM | Type-1_BR.RM.like.systems | <-RNase<-?\|\|tRNA-><-EVE<-REase<-TUDOR+SF1-SF2-Helicase+iREase+CTD+REase-p<-Cluster35_66clades<-DUF499*<-METHYLASE<-?<-ABC-ATPase+SbcC | 1128 | Archaea>Euryarchaeota>Thermococci>Thermococcales>Thermococcaceae | GCF_001647085.1 | Thermococcus_piezophilus | A7C91_RS04480 | 826311..829697 | 1712654 | ATP-binding-protein-[Thermococcus-piezophilus] |
| HIH73327.1 | AAA+ATPase+wHTH+Alpha+Beta-domain+wHTH+RRM | Type-1_BR.RM.like.systems | SF1-SF2-Helicase-><-EVE\|\|METHYLASE->DUF499*->Cluster35_66clades->TUDOR+SF1-SF2-Helicase+iREase+CTD+REase-><-?<-PIN<-PADR-HTH | 1125 | Archaea>Euryarchaeota>Thermococci>Thermococcales>Thermococcaceae | GCA_013330365.1 | Thermococcaceae_archaeon | HA300_08290 | 15719..19096 | 2666349 | DUF499-domain-containing-protein-[Thermococcaceae-archaeon] |
| WP_058946124.1 | AAA+ATPase+wHTH+Alpha+Beta-domain+wHTH+RRM | Type-1_BR.RM.like.systems | Phage_integrase-><-PADR-HTH<-EVE\|\|?->METHYLASE->DUF499*->Cluster35_66clades->TUDOR+SF1-SF2-Helicase+iREase+CTD+REase->?->?->EVE-> | 1128 | Archaea>Euryarchaeota>Thermococci>Thermococcales>Thermococcaceae | GCF_001484685.1 | Thermococcus_sp._2319x1 | ADU37_RS02440 | 442827..446213 | 1674923 | ATP-binding-protein-[Thermococcus-sp.-2319x1] |
| WP_068577206.1 | AAA+ATPase+wHTH+Alpha+Beta-domain+wHTH+RRM | Type-1_BR.RM.like.systems | <-McrC-NTD+REase<-?<-TUDOR+SF1-SF2-Helicase+iREase+CTD+REase<-Cluster35_66clades<-DUF499*<-?<-AbiEii<-CRP-HTH<-?<-METHYLASE | 1129 | Archaea>Euryarchaeota>Thermococci>Thermococcales>Thermococcaceae | GCF_002214605.1 | Thermococcus_chitonophagus | A3L04_RS08565 | 1523280..1526669 | 54262 | DUF499-domain-containing-protein-[Thermococcus-chitonophagus] |
| WP_062389038.1 | AAA+ATPase+wHTH+Alpha+Beta-domain+wHTH+RRM | Type-1_BR.RM.like.systems | <-EVE<-?<-?<-TUDOR+SF1-SF2-Helicase+iREase+CTD+REase<-Cluster35_66clades<-DUF499*<-METHYLASE<-?\|\|EVE->PADR-HTH->PIN-> | 1128 | Archaea>Euryarchaeota>Thermococci>Thermococcales>Thermococcaceae | GCF_001592435.1 | Thermococcus_peptonophilus | A0127_RS05640 | 1032348..1035734 | 53952 | ATP-binding-protein-[Thermococcus-peptonophilus] |
| WP_167904334.1 | AAA+ATPase+wHTH+Alpha+Beta-domain+wHTH+RRM | Type-1_BR.RM.like.systems | <-EVE<-?<-?<-TUDOR+SF1-SF2-Helicase+iREase+CTD+REase<-Cluster35_66clades<-DUF499*<-METHYLASE\|\|PADR-HTH->PIN-> | 1128 | Archaea>Euryarchaeota>Thermococci>Thermococcales>Thermococcaceae | GCF_012027495.1 | Thermococcus_sp._JdF3 | E3E42_RS09505 | 200796..204182 | 1638258 | ATP-binding-protein-[Thermococcus-sp.-JdF3] |
| WP_167899657.1 | AAA+ATPase+wHTH+Alpha+Beta-domain+wHTH+RRM | Type-1_BR.RM.like.systems | <-REase<-EVE\|\|?->METHYLASE->DUF499*->Cluster35_66clades->TUDOR+SF1-SF2-Helicase+iREase+CTD+REase->?->?->EVE-> | 1139 | Archaea>Euryarchaeota>Thermococci>Thermococcales>Thermococcaceae | GCF_012027395.1 | Thermococcus_sp._LS1 | E3E26_RS01870 | 341097..344516 | 1638259 | DUF499-domain-containing-protein-[Thermococcus-sp.-LS1] |
| MCD6143899.1 | AAA+ATPase+wHTH+Alpha+Beta-domain+wHTH+RRM | Type-1_BR.RM.like.systems | <-TUDOR+SF1-SF2-Helicase+iREase+CTD+REase<-Cluster35_66clades<-DUF499*<-METHYLASE\|\|?->?->DNA-glycosylase-> | 1121 | Archaea>Euryarchaeota>Thermococci>Thermococcales>Thermococcaceae | GCA_021159205.1 | Thermococcus_sp. | J7J04_06235 | 3390..6755 | 35749 | DUF499-domain-containing-protein-[Thermococcus-sp.] |
| WP_240912577.1 | AAA+ATPase+wHTH+Alpha+Beta-domain+wHTH+RRM | Type-1_BR.RM.like.systems | <-TUDOR+SF1-SF2-Helicase+iREase+CTD+REase<-Cluster35_66clades<-DUF499*<-METHYLASE\|\|EVE->?-><-?\|\|?-><-MNS+HTH | 1116 | Archaea>Euryarchaeota>Thermococci>Thermococcales>Thermococcaceae | GCF_012027375.1 | Thermococcus_sp._LS2 | E3E24_RS11855 | 7119..10469 | 1638260 | DUF499-domain-containing-protein-[Thermococcus-sp.-LS2] |
| NJE13749.1 | AAA+ATPase+wHTH+Alpha+Beta-domain+wHTH+RRM | Type-1_BR.RM.like.systems | <-TUDOR+SF1-SF2-Helicase+iREase+CTD+REase<-Cluster35_66clades<-DUF499*<-METHYLASE\|\|EVE->?-><-?<-MNS+HTH | 1114 | Archaea>Euryarchaeota>Thermococci>Thermococcales>Thermococcaceae | GCA_012027375.1 | Thermococcus_sp._LS2 | E3E24_11715 | 7119..10463 | 1638260 | DUF499-domain-containing-protein-[Thermococcus-sp.-LS2] |
| WP_088863529.1 | AAA+ATPase+wHTH+Alpha+Beta-domain+wHTH+RRM | Type-1_BR.RM.like.systems | <-EVE\|\|METHYLASE->DUF499*->Cluster35_66clades->TUDOR+SF1-SF2-Helicase+iREase+CTD+REase-> | 1129 | Archaea>Euryarchaeota>Thermococci>Thermococcales>Thermococcaceae | GCF_002214365.1 | Thermococcus_celer | A3L02_RS08655 | 1592921..1596310 | 2264 | ATP-binding-protein-[Thermococcus-celer] |
| MCD6144065.1 | AAA+ATPase+wHTH+Alpha+Beta-domain+wHTH+RRM | Type-1_BR.RM.like.systems | <-HEPN\|\|METHYLASE->DUF499*->?->TUDOR+SF1-SF2-Helicase+iREase+CTD+REase-><-DNA-glycosylase | 1126 | Archaea>Euryarchaeota>Thermococci>Thermococcales>Thermococcaceae | GCA_021159205.1 | Thermococcus_sp. | J7J04_07155 | 4954..8334 | 35749 | DUF499-domain-containing-protein-[Thermococcus-sp.] |
| WP_015859240.1 | AAA+ATPase+wHTH+Alpha+Beta-domain+wHTH+RRM | Type-1_BR.RM.like.systems | GDResolvase+HIN-HTH->?->TUDOR+SF1-SF2-Helicase+iREase+CTD+REase->METHYLASE-><-?\|\|MNS+HTH->DUF499*->?->?->SF1-SF2-Helicase-><-tRNA\|\|TOPRIM->Phage_int_SAM_1+Phage_integrase-> | 882 | Archaea>Euryarchaeota>Thermococci>Thermococcales>Thermococcaceae | GCF_000022365.1 | Thermococcus_gammatolerans | TGAM_RS08235 | 1533229..1535877 | 187878 | DUF499-domain-containing-protein-[Thermococcus-gammatolerans] |
| WP_139681370.1 | AAA+ATPase+wHTH+Alpha+Beta-domain+wHTH+RRM | Type-1_BR.RM.like.systems | <-GDResolvase+HIN-HTH\|\|TUDOR+SF1-SF2-Helicase+iREase+CTD+REase->METHYLASE-><-PIN<-?\|\|?->DUF499*->?->?->SF1-SF2-Helicase-><-tRNA\|\|?->REase-> | 883 | Archaea>Euryarchaeota>Thermococci>Thermococcales>Thermococcaceae | GCF_006274605.1 | Thermococcus_indicus | FH039_RS11140 | 2049351..2052002 | 2586643 | DUF499-domain-containing-protein-[Thermococcus-indicus] |
| WP_010477366.1 | AAA+ATPase+wHTH+Alpha+Beta-domain+wHTH+RRM | Type-1_BR.RM.like.systems | <-Phage_int_SAM_1+Phage_integrase<-TOPRIM\|\|tRNA-><-DUF499*\|\|?->PIN-><-METHYLASE<-TUDOR+SF1-SF2-Helicase+iREase+CTD+REase | 887 | Archaea>Euryarchaeota>Thermococci>Thermococcales>Thermococcaceae | GCF_000258515.1 | Thermococcus_zilligii | TZI_RS0101400 | 259596..262259 | 54076 | DUF499-domain-containing-protein-[Thermococcus-zilligii] |
| WP_048151392.1 | AAA+ATPase+wHTH+Alpha+Beta-domain+wHTH+RRM | Type-1_BR.RM.like.systems | TUDOR+SF1-SF2-Helicase+iREase+CTD+REase->METHYLASE->MNS+HTH->?->?->DUF499*-><-?<-?\|\|REase-><-ORC-CDC6+HTH\|\|?-><-HTH_36 | 878 | Archaea>Euryarchaeota>Thermococci>Thermococcales>Thermococcaceae | GCF_000966265.1 | Palaeococcus_ferrophilus | PFER_RS09170 | 22993..25629 | 83868 | DUF499-domain-containing-protein-[Palaeococcus-ferrophilus] |
| WP_167910221.1 | AAA+ATPase+wHTH+Alpha+Beta-domain+wHTH+RRM | Type-1_BR.RM.like.systems | <-PadR-HTH\|\|?->Phage_integrase->?->?->?-><-DUF499*<-?\|\|?->PIN-><-METHYLASE<-?<-TUDOR+SF1-SF2-Helicase+iREase+CTD+REase | 885 | Archaea>Euryarchaeota>Thermococci>Thermococcales>Thermococcaceae | GCF_012027595.1 | Thermococcus_sp._Bubb.Bath | E3E29_RS06900 | 296042..298699 | 1638242 | DUF499-domain-containing-protein-[Thermococcus-sp.-Bubb.Bath] |
| WP_167893361.1 | AAA+ATPase+wHTH+Alpha+Beta-domain+wHTH+RRM | Type-1_BR.RM.like.systems | <-REase<-?\|\|tRNA-><-SF1-SF2-Helicase<-?<-?<-DUF499*<-?\|\|?->PIN-><-METHYLASE<-?<-TUDOR+SF1-SF2-Helicase+iREase+CTD+REase | 880 | Archaea>Euryarchaeota>Thermococci>Thermococcales>Thermococcaceae | GCF_012027305.1 | Thermococcus_sp._MAR1 | E3E25_RS11215 | 186255..188897 | 1638263 | DUF499-domain-containing-protein-[Thermococcus-sp.-MAR1] |
| HIP74717.1 | AAA+ATPase+wHTH+Alpha+Beta-domain+wHTH+RRM | Type-1_BR.RM.like.systems | ZnR->tRNA-><-DUF499*<-METHYLASE-p | 867 | Archaea>Euryarchaeota>Thermococci>Thermococcales>Thermococcaceae | GCA_015662375.1 | Thermococcus_paralvinellae | EYH13_00890 | 4304..6907 | 582419 | ATP-binding-protein-[Thermococcus-paralvinellae] |
| MCX6666484.1 | AAA+ATPase+wHTH+Alpha+Beta-domain+wHTH+RRM | Type-1_BR.RM.like.systems | TUDOR+SF1-SF2-Helicase+iREase+CTD+REase-p->METHYLASE->DUF499*-><-?<-?<-?<-REase\|\|RNase-> | 921 | Archaea>Euryarchaeota>unclassified_Euryarchaeota | GCA_026394495.1 | Euryarchaeota_archaeon | NT038_10595 | 4419..7184 | 2026739 | MAG:-DUF499-domain-containing-protein-[Euryarchaeota-archaeon] |
| MA052903.1 | AAA+ATPase+wHTH+Alpha+Beta-domain+wHTH+RRM | Type-1_BR.RM.like.systems | <-HKD-DNase\|\|METHYLASE->DUF499*-> | 948 | Archaea>Euryarchaeota>unclassified_Euryarchaeota | GCA_002693705.1 | Euryarchaeota_archaeon | CL984_00015 | 3819..6665 | 2026739 | hypothetical-protein-[Euryarchaeota-archaeon] |
| MBU4220744.1 | AAA+ATPase+wHTH+Alpha+Beta-domain+wHTH+RRM | Type-1_BR.RM.like.systems | <-HEPN<-PolB\|\|?-><-DUF499*<-TOPRIM<-OLD-ABC<-METHYLASE<-?<-SbcC<-METHYLASE-p | 846 | Archaea>Euryarchaeota>unclassified_Euryarchaeota | GCA_018898145.1 | Euryarchaeota_archaeon | KKA10_03835 | 1706..4246 | 2026739 | DUF499-domain-containing-protein-[Euryarchaeota-archaeon] |
| MCW4015124.1 | AAA+ATPase+wHTH+Alpha+Beta-domain+wHTH+RRM | Type-1_BR.RM.like.systems | <-DUF499*<-METHYLASE<-TUDOR+SF1-SF2-Helicase+iREase+CTD+REase-p | 667 | Archaea>TACK_group>Candidatus_Bathyarchaeota>Candidatus_Bathyarchaeia>Candidatus_Bathyarchaeales | GCA_026014495.1 | Candidatus_Bathyarchaeum_sp. | NWF06_02025 | 1..2000 | 3072974 | MAG:-DUF499-domain-containing-protein,-partial-[Candidatus-Bathyarchaeota-archaeon] |
| MCW4008815.1 | AAA+ATPase+wHTH+Alpha+Beta-domain+wHTH+RRM | Type-1_BR.RM.like.systems | <-TUDOR+SF1-SF2-Helicase+iREase+CTD+REase<-Cluster35_66clades<-DUF499*<-METHYLASE<-?\|\|N6_N4_MTase-><-?<-VirD4-TraG+HerA | 1119 | Archaea>TACK_group>Candidatus_Bathyarchaeota>unclassified_Candidatus_Bathyarchaeota | GCA_026014545.1 | Candidatus_Bathyarchaeota_archaeon | NWF09_09045 | 2664..6023 | 2026714 | MAG:-ATP-binding-protein-[Candidatus-Bathyarchaeota-archaeon] |
| RLI36047.1 | AAA+ATPase+wHTH+Alpha+Beta-domain+wHTH+RRM | Type-1_BR.RM.like.systems | <-TUDOR+SF1-SF2-Helicase+iREase+CTD+REase-p<-Cluster35_66clades<-DUF499*<-METHYLASE-p | 1121 | Archaea>TACK_group>Candidatus_Bathyarchaeota>unclassified_Candidatus_Bathyarchaeota | GCA_003662645.1 | Candidatus_Bathyarchaeota_archaeon | DR055_04280 | 1001..4366 | 2026714 | hypothetical-protein-DR055_04280-[Candidatus-Bathyarchaeota-archaeon] |
| RLI28985.1 | AAA+ATPase+wHTH+Alpha+Beta-domain+wHTH+RRM | Type-1_BR.RM.like.systems | METHYLASE->DUF499*->Cluster35_66clades->TUDOR+SF1-SF2-Helicase+iREase+CTD+REase-> | 1110 | Archaea>TACK_group>Candidatus_Bathyarchaeota>unclassified_Candidatus_Bathyarchaeota | GCA_003662445.1 | Candidatus_Bathyarchaeota_archaeon | DR058_00900 | 2294..5626 | 2026714 | hypothetical-protein-DR058_00900-[Candidatus-Bathyarchaeota-archaeon] |
| RLG99044.1 | AAA+ATPase+wHTH+Alpha+Beta-domain+wHTH+RRM | Type-1_BR.RM.like.systems | DUF499*->Cluster35_66clades->TUDOR+SF1-SF2-Helicase+iREase+CTD+REase-> | 386 | Archaea>TACK_group>Candidatus_Bathyarchaeota>unclassified_Candidatus_Bathyarchaeota | GCA_003662065.1 | Candidatus_Bathyarchaeota_archaeon | DR019_03370 | 1..1161 | 2026714 | hypothetical-protein-DR019_03370,-partial-[Candidatus-Bathyarchaeota-archaeon] |
| HDO72515.1 | AAA+ATPase+wHTH+Alpha+Beta-domain+wHTH+RRM | Type-1_BR.RM.like.systems | DUF499*->Cluster35_66clades->TUDOR+SF1-SF2-Helicase+iREase+CTD+REase-> | 520 | Archaea>TACK_group>Candidatus_Bathyarchaeota>unclassified_Candidatus_Bathyarchaeota | GCA_011044055.1 | Candidatus_Bathyarchaeota_archaeon | ENG10_04435 | 1..1564 | 2026714 | DUF499-domain-containing-protein-[Candidatus-Bathyarchaeota-archaeon] |
| MCD6241711.1 | AAA+ATPase+wHTH+Alpha+Beta-domain+wHTH+RRM | Type-1_BR.RM.like.systems | <-OLD-ABC+TOPRIM<-?<-PIWI<-?<-?\|\|?-><-DUF499*<-METHYLASE<-REase<-TUDOR+SF1-SF2-Helicase+iREase+CTD+REase | 896 | Archaea>TACK_group>Candidatus_Bathyarchaeota>unclassified_Candidatus_Bathyarchaeota | GCA_021158145.1 | Candidatus_Bathyarchaeota_archaeon | J7K27_09445 | 7902..10592 | 2026714 | ATP-binding-protein-[Candidatus-Bathyarchaeota-archaeon] |
| OYT64646.1 | AAA+ATPase+wHTH+Alpha+Beta-domain+wHTH+RRM | Type-1_BR.RM.like.systems | <-SbcC<-DUF499*<-PIN<-SbcC<-METHYLASE-p<-HEPN<-PolB | 851 | Archaea>TACK_group>Candidatus_Bathyarchaeota>unclassified_Candidatus_Bathyarchaeota | GCA_002255045.1 | Candidatus_Bathyarchaeota_archaeon_ex4484_205 | B6U74_05075 | 1011..3566 | 2012510 | AAA-family-ATPase-[Candidatus-Bathyarchaeota-archaeon-ex4484_205] |
| MBE3116226.1 | AAA+ATPase+wHTH+Alpha+Beta-domain+wHTH+RRM | Type-1_BR.RM.like.systems | <-DUF499*<-METHYLASE<-TUDOR+SF1-SF2-Helicase+iREase+CTD+REase-p | 468 | Archaea>TACK_group>Candidatus_Bathyarchaeota>unclassified_Candidatus_Bathyarchaeota | GCA_014894195.1 | Candidatus_Bathyarchaeota_archaeon | IMZ68_03400 | 1..1405 | 2026714 | ATP-binding-protein-[Candidatus-Bathyarchaeota-archaeon] |
| MBO3840365.1 | AAA+ATPase+wHTH+Alpha+Beta-domain+wHTH+RRM | Type-1_BR.RM.like.systems | METHYLASE->DUF499*->Cluster35_66clades->TUDOR+SF1-SF2-Helicase+iREase+CTD+REase-> | 1126 | Archaea>TACK_group>Candidatus_Brockarchaeota>unclassified_Candidatus_Brockarchaeota | GCA_017601185.1 | Candidatus_Brockarchaeota_archaeon | JTT17_06060 | 6478..9858 | 2815303 | DUF499-domain-containing-protein-[Candidatus-Brockarchaeota-archaeon] |
| MBO3798316.1 | AAA+ATPase+wHTH+Alpha+Beta-domain+wHTH+RRM | Type-1_BR.RM.like.systems | <-EVE<-HEPN<-DUF499*<-METHYLASE<-TUDOR+SF1-SF2-Helicase+iREase+CTD+REase<-URI-DNase+RAMA\|\|?->?->URI-DNase-> | 887 | Archaea>TACK_group>Candidatus_Brockarchaeota>unclassified_Candidatus_Brockarchaeota | GCA_017601225.1 | Candidatus_Brockarchaeota_archaeon | JTT13_05580 | 80557..83220 | 2815303 | ATP-binding-protein-[Candidatus-Brockarchaeota-archaeon] |
| WP_125670531.1 | AAA+ATPase+wHTH+Alpha+Beta-domain+wHTH+RRM | Type-1_BR.RM.like.systems | TUDOR+SF1-SF2-Helicase+iREase+CTD+REase->METHYLASE->DUF499*-> | 845 | Archaea>TACK_group>Candidatus_Korarchaeota>Candidatus_Methanodesulfokores | GCF_003947435.1 | Candidatus_Methanodesulfokores_washburnensis | D6D85_RS02645 | 18281..20818 | 2478471 | ATP-binding-protein-[Candidatus-Methanodesulfokores-washburnensis] |
| WP_125740982.1 | AAA+ATPase+wHTH+Alpha+Beta-domain+wHTH+RRM | Type-1_BR.RM.like.systems | SF1-SF2-helicase+DUF1998-><-?<-TUDOR+SF1-SF2-Helicase+iREase+CTD+REase<-DUF499*<-METHYLASE-p<-METHYLASE<-RNase+PIN | 1120 | Archaea>TACK_group>Candidatus_Korarchaeota>Korarchaeum | GCF_003948265.1 | Candidatus_Korarchaeum_cryptofilum | D9Q81_RS02265 | 14543..17905 | 498846 | ATP-binding-protein-[Candidatus-Korarchaeum-cryptofilum] |
| WP_012309349.1 | AAA+ATPase+wHTH+Alpha+Beta-domain+wHTH+RRM | Type-1_BR.RM.like.systems | SF1-SF2-helicase+DUF1998-><-?<-TUDOR+SF1-SF2-Helicase+iREase+CTD+REase<-DUF499*<-METHYLASE<-RNase+PIN\|\|EVE-> | 1120 | Archaea>TACK_group>Candidatus_Korarchaeota>Korarchaeum | GCF_000019605.1 | Candidatus_Korarchaeum_cryptofilum | KCR_RS03625 | 719792..723154 | 498846 | DUF499-domain-containing-protein-[Candidatus-Korarchaeum-cryptofilum] |
| WP_125740641.1 | AAA+ATPase+wHTH+Alpha+Beta-domain+wHTH+RRM | Type-1_BR.RM.like.systems | METHYLASE-p->DUF499*-> | 847 | Archaea>TACK_group>Candidatus_Korarchaeota>Korarchaeum | GCF_003948265.1 | Candidatus_Korarchaeum_cryptofilum | D9Q81_RS01175 | 1850..4393 | 498846 | ATP-binding-protein-[Candidatus-Korarchaeum-cryptofilum] |
| MCL4551940.1 | AAA+ATPase+wHTH+Alpha+Beta-domain+wHTH+RRM | Type-1_BR.RM.like.systems | XIS-HTH->OLD-ABC->HNH->METHYLASE->DUF499*->DUF499-p->SF1-SF2-Helicase+REase-> | 336 | Archaea>TACK_group>Candidatus_Marsarchaeota>unclassified_Candidatus_Marsarchaeota | GCA_023381795.1 | Candidatus_Marsarchaeota_archaeon | M1305_00065 | 10946..11954 | 2823144 | DUF499-domain-containing-protein-[Candidatus-Marsarchaeota-archaeon] |
| MCS7097590.1 | AAA+ATPase+wHTH+Alpha+Beta-domain+wHTH+RRM | Type-1_BR.RM.like.systems | <-SF1-SF2-Helicase\|\|?-><-DNA-glycosylase<-?<-TUDOR+SF1-SF2-Helicase+iREase+CTD+REase<-Cluster35_66clades<-DUF499*<-METHYLASE | 1122 | Archaea>TACK_group>Candidatus_Verstraetearchaeota>Candidatus_Methanomethylicia>Candidatus_Methanomethylicales | GCA_025057955.1 | Candidatus_Methanomethyliaceae_archaeon | NZ922_01240 | 95023..98391 | 2282146 | ATP-binding-protein-[Candidatus-Methanomethyliaceae-archaeon] |
| MCQ5340613.1 | AAA+ATPase+wHTH+Alpha+Beta-domain+wHTH+RRM | Type-1_BR.RM.like.systems | ZnR->METHYLASE->DUF499*->Cluster35_66clades->TUDOR+SF1-SF2-Helicase+iREase+CTD+REase-> | 1137 | Archaea>TACK_group>Candidatus_Verstraetearchaeota>Candidatus_Methanomethylicia>unclassified_Candidatus_Methanomethylicia | - | Candidatus_Methanomethylicia_archaeon | N0483_02750 | 25514..28927 | 2268187 | ATP-binding-protein-[Candidatus-Methanomethylicia-archaeon] |
| MCC6018684.1 | AAA+ATPase+wHTH+Alpha+Beta-domain+wHTH+RRM | Type-1_BR.RM.like.systems | <-MORC+DUF3684+REase<-SF1-SF2-Helicase\|\|METHYLASE->DUF499*->Cluster35_66clades->TUDOR+SF1-SF2-Helicase+iREase+CTD+REase->?->TPR+TPR-> | 1145 | Archaea>TACK_group>Candidatus_Verstraetearchaeota>unclassified_Candidatus_Verstraetearchaeota | GCA_020833005.1 | Candidatus_Verstraetearchaeota_archaeon | LM601_06620 | 11919..15356 | 2250257 | ATP-binding-protein-[Candidatus-Verstraetearchaeota-archaeon] |
| MCC6013990.1 | AAA+ATPase+wHTH+Alpha+Beta-domain+wHTH+RRM | Type-1_BR.RM.like.systems | <-TUDOR+SF1-SF2-Helicase+iREase+CTD+REase<-Cluster35_66clades<-DUF499*<-METHYLASE-p | 1137 | Archaea>TACK_group>Candidatus_Verstraetearchaeota>unclassified_Candidatus_Verstraetearchaeota | GCA_020832905.1 | Candidatus_Verstraetearchaeota_archaeon | LM593_06460 | 4830..8243 | 2250257 | ATP-binding-protein-[Candidatus-Verstraetearchaeota-archaeon] |
| MCC6018921.1 | AAA+ATPase+wHTH+Alpha+Beta-domain+wHTH+RRM | Type-1_BR.RM.like.systems | METHYLASE->DUF499*->Cluster35_66clades->TUDOR+SF1-SF2-Helicase+iREase+CTD->?-><-REase | 1124 | Archaea>TACK_group>Candidatus_Verstraetearchaeota>unclassified_Candidatus_Verstraetearchaeota | GCA_020833005.1 | Candidatus_Verstraetearchaeota_archaeon | LM601_07815 | 5184..8558 | 2250257 | ATP-binding-protein-[Candidatus-Verstraetearchaeota-archaeon] |
| MXX20696.1 | AAA+ATPase+wHTH+Alpha+Beta-domain+wHTH+RRM | Type-1_BR.RM.like.systems | PIN->Schlafen+HATPase+HTH->DUF499*->METHYLASE->SF1-SF2-Helicase+iREase+CTD+REase-> | 822 | Archaea>TACK_group>Nitrososphaerota>Cenarchaeales>Cenarchaeaceae | GCA_009836545.1 | Cenarchaeum_sp._SB0667_bin_13 | F4Z46_02780 | 8230..10698 | 2604878 | ATP-binding-protein-[Cenarchaeum-sp.-SB0667_bin_13] |
| MYG32491.1 | AAA+ATPase+wHTH+Alpha+Beta-domain+wHTH+RRM | Type-1_BR.RM.like.systems | BirA-HTH->?->PIN->DUF499*->METHYLASE->SF1-SF2-Helicase+iREase+CTD+REase-> | 806 | Archaea>TACK_group>Nitrososphaerota>Cenarchaeales>Cenarchaeaceae | GCA_009842575.1 | Cenarchaeum_sp._SB0677_bin_16 | F4202_00565 | 1815..4235 | 2604883 | ATP-binding-protein-[Cenarchaeum-sp.-SB0677_bin_16] |
| MYG32966.1 | AAA+ATPase+wHTH+Alpha+Beta-domain+wHTH+RRM | Type-1_BR.RM.like.systems | PIN->DUF499*->METHYLASE->SF1-SF2-Helicase+iREase+CTD+REase-> | 820 | Archaea>TACK_group>Nitrososphaerota>Cenarchaeales>Cenarchaeaceae | GCA_009842575.1 | Cenarchaeum_sp._SB0677_bin_16 | F4202_03070 | 8355..10817 | 2604883 | ATP-binding-protein-[Cenarchaeum-sp.-SB0677_bin_16] |
| MXY61028.1 | AAA+ATPase+wHTH+Alpha+Beta-domain+wHTH+RRM | Type-1_BR.RM.like.systems | PIN->DUF499*->METHYLASE->SF1-SF2-Helicase+iREase+CTD+REase-> | 820 | Archaea>TACK_group>Nitrososphaerota>Cenarchaeales>Cenarchaeaceae | GCA_009837825.1 | Cenarchaeum_sp._SB0665_bin_23 | F4Y82_02795 | 8011..10473 | 2604876 | ATP-binding-protein-[Cenarchaeum-sp.-SB0665_bin_23] |
| MYG33577.1 | AAA+ATPase+wHTH+Alpha+Beta-domain+wHTH+RRM | Type-1_BR.RM.like.systems | <-METHYLASE<-DUF499*\|\|ACYC-> | 841 | Archaea>TACK_group>Nitrososphaerota>Cenarchaeales>Cenarchaeaceae | GCA_009842575.1 | Cenarchaeum_sp._SB0677_bin_16 | F4202_06245 | 4059..6584 | 2604883 | DUF499-domain-containing-protein-[Cenarchaeum-sp.-SB0677_bin_16] |
| MYB46238.1 | AAA+ATPase+wHTH+Alpha+Beta-domain+wHTH+RRM | Type-1_BR.RM.like.systems | <-ACYC\|\|DUF499*->METHYLASE-> | 841 | Archaea>TACK_group>Nitrososphaerota>Cenarchaeales>Cenarchaeaceae | GCA_009840065.1 | Cenarchaeum_sp._SB0662_bin_33 | F4X71_00655 | 8327..10852 | 2604873 | DUF499-domain-containing-protein-[Cenarchaeum-sp.-SB0662_bin_33] |
| MXZ93207.1 | AAA+ATPase+wHTH+Alpha+Beta-domain+wHTH+RRM | Type-1_BR.RM.like.systems | <-ACYC\|\|DUF499*->METHYLASE-> | 841 | Archaea>TACK_group>Nitrososphaerota>Cenarchaeales>Cenarchaeaceae | GCA_009837245.1 | Cenarchaeum_sp._SB0666_bin_15 | F4Z11_01710 | 8361..10886 | 2604877 | DUF499-domain-containing-protein-[Cenarchaeum-sp.-SB0666_bin_15] |
| WP_174448675.1 | AAA+ATPase+wHTH+Alpha+Beta-domain+wHTH+RRM | Type-1_BR.RM.like.systems | <-N6_N4_MTase<-MazE_antitoxin\|\|VirD4-TraG+HerA-><-?<-?<-?<-DUF499*<-METHYLASE<-SF1-SF2-Helicase+iREase+CTD+REase\|\|?->REase->VirD4-TraG+HerA-> | 988 | Archaea>TACK_group>Nitrososphaerota>Conexivisphaeria>Conexivisphaerales | GCF_013340765.1 | Conexivisphaera_calida | NAS2_RS05260 | 1016516..1019482 | 1874277 | DUF499-domain-containing-protein-[Conexivisphaera-calida] |
| KFM20182.1 | AAA+ATPase+wHTH+Alpha+Beta-domain+wHTH+RRM | Type-1_BR.RM.like.systems | <-HEPN<-DUF499*<-METHYLASE<-TUDOR+SF1-SF2-Helicase+iREase+CTD+REase<-?\|\|?-><-REase+DUF5655 | 852 | Archaea>TACK_group>Nitrososphaerota>Nitrososphaerota_incertae_sedis>Marine_Group_I | GCA_000746685.1 | Marine_Group_I_thaumarchaeote_SCGC_AAA799-P11 | AAA799P11_00258 | 97012..99570 | 1502295 | ATPase-AAA-protein-[Marine-Group-I-thaumarchaeote-SCGC-AAA799-P11] |
| KEQ56705.1 | AAA+ATPase+wHTH+Alpha+Beta-domain+wHTH+RRM | Type-1_BR.RM.like.systems | TUDOR+SF1-SF2-Helicase+iREase+CTD+REase->METHYLASE->DUF499*->HEPN-> | 855 | Archaea>TACK_group>Nitrososphaerota>Nitrososphaerota_incertae_sedis>Marine_Group_I | GCA_000722915.1 | Marine_Group_I_thaumarchaeote_SCGC_AAA799-N04 | AAA799N04_00839 | 23778..26345 | 1502293 | ATPase-AAA-protein-[Marine-Group-I-thaumarchaeote-SCGC-AAA799-N04] |
| MDE1878099.1 | AAA+ATPase+wHTH+Alpha+Beta-domain+wHTH+RRM | Type-1_BR.RM.like.systems | SF1-SF2-Helicase+REase->?->EVE-><-DUF499*<-METHYLASE<-TUDOR+SF1-SF2-Helicase+iREase+CTD+REase | 916 | Archaea>TACK_group>Nitrososphaerota>unclassified_Nitrososphaerota | GCA_028866635.1 | Nitrososphaerota_archaeon | KGI07_06160 | 3651..6401 | 2026795 | MAG:-ATP-binding-protein-[Thaumarchaeota-archaeon] |
| MCC6023568.1 | AAA+ATPase+wHTH+Alpha+Beta-domain+wHTH+RRM | Type-1_BR.RM.like.systems | <-TUDOR+SF1-SF2-Helicase+iREase+CTD+REase<-Cluster35_66clades<-DUF499*<-METHYLASE | 1143 | Archaea>TACK_group>Nitrososphaerota>unclassified_Nitrososphaerota | GCA_020833125.1 | Nitrososphaerota_archaeon | LM600_02580 | 6412..9843 | 2026795 | ATP-binding-protein-[Thaumarchaeota-archaeon] |
| MCC6023259.1 | AAA+ATPase+wHTH+Alpha+Beta-domain+wHTH+RRM | Type-1_BR.RM.like.systems | <-TUDOR+SF1-SF2-Helicase+iREase+CTD+REase<-Cluster35_66clades<-DUF499*<-METHYLASE | 1133 | Archaea>TACK_group>Nitrososphaerota>unclassified_Nitrososphaerota | GCA_020833125.1 | Nitrososphaerota_archaeon | LM600_00990 | 12204..15605 | 2026795 | ATP-binding-protein-[Thaumarchaeota-archaeon] |
| MCC6024181.1 | AAA+ATPase+wHTH+Alpha+Beta-domain+wHTH+RRM | Type-1_BR.RM.like.systems | <-TUDOR+SF1-SF2-Helicase+iREase+CTD+REase-p<-Cluster35_66clades<-DUF499* | 1033 | Archaea>TACK_group>Nitrososphaerota>unclassified_Nitrososphaerota | GCA_020833125.1 | Nitrososphaerota_archaeon | LM600_05715 | 1238..4341 | 2026795 | ATP-binding-protein-[Thaumarchaeota-archaeon] |
| MCC6023551.1 | AAA+ATPase+wHTH+Alpha+Beta-domain+wHTH+RRM | Type-1_BR.RM.like.systems | EVE-><-SF1-SF2-Helicase+REase<-Cluster35_66clades<-DUF499*<-METHYLASE | 1124 | Archaea>TACK_group>Nitrososphaerota>unclassified_Nitrososphaerota | GCA_020833125.1 | Nitrososphaerota_archaeon | LM600_02495 | 4852..8226 | 2026795 | ATP-binding-protein-[Thaumarchaeota-archaeon] |
| RLG03319.1 | AAA+ATPase+wHTH+Alpha+Beta-domain+wHTH+RRM | Type-1_BR.RM.like.systems | <-DUF499<-DUF499*<-METHYLASE-p | 526 | Archaea>TACK_group>Nitrososphaerota>unclassified_Nitrososphaerota | GCA_003660695.1 | Nitrososphaerota_archaeon | DRN61_05000 | 1024..2604 | 2026795 | hypothetical-protein-DRN61_05000-[Nitrososphaerota-archaeon] |
| RLG03318.1 | AAA+ATPase+wHTH+Alpha+Beta-domain+wHTH+RRM | Type-1_BR.RM.like.systems | <-DUF499*<-DUF499<-METHYLASE-p | 274 | Archaea>TACK_group>Nitrososphaerota>unclassified_Nitrososphaerota | GCA_003660695.1 | Nitrososphaerota_archaeon | DRN61_04995 | 1..822 | 2026795 | hypothetical-protein-DRN61_04995,-partial-[Nitrososphaerota-archaeon] |
| MBI1658876.1 | AAA+ATPase+wHTH+Alpha+Beta-domain+wHTH+RRM | Type-1_BR.RM.like.systems | <-METHYLASE<-DUF499*<-SF1-SF2-Helicase+iREase+CTD+REase | 811 | Archaea>TACK_group>Nitrososphaerota>unclassified_Nitrososphaerota | GCA_016126025.1 | Nitrososphaerota_archaeon | IS632_08940 | 4229..6664 | 2026795 | ATP-binding-protein-[Nitrososphaerota-archaeon] |
| MBT6647711.1 | AAA+ATPase+wHTH+Alpha+Beta-domain+wHTH+RRM | Type-1_BR.RM.like.systems | <-HEPN<-DUF499*<-METHYLASE-p\|\|METHYLASE-p-> | 834 | Archaea>TACK_group>Nitrososphaerota>unclassified_Nitrososphaerota | GCA_018697615.1 | Nitrososphaerota_archaeon | HOB51_09415 | 855..3359 | 2026795 | ATP-binding-protein-[Thaumarchaeota-archaeon] |
| MCH7560423.1 | AAA+ATPase+wHTH+Alpha+Beta-domain+wHTH+RRM | Type-1_BR.RM.like.systems | <-HEPN<-DUF499*<-METHYLASE | 852 | Archaea>TACK_group>Nitrososphaerota>unclassified_Nitrososphaerota | GCA_022561135.1 | Nitrososphaerota_archaeon | IIC67_03480 | 947..3505 | 2026795 | ATP-binding-protein-[Thaumarchaeota-archaeon] |
| NAZ13696.1 | AAA+ATPase+wHTH+Alpha+Beta-domain+wHTH+RRM | Type-1_BR.RM.like.systems | ParA-Soj-PloopNTPase->ParA-Soj-PloopNTPase->?->METHYLASE->DUF499*->Cluster35_66clades->TUDOR+SF1-SF2-Helicase+iREase+CTD-><-DUF2791 | 1122 | Archaea>TACK_group>Thermoproteota>Thermoprotei>Desulfurococcales | GCA_009903825.1 | Desulfurococcales_archaeon | GU359_06060 | 7964..11332 | 2480821 | DUF499-domain-containing-protein-[Desulfurococcales-archaeon] |
| ADM27560.1 | AAA+ATPase+wHTH+Alpha+Beta-domain+wHTH+RRM | Type-1_BR.RM.like.systems | <-PRTase+PRTase\|\|N6_MTase-><-?<-?<-TUDOR+SF1-SF2-Helicase+iREase+CTD+REase<-Cluster35_66clades<-DUF499*<-METHYLASE\|\|EVE-> | 1146 | Archaea>TACK_group>Thermoproteota>Thermoprotei>Desulfurococcales | GCA_000145985.1 | Ignisphaera_aggregans_DSM_17230 | Igag_0730 | 682542..685982 | 583356 | ATPase-[Ignisphaera-aggregans-DSM-17230] |
| WP_075049293.1 | AAA+ATPase+wHTH+Alpha+Beta-domain+wHTH+RRM | Type-1_BR.RM.like.systems | MORC-><-?<-Radical-SAM<-?\|\|?-><-METHYLASE\|\|DUF499*->Cluster35_66clades->TUDOR+SF1-SF2-Helicase+iREase+CTD+REase-> | 1085 | Archaea>TACK_group>Thermoproteota>Thermoprotei>Desulfurococcales | GCF_001481685.1 | Ignicoccus_islandicus | EYM_RS01160 | 199793..203050 | 54259 | DUF499-domain-containing-protein-[Ignicoccus-islandicus] |
| WP_058370865.1 | AAA+ATPase+wHTH+Alpha+Beta-domain+wHTH+RRM | Type-1_BR.RM.like.systems | <-ABC-ATPase+SbcC<-?\|\|METHYLASE-p-><-DUF499*<-METHYLASE\|\|REase->SF1-SF2-Helicase-><-ABC-ATPase+SbcC<-?<-REase | 1039 | Archaea>TACK_group>Thermoproteota>Thermoprotei>Desulfurococcales | GCF_001462395.1 | Pyrodictium_occultum | CF15_RS05340 | 1008872..1011991 | 2309 | DUF499-domain-containing-protein-[Pyrodictium-occultum] |
| WP_042667294.1 | AAA+ATPase+wHTH+Alpha+Beta-domain+wHTH+RRM | Type-1_BR.RM.like.systems | <-TUDOR+SF1-SF2-Helicase+iREase+CTD+REase<-Cluster35_66clades<-DUF499*<-METHYLASE\|\|?->?-><-Methylase | 1140 | Archaea>TACK_group>Thermoproteota>Thermoprotei>Desulfurococcales | GCF_000513855.1 | Desulfurococcus_amylolyticus | SPHMEL_RS02945 | 547501..550923 | 94694 | DUF499-domain-containing-protein-[Desulfurococcus-amylolyticus] |
| WP_013142539.1 | AAA+ATPase+wHTH+Alpha+Beta-domain+wHTH+RRM | Type-1_BR.RM.like.systems | URI-DNase->?-><-REase<-?<-?<-DUF499*<-METHYLASE\|\|Methyltransf->METHYLASE-p-><-SF1-SF2-Helicase<-REase | 1139 | Archaea>TACK_group>Thermoproteota>Thermoprotei>Desulfurococcales | GCF_000092465.1 | Staphylothermus_hellenicus | SHELL_RS01010 | 174708..178127 | 84599 | ATP-binding-protein-[Staphylothermus-hellenicus] |
| WP_011838873.1 | AAA+ATPase+wHTH+Alpha+Beta-domain+wHTH+RRM | Type-1_BR.RM.like.systems | Phage_integrase-><-?<-?<-REase<-?<-?<-DUF499*<-METHYLASE\|\|METHYLASE-p-><-SF1-SF2-Helicase<-REase | 1140 | Archaea>TACK_group>Thermoproteota>Thermoprotei>Desulfurococcales | GCF_000015945.1 | Staphylothermus_marinus | SMAR_RS02915 | 576255..579677 | 2280 | ATP-binding-protein-[Staphylothermus-marinus] |
| MCD6301539.1 | AAA+ATPase+wHTH+Alpha+Beta-domain+wHTH+RRM | Type-1_BR.RM.like.systems | METHYLASE-p->DUF499*->Cluster35_66clades->TUDOR+SF1-SF2-Helicase+iREase+CTD+REase-> | 1134 | Archaea>TACK_group>Thermoproteota>Thermoprotei>Desulfurococcales | GCA_021159785.1 | Staphylothermus_sp. | J7L82_05660 | 1324..4728 | 2820299 | DUF499-domain-containing-protein-[Staphylothermus-sp.] |
| WP_048163335.1 | AAA+ATPase+wHTH+Alpha+Beta-domain+wHTH+RRM | Type-1_BR.RM.like.systems | ABC-ATPase+SbcC->METHYLASE->DUF499*->?->SF1-SF2-Helicase+REase->HHH+ENDO3+NUDIX-> | 1244 | Archaea>TACK_group>Thermoproteota>Thermoprotei>Desulfurococcales | GCF_000264495.1 | Thermogladius_calderae | TCELL_RS06135 | 1136275..1140009 | 1200300 | DUF499-domain-containing-protein-[Thermogladius-calderae] |
| WP_013129635.1 | AAA+ATPase+wHTH+Alpha+Beta-domain+wHTH+RRM | Type-1_BR.RM.like.systems | <-SF1-SF2-Helicase+REase<-?<-DUF499*<-METHYLASE\|\|?->MerR-HTH+GDResolvase->ZnR->HNH-> | 1246 | Archaea>TACK_group>Thermoproteota>Thermoprotei>Desulfurococcales | GCF_000092185.1 | Thermosphaera_aggregans | TAGG_RS03820 | 727041..730781 | 54254 | DUF499-domain-containing-protein-[Thermosphaera-aggregans] |
| AFK51621.1 | AAA+ATPase+wHTH+Alpha+Beta-domain+wHTH+RRM | Type-1_BR.RM.like.systems | ABC-ATPase+SbcC->?->METHYLASE->DUF499*->?->SF1-SF2-Helicase+REase->HHH+ENDO3+NUDIX-> | 1233 | Archaea>TACK_group>Thermoproteota>Thermoprotei>Desulfurococcales | GCA_000264495.1 | Thermogladius_calderae_1633 | TCELL_1198 | 1136308..1140009 | 1184251 | hypothetical-protein-TCELL_1198-[Thermogladius-calderae-1633] |
| MCE4610565.1 | AAA+ATPase+wHTH+Alpha+Beta-domain+wHTH+RRM | Type-1_BR.RM.like.systems | TOPRIM-><-?<-tRNA<-?<-sGTP<-?\|\|?-><-DUF499*<-METHYLASE\|\|REase->SF1-SF2-Helicase->METHYLASE-p-> | 1071 | Archaea>TACK_group>Thermoproteota>Thermoprotei>Desulfurococcales | GCA_015523465.1 | Desulfurococcales_archaeon | F7B17_01170 | 7104..10319 | 2480821 | ATP-binding-protein-[Desulfurococcales-archaeon] |
| WP_055408547.1 | AAA+ATPase+wHTH+Alpha+Beta-domain+wHTH+RRM | Type-1_BR.RM.like.systems | <-METHYLASE-p<-REase<-SF1-SF2-Helicase<-REase\|\|METHYLASE->DUF499*-><-?<-HNH | 1047 | Archaea>TACK_group>Thermoproteota>Thermoprotei>Desulfurococcales | GCF_001412615.1 | Pyrodictium_delaneyi | Pyrde_RS04390 | 857202..860345 | 1273541 | DUF499-domain-containing-protein-[Pyrodictium-delaneyi] |
| WP_010865722.1 | AAA+ATPase+wHTH+Alpha+Beta-domain+wHTH+RRM | Type-1_BR.RM.like.systems | <-TUDOR+SF1-SF2-Helicase+iREase+CTD+REase<-?<-DUF499*<-METHYLASE<-tRNA<-tRNA<-TraJ-RHH | 1188 | Archaea>TACK_group>Thermoproteota>Thermoprotei>Desulfurococcales | GCF_000011125.1 | Aeropyrum_pernix | APE_RS01525 | 288073..291639 | 56636 | DUF499-domain-containing-protein-[Aeropyrum-pernix] |
| WP_276815000.1 | AAA+ATPase+wHTH+Alpha+Beta-domain+wHTH+RRM | Type-1_BR.RM.like.systems | <-TUDOR+SF1-SF2-Helicase+iREase+CTD+REase<-Cluster35_66clades<-DUF499* | 1060 | Archaea>TACK_group>Thermoproteota>Thermoprotei>Desulfurococcales | GCF_023256035.1 | Desulfurococcus_amylolyticus | P6716_RS06675 | 21797..24977 | 94694 | DUF499-domain-containing-protein,-partial-[Desulfurococcus-amylolyticus] |
| MCE4605485.1 | AAA+ATPase+wHTH+Alpha+Beta-domain+wHTH+RRM | Type-1_BR.RM.like.systems | REase-><-DUF499*<-METHYLASE\|\|REase->SF1-SF2-Helicase-><-?<-?\|\|N6_N4_MTase-> | 1031 | Archaea>TACK_group>Thermoproteota>Thermoprotei>Desulfurococcales | GCA_015521665.1 | Desulfurococcales_archaeon | F7C08_03020 | 104683..107778 | 2480821 | DUF499-domain-containing-protein-[Desulfurococcales-archaeon] |
| MCE4602804.1 | AAA+ATPase+wHTH+Alpha+Beta-domain+wHTH+RRM | Type-1_BR.RM.like.systems | <-N6_N4_MTase\|\|?->?-><-SF1-SF2-Helicase<-REase\|\|METHYLASE->DUF499*-> | 996 | Archaea>TACK_group>Thermoproteota>Thermoprotei>Desulfurococcales | GCA_015523445.1 | Desulfurococcales_archaeon | F7B18_06450 | 10084..13072 | 2480821 | DUF499-domain-containing-protein-[Desulfurococcales-archaeon] |
| MCE4606659.1 | AAA+ATPase+wHTH+Alpha+Beta-domain+wHTH+RRM | Type-1_BR.RM.like.systems | <-TUDOR+SF1-SF2-Helicase+iREase+CTD+REase<-?<-DUF499*<-METHYLASE | 1190 | Archaea>TACK_group>Thermoproteota>Thermoprotei>Desulfurococcales | GCA_015522645.1 | Desulfurococcales_archaeon | F7B59_04910 | 5670..9242 | 2480821 | ATP-binding-protein-[Desulfurococcales-archaeon] |
| MCC6010980.1 | AAA+ATPase+wHTH+Alpha+Beta-domain+wHTH+RRM | Type-1_BR.RM.like.systems | <-REase<-SF1-SF2-Helicase+REase<-?<-DUF499*<-METHYLASE-p | 1248 | Archaea>TACK_group>Thermoproteota>Thermoprotei>Desulfurococcales | GCA_020832865.1 | Desulfurococcaceae_archaeon | LM556_01090 | 5254..9000 | 2184738 | ATP-binding-protein-[Desulfurococcaceae-archaeon] |
| NAZ12665.1 | AAA+ATPase+wHTH+Alpha+Beta-domain+wHTH+RRM | Type-1_BR.RM.like.systems | <-SF1-SF2-Helicase+REase<-?<-DUF499*<-METHYLASE | 1252 | Archaea>TACK_group>Thermoproteota>Thermoprotei>Desulfurococcales | GCA_009903825.1 | Desulfurococcales_archaeon | GU359_00740 | 3566..7324 | 2480821 | DUF499-domain-containing-protein-[Desulfurococcales-archaeon] |
| MCE4612152.1 | AAA+ATPase+wHTH+Alpha+Beta-domain+wHTH+RRM | Type-1_BR.RM.like.systems | <-DUF499*<-METHYLASE-p\|\|?->?->?->?->PadR-HTH-> | 862 | Archaea>TACK_group>Thermoproteota>Thermoprotei>Desulfurococcales | GCA_015523465.1 | Desulfurococcales_archaeon | F7B17_09305 | 31931..34519 | 2480821 | ATP-binding-protein-[Desulfurococcales-archaeon] |
| MCI4462011.1 | AAA+ATPase+wHTH+Alpha+Beta-domain+wHTH+RRM | Type-1_BR.RM.like.systems | <-DUF499-p<-DUF499*<-METHYLASE-p | 467 | Archaea>TACK_group>Thermoproteota>Thermoprotei>Desulfurococcales | GCA_022759365.1 | Thermogladius_sp. | JHC12_03500 | 2242..3641 | 2023064 | DUF499-domain-containing-protein-[Thermogladius-sp.] |
| PWV37728.1 | AAA+ATPase+wHTH+Alpha+Beta-domain+wHTH+RRM | Type-1_BR.RM.like.systems | <-N6_N4_MTase\|\|TUDOR+SF1-SF2-Helicase+iREase+CTD+REase->METHYLASE->DUF499*-><-?\|\|?->ZnR-> | 863 | Archaea>TACK_group>Thermoproteota>Thermoprotei>Desulfurococcales | GCA_003177295.1 | Desulfurococcaceae_archaeon | DJ555_00765 | 8666..11257 | 2184738 | AAA-family-ATPase-[Desulfurococcaceae-archaeon] |
| WP_228009360.1 | AAA+ATPase+wHTH+Alpha+Beta-domain+wHTH+RRM | Type-1_BR.RM.like.systems | HEPN->?->PIN->?->METHYLASE->DUF499*->Cluster35_66clades->TUDOR+SF1-SF2-Helicase+iREase+CTD+REase-><-?\|\|PIN-> | 1042 | Archaea>TACK_group>Thermoproteota>Thermoprotei>Fervidicoccales | - | Fervidicoccus_fontis | IOK49_RS01845 | 332374..335502 | 683846 | DUF499-domain-containing-protein-[Fervidicoccus-fontis] |
| MBE9390828.1 | AAA+ATPase+wHTH+Alpha+Beta-domain+wHTH+RRM | Type-1_BR.RM.like.systems | HEPN->?->PIN->?->METHYLASE->DUF499*->Cluster35_66clades->TUDOR+SF1-SF2-Helicase+iREase+CTD+REase-><-?\|\|PIN-> | 1144 | Archaea>TACK_group>Thermoproteota>Thermoprotei>Fervidicoccales | GCA_015209645.1 | Fervidicoccus_fontis | IOK49_01845 | 332068..335502 | 683846 | DUF499-domain-containing-protein-[Fervidicoccus-fontis] |
| PMB78266.1 | AAA+ATPase+wHTH+Alpha+Beta-domain+wHTH+RRM | Type-1_BR.RM.like.systems | TUDOR+SF1-SF2-Helicase+iREase+CTD+REase->METHYLASE-p->HEPN->NTP_transf_2->PolB->HEPN->DUF499*-> | 859 | Archaea>TACK_group>Thermoproteota>Thermoprotei>Fervidicoccales | GCA_002871295.1 | Fervidicoccus_fontis | C0177_00595 | 9295..11874 | 683846 | AAA-family-ATPase-[Fervidicoccus-fontis] |
| WP_229570012.1 | AAA+ATPase+wHTH+Alpha+Beta-domain+wHTH+RRM | Type-1_BR.RM.like.systems | <-ZnR+Methylase<-?<-TUDOR+SF1-SF2-Helicase+iREase+CTD+REase<-Cluster35_66clades<-DUF499*<-METHYLASE\|\|?->SF1-SF2-Helicase->HKD-DNase-><-MethyltransfD12\|\|Zn_Tnp_IS1+HTH_Tnp_IS1+DDE_Tnp_IS1-> | 1094 | Archaea>TACK_group>Thermoproteota>Thermoprotei>Sulfolobales | GCF_020886315.1 | Saccharolobus_caldissimus | SACC_RS13230 | 2411160..2414444 | 1702097 | DUF499-domain-containing-protein-[Saccharolobus-caldissimus] |
| WP_153425346.1 | AAA+ATPase+wHTH+Alpha+Beta-domain+wHTH+RRM | Type-1_BR.RM.like.systems | SF1-SF2-Helicase->HKD-DNase-><-MethyltransfD12\|\|METHYLASE-p->METHYLASE->DUF499*->Cluster35_66clades->TUDOR+SF1-SF2-Helicase+iREase+CTD+REase->?-><-ZnR+Methylase | 1086 | Archaea>TACK_group>Thermoproteota>Thermoprotei>Sulfolobales | GCF_009602405.1 | Sulfolobus_sp._E11-6 | GFS33_RS10400 | 1934011..1937271 | 2663020 | DUF499-domain-containing-protein-[Sulfolobus-sp.-E11-6] |
| WP_110271302.1 | AAA+ATPase+wHTH+Alpha+Beta-domain+wHTH+RRM | Type-1_BR.RM.like.systems | HEPN-><-?<-ZnR+Methylase<-?<-TUDOR+SF1-SF2-Helicase+iREase+CTD+REase<-Cluster35_66clades<-DUF499*<-METHYLASE<-?<-REase+SF1-SF2-Helicase\|\|?->MethyltransfD12-><-HKD-DNase | 1095 | Archaea>TACK_group>Thermoproteota>Thermoprotei>Sulfolobales | GCF_003201835.2 | Acidianus_brierleyi | DFR85_RS29145 | 2432274..2435561 | 41673 | DUF499-domain-containing-protein-[Acidianus-brierleyi] |
| WP_012715695.1 | AAA+ATPase+wHTH+Alpha+Beta-domain+wHTH+RRM | Type-1_BR.RM.like.systems | MethyltransfD12-><-HKD-DNase<-SF1-SF2-Helicase<-?\|\|METHYLASE->DUF499*->Cluster35_66clades->TUDOR+SF1-SF2-Helicase+iREase+CTD+REase->?->ZnR+Methylase-><-?\|\|MarR-HTH-> | 1094 | Archaea>TACK_group>Thermoproteota>Thermoprotei>Sulfolobales | GCF_000022465.1 | Sulfolobus_islandicus | YG5714_RS02790 | 524842..528126 | 43080 | DUF499-domain-containing-protein-[Sulfolobus-islandicus] |
| MCX8185269.1 | AAA+ATPase+wHTH+Alpha+Beta-domain+wHTH+RRM | Type-1_BR.RM.like.systems | ABhydrolase-><-MetJarc-HTH\|\|N6_MTase-p->METHYLASE->DUF499*->Cluster35_66clades->TUDOR+SF1-SF2-Helicase+iREase+CTD+REase-><-tRNA<-Aminotran_3<-?<-HHH+ENDO3+NUDIX | 1123 | Archaea>TACK_group>Thermoproteota>Thermoprotei>Sulfolobales | GCA_026413885.1 | Sulfolobales_archaeon | N3E43_05930 | 38123..41494 | 2268200 | MAG:-ATP-binding-protein-[Sulfolobales-archaeon] |
| WP_156014062.1 | AAA+ATPase+wHTH+Alpha+Beta-domain+wHTH+RRM | Type-1_BR.RM.like.systems | <-ZnR+Methylase<-?<-TUDOR+SF1-SF2-Helicase+iREase+CTD+REase<-Cluster35_66clades<-DUF499*<-METHYLASE\|\|METHYLASE-p->METHYLASE-p->METHYLASE-p->METHYLASE-p-> | 1094 | Archaea>TACK_group>Thermoproteota>Thermoprotei>Sulfolobales | GCF_009729055.1 | Sulfurisphaera_ohwakuensis | D1869_RS04270 | 715568..718852 | 69656 | DUF499-domain-containing-protein-[Sulfurisphaera-ohwakuensis] |
| WP_168064839.1 | AAA+ATPase+wHTH+Alpha+Beta-domain+wHTH+RRM | Type-1_BR.RM.like.systems | REase-><-?<-METHYLASE-p<-METHYLASE-p\|\|METHYLASE->DUF499*->Cluster35_66clades->TUDOR+SF1-SF2-Helicase+iREase+CTD+REase->?-><-ZnR+Methylase | 1091 | Archaea>TACK_group>Thermoproteota>Thermoprotei>Sulfolobales | GCF_012222305.1 | Sulfolobus_sp._S-194 | EWF20_RS06040 | 1155575..1158850 | 2512240 | DUF499-domain-containing-protein-[Sulfolobus-sp.-S-194] |
| WP_218260487.1 | AAA+ATPase+wHTH+Alpha+Beta-domain+wHTH+RRM | Type-1_BR.RM.like.systems | Transposase->ZnR+Methylase-><-?<-TUDOR+SF1-SF2-Helicase+iREase+CTD+REase<-Cluster35_66clades<-DUF499*<-METHYLASE\|\|METHYLASE-><-?\|\|?-><-?\|\|ABC-ATPase+SbcC-> | 1093 | Archaea>TACK_group>Thermoproteota>Thermoprotei>Sulfolobales | GCF_019175325.1 | Saccharolobus_shibatae | J5U21_RS08220 | 1462926..1466207 | 2286 | DUF499-domain-containing-protein-[Saccharolobus-shibatae] |
| WP_206760616.1 | AAA+ATPase+wHTH+Alpha+Beta-domain+wHTH+RRM | Type-1_BR.RM.like.systems | <-ABC-ATPase+SbcC<-ZnR+Methylase<-?<-TUDOR+SF1-SF2-Helicase+iREase+CTD+REase<-Cluster35_66clades<-DUF499*<-METHYLASE\|\|<-HEPN\|\|HEPN-> | 1098 | Archaea>TACK_group>Thermoproteota>Thermoprotei>Sulfolobales | GCF_000189555.1 | Sulfolobus_islandicus | SIRE_RS03030 | 568473..571769 | 43080 | DUF499-domain-containing-protein-[Sulfolobus-islandicus] |
| WP_221290325.1 | AAA+ATPase+wHTH+Alpha+Beta-domain+wHTH+RRM | Type-1_BR.RM.like.systems | <-METHYLASE<-METHYLASE\|\|METHYLASE->DUF499*->Cluster35_66clades->TUDOR+SF1-SF2-Helicase+iREase+CTD+REase->?-><-ZnR+Methylase\|\|SbcC-> | 1093 | Archaea>TACK_group>Thermoproteota>Thermoprotei>Sulfolobales | GCF_019704315.1 | Stygiolobus_caldivivus | KN1_RS06645 | 1407029..1410310 | 2824673 | DUF499-domain-containing-protein-[Stygiolobus-caldivivus] |
| WP_218257720.1 | AAA+ATPase+wHTH+Alpha+Beta-domain+wHTH+RRM | Type-1_BR.RM.like.systems | ZnR+Methylase-><-?<-TUDOR+SF1-SF2-Helicase+iREase+CTD+REase<-Cluster35_66clades<-DUF499*<-METHYLASE\|\|METHYLASE->HD-><-?\|\|?-><-Transposase | 1093 | Archaea>TACK_group>Thermoproteota>Thermoprotei>Sulfolobales | GCF_019175305.1 | Saccharolobus_shibatae | J5U22_RS07505 | 1396882..1400163 | 2286 | DUF499-domain-containing-protein-[Saccharolobus-shibatae] |
| WP_012713458.1 | AAA+ATPase+wHTH+Alpha+Beta-domain+wHTH+RRM | Type-1_BR.RM.like.systems | <-MarR-HTH\|\|?-><-ZnR+Methylase<-?<-TUDOR+SF1-SF2-Helicase+iREase+CTD+REase<-Cluster35_66clades<-DUF499*<-METHYLASE\|\|?->?-><-PADR-HTH\|\|HEPN-> | 1094 | Archaea>TACK_group>Thermoproteota>Thermoprotei>Sulfolobales | GCF_000022385.1 | Sulfolobus_islandicus | LS215_RS05230 | 976652..979936 | 43080 | DUF499-domain-containing-protein-[Sulfolobus-islandicus] |
| MUM65787.1 | AAA+ATPase+wHTH+Alpha+Beta-domain+wHTH+RRM | Type-1_BR.RM.like.systems | <-SF1-SF2-Helicase<-?<-ZnR+Methylase<-?<-TUDOR+SF1-SF2-Helicase+iREase+CTD+REase<-Cluster35_66clades<-DUF499*<-METHYLASE\|\|?-><-?<-?<-?\|\|ZnR-> | 1093 | Archaea>TACK_group>Thermoproteota>Thermoprotei>Sulfolobales | GCA_009729545.1 | Acidianus_infernus | D1867_11170 | 1971916..1975197 | 12915 | DUF499-domain-containing-protein-[Acidianus-infernus] |
| ADX84676.1 | AAA+ATPase+wHTH+Alpha+Beta-domain+wHTH+RRM | Type-1_BR.RM.like.systems | <-ABC-ATPase+SbcC<-ZnR+Methylase<-?<-TUDOR+SF1-SF2-Helicase+iREase+CTD+REase<-Cluster35_66clades<-DUF499*<-METHYLASE\|\|?->HEPN-> | 1094 | Archaea>TACK_group>Thermoproteota>Thermoprotei>Sulfolobales | GCA_000189555.1 | Sulfolobus_islandicus_REY15A | SiRe_0593 | 568473..571757 | 930945 | ATPase-(AAA+-superfamily)-like-protein-[Sulfolobus-islandicus-REY15A] |
| WP_016729821.1 | AAA+ATPase+wHTH+Alpha+Beta-domain+wHTH+RRM | Type-1_BR.RM.like.systems | PADR-HTH->?->?-><-?\|\|METHYLASE->DUF499*->Cluster35_66clades->TUDOR+SF1-SF2-Helicase+iREase+CTD+REase->?->ZnR+Methylase-><-?\|\|MarR-HTH-> | 1094 | Archaea>TACK_group>Thermoproteota>Thermoprotei>Sulfolobales | GCF_000245095.1 | Sulfolobus_islandicus | M162_RS0103490 | 602574..605858 | 43080 | DUF499-domain-containing-protein-[Sulfolobus-islandicus] |

| Protein-ID | DUF499-architecture | System | Gene-Neighbourhoods | Length | Taxonomy | Assembly-ID | Strain | Locus-Tag | Genomic-coordinates | TaxID | Fasta-defline |
|---|---|---|---|---|---|---|---|---|---|---|---|
| QIW23757.1 | AAA+ATPase+wHTH+Alpha+Beta-domain+wHTH+RRM | Type-1_BR.RM.like.systems | <-CARF+HTH\|\|REase-><-?\|\|METHYLASE->DUF499*->Cluster35_66clades->TUDOR+SF1-SF2-Helicase+iREase+CTD+REase->?-><-ZnR+Methylase | 1091 | Archaea>TACK_group>Thermoproteota>Thermoprotei>Sulfolobales | GCA_012222305.1 | Sulfolobus_sp._S-194 | EWF20_06020 | 1155575..1158850 | 2512240 | DUF499-domain-containing-protein-[Sulfolobus-sp.-S-194] |
| WP_240872372.1 | AAA+ATPase+wHTH+Alpha+Beta-domain+wHTH+RRM | Type-1_BR.RM.like.systems | <-SF1-SF2-Helicase<-?<-ZnR+Methylase<-?<-TUDOR+SF1-SF2-Helicase+iREase+CTD+REase<-Cluster35_66clades<-DUF499*<-METHYLASE | 1039 | Archaea>TACK_group>Thermoproteota>Thermoprotei>Sulfolobales | GCF_009729545.1 | Acidianus_infernus | D1867_RS11160 | 1971916..1975035 | 12915 | DUF499-domain-containing-protein-[Acidianus-infernus] |
| EHP69856.1 | AAA+ATPase+wHTH+Alpha+Beta-domain+wHTH+RRM | Type-1_BR.RM.like.systems | <-ORC-CDC6+REase<-?\|\|?->PIN->?->Transposase-><-DUF499*<-DUF499-p<-METHYLASE<-?<-REase+SF1-SF2-Helicase\|\|?->MethyltransfD12-> | 129 | Archaea>TACK_group>Thermoproteota>Thermoprotei>Sulfolobales | GCA_000243315.1 | Metallosphaera_yellowstonensis_MK1 | MetMK1DRAFT_00003580 | 133014..133403 | 671065 | putative-ATPase-(AAA+-superfamily)-[Metallosphaera-yellowstonensis-MK1] |
| WP_156006737.1 | AAA+ATPase+wHTH+Alpha+Beta-domain+wHTH+RRM | Type-1_BR.RM.like.systems | SF1-SF2-Helicase-p-><-?<-TUDOR+SF1-SF2-Helicase+iREase+CTD+REase<-Cluster35_66clades<-DUF499*<-METHYLASE<-?\|\|?-><-ZnR | 1143 | Archaea>TACK_group>Thermoproteota>Thermoprotei>Sulfolobales | GCF_009729035.1 | Stygiolobus_azoricus | D1868_RS06600 | 1190178..1193609 | 41675 | DUF499-domain-containing-protein-[Stygiolobus-azoricus] |
| WP_016730556.1 | AAA+ATPase+wHTH+Alpha+Beta-domain+wHTH+RRM | Type-1_BR.RM.like.systems | PADR-HTH->?->\|\|METHYLASE->DUF499*->Cluster35_66clades->TUDOR+SF1-SF2-Helicase+iREase+CTD+REase->?->?->?->ZnR+Methylase-> | 1092 | Archaea>TACK_group>Thermoproteota>Thermoprotei>Sulfolobales | GCF_000245115.1 | Sulfolobus_islandicus | M1622_RS0103355 | 558017..561295 | 43080 | DUF499-domain-containing-protein-[Sulfolobus-islandicus] |
| PVU69402.1 | AAA+ATPase+wHTH+Alpha+Beta-domain+wHTH+RRM | Type-1_BR.RM.like.systems | DUF499*->Cluster35_66clades->TUDOR+SF1-SF2-Helicase+iREase+CTD+REase->?-><-HEPN<-PolB-NTase\|\|NTP_transf-> | 740 | Archaea>TACK_group>Thermoproteota>Thermoprotei>Sulfolobales | GCA_003086575.1 | Sulfolobales_archaeon_SCGC_AB-777_K09 | DDW04_00535 | 1..2225 | 2005260 | AAA-family-ATPase,-partial-[Sulfolobales-archaeon-SCGC-AB-777_K09] |
| MCI4456204.1 | AAA+ATPase+wHTH+Alpha+Beta-domain+wHTH+RRM | Type-1_BR.RM.like.systems | ZnR+Methylase-><-?<-TUDOR+SF1-SF2-Helicase+iREase+CTD+REase<-Cluster35_66clades<-DUF499*<-METHYLASE-p | 1102 | Archaea>TACK_group>Thermoproteota>Thermoprotei>Sulfolobales | GCA_022759265.1 | Sulfolobus_sp. | JHC23_05755 | 6447..9755 | 2288 | DUF499-domain-containing-protein-[Sulfolobus-sp.] |
| WP_009070048.1 | AAA+ATPase+wHTH+Alpha+Beta-domain+wHTH+RRM | Type-1_BR.RM.like.systems | <-ORC-CDC6+REase\|\|?->?->?->PIN-><-DUF499*<-METHYLASE<-?<-REase+SF1-SF2-Helicase\|\|?->MethyltransfD12-><-HKD-DNase | 137 | Archaea>TACK_group>Thermoproteota>Thermoprotei>Sulfolobales | GCF_000243315.1 | Metallosphaera_yellowstonensis | METMK1DRAFT_RS18680 | 133361..133774 | 1111107 | DUF499-domain-containing-protein-[Metallosphaera-yellowstonensis] |
| MCG2871562.1 | AAA+ATPase+wHTH+Alpha+Beta-domain+wHTH+RRM | Type-1_BR.RM.like.systems | ZnR+Methylase-><-?<-TUDOR+SF1-SF2-Helicase+iREase+CTD+REase<-Cluster35_66clades<-DUF499*<-METHYLASE | 1107 | Archaea>TACK_group>Thermoproteota>Thermoprotei>Sulfolobales | GCA_022014915.1 | Sulfolobales_archaeon | L7H13_04865 | 6840..10163 | 2268200 | ATP-binding-protein-[Sulfolobales-archaeon] |
| MCX8186953.1 | AAA+ATPase+wHTH+Alpha+Beta-domain+wHTH+RRM | Type-1_BR.RM.like.systems | <-TUDOR+SF1-SF2-Helicase+iREase+CTD+REase<-Cluster35_66clades<-DUF499*<-METHYLASE-p | 1123 | Archaea>TACK_group>Thermoproteota>Thermoprotei>Sulfolobales | GCA_026413615.1 | Sulfolobales_archaeon | N3G48_07625 | 3009..6380 | 2268200 | MAG:-ATP-binding-protein-[Sulfolobales-archaeon] |
| MCG2871663.1 | AAA+ATPase+wHTH+Alpha+Beta-domain+wHTH+RRM | Type-1_BR.RM.like.systems | <-TUDOR+SF1-SF2-Helicase+iREase+CTD+REase<-Cluster35_66clades<-DUF499*<-METHYLASE | 1094 | Archaea>TACK_group>Thermoproteota>Thermoprotei>Sulfolobales | GCA_022014915.1 | Sulfolobales_archaeon | L7H13_05415 | 4741..8025 | 2268200 | ATP-binding-protein-[Sulfolobales-archaeon] |
| WP_198968357.1 | AAA+ATPase+wHTH+Alpha+Beta-domain+wHTH+RRM | Type-1_BR.RM.like.systems | DUF499*->Cluster35_66clades->TUDOR+SF1-SF2-Helicase+iREase+CTD+REase-p-> | 366 | Archaea>TACK_group>Thermoproteota>Thermoprotei>Sulfolobales | GCF_016415645.1 | Sulfolobus_islandicus | IEQ43_RS13155 | 1..1101 | 43080 | DUF499-domain-containing-protein,-partial-[Sulfolobus-islandicus] |
| MCQ4448867.1 | AAA+ATPase+wHTH+Alpha+Beta-domain+wHTH+RRM | Type-1_BR.RM.like.systems | DUF499*->Cluster35_66clades->TUDOR+SF1-SF2-Helicase+iREase+CTD+REase-p-> | 918 | Archaea>TACK_group>Thermoproteota>Thermoprotei>Sulfolobales | GCA_024451005.1 | Sulfolobales_archaeon | NO076_06470 | 1..2757 | 2268200 | ATP-binding-protein-[Sulfolobales-archaeon] |
| MCQ4345626.1 | AAA+ATPase+wHTH+Alpha+Beta-domain+wHTH+RRM | Type-1_BR.RM.like.systems | <-TUDOR+SF1-SF2-Helicase+iREase+CTD+REase-p<-Cluster35_66clades<-DUF499* | 769 | Archaea>TACK_group>Thermoproteota>Thermoprotei>Sulfolobales | GCA_024451075.1 | Sulfolobales_archaeon | NO086_05565 | 1164..3473 | 2268200 | ATP-binding-protein-[Sulfolobales-archaeon] |
| BCU66881.1 | AAA+ATPase+wHTH+Alpha+Beta-domain+wHTH+RRM | Type-1_BR.RM.like.systems | MarR-HTH-><-?\|\|?-><-METHYLASE\|\|DUF499*->?-><-SF1-SF2-Helicase+iREase+CTD+REase | 1008 | Archaea>TACK_group>Thermoproteota>Thermoprotei>Sulfolobales | GCA_019704295.1 | Sulfolobales_archaeon_HS-7 | HS7_03180 | 325048..328074 | 2823152 | hypothetical-protein-HS7_03180-[Sulfolobales-archaeon-HS-7] |
| WP_277346522.1 | AAA+ATPase+wHTH+Alpha+Beta-domain+wHTH+RRM | Type-1_BR.RM.like.systems | ZnR->CARF->METHYLASE->DUF499*->Cluster35_66clades-> | 969 | Archaea>TACK_group>Thermoproteota>Thermoprotei>Sulfolobales | GCF_003201765.2 | Acidianus_sulfidivorans | DFR86_RS23580 | 2261261..2264170 | 312539 | DUF499-domain-containing-protein-[Acidianus-sulfidivorans] |
| AWR98153.1 | AAA+ATPase+wHTH+Alpha+Beta-domain+wHTH+RRM | Type-1_BR.RM.like.systems | CARF->METHYLASE->DUF499*->Cluster35_66clades-> | 1046 | Archaea>TACK_group>Thermoproteota>Thermoprotei>Sulfolobales | GCA_003201765.2 | Acidianus_sulfidivorans_JP7 | DFR86_11810 | 2261030..2264170 | 619593 | DUF499-domain-containing-protein-[Acidianus-sulfidivorans-JP7] |
| MCQ4448546.1 | AAA+ATPase+wHTH+Alpha+Beta-domain+wHTH+RRM | Type-1_BR.RM.like.systems | <-Cluster35_66clades<-DUF499*<-METHYLASE | 1086 | Archaea>TACK_group>Thermoproteota>Thermoprotei>Sulfolobales | GCA_024451005.1 | Sulfolobales_archaeon | NO076_04785 | 187..3447 | 2268200 | ATP-binding-protein-[Sulfolobales-archaeon] |
| WP_052883987.1 | AAA+ATPase+wHTH+Alpha+Beta-domain+wHTH+RRM | Type-1_BR.RM.like.systems | Radical-SAM-><-?<-TUDOR+SF1-SF2-Helicase+iREase+CTD+REase<-Cluster35_66clades<-DUF499*<-METHYLASE<-?\|\|?-><-?<-SF1-SF2-Helicase | 1064 | Archaea>TACK_group>Thermoproteota>Thermoprotei>Thermofilales | GCF_000993805.1 | Infirmifilum_uzonense | MA03_RS03695 | 730480..733674 | 1550241 | DUF499-domain-containing-protein-[Infirmifilum-uzonense] |
| WP_020963485.1 | AAA+ATPase+wHTH+Alpha+Beta-domain+wHTH+RRM | Type-1_BR.RM.like.systems | <-HEPN<-EVE<-?\|\|METHYLASE->DUF499*->Cluster35_66clades->TUDOR+SF1-SF2-Helicase+iREase+CTD+REase-> | 1066 | Archaea>TACK_group>Thermoproteota>Thermoprotei>Thermofilales | GCF_000446015.1 | Thermofilum_adornatum | N186_RS08890 | 1646491..1649691 | 1365176 | DUF499-domain-containing-protein-[Thermofilum-adornatum] |
| MCI4409924.1 | AAA+ATPase+wHTH+Alpha+Beta-domain+wHTH+RRM | Type-1_BR.RM.like.systems | METHYLASE-p->DUF499*->DUF499->Cluster35_66clades->TUDOR+SF1-SF2-Helicase+iREase+CTD+REase-> | 298 | Archaea>TACK_group>Thermoproteota>Thermoprotei>Thermofilales | GCA_022759215.1 | Thermofilum_sp. | JHC26_12595 | 249..1144 | 1961369 | ATP-binding-protein-[Thermofilum-sp.] |
| MCC6050142.1 | AAA+ATPase+wHTH+Alpha+Beta-domain+wHTH+RRM | Type-1_BR.RM.like.systems | <-TUDOR+SF1-SF2-Helicase+iREase+CTD+REase-p<-Cluster35_66clades<-?<-DUF499*<-METHYLASE-p | 947 | Archaea>TACK_group>Thermoproteota>Thermoprotei>Thermofilales | GCA_020833305.1 | Thermofilum_sp. | LM580_05520 | 1720..4563 | 1961369 | ATP-binding-protein-[Thermofilum-sp.] |
| MCI4407789.1 | AAA+ATPase+wHTH+Alpha+Beta-domain+wHTH+RRM | Type-1_BR.RM.like.systems | <-TUDOR+SF1-SF2-Helicase+iREase+CTD+REase<-Cluster35_66clades<-DUF499*<-METHYLASE | 1094 | Archaea>TACK_group>Thermoproteota>Thermoprotei>Thermofilales | GCA_022759215.1 | Thermofilum_sp. | JHC26_01765 | 3825..7109 | 1961369 | DUF499-domain-containing-protein-[Thermofilum-sp.] |
| MCS7105253.1 | AAA+ATPase+wHTH+Alpha+Beta-domain+wHTH+RRM | Type-1_BR.RM.like.systems | REase->?-><-?\|\|PIN-><-?<-TUDOR+SF1-SF2-Helicase+iREase+CTD+REase<-DUF499*<-METHYLASE | 1067 | Archaea>TACK_group>Thermoproteota>Thermoprotei>Thermofilales | GCA_025057995.1 | Thermofilaceae_archaeon | NZ954_06790 | 22675..25878 | 2268203 | ATP-binding-protein-[Thermofilaceae-archaeon] |
| MCC6004216.1 | AAA+ATPase+wHTH+Alpha+Beta-domain+wHTH+RRM | Type-1_BR.RM.like.systems | <-TUDOR+SF1-SF2-Helicase+iREase+CTD+REase<-Cluster35_66clades<-DUF499* | 680 | Archaea>TACK_group>Thermoproteota>Thermoprotei>Thermofilales | GCA_020832965.1 | Thermofilum_sp. | LM590_07730 | 9082..11126 | 1961369 | ATP-binding-protein-[Thermofilum-sp.] |
| MCC6003567.1 | AAA+ATPase+wHTH+Alpha+Beta-domain+wHTH+RRM | Type-1_BR.RM.like.systems | <-DUF499*<-METHYLASE\|\|?->?-><-EVE<-ZnR | 407 | Archaea>TACK_group>Thermoproteota>Thermoprotei>Thermofilales | GCA_020832965.1 | Thermofilum_sp. | LM590_04430 | 1..1222 | 1961369 | DUF499-domain-containing-protein-[Thermofilum-sp.] |
| MCC6065425.1 | AAA+ATPase+wHTH+Alpha+Beta-domain+wHTH+RRM | Type-1_BR.RM.like.systems | <-DUF499*<-METHYLASE\|\|EVE-> | 185 | Archaea>TACK_group>Thermoproteota>Thermoprotei>Thermofilales | GCA_020833625.1 | Thermofilum_sp. | LM576_05525 | 1..554 | 1961369 | DUF499-domain-containing-protein-[Thermofilum-sp.] |
| KU092820.1 | AAA+ATPase+wHTH+Alpha+Beta-domain+wHTH+RRM | Type-1_BR.RM.like.systems | <-ABC-ATPase+SbcC\|\|?->Radical-SAM->tRNA->?->?->METHYLASE->DUF499*->SF1-SF2-Helicase+REase-><-?<-ZnR<-HKD-DNase<-SF1-SF2-Helicase<-METHYLASE | 1052 | Archaea>TACK_group>Thermoproteota>Thermoprotei>Thermoproteales | GCA_001516585.1 | Thermocladium_sp._ECH_B | AT710_02110 | 21625..24783 | 1714261 | MAG:-hypothetical-protein-AT710_02110-[Thermocladium-sp.-ECH_B] |
| WP_013680508.1 | AAA+ATPase+wHTH+Alpha+Beta-domain+wHTH+RRM | Type-1_BR.RM.like.systems | <-MORC+DUF3684+REase\|\|?-><-APATPase+TPR\|\|<-PIWI\|\|METHYLASE->DUF499*->?->SF1-SF2-Helicase+REase-><-ZnR<-?<-?\|\|HEPN-> | 1023 | Archaea>TACK_group>Thermoproteota>Thermoprotei>Thermoproteales | GCF_000193375.1 | Thermoproteus_uzoniensis | TUZN_RS08265 | 1481435..1484506 | 184117 | DUF499-domain-containing-protein-[Thermoproteus-uzoniensis] |
| WP_014290093.1 | AAA+ATPase+wHTH+Alpha+Beta-domain+wHTH+RRM | Type-1_BR.RM.like.systems | <-SF1-SF2-Helicase\|\|ZnR-><-SF1-SF2-Helicase+REase<-?<-DUF499*<-METHYLASE\|\|PIWI->\|\|?->APATPase+TPR-> | 1023 | Archaea>TACK_group>Thermoproteota>Thermoprotei>Thermoproteales | GCF_000234805.1 | Pyrobaculum_ferrireducens | P186_RS13545 | 2438033..2441104 | 1104324 | DUF499-domain-containing-protein-[Pyrobaculum-ferrireducens] |
| WP_014287586.1 | AAA+ATPase+wHTH+Alpha+Beta-domain+wHTH+RRM | Type-1_BR.RM.like.systems | <-DUF499*<-METHYLASE-p<-METHYLASE-p<-SF1-SF2-Helicase+iREase+CTD+REase\|\|?->\|\|?->APATPase+TPR-> | 912 | Archaea>TACK_group>Thermoproteota>Thermoprotei>Thermoproteales | GCF_000234805.1 | Pyrobaculum_ferrireducens | P186_RS01365 | 250583..253321 | 1104324 | DUF499-domain-containing-protein-[Pyrobaculum-ferrireducens] |
| NAZ33307.1 | AAA+ATPase+wHTH+Alpha+Beta-domain+wHTH+RRM | Type-1_BR.RM.like.systems | METHYLASE->DUF499*->?->SF1-SF2-Helicase+REase-> | 1024 | Archaea>TACK_group>Thermoproteota>Thermoprotei>Thermoproteales | GCA_009903905.1 | Pyrobaculum_sp. | GU356_03295 | 3707..6781 | 2004705 | DUF499-domain-containing-protein-[Pyrobaculum-sp.] |
| NAZ33684.1 | AAA+ATPase+wHTH+Alpha+Beta-domain+wHTH+RRM | Type-1_BR.RM.like.systems | SF1-SF2-Helicase->DUF499*-><-METHYLASE-p | 923 | Archaea>TACK_group>Thermoproteota>Thermoprotei>Thermoproteales | GCA_009903905.1 | Pyrobaculum_sp. | GU356_05300 | 5117..7888 | 2004705 | DUF499-domain-containing-protein-[Pyrobaculum-sp.] |
| RLG82277.1 | AAA+ATPase+wHTH+Alpha+Beta-domain+wHTH+RRM | Type-1_BR.RM.like.systems | EVE->METHYLASE->DUF499*->Cluster35_66clades->TUDOR+SF1-SF2-Helicase+iREase+CTD+REase-> | 1127 | Archaea>TACK_group>Thermoproteota>Thermoprotei>unclassified_Thermoprotei | GCA_003661845.1 | Thermoprotei_archaeon | DRO09_00675 | 3464..6847 | 2250277 | MAG:-hypothetical-protein-DRO09_00675-[Thermoprotei-archaeon] |
| RLE92416.1 | AAA+ATPase+wHTH+Alpha+Beta-domain+wHTH+RRM | Type-1_BR.RM.like.systems | <-SF1-SF2-Helicase<-?<-TUDOR+SF1-SF2-Helicase+iREase+CTD+REase<-?<-DUF499*<-METHYLASE | 1186 | Archaea>TACK_group>Thermoproteota>Thermoprotei>unclassified_Thermoprotei | GCA_003649445.1 | Thermoprotei_archaeon | DRN04_10370 | 7322..10882 | 2250277 | MAG:-hypothetical-protein-DRN04_10370-[Thermoprotei-archaeon] |
| RLE92006.1 | AAA+ATPase+wHTH+Alpha+Beta-domain+wHTH+RRM | Type-1_BR.RM.like.systems | <-TUDOR+SF1-SF2-Helicase+iREase+CTD+REase-p<-?<-DUF499*<-METHYLASE<-?\|\|EVE-> | 1179 | Archaea>TACK_group>Thermoproteota>Thermoprotei>unclassified_Thermoprotei | GCA_003649445.1 | Thermoprotei_archaeon | DRN04_11430 | 1781..5320 | 2250277 | MAG:-hypothetical-protein-DRN04_11430-[Thermoprotei-archaeon] |
| RLF11833.1 | AAA+ATPase+wHTH+Alpha+Beta-domain+wHTH+RRM | Type-1_BR.RM.like.systems | <-TUDOR+SF1-SF2-Helicase+iREase+CTD+REase<-?<-DUF499*<-METHYLASE-p | 1188 | Archaea>TACK_group>Thermoproteota>Thermoprotei>unclassified_Thermoprotei | GCA_003650845.1 | Thermoprotei_archaeon | DRJ62_03175 | 5160..8726 | 2250277 | MAG:-hypothetical-protein-DRJ62_03175-[Thermoprotei-archaeon] |
| RLF01937.1 | AAA+ATPase+wHTH+Alpha+Beta-domain+wHTH+RRM | Type-1_BR.RM.like.systems | METHYLASE-p->DUF499*->BrxL-> | 1076 | Archaea>TACK_group>Thermoproteota>Thermoprotei>unclassified_Thermoprotei | GCA_003649515.1 | Thermoprotei_archaeon | DRJ59_04920 | 2032..5262 | 2250277 | MAG:-hypothetical-protein-DRJ59_04920-[Thermoprotei-archaeon] |
| NPA85109.1 | AAA+ATPase+wHTH+Alpha+Beta-domain+wHTH+RRM | Type-1_BR.RM.like.systems | <-METHYLASE\|\|DUF499*->Cluster35_66clades->TUDOR+SF1-SF2-Helicase+iREase+CTD+REase-><-?<-OLD-ABC+TOPRIM | 1124 | Archaea>TACK_group>Thermoproteota>unclassified_Thermoproteota | GCA_013154085.1 | Thermoproteota_archaeon | GXO07_03785 | 3198..6572 | 2056631 | DUF499-domain-containing-protein-[Thermoproteota-archaeon] |
| NPA70922.1 | AAA+ATPase+wHTH+Alpha+Beta-domain+wHTH+RRM | Type-1_BR.RM.like.systems | <-TUDOR+SF1-SF2-Helicase+iREase+CTD+REase-p<-Cluster35_66clades<-DUF499* | 1151 | Archaea>TACK_group>Thermoproteota>unclassified_Thermoproteota | GCA_013153895.1 | Thermoproteota_archaeon | GXO26_08940 | 1756..5211 | 2056631 | DUF499-domain-containing-protein-[Crenarchaeota-archaeon] |
| NPA84494.1 | AAA+ATPase+wHTH+Alpha+Beta-domain+wHTH+RRM | Type-1_BR.RM.like.systems | <-SF1-SF2-Helicase<-REase\|\|METHYLASE->DUF499*-> | 785 | Archaea>TACK_group>Thermoproteota>unclassified_Thermoproteota | GCA_013154085.1 | Thermoproteota_archaeon | GXO07_00630 | 7378..9733 | 2056631 | DUF499-domain-containing-protein-[Crenarchaeota-archaeon] |
| MBT7567302.1 | AAA+ATPase+wHTH+Alpha+Beta-domain+wHTH+RRM | Type-1_BR.RM.like.systems | Phage_integrase->?-><-?<-?<-?<-DUF499*<-METHYLASE-p | 942 | Archaea>unclassified_Archaea | GCA_018674515.1 | archaeon | HN630_00165 | 7617..10445 | 1906665 | ATP-binding-protein-[archaeon] |
| HDZ17009.1 | AAA+ATPase+wHTH+Alpha+Beta-domain+wHTH+RRM | Type-1_BR.RM.like.systems | SF1-SF2-Helicase+REase->METHYLASE->DUF499*-> | 949 | Archaea>unclassified_Archaea | GCA_011050315.1 | archaeon | ENH75_01880 | 4806..7655 | 1906665 | ATP-binding-protein-[archaeon] |
| MBA2524268.1 | AAA+ATPase+wHTH+Alpha+Beta-domain+wHTH+RRM | Type-1_BR.RM.like.systems | REase->METHYLASE->REase->DUF499*-><-?<-N6_N4_MTase | 930 | Bacteria>Acidobacteriota>Blastocatellia>Blastocatellales>Pyrinomonadaceae | GCA_013696625.1 | Pyrinomonadaceae_bacterium | HOV18_00595 | 5309..8101 | 2283092 | ATP-binding-protein-[Pyrinomonadaceae-bacterium] |
| WP_211197588.1 | AAA+ATPase+wHTH+Alpha+Beta-domain+wHTH+RRM | Type-1_BR.RM.like.systems | <-DUF499*<-METHYLASE<-TUDOR+SF1-SF2-Helicase+iREase+CTD+REase\|\|?->?-><-?<-ABhydrolase | 940 | Bacteria>Acidobacteriota>Blastocatellia>Blastocatellales>Pyrinomonadaceae | GCF_000820845.2 | Pyrinomonas_methylaliphatogenes | PYK22_RS02075 | 98020..100842 | 454194 | DUF499-domain-containing-protein-[Pyrinomonas-methylaliphatogenes] |
| CDM64418.1 | AAA+ATPase+wHTH+Alpha+Beta-domain+wHTH+RRM | Type-1_BR.RM.like.systems | <-DUF499*<-METHYLASE<-TUDOR+SF1-SF2-Helicase+iREase+CTD+REase\|\|?->?-><-?<-ABhydrolase | 922 | Bacteria>Acidobacteriota>Blastocatellia>Blastocatellales>Pyrinomonadaceae | GCA_000820845.2 | Pyrinomonas_methylaliphatogenes | PYK22_00412 | 98020..100788 | 454194 | predicted-ATPase-(AAA+-superfamily)-[Pyrinomonas-methylaliphatogenes] |
| WP_211425996.1 | AAA+ATPase+wHTH+Alpha+Beta-domain+wHTH+RRM | Type-1_BR.RM.like.systems | SAM-methylase-><-?<-?\|\|TUDOR+SF1-SF2-Helicase+iREase+CTD+REase->PIN->DUF499*->sGTP-> | 933 | Bacteria>Acidobacteriota>Blastocatellia>Chloracidobacterium | GCF_018304805.1 | Chloracidobacterium_aggregatum | J8C02_RS10960 | 2617767..2620568 | 2851959 | DUF499-domain-containing-protein-[Chloracidobacterium-aggregatum] |
| WP_014100413.1 | AAA+ATPase+wHTH+Alpha+Beta-domain+wHTH+RRM | Type-1_BR.RM.like.systems | <-SIGMA-HTH\|\|?-><-DUF499*\|\|Y1_Tnp->OrfB_IS605+OrfB_Zn_ribbon-><-HEPN<-HEPN<-PolB-NTase<-METHYLASE | 876 | Bacteria>Acidobacteriota>Blastocatellia>Chloracidobacterium | GCF_000226295.1 | Chloracidobacterium_thermophilum | CABTHER_RS09470 | 2292101..2294731 | 458033 | DUF499-domain-containing-protein-[Chloracidobacterium-thermophilum] |
| OYT70848.1 | AAA+ATPase+wHTH+Alpha+Beta-domain+wHTH+RRM | Type-1_BR.RM.like.systems | <-sGTP\|\|<-?<-DUF499*<-METHYLASE-p | 948 | Bacteria>Acidobacteriota>Blastocatellia>Chloracidobacterium>unclassified_Chloracidobacterium | GCA_002254845.1 | Chloracidobacterium_sp._CP2_5A | CFK52_09955 | 37798..40644 | 2012633 | MAG:-hypothetical-protein-CFK52_09955-[Chloracidobacterium-sp.-CP2_5A] |
| WP_211431750.1 | AAA+ATPase+wHTH+Alpha+Beta-domain+wHTH+RRM | Type-1_BR.RM.like.systems | <-SIGMA-HTH\|\|?-><-DUF499*\|\|Y1_Tnp->OrfB_IS605+OrfB_Zn_ribbon-><-HEPN<-PolB-NTase<-METHYLASE<-TUDOR+SF1-SF2-Helicase+iREase+CTD+REase | 876 | Bacteria>Acidobacteriota>Blastocatellia>Chloracidobacterium>unclassified_Chloracidobacterium | GCF_018304685.1 | Chloracidobacterium_sp._D | J8C01_RS09520 | 2253540..2256170 | 2821536 | DUF499-domain-containing-protein-[Chloracidobacterium-sp.-D] |
| MCS6804914.1 | AAA+ATPase+wHTH+Alpha+Beta-domain+wHTH+RRM | Type-1_BR.RM.like.systems | <-NTP_transf<-?\|\|?->BetaPropeller+ABhydrolase->?-><-DUF499*<-?<-?<-?<-METHYLASE-p | 970 | Bacteria>Acidobacteriota>Blastocatellia>unclassified_Blastocatellia | GCA_025054955.1 | Blastocatellia_bacterium | NZ823_07190 | 35424..38336 | 2052146 | DUF499-domain-containing-protein-[Blastocatellia-bacterium] |
| MCS7158686.1 | AAA+ATPase+wHTH+Alpha+Beta-domain+wHTH+RRM | Type-1_BR.RM.like.systems | <-DUF499*<-METHYLASE-p<-METHYLASE-p<-METHYLASE-p | 782 | Bacteria>Acidobacteriota>Blastocatellia>unclassified_Blastocatellia | GCA_025060035.1 | Blastocatellia_bacterium | NOA16_13315 | 103..2451 | 2052146 | DUF499-domain-containing-protein-[Blastocatellia-bacterium] |
| MCS6884392.1 | AAA+ATPase+wHTH+Alpha+Beta-domain+wHTH+RRM | Type-1_BR.RM.like.systems | METHYLASE->DUF499*->?->?->?-><-?<-PfkB | 877 | Bacteria>Acidobacteriota>Blastocatellia>unclassified_Blastocatellia | GCA_025057015.1 | Blastocatellia_bacterium | NZ558_03095 | 22764..25397 | 2052146 | ATP-binding-protein-[Blastocatellia-bacterium] |
| MBP1626439.1 | AAA+ATPase+wHTH+Alpha+Beta-domain+wHTH+RRM | Type-1_BR.RM.like.systems | HhH-RADC+JAB-><-PHP+SbcC<-DUF499*<-METHYLASE<-HEPN<-TUDOR+SF1-SF2-Helicase+iREase+CTD+REase<-HTH\|\|XIS-HTH-> | 952 | Bacteria>Acidobacteriota>Holophagae>Holophagales>Holophagaceae | GCA_017859955.1 | Holophagaceae_bacterium | H6Q00_914 | 10441..13299 | 2052165 | hypothetical-protein-[Holophagaceae-bacterium] |
| MCL6546792.1 | AAA+ATPase+wHTH+Alpha+Beta-domain+wHTH+RRM | Type-1_BR.RM.like.systems | <-DUF499*<-?<-HicB_lk_antitox<-METHYLASE-p | 921 | Bacteria>Acidobacteriota>Terriglobia>Bryobacterales>Bryobacteraceae | GCA_023511855.1 | Bryobacteraceae_bacterium | K6T61_16365 | 960..3725 | 2212468 | ATP-binding-protein-[Bryobacteraceae-bacterium] |
| MCL6508857.1 | AAA+ATPase+wHTH+Alpha+Beta-domain+wHTH+RRM | Type-1_BR.RM.like.systems | <-DUF499*<-HTH<-BrnT_toxin<-METHYLASE-p | 227 | Bacteria>Acidobacteriota>Terriglobia>Bryobacterales>Bryobacteraceae | GCA_023511775.1 | Bryobacteraceae_bacterium | K6T59_17750 | 1..681 | 2212468 | DUF499-domain-containing-protein-[Bryobacteraceae-bacterium] |
| MCZ2075917.1 | AAA+ATPase+wHTH+Alpha+Beta-domain+wHTH+RRM | Type-1_BR.RM.like.systems | <-MerR-HTH<-SF1-SF2-helicase+RdgC+Beta-barrel+REase<-DUF499*<-?<-RNaseH<-METHYLASE<-TUDOR+SF1-SF2-Helicase+iREase+CTD+REase<-XIS-HTH | 925 | Bacteria>Acidobacteriota>Terriglobia>Bryobacterales>unclassified_Bryobacterales | GCA_026928105.1 | Bryobacterales_bacterium | LC130_13070 | 122671..125448 | 2026791 | MAG:-DUF499-domain-containing-protein-[Bryobacterales-bacterium] |
| MCC7341998.1 | AAA+ATPase+wHTH+Alpha+Beta-domain+wHTH+RRM | Type-1_BR.RM.like.systems | <-DUF499*<-?<-RNaseH<-METHYLASE<-TUDOR+SF1-SF2-Helicase+iREase+CTD+REase<-AbiEii<-AbiEi_4 | 926 | Bacteria>Acidobacteriota>Terriglobia>Bryobacterales>unclassified_Bryobacterales | GCA_020853155.1 | Bryobacterales_bacterium | IT170_13025 | 317..3097 | 2026791 | ATP-binding-protein-[Bryobacterales-bacterium] |
| MCC6263216.1 | AAA+ATPase+wHTH+Alpha+Beta-domain+wHTH+RRM | Type-1_BR.RM.like.systems | <-DUF499*<-Schlafen+HATPase+HTH<-METHYLASE<-TUDOR+SF1-SF2-Helicase+iREase+CTD+REase | 926 | Bacteria>Acidobacteriota>Terriglobia>Bryobacterales>unclassified_Bryobacterales | GCA_020847315.1 | Bryobacterales_bacterium | IT169_06530 | 310..3090 | 2026791 | ATP-binding-protein-[Bryobacterales-bacterium] |
| MCU1227020.1 | AAA+ATPase+wHTH+Alpha+Beta-domain+wHTH+RRM | Type-1_BR.RM.like.systems | <-DUF499*<-METHYLASE-p<-?<-?<-CHTH<-Phage_integrase<-Phage_integrase | 740 | Bacteria>Acidobacteriota>Terriglobia>Terriglobales>Acidobacteriaceae | GCA_025462645.1 | Edaphobacter_sp. | JWQ42_5113 | 5314..7536 | 1934404 | hypothetical-protein-[Edaphobacter-sp.] |
| MCU1227032.1 | AAA+ATPase+wHTH+Alpha+Beta-domain+wHTH+RRM | Type-1_BR.RM.like.systems | <-ParB+HTH\|\|?->?->DUF499*-><-MPTase<-HTH_3\|\|?->HTH+UbE1->?->RnaseHII-> | 328 | Bacteria>Acidobacteriota>Terriglobia>Terriglobales>Acidobacteriaceae | GCA_025462645.1 | Edaphobacter_sp. | JWQ42_5125 | 3749..4735 | 1934404 | hypothetical-protein-[Edaphobacter-sp.] |
| MBZ5523276.1 | AAA+ATPase+wHTH+Alpha+Beta-domain+wHTH+RRM | Type-1_BR.RM.like.systems | <-ABhydrolase<-?\|\|ParB+HNH->METHYLASE-><-HEPN\|\|DUF499*-><-PIN<-?\|\|MazE_antitoxin->PemK_toxin->?->PIN-> | 924 | Bacteria>Acidobacteriota>Terriglobia>unclassified_Terriglobia | GCA_020072595.1 | Terriglobia_bacterium | LAP21_13650 | 15770..18544 | 2081523 | DUF499-domain-containing-protein-[Acidobacteriia-bacterium] |
| MBZ5536629.1 | AAA+ATPase+wHTH+Alpha+Beta-domain+wHTH+RRM | Type-1_BR.RM.like.systems | TUDOR+SF1-SF2-Helicase+iREase+CTD+REase->DOC+HTH->?->METHYLASE->REase->DUF499*-><-?<-?<-?<-?<-TPR+TPR<-NTP_transf_3 | 926 | Bacteria>Acidobacteriota>Terriglobia>unclassified_Terriglobia | GCA_020072965.1 | Terriglobia_bacterium | LAO31_11795 | 35949..38729 | 2081523 | DUF499-domain-containing-protein-[Acidobacteriia-bacterium] |
| MBZ5490311.1 | AAA+ATPase+wHTH+Alpha+Beta-domain+wHTH+RRM | Type-1_BR.RM.like.systems | <-DUF499*<-REase<-METHYLASE<-OLD-ABC<-TUDOR+SF1-SF2-Helicase+iREase+CTD+REase | 921 | Bacteria>Acidobacteriota>Terriglobia>unclassified_Terriglobia | GCA_020072645.1 | Terriglobia_bacterium | LAO76_05190 | 365622..368387 | 2081523 | DUF499-domain-containing-protein-[Acidobacteriia-bacterium] |
| MCL6564987.1 | AAA+ATPase+wHTH+Alpha+Beta-domain+wHTH+RRM | Type-1_BR.RM.like.systems | TUDOR+SF1-SF2-Helicase+iREase+CTD+REase->?->METHYLASE->HEPN->DUF499*-><-?\|\|?-><-?<-HTH_17 | 920 | Bacteria>Acidobacteriota>Terriglobia>unclassified_Terriglobia | GCA_023512135.1 | Terriglobia_bacterium | K6U09_01035 | 242502..245264 | 2081523 | DUF499-domain-containing-protein-[Acidobacteriia-bacterium] |
| MBZ5590550.1 | AAA+ATPase+wHTH+Alpha+Beta-domain+wHTH+RRM | Type-1_BR.RM.like.systems | METHYLASE-p->DOC+HTH->DUF499*-> | 774 | Bacteria>Acidobacteriota>Terriglobia>unclassified_Terriglobia | GCA_020073245.1 | Terriglobia_bacterium | LAO05_18525 | 1697..4018 | 2081523 | DUF499-domain-containing-protein-[Acidobacteriia-bacterium] |
| MBP7799241.1 | AAA+ATPase+wHTH+Alpha+Beta-domain+wHTH+RRM | Type-1_BR.RM.like.systems | OLD-ABC+TOPRIM->METHYLASE->OLD-ABC->TOPRIM->?->?->DUF499*-> | 919 | Bacteria>Acidobacteriota>Thermoanaerobaculia>Thermoanaerobaculales>Thermoanaerobaculaceae | GCA_018000955.1 | Thermoanaerobaculaceae_bacterium | KA072_11485 | 15680..18439 | 2699758 | ATP-binding-protein-[Thermoanaerobaculaceae-bacterium] |
| MCU0292531.1 | AAA+ATPase+wHTH+Alpha+Beta-domain+wHTH+RRM | Type-1_BR.RM.like.systems | <-PIN<-?\|\|?->?-><-DUF499-p<-DUF499*<-TOPRIM<-OLD-ABC<-METHYLASE | 869 | Bacteria>Acidobacteriota>Thermoanaerobaculia>Thermoanaerobaculales>Thermoanaerobaculaceae | GCA_025456225.1 | Thermoanaerobaculaceae_bacterium | MUF10_11190 | 5913..8522 | 2699758 | DUF499-domain-containing-protein-[Thermoanaerobaculaceae-bacterium] |
| QQS46399.1 | AAA+ATPase+wHTH+Alpha+Beta-domain+wHTH+RRM | Type-1_BR.RM.like.systems | TUDOR+SF1-SF2-Helicase+iREase+CTD+REase->Phd-YefM->PIN->METHYLASE->?->DUF499*->Phd-YefM-> | 931 | Bacteria>Acidobacteriota>unclassified_Acidobacteriota | GCA_016700195.1 | Acidobacteriota_bacterium | IPM66_21230 | 5343713..5346508 | 1978231 | ATP-binding-protein-[Acidobacteria-bacterium] |
| MBI3940688.1 | AAA+ATPase+wHTH+Alpha+Beta-domain+wHTH+RRM | Type-1_BR.RM.like.systems | <-CarboxypepD-reg-IG\|\|?-><-CarboxypepD-reg-IG<-DUF499* | 592 | Bacteria>Acidobacteriota>unclassified_Acidobacteriota | GCA_016199665.1 | Acidobacteriota_bacterium | HY315_07615 | 13474..15253 | 1978231 | AAA+-family-ATPase-[Acidobacteriota-bacterium] |
| MCB9399522.1 | AAA+ATPase+wHTH+Alpha+Beta-domain+wHTH+RRM | Type-1_BR.RM.like.systems | Pkinase+RNA-Helicase-><-?<-DUF499*\|\|TPR->?-><-?<-ABC-ATPase+SbcC | 1390 | Bacteria>Acidobacteriota>unclassified_Acidobacteriota | GCA_020637335.1 | Acidobacteriota_bacterium | H6510_17050 | 81338..85510 | 1978231 | hypothetical-protein-[Acidobacteria-bacterium] |
| MBN1568514.1 | AAA+ATPase+wHTH+Alpha+Beta-domain+wHTH+RRM | Type-1_BR.RM.like.systems | ABC-ATPase+SbcC->ABC-ATPase+SbcC->ABC-ATPase+SbcC->TUDOR+SF1-SF2-Helicase+iREase+CTD+REase->METHYLASE->REase->DUF499*->?->?->?->HEPN->OLD-ABC+TOPRIM->SF1-SF2-Helicase-> | 971 | Bacteria>Acidobacteriota>unclassified_Acidobacteriota | GCA_016929335.1 | Acidobacteriota_bacterium | JXA73_11765 | 13094..16009 | 1978231 | ATP-binding-protein-[Acidobacteria-bacterium] |
| MBV8855354.1 | AAA+ATPase+wHTH+Alpha+Beta-domain+wHTH+RRM | Type-1_BR.RM.like.systems | <-HicB<-?<-?\|\|HicB->SF1-SF2-Helicase->REase-><-DUF499*<-METHYLASE<-TUDOR+SF1-SF2-Helicase+iREase+CTD+REase<-?\|\|HTH-><-?\|\|HTH_3-> | 949 | Bacteria>Acidobacteriota>unclassified_Acidobacteriota | GCA_019242475.1 | Acidobacteriota_bacterium | JOZ02_00185 | 45350..48199 | 1978231 | ATP-binding-protein-[Acidobacteria-bacterium] |
| MCY3932061.1 | AAA+ATPase+wHTH+Alpha+Beta-domain+wHTH+RRM | Type-1_BR.RM.like.systems | TUDOR+SF1-SF2-Helicase+iREase+CTD+REase->ParB+HNH->Schlafen->?->METHYLASE-p->METHYLASE->DUF499*-><-?<-?\|\|Peptidase_M28-> | 937 | Bacteria>Acidobacteriota>unclassified_Acidobacteriota | GCA_026707325.1 | Acidobacteriota_bacterium | OXH70_10095 | 67644..70457 | 1978231 | MAG:-DUF499-domain-containing-protein-[Acidobacteria-bacterium] |
| PYV70338.1 | AAA+ATPase+wHTH+Alpha+Beta-domain+wHTH+RRM | Type-1_BR.RM.like.systems | REase+35exo-><-?\|\|TUDOR+SF1-SF2-Helicase+iREase+CTD+REase-p->METHYLASE->METHYLASE-p->REase->DUF499*-> | 691 | Bacteria>Acidobacteriota>unclassified_Acidobacteriota | GCA_003225375.1 | Acidobacteriota_bacterium | DMG97_19575 | 8894..10968 | 1978231 | AAA-family-ATPase,-partial-[Acidobacteriota-bacterium] |
| MDD8033825.1 | AAA+ATPase+wHTH+Alpha+Beta-domain+wHTH+RRM | Type-1_BR.RM.like.systems | <-Methyltransf_25<-REase<-?<-?<-DUF499*<-METHYLASE<-SbcC<-?<-?<-TUDOR+SF1-SF2-Helicase+iREase+CTD+REase<-XIS-HTH | 929 | Bacteria>Acidobacteriota>unclassified_Acidobacteriota | GCA_028749145.1 | Acidobacteriota_bacterium | PHF93_08405 | 53608..56397 | 1978231 | MAG:-DUF499-domain-containing-protein-[Acidobacteriota-bacterium] |
| MBI4730026.1 | AAA+ATPase+wHTH+Alpha+Beta-domain+wHTH+RRM | Type-1_BR.RM.like.systems | TUDOR+SF1-SF2-Helicase+iREase+CTD+REase->?->HEPN+REase->METHYLASE->?->HEPN->DUF499*->?->?-><-?<-sGTP<-?<-SAM-methylase | 918 | Bacteria>Acidobacteriota>unclassified_Acidobacteriota | GCA_016208485.1 | Acidobacteriota_bacterium | HY775_11110 | 79906..82662 | 1978231 | DUF499-domain-containing-protein-[Acidobacteria-bacterium] |
| MDA2929041.1 | AAA+ATPase+wHTH+Alpha+Beta-domain+wHTH+RRM | Type-1_BR.RM.like.systems | JAB->GDResolvase+HTH->HicB+ZnR->?->TUDOR+SF1-SF2-Helicase+iREase+CTD+REase->METHYLASE->DUF499*->HEPN-> | 981 | Bacteria>Acidobacteriota>unclassified_Acidobacteriota | GCA_027717865.1 | Acidobacteria_bacterium_AH-259-O06 | MYX84_03670 | 12694..15639 | 2935772 | DUF499-domain-containing-protein-[Acidobacteria-bacterium-AH-259-O06] |
| MBI3493153.1 | AAA+ATPase+wHTH+Alpha+Beta-domain+wHTH+RRM | Type-1_BR.RM.like.systems | TUDOR+SF1-SF2-Helicase+iREase+CTD+REase->METHYLASE->DUF433->HicB_lk_antitox->HicA_toxin->DUF499*-> | 593 | Bacteria>Acidobacteriota>unclassified_Acidobacteriota | GCA_016195295.1 | Acidobacteriota_bacterium | HY047_15445 | 8948..10727 | 1978231 | ATP-binding-protein-[Acidobacteriota-bacterium] |
| MCY4659240.1 | AAA+ATPase+wHTH+Alpha+Beta-domain+wHTH+RRM | Type-1_BR.RM.like.systems | <-ORC-CDC6+REase<-DUF499*<-PIN<-OLD-ABC<-METHYLASE<-ParB+HNH+Schlafen<-TOPRIM<-OLD-ABC | 941 | Bacteria>Acidobacteriota>unclassified_Acidobacteriota | GCA_026713985.1 | Acidobacteriota_bacterium | OXF93_05425 | 1121..3946 | 1978231 | MAG:-DUF499-domain-containing-protein-[Acidobacteria-bacterium] |
| MXX86607.1 | AAA+ATPase+wHTH+Alpha+Beta-domain+wHTH+RRM | Type-1_BR.RM.like.systems | TUDOR+SF1-SF2-Helicase+iREase+CTD+REase->PolB->HEPN->METHYLASE->DUF499*-><-?<-?<-?<-?\|\|PP_kinase+HKD-DNase-> | 922 | Bacteria>Acidobacteriota>unclassified_Acidobacteriota | GCA_009838025.1 | Acidobacteriota_bacterium | F4Y71_09150 | 13575..16343 | 1978231 | ATP-binding-protein-[Acidobacteria-bacterium] |
| MDE2925620.1 | AAA+ATPase+wHTH+Alpha+Beta-domain+wHTH+RRM | Type-1_BR.RM.like.systems | METHYLASE->DUF499*->Arm-DNA-bind_3+Phage_integrase-><-?<-HTH\|\|DNA_methylase-> | 933 | Bacteria>Acidobacteriota>unclassified_Acidobacteriota | GCA_028874215.1 | Acidobacteriota_bacterium | OXT71_04395 | 1686..4487 | 1978231 | MAG:-DUF499-domain-containing-protein-[Acidobacteriota-bacterium] |
| MXY24404.1 | AAA+ATPase+wHTH+Alpha+Beta-domain+wHTH+RRM | Type-1_BR.RM.like.systems | TUDOR+SF1-SF2-Helicase+iREase+CTD+REase->SIR2+SIR2->METHYLASE->DUF499*-><-HEPN\|\|ParB+HNH-> | 934 | Bacteria>Acidobacteriota>unclassified_Acidobacteriota | GCA_009838525.1 | Acidobacteriota_bacterium | F4Y45_07765 | 419511..422315 | 1978231 | ATP-binding-protein-[Acidobacteria-bacterium] |
| MDE2979142.1 | AAA+ATPase+wHTH+Alpha+Beta-domain+wHTH+RRM | Type-1_BR.RM.like.systems | ParB+HNH+Schlafen->PolB->METHYLASE->?->OLD-ABC->TOPRIM->DUF499*->XIS-HTH+RVT-> | 923 | Bacteria>Acidobacteriota>unclassified_Acidobacteriota | GCA_028873735.1 | Acidobacteriota_bacterium | OXU63_16720 | 13221..15992 | 1978231 | MAG:-DUF499-domain-containing-protein-[Acidobacteriota-bacterium] |
| HBL27432.1 | AAA+ATPase+wHTH+Alpha+Beta-domain+wHTH+RRM | Type-1_BR.RM.like.systems | <-DUF499*<-?<-DUF499-p<-REase<-REase<-METHYLASE<-TUDOR+SF1-SF2-Helicase+iREase+CTD+REase | 720 | Bacteria>Acidobacteriota>unclassified_Acidobacteriota | GCA_003504885.1 | Acidobacteriota_bacterium | DD490_11410 | 1..2162 | 1978231 | AAA-family-ATPase-[Acidobacteria-bacterium] |
| MCI0388707.1 | AAA+ATPase+wHTH+Alpha+Beta-domain+wHTH+RRM | Type-1_BR.RM.like.systems | TUDOR+SF1-SF2-Helicase+iREase+CTD+REase->?->?-><-?\|\|METHYLASE->?->DUF499*->?-><-?<-?<-?<-RelE-ParE<-DUF433 | 943 | Bacteria>Acidobacteriota>unclassified_Acidobacteriota | GCA_022614935.1 | Acidobacteriota_bacterium | MOB07_08065 | 27612..30443 | 1978231 | DUF499-domain-containing-protein-[Acidobacteria-bacterium] |
| MYN64243.1 | AAA+ATPase+wHTH+Alpha+Beta-domain+wHTH+RRM | Type-1_BR.RM.like.systems | OLD-ABC->PIN->OLD-ABC->TOPRIM->MetJarc-HTH->PIN->DUF499*-><-?\|\|?->?->PIN-> | 935 | Bacteria>Acidobacteriota>unclassified_Acidobacteriota | GCA_009861545.1 | Acidobacteriota_bacterium | F4X11_04340 | 45133..47940 | 1978231 | ATP-binding-protein-[Acidobacteria-bacterium] |
| MCY3844246.1 | AAA+ATPase+wHTH+Alpha+Beta-domain+wHTH+RRM | Type-1_BR.RM.like.systems | <-PRTase+PRTase<-?<-DUF499*<-METHYLASE<-TUDOR+SF1-SF2-Helicase+iREase+CTD+REase | 734 | Bacteria>Acidobacteriota>unclassified_Acidobacteriota | GCA_026706755.1 | Acidobacteriota_bacterium | OXH69_11995 | 7370..9574 | 1978231 | MAG:-DUF499-domain-containing-protein-[Acidobacteria-bacterium] |
| MBI3951052.1 | AAA+ATPase+wHTH+Alpha+Beta-domain+wHTH+RRM | Type-1_BR.RM.like.systems | Schlafen+HATPase+HTH->MNS->HEPN->METHYLASE->?->HicA_toxin->DUF499*-> | 965 | Bacteria>Acidobacteriota>unclassified_Acidobacteriota | GCA_016199695.1 | Acidobacteriota_bacterium | HY314_11430 | 12465..15362 | 1978231 | ATP-binding-protein-[Acidobacteria-bacterium] |
| MXW39171.1 | AAA+ATPase+wHTH+Alpha+Beta-domain+wHTH+RRM | Type-1_BR.RM.like.systems | <-DUF499*<-PIN<-DUF433<-DUF433<-METHYLASE<-HEPN | 922 | Bacteria>Acidobacteriota>unclassified_Acidobacteriota | GCA_009836105.1 | Acidobacteriota_bacterium | F4Z65_13025 | 3012..5780 | 1978231 | DUF499-domain-containing-protein-[Acidobacteria-bacterium] |
| MBI2833279.1 | AAA+ATPase+wHTH+Alpha+Beta-domain+wHTH+RRM | Type-1_BR.RM.like.systems | TOPRIM->OLD-ABC->TOPRIM->REase->?->?->DUF499*->sGTP-> | 917 | Bacteria>Acidobacteriota>unclassified_Acidobacteriota | GCA_016190995.1 | Acidobacteriota_bacterium | HYX76_02505 | 4163..6916 | 1978231 | ATP-binding-protein-[Acidobacteria-bacterium] |
| MCY4077553.1 | AAA+ATPase+wHTH+Alpha+Beta-domain+wHTH+RRM | Type-1_BR.RM.like.systems | <-Radical-SAM<-?\|\|?->?-><-OLD-ABC+TOPRIM<-DUF499*<-METHYLASE-p | 876 | Bacteria>Acidobacteriota>unclassified_Acidobacteriota | GCA_026707615.1 | Acidobacteriota_bacterium | OXH04_19220 | 9433..12063 | 1978231 | MAG:-DUF499-domain-containing-protein-[Acidobacteria-bacterium] |
| MCI0723926.1 | AAA+ATPase+wHTH+Alpha+Beta-domain+wHTH+RRM | Type-1_BR.RM.like.systems | <-MPTase<-HTH_3<-?<-DUF499*\|\|HEPN-><-?<-METHYLASE | 918 | Bacteria>Acidobacteriota>unclassified_Acidobacteriota | GCA_022616615.1 | Acidobacteriota_bacterium | L0338_33955 | 3841..6597 | 1978231 | DUF499-domain-containing-protein-[Acidobacteria-bacterium] |
| MYD85224.1 | AAA+ATPase+wHTH+Alpha+Beta-domain+wHTH+RRM | Type-1_BR.RM.like.systems | <-REase+PNTD1+PNTD2\|\|?-><-DUF499* | 819 | Bacteria>Acidobacteriota>unclassified_Acidobacteriota | GCA_009839145.1 | Acidobacteriota_bacterium | F4Y14_03015 | 2592..5051 | 1978231 | DUF499-domain-containing-protein-[Acidobacteria-bacterium] |
| MBM3746695.1 | AAA+ATPase+wHTH+Alpha+Beta-domain+wHTH+RRM | Type-1_BR.RM.like.systems | <-DUF499*<-?<-RNaseH<-METHYLASE-p | 914 | Bacteria>Acidobacteriota>unclassified_Acidobacteriota | GCA_016871095.1 | Acidobacteriota_bacterium | FJW34_12945 | 2237..4981 | 1978231 | ATP-binding-protein-[Acidobacteria-bacterium] |
| RMG44917.1 | AAA+ATPase+wHTH+Alpha+Beta-domain+wHTH+RRM | Type-1_BR.RM.like.systems | METHYLASE-p->REase->DUF499*-> | 649 | Bacteria>Acidobacteriota>unclassified_Acidobacteriota | GCA_003696865.1 | Acidobacteriota_bacterium | D6723_18895 | 1491..3439 | 1978231 | ATP-binding-protein,-partial-[Acidobacteriota-bacterium] |
| MDE2906585.1 | AAA+ATPase+wHTH+Alpha+Beta-domain+wHTH+RRM | Type-1_BR.RM.like.systems | <-DUF499*<-METHYLASE<-REase | 963 | Bacteria>Acidobacteriota>unclassified_Acidobacteriota | GCA_028874255.1 | Acidobacteriota_bacterium | OXQ28_10940 | 1..2891 | 1978231 | MAG:-DUF499-domain-containing-protein,-partial-[Acidobacteriota-bacterium] |
| MCC6064135.1 | AAA+ATPase+wHTH+Alpha+Beta-domain+wHTH+RRM | Type-1_BR.RM.like.systems | TUDOR+SF1-SF2-Helicase+iREase+CTD+REase-p->METHYLASE-><-?\|\|METHYLASE-p->DUF499*->?-><-Cysteine_desulfurase<-ParA-Soj-PloopNTPase\|\|?->?->35exo-> | 860 | Bacteria>Aquificota>Aquificae>Aquificales>Aquificaceae | GCA_020833645.1 | Thermocrinis_sp. | LM570_06395 | 4147..6729 | 2024383 | DUF499-domain-containing-protein-[Thermocrinis-sp.] |
| WP_012992626.1 | AAA+ATPase+wHTH+Alpha+Beta-domain+wHTH+RRM | Type-1_BR.RM.like.systems | Peptidase-><-?<-?<-tRNA\|\|TUDOR+SF1-SF2-Helicase+iREase+CTD+REase->METHYLASE->DUF499*->?->?->?->?->?->Phage_integrase-> | 851 | Bacteria>Aquificota>Aquificae>Aquificales>Aquificaceae | GCF_000025605.1 | Thermocrinis_albus | THAL_RS08090 | 1486008..1488563 | 136094 | ATP-binding-protein-[Thermocrinis-albus] |
| MCI4458224.1 | AAA+ATPase+wHTH+Alpha+Beta-domain+wHTH+RRM | Type-1_BR.RM.like.systems | <-35exo\|\|ParA-Soj-PloopNTPase->Cysteine_desulfurase-><-?<-DUF499*<-METHYLASE-p | 860 | Bacteria>Aquificota>Aquificae>Aquificales>Aquificaceae | GCA_022759285.1 | Thermocrinis_sp. | JHC21_01225 | 3196..5778 | 2024383 | ATP-binding-protein-[Thermocrinis-sp.] |
| MCY0866314.1 | AAA+ATPase+wHTH+Alpha+Beta-domain+wHTH+RRM | Type-1_BR.RM.like.systems | TUDOR+SF1-SF2-Helicase+iREase+CTD+REase->METHYLASE->DUF499-p->DUF499*-><-?<-?<-?<-?<-?<-OrfB_Zn_ribbon | 691 | Bacteria>Aquificota>Aquificae>Aquificales>Aquificaceae | GCA_026627465.1 | Aquificaceae_bacterium | OWQ49_01975 | 39953..42028 | 2053503 | MAG:-DUF499-domain-containing-protein-[Aquificaceae-bacterium] |
| HCO39566.1 | AAA+ATPase+wHTH+Alpha+Beta-domain+wHTH+RRM | Type-1_BR.RM.like.systems | METHYLASE-p->Polbeta->PolB->METHYLASE-p->DUF499*-> | 317 | Bacteria>Aquificota>Aquificae>Aquificales>Aquificaceae | GCA_003534055.1 | Aquificaceae_bacterium | DIT36_06355 | 2216..3168 | 2053503 | AAA-family-ATPase-[Aquificaceae-bacterium] |
| RUM61916.1 | AAA+ATPase+wHTH+Alpha+Beta-domain+wHTH+RRM | Type-1_BR.RM.like.systems | METHYLASE-p->MazE_antitoxin->PIN->DUF499*-> | 858 | Bacteria>Aquificota>Aquificae>Aquificales>Hydrogenothermaceae | GCA_003978555.1 | Persephonella_sp. | DSY59_01055 | 1949..4525 | 2060922 | MAG:-AAA-family-ATPase-[Persephonella-sp.] |
| NHZ71967.1 | AAA+ATPase+wHTH+Alpha+Beta-domain+wHTH+RRM | Type-1_BR.RM.like.systems | <-DUF499*<-METHYLASE-p<-METHYLASE-p | 696 | Bacteria>Aquificota>Aquificae>Aquificales>unclassified_Aquificales | GCA_011682295.1 | Aquificales_bacterium | GWP17_02645 | 1..2088 | 2109902 | DUF499-domain-containing-protein-[Aquificales-bacterium] |
| QWK13922.1 | AAA+ATPase+wHTH+Alpha+Beta-domain+wHTH+RRM | Type-1_BR.RM.like.systems | METHYLASE-p->?->?->?->?->?-><-DUF499*<-METHYLASE<-TUDOR+SF1-SF2-Helicase+iREase+CTD+REase\|\|?->?->sGTP-> | 851 | Bacteria>Aquificota>unclassified_Aquificota | GCA_018771605.1 | Aquificota_bacterium | KNN14_04835 | 850471..853026 | 2202151 | MAG:-DUF499-domain-containing-protein-[Aquificae-bacterium] |
| MBN2396704.1 | AAA+ATPase+wHTH+Alpha+Beta-domain+wHTH+RRM | Type-1_BR.RM.like.systems | <-DUF499*<-METHYLASE<-TUDOR+SF1-SF2-Helicase+iREase+CTD+REase\|\|PAIREDC-HTH-> | 822 | Bacteria>Atribacterota>Atribacteria>unclassified_Atribacteria | GCA_016934875.1 | Candidatus_Atribacteria_bacterium | JXC36_09655 | 1..2468 | 2053509 | ATP-binding-protein-[Candidatus-Atribacteria-bacterium] |
| MBC7217419.1 | AAA+ATPase+wHTH+Alpha+Beta-domain+wHTH+RRM | Type-1_BR.RM.like.systems | Schlafen+HATPase+HTH->Schlafen->METHYLASE->DUF499*-> | 870 | Bacteria>Atribacterota>Atribacterota_incertae_sedis>Candidatus_Caldatribacterium>unclassified_Candidatus_Caldatribacterium | GCA_014359405.1 | Candidatus_Caldatribacterium_sp. | H5U36_04510 | 5319..7931 | 2282143 | ATP-binding-protein-[Candidatus-Caldatribacterium-sp.] |
| MCS7241463.1 | AAA+ATPase+wHTH+Alpha+Beta-domain+wHTH+RRM | Type-1_BR.RM.like.systems | <-DUF499*<-PolB<-METHYLASE-p | 879 | Bacteria>Atribacterota>Atribacterota_incertae_sedis>Candidatus_Caldatribacterium>unclassified_Candidatus_Caldatribacterium | GCA_025060675.1 | Candidatus_Caldatribacterium_sp. | NZ706_02105 | 4564..7203 | 2282143 | DUF499-domain-containing-protein-[Candidatus-Caldatribacterium-sp.] |
| PKP59483.1 | AAA+ATPase+wHTH+Alpha+Beta-domain+wHTH+RRM | Type-1_BR.RM.like.systems | METHYLASE->DUF499*->HEPN->RNase-> | 876 | Bacteria>Atribacterota>unclassified_Atribacterota | GCA_002841055.1 | Candidatus_Atribacteria_bacterium_HGW-Atribacteria-1 | CVT91_06625 | 3132..5762 | 2013676 | AAA-family-ATPase-[Candidatus-Atribacteria-bacterium-HGW-Atribacteria-1] |
| MBN1482772.1 | AAA+ATPase+wHTH+Alpha+Beta-domain+wHTH+RRM | Type-1_BR.RM.like.systems | TUDOR+SF1-SF2-Helicase+iREase+CTD+REase->METHYLASE->DUF499*->?->?->?->REase->REase-> | 982 | Bacteria>Bacteria_incertae_sedis>Bacteria_candidate_phyla>candidate_division_KSB1>unclassified_candidate_division_KSB1 | GCA_016928775.1 | candidate_division_KSB1_bacterium | JXA70_21040 | 9299..12247 | 2172550 | DUF499-domain-containing-protein-[candidate-division-KSB1-bacterium] |
| MDD5559009.1 | AAA+ATPase+wHTH+Alpha+Beta-domain+wHTH+RRM | Type-1_BR.RM.like.systems | HD->DUF4065->METHYLASE-p->OLD-ABC->TOPRIM->METHYLASE->DUF499*-><-?<-REase<-?<-?<-?<-CopG_antitoxin | 950 | Bacteria>Bacteria_incertae_sedis>Bacteria_candidate_phyla>candidate_division_NC10>Methylomirabilis | GCA_028716865.1 | Candidatus_Methylomirabilis_sp. | PHV01_05045 | 55831..58683 | 2032687 | MAG:-DUF499-domain-containing-protein-[Candidatus-Methylomirabilis-sp.] |
| MBZ0160824.1 | AAA+ATPase+wHTH+Alpha+Beta-domain+wHTH+RRM | Type-1_BR.RM.like.systems | <-DUF499*<-PIN<-DUF433<-REase<-HTH<-BrnT_toxin<-METHYLASE | 920 | Bacteria>Bacteria_incertae_sedis>Bacteria_candidate_phyla>candidate_division_NC10>Methylomirabilis | GCA_019912425.1 | Candidatus_Methylomirabilis_sp. | K8G79_11930 | 646..3408 | 2032687 | DUF499-domain-containing-protein-[bacterium] |
| NJD68050.1 | AAA+ATPase+wHTH+Alpha+Beta-domain+wHTH+RRM | Type-1_BR.RM.like.systems | DUF499*->?->?-><-?<-REase<-?<-CopG_antitoxin | 854 | Bacteria>Bacteria_incertae_sedis>Bacteria_candidate_phyla>candidate_division_NC10>unclassified_candidate_division_NC10 | GCA_012026875.1 | candidate_division_NC10_bacterium | FIA90_05185 | 1..2567 | 2072417 | ATP-binding-protein-[candidate-division-NC10-bacterium] |
| PWB47222.1 | AAA+ATPase+wHTH+Alpha+Beta-domain+wHTH+RRM | Type-1_BR.RM.like.systems | <-GDResolvase+Recombinase+ZnR<-GDResolvase+Recombinase+ZnR<-GDResolvase<-?<-?<-?<-DUF499*<-?<-METHYLASE<-Peptidase-S8<-Classical-AAA<-wHTH+REase<-TUDOR+SF1-SF2-Helicase+iREase+CTD+REase | 919 | Bacteria>Bacteria_incertae_sedis>Bacteria_candidate_phyla>candidate_division_NC10>unclassified_candidate_division_NC10 | GCA_003104975.1 | candidate_division_NC10_bacterium | C3F12_04390 | 145776..148535 | 2072417 | AAA-family-ATPase-[candidate-division-NC10-bacterium] |
| NJD67784.1 | AAA+ATPase+wHTH+Alpha+Beta-domain+wHTH+RRM | Type-1_BR.RM.like.systems | <-GDResolvase+Recombinase+ZnR<-GDResolvase+Recombinase+ZnR<-GDResolvase<-?<-?<-?<-DUF499*<-?<-METHYLASE<-Peptidase-S8<-Classical-AAA<-wHTH+REase<-TUDOR+SF1-SF2-Helicase+iREase+CTD+REase | 919 | Bacteria>Bacteria_incertae_sedis>Bacteria_candidate_phyla>candidate_division_NC10>unclassified_candidate_division_NC10 | GCA_012026875.1 | candidate_division_NC10_bacterium | FIA90_03770 | 15454..18213 | 2072417 | ATP-binding-protein-[candidate-division-NC10-bacterium] |
| MDE2058207.1 | AAA+ATPase+wHTH+Alpha+Beta-domain+wHTH+RRM | Type-1_BR.RM.like.systems | OLD-ABC->TOPRIM->METHYLASE->BrnT_toxin->HTH->REase->DUF499*->?->HTH+PolB->HEPN-> | 931 | Bacteria>Bacteria_incertae_sedis>Bacteria_candidate_phyla>candidate_division_NC10>unclassified_candidate_division_NC10 | GCA_028864815.1 | candidate_division_NC10_bacterium | KGJ27_00545 | 34245..37040 | 2072417 | MAG:-ATP-binding-protein-[candidate-division-NC10-bacterium] |
| MBI4536775.1 | AAA+ATPase+wHTH+Alpha+Beta-domain+wHTH+RRM | Type-1_BR.RM.like.systems | <-REase<-sGTP<-DUF499*<-?<-TOPRIM<-OLD-ABC<-METHYLASE-p<-?<-METHYLASE | 938 | Bacteria>Bacteria_incertae_sedis>Bacteria_candidate_phyla>candidate_division_NC10>unclassified_candidate_division_NC10 | GCA_016209735.1 | candidate_division_NC10_bacterium | HY712_02325 | 61380..64196 | 2072417 | ATP-binding-protein-[candidate-division-NC10-bacterium] |
| MCX6842949.1 | AAA+ATPase+wHTH+Alpha+Beta-domain+wHTH+RRM | Type-1_BR.RM.like.systems | <-PIN<-?<-DUF499*<-METHYLASE<-TOPRIM<-OLD-ABC<-REase<-TUDOR+SF1-SF2-Helicase+iREase+CTD+REase | 912 | Bacteria>Bacteria_incertae_sedis>Bacteria_candidate_phyla>candidate_division_WOR-3 | GCA_026397595.1 | candidate_division_WOR-3_bacterium | NTX53_11780 | 2277..5015 | 2052148 | MAG:-DUF499-domain-containing-protein-[candidate-division-WOR-3-bacterium] |
| RLE11804.1 | AAA+ATPase+wHTH+Alpha+Beta-domain+wHTH+RRM | Type-1_BR.RM.like.systems | <-HEPN<-PolB<-DUF499<-DUF499*<-DUF499-p<-XIS-HTH | 105 | Bacteria>Bacteria_incertae_sedis>Bacteria_candidate_phyla>Candidatus_Aerophobetes>unclassified_Candidatus_Aerophobetes | GCA_003648845.1 | Candidatus_Aerophobetes_bacterium | DRJ04_07360 | 1376..1693 | 2030807 | hypothetical-protein-DRJ04_07360-[Candidatus-Aerophobetes-bacterium] |
| RLE11803.1 | AAA+ATPase+wHTH+Alpha+Beta-domain+wHTH+RRM | Type-1_BR.RM.like.systems | <-HEPN<-PolB<-DUF499*<-DUF499<-DUF499-p<-XIS-HTH | 147 | Bacteria>Bacteria_incertae_sedis>Bacteria_candidate_phyla>Candidatus_Aerophobetes>unclassified_Candidatus_Aerophobetes | GCA_003648845.1 | Candidatus_Aerophobetes_bacterium | DRJ04_07355 | 996..1439 | 2030807 | hypothetical-protein-DRJ04_07355-[Candidatus-Aerophobetes-bacterium] |
| MCD6472211.1 | AAA+ATPase+wHTH+Alpha+Beta-domain+wHTH+RRM | Type-1_BR.RM.like.systems | <-SF1-SF2-Helicase+wHTH+PRTase<-?<-HicB_lk_antitox<-DUF499*<-METHYLASE<-TUDOR+SF1-SF2-Helicase+iREase+CTD+REase<-PolB<-?<-Calcineurin<-PIN | 847 | Bacteria>Bacteria_incertae_sedis>Bacteria_candidate_phyla>Candidatus_Aerophobetes>unclassified_Candidatus_Aerophobetes | GCA_021163155.1 | Candidatus_Aerophobetes_bacterium | J7K55_03705 | 5998..8541 | 2030807 | ATP-binding-protein-[Candidatus-Aerophobetes-bacterium] |
| MCD6083667.1 | AAA+ATPase+wHTH+Alpha+Beta-domain+wHTH+RRM | Type-1_BR.RM.like.systems | TUDOR+SF1-SF2-Helicase+iREase+CTD+REase->BirA-HTH->METHYLASE->DUF499*-> | 836 | Bacteria>Bacteria_incertae_sedis>Bacteria_candidate_phyla>Candidatus_Aerophobetes>unclassified_Candidatus_Aerophobetes | GCA_021158705.1 | Candidatus_Aerophobetes_bacterium | J7J59_06145 | 7153..9663 | 2030807 | ATP-binding-protein-[Candidatus-Aerophobetes-bacterium] |
| MCD6257887.1 | AAA+ATPase+wHTH+Alpha+Beta-domain+wHTH+RRM | Type-1_BR.RM.like.systems | METHYLASE-p->PolB->HEPN->DUF499*-> | 846 | Bacteria>Bacteria_incertae_sedis>Bacteria_candidate_phyla>Candidatus_Aerophobetes>unclassified_Candidatus_Aerophobetes | GCA_021163905.1 | Candidatus_Aerophobetes_bacterium | J7J45_07510 | 2315..4855 | 2030807 | ATP-binding-protein-[Candidatus-Aerophobetes-bacterium] |
| MBC7362513.1 | AAA+ATPase+wHTH+Alpha+Beta-domain+wHTH+RRM | Type-1_BR.RM.like.systems | <-HEPN<-HEPN<-REase<-?<-MerR-HTH<-SF1-SF2-Helicase<-DUF499*<-METHYLASE<-TUDOR+SF1-SF2-Helicase+iREase+CTD+REase<-?<-HTH+MPTase\|\|Peptidase-> | 847 | Bacteria>Bacteria_incertae_sedis>Bacteria_candidate_phyla>Candidatus_Aminicenantes>unclassified_Candidatus_Aminicenantes | GCA_014360015.1 | Candidatus_Aminicenantes_bacterium | H5U06_09580 | 13628..16171 | 2052149 | ATP-binding-protein-[Candidatus-Aminicenantes-bacterium] |
| NQT80771.1 | AAA+ATPase+wHTH+Alpha+Beta-domain+wHTH+RRM | Type-1_BR.RM.like.systems | <-URI-DNase<-HEPN<-REase<-McrC-NTD+REase<-?<-REase<-DUF499*<-METHYLASE<-TUDOR+SF1-SF2-Helicase+iREase+CTD+REase | 981 | Bacteria>Bacteria_incertae_sedis>Bacteria_candidate_phyla>Candidatus_Aminicenantes>unclassified_Candidatus_Aminicenantes | GCA_013203055.1 | Candidatus_Aminicenantes_bacterium | HQ555_10335 | 19157..22102 | 2052149 | ATP-binding-protein-[Candidatus-Aminicenantes-bacterium] |

| Protein-ID | DUF499-architecture | System | Gene-Neighbourhoods | Length | Taxonomy | Assembly-ID | Strain | Locus-Tag | Genomic-coordinates | TaxID | Fasta-defline |
|---|---|---|---|---|---|---|---|---|---|---|---|
| NIM17954.1 | AAA+ATPase+wHTH+Alpha+Beta-domain+wHTH+RRM | Type-1_BR.RM.like.systems | PIN->?->?->RADICAL-SAM-><-MNS+REase<-DUF499*<-PIN<-MazE_antitoxin<-METHYLASE-p | 846 | Bacteria>Bacteria_incertae_sedis>Bacteria_candidate_phyla>Candidatus_Aminicenantes>unclassified_Candidatus_Aminicenantes | GCA_011771225.1 | Candidatus_Aminicenantes_bacterium | GT081_38750 | 19109..21649 | 2052149 | DUF499-domain-containing-protein-[Candidatus-Aminicenantes-bacterium] |
| HDP95525.1 | AAA+ATPase+wHTH+Alpha+Beta-domain+wHTH+RRM | Type-1_BR.RM.like.systems | <-DUF499*<-REase<-?<-METHYLASE<-DOC+HTH<-TUDOR+SF1-SF2-Helicase+iREase+CTD+REase | 933 | Bacteria>Bacteria_incertae_sedis>Bacteria_candidate_phyla>Candidatus_Aminicenantes>unclassified_Candidatus_Aminicenantes | GCA_011049425.1 | Candidatus_Aminicenantes_bacterium | ENN40_09225 | 209..3010 | 2052149 | ATP-binding-protein-[Candidatus-Aminicenantes-bacterium] |
| MCX6557429.1 | AAA+ATPase+wHTH+Alpha+Beta-domain+wHTH+RRM | Type-1_BR.RM.like.systems | Schlafen+HATPase+HTH->METHYLASE->TIR->TIR->?->DUF499*-> | 896 | Bacteria>Bacteria_incertae_sedis>Bacteria_candidate_phyla>Candidatus_Aminicenantes>unclassified_Candidatus_Aminicenantes | GCA_026393795.1 | Candidatus_Aminicenantes_bacterium | NTW95_08395 | 10594..13284 | 2052149 | MAG:-DUF499-domain-containing-protein-[Candidatus-Aminicenantes-bacterium] |
| MCX7973587.1 | AAA+ATPase+wHTH+Alpha+Beta-domain+wHTH+RRM | Type-1_BR.RM.like.systems | METHYLASE-p->METHYLASE-p->DUF499*-><-REase | 845 | Bacteria>Bacteria_incertae_sedis>Bacteria_candidate_phyla>Candidatus_Aminicenantes>unclassified_Candidatus_Aminicenantes | GCA_026416195.1 | Candidatus_Aminicenantes_bacterium | N3B16_03705 | 1453..3990 | 2052149 | MAG:-DUF499-domain-containing-protein-[Candidatus-Aminicenantes-bacterium] |
| MBE0461501.1 | AAA+ATPase+wHTH+Alpha+Beta-domain+wHTH+RRM | Type-1_BR.RM.like.systems | <-HEPN<-ParB+HNH<-DUF499*<-METHYLASE-p | 878 | Bacteria>Bacteria_incertae_sedis>Bacteria_candidate_phyla>Candidatus_Aminicenantes>unclassified_Candidatus_Aminicenantes | GCA_014859315.1 | Candidatus_Aminicenantes_bacterium | IBX60_07695 | 3318..5954 | 2052149 | ATP-binding-protein-[Candidatus-Aminicenantes-bacterium] |
| MBC7349956.1 | AAA+ATPase+wHTH+Alpha+Beta-domain+wHTH+RRM | Type-1_BR.RM.like.systems | METHYLASE->DUF499*->SF1-SF2-Helicase-> | 837 | Bacteria>Bacteria_incertae_sedis>Bacteria_candidate_phyla>Candidatus_Aminicenantes>unclassified_Candidatus_Aminicenantes | GCA_014360005.1 | Candidatus_Aminicenantes_bacterium | H5U05_08285 | 2604..5117 | 2052149 | ATP-binding-protein-[Candidatus-Aminicenantes-bacterium] |
| PMP94108.1 | AAA+ATPase+wHTH+Alpha+Beta-domain+wHTH+RRM | Type-1_BR.RM.like.systems | METHYLASE-p->DUF499*-><-?<-?<-BirA-HTH | 845 | Bacteria>Bacteria_incertae_sedis>Bacteria_candidate_phyla>Candidatus_Aminicenantes>unclassified_Candidatus_Aminicenantes | GCA_002877915.1 | Candidatus_Aminicenantes_bacterium | C0168_10035 | 1036..3573 | 2052149 | AAA-family-ATPase-[Candidatus-Aminicenantes-bacterium] |
| GIW40409.1 | AAA+ATPase+wHTH+Alpha+Beta-domain+wHTH+RRM | Type-1_BR.RM.like.systems | <-Phage_int_SAM_1+Phage_integrase<-DUF499*<-?<-METHYLASE-p<-METHYLASE-p<-?<-TUDOR+SF1-SF2-Helicase+iREase+CTD+REase | 921 | Bacteria>Bacteria_incertae_sedis>Bacteria_candidate_phyla>Candidatus_Binatota>Candidatus_Binatia | GCA_026004195.1 | Candidatus_Binatia_bacterium | KatS3mg076_0986 | 1066498..1069263 | 2838779 | MAG:-hypothetical-protein-KatS3mg076_0986-[Candidatus-Binatia-bacterium] |
| GIW42343.1 | AAA+ATPase+wHTH+Alpha+Beta-domain+wHTH+RRM | Type-1_BR.RM.like.systems | HicB->PIN->tRNA-><-DUF499-p<-DUF499*<-HEPN<-PolB<-DUF499-p<-HEPN+HEPN<-RAMPs | 391 | Bacteria>Bacteria_incertae_sedis>Bacteria_candidate_phyla>Candidatus_Binatota>Candidatus_Binatia | GCA_026004195.1 | Candidatus_Binatia_bacterium | KatS3mg076_2920 | 430463..431638 | 2838779 | MAG:-hypothetical-protein-KatS3mg076_2920-[Candidatus-Binatia-bacterium] |
| MBF6571348.1 | AAA+ATPase+wHTH+Alpha+Beta-domain+wHTH+RRM | Type-1_BR.RM.like.systems | METHYLASE->?->XIS-HTH->?->PIN->DUF499*-> | 924 | Bacteria>Bacteria_incertae_sedis>Bacteria_candidate_phyla>Candidatus_Binatota>Candidatus_Binatia | GCA_015478625.1 | Candidatus_Binataceae_bacterium | IVW54_21020 | 19322..22096 | 2785449 | ATP-binding-protein-[Candidatus-Binataceae-bacterium] |
| MBF6571679.1 | AAA+ATPase+wHTH+Alpha+Beta-domain+wHTH+RRM | Type-1_BR.RM.like.systems | <-DUF499*<-DUF499-p<-PIN<-DUF433<-?<-METHYLASE | 905 | Bacteria>Bacteria_incertae_sedis>Bacteria_candidate_phyla>Candidatus_Binatota>Candidatus_Binatia | GCA_015478625.1 | Candidatus_Binataceae_bacterium | IVW54_22790 | 1..2714 | 2785449 | ATP-binding-protein-[Candidatus-Binataceae-bacterium] |
| QAA76831.1 | AAA+ATPase+wHTH+Alpha+Beta-domain+wHTH+RRM | Type-1_BR.RM.like.systems | METHYLASE-p->TUDOR+SF1-SF2-Helicase+iREase+CTD+REase->?->SbcC->METHYLASE->REase->DUF499*->DEDD_Tnp_IS110+Transposase_20->Glyco_transf_20-><-ABC-ATPase+SbcC<-ABC-ATPase+SbcC | 920 | Bacteria>Bacteria_incertae_sedis>Bacteria_candidate_phyla>Candidatus_Bipolaricaulota>Bipolaricaulis | GCA_004102645.1 | Candidatus_Bipolaricaulis_sibiricus | BIP78_1065 | 1085101..1087863 | 2501609 | hypothetical-protein-BIP78_1065-[Candidatus-Bipolaricaulis-sibiricus] |
| MDD5453217.1 | AAA+ATPase+wHTH+Alpha+Beta-domain+wHTH+RRM | Type-1_BR.RM.like.systems | SF1-SF2-Helicase-p->PIN->ParB+HNH+Schlafen->METHYLASE->PolB->HEPN->DUF499*-><-ABC-ATPase | 932 | Bacteria>Bacteria_incertae_sedis>Bacteria_candidate_phyla>Candidatus_Bipolaricaulota>Bipolaricaulis | GCA_028715685.1 | Candidatus_Bipolaricaulis_sp. | PHZ21_02925 | 36651..39449 | 2814139 | MAG:-DUF499-domain-containing-protein-[Candidatus-Bipolaricaulis-sp.] |
| MCD6320860.1 | AAA+ATPase+wHTH+Alpha+Beta-domain+wHTH+RRM | Type-1_BR.RM.like.systems | <-DUF499*<-METHYLASE-p<-HEPN<-PolB<-METHYLASE-p<-REase<-TUDOR+SF1-SF2-Helicase+iREase+CTD+REase | 867 | Bacteria>Bacteria_incertae_sedis>Bacteria_candidate_phyla>Candidatus_Bipolaricaulota>unclassified_Candidatus_Bipolaricaulota | GCA_021160365.1 | Candidatus_Bipolaricaulota_bacterium | J7L84_01245 | 3909..6512 | 2306012 | DUF499-domain-containing-protein-[Candidatus-Bipolaricaulota-bacterium] |
| MBC7170403.1 | AAA+ATPase+wHTH+Alpha+Beta-domain+wHTH+RRM | Type-1_BR.RM.like.systems | ABC-ATPase+SbcC->ABC-ATPase+SbcC-><-Glyco_transf_20<-DUF499*<-REase<-METHYLASE | 920 | Bacteria>Bacteria_incertae_sedis>Bacteria_candidate_phyla>Candidatus_Bipolaricaulota>unclassified_Candidatus_Bipolaricaulota | GCA_014361015.1 | Candidatus_Bipolaricaulota_bacterium | H5T54_05700 | 11895..14657 | 2306012 | ATP-binding-protein-[Candidatus-Bipolaricaulota-bacterium] |
| MCF7889851.1 | AAA+ATPase+wHTH+Alpha+Beta-domain+wHTH+RRM | Type-1_BR.RM.like.systems | TUDOR+SF1-SF2-Helicase+iREase+CTD+REase->METHYLASE->DUF499*->ParB+HNH->XIS-HTH-> | 911 | Bacteria>Bacteria_incertae_sedis>Bacteria_candidate_phyla>Candidatus_Bipolaricaulota>unclassified_Candidatus_Bipolaricaulota | GCA_021735245.1 | Candidatus_Bipolaricaulota_bacterium | K9M78_01380 | 69042..71777 | 2306012 | DUF499-domain-containing-protein-[Candidatus-Bipolaricaulota-bacterium] |
| MCD6541617.1 | AAA+ATPase+wHTH+Alpha+Beta-domain+wHTH+RRM | Type-1_BR.RM.like.systems | ABC-ATPase+SbcC->?-><-DUF499*<-?<-PolB<-METHYLASE<-PIN<-DUF433 | 885 | Bacteria>Bacteria_incertae_sedis>Bacteria_candidate_phyla>Candidatus_Bipolaricaulota>unclassified_Candidatus_Bipolaricaulota | GCA_021163465.1 | Candidatus_Bipolaricaulota_bacterium | J7K76_07320 | 5334..7991 | 2306012 | ATP-binding-protein-[Candidatus-Bipolaricaulota-bacterium] |
| KUK26300.1 | AAA+ATPase+wHTH+Alpha+Beta-domain+wHTH+RRM | Type-1_BR.RM.like.systems | <-DUF499*<-HEPN<-PolB<-METHYLASE<-TUDOR+SF1-SF2-Helicase+iREase+CTD+REase | 889 | Bacteria>Bacteria_incertae_sedis>Bacteria_candidate_phyla>Candidatus_Bipolaricaulota>unclassified_Candidatus_Bipolaricaulota | GCA_001508155.1 | Acetothermia_bacterium_64_32 | XD60_1536 | 19316..21985 | 1635286 | Uncharacterized-protein-XD60_1536-[Acetothermia-bacterium-64_32] |
| RLE33859.1 | AAA+ATPase+wHTH+Alpha+Beta-domain+wHTH+RRM | Type-1_BR.RM.like.systems | <-DUF499*<-HEPN<-PolB<-METHYLASE-p | 930 | Bacteria>Bacteria_incertae_sedis>Bacteria_candidate_phyla>Candidatus_Bipolaricaulota>unclassified_Candidatus_Bipolaricaulota | GCA_003650505.1 | Candidatus_Acetothermia_bacterium | DRJ24_04700 | 120..2912 | 2053493 | hypothetical-protein-DRJ24_04700-[Candidatus-Acetothermia-bacterium] |
| HAZ27921.1 | AAA+ATPase+wHTH+Alpha+Beta-domain+wHTH+RRM | Type-1_BR.RM.like.systems | METHYLASE-p->PolB->HEPN->DUF499*-> | 742 | Bacteria>Bacteria_incertae_sedis>Bacteria_candidate_phyla>Candidatus_Bipolaricaulota>unclassified_Candidatus_Bipolaricaulota | GCA_003485525.1 | Candidatus_Acetothermia_bacterium | DCY67_04830 | 1017..3242 | 2053493 | AAA-family-ATPase-[Candidatus-Acetothermia-bacterium] |
| MCD6129002.1 | AAA+ATPase+wHTH+Alpha+Beta-domain+wHTH+RRM | Type-1_BR.RM.like.systems | METHYLASE-p->DUF499*-><-?\|\|?->?->?->DUF433-> | 1039 | Bacteria>Bacteria_incertae_sedis>Bacteria_candidate_phyla>Candidatus_Bipolaricaulota>unclassified_Candidatus_Bipolaricaulota | GCA_021159405.1 | Candidatus_Bipolaricaulota_bacterium | J7J35_04765 | 861..3980 | 2306012 | ATP-binding-protein-[Candidatus-Bipolaricaulota-bacterium] |
| MBC7221132.1 | AAA+ATPase+wHTH+Alpha+Beta-domain+wHTH+RRM | Type-1_BR.RM.like.systems | METHYLASE->DUF499*-> | 921 | Bacteria>Bacteria_incertae_sedis>Bacteria_candidate_phyla>Candidatus_Bipolaricaulota>unclassified_Candidatus_Bipolaricaulota | GCA_014360985.1 | Candidatus_Bipolaricaulota_bacterium | H5T55_06690 | 2961..5726 | 2306012 | ATP-binding-protein-[Candidatus-Bipolaricaulota-bacterium] |
| MCS7213831.1 | AAA+ATPase+wHTH+Alpha+Beta-domain+wHTH+RRM | Type-1_BR.RM.like.systems | METHYLASE-p->DUF499*->DUF499-p-><-Transposase_31+Yae1_N<-Transposase_31+Yae1_N | 416 | Bacteria>Bacteria_incertae_sedis>Bacteria_candidate_phyla>Candidatus_Calescamantes>Calescibacterium | GCA_025060505.1 | Candidatus_Calescibacterium_sp. | NZ927_06390 | 1425..2675 | 2972667 | DUF499-domain-containing-protein-[Candidatus-Calescibacterium-sp.] |
| MCC6650376.1 | AAA+ATPase+wHTH+Alpha+Beta-domain+wHTH+RRM | Type-1_BR.RM.like.systems | HTH_Tnp_1->HTH_29+rve+rve_3-><-HerA/FtsK<-?<-DUF499*<-METHYLASE<-TUDOR+SF1-SF2-Helicase+iREase+CTD+REase | 925 | Bacteria>Bacteria_incertae_sedis>Bacteria_candidate_phyla>Candidatus_Eisenbacteria>unclassified_Candidatus_Eisenbacteria | GCA_020849095.1 | Candidatus_Eisenbacteria_bacterium | IT348_04415 | 7803..10580 | 2212470 | ATP-binding-protein-[Candidatus-Eisenbacteria-bacterium] |
| MBI5169750.1 | AAA+ATPase+wHTH+Alpha+Beta-domain+wHTH+RRM | Type-1_BR.RM.like.systems | DUF499*->ABhydrolase-><-?\|\|ABhydrolase-><-?<-?<-ABC-ATPase+SbcC | 918 | Bacteria>Bacteria_incertae_sedis>Bacteria_candidate_phyla>Candidatus_Eisenbacteria>unclassified_Candidatus_Eisenbacteria | GCA_016215625.1 | Candidatus_Eisenbacteria_bacterium | HZA61_09705 | 1113..3869 | 2212470 | ATP-binding-protein-[Candidatus-Eisenbacteria-bacterium] |
| MBP6875772.1 | AAA+ATPase+wHTH+Alpha+Beta-domain+wHTH+RRM | Type-1_BR.RM.like.systems | HerA/FtsK->?->METHYLASE->REase->DUF499*-><-ParB | 924 | Bacteria>Bacteria_incertae_sedis>Bacteria_candidate_phyla>Candidatus_Eisenbacteria>unclassified_Candidatus_Eisenbacteria | GCA_017991665.1 | Candidatus_Eisenbacteria_bacterium | KBD56_06850 | 148901..151675 | 2212470 | DUF499-domain-containing-protein-[Candidatus-Eisenbacteria-bacterium] |
| WP_259099629.1 | AAA+ATPase+wHTH+Alpha+Beta-domain+wHTH+RRM | Type-1_BR.RM.like.systems | TUDOR+SF1-SF2-Helicase+iREase+CTD+REase->Schlafen+HATPase+HTH->METHYLASE->DUF499*->?->?->PfkB-> | 880 | Bacteria>Bacteria_incertae_sedis>Bacteria_candidate_phyla>Candidatus_Fervidibacteria>Candidatus_Fervidibacter | GCF_030520105.1 | Candidatus_Fervidibacter_sacchari | M2350_RS13990 | 282876..285518 | 1448929 | DUF499-domain-containing-protein-[Candidatus-Fervidibacter-sacchari] |
| WP_259102400.1 | AAA+ATPase+wHTH+Alpha+Beta-domain+wHTH+RRM | Type-1_BR.RM.like.systems | TUDOR+SF1-SF2-Helicase+iREase+CTD+REase->?->METHYLASE->?->DUF499*-><-NACHT | 954 | Bacteria>Bacteria_incertae_sedis>Bacteria_candidate_phyla>Candidatus_Fervidibacteria>Candidatus_Fervidibacter | GCF_030520105.1 | Candidatus_Fervidibacter_sacchari | M2350_RS18330 | 29934..32798 | 1448929 | DUF499-domain-containing-protein-[Candidatus-Fervidibacter-sacchari] |
| OGG53870.1 | AAA+ATPase+wHTH+Alpha+Beta-domain+wHTH+RRM | Type-1_BR.RM.like.systems | METHYLASE->?->PIN->DUF499*->?->?->?->NTP_transf-> | 943 | Bacteria>Bacteria_incertae_sedis>Bacteria_candidate_phyla>Candidatus_Handelsmanbacteria | GCA_001781055.1 | Candidatus_Handelsmanbacteria_bacterium_RIFCSPLOWO2_12_FULL_64_10 | A3F84_03685 | 4071..6902 | 1817868 | hypothetical-protein-A3F84_03685-[Candidatus-Handelsmanbacteria-bacterium-RIFCSPLOWO2_12_FULL_64_10] |
| RCK78428.1 | AAA+ATPase+wHTH+Alpha+Beta-domain+wHTH+RRM | Type-1_BR.RM.like.systems | PP_kinase+HKD-DNase-><-DUF499*<-BrnA_antitoxin<-METHYLASE-p | 937 | Bacteria>Bacteria_incertae_sedis>Bacteria_candidate_phyla>Candidatus_Ozemobacteria>Candidatus_Ozemobacterales | GCA_003327425.1 | Candidatus_Ozemobacter_sibiricus | OZSIB_1530 | 101175..103988 | 2268124 | putative-ATPase-(AAA+-superfamily)-[Candidatus-Ozemobacter-sibiricus] |
| MDE0688672.1 | AAA+ATPase+wHTH+Alpha+Beta-domain+wHTH+RRM | Type-1_BR.RM.like.systems | <-PspA+ParA-Soj-PloopNTPase<-?<-?<-DUF499*<-METHYLASE<-?<-HEPN<-TUDOR+SF1-SF2-Helicase+iREase+CTD+REase\|\|OrfB_IS605+OrfB_Zn_ribbon-> | 937 | Bacteria>Bacteria_incertae_sedis>Bacteria_candidate_phyla>Candidatus_Poribacteria>unclassified_Candidatus_Poribacteria | GCA_028824525.1 | Candidatus_Poribacteria_bacterium | OXI61_10940 | 39625..42438 | 2026781 | MAG:-DUF499-domain-containing-protein-[Candidatus-Poribacteria-bacterium] |
| MDE0634775.1 | AAA+ATPase+wHTH+Alpha+Beta-domain+wHTH+RRM | Type-1_BR.RM.like.systems | TUDOR+SF1-SF2-Helicase+iREase+CTD+REase->SMODS->HNH+SAVED->HEPN+REase->?->METHYLASE->DUF499*->?->TPR+TPR->SbcC-> | 935 | Bacteria>Bacteria_incertae_sedis>Bacteria_candidate_phyla>Candidatus_Poribacteria>unclassified_Candidatus_Poribacteria | GCA_028824125.1 | Candidatus_Poribacteria_bacterium | OXI43_02830 | 28592..31399 | 2026781 | MAG:-DUF499-domain-containing-protein-[Candidatus-Poribacteria-bacterium] |
| MYA68863.1 | AAA+ATPase+wHTH+Alpha+Beta-domain+wHTH+RRM | Type-1_BR.RM.like.systems | TUDOR+SF1-SF2-Helicase+iREase+CTD+REase-p->?->METHYLASE->?->DUF499*-> | 937 | Bacteria>Bacteria_incertae_sedis>Bacteria_candidate_phyla>Candidatus_Poribacteria>unclassified_Candidatus_Poribacteria | GCA_009839085.1 | Candidatus_Poribacteria_bacterium | F4Y19_01975 | 9618..12431 | 2026781 | ATP-binding-protein-[Candidatus-Poribacteria-bacterium] |
| MXY28219.1 | AAA+ATPase+wHTH+Alpha+Beta-domain+wHTH+RRM | Type-1_BR.RM.like.systems | DUF499*->REase+HTH->REase->METHYLASE->URI-DNase-> | 895 | Bacteria>Bacteria_incertae_sedis>Bacteria_candidate_phyla>Candidatus_Poribacteria>unclassified_Candidatus_Poribacteria | GCA_009838315.1 | Candidatus_Poribacteria_bacterium | F4Y59_08690 | 1..2690 | 2026781 | ATP-binding-protein-[Candidatus-Poribacteria-bacterium] |
| MYK19441.1 | AAA+ATPase+wHTH+Alpha+Beta-domain+wHTH+RRM | Type-1_BR.RM.like.systems | <-DUF499*<-HNH<-?<-METHYLASE | 934 | Bacteria>Bacteria_incertae_sedis>Bacteria_candidate_phyla>Candidatus_Poribacteria>unclassified_Candidatus_Poribacteria | GCA_009845505.1 | Candidatus_Poribacteria_bacterium | F4O55_14930 | 451..3255 | 2026781 | ATP-binding-protein-[Candidatus-Poribacteria-bacterium] |
| MXV82112.1 | AAA+ATPase+wHTH+Alpha+Beta-domain+wHTH+RRM | Type-1_BR.RM.like.systems | TPR->?->?->METHYLASE->DUF499*-> | 937 | Bacteria>Bacteria_incertae_sedis>Bacteria_candidate_phyla>Candidatus_Poribacteria>unclassified_Candidatus_Poribacteria | GCA_009835485.1 | Candidatus_Poribacteria_bacterium | F4Z98_01965 | 5329..8142 | 2026781 | ATP-binding-protein-[Candidatus-Poribacteria-bacterium] |
| MDE0555374.1 | AAA+ATPase+wHTH+Alpha+Beta-domain+wHTH+RRM | Type-1_BR.RM.like.systems | <-DUF499*<-HNH<-?<-METHYLASE | 546 | Bacteria>Bacteria_incertae_sedis>Bacteria_candidate_phyla>Candidatus_Poribacteria>unclassified_Candidatus_Poribacteria | GCA_028824855.1 | Candidatus_Poribacteria_bacterium | OXI24_14220 | 20..1660 | 2026781 | MAG:-DUF499-domain-containing-protein-[Candidatus-Poribacteria-bacterium] |
| MCG9128542.1 | AAA+ATPase+wHTH+Alpha+Beta-domain+wHTH+RRM | Type-1_BR.RM.like.systems | STAS-><-?\|\|?->Glyco_transf_28->?-><-DUF499*<-METHYLASE<-TUDOR+SF1-SF2-Helicase+iREase+CTD+REase | 936 | Bacteria>Bacteria_incertae_sedis>Bacteria_candidate_phyla>Candidatus_Poribacteria>unclassified_Candidatus_Poribacteria | GCA_022394755.1 | Candidatus_Poribacteria_bacterium | JT359_13160 | 38351..41161 | 2026781 | ATP-binding-protein-[Candidatus-Poribacteria-bacterium] |
| MCD6507091.1 | AAA+ATPase+wHTH+Alpha+Beta-domain+wHTH+RRM | Type-1_BR.RM.like.systems | TUDOR+SF1-SF2-Helicase+iREase+CTD+REase->Schlafen->BirA-HTH->METHYLASE->DUF499*-> | 864 | Bacteria>Bacteria_incertae_sedis>Bacteria_candidate_phyla>Candidatus_Poribacteria>unclassified_Candidatus_Poribacteria | GCA_021162805.1 | Candidatus_Poribacteria_bacterium | J7M22_10765 | 25392..27986 | 2026781 | ATP-binding-protein-[Candidatus-Poribacteria-bacterium] |
| MAE19516.1 | AAA+ATPase+wHTH+Alpha+Beta-domain+wHTH+RRM | Type-1_BR.RM.like.systems | ABhydrolase-><-?<-?<-Aminotran_4<-ACET<-DUF499*<-METHYLASE<-REase | 922 | Bacteria>Bacteria_incertae_sedis>Bacteria_candidate_phyla>Candidatus_Poribacteria>unclassified_Candidatus_Poribacteria | GCA_002686625.1 | Candidatus_Poribacteria_bacterium | CMK12_11405 | 49414..52182 | 2026781 | AAA-family-ATPase-[Candidatus-Poribacteria-bacterium] |
| MBM3210709.1 | AAA+ATPase+wHTH+Alpha+Beta-domain+wHTH+RRM | Type-1_BR.RM.like.systems | <-XIS-HTH<-DUF499*<-PIN<-?<-METHYLASE<-TUDOR+SF1-SF2-Helicase+iREase+CTD+REase-p | 903 | Bacteria>Bacteria_incertae_sedis>Bacteria_candidate_phyla>Candidatus_Poribacteria>unclassified_Candidatus_Poribacteria | GCA_016866835.1 | Candidatus_Poribacteria_bacterium | FJZ33_00700 | 511..3222 | 2026781 | DUF499-domain-containing-protein-[Candidatus-Poribacteria-bacterium] |
| MYB96235.1 | AAA+ATPase+wHTH+Alpha+Beta-domain+wHTH+RRM | Type-1_BR.RM.like.systems | STAS-><-?\|\|?->Glyco_transf_28->?-><-DUF499*<-METHYLASE | 928 | Bacteria>Bacteria_incertae_sedis>Bacteria_candidate_phyla>Candidatus_Poribacteria>unclassified_Candidatus_Poribacteria | GCA_009838805.1 | Candidatus_Poribacteria_bacterium | F4Y43_20500 | 7286..10072 | 2026781 | ATP-binding-protein-[Candidatus-Poribacteria-bacterium] |
| MCZ6679540.1 | AAA+ATPase+wHTH+Alpha+Beta-domain+wHTH+RRM | Type-1_BR.RM.like.systems | <-DUF499-p<-DUF499*<-?<-?<-?<-METHYLASE-p<-METHYLASE-p | 746 | Bacteria>Bacteria_incertae_sedis>Bacteria_candidate_phyla>Candidatus_Poribacteria>unclassified_Candidatus_Poribacteria | GCA_027296925.1 | Candidatus_Poribacteria_bacterium | O7E52_20100 | 775..3015 | 2026781 | MAG:-DUF499-domain-containing-protein-[Candidatus-Poribacteria-bacterium] |
| MCY3548948.1 | AAA+ATPase+wHTH+Alpha+Beta-domain+wHTH+RRM | Type-1_BR.RM.like.systems | STAS-><-?\|\|?->Glyco_transf_28->?-><-DUF499*<-METHYLASE | 938 | Bacteria>Bacteria_incertae_sedis>Bacteria_candidate_phyla>Candidatus_Poribacteria>unclassified_Candidatus_Poribacteria | GCA_026702755.1 | Candidatus_Poribacteria_bacterium | OXH39_00700 | 158899..161715 | 2026781 | MAG:-DUF499-domain-containing-protein-[Candidatus-Poribacteria-bacterium] |
| MXV84967.1 | AAA+ATPase+wHTH+Alpha+Beta-domain+wHTH+RRM | Type-1_BR.RM.like.systems | <-DUF499*<-METHYLASE<-METHYLASE-p | 948 | Bacteria>Bacteria_incertae_sedis>Bacteria_candidate_phyla>Candidatus_Poribacteria>unclassified_Candidatus_Poribacteria | GCA_009835485.1 | Candidatus_Poribacteria_bacterium | F4Z98_16545 | 1006..3852 | 2026781 | DUF499-domain-containing-protein-[Candidatus-Poribacteria-bacterium] |
| MYB00594.1 | AAA+ATPase+wHTH+Alpha+Beta-domain+wHTH+RRM | Type-1_BR.RM.like.systems | URI-DNase->METHYLASE->DUF499*-> | 473 | Bacteria>Bacteria_incertae_sedis>Bacteria_candidate_phyla>Candidatus_Poribacteria>unclassified_Candidatus_Poribacteria | GCA_009839665.1 | Candidatus_Poribacteria_bacterium | F4X90_13090 | 6208..7628 | 2026781 | ATP-binding-protein-[Candidatus-Poribacteria-bacterium] |
| MCH8296057.1 | AAA+ATPase+wHTH+Alpha+Beta-domain+wHTH+RRM | Type-1_BR.RM.like.systems | HEPN->PolB->HEPN->HNH->HEPN->DUF499*-> | 234 | Bacteria>Bacteria_incertae_sedis>Bacteria_candidate_phyla>Candidatus_Poribacteria>unclassified_Candidatus_Poribacteria | GCA_022567695.1 | Candidatus_Poribacteria_bacterium | IH992_33685 | 3295..3996 | 2026781 | ATP-binding-protein-[Candidatus-Poribacteria-bacterium] |
| MBF0502329.1 | AAA+ATPase+wHTH+Alpha+Beta-domain+wHTH+RRM | Type-1_BR.RM.like.systems | TUDOR+SF1-SF2-Helicase+iREase+CTD+REase->?->?->MazF->METHYLASE->?->DUF499*-> | 955 | Bacteria>Bacteria_incertae_sedis>Bacteria_candidate_phyla>Candidatus_Riflebacteria>unclassified_Candidatus_Riflebacteria | GCA_015233155.1 | Candidatus_Riflebacteria_bacterium | HQM09_19470 | 31087..33954 | 2053586 | ATP-binding-protein-[Candidatus-Riflebacteria-bacterium] |
| MBI3629558.1 | AAA+ATPase+wHTH+Alpha+Beta-domain+wHTH+RRM | Type-1_BR.RM.like.systems | DUF499*-><-?\|\|?-><-tRNA\|\|?->?->?->METHYLASE-p-> | 612 | Bacteria>Bacteria_incertae_sedis>Bacteria_candidate_phyla>Candidatus_Rokubacteria>unclassified_Candidatus_Rokubacteria | GCA_016201575.1 | Candidatus_Rokubacteria_bacterium | HY217_08280 | 1..1841 | 2053607 | AAA+-family-ATPase-[Candidatus-Rokubacteria-bacterium] |
| MBI2198940.1 | AAA+ATPase+wHTH+Alpha+Beta-domain+wHTH+RRM | Type-1_BR.RM.like.systems | <-ACET<-SF1-SF2-Helicase+iREase+CTD<-?<-PIN\|\|?-><-DUF499*<-ACET | 539 | Bacteria>Bacteria_incertae_sedis>Bacteria_candidate_phyla>Candidatus_Rokubacteria>unclassified_Candidatus_Rokubacteria | GCA_016184305.1 | Candidatus_Rokubacteria_bacterium | HYU42_10145 | 6805..8424 | 2053607 | hypothetical-protein-[Candidatus-Rokubacteria-bacterium] |
| MBI4609709.1 | AAA+ATPase+wHTH+Alpha+Beta-domain+wHTH+RRM | Type-1_BR.RM.like.systems | PIN-><-?\|\|?->?->PhdYeFM_antitox->PIN-><-DUF499*<-HTH<-BrnT_toxin<-METHYLASE-p<-DDE_Tnp_1<-METHYLASE-p\|\|PAIREDC-HTH+HIN-HTH-> | 919 | Bacteria>Bacteria_incertae_sedis>Bacteria_candidate_phyla>Candidatus_Rokubacteria>unclassified_Candidatus_Rokubacteria | GCA_016209385.1 | Candidatus_Rokubacteria_bacterium | HY726_11955 | 6098..8857 | 2053607 | ATP-binding-protein-[Candidatus-Rokubacteria-bacterium] |
| MBI2198602.1 | AAA+ATPase+wHTH+Alpha+Beta-domain+wHTH+RRM | Type-1_BR.RM.like.systems | <-TetR-HTH\|\|?->TetR-HTH->ABhydrolase-><-Radical-SAM\|\|<-DUF499* | 914 | Bacteria>Bacteria_incertae_sedis>Bacteria_candidate_phyla>Candidatus_Rokubacteria>unclassified_Candidatus_Rokubacteria | GCA_016184305.1 | Candidatus_Rokubacteria_bacterium | HYU42_08400 | 18178..20922 | 2053607 | ATP-binding-protein-[Candidatus-Rokubacteria-bacterium] |
| HAM55582.1 | AAA+ATPase+wHTH+Alpha+Beta-domain+wHTH+RRM | Type-1_BR.RM.like.systems | <-DUF499-p<-DUF499*<-METHYLASE<-AAA-ATPase+REase | 911 | Bacteria>Bacteria_incertae_sedis>Bacteria_candidate_phyla>Candidatus_Rokubacteria>unclassified_Candidatus_Rokubacteria | GCA_003452995.1 | Candidatus_Rokubacteria_bacterium | DCQ64_09305 | 854..3589 | 2053607 | hypothetical-protein-[Candidatus-Rokubacteria-bacterium] |
| MBI2567004.1 | AAA+ATPase+wHTH+Alpha+Beta-domain+wHTH+RRM | Type-1_BR.RM.like.systems | <-REase\|\|XIS-HTH->HhH-RADC+JAB->TUDOR+SF1-SF2-Helicase+iREase+CTD+REase->METHYLASE-p->METHYLASE->DUF499*->OLD-ABC->TOPRIM->?->McrC-NTD+REase-> | 962 | Bacteria>Bacteria_incertae_sedis>Bacteria_candidate_phyla>Candidatus_Schekmanbacteria>unclassified_Candidatus_Schekmanbacteria | GCA_016188575.1 | Candidatus_Schekmanbacteria_bacterium | HYV63_08250 | 57213..60101 | 2419984 | ATP-binding-protein-[Candidatus-Schekmanbacteria-bacterium] |
| MBI2567876.1 | AAA+ATPase+wHTH+Alpha+Beta-domain+wHTH+RRM | Type-1_BR.RM.like.systems | TUDOR+SF1-SF2-Helicase+iREase+CTD+REase->PolB->HEPN->?->DUF499*-> | 808 | Bacteria>Bacteria_incertae_sedis>Bacteria_candidate_phyla>Candidatus_Schekmanbacteria>unclassified_Candidatus_Schekmanbacteria | GCA_016188575.1 | Candidatus_Schekmanbacteria_bacterium | HYV63_12665 | 36519..38945 | 2419984 | DUF499-domain-containing-protein-[Candidatus-Schekmanbacteria-bacterium] |
| MCX7047270.1 | AAA+ATPase+wHTH+Alpha+Beta-domain+wHTH+RRM | Type-1_BR.RM.like.systems | Radical-SAM->tRNA-><-DUF499<-DUF499*<-Schlafen+HATPase+HTH<-?<-METHYLASE<-Schlafen+HATPase+HTH<-?<-TUDOR+SF1-SF2-Helicase+iREase+CTD+REase | 330 | Bacteria>Bacteria_incertae_sedis>Bacteria_candidate_phyla>Candidatus_Sumerlaeota>unclassified_Candidatus_Sumerlaeota | GCA_026398575.1 | Candidatus_Sumerlaeota_bacterium | NTX50_17510 | 7797..8789 | 2107588 | MAG:-DUF499-domain-containing-protein-[Candidatus-Sumerlaeota-bacterium] |
| MCX7047269.1 | AAA+ATPase+wHTH+Alpha+Beta-domain+wHTH+RRM | Type-1_BR.RM.like.systems | DDE_Tnp_1-><-?<-?<-?<-tRNA\|\|?->Radical-SAM->tRNA-><-DUF499*<-DUF499<-Schlafen+HATPase+HTH<-?<-METHYLASE<-Schlafen+HATPase+HTH | 616 | Bacteria>Bacteria_incertae_sedis>Bacteria_candidate_phyla>Candidatus_Sumerlaeota>unclassified_Candidatus_Sumerlaeota | GCA_026398575.1 | Candidatus_Sumerlaeota_bacterium | NTX50_17505 | 5846..7696 | 2107588 | MAG:-DUF499-domain-containing-protein-[Candidatus-Sumerlaeota-bacterium] |
| MCX7011324.1 | AAA+ATPase+wHTH+Alpha+Beta-domain+wHTH+RRM | Type-1_BR.RM.like.systems | <-DUF499-p<-DUF499*<-?<-?<-TUDOR+SF1-SF2-Helicase+iREase+CTD+REase-p | 320 | Bacteria>Bacteria_incertae_sedis>Bacteria_candidate_phyla>Candidatus_Sumerlaeota>unclassified_Candidatus_Sumerlaeota | GCA_026398515.1 | Candidatus_Sumerlaeota_bacterium | NTW86_01935 | 1754..2716 | 2107588 | MAG:-DUF499-domain-containing-protein-[Candidatus-Sumerlaeota-bacterium] |
| MBM3272924.1 | AAA+ATPase+wHTH+Alpha+Beta-domain+wHTH+RRM | Type-1_BR.RM.like.systems | HIN-HTH->TUDOR+SF1-SF2-Helicase+iREase+CTD+REase->HEPN->METHYLASE->DUF499*-> | 220 | Bacteria>Bacteria_incertae_sedis>Bacteria_candidate_phyla>Patescibacteria_group>Parcubacteria_group | GCA_016867345.1 | Candidatus_Kaiserbacteria_bacterium | FJY94_06720 | 8221..8881 | 2030810 | ATP-binding-protein-[Candidatus-Kaiserbacteria-bacterium] |
| MDE2100040.1 | AAA+ATPase+wHTH+Alpha+Beta-domain+wHTH+RRM | Type-1_BR.RM.like.systems | HTH_26->Phage_integrase->TUDOR+SF1-SF2-Helicase+iREase+CTD+REase->METHYLASE->REase->DUF499*-> | 970 | Bacteria>Bacteria_incertae_sedis>Bacteria_candidate_phyla>Patescibacteria_group>unclassified_Patescibacteria_group | GCA_028872085.1 | Patescibacteria_group_bacterium | KGL39_22490 | 13393..16305 | 2052139 | MAG:-ATP-binding-protein-[Patescibacteria-group-bacterium] |
| MCP6718634.1 | AAA+ATPase+wHTH+Alpha+Beta-domain+wHTH+RRM | Type-1_BR.RM.like.systems | TUDOR+SF1-SF2-Helicase+iREase+CTD+REase->METHYLASE->DUF499*->wHTH+REase-><-?<-?\|\|?-><-ZnR | 947 | Bacteria>Bacteria_incertae_sedis>Bacteria_candidate_phyla>Patescibacteria_group>unclassified_Patescibacteria_group | GCA_024239015.1 | Patescibacteria_group_bacterium | KJI71_00195 | 43190..46033 | 2052139 | hypothetical-protein-[Patescibacteria-group-bacterium] |
| MBU2264520.1 | AAA+ATPase+wHTH+Alpha+Beta-domain+wHTH+RRM | Type-1_BR.RM.like.systems | METHYLASE-p->REase->PolB->HEPN->DUF499*-> | 810 | Bacteria>Bacteria_incertae_sedis>Bacteria_candidate_phyla>Patescibacteria_group>unclassified_Patescibacteria_group | GCA_018827685.1 | Patescibacteria_group_bacterium | KJ797_04325 | 2908..5337 | 2052139 | DUF499-domain-containing-protein-[Patescibacteria-group-bacterium] |
| MBL6992047.1 | AAA+ATPase+wHTH+Alpha+Beta-domain+wHTH+RRM | Type-1_BR.RM.like.systems | <-DUF499*<-XIS-HTH<-?\|\|REase+35exo-> | 433 | Bacteria>Bdellovibrionota>Bacteriovoracia>Bacteriovoracales>Bacteriovoracaceae | GCA_016784965.1 | Bacteriovoracaceae_bacterium | ISR65_19850 | 121..1422 | 2081706 | hypothetical-protein-[Bacteriovoracaceae-bacterium] |
| MCC7404109.1 | AAA+ATPase+wHTH+Alpha+Beta-domain+wHTH+RRM | Type-1_BR.RM.like.systems | XIS-HTH->TUDOR+SF1-SF2-Helicase+iREase+CTD+REase->METHYLASE->DUF499*->?->?->Radical-SAM->SF1-SF2-Helicase-> | 947 | Bacteria>Bdellovibrionota>Bdellovibrionia>Bdellovibrionales>unclassified_Bdellovibrionales | GCA_020850475.1 | Bdellovibrionales_bacterium | IT288_06880 | 16387..19230 | 2053517 | ATP-binding-protein-[Bdellovibrionales-bacterium] |
| MBS1963094.1 | AAA+ATPase+wHTH+Alpha+Beta-domain+wHTH+RRM | Type-1_BR.RM.like.systems | <-HISKIN+MORC<-?<-?\|\|XIS-HTH->TUDOR+SF1-SF2-Helicase+iREase+CTD+REase->METHYLASE->DUF499*->?->?->Radical-SAM-> | 948 | Bacteria>Bdellovibrionota>Bdellovibrionia>Bdellovibrionales>unclassified_Bdellovibrionales | GCA_018266355.1 | Bdellovibrionales_bacterium | JST04_12840 | 180746..183592 | 2053517 | ATP-binding-protein-[Bdellovibrionales-bacterium] |
| MBK8201826.1 | AAA+ATPase+wHTH+Alpha+Beta-domain+wHTH+RRM | Type-1_BR.RM.like.systems | <-HTH\|\|HTH->?->?->HEPN+DUF499->DUF499*->MarR-HTH-> | 741 | Bacteria>Bdellovibrionota>Bdellovibrionia>Bdellovibrionales>unclassified_Bdellovibrionales | GCA_016712575.1 | Bdellovibrionales_bacterium | IPK68_05760 | 73468..75693 | 2053517 | ATP-binding-protein-[Bdellovibrionales-bacterium] |
| MCR4433941.1 | AAA+ATPase+wHTH+Alpha+Beta-domain+wHTH+RRM | Type-1_BR.RM.like.systems | TUDOR+SF1-SF2-Helicase+iREase+CTD+REase->?->METHYLASE->REase->DUF499*-> | 924 | Bacteria>Caldisericota/Cryosericota_group>Caldisericota>unclassified_Caldisericota | GCA_024655865.1 | Caldisericota_bacterium | NUV70_07825 | 12514..15288 | 2052147 | DUF499-domain-containing-protein-[Caldisericota-bacterium] |
| KAA3660235.1 | AAA+ATPase+wHTH+Alpha+Beta-domain+wHTH+RRM | Type-1_BR.RM.like.systems | BirA-HTH->?-><-DUF499*<-METHYLASE<-TUDOR+SF1-SF2-Helicase+iREase+CTD+REase-p | 852 | Bacteria>Calditrichota>unclassified_Calditrichota | GCA_008501545.1 | Calditrichota_bacterium | DWQ10_07140 | 2521..5079 | 2212469 | DUF499-domain-containing-protein-[Calditrichaeota-bacterium] |
| GAK51397.1 | AAA+ATPase+wHTH+Alpha+Beta-domain+wHTH+RRM | Type-1_BR.RM.like.systems | ABhydrolase->?-><-?<-HEPN<-?<-?<-DUF499*\|\|?->PRTase->?->PIN-><-ABC-ATPase+SbcC | 837 | Bacteria>Candidatus_Moduliflexota>Candidatus_Moduliflexia>Candidatus_Moduliflexales>Candidatus_Moduliflexaceae | GCA_000739515.1 | Candidatus_Moduliflexus_flocculans | U14_02641 | 303019..305532 | 1499966 | ATPase-(AAA+-superfamily)-like-protein-[Candidatus-Moduliflexus-flocculans] |
| GAK55088.1 | AAA+ATPase+wHTH+Alpha+Beta-domain+wHTH+RRM | Type-1_BR.RM.like.systems | CopG_antitoxin->PolB->METHYLASE->METHYLASE-p->PolB->?->DUF499*-> | 855 | Bacteria>Candidatus_Moduliflexota>Candidatus_Vecturitrichia>Candidatus_Vecturitrichales>Candidatus_Vecturitrichaceae | GCA_000739535.1 | Candidatus_Vecturithrix_granuli | U27_01919 | 416155..418722 | 1499967 | ATPase-(AAA+-superfamily)-like-protein-[Candidatus-Vecturithrix-granuli] |
| GAK56408.1 | AAA+ATPase+wHTH+Alpha+Beta-domain+wHTH+RRM | Type-1_BR.RM.like.systems | TUDOR+SF1-SF2-Helicase+iREase+CTD+REase->?->PIN->METHYLASE->DUF499*->?->?-><-?<-?<-REase | 1005 | Bacteria>Candidatus_Moduliflexota>Candidatus_Vecturitrichia>Candidatus_Vecturitrichales>Candidatus_Vecturitrichaceae | GCA_000739535.1 | Candidatus_Vecturithrix_granuli | U27_03370 | 807271..810288 | 1499967 | ATPase-(AAA+-superfamily)-like-protein-[Candidatus-Vecturithrix-granuli] |
| GAK55935.1 | AAA+ATPase+wHTH+Alpha+Beta-domain+wHTH+RRM | Type-1_BR.RM.like.systems | <-DUF499*\|\|?->?->?->ABC-ATPase+SbcC->ABC-ATPase+SbcC-> | 1007 | Bacteria>Candidatus_Moduliflexota>Candidatus_Vecturitrichia>Candidatus_Vecturitrichales>Candidatus_Vecturitrichaceae | GCA_000739535.1 | Candidatus_Vecturithrix_granuli | U27_02896 | 302260..305283 | 1499967 | ATPase-(AAA+-superfamily)-like-protein-[Candidatus-Vecturithrix-granuli] |
| NPV88406.1 | AAA+ATPase+wHTH+Alpha+Beta-domain+wHTH+RRM | Type-1_BR.RM.like.systems | TUDOR+SF1-SF2-Helicase+iREase+CTD+REase->METHYLASE->DUF499*->ParB+HNH->HEPN+Mrr_N->HEPN+Mrr_N-> | 865 | Bacteria>Coprothermobacterota>Coprothermobacteria>Coprothermobacterales>Coprothermobacteraceae | GCA_013178255.1 | Coprothermobacteraceae_bacterium | HPY42_02620 | 513240..515837 | 2736298 | ATP-binding-protein-[Coprothermobacteraceae-bacterium] |
| MCX5970039.1 | AAA+ATPase+wHTH+Alpha+Beta-domain+wHTH+RRM | Type-1_BR.RM.like.systems | DUF499*->?->?->?->ABC-ATPase+SbcC-><-Peptidase_M20\|\|Mbetalac-> | 962 | Bacteria>Coprothermobacterota>unclassified_Coprothermobacterota | GCA_026389425.1 | Coprothermobacterota_bacterium | NTV14_00745 | 1..2889 | 2874057 | MAG:-DUF499-domain-containing-protein,-partial-[Coprothermobacterota-bacterium] |
| WP_237889222.1 | AAA+ATPase+wHTH+Alpha+Beta-domain+wHTH+RRM | Type-1_BR.RM.like.systems | TUDOR+SF1-SF2-Helicase+iREase+CTD+REase->Schlafen+HATPase+HTH->METHYLASE->DUF499*->?->?->tRNA->ABC-ATPase+SbcC-> | 970 | Bacteria>delta/epsilon_subdivisions>Deltaproteobacteria>Deltaproteobacteria_incertae_sedis>Dissulfurimicrobium | GCF_022026155.1 | Dissulfurimicrobium_hydrothermale | LGS26_RS03280 | 705149..708061 | 1750598 | DUF499-domain-containing-protein-[Dissulfurimicrobium-hydrothermale] |
| OGR05786.1 | AAA+ATPase+wHTH+Alpha+Beta-domain+wHTH+RRM | Type-1_BR.RM.like.systems | METHYLASE->DUF499*->?->Schlafen+HATPase+HTH-> | 944 | Bacteria>delta/epsilon_subdivisions>Deltaproteobacteria>unclassified_Deltaproteobacteria | GCA_001799225.1 | Deltaproteobacteria_bacterium_RIFOXYD12_FULL_50_9 | A2511_04565 | 2695..5529 | 1797902 | hypothetical-protein-A2511_04565-[Deltaproteobacteria-bacterium-RIFOXYD12_FULL_50_9] |
| OIP30648.1 | AAA+ATPase+wHTH+Alpha+Beta-domain+wHTH+RRM | Type-1_BR.RM.like.systems | TUDOR+SF1-SF2-Helicase+iREase+CTD+REase-p->?->?-><-DNA-glycosylase<-DUF499*<-METHYLASE<-TUDOR+SF1-SF2-Helicase+iREase+CTD+REase-p<-?<-ParB+METHYLASE<-HKD-DNase | 935 | Bacteria>delta/epsilon_subdivisions>Deltaproteobacteria>unclassified_Deltaproteobacteria | GCA_001873925.1 | Deltaproteobacteria_bacterium_CG2_30_63_29 | AUK47_24645 | 4007..6814 | 1805126 | AAA-family-ATPase-[Deltaproteobacteria-bacterium-CG2_30_63_29] |
| OGP57634.1 | AAA+ATPase+wHTH+Alpha+Beta-domain+wHTH+RRM | Type-1_BR.RM.like.systems | <-ABC-ATPase+SbcC<-ABC-ATPase+SbcC<-?<-?<-RADICAL-SAM<-RADICAL-SAM<-DUF499*<-METHYLASE\|\|?-><-TUDOR+SF1-SF2-Helicase+iREase+CTD+REase | 942 | Bacteria>delta/epsilon_subdivisions>Deltaproteobacteria>unclassified_Deltaproteobacteria | GCA_001797815.1 | Deltaproteobacteria_bacterium_RBG_13_61_14 | A2V67_09195 | 7345..10173 | 1797834 | AAA-family-ATPase-[Deltaproteobacteria-bacterium-RBG_13_61_14] |
| PKN67833.1 | AAA+ATPase+wHTH+Alpha+Beta-domain+wHTH+RRM | Type-1_BR.RM.like.systems | <-SF1-SF2-Helicase<-?<-?<-?<-?<-DUF499*<-METHYLASE<-DrHyd+BirA-HTH<-TUDOR+SF1-SF2-Helicase+iREase+CTD+REase<-SbcC+SbcC<-Calcineurin | 970 | Bacteria>delta/epsilon_subdivisions>Deltaproteobacteria>unclassified_Deltaproteobacteria | GCA_002840535.1 | Deltaproteobacteria_bacterium_HGW-Deltaproteobacteria-15 | CVU57_01060 | 27172..30084 | 2013742 | AAA-family-ATPase-[Deltaproteobacteria-bacterium-HGW-Deltaproteobacteria-15] |
| OGP05895.1 | AAA+ATPase+wHTH+Alpha+Beta-domain+wHTH+RRM | Type-1_BR.RM.like.systems | ParA-Soj-PloopNTPase->Peptidase->TUDOR+SF1-SF2-Helicase+iREase+CTD+REase->wHTH+REase->METHYLASE->DUF499*-> | 918 | Bacteria>delta/epsilon_subdivisions>Deltaproteobacteria>unclassified_Deltaproteobacteria | GCA_001795495.1 | Deltaproteobacteria_bacterium_GWA2_42_85 | A2056_04335 | 11127..13883 | 1797803 | AAA-family-ATPase-[Deltaproteobacteria-bacterium-GWA2_42_85] |
| PKN69055.1 | AAA+ATPase+wHTH+Alpha+Beta-domain+wHTH+RRM | Type-1_BR.RM.like.systems | <-HKD-DNase<-?<-?<-?<-?<-DUF499*<-METHYLASE<-TUDOR+SF1-SF2-Helicase+iREase+CTD+REase<-HEPN<-REase | 913 | Bacteria>delta/epsilon_subdivisions>Deltaproteobacteria>unclassified_Deltaproteobacteria | GCA_002840565.1 | Deltaproteobacteria_bacterium_HGW-Deltaproteobacteria-12 | CVU54_12145 | 60436..63177 | 2013739 | hypothetical-protein-CVU54_12145-[Deltaproteobacteria-bacterium-HGW-Deltaproteobacteria-12] |
| OQB75596.1 | AAA+ATPase+wHTH+Alpha+Beta-domain+wHTH+RRM | Type-1_BR.RM.like.systems | <-DUF499*<-DrHyd+BirA-HTH<-METHYLASE<-TUDOR+SF1-SF2-Helicase+iREase+CTD+REase | 981 | Bacteria>delta/epsilon_subdivisions>Deltaproteobacteria>unclassified_Deltaproteobacteria | GCA_002070575.1 | Deltaproteobacteria_bacterium_ADurb.Bin135 | BWX92_02548 | 501..3446 | 1852867 | hypothetical-protein-BWX92_02548-[Deltaproteobacteria-bacterium-ADurb.Bin135] |
| PIV20448.1 | AAA+ATPase+wHTH+Alpha+Beta-domain+wHTH+RRM | Type-1_BR.RM.like.systems | MazE_antitoxin->PIN->?->METHYLASE->PolB->HEPN->DUF499*-> | 921 | Bacteria>delta/epsilon_subdivisions>Deltaproteobacteria>unclassified_Deltaproteobacteria | GCA_002780715.1 | Deltaproteobacteria_bacterium_CG03_land_8_20_14_0_80_45_14 | COS40_12540 | 8362..11127 | 1973969 | AAA-family-ATPase-[Deltaproteobacteria-bacterium-CG03_land_8_20_14_0_80_45_14] |
| PMQ01329.1 | AAA+ATPase+wHTH+Alpha+Beta-domain+wHTH+RRM | Type-1_BR.RM.like.systems | <-ABC-ATPase+SbcC\|\|tRNA-><-?\|\|?-><-DUF499*<-METHYLASE<-TUDOR+SF1-SF2-Helicase+iREase+CTD+REase | 845 | Bacteria>Dictyoglomota>Dictyoglomia>Dictyoglomales>Dictyoglomaceae | GCA_002878375.1 | Dictyoglomus_sp._NZ13-RE01 | CBR30_06655 | 56305..58842 | 2010949 | MAG:-AAA-family-ATPase-[Dictyoglomus-sp.-NZ13-RE01] |
| MBN4059261.1 | AAA+ATPase+wHTH+Alpha+Beta-domain+wHTH+RRM | Type-1_BR.RM.like.systems | <-zf-IS66+DDE_Tnp_IS66\|\|METHYLASE-p->DUF499*-> | 305 | Bacteria>Elusimicrobiota>Endomicrobiia>Endomicrobiales>Endomicrobiaceae | GCA_017113195.1 | Endomicrobium_sp._AH-315-J14 | JYT22_01255 | 2291..3205 | 2814560 | ATP-binding-protein-[Endomicrobium-sp.-AH-315-J14] |
| MBI4801733.1 | AAA+ATPase+wHTH+Alpha+Beta-domain+wHTH+RRM | Type-1_BR.RM.like.systems | DUF499*->?->HKD-DNase->?->?->?->Glyoxalase-> | 973 | Bacteria>Elusimicrobiota>unclassified_Elusimicrobiota | GCA_016208085.1 | Elusimicrobiota_bacterium | HY796_04325 | 1379..4300 | 2030800 | ATP-binding-protein-[Elusimicrobia-bacterium] |
| MBI4669491.1 | AAA+ATPase+wHTH+Alpha+Beta-domain+wHTH+RRM | Type-1_BR.RM.like.systems | METHYLASE-p->?->DUF1828+REase->DUF499*-><-SIGMA-HTH+SIGMA-HTH<-ZnR+TOPRIM+DnaB<-?<-?<-Radical-SAM+TRAM<-Peptidase_M22 | 930 | Bacteria>Elusimicrobiota>unclassified_Elusimicrobiota | GCA_016209025.1 | Elusimicrobiota_bacterium | HY747_09985 | 2681..5473 | 2030800 | ATP-binding-protein-[Elusimicrobia-bacterium] |
| MDE2039591.1 | AAA+ATPase+wHTH+Alpha+Beta-domain+wHTH+RRM | Type-1_BR.RM.like.systems | <-MerR-HTH<-?<-?<-?<-ASCH<-ACET+PIN+ACET<-DUF499*<-METHYLASE<-TUDOR+SF1-SF2-Helicase+iREase+CTD+REase\|\|?-><-JAB\|\|PadR-HTH-> | 920 | Bacteria>Elusimicrobiota>unclassified_Elusimicrobiota | GCA_028865065.1 | Elusimicrobiota_bacterium | KGJ45_06180 | 87247..90009 | 2030800 | MAG:-ATP-binding-protein-[Elusimicrobia-bacterium] |
| MBI4676018.1 | AAA+ATPase+wHTH+Alpha+Beta-domain+wHTH+RRM | Type-1_BR.RM.like.systems | <-Peptidase<-DUF499*<-TOPRIM<-OLD-ABC<-TOPRIM<-OLD-ABC<-METHYLASE<-ORC-CDC6 | 925 | Bacteria>Elusimicrobiota>unclassified_Elusimicrobiota | GCA_016209035.1 | Elusimicrobiota_bacterium | HY748_00360 | 72321..75098 | 2030800 | ATP-binding-protein-[Elusimicrobia-bacterium] |
| MBI5244845.1 | AAA+ATPase+wHTH+Alpha+Beta-domain+wHTH+RRM | Type-1_BR.RM.like.systems | <-DUF499*<-HEPN<-PolB<-METHYLASE | 944 | Bacteria>Elusimicrobiota>unclassified_Elusimicrobiota | GCA_016212885.1 | Elusimicrobiota_bacterium | HY922_14340 | 21906..24740 | 2030800 | ATP-binding-protein-[Elusimicrobia-bacterium] |
| MCZ2341133.1 | AAA+ATPase+wHTH+Alpha+Beta-domain+wHTH+RRM | Type-1_BR.RM.like.systems | TUDOR+SF1-SF2-Helicase+iREase+CTD+REase->?->METHYLASE->Phd-YefM->PIN->DUF499*-> | 944 | Bacteria>FCB_group>Bacteroidota/Chlorobiota_group>Bacteroidota>Bacteroidia | GCA_026931725.1 | Bacteroidales_bacterium | LC104_04965 | 19853..22687 | 2030927 | MAG:-DUF499-domain-containing-protein-[Bacteroidales-bacterium] |
| MCS2505373.1 | AAA+ATPase+wHTH+Alpha+Beta-domain+wHTH+RRM | Type-1_BR.RM.like.systems | TUDOR+SF1-SF2-Helicase+iREase+CTD+REase-p->METHYLASE-p->METHYLASE-p->HEPN-p->HEPN+DUF499->DUF499*-><-?<-HicB<-TOPRIM<-SbcC\|\|BirA-HTH-><-Phage_int_SAM_5+Phage_integrase | 640 | Bacteria>FCB_group>Bacteroidota/Chlorobiota_group>Bacteroidota>Bacteroidia | GCA_024788865.1 | Phocaeicola_vulgatus | NXW48_05310 | 1160902..1162824 | 821 | ATP-binding-protein-[Phocaeicola-vulgatus] |
| UVP11344.1 | AAA+ATPase+wHTH+Alpha+Beta-domain+wHTH+RRM | Type-1_BR.RM.like.systems | TUDOR+SF1-SF2-Helicase+iREase+CTD+REase-p->zf-ISL3+DDE_Tnp_ISL3->TUDOR+SF1-SF2-Helicase+iREase+CTD+REase-p->METHYLASE->METHYLASE-p->HEPN+DUF499-p->DUF499*-> | 689 | Bacteria>FCB_group>Bacteroidota/Chlorobiota_group>Bacteroidota>Bacteroidia | GCA_024758285.1 | Bacteroides_ovatus | NXW52_28300 | 6817403..6819472 | 28116 | ATP-binding-protein-[Bacteroides-ovatus] |
| MCS2524521.1 | AAA+ATPase+wHTH+Alpha+Beta-domain+wHTH+RRM | Type-1_BR.RM.like.systems | Peptidase-><-?<-?<-?<-DUF499-p<-DUF499*<-HEPN-p<-METHYLASE-p<-METHYLASE<-TUDOR+SF1-SF2-Helicase+iREase+CTD+REase<-TUDOR+SF1-SF2-Helicase+iREase+CTD+REase-p | 704 | Bacteria>FCB_group>Bacteroidota/Chlorobiota_group>Bacteroidota>Bacteroidia | GCA_024789005.1 | Bacteroides_ovatus | NXV02_14125 | 3438684..3440798 | 28116 | DUF499-domain-containing-protein-[Bacteroides-ovatus] |
| GIV23748.1 | AAA+ATPase+wHTH+Alpha+Beta-domain+wHTH+RRM | Type-1_BR.RM.like.systems | TUDOR+SF1-SF2-Helicase+iREase+CTD+REase->Schlafen+HATPase+HTH->METHYLASE->DUF499*->DUF499-p->?->?->TUDOR+SF1-SF2-Helicase+iREase+CTD->?->METHYLASE-p-> | 393 | Bacteria>FCB_group>Bacteroidota/Chlorobiota_group>Bacteroidota>Bacteroidia | GCA_026003215.1 | Bacteroidia_bacterium | KatS3mg025_1407 | 1616391..1617572 | 2044936 | MAG:-hypothetical-protein-KatS3mg025_1407-[Bacteroidia-bacterium] |
| MBN1340543.1 | AAA+ATPase+wHTH+Alpha+Beta-domain+wHTH+RRM | Type-1_BR.RM.like.systems | TUDOR+SF1-SF2-Helicase+iREase+CTD+REase->BirA-HTH->METHYLASE->wHTH+REase->DUF499*-><-?<-tRNA\|\|sGTP->sGTP-> | 974 | Bacteria>FCB_group>Bacteroidota/Chlorobiota_group>Bacteroidota>Bacteroidia | GCA_016927695.1 | Bacteroidales_bacterium | JXA03_14535 | 29128..32052 | 2030927 | ATP-binding-protein-[Bacteroidales-bacterium] |
| EEO54231.1 | AAA+ATPase+wHTH+Alpha+Beta-domain+wHTH+RRM | Type-1_BR.RM.like.systems | DEDD_Tnp_IS110-><-?\|\|HTH_3->TUDOR+SF1-SF2-Helicase+iREase+CTD+REase->METHYLASE-p->METHYLASE->DUF499*-> | 961 | Bacteria>FCB_group>Bacteroidota/Chlorobiota_group>Bacteroidota>Bacteroidia | GCA_000157055.1 | Bacteroides_sp._2_2_4 | BSCG_01156 | 592841..595726 | 469590 | hypothetical-protein-BSCG_01156-[Bacteroides-sp.-2_2_4] |
| MBR2245017.1 | AAA+ATPase+wHTH+Alpha+Beta-domain+wHTH+RRM | Type-1_BR.RM.like.systems | <-OLD-ABC\|\|TUDOR+SF1-SF2-Helicase+iREase+CTD+REase->METHYLASE->DUF499*-><-HTH_3<-?\|\|wHTH+AbiEi->AbiEii-> | 933 | Bacteria>FCB_group>Bacteroidota/Chlorobiota_group>Bacteroidota>Bacteroidia | GCA_017475205.1 | Prevotella_sp. | IJ888_09875 | 11611..14412 | 59823 | ATP-binding-protein-[Prevotella-sp.] |
| MBR3480399.1 | AAA+ATPase+wHTH+Alpha+Beta-domain+wHTH+RRM | Type-1_BR.RM.like.systems | Glyco_hydro_43+CBM_6->?->?->TUDOR+SF1-SF2-Helicase+iREase+CTD+REase->METHYLASE->DUF499*-><-35exo | 929 | Bacteria>FCB_group>Bacteroidota/Chlorobiota_group>Bacteroidota>Bacteroidia | GCA_017509035.1 | Prevotella_sp. | IKH32_06860 | 45858..48647 | 59823 | ATP-binding-protein-[Prevotella-sp.] |
| WP_197018884.1 | AAA+ATPase+wHTH+Alpha+Beta-domain+wHTH+RRM | Type-1_BR.RM.like.systems | DDE_Tnp_1->TUDOR+SF1-SF2-Helicase+iREase+CTD+REase->METHYLASE->DUF499*-><-?\|\|?->HTH->?-><-RelE-ParE | 931 | Bacteria>FCB_group>Bacteroidota/Chlorobiota_group>Bacteroidota>Bacteroidia | GCF_000619985.1 | Prevotella_sp._P6B4 | T491_RS16740 | 7401..10196 | 1410614 | DUF499-domain-containing-protein-[Prevotella-sp.-P6B4] |
| MBR6189438.1 | AAA+ATPase+wHTH+Alpha+Beta-domain+wHTH+RRM | Type-1_BR.RM.like.systems | 35exo-><-DUF499*<-METHYLASE<-TUDOR+SF1-SF2-Helicase+iREase+CTD+REase<-?<-?<-Glyco_hydro_43+CBM_6 | 929 | Bacteria>FCB_group>Bacteroidota/Chlorobiota_group>Bacteroidota>Bacteroidia | GCA_017540805.1 | Prevotella_sp. | IKQ59_10915 | 405..3194 | 59823 | ATP-binding-protein-[Prevotella-sp.] |
| MBR2238254.1 | AAA+ATPase+wHTH+Alpha+Beta-domain+wHTH+RRM | Type-1_BR.RM.like.systems | Glyco_hydro_43+CBM_6->?->?->TUDOR+SF1-SF2-Helicase+iREase+CTD+REase->METHYLASE->DUF499*-><-35exo | 929 | Bacteria>FCB_group>Bacteroidota/Chlorobiota_group>Bacteroidota>Bacteroidia | GCA_017475265.1 | Prevotella_sp. | IJ887_10335 | 38305..41094 | 59823 | ATP-binding-protein-[Prevotella-sp.] |
| GAP68724.1 | AAA+ATPase+wHTH+Alpha+Beta-domain+wHTH+RRM | Type-1_BR.RM.like.systems | <-PAIREDC-HTH<-?<-?\|\|?->TUDOR+SF1-SF2-Helicase+iREase+CTD+REase->METHYLASE->DUF499*->HKD-DNase-> | 974 | Bacteria>FCB_group>Bacteroidota/Chlorobiota_group>Bacteroidota>Bacteroidia | GCA_001270045.2 | Bacteroidales_bacterium_6E | BA6E_104172 | 195307..198231 | 1688776 | hypothetical-protein-BA6E_104172-[Bacteroidales-bacterium-6E] |
| MBI4648362.1 | AAA+ATPase+wHTH+Alpha+Beta-domain+wHTH+RRM | Type-1_BR.RM.like.systems | TUDOR+SF1-SF2-Helicase+iREase+CTD+REase->?->METHYLASE->DUF499*->DUF499-p->?-><-wHTH+REase | 388 | Bacteria>FCB_group>Bacteroidota/Chlorobiota_group>Bacteroidota>Bacteroidia | GCA_016209245.1 | Bacteroidia_bacterium | HY738_17700 | 5747..6913 | 2044936 | ATP-binding-protein-[Bacteroidia-bacterium] |
| WP_074760634.1 | AAA+ATPase+wHTH+Alpha+Beta-domain+wHTH+RRM | Type-1_BR.RM.like.systems | Schlafen+HATPase+HTH->TUDOR+SF1-SF2-Helicase+iREase+CTD+REase->METHYLASE->DUF499*-> | 930 | Bacteria>FCB_group>Bacteroidota/Chlorobiota_group>Bacteroidota>Bacteroidia | GCF_900107775.1 | Xylanibacter_ruminicola | BLU97_RS05645 | 384764..387556 | 839 | DUF499-domain-containing-protein-[Prevotella-ruminicola] |
| MBK9661552.1 | AAA+ATPase+wHTH+Alpha+Beta-domain+wHTH+RRM | Type-1_BR.RM.like.systems | <-Integrase_1+Phage_integrase\|\|?->DUF488-1-><-?<-DUF499*<-wHTH+REase<-METHYLASE<-HEPN<-TUDOR+SF1-SF2-Helicase+iREase+CTD+REase<-HTH+WYL\|\|RnaseHII-> | 943 | Bacteria>FCB_group>Bacteroidota/Chlorobiota_group>Bacteroidota>Chitinophagia | GCA_016717645.1 | Chitinophagaceae_bacterium | IPO68_16915 | 5414..8245 | 1869212 | ATP-binding-protein-[Chitinophagaceae-bacterium] |
| MBK9465916.1 | AAA+ATPase+wHTH+Alpha+Beta-domain+wHTH+RRM | Type-1_BR.RM.like.systems | ABhydrolase-><-?<-Integrase_1+Phage_integrase\|\|?->DUF488-1-><-?<-DUF499*<-wHTH+REase<-HEPN<-HTH+WYL\|\|RnaseHII-> | 943 | Bacteria>FCB_group>Bacteroidota/Chlorobiota_group>Bacteroidota>Chitinophagia | GCA_016719015.1 | Chitinophagaceae_bacterium | IPN56_14270 | 8966..11797 | 1869212 | ATP-binding-protein-[Chitinophagaceae-bacterium] |
| MBX7108627.1 | AAA+ATPase+wHTH+Alpha+Beta-domain+wHTH+RRM | Type-1_BR.RM.like.systems | HTH->TUDOR+SF1-SF2-Helicase+iREase+CTD+REase->METHYLASE->ParB+HNH->OLD-ABC->?->DUF499*->?->PIWI-> | 925 | Bacteria>FCB_group>Bacteroidota/Chlorobiota_group>Bacteroidota>Chitinophagia | GCA_019694975.1 | Chitinophagales_bacterium | K1X61_08290 | 480306..483083 | 2448779 | DUF499-domain-containing-protein-[Chitinophagales-bacterium] |
| WP_207513720.1 | AAA+ATPase+wHTH+Alpha+Beta-domain+wHTH+RRM | Type-1_BR.RM.like.systems | Glyco_hydro_43+CBM_6->TUDOR+SF1-SF2-Helicase+iREase+CTD+REase->METHYLASE->DUF499*-> | 930 | Bacteria>FCB_group>Bacteroidota/Chlorobiota_group>Bacteroidota>Chitinophagia | GCF_017356035.1 | Longitalea_luteola | JUB15_RS17135 | 322919..325711 | 2812563 | DUF499-domain-containing-protein-[Longitalea-luteola] |
| MBP8114174.1 | AAA+ATPase+wHTH+Alpha+Beta-domain+wHTH+RRM | Type-1_BR.RM.like.systems | HTH->TUDOR+SF1-SF2-Helicase+iREase+CTD+REase->METHYLASE->ParB+HNH->OLD-ABC->?->DUF499*-> | 925 | Bacteria>FCB_group>Bacteroidota/Chlorobiota_group>Bacteroidota>Chitinophagia | GCA_018005575.1 | Chitinophagaceae_bacterium | KAY50_02395 | 14241..17018 | 1869212 | ATP-binding-protein-[Chitinophagaceae-bacterium] |
| NCU04700.1 | AAA+ATPase+wHTH+Alpha+Beta-domain+wHTH+RRM | Type-1_BR.RM.like.systems | wHTH+AbiEi->AbiEii->METHYLASE->ParB+HNH->OLD-ABC->?->DUF499*-> | 237 | Bacteria>FCB_group>Bacteroidota/Chlorobiota_group>Bacteroidota>Chitinophagia | GCA_009996555.1 | Chitinophagaceae_bacterium | GXC73_12005 | 16910..17621 | 1869212 | ATP-binding-protein-[Chitinophagaceae-bacterium] |
| WP_135533129.1 | AAA+ATPase+wHTH+Alpha+Beta-domain+wHTH+RRM | Type-1_BR.RM.like.systems | Phage_integrase-><-?\|\|?->TUDOR+SF1-SF2-Helicase+iREase+CTD+REase->METHYLASE->DUF499*->?->?-><-?<-DOC<-ZnR | 953 | Bacteria>FCB_group>Bacteroidota/Chlorobiota_group>Bacteroidota>Cytophagia | GCF_004745865.1 | Hymenobacter_wooponensis | EU557_RS24810 | 69166..72027 | 1525360 | DUF499-domain-containing-protein-[Hymenobacter-wooponensis] |
| WP_152126729.1 | AAA+ATPase+wHTH+Alpha+Beta-domain+wHTH+RRM | Type-1_BR.RM.like.systems | <-REase+PNTD1+PNTD2<-?\|\|BirA-HTH->AbiEii->TUDOR+SF1-SF2-Helicase+iREase+CTD+REase->METHYLASE->DUF499*->?-><-?<-MobL+ZnR+TOPRIM<-?\|\|?->HTH-> | 987 | Bacteria>FCB_group>Bacteroidota/Chlorobiota_group>Bacteroidota>Cytophagia | GCF_009184635.1 | Rudanella_paleaurantiibacter | F5984_RS23815 | 78854..81817 | 2614655 | DUF499-domain-containing-protein-[Rudanella-paleaurantiibacter] |
| WP_086597021.1 | AAA+ATPase+wHTH+Alpha+Beta-domain+wHTH+RRM | Type-1_BR.RM.like.systems | TUDOR+SF1-SF2-Helicase+iREase+CTD+REase->ParB+HNH->METHYLASE->DUF499*-><-?\|\|?->Peptidase_S74-> | 972 | Bacteria>FCB_group>Bacteroidota/Chlorobiota_group>Bacteroidota>Cytophagia | GCF_002154225.1 | Hymenobacter_crusticola | BXP70_RS25835 | 30420..33338 | 1770526 | DUF499-domain-containing-protein-[Hymenobacter-crusticola] |
| MCB0808336.1 | AAA+ATPase+wHTH+Alpha+Beta-domain+wHTH+RRM | Type-1_BR.RM.like.systems | <-DUF499*<-METHYLASE<-HNH<-OLD-ABC<-TUDOR+SF1-SF2-Helicase+iREase+CTD+REase | 759 | Bacteria>FCB_group>Bacteroidota/Chlorobiota_group>Bacteroidota>Flavobacteriia | GCA_020438005.1 | Flavobacteriales_bacterium | KDB96_03570 | 1..2279 | 2021391 | ATP-binding-protein-[Flavobacteriales-bacterium] |
| MCX7650091.1 | AAA+ATPase+wHTH+Alpha+Beta-domain+wHTH+RRM | Type-1_BR.RM.like.systems | <-DUF499*<-AAA-ATPase+REase<-METHYLASE-p | 926 | Bacteria>FCB_group>Bacteroidota/Chlorobiota_group>Bacteroidota>Flavobacteriia | GCA_026419215.1 | Flavobacteriales_bacterium | N2050_06005 | 2592..5372 | 2021391 | MAG:-DUF499-domain-containing-protein-[Flavobacteriales-bacterium] |
| MBK7304321.1 | AAA+ATPase+wHTH+Alpha+Beta-domain+wHTH+RRM | Type-1_BR.RM.like.systems | HTH->METHYLASE-p->METHYLASE-p->METHYLASE-p->METHYLASE-p->METHYLASE-p->DUF499*->DUF499-p->?-><-?<-?\|\|?->HTH_3-> | 390 | Bacteria>FCB_group>Bacteroidota/Chlorobiota_group>Bacteroidota>Saprospiria | GCA_016705745.1 | Candidatus_Vicinibacter_affinis | IPI90_14000 | 15462..16634 | 2982026 | ATP-binding-protein-[Saprospiraceae-bacterium] |
| MBK9716010.1 | AAA+ATPase+wHTH+Alpha+Beta-domain+wHTH+RRM | Type-1_BR.RM.like.systems | SIR2->HTH->TUDOR+SF1-SF2-Helicase+iREase+CTD+REase->METHYLASE->METHYLASE-p->DUF499*->DUF499-p->?-><-?<-?\|\|HTH_3-> | 749 | Bacteria>FCB_group>Bacteroidota/Chlorobiota_group>Bacteroidota>Saprospiria | GCA_016717245.1 | Candidatus_Defluviibacterium_haderslevense | IPO85_00510 | 27995..30244 | 2981993 | ATP-binding-protein-[Saprospiraceae-bacterium] |
| WP_068215357.1 | AAA+ATPase+wHTH+Alpha+Beta-domain+wHTH+RRM | Type-1_BR.RM.like.systems | <-TPR+TPR\|\|?->TUDOR+SF1-SF2-Helicase+iREase+CTD+REase->METHYLASE->DUF499*->SF1-SF2-Helicase->?->MACRODOMAIN-> | 933 | Bacteria>FCB_group>Bacteroidota/Chlorobiota_group>Bacteroidota>Saprospiria | GCF_001625015.1 | Lewinella_sp._4G2 | A3850_RS07910 | 1744205..1747006 | 1803372 | DUF499-domain-containing-protein-[Lewinella-sp.-4G2] |
| MBK8405311.1 | AAA+ATPase+wHTH+Alpha+Beta-domain+wHTH+RRM | Type-1_BR.RM.like.systems | SIR2->HTH->METHYLASE-p->METHYLASE-p->METHYLASE-p->DUF499*->DUF499-p-> | 284 | Bacteria>FCB_group>Bacteroidota/Chlorobiota_group>Bacteroidota>Saprospiria | GCA_016711565.1 | Candidatus_Vicinibacter_affinis | IPL25_14925 | 8413..9267 | 2982026 | ATP-binding-protein-[Saprospiraceae-bacterium] |
| MBK6701489.1 | AAA+ATPase+wHTH+Alpha+Beta-domain+wHTH+RRM | Type-1_BR.RM.like.systems | <-DUF499-p<-DUF499*<-METHYLASE<-PIN<-PIN<-HTH<-HTH<-GDResolvase+Recombinase+ZnR | 823 | Bacteria>FCB_group>Bacteroidota/Chlorobiota_group>Bacteroidota>Saprospiria | GCA_016703345.1 | Candidatus_Defluviibacterium_haderslevense | IPG55_16710 | 1974881..1977352 | 2981993 | ATP-binding-protein-[Saprospiraceae-bacterium] |
| MBK8644365.1 | AAA+ATPase+wHTH+Alpha+Beta-domain+wHTH+RRM | Type-1_BR.RM.like.systems | <-HTH_3<-?\|\|?->?-><-?<-DUF499*<-METHYLASE<-METHYLASE-p<-PIN<-TUDOR+SF1-SF2-Helicase+iREase+CTD+REase<-HTH<-SIR2 | 923 | Bacteria>FCB_group>Bacteroidota/Chlorobiota_group>Bacteroidota>Saprospiria | GCA_016715705.1 | Candidatus_Vicinibacter_affinis | IPN15_19785 | 428646..431417 | 2982026 | ATP-binding-protein-[Saprospiraceae-bacterium] |
| MBK7697113.1 | AAA+ATPase+wHTH+Alpha+Beta-domain+wHTH+RRM | Type-1_BR.RM.like.systems | PIN->PIN->METHYLASE-p->METHYLASE-p->?->METHYLASE-p->DUF499*->DUF499-p->?-><-?<-?\|\|?->HTH_3-> | 749 | Bacteria>FCB_group>Bacteroidota/Chlorobiota_group>Bacteroidota>Saprospiria | GCA_016706925.1 | Candidatus_Vicinibacter_affinis | IPI30_23260 | 68066..70315 | 2982026 | ATP-binding-protein-[Saprospiraceae-bacterium] |
| MBN1449251.1 | AAA+ATPase+wHTH+Alpha+Beta-domain+wHTH+RRM | Type-1_BR.RM.like.systems | <-DUF499*<-ParB+HNH<-METHYLASE | 939 | Bacteria>FCB_group>Bacteroidota/Chlorobiota_group>Bacteroidota>unclassified_Bacteroidota | GCA_016928515.1 | Bacteroidota_bacterium | JXA28_15095 | 364..3183 | 1898104 | ATP-binding-protein-[Bacteroidetes-bacterium] |
| BCX00053.1 | AAA+ATPase+wHTH+Alpha+Beta-domain+wHTH+RRM | Type-1_BR.RM.like.systems | <-35exo\|\|?-><-Methyltransf_4<-?<-ABhydrolase<-DUF499*<-METHYLASE<-Schlafen+HATPase+HTH<-TUDOR+SF1-SF2-Helicase+iREase+CTD+REase<-?<-?<-Endonuclease_5 | 878 | Bacteria>FCB_group>Bacteroidota/Chlorobiota_group>Bacteroidota>unclassified_Bacteroidota | GCA_025998875.1 | Bacteroidota_bacterium | KatS3mg041_0099 | 106704..109340 | 1898104 | MAG:-hypothetical-protein-KatS3mg041_0099-[Bacteroidota-bacterium] |

| Protein-ID | DUF499-architecture | System | Gene-Neighbourhoods | Length | Taxonomy | Assembly-ID | Strain | Locus-Tag | Genomic-coordinates | TaxID | Fasta-defline |
|---|---|---|---|---|---|---|---|---|---|---|---|
| MDE2644924.1 | AAA+ATPase+wHTH+Alpha+Beta-domain+wHTH+RRM | Type-1_BR.RM.like.systems | <-NTP_transferase<-REase+NACHT\|\|TUDOR+SF1-SF2-Helicase+iREase+CTD+REase->SF1-SF2-Helicase->METHYLASE->DUF499*->?-><-?\|\|?-><-?<-?<-PIN | 933 | Bacteria>FCB_group>Bacteroidota/Chlorobiota_group>Bacteroidota>unclassified_Bacteroidota | GCA_028875465.1 | Bacteroidota_bacterium | OXI05_03670 | 47472..50273 | 1898104 | MAG:-DUF499-domain-containing-protein-[Bacteroidota-bacterium] |
| MDE2646784.1 | AAA+ATPase+wHTH+Alpha+Beta-domain+wHTH+RRM | Type-1_BR.RM.like.systems | ABhydrolase-><-DUF499*<-PIN<-OLD-ABC<-METHYLASE<-METHYLASE-p<-wHTH+REase<-TUDOR+SF1-SF2-Helicase+iREase+CTD+REase | 924 | Bacteria>FCB_group>Bacteroidota/Chlorobiota_group>Bacteroidota>unclassified_Bacteroidota | GCA_028875465.1 | Bacteroidota_bacterium | OXI05_13240 | 2968..5742 | 1898104 | MAG:-DUF499-domain-containing-protein-[Bacteroidota-bacterium] |
| MCL5028412.1 | AAA+ATPase+wHTH+Alpha+Beta-domain+wHTH+RRM | Type-1_BR.RM.like.systems | HTH_3->RVT-><-?<-?<-?<-HISKIN<-DUF499*<-METHYLASE<-TIR+PNTD2<-?<-DUF488-l<-TUDOR+SF1-SF2-Helicase+iREase+CTD+REase<-AbiEii | 919 | Bacteria>FCB_group>Bacteroidota/Chlorobiota_group>Bacteroidota>unclassified_Bacteroidota | GCA_023475265.1 | Bacteroidota_bacterium | M1480_05255 | 11247..14006 | 1898104 | DUF499-domain-containing-protein-[Bacteroidetes-bacterium] |
| MCY3594931.1 | AAA+ATPase+wHTH+Alpha+Beta-domain+wHTH+RRM | Type-1_BR.RM.like.systems | Peptidase_S41-><-?<-DUF499*<-REase<-PIN<-OLD-ABC<-METHYLASE<-TUDOR+SF1-SF2-Helicase+iREase+CTD+REase | 913 | Bacteria>FCB_group>Bacteroidota/Chlorobiota_group>Bacteroidota>unclassified_Bacteroidota | GCA_026705205.1 | Bacteroidota_bacterium | OXH01_06810 | 22578..25319 | 1898104 | MAG:-DUF499-domain-containing-protein-[Bacteroidetes-bacterium] |
| TAL70497.1 | AAA+ATPase+wHTH+Alpha+Beta-domain+wHTH+RRM | Type-1_BR.RM.like.systems | <-XIS-HTH<-?\|\|TUDOR+SF1-SF2-Helicase+iREase+CTD+REase->METHYLASE->?->?->DUF499*->?-><-?<-?<-TPR+TPR<-?<-SIGMA-HTH | 980 | Bacteria>FCB_group>Bacteroidota/Chlorobiota_group>Bacteroidota>unclassified_Bacteroidota | GCA_004322215.1 | Bacteroidota_bacterium | EPN82_02485 | 152663..155605 | 1898104 | ATP-binding-protein-[Bacteroidetes-bacterium] |
| MBS1776511.1 | AAA+ATPase+wHTH+Alpha+Beta-domain+wHTH+RRM | Type-1_BR.RM.like.systems | XIS-HTH->?->HTH->TUDOR+SF1-SF2-Helicase+iREase+CTD+REase->METHYLASE->DUF499*->PIWI->ParB+HNH-> | 925 | Bacteria>FCB_group>Bacteroidota/Chlorobiota_group>Bacteroidota>unclassified_Bacteroidota | GCA_018267925.1 | Bacteroidota_bacterium | JSS64_09560 | 68237..71014 | 1898104 | ATP-binding-protein-[Bacteroidetes-bacterium] |
| MCY3630037.1 | AAA+ATPase+wHTH+Alpha+Beta-domain+wHTH+RRM | Type-1_BR.RM.like.systems | <-DUF499*<-PIN<-OLD-ABC<-METHYLASE<-wHTH+REase<-TUDOR+SF1-SF2-Helicase+iREase+CTD+REase | 884 | Bacteria>FCB_group>Bacteroidota/Chlorobiota_group>Bacteroidota>unclassified_Bacteroidota | GCA_026705375.1 | Bacteroidota_bacterium | OXG94_08505 | 1..2653 | 1898104 | MAG:-DUF499-domain-containing-protein,-partial-[Bacteroidetes-bacterium] |
| MCY4170340.1 | AAA+ATPase+wHTH+Alpha+Beta-domain+wHTH+RRM | Type-1_BR.RM.like.systems | OLD-ABC-><-DUF499*<-REase<-?<-METHYLASE<-TUDOR+SF1-SF2-Helicase+iREase+CTD+REase | 912 | Bacteria>FCB_group>Bacteroidota/Chlorobiota_group>Bacteroidota>unclassified_Bacteroidota | GCA_026709065.1 | Bacteroidota_bacterium | OXF08_01935 | 1316..4054 | 1898104 | MAG:-DUF499-domain-containing-protein-[Bacteroidetes-bacterium] |
| MCX6246558.1 | AAA+ATPase+wHTH+Alpha+Beta-domain+wHTH+RRM | Type-1_BR.RM.like.systems | TUDOR+SF1-SF2-Helicase+iREase+CTD+REase->METHYLASE->DUF499*-><-?<-Peptidase_C47 | 857 | Bacteria>FCB_group>Bacteroidota/Chlorobiota_group>Bacteroidota>unclassified_Bacteroidota | GCA_026391795.1 | Bacteroidota_bacterium | NTW10_02390 | 79074..81647 | 1898104 | MAG:-DUF499-domain-containing-protein-[Bacteroidetes-bacterium] |
| MQY79011.1 | AAA+ATPase+wHTH+Alpha+Beta-domain+wHTH+RRM | Type-1_BR.RM.like.systems | TUDOR+SF1-SF2-Helicase+iREase+CTD+REase->Schlafen+HATPase+HTH->METHYLASE->?->DUF499*-> | 491 | Bacteria>FCB_group>Bacteroidota/Chlorobiota_group>Bacteroidota>unclassified_Bacteroidota | GCA_009619225.1 | Bacteroidota_bacterium | GH151_07430 | 8478..9950 | 1898104 | DUF499-domain-containing-protein-[Bacteroidota-bacterium] |
| MCB2218983.1 | AAA+ATPase+wHTH+Alpha+Beta-domain+wHTH+RRM | Type-1_BR.RM.like.systems | <-DUF499*<-METHYLASE<-TUDOR+SF1-SF2-Helicase+iREase+CTD+REase<-?<-?<-?<-XIS-HTH | 860 | Bacteria>FCB_group>Bacteroidota/Chlorobiota_group>Bacteroidota>unclassified_Bacteroidota | GCA_020444515.1 | Bacteroidota_bacterium | KQI35_01220 | 325627..328209 | 1898104 | DUF499-domain-containing-protein-[Bacteroidetes-bacterium] |
| MCY4234690.1 | AAA+ATPase+wHTH+Alpha+Beta-domain+wHTH+RRM | Type-1_BR.RM.like.systems | <-METHYLASE-p<-?<-?<-?<-?<-Peptidase<-DUF499*\|\|?-><-PIN<-OLD-ABC | 893 | Bacteria>FCB_group>Bacteroidota/Chlorobiota_group>Bacteroidota>unclassified_Bacteroidota | GCA_026709525.1 | Bacteroidota_bacterium | OXE59_13250 | 15773..18454 | 1898104 | MAG:-DUF499-domain-containing-protein-[Bacteroidetes-bacterium] |
| MBM3454155.1 | AAA+ATPase+wHTH+Alpha+Beta-domain+wHTH+RRM | Type-1_BR.RM.like.systems | TUDOR+SF1-SF2-Helicase+iREase+CTD+REase->METHYLASE->DUF499*-> | 974 | Bacteria>FCB_group>Bacteroidota/Chlorobiota_group>Bacteroidota>unclassified_Bacteroidota | GCA_016868905.1 | Bacteroidota_bacterium | FJX80_03285 | 10645..13569 | 1898104 | ATP-binding-protein-[Bacteroidetes-bacterium] |
| MCY4001546.1 | AAA+ATPase+wHTH+Alpha+Beta-domain+wHTH+RRM | Type-1_BR.RM.like.systems | wHTH+REase->METHYLASE->OLD-ABC->PIN->DUF499*-> | 903 | Bacteria>FCB_group>Bacteroidota/Chlorobiota_group>Bacteroidota>unclassified_Bacteroidota | GCA_026708685.1 | Bacteroidota_bacterium | OXF84_12180 | 5800..8511 | 1898104 | MAG:-DUF499-domain-containing-protein-[Bacteroidetes-bacterium] |
| MCY3614378.1 | AAA+ATPase+wHTH+Alpha+Beta-domain+wHTH+RRM | Type-1_BR.RM.like.systems | SWACOS-><-DUF499*<-PIN<-OLD-ABC<-METHYLASE-p | 924 | Bacteria>FCB_group>Bacteroidota/Chlorobiota_group>Bacteroidota>unclassified_Bacteroidota | GCA_026705365.1 | Bacteroidota_bacterium | OXH03_04690 | 2086..4860 | 1898104 | MAG:-DUF499-domain-containing-protein-[Bacteroidetes-bacterium] |
| GIV55103.1 | AAA+ATPase+wHTH+Alpha+Beta-domain+wHTH+RRM | Type-1_BR.RM.like.systems | TUDOR+SF1-SF2-Helicase+iREase+CTD+REase->Schlafen->BirA-HTH->METHYLASE-p->METHYLASE-p->DUF499*->ParB+HNH-> | 875 | Bacteria>FCB_group>Bacteroidota/Chlorobiota_group>Candidatus_Kapabacteria>unclassified_Candidatus_Kapabacteria | GCA_026003475.1 | Candidatus_Kapabacteria_bacterium | KatS3mg039_1621 | 1981096..1983723 | 2447898 | MAG:-hypothetical-protein-KatS3mg039_1621-[Candidatus-Kapabacteria-bacterium] |
| WP_011744524.1 | AAA+ATPase+wHTH+Alpha+Beta-domain+wHTH+RRM | Type-1_BR.RM.like.systems | TUDOR+SF1-SF2-Helicase+iREase+CTD+REase->?->METHYLASE->DUF499*->NACHT->?->ArdCN+MPTase-><-DOC+DOC\|\|BirA-HTH->HIN-HTH-> | 941 | Bacteria>FCB_group>Bacteroidota/Chlorobiota_group>Chlorobiota>Chlorobiia | GCF_000015125.1 | Chlorobium_phaeobacteroides | CPHA266_RS03310 | 722346..725171 | 1096 | DUF499-domain-containing-protein-[Chlorobium-phaeobacteroides] |
| MBZ4218777.1 | AAA+ATPase+wHTH+Alpha+Beta-domain+wHTH+RRM | Type-1_BR.RM.like.systems | <-ZDNA-HTH<-?<-ArdCN+MPTase<-TIR<-?<-DUF499*<-METHYLASE<-?<-TUDOR+SF1-SF2-Helicase+iREase+CTD+REase | 942 | Bacteria>FCB_group>Bacteroidota/Chlorobiota_group>Chlorobiota>Chlorobiia | GCA_020035305.1 | Chlorobium_sp. | LAC69_01110 | 16878..19706 | 1095 | DUF499-domain-containing-protein-[Chlorobium-sp.] |
| MBL7989353.1 | AAA+ATPase+wHTH+Alpha+Beta-domain+wHTH+RRM | Type-1_BR.RM.like.systems | <-REase<-?<-?<-?\|\|ORC-CDC6+REase-><-DUF499* | 945 | Bacteria>FCB_group>Bacteroidota/Chlorobiota_group>Chlorobiota>unclassified_Chlorobiota | GCA_016788565.1 | Chlorobiota_bacterium | JNJ94_14925 | 66381..69218 | 2268192 | ATP-binding-protein-[Chlorobi-bacterium] |
| KXB97080.1 | AAA+ATPase+wHTH+Alpha+Beta-domain+wHTH+RRM | Type-1_BR.RM.like.systems | TUDOR+SF1-SF2-Helicase+iREase+CTD+REase->REase->?->METHYLASE->DUF499*-> | 876 | Bacteria>FCB_group>Bacteroidota/Chlorobiota_group>Chlorobiota>unclassified_Chlorobiota | GCA_001302225.1 | Chlorobi_bacterium_NICIL-2 | AA908_08510 | 7880..10510 | 1662223 | hypothetical-protein-AA908_08510-[Chlorobi-bacterium-NICIL-2] |
| MBV6446375.1 | AAA+ATPase+wHTH+Alpha+Beta-domain+wHTH+RRM | Type-1_BR.RM.like.systems | HTH->VirB11+VirB4-FtsK->?->?->?-><-?\|\|DUF499*-> | 771 | Bacteria>FCB_group>Bacteroidota/Chlorobiota_group>Ignavibacteriota>Ignavibacteria | GCA_019187245.1 | Ignavibacteriaceae_bacterium | IFNCLDLE_02673 | 21016..23331 | 2651163 | hypothetical-protein-[Ignavibacteriaceae-bacterium] |
| MCW8961935.1 | AAA+ATPase+wHTH+Alpha+Beta-domain+wHTH+RRM | Type-1_BR.RM.like.systems | METHYLASE-p->HEPN->DUF499*-> | 543 | Bacteria>FCB_group>Bacteroidota/Chlorobiota_group>Ignavibacteriota>Ignavibacteria | GCA_026196095.1 | Ignavibacteriaceae_bacterium | OQK29_10260 | 1268..2896 | 2651163 | MAG:-DUF499-domain-containing-protein,-partial-[Ignavibacteriaceae-bacterium] |
| BDQ01547.1 | AAA+ATPase+wHTH+Alpha+Beta-domain+wHTH+RRM | Type-1_BR.RM.like.systems | TPR+TPR->SF1-SF2-Helicase->NERD+SF1-SF2-Helicase->TUDOR+SF1-SF2-Helicase+iREase+CTD+REase->METHYLASE-p->METHYLASE-p->DUF499*->HKD-DNase->?->?->?->SIR2->HEPN-> | 848 | Bacteria>FCB_group>Bacteroidota/Chlorobiota_group>Ignavibacteriota>Ignavibacteria | GCA_025998815.1 | Ignavibacterium_sp. | KatS3mg037_0122 | 160550..163096 | 2651167 | MAG:-hypothetical-protein-KatS3mg037_0122-[Ignavibacterium-sp.] |
| MCR4416749.1 | AAA+ATPase+wHTH+Alpha+Beta-domain+wHTH+RRM | Type-1_BR.RM.like.systems | Cas_Cmr3->?->?->?->?->?-><-DUF499*<-HEPN<-PolB<-METHYLASE<-TUDOR+SF1-SF2-Helicase+iREase+CTD+REase<-tRNA\|\|?->Peptidase_M16+Peptidase_M16_C-> | 847 | Bacteria>FCB_group>Bacteroidota/Chlorobiota_group>Ignavibacteriota>Ignavibacteria | GCA_024653255.1 | Ignavibacteria_bacterium | NUV92_03270 | 797868..800411 | 2053306 | DUF499-domain-containing-protein-[Ignavibacteria-bacterium] |
| MCX6133013.1 | AAA+ATPase+wHTH+Alpha+Beta-domain+wHTH+RRM | Type-1_BR.RM.like.systems | <-DUF932<-?\|\|TUDOR+SF1-SF2-Helicase+iREase+CTD+REase->METHYLASE->REase->DUF499*->?->DUF488-l-> | 972 | Bacteria>FCB_group>Bacteroidota/Chlorobiota_group>Ignavibacteriota>Ignavibacteria | GCA_026390595.1 | Ignavibacteriales_bacterium | NTU47_04275 | 82490..85408 | 2049428 | MAG:-DUF499-domain-containing-protein-[Ignavibacteriales-bacterium] |
| WP_014560100.1 | AAA+ATPase+wHTH+Alpha+Beta-domain+wHTH+RRM | Type-1_BR.RM.like.systems | HNH->?->tRNA->tRNA-><-?<-DOC+HTH<-DUF499*<-AbiEii<-BirA-HTH<-METHYLASE<-TUDOR+SF1-SF2-Helicase+iREase+CTD+REase | 849 | Bacteria>FCB_group>Bacteroidota/Chlorobiota_group>Ignavibacteriota>Ignavibacteria | GCF_000258405.1 | Ignavibacterium_album | IALB_RS06070 | 1399560..1402109 | 591197 | DUF499-domain-containing-protein-[Ignavibacterium-album] |
| MCZ7603144.1 | AAA+ATPase+wHTH+Alpha+Beta-domain+wHTH+RRM | Type-1_BR.RM.like.systems | TUDOR+SF1-SF2-Helicase+iREase+CTD+REase->METHYLASE->DUF499*-><-?<-UbE1+JAB<-?\|\|?-><-RVT | 843 | Bacteria>FCB_group>Bacteroidota/Chlorobiota_group>Ignavibacteriota>Ignavibacteria | GCA_027493725.1 | Melioribacteraceae_bacterium | M5R37_09760 | 2173982..2176513 | 2614255 | MAG:-DUF499-domain-containing-protein-[Melioribacteraceae-bacterium] |
| MCX7909406.1 | AAA+ATPase+wHTH+Alpha+Beta-domain+wHTH+RRM | Type-1_BR.RM.like.systems | METHYLASE-p->DUF499*->?-><-?<-SF1-SF2-Helicase | 884 | Bacteria>FCB_group>Bacteroidota/Chlorobiota_group>Ignavibacteriota>Ignavibacteria | GCA_026416955.1 | Ignavibacteria_bacterium | N2560_07820 | 715..3369 | 2053306 | MAG:-DUF499-domain-containing-protein-[Ignavibacteria-bacterium] |
| MBI3194966.1 | AAA+ATPase+wHTH+Alpha+Beta-domain+wHTH+RRM | Type-1_BR.RM.like.systems | <-DUF499*<-AbiEii<-BirA-HTH<-METHYLASE<-METHYLASE-p<-TUDOR+SF1-SF2-Helicase+iREase+CTD+REase-p | 312 | Bacteria>FCB_group>Bacteroidota/Chlorobiota_group>Ignavibacteriota>unclassified_Ignavibacteriota | GCA_016191715.1 | Ignavibacteriota_bacterium | HYZ34_10965 | 1..937 | 2026749 | ATP-binding-protein-[Ignavibacteriota-bacterium] |
| MBI5470879.1 | AAA+ATPase+wHTH+Alpha+Beta-domain+wHTH+RRM | Type-1_BR.RM.like.systems | <-DUF499*<-METHYLASE<-JAB<-UbE1+JAB<-SMODS<-REase+SAVED<-TUDOR+SF1-SF2-Helicase+iREase+CTD+REase | 600 | Bacteria>FCB_group>Bacteroidota/Chlorobiota_group>Ignavibacteriota>unclassified_Ignavibacteriota | GCA_016218065.1 | Ignavibacteriota_bacterium | HY961_00895 | 1..1799 | 2026749 | ATP-binding-protein-[Ignavibacteriota-bacterium] |
| WP_178139398.1 | AAA+ATPase+wHTH+Alpha+Beta-domain+wHTH+RRM | Type-1_BR.RM.like.systems | <-HDRnase+SF1-SF2-Helicase<-35exo<-Peptidase_M20\|\|?->?->tRNA-><-TUDOR+SF1-SF2-Helicase+iREase+CTD+REase<-DUF499*<-MORC+DUF3684+REase<-METHYLASE<-35exo | 1042 | Bacteria>FCB_group>Bacteroidota/Chlorobiota_group>Rhodothermota>Rhodothermia | GCF_900142415.1 | Rhodothermus_profundi | BUA15_RS08290 | 58182..61310 | 633813 | DUF499-domain-containing-protein-[Rhodothermus-profundi] |
| SHK67770.1 | AAA+ATPase+wHTH+Alpha+Beta-domain+wHTH+RRM | Type-1_BR.RM.like.systems | <-HDRnase+SF1-SF2-Helicase<-35exo<-Peptidase_M20\|\|?->?->tRNA-><-TUDOR+SF1-SF2-Helicase+iREase+CTD+REase<-DUF499*<-MORC+DUF3684+REase<-METHYLASE<-35exo | 1047 | Bacteria>FCB_group>Bacteroidota/Chlorobiota_group>Rhodothermota>Rhodothermia | GCA_900142415.1 | Rhodothermus_profundi | SAMN04488087_1677 | 58182..61325 | 633813 | Predicted-ATPase,-AAA+-superfamily-[Rhodothermus-profundi] |
| WP_259054315.1 | AAA+ATPase+wHTH+Alpha+Beta-domain+wHTH+RRM | Type-1_BR.RM.like.systems | TUDOR+SF1-SF2-Helicase+iREase+CTD+REase->METHYLASE->SbcC->?->DUF499*->?->McrC-NTD+REase->?-><-MULE-TRANSPOSASE<-REase | 921 | Bacteria>FCB_group>Bacteroidota/Chlorobiota_group>Rhodothermota>Rhodothermia | GCF_024809575.1 | Salinibacter_ruber | GGP67_RS06675 | 1589032..1591797 | 146919 | DUF499-domain-containing-protein-[Salinibacter-ruber] |
| MYH11281.1 | AAA+ATPase+wHTH+Alpha+Beta-domain+wHTH+RRM | Type-1_BR.RM.like.systems | TUDOR+SF1-SF2-Helicase+iREase+CTD+REase->wHTH+REase->METHYLASE->OLD-ABC->PIN->DUF499*-><-ABhydrolase | 924 | Bacteria>FCB_group>Bacteroidota/Chlorobiota_group>Rhodothermota>Rhodothermia | GCA_009842805.1 | Rhodothermaceae_bacterium | F4191_00350 | 10395..13169 | 2026787 | ATP-binding-protein-[Rhodothermaceae-bacterium] |
| MXZ58767.1 | AAA+ATPase+wHTH+Alpha+Beta-domain+wHTH+RRM | Type-1_BR.RM.like.systems | ABhydrolase-><-DUF499*<-PIN<-OLD-ABC<-METHYLASE<-wHTH+REase<-TUDOR+SF1-SF2-Helicase+iREase+CTD+REase | 924 | Bacteria>FCB_group>Bacteroidota/Chlorobiota_group>Rhodothermota>Rhodothermia | GCA_009837125.1 | Rhodothermaceae_bacterium | F4Z16_09865 | 2884..5658 | 2026787 | ATP-binding-protein-[Rhodothermaceae-bacterium] |
| MXX97465.1 | AAA+ATPase+wHTH+Alpha+Beta-domain+wHTH+RRM | Type-1_BR.RM.like.systems | TUDOR+SF1-SF2-Helicase+iREase+CTD+REase->wHTH+REase->METHYLASE->OLD-ABC->PIN->DUF499*-><-ABhydrolase | 907 | Bacteria>FCB_group>Bacteroidota/Chlorobiota_group>Rhodothermota>Rhodothermia | GCA_009838195.1 | Rhodothermaceae_bacterium | F4Y64_07620 | 10395..13118 | 2026787 | ATP-binding-protein-[Rhodothermaceae-bacterium] |
| MYC03829.1 | AAA+ATPase+wHTH+Alpha+Beta-domain+wHTH+RRM | Type-1_BR.RM.like.systems | ABhydrolase-><-DUF499*<-PIN<-OLD-ABC<-wHTH+REase<-TUDOR+SF1-SF2-Helicase+iREase+CTD+REase | 907 | Bacteria>FCB_group>Bacteroidota/Chlorobiota_group>Rhodothermota>Rhodothermia | GCA_009840265.1 | Rhodothermaceae_bacterium | F4X61_04285 | 3864..6587 | 2026787 | ATP-binding-protein-[Rhodothermaceae-bacterium] |
| MXW15222.1 | AAA+ATPase+wHTH+Alpha+Beta-domain+wHTH+RRM | Type-1_BR.RM.like.systems | TUDOR+SF1-SF2-Helicase+iREase+CTD+REase->wHTH+REase->OLD-ABC->PIN->DUF499*-><-ABhydrolase | 924 | Bacteria>FCB_group>Bacteroidota/Chlorobiota_group>Rhodothermota>Rhodothermia | GCA_009835915.1 | Rhodothermaceae_bacterium | F4Z76_09955 | 11969..14743 | 2026787 | ATP-binding-protein-[Rhodothermaceae-bacterium] |
| WP_183956978.1 | AAA+ATPase+wHTH+Alpha+Beta-domain+wHTH+RRM | Type-1_BR.RM.like.systems | <-SF1-SF2-Helicase\|\|?->?->TUDOR+SF1-SF2-Helicase+iREase+CTD+REase->METHYLASE->DUF499*-> | 917 | Bacteria>FCB_group>Bacteroidota/Chlorobiota_group>Rhodothermota>Rhodothermia | GCF_014197195.1 | Salinibacter_ruber | GGQ18_RS08915 | 173852..176605 | 146919 | DUF499-domain-containing-protein-[Salinibacter-ruber] |
| MCP4581764.1 | AAA+ATPase+wHTH+Alpha+Beta-domain+wHTH+RRM | Type-1_BR.RM.like.systems | HEPN->?->DOC+DOC->TUDOR+SF1-SF2-Helicase+iREase+CTD+REase->METHYLASE->DUF499*->?-><-ABC-ATPase<-ABC-ATPase+SbcC<-ABC-ATPase+SbcC<-?<-METHYLASE | 978 | Bacteria>FCB_group>candidate_division_Zixibacteria>unclassified_candidate_division_Zixibacteria | GCA_024231855.1 | candidate_division_Zixibacteria_bacterium | GY839_09100 | 16694..19630 | 2053527 | ATP-binding-protein-[candidate-division-Zixibacteria-bacterium] |
| HCM14507.1 | AAA+ATPase+wHTH+Alpha+Beta-domain+wHTH+RRM | Type-1_BR.RM.like.systems | HIN-HTH->?->METHYLASE-p->MarR-HTH->?->DUF499-p->DUF499*->MORC+DUF3684+REase->?->?->?->?->Peptidase-> | 736 | Bacteria>FCB_group>Candidatus_Cloacimonadota>Candidatus_Cloacimonadia>Candidatus_Cloacimonadales | GCA_003527465.1 | Candidatus_Cloacimonas_sp. | DHW79_00910 | 17568..19778 | 2053534 | AAA-family-ATPase-[Candidatus-Cloacimonas-sp.] |
| NLH93351.1 | AAA+ATPase+wHTH+Alpha+Beta-domain+wHTH+RRM | Type-1_BR.RM.like.systems | HEPN->TUDOR+SF1-SF2-Helicase+iREase+CTD+REase->ZDNA-HTH->METHYLASE->DUF499*->?->SF1-SF2-Helicase-><-DUF772+DDE_Tnp_1 | 945 | Bacteria>FCB_group>Candidatus_Cloacimonadota>unclassified_Candidatus_Cloacimonadota | GCA_012516855.1 | Candidatus_Cloacimonadota_bacterium | GX466_03915 | 18260..21097 | 2030808 | ATP-binding-protein-[Candidatus-Cloacimonetes-bacterium] |
| PKN79833.1 | AAA+ATPase+wHTH+Alpha+Beta-domain+wHTH+RRM | Type-1_BR.RM.like.systems | <-ParB+HNH-p<-REase<-NERD<-?<-DUF499*<-METHYLASE<-TUDOR+SF1-SF2-Helicase+iREase+CTD+REase | 946 | Bacteria>FCB_group>Candidatus_Cloacimonadota>unclassified_Candidatus_Cloacimonadota | GCA_002841975.1 | Candidatus_Cloacimonetes_bacterium_HGW-Cloacimonetes-1 | CVU48_04095 | 13888..16728 | 2013733 | hypothetical-protein-CVU48_04095-[Candidatus-Cloacimonetes-bacterium-HGW-Cloacimonetes-1] |
| MCK9584245.1 | AAA+ATPase+wHTH+Alpha+Beta-domain+wHTH+RRM | Type-1_BR.RM.like.systems | DUF499*->OLD-ABC->REase->REase+nSTAND3-><-SF1-SF2-Helicase | 677 | Bacteria>FCB_group>Candidatus_Cloacimonadota>unclassified_Candidatus_Cloacimonadota | GCA_023228125.1 | Candidatus_Cloacimonadota_bacterium | MOR69_04960 | 1..2035 | 2030808 | DUF499-domain-containing-protein-[Candidatus-Cloacimonadota-bacterium] |
| MBW7876631.1 | AAA+ATPase+wHTH+Alpha+Beta-domain+wHTH+RRM | Type-1_BR.RM.like.systems | <-DUF499*<-OLD-ABC<-?<-METHYLASE<-JAB<-UbE1+JAB<-SMODS | 944 | Bacteria>FCB_group>Candidatus_Cloacimonadota>unclassified_Candidatus_Cloacimonadota | GCA_019454875.1 | Candidatus_Cloacimonadota_bacterium | H3C47_11655 | 180..3014 | 2030808 | ATP-binding-protein-[Candidatus-Cloacimonetes-bacterium] |
| MCB5288062.1 | AAA+ATPase+wHTH+Alpha+Beta-domain+wHTH+RRM | Type-1_BR.RM.like.systems | <-SbcC<-DUF499*<-Schlafen+HATPase+HTH<-METHYLASE | 940 | Bacteria>FCB_group>Candidatus_Cloacimonadota>unclassified_Candidatus_Cloacimonadota | GCA_020532145.1 | Candidatus_Cloacimonadota_bacterium | LHW64_09695 | 2385..5207 | 2030808 | DUF499-domain-containing-protein-[Candidatus-Cloacimonetes-bacterium] |
| MCB5253275.1 | AAA+ATPase+wHTH+Alpha+Beta-domain+wHTH+RRM | Type-1_BR.RM.like.systems | SF1-SF2-Helicase-><-REase<-NERD<-DUF499* | 944 | Bacteria>FCB_group>Candidatus_Cloacimonadota>unclassified_Candidatus_Cloacimonadota | GCA_020532345.1 | Candidatus_Cloacimonadota_bacterium | LHW51_09920 | 5656..8490 | 2030808 | DUF499-domain-containing-protein-[Candidatus-Cloacimonetes-bacterium] |
| MCB5262479.1 | AAA+ATPase+wHTH+Alpha+Beta-domain+wHTH+RRM | Type-1_BR.RM.like.systems | <-DUF499*<-Schlafen+HATPase+HTH<-METHYLASE-p | 826 | Bacteria>FCB_group>Candidatus_Cloacimonadota>unclassified_Candidatus_Cloacimonadota | GCA_020532425.1 | Candidatus_Cloacimonadota_bacterium | LHW53_07100 | 1..2478 | 2030808 | DUF499-domain-containing-protein-[Candidatus-Cloacimonetes-bacterium] |
| MCB5246762.1 | AAA+ATPase+wHTH+Alpha+Beta-domain+wHTH+RRM | Type-1_BR.RM.like.systems | METHYLASE->OLD-ABC->REase->DUF499*-> | 957 | Bacteria>FCB_group>Candidatus_Cloacimonadota>unclassified_Candidatus_Cloacimonadota | GCA_020532215.1 | Candidatus_Cloacimonadota_bacterium | LHW57_01885 | 3263..6136 | 2030808 | DUF499-domain-containing-protein-[Candidatus-Cloacimonetes-bacterium] |
| MCK9335735.1 | AAA+ATPase+wHTH+Alpha+Beta-domain+wHTH+RRM | Type-1_BR.RM.like.systems | METHYLASE-p->BirA-HTH->DUF499*-> | 836 | Bacteria>FCB_group>Candidatus_Cloacimonadota>unclassified_Candidatus_Cloacimonadota | GCA_023230025.1 | Candidatus_Cloacimonadota_bacterium | MOQ99_10545 | 1838..4346 | 2030808 | DUF499-domain-containing-protein-[Candidatus-Cloacimonetes-bacterium] |
| MCB5268142.1 | AAA+ATPase+wHTH+Alpha+Beta-domain+wHTH+RRM | Type-1_BR.RM.like.systems | <-DUF499*<-BirA-HTH<-METHYLASE-p | 828 | Bacteria>FCB_group>Candidatus_Cloacimonadota>unclassified_Candidatus_Cloacimonadota | GCA_020532165.1 | Candidatus_Cloacimonadota_bacterium | LHW46_08560 | 1..2483 | 2030808 | DUF499-domain-containing-protein-[Candidatus-Cloacimonetes-bacterium] |
| MBC8384565.1 | AAA+ATPase+wHTH+Alpha+Beta-domain+wHTH+RRM | Type-1_BR.RM.like.systems | <-Radical-SAM<-?\|\|zf-ISL3+HTH_Tnp_ISL3+DDE_Tnp_ISL3-><-DUF499* | 861 | Bacteria>FCB_group>Candidatus_Cloacimonadota>unclassified_Candidatus_Cloacimonadota | GCA_014381995.1 | Candidatus_Cloacimonadota_bacterium | H8E57_03485 | 3984..6569 | 2030808 | ATP-binding-protein-[Candidatus-Cloacimonetes-bacterium] |
| MCA1901348.1 | AAA+ATPase+wHTH+Alpha+Beta-domain+wHTH+RRM | Type-1_BR.RM.like.systems | <-DUF499*<-METHYLASE<-TUDOR+SF1-SF2-Helicase+iREase+CTD+REase-p | 845 | Bacteria>FCB_group>Candidatus_Hydrogenedentes>Hydrogenedentia>unclassified_Candidatus_Hydrogenedens | GCA_020216745.1 | Candidatus_Hydrogenedens_sp. | LDL53_03900 | 3377..5914 | 2654184 | DUF499-domain-containing-protein-[Candidatus-Hydrogenedens-sp.] |
| MBP7427878.1 | AAA+ATPase+wHTH+Alpha+Beta-domain+wHTH+RRM | Type-1_BR.RM.like.systems | ABC-ATPase+SbcC->ABC-ATPase+SbcC->ABC-ATPase+SbcC-><-DUF499*<-?<-RNaseH<-METHYLASE<-?<-TUDOR+SF1-SF2-Helicase+iREase+CTD+REase<-ZnR+TOPRIM+primpol+HTH | 985 | Bacteria>FCB_group>Candidatus_Hydrogenedentes>unclassified_Candidatus_Hydrogenedentes | GCA_017997415.1 | Candidatus_Hydrogenedentes_bacterium | KBC05_00520 | 30442..33399 | 2030809 | ATP-binding-protein-[Candidatus-Hydrogenedentes-bacterium] |
| MBW7864504.1 | AAA+ATPase+wHTH+Alpha+Beta-domain+wHTH+RRM | Type-1_BR.RM.like.systems | PolB->HEPN->HEPN->METHYLASE->?->?->DUF499*->?->?->XIS-HTH->?->Primpol+PriCT1->HTH-> | 944 | Bacteria>FCB_group>Candidatus_Hydrogenedentes>unclassified_Candidatus_Hydrogenedentes | GCA_019455225.1 | Candidatus_Hydrogenedentes_bacterium | H3C30_08835 | 18999..21833 | 2030809 | ATP-binding-protein-[Candidatus-Hydrogenedentes-bacterium] |
| MBI5093542.1 | AAA+ATPase+wHTH+Alpha+Beta-domain+wHTH+RRM | Type-1_BR.RM.like.systems | DUF4065->HEPN->REase->METHYLASE-p->METHYLASE->?->DUF499*->?->?->HISKIN+MORC->REC-> | 944 | Bacteria>FCB_group>Candidatus_Hydrogenedentes>unclassified_Candidatus_Hydrogenedentes | GCA_016215105.1 | Candidatus_Hydrogenedentes_bacterium | HZB26_14005 | 11856..14690 | 2030809 | ATP-binding-protein-[Candidatus-Hydrogenedentes-bacterium] |
| NLV46598.1 | AAA+ATPase+wHTH+Alpha+Beta-domain+wHTH+RRM | Type-1_BR.RM.like.systems | <-DUF499*<-?<-METHYLASE<-?<-?<-ZDNA-HTH<-TUDOR+SF1-SF2-Helicase+iREase+CTD+REase | 985 | Bacteria>FCB_group>Candidatus_Hydrogenedentes>unclassified_Candidatus_Hydrogenedentes | GCA_012730105.1 | Candidatus_Hydrogenedentes_bacterium | GXY07_19110 | 217..3174 | 2030809 | ATP-binding-protein-[Candidatus-Hydrogenedentes-bacterium] |
| WP_072150613.1 | AAA+ATPase+wHTH+Alpha+Beta-domain+wHTH+RRM | Type-1_BR.RM.like.systems | TUDOR+SF1-SF2-Helicase+iREase+CTD+REase-p->METHYLASE->DUF499*-><-tRNA\|\|?->PIN-> | 852 | Bacteria>FCB_group>Candidatus_Kryptonia>Kryptobacter | GCF_001458495.1 | Candidatus_Kryptobacter_tengchongensis | AT691_RS06200 | 4768..7326 | 1643429 | AAA-family-ATPase-[Candidatus-Kryptobacter-tengchongensis] |
| WP_140945780.1 | AAA+ATPase+wHTH+Alpha+Beta-domain+wHTH+RRM | Type-1_BR.RM.like.systems | DUF499*-><-?<-?<-?<-?<-ABC-ATPase+SbcC | 837 | Bacteria>FCB_group>Candidatus_Kryptonia>Thermokryptus | GCF_900070205.1 | Candidatus_Thermokryptus_mobilis | FKZ43_RS10140 | 610..3123 | 1643428 | DUF499-domain-containing-protein-[Candidatus-Thermokryptus-mobilis] |
| MCD6099133.1 | AAA+ATPase+wHTH+Alpha+Beta-domain+wHTH+RRM | Type-1_BR.RM.like.systems | REase->DUF499*->OBFOLD-NUCLEASE->SIR2+SIR2->PIWI-> | 932 | Bacteria>FCB_group>Candidatus_Marinimicrobia>unclassified_Candidatus_Marinimicrobia | GCA_021158485.1 | Candidatus_Marinimicrobia_bacterium | J7K33_00965 | 724..3522 | 2026760 | ATP-binding-protein-[Candidatus-Marinimicrobia-bacterium] |
| MCD6101177.1 | AAA+ATPase+wHTH+Alpha+Beta-domain+wHTH+RRM | Type-1_BR.RM.like.systems | TUDOR+SF1-SF2-Helicase+iREase+CTD+REase->METHYLASE->DUF499*-> | 915 | Bacteria>FCB_group>Candidatus_Marinimicrobia>unclassified_Candidatus_Marinimicrobia | GCA_021158485.1 | Candidatus_Marinimicrobia_bacterium | J7K33_11430 | 6522..9269 | 2026760 | ATP-binding-protein-[Candidatus-Marinimicrobia-bacterium] |
| MBO8151135.1 | AAA+ATPase+wHTH+Alpha+Beta-domain+wHTH+RRM | Type-1_BR.RM.like.systems | TUDOR+SF1-SF2-Helicase+iREase+CTD+REase->?->PIN->DUF499*-> | 854 | Bacteria>FCB_group>Candidatus_Marinimicrobia>unclassified_Candidatus_Marinimicrobia | GCA_017656455.1 | Candidatus_Marinimicrobia_bacterium | HOZ30_01400 | 332624..335188 | 2026760 | ATP-binding-protein-[Candidatus-Marinimicrobia-bacterium] |
| MBN1128503.1 | AAA+ATPase+wHTH+Alpha+Beta-domain+wHTH+RRM | Type-1_BR.RM.like.systems | <-DUF499*<-URI-DNase<-METHYLASE-p | 944 | Bacteria>FCB_group>Fibrobacterota>Chitinispirillia>Chitinispirillales | GCA_016926315.1 | Chitinispirillaceae_bacterium | JXA71_05930 | 3630..6464 | 2813024 | ATP-binding-protein-[Chitinispirillaceae-bacterium] |
| MBN1601227.1 | AAA+ATPase+wHTH+Alpha+Beta-domain+wHTH+RRM | Type-1_BR.RM.like.systems | <-PIN<-OLD-ABC<-?<-?<-DUF499*<-wHTH+REase<-?<-METHYLASE<-TUDOR+SF1-SF2-Helicase+iREase+CTD+REase<-REase | 922 | Bacteria>FCB_group>Fibrobacterota>Chitinispirillia>Chitinispirillales | GCA_016929595.1 | Chitinispirillaceae_bacterium | JW915_06440 | 4163..6931 | 2813024 | ATP-binding-protein-[Chitinispirillaceae-bacterium] |
| MBR2089170.1 | AAA+ATPase+wHTH+Alpha+Beta-domain+wHTH+RRM | Type-1_BR.RM.like.systems | <-DUF4417<-PIN<-HTH+MPTase<-?<-?<-HEPN<-DUF499*<-METHYLASE<-TUDOR+SF1-SF2-Helicase+iREase+CTD+REase-p | 933 | Bacteria>FCB_group>Fibrobacterota>Fibrobacteria>Fibrobacterales | - | Fibrobacter_sp. | IJ905_02095 | 11314..14115 | 35828 | ATP-binding-protein-[Fibrobacter-sp.] |
| MBR1746391.1 | AAA+ATPase+wHTH+Alpha+Beta-domain+wHTH+RRM | Type-1_BR.RM.like.systems | <-DUF499*<-METHYLASE<-TUDOR+SF1-SF2-Helicase+iREase+CTD+REase-p | 850 | Bacteria>FCB_group>Fibrobacterota>Fibrobacteria>Fibrobacterales | GCA_017476345.1 | Fibrobacter_sp. | IJ734_10590 | 1..2551 | 35828 | ATP-binding-protein-[Fibrobacter-sp.] |
| MYC87147.1 | AAA+ATPase+wHTH+Alpha+Beta-domain+wHTH+RRM | Type-1_BR.RM.like.systems | TUDOR+SF1-SF2-Helicase+iREase+CTD+REase->OLD-ABC-p->OLD-ABC-p->REase->DUF499*->?-><-?<-?<-?<-SF1-SF2-Helicase<-HKD-DNase<-DUF6361 | 922 | Bacteria>FCB_group>Gemmatimonadota>Gemmatimonadetes>Candidatus_Palauibacterales | GCA_009841015.1 | Candidatus_Palauibacter_denitrificans | F4X22_02775 | 10288..13056 | 3056734 | ATP-binding-protein-[Gemmatimonadales-bacterium] |
| WP_284352185.1 | AAA+ATPase+wHTH+Alpha+Beta-domain+wHTH+RRM | Type-1_BR.RM.like.systems | TUDOR+SF1-SF2-Helicase+iREase+CTD+REase->METHYLASE->DUF499*->?->\|\|?-><-ParB+Schlafen\|\|?->?->WYL-> | 940 | Bacteria>FCB_group>Gemmatimonadota>Gemmatimonadetes>Gemmatimonadales | GCF_030159095.1 | Roseisolibacter_agri | rosag_RS21265 | 15472..18294 | 2014610 | DUF499-domain-containing-protein-[Roseisolibacter-agri] |
| MXW55543.1 | AAA+ATPase+wHTH+Alpha+Beta-domain+wHTH+RRM | Type-1_BR.RM.like.systems | <-DUF499*<-HEPN-p<-METHYLASE-p | 977 | Bacteria>FCB_group>Gemmatimonadota>Gemmatimonadetes>Gemmatimonadales | GCA_009836275.1 | Gemmatimonadales_bacterium | F4Z59_02325 | 190..3123 | 2448054 | ATP-binding-protein-[Gemmatimonadales-bacterium] |
| MYA34704.1 | AAA+ATPase+wHTH+Alpha+Beta-domain+wHTH+RRM | Type-1_BR.RM.like.systems | HEPN->TUDOR+SF1-SF2-Helicase+iREase+CTD+REase->METHYLASE->OLD-ABC->TOPRIM->ParB+HNH+Schlafen->DUF499*-><-?\|\|HTH->MPTase->?->SMODS-><-ABhydrolase | 928 | Bacteria>FCB_group>Gemmatimonadota>Gemmatimonadetes>Gemmatimonadales | GCA_009838765.1 | Gemmatimonadales_bacterium | F4Y33_13800 | 70561..73347 | 2448054 | ATP-binding-protein-[Gemmatimonadales-bacterium] |
| MQA92114.1 | AAA+ATPase+wHTH+Alpha+Beta-domain+wHTH+RRM | Type-1_BR.RM.like.systems | NTP_transf-><-?<-?<-MPTase\|\|HTH+UbE1-><-DUF499* | 920 | Bacteria>FCB_group>Gemmatimonadota>Gemmatimonadetes>Gemmatimonadales | GCA_009380025.1 | Gemmatimonas_sp. | GEU90_18125 | 8154..10916 | 1962908 | DUF499-domain-containing-protein-[Gemmatimonas-sp.] |
| MXX77582.1 | AAA+ATPase+wHTH+Alpha+Beta-domain+wHTH+RRM | Type-1_BR.RM.like.systems | <-DUF499*<-HicB_lk_antitox<-METHYLASE-p | 649 | Bacteria>FCB_group>Gemmatimonadota>Gemmatimonadetes>Gemmatimonadales | GCA_009836785.1 | Gemmatimonadales_bacterium | F4Z33_01020 | 1..1948 | 2448054 | ATP-binding-protein-[Gemmatimonadales-bacterium] |
| MXW55615.1 | AAA+ATPase+wHTH+Alpha+Beta-domain+wHTH+RRM | Type-1_BR.RM.like.systems | METHYLASE->HicB_lk_antitox->DUF499*-> | 427 | Bacteria>FCB_group>Gemmatimonadota>Gemmatimonadetes>Gemmatimonadales | GCA_009836275.1 | Gemmatimonadales_bacterium | F4Z59_02700 | 2593..3872 | 2448054 | ATP-binding-protein-[Gemmatimonadales-bacterium] |
| MDE2981419.1 | AAA+ATPase+wHTH+Alpha+Beta-domain+wHTH+RRM | Type-1_BR.RM.like.systems | ZDNA-HTH->SF1-SF2-Helicase->Schlafen+HATPase+HTH-><-TPR\|\|METHYLASE->REase->DUF499*->?->URI-DNase->HEPN-> | 922 | Bacteria>FCB_group>Gemmatimonadota>unclassified_Gemmatimonadota | GCA_028873815.1 | Gemmatimonadota_bacterium | OXU74_09510 | 42276..45044 | 2026742 | MAG:-DUF499-domain-containing-protein-[Gemmatimonadota-bacterium] |
| MYG34810.1 | AAA+ATPase+wHTH+Alpha+Beta-domain+wHTH+RRM | Type-1_BR.RM.like.systems | <-XIS-HTH\|\|LexA-protease->TUDOR+SF1-SF2-Helicase+iREase+CTD+REase->METHYLASE->DUF499*->Schlafen+HATPase+HTH-> | 963 | Bacteria>FCB_group>Gemmatimonadota>unclassified_Gemmatimonadota | GCA_009842515.1 | Gemmatimonadota_bacterium | F4205_04900 | 8108..10999 | 2026742 | ATP-binding-protein-[Gemmatimonadetes-bacterium] |
| MYB06457.1 | AAA+ATPase+wHTH+Alpha+Beta-domain+wHTH+RRM | Type-1_BR.RM.like.systems | McrC-NTD+REase-><-?\|\|?->?-><-DUF499*<-METHYLASE<-SF1-SF2-Helicase-p<-TUDOR+SF1-SF2-Helicase+iREase+CTD+REase | 946 | Bacteria>FCB_group>Gemmatimonadota>unclassified_Gemmatimonadota | GCA_009838975.1 | Gemmatimonadota_bacterium | F4Y24_08895 | 37066..39906 | 2026742 | ATP-binding-protein-[Gemmatimonadetes-bacterium] |
| MCY4604211.1 | AAA+ATPase+wHTH+Alpha+Beta-domain+wHTH+RRM | Type-1_BR.RM.like.systems | METHYLASE->METHYLASE-p->DUF433->PIN->METHYLASE-p->REase->DUF499*-> | 922 | Bacteria>FCB_group>Gemmatimonadota>unclassified_Gemmatimonadota | GCA_026713655.1 | Gemmatimonadota_bacterium | OXE49_08275 | 9321..12089 | 2026742 | MAG:-DUF499-domain-containing-protein-[Gemmatimonadetes-bacterium] |
| MDE2763067.1 | AAA+ATPase+wHTH+Alpha+Beta-domain+wHTH+RRM | Type-1_BR.RM.like.systems | <-HTH<-?\|\|?-><-?\|\|<-ORC-CDC6+REase<-?<-DUF499*<-MNS+ZDNA-HTH<-DDE_Tnp_1 | 934 | Bacteria>FCB_group>Gemmatimonadota>unclassified_Gemmatimonadota | GCA_028874985.1 | Gemmatimonadota_bacterium | OXX74_09825 | 20104..22908 | 2026742 | MAG:-DUF499-domain-containing-protein-[Gemmatimonadota-bacterium] |
| MDE2973202.1 | AAA+ATPase+wHTH+Alpha+Beta-domain+wHTH+RRM | Type-1_BR.RM.like.systems | TUDOR+SF1-SF2-Helicase+iREase+CTD+REase->wHTH+REase->METHYLASE->DUF499*-> | 933 | Bacteria>FCB_group>Gemmatimonadota>unclassified_Gemmatimonadota | GCA_028873825.1 | Gemmatimonadota_bacterium | OXU64_00560 | 8847..11648 | 2026742 | MAG:-DUF499-domain-containing-protein-[Gemmatimonadota-bacterium] |
| MYE70563.1 | AAA+ATPase+wHTH+Alpha+Beta-domain+wHTH+RRM | Type-1_BR.RM.like.systems | METHYLASE->METHYLASE-p->REase->DUF499*-> | 400 | Bacteria>FCB_group>Gemmatimonadota>unclassified_Gemmatimonadota | GCA_009841855.1 | Gemmatimonadota_bacterium | F4237_11010 | 2710..3911 | 2026742 | ATP-binding-protein-[Gemmatimonadota-bacterium] |
| MYB67636.1 | AAA+ATPase+wHTH+Alpha+Beta-domain+wHTH+RRM | Type-1_BR.RM.like.systems | <-DUF499*<-TOPRIM<-OLD-ABC<-HEPN<-PolB<-?<-TraJ-RHH | 924 | Bacteria>FCB_group>Gemmatimonadota>unclassified_Gemmatimonadota | GCA_009840005.1 | Gemmatimonadota_bacterium | F4X75_03955 | 7160..9934 | 2026742 | ATP-binding-protein-[Gemmatimonadetes-bacterium] |
| MBA4158336.1 | AAA+ATPase+wHTH+Alpha+Beta-domain+wHTH+RRM | Type-1_BR.RM.like.systems | HEPN->DUF499*->SF1-SF2-Helicase+wHTH+PRTase->?->?->XIS-HTH-> | 907 | Bacteria>FCB_group>Gemmatimonadota>unclassified_Gemmatimonadota | GCA_013820915.1 | Gemmatimonadota_bacterium | HOX65_12770 | 579..3302 | 2026742 | ATP-binding-protein-[Gemmatimonadetes-bacterium] |
| MDE2874623.1 | AAA+ATPase+wHTH+Alpha+Beta-domain+wHTH+RRM | Type-1_BR.RM.like.systems | <-DUF499*<-METHYLASE<-HKD-DNase+SF1-SF2-Helicase | 923 | Bacteria>FCB_group>Gemmatimonadota>unclassified_Gemmatimonadota | GCA_028874445.1 | Gemmatimonadota_bacterium | OXQ93_04215 | 979..3750 | 2026742 | MAG:-DUF499-domain-containing-protein-[Gemmatimonadota-bacterium] |
| MYE95017.1 | AAA+ATPase+wHTH+Alpha+Beta-domain+wHTH+RRM | Type-1_BR.RM.like.systems | <-DUF499*<-METHYLASE<-REase<-DNA_methylase<-HNH | 358 | Bacteria>FCB_group>Gemmatimonadota>unclassified_Gemmatimonadota | GCA_009841825.1 | Gemmatimonadota_bacterium | F4238_16835 | 1..1075 | 2026742 | ATP-binding-protein-[Gemmatimonadota-bacterium] |
| MCY3788288.1 | AAA+ATPase+wHTH+Alpha+Beta-domain+wHTH+RRM | Type-1_BR.RM.like.systems | REase->wHTH+REase->METHYLASE->REase->DUF499*-> | 918 | Bacteria>FCB_group>Gemmatimonadota>unclassified_Gemmatimonadota | GCA_026706505.1 | Gemmatimonadota_bacterium | OXH63_05815 | 5478..8234 | 2026742 | MAG:-DUF499-domain-containing-protein-[Gemmatimonadetes-bacterium] |
| MYA24522.1 | AAA+ATPase+wHTH+Alpha+Beta-domain+wHTH+RRM | Type-1_BR.RM.like.systems | <-DUF499*<-TOPRIM | 924 | Bacteria>FCB_group>Gemmatimonadota>unclassified_Gemmatimonadota | GCA_009836855.1 | Gemmatimonadota_bacterium | F4Z30_17850 | 5045..7819 | 2026742 | ATP-binding-protein-[Gemmatimonadetes-bacterium] |
| MDE2723058.1 | AAA+ATPase+wHTH+Alpha+Beta-domain+wHTH+RRM | Type-1_BR.RM.like.systems | <-DUF499*<-TOPRIM<-OLD-ABC<-HTH_3<-METHYLASE<-TOPRIM | 656 | Bacteria>FCB_group>Gemmatimonadota>unclassified_Gemmatimonadota | GCA_028875455.1 | Gemmatimonadota_bacterium | OXI59_06750 | 1..1968 | 2026742 | MAG:-DUF499-domain-containing-protein,-partial-[Gemmatimonadota-bacterium] |
| WP_171805995.1 | AAA+ATPase+wHTH+Alpha+Beta-domain+wHTH+RRM | Type-1_BR.RM.like.systems | TUDOR+SF1-SF2-Helicase+iREase+CTD+REase->OLD-ABC->HNH->METHYLASE->DUF499*->DNA-glycosylase->CHTH->?->LZ_Tnp_IS66+zf-IS66+DDE_Tnp_IS66+DDE_Tnp_IS66_C->DUF4096+DDE_Tnp_1+DDE_Tnp_1_2-> | 942 | Bacteria>Myxococcota>Myxococcia>Myxococcales>Cystobacterineae | GCF_013155195.1 | Corallococcus_exiguus | HPP06_RS00265 | 63781..66609 | 83462 | DUF499-domain-containing-protein-[Corallococcus-exiguus] |
| WP_174489000.1 | AAA+ATPase+wHTH+Alpha+Beta-domain+wHTH+RRM | Type-1_BR.RM.like.systems | <-METHYLASE+DNA_methylase\|\|tRNA-><-DNA-glycosylase<-?<-Pkinase<-REase<-DUF499*<-METHYLASE<-HNH<-OLD-ABC<-TUDOR+SF1-SF2-Helicase+iREase+CTD+REase<-XIS-HTH<-PAIREDC-HTH | 943 | Bacteria>Myxococcota>Myxococcia>Myxococcales>Cystobacterineae | GCF_013336725.1 | Myxococcus_sp._CA040A | HUA75_RS36310 | 164891..167722 | 2741738 | DUF499-domain-containing-protein-[Myxococcus-sp.-CA040A] |
| WP_121761665.1 | AAA+ATPase+wHTH+Alpha+Beta-domain+wHTH+RRM | Type-1_BR.RM.like.systems | TUDOR+SF1-SF2-Helicase+iREase+CTD+REase->METHYLASE->DUF499*->REase-> | 947 | Bacteria>Myxococcota>Myxococcia>Myxococcales>Cystobacterineae | GCF_003668955.1 | Corallococcus_sp._AB038B | D7Y04_RS31655 | 364596..367439 | 2316718 | DUF499-domain-containing-protein-[Corallococcus-sp.-AB038B] |
| WP_012098801.1 | AAA+ATPase+wHTH+Alpha+Beta-domain+wHTH+RRM | Type-1_BR.RM.like.systems | <-Phage_integrase<-XIS-HTH<-?<-wHTH+SAD+HNH<-HKD-DNase+SF1-SF2-Helicase+TRD<-DUF499*<-REase<-METHYLASE<-METHYLASE<-TUDOR+SF1-SF2-Helicase+iREase+CTD+REase\|\|DUF6361->HKD-DNase-> | 932 | Bacteria>Myxococcota>Myxococcia>Myxococcales>Cystobacterineae | GCF_000017505.1 | Anaeromyxobacter_sp._Fw109-5 | ANAE109_RS20460 | 4648491..4651289 | 404589 | DUF499-domain-containing-protein-[Anaeromyxobacter-sp.-Fw109-5] |
| MCG5052821.1 | AAA+ATPase+wHTH+Alpha+Beta-domain+wHTH+RRM | Type-1_BR.RM.like.systems | METHYLASE-p->DUF499-p->TUDOR+SF1-SF2-Helicase+iREase+CTD+REase->PIN->METHYLASE->METHYLASE-p->DUF499*->?-><-DDE_Tnp_ISAZ013<-?\|\|?->SIGMA-HTH-> | 941 | Bacteria>Myxococcota>Myxococcia>Myxococcales>unclassified_Myxococcales | GCA_022184915.1 | Myxococcales_bacterium | KA712_07655 | 10615..13440 | 2026763 | DUF499-domain-containing-protein-[Myxococcales-bacterium] |
| MBK6464838.1 | AAA+ATPase+wHTH+Alpha+Beta-domain+wHTH+RRM | Type-1_BR.RM.like.systems | <-NTP_transferase+MannoseP_isomer\|\|?-><-Peptidase_M4+Peptidase_M4_C<-?<-?<-?<-DUF499*<-?\|\|REase-><-METHYLASE-p | 959 | Bacteria>Myxococcota>Myxococcia>Myxococcales>unclassified_Myxococcales | GCA_016704525.1 | Myxococcales_bacterium | IPF92_28125 | 158068..160947 | 2026763 | ATP-binding-protein-[Myxococcales-bacterium] |
| MBL0196370.1 | AAA+ATPase+wHTH+Alpha+Beta-domain+wHTH+RRM | Type-1_BR.RM.like.systems | <-Peptidase_M4+Peptidase_M4_C<-?<-?<-?<-?<-DUF499*<-?<-?\|\|REase-p-> | 959 | Bacteria>Myxococcota>Myxococcia>Myxococcales>unclassified_Myxococcales | GCA_016720545.1 | Myxococcales_bacterium | IPQ09_19515 | 186242..189121 | 2026763 | ATP-binding-protein-[Myxococcales-bacterium] |
| MBI3184665.1 | AAA+ATPase+wHTH+Alpha+Beta-domain+wHTH+RRM | Type-1_BR.RM.like.systems | <-HAD<-?\|\|XIS-HTH->TUDOR+SF1-SF2-Helicase+iREase+CTD+REase->METHYLASE->REase->DUF499*->?->Radical-SAM->ParB->?->XIS-HTH-><-REase | 930 | Bacteria>Myxococcota>Myxococcia>Myxococcales>unclassified_Myxococcales | GCA_016191815.1 | Myxococcales_bacterium | HYZ28_21220 | 50523..53315 | 2026763 | ATP-binding-protein-[Myxococcales-bacterium] |
| MBL9006089.1 | AAA+ATPase+wHTH+Alpha+Beta-domain+wHTH+RRM | Type-1_BR.RM.like.systems | XIS-HTH->TUDOR+SF1-SF2-Helicase+iREase+CTD+REase->OLD-ABC->METHYLASE->METHYLASE->REase->DUF499*-> | 928 | Bacteria>Myxococcota>Myxococcia>Myxococcales>unclassified_Myxococcales | GCA_016794505.1 | Myxococcales_bacterium | JNJ46_17680 | 14814..17600 | 2026763 | ATP-binding-protein-[Myxococcales-bacterium] |
| MBF0104316.1 | AAA+ATPase+wHTH+Alpha+Beta-domain+wHTH+RRM | Type-1_BR.RM.like.systems | <-Radical-SAM\|\|METHYLASE->Methyltrans_RNA-><-TPR+TPR<-TPR+TPR<-?<-DUF499*<-Schlafen+HATPase+HTH<-METHYLASE<-METHYLASE-p<-TUDOR+SF1-SF2-Helicase+iREase+CTD+REase-p<-TPR+TPR | 375 | Bacteria>Myxococcota>Myxococcia>unclassified_Myxococcia | GCA_015228875.1 | Deltaproteobacteria_bacterium | HQM16_03205 | 33942..35069 | 2026735 | ATP-binding-protein-[Deltaproteobacteria-bacterium] |
| MBN2361667.1 | AAA+ATPase+wHTH+Alpha+Beta-domain+wHTH+RRM | Type-1_BR.RM.like.systems | TUDOR+SF1-SF2-Helicase+iREase+CTD+REase->PIN->METHYLASE->DUF499*->Schlafen->EVE-> | 935 | Bacteria>Myxococcota>Myxococcia>unclassified_Myxococcia | GCA_016934455.1 | Deltaproteobacteria_bacterium | JXR83_19595 | 9461..12268 | 2026735 | ATP-binding-protein-[Deltaproteobacteria-bacterium] |
| MBT6612314.1 | AAA+ATPase+wHTH+Alpha+Beta-domain+wHTH+RRM | Type-1_BR.RM.like.systems | <-DUF499*<-METHYLASE<-TUDOR+SF1-SF2-Helicase+iREase+CTD+REase | 423 | Bacteria>Myxococcota>Myxococcia>unclassified_Myxococcia | GCA_018697355.1 | Deltaproteobacteria_bacterium | HOB38_09410 | 1..1269 | 2026735 | ATP-binding-protein-[Deltaproteobacteria-bacterium] |
| MBT4265466.1 | AAA+ATPase+wHTH+Alpha+Beta-domain+wHTH+RRM | Type-1_BR.RM.like.systems | <-DUF499*<-METHYLASE<-TUDOR+SF1-SF2-Helicase+iREase+CTD+REase | 939 | Bacteria>Myxococcota>Myxococcia>unclassified_Myxococcia | GCA_018651965.1 | Deltaproteobacteria_bacterium | HOD85_14800 | 1158..3977 | 2026735 | ATP-binding-protein-[Deltaproteobacteria-bacterium] |
| MBT4088220.1 | AAA+ATPase+wHTH+Alpha+Beta-domain+wHTH+RRM | Type-1_BR.RM.like.systems | <-DUF499*<-METHYLASE<-TUDOR+SF1-SF2-Helicase+iREase+CTD+REase | 438 | Bacteria>Myxococcota>Myxococcia>unclassified_Myxococcia | GCA_018651115.1 | Deltaproteobacteria_bacterium | HOE30_06990 | 1..1314 | 2026735 | ATP-binding-protein-[Deltaproteobacteria-bacterium] |
| MCL5406608.1 | AAA+ATPase+wHTH+Alpha+Beta-domain+wHTH+RRM | Type-1_BR.RM.like.systems | DUF499*->REase->?->?-><-?<-?<-GDResolvase+Recombinase+ZnR | 564 | Bacteria>Myxococcota>Myxococcia>unclassified_Myxococcia | GCA_023484975.1 | Deltaproteobacteria_bacterium | M1398_07815 | 1..1696 | 2026735 | DUF499-domain-containing-protein-[Deltaproteobacteria-bacterium] |
| MCL5406869.1 | AAA+ATPase+wHTH+Alpha+Beta-domain+wHTH+RRM | Type-1_BR.RM.like.systems | METHYLASE->DUF499*->?-><-McrC-NTD+REase | 939 | Bacteria>Myxococcota>Myxococcia>unclassified_Myxococcia | GCA_023484975.1 | Deltaproteobacteria_bacterium | M1398_09200 | 2598..5417 | 2026735 | DUF499-domain-containing-protein-[Deltaproteobacteria-bacterium] |
| RLB11002.1 | AAA+ATPase+wHTH+Alpha+Beta-domain+wHTH+RRM | Type-1_BR.RM.like.systems | <-TUDOR+SF1-SF2-Helicase+iREase+CTD+REase-p<-Cluster35_66clades<-DUF499*<-METHYLASE-p | 1107 | Bacteria>Myxococcota>Myxococcia>unclassified_Myxococcia | GCA_003646755.1 | Deltaproteobacteria_bacterium | DRG27_02045 | 1419..4742 | 2026735 | hypothetical-protein-DRG27_02045-[Deltaproteobacteria-bacterium] |
| MCD6255997.1 | AAA+ATPase+wHTH+Alpha+Beta-domain+wHTH+RRM | Type-1_BR.RM.like.systems | DUF499->DUF499*->Cluster35_66clades->TUDOR+SF1-SF2-Helicase+iREase+CTD+REase-> | 839 | Bacteria>Myxococcota>Myxococcia>unclassified_Myxococcia | GCA_021163945.1 | Deltaproteobacteria_bacterium | J7J44_06980 | 851..3370 | 2026735 | DUF499-domain-containing-protein-[Deltaproteobacteria-bacterium] |
| MCD6255996.1 | AAA+ATPase+wHTH+Alpha+Beta-domain+wHTH+RRM | Type-1_BR.RM.like.systems | DUF499*->DUF499->Cluster35_66clades->TUDOR+SF1-SF2-Helicase+iREase+CTD+REase-> | 286 | Bacteria>Myxococcota>Myxococcia>unclassified_Myxococcia | GCA_021163945.1 | Deltaproteobacteria_bacterium | J7J44_06975 | 1..861 | 2026735 | ATP-binding-protein-[Deltaproteobacteria-bacterium] |
| RLB78014.1 | AAA+ATPase+wHTH+Alpha+Beta-domain+wHTH+RRM | Type-1_BR.RM.like.systems | <-HKD-DNase<-BirA-HTH<-?<-Phage_integrase\|\|DUF499*-> | 295 | Bacteria>Myxococcota>Myxococcia>unclassified_Myxococcia | GCA_003647245.1 | Deltaproteobacteria_bacterium | DRH24_15195 | 3532..4418 | 2026735 | hypothetical-protein-DRH24_15195,-partial-[Deltaproteobacteria-bacterium] |
| MBN1102533.1 | AAA+ATPase+wHTH+Alpha+Beta-domain+wHTH+RRM | Type-1_BR.RM.like.systems | <-DUF499*\|\|?->?->?-><-?\|\|NTP_transferase+MannoseP_isomer-> | 676 | Bacteria>Myxococcota>Myxococcia>unclassified_Myxococcia | GCA_016926135.1 | Deltaproteobacteria_bacterium | JXL84_03865 | 1..2027 | 2026735 | hypothetical-protein-[Deltaproteobacteria-bacterium] |
| NLN59460.1 | AAA+ATPase+wHTH+Alpha+Beta-domain+wHTH+RRM | Type-1_BR.RM.like.systems | <-SF1-SF2-helicase<-?<-PAIREDC-HTH+HIN-HTH<-Methyltransf_25<-?<-?<-DUF499*<-METHYLASE<-TOPRIM<-OLD-ABC<-DOC+HTH<-TUDOR+SF1-SF2-Helicase+iREase+CTD+REase<-XIS-HTH | 920 | Bacteria>Myxococcota>Myxococcia>unclassified_Myxococcia | GCA_012522415.1 | Deltaproteobacteria_bacterium | GX147_01890 | 29834..32596 | 2026735 | ATP-binding-protein-[Deltaproteobacteria-bacterium] |
| MCC6812134.1 | AAA+ATPase+wHTH+Alpha+Beta-domain+wHTH+RRM | Type-1_BR.RM.like.systems | <-DNA-glycosylase<-Peptidase-S8<-Classical-AAA<-McrC-NTD+REase<-?<-?<-DUF499*<-METHYLASE<-TUDOR+SF1-SF2-Helicase+iREase+CTD+REase<-HhH-RADC+JAB<-?<-?\|\|REase-> | 954 | Bacteria>Myxococcota>Myxococcia>unclassified_Myxococcia | GCA_020848545.1 | Deltaproteobacteria_bacterium | IT381_32235 | 12252..15116 | 2026735 | DUF499-domain-containing-protein-[Deltaproteobacteria-bacterium] |
| MBI5446667.1 | AAA+ATPase+wHTH+Alpha+Beta-domain+wHTH+RRM | Type-1_BR.RM.like.systems | <-Peptidase_S8<-Classical-AAA<-DUF499*<-CASPASE<-METHYLASE<-SF1-SF2-Helicase<-TUDOR+SF1-SF2-Helicase+iREase+CTD+REase<-XIS-HTH | 949 | Bacteria>Myxococcota>Myxococcia>unclassified_Myxococcia | GCA_016218905.1 | Deltaproteobacteria_bacterium | HY900_36290 | 8105..10954 | 2026735 | DUF499-domain-containing-protein-[Deltaproteobacteria-bacterium] |
| MBI4700963.1 | AAA+ATPase+wHTH+Alpha+Beta-domain+wHTH+RRM | Type-1_BR.RM.like.systems | PemK_toxin-><-AbiEi_1<-?<-TPR+TPR<-?<-?<-DUF499*<-RNaseH<-METHYLASE<-TUDOR+SF1-SF2-Helicase+iREase+CTD+REase<-?\|\|<-tRNA<-?<-SF1-SF2-Helicase | 954 | Bacteria>Myxococcota>Myxococcia>unclassified_Myxococcia | GCA_016209145.1 | Deltaproteobacteria_bacterium | HY744_07350 | 22677..25541 | 2026735 | ATP-binding-protein-[Deltaproteobacteria-bacterium] |
| PXF54128.1 | AAA+ATPase+wHTH+Alpha+Beta-domain+wHTH+RRM | Type-1_BR.RM.like.systems | Phage_int_SAM_1+Phage_integrase-><-SbcC+SbcC<-Calcineurin<-?<-?<-DUF499*<-Schlafen+HATPase+HTH<-METHYLASE-p<-TOPRIM<-OLD-ABC | 1011 | Bacteria>Myxococcota>Myxococcia>unclassified_Myxococcia | GCA_003194485.1 | Deltaproteobacteria_bacterium | C4B57_08240 | 17848..20883 | 2026735 | AAA-family-ATPase-[Deltaproteobacteria-bacterium] |
| MBI4516470.1 | AAA+ATPase+wHTH+Alpha+Beta-domain+wHTH+RRM | Type-1_BR.RM.like.systems | REase->Y1_Tnp->DUF499*->Terminase->?->?->TOPRIM+SF1-SF2-Helicase+HTH-><-?\|\|SF1-SF2-Helicase-p-> | 971 | Bacteria>Myxococcota>Myxococcia>unclassified_Myxococcia | GCA_016210005.1 | Deltaproteobacteria_bacterium | HY699_11720 | 1180..4095 | 2026735 | ATP-binding-protein-[Deltaproteobacteria-bacterium] |
| GIW47700.1 | AAA+ATPase+wHTH+Alpha+Beta-domain+wHTH+RRM | Type-1_BR.RM.like.systems | <-HEPN<-PolB\|\|?-><-DUF499*<-HEPN<-PolB<-METHYLASE<-Schlafen+HATPase+HTH<-TUDOR+SF1-SF2-Helicase+iREase+CTD+REase | 857 | Bacteria>Myxococcota>Myxococcia>unclassified_Myxococcia | GCA_026004235.1 | Deltaproteobacteria_bacterium | KatS3mg078_1577 | 289556..292129 | 2026735 | MAG:-hypothetical-protein-KatS3mg078_1577-[Deltaproteobacteria-bacterium] |
| MBI4814845.1 | AAA+ATPase+wHTH+Alpha+Beta-domain+wHTH+RRM | Type-1_BR.RM.like.systems | <-DNA-glycosylase\|\|TUDOR+SF1-SF2-Helicase+iREase+CTD+REase->METHYLASE->?->PIN->METHYLASE->DUF499*->REase+NACHT-> | 920 | Bacteria>Myxococcota>Myxococcia>unclassified_Myxococcia | GCA_016208175.1 | Deltaproteobacteria_bacterium | HY791_01225 | 36261..39023 | 2026735 | ATP-binding-protein-[Deltaproteobacteria-bacterium] |
| RLC12751.1 | AAA+ATPase+wHTH+Alpha+Beta-domain+wHTH+RRM | Type-1_BR.RM.like.systems | <-DUF499*<-Schlafen+HATPase+HTH<-METHYLASE-p<-METHYLASE<-MNS<-MACRODOMAIN<-TUDOR+SF1-SF2-Helicase+iREase+CTD+REase | 645 | Bacteria>Myxococcota>Myxococcia>unclassified_Myxococcia | GCA_003647435.1 | Deltaproteobacteria_bacterium | DRH43_00745 | 1..1934 | 2026735 | AAA-family-ATPase,-partial-[Deltaproteobacteria-bacterium] |
| MBN1381305.1 | AAA+ATPase+wHTH+Alpha+Beta-domain+wHTH+RRM | Type-1_BR.RM.like.systems | <-ABhydrolase\|\|?-><-?<-?\|\|BirA-HTH-><-DUF499*<-?<-HEPN<-PolB<-HEPN<-Schlafen+HATPase+HTH<-DOC+HTH | 916 | Bacteria>Myxococcota>Myxococcia>unclassified_Myxococcia | GCA_016928025.1 | Deltaproteobacteria_bacterium | JXA41_06500 | 6371..9121 | 2026735 | ATP-binding-protein-[Deltaproteobacteria-bacterium] |
| MDD9854174.1 | AAA+ATPase+wHTH+Alpha+Beta-domain+wHTH+RRM | Type-1_BR.RM.like.systems | <-DUF499*<-OLD-ABC<-METHYLASE<-TPR+TPR<-Schlafen+HATPase+HTH<-TUDOR+SF1-SF2-Helicase+iREase+CTD+REase | 946 | Bacteria>Myxococcota>Myxococcia>unclassified_Myxococcia | GCA_028817165.1 | Deltaproteobacteria_bacterium | OXU78_09525 | 86..2926 | 2026735 | MAG:-DUF499-domain-containing-protein-[Deltaproteobacteria-bacterium] |
| MBM4305128.1 | AAA+ATPase+wHTH+Alpha+Beta-domain+wHTH+RRM | Type-1_BR.RM.like.systems | JAB->?->?->DUF932-><-REase<-?<-DUF499*<-?<-TUDOR+SF1-SF2-Helicase+iREase+CTD+REase<-HTH+WYL<-?<-?\|\|REase-> | 982 | Bacteria>Myxococcota>Myxococcia>unclassified_Myxococcia | GCA_016875055.1 | Deltaproteobacteria_bacterium | FJ123_00175 | 29657..32605 | 2026735 | DUF499-domain-containing-protein-[Deltaproteobacteria-bacterium] |
| MBN1842594.1 | AAA+ATPase+wHTH+Alpha+Beta-domain+wHTH+RRM | Type-1_BR.RM.like.systems | <-ParB+HNH<-?<-EVE<-DUF499*<-METHYLASE<-HerA/FtsK<-?<-TUDOR+SF1-SF2-Helicase+iREase+CTD+REase | 917 | Bacteria>Myxococcota>Myxococcia>unclassified_Myxococcia | GCA_016930875.1 | Deltaproteobacteria_bacterium | JW883_09985 | 3841..6594 | 2026735 | ATP-binding-protein-[Deltaproteobacteria-bacterium] |
| MBI3799269.1 | AAA+ATPase+wHTH+Alpha+Beta-domain+wHTH+RRM | Type-1_BR.RM.like.systems | METHYLASE->DUF1828+REase->METHYLASE-p->DUF433->DUF433->PIN->DUF499*-> | 939 | Bacteria>Myxococcota>Myxococcia>unclassified_Myxococcia | GCA_016200595.1 | Deltaproteobacteria_bacterium | HY268_20180 | 14299..17118 | 2026735 | ATP-binding-protein-[Deltaproteobacteria-bacterium] |
| GIW46171.1 | AAA+ATPase+wHTH+Alpha+Beta-domain+wHTH+RRM | Type-1_BR.RM.like.systems | PolB->METHYLASE-p->METHYLASE-p->PolB->?->?->DUF499*->?-><-?<-?\|\|?->NAD_binding_2+6PGD-> | 872 | Bacteria>Myxococcota>Myxococcia>unclassified_Myxococcia | GCA_026004235.1 | Deltaproteobacteria_bacterium | KatS3mg078_0048 | 41535..44153 | 2026735 | MAG:-hypothetical-protein-KatS3mg078_0048-[Deltaproteobacteria-bacterium] |
| MBW1928071.1 | AAA+ATPase+wHTH+Alpha+Beta-domain+wHTH+RRM | Type-1_BR.RM.like.systems | SIR2+SIR2->TUDOR+SF1-SF2-Helicase+iREase+CTD+REase->HD->DUF4065->METHYLASE->?->DUF499*-> | 975 | Bacteria>Myxococcota>Myxococcia>unclassified_Myxococcia | GCA_019307815.1 | Deltaproteobacteria_bacterium | JRJ13_02825 | 203194..206121 | 2026735 | DUF499-domain-containing-protein-[Deltaproteobacteria-bacterium] |
| MBC8466666.1 | AAA+ATPase+wHTH+Alpha+Beta-domain+wHTH+RRM | Type-1_BR.RM.like.systems | <-DUF499*<-REase<-METHYLASE<-?<-DrHyd+BirA-HTH<-TUDOR+SF1-SF2-Helicase+iREase+CTD+REase | 918 | Bacteria>Myxococcota>Myxococcia>unclassified_Myxococcia | GCA_014384065.1 | Deltaproteobacteria_bacterium | H8D55_02390 | 3412..6168 | 2026735 | ATP-binding-protein-[Deltaproteobacteria-bacterium] |

| Protein-ID | DUF499-architecture | System | Gene-Neighbourhoods | Length | Taxonomy | Assembly-ID | Strain | Locus-Tag | Genomic-coordinates | TaxID | Fasta-defline |
|---|---|---|---|---|---|---|---|---|---|---|---|
| MCD6198158.1 | AAA+ATPase+wHTH+Alpha+Beta-domain+wHTH+RRM | Type-1_BR.RM.like.systems | <-DUF499*<-OLD-ABC+TOPRIM<-MACRODOMAIN<-?<-TUDOR+SF1-SF2-Helicase+iREase+CTD+REase<-XIS-HTH | 992 | Bacteria>Myxococcota>Myxococcia>unclassified_Myxococcia | GCA_021159445.1 | Deltaproteobacteria_bacterium | J7K15_06235 | 2443..5421 | 2026735 | ATP-binding-protein-[Deltaproteobacteria-bacterium] |
| MBM4341420.1 | AAA+ATPase+wHTH+Alpha+Beta-domain+wHTH+RRM | Type-1_BR.RM.like.systems | <-DUF499*<-Schlafen+HATPase+HTH<-METHYLASE<-SIR2<-TUDOR+SF1-SF2-Helicase+iREase+CTD+REase | 454 | Bacteria>Myxococcota>Myxococcia>unclassified_Myxococcia | GCA_016875195.1 | Deltaproteobacteria_bacterium | FJ110_17985 | 1..1363 | 2026735 | ATP-binding-protein-[Deltaproteobacteria-bacterium] |
| MBP7321436.1 | AAA+ATPase+wHTH+Alpha+Beta-domain+wHTH+RRM | Type-1_BR.RM.like.systems | <-ZnR\|\|?-><-SF1-SF2-Helicase<-?<-RVT<-?<-DUF499*<-?<-METHYLASE-p<-METHYLASE | 924 | Bacteria>Myxococcota>Myxococcia>unclassified_Myxococcia | GCA_017996625.1 | Deltaproteobacteria_bacterium | KBA65_00615 | 58214..60988 | 2026735 | ATP-binding-protein-[Deltaproteobacteria-bacterium] |
| MBM4288095.1 | AAA+ATPase+wHTH+Alpha+Beta-domain+wHTH+RRM | Type-1_BR.RM.like.systems | TUDOR+SF1-SF2-Helicase+iREase+CTD+REase->METHYLASE->HicB->HicA_toxin->DUF499*-> | 914 | Bacteria>Myxococcota>Myxococcia>unclassified_Myxococcia | GCA_016874895.1 | Deltaproteobacteria_bacterium | FJ135_08115 | 7709..10453 | 2026735 | ATP-binding-protein-[Deltaproteobacteria-bacterium] |
| MBW1911012.1 | AAA+ATPase+wHTH+Alpha+Beta-domain+wHTH+RRM | Type-1_BR.RM.like.systems | Radical-SAM-><-DUF499*<-PIN<-?<-DUF433<-METHYLASE<-TOPRIM<-OLD-ABC-p | 941 | Bacteria>Myxococcota>Myxococcia>unclassified_Myxococcia | GCA_019307765.1 | Deltaproteobacteria_bacterium | JRJ11_15995 | 905..3730 | 2026735 | ATP-binding-protein-[Deltaproteobacteria-bacterium] |
| MCX5850425.1 | AAA+ATPase+wHTH+Alpha+Beta-domain+wHTH+RRM | Type-1_BR.RM.like.systems | TUDOR+SF1-SF2-Helicase+iREase+CTD+REase-p->TUDOR+SF1-SF2-Helicase+iREase+CTD+REase-p->?->?->DUF499*-> | 831 | Bacteria>Myxococcota>Myxococcia>unclassified_Myxococcia | GCA_026388535.1 | Deltaproteobacteria_bacterium | NTW65_13435 | 11180..13675 | 2026735 | MAG:-DUF499-domain-containing-protein-[Deltaproteobacteria-bacterium] |
| MCL5125416.1 | AAA+ATPase+wHTH+Alpha+Beta-domain+wHTH+RRM | Type-1_BR.RM.like.systems | <-DUF499*<-METHYLASE-p<-METHYLASE<-?<-?<-TUDOR+SF1-SF2-Helicase+iREase+CTD+REase | 851 | Bacteria>Myxococcota>Myxococcia>unclassified_Myxococcia | GCA_023474025.1 | Deltaproteobacteria_bacterium | M1511_13160 | 645..3200 | 2026735 | DUF499-domain-containing-protein-[Deltaproteobacteria-bacterium] |
| MBI4706258.1 | AAA+ATPase+wHTH+Alpha+Beta-domain+wHTH+RRM | Type-1_BR.RM.like.systems | <-DUF499*<-?<-SbcC<-OLD-ABC+TOPRIM<-METHYLASE<-TOPRIM<-OLD-ABC | 925 | Bacteria>Myxococcota>Myxococcia>unclassified_Myxococcia | GCA_016209145.1 | Deltaproteobacteria_bacterium | HY744_34610 | 6205..8982 | 2026735 | ATP-binding-protein-[Deltaproteobacteria-bacterium] |
| RLB21509.1 | AAA+ATPase+wHTH+Alpha+Beta-domain+wHTH+RRM | Type-1_BR.RM.like.systems | <-PolB<-Calcineurin<-DUF499*<-DUF1828+REase<-?<-METHYLASE-p | 920 | Bacteria>Myxococcota>Myxococcia>unclassified_Myxococcia | GCA_003646865.1 | Deltaproteobacteria_bacterium | DRG76_08830 | 848..3610 | 2026735 | AAA-family-ATPase-[Deltaproteobacteria-bacterium] |
| MBW1732442.1 | AAA+ATPase+wHTH+Alpha+Beta-domain+wHTH+RRM | Type-1_BR.RM.like.systems | METHYLASE-p->?->DUF1828+REase->DUF499*->Calcineurin->PolB-> | 920 | Bacteria>Myxococcota>Myxococcia>unclassified_Myxococcia | GCA_019306325.1 | Deltaproteobacteria_bacterium | JRJ75_16450 | 1746..4508 | 2026735 | ATP-binding-protein-[Deltaproteobacteria-bacterium] |
| MBI4527351.1 | AAA+ATPase+wHTH+Alpha+Beta-domain+wHTH+RRM | Type-1_BR.RM.like.systems | Schlafen+HATPase+HTH->METHYLASE->?->PIN->HicB_lk_antitox->?->DUF499*-> | 920 | Bacteria>Myxococcota>Myxococcia>unclassified_Myxococcia | GCA_016210115.1 | Deltaproteobacteria_bacterium | HY695_26450 | 25107..27869 | 2026735 | ATP-binding-protein-[Deltaproteobacteria-bacterium] |
| MBI2179305.1 | AAA+ATPase+wHTH+Alpha+Beta-domain+wHTH+RRM | Type-1_BR.RM.like.systems | METHYLASE-p->DUF433->PIN->PIN->DUF499*->?-><-?\|\|?-><-Glyoxalase | 937 | Bacteria>Myxococcota>Myxococcia>unclassified_Myxococcia | GCA_016179825.1 | Deltaproteobacteria_bacterium | HYU31_00605 | 1817..4630 | 2026735 | ATP-binding-protein-[Deltaproteobacteria-bacterium] |
| PXF56405.1 | AAA+ATPase+wHTH+Alpha+Beta-domain+wHTH+RRM | Type-1_BR.RM.like.systems | METHYLASE->?->HicB->HicA->DUF499*->?->PRTase-><-?<-?<-HISKIN+MORC | 984 | Bacteria>Myxococcota>Myxococcia>unclassified_Myxococcia | GCA_003194495.1 | Deltaproteobacteria_bacterium | C4B58_13545 | 3402..6356 | 2026735 | AAA-family-ATPase-[Deltaproteobacteria-bacterium] |
| MBS3919360.1 | AAA+ATPase+wHTH+Alpha+Beta-domain+wHTH+RRM | Type-1_BR.RM.like.systems | SF1-SF2-Helicase->METHYLASE->DUF433->PIN->DUF499*-> | 921 | Bacteria>Myxococcota>Myxococcia>unclassified_Myxococcia | GCA_018333535.1 | Deltaproteobacteria_bacterium | KG012_10765 | 28833..31598 | 2026735 | ATP-binding-protein-[Deltaproteobacteria-bacterium] |
| RLA85517.1 | AAA+ATPase+wHTH+Alpha+Beta-domain+wHTH+RRM | Type-1_BR.RM.like.systems | TUDOR+SF1-SF2-Helicase+iREase+CTD+REase->METHYLASE-p->DUF499*-> | 474 | Bacteria>Myxococcota>Myxococcia>unclassified_Myxococcia | GCA_003646585.1 | Deltaproteobacteria_bacterium | DRG31_02795 | 5720..7142 | 2026735 | AAA-family-ATPase,-partial-[Deltaproteobacteria-bacterium] |
| MBW2149119.1 | AAA+ATPase+wHTH+Alpha+Beta-domain+wHTH+RRM | Type-1_BR.RM.like.systems | METHYLASE-p->METHYLASE-p->DUF433->PIN->DUF499*-> | 939 | Bacteria>Myxococcota>Myxococcia>unclassified_Myxococcia | GCA_019309515.1 | Deltaproteobacteria_bacterium | JRI22_19100 | 2839..5658 | 2026735 | ATP-binding-protein-[Deltaproteobacteria-bacterium] |
| MBW1922689.1 | AAA+ATPase+wHTH+Alpha+Beta-domain+wHTH+RRM | Type-1_BR.RM.like.systems | METHYLASE->OLD-ABC->TOPRIM->DUF499*->EVE-><-HIN-HTH | 965 | Bacteria>Myxococcota>Myxococcia>unclassified_Myxococcia | GCA_019307945.1 | Deltaproteobacteria_bacterium | JRJ35_04385 | 5613..8510 | 2026735 | DUF499-domain-containing-protein-[Deltaproteobacteria-bacterium] |
| MBW2309273.1 | AAA+ATPase+wHTH+Alpha+Beta-domain+wHTH+RRM | Type-1_BR.RM.like.systems | <-DUF499*<-PIN<-DUF433<-METHYLASE-p<-METHYLASE | 908 | Bacteria>Myxococcota>Myxococcia>unclassified_Myxococcia | GCA_019310535.1 | Deltaproteobacteria_bacterium | JRG73_20315 | 275..3001 | 2026735 | ATP-binding-protein-[Deltaproteobacteria-bacterium] |
| MBN2034424.1 | AAA+ATPase+wHTH+Alpha+Beta-domain+wHTH+RRM | Type-1_BR.RM.like.systems | TUDOR+SF1-SF2-Helicase+iREase+CTD+REase->METHYLASE->?->DUF499*-> | 980 | Bacteria>Myxococcota>Myxococcia>unclassified_Myxococcia | GCA_016932115.1 | Deltaproteobacteria_bacterium | JW836_14215 | 7336..10278 | 2026735 | ATP-binding-protein-[Deltaproteobacteria-bacterium] |
| MBK06773.1 | AAA+ATPase+wHTH+Alpha+Beta-domain+wHTH+RRM | Type-1_BR.RM.like.systems | <-TPR\|\|LON+AAA->TetR-HTH->?->?->?->DUF499*-><-HDRnase | 951 | Bacteria>Myxococcota>Myxococcia>unclassified_Myxococcia | GCA_002720785.1 | Deltaproteobacteria_bacterium | CL932_18625 | 20896..23751 | 2026735 | hypothetical-protein-[Deltaproteobacteria-bacterium] |
| MBM4285998.1 | AAA+ATPase+wHTH+Alpha+Beta-domain+wHTH+RRM | Type-1_BR.RM.like.systems | <-DUF499*<-?<-HicB<-TUDOR+SF1-SF2-Helicase+iREase+CTD+REase | 915 | Bacteria>Myxococcota>Myxococcia>unclassified_Myxococcia | GCA_016874975.1 | Deltaproteobacteria_bacterium | FJ128_12245 | 624..3371 | 2026735 | ATP-binding-protein-[Deltaproteobacteria-bacterium] |
| MDD9827665.1 | AAA+ATPase+wHTH+Alpha+Beta-domain+wHTH+RRM | Type-1_BR.RM.like.systems | <-LON-AAA<-?\|\|?->?->?-><-DUF499*<-OLD-ABC<-METHYLASE-p | 946 | Bacteria>Myxococcota>Myxococcia>unclassified_Myxococcia | GCA_028816715.1 | Deltaproteobacteria_bacterium | OXU92_04690 | 14260..17100 | 2026735 | MAG:-DUF499-domain-containing-protein-[Deltaproteobacteria-bacterium] |
| MBW2135777.1 | AAA+ATPase+wHTH+Alpha+Beta-domain+wHTH+RRM | Type-1_BR.RM.like.systems | METHYLASE-p->PolB->HEPN->DUF499*-> | 916 | Bacteria>Myxococcota>Myxococcia>unclassified_Myxococcia | GCA_019309545.1 | Deltaproteobacteria_bacterium | JRG72_11235 | 2815..5565 | 2026735 | ATP-binding-protein-[Deltaproteobacteria-bacterium] |
| MBI2988835.1 | AAA+ATPase+wHTH+Alpha+Beta-domain+wHTH+RRM | Type-1_BR.RM.like.systems | <-sGTP<-?<-?<-?<-?<-DUF499*<-METHYLASE<-?<-REase | 931 | Bacteria>Myxococcota>Myxococcia>unclassified_Myxococcia | GCA_016193465.1 | Deltaproteobacteria_bacterium | HYY45_18895 | 27398..30193 | 2026735 | DUF499-domain-containing-protein-[Deltaproteobacteria-bacterium] |
| MBC8458333.1 | AAA+ATPase+wHTH+Alpha+Beta-domain+wHTH+RRM | Type-1_BR.RM.like.systems | METHYLASE->PolB->HEPN->DUF499*-> | 802 | Bacteria>Myxococcota>Myxococcia>unclassified_Myxococcia | GCA_014383825.1 | Deltaproteobacteria_bacterium | H8D67_10095 | 2315..4719 | 2026735 | ATP-binding-protein-[Deltaproteobacteria-bacterium] |
| RLC08398.1 | AAA+ATPase+wHTH+Alpha+Beta-domain+wHTH+RRM | Type-1_BR.RM.like.systems | <-DUF499*<-TOPRIM<-OLD-ABC<-METHYLASE-p | 353 | Bacteria>Myxococcota>Myxococcia>unclassified_Myxococcia | GCA_003647435.1 | Deltaproteobacteria_bacterium | DRH43_10115 | 1..1059 | 2026735 | AAA-family-ATPase,-partial-[Deltaproteobacteria-bacterium] |
| RLC09317.1 | AAA+ATPase+wHTH+Alpha+Beta-domain+wHTH+RRM | Type-1_BR.RM.like.systems | <-DUF499*<-REase<-HEPN<-METHYLASE-p | 517 | Bacteria>Myxococcota>Myxococcia>unclassified_Myxococcia | GCA_003647435.1 | Deltaproteobacteria_bacterium | DRH43_08475 | 1..1550 | 2026735 | AAA-family-ATPase,-partial-[Deltaproteobacteria-bacterium] |
| MBW1926153.1 | AAA+ATPase+wHTH+Alpha+Beta-domain+wHTH+RRM | Type-1_BR.RM.like.systems | <-DUF499*<-DUF1828+REase<-?<-METHYLASE | 912 | Bacteria>Myxococcota>Myxococcia>unclassified_Myxococcia | GCA_019307845.1 | Deltaproteobacteria_bacterium | JRI35_02965 | 2089..4827 | 2026735 | ATP-binding-protein-[Deltaproteobacteria-bacterium] |
| RLC30199.1 | AAA+ATPase+wHTH+Alpha+Beta-domain+wHTH+RRM | Type-1_BR.RM.like.systems | METHYLASE-p->?->HicB->HicA->DUF499*-> | 676 | Bacteria>Myxococcota>Myxococcia>unclassified_Myxococcia | GCA_003647535.1 | Deltaproteobacteria_bacterium | DRH37_05905 | 2759..4788 | 2026735 | AAA-family-ATPase,-partial-[Deltaproteobacteria-bacterium] |
| MBC8460283.1 | AAA+ATPase+wHTH+Alpha+Beta-domain+wHTH+RRM | Type-1_BR.RM.like.systems | HEPN+REase->METHYLASE->DUF499*-> | 926 | Bacteria>Myxococcota>Myxococcia>unclassified_Myxococcia | GCA_014383825.1 | Deltaproteobacteria_bacterium | H8D67_20045 | 4060..6840 | 2026735 | ATP-binding-protein-[Deltaproteobacteria-bacterium] |
| MBW2309256.1 | AAA+ATPase+wHTH+Alpha+Beta-domain+wHTH+RRM | Type-1_BR.RM.like.systems | METHYLASE-p->HEPN->DUF499*-> | 920 | Bacteria>Myxococcota>Myxococcia>unclassified_Myxococcia | GCA_019310535.1 | Deltaproteobacteria_bacterium | JRG73_20230 | 2438..5200 | 2026735 | ATP-binding-protein-[Deltaproteobacteria-bacterium] |
| MBI4528434.1 | AAA+ATPase+wHTH+Alpha+Beta-domain+wHTH+RRM | Type-1_BR.RM.like.systems | TPR+TPR-><-DUF499*<-METHYLASE | 957 | Bacteria>Myxococcota>Myxococcia>unclassified_Myxococcia | GCA_016210115.1 | Deltaproteobacteria_bacterium | HY695_31965 | 1335..4208 | 2026735 | ATP-binding-protein-[Deltaproteobacteria-bacterium] |
| MBC8392014.1 | AAA+ATPase+wHTH+Alpha+Beta-domain+wHTH+RRM | Type-1_BR.RM.like.systems | METHYLASE->REase->DUF499*-> | 427 | Bacteria>Myxococcota>Myxococcia>unclassified_Myxococcia | GCA_014382765.1 | Deltaproteobacteria_bacterium | H8E17_05550 | 4033..5315 | 2026735 | ATP-binding-protein-[Deltaproteobacteria-bacterium] |
| MBV8356666.1 | AAA+ATPase+wHTH+Alpha+Beta-domain+wHTH+RRM | Type-1_BR.RM.like.systems | <-DUF499*<-DUF1828+REase | 925 | Bacteria>Myxococcota>Myxococcia>unclassified_Myxococcia | GCA_019235735.1 | Deltaproteobacteria_bacterium | JO189_01845 | 1414..4191 | 2026735 | ATP-binding-protein-[Deltaproteobacteria-bacterium] |
| MBW1705706.1 | AAA+ATPase+wHTH+Alpha+Beta-domain+wHTH+RRM | Type-1_BR.RM.like.systems | <-SMS-KaiC<-RelE-ParE<-MetJarc-HTH\|\|TUDOR+SF1-SF2-Helicase+iREase+CTD+REase->wHTH+REase->METHYLASE->DUF499*-> | 670 | Bacteria>Myxococcota>Myxococcia>unclassified_Myxococcia | GCA_019306165.1 | Deltaproteobacteria_bacterium | JRJ86_11145 | 11472..13481 | 2026735 | ATP-binding-protein-[Deltaproteobacteria-bacterium] |
| MBM4274058.1 | AAA+ATPase+wHTH+Alpha+Beta-domain+wHTH+RRM | Type-1_BR.RM.like.systems | TUDOR+SF1-SF2-Helicase+iREase+CTD+REase->HicB->HicA_toxin->DUF499*->DUF499-p->TUDOR+SF1-SF2-Helicase+iREase+CTD-> | 422 | Bacteria>Myxococcota>Myxococcia>unclassified_Myxococcia | GCA_016874915.1 | Deltaproteobacteria_bacterium | FJ134_06320 | 6113..7381 | 2026735 | ATP-binding-protein-[Deltaproteobacteria-bacterium] |
| MCX5910053.1 | AAA+ATPase+wHTH+Alpha+Beta-domain+wHTH+RRM | Type-1_BR.RM.like.systems | OLD-ABC-p->TOPRIM->DUF499-p->DUF499*->DUF499-p->AbiEii-> | 258 | Bacteria>Myxococcota>Myxococcia>unclassified_Myxococcia | GCA_026388955.1 | Deltaproteobacteria_bacterium | NTY64_23470 | 1666..2438 | 2026735 | MAG:-DUF499-domain-containing-protein,-partial-[Deltaproteobacteria-bacterium] |
| MCX5909644.1 | AAA+ATPase+wHTH+Alpha+Beta-domain+wHTH+RRM | Type-1_BR.RM.like.systems | METHYLASE-p->METHYLASE-p->METHYLASE-p->DUF499*-> | 339 | Bacteria>Myxococcota>Myxococcia>unclassified_Myxococcia | GCA_026388955.1 | Deltaproteobacteria_bacterium | NTY64_21345 | 2877..3893 | 2026735 | MAG:-DUF499-domain-containing-protein,-partial-[Deltaproteobacteria-bacterium] |
| MBW2050568.1 | AAA+ATPase+wHTH+Alpha+Beta-domain+wHTH+RRM | Type-1_BR.RM.like.systems | <-DUF499-p<-DUF499*<-METHYLASE-p | 639 | Bacteria>Myxococcota>Myxococcia>unclassified_Myxococcia | GCA_019308945.1 | Deltaproteobacteria_bacterium | JRJ09_18885 | 1236..3155 | 2026735 | ATP-binding-protein-[Deltaproteobacteria-bacterium] |
| MBK8259833.1 | AAA+ATPase+wHTH+Alpha+Beta-domain+wHTH+RRM | Type-1_BR.RM.like.systems | TUDOR+SF1-SF2-Helicase+iREase+CTD+REase->METHYLASE->DUF499*-><-?<-HTH_Tnp_1+HTH_21+rve+rve_2\|\|?->CR-ATPase9+REase-> | 929 | Bacteria>Myxococcota>Polyangia>Nannocystales>Nannocystaceae | GCA_016712315.1 | Nannocystis_sp. | IPK80_00625 | 22848..25637 | 1962667 | ATP-binding-protein-[Nannocystis-sp.] |
| MBX3230656.1 | AAA+ATPase+wHTH+Alpha+Beta-domain+wHTH+RRM | Type-1_BR.RM.like.systems | <-Cas_Cmr5<-?<-Cas_Cmr3<-?<-?<-DUF499*<-?<-METHYLASE<-SIGMA-HTH<-TUDOR+SF1-SF2-Helicase+iREase+CTD+REase<-DUF4236+TPR | 942 | Bacteria>Myxococcota>Polyangia>Polyangiales>Labilitrichaceae | GCA_019637155.1 | Labilithrix_sp. | KF837_25280 | 104430..107258 | 1971228 | ATP-binding-protein-[Labilithrix-sp.] |
| KYF87240.1 | AAA+ATPase+wHTH+Alpha+Beta-domain+wHTH+RRM | Type-1_BR.RM.like.systems | <-DUF499*<-REase<-OLD-ABC<-METHYLASE<-HNH<-OLD-ABC<-TUDOR+SF1-SF2-Helicase+iREase+CTD+REase | 954 | Bacteria>Myxococcota>Polyangia>Polyangiales>Polyangiaceae | GCA_001589185.1 | Sorangium_cellulosum | BE17_46125 | 4063..6927 | 56 | hypothetical-protein-BE17_46125-[Sorangium-cellulosum] |
| NUQ73302.1 | AAA+ATPase+wHTH+Alpha+Beta-domain+wHTH+RRM | Type-1_BR.RM.like.systems | TUDOR+SF1-SF2-Helicase+iREase+CTD+REase->METHYLASE->DUF499*->Pkinase-> | 948 | Bacteria>Myxococcota>Polyangia>Polyangiales>Polyangiaceae | GCA_013360665.1 | Polyangiaceae_bacterium | HUU21_07085 | 10269..13115 | 2268199 | ATP-binding-protein-[Polyangiaceae-bacterium] |
| WP_050434136.1 | AAA+ATPase+wHTH+Alpha+Beta-domain+wHTH+RRM | Type-1_BR.RM.like.systems | <-SF1-SF2-Helicase<-PAIREDC-HTH<-DNA-glycosylase<-DUF499*<-REase<-METHYLASE<-BirA-HTH<-METHYLASE<-OLD-ABC<-TUDOR+SF1-SF2-Helicase+iREase+CTD+REase | 936 | Bacteria>Myxococcota>Polyangia>Polyangiales>Polyangiaceae | GCF_001189295.1 | Chondromyces_crocatus | CMC5_RS33060 | 9153103..9155913 | 52 | DUF499-domain-containing-protein-[Chondromyces-crocatus] |
| MCS6856320.1 | AAA+ATPase+wHTH+Alpha+Beta-domain+wHTH+RRM | Type-1_BR.RM.like.systems | <-DUF499*<-METHYLASE<-TUDOR+SF1-SF2-Helicase+iREase+CTD+REase<-?\|\|ABhydrolase-> | 902 | Bacteria>Myxococcota>Polyangia>Polyangiales>Sandaracinaceae | GCA_025056955.1 | Sandaracinaceae_bacterium | NZM37_01275 | 4955..7663 | 2510337 | DUF499-domain-containing-protein-[Sandaracinaceae-bacterium] |
| MCZ7618403.1 | AAA+ATPase+wHTH+Alpha+Beta-domain+wHTH+RRM | Type-1_BR.RM.like.systems | <-SF1-SF2-Helicase\|\|?->KAP-NTPase->?->PAPS_reduct->?-><-DUF499*<-METHYLASE\|\|DDE_Tnp_1_4-><-METHYLASE-p\|\|REase->JAB->ArdCN+MPTase-> | 931 | Bacteria>Myxococcota>unclassified_Myxococcota | GCA_027493655.1 | Myxococcota_bacterium | M5U32_08850 | 1982126..1984921 | 2818507 | MAG:-ATP-binding-protein-[Myxococcota-bacterium] |
| MBU0551512.1 | AAA+ATPase+wHTH+Alpha+Beta-domain+wHTH+RRM | Type-1_BR.RM.like.systems | TUDOR+SF1-SF2-Helicase+iREase+CTD+REase-p->HTH+MPTase->PIN->METHYLASE->OLD-ABC->REase->DUF499*-><-?<-?<-RecR+TOPRIM | 942 | Bacteria>Myxococcota>unclassified_Myxococcota | GCA_018813065.1 | Myxococcota_bacterium | KJ940_08450 | 6900..9728 | 2818507 | DUF499-domain-containing-protein-[Myxococcota-bacterium] |
| MBU1411233.1 | AAA+ATPase+wHTH+Alpha+Beta-domain+wHTH+RRM | Type-1_BR.RM.like.systems | TUDOR+SF1-SF2-Helicase+iREase+CTD+REase->METHYLASE->DUF499*-> | 746 | Bacteria>Myxococcota>unclassified_Myxococcota | GCA_018820615.1 | Myxococcota_bacterium | KKC22_06895 | 4846..7084 | 2818507 | DUF499-domain-containing-protein-[Myxococcota-bacterium] |
| MCS6797210.1 | AAA+ATPase+wHTH+Alpha+Beta-domain+wHTH+RRM | Type-1_BR.RM.like.systems | Peptidase->?-><-?<-?<-DUF499*<-METHYLASE-p<-METHYLASE<-?<-?<-TOPRIM<-OLD-ABC | 1007 | Bacteria>Myxococcota>unclassified_Myxococcota | GCA_025054695.1 | Myxococcota_bacterium | NZ898_01550 | 47913..50936 | 2818507 | DUF499-domain-containing-protein-[Myxococcota-bacterium] |
| MBI3326464.1 | AAA+ATPase+wHTH+Alpha+Beta-domain+wHTH+RRM | Type-1_BR.RM.like.systems | <-METHYLASE-p<-DUF499<-DUF499* | 233 | Bacteria>Nitrospinae/Tectomicrobia_group>Nitrospinota>unclassified_Nitrospinota | GCA_016196645.1 | Nitrospinota_bacterium | HYZ81_07165 | 2240..2941 | 2026769 | hypothetical-protein-[Nitrospinota-bacterium] |
| MBI3326463.1 | AAA+ATPase+wHTH+Alpha+Beta-domain+wHTH+RRM | Type-1_BR.RM.like.systems | <-METHYLASE-p<-DUF499*<-DUF499 | 624 | Bacteria>Nitrospinae/Tectomicrobia_group>Nitrospinota>unclassified_Nitrospinota | GCA_016196645.1 | Nitrospinota_bacterium | HYZ81_07160 | 350..2224 | 2026769 | hypothetical-protein-[Nitrospinota-bacterium] |
| MDE0331736.1 | AAA+ATPase+wHTH+Alpha+Beta-domain+wHTH+RRM | Type-1_BR.RM.like.systems | <-OLD-ABC+TOPRIM<-DUF499*<-METHYLASE<-TUDOR+SF1-SF2-Helicase+iREase+CTD+REase | 967 | Bacteria>Nitrospinae/Tectomicrobia_group>Nitrospinota>unclassified_Nitrospinota | GCA_028820795.1 | Nitrospinota_bacterium | OXL41_07695 | 2738..5641 | 2026769 | MAG:-DUF499-domain-containing-protein-[Nitrospinae-bacterium] |
| MBS0175039.1 | AAA+ATPase+wHTH+Alpha+Beta-domain+wHTH+RRM | Type-1_BR.RM.like.systems | HTH+WYL->TUDOR+SF1-SF2-Helicase+iREase+CTD+REase->HEPN->METHYLASE->DUF499*->?->?->?-><-Phage_lysozyme | 941 | Bacteria>Nitrospirota>Nitrospiria>Nitrospirales>Nitrospiraceae | GCA_018242785.1 | Nitrospira_sp. | JSR64_13460 | 8684..11509 | 70125 | ATP-binding-protein-[Nitrospira-sp.] |
| MCP9448661.1 | AAA+ATPase+wHTH+Alpha+Beta-domain+wHTH+RRM | Type-1_BR.RM.like.systems | sGTP->?->?->?->?->?-><-DUF499*<-HEPN<-ORC-CDC6+REase<-METHYLASE-p\|\|?-><-Radical-SAM | 987 | Bacteria>Nitrospirota>Nitrospiria>Nitrospirales>Nitrospiraceae | GCA_024331085.1 | Nitrospira_sp. | NNA21_01185 | 6072..9035 | 70125 | DUF499-domain-containing-protein-[Nitrospira-sp.] |
| MDA8112640.1 | AAA+ATPase+wHTH+Alpha+Beta-domain+wHTH+RRM | Type-1_BR.RM.like.systems | <-ABC-ATPase+SbcC<-?\|\|ADH_N+ADH_zinc_N->ABhydrolase->?-><-DUF499* | 798 | Bacteria>Nitrospirota>Nitrospiria>Nitrospirales>Nitrospiraceae | GCA_028064555.1 | Nitrospiraceae_bacterium | MOT83_09320 | 8500..10896 | 2026770 | MAG:-DUF499-domain-containing-protein-[Nitrospiraceae-bacterium] |
| MBS0166207.1 | AAA+ATPase+wHTH+Alpha+Beta-domain+wHTH+RRM | Type-1_BR.RM.like.systems | <-TrwC+SF1-SF2-helicase+TOPRIM<-VirD4-TraG+VirD4-FtsK<-?<-?<-?<-DUF499*<-PIN<-?<-METHYLASE<-HicA_toxin<-HicB_lk_antitox<-TUDOR+SF1-SF2-Helicase+iREase+CTD+REase | 920 | Bacteria>Nitrospirota>Nitrospiria>Nitrospirales>Nitrospiraceae | GCA_018242765.1 | Nitrospira_sp. | JSR29_09005 | 53219..55981 | 70125 | ATP-binding-protein-[Nitrospira-sp.] |
| MCP9446717.1 | AAA+ATPase+wHTH+Alpha+Beta-domain+wHTH+RRM | Type-1_BR.RM.like.systems | <-REase<-?<-?<-?<-N6_N4_MTase<-?<-DUF499*<-REase<-METHYLASE<-Schlafen+HATPase+HTH<-TUDOR+SF1-SF2-Helicase+iREase+CTD+REase | 941 | Bacteria>Nitrospirota>Nitrospiria>Nitrospirales>Nitrospiraceae | GCA_024330945.1 | Nitrospira_sp. | NNA22_03990 | 9048..11873 | 70125 | DUF499-domain-containing-protein-[Nitrospira-sp.] |
| MCP9443508.1 | AAA+ATPase+wHTH+Alpha+Beta-domain+wHTH+RRM | Type-1_BR.RM.like.systems | <-Peptidase_S8<-Classical-AAA<-DUF499*<-METHYLASE<-?<-?<-TUDOR+SF1-SF2-Helicase+iREase+CTD+REase<-XIS-HTH | 970 | Bacteria>Nitrospirota>Nitrospiria>Nitrospirales>Nitrospiraceae | GCA_024330965.1 | Nitrospira_sp. | NNA20_13090 | 5814..8726 | 70125 | DUF499-domain-containing-protein-[Nitrospira-sp.] |
| MDC8448550.1 | AAA+ATPase+wHTH+Alpha+Beta-domain+wHTH+RRM | Type-1_BR.RM.like.systems | rve+rve_3->?-><-DUF499*<-PIN<-?<-METHYLASE-p<-METHYLASE-p<-HicA_toxin<-HicB_lk_antitox | 922 | Bacteria>Nitrospirota>Nitrospiria>Nitrospirales>Nitrospiraceae | GCA_028580895.1 | Nitrospira_sp. | LV473_09340 | 1848..4616 | 70125 | MAG:-DUF499-domain-containing-protein-[Nitrospira-sp.] |
| WP_213042468.1 | AAA+ATPase+wHTH+Alpha+Beta-domain+wHTH+RRM | Type-1_BR.RM.like.systems | LuxR-HTH-><-?<-?<-DUF499*<-METHYLASE<-HKD-DNase<-TUDOR+SF1-SF2-Helicase+iREase+CTD+REase\|\|XIS-HTH-> | 914 | Bacteria>Nitrospirota>Nitrospiria>Nitrospirales>Nitrospiraceae | GCF_905220995.1 | Nitrospira_defluvii | KJA79_RS12930 | 291076..293820 | 330214 | DUF499-domain-containing-protein-[Nitrospira-defluvii] |
| HBI24139.1 | AAA+ATPase+wHTH+Alpha+Beta-domain+wHTH+RRM | Type-1_BR.RM.like.systems | TUDOR+SF1-SF2-Helicase+iREase+CTD+REase-p->METHYLASE->PolB->?->PolB->HEPN->DUF499*->?->?->REase-p-> | 849 | Bacteria>Nitrospirota>Nitrospiria>Nitrospirales>Nitrospiraceae | GCA_003508715.1 | Nitrospiraceae_bacterium | DDX84_08070 | 5508..8057 | 2026770 | AAA-family-ATPase-[Nitrospiraceae-bacterium] |
| MCP9469842.1 | AAA+ATPase+wHTH+Alpha+Beta-domain+wHTH+RRM | Type-1_BR.RM.like.systems | <-Glyoxalase\|\|?->HTH->?-><-DUF499*<-HEPN<-PolB<-METHYLASE<-?<-TUDOR+SF1-SF2-Helicase+iREase+CTD+REase | 919 | Bacteria>Nitrospirota>Nitrospiria>Nitrospirales>Nitrospiraceae | GCA_024331125.1 | Nitrospira_sp. | NNA31_07565 | 179179..181938 | 70125 | DUF499-domain-containing-protein-[Nitrospira-sp.] |
| MBS0156596.1 | AAA+ATPase+wHTH+Alpha+Beta-domain+wHTH+RRM | Type-1_BR.RM.like.systems | TraJ-RHH->?->TUDOR+SF1-SF2-Helicase+iREase+CTD+REase->METHYLASE->DUF499*->?->?->?->?-><-PIWI<-TIR+PNTD2 | 920 | Bacteria>Nitrospirota>Nitrospiria>Nitrospirales>Nitrospiraceae | GCA_018242825.1 | Nitrospira_sp. | JSS38_18585 | 9725..12487 | 70125 | ATP-binding-protein-[Nitrospira-sp.] |
| MDD9860639.1 | AAA+ATPase+wHTH+Alpha+Beta-domain+wHTH+RRM | Type-1_BR.RM.like.systems | SPOUT-><-LuxR-HTH\|\|?->?->?->?-><-DUF499*<-PIN<-OLD-ABC<-METHYLASE-p | 939 | Bacteria>Nitrospirota>Nitrospiria>Nitrospirales>Nitrospiraceae | GCA_028817215.1 | Nitrospira_sp. | OXU40_09370 | 15116..17935 | 70125 | MAG:-DUF499-domain-containing-protein-[Nitrospira-sp.] |
| MCW5796793.1 | AAA+ATPase+wHTH+Alpha+Beta-domain+wHTH+RRM | Type-1_BR.RM.like.systems | DUF4096+DDE_Tnp_1+DDE_Tnp->?-><-DUF499*<-PIN<-?<-?<-METHYLASE | 852 | Bacteria>Nitrospirota>Nitrospiria>Nitrospirales>Nitrospiraceae | GCA_026128925.1 | Nitrospira_sp. | KIS97_20860 | 1483..4041 | 70125 | MAG:-DUF499-domain-containing-protein-[Nitrospira-sp.] |
| MCP9450544.1 | AAA+ATPase+wHTH+Alpha+Beta-domain+wHTH+RRM | Type-1_BR.RM.like.systems | METHYLASE->?->DUF499*->REase+N6_MTase-> | 918 | Bacteria>Nitrospirota>Nitrospiria>Nitrospirales>Nitrospiraceae | GCA_024331085.1 | Nitrospira_sp. | NNA21_10860 | 3927..6683 | 70125 | DUF499-domain-containing-protein-[Nitrospira-sp.] |
| NGZ60735.1 | AAA+ATPase+wHTH+Alpha+Beta-domain+wHTH+RRM | Type-1_BR.RM.like.systems | <-DUF499*<-HEPN<-PolB<-METHYLASE-p | 916 | Bacteria>Nitrospirota>Nitrospiria>Nitrospirales>Nitrospiraceae | GCA_011090395.1 | Nitrospira_sp._LK265 | CV081_09575 | 239..2989 | 2039810 | AAA-family-ATPase-[Nitrospira-sp.-LK265] |
| MDD9819374.1 | AAA+ATPase+wHTH+Alpha+Beta-domain+wHTH+RRM | Type-1_BR.RM.like.systems | <-DUF499*<-PIN<-OLD-ABC<-METHYLASE-p | 939 | Bacteria>Nitrospirota>Nitrospiria>Nitrospirales>Nitrospiraceae | GCA_028817545.1 | Nitrospira_sp. | OXR07_00640 | 2590..5409 | 70125 | MAG:-DUF499-domain-containing-protein-[Nitrospira-sp.] |
| MCS7214945.1 | AAA+ATPase+wHTH+Alpha+Beta-domain+wHTH+RRM | Type-1_BR.RM.like.systems | <-GDResolvase+Recombinase+ZnR<-HDRnase\|\|HTH_26->TUDOR+SF1-SF2-Helicase+iREase+CTD+REase->METHYLASE->DUF499*->?->REase->?->Phage_integrase-><-?<-GDResolvase+Recombinase+ZnR | 911 | Bacteria>Nitrospirota>Thermodesulfovibrionia>Thermodesulfovibrionales>Thermodesulfovibrionaceae | GCA_025060625.1 | Thermodesulfovibrio_sp. | NZ826_02170 | 14884..17619 | 2067987 | ATP-binding-protein-[Thermodesulfovibrio-sp.] |
| WP_059175576.1 | AAA+ATPase+wHTH+Alpha+Beta-domain+wHTH+RRM | Type-1_BR.RM.like.systems | REase->TUDOR+SF1-SF2-Helicase+iREase+CTD+REase->METHYLASE->PolB->HEPN->METHYLASE-p->DUF499*-><-?<-?\|\|SAM-methylase-> | 842 | Bacteria>Nitrospirota>Thermodesulfovibrionia>Thermodesulfovibrionales>Thermodesulfovibrionaceae | GCF_001514535.1 | Thermodesulfovibrio_aggregans | TAGGR_RS01355 | 271390..273918 | 86166 | DUF499-domain-containing-protein-[Thermodesulfovibrio-aggregans] |
| WP_281755187.1 | AAA+ATPase+wHTH+Alpha+Beta-domain+wHTH+RRM | Type-1_BR.RM.like.systems | <-Phage_integrase<-XIS-HTH<-?\|\|?-><-DUF499*<-METHYLASE<-PIN<-?<-TUDOR+SF1-SF2-Helicase+iREase+CTD+REase | 846 | Bacteria>Nitrospirota>Thermodesulfovibrionia>Thermodesulfovibrionales>Thermodesulfovibrionaceae | GCF_027925485.1 | Thermodesulfovibrio_islandicus | QMF67_RS07335 | 1419604..1422144 | 53423 | DUF499-domain-containing-protein-[Thermodesulfovibrio-islandicus] |
| PMP69410.1 | AAA+ATPase+wHTH+Alpha+Beta-domain+wHTH+RRM | Type-1_BR.RM.like.systems | METHYLASE-p->DUF499*-><-tRNA\|\|?->SPOUT-> | 843 | Bacteria>Nitrospirota>Thermodesulfovibrionia>Thermodesulfovibrionales>Thermodesulfovibrionaceae | GCA_002878055.1 | Thermodesulfovibrio_aggregans | C0186_06805 | 924..3455 | 86166 | AAA-family-ATPase-[Thermodesulfovibrio-aggregans] |
| MCX8028203.1 | AAA+ATPase+wHTH+Alpha+Beta-domain+wHTH+RRM | Type-1_BR.RM.like.systems | PIN->MerR-HTH->?-><-DUF499*<-METHYLASE-p | 843 | Bacteria>Nitrospirota>Thermodesulfovibrionia>Thermodesulfovibrionales>unclassified_Thermodesulfovibrionales | GCA_026415695.1 | Thermodesulfovibrionales_bacterium | N3A62_10190 | 4116..6647 | 2814272 | MAG:-DUF499-domain-containing-protein-[Thermodesulfovibrionales-bacterium] |
| MBI5634975.1 | AAA+ATPase+wHTH+Alpha+Beta-domain+wHTH+RRM | Type-1_BR.RM.like.systems | TUDOR+SF1-SF2-Helicase+iREase+CTD+REase->?->METHYLASE->HEPN->DUF499*->HTH+MPTase->?->?-><-Schlafen+HATPase+HTH<-MqsA_antitoxin+HTH_3\|\|REase-> | 927 | Bacteria>Nitrospirota>unclassified_Nitrospirota | GCA_016217675.1 | Nitrospirota_bacterium | HZA15_16020 | 27832..30615 | 2026887 | ATP-binding-protein-[Nitrospirae-bacterium] |
| MBI2060378.1 | AAA+ATPase+wHTH+Alpha+Beta-domain+wHTH+RRM | Type-1_BR.RM.like.systems | PHP+SbcC->METHYLASE->?->DUF1828+REase->?->DUF499*-><-PAIREDC-HTH<-?<-?<-ZnR+TOPRIM+D5ATPase<-?<-XIS-HTH | 918 | Bacteria>Nitrospirota>unclassified_Nitrospirota | GCA_016180645.1 | Nitrospirota_bacterium | HYT87_11470 | 28661..31417 | 2026887 | ATP-binding-protein-[Nitrospirae-bacterium] |
| MBI5740568.1 | AAA+ATPase+wHTH+Alpha+Beta-domain+wHTH+RRM | Type-1_BR.RM.like.systems | SbcC->METHYLASE->PolB->HEPN->DUF499*->?->?->?-><-Peptidase_U32+Peptidase_U32_C | 919 | Bacteria>Nitrospirota>unclassified_Nitrospirota | GCA_016234115.1 | Nitrospirota_bacterium | HZA16_07595 | 59463..62222 | 2026887 | ATP-binding-protein-[Nitrospirae-bacterium] |
| PIV85847.1 | AAA+ATPase+wHTH+Alpha+Beta-domain+wHTH+RRM | Type-1_BR.RM.like.systems | <-DUF499*<-REase<-REase<-DUF1828+REase<-?<-METHYLASE-p | 916 | Bacteria>Nitrospirota>unclassified_Nitrospirota | GCA_002781445.1 | Nitrospirae_bacterium_CG17_big_fil_post_rev_8_21_14_2_50_50_9 | COW52_00060 | 1390..4140 | 1974070 | AAA-family-ATPase,-partial-[Nitrospirae-bacterium-CG17_big_fil_post_rev_8_21_14_2_50_50_9] |
| OIP61731.1 | AAA+ATPase+wHTH+Alpha+Beta-domain+wHTH+RRM | Type-1_BR.RM.like.systems | <-DUF499*<-REase<-REase<-DUF1828+REase<-?<-METHYLASE-p | 918 | Bacteria>Nitrospirota>unclassified_Nitrospirota | GCA_001873285.1 | Nitrospirae_bacterium_CG2_30_53_67 | AUK29_09775 | 1402..4158 | 1805278 | AAA-family-ATPase-[Nitrospirae-bacterium-CG2_30_53_67] |
| RMH36427.1 | AAA+ATPase+wHTH+Alpha+Beta-domain+wHTH+RRM | Type-1_BR.RM.like.systems | <-DUF499*<-PIN<-DUF433<-REase<-HTH | 933 | Bacteria>Nitrospirota>unclassified_Nitrospirota | GCA_003695915.1 | Nitrospirota_bacterium | D6690_05850 | 2045..4846 | 2026887 | ATP-binding-protein-[Nitrospirae-bacterium] |
| WP_215870431.1 | AAA+ATPase+wHTH+Alpha+Beta-domain+wHTH+RRM | Type-1_BR.RM.like.systems | <-DUF499*<-METHYLASE<-PIN<-TUDOR+SF1-SF2-Helicase+iREase+CTD+REase | 949 | Bacteria>Pseudomonadota>Acidithiobacillia>Acidithiobacillales>Acidithiobacillaceae | GCF_018853695.1 | Acidithiobacillus_caldus | HAQ02_RS11835 | 505..3354 | 33059 | DUF499-domain-containing-protein-[Acidithiobacillus-caldus] |
| MBU2768209.1 | AAA+ATPase+wHTH+Alpha+Beta-domain+wHTH+RRM | Type-1_BR.RM.like.systems | <-DUF499*<-REase<-OLD-ABC<-wHTH+REase<-METHYLASE<-HNH<-OLD-ABC | 542 | Bacteria>Pseudomonadota>Acidithiobacillia>Acidithiobacillales>Acidithiobacillaceae | GCF_018853935.1 | Acidithiobacillus_ferrivorans | HAP94_19055 | 1..1626 | 160808 | ATP-binding-protein-[Acidithiobacillus-ferrivorans] |
| WP_198926060.1 | AAA+ATPase+wHTH+Alpha+Beta-domain+wHTH+RRM | Type-1_BR.RM.like.systems | <-DUF499-p<-DUF499*<-wHTH+REase | 460 | Bacteria>Pseudomonadota>Acidithiobacillia>Acidithiobacillales>Acidithiobacillaceae | GCF_001705725.1 | Acidithiobacillus_thiooxidans | A6M27_RS22710 | 4222..5604 | 930 | DUF499-domain-containing-protein-[Acidithiobacillus-thiooxidans] |
| WP_056620958.1 | AAA+ATPase+wHTH+Alpha+Beta-domain+wHTH+RRM | Type-1_BR.RM.like.systems | HTH+WYL->REase+35exo+Beta-Barrel+SF1-SF2-Helicase->TUDOR+SF1-SF2-Helicase+iREase+CTD->?->TUDOR+SF1-SF2-Helicase+iREase+CTD+REase->METHYLASE->DUF499*-><-?<-35exo | 939 | Bacteria>Pseudomonadota>Alphaproteobacteria>Caulobacterales>Caulobacteraceae | GCF_001426805.1 | Brevundimonas_sp._Root1423 | ASD25_RS23505 | 377520..380339 | 1736462 | DUF499-domain-containing-protein-[Brevundimonas-sp.-Root1423] |
| WP_244532343.1 | AAA+ATPase+wHTH+Alpha+Beta-domain+wHTH+RRM | Type-1_BR.RM.like.systems | METHYLASE->HEPN->DUF499*-><-?\|\|MODE-HTH+PBPII->\|\|?->?->HTH_24->ZnR-> | 752 | Bacteria>Pseudomonadota>Alphaproteobacteria>Hyphomicrobiales>Beijerinckiaceae | GCF_900114285.1 | Methylocapsa_palsarum | BM177_RS13845 | 3654..5912 | 1612308 | DUF499-domain-containing-protein-[Methylocapsa-palsarum] |
| SFK63485.1 | AAA+ATPase+wHTH+Alpha+Beta-domain+wHTH+RRM | Type-1_BR.RM.like.systems | METHYLASE->DUF499*->DUF499-p-><-?\|\|MODE-HTH+PBPII->?->?->HTH_24-> | 896 | Bacteria>Pseudomonadota>Alphaproteobacteria>Hyphomicrobiales>Beijerinckiaceae | GCF_900114285.1 | Methylocapsa_palsarum | SAMN05444581_1132 | 3222..5912 | 1612308 | Predicted-ATPase,-AAA+-superfamily-[Methylocapsa-palsarum] |
| MBX9865205.1 | AAA+ATPase+wHTH+Alpha+Beta-domain+wHTH+RRM | Type-1_BR.RM.like.systems | <-TIR<-?<-?<-?<-DUF499*<-METHYLASE<-TUDOR+SF1-SF2-Helicase+iREase+CTD+REase<-HTH_Tnp<-HTH+WYL\|\|AbiEi_4->AbiEii-> | 948 | Bacteria>Pseudomonadota>Alphaproteobacteria>Hyphomicrobiales>Hyphomicrobiaceae | GCA_019744875.1 | Hyphomicrobium_sp. | K2Y42_20880 | 17553..20399 | 82 | DUF499-domain-containing-protein-[Hyphomicrobium-sp.] |
| WP_147850359.1 | AAA+ATPase+wHTH+Alpha+Beta-domain+wHTH+RRM | Type-1_BR.RM.like.systems | TUDOR+SF1-SF2-Helicase+iREase+CTD+REase-p->HhH-RADC+JAB->TUDOR+SF1-SF2-Helicase+iREase+CTD+REase->METHYLASE->DUF499*->?->McrC-NTD+REase-><-REase<-Phage_integrase\|\|HNH-><-DDE_Tnp_ISAZ013 | 952 | Bacteria>Pseudomonadota>Alphaproteobacteria>Hyphomicrobiales>Hyphomicrobiales_incertae_sedis | GCF_008039615.1 | Vineibacter_terrae | FHP25_RS28385 | 20762..23620 | 2586908 | DUF499-domain-containing-protein-[Vineibacter-terrae] |
| WP_244479882.1 | AAA+ATPase+wHTH+Alpha+Beta-domain+wHTH+RRM | Type-1_BR.RM.like.systems | TUDOR+SF1-SF2-Helicase+iREase+CTD+REase->OLD-ABC->HNH->METHYLASE->DUF499*->GDResolvase+HTH->ParB+HNH-><-?\|\|HTH_3-><-?\|\|REase-> | 950 | Bacteria>Pseudomonadota>Alphaproteobacteria>Hyphomicrobiales>Methylobacteriaceae | GCF_001426045.1 | Methylobacterium_sp._Leaf94 | ASG63_RS11585 | 13925..16777 | 1736250 | ATP-binding-protein-[Methylobacterium-sp.-Leaf94] |
| KQU25484.1 | AAA+ATPase+wHTH+Alpha+Beta-domain+wHTH+RRM | Type-1_BR.RM.like.systems | REase-><-?\|\|?->TUDOR+SF1-SF2-Helicase+iREase+CTD+REase->OLD-ABC-><-?\|\|DUF499*->GDResolvase+HTH-><-?\|\|HTH_3-><-?\|\|REase-> | 944 | Bacteria>Pseudomonadota>Alphaproteobacteria>Hyphomicrobiales>Methylobacteriaceae | GCA_001426045.1 | Methylobacterium_sp._Leaf94 | ASG63_20450 | 13943..16777 | 1736250 | hypothetical-protein-ASG63_20450-[Methylobacterium-sp.-Leaf94] |
| WP_047191423.1 | AAA+ATPase+wHTH+Alpha+Beta-domain+wHTH+RRM | Type-1_BR.RM.like.systems | Phage_integrase->HTH_3->TUDOR+SF1-SF2-Helicase+iREase+CTD+REase->METHYLASE->DUF499*->?->REase+SAVED-> | 936 | Bacteria>Pseudomonadota>Alphaproteobacteria>Hyphomicrobiales>Methylobacteriaceae | GCF_001017175.1 | Microvirga_vignae | AA309_RS23290 | 25634..28444 | 1225564 | ATP-binding-protein-[Microvirga-vignae] |
| WP_262028302.1 | AAA+ATPase+wHTH+Alpha+Beta-domain+wHTH+RRM | Type-1_BR.RM.like.systems | HTH_3->TUDOR+SF1-SF2-Helicase+iREase+CTD+REase->METHYLASE->DUF499*-> | 939 | Bacteria>Pseudomonadota>Alphaproteobacteria>Hyphomicrobiales>Methylobacteriaceae | GCF_024519725.1 | Microvirga_sp._Mcv34 | OC446_RS06820 | 11099..13918 | 2926016 | ATP-binding-protein-[Microvirga-sp.-Mcv34] |
| WP_203270238.1 | AAA+ATPase+wHTH+Alpha+Beta-domain+wHTH+RRM | Type-1_BR.RM.like.systems | Methyltransf_25-><-?<-?<-DUF499*<-METHYLASE<-TUDOR+SF1-SF2-Helicase+iREase+CTD+REase | 1004 | Bacteria>Pseudomonadota>Alphaproteobacteria>Hyphomicrobiales>Methylobacteriaceae | GCF_016811235.1 | Microvirga_arabica | JQC81_RS03035 | 153789..156803 | 1128671 | DUF499-domain-containing-protein-[Microvirga-arabica] |
| WP_246744939.1 | AAA+ATPase+wHTH+Alpha+Beta-domain+wHTH+RRM | Type-1_BR.RM.like.systems | CHASE2+ACYC->?->\|\|DUF499*-><-?<-?\|\|?->FAD-NAD-dep-oxidoreductase-> | 513 | Bacteria>Pseudomonadota>Alphaproteobacteria>Hyphomicrobiales>Methylocystaceae | GCF_016745395.1 | Methylocystis_sp._Sn-Cys | JJT62_RS15690 | 194164..195705 | 1701263 | DUF499-domain-containing-protein-[Methylocystis-sp.-Sn-Cys] |
| GAJ31429.1 | AAA+ATPase+wHTH+Alpha+Beta-domain+wHTH+RRM | Type-1_BR.RM.like.systems | ZnR+TOPRIM->TOPRIM+SF1-SF2-Helicase+HTH->?->?->REase->REase-><-DUF499*<-METHYLASE<-ASCH<-TUDOR+SF1-SF2-Helicase+iREase+CTD+REase<-?<-MULE-TRANSPOSASE<-tRNA<-HISKIN+MORC | 927 | Bacteria>Pseudomonadota>Alphaproteobacteria>Hyphomicrobiales>Nitrobacteraceae | GCA_000617845.2 | Bradyrhizobium_sp._DOA9 | BDOA9_0106080 | 657043..659826 | 1126627 | hypothetical-protein-BDOA9_0106080-[Bradyrhizobium-sp.-DOA9] |
| RXG91618.1 | AAA+ATPase+wHTH+Alpha+Beta-domain+wHTH+RRM | Type-1_BR.RM.like.systems | Methylase->Radical-SAM->Phage_integrase->TUDOR+SF1-SF2-Helicase+iREase+CTD+REase->METHYLASE->DUF499*->?-><-PIN<-?<-?\|\|?-><-Phage_integrase | 946 | Bacteria>Pseudomonadota>Alphaproteobacteria>Hyphomicrobiales>Nitrobacteraceae | GCA_004114465.1 | Bradyrhizobium_zhanjiangense | EAS62_24370 | 123719..126559 | 1325107 | ATP-binding-protein-[Bradyrhizobium-zhanjiangense] |
| WP_245509281.1 | AAA+ATPase+wHTH+Alpha+Beta-domain+wHTH+RRM | Type-1_BR.RM.like.systems | Methylase->Radical-SAM->Phage_integrase->?->TUDOR+SF1-SF2-Helicase+iREase+CTD+REase->METHYLASE->DUF499*->?-><-PIN<-?<-?<-?<-Phage_integrase | 940 | Bacteria>Pseudomonadota>Alphaproteobacteria>Hyphomicrobiales>Nitrobacteraceae | GCF_004114465.1 | Bradyrhizobium_zhanjiangense | EAS62_RS24260 | 123737..126559 | 1325107 | ATP-binding-protein-[Bradyrhizobium-zhanjiangense] |
| WP_166097056.1 | AAA+ATPase+wHTH+Alpha+Beta-domain+wHTH+RRM | Type-1_BR.RM.like.systems | MerR-HTH->tRNA->TUDOR+SF1-SF2-Helicase+iREase+CTD+REase->OLD-ABC->HNH->METHYLASE->DUF499*->DUF433->PIN-><-Phage_integrase | 945 | Bacteria>Pseudomonadota>Alphaproteobacteria>Hyphomicrobiales>Nitrobacteraceae | GCF_011516635.5 | Bradyrhizobium_barranii | HAP41_RS22275 | 4684908..4687745 | 2992140 | ATP-binding-protein-[Bradyrhizobium-barranii] |
| BCF44179.1 | AAA+ATPase+wHTH+Alpha+Beta-domain+wHTH+RRM | Type-1_BR.RM.like.systems | TUDOR+SF1-SF2-Helicase+iREase+CTD+REase->METHYLASE->DUF499*->?->?-><-?<-?<-ParB | 947 | Bacteria>Pseudomonadota>Alphaproteobacteria>Hyphomicrobiales>Nitrobacteraceae | - | Bradyrhizobium_diazoefficiens | XF16B_46690 | 4996597..4999440 | 1355477 | hypothetical-protein-XF16B_46690-[Bradyrhizobium-diazoefficiens] |
| WP_257787586.1 | AAA+ATPase+wHTH+Alpha+Beta-domain+wHTH+RRM | Type-1_BR.RM.like.systems | <-AbiEii<-AbiEi_4\|\|CHTH->?-><-?\|\|REase-><-DUF499* | 841 | Bacteria>Pseudomonadota>Alphaproteobacteria>Hyphomicrobiales>Nitrobacteraceae | GCF_000648595.1 | Bradyrhizobium_diazoefficiens | QIU08_RS38570 | 94057..96582 | 1355477 | DUF499-domain-containing-protein-[Bradyrhizobium-diazoefficiens] |
| WP_247337805.1 | AAA+ATPase+wHTH+Alpha+Beta-domain+wHTH+RRM | Type-1_BR.RM.like.systems | SF1-SF2-Helicase+REase->SF1-SF2-Helicase+REase->?->HKD-DNase+HKD-DNase-><-McrC-NTD+REase<-?<-DUF499*<-METHYLASE<-TUDOR+SF1-SF2-Helicase+iREase+CTD+REase<-?<-XIS-HTH<-Phage_integrase | 960 | Bacteria>Pseudomonadota>Alphaproteobacteria>Hyphomicrobiales>Nitrobacteraceae | GCF_023100665.1 | unclassified_Bradyrhizobium | IVB54_RS34195 | 7122517..7125399 | 2631580 | MULTISPECIES:-DUF499-domain-containing-protein-[unclassified-Bradyrhizobium] |
| WP_271594377.1 | AAA+ATPase+wHTH+Alpha+Beta-domain+wHTH+RRM | Type-1_BR.RM.like.systems | SF1-SF2-Helicase+REase->SF1-SF2-Helicase+REase->?-><-McrC-NTD+REase<-?<-DUF499*<-METHYLASE<-TUDOR+SF1-SF2-Helicase+iREase+CTD+REase<-XIS-HTH<-Phage_integrase | 960 | Bacteria>Pseudomonadota>Alphaproteobacteria>Hyphomicrobiales>Nitrobacteraceae | GCF_028128835.1 | Bradyrhizobium_sp._CCBAU_65884 | XI03_RS32350 | 114486..117368 | 722477 | DUF499-domain-containing-protein-[Bradyrhizobium-sp.-CCBAU-65884] |
| WP_232674387.1 | AAA+ATPase+wHTH+Alpha+Beta-domain+wHTH+RRM | Type-1_BR.RM.like.systems | Phage_integrase->XIS-HTH->?->TUDOR+SF1-SF2-Helicase+iREase+CTD+REase->METHYLASE->DUF499*->?->McrC-NTD+REase-><-?<-SF1-SF2-Helicase+REase<-SF1-SF2-Helicase+REase | 960 | Bacteria>Pseudomonadota>Alphaproteobacteria>Hyphomicrobiales>Nitrobacteraceae | - | Bradyrhizobium_japonicum | LUG63_RS19105 | 362909..365791 | 375 | DUF499-domain-containing-protein-[Bradyrhizobium-japonicum] |
| WP_038382719.1 | AAA+ATPase+wHTH+Alpha+Beta-domain+wHTH+RRM | Type-1_BR.RM.like.systems | HTH_3-><-?\|\|TUDOR+SF1-SF2-Helicase+iREase+CTD+REase->METHYLASE->DUF499*->Phage_integrase-><-?\|\|GDResolvase+PPP1-HTH->ParB->ParB->HEPN-> | 928 | Bacteria>Pseudomonadota>Alphaproteobacteria>Hyphomicrobiales>Nitrobacteraceae | GCF_000261505.1 | Bradyrhizobium_elkanii | IC57_RS05230 | 16881..19667 | 29448 | DUF499-domain-containing-protein-[Bradyrhizobium-elkanii] |
| WP_116622075.1 | AAA+ATPase+wHTH+Alpha+Beta-domain+wHTH+RRM | Type-1_BR.RM.like.systems | HTH_3->TUDOR+SF1-SF2-Helicase+iREase+CTD+REase->OLD-ABC->HNH->ASCH->METHYLASE->DUF499*-><-?<-TraD\|\|MobA_MobL+SF1-SF2-helicase->XIS-HTH->?->PhdYeFM_antitox-> | 945 | Bacteria>Pseudomonadota>Alphaproteobacteria>Hyphomicrobiales>Phyllobacteriaceae | GCF_003403035.1 | Mesorhizobium_denitrificans | DY251_RS01610 | 349492..352329 | 2294114 | DUF499-domain-containing-protein-[Mesorhizobium-denitrificans] |
| WP_280515628.1 | AAA+ATPase+wHTH+Alpha+Beta-domain+wHTH+RRM | Type-1_BR.RM.like.systems | <-HTH_Tnp_1+HTH_21+rve+rve_3\|\|?->?->METHYLASE-p->METHYLASE->HEPN+DUF499->DUF499*->ParB+HTH->HTH_3->ParB+HNH-p->ParB+HNH-p->URI-DNase-> | 678 | Bacteria>Pseudomonadota>Alphaproteobacteria>Hyphomicrobiales>Phyllobacteriaceae | - | Mesorhizobium_sediminum | HK436_RS32255 | 845119..847155 | 1849104 | DUF499-domain-containing-protein-[Mesorhizobium-sediminum] |
| WP_140734688.1 | AAA+ATPase+wHTH+Alpha+Beta-domain+wHTH+RRM | Type-1_BR.RM.like.systems | Phage_terminase->TUDOR+SF1-SF2-Helicase+iREase+CTD+REase->OLD-ABC->HNH->METHYLASE->DUF499*->?->\|\|?-><-?<-?<-?\|\|Phage_integrase-> | 942 | Bacteria>Pseudomonadota>Alphaproteobacteria>Hyphomicrobiales>Phyllobacteriaceae | GCF_006442905.1 | Mesorhizobium_sp._B2-4-9 | FJ945_RS19675 | 32509..35337 | 2589940 | ATP-binding-protein-[Mesorhizobium-sp.-B2-4-9] |
| RWB26422.1 | AAA+ATPase+wHTH+Alpha+Beta-domain+wHTH+RRM | Type-1_BR.RM.like.systems | <-GDResolvase+Recombinase<-?\|\|?->N6_MTase->REase-><-DUF499*<-METHYLASE-p | 944 | Bacteria>Pseudomonadota>Alphaproteobacteria>Hyphomicrobiales>Phyllobacteriaceae | GCA_004016585.1 | Mesorhizobium_sp. | EOQ43_30895 | 29681..32515 | 1871066 | ATP-binding-protein-[Mesorhizobium-sp.] |
| WP_216964241.1 | AAA+ATPase+wHTH+Alpha+Beta-domain+wHTH+RRM | Type-1_BR.RM.like.systems | Phage_AlpA->XIS-HTH->Phage_integrase->GDResolvase+Recombinase+ZnR->TUDOR+SF1-SF2-Helicase+iREase+CTD+REase->METHYLASE->DUF499*-> | 942 | Bacteria>Pseudomonadota>Alphaproteobacteria>Hyphomicrobiales>Reyranellaceae | GCF_019039055.1 | Reyranella_sp._MMS21-HV4-11 | KQ910_RS19035 | 150193..153021 | 2849149 | ATP-binding-protein-[Reyranella-sp.-MMS21-HV4-11] |
| WP_164056349.1 | AAA+ATPase+wHTH+Alpha+Beta-domain+wHTH+RRM | Type-1_BR.RM.like.systems | Arm-DNA-bind_3+Phage_integrase-><-XIS-HTH<-MobA_MobL+SF1-SF2-helicase\|\|TraD->?-><-DUF499*<-METHYLASE<-HNH<-OLD-ABC<-TUDOR+SF1-SF2-Helicase+iREase+CTD+REase<-HTH_3<-HigB | 944 | Bacteria>Pseudomonadota>Alphaproteobacteria>Hyphomicrobiales>Rhizobiaceae | GCF_010669145.1 | Rhizobium_oryzihabitans | G3A56_RS10085 | 1920588..1923422 | 2267833 | ATP-binding-protein-[Rhizobium-oryzihabitans] |
| WP_200985559.1 | AAA+ATPase+wHTH+Alpha+Beta-domain+wHTH+RRM | Type-1_BR.RM.like.systems | <-VirD4-TraG+VirD4-FtsK<-?<-Classical-AAA<-PIN+OST-HTH\|\|MobA_MobL+SF1-SF2-helicase-><-DUF499*<-METHYLASE<-HNH<-OLD-ABC<-TUDOR+SF1-SF2-Helicase+iREase+CTD+REase<-HTH_3 | 944 | Bacteria>Pseudomonadota>Alphaproteobacteria>Hyphomicrobiales>Rhizobiaceae | - | Rhizobium_rhizogenes | JL018_RS00080 | 14669..17503 | 359 | ATP-binding-protein-[Agrobacterium-rhizogenes] |
| WP_242223016.1 | AAA+ATPase+wHTH+Alpha+Beta-domain+wHTH+RRM | Type-1_BR.RM.like.systems | HigB->HTH_3->TUDOR+SF1-SF2-Helicase+iREase+CTD+REase->OLD-ABC->HNH->METHYLASE->DUF499*-><-?<-TraD\|\|MobA_MobL+SF1-SF2-helicase-><-?\|\|XIS-HTH-> | 946 | Bacteria>Pseudomonadota>Alphaproteobacteria>Hyphomicrobiales>Rhizobiaceae | GCF_022682305.1 | Shinella_zoogloeoides | MOE34_RS09030 | 1765134..1767974 | 352475 | ATP-binding-protein-[Shinella-zoogloeoides] |
| WP_281435123.1 | AAA+ATPase+wHTH+Alpha+Beta-domain+wHTH+RRM | Type-1_BR.RM.like.systems | <-XIS-HTH\|\|HTH_3->TUDOR+SF1-SF2-Helicase+iREase+CTD+REase->OLD-ABC->HNH->METHYLASE->DUF499*->MODE-HTH+PBPII-><-ParG_RHH | 943 | Bacteria>Pseudomonadota>Alphaproteobacteria>Hyphomicrobiales>Rhizobiaceae | GCF_022968395.1 | Rhizobium_lemnae | MUU53_RS12880 | 44328..47159 | 1214924 | ATP-binding-protein-[Rhizobium-lemnae] |
| WP_247261244.1 | AAA+ATPase+wHTH+Alpha+Beta-domain+wHTH+RRM | Type-1_BR.RM.like.systems | <-XIS-HTH\|\|HTH_3->TUDOR+SF1-SF2-Helicase+iREase+CTD+REase->OLD-ABC->HNH->METHYLASE->DUF499*->MODE-HTH+PBPII-><-ParG_RHH | 937 | Bacteria>Pseudomonadota>Alphaproteobacteria>Hyphomicrobiales>Rhizobiaceae | - | Rhizobium_lemnae | MUU53_12875 | 44346..47159 | 1214924 | ATP-binding-protein-[Rhizobium-lemnae] |
| MCJ8508804.1 | AAA+ATPase+wHTH+Alpha+Beta-domain+wHTH+RRM | Type-1_BR.RM.like.systems | <-XIS-HTH\|\|HTH_3->TUDOR+SF1-SF2-Helicase+iREase+CTD+REase->OLD-ABC->HNH->METHYLASE->DUF499*->MODE-HTH+PBPII-><-ParG_RHH | 937 | Bacteria>Pseudomonadota>Alphaproteobacteria>Hyphomicrobiales>Rhizobiaceae | GCA_022968395.1 | Rhizobium_lemnae | MUU53_12875 | 44346..47159 | 1214924 | ATP-binding-protein-[Rhizobium-lemnae] |
| WP_069615760.1 | AAA+ATPase+wHTH+Alpha+Beta-domain+wHTH+RRM | Type-1_BR.RM.like.systems | HTH_3->TUDOR+SF1-SF2-Helicase+iREase+CTD+REase->OLD-ABC->HNH->METHYLASE->DUF499*-><-?<-N6_MTase<-HTH<-?\|\|?->McrC-NTD+REase-> | 945 | Bacteria>Pseudomonadota>Alphaproteobacteria>Hyphomicrobiales>Rhizobiaceae | GCF_001729315.1 | Rhizobium_sp._YK2 | A9Z06_RS30975 | 24492..27329 | 1860096 | ATP-binding-protein-[Rhizobium-sp.-YK2] |
| WP_012755925.1 | AAA+ATPase+wHTH+Alpha+Beta-domain+wHTH+RRM | Type-1_BR.RM.like.systems | Phage_integrase->?->?->OLD-ABC+TOPRIM->?-><-DUF499*<-METHYLASE<-HNH<-OLD-ABC<-TUDOR+SF1-SF2-Helicase+iREase+CTD+REase<-HerA/FtsK | 945 | Bacteria>Pseudomonadota>Alphaproteobacteria>Hyphomicrobiales>Rhizobiaceae | GCF_000023185.1 | Rhizobium_leguminosarum | RLEG_RS01005 | 199729..202566 | 384 | ATP-binding-protein-[Rhizobium-leguminosarum] |
| WP_164568970.1 | AAA+ATPase+wHTH+Alpha+Beta-domain+wHTH+RRM | Type-1_BR.RM.like.systems | XIS-HTH->TUDOR+SF1-SF2-Helicase+iREase+CTD+REase->OLD-ABC->HNH->METHYLASE->DUF499*-><-?<-Radical-SAM | 943 | Bacteria>Pseudomonadota>Alphaproteobacteria>Hyphomicrobiales>Rhizobiaceae | GCF_010668765.1 | Rhizobium_ruizarguesonis | GUK21_31970 | 69157..71988 | 2081791 | ATP-binding-protein-[Rhizobium-leguminosarum] |
| NEJ60793.1 | AAA+ATPase+wHTH+Alpha+Beta-domain+wHTH+RRM | Type-1_BR.RM.like.systems | XIS-HTH->TUDOR+SF1-SF2-Helicase+iREase+CTD+REase->OLD-ABC->HNH->METHYLASE->DUF499*-><-?<-Radical-SAM | 943 | Bacteria>Pseudomonadota>Alphaproteobacteria>Hyphomicrobiales>Rhizobiaceae | GCA_010668765.1 | Rhizobium_ruizarguesonis | GUK21_31970 | 69157..71988 | 2081791 | DUF499-domain-containing-protein-[Rhizobium-leguminosarum] |
| WP_010968623.1 | AAA+ATPase+wHTH+Alpha+Beta-domain+wHTH+RRM | Type-1_BR.RM.like.systems | HTH_3->TUDOR+SF1-SF2-Helicase+iREase+CTD+REase->OLD-ABC->HNH->METHYLASE->DUF499*-><-?<-REase<-HTH_3 | 942 | Bacteria>Pseudomonadota>Alphaproteobacteria>Hyphomicrobiales>Rhizobiaceae | GCF_000006965.1 | Sinorhizobium_meliloti | J3S94_RS16455 | 3478625..3481453 | 382 | ATP-binding-protein-[Sinorhizobium-meliloti] |
| WP_130838769.1 | AAA+ATPase+wHTH+Alpha+Beta-domain+wHTH+RRM | Type-1_BR.RM.like.systems | <-REase\|\|TUDOR+SF1-SF2-Helicase+iREase+CTD+REase->OLD-ABC->HNH->METHYLASE->DUF499*-> | 944 | Bacteria>Pseudomonadota>Alphaproteobacteria>Hyphomicrobiales>Rhizobiaceae | GCF_004305895.1 | Rhizobium_ruizarguesonis | HA464_RS06765 | 1375987..1378821 | 2081791 | ATP-binding-protein-[Rhizobium-leguminosarum] |
| WP_248132970.1 | AAA+ATPase+wHTH+Alpha+Beta-domain+wHTH+RRM | Type-1_BR.RM.like.systems | <-DUF499*<-METHYLASE<-HNH<-OLD-ABC<-TUDOR+SF1-SF2-Helicase+iREase+CTD+REase<-HTH_3 | 506 | Bacteria>Pseudomonadota>Alphaproteobacteria>Hyphomicrobiales>Rhizobiaceae | GCF_023119065.1 | Sinorhizobium_meliloti | J3S92_RS00005 | 1..1518 | 382 | DUF499-domain-containing-protein,-partial-[Sinorhizobium-meliloti] |
| WP_281418752.1 | AAA+ATPase+wHTH+Alpha+Beta-domain+wHTH+RRM | Type-1_BR.RM.like.systems | <-DUF499*\|\|?->?->?->?-><-GDResolvase+HIN-HTH<-GDResolvase+HIN-HTH | 240 | Bacteria>Pseudomonadota>Alphaproteobacteria>Hyphomicrobiales>Rhizobiaceae | GCF_014202175.1 | Rhizobium_rosettiformans | HNR26_RS22135 | 295..1017 | 1368430 | DUF499-domain-containing-protein-[Rhizobium-rosettiformans] |
| WP_246035161.1 | AAA+ATPase+wHTH+Alpha+Beta-domain+wHTH+RRM | Type-1_BR.RM.like.systems | <-DUF499*\|\|?->?->?->?-><-GDResolvase+HIN-HTH<-DUF4158+DDE_Tnp_Tn3 | 249 | Bacteria>Pseudomonadota>Alphaproteobacteria>Hyphomicrobiales>Rhizobiaceae | GCF_004912135.1 | Rhizobium_rosettiformans | FAA86_RS21860 | 201..950 | 1368430 | DUF499-domain-containing-protein-[Rhizobium-rosettiformans] |
| HCX68724.1 | AAA+ATPase+wHTH+Alpha+Beta-domain+wHTH+RRM | Type-1_BR.RM.like.systems | TUDOR+SF1-SF2-Helicase+iREase+CTD+REase->?->METHYLASE->HEPN->DUF499*-> | 525 | Bacteria>Pseudomonadota>Alphaproteobacteria>Hyphomicrobiales>Rhodobiaceae | GCA_003538155.1 | Rhodobiaceae_bacterium | DHK64_14660 | 8558..10133 | 2026785 | AAA-family-ATPase-[Rhodobiaceae-bacterium] |
| SDR63348.1 | AAA+ATPase+wHTH+Alpha+Beta-domain+wHTH+RRM | Type-1_BR.RM.like.systems | Phage_int_SAM_1+Phage_integrase->REase-><-?<-REase+35exo+Beta-Barrel+SF1-SF2-Helicase<-DUF499*<-METHYLASE<-HNH<-OLD-ABC<-TUDOR+SF1-SF2-Helicase+iREase+CTD+REase | 942 | Bacteria>Pseudomonadota>Alphaproteobacteria>Hyphomicrobiales>unclassified_Hyphomicrobiales | GCA_900103895.1 | Rhizobiales_bacterium_GAS113 | SAMN05519103_08517 | 30774..33602 | 1884352 | hypothetical-protein-SAMN05519103_08517-[Rhizobiales-bacterium-GAS113] |

| Protein-ID | DUF499-architecture | System | Gene-Neighbourhoods | Length | Taxonomy | Assembly-ID | Strain | Locus-Tag | Genomic-coordinates | TaxID | Fasta-defline |
|---|---|---|---|---|---|---|---|---|---|---|---|
| MCU0732818.1 | AAA+ATPase+wHTH+Alpha+Beta-domain+wHTH+RRM | Type-1_BR.RM.like.systems | METHYLASE-p->DUF499*->DUF499-p-><-?<-TraD\|\|MobA_MobL+SF1-SF2-helicase-> | 672 | Bacteria>Pseudomonadota>Alphaproteobacteria>Hyphomonadales>Hyphomonadaceae | GCA_025459075.1 | Hyphomonas_sp. | MUE84_14685 | 1072..3090 | 87 | ATP-binding-protein-[Hyphomonas-sp.] |
| WP_011714306.1 | AAA+ATPase+wHTH+Alpha+Beta-domain+wHTH+RRM | Type-1_BR.RM.like.systems | <-HISKIN+MORC<-SbcC\|\|HTH+WYL->REase+35exo+Beta-Barrel+SF1-SF2-Helicase->TUDOR+SF1-SF2-Helicase+iREase+CTD+REase->METHYLASE->DUF499*->\|\|Y1_Tnp-><-DDE_5 | 948 | Bacteria>Pseudomonadota>Alphaproteobacteria>Magnetococcales>Magnetococcaceae | GCF_000014865.1 | Magnetococcus_marinus | MMC1_RS13905 | 3444637..3447483 | 1124597 | DUF499-domain-containing-protein-[Magnetococcus-marinus] |
| MBF0185782.1 | AAA+ATPase+wHTH+Alpha+Beta-domain+wHTH+RRM | Type-1_BR.RM.like.systems | REase+35exo+Beta-Barrel+SF1-SF2-Helicase->TUDOR+SF1-SF2-Helicase+iREase+CTD+REase->HEPN->METHYLASE->?->PIN->DUF499*-> | 943 | Bacteria>Pseudomonadota>Alphaproteobacteria>Magnetococcales>unclassified_Magnetococcales | GCA_015231215.1 | Magnetococcales_bacterium | HQM06_15545 | 19234..22065 | 2026759 | ATP-binding-protein-[Magnetococcales-bacterium] |
| MBF0340191.1 | AAA+ATPase+wHTH+Alpha+Beta-domain+wHTH+RRM | Type-1_BR.RM.like.systems | <-DUF499*<-PIN<-?<-HEPN<-TUDOR+SF1-SF2-Helicase+iREase+CTD+REase<-HTH+WYL\|\|REase+SAVED-> | 705 | Bacteria>Pseudomonadota>Alphaproteobacteria>Magnetococcales>unclassified_Magnetococcales | GCA_015232245.1 | Magnetococcales_bacterium | HQL95_04415 | 72..2189 | 2026759 | ATP-binding-protein-[Magnetococcales-bacterium] |
| MBF0214027.1 | AAA+ATPase+wHTH+Alpha+Beta-domain+wHTH+RRM | Type-1_BR.RM.like.systems | TUDOR+SF1-SF2-Helicase+iREase+CTD+REase->PIN->METHYLASE->?->PIN->DUF499*-> | 946 | Bacteria>Pseudomonadota>Alphaproteobacteria>Magnetococcales>unclassified_Magnetococcales | GCA_015231295.1 | Magnetococcales_bacterium | HQM00_10780 | 8228..11068 | 2026759 | ATP-binding-protein-[Magnetococcales-bacterium] |
| MBF0177023.1 | AAA+ATPase+wHTH+Alpha+Beta-domain+wHTH+RRM | Type-1_BR.RM.like.systems | SAM-methylase-><-?<-?<-?<-SF1-SF2-Helicase+HRDC<-DUF499*<-REase<-OLD-ABC<-METHYLASE | 950 | Bacteria>Pseudomonadota>Alphaproteobacteria>Magnetococcales>unclassified_Magnetococcales | GCA_015231175.1 | Magnetococcales_bacterium | HQL63_09275 | 22537..25389 | 2026759 | ATP-binding-protein-[Magnetococcales-bacterium] |
| MBF0143013.1 | AAA+ATPase+wHTH+Alpha+Beta-domain+wHTH+RRM | Type-1_BR.RM.like.systems | TUDOR+SF1-SF2-Helicase+iREase+CTD+REase->PIN->METHYLASE->?->PIN->DUF499*-> | 945 | Bacteria>Pseudomonadota>Alphaproteobacteria>Magnetococcales>unclassified_Magnetococcales | GCA_015229005.1 | Magnetococcales_bacterium | HQL59_06090 | 11314..14151 | 2026759 | ATP-binding-protein-[Magnetococcales-bacterium] |
| MBF0138066.1 | AAA+ATPase+wHTH+Alpha+Beta-domain+wHTH+RRM | Type-1_BR.RM.like.systems | OLD-ABC->?->DUF499*->?->?->Radical-SAM+TRAM->PIN+PhoH-> | 961 | Bacteria>Pseudomonadota>Alphaproteobacteria>Magnetococcales>unclassified_Magnetococcales | GCA_015228935.1 | Magnetococcales_bacterium | HQL65_17685 | 1441..4326 | 2026759 | DUF499-domain-containing-protein-[Magnetococcales-bacterium] |
| MBF0416113.1 | AAA+ATPase+wHTH+Alpha+Beta-domain+wHTH+RRM | Type-1_BR.RM.like.systems | <-TOPRIM<-?<-REase+HTH<-DUF499*<-METHYLASE-p | 927 | Bacteria>Pseudomonadota>Alphaproteobacteria>Magnetococcales>unclassified_Magnetococcales | - | Magnetococcales_bacterium | HQL79_10150 | 22485..25268 | 2026759 | ATP-binding-protein-[Magnetococcales-bacterium] |
| MBF8274722.1 | AAA+ATPase+wHTH+Alpha+Beta-domain+wHTH+RRM | Type-1_BR.RM.like.systems | <-REase+HTH<-DUF499*<-METHYLASE | 935 | Bacteria>Pseudomonadota>Alphaproteobacteria>Magnetococcales>unclassified_Magnetococcales | GCA_015660855.1 | Magnetococcales_bacterium | HW380_3827 | 1830..4637 | 2026759 | hypothetical-protein-[Magnetococcales-bacterium] |
| MBF0182290.1 | AAA+ATPase+wHTH+Alpha+Beta-domain+wHTH+RRM | Type-1_BR.RM.like.systems | METHYLASE->?->MPTase->DUF499*-><-Mbetalac<-TPR+TPR | 947 | Bacteria>Pseudomonadota>Alphaproteobacteria>Magnetococcales>unclassified_Magnetococcales | GCA_015231235.1 | Magnetococcales_bacterium | HQM03_19925 | 5811..8654 | 2026759 | ATP-binding-protein-[Magnetococcales-bacterium] |
| MBF0182505.1 | AAA+ATPase+wHTH+Alpha+Beta-domain+wHTH+RRM | Type-1_BR.RM.like.systems | <-DUF499*<-MPTase<-?<-METHYLASE\|\|DUF433-> | 449 | Bacteria>Pseudomonadota>Alphaproteobacteria>Magnetococcales>unclassified_Magnetococcales | GCA_015231235.1 | Magnetococcales_bacterium | HQM03_21010 | 25..1374 | 2026759 | ATP-binding-protein-[Magnetococcales-bacterium] |
| WP_013654352.1 | AAA+ATPase+wHTH+Alpha+Beta-domain+wHTH+RRM | Type-1_BR.RM.like.systems | <-PIN<-XIS-HTH\|\|?-><-ArdCN+MPTase<-HEPN<-DUF499*<-METHYLASE<-HNH<-OLD-ABC<-TUDOR+SF1-SF2-Helicase+iREase+CTD+REase<-HTH_3<-HigB | 944 | Bacteria>Pseudomonadota>Alphaproteobacteria>Rhodobacterales>Paracoccaceae | GCF_000192745.1 | Polymorphum_gilvum | SL003B_RS18265 | 3920112..3922946 | 991904 | DUF499-domain-containing-protein-[Polymorphum-gilvum] |
| WP_221828739.1 | AAA+ATPase+wHTH+Alpha+Beta-domain+wHTH+RRM | Type-1_BR.RM.like.systems | ParG_RHH-><-TetR-HTH\|\|MODE-HTH+PBPII-><-?<-DUF499*\|\|VirB2->?->FtsK+VirB4-FtsK-> | 867 | Bacteria>Pseudomonadota>Alphaproteobacteria>Rhodobacterales>Paracoccaceae | GCF_019748455.1 | Paracoccus_yeei | H7K23_RS03540 | 14446..17049 | 147645 | DUF499-domain-containing-protein-[Paracoccus-yeei] |
| MYA89893.1 | AAA+ATPase+wHTH+Alpha+Beta-domain+wHTH+RRM | Type-1_BR.RM.like.systems | Arm-DNA-bind_3+Phage_int_SAM_3+Phage_integrase->Arm-DNA-bind_3+Phage_int_SAM_3+Phage_integrase-><-?<-DUF499*<-PIN<-DUF433<-METHYLASE-p | 926 | Bacteria>Pseudomonadota>Alphaproteobacteria>Rhodobacterales>Paracoccaceae | GCA_009839535.1 | Boseongicola_sp._SB0662_bin_57 | F4X97_15820 | 7656..10436 | 2604843 | ATP-binding-protein-[Boseongicola-sp.-SB0662_bin_57] |
| MBO6086676.1 | AAA+ATPase+wHTH+Alpha+Beta-domain+wHTH+RRM | Type-1_BR.RM.like.systems | DUF499*->FAD-NAD-dep-oxidoreductase->?->?->SF1-SF2-Helicase-p->HKD-DNase+SF1-SF2-Helicase-p-> | 48 | Bacteria>Pseudomonadota>Alphaproteobacteria>Rhodospirillales>Acetobacteraceae | GCA_017633635.1 | Acetobacter_sp. | J6P19_09845 | 1732..1878 | 440 | hypothetical-protein-[Acetobacter-sp.] |
| WP_253545856.1 | AAA+ATPase+wHTH+Alpha+Beta-domain+wHTH+RRM | Type-1_BR.RM.like.systems | <-HTH<-ArsR-HTH+YTCD-HTH\|\|?->GDResolvase+HIN-HTH->DUF499*->METHYLASE-p-><-HTH | 912 | Bacteria>Pseudomonadota>Alphaproteobacteria>Rhodospirillales>Acetobacteraceae | GCF_024158445.1 | Acetobacter_lovaniensis | NKW44_RS12945 | 30888..33626 | 104100 | DUF499-domain-containing-protein-[Acetobacter-lovaniensis] |
| TXH33435.1 | AAA+ATPase+wHTH+Alpha+Beta-domain+wHTH+RRM | Type-1_BR.RM.like.systems | HTH_3->TUDOR+SF1-SF2-Helicase+iREase+CTD+REase->OLD-ABC->HNH->ASCH->METHYLASE->DUF499*-><-?<-TraD\|\|MobA_MobL+SF1-SF2-helicase-> | 945 | Bacteria>Pseudomonadota>Alphaproteobacteria>Rhodospirillales>Rhodospirillaceae | GCA_008012315.1 | Rhodospirillaceae_bacterium | E6Q98_22210 | 66565..69402 | 1898112 | DUF499-domain-containing-protein-[Rhodospirillaceae-bacterium] |
| MBT3398604.1 | AAA+ATPase+wHTH+Alpha+Beta-domain+wHTH+RRM | Type-1_BR.RM.like.systems | REase->METHYLASE->DUF499*->HEPN->Methylase-><-Radical-SAM<-?<-Peptidase_M15_4 | 922 | Bacteria>Pseudomonadota>Alphaproteobacteria>Rhodospirillales>Rhodospirillaceae | GCA_018648085.1 | Rhodospirillaceae_bacterium | HN420_01275 | 4129..6897 | 1898112 | ATP-binding-protein-[Rhodospirillaceae-bacterium] |
| MCZ0954488.1 | AAA+ATPase+wHTH+Alpha+Beta-domain+wHTH+RRM | Type-1_BR.RM.like.systems | <-DUF499*<-REase<-METHYLASE | 920 | Bacteria>Pseudomonadota>Alphaproteobacteria>Rhodospirillales>Rhodospirillaceae | GCA_026914105.1 | Rhodospirillaceae_bacterium | DXJ56_18105 | 195..2957 | 1898112 | MAG:-DUF499-domain-containing-protein-[Rhodospirillaceae-bacterium] |
| KAF0112375.1 | AAA+ATPase+wHTH+Alpha+Beta-domain+wHTH+RRM | Type-1_BR.RM.like.systems | <-DDE_Tnp_1_2<-?\|\|DUF499*->?-><-?<-?<-Arm-DNA-bind_3+Phage_integrase | 756 | Bacteria>Pseudomonadota>Alphaproteobacteria>Rhodospirillales>Rhodospirillaceae | GCA_009773045.1 | Rhodospirillaceae_bacterium | FD149_2445 | 759..3029 | 1898112 | hypothetical-protein-FD149_2445-[Rhodospirillaceae-bacterium] |
| MDA8051502.1 | AAA+ATPase+wHTH+Alpha+Beta-domain+wHTH+RRM | Type-1_BR.RM.like.systems | TUDOR+SF1-SF2-Helicase+iREase+CTD+REase->OLD-ABC->TOPRIM->METHYLASE->DUF499*-><-?\|\|?->Arm-DNA-bind_3+Phage_int_SAM_3+Phage_integrase->?->ZnR+TOPRIM->D5N+D5ATPase-> | 933 | Bacteria>Pseudomonadota>Alphaproteobacteria>Rhodospirillales>unclassified_Rhodospirillales | GCA_028063655.1 | Rhodospirillales_bacterium | M0002_16125 | 8854..11655 | 2026786 | MAG:-DUF499-domain-containing-protein-[Rhodospirillales-bacterium] |
| TVR84793.1 | AAA+ATPase+wHTH+Alpha+Beta-domain+wHTH+RRM | Type-1_BR.RM.like.systems | NTP_transferase-><-DDE_Tnp_1\|\|PhdYeFM_antitox->TUDOR+SF1-SF2-Helicase+iREase+CTD+REase->METHYLASE->?->DUF499*->?->HerA/FtsK->?-><-REase+RNase | 920 | Bacteria>Pseudomonadota>Alphaproteobacteria>Rhodospirillales>unclassified_Rhodospirillales | GCA_007695045.1 | Rhodospirillales_bacterium | EA405_00800 | 51145..53907 | 2026786 | ATP-binding-protein-[Rhodospirillales-bacterium] |
| WP_066559804.1 | AAA+ATPase+wHTH+Alpha+Beta-domain+wHTH+RRM | Type-1_BR.RM.like.systems | CopG/RHH->TUDOR+SF1-SF2-Helicase+iREase+CTD+REase->OLD-ABC->HNH->ASCH->METHYLASE->DUF499*-><-?\|\|LexA-protease-><-wHTH+SAD+HNH<-TrwC+SF1-SF2-helicase<-VirD4-TraG+VirD4-FtsK<-CopG/RHH | 940 | Bacteria>Pseudomonadota>Alphaproteobacteria>Sphingomonadales>Erythrobacteraceae | GCF_001634625.1 | Croceicoccus_bisphenolivorans | A7A77_RS14325 | 44266..47088 | 1783232 | ATP-binding-protein-[Croceicoccus-bisphenolivorans] |
| MBL8657365.1 | AAA+ATPase+wHTH+Alpha+Beta-domain+wHTH+RRM | Type-1_BR.RM.like.systems | <-DUF499*<-METHYLASE<-ASCH<-HNH<-OLD-ABC<-TUDOR+SF1-SF2-Helicase+iREase+CTD+REase<-HTH_3 | 944 | Bacteria>Pseudomonadota>Alphaproteobacteria>Sphingomonadales>Erythrobacteraceae | GCA_016793865.1 | Altererythrobacter_sp. | JNJ92_08435 | 1147..3981 | 1872480 | DUF499-domain-containing-protein-[Altererythrobacter-sp.] |
| WP_159367773.1 | AAA+ATPase+wHTH+Alpha+Beta-domain+wHTH+RRM | Type-1_BR.RM.like.systems | <-DDE_Tnp_IS240<-?<-LZ_Tnp_IS66+zf-IS66+DDE_Tnp_IS66+DDE_Tnp_IS66_C<-?<-HIN-HTH<-PAIREDC-HTH+HIN-HTH<-DUF499*<-METHYLASE<-HNH<-OLD-ABC<-TUDOR+SF1-SF2-Helicase+iREase+CTD+REase<-CopG/RHH<-HTH+WYL | 940 | Bacteria>Pseudomonadota>Alphaproteobacteria>Sphingomonadales>Sphingomonadaceae | GCF_009856545.1 | Sphingobium_yanoikuyae | GS397_RS23435 | 4919345..4922167 | 13690 | ATP-binding-protein-[Sphingobium-yanoikuyae] |
| WP_168853122.1 | AAA+ATPase+wHTH+Alpha+Beta-domain+wHTH+RRM | Type-1_BR.RM.like.systems | CopG/RHH->VirD4-TraG+VirD4-FtsK->TrwC+SF1-SF2-helicase->wHTH(SBD)+HNH-><-LexA-protease\|\|?-><-DUF499*<-METHYLASE<-HNH<-OLD-ABC<-TUDOR+SF1-SF2-Helicase+iREase+CTD+REase<-CopG/RHH<-HTH+WYL | 940 | Bacteria>Pseudomonadota>Alphaproteobacteria>Sphingomonadales>Sphingomonadaceae | GCF_012641335.1 | Novosphingobium_sp._ERN07 | HGI47_RS12585 | 95856..98678 | 2726187 | ATP-binding-protein-[Novosphingobium-sp.-ERN07] |
| WP_130751563.1 | AAA+ATPase+wHTH+Alpha+Beta-domain+wHTH+RRM | Type-1_BR.RM.like.systems | CopG/RHH->VirD4-TraG+VirD4-FtsK->TrwC+SF1-SF2-helicase->wHTH(SBD)+HNH-><-LexA-protease\|\|?-><-DUF499*<-METHYLASE<-HNH<-OLD-ABC<-TUDOR+SF1-SF2-Helicase+iREase+CTD+REase<-CopG/RHH<-HTH+WYL | 940 | Bacteria>Pseudomonadota>Alphaproteobacteria>Sphingomonadales>Sphingomonadaceae | GCF_004305355.1 | Sphingobium_xenophagum | MBESOW_RS00075 | 12724..15546 | 121428 | ATP-binding-protein-[Sphingobium-xenophagum] |
| WP_099185493.1 | AAA+ATPase+wHTH+Alpha+Beta-domain+wHTH+RRM | Type-1_BR.RM.like.systems | HTH+WYL->CopG/RHH->TUDOR+SF1-SF2-Helicase+iREase+CTD+REase->OLD-ABC->HNH->METHYLASE->DUF499*-><-?\|\|LexA-protease-><-wHTH(SBD)+HNH<-TrwC+SF1-SF2-helicase<-VirD4-TraG+VirD4-FtsK<-CopG/RHH | 940 | Bacteria>Pseudomonadota>Alphaproteobacteria>Sphingomonadales>Sphingomonadaceae | GCF_002723855.1 | Sphingobium | SFOMI_RS05000 | 866506..869328 | 165695 | ATP-binding-protein-[Sphingobium-fuliginis] |
| GAY20452.1 | AAA+ATPase+wHTH+Alpha+Beta-domain+wHTH+RRM | Type-1_BR.RM.like.systems | HTH+WYL->TUDOR+SF1-SF2-Helicase+iREase+CTD+REase->OLD-ABC-><-?\|\|METHYLASE-><-?\|\|DUF499*->LexA-protease-><-wHTH+SAD+HNH<-TrwC+SF1-SF2-helicase<-VirD4-TraG+VirD4-FtsK | 851 | Bacteria>Pseudomonadota>Alphaproteobacteria>Sphingomonadales>Sphingomonadaceae | GCA_002723855.1 | Sphingobium_fuliginis | SFOMI_0976 | 866773..869328 | 336203 | predicted-ATPase-[Sphingobium-fuliginis] |
| WP_260153764.1 | AAA+ATPase+wHTH+Alpha+Beta-domain+wHTH+RRM | Type-1_BR.RM.like.systems | TUDOR+SF1-SF2-Helicase+iREase+CTD+REase->OLD-ABC->HNH->METHYLASE->DUF499*-> | 822 | Bacteria>Pseudomonadota>Alphaproteobacteria>Sphingomonadales>Sphingomonadaceae | - | Sphingomonas_desiccabilis | GGQ81_RS01635 | 357438..359906 | 429134 | DUF499-domain-containing-protein-[Sphingomonas-desiccabilis] |
| WP_206364336.1 | AAA+ATPase+wHTH+Alpha+Beta-domain+wHTH+RRM | Type-1_BR.RM.like.systems | TUDOR+SF1-SF2-Helicase+iREase+CTD+REase->OLD-ABC->HNH->METHYLASE->DUF499*-> | 934 | Bacteria>Pseudomonadota>Alphaproteobacteria>Sphingomonadales>Sphingomonadaceae | GCF_014196135.1 | Sphingomonas_desiccabilis | EO081_RS01620 | 357134..359938 | 429134 | ATP-binding-protein-[Sphingomonas-desiccabilis] |
| RXZ34416.1 | AAA+ATPase+wHTH+Alpha+Beta-domain+wHTH+RRM | Type-1_BR.RM.like.systems | TUDOR+SF1-SF2-Helicase+iREase+CTD+REase->OLD-ABC->HNH->METHYLASE->DUF499*-> | 940 | Bacteria>Pseudomonadota>Alphaproteobacteria>Sphingomonadales>Sphingomonadaceae | GCA_004135605.1 | Sphingomonas_desiccabilis | EO081_01620 | 357116..359938 | 429134 | DUF499-domain-containing-protein-[Sphingomonas-desiccabilis] |
| PXW67391.1 | AAA+ATPase+wHTH+Alpha+Beta-domain+wHTH+RRM | Type-1_BR.RM.like.systems | <-DUF499*<-HTH+WYL<-SbcC+SbcC<-Calcineurin\|\|ArdCN+MPTase->?->ParB+HTH-> | 56 | Bacteria>Pseudomonadota>Alphaproteobacteria>Sphingomonadales>Sphingomonadaceae | GCA_003201955.1 | Blastomonas_natatoria | C7451_1301 | 53..223 | 34015 | hypothetical-protein-C7451_1301-[Blastomonas-natatoria] |
| WP_037491503.1 | AAA+ATPase+wHTH+Alpha+Beta-domain+wHTH+RRM | Type-1_BR.RM.like.systems | <-ArdCN+MPTase\|\|Calcineurin->SbcC+SbcC->HTH+WYL->DUF499*-> | 210 | Bacteria>Pseudomonadota>Alphaproteobacteria>Sphingomonadales>Sphingomonadaceae | GCF_000382885.1 | Sphingobium_indicum | EBBID32_RS17205 | 7510..8139 | 332055 | ATP-binding-protein,-partial-[Sphingobium-indicum] |
| MDD3182285.1 | AAA+ATPase+wHTH+Alpha+Beta-domain+wHTH+RRM | Type-1_BR.RM.like.systems | <-ACET\|\|Peptidase_M15->?-><-SF1-SF2-Helicase<-OLD-ABC+TOPRIM<-DUF499*<-METHYLASE<-PIN<-TUDOR+SF1-SF2-Helicase+iREase+CTD+REase<-HTH+WYL | 940 | Bacteria>Pseudomonadota>Alphaproteobacteria>unclassified_Alphaproteobacteria | GCA_028691835.1 | Alphaproteobacteria_bacterium | PHD48_05740 | 6780..9602 | 1913988 | MAG:-DUF499-domain-containing-protein-[Alphaproteobacteria-bacterium] |
| EDP62636.1 | AAA+ATPase+wHTH+Alpha+Beta-domain+wHTH+RRM | Type-1_BR.RM.like.systems | <-GST_N+GST_C\|\|PADR-HTH->?-><-?\|\|?-><-DUF499*<-METHYLASE-p<-TUDOR+SF1-SF2-Helicase+iREase+CTD+REase<-HTH+WYL<-?\|\|DDE_Tnp_1_4-> | 744 | Bacteria>Pseudomonadota>Alphaproteobacteria>unclassified_Alphaproteobacteria | GCA_000171835.1 | alpha_proteobacterium_BAL199 | BAL199_22547 | 18891..21125 | 331869 | hypothetical-protein-BAL199_22547-[alpha-proteobacterium-BAL199] |
| MCB9682177.1 | AAA+ATPase+wHTH+Alpha+Beta-domain+wHTH+RRM | Type-1_BR.RM.like.systems | <-REase<-?\|\|TUDOR+SF1-SF2-Helicase+iREase+CTD+REase->METHYLASE->OLD-ABC->REase->DUF499*-><-CNMP+TIR<-ACYC+ACYC | 951 | Bacteria>Pseudomonadota>Alphaproteobacteria>unclassified_Alphaproteobacteria | GCA_020632915.1 | Alphaproteobacteria_bacterium | H6733_11990 | 40994..43849 | 1913988 | ATP-binding-protein-[Alphaproteobacteria-bacterium] |
| MDD3183268.1 | AAA+ATPase+wHTH+Alpha+Beta-domain+wHTH+RRM | Type-1_BR.RM.like.systems | <-DUF499*<-METHYLASE<-PIN<-TUDOR+SF1-SF2-Helicase+iREase+CTD+REase<-HTH+WYL | 795 | Bacteria>Pseudomonadota>Alphaproteobacteria>unclassified_Alphaproteobacteria | GCA_028691835.1 | Alphaproteobacteria_bacterium | PHD48_10775 | 1..2386 | 1913988 | MAG:-DUF499-domain-containing-protein,-partial-[Alphaproteobacteria-bacterium] |
| MBU1213428.1 | AAA+ATPase+wHTH+Alpha+Beta-domain+wHTH+RRM | Type-1_BR.RM.like.systems | <-DUF499*<-?<-?<-REase<-DNA_methylase<-?<-PIN | 419 | Bacteria>Pseudomonadota>Alphaproteobacteria>unclassified_Alphaproteobacteria | GCA_018819265.1 | Alphaproteobacteria_bacterium | KJ587_19500 | 798..2057 | 1913988 | DUF499-domain-containing-protein-[Alphaproteobacteria-bacterium] |
| MBV9199724.1 | AAA+ATPase+wHTH+Alpha+Beta-domain+wHTH+RRM | Type-1_BR.RM.like.systems | TUDOR+SF1-SF2-Helicase+iREase+CTD+REase-p->METHYLASE->?->DUF1828+REase->DUF499*->?->XIS-HTH->?->Primpol+PriCT1->DnaB-> | 917 | Bacteria>Pseudomonadota>Alphaproteobacteria>unclassified_Alphaproteobacteria | GCA_019246595.1 | Alphaproteobacteria_bacterium | JOY83_08345 | 6016..8769 | 1913988 | ATP-binding-protein-[Alphaproteobacteria-bacterium] |
| RMF09073.1 | AAA+ATPase+wHTH+Alpha+Beta-domain+wHTH+RRM | Type-1_BR.RM.like.systems | SF1-SF2-Helicase->TUDOR+SF1-SF2-Helicase+iREase+CTD+REase->METHYLASE->DUF499*-> | 923 | Bacteria>Pseudomonadota>Alphaproteobacteria>unclassified_Alphaproteobacteria | GCA_003694925.1 | Alphaproteobacteria_bacterium | D6773_02065 | 8427..11198 | 1913988 | ATP-binding-protein-[Alphaproteobacteria-bacterium] |
| MDA8001255.1 | AAA+ATPase+wHTH+Alpha+Beta-domain+wHTH+RRM | Type-1_BR.RM.like.systems | <-DUF499*<-DUF1828+REase<-?<-METHYLASE | 799 | Bacteria>Pseudomonadota>Alphaproteobacteria>unclassified_Alphaproteobacteria | GCA_028066975.1 | Alphaproteobacteria_bacterium | MPL62_08195 | 1..2396 | 1913988 | MAG:-DUF499-domain-containing-protein,-partial-[Alphaproteobacteria-bacterium] |
| MBL8396550.1 | AAA+ATPase+wHTH+Alpha+Beta-domain+wHTH+RRM | Type-1_BR.RM.like.systems | HTH+WYL->TUDOR+SF1-SF2-Helicase+iREase+CTD+REase->HEPN->METHYLASE->wHTH+REase->DUF499*->DDE_5+DDE_Tnp_1->?-><-MORC+DNA_gyraseB+TOPRIM | 939 | Bacteria>Pseudomonadota>Betaproteobacteria>Betaproteobacteria_incertae_sedis>Candidatus_Accumulibacter | GCA_016789495.1 | Accumulibacter_sp. | JNL84_00165 | 43696..46515 | 2053492 | ATP-binding-protein-[Candidatus-Accumulibacter-sp.] |
| WP_248595613.1 | AAA+ATPase+wHTH+Alpha+Beta-domain+wHTH+RRM | Type-1_BR.RM.like.systems | RelE-ParE->Phage_int_SAM_4+Phage_integrase->?->OLD-ABC->HNH->METHYLASE->DUF499*->DUF499-p->Phage_int_SAM_4+Phage_integrase-> | 644 | Bacteria>Pseudomonadota>Betaproteobacteria>Betaproteobacteria_incertae_sedis>Candidatus_Accumulibacter | GCF_012939955.1 | Candidatus_Accumulibacter_phosphatis | E4Q23_RS20285 | 20006..21940 | 327160 | DUF499-domain-containing-protein-[Candidatus-Accumulibacter-phosphatis] |
| MCM8600113.1 | AAA+ATPase+wHTH+Alpha+Beta-domain+wHTH+RRM | Type-1_BR.RM.like.systems | <-SbcC<-DUF499*<-Schlafen+HATPase+HTH<-METHYLASE<-METHYLASE<-HEPN<-TUDOR+SF1-SF2-Helicase+iREase+CTD+REase<-HTH_Tnp | 949 | Bacteria>Pseudomonadota>Betaproteobacteria>Betaproteobacteria_incertae_sedis>Candidatus_Accumulibacter | GCA_023806585.1 | Accumulibacter_sp. | NFW04_15895 | 29772..32621 | 2053492 | DUF499-domain-containing-protein-[Candidatus-Accumulibacter-sp.] |
| QKS27499.1 | AAA+ATPase+wHTH+Alpha+Beta-domain+wHTH+RRM | Type-1_BR.RM.like.systems | RNase->?-><-?<-?<-?<-DUF499*<-METHYLASE<-TUDOR+SF1-SF2-Helicase+iREase+CTD+REase<-HTH_Tnp<-HTH+WYL\|\|HIN-HTH-> | 948 | Bacteria>Pseudomonadota>Betaproteobacteria>Betaproteobacteria_incertae_sedis>Candidatus_Accumulibacter | GCA_013347225.1 | Candidatus_Accumulibacter_similis | HT579_00075 | 12929..15775 | 2954388 | MAG:-ATP-binding-protein-[Candidatus-Accumulibacter-similis] |
| SBT04363.1 | AAA+ATPase+wHTH+Alpha+Beta-domain+wHTH+RRM | Type-1_BR.RM.like.systems | TP_methylase-><-DUF499-p<-DUF499*<-REase<-OLD-ABC<-METHYLASE-p | 671 | Bacteria>Pseudomonadota>Betaproteobacteria>Betaproteobacteria_incertae_sedis>Candidatus_Accumulibacter | GCA_900089955.1 | Candidatus_Accumulibacter_aalborgensis | ACCAA_130242 | 226085..228100 | 1860102 | conserved-hypothetical-protein-[Candidatus-Accumulibacter-aalborgensis] |
| MCM8638973.1 | AAA+ATPase+wHTH+Alpha+Beta-domain+wHTH+RRM | Type-1_BR.RM.like.systems | METHYLASE-p->PIN->DUF499*-> | 937 | Bacteria>Pseudomonadota>Betaproteobacteria>Betaproteobacteria_incertae_sedis>Candidatus_Accumulibacter | GCA_023806635.1 | Accumulibacter_sp. | NFW30_04525 | 1720..4533 | 2053492 | DUF499-domain-containing-protein-[Candidatus-Accumulibacter-sp.] |
| KFB70630.1 | AAA+ATPase+wHTH+Alpha+Beta-domain+wHTH+RRM | Type-1_BR.RM.like.systems | <-DUF499-p<-DUF499*<-Schlafen+HATPase+HTH<-Schlafen<-AAA_21 | 484 | Bacteria>Pseudomonadota>Betaproteobacteria>Betaproteobacteria_incertae_sedis>Candidatus_Accumulibacter | GCA_000585035.2 | Candidatus_Accumulibacter_phosphatis | AW09_004270 | 1814..3268 | 327160 | MAG:-hypothetical-protein-AW09_004270-[Candidatus-Accumulibacter-phosphatis] |
| MCX8099543.1 | AAA+ATPase+wHTH+Alpha+Beta-domain+wHTH+RRM | Type-1_BR.RM.like.systems | <-SF1-SF2-Helicase+DUF1998<-ORC-CDC6+REase<-?<-DUF499*<-PIN<-HTH<-METHYLASE<-TOPRIM<-OLD-ABC<-TUDOR+SF1-SF2-Helicase+iREase+CTD+REase | 918 | Bacteria>Pseudomonadota>Betaproteobacteria>Betaproteobacteria_incertae_sedis>Casimicrobiaceae | GCA_026414825.1 | Casimicrobiaceae_bacterium | N3F11_11190 | 5518..8274 | 2906524 | MAG:-DUF499-domain-containing-protein-[Casimicrobiaceae-bacterium] |
| WP_206111008.1 | AAA+ATPase+wHTH+Alpha+Beta-domain+wHTH+RRM | Type-1_BR.RM.like.systems | HTH+WYL->HTH_Tnp->TUDOR+SF1-SF2-Helicase+iREase+CTD+REase->PIN->HEPN->METHYLASE->DUF499*->TIR->SLATT->SMODS->Methylase-><-Radical-SAM<-Phage_lysozyme | 955 | Bacteria>Pseudomonadota>Betaproteobacteria>Burkholderiales | GCF_017150735.1 | Burkholderiaceae | KTD10_RS22775 | 57014..59881 | 119060 | MULTISPECIES:-DUF499-domain-containing-protein-[Burkholderiaceae] |
| WP_033446622.1 | AAA+ATPase+wHTH+Alpha+Beta-domain+wHTH+RRM | Type-1_BR.RM.like.systems | TetR-HTH->?->MarR-HTH->Phage_int_SAM_2+Integrase_1-><-DUF499*<-Schlafen+HATPase+HTH<-METHYLASE<-METHYLASE<-HEPN<-TUDOR+SF1-SF2-Helicase+iREase+CTD+REase<-HTH+WYL | 947 | Bacteria>Pseudomonadota>Betaproteobacteria>Burkholderiales>Alcaligenaceae | GCF_000690475.1 | Bordetella_bronchiseptica | EL288_RS19650 | 4193914..4196757 | 518 | DUF499-domain-containing-protein-[Bordetella-bronchiseptica] |
| WP_160982338.1 | AAA+ATPase+wHTH+Alpha+Beta-domain+wHTH+RRM | Type-1_BR.RM.like.systems | METHYLASE->METHYLASE->SF1-SF2-Helicase-><-?\|\|?-><-DUF499*<-?<-METHYLASE<-TUDOR+SF1-SF2-Helicase+iREase+CTD+REase<-HTH+WYL<-?\|\|Tn7_Tnp_TnsA_N+Tn7_Tnp_TnsA_C-> | 954 | Bacteria>Pseudomonadota>Betaproteobacteria>Burkholderiales>Alcaligenaceae | - | Pusillimonas_sp._TS35 | GSY71_RS17660 | 19469..22333 | 2695269 | DUF499-domain-containing-protein-[Pusillimonas-sp.-TS35] |
| WP_259661626.1 | AAA+ATPase+wHTH+Alpha+Beta-domain+wHTH+RRM | Type-1_BR.RM.like.systems | TUDOR+SF1-SF2-Helicase+iREase+CTD+REase->METHYLASE->DUF932->DUF499*-><-wHTH+HNH\|\|SAM-methylase-><-?<-?\|\|NERD+SF1-SF2-Helicase-> | 940 | Bacteria>Pseudomonadota>Betaproteobacteria>Burkholderiales>Alcaligenaceae | GCF_019923725.1 | Zwartia_hollandica | KZZ10_RS11225 | 114474..117296 | 324606 | DUF499-domain-containing-protein-[Zwartia-hollandica] |
| MBZ1351211.1 | AAA+ATPase+wHTH+Alpha+Beta-domain+wHTH+RRM | Type-1_BR.RM.like.systems | TUDOR+SF1-SF2-Helicase+iREase+CTD+REase->METHYLASE->DUF932->DUF499*-><-wHTH+HNH\|\|SAM-methylase-><-?<-?\|\|NERD+SF1-SF2-Helicase-> | 940 | Bacteria>Pseudomonadota>Betaproteobacteria>Burkholderiales>Alcaligenaceae | GCA_019923725.1 | Zwartia_hollandica | KZZ10_11185 | 114474..117296 | 324606 | DUF499-domain-containing-protein-[Alcaligenaceae-bacterium-LF4-65] |
| WP_226954710.1 | AAA+ATPase+wHTH+Alpha+Beta-domain+wHTH+RRM | Type-1_BR.RM.like.systems | <-GNTR-HTH<-?\|\|?-><-DUF499*<-METHYLASE<-TUDOR+SF1-SF2-Helicase+iREase+CTD+REase<-HTH+WYL\|\|SbcC-><-?<-Phage_integrase | 952 | Bacteria>Pseudomonadota>Betaproteobacteria>Burkholderiales>Alcaligenaceae | GCF_020539505.1 | Mesopusillimonas_faecipullorum | H0484_RS11140 | 214757..217615 | 2755040 | DUF499-domain-containing-protein-[Pusillimonas-faecipullorum] |
| KCV30144.1 | AAA+ATPase+wHTH+Alpha+Beta-domain+wHTH+RRM | Type-1_BR.RM.like.systems | <-DUF499*<-XIS-HTH<-Phage_int_SAM_2+Integrase_1<-MarR-HTH | 118 | Bacteria>Pseudomonadota>Betaproteobacteria>Burkholderiales>Alcaligenaceae | GCA_000662075.1 | Bordetella_bronchiseptica_00-P-2730 | L489_1003 | 1..354 | 1331198 | hypothetical-protein-L489_1003,-partial-[Bordetella-bronchiseptica-00-P-2730] |
| WP_246436895.1 | AAA+ATPase+wHTH+Alpha+Beta-domain+wHTH+RRM | Type-1_BR.RM.like.systems | <-HISKIN+MORC<-?<-?\|\|?-><-DUF499*<-ParB+HTH<-?\|\|HTH_19->XIS-HTH-><-Arm-DNA-bind_3+Phage_integrase | 127 | Bacteria>Pseudomonadota>Betaproteobacteria>Burkholderiales>Alcaligenaceae | GCF_013416525.1 | Parapusillimonas_granuli | H0A72_RS07635 | 4988..5371 | 380911 | hypothetical-protein-[Parapusillimonas-granuli] |
| TAL53973.1 | AAA+ATPase+wHTH+Alpha+Beta-domain+wHTH+RRM | Type-1_BR.RM.like.systems | HTH->STYKinase-like-><-DUF2357+REase<-Preader7+McrB-NTD+McrB-AAA\|\|HTH->STYKinase-like-><-DUF499*<-METHYLASE<-HEPN<-HEPN<-TUDOR+SF1-SF2-Helicase+iREase+CTD+REase | 966 | Bacteria>Pseudomonadota>Betaproteobacteria>Burkholderiales>Burkholderiaceae | GCA_004297875.1 | Pandoraea_sp. | EPN80_13305 | 57479..60379 | 1883445 | MAG:-ATP-binding-protein-[Pandoraea-sp.] |
| WP_090542133.1 | AAA+ATPase+wHTH+Alpha+Beta-domain+wHTH+RRM | Type-1_BR.RM.like.systems | <-HIN-HTH\|\|<-?\|\|HTH+WYL->TUDOR+SF1-SF2-Helicase+iREase+CTD+REase->METHYLASE->DUF499*-><-CHTH<-RelE<-Arm-DNA-bind_3+Phage_integrase+Phage_integr_3 | 948 | Bacteria>Pseudomonadota>Betaproteobacteria>Burkholderiales>Burkholderiaceae | GCF_900104845.1 | Paraburkholderia_caballeronis | BLV92_RS03095 | 696786..699632 | 416943 | DUF499-domain-containing-protein-[Paraburkholderia-caballeronis] |
| WP_045141103.1 | AAA+ATPase+wHTH+Alpha+Beta-domain+wHTH+RRM | Type-1_BR.RM.like.systems | MORC-><-?\|\|HTH+WYL->TUDOR+SF1-SF2-Helicase+iREase+CTD+REase->METHYLASE->DUF499*-><-CHTH<-RelE<-Arm-DNA-bind_3+Phage_integrase+Phage_integr_3\|\|tRNA->?->N6_N4_MTase-> | 948 | Bacteria>Pseudomonadota>Betaproteobacteria>Burkholderiales>Burkholderiaceae | GCF_001321165.1 | Burkholderia_pseudomallei | AP538_RS23015 | 37621..40467 | 28450 | DUF499-domain-containing-protein-[Burkholderia-pseudomallei] |
| WP_217096864.1 | AAA+ATPase+wHTH+Alpha+Beta-domain+wHTH+RRM | Type-1_BR.RM.like.systems | <-ACET\|\|?->?->MarR-HTH->Phage_int_SAM_2+Integrase_1-><-DUF499*<-Schlafen+HATPase+HTH<-METHYLASE<-METHYLASE<-HNH<-OLD-ABC<-TUDOR+SF1-SF2-Helicase+iREase+CTD+REase | 947 | Bacteria>Pseudomonadota>Betaproteobacteria>Burkholderiales>Burkholderiaceae | GCF_019041095.1 | Burkholderia_multivorans | KTE23_RS28415 | 22880..25723 | 87883 | DUF499-domain-containing-protein-[Burkholderia-multivorans] |
| WP_060088428.1 | AAA+ATPase+wHTH+Alpha+Beta-domain+wHTH+RRM | Type-1_BR.RM.like.systems | MORC-><-?\|\|HTH+WYL->TUDOR+SF1-SF2-Helicase+iREase+CTD+REase->METHYLASE->DUF499*-><-CHTH<-RelE<-Arm-DNA-bind_3+Phage_integrase+Phage_integr_3\|\|PIN-> | 947 | Bacteria>Pseudomonadota>Betaproteobacteria>Burkholderiales>Burkholderiaceae | GCF_001529865.1 | Burkholderia_ubonensis | WK66_RS31340 | 90789..93632 | 101571 | DUF499-domain-containing-protein-[Burkholderia-ubonensis] |
| WP_257803454.1 | AAA+ATPase+wHTH+Alpha+Beta-domain+wHTH+RRM | Type-1_BR.RM.like.systems | <-HIN-HTH<-?\|\|HTH+WYL->TUDOR+SF1-SF2-Helicase+iREase+CTD+REase->METHYLASE->DUF499*-><-CHTH<-RelE<-Arm-DNA-bind_3+Phage_integrase+Phage_integr_3 | 948 | Bacteria>Pseudomonadota>Betaproteobacteria>Burkholderiales>Burkholderiaceae | GCF_024664235.1 | Burkholderia_glumae | NIX97_RS01775 | 371737..374583 | 337 | DUF499-domain-containing-protein-[Burkholderia-glumae] |
| WP_065497253.1 | AAA+ATPase+wHTH+Alpha+Beta-domain+wHTH+RRM | Type-1_BR.RM.like.systems | TUDOR+SF1-SF2-Helicase+iREase+CTD+REase-p->TUDOR+SF1-SF2-Helicase+iREase+CTD+REase->PIN->METHYLASE->METHYLASE->Schlafen+HATPase+HTH->DUF499*->HEPN-> | 955 | Bacteria>Pseudomonadota>Betaproteobacteria>Burkholderiales>Burkholderiaceae | GCF_001685525.1 | Burkholderia_sp._CCA53 | BCR55_RS29355 | 26270..29137 | 1776288 | DUF499-domain-containing-protein-[Burkholderia-sp.-CCA53] |
| WP_116311964.1 | AAA+ATPase+wHTH+Alpha+Beta-domain+wHTH+RRM | Type-1_BR.RM.like.systems | <-RNaseP<-?\|\|?->?->MORC+DNA_gyraseB+TOPRIM-><-REase<-DUF499*<-METHYLASE<-TUDOR+SF1-SF2-Helicase+iREase+CTD+REase<-HTH+WYL\|\|STYKinase-like->?->DUF6361-> | 963 | Bacteria>Pseudomonadota>Betaproteobacteria>Burkholderiales>Burkholderiaceae | GCF_900249825.1 | Cupriavidus_taiwanensis | CBM2633_RS00535 | 131730..134621 | 164546 | DUF499-domain-containing-protein-[Cupriavidus-taiwanensis] |
| WP_064572931.1 | AAA+ATPase+wHTH+Alpha+Beta-domain+wHTH+RRM | Type-1_BR.RM.like.systems | MODE-HTH+PBPII-><-?\|\|HTH+WYL->HTH_Tnp->TUDOR+SF1-SF2-Helicase+iREase+CTD+REase->METHYLASE->DUF499*->SIR2+STALD->TIR->TIR->\|\|<-HAD<-METHYLASE-p | 951 | Bacteria>Pseudomonadota>Betaproteobacteria>Burkholderiales>Burkholderiaceae | GCF_001658125.1 | Cupriavidus_gilardii | JZ4iCDA_RS02255 | 517357..520212 | 82541 | DUF499-domain-containing-protein-[Cupriavidus-gilardii] |
| WP_211950277.1 | AAA+ATPase+wHTH+Alpha+Beta-domain+wHTH+RRM | Type-1_BR.RM.like.systems | <-STYKinase-like\|\|HTH+WYL->TUDOR+SF1-SF2-Helicase+iREase+CTD+REase->HEPN->METHYLASE->DUF499*->?->HEPN+CBS->?-><-?<-?<-Phage_int_SAM_1+Phage_integrase | 966 | Bacteria>Pseudomonadota>Betaproteobacteria>Burkholderiales>Burkholderiaceae | GCF_905397275.1 | Cupriavidus_yeonceonensis | KHD22_RS27195 | 69520..72420 | 1462994 | DUF499-domain-containing-protein-[Cupriavidus-yeonceonensis] |
| WP_204383624.1 | AAA+ATPase+wHTH+Alpha+Beta-domain+wHTH+RRM | Type-1_BR.RM.like.systems | MORC-><-?\|\|HTH+WYL->TUDOR+SF1-SF2-Helicase+iREase+CTD+REase->METHYLASE->DUF499*-><-CHTH<-RelE<-Arm-DNA-bind_3+Phage_integrase+Phage_integr_3 | 944 | Bacteria>Pseudomonadota>Betaproteobacteria>Burkholderiales>Burkholderiaceae | GCF_016899425.1 | Burkholderia_sp._MS389 | GJG85_RS02305 | 491304..494138 | 2811789 | DUF499-domain-containing-protein-[Burkholderia-sp.-MS389] |
| WP_222267509.1 | AAA+ATPase+wHTH+Alpha+Beta-domain+wHTH+RRM | Type-1_BR.RM.like.systems | <-MAD-NTD+HNH<-?\|\|HTH_3-><-Peptidase_C70<-?<-DUF499*<-METHYLASE<-HEPN<-TUDOR+SF1-SF2-Helicase+iREase+CTD+REase<-HTH+WYL\|\|REase->Resolvase-> | 943 | Bacteria>Pseudomonadota>Betaproteobacteria>Burkholderiales>Burkholderiaceae | GCF_020419455.1 | Burkholderia_arboris | LGM95_RS31475 | 204448..207279 | 488730 | DUF499-domain-containing-protein-[Burkholderia-arboris] |
| WP_186166579.1 | AAA+ATPase+wHTH+Alpha+Beta-domain+wHTH+RRM | Type-1_BR.RM.like.systems | Arm-DNA-bind_3+Phage_integrase+Phage_integr_3->CHTH-><-DUF499*<-METHYLASE<-TUDOR+SF1-SF2-Helicase+iREase+CTD+REase<-HTH+WYL\|\|?-><-MORC | 944 | Bacteria>Pseudomonadota>Betaproteobacteria>Burkholderiales>Burkholderiaceae | GCF_902831855.1 | Burkholderia | H9E02_RS31470 | 59730..62564 | 32008 | DUF499-domain-containing-protein-[Burkholderia-gladioli] |
| WP_186144236.1 | AAA+ATPase+wHTH+Alpha+Beta-domain+wHTH+RRM | Type-1_BR.RM.like.systems | HTH+WYL->TIR+PNTD2->PIWI->TUDOR+SF1-SF2-Helicase+iREase+CTD+REase->HEPN->METHYLASE->DUF499*->REase-><-tRNA\|\|?->?-><-?\|\|MODE-HTH+PBPII-><-TOPRIM+ZnR | 955 | Bacteria>Pseudomonadota>Betaproteobacteria>Burkholderiales>Burkholderiaceae | GCF_902831945.1 | Burkholderia_gladioli | H9E34_RS32225 | 51067..53934 | 28095 | DUF499-domain-containing-protein-[Burkholderia-gladioli] |
| WP_238209455.1 | AAA+ATPase+wHTH+Alpha+Beta-domain+wHTH+RRM | Type-1_BR.RM.like.systems | wHTH+SAD+HNH-><-AraC-HTH+SHS2+SHS2\|\|?->SF1-SF2-Helicase-><-?\|\|DUF488-u-><-DUF499*\|\|?-><-METHYLASE<-TUDOR+SF1-SF2-Helicase+iREase+CTD+REase | 797 | Bacteria>Pseudomonadota>Betaproteobacteria>Burkholderiales>Burkholderiaceae | - | Caballeronia_novacaledonica | CBA19CS42_00990 | 207479..209872 | 1544861 | DUF499-domain-containing-protein-[Caballeronia-novacaledonica] |
| WP_060256345.1 | AAA+ATPase+wHTH+Alpha+Beta-domain+wHTH+RRM | Type-1_BR.RM.like.systems | <-RNaseP<-?\|\|?->?->MORC+DNA_gyraseB+TOPRIM-><-?<-DUF499*<-METHYLASE<-TUDOR+SF1-SF2-Helicase+iREase+CTD+REase<-HTH+WYL\|\|ZDNA-HTH-><-ASCH<-GNAT | 947 | Bacteria>Pseudomonadota>Betaproteobacteria>Burkholderiales>Burkholderiaceae | GCF_001533365.1 | Burkholderia_ubonensis | WL76_RS09405 | 506654..509497 | 101571 | DUF499-domain-containing-protein-[Burkholderia-ubonensis] |
| GJH23036.1 | AAA+ATPase+wHTH+Alpha+Beta-domain+wHTH+RRM | Type-1_BR.RM.like.systems | wHTH+SAD+HNH-><-AraC-HTH+SHS2+SHS2\|\|?->SF1-SF2-Helicase-><-?\|\|DUF488-u-><-DUF499*\|\|?-><-METHYLASE<-TUDOR+SF1-SF2-Helicase+iREase+CTD+REase | 797 | Bacteria>Pseudomonadota>Betaproteobacteria>Burkholderiales>Burkholderiaceae | GCA_022179885.1 | Caballeronia_novacaledonica | CBA19CS42_00990 | 207479..209872 | 1544861 | DUF499-domain-containing-protein-[Caballeronia-novacaledonica] |
| GJH07681.1 | AAA+ATPase+wHTH+Alpha+Beta-domain+wHTH+RRM | Type-1_BR.RM.like.systems | TUDOR+SF1-SF2-Helicase+iREase+CTD+REase->METHYLASE-><-?\|\|DUF499*-><-DUF488-u\|\|?-><-SF1-SF2-Helicase<-?\|\|AraC-HTH+SHS2+SHS2-><-wHTH+SAD+HNH | 744 | Bacteria>Pseudomonadota>Betaproteobacteria>Burkholderiales>Burkholderiaceae | GCA_022179845.1 | Caballeronia_novacaledonica | CBA19CS11_02605 | 567311..569545 | 1544861 | DUF499-domain-containing-protein-[Caballeronia-novacaledonica] |
| WP_273654900.1 | AAA+ATPase+wHTH+Alpha+Beta-domain+wHTH+RRM | Type-1_BR.RM.like.systems | wHTH+SAD+HNH-><-AraC-HTH+SHS2+SHS2\|\|?->SF1-SF2-Helicase-><-?\|\|DUF488-u-><-DUF499*<-METHYLASE<-TUDOR+SF1-SF2-Helicase+iREase+CTD+REase | 938 | Bacteria>Pseudomonadota>Betaproteobacteria>Burkholderiales>Burkholderiaceae | GCF_022179885.1 | Caballeronia_novacaledonica | cba_RS01005 | 207479..210295 | 1544861 | DUF499-domain-containing-protein-[Caballeronia-novacaledonica] |
| WP_238285683.1 | AAA+ATPase+wHTH+Alpha+Beta-domain+wHTH+RRM | Type-1_BR.RM.like.systems | TUDOR+SF1-SF2-Helicase+iREase+CTD+REase->METHYLASE->DUF499*-><-DUF488-u\|\|?-><-SF1-SF2-Helicase<-?\|\|AraC-HTH+SHS2+SHS2-><-wHTH+SAD+HNH | 938 | Bacteria>Pseudomonadota>Betaproteobacteria>Burkholderiales>Burkholderiaceae | - | Caballeronia_novacaledonica | MFF65_RS02620 | 566729..569545 | 1544861 | DUF499-domain-containing-protein-[Caballeronia-novacaledonica] |
| WP_102632169.1 | AAA+ATPase+wHTH+Alpha+Beta-domain+wHTH+RRM | Type-1_BR.RM.like.systems | HTH-><-Peptidase_C70<-?<-DUF499*<-METHYLASE<-HEPN<-TUDOR+SF1-SF2-Helicase+iREase+CTD+REase<-HTH+WYL\|\|Resolvase-><-Peptidase_S8 | 943 | Bacteria>Pseudomonadota>Betaproteobacteria>Burkholderiales>Burkholderiaceae | GCF_902859775.1 | Paraburkholderia_rhynchosiae | HV734_RS08115 | 141591..144422 | 487049 | DUF499-domain-containing-protein-[Paraburkholderia-rhynchosiae] |
| WP_211616585.1 | AAA+ATPase+wHTH+Alpha+Beta-domain+wHTH+RRM | Type-1_BR.RM.like.systems | <-ArdB\|\|ParB+HTH-><-?\|\|MobA_MobL+SF1-SF2-helicase-><-?<-DUF499*<-METHYLASE<-TUDOR+SF1-SF2-Helicase+iREase+CTD+REase | 965 | Bacteria>Pseudomonadota>Betaproteobacteria>Burkholderiales>Burkholderiaceae | GCF_905220975.1 | Paraburkholderia_haematera | KHD23_RS31730 | 44872..47769 | 2793077 | DUF499-domain-containing-protein-[Paraburkholderia-haematera] |
| WP_175712494.1 | AAA+ATPase+wHTH+Alpha+Beta-domain+wHTH+RRM | Type-1_BR.RM.like.systems | <-DUF499*<-METHYLASE<-?<-TUDOR+SF1-SF2-Helicase+iREase+CTD+REase<-HTH+WYL\|\|HTH_31->rve+rve_3+Mu-transpos_C-> | 955 | Bacteria>Pseudomonadota>Betaproteobacteria>Burkholderiales>Burkholderiaceae | GCF_902829595.1 | Burkholderia_ambifaria | HWW29_RS33350 | 873..3740 | 152480 | DUF499-domain-containing-protein-[Burkholderia-ambifaria] |
| WP_150698381.1 | AAA+ATPase+wHTH+Alpha+Beta-domain+wHTH+RRM | Type-1_BR.RM.like.systems | HTH->GDResolvase+HIN-HTH->URI-DNase-><-DUF499*<-METHYLASE<-HEPN<-TUDOR+SF1-SF2-Helicase+iREase+CTD+REase | 944 | Bacteria>Pseudomonadota>Betaproteobacteria>Burkholderiales>Burkholderiaceae | GCF_902459695.1 | Pandoraea_terrae | PTE30175_RS17675 | 118123..120957 | 1537710 | DUF499-domain-containing-protein-[Pandoraea-terrae] |
| WP_251118741.1 | AAA+ATPase+wHTH+Alpha+Beta-domain+wHTH+RRM | Type-1_BR.RM.like.systems | ZnR+TOPRIM+D5ATPase->HTH+WYL->TUDOR+SF1-SF2-Helicase+iREase+CTD+REase->METHYLASE->DUF499*-><-CHTH<-RelE | 948 | Bacteria>Pseudomonadota>Betaproteobacteria>Burkholderiales>Burkholderiaceae | GCF_023702455.1 | Burkholderia_glumae | M8545_RS05295 | 854265..857111 | 337 | DUF499-domain-containing-protein-[Burkholderia-glumae] |
| WP_069222598.1 | AAA+ATPase+wHTH+Alpha+Beta-domain+wHTH+RRM | Type-1_BR.RM.like.systems | <-SIR2<-?\|\|?->?->?-><-HIN-HTH\|\|<-DUF499*<-METHYLASE<-TUDOR+SF1-SF2-Helicase+iREase+CTD+REase<-HTH+WYL\|\|?-><-ParB<-ParB | 951 | Bacteria>Pseudomonadota>Betaproteobacteria>Burkholderiales>Burkholderiaceae | GCF_001718495.1 | Burkholderia_vietnamiensis | WJ02_RS14585 | 3208702..3211557 | 60552 | DUF499-domain-containing-protein-[Burkholderia-vietnamiensis] |
| WP_013088136.1 | AAA+ATPase+wHTH+Alpha+Beta-domain+wHTH+RRM | Type-1_BR.RM.like.systems | Phage_integrase->TUDOR+SF1-SF2-Helicase+iREase+CTD+REase->METHYLASE->DUF499*-><-?<-STYKinase-like<-TPR | 967 | Bacteria>Pseudomonadota>Betaproteobacteria>Burkholderiales>Burkholderiaceae | GCF_000092885.1 | Paraburkholderia_atlantica | BC1002_RS00635 | 142945..145848 | 2654982 | DUF499-domain-containing-protein-[Paraburkholderia-atlantica] |
| KAK43608.1 | AAA+ATPase+wHTH+Alpha+Beta-domain+wHTH+RRM | Type-1_BR.RM.like.systems | METHYLASE-p->DUF499*-><-DUF488-u\|\|?-><-SF1-SF2-Helicase<-?\|\|AraC-HTH+SHS2+SHS2-><-MORC+DNA_gyraseB+TOPRIM | 938 | Bacteria>Pseudomonadota>Betaproteobacteria>Burkholderiales>Burkholderiaceae | GCA_000648925.1 | Caballeronia_jiangsuensis | BG58_31250 | 6996..9812 | 1458357 | hypothetical-protein-BG58_31250-[Caballeronia-jiangsuensis] |
| EIF53189.1 | AAA+ATPase+wHTH+Alpha+Beta-domain+wHTH+RRM | Type-1_BR.RM.like.systems | <-DUF932<-?\|\|HTH+WYL->TUDOR+SF1-SF2-Helicase+iREase+CTD+REase->METHYLASE->DUF499*-><-?<-?<-tRNA\|\|?->?-><-?\|\|MODE-HTH+PBPII-> | 929 | Bacteria>Pseudomonadota>Betaproteobacteria>Burkholderiales>Burkholderiaceae | GCA_000259755.1 | Burkholderia_pseudomallei_1258a | BP1258A_5453 | 40076..42865 | 1086032 | hypothetical-protein-BP1258A_5453-[Burkholderia-pseudomallei-1258a] |
| WP_266261887.1 | AAA+ATPase+wHTH+Alpha+Beta-domain+wHTH+RRM | Type-1_BR.RM.like.systems | HKD-DNase->?->?->?-><-?<-DUF499*<-METHYLASE<-TUDOR+SF1-SF2-Helicase+iREase+CTD+REase\|\|RelB->RelE-ParE-> | 959 | Bacteria>Pseudomonadota>Betaproteobacteria>Burkholderiales>Burkholderiaceae | GCF_030464835.1 | Paraburkholderia | MOD64_RS41395 | 20884..23763 | 1822464 | DUF499-domain-containing-protein-[Paraburkholderia-madseniana] |
| SAL19074.1 | AAA+ATPase+wHTH+Alpha+Beta-domain+wHTH+RRM | Type-1_BR.RM.like.systems | HNH->?-><-RNase<-DUF499*<-METHYLASE<-?<-TUDOR+SF1-SF2-Helicase+iREase+CTD+REase\|\|?->Glyoxalase-> | 968 | Bacteria>Pseudomonadota>Betaproteobacteria>Burkholderiales>Burkholderiaceae | GCA_001544655.2 | Caballeronia_turbans | AWB73_01129 | 180728..183634 | 1777134 | hypothetical-protein-AWB73_01129-[Caballeronia-turbans] |
| WP_151072628.1 | AAA+ATPase+wHTH+Alpha+Beta-domain+wHTH+RRM | Type-1_BR.RM.like.systems | <-RVT<-DUF499*<-METHYLASE<-TUDOR+SF1-SF2-Helicase+iREase+CTD+REase<-HTH+WYL\|\|?->DUF932-> | 967 | Bacteria>Pseudomonadota>Betaproteobacteria>Burkholderiales>Burkholderiaceae | GCF_008807855.1 | Cupriavidus_oxalaticus | D2917_RS28525 | 3422973..3425876 | 96344 | DUF499-domain-containing-protein-[Cupriavidus-oxalaticus] |
| WP_104766975.1 | AAA+ATPase+wHTH+Alpha+Beta-domain+wHTH+RRM | Type-1_BR.RM.like.systems | <-DUF499*<-METHYLASE<-TUDOR+SF1-SF2-Helicase+iREase+CTD+REase<-HTH+WYL<-?<-ParB<-ParB | 951 | Bacteria>Pseudomonadota>Betaproteobacteria>Burkholderiales>Burkholderiaceae | GCF_002942785.2 | Burkholderia_cepacia | C5075_RS18605 | 865..3720 | 292 | DUF499-domain-containing-protein-[Burkholderia-cepacia] |
| WP_264254228.1 | AAA+ATPase+wHTH+Alpha+Beta-domain+wHTH+RRM | Type-1_BR.RM.like.systems | <-DUF499*<-METHYLASE<-TUDOR+SF1-SF2-Helicase+iREase+CTD+REase<-HTH+WYL\|\|?-><-MORC | 944 | Bacteria>Pseudomonadota>Betaproteobacteria>Burkholderiales>Burkholderiaceae | GCF_025848075.1 | Burkholderia_pseudomallei | DIV61_RS24740 | 25495..28329 | 28450 | DUF499-domain-containing-protein-[Burkholderia-pseudomallei] |
| WP_076897581.1 | AAA+ATPase+wHTH+Alpha+Beta-domain+wHTH+RRM | Type-1_BR.RM.like.systems | <-DUF499*<-METHYLASE<-TUDOR+SF1-SF2-Helicase+iREase+CTD+REase<-HTH+WYL\|\|?-><-MORC | 944 | Bacteria>Pseudomonadota>Betaproteobacteria>Burkholderiales>Burkholderiaceae | GCF_001976675.1 | Burkholderia_pseudomallei | GPJ57_RS06820 | 53492..56326 | 28450 | DUF499-domain-containing-protein-[Burkholderia-pseudomallei] |
| WP_038716142.1 | AAA+ATPase+wHTH+Alpha+Beta-domain+wHTH+RRM | Type-1_BR.RM.like.systems | MORC-><-?\|\|HTH+WYL->TUDOR+SF1-SF2-Helicase+iREase+CTD+REase->METHYLASE->DUF499*-> | 944 | Bacteria>Pseudomonadota>Betaproteobacteria>Burkholderiales>Burkholderiaceae | GCF_000259775.1 | Burkholderia_pseudomallei | BP1258B_RS28565 | 40035..42869 | 28450 | DUF499-domain-containing-protein-[Burkholderia-pseudomallei] |
| WP_106351229.1 | AAA+ATPase+wHTH+Alpha+Beta-domain+wHTH+RRM | Type-1_BR.RM.like.systems | <-DUF499*<-METHYLASE<-?<-TUDOR+SF1-SF2-Helicase+iREase+CTD+REase<-?\|\|?->HTH+WYL-> | 962 | Bacteria>Pseudomonadota>Betaproteobacteria>Burkholderiales>Burkholderiaceae | GCF_003002115.1 | Paraburkholderia_fungorum | BX589_RS00015 | 1094..3982 | 134537 | DUF499-domain-containing-protein-[Paraburkholderia-insulsa] |
| WP_028211840.1 | AAA+ATPase+wHTH+Alpha+Beta-domain+wHTH+RRM | Type-1_BR.RM.like.systems | <-DUF499*<-METHYLASE<-TUDOR+SF1-SF2-Helicase+iREase+CTD+REase\|\|HEPN-p-> | 939 | Bacteria>Pseudomonadota>Betaproteobacteria>Burkholderiales>Burkholderiaceae | GCF_000472845.1 | Paraburkholderia_mimosarum | A19S_RS0115645 | 1157..3976 | 312026 | DUF499-domain-containing-protein-[Paraburkholderia-mimosarum] |
| WP_144832727.1 | AAA+ATPase+wHTH+Alpha+Beta-domain+wHTH+RRM | Type-1_BR.RM.like.systems | HTH_3->Radical-SAM-><-Methylase<-SMODS<-SLATT<-TIR<-DUF499* | 950 | Bacteria>Pseudomonadota>Betaproteobacteria>Burkholderiales>Burkholderiaceae | GCF_007829435.1 | Cupriavidus_gilardii | L602_RS14240 | 51207..54059 | 82541 | DUF499-domain-containing-protein-[Cupriavidus-gilardii] |
| WP_254661892.1 | AAA+ATPase+wHTH+Alpha+Beta-domain+wHTH+RRM | Type-1_BR.RM.like.systems | PIN->METHYLASE->AAA_21->Schlafen+HATPase+HTH->STYKIN-><-?\|\|DUF499*-> | 314 | Bacteria>Pseudomonadota>Betaproteobacteria>Burkholderiales>Burkholderiaceae | GCF_001685525.1 | Burkholderia_sp._CCA53 | BCR55_RS29865 | 130717..131658 | 1776288 | DUF499-domain-containing-protein,-partial-[Burkholderia-sp.-CCA53] |
| MBX9834804.1 | AAA+ATPase+wHTH+Alpha+Beta-domain+wHTH+RRM | Type-1_BR.RM.like.systems | METHYLASE->HEPN->DUF499*->?->?-><-SAM-methylase | 945 | Bacteria>Pseudomonadota>Betaproteobacteria>Burkholderiales>Burkholderiaceae | GCA_019746655.1 | Burkholderiaceae_bacterium | K2X65_01340 | 3559..6396 | 2030806 | DUF499-domain-containing-protein-[Burkholderiaceae-bacterium] |
| MBP6559243.1 | AAA+ATPase+wHTH+Alpha+Beta-domain+wHTH+RRM | Type-1_BR.RM.like.systems | <-DUF499*<-REase<-OLD-ABC<-METHYLASE-p | 945 | Bacteria>Pseudomonadota>Betaproteobacteria>Burkholderiales>Burkholderiaceae | GCA_017989415.1 | Burkholderiaceae_bacterium | KA212_02995 | 512..3349 | 2030806 | ATP-binding-protein-[Burkholderiaceae-bacterium] |
| WP_059939474.1 | AAA+ATPase+wHTH+Alpha+Beta-domain+wHTH+RRM | Type-1_BR.RM.like.systems | Phage_AlpA->DnaB->?->TraJ-RHH->RHH->XIS-HTH->DUF499*-> | 315 | Bacteria>Pseudomonadota>Betaproteobacteria>Burkholderiales>Burkholderiaceae | GCF_001528605.1 | Burkholderia_multivorans | WK33_RS15135 | 116003..116950 | 87883 | hypothetical-protein-[Burkholderia-multivorans] |
| WP_027802422.1 | AAA+ATPase+wHTH+Alpha+Beta-domain+wHTH+RRM | Type-1_BR.RM.like.systems | Arm-DNA-bind_3+Phage_int_SAM_3+Phage_integrase->Phage_AlpA->?->RHH_1->TUDOR+SF1-SF2-Helicase+iREase+CTD+REase->METHYLASE->DUF499*-><-?<-?\|\|?->?->REase+SPF-cons-> | 930 | Bacteria>Pseudomonadota>Betaproteobacteria>Burkholderiales>Burkholderiaceae | GCF_000472525.1 | Paraburkholderia_dilworthii | F759_RS0125900 | 42632..45424 | 948106 | DUF499-domain-containing-protein-[Paraburkholderia-dilworthii] |
| WP_008352254.1 | AAA+ATPase+wHTH+Alpha+Beta-domain+wHTH+RRM | Type-1_BR.RM.like.systems | HIN-HTH-><-?<-DUF932\|\|TUDOR+SF1-SF2-Helicase+iREase+CTD+REase->METHYLASE->DUF499*->\|\|HIN-HTH->?->?-><-?\|\|Glyoxalase-> | 929 | Bacteria>Pseudomonadota>Betaproteobacteria>Burkholderiales>Burkholderiaceae | GCF_022879815.1 | Caballeronia_zhejiangensis | LDZ23_RS14465 | 3017077..3019866 | 871203 | DUF499-domain-containing-protein-[Caballeronia-zhejiangensis] |
| WP_217193651.1 | AAA+ATPase+wHTH+Alpha+Beta-domain+wHTH+RRM | Type-1_BR.RM.like.systems | RHH_1->TUDOR+SF1-SF2-Helicase+iREase+CTD+REase->METHYLASE->DUF499*-><-?<-?<-ParB+HNH\|\|HTH+WYL-> | 929 | Bacteria>Pseudomonadota>Betaproteobacteria>Burkholderiales>Burkholderiaceae | GCF_019040475.1 | Burkholderia_multivorans | KTE35_RS28955 | 6892..9681 | 87883 | DUF499-domain-containing-protein-[Burkholderia-multivorans] |
| WP_115688423.1 | AAA+ATPase+wHTH+Alpha+Beta-domain+wHTH+RRM | Type-1_BR.RM.like.systems | <-DUF499*<-METHYLASE<-TUDOR+SF1-SF2-Helicase+iREase+CTD+REase<-RHH_1\|\|?->SF1-SF2-Helicase-> | 928 | Bacteria>Pseudomonadota>Betaproteobacteria>Burkholderiales>Burkholderiaceae | GCF_900249765.1 | Cupriavidus_taiwanensis | CBM2634_RS16190 | 3449898..3452684 | 164546 | DUF499-domain-containing-protein-[Cupriavidus-taiwanensis] |
| MBE2264050.1 | AAA+ATPase+wHTH+Alpha+Beta-domain+wHTH+RRM | Type-1_BR.RM.like.systems | <-DUF499*\|\|DUF433->PIN-><-REase | 760 | Bacteria>Pseudomonadota>Betaproteobacteria>Burkholderiales>Burkholderiaceae | GCA_014879655.1 | Burkholderiaceae_bacterium | IAE92_14980 | 1..2282 | 2030806 | ATP-binding-protein-[Burkholderiaceae-bacterium] |
| WP_132962623.1 | AAA+ATPase+wHTH+Alpha+Beta-domain+wHTH+RRM | Type-1_BR.RM.like.systems | GST_N+GST_C->tRNA->tRNA-><-DUF499*<-REase<-OLD-ABC<-METHYLASE<-PIN<-TUDOR+SF1-SF2-Helicase+iREase+CTD+REase\|\|Peptidase_M24-> | 949 | Bacteria>Pseudomonadota>Betaproteobacteria>Burkholderiales>Burkholderiales_genera_incertae_sedis | GCF_004342625.1 | Tepidimonas_ignava | EDC36_RS07390 | 30173..33022 | 114249 | DUF499-domain-containing-protein-[Tepidimonas-ignava] |
| WP_143969665.1 | AAA+ATPase+wHTH+Alpha+Beta-domain+wHTH+RRM | Type-1_BR.RM.like.systems | HTH+WYL->TUDOR+SF1-SF2-Helicase+iREase+CTD+REase->PIN->METHYLASE->OLD-ABC->TOPRIM->DUF499*->HEPN-> | 952 | Bacteria>Pseudomonadota>Betaproteobacteria>Burkholderiales>Burkholderiales_genera_incertae_sedis | GCF_007556755.1 | Tepidimonas_fonticaldi | Tfont_RS12360 | 12901..15759 | 1101373 | DUF499-domain-containing-protein-[Tepidimonas-fonticaldi] |
| WP_043699941.1 | AAA+ATPase+wHTH+Alpha+Beta-domain+wHTH+RRM | Type-1_BR.RM.like.systems | PIN->METHYLASE->OLD-ABC->TOPRIM->METHYLASE->Schlafen+HATPase+HTH->DUF499*-><-?<-?<-MORC<-?<-VirB11+MoxR-AAA | 945 | Bacteria>Pseudomonadota>Betaproteobacteria>Burkholderiales>Burkholderiales_genera_incertae_sedis | GCF_020162115.1 | Tepidimonas_taiwanensis | PL21_RS04270 | 89059..91896 | 307486 | DUF499-domain-containing-protein-[Tepidimonas-taiwanensis] |
| WP_166837252.1 | AAA+ATPase+wHTH+Alpha+Beta-domain+wHTH+RRM | Type-1_BR.RM.like.systems | TUDOR+SF1-SF2-Helicase+iREase+CTD+REase->ParB+HNH->PIN->METHYLASE->DUF499*-> | 936 | Bacteria>Pseudomonadota>Betaproteobacteria>Burkholderiales>Burkholderiales_genera_incertae_sedis | GCF_011742065.1 | Aquabacterium_sp._A08 | HA630_RS20340 | 28723..31533 | 2718532 | DUF499-domain-containing-protein-[Aquabacterium-sp.-A08] |
| WP_143969782.1 | AAA+ATPase+wHTH+Alpha+Beta-domain+wHTH+RRM | Type-1_BR.RM.like.systems | <-DUF772+DDE_Tnp_1\|\|<-DUF499*<-REase<-METHYLASE<-TOPRIM<-OLD-ABC<-TUDOR+SF1-SF2-Helicase+iREase+CTD+REase<-XIS-HTH | 948 | Bacteria>Pseudomonadota>Betaproteobacteria>Burkholderiales>Burkholderiales_genera_incertae_sedis | GCF_007556755.1 | Tepidimonas_fonticaldi | Tfont_RS13255 | 1599..4445 | 1101373 | DUF499-domain-containing-protein-[Tepidimonas-fonticaldi] |
| WP_180684522.1 | AAA+ATPase+wHTH+Alpha+Beta-domain+wHTH+RRM | Type-1_BR.RM.like.systems | XIS-HTH->TUDOR+SF1-SF2-Helicase+iREase+CTD+REase->OLD-ABC->METHYLASE->REase->DUF499*-><-Phage_lysozyme | 939 | Bacteria>Pseudomonadota>Betaproteobacteria>Burkholderiales>Burkholderiales_genera_incertae_sedis | GCF_013432195.1 | Tepidicella_baoligensis | HZS63_RS13535 | 8909..11728 | 2707016 | DUF499-domain-containing-protein-[Tepidicella-baoligensis] |
| WP_139702092.1 | AAA+ATPase+wHTH+Alpha+Beta-domain+wHTH+RRM | Type-1_BR.RM.like.systems | Arm-DNA-bind_3+Phage_integrase-><-HEPN+CBS\|\|REase->?-><-MORC+HisKin<-?<-DUF499*<-METHYLASE<-TUDOR+SF1-SF2-Helicase+iREase+CTD+REase<-HTH+WYL\|\|OLD-ABC+TOPRIM->SF1-SF2-Helicase->JAB-> | 961 | Bacteria>Pseudomonadota>Betaproteobacteria>Burkholderiales>Comamonadaceae | GCF_006323185.2 | Variovorax_sp._KBS0712 | FFI97_RS04005 | 912155..915040 | 2578111 | DUF499-domain-containing-protein-[Variovorax-sp.-KBS0712] |
| WP_252101816.1 | AAA+ATPase+wHTH+Alpha+Beta-domain+wHTH+RRM | Type-1_BR.RM.like.systems | TUDOR+SF1-SF2-Helicase+iREase+CTD+REase->Schlafen+HATPase+HTH->?->METHYLASE->OLD-ABC->REase->DUF499*-><-?<-?<-Phage_integrase+Phage_integrase<-?<-HIN-HTH<-Phage_integrase+Phage_integrase | 945 | Bacteria>Pseudomonadota>Betaproteobacteria>Burkholderiales>Comamonadaceae | - | [Acidovorax]_ebreus | MRB47_RS02565 | 530947..533784 | 721785 | DUF499-domain-containing-protein-[[Acidovorax] |
| WP_088958124.1 | AAA+ATPase+wHTH+Alpha+Beta-domain+wHTH+RRM | Type-1_BR.RM.like.systems | ParB->HTH+WYL->HTH_Tnp->TUDOR+SF1-SF2-Helicase+iREase+CTD+REase->HEPN->METHYLASE->DUF499*->TIR->SLATT->SMODS->Methylase-><-Radical-SAM<-HTH_3 | 951 | Bacteria>Pseudomonadota>Betaproteobacteria>Burkholderiales>Comamonadaceae | GCF_900090195.1 | Variovorax_sp._HW608 | VAR608DRAFT_RS34345 | 7257646..7260501 | 1034889 | DUF499-domain-containing-protein-[Variovorax-sp.-HW608] |
| WP_011810371.1 | AAA+ATPase+wHTH+Alpha+Beta-domain+wHTH+RRM | Type-1_BR.RM.like.systems | <-S1+RNase+2C<-?\|\|?-><-SPOUT_MTase<-?\|\|?-><-DUF499*<-Schlafen+HATPase+HTH<-METHYLASE<-METHYLASE<-DUF4096+DDE_Tnp_1+DDE_Tnp_1_2<-PAIREDC-HTH+HIN-HTH<-HAMP+HISKIN | 936 | Bacteria>Pseudomonadota>Betaproteobacteria>Burkholderiales>Comamonadaceae | GCF_000015565.1 | Verminephrobacter_eiseniae | VEIS_RS12785 | 2967373..2970183 | 364317 | DUF499-domain-containing-protein-[Verminephrobacter-eiseniae] |
| WP_095541484.1 | AAA+ATPase+wHTH+Alpha+Beta-domain+wHTH+RRM | Type-1_BR.RM.like.systems | METHYLASE-p->?->HAD->Arm-DNA-bind_3->?-><-DUF499*<-ParB+HNH<-METHYLASE<-HEPN<-TUDOR+SF1-SF2-Helicase+iREase+CTD+REase<-HTH_Tnp<-HTH+WYL | 948 | Bacteria>Pseudomonadota>Betaproteobacteria>Burkholderiales>Comamonadaceae | GCF_002285255.1 | Vandammella_animalimorsus | CLI93_RS07370 | 83443..86289 | 2029117 | DUF499-domain-containing-protein-[Vandammella-animalimorsus] |
| WP_055397337.1 | AAA+ATPase+wHTH+Alpha+Beta-domain+wHTH+RRM | Type-1_BR.RM.like.systems | <-ArdCN+MPTase<-?<-?<-MobA_MobL+SF1-SF2-helicase\|\|?-><-?<-DUF499*<-REase<-OLD-ABC<-METHYLASE<-HNH<-OLD-ABC<-TUDOR+SF1-SF2-Helicase+iREase+CTD+REase | 945 | Bacteria>Pseudomonadota>Betaproteobacteria>Burkholderiales>Comamonadaceae | GCF_017355385.1 | Acidovorax | AE621_RS12465 | 7782..10619 | 12916 | MULTISPECIES:-DUF499-domain-containing-protein-[Acidovorax] |
| MBX3657558.1 | AAA+ATPase+wHTH+Alpha+Beta-domain+wHTH+RRM | Type-1_BR.RM.like.systems | HTH+WYL->TUDOR+SF1-SF2-Helicase+iREase+CTD+REase->HEPN->METHYLASE->?->?->DUF499*->SbcC->HIN-HTH->?-><-?<-Phage_lysozyme<-Phage_holin | 951 | Bacteria>Pseudomonadota>Betaproteobacteria>Burkholderiales>Comamonadaceae | GCA_019635415.1 | Ramlibacter_sp. | KF740_03900 | 804820..807675 | 1917967 | ATP-binding-protein-[Ramlibacter-sp.] |
| WP_210930504.1 | AAA+ATPase+wHTH+Alpha+Beta-domain+wHTH+RRM | Type-1_BR.RM.like.systems | HTH->?->Phage_integrase->STYKinase-like-><-?\|\|DDE_Tnp_1_5+DDE_Tnp_1-><-DUF499*<-REase<-OLD-ABC<-HEPN<-METHYLASE<-HNH<-OLD-ABC | 945 | Bacteria>Pseudomonadota>Betaproteobacteria>Burkholderiales>Comamonadaceae | GCF_018064205.1 | Comamonas_aquatica | KAQ61_RS03430 | 737208..740045 | 225991 | DUF499-domain-containing-protein-[Comamonas-aquatica] |
| WP_127807271.1 | AAA+ATPase+wHTH+Alpha+Beta-domain+wHTH+RRM | Type-1_BR.RM.like.systems | TUDOR+SF1-SF2-Helicase+iREase+CTD+REase->PIN->HEPN->METHYLASE->?->?->DUF499*->TIR->?-><-Radical-SAM<-HTH_3<-?<-Phage_lysozyme | 950 | Bacteria>Pseudomonadota>Betaproteobacteria>Burkholderiales>Comamonadaceae | GCF_004006415.1 | Hydrogenophaga_sp._NH-16 | DD542_RS03485 | 729405..732257 | 2184519 | DUF499-domain-containing-protein-[Hydrogenophaga-sp.-NH-16] |
| WP_042427102.1 | AAA+ATPase+wHTH+Alpha+Beta-domain+wHTH+RRM | Type-1_BR.RM.like.systems | Phage_lysozyme->?->HTH_3->Radical-SAM-><-?<-TIR<-DUF499*<-?<-?<-METHYLASE<-HEPN<-PIN<-TUDOR+SF1-SF2-Helicase+iREase+CTD+REase | 951 | Bacteria>Pseudomonadota>Betaproteobacteria>Burkholderiales>Comamonadaceae | GCF_000739995.1 | Comamonas_granuli | CGR01S_RS11705 | 67732..70587 | 290309 | DUF499-domain-containing-protein-[Comamonas-granuli] |
| TFI45392.1 | AAA+ATPase+wHTH+Alpha+Beta-domain+wHTH+RRM | Type-1_BR.RM.like.systems | <-MobA_MobL+SF1-SF2-helicase\|\|?-><-?<-DUF499*<-REase<-OLD-ABC<-METHYLASE<-PIN<-TUDOR+SF1-SF2-Helicase+iREase+CTD+REase | 945 | Bacteria>Pseudomonadota>Betaproteobacteria>Burkholderiales>Comamonadaceae | - | Diaphorobacter_sp._DS2 | E4093_19980 | 3252..6089 | 1302548 | ATP-binding-protein-[Diaphorobacter-sp.-DS2] |
| MDD2610199.1 | AAA+ATPase+wHTH+Alpha+Beta-domain+wHTH+RRM | Type-1_BR.RM.like.systems | TUDOR+SF1-SF2-Helicase+iREase+CTD+REase->OLD-ABC->HNH->METHYLASE-p->METHYLASE->?->DUF499*->?-><-?\|\|<-JAB<-UbE1+JAB<-SMODS | 951 | Bacteria>Pseudomonadota>Betaproteobacteria>Burkholderiales>Comamonadaceae | GCA_028686475.1 | Giesbergeria_sp. | PHX60_11025 | 19718..22573 | 2818473 | MAG:-DUF499-domain-containing-protein-[Giesbergeria-sp.] |

| Protein-ID | DUF499-architecture | System | Gene-Neighbourhoods | Length | Taxonomy | Assembly-ID | Strain | Locus-Tag | Genomic-coordinates | TaxID | Fasta-defline |
|---|---|---|---|---|---|---|---|---|---|---|---|
| WP_183017341.1 | AAA+ATPase+wHTH+Alpha+Beta-domain+wHTH+RRM | Type-1_BR.RM.like.systems | Phage_integrase-><-MORC+HisKin<-REase<-?<-?<-DUF499*<-METHYLASE<-TUDOR+SF1-SF2-Helicase+iREase+CTD+REase<-HTH+WYL | 955 | Bacteria>Pseudomonadota>Betaproteobacteria>Burkholderiales>Comamonadaceae | GCF_014170385.1 | Delftia_sp._UME58 | A9976_RS00455 | 77977..80844 | 1862322 | DUF499-domain-containing-protein-[Delftia-sp.-UME58] |
| NMM11921.1 | AAA+ATPase+wHTH+Alpha+Beta-domain+wHTH+RRM | Type-1_BR.RM.like.systems | <-HEPN<-REase<-?<-?<-DUF499*<-METHYLASE<-HEPN<-TUDOR+SF1-SF2-Helicase+iREase+CTD+REase<-HTH+WYL\|\|STYKinase-like-> | 961 | Bacteria>Pseudomonadota>Betaproteobacteria>Burkholderiales>Comamonadaceae | GCA_012927445.1 | Rhodoferax_sp. | HHJ17_00055 | 10474..13359 | 50421 | ATP-binding-protein-[Rhodoferax-sp.] |
| WP_265289361.1 | AAA+ATPase+wHTH+Alpha+Beta-domain+wHTH+RRM | Type-1_BR.RM.like.systems | METHYLASE-p->METHYLASE-><-?\|\|METHYLASE->Schlafen+HATPase+HTH->Schlafen+HATPase+HTH->DUF499*-><-?\|\|?->SPOUT_MTase->?->S1+RNase+2C-> | 942 | Bacteria>Pseudomonadota>Betaproteobacteria>Burkholderiales>Comamonadaceae | GCF_026127525.1 | Verminephrobacter_eiseniae | D5047_RS20580 | 4396867..4399695 | 364317 | DUF499-domain-containing-protein-[Verminephrobacter-eiseniae] |
| WP_012839779.1 | AAA+ATPase+wHTH+Alpha+Beta-domain+wHTH+RRM | Type-1_BR.RM.like.systems | <-MORC+HisKin<-REase<-?<-DUF499*<-METHYLASE<-HEPN<-TUDOR+SF1-SF2-Helicase+iREase+CTD+REase<-HTH+WYL | 955 | Bacteria>Pseudomonadota>Betaproteobacteria>Burkholderiales>Comamonadaceae | GCF_000093145.3 | Comamonas_thiooxydans | HNB49_RS10950 | 10966..13833 | 363952 | DUF499-domain-containing-protein-[Comamonas-thiooxydans] |
| WP_142812147.1 | AAA+ATPase+wHTH+Alpha+Beta-domain+wHTH+RRM | Type-1_BR.RM.like.systems | <-PIN\|\|?-><-?<-DUF499*<-METHYLASE<-TUDOR+SF1-SF2-Helicase+iREase+CTD+REase<-HTH+WYL\|\|STYKinase-like-> | 960 | Bacteria>Pseudomonadota>Betaproteobacteria>Burkholderiales>Comamonadaceae | GCF_006974105.1 | Rhodoferax_aquaticus | EXZ61_RS12885 | 2804871..2807753 | 2527691 | DUF499-domain-containing-protein-[Rhodoferax-aquaticus] |
| WP_045536717.1 | AAA+ATPase+wHTH+Alpha+Beta-domain+wHTH+RRM | Type-1_BR.RM.like.systems | TUDOR+SF1-SF2-Helicase+iREase+CTD+REase->PIN->HEPN->METHYLASE-p->METHYLASE->ParB+HNH->DUF499*-> | 950 | Bacteria>Pseudomonadota>Betaproteobacteria>Burkholderiales>Comamonadaceae | GCF_000828915.1 | Serpentinimonas_maccroryi | SMCB_RS09950 | 2162226..2165078 | 1458426 | DUF499-domain-containing-protein-[Serpentinimonas-maccroryi] |
| WP_210568820.1 | AAA+ATPase+wHTH+Alpha+Beta-domain+wHTH+RRM | Type-1_BR.RM.like.systems | TUDOR+SF1-SF2-Helicase+iREase+CTD+REase->OLD-ABC->HNH->METHYLASE->OLD-ABC->REase->DUF499*->?->?->?-><-?\|\|SPOUT-> | 945 | Bacteria>Pseudomonadota>Betaproteobacteria>Burkholderiales>Comamonadaceae | GCF_017939685.1 | Acidovorax_sp._JG5 | J8G26_RS10065 | 96032..98869 | 2822718 | DUF499-domain-containing-protein-[Acidovorax-sp.-JG5] |
| WP_087746901.1 | AAA+ATPase+wHTH+Alpha+Beta-domain+wHTH+RRM | Type-1_BR.RM.like.systems | <-SPOUT\|\|?-><-?<-?<-DUF499*<-REase<-OLD-ABC<-METHYLASE<-HNH<-OLD-ABC<-TUDOR+SF1-SF2-Helicase+iREase+CTD+REase | 945 | Bacteria>Pseudomonadota>Betaproteobacteria>Burkholderiales>Comamonadaceae | GCF_002176815.1 | unclassified_Acidovorax | CCX87_RS13165 | 2797139..2799976 | 2684926 | MULTISPECIES:-DUF499-domain-containing-protein-[unclassified-Acidovorax] |
| MCW5668677.1 | AAA+ATPase+wHTH+Alpha+Beta-domain+wHTH+RRM | Type-1_BR.RM.like.systems | TUDOR+SF1-SF2-Helicase+iREase+CTD+REase->OLD-ABC->HNH->METHYLASE->?->?->DUF499*-><-HTH_3<-?<-HD\|\|NUDIX-> | 937 | Bacteria>Pseudomonadota>Betaproteobacteria>Burkholderiales>Comamonadaceae | GCA_026125825.1 | Hydrogenophaga_sp. | KIT86_03380 | 51119..53932 | 1904254 | MAG:-ATP-binding-protein-[Hydrogenophaga-sp.] |
| WP_265654212.1 | AAA+ATPase+wHTH+Alpha+Beta-domain+wHTH+RRM | Type-1_BR.RM.like.systems | Peptidase_C26->?->?-><-?<-OLD-ABC+TOPRIM<-DUF499*<-METHYLASE<-HEPN<-TUDOR+SF1-SF2-Helicase+iREase+CTD+REase | 951 | Bacteria>Pseudomonadota>Betaproteobacteria>Burkholderiales>Comamonadaceae | GCF_026192715.1 | Verminephrobacter_aporrectodeae | D8B27_RS10360 | 21500..24355 | 1110389 | DUF499-domain-containing-protein-[Verminephrobacter-aporrectodeae] |
| WP_265282594.1 | AAA+ATPase+wHTH+Alpha+Beta-domain+wHTH+RRM | Type-1_BR.RM.like.systems | TUDOR+SF1-SF2-Helicase+iREase+CTD+REase->HEPN->METHYLASE->DUF499*->OLD-ABC+TOPRIM->?->?-><-?<-?<-Peptidase_C26 | 951 | Bacteria>Pseudomonadota>Betaproteobacteria>Burkholderiales>Comamonadaceae | GCF_026127365.1 | Verminephrobacter_aporrectodeae | D5039_RS14590 | 1238377..1241232 | 1110389 | DUF499-domain-containing-protein-[Verminephrobacter-aporrectodeae] |
| WP_209423136.1 | AAA+ATPase+wHTH+Alpha+Beta-domain+wHTH+RRM | Type-1_BR.RM.like.systems | SF1-SF2-Helicase+DUF1998->?-><-?<-?<-?\|\|?-><-DUF499*<-REase<-OLD-ABC<-TOPRIM<-OLD-ABC<-METHYLASE-p<-METHYLASE | 943 | Bacteria>Pseudomonadota>Betaproteobacteria>Burkholderiales>Comamonadaceae | GCF_021249385.1 | Melaminivora_jejuensis | LVC68_RS01445 | 284242..287073 | 1267217 | DUF499-domain-containing-protein-[Melaminivora-jejuensis] |
| WP_265259651.1 | AAA+ATPase+wHTH+Alpha+Beta-domain+wHTH+RRM | Type-1_BR.RM.like.systems | METHYLASE->METHYLASE->Schlafen+HATPase+HTH->DUF499*-><-?\|\|?->SPOUT_MTase-><-?\|\|?->S1+RNase+2C-> | 936 | Bacteria>Pseudomonadota>Betaproteobacteria>Burkholderiales>Comamonadaceae | GCF_026127445.1 | Verminephrobacter_eiseniae | D8B29_RS12175 | 5874..8684 | 364317 | DUF499-domain-containing-protein-[Verminephrobacter-eiseniae] |
| WP_265302252.1 | AAA+ATPase+wHTH+Alpha+Beta-domain+wHTH+RRM | Type-1_BR.RM.like.systems | <-Phage_integrase\|\|?->DUF4096+DDE_Tnp_1+DDE_Tnp_1_2->?->?->?->DUF499*-><-?<-?\|\|?->SPOUT_MTase-> | 936 | Bacteria>Pseudomonadota>Betaproteobacteria>Burkholderiales>Comamonadaceae | GCF_026127465.1 | Verminephrobacter_eiseniae | D5045_RS08515 | 1712290..1715100 | 364317 | DUF499-domain-containing-protein-[Verminephrobacter-eiseniae] |
| WP_086929120.1 | AAA+ATPase+wHTH+Alpha+Beta-domain+wHTH+RRM | Type-1_BR.RM.like.systems | TUDOR+SF1-SF2-Helicase+iREase+CTD+REase->METHYLASE->DUF499*->?->OLD-ABC+TOPRIM-> | 970 | Bacteria>Pseudomonadota>Betaproteobacteria>Burkholderiales>Comamonadaceae | GCF_002157185.1 | Acidovorax_carolinensis | CBP35_RS20910 | 151033..153945 | 553814 | DUF499-domain-containing-protein-[Acidovorax-carolinensis] |
| NMM07407.1 | AAA+ATPase+wHTH+Alpha+Beta-domain+wHTH+RRM | Type-1_BR.RM.like.systems | TUDOR+SF1-SF2-Helicase+iREase+CTD+REase->HEPN->METHYLASE->DUF499*-> | 948 | Bacteria>Pseudomonadota>Betaproteobacteria>Burkholderiales>Comamonadaceae | GCA_012927455.1 | Polaromonas_sp. | HHJ18_14460 | 504043..506889 | 1869339 | ATP-binding-protein-[Polaromonas-sp.] |
| MBK6748240.1 | AAA+ATPase+wHTH+Alpha+Beta-domain+wHTH+RRM | Type-1_BR.RM.like.systems | <-ABhydrolase\|\|<-?<-HEPN\|\|wHTH+REase->DUF499*-><-tRNA<-?<-?\|\|?->MoxR-AAA-> | 942 | Bacteria>Pseudomonadota>Betaproteobacteria>Burkholderiales>Comamonadaceae | GCA_016703145.1 | Ottowia_sp. | IPG65_18050 | 1102212..1105040 | 1898956 | ATP-binding-protein-[Ottowia-sp.] |
| MBK6612629.1 | AAA+ATPase+wHTH+Alpha+Beta-domain+wHTH+RRM | Type-1_BR.RM.like.systems | <-RHH<-MULE-TRANSPOSASE<-?<-?<-DUF499*\|\|HEPN->?->\|\|ABhydrolase-> | 942 | Bacteria>Pseudomonadota>Betaproteobacteria>Burkholderiales>Comamonadaceae | GCA_016703855.1 | Ottowia_sp. | IPG27_00540 | 122634..125462 | 1898956 | ATP-binding-protein-[Ottowia-sp.] |
| MCB1982133.1 | AAA+ATPase+wHTH+Alpha+Beta-domain+wHTH+RRM | Type-1_BR.RM.like.systems | <-DUF499*<-HEPN<-HEPN-p<-HEPN<-PolB<-PIN<-OLD-ABC | 869 | Bacteria>Pseudomonadota>Betaproteobacteria>Burkholderiales>Comamonadaceae | GCA_020446185.1 | Rhodoferax_sp. | KDF63_09470 | 259..2868 | 50421 | ATP-binding-protein-[Rhodoferax-sp.] |
| MBP7545005.1 | AAA+ATPase+wHTH+Alpha+Beta-domain+wHTH+RRM | Type-1_BR.RM.like.systems | <-DUF499*<-REase<-OLD-ABC<-METHYLASE-p | 949 | Bacteria>Pseudomonadota>Betaproteobacteria>Burkholderiales>Comamonadaceae | GCA_017998835.1 | Acidovorax_sp. | KA749_05805 | 238..3087 | 1872122 | ATP-binding-protein-[Acidovorax-sp.] |
| UUZ52278.1 | AAA+ATPase+wHTH+Alpha+Beta-domain+wHTH+RRM | Type-1_BR.RM.like.systems | SF1-SF2-Helicase-p->?-><-REase<-DUF499-p<-DUF499-p<-DUF499*<-METHYLASE-p<-METHYLASE-p<-HEPN<-TUDOR+SF1-SF2-Helicase+iREase+CTD+REase-p<-TUDOR+SF1-SF2-Helicase+iREase+CTD+REase<-TUDOR+SF1-SF2-Helicase+iREase+CTD+REase-p | 325 | Bacteria>Pseudomonadota>Betaproteobacteria>Burkholderiales>Oxalobacteraceae | GCA_024703915.1 | Massilia_sp._H-1 | LP419_21985 | 3096112..3097089 | 2895566 | DUF499-domain-containing-protein-[Massilia-sp.-H-1] |
| WP_012080583.1 | AAA+ATPase+wHTH+Alpha+Beta-domain+wHTH+RRM | Type-1_BR.RM.like.systems | Phage_lysozyme-><-TIR<-TIR<-SIR2+STALD<-DUF499*<-METHYLASE<-TUDOR+SF1-SF2-Helicase+iREase+CTD+REase<-HTH+WYL<-?<-HTH<-GDResolvase+Recombinase+ZnR | 950 | Bacteria>Pseudomonadota>Betaproteobacteria>Burkholderiales>Oxalobacteraceae | GCF_000013625.1 | Janthinobacterium_sp._Marseille | MMA_RS14185 | 3073154..3076006 | 375286 | DUF499-domain-containing-protein-[Janthinobacterium-sp.-Marseille] |
| WP_281913453.1 | AAA+ATPase+wHTH+Alpha+Beta-domain+wHTH+RRM | Type-1_BR.RM.like.systems | <-XIS-HTH\|\|?->?->HTH+WYL->TUDOR+SF1-SF2-Helicase+iREase+CTD+REase->METHYLASE->DUF499*->?-><-HHH_4+SF1-SF2-helicase<-?<-?<-?<-Phage_integrase | 963 | Bacteria>Pseudomonadota>Betaproteobacteria>Burkholderiales>Oxalobacteraceae | GCF_027923905.1 | Massilia_varians | MasN3_RS08140 | 1772943..1775834 | 457921 | DUF499-domain-containing-protein-[Massilia-varians] |
| WP_170204580.1 | AAA+ATPase+wHTH+Alpha+Beta-domain+wHTH+RRM | Type-1_BR.RM.like.systems | <-RVT\|\|HTH_Tnp_1+HTH_21+rve+rve_3-><-?<-?<-DUF499*<-METHYLASE<-HEPN<-TUDOR+SF1-SF2-Helicase+iREase+CTD+REase\|\|STYKinase-like-> | 956 | Bacteria>Pseudomonadota>Betaproteobacteria>Burkholderiales>Oxalobacteraceae | GCF_012849555.1 | Massilia_forsythiae | HH212_RS22790 | 5438600..5441470 | 2728020 | DUF499-domain-containing-protein-[Massilia-forsythiae] |
| WP_182994400.1 | AAA+ATPase+wHTH+Alpha+Beta-domain+wHTH+RRM | Type-1_BR.RM.like.systems | HTH+WYL->TUDOR+SF1-SF2-Helicase+iREase+CTD+REase->OLD-ABC->HNH->METHYLASE->DUF499*-><-?\|\|?-><-?<-Arm-DNA-bind_3+Phage_integrase | 941 | Bacteria>Pseudomonadota>Betaproteobacteria>Burkholderiales>Oxalobacteraceae | GCF_014171515.1 | Herbaspirillum_frisingense | G5S34_RS08630 | 1936607..1939432 | 92645 | DUF499-domain-containing-protein-[Herbaspirillum-frisingense] |
| UUZ46680.1 | AAA+ATPase+wHTH+Alpha+Beta-domain+wHTH+RRM | Type-1_BR.RM.like.systems | SF1-SF2-Helicase-p->?->?-><-REase<-DUF499-p<-DUF499-p<-DUF499*<-METHYLASE-p<-METHYLASE-p<-METHYLASE-p<-HEPN\|\|?->RNaseP-> | 339 | Bacteria>Pseudomonadota>Betaproteobacteria>Burkholderiales>Oxalobacteraceae | GCA_024703935.1 | Massilia_sp._B-10 | LP420_22520 | 3094423..3095442 | 2895567 | DUF499-domain-containing-protein-[Massilia-sp.-B-10] |
| WP_072453399.1 | AAA+ATPase+wHTH+Alpha+Beta-domain+wHTH+RRM | Type-1_BR.RM.like.systems | <-STYKinase-like\|\|HTH+WYL->TUDOR+SF1-SF2-Helicase+iREase+CTD+REase->HEPN->METHYLASE->DUF499*->REase-> | 961 | Bacteria>Pseudomonadota>Betaproteobacteria>Burkholderiales>Oxalobacteraceae | GCF_900119665.1 | Janthinobacterium_lividum | BT188_RS05025 | 1132446..1135331 | 29581 | DUF499-domain-containing-protein-[Janthinobacterium-lividum] |
| WP_027868039.1 | AAA+ATPase+wHTH+Alpha+Beta-domain+wHTH+RRM | Type-1_BR.RM.like.systems | SF1-SF2-Helicase-><-DUF499*<-METHYLASE<-TUDOR+SF1-SF2-Helicase+iREase+CTD+REase<-HTH+WYL\|\|HIN-HTH-> | 963 | Bacteria>Pseudomonadota>Betaproteobacteria>Burkholderiales>Oxalobacteraceae | GCF_000427785.1 | Massilia_alkalitolerans | YWC_RS0126390 | 1159..4050 | 286638 | DUF499-domain-containing-protein-[Massilia-alkalitolerans] |
| MCM1512954.1 | AAA+ATPase+wHTH+Alpha+Beta-domain+wHTH+RRM | Type-1_BR.RM.like.systems | HTH_3->?->?->?->TUDOR+SF1-SF2-Helicase+iREase+CTD+REase->METHYLASE->DUF499*->OLD-ABC->HNH-><-?\|\|MazE_antitoxin-> | 937 | Bacteria>Pseudomonadota>Betaproteobacteria>Burkholderiales>Oxalobacteraceae | GCA_023667225.1 | Oxalobacter_formigenes | NC112_07605 | 9124..11937 | 847 | DUF499-domain-containing-protein-[Oxalobacter-formigenes] |
| RYF19713.1 | AAA+ATPase+wHTH+Alpha+Beta-domain+wHTH+RRM | Type-1_BR.RM.like.systems | MerR-HTH->GDResolvase+HTH-><-?\|\|HTH_3->DUF499*-> | 465 | Bacteria>Pseudomonadota>Betaproteobacteria>Burkholderiales>Oxalobacteraceae | GCA_004144765.1 | Oxalobacteraceae_bacterium | EOO77_09540 | 2015..3409 | 1871070 | DUF499-domain-containing-protein,-partial-[Oxalobacteraceae-bacterium] |
| RZL03828.1 | AAA+ATPase+wHTH+Alpha+Beta-domain+wHTH+RRM | Type-1_BR.RM.like.systems | HTH+WYL->TIR+PNTD2->PIWI->TUDOR+SF1-SF2-Helicase+iREase+CTD+REase->HEPN->METHYLASE->DUF499*->?->KAP-NTPase->?->PAPS_reduct-> | 956 | Bacteria>Pseudomonadota>Betaproteobacteria>Burkholderiales>Sphaerotilaceae | GCA_004211355.1 | Rubrivivax_sp. | EOP36_03025 | 28587..31457 | 50259 | MAG:-ATP-binding-protein-[Rubrivivax-sp.] |
| WP_066335883.1 | AAA+ATPase+wHTH+Alpha+Beta-domain+wHTH+RRM | Type-1_BR.RM.like.systems | <-RNase+SF1-SF2-Helicase<-?<-?<-MACRODOMAIN<-DUF499*<-REase<-OLD-ABC<-METHYLASE<-?<-TUDOR+SF1-SF2-Helicase+iREase+CTD+REase | 939 | Bacteria>Pseudomonadota>Betaproteobacteria>Burkholderiales>Sphaerotilaceae | GCF_001571085.1 | Azohydromonas_lata | ALA01S_RS10810 | 44580..47399 | 45677 | DUF499-domain-containing-protein-[Azohydromonas-lata] |
| NRT58384.1 | AAA+ATPase+wHTH+Alpha+Beta-domain+wHTH+RRM | Type-1_BR.RM.like.systems | HAD->?-><-rve_3<-ParB+HTH<-?\|\|?-><-DUF499*<-TOPRIM<-OLD-ABC<-METHYLASE<-HNH<-OLD-ABC<-TUDOR+SF1-SF2-Helicase+iREase+CTD+REase | 953 | Bacteria>Pseudomonadota>Betaproteobacteria>Burkholderiales>Sphaerotilaceae | GCA_013294065.1 | Leptothrix_sp._C29 | HNQ01_004152 | 18472..21333 | 2735897 | hypothetical-protein-[Leptothrix-sp.-C29] |
| WP_273473498.1 | AAA+ATPase+wHTH+Alpha+Beta-domain+wHTH+RRM | Type-1_BR.RM.like.systems | <-zf-IS66+DDE_Tnp_IS66<-DDE_Tnp_1_assoc+DDE_Tnp_1<-ParB+HTH<-?\|\|?-><-DUF499*<-TOPRIM<-OLD-ABC<-METHYLASE<-?<-PIN<-OLD-ABC | 955 | Bacteria>Pseudomonadota>Betaproteobacteria>Burkholderiales>Sphaerotilaceae | GCF_023150255.1 | Sphaerotilus_sulfidivorans | L6Q74_RS20510 | 5829..8696 | 639200 | DUF499-domain-containing-protein-[Sphaerotilus-sulfidivorans] |
| MCK6404267.1 | AAA+ATPase+wHTH+Alpha+Beta-domain+wHTH+RRM | Type-1_BR.RM.like.systems | <-zf-IS66+DDE_Tnp_IS66<-DDE_Tnp_1_assoc+DDE_Tnp_1<-ParB+HTH<-?\|\|?-><-DUF499*<-TOPRIM<-OLD-ABC<-METHYLASE<-?<-PIN<-OLD-ABC | 955 | Bacteria>Pseudomonadota>Betaproteobacteria>Burkholderiales>Sphaerotilaceae | GCA_023150255.1 | Sphaerotilus_sulfidivorans | L6Q74_20550 | 5829..8696 | 639200 | DUF499-domain-containing-protein-[Sphaerotilus-sulfidivorans] |
| WP_245558353.1 | AAA+ATPase+wHTH+Alpha+Beta-domain+wHTH+RRM | Type-1_BR.RM.like.systems | RNase->?->?->Phage_lysozyme-><-TIR<-DUF499*<-?<-?<-METHYLASE<-PIN<-TUDOR+SF1-SF2-Helicase+iREase+CTD+REase<-HTH_Tnp | 934 | Bacteria>Pseudomonadota>Betaproteobacteria>Burkholderiales>Sphaerotilaceae | GCF_000381125.1 | Caldimonas_manganoxidans | F615_RS0111630 | 4122..6926 | 196015 | DUF499-domain-containing-protein-[Caldimonas-manganoxidans] |
| WP_173807403.1 | AAA+ATPase+wHTH+Alpha+Beta-domain+wHTH+RRM | Type-1_BR.RM.like.systems | HAD->?-><-ParB+HTH<-?\|\|?-><-DUF499*<-TOPRIM<-OLD-ABC<-METHYLASE<-HNH<-OLD-ABC<-TUDOR+SF1-SF2-Helicase+iREase+CTD+REase | 953 | Bacteria>Pseudomonadota>Betaproteobacteria>Burkholderiales>Sphaerotilaceae | GCF_013294065.1 | Leptothrix_sp._C29 | HNQ01_RS20655 | 18472..21333 | 2735897 | DUF499-domain-containing-protein-[Leptothrix-sp.-C29] |
| WP_110402335.1 | AAA+ATPase+wHTH+Alpha+Beta-domain+wHTH+RRM | Type-1_BR.RM.like.systems | <-ABhydrolase\|\|?->?-><-?<-DUF499*<-REase<-OLD-ABC<-METHYLASE<-HNH<-OLD-ABC<-TUDOR+SF1-SF2-Helicase+iREase+CTD+REase | 958 | Bacteria>Pseudomonadota>Betaproteobacteria>Burkholderiales>Sphaerotilaceae | - | Sphaerotilus_hippei | C7444_RS18460 | 22920..25796 | 744406 | DUF499-domain-containing-protein-[Sphaerotilus-hippei] |
| WP_140628337.1 | AAA+ATPase+wHTH+Alpha+Beta-domain+wHTH+RRM | Type-1_BR.RM.like.systems | PIN+OST-HTH-><-?\|\|HTH_3-><-?<-?<-?<-DUF499*<-?<-?<-METHYLASE<-HNH<-OLD-ABC<-TUDOR+SF1-SF2-Helicase+iREase+CTD+REase | 936 | Bacteria>Pseudomonadota>Betaproteobacteria>Burkholderiales>Sphaerotilaceae | GCF_006386545.1 | Methylibium_rhizosphaerae | E8D38_RS09385 | 117386..120196 | 2570323 | DUF499-domain-containing-protein-[Pseudorivibacter-rhizosphaerae] |
| WP_157271680.1 | AAA+ATPase+wHTH+Alpha+Beta-domain+wHTH+RRM | Type-1_BR.RM.like.systems | <-MACRODOMAIN<-DUF499*<-ParB+HNH<-METHYLASE<-TUDOR+SF1-SF2-Helicase+iREase+CTD+REase | 943 | Bacteria>Pseudomonadota>Betaproteobacteria>Burkholderiales>Sphaerotilaceae | GCF_009760915.1 | Azohydromonas_aeria | FJ694_RS32510 | 1271..4102 | 2590212 | DUF499-domain-containing-protein-[Azohydromonas-aeria] |
| WP_281915166.1 | AAA+ATPase+wHTH+Alpha+Beta-domain+wHTH+RRM | Type-1_BR.RM.like.systems | Phage_lysozyme-><-HEPN<-DUF499*<-Schlafen+HATPase+HTH<-METHYLASE<-METHYLASE | 955 | Bacteria>Pseudomonadota>Betaproteobacteria>Burkholderiales>Sphaerotilaceae | GCF_937957785.1 | Caldimonas_thermodepolymerans | QMG53_RS04175 | 1459..4326 | 215580 | DUF499-domain-containing-protein-[Caldimonas-thermodepolymerans] |
| MBP7519657.1 | AAA+ATPase+wHTH+Alpha+Beta-domain+wHTH+RRM | Type-1_BR.RM.like.systems | <-METHYLASE-p\|\|?->?-><-DUF499* | 649 | Bacteria>Pseudomonadota>Betaproteobacteria>Burkholderiales>Sphaerotilaceae | GCA_017998205.1 | Leptothrix_sp._(in:_b-proteobacteria) | KA896_04365 | 29319..31268 | 34030 | ATP-binding-protein-[Leptothrix-sp.-(in:-b-proteobacteria)] |
| MCA3215581.1 | AAA+ATPase+wHTH+Alpha+Beta-domain+wHTH+RRM | Type-1_BR.RM.like.systems | <-MetJarc-HTH\|\|HTH+WYL->HTH_Tnp->TUDOR+SF1-SF2-Helicase+iREase+CTD+REase->HEPN->METHYLASE->DUF499*->?->Methylase-><-Radical-SAM<-HTH_3 | 948 | Bacteria>Pseudomonadota>Betaproteobacteria>Burkholderiales>unclassified_Burkholderiales | GCA_020248845.1 | Burkholderiales_bacterium | ING39_09185 | 33455..36301 | 1891238 | ATP-binding-protein-[Burkholderiales-bacterium] |
| MCZ2134305.1 | AAA+ATPase+wHTH+Alpha+Beta-domain+wHTH+RRM | Type-1_BR.RM.like.systems | <-RAMA<-?<-?<-HTH_21+rve+rve_3<-HIN-HTH<-?<-DUF499*<-wHTH+REase<-METHYLASE<-HEPN<-PIN<-TUDOR+SF1-SF2-Helicase+iREase+CTD+REase\|\|HIN-HTH-> | 956 | Bacteria>Pseudomonadota>Betaproteobacteria>Burkholderiales>unclassified_Burkholderiales | GCA_026932525.1 | Burkholderiales_bacterium | LC098_02580 | 42253..45123 | 1891238 | MAG:-DUF499-domain-containing-protein-[Burkholderiales-bacterium] |
| NNM65793.1 | AAA+ATPase+wHTH+Alpha+Beta-domain+wHTH+RRM | Type-1_BR.RM.like.systems | HTH+WYL->TUDOR+SF1-SF2-Helicase+iREase+CTD+REase->METHYLASE->DUF499*->OLD-ABC+TOPRIM->SF1-SF2-Helicase->?->OLD-ABC+TOPRIM-> | 971 | Bacteria>Pseudomonadota>Betaproteobacteria>Burkholderiales>unclassified_Burkholderiales | GCA_013044035.1 | Burkholderiales_bacterium | HKL99_14520 | 53445..56360 | 1891238 | ATP-binding-protein-[Burkholderiales-bacterium] |
| MBX9611601.1 | AAA+ATPase+wHTH+Alpha+Beta-domain+wHTH+RRM | Type-1_BR.RM.like.systems | TUDOR+SF1-SF2-Helicase+iREase+CTD+REase->HEPN->METHYLASE->DUF499*-><-?<-?<-SF1-SF2-Helicase+HRDC | 953 | Bacteria>Pseudomonadota>Betaproteobacteria>Burkholderiales>unclassified_Burkholderiales | GCA_019747995.1 | Burkholderiales_bacterium | K2X51_08260 | 22656..25517 | 1891238 | DUF499-domain-containing-protein-[Burkholderiales-bacterium] |
| MCZ2407521.1 | AAA+ATPase+wHTH+Alpha+Beta-domain+wHTH+RRM | Type-1_BR.RM.like.systems | Phage_integrase+Phage_integrase->?-><-DUF499*<-REase<-OLD-ABC<-METHYLASE | 945 | Bacteria>Pseudomonadota>Betaproteobacteria>Burkholderiales>unclassified_Burkholderiales | GCA_026931285.1 | Burkholderiales_bacterium | LC097_12425 | 115108..117945 | 1891238 | MAG:-DUF499-domain-containing-protein-[Burkholderiales-bacterium] |
| MCA3175663.1 | AAA+ATPase+wHTH+Alpha+Beta-domain+wHTH+RRM | Type-1_BR.RM.like.systems | METHYLASE-p->wHTH+REase->DUF499*-><-tRNA\|\|?->?->Methyltransf_4-> | 941 | Bacteria>Pseudomonadota>Betaproteobacteria>Burkholderiales>unclassified_Burkholderiales | GCA_020248585.1 | Burkholderiales_bacterium | ING36_09020 | 1791..4616 | 1891238 | ATP-binding-protein-[Burkholderiales-bacterium] |
| MBK6716517.1 | AAA+ATPase+wHTH+Alpha+Beta-domain+wHTH+RRM | Type-1_BR.RM.like.systems | METHYLASE-p->OLD-ABC->REase->DUF499*-><-?<-?\|\|METHYLASE-p-> | 948 | Bacteria>Pseudomonadota>Betaproteobacteria>Burkholderiales>unclassified_Burkholderiales | GCA_016703325.1 | Burkholderiales_bacterium | IPG57_15845 | 2519..5365 | 1891238 | ATP-binding-protein-[Burkholderiales-bacterium] |
| QKE41939.1 | AAA+ATPase+wHTH+Alpha+Beta-domain+wHTH+RRM | Type-1_BR.RM.like.systems | DDE_Tnp_1_assoc+DDE_Tnp_1->HTH+WYL->TUDOR+SF1-SF2-Helicase+iREase+CTD+REase->OLD-ABC->HNH->wHTH+REase->DUF499*->DDE_Tnp_ISAZ013-><-?<-Hydrolase_3 | 942 | Bacteria>Pseudomonadota>Betaproteobacteria>Ferrovales>Ferrovaceae | GCA_013181415.1 | Ferrovum_myxofaciens | HO274_11940 | 2423876..2426704 | 416213 | ATP-binding-protein-[Ferrovum-myxofaciens] |
| WP_273119931.1 | AAA+ATPase+wHTH+Alpha+Beta-domain+wHTH+RRM | Type-1_BR.RM.like.systems | TUDOR+SF1-SF2-Helicase+iREase+CTD+REase->OLD-ABC->HNH->METHYLASE-p->METHYLASE-p->wHTH+REase->DUF499*->DDE_Tnp_ISAZ013-><-?<-?<-Hydrolase_3 | 942 | Bacteria>Pseudomonadota>Betaproteobacteria>Ferrovales>Ferrovaceae | GCF_013181415.1 | Ferrovum_myxofaciens | HO274_RS11980 | 2423876..2426704 | 416213 | DUF499-domain-containing-protein-[Ferrovum-myxofaciens] |
| WP_031597390.1 | AAA+ATPase+wHTH+Alpha+Beta-domain+wHTH+RRM | Type-1_BR.RM.like.systems | HTH+WYL->TUDOR+SF1-SF2-Helicase+iREase+CTD+REase->OLD-ABC->HNH->METHYLASE->DUF499*->?->?->?-><-?<-?<-DUF2163+Phage_BR0599 | 944 | Bacteria>Pseudomonadota>Betaproteobacteria>Ferrovales>Ferrovaceae | GCF_000735045.1 | Ferrovum_myxofaciens | IV91_RS0107820 | 14389..17223 | 416213 | DUF499-domain-containing-protein-[Ferrovum-myxofaciens] |
| NNM68915.1 | AAA+ATPase+wHTH+Alpha+Beta-domain+wHTH+RRM | Type-1_BR.RM.like.systems | METHYLASE-p->wHTH+REase->DUF499*-> | 522 | Bacteria>Pseudomonadota>Betaproteobacteria>Nitrosomonadales>Gallionellaceae | GCA_013043965.1 | Gallionella_sp. | HKM00_02900 | 1745..3312 | 2052161 | ATP-binding-protein-[Gallionella-sp.] |
| WP_090702677.1 | AAA+ATPase+wHTH+Alpha+Beta-domain+wHTH+RRM | Type-1_BR.RM.like.systems | rve_3-><-?<-HerA/FtsK<-?<-DUF499*<-METHYLASE<-TUDOR+SF1-SF2-Helicase+iREase+CTD+REase<-XIS-HTH\|\|REase+RNase-><-tRNA<-Arm-DNA-bind_2+Phage_int_SAM_3+Phage_integrase | 942 | Bacteria>Pseudomonadota>Betaproteobacteria>Nitrosomonadales>Nitrosomonadaceae | GCF_900114305.1 | Nitrosomonas_aestuarii | BM096_RS15665 | 3734..6562 | 52441 | DUF499-domain-containing-protein-[Nitrosomonas-aestuarii] |
| TXI39738.1 | AAA+ATPase+wHTH+Alpha+Beta-domain+wHTH+RRM | Type-1_BR.RM.like.systems | <-VirB11+VirB4-FtsK<-HTH\|\|ParB+HTH->?->TUDOR+SF1-SF2-Helicase+iREase+CTD+REase->METHYLASE->DUF499*-><-JAB<-UbE1+JAB<-SMODS<-?\|\|HTH+WYL-> | 930 | Bacteria>Pseudomonadota>Betaproteobacteria>Nitrosomonadales>Nitrosomonadaceae | GCA_008015645.1 | Nitrosomonas_sp. | E6Q59_04335 | 20804..23596 | 42353 | ATP-binding-protein-[Nitrosomonas-sp.] |
| WP_090700444.1 | AAA+ATPase+wHTH+Alpha+Beta-domain+wHTH+RRM | Type-1_BR.RM.like.systems | Y1_Tnp->?->?-><-DUF499*<-METHYLASE<-TUDOR+SF1-SF2-Helicase+iREase+CTD+REase<-?<-?<-?<-ZnR+TOPRIM+primpol+HTH | 946 | Bacteria>Pseudomonadota>Betaproteobacteria>Nitrosomonadales>Nitrosomonadaceae | GCF_900114305.1 | Nitrosomonas_aestuarii | BM096_RS10660 | 61898..64738 | 52441 | DUF499-domain-containing-protein-[Nitrosomonas-aestuarii] |
| MBX3631056.1 | AAA+ATPase+wHTH+Alpha+Beta-domain+wHTH+RRM | Type-1_BR.RM.like.systems | <-DUF499*<-METHYLASE<-HNH<-OLD-ABC<-TUDOR+SF1-SF2-Helicase+iREase+CTD+REase<-REase+RNase | 949 | Bacteria>Pseudomonadota>Betaproteobacteria>Nitrosomonadales>Nitrosomonadaceae | GCA_019635215.1 | Nitrosomonas_sp. | KF908_14345 | 13134..15983 | 42353 | ATP-binding-protein-[Nitrosomonas-sp.] |
| MCW5608748.1 | AAA+ATPase+wHTH+Alpha+Beta-domain+wHTH+RRM | Type-1_BR.RM.like.systems | HNH->METHYLASE->DUF499*-> | 949 | Bacteria>Pseudomonadota>Betaproteobacteria>Nitrosomonadales>Nitrosomonadaceae | GCA_026125505.1 | Nitrosomonas_sp. | KIT26_11955 | 3990..6839 | 42353 | MAG:-ATP-binding-protein-[Nitrosomonas-sp.] |
| MCC7313057.1 | AAA+ATPase+wHTH+Alpha+Beta-domain+wHTH+RRM | Type-1_BR.RM.like.systems | TUDOR+SF1-SF2-Helicase+iREase+CTD+REase->PRTase+PRTase->?->METHYLASE->DUF499*-> | 937 | Bacteria>Pseudomonadota>Betaproteobacteria>Nitrosomonadales>Sterolibacteriaceae | GCA_020853215.1 | Sulfuritalea_sp. | IT510_17610 | 8921..11734 | 2480090 | ATP-binding-protein-[Sulfuritalea-sp.] |
| ODU50877.1 | AAA+ATPase+wHTH+Alpha+Beta-domain+wHTH+RRM | Type-1_BR.RM.like.systems | <-HAD<-?\|\|?-><-?<-?\|\|?-><-DUF499*<-?<-METHYLASE<-?<-Schlafen+HATPase+HTH<-TUDOR+SF1-SF2-Helicase+iREase+CTD+REase | 937 | Bacteria>Pseudomonadota>Betaproteobacteria>Nitrosomonadales>Thiobacillaceae | GCA_001725135.1 | Thiobacillus_sp._SCN_63-374 | ABS92_01640 | 11556..14369 | 1660149 | MAG:-hypothetical-protein-ABS92_01640-[Thiobacillus-sp.-SCN-63-374] |
| MBP6189859.1 | AAA+ATPase+wHTH+Alpha+Beta-domain+wHTH+RRM | Type-1_BR.RM.like.systems | DUF488-1-><-?<-DUF499*<-wHTH+REase<-METHYLASE<-HEPN | 943 | Bacteria>Pseudomonadota>Betaproteobacteria>Rhodocyclales>Azonexaceae | GCA_017985755.1 | Azonexus_sp. | KA440_12990 | 2046..4877 | 1872668 | ATP-binding-protein-[Azonexus-sp.] |
| WP_169202394.1 | AAA+ATPase+wHTH+Alpha+Beta-domain+wHTH+RRM | Type-1_BR.RM.like.systems | TUDOR+SF1-SF2-Helicase+iREase+CTD+REase->?->?-><-PAIREDC-HTH+HIN-HTH\|\|METHYLASE->HEPN->wHTH+REase->DUF499*-><-?<-?<-NADAR<-?<-NUDIX | 941 | Bacteria>Pseudomonadota>Betaproteobacteria>Rhodocyclales>Rhodocyclaceae | GCF_017894365.1 | Aromatoleum_bremense | GPA24_RS09420 | 37549..40374 | 76115 | DUF499-domain-containing-protein-[Aromatoleum-bremense] |
| MCA3105992.1 | AAA+ATPase+wHTH+Alpha+Beta-domain+wHTH+RRM | Type-1_BR.RM.like.systems | Thioredoxin->TraH->?-><-TIR<-DUF499*<-METHYLASE<-TUDOR+SF1-SF2-Helicase+iREase+CTD+REase<-HTH_Tnp<-HTH+WYL\|\|?->SMODS-> | 951 | Bacteria>Pseudomonadota>Betaproteobacteria>Rhodocyclales>Rhodocyclaceae | GCA_020248145.1 | Rhodocyclaceae_bacterium | IOD09_08055 | 7409..10264 | 1898103 | ATP-binding-protein-[Rhodocyclaceae-bacterium] |
| MBX3668836.1 | AAA+ATPase+wHTH+Alpha+Beta-domain+wHTH+RRM | Type-1_BR.RM.like.systems | HTH_Tnp->TUDOR+SF1-SF2-Helicase+iREase+CTD+REase->OLD-ABC->HNH->METHYLASE->DUF499*->?-><-Phage_lysozyme<-PIN<-?\|\|?->HTH-> | 945 | Bacteria>Pseudomonadota>Betaproteobacteria>Rhodocyclales>Rhodocyclaceae | GCA_019635235.1 | Rhodocyclaceae_bacterium | KF778_10575 | 185865..188702 | 1898103 | ATP-binding-protein-[Rhodocyclaceae-bacterium] |
| WP_169202998.1 | AAA+ATPase+wHTH+Alpha+Beta-domain+wHTH+RRM | Type-1_BR.RM.like.systems | <-DUF499*<-REase<-OLD-ABC<-METHYLASE<-HNH<-OLD-ABC<-TUDOR+SF1-SF2-Helicase+iREase+CTD+REase | 951 | Bacteria>Pseudomonadota>Betaproteobacteria>Rhodocyclales>Rhodocyclaceae | GCF_017894365.1 | Aromatoleum_bremense | GPA24_RS12870 | 12346..15201 | 76115 | DUF499-domain-containing-protein-[Aromatoleum-bremense] |
| MCX8086911.1 | AAA+ATPase+wHTH+Alpha+Beta-domain+wHTH+RRM | Type-1_BR.RM.like.systems | <-Phage_integrase\|\|tRNA-><-DUF499*<-Schlafen+HATPase+HTH<-METHYLASE<-TOPRIM<-OLD-ABC<-METHYLASE | 943 | Bacteria>Pseudomonadota>Betaproteobacteria>Rhodocyclales>Rhodocyclaceae | GCA_026415155.1 | Rhodocyclaceae_bacterium | N3C63_08430 | 24540..27371 | 1898103 | MAG:-DUF499-domain-containing-protein-[Rhodocyclaceae-bacterium] |
| MBQ7625733.1 | AAA+ATPase+wHTH+Alpha+Beta-domain+wHTH+RRM | Type-1_BR.RM.like.systems | METHYLASE-p->ParB+HNH->DUF499*-><-REase\|\|TP_methylase->?->ncRNA->?->Methyltransf_5-> | 945 | Bacteria>Pseudomonadota>Betaproteobacteria>Rhodocyclales>Rhodocyclaceae | GCA_017444505.1 | Rhodocyclaceae_bacterium | IJS87_01550 | 2639..5476 | 1898103 | ATP-binding-protein-[Rhodocyclaceae-bacterium] |
| MCX7164773.1 | AAA+ATPase+wHTH+Alpha+Beta-domain+wHTH+RRM | Type-1_BR.RM.like.systems | Arm-DNA-bind_3+Phage_integrase->XIS-HTH->Peptidase_C39-><-?\|\|HTH_3->RVT-><-DUF499*<-METHYLASE<-TUDOR+SF1-SF2-Helicase+iREase+CTD+REase\|\|?-><-?<-Trypsin<-MORC+WLM | 940 | Bacteria>Pseudomonadota>Betaproteobacteria>Rhodocyclales>unclassified_Rhodocyclales | GCA_026399475.1 | Rhodocyclales_bacterium | NTV11_00675 | 10515..13337 | 2072421 | MAG:-DUF499-domain-containing-protein-[Rhodocyclales-bacterium] |
| MBI5781383.1 | AAA+ATPase+wHTH+Alpha+Beta-domain+wHTH+RRM | Type-1_BR.RM.like.systems | OrfB_IS605+OrfB_Zn_ribbon->DUF499*-><-?<-?<-ParA-Soj-PloopNTPase | 947 | Bacteria>Pseudomonadota>Betaproteobacteria>Rhodocyclales>unclassified_Rhodocyclales | GCA_016235745.1 | Rhodocyclales_bacterium | HZA65_06945 | 891..3734 | 2072421 | ATP-binding-protein-[Rhodocyclales-bacterium] |
| WP_002936068.1 | AAA+ATPase+wHTH+Alpha+Beta-domain+wHTH+RRM | Type-1_BR.RM.like.systems | <-RNaseP<-?\|\|?->?->MORC+DNA_gyraseB+TOPRIM->?-><-DUF499*<-wHTH+REase<-METHYLASE<-HEPN<-ParB+HNH<-TUDOR+SF1-SF2-Helicase+iREase+CTD+REase | 941 | Bacteria>Pseudomonadota>Betaproteobacteria>Rhodocyclales>Zoogloeaceae | GCF_000310125.2 | Thauera_sp._27 | B447_RS02400 | 104287..107112 | 305700 | DUF499-domain-containing-protein-[Thauera-sp.-27] |
| WP_153168444.1 | AAA+ATPase+wHTH+Alpha+Beta-domain+wHTH+RRM | Type-1_BR.RM.like.systems | <-Radical-SAM+TRAM<-?\|\|tRNA->?->?->?-><-?<-DUF499*<-wHTH+REase<-REase<-OLD-ABC<-METHYLASE<-HEPN<-TUDOR+SF1-SF2-Helicase+iREase+CTD+REase | 941 | Bacteria>Pseudomonadota>Betaproteobacteria>Rhodocyclales>Zoogloeaceae | GCF_009469595.2 | Thauera_sp._2A1 | GH664_RS12700 | 85060..87885 | 2570191 | DUF499-domain-containing-protein-[Thauera-sp.-2A1] |
| WP_212351537.1 | AAA+ATPase+wHTH+Alpha+Beta-domain+wHTH+RRM | Type-1_BR.RM.like.systems | <-Phage_integrase\|\|HTH+WYL->TUDOR+SF1-SF2-Helicase+iREase+CTD+REase->OLD-ABC->HNH->METHYLASE->DUF499*->?-><-?<-RelA | 954 | Bacteria>Pseudomonadota>Betaproteobacteria>Rhodocyclales>Zoogloeaceae | GCF_018122735.1 | Azoarcus_sp._L1K30 | J5J83_RS19190 | 4181146..4184010 | 2820277 | DUF499-domain-containing-protein-[Azoarcus-sp.-L1K30] |
| WP_159691715.1 | AAA+ATPase+wHTH+Alpha+Beta-domain+wHTH+RRM | Type-1_BR.RM.like.systems | MODE-HTH+PBPII-><-TPR<-DUF499*<-REase<-OLD-ABC<-METHYLASE<-HNH<-OLD-ABC<-TUDOR+SF1-SF2-Helicase+iREase+CTD+REase | 943 | Bacteria>Pseudomonadota>Betaproteobacteria>Rhodocyclales>Zoogloeaceae | GCF_009800965.1 | Cognatazoarcus_halotolerans | GQB03_RS16405 | 71425..74256 | 2686016 | DUF499-domain-containing-protein-[Cognatazoarcus-halotolerans] |
| WP_168986546.1 | AAA+ATPase+wHTH+Alpha+Beta-domain+wHTH+RRM | Type-1_BR.RM.like.systems | Phage_integrase-><-DUF499*<-?<-?<-ParB+HNH<-METHYLASE<-PIN<-TUDOR+SF1-SF2-Helicase+iREase+CTD+REase | 944 | Bacteria>Pseudomonadota>Betaproteobacteria>Rhodocyclales>Zoogloeaceae | GCF_012910885.1 | Azoarcus_taiwanensis | GPA21_RS02025 | 735..3569 | 666964 | DUF499-domain-containing-protein-[Azoarcus-taiwanensis] |
| TMW75521.1 | AAA+ATPase+wHTH+Alpha+Beta-domain+wHTH+RRM | Type-1_BR.RM.like.systems | <-DUF499*<-METHYLASE<-HNH<-OLD-ABC<-DOC+HTH<-TUDOR+SF1-SF2-Helicase+iREase+CTD+REase\|\|HTH_31-> | 622 | Bacteria>Pseudomonadota>Betaproteobacteria>Rhodocyclales>Zoogloeaceae | GCA_006007845.1 | Thauera_sp._UPWRP | FG147_05165 | 1..1865 | 2488794 | ATP-binding-protein,-partial-[Thauera-sp.-UPWRP] |
| WP_191658157.1 | AAA+ATPase+wHTH+Alpha+Beta-domain+wHTH+RRM | Type-1_BR.RM.like.systems | <-DUF499*<-wHTH+REase<-HEPN<-METHYLASE<-?<-ParB+HNH<-TUDOR+SF1-SF2-Helicase+iREase+CTD+REase | 944 | Bacteria>Pseudomonadota>Betaproteobacteria>Rhodocyclales>Zoogloeaceae | - | Azoarcus_sp._Aa7 | AZOA_12030 | 1309..4143 | 2303328 | DUF499-domain-containing-protein-[Azoarcus-sp.-Aa7] |
| MBD5801779.1 | AAA+ATPase+wHTH+Alpha+Beta-domain+wHTH+RRM | Type-1_BR.RM.like.systems | <-DUF499*<-wHTH+REase<-HEPN<-METHYLASE<-?<-ParB+HNH<-TUDOR+SF1-SF2-Helicase+iREase+CTD+REase | 944 | Bacteria>Pseudomonadota>Betaproteobacteria>Rhodocyclales>Zoogloeaceae | GCA_014822185.1 | Azoarcus_sp._Aa7 | AZOA_12030 | 1309..4143 | 2303328 | hypothetical-protein-[Azoarcus-sp.-Aa7] |
| WP_148581647.1 | AAA+ATPase+wHTH+Alpha+Beta-domain+wHTH+RRM | Type-1_BR.RM.like.systems | DUF772+DDE_Tnp_1->METHYLASE->?->?->DUF499*-><-?\|\|HTH_3-><-DDE_Tnp_1_assoc+DDE_Tnp_1 | 946 | Bacteria>Pseudomonadota>Betaproteobacteria>Rhodocyclales>Zoogloeaceae | GCF_008107625.1 | Zoogloea_oleivorans | ETQ85_RS24610 | 14525..17365 | 1552750 | DUF499-domain-containing-protein-[Zoogloea-oleivorans] |
| MCP5226772.1 | AAA+ATPase+wHTH+Alpha+Beta-domain+wHTH+RRM | Type-1_BR.RM.like.systems | wHTH+REase->?->ParB+HNH->?->?->DUF499*->TPR-><-MORC+DNA_gyraseB+TOPRIM<-?<-?\|\|?->RNaseP-> | 941 | Bacteria>Pseudomonadota>Betaproteobacteria>Rhodocyclales>Zoogloeaceae | GCA_024235615.1 | Thauera_sp. | H6946_16890 | 24788..27613 | 1905334 | ATP-binding-protein-[Thauera-sp.] |
| MDD3354816.1 | AAA+ATPase+wHTH+Alpha+Beta-domain+wHTH+RRM | Type-1_BR.RM.like.systems | PIN-><-DUF499*<-REase<-OLD-ABC<-METHYLASE | 948 | Bacteria>Pseudomonadota>Betaproteobacteria>Rhodocyclales>Zoogloeaceae | GCA_028693495.1 | Zoogloea_sp. | PHD22_14435 | 4878..7724 | 49181 | MAG:-DUF499-domain-containing-protein-[Zoogloea-sp.] |
| GHU44749.1 | AAA+ATPase+wHTH+Alpha+Beta-domain+wHTH+RRM | Type-1_BR.RM.like.systems | <-Aminotran_1_2<-DUF499*<-Schlafen+HATPase+HTH<-METHYLASE<-METHYLASE<-PIN<-TUDOR+SF1-SF2-Helicase+iREase+CTD+REase\|\|Aminotran_3-> | 945 | Bacteria>Pseudomonadota>Betaproteobacteria>unclassified_Betaproteobacteria | GCA_025997095.1 | Betaproteobacteria_bacterium | AGMMS50289_13810 | 2255..5092 | 1891241 | hypothetical-protein-AGMMS50289_13810-[Betaproteobacteria-bacterium] |
| MDA8417189.1 | AAA+ATPase+wHTH+Alpha+Beta-domain+wHTH+RRM | Type-1_BR.RM.like.systems | TUDOR+SF1-SF2-Helicase+iREase+CTD+REase->RVT->OLD-ABC->HNH->METHYLASE->wHTH+REase->DUF499*-><-?\|\|?->?->?-><-Response_reg+LuxR-HTH | 946 | Bacteria>Pseudomonadota>Betaproteobacteria>unclassified_Betaproteobacteria | GCA_028069095.1 | Betaproteobacteria_bacterium | MOZ78_09075 | 11618..14458 | 1891241 | MAG:-DUF499-domain-containing-protein-[Betaproteobacteria-bacterium] |
| MDA8257135.1 | AAA+ATPase+wHTH+Alpha+Beta-domain+wHTH+RRM | Type-1_BR.RM.like.systems | HTH+WYL->TUDOR+SF1-SF2-Helicase+iREase+CTD+REase->METHYLASE->DUF499*->?->?->?-><-?<-?<-Phage_lysozyme | 950 | Bacteria>Pseudomonadota>Betaproteobacteria>unclassified_Betaproteobacteria | GCA_028065725.1 | Betaproteobacteria_bacterium | MOZ99_16155 | 7897..10749 | 1891241 | MAG:-DUF499-domain-containing-protein-[Betaproteobacteria-bacterium] |
| GHT86461.1 | AAA+ATPase+wHTH+Alpha+Beta-domain+wHTH+RRM | Type-1_BR.RM.like.systems | ABC-ATPase+SbcC->?->?-><-DUF499*<-?<-?<-METHYLASE | 951 | Bacteria>Pseudomonadota>Betaproteobacteria>unclassified_Betaproteobacteria | GCA_025996925.1 | Betaproteobacteria_bacterium | AGMMS49543_19740 | 10642..13497 | 1891241 | hypothetical-protein-AGMMS49543_19740-[Betaproteobacteria-bacterium] |
| MSQ72376.1 | AAA+ATPase+wHTH+Alpha+Beta-domain+wHTH+RRM | Type-1_BR.RM.like.systems | ABC-ATPase+SbcC->?->?->?->Phd-YefM->PIN->DUF499*-><-DOC+HTH<-HTH<-RelE-ParE\|\|?-><-BirA-HTH | 104 | Bacteria>Pseudomonadota>Betaproteobacteria>unclassified_Betaproteobacteria | GCA_009693435.1 | Betaproteobacteria_bacterium | EXR27_13930 | 25540..25854 | 1891241 | hypothetical-protein-[Betaproteobacteria-bacterium] |
| OGA55362.1 | AAA+ATPase+wHTH+Alpha+Beta-domain+wHTH+RRM | Type-1_BR.RM.like.systems | <-DUF499*<-REase<-METHYLASE<-Schlafen+HATPase+HTH<-TUDOR+SF1-SF2-Helicase+iREase+CTD+REase | 941 | Bacteria>Pseudomonadota>Betaproteobacteria>unclassified_Betaproteobacteria | GCA_001772005.1 | Betaproteobacteria_bacterium_RIFCSPLOWO2_12_FULL_62_58 | A3F74_07155 | 338..3163 | 1797497 | AAA-family-ATPase-[Betaproteobacteria-bacterium-RIFCSPLOWO2_12_FULL_62_58] |
| MBI5922415.1 | AAA+ATPase+wHTH+Alpha+Beta-domain+wHTH+RRM | Type-1_BR.RM.like.systems | <-DUF499*<-REase<-METHYLASE<-?<-Schlafen+HATPase+HTH\|\|URI-DNase-> | 933 | Bacteria>Pseudomonadota>Betaproteobacteria>unclassified_Betaproteobacteria | GCA_016234745.1 | Betaproteobacteria_bacterium | HY847_12340 | 39471..42272 | 1891241 | ATP-binding-protein-[Betaproteobacteria-bacterium] |
| MBI3917901.1 | AAA+ATPase+wHTH+Alpha+Beta-domain+wHTH+RRM | Type-1_BR.RM.like.systems | OLD-ABC->PHP+SbcC->Schlafen+HATPase+HTH->METHYLASE->OLD-ABC->TOPRIM->DUF499*-> | 927 | Bacteria>Pseudomonadota>Betaproteobacteria>unclassified_Betaproteobacteria | GCA_016199545.1 | Betaproteobacteria_bacterium | HY322_12940 | 28133..30916 | 1891241 | ATP-binding-protein-[Betaproteobacteria-bacterium] |
| OGA43321.1 | AAA+ATPase+wHTH+Alpha+Beta-domain+wHTH+RRM | Type-1_BR.RM.like.systems | METHYLASE->PolB->HEPN->REase->DUF499*-> | 927 | Bacteria>Pseudomonadota>Betaproteobacteria>unclassified_Betaproteobacteria | GCA_001771985.1 | Betaproteobacteria_bacterium_RIFCSPLOWO2_12_FULL_62_13 | A3G24_09290 | 4491..7274 | 1797495 | AAA-family-ATPase-[Betaproteobacteria-bacterium-RIFCSPLOWO2_12_FULL_62_13] |
| WP_021314752.1 | AAA+ATPase+wHTH+Alpha+Beta-domain+wHTH+RRM | Type-1_BR.RM.like.systems | XIS-HTH->ZnR->HIN-HTH->TUDOR+SF1-SF2-Helicase+iREase+CTD+REase->SbcC->METHYLASE->DUF499*->RVT-><-LZ_Tnp_IS66+zf-IS66+DDE_Tnp_IS66+DDE_Tnp_IS66_C<-?<-?<-HIN-HTH\|\|DEDD_Tnp_IS110+Transposase_20-> | 947 | Bacteria>Pseudomonadota>Gammaproteobacteria | GCF_902159515.1 | Enterobacterales | OI978_RS10445 | 2192987..2195830 | 91347 | MULTISPECIES:-DUF499-domain-containing-protein-[Enterobacterales] |
| WP_096459285.1 | AAA+ATPase+wHTH+Alpha+Beta-domain+wHTH+RRM | Type-1_BR.RM.like.systems | TUDOR+SF1-SF2-Helicase+iREase+CTD+REase->OLD-ABC->METHYLASE->REase->DUF499*->?->?->?->?->?-><-HTH_3 | 943 | Bacteria>Pseudomonadota>Gammaproteobacteria>Acidiferrobacterales>Acidiferrobacteraceae | GCF_002355415.1 | Sulfurifustis_variabilis | SVA_RS04285 | 892754..895585 | 1675686 | DUF499-domain-containing-protein-[Sulfurifustis-variabilis] |
| WP_257709589.1 | AAA+ATPase+wHTH+Alpha+Beta-domain+wHTH+RRM | Type-1_BR.RM.like.systems | TUDOR+SF1-SF2-Helicase+iREase+CTD+REase->OLD-ABC->HNH->METHYLASE->OLD-ABC->TOPRIM->DUF499*-><-RlpA<-CbeA_antitoxin<-JAB<-?\|\|<-Arm-DNA-bind_3+Phage_integrase | 948 | Bacteria>Pseudomonadota>Gammaproteobacteria>Aeromonadales>Aeromonadaceae | GCF_024649925.1 | Aeromonas_caviae | NUK49_RS00060 | 12274..15120 | 648 | DUF499-domain-containing-protein-[Aeromonas-caviae] |
| WP_204794524.1 | AAA+ATPase+wHTH+Alpha+Beta-domain+wHTH+RRM | Type-1_BR.RM.like.systems | Phage_integrase-><-?<-DDE_Tnp_1_assoc+DDE_Tnp_1<-?<-?\|\|?<-DUF499*<-METHYLASE<-TUDOR+SF1-SF2-Helicase+iREase+CTD+REase<-?\|\|MODE-HTH+PBPII-> | 955 | Bacteria>Pseudomonadota>Gammaproteobacteria>Aeromonadales>Aeromonadaceae | GCF_016907015.1 | Oceanisphaera_litoralis | HNR62_RS02530 | 74478..77345 | 225144 | DUF499-domain-containing-protein-[Oceanisphaera-litoralis] |
| WP_223918005.1 | AAA+ATPase+wHTH+Alpha+Beta-domain+wHTH+RRM | Type-1_BR.RM.like.systems | TUDOR+SF1-SF2-Helicase+iREase+CTD+REase->OLD-ABC->HNH->METHYLASE->OLD-ABC->TOPRIM->DUF499*-><-RlpA<-CbeA_antitoxin<-JAB | 948 | Bacteria>Pseudomonadota>Gammaproteobacteria>Aeromonadales>Aeromonadaceae | GCF_019971655.1 | Aeromonas_caviae | LDN21_RS02455 | 71531..74377 | 648 | DUF499-domain-containing-protein-[Aeromonas-caviae] |
| WP_216991887.1 | AAA+ATPase+wHTH+Alpha+Beta-domain+wHTH+RRM | Type-1_BR.RM.like.systems | TUDOR+SF1-SF2-Helicase+iREase+CTD+REase->OLD-ABC->HNH->METHYLASE->OLD-ABC->TOPRIM->DUF499*-><-RlpA<-CbeA_antitoxin<-JAB | 948 | Bacteria>Pseudomonadota>Gammaproteobacteria>Aeromonadales>Aeromonadaceae | GCF_019048725.1 | Aeromonas_sp._FDAARGOS_1407 | I6L37_RS11490 | 2395827..2398673 | 2778056 | DUF499-domain-containing-protein-[Aeromonas-sp.-FDAARGOS-1407] |
| WP_201871420.1 | AAA+ATPase+wHTH+Alpha+Beta-domain+wHTH+RRM | Type-1_BR.RM.like.systems | JAB->CbeA_antitoxin->RlpA-><-DUF499*<-TOPRIM<-OLD-ABC<-METHYLASE<-HNH<-OLD-ABC<-TUDOR+SF1-SF2-Helicase+iREase+CTD+REase | 948 | Bacteria>Pseudomonadota>Gammaproteobacteria>Aeromonadales>Aeromonadaceae | GCF_016728905.1 | Aeromonas_caviae | JD551_RS01070 | 83459..86305 | 648 | DUF499-domain-containing-protein-[Aeromonas-caviae] |
| WP_103858986.1 | AAA+ATPase+wHTH+Alpha+Beta-domain+wHTH+RRM | Type-1_BR.RM.like.systems | JAB->CbeA_antitoxin->RlpA-><-DUF499*<-TOPRIM<-OLD-ABC<-METHYLASE<-HNH<-OLD-ABC<-TUDOR+SF1-SF2-Helicase+iREase+CTD+REase | 948 | Bacteria>Pseudomonadota>Gammaproteobacteria>Aeromonadales>Aeromonadaceae | GCF_002919855.1 | Aeromonas_sp._ASNIH8 | C3418_RS17745 | 3201..6047 | 1920113 | DUF499-domain-containing-protein-[Aeromonas-sp.-ASNIH8] |
| WP_223954701.1 | AAA+ATPase+wHTH+Alpha+Beta-domain+wHTH+RRM | Type-1_BR.RM.like.systems | JAB->CbeA_antitoxin->RlpA-><-DUF499*<-REase<-OLD-ABC<-METHYLASE<-HNH<-OLD-ABC<-TUDOR+SF1-SF2-Helicase+iREase+CTD+REase | 946 | Bacteria>Pseudomonadota>Gammaproteobacteria>Aeromonadales>Aeromonadaceae | GCF_019972775.1 | Aeromonas_caviae | LDN68_RS17105 | 3642842..3645682 | 648 | DUF499-domain-containing-protein-[Aeromonas-caviae] |
| WP_219296379.1 | AAA+ATPase+wHTH+Alpha+Beta-domain+wHTH+RRM | Type-1_BR.RM.like.systems | JAB->RlpA-><-HTH\|\|?-><-DUF499*<-TOPRIM<-OLD-ABC<-METHYLASE<-HNH<-OLD-ABC<-TUDOR+SF1-SF2-Helicase+iREase+CTD+REase | 952 | Bacteria>Pseudomonadota>Gammaproteobacteria>Aeromonadales>Aeromonadaceae | GCF_019348625.1 | Aeromonas_veronii | GL270_RS12655 | 107539..110397 | 654 | DUF499-domain-containing-protein-[Aeromonas-veronii] |
| WP_264787584.1 | AAA+ATPase+wHTH+Alpha+Beta-domain+wHTH+RRM | Type-1_BR.RM.like.systems | HTH+WYL->TUDOR+SF1-SF2-Helicase+iREase+CTD+REase->?->METHYLASE->OLD-ABC->TOPRIM->DUF499*-><-RlpA<-?<-JAB<-?<-?<-ParB | 945 | Bacteria>Pseudomonadota>Gammaproteobacteria>Aeromonadales>Aeromonadaceae | GCF_026000055.1 | Aeromonas_caviae | OPI30_RS08280 | 1646827..1649664 | 648 | DUF499-domain-containing-protein-[Aeromonas-caviae] |
| WP_244812704.1 | AAA+ATPase+wHTH+Alpha+Beta-domain+wHTH+RRM | Type-1_BR.RM.like.systems | METHYLASE-p->OLD-ABC->TOPRIM->DUF499*-><-RlpA<-CbeA_antitoxin<-JAB<-?\|\|<-Arm-DNA-bind_3+Phage_integrase | 948 | Bacteria>Pseudomonadota>Gammaproteobacteria>Aeromonadales>Aeromonadaceae | GCF_022869435.1 | Aeromonas | MUW49_RS10495 | 3086..5932 | 642 | DUF499-domain-containing-protein-[Aeromonas-sp.-LsrichE-8G] |
| WP_064338617.1 | AAA+ATPase+wHTH+Alpha+Beta-domain+wHTH+RRM | Type-1_BR.RM.like.systems | JAB->RlpA-><-HTH\|\|?-><-DUF499*<-TOPRIM<-OLD-ABC<-DDE_Tnp_1_assoc+DDE_Tnp_1<-OLD-ABC-p<-METHYLASE<-HNH | 954 | Bacteria>Pseudomonadota>Gammaproteobacteria>Aeromonadales>Aeromonadaceae | GCF_001634325.1 | Aeromonas_veronii | WM43_RS07825 | 1773937..1776801 | 654 | DUF499-domain-containing-protein-[Aeromonas-veronii] |
| WP_064341063.1 | AAA+ATPase+wHTH+Alpha+Beta-domain+wHTH+RRM | Type-1_BR.RM.like.systems | <-HTH\|\|?-><-HIN-HTH\|\|?->?->DDE_Tnp_1-><-DUF499*<-TOPRIM<-OLD-ABC<-OLD-ABC-p<-METHYLASE<-METHYLASE-p<-HNH | 954 | Bacteria>Pseudomonadota>Gammaproteobacteria>Aeromonadales>Aeromonadaceae | GCF_014169835.1 | Aeromonas_veronii | QBS04_RS15305 | 3271066..3273930 | 654 | DUF499-domain-containing-protein-[Aeromonas-veronii] |
| WP_270793588.1 | AAA+ATPase+wHTH+Alpha+Beta-domain+wHTH+RRM | Type-1_BR.RM.like.systems | TUDOR+SF1-SF2-Helicase+iREase+CTD+REase->?->METHYLASE->OLD-ABC->TOPRIM->DUF499*-><-RlpA<-?<-JAB | 945 | Bacteria>Pseudomonadota>Gammaproteobacteria>Aeromonadales>Aeromonadaceae | GCF_027705385.1 | unclassified_Aeromonas | OKC33_RS16320 | 9044..11881 | 257493 | MULTISPECIES:-DUF499-domain-containing-protein-[unclassified-Aeromonas] |
| WP_235462717.1 | AAA+ATPase+wHTH+Alpha+Beta-domain+wHTH+RRM | Type-1_BR.RM.like.systems | TUDOR+SF1-SF2-Helicase+iREase+CTD+REase->OLD-ABC->HNH->METHYLASE->OLD-ABC->TOPRIM->DUF499*-> | 897 | Bacteria>Pseudomonadota>Gammaproteobacteria>Aeromonadales>Aeromonadaceae | GCF_021538905.1 | Aeromonas_australiensis | EHZ86_RS11610 | 15978..18671 | 1114880 | DUF499-domain-containing-protein-[Aeromonas-australiensis] |
| MCF3097928.1 | AAA+ATPase+wHTH+Alpha+Beta-domain+wHTH+RRM | Type-1_BR.RM.like.systems | TUDOR+SF1-SF2-Helicase+iREase+CTD+REase->HNH->METHYLASE->OLD-ABC->TOPRIM->DUF499*-> | 954 | Bacteria>Pseudomonadota>Gammaproteobacteria>Aeromonadales>Aeromonadaceae | GCA_021538905.1 | Aeromonas_australiensis | EHZ86_11640 | 15807..18671 | 1114880 | ATP-binding-protein-[Aeromonas-australiensis] |
| QWL60397.1 | AAA+ATPase+wHTH+Alpha+Beta-domain+wHTH+RRM | Type-1_BR.RM.like.systems | URI-DNase-><-PAIREDC-HTH\|\|?->OLD-ABC->HNH->METHYLASE->DUF499*-><-?\|\|HTH-><-RlpA<-JAB | 954 | Bacteria>Pseudomonadota>Gammaproteobacteria>Aeromonadales>Aeromonadaceae | GCA_018802265.1 | Aeromonas_jandaei | HQ400_01350 | 291564..294428 | 650 | ATP-binding-protein-[Aeromonas-jandaei] |
| WP_270668501.1 | AAA+ATPase+wHTH+Alpha+Beta-domain+wHTH+RRM | Type-1_BR.RM.like.systems | <-DUF499*<-METHYLASE<-HNH<-OLD-ABC<-?<-TUDOR+SF1-SF2-Helicase+iREase+CTD+REase | 954 | Bacteria>Pseudomonadota>Gammaproteobacteria>Aeromonadales>Aeromonadaceae | GCF_027705205.1 | unclassified_Aeromonas | OKB91_RS16400 | 106..2970 | 257493 | MULTISPECIES:-DUF499-domain-containing-protein-[unclassified-Aeromonas] |
| WP_020742688.1 | AAA+ATPase+wHTH+Alpha+Beta-domain+wHTH+RRM | Type-1_BR.RM.like.systems | <-METHYLASE<-Methyltransf_4<-ABhydrolase\|\|HHH+ENDO3+NUDIX->?->tRNA-><-Schlafen<-DUF499*<-METHYLASE-p\|\|<-PAIREDC-HTH\|\|?->Resolvase->HTH_28+rve+Mu-transpos_C->GspA-ATPase-> | 939 | Bacteria>Pseudomonadota>Gammaproteobacteria>Alteromonadales>Alteromonadaceae | GCF_000439515.1 | Alteromonas_mediterranea | I876_RS01280 | 288971..291790 | 314275 | DUF499-domain-containing-protein-[Alteromonas-mediterranea] |
| WP_250655185.1 | AAA+ATPase+wHTH+Alpha+Beta-domain+wHTH+RRM | Type-1_BR.RM.like.systems | HTH+WYL->TUDOR+SF1-SF2-Helicase+iREase+CTD+REase->OLD-ABC->HNH->TUDOR+SF1-SF2-Helicase+iREase+CTD+REase-p->METHYLASE->DUF499*->SMODS->?-><-RlpA<-Phage_lysozyme | 945 | Bacteria>Pseudomonadota>Gammaproteobacteria>Alteromonadales>Alteromonadaceae | GCF_023650515.1 | Alkalimarinus_coralli | MY525_RS07060 | 1557790..1560627 | 2935863 | MULTISPECIES:-DUF499-domain-containing-protein-[unclassified-Alkalimarinus] |
| WP_101201433.1 | AAA+ATPase+wHTH+Alpha+Beta-domain+wHTH+RRM | Type-1_BR.RM.like.systems | HTH+WYL->TUDOR+SF1-SF2-Helicase+iREase+CTD+REase->OLD-ABC->HNH->METHYLASE->DUF499*-><-RlpA<-?\|\|?->Phage_integrase-> | 944 | Bacteria>Pseudomonadota>Gammaproteobacteria>Alteromonadales>Alteromonadaceae | GCF_002835935.1 | Paraglaciecola_sp._MB-3u-78 | CXF95_RS26560 | 32531..35365 | 2058332 | DUF499-domain-containing-protein-[Paraglaciecola-sp.-MB-3u-78] |
| APD87299.1 | AAA+ATPase+wHTH+Alpha+Beta-domain+wHTH+RRM | Type-1_BR.RM.like.systems | <-DUF499*<-METHYLASE<-TUDOR+SF1-SF2-Helicase+iREase+CTD+REase<-HTH+WYL\|\|?->REase->PIN+OST-HTH-> | 938 | Bacteria>Pseudomonadota>Gammaproteobacteria>Alteromonadales>Alteromonadaceae | GCA_001886375.1 | Alteromonas_sp._Mex14 | BM527_15040 | 3539852..3542668 | 1917157 | hypothetical-protein-BM527_15040-[Alteromonas-sp.-Mex14] |
| WP_087504467.1 | AAA+ATPase+wHTH+Alpha+Beta-domain+wHTH+RRM | Type-1_BR.RM.like.systems | <-Phage_integrase\|\|HTH+WYL->TUDOR+SF1-SF2-Helicase+iREase+CTD+REase->METHYLASE->OLD-ABC->REase->DUF499*-><-?<-?<-?<-?<-?\|\|REase-> | 940 | Bacteria>Pseudomonadota>Gammaproteobacteria>Alteromonadales>Echinimonadaceae | GCF_014636195.1 | Neiella_marina | CBE68_RS02320 | 109774..112596 | 508461 | DUF499-domain-containing-protein-[Neiella-marina] |
| KTG26834.1 | AAA+ATPase+wHTH+Alpha+Beta-domain+wHTH+RRM | Type-1_BR.RM.like.systems | Arm-DNA-bind_3+Phage_integrase-><-Phage_AlpA<-ZnR+TOPRIM+primpol+HTH<-XIS-HTH\|\|MobI-><-DUF499*<-REase<-OLD-ABC<-METHYLASE<-OLD-ABC<-TUDOR+SF1-SF2-Helicase+iREase+CTD+REase<-HTH+WYL | 953 | Bacteria>Pseudomonadota>Gammaproteobacteria>Alteromonadales>Idiomarinaceae | GCA_001469885.1 | Idiomarina_sp._H105 | AUR68_16335 | 105470..108331 | 1766622 | hypothetical-protein-AUR68_16335-[Idiomarina-sp.-H105] |
| WP_219013226.1 | AAA+ATPase+wHTH+Alpha+Beta-domain+wHTH+RRM | Type-1_BR.RM.like.systems | HTH_3->TUDOR+SF1-SF2-Helicase+iREase+CTD+REase->OLD-ABC->HNH->METHYLASE->DUF499*-><-RlpA<-Arm-DNA-bind_3+Phage_integrase<-?\|\|RNase->PRTase-> | 943 | Bacteria>Pseudomonadota>Gammaproteobacteria>Alteromonadales>Shewanellaceae | GCF_019303275.1 | Shewanella_algae | KOE54_RS14875 | 19508..22339 | 38313 | DUF499-domain-containing-protein-[Shewanella-algae] |
| WP_186265255.1 | AAA+ATPase+wHTH+Alpha+Beta-domain+wHTH+RRM | Type-1_BR.RM.like.systems | HTH_3->TUDOR+SF1-SF2-Helicase+iREase+CTD+REase->OLD-ABC->HNH->METHYLASE->DUF499*-><-RlpA<-Arm-DNA-bind_3+Phage_integrase<-?\|\|RNase->PRTase-> | 943 | Bacteria>Pseudomonadota>Gammaproteobacteria>Alteromonadales>Shewanellaceae | GCF_014263185.1 | Shewanella_algae | HU689_RS02010 | 444450..447281 | 38313 | DUF499-domain-containing-protein-[Shewanella-algae] |
| BCV35016.1 | AAA+ATPase+wHTH+Alpha+Beta-domain+wHTH+RRM | Type-1_BR.RM.like.systems | <-PRTase<-RNase\|\|?->Arm-DNA-bind_3+Phage_integrase->RlpA-><-DUF499*<-METHYLASE<-HNH<-OLD-ABC<-TUDOR+SF1-SF2-Helicase+iREase+CTD+REase<-HTH_3 | 943 | Bacteria>Pseudomonadota>Gammaproteobacteria>Alteromonadales>Shewanellaceae | GCA_019670565.1 | Shewanella_chilikensis | TUM17377_03440 | 374214..377045 | 558541 | hypothetical-protein-TUM17377_03440-[Shewanella-chilikensis] |
| WP_115016378.1 | AAA+ATPase+wHTH+Alpha+Beta-domain+wHTH+RRM | Type-1_BR.RM.like.systems | <-JAB\|\|XIS-HTH->REase-p->REase->RlpA-><-DUF499*<-METHYLASE<-HNH<-OLD-ABC<-TUDOR+SF1-SF2-Helicase+iREase+CTD+REase\|\|?->DDE_Tnp_1_3-> | 943 | Bacteria>Pseudomonadota>Gammaproteobacteria>Alteromonadales>Shewanellaceae | GCF_019266125.1 | Shewanella | DX079_RS11245 | 2521097..2523928 | 22 | MULTISPECIES:-DUF499-domain-containing-protein-[Shewanella] |
| WP_079541822.1 | AAA+ATPase+wHTH+Alpha+Beta-domain+wHTH+RRM | Type-1_BR.RM.like.systems | TUDOR+SF1-SF2-Helicase+iREase+CTD+REase->OLD-ABC->HNH->METHYLASE->OLD-ABC->TOPRIM->DUF499*-> | 937 | Bacteria>Pseudomonadota>Gammaproteobacteria>Cardiobacteriales>Cardiobacteriaceae | GCF_900079085.1 | Cardiobacterium_hominis | CHUV0807_RS10775 | 37520..40333 | 2718 | DUF499-domain-containing-protein-[Cardiobacterium-hominis] |
| WP_182173095.1 | AAA+ATPase+wHTH+Alpha+Beta-domain+wHTH+RRM | Type-1_BR.RM.like.systems | Phage_integrase-><-Phage_AlpA<-ZnR+TOPRIM+primpol+HTH<-XIS-HTH\|\|MobI-><-DUF499*<-Schlafen+HATPase+HTH<-METHYLASE<-METHYLASE<-SbcC+SbcC<-TUDOR+SF1-SF2-Helicase+iREase+CTD+REase<-HTH+WYL | 951 | Bacteria>Pseudomonadota>Gammaproteobacteria>Cellvibrionales>Halieaceae | GCF_014077625.1 | Sediminihaliea_albiluteola | H2508_RS10780 | 844621..847476 | 2758564 | DUF499-domain-containing-protein-[Sediminihaliea-albiluteola] |
| WP_135445665.1 | AAA+ATPase+wHTH+Alpha+Beta-domain+wHTH+RRM | Type-1_BR.RM.like.systems | METHYLASE->?->?->OLD-ABC->TOPRIM->DUF499*->REase->MORC+DUF3684+REase-><-SF1-SF2-Helicase+REase<-SF1-SF2-Helicase+REase\|\|?->HKD-DNase+HKD-DNase-> | 947 | Bacteria>Pseudomonadota>Gammaproteobacteria>Cellvibrionales>Halieaceae | GCF_004745945.1 | Haliea_sp._SAOS-164 | E4634_RS16035 | 159390..162233 | 2562682 | DUF499-domain-containing-protein-[Haliea-sp.-SAOS-164] |
| WP_230198391.1 | AAA+ATPase+wHTH+Alpha+Beta-domain+wHTH+RRM | Type-1_BR.RM.like.systems | Phage_integrase->?->?-><-?<-DUF499*<-METHYLASE<-TUDOR+SF1-SF2-Helicase+iREase+CTD+REase | 965 | Bacteria>Pseudomonadota>Gammaproteobacteria>Cellvibrionales>Halieaceae | GCF_000158175.1 | Luminiphilus_syltensis | NOR51B_RS03435 | 755103..758000 | 1341119 | DUF499-domain-containing-protein-[Luminiphilus-syltensis] |
| EED36300.1 | AAA+ATPase+wHTH+Alpha+Beta-domain+wHTH+RRM | Type-1_BR.RM.like.systems | Phage_integrase->?->?-><-?<-DUF499*<-METHYLASE<-TUDOR+SF1-SF2-Helicase+iREase+CTD+REase | 950 | Bacteria>Pseudomonadota>Gammaproteobacteria>Cellvibrionales>Halieaceae | GCA_000158175.1 | Luminiphilus_syltensis_NOR5-1B | NOR51B_2250 | 755103..757955 | 565045 | hypothetical-protein-NOR51B_2250-[Luminiphilus-syltensis-NOR5-1B] |
| WP_237067886.1 | AAA+ATPase+wHTH+Alpha+Beta-domain+wHTH+RRM | Type-1_BR.RM.like.systems | TOPRIM-><-?\|\|HTH+WYL->TUDOR+SF1-SF2-Helicase+iREase+CTD+REase->METHYLASE->DUF499*->?->?->?->?->?-><-ADH_N+ADH_zinc_N | 947 | Bacteria>Pseudomonadota>Gammaproteobacteria>Cellvibrionales>Microbulbiferaceae | GCF_021729075.1 | Microbulbifer_guangxiensis | LVD12_RS11675 | 459683..462526 | 2904249 | DUF499-domain-containing-protein-[Microbulbifer-sp.-L3] |
| WP_266066709.1 | AAA+ATPase+wHTH+Alpha+Beta-domain+wHTH+RRM | Type-1_BR.RM.like.systems | HTH+WYL->TUDOR+SF1-SF2-Helicase+iREase+CTD+REase->METHYLASE->DUF499*-> | 945 | Bacteria>Pseudomonadota>Gammaproteobacteria>Cellvibrionales>Microbulbiferaceae | GCF_026241635.1 | Microbulbifer_thermotolerans | OQJ68_RS15995 | 7461..10298 | 252514 | DUF499-domain-containing-protein-[Microbulbifer-thermotolerans] |

| Protein-ID | DUF499-architecture | System | Gene-Neighbourhoods | Length | Taxonomy | Assembly-ID | Strain | Locus-Tag | Genomic-coordinates | TaxID | Fasta-defline |
|---|---|---|---|---|---|---|---|---|---|---|---|
| MAT52642.1 | AAA+ATPase+wHTH+Alpha+Beta-domain+wHTH+RRM | Type-1_BR.RM.like.systems | <-KAP-NTPase\|\|HTH+WYL->TUDOR+SF1-SF2-Helicase+iREase+CTD+REase->METHYLASE->METHYLASE->Schlafen+HATPase+HTH->DUF499*-><-?<-TOPRIM<-XIS-HTH<-?<-Arm-DNA-bind_3+Phage_int_SAM_3+Phage_integrase | 949 | Bacteria>Pseudomonadota>Gammaproteobacteria>Cellvibrionales>Porticoccaceae | GCA_002702725.1 | Porticoccaceae_bacterium | CMK32_15795 | 17542..20391 | 2026782 | hypothetical-protein-[Porticoccaceae-bacterium] |
| WP_272991604.1 | AAA+ATPase+wHTH+Alpha+Beta-domain+wHTH+RRM | Type-1_BR.RM.like.systems | <-PAPS_reduct<-?<-KAP-NTPase\|\|HTH+WYL->TUDOR+SF1-SF2-Helicase+iREase+CTD+REase->METHYLASE->DUF499*->?-><-TraG_N<-?<-TraH | 949 | Bacteria>Pseudomonadota>Gammaproteobacteria>Cellvibrionales>Porticoccaceae | GCF_002364235.1 | Porticoccus_hydrocarbonoclasticus | B2U98_RS00265 | 37905..40754 | 1073414 | DUF499-domain-containing-protein-[Porticoccus-hydrocarbonoclasticus] |
| MBG58119.1 | AAA+ATPase+wHTH+Alpha+Beta-domain+wHTH+RRM | Type-1_BR.RM.like.systems | TraH->?->TraG_N-><-?<-DUF499*<-METHYLASE<-TUDOR+SF1-SF2-Helicase+iREase+CTD+REase<-HTH+WYL\|\|HNH->KAP-NTPase-> | 949 | Bacteria>Pseudomonadota>Gammaproteobacteria>Cellvibrionales>Porticoccaceae | GCA_002715835.1 | Porticoccus_sp. | CMK46_07505 | 60449..63298 | 2024853 | hypothetical-protein-[Porticoccus-sp.] |
| MDA8652393.1 | AAA+ATPase+wHTH+Alpha+Beta-domain+wHTH+RRM | Type-1_BR.RM.like.systems | <-DUF499*<-METHYLASE<-TUDOR+SF1-SF2-Helicase+iREase+CTD+REase-p | 942 | Bacteria>Pseudomonadota>Gammaproteobacteria>Cellvibrionales>Porticoccaceae | GCA_028099245.1 | Porticoccaceae_bacterium | N9L75_07475 | 196..3024 | 2026782 | DUF499-domain-containing-protein-[Porticoccaceae-bacterium] |
| MCK9506376.1 | AAA+ATPase+wHTH+Alpha+Beta-domain+wHTH+RRM | Type-1_BR.RM.like.systems | OLD-ABC->REase->DUF499*->Classical-AAA-><-?<-DUF932 | 951 | Bacteria>Pseudomonadota>Gammaproteobacteria>Cellvibrionales>Porticoccaceae | GCA_023228865.1 | Porticoccaceae_bacterium | M0Q95_19625 | 878..3733 | 2026782 | DUF499-domain-containing-protein-[Porticoccaceae-bacterium] |
| WP_273009210.1 | AAA+ATPase+wHTH+Alpha+Beta-domain+wHTH+RRM | Type-1_BR.RM.like.systems | <-DUF499*<-METHYLASE<-PIN<-TUDOR+SF1-SF2-Helicase+iREase+CTD+REase | 943 | Bacteria>Pseudomonadota>Gammaproteobacteria>Cellvibrionales>Spongiibacteraceae | GCF_002435105.1 | Spongiibacter_tropicus | B3U69_RS04085 | 395..3226 | 454602 | DUF499-domain-containing-protein-[Spongiibacter-tropicus] |
| WP_213378539.1 | AAA+ATPase+wHTH+Alpha+Beta-domain+wHTH+RRM | Type-1_BR.RM.like.systems | MerR-HTH+GDResolvase->HTH_OrfB_IS605+OrfB_IS605+OrfB_Zn_ribbon->?-><-DUF499*<-REase<-OLD-ABC<-METHYLASE<-TUDOR+SF1-SF2-Helicase+iREase+CTD+REase<-HTH+WYL\|\|HTH_OrfB_IS605+OrfB_IS605+OrfB_Zn_ribbon-> | 943 | Bacteria>Pseudomonadota>Gammaproteobacteria>Chromatiales>Chromatiaceae | GCF_018409545.1 | Allochromatium_tepidum | Atep_RS10840 | 2243146..2245977 | 553982 | DUF499-domain-containing-protein-[Allochromatium-tepidum] |
| WP_244860024.1 | AAA+ATPase+wHTH+Alpha+Beta-domain+wHTH+RRM | Type-1_BR.RM.like.systems | TraH->?->TraG_N-><-?<-DUF499*<-METHYLASE<-TUDOR+SF1-SF2-Helicase+iREase+CTD+REase\|\|?->DDE_3-><-TUDOR+SF1-SF2-Helicase+iREase+CTD+REase-p<-HTH+WYL | 964 | Bacteria>Pseudomonadota>Gammaproteobacteria>Chromatiales>Chromatiaceae | GCF_000012805.1 | Nitrosococcus_oceani | NOC_RS03615 | 703535..706429 | 1229 | DUF499-domain-containing-protein-[Nitrosococcus-oceani] |
| WP_036499483.1 | AAA+ATPase+wHTH+Alpha+Beta-domain+wHTH+RRM | Type-1_BR.RM.like.systems | <-BrnA_antitoxin<-BrnT_toxin<-GDResolvase+HIN-HTH\|\|DDE_3-><-DUF499*<-METHYLASE-p<-METHYLASE-p<-TUDOR+SF1-SF2-Helicase+iREase+CTD+REase | 942 | Bacteria>Pseudomonadota>Gammaproteobacteria>Chromatiales>Chromatiaceae | GCF_000740735.1 | Nitrosococcus_oceani | HW44_RS03170 | 39038..41866 | 1229 | DUF499-domain-containing-protein-[Nitrosococcus-oceani] |
| WP_200208561.1 | AAA+ATPase+wHTH+Alpha+Beta-domain+wHTH+RRM | Type-1_BR.RM.like.systems | RHH->TUDOR+SF1-SF2-Helicase+iREase+CTD+REase->METHYLASE->RelE-ParE->HTH_3->DUF499*->?->HD-><-?\|\|BrnA_antitoxin->DnaB+SMS-KaiC-> | 941 | Bacteria>Pseudomonadota>Gammaproteobacteria>Chromatiales>Chromatiaceae | - | Chromatium_weissei | CK009_11850 | 40098..42923 | 626372 | ATP-binding-protein-[Chromatium-weissei] |
| MBK1695425.1 | AAA+ATPase+wHTH+Alpha+Beta-domain+wHTH+RRM | Type-1_BR.RM.like.systems | RHH->TUDOR+SF1-SF2-Helicase+iREase+CTD+REase->METHYLASE->RelE-ParE->HTH_3->DUF499*->?->HD-><-?\|\|BrnA_antitoxin->DnaB+SMS-KaiC-> | 941 | Bacteria>Pseudomonadota>Gammaproteobacteria>Chromatiales>Chromatiaceae | GCA_016583765.1 | Chromatium_weissei | CK009_11850 | 40098..42923 | 626372 | hypothetical-protein-[Chromatium-weissei] |
| WP_093036865.1 | AAA+ATPase+wHTH+Alpha+Beta-domain+wHTH+RRM | Type-1_BR.RM.like.systems | <-wHTH+REase<-?<-DUF499*<-METHYLASE<-TUDOR+SF1-SF2-Helicase+iREase+CTD+REase<-?<-?<-MobL+ZnR+TOPRIM\|\|Phage_integrase-> | 940 | Bacteria>Pseudomonadota>Gammaproteobacteria>Chromatiales>Chromatiaceae | GCF_900106925.1 | Thiocapsa_roseopersicina | BLV04_RS20440 | 8152..10974 | 1058 | ATP-binding-protein-[Thiocapsa-roseopersicina] |
| WP_232410038.1 | AAA+ATPase+wHTH+Alpha+Beta-domain+wHTH+RRM | Type-1_BR.RM.like.systems | DDE_Tnp_1->METHYLASE-p->PolB->HEPN->HEPN-><-MULE-TRANSPOSASE\|\|DUF499*-><-DDE_Tnp_1\|\|DUF499-p-> | 794 | Bacteria>Pseudomonadota>Gammaproteobacteria>Chromatiales>Chromatiaceae | GCF_021228795.1 | Thiocapsa_bogorovii | LT988_RS10220 | 2255421..2257805 | 521689 | DUF499-domain-containing-protein-[Thiocapsa-bogorovii] |
| WP_002813791.1 | AAA+ATPase+wHTH+Alpha+Beta-domain+wHTH+RRM | Type-1_BR.RM.like.systems | TraH->?->TraG_N-><-?<-DUF499*<-METHYLASE<-TUDOR+SF1-SF2-Helicase+iREase+CTD+REase | 949 | Bacteria>Pseudomonadota>Gammaproteobacteria>Chromatiales>Chromatiaceae | - | Nitrosococcus_oceani | IB75_RS03280 | 66265..69114 | 1229 | DUF499-domain-containing-protein-[Nitrosococcus-oceani] |
| MTW22277.1 | AAA+ATPase+wHTH+Alpha+Beta-domain+wHTH+RRM | Type-1_BR.RM.like.systems | TUDOR+SF1-SF2-Helicase+iREase+CTD+REase->OLD-ABC->HNH->OLD-ABC->TOPRIM->DUF499*-> | 885 | Bacteria>Pseudomonadota>Gammaproteobacteria>Chromatiales>Chromatiaceae | GCA_009720725.1 | Allochromatium_palmeri | GJ668_14455 | 38860..41517 | 231048 | DUF499-domain-containing-protein-[Allochromatium-palmeri] |
| WP_150093824.1 | AAA+ATPase+wHTH+Alpha+Beta-domain+wHTH+RRM | Type-1_BR.RM.like.systems | <-DUF499*<-HTH<-?<-METHYLASE<-TUDOR+SF1-SF2-Helicase+iREase+CTD+REase | 516 | Bacteria>Pseudomonadota>Gammaproteobacteria>Chromatiales>Chromatiaceae | GCF_008632335.1 | Thiohalocapsa_marina | F2Q65_RS12855 | 1..1547 | 424902 | DUF499-domain-containing-protein,-partial-[Thiohalocapsa-marina] |
| WP_155450846.1 | AAA+ATPase+wHTH+Alpha+Beta-domain+wHTH+RRM | Type-1_BR.RM.like.systems | OLD-ABC->HNH->METHYLASE-p->METHYLASE-p->OLD-ABC->TOPRIM->DUF499*-> | 883 | Bacteria>Pseudomonadota>Gammaproteobacteria>Chromatiales>Chromatiaceae | GCF_009720725.1 | Allochromatium_palmeri | GJ668_RS14460 | 38866..41517 | 231048 | DUF499-domain-containing-protein-[Allochromatium-palmeri] |
| WP_020507360.1 | AAA+ATPase+wHTH+Alpha+Beta-domain+wHTH+RRM | Type-1_BR.RM.like.systems | <-McrC-NTD+REase<-?<-DUF499*<-wHTH+REase<-METHYLASE<-TOPRIM<-OLD-ABC<-METHYLASE-p<-TUDOR+SF1-SF2-Helicase+iREase+CTD+REase | 961 | Bacteria>Pseudomonadota>Gammaproteobacteria>Chromatiales>Chromatiaceae | GCF_000379525.1 | Lamprocystis_purpurea | A390_RS0124805 | 120589..123474 | 61598 | DUF499-domain-containing-protein-[Lamprocystis-purpurea] |
| WP_020506626.1 | AAA+ATPase+wHTH+Alpha+Beta-domain+wHTH+RRM | Type-1_BR.RM.like.systems | HTH->OLD-ABC->METHYLASE->?->DUF433->DUF433->DUF499*-><-Mbetalac | 976 | Bacteria>Pseudomonadota>Gammaproteobacteria>Chromatiales>Chromatiaceae | GCF_000379525.1 | Lamprocystis_purpurea | A390_RS0120930 | 29422..32352 | 61598 | DUF499-domain-containing-protein-[Lamprocystis-purpurea] |
| WP_100921859.1 | AAA+ATPase+wHTH+Alpha+Beta-domain+wHTH+RRM | Type-1_BR.RM.like.systems | <-DUF499*<-DUF433<-?<-METHYLASE<-TUDOR+SF1-SF2-Helicase+iREase+CTD+REase | 981 | Bacteria>Pseudomonadota>Gammaproteobacteria>Chromatiales>Chromatiaceae | GCF_002813775.1 | Candidatus_Thiodictyon_syntrophicum | THSYN_RS26980 | 6283771..6286716 | 1166950 | DUF499-domain-containing-protein-[Candidatus-Thiodictyon-syntrophicum] |
| MBV5272245.1 | AAA+ATPase+wHTH+Alpha+Beta-domain+wHTH+RRM | Type-1_BR.RM.like.systems | REase->wHTH+REase->DUF499*->?->McrC-NTD+REase-> | 961 | Bacteria>Pseudomonadota>Gammaproteobacteria>Chromatiales>Chromatiaceae | GCA_019163055.1 | Lamprocystis_purpurea | JZU52_00945 | 1014..3899 | 61598 | ATP-binding-protein-[Lamprocystis-purpurea] |
| WP_023412668.1 | AAA+ATPase+wHTH+Alpha+Beta-domain+wHTH+RRM | Type-1_BR.RM.like.systems | METHYLASE->DUF499*->?->McrC-NTD+REase-> | 963 | Bacteria>Pseudomonadota>Gammaproteobacteria>Chromatiales>Chromatiaceae | - | Thiohalocapsa_sp._PB-PSB1 | N838_12705 | 2059..4950 | 1385625 | DUF499-domain-containing-protein-[Thiohalocapsa-sp.-PB-PSB1] |
| ESQ14149.1 | AAA+ATPase+wHTH+Alpha+Beta-domain+wHTH+RRM | Type-1_BR.RM.like.systems | METHYLASE->DUF499*->?->McrC-NTD+REase-> | 963 | Bacteria>Pseudomonadota>Gammaproteobacteria>Chromatiales>Chromatiaceae | GCA_000495615.1 | Thiohalocapsa_sp._PB-PSB1 | N838_12705 | 2059..4950 | 1385625 | MAG:-hypothetical-protein-N838_12705-[Thiohalocapsa-sp.-PB-PSB1] |
| HBG95093.1 | AAA+ATPase+wHTH+Alpha+Beta-domain+wHTH+RRM | Type-1_BR.RM.like.systems | <-DUF499-p<-DUF499*<-METHYLASE | 558 | Bacteria>Pseudomonadota>Gammaproteobacteria>Chromatiales>Chromatiaceae | GCA_003499475.1 | Chromatiaceae_bacterium | DDY14_07165 | 3311..4984 | 2049432 | hypothetical-protein-[Chromatiaceae-bacterium] |
| WP_019627826.1 | AAA+ATPase+wHTH+Alpha+Beta-domain+wHTH+RRM | Type-1_BR.RM.like.systems | Arm-DNA-bind_2+Phage_int_SAM_3+Phage_integrase->HDRnase->ZnR->TUDOR+SF1-SF2-Helicase+iREase+CTD+REase->PIN->METHYLASE->DUF499*->?->SF1-SF2-Helicase->?-><-?<-Phage_integrase | 951 | Bacteria>Pseudomonadota>Gammaproteobacteria>Chromatiales>Ectothiorhodospiraceae | GCF_000381845.1 | Thioalkalivibrio_sp._AKL10 | G333_RS0104940 | 1030421..1033276 | 1158158 | DUF499-domain-containing-protein-[Thioalkalivibrio-sp.-AKL10] |
| WP_025282662.1 | AAA+ATPase+wHTH+Alpha+Beta-domain+wHTH+RRM | Type-1_BR.RM.like.systems | TraH->?->TraG_N-><-?\|\|AbiEi_4-><-DUF499*<-REase<-OLD-ABC<-METHYLASE<-?<-TUDOR+SF1-SF2-Helicase+iREase+CTD+REase\|\|TPR+TPR-> | 956 | Bacteria>Pseudomonadota>Gammaproteobacteria>Chromatiales>Ectothiorhodospiraceae | GCF_000583815.1 | Ectothiorhodospira_haloalkaliphila | ECTHA_RS0102365 | 117439..120309 | 421628 | DUF499-domain-containing-protein-[Ectothiorhodospira-haloalkaliphila] |
| WP_240480980.1 | AAA+ATPase+wHTH+Alpha+Beta-domain+wHTH+RRM | Type-1_BR.RM.like.systems | <-HTH_Tnp_1+HTH_21+rve\|\|EVE-><-Y1_Tnp\|\|HTH_OrfB_IS605+OrfB_IS605+OrfB_Zn_ribbon->?->DUF499*->?-><-AbiEi_4\|\|?-><-?<-TraG_N | 924 | Bacteria>Pseudomonadota>Gammaproteobacteria>Chromatiales>Ectothiorhodospiraceae | GCF_001632845.1 | Ectothiorhodospira_sp._BSL-9 | ECTOBSL9_RS12585 | 2708145..2710919 | 1442136 | DUF499-domain-containing-protein-[Ectothiorhodospira-sp.-BSL-9] |
| WP_090202860.1 | AAA+ATPase+wHTH+Alpha+Beta-domain+wHTH+RRM | Type-1_BR.RM.like.systems | ABC-ATPase+SbcC->?-><-?<-?<-?<-PUA+METHYLASE<-DUF499*<-METHYLASE<-TUDOR+SF1-SF2-Helicase+iREase+CTD+REase\|\|?->?-><-?<-METHYLASE-p | 954 | Bacteria>Pseudomonadota>Gammaproteobacteria>Chromatiales>Ectothiorhodospiraceae | GCF_900110965.1 | Ectothiorhodospira_magna | BMX62_RS01320 | 108182..111046 | 867345 | DUF499-domain-containing-protein-[Ectothiorhodospira-magna] |
| TQE99812.1 | AAA+ATPase+wHTH+Alpha+Beta-domain+wHTH+RRM | Type-1_BR.RM.like.systems | HEPN->ParA-Soj-PloopNTPase->ORC-CDC6->METHYLASE->OLD-ABC->REase->DUF499*->?->UbE1+JAB->?->Schlafen-> | 955 | Bacteria>Pseudomonadota>Gammaproteobacteria>Chromatiales>Ectothiorhodospiraceae | GCA_006569195.1 | Spiribacter_salinus | FKY71_06670 | 12316..15183 | 1335746 | ATP-binding-protein-[Spiribacter-salinus] |
| ANB02986.1 | AAA+ATPase+wHTH+Alpha+Beta-domain+wHTH+RRM | Type-1_BR.RM.like.systems | <-HTH_21+rve+rve_3<-?\|\|EVE-><-Y1_Tnp\|\|?->DUF499*->?-><-AbiEi_4\|\|?-><-?<-?<-GspA-ATPase | 956 | Bacteria>Pseudomonadota>Gammaproteobacteria>Chromatiales>Ectothiorhodospiraceae | GCA_001632845.1 | Ectothiorhodospira_sp._BSL-9 | ECTOBSL9_2519 | 2708049..2710919 | 1442136 | hypothetical-protein-ECTOBSL9_2519-[Ectothiorhodospira-sp.-BSL-9] |
| TQE98724.1 | AAA+ATPase+wHTH+Alpha+Beta-domain+wHTH+RRM | Type-1_BR.RM.like.systems | ParB+HNH->DUF499*-><-?<-REase+35exo<-HTH+WYL | 939 | Bacteria>Pseudomonadota>Gammaproteobacteria>Chromatiales>Ectothiorhodospiraceae | GCA_006569195.1 | Spiribacter_salinus | FKY71_12295 | 713..3532 | 1335746 | ATP-binding-protein-[Spiribacter-salinus] |
| MCG7866473.1 | AAA+ATPase+wHTH+Alpha+Beta-domain+wHTH+RRM | Type-1_BR.RM.like.systems | <-KAP-NTPase<-HNH\|\|HTH+WYL->TUDOR+SF1-SF2-Helicase+iREase+CTD+REase->METHYLASE->DUF499*->?->?-><-?<-?<-?<-N6_MTase-p | 944 | Bacteria>Pseudomonadota>Gammaproteobacteria>Chromatiales>Sedimenticolaceae | GCA_016842585.1 | Candidatus_Thiodiazotropha_taylori | JAY74_08885 | 46162..48996 | 2792791 | DUF499-domain-containing-protein-[Candidatus-Thiodiazotropha-taylori] |
| WP_114191418.1 | AAA+ATPase+wHTH+Alpha+Beta-domain+wHTH+RRM | Type-1_BR.RM.like.systems | ParB+HNH->ParB+HNH->?->LZ_Tnp_IS66+zf-IS66+DDE_Tnp_IS66+DDE_Tnp_IS66_C->RlpA-><-DUF499*<-METHYLASE<-SbcC<-TUDOR+SF1-SF2-Helicase+iREase+CTD+REase<-?<-?<-XIS-HTH | 947 | Bacteria>Pseudomonadota>Gammaproteobacteria>Enterobacterales | GCF_013317085.1 | Enterobacteriaceae | MS6194_RS10935 | 1850138..1852981 | 543 | MULTISPECIES:-DUF499-domain-containing-protein-[Enterobacteriaceae] |
| WP_134906061.1 | AAA+ATPase+wHTH+Alpha+Beta-domain+wHTH+RRM | Type-1_BR.RM.like.systems | Arm-DNA-bind_3+Phage_integrase->RlpA-><-DUF499*<-REase<-OLD-ABC<-METHYLASE<-SbcC<-TUDOR+SF1-SF2-Helicase+iREase+CTD+REase<-HIN-HTH | 946 | Bacteria>Pseudomonadota>Gammaproteobacteria>Enterobacterales | GCF_004801595.1 | Enterobacteriaceae | I8Q62_RS26385 | 2639..5479 | 543 | MULTISPECIES:-DUF499-domain-containing-protein-[Enterobacteriaceae] |
| WP_220756270.1 | AAA+ATPase+wHTH+Alpha+Beta-domain+wHTH+RRM | Type-1_BR.RM.like.systems | TUDOR+SF1-SF2-Helicase+iREase+CTD+REase->OLD-ABC->HNH->METHYLASE->DUF499*-><-RlpA<-RVT+HNH<-?<-ParB+HNH | 947 | Bacteria>Pseudomonadota>Gammaproteobacteria>Enterobacterales | GCF_019655755.1 | Enterobacteriaceae | K5R72_RS10955 | 43358..46201 | 543 | DUF499-domain-containing-protein-[Escherichia-coli] |
| WP_008325059.1 | AAA+ATPase+wHTH+Alpha+Beta-domain+wHTH+RRM | Type-1_BR.RM.like.systems | TUDOR+SF1-SF2-Helicase+iREase+CTD+REase->OLD-ABC->HNH->METHYLASE->DUF499*-><-RlpA | 947 | Bacteria>Pseudomonadota>Gammaproteobacteria>Enterobacterales | GCF_000277565.1 | Enterobacteriaceae | B397_RS01195 | 10694..13537 | 543 | MULTISPECIES:-DUF499-domain-containing-protein-[Enterobacteriaceae] |
| WP_236874409.1 | AAA+ATPase+wHTH+Alpha+Beta-domain+wHTH+RRM | Type-1_BR.RM.like.systems | DEDD_Tnp_IS110+Transposase_20-><-LZ_Tnp_IS66+zf-IS66+DDE_Tnp_IS66+DDE_Tnp_IS66_C<-?<-HIN-HTH\|\|RlpA-><-DUF499*<-METHYLASE<-SbcC<-TUDOR+SF1-SF2-Helicase+iREase+CTD+REase<-HIN-HTH<-ZnR<-XIS-HTH | 947 | Bacteria>Pseudomonadota>Gammaproteobacteria>Enterobacterales>Enterobacteriaceae | GCF_021729115.1 | Citrobacter_portucalensis | LXK91_RS21210 | 4411655..4414498 | 1639133 | DUF499-domain-containing-protein-[Citrobacter-portucalensis] |
| MCS6087896.1 | AAA+ATPase+wHTH+Alpha+Beta-domain+wHTH+RRM | Type-1_BR.RM.like.systems | HEPN-><-AbiEii<-AbiEi_4\|\|?->RlpA-><-DUF499-p<-DUF499*<-DUF499-p<-METHYLASE-p<-METHYLASE-p<-METHYLASE-p<-TUDOR+SF1-SF2-Helicase+iREase+CTD+REase-p<-TUDOR+SF1-SF2-Helicase+iREase+CTD+REase | 303 | Bacteria>Pseudomonadota>Gammaproteobacteria>Enterobacterales>Enterobacteriaceae | GCA_024988895.1 | Klebsiella_pneumoniae_subsp._pneumoniae | LNO03_00250 | 50022..50933 | 72407 | hypothetical-protein-[Klebsiella-pneumoniae-subsp.-pneumoniae] |
| WP_186588561.1 | AAA+ATPase+wHTH+Alpha+Beta-domain+wHTH+RRM | Type-1_BR.RM.like.systems | HTH_3->\|\|?->TUDOR+SF1-SF2-Helicase+iREase+CTD+REase->SbcC->SbcC->METHYLASE->DUF499*-><-RlpA\|\|<-DnaB<-XIS-HTH<-Arm-DNA-bind_3+Phage_integrase<-Phage_integrase+Phage_integrase | 947 | Bacteria>Pseudomonadota>Gammaproteobacteria>Enterobacterales>Enterobacteriaceae | GCF_026163855.1 | Klebsiella_pneumoniae | ORK99_RS12855 | 13102..15945 | 573 | DUF499-domain-containing-protein-[Klebsiella-pneumoniae] |
| WP_177331926.1 | AAA+ATPase+wHTH+Alpha+Beta-domain+wHTH+RRM | Type-1_BR.RM.like.systems | HTH_3->\|\|?->TUDOR+SF1-SF2-Helicase+iREase+CTD+REase->SbcC->METHYLASE->DUF499*-><-RlpA\|\|<-DnaB<-XIS-HTH<-Arm-DNA-bind_3+Phage_integrase<-Phage_integrase+Phage_integrase | 947 | Bacteria>Pseudomonadota>Gammaproteobacteria>Enterobacterales>Enterobacteriaceae | GCF_902807095.1 | Klebsiella_michiganensis | HUW14_RS12480 | 31267..34110 | 1134687 | DUF499-domain-containing-protein-[Klebsiella-michiganensis] |
| MCQ9222683.1 | AAA+ATPase+wHTH+Alpha+Beta-domain+wHTH+RRM | Type-1_BR.RM.like.systems | RlpA-><-DUF499*<-DUF499*<-METHYLASE<-TUDOR+SF1-SF2-Helicase+iREase+CTD+REase<-TUDOR+SF1-SF2-Helicase+iREase+CTD+REase-p<-TUDOR+SF1-SF2-Helicase+iREase+CTD+REase-p | 208 | Bacteria>Pseudomonadota>Gammaproteobacteria>Enterobacterales>Enterobacteriaceae | GCA_024580235.1 | Klebsiella_pneumoniae | NQ122_29740 | 3455..4081 | 573 | DUF499-domain-containing-protein-[Klebsiella-pneumoniae] |
| MCQ9222682.1 | AAA+ATPase+wHTH+Alpha+Beta-domain+wHTH+RRM | Type-1_BR.RM.like.systems | RlpA-><-DUF499<-DUF499*<-METHYLASE<-TUDOR+SF1-SF2-Helicase+iREase+CTD+REase<-TUDOR+SF1-SF2-Helicase+iREase+CTD+REase-p<-TUDOR+SF1-SF2-Helicase+iREase+CTD+REase-p | 743 | Bacteria>Pseudomonadota>Gammaproteobacteria>Enterobacterales>Enterobacteriaceae | GCA_024580235.1 | Klebsiella_pneumoniae | NQ122_29735 | 1255..3486 | 573 | hypothetical-protein-[Klebsiella-pneumoniae] |
| WP_172615335.1 | AAA+ATPase+wHTH+Alpha+Beta-domain+wHTH+RRM | Type-1_BR.RM.like.systems | <-nSTAND1+sGTP\|\|<-HTH_Tnp_1+HTH_21+rve+rve_2\|\|HIN-HTH->?-><-?<-DUF499*<-METHYLASE<-SbcC<-TUDOR+SF1-SF2-Helicase+iREase+CTD+REase<-?\|\|SF1-SF2-Helicase-><-DUF772+DDE_Tnp_1 | 947 | Bacteria>Pseudomonadota>Gammaproteobacteria>Enterobacterales>Enterobacteriaceae | GCF_013201175.1 | Escherichia_coli | HP434_RS13950 | 2903325..2906168 | 562 | DUF499-domain-containing-protein-[Escherichia-coli] |
| WP_265179947.1 | AAA+ATPase+wHTH+Alpha+Beta-domain+wHTH+RRM | Type-1_BR.RM.like.systems | TUDOR+SF1-SF2-Helicase+iREase+CTD+REase-p->TUDOR+SF1-SF2-Helicase+iREase+CTD+REase->SbcC->METHYLASE-p->METHYLASE->DUF499-p->DUF499*-><-RlpA<-?<-tRNA\|\|?->?->ADH_N+ADH_zinc_N-> | 787 | Bacteria>Pseudomonadota>Gammaproteobacteria>Enterobacterales>Enterobacteriaceae | - | Klebsiella_pneumoniae | LH689_RS19250 | 815009..817372 | 573 | DUF499-domain-containing-protein-[Klebsiella-pneumoniae] |
| WP_234573269.1 | AAA+ATPase+wHTH+Alpha+Beta-domain+wHTH+RRM | Type-1_BR.RM.like.systems | TUDOR+SF1-SF2-Helicase+iREase+CTD+REase-p->TUDOR+SF1-SF2-Helicase+iREase+CTD+REase->SbcC->METHYLASE-p->METHYLASE->DUF499-p->DUF499*-><-RlpA<-?<-tRNA\|\|?->?->ADH_N+ADH_zinc_N-> | 779 | Bacteria>Pseudomonadota>Gammaproteobacteria>Enterobacterales>Enterobacteriaceae | GCF_021461365.1 | Klebsiella_pneumoniae | LH689_19265 | 815033..817372 | 573 | hypothetical-protein-[Klebsiella-pneumoniae] |
| MCF1375490.1 | AAA+ATPase+wHTH+Alpha+Beta-domain+wHTH+RRM | Type-1_BR.RM.like.systems | TUDOR+SF1-SF2-Helicase+iREase+CTD+REase-p->TUDOR+SF1-SF2-Helicase+iREase+CTD+REase->SbcC->METHYLASE-p->METHYLASE->DUF499-p->DUF499*-><-RlpA<-?<-tRNA\|\|?->?->ADH_N+ADH_zinc_N-> | 779 | Bacteria>Pseudomonadota>Gammaproteobacteria>Enterobacterales>Enterobacteriaceae | GCA_021461365.1 | Klebsiella_pneumoniae | LH689_19265 | 815033..817372 | 573 | hypothetical-protein-[Klebsiella-pneumoniae] |
| MBC4043610.1 | AAA+ATPase+wHTH+Alpha+Beta-domain+wHTH+RRM | Type-1_BR.RM.like.systems | Phage_integrase+Phage_integrase->Arm-DNA-bind_3+Phage_integrase->XIS-HTH->DnaB->\|\|RlpA-><-DUF499*<-METHYLASE<-TUDOR+SF1-SF2-Helicase+iREase+CTD+REase<-?\|\|<-HTH_3 | 947 | Bacteria>Pseudomonadota>Gammaproteobacteria>Enterobacterales>Enterobacteriaceae | GCA_014283365.1 | Klebsiella_pneumoniae | H8M87_14600 | 163621..166464 | 573 | ATP-binding-protein-[Klebsiella-pneumoniae] |
| MBY5090623.1 | AAA+ATPase+wHTH+Alpha+Beta-domain+wHTH+RRM | Type-1_BR.RM.like.systems | <-FtsK+HerA+A32<-ADH_N+ADH_zinc_N<-SAM-methylase<-wHTH+HNH\|\|tRNA->?->RlpA-><-DUF499*<-METHYLASE<-TUDOR+SF1-SF2-Helicase+iREase+CTD+REase<-?\|\|?->MAD-NTD+HNH-><-SymE_toxin | 949 | Bacteria>Pseudomonadota>Gammaproteobacteria>Enterobacterales>Enterobacteriaceae | GCA_019795045.1 | Citrobacter_freundii | K4X46_07840 | 164380..167229 | 546 | DUF499-domain-containing-protein-[Citrobacter-freundii] |
| WP_219173436.1 | AAA+ATPase+wHTH+Alpha+Beta-domain+wHTH+RRM | Type-1_BR.RM.like.systems | HIN-HTH->TUDOR+SF1-SF2-Helicase+iREase+CTD+REase->SbcC->METHYLASE->OLD-ABC->REase->DUF499*-><-RlpA<-Arm-DNA-bind_3+Phage_integrase\|\|HIN-HTH->\|\|?->?->ParB+HNH-> | 946 | Bacteria>Pseudomonadota>Gammaproteobacteria>Enterobacterales>Enterobacteriaceae | GCF_019334445.1 | Klebsiella/Raoultella_group | LVJ43_RS03725 | 746872..749712 | 2890311 | DUF499-domain-containing-protein-[Raoultella-ornithinolytica] |
| WP_121585419.1 | AAA+ATPase+wHTH+Alpha+Beta-domain+wHTH+RRM | Type-1_BR.RM.like.systems | <-ParB+BAM<-XIS-HTH<-?\|\|wHTH(SBD)+HNH->TUDOR+SF1-SF2-Helicase+iREase+CTD+REase->METHYLASE->DUF499*-><-RlpA<-?\|\|AbiEi_4->AbiEii-><-wHTH(SBD)+HNH\|\|MAD-NTD+HNH-> | 941 | Bacteria>Pseudomonadota>Gammaproteobacteria>Enterobacterales>Enterobacteriaceae | GCF_003665575.1 | Citrobacter | CUC49_RS22185 | 4547789..4550614 | 544 | MULTISPECIES:-DUF499-domain-containing-protein-[Citrobacter] |
| EFO1856148.1 | AAA+ATPase+wHTH+Alpha+Beta-domain+wHTH+RRM | Type-1_BR.RM.like.systems | TUDOR+SF1-SF2-Helicase+iREase+CTD+REase->OLD-ABC->HNH->METHYLASE->DUF499*-><-HTH_Tnp_1+HTH_21+rve+rve_2<-HTH_Tnp_1+HTH_21+rve+rve_2\|\|<-?<-?\|\|XIS-HTH->DUF3987-> | 947 | Bacteria>Pseudomonadota>Gammaproteobacteria>Enterobacterales>Enterobacteriaceae | GCA_013065125.1 | Escherichia_coli | DUZ37_17320 | 27712..30555 | 562 | DUF499-domain-containing-protein-[Escherichia-coli] |
| WP_251853646.1 | AAA+ATPase+wHTH+Alpha+Beta-domain+wHTH+RRM | Type-1_BR.RM.like.systems | <-SymE_toxin\|\|wHTH(SBD)+HNH-><-AbiEii<-AbiEi_4\|\|?->RlpA-><-DUF499*<-METHYLASE<-TUDOR+SF1-SF2-Helicase+iREase+CTD+REase<-wHTH(SBD)+HNH\|\|?->XIS-HTH->ParB+BAM-> | 941 | Bacteria>Pseudomonadota>Gammaproteobacteria>Enterobacterales>Enterobacteriaceae | GCF_903935915.1 | Enterobacter_cloacae | NGL52_RS07590 | 54377..57202 | 550 | DUF499-domain-containing-protein-[Enterobacter-cloacae] |
| WP_155022959.1 | AAA+ATPase+wHTH+Alpha+Beta-domain+wHTH+RRM | Type-1_BR.RM.like.systems | <-ParB+BAM<-XIS-HTH<-?\|\|wHTH(SBD)+HNH->TUDOR+SF1-SF2-Helicase+iREase+CTD+REase->METHYLASE->DUF499*-><-RlpA<-?\|\|AbiEi_4->AbiEii-><-wHTH(SBD)+HNH\|\|SymE_toxin-> | 941 | Bacteria>Pseudomonadota>Gammaproteobacteria>Enterobacterales>Enterobacteriaceae | GCF_015704165.1 | Enterobacter_hormaechei | I4P33_RS13240 | 224794..227619 | 158836 | MULTISPECIES:-DUF499-domain-containing-protein-[Enterobacter-cloacae-complex] |
| WP_101977492.1 | AAA+ATPase+wHTH+Alpha+Beta-domain+wHTH+RRM | Type-1_BR.RM.like.systems | Arm-DNA-bind_3+Phage_integrase->XIS-HTH->DnaB->\|\|RlpA-><-DUF499*<-METHYLASE<-SbcC<-SbcC<-TUDOR+SF1-SF2-Helicase+iREase+CTD+REase<-?<-PAIREDC-HTH+HIN-HTH | 951 | Bacteria>Pseudomonadota>Gammaproteobacteria>Enterobacterales>Enterobacteriaceae | GCF_002851295.1 | Klebsiella_quasipneumoniae | B6J05_RS15930 | 4794..7649 | 1463165 | DUF499-domain-containing-protein-[Klebsiella-quasipneumoniae] |
| WP_191238898.1 | AAA+ATPase+wHTH+Alpha+Beta-domain+wHTH+RRM | Type-1_BR.RM.like.systems | <-MAD-NTD+HNH<-?\|\|?->TUDOR+SF1-SF2-Helicase+iREase+CTD+REase->SbcC->METHYLASE->DUF499*-><-RlpA<-tRNA\|\|wHTH+HNH->SAM-methylase->ADH_N+ADH_zinc_N->FtsK+HerA+A32-> | 949 | Bacteria>Pseudomonadota>Gammaproteobacteria>Enterobacterales>Enterobacteriaceae | GCF_030332975.1 | unclassified_Citrobacter | OGY02_RS18160 | 260747..263596 | 2644389 | DUF499-domain-containing-protein-[Citrobacter-freundii] |
| WP_054625948.1 | AAA+ATPase+wHTH+Alpha+Beta-domain+wHTH+RRM | Type-1_BR.RM.like.systems | ParB+HNH->ParB+HNH->?->LZ_Tnp_IS66+zf-IS66+DDE_Tnp_IS66+DDE_Tnp_IS66_C->RlpA-><-DUF499*<-METHYLASE<-HNH<-OLD-ABC<-TUDOR+SF1-SF2-Helicase+iREase+CTD+REase | 947 | Bacteria>Pseudomonadota>Gammaproteobacteria>Enterobacterales>Enterobacteriaceae | GCF_018085225.1 | Escherichia_coli | J7886_RS18855 | 3900823..3903666 | 562 | DUF499-domain-containing-protein-[Escherichia-coli] |
| WP_048233987.1 | AAA+ATPase+wHTH+Alpha+Beta-domain+wHTH+RRM | Type-1_BR.RM.like.systems | SymE_toxin->wHTH(SBD)+HNH-><-AbiEii<-AbiEi_4\|\|?->RlpA-><-DUF499*<-METHYLASE<-TUDOR+SF1-SF2-Helicase+iREase+CTD+REase<-wHTH(SBD)+HNH\|\|?->XIS-HTH->ParB+BAM-> | 941 | Bacteria>Pseudomonadota>Gammaproteobacteria>Enterobacterales>Enterobacteriaceae | GCF_001034485.1 | Citrobacter_sp._MGH104 | SK34_RS19955 | 343965..346790 | 1686379 | DUF499-domain-containing-protein-[Citrobacter-sp.-MGH104] |
| EHD5776078.1 | AAA+ATPase+wHTH+Alpha+Beta-domain+wHTH+RRM | Type-1_BR.RM.like.systems | <-ADH_N+ADH_zinc_N\|\|tRNA->HTH_Tnp_1+HTH_21+rve+rve_2->RlpA-><-DUF499*<-METHYLASE<-TUDOR+SF1-SF2-Helicase+iREase+CTD+REase<-HTH_21+rve+rve_3<-HIN-HTH | 941 | Bacteria>Pseudomonadota>Gammaproteobacteria>Enterobacterales>Enterobacteriaceae | GCA_017158295.1 | Escherichia_coli | JQY67_002368 | 23024..25849 | 562 | ATP-binding-protein-[Escherichia-coli] |
| WP_227818485.1 | AAA+ATPase+wHTH+Alpha+Beta-domain+wHTH+RRM | Type-1_BR.RM.like.systems | <-AbiEii<-AbiEi_4\|\|?->RlpA-><-DUF499-p<-DDE_Tnp_1_5+DDE_Tnp_1<-DUF499*<-METHYLASE<-HIN-HTH<-METHYLASE-p<-TUDOR+SF1-SF2-Helicase+iREase+CTD+REase | 723 | Bacteria>Pseudomonadota>Gammaproteobacteria>Enterobacterales>Enterobacteriaceae | GCF_012113635.1 | Klebsiella_pneumoniae | HC649_RS22790 | 4559312..4561483 | 573 | DUF499-domain-containing-protein-[Klebsiella-pneumoniae] |
| HCI7559692.1 | AAA+ATPase+wHTH+Alpha+Beta-domain+wHTH+RRM | Type-1_BR.RM.like.systems | <-HIN-HTH\|\|?->TUDOR+SF1-SF2-Helicase+iREase+CTD+REase->OLD-ABC->HNH->METHYLASE->DUF499*-><-RlpA<-?<-Arm-DNA-bind_3+Phage_integrase<-tRNA\|\|?->?->ADH_N+ADH_zinc_N-> | 947 | Bacteria>Pseudomonadota>Gammaproteobacteria>Enterobacterales>Enterobacteriaceae | GCA_024446695.1 | Klebsiella_pneumoniae | NPW42_004570 | 42168..45011 | 573 | ATP-binding-protein-[Klebsiella-pneumoniae] |
| WP_101970653.1 | AAA+ATPase+wHTH+Alpha+Beta-domain+wHTH+RRM | Type-1_BR.RM.like.systems | Arm-DNA-bind_3+Phage_integrase->?->?->\|\|TUDOR+SF1-SF2-Helicase+iREase+CTD+REase->METHYLASE->DUF499*-><-RlpA<-?\|\|AbiEi_4->AbiEii->HIN-HTH-><-HEPN | 941 | Bacteria>Pseudomonadota>Gammaproteobacteria>Enterobacterales>Enterobacteriaceae | GCF_900774085.1 | Klebsiella_pneumoniae | E7T50_RS20015 | 299802..302627 | 573 | DUF499-domain-containing-protein-[Klebsiella-pneumoniae] |
| WP_048337983.1 | AAA+ATPase+wHTH+Alpha+Beta-domain+wHTH+RRM | Type-1_BR.RM.like.systems | HEPN-><-HIN-HTH<-AbiEii<-AbiEi_4\|\|?->RlpA-><-DUF499*<-METHYLASE<-TUDOR+SF1-SF2-Helicase+iREase+CTD+REase\|\|<-?<-?<-Arm-DNA-bind_3+Phage_integrase | 941 | Bacteria>Pseudomonadota>Gammaproteobacteria>Enterobacterales>Enterobacteriaceae | GCF_004313005.1 | Klebsiella_pneumoniae | CCZ31_RS16900 | 43841..46666 | 573 | DUF499-domain-containing-protein-[Klebsiella-pneumoniae] |
| MCJ4335970.1 | AAA+ATPase+wHTH+Alpha+Beta-domain+wHTH+RRM | Type-1_BR.RM.like.systems | HEPN-><-HIN-HTH<-AbiEii<-AbiEi_4\|\|?->RlpA-><-DUF499*<-METHYLASE<-TUDOR+SF1-SF2-Helicase+iREase+CTD+REase\|\|<-?<-?<-Arm-DNA-bind_3+Phage_integrase | 941 | Bacteria>Pseudomonadota>Gammaproteobacteria>Enterobacterales>Enterobacteriaceae | GCA_022853825.1 | Klebsiella_pneumoniae | LNV93_18595 | 43805..46630 | 573 | DUF499-domain-containing-protein-[Klebsiella-pneumoniae] |
| WP_080831491.1 | AAA+ATPase+wHTH+Alpha+Beta-domain+wHTH+RRM | Type-1_BR.RM.like.systems | Phage_integrase->?->?->\|\|TUDOR+SF1-SF2-Helicase+iREase+CTD+REase->METHYLASE->DUF499*-><-RlpA<-?\|\|AbiEi_4->AbiEii-><-wHTH+SAD+HNH\|\|MAD-NTD+HNH-> | 941 | Bacteria>Pseudomonadota>Gammaproteobacteria>Enterobacterales>Enterobacteriaceae | GCF_900173235.1 | Klebsiella_pneumoniae | CI699_RS15160 | 12102..14927 | 573 | DUF499-domain-containing-protein-[Klebsiella-pneumoniae] |
| WP_025368507.1 | AAA+ATPase+wHTH+Alpha+Beta-domain+wHTH+RRM | Type-1_BR.RM.like.systems | RlpA-><-DUF499*<-METHYLASE<-SbcC<-TUDOR+SF1-SF2-Helicase+iREase+CTD+REase<-TUDOR+SF1-SF2-Helicase+iREase+CTD+REase-p<-?\|\|<-McrC-NTD+REase | 947 | Bacteria>Pseudomonadota>Gammaproteobacteria>Enterobacterales>Enterobacteriaceae | GCF_900775085.1 | Klebsiella_pneumoniae | CQR95_RS27300 | 2719..5562 | 573 | DUF499-domain-containing-protein-[Klebsiella-pneumoniae] |
| HAU5791339.1 | AAA+ATPase+wHTH+Alpha+Beta-domain+wHTH+RRM | Type-1_BR.RM.like.systems | Phage_ASH->?->?->TOPRIM+DNAG+D5N+D5ATPase+P4PRIMASE-HTH->?->RlpA-><-DUF499*<-METHYLASE<-HNH<-OLD-ABC<-TUDOR+SF1-SF2-Helicase+iREase+CTD+REase | 947 | Bacteria>Pseudomonadota>Gammaproteobacteria>Enterobacterales>Enterobacteriaceae | GCA_016076255.1 | Citrobacter_freundii | JD367_15405 | 174239..177082 | 546 | ATP-binding-protein-[Citrobacter-freundii] |
| WP_086872901.1 | AAA+ATPase+wHTH+Alpha+Beta-domain+wHTH+RRM | Type-1_BR.RM.like.systems | TUDOR+SF1-SF2-Helicase+iREase+CTD+REase->OLD-ABC->HNH->METHYLASE->DUF499*-><-RlpA<-Arm-DNA-bind_3+Phage_integrase<-tRNA\|\|REC+LuxR-HTH-> | 953 | Bacteria>Pseudomonadota>Gammaproteobacteria>Enterobacterales>Enterobacteriaceae | GCF_900184035.1 | Kosakonia_pseudosacchari | BN2591_RS15655 | 26880..29741 | 1646340 | DUF499-domain-containing-protein-[Kosakonia-pseudosacchari] |
| WP_256633221.1 | AAA+ATPase+wHTH+Alpha+Beta-domain+wHTH+RRM | Type-1_BR.RM.like.systems | <-FtsK+HerA+A32<-ADH_N+ADH_zinc_N\|\|tRNA->RlpA-><-DUF499-p<-DUF499*<-METHYLASE-p<-METHYLASE-p\|\|Zn_Tnp_IS1+HTH_Tnp_IS1+DDE_Tnp_IS1-> | 847 | Bacteria>Pseudomonadota>Gammaproteobacteria>Enterobacterales>Enterobacteriaceae | GCF_024502275.1 | Escherichia_coli | MRO39_RS14565 | 64937..67480 | 562 | DUF499-domain-containing-protein-[Escherichia-coli] |
| WP_232585745.1 | AAA+ATPase+wHTH+Alpha+Beta-domain+wHTH+RRM | Type-1_BR.RM.like.systems | <-Phage_integrase\|\|?->?->?->\|\|wHTH+SAD+HNH-><-Zn_Tnp_IS1+HTH_Tnp_IS1+DDE_Tnp_IS1\|\|DUF499*-><-RlpA<-tRNA\|\|ADH_N+ADH_zinc_N->FtsK+HerA+A32-> | 810 | Bacteria>Pseudomonadota>Gammaproteobacteria>Enterobacterales>Enterobacteriaceae | GCF_021229275.1 | Escherichia_coli | LUR14_19180 | 3924175..3926607 | 562 | DUF499-domain-containing-protein-[Escherichia-coli] |
| HAW1058251.1 | AAA+ATPase+wHTH+Alpha+Beta-domain+wHTH+RRM | Type-1_BR.RM.like.systems | <-nSTAND1+sGTP\|\|?->TUDOR+SF1-SF2-Helicase+iREase+CTD+REase->OLD-ABC->HNH->METHYLASE->DUF499*-><-DEDD_Tnp_IS110+Transposase_20 | 947 | Bacteria>Pseudomonadota>Gammaproteobacteria>Enterobacterales>Enterobacteriaceae | GCA_016672425.1 | Escherichia_coli | JLK83_004317 | 23199..26042 | 562 | ATP-binding-protein-[Escherichia-coli] |
| WP_061359526.1 | AAA+ATPase+wHTH+Alpha+Beta-domain+wHTH+RRM | Type-1_BR.RM.like.systems | <-JetD<-?\|\|?->TUDOR+SF1-SF2-Helicase+iREase+CTD+REase->SbcC->METHYLASE->DUF499*-><-RlpA<-tRNA\|\|ADH_N+ADH_zinc_N->FtsK+HerA+A32-> | 947 | Bacteria>Pseudomonadota>Gammaproteobacteria>Enterobacterales>Enterobacteriaceae | GCF_030295075.1 | Escherichia_coli | NLZ09_RS19755 | 4038306..4041149 | 562 | DUF499-domain-containing-protein-[Escherichia-coli] |
| WP_247754246.1 | AAA+ATPase+wHTH+Alpha+Beta-domain+wHTH+RRM | Type-1_BR.RM.like.systems | XIS-HTH->?->?->TUDOR+SF1-SF2-Helicase+iREase+CTD+REase->SbcC->METHYLASE->DUF499*-><-RlpA<-Arm-DNA-bind_3+Phage_integrase | 957 | Bacteria>Pseudomonadota>Gammaproteobacteria>Enterobacterales>Enterobacteriaceae | GCF_013748855.1 | Citrobacter_freundii | HV201_RS07875 | 1647485..1650358 | 546 | DUF499-domain-containing-protein-[Citrobacter-freundii] |
| WP_097584906.1 | AAA+ATPase+wHTH+Alpha+Beta-domain+wHTH+RRM | Type-1_BR.RM.like.systems | <-REC+LuxR-HTH<-SAM-methylase<-wHTH(SBD)+HNH\|\|tRNA->RlpA-><-DUF499*<-METHYLASE<-SbcC<-TUDOR+SF1-SF2-Helicase+iREase+CTD+REase | 947 | Bacteria>Pseudomonadota>Gammaproteobacteria>Enterobacterales>Enterobacteriaceae | GCF_002532355.1 | Cronobacter_sakazakii | FC600_RS12565 | 210517..213360 | 28141 | DUF499-domain-containing-protein-[Cronobacter-sakazakii] |
| WP_086660584.1 | AAA+ATPase+wHTH+Alpha+Beta-domain+wHTH+RRM | Type-1_BR.RM.like.systems | <-nSTAND1+sGTP\|\|?->TUDOR+SF1-SF2-Helicase+iREase+CTD+REase->OLD-ABC->HNH->METHYLASE->DUF499*->\|\|?->CbeA_antitoxin->?-><-tRNA<-Aldedh | 947 | Bacteria>Pseudomonadota>Gammaproteobacteria>Enterobacterales>Enterobacteriaceae | GCF_014216435.1 | Escherichia_coli | GVI19_RS16185 | 90398..93241 | 562 | DUF499-domain-containing-protein-[Escherichia-coli] |
| WP_047363208.1 | AAA+ATPase+wHTH+Alpha+Beta-domain+wHTH+RRM | Type-1_BR.RM.like.systems | XIS-HTH->?->?->TUDOR+SF1-SF2-Helicase+iREase+CTD+REase->SbcC->METHYLASE->DUF499*-><-RlpA<-Arm-DNA-bind_3+Phage_integrase | 947 | Bacteria>Pseudomonadota>Gammaproteobacteria>Enterobacterales>Enterobacteriaceae | GCF_001022495.2 | Enterobacter_kobei | ABR31_RS11720 | 10560..13403 | 208224 | DUF499-domain-containing-protein-[Enterobacter-kobei] |
| WP_277745132.1 | AAA+ATPase+wHTH+Alpha+Beta-domain+wHTH+RRM | Type-1_BR.RM.like.systems | McrC-NTD+REase->\|\|?->TUDOR+SF1-SF2-Helicase+iREase+CTD+REase->SbcC->METHYLASE->DUF499*-><-RlpA<-?<-tRNA\|\|?->?->ADH_N+ADH_zinc_N-> | 947 | Bacteria>Pseudomonadota>Gammaproteobacteria>Enterobacterales>Enterobacteriaceae | GCF_029541825.1 | Klebsiella_pneumoniae | LVQ70_RS22825 | 4647607..4650450 | 573 | DUF499-domain-containing-protein-[Klebsiella-pneumoniae] |
| WP_256043565.1 | AAA+ATPase+wHTH+Alpha+Beta-domain+wHTH+RRM | Type-1_BR.RM.like.systems | Arm-DNA-bind_3+Phage_integrase->RlpA-><-DUF499*<-METHYLASE<-HNH<-OLD-ABC<-TUDOR+SF1-SF2-Helicase+iREase+CTD+REase | 947 | Bacteria>Pseudomonadota>Gammaproteobacteria>Enterobacterales>Enterobacteriaceae | GCF_024451305.1 | Enterobacter_kobei | NOX26_RS15660 | 3292884..3295727 | 208224 | DUF499-domain-containing-protein-[Enterobacter-kobei] |
| WP_222563661.1 | AAA+ATPase+wHTH+Alpha+Beta-domain+wHTH+RRM | Type-1_BR.RM.like.systems | Arm-DNA-bind_3+Phage_integrase->RlpA-><-DUF499*<-METHYLASE<-HNH<-OLD-ABC<-TUDOR+SF1-SF2-Helicase+iREase+CTD+REase | 947 | Bacteria>Pseudomonadota>Gammaproteobacteria>Enterobacterales>Enterobacteriaceae | GCF_019803045.1 | Citrobacter_farmeri | Cf24236_RS16360 | 3446361..3449204 | 67824 | MULTISPECIES:-DUF499-domain-containing-protein-[Enterobacteriaceae] |
| MCQ4415112.1 | AAA+ATPase+wHTH+Alpha+Beta-domain+wHTH+RRM | Type-1_BR.RM.like.systems | Arm-DNA-bind_3+Phage_integrase->RlpA-><-DUF499*<-METHYLASE<-HNH<-OLD-ABC<-TUDOR+SF1-SF2-Helicase+iREase+CTD+REase | 947 | Bacteria>Pseudomonadota>Gammaproteobacteria>Enterobacterales>Enterobacteriaceae | GCA_024451305.1 | Enterobacter_kobei | NOX26_15660 | 3292884..3295727 | 208224 | DUF499-domain-containing-protein-[Enterobacter-kobei] |
| HBV3413501.1 | AAA+ATPase+wHTH+Alpha+Beta-domain+wHTH+RRM | Type-1_BR.RM.like.systems | RlpA-><-DUF499*<-METHYLASE<-TUDOR+SF1-SF2-Helicase+iREase+CTD+REase\|\|HIN-HTH-><-?<-?<-Arm-DNA-bind_3+Phage_integrase | 941 | Bacteria>Pseudomonadota>Gammaproteobacteria>Enterobacterales>Enterobacteriaceae | GCA_022140105.1 | Klebsiella_pneumoniae | MDU10_005213 | 758..3583 | 573 | ATP-binding-protein-[Klebsiella-pneumoniae] |
| HBR2941452.1 | AAA+ATPase+wHTH+Alpha+Beta-domain+wHTH+RRM | Type-1_BR.RM.like.systems | RlpA-><-DUF499*<-METHYLASE<-TUDOR+SF1-SF2-Helicase+iREase+CTD+REase\|\|HIN-HTH-><-?<-?<-Arm-DNA-bind_3+Phage_integrase | 941 | Bacteria>Pseudomonadota>Gammaproteobacteria>Enterobacterales>Enterobacteriaceae | GCA_021919485.1 | Klebsiella_pneumoniae | L9R10_004957 | 796..3621 | 573 | ATP-binding-protein-[Klebsiella-pneumoniae] |
| MBA8327618.1 | AAA+ATPase+wHTH+Alpha+Beta-domain+wHTH+RRM | Type-1_BR.RM.like.systems | Phage_ASH->?->?->TOPRIM+DNAG+D5N+D5ATPase+P4PRIMASE-HTH->?->RlpA-><-DUF499*<-DUF772+DDE_Tnp_1\|\|?->HIN-HTH-><-SymE_toxin | 951 | Bacteria>Pseudomonadota>Gammaproteobacteria>Enterobacterales>Enterobacteriaceae | GCA_014105425.1 | Citrobacter_freundii | HVX14_08920 | 1881381..1884236 | 546 | ATP-binding-protein-[Citrobacter-freundii] |
| WP_283999135.1 | AAA+ATPase+wHTH+Alpha+Beta-domain+wHTH+RRM | Type-1_BR.RM.like.systems | <-JAB<-nSTAND1+sGTP\|\|?->TUDOR+SF1-SF2-Helicase+iREase+CTD+REase->SbcC->METHYLASE->DUF499*-><-RlpA<-?<-?<-?<-Phage_AlpA | 951 | Bacteria>Pseudomonadota>Gammaproteobacteria>Enterobacterales>Enterobacteriaceae | GCF_030126265.1 | Cronobacter_malonaticus | QMS61_RS01615 | 345961..348816 | 413503 | DUF499-domain-containing-protein-[Cronobacter-malonaticus] |
| HBU8410481.1 | AAA+ATPase+wHTH+Alpha+Beta-domain+wHTH+RRM | Type-1_BR.RM.like.systems | HEPN-><-HIN-HTH<-AbiEii<-AbiEi_4\|\|?->RlpA-><-DUF499*<-HIN-HTH<-TUDOR+SF1-SF2-Helicase+iREase+CTD+REase\|\|RVT-> | 912 | Bacteria>Pseudomonadota>Gammaproteobacteria>Enterobacterales>Enterobacteriaceae | GCA_022109915.1 | Klebsiella_pneumoniae | MDC63_004746 | 22723..25461 | 573 | ATP-binding-protein-[Klebsiella-pneumoniae] |
| WP_208595561.1 | AAA+ATPase+wHTH+Alpha+Beta-domain+wHTH+RRM | Type-1_BR.RM.like.systems | wHTH+SAD+HNH-><-?\|\|METHYLASE-p->METHYLASE-p->DUF499*-><-RlpA<-tRNA\|\|ADH_N+ADH_zinc_N->FtsK+HerA+A32-><-?<-?\|\|Peptidase-> | 947 | Bacteria>Pseudomonadota>Gammaproteobacteria>Enterobacterales>Enterobacteriaceae | GCF_017601385.1 | Escherichia_coli | J5660_RS20445 | 4118728..4121571 | 562 | DUF499-domain-containing-protein-[Escherichia-coli] |
| HDK5497440.1 | AAA+ATPase+wHTH+Alpha+Beta-domain+wHTH+RRM | Type-1_BR.RM.like.systems | Arm-DNA-bind_3+Phage_integrase->?->?-><-HIN-HTH\|\|TUDOR+SF1-SF2-Helicase+iREase+CTD+REase->METHYLASE->DUF499*-> | 349 | Bacteria>Pseudomonadota>Gammaproteobacteria>Enterobacterales>Enterobacteriaceae | GCA_028561575.1 | Klebsiella_pneumoniae | PTA52_005037 | 31576..32623 | 573 | TPA:-ATP-binding-protein,-partial-[Klebsiella-pneumoniae] |
| WP_229532552.1 | AAA+ATPase+wHTH+Alpha+Beta-domain+wHTH+RRM | Type-1_BR.RM.like.systems | ACET-><-Phage_integrase\|\|RlpA-><-DUF499*<-METHYLASE<-HNH<-?<-OLD-ABC<-TUDOR+SF1-SF2-Helicase+iREase+CTD+REase | 947 | Bacteria>Pseudomonadota>Gammaproteobacteria>Enterobacterales>Enterobacteriaceae | GCF_020577415.1 | Klebsiella_variicola | KAG03_RS12050 | 331978..334821 | 244366 | DUF499-domain-containing-protein-[Klebsiella-variicola] |
| HAV2024006.1 | AAA+ATPase+wHTH+Alpha+Beta-domain+wHTH+RRM | Type-1_BR.RM.like.systems | <-ParB+BAM<-XIS-HTH<-?\|\|wHTH(SBD)+HNH->TUDOR+SF1-SF2-Helicase+iREase+CTD+REase->METHYLASE->DUF499*-><-RlpA | 941 | Bacteria>Pseudomonadota>Gammaproteobacteria>Enterobacterales>Enterobacteriaceae | GCA_016444345.1 | Citrobacter_koseri | JHJ49_004194 | 16997..19822 | 545 | ATP-binding-protein-[Citrobacter-koseri] |
| WP_061423982.1 | AAA+ATPase+wHTH+Alpha+Beta-domain+wHTH+RRM | Type-1_BR.RM.like.systems | wHTH(SBD)+HNH->TUDOR+SF1-SF2-Helicase+iREase+CTD+REase->METHYLASE->DUF499*-><-RlpA<-?\|\|AbiEi_4->AbiEii-> | 941 | Bacteria>Pseudomonadota>Gammaproteobacteria>Enterobacterales>Enterobacteriaceae | GCF_004194635.3 | Salmonella_enterica | DBZ85_RS25390 | 353324..356149 | 28901 | DUF499-domain-containing-protein-[Salmonella-enterica] |
| WP_080321606.1 | AAA+ATPase+wHTH+Alpha+Beta-domain+wHTH+RRM | Type-1_BR.RM.like.systems | HTH_3->?-><-Phage_integrase\|\|RlpA-><-DUF499*<-METHYLASE<-SbcC<-TUDOR+SF1-SF2-Helicase+iREase+CTD+REase | 947 | Bacteria>Pseudomonadota>Gammaproteobacteria>Enterobacterales>Enterobacteriaceae | GCF_002071765.2 | Cronobacter_sakazakii | B7T09_RS01300 | 242227..245070 | 28141 | DUF499-domain-containing-protein-[Cronobacter-sakazakii] |
| WP_262806643.1 | AAA+ATPase+wHTH+Alpha+Beta-domain+wHTH+RRM | Type-1_BR.RM.like.systems | <-IclR-HTH<-?<-?\|\|?->?->?->tRNA-><-DUF499*<-METHYLASE<-TUDOR+SF1-SF2-Helicase+iREase+CTD+REase<-SAM-methylase<-wHTH+HNH | 947 | Bacteria>Pseudomonadota>Gammaproteobacteria>Enterobacterales>Enterobacteriaceae | GCF_025535395.1 | Enterobacter_bugandensis | KWH72_RS14435 | 121926..124769 | 881260 | DUF499-domain-containing-protein-[Enterobacter-bugandensis] |
| MCM7403168.1 | AAA+ATPase+wHTH+Alpha+Beta-domain+wHTH+RRM | Type-1_BR.RM.like.systems | RlpA-><-DUF499*<-METHYLASE<-SbcC<-TUDOR+SF1-SF2-Helicase+iREase+CTD+REase<-?\|\|ParB->MethyltransfD12-> | 951 | Bacteria>Pseudomonadota>Gammaproteobacteria>Enterobacterales>Enterobacteriaceae | GCA_023751075.1 | Enterobacter_hormaechei | M8R60_15845 | 109739..112594 | 158836 | DUF499-domain-containing-protein-[Enterobacter-hormaechei] |
| WP_193132801.1 | AAA+ATPase+wHTH+Alpha+Beta-domain+wHTH+RRM | Type-1_BR.RM.like.systems | HTH->?->HEPN->RlpA-><-DUF499*<-METHYLASE<-HNH<-OLD-ABC<-TUDOR+SF1-SF2-Helicase+iREase+CTD+REase<-?\|\|ParB-> | 942 | Bacteria>Pseudomonadota>Gammaproteobacteria>Enterobacterales>Enterobacteriaceae | GCF_032369155.1 | Enterobacter_hormaechei | CXM24_RS20100 | 4287724..4290552 | 158836 | DUF499-domain-containing-protein-[Enterobacter-hormaechei] |
| WP_282376200.1 | AAA+ATPase+wHTH+Alpha+Beta-domain+wHTH+RRM | Type-1_BR.RM.like.systems | Arm-DNA-bind_3+Phage_integrase-><-DUF499*<-METHYLASE<-TUDOR+SF1-SF2-Helicase+iREase+CTD+REase | 949 | Bacteria>Pseudomonadota>Gammaproteobacteria>Enterobacterales>Enterobacteriaceae | GCF_029960465.1 | Enterobacter_kobei | QK340_RS16355 | 158507..161356 | 208224 | DUF499-domain-containing-protein-[Enterobacter-kobei] |
| AHE46601.1 | AAA+ATPase+wHTH+Alpha+Beta-domain+wHTH+RRM | Type-1_BR.RM.like.systems | <-HIN-HTH<-HIN-HTH\|\|?->TUDOR+SF1-SF2-Helicase+iREase+CTD+REase->SbcC->METHYLASE->DUF499*-><-RlpA | 963 | Bacteria>Pseudomonadota>Gammaproteobacteria>Enterobacterales>Enterobacteriaceae | GCA_000512165.1 | Klebsiella_pneumoniae_subsp._pneumoniae_Kp13 | KP13_01335 | 4689607..4692498 | 1123862 | hypothetical-protein-KP13_01335-[Klebsiella-pneumoniae-subsp.-pneumoniae-Kp13] |
| WP_070182596.1 | AAA+ATPase+wHTH+Alpha+Beta-domain+wHTH+RRM | Type-1_BR.RM.like.systems | RlpA-><-DUF499*<-METHYLASE<-SbcC<-TUDOR+SF1-SF2-Helicase+iREase+CTD+REase<-?\|\|<-McrC-NTD+REase | 947 | Bacteria>Pseudomonadota>Gammaproteobacteria>Enterobacterales>Enterobacteriaceae | GCF_001756835.2 | Klebsiella_pneumoniae | BA903_RS22470 | 906..3749 | 573 | DUF499-domain-containing-protein-[Klebsiella-pneumoniae] |
| WP_181047111.1 | AAA+ATPase+wHTH+Alpha+Beta-domain+wHTH+RRM | Type-1_BR.RM.like.systems | METHYLASE-p->METHYLASE-p->DUF499*-><-RlpA<-tRNA\|\|ADH_N+ADH_zinc_N->FtsK+HerA+A32-><-?<-?\|\|Peptidase-> | 947 | Bacteria>Pseudomonadota>Gammaproteobacteria>Enterobacterales>Enterobacteriaceae | GCF_022548015.1 | Escherichia_coli | EFK04_RS21050 | 1469..4312 | 562 | DUF499-domain-containing-protein-[Escherichia-coli] |
| WP_249920526.1 | AAA+ATPase+wHTH+Alpha+Beta-domain+wHTH+RRM | Type-1_BR.RM.like.systems | METHYLASE-p->METHYLASE-p->DUF499-p->DUF499*-><-RlpA<-tRNA\|\|ADH_N+ADH_zinc_N->FtsK+HerA+A32-> | 802 | Bacteria>Pseudomonadota>Gammaproteobacteria>Enterobacterales>Enterobacteriaceae | GCF_003302135.1 | Escherichia_coli | DUM19_RS12745 | 2364..4772 | 562 | DUF499-domain-containing-protein-[Escherichia-coli] |
| WP_196359062.1 | AAA+ATPase+wHTH+Alpha+Beta-domain+wHTH+RRM | Type-1_BR.RM.like.systems | RlpA-><-DUF499*<-METHYLASE<-HNH<-OLD-ABC<-TUDOR+SF1-SF2-Helicase+iREase+CTD+REase<-?\|\|ParB-> | 947 | Bacteria>Pseudomonadota>Gammaproteobacteria>Enterobacterales>Enterobacteriaceae | GCF_015701305.1 | Enterobacter_hormaechei | I4P56_RS16415 | 165030..167873 | 158836 | DUF499-domain-containing-protein-[Enterobacter-hormaechei] |
| EGQ7342643.1 | AAA+ATPase+wHTH+Alpha+Beta-domain+wHTH+RRM | Type-1_BR.RM.like.systems | RlpA-><-DUF499*<-METHYLASE<-HNH<-OLD-ABC<-TUDOR+SF1-SF2-Helicase+iREase+CTD+REase<-?\|\|ParB-> | 947 | Bacteria>Pseudomonadota>Gammaproteobacteria>Enterobacterales>Enterobacteriaceae | GCA_015730265.1 | Enterobacter_cloacae | I6C89_001957 | 175814..178657 | 550 | ATP-binding-protein-[Enterobacter-cloacae] |
| WP_096194529.1 | AAA+ATPase+wHTH+Alpha+Beta-domain+wHTH+RRM | Type-1_BR.RM.like.systems | PfkB->?->?-><-DUF499*<-METHYLASE<-SbcC<-TUDOR+SF1-SF2-Helicase+iREase+CTD+REase<-?<-?\|\|HIN-HTH-> | 947 | Bacteria>Pseudomonadota>Gammaproteobacteria>Enterobacterales>Enterobacteriaceae | GCF_018279125.1 | Escherichia_coli | NBV25_RS22590 | 5405..8248 | 562 | DUF499-domain-containing-protein-[Escherichia-coli] |
| WP_094392449.1 | AAA+ATPase+wHTH+Alpha+Beta-domain+wHTH+RRM | Type-1_BR.RM.like.systems | HIN-HTH->\|\|?->TUDOR+SF1-SF2-Helicase+iREase+CTD+REase->SbcC->METHYLASE->DUF499*-><-RVT | 947 | Bacteria>Pseudomonadota>Gammaproteobacteria>Enterobacterales>Enterobacteriaceae | GCF_902703155.1 | Escherichia_coli | HSY94_RS20435 | 10437..13280 | 562 | DUF499-domain-containing-protein-[Escherichia-coli] |
| WP_262790701.1 | AAA+ATPase+wHTH+Alpha+Beta-domain+wHTH+RRM | Type-1_BR.RM.like.systems | RlpA-><-DUF499*<-METHYLASE<-SbcC<-TUDOR+SF1-SF2-Helicase+iREase+CTD+REase<-?<-?\|\|ParB-> | 951 | Bacteria>Pseudomonadota>Gammaproteobacteria>Enterobacterales>Enterobacteriaceae | GCF_025536435.1 | Enterobacter_quasiroggenkampii | KW818_RS19435 | 30782..33637 | 2497436 | DUF499-domain-containing-protein-[Enterobacter-quasiroggenkampii] |
| WP_224381981.1 | AAA+ATPase+wHTH+Alpha+Beta-domain+wHTH+RRM | Type-1_BR.RM.like.systems | TUDOR+SF1-SF2-Helicase+iREase+CTD+REase->SbcC->METHYLASE->OLD-ABC->REase->DUF499*-> | 350 | Bacteria>Pseudomonadota>Gammaproteobacteria>Enterobacterales>Enterobacteriaceae | GCF_020118495.1 | Klebsiella_michiganensis | FMK66_RS24360 | 60161..61211 | 1134687 | DUF499-domain-containing-protein,-partial-[Klebsiella-michiganensis] |
| WP_186934038.1 | AAA+ATPase+wHTH+Alpha+Beta-domain+wHTH+RRM | Type-1_BR.RM.like.systems | <-DUF499*<-REase<-OLD-ABC<-METHYLASE<-SbcC<-TUDOR+SF1-SF2-Helicase+iREase+CTD+REase | 946 | Bacteria>Pseudomonadota>Gammaproteobacteria>Enterobacterales>Enterobacteriaceae | GCF_014285945.1 | Klebsiella | H8M57_RS27310 | 138..2978 | 570 | MULTISPECIES:-DUF499-domain-containing-protein-[Klebsiella] |
| HBC9812127.1 | AAA+ATPase+wHTH+Alpha+Beta-domain+wHTH+RRM | Type-1_BR.RM.like.systems | TUDOR+SF1-SF2-Helicase+iREase+CTD+REase-p->OLD-ABC->HNH->METHYLASE->DUF499*-><-RlpA | 951 | Bacteria>Pseudomonadota>Gammaproteobacteria>Enterobacterales>Enterobacteriaceae | GCA_018195115.1 | Escherichia_coli | KFP54_004194 | 5317..8172 | 562 | ATP-binding-protein-[Escherichia-coli] |
| WP_214577408.1 | AAA+ATPase+wHTH+Alpha+Beta-domain+wHTH+RRM | Type-1_BR.RM.like.systems | RlpA-><-DUF499*<-METHYLASE<-HNH<-OLD-ABC<-TUDOR+SF1-SF2-Helicase+iREase+CTD+REase | 947 | Bacteria>Pseudomonadota>Gammaproteobacteria>Enterobacterales>Enterobacteriaceae | GCF_018597265.1 | Enterobacter_dykesii | KK116_RS19460 | 158324..161167 | 2797506 | DUF499-domain-containing-protein-[Enterobacter-dykesii] |
| WP_048997346.1 | AAA+ATPase+wHTH+Alpha+Beta-domain+wHTH+RRM | Type-1_BR.RM.like.systems | RlpA-><-DUF499*<-METHYLASE<-HNH<-OLD-ABC<-TUDOR+SF1-SF2-Helicase+iREase+CTD+REase | 947 | Bacteria>Pseudomonadota>Gammaproteobacteria>Enterobacterales>Enterobacteriaceae | GCF_021869685.1 | Enterobacter | L3068_RS02450 | 520532..523375 | 547 | MULTISPECIES:-DUF499-domain-containing-protein-[Enterobacter] |
| WP_045284468.1 | AAA+ATPase+wHTH+Alpha+Beta-domain+wHTH+RRM | Type-1_BR.RM.like.systems | RlpA-><-DUF499*<-METHYLASE<-HNH<-OLD-ABC<-TUDOR+SF1-SF2-Helicase+iREase+CTD+REase | 947 | Bacteria>Pseudomonadota>Gammaproteobacteria>Enterobacterales>Enterobacteriaceae | GCF_000958085.1 | Enterobacter_sichuanensis | SS37_RS01295 | 119849..122692 | 2071710 | DUF499-domain-containing-protein-[Enterobacter-sichuanensis] |
| HBW9592181.1 | AAA+ATPase+wHTH+Alpha+Beta-domain+wHTH+RRM | Type-1_BR.RM.like.systems | TUDOR+SF1-SF2-Helicase+iREase+CTD+REase->OLD-ABC->HNH->METHYLASE->DUF499*-><-RlpA | 953 | Bacteria>Pseudomonadota>Gammaproteobacteria>Enterobacterales>Enterobacteriaceae | GCA_022207645.1 | Klebsiella_pneumoniae | MFS54_26490 | 9564..12425 | 573 | DUF499-domain-containing-protein-[Klebsiella-pneumoniae] |
| HAU4285776.1 | AAA+ATPase+wHTH+Alpha+Beta-domain+wHTH+RRM | Type-1_BR.RM.like.systems | TUDOR+SF1-SF2-Helicase+iREase+CTD+REase->OLD-ABC->HNH->METHYLASE->DUF499*-><-RlpA | 947 | Bacteria>Pseudomonadota>Gammaproteobacteria>Enterobacterales>Enterobacteriaceae | GCA_016065535.1 | Citrobacter_freundii | F6W46_24070 | 10411..13254 | 546 | ATP-binding-protein-[Citrobacter-freundii] |
| WP_261783471.1 | AAA+ATPase+wHTH+Alpha+Beta-domain+wHTH+RRM | Type-1_BR.RM.like.systems | METHYLASE-p->DUF499*-><-RlpA<-tRNA\|\|ADH_N+ADH_zinc_N->FtsK+HerA+A32-><-?<-?\|\|Peptidase-> | 927 | Bacteria>Pseudomonadota>Gammaproteobacteria>Enterobacterales>Enterobacteriaceae | GCF_019841165.1 | Escherichia_coli | K6725_RS09080 | 1342..4125 | 562 | DUF499-domain-containing-protein-[Escherichia-coli] |
| MCS0753638.1 | AAA+ATPase+wHTH+Alpha+Beta-domain+wHTH+RRM | Type-1_BR.RM.like.systems | <-FtsK+HerA+A32<-ADH_N+ADH_zinc_N\|\|tRNA->RlpA-><-DUF499*<-METHYLASE-p<-METHYLASE-p | 971 | Bacteria>Pseudomonadota>Gammaproteobacteria>Enterobacterales>Enterobacteriaceae | GCA_024753945.1 | Escherichia_coli | NXZ26_22935 | 12447..15362 | 562 | DUF499-domain-containing-protein-[Escherichia-coli] |
| WP_249548439.1 | AAA+ATPase+wHTH+Alpha+Beta-domain+wHTH+RRM | Type-1_BR.RM.like.systems | <-FtsK+HerA+A32<-ADH_N+ADH_zinc_N\|\|tRNA->RlpA-><-DUF499-p<-DUF499*<-METHYLASE-p | 802 | Bacteria>Pseudomonadota>Gammaproteobacteria>Enterobacterales>Enterobacteriaceae | GCF_002464325.1 | Escherichia_coli | AS924_RS15765 | 55179..57587 | 562 | DUF499-domain-containing-protein-[Escherichia-coli] |
| WP_213201037.1 | AAA+ATPase+wHTH+Alpha+Beta-domain+wHTH+RRM | Type-1_BR.RM.like.systems | TUDOR+SF1-SF2-Helicase+iREase+CTD+REase->SbcC->METHYLASE->DUF499*-> | 330 | Bacteria>Pseudomonadota>Gammaproteobacteria>Enterobacterales>Enterobacteriaceae | GCF_018348455.1 | Cronobacter_sakazakii | KHP24_RS19350 | 7698..8687 | 28141 | DUF499-domain-containing-protein,-partial-[Cronobacter-sakazakii] |
| WP_101978990.1 | AAA+ATPase+wHTH+Alpha+Beta-domain+wHTH+RRM | Type-1_BR.RM.like.systems | TUDOR+SF1-SF2-Helicase+iREase+CTD+REase-p->METHYLASE->DUF499*-> | 892 | Bacteria>Pseudomonadota>Gammaproteobacteria>Enterobacterales>Enterobacteriaceae | GCF_002855915.1 | Klebsiella_pneumoniae | CWN66_RS19170 | 3813..6489 | 573 | DUF499-domain-containing-protein,-partial-[Klebsiella-pneumoniae] |
| PLN33546.1 | AAA+ATPase+wHTH+Alpha+Beta-domain+wHTH+RRM | Type-1_BR.RM.like.systems | TUDOR+SF1-SF2-Helicase+iREase+CTD+REase-p->METHYLASE->DUF499*-> | 905 | Bacteria>Pseudomonadota>Gammaproteobacteria>Enterobacterales>Enterobacteriaceae | GCA_002856755.1 | Klebsiella_pneumoniae | CWM91_06635 | 4593..7307 | 573 | hypothetical-protein-CWM91_06635,-partial-[Klebsiella-pneumoniae] |
| HCF8312688.1 | AAA+ATPase+wHTH+Alpha+Beta-domain+wHTH+RRM | Type-1_BR.RM.like.systems | TUDOR+SF1-SF2-Helicase+iREase+CTD+REase-p->METHYLASE->DUF499*-> | 887 | Bacteria>Pseudomonadota>Gammaproteobacteria>Enterobacterales>Enterobacteriaceae | GCA_023979705.1 | Klebsiella_pneumoniae | NIV18_005199 | 3718..6380 | 573 | ATP-binding-protein-[Klebsiella-pneumoniae] |
| HAL1866777.1 | AAA+ATPase+wHTH+Alpha+Beta-domain+wHTH+RRM | Type-1_BR.RM.like.systems | DUF499*-><-?<-tRNA\|\|ADH_N+ADH_zinc_N->FtsK+HerA+A32-><-?<-?\|\|Peptidase-> | 813 | Bacteria>Pseudomonadota>Gammaproteobacteria>Enterobacterales>Enterobacteriaceae | GCA_014081005.1 | Escherichia_coli | H5W94_002966 | 1..2442 | 562 | ATP-binding-protein-[Escherichia-coli] |
| MBS4494223.1 | AAA+ATPase+wHTH+Alpha+Beta-domain+wHTH+RRM | Type-1_BR.RM.like.systems | DUF499*-><-RlpA<-?<-tRNA\|\|wHTH+HNH->SAM-methylase->REC+LuxR-HTH-> | 613 | Bacteria>Pseudomonadota>Gammaproteobacteria>Enterobacterales>Enterobacteriaceae | GCA_018348455.1 | Cronobacter_sakazakii | KHP24_14120 | 1..1843 | 28141 | ATP-binding-protein-[Cronobacter-sakazakii] |
| WP_047625607.1 | AAA+ATPase+wHTH+Alpha+Beta-domain+wHTH+RRM | Type-1_BR.RM.like.systems | METHYLASE-p->DUF499*-><-Arm-DNA-bind_3+Phage_integrase | 951 | Bacteria>Pseudomonadota>Gammaproteobacteria>Enterobacterales>Enterobacteriaceae | GCF_000798855.1 | Enterobacter_sp._ZOR0014 | L977_RS08775 | 1412..4267 | 1339232 | DUF499-domain-containing-protein-[Enterobacter-sp.-ZOR0014] |
| HDK5497304.1 | AAA+ATPase+wHTH+Alpha+Beta-domain+wHTH+RRM | Type-1_BR.RM.like.systems | HEPN-><-HIN-HTH<-AbiEii<-AbiEi_4\|\|?->RlpA-><-DUF499* | 593 | Bacteria>Pseudomonadota>Gammaproteobacteria>Enterobacterales>Enterobacteriaceae | GCA_028561575.1 | Klebsiella_pneumoniae | PTA52_004894 | 43712..45495 | 573 | TPA:-ATP-binding-protein,-partial-[Klebsiella-pneumoniae] |
| EHE2763427.1 | AAA+ATPase+wHTH+Alpha+Beta-domain+wHTH+RRM | Type-1_BR.RM.like.systems | DUF499*-><-?<-tRNA\|\|ADH_N+ADH_zinc_N-><-?<-?\|\|Peptidase-> | 947 | Bacteria>Pseudomonadota>Gammaproteobacteria>Enterobacterales>Enterobacteriaceae | GCA_017227475.1 | Escherichia_coli | JND72_004217 | 1490..4333 | 562 | ATP-binding-protein-[Escherichia-coli] |
| HBV3519256.1 | AAA+ATPase+wHTH+Alpha+Beta-domain+wHTH+RRM | Type-1_BR.RM.like.systems | <-DUF772+DDE_Tnp_1\|\|METHYLASE->DUF499*-> | 759 | Bacteria>Pseudomonadota>Gammaproteobacteria>Enterobacterales>Enterobacteriaceae | GCA_022140685.1 | Klebsiella_pneumoniae | MDU84_005291 | 4660..6937 | 573 | ATP-binding-protein-[Klebsiella-pneumoniae] |
| WP_279936012.1 | AAA+ATPase+wHTH+Alpha+Beta-domain+wHTH+RRM | Type-1_BR.RM.like.systems | <-DUF499*<-METHYLASE<-HNH | 951 | Bacteria>Pseudomonadota>Gammaproteobacteria>Enterobacterales>Enterobacteriaceae | GCF_029837515.1 | Klebsiella_oxytoca | N7642_RS28020 | 124..2979 | 571 | DUF499-domain-containing-protein-[Klebsiella-oxytoca] |

| Protein-ID | DUF499-architecture | System | Gene-Neighbourhoods | Length | Taxonomy | Assembly-ID | Strain | Locus-Tag | Genomic-coordinates | TaxID | Fasta-defline |
|---|---|---|---|---|---|---|---|---|---|---|---|
| WP_024146115.1 | AAA+ATPase+wHTH+Alpha+Beta-domain+wHTH+RRM | Type-1_BR.RM.like.systems | TraG_N->MULE-TRANSPOSASE->MORC+HISKIN->?-><-PAPS_reduct<-DUF499*<-DUF4007<-HTH_Tnp_1+HTH_21+rve+rve_2\|\|ArdCN+MPTase->HDRnase->Phage_int_SAM_5+Phage_integrase-><-tRNA\|\|ADH_N+ADH_zinc_N-> | 1107 | Bacteria>Pseudomonadota>Gammaproteobacteria>Enterobacterales>Enterobacteriaceae | GCF_000486445.2 | Salmonella_enterica | SEESL791_RS19985 | 4155736..4159059 | 28901 | ATP-binding-protein-[Salmonella-enterica] |
| WP_201435379.1 | AAA+ATPase+wHTH+Alpha+Beta-domain+wHTH+RRM | Type-1_BR.RM.like.systems | <-DUF1176\|\|REase->SymE_toxin->Glyoxalase->\|\|DUF4007->DUF499*->PAPS_reduct-><-?<-MORC+HISKIN<-Arm-DNA-bind_3+Phage_integrase<-tRNA\|\|ADH_N+ADH_zinc_N-> | 989 | Bacteria>Pseudomonadota>Gammaproteobacteria>Enterobacterales>Enterobacteriaceae | - | Klebsiella_pasteurii | LVK93_004870 | 21596..24565 | 2587529 | ATP-binding-protein-[Klebsiella-pasteurii] |
| WP_023339500.1 | AAA+ATPase+wHTH+Alpha+Beta-domain+wHTH+RRM | Type-1_BR.RM.like.systems | Glyco_hydro_43+CBM_6-><-MAD-NTD+HNH\|\|MORC+HISKIN->?-><-PAPS_reduct<-DUF499*<-DUF4007\|\|ArdCN+MPTase->HDRnase->Phage_int_SAM_5+Phage_integrase-> | 1107 | Bacteria>Pseudomonadota>Gammaproteobacteria>Enterobacterales>Enterobacteriaceae | GCF_901420855.1 | Klebsiella_variicola | FQT79_RS24945 | 4878943..4882266 | 244366 | hypothetical-protein-[Klebsiella-variicola] |
| HCB1226982.1 | AAA+ATPase+wHTH+Alpha+Beta-domain+wHTH+RRM | Type-1_BR.RM.like.systems | <-NACHT+SF1-SF2-Helicase+HKD-DNase\|\|HIN-HTH->HTH_21+rve+rve_3->DUF4007->DUF499*->PAPS_reduct-><-?<-MORC+HISKIN<-tRNA\|\|?->?->ADH_N+ADH_zinc_N-> | 989 | Bacteria>Pseudomonadota>Gammaproteobacteria>Enterobacterales>Enterobacteriaceae | GCA_023033095.1 | Klebsiella_variicola_subsp._variicola | MYN71_004237 | 37585..40554 | 2590157 | ATP-binding-protein-[Klebsiella-variicola-subsp.-variicola] |
| EHO3141308.1 | AAA+ATPase+wHTH+Alpha+Beta-domain+wHTH+RRM | Type-1_BR.RM.like.systems | <-FtsK+HerA+A32\|\|<-ADH_N+ADH_zinc_N\|\|tRNA-><-HDRnase<-ArdCN+MPTase<-?\|\|DUF4007->DUF499*->PAPS_reduct-><-?<-MORC+HISKIN<-TraG_N<-?<-TraH | 986 | Bacteria>Pseudomonadota>Gammaproteobacteria>Enterobacterales>Enterobacteriaceae | GCA_018501385.1 | Salmonella_enterica | KK458_002739 | 251186..254146 | 28901 | ATP-binding-protein-[Salmonella-enterica] |
| EDR5086716.1 | AAA+ATPase+wHTH+Alpha+Beta-domain+wHTH+RRM | Type-1_BR.RM.like.systems | MORC+HISKIN->?-><-PAPS_reduct<-DUF499*<-DUF4007\|\|?->ArdCN+MPTase->NUDIX->Phage_int_SAM_5+Phage_integrase-><-tRNA\|\|ADH_N+ADH_zinc_N-> | 1107 | Bacteria>Pseudomonadota>Gammaproteobacteria>Enterobacterales>Enterobacteriaceae | GCA_010449355.1 | Salmonella_enterica_subsp._enterica_serovar_Kingston | GRZ33_002994 | 5595..8918 | 2161663 | ATP-binding-protein-[Salmonella-enterica-subsp.-enterica] |
| HCF8019186.1 | AAA+ATPase+wHTH+Alpha+Beta-domain+wHTH+RRM | Type-1_BR.RM.like.systems | TraH->?->TraG_N->MORC+HISKIN->?-><-PAPS_reduct<-DUF499*<-DUF4007\|\|?->ArdCN+MPTase->HDRnase->Phage_int_SAM_5+Phage_integrase-> | 986 | Bacteria>Pseudomonadota>Gammaproteobacteria>Enterobacterales>Enterobacteriaceae | GCA_023978205.1 | Klebsiella_aerogenes | NIU39_003418 | 64348..67308 | 548 | ATP-binding-protein-[Klebsiella-aerogenes] |
| HBQ8474228.1 | AAA+ATPase+wHTH+Alpha+Beta-domain+wHTH+RRM | Type-1_BR.RM.like.systems | TraH->?->TraG_N->MORC+HISKIN->?-><-PAPS_reduct<-DUF499*<-DUF4007\|\|?->ArdCN+MPTase->HDRnase->Phage_int_SAM_5+Phage_integrase-> | 986 | Bacteria>Pseudomonadota>Gammaproteobacteria>Enterobacterales>Enterobacteriaceae | GCA_021895595.1 | Klebsiella_pneumoniae | L9A49_004132 | 61808..64768 | 573 | ATP-binding-protein-[Klebsiella-pneumoniae] |
| BBV79047.1 | AAA+ATPase+wHTH+Alpha+Beta-domain+wHTH+RRM | Type-1_BR.RM.like.systems | <-DUF772+DDE_Tnp_1<-HTH<-RelE-ParE<-DDE_Tnp_IS240\|\|XIS-HTH->DUF499*->DDE_Tnp_IS240-><-?<-?<-?<-NTP_transferase | 88 | Bacteria>Pseudomonadota>Gammaproteobacteria>Enterobacterales>Enterobacteriaceae | GCA_015135635.1 | Raoultella_planticola | STW0522RA056_P10600 | 45634..45900 | 575 | hypothetical-protein-STW0522RA056_P10600-[Raoultella-planticola] |
| EFM2416768.1 | AAA+ATPase+wHTH+Alpha+Beta-domain+wHTH+RRM | Type-1_BR.RM.like.systems | RVT-><-zf-ISL3+HTH_Tnp_ISL3+DDE_Tnp_ISL3<-?\|\|ArsR-HTH+YTCD-HTH->HTH_Tnp->XIS-HTH->DUF499*-> | 156 | Bacteria>Pseudomonadota>Gammaproteobacteria>Enterobacterales>Enterobacteriaceae | GCA_012855175.1 | Escherichia_coli | HCP85_004433 | 17961..18429 | 562 | hypothetical-protein-[Escherichia-coli] |
| MBJ9237166.1 | AAA+ATPase+wHTH+Alpha+Beta-domain+wHTH+RRM | Type-1_BR.RM.like.systems | ABC-ATPase+SbcC->?->?->DUF4007->DUF499*->PAPS_reduct-><-?<-MORC+HISKIN\|\|ArdCN+MPTase-> | 989 | Bacteria>Pseudomonadota>Gammaproteobacteria>Enterobacterales>Enterobacteriaceae | GCA_016504745.1 | Citrobacter_koseri | GHV37_21265 | 31592..34561 | 545 | ATP-binding-protein-[Citrobacter-koseri] |
| EIN2284072.1 | AAA+ATPase+wHTH+Alpha+Beta-domain+wHTH+RRM | Type-1_BR.RM.like.systems | <-DUF499*<-XIS-HTH<-HTH_Tnp<-tRNA\|\|ADH_N+ADH_zinc_N->FtsK+HerA+A32-> | 144 | Bacteria>Pseudomonadota>Gammaproteobacteria>Enterobacterales>Enterobacteriaceae | GCA_020874335.1 | Escherichia_coli | LOP93_004303 | 1..432 | 562 | hypothetical-protein-[Escherichia-coli] |
| WP_244570774.1 | AAA+ATPase+wHTH+Alpha+Beta-domain+wHTH+RRM | Type-1_BR.RM.like.systems | XIS-HTH->XIS-HTH->DUF499*-><-?<-HTH_Tnp_1+HTH_21+rve+rve_2 | 161 | Bacteria>Pseudomonadota>Gammaproteobacteria>Enterobacterales>Enterobacteriaceae | GCF_963281085.1 | Escherichia_coli | ATA54_RS24775 | 2135..2620 | 562 | hypothetical-protein-[Escherichia-coli] |
| WP_198924247.1 | AAA+ATPase+wHTH+Alpha+Beta-domain+wHTH+RRM | Type-1_BR.RM.like.systems | MORC+HISKIN->?-><-PAPS_reduct<-DUF499*<-DUF4007 | 986 | Bacteria>Pseudomonadota>Gammaproteobacteria>Enterobacterales>Enterobacteriaceae | - | Enterobacter_hormaechei | OMS03_001576 | 5477..8437 | 158836 | ATP-binding-protein-[Enterobacter-hormaechei] |
| HAM2531449.1 | AAA+ATPase+wHTH+Alpha+Beta-domain+wHTH+RRM | Type-1_BR.RM.like.systems | <-DUF499*<-XIS-HTH<-XIS-HTH<-?<-?<-DUF3987 | 235 | Bacteria>Pseudomonadota>Gammaproteobacteria>Enterobacterales>Enterobacteriaceae | GCA_014365645.1 | Escherichia_coli | IBD21_004826 | 1..706 | 562 | hypothetical-protein-[Escherichia-coli] |
| WP_264228023.1 | AAA+ATPase+wHTH+Alpha+Beta-domain+wHTH+RRM | Type-1_BR.RM.like.systems | SbcC+SbcC->?->RlpA-><-DUF499*<-METHYLASE<-SbcC<-TUDOR+SF1-SF2-Helicase+iREase+CTD+REase<-?\|\|nSTAND1+sGTP->HTH+WYL-> | 951 | Bacteria>Pseudomonadota>Gammaproteobacteria>Enterobacterales>Erwiniaceae | GCF_025882065.1 | Pantoea_ananatis | NB704_RS06295 | 1388583..1391438 | 553 | DUF499-domain-containing-protein-[Pantoea-ananatis] |
| PPS64671.1 | AAA+ATPase+wHTH+Alpha+Beta-domain+wHTH+RRM | Type-1_BR.RM.like.systems | <-ABC-ATPase+SbcC<-?<-?\|\|RlpA-><-DUF499*<-METHYLASE | 951 | Bacteria>Pseudomonadota>Gammaproteobacteria>Enterobacterales>Erwiniaceae | GCA_002939445.1 | Pantoea_sp._BRM17 | CRX72_02445 | 37760..40615 | 1343867 | hypothetical-protein-CRX72_02445-[Pantoea-sp.-BRM17] |
| WP_077999168.1 | AAA+ATPase+wHTH+Alpha+Beta-domain+wHTH+RRM | Type-1_BR.RM.like.systems | JAB->RlpA-><-DUF499*<-REase<-OLD-ABC<-METHYLASE<-HNH<-OLD-ABC<-TUDOR+SF1-SF2-Helicase+iREase+CTD+REase | 949 | Bacteria>Pseudomonadota>Gammaproteobacteria>Enterobacterales>Hafniaceae | GCF_002006945.1 | Edwardsiella_tarda | BTN58_RS02535 | 130022..132871 | 636 | DUF499-domain-containing-protein-[Edwardsiella-tarda] |
| WP_196717876.1 | AAA+ATPase+wHTH+Alpha+Beta-domain+wHTH+RRM | Type-1_BR.RM.like.systems | Phage_AlpA->TOPRIM->?->\|\|RlpA-><-?<-SMODS<-DUF499*<-METHYLASE<-TUDOR+SF1-SF2-Helicase+iREase+CTD+REase<-HTH+WYL\|\|nSTAND1+sGTP-> | 947 | Bacteria>Pseudomonadota>Gammaproteobacteria>Enterobacterales>Morganellaceae | GCF_015739395.1 | Providencia_stuartii | I5E15_RS14725 | 67397..70240 | 588 | DUF499-domain-containing-protein-[Providencia-stuartii] |
| HCZ8677551.1 | AAA+ATPase+wHTH+Alpha+Beta-domain+wHTH+RRM | Type-1_BR.RM.like.systems | <-DUF499*<-XIS-HTH<-HTH_Tnp<-XIS-HTH<-?\|\|CopG/RHH->nSTAND1+sGTP-> | 587 | Bacteria>Pseudomonadota>Gammaproteobacteria>Enterobacterales>Morganellaceae | GCA_026957575.1 | Proteus_mirabilis | O4E69_003747 | 1..1762 | 584 | TPA:-hypothetical-protein-O4E69_003747,-partial-[Proteus-mirabilis] |
| WP_205545050.1 | AAA+ATPase+wHTH+Alpha+Beta-domain+wHTH+RRM | Type-1_BR.RM.like.systems | <-PAIREDC-HTH+HIN-HTH\|\|?->TUDOR+SF1-SF2-Helicase+iREase+CTD+REase->OLD-ABC->HNH->METHYLASE->DUF499*-><-?<-HNH<-?<-DHTH+WYL\|\|nSTAND1+sGTP->HTH+WYL-> | 951 | Bacteria>Pseudomonadota>Gammaproteobacteria>Enterobacterales>Pectobacteriaceae | GCF_016949795.1 | Pectobacterium_brasiliense | H4F47_RS10595 | 47611..50466 | 180957 | DUF499-domain-containing-protein-[Pectobacterium-brasiliense] |
| WP_115183101.1 | AAA+ATPase+wHTH+Alpha+Beta-domain+wHTH+RRM | Type-1_BR.RM.like.systems | <-nSTAND1+sGTP\|\|?->TUDOR+SF1-SF2-Helicase+iREase+CTD+REase->OLD-ABC->HNH->METHYLASE->DUF499*-><-RlpA<-SF1-SF2-Helicase<-?<-GDResolvase+SIGMA-HTH<-?<-XIS-HTH | 951 | Bacteria>Pseudomonadota>Gammaproteobacteria>Enterobacterales>Yersiniaceae | GCF_946228205.1 | Serratia | BSR03_RS13425 | 143107..145962 | 613 | MULTISPECIES:-DUF499-domain-containing-protein-[Serratia] |
| CAI1753142.1 | AAA+ATPase+wHTH+Alpha+Beta-domain+wHTH+RRM | Type-1_BR.RM.like.systems | TUDOR+SF1-SF2-Helicase+iREase+CTD+REase->OLD-ABC->HNH->METHYLASE->DUF499*-><-RlpA<-SF1-SF2-Helicase<-?<-GDResolvase+SIGMA-HTH<-?<-XIS-HTH | 975 | Bacteria>Pseudomonadota>Gammaproteobacteria>Enterobacterales>Yersiniaceae | GCA_946228205.1 | Serratia_quinivorans | SAMEA4857275_02169 | 318450..321377 | 137545 | Protein-of-uncharacterised-function-(DUF499)-[Serratia-quinivorans] |
| WP_047727733.1 | AAA+ATPase+wHTH+Alpha+Beta-domain+wHTH+RRM | Type-1_BR.RM.like.systems | Arm-DNA-bind_3+Phage_int_SAM_3+Phage_integrase-><-?<-?<-DUF499*<-METHYLASE<-HEPN<-TUDOR+SF1-SF2-Helicase+iREase+CTD+REase<-HTH+WYL | 942 | Bacteria>Pseudomonadota>Gammaproteobacteria>Enterobacterales>Yersiniaceae | GCF_001022215.1 | Serratia | DKC05_RS04355 | 483774..486602 | 613 | MULTISPECIES:-DUF499-domain-containing-protein-[Serratia] |
| SUI50248.1 | AAA+ATPase+wHTH+Alpha+Beta-domain+wHTH+RRM | Type-1_BR.RM.like.systems | GDResolvase+SIGMA-HTH->?->SF1-SF2-Helicase->RlpA-><-DUF499*<-METHYLASE<-HNH<-OLD-ABC<-TUDOR+SF1-SF2-Helicase+iREase+CTD+REase | 975 | Bacteria>Pseudomonadota>Gammaproteobacteria>Enterobacterales>Yersiniaceae | GCA_900457075.1 | Serratia_quinivorans | NCTC11544_01113 | 1025353..1028280 | 137545 | Protein-of-uncharacterised-function-(DUF499)-[Serratia-quinivorans] |
| HDL6629710.1 | AAA+ATPase+wHTH+Alpha+Beta-domain+wHTH+RRM | Type-1_BR.RM.like.systems | DOC-><-?\|\|Phage_AlpA->?->?->RlpA-><-DUF499*<-METHYLASE<-TUDOR+SF1-SF2-Helicase+iREase+CTD+REase<-?\|\|JAB->nSTAND1+sGTP->HTH+WYL-> | 951 | Bacteria>Pseudomonadota>Gammaproteobacteria>Enterobacterales>Yersiniaceae | GCA_028833845.1 | Yersinia_enterocolitica | PXJ35_002272 | 241778..244633 | 630 | TPA:-ATP-binding-protein-[Yersinia-enterocolitica] |
| BAL53124.1 | AAA+ATPase+wHTH+Alpha+Beta-domain+wHTH+RRM | Type-1_BR.RM.like.systems | TUDOR+SF1-SF2-Helicase+iREase+CTD+REase->OLD-ABC->TOPRIM->METHYLASE->?->?->DUF499*-><-?<-?<-tRNA\|\|?-><-?<-TRAM+Methyltransf_31 | 967 | Bacteria>Pseudomonadota>Gammaproteobacteria>environmental_samples | - | uncultured_Gammaproteobacteria_bacterium | HGMM_F04D09C21 | 20323..23226 | 86473 | AAA-family-ATPase-[uncultured-Gammaproteobacteria-bacterium] |
| MBK8754773.1 | AAA+ATPase+wHTH+Alpha+Beta-domain+wHTH+RRM | Type-1_BR.RM.like.systems | METHYLASE->OLD-ABC->TOPRIM->?->PIN->DUF499*->?-><-?<-CASPASE<-TIR<-?<-ADH_N+ADH_zinc_N | 946 | Bacteria>Pseudomonadota>Gammaproteobacteria>Gammaproteobacteria_incertae_sedis>Candidatus_Competibacteraceae | GCA_016710535.1 | Candidatus_Competibacteraceae_bacterium | IPL99_25425 | 33622..36462 | 2053538 | ATP-binding-protein-[Candidatus-Competibacteraceae-bacterium] |
| RUQ36603.1 | AAA+ATPase+wHTH+Alpha+Beta-domain+wHTH+RRM | Type-1_BR.RM.like.systems | ParB+HNH->DUF499*-><-ABhydrolase+ABhydrolase<-?<-tRNA<-?<-HHH+ENDO3+NUDIX | 937 | Bacteria>Pseudomonadota>Gammaproteobacteria>Gammaproteobacteria_incertae_sedis>Candidatus_Competibacteraceae | GCA_003989065.1 | Candidatus_Competibacteraceae_bacterium | EKK71_07765 | 2719..5532 | 2053538 | ATP-binding-protein-[Candidatus-Competibacteraceae-bacterium] |
| MCC8994173.1 | AAA+ATPase+wHTH+Alpha+Beta-domain+wHTH+RRM | Type-1_BR.RM.like.systems | <-HTH\|\|CHTH->?->?-><-DUF499*<-HTH<-HigB_toxin\|\|?->?-><-METHYLASE<-DOC+HTH | 946 | Bacteria>Pseudomonadota>Gammaproteobacteria>Gammaproteobacteria_incertae_sedis>Candidatus_Competibacteraceae | GCA_020890935.1 | Candidatus_Contendobacter_sp. | LM550_10905 | 15829..18669 | 2651161 | DUF499-domain-containing-protein-[Candidatus-Contendobacter-sp.] |
| MBL8253032.1 | AAA+ATPase+wHTH+Alpha+Beta-domain+wHTH+RRM | Type-1_BR.RM.like.systems | <-DUF499*<-PIN<-DUF433<-METHYLASE<-TUDOR+SF1-SF2-Helicase+iREase+CTD+REase | 937 | Bacteria>Pseudomonadota>Gammaproteobacteria>Gammaproteobacteria_incertae_sedis>Candidatus_Competibacteraceae | GCA_016789525.1 | Candidatus_Competibacter_sp. | JNJ76_05450 | 1..2812 | 2800415 | ATP-binding-protein-[Candidatus-Competibacter-sp.] |
| PLY13647.1 | AAA+ATPase+wHTH+Alpha+Beta-domain+wHTH+RRM | Type-1_BR.RM.like.systems | <-SIR2+STALD<-TIR\|\|HTH+WYL->TUDOR+SF1-SF2-Helicase+iREase+CTD+REase->METHYLASE->DUF499*-> | 949 | Bacteria>Pseudomonadota>Gammaproteobacteria>Gammaproteobacteria_incertae_sedis>Sedimenticola | GCA_002869585.1 | Sedimenticola_sp. | CO631_13485 | 9569..12418 | 1940285 | MAG:-hypothetical-protein-CO631_13485-[Sedimenticola-sp.] |
| WP_058474609.1 | AAA+ATPase+wHTH+Alpha+Beta-domain+wHTH+RRM | Type-1_BR.RM.like.systems | HTH_60->XIS-HTH->TUDOR+SF1-SF2-Helicase+iREase+CTD+REase->TUDOR+SF1-SF2-Helicase+iREase+CTD+REase-p->METHYLASE->DUF499*->SIR2+STALD->TIR-><-HTH+LexA-protease\|\|?->Phage_lambda-> | 948 | Bacteria>Pseudomonadota>Gammaproteobacteria>Legionellales>Legionellaceae | GCF_900640005.1 | Legionella_quateirensis | DXZ53_RS13215 | 3230721..3233567 | 45072 | DUF499-domain-containing-protein-[Legionella-quateirensis] |
| WP_106141259.1 | AAA+ATPase+wHTH+Alpha+Beta-domain+wHTH+RRM | Type-1_BR.RM.like.systems | HTH_60->XIS-HTH->TUDOR+SF1-SF2-Helicase+iREase+CTD+REase->METHYLASE-p->METHYLASE->DUF499*-><-HTH+LexA-protease\|\|?->Phage_lambda->?->VirB11+VirB4-FtsK-> | 948 | Bacteria>Pseudomonadota>Gammaproteobacteria>Legionellales>Legionellaceae | GCF_003004175.1 | Legionella_pneumophila | CA922_RS12145 | 2667750..2670596 | 446 | DUF499-domain-containing-protein-[Legionella-pneumophila] |
| WP_269562751.1 | AAA+ATPase+wHTH+Alpha+Beta-domain+wHTH+RRM | Type-1_BR.RM.like.systems | <-VirB11+VirB4-FtsK<-?<-Phage_lambda<-?\|\|HTH+LexA-protease-><-DUF499*<-METHYLASE<-TUDOR+SF1-SF2-Helicase+iREase+CTD+REase<-XIS-HTH<-HTH_60 | 954 | Bacteria>Pseudomonadota>Gammaproteobacteria>Legionellales>Legionellaceae | GCF_027273505.1 | Legionella_pneumophila | O6C85_RS13950 | 27428..30292 | 446 | DUF499-domain-containing-protein-[Legionella-pneumophila] |
| WP_202967569.1 | AAA+ATPase+wHTH+Alpha+Beta-domain+wHTH+RRM | Type-1_BR.RM.like.systems | <-VirB11+VirB4-FtsK<-?<-Phage_lambda<-?\|\|HTH+LexA-protease-><-DUF499*<-METHYLASE<-TUDOR+SF1-SF2-Helicase+iREase+CTD+REase<-XIS-HTH<-HTH_60 | 894 | Bacteria>Pseudomonadota>Gammaproteobacteria>Legionellales>Legionellaceae | GCF_900640205.1 | Legionella_jamestowniensis | BM259_RS00525 | 103607..106291 | 455 | DUF499-domain-containing-protein-[Legionella-jamestowniensis] |
| WP_058450411.1 | AAA+ATPase+wHTH+Alpha+Beta-domain+wHTH+RRM | Type-1_BR.RM.like.systems | <-VirB11+VirB4-FtsK<-?<-Phage_lambda<-?\|\|HTH+LexA-protease-><-DUF499*<-METHYLASE<-TUDOR+SF1-SF2-Helicase+iREase+CTD+REase<-XIS-HTH<-HTH_60 | 951 | Bacteria>Pseudomonadota>Gammaproteobacteria>Legionellales>Legionellaceae | GCF_019974095.1 | Legionella_pneumophila | A1308_RS13960 | 33285..36140 | 446 | DUF499-domain-containing-protein-[Legionella-pneumophila] |
| WP_042234453.1 | AAA+ATPase+wHTH+Alpha+Beta-domain+wHTH+RRM | Type-1_BR.RM.like.systems | HTH_60->XIS-HTH->TUDOR+SF1-SF2-Helicase+iREase+CTD+REase->METHYLASE->DUF499*-><-HTH+LexA-protease\|\|?->Phage_lambda->?->VirB11+VirB4-FtsK-> | 951 | Bacteria>Pseudomonadota>Gammaproteobacteria>Legionellales>Legionellaceae | GCF_000694995.1 | Legionella_pneumophila | A0W40_RS11710 | 109101..111956 | 446 | DUF499-domain-containing-protein-[Legionella-pneumophila] |
| HAU0909273.1 | AAA+ATPase+wHTH+Alpha+Beta-domain+wHTH+RRM | Type-1_BR.RM.like.systems | <-VirB11+VirB4-FtsK<-?<-Phage_lambda<-?\|\|HTH+LexA-protease-><-DUF499*<-METHYLASE<-TUDOR+SF1-SF2-Helicase+iREase+CTD+REase<-XIS-HTH<-HTH_60 | 951 | Bacteria>Pseudomonadota>Gammaproteobacteria>Legionellales>Legionellaceae | GCA_015983645.1 | Legionella_pneumophila | JBH59_05555 | 25593..28448 | 446 | ATP-binding-protein-[Legionella-pneumophila] |
| WP_112220365.1 | AAA+ATPase+wHTH+Alpha+Beta-domain+wHTH+RRM | Type-1_BR.RM.like.systems | <-Phage_lambda<-?\|\|HTH+LexA-protease-><-TIR<-SIR2+STALD<-DUF499*<-METHYLASE<-TUDOR+SF1-SF2-Helicase+iREase+CTD+REase<-XIS-HTH<-HTH_60 | 948 | Bacteria>Pseudomonadota>Gammaproteobacteria>Legionellales>Legionellaceae | GCF_003265995.1 | Legionella_quinlivanii | B1207_RS12955 | 55194..58040 | 45073 | DUF499-domain-containing-protein-[Legionella-quinlivanii] |
| WP_115710943.1 | AAA+ATPase+wHTH+Alpha+Beta-domain+wHTH+RRM | Type-1_BR.RM.like.systems | <-ACET<-?\|\|?->HTH_60-><-DUF499*<-METHYLASE<-TUDOR+SF1-SF2-Helicase+iREase+CTD+REase<-HTH+LexA-protease\|\|?->Phage_lambda-> | 939 | Bacteria>Pseudomonadota>Gammaproteobacteria>Legionellales>Legionellaceae | GCF_900492725.1 | Legionella_sainthelensi | DY395_RS14085 | 98851..101670 | 28087 | DUF499-domain-containing-protein-[Legionella-sainthelensi] |
| WP_064023787.1 | AAA+ATPase+wHTH+Alpha+Beta-domain+wHTH+RRM | Type-1_BR.RM.like.systems | PIN-><-?<-RelE-ParE<-TraJ-RHH\|\|Phage_int_SAM_1+Phage_integrase-><-HEPN<-DUF499*<-METHYLASE<-TUDOR+SF1-SF2-Helicase+iREase+CTD+REase<-RHH<-HIN-HTH | 936 | Bacteria>Pseudomonadota>Gammaproteobacteria>Methylococcales>Methylococcaceae | GCF_001644685.1 | Methylomonas | A1356_RS16225 | 62920..65730 | 416 | MULTISPECIES:-DUF499-domain-containing-protein-[Methylomonas] |
| WP_017365774.1 | AAA+ATPase+wHTH+Alpha+Beta-domain+wHTH+RRM | Type-1_BR.RM.like.systems | XIS-HTH->?->?->?->?-><-TIR<-DUF499*<-Schlafen+HATPase+HTH<-METHYLASE<-PemK_toxin<-PrlF_antitoxin<-TUDOR+SF1-SF2-Helicase+iREase+CTD+REase | 942 | Bacteria>Pseudomonadota>Gammaproteobacteria>Methylococcales>Methylococcaceae | GCF_000297615.1 | Methylococcus_capsulatus | QMX71_RS07545 | 1581839..1584667 | 414 | DUF499-domain-containing-protein-[Methylococcus-capsulatus] |
| WP_218796623.1 | AAA+ATPase+wHTH+Alpha+Beta-domain+wHTH+RRM | Type-1_BR.RM.like.systems | METHYLASE-p-><-?\|\|TUDOR+SF1-SF2-Helicase+iREase+CTD+REase->PrlF_antitoxin->PemK_toxin->METHYLASE->DUF499*->TIR-><-?\|\|XIS-HTH-><-TIR | 948 | Bacteria>Pseudomonadota>Gammaproteobacteria>Methylococcales>Methylococcaceae | GCF_019285555.1 | Methylococcus_capsulatus | KW114_RS08265 | 1823970..1826816 | 414 | DUF499-domain-containing-protein-[Methylococcus-capsulatus] |
| WP_010962200.1 | AAA+ATPase+wHTH+Alpha+Beta-domain+wHTH+RRM | Type-1_BR.RM.like.systems | TUDOR+SF1-SF2-Helicase+iREase+CTD+REase->PrlF_antitoxin->PemK_toxin->METHYLASE->METHYLASE->Schlafen+HATPase+HTH->DUF499*->TIR-> | 942 | Bacteria>Pseudomonadota>Gammaproteobacteria>Methylococcales>Methylococcaceae | GCF_000008325.1 | Methylococcus_capsulatus | MCA_RS14770 | 3194105..3196933 | 414 | DUF499-domain-containing-protein-[Methylococcus-capsulatus] |
| WP_020565890.1 | AAA+ATPase+wHTH+Alpha+Beta-domain+wHTH+RRM | Type-1_BR.RM.like.systems | <-REase\|\|MetJarc-HTH->RelE-ParE->TUDOR+SF1-SF2-Helicase+iREase+CTD+REase->METHYLASE->DUF499*->?->HerA/FtsK-><-?<-SMODS | 932 | Bacteria>Pseudomonadota>Gammaproteobacteria>Methylococcales>Methylococcaceae | GCF_000372865.1 | Methylosarcina_fibrata | A3OW_RS0123395 | 11822..14620 | 105972 | DUF499-domain-containing-protein-[Methylosarcina-fibrata] |
| WP_218800673.1 | AAA+ATPase+wHTH+Alpha+Beta-domain+wHTH+RRM | Type-1_BR.RM.like.systems | <-TIR<-DUF499*<-REase<-OLD-ABC<-METHYLASE<-PIN<-TUDOR+SF1-SF2-Helicase+iREase+CTD+REase<-HTH_Tnp | 942 | Bacteria>Pseudomonadota>Gammaproteobacteria>Methylococcales>Methylococcaceae | GCF_019285535.1 | Methylococcus_capsulatus | M3M30_RS10970 | 2380003..2382831 | 414 | DUF499-domain-containing-protein-[Methylococcus-capsulatus] |
| WP_064041552.1 | AAA+ATPase+wHTH+Alpha+Beta-domain+wHTH+RRM | Type-1_BR.RM.like.systems | DUF499*->?-><-LZ_Tnp_IS66+zf-IS66+DDE_Tnp_IS66+DDE_Tnp_IS66_C<-?<-HIN-HTH | 937 | Bacteria>Pseudomonadota>Gammaproteobacteria>Methylococcales>Methylococcaceae | GCF_001644135.1 | Methylomonas_koyamae | A1507_RS15520 | 699..3512 | 702114 | DUF499-domain-containing-protein-[Methylomonas-koyamae] |
| NOT11404.1 | AAA+ATPase+wHTH+Alpha+Beta-domain+wHTH+RRM | Type-1_BR.RM.like.systems | PemK_toxin->OLD-ABC->TOPRIM->METHYLASE->OLD-ABC->TOPRIM->DUF499*-> | 939 | Bacteria>Pseudomonadota>Gammaproteobacteria>Methylococcales>Methylococcaceae | GCA_013140705.1 | Methylococcaceae_bacterium | HOP23_06170 | 47500..50319 | 1933926 | ATP-binding-protein-[Methylococcaceae-bacterium] |
| WP_282213184.1 | AAA+ATPase+wHTH+Alpha+Beta-domain+wHTH+RRM | Type-1_BR.RM.like.systems | HHH_5+SF1-SF2-helicase->XIS-HTH->TUDOR+SF1-SF2-Helicase+iREase+CTD+REase->REase->METHYLASE->REase->DUF499*->Methylase-><-Radical-SAM<-HTH_3 | 939 | Bacteria>Pseudomonadota>Gammaproteobacteria>Methylococcales>Methylococcaceae | GCF_949769275.1 | Methylococcus_capsulatus | QMX71_RS08995 | 1893622..1896441 | 414 | DUF499-domain-containing-protein-[Methylococcus-capsulatus] |
| WP_218796637.1 | AAA+ATPase+wHTH+Alpha+Beta-domain+wHTH+RRM | Type-1_BR.RM.like.systems | XIS-HTH->TUDOR+SF1-SF2-Helicase+iREase+CTD+REase->OLD-ABC->TOPRIM->METHYLASE->REase->DUF499*->MORC+REase->HEPN-><-Phage_lysozyme<-?<-?<-RNase | 939 | Bacteria>Pseudomonadota>Gammaproteobacteria>Methylococcales>Methylococcaceae | GCF_019285555.1 | Methylococcus_capsulatus | KW114_RS08340 | 1841753..1844572 | 414 | DUF499-domain-containing-protein-[Methylococcus-capsulatus] |
| WP_218797031.1 | AAA+ATPase+wHTH+Alpha+Beta-domain+wHTH+RRM | Type-1_BR.RM.like.systems | HHH_5+SF1-SF2-helicase->XIS-HTH->TUDOR+SF1-SF2-Helicase+iREase+CTD+REase->TIR->METHYLASE->REase->DUF499*->Methylase-><-Radical-SAM<-HTH_3 | 939 | Bacteria>Pseudomonadota>Gammaproteobacteria>Methylococcales>Methylococcaceae | GCF_019285555.1 | Methylococcus_capsulatus | KW114_RS10930 | 2381240..2384059 | 414 | DUF499-domain-containing-protein-[Methylococcus-capsulatus] |
| WP_156881498.1 | AAA+ATPase+wHTH+Alpha+Beta-domain+wHTH+RRM | Type-1_BR.RM.like.systems | <-SLATT\|\|XIS-HTH->TUDOR+SF1-SF2-Helicase+iREase+CTD+REase->OLD-ABC->METHYLASE->REase->DUF499*->?-><-Radical-SAM<-HTH_3<-Phage_lysozyme | 939 | Bacteria>Pseudomonadota>Gammaproteobacteria>Methylococcales>Methylococcaceae | GCF_017832375.1 | Methylocaldum_sp._RMAD-M | JOD69_RS15380 | 37425..40244 | 2806557 | DUF499-domain-containing-protein-[Methylocaldum-sp.-RMAD-M] |
| PZN71299.1 | AAA+ATPase+wHTH+Alpha+Beta-domain+wHTH+RRM | Type-1_BR.RM.like.systems | <-DUF499*<-PIN<-METHYLASE<-HNH<-OLD-ABC<-PemK_toxin<-PrlF_antitoxin | 955 | Bacteria>Pseudomonadota>Gammaproteobacteria>Methylococcales>Methylococcales_incertae_sedis | GCA_003242955.1 | Candidatus_Methylumidiphilus_alinenensis | DM484_26720 | 13367..16234 | 2202197 | hypothetical-protein-DM484_26720-[Candidatus-Methylumidiphilus-alinenensis] |
| WP_022949219.1 | AAA+ATPase+wHTH+Alpha+Beta-domain+wHTH+RRM | Type-1_BR.RM.like.systems | LON+AAA->TUDOR+SF1-SF2-Helicase+iREase+CTD+REase->METHYLASE->OLD-ABC->REase->DUF499*->?->RADICAL-SAM->Glyco_transf_8-> | 952 | Bacteria>Pseudomonadota>Gammaproteobacteria>Methylococcales>Methylothermaceae | GCF_000421465.1 | Methylohalobius_crimeensis | HO35_RS0111990 | 2436080..2438938 | 244365 | DUF499-domain-containing-protein-[Methylohalobius-crimeensis] |
| WP_194364110.1 | AAA+ATPase+wHTH+Alpha+Beta-domain+wHTH+RRM | Type-1_BR.RM.like.systems | ParB+HNH->ParB+HNH->TUDOR+SF1-SF2-Helicase+iREase+CTD+REase->METHYLASE->DUF499*-><-?<-?<-Arm-DNA-bind_2+Phage_int_SAM_3+Phage_integrase<-?\|\|HTH_26-> | 949 | Bacteria>Pseudomonadota>Gammaproteobacteria>Moraxellales>Moraxellaceae | GCF_015277735.1 | Psychrobacter_sp._NG254 | IP510_RS07350 | 481008..483857 | 2782003 | DUF499-domain-containing-protein-[Psychrobacter-sp.-NG254] |
| MAE39903.1 | AAA+ATPase+wHTH+Alpha+Beta-domain+wHTH+RRM | Type-1_BR.RM.like.systems | HTH_26->TUDOR+SF1-SF2-Helicase+iREase+CTD+REase->METHYLASE->OLD-ABC->REase->DUF499*->?->?-><-Phage_integrase<-Phage_integrase | 941 | Bacteria>Pseudomonadota>Gammaproteobacteria>Moraxellales>Moraxellaceae | GCA_002686735.1 | Psychrobacter_sp. | CML07_03075 | 11656..14481 | 56811 | hypothetical-protein-[Psychrobacter-sp.] |
| WP_107866351.1 | AAA+ATPase+wHTH+Alpha+Beta-domain+wHTH+RRM | Type-1_BR.RM.like.systems | METHYLASE->OLD-ABC->REase->DUF499*-><-ParB+HNH<-?<-ParB | 941 | Bacteria>Pseudomonadota>Gammaproteobacteria>Moraxellales>Moraxellaceae | GCF_003051055.1 | Agitococcus_lubricus | C8N29_RS12225 | 4668..7493 | 1077255 | DUF499-domain-containing-protein-[Agitococcus-lubricus] |
| MCC6374735.1 | AAA+ATPase+wHTH+Alpha+Beta-domain+wHTH+RRM | Type-1_BR.RM.like.systems | <-DUF499*<-REase<-OLD-ABC<-METHYLASE-p | 878 | Bacteria>Pseudomonadota>Gammaproteobacteria>Moraxellales>Moraxellaceae | GCA_020846355.1 | Moraxellaceae_bacterium | IT465_07990 | 1..2635 | 1889775 | ATP-binding-protein-[Moraxellaceae-bacterium] |
| WP_142947823.1 | AAA+ATPase+wHTH+Alpha+Beta-domain+wHTH+RRM | Type-1_BR.RM.like.systems | Arm-DNA-bind_3+Phage_integrase-><-Phage_AlpA<-ZnR+TOPRIM+primpol+HTH<-XIS-HTH\|\|MobI-><-DUF499*<-Schlafen+HATPase+HTH<-METHYLASE<-METHYLASE<-HNH<-OLD-ABC<-TUDOR+SF1-SF2-Helicase+iREase+CTD+REase | 953 | Bacteria>Pseudomonadota>Gammaproteobacteria>Oceanospirillales>Alcanivoracaceae | - | Alcanivorax_sp._DSM_26295 | FN816_RS03075 | 107194..110055 | 1938738 | DUF499-domain-containing-protein-[Alcanivorax-sp.-DSM-26295] |
| WP_246952339.1 | AAA+ATPase+wHTH+Alpha+Beta-domain+wHTH+RRM | Type-1_BR.RM.like.systems | HTH+WYL->TUDOR+SF1-SF2-Helicase+iREase+CTD+REase->PIN->METHYLASE->DUF499*->SIR2-><-Phage_lysozyme<-?<-SF1-SF2-Helicase<-Schlafen+HATPase+HTH<-TRD+TRD | 947 | Bacteria>Pseudomonadota>Gammaproteobacteria>Oceanospirillales>Alcanivoracaceae | GCF_023051175.1 | Alcanivorax_quisquiliarum | MU846_10190 | 134529..137372 | 2933565 | DUF499-domain-containing-protein-[Alcanivorax-sp.-CY1518] |
| MCK0538078.1 | AAA+ATPase+wHTH+Alpha+Beta-domain+wHTH+RRM | Type-1_BR.RM.like.systems | HTH+WYL->TUDOR+SF1-SF2-Helicase+iREase+CTD+REase->PIN->METHYLASE->DUF499*->SIR2-><-Phage_lysozyme<-?<-SF1-SF2-Helicase<-Schlafen+HATPase+HTH<-TRD+TRD | 947 | Bacteria>Pseudomonadota>Gammaproteobacteria>Oceanospirillales>Alcanivoracaceae | GCA_023051175.1 | Alcanivorax_quisquiliarum | MU846_10190 | 134529..137372 | 2933565 | DUF499-domain-containing-protein-[Alcanivorax-sp.-CY1518] |
| WP_277673134.1 | AAA+ATPase+wHTH+Alpha+Beta-domain+wHTH+RRM | Type-1_BR.RM.like.systems | <-DUF499*<-METHYLASE<-TUDOR+SF1-SF2-Helicase+iREase+CTD+REase<-HTH+WYL\|\|KAP-NTPase->?->PAPS_reduct-> | 949 | Bacteria>Pseudomonadota>Gammaproteobacteria>Oceanospirillales>Alcanivoracaceae | GCF_029210245.1 | Alcanivorax_jadensis | P1U79_RS15395 | 46..2895 | 64988 | DUF499-domain-containing-protein-[Alcanivorax-jadensis] |
| WP_161384178.1 | AAA+ATPase+wHTH+Alpha+Beta-domain+wHTH+RRM | Type-1_BR.RM.like.systems | HTH+WYL->TUDOR+SF1-SF2-Helicase+iREase+CTD+REase->METHYLASE->DUF499*->SIR2-><-Phage_lysozyme | 947 | Bacteria>Pseudomonadota>Gammaproteobacteria>Oceanospirillales>Alcanivoracaceae | - | Alcanivorax_sp._ZXX171 | GN155_RS08585 | 8972..11815 | 2695858 | DUF499-domain-containing-protein-[Alcanivorax-sp.-ZXX171] |
| WP_233917946.1 | AAA+ATPase+wHTH+Alpha+Beta-domain+wHTH+RRM | Type-1_BR.RM.like.systems | Phage_lysozyme-><-?<-SMODS<-DUF499*<-METHYLASE | 947 | Bacteria>Pseudomonadota>Gammaproteobacteria>Oceanospirillales>Alcanivoracaceae | GCF_021395115.1 | Alloalcanivorax_xenomutans | LZG34_RS15945 | 141276..144119 | 1094342 | DUF499-domain-containing-protein-[Alcanivorax-xenomutans] |
| MCP5162750.1 | AAA+ATPase+wHTH+Alpha+Beta-domain+wHTH+RRM | Type-1_BR.RM.like.systems | <-ABC-ATPase+SbcC<-?<-DUF499*<-METHYLASE<-HNH<-OLD-ABC<-TUDOR+SF1-SF2-Helicase+iREase+CTD+REase<-HTH+WYL | 944 | Bacteria>Pseudomonadota>Gammaproteobacteria>Oceanospirillales>Hahellaceae | GCA_024234505.1 | Hahellaceae_bacterium | H7A00_13970 | 74140..76974 | 2048971 | ATP-binding-protein-[Hahellaceae-bacterium] |
| WP_231663150.1 | AAA+ATPase+wHTH+Alpha+Beta-domain+wHTH+RRM | Type-1_BR.RM.like.systems | HTH+WYL->TUDOR+SF1-SF2-Helicase+iREase+CTD+REase->METHYLASE->OLD-ABC->REase->ParB+HNH->DUF499*->Primpol+D5N+D5ATPase-><-Arm-DNA-bind_3+Phage_integrase\|\|?->FAD-NAD-dep-oxidoreductase-> | 957 | Bacteria>Pseudomonadota>Gammaproteobacteria>Oceanospirillales>Halomonadaceae | GCF_021151495.1 | Halomonas_sp._IOP_31 | K7I12_RS04335 | 15378..18251 | 2876584 | DUF499-domain-containing-protein-[Halomonas-sp.-IOP_31] |
| WP_284453091.1 | AAA+ATPase+wHTH+Alpha+Beta-domain+wHTH+RRM | Type-1_BR.RM.like.systems | Arm-DNA-bind_3+Phage_integrase-><-Phage_AlpA<-ZnR+TOPRIM+primpol+HTH<-XIS-HTH\|\|MobI-><-DUF499*<-REase<-OLD-ABC<-METHYLASE<-HNH<-OLD-ABC<-TUDOR+SF1-SF2-Helicase+iREase+CTD+REase | 953 | Bacteria>Pseudomonadota>Gammaproteobacteria>Oceanospirillales>Halomonadaceae | GCF_030145725.1 | Halomonas_meridiana | KYX60_RS01235 | 106598..109459 | 29570 | DUF499-domain-containing-protein-[Halomonas-meridiana] |
| WP_137094585.1 | AAA+ATPase+wHTH+Alpha+Beta-domain+wHTH+RRM | Type-1_BR.RM.like.systems | TUDOR+SF1-SF2-Helicase+iREase+CTD+REase->OLD-ABC->HNH->METHYLASE->OLD-ABC->REase->DUF499*-><-MobI\|\|XIS-HTH->ZnR+TOPRIM+primpol+HTH->Phage_AlpA-><-Arm-DNA-bind_3+Phage_integrase | 953 | Bacteria>Pseudomonadota>Gammaproteobacteria>Oceanospirillales>Halomonadaceae | GCF_028580285.1 | Halomonas | E8Q34_RS13770 | 13695..16556 | 2745 | DUF499-domain-containing-protein-[Halomonas-sp.-15WGF] |
| WP_159342933.1 | AAA+ATPase+wHTH+Alpha+Beta-domain+wHTH+RRM | Type-1_BR.RM.like.systems | zf-ISL3+HTH_Tnp_ISL3+DDE_Tnp_ISL3->?->REase->?-><-HTH_Tnp_1+HTH_21+rve+rve_3\|\|TPR+TPR+TPR+TPR+TPR+TPR+TPR+TPR-><-DUF499*<-McrC-NTD+REase<-EVE+McrB-AAA<-REase<-OLD-ABC<-METHYLASE<-HNH | 958 | Bacteria>Pseudomonadota>Gammaproteobacteria>Oceanospirillales>Halomonadaceae | GCF_009846525.1 | Halomonas_meridiana | CTT34_RS13795 | 2879494..2882370 | 29570 | DUF499-domain-containing-protein-[Halomonas-meridiana] |
| WP_253027667.1 | AAA+ATPase+wHTH+Alpha+Beta-domain+wHTH+RRM | Type-1_BR.RM.like.systems | Phage_integrase-><-Phage_AlpA<-ZnR+TOPRIM+primpol+HTH<-XIS-HTH\|\|MobI-><-DUF499*<-REase<-OLD-ABC<-METHYLASE<-OLD-ABC<-TUDOR+SF1-SF2-Helicase+iREase+CTD+REase<-HTH+WYL | 953 | Bacteria>Pseudomonadota>Gammaproteobacteria>Oceanospirillales>Halomonadaceae | GCF_024124725.1 | Halomonas_sp._Ps84H-12 | NH344_RS06635 | 49336..52197 | 2954501 | DUF499-domain-containing-protein-[Halomonas-sp.-Ps84H-12] |
| WP_097789710.1 | AAA+ATPase+wHTH+Alpha+Beta-domain+wHTH+RRM | Type-1_BR.RM.like.systems | Peptidase_M20->Arm-DNA-bind_3+Phage_integrase->TPR+TPR-><-?<-DUF499*<-McrC-NTD+REase<-MAD-NTD+EVE+McrB-AAA<-REase<-OLD-ABC<-HEPN+CBS<-METHYLASE | 957 | Bacteria>Pseudomonadota>Gammaproteobacteria>Oceanospirillales>Halomonadaceae | GCF_002549795.1 | Halomonas_beimenensis | BEI_RS11820 | 2512898..2515771 | 475662 | DUF499-domain-containing-protein-[Halomonas-beimenensis] |
| WP_239285448.1 | AAA+ATPase+wHTH+Alpha+Beta-domain+wHTH+RRM | Type-1_BR.RM.like.systems | TUDOR+SF1-SF2-Helicase+iREase+CTD+REase->OLD-ABC->HNH->METHYLASE->ParB+HNH->DUF499*->HTH_Tnp_1+HTH_21+rve+rve_3->?->?->AbiEi-><-CHTH<-RelE | 962 | Bacteria>Pseudomonadota>Gammaproteobacteria>Oceanospirillales>Halomonadaceae | GCF_022348135.1 | unclassified_Halomonas | MHM85_RS05495 | 207037..209925 | 2609666 | MULTISPECIES:-DUF499-domain-containing-protein-[unclassified-Halomonas] |
| BBI71108.2 | AAA+ATPase+wHTH+Alpha+Beta-domain+wHTH+RRM | Type-1_BR.RM.like.systems | Phage_integrase-><-Phage_AlpA<-TOPRIM+primpol<-?<-XIS-HTH<-DUF499-p<-DUF499*<-DUF499-p<-REase<-OLD-ABC<-?<-METHYLASE-p<-METHYLASE-p | 386 | Bacteria>Pseudomonadota>Gammaproteobacteria>Oceanospirillales>Halomonadaceae | GCA_007163885.2 | Halomonas_axialensis | HAALTHF_10150n | 665831..666991 | 115555 | hypothetical-protein-HAALTHF_10150n-[Halomonas-axialensis] |
| WP_044628625.1 | AAA+ATPase+wHTH+Alpha+Beta-domain+wHTH+RRM | Type-1_BR.RM.like.systems | METHYLASE->OLD-ABC->REase->DUF499*-><-MobI\|\|XIS-HTH->ZnR+TOPRIM+primpol+HTH->Phage_AlpA-><-Arm-DNA-bind_3+Phage_integrase | 953 | Bacteria>Pseudomonadota>Gammaproteobacteria>Oceanospirillales>Halomonadaceae | GCF_011398715.1 | Halomonas | QWE69_RS13020 | 3280..6141 | 2745 | DUF499-domain-containing-protein-[Halomonas] |
| WP_239295914.1 | AAA+ATPase+wHTH+Alpha+Beta-domain+wHTH+RRM | Type-1_BR.RM.like.systems | <-HTH_Tnp_1+HTH_21+rve+rve_2<-?<-DUF499*<-?<-REase<-OLD-ABC<-METHYLASE<-?<-TUDOR+SF1-SF2-Helicase+iREase+CTD+REase | 958 | Bacteria>Pseudomonadota>Gammaproteobacteria>Oceanospirillales>Halomonadaceae | GCF_022348215.1 | unclassified_Halomonas | MHM38_RS03720 | 8315..11191 | 2609666 | MULTISPECIES:-DUF499-domain-containing-protein-[unclassified-Halomonas] |
| WP_030073674.1 | AAA+ATPase+wHTH+Alpha+Beta-domain+wHTH+RRM | Type-1_BR.RM.like.systems | RelE->CHTH-><-AbiEi\|\|?-><-DUF499*<-McrC-NTD+REase<-MAD-NTD+EVE+McrB-AAA<-OLD-ABC<-?<-METHYLASE<-HNH | 961 | Bacteria>Pseudomonadota>Gammaproteobacteria>Oceanospirillales>Halomonadaceae | GCF_000712975.1 | Halomonas_alkaliantarctica | CR46_RS0113290 | 253918..256803 | 232346 | DUF499-domain-containing-protein-[Halomonas-alkaliantarctica] |
| WP_253017818.1 | AAA+ATPase+wHTH+Alpha+Beta-domain+wHTH+RRM | Type-1_BR.RM.like.systems | HNH->METHYLASE->OLD-ABC->REase->EVE+McrB-AAA->McrC-NTD+REase->DUF499*->?->AbiEi-><-CHTH<-RelE | 961 | Bacteria>Pseudomonadota>Gammaproteobacteria>Oceanospirillales>Halomonadaceae | GCF_024124605.1 | Halomonas_sp._CnH100-B | NH398_RS08385 | 365976..368861 | 2954490 | DUF499-domain-containing-protein-[Halomonas-sp.-CnH100-B] |
| WP_197030572.1 | AAA+ATPase+wHTH+Alpha+Beta-domain+wHTH+RRM | Type-1_BR.RM.like.systems | OLD-ABC->REase->MAD-NTD+EVE+McrB-AAA->?->McrC-NTD+REase->?->DUF499*->DUF499->?-><-?\|\|?->REase-> | 264 | Bacteria>Pseudomonadota>Gammaproteobacteria>Oceanospirillales>Halomonadaceae | GCF_000568385.1 | Halomonas_sp._BC04 | Q427_RS35225 | 32151..32945 | 1403540 | DUF499-domain-containing-protein-[Halomonas-sp.-BC04] |
| EWH01940.1 | AAA+ATPase+wHTH+Alpha+Beta-domain+wHTH+RRM | Type-1_BR.RM.like.systems | REase->MAD-NTD+EVE+McrB-AAA->?->McrC-NTD+REase->?->DUF499-p->DUF499*->?->?-><-?\|\|?->REase-> | 676 | Bacteria>Pseudomonadota>Gammaproteobacteria>Oceanospirillales>Halomonadaceae | GCA_000568385.1 | Halomonas_sp._BC04 | Q427_11510 | 32984..35014 | 1403540 | hypothetical-protein-Q427_11510-[Halomonas-sp.-BC04] |
| WP_234246274.1 | AAA+ATPase+wHTH+Alpha+Beta-domain+wHTH+RRM | Type-1_BR.RM.like.systems | METHYLASE->METHYLASE->Schlafen+HATPase+HTH->HEPN+CBS->OLD-ABC->REase->DUF499*-> | 957 | Bacteria>Pseudomonadota>Gammaproteobacteria>Oceanospirillales>Halomonadaceae | GCF_021404345.1 | Halomonas_desiderata | HOP57_RS15495 | 85349..88222 | 52021 | DUF499-domain-containing-protein-[Halomonas-desiderata] |
| WP_269321216.1 | AAA+ATPase+wHTH+Alpha+Beta-domain+wHTH+RRM | Type-1_BR.RM.like.systems | REase->MAD-NTD+EVE+McrB-AAA->?->McrC-NTD+REase->?->DUF499->DUF499*->?-><-?\|\|?->REase-> | 680 | Bacteria>Pseudomonadota>Gammaproteobacteria>Oceanospirillales>Halomonadaceae | GCF_000568385.1 | Halomonas_sp._BC04 | Q427_RS09405 | 32972..35014 | 1403540 | DUF499-domain-containing-protein-[Halomonas-sp.-BC04] |
| MCD6438846.1 | AAA+ATPase+wHTH+Alpha+Beta-domain+wHTH+RRM | Type-1_BR.RM.like.systems | HNH->METHYLASE->OLD-ABC->REase->EVE+McrB-AAA->McrC-NTD+REase->DUF499*-> | 282 | Bacteria>Pseudomonadota>Gammaproteobacteria>Oceanospirillales>Halomonadaceae | GCA_021161325.1 | Halomonas_sp. | J7L56_11360 | 10325..11171 | 1486246 | ATP-binding-protein-[Halomonas-sp.] |
| WP_161430580.1 | AAA+ATPase+wHTH+Alpha+Beta-domain+wHTH+RRM | Type-1_BR.RM.like.systems | SbcC->?-><-HTH_29+rve+rve_3<-PAIREDC-HTH<-PAPS_reduct<-DUF499*<-DUF4007<-Cysteine_desulfurase<-?<-ParB+HNH\|\|?->HEPN-> | 1106 | Bacteria>Pseudomonadota>Gammaproteobacteria>Oceanospirillales>Halomonadaceae | GCF_009902005.1 | Halomonas_alimentaria | GRB96_RS03575 | 733928..737248 | 147248 | hypothetical-protein-[Halomonas-alimentaria] |
| WP_072841218.1 | AAA+ATPase+wHTH+Alpha+Beta-domain+wHTH+RRM | Type-1_BR.RM.like.systems | TUDOR+SF1-SF2-Helicase+iREase+CTD+REase->OLD-ABC->HNH->METHYLASE->OLD-ABC->REase->DUF499*->SMODS->?-><-RlpA\|\|?-><-HTH_26 | 942 | Bacteria>Pseudomonadota>Gammaproteobacteria>Oceanospirillales>Oceanospirillaceae | GCF_900129155.1 | Marinomonas_polaris | BUA59_RS18830 | 76459..79287 | 293552 | DUF499-domain-containing-protein-[Marinomonas-polaris] |
| MDB9958356.1 | AAA+ATPase+wHTH+Alpha+Beta-domain+wHTH+RRM | Type-1_BR.RM.like.systems | <-DUF499*<-METHYLASE<-HEPN<-TUDOR+SF1-SF2-Helicase+iREase+CTD+REase | 942 | Bacteria>Pseudomonadota>Gammaproteobacteria>Oceanospirillales>Oceanospirillaceae | GCA_028340495.1 | Oceanospirillaceae_bacterium | OAD42_04655 | 239..3067 | 1899355 | DUF499-domain-containing-protein-[Oceanospirillaceae-bacterium] |
| MBE0508133.1 | AAA+ATPase+wHTH+Alpha+Beta-domain+wHTH+RRM | Type-1_BR.RM.like.systems | <-DUF499*<-ParB+HNH<-METHYLASE | 954 | Bacteria>Pseudomonadota>Gammaproteobacteria>Oceanospirillales>Oceanospirillaceae | GCA_014859565.1 | Marinospirillum_sp. | IBX50_15700 | 354..3218 | 2183934 | ATP-binding-protein-[Marinospirillum-sp.] |
| WP_116685595.1 | AAA+ATPase+wHTH+Alpha+Beta-domain+wHTH+RRM | Type-1_BR.RM.like.systems | HTH+WYL->TUDOR+SF1-SF2-Helicase+iREase+CTD+REase->OLD-ABC->HNH->METHYLASE->DUF499*-><-?<-?\|\|?->TIR-> | 935 | Bacteria>Pseudomonadota>Gammaproteobacteria>Oceanospirillales>Oceanospirillales_incertae_sedis | GCF_003097315.1 | Pelagibaculum_spongiae | DC094_RS03025 | 690947..693754 | 2080658 | DUF499-domain-containing-protein-[Pelagibaculum-spongiae] |
| WP_253043929.1 | AAA+ATPase+wHTH+Alpha+Beta-domain+wHTH+RRM | Type-1_BR.RM.like.systems | <-DUF499*<-METHYLASE<-TUDOR+SF1-SF2-Helicase+iREase+CTD+REase<-HTH+WYL\|\|JetA->JetB->SbcC-> | 951 | Bacteria>Pseudomonadota>Gammaproteobacteria>Oceanospirillales>Pleioneaceae | GCF_024124665.1 | Pleionea_sp._CnH1-48 | NH399_RS24165 | 522..3377 | 2954494 | DUF499-domain-containing-protein-[Pleionea-sp.-CnH1-48] |
| WP_200203476.1 | AAA+ATPase+wHTH+Alpha+Beta-domain+wHTH+RRM | Type-1_BR.RM.like.systems | Arm-DNA-bind_3+Phage_integrase-><-Phage_AlpA<-ZnR+TOPRIM+primpol+HTH<-?<-XIS-HTH\|\|<-DUF499*<-Schlafen+HATPase+HTH<-METHYLASE<-METHYLASE<-?<-Schlafen+HATPase+HTH<-TUDOR+SF1-SF2-Helicase+iREase+CTD+REase | 951 | Bacteria>Pseudomonadota>Gammaproteobacteria>Pseudomonadales>Marinobacteraceae | GCF_016597775.1 | Marinobacter_sp._1-3A | FE848_RS12575 | 132618..135473 | 2582920 | DUF499-domain-containing-protein-[Marinobacter-sp.-1-3A] |
| WP_230713447.1 | AAA+ATPase+wHTH+Alpha+Beta-domain+wHTH+RRM | Type-1_BR.RM.like.systems | Phage_integrase-><-Phage_AlpA<-ZnR+TOPRIM+primpol+HTH<-XIS-HTH\|\|MobI-><-DUF499*<-REase<-OLD-ABC<-METHYLASE<-OLD-ABC<-TUDOR+SF1-SF2-Helicase+iREase+CTD+REase<-HTH+WYL | 953 | Bacteria>Pseudomonadota>Gammaproteobacteria>Pseudomonadales>Marinobacteraceae | GCF_021021515.1 | Marinobacter_shengliensis | K7I15_RS06080 | 103063..105924 | 1389223 | DUF499-domain-containing-protein-[Marinobacter-shengliensis] |
| WP_091985110.1 | AAA+ATPase+wHTH+Alpha+Beta-domain+wHTH+RRM | Type-1_BR.RM.like.systems | TUDOR+SF1-SF2-Helicase+iREase+CTD+REase->OLD-ABC->HNH->METHYLASE->MAD-NTD+EVE+McrB-AAA->McrC-NTD+REase->DUF499*->SAM-methylase->HTH+HNH-><-?<-CHTH | 947 | Bacteria>Pseudomonadota>Gammaproteobacteria>Pseudomonadales>Marinobacteraceae | GCF_900115175.1 | Marinobacter_gudaonensis | BM344_RS01255 | 264147..266990 | 375760 | DUF499-domain-containing-protein-[Marinobacter-gudaonensis] |
| WP_098418633.1 | AAA+ATPase+wHTH+Alpha+Beta-domain+wHTH+RRM | Type-1_BR.RM.like.systems | XIS-HTH->TUDOR+SF1-SF2-Helicase+iREase+CTD+REase->OLD-ABC->HNH->METHYLASE->ParB+HNH->DUF499*->GDResolvase+HIN-HTH-><-?<-REase<-HNH<-?\|\|HIN-HTH-> | 946 | Bacteria>Pseudomonadota>Gammaproteobacteria>Pseudomonadales>Marinobacteraceae | GCF_002563885.1 | Marinobacter_sp._LV10MA510-1 | ATI45_RS05630 | 1155568..1158408 | 1415567 | DUF499-domain-containing-protein-[Marinobacter-sp.-LV10MA510-1] |
| AKV97706.1 | AAA+ATPase+wHTH+Alpha+Beta-domain+wHTH+RRM | Type-1_BR.RM.like.systems | ParB->?->REase->?->DDE_Tnp_1_4->VirB4-FtsK+HerA-><-DUF499*<-METHYLASE-p\|\|?-><-HNH<-OLD-ABC<-TUDOR+SF1-SF2-Helicase+iREase+CTD+REase<-XIS-HTH | 951 | Bacteria>Pseudomonadota>Gammaproteobacteria>Pseudomonadales>Marinobacteraceae | GCA_001266795.1 | Marinobacter_sp._CP1 | ACP86_16945 | 3679225..3682080 | 1671721 | hypothetical-protein-ACP86_16945-[Marinobacter-sp.-CP1] |
| WP_200905073.1 | AAA+ATPase+wHTH+Alpha+Beta-domain+wHTH+RRM | Type-1_BR.RM.like.systems | REase->?->DDE_Tnp_1_4->?->VirB4-FtsK+HerA-><-DUF499*<-METHYLASE-p\|\|?-><-METHYLASE<-HNH<-OLD-ABC<-TUDOR+SF1-SF2-Helicase+iREase+CTD+REase | 935 | Bacteria>Pseudomonadota>Gammaproteobacteria>Pseudomonadales>Marinobacteraceae | GCF_001266795.1 | Marinobacter_sp._CP1 | ACP86_RS16950 | 3679225..3682032 | 1671721 | DUF499-domain-containing-protein-[Marinobacter-sp.-CP1] |
| WP_098424145.1 | AAA+ATPase+wHTH+Alpha+Beta-domain+wHTH+RRM | Type-1_BR.RM.like.systems | XIS-HTH->TUDOR+SF1-SF2-Helicase+iREase+CTD+REase->OLD-ABC->HNH->METHYLASE->ParB+HNH->DUF499*->GDResolvase+HIN-HTH-><-?<-REase<-HNH | 946 | Bacteria>Pseudomonadota>Gammaproteobacteria>Pseudomonadales>Marinobacteraceae | GCF_002563815.1 | Marinobacter_sp._LV10R520-4 | ATG98_RS19590 | 4036541..4039381 | 1761796 | DUF499-domain-containing-protein-[Marinobacter-sp.-LV10R520-4] |
| WP_230706347.1 | AAA+ATPase+wHTH+Alpha+Beta-domain+wHTH+RRM | Type-1_BR.RM.like.systems | Phage_integrase-><-Phage_AlpA<-ZnR+TOPRIM+primpol+HTH<-XIS-HTH\|\|MobI-><-DUF499*<-REase<-OLD-ABC\|\|\|\|?-><-?<-?<-Resolvase | 953 | Bacteria>Pseudomonadota>Gammaproteobacteria>Pseudomonadales>Marinobacteraceae | GCF_021021495.1 | Marinobacter_adhaerens | K7I10_RS06135 | 1373639..1376500 | 1033846 | DUF499-domain-containing-protein-[Marinobacter-adhaerens] |
| WP_199457594.1 | AAA+ATPase+wHTH+Alpha+Beta-domain+wHTH+RRM | Type-1_BR.RM.like.systems | <-DUF499*<-ParB+HNH<-METHYLASE | 953 | Bacteria>Pseudomonadota>Gammaproteobacteria>Pseudomonadales>Marinobacteraceae | GCF_016463955.1 | unclassified_Marinobacter | JIF54_RS10850 | 749..3610 | 83889 | MULTISPECIES:-DUF499-domain-containing-protein-[unclassified-Marinobacter] |
| WP_192907661.1 | AAA+ATPase+wHTH+Alpha+Beta-domain+wHTH+RRM | Type-1_BR.RM.like.systems | TUDOR+SF1-SF2-Helicase+iREase+CTD+REase->OLD-ABC->HNH->METHYLASE->HEPN->?->DUF499*-><-RlpA\|\|SF1-SF2-helicase+RdgC+Beta-barrel+REase-><-REase<-Phage_int_SAM<-Integrase_1<-Arm-DNA-bind_3+Phage_integrase | 941 | Bacteria>Pseudomonadota>Gammaproteobacteria>Pseudomonadales>Pseudomonadaceae | GCF_014883875.1 | Pseudomonas_taiwanensis | ICN73_RS10045 | 2229206..2232031 | 470150 | DUF499-domain-containing-protein-[Pseudomonas-taiwanensis] |
| WP_280091279.1 | AAA+ATPase+wHTH+Alpha+Beta-domain+wHTH+RRM | Type-1_BR.RM.like.systems | Arm-DNA-bind_3+Phage_integrase->Phage_int_SAM-><-HNH+SAVED<-SF1-SF2-helicase+RdgC+Beta-barrel+REase\|\|RlpA-><-DUF499*<-HEPN<-METHYLASE<-HNH<-OLD-ABC<-TUDOR+SF1-SF2-Helicase+iREase+CTD+REase | 941 | Bacteria>Pseudomonadota>Gammaproteobacteria>Pseudomonadales>Pseudomonadaceae | GCF_947311875.1 | Pseudomonas | QME24_RS13335 | 66050..68875 | 286 | DUF499-domain-containing-protein-[Pseudomonas-juntendi] |
| MBG7398186.1 | AAA+ATPase+wHTH+Alpha+Beta-domain+wHTH+RRM | Type-1_BR.RM.like.systems | SF1-SF2-Helicase-><-SF1-SF2-helicase+RdgC+Beta-barrel+REase<-35exo+SF1-SF2-Helicase\|\|ABhydrolase->?->RlpA-><-DUF499*<-REase<-OLD-ABC<-METHYLASE<-HEPN<-TUDOR+SF1-SF2-Helicase+iREase+CTD+REase | 942 | Bacteria>Pseudomonadota>Gammaproteobacteria>Pseudomonadales>Pseudomonadaceae | GCA_015766545.1 | Pseudomonas_aeruginosa | GHU66_27420 | 38429..41257 | 287 | ATP-binding-protein-[Pseudomonas-aeruginosa] |
| WP_126634673.1 | AAA+ATPase+wHTH+Alpha+Beta-domain+wHTH+RRM | Type-1_BR.RM.like.systems | <-DDE_Tnp_1\|\|?-><-LZ_Tnp_IS66+zf-IS66+DDE_Tnp_IS66+DDE_Tnp_IS66_C<-?<-HIN-HTH\|\|RlpA-><-DUF499*<-?<-SF1-SF2-Helicase+wHTH+PRTase<-METHYLASE<-HNH<-OLD-ABC<-TUDOR+SF1-SF2-Helicase+iREase+CTD+REase | 943 | Bacteria>Pseudomonadota>Gammaproteobacteria>Pseudomonadales>Pseudomonadaceae | GCF_003970625.1 | Pseudomonas_aeruginosa | E5D53_RS31320 | 11116..13947 | 287 | DUF499-domain-containing-protein-[Pseudomonas-aeruginosa] |
| WP_125867554.1 | AAA+ATPase+wHTH+Alpha+Beta-domain+wHTH+RRM | Type-1_BR.RM.like.systems | SbcC->MORC+DNA_gyraseB+TOPRIM->TUDOR+SF1-SF2-Helicase+iREase+CTD+REase->PIN->METHYLASE->ParB+HNH->DUF499*->HIN-HTH->?->LZ_Tnp_IS66+zf-IS66+DDE_Tnp_IS66+DDE_Tnp_IS66_C-><-?<-?<-Peptidase_S8 | 946 | Bacteria>Pseudomonadota>Gammaproteobacteria>Pseudomonadales>Pseudomonadaceae | - | Stutzerimonas_stutzeri | EGI99_RS04500 | 994408..997248 | 316 | DUF499-domain-containing-protein-[Stutzerimonas-stutzeri] |
| WP_046835095.1 | AAA+ATPase+wHTH+Alpha+Beta-domain+wHTH+RRM | Type-1_BR.RM.like.systems | TUDOR+SF1-SF2-Helicase+iREase+CTD+REase->PIN->METHYLASE-p->DUF772+DDE_Tnp_1->METHYLASE->ParB+HNH->DUF499*-><-RlpA<-?\|\|CHTH-><-LZ_Tnp_IS66+zf-IS66+DDE_Tnp_IS66+DDE_Tnp_IS66_C | 944 | Bacteria>Pseudomonadota>Gammaproteobacteria>Pseudomonadales>Pseudomonadaceae | GCF_030252675.1 | Pseudomonas_amygdali | Q0021_RS27770 | 6030244..6033078 | 47877 | DUF499-domain-containing-protein-[Pseudomonas-amygdali] |
| WP_122622773.1 | AAA+ATPase+wHTH+Alpha+Beta-domain+wHTH+RRM | Type-1_BR.RM.like.systems | <-ParB+HNH\|\|HTH+WYL->TUDOR+SF1-SF2-Helicase+iREase+CTD+REase->OLD-ABC->HNH->METHYLASE->DUF499*-><-RlpA<-Phage_int_SAM<-?<-TOPRIM<-Mbetalac<-Arm-DNA-bind_3+Phage_integrase | 942 | Bacteria>Pseudomonadota>Gammaproteobacteria>Pseudomonadales>Pseudomonadaceae | - | Pseudomonas_viridiflava | EEX91_RS04620 | 101897..104725 | 33069 | DUF499-domain-containing-protein-[Pseudomonas-viridiflava] |
| WP_272438616.1 | AAA+ATPase+wHTH+Alpha+Beta-domain+wHTH+RRM | Type-1_BR.RM.like.systems | HKD-DNase->SF1-SF2-Helicase-><-SF1-SF2-helicase+RdgC+Beta-barrel+REase\|\|?->?->RlpA-><-DUF499*<-REase<-OLD-ABC<-METHYLASE<-HEPN<-TUDOR+SF1-SF2-Helicase+iREase+CTD+REase | 942 | Bacteria>Pseudomonadota>Gammaproteobacteria>Pseudomonadales>Pseudomonadaceae | GCF_028435625.1 | Pseudomonas_aeruginosa | KGZ09_RS30315 | 38429..41257 | 287 | DUF499-domain-containing-protein-[Pseudomonas-aeruginosa] |
| WP_049308056.1 | AAA+ATPase+wHTH+Alpha+Beta-domain+wHTH+RRM | Type-1_BR.RM.like.systems | TUDOR+SF1-SF2-Helicase+iREase+CTD+REase->HEPN->METHYLASE->OLD-ABC->REase->DUF499*-><-RlpA<-?<-?\|\|SF1-SF2-helicase+RdgC+Beta-barrel+REase-><-SF1-SF2-Helicase<-HKD-DNase | 942 | Bacteria>Pseudomonadota>Gammaproteobacteria>Pseudomonadales>Pseudomonadaceae | GCF_001061475.1 | Pseudomonas_aeruginosa | ADM76_RS18680 | 17855..20683 | 287 | DUF499-domain-containing-protein-[Pseudomonas-aeruginosa] |
| WP_128722906.1 | AAA+ATPase+wHTH+Alpha+Beta-domain+wHTH+RRM | Type-1_BR.RM.like.systems | TUDOR+SF1-SF2-Helicase+iREase+CTD+REase->OLD-ABC->HNH->METHYLASE->SF1-SF2-Helicase+wHTH+PRTase->?->DUF499*-><-RlpA<-SymE_toxin<-?<-ZNR+Pkinase+SF1-SF2-Helicase+REase | 942 | Bacteria>Pseudomonadota>Gammaproteobacteria>Pseudomonadales>Pseudomonadaceae | GCF_029954295.1 | Pseudomonas_aeruginosa | OLM70_RS05770 | 1186505..1189333 | 287 | DUF499-domain-containing-protein-[Pseudomonas-aeruginosa] |
| WP_033888172.1 | AAA+ATPase+wHTH+Alpha+Beta-domain+wHTH+RRM | Type-1_BR.RM.like.systems | TUDOR+SF1-SF2-Helicase+iREase+CTD+REase->OLD-ABC->HNH->METHYLASE->SF1-SF2-Helicase+wHTH+PRTase->?->DUF499*-><-RlpA<-SymE_toxin<-?<-ZNR+Pkinase+SF1-SF2-Helicase+REase | 942 | Bacteria>Pseudomonadota>Gammaproteobacteria>Pseudomonadales>Pseudomonadaceae | GCF_000506025.1 | Pseudomonas_aeruginosa | HB728_RS10975 | 2312198..2315026 | 287 | DUF499-domain-containing-protein-[Pseudomonas-aeruginosa] |
| WP_226045701.1 | AAA+ATPase+wHTH+Alpha+Beta-domain+wHTH+RRM | Type-1_BR.RM.like.systems | SbcC->MORC+DNA_gyraseB+TOPRIM->TUDOR+SF1-SF2-Helicase+iREase+CTD+REase->PIN->METHYLASE->ParB+HNH->DUF499*-><-RlpA<-?<-Peptidase_S8<-Classical-AAA<-ParB<-ParB | 946 | Bacteria>Pseudomonadota>Gammaproteobacteria>Pseudomonadales>Pseudomonadaceae | GCF_020401845.1 | Pseudomonas_sp._HS-18 | LDJ84_RS00045 | 14958..17798 | 2879114 | DUF499-domain-containing-protein-[Pseudomonas-sp.-HS-18] |
| WP_213582008.1 | AAA+ATPase+wHTH+Alpha+Beta-domain+wHTH+RRM | Type-1_BR.RM.like.systems | ParB-><-HAD\|\|Classical-AAA->Peptidase_S8-><-?\|\|RlpA-><-DUF499*<-ParB+HNH<-METHYLASE<-TUDOR+SF1-SF2-Helicase+iREase+CTD+REase\|\|?-><-MORC+DNA_gyraseB+TOPRIM<-SbcC | 946 | Bacteria>Pseudomonadota>Gammaproteobacteria>Pseudomonadales>Pseudomonadaceae | GCF_018383465.1 | Pseudomonas_syringae | KID98_RS09080 | 2027689..2030529 | 317 | DUF499-domain-containing-protein-[Pseudomonas-syringae] |
| WP_192069117.1 | AAA+ATPase+wHTH+Alpha+Beta-domain+wHTH+RRM | Type-1_BR.RM.like.systems | Integrase_1->Phage_int_SAM->N6_N4_MTase->REase+SF1-SF2-Helicase->RlpA-><-DUF499*<-REase<-OLD-ABC<-METHYLASE<-TUDOR+SF1-SF2-Helicase+iREase+CTD+REase\|\|?-><-HTH_3 | 941 | Bacteria>Pseudomonadota>Gammaproteobacteria>Pseudomonadales>Pseudomonadaceae | GCF_014842375.1 | Pseudomonas_coleopterorum | IFT40_RS20700 | 77168..79993 | 1605838 | DUF499-domain-containing-protein-[Pseudomonas-coleopterorum] |
| WP_143508093.1 | AAA+ATPase+wHTH+Alpha+Beta-domain+wHTH+RRM | Type-1_BR.RM.like.systems | SbcC->MORC+DNA_gyraseB+TOPRIM->TUDOR+SF1-SF2-Helicase+iREase+CTD+REase->PIN->METHYLASE->ParB+HNH->DUF499*-><-RlpA<-?<-Peptidase_S8<-Classical-AAA<-ParB<-ParB | 946 | Bacteria>Pseudomonadota>Gammaproteobacteria>Pseudomonadales>Pseudomonadaceae | GCF_007049885.1 | Pseudomonas_mendocina | EQ828_RS12690 | 220581..223421 | 300 | DUF499-domain-containing-protein-[Pseudomonas-mendocina] |
| WP_129484085.1 | AAA+ATPase+wHTH+Alpha+Beta-domain+wHTH+RRM | Type-1_BR.RM.like.systems | SbcC->MORC+DNA_gyraseB+TOPRIM->TUDOR+SF1-SF2-Helicase+iREase+CTD+REase->PIN->METHYLASE->ParB+HNH->DUF499*-><-RlpA<-?<-Peptidase_S8<-Classical-AAA<-ParB<-ParB | 946 | Bacteria>Pseudomonadota>Gammaproteobacteria>Pseudomonadales>Pseudomonadaceae | GCF_021495345.1 | Pseudomonas_hydrolytica | L1F06_RS00045 | 14914..17754 | 2493633 | DUF499-domain-containing-protein-[Pseudomonas-hydrolytica] |
| MTI93108.1 | AAA+ATPase+wHTH+Alpha+Beta-domain+wHTH+RRM | Type-1_BR.RM.like.systems | ParB->ParB->Classical-AAA->Peptidase_S8->?->RlpA-><-DUF499*<-ParB+HNH<-METHYLASE<-PIN<-TUDOR+SF1-SF2-Helicase+iREase+CTD+REase<-MORC+DNA_gyraseB+TOPRIM<-SbcC | 946 | Bacteria>Pseudomonadota>Gammaproteobacteria>Pseudomonadales>Pseudomonadaceae | GCA_009711815.1 | Stutzerimonas_stutzeri | FH750_17245 | 14123..16963 | 316 | ATP-binding-protein-[Stutzerimonas-stutzeri] |
| MBG3917698.1 | AAA+ATPase+wHTH+Alpha+Beta-domain+wHTH+RRM | Type-1_BR.RM.like.systems | SbcC->MORC+DNA_gyraseB+TOPRIM->TUDOR+SF1-SF2-Helicase+iREase+CTD+REase->PIN->METHYLASE->ParB+HNH->DUF499*-><-RlpA<-?<-Peptidase_S8<-Classical-AAA<-ParB<-ParB | 946 | Bacteria>Pseudomonadota>Gammaproteobacteria>Pseudomonadales>Pseudomonadaceae | GCA_015732935.1 | Pseudomonas_aeruginosa | I4931_21710 | 92367..95207 | 287 | ATP-binding-protein-[Pseudomonas-aeruginosa] |
| WP_232392569.1 | AAA+ATPase+wHTH+Alpha+Beta-domain+wHTH+RRM | Type-1_BR.RM.like.systems | DUF932->?->ParB->JAB->RlpA-><-DUF499*<-METHYLASE<-HNH<-OLD-ABC<-TUDOR+SF1-SF2-Helicase+iREase+CTD+REase-p<-TUDOR+SF1-SF2-Helicase+iREase+CTD+REase<-HTH+WYL | 921 | Bacteria>Pseudomonadota>Gammaproteobacteria>Pseudomonadales>Pseudomonadaceae | GCF_021228675.1 | Pseudomonas_sp._NIBR-H-19 | LS633_RS06540 | 1376164..1378929 | 2901380 | DUF499-domain-containing-protein-[Pseudomonas-sp.-NIBR-H-19] |
| MCF6804930.1 | AAA+ATPase+wHTH+Alpha+Beta-domain+wHTH+RRM | Type-1_BR.RM.like.systems | ParB->Classical-AAA->Peptidase_S8->?->?->RlpA-><-DUF499*<-ParB+HNH<-METHYLASE<-PIN<-TUDOR+SF1-SF2-Helicase+iREase+CTD+REase<-MORC+DNA_gyraseB+TOPRIM<-SbcC | 889 | Bacteria>Pseudomonadota>Gammaproteobacteria>Pseudomonadales>Pseudomonadaceae | GCA_021713635.1 | Stutzerimonas_stutzeri | L3X13_09150 | 18126..20795 | 316 | DUF499-domain-containing-protein-[Stutzerimonas-stutzeri] |
| MCF6781034.1 | AAA+ATPase+wHTH+Alpha+Beta-domain+wHTH+RRM | Type-1_BR.RM.like.systems | ParB->Classical-AAA->Peptidase_S8->?->?->RlpA-><-DUF499*<-ParB+HNH<-METHYLASE<-PIN<-TUDOR+SF1-SF2-Helicase+iREase+CTD+REase<-MORC+DNA_gyraseB+TOPRIM<-SbcC | 946 | Bacteria>Pseudomonadota>Gammaproteobacteria>Pseudomonadales>Pseudomonadaceae | GCA_021713665.1 | Stutzerimonas_stutzeri | L3X17_04510 | 18122..20962 | 316 | DUF499-domain-containing-protein-[Stutzerimonas-stutzeri] |
| HIQ43647.1 | AAA+ATPase+wHTH+Alpha+Beta-domain+wHTH+RRM | Type-1_BR.RM.like.systems | ParB->ParB-><-MORC+HisKin\|\|<-RVT\|\|?->RlpA-><-DUF499*<-ParB+HNH<-METHYLASE<-PIN<-TUDOR+SF1-SF2-Helicase+iREase+CTD+REase<-MORC+DNA_gyraseB+TOPRIM<-SbcC | 946 | Bacteria>Pseudomonadota>Gammaproteobacteria>Pseudomonadales>Pseudomonadaceae | GCA_015661735.1 | Pseudomonas_oleovorans | EYH47_12960 | 31988..34828 | 301 | ATP-binding-protein-[Pseudomonas-oleovorans] |
| WP_213918359.1 | AAA+ATPase+wHTH+Alpha+Beta-domain+wHTH+RRM | Type-1_BR.RM.like.systems | <-rve_3+Mu-transpos_C<-REase\|\|ParB->RlpA-><-DUF499*<-HEPN<-METHYLASE<-PIN<-TUDOR+SF1-SF2-Helicase+iREase+CTD+REase<-MORC+DNA_gyraseB+TOPRIM<-SbcC | 946 | Bacteria>Pseudomonadota>Gammaproteobacteria>Pseudomonadales>Pseudomonadaceae | GCF_018457005.1 | Stutzerimonas_stutzeri | JR047_RS05385 | 4622..7462 | 316 | DUF499-domain-containing-protein-[Stutzerimonas-stutzeri] |
| WP_085592307.1 | AAA+ATPase+wHTH+Alpha+Beta-domain+wHTH+RRM | Type-1_BR.RM.like.systems | MerR-HTH->tRNA->TUDOR+SF1-SF2-Helicase+iREase+CTD+REase->METHYLASE->OLD-ABC->REase->DUF499*-><-RlpA<-REase+SF1-SF2-Helicase<-N6_N4_MTase\|\|MAD-NTD+HNH-> | 941 | Bacteria>Pseudomonadota>Gammaproteobacteria>Pseudomonadales>Pseudomonadaceae | GCF_002113255.1 | Pseudomonas_sp._B14(2017) | CAG34_RS21645 | 108391..111216 | 1981745 | DUF499-domain-containing-protein-[Pseudomonas-sp.-B14(2017)] |
| WP_020308343.1 | AAA+ATPase+wHTH+Alpha+Beta-domain+wHTH+RRM | Type-1_BR.RM.like.systems | <-RelE-ParE<-RelB<-PIN+OST-HTH<-?\|\|?->RlpA-><-DUF499*<-HEPN<-METHYLASE<-PIN<-TUDOR+SF1-SF2-Helicase+iREase+CTD+REase<-MORC+DNA_gyraseB+TOPRIM<-SbcC | 946 | Bacteria>Pseudomonadota>Gammaproteobacteria>Pseudomonadales>Pseudomonadaceae | - | Stutzerimonas_stutzeri | B382_19940 | 6557..9397 | 316 | DUF499-domain-containing-protein-[Pseudomonas-stutzeri] |
| EPL60615.1 | AAA+ATPase+wHTH+Alpha+Beta-domain+wHTH+RRM | Type-1_BR.RM.like.systems | <-RelE-ParE<-RelB<-PIN+OST-HTH<-?\|\|?->RlpA-><-DUF499*<-HEPN<-METHYLASE<-PIN<-TUDOR+SF1-SF2-Helicase+iREase+CTD+REase<-MORC+DNA_gyraseB+TOPRIM<-SbcC | 946 | Bacteria>Pseudomonadota>Gammaproteobacteria>Pseudomonadales>Pseudomonadaceae | GCA_000416345.1 | Stutzerimonas_stutzeri_B1SMN1 | B382_19940 | 6557..9397 | 1212549 | hypothetical-protein-B382_19940-[Stutzerimonas-stutzeri-B1SMN1] |
| WP_236481327.1 | AAA+ATPase+wHTH+Alpha+Beta-domain+wHTH+RRM | Type-1_BR.RM.like.systems | <-NERD+SF1-SF2-Helicase\|\|RlpA-><-DUF499*<-ParB+HNH<-METHYLASE<-HNH<-OLD-ABC<-TUDOR+SF1-SF2-Helicase+iREase+CTD+REase<-MORC+DNA_gyraseB+TOPRIM | 946 | Bacteria>Pseudomonadota>Gammaproteobacteria>Pseudomonadales>Pseudomonadaceae | - | Pseudomonas_syringae | GIW69_RS06575 | 2105..4945 | 317 | DUF499-domain-containing-protein-[Pseudomonas-syringae] |

| Protein-ID | DUF499-architecture | System | Gene-Neighbourhoods | Length | Taxonomy | Assembly-ID | Strain | Locus-Tag | Genomic-coordinates | TaxID | Fasta-defline |
|---|---|---|---|---|---|---|---|---|---|---|---|
| WP_110599711.1 | AAA+ATPase+wHTH+Alpha+Beta-domain+wHTH+RRM | Type-1_BR.RM.like.systems | ParB-><-PIN<-HTH+MPTase\|\|?->?->RlpA-><-DUF499*<-ParB+HNH<-METHYLASE<-PIN<-TUDOR+SF1-SF2-Helicase+iREase+CTD+REase<-MORC+DNA_gyraseB+TOPRIM<-SbcC | 945 | Bacteria>Pseudomonadota>Gammaproteobacteria>Pseudomonadales>Pseudomonadaceae | - | Pseudomonas_koreensis | DMX04_RS04460 | 134660..137497 | 198620 | DUF499-domain-containing-protein-[Pseudomonas-koreensis] |
| WP_033983017.1 | AAA+ATPase+wHTH+Alpha+Beta-domain+wHTH+RRM | Type-1_BR.RM.like.systems | TUDOR+SF1-SF2-Helicase+iREase+CTD+REase->OLD-ABC->HNH->METHYLASE->SF1-SF2-Helicase+wHTH+PRTase->?->DUF499*-><-RlpA<-?<-JAB<-ParB<-?\|\|MazE_antitoxin-> | 943 | Bacteria>Pseudomonadota>Gammaproteobacteria>Pseudomonadales>Pseudomonadaceae | GCF_000789805.1 | Pseudomonas_aeruginosa | NR66_RS02500 | 146772..149603 | 287 | DUF499-domain-containing-protein-[Pseudomonas-aeruginosa] |
| WP_276226492.1 | AAA+ATPase+wHTH+Alpha+Beta-domain+wHTH+RRM | Type-1_BR.RM.like.systems | TUDOR+SF1-SF2-Helicase+iREase+CTD+REase->OLD-ABC->HNH->METHYLASE->SF1-SF2-Helicase+wHTH+PRTase->?->DUF499*-><-RlpA\|\|<-METHYLASE\|\|HTH->SbcC-> | 939 | Bacteria>Pseudomonadota>Gammaproteobacteria>Pseudomonadales>Pseudomonadaceae | GCF_029335315.1 | Pseudomonas_citronellolis | P3W75_RS06225 | 49283..52102 | 53408 | DUF499-domain-containing-protein-[Pseudomonas-citronellolis] |
| WP_200582036.1 | AAA+ATPase+wHTH+Alpha+Beta-domain+wHTH+RRM | Type-1_BR.RM.like.systems | TUDOR+SF1-SF2-Helicase+iREase+CTD+REase->HEPN->METHYLASE->OLD-ABC->REase->DUF499*-><-RlpA<-?<-SymE_toxin<-?<-ZNR+Pkinase+SF1-SF2-Helicase+iREase | 943 | Bacteria>Pseudomonadota>Gammaproteobacteria>Pseudomonadales>Pseudomonadaceae | GCF_016622185.1 | Pseudomonas_aeruginosa | GFM12_RS08425 | 1738931..1741762 | 287 | DUF499-domain-containing-protein-[Pseudomonas-aeruginosa] |
| WP_033975677.1 | AAA+ATPase+wHTH+Alpha+Beta-domain+wHTH+RRM | Type-1_BR.RM.like.systems | TUDOR+SF1-SF2-Helicase+iREase+CTD+REase->HEPN->METHYLASE->OLD-ABC->REase->DUF499*-><-RlpA<-?<-SymE_toxin<-?<-ZNR+Pkinase+SF1-SF2-Helicase+iREase | 943 | Bacteria>Pseudomonadota>Gammaproteobacteria>Pseudomonadales>Pseudomonadaceae | GCF_022569915.1 | Pseudomonas_aeruginosa | CGU39_RS16575 | 614189..617020 | 287 | DUF499-domain-containing-protein-[Pseudomonas-aeruginosa] |
| WP_279641602.1 | AAA+ATPase+wHTH+Alpha+Beta-domain+wHTH+RRM | Type-1_BR.RM.like.systems | SbcC->MORC+DNA_gyraseB+TOPRIM->TUDOR+SF1-SF2-Helicase+iREase+CTD+REase->PIN->METHYLASE->ParB+HNH->DUF499*->?-><-RlpA<-?\|\|RVT-><-ParB<-ParB | 946 | Bacteria>Pseudomonadota>Gammaproteobacteria>Pseudomonadales>Pseudomonadaceae | GCF_029839225.1 | Stutzerimonas_stutzeri | N5C32_RS13925 | 184183..187023 | 316 | DUF499-domain-containing-protein-[Stutzerimonas-stutzeri] |
| WP_120125136.1 | AAA+ATPase+wHTH+Alpha+Beta-domain+wHTH+RRM | Type-1_BR.RM.like.systems | <-MODE-HTH+PBPII\|\|?->HTH+WYL->TUDOR+SF1-SF2-Helicase+iREase+CTD+REase->METHYLASE->DUF499*->?->RlpA-><-Phage_int_SAM<-Integrase_1<-?<-TOPRIM | 962 | Bacteria>Pseudomonadota>Gammaproteobacteria>Pseudomonadales>Pseudomonadaceae | GCF_003605735.1 | Pseudomonas_sp._LS-2 | D3M70_RS07205 | 26051..28939 | 2315859 | DUF499-domain-containing-protein-[Pseudomonas-sp.-LS-2] |
| WP_232896897.1 | AAA+ATPase+wHTH+Alpha+Beta-domain+wHTH+RRM | Type-1_BR.RM.like.systems | DUF932->?->ParB->JAB-><-?\|\|RlpA-><-DUF499*<-?<-SF1-SF2-Helicase+wHTH+PRTase<-METHYLASE<-HNH<-OLD-ABC<-TUDOR+SF1-SF2-Helicase+iREase+CTD+REase | 928 | Bacteria>Pseudomonadota>Gammaproteobacteria>Pseudomonadales>Pseudomonadaceae | GCF_021283465.2 | Pseudomonas | LU687_RS25005 | 5299214..5302000 | 286 | MULTISPECIES:-DUF499-domain-containing-protein-[Pseudomonas] |
| WP_015271883.1 | AAA+ATPase+wHTH+Alpha+Beta-domain+wHTH+RRM | Type-1_BR.RM.like.systems | DUF932->?->ParB->JAB-><-?\|\|RlpA-><-DUF499*<-?<-SF1-SF2-Helicase+wHTH+PRTase<-METHYLASE<-HNH<-OLD-ABC<-TUDOR+SF1-SF2-Helicase+iREase+CTD+REase | 943 | Bacteria>Pseudomonadota>Gammaproteobacteria>Pseudomonadales>Pseudomonadaceae | GCF_000325725.1 | Pseudomonas | LU639_RS23750 | 5103653..5106484 | 286 | MULTISPECIES:-DUF499-domain-containing-protein-[Pseudomonas] |
| WP_125852814.1 | AAA+ATPase+wHTH+Alpha+Beta-domain+wHTH+RRM | Type-1_BR.RM.like.systems | SbcC->MORC+DNA_gyraseB+TOPRIM->TUDOR+SF1-SF2-Helicase+iREase+CTD+REase->PIN->METHYLASE->ParB+HNH->DUF499*-><-RlpA<-ParB\|\|REase-> | 946 | Bacteria>Pseudomonadota>Gammaproteobacteria>Pseudomonadales>Pseudomonadaceae | GCF_003936145.1 | Stutzerimonas_stutzeri | EGJ42_RS05720 | 52381..55221 | 316 | DUF499-domain-containing-protein-[Stutzerimonas-stutzeri] |
| WP_104882974.1 | AAA+ATPase+wHTH+Alpha+Beta-domain+wHTH+RRM | Type-1_BR.RM.like.systems | DUF932->?->ParB->JAB->RlpA-><-DUF499*<-?<-SF1-SF2-Helicase+wHTH+PRTase<-METHYLASE<-HNH<-OLD-ABC<-TUDOR+SF1-SF2-Helicase+iREase+CTD+REase | 943 | Bacteria>Pseudomonadota>Gammaproteobacteria>Pseudomonadales>Pseudomonadaceae | GCF_002947915.1 | Pseudomonas_sp._SWI44 | C4Q26_RS08410 | 1754987..1757818 | 2083053 | DUF499-domain-containing-protein-[Pseudomonas-sp.-SWI44] |
| WP_285534180.1 | AAA+ATPase+wHTH+Alpha+Beta-domain+wHTH+RRM | Type-1_BR.RM.like.systems | ZnR+ZnR-><-HTH_3\|\|?->?->RlpA-><-DUF499*<-?<-SF1-SF2-Helicase+wHTH+PRTase<-METHYLASE<-HNH<-OLD-ABC<-TUDOR+SF1-SF2-Helicase+iREase+CTD+REase | 942 | Bacteria>Pseudomonadota>Gammaproteobacteria>Pseudomonadales>Pseudomonadaceae | GCF_030269585.1 | Stutzerimonas_stutzeri | QSJ82_RS21895 | 7754..10582 | 316 | DUF499-domain-containing-protein-[Stutzerimonas-stutzeri] |
| WP_219806875.1 | AAA+ATPase+wHTH+Alpha+Beta-domain+wHTH+RRM | Type-1_BR.RM.like.systems | <-MazE_antitoxin\|\|?->ParB->JAB->?->RlpA-><-DUF499*<-REase<-OLD-ABC<-METHYLASE<-HEPN<-TUDOR+SF1-SF2-Helicase+iREase+CTD+REase<-Peptidase | 943 | Bacteria>Pseudomonadota>Gammaproteobacteria>Pseudomonadales>Pseudomonadaceae | GCF_019429735.1 | Pseudomonas_aeruginosa | KZ843_RS17265 | 5657..8488 | 287 | DUF499-domain-containing-protein-[Pseudomonas-aeruginosa] |
| WP_031691594.1 | AAA+ATPase+wHTH+Alpha+Beta-domain+wHTH+RRM | Type-1_BR.RM.like.systems | ZnR+ZnR-><-HTH_3\|\|?->?->RlpA-><-DUF499*<-?<-SF1-SF2-Helicase+wHTH+PRTase<-METHYLASE<-HNH<-OLD-ABC<-TUDOR+SF1-SF2-Helicase+iREase+CTD+REase | 954 | Bacteria>Pseudomonadota>Gammaproteobacteria>Pseudomonadales>Pseudomonadaceae | GCF_000629125.1 | Pseudomonas_aeruginosa | AJ70_RS18015 | 30417..33281 | 287 | DUF499-domain-containing-protein-[Pseudomonas-aeruginosa] |
| WCY08688.1 | AAA+ATPase+wHTH+Alpha+Beta-domain+wHTH+RRM | Type-1_BR.RM.like.systems | TUDOR+SF1-SF2-Helicase+iREase+CTD+REase->OLD-ABC->HNH->SF1-SF2-Helicase+wHTH+PRTase->?->DUF499*-><-RlpA<-?<-JAB<-ParB<-?\|\|MazE_antitoxin-> | 943 | Bacteria>Pseudomonadota>Gammaproteobacteria>Pseudomonadales>Pseudomonadaceae | GCA_028609905.1 | Pseudomonas_aeruginosa | KK229_11705 | 2446793..2449624 | 287 | DUF499-domain-containing-protein-[Pseudomonas-aeruginosa] |
| MBI8725677.1 | AAA+ATPase+wHTH+Alpha+Beta-domain+wHTH+RRM | Type-1_BR.RM.like.systems | TUDOR+SF1-SF2-Helicase+iREase+CTD+REase->OLD-ABC->HNH->METHYLASE->SF1-SF2-Helicase+wHTH+PRTase->?->DUF499*-><-RlpA<-?<-?\|\|HTH_3-><-ZnR+ZnR | 952 | Bacteria>Pseudomonadota>Gammaproteobacteria>Pseudomonadales>Pseudomonadaceae | GCA_016342885.1 | Pseudomonas_aeruginosa | JEZ69_03815 | 204270..207128 | 287 | ATP-binding-protein-[Pseudomonas-aeruginosa] |
| WP_064616121.1 | AAA+ATPase+wHTH+Alpha+Beta-domain+wHTH+RRM | Type-1_BR.RM.like.systems | HTH+WYL->TUDOR+SF1-SF2-Helicase+iREase+CTD+REase->OLD-ABC->HNH->METHYLASE->DUF499*-><-RlpA<-Phage_int_SAM<-Integrase_1<-?<-TOPRIM | 931 | Bacteria>Pseudomonadota>Gammaproteobacteria>Pseudomonadales>Pseudomonadaceae | GCF_001655615.1 | Pseudomonas_sp._GR_6-02 | PGR6_RS03540 | 827904..830699 | 1659194 | DUF499-domain-containing-protein-[Pseudomonas-sp.-GR-6-02] |
| WP_283258799.1 | AAA+ATPase+wHTH+Alpha+Beta-domain+wHTH+RRM | Type-1_BR.RM.like.systems | HIN-HTH->?->RlpA-><-DUF499*<-ParB+HNH<-METHYLASE<-PIN<-TUDOR+SF1-SF2-Helicase+iREase+CTD+REase<-MORC+DNA_gyraseB+TOPRIM<-SbcC | 946 | Bacteria>Pseudomonadota>Gammaproteobacteria>Pseudomonadales>Pseudomonadaceae | GCF_030056015.1 | Stutzerimonas_stutzeri | QM290_RS03915 | 3700..6540 | 316 | DUF499-domain-containing-protein-[Stutzerimonas-stutzeri] |
| WP_279947325.1 | AAA+ATPase+wHTH+Alpha+Beta-domain+wHTH+RRM | Type-1_BR.RM.like.systems | DUF932->?->ParB->JAB->?->RlpA-><-DUF499*<-?<-SF1-SF2-Helicase+wHTH+PRTase<-METHYLASE<-HEPN<-TUDOR+SF1-SF2-Helicase+iREase+CTD+REase | 944 | Bacteria>Pseudomonadota>Gammaproteobacteria>Pseudomonadales>Pseudomonadaceae | GCF_029843075.1 | Stutzerimonas_stutzeri | N5E83_RS18215 | 56624..59458 | 316 | DUF499-domain-containing-protein-[Stutzerimonas-stutzeri] |
| WP_270067442.1 | AAA+ATPase+wHTH+Alpha+Beta-domain+wHTH+RRM | Type-1_BR.RM.like.systems | TUDOR+SF1-SF2-Helicase+iREase+CTD+REase->HEPN->METHYLASE->SF1-SF2-Helicase+wHTH+PRTase->?->DUF499*-><-RlpA<-?<-JAB<-ParB<-?<-DUF932 | 944 | Bacteria>Pseudomonadota>Gammaproteobacteria>Pseudomonadales>Pseudomonadaceae | GCF_027594645.1 | Stutzerimonas_stutzeri | N7388_RS04725 | 1052728..1055562 | 316 | DUF499-domain-containing-protein-[Stutzerimonas-stutzeri] |
| WP_003246716.1 | AAA+ATPase+wHTH+Alpha+Beta-domain+wHTH+RRM | Type-1_BR.RM.like.systems | SbcC->MORC+DNA_gyraseB+TOPRIM->TUDOR+SF1-SF2-Helicase+iREase+CTD+REase->PIN->METHYLASE->ParB+HNH->DUF499*-><-RlpA<-?<-?\|\|RVT-> | 946 | Bacteria>Pseudomonadota>Gammaproteobacteria>Pseudomonadales>Pseudomonadaceae | - | Pseudomonas_mendocina | A471_RS19815 | 19321..22161 | 300 | DUF499-domain-containing-protein-[Pseudomonas-mendocina] |
| WP_093986953.1 | AAA+ATPase+wHTH+Alpha+Beta-domain+wHTH+RRM | Type-1_BR.RM.like.systems | TUDOR+SF1-SF2-Helicase+iREase+CTD+REase->OLD-ABC->HNH->METHYLASE->SF1-SF2-Helicase+wHTH+PRTase->?->DUF499*-><-RlpA<-JAB<-ParB | 943 | Bacteria>Pseudomonadota>Gammaproteobacteria>Pseudomonadales>Pseudomonadaceae | GCF_014635625.1 | Pseudomonas_fluvialis | CFY91_RS13770 | 27997..30828 | 1793966 | DUF499-domain-containing-protein-[Pseudomonas-fluvialis] |
| MCO2346212.1 | AAA+ATPase+wHTH+Alpha+Beta-domain+wHTH+RRM | Type-1_BR.RM.like.systems | TUDOR+SF1-SF2-Helicase+iREase+CTD+REase-><-?\|\|METHYLASE->OLD-ABC->REase->DUF499*-><-RlpA<-?\|\|35exo+SF1-SF2-Helicase-><-?<-HIN-HTH | 942 | Bacteria>Pseudomonadota>Gammaproteobacteria>Pseudomonadales>Pseudomonadaceae | GCA_023907105.1 | Pseudomonas_aeruginosa | FA362_14075 | 17744..20572 | 287 | ATP-binding-protein-[Pseudomonas-aeruginosa] |
| WP_054902927.1 | AAA+ATPase+wHTH+Alpha+Beta-domain+wHTH+RRM | Type-1_BR.RM.like.systems | DUF932->?->ParB->JAB->?->RlpA-><-DUF499*<-REase<-OLD-ABC<-METHYLASE<-HEPN<-TUDOR+SF1-SF2-Helicase+iREase+CTD+REase\|\|HTH+WYL-> | 944 | Bacteria>Pseudomonadota>Gammaproteobacteria>Pseudomonadales>Pseudomonadaceae | GCF_001320525.1 | Pseudomonas_sp._NBRC_111131 | APH34_RS22145 | 20096..22930 | 1661046 | DUF499-domain-containing-protein-[Pseudomonas-sp.-NBRC-111131] |
| HCK4430924.1 | AAA+ATPase+wHTH+Alpha+Beta-domain+wHTH+RRM | Type-1_BR.RM.like.systems | TUDOR+SF1-SF2-Helicase+iREase+CTD+REase->HEPN->METHYLASE->OLD-ABC->TOPRIM->DUF499*-><-RlpA<-?<-JAB<-ParB<-?\|\|MazE_antitoxin-> | 942 | Bacteria>Pseudomonadota>Gammaproteobacteria>Pseudomonadales>Pseudomonadaceae | GCA_024830635.1 | Pseudomonas_aeruginosa | NOG72_005469 | 46146..48974 | 287 | ATP-binding-protein-[Pseudomonas-aeruginosa] |
| WP_201266774.1 | AAA+ATPase+wHTH+Alpha+Beta-domain+wHTH+RRM | Type-1_BR.RM.like.systems | ZNR+Pkinase+SF1-SF2-Helicase+REase->?->SymE_toxin->RlpA-><-DUF499*<-SF1-SF2-Helicase+wHTH+PRTase<-METHYLASE\|\|HIN-HTH-> | 601 | Bacteria>Pseudomonadota>Gammaproteobacteria>Pseudomonadales>Pseudomonadaceae | GCF_003834385.1 | Pseudomonas_aeruginosa | IPC1227_RS29350 | 50900..52705 | 287 | ATP-binding-protein-[Pseudomonas-aeruginosa] |
| WP_054093143.1 | AAA+ATPase+wHTH+Alpha+Beta-domain+wHTH+RRM | Type-1_BR.RM.like.systems | <-PIN<-MazE_antitoxin\|\|?->ParB->JAB->?-><-DUF499*<-METHYLASE<-TUDOR+SF1-SF2-Helicase+iREase+CTD+REase<-HTH+WYL<-?<-tRNA<-MerR-HTH | 943 | Bacteria>Pseudomonadota>Gammaproteobacteria>Pseudomonadales>Pseudomonadaceae | - | Stutzerimonas_degradans | NHN96_001854 | 574742..577573 | 2968968 | DUF499-domain-containing-protein-[Pseudomonas-stutzeri] |
| HCE6219529.1 | AAA+ATPase+wHTH+Alpha+Beta-domain+wHTH+RRM | Type-1_BR.RM.like.systems | <-PIN<-MazE_antitoxin\|\|?->ParB->JAB->?-><-DUF499*<-METHYLASE<-TUDOR+SF1-SF2-Helicase+iREase+CTD+REase<-HTH+WYL<-?<-tRNA<-MerR-HTH | 943 | Bacteria>Pseudomonadota>Gammaproteobacteria>Pseudomonadales>Pseudomonadaceae | GCA_023915785.1 | Pseudomonas_aeruginosa | NHN96_001854 | 574742..577573 | 287 | ATP-binding-protein-[Pseudomonas-aeruginosa] |
| WP_013982993.1 | AAA+ATPase+wHTH+Alpha+Beta-domain+wHTH+RRM | Type-1_BR.RM.like.systems | <-PIN<-MazE_antitoxin\|\|?->ParB->JAB->?-><-DUF499*<-METHYLASE<-TUDOR+SF1-SF2-Helicase+iREase+CTD+REase<-HTH+WYL<-?<-HIN-HTH | 943 | Bacteria>Pseudomonadota>Gammaproteobacteria>Pseudomonadales>Pseudomonadaceae | GCF_000219605.1 | Stutzerimonas_stutzeri | I6G32_RS01545 | 359408..362239 | 316 | DUF499-domain-containing-protein-[Stutzerimonas-stutzeri] |
| WP_090941615.1 | AAA+ATPase+wHTH+Alpha+Beta-domain+wHTH+RRM | Type-1_BR.RM.like.systems | TUDOR+SF1-SF2-Helicase+iREase+CTD+REase->OLD-ABC->HNH->METHYLASE->OLD-ABC->REase->DUF499*-><-RlpA<-?<-?<-ZnR+TOPRIM+D5ATPase | 929 | Bacteria>Pseudomonadota>Gammaproteobacteria>Pseudomonadales>Pseudomonadaceae | GCF_900114395.1 | Azotobacter_beijerinckii | BM240_RS15650 | 17867..20656 | 170623 | DUF499-domain-containing-protein-[Azotobacter-beijerinckii] |
| WP_232885556.1 | AAA+ATPase+wHTH+Alpha+Beta-domain+wHTH+RRM | Type-1_BR.RM.like.systems | <-HTH+WYL\|\|?->TUDOR+SF1-SF2-Helicase+iREase+CTD+REase->OLD-ABC->HNH->METHYLASE->DUF499*-><-RlpA\|\|?-><-JAB<-ParB<-?<-DUF932 | 943 | Bacteria>Pseudomonadota>Gammaproteobacteria>Pseudomonadales>Pseudomonadaceae | GCF_021282765.1 | Pseudomonas | QK328_RS05450 | 308623..311454 | 286 | DUF499-domain-containing-protein-[Pseudomonas-sp.-NMI795_08] |
| WP_124078188.1 | AAA+ATPase+wHTH+Alpha+Beta-domain+wHTH+RRM | Type-1_BR.RM.like.systems | ZnR+ZnR-><-HTH_3\|\|?->?->RlpA-><-DUF499*<-TOPRIM<-OLD-ABC<-METHYLASE<-HNH<-OLD-ABC<-TUDOR+SF1-SF2-Helicase+iREase+CTD+REase | 943 | Bacteria>Pseudomonadota>Gammaproteobacteria>Pseudomonadales>Pseudomonadaceae | GCF_033140865.1 | Pseudomonas_aeruginosa | IPC1239_RS13835 | 644387..647218 | 287 | DUF499-domain-containing-protein-[Pseudomonas-aeruginosa] |
| WP_103518020.1 | AAA+ATPase+wHTH+Alpha+Beta-domain+wHTH+RRM | Type-1_BR.RM.like.systems | <-HTH+WYL\|\|?->TUDOR+SF1-SF2-Helicase+iREase+CTD+REase->OLD-ABC->HNH->METHYLASE->DUF499*-><-RlpA\|\|?-><-JAB<-ParB<-?<-DUF932 | 942 | Bacteria>Pseudomonadota>Gammaproteobacteria>Pseudomonadales>Pseudomonadaceae | GCF_013305665.1 | Pseudomonas | CW309_RS13950 | 90223..93051 | 286 | MULTISPECIES:-DUF499-domain-containing-protein-[Pseudomonas] |
| WP_146538006.1 | AAA+ATPase+wHTH+Alpha+Beta-domain+wHTH+RRM | Type-1_BR.RM.like.systems | TUDOR+SF1-SF2-Helicase+iREase+CTD+REase->OLD-ABC->HNH->METHYLASE->SF1-SF2-Helicase+wHTH+PRTase->?->DUF499*-><-RlpA | 942 | Bacteria>Pseudomonadota>Gammaproteobacteria>Pseudomonadales>Pseudomonadaceae | GCF_007896855.1 | Pseudomonas_aeruginosa | FQA91_RS32330 | 37996..40824 | 287 | DUF499-domain-containing-protein-[Pseudomonas-aeruginosa] |
| WP_135265733.1 | AAA+ATPase+wHTH+Alpha+Beta-domain+wHTH+RRM | Type-1_BR.RM.like.systems | TUDOR+SF1-SF2-Helicase+iREase+CTD+REase->OLD-ABC->HNH->METHYLASE->SF1-SF2-Helicase+wHTH+PRTase->?->DUF499*-><-RlpA | 943 | Bacteria>Pseudomonadota>Gammaproteobacteria>Pseudomonadales>Pseudomonadaceae | GCF_004683755.1 | Stutzerimonas_stutzeri | AK6_RS01615 | 334422..337253 | 316 | DUF499-domain-containing-protein-[Stutzerimonas-stutzeri] |
| WP_131349395.1 | AAA+ATPase+wHTH+Alpha+Beta-domain+wHTH+RRM | Type-1_BR.RM.like.systems | TUDOR+SF1-SF2-Helicase+iREase+CTD+REase->OLD-ABC->HNH->METHYLASE->OLD-ABC->REase->DUF499*-><-RlpA<-TPR<-?<-?\|\|?-><-Mbetalac | 929 | Bacteria>Pseudomonadota>Gammaproteobacteria>Pseudomonadales>Pseudomonadaceae | GCF_004327885.1 | Azotobacter_chroococcum | E0E52_RS08250 | 59995..62784 | 353 | DUF499-domain-containing-protein-[Azotobacter-chroococcum] |
| WP_063540911.1 | AAA+ATPase+wHTH+Alpha+Beta-domain+wHTH+RRM | Type-1_BR.RM.like.systems | MarR-HTH->XIS-HTH->?->HTH+WYL->TUDOR+SF1-SF2-Helicase+iREase+CTD+REase->METHYLASE->DUF499*-><-?<-JAB<-ParB<-?<-DUF932 | 941 | Bacteria>Pseudomonadota>Gammaproteobacteria>Pseudomonadales>Pseudomonadaceae | GCF_001635435.1 | Stutzerimonas_frequens | A3710_RS09070 | 360677..363502 | 2968969 | DUF499-domain-containing-protein-[Stutzerimonas-frequens] |
| WP_031643596.1 | AAA+ATPase+wHTH+Alpha+Beta-domain+wHTH+RRM | Type-1_BR.RM.like.systems | TUDOR+SF1-SF2-Helicase+iREase+CTD+REase->HEPN->METHYLASE->OLD-ABC->REase->DUF499*-><-RlpA<-?<-ABhydrolase<-?<-?\|\|HTH_3-> | 950 | Bacteria>Pseudomonadota>Gammaproteobacteria>Pseudomonadales>Pseudomonadaceae | GCF_000629305.1 | Pseudomonas_aeruginosa | AJ61_RS31490 | 12829..15681 | 287 | DUF499-domain-containing-protein-[Pseudomonas-aeruginosa] |
| WP_023835622.1 | AAA+ATPase+wHTH+Alpha+Beta-domain+wHTH+RRM | Type-1_BR.RM.like.systems | TUDOR+SF1-SF2-Helicase+iREase+CTD+REase->HEPN->METHYLASE->OLD-ABC->REase->DUF499*-><-RlpA<-?<-JAB<-ParB<-?\|\|HIN-HTH-> | 944 | Bacteria>Pseudomonadota>Gammaproteobacteria>Pseudomonadales>Pseudomonadaceae | GCF_000503175.1 | Pseudomonas_aeruginosa | CF344_RS23300 | 12270..15104 | 287 | DUF499-domain-containing-protein-[Pseudomonas-aeruginosa] |
| WP_266022655.1 | AAA+ATPase+wHTH+Alpha+Beta-domain+wHTH+RRM | Type-1_BR.RM.like.systems | TUDOR+SF1-SF2-Helicase+iREase+CTD+REase->HEPN->METHYLASE->OLD-ABC->REase->DUF499*-><-RlpA<-?<-JAB<-ParB<-?<-DUF932 | 948 | Bacteria>Pseudomonadota>Gammaproteobacteria>Pseudomonadales>Pseudomonadaceae | GCF_026241065.1 | unclassified_Pseudomonas | OO409_RS22205 | 32872..35718 | 196821 | MULTISPECIES:-DUF499-domain-containing-protein-[unclassified-Pseudomonas] |
| WP_223119133.1 | AAA+ATPase+wHTH+Alpha+Beta-domain+wHTH+RRM | Type-1_BR.RM.like.systems | ZnR+ZnR-><-HTH_3\|\|?->?->RlpA-><-DUF499*<-HEPN<-METHYLASE<-?<-Schlafen+HATPase+HTH<-TUDOR+SF1-SF2-Helicase+iREase+CTD+REase | 942 | Bacteria>Pseudomonadota>Gammaproteobacteria>Pseudomonadales>Pseudomonadaceae | GCF_020036175.1 | Pseudomonas_aeruginosa | NFY03_RS10210 | 2202477..2205305 | 287 | DUF499-domain-containing-protein-[Pseudomonas-aeruginosa] |
| WP_198580799.1 | AAA+ATPase+wHTH+Alpha+Beta-domain+wHTH+RRM | Type-1_BR.RM.like.systems | TUDOR+SF1-SF2-Helicase+iREase+CTD+REase->HEPN->METHYLASE->OLD-ABC->REase->DUF499*-><-RlpA<-?<-JAB<-ParB<-?<-DUF932 | 944 | Bacteria>Pseudomonadota>Gammaproteobacteria>Pseudomonadales>Pseudomonadaceae | GCF_016134915.1 | Pseudomonas_aeruginosa | QMY34_RS12045 | 381175..384009 | 287 | DUF499-domain-containing-protein-[Pseudomonas-aeruginosa] |
| WP_154355052.1 | AAA+ATPase+wHTH+Alpha+Beta-domain+wHTH+RRM | Type-1_BR.RM.like.systems | TUDOR+SF1-SF2-Helicase+iREase+CTD+REase->HEPN->METHYLASE->OLD-ABC->REase->DUF499*-><-RlpA<-?<-JAB<-ParB<-?<-DUF932 | 942 | Bacteria>Pseudomonadota>Gammaproteobacteria>Pseudomonadales>Pseudomonadaceae | GCF_009676785.1 | Pseudomonas_aeruginosa | GJV02_RS03270 | 700187..703015 | 287 | DUF499-domain-containing-protein-[Pseudomonas-aeruginosa] |
| WP_150035379.1 | AAA+ATPase+wHTH+Alpha+Beta-domain+wHTH+RRM | Type-1_BR.RM.like.systems | TUDOR+SF1-SF2-Helicase+iREase+CTD+REase->Schlafen+HATPase+HTH->?->METHYLASE->HEPN->DUF499*-><-RlpA<-?<-?\|\|HTH_3-><-ZnR+ZnR | 942 | Bacteria>Pseudomonadota>Gammaproteobacteria>Pseudomonadales>Pseudomonadaceae | GCF_022627285.1 | Pseudomonas_aeruginosa | JNB05_RS00615 | 132976..135804 | 287 | DUF499-domain-containing-protein-[Pseudomonas-aeruginosa] |
| WP_132674023.1 | AAA+ATPase+wHTH+Alpha+Beta-domain+wHTH+RRM | Type-1_BR.RM.like.systems | DUF932->?->ParB->JAB->?->RlpA-><-DUF499*<-REase<-OLD-ABC<-METHYLASE<-HEPN<-TUDOR+SF1-SF2-Helicase+iREase+CTD+REase | 944 | Bacteria>Pseudomonadota>Gammaproteobacteria>Pseudomonadales>Pseudomonadaceae | - | Pseudomonas_aeruginosa | EQ623_RS00180 | 30601..33435 | 287 | DUF499-domain-containing-protein-[Pseudomonas-aeruginosa] |
| WP_074197337.1 | AAA+ATPase+wHTH+Alpha+Beta-domain+wHTH+RRM | Type-1_BR.RM.like.systems | DUF932->?->ParB->JAB->?->RlpA-><-DUF499*<-REase<-OLD-ABC<-METHYLASE<-HEPN<-TUDOR+SF1-SF2-Helicase+iREase+CTD+REase | 946 | Bacteria>Pseudomonadota>Gammaproteobacteria>Pseudomonadales>Pseudomonadaceae | GCF_001909445.1 | Pseudomonas_aeruginosa | BH596_RS15220 | 11473..14313 | 287 | DUF499-domain-containing-protein-[Pseudomonas-aeruginosa] |
| WP_033979009.1 | AAA+ATPase+wHTH+Alpha+Beta-domain+wHTH+RRM | Type-1_BR.RM.like.systems | ZnR+ZnR-><-HTH_3\|\|?->?->RlpA-><-DUF499*<-HEPN<-METHYLASE<-?<-Schlafen+HATPase+HTH<-TUDOR+SF1-SF2-Helicase+iREase+CTD+REase | 943 | Bacteria>Pseudomonadota>Gammaproteobacteria>Pseudomonadales>Pseudomonadaceae | - | Pseudomonas_aeruginosa | CJT50_RS28650 | 19614..22445 | 287 | DUF499-domain-containing-protein-[Pseudomonas-aeruginosa] |
| MCO2926705.1 | AAA+ATPase+wHTH+Alpha+Beta-domain+wHTH+RRM | Type-1_BR.RM.like.systems | TUDOR+SF1-SF2-Helicase+iREase+CTD+REase->Schlafen+HATPase+HTH->?->METHYLASE->HEPN->DUF499*-><-RlpA<-?<-?\|\|HTH_3-><-ZnR+ZnR | 943 | Bacteria>Pseudomonadota>Gammaproteobacteria>Pseudomonadales>Pseudomonadaceae | GCA_023909145.1 | Pseudomonas_aeruginosa | FA260_26715 | 18266..21097 | 287 | ATP-binding-protein-[Pseudomonas-aeruginosa] |
| WP_151883645.1 | AAA+ATPase+wHTH+Alpha+Beta-domain+wHTH+RRM | Type-1_BR.RM.like.systems | TUDOR+SF1-SF2-Helicase+iREase+CTD+REase->OLD-ABC->HNH->METHYLASE->DUF499*-><-RlpA\|\|?-><-JAB<-ParB<-?<-DUF932 | 943 | Bacteria>Pseudomonadota>Gammaproteobacteria>Pseudomonadales>Pseudomonadaceae | GCF_009012945.1 | Pseudomonas | GB470_RS16305 | 7805..10636 | 286 | MULTISPECIES:-DUF499-domain-containing-protein-[Pseudomonas] |
| WP_033982644.1 | AAA+ATPase+wHTH+Alpha+Beta-domain+wHTH+RRM | Type-1_BR.RM.like.systems | ABhydrolase->?->RlpA-><-DUF499*<-REase<-OLD-ABC<-METHYLASE<-HEPN<-TUDOR+SF1-SF2-Helicase+iREase+CTD+REase | 942 | Bacteria>Pseudomonadota>Gammaproteobacteria>Pseudomonadales>Pseudomonadaceae | GCF_900707645.1 | Pseudomonas_aeruginosa | BUH55_RS18155 | 7471..10299 | 287 | DUF499-domain-containing-protein-[Pseudomonas-aeruginosa] |
| HCI3160560.1 | AAA+ATPase+wHTH+Alpha+Beta-domain+wHTH+RRM | Type-1_BR.RM.like.systems | ABhydrolase->?->RlpA-><-DUF499*<-REase<-OLD-ABC<-METHYLASE<-HEPN<-TUDOR+SF1-SF2-Helicase+iREase+CTD+REase | 942 | Bacteria>Pseudomonadota>Gammaproteobacteria>Pseudomonadales>Pseudomonadaceae | GCA_024356045.1 | Pseudomonas_aeruginosa | NOV63_004506 | 69244..72072 | 287 | ATP-binding-protein-[Pseudomonas-aeruginosa] |
| HBO5114778.1 | AAA+ATPase+wHTH+Alpha+Beta-domain+wHTH+RRM | Type-1_BR.RM.like.systems | TUDOR+SF1-SF2-Helicase+iREase+CTD+REase->HEPN->METHYLASE->OLD-ABC->REase->DUF499*-><-RlpA<-?<-ABhydrolase | 946 | Bacteria>Pseudomonadota>Gammaproteobacteria>Pseudomonadales>Pseudomonadaceae | GCA_021662155.1 | Pseudomonas_aeruginosa | L4W37_005387 | 12730..15570 | 287 | ATP-binding-protein-[Pseudomonas-aeruginosa] |
| WP_213870614.1 | AAA+ATPase+wHTH+Alpha+Beta-domain+wHTH+RRM | Type-1_BR.RM.like.systems | JAB->?-><-HTH_3\|\|?->?->RlpA-><-DUF499*<-HEPN<-METHYLASE<-HNH<-OLD-ABC<-TUDOR+SF1-SF2-Helicase+iREase+CTD+REase | 952 | Bacteria>Pseudomonadota>Gammaproteobacteria>Pseudomonadales>Pseudomonadaceae | GCF_029201245.1 | Pseudomonas_aeruginosa | KKT79_RS19565 | 32215..35073 | 287 | DUF499-domain-containing-protein-[Pseudomonas-aeruginosa] |
| WP_090734260.1 | AAA+ATPase+wHTH+Alpha+Beta-domain+wHTH+RRM | Type-1_BR.RM.like.systems | TUDOR+SF1-SF2-Helicase+iREase+CTD+REase->?->METHYLASE->OLD-ABC->REase->DUF499*-><-RlpA<-?\|\|PAIREDC-HTH-> | 929 | Bacteria>Pseudomonadota>Gammaproteobacteria>Pseudomonadales>Pseudomonadaceae | GCF_900108885.1 | Azotobacter_beijerinckii | BMW56_RS18070 | 16681..19470 | 170623 | DUF499-domain-containing-protein-[Azotobacter-beijerinckii] |
| WP_165892256.1 | AAA+ATPase+wHTH+Alpha+Beta-domain+wHTH+RRM | Type-1_BR.RM.like.systems | TPR->RlpA-><-DUF499*<-ParB+HNH<-METHYLASE<-HNH<-OLD-ABC<-TUDOR+SF1-SF2-Helicase+iREase+CTD+REase | 933 | Bacteria>Pseudomonadota>Gammaproteobacteria>Pseudomonadales>Pseudomonadaceae | GCF_011470035.1 | Azotobacter_chroococcum | HA520_RS07840 | 19913..22714 | 353 | DUF499-domain-containing-protein-[Azotobacter-chroococcum] |
| WP_128472827.1 | AAA+ATPase+wHTH+Alpha+Beta-domain+wHTH+RRM | Type-1_BR.RM.like.systems | RlpA-><-DUF499*<-TOPRIM<-OLD-ABC<-METHYLASE<-HEPN<-TUDOR+SF1-SF2-Helicase+iREase+CTD+REase | 943 | Bacteria>Pseudomonadota>Gammaproteobacteria>Pseudomonadales>Pseudomonadaceae | GCF_004053955.1 | Pseudomonas_aeruginosa | EQH74_RS32310 | 490..3321 | 287 | DUF499-domain-containing-protein-[Pseudomonas-aeruginosa] |
| HCL3757805.1 | AAA+ATPase+wHTH+Alpha+Beta-domain+wHTH+RRM | Type-1_BR.RM.like.systems | TUDOR+SF1-SF2-Helicase+iREase+CTD+REase->OLD-ABC->HNH->METHYLASE->OLD-ABC->TOPRIM->DUF499*-> | 667 | Bacteria>Pseudomonadota>Gammaproteobacteria>Pseudomonadales>Pseudomonadaceae | GCA_025003865.1 | Pseudomonas_aeruginosa | N2B17_005733 | 10859..12860 | 287 | ATP-binding-protein-[Pseudomonas-aeruginosa] |
| WP_262384438.1 | AAA+ATPase+wHTH+Alpha+Beta-domain+wHTH+RRM | Type-1_BR.RM.like.systems | PIN->METHYLASE->METHYLASE-p->ParB+HNH->DUF499*-><-?\|\|HTH+MPTase->PIN->HAD->HAD-><-ParB | 415 | Bacteria>Pseudomonadota>Gammaproteobacteria>Pseudomonadales>Pseudomonadaceae | GCF_012986835.1 | Pseudomonas_fragi | HBN88_RS02020 | 44440..45687 | 296 | DUF499-domain-containing-protein-[Pseudomonas-fragi] |
| MCR7692359.1 | AAA+ATPase+wHTH+Alpha+Beta-domain+wHTH+RRM | Type-1_BR.RM.like.systems | TUDOR+SF1-SF2-Helicase+iREase+CTD+REase->HEPN->METHYLASE->OLD-ABC->REase->DUF499*-> | 417 | Bacteria>Pseudomonadota>Gammaproteobacteria>Pseudomonadales>Pseudomonadaceae | GCA_024717035.1 | Pseudomonas_aeruginosa | NTA33_29240 | 17686..18936 | 287 | DUF499-domain-containing-protein-[Pseudomonas-aeruginosa] |
| WP_003291474.1 | AAA+ATPase+wHTH+Alpha+Beta-domain+wHTH+RRM | Type-1_BR.RM.like.systems | HTH+WYL->TUDOR+SF1-SF2-Helicase+iREase+CTD+REase-><-?\|\|METHYLASE-p->DUF499*-><-?<-JAB | 943 | Bacteria>Pseudomonadota>Gammaproteobacteria>Pseudomonadales>Pseudomonadaceae | - | Stutzerimonas_stutzeri | YO5_00765 | 7551..10382 | 316 | DUF499-domain-containing-protein-[Stutzerimonas-stutzeri] |
| EIK51241.1 | AAA+ATPase+wHTH+Alpha+Beta-domain+wHTH+RRM | Type-1_BR.RM.like.systems | HTH+WYL->TUDOR+SF1-SF2-Helicase+iREase+CTD+REase-><-?\|\|METHYLASE-p->DUF499*-><-?<-JAB | 943 | Bacteria>Pseudomonadota>Gammaproteobacteria>Pseudomonadales>Pseudomonadaceae | GCA_000263395.1 | Stutzerimonas_stutzeri_TS44 | YO5_00765 | 7551..10382 | 477228 | hypothetical-protein-YO5_00765-[Stutzerimonas-stutzeri-TS44] |
| WP_188258395.1 | AAA+ATPase+wHTH+Alpha+Beta-domain+wHTH+RRM | Type-1_BR.RM.like.systems | DUF932->?->ParB->JAB->?->RlpA-><-DUF499*<-?<-SF1-SF2-Helicase+wHTH+PRTase<-METHYLASE-p | 943 | Bacteria>Pseudomonadota>Gammaproteobacteria>Pseudomonadales>Pseudomonadaceae | GCF_014596275.1 | Pseudomonas_aeruginosa | ID863_RS29425 | 26310..29141 | 287 | DUF499-domain-containing-protein-[Pseudomonas-aeruginosa] |
| WP_130888416.1 | AAA+ATPase+wHTH+Alpha+Beta-domain+wHTH+RRM | Type-1_BR.RM.like.systems | <-HTH_3\|\|?->?-><-RlpA<-DUF499*<-METHYLASE<-TUDOR+SF1-SF2-Helicase+iREase+CTD+REase | 960 | Bacteria>Pseudomonadota>Gammaproteobacteria>Pseudomonadales>Pseudomonadaceae | GCF_023241715.1 | Pseudomonas | M1B34_RS15405 | 61298..64180 | 286 | DUF499-domain-containing-protein-[Pseudomonas-sp.-Sample_21] |
| WP_037052364.1 | AAA+ATPase+wHTH+Alpha+Beta-domain+wHTH+RRM | Type-1_BR.RM.like.systems | ParB->Classical-AAA->Peptidase_S8->?->?->RlpA-><-DUF499*<-ParB+HNH | 946 | Bacteria>Pseudomonadota>Gammaproteobacteria>Pseudomonadales>Pseudomonadaceae | - | Pseudomonas_oleovorans | GL31_RS12560 | 122696..125536 | 301 | DUF499-domain-containing-protein-[Pseudomonas-oleovorans] |
| WP_161612279.1 | AAA+ATPase+wHTH+Alpha+Beta-domain+wHTH+RRM | Type-1_BR.RM.like.systems | METHYLASE->ParB+HNH->DUF499*->?-><-RlpA<-?\|\|RVT-><-ParB<-ParB | 946 | Bacteria>Pseudomonadota>Gammaproteobacteria>Pseudomonadales>Pseudomonadaceae | - | Stutzerimonas_stutzeri | GXB82_RS15495 | 4443..7283 | 316 | DUF499-domain-containing-protein-[Stutzerimonas-stutzeri] |
| HCF1493245.1 | AAA+ATPase+wHTH+Alpha+Beta-domain+wHTH+RRM | Type-1_BR.RM.like.systems | <-MORC+HisKin<-RVT\|\|?->?->RlpA-><-?<-DUF499*<-HEPN<-METHYLASE-p | 948 | Bacteria>Pseudomonadota>Gammaproteobacteria>Pseudomonadales>Pseudomonadaceae | GCA_023941105.1 | Pseudomonas_aeruginosa | NH848_005990 | 4619..7465 | 287 | ATP-binding-protein-[Pseudomonas-aeruginosa] |
| WP_049271643.1 | AAA+ATPase+wHTH+Alpha+Beta-domain+wHTH+RRM | Type-1_BR.RM.like.systems | ParB->JAB->?->RlpA-><-DUF499*<-REase<-OLD-ABC<-METHYLASE | 943 | Bacteria>Pseudomonadota>Gammaproteobacteria>Pseudomonadales>Pseudomonadaceae | GCF_001077135.1 | Pseudomonas_aeruginosa | ADL79_RS26490 | 2020..4851 | 287 | DUF499-domain-containing-protein-[Pseudomonas-aeruginosa] |
| TXH86520.1 | AAA+ATPase+wHTH+Alpha+Beta-domain+wHTH+RRM | Type-1_BR.RM.like.systems | JAB->RlpA-><-DUF499*<-TOPRIM<-OLD-ABC<-METHYLASE | 942 | Bacteria>Pseudomonadota>Gammaproteobacteria>Pseudomonadales>Pseudomonadaceae | GCA_008015405.1 | Pseudomonas_sp. | E6Q71_06445 | 770..3598 | 306 | ATP-binding-protein-[Pseudomonas-sp.] |
| WP_079328502.1 | AAA+ATPase+wHTH+Alpha+Beta-domain+wHTH+RRM | Type-1_BR.RM.like.systems | ParB+HNH->DUF499*-><-RlpA<-?<-?\|\|RVT->RelB-><-ParB | 946 | Bacteria>Pseudomonadota>Gammaproteobacteria>Pseudomonadales>Pseudomonadaceae | GCF_002027175.1 | Stutzerimonas_stutzeri | B2J73_RS08375 | 2891..5731 | 316 | DUF499-domain-containing-protein-[Stutzerimonas-stutzeri] |
| WP_279640682.1 | AAA+ATPase+wHTH+Alpha+Beta-domain+wHTH+RRM | Type-1_BR.RM.like.systems | HEPN->DUF499*-><-RlpA<-?<-?\|\|?->PIN+OST-HTH->RelB-> | 946 | Bacteria>Pseudomonadota>Gammaproteobacteria>Pseudomonadales>Pseudomonadaceae | GCF_029835615.1 | Stutzerimonas_stutzeri | N7363_RS16470 | 1848..4688 | 316 | DUF499-domain-containing-protein-[Stutzerimonas-stutzeri] |
| WP_263875167.1 | AAA+ATPase+wHTH+Alpha+Beta-domain+wHTH+RRM | Type-1_BR.RM.like.systems | DUF499*-><-RlpA<-?<-JAB<-ParB<-?<-DUF932 | 749 | Bacteria>Pseudomonadota>Gammaproteobacteria>Pseudomonadales>Pseudomonadaceae | GCF_025799375.1 | Pseudomonas_aeruginosa | OFL37_RS34015 | 83..2332 | 287 | DUF499-domain-containing-protein-[Pseudomonas-aeruginosa] |
| MCO2182137.1 | AAA+ATPase+wHTH+Alpha+Beta-domain+wHTH+RRM | Type-1_BR.RM.like.systems | SymE_toxin->RlpA-><-DUF499* | 927 | Bacteria>Pseudomonadota>Gammaproteobacteria>Pseudomonadales>Pseudomonadaceae | GCA_023906505.1 | Pseudomonas_aeruginosa | FA392_27080 | 1021..3806 | 287 | ATP-binding-protein-[Pseudomonas-aeruginosa] |
| UPL05438.1 | AAA+ATPase+wHTH+Alpha+Beta-domain+wHTH+RRM | Type-1_BR.RM.like.systems | JAB->DUF4007->DUF499*->PAPS_reduct->HNH-><-?<-MORC+HISKIN | 1110 | Bacteria>Pseudomonadota>Gammaproteobacteria>Pseudomonadales>Pseudomonadaceae | GCA_023101365.1 | Pseudomonas_sp._IsoF | PisoF_01086 | 1193586..1196918 | 2821559 | ATP-binding-protein-[Pseudomonas-sp.-IsoF] |
| MCG8614350.1 | AAA+ATPase+wHTH+Alpha+Beta-domain+wHTH+RRM | Type-1_BR.RM.like.systems | HISKIN-><-?\|\|?->Phage_int_SAM_4-><-?\|\|?-><-DUF499*<-TOPRIM<-OLD-ABC<-METHYLASE<-TUDOR+SF1-SF2-Helicase+iREase+CTD+REase-p | 947 | Bacteria>Pseudomonadota>Gammaproteobacteria>Pseudomonadales>unclassified_Pseudomonadales | GCA_022361325.1 | Pseudomonadales_bacterium | MI864_27885 | 9196..12039 | 1891229 | DUF499-domain-containing-protein-[Pseudomonadales-bacterium] |
| MCP5147023.1 | AAA+ATPase+wHTH+Alpha+Beta-domain+wHTH+RRM | Type-1_BR.RM.like.systems | GNAT->?->?-><-?<-DUF499*<-METHYLASE<-TUDOR+SF1-SF2-Helicase+iREase+CTD+REase<-RHH<-?<-?<-ArdCN+MPTase | 933 | Bacteria>Pseudomonadota>Gammaproteobacteria>Pseudomonadales>unclassified_Pseudomonadales | GCA_024234795.1 | Pseudomonadales_bacterium | H6986_02770 | 290054..292855 | 1891229 | ATP-binding-protein-[Pseudomonadales-bacterium] |
| MCB1656848.1 | AAA+ATPase+wHTH+Alpha+Beta-domain+wHTH+RRM | Type-1_BR.RM.like.systems | <-HIN-HTH\|\|ParB-><-DUF499*<-ParB+HNH<-METHYLASE | 939 | Bacteria>Pseudomonadota>Gammaproteobacteria>Pseudomonadales>unclassified_Pseudomonadales | GCA_020443205.1 | Pseudomonadales_bacterium | KDI39_01315 | 4825..7644 | 1891229 | ATP-binding-protein-[Pseudomonadales-bacterium] |
| MAS10374.1 | AAA+ATPase+wHTH+Alpha+Beta-domain+wHTH+RRM | Type-1_BR.RM.like.systems | <-HhH-RADC+JAB\|\|?->?->?-><-DUF499*<-TOPRIM<-OLD-ABC<-METHYLASE<-PIN<-TUDOR+SF1-SF2-Helicase+iREase+CTD+REase<-JetD | 942 | Bacteria>Pseudomonadota>Gammaproteobacteria>Salinisphaerales>Salinisphaeraceae | GCA_002702145.1 | Salinisphaera_sp. | CMN26_09985 | 5272..8100 | 1914330 | hypothetical-protein-[Salinisphaera-sp.] |
| WP_203168599.1 | AAA+ATPase+wHTH+Alpha+Beta-domain+wHTH+RRM | Type-1_BR.RM.like.systems | SF1-SF2-Helicase+DUF1998->SF1-SF2-Helicase+DUF1998->?-><-?<-DUF499*<-REase<-METHYLASE<-TUDOR+SF1-SF2-Helicase+iREase+CTD+REase<-XIS-HTH | 913 | Bacteria>Pseudomonadota>Gammaproteobacteria>Steroidobacterales>Steroidobacteraceae | GCF_016801985.1 | Steroidobacter_gossypii | JM946_RS17390 | 830297..833038 | 2805490 | DUF499-domain-containing-protein-[Steroidobacter-gossypii] |
| WP_054965897.1 | AAA+ATPase+wHTH+Alpha+Beta-domain+wHTH+RRM | Type-1_BR.RM.like.systems | <-REase<-?<-?\|\|TUDOR+SF1-SF2-Helicase+iREase+CTD+REase->METHYLASE->DUF499*->ArdCN+MPTase->DUF6361->HKD-DNase->SF1-SF2-Helicase-> | 947 | Bacteria>Pseudomonadota>Gammaproteobacteria>Thiohalorhabdales>Thiohalorhabdaceae | GCF_900101365.1 | Thiohalorhabdus_denitrificans | BLP36_RS04835 | 12631..15474 | 381306 | DUF499-domain-containing-protein-[Thiohalorhabdus-denitrificans] |
| WP_029406945.1 | AAA+ATPase+wHTH+Alpha+Beta-domain+wHTH+RRM | Type-1_BR.RM.like.systems | Phage_integrase-><-GDResolvase+HIN-HTH<-DUF499*<-METHYLASE<-HNH<-OLD-ABC<-TUDOR+SF1-SF2-Helicase+iREase+CTD+REase<-?<-HTH | 941 | Bacteria>Pseudomonadota>Gammaproteobacteria>Thiotrichales>Piscirickettsiaceae | GCF_000702325.1 | Thiomicrorhabdus_sp._Milos-T2 | BS34_RS0102315 | 537290..540115 | 90814 | DUF499-domain-containing-protein-[Thiomicrorhabdus-sp.-Milos-T2] |
| WP_020558458.1 | AAA+ATPase+wHTH+Alpha+Beta-domain+wHTH+RRM | Type-1_BR.RM.like.systems | 35exo->?->?->?-><-DUF499*<-METHYLASE<-HNH<-OLD-ABC<-TUDOR+SF1-SF2-Helicase+iREase+CTD+REase<-SF1-SF2-Helicase | 942 | Bacteria>Pseudomonadota>Gammaproteobacteria>Thiotrichales>Thiofilaceae | GCF_000380185.1 | Thiofilum_flexile | A3IK_RS0102855 | 95361..98189 | 125627 | DUF499-domain-containing-protein-[Thiofilum-flexile] |
| WP_103920297.1 | AAA+ATPase+wHTH+Alpha+Beta-domain+wHTH+RRM | Type-1_BR.RM.like.systems | TUDOR+SF1-SF2-Helicase+iREase+CTD+REase->OLD-ABC->HNH->METHYLASE->DUF499*->?->?->ParA-Soj-PloopNTPase-> | 934 | Bacteria>Pseudomonadota>Gammaproteobacteria>Thiotrichales>Thiotrichaceae | GCF_900099115.1 | Candidatus_Venteria_ishoeyi | MBHS_RS13590 | 35251..38055 | 1899563 | DUF499-domain-containing-protein-[Candidatus-Venteria-ishoeyi] |
| SEH06532.1 | AAA+ATPase+wHTH+Alpha+Beta-domain+wHTH+RRM | Type-1_BR.RM.like.systems | TUDOR+SF1-SF2-Helicase+iREase+CTD+REase->OLD-ABC->HNH->METHYLASE->DUF499*->?->?->?->ParA-Soj-PloopNTPase-> | 934 | Bacteria>Pseudomonadota>Gammaproteobacteria>Thiotrichales>Thiotrichaceae | GCA_900099115.2 | Candidatus_Venteria_ishoeyi | MBHS_02395 | 35251..38055 | 1899563 | Uncharacterised-protein-[Thiotrichales-bacterium-HS_08] |
| WP_109837975.1 | AAA+ATPase+wHTH+Alpha+Beta-domain+wHTH+RRM | Type-1_BR.RM.like.systems | <-ORC-CDC6+REase\|\|TUDOR+SF1-SF2-Helicase+iREase+CTD+REase->OLD-ABC->HNH->METHYLASE->DUF499*-> | 944 | Bacteria>Pseudomonadota>Gammaproteobacteria>Thiotrichales>Thiotrichaceae | GCF_003172875.1 | Leucothrix_pacifica | DKW60_RS12435 | 56874..59708 | 1247513 | DUF499-domain-containing-protein-[Leucothrix-pacifica] |
| WP_078924154.1 | AAA+ATPase+wHTH+Alpha+Beta-domain+wHTH+RRM | Type-1_BR.RM.like.systems | TUDOR+SF1-SF2-Helicase+iREase+CTD+REase->OLD-ABC->HNH->METHYLASE->DUF499*-> | 937 | Bacteria>Pseudomonadota>Gammaproteobacteria>Thiotrichales>Thiotrichaceae | GCF_900167255.1 | Thiothrix_eikelboomii | B5D50_RS18550 | 28151..30964 | 92487 | DUF499-domain-containing-protein-[Thiothrix-eikelboomii] |
| MCP5421179.1 | AAA+ATPase+wHTH+Alpha+Beta-domain+wHTH+RRM | Type-1_BR.RM.like.systems | GDResolvase+HIN-HTH->?->TUDOR+SF1-SF2-Helicase+iREase+CTD+REase->METHYLASE->ParB+HNH->DUF499*->wHTH+REase->SbcC-><-?\|\|BrnT_toxin->BrnA_antitoxin-> | 941 | Bacteria>Pseudomonadota>Gammaproteobacteria>unclassified_Gammaproteobacteria | GCA_024235155.1 | Gammaproteobacteria_bacterium | H6969_11925 | 8981..11806 | 1913989 | ATP-binding-protein-[Gammaproteobacteria-bacterium] |
| MBK9130403.1 | AAA+ATPase+wHTH+Alpha+Beta-domain+wHTH+RRM | Type-1_BR.RM.like.systems | TraH->?->TraG_N-><-?<-DUF499*<-METHYLASE<-TUDOR+SF1-SF2-Helicase+iREase+CTD+REase<-HTH+WYL\|\|KAP-NTPase->?->PAPS_reduct-> | 949 | Bacteria>Pseudomonadota>Gammaproteobacteria>unclassified_Gammaproteobacteria | GCA_016716465.1 | Gammaproteobacteria_bacterium | IPM20_01990 | 416080..418929 | 1913989 | ATP-binding-protein-[Gammaproteobacteria-bacterium] |
| TAK54048.1 | AAA+ATPase+wHTH+Alpha+Beta-domain+wHTH+RRM | Type-1_BR.RM.like.systems | <-HTH<-HigB_toxin<-?<-DUF499*<-METHYLASE<-TUDOR+SF1-SF2-Helicase+iREase+CTD+REase<-HTH+WYL\|\|?->SMODS->UbE1+JAB-> | 939 | Bacteria>Pseudomonadota>Gammaproteobacteria>unclassified_Gammaproteobacteria | GCA_004297335.1 | Gammaproteobacteria_bacterium | EPO25_08575 | 7906..10725 | 1913989 | ATP-binding-protein-[Gammaproteobacteria-bacterium] |
| MCY4533098.1 | AAA+ATPase+wHTH+Alpha+Beta-domain+wHTH+RRM | Type-1_BR.RM.like.systems | HTH+WYL->TUDOR+SF1-SF2-Helicase+iREase+CTD+REase->PIN->METHYLASE->DUF499*->?-><-Peptidase_S8<-Classical-AAA | 931 | Bacteria>Pseudomonadota>Gammaproteobacteria>unclassified_Gammaproteobacteria | GCA_026713185.1 | Gammaproteobacteria_bacterium | OXC84_12365 | 15998..18793 | 1913989 | MAG:-DUF499-domain-containing-protein-[Gammaproteobacteria-bacterium] |
| NLN57017.1 | AAA+ATPase+wHTH+Alpha+Beta-domain+wHTH+RRM | Type-1_BR.RM.like.systems | ParB-><-DOC<-?\|\|TUDOR+SF1-SF2-Helicase+iREase+CTD+REase->METHYLASE->DUF499*-> | 947 | Bacteria>Pseudomonadota>Gammaproteobacteria>unclassified_Gammaproteobacteria | GCA_012522365.1 | Gammaproteobacteria_bacterium | GX151_03495 | 14614..17457 | 1913989 | DUF499-domain-containing-protein-[Gammaproteobacteria-bacterium] |
| MDA8120038.1 | AAA+ATPase+wHTH+Alpha+Beta-domain+wHTH+RRM | Type-1_BR.RM.like.systems | <-DUF499*<-METHYLASE<-TUDOR+SF1-SF2-Helicase+iREase+CTD+REase | 950 | Bacteria>Pseudomonadota>Gammaproteobacteria>unclassified_Gammaproteobacteria | GCA_028064935.1 | Gammaproteobacteria_bacterium | MOZ85_08355 | 3..2855 | 1913989 | MAG:-DUF499-domain-containing-protein-[Gammaproteobacteria-bacterium] |
| PJA03750.1 | AAA+ATPase+wHTH+Alpha+Beta-domain+wHTH+RRM | Type-1_BR.RM.like.systems | DUF499*->?->?->Phage_int_SAM_2+Integrase_1->XIS-HTH->ABhydrolase-> | 667 | Bacteria>Pseudomonadota>Gammaproteobacteria>unclassified_Gammaproteobacteria | GCA_002793195.1 | Gammaproteobacteria_bacterium_CG_4_10_14_0_2_um_filter_38_22 | COX72_03525 | 88..2091 | 1974020 | hypothetical-protein-COX72_03525-[Gammaproteobacteria-bacterium-CG_4_10_14_0_2_um_filter_38_22] |
| PIZ04909.1 | AAA+ATPase+wHTH+Alpha+Beta-domain+wHTH+RRM | Type-1_BR.RM.like.systems | DUF499*->?->?->Phage_int_SAM_2+Integrase_1->XIS-HTH->ABhydrolase-> | 678 | Bacteria>Pseudomonadota>Gammaproteobacteria>unclassified_Gammaproteobacteria | GCA_002785485.1 | Gammaproteobacteria_bacterium_CG_4_10_14_0_8_um_filter_38_16 | COY58_01735 | 1..2037 | 1974021 | hypothetical-protein-COY58_01735,-partial-[Gammaproteobacteria-bacterium-CG_4_10_14_0_8_um_filter_38_16] |
| RTZ75007.1 | AAA+ATPase+wHTH+Alpha+Beta-domain+wHTH+RRM | Type-1_BR.RM.like.systems | <-NERD+SF1-SF2-Helicase<-DUF499*<-REase<-OLD-ABC<-METHYLASE-p | 943 | Bacteria>Pseudomonadota>Gammaproteobacteria>unclassified_Gammaproteobacteria | GCA_003973075.1 | Gammaproteobacteria_bacterium | DSZ00_02995 | 2492..5323 | 1913989 | hypothetical-protein-DSZ00_02995-[Gammaproteobacteria-bacterium] |
| MBU2039164.1 | AAA+ATPase+wHTH+Alpha+Beta-domain+wHTH+RRM | Type-1_BR.RM.like.systems | DUF499*->SIR2+STALD->TIR-><-Phage_lysozyme | 945 | Bacteria>Pseudomonadota>Gammaproteobacteria>unclassified_Gammaproteobacteria | GCA_018826465.1 | Gammaproteobacteria_bacterium | KKH95_08475 | 3..2840 | 1913989 | DUF499-domain-containing-protein-[Gammaproteobacteria-bacterium] |
| HFD86238.1 | AAA+ATPase+wHTH+Alpha+Beta-domain+wHTH+RRM | Type-1_BR.RM.like.systems | <-DUF499*<-REase<-OLD-ABC<-METHYLASE-p | 692 | Bacteria>Pseudomonadota>Gammaproteobacteria>unclassified_Gammaproteobacteria | GCA_011322685.1 | Gammaproteobacteria_bacterium | ENJ35_01010 | 1..2075 | 1913989 | ATP-binding-protein-[Gammaproteobacteria-bacterium] |
| TNF00690.1 | AAA+ATPase+wHTH+Alpha+Beta-domain+wHTH+RRM | Type-1_BR.RM.like.systems | METHYLASE-p->DUF499*->?-><-TraG_N | 949 | Bacteria>Pseudomonadota>Gammaproteobacteria>unclassified_Gammaproteobacteria | GCA_006226625.1 | Gammaproteobacteria_bacterium | EP328_00985 | 580..3429 | 1913989 | ATP-binding-protein-[Gammaproteobacteria-bacterium] |
| MCL4555400.1 | AAA+ATPase+wHTH+Alpha+Beta-domain+wHTH+RRM | Type-1_BR.RM.like.systems | <-DUF499*<-?<-METHYLASE<-HNH | 513 | Bacteria>Pseudomonadota>Gammaproteobacteria>unclassified_Gammaproteobacteria | GCA_023382035.1 | Gammaproteobacteria_bacterium | M1572_00725 | 1..1540 | 1913989 | DUF499-domain-containing-protein-[Gammaproteobacteria-bacterium] |
| MCP5127473.1 | AAA+ATPase+wHTH+Alpha+Beta-domain+wHTH+RRM | Type-1_BR.RM.like.systems | 35exo->DOC+HTH->TUDOR+SF1-SF2-Helicase+iREase+CTD+REase->Schlafen+HATPase+HTH->METHYLASE->DUF499*->METHYLASE->XIS-HTH+RVT-><-MobL+ZnR+TOPRIM<-DOC<-MazE_antitoxin<-Phage_int_SAM_4+Phage_integrase | 986 | Bacteria>Pseudomonadota>Gammaproteobacteria>unclassified_Gammaproteobacteria | GCA_024235095.1 | Gammaproteobacteria_bacterium | H6973_18120 | 278648..281608 | 1913989 | ATP-binding-protein-[Gammaproteobacteria-bacterium] |
| MDE0453907.1 | AAA+ATPase+wHTH+Alpha+Beta-domain+wHTH+RRM | Type-1_BR.RM.like.systems | <-HEPN+REase<-Schlafen+HATPase+HTH<-Schlafen<-DUF499*<-METHYLASE<-TUDOR+SF1-SF2-Helicase<-OLD-ABC+TOPRIM<-TUDOR+SF1-SF2-Helicase+iREase+CTD+REase<-AbiEii<-AbiEi_4 | 967 | Bacteria>Pseudomonadota>Gammaproteobacteria>unclassified_Gammaproteobacteria | GCA_028821145.1 | Gammaproteobacteria_bacterium | OXJ63_01205 | 18382..21285 | 1913989 | MAG:-DUF499-domain-containing-protein-[Gammaproteobacteria-bacterium] |
| MXY51432.1 | AAA+ATPase+wHTH+Alpha+Beta-domain+wHTH+RRM | Type-1_BR.RM.like.systems | <-PhdYeFM_antitox\|\|TUDOR+SF1-SF2-Helicase+iREase+CTD+REase->OLD-ABC+TOPRIM->SF1-SF2-Helicase->METHYLASE-p->METHYLASE->DUF499*->TIR-> | 968 | Bacteria>Pseudomonadota>Gammaproteobacteria>unclassified_Gammaproteobacteria | GCA_009837725.1 | Gammaproteobacteria_bacterium | F4Y86_02730 | 44418..47324 | 1913989 | ATP-binding-protein-[Gammaproteobacteria-bacterium] |
| MCY4199355.1 | AAA+ATPase+wHTH+Alpha+Beta-domain+wHTH+RRM | Type-1_BR.RM.like.systems | AbiEi_1->?-><-HEPN<-?<-TPR+TPR+PIN+PIN<-DUF499*<-PIN<-DUF433<-METHYLASE<-?<-TUDOR+SF1-SF2-Helicase+iREase+CTD+REase\|\|RVT-> | 924 | Bacteria>Pseudomonadota>Gammaproteobacteria>unclassified_Gammaproteobacteria | GCA_026709225.1 | Gammaproteobacteria_bacterium | OXF31_05975 | 33360..36134 | 1913989 | MAG:-DUF499-domain-containing-protein-[Gammaproteobacteria-bacterium] |
| MDE0129995.1 | AAA+ATPase+wHTH+Alpha+Beta-domain+wHTH+RRM | Type-1_BR.RM.like.systems | <-SF1-SF2-Helicase<-OLD-ABC+TOPRIM<-Schlafen+HATPase+HTH<-DUF499*<-METHYLASE<-TUDOR+SF1-SF2-Helicase+iREase+CTD+REase<-?<-OLD-ABC+TOPRIM | 968 | Bacteria>Pseudomonadota>Gammaproteobacteria>unclassified_Gammaproteobacteria | GCA_028819355.1 | Gammaproteobacteria_bacterium | OXQ86_10575 | 58066..60972 | 1913989 | MAG:-DUF499-domain-containing-protein-[Gammaproteobacteria-bacterium] |
| MCY4342454.1 | AAA+ATPase+wHTH+Alpha+Beta-domain+wHTH+RRM | Type-1_BR.RM.like.systems | ORC-CDC6+REase-><-DUF499*<-PIN<-DUF433<-METHYLASE-p<-METHYLASE<-Peptidase_S8<-Classical-AAA | 924 | Bacteria>Pseudomonadota>Gammaproteobacteria>unclassified_Gammaproteobacteria | GCA_026710215.1 | Gammaproteobacteria_bacterium | OXE83_02620 | 2601..5375 | 1913989 | MAG:-DUF499-domain-containing-protein-[Gammaproteobacteria-bacterium] |
| MCY3812415.1 | AAA+ATPase+wHTH+Alpha+Beta-domain+wHTH+RRM | Type-1_BR.RM.like.systems | <-DUF499*<-BrnA_antitoxin<-BrnT_toxin<-METHYLASE<-?<-METHYLASE-p<-TUDOR+SF1-SF2-Helicase+iREase+CTD+REase | 916 | Bacteria>Pseudomonadota>Gammaproteobacteria>unclassified_Gammaproteobacteria | GCA_026706385.1 | Gammaproteobacteria_bacterium | OXH15_11520 | 652..3402 | 1913989 | MAG:-DUF499-domain-containing-protein-[Gammaproteobacteria-bacterium] |
| MCY4340924.1 | AAA+ATPase+wHTH+Alpha+Beta-domain+wHTH+RRM | Type-1_BR.RM.like.systems | LD-peptidase-><-DUF499*<-PIN<-DUF433<-METHYLASE<-?<-?<-TUDOR+SF1-SF2-Helicase+iREase+CTD+REase | 954 | Bacteria>Pseudomonadota>Gammaproteobacteria>unclassified_Gammaproteobacteria | GCA_026710305.1 | Gammaproteobacteria_bacterium | OXE48_06110 | 63766..66630 | 1913989 | MAG:-DUF499-domain-containing-protein-[Gammaproteobacteria-bacterium] |
| MCY4255936.1 | AAA+ATPase+wHTH+Alpha+Beta-domain+wHTH+RRM | Type-1_BR.RM.like.systems | TUDOR+SF1-SF2-Helicase+iREase+CTD+REase->?->?->METHYLASE->DUF433->PIN->DUF499*-><-LD-peptidase | 953 | Bacteria>Pseudomonadota>Gammaproteobacteria>unclassified_Gammaproteobacteria | GCA_026709685.1 | Gammaproteobacteria_bacterium | OXE51_07470 | 8826..11687 | 1913989 | MAG:-DUF499-domain-containing-protein-[Gammaproteobacteria-bacterium] |
| MCY4164747.1 | AAA+ATPase+wHTH+Alpha+Beta-domain+wHTH+RRM | Type-1_BR.RM.like.systems | TUDOR+SF1-SF2-Helicase+iREase+CTD+REase->?->?->METHYLASE->DUF433->PIN->DUF499*-><-LD-peptidase | 953 | Bacteria>Pseudomonadota>Gammaproteobacteria>unclassified_Gammaproteobacteria | GCA_026709045.1 | Gammaproteobacteria_bacterium | OXF03_01150 | 156657..159518 | 1913989 | MAG:-DUF499-domain-containing-protein-[Gammaproteobacteria-bacterium] |
| MCY4157213.1 | AAA+ATPase+wHTH+Alpha+Beta-domain+wHTH+RRM | Type-1_BR.RM.like.systems | TUDOR+SF1-SF2-Helicase+iREase+CTD+REase->?->?->METHYLASE->DUF433->PIN->DUF499*-><-LD-peptidase | 954 | Bacteria>Pseudomonadota>Gammaproteobacteria>unclassified_Gammaproteobacteria | GCA_026708965.1 | Gammaproteobacteria_bacterium | OXF66_07255 | 51365..54229 | 1913989 | MAG:-DUF499-domain-containing-protein-[Gammaproteobacteria-bacterium] |
| MDE0157838.1 | AAA+ATPase+wHTH+Alpha+Beta-domain+wHTH+RRM | Type-1_BR.RM.like.systems | TUDOR+SF1-SF2-Helicase+iREase+CTD+REase->N6_N4_MTase->?->?->METHYLASE->REase->DUF499*-> | 926 | Bacteria>Pseudomonadota>Gammaproteobacteria>unclassified_Gammaproteobacteria | GCA_028819495.1 | Gammaproteobacteria_bacterium | OXS28_19845 | 41740..44520 | 1913989 | MAG:-DUF499-domain-containing-protein-[Gammaproteobacteria-bacterium] |
| MDD9824620.1 | AAA+ATPase+wHTH+Alpha+Beta-domain+wHTH+RRM | Type-1_BR.RM.like.systems | <-DUF499*<-PIN<-OLD-ABC<-OLD-ABC+TOPRIM<-METHYLASE<-TPR+TPR | 928 | Bacteria>Pseudomonadota>Gammaproteobacteria>unclassified_Gammaproteobacteria | GCA_028817255.1 | Gammaproteobacteria_bacterium | OXU43_05565 | 147..2933 | 1913989 | MAG:-DUF499-domain-containing-protein-[Gammaproteobacteria-bacterium] |
| MYA16811.1 | AAA+ATPase+wHTH+Alpha+Beta-domain+wHTH+RRM | Type-1_BR.RM.like.systems | OLD-ABC-p->OLD-ABC-p->REase->DUF499*->URI-DNase-> | 922 | Bacteria>Pseudomonadota>Gammaproteobacteria>unclassified_Gammaproteobacteria | GCA_009836885.1 | Gammaproteobacteria_bacterium | F4Z28_08615 | 8657..11425 | 1913989 | ATP-binding-protein-[Gammaproteobacteria-bacterium] |
| MXW54998.1 | AAA+ATPase+wHTH+Alpha+Beta-domain+wHTH+RRM | Type-1_BR.RM.like.systems | <-METHYLASE-p<-DUF499*\|\|SF1-SF2-Helicase+iREase+CTD+REase-> | 809 | Bacteria>Pseudomonadota>Gammaproteobacteria>unclassified_Gammaproteobacteria | GCA_009836115.1 | Gammaproteobacteria_bacterium | F4Z66_13760 | 292..2721 | 1913989 | ATP-binding-protein-[Gammaproteobacteria-bacterium] |
| MDD9822476.1 | AAA+ATPase+wHTH+Alpha+Beta-domain+wHTH+RRM | Type-1_BR.RM.like.systems | METHYLASE-p->OLD-ABC+TOPRIM->DUF499*-> | 928 | Bacteria>Pseudomonadota>Gammaproteobacteria>unclassified_Gammaproteobacteria | GCA_028816685.1 | Gammaproteobacteria_bacterium | OXU98_06290 | 2678..5464 | 1913989 | MAG:-DUF499-domain-containing-protein-[Gammaproteobacteria-bacterium] |
| MDE0157713.1 | AAA+ATPase+wHTH+Alpha+Beta-domain+wHTH+RRM | Type-1_BR.RM.like.systems | OLD-ABC->TOPRIM->METHYLASE-p->?->?->DUF499*-> | 952 | Bacteria>Pseudomonadota>Gammaproteobacteria>unclassified_Gammaproteobacteria | GCA_028819495.1 | Gammaproteobacteria_bacterium | OXS28_19190 | 7492..10350 | 1913989 | MAG:-DUF499-domain-containing-protein-[Gammaproteobacteria-bacterium] |
| MDE0512489.1 | AAA+ATPase+wHTH+Alpha+Beta-domain+wHTH+RRM | Type-1_BR.RM.like.systems | <-DUF499*<-?<-?<-METHYLASE<-TOPRIM<-OLD-ABC | 952 | Bacteria>Pseudomonadota>Gammaproteobacteria>unclassified_Gammaproteobacteria | GCA_028823865.1 | Gammaproteobacteria_bacterium | OXI88_11955 | 1695..4553 | 1913989 | MAG:-DUF499-domain-containing-protein-[Gammaproteobacteria-bacterium] |
| MYD00428.1 | AAA+ATPase+wHTH+Alpha+Beta-domain+wHTH+RRM | Type-1_BR.RM.like.systems | <-HerA/FtsK<-?<-DUF499*<-METHYLASE | 933 | Bacteria>Pseudomonadota>Gammaproteobacteria>unclassified_Gammaproteobacteria | GCA_009841185.1 | Gammaproteobacteria_bacterium | F4X13_14340 | 42575..45376 | 1913989 | ATP-binding-protein-[Gammaproteobacteria-bacterium] |
| MDE0285439.1 | AAA+ATPase+wHTH+Alpha+Beta-domain+wHTH+RRM | Type-1_BR.RM.like.systems | <-DUF499*<-REase<-METHYLASE-p | 924 | Bacteria>Pseudomonadota>Gammaproteobacteria>unclassified_Gammaproteobacteria | GCA_028820555.1 | Gammaproteobacteria_bacterium | OXN26_12940 | 1577..4351 | 1913989 | MAG:-DUF499-domain-containing-protein-[Gammaproteobacteria-bacterium] |
| MCP5135119.1 | AAA+ATPase+wHTH+Alpha+Beta-domain+wHTH+RRM | Type-1_BR.RM.like.systems | TUDOR+SF1-SF2-Helicase+iREase+CTD+REase->METHYLASE->DDE_Tnp_1->?->?->DUF499*->?->DUF499-p->DUF499-p-><-RVT+HNH<-GDResolvase+Recombinase+ZnR | 428 | Bacteria>Pseudomonadota>Gammaproteobacteria>unclassified_Gammaproteobacteria | GCA_024234995.1 | Gammaproteobacteria_bacterium | H6976_16380 | 20781..22067 | 1913989 | ATP-binding-protein-[Gammaproteobacteria-bacterium] |
| MXY51727.1 | AAA+ATPase+wHTH+Alpha+Beta-domain+wHTH+RRM | Type-1_BR.RM.like.systems | <-DUF499*<-METHYLASE<-METHYLASE-p<-HEPN<-DNA_methylase<-MORC+REase<-TUDOR+SF1-SF2-Helicase+iREase+CTD+REase | 951 | Bacteria>Pseudomonadota>Gammaproteobacteria>unclassified_Gammaproteobacteria | GCA_009837725.1 | Gammaproteobacteria_bacterium | F4Y86_04245 | 1..2853 | 1913989 | ATP-binding-protein-[Gammaproteobacteria-bacterium] |
| MYA16109.1 | AAA+ATPase+wHTH+Alpha+Beta-domain+wHTH+RRM | Type-1_BR.RM.like.systems | <-NERD<-DUF499*<-METHYLASE<-TUDOR+SF1-SF2-Helicase+iREase+CTD+REase | 1006 | Bacteria>Pseudomonadota>Gammaproteobacteria>unclassified_Gammaproteobacteria | GCA_009836885.1 | Gammaproteobacteria_bacterium | F4Z28_04740 | 2858..5878 | 1913989 | DUF499-domain-containing-protein-[Gammaproteobacteria-bacterium] |
| WP_062660547.1 | AAA+ATPase+wHTH+Alpha+Beta-domain+wHTH+RRM | Type-1_BR.RM.like.systems | HTH+WYL->TUDOR+SF1-SF2-Helicase+iREase+CTD+REase->OLD-ABC->HNH->TUDOR+SF1-SF2-Helicase+iREase+CTD+REase-p->METHYLASE->DUF499*-><-RlpA<-?\|\|?->?-><-HNH<-OBFOLD-NUCLEASE | 947 | Bacteria>Pseudomonadota>Gammaproteobacteria>Vibrionales>Vibrionaceae | GCF_900055185.1 | Grimontia_celer | GCE9029_RS00270 | 67941..70784 | 1796497 | DUF499-domain-containing-protein-[Grimontia-celer] |
| WP_248386405.1 | AAA+ATPase+wHTH+Alpha+Beta-domain+wHTH+RRM | Type-1_BR.RM.like.systems | <-HTH\|\|SF1-SF2-Helicase->?-><-Resolvase\|\|TUDOR+SF1-SF2-Helicase+iREase+CTD+REase->METHYLASE->DUF499*-><-RlpA<-?<-Arm-DNA-bind_3+Phage_integrase | 954 | Bacteria>Pseudomonadota>Gammaproteobacteria>Vibrionales>Vibrionaceae | GCF_023206395.1 | Vibrio_sp._ED004 | ITG10_RS11530 | 2564414..2567278 | 2785124 | DUF499-domain-containing-protein-[Vibrio-sp.-ED004] |
| HDI3197646.1 | AAA+ATPase+wHTH+Alpha+Beta-domain+wHTH+RRM | Type-1_BR.RM.like.systems | <-VirD4-TraG+VirB4-FtsK<-HDRNase\|\|RlpA-><-?<-SMODS<-DUF499*<-METHYLASE-p<-METHYLASE<-TUDOR+SF1-SF2-Helicase+iREase+CTD+REase<-HTH+WYL<-?<-35exo | 947 | Bacteria>Pseudomonadota>Gammaproteobacteria>Vibrionales>Vibrionaceae | GCA_028119985.1 | Vibrio_cholerae | PMB20_000302 | 85117..87960 | 666 | TPA:-ATP-binding-protein-[Vibrio-cholerae] |
| WP_254620468.1 | AAA+ATPase+wHTH+Alpha+Beta-domain+wHTH+RRM | Type-1_BR.RM.like.systems | Arm-DNA-bind_3+Phage_integrase->RlpA-><-DUF499*<-METHYLASE<-HNH<-OLD-ABC<-TUDOR+SF1-SF2-Helicase+iREase+CTD+REase<-HTH+WYL<-XIS-HTH | 856 | Bacteria>Pseudomonadota>Gammaproteobacteria>Vibrionales>Vibrionaceae | - | Vibrio_cholerae | KVS75_RS02420 | 551801..554371 | 666 | DUF499-domain-containing-protein-[Vibrio-cholerae] |

| Protein-ID | DUF499-architecture | System | Gene-Neighbourhoods | Length | Taxonomy | Assembly-ID | Strain | Locus-Tag | Genomic-coordinates | TaxID | Fasta-defline |
|---|---|---|---|---|---|---|---|---|---|---|---|
| WP_213964958.1 | AAA+ATPase+wHTH+Alpha+Beta-domain+wHTH+RRM | Type-1_BR.RM.like.systems | XIS-HTH->HTH+WYL->TUDOR+SF1-SF2-Helicase+iREase+CTD+REase->OLD-ABC->HNH->METHYLASE->DUF499*-><-RlpA<-Arm-DNA-bind_3+Phage_integrase | 940 | Bacteria>Pseudomonadota>Gammaproteobacteria>Vibrionales>Vibrionaceae | GCF_018457925.1 | Vibrio_alginolyticus | J4H38_RS12950 | 94691..97513 | 663 | DUF499-domain-containing-protein-[Vibrio-alginolyticus] |
| HCH6462476.1 | AAA+ATPase+wHTH+Alpha+Beta-domain+wHTH+RRM | Type-1_BR.RM.like.systems | XIS-HTH->HTH+WYL->TUDOR+SF1-SF2-Helicase+iREase+CTD+REase->OLD-ABC->HNH->METHYLASE->DUF499*-><-RlpA<-Arm-DNA-bind_3+Phage_integrase | 940 | Bacteria>Pseudomonadota>Gammaproteobacteria>Vibrionales>Vibrionaceae | GCA_024102875.1 | Vibrio_parahaemolyticus | NK401_003629 | 41421..44243 | 670 | ATP-binding-protein-[Vibrio-parahaemolyticus] |
| EJR0961152.1 | AAA+ATPase+wHTH+Alpha+Beta-domain+wHTH+RRM | Type-1_BR.RM.like.systems | Arm-DNA-bind_3+Phage_integrase->RlpA-><-DUF499*<-METHYLASE<-HNH<-OLD-ABC<-TUDOR+SF1-SF2-Helicase+iREase+CTD+REase<-HTH+WYL<-XIS-HTH | 940 | Bacteria>Pseudomonadota>Gammaproteobacteria>Vibrionales>Vibrionaceae | GCA_024870505.1 | Vibrio_parahaemolyticus | NYK10_002560 | 278932..281754 | 670 | ATP-binding-protein-[Vibrio-parahaemolyticus] |
| EJL6982897.1 | AAA+ATPase+wHTH+Alpha+Beta-domain+wHTH+RRM | Type-1_BR.RM.like.systems | XIS-HTH->HTH+WYL->TUDOR+SF1-SF2-Helicase+iREase+CTD+REase->OLD-ABC->HNH->METHYLASE->DUF499*-><-RlpA<-Arm-DNA-bind_3+Phage_integrase | 915 | Bacteria>Pseudomonadota>Gammaproteobacteria>Vibrionales>Vibrionaceae | GCA_024219855.1 | Vibrio_cholerae | NMT40_002449 | 120290..123037 | 666 | ATP-binding-protein-[Vibrio-cholerae] |
| TBT35889.1 | AAA+ATPase+wHTH+Alpha+Beta-domain+wHTH+RRM | Type-1_BR.RM.like.systems | Arm-DNA-bind_3+Phage_integrase->RlpA-><-DUF499*<-TUDOR+SF1-SF2-Helicase+iREase+CTD+REase-p<-HNH<-OLD-ABC<-HTH+WYL<-XIS-HTH\|\|JAB-> | 940 | Bacteria>Pseudomonadota>Gammaproteobacteria>Vibrionales>Vibrionaceae | GCA_004324405.1 | Vibrio_parahaemolyticus | D5E85_08495 | 430725..433547 | 670 | ATP-binding-protein-[Vibrio-parahaemolyticus] |
| MBM4953950.1 | AAA+ATPase+wHTH+Alpha+Beta-domain+wHTH+RRM | Type-1_BR.RM.like.systems | Arm-DNA-bind_3+Phage_integrase->RlpA-><-DUF499*<-METHYLASE<-TUDOR+SF1-SF2-Helicase+iREase+CTD+REase<-HTH+WYL<-XIS-HTH<-?\|\|JAB-> | 947 | Bacteria>Pseudomonadota>Gammaproteobacteria>Vibrionales>Vibrionaceae | GCA_016876875.1 | Vibrio_parahaemolyticus | HYO25_15375 | 99251..102094 | 670 | ATP-binding-protein-[Vibrio-parahaemolyticus] |
| WP_206975067.1 | AAA+ATPase+wHTH+Alpha+Beta-domain+wHTH+RRM | Type-1_BR.RM.like.systems | 35exo->?->HTH+WYL->TUDOR+SF1-SF2-Helicase+iREase+CTD+REase->METHYLASE->DUF499*-><-RlpA\|\|HDRNase->VirD4-TraG+VirB4-FtsK-> | 947 | Bacteria>Pseudomonadota>Gammaproteobacteria>Vibrionales>Vibrionaceae | GCF_017313825.1 | Vibrio_parahaemolyticus | J0689_RS06960 | 11079..13922 | 670 | DUF499-domain-containing-protein-[Vibrio-parahaemolyticus] |
| WP_102552172.1 | AAA+ATPase+wHTH+Alpha+Beta-domain+wHTH+RRM | Type-1_BR.RM.like.systems | Arm-DNA-bind_3+Phage_integrase->?->RlpA-><-DUF499*<-METHYLASE<-METHYLASE-p<-TUDOR+SF1-SF2-Helicase+iREase+CTD+REase | 960 | Bacteria>Pseudomonadota>Gammaproteobacteria>Vibrionales>Vibrionaceae | GCF_002875995.1 | Vibrio_splendidus | BCT54_RS08470 | 12202..15084 | 29497 | DUF499-domain-containing-protein-[Vibrio-splendidus] |
| HDM8159538.1 | AAA+ATPase+wHTH+Alpha+Beta-domain+wHTH+RRM | Type-1_BR.RM.like.systems | SbcC+SbcC->?-><-?<-?\|\|?->Phage_integrase-><-DUF499*<-METHYLASE<-TUDOR+SF1-SF2-Helicase+iREase+CTD+REase\|\|Resolvase-> | 960 | Bacteria>Pseudomonadota>Gammaproteobacteria>Vibrionales>Vibrionaceae | GCA_028991035.1 | Vibrio_harveyi | POE13_002783 | 107272..110154 | 669 | TPA:-ATP-binding-protein-[Vibrio-harveyi] |
| WP_077459495.1 | AAA+ATPase+wHTH+Alpha+Beta-domain+wHTH+RRM | Type-1_BR.RM.like.systems | METHYLASE->DUF499*->?-><-Arm-DNA-bind_3+Phage_integrase<-?\|\|RNase->PRTase-><-TetR-HTH | 949 | Bacteria>Pseudomonadota>Gammaproteobacteria>Vibrionales>Vibrionaceae | GCF_001995805.1 | Salinivibrio_kushneri | BZG09_RS15020 | 2338..5187 | 1908198 | DUF499-domain-containing-protein-[Salinivibrio-kushneri] |
| WP_017630403.1 | AAA+ATPase+wHTH+Alpha+Beta-domain+wHTH+RRM | Type-1_BR.RM.like.systems | <-DUF499*<-METHYLASE-p<-METHYLASE<-TUDOR+SF1-SF2-Helicase+iREase+CTD+REase | 880 | Bacteria>Pseudomonadota>Gammaproteobacteria>Vibrionales>Vibrionaceae | GCF_000316985.1 | Vibrio_sp._624788 | C658_RS25835 | 1..2642 | 1234362 | DUF499-domain-containing-protein,-partial-[Vibrio-sp.-624788] |
| WP_077605710.1 | AAA+ATPase+wHTH+Alpha+Beta-domain+wHTH+RRM | Type-1_BR.RM.like.systems | <-HIN-HTH\|\|METHYLASE->DUF499*->?-><-Arm-DNA-bind_3+Phage_integrase | 949 | Bacteria>Pseudomonadota>Gammaproteobacteria>Vibrionales>Vibrionaceae | GCF_001995785.1 | Salinivibrio_kushneri | BZG06_RS14975 | 2300..5149 | 1908198 | DUF499-domain-containing-protein-[Salinivibrio-kushneri] |
| WP_043696850.1 | AAA+ATPase+wHTH+Alpha+Beta-domain+wHTH+RRM | Type-1_BR.RM.like.systems | HTH+WYL->TUDOR+SF1-SF2-Helicase+iREase+CTD+REase->SbcC+SbcC->METHYLASE->METHYLASE->Schlafen+HATPase+HTH->DUF499*-><-Phage_int_SAM_2+Integrase_1<-MarR-HTH\|\|?-><-?<-MODE-HTH+PBPII\|\|ADH_N+ADH_zinc_N-> | 947 | Bacteria>Pseudomonadota>Gammaproteobacteria>Xanthomonadales>Rhodanobacteraceae | GCF_000800155.1 | Luteibacter_sp._9133 | FA87_RS19045 | 4371799..4374642 | 1500891 | DUF499-domain-containing-protein-[Luteibacter-sp.-9133] |
| WP_091239202.1 | AAA+ATPase+wHTH+Alpha+Beta-domain+wHTH+RRM | Type-1_BR.RM.like.systems | Phage_holin->Phage_lysozyme->?->HTH_3->Radical-SAM-><-?<-DUF499*<-ParB+HNH<-METHYLASE<-HEPN<-TUDOR+SF1-SF2-Helicase+iREase+CTD+REase<-HTH_Tnp<-HTH+WYL | 949 | Bacteria>Pseudomonadota>Gammaproteobacteria>Xanthomonadales>Rhodanobacteraceae | GCF_900101825.1 | Aquimonas_voraii | BLP83_RS04255 | 981108..983957 | 265719 | DUF499-domain-containing-protein-[Aquimonas-voraii] |
| WP_008437332.1 | AAA+ATPase+wHTH+Alpha+Beta-domain+wHTH+RRM | Type-1_BR.RM.like.systems | <-AbiEi_4\|\|?->?->XIS-HTH->XIS-HTH->Phage_int_SAM_1+Phage_integrase-><-DUF499*<-?<-METHYLASE<-?<-PIN<-TUDOR+SF1-SF2-Helicase+iREase+CTD+REase | 933 | Bacteria>Pseudomonadota>Gammaproteobacteria>Xanthomonadales>Rhodanobacteraceae | GCF_000264375.1 | Rhodanobacter_thiooxydans | RHOFW104T7_RS11960 | 24308..27109 | 416169 | DUF499-domain-containing-protein-[Rhodanobacter-thiooxydans] |
| WP_243041017.1 | AAA+ATPase+wHTH+Alpha+Beta-domain+wHTH+RRM | Type-1_BR.RM.like.systems | HTH+WYL->HTH_Tnp->TUDOR+SF1-SF2-Helicase+iREase+CTD+REase->PIN->METHYLASE->DUF499*->SbcC-><-Phage_lysozyme<-?<-?<-RNase | 955 | Bacteria>Pseudomonadota>Gammaproteobacteria>Xanthomonadales>Rhodanobacteraceae | GCF_022760995.1 | Dyella_sedimenti | MJ605_RS08555 | 206646..209513 | 2919947 | DUF499-domain-containing-protein-[Dyella-sp.-NS14] |
| TAN08689.1 | AAA+ATPase+wHTH+Alpha+Beta-domain+wHTH+RRM | Type-1_BR.RM.like.systems | <-STYKinase-like<-HTH\|\|OLD-ABC+TOPRIM->METHYLASE->METHYLASE->Schlafen+HATPase+HTH->DUF499*-><-BirA-HTH<-?<-Phage_integrase<-?<-ZnR | 934 | Bacteria>Pseudomonadota>Gammaproteobacteria>Xanthomonadales>Rhodanobacteraceae | GCA_004299555.1 | Rhodanobacteraceae_bacterium | EPN38_01845 | 144698..147502 | 2052182 | ATP-binding-protein-[Rhodanobacteraceae-bacterium] |
| MCC7256721.1 | AAA+ATPase+wHTH+Alpha+Beta-domain+wHTH+RRM | Type-1_BR.RM.like.systems | <-KAP-NTPase<-?<-DUF499*<-KAP-NTPase<-METHYLASE<-HNH<-OLD-ABC<-TUDOR+SF1-SF2-Helicase+iREase+CTD+REase<-PIWI | 930 | Bacteria>Pseudomonadota>Gammaproteobacteria>Xanthomonadales>Rhodanobacteraceae | GCA_020852005.1 | Dokdonella_sp. | IT467_12405 | 2246..5038 | 2291710 | ATP-binding-protein-[Dokdonella-sp.] |
| WHZ20488.1 | AAA+ATPase+wHTH+Alpha+Beta-domain+wHTH+RRM | Type-1_BR.RM.like.systems | <-XIS-HTH<-?<-Arm-DNA-bind_3+Phage_integrase<-?<-?<-DUF499*<-HTH<-HigB_toxin<-?<-METHYLASE<-PIN<-PemK_toxin | 932 | Bacteria>Pseudomonadota>Gammaproteobacteria>Xanthomonadales>Rhodanobacteraceae | GCA_030123585.1 | Rhodanobacteraceae_bacterium | OJF55_002637 | 2854033..2856831 | 2052182 | MAG:-putative-ATPase-(AAA+-superfamily)-[Rhodanobacteraceae-bacterium] |
| WP_182517583.1 | AAA+ATPase+wHTH+Alpha+Beta-domain+wHTH+RRM | Type-1_BR.RM.like.systems | XIS-HTH->TUDOR+SF1-SF2-Helicase+iREase+CTD+REase->HEPN->METHYLASE->OLD-ABC->TOPRIM->DUF499*-> | 930 | Bacteria>Pseudomonadota>Gammaproteobacteria>Xanthomonadales>Rhodanobacteraceae | GCF_014138295.1 | Dokdonella_fugitiva | FHW13_RS05515 | 304049..306841 | 328517 | DUF499-domain-containing-protein-[Dokdonella-fugitiva] |
| MCK9487878.1 | AAA+ATPase+wHTH+Alpha+Beta-domain+wHTH+RRM | Type-1_BR.RM.like.systems | <-SIGMA-HTH<-OLD-ABC<-?<-?<-?<-DUF499*<-REase<-OLD-ABC<-METHYLASE<-HEPN<-PIN<-PemK_toxin | 944 | Bacteria>Pseudomonadota>Gammaproteobacteria>Xanthomonadales>unclassified_Xanthomonadales | GCA_023228845.1 | Xanthomonadales_bacterium | M0Q42_00470 | 105333..108167 | 2006849 | DUF499-domain-containing-protein-[Xanthomonadales-bacterium] |
| MCU0753612.1 | AAA+ATPase+wHTH+Alpha+Beta-domain+wHTH+RRM | Type-1_BR.RM.like.systems | <-ArsR-HTH\|\|?->?->?->?-><-DUF499*<-Schlafen+HATPase+HTH<-ParB+HNH<-REase<-OLD-ABC<-METHYLASE<-HNH | 955 | Bacteria>Pseudomonadota>Gammaproteobacteria>Xanthomonadales>unclassified_Xanthomonadales | GCA_025458445.1 | Xanthomonadales_bacterium | MUE46_00615 | 13163..16030 | 2006849 | DUF499-domain-containing-protein-[Xanthomonadales-bacterium] |
| WP_246149960.1 | AAA+ATPase+wHTH+Alpha+Beta-domain+wHTH+RRM | Type-1_BR.RM.like.systems | HTH+WYL->HTH_Tnp->TUDOR+SF1-SF2-Helicase+iREase+CTD+REase->PIN->METHYLASE->DUF499*->SIR2+STALD->TIR->Methylase-><-Radical-SAM<-HTH_3<-Phage_lysozyme | 959 | Bacteria>Pseudomonadota>Gammaproteobacteria>Xanthomonadales>Xanthomonadaceae | GCF_008386465.1 | Arenimonas_fontis | F0415_RS12225 | 12028..14907 | 2608255 | DUF499-domain-containing-protein-[Arenimonas-fontis] |
| KAA2283481.1 | AAA+ATPase+wHTH+Alpha+Beta-domain+wHTH+RRM | Type-1_BR.RM.like.systems | HTH+WYL->HTH_Tnp->TUDOR+SF1-SF2-Helicase+iREase+CTD+REase->PIN->METHYLASE->DUF499*->SIR2+STALD->TIR->Methylase-><-Radical-SAM<-HTH_3<-Phage_lysozyme | 955 | Bacteria>Pseudomonadota>Gammaproteobacteria>Xanthomonadales>Xanthomonadaceae | GCA_008386465.1 | Arenimonas_fontis | F0415_12220 | 12040..14907 | 2608255 | ATP-binding-protein-[Arenimonas-fontis] |
| WP_194929585.1 | AAA+ATPase+wHTH+Alpha+Beta-domain+wHTH+RRM | Type-1_BR.RM.like.systems | DUF3987-><-?<-REase+35exo+Beta-Barrel+SF1-SF2-Helicase<-DUF499*<-METHYLASE<-TUDOR+SF1-SF2-Helicase+iREase+CTD+REase<-HTH_Tnp<-HTH+WYL<-JAB | 951 | Bacteria>Pseudomonadota>Gammaproteobacteria>Xanthomonadales>Xanthomonadaceae | GCF_015453285.1 | Lysobacter_niastensis | IU514_RS02875 | 30934..33789 | 380629 | DUF499-domain-containing-protein-[Lysobacter-niastensis] |
| WP_055771167.1 | AAA+ATPase+wHTH+Alpha+Beta-domain+wHTH+RRM | Type-1_BR.RM.like.systems | <-D5N+D5ATPase<-?<-?<-?<-?<-DUF499*<-ParB+HNH<-?<-METHYLASE<-PIN<-TUDOR+SF1-SF2-Helicase+iREase+CTD+REase<-Phage_integrase | 929 | Bacteria>Pseudomonadota>Gammaproteobacteria>Xanthomonadales>Xanthomonadaceae | GCF_001422025.1 | Stenotrophomonas_sp._Leaf70 | ASF01_RS13960 | 54995..57784 | 1736233 | DUF499-domain-containing-protein-[Stenotrophomonas-sp.-Leaf70] |
| MBH2008931.1 | AAA+ATPase+wHTH+Alpha+Beta-domain+wHTH+RRM | Type-1_BR.RM.like.systems | <-SF1-SF2-Helicase<-OLD-ABC+TOPRIM<-DUF499*<-METHYLASE<-HEPN<-TUDOR+SF1-SF2-Helicase+iREase+CTD+REase<-?\|\|DDE_Tnp_1_2-> | 969 | Bacteria>Pseudomonadota>Gammaproteobacteria>Xanthomonadales>Xanthomonadaceae | GCA_015999665.1 | Xanthomonadaceae_bacterium | I8H71_04430 | 4236..7145 | 1926873 | ATP-binding-protein-[Xanthomonadaceae-bacterium] |
| MTI74440.1 | AAA+ATPase+wHTH+Alpha+Beta-domain+wHTH+RRM | Type-1_BR.RM.like.systems | <-JAB<-?<-D5N+D5ATPase<-?<-?<-?<-DUF499*<-?<-?<-TOPRIM<-OLD-ABC<-TUDOR+SF1-SF2-Helicase+iREase+CTD+REase<-Phage_integrase | 931 | Bacteria>Pseudomonadota>Gammaproteobacteria>Xanthomonadales>Xanthomonadaceae | GCA_009711865.1 | Stenotrophomonas_sp. | FH747_12410 | 103914..106709 | 69392 | ATP-binding-protein-[Stenotrophomonas-sp.] |
| WP_049462969.1 | AAA+ATPase+wHTH+Alpha+Beta-domain+wHTH+RRM | Type-1_BR.RM.like.systems | HTH_26->TUDOR+SF1-SF2-Helicase+iREase+CTD+REase->OLD-ABC->HNH->METHYLASE->ParB+HNH->DUF499*->?->?->?->?->TPR-> | 930 | Bacteria>Pseudomonadota>Gammaproteobacteria>Xanthomonadales>Xanthomonadaceae | GCF_001075485.1 | Stenotrophomonas_maltophilia | ADM82_RS13495 | 40138..42930 | 40324 | DUF499-domain-containing-protein-[Stenotrophomonas-maltophilia] |
| PJK14600.1 | AAA+ATPase+wHTH+Alpha+Beta-domain+wHTH+RRM | Type-1_BR.RM.like.systems | <-DUF499*<-ParB+HNH<-METHYLASE<-HNH<-OLD-ABC<-TUDOR+SF1-SF2-Helicase+iREase+CTD+REase | 945 | Bacteria>Pseudomonadota>Gammaproteobacteria>Xanthomonadales>Xanthomonadaceae | GCA_002798295.1 | Xanthomonadaceae_bacterium_NML07-0707 | CO613_05135 | 84524..87361 | 2032574 | hypothetical-protein-CO613_05135-[Xanthomonadaceae-bacterium-NML07-0707] |
| WP_161614005.1 | AAA+ATPase+wHTH+Alpha+Beta-domain+wHTH+RRM | Type-1_BR.RM.like.systems | TUDOR+SF1-SF2-Helicase+iREase+CTD+REase->HEPN->METHYLASE->?->?->?->DUF499*->?->?->?->?->?->D5N+D5ATPase-> | 929 | Bacteria>Pseudomonadota>Gammaproteobacteria>Xanthomonadales>Xanthomonadaceae | GCF_009996725.1 | Stenotrophomonas_acidaminiphila | GXB84_RS06675 | 54604..57393 | 128780 | DUF499-domain-containing-protein-[Stenotrophomonas-acidaminiphila] |
| WP_106991537.1 | AAA+ATPase+wHTH+Alpha+Beta-domain+wHTH+RRM | Type-1_BR.RM.like.systems | <-DUF499*<-?<-METHYLASE<-HEPN<-PIN<-TUDOR+SF1-SF2-Helicase+iREase+CTD+REase<-HTH_26 | 930 | Bacteria>Pseudomonadota>Gammaproteobacteria>Xanthomonadales>Xanthomonadaceae | GCF_003024235.1 | Arenimonas_caeni | C6N40_RS13415 | 668..3460 | 2058085 | DUF499-domain-containing-protein-[Arenimonas-caeni] |
| PZO64463.1 | AAA+ATPase+wHTH+Alpha+Beta-domain+wHTH+RRM | Type-1_BR.RM.like.systems | <-DUF499*<-TOPRIM<-OLD-ABC<-METHYLASE<-PIN<-TUDOR+SF1-SF2-Helicase+iREase+CTD+REase | 932 | Bacteria>Pseudomonadota>Gammaproteobacteria>Xanthomonadales>Xanthomonadaceae | GCA_003240695.1 | Pseudoxanthomonas_suwonensis | DI635_05295 | 731..3529 | 314722 | hypothetical-protein-DI635_05295-[Pseudoxanthomonas-suwonensis] |
| ODS64316.1 | AAA+ATPase+wHTH+Alpha+Beta-domain+wHTH+RRM | Type-1_BR.RM.like.systems | <-HTH_Tnp<-?\|\|?->TUDOR+SF1-SF2-Helicase+iREase+CTD+REase->DOC+HTH->?->DUF499*->HIN-HTH-> | 937 | Bacteria>Pseudomonadota>Gammaproteobacteria>Xanthomonadales>Xanthomonadaceae | GCA_001724815.1 | Arenimonas_sp._SCN_70-307 | ABS41_03390 | 88987..91800 | 1660089 | MAG:-hypothetical-protein-ABS41_03390-[Arenimonas-sp.-SCN-70-307] |
| MBS3959804.1 | AAA+ATPase+wHTH+Alpha+Beta-domain+wHTH+RRM | Type-1_BR.RM.like.systems | TUDOR+SF1-SF2-Helicase+iREase+CTD+REase-p->URI-DNase+RAMA->METHYLASE->DUF499*-> | 947 | Bacteria>Pseudomonadota>Gammaproteobacteria>Xanthomonadales>Xanthomonadaceae | GCA_018333835.1 | Xanthomonadaceae_bacterium | KGZ52_10435 | 4737..7580 | 1926873 | ATP-binding-protein-[Xanthomonadaceae-bacterium] |
| MBP7655321.1 | AAA+ATPase+wHTH+Alpha+Beta-domain+wHTH+RRM | Type-1_BR.RM.like.systems | HEPN->METHYLASE->?->?->DUF499*->TIR->Methylase-><-Radical-SAM<-HTH_3<-?<-Phage_lysozyme | 950 | Bacteria>Pseudomonadota>Gammaproteobacteria>Xanthomonadales>Xanthomonadaceae | GCA_017999045.1 | Pseudoxanthomonas_sp. | KA742_02095 | 7019..9871 | 1871049 | ATP-binding-protein-[Pseudoxanthomonas-sp.] |
| WP_034214097.1 | AAA+ATPase+wHTH+Alpha+Beta-domain+wHTH+RRM | Type-1_BR.RM.like.systems | <-DUF499*<-?<-METHYLASE<-PIN<-DOC+HTH<-TUDOR+SF1-SF2-Helicase+iREase+CTD+REase | 933 | Bacteria>Pseudomonadota>Gammaproteobacteria>Xanthomonadales>Xanthomonadaceae | GCF_000747155.1 | Arenimonas_metalli | N787_RS11310 | 49441..52242 | 948077 | DUF499-domain-containing-protein-[Arenimonas-metalli] |
| WP_053505566.1 | AAA+ATPase+wHTH+Alpha+Beta-domain+wHTH+RRM | Type-1_BR.RM.like.systems | HEPN+TPR+RhoN->HicA_toxin->METHYLASE->?->?->ParB+HNH->DUF499*->?->NACHT-> | 930 | Bacteria>Pseudomonadota>Gammaproteobacteria>Xanthomonadales>Xanthomonadaceae | - | Stenotrophomonas_maltophilia | ABW45_RS08210 | 26513..29305 | 40324 | DUF499-domain-containing-protein-[Stenotrophomonas-maltophilia] |
| GIX36093.1 | AAA+ATPase+wHTH+Alpha+Beta-domain+wHTH+RRM | Type-1_BR.RM.like.systems | METHYLASE-p->METHYLASE-p->OLD-ABC->REase->DUF499*-><-?<-?<-?<-?<-?<-Radical-SAM | 950 | Bacteria>Pseudomonadota>Gammaproteobacteria>Xanthomonadales>Xanthomonadaceae | GCA_026005075.1 | Xanthomonadaceae_bacterium | KatS3mg126_1872 | 4497..7349 | 1926873 | MAG:-hypothetical-protein-KatS3mg126_1872-[Xanthomonadaceae-bacterium] |
| WP_144815145.1 | AAA+ATPase+wHTH+Alpha+Beta-domain+wHTH+RRM | Type-1_BR.RM.like.systems | METHYLASE-p->?->ParB+HNH->DUF499*->?->?->?->?->?->D5N+D5ATPase-> | 930 | Bacteria>Pseudomonadota>Gammaproteobacteria>Xanthomonadales>Xanthomonadaceae | GCF_007830115.1 | Lysobacter_ruishenii | IP93_RS09870 | 5117..7909 | 686800 | DUF499-domain-containing-protein-[Lysobacter-ruishenii] |
| MBU6247856.1 | AAA+ATPase+wHTH+Alpha+Beta-domain+wHTH+RRM | Type-1_BR.RM.like.systems | <-DUF499*<-ParB+HNH<-?<-HEPN+CBS<-METHYLASE<-PIN | 934 | Bacteria>Pseudomonadota>Gammaproteobacteria>Xanthomonadales>Xanthomonadaceae | GCA_018970895.1 | Xanthomonadaceae_bacterium | KGN77_08870 | 9374..12178 | 1926873 | DUF499-domain-containing-protein-[Xanthomonadaceae-bacterium] |
| WP_169116186.1 | AAA+ATPase+wHTH+Alpha+Beta-domain+wHTH+RRM | Type-1_BR.RM.like.systems | HEPN->PolB->TUDOR+SF1-SF2-Helicase+iREase+CTD+REase->PIN->METHYLASE->DUF499*->?-><-?<-RuvC<-?\|\|SF1-SF2-Helicase-> | 948 | Bacteria>Pseudomonadota>Hydrogenophilia>Hydrogenophilales>Hydrogenophilaceae | GCF_012911495.1 | Tepidiphilus_baoligensis | GV368_RS08285 | 50966..53812 | 2698687 | DUF499-domain-containing-protein-[Tepidiphilus-baoligensis] |
| WP_219972353.1 | AAA+ATPase+wHTH+Alpha+Beta-domain+wHTH+RRM | Type-1_BR.RM.like.systems | MODE-HTH+PBPII->?-><-tRNA\|\|?->?-><-?<-DUF499*<-?<-METHYLASE<-PIN<-TUDOR+SF1-SF2-Helicase+iREase+CTD+REase | 955 | Bacteria>Pseudomonadota>Hydrogenophilia>Hydrogenophilales>Hydrogenophilaceae | GCF_019448605.1 | Hydrogenophilus_thermoluteolus | K1I42_RS04670 | 179917..182784 | 297 | DUF499-domain-containing-protein-[Hydrogenophilus-thermoluteolus] |
| WP_028873821.1 | AAA+ATPase+wHTH+Alpha+Beta-domain+wHTH+RRM | Type-1_BR.RM.like.systems | DDE_Tnp_1_assoc+DDE_Tnp_1->PIN->HEPN->METHYLASE->?->DUF499*->?->PIN->?-><-?<-RuvC | 947 | Bacteria>Pseudomonadota>Hydrogenophilia>Hydrogenophilales>Hydrogenophilaceae | GCF_000425565.1 | Tepidiphilus_margaritifer | H607_RS0100570 | 5909..8752 | 203471 | DUF499-domain-containing-protein-[Tepidiphilus-margaritifer] |
| MBS0576259.1 | AAA+ATPase+wHTH+Alpha+Beta-domain+wHTH+RRM | Type-1_BR.RM.like.systems | RNase->?->?->Phage_lysozyme-><-SbcC<-DUF499*<-METHYLASE<-HEPN<-PIN<-TUDOR+SF1-SF2-Helicase+iREase+CTD+REase<-HTH_Tnp<-HTH+WYL | 955 | Bacteria>Pseudomonadota>unclassified_Pseudomonadota | GCA_018240435.1 | Pseudomonadota_bacterium | JSS45_07535 | 37839..40706 | 1977087 | ATP-binding-protein-[Proteobacteria-bacterium] |
| MBU4261625.1 | AAA+ATPase+wHTH+Alpha+Beta-domain+wHTH+RRM | Type-1_BR.RM.like.systems | <-DUF499*<-METHYLASE<-OLD-ABC<-TUDOR+SF1-SF2-Helicase+iREase+CTD+REase<-NERD<-Phage_integrase\|\|MetJarc-HTH-> | 944 | Bacteria>Pseudomonadota>unclassified_Pseudomonadota | GCA_018897275.1 | Pseudomonadota_bacterium | KKC76_07060 | 397..3231 | 1977087 | DUF499-domain-containing-protein-[Proteobacteria-bacterium] |
| NBX75138.1 | AAA+ATPase+wHTH+Alpha+Beta-domain+wHTH+RRM | Type-1_BR.RM.like.systems | <-HTH\|\|Calcineurin->?->HTH+WYL->TUDOR+SF1-SF2-Helicase+iREase+CTD+REase->METHYLASE->DUF499*-><-?<-Peptidase | 941 | Bacteria>Pseudomonadota>unclassified_Pseudomonadota | GCA_009927005.1 | Pseudomonadota_bacterium | EBQ92_01110 | 10725..13550 | 1977087 | ATP-binding-protein-[Proteobacteria-bacterium] |
| MBS0558266.1 | AAA+ATPase+wHTH+Alpha+Beta-domain+wHTH+RRM | Type-1_BR.RM.like.systems | <-DUF499*<-?<-METHYLASE<-HicA_toxin<-PemK_toxin<-PrlF_antitoxin<-TUDOR+SF1-SF2-Helicase+iREase+CTD+REase | 923 | Bacteria>Pseudomonadota>unclassified_Pseudomonadota | GCA_018240565.1 | Pseudomonadota_bacterium | JSR27_12745 | 54620..57391 | 1977087 | ATP-binding-protein-[Proteobacteria-bacterium] |
| MBU0995376.1 | AAA+ATPase+wHTH+Alpha+Beta-domain+wHTH+RRM | Type-1_BR.RM.like.systems | ParB-><-?\|\|HTH->MPTase->ZnR-><-?<-DUF499*<-METHYLASE<-TUDOR+SF1-SF2-Helicase+iREase+CTD+REase<-XIS-HTH | 940 | Bacteria>Pseudomonadota>unclassified_Pseudomonadota | GCA_018816735.1 | Pseudomonadota_bacterium | KJ737_23005 | 22676..25498 | 1977087 | DUF499-domain-containing-protein-[Proteobacteria-bacterium] |
| MDE3269952.1 | AAA+ATPase+wHTH+Alpha+Beta-domain+wHTH+RRM | Type-1_BR.RM.like.systems | REase-><-TUDOR+SF1-SF2-Helicase+iREase+CTD+REase<-?<-DUF499*<-PIN<-MPTase<-HTH<-?<-METHYLASE | 917 | Bacteria>Pseudomonadota>unclassified_Pseudomonadota | GCA_028877195.1 | Pseudomonadota_bacterium | OYH77_06680 | 5373..8126 | 1977087 | MAG:-hypothetical-protein-OYH77_06680-[Pseudomonadota-bacterium] |
| MBU0673024.1 | AAA+ATPase+wHTH+Alpha+Beta-domain+wHTH+RRM | Type-1_BR.RM.like.systems | TUDOR+SF1-SF2-Helicase+iREase+CTD+REase->PIN->?->METHYLASE->OLD-ABC->REase->DUF499*-> | 943 | Bacteria>Pseudomonadota>unclassified_Pseudomonadota | GCA_018814305.1 | Pseudomonadota_bacterium | KJ950_00090 | 19515..22346 | 1977087 | DUF499-domain-containing-protein-[Proteobacteria-bacterium] |
| MBS0225815.1 | AAA+ATPase+wHTH+Alpha+Beta-domain+wHTH+RRM | Type-1_BR.RM.like.systems | ACET->?-><-?\|\|HEPN-><-DUF499*<-?<-?<-PIN<-TUDOR+SF1-SF2-Helicase+iREase+CTD+REase | 929 | Bacteria>Pseudomonadota>unclassified_Pseudomonadota | GCA_018242465.1 | Pseudomonadota_bacterium | JSS25_05685 | 37623..40412 | 1977087 | ATP-binding-protein-[Proteobacteria-bacterium] |
| NCC05043.1 | AAA+ATPase+wHTH+Alpha+Beta-domain+wHTH+RRM | Type-1_BR.RM.like.systems | TUDOR+SF1-SF2-Helicase+iREase+CTD+REase->METHYLASE->DUF499*-> | 621 | Bacteria>Pseudomonadota>unclassified_Pseudomonadota | GCA_009929575.1 | Pseudomonadota_bacterium | EOM37_13610 | 6622..8484 | 1977087 | ATP-binding-protein-[Pseudomonadota-bacterium] |
| MCL1861026.1 | AAA+ATPase+wHTH+Alpha+Beta-domain+wHTH+RRM | Type-1_BR.RM.like.systems | PIN->METHYLASE->?->DUF499*-><-?<-?<-ZnR+TOPRIM+D5ATPase<-?<-HTH_17 | 943 | Bacteria>Pseudomonadota>unclassified_Pseudomonadota | GCA_009778425.1 | Pseudomonadota_bacterium | FWG52_05760 | 4102..6933 | 1977087 | DUF499-domain-containing-protein-[Proteobacteria-bacterium] |
| MDA8349445.1 | AAA+ATPase+wHTH+Alpha+Beta-domain+wHTH+RRM | Type-1_BR.RM.like.systems | METHYLASE-p->DUF499*->REase+nSTAND1-> | 941 | Bacteria>Pseudomonadota>unclassified_Pseudomonadota | GCA_028067895.1 | Pseudomonadota_bacterium | M0038_11690 | 747..3572 | 1977087 | MAG:-DUF499-domain-containing-protein-[Pseudomonadota-bacterium] |
| MBU1696973.1 | AAA+ATPase+wHTH+Alpha+Beta-domain+wHTH+RRM | Type-1_BR.RM.like.systems | <-DUF499*<-XIS-HTH\|\|HTH_3->STYKinase-like-> | 134 | Bacteria>Pseudomonadota>unclassified_Pseudomonadota | GCA_018823085.1 | Pseudomonadota_bacterium | KKD21_08020 | 95..499 | 1977087 | hypothetical-protein-[Pseudomonadota-bacterium] |
| MCG8418451.1 | AAA+ATPase+wHTH+Alpha+Beta-domain+wHTH+RRM | Type-1_BR.RM.like.systems | <-DUF499*<-REase<-METHYLASE<-METHYLASE<-TUDOR+SF1-SF2-Helicase+iREase+CTD+REase\|\|Phage_integrase-> | 934 | Bacteria>Pseudomonadota>unclassified_Pseudomonadota | GCA_022360315.1 | Pseudomonadota_bacterium | MJE77_10965 | 1001..3805 | 1977087 | DUF499-domain-containing-protein-[Proteobacteria-bacterium] |
| MBU2056000.1 | AAA+ATPase+wHTH+Alpha+Beta-domain+wHTH+RRM | Type-1_BR.RM.like.systems | wHTH+REase->SF1-SF2-Helicase->METHYLASE->REase->PolB->HEPN->DUF499*-><-?<-?<-HTH | 950 | Bacteria>Pseudomonadota>unclassified_Pseudomonadota | GCA_018825885.1 | Pseudomonadota_bacterium | KKC25_13705 | 11854..14706 | 1977087 | DUF499-domain-containing-protein-[Proteobacteria-bacterium] |
| MCX7113477.1 | AAA+ATPase+wHTH+Alpha+Beta-domain+wHTH+RRM | Type-1_BR.RM.like.systems | <-DUF499*<-?<-METHYLASE<-?<-ORC-CDC6+REase<-TUDOR+SF1-SF2-Helicase+iREase+CTD+REase | 938 | Bacteria>Pseudomonadota>unclassified_Pseudomonadota | GCA_026399135.1 | Pseudomonadota_bacterium | NTX45_25930 | 1011..3827 | 1977087 | MAG:-DUF499-domain-containing-protein-[Proteobacteria-bacterium] |
| MBU1184930.1 | AAA+ATPase+wHTH+Alpha+Beta-domain+wHTH+RRM | Type-1_BR.RM.like.systems | Synaptojanin->?->SF1-SF2-Helicase->tRNA-><-DUF499*<-METHYLASE | 920 | Bacteria>Pseudomonadota>unclassified_Pseudomonadota | GCA_018819165.1 | Pseudomonadota_bacterium | KJ663_07105 | 30478..33240 | 1977087 | DUF499-domain-containing-protein-[Proteobacteria-bacterium] |
| MBU0652986.1 | AAA+ATPase+wHTH+Alpha+Beta-domain+wHTH+RRM | Type-1_BR.RM.like.systems | AAA-ATPase+REase->DUF499*-> | 325 | Bacteria>Pseudomonadota>unclassified_Pseudomonadota | GCA_018814185.1 | Pseudomonadota_bacterium | KKG96_08920 | 1165..2139 | 1977087 | DUF499-domain-containing-protein-[Pseudomonadota-bacterium] |
| OGW72082.1 | AAA+ATPase+wHTH+Alpha+Beta-domain+wHTH+RRM | Type-1_BR.RM.like.systems | ParB+HNH->REase->TUDOR+SF1-SF2-Helicase+iREase+CTD+REase->?->METHYLASE->DUF499-p->DUF499*->REase-><-DNA_methylase\|\|MORC+HISKIN->REC-><-Phage_integrase+ZnR | 616 | Bacteria>PVC_group>Candidatus_Omnitrophota>unclassified_Candidatus_Omnitrophota | GCA_001803975.1 | Omnitrophica_bacterium_GWA2_41_15 | A2047_01995 | 16552..18402 | 1801814 | MAG:-hypothetical-protein-A2047_01995-[Omnitrophica-bacterium-GWA2_41_15] |
| OQX87553.1 | AAA+ATPase+wHTH+Alpha+Beta-domain+wHTH+RRM | Type-1_BR.RM.like.systems | <-TPR<-Radical-SAM+TRAM<-?<-?<-?<-DUF499*\|\|?-><-?<-METHYLASE<-?<-?<-TUDOR+SF1-SF2-Helicase+iREase+CTD+REase | 872 | Bacteria>PVC_group>Candidatus_Omnitrophota>unclassified_Candidatus_Omnitrophota | GCA_002085345.1 | Candidatus_Omnitrophica_bacterium_4484_70.2 | B6D55_03070 | 22571..25189 | 1970783 | hypothetical-protein-B6D55_03070-[Candidatus-Omnitrophica-bacterium-4484_70.2] |
| MBI3319953.1 | AAA+ATPase+wHTH+Alpha+Beta-domain+wHTH+RRM | Type-1_BR.RM.like.systems | <-ORC-CDC6+REase<-?<-?<-MqsA_antitoxin+HTH_3<-?<-DUF499*<-HEPN<-PolB<-METHYLASE | 925 | Bacteria>PVC_group>Candidatus_Omnitrophota>unclassified_Candidatus_Omnitrophota | GCA_016196435.1 | Candidatus_Omnitrophota_bacterium | HYZ89_05150 | 6165..8942 | 2035772 | ATP-binding-protein-[Candidatus-Omnitrophica-bacterium] |
| MCM8768828.1 | AAA+ATPase+wHTH+Alpha+Beta-domain+wHTH+RRM | Type-1_BR.RM.like.systems | PolB->HEPN->DUF499*-><-?<-?<-?<-?<-Peptidase_M55<-Radical-SAM | 875 | Bacteria>PVC_group>Candidatus_Omnitrophota>unclassified_Candidatus_Omnitrophota | GCA_023818865.1 | Candidatus_Omnitrophota_bacterium | NC911_03985 | 646..3273 | 2035772 | DUF499-domain-containing-protein-[Candidatus-Omnitrophica-bacterium] |
| MBQ9726423.1 | AAA+ATPase+wHTH+Alpha+Beta-domain+wHTH+RRM | Type-1_BR.RM.like.systems | VirB11+VirB4-FtsK->BirA-HTH->TUDOR+SF1-SF2-Helicase+iREase+CTD+REase->METHYLASE->DUF499*->DUF4417->?->?->?->HTH+MPTase-> | 933 | Bacteria>PVC_group>Kiritimatiellota>Kiritimatiellia>unclassified_Kiritimatiellia | GCA_017457625.1 | Kiritimatiellia_bacterium | IJV65_02840 | 14824..17625 | 2212473 | ATP-binding-protein-[Kiritimatiellae-bacterium] |
| MBQ7190578.1 | AAA+ATPase+wHTH+Alpha+Beta-domain+wHTH+RRM | Type-1_BR.RM.like.systems | TUDOR+SF1-SF2-Helicase+iREase+CTD+REase-p->METHYLASE->DUF499*->?->ORC-CDC6+REase-> | 952 | Bacteria>PVC_group>Kiritimatiellota>Kiritimatiellia>unclassified_Kiritimatiellia | GCA_017450695.1 | Kiritimatiellia_bacterium | IJR99_14315 | 4122..6980 | 2212473 | ATP-binding-protein-[Kiritimatiellae-bacterium] |
| MCZ7591035.1 | AAA+ATPase+wHTH+Alpha+Beta-domain+wHTH+RRM | Type-1_BR.RM.like.systems | <-PAIREDC-HTH\|\|?-><-DUF499*<-?<-PIWI<-?<-METHYLASE-p<-METHYLASE<-HEPN | 493 | Bacteria>PVC_group>Kiritimatiellota>Kiritimatiellia>unclassified_Kiritimatiellia | GCA_027493775.1 | Kiritimatiellia_bacterium | M5U15_02285 | 473098..474579 | 2212473 | MAG:-DUF499-domain-containing-protein-[Kiritimatiellae-bacterium] |
| WP_168963703.1 | AAA+ATPase+wHTH+Alpha+Beta-domain+wHTH+RRM | Type-1_BR.RM.like.systems | XIS-HTH-><-?<-Phage_integrase\|\|ORC-CDC6+REase->?-><-?<-DUF499*<-NERD<-OLD-ABC<-METHYLASE<-HNH<-OLD-ABC<-TUDOR+SF1-SF2-Helicase+iREase+CTD+REase | 935 | Bacteria>PVC_group>Lentisphaerota>Lentisphaeria>Victivallales | GCF_012843305.1 | Victivallis_vadensis | HF882_RS18870 | 10950..13757 | 172901 | DUF499-domain-containing-protein-[Victivallis-vadensis] |
| MBR6471930.1 | AAA+ATPase+wHTH+Alpha+Beta-domain+wHTH+RRM | Type-1_BR.RM.like.systems | TUDOR+SF1-SF2-Helicase+iREase+CTD+REase->METHYLASE->OLD-ABC->NERD->?->DUF499*-> | 933 | Bacteria>PVC_group>Lentisphaerota>Lentisphaeria>Victivallales | GCA_017538465.1 | Victivallales_bacterium | IKS83_09045 | 14301..17102 | 2212475 | ATP-binding-protein-[Victivallales-bacterium] |
| MBQ6473657.1 | AAA+ATPase+wHTH+Alpha+Beta-domain+wHTH+RRM | Type-1_BR.RM.like.systems | Schlafen+HATPase+HTH->OLD-ABC->HNH->METHYLASE->OLD-ABC->NERD->DUF499*-> | 944 | Bacteria>PVC_group>Lentisphaerota>Lentisphaeria>Victivallales | GCA_017431185.1 | Victivallales_bacterium | IJJ33_16845 | 18968..21802 | 2212475 | ATP-binding-protein-[Victivallales-bacterium] |
| MBR6058788.1 | AAA+ATPase+wHTH+Alpha+Beta-domain+wHTH+RRM | Type-1_BR.RM.like.systems | METHYLASE->OLD-ABC->NERD->OLD-ABC->NERD->DUF499*-><-?<-?<-?<-?\|\|?-><-Mbetalac | 929 | Bacteria>PVC_group>Lentisphaerota>Lentisphaeria>Victivallales | GCA_017542865.1 | Victivallales_bacterium | IKP58_11525 | 37212..40001 | 2212475 | ATP-binding-protein-[Victivallales-bacterium] |
| OGV65464.1 | AAA+ATPase+wHTH+Alpha+Beta-domain+wHTH+RRM | Type-1_BR.RM.like.systems | BrnT_toxin->CopG_antitoxin->?-><-?<-?<-?<-DUF499*<-?<-METHYLASE<-?<-HEPN<-TUDOR+SF1-SF2-Helicase+iREase+CTD+REase<-HEPN | 939 | Bacteria>PVC_group>Lentisphaerota>unclassified_Lentisphaerota | GCA_001803235.1 | Lentisphaerae_bacterium_RIFOXYA12_64_32 | A3K18_32130 | 5290..8109 | 1798577 | MAG:-hypothetical-protein-A3K18_32130-[Lentisphaerae-bacterium-RIFOXYA12_64_32] |
| MBE6357833.1 | AAA+ATPase+wHTH+Alpha+Beta-domain+wHTH+RRM | Type-1_BR.RM.like.systems | ParB-HTH->?-><-?<-?<-DUF499*<-REase<-OLD-ABC<-METHYLASE<-HNH<-OLD-ABC<-TUDOR+SF1-SF2-Helicase+iREase+CTD+REase | 938 | Bacteria>PVC_group>Lentisphaerota>unclassified_Lentisphaerota | GCA_015061525.1 | Lentisphaerota_bacterium | E7057_01125 | 3177..5993 | 1932692 | ATP-binding-protein-[Lentisphaerae-bacterium] |
| OQC12437.1 | AAA+ATPase+wHTH+Alpha+Beta-domain+wHTH+RRM | Type-1_BR.RM.like.systems | TUDOR+SF1-SF2-Helicase+iREase+CTD+REase->MqsA_antitoxin+HTH_3->?->METHYLASE->OLD-ABC->TOPRIM->DUF499*-><-?<-?<-ZnR<-35exo+SF1-SF2-Helicase | 976 | Bacteria>PVC_group>Lentisphaerota>unclassified_Lentisphaerota | GCA_002070945.1 | Lentisphaerae_bacterium_ADurb.Bin082 | BWX73_02930 | 53685..56615 | 1852899 | MAG:-hypothetical-protein-BWX73_02930-[Lentisphaerae-bacterium-ADurb.Bin082] |
| NLG01020.1 | AAA+ATPase+wHTH+Alpha+Beta-domain+wHTH+RRM | Type-1_BR.RM.like.systems | <-DOC+HTH<-?<-DUF499*<-?<-TIR<-TIR<-METHYLASE<-REase<-SF1-SF2-Helicase | 925 | Bacteria>PVC_group>Lentisphaerota>unclassified_Lentisphaerota | GCA_012515415.1 | Lentisphaerota_bacterium | GX565_12840 | 2112..4889 | 1932692 | ATP-binding-protein-[Lentisphaerae-bacterium] |
| NLN03021.1 | AAA+ATPase+wHTH+Alpha+Beta-domain+wHTH+RRM | Type-1_BR.RM.like.systems | <-DUF499*<-REase<-METHYLASE-p | 437 | Bacteria>PVC_group>Lentisphaerota>unclassified_Lentisphaerota | GCA_012521875.1 | Lentisphaerota_bacterium | GX174_14140 | 1..1313 | 1932692 | ATP-binding-protein-[Lentisphaerota-bacterium] |
| NLF19752.1 | AAA+ATPase+wHTH+Alpha+Beta-domain+wHTH+RRM | Type-1_BR.RM.like.systems | <-DUF499*<-wHTH+REase<-METHYLASE<-?<-?<-?<-Schlafen+HATPase+HTH | 905 | Bacteria>PVC_group>Lentisphaerota>unclassified_Lentisphaerota | GCA_012514865.1 | Lentisphaerota_bacterium | GX595_21175 | 1..2716 | 1932692 | ATP-binding-protein-[Lentisphaerae-bacterium] |
| NLE53431.1 | AAA+ATPase+wHTH+Alpha+Beta-domain+wHTH+RRM | Type-1_BR.RM.like.systems | METHYLASE-p->Schlafen+HATPase+HTH->DUF499*-> | 656 | Bacteria>PVC_group>Lentisphaerota>unclassified_Lentisphaerota | GCA_012514365.1 | Lentisphaerota_bacterium | GX617_00715 | 3231..5197 | 1932692 | ATP-binding-protein-[Lentisphaerota-bacterium] |
| KAA3596712.1 | AAA+ATPase+wHTH+Alpha+Beta-domain+wHTH+RRM | Type-1_BR.RM.like.systems | METHYLASE-p->?->HicB->HicA->DUF499*-><-?\|\|zf-ISL3+HTH_Tnp_ISL3+DDE_Tnp_ISL3-> | 981 | Bacteria>PVC_group>Planctomycetota>Candidatus_Brocadiia>Candidatus_Brocadiales | GCA_008501815.1 | Candidatus_Scalindua_sp. | DWQ00_14770 | 3047..5992 | 2268189 | MAG:-DUF499-domain-containing-protein-[Candidatus-Scalindua-sp.] |
| TVL98635.1 | AAA+ATPase+wHTH+Alpha+Beta-domain+wHTH+RRM | Type-1_BR.RM.like.systems | TUDOR+SF1-SF2-Helicase+iREase+CTD+REase->OLD-ABC->METHYLASE->REase->DUF499*-> | 981 | Bacteria>PVC_group>Planctomycetota>Candidatus_Brocadiia>Candidatus_Brocadiales | GCA_007618135.1 | Candidatus_Brocadia_sp._WS118 | CV087_21015 | 12418..15363 | 2040625 | MAG:-AAA-family-ATPase-[Candidatus-Brocadia-sp.-WS118] |
| WP_052562727.1 | AAA+ATPase+wHTH+Alpha+Beta-domain+wHTH+RRM | Type-1_BR.RM.like.systems | TUDOR+SF1-SF2-Helicase+iREase+CTD+REase->?->?->PIN->METHYLASE->DUF433->DUF499*->?->?-><-?\|\|DUF1264-> | 925 | Bacteria>PVC_group>Planctomycetota>Candidatus_Brocadiia>Candidatus_Brocadiales | GCF_000949635.1 | Brocadia | FBQ80_RS01160 | 33537..36314 | 380240 | DUF499-domain-containing-protein-[Candidatus-Brocadia-sinica] |
| KXK32838.1 | AAA+ATPase+wHTH+Alpha+Beta-domain+wHTH+RRM | Type-1_BR.RM.like.systems | <-DUF1264\|\|?-><-?<-?<-DUF499*<-DUF433<-METHYLASE-p<-METHYLASE<-PIN | 925 | Bacteria>PVC_group>Planctomycetota>Candidatus_Brocadiia>Candidatus_Brocadiales | GCA_001567345.1 | Candidatus_Brocadia_sinica | UZ01_00260 | 122708..125485 | 795830 | hypothetical-protein-UZ01_00260-[Candidatus-Brocadia-sinica] |
| UJS18027.1 | AAA+ATPase+wHTH+Alpha+Beta-domain+wHTH+RRM | Type-1_BR.RM.like.systems | Aldedh->TUDOR+SF1-SF2-Helicase+iREase+CTD+REase->METHYLASE->ORC-CDC6+REase->DUF499*->DUF499-p->SLATT-><-?<-URI-DNase+HhH-RADC | 841 | Bacteria>PVC_group>Planctomycetota>Candidatus_Brocadiia>Candidatus_Brocadiales | GCA_021650895.1 | Candidatus_Jettenia_sp. | L3J17_02955 | 714724..717249 | 2170669 | MAG:-DUF499-domain-containing-protein-[Candidatus-Jettenia-sp.] |
| MBX3409892.1 | AAA+ATPase+wHTH+Alpha+Beta-domain+wHTH+RRM | Type-1_BR.RM.like.systems | RAMA->GDResolvase+Recombinase+ZnR->ParB-><-?<-Radical-SAM<-DUF499*<-METHYLASE<-HNH<-OLD-ABC<-TUDOR+SF1-SF2-Helicase+iREase+CTD+REase\|\|HTH->MPTase-> | 947 | Bacteria>PVC_group>Planctomycetota>Phycisphaerae>Phycisphaerales | GCA_019636575.1 | Phycisphaeraceae_bacterium | KF859_08400 | 137437..140280 | 2026777 | ATP-binding-protein-[Phycisphaeraceae-bacterium] |
| MBX3408154.1 | AAA+ATPase+wHTH+Alpha+Beta-domain+wHTH+RRM | Type-1_BR.RM.like.systems | RAMA->GDResolvase+Recombinase+ZnR->ParB-><-?<-Radical-SAM<-DUF499*<-METHYLASE<-HNH<-OLD-ABC<-TUDOR+SF1-SF2-Helicase+iREase+CTD+REase | 858 | Bacteria>PVC_group>Planctomycetota>Phycisphaerae>Phycisphaerales | GCA_019636415.1 | Phycisphaeraceae_bacterium | KF869_15485 | 6654..9230 | 2026777 | ATP-binding-protein-[Phycisphaeraceae-bacterium] |
| MBX3387703.1 | AAA+ATPase+wHTH+Alpha+Beta-domain+wHTH+RRM | Type-1_BR.RM.like.systems | <-DUF499*<-METHYLASE<-TUDOR+SF1-SF2-Helicase+iREase+CTD+REase\|\|HTH->MPTase->N6_N4_MTase-> | 599 | Bacteria>PVC_group>Planctomycetota>Phycisphaerae>Phycisphaerales | GCA_019636535.1 | Phycisphaeraceae_bacterium | KF768_14130 | 1..1798 | 2026777 | ATP-binding-protein-[Phycisphaeraceae-bacterium] |
| TVQ33187.1 | AAA+ATPase+wHTH+Alpha+Beta-domain+wHTH+RRM | Type-1_BR.RM.like.systems | FURR-HTH->tRNA->?->RAMA->GDResolvase+Recombinase+ZnR->ParB-><-DUF499* | 951 | Bacteria>PVC_group>Planctomycetota>Phycisphaerae>Phycisphaerales | GCA_007693965.1 | Phycisphaeraceae_bacterium | EA376_03820 | 175165..178020 | 2026777 | MAG:-ATP-binding-protein-[Phycisphaeraceae-bacterium] |
| MBX3387113.1 | AAA+ATPase+wHTH+Alpha+Beta-domain+wHTH+RRM | Type-1_BR.RM.like.systems | DUF499*->Radical-SAM->?->?-><-GDResolvase+Recombinase+ZnR<-RAMA | 880 | Bacteria>PVC_group>Planctomycetota>Phycisphaerae>Phycisphaerales | GCA_019636535.1 | Phycisphaeraceae_bacterium | KF768_11125 | 1..2643 | 2026777 | ATP-binding-protein-[Phycisphaeraceae-bacterium] |
| MBX3408175.1 | AAA+ATPase+wHTH+Alpha+Beta-domain+wHTH+RRM | Type-1_BR.RM.like.systems | <-NADAR<-?<-?<-Y2_Tnp\|\|?-><-Phage_integrase<-DUF499* | 917 | Bacteria>PVC_group>Planctomycetota>Phycisphaerae>Phycisphaerales | GCA_019636415.1 | Phycisphaeraceae_bacterium | KF869_15590 | 13486..16239 | 2026777 | ATP-binding-protein-[Phycisphaeraceae-bacterium] |
| MCL4742828.1 | AAA+ATPase+wHTH+Alpha+Beta-domain+wHTH+RRM | Type-1_BR.RM.like.systems | <-N6_N4_MTase<-MPTase<-HTH<-DUF499*<-METHYLASE<-TUDOR+SF1-SF2-Helicase+iREase+CTD+REase<-ParB<-GDResolvase+Recombinase+ZnR<-RAMA\|\|HNH-> | 949 | Bacteria>PVC_group>Planctomycetota>Phycisphaerae>Phycisphaerales | GCA_023384195.1 | Phycisphaerales_bacterium | KJZ54_11580 | 4360..7209 | 2052180 | DUF499-domain-containing-protein-[Phycisphaerales-bacterium] |
| NOT00635.1 | AAA+ATPase+wHTH+Alpha+Beta-domain+wHTH+RRM | Type-1_BR.RM.like.systems | <-DUF499*<-DOC+HTH<-METHYLASE<-TUDOR+SF1-SF2-Helicase+iREase+CTD+REase\|\|?-><-TUDOR+SF1-SF2-Helicase+iREase+CTD+REase-p<-GDResolvase+Recombinase+ZnR | 953 | Bacteria>PVC_group>Planctomycetota>Phycisphaerae>Phycisphaerales | GCA_013140575.1 | Phycisphaerales_bacterium | HOP29_08410 | 245..3106 | 2052180 | DUF499-domain-containing-protein-[Phycisphaerales-bacterium] |
| MBK8915085.1 | AAA+ATPase+wHTH+Alpha+Beta-domain+wHTH+RRM | Type-1_BR.RM.like.systems | HIN-HTH->?-><-zf-IS66+DDE_Tnp_IS66\|\|?->?-><-DUF499*<-?<-METHYLASE<-HKD-DNase<-TUDOR+SF1-SF2-Helicase+iREase+CTD+REase | 943 | Bacteria>PVC_group>Planctomycetota>Phycisphaerae>Phycisphaerales | GCA_016710275.1 | Phycisphaerales_bacterium | IPM64_10895 | 2678666..2681497 | 2052180 | ATP-binding-protein-[Phycisphaerales-bacterium] |
| HAU37798.1 | AAA+ATPase+wHTH+Alpha+Beta-domain+wHTH+RRM | Type-1_BR.RM.like.systems | <-AbiEi_4<-LPD36+METHYLASE+SF1-SF2-Helicase<-?\|\|?-><-PIWI<-TIR+PNTD2<-DUF499*<-?<-METHYLASE | 924 | Bacteria>PVC_group>Planctomycetota>Phycisphaerae>Phycisphaerales | GCA_003485185.1 | Phycisphaerales_bacterium | DCX07_08790 | 42407..45181 | 2052180 | AAA-family-ATPase-[Phycisphaerales-bacterium] |
| UCC29403.1 | AAA+ATPase+wHTH+Alpha+Beta-domain+wHTH+RRM | Type-1_BR.RM.like.systems | <-Pkinase<-SIGMA-HTH<-?\|\|ORC-CDC6+REase-><-?<-DUF499*\|\|?->?->MPTase->?->?->MFS_1-> | 503 | Bacteria>PVC_group>Planctomycetota>Phycisphaerae>Phycisphaerales | GCA_020344555.1 | Phycisphaerales_bacterium | JSU86_14520 | 3569391..3570900 | 2052180 | MAG:-DUF499-domain-containing-protein,-partial-[Phycisphaerales-bacterium] |
| MBN1125969.1 | AAA+ATPase+wHTH+Alpha+Beta-domain+wHTH+RRM | Type-1_BR.RM.like.systems | Phage_integrase-><-?<-DUF499*<-REase<-METHYLASE<-SF1-SF2-Helicase<-TUDOR+SF1-SF2-Helicase+iREase+CTD+REase\|\|?->PAPS_reduct-> | 912 | Bacteria>PVC_group>Planctomycetota>Phycisphaerae>Sedimentisphaerales | GCA_016926155.1 | Sedimentisphaerales_bacterium | JXA82_13255 | 1576..4314 | 2813034 | ATP-binding-protein-[Sedimentisphaerales-bacterium] |
| MBN2133793.1 | AAA+ATPase+wHTH+Alpha+Beta-domain+wHTH+RRM | Type-1_BR.RM.like.systems | <-XIS-HTH<-DUF499*<-?<-?<-METHYLASE<-SF1-SF2-Helicase<-TUDOR+SF1-SF2-Helicase+iREase+CTD+REase | 956 | Bacteria>PVC_group>Planctomycetota>Phycisphaerae>Sedimentisphaerales | GCA_016932895.1 | Sedimentisphaerales_bacterium | JW741_30100 | 1712..4582 | 2813034 | ATP-binding-protein-[Sedimentisphaerales-bacterium] |
| MBN1509225.1 | AAA+ATPase+wHTH+Alpha+Beta-domain+wHTH+RRM | Type-1_BR.RM.like.systems | <-PAPS_reduct<-?<-?<-KAP-NTPase<-DUF499*<-HTH<-REase<-METHYLASE | 922 | Bacteria>PVC_group>Planctomycetota>Phycisphaerae>Sedimentisphaerales | GCA_016928975.1 | Sedimentisphaerales_bacterium | JW955_20435 | 4212..6980 | 2813034 | ATP-binding-protein-[Sedimentisphaerales-bacterium] |
| MBN2561533.1 | AAA+ATPase+wHTH+Alpha+Beta-domain+wHTH+RRM | Type-1_BR.RM.like.systems | <-METHYLASE-p<-Radical-SAM<-?<-?<-DUF499*<-AbiGi<-METHYLASE<-SF1-SF2-Helicase<-TUDOR+SF1-SF2-Helicase+iREase+CTD+REase<-XIS-HTH\|\|XIS-HTH-> | 994 | Bacteria>PVC_group>Planctomycetota>Phycisphaerae>unclassified_Phycisphaerae | GCA_016935885.1 | Phycisphaerae_bacterium | JXQ75_11445 | 19066..22050 | 2026778 | ATP-binding-protein-[Phycisphaerae-bacterium] |
| MAO24327.1 | AAA+ATPase+wHTH+Alpha+Beta-domain+wHTH+RRM | Type-1_BR.RM.like.systems | TUDOR+SF1-SF2-Helicase+iREase+CTD+REase->TIR->METHYLASE->REase->DUF499*-><-GDResolvase+Recombinase+ZnR<-RAMA | 991 | Bacteria>PVC_group>Planctomycetota>Phycisphaerae>unclassified_Phycisphaerae | GCA_002696165.1 | Phycisphaerae_bacterium | CMJ25_26550 | 12971..15946 | 2026778 | AAA-family-ATPase-[Phycisphaerae-bacterium] |
| GJQ27840.1 | AAA+ATPase+wHTH+Alpha+Beta-domain+wHTH+RRM | Type-1_BR.RM.like.systems | <-N6_N4_MTase<-MPTase<-HTH<-DUF499*<-?<-METHYLASE<-Schlafen+HATPase+HTH\|\|?-><-?<-TUDOR+SF1-SF2-Helicase+iREase+CTD+REase | 926 | Bacteria>PVC_group>Planctomycetota>Phycisphaerae>unclassified_Phycisphaerae | GCA_022180675.1 | Phycisphaerae_bacterium | HBSAPP02_28720 | 66529..69309 | 2026778 | hypothetical-protein-HBSAPP02_28720-[Phycisphaerae-bacterium] |
| MCK4341360.1 | AAA+ATPase+wHTH+Alpha+Beta-domain+wHTH+RRM | Type-1_BR.RM.like.systems | HTH_3-><-GDResolvase+Recombinase+ZnR<-?<-DUF499*<-PIN<-?<-?<-METHYLASE-p | 947 | Bacteria>PVC_group>Planctomycetota>Phycisphaerae>unclassified_Phycisphaerae | GCA_023128695.1 | Phycisphaerae_bacterium | KAY37_06525 | 2745..5588 | 2026778 | ATP-binding-protein-[Phycisphaerae-bacterium] |
| NLX57996.1 | AAA+ATPase+wHTH+Alpha+Beta-domain+wHTH+RRM | Type-1_BR.RM.like.systems | TUDOR+SF1-SF2-Helicase+iREase+CTD+REase->HEPN->?->METHYLASE->DUF499*-> | 975 | Bacteria>PVC_group>Planctomycetota>Phycisphaerae>unclassified_Phycisphaerae | - | Phycisphaerae_bacterium | GXY74_02785 | 9938..12865 | 2026778 | ATP-binding-protein-[Phycisphaerae-bacterium] |
| MCJ7543598.1 | AAA+ATPase+wHTH+Alpha+Beta-domain+wHTH+RRM | Type-1_BR.RM.like.systems | <-DUF499*<-?<-HKD-DNase<-?<-METHYLASE | 893 | Bacteria>PVC_group>Planctomycetota>Phycisphaerae>unclassified_Phycisphaerae | GCA_022866005.1 | Phycisphaerae_bacterium | MUP47_03365 | 1912..4593 | 2026778 | ATP-binding-protein-[Phycisphaerae-bacterium] |
| MDB5306879.1 | AAA+ATPase+wHTH+Alpha+Beta-domain+wHTH+RRM | Type-1_BR.RM.like.systems | <-DUF499*<-PIN<-Phd-YefM<-METHYLASE<-TUDOR+SF1-SF2-Helicase+iREase+CTD+REase-p<-TUDOR+SF1-SF2-Helicase+iREase+CTD+REase | 207 | Bacteria>PVC_group>Planctomycetota>Planctomycetia>Gemmatales | GCA_028288765.1 | Gemmataceae_bacterium | JWO38_1081 | 91..714 | 2052164 | MAG:-hypothetical-protein-JWO38_1081-[Gemmataceae-bacterium] |
| GIW84013.1 | AAA+ATPase+wHTH+Alpha+Beta-domain+wHTH+RRM | Type-1_BR.RM.like.systems | GDResolvase+Recombinase+ZnR->ParB->TUDOR+SF1-SF2-Helicase+iREase+CTD+REase->METHYLASE->DUF433->PIN->DUF499*->HTH->MPTase-><-DUF4096+DDE_Tnp_1+DDE_Tnp | 924 | Bacteria>PVC_group>Planctomycetota>Planctomycetia>Gemmatales | GCA_026004695.1 | Gemmataceae_bacterium | KatS3mg106_526 | 15876..18650 | 2052164 | MAG:-hypothetical-protein-KatS3mg106_526-[Gemmataceae-bacterium] |
| WP_210660196.1 | AAA+ATPase+wHTH+Alpha+Beta-domain+wHTH+RRM | Type-1_BR.RM.like.systems | <-PadR-HTH\|\|?->?-><-?<-DUF499*<-REase<-METHYLASE<-DUF4065<-HD<-TUDOR+SF1-SF2-Helicase+iREase+CTD+REase<-ZnR+TOPRIM+D5ATPase | 985 | Bacteria>PVC_group>Planctomycetota>Planctomycetia>Gemmatales | GCF_017939745.1 | Gemmata_palustris | J8F10_RS30110 | 7260785..7263742 | 2822762 | DUF499-domain-containing-protein-[Gemmata-palustris] |
| WP_171472664.1 | AAA+ATPase+wHTH+Alpha+Beta-domain+wHTH+RRM | Type-1_BR.RM.like.systems | <-ParB<-HTH\|\|TUDOR+SF1-SF2-Helicase+iREase+CTD+REase->METHYLASE->DUF499*->?->?-><-?<-Phage_integrase<-?<-ZnR+TOPRIM+D5ATPase | 941 | Bacteria>PVC_group>Planctomycetota>Planctomycetia>Gemmatales | GCF_013128195.2 | Frigoriglobus_tundricola | FTUN_RS21585 | 5224957..5227782 | 2774151 | DUF499-domain-containing-protein-[Frigoriglobus-tundricola] |
| MCI0639191.1 | AAA+ATPase+wHTH+Alpha+Beta-domain+wHTH+RRM | Type-1_BR.RM.like.systems | TUDOR+SF1-SF2-Helicase+iREase+CTD+REase-p->TUDOR+SF1-SF2-Helicase+iREase+CTD+REase->METHYLASE->DUF499*-> | 949 | Bacteria>PVC_group>Planctomycetota>Planctomycetia>Gemmatales | GCA_022616245.1 | Gemmataceae_bacterium | LOY70_08955 | 7374..10223 | 2052164 | DUF499-domain-containing-protein-[Gemmataceae-bacterium] |
| MSR55922.1 | AAA+ATPase+wHTH+Alpha+Beta-domain+wHTH+RRM | Type-1_BR.RM.like.systems | <-DUF499*<-REase<-PIN<-DUF433<-METHYLASE-p | 984 | Bacteria>PVC_group>Planctomycetota>Planctomycetia>Gemmatales | GCA_009691845.1 | Gemmataceae_bacterium | EXS09_22015 | 769..3723 | 2052164 | ATP-binding-protein-[Gemmataceae-bacterium] |
| MCI0378738.1 | AAA+ATPase+wHTH+Alpha+Beta-domain+wHTH+RRM | Type-1_BR.RM.like.systems | <-DUF499*<-METHYLASE<-REase | 207 | Bacteria>PVC_group>Planctomycetota>Planctomycetia>Gemmatales | GCA_022614915.1 | Gemmataceae_bacterium | LO215_14110 | 1..621 | 2052164 | DUF499-domain-containing-protein-[Gemmataceae-bacterium] |
| MCS7166809.1 | AAA+ATPase+wHTH+Alpha+Beta-domain+wHTH+RRM | Type-1_BR.RM.like.systems | METHYLASE-p->DUF499*-> | 879 | Bacteria>PVC_group>Planctomycetota>Planctomycetia>Gemmatales | GCA_025060195.1 | Gemmatales_bacterium | NZ914_04165 | 501..3140 | 2838789 | DUF499-domain-containing-protein-[Gemmatales-bacterium] |
| WP_013566219.1 | AAA+ATPase+wHTH+Alpha+Beta-domain+wHTH+RRM | Type-1_BR.RM.like.systems | <-NurA<-DUF4338<-?\|\|?-><-REase<-?<-DUF499*<-PIN<-DUF433<-METHYLASE<-TUDOR+SF1-SF2-Helicase+iREase+CTD+REase<-SF1-SF2-Helicase-p<-METHYLASE | 959 | Bacteria>PVC_group>Planctomycetota>Planctomycetia>Isosphaerales | GCF_000186345.1 | Isosphaera_pallida | ISOP_RS17995 | 4934004..4936883 | 128 | ATP-binding-protein-[Isosphaera-pallida] |
| WP_076345234.1 | AAA+ATPase+wHTH+Alpha+Beta-domain+wHTH+RRM | Type-1_BR.RM.like.systems | Glyoxalase->?->?->?->?->tRNA-><-XIS-HTH<-DUF499*<-REase<-OLD-ABC<-METHYLASE<-TUDOR+SF1-SF2-Helicase+iREase+CTD+REase<-ParB | 941 | Bacteria>PVC_group>Planctomycetota>Planctomycetia>Isosphaerales | GCF_001956985.1 | Paludisphaera_borealis | BSF38_RS09985 | 2556227..2559052 | 1387353 | DUF499-domain-containing-protein-[Paludisphaera-borealis] |
| WP_250846938.1 | AAA+ATPase+wHTH+Alpha+Beta-domain+wHTH+RRM | Type-1_BR.RM.like.systems | <-DUF499*<-REase<-OLD-ABC<-METHYLASE<-TUDOR+SF1-SF2-Helicase+iREase+CTD+REase | 939 | Bacteria>PVC_group>Planctomycetota>Planctomycetia>Isosphaerales | GCF_011064615.1 | Aquisphaera_insulae | G6054_RS16920 | 61801..64620 | 2712864 | DUF499-domain-containing-protein-[Aquisphaera-sp.-JC669] |
| WP_148590534.1 | AAA+ATPase+wHTH+Alpha+Beta-domain+wHTH+RRM | Type-1_BR.RM.like.systems | Phd-YefM->HTH_37-><-?<-?<-DUF499*<-?<-METHYLASE<-OLD-ABC<-TUDOR+SF1-SF2-Helicase+iREase+CTD+REase | 981 | Bacteria>PVC_group>Planctomycetota>Planctomycetia>Isosphaerales | GCF_008087625.1 | Aquisphaera_giovannonii | OJF2_RS01285 | 356690..359635 | 406548 | DUF499-domain-containing-protein-[Aquisphaera-giovannonii] |
| MBV8611623.1 | AAA+ATPase+wHTH+Alpha+Beta-domain+wHTH+RRM | Type-1_BR.RM.like.systems | METHYLASE-p->DUF433->PIN->HTH->DUF499*-> | 454 | Bacteria>PVC_group>Planctomycetota>Planctomycetia>Isosphaerales | GCA_019233025.1 | Singulisphaera_sp. | JO034_29790 | 1731..3093 | 1906241 | ATP-binding-protein-[Singulisphaera-sp.] |
| WP_198953633.1 | AAA+ATPase+wHTH+Alpha+Beta-domain+wHTH+RRM | Type-1_BR.RM.like.systems | <-wHTH(SBD)+HNH<-URI-DNase+RAMA<-REase<-EVE<-REase<-DUF499*<-HNH<-OLD-ABC<-METHYLASE<-Peptidase_S8<-Classical-AAA<-TUDOR+SF1-SF2-Helicase+iREase+CTD+REase | 951 | Bacteria>PVC_group>Planctomycetota>Planctomycetia>Pirellulales | GCF_002251265.1 | Rhodopirellula_sp._MGV | CGZ80_RS05865 | 9695..12550 | 2023130 | DUF499-domain-containing-protein-[Rhodopirellula-sp.-MGV] |
| OYP37234.1 | AAA+ATPase+wHTH+Alpha+Beta-domain+wHTH+RRM | Type-1_BR.RM.like.systems | <-wHTH(SBD)+HNH<-URI-DNase+RAMA<-REase<-EVE<-REase<-DUF499*<-HNH<-OLD-ABC<-METHYLASE<-Peptidase_S8<-Classical-AAA<-TUDOR+SF1-SF2-Helicase+iREase+CTD+REase | 948 | Bacteria>PVC_group>Planctomycetota>Planctomycetia>Pirellulales | GCF_002251265.1 | Rhodopirellula_sp._MGV | CGZ80_05855 | 9695..12541 | 2023130 | hypothetical-protein-CGZ80_05855-[Rhodopirellula-sp.-MGV] |
| WP_230262563.1 | AAA+ATPase+wHTH+Alpha+Beta-domain+wHTH+RRM | Type-1_BR.RM.like.systems | <-RAMA<-URI-DNase+RAMA<-REase<-EVE<-REase<-DUF499*<-REase<-OLD-ABC<-METHYLASE<-Peptidase_S8<-Classical-AAA<-TUDOR+SF1-SF2-Helicase+iREase+CTD+REase | 950 | Bacteria>PVC_group>Planctomycetota>Planctomycetia>Pirellulales | GCF_020966635.1 | Stieleria_sp._JC731 | LOC67_RS10680 | 1300044..1302896 | 2894195 | DUF499-domain-containing-protein-[Stieleria-sp.-JC731] |
| WP_224435288.1 | AAA+ATPase+wHTH+Alpha+Beta-domain+wHTH+RRM | Type-1_BR.RM.like.systems | <-NERD+Pkinase+Pkinase+nSTAND3<-HKD-DNase<-SbcC<-DUF1679<-SIR2<-DUF499*<-METHYLASE<-TUDOR+SF1-SF2-Helicase+iREase+CTD+REase<-XIS-HTH | 947 | Bacteria>PVC_group>Planctomycetota>Planctomycetia>Pirellulales | GCF_020150455.1 | Bremerella_volcania | LA756_RS13690 | 3459073..3461916 | 2527984 | DUF499-domain-containing-protein-[Bremerella-volcania] |
| MCE2752488.1 | AAA+ATPase+wHTH+Alpha+Beta-domain+wHTH+RRM | Type-1_BR.RM.like.systems | Methyltransf_21->TUDOR+SF1-SF2-Helicase+iREase+CTD+REase->?->?->METHYLASE->ParB+HNH->DUF499*->REase->ParB+HNH->?->URI-DNase-> | 957 | Bacteria>PVC_group>Planctomycetota>Planctomycetia>Pirellulales | GCA_021299695.1 | Pirellula_sp. | LW720_11440 | 12221..15094 | 117 | ATP-binding-protein-[Pirellula-sp.] |
| MCF7962597.1 | AAA+ATPase+wHTH+Alpha+Beta-domain+wHTH+RRM | Type-1_BR.RM.like.systems | <-Pkinase\|\|Phage_int_SAM_4+Phage_integrase-><-OLD-ABC+TOPRIM<-?<-DUF499*<-METHYLASE | 956 | Bacteria>PVC_group>Planctomycetota>Planctomycetia>Pirellulales | GCA_021736575.1 | Pirellula_sp. | K9M08_17805 | 7810..10680 | 117 | ATP-binding-protein-[Pirellula-sp.] |
| WP_146679086.1 | AAA+ATPase+wHTH+Alpha+Beta-domain+wHTH+RRM | Type-1_BR.RM.like.systems | XIS-HTH->TUDOR+SF1-SF2-Helicase+iREase+CTD+REase->HEPN->METHYLASE->METHYLASE->REase->DUF499*->REase+ParB->RVT->?->RAMA->GDResolvase+Recombinase+ZnR->GDResolvase+Recombinase+ZnR-> | 981 | Bacteria>PVC_group>Planctomycetota>Planctomycetia>Pirellulales | GCF_001610875.1 | Pirellula_sp._SH-Sr6A | VN12_RS23070 | 5951575..5954520 | 1632865 | DUF499-domain-containing-protein-[Pirellula-sp.-SH-Sr6A] |
| WP_146436433.1 | AAA+ATPase+wHTH+Alpha+Beta-domain+wHTH+RRM | Type-1_BR.RM.like.systems | ST-Hiskin->tRNA-><-DUF499*<-REase<-METHYLASE<-PRTase+PRTase<-TUDOR+SF1-SF2-Helicase+iREase+CTD+REase<-XIS-HTH<-ParB | 983 | Bacteria>PVC_group>Planctomycetota>Planctomycetia>Pirellulales | GCF_007859755.1 | Blastopirellula_retiformator | Enr8_RS23480 | 388216..391167 | 2527970 | DUF499-domain-containing-protein-[Blastopirellula-retiformator] |

| Protein-ID | DUF499-architecture | System | Gene-Neighbourhoods | Length | Taxonomy | Assembly-ID | Strain | Locus-Tag | Genomic-coordinates | TaxID | Fasta-defline |
|---|---|---|---|---|---|---|---|---|---|---|---|
| NLF73683.1 | AAA+ATPase+wHTH+Alpha+Beta-domain+wHTH+RRM | Type-1_BR.RM.like.systems | HD->Peptidase_M48->TUDOR+SF1-SF2-Helicase+iREase+CTD+REase->Schlafen+HATPase+HTH->METHYLASE->?->DUF499*->?->?->Primpol->Phage_integrase-> | 979 | Bacteria>PVC_group>Planctomycetota>Planctomycetia>Pirellulales | GCA_012515235.1 | Candidatus_Anammoximicrobium_sp. | GX575_31980 | 17079..20018 | 2699742 | ATP-binding-protein-[Candidatus-Anammoximicrobium-sp.] |
| NLX98932.1 | AAA+ATPase+wHTH+Alpha+Beta-domain+wHTH+RRM | Type-1_BR.RM.like.systems | TUDOR+SF1-SF2-Helicase+iREase+CTD+REase->ParB+HNH->?->METHYLASE->REase->DUF499*->REase->MORC+REase-><-DNA_methylase<-HTH | 983 | Bacteria>PVC_group>Planctomycetota>Planctomycetia>Pirellulales | GCA_012728855.1 | Rhodopirellula_sp. | GXY83_22530 | 8787..11738 | 2024855 | ATP-binding-protein-[Rhodopirellula-sp.] |
| MBN1909337.1 | AAA+ATPase+wHTH+Alpha+Beta-domain+wHTH+RRM | Type-1_BR.RM.like.systems | TUDOR+SF1-SF2-Helicase+iREase+CTD+REase->REase->METHYLASE->?->?->HTH->DUF499*-><-PIN<-HTH+MPTase<-PAPS_reduct<-?\|\|HEPN->HTH-> | 936 | Bacteria>PVC_group>Planctomycetota>Planctomycetia>Pirellulales | GCA_016931075.1 | Pirellulales_bacterium | JW818_06335 | 16839..19649 | 2715253 | ATP-binding-protein-[Pirellulales-bacterium] |
| MBX9790027.1 | AAA+ATPase+wHTH+Alpha+Beta-domain+wHTH+RRM | Type-1_BR.RM.like.systems | Classical-AAA->Peptidase_S8->TUDOR+SF1-SF2-Helicase+iREase+CTD+REase->REase->METHYLASE->?->DUF499*-><-SF1-SF2-Helicase | 926 | Bacteria>PVC_group>Planctomycetota>Planctomycetia>Pirellulales | GCA_019745975.1 | Pirellulales_bacterium | K2Y37_14000 | 59028..61808 | 2715253 | DUF499-domain-containing-protein-[Pirellulales-bacterium] |
| WP_262025600.1 | AAA+ATPase+wHTH+Alpha+Beta-domain+wHTH+RRM | Type-1_BR.RM.like.systems | TUDOR+SF1-SF2-Helicase+iREase+CTD+REase->OLD-ABC->?->HEPN->METHYLASE->?->DUF499*->?->?->?->?->HKD-DNase->SF1-SF2-Helicase-> | 962 | Bacteria>PVC_group>Planctomycetota>Planctomycetia>Pirellulales | GCF_024584705.1 | Blastopirellula_sp._J2-11 | M4951_RS06120 | 1475319..1478207 | 2943192 | DUF499-domain-containing-protein-[Blastopirellula-sp.-J2-11] |
| MBN2290917.1 | AAA+ATPase+wHTH+Alpha+Beta-domain+wHTH+RRM | Type-1_BR.RM.like.systems | METHYLASE->?->HTH->DUF499*->?->XIS-HTH->ZnR+TOPRIM+D5ATPase->?->?->HTH_26-> | 932 | Bacteria>PVC_group>Planctomycetota>Planctomycetia>Pirellulales | GCA_016934075.1 | Pirellulales_bacterium | JXM70_00745 | 4056..6854 | 2715253 | ATP-binding-protein-[Pirellulales-bacterium] |
| GIW94553.1 | AAA+ATPase+wHTH+Alpha+Beta-domain+wHTH+RRM | Type-1_BR.RM.like.systems | <-DUF499*<-REase<-?<-TOPRIM<-OLD-ABC<-METHYLASE-p<-METHYLASE-p | 938 | Bacteria>PVC_group>Planctomycetota>Planctomycetia>Pirellulales | GCA_026004775.1 | Pirellulaceae_bacterium | KatS3mg110_2594 | 12784..15600 | 2699754 | MAG:-hypothetical-protein-KatS3mg110_2594-[Pirellulaceae-bacterium] |
| MCL6502992.1 | AAA+ATPase+wHTH+Alpha+Beta-domain+wHTH+RRM | Type-1_BR.RM.like.systems | <-OBFOLD-NUCLEASE<-DUF499*<-?<-Schlafen+HATPase+HTH<-METHYLASE-p | 973 | Bacteria>PVC_group>Planctomycetota>Planctomycetia>Pirellulales | GCA_023511675.1 | Pirellulales_bacterium | K6T86_09935 | 15292..18213 | 2715253 | DUF499-domain-containing-protein-[Pirellulales-bacterium] |
| NLX99940.1 | AAA+ATPase+wHTH+Alpha+Beta-domain+wHTH+RRM | Type-1_BR.RM.like.systems | <-DUF499*<-PIN<-?<-REase<-?<-METHYLASE<-REase | 981 | Bacteria>PVC_group>Planctomycetota>Planctomycetia>Pirellulales | GCA_012728855.1 | Rhodopirellula_sp. | GXY83_27900 | 2887..5832 | 2024855 | ATP-binding-protein-[Rhodopirellula-sp.] |
| NLF68873.1 | AAA+ATPase+wHTH+Alpha+Beta-domain+wHTH+RRM | Type-1_BR.RM.like.systems | ParB->ParB-><-HEPN<-PIWI<-REase+PNTD1+PNTD2<-?<-DUF499* | 982 | Bacteria>PVC_group>Planctomycetota>Planctomycetia>Pirellulales | GCA_012515235.1 | Candidatus_Anammoximicrobium_sp. | GX575_07440 | 18006..20954 | 2699742 | ATP-binding-protein-[Candidatus-Anammoximicrobium-sp.] |
| MCS7305004.1 | AAA+ATPase+wHTH+Alpha+Beta-domain+wHTH+RRM | Type-1_BR.RM.like.systems | <-ABC-ATPase+SbcC<-?<-?<-?<-?<-Peptidase\|\|DUF499*->?-><-tRNA<-?<-GNAT | 844 | Bacteria>PVC_group>Planctomycetota>Planctomycetia>Pirellulales | GCA_025062105.1 | Thermoguttaceae_bacterium | NZ602_07845 | 39030..41564 | 2691551 | DUF499-domain-containing-protein-[Thermoguttaceae-bacterium] |
| MCS7237814.1 | AAA+ATPase+wHTH+Alpha+Beta-domain+wHTH+RRM | Type-1_BR.RM.like.systems | sGTP-><-?\|\|?-><-?<-?<-?<-DUF499*<-METHYLASE-p<-?<-PolB | 882 | Bacteria>PVC_group>Planctomycetota>Planctomycetia>Pirellulales | GCA_025060325.1 | Thermoguttaceae_bacterium | NZ899_06035 | 153556..156204 | 2691551 | DUF499-domain-containing-protein-[Thermoguttaceae-bacterium] |
| NMC20499.1 | AAA+ATPase+wHTH+Alpha+Beta-domain+wHTH+RRM | Type-1_BR.RM.like.systems | <-DUF499*<-PIN<-DUF433<-METHYLASE-p | 919 | Bacteria>PVC_group>Planctomycetota>Planctomycetia>Pirellulales | GCA_012798155.1 | Thermogutta_sp. | GYA33_08775 | 1111..3870 | 1962930 | ATP-binding-protein-[Thermogutta-sp.] |
| WP_146679186.1 | AAA+ATPase+wHTH+Alpha+Beta-domain+wHTH+RRM | Type-1_BR.RM.like.systems | TUDOR+SF1-SF2-Helicase+iREase+CTD+REase-><-DUF772+DDE_Tnp_1+DDE_Tnp_1_6\|\|?->?->METHYLASE->DUF499*->DDE_Tnp_1_4->DUF499-> | 338 | Bacteria>PVC_group>Planctomycetota>Planctomycetia>Pirellulales | GCF_001610875.1 | Pirellula_sp._SH-Sr6A | VN12_RS23345 | 6012920..6013936 | 1632865 | DUF499-domain-containing-protein-[Pirellula-sp.-SH-Sr6A] |
| MBX3448931.1 | AAA+ATPase+wHTH+Alpha+Beta-domain+wHTH+RRM | Type-1_BR.RM.like.systems | TUDOR+SF1-SF2-Helicase+iREase+CTD+REase->OLD-ABC->HNH->METHYLASE->OLD-ABC->REase->DUF499*->?->MORC->?->?-><-wHTH(SBD)+HNH<-DNA_methylase | 948 | Bacteria>PVC_group>Planctomycetota>Planctomycetia>Planctomycetales | GCA_019636285.1 | Planctomycetaceae_bacterium | KF777_05185 | 10482..13328 | 2026779 | ATP-binding-protein-[Planctomycetaceae-bacterium] |
| MCA9232427.1 | AAA+ATPase+wHTH+Alpha+Beta-domain+wHTH+RRM | Type-1_BR.RM.like.systems | <-ParB+HNH<-DUF499*<-METHYLASE<-TOPRIM<-OLD-ABC<-TUDOR+SF1-SF2-Helicase+iREase+CTD+REase-p | 953 | Bacteria>PVC_group>Planctomycetota>Planctomycetia>Planctomycetales | GCA_020427155.1 | Planctomycetales_bacterium | KDA57_17390 | 2105..4966 | 2053591 | ATP-binding-protein-[Planctomycetales-bacterium] |
| MCA9013093.1 | AAA+ATPase+wHTH+Alpha+Beta-domain+wHTH+RRM | Type-1_BR.RM.like.systems | <-DUF499*<-PIN<-?<-METHYLASE<-METHYLASE-p<-TUDOR+SF1-SF2-Helicase+iREase+CTD+REase | 944 | Bacteria>PVC_group>Planctomycetota>Planctomycetia>Planctomycetales | GCA_020426105.1 | Planctomycetaceae_bacterium | KDB01_25255 | 1526..4360 | 2026779 | ATP-binding-protein-[Planctomycetaceae-bacterium] |
| MDB5334583.1 | AAA+ATPase+wHTH+Alpha+Beta-domain+wHTH+RRM | Type-1_BR.RM.like.systems | TUDOR+SF1-SF2-Helicase+iREase+CTD+REase->OLD-ABC->HNH->METHYLASE->DUF499*-> | 942 | Bacteria>PVC_group>Planctomycetota>Planctomycetia>Planctomycetales | GCA_028291605.1 | Planctomycetaceae_bacterium | JWN70_202 | 14767..17595 | 2026779 | MAG:-hypothetical-protein-JWN70_202-[Planctomycetaceae-bacterium] |
| MCX7388316.1 | AAA+ATPase+wHTH+Alpha+Beta-domain+wHTH+RRM | Type-1_BR.RM.like.systems | <-NUDIX<-?\|\|?-><-MPTase<-HTH<-DUF499*<-DUF499-p\|\|DDE_Tnp_1-> | 503 | Bacteria>PVC_group>Planctomycetota>Planctomycetia>Planctomycetales | GCA_026400875.1 | Planctomycetales_bacterium | NTX48_16755 | 35618..37129 | 2053591 | MAG:-DUF499-domain-containing-protein-[Planctomycetales-bacterium] |
| HBI42788.1 | AAA+ATPase+wHTH+Alpha+Beta-domain+wHTH+RRM | Type-1_BR.RM.like.systems | SIGMA-HTH->?-><-DUF499*<-REase<-OLD-ABC<-METHYLASE | 939 | Bacteria>PVC_group>Planctomycetota>Planctomycetia>Planctomycetales | GCA_003508275.1 | Planctomycetales_bacterium | DDY78_08010 | 2578..5397 | 2053591 | hypothetical-protein-[Planctomycetales-bacterium] |
| MCX7393563.1 | AAA+ATPase+wHTH+Alpha+Beta-domain+wHTH+RRM | Type-1_BR.RM.like.systems | CRP-HTH->?-><-SbcC<-DUF499*<-METHYLASE-p | 951 | Bacteria>PVC_group>Planctomycetota>Planctomycetia>Planctomycetales | GCA_026400915.1 | Planctomycetales_bacterium | NTW75_05495 | 276726..279581 | 2053591 | MAG:-ATP-binding-protein-[Planctomycetales-bacterium] |
| MCA9239533.1 | AAA+ATPase+wHTH+Alpha+Beta-domain+wHTH+RRM | Type-1_BR.RM.like.systems | <-SbcC<-DUF499*<-METHYLASE-p | 956 | Bacteria>PVC_group>Planctomycetota>Planctomycetia>Planctomycetales | GCA_020427175.1 | Planctomycetales_bacterium | KDA37_05015 | 2149..5019 | 2053591 | ATP-binding-protein-[Planctomycetales-bacterium] |
| MBL8874046.1 | AAA+ATPase+wHTH+Alpha+Beta-domain+wHTH+RRM | Type-1_BR.RM.like.systems | <-TOPRIM<-SbcC<-?\|\|XIS-HTH->DUF499*->?->RAMA->Glyoxalase->?->?->Trypsin-> | 156 | Bacteria>PVC_group>Planctomycetota>Planctomycetia>Planctomycetales | GCA_016792705.1 | Planctomycetaceae_bacterium | JNK90_30005 | 20016..20486 | 2026779 | hypothetical-protein-[Planctomycetaceae-bacterium] |
| WP_145201095.1 | AAA+ATPase+wHTH+Alpha+Beta-domain+wHTH+RRM | Type-1_BR.RM.like.systems | <-JAB+UbE1<-?<-?<-MPTase<-?<-SIGMA-HTH<-DUF499*<-REase<-?<-METHYLASE<-OLD-ABC<-TUDOR+SF1-SF2-Helicase+iREase+CTD+REase<-XIS-HTH | 984 | Bacteria>PVC_group>Planctomycetota>Planctomycetia>Planctomycetales | GCF_007744255.1 | Thalassoglobus_polymorphus | Mal48_RS14940 | 4184383..4187337 | 2527994 | DUF499-domain-containing-protein-[Thalassoglobus-polymorphus] |
| MCB9950010.1 | AAA+ATPase+wHTH+Alpha+Beta-domain+wHTH+RRM | Type-1_BR.RM.like.systems | <-JAB+UbE1<-?<-?<-MPTase<-?<-SIGMA-HTH<-DUF499*<-REase<-?<-SAM-methylase<-TUDOR+SF1-SF2-Helicase+iREase+CTD+REase<-XIS-HTH | 984 | Bacteria>PVC_group>Planctomycetota>Planctomycetia>Planctomycetales | GCA_020638955.1 | Planctomycetaceae_bacterium | H6824_03425 | 868726..871680 | 2026779 | ATP-binding-protein-[Planctomycetaceae-bacterium] |
| QDT36019.1 | AAA+ATPase+wHTH+Alpha+Beta-domain+wHTH+RRM | Type-1_BR.RM.like.systems | <-DUF499*<-?<-METHYLASE<-ParB+HNH<-TUDOR+SF1-SF2-Helicase+iREase+CTD+REase<-GDResolvase+Recombinase+ZnR | 961 | Bacteria>PVC_group>Planctomycetota>Planctomycetia>Planctomycetales | GCA_007744515.1 | Stratiformator_vulcanicus | Pan189_03740 | 483899..486784 | 2527980 | hypothetical-protein-Pan189_03740-[Planctomycetes-bacterium-Pan189] |
| HBN75528.1 | AAA+ATPase+wHTH+Alpha+Beta-domain+wHTH+RRM | Type-1_BR.RM.like.systems | XIS-HTH->?->?->?->HKD-DNase->DUF499*->SIGMA-HTH->?->MPTase->?->?->JAB+UbE1-> | 984 | Bacteria>PVC_group>Planctomycetota>Planctomycetia>Planctomycetales | GCA_003503535.1 | Planctomycetaceae_bacterium | DD473_06870 | 45061..48015 | 2026779 | AAA-family-ATPase-[Planctomycetaceae-bacterium] |
| MCA9067325.1 | AAA+ATPase+wHTH+Alpha+Beta-domain+wHTH+RRM | Type-1_BR.RM.like.systems | <-DUF499*<-?<-REase<-REase | 982 | Bacteria>PVC_group>Planctomycetota>Planctomycetia>Planctomycetales | GCA_020426385.1 | Planctomycetaceae_bacterium | KDA84_00275 | 118..3066 | 2026779 | ATP-binding-protein-[Planctomycetaceae-bacterium] |
| MDA8378650.1 | AAA+ATPase+wHTH+Alpha+Beta-domain+wHTH+RRM | Type-1_BR.RM.like.systems | <-35exo\|\|?->TUDOR+SF1-SF2-Helicase+iREase+CTD+REase->HEPN->METHYLASE->wHTH+REase->DUF499*-> | 945 | Bacteria>PVC_group>Planctomycetota>Planctomycetia>unclassified_Planctomycetia | GCA_028068345.1 | Planctomycetia_bacterium | MOZ50_16785 | 11833..14670 | 2052181 | MAG:-DUF499-domain-containing-protein-[Planctomycetia-bacterium] |
| MBI2825945.1 | AAA+ATPase+wHTH+Alpha+Beta-domain+wHTH+RRM | Type-1_BR.RM.like.systems | <-REase<-?<-?<-?<-DNA_methylase<-DUF499*<-HTH<-REase<-RNaseH<-METHYLASE<-METHYLASE-p | 930 | Bacteria>PVC_group>Planctomycetota>Planctomycetia>unclassified_Planctomycetia | GCA_016191115.1 | Planctomycetia_bacterium | HYX69_14775 | 236780..239572 | 2052181 | ATP-binding-protein-[Planctomycetia-bacterium] |
| MBE7443980.1 | AAA+ATPase+wHTH+Alpha+Beta-domain+wHTH+RRM | Type-1_BR.RM.like.systems | METHYLASE->?->DUF433->?->?->DUF499*->?-><-RADICAL-SAM | 844 | Bacteria>PVC_group>Planctomycetota>Planctomycetia>unclassified_Planctomycetia | GCA_015075005.1 | Planctomycetia_bacterium | HS132_01435 | 268107..270641 | 2052181 | ATP-binding-protein-[Planctomycetia-bacterium] |
| MCY3008131.1 | AAA+ATPase+wHTH+Alpha+Beta-domain+wHTH+RRM | Type-1_BR.RM.like.systems | DUF4096+DDE_Tnp_1+DDE_Tnp_1_2-><-PAPS_reduct<-?<-DUF499*<-METHYLASE<-TUDOR+SF1-SF2-Helicase+iREase+CTD+REase<-XIS-HTH\|\|RAMA->GDResolvase+Recombinase+ZnR->HTH-> | 956 | Bacteria>PVC_group>Planctomycetota>unclassified_Planctomycetota | GCA_026692845.1 | Planctomycetota_bacterium | NTV29_19435 | 12132..15002 | 2026780 | MAG:-ATP-binding-protein-[Planctomycetota-bacterium] |
| MCY2978566.1 | AAA+ATPase+wHTH+Alpha+Beta-domain+wHTH+RRM | Type-1_BR.RM.like.systems | <-DUF499*<-REase<-OLD-ABC<-METHYLASE<-TUDOR+SF1-SF2-Helicase+iREase+CTD+REase<-XIS-HTH | 968 | Bacteria>PVC_group>Planctomycetota>unclassified_Planctomycetota | GCA_026692945.1 | Planctomycetota_bacterium | NTU79_07885 | 10024..12930 | 2026780 | MAG:-ATP-binding-protein-[Planctomycetota-bacterium] |
| MBA3683762.1 | AAA+ATPase+wHTH+Alpha+Beta-domain+wHTH+RRM | Type-1_BR.RM.like.systems | <-DUF499*<-METHYLASE<-PolB<-HEPN<-TUDOR+SF1-SF2-Helicase+iREase+CTD+REase<-tRNA<-tRNA\|\|PRTase-> | 945 | Bacteria>PVC_group>Planctomycetota>unclassified_Planctomycetota | GCA_013816645.1 | Planctomycetota_bacterium | HOW72_00725 | 4185..7022 | 2026780 | ATP-binding-protein-[Planctomycetota-bacterium] |
| RLS82192.1 | AAA+ATPase+wHTH+Alpha+Beta-domain+wHTH+RRM | Type-1_BR.RM.like.systems | <-DUF499*<-?<-METHYLASE<-TUDOR+SF1-SF2-Helicase+iREase+CTD+REase-p | 216 | Bacteria>PVC_group>Planctomycetota>unclassified_Planctomycetota | GCA_003669255.1 | Planctomycetota_bacterium | DWI04_05200 | 1..647 | 2026780 | MAG:-ATP-binding-protein,-partial-[Planctomycetota-bacterium] |
| MCY2995756.1 | AAA+ATPase+wHTH+Alpha+Beta-domain+wHTH+RRM | Type-1_BR.RM.like.systems | TUDOR+SF1-SF2-Helicase+iREase+CTD+REase->?->METHYLASE->DUF499*-> | 947 | Bacteria>PVC_group>Planctomycetota>unclassified_Planctomycetota | GCA_026692885.1 | Planctomycetota_bacterium | NTY19_49095 | 11248..14091 | 2026780 | MAG:-ATP-binding-protein-[Planctomycetota-bacterium] |
| MCA8917196.1 | AAA+ATPase+wHTH+Alpha+Beta-domain+wHTH+RRM | Type-1_BR.RM.like.systems | XIS-HTH->?-><-DUF499*<-REase<-OLD-ABC<-METHYLASE | 951 | Bacteria>PVC_group>Planctomycetota>unclassified_Planctomycetota | GCA_020425605.1 | Planctomycetota_bacterium | KDB90_17530 | 3869..6724 | 2026780 | ATP-binding-protein-[Planctomycetota-bacterium] |
| MCX5685413.1 | AAA+ATPase+wHTH+Alpha+Beta-domain+wHTH+RRM | Type-1_BR.RM.like.systems | <-METHYLASE-p<-?<-DUF499*<-XIS-HTH<-PIWI<-REase+PNTD1+PNTD2 | 1212 | Bacteria>PVC_group>Planctomycetota>unclassified_Planctomycetota | GCA_026386975.1 | Planctomycetota_bacterium | NT049_17285 | 1491..5129 | 2026780 | MAG:-hypothetical-protein-NT049_17285-[Planctomycetota-bacterium] |
| MCE9554597.1 | AAA+ATPase+wHTH+Alpha+Beta-domain+wHTH+RRM | Type-1_BR.RM.like.systems | SMS-KaiC->?->GDResolvase->?-><-SF1-SF2-Helicase-p<-SF1-SF2-Helicase<-DUF499*<-wHTH+REase<-METHYLASE<-DUF4065<-HD<-TUDOR+SF1-SF2-Helicase+iREase+CTD+REase | 950 | Bacteria>PVC_group>Planctomycetota>unclassified_Planctomycetota | GCA_021413705.1 | Planctomycetota_bacterium | K8T91_14665 | 30922..33774 | 2026780 | DUF499-domain-containing-protein-[Planctomycetota-bacterium] |
| MBE7491676.1 | AAA+ATPase+wHTH+Alpha+Beta-domain+wHTH+RRM | Type-1_BR.RM.like.systems | PAIREDC-HTH+HIN-HTH->?-><-DDE_Tnp_1+DDE_Tnp_1_2<-DUF4096<-?<-DUF499*<-REase<-METHYLASE<-OLD-ABC<-OLD-ABC-p<-TUDOR+SF1-SF2-Helicase+iREase+CTD+REase | 974 | Bacteria>PVC_group>Planctomycetota>unclassified_Planctomycetota | GCA_015075515.1 | Planctomycetota_bacterium | HS108_07990 | 570856..573780 | 2026780 | ATP-binding-protein-[Planctomycetota-bacterium] |
| MBE3098083.1 | AAA+ATPase+wHTH+Alpha+Beta-domain+wHTH+RRM | Type-1_BR.RM.like.systems | <-XIS-HTH\|\|TUDOR+SF1-SF2-Helicase+iREase+CTD+REase->?->METHYLASE->?->DUF499*-><-Arm-DNA-bind_3+Phage_integrase<-Phage_int_SAM_4+Phage_integrase<-?<-?<-HTH_36 | 953 | Bacteria>PVC_group>Planctomycetota>unclassified_Planctomycetota | GCA_014894695.1 | Planctomycetota_bacterium | IMZ44_13275 | 12925..15786 | 2026780 | ATP-binding-protein-[Planctomycetota-bacterium] |
| MBI2193014.1 | AAA+ATPase+wHTH+Alpha+Beta-domain+wHTH+RRM | Type-1_BR.RM.like.systems | TUDOR+SF1-SF2-Helicase+iREase+CTD+REase->Schlafen+HATPase+HTH->METHYLASE->METHYLASE->Y1_Tnp->DUF433->DUF499*->ABhydrolase-><-TPR<-SIGMA-HTH | 946 | Bacteria>PVC_group>Planctomycetota>unclassified_Planctomycetota | GCA_016179685.1 | Planctomycetota_bacterium | HYU36_13630 | 54036..56876 | 2026780 | ATP-binding-protein-[Planctomycetota-bacterium] |
| MBI3818879.1 | AAA+ATPase+wHTH+Alpha+Beta-domain+wHTH+RRM | Type-1_BR.RM.like.systems | <-PIN<-HTH+MPTase<-?<-?\|\|TUDOR+SF1-SF2-Helicase+iREase+CTD+REase->METHYLASE->DUF499*->REase+nSTAND1-><-ParB<-Peptidase<-ABC-ATPase+SbcC | 945 | Bacteria>PVC_group>Planctomycetota>unclassified_Planctomycetota | GCA_016200225.1 | Planctomycetota_bacterium | HY286_09340 | 45594..48431 | 2026780 | ATP-binding-protein-[Planctomycetota-bacterium] |
| MCH8343169.1 | AAA+ATPase+wHTH+Alpha+Beta-domain+wHTH+RRM | Type-1_BR.RM.like.systems | <-METHYLASE\|\|TUDOR+SF1-SF2-Helicase+iREase+CTD+REase->METHYLASE->DUF499*->REase->?->XIS-HTH->ZnR+TOPRIM+D5ATPase->XIS-HTH->HTH-> | 982 | Bacteria>PVC_group>Planctomycetota>unclassified_Planctomycetota | GCA_022568135.1 | Planctomycetota_bacterium | IH983_04190 | 73866..76814 | 2026780 | ATP-binding-protein-[Planctomycetota-bacterium] |
| MCE9567305.1 | AAA+ATPase+wHTH+Alpha+Beta-domain+wHTH+RRM | Type-1_BR.RM.like.systems | <-SF1-SF2-Helicase+REase<-?<-SIGMA-HTH\|\|ParB+HNH->METHYLASE->REase->DUF499*->ParB+HNH->ParB+HNH-> | 986 | Bacteria>PVC_group>Planctomycetota>unclassified_Planctomycetota | GCA_021413735.1 | Planctomycetota_bacterium | K8U57_35335 | 37179..40139 | 2026780 | DUF499-domain-containing-protein-[Planctomycetota-bacterium] |
| MBI5364695.1 | AAA+ATPase+wHTH+Alpha+Beta-domain+wHTH+RRM | Type-1_BR.RM.like.systems | Phage_integrase->TUDOR+SF1-SF2-Helicase+iREase+CTD+REase->OLD-ABC->METHYLASE->REase->DUF499*->DNA-glycosylase-> | 936 | Bacteria>PVC_group>Planctomycetota>unclassified_Planctomycetota | GCA_016220035.1 | Planctomycetota_bacterium | HZA53_16080 | 12813..15623 | 2026780 | ATP-binding-protein-[Planctomycetota-bacterium] |
| MBN8597233.1 | AAA+ATPase+wHTH+Alpha+Beta-domain+wHTH+RRM | Type-1_BR.RM.like.systems | TUDOR+SF1-SF2-Helicase+iREase+CTD+REase->?->?->METHYLASE->?->REase->DUF499*-><-?<-?\|\|?->?->RAMA->GDResolvase+Recombinase+ZnR-> | 987 | Bacteria>PVC_group>Planctomycetota>unclassified_Planctomycetota | GCA_017303555.1 | Planctomycetota_bacterium | JOL78_06125 | 35545..38508 | 2026780 | ATP-binding-protein-[Planctomycetota-bacterium] |
| MBI5763662.1 | AAA+ATPase+wHTH+Alpha+Beta-domain+wHTH+RRM | Type-1_BR.RM.like.systems | <-ParB+N6_N4_MTase<-?<-ZnR+TOPRIM+D5ATPase<-?<-?<-HIN-HTH<-DUF499*<-REase<-?<-METHYLASE<-?\|\|?-><-TUDOR+SF1-SF2-Helicase+iREase+CTD+REase | 921 | Bacteria>PVC_group>Planctomycetota>unclassified_Planctomycetota | GCA_016235815.1 | Planctomycetota_bacterium | HZA51_09085 | 13223..15988 | 2026780 | ATP-binding-protein-[Planctomycetota-bacterium] |
| RIK66096.1 | AAA+ATPase+wHTH+Alpha+Beta-domain+wHTH+RRM | Type-1_BR.RM.like.systems | TUDOR+SF1-SF2-Helicase+iREase+CTD+REase-><-?\|\|?->METHYLASE->MORC->REase->DUF499*-><-?\|\|HTH_3-><-?<-?\|\|HTH_3->MPTase-> | 974 | Bacteria>PVC_group>Planctomycetota>unclassified_Planctomycetota | GCA_003576905.1 | Planctomycetota_bacterium | DCC65_10760 | 38314..41238 | 2026780 | MAG:-AAA-family-ATPase-[Planctomycetota-bacterium] |
| MBI3271453.1 | AAA+ATPase+wHTH+Alpha+Beta-domain+wHTH+RRM | Type-1_BR.RM.like.systems | <-Phage_int_SAM_1+Phage_integrase<-?<-?<-ParB+HNH<-?<-DUF499*<-PIN<-?<-?<-METHYLASE<-HEPN<-TUDOR+SF1-SF2-Helicase+iREase+CTD+REase | 953 | Bacteria>PVC_group>Planctomycetota>unclassified_Planctomycetota | GCA_016197205.1 | Planctomycetota_bacterium | HYZ53_20855 | 61698..64559 | 2026780 | ATP-binding-protein-[Planctomycetota-bacterium] |
| MCY2928835.1 | AAA+ATPase+wHTH+Alpha+Beta-domain+wHTH+RRM | Type-1_BR.RM.like.systems | HTH_3->MPTase->?->?->?->?-><-DUF499*<-REase<-METHYLASE<-?<-?<-TUDOR+SF1-SF2-Helicase+iREase+CTD+REase\|\|ParB+HNH-> | 970 | Bacteria>PVC_group>Planctomycetota>unclassified_Planctomycetota | GCA_026693065.1 | Planctomycetota_bacterium | NTV86_04950 | 16719..19631 | 2026780 | MAG:-DUF499-domain-containing-protein-[Planctomycetota-bacterium] |
| MBL8860740.1 | AAA+ATPase+wHTH+Alpha+Beta-domain+wHTH+RRM | Type-1_BR.RM.like.systems | TUDOR+SF1-SF2-Helicase+iREase+CTD+REase->OLD-ABC+TOPRIM->METHYLASE->DUF499*->DNA-glycosylase->?->?-><-?<-TPR+TPR\|\|Phage_integrase-> | 888 | Bacteria>PVC_group>Planctomycetota>unclassified_Planctomycetota | GCA_016795205.1 | Planctomycetota_bacterium | JNK02_01915 | 8775..11441 | 2026780 | DUF499-domain-containing-protein-[Planctomycetota-bacterium] |
| GIW70817.1 | AAA+ATPase+wHTH+Alpha+Beta-domain+wHTH+RRM | Type-1_BR.RM.like.systems | <-Glyco_hydro_57+DUF3536<-?<-DUF499*<-REase<-REase<-TOPRIM<-OLD-ABC<-METHYLASE-p<-METHYLASE-p | 942 | Bacteria>PVC_group>Planctomycetota>unclassified_Planctomycetota | GCA_026004615.1 | Planctomycetota_bacterium | KatS3mg102_0359 | 449544..452372 | 2026780 | MAG:-hypothetical-protein-KatS3mg102_0359-[Planctomycetota-bacterium] |
| MBI2191311.1 | AAA+ATPase+wHTH+Alpha+Beta-domain+wHTH+RRM | Type-1_BR.RM.like.systems | <-DUF499*<-REase<-METHYLASE<-HEPN<-HEPN<-Schlafen+HATPase+HTH\|\|DEDD_Tnp_IS110+Transposase_20-> | 926 | Bacteria>PVC_group>Planctomycetota>unclassified_Planctomycetota | GCA_016179685.1 | Planctomycetota_bacterium | HYU36_04935 | 72260..75040 | 2026780 | ATP-binding-protein-[Planctomycetota-bacterium] |
| MBI2806086.1 | AAA+ATPase+wHTH+Alpha+Beta-domain+wHTH+RRM | Type-1_BR.RM.like.systems | <-ParB+HNH<-?<-PIWI<-?<-DUF499*<-HTH<-METHYLASE<-REase<-HEPN<-TUDOR+SF1-SF2-Helicase+iREase+CTD+REase\|\|HTH-> | 936 | Bacteria>PVC_group>Planctomycetota>unclassified_Planctomycetota | GCA_016191165.1 | Planctomycetota_bacterium | HYX68_13985 | 123843..126653 | 2026780 | ATP-binding-protein-[Planctomycetota-bacterium] |
| MBI4584119.1 | AAA+ATPase+wHTH+Alpha+Beta-domain+wHTH+RRM | Type-1_BR.RM.like.systems | Schlafen+HATPase+HTH->?->METHYLASE->REase->OLD-ABC->TOPRIM->DUF499*->HicB_lk_antitox->HicA_toxin-> | 920 | Bacteria>PVC_group>Planctomycetota>unclassified_Planctomycetota | GCA_016209605.1 | Planctomycetota_bacterium | HY717_08860 | 13681..16443 | 2026780 | ATP-binding-protein-[Planctomycetota-bacterium] |
| MBI1826653.1 | AAA+ATPase+wHTH+Alpha+Beta-domain+wHTH+RRM | Type-1_BR.RM.like.systems | <-DUF499*<-TOPRIM<-OLD-ABC<-METHYLASE<-PHP+SbcC<-TUDOR+SF1-SF2-Helicase+iREase+CTD+REase<-ArdCN+MPTase | 948 | Bacteria>PVC_group>Planctomycetota>unclassified_Planctomycetota | GCA_016178855.1 | Planctomycetota_bacterium | HYR83_09750 | 14494..17340 | 2026780 | ATP-binding-protein-[Planctomycetota-bacterium] |
| MBM4070796.1 | AAA+ATPase+wHTH+Alpha+Beta-domain+wHTH+RRM | Type-1_BR.RM.like.systems | <-REase\|\|?->wHTH+HNH-><-ParB+Schlafen<-?<-DUF499*<-METHYLASE<-TUDOR+SF1-SF2-Helicase+iREase+CTD+REase | 938 | Bacteria>PVC_group>Planctomycetota>unclassified_Planctomycetota | GCA_016873095.1 | Planctomycetota_bacterium | FJ271_17850 | 19856..22672 | 2026780 | ATP-binding-protein-[Planctomycetota-bacterium] |
| MBI1914948.1 | AAA+ATPase+wHTH+Alpha+Beta-domain+wHTH+RRM | Type-1_BR.RM.like.systems | ABC-ATPase-><-DUF499*<-REase<-METHYLASE<-?<-DUF4065<-?<-HIN-HTH | 974 | Bacteria>PVC_group>Planctomycetota>unclassified_Planctomycetota | GCA_016177925.1 | Planctomycetota_bacterium | HYS12_09465 | 5117..8041 | 2026780 | DUF499-domain-containing-protein-[Planctomycetota-bacterium] |
| MCY2991648.1 | AAA+ATPase+wHTH+Alpha+Beta-domain+wHTH+RRM | Type-1_BR.RM.like.systems | <-DUF499*<-PIN<-DUF433<-DUF433<-METHYLASE<-REase<-TUDOR+SF1-SF2-Helicase+iREase+CTD+REase | 983 | Bacteria>PVC_group>Planctomycetota>unclassified_Planctomycetota | GCA_026692885.1 | Planctomycetota_bacterium | NTY19_27810 | 2128..5079 | 2026780 | MAG:-DUF499-domain-containing-protein-[Planctomycetota-bacterium] |
| MBM4038866.1 | AAA+ATPase+wHTH+Alpha+Beta-domain+wHTH+RRM | Type-1_BR.RM.like.systems | <-DUF499*<-?<-METHYLASE-p<-METHYLASE<-Schlafen+HATPase+HTH<-TUDOR+SF1-SF2-Helicase+iREase+CTD+REase | 938 | Bacteria>PVC_group>Planctomycetota>unclassified_Planctomycetota | GCA_016872855.1 | Planctomycetota_bacterium | FJ290_10155 | 3372..6188 | 2026780 | ATP-binding-protein-[Planctomycetota-bacterium] |
| MCX5669700.1 | AAA+ATPase+wHTH+Alpha+Beta-domain+wHTH+RRM | Type-1_BR.RM.like.systems | TUDOR+SF1-SF2-Helicase+iREase+CTD+REase->OLD-ABC+TOPRIM->METHYLASE->PolB->HEPN->DUF499*-> | 924 | Bacteria>PVC_group>Planctomycetota>unclassified_Planctomycetota | GCA_026387055.1 | Planctomycetota_bacterium | NTU94_00045 | 8049..10823 | 2026780 | MAG:-DUF499-domain-containing-protein-[Planctomycetota-bacterium] |
| RME04985.1 | AAA+ATPase+wHTH+Alpha+Beta-domain+wHTH+RRM | Type-1_BR.RM.like.systems | <-HISKIN\|\|?->?->MoxR-AAA-><-?<-DUF499*<-Peptidase_M22<-Peptidase_S41 | 867 | Bacteria>PVC_group>Planctomycetota>unclassified_Planctomycetota | GCA_003694635.1 | Planctomycetota_bacterium | D6805_01125 | 7249..9852 | 2026780 | MAG:-ATP-binding-protein-[Planctomycetota-bacterium] |
| MBN8644244.1 | AAA+ATPase+wHTH+Alpha+Beta-domain+wHTH+RRM | Type-1_BR.RM.like.systems | <-McrC-NTD+REase<-?<-DUF499*<-REase<-METHYLASE | 978 | Bacteria>PVC_group>Planctomycetota>unclassified_Planctomycetota | GCA_017303565.1 | Planctomycetota_bacterium | JOL61_03265 | 4226..7162 | 2026780 | ATP-binding-protein-[Planctomycetota-bacterium] |
| MBI2930414.1 | AAA+ATPase+wHTH+Alpha+Beta-domain+wHTH+RRM | Type-1_BR.RM.like.systems | <-DUF499*<-METHYLASE<-REase<-TUDOR+SF1-SF2-Helicase+iREase+CTD+REase-p | 940 | Bacteria>PVC_group>Planctomycetota>unclassified_Planctomycetota | GCA_016190155.1 | Planctomycetota_bacterium | HYY16_02075 | 259..3081 | 2026780 | ATP-binding-protein-[Planctomycetota-bacterium] |
| MCY2987752.1 | AAA+ATPase+wHTH+Alpha+Beta-domain+wHTH+RRM | Type-1_BR.RM.like.systems | TUDOR+SF1-SF2-Helicase+iREase+CTD+REase->METHYLASE->RNase->?->DUF499*-> | 963 | Bacteria>PVC_group>Planctomycetota>unclassified_Planctomycetota | GCA_026692885.1 | Planctomycetota_bacterium | NTY19_07800 | 12759..15650 | 2026780 | MAG:-DUF499-domain-containing-protein-[Planctomycetota-bacterium] |
| RKY15801.1 | AAA+ATPase+wHTH+Alpha+Beta-domain+wHTH+RRM | Type-1_BR.RM.like.systems | METHYLASE->DUF499*-><-?<-Radical-SAM\|\|Polbeta->HEPN-> | 928 | Bacteria>PVC_group>Planctomycetota>unclassified_Planctomycetota | GCA_003644235.1 | Planctomycetota_bacterium | DRP82_00780 | 2911..5697 | 2026780 | MAG:-AAA-family-ATPase-[Planctomycetota-bacterium] |
| MBI3467145.1 | AAA+ATPase+wHTH+Alpha+Beta-domain+wHTH+RRM | Type-1_BR.RM.like.systems | <-DUF499*<-HTH<-TIR<-METHYLASE<-REase<-?<-REase | 942 | Bacteria>PVC_group>Planctomycetota>unclassified_Planctomycetota | GCA_016196255.1 | Planctomycetota_bacterium | HY000_29375 | 4772..7600 | 2026780 | ATP-binding-protein-[Planctomycetota-bacterium] |
| MBP7751892.1 | AAA+ATPase+wHTH+Alpha+Beta-domain+wHTH+RRM | Type-1_BR.RM.like.systems | <-MoxR-AAA<-?\|\|?->?->?-><-DUF499*<-REase<-METHYLASE-p | 932 | Bacteria>PVC_group>Planctomycetota>unclassified_Planctomycetota | GCA_018000135.1 | Planctomycetota_bacterium | KA323_14485 | 46655..49453 | 2026780 | ATP-binding-protein-[Planctomycetota-bacterium] |
| MCH7685161.1 | AAA+ATPase+wHTH+Alpha+Beta-domain+wHTH+RRM | Type-1_BR.RM.like.systems | METHYLASE-p->?->DUF433->PIN->HTH->DUF499*-> | 617 | Bacteria>PVC_group>Planctomycetota>unclassified_Planctomycetota | GCA_022562015.1 | Planctomycetota_bacterium | IH899_00515 | 3772..5625 | 2026780 | ATP-binding-protein-[Planctomycetota-bacterium] |
| MCH8852384.1 | AAA+ATPase+wHTH+Alpha+Beta-domain+wHTH+RRM | Type-1_BR.RM.like.systems | <-DUF499*<-HTH<-?<-RNaseH<-?<-METHYLASE-p | 775 | Bacteria>PVC_group>Planctomycetota>unclassified_Planctomycetota | GCA_022574515.1 | Planctomycetota_bacterium | IID41_06975 | 1..2325 | 2026780 | ATP-binding-protein-[Planctomycetota-bacterium] |
| NMD35173.1 | AAA+ATPase+wHTH+Alpha+Beta-domain+wHTH+RRM | Type-1_BR.RM.like.systems | <-DUF499*<-REase<-METHYLASE<-OLD-ABC+TOPRIM | 932 | Bacteria>PVC_group>Planctomycetota>unclassified_Planctomycetota | GCA_012797405.1 | Planctomycetota_bacterium | GYA73_04750 | 409..3207 | 2026780 | ATP-binding-protein-[Planctomycetota-bacterium] |
| MCH8840527.1 | AAA+ATPase+wHTH+Alpha+Beta-domain+wHTH+RRM | Type-1_BR.RM.like.systems | <-DUF499*<-DUF499-p<-?<-METHYLASE-p | 442 | Bacteria>PVC_group>Planctomycetota>unclassified_Planctomycetota | GCA_022574455.1 | Planctomycetota_bacterium | IH831_07595 | 1..1329 | 2026780 | DUF499-domain-containing-protein-[Planctomycetota-bacterium] |
| NQW48683.1 | AAA+ATPase+wHTH+Alpha+Beta-domain+wHTH+RRM | Type-1_BR.RM.like.systems | TUDOR+SF1-SF2-Helicase+iREase+CTD+REase->SF1-SF2-Helicase->METHYLASE->OLD-ABC->TOPRIM->DUF499*-> | 332 | Bacteria>PVC_group>Planctomycetota>unclassified_Planctomycetota | GCA_013205375.1 | Planctomycetota_bacterium | HQ464_13060 | 11784..12781 | 2026780 | ATP-binding-protein-[Planctomycetota-bacterium] |
| MBM4083197.1 | AAA+ATPase+wHTH+Alpha+Beta-domain+wHTH+RRM | Type-1_BR.RM.like.systems | <-DUF499*<-TOPRIM<-OLD-ABC<-METHYLASE<-BirA-HTH | 878 | Bacteria>PVC_group>Planctomycetota>unclassified_Planctomycetota | GCA_016873055.1 | Planctomycetota_bacterium | FJ272_00230 | 1..2634 | 2026780 | ATP-binding-protein-[Planctomycetota-bacterium] |
| MBF0245070.1 | AAA+ATPase+wHTH+Alpha+Beta-domain+wHTH+RRM | Type-1_BR.RM.like.systems | METHYLASE->URI-DNase->DUF499*-> | 775 | Bacteria>PVC_group>Planctomycetota>unclassified_Planctomycetota | GCA_015231495.1 | Planctomycetota_bacterium | HQL31_07330 | 2271..4595 | 2026780 | ATP-binding-protein-[Planctomycetota-bacterium] |
| MBM4022804.1 | AAA+ATPase+wHTH+Alpha+Beta-domain+wHTH+RRM | Type-1_BR.RM.like.systems | <-DUF499*<-PIN<-METHYLASE-p | 620 | Bacteria>PVC_group>Planctomycetota>unclassified_Planctomycetota | GCA_016872955.1 | Planctomycetota_bacterium | FJ284_11300 | 1..1859 | 2026780 | ATP-binding-protein-[Planctomycetota-bacterium] |
| MCS7063069.1 | AAA+ATPase+wHTH+Alpha+Beta-domain+wHTH+RRM | Type-1_BR.RM.like.systems | <-DUF499*<-METHYLASE-p | 884 | Bacteria>PVC_group>Verrucomicrobiota>Methylacidiphilae>Methylacidiphilales | GCA_025056655.1 | Candidatus_Methylacidiphilales_bacterium | NZM04_03315 | 528..3182 | 2720631 | DUF499-domain-containing-protein-[Methylacidiphilales-bacterium] |
| PTY07976.1 | AAA+ATPase+wHTH+Alpha+Beta-domain+wHTH+RRM | Type-1_BR.RM.like.systems | TUDOR+SF1-SF2-Helicase+iREase+CTD+REase->?->METHYLASE->?->DUF499*-> | 938 | Bacteria>PVC_group>Verrucomicrobiota>Opitutae>Opitutales | GCA_003054665.1 | Opitutaceae_bacterium_EW11 | DB347_07080 | 585687..588503 | 2161865 | AAA-family-ATPase-[Opitutaceae-bacterium-EW11] |
| MBE2203055.1 | AAA+ATPase+wHTH+Alpha+Beta-domain+wHTH+RRM | Type-1_BR.RM.like.systems | <-JAB\|\|?-><-REase<-?<-?<-MORC+HisKin+REase<-DUF499*<-METHYLASE<-TOPRIM<-OLD-ABC<-TUDOR+SF1-SF2-Helicase+iREase+CTD+REase | 938 | Bacteria>PVC_group>Verrucomicrobiota>Spartobacteria>Chthoniobacterales | GCA_014879575.1 | Chthoniobacterales_bacterium | IAE94_01770 | 19431..22247 | 2201154 | ATP-binding-protein-[Chthoniobacterales-bacterium] |
| MBE2180625.1 | AAA+ATPase+wHTH+Alpha+Beta-domain+wHTH+RRM | Type-1_BR.RM.like.systems | <-DUF499*<-DOC+HTH<-METHYLASE<-TUDOR+SF1-SF2-Helicase+iREase+CTD+REase<-?<-HISKIN+MORC<-SbcC | 938 | Bacteria>PVC_group>Verrucomicrobiota>Spartobacteria>Chthoniobacterales | GCA_014879535.1 | Chthoniobacterales_bacterium | IAE97_09165 | 18976..21792 | 2201154 | ATP-binding-protein-[Chthoniobacterales-bacterium] |
| MCX6879289.1 | AAA+ATPase+wHTH+Alpha+Beta-domain+wHTH+RRM | Type-1_BR.RM.like.systems | <-HISKIN<-DUF499*<-REase<-OLD-ABC<-METHYLASE<-HNH<-OLD-ABC<-TUDOR+SF1-SF2-Helicase+iREase+CTD+REase | 941 | Bacteria>PVC_group>Verrucomicrobiota>unclassified_Verrucomicrobiota | GCA_026397725.1 | Verrucomicrobiota_bacterium | NTW21_36620 | 9837..12662 | 2026799 | MAG:-DUF499-domain-containing-protein-[Verrucomicrobia-bacterium] |
| MCX6953048.1 | AAA+ATPase+wHTH+Alpha+Beta-domain+wHTH+RRM | Type-1_BR.RM.like.systems | TUDOR+SF1-SF2-Helicase+iREase+CTD+REase->OLD-ABC->HNH->METHYLASE->REase->DUF499*->?->?->?->ABhydrolase-> | 941 | Bacteria>PVC_group>Verrucomicrobiota>unclassified_Verrucomicrobiota | GCA_026398065.1 | Verrucomicrobiota_bacterium | NTV51_12915 | 8536..11361 | 2026799 | MAG:-DUF499-domain-containing-protein-[Verrucomicrobia-bacterium] |
| MBI5767835.1 | AAA+ATPase+wHTH+Alpha+Beta-domain+wHTH+RRM | Type-1_BR.RM.like.systems | HTH->TUDOR+SF1-SF2-Helicase+iREase+CTD+REase->OLD-ABC->HNH->METHYLASE->DUF499*->Radical-SAM-> | 944 | Bacteria>PVC_group>Verrucomicrobiota>unclassified_Verrucomicrobiota | GCA_016235555.1 | Verrucomicrobiota_bacterium | HZA93_08560 | 101523..104357 | 2026799 | ATP-binding-protein-[Verrucomicrobia-bacterium] |
| MCX6966137.1 | AAA+ATPase+wHTH+Alpha+Beta-domain+wHTH+RRM | Type-1_BR.RM.like.systems | TUDOR+SF1-SF2-Helicase+iREase+CTD+REase->OLD-ABC->TOPRIM->METHYLASE->?->DUF499*->?->?-><-Peptidase | 941 | Bacteria>PVC_group>Verrucomicrobiota>unclassified_Verrucomicrobiota | GCA_026398255.1 | Verrucomicrobiota_bacterium | NTW41_12430 | 37940..40765 | 2026799 | MAG:-DUF499-domain-containing-protein-[Verrucomicrobia-bacterium] |
| MBM3833977.1 | AAA+ATPase+wHTH+Alpha+Beta-domain+wHTH+RRM | Type-1_BR.RM.like.systems | <-SF1-SF2-Helicase+iREase+CTD<-METHYLASE-p<-ZnR+ZnR<-DUF499* | 350 | Bacteria>PVC_group>Verrucomicrobiota>unclassified_Verrucomicrobiota | GCA_016871515.1 | Verrucomicrobiota_bacterium | FJ403_12050 | 33379..34431 | 2026799 | hypothetical-protein-[Verrucomicrobiota-bacterium] |
| TAE90777.1 | AAA+ATPase+wHTH+Alpha+Beta-domain+wHTH+RRM | Type-1_BR.RM.like.systems | <-DUF499*<-REase<-METHYLASE<-HNH<-OLD-ABC<-TOPRIM<-OLD-ABC | 939 | Bacteria>PVC_group>Verrucomicrobiota>unclassified_Verrucomicrobiota | GCA_004293285.1 | Verrucomicrobiota_bacterium | EAZ81_08635 | 280..3099 | 2026799 | ATP-binding-protein-[Verrucomicrobia-bacterium] |
| MCL4785388.1 | AAA+ATPase+wHTH+Alpha+Beta-domain+wHTH+RRM | Type-1_BR.RM.like.systems | <-URI-DNase<-?<-?\|\|ABhydrolase+ABhydrolase->HHH_5+SF1-SF2-helicase-><-ASCH<-DUF499*<-?<-METHYLASE<-?<-TUDOR+SF1-SF2-Helicase+iREase+CTD+REase<-PIN<-HTH+MPTase | 915 | Bacteria>PVC_group>Verrucomicrobiota>unclassified_Verrucomicrobiota | GCA_023383755.1 | Verrucomicrobiota_bacterium | KJ070_01140 | 19812..22559 | 2026799 | DUF499-domain-containing-protein-[Verrucomicrobia-bacterium] |
| MBI3850446.1 | AAA+ATPase+wHTH+Alpha+Beta-domain+wHTH+RRM | Type-1_BR.RM.like.systems | <-ACET+PIN+ACET\|\|TUDOR+SF1-SF2-Helicase+iREase+CTD+REase->?->SF1-SF2-Helicase->?->METHYLASE->DUF499*->HEPN->?->REase+HTH->NTP_transf->?-><-35exo | 961 | Bacteria>PVC_group>Verrucomicrobiota>unclassified_Verrucomicrobiota | GCA_016200005.1 | Verrucomicrobiota_bacterium | HY298_09175 | 22264..25149 | 2026799 | DUF499-domain-containing-protein-[Verrucomicrobia-bacterium] |
| OQC32217.1 | AAA+ATPase+wHTH+Alpha+Beta-domain+wHTH+RRM | Type-1_BR.RM.like.systems | DDE_Tnp_IS66+DDE_Tnp_IS66_C-><-?<-Radical-SAM<-?<-DUF499*<-?<-?<-METHYLASE<-HEPN<-ParB+HNH<-TUDOR+SF1-SF2-Helicase+iREase+CTD+REase | 945 | Bacteria>PVC_group>Verrucomicrobiota>unclassified_Verrucomicrobiota | GCA_002071025.1 | Verrucomicrobia_bacterium_ADurb.Bin070 | BWX70_00665 | 3737..6574 | 1852927 | hypothetical-protein-BWX70_00665-[Verrucomicrobia-bacterium-ADurb.Bin070] |
| PWU13198.1 | AAA+ATPase+wHTH+Alpha+Beta-domain+wHTH+RRM | Type-1_BR.RM.like.systems | TUDOR+SF1-SF2-Helicase+iREase+CTD+REase->METHYLASE->REase->OLD-ABC->TOPRIM->DUF499*->?->?->?->?->Cluster1029_5clades-> | 928 | Bacteria>PVC_group>Verrucomicrobiota>unclassified_Verrucomicrobiota | GCA_003176045.1 | Verrucomicrobiota_bacterium | C5B50_19760 | 19033..21819 | 2026799 | AAA-family-ATPase-[Verrucomicrobia-bacterium] |
| NLH75306.1 | AAA+ATPase+wHTH+Alpha+Beta-domain+wHTH+RRM | Type-1_BR.RM.like.systems | AbiEi_4->TUDOR+SF1-SF2-Helicase+iREase+CTD+REase->SF1-SF2-Helicase->METHYLASE->?->DUF499*->?->?->?->?->HEPN->HEPN-> | 915 | Bacteria>PVC_group>Verrucomicrobiota>unclassified_Verrucomicrobiota | GCA_012517065.1 | Verrucomicrobiota_bacterium | GX456_19860 | 11784..14531 | 2026799 | ATP-binding-protein-[Verrucomicrobia-bacterium] |
| MBU4201307.1 | AAA+ATPase+wHTH+Alpha+Beta-domain+wHTH+RRM | Type-1_BR.RM.like.systems | <-DUF499*<-METHYLASE<-TOPRIM<-OLD-ABC<-?<-TUDOR+SF1-SF2-Helicase+iREase+CTD+REase<-ORC-CDC6+REase | 941 | Bacteria>PVC_group>Verrucomicrobiota>unclassified_Verrucomicrobiota | GCA_018898455.1 | Verrucomicrobiota_bacterium | KJ919_15475 | 11057..13882 | 2026799 | DUF499-domain-containing-protein-[Verrucomicrobia-bacterium] |
| MCX6880519.1 | AAA+ATPase+wHTH+Alpha+Beta-domain+wHTH+RRM | Type-1_BR.RM.like.systems | <-DUF499*<-REase<-SPOUT<-METHYLASE<-DrHyd+BirA-HTH<-REase | 689 | Bacteria>PVC_group>Verrucomicrobiota>unclassified_Verrucomicrobiota | GCA_026397725.1 | Verrucomicrobiota_bacterium | NTW21_42945 | 1..2067 | 2026799 | MAG:-DUF499-domain-containing-protein,-partial-[Verrucomicrobia-bacterium] |
| MBU6399160.1 | AAA+ATPase+wHTH+Alpha+Beta-domain+wHTH+RRM | Type-1_BR.RM.like.systems | METHYLASE-p->OLD-ABC->TOPRIM->DUF499*->MORC+HisKin-><-SIGMA-HTH | 908 | Bacteria>PVC_group>Verrucomicrobiota>unclassified_Verrucomicrobiota | GCA_018969165.1 | Verrucomicrobiota_bacterium | KGS61_02480 | 2710..5436 | 2026799 | DUF499-domain-containing-protein-[Verrucomicrobia-bacterium] |
| MCU0784170.1 | AAA+ATPase+wHTH+Alpha+Beta-domain+wHTH+RRM | Type-1_BR.RM.like.systems | REase->?->METHYLASE->ORC-CDC6+REase->DUF499*->HTH_31->Peptidase-> | 935 | Bacteria>PVC_group>Verrucomicrobiota>unclassified_Verrucomicrobiota | GCA_025457745.1 | Verrucomicrobiota_bacterium | MUF81_09035 | 5492..8299 | 2026799 | DUF499-domain-containing-protein-[Verrucomicrobiota-bacterium] |
| MBI2947997.1 | AAA+ATPase+wHTH+Alpha+Beta-domain+wHTH+RRM | Type-1_BR.RM.like.systems | METHYLASE-p->AAA-ATPase+REase->?->PIN->HicB->DUF499*-> | 264 | Bacteria>PVC_group>Verrucomicrobiota>unclassified_Verrucomicrobiota | GCA_016190065.1 | Verrucomicrobiota_bacterium | HYY23_10135 | 3501..4294 | 2026799 | ATP-binding-protein-[Verrucomicrobiota-bacterium] |
| MBM3880410.1 | AAA+ATPase+wHTH+Alpha+Beta-domain+wHTH+RRM | Type-1_BR.RM.like.systems | HEPN->?->?->METHYLASE->ZDNA-HTH->REase->DUF499*->?-><-PIN<-?<-?<-?<-PIN | 967 | Bacteria>PVC_group>Verrucomicrobiota>unclassified_Verrucomicrobiota | GCA_016871715.1 | Verrucomicrobiota_bacterium | FJ387_11950 | 21225..24128 | 2026799 | ATP-binding-protein-[Verrucomicrobia-bacterium] |
| MCX6969348.1 | AAA+ATPase+wHTH+Alpha+Beta-domain+wHTH+RRM | Type-1_BR.RM.like.systems | DUF499*->?->?->DNA_pol3_delta+DNA_pol3_gamma3-><-TPR+TPR | 938 | Bacteria>PVC_group>Verrucomicrobiota>unclassified_Verrucomicrobiota | GCA_026398275.1 | Verrucomicrobiota_bacterium | NTV93_04230 | 321..3137 | 2026799 | MAG:-DUF499-domain-containing-protein-[Verrucomicrobia-bacterium] |
| RME92999.1 | AAA+ATPase+wHTH+Alpha+Beta-domain+wHTH+RRM | Type-1_BR.RM.like.systems | DUF499*->?->TUDOR+SF1-SF2-Helicase+iREase+CTD+REase-p->?->OLD-ABC->TOPRIM-> | 774 | Bacteria>PVC_group>Verrucomicrobiota>unclassified_Verrucomicrobiota | GCA_003695015.1 | Verrucomicrobiota_bacterium | D6766_09020 | 1..2326 | 2026799 | DUF499-domain-containing-protein,-partial-[Verrucomicrobia-bacterium] |
| MBI2925504.1 | AAA+ATPase+wHTH+Alpha+Beta-domain+wHTH+RRM | Type-1_BR.RM.like.systems | ZnR->?->?->?-><-DUF499*<-REase<-METHYLASE-p | 941 | Bacteria>PVC_group>Verrucomicrobiota>unclassified_Verrucomicrobiota | GCA_016190045.1 | Verrucomicrobiota_bacterium | HYY24_07350 | 8775..11600 | 2026799 | ATP-binding-protein-[Verrucomicrobia-bacterium] |
| PYL05758.1 | AAA+ATPase+wHTH+Alpha+Beta-domain+wHTH+RRM | Type-1_BR.RM.like.systems | <-DUF499<-DUF499*<-?<-?<-METHYLASE-p<-METHYLASE-p | 253 | Bacteria>PVC_group>Verrucomicrobiota>unclassified_Verrucomicrobiota | GCA_003219695.1 | Verrucomicrobiota_bacterium | DME33_15670 | 11312..12073 | 2026799 | hypothetical-protein-DME33_15670-[Verrucomicrobiota-bacterium] |
| PYL05757.1 | AAA+ATPase+wHTH+Alpha+Beta-domain+wHTH+RRM | Type-1_BR.RM.like.systems | <-DUF499*<-DUF499<-?<-?<-METHYLASE-p<-METHYLASE-p | 685 | Bacteria>PVC_group>Verrucomicrobiota>unclassified_Verrucomicrobiota | GCA_003219695.1 | Verrucomicrobiota_bacterium | DME33_15665 | 9325..11382 | 2026799 | hypothetical-protein-DME33_15665-[Verrucomicrobiota-bacterium] |
| MBI2926623.1 | AAA+ATPase+wHTH+Alpha+Beta-domain+wHTH+RRM | Type-1_BR.RM.like.systems | METHYLASE->?->?->MazF->DUF499*->?->RADICAL-SAM-> | 992 | Bacteria>PVC_group>Verrucomicrobiota>unclassified_Verrucomicrobiota | GCA_016190045.1 | Verrucomicrobiota_bacterium | HYY24_13085 | 4053..7031 | 2026799 | ATP-binding-protein-[Verrucomicrobia-bacterium] |
| MBI2924950.1 | AAA+ATPase+wHTH+Alpha+Beta-domain+wHTH+RRM | Type-1_BR.RM.like.systems | METHYLASE-p->DUF499*->?->?->?->PIN-><-?<-REase | 964 | Bacteria>PVC_group>Verrucomicrobiota>unclassified_Verrucomicrobiota | GCA_016190045.1 | Verrucomicrobiota_bacterium | HYY24_04500 | 279..3173 | 2026799 | DUF499-domain-containing-protein-[Verrucomicrobia-bacterium] |
| WP_165107642.1 | AAA+ATPase+wHTH+Alpha+Beta-domain+wHTH+RRM | Type-1_BR.RM.like.systems | Cluster1029_5clades-><-?\|\|?-><-SF1-SF2-Helicase\|\|?-><-DUF499*<-REase<-HTH<-REase<-METHYLASE<-?<-TUDOR+SF1-SF2-Helicase+iREase+CTD+REase | 925 | Bacteria>PVC_group>Verrucomicrobiota>Verrucomicrobiae>Limisphaerales | GCF_011044775.1 | Limisphaera_ngatamarikiensis | G4L39_RS09160 | 324178..326955 | 1324935 | DUF499-domain-containing-protein-[Limisphaera-ngatamarikiensis] |
| WP_165106894.1 | AAA+ATPase+wHTH+Alpha+Beta-domain+wHTH+RRM | Type-1_BR.RM.like.systems | Cluster1029_5clades->?-><-DUF499*<-DUF488-1<-METHYLASE<-MNS<-TOPRIM<-OLD-ABC<-TUDOR+SF1-SF2-Helicase+iREase+CTD+REase | 963 | Bacteria>PVC_group>Verrucomicrobiota>Verrucomicrobiae>Limisphaerales | GCF_011044775.1 | Limisphaera_ngatamarikiensis | G4L39_RS06720 | 76974..79865 | 1324935 | DUF499-domain-containing-protein-[Limisphaera-ngatamarikiensis] |
| MCC6819872.1 | AAA+ATPase+wHTH+Alpha+Beta-domain+wHTH+RRM | Type-1_BR.RM.like.systems | GDResolvase+PPP1-HTH->ParB->ParB-><-DUF499*<-MACRODOMAIN | 924 | Bacteria>PVC_group>Verrucomicrobiota>Verrucomicrobiae>Limisphaerales | GCA_020848635.1 | Limisphaerales_bacterium | IT579_04000 | 2922..5696 | 2052186 | ATP-binding-protein-[Verrucomicrobia-subdivision-3-bacterium] |
| HAO79056.1 | AAA+ATPase+wHTH+Alpha+Beta-domain+wHTH+RRM | Type-1_BR.RM.like.systems | METHYLASE-p->RNaseH->?->DUF499*-><-HNH\|\|TPR+TPR-> | 924 | Bacteria>PVC_group>Verrucomicrobiota>Verrucomicrobiae>Limisphaerales | GCA_003455565.1 | Limisphaerales_bacterium | DCQ92_08790 | 3206..5980 | 2052186 | AAA-family-ATPase-[Verrucomicrobia-subdivision-3-bacterium] |
| MBQ7024153.1 | AAA+ATPase+wHTH+Alpha+Beta-domain+wHTH+RRM | Type-1_BR.RM.like.systems | <-DUF499*<-METHYLASE<-TUDOR+SF1-SF2-Helicase+iREase+CTD+REase | 954 | Bacteria>PVC_group>Verrucomicrobiota>Verrucomicrobiae>Verrucomicrobiales | GCA_017442145.1 | Akkermansia_sp. | IJN29_11255 | 160..3024 | 1872421 | ATP-binding-protein-[Akkermansia-sp.] |
| MBE7497142.1 | AAA+ATPase+wHTH+Alpha+Beta-domain+wHTH+RRM | Type-1_BR.RM.like.systems | HKD-DNase->ParB+HNH->METHYLASE->?->nSTAND1+sGTP->REase->DUF499*->tRNA-><-?\|\|?-><-NTP_transf | 920 | Bacteria>PVC_group>Verrucomicrobiota>Verrucomicrobiae>Verrucomicrobiales | GCA_015075365.1 | Verrucomicrobiaceae_bacterium | HS117_19545 | 529489..532251 | 2026800 | ATP-binding-protein-[Verrucomicrobiaceae-bacterium] |
| WP_155996382.1 | AAA+ATPase+wHTH+Alpha+Beta-domain+wHTH+RRM | Type-1_BR.RM.like.systems | <-ParA-Soj-PloopNTPase\|\|?-><-DUF499*<-?<-?<-?<-METHYLASE<-?\|\|DDE_3-> | 966 | Bacteria>PVC_group>Verrucomicrobiota>Verrucomicrobiae>Verrucomicrobiales | GCF_000379365.1 | Verrucomicrobium_sp._3C | A37A_RS0100780 | 120354..123254 | 1134055 | DUF499-domain-containing-protein-[Verrucomicrobium-sp.-3C] |
| MCI0534235.1 | AAA+ATPase+wHTH+Alpha+Beta-domain+wHTH+RRM | Type-1_BR.RM.like.systems | <-DUF499*<-DOC+HTH<-DOC+HTH<-METHYLASE-p | 957 | Bacteria>PVC_group>Verrucomicrobiota>Verrucomicrobiae>Verrucomicrobiales | GCA_022615925.1 | Verrucomicrobiales_bacterium | LOZ50_03300 | 263..3136 | 2026801 | ATP-binding-protein-[Verrucomicrobiales-bacterium] |
| MBE7503149.1 | AAA+ATPase+wHTH+Alpha+Beta-domain+wHTH+RRM | Type-1_BR.RM.like.systems | wHTH+REase->?->?->METHYLASE->OLD-ABC->TOPRIM->DUF499*->DUF499-> | 393 | Bacteria>PVC_group>Verrucomicrobiota>Verrucomicrobiae>Verrucomicrobiales | GCA_015075445.1 | Verrucomicrobiales_bacterium | HS113_23265 | 55757..56938 | 2026801 | ATP-binding-protein-[Verrucomicrobiales-bacterium] |
| WP_142661086.1 | AAA+ATPase+wHTH+Alpha+Beta-domain+wHTH+RRM | Type-1_BR.RM.like.systems | Schlafen+HATPase+HTH->METHYLASE->DUF499*-> | 945 | Bacteria>PVC_group>Verrucomicrobiota>Verrucomicrobiota_incertae_sedis>Methylacidimicrobium | GCF_902143375.2 | Methylacidimicrobium_tartarophylax | MAMT_RS11075 | 5444..8281 | 1041768 | DUF499-domain-containing-protein-[Methylacidimicrobium-tartarophylax] |
| MBK9498689.1 | AAA+ATPase+wHTH+Alpha+Beta-domain+wHTH+RRM | Type-1_BR.RM.like.systems | METHYLASE->ParB+HNH->OLD-ABC->?->?->DUF499*-><-?\|\|?-><-STYKIN<-HKD-DNase | 972 | Bacteria>Spirochaetota>Spirochaetia>Leptospirales>Leptospiraceae | GCA_016718635.1 | Leptospiraceae_bacterium | IPO06_04755 | 70687..73605 | 2026757 | ATP-binding-protein-[Leptospiraceae-bacterium] |
| MBP6738374.1 | AAA+ATPase+wHTH+Alpha+Beta-domain+wHTH+RRM | Type-1_BR.RM.like.systems | <-Phage_integrase<-DUF499*<-?<-OLD-ABC<-ParB+HNH<-METHYLASE | 972 | Bacteria>Spirochaetota>Spirochaetia>Leptospirales>Leptospiraceae | GCA_017990775.1 | Leptospiraceae_bacterium | KA146_00215 | 1347..4265 | 2026757 | ATP-binding-protein-[Leptospiraceae-bacterium] |
| MCX7997938.1 | AAA+ATPase+wHTH+Alpha+Beta-domain+wHTH+RRM | Type-1_BR.RM.like.systems | <-DUF499*<-HEPN<-PolB<-METHYLASE-p | 838 | Bacteria>Spirochaetota>Spirochaetia>Leptospirales>Leptospiraceae | GCA_026415905.1 | Leptospiraceae_bacterium | N3A69_03160 | 811..3327 | 2026757 | MAG:-hypothetical-protein-N3A69_03160-[Leptospiraceae-bacterium] |
| MCB1144525.1 | AAA+ATPase+wHTH+Alpha+Beta-domain+wHTH+RRM | Type-1_BR.RM.like.systems | <-DUF499*<-?<-?<-OLD-ABC<-ParB+HNH<-METHYLASE | 972 | Bacteria>Spirochaetota>Spirochaetia>Leptospirales>Leptospiraceae | GCA_020440865.1 | Leptospiraceae_bacterium | KDK54_19880 | 628..3546 | 2026757 | ATP-binding-protein-[Leptospiraceae-bacterium] |
| MDD4082191.1 | AAA+ATPase+wHTH+Alpha+Beta-domain+wHTH+RRM | Type-1_BR.RM.like.systems | <-SF1-SF2-Helicase-p<-HEPN<-DUF499*<-METHYLASE-p<-METHYLASE<-TUDOR+SF1-SF2-Helicase+iREase+CTD+REase | 924 | Bacteria>Spirochaetota>Spirochaetia>Spirochaetales>Sphaerochaetaceae | GCA_028701295.1 | Sphaerochaetaceae_bacterium | PHD05_02270 | 2146..4920 | 2791017 | MAG:-DUF499-domain-containing-protein-[Sphaerochaetaceae-bacterium] |
| HCO48204.1 | AAA+ATPase+wHTH+Alpha+Beta-domain+wHTH+RRM | Type-1_BR.RM.like.systems | <-DUF499-p<-DUF499*<-METHYLASE | 631 | Bacteria>Spirochaetota>Spirochaetia>Spirochaetales>Spirochaetaceae | GCA_003533845.1 | Spirochaetaceae_bacterium | DIT55_01685 | 6809..8700 | 1898206 | hypothetical-protein-[Spirochaetaceae-bacterium] |
| MCL2137974.1 | AAA+ATPase+wHTH+Alpha+Beta-domain+wHTH+RRM | Type-1_BR.RM.like.systems | TUDOR+SF1-SF2-Helicase+iREase+CTD+REase->SbcC->?->METHYLASE->MetJarc-HTH->PIN->DUF499*-> | 821 | Bacteria>Spirochaetota>Spirochaetia>Spirochaetales>Treponemataceae | GCA_009776715.1 | Treponema_sp. | FWH41_00405 | 14879..17342 | 166 | DUF499-domain-containing-protein-[Treponema-sp.] |
| WP_010252910.1 | AAA+ATPase+wHTH+Alpha+Beta-domain+wHTH+RRM | Type-1_BR.RM.like.systems | <-wHTH+REase<-DUF499*<-METHYLASE-p<-METHYLASE<-TUDOR+SF1-SF2-Helicase+iREase+CTD+REase\|\|?->ORC-CDC6+REase->Phage_integrase-> | 929 | Bacteria>Spirochaetota>Spirochaetia>Spirochaetales>Treponemataceae | GCF_000297095.1 | Treponema_primitia | TPRIMZ1_RS0100035 | 3238..6027 | 88058 | DUF499-domain-containing-protein-[Treponema-primitia] |
| MDD6486202.1 | AAA+ATPase+wHTH+Alpha+Beta-domain+wHTH+RRM | Type-1_BR.RM.like.systems | <-HTH\|\|TUDOR+SF1-SF2-Helicase+iREase+CTD+REase->METHYLASE->APATPase+iSTAND3->Schlafen+HATPase+HTH->DUF499*->HKD-DNase+HKD-DNase->?->HicB_lk_antitox->tRNA->tRNA->?->Mbetalac-> | 948 | Bacteria>Spirochaetota>Spirochaetia>Spirochaetales>unclassified_Spirochaetales | GCA_029227075.1 | Spirochaetales_bacterium | PUF61_04585 | 321927..324773 | 2026792 | MAG:-DUF499-domain-containing-protein-[Spirochaetales-bacterium] |
| MBN2536457.1 | AAA+ATPase+wHTH+Alpha+Beta-domain+wHTH+RRM | Type-1_BR.RM.like.systems | <-DUF499*<-METHYLASE-p<-METHYLASE<-TUDOR+SF1-SF2-Helicase+iREase+CTD+REase-p | 981 | Bacteria>Spirochaetota>Spirochaetia>Spirochaetales>unclassified_Spirochaetales | GCA_016935735.1 | Spirochaetales_bacterium | JXB88_26470 | 255..3200 | 2026792 | ATP-binding-protein-[Spirochaetales-bacterium] |

| Protein-ID | DUF499-architecture | System | Gene-Neighbourhoods | Length | Taxonomy | Assembly-ID | Strain | Locus-Tag | Genomic-coordinates | TaxID | Fasta-defline |
|---|---|---|---|---|---|---|---|---|---|---|---|
| MCX8124816.1 | AAA+ATPase+wHTH+Alpha+Beta-domain+wHTH+RRM | Type-1_BR.RM.like.systems | <-DUF499*<-METHYLASE<-TUDOR+SF1-SF2-Helicase+iREase+CTD+REase\|\|?->?->PRTase-> | 824 | Bacteria>Spirochaetota>unclassified_Spirochaetota | GCA_026414805.1 | Spirochaetota_bacterium | N3F66_11750 | 1..2471 | 2202144 | MAG:-AAA-family-ATPase,-partial-[Spirochaetes-bacterium] |
| MBQ7577078.1 | AAA+ATPase+wHTH+Alpha+Beta-domain+wHTH+RRM | Type-1_BR.RM.like.systems | METHYLASE-p->DUF499*->?->GspA-ATPase-><-?<-ABC-ATPase+SbcC | 919 | Bacteria>Synergistota>Synergistia>Synergistales>Synergistaceae | GCA_017443945.1 | Synergistaceae_bacterium | IJT21_02305 | 1187..3946 | 2053624 | ATP-binding-protein-[Synergistaceae-bacterium] |
| MBQ9433643.1 | AAA+ATPase+wHTH+Alpha+Beta-domain+wHTH+RRM | Type-1_BR.RM.like.systems | REase-><-?<-?<-?<-?\|\|?-><-DUF499*<-METHYLASE | 916 | Bacteria>Synergistota>Synergistia>Synergistales>Synergistaceae | GCA_017455085.1 | Synergistaceae_bacterium | IJU26_05445 | 12618..15368 | 2053624 | ATP-binding-protein-[Synergistaceae-bacterium] |
| MBV5322918.1 | AAA+ATPase+wHTH+Alpha+Beta-domain+wHTH+RRM | Type-1_BR.RM.like.systems | HEPN->OLD-ABC->HNH->METHYLASE->METHYLASE->HEPN->DUF499*-> | 872 | Bacteria>Terrabacteria_group>Actinomycetota>Acidimicrobiia>Acidimicrobiales | GCA_019162935.1 | Ilumatobacteraceae_bacterium | JZU60_03690 | 12690..15310 | 2497619 | ATP-binding-protein-[Ilumatobacteraceae-bacterium] |
| MCB0995949.1 | AAA+ATPase+wHTH+Alpha+Beta-domain+wHTH+RRM | Type-1_BR.RM.like.systems | NERD->DUF499-p->?-><-NTD+SF1-SF2-Helicase<-DUF499* | 793 | Bacteria>Terrabacteria_group>Actinomycetota>Acidimicrobiia>Acidimicrobiales | GCA_020439465.1 | Acidimicrobiales_bacterium | KDB21_12705 | 17689..20072 | 2201156 | ATP-binding-protein-[Acidimicrobiales-bacterium] |
| MDE0160529.1 | AAA+ATPase+wHTH+Alpha+Beta-domain+wHTH+RRM | Type-1_BR.RM.like.systems | TUDOR+SF1-SF2-Helicase+iREase+CTD+REase->DUF433->PIN->METHYLASE->DUF433->PIN->DUF499*->?-><-tRNA\|\|?->?->FAD-NAD-dep-oxidoreductase-><-HEPN-p | 923 | Bacteria>Terrabacteria_group>Actinomycetota>Acidimicrobiia>Acidimicrobiales | GCA_028819505.1 | Acidimicrobiaceae_bacterium | OXL98_02215 | 9144..11915 | 2024894 | MAG:-DUF499-domain-containing-protein-[Acidimicrobiaceae-bacterium] |
| MCY4424448.1 | AAA+ATPase+wHTH+Alpha+Beta-domain+wHTH+RRM | Type-1_BR.RM.like.systems | <-SIR2+SIR2\|\|TUDOR+SF1-SF2-Helicase+iREase+CTD+REase->METHYLASE->DUF499*->MORC+REase-><-MNS+HTH<-?\|\|?->?->ABC-ATPase+SbcC-> | 924 | Bacteria>Terrabacteria_group>Actinomycetota>Acidimicrobiia>Acidimicrobiales | GCA_026710865.1 | Acidimicrobiaceae_bacterium | OXC06_15410 | 11854..14628 | 2024894 | MAG:-DUF499-domain-containing-protein-[Acidimicrobiaceae-bacterium] |
| MYH43296.1 | AAA+ATPase+wHTH+Alpha+Beta-domain+wHTH+RRM | Type-1_BR.RM.like.systems | HNH->?->?->TUDOR+SF1-SF2-Helicase+iREase+CTD+REase->METHYLASE->DUF499*-><-?<-HEPN<-?\|\|?->Aminotran_1_2->ACET+PIN+ACET-> | 923 | Bacteria>Terrabacteria_group>Actinomycetota>Acidimicrobiia>Acidimicrobiales | GCA_009843995.1 | Acidimicrobiaceae_bacterium | F4131_06280 | 9069..11840 | 2024894 | ATP-binding-protein-[Acidimicrobiaceae-bacterium] |
| MDE0606020.1 | AAA+ATPase+wHTH+Alpha+Beta-domain+wHTH+RRM | Type-1_BR.RM.like.systems | MAD-NTD+HNH-><-MORC+REase<-DNA_methylase<-DUF499*<-METHYLASE<-REase<-TUDOR+SF1-SF2-Helicase+iREase+CTD+REase\|\|DUF222+HNH-><-PIN | 983 | Bacteria>Terrabacteria_group>Actinomycetota>Acidimicrobiia>Acidimicrobiales | GCA_028824095.1 | Acidimicrobiaceae_bacterium | OXH78_02565 | 4211..7162 | 2024894 | MAG:-DUF499-domain-containing-protein-[Acidimicrobiaceae-bacterium] |
| MDE0664813.1 | AAA+ATPase+wHTH+Alpha+Beta-domain+wHTH+RRM | Type-1_BR.RM.like.systems | HIN-HTH->?->TUDOR+SF1-SF2-Helicase+iREase+CTD+REase->METHYLASE->DUF499*->?->PIN-><-tRNA<-DDE_Tnp_1_4<-Phage_int_SAM_1<-Phage_int_SAM_1 | 991 | Bacteria>Terrabacteria_group>Actinomycetota>Acidimicrobiia>Acidimicrobiales | GCA_028824455.1 | Acidimicrobiaceae_bacterium | OXH67_04395 | 41245..44220 | 2024894 | MAG:-DUF499-domain-containing-protein-[Acidimicrobiaceae-bacterium] |
| MYD34460.1 | AAA+ATPase+wHTH+Alpha+Beta-domain+wHTH+RRM | Type-1_BR.RM.like.systems | NTP_transf-><-DUF499*<-METHYLASE<-REase<-TUDOR+SF1-SF2-Helicase+iREase+CTD+REase\|\|?-><-Cysteine_desulfurase\|\|Aminotran_3-> | 970 | Bacteria>Terrabacteria_group>Actinomycetota>Acidimicrobiia>Acidimicrobiales | GCA_009841525.1 | Acidimicrobiales_bacterium | F4W98_11365 | 6251..9163 | 2201156 | ATP-binding-protein-[Acidimicrobiales-bacterium] |
| MCY4516058.1 | AAA+ATPase+wHTH+Alpha+Beta-domain+wHTH+RRM | Type-1_BR.RM.like.systems | <-Aminotran_3\|\|Cysteine_desulfurase-><-?\|\|TUDOR+SF1-SF2-Helicase+iREase+CTD+REase->REase->METHYLASE->DUF499*-><-NTP_transf | 966 | Bacteria>Terrabacteria_group>Actinomycetota>Acidimicrobiia>Acidimicrobiales | GCA_026713105.1 | Acidimicrobiaceae_bacterium | OXB99_02335 | 12101..15001 | 2024894 | MAG:-DUF499-domain-containing-protein-[Acidimicrobiaceae-bacterium] |
| MDE0133758.1 | AAA+ATPase+wHTH+Alpha+Beta-domain+wHTH+RRM | Type-1_BR.RM.like.systems | <-Aminotran_3\|\|Cysteine_desulfurase->?->TUDOR+SF1-SF2-Helicase+iREase+CTD+REase->REase->METHYLASE->DUF499*-><-NTP_transf | 970 | Bacteria>Terrabacteria_group>Actinomycetota>Acidimicrobiia>Acidimicrobiales | GCA_028819405.1 | Acidimicrobiaceae_bacterium | OXM54_02860 | 72167..75079 | 2024894 | MAG:-DUF499-domain-containing-protein-[Acidimicrobiaceae-bacterium] |
| MXW60960.1 | AAA+ATPase+wHTH+Alpha+Beta-domain+wHTH+RRM | Type-1_BR.RM.like.systems | <-DUF222+HNH\|\|TUDOR+SF1-SF2-Helicase+iREase+CTD+REase->METHYLASE->DUF499*->DNA_methylase->MORC+REase-><-MAD-NTD+HNH | 983 | Bacteria>Terrabacteria_group>Actinomycetota>Acidimicrobiia>Acidimicrobiales | GCA_009835385.1 | Acidimicrobiaceae_bacterium | F4003_04135 | 32450..35401 | 2024894 | DUF499-domain-containing-protein-[Acidimicrobiaceae-bacterium] |
| MXZ98307.1 | AAA+ATPase+wHTH+Alpha+Beta-domain+wHTH+RRM | Type-1_BR.RM.like.systems | <-DDE_3<-DUF499-p<-DUF499*<-METHYLASE-p<-METHYLASE-p\|\|Tox-RelE->HTH_3-><-TUDOR+SF1-SF2-Helicase+iREase+CTD+REase | 711 | Bacteria>Terrabacteria_group>Actinomycetota>Acidimicrobiia>Acidimicrobiales | GCA_009836995.1 | Acidimicrobiaceae_bacterium | F4Z23_04625 | 8716..10851 | 2024894 | ATP-binding-protein-[Acidimicrobiaceae-bacterium] |
| MDE0515901.1 | AAA+ATPase+wHTH+Alpha+Beta-domain+wHTH+RRM | Type-1_BR.RM.like.systems | ZnR+TOPRIM+DnaB->tRNA->TUDOR+SF1-SF2-Helicase+iREase+CTD+REase->METHYLASE->DUF499*-><-TPR<-?\|\|HEPN->?-><-?\|\|REase+NACHT-> | 974 | Bacteria>Terrabacteria_group>Actinomycetota>Acidimicrobiia>Acidimicrobiales | GCA_028824915.1 | Acidimicrobiaceae_bacterium | OXH43_06180 | 15501..18425 | 2024894 | MAG:-DUF499-domain-containing-protein-[Acidimicrobiaceae-bacterium] |
| MDE0136684.1 | AAA+ATPase+wHTH+Alpha+Beta-domain+wHTH+RRM | Type-1_BR.RM.like.systems | DUF4527-><-HTH+MPTase<-RelE-ParE\|\|SIR2+SIR2->?-><-DUF499*<-METHYLASE<-TUDOR+SF1-SF2-Helicase+iREase+CTD+REase | 991 | Bacteria>Terrabacteria_group>Actinomycetota>Acidimicrobiia>Acidimicrobiales | GCA_028819405.1 | Acidimicrobiaceae_bacterium | OXM54_17820 | 6064..9039 | 2024894 | MAG:-DUF499-domain-containing-protein-[Acidimicrobiaceae-bacterium] |
| MCY4294037.1 | AAA+ATPase+wHTH+Alpha+Beta-domain+wHTH+RRM | Type-1_BR.RM.like.systems | <-DUF499*<-METHYLASE-p<-METHYLASE-p<-HTH_Tnp+HTH_Tnp<-RelE-ParE<-TUDOR+SF1-SF2-Helicase+iREase+CTD+REase | 605 | Bacteria>Terrabacteria_group>Actinomycetota>Acidimicrobiia>Acidimicrobiales | GCA_026710015.1 | Acidimicrobiaceae_bacterium | OXC58_04270 | 1..1814 | 2024894 | MAG:-DUF499-domain-containing-protein,-partial-[Acidimicrobiaceae-bacterium] |
| MCY4174746.1 | AAA+ATPase+wHTH+Alpha+Beta-domain+wHTH+RRM | Type-1_BR.RM.like.systems | <-DUF499*<-METHYLASE-p<-METHYLASE-p<-HTH_Tnp+HTH_Tnp<-RelE-ParE<-TUDOR+SF1-SF2-Helicase+iREase+CTD+REase | 976 | Bacteria>Terrabacteria_group>Actinomycetota>Acidimicrobiia>Acidimicrobiales | GCA_026709125.1 | Acidimicrobiaceae_bacterium | OXD37_00225 | 22754..25684 | 2024894 | MAG:-DUF499-domain-containing-protein-[Acidimicrobiaceae-bacterium] |
| MCY4068979.1 | AAA+ATPase+wHTH+Alpha+Beta-domain+wHTH+RRM | Type-1_BR.RM.like.systems | TUDOR+SF1-SF2-Helicase+iREase+CTD+REase->RelE-ParE->HTH_Tnp+HTH_Tnp->METHYLASE-p->METHYLASE-p->DUF499*-> | 976 | Bacteria>Terrabacteria_group>Actinomycetota>Acidimicrobiia>Acidimicrobiales | GCA_026708645.1 | Acidimicrobiaceae_bacterium | OXE79_07740 | 35900..38830 | 2024894 | MAG:-DUF499-domain-containing-protein-[Acidimicrobiaceae-bacterium] |
| MCY3585837.1 | AAA+ATPase+wHTH+Alpha+Beta-domain+wHTH+RRM | Type-1_BR.RM.like.systems | ABC-ATPase-><-?<-RelA\|\|TUDOR+SF1-SF2-Helicase+iREase+CTD+REase->METHYLASE->DUF499*->MethyltransfD12-><-?<-ParB | 980 | Bacteria>Terrabacteria_group>Actinomycetota>Acidimicrobiia>Acidimicrobiales | GCA_026705175.1 | Acidimicrobiaceae_bacterium | OXG76_09110 | 500669..503611 | 2024894 | MAG:-DUF499-domain-containing-protein-[Acidimicrobiaceae-bacterium] |
| MXW97851.1 | AAA+ATPase+wHTH+Alpha+Beta-domain+wHTH+RRM | Type-1_BR.RM.like.systems | TUDOR+SF1-SF2-Helicase+iREase+CTD+REase->RelE-ParE->HTH_Tnp+HTH_Tnp->METHYLASE->DUF499*->OLD-ABC->TOPRIM-> | 977 | Bacteria>Terrabacteria_group>Actinomycetota>Acidimicrobiia>Acidimicrobiales | GCA_009835735.1 | Acidimicrobiaceae_bacterium | F4Z89_04460 | 7235..10168 | 2024894 | ATP-binding-protein-[Acidimicrobiaceae-bacterium] |
| MYJ80998.1 | AAA+ATPase+wHTH+Alpha+Beta-domain+wHTH+RRM | Type-1_BR.RM.like.systems | <-DDE_3<-DUF499*<-METHYLASE-p<-METHYLASE-p\|\|Tox-RelE->HTH_3-><-TUDOR+SF1-SF2-Helicase+iREase+CTD+REase | 994 | Bacteria>Terrabacteria_group>Actinomycetota>Acidimicrobiia>Acidimicrobiales | GCA_009845665.1 | Acidimicrobiaceae_bacterium | F4046_04140 | 2126..5110 | 2024894 | ATP-binding-protein-[Acidimicrobiaceae-bacterium] |
| MXW95144.1 | AAA+ATPase+wHTH+Alpha+Beta-domain+wHTH+RRM | Type-1_BR.RM.like.systems | TUDOR+SF1-SF2-Helicase+iREase+CTD+REase-><-HTH_3<-Tox-RelE\|\|METHYLASE->DUF499*->DDE_3-><-?<-?<-?<-?<-sGTP | 994 | Bacteria>Terrabacteria_group>Actinomycetota>Acidimicrobiia>Acidimicrobiales | GCA_009836165.1 | Acidimicrobiaceae_bacterium | F4Z64_06505 | 82291..85275 | 2024894 | ATP-binding-protein-[Acidimicrobiaceae-bacterium] |
| MDE0161736.1 | AAA+ATPase+wHTH+Alpha+Beta-domain+wHTH+RRM | Type-1_BR.RM.like.systems | <-AbiEii<-?\|\|?->?-><-DUF499*<-METHYLASE<-TUDOR+SF1-SF2-Helicase+iREase+CTD+REase<-tRNA<-ZnR+TOPRIM+DnaB | 974 | Bacteria>Terrabacteria_group>Actinomycetota>Acidimicrobiia>Acidimicrobiales | GCA_028819505.1 | Acidimicrobiaceae_bacterium | OXL98_08335 | 5758..8682 | 2024894 | MAG:-DUF499-domain-containing-protein-[Acidimicrobiaceae-bacterium] |
| MCY4424295.1 | AAA+ATPase+wHTH+Alpha+Beta-domain+wHTH+RRM | Type-1_BR.RM.like.systems | METHYLASE-p->METHYLASE-p->DUF499*->RelE-ParE->HTH_Tnp+HTH_Tnp-><-?<-?<-?<-REase-p | 983 | Bacteria>Terrabacteria_group>Actinomycetota>Acidimicrobiia>Acidimicrobiales | GCA_026710865.1 | Acidimicrobiaceae_bacterium | OXC06_14640 | 4154..7105 | 2024894 | MAG:-DUF499-domain-containing-protein-[Acidimicrobiaceae-bacterium] |
| MXV86625.1 | AAA+ATPase+wHTH+Alpha+Beta-domain+wHTH+RRM | Type-1_BR.RM.like.systems | ABC-ATPase-><-?<-?\|\|TUDOR+SF1-SF2-Helicase+iREase+CTD+REase->METHYLASE->DUF499*-><-MACRODOMAIN | 980 | Bacteria>Terrabacteria_group>Actinomycetota>Acidimicrobiia>Acidimicrobiales | GCA_009835365.1 | Acidimicrobiales_bacterium | F4005_03760 | 18560..21502 | 2201156 | DUF499-domain-containing-protein-[Acidimicrobiales-bacterium] |
| MYH42421.1 | AAA+ATPase+wHTH+Alpha+Beta-domain+wHTH+RRM | Type-1_BR.RM.like.systems | <-DDE_3<-DUF499*<-METHYLASE\|\|Tox-RelE->HTH_3-><-TUDOR+SF1-SF2-Helicase+iREase+CTD+REase | 994 | Bacteria>Terrabacteria_group>Actinomycetota>Acidimicrobiia>Acidimicrobiales | GCA_009843995.1 | Acidimicrobiaceae_bacterium | F4131_01795 | 8591..11575 | 2024894 | ATP-binding-protein-[Acidimicrobiaceae-bacterium] |
| MDE0319207.1 | AAA+ATPase+wHTH+Alpha+Beta-domain+wHTH+RRM | Type-1_BR.RM.like.systems | MACRODOMAIN-><-DUF499*<-METHYLASE<-TUDOR+SF1-SF2-Helicase+iREase+CTD+REase\|\|RelA-> | 980 | Bacteria>Terrabacteria_group>Actinomycetota>Acidimicrobiia>Acidimicrobiales | GCA_028820815.1 | Acidimicrobiaceae_bacterium | OXI97_04940 | 6938..9880 | 2024894 | MAG:-DUF499-domain-containing-protein-[Acidimicrobiaceae-bacterium] |
| MXZ14251.1 | AAA+ATPase+wHTH+Alpha+Beta-domain+wHTH+RRM | Type-1_BR.RM.like.systems | NTP_transf-><-DUF499*<-METHYLASE<-REase<-TUDOR+SF1-SF2-Helicase+iREase+CTD+REase | 970 | Bacteria>Terrabacteria_group>Actinomycetota>Acidimicrobiia>Acidimicrobiales | GCA_009837945.1 | Acidimicrobiales_bacterium | F4Y76_01880 | 8322..11234 | 2201156 | ATP-binding-protein-[Acidimicrobiales-bacterium] |
| MCY3851371.1 | AAA+ATPase+wHTH+Alpha+Beta-domain+wHTH+RRM | Type-1_BR.RM.like.systems | TUDOR+SF1-SF2-Helicase+iREase+CTD+REase->METHYLASE->DUF499*-> | 1006 | Bacteria>Terrabacteria_group>Actinomycetota>Acidimicrobiia>Acidimicrobiales | GCA_026706815.1 | Acidimicrobiaceae_bacterium | OXF75_11330 | 7755..10775 | 2024894 | MAG:-PASTA-domain-containing-protein-[Acidimicrobiaceae-bacterium] |
| MXY02283.1 | AAA+ATPase+wHTH+Alpha+Beta-domain+wHTH+RRM | Type-1_BR.RM.like.systems | NTP_transf-><-DUF499*<-METHYLASE<-REase | 970 | Bacteria>Terrabacteria_group>Actinomycetota>Acidimicrobiia>Acidimicrobiales | GCA_009838155.1 | Acidimicrobiales_bacterium | F4Y66_05500 | 5633..8545 | 2201156 | DUF499-domain-containing-protein-[Acidimicrobiales-bacterium] |
| MXX43802.1 | AAA+ATPase+wHTH+Alpha+Beta-domain+wHTH+RRM | Type-1_BR.RM.like.systems | NTP_transf-><-DUF499*<-METHYLASE<-REase | 970 | Bacteria>Terrabacteria_group>Actinomycetota>Acidimicrobiia>Acidimicrobiales | GCA_009836375.1 | Acidimicrobiales_bacterium | F4Z53_12210 | 6244..9156 | 2201156 | ATP-binding-protein-[Acidimicrobiales-bacterium] |
| MXV87683.1 | AAA+ATPase+wHTH+Alpha+Beta-domain+wHTH+RRM | Type-1_BR.RM.like.systems | NTP_transf-><-DUF499*<-METHYLASE<-REase | 970 | Bacteria>Terrabacteria_group>Actinomycetota>Acidimicrobiia>Acidimicrobiales | GCA_009835365.1 | Acidimicrobiales_bacterium | F4005_09195 | 34597..37509 | 2201156 | ATP-binding-protein-[Acidimicrobiales-bacterium] |
| MDE0498678.1 | AAA+ATPase+wHTH+Alpha+Beta-domain+wHTH+RRM | Type-1_BR.RM.like.systems | NTP_transf-><-DUF499*<-METHYLASE<-REase | 970 | Bacteria>Terrabacteria_group>Actinomycetota>Acidimicrobiia>Acidimicrobiales | GCA_028823965.1 | Acidimicrobiaceae_bacterium | OXH86_15120 | 56164..59076 | 2024894 | MAG:-DUF499-domain-containing-protein-[Acidimicrobiaceae-bacterium] |
| MDE0319984.1 | AAA+ATPase+wHTH+Alpha+Beta-domain+wHTH+RRM | Type-1_BR.RM.like.systems | NTP_transf-><-DUF499*<-METHYLASE<-REase | 970 | Bacteria>Terrabacteria_group>Actinomycetota>Acidimicrobiia>Acidimicrobiales | GCA_028820815.1 | Acidimicrobiaceae_bacterium | OXI97_08880 | 1201..4113 | 2024894 | MAG:-DUF499-domain-containing-protein-[Acidimicrobiaceae-bacterium] |
| MCY3894574.1 | AAA+ATPase+wHTH+Alpha+Beta-domain+wHTH+RRM | Type-1_BR.RM.like.systems | REase->METHYLASE->DUF499*-><-NTP_transf | 970 | Bacteria>Terrabacteria_group>Actinomycetota>Acidimicrobiia>Acidimicrobiales | GCA_026707105.1 | Acidimicrobiaceae_bacterium | OXF65_14910 | 3656..6568 | 2024894 | MAG:-DUF499-domain-containing-protein-[Acidimicrobiaceae-bacterium] |
| MXZ30355.1 | AAA+ATPase+wHTH+Alpha+Beta-domain+wHTH+RRM | Type-1_BR.RM.like.systems | ORC-CDC6+REase-><-PIN<-MPTase<-PIN\|\|METHYLASE-><-?\|\|DUF499*-> | 778 | Bacteria>Terrabacteria_group>Actinomycetota>Acidimicrobiia>unclassified_Acidimicrobiia | GCA_009837005.1 | Acidimicrobiia_bacterium | F4Z22_05835 | 49323..51659 | 2080302 | ATP-binding-protein-[Acidimicrobiia-bacterium] |
| MXZ84910.1 | AAA+ATPase+wHTH+Alpha+Beta-domain+wHTH+RRM | Type-1_BR.RM.like.systems | <-HEPN\|\|TUDOR+SF1-SF2-Helicase+iREase+CTD+REase->METHYLASE->DUF499*-><-?<-?<-?<-?<-PIN<-RHH_1 | 933 | Bacteria>Terrabacteria_group>Actinomycetota>Acidimicrobiia>unclassified_Acidimicrobiia | GCA_009837425.1 | Acidimicrobiia_bacterium | F4Z02_04560 | 13074..15875 | 2080302 | ATP-binding-protein-[Acidimicrobiia-bacterium] |
| MCQ3809930.1 | AAA+ATPase+wHTH+Alpha+Beta-domain+wHTH+RRM | Type-1_BR.RM.like.systems | RHH_1->?-><-TIR<-?<-DUF499*<-METHYLASE<-TUDOR+SF1-SF2-Helicase+iREase+CTD+REase\|\|HEPN-> | 933 | Bacteria>Terrabacteria_group>Actinomycetota>Acidimicrobiia>unclassified_Acidimicrobiia | GCA_024421205.1 | Acidimicrobiia_bacterium | KTV68_05170 | 18978..21779 | 2080302 | DUF499-domain-containing-protein-[Acidimicrobiia-bacterium] |
| MYB43816.1 | AAA+ATPase+wHTH+Alpha+Beta-domain+wHTH+RRM | Type-1_BR.RM.like.systems | TUDOR+SF1-SF2-Helicase+iREase+CTD+REase->METHYLASE-p->Zn_Tnp_IS1595+DDE_Tnp_IS1595->N6_N4_MTase->?->METHYLASE-p->DUF499*->?-><-?<-Classical-AAA<-?<-SF1-SF2-Helicase+wHTH+PRTase | 962 | Bacteria>Terrabacteria_group>Actinomycetota>Acidimicrobiia>unclassified_Acidimicrobiia | GCA_009839965.1 | Acidimicrobiia_bacterium | F4X74_02590 | 20780..23668 | 2080302 | DUF499-domain-containing-protein-[Acidimicrobiia-bacterium] |
| MCZ7536063.1 | AAA+ATPase+wHTH+Alpha+Beta-domain+wHTH+RRM | Type-1_BR.RM.like.systems | <-DUF499<-DUF499<-DUF499*<-METHYLASE-p<-METHYLASE-p<-REase<-?<-HKD-DNase+SF1-SF2-Helicase-p | 207 | Bacteria>Terrabacteria_group>Actinomycetota>Acidimicrobiia>unclassified_Acidimicrobiia | GCA_027494135.1 | Acidimicrobiia_bacterium | M5T61_09145 | 13163..13786 | 2080302 | MAG:-hypothetical-protein-M5T61_09145-[Acidimicrobiia-bacterium] |
| MCZ7536062.1 | AAA+ATPase+wHTH+Alpha+Beta-domain+wHTH+RRM | Type-1_BR.RM.like.systems | <-DUF499<-DUF499*<-DUF499<-METHYLASE-p<-METHYLASE-p<-REase<-?<-HKD-DNase+SF1-SF2-Helicase-p | 184 | Bacteria>Terrabacteria_group>Actinomycetota>Acidimicrobiia>unclassified_Acidimicrobiia | GCA_027494135.1 | Acidimicrobiia_bacterium | M5T61_09140 | 12615..13169 | 2080302 | MAG:-hypothetical-protein-M5T61_09140-[Acidimicrobiia-bacterium] |
| MXY74133.1 | AAA+ATPase+wHTH+Alpha+Beta-domain+wHTH+RRM | Type-1_BR.RM.like.systems | METHYLASE-p-><-?<-tRNA\|\|TUDOR+SF1-SF2-Helicase+iREase+CTD+REase->METHYLASE->DUF499*-> | 260 | Bacteria>Terrabacteria_group>Actinomycetota>Acidimicrobiia>unclassified_Acidimicrobiia | GCA_009837815.1 | Acidimicrobiia_bacterium | F4Y83_04300 | 12600..13380 | 2080302 | AAA-family-ATPase-[Acidimicrobiia-bacterium] |
| MBP8180265.1 | AAA+ATPase+wHTH+Alpha+Beta-domain+wHTH+RRM | Type-1_BR.RM.like.systems | <-TrwC+SF1-SF2-helicase+TOPRIM\|\|?->TUDOR+SF1-SF2-Helicase+iREase+CTD+REase->METHYLASE->DUF499*-> | 988 | Bacteria>Terrabacteria_group>Actinomycetota>Acidimicrobiia>unclassified_Acidimicrobiia | GCA_018006325.1 | Acidimicrobiia_bacterium | KAZ88_04660 | 11065..14031 | 2080302 | DUF499-domain-containing-protein-[Acidimicrobiia-bacterium] |
| MCZ7536061.1 | AAA+ATPase+wHTH+Alpha+Beta-domain+wHTH+RRM | Type-1_BR.RM.like.systems | <-DUF499*<-DUF499<-DUF499<-METHYLASE-p<-METHYLASE-p<-REase | 300 | Bacteria>Terrabacteria_group>Actinomycetota>Acidimicrobiia>unclassified_Acidimicrobiia | GCA_027494135.1 | Acidimicrobiia_bacterium | M5T61_09135 | 11348..12250 | 2080302 | MAG:-hypothetical-protein-M5T61_09135-[Acidimicrobiia-bacterium] |
| MYB24421.1 | AAA+ATPase+wHTH+Alpha+Beta-domain+wHTH+RRM | Type-1_BR.RM.like.systems | TUDOR+SF1-SF2-Helicase+iREase+CTD+REase->REase->?->?->METHYLASE->DUF499*-> | 1025 | Bacteria>Terrabacteria_group>Actinomycetota>Acidimicrobiia>unclassified_Acidimicrobiia | GCA_009840735.1 | Acidimicrobiia_bacterium | F4X37_04860 | 55279..58356 | 2080302 | DUF499-domain-containing-protein-[Acidimicrobiia-bacterium] |
| MCL6550890.1 | AAA+ATPase+wHTH+Alpha+Beta-domain+wHTH+RRM | Type-1_BR.RM.like.systems | DUF499*->Phage_integrase-><-?<-?<-?<-?<-ZnR+TOPRIM+D5ATPase | 626 | Bacteria>Terrabacteria_group>Actinomycetota>Actinomycetes>Acidothermales | GCA_023512095.1 | Acidothermus_cellulolyticus | K6T37_07780 | 1..1881 | 28049 | ATP-binding-protein-[Acidothermus-cellulolyticus] |
| WP_252863026.1 | AAA+ATPase+wHTH+Alpha+Beta-domain+wHTH+RRM | Type-1_BR.RM.like.systems | <-METHYLASE-p<-REase<-HTH+SF1-SF2-Helicase+REase<-REase<-MORC<-DUF499*<-HEPN<-METHYLASE\|\|DNA_methylase-> | 912 | Bacteria>Terrabacteria_group>Actinomycetota>Actinomycetes>Actinomycetales | - | Actinomyces_oris | BKH30_RS08650 | 21386..24124 | 544580 | DUF499-domain-containing-protein-[Actinomyces-oris] |
| WP_143180508.1 | AAA+ATPase+wHTH+Alpha+Beta-domain+wHTH+RRM | Type-1_BR.RM.like.systems | DUF499*->SbcC->SbcC-><-DNA_methylase\|\|<-DNA_methylase | 702 | Bacteria>Terrabacteria_group>Actinomycetota>Actinomycetes>Actinomycetales | GCF_001907295.1 | Bowdeniella_nasicola | BSZ39_RS11170 | 1..2109 | 208480 | DUF499-domain-containing-protein,-partial-[Bowdeniella-nasicola] |
| WP_284940342.1 | AAA+ATPase+wHTH+Alpha+Beta-domain+wHTH+RRM | Type-1_BR.RM.like.systems | HEPN-><-HEPN<-ParB+HNH<-DUF499*<-TraJ-RHH | 797 | Bacteria>Terrabacteria_group>Actinomycetota>Actinomycetes>Bifidobacteriales | GCF_030250405.1 | Bifidobacterium_catenulatum | K1513_RS02315 | 519682..522075 | 1686 | SNF2-related-protein-[Bifidobacterium-catenulatum] |
| WP_073264319.1 | AAA+ATPase+wHTH+Alpha+Beta-domain+wHTH+RRM | Type-1_BR.RM.like.systems | GDResolvase-><-DUF499*<-METHYLASE<-TUDOR+SF1-SF2-Helicase+iREase+CTD+REase\|\|?-><-?<-HTH+HKD-DNase\|\|GNTR-HTH-> | 992 | Bacteria>Terrabacteria_group>Actinomycetota>Actinomycetes>Cryptosporangiales | GCF_900143005.1 | Cryptosporangium_aurantiacum | BUB75_RS37295 | 3783..6761 | 134849 | DUF499-domain-containing-protein-[Cryptosporangium-aurantiacum] |
| WP_264763297.1 | AAA+ATPase+wHTH+Alpha+Beta-domain+wHTH+RRM | Type-1_BR.RM.like.systems | <-TetR-HTH\|\|?-><-Phage_integrase<-DUF499*<-METHYLASE<-TUDOR+SF1-SF2-Helicase+iREase+CTD+REase | 889 | Bacteria>Terrabacteria_group>Actinomycetota>Actinomycetes>Frankiales | - | Frankia_gtarii | NBZ83_RS14085 | 13561..16230 | 2950102 | ATP-binding-protein-[Frankia-gtarii] |
| WP_054571357.1 | AAA+ATPase+wHTH+Alpha+Beta-domain+wHTH+RRM | Type-1_BR.RM.like.systems | <-METHYLASE-p\|\|?->TUDOR+SF1-SF2-Helicase+iREase+CTD+REase->METHYLASE->DUF499*-> | 954 | Bacteria>Terrabacteria_group>Actinomycetota>Actinomycetes>Frankiales | GCF_001306465.1 | Frankia_sp._R43 | ACG83_RS39405 | 169532..172396 | 269536 | DUF499-domain-containing-protein-[Frankia-sp.-R43] |
| MCU1612622.1 | AAA+ATPase+wHTH+Alpha+Beta-domain+wHTH+RRM | Type-1_BR.RM.like.systems | <-DUF499*<-METHYLASE<-TUDOR+SF1-SF2-Helicase+iREase+CTD+REase\|\|?-><-HTH+MPTase\|\|HTH_17-> | 979 | Bacteria>Terrabacteria_group>Actinomycetota>Actinomycetes>Frankiales | GCA_025465155.1 | Frankiales_bacterium | JWD98_162 | 7300..10239 | 1909291 | putative-ATPase-superfamily-[Frankiales-bacterium] |
| WP_158716411.1 | AAA+ATPase+wHTH+Alpha+Beta-domain+wHTH+RRM | Type-1_BR.RM.like.systems | DUF499*->MORC->REase->HTH+SF1-SF2-Helicase+REase->SIGMA-HTH->HTH_Tnp_1+HTH_21+rve+rve_3-> | 941 | Bacteria>Terrabacteria_group>Actinomycetota>Actinomycetes>Geodermatophilales | GCF_900292035.1 | Blastococcus_sp._Marseille-P5729 | DAA40_RS12375 | 801548..804373 | 2086582 | DUF499-domain-containing-protein-[Blastococcus-sp.-Marseille-P5729] |
| WP_139263554.1 | AAA+ATPase+wHTH+Alpha+Beta-domain+wHTH+RRM | Type-1_BR.RM.like.systems | DnaB->XIS-HTH-><-?<-REase\|\|TUDOR+SF1-SF2-Helicase+iREase+CTD+REase->METHYLASE->DUF499*-><-?<-HEPN | 975 | Bacteria>Terrabacteria_group>Actinomycetota>Actinomycetes>Geodermatophilales | GCF_900107105.1 | Geodermatophilus_africanus | BLW63_RS26160 | 226185..229112 | 1137993 | DUF499-domain-containing-protein-[Geodermatophilus-africanus] |
| WP_255137076.1 | AAA+ATPase+wHTH+Alpha+Beta-domain+wHTH+RRM | Type-1_BR.RM.like.systems | <-ESAT<-ESAT\|\|?-><-?<-?\|\|?-><-DUF499*<-METHYLASE<-TUDOR+SF1-SF2-Helicase+iREase+CTD+REase<-ZDNA-HTH | 695 | Bacteria>Terrabacteria_group>Actinomycetota>Actinomycetes>Kitasatosporales | GCF_024349085.1 | Streptomyces_somaliensis | LUX12_RS04145 | 951894..953981 | 78355 | DUF499-domain-containing-protein-[Streptomyces-somaliensis] |
| WP_189449183.1 | AAA+ATPase+wHTH+Alpha+Beta-domain+wHTH+RRM | Type-1_BR.RM.like.systems | VirD4-TraG+VirD4-FtsK->DNA_methylase-><-HNH<-ABC-ATPase<-DUF499*<-METHYLASE<-TUDOR+SF1-SF2-Helicase+iREase+CTD+REase<-REase+DUF5655<-TRAM+Methyltransf_31 | 922 | Bacteria>Terrabacteria_group>Actinomycetota>Actinomycetes>Kitasatosporales | GCF_014648775.1 | Streptomyces_abikoensis | IE096_RS06315 | 1448410..1451178 | 97398 | DUF499-domain-containing-protein-[Streptomyces-abikoensis] |
| WP_190197152.1 | AAA+ATPase+wHTH+Alpha+Beta-domain+wHTH+RRM | Type-1_BR.RM.like.systems | ABhydrolase+ABhydrolase+ABhydrolase->Peptidase->?-><-HNH<-ABC-ATPase<-HEPN<-DUF499*<-METHYLASE<-TUDOR+SF1-SF2-Helicase+iREase+CTD+REase\|\|HIN-HTH-> | 910 | Bacteria>Terrabacteria_group>Actinomycetota>Actinomycetes>Kitasatosporales | GCF_014651075.1 | Streptomyces_djakartensis | IE212_RS08750 | 190992..193724 | 68193 | DUF499-domain-containing-protein-[Streptomyces-djakartensis] |
| WP_019064605.1 | AAA+ATPase+wHTH+Alpha+Beta-domain+wHTH+RRM | Type-1_BR.RM.like.systems | <-ABC-ATPase<-PIN<-?<-?<-ABhydrolase+ABhydrolase+ABhydrolase\|\|?-><-DUF499*<-METHYLASE<-TUDOR+SF1-SF2-Helicase+iREase+CTD+REase<-?\|\|REase+DUF5655-> | 905 | Bacteria>Terrabacteria_group>Actinomycetota>Actinomycetes>Kitasatosporales | GCF_000367365.1 | Streptomyces_prunicolor | SPRO2S_RS50025 | 43497..46214 | 67348 | DUF499-domain-containing-protein-[Streptomyces-prunicolor] |
| WP_160312248.1 | AAA+ATPase+wHTH+Alpha+Beta-domain+wHTH+RRM | Type-1_BR.RM.like.systems | ABC-ATPase+SbcC->?-><-MODE-HTH+PBPII\|\|?->Glyoxalase->?-><-DUF499*<-METHYLASE<-TUDOR+SF1-SF2-Helicase+iREase+CTD+REase\|\|ABC-ATPase->HNH-> | 912 | Bacteria>Terrabacteria_group>Actinomycetota>Actinomycetes>Kitasatosporales | GCF_000787755.1 | Streptacidiphilus_albus | BS75_RS18810 | 4369420..4372158 | 105425 | DUF499-domain-containing-protein-[Streptacidiphilus-albus] |
| TDU05561.1 | AAA+ATPase+wHTH+Alpha+Beta-domain+wHTH+RRM | Type-1_BR.RM.like.systems | <-DDE_Tnp_1<-Schlafen+HATPase+HTH\|\|TUDOR+SF1-SF2-Helicase+iREase+CTD+REase->METHYLASE->DUF499*-><-HAD\|\|HTH_31->?->RADICAL-SAM-> | 960 | Bacteria>Terrabacteria_group>Actinomycetota>Actinomycetes>Kitasatosporales | GCA_004365705.1 | Streptomyces_sp._846.5 | EDD99_4083 | 4433097..4435979 | 2485156 | uncharacterized-protein-DUF499-[Streptomyces-sp.-846.5] |
| WP_037620150.1 | AAA+ATPase+wHTH+Alpha+Beta-domain+wHTH+RRM | Type-1_BR.RM.like.systems | REase->Peptidase-S8+Subtilisin->?->?->TUDOR+SF1-SF2-Helicase+iREase+CTD+REase->METHYLASE->DUF499*->HEPN->ABC-ATPase->HNH-> | 910 | Bacteria>Terrabacteria_group>Actinomycetota>Actinomycetes>Kitasatosporales | GCF_000725495.1 | Streptomyces_aureus | IO36_RS15000 | 183930..186662 | 193461 | DUF499-domain-containing-protein-[Streptomyces-aureus] |
| WP_176189011.1 | AAA+ATPase+wHTH+Alpha+Beta-domain+wHTH+RRM | Type-1_BR.RM.like.systems | <-Radical-SAM<-?\|\|?->TUDOR+SF1-SF2-Helicase+iREase+CTD+REase->METHYLASE->DUF499*-><-HTH_21+rve+rve_3<-SIGMA-HTH | 930 | Bacteria>Terrabacteria_group>Actinomycetota>Actinomycetes>Kitasatosporales | GCF_013364235.1 | Kitasatospora_sp._NA04385 | HUT16_RS17010 | 3826130..3828922 | 2742135 | DUF499-domain-containing-protein-[Kitasatospora-sp.-NA04385] |
| WP_254370730.1 | AAA+ATPase+wHTH+Alpha+Beta-domain+wHTH+RRM | Type-1_BR.RM.like.systems | Methyltransf->?-><-?\|\|?-><-DUF499*<-METHYLASE<-TUDOR+SF1-SF2-Helicase+iREase+CTD+REase\|\|Schlafen+HATPase+HTH->?-><-?\|\|ABhydrolase-> | 953 | Bacteria>Terrabacteria_group>Actinomycetota>Actinomycetes>Kitasatosporales | GCF_024213665.1 | Streptomyces_sp._A3M-1-3 | NLX86_07020 | 25922..28783 | 2962044 | DUF499-domain-containing-protein-[Streptomyces-sp.-A3M-1-3] |
| WP_030324200.1 | AAA+ATPase+wHTH+Alpha+Beta-domain+wHTH+RRM | Type-1_BR.RM.like.systems | wHTH+SAD+HNH->MetJarc-HTH->?-><-Schlafen+HATPase+HTH\|\|TUDOR+SF1-SF2-Helicase+iREase+CTD+REase->METHYLASE->DUF499*-> | 931 | Bacteria>Terrabacteria_group>Actinomycetota>Actinomycetes>Kitasatosporales | GCF_000719505.1 | Streptomyces_sp._NRRL_B-3229 | IF54_RS0129580 | 225412..228207 | 1463836 | DUF499-domain-containing-protein-[Streptomyces-sp.-NRRL-B-3229] |
| AWN29968.1 | AAA+ATPase+wHTH+Alpha+Beta-domain+wHTH+RRM | Type-1_BR.RM.like.systems | REase+DUF5655-><-?\|\|?->\|\|TUDOR+SF1-SF2-Helicase+iREase+CTD+REase->METHYLASE->DUF499*->ABC-ATPase-><-JetD<-?<-?<-?<-DnaB+DnaB | 917 | Bacteria>Terrabacteria_group>Actinomycetota>Actinomycetes>Kitasatosporales | GCA_003173275.1 | Streptomyces_sp._NEAU-S7GS2 | DKG71_30770 | 7152841..7155594 | 2202000 | hypothetical-protein-DKG71_30770-[Streptomyces-sp.-NEAU-S7GS2] |
| WP_166682464.1 | AAA+ATPase+wHTH+Alpha+Beta-domain+wHTH+RRM | Type-1_BR.RM.like.systems | <-Schlafen+HATPase+HTH\|\|TUDOR+SF1-SF2-Helicase+iREase+CTD+REase->METHYLASE->DUF499*-><-HAD\|\|HTH_31->?->?->RADICAL-SAM-> | 955 | Bacteria>Terrabacteria_group>Actinomycetota>Actinomycetes>Kitasatosporales | GCF_004365705.1 | Streptomyces_sp._846.5 | EDD99_RS20240 | 4433112..4435979 | 2485156 | DUF499-domain-containing-protein-[Streptomyces-sp.-846.5] |
| WP_239157864.1 | AAA+ATPase+wHTH+Alpha+Beta-domain+wHTH+RRM | Type-1_BR.RM.like.systems | VapB_antitoxin-><-DUF499*<-METHYLASE<-TUDOR+SF1-SF2-Helicase+iREase+CTD+REase\|\|Schlafen+HATPase+HTH->REase-> | 850 | Bacteria>Terrabacteria_group>Actinomycetota>Actinomycetes>Kitasatosporales | GCF_010548375.1 | Streptomyces_sp._SID13726 | G3I78_RS15960 | 365366..367918 | 2706058 | DUF499-domain-containing-protein-[Streptomyces-sp.-SID13726] |
| NEB00544.1 | AAA+ATPase+wHTH+Alpha+Beta-domain+wHTH+RRM | Type-1_BR.RM.like.systems | VapB_antitoxin->PIN-><-DUF499*<-METHYLASE<-TUDOR+SF1-SF2-Helicase+iREase+CTD+REase\|\|Schlafen+HATPase+HTH-> | 931 | Bacteria>Terrabacteria_group>Actinomycetota>Actinomycetes>Kitasatosporales | GCA_010548375.1 | Streptomyces_sp._SID13726 | G3I78_15960 | 365366..368161 | 2706058 | ATP-binding-protein-[Streptomyces-sp.-SID13726] |
| WP_162596947.1 | AAA+ATPase+wHTH+Alpha+Beta-domain+wHTH+RRM | Type-1_BR.RM.like.systems | REase+DUF5655-><-?\|\|?->\|\|TUDOR+SF1-SF2-Helicase+iREase+CTD+REase->METHYLASE->DUF499*->ABC-ATPase->HNH-><-JetD | 910 | Bacteria>Terrabacteria_group>Actinomycetota>Actinomycetes>Kitasatosporales | GCF_003173275.1 | Streptomyces_sp._NEAU-S7GS2 | DKG71_RS30770 | 7152862..7155594 | 2202000 | DUF499-domain-containing-protein-[Streptomyces-sp.-NEAU-S7GS2] |
| WP_086599862.1 | AAA+ATPase+wHTH+Alpha+Beta-domain+wHTH+RRM | Type-1_BR.RM.like.systems | HTH-><-HEPN\|\|?->DndB->TUDOR+SF1-SF2-Helicase+iREase+CTD+REase->METHYLASE->DUF499*->ABC-ATPase->HNH-> | 909 | Bacteria>Terrabacteria_group>Actinomycetota>Actinomycetes>Kitasatosporales | GCF_002148965.1 | Streptomyces_swartbergensis | CA983_RS06125 | 21814..24543 | 487165 | DUF499-domain-containing-protein-[Streptomyces-swartbergensis] |
| NUQ96612.1 | AAA+ATPase+wHTH+Alpha+Beta-domain+wHTH+RRM | Type-1_BR.RM.like.systems | <-HNH<-ABC-ATPase<-HEPN<-DUF499*<-METHYLASE<-TUDOR+SF1-SF2-Helicase+iREase+CTD+REase\|\|?->?-><-REase | 910 | Bacteria>Terrabacteria_group>Actinomycetota>Actinomycetes>Kitasatosporales | GCA_013361825.1 | Streptomyces_sp. | HOY79_08625 | 38052..40784 | 1931 | ATP-binding-protein-[Streptomyces-sp.] |
| WP_263976853.1 | AAA+ATPase+wHTH+Alpha+Beta-domain+wHTH+RRM | Type-1_BR.RM.like.systems | <-DUF499*<-TUDOR+SF1-SF2-Helicase+iREase+CTD+REase\|\|Schlafen+HATPase+HTH-><-?<-HTH_3\|\|ST-Hiskin-> | 851 | Bacteria>Terrabacteria_group>Actinomycetota>Actinomycetes>Kitasatosporales | - | Streptomyces_sp._NK08204 | K7G59_RS27400 | 90033..92588 | 2873260 | DUF499-domain-containing-protein-[Streptomyces-sp.-NK08204] |
| WP_256362104.1 | AAA+ATPase+wHTH+Alpha+Beta-domain+wHTH+RRM | Type-1_BR.RM.like.systems | HTH+DUF5753->?-><-REase<-?<-Schlafen+HATPase+HTH\|\|TUDOR+SF1-SF2-Helicase+iREase+CTD+REase->DUF499*-> | 811 | Bacteria>Terrabacteria_group>Actinomycetota>Actinomycetes>Kitasatosporales | - | Streptomyces_sp._TRM70350 | KWI83_RS10510 | 186214..188649 | 2856165 | DUF499-domain-containing-protein-[Streptomyces-sp.-TRM70350] |
| WP_147255375.1 | AAA+ATPase+wHTH+Alpha+Beta-domain+wHTH+RRM | Type-1_BR.RM.like.systems | <-HEPN\|\|TUDOR+SF1-SF2-Helicase+iREase+CTD+REase->METHYLASE->DUF499*->ABC-ATPase->HNH-> | 940 | Bacteria>Terrabacteria_group>Actinomycetota>Actinomycetes>Kitasatosporales | GCF_003311645.1 | Streptomyces_sp._PT12 | DEH69_RS03700 | 8734..11556 | 1510197 | DUF499-domain-containing-protein-[Streptomyces-sp.-PT12] |
| RBM22970.1 | AAA+ATPase+wHTH+Alpha+Beta-domain+wHTH+RRM | Type-1_BR.RM.like.systems | <-HEPN\|\|TUDOR+SF1-SF2-Helicase+iREase+CTD+REase->METHYLASE->DUF499*->ABC-ATPase->HNH-> | 921 | Bacteria>Terrabacteria_group>Actinomycetota>Actinomycetes>Kitasatosporales | GCA_003311645.1 | Streptomyces_sp._PT12 | DEH69_03700 | 8791..11556 | 1510197 | hypothetical-protein-DEH69_03700-[Streptomyces-sp.-PT12] |
| WP_280886836.1 | AAA+ATPase+wHTH+Alpha+Beta-domain+wHTH+RRM | Type-1_BR.RM.like.systems | <-wHTH+SAD+HNH\|\|?->?->Methyltransf->?-><-?<-DUF499*<-METHYLASE-p<-tRNA<-?<-Calcineurin | 931 | Bacteria>Terrabacteria_group>Actinomycetota>Actinomycetes>Kitasatosporales | GCF_029893585.1 | Streptomyces_sp._LBL | M2271_RS10380 | 53258..56053 | 2940562 | DUF499-domain-containing-protein-[Streptomyces-sp.-LBL] |
| WP_218514040.1 | AAA+ATPase+wHTH+Alpha+Beta-domain+wHTH+RRM | Type-1_BR.RM.like.systems | <-REase<-?<-Schlafen+HATPase+HTH\|\|TUDOR+SF1-SF2-Helicase+iREase+CTD+REase->METHYLASE->DUF499*-> | 931 | Bacteria>Terrabacteria_group>Actinomycetota>Actinomycetes>Kitasatosporales | GCF_019219825.1 | Streptomyces_sp._TRM70350 | KWI83_10510 | 185854..188649 | 2856165 | DUF499-domain-containing-protein-[Streptomyces-sp.-TRM70350] |
| MBV7696079.1 | AAA+ATPase+wHTH+Alpha+Beta-domain+wHTH+RRM | Type-1_BR.RM.like.systems | <-REase<-?<-Schlafen+HATPase+HTH\|\|TUDOR+SF1-SF2-Helicase+iREase+CTD+REase->METHYLASE->DUF499*-> | 931 | Bacteria>Terrabacteria_group>Actinomycetota>Actinomycetes>Kitasatosporales | GCA_019219825.1 | Streptomyces_sp._TRM70350 | KWI83_10510 | 185854..188649 | 2856165 | DUF499-domain-containing-protein-[Streptomyces-sp.-TRM70350] |
| WP_221752200.1 | AAA+ATPase+wHTH+Alpha+Beta-domain+wHTH+RRM | Type-1_BR.RM.like.systems | <-Schlafen+HATPase+HTH\|\|TUDOR+SF1-SF2-Helicase+iREase+CTD+REase->METHYLASE->DUF499*-> | 932 | Bacteria>Terrabacteria_group>Actinomycetota>Actinomycetes>Kitasatosporales | GCF_019733235.1 | Streptomyces_sp._TRM72054 | K4749_28635 | 8261..11059 | 2870562 | DUF499-domain-containing-protein-[Streptomyces-sp.-TRM72054] |
| MBX9397445.1 | AAA+ATPase+wHTH+Alpha+Beta-domain+wHTH+RRM | Type-1_BR.RM.like.systems | <-Schlafen+HATPase+HTH\|\|TUDOR+SF1-SF2-Helicase+iREase+CTD+REase->METHYLASE->DUF499*-> | 932 | Bacteria>Terrabacteria_group>Actinomycetota>Actinomycetes>Kitasatosporales | GCA_019733235.1 | Streptomyces_sp._TRM72054 | K4749_28635 | 8261..11059 | 2870562 | DUF499-domain-containing-protein-[Streptomyces-sp.-TRM72054] |
| WP_260463288.1 | AAA+ATPase+wHTH+Alpha+Beta-domain+wHTH+RRM | Type-1_BR.RM.like.systems | <-REase<-Schlafen+HATPase+HTH\|\|TUDOR+SF1-SF2-Helicase+iREase+CTD+REase->DUF499*-> | 851 | Bacteria>Terrabacteria_group>Actinomycetota>Actinomycetes>Kitasatosporales | - | Streptomyces_sp._TRM72054 | K4749_28650 | 8504..11059 | 2870562 | DUF499-domain-containing-protein-[Streptomyces-sp.-TRM72054] |
| WP_030589947.1 | AAA+ATPase+wHTH+Alpha+Beta-domain+wHTH+RRM | Type-1_BR.RM.like.systems | <-HTH_28+rve+Mu-transpos_C<-REase\|\|DUF4096+DDE_Tnp_1+DDE_Tnp_1_2->MNS+ZDNA-HTH->TUDOR+SF1-SF2-Helicase+iREase+CTD+REase->METHYLASE->DUF499*->?-><-TetR-HTH<-Phage_integrase<-Phage_int_SAM_4+Phage_integrase<-DDE_5 | 1000 | Bacteria>Terrabacteria_group>Actinomycetota>Actinomycetes>Kitasatosporales | GCF_000717105.1 | Streptomyces_anulatus | IG90_RS0127240 | 12878..15880 | 1892 | DUF499-domain-containing-protein-[Streptomyces-anulatus] |
| WP_123556145.1 | AAA+ATPase+wHTH+Alpha+Beta-domain+wHTH+RRM | Type-1_BR.RM.like.systems | Response_reg+LuxR-HTH->?-><-pPIWI+MID+RNaseH<-pPIWI+SF1-SF2-Helicase<-pPIWI+pPIWI<-HTH_3<-DUF499*<-METHYLASE<-TUDOR+SF1-SF2-Helicase+iREase+CTD+REase<-MNS+ZDNA-HTH<-Phage_integrase<-HTH\|\|XIS-HTH-> | 1001 | Bacteria>Terrabacteria_group>Actinomycetota>Actinomycetes>Kitasatosporales | GCF_003751605.1 | Kitasatospora_cineracea | EDD39_RS14405 | 3166835..3169840 | 88074 | DUF499-domain-containing-protein-[Kitasatospora-cineracea] |
| WP_030897848.1 | AAA+ATPase+wHTH+Alpha+Beta-domain+wHTH+RRM | Type-1_BR.RM.like.systems | <-pPIWI+MID+RNaseH<-pPIWI+SF1-SF2-Helicase<-pPIWI+pPIWI<-HTH_3<-DUF499*<-METHYLASE<-TUDOR+SF1-SF2-Helicase+iREase+CTD+REase\|\|REase+nSTAND3->?-><-Aminotran_1_2\|\|METHYLASE-p-> | 999 | Bacteria>Terrabacteria_group>Actinomycetota>Actinomycetes>Kitasatosporales | GCF_000722015.1 | Streptomyces_sp._NRRL_S-474 | IH12_RS0119650 | 193207..196206 | 1463909 | DUF499-domain-containing-protein-[Streptomyces-sp.-NRRL-S-474] |
| WP_030380972.1 | AAA+ATPase+wHTH+Alpha+Beta-domain+wHTH+RRM | Type-1_BR.RM.like.systems | <-DDE_5\|\|MNS+ZDNA-HTH->TUDOR+SF1-SF2-Helicase+iREase+CTD+REase->METHYLASE->DUF499*-><-TetR-HTH<-Phage_integrase<-Phage_int_SAM_4+Phage_integrase\|\|MULE-TRANSPOSASE-> | 1000 | Bacteria>Terrabacteria_group>Actinomycetota>Actinomycetes>Kitasatosporales | GCF_000716615.1 | unclassified_Streptomyces | IO29_RS21280 | 56589..59591 | 2593676 | MULTISPECIES:-DUF499-domain-containing-protein-[unclassified-Streptomyces] |
| WP_062189925.1 | AAA+ATPase+wHTH+Alpha+Beta-domain+wHTH+RRM | Type-1_BR.RM.like.systems | URI-DNase->DDE_5->Phage_int_SAM_4+Phage_integrase->Phage_integrase->TetR-HTH-><-?<-DUF499*<-METHYLASE<-TUDOR+SF1-SF2-Helicase+iREase+CTD+REase<-MNS+ZDNA-HTH\|\|DDE_5-> | 1000 | Bacteria>Terrabacteria_group>Actinomycetota>Actinomycetes>Kitasatosporales | GCF_001469455.2 | Streptomyces_sp._AVP053U2 | APS67_RS29825 | 26807..29809 | 1737066 | DUF499-domain-containing-protein-[Streptomyces-sp.-AVP053U2] |
| ODA69735.1 | AAA+ATPase+wHTH+Alpha+Beta-domain+wHTH+RRM | Type-1_BR.RM.like.systems | URI-DNase->DDE_5->Phage_int_SAM_4+Phage_integrase->Phage_integrase->TetR-HTH-><-?<-DUF499*<-METHYLASE<-TUDOR+SF1-SF2-Helicase+iREase+CTD+REase<-MNS+ZDNA-HTH\|\|?->DDE_5-> | 1000 | Bacteria>Terrabacteria_group>Actinomycetota>Actinomycetes>Kitasatosporales | GCA_001469455.2 | Streptomyces_sp._AVP053U2 | APS67_006117 | 26807..29809 | 1737066 | hypothetical-protein-APS67_006117-[Streptomyces-sp.-AVP053U2] |
| WP_093745813.1 | AAA+ATPase+wHTH+Alpha+Beta-domain+wHTH+RRM | Type-1_BR.RM.like.systems | HTH_3->NUDIX->TUDOR+SF1-SF2-Helicase+iREase+CTD+REase->METHYLASE->DUF499*-><-?\|\|Methyltransf->REase+N6_MTase-><-REase+SF1-SF2-Helicase\|\|PIN-> | 993 | Bacteria>Terrabacteria_group>Actinomycetota>Actinomycetes>Kitasatosporales | GCF_900105465.1 | Streptomyces_sp._PAN_FS17 | BLW73_RS14605 | 3279915..3282896 | 1855351 | DUF499-domain-containing-protein-[Streptomyces-sp.-PAN_FS17] |
| WP_283466091.1 | AAA+ATPase+wHTH+Alpha+Beta-domain+wHTH+RRM | Type-1_BR.RM.like.systems | <-AdoMet_Synthase<-Pkinase\|\|HTH_31->NUDIX->?->?-><-DUF499*<-METHYLASE<-TUDOR+SF1-SF2-Helicase+iREase+CTD+REase\|\|<-?<-SF1-SF2-Helicase<-RelE-ParE<-PhdYeFM_antitox | 999 | Bacteria>Terrabacteria_group>Actinomycetota>Actinomycetes>Kitasatosporales | GCF_030063325.1 | unclassified_Streptomyces | QMK19_RS02210 | 466319..469318 | 2593676 | MULTISPECIES:-DUF499-domain-containing-protein-[unclassified-Streptomyces] |
| WP_169439034.1 | AAA+ATPase+wHTH+Alpha+Beta-domain+wHTH+RRM | Type-1_BR.RM.like.systems | <-DDE_Tnp_1_4<-Phage_int_SAM_1<-?<-?\|\|TUDOR+SF1-SF2-Helicase+iREase+CTD+REase->METHYLASE->DUF499*->?->HTH+MPTase-><-HTH_3\|\|?-><-IclR-HTH<-Peptidase_M6+Peptidase_M7 | 996 | Bacteria>Terrabacteria_group>Actinomycetota>Actinomycetes>Kitasatosporales | GCF_012933395.1 | Streptomyces_sp._GMY02 | HG826_RS27775 | 120488..123478 | 1333528 | DUF499-domain-containing-protein-[Streptomyces-sp.-GMY02] |
| WP_234466616.1 | AAA+ATPase+wHTH+Alpha+Beta-domain+wHTH+RRM | Type-1_BR.RM.like.systems | HEPN->MNS+ZDNA-HTH->TUDOR+SF1-SF2-Helicase+iREase+CTD+REase->METHYLASE->DUF499*-><-TetR-HTH<-Phage_integrase<-Phage_int_SAM_4+Phage_integrase<-DDE_5 | 989 | Bacteria>Terrabacteria_group>Actinomycetota>Actinomycetes>Kitasatosporales | - | Streptomyces_sp._MBT60 | JHN56_RS23145 | 10506..13475 | 2800409 | DUF499-domain-containing-protein-[Streptomyces-sp.-MBT60] |
| MBK3545942.1 | AAA+ATPase+wHTH+Alpha+Beta-domain+wHTH+RRM | Type-1_BR.RM.like.systems | HEPN->MNS+ZDNA-HTH->TUDOR+SF1-SF2-Helicase+iREase+CTD+REase->METHYLASE->DUF499*-><-TetR-HTH<-Phage_integrase<-Phage_int_SAM_4+Phage_integrase<-DDE_5 | 971 | Bacteria>Terrabacteria_group>Actinomycetota>Actinomycetes>Kitasatosporales | GCA_016612665.1 | Streptomyces_sp._MBT60 | JHN56_23135 | 10560..13475 | 2800409 | ATP-binding-protein-[Streptomyces-sp.-MBT60] |
| WP_208865670.1 | AAA+ATPase+wHTH+Alpha+Beta-domain+wHTH+RRM | Type-1_BR.RM.like.systems | <-HTH+WYL+WYL-C\|\|?->?->wHTH+SAD+HNH->?->?-><-DUF499*<-METHYLASE<-TUDOR+SF1-SF2-Helicase+iREase+CTD+REase<-DEDD_Tnp_IS110+Transposase_20<-?\|\|GNTR-HTH->ST-Hiskin-> | 984 | Bacteria>Terrabacteria_group>Actinomycetota>Actinomycetes>Kitasatosporales | GCF_000158955.1 | Streptomyces_viridochromogenes | SSQG_RS26535 | 5986550..5989504 | 1938 | DUF499-domain-containing-protein-[Streptomyces-viridochromogenes] |
| EFL34855.1 | AAA+ATPase+wHTH+Alpha+Beta-domain+wHTH+RRM | Type-1_BR.RM.like.systems | DNA-glycosylase->?->wHTH+SAD+HNH->?->?-><-DUF499*<-METHYLASE<-TUDOR+SF1-SF2-Helicase+iREase+CTD+REase<-DEDD_Tnp_IS110+Transposase_20<-?\|\|GNTR-HTH->ST-Hiskin-> | 996 | Bacteria>Terrabacteria_group>Actinomycetota>Actinomycetes>Kitasatosporales | GCA_000158955.1 | Streptomyces_viridochromogenes_DSM_40736 | SSQG_05373 | 5986550..5989540 | 591159 | AAA-family-ATPase-[Streptomyces-viridochromogenes-DSM-40736] |
| WP_127354347.1 | AAA+ATPase+wHTH+Alpha+Beta-domain+wHTH+RRM | Type-1_BR.RM.like.systems | zf-IS66+DDE_Tnp_IS66-><-SbcC\|\|?-><-?<-HTH_36\|\|<-?<-DUF499*<-METHYLASE<-TUDOR+SF1-SF2-Helicase+iREase+CTD+REase<-GNTR-HTH-p\|\|?-><-REase+ParB<-DDE_Tnp_1 | 991 | Bacteria>Terrabacteria_group>Actinomycetota>Actinomycetes>Kitasatosporales | GCF_003999195.1 | Actinacidiphila_soli | EFK10_RS03775 | 22787..25762 | 2487275 | DUF499-domain-containing-protein-[Streptomyces-sp.-LAM7114] |
| WP_030261137.1 | AAA+ATPase+wHTH+Alpha+Beta-domain+wHTH+RRM | Type-1_BR.RM.like.systems | TraJ-RHH->PIN-><-?<-?<-?<-?<-DUF499*<-METHYLASE<-TUDOR+SF1-SF2-Helicase+iREase+CTD+REase<-ACET+ACET<-DEDD_Tnp_IS110+Transposase_20\|\|Phage_integrase-> | 977 | Bacteria>Terrabacteria_group>Actinomycetota>Actinomycetes>Kitasatosporales | GCF_000719095.1 | Streptacidiphilus_jeojiense | IF29_RS0129245 | 48184..51117 | 436229 | hypothetical-protein-[Streptacidiphilus-jeojiense] |
| WP_058852360.1 | AAA+ATPase+wHTH+Alpha+Beta-domain+wHTH+RRM | Type-1_BR.RM.like.systems | <-Pkinase<-AdoMet_Synthase<-Pkinase\|\|?->?-><-DUF499*<-METHYLASE<-TUDOR+SF1-SF2-Helicase+iREase+CTD+REase\|\|35exo-><-?\|\|REase->PhdYeFM_antitox-> | 999 | Bacteria>Terrabacteria_group>Actinomycetota>Actinomycetes>Kitasatosporales | GCF_001482415.1 | Streptomyces_silvensis | AT728_RS38090 | 13270..16269 | 1765722 | DUF499-domain-containing-protein-[Streptomyces-silvensis] |
| WP_228917173.1 | AAA+ATPase+wHTH+Alpha+Beta-domain+wHTH+RRM | Type-1_BR.RM.like.systems | DEDD_Tnp_IS110+Transposase_20-><-?<-REase\|\|HNH-><-DNA_methylase<-DUF499*<-METHYLASE<-TUDOR+SF1-SF2-Helicase+iREase+CTD+REase<-PADR-HTH | 991 | Bacteria>Terrabacteria_group>Actinomycetota>Actinomycetes>Kitasatosporales | GCF_020873815.1 | Streptomyces_sp._DH20 | LPA43_RS09485 | 100759..103734 | 2857009 | DUF499-domain-containing-protein-[Streptomyces-sp.-DH20] |
| WP_275448697.1 | AAA+ATPase+wHTH+Alpha+Beta-domain+wHTH+RRM | Type-1_BR.RM.like.systems | <-Phage_integrase\|\|?->Peptidase-M15->MNS+ZDNA-HTH->TUDOR+SF1-SF2-Helicase+iREase+CTD+REase->METHYLASE->DUF499*->?-><-?<-HTH_3\|\|NUDIX-> | 989 | Bacteria>Terrabacteria_group>Actinomycetota>Actinomycetes>Kitasatosporales | - | Streptomyces_sp._OfavH-34-F | MHW47_RS00605 | 125878..128847 | 2917760 | DUF499-domain-containing-protein-[Streptomyces-sp.-OfavH-34-F] |
| WP_239220568.1 | AAA+ATPase+wHTH+Alpha+Beta-domain+wHTH+RRM | Type-1_BR.RM.like.systems | <-Phage_integrase\|\|?->Peptidase-M15->MNS+ZDNA-HTH->TUDOR+SF1-SF2-Helicase+iREase+CTD+REase->METHYLASE->DUF499*->?-><-?<-HTH_3\|\|NUDIX-> | 1000 | Bacteria>Terrabacteria_group>Actinomycetota>Actinomycetes>Kitasatosporales | GCF_022347745.1 | Streptomyces_sp._OfavH-34-F | MHW47_00605 | 125845..128847 | 2917760 | DUF499-domain-containing-protein-[Streptomyces-sp.-OfavH-34-F] |
| WP_030918063.1 | AAA+ATPase+wHTH+Alpha+Beta-domain+wHTH+RRM | Type-1_BR.RM.like.systems | ABhydrolase-><-?<-?<-?\|\|TUDOR+SF1-SF2-Helicase+iREase+CTD+REase->METHYLASE->DUF499*-><-?\|\|HEPN->GNTR-HTH->HDRnase-><-NUDIX<-HTH+HKD-DNase | 991 | Bacteria>Terrabacteria_group>Actinomycetota>Actinomycetes>Kitasatosporales | GCF_000721715.1 | Streptomyces_sp._NRRL_B-24720 | IH49_RS0107280 | 93522..96497 | 1476876 | DUF499-domain-containing-protein-[Streptomyces-sp.-NRRL-B-24720] |
| MCG7522956.1 | AAA+ATPase+wHTH+Alpha+Beta-domain+wHTH+RRM | Type-1_BR.RM.like.systems | <-Phage_integrase\|\|?->Peptidase-M15->MNS+ZDNA-HTH->TUDOR+SF1-SF2-Helicase+iREase+CTD+REase->METHYLASE->DUF499*->?-><-?<-HTH_3\|\|NUDIX-> | 1000 | Bacteria>Terrabacteria_group>Actinomycetota>Actinomycetes>Kitasatosporales | GCA_022347745.1 | Streptomyces_sp._OfavH-34-F | MHW47_00605 | 125845..128847 | 2917760 | DUF499-domain-containing-protein-[Streptomyces-sp.-OfavH-34-F] |
| WP_109179522.1 | AAA+ATPase+wHTH+Alpha+Beta-domain+wHTH+RRM | Type-1_BR.RM.like.systems | <-Radical-SAM<-?\|\|HTH+HKD-DNase->NUDIX-><-HDRnase<-DUF499*<-METHYLASE<-TUDOR+SF1-SF2-Helicase+iREase+CTD+REase<-Phage_terminase | 999 | Bacteria>Terrabacteria_group>Actinomycetota>Actinomycetes>Kitasatosporales | GCF_003116755.1 | Streptomyces_sp._CS131 | DBP20_RS19500 | 4282075..4285074 | 2162711 | DUF499-domain-containing-protein-[Streptomyces-sp.-CS131] |
| WP_268985962.1 | AAA+ATPase+wHTH+Alpha+Beta-domain+wHTH+RRM | Type-1_BR.RM.like.systems | N6_N4_MTase->?-><-REase+DUF5655\|\|TUDOR+SF1-SF2-Helicase+iREase+CTD+REase->METHYLASE->DUF499*-><-?<-?<-?\|\|HTH+DSBH->tRNA-><-?\|\|<-Methyltransf_4 | 981 | Bacteria>Terrabacteria_group>Actinomycetota>Actinomycetes>Kitasatosporales | - | Streptomyces_sp._CC210A | L6E18_RS02680 | 590724..593669 | 2898184 | DUF499-domain-containing-protein-[Streptomyces-sp.-CC210A] |
| WP_187821572.1 | AAA+ATPase+wHTH+Alpha+Beta-domain+wHTH+RRM | Type-1_BR.RM.like.systems | HTH_31->?->HTH-><-?\|\|TUDOR+SF1-SF2-Helicase+iREase+CTD+REase->METHYLASE->DUF499*-><-ParB\|\|JetA->JetB->?->?-><-Primpol+D5N+D5ATPase | 979 | Bacteria>Terrabacteria_group>Actinomycetota>Actinomycetes>Kitasatosporales | GCF_014495685.1 | Streptomyces_sp._TRM68367 | H8R17_RS03215 | 77833..80772 | 2758415 | DUF499-domain-containing-protein-[Streptomyces-sp.-TRM68367] |
| WP_100598678.1 | AAA+ATPase+wHTH+Alpha+Beta-domain+wHTH+RRM | Type-1_BR.RM.like.systems | NACHT-><-HTH_21+rve+rve_3\|\|?->HTH->TUDOR+SF1-SF2-Helicase+iREase+CTD+REase->METHYLASE->DUF499*-><-?<-?<-?\|\|REase->HEPN->HEPN-> | 976 | Bacteria>Terrabacteria_group>Actinomycetota>Actinomycetes>Kitasatosporales | GCF_002803065.1 | Streptomyces_sp._CB01635 | CG723_RS41180 | 54423..57353 | 2020326 | hypothetical-protein-[Streptomyces-sp.-CB01635] |
| WP_285367923.1 | AAA+ATPase+wHTH+Alpha+Beta-domain+wHTH+RRM | Type-1_BR.RM.like.systems | ParB-><-?<-?<-?\|\|TUDOR+SF1-SF2-Helicase+iREase+CTD+REase->METHYLASE->DUF499*->?-><-SF1-SF2-Helicase<-PIN\|\|?->tRNA->Peptidase_M20->ACET-> | 965 | Bacteria>Terrabacteria_group>Actinomycetota>Actinomycetes>Kitasatosporales | - | Streptomyces_sp._CWNU-1 | NBG84_RS27845 | 168103..171000 | 2897419 | DUF499-domain-containing-protein-[Streptomyces-sp.-CWNU-1] |
| WP_250922371.1 | AAA+ATPase+wHTH+Alpha+Beta-domain+wHTH+RRM | Type-1_BR.RM.like.systems | ParB-><-?<-?<-?\|\|TUDOR+SF1-SF2-Helicase+iREase+CTD+REase->METHYLASE->DUF499*->?-><-SF1-SF2-Helicase<-PIN\|\|?->tRNA->Peptidase_M20->ACET-> | 984 | Bacteria>Terrabacteria_group>Actinomycetota>Actinomycetes>Kitasatosporales | GCF_023701525.1 | Streptomyces_sp._CWNU-1 | NBG84_27855 | 168046..171000 | 2897419 | DUF499-domain-containing-protein-[Streptomyces-sp.-CWNU-1] |
| MCM2392056.1 | AAA+ATPase+wHTH+Alpha+Beta-domain+wHTH+RRM | Type-1_BR.RM.like.systems | ParB-><-?<-?<-?\|\|TUDOR+SF1-SF2-Helicase+iREase+CTD+REase->METHYLASE->DUF499*->?-><-SF1-SF2-Helicase<-PIN\|\|?->tRNA->Peptidase_M20->ACET-> | 984 | Bacteria>Terrabacteria_group>Actinomycetota>Actinomycetes>Kitasatosporales | GCA_023701525.1 | Streptomyces_sp._CWNU-1 | NBG84_27855 | 168046..171000 | 2897419 | DUF499-domain-containing-protein-[Streptomyces-sp.-CWNU-1] |
| GHG49832.1 | AAA+ATPase+wHTH+Alpha+Beta-domain+wHTH+RRM | Type-1_BR.RM.like.systems | TUDOR+SF1-SF2-Helicase+iREase+CTD+REase->METHYLASE->DUF499->DUF499*-><-pPIWI+MID+RNaseH<-pPIWI<-pPIWI+pPIWI\|\|?-><-?<-HD | 490 | Bacteria>Terrabacteria_group>Actinomycetota>Actinomycetes>Kitasatosporales | GCA_014654935.1 | Streptomyces_capoamus | GCM10018980_31170 | 125555..127027 | 68183 | hypothetical-protein-GCM10018980_31170-[Streptomyces-capoamus] |
| WP_239081086.1 | AAA+ATPase+wHTH+Alpha+Beta-domain+wHTH+RRM | Type-1_BR.RM.like.systems | ABhydrolase->TetR-HTH->\|\|TUDOR+SF1-SF2-Helicase+iREase+CTD+REase->METHYLASE->DUF499*->?->HEPN-><-HEPN<-?\|\|TetR-HTH-> | 848 | Bacteria>Terrabacteria_group>Actinomycetota>Actinomycetes>Kitasatosporales | GCF_010548385.1 | Streptomyces_coelicoflavus | G3I32_RS20995 | 9819..12365 | 285562 | DUF499-domain-containing-protein-[Streptomyces-coelicoflavus] |
| NEB11285.1 | AAA+ATPase+wHTH+Alpha+Beta-domain+wHTH+RRM | Type-1_BR.RM.like.systems | ABhydrolase->TetR-HTH-><-?\|\|TUDOR+SF1-SF2-Helicase+iREase+CTD+REase->METHYLASE->DUF499*->?->HEPN-><-HEPN<-?\|\|TetR-HTH-> | 998 | Bacteria>Terrabacteria_group>Actinomycetota>Actinomycetes>Kitasatosporales | GCA_010548385.1 | Streptomyces_coelicoflavus | G3I32_21005 | 9369..12365 | 285562 | DUF499-domain-containing-protein-[Streptomyces-coelicoflavus] |
| WP_215040121.1 | AAA+ATPase+wHTH+Alpha+Beta-domain+wHTH+RRM | Type-1_BR.RM.like.systems | METHYLASE-p->DUF499*->DUF499-p-><-pPIWI+MID+RNaseH<-pPIWI+SF1-SF2-Helicase<-pPIWI+pPIWI<-HTH_3 | 831 | Bacteria>Terrabacteria_group>Actinomycetota>Actinomycetes>Kitasatosporales | GCF_018614515.1 | Streptomyces_sp._ISL-111 | J7E90_RS31900 | 2399..4894 | 2819175 | DUF499-domain-containing-protein-[Streptomyces-sp.-ISL-111] |
| WP_049564944.1 | AAA+ATPase+wHTH+Alpha+Beta-domain+wHTH+RRM | Type-1_BR.RM.like.systems | <-SF1-SF2-Helicase\|\|?->\|\|?-><-DUF499*<-METHYLASE<-TUDOR+SF1-SF2-Helicase+iREase+CTD+REase<-Cysteine_desulfurase\|\|?-><-ACET | 984 | Bacteria>Terrabacteria_group>Actinomycetota>Actinomycetes>Kitasatosporales | GCF_001083795.1 | Streptomyces_sp._SBT349 | ADY01_RS02885 | 8043..10997 | 1580539 | DUF499-domain-containing-protein-[Streptomyces-sp.-SBT349] |
| WP_266941289.1 | AAA+ATPase+wHTH+Alpha+Beta-domain+wHTH+RRM | Type-1_BR.RM.like.systems | KAP-NTPase->?-><-?\|\|HTH_Tnp_4+DDE_Tnp_4->TUDOR+SF1-SF2-Helicase+iREase+CTD+REase->METHYLASE->DUF499*->DUF4096->?->?-><-?<-HTH | 998 | Bacteria>Terrabacteria_group>Actinomycetota>Actinomycetes>Kitasatosporales | GCF_026342745.1 | Streptomyces_mirabilis | OG901_RS30250 | 6669570..6672566 | 68239 | DUF499-domain-containing-protein-[Streptomyces-mirabilis] |
| WP_127832767.1 | AAA+ATPase+wHTH+Alpha+Beta-domain+wHTH+RRM | Type-1_BR.RM.like.systems | SIGMA-HTH->?-><-?\|\|TUDOR+SF1-SF2-Helicase+iREase+CTD+REase->METHYLASE->DUF499*->?->?->REase+SAVED-><-?\|\|SMODS->UbE1+JAB-> | 992 | Bacteria>Terrabacteria_group>Actinomycetota>Actinomycetes>Kitasatosporales | GCF_004005495.1 | Streptomyces_sp._San01 | EOT10_RS36885 | 33671..36649 | 2494256 | DUF499-domain-containing-protein-[Streptomyces-sp.-San01] |
| WP_229899785.1 | AAA+ATPase+wHTH+Alpha+Beta-domain+wHTH+RRM | Type-1_BR.RM.like.systems | TUDOR+SF1-SF2-Helicase+iREase+CTD+REase->METHYLASE->DUF499*-><-pPIWI+MID+RNaseH<-pPIWI<-pPIWI+pPIWI\|\|?-><-?<-HD | 459 | Bacteria>Terrabacteria_group>Actinomycetota>Actinomycetes>Kitasatosporales | GCF_014654935.1 | Streptomyces_capoamus | IE312_RS40150 | 123991..125367 | 68183 | DUF499-domain-containing-protein,-partial-[Streptomyces-capoamus] |
| WP_189482703.1 | AAA+ATPase+wHTH+Alpha+Beta-domain+wHTH+RRM | Type-1_BR.RM.like.systems | TUDOR+SF1-SF2-Helicase+iREase+CTD+REase->METHYLASE->DUF499*->?->?-><-DEDD_Tnp_IS110+Transposase_20<-?<-?\|\|HTH+DUF5753-> | 993 | Bacteria>Terrabacteria_group>Actinomycetota>Actinomycetes>Kitasatosporales | GCF_014649895.1 | Streptomyces_rubiginosus | IE146_RS30935 | 8308..11289 | 1929 | DUF499-domain-containing-protein-[Streptomyces-rubiginosus] |
| GHG49827.1 | AAA+ATPase+wHTH+Alpha+Beta-domain+wHTH+RRM | Type-1_BR.RM.like.systems | TUDOR+SF1-SF2-Helicase+iREase+CTD+REase->METHYLASE->DUF499*->DUF499-><-pPIWI+MID+RNaseH<-pPIWI<-pPIWI+pPIWI | 487 | Bacteria>Terrabacteria_group>Actinomycetota>Actinomycetes>Kitasatosporales | GCA_014654935.1 | Streptomyces_capoamus | GCM10018980_31160 | 123991..125454 | 68183 | hypothetical-protein-GCM10018980_31160-[Streptomyces-capoamus] |
| WP_030819391.1 | AAA+ATPase+wHTH+Alpha+Beta-domain+wHTH+RRM | Type-1_BR.RM.like.systems | <-METHYLASE-p<-?<-?\|\|MNS+ZDNA-HTH->TUDOR+SF1-SF2-Helicase+iREase+CTD+REase->METHYLASE->DUF499*->?-><-DDE_5\|\|?-><-HTH_3 | 1000 | Bacteria>Terrabacteria_group>Actinomycetota>Actinomycetes>Kitasatosporales | GCF_000718375.1 | Streptomyces | ADK41_RS01085 | 230582..233584 | 1883 | MULTISPECIES:-DUF499-domain-containing-protein-[Streptomyces] |
| GGW17888.1 | AAA+ATPase+wHTH+Alpha+Beta-domain+wHTH+RRM | Type-1_BR.RM.like.systems | HD->?-><-?\|\|pPIWI+pPIWI->pPIWI->pPIWI+MID+RNaseH-><-DUF499*<-METHYLASE<-TUDOR+SF1-SF2-Helicase+iREase+CTD+REase | 999 | Bacteria>Terrabacteria_group>Actinomycetota>Actinomycetes>Kitasatosporales | GCA_014649295.1 | Streptomyces_libani_subsp._rufus | GCM10010501_40440 | 198564..201563 | 249582 | hypothetical-protein-GCM10010501_40440-[Streptomyces-libani-subsp.-rufus] |
| MXQ57531.1 | AAA+ATPase+wHTH+Alpha+Beta-domain+wHTH+RRM | Type-1_BR.RM.like.systems | <-FtsK<-?\|\|GNTR-HTH->TUDOR+SF1-SF2-Helicase+iREase+CTD+REase->METHYLASE->DUF499*-><-?<-?<-?\|\|HTH_3-><-Pkinase<-PIN | 996 | Bacteria>Terrabacteria_group>Actinomycetota>Actinomycetes>Kitasatosporales | GCA_009831155.1 | Streptomyces_sp._XHT-2 | GSE51_07935 | 14550..17540 | 2692621 | DUF499-domain-containing-protein-[Streptomyces-sp.-XHT-2] |

| Protein-ID | DUF499-architecture | System | Gene-Neighbourhoods | Length | Taxonomy | Assembly-ID | Strain | Locus-Tag | Genomic-coordinates | TaxID | Fasta-defline |
|---|---|---|---|---|---|---|---|---|---|---|---|
| WP_148710267.1 | AAA+ATPase+wHTH+Alpha+Beta-domain+wHTH+RRM | Type-1_BR.RM.like.systems | <-ABhydrolase<-?<-LuxR-HTH\|\|?-><-DUF499*<-METHYLASE<-TUDOR+SF1-SF2-Helicase+iREase+CTD+REase<-NUDIX<-HTH_31 | 984 | Bacteria>Terrabacteria_group>Actinomycetota>Actinomycetes>Kitasatosporales | GCF_003634815.1 | Streptomyces_sp._3211.6 | BX286_RS12800 | 2784150..2787104 | 1938845 | DUF499-domain-containing-protein-[Streptomyces-sp.-3211.6] |
| WP_123995653.1 | AAA+ATPase+wHTH+Alpha+Beta-domain+wHTH+RRM | Type-1_BR.RM.like.systems | <-ABhydrolase<-?<-LuxR-HTH\|\|?-><-DUF499*<-METHYLASE<-TUDOR+SF1-SF2-Helicase+iREase+CTD+REase<-NUDIX<-HTH_31 | 984 | Bacteria>Terrabacteria_group>Actinomycetota>Actinomycetes>Kitasatosporales | GCF_002028425.1 | Streptomyces | B4U46_RS11765 | 2605267..2608221 | 1883 | MULTISPECIES:-DUF499-domain-containing-protein-[Streptomyces] |
| WP_067368937.1 | AAA+ATPase+wHTH+Alpha+Beta-domain+wHTH+RRM | Type-1_BR.RM.like.systems | HTH+DSBH->?->?-><-?\|\|TUDOR+SF1-SF2-Helicase+iREase+CTD+REase->METHYLASE->DUF499*-><-?<-SbcC<-?<-?<-ST-Hiskin<-NUDIX | 991 | Bacteria>Terrabacteria_group>Actinomycetota>Actinomycetes>Kitasatosporales | GCF_002335465.1 | Streptomyces_olivochromogenes | AQJ27_RS17200 | 471830..474805 | 1963 | DUF499-domain-containing-protein-[Streptomyces-olivochromogenes] |
| KOT46903.1 | AAA+ATPase+wHTH+Alpha+Beta-domain+wHTH+RRM | Type-1_BR.RM.like.systems | <-METHYLASE-p<-?<-?\|\|MNS+ZDNA-HTH->TUDOR+SF1-SF2-Helicase+iREase+CTD+REase->METHYLASE->DUF499*->?->?-><-HTH_3 | 989 | Bacteria>Terrabacteria_group>Actinomycetota>Actinomycetes>Kitasatosporales | GCA_001279005.1 | Streptomyces_caelestis | ADK41_01090 | 230615..233584 | 36816 | ATPase-AAA-[Streptomyces-caelestis] |
| MCZ1001153.1 | AAA+ATPase+wHTH+Alpha+Beta-domain+wHTH+RRM | Type-1_BR.RM.like.systems | Phage_cap_E->?->?->?->TUDOR+SF1-SF2-Helicase+iREase+CTD+REase->METHYLASE->DUF499*-><-?<-HTH_3\|\|?->HISKIN-> | 999 | Bacteria>Terrabacteria_group>Actinomycetota>Actinomycetes>Kitasatosporales | GCA_026913145.1 | Streptomyces_mirabilis | O1M63_28865 | 330805..333804 | 68239 | DUF499-domain-containing-protein-[Streptomyces-mirabilis] |
| WP_242328934.1 | AAA+ATPase+wHTH+Alpha+Beta-domain+wHTH+RRM | Type-1_BR.RM.like.systems | Phage_terminase->TUDOR+SF1-SF2-Helicase+iREase+CTD+REase->METHYLASE->DUF499*->?->HNH->?->?->REase-> | 999 | Bacteria>Terrabacteria_group>Actinomycetota>Actinomycetes>Kitasatosporales | GCF_022647665.1 | Streptomyces_formicae | J4032_RS01965 | 417762..420761 | 1616117 | DUF499-domain-containing-protein-[Streptomyces-formicae] |
| WP_114531710.1 | AAA+ATPase+wHTH+Alpha+Beta-domain+wHTH+RRM | Type-1_BR.RM.like.systems | <-HTH_20<-ABC-ATPase+SbcC<-?\|\|TUDOR+SF1-SF2-Helicase+iREase+CTD+REase->METHYLASE->DUF499*-> | 998 | Bacteria>Terrabacteria_group>Actinomycetota>Actinomycetes>Kitasatosporales | GCF_003344905.1 | Streptomyces_parvulus | DVZ84_RS28705 | 98854..101850 | 146923 | DUF499-domain-containing-protein-[Streptomyces-parvulus] |
| RDD85728.1 | AAA+ATPase+wHTH+Alpha+Beta-domain+wHTH+RRM | Type-1_BR.RM.like.systems | <-HTH_20<-ABC-ATPase+SbcC<-?\|\|TUDOR+SF1-SF2-Helicase+iREase+CTD+REase->METHYLASE->DUF499*-> | 998 | Bacteria>Terrabacteria_group>Actinomycetota>Actinomycetes>Kitasatosporales | GCA_003344905.1 | Streptomyces_parvulus | DVZ84_28715 | 98854..101850 | 146923 | DUF499-domain-containing-protein-[Streptomyces-parvulus] |
| WP_206961363.1 | AAA+ATPase+wHTH+Alpha+Beta-domain+wHTH+RRM | Type-1_BR.RM.like.systems | <-TraJ-RHH<-?<-?<-?\|\|HTH->XIS-HTH-><-DUF499*<-METHYLASE<-TUDOR+SF1-SF2-Helicase+iREase+CTD+REase | 968 | Bacteria>Terrabacteria_group>Actinomycetota>Actinomycetes>Kitasatosporales | GCF_017315755.1 | Streptomyces_beijiangensis | J0695_RS09060 | 13885..16791 | 163361 | DUF499-domain-containing-protein-[Streptomyces-beijiangensis] |
| WP_043224560.1 | AAA+ATPase+wHTH+Alpha+Beta-domain+wHTH+RRM | Type-1_BR.RM.like.systems | <-NUDIX<-?<-?\|\|MNS+ZDNA-HTH->TUDOR+SF1-SF2-Helicase+iREase+CTD+REase->METHYLASE->DUF499*->?-><-PIN | 1000 | Bacteria>Terrabacteria_group>Actinomycetota>Actinomycetes>Kitasatosporales | GCF_000721515.1 | Streptomyces_sp._NRRL_F-5193 | IH76_RS14460 | 112777..115779 | 1463860 | DUF499-domain-containing-protein-[Streptomyces-sp.-NRRL-F-5193] |
| WP_277438204.1 | AAA+ATPase+wHTH+Alpha+Beta-domain+wHTH+RRM | Type-1_BR.RM.like.systems | <-DUF499*<-METHYLASE<-TUDOR+SF1-SF2-Helicase+iREase+CTD+REase<-MNS+ZDNA-HTH<-METHYLASE-p | 997 | Bacteria>Terrabacteria_group>Actinomycetota>Actinomycetes>Kitasatosporales | GCF_029504575.1 | Streptomyces_sp._SPB162 | M2266_RS12075 | 2617854..2620847 | 2940560 | DUF499-domain-containing-protein-[Streptomyces-sp.-SPB162] |
| WP_234477546.1 | AAA+ATPase+wHTH+Alpha+Beta-domain+wHTH+RRM | Type-1_BR.RM.like.systems | TUDOR+SF1-SF2-Helicase+iREase+CTD+REase->METHYLASE->DUF499*->?->?->Phage_cap_E->?->?->METHYLASE-p-> | 963 | Bacteria>Terrabacteria_group>Actinomycetota>Actinomycetes>Kitasatosporales | - | Streptomyces_sp._MBT65 | JHN63_RS22530 | 17243..20134 | 1488395 | DUF499-domain-containing-protein-[Streptomyces-sp.-MBT65] |
| MBK3576532.1 | AAA+ATPase+wHTH+Alpha+Beta-domain+wHTH+RRM | Type-1_BR.RM.like.systems | TUDOR+SF1-SF2-Helicase+iREase+CTD+REase->METHYLASE->DUF499*->?->?->Phage_cap_E->?->?->METHYLASE-p-> | 982 | Bacteria>Terrabacteria_group>Actinomycetota>Actinomycetes>Kitasatosporales | GCA_016612545.1 | Streptomyces_sp._MBT65 | JHN63_22535 | 17186..20134 | 1488395 | DUF499-domain-containing-protein-[Streptomyces-sp.-MBT65] |
| WP_053212854.1 | AAA+ATPase+wHTH+Alpha+Beta-domain+wHTH+RRM | Type-1_BR.RM.like.systems | HD->?-><-?\|\|pPIWI+pPIWI->pPIWI->pPIWI+MID+RNaseH-><-DUF499*<-METHYLASE-p | 999 | Bacteria>Terrabacteria_group>Actinomycetota>Actinomycetes>Kitasatosporales | GCF_001270635.1 | Streptomyces_antibioticus | ADK77_RS41100 | 32734..35733 | 1890 | DUF499-domain-containing-protein-[Streptomyces-antibioticus] |
| WP_096624439.1 | AAA+ATPase+wHTH+Alpha+Beta-domain+wHTH+RRM | Type-1_BR.RM.like.systems | <-DUF499*<-METHYLASE<-TUDOR+SF1-SF2-Helicase+iREase+CTD+REase<-MNS+ZDNA-HTH | 1000 | Bacteria>Terrabacteria_group>Actinomycetota>Actinomycetes>Kitasatosporales | - | Streptomyces_sp._WZ.A104 | CIB93_RS04970 | 106791..109793 | 2023771 | DUF499-domain-containing-protein-[Streptomyces-sp.-WZ.A104] |
| PCG87192.1 | AAA+ATPase+wHTH+Alpha+Beta-domain+wHTH+RRM | Type-1_BR.RM.like.systems | <-DUF499*<-METHYLASE<-TUDOR+SF1-SF2-Helicase+iREase+CTD+REase<-MNS+ZDNA-HTH | 1000 | Bacteria>Terrabacteria_group>Actinomycetota>Actinomycetes>Kitasatosporales | - | Streptomyces_sp._WZ.A104 | CIB93_04970 | 106791..109793 | 2023771 | AAA-family-ATPase-[Streptomyces-sp.-WZ.A104] |
| WP_055694199.1 | AAA+ATPase+wHTH+Alpha+Beta-domain+wHTH+RRM | Type-1_BR.RM.like.systems | <-DUF499*<-METHYLASE<-TUDOR+SF1-SF2-Helicase+iREase+CTD+REase<-ACET | 998 | Bacteria>Terrabacteria_group>Actinomycetota>Actinomycetes>Kitasatosporales | GCF_001419695.1 | Streptomyces_prasinopilosus | AOK18_RS10915 | 1183..4179 | 67344 | DUF499-domain-containing-protein-[Streptomyces-prasinopilosus] |
| WP_075971379.1 | AAA+ATPase+wHTH+Alpha+Beta-domain+wHTH+RRM | Type-1_BR.RM.like.systems | HTH_3->?->HTH_3->?-><-DUF499*<-METHYLASE | 977 | Bacteria>Terrabacteria_group>Actinomycetota>Actinomycetes>Kitasatosporales | GCF_001941845.1 | Streptomyces_sp._TN58 | BSL84_RS27585 | 6105920..6108853 | 234612 | DUF499-domain-containing-protein-[Streptomyces-sp.-TN58] |
| WBM79848.1 | AAA+ATPase+wHTH+Alpha+Beta-domain+wHTH+RRM | Type-1_BR.RM.like.systems | XIS-HTH->TUDOR+SF1-SF2-Helicase+iREase+CTD+REase-p->TUDOR+SF1-SF2-Helicase+iREase+CTD+REase->METHYLASE->ParB+HNH->HEPN->DUF499*->DUF499-p->DUF499-p-><-?\|\|REase->REase-> | 285 | Bacteria>Terrabacteria_group>Actinomycetota>Actinomycetes>Micrococcales | GCA_027594785.1 | Cryobacterium_breve | KIV56_17020 | 3683119..3683976 | 1259258 | DUF499-domain-containing-protein-[Cryobacterium-breve] |
| WP_281534461.1 | AAA+ATPase+wHTH+Alpha+Beta-domain+wHTH+RRM | Type-1_BR.RM.like.systems | TUDOR+SF1-SF2-Helicase+iREase+CTD+REase-p->TUDOR+SF1-SF2-Helicase+iREase+CTD+REase->METHYLASE->ParB+HNH->HEPN->DUF499-p->DUF499*->DUF499-p-><-?\|\|REase->?->?-><-RelA | 430 | Bacteria>Terrabacteria_group>Actinomycetota>Actinomycetes>Micrococcales | GCF_027594785.1 | Cryobacterium_breve | KIV56_RS17895 | 3683885..3685177 | 1259258 | DUF499-domain-containing-protein-[Cryobacterium-breve] |
| WP_262391126.1 | AAA+ATPase+wHTH+Alpha+Beta-domain+wHTH+RRM | Type-1_BR.RM.like.systems | Peptidase_S66+Peptidase_S66C-><-HTH+SF1-SF2-Helicase+REase<-REase<-MORC<-MORC<-DUF499*<-HEPN<-METHYLASE<-DNA_methylase\|\|REase-> | 679 | Bacteria>Terrabacteria_group>Actinomycetota>Actinomycetes>Micrococcales | GCF_003071525.2 | Micrococcus_luteus | CO205_RS12620 | 704510..706549 | 1270 | DUF499-domain-containing-protein-[Micrococcus-luteus] |
| MDD0856750.1 | AAA+ATPase+wHTH+Alpha+Beta-domain+wHTH+RRM | Type-1_BR.RM.like.systems | ABC-ATPase+SbcC-><-?<-TrwC+SF1-SF2-Helicase<-?<-?\|\|<-DUF499-p<-DUF499*<-DUF499-p<-HEPN<-METHYLASE<-?<-METHYLASE-p | 458 | Bacteria>Terrabacteria_group>Actinomycetota>Actinomycetes>Micrococcales | GCA_024137885.2 | Arthrobacter_alpinus | NHF46_00925 | 169502..170878 | 656366 | DUF499-domain-containing-protein-[Arthrobacter-alpinus] |
| WP_047688630.1 | AAA+ATPase+wHTH+Alpha+Beta-domain+wHTH+RRM | Type-1_BR.RM.like.systems | Glyoxalase-><-?<-HTH+SF1-SF2-Helicase+REase<-REase<-MORC<-DUF499* | 699 | Bacteria>Terrabacteria_group>Actinomycetota>Actinomycetes>Micrococcales | GCF_000800955.1 | Kocuria_sp._ZOR0020 | L979_RS02200 | 64471..66571 | 1339234 | DUF499-domain-containing-protein,-partial-[Kocuria-sp.-ZOR0020] |
| MCL1800470.1 | AAA+ATPase+wHTH+Alpha+Beta-domain+wHTH+RRM | Type-1_BR.RM.like.systems | ACET-><-?<-?<-DUF499*\|\|?->HTH_21+rve+rve_3->METHYLASE-p-> | 751 | Bacteria>Terrabacteria_group>Actinomycetota>Actinomycetes>Micrococcales | GCA_009782595.1 | Promicromonosporaceae_bacterium | FWG25_03780 | 2310..4565 | 2304674 | DUF499-domain-containing-protein-[Promicromonosporaceae-bacterium] |
| WP_202978393.1 | AAA+ATPase+wHTH+Alpha+Beta-domain+wHTH+RRM | Type-1_BR.RM.like.systems | <-Phage_int_SAM_1+Phage_integrase<-XIS-HTH\|\|TrwC+SF1-SF2-helicase+TOPRIM->XIS-HTH+RVT-><-SbcC+SbcC<-DUF499*<-METHYLASE<-TUDOR+SF1-SF2-Helicase+iREase+CTD+REase\|\|?->?-><-?<-ABhydrolase+ABhydrolase | 979 | Bacteria>Terrabacteria_group>Actinomycetota>Actinomycetes>Micrococcales | GCF_006337125.1 | Serinicoccus_chungangensis | FHD63_RS00150 | 33683..36622 | 767452 | DUF499-domain-containing-protein-[Serinicoccus-chungangensis] |
| MBE4717295.1 | AAA+ATPase+wHTH+Alpha+Beta-domain+wHTH+RRM | Type-1_BR.RM.like.systems | DDE_Tnp_1_4-><-DUF499*<-METHYLASE<-TUDOR+SF1-SF2-Helicase+iREase+CTD+REase<-HKD-DNase+SF1-SF2-Helicase+TRD\|\|MACRODOMAIN+SIR2-><-?<-Aminotran_1_2 | 948 | Bacteria>Terrabacteria_group>Actinomycetota>Actinomycetes>Micrococcales | GCA_014926485.1 | Pseudarthrobacter_sp._AB1 | DAD99_03775 | 802..3648 | 2138309 | AAA-family-ATPase-[Pseudarthrobacter-sp.-AB1] |
| WP_227007644.1 | AAA+ATPase+wHTH+Alpha+Beta-domain+wHTH+RRM | Type-1_BR.RM.like.systems | DDE_Tnp_1_4-><-DUF499*<-METHYLASE<-TUDOR+SF1-SF2-Helicase+iREase+CTD+REase<-HKD-DNase+SF1-SF2-Helicase+TRD\|\|MACRODOMAIN+SIR2-> | 796 | Bacteria>Terrabacteria_group>Actinomycetota>Actinomycetes>Micrococcales | - | Pseudarthrobacter_sp._AB1 | DAD99_RS03770 | 802..3192 | 2138309 | DUF499-domain-containing-protein-[Pseudarthrobacter-sp.-AB1] |
| WP_226963983.1 | AAA+ATPase+wHTH+Alpha+Beta-domain+wHTH+RRM | Type-1_BR.RM.like.systems | VirB4-FtsK->?->TUDOR+SF1-SF2-Helicase+iREase+CTD+REase->METHYLASE->DUF499*->?->VirD4-TraG+VirD4-FtsK->?->?-><-TrwC+SF1-SF2-helicase+TOPRIM | 884 | Bacteria>Terrabacteria_group>Actinomycetota>Actinomycetes>Micrococcales | GCF_009184895.1 | Tetrasphaera_sp._F2B08 | GA707_RS19590 | 53747..56401 | 2653936 | DUF499-domain-containing-protein-[Tetrasphaera-sp.-F2B08] |
| KAB7740356.1 | AAA+ATPase+wHTH+Alpha+Beta-domain+wHTH+RRM | Type-1_BR.RM.like.systems | VirB4-FtsK->?->TUDOR+SF1-SF2-Helicase+iREase+CTD+REase->DUF499*->?->VirD4-TraG+VirD4-FtsK->?->?-><-TrwC+SF1-SF2-helicase+TOPRIM | 976 | Bacteria>Terrabacteria_group>Actinomycetota>Actinomycetes>Micrococcales | GCA_009184895.1 | Tetrasphaera_sp._F2B08 | GA707_19585 | 53471..56401 | 2653936 | DUF499-domain-containing-protein-[Tetrasphaera-sp.-F2B08] |
| MBT2248359.1 | AAA+ATPase+wHTH+Alpha+Beta-domain+wHTH+RRM | Type-1_BR.RM.like.systems | HEPN->?->HKD-DNase+SF1-SF2-Helicase+TRD->TUDOR+SF1-SF2-Helicase+iREase+CTD+REase->METHYLASE->DUF499*-><-PfkB\|\|?->?->?-><-HAD | 981 | Bacteria>Terrabacteria_group>Actinomycetota>Actinomycetes>Micrococcales | GCA_018604465.1 | Arthrobacter_sp._BHU_FT2 | JHV56_06580 | 37589..40534 | 2800388 | DUF499-domain-containing-protein-[Arthrobacter-sp.-BHU-FT2] |
| WP_218122143.1 | AAA+ATPase+wHTH+Alpha+Beta-domain+wHTH+RRM | Type-1_BR.RM.like.systems | HNH->\|\|?-><-?<-?<-MPTase<-DUF499*<-METHYLASE<-TUDOR+SF1-SF2-Helicase+iREase+CTD+REase<-ParB+HNH\|\|?-><-?\|\|METHYLASE-> | 977 | Bacteria>Terrabacteria_group>Actinomycetota>Actinomycetes>Micrococcales | GCF_900102895.1 | Arthrobacter_sp._ov407 | BLS42_RS19705 | 16403..19336 | 1761748 | DUF499-domain-containing-protein-[Arthrobacter-sp.-ov407] |
| WP_072313384.1 | AAA+ATPase+wHTH+Alpha+Beta-domain+wHTH+RRM | Type-1_BR.RM.like.systems | XIS-HTH-><-SIR2<-?<-?\|\|TUDOR+SF1-SF2-Helicase+iREase+CTD+REase->METHYLASE->DUF499*->?-><-?\|\|?->\|\|RNase-> | 986 | Bacteria>Terrabacteria_group>Actinomycetota>Actinomycetes>Micrococcales | GCF_900117395.1 | Agrococcus_sp._Marseille-P2731 | BQ6202_RS05015 | 226692..229652 | 1841862 | DUF499-domain-containing-protein-[Agrococcus-sp.-Marseille-P2731] |
| WP_111008500.1 | AAA+ATPase+wHTH+Alpha+Beta-domain+wHTH+RRM | Type-1_BR.RM.like.systems | McrC-NTD+REase->TUDOR+SF1-SF2-Helicase+iREase+CTD+REase->METHYLASE->DUF499*-> | 970 | Bacteria>Terrabacteria_group>Actinomycetota>Actinomycetes>Micrococcales | GCF_003234565.1 | Curtobacterium_sp._MCBD17_013 | DEI81_RS00160 | 36382..39294 | 2175668 | DUF499-domain-containing-protein-[Curtobacterium-sp.-MCBD17_013] |
| WP_134511046.1 | AAA+ATPase+wHTH+Alpha+Beta-domain+wHTH+RRM | Type-1_BR.RM.like.systems | rve_3-><-?<-DUF499*<-METHYLASE<-TUDOR+SF1-SF2-Helicase+iREase+CTD+REase | 977 | Bacteria>Terrabacteria_group>Actinomycetota>Actinomycetes>Micrococcales | GCF_004402815.1 | Cryobacterium_sp._Hz7 | E3N86_RS00015 | 936..3869 | 1259166 | DUF499-domain-containing-protein-[Cryobacterium-sp.-Hz7] |
| WP_281444405.1 | AAA+ATPase+wHTH+Alpha+Beta-domain+wHTH+RRM | Type-1_BR.RM.like.systems | TUDOR+SF1-SF2-Helicase+iREase+CTD+REase->METHYLASE->DUF499*-> | 985 | Bacteria>Terrabacteria_group>Actinomycetota>Actinomycetes>Micrococcales | GCF_029952465.1 | Paenarthrobacter_nicotinovorans | PJL18_RS02445 | 12304..15261 | 29320 | DUF499-domain-containing-protein-[Paenarthrobacter-nicotinovorans] |
| WP_262285937.1 | AAA+ATPase+wHTH+Alpha+Beta-domain+wHTH+RRM | Type-1_BR.RM.like.systems | TUDOR+SF1-SF2-Helicase+iREase+CTD+REase->METHYLASE-><-?\|\|DUF499*-><-?<-?<-RAMA | 936 | Bacteria>Terrabacteria_group>Actinomycetota>Actinomycetes>Micromonosporales | GCF_025460765.1 | Micromonospora_sp._MA102 | NFX44_RS24460 | 45695..48505 | 2952755 | DUF499-domain-containing-protein-[Micromonospora-sp.-MA102] |
| WP_208869683.1 | AAA+ATPase+wHTH+Alpha+Beta-domain+wHTH+RRM | Type-1_BR.RM.like.systems | PAIREDC-HTH->rve_3-><-?<-DUF499<-DDE_Tnp_1_assoc+DDE_Tnp_1<-DUF499*<-METHYLASE<-TUDOR+SF1-SF2-Helicase+iREase+CTD+REase\|\|GDResolvase+Recombinase+ZnR-> | 494 | Bacteria>Terrabacteria_group>Actinomycetota>Actinomycetes>Micromonosporales | GCF_900143515.1 | Micromonospora_cremea | BUS84_RS16705 | 733392..734876 | 709881 | DUF499-domain-containing-protein-[Micromonospora-cremea] |
| WP_074313671.1 | AAA+ATPase+wHTH+Alpha+Beta-domain+wHTH+RRM | Type-1_BR.RM.like.systems | PAIREDC-HTH->rve_3-><-?<-DUF499<-DDE_Tnp_1_assoc+DDE_Tnp_1<-DUF499<-METHYLASE<-TUDOR+SF1-SF2-Helicase+iREase+CTD+REase\|\|GDResolvase+Recombinase+ZnR-> | 493 | Bacteria>Terrabacteria_group>Actinomycetota>Actinomycetes>Micromonosporales | GCF_900143515.1 | Micromonospora_cremea | BUS84_RS16695 | 730523..732004 | 709881 | hypothetical-protein-[Micromonospora-cremea] |
| SIN14395.1 | AAA+ATPase+wHTH+Alpha+Beta-domain+wHTH+RRM | Type-1_BR.RM.like.systems | PAIREDC-HTH-><-?<-DUF499-p<-DDE_Tnp_1_assoc+DDE_Tnp_1<-DUF499*<-METHYLASE<-TUDOR+SF1-SF2-Helicase+iREase+CTD+REase\|\|?->GDResolvase+Recombinase+ZnR-> | 497 | Bacteria>Terrabacteria_group>Actinomycetota>Actinomycetes>Micromonosporales | GCA_900143515.1 | Micromonospora_cremea | SAMN04489832_3406 | 733392..734885 | 709881 | Protein-of-unknown-function-[Micromonospora-cremea] |
| WP_275979722.1 | AAA+ATPase+wHTH+Alpha+Beta-domain+wHTH+RRM | Type-1_BR.RM.like.systems | JetB->SbcC->?->?->TUDOR+SF1-SF2-Helicase+iREase+CTD+REase->METHYLASE->DUF499*->SIGMA-HTH+CR-ATPase9+Schlafen-><-?\|\|ACET+ACET-><-HTH_3\|\|?->HTH+HKD-DNase-> | 803 | Bacteria>Terrabacteria_group>Actinomycetota>Actinomycetes>Micromonosporales | - | Hamadaea_flava | HDA40_RS34820 | 7447529..7449940 | 1742688 | DUF499-domain-containing-protein-[Hamadaea-flava] |
| WP_204014264.1 | AAA+ATPase+wHTH+Alpha+Beta-domain+wHTH+RRM | Type-1_BR.RM.like.systems | <-TPR\|\|?-><-?\|\|TUDOR+SF1-SF2-Helicase+iREase+CTD+REase->METHYLASE->DUF499*-><-REase<-SF1-SF2-Helicase<-SbcC\|\|ACET->?->NUDIX-> | 981 | Bacteria>Terrabacteria_group>Actinomycetota>Actinomycetes>Micromonosporales | GCF_016863635.1 | Virgisporangium_aurantiacum | Vau01_RS61330 | 13451..16396 | 175570 | DUF499-domain-containing-protein-[Virgisporangium-aurantiacum] |
| MCP2328441.1 | AAA+ATPase+wHTH+Alpha+Beta-domain+wHTH+RRM | Type-1_BR.RM.like.systems | TUDOR+SF1-SF2-Helicase+iREase+CTD+REase->METHYLASE->DUF499*->SIGMA-HTH+CR-ATPase9+Schlafen-><-?\|\|ACET+ACET-><-HTH_3 | 944 | Bacteria>Terrabacteria_group>Actinomycetota>Actinomycetes>Micromonosporales | GCA_024172085.1 | Hamadaea_flava | HDA40_006948 | 7447106..7449940 | 1742688 | hypothetical-protein-[Hamadaea-flava] |
| WP_119663874.1 | AAA+ATPase+wHTH+Alpha+Beta-domain+wHTH+RRM | Type-1_BR.RM.like.systems | <-DUF499*<-METHYLASE<-TUDOR+SF1-SF2-Helicase+iREase+CTD+REase<-ZnR<-GDresolvase<-ParB+HTH\|\|Schlafen+HATPase+HTH-> | 581 | Bacteria>Terrabacteria_group>Actinomycetota>Actinomycetes>Micromonosporales | GCF_003581605.1 | Micromonospora_endophytica | D3H59_RS20815 | 1..1744 | 515350 | DUF499-domain-containing-protein,-partial-[Micromonospora-endophytica] |
| WP_111242810.1 | AAA+ATPase+wHTH+Alpha+Beta-domain+wHTH+RRM | Type-1_BR.RM.like.systems | <-Schlafen+HATPase+HTH\|\|ParB+HTH->GDresolvase->ZnR->TUDOR+SF1-SF2-Helicase+iREase+CTD+REase->METHYLASE->DUF499*-> | 978 | Bacteria>Terrabacteria_group>Actinomycetota>Actinomycetes>Micromonosporales | GCF_018326245.1 | Micromonospora_endophytica | Jiend_RS24020 | 5144956..5147892 | 515350 | DUF499-domain-containing-protein-[Micromonospora-endophytica-(Xie-et-al.-2001)-Li-et-al.-2019] |
| WP_240955535.1 | AAA+ATPase+wHTH+Alpha+Beta-domain+wHTH+RRM | Type-1_BR.RM.like.systems | ParB+HTH->GDresolvase->ZnR->TUDOR+SF1-SF2-Helicase+iREase+CTD+REase->METHYLASE->DUF499*-> | 805 | Bacteria>Terrabacteria_group>Actinomycetota>Actinomycetes>Micromonosporales | GCF_012726175.1 | Micromonospora_sp._HNM0581 | HCA58_RS04925 | 452745..455162 | 2716341 | DUF499-domain-containing-protein-[Micromonospora-sp.-HNM0581] |
| NLU77749.1 | AAA+ATPase+wHTH+Alpha+Beta-domain+wHTH+RRM | Type-1_BR.RM.like.systems | ParB+HTH->GDresolvase->ZnR->TUDOR+SF1-SF2-Helicase+iREase+CTD+REase->METHYLASE->DUF499*-> | 978 | Bacteria>Terrabacteria_group>Actinomycetota>Actinomycetes>Micromonosporales | GCA_012726175.1 | Micromonospora_sp._HNM0581 | HCA58_04925 | 452226..455162 | 2716341 | DUF499-domain-containing-protein-[Micromonospora-sp.-HNM0581] |
| WP_207783825.1 | AAA+ATPase+wHTH+Alpha+Beta-domain+wHTH+RRM | Type-1_BR.RM.like.systems | <-HTH_31<-?\|\|?-><-DUF499*<-METHYLASE<-TUDOR+SF1-SF2-Helicase+iREase+CTD+REase<-REase<-?<-?\|\|REase-> | 984 | Bacteria>Terrabacteria_group>Actinomycetota>Actinomycetes>Micromonosporales | GCF_003176735.1 | Micromonospora_globispora | DLJ46_RS10730 | 15153..18107 | 1450148 | DUF499-domain-containing-protein-[Micromonospora-globispora] |
| PWU48923.1 | AAA+ATPase+wHTH+Alpha+Beta-domain+wHTH+RRM | Type-1_BR.RM.like.systems | <-HTH_31<-?<-DUF499*<-METHYLASE<-TUDOR+SF1-SF2-Helicase+iREase+CTD+REase<-REase<-?\|\|REase-> | 962 | Bacteria>Terrabacteria_group>Actinomycetota>Actinomycetes>Micromonosporales | GCA_003176735.1 | Micromonospora_globispora | DLJ46_10740 | 15153..18041 | 1450148 | AAA-family-ATPase-[Micromonospora-globispora] |
| WP_255544576.1 | AAA+ATPase+wHTH+Alpha+Beta-domain+wHTH+RRM | Type-1_BR.RM.like.systems | DUF499*-><-DEDD_Tnp_IS110+Transposase_20<-?<-?\|\|HTH+DUF5753-> | 812 | Bacteria>Terrabacteria_group>Actinomycetota>Actinomycetes>Micromonosporales | - | Micromonospora_sp._ATA51 | JNW87_RS31265 | 566..3004 | 2806098 | DUF499-domain-containing-protein-[Micromonospora-sp.-ATA51] |
| WP_238454909.1 | AAA+ATPase+wHTH+Alpha+Beta-domain+wHTH+RRM | Type-1_BR.RM.like.systems | DUF499*-><-DEDD_Tnp_IS110+Transposase_20<-?<-?\|\|HTH+DUF5753-> | 988 | Bacteria>Terrabacteria_group>Actinomycetota>Actinomycetes>Micromonosporales | GCF_016802855.1 | Micromonospora_sp._ATA51 | JNW87_30660 | 38..3004 | 2806098 | DUF499-domain-containing-protein-[Micromonospora-sp.-ATA51] |
| MBM0229195.1 | AAA+ATPase+wHTH+Alpha+Beta-domain+wHTH+RRM | Type-1_BR.RM.like.systems | DUF499*-><-DEDD_Tnp_IS110+Transposase_20<-?<-?\|\|HTH+DUF5753-> | 988 | Bacteria>Terrabacteria_group>Actinomycetota>Actinomycetes>Micromonosporales | GCF_016802855.1 | Micromonospora_sp._ATA51 | JNW87_30660 | 38..3004 | 2806098 | DUF499-domain-containing-protein-[Micromonospora-sp.-ATA51] |
| WP_267325377.1 | AAA+ATPase+wHTH+Alpha+Beta-domain+wHTH+RRM | Type-1_BR.RM.like.systems | TUDOR+SF1-SF2-Helicase+iREase+CTD+REase-p->METHYLASE-><-GspA-ATPase<-HTH_28+rve+Mu-transpos_C<-HIN-HTH\|\|DUF499*-><-?<-?\|\|VapB_antitoxin->PIN-><-JetD | 663 | Bacteria>Terrabacteria_group>Actinomycetota>Actinomycetes>Mycobacteriales | - | Mycolicibacterium_mucogenicum | OS121_RS17040 | 9178..11169 | 56689 | DUF499-domain-containing-protein-[Mycolicibacterium-mucogenicum] |
| WP_267324306.1 | AAA+ATPase+wHTH+Alpha+Beta-domain+wHTH+RRM | Type-1_BR.RM.like.systems | <-DUF499*<-zf-ISL3+DDE_Tnp_ISL3<-HEPN<-METHYLASE<-TUDOR+SF1-SF2-Helicase+iREase+CTD+REase-p | 810 | Bacteria>Terrabacteria_group>Actinomycetota>Actinomycetes>Mycobacteriales | GCF_026501155.1 | Mycolicibacterium_mucogenicum | OS121_RS10005 | 2088715..2091147 | 56689 | DUF499-domain-containing-protein-[Mycolicibacterium-mucogenicum] |
| CCK55004.1 | AAA+ATPase+wHTH+Alpha+Beta-domain+wHTH+RRM | Type-1_BR.RM.like.systems | STYKinase->HIN-HTH->?-><-?<-REase<-DUF499*<-HEPN<-?<-?<-HIN-HTH<-HIN-HTH<-METHYLASE-p | 792 | Bacteria>Terrabacteria_group>Actinomycetota>Actinomycetes>Mycobacteriales | GCA_000328845.1 | Mycobacterium_canettii_CIPT_140070008 | BN43_30077 | 1164706..1167084 | 1205675 | Conserved-protein-of-unknown-function-[Mycobacterium-canettii-CIPT-140070008] |
| WP_231390207.1 | AAA+ATPase+wHTH+Alpha+Beta-domain+wHTH+RRM | Type-1_BR.RM.like.systems | HTH_31-><-DUF499*<-METHYLASE\|\|TetR-HTH-> | 984 | Bacteria>Terrabacteria_group>Actinomycetota>Actinomycetes>Mycobacteriales | GCF_000482385.1 | Nocardia_sp._CNY236 | K940_RS26625 | 381081..384035 | 1169152 | DUF499-domain-containing-protein-[Nocardia-sp.-CNY236] |
| WP_162113231.1 | AAA+ATPase+wHTH+Alpha+Beta-domain+wHTH+RRM | Type-1_BR.RM.like.systems | DUF499*->REase-><-GDResolvase+HIN-HTH | 883 | Bacteria>Terrabacteria_group>Actinomycetota>Actinomycetes>Mycobacteriales | GCF_010093495.1 | Mycolicibacter_kumamotonensis | GWR20_RS22070 | 1..2652 | 354243 | DUF499-domain-containing-protein,-partial-[Mycolicibacter-kumamotonensis] |
| WP_157762129.1 | AAA+ATPase+wHTH+Alpha+Beta-domain+wHTH+RRM | Type-1_BR.RM.like.systems | BirA-HTH-><-Schlafen+HATPase+HTH\|\|TUDOR+SF1-SF2-Helicase+iREase+CTD+REase->METHYLASE->DUF499*-><-?<-tRNA\|\|CR-Kinase-><-TetR-HTH | 911 | Bacteria>Terrabacteria_group>Actinomycetota>Actinomycetes>Mycobacteriales | GCF_001613325.1 | Nocardia_yamanashiensis | NY1_RS09400 | 81283..84018 | 209247 | DUF499-domain-containing-protein-[Nocardia-yamanashiensis] |
| WP_084987643.1 | AAA+ATPase+wHTH+Alpha+Beta-domain+wHTH+RRM | Type-1_BR.RM.like.systems | <-XIS-HTH<-?\|\|ParA+HTH->TUDOR+SF1-SF2-Helicase+iREase+CTD+REase->METHYLASE->DUF499*-><-TetR-HTH<-Phage_integrase+Phage_integrase<-Phage_integrase+Phage_integrase<-Phage_integrase<-LON-AAA<-REase | 999 | Bacteria>Terrabacteria_group>Actinomycetota>Actinomycetes>Mycobacteriales | GCF_002095295.1 | Prescottella_equi | A5905_RS18185 | 281118..284117 | 43767 | DUF499-domain-containing-protein-[Prescottella-equi] |
| WP_064917862.1 | AAA+ATPase+wHTH+Alpha+Beta-domain+wHTH+RRM | Type-1_BR.RM.like.systems | VirD4-TraG+VirD4-FtsK->BirA-HTH->?->REase-><-HTH_Tnp_1+HTH_21+rve+rve_2\|\|?-><-DUF499*<-METHYLASE<-TUDOR+SF1-SF2-Helicase+iREase+CTD+REase<-TrwC+SF1-SF2-helicase+TOPRIM\|\|HTH+ZnR->HNH+HTH-> | 988 | Bacteria>Terrabacteria_group>Actinomycetota>Actinomycetes>Mycobacteriales | GCF_001667305.1 | Mycolicibacterium_conceptionense | A5718_RS25135 | 34961..37927 | 451644 | DUF499-domain-containing-protein-[Mycolicibacterium-conceptionense] |
| WP_257017806.1 | AAA+ATPase+wHTH+Alpha+Beta-domain+wHTH+RRM | Type-1_BR.RM.like.systems | SF1-SF2-Helicase+REase->SF1-SF2-helicase-><-?<-?\|\|TUDOR+SF1-SF2-Helicase+iREase+CTD+REase->METHYLASE->DUF499*-><-wHTH(SBD)+HNH<-REase+SF1-SF2-Helicase<-MazG | 1022 | Bacteria>Terrabacteria_group>Actinomycetota>Actinomycetes>Mycobacteriales | GCF_900215125.1 | unclassified_Rhodococcus_(in:_high_G+C_Gram-positive_bacteria) | C8K38_RS25080 | 92076..95144 | 192944 | MULTISPECIES:-DUF499-domain-containing-protein-[unclassified-Rhodococcus-(in:-high-G+C-Gram-positive-bacteria)] |
| PTR36692.1 | AAA+ATPase+wHTH+Alpha+Beta-domain+wHTH+RRM | Type-1_BR.RM.like.systems | SF1-SF2-Helicase+REase->SF1-SF2-helicase-><-?<-?\|\|TUDOR+SF1-SF2-Helicase+iREase+CTD+REase->METHYLASE->DUF499*->?-><-wHTH+SAD+HNH<-REase+SF1-SF2-Helicase<-MazG | 1022 | Bacteria>Terrabacteria_group>Actinomycetota>Actinomycetes>Mycobacteriales | GCA_003050885.1 | Rhodococcus_sp._OK611 | C8K38_12385 | 92076..95144 | 2135731 | uncharacterized-protein-DUF499-[Rhodococcus-sp.-OK611] |
| WP_235881881.1 | AAA+ATPase+wHTH+Alpha+Beta-domain+wHTH+RRM | Type-1_BR.RM.like.systems | METHYLASE->SF1-SF2-Helicase-><-?<-TIR\|\|TUDOR+SF1-SF2-Helicase+iREase+CTD+REase->METHYLASE->DUF499*-><-UDG\|\|?-><-SIGMA-HTH\|\|?-><-METHYLASE-p<-RADICAL-SAM | 1001 | Bacteria>Terrabacteria_group>Actinomycetota>Actinomycetes>Mycobacteriales | GCF_021559935.1 | Mycolicibacterium_vanbaalenii | HZU38_RS30730 | 104516..107521 | 110539 | DUF499-domain-containing-protein-[Mycolicibacterium-vanbaalenii] |
| WP_163797615.1 | AAA+ATPase+wHTH+Alpha+Beta-domain+wHTH+RRM | Type-1_BR.RM.like.systems | HTH+ZnR->?->?-><-URI-DNase\|\|TUDOR+SF1-SF2-Helicase+iREase+CTD+REase->METHYLASE->DUF499*->NERD+SF1-SF2-Helicase->Mbetalac->MORC+DUF3684+REase-> | 1001 | Bacteria>Terrabacteria_group>Actinomycetota>Actinomycetes>Mycobacteriales | GCF_010731735.1 | Mycolicibacterium_sediminis | G6N64_RS14060 | 2936388..2939393 | 1286180 | DUF499-domain-containing-protein-[Mycolicibacterium-sediminis] |
| WP_229583267.1 | AAA+ATPase+wHTH+Alpha+Beta-domain+wHTH+RRM | Type-1_BR.RM.like.systems | <-HIN-HTH\|\|RAMA-><-DUF499*<-METHYLASE\|\|MULE-TRANSPOSASE-><-REase\|\|MULE-TRANSPOSASE-><-TUDOR+SF1-SF2-Helicase+iREase+CTD+REase<-ParA+HTH | 1005 | Bacteria>Terrabacteria_group>Actinomycetota>Actinomycetes>Mycobacteriales | GCF_020637575.2 | Rhodococcus_rhodochrous | KUM34_RS27995 | 528724..531741 | 1829 | DUF499-domain-containing-protein-[Rhodococcus-rhodochrous] |
| WP_209449044.1 | AAA+ATPase+wHTH+Alpha+Beta-domain+wHTH+RRM | Type-1_BR.RM.like.systems | <-ParB+HNH\|\|?-><-HNH<-?<-DUF499*<-METHYLASE<-TUDOR+SF1-SF2-Helicase+iREase+CTD+REase\|\|<-?<-HNH+HTH<-HTH+ZnR\|\|TrwC+SF1-SF2-helicase+TOPRIM-> | 1011 | Bacteria>Terrabacteria_group>Actinomycetota>Actinomycetes>Mycobacteriales | GCF_017795725.1 | Rhodococcus_qingshengii | HRR33_RS34870 | 13815..16850 | 334542 | DUF499-domain-containing-protein-[Rhodococcus-qingshengii] |
| WP_099698437.1 | AAA+ATPase+wHTH+Alpha+Beta-domain+wHTH+RRM | Type-1_BR.RM.like.systems | <-DUF4236+TPR<-REase\|\|?->TetR-HTH->TUDOR+SF1-SF2-Helicase+iREase+CTD+REase->METHYLASE->DUF499*->Phage_integrase-><-?<-HIN-HTH | 1005 | Bacteria>Terrabacteria_group>Actinomycetota>Actinomycetes>Mycobacteriales | GCF_002744595.1 | Rhodococcus_qingshengii | CHR55_RS26310 | 49083..52100 | 334542 | DUF499-domain-containing-protein-[Rhodococcus-qingshengii] |
| ETA92372.1 | AAA+ATPase+wHTH+Alpha+Beta-domain+wHTH+RRM | Type-1_BR.RM.like.systems | <-HIN-HTH\|\|?->?-><-TetR-HTH\|\|?-><-DUF499*<-METHYLASE<-TUDOR+SF1-SF2-Helicase+iREase+CTD+REase\|\|DNA-glycosylase->?->?-><-ABhydrolase | 1000 | Bacteria>Terrabacteria_group>Actinomycetota>Actinomycetes>Mycobacteriales | GCA_000504805.1 | Mycobacterium_avium_10-5581 | O982_23635 | 4803..7805 | 1392000 | ATPase-AAA-[Mycobacterium-avium-10-5581] |
| WP_157574125.1 | AAA+ATPase+wHTH+Alpha+Beta-domain+wHTH+RRM | Type-1_BR.RM.like.systems | <-RADICAL-SAM<-Radical-SAM<-?\|\|HIN-HTH->TUDOR+SF1-SF2-Helicase+iREase+CTD+REase->METHYLASE->DUF499*-><-?<-?\|\|?->?-><-?\|\|HDRnase-> | 979 | Bacteria>Terrabacteria_group>Actinomycetota>Actinomycetes>Mycobacteriales | GCF_001613145.1 | Nocardia_jejuensis | NJ1_RS21185 | 183794..186733 | 328049 | DUF499-domain-containing-protein-[Nocardia-jejuensis] |
| WP_217580599.1 | AAA+ATPase+wHTH+Alpha+Beta-domain+wHTH+RRM | Type-1_BR.RM.like.systems | ParA+HTH->TUDOR+SF1-SF2-Helicase+iREase+CTD+REase->METHYLASE->DUF499*-><-RAMA<-?<-?\|\|?->?-><-DEDD_Tnp_IS110+Transposase_20 | 1005 | Bacteria>Terrabacteria_group>Actinomycetota>Actinomycetes>Mycobacteriales | GCF_019095205.1 | Rhodococcus_pyridinivorans | HBA53_RS24680 | 43300..46317 | 103816 | DUF499-domain-containing-protein-[Rhodococcus-pyridinivorans] |
| WP_283569455.1 | AAA+ATPase+wHTH+Alpha+Beta-domain+wHTH+RRM | Type-1_BR.RM.like.systems | <-rve+rve_3<-HIN-HTH\|\|?->PemK_toxin->TUDOR+SF1-SF2-Helicase+iREase+CTD+REase->METHYLASE->DUF499*->REase+TPR-> | 982 | Bacteria>Terrabacteria_group>Actinomycetota>Actinomycetes>Mycobacteriales | GCF_030063715.1 | Rhodococcus_qingshengii | QNA24_RS28365 | 79281..82229 | 334542 | DUF499-domain-containing-protein-[Rhodococcus-qingshengii] |
| WP_152965793.1 | AAA+ATPase+wHTH+Alpha+Beta-domain+wHTH+RRM | Type-1_BR.RM.like.systems | <-DUF499*<-METHYLASE<-TUDOR+SF1-SF2-Helicase+iREase+CTD+REase\|\|DEDD_Tnp_IS110+Transposase_20-><-?\|\|DUF4192-> | 906 | Bacteria>Terrabacteria_group>Actinomycetota>Actinomycetes>Mycobacteriales | GCF_001242945.1 | Rhodococcus_sp._311R | ALC46_RS08430 | 1..2719 | 1617904 | DUF499-domain-containing-protein,-partial-[Rhodococcus-sp.-311R] |
| WP_006243978.1 | AAA+ATPase+wHTH+Alpha+Beta-domain+wHTH+RRM | Type-1_BR.RM.like.systems | DOC+DOC-><-URI-DNase<-?\|\|TUDOR+SF1-SF2-Helicase+iREase+CTD+REase->METHYLASE->DUF499*-><-UDG\|\|?-><-?<-SIGMA-HTH | 1003 | Bacteria>Terrabacteria_group>Actinomycetota>Actinomycetes>Mycobacteriales | GCF_000243415.2 | Mycolicibacterium_tusciae | MYCTUDRAFT_RS0201730 | 175659..178670 | 75922 | DUF499-domain-containing-protein-[Mycolicibacterium-tusciae] |
| WP_094724089.1 | AAA+ATPase+wHTH+Alpha+Beta-domain+wHTH+RRM | Type-1_BR.RM.like.systems | Phage_integrase->?->TetR-HTH->ParA+HTH->TUDOR+SF1-SF2-Helicase+iREase+CTD+REase->METHYLASE->DUF499*-> | 999 | Bacteria>Terrabacteria_group>Actinomycetota>Actinomycetes>Mycobacteriales | GCF_002258205.1 | Rhodococcus_sp._06-621-2 | CH267_RS06730 | 238159..241158 | 2022500 | DUF499-domain-containing-protein-[Rhodococcus-sp.-06-621-2] |
| OXM18093.1 | AAA+ATPase+wHTH+Alpha+Beta-domain+wHTH+RRM | Type-1_BR.RM.like.systems | MULE-TRANSPOSASE-><-?\|\|?->?-><-HNH<-?<-DUF499*<-?\|\|DUF4192->DUF4192-><-?<-HNH+HTH<-HTH+ZnR | 1011 | Bacteria>Terrabacteria_group>Actinomycetota>Actinomycetes>Mycobacteriales | GCA_002233715.1 | Rhodococcus_erythropolis | CBI33_27980 | 22488..25523 | 1833 | AAA-family-ATPase-[Rhodococcus-erythropolis] |
| WP_225329404.1 | AAA+ATPase+wHTH+Alpha+Beta-domain+wHTH+RRM | Type-1_BR.RM.like.systems | ArsR-HTH->?->ACET->?-><-?<-ArsR-HTH<-DUF499*<-METHYLASE<-TUDOR+SF1-SF2-Helicase+iREase+CTD+REase | 1001 | Bacteria>Terrabacteria_group>Actinomycetota>Actinomycetes>Mycobacteriales | GCF_020217485.1 | unclassified_Mycobacterium | JF781_RS21255 | 9007..12012 | 2642494 | MULTISPECIES:-DUF499-domain-containing-protein-[unclassified-Mycobacterium] |
| WP_070355366.1 | AAA+ATPase+wHTH+Alpha+Beta-domain+wHTH+RRM | Type-1_BR.RM.like.systems | <-URI-DNase<-?\|\|TUDOR+SF1-SF2-Helicase+iREase+CTD+REase->METHYLASE->DUF499*-><-UDG<-?\|\|?-><-HNH | 1003 | Bacteria>Terrabacteria_group>Actinomycetota>Actinomycetes>Mycobacteriales | GCF_001766635.1 | Mycobacterium_grossiae | BEL07_RS22915 | 16391..19402 | 1552759 | DUF499-domain-containing-protein-[Mycobacterium-grossiae] |
| WP_015298134.1 | AAA+ATPase+wHTH+Alpha+Beta-domain+wHTH+RRM | Type-1_BR.RM.like.systems | HNH->?->?->DNA-glycosylase->?->UDG-><-DUF499*<-METHYLASE<-TUDOR+SF1-SF2-Helicase+iREase+CTD+REase | 1001 | Bacteria>Terrabacteria_group>Actinomycetota>Actinomycetes>Mycobacteriales | GCF_000328565.1 | Mycobacterium_sp._JS623 | MYCSM_RS34275 | 9437..12442 | 212767 | DUF499-domain-containing-protein-[Mycobacterium-sp.-JS623] |
| WP_260938236.1 | AAA+ATPase+wHTH+Alpha+Beta-domain+wHTH+RRM | Type-1_BR.RM.like.systems | METHYLASE-p->DUF499*->HTH_Tnp_1+HTH_21+rve+rve_3->DUF499-p->?->TPR+TPR-> | 651 | Bacteria>Terrabacteria_group>Actinomycetota>Actinomycetes>Mycobacteriales | GCF_025331195.1 | Mycolicibacterium_llatzerense | A7G45_RS14500 | 2489..4444 | 280871 | DUF499-domain-containing-protein-[Mycolicibacterium-llatzerense] |
| WP_155903314.1 | AAA+ATPase+wHTH+Alpha+Beta-domain+wHTH+RRM | Type-1_BR.RM.like.systems | <-DUF499*<-METHYLASE<-TUDOR+SF1-SF2-Helicase+iREase+CTD+REase<-PAIREDC-HTH\|\|?->HicB-> | 1015 | Bacteria>Terrabacteria_group>Actinomycetota>Actinomycetes>Mycobacteriales | GCF_009729075.1 | Mycolicibacterium_sp._CBMA_226 | FZI94_RS22285 | 57486..60533 | 2606611 | DUF499-domain-containing-protein-[Mycolicibacterium-sp.-CBMA-226] |
| WP_085292636.1 | AAA+ATPase+wHTH+Alpha+Beta-domain+wHTH+RRM | Type-1_BR.RM.like.systems | <-DNA-glycosylase\|\|TUDOR+SF1-SF2-Helicase+iREase+CTD+REase->METHYLASE->DUF499*-> | 999 | Bacteria>Terrabacteria_group>Actinomycetota>Actinomycetes>Mycobacteriales | GCF_002104765.1 | Mycolicibacterium_vulneris | B8W69_RS26320 | 11897..14896 | 547163 | DUF499-domain-containing-protein-[Mycobacterium-vulneris] |
| WP_156888996.1 | AAA+ATPase+wHTH+Alpha+Beta-domain+wHTH+RRM | Type-1_BR.RM.like.systems | ACET-><-?<-?<-?\|\|?->ACET-><-DUF499*<-MULE-TRANSPOSASE\|\|Phage_integrase-> | 672 | Bacteria>Terrabacteria_group>Actinomycetota>Actinomycetes>Mycobacteriales | GCF_002079265.1 | Rhodococcus_sp._BH4 | A0W34_RS32325 | 630252..632270 | 1807790 | DUF499-domain-containing-protein-[Rhodococcus-sp.-BH4] |
| WP_007295669.1 | AAA+ATPase+wHTH+Alpha+Beta-domain+wHTH+RRM | Type-1_BR.RM.like.systems | <-DUF499*<-METHYLASE<-TUDOR+SF1-SF2-Helicase+iREase+CTD+REase<-PemK_toxin | 981 | Bacteria>Terrabacteria_group>Actinomycetota>Actinomycetes>Mycobacteriales | GCF_000260815.1 | Rhodococcus | W59_RS01160 | 2200..5145 | 1827 | MULTISPECIES:-DUF499-domain-containing-protein-[Rhodococcus] |
| WP_181699039.1 | AAA+ATPase+wHTH+Alpha+Beta-domain+wHTH+RRM | Type-1_BR.RM.like.systems | <-DUF932<-?<-?\|\|TUDOR+SF1-SF2-Helicase+iREase+CTD+REase->METHYLASE->DUF499*-> | 997 | Bacteria>Terrabacteria_group>Actinomycetota>Actinomycetes>Mycobacteriales | GCF_013839555.1 | Nocardia_sp._GTS18 | HYG95_RS17445 | 77605..80598 | 1778064 | DUF499-domain-containing-protein-[Nocardia-sp.-GTS18] |
| ARE38116.1 | AAA+ATPase+wHTH+Alpha+Beta-domain+wHTH+RRM | Type-1_BR.RM.like.systems | ACET-><-?<-?<-?\|\|<-?<-DUF499*<-MULE-TRANSPOSASE\|\|Phage_integrase-> | 633 | Bacteria>Terrabacteria_group>Actinomycetota>Actinomycetes>Mycobacteriales | GCF_002079265.1 | Rhodococcus_sp._BH4 | A0W34_32100 | 630252..632153 | 1807790 | hypothetical-protein-A0W34_32100-[Rhodococcus-sp.-BH4] |
| SBS79264.1 | AAA+ATPase+wHTH+Alpha+Beta-domain+wHTH+RRM | Type-1_BR.RM.like.systems | TUDOR+SF1-SF2-Helicase+iREase+CTD+REase->REase->METHYLASE->DUF499*-> | 986 | Bacteria>Terrabacteria_group>Actinomycetota>Actinomycetes>Mycobacteriales | GCA_900078375.1 | uncultured_Mycobacterium_sp. | MHPYR_700004 | 5438..8398 | 171292 | AAA-family-ATPase-[uncultured-Mycobacterium-sp.] |
| EME19358.1 | AAA+ATPase+wHTH+Alpha+Beta-domain+wHTH+RRM | Type-1_BR.RM.like.systems | DUF499*-><-HTH_21+rve+rve_3<-HIN-HTH<-HEPN | 947 | Bacteria>Terrabacteria_group>Actinomycetota>Actinomycetes>Mycobacteriales | GCA_000341815.1 | Rhodococcus_qingshengii_BKS_20-40 | G418_17670 | 1..2845 | 1278077 | ATPase-AAA,-partial-[Rhodococcus-qingshengii-BKS-20-40] |
| WP_132399704.1 | AAA+ATPase+wHTH+Alpha+Beta-domain+wHTH+RRM | Type-1_BR.RM.like.systems | <-pPIWI+MID+RNaseH<-pPIWI+SF1-SF2-Helicase<-pPIWI+pPIWI<-HTH_3\|\|TUDOR+SF1-SF2-Helicase+iREase+CTD+REase->METHYLASE->DUF499*-><-?\|\|Lsr2-><-PAIREDC-HTH\|\|HTH_3-><-ABhydrolase\|\|METHYLASE-p-> | 992 | Bacteria>Terrabacteria_group>Actinomycetota>Actinomycetes>Propionibacteriales | GCF_004348455.1 | Kribbella_albertanoniae | E1261_RS00055 | 18250..21228 | 1266829 | DUF499-domain-containing-protein-[Kribbella-albertanoniae] |
| WP_056540362.1 | AAA+ATPase+wHTH+Alpha+Beta-domain+wHTH+RRM | Type-1_BR.RM.like.systems | TUDOR+SF1-SF2-Helicase+iREase+CTD+REase->METHYLASE->DUF499*->TrwC+SF1-SF2-helicase->XIS-HTH->XIS-HTH->XIS-HTH-> | 999 | Bacteria>Terrabacteria_group>Actinomycetota>Actinomycetes>Propionibacteriales | GCF_001424755.1 | Marmoricola_sp._Leaf446 | ASG49_RS08760 | 147499..150498 | 1736379 | DUF499-domain-containing-protein-[Marmoricola-sp.-Leaf446] |
| WP_243852381.1 | AAA+ATPase+wHTH+Alpha+Beta-domain+wHTH+RRM | Type-1_BR.RM.like.systems | NTP_transferase->NUDIX->?->Primpol->?-><-DUF499*<-METHYLASE<-TUDOR+SF1-SF2-Helicase+iREase+CTD+REase\|\|?->Radical-SAM->Radical-SAM->Peptidase_M20-> | 850 | Bacteria>Terrabacteria_group>Actinomycetota>Actinomycetes>Pseudonocardiales | - | Saccharomonospora_amisosensis | FHU38_RS04090 | 814094..816646 | 1128677 | DUF499-domain-containing-protein-[Saccharomonospora-amisosensis] |
| NIJ10487.1 | AAA+ATPase+wHTH+Alpha+Beta-domain+wHTH+RRM | Type-1_BR.RM.like.systems | NTP_transferase->NUDIX->?->Primpol->?->?-><-DUF499*<-METHYLASE<-TUDOR+SF1-SF2-Helicase+iREase+CTD+REase\|\|?->?->Radical-SAM->Radical-SAM-> | 950 | Bacteria>Terrabacteria_group>Actinomycetota>Actinomycetes>Pseudonocardiales | GCA_011761185.1 | Saccharomonospora_amisosensis | FHU38_000831 | 814094..816946 | 1128677 | hypothetical-protein-[Saccharomonospora-amisosensis] |
| WP_192909784.1 | AAA+ATPase+wHTH+Alpha+Beta-domain+wHTH+RRM | Type-1_BR.RM.like.systems | HTH+DUF5753->DUF397->TUDOR+SF1-SF2-Helicase+iREase+CTD+REase->METHYLASE->DUF499*-><-SbcC | 951 | Bacteria>Terrabacteria_group>Actinomycetota>Actinomycetes>Pseudonocardiales | GCF_005405885.1 | Gandjariella_thermophila | FFT21_RS27215 | 8693..11548 | 1931992 | DUF499-domain-containing-protein-[Gandjariella-thermophila] |
| WP_255481987.1 | AAA+ATPase+wHTH+Alpha+Beta-domain+wHTH+RRM | Type-1_BR.RM.like.systems | <-REase+SF1-SF2-Helicase\|\|REase+SF1-SF2-Helicase-><-?\|\|LuxR-HTH-><-DUF499*<-METHYLASE<-TUDOR+SF1-SF2-Helicase+iREase+CTD+REase\|\|tRNA-><-?<-SF1-SF2-Helicase<-Phage_integrase<-Phage_integrase | 802 | Bacteria>Terrabacteria_group>Actinomycetota>Actinomycetes>Pseudonocardiales | GCF_009910495.1 | Amycolatopsis_sp._SID8362 | GTY80_RS34290 | 8314..10722 | 2690346 | DUF499-domain-containing-protein-[Amycolatopsis-sp.-SID8362] |
| NBH08296.1 | AAA+ATPase+wHTH+Alpha+Beta-domain+wHTH+RRM | Type-1_BR.RM.like.systems | <-REase+SF1-SF2-Helicase\|\|REase+SF1-SF2-Helicase-><-?\|\|LuxR-HTH-><-DUF499*<-METHYLASE<-TUDOR+SF1-SF2-Helicase+iREase+CTD+REase\|\|tRNA-><-?<-SF1-SF2-Helicase<-Phage_integrase<-Phage_integrase | 978 | Bacteria>Terrabacteria_group>Actinomycetota>Actinomycetes>Pseudonocardiales | GCF_009910495.1 | Amycolatopsis_sp._SID8362 | GTY80_34300 | 8314..11250 | 2690346 | DUF499-domain-containing-protein-[Amycolatopsis-sp.-SID8362] |
| TQM11049.1 | AAA+ATPase+wHTH+Alpha+Beta-domain+wHTH+RRM | Type-1_BR.RM.like.systems | PAIREDC-HTH->\|\|HIN-HTH->?->?->TPR+TPR+TPR+TPR-><-DUF499*<-METHYLASE<-TUDOR+SF1-SF2-Helicase+iREase+CTD+REase\|\|?-><-PAIREDC-HTH<-Mbetalac | 968 | Bacteria>Terrabacteria_group>Actinomycetota>Actinomycetes>Pseudonocardiales | GCA_006716445.1 | Pseudonocardia_kunmingensis | FB558_3587 | 391911..394817 | 630975 | uncharacterized-protein-DUF499-[Pseudonocardia-kunmingensis] |
| WP_244326026.1 | AAA+ATPase+wHTH+Alpha+Beta-domain+wHTH+RRM | Type-1_BR.RM.like.systems | <-ABhydrolase<-ABhydrolase\|\|TetR-HTH->ParB+HNH-p-><-?<-DUF499*<-METHYLASE<-TUDOR+SF1-SF2-Helicase+iREase+CTD+REase\|\|?->?-><-TrwC+SF1-SF2-helicase+TOPRIM | 798 | Bacteria>Terrabacteria_group>Actinomycetota>Actinomycetes>Pseudonocardiales | - | Pseudonocardia_alni | GSF26_RS03555 | 63253..65649 | 33907 | DUF499-domain-containing-protein-[Pseudonocardia-alni] |
| MBO4236828.1 | AAA+ATPase+wHTH+Alpha+Beta-domain+wHTH+RRM | Type-1_BR.RM.like.systems | <-ABhydrolase<-ABhydrolase\|\|TetR-HTH->ParB+HNH-><-?<-DUF499*<-METHYLASE<-TUDOR+SF1-SF2-Helicase+iREase+CTD+REase\|\|?->?-><-TrwC+SF1-SF2-helicase+TOPRIM | 970 | Bacteria>Terrabacteria_group>Actinomycetota>Actinomycetes>Pseudonocardiales | GCA_017615005.1 | Pseudonocardia_alni | GSF26_03560 | 63253..66165 | 33907 | DUF499-domain-containing-protein-[Pseudonocardia-alni] |
| WP_246106684.1 | AAA+ATPase+wHTH+Alpha+Beta-domain+wHTH+RRM | Type-1_BR.RM.like.systems | TPR+TPR+TPR+TPR-><-DDE_3<-?<-?<-DUF499*<-METHYLASE<-TUDOR+SF1-SF2-Helicase+iREase+CTD+REase<-?\|\|?-><-PAIREDC-HTH<-PAIREDC-HTH | 796 | Bacteria>Terrabacteria_group>Actinomycetota>Actinomycetes>Pseudonocardiales | GCF_006716445.1 | Pseudonocardia_kunmingensis | FB558_RS17695 | 391911..394301 | 630975 | DUF499-domain-containing-protein-[Pseudonocardia-kunmingensis] |
| WP_196777628.1 | AAA+ATPase+wHTH+Alpha+Beta-domain+wHTH+RRM | Type-1_BR.RM.like.systems | HKD-DNase+HKD-DNase-><-REase+N6_MTase<-TRD+TRD<-?<-DUF499*<-METHYLASE<-TUDOR+SF1-SF2-Helicase+iREase+CTD+REase | 981 | Bacteria>Terrabacteria_group>Actinomycetota>Actinomycetes>Pseudonocardiales | GCF_000955955.1 | Lentzea_aerocolonigenes | UK23_RS06015 | 52073..55018 | 68170 | DUF499-domain-containing-protein-[Lentzea-aerocolonigenes] |
| WP_255520953.1 | AAA+ATPase+wHTH+Alpha+Beta-domain+wHTH+RRM | Type-1_BR.RM.like.systems | <-Recombinase\|\|TUDOR+SF1-SF2-Helicase+iREase+CTD+REase->METHYLASE->DUF499*-><-GDResolvase<-?<-?\|\|NUDIX-><-NUDIX | 846 | Bacteria>Terrabacteria_group>Actinomycetota>Actinomycetes>Pseudonocardiales | - | Saccharopolyspora_sp._HNM0986 | INP57_RS24035 | 230963..233503 | 2781241 | DUF499-domain-containing-protein-[Saccharopolyspora-sp.-HNM0986] |
| MBV9161262.1 | AAA+ATPase+wHTH+Alpha+Beta-domain+wHTH+RRM | Type-1_BR.RM.like.systems | NUDIX->ABhydrolase->TUDOR+SF1-SF2-Helicase+iREase+CTD+REase->METHYLASE->DUF499*->tRNA-><-Phage_integrase | 983 | Bacteria>Terrabacteria_group>Actinomycetota>Actinomycetes>Pseudonocardiales | GCA_019246555.1 | Pseudonocardiales_bacterium | JO281_06860 | 19901..22852 | 2201155 | DUF499-domain-containing-protein-[Pseudonocardiales-bacterium] |
| MBK0869891.1 | AAA+ATPase+wHTH+Alpha+Beta-domain+wHTH+RRM | Type-1_BR.RM.like.systems | <-Recombinase\|\|TUDOR+SF1-SF2-Helicase+iREase+CTD+REase->METHYLASE->DUF499*-><-?<-?\|\|NUDIX-><-NUDIX | 992 | Bacteria>Terrabacteria_group>Actinomycetota>Actinomycetes>Pseudonocardiales | GCA_016526145.1 | Saccharopolyspora_sp._HNM0986 | INP57_24045 | 230525..233503 | 2781241 | ATP-binding-protein-[Saccharopolyspora-sp.-HNM0986] |
| KJK51672.1 | AAA+ATPase+wHTH+Alpha+Beta-domain+wHTH+RRM | Type-1_BR.RM.like.systems | HKD-DNase+HKD-DNase-><-?\|\|?-><-?<-DUF499*<-METHYLASE<-TUDOR+SF1-SF2-Helicase+iREase+CTD+REase | 991 | Bacteria>Terrabacteria_group>Actinomycetota>Actinomycetes>Pseudonocardiales | GCA_000955955.1 | Lentzea_aerocolonigenes | UK23_06010 | 52073..55048 | 68170 | ATPase-AAA-[Lentzea-aerocolonigenes] |
| WP_239071825.1 | AAA+ATPase+wHTH+Alpha+Beta-domain+wHTH+RRM | Type-1_BR.RM.like.systems | Phage_integrase->?-><-?\|\|?-><-?<-DUF499*<-METHYLASE<-TUDOR+SF1-SF2-Helicase+iREase+CTD+REase | 784 | Bacteria>Terrabacteria_group>Actinomycetota>Actinomycetes>Pseudonocardiales | GCF_009910495.1 | Amycolatopsis_sp._SID8362 | GTY80_RS44855 | 28444..30798 | 2690346 | DUF499-domain-containing-protein-[Amycolatopsis-sp.-SID8362] |
| NBH10357.1 | AAA+ATPase+wHTH+Alpha+Beta-domain+wHTH+RRM | Type-1_BR.RM.like.systems | Phage_integrase->?-><-?\|\|?-><-DUF499*<-METHYLASE<-TUDOR+SF1-SF2-Helicase+iREase+CTD+REase | 956 | Bacteria>Terrabacteria_group>Actinomycetota>Actinomycetes>Pseudonocardiales | - | Amycolatopsis_sp._SID8362 | GTY80_44850 | 28444..31314 | 2690346 | DUF499-domain-containing-protein-[Amycolatopsis-sp.-SID8362] |
| WP_116051372.1 | AAA+ATPase+wHTH+Alpha+Beta-domain+wHTH+RRM | Type-1_BR.RM.like.systems | <-DUF499*<-METHYLASE<-TUDOR+SF1-SF2-Helicase+iREase+CTD+REase<-?<-?\|\|?->HEPN-> | 994 | Bacteria>Terrabacteria_group>Actinomycetota>Actinomycetes>Pseudonocardiales | GCF_003385185.1 | Pseudonocardiaceae | DOC15_RS29340 | 1670..4654 | 2070 | MULTISPECIES:-DUF499-domain-containing-protein-[Pseudonocardiaceae] |
| WP_131878611.1 | AAA+ATPase+wHTH+Alpha+Beta-domain+wHTH+RRM | Type-1_BR.RM.like.systems | ST-Hiskin->?->TraJ-RHH->MobL->?->XIS-HTH-><-DUF499*<-METHYLASE<-TUDOR+SF1-SF2-Helicase+iREase+CTD+REase<-?\|\|REase+nSTAND3-> | 938 | Bacteria>Terrabacteria_group>Actinomycetota>Actinomycetes>Streptosporangiales | GCF_004348515.1 | Actinomadura_sp._KC345 | E1281_RS15270 | 16620..19436 | 2530371 | DUF499-domain-containing-protein-[Actinomadura-sp.-KC345] |
| WP_238498630.1 | AAA+ATPase+wHTH+Alpha+Beta-domain+wHTH+RRM | Type-1_BR.RM.like.systems | <-TetR-HTH<-tRNA<-?<-DUF499*<-METHYLASE<-TUDOR+SF1-SF2-Helicase+iREase+CTD+REase<-HEPN<-XIS-HTH+RVT<-?<-HTH+DUF5753 | 939 | Bacteria>Terrabacteria_group>Actinomycetota>Actinomycetes>Streptosporangiales | - | Streptosporangium_sp._KLBMP_9127 | MBA17_23095 | 277278..280097 | 2916433 | DUF499-domain-containing-protein-[Streptosporangium-sp.-KLBMP-9127] |
| WP_285915486.1 | AAA+ATPase+wHTH+Alpha+Beta-domain+wHTH+RRM | Type-1_BR.RM.like.systems | TraJ-RHH->MobL->?->XIS-HTH->TUDOR+SF1-SF2-Helicase+iREase+CTD+REase->METHYLASE->DUF499*-><-?<-tRNA<-Calcineurin\|\|?-><-?\|\|?-><-ParA-Soj-PloopNTPase | 997 | Bacteria>Terrabacteria_group>Actinomycetota>Actinomycetes>Streptosporangiales | GCF_030283205.1 | Actinomadura_sp._OS1-43 | QP089_RS28315 | 18384..21377 | 604315 | DUF499-domain-containing-protein-[Actinomadura-sp.-OS1-43] |
| WP_142566095.1 | AAA+ATPase+wHTH+Alpha+Beta-domain+wHTH+RRM | Type-1_BR.RM.like.systems | PAIREDC-HTH-><-tRNA\|\|TUDOR+SF1-SF2-Helicase+iREase+CTD+REase->METHYLASE->DUF499*->HEPN-><-HTH+HKD-DNase\|\|?->GNTR-HTH->HDRnase-> | 976 | Bacteria>Terrabacteria_group>Actinomycetota>Actinomycetes>Streptosporangiales | GCF_006874435.1 | Microbispora_sp._KK1-11 | FLW16_RS39395 | 9114..12044 | 2053005 | DUF499-domain-containing-protein-[Microbispora-sp.-KK1-11] |
| WP_246641040.1 | AAA+ATPase+wHTH+Alpha+Beta-domain+wHTH+RRM | Type-1_BR.RM.like.systems | HTH+TPR+TPR->?-><-Phage_integrase\|\|?-><-DUF499*<-METHYLASE<-TUDOR+SF1-SF2-Helicase+iREase+CTD+REase<-MNS+ZDNA-HTH\|\|?->?-><-HIN-HTH | 849 | Bacteria>Terrabacteria_group>Actinomycetota>Actinomycetes>Streptosporangiales | - | Nonomuraea_harbinensis | KME79_RS36625 | 18799..21348 | 1286938 | DUF499-domain-containing-protein-[Nonomuraea-harbinensis] |
| WP_145851069.1 | AAA+ATPase+wHTH+Alpha+Beta-domain+wHTH+RRM | Type-1_BR.RM.like.systems | Phage_int_SAM_1+Phage_integrase->?->TUDOR+SF1-SF2-Helicase+iREase+CTD+REase->METHYLASE->DUF499*-><-?<-?<-?<-?<-SF1-SF2-Helicase | 986 | Bacteria>Terrabacteria_group>Actinomycetota>Actinomycetes>Streptosporangiales | GCF_007827045.1 | Trebonia_kvetii | EAS64_RS02465 | 593130..596090 | 2480626 | DUF499-domain-containing-protein-[Trebonia-kvetii] |
| WP_165975551.1 | AAA+ATPase+wHTH+Alpha+Beta-domain+wHTH+RRM | Type-1_BR.RM.like.systems | <-DUF499*<-METHYLASE<-TUDOR+SF1-SF2-Helicase+iREase+CTD+REase\|\|<-pPIWI+MID+RNaseH<-pPIWI<-pPIWI+pPIWI<-REase | 848 | Bacteria>Terrabacteria_group>Actinomycetota>Actinomycetes>Streptosporangiales | GCF_004349215.1 | Actinomadura_rubrisoli | E1298_RS19365 | 1..2546 | 2530368 | DUF499-domain-containing-protein,-partial-[Actinomadura-rubrisoli] |
| TDD84857.1 | AAA+ATPase+wHTH+Alpha+Beta-domain+wHTH+RRM | Type-1_BR.RM.like.systems | <-DUF499*<-TUDOR+SF1-SF2-Helicase+iREase+CTD+REase<-pPIWI+MID+RNaseH<-pPIWI<-pPIWI+pPIWI<-REase\|\|Pkinase-> | 918 | Bacteria>Terrabacteria_group>Actinomycetota>Actinomycetes>Streptosporangiales | GCA_004349215.1 | Actinomadura_rubrisoli | E1298_19375 | 1..2756 | 2530368 | DUF499-domain-containing-protein,-partial-[Actinomadura-rubrisoli] |
| WP_246641376.1 | AAA+ATPase+wHTH+Alpha+Beta-domain+wHTH+RRM | Type-1_BR.RM.like.systems | <-DUF499*<-METHYLASE<-TUDOR+SF1-SF2-Helicase+iREase+CTD+REase\|\|?-><-GNTR-HTH-p\|\|<-GNTR-HTH-p | 842 | Bacteria>Terrabacteria_group>Actinomycetota>Actinomycetes>Streptosporangiales | - | Nonomuraea_harbinensis | KME79_RS43360 | 1787..4315 | 1286938 | DUF499-domain-containing-protein-[Nonomuraea-harbinensis] |
| WP_236004105.1 | AAA+ATPase+wHTH+Alpha+Beta-domain+wHTH+RRM | Type-1_BR.RM.like.systems | <-CHTH<-?<-DUF499*<-METHYLASE<-TUDOR+SF1-SF2-Helicase+iREase+CTD+REase<-NUDIX<-GNTR-HTH-p | 845 | Bacteria>Terrabacteria_group>Actinomycetota>Actinomycetes>Streptosporangiales | GCF_013283785.1 | Nonomuraea_antri | HII36_RS32990 | 6504..9041 | 2730852 | DUF499-domain-containing-protein-[Nonomuraea-antri] |
| NRQ36622.1 | AAA+ATPase+wHTH+Alpha+Beta-domain+wHTH+RRM | Type-1_BR.RM.like.systems | <-CHTH<-?<-DUF499*<-METHYLASE<-TUDOR+SF1-SF2-Helicase+iREase+CTD+REase<-NUDIX<-GNTR-HTH-p | 995 | Bacteria>Terrabacteria_group>Actinomycetota>Actinomycetes>Streptosporangiales | GCF_013283785.1 | Nonomuraea_antri | HII36_33020 | 6504..9491 | 2730852 | ATP-binding-protein-[Nonomuraea-antri] |
| WP_067453152.1 | AAA+ATPase+wHTH+Alpha+Beta-domain+wHTH+RRM | Type-1_BR.RM.like.systems | <-DDE_5+DDE_Tnp_1<-?\|\|TUDOR+SF1-SF2-Helicase+iREase+CTD+REase->METHYLASE->DUF499*-> | 976 | Bacteria>Terrabacteria_group>Actinomycetota>Actinomycetes>Streptosporangiales | GCF_001552215.1 | Actinomadura_macra | AMA02S_RS06615 | 390026..392956 | 46164 | DUF499-domain-containing-protein-[Actinomadura-macra] |
| WP_132180042.1 | AAA+ATPase+wHTH+Alpha+Beta-domain+wHTH+RRM | Type-1_BR.RM.like.systems | <-VirB4-FtsK<-?<-?<-?<-?\|\|METHYLASE->DUF499*-> | 872 | Bacteria>Terrabacteria_group>Actinomycetota>Actinomycetes>Streptosporangiales | GCF_004348535.1 | Actinomadura_sp._KC216 | E1264_RS33680 | 7949..10563 | 2530370 | DUF499-domain-containing-protein,-partial-[Actinomadura-sp.-KC216] |
| WP_147945511.1 | AAA+ATPase+wHTH+Alpha+Beta-domain+wHTH+RRM | Type-1_BR.RM.like.systems | <-HEPN<-DUF499*<-METHYLASE | 976 | Bacteria>Terrabacteria_group>Actinomycetota>Actinomycetes>Streptosporangiales | GCF_008033685.1 | Microbispora_sp._CSR-4 | FLV34_RS38505 | 1003..3933 | 2592813 | DUF499-domain-containing-protein-[Microbispora-sp.-CSR-4] |
| MCL8207108.1 | AAA+ATPase+wHTH+Alpha+Beta-domain+wHTH+RRM | Type-1_BR.RM.like.systems | <-Arm-DNA-bind_4+Phage_int_SAM_3+Phage_integrase<-?<-?\|\|?->TUDOR+SF1-SF2-Helicase+iREase+CTD+REase->METHYLASE->DUF499*-><-ParB+N6_N4_MTase<-Phage_integrase<-?<-TOPRIM+ZnR | 959 | Bacteria>Terrabacteria_group>Actinomycetota>Actinomycetes>unclassified_Actinomycetes | GCA_023565505.1 | Actinomycetes_bacterium | K6V97_03400 | 11854..14733 | 1883427 | DUF499-domain-containing-protein-[Actinomycetia-bacterium] |
| MCY4088355.1 | AAA+ATPase+wHTH+Alpha+Beta-domain+wHTH+RRM | Type-1_BR.RM.like.systems | HTH+UbE1->?-><-?\|\|TUDOR+SF1-SF2-Helicase+iREase+CTD+REase->REase->METHYLASE->DUF499*-><-SF1-SF2-Helicase<-ParB+HNH\|\|DUF4065-><-HD\|\|Peptidase-> | 929 | Bacteria>Terrabacteria_group>Actinomycetota>Actinomycetes>unclassified_Actinomycetes | GCA_026708005.1 | Actinomycetes_bacterium | OXG37_16080 | 16647..19436 | 1883427 | MAG:-DUF499-domain-containing-protein-[Actinomycetia-bacterium] |
| MCL8207344.1 | AAA+ATPase+wHTH+Alpha+Beta-domain+wHTH+RRM | Type-1_BR.RM.like.systems | ABC-ATPase+SbcC->ABC-ATPase+SbcC->ABC-ATPase->?->?->ABhydrolase-><-DUF499*<-METHYLASE<-TUDOR+SF1-SF2-Helicase+iREase+CTD+REase | 953 | Bacteria>Terrabacteria_group>Actinomycetota>Actinomycetes>unclassified_Actinomycetes | GCA_023565505.1 | Actinomycetes_bacterium | K6V97_04620 | 84887..87748 | 1883427 | DUF499-domain-containing-protein-[Actinomycetia-bacterium] |
| WP_176235274.1 | AAA+ATPase+wHTH+Alpha+Beta-domain+wHTH+RRM | Type-1_BR.RM.like.systems | <-DUF499*<-XIS-HTH\|\|tRNA->AbiEi_1->AbiEii-><-Polbeta | 871 | Bacteria>Terrabacteria_group>Actinomycetota>Actinomycetota_incertae_sedis>Candidatus_Hakubellales | GCF_013281975.1 | Candidatus_Hakubella_thermalkaliphila | HW443_RS03710 | 631..3246 | 2754717 | hypothetical-protein-[Candidatus-Hakubanella-thermoalkaliphilus] |
| WP_176237819.1 | AAA+ATPase+wHTH+Alpha+Beta-domain+wHTH+RRM | Type-1_BR.RM.like.systems | TUDOR+SF1-SF2-Helicase+iREase+CTD+REase-p->TUDOR+SF1-SF2-Helicase+iREase+CTD+REase->METHYLASE->PHP+SbcC->Schlafen+HATPase+HTH->DUF499*->?-><-Peptidase_M29\|\|?->?->DDE_Tnp_1-> | 867 | Bacteria>Terrabacteria_group>Actinomycetota>Actinomycetota_incertae_sedis>Candidatus_Hakubellales | - | Candidatus_Hakubella_thermalkaliphila | HKBW3S34_00093 | 39810..42413 | 2754717 | DUF499-domain-containing-protein-[Candidatus-Hakubanella-thermoalkaliphilus] |
| GFP29174.1 | AAA+ATPase+wHTH+Alpha+Beta-domain+wHTH+RRM | Type-1_BR.RM.like.systems | TUDOR+SF1-SF2-Helicase+iREase+CTD+REase-p->TUDOR+SF1-SF2-Helicase+iREase+CTD+REase->METHYLASE->PHP+SbcC->Schlafen+HATPase+HTH->DUF499*->?-><-Peptidase_M29\|\|?->?->DDE_Tnp_1-> | 867 | Bacteria>Terrabacteria_group>Actinomycetota>Actinomycetota_incertae_sedis>Candidatus_Hakubellales | GCA_013281975.1 | Candidatus_Hakubella_thermalkaliphila | HKBW3S34_00093 | 39810..42413 | 2754717 | hypothetical-protein-HKBW3S34_00093-[Candidatus-Hakubanella-thermoalkaliphilus] |

| Protein-ID | DUF499-architecture | System | Gene-Neighbourhoods | Length | Taxonomy | Assembly-ID | Strain | Locus-Tag | Genomic-coordinates | TaxID | Fasta-defline |
|---|---|---|---|---|---|---|---|---|---|---|---|
| WP_258190433.1 | AAA+ATPase+wHTH+Alpha+Beta-domain+wHTH+RRM | Type-1_BR.RM.like.systems | TUDOR+SF1-SF2-Helicase+iREase+CTD+REase-p->TUDOR+SF1-SF2-Helicase+iREase+CTD+REase->METHYLASE->PHP+SbcC->Schlafen+HATPase+HTH->DUF499*->?-><-Peptidase_M29 | 869 | Bacteria>Terrabacteria_group>Actinomycetota>Actinomycetota_incertae_sedis>Candidatus_Hakubellales | GCF_013281975.1 | Candidatus_Hakubella_thermalkaliphila | HW443_RS00535 | 39804..42413 | 2754717 | DUF499-domain-containing-protein-[Candidatus-Hakubanella-thermoalkaliphilus] |
| WP_176229285.1 | AAA+ATPase+wHTH+Alpha+Beta-domain+wHTH+RRM | Type-1_BR.RM.like.systems | Peptidase_M29-><-PemK_toxin<-?<-DUF499*<-Schlafen+HATPase+HTH<-PHP+SbcC<-METHYLASE<-TUDOR+SF1-SF2-Helicase+iREase+CTD+REase | 867 | Bacteria>Terrabacteria_group>Actinomycetota>Actinomycetota_incertae_sedis>Candidatus_Hakubellales | GCF_013282015.1 | Candidatus_Hakubella_thermalkaliphila | HW437_RS01600 | 1284..3887 | 2754717 | DUF499-domain-containing-protein-[Candidatus-Hakubanella-thermoalkaliphilus] |
| GFP42119.1 | AAA+ATPase+wHTH+Alpha+Beta-domain+wHTH+RRM | Type-1_BR.RM.like.systems | <-DUF499*<-Schlafen+HATPase+HTH<-PHP+SbcC<-METHYLASE<-METHYLASE-p<-TUDOR+SF1-SF2-Helicase+iREase+CTD+REase-p | 603 | Bacteria>Terrabacteria_group>Actinomycetota>Actinomycetota_incertae_sedis>Candidatus_Hakubellales | GCA_013282075.1 | Candidatus_Hakubella_thermalkaliphila | HKBW3C_01245 | 1..1809 | 2754717 | hypothetical-protein-HKBW3C_01245,-partial-[Candidatus-Hakubanella-thermoalkaliphilus] |
| WP_176233419.1 | AAA+ATPase+wHTH+Alpha+Beta-domain+wHTH+RRM | Type-1_BR.RM.like.systems | DDE_Tnp_1->Phage_int_SAM_1-><-?<-sGTP<-?<-?<-DUF499*<-TOPRIM<-OLD-ABC<-METHYLASE-p | 683 | Bacteria>Terrabacteria_group>Actinomycetota>Actinomycetota_incertae_sedis>Candidatus_Hakubellales | GCF_013281955.1 | Candidatus_Hakubella_thermalkaliphila | HW440_RS05440 | 5834..7885 | 2754717 | DUF499-domain-containing-protein-[Candidatus-Hakubanella-thermoalkaliphilus] |
| WP_219852434.1 | AAA+ATPase+wHTH+Alpha+Beta-domain+wHTH+RRM | Type-1_BR.RM.like.systems | REase+HTH->NTP_transf->TUDOR+SF1-SF2-Helicase+iREase+CTD+REase->METHYLASE->DUF499*-> | 342 | Bacteria>Terrabacteria_group>Actinomycetota>Actinomycetota_incertae_sedis>Candidatus_Hakubellales | GCF_013281895.1 | Candidatus_Hakubella_thermalkaliphila | HW426_RS02085 | 12413..13438 | 2754717 | DUF499-domain-containing-protein,-partial-[Candidatus-Hakubanella-thermoalkaliphilus] |
| WP_176229130.1 | AAA+ATPase+wHTH+Alpha+Beta-domain+wHTH+RRM | Type-1_BR.RM.like.systems | REase+HTH->NTP_transf->TUDOR+SF1-SF2-Helicase+iREase+CTD+REase->METHYLASE->DUF499*-> | 989 | Bacteria>Terrabacteria_group>Actinomycetota>Actinomycetota_incertae_sedis>Candidatus_Hakubellales | GCF_013282015.1 | Candidatus_Hakubella_thermalkaliphila | HW437_RS00625 | 20895..23864 | 2754717 | DUF499-domain-containing-protein-[Candidatus-Hakubanella-thermoalkaliphilus] |
| GFP21179.1 | AAA+ATPase+wHTH+Alpha+Beta-domain+wHTH+RRM | Type-1_BR.RM.like.systems | REase+HTH->NTP_transf->TUDOR+SF1-SF2-Helicase+iREase+CTD+REase->METHYLASE->DUF499*-> | 345 | Bacteria>Terrabacteria_group>Actinomycetota>Actinomycetota_incertae_sedis>Candidatus_Hakubellales | GCA_013281895.1 | Candidatus_Hakubella_thermalkaliphila | HKBW3S06_00405 | 12404..13438 | 2754717 | hypothetical-protein-HKBW3S06_00405,-partial-[Candidatus-Hakubanella-thermoalkaliphilus] |
| GFP41713.1 | AAA+ATPase+wHTH+Alpha+Beta-domain+wHTH+RRM | Type-1_BR.RM.like.systems | <-DUF499*<-METHYLASE<-TUDOR+SF1-SF2-Helicase+iREase+CTD+REase<-NTP_transf<-HTH | 989 | Bacteria>Terrabacteria_group>Actinomycetota>Actinomycetota_incertae_sedis>Candidatus_Hakubellales | GCA_013282075.1 | Candidatus_Hakubella_thermalkaliphila | HKBW3C_00839 | 2157..5126 | 2754717 | hypothetical-protein-HKBW3C_00839-[Candidatus-Hakubanella-thermoalkaliphilus] |
| WP_176233286.1 | AAA+ATPase+wHTH+Alpha+Beta-domain+wHTH+RRM | Type-1_BR.RM.like.systems | METHYLASE->PolB->HEPN->DUF499*-><-tRNA\|\|?->NTP_transferase->?->?-><-ACET | 876 | Bacteria>Terrabacteria_group>Actinomycetota>Actinomycetota_incertae_sedis>Candidatus_Hakubellales | GCF_013281955.1 | Candidatus_Hakubella_thermalkaliphila | HW440_RS04150 | 3204..5834 | 2754717 | DUF499-domain-containing-protein-[Candidatus-Hakubanella-thermoalkaliphilus] |
| GFP42415.1 | AAA+ATPase+wHTH+Alpha+Beta-domain+wHTH+RRM | Type-1_BR.RM.like.systems | DDE_Tnp_1-><-?\|\|Phage_int_SAM_1-><-?<-sGTP<-?<-DUF499*<-TOPRIM | 683 | Bacteria>Terrabacteria_group>Actinomycetota>Actinomycetota_incertae_sedis>Candidatus_Hakubellales | GCA_013282075.1 | Candidatus_Hakubella_thermalkaliphila | HKBW3C_01541 | 5550..7601 | 2754717 | hypothetical-protein-HKBW3C_01541-[Candidatus-Hakubanella-thermoalkaliphilus] |
| GFP32491.1 | AAA+ATPase+wHTH+Alpha+Beta-domain+wHTH+RRM | Type-1_BR.RM.like.systems | <-DUF499*<-HEPN<-PolB<-METHYLASE | 577 | Bacteria>Terrabacteria_group>Actinomycetota>Actinomycetota_incertae_sedis>Candidatus_Hakubellales | GCA_013281995.1 | Candidatus_Hakubella_thermalkaliphila | HKBW3S42_00795 | 50..1783 | 2754717 | hypothetical-protein-HKBW3S42_00795-[Candidatus-Hakubanella-thermoalkaliphilus] |
| GAV31939.1 | AAA+ATPase+wHTH+Alpha+Beta-domain+wHTH+RRM | Type-1_BR.RM.like.systems | PspA+PIN->?-><-DUF499*<-?<-MqsA_antitoxin+HTH_3<-METHYLASE<-ParB+HNH<-TUDOR+SF1-SF2-Helicase+iREase+CTD+REase | 950 | Bacteria>Terrabacteria_group>Actinomycetota>Coriobacteriia>Coriobacteriales | GCA_001950395.1 | Coriobacteriaceae_bacterium_EMTCatB1 | emb_1c0561 | 1370215..1373067 | 1927122 | predicted-ATPase-[Coriobacteriaceae-bacterium-EMTCatB1] |
| WP_259314362.1 | AAA+ATPase+wHTH+Alpha+Beta-domain+wHTH+RRM | Type-1_BR.RM.like.systems | PIN->tRNA->TUDOR+SF1-SF2-Helicase+iREase+CTD+REase->METHYLASE->DUF499*->?-><-?\|\|XIS-HTH->?->?->Phage_int_SAM_1+Phage_integrase-> | 940 | Bacteria>Terrabacteria_group>Actinomycetota>Thermoleophilia>Solirubrobacterales | GCF_021172045.1 | Capillimicrobium_parvum | DSM104329_RS05360 | 1087018..1089840 | 2884022 | DUF499-domain-containing-protein-[Capillimicrobium-parvum] |
| UGS34698.1 | AAA+ATPase+wHTH+Alpha+Beta-domain+wHTH+RRM | Type-1_BR.RM.like.systems | PIN->tRNA->TUDOR+SF1-SF2-Helicase+iREase+CTD+REase->METHYLASE->DUF499*-><-?\|\|?->?->Phage_int_SAM_1+Phage_integrase-><-HTH_3 | 940 | Bacteria>Terrabacteria_group>Actinomycetota>Thermoleophilia>Solirubrobacterales | GCF_021172045.1 | Capillimicrobium_parvum | DSM104329_01080 | 1087018..1089840 | 2884022 | hypothetical-protein-DSM104329_01080-[Solirubrobacterales-bacterium-0166_1] |
| WP_270039609.1 | AAA+ATPase+wHTH+Alpha+Beta-domain+wHTH+RRM | Type-1_BR.RM.like.systems | SAD+HNH->?->HISKIN->rve_3-><-?<-ABC-ATPase<-DUF499*<-METHYLASE<-TUDOR+SF1-SF2-Helicase+iREase+CTD+REase<-?\|\|HTH+HKD-DNase->HDRnase-> | 981 | Bacteria>Terrabacteria_group>Actinomycetota>Thermoleophilia>Solirubrobacterales | GCF_027587205.1 | Solirubrobacter_ginsenosidimutans | OM076_RS10245 | 130383..133328 | 490573 | DUF499-domain-containing-protein-[Solirubrobacter-ginsenosidimutans] |
| UUY05125.1 | AAA+ATPase+wHTH+Alpha+Beta-domain+wHTH+RRM | Type-1_BR.RM.like.systems | ParA-Soj-PloopNTPase->?->VirD4-TraG+HerA->?->?->DUF499*-> | 590 | Bacteria>Terrabacteria_group>Actinomycetota>Thermoleophilia>Solirubrobacterales | GCA_024666565.1 | Parviterribacteraceae_bacterium_J379 | LRS13_06240 | 1274832..1276604 | 2898438 | hypothetical-protein-LRS13_06240-[Parviterribacteraceae-bacterium-J379] |
| MBA3420890.1 | AAA+ATPase+wHTH+Alpha+Beta-domain+wHTH+RRM | Type-1_BR.RM.like.systems | <-DUF499*<-?<-?<-?<-?<-METHYLASE-p | 938 | Bacteria>Terrabacteria_group>Actinomycetota>Thermoleophilia>Thermoleophilales | GCA_013812065.1 | Thermoleophilaceae_bacterium | HOU12_03190 | 10493..13309 | 2732252 | ATP-binding-protein-[Thermoleophilaceae-bacterium] |
| MBE3095474.1 | AAA+ATPase+wHTH+Alpha+Beta-domain+wHTH+RRM | Type-1_BR.RM.like.systems | TUDOR+SF1-SF2-Helicase+iREase+CTD+REase->METHYLASE->DUF499*-> | 181 | Bacteria>Terrabacteria_group>Actinomycetota>unclassified_Actinomycetota | GCA_014894515.1 | Actinomycetota_bacterium | IMZ52_10625 | 4914..5457 | 2900548 | ATP-binding-protein-[Actinomycetia-bacterium] |
| MCE7883902.1 | AAA+ATPase+wHTH+Alpha+Beta-domain+wHTH+RRM | Type-1_BR.RM.like.systems | <-REase+SF1-SF2-Helicase<-DUF499*<-DUF499-p\|\|XIS-HTH-> | 594 | Bacteria>Terrabacteria_group>Actinomycetota>unclassified_Actinomycetota | GCA_021404985.1 | Actinobacteria_bacterium_ATB1 | DYH08_08590 | 2755..4539 | 2293627 | ATP-binding-protein-[Actinobacteria-bacterium-ATB1] |
| MBU2110960.1 | AAA+ATPase+wHTH+Alpha+Beta-domain+wHTH+RRM | Type-1_BR.RM.like.systems | DUF499*->MORC->REase->HTH+SF1-SF2-Helicase+REase-> | 991 | Bacteria>Terrabacteria_group>Actinomycetota>unclassified_Actinomycetota | GCA_018826625.1 | Actinomycetota_bacterium | KKE65_04805 | 1..2976 | 2900548 | DUF499-domain-containing-protein-[Actinomycetia-bacterium] |
| MDA8315608.1 | AAA+ATPase+wHTH+Alpha+Beta-domain+wHTH+RRM | Type-1_BR.RM.like.systems | DUF499*->DUF499-p->DDE_Tnp_1_assoc+DDE_Tnp_1-> | 659 | Bacteria>Terrabacteria_group>Actinomycetota>unclassified_Actinomycetota | GCA_028067765.1 | Actinomycetota_bacterium | M0010_10610 | 1..1978 | 2900548 | MAG:-DUF499-domain-containing-protein,-partial-[Actinomycetota-bacterium] |
| MDA8281345.1 | AAA+ATPase+wHTH+Alpha+Beta-domain+wHTH+RRM | Type-1_BR.RM.like.systems | Phage_integrase->PIN-><-XIS-HTH\|\|TUDOR+SF1-SF2-Helicase+iREase+CTD+REase->ParB+HNH->METHYLASE->DUF499*->DUF4527->?->?->HTH-> | 939 | Bacteria>Terrabacteria_group>Actinomycetota>unclassified_Actinomycetota | GCA_028067595.1 | Actinomycetota_bacterium | MOZ63_13180 | 23767..26586 | 2900548 | MAG:-DUF499-domain-containing-protein-[Actinomycetota-bacterium] |
| MDA8047538.1 | AAA+ATPase+wHTH+Alpha+Beta-domain+wHTH+RRM | Type-1_BR.RM.like.systems | TUDOR+SF1-SF2-Helicase+iREase+CTD+REase->METHYLASE->DUF499*->OLD-ABC+TOPRIM-><-PIWI<-REase+PNTD1+PNTD2\|\|HicB-> | 941 | Bacteria>Terrabacteria_group>Actinomycetota>unclassified_Actinomycetota | GCA_028063615.1 | Actinomycetota_bacterium | MOZ30_20285 | 6559..9384 | 2900548 | MAG:-DUF499-domain-containing-protein-[Actinomycetota-bacterium] |
| MCL5942545.1 | AAA+ATPase+wHTH+Alpha+Beta-domain+wHTH+RRM | Type-1_BR.RM.like.systems | HISKIN+MORC-><-SIGMA-HTH<-?\|\|DDE_Tnp_1->DDE_Tnp_1-><-DUF499* | 979 | Bacteria>Terrabacteria_group>Actinomycetota>unclassified_Actinomycetota | GCA_023488925.1 | Actinomycetota_bacterium | M1325_03350 | 4054..6993 | 2900548 | DUF499-domain-containing-protein-[Actinobacteria-bacterium] |
| MBE3129082.1 | AAA+ATPase+wHTH+Alpha+Beta-domain+wHTH+RRM | Type-1_BR.RM.like.systems | <-DUF499*<-METHYLASE-p<-TUDOR+SF1-SF2-Helicase+iREase+CTD+REase | 311 | Bacteria>Terrabacteria_group>Actinomycetota>unclassified_Actinomycetota | GCA_014894335.1 | Actinomycetota_bacterium | IMZ60_05330 | 1..934 | 2900548 | ATP-binding-protein-[Actinomycetia-bacterium] |
| MCA1701756.1 | AAA+ATPase+wHTH+Alpha+Beta-domain+wHTH+RRM | Type-1_BR.RM.like.systems | <-DUF499*<-METHYLASE<-TUDOR+SF1-SF2-Helicase+iREase+CTD+REase-p | 799 | Bacteria>Terrabacteria_group>Actinomycetota>unclassified_Actinomycetota | GCA_020200655.1 | Actinomycetota_bacterium | LC808_00175 | 1..2398 | 2900548 | DUF499-domain-containing-protein-[Actinomycetia-bacterium] |
| MCL5942956.1 | AAA+ATPase+wHTH+Alpha+Beta-domain+wHTH+RRM | Type-1_BR.RM.like.systems | <-DUF499*<-METHYLASE-p | 957 | Bacteria>Terrabacteria_group>Actinomycetota>unclassified_Actinomycetota | GCA_023488925.1 | Actinomycetota_bacterium | M1325_05520 | 7262..10135 | 2900548 | DUF499-domain-containing-protein-[Actinobacteria-bacterium] |
| MDA3014853.1 | AAA+ATPase+wHTH+Alpha+Beta-domain+wHTH+RRM | Type-1_BR.RM.like.systems | <-rve+rve_3+Mu-transpos_C<-?\|\|?->TUDOR+SF1-SF2-Helicase+iREase+CTD+REase->METHYLASE->DUF499*->?-><-?<-?\|\|?-><-?\|\|ABC-ATPase+SbcC-> | 979 | Bacteria>Terrabacteria_group>Actinomycetota>unclassified_Actinomycetota | GCA_027730025.1 | Actinomycetota_bacterium | O3A89_04725 | 12267..15206 | 2900548 | MAG:-DUF499-domain-containing-protein-[Actinomycetota-bacterium] |
| WP_075162425.1 | AAA+ATPase+wHTH+Alpha+Beta-domain+wHTH+RRM | Type-1_BR.RM.like.systems | <-REase<-?\|\|HAD-><-?<-?\|\|?-><-DUF499*<-METHYLASE-p<-METHYLASE-p<-HNH<-SMS-KaiC<-TUDOR+SF1-SF2-Helicase+iREase+CTD+REase | 937 | Bacteria>Terrabacteria_group>Armatimonadota>Chthonomonadetes>Chthonomonadales | GCF_001457975.1 | Chthonomonas_calidirosea | CP488_RS03500 | 839037..841850 | 454171 | DUF499-domain-containing-protein-[Chthonomonas-calidirosea] |
| WP_075165136.1 | AAA+ATPase+wHTH+Alpha+Beta-domain+wHTH+RRM | Type-1_BR.RM.like.systems | <-REase<-?\|\|HAD-><-?<-?\|\|?-><-DUF499*<-METHYLASE-p<-METHYLASE<-TUDOR+SF1-SF2-Helicase+iREase+CTD+REase<-?<-?<-tRNA\|\|MORC-> | 937 | Bacteria>Terrabacteria_group>Armatimonadota>Chthonomonadetes>Chthonomonadales | GCF_001457855.1 | Chthonomonas_calidirosea | CWRG_RS03480 | 839017..841830 | 454171 | DUF499-domain-containing-protein-[Chthonomonas-calidirosea] |
| WP_016481846.1 | AAA+ATPase+wHTH+Alpha+Beta-domain+wHTH+RRM | Type-1_BR.RM.like.systems | TUDOR+SF1-SF2-Helicase+iREase+CTD+REase->SMS-KaiC->HNH->METHYLASE->DUF499*-><-?\|\|?->?-><-HAD\|\|?->REase-> | 937 | Bacteria>Terrabacteria_group>Armatimonadota>Chthonomonadetes>Chthonomonadales | GCF_000427095.1 | Chthonomonas_calidirosea | CCALI_RS02245 | 511927..514740 | 454171 | DUF499-domain-containing-protein-[Chthonomonas-calidirosea] |
| NSW79111.1 | AAA+ATPase+wHTH+Alpha+Beta-domain+wHTH+RRM | Type-1_BR.RM.like.systems | <-DUF499*<-METHYLASE<-TUDOR+SF1-SF2-Helicase+iREase+CTD+REase | 866 | Bacteria>Terrabacteria_group>Armatimonadota>Chthonomonadetes>unclassified_Chthonomonadetes | GCA_013314755.1 | Chthonomonadetes_bacterium | HPY54_08770 | 75702..78302 | 2282152 | ATP-binding-protein-[Chthonomonadetes-bacterium] |
| MCC7491466.1 | AAA+ATPase+wHTH+Alpha+Beta-domain+wHTH+RRM | Type-1_BR.RM.like.systems | METHYLASE-p->?->?->?->?->?->DUF499*-><-?<-ABC-ATPase+SbcC<-?\|\|N6_N4_MTase-> | 1453 | Bacteria>Terrabacteria_group>Armatimonadota>Fimbriimonadia>Fimbriimonadales | GCA_020850355.1 | Fimbriimonadaceae_bacterium | IT204_03925 | 23622..27983 | 2682144 | hypothetical-protein-[Fimbriimonadaceae-bacterium] |
| GIV04311.1 | AAA+ATPase+wHTH+Alpha+Beta-domain+wHTH+RRM | Type-1_BR.RM.like.systems | <-DUF499*<-HTH<-METHYLASE<-OLD-ABC+TOPRIM<-SF1-SF2-Helicase-p<-Schlafen+HATPase+HTH<-TUDOR+SF1-SF2-Helicase+iREase+CTD+REase | 944 | Bacteria>Terrabacteria_group>Armatimonadota>Fimbriimonadia>Fimbriimonadales | GCA_026003055.1 | Fimbriimonadales_bacterium | KatS3mg015_3141 | 322..3156 | 2682145 | MAG:-hypothetical-protein-KatS3mg015_3141-[Fimbriimonadales-bacterium] |
| GIV02394.1 | AAA+ATPase+wHTH+Alpha+Beta-domain+wHTH+RRM | Type-1_BR.RM.like.systems | <-URI-DNase+RAMA<-?<-HEPN+REase<-?<-REase<-Schlafen+HATPase+HTH<-DUF499*<-REase<-?<-METHYLASE-p\|\|?-><-?<-METHYLASE | 920 | Bacteria>Terrabacteria_group>Armatimonadota>Fimbriimonadia>Fimbriimonadales | GCA_026003055.1 | Fimbriimonadales_bacterium | KatS3mg015_1224 | 1249734..1252496 | 2682145 | MAG:-hypothetical-protein-KatS3mg015_1224-[Fimbriimonadales-bacterium] |
| MCS7301717.1 | AAA+ATPase+wHTH+Alpha+Beta-domain+wHTH+RRM | Type-1_BR.RM.like.systems | <-TUDOR+SF1-SF2-Helicase+iREase+CTD+REase-p<-DUF499*<-HEPN<-PolB<-HEPN<-PolB<-METHYLASE-p | 950 | Bacteria>Terrabacteria_group>Armatimonadota>Fimbriimonadia>Fimbriimonadales | GCA_025062205.1 | Fimbriimonadales_bacterium | NZ556_09225 | 1592..4444 | 2682145 | DUF499-domain-containing-protein-[Fimbriimonadales-bacterium] |
| MCX7689026.1 | AAA+ATPase+wHTH+Alpha+Beta-domain+wHTH+RRM | Type-1_BR.RM.like.systems | TUDOR+SF1-SF2-Helicase+iREase+CTD+REase-p->Schlafen+HATPase+HTH->METHYLASE->METHYLASE-p->DUF499*-> | 872 | Bacteria>Terrabacteria_group>Armatimonadota>Fimbriimonadia>Fimbriimonadales | GCA_026418845.1 | Fimbriimonadales_bacterium | N2045_13770 | 4406..7024 | 2682145 | MAG:-DUF499-domain-containing-protein-[Fimbriimonadales-bacterium] |
| GIV14563.1 | AAA+ATPase+wHTH+Alpha+Beta-domain+wHTH+RRM | Type-1_BR.RM.like.systems | Cluster1029_5clades-><-TUDOR+SF1-SF2-Helicase+iREase+CTD+REase\|\|?-><-?<-DUF499* | 798 | Bacteria>Terrabacteria_group>Armatimonadota>Fimbriimonadia>Fimbriimonadales | GCA_026003155.1 | Fimbriimonadales_bacterium | KatS3mg021_2845 | 184694..187090 | 2682145 | MAG:-hypothetical-protein-KatS3mg021_2845-[Fimbriimonadales-bacterium] |
| MBM3465143.1 | AAA+ATPase+wHTH+Alpha+Beta-domain+wHTH+RRM | Type-1_BR.RM.like.systems | DUF499*->Arm-DNA-bind_3+Phage_int_SAM_3+Phage_integrase-> | 700 | Bacteria>Terrabacteria_group>Armatimonadota>unclassified_Armatimonadota | GCA_016868975.1 | Armatimonadota_bacterium | FJX76_23850 | 1..2104 | 2033014 | ATP-binding-protein-[Armatimonadetes-bacterium] |
| MDE2206237.1 | AAA+ATPase+wHTH+Alpha+Beta-domain+wHTH+RRM | Type-1_BR.RM.like.systems | DUF499*->HEPN+REase->HTH->MPTase->?->HTH+UbE1->MarR-HTH-> | 600 | Bacteria>Terrabacteria_group>Armatimonadota>unclassified_Armatimonadota | GCA_028871005.1 | Armatimonadota_bacterium | KGK12_06315 | 1..1805 | 2033014 | MAG:-ATP-binding-protein,-partial-[Armatimonadetes-bacterium] |
| MBM3494074.1 | AAA+ATPase+wHTH+Alpha+Beta-domain+wHTH+RRM | Type-1_BR.RM.like.systems | <-REase+DUF4268<-DUF499* | 989 | Bacteria>Terrabacteria_group>Armatimonadota>unclassified_Armatimonadota | GCA_016869115.1 | Armatimonadota_bacterium | FJX72_07105 | 2132..5102 | 2033014 | ATP-binding-protein-[Armatimonadetes-bacterium] |
| MCC6485536.1 | AAA+ATPase+wHTH+Alpha+Beta-domain+wHTH+RRM | Type-1_BR.RM.like.systems | <-ABC-ATPase+SbcC<-ABC-ATPase+SbcC<-?\|\|Phage_int_SAM_4+Phage_integrase->TUDOR+SF1-SF2-Helicase+iREase+CTD+REase->METHYLASE->DUF499*->MNS+HTH+REase-> | 923 | Bacteria>Terrabacteria_group>Armatimonadota>unclassified_Armatimonadota | GCA_020846715.1 | Armatimonadota_bacterium | IT209_11875 | 12135..14906 | 2033014 | ATP-binding-protein-[Armatimonadetes-bacterium] |
| GIV15264.1 | AAA+ATPase+wHTH+Alpha+Beta-domain+wHTH+RRM | Type-1_BR.RM.like.systems | TUDOR+SF1-SF2-Helicase+iREase+CTD+REase->?->Schlafen+HATPase+HTH->Schlafen+HATPase+HTH->METHYLASE-p->METHYLASE-p->DUF499*-> | 873 | Bacteria>Terrabacteria_group>Armatimonadota>unclassified_Armatimonadota | GCA_026003175.1 | Armatimonadota_bacterium | KatS3mg022_0699 | 793472..796093 | 2033014 | MAG:-hypothetical-protein-KatS3mg022_0699-[Armatimonadota-bacterium] |
| NIM68640.1 | AAA+ATPase+wHTH+Alpha+Beta-domain+wHTH+RRM | Type-1_BR.RM.like.systems | TUDOR+SF1-SF2-Helicase+iREase+CTD+REase->NTP_transf->REase->METHYLASE->DUF499*->DNA-glycosylase-> | 918 | Bacteria>Terrabacteria_group>Armatimonadota>unclassified_Armatimonadota | GCA_011771675.1 | Armatimonadota_bacterium | GTO55_11015 | 8511..11267 | 2033014 | DUF499-domain-containing-protein-[Armatimonadetes-bacterium] |
| MCS7225146.1 | AAA+ATPase+wHTH+Alpha+Beta-domain+wHTH+RRM | Type-1_BR.RM.like.systems | <-DUF499*<-METHYLASE<-?<-TUDOR+SF1-SF2-Helicase+iREase+CTD+REase\|\|HHH+ENDO3+NUDIX->HDRnase-> | 870 | Bacteria>Terrabacteria_group>Armatimonadota>unclassified_Armatimonadota | GCA_025057575.1 | Armatimonadota_bacterium | NZ959_11445 | 842..3454 | 2033014 | DUF499-domain-containing-protein-[Armatimonadota-bacterium] |
| PIU67978.1 | AAA+ATPase+wHTH+Alpha+Beta-domain+wHTH+RRM | Type-1_BR.RM.like.systems | HEPN-p->METHYLASE->DUF499*->?-><-?<-?<-SF1-SF2-Helicase<-Schlafen+HATPase+HTH | 944 | Bacteria>Terrabacteria_group>Armatimonadota>unclassified_Armatimonadota | GCA_002779575.1 | Armatimonadetes_bacterium_CG07_land_8_20_14_0_80_40_9 | COS84_02435 | 3211..6045 | 1973916 | AAA-family-ATPase-[Armatimonadetes-bacterium-CG07_land_8_20_14_0_80_40_9] |
| NIM06025.1 | AAA+ATPase+wHTH+Alpha+Beta-domain+wHTH+RRM | Type-1_BR.RM.like.systems | <-DNA-glycosylase<-DUF499*<-REase<-METHYLASE-p | 976 | Bacteria>Terrabacteria_group>Armatimonadota>unclassified_Armatimonadota | GCA_011771275.1 | Armatimonadota_bacterium | GTO78_10545 | 837..3767 | 2033014 | DUF499-domain-containing-protein-[Armatimonadetes-bacterium] |
| MCL6533304.1 | AAA+ATPase+wHTH+Alpha+Beta-domain+wHTH+RRM | Type-1_BR.RM.like.systems | TOPRIM+ZnR->METHYLASE->DUF499*->TUDOR+SF1-SF2-Helicase+iREase+CTD+REase-p-> | 880 | Bacteria>Terrabacteria_group>Armatimonadota>unclassified_Armatimonadota | GCA_023511975.1 | Armatimonadota_bacterium | K6U12_08540 | 25790..28432 | 2033014 | DUF499-domain-containing-protein-[Armatimonadetes-bacterium] |
| NLB94469.1 | AAA+ATPase+wHTH+Alpha+Beta-domain+wHTH+RRM | Type-1_BR.RM.like.systems | REase->HEPN->METHYLASE->DUF499*->DNA-glycosylase-> | 991 | Bacteria>Terrabacteria_group>Armatimonadota>unclassified_Armatimonadota | GCA_012512265.1 | Armatimonadota_bacterium | GX785_02050 | 5194..8169 | 2033014 | DUF499-domain-containing-protein-[Armatimonadetes-bacterium] |
| MCA1947578.1 | AAA+ATPase+wHTH+Alpha+Beta-domain+wHTH+RRM | Type-1_BR.RM.like.systems | URI-DNase-><-DUF499*<-HEPN<-PolB<-METHYLASE | 919 | Bacteria>Terrabacteria_group>Armatimonadota>unclassified_Armatimonadota | GCA_020217215.1 | Armatimonadota_bacterium | LDL55_05595 | 1010..3769 | 2033014 | DUF499-domain-containing-protein-[Armatimonadetes-bacterium] |
| MCX7968313.1 | AAA+ATPase+wHTH+Alpha+Beta-domain+wHTH+RRM | Type-1_BR.RM.like.systems | TUDOR+SF1-SF2-Helicase+iREase+CTD+REase->DUF499*-> | 1026 | Bacteria>Terrabacteria_group>Armatimonadota>unclassified_Armatimonadota | GCA_026416275.1 | Armatimonadota_bacterium | N3B10_07465 | 4197..7277 | 2033014 | MAG:-DUF499-domain-containing-protein-[Armatimonadetes-bacterium] |
| MCL6520426.1 | AAA+ATPase+wHTH+Alpha+Beta-domain+wHTH+RRM | Type-1_BR.RM.like.systems | METHYLASE-p->DUF499*-> | 875 | Bacteria>Terrabacteria_group>Armatimonadota>unclassified_Armatimonadota | GCA_023511795.1 | Armatimonadota_bacterium | K6T99_11405 | 1472..4099 | 2033014 | DUF499-domain-containing-protein-[Armatimonadetes-bacterium] |
| MCX6379001.1 | AAA+ATPase+wHTH+Alpha+Beta-domain+wHTH+RRM | Type-1_BR.RM.like.systems | <-DUF499*<-Schlafen+HATPase+HTH<-METHYLASE<-MORC+DUF3684+REase<-TUDOR+SF1-SF2-Helicase+iREase+CTD+REase<-?\|\|XIS-HTH-> | 625 | Bacteria>Terrabacteria_group>Armatimonadota>unclassified_Armatimonadota | GCA_026392135.1 | Armatimonadota_bacterium | NTO23_05910 | 1..1875 | 2033014 | MAG:-DUF499-domain-containing-protein,-partial-[Armatimonadetes-bacterium] |
| PIY44270.1 | AAA+ATPase+wHTH+Alpha+Beta-domain+wHTH+RRM | Type-1_BR.RM.like.systems | METHYLASE->DUF433->PIN->DUF499*-> | 318 | Bacteria>Terrabacteria_group>Armatimonadota>unclassified_Armatimonadota | GCA_002784925.1 | Armatimonadetes_bacterium_CG_4_10_14_3_um_filter_59_10 | COZ05_08660 | 3346..4299 | 1973909 | hypothetical-protein-COZ05_08660,-partial-[Armatimonadetes-bacterium-CG_4_10_14_3_um_filter_59_10] |
| WP_237700092.1 | AAA+ATPase+wHTH+Alpha+Beta-domain+wHTH+RRM | Type-1_BR.RM.like.systems | TUDOR+SF1-SF2-Helicase+iREase+CTD+REase->METHYLASE-p->DUF4277+DDE_Tnp_1->METHYLASE->DUF499*-><-?<-HIN-HTH<-DUF772+DDE_Tnp_1+DDE_Tnp_1_6<-?\|\|?->GNTR-HTH-p-> | 508 | Bacteria>Terrabacteria_group>Bacillota>Bacilli>Bacillales | GCF_000219875.1 | Alicyclobacillus_acidocaldarius | TC41_RS05325 | 1151814..1153340 | 405212 | DUF499-domain-containing-protein-[Alicyclobacillus-acidocaldarius] |
| WP_141336163.1 | AAA+ATPase+wHTH+Alpha+Beta-domain+wHTH+RRM | Type-1_BR.RM.like.systems | <-TPR<-DUF2357+REase<-?<-?<-?<-DUF499*<-METHYLASE<-TUDOR+SF1-SF2-Helicase+iREase+CTD+REase<-?<-SF1-SF2-Helicase+REase<-Exonuc_V_gamma+REase | 951 | Bacteria>Terrabacteria_group>Bacillota>Bacilli>Bacillales | GCF_006519665.1 | Paenibacillus_sp._tmac-D7 | FKP33_RS19830 | 6433..9288 | 2591462 | DUF499-domain-containing-protein-[Paenibacillus-sp.-tmac-D7] |
| WP_119798229.1 | AAA+ATPase+wHTH+Alpha+Beta-domain+wHTH+RRM | Type-1_BR.RM.like.systems | <-Phage_integrase<-REase<-DUF499*<-METHYLASE<-TUDOR+SF1-SF2-Helicase+iREase+CTD+REase\|\|TraJ-RHH-><-?<-HerA/FtsK | 952 | Bacteria>Terrabacteria_group>Bacillota>Bacilli>Bacillales | GCF_003591485.1 | Paenibacillus_sp._1011MAR3C5 | D3P07_RS00960 | 194996..197854 | 1675787 | DUF499-domain-containing-protein-[Paenibacillus-sp.-1011MAR3C5] |
| WP_218917114.1 | AAA+ATPase+wHTH+Alpha+Beta-domain+wHTH+RRM | Type-1_BR.RM.like.systems | Cysteine_desulfurase->?->tRNA->tRNA->tRNA->TUDOR+SF1-SF2-Helicase+iREase+CTD+REase->METHYLASE->DUF499*->?->HerA/FtsK-><-DDE_Tnp_1 | 961 | Bacteria>Terrabacteria_group>Bacillota>Bacilli>Bacillales | GCF_000024285.1 | Alicyclobacillus_acidocaldarius | AACI_RS06130 | 1286262..1289147 | 405212 | DUF499-domain-containing-protein-[Alicyclobacillus-acidocaldarius] |
| ACV58265.1 | AAA+ATPase+wHTH+Alpha+Beta-domain+wHTH+RRM | Type-1_BR.RM.like.systems | Cysteine_desulfurase->?->tRNA->tRNA->tRNA->TUDOR+SF1-SF2-Helicase+iREase+CTD+REase->METHYLASE->DUF499*->?->HerA/FtsK-><-DDE_Tnp_1 | 959 | Bacteria>Terrabacteria_group>Bacillota>Bacilli>Bacillales | GCA_000024285.1 | Alicyclobacillus_acidocaldarius_subsp._acidocaldarius_DSM_446 | Aaci_1236 | 1286268..1289147 | 521098 | conserved-hypothetical-protein-[Alicyclobacillus-acidocaldarius-subsp.-acidocaldarius-DSM-446] |
| WP_082451901.1 | AAA+ATPase+wHTH+Alpha+Beta-domain+wHTH+RRM | Type-1_BR.RM.like.systems | <-REase<-DUF499*<-METHYLASE<-TUDOR+SF1-SF2-Helicase+iREase+CTD+REase<-?<-?<-?\|\|HTH_3-> | 838 | Bacteria>Terrabacteria_group>Bacillota>Bacilli>Bacillales | GCF_902375745.1 | Paenibacillus_ihuae | FY521_RS18285 | 3939778..3942291 | 1232431 | DUF499-domain-containing-protein,-partial-[Paenibacillus-ihuae] |
| WP_269447895.1 | AAA+ATPase+wHTH+Alpha+Beta-domain+wHTH+RRM | Type-1_BR.RM.like.systems | <-GNTR-HTH-p<-?<-?<-?<-?<-SMODS<-DUF499*<-METHYLASE | 902 | Bacteria>Terrabacteria_group>Bacillota>Bacilli>Bacillales | - | Alicyclobacillus_mali_(ex_Roth_et_al._2021) | AMA01_RS08555 | 91717..94425 | 1123961 | DUF499-domain-containing-protein-[Alicyclobacillus-mali-(ex-Roth-et-al.-2021)] |
| HHW19260.1 | AAA+ATPase+wHTH+Alpha+Beta-domain+wHTH+RRM | Type-1_BR.RM.like.systems | TUDOR+SF1-SF2-Helicase+iREase+CTD+REase->SbcC->METHYLASE->REase->DUF499*->DUF433->PIN-> | 932 | Bacteria>Terrabacteria_group>Bacillota>Candidatus_Fermentithermobacillia>Candidatus_Fermentithermobacillales | GCA_012840505.1 | Candidatus_Fermentithermobacillaceae_bacterium | GXX30_10280 | 57551..60349 | 3085339 | ATP-binding-protein-[Firmicutes-bacterium] |
| WP_013432067.1 | AAA+ATPase+wHTH+Alpha+Beta-domain+wHTH+RRM | Type-1_BR.RM.like.systems | MazE_antitoxin->?->\|\|REase+NACHT->REase+NACHT->TUDOR+SF1-SF2-Helicase+iREase+CTD+REase->METHYLASE->DUF499*->ParB+HNH-p->DDE_Tnp_1->ParB+HNH-p-> | 913 | Bacteria>Terrabacteria_group>Bacillota>Clostridia>Caldicellulosiruptorales | GCF_000166695.1 | Caldicellulosiruptor_acetigenus | CALKR_RS03530 | 770817..773558 | 301953 | DUF499-domain-containing-protein-[Caldicellulosiruptor-acetigenus] |
| NSW83680.1 | AAA+ATPase+wHTH+Alpha+Beta-domain+wHTH+RRM | Type-1_BR.RM.like.systems | <-DDE_Tnp\|\|HIN-HTH-><-DDE_Tnp_1<-NACHT+REase<-DUF499*<-METHYLASE<-TUDOR+SF1-SF2-Helicase+iREase+CTD+REase<-PrlF_antitoxin | 905 | Bacteria>Terrabacteria_group>Bacillota>Clostridia>Eubacteriales | GCA_013314835.1 | Syntrophothermus_sp. | HPY90_10510 | 22691..25408 | 2736299 | ATP-binding-protein-[Syntrophothermus-sp.] |
| MDD2509910.1 | AAA+ATPase+wHTH+Alpha+Beta-domain+wHTH+RRM | Type-1_BR.RM.like.systems | SF1-SF2-Helicase+REase->?->TUDOR+SF1-SF2-Helicase+iREase+CTD+REase->METHYLASE->RNase->DUF499*->?->?->?->ZDNA-HTH-><-?\|\|HTH+MPTase-> | 977 | Bacteria>Terrabacteria_group>Bacillota>Clostridia>Eubacteriales | GCA_028685375.1 | Syntrophomonas_sp. | PHG94_02110 | 14056..16989 | 2053627 | MAG:-DUF499-domain-containing-protein-[Syntrophomonas-sp.] |
| BAF59044.1 | AAA+ATPase+wHTH+Alpha+Beta-domain+wHTH+RRM | Type-1_BR.RM.like.systems | TUDOR+SF1-SF2-Helicase+iREase+CTD+REase->?->METHYLASE->HEPN->HEPN+Mrr_N->DUF499*-><-?\|\|?->?->DNA_pol3_delta+DNA_pol3_gamma3-> | 885 | Bacteria>Terrabacteria_group>Bacillota>Clostridia>Eubacteriales | GCA_000010565.1 | Pelotomaculum_thermopropionicum_SI | PTH_0863 | 877158..879815 | 370438 | predicted-ATPase-[Pelotomaculum-thermopropionicum-SI] |
| WP_012301643.1 | AAA+ATPase+wHTH+Alpha+Beta-domain+wHTH+RRM | Type-1_BR.RM.like.systems | REase->TUDOR+SF1-SF2-Helicase+iREase+CTD+REase->?->OLD-ABC+TOPRIM->METHYLASE->REase->DUF499*->RNase->?->?->?->HicB_lk_antitox-> | 924 | Bacteria>Terrabacteria_group>Bacillota>Clostridia>Eubacteriales | GCF_000018425.1 | Candidatus_Desulforudis_audaxviator | Daudx_RS02685 | 550555..553329 | 471827 | DUF499-domain-containing-protein-[Candidatus-Desulforudis-audaxviator] |
| WP_027356905.1 | AAA+ATPase+wHTH+Alpha+Beta-domain+wHTH+RRM | Type-1_BR.RM.like.systems | TUDOR+SF1-SF2-Helicase+iREase+CTD+REase->METHYLASE->DUF499*-><-Polbeta<-PIN<-PhdYeFM_antitox\|\|?->HIN-HTH-> | 896 | Bacteria>Terrabacteria_group>Bacillota>Clostridia>Eubacteriales | GCF_000686645.1 | Desulfofundulus_thermocisternus | BR83_RS0109605 | 50288..52978 | 42471 | DUF499-domain-containing-protein-[Desulfofundulus-thermocisternus] |
| HHW42091.1 | AAA+ATPase+wHTH+Alpha+Beta-domain+wHTH+RRM | Type-1_BR.RM.like.systems | <-Phage_integrase\|\|?->TUDOR+SF1-SF2-Helicase+iREase+CTD+REase->METHYLASE->DUF499*->?->?-><-?\|\|?->?-><-LD-peptidase | 911 | Bacteria>Terrabacteria_group>Bacillota>Clostridia>Eubacteriales | GCA_012840385.1 | Syntrophomonadaceae_bacterium | GXX19_13315 | 8270..11005 | 2093811 | ATP-binding-protein-[Syntrophomonadaceae-bacterium] |
| WP_204488211.1 | AAA+ATPase+wHTH+Alpha+Beta-domain+wHTH+RRM | Type-1_BR.RM.like.systems | <-HAMP+HISKIN<-?\|\|?->tRNA->NUDIX->TUDOR+SF1-SF2-Helicase+iREase+CTD+REase->METHYLASE->DUF499*->?->?->?->?->?-><-HTH+DSBH | 906 | Bacteria>Terrabacteria_group>Bacillota>Clostridia>Eubacteriales | GCF_016908415.1 | Caldicoprobacter_guelmensis | JOD02_RS06990 | 180683..183403 | 1170224 | DUF499-domain-containing-protein-[Caldicoprobacter-guelmensis] |
| WP_011641394.1 | AAA+ATPase+wHTH+Alpha+Beta-domain+wHTH+RRM | Type-1_BR.RM.like.systems | DDE_Tnp_1-><-?<-MPTase<-HTH<-?<-?<-DUF499*<-?<-METHYLASE<-TUDOR+SF1-SF2-Helicase+iREase+CTD+REase | 982 | Bacteria>Terrabacteria_group>Bacillota>Clostridia>Eubacteriales | GCF_000014725.1 | Syntrophomonas_wolfei | SWOL_RS10375 | 2293555..2296503 | 863 | DUF499-domain-containing-protein-[Syntrophomonas-wolfei] |
| BAF60584.1 | AAA+ATPase+wHTH+Alpha+Beta-domain+wHTH+RRM | Type-1_BR.RM.like.systems | DDE_Tnp_IS66+DDE_Tnp_IS66_C-><-?<-?<-?<-?<-?<-DUF499*<-METHYLASE\|\|PAIREDC-HTH->DDE_5-> | 884 | Bacteria>Terrabacteria_group>Bacillota>Clostridia>Eubacteriales | GCA_000010565.1 | Pelotomaculum_thermopropionicum_SI | PTH_2403 | 2504109..2506763 | 370438 | predicted-ATPase-[Pelotomaculum-thermopropionicum-SI] |
| MBO8129694.1 | AAA+ATPase+wHTH+Alpha+Beta-domain+wHTH+RRM | Type-1_BR.RM.like.systems | TUDOR+SF1-SF2-Helicase+iREase+CTD+REase->ParB+HNH->METHYLASE->DUF499*-> | 984 | Bacteria>Terrabacteria_group>Bacillota>Clostridia>Eubacteriales | GCA_017656185.1 | Peptococcaceae_bacterium | HOZ39_10965 | 8170..11124 | 2052179 | DUF499-domain-containing-protein-[Peptococcaceae-bacterium] |
| NPV74526.1 | AAA+ATPase+wHTH+Alpha+Beta-domain+wHTH+RRM | Type-1_BR.RM.like.systems | TUDOR+SF1-SF2-Helicase+iREase+CTD+REase->?->METHYLASE->DUF499*-> | 885 | Bacteria>Terrabacteria_group>Bacillota>Clostridia>Eubacteriales | GCA_013178385.1 | Pelotomaculum_sp. | HPY89_12310 | 13139..15796 | 2053590 | DUF499-domain-containing-protein-[Pelotomaculum-sp.] |
| NLU51029.1 | AAA+ATPase+wHTH+Alpha+Beta-domain+wHTH+RRM | Type-1_BR.RM.like.systems | SF1-SF2-Helicase->?->?-><-DUF499*<-METHYLASE | 977 | Bacteria>Terrabacteria_group>Bacillota>Clostridia>Eubacteriales | GCA_012719535.1 | Syntrophomonadaceae_bacterium | GXX09_11645 | 5429..8362 | 2093811 | ATP-binding-protein-[Syntrophomonadaceae-bacterium] |
| MDD3890320.1 | AAA+ATPase+wHTH+Alpha+Beta-domain+wHTH+RRM | Type-1_BR.RM.like.systems | <-DUF499-p<-DUF499*<-?<-METHYLASE | 430 | Bacteria>Terrabacteria_group>Bacillota>Clostridia>Eubacteriales | GCA_028699095.1 | Syntrophomonadaceae_bacterium | PHR65_10455 | 753..2045 | 2093811 | MAG:-DUF499-domain-containing-protein-[Syntrophomonadaceae-bacterium] |
| NLE27663.1 | AAA+ATPase+wHTH+Alpha+Beta-domain+wHTH+RRM | Type-1_BR.RM.like.systems | TPR+TPR->?-><-?<-?<-?<-DUF499*<-METHYLASE | 988 | Bacteria>Terrabacteria_group>Bacillota>Clostridia>Eubacteriales | GCA_012514215.1 | Clostridiaceae_bacterium | GX625_20470 | 9503..12469 | 1898204 | ATP-binding-protein-[Clostridiaceae-bacterium] |
| HCW51868.1 | AAA+ATPase+wHTH+Alpha+Beta-domain+wHTH+RRM | Type-1_BR.RM.like.systems | <-DUF499*<-METHYLASE<-TUDOR+SF1-SF2-Helicase+iREase+CTD+REase | 936 | Bacteria>Terrabacteria_group>Bacillota>Clostridia>Eubacteriales | GCA_003541445.1 | Clostridiales_bacterium | DGR79_07395 | 1..2808 | 1898207 | hypothetical-protein-[Clostridiales-bacterium] |
| WP_169100840.1 | AAA+ATPase+wHTH+Alpha+Beta-domain+wHTH+RRM | Type-1_BR.RM.like.systems | LuxR-HTH->?->?->?-><-?<-DUF499-p<-DUF499*<-METHYLASE-p | 856 | Bacteria>Terrabacteria_group>Bacillota>Clostridia>Eubacteriales | GCF_012933365.1 | Sulfobacillus_harzensis | HIJ39_RS14250 | 21480..24050 | 2729629 | DUF499-domain-containing-protein-[Sulfobacillus-sp.-DSM-109850] |
| WP_054935692.1 | AAA+ATPase+wHTH+Alpha+Beta-domain+wHTH+RRM | Type-1_BR.RM.like.systems | <-HIN-HTH<-?<-?<-?<-SLATT<-SMODS<-DUF499*<-METHYLASE<-TUDOR+SF1-SF2-Helicase+iREase+CTD+REase<-Phage_int_SAM_4+Phage_integrase\|\|?->REase-> | 951 | Bacteria>Terrabacteria_group>Bacillota>Clostridia>Moorellales | - | Moorella_glycerini | 487 | 3912..6767 | 55779 | DUF499-domain-containing-protein-[Moorella-glycerini] |
| CEP66429.1 | AAA+ATPase+wHTH+Alpha+Beta-domain+wHTH+RRM | Type-1_BR.RM.like.systems | <-HIN-HTH<-?<-?<-?<-SLATT<-SMODS<-DUF499*<-METHYLASE<-TUDOR+SF1-SF2-Helicase+iREase+CTD+REase<-Phage_int_SAM_4+Phage_integrase\|\|?->REase-> | 951 | Bacteria>Terrabacteria_group>Bacillota>Clostridia>Moorellales | GCA_001373375.1 | Moorella_glycerini | 487 | 3912..6767 | 55779 | Protein-of-unknown-function-DUF499-[Moorella-glycerini] |
| WP_088552559.1 | AAA+ATPase+wHTH+Alpha+Beta-domain+wHTH+RRM | Type-1_BR.RM.like.systems | HKD-DNase+SF1-SF2-Helicase+TRD->?->?->TUDOR+SF1-SF2-Helicase+iREase+CTD+REase->METHYLASE->DUF499*->ParB+HNH->?->PIN-><-35exo | 850 | Bacteria>Terrabacteria_group>Bacillota>Clostridia>Moorellales | GCF_002207765.1 | Calderihabitans_maritimus | KKC1_RS00515 | 36542..39094 | 1246530 | DUF499-domain-containing-protein-[Calderihabitans-maritimus] |
| WP_227763399.1 | AAA+ATPase+wHTH+Alpha+Beta-domain+wHTH+RRM | Type-1_BR.RM.like.systems | <-ArsR-HTH<-?<-?<-ArsR-HTH<-?<-NACHT+REase<-DUF499*<-METHYLASE<-TUDOR+SF1-SF2-Helicase+iREase+CTD+REase<-PrlF_antitoxin | 905 | Bacteria>Terrabacteria_group>Bacillota>Clostridia>Moorellales | GCF_004353525.1 | Zhaonella_formicivorans | EYS13_RS12840 | 2618315..2621032 | 2528593 | DUF499-domain-containing-protein-[Zhaonella-formicivorans] |
| WP_196768295.1 | AAA+ATPase+wHTH+Alpha+Beta-domain+wHTH+RRM | Type-1_BR.RM.like.systems | TUDOR+SF1-SF2-Helicase+iREase+CTD+REase->OLD-ABC->METHYLASE->DUF499*-><-?<-?\|\|MazE_antitoxin->ParB+HNH->URI-DNase-> | 989 | Bacteria>Terrabacteria_group>Bacillota>Clostridia>Moorellales | GCF_001874605.1 | Moorella_thermoacetica | MTJW_RS02925 | 594709..597678 | 1525 | DUF499-domain-containing-protein-[Moorella-thermoacetica] |
| WP_106007145.1 | AAA+ATPase+wHTH+Alpha+Beta-domain+wHTH+RRM | Type-1_BR.RM.like.systems | <-DUF499*<-METHYLASE<-TUDOR+SF1-SF2-Helicase+iREase+CTD+REase<-Phage_int_SAM_4+Phage_integrase\|\|?->REase-> | 857 | Bacteria>Terrabacteria_group>Bacillota>Clostridia>Moorellales | GCF_002995805.1 | Moorella_stamsii | MOST_RS10850 | 1..2573 | 1266720 | DUF499-domain-containing-protein,-partial-[Moorella-stamsii] |
| WP_015050308.1 | AAA+ATPase+wHTH+Alpha+Beta-domain+wHTH+RRM | Type-1_BR.RM.like.systems | <-MPTase<-XIS-HTH<-?<-DUF499*<-METHYLASE<-TUDOR+SF1-SF2-Helicase+iREase+CTD+REase | 909 | Bacteria>Terrabacteria_group>Bacillota>Clostridia>Thermoanaerobacterales | GCF_000305935.1 | Thermacetogenium_phaeum | THEPH_RS0106005 | 1147808..1150537 | 85874 | DUF499-domain-containing-protein-[Thermacetogenium-phaeum] |
| KUK37245.1 | AAA+ATPase+wHTH+Alpha+Beta-domain+wHTH+RRM | Type-1_BR.RM.like.systems | <-MPTase<-XIS-HTH<-?<-DUF499*<-METHYLASE<-TUDOR+SF1-SF2-Helicase+iREase+CTD+REase | 909 | Bacteria>Terrabacteria_group>Bacillota>Clostridia>Thermoanaerobacterales | GCA_001508215.1 | Thermacetogenium_phaeum | XD66_0054 | 8673..11402 | 85874 | ATPase-[Thermacetogenium-phaeum] |
| HHW04285.1 | AAA+ATPase+wHTH+Alpha+Beta-domain+wHTH+RRM | Type-1_BR.RM.like.systems | REase->METHYLASE->DUF499*-> | 910 | Bacteria>Terrabacteria_group>Bacillota>Clostridia>Thermoanaerobacterales | GCA_012840135.1 | Thermoanaerobacterales_bacterium | GXX35_16055 | 4069..6801 | 2304039 | ATP-binding-protein-[Thermoanaerobacterales-bacterium] |
| MCG0276507.1 | AAA+ATPase+wHTH+Alpha+Beta-domain+wHTH+RRM | Type-1_BR.RM.like.systems | <-DUF499*<-METHYLASE<-TUDOR+SF1-SF2-Helicase+iREase+CTD+REase | 905 | Bacteria>Terrabacteria_group>Bacillota>Clostridia>Thermosediminibacterales | GCA_021890755.1 | Thermosediminibacteraceae_bacterium | L5655_10185 | 1667..4384 | 2885163 | DUF499-domain-containing-protein-[Thermosediminibacteraceae-bacterium] |
| MBS3948798.1 | AAA+ATPase+wHTH+Alpha+Beta-domain+wHTH+RRM | Type-1_BR.RM.like.systems | <-SF1-SF2-Helicase<-Schlafen+HATPase+HTH<-?<-RelE-ParE<-Phd-YefM<-DUF499* | 711 | Bacteria>Terrabacteria_group>Bacillota>Dethiobacteria>Dethiobacterales | GCA_018333395.1 | Dethiobacter_sp. | KGZ57_10990 | 89378..91513 | 2093370 | DUF499-domain-containing-protein-[Dethiobacter-sp.] |
| WP_278674830.1 | AAA+ATPase+wHTH+Alpha+Beta-domain+wHTH+RRM | Type-1_BR.RM.like.systems | <-REase<-?<-?<-SIGMA-HTH<-?<-HTH_3<-DUF499*<-METHYLASE<-METHYLASE | 865 | Bacteria>Terrabacteria_group>Bacillota>Negativicutes>Acidaminococcales | - | Acidaminococcus_fermentans | QDI05_RS06240 | 10767..13364 | 905 | DUF499-domain-containing-protein-[Acidaminococcus-fermentans] |
| MBE8949964.1 | AAA+ATPase+wHTH+Alpha+Beta-domain+wHTH+RRM | Type-1_BR.RM.like.systems | ABC-ATPase+SbcC->?->?->ABC-ATPase+SbcC-><-?\|\|?-><-DUF499*<-METHYLASE<-?<-TUDOR+SF1-SF2-Helicase+iREase+CTD+REase\|\|?-><-ABC-ATPase+SbcC | 915 | Bacteria>Terrabacteria_group>Bacillota>Negativicutes>Negativicutes_incertae_sedis | GCA_015206825.1 | Quinella_sp._3Q1 | SR3Q1_05115 | 63786..66533 | 1958791 | ATP-binding-protein-[Quinella-sp.-3Q1] |
| MCR5834671.1 | AAA+ATPase+wHTH+Alpha+Beta-domain+wHTH+RRM | Type-1_BR.RM.like.systems | TUDOR+SF1-SF2-Helicase+iREase+CTD+REase->METHYLASE->DUF499*-><-?<-RADICAL-SAM<-ABC-ATPase+SbcC\|\|?->PRTase->SbcC+SbcC-> | 931 | Bacteria>Terrabacteria_group>Bacillota>Negativicutes>Selenomonadales | GCA_024698045.1 | Selenomonadaceae_bacterium | K6G55_08515 | 16320..19115 | 2652289 | DUF499-domain-containing-protein-[Selenomonadaceae-bacterium] |
| MBR1646577.1 | AAA+ATPase+wHTH+Alpha+Beta-domain+wHTH+RRM | Type-1_BR.RM.like.systems | <-REase\|\|ncRNA->?->?->?->SF1-SF2-Helicase+HNH-><-DUF499*<-METHYLASE<-?<-TUDOR+SF1-SF2-Helicase+iREase+CTD+REase\|\|?->METHYLASE-> | 911 | Bacteria>Terrabacteria_group>Bacillota>Negativicutes>Selenomonadales | GCA_017470715.1 | Selenomonadaceae_bacterium | IJ685_07350 | 8677..11412 | 2652289 | ATP-binding-protein-[Selenomonadaceae-bacterium] |
| MBR1645373.1 | AAA+ATPase+wHTH+Alpha+Beta-domain+wHTH+RRM | Type-1_BR.RM.like.systems | <-CopG_antitoxin\|\|?-><-GDResolvase+Recombinase+ZnR\|\|Transposase_31-><-DUF499*<-METHYLASE\|\|DnaB+SMS-KaiC-> | 913 | Bacteria>Terrabacteria_group>Bacillota>Negativicutes>Selenomonadales | GCA_017470715.1 | Selenomonadaceae_bacterium | IJ685_01220 | 10518..13259 | 2652289 | ATP-binding-protein-[Selenomonadaceae-bacterium] |
| MBR6888673.1 | AAA+ATPase+wHTH+Alpha+Beta-domain+wHTH+RRM | Type-1_BR.RM.like.systems | ABC-ATPase+SbcC->?->?->ABC-ATPase+SbcC-><-?\|\|?-><-DUF499*<-METHYLASE | 915 | Bacteria>Terrabacteria_group>Bacillota>Negativicutes>Selenomonadales | GCA_017529935.1 | Selenomonadaceae_bacterium | IKN16_09550 | 12326..15073 | 2652289 | ATP-binding-protein-[Selenomonadaceae-bacterium] |
| MBR3747426.1 | AAA+ATPase+wHTH+Alpha+Beta-domain+wHTH+RRM | Type-1_BR.RM.like.systems | METHYLASE-p->DUF499*->DUF499-p-> | 478 | Bacteria>Terrabacteria_group>Bacillota>Negativicutes>Selenomonadales | GCA_017529545.1 | Selenomonadaceae_bacterium | IKN27_10770 | 501..1937 | 2652289 | ATP-binding-protein-[Selenomonadaceae-bacterium] |
| NCB76905.1 | AAA+ATPase+wHTH+Alpha+Beta-domain+wHTH+RRM | Type-1_BR.RM.like.systems | Ntox49->?->Resolvase->Recombinase+ZnR->?->PIN+OST-HTH-><-DUF499*<-TOPRIM<-OLD-ABC<-METHYLASE<-TUDOR+SF1-SF2-Helicase+iREase+CTD+REase<-?\|\|NACHT-> | 964 | Bacteria>Terrabacteria_group>Bacillota>Negativicutes>unclassified_Negativicutes | GCA_009929485.1 | Negativicutes_bacterium | EOM42_07415 | 5128..8022 | 2499140 | ATP-binding-protein-[Negativicutes-bacterium] |
| NLG79149.1 | AAA+ATPase+wHTH+Alpha+Beta-domain+wHTH+RRM | Type-1_BR.RM.like.systems | <-XIS-HTH+RVT\|\|?->?-><-REase<-DUF499*<-METHYLASE<-TUDOR+SF1-SF2-Helicase+iREase+CTD+REase | 946 | Bacteria>Terrabacteria_group>Bacillota>unclassified_Bacillota | GCA_012516125.1 | Bacillota_bacterium | GX492_04905 | 11356..14196 | 1879010 | ATP-binding-protein-[Firmicutes-bacterium] |
| NPV53644.1 | AAA+ATPase+wHTH+Alpha+Beta-domain+wHTH+RRM | Type-1_BR.RM.like.systems | ParB+HNH->?->TUDOR+SF1-SF2-Helicase+iREase+CTD+REase->METHYLASE->DUF499*->ACET+PIN+ACET->ASCH->HEPN-><-?<-Phage_integrase | 897 | Bacteria>Terrabacteria_group>Bacillota>unclassified_Bacillota | GCA_013178125.1 | Bacillota_bacterium | HPY71_08975 | 9373..12066 | 1879010 | ATP-binding-protein-[Firmicutes-bacterium] |
| HHV78173.1 | AAA+ATPase+wHTH+Alpha+Beta-domain+wHTH+RRM | Type-1_BR.RM.like.systems | DNA-glycosylase->REase->PrlF_antitoxin->TUDOR+SF1-SF2-Helicase+iREase+CTD+REase->METHYLASE->DUF499*->NACHT+REase-> | 906 | Bacteria>Terrabacteria_group>Bacillota>unclassified_Bacillota | GCA_012840015.1 | Bacillota_bacterium | GXX40_00925 | 106213..108933 | 1879010 | ATP-binding-protein-[Firmicutes-bacterium] |
| NPV45159.1 | AAA+ATPase+wHTH+Alpha+Beta-domain+wHTH+RRM | Type-1_BR.RM.like.systems | REase->?->HEPN-><-KAP-NTPase\|\|TUDOR+SF1-SF2-Helicase+iREase+CTD+REase->METHYLASE->DUF499*-> | 851 | Bacteria>Terrabacteria_group>Bacillota>unclassified_Bacillota | GCA_013178105.1 | Bacillota_bacterium | HPY70_14490 | 186371..188926 | 1879010 | ATP-binding-protein-[Firmicutes-bacterium] |
| NLI69745.1 | AAA+ATPase+wHTH+Alpha+Beta-domain+wHTH+RRM | Type-1_BR.RM.like.systems | TUDOR+SF1-SF2-Helicase+iREase+CTD+REase->?->HerA/FtsK->METHYLASE->DUF499*->HIN-HTH->?->?->HTH-> | 929 | Bacteria>Terrabacteria_group>Bacillota>unclassified_Bacillota | GCA_012518215.1 | Bacillota_bacterium | GX364_02625 | 10971..13760 | 1879010 | ATP-binding-protein-[Firmicutes-bacterium] |
| MBC7083917.1 | AAA+ATPase+wHTH+Alpha+Beta-domain+wHTH+RRM | Type-1_BR.RM.like.systems | <-DUF499*<-REase<-METHYLASE<-REase<-TUDOR+SF1-SF2-Helicase+iREase+CTD+REase | 907 | Bacteria>Terrabacteria_group>Bacillota>unclassified_Bacillota | GCA_014360195.1 | Bacillota_bacterium | H5T95_10620 | 51..2774 | 1879010 | ATP-binding-protein-[Firmicutes-bacterium] |
| MCL6474638.1 | AAA+ATPase+wHTH+Alpha+Beta-domain+wHTH+RRM | Type-1_BR.RM.like.systems | PIN->DUF499*->TUDOR+SF1-SF2-Helicase+iREase+CTD+REase->?-><-REase | 879 | Bacteria>Terrabacteria_group>Bacillota>unclassified_Bacillota | GCA_023511455.1 | Bacillota_bacterium | K6U75_06255 | 18500..21139 | 1879010 | DUF499-domain-containing-protein-[Firmicutes-bacterium] |
| MBT9164708.1 | AAA+ATPase+wHTH+Alpha+Beta-domain+wHTH+RRM | Type-1_BR.RM.like.systems | <-DUF499*<-TOPRIM<-OLD-ABC<-METHYLASE | 873 | Bacteria>Terrabacteria_group>Candidatus_Lithacetigenota>Candidatus_Lithacetigena | GCA_018725485.1 | Candidatus_Lithacetigena_glycinireducens | DDT23_00712 | 1258..3879 | 2986055 | hypothetical-protein-[candidate-division-WS2-bacterium] |
| MBT9144502.1 | AAA+ATPase+wHTH+Alpha+Beta-domain+wHTH+RRM | Type-1_BR.RM.like.systems | <-Phage_int_SAM_1<-?<-?<-REase<-?<-DUF499* | 806 | Bacteria>Terrabacteria_group>Candidatus_Lithacetigenota>Candidatus_Psychracetigena | GCA_018725125.1 | Candidatus_Psychracetigena_formicireducens | DDT42_00343 | 27775..30195 | 2986056 | hypothetical-protein-[candidate-division-WS2-bacterium] |
| MCA9874333.1 | AAA+ATPase+wHTH+Alpha+Beta-domain+wHTH+RRM | Type-1_BR.RM.like.systems | SIR2+nSTAND1+FGS->SIR2+CASPASE->?->nSTAND1-><-HKD-DNase+HKD-DNase<-?<-DUF499* | 787 | Bacteria>Terrabacteria_group>Chloroflexota>Anaerolineae>Anaerolineales | GCA_020432445.1 | Anaerolineales_bacterium | KC441_11780 | 12847..15212 | 2073117 | ATP-binding-protein-[Anaerolineales-bacterium] |
| MCL4824052.1 | AAA+ATPase+wHTH+Alpha+Beta-domain+wHTH+RRM | Type-1_BR.RM.like.systems | METHYLASE-p->METHYLASE-p->DUF499*-> | 400 | Bacteria>Terrabacteria_group>Chloroflexota>Anaerolineae>Anaerolineales | GCA_023384535.1 | Anaerolineales_bacterium | KJZ57_07515 | 1228..2427 | 2073117 | DUF499-domain-containing-protein-[Anaerolineales-bacterium] |
| MCL4271579.1 | AAA+ATPase+wHTH+Alpha+Beta-domain+wHTH+RRM | Type-1_BR.RM.like.systems | DUF499*->REase+PNTD1+PNTD2->PIWI-> | 976 | Bacteria>Terrabacteria_group>Chloroflexota>Anaerolineae>Anaerolineales | GCA_023382575.1 | Anaerolineales_bacterium | KJZ72_18620 | 1..2933 | 2073117 | DUF499-domain-containing-protein-[Anaerolineales-bacterium] |
| WP_075063424.1 | AAA+ATPase+wHTH+Alpha+Beta-domain+wHTH+RRM | Type-1_BR.RM.like.systems | LuxR-HTH->?->SF1-SF2-Helicase+iREase+CTD+REase->METHYLASE->DUF499*->REase+SF1-SF2-Helicase->N6_N4_MTase-><-SF1-SF2-Helicase | 1097 | Bacteria>Terrabacteria_group>Chloroflexota>Anaerolineae>Anaerolineales | GCF_001306115.1 | Ornatilinea_apprima | ADN00_RS13000 | 57976..61269 | 1134406 | DUF499-domain-containing-protein-[Ornatilinea-apprima] |
| MBN1306185.1 | AAA+ATPase+wHTH+Alpha+Beta-domain+wHTH+RRM | Type-1_BR.RM.like.systems | AbiEi_4->?->URI-DNase+RAMA->TUDOR+SF1-SF2-Helicase+iREase+CTD+REase->METHYLASE->DUF499*->?->?-><-PIN<-?<-AbiEii | 1220 | Bacteria>Terrabacteria_group>Chloroflexota>Anaerolineae>Anaerolineales | GCA_016927315.1 | Anaerolineales_bacterium | JXA13_17245 | 40765..44427 | 2073117 | ATP-binding-protein-[Anaerolineales-bacterium] |
| MAU11782.1 | AAA+ATPase+wHTH+Alpha+Beta-domain+wHTH+RRM | Type-1_BR.RM.like.systems | <-VirD4-TraG+VirD4-FtsK<-?<-?<-MetJarc-HTH\|\|TUDOR+SF1-SF2-Helicase+iREase+CTD+REase->METHYLASE->DUF499*-> | 1108 | Bacteria>Terrabacteria_group>Chloroflexota>Anaerolineae>Anaerolineales | GCA_002699585.1 | Anaerolineaceae_bacterium | CL607_18295 | 54905..58231 | 2024896 | hypothetical-protein-[Anaerolineaceae-bacterium] |
| MBK8904525.1 | AAA+ATPase+wHTH+Alpha+Beta-domain+wHTH+RRM | Type-1_BR.RM.like.systems | <-HhH-RADC+JAB<-?<-?<-DDE_5<-DUF499*\|\|?-><-?<-?<-REase<-METHYLASE-p<-HTH_Tnp_4 | 948 | Bacteria>Terrabacteria_group>Chloroflexota>Anaerolineae>Anaerolineales | GCA_016710325.1 | Anaerolineaceae_bacterium | IPM53_25325 | 3764417..3767263 | 2024896 | ATP-binding-protein-[Anaerolineaceae-bacterium] |
| MBE7469119.1 | AAA+ATPase+wHTH+Alpha+Beta-domain+wHTH+RRM | Type-1_BR.RM.like.systems | METHYLASE-p->?->?->?-><-DUF499*<-REase<-METHYLASE<-HEPN<-PolB<-HEPN<-Polbeta | 906 | Bacteria>Terrabacteria_group>Chloroflexota>Anaerolineae>Anaerolineales | GCA_015075405.1 | Anaerolineales_bacterium | HS114_08270 | 1909271..1911991 | 2073117 | ATP-binding-protein-[Anaerolineales-bacterium] |
| MBE2269142.1 | AAA+ATPase+wHTH+Alpha+Beta-domain+wHTH+RRM | Type-1_BR.RM.like.systems | <-DUF499*<-METHYLASE<-PIN<-DUF433<-TUDOR+SF1-SF2-Helicase+iREase+CTD+REase | 1106 | Bacteria>Terrabacteria_group>Chloroflexota>Anaerolineae>Anaerolineales | GCA_014879865.1 | Anaerolinea_sp. | IAE80_13000 | 195..3515 | 1872519 | ATP-binding-protein-[Anaerolinea-sp.] |
| MCS7249243.1 | AAA+ATPase+wHTH+Alpha+Beta-domain+wHTH+RRM | Type-1_BR.RM.like.systems | METHYLASE-p->DUF499*->?->?->ABC-ATPase+SbcC->ABC-ATPase+SbcC-> | 883 | Bacteria>Terrabacteria_group>Chloroflexota>Anaerolineae>Anaerolineales | GCA_025057645.1 | Anaerolineales_bacterium | NZ840_13535 | 969..3620 | 2073117 | DUF499-domain-containing-protein-[Anaerolineales-bacterium] |
| MBN1316514.1 | AAA+ATPase+wHTH+Alpha+Beta-domain+wHTH+RRM | Type-1_BR.RM.like.systems | <-DUF499*<-METHYLASE<-?<-PAPS_reduct<-?<-?<-REase-p | 970 | Bacteria>Terrabacteria_group>Chloroflexota>Anaerolineae>Anaerolineales | GCA_016927355.1 | Anaerolineales_bacterium | JXA42_13645 | 1..2912 | 2073117 | ATP-binding-protein-[Anaerolineales-bacterium] |
| OQY42893.1 | AAA+ATPase+wHTH+Alpha+Beta-domain+wHTH+RRM | Type-1_BR.RM.like.systems | <-DUF499*<-?<-OLD-ABC<-METHYLASE | 480 | Bacteria>Terrabacteria_group>Chloroflexota>Anaerolineae>Anaerolineales | GCA_002084875.1 | Anaerolineaceae_bacterium_4572_78 | B6242_15635 | 321..1763 | 1972460 | hypothetical-protein-B6242_15635-[Anaerolineaceae-bacterium-4572_78] |
| MBN1317038.1 | AAA+ATPase+wHTH+Alpha+Beta-domain+wHTH+RRM | Type-1_BR.RM.like.systems | METHYLASE->OLD-ABC->TOPRIM->?->SWACOS->DUF499*-> | 449 | Bacteria>Terrabacteria_group>Chloroflexota>Anaerolineae>Anaerolineales | GCA_016927355.1 | Anaerolineales_bacterium | JXA42_16270 | 39202..40547 | 2073117 | ATP-binding-protein-[Anaerolineales-bacterium] |
| MBK9093501.1 | AAA+ATPase+wHTH+Alpha+Beta-domain+wHTH+RRM | Type-1_BR.RM.like.systems | TUDOR+SF1-SF2-Helicase+iREase+CTD+REase->OLD-ABC->HNH->METHYLASE->OLD-ABC->REase->DUF499*->?->ParB+HNH->?->REase-p-> | 943 | Bacteria>Terrabacteria_group>Chloroflexota>Anaerolineae>unclassified_Anaerolineae | GCA_016719785.1 | Anaerolineae_bacterium | IPM84_12145 | 210722..213553 | 2052143 | ATP-binding-protein-[Anaerolineae-bacterium] |
| MBN1429503.1 | AAA+ATPase+wHTH+Alpha+Beta-domain+wHTH+RRM | Type-1_BR.RM.like.systems | <-ZnR<-SIGMA-HTH\|\|ABC-ATPase+SbcC->?-><-?<-DUF499* | 601 | Bacteria>Terrabacteria_group>Chloroflexota>Anaerolineae>unclassified_Anaerolineae | GCA_016928235.1 | Anaerolineae_bacterium | JXB07_14105 | 7224..9030 | 2052143 | ATP-binding-protein-[Anaerolineae-bacterium] |
| MBK9231664.1 | AAA+ATPase+wHTH+Alpha+Beta-domain+wHTH+RRM | Type-1_BR.RM.like.systems | <-ABhydrolase<-?<-DUF499*<-HEPN | 801 | Bacteria>Terrabacteria_group>Chloroflexota>Anaerolineae>unclassified_Anaerolineae | GCA_016718485.1 | Anaerolineae_bacterium | IPO15_12650 | 109647..112052 | 2052143 | ATP-binding-protein-[Anaerolineae-bacterium] |
| MBK6433234.1 | AAA+ATPase+wHTH+Alpha+Beta-domain+wHTH+RRM | Type-1_BR.RM.like.systems | <-DUF499*<-REase<-OLD-ABC<-HNH | 943 | Bacteria>Terrabacteria_group>Chloroflexota>Anaerolineae>unclassified_Anaerolineae | GCA_016704685.1 | Anaerolineae_bacterium | IPF85_21385 | 32175..35006 | 2052143 | ATP-binding-protein-[Anaerolineae-bacterium] |
| MBN1874410.1 | AAA+ATPase+wHTH+Alpha+Beta-domain+wHTH+RRM | Type-1_BR.RM.like.systems | XIS-HTH->?->?->DUF499*->?->Peptidase->METHYLASE-p-> | 1323 | Bacteria>Terrabacteria_group>Chloroflexota>Anaerolineae>unclassified_Anaerolineae | GCA_016931185.1 | Anaerolineae_bacterium | JXA33_09275 | 6301..10272 | 2052143 | hypothetical-protein-[Anaerolineae-bacterium] |

| Protein-ID | DUF499-architecture | System | Gene-Neighbourhoods | Length | Taxonomy | Assembly-ID | Strain | Locus-Tag | Genomic-coordinates | TaxID | Fasta-defline |
|---|---|---|---|---|---|---|---|---|---|---|---|
| MCZ7543294.1 | AAA+ATPase+wHTH+Alpha+Beta-domain+wHTH+RRM | Type-1_BR.RM.like.systems | METHYLASE-p->?->METHYLASE-p->METHYLASE-p->?->DUF499->DUF499*->PAIREDC-HTH->DDE_3->?->SF1-SF2-Helicase-p->SF1-SF2-Helicase-p->SF1-SF2-Helicase-p-> | 394 | Bacteria>Terrabacteria_group>Chloroflexota>Anaerolineae>unclassified_Anaerolineae | GCA_027494055.1 | Anaerolineae_bacterium | M5R40_07020 | 10938..12122 | 2052143 | MAG:-DUF499-domain-containing-protein-[Anaerolineae-bacterium] |
| NPV78215.1 | AAA+ATPase+wHTH+Alpha+Beta-domain+wHTH+RRM | Type-1_BR.RM.like.systems | LuxR-HTH->?->XIS-HTH->TUDOR+SF1-SF2-Helicase+iREase+CTD+REase->SF1-SF2-Helicase->METHYLASE->DUF499*->?-><-SF1-SF2-Helicase<-SF1-SF2-Helicase-p<-?\|\|MerR-HTH-> | 945 | Bacteria>Terrabacteria_group>Chloroflexota>Anaerolineae>unclassified_Anaerolineae | GCA_013178185.1 | Anaerolineae_bacterium | HPY59_17805 | 13533..16370 | 2052143 | ATP-binding-protein-[Anaerolineae-bacterium] |
| MBK9232391.1 | AAA+ATPase+wHTH+Alpha+Beta-domain+wHTH+RRM | Type-1_BR.RM.like.systems | SF1-SF2-Helicase+iREase+CTD+REase-p->REase->METHYLASE-p->METHYLASE-p->DUF499-p->DUF499*->DUF499-p->REase+DUF4268->REase-><-?<-?<-NTP_transferase | 403 | Bacteria>Terrabacteria_group>Chloroflexota>Anaerolineae>unclassified_Anaerolineae | GCA_016718485.1 | Anaerolineae_bacterium | IPO15_16470 | 5943..7154 | 2052143 | DUF499-domain-containing-protein-[Anaerolineae-bacterium] |
| MCZ7543293.1 | AAA+ATPase+wHTH+Alpha+Beta-domain+wHTH+RRM | Type-1_BR.RM.like.systems | METHYLASE-p->?->METHYLASE-p->METHYLASE-p->?->DUF499*->DUF499->PAIREDC-HTH->DDE_3->?->SF1-SF2-Helicase-p->SF1-SF2-Helicase-p-> | 241 | Bacteria>Terrabacteria_group>Chloroflexota>Anaerolineae>unclassified_Anaerolineae | GCA_027494055.1 | Anaerolineae_bacterium | M5R40_07015 | 10253..10978 | 2052143 | MAG:-DUF499-domain-containing-protein-[Anaerolineae-bacterium] |
| MBX3056674.1 | AAA+ATPase+wHTH+Alpha+Beta-domain+wHTH+RRM | Type-1_BR.RM.like.systems | Response_reg+LuxR-HTH->Phage_integrase-><-?<-HKD-DNase+HKD-DNase<-?<-DUF499*<-DUF433<-REase\|\|?-><-?<-METHYLASE | 932 | Bacteria>Terrabacteria_group>Chloroflexota>Anaerolineae>unclassified_Anaerolineae | GCA_019637635.1 | Anaerolineae_bacterium | KF770_09400 | 8633..11431 | 2052143 | ATP-binding-protein-[Anaerolineae-bacterium] |
| MCL4264803.1 | AAA+ATPase+wHTH+Alpha+Beta-domain+wHTH+RRM | Type-1_BR.RM.like.systems | TUDOR+SF1-SF2-Helicase+iREase+CTD+REase->?-><-DUF4096+DDE_Tnp_1+DDE_Tnp_1_2\|\|?->PIN->METHYLASE->DUF499*->?->?->?->XIS-HTH-><-?\|\|HhH-RADC+JAB-> | 1222 | Bacteria>Terrabacteria_group>Chloroflexota>Anaerolineae>unclassified_Anaerolineae | GCA_023382655.1 | Anaerolineae_bacterium | KJ069_16390 | 143359..147027 | 2052143 | DUF499-domain-containing-protein-[Anaerolineae-bacterium] |
| MCB0190758.1 | AAA+ATPase+wHTH+Alpha+Beta-domain+wHTH+RRM | Type-1_BR.RM.like.systems | Mu-transpos_C->TnsC-ATPase->?->?->?-><-?<-DUF499*<-CR-ATPase9<-CR-ATPase9<-METHYLASE<-?<-?<-SF1-SF2-Helicase+HRDC | 1216 | Bacteria>Terrabacteria_group>Chloroflexota>Anaerolineae>unclassified_Anaerolineae | GCA_020433805.1 | Anaerolineae_bacterium | KDJ65_02340 | 22828..26478 | 2052143 | ATP-binding-protein-[Anaerolineae-bacterium] |
| MCG3142365.1 | AAA+ATPase+wHTH+Alpha+Beta-domain+wHTH+RRM | Type-1_BR.RM.like.systems | METHYLASE->REase->DUF499*-><-?<-?<-?\|\|?->Phage_int_SAM_1+Phage_integrase->Phage_integrase-> | 937 | Bacteria>Terrabacteria_group>Chloroflexota>Anaerolineae>unclassified_Anaerolineae | GCA_022072245.1 | Anaerolineae_bacterium | HDKAJFGB_03811 | 2691..5504 | 2052143 | hypothetical-protein-[Anaerolineae-bacterium] |
| MBN8655123.1 | AAA+ATPase+wHTH+Alpha+Beta-domain+wHTH+RRM | Type-1_BR.RM.like.systems | PIN->REase+NACHT->TUDOR+SF1-SF2-Helicase+iREase+CTD+REase->METHYLASE->HEPN->?->DUF499*-><-SF1-SF2-Helicase | 853 | Bacteria>Terrabacteria_group>Chloroflexota>Anaerolineae>unclassified_Anaerolineae | GCA_017303735.1 | Anaerolineae_bacterium | JOM11_05235 | 15902..18463 | 2052143 | DUF499-domain-containing-protein-[Anaerolineae-bacterium] |
| MBN2002245.1 | AAA+ATPase+wHTH+Alpha+Beta-domain+wHTH+RRM | Type-1_BR.RM.like.systems | STYKIN->REase->Trypsin->REase->SF1-SF2-Helicase+iREase+CTD+REase->METHYLASE->DUF499*->?-><-URI-DNase | 1003 | Bacteria>Terrabacteria_group>Chloroflexota>Anaerolineae>unclassified_Anaerolineae | GCA_016931835.1 | Anaerolineae_bacterium | JXA21_02715 | 14285..17296 | 2052143 | ATP-binding-protein-[Anaerolineae-bacterium] |
| MCQ3973997.1 | AAA+ATPase+wHTH+Alpha+Beta-domain+wHTH+RRM | Type-1_BR.RM.like.systems | PIN->CHTH->METHYLASE-p->?->METHYLASE->CR-ATPase9->DUF499*->?->HEPN-><-?<-Phage_integrase | 1225 | Bacteria>Terrabacteria_group>Chloroflexota>Anaerolineae>unclassified_Anaerolineae | GCA_024434315.1 | Anaerolineae_bacterium | DPW09_11170 | 46079..49756 | 2052143 | hypothetical-protein-[Anaerolineae-bacterium] |
| MBL7062619.1 | AAA+ATPase+wHTH+Alpha+Beta-domain+wHTH+RRM | Type-1_BR.RM.like.systems | MazE_antitoxin->PIN->?->AbiEii->METHYLASE->REase->DUF499*-><-?<-ABC-ATPase+SbcC | 858 | Bacteria>Terrabacteria_group>Chloroflexota>Anaerolineae>unclassified_Anaerolineae | GCA_016783285.1 | Anaerolineae_bacterium | ISS49_00240 | 51879..54455 | 2052143 | ATP-binding-protein-[Anaerolineae-bacterium] |
| MCB0169348.1 | AAA+ATPase+wHTH+Alpha+Beta-domain+wHTH+RRM | Type-1_BR.RM.like.systems | Zn_Tnp_IS1595+DDE_Tnp_IS1595->Zn_Tnp_IS1595+DDE_Tnp_IS1595->DUF499*-> | 1118 | Bacteria>Terrabacteria_group>Chloroflexota>Anaerolineae>unclassified_Anaerolineae | GCA_020433815.1 | Anaerolineae_bacterium | KDJ97_02250 | 15974..19330 | 2052143 | ATP-binding-protein-[Anaerolineae-bacterium] |
| MBN1139287.1 | AAA+ATPase+wHTH+Alpha+Beta-domain+wHTH+RRM | Type-1_BR.RM.like.systems | MazE_antitoxin->PIN-><-?<-DUF433<-DUF499*\|\|?->PIN-><-ParB+HNH<-METHYLASE | 1237 | Bacteria>Terrabacteria_group>Chloroflexota>Anaerolineae>unclassified_Anaerolineae | GCA_016926595.1 | Anaerolineae_bacterium | JXM73_22110 | 53334..57047 | 2052143 | ATP-binding-protein-[Anaerolineae-bacterium] |
| CAG1014975.1 | AAA+ATPase+wHTH+Alpha+Beta-domain+wHTH+RRM | Type-1_BR.RM.like.systems | PHP+SbcC->Schlafen+HATPase+HTH->METHYLASE-p->?->METHYLASE->HEPN->DUF499*-> | 923 | Bacteria>Terrabacteria_group>Chloroflexota>Anaerolineae>unclassified_Anaerolineae | GCA_905339325.1 | Anaerolineae_bacterium | ANRL4_05364 | 17229..20000 | 2052143 | hypothetical-protein-ANRL4_05364-[Anaerolineae-bacterium] |
| MCB0164247.1 | AAA+ATPase+wHTH+Alpha+Beta-domain+wHTH+RRM | Type-1_BR.RM.like.systems | <-HKD-DNase<-REase\|\|?-><-DUF499*<-?<-METHYLASE<-CHTH<-REase<-?<-REase | 1226 | Bacteria>Terrabacteria_group>Chloroflexota>Anaerolineae>unclassified_Anaerolineae | GCA_020433785.1 | Anaerolineae_bacterium | KDI79_08485 | 9055..12735 | 2052143 | ATP-binding-protein-[Anaerolineae-bacterium] |
| MCL4300573.1 | AAA+ATPase+wHTH+Alpha+Beta-domain+wHTH+RRM | Type-1_BR.RM.like.systems | TUDOR+SF1-SF2-Helicase+iREase+CTD+REase->?->?->METHYLASE->REase->?->DUF499*-><-?\|\|?->?->?-><-HNH | 929 | Bacteria>Terrabacteria_group>Chloroflexota>Anaerolineae>unclassified_Anaerolineae | GCA_023382535.1 | Anaerolineae_bacterium | KJ077_32855 | 82827..85616 | 2052143 | DUF499-domain-containing-protein-[Anaerolineae-bacterium] |
| MBN1262538.1 | AAA+ATPase+wHTH+Alpha+Beta-domain+wHTH+RRM | Type-1_BR.RM.like.systems | Phage_int_SAM_4+Phage_integrase-><-DUF499*<-PAIREDC-HTH | 868 | Bacteria>Terrabacteria_group>Chloroflexota>Anaerolineae>unclassified_Anaerolineae | GCA_016926915.1 | Anaerolineae_bacterium | JXB35_17790 | 1172..3778 | 2052143 | ATP-binding-protein-[Anaerolineae-bacterium] |
| MCD6553420.1 | AAA+ATPase+wHTH+Alpha+Beta-domain+wHTH+RRM | Type-1_BR.RM.like.systems | TUDOR+SF1-SF2-Helicase+iREase+CTD+REase-p->?->AbiEii->METHYLASE-><-?\|\|DUF499*-> | 683 | Bacteria>Terrabacteria_group>Chloroflexota>Anaerolineae>unclassified_Anaerolineae | GCA_021162125.1 | Anaerolineae_bacterium | J7M16_05360 | 6277..8326 | 2052143 | DUF499-domain-containing-protein-[Anaerolineae-bacterium] |
| MBZ0275669.1 | AAA+ATPase+wHTH+Alpha+Beta-domain+wHTH+RRM | Type-1_BR.RM.like.systems | SF1-SF2-Helicase+iREase+CTD+REase->METHYLASE->DUF499*->REase-> | 1097 | Bacteria>Terrabacteria_group>Chloroflexota>Anaerolineae>unclassified_Anaerolineae | GCA_019912915.1 | Anaerolineae_bacterium | K8I60_05975 | 6820..10113 | 2052143 | DUF499-domain-containing-protein-[Anaerolineae-bacterium] |
| MBN1920387.1 | AAA+ATPase+wHTH+Alpha+Beta-domain+wHTH+RRM | Type-1_BR.RM.like.systems | DUF433->?->?->OLD-ABC+TOPRIM->METHYLASE->?->DUF499*-> | 909 | Bacteria>Terrabacteria_group>Chloroflexota>Anaerolineae>unclassified_Anaerolineae | GCA_016931315.1 | Anaerolineae_bacterium | JW892_04030 | 10544..13273 | 2052143 | ATP-binding-protein-[Anaerolineae-bacterium] |
| MCS7282892.1 | AAA+ATPase+wHTH+Alpha+Beta-domain+wHTH+RRM | Type-1_BR.RM.like.systems | METHYLASE-p->DUF499*-><-PIN<-PhdYeFM_antitox | 1035 | Bacteria>Terrabacteria_group>Chloroflexota>Anaerolineae>unclassified_Anaerolineae | GCA_025062415.1 | Anaerolineae_bacterium | NZ769_06035 | 1264..4371 | 2052143 | DUF499-domain-containing-protein-[Anaerolineae-bacterium] |
| MBN1287263.1 | AAA+ATPase+wHTH+Alpha+Beta-domain+wHTH+RRM | Type-1_BR.RM.like.systems | <-DUF499*<-TPR+TPR+TPR<-REase<-METHYLASE-p | 1083 | Bacteria>Terrabacteria_group>Chloroflexota>Anaerolineae>unclassified_Anaerolineae | GCA_016927535.1 | Anaerolineae_bacterium | JXB47_17825 | 1830..5081 | 2052143 | ATP-binding-protein-[Anaerolineae-bacterium] |
| MBN1921823.1 | AAA+ATPase+wHTH+Alpha+Beta-domain+wHTH+RRM | Type-1_BR.RM.like.systems | REase->DUF499*->McrC-NTD+REase-> | 924 | Bacteria>Terrabacteria_group>Chloroflexota>Anaerolineae>unclassified_Anaerolineae | GCA_016931315.1 | Anaerolineae_bacterium | JW892_11295 | 1846..4620 | 2052143 | ATP-binding-protein-[Anaerolineae-bacterium] |
| MCD6553270.1 | AAA+ATPase+wHTH+Alpha+Beta-domain+wHTH+RRM | Type-1_BR.RM.like.systems | <-DUF499*<-REase<-TPR+TPR+TPR<-?<-?<-METHYLASE | 901 | Bacteria>Terrabacteria_group>Chloroflexota>Anaerolineae>unclassified_Anaerolineae | GCA_021162125.1 | Anaerolineae_bacterium | J7M16_04575 | 6228..8933 | 2052143 | ATP-binding-protein-[Anaerolineae-bacterium] |
| MBN2393686.1 | AAA+ATPase+wHTH+Alpha+Beta-domain+wHTH+RRM | Type-1_BR.RM.like.systems | <-DUF499*<-REase<-?<-?<-METHYLASE-p | 955 | Bacteria>Terrabacteria_group>Chloroflexota>Anaerolineae>unclassified_Anaerolineae | GCA_016934645.1 | Anaerolineae_bacterium | JXR84_23340 | 10232..13099 | 2052143 | DUF499-domain-containing-protein-[Anaerolineae-bacterium] |
| PJH76238.1 | AAA+ATPase+wHTH+Alpha+Beta-domain+wHTH+RRM | Type-1_BR.RM.like.systems | DUF499*->?->ORC-CDC6+REase-> | 750 | Bacteria>Terrabacteria_group>Chloroflexota>Anaerolineae>unclassified_Anaerolineae | GCA_002796565.1 | Anaerolineae_bacterium_CG_4_9_14_0_8_um_filter_58_9 | CO064_02300 | 1..2255 | 1973903 | MAG:-AAA-family-ATPase,-partial-[Anaerolineae-bacterium-CG_4_9_14_0_8_um_filter_58_9] |
| MBN1956153.1 | AAA+ATPase+wHTH+Alpha+Beta-domain+wHTH+RRM | Type-1_BR.RM.like.systems | <-DUF499*\|\|?->?-><-METHYLASE-p | 1091 | Bacteria>Terrabacteria_group>Chloroflexota>Anaerolineae>unclassified_Anaerolineae | GCA_016931695.1 | Anaerolineae_bacterium | JW900_14040 | 661..3936 | 2052143 | ATP-binding-protein-[Anaerolineae-bacterium] |
| MBN1814375.1 | AAA+ATPase+wHTH+Alpha+Beta-domain+wHTH+RRM | Type-1_BR.RM.like.systems | DDE_3-><-REase<-DUF499* | 1242 | Bacteria>Terrabacteria_group>Chloroflexota>Anaerolineae>unclassified_Anaerolineae | GCA_016930605.1 | Anaerolineae_bacterium | JXA14_21225 | 14526..18254 | 2052143 | ATP-binding-protein-[Anaerolineae-bacterium] |
| MBN2007579.1 | AAA+ATPase+wHTH+Alpha+Beta-domain+wHTH+RRM | Type-1_BR.RM.like.systems | TUDOR+SF1-SF2-Helicase+iREase+CTD+REase->REase->METHYLASE->OLD-ABC->TOPRIM->DUF499*-> | 737 | Bacteria>Terrabacteria_group>Chloroflexota>Anaerolineae>unclassified_Anaerolineae | GCA_016931835.1 | Anaerolineae_bacterium | JXA21_29810 | 7893..10102 | 2052143 | ATP-binding-protein-[Anaerolineae-bacterium] |
| MCZ7573949.1 | AAA+ATPase+wHTH+Alpha+Beta-domain+wHTH+RRM | Type-1_BR.RM.like.systems | <-Pkinase<-?<-?<-SIGMA-HTH<-?<-DUF499*<-HTH<-PolB<-METHYLASE<-?<-PolB<-REase | 933 | Bacteria>Terrabacteria_group>Chloroflexota>Ardenticatenia>Ardenticatenales | GCA_027493995.1 | Ardenticatenaceae_bacterium | M5U01_35810 | 8592301..8595102 | 2614247 | MAG:-DUF499-domain-containing-protein-[Ardenticatenaceae-bacterium] |
| MDE0464749.1 | AAA+ATPase+wHTH+Alpha+Beta-domain+wHTH+RRM | Type-1_BR.RM.like.systems | METHYLASE->PolB->HEPN->OLD-ABC->TOPRIM->HEPN->DUF499*->?->?-><-?\|\|MazE_antitoxin-><-DSBH | 775 | Bacteria>Terrabacteria_group>Chloroflexota>Caldilineae>Caldilineales | GCA_028821465.1 | Caldilineaceae_bacterium | OXH93_20190 | 7230..9557 | 2614250 | MAG:-DUF499-domain-containing-protein-[Caldilineaceae-bacterium] |
| MDE0312852.1 | AAA+ATPase+wHTH+Alpha+Beta-domain+wHTH+RRM | Type-1_BR.RM.like.systems | HTH_3->MPTase->TUDOR+SF1-SF2-Helicase+iREase+CTD+REase->OLD-ABC+TOPRIM->METHYLASE->DUF499*-><-?\|\|?->BrnA_antitoxin->?->DOC-> | 928 | Bacteria>Terrabacteria_group>Chloroflexota>Caldilineae>Caldilineales | GCA_028820765.1 | Caldilineaceae_bacterium | OXI52_11355 | 17311..20097 | 2614250 | MAG:-DUF499-domain-containing-protein-[Caldilineaceae-bacterium] |
| MYD91965.1 | AAA+ATPase+wHTH+Alpha+Beta-domain+wHTH+RRM | Type-1_BR.RM.like.systems | 35exo+TPR->tRNA->TUDOR+SF1-SF2-Helicase+iREase+CTD+REase->METHYLASE->DUF499*->OLD-ABC+TOPRIM->\|\|ORC-CDC6+REase-> | 967 | Bacteria>Terrabacteria_group>Chloroflexota>Caldilineae>Caldilineales | GCA_009839295.1 | Caldilineaceae_bacterium_SB0662_bin_9 | F4Y08_16820 | 82330..85233 | 2605258 | ATP-binding-protein-[Caldilineaceae-bacterium-SB0662_bin_9] |
| MYC94583.1 | AAA+ATPase+wHTH+Alpha+Beta-domain+wHTH+RRM | Type-1_BR.RM.like.systems | REase+SAVED->?->TUDOR+SF1-SF2-Helicase+iREase+CTD+REase->METHYLASE->DUF499*-> | 963 | Bacteria>Terrabacteria_group>Chloroflexota>Caldilineae>Caldilineales | GCA_009841235.1 | Caldilineaceae_bacterium_SB0661_bin_32 | F4X14_06395 | 119418..122309 | 2605255 | ATP-binding-protein-[Caldilineaceae-bacterium-SB0661_bin_32] |
| MDE0143354.1 | AAA+ATPase+wHTH+Alpha+Beta-domain+wHTH+RRM | Type-1_BR.RM.like.systems | <-DUF499*<-METHYLASE<-OLD-ABC+TOPRIM<-TUDOR+SF1-SF2-Helicase+iREase+CTD+REase | 926 | Bacteria>Terrabacteria_group>Chloroflexota>Caldilineae>Caldilineales | GCA_028819395.1 | Caldilineaceae_bacterium | OXN19_17910 | 956..3736 | 2614250 | MAG:-DUF499-domain-containing-protein-[Caldilineaceae-bacterium] |
| GIV69804.1 | AAA+ATPase+wHTH+Alpha+Beta-domain+wHTH+RRM | Type-1_BR.RM.like.systems | Schlafen+HATPase+HTH->METHYLASE->REase-><-?\|\|DUF499*-> | 854 | Bacteria>Terrabacteria_group>Chloroflexota>Caldilineae>Caldilineales | GCA_026003635.1 | Caldilinea_sp. | KatS3mg048_2666 | 137004..139568 | 2293560 | MAG:-hypothetical-protein-KatS3mg048_2666-[Caldilinea-sp.] |
| MYC61956.1 | AAA+ATPase+wHTH+Alpha+Beta-domain+wHTH+RRM | Type-1_BR.RM.like.systems | <-DUF499*<-METHYLASE-p<-zf-IS66+DDE_Tnp_IS66<-METHYLASE<-REase | 218 | Bacteria>Terrabacteria_group>Chloroflexota>Caldilineae>Caldilineales | GCA_009841145.1 | Caldilineaceae_bacterium_SB0661_bin_34 | F4X16_03815 | 38..694 | 2605256 | hypothetical-protein-[Caldilineaceae-bacterium-SB0661_bin_34] |
| HEY65250.1 | AAA+ATPase+wHTH+Alpha+Beta-domain+wHTH+RRM | Type-1_BR.RM.like.systems | METHYLASE->METHYLASE-p->DUF499*->?->?->?->?->?-><-Radical-SAM | 859 | Bacteria>Terrabacteria_group>Chloroflexota>Caldilineae>unclassified_Caldilineae | GCA_011176545.1 | Caldilineae_bacterium | G4D02_11835 | 3040..5619 | 2420332 | ATP-binding-protein-[Caldilineae-bacterium] |
| BCX03099.1 | AAA+ATPase+wHTH+Alpha+Beta-domain+wHTH+RRM | Type-1_BR.RM.like.systems | Glyco_transf_28->TUDOR+SF1-SF2-Helicase+iREase+CTD+REase->METHYLASE-p->METHYLASE-p-><-DDE_Tnp_1\|\|METHYLASE-p->DUF499*->Peptidase_M50->Peptidase_S41-><-Peptidase_M20 | 803 | Bacteria>Terrabacteria_group>Chloroflexota>Candidatus_Thermofonsia>Candidatus_Thermofonsia_Clade_3 | GCA_025998955.1 | Candidatus_Roseilinea_sp. | KatS3mg053_1037 | 1172091..1174502 | 2838777 | MAG:-hypothetical-protein-KatS3mg053_1037-[Candidatus-Roseilinea-sp.] |
| WP_097653976.1 | AAA+ATPase+wHTH+Alpha+Beta-domain+wHTH+RRM | Type-1_BR.RM.like.systems | <-RelE-ParE<-?<-?\|\|TUDOR+SF1-SF2-Helicase+iREase+CTD+REase->METHYLASE->DUF499*->CASPASE+APATPase+TPR+TPR+TPR->?->?-><-Glyoxalase | 952 | Bacteria>Terrabacteria_group>Chloroflexota>Chloroflexia>Chloroflexales | GCF_002532075.1 | Candidatus_Chloroploca_asiatica | A9Q02_RS15395 | 14881..17739 | 1506545 | ATP-binding-protein-[Candidatus-Chloroploca-asiatica] |
| WP_129677650.1 | AAA+ATPase+wHTH+Alpha+Beta-domain+wHTH+RRM | Type-1_BR.RM.like.systems | TUDOR+SF1-SF2-Helicase+iREase+CTD+REase->METHYLASE->DUF499*->CASPASE+APATPase+TPR+TPR+TPR->?->?->PIN->REase-> | 952 | Bacteria>Terrabacteria_group>Chloroflexota>Chloroflexia>Chloroflexales | GCF_004138165.1 | Candidatus_Chloroploca_sp._Khr17 | EKD05_RS23205 | 8249..11107 | 2496869 | ATP-binding-protein-[Candidatus-Chloroploca-sp.-Khr17] |
| MBO9336877.1 | AAA+ATPase+wHTH+Alpha+Beta-domain+wHTH+RRM | Type-1_BR.RM.like.systems | DUF499*->TUDOR+SF1-SF2-Helicase+iREase+CTD+REase-> | 610 | Bacteria>Terrabacteria_group>Chloroflexota>Chloroflexia>Chloroflexales | GCA_017743555.1 | Roseiflexus_sp. | J7455_19810 | 1..1833 | 2562120 | ATP-binding-protein-[Roseiflexus-sp.] |
| WP_012120125.1 | AAA+ATPase+wHTH+Alpha+Beta-domain+wHTH+RRM | Type-1_BR.RM.like.systems | <-Peptidase_C14<-Peptidase_C14<-DUF499*<-Schlafen+HATPase+HTH<-PHP+SbcC+PspA+SbcC<-METHYLASE<-TUDOR+SF1-SF2-Helicase+iREase+CTD+REase<-AbiEii<-AbiEi_4 | 854 | Bacteria>Terrabacteria_group>Chloroflexota>Chloroflexia>Chloroflexales | GCF_000017805.1 | Roseiflexus_castenholzii | RCAS_RS08215 | 2067227..2069791 | 120962 | DUF499-domain-containing-protein-[Roseiflexus-castenholzii] |
| WP_011955032.1 | AAA+ATPase+wHTH+Alpha+Beta-domain+wHTH+RRM | Type-1_BR.RM.like.systems | MethyltransfD12-><-?<-?\|\|MODE-HTH+PBPII-><-DUF499*<-REase<-METHYLASE<-TUDOR+SF1-SF2-Helicase+iREase+CTD+REase<-DDE_Tnp_1<-DDE_Tnp_1 | 865 | Bacteria>Terrabacteria_group>Chloroflexota>Chloroflexia>Chloroflexales | GCF_000016665.1 | Roseiflexus_sp._RS-1 | ROSERS_RS01210 | 293633..296230 | 357808 | DUF499-domain-containing-protein-[Roseiflexus-sp.-RS-1] |
| MBK9940845.1 | AAA+ATPase+wHTH+Alpha+Beta-domain+wHTH+RRM | Type-1_BR.RM.like.systems | WYL->TUDOR+SF1-SF2-Helicase+iREase+CTD+REase->?->METHYLASE->HISKIN->METHYLASE->DUF499*->?->?->SF1-SF2-Helicase-><-?\|\|DDE_Tnp_1-> | 1103 | Bacteria>Terrabacteria_group>Chloroflexota>Chloroflexia>Chloroflexales | - | Kouleothrix_sp. | IPP13_04390 | 1113074..1116385 | 2779161 | ATP-binding-protein-[Kouleothrix-sp.] |
| WP_097654147.1 | AAA+ATPase+wHTH+Alpha+Beta-domain+wHTH+RRM | Type-1_BR.RM.like.systems | TUDOR+SF1-SF2-Helicase+iREase+CTD+REase-p->TUDOR+SF1-SF2-Helicase+iREase+CTD+REase->?->METHYLASE->DUF499*->?->?->?->?->HEPN-p-> | 1005 | Bacteria>Terrabacteria_group>Chloroflexota>Chloroflexia>Chloroflexales | GCF_002532075.1 | Candidatus_Chloroploca_asiatica | A9Q02_RS16295 | 22212..25229 | 1506545 | DUF499-domain-containing-protein-[Candidatus-Chloroploca-asiatica] |
| WP_165360624.1 | AAA+ATPase+wHTH+Alpha+Beta-domain+wHTH+RRM | Type-1_BR.RM.like.systems | TRAM+Methyltransf_31-><-?<-?<-?<-DUF499*<-METHYLASE<-?<-?<-TUDOR+SF1-SF2-Helicase+iREase+CTD+REase | 1030 | Bacteria>Terrabacteria_group>Chloroflexota>Chloroflexia>Chloroflexales | GCF_004138165.1 | Candidatus_Chloroploca_sp._Khr17 | EKD05_RS07125 | 87778..90870 | 2496869 | DUF499-domain-containing-protein-[Candidatus-Chloroploca-sp.-Khr17] |
| MCS6840106.1 | AAA+ATPase+wHTH+Alpha+Beta-domain+wHTH+RRM | Type-1_BR.RM.like.systems | METHYLASE-p->SbcC->?->DUF499*-><-?<-?<-GDResolvase+Recombinase+ZnR\|\|HTH-> | 932 | Bacteria>Terrabacteria_group>Chloroflexota>Chloroflexia>Chloroflexales | GCA_025054995.1 | Roseiflexus_sp. | NZ701_04935 | 3430..6228 | 2562120 | DUF499-domain-containing-protein-[Roseiflexus-sp.] |
| MCS7288799.1 | AAA+ATPase+wHTH+Alpha+Beta-domain+wHTH+RRM | Type-1_BR.RM.like.systems | METHYLASE-p->REase->DUF499*-><-TPR+TPR<-?<-STYKinase<-ZnR | 857 | Bacteria>Terrabacteria_group>Chloroflexota>Chloroflexia>Chloroflexales | GCA_025062325.1 | Roseiflexus_sp. | NZ699_06675 | 1458..4031 | 2562120 | DUF499-domain-containing-protein-[Roseiflexus-sp.] |
| NJL06255.1 | AAA+ATPase+wHTH+Alpha+Beta-domain+wHTH+RRM | Type-1_BR.RM.like.systems | <-RNase\|\|?->TUDOR+SF1-SF2-Helicase+iREase+CTD+REase->METHYLASE->DUF499*-> | 1017 | Bacteria>Terrabacteria_group>Chloroflexota>Chloroflexia>Chloroflexales | GCA_012031225.1 | Chloroflexaceae_bacterium | HC911_15390 | 8460..11513 | 2562117 | ATP-binding-protein-[Chloroflexaceae-bacterium] |
| MBO9389921.1 | AAA+ATPase+wHTH+Alpha+Beta-domain+wHTH+RRM | Type-1_BR.RM.like.systems | DUF499*-><-MODE-HTH+PBPII\|\|?-><-MethyltransfD12 | 865 | Bacteria>Terrabacteria_group>Chloroflexota>Chloroflexia>Chloroflexales | GCA_017743475.1 | Roseiflexus_sp. | J7482_14180 | 236..2833 | 2562120 | ATP-binding-protein-[Roseiflexus-sp.] |
| MDD5510539.1 | AAA+ATPase+wHTH+Alpha+Beta-domain+wHTH+RRM | Type-1_BR.RM.like.systems | <-NERD<-DUF499*<-METHYLASE<-TUDOR+SF1-SF2-Helicase+iREase+CTD+REase | 849 | Bacteria>Terrabacteria_group>Chloroflexota>Dehalococcoidia>Dehalococcoidales | GCA_028716225.1 | Dehalococcoidales_bacterium | PHI12_07010 | 4311..6860 | 2026733 | MAG:-DUF499-domain-containing-protein-[Dehalococcoidales-bacterium] |
| MCL0075718.1 | AAA+ATPase+wHTH+Alpha+Beta-domain+wHTH+RRM | Type-1_BR.RM.like.systems | SF1-SF2-Helicase+DUF1998->?->SLATT->HicB->TUDOR+SF1-SF2-Helicase+iREase+CTD+REase->METHYLASE->DUF499*->REase->Phage_int_SAM_4+Phage_integrase->?->HEPN-> | 847 | Bacteria>Terrabacteria_group>Chloroflexota>Dehalococcoidia>unclassified_Dehalococcoidia | GCA_023271455.1 | Dehalococcoidia_bacterium | M1O17_02415 | 8590..11133 | 2026734 | DUF499-domain-containing-protein-[Dehalococcoidia-bacterium] |
| MCL0064885.1 | AAA+ATPase+wHTH+Alpha+Beta-domain+wHTH+RRM | Type-1_BR.RM.like.systems | <-TPR+TPR+TPR+TPR+TPR+TPR+TPR+TPR<-?<-DUF499*<-REase<-REase<-OLD-ABC<-METHYLASE | 925 | Bacteria>Terrabacteria_group>Chloroflexota>Dehalococcoidia>unclassified_Dehalococcoidia | GCA_023271695.1 | Dehalococcoidia_bacterium | M1N84_03205 | 1953..4730 | 2026734 | DUF499-domain-containing-protein-[Dehalococcoidia-bacterium] |
| MYH67239.1 | AAA+ATPase+wHTH+Alpha+Beta-domain+wHTH+RRM | Type-1_BR.RM.like.systems | <-REase+PNTD1+PNTD2\|\|?-><-DUF499*<-DUF499-p<-?<-TUDOR+SF1-SF2-Helicase+iREase+CTD+REase\|\|?-><-PIN<-DUF433 | 780 | Bacteria>Terrabacteria_group>Chloroflexota>Dehalococcoidia>unclassified_Dehalococcoidia | GCA_009843575.1 | Dehalococcoidia_bacterium | F4152_01280 | 9438..11780 | 2026734 | DUF499-domain-containing-protein-[Dehalococcoidia-bacterium] |
| MCD6599418.1 | AAA+ATPase+wHTH+Alpha+Beta-domain+wHTH+RRM | Type-1_BR.RM.like.systems | TUDOR+SF1-SF2-Helicase+iREase+CTD+REase->METHYLASE->DUF499*-><-Methyltransf_25<-SF1-SF2-Helicase | 847 | Bacteria>Terrabacteria_group>Chloroflexota>Dehalococcoidia>unclassified_Dehalococcoidia | GCA_021162245.1 | Dehalococcoidia_bacterium | J7L19_02445 | 13492..16035 | 2026734 | ATP-binding-protein-[Dehalococcoidia-bacterium] |
| MYD51869.1 | AAA+ATPase+wHTH+Alpha+Beta-domain+wHTH+RRM | Type-1_BR.RM.like.systems | TUDOR+SF1-SF2-Helicase+iREase+CTD+REase->PolB->HEPN->METHYLASE->DUF433->PIN->DUF499*-> | 928 | Bacteria>Terrabacteria_group>Chloroflexota>Dehalococcoidia>unclassified_Dehalococcoidia | GCA_009841605.1 | Dehalococcoidia_bacterium | F4W93_10360 | 50124..52910 | 2026734 | ATP-binding-protein-[Dehalococcoidia-bacterium] |
| MBK6562039.1 | AAA+ATPase+wHTH+Alpha+Beta-domain+wHTH+RRM | Type-1_BR.RM.like.systems | <-REase<-REase+DUF4268<-DUF499-p<-?<-DUF499*<-DUF499-p<-METHYLASE-p | 582 | Bacteria>Terrabacteria_group>Chloroflexota>Dehalococcoidia>unclassified_Dehalococcoidia | GCA_016704105.1 | Dehalococcoidia_bacterium | IPG14_10980 | 2567..4315 | 2026734 | DUF499-domain-containing-protein-[Dehalococcoidia-bacterium] |
| MCL0092366.1 | AAA+ATPase+wHTH+Alpha+Beta-domain+wHTH+RRM | Type-1_BR.RM.like.systems | PemK_toxin-><-DUF499*<-METHYLASE<-TUDOR+SF1-SF2-Helicase+iREase+CTD+REase | 868 | Bacteria>Terrabacteria_group>Chloroflexota>Dehalococcoidia>unclassified_Dehalococcoidia | GCA_023271105.1 | Dehalococcoidia_bacterium | M1N92_01640 | 7241..9847 | 2026734 | DUF499-domain-containing-protein-[Dehalococcoidia-bacterium] |
| MCL0078596.1 | AAA+ATPase+wHTH+Alpha+Beta-domain+wHTH+RRM | Type-1_BR.RM.like.systems | PemK_toxin-><-DUF499*<-METHYLASE<-TUDOR+SF1-SF2-Helicase+iREase+CTD+REase | 759 | Bacteria>Terrabacteria_group>Chloroflexota>Dehalococcoidia>unclassified_Dehalococcoidia | GCA_023271435.1 | Dehalococcoidia_bacterium | M1O56_02885 | 6086..8365 | 2026734 | ATP-binding-protein-[Dehalococcoidia-bacterium] |
| HAZ31511.1 | AAA+ATPase+wHTH+Alpha+Beta-domain+wHTH+RRM | Type-1_BR.RM.like.systems | <-ABC-ATPase+SbcC<-DUF499*<-METHYLASE-p | 847 | Bacteria>Terrabacteria_group>Chloroflexota>Dehalococcoidia>unclassified_Dehalococcoidia | GCA_003485725.1 | Dehalococcoidia_bacterium | DCY61_02240 | 3680..6223 | 2026734 | AAA-family-ATPase-[Dehalococcoidia-bacterium] |
| MBA2393113.1 | AAA+ATPase+wHTH+Alpha+Beta-domain+wHTH+RRM | Type-1_BR.RM.like.systems | <-ABC-ATPase+SbcC<-ABC-ATPase+SbcC<-MarR-HTH\|\|METHYLASE->DUF499*-><-ParA-Soj-PloopNTPase | 997 | Bacteria>Terrabacteria_group>Chloroflexota>Ktedonobacteria>Ktedonobacterales | GCA_013695595.1 | Ktedonobacteraceae_bacterium | HOV70_10255 | 7422..10415 | 2740538 | ATP-binding-protein-[Ktedonobacteraceae-bacterium] |
| WP_200808050.1 | AAA+ATPase+wHTH+Alpha+Beta-domain+wHTH+RRM | Type-1_BR.RM.like.systems | <-ABC-ATPase+SbcC<-?<-?<-?<-SIGMA-HTH<-DUF499*<-METHYLASE<-TOPRIM<-OLD-ABC<-TUDOR+SF1-SF2-Helicase+iREase+CTD+REase<-?<-SF1-SF2-helicase+DUF1998 | 1109 | Bacteria>Terrabacteria_group>Chloroflexota>Thermoflexia>Thermoflexales | GCF_900187885.1 | Thermoflexus_hugenholtzii | CFB18_RS02785 | 14938..18267 | 1495650 | DUF499-domain-containing-protein-[Thermoflexus-hugenholtzii] |
| MBN1890000.1 | AAA+ATPase+wHTH+Alpha+Beta-domain+wHTH+RRM | Type-1_BR.RM.like.systems | REase->DUF499*-><-?\|\|?->ABhydrolase->DOC+HTH-> | 987 | Bacteria>Terrabacteria_group>Chloroflexota>Thermoflexia>Thermoflexales | GCA_016931085.1 | Thermoflexales_bacterium | JW850_18535 | 987..3950 | 2497639 | ATP-binding-protein-[Thermoflexales-bacterium] |
| MCS7088122.1 | AAA+ATPase+wHTH+Alpha+Beta-domain+wHTH+RRM | Type-1_BR.RM.like.systems | Radical-SAM->?->?-><-?<-?<-tRNA\|\|?-><-DUF499*<-METHYLASE | 872 | Bacteria>Terrabacteria_group>Chloroflexota>Thermoflexia>Thermoflexales | GCA_025057935.1 | Thermoflexales_bacterium | NZL91_05445 | 28674..31292 | 2497639 | DUF499-domain-containing-protein-[Thermoflexales-bacterium] |
| MBN1889079.1 | AAA+ATPase+wHTH+Alpha+Beta-domain+wHTH+RRM | Type-1_BR.RM.like.systems | <-SF1-SF2-Helicase+iREase+CTD+REase<-DUF499* | 1100 | Bacteria>Terrabacteria_group>Chloroflexota>Thermoflexia>Thermoflexales | GCA_016931085.1 | Thermoflexales_bacterium | JW850_13880 | 3323..6625 | 2497639 | ATP-binding-protein-[Thermoflexales-bacterium] |
| MCL5961065.1 | AAA+ATPase+wHTH+Alpha+Beta-domain+wHTH+RRM | Type-1_BR.RM.like.systems | TUDOR+SF1-SF2-Helicase+iREase+CTD+REase->Schlafen+HATPase+HTH->METHYLASE->OLD-ABC->REase->DUF499*->MerR-HTH->?->REase->IclR-HTH-> | 939 | Bacteria>Terrabacteria_group>Chloroflexota>unclassified_Chloroflexota | GCA_023489135.1 | Chloroflexota_bacterium | M1358_17465 | 13442..16261 | 2026724 | ATP-binding-protein-[Chloroflexi-bacterium] |
| MBK7919049.1 | AAA+ATPase+wHTH+Alpha+Beta-domain+wHTH+RRM | Type-1_BR.RM.like.systems | <-DDE_Tnp_IS1\|\|METHYLASE-p->?->?->HEPN-p->HEPN+DUF499->DUF499*-><-DDE_5<-DDE_5\|\|?-><-DDE_5+DDE_Tnp_2 | 452 | Bacteria>Terrabacteria_group>Chloroflexota>unclassified_Chloroflexota | GCA_016713795.1 | Chloroflexota_bacterium | IPJ94_22865 | 31366..32724 | 2026724 | hypothetical-protein-[Chloroflexota-bacterium] |
| MBI4322294.1 | AAA+ATPase+wHTH+Alpha+Beta-domain+wHTH+RRM | Type-1_BR.RM.like.systems | TUDOR+SF1-SF2-Helicase+iREase+CTD+REase->Schlafen+HATPase+HTH->?->METHYLASE->OLD-ABC->REase->DUF499*-> | 938 | Bacteria>Terrabacteria_group>Chloroflexota>unclassified_Chloroflexota | GCA_016210495.1 | Chloroflexota_bacterium | HY675_27690 | 51598..54414 | 2026724 | ATP-binding-protein-[Chloroflexi-bacterium] |
| MCZ7670577.1 | AAA+ATPase+wHTH+Alpha+Beta-domain+wHTH+RRM | Type-1_BR.RM.like.systems | Peptidase_C26->?->tRNA-><-?<-?<-DUF499-p<-DUF499*<-HEPN<-?<-METHYLASE-p\|\|?-><-METHYLASE-p<-METHYLASE-p | 431 | Bacteria>Terrabacteria_group>Chloroflexota>unclassified_Chloroflexota | GCA_027493515.1 | Chloroflexota_bacterium | M5U34_27215 | 48065..49360 | 2026724 | MAG:-DUF499-domain-containing-protein-[Chloroflexi-bacterium] |
| MCY4113251.1 | AAA+ATPase+wHTH+Alpha+Beta-domain+wHTH+RRM | Type-1_BR.RM.like.systems | <-METHYLASE-p<-?<-Radical-SAM\|\|?-><-?<-?\|\|DUF499*->METHYLASE->URI-DNase->?->?->?->ABC-ATPase+SbcC-> | 862 | Bacteria>Terrabacteria_group>Chloroflexota>unclassified_Chloroflexota | GCA_026708035.1 | Chloroflexota_bacterium | OXG33_04830 | 101728..104316 | 2026724 | MAG:-DUF499-domain-containing-protein-[Chloroflexi-bacterium] |
| PZR94485.1 | AAA+ATPase+wHTH+Alpha+Beta-domain+wHTH+RRM | Type-1_BR.RM.like.systems | TUDOR+SF1-SF2-Helicase+iREase+CTD+REase->METHYLASE->METHYLASE-p->DUF499*-> | 951 | Bacteria>Terrabacteria_group>Chloroflexota>unclassified_Chloroflexota | GCA_003243865.1 | Chloroflexota_bacterium | DLM69_12410 | 11972..14827 | 2026724 | hypothetical-protein-DLM69_12410-[Chloroflexi-bacterium] |
| GIW20843.1 | AAA+ATPase+wHTH+Alpha+Beta-domain+wHTH+RRM | Type-1_BR.RM.like.systems | HEPN->DUF499*->DUF499-p->?->PemK_toxin-><-?<-MULE-TRANSPOSASE<-MULE-TRANSPOSASE | 667 | Bacteria>Terrabacteria_group>Chloroflexota>unclassified_Chloroflexota | GCA_026004015.1 | Chloroflexota_bacterium | KatS3mg065_1139 | 2064..4067 | 2026724 | MAG:-hypothetical-protein-KatS3mg065_1139-[Chloroflexota-bacterium] |
| MBM2811009.1 | AAA+ATPase+wHTH+Alpha+Beta-domain+wHTH+RRM | Type-1_BR.RM.like.systems | METHYLASE-p-><-?\|\|?->HicB_lk_antitox->HicA_toxin->HEPN+DUF499->DUF499*-> | 696 | Bacteria>Terrabacteria_group>Chloroflexota>unclassified_Chloroflexota | - | Chloroflexota_bacterium | HW416_1768 | 4823..6913 | 2026724 | ATPase-superfamily-[Chloroflexota-bacterium] |
| MDA1330708.1 | AAA+ATPase+wHTH+Alpha+Beta-domain+wHTH+RRM | Type-1_BR.RM.like.systems | <-DUF499*<-METHYLASE<-TUDOR+SF1-SF2-Helicase+iREase+CTD+REase-p | 949 | Bacteria>Terrabacteria_group>Chloroflexota>unclassified_Chloroflexota | GCA_027624805.1 | Chloroflexota_bacterium | O3B43_06560 | 287..3136 | 2026724 | MAG:-DUF499-domain-containing-protein-[Chloroflexi-bacterium] |
| MCY4538911.1 | AAA+ATPase+wHTH+Alpha+Beta-domain+wHTH+RRM | Type-1_BR.RM.like.systems | <-SF1-SF2-Helicase<-N6_N4_MTase\|\|DUF499*-><-GNTR-HTH | 633 | Bacteria>Terrabacteria_group>Chloroflexota>unclassified_Chloroflexota | GCA_026713245.1 | Chloroflexota_bacterium | OXE52_11890 | 5551..7452 | 2026724 | MAG:-DUF499-domain-containing-protein-[Chloroflexi-bacterium] |
| OQA96160.1 | AAA+ATPase+wHTH+Alpha+Beta-domain+wHTH+RRM | Type-1_BR.RM.like.systems | <-RAMPs<-?<-Phage_int_SAM_1+Phage_integrase<-DUF499* | 781 | Bacteria>Terrabacteria_group>Chloroflexota>unclassified_Chloroflexota | GCA_002069935.1 | Chloroflexi_bacterium_ADurb.Bin222 | BWY25_02290 | 3784..6129 | 1852858 | hypothetical-protein-BWY25_02290-[Chloroflexi-bacterium-ADurb.Bin222] |
| NMC03209.1 | AAA+ATPase+wHTH+Alpha+Beta-domain+wHTH+RRM | Type-1_BR.RM.like.systems | DUF499*->Phage_int_SAM_1+Phage_integrase-> | 614 | Bacteria>Terrabacteria_group>Chloroflexota>unclassified_Chloroflexota | GCA_012798275.1 | Chloroflexota_bacterium | GYA30_12650 | 1..1846 | 2026724 | ATP-binding-protein-[Chloroflexota-bacterium] |
| MCH7607819.1 | AAA+ATPase+wHTH+Alpha+Beta-domain+wHTH+RRM | Type-1_BR.RM.like.systems | Phage_int_SAM_4+Phage_integrase->?-><-?<-DUF499* | 531 | Bacteria>Terrabacteria_group>Chloroflexota>unclassified_Chloroflexota | GCA_022561455.1 | Chloroflexota_bacterium | IIC94_04545 | 15177..16772 | 2026724 | ATP-binding-protein-[Chloroflexota-bacterium] |
| MBU6289535.1 | AAA+ATPase+wHTH+Alpha+Beta-domain+wHTH+RRM | Type-1_BR.RM.like.systems | METHYLASE-p-><-DUF2961<-DUF499* | 692 | Bacteria>Terrabacteria_group>Chloroflexota>unclassified_Chloroflexota | GCA_018970655.1 | Chloroflexota_bacterium | KGS10_15355 | 6260..8340 | 2026724 | DUF499-domain-containing-protein-[Chloroflexota-bacterium] |
| MBK8032220.1 | AAA+ATPase+wHTH+Alpha+Beta-domain+wHTH+RRM | Type-1_BR.RM.like.systems | TUDOR+SF1-SF2-Helicase+iREase+CTD+REase->TIR+PNTD2->METHYLASE->METHYLASE-p->TIR+PNTD2->TIR+PNTD2->DUF499*->DUF499-p->DUF499-p->REase->REase+DUF4268->?->REase-> | 511 | Bacteria>Terrabacteria_group>Chloroflexota>unclassified_Chloroflexota | GCA_016713355.1 | Chloroflexota_bacterium | IPK17_22630 | 29610..31145 | 2026724 | ATP-binding-protein-[Chloroflexota-bacterium] |
| MCL5950399.1 | AAA+ATPase+wHTH+Alpha+Beta-domain+wHTH+RRM | Type-1_BR.RM.like.systems | <-ParB+HNH<-REase<-REase<-PIWI<-DUF499*<-REase<-METHYLASE<-MarR-HTH<-HKD-DNase<-HKD-DNase<-TUDOR+SF1-SF2-Helicase+iREase+CTD+REase | 935 | Bacteria>Terrabacteria_group>Chloroflexota>unclassified_Chloroflexota | GCA_023489485.1 | Chloroflexota_bacterium | M1132_01520 | 20661..23468 | 2026724 | DUF499-domain-containing-protein-[Chloroflexi-bacterium] |
| MCX6032091.1 | AAA+ATPase+wHTH+Alpha+Beta-domain+wHTH+RRM | Type-1_BR.RM.like.systems | <-SF1-SF2-Helicase+REase<-Exonuc_V_gamma+REase<-SF1-SF2-Helicase+DUF1998<-?<-?<-DUF499*<-?<-?<-METHYLASE<-REase<-TUDOR+SF1-SF2-Helicase+iREase+CTD+REase | 1118 | Bacteria>Terrabacteria_group>Chloroflexota>unclassified_Chloroflexota | GCA_026389845.1 | Chloroflexota_bacterium | NT169_22690 | 15055..18411 | 2026724 | MAG:-DUF499-domain-containing-protein-[Chloroflexi-bacterium] |
| MBI4322283.1 | AAA+ATPase+wHTH+Alpha+Beta-domain+wHTH+RRM | Type-1_BR.RM.like.systems | TUDOR+SF1-SF2-Helicase+iREase+CTD+REase->HD->DUF4065->METHYLASE-p->REase->METHYLASE->DUF499*->?-><-?<-?<-?\|\|TUDOR+SF1-SF2-Helicase+iREase+CTD+REase->Schlafen+HATPase+HTH-> | 979 | Bacteria>Terrabacteria_group>Chloroflexota>unclassified_Chloroflexota | GCA_016210495.1 | Chloroflexota_bacterium | HY675_27635 | 34699..37638 | 2026724 | ATP-binding-protein-[Chloroflexi-bacterium] |
| MCY3783979.1 | AAA+ATPase+wHTH+Alpha+Beta-domain+wHTH+RRM | Type-1_BR.RM.like.systems | MORC->?->?->?-><-?<-DUF499*<-METHYLASE<-ParB+HNH+Schlafen<-SF1-SF2-Helicase-p<-VirB4-FtsK+HerA<-?<-TUDOR+SF1-SF2-Helicase+iREase+CTD+REase | 920 | Bacteria>Terrabacteria_group>Chloroflexota>unclassified_Chloroflexota | GCA_026706485.1 | Chloroflexota_bacterium | OXG79_09360 | 384253..387015 | 2026724 | MAG:-DUF499-domain-containing-protein-[Chloroflexi-bacterium] |
| MBK9748864.1 | AAA+ATPase+wHTH+Alpha+Beta-domain+wHTH+RRM | Type-1_BR.RM.like.systems | <-VirD4-TraG+VirD4-FtsK<-?<-?<-MetJarc-HTH<-DUF499*<-METHYLASE<-PIN<-DUF433<-TUDOR+SF1-SF2-Helicase+iREase+CTD+REase\|\|?-><-PIWI | 1104 | Bacteria>Terrabacteria_group>Chloroflexota>unclassified_Chloroflexota | GCA_016717205.1 | Chloroflexota_bacterium | IPO91_19080 | 75969..79283 | 2026724 | ATP-binding-protein-[Chloroflexi-bacterium] |
| MCL4543926.1 | AAA+ATPase+wHTH+Alpha+Beta-domain+wHTH+RRM | Type-1_BR.RM.like.systems | TUDOR+SF1-SF2-Helicase+iREase+CTD+REase->METHYLASE->PolB->HEPN->DUF499*->TraJ-RHH->PIN->DDE_Tnp_1_3-> | 860 | Bacteria>Terrabacteria_group>Chloroflexota>unclassified_Chloroflexota | GCA_023381895.1 | Chloroflexota_bacterium | M1118_04890 | 7735..10317 | 2026724 | DUF499-domain-containing-protein-[Chloroflexi-bacterium] |
| MCY3958689.1 | AAA+ATPase+wHTH+Alpha+Beta-domain+wHTH+RRM | Type-1_BR.RM.like.systems | SF1-SF2-Helicase-p->Peptidase->?-><-DUF499*<-PIN<-DUF433<-METHYLASE<-?<-TUDOR+SF1-SF2-Helicase+iREase+CTD+REase | 926 | Bacteria>Terrabacteria_group>Chloroflexota>unclassified_Chloroflexota | GCA_026707585.1 | Chloroflexota_bacterium | OXG65_10380 | 10889..13669 | 2026724 | MAG:-DUF499-domain-containing-protein-[Chloroflexi-bacterium] |
| MBM3127627.1 | AAA+ATPase+wHTH+Alpha+Beta-domain+wHTH+RRM | Type-1_BR.RM.like.systems | Pkinase->?->?-><-SF1-SF2-Helicase+DUF1998<-?<-DUF499*<-REase<-TIR<-METHYLASE<-?<-REase+ParB<-RNase | 922 | Bacteria>Terrabacteria_group>Chloroflexota>unclassified_Chloroflexota | GCA_016865935.1 | Chloroflexota_bacterium | FJ009_03225 | 26548..29316 | 2026724 | ATP-binding-protein-[Chloroflexi-bacterium] |
| GIW20263.1 | AAA+ATPase+wHTH+Alpha+Beta-domain+wHTH+RRM | Type-1_BR.RM.like.systems | TUDOR+SF1-SF2-Helicase+iREase+CTD+REase->PIN->METHYLASE->?->DUF499*-><-?\|\|?-><-tRNA<-Peptidase<-tRNA<-tRNA<-tRNA<-REase | 906 | Bacteria>Terrabacteria_group>Chloroflexota>unclassified_Chloroflexota | GCA_026004015.1 | Chloroflexota_bacterium | KatS3mg065_0559 | 209249..211969 | 2026724 | MAG:-hypothetical-protein-KatS3mg065_0559-[Chloroflexota-bacterium] |
| MCY4614892.1 | AAA+ATPase+wHTH+Alpha+Beta-domain+wHTH+RRM | Type-1_BR.RM.like.systems | TUDOR+SF1-SF2-Helicase+iREase+CTD+REase->METHYLASE->?->DUF499*-><-HicB<-PadR-HTH<-VirD4-TraG+HerA | 930 | Bacteria>Terrabacteria_group>Chloroflexota>unclassified_Chloroflexota | GCA_026713725.1 | Chloroflexota_bacterium | OXC71_00655 | 12945..15737 | 2026724 | MAG:-DUF499-domain-containing-protein-[Chloroflexi-bacterium] |
| MDE2669829.1 | AAA+ATPase+wHTH+Alpha+Beta-domain+wHTH+RRM | Type-1_BR.RM.like.systems | <-REase+NACHT\|\|TUDOR+SF1-SF2-Helicase+iREase+CTD+REase->METHYLASE->DUF433->PIN->DUF499*-> | 918 | Bacteria>Terrabacteria_group>Chloroflexota>unclassified_Chloroflexota | GCA_028875515.1 | Chloroflexota_bacterium | OXI51_09280 | 52276..55032 | 2026724 | MAG:-DUF499-domain-containing-protein-[Chloroflexota-bacterium] |
| MBM3143888.1 | AAA+ATPase+wHTH+Alpha+Beta-domain+wHTH+RRM | Type-1_BR.RM.like.systems | TUDOR+SF1-SF2-Helicase+iREase+CTD+REase->HEPN->METHYLASE->DUF499*->?->PIN-><-?\|\|?->Peptidase_M24-> | 927 | Bacteria>Terrabacteria_group>Chloroflexota>unclassified_Chloroflexota | GCA_016865865.1 | Chloroflexota_bacterium | FJ010_02740 | 17191..19974 | 2026724 | ATP-binding-protein-[Chloroflexi-bacterium] |
| HCG01542.1 | AAA+ATPase+wHTH+Alpha+Beta-domain+wHTH+RRM | Type-1_BR.RM.like.systems | <-Polbeta<-?<-DUF499*<-METHYLASE<-TUDOR+SF1-SF2-Helicase+iREase+CTD+REase<-SF1-SF2-Helicase | 935 | Bacteria>Terrabacteria_group>Chloroflexota>unclassified_Chloroflexota | GCA_003530725.1 | Chloroflexota_bacterium | DEV93_13470 | 2544..5351 | 2026724 | hypothetical-protein-[Chloroflexi-bacterium] |
| MCX6033329.1 | AAA+ATPase+wHTH+Alpha+Beta-domain+wHTH+RRM | Type-1_BR.RM.like.systems | REase->?->METHYLASE->ParB+HNH->APATPase+iSTAND3->DUF499*->STYKIN-> | 993 | Bacteria>Terrabacteria_group>Chloroflexota>unclassified_Chloroflexota | GCA_026389845.1 | Chloroflexota_bacterium | NT169_29110 | 10350..13331 | 2026724 | MAG:-DUF499-domain-containing-protein-[Chloroflexi-bacterium] |
| MBI3749115.1 | AAA+ATPase+wHTH+Alpha+Beta-domain+wHTH+RRM | Type-1_BR.RM.like.systems | <-Primpol+PriCT1<-?<-?<-?\|\|TUDOR+SF1-SF2-Helicase+iREase+CTD+REase->METHYLASE->DUF499*-> | 981 | Bacteria>Terrabacteria_group>Chloroflexota>unclassified_Chloroflexota | GCA_016200745.1 | Chloroflexota_bacterium | HY262_09770 | 11576..14521 | 2026724 | ATP-binding-protein-[Chloroflexi-bacterium] |
| RIK37466.1 | AAA+ATPase+wHTH+Alpha+Beta-domain+wHTH+RRM | Type-1_BR.RM.like.systems | TUDOR+SF1-SF2-Helicase+iREase+CTD+REase->?->HicB_lk_antitox->METHYLASE->DUF499*-> | 959 | Bacteria>Terrabacteria_group>Chloroflexota>unclassified_Chloroflexota | GCA_003577005.1 | Chloroflexota_bacterium | DCC55_24215 | 9691..12570 | 2026724 | AAA-family-ATPase-[Chloroflexi-bacterium] |
| MCL4466049.1 | AAA+ATPase+wHTH+Alpha+Beta-domain+wHTH+RRM | Type-1_BR.RM.like.systems | Phage_integrase-><-DUF499*<-OLD-ABC+TOPRIM<-?<-METHYLASE<-METHYLASE<-?<-SPOUT | 980 | Bacteria>Terrabacteria_group>Chloroflexota>unclassified_Chloroflexota | GCA_023382155.1 | Chloroflexota_bacterium | M1389_08480 | 1506..4448 | 2026724 | DUF499-domain-containing-protein-[Chloroflexi-bacterium] |
| MDE2823077.1 | AAA+ATPase+wHTH+Alpha+Beta-domain+wHTH+RRM | Type-1_BR.RM.like.systems | Schlafen+HATPase+HTH->SF1-SF2-Helicase->METHYLASE->DUF499*-><-PIN | 949 | Bacteria>Terrabacteria_group>Chloroflexota>unclassified_Chloroflexota | GCA_028874805.1 | Chloroflexota_bacterium | OXK79_06190 | 6326..9175 | 2026724 | MAG:-DUF499-domain-containing-protein-[Chloroflexota-bacterium] |
| MBI2917634.1 | AAA+ATPase+wHTH+Alpha+Beta-domain+wHTH+RRM | Type-1_BR.RM.like.systems | <-URI-DNase<-?\|\|?->?->?->?-><-DUF499*<-METHYLASE<-TUDOR+SF1-SF2-Helicase+iREase+CTD+REase | 935 | Bacteria>Terrabacteria_group>Chloroflexota>unclassified_Chloroflexota | GCA_016190505.1 | Chloroflexota_bacterium | HYY01_06525 | 30004..32811 | 2026724 | ATP-binding-protein-[Chloroflexi-bacterium] |
| HAF11537.1 | AAA+ATPase+wHTH+Alpha+Beta-domain+wHTH+RRM | Type-1_BR.RM.like.systems | <-HTH<-SIGMA-HTH<-Peptidase-S8<-Classical-AAA<-?<-?\|\|DUF499*-> | 834 | Bacteria>Terrabacteria_group>Chloroflexota>unclassified_Chloroflexota | GCA_003446795.1 | Chloroflexota_bacterium | DCK98_15855 | 19417..21921 | 2026724 | AAA-family-ATPase-[Chloroflexi-bacterium] |
| MBC8450530.1 | AAA+ATPase+wHTH+Alpha+Beta-domain+wHTH+RRM | Type-1_BR.RM.like.systems | <-HAD<-?\|\|METHYLASE->REase->DUF499*->ABhydrolase-> | 946 | Bacteria>Terrabacteria_group>Chloroflexota>unclassified_Chloroflexota | GCA_014383605.1 | Chloroflexota_bacterium | H8D78_22580 | 11778..14618 | 2026724 | ATP-binding-protein-[Chloroflexi-bacterium] |
| MBU0491443.1 | AAA+ATPase+wHTH+Alpha+Beta-domain+wHTH+RRM | Type-1_BR.RM.like.systems | METHYLASE->?->?->?-><-?<-RNase\|\|DUF499*->?->Peptidase_C14-> | 1092 | Bacteria>Terrabacteria_group>Chloroflexota>unclassified_Chloroflexota | GCA_018812555.1 | Chloroflexota_bacterium | KKB13_06300 | 10013..13291 | 2026724 | DUF499-domain-containing-protein-[Chloroflexi-bacterium] |
| NBD36354.1 | AAA+ATPase+wHTH+Alpha+Beta-domain+wHTH+RRM | Type-1_BR.RM.like.systems | <-Mbetalac<-?<-DUF499*<-TPR+TPR+TPR+TPR\|\|?->PIN-> | 629 | Bacteria>Terrabacteria_group>Chloroflexota>unclassified_Chloroflexota | GCA_009908585.1 | Chloroflexota_bacterium | GVY30_10225 | 3115..5004 | 2026724 | DUF499-domain-containing-protein-[Chloroflexota-bacterium] |
| MDE3090299.1 | AAA+ATPase+wHTH+Alpha+Beta-domain+wHTH+RRM | Type-1_BR.RM.like.systems | <-DUF499*<-HEPN<-PolB<-PhdYeFM_antitox | 499 | Bacteria>Terrabacteria_group>Chloroflexota>unclassified_Chloroflexota | GCA_028873035.1 | Chloroflexota_bacterium | KGJ80_13035 | 1..1497 | 2026724 | MAG:-ATP-binding-protein,-partial-[Chloroflexota-bacterium] |
| MBU1747544.1 | AAA+ATPase+wHTH+Alpha+Beta-domain+wHTH+RRM | Type-1_BR.RM.like.systems | <-DUF499*<-REase<-?<-METHYLASE | 437 | Bacteria>Terrabacteria_group>Chloroflexota>unclassified_Chloroflexota | GCA_018823415.1 | Chloroflexota_bacterium | KKA73_07640 | 1..1312 | 2026724 | DUF499-domain-containing-protein-[Chloroflexota-bacterium] |
| MBO9368960.1 | AAA+ATPase+wHTH+Alpha+Beta-domain+wHTH+RRM | Type-1_BR.RM.like.systems | DUF499*->HKD-DNase+SF1-SF2-Helicase-> | 856 | Bacteria>Terrabacteria_group>Chloroflexota>unclassified_Chloroflexota | GCA_017743695.1 | Chloroflexota_bacterium | J7555_09500 | 1..2571 | 2026724 | ATP-binding-protein-[Chloroflexi-bacterium] |
| MCE7937788.1 | AAA+ATPase+wHTH+Alpha+Beta-domain+wHTH+RRM | Type-1_BR.RM.like.systems | <-DUF499*<-TOPRIM<-OLD-ABC<-METHYLASE-p | 923 | Bacteria>Terrabacteria_group>Chloroflexota>unclassified_Chloroflexota | GCA_021404695.1 | Chloroflexi_bacterium_CFX6 | DYG90_04145 | 392..3163 | 2293710 | ATP-binding-protein-[Chloroflexi-bacterium-CFX6] |
| MBI4630530.1 | AAA+ATPase+wHTH+Alpha+Beta-domain+wHTH+RRM | Type-1_BR.RM.like.systems | METHYLASE-p->?->DUF499*->HAMP+ACYC-> | 1116 | Bacteria>Terrabacteria_group>Chloroflexota>unclassified_Chloroflexota | GCA_016209165.1 | Chloroflexota_bacterium | HY740_02235 | 3036..6386 | 2026724 | ATP-binding-protein-[Chloroflexi-bacterium] |
| MBI4329820.1 | AAA+ATPase+wHTH+Alpha+Beta-domain+wHTH+RRM | Type-1_BR.RM.like.systems | <-DUF499*<-MACRODOMAIN<-?<-REase | 957 | Bacteria>Terrabacteria_group>Chloroflexota>unclassified_Chloroflexota | GCA_016210565.1 | Chloroflexota_bacterium | HY673_00895 | 861..3734 | 2026724 | DUF499-domain-containing-protein-[Chloroflexi-bacterium] |
| MBU1750077.1 | AAA+ATPase+wHTH+Alpha+Beta-domain+wHTH+RRM | Type-1_BR.RM.like.systems | METHYLASE->?->?->?-><-?<-RNase\|\|DUF499*-> | 575 | Bacteria>Terrabacteria_group>Chloroflexota>unclassified_Chloroflexota | GCA_018823415.1 | Chloroflexota_bacterium | KKA73_20530 | 10013..11738 | 2026724 | DUF499-domain-containing-protein-[Chloroflexota-bacterium] |
| MBC8450556.1 | AAA+ATPase+wHTH+Alpha+Beta-domain+wHTH+RRM | Type-1_BR.RM.like.systems | METHYLASE-p->REase->DUF499*-> | 874 | Bacteria>Terrabacteria_group>Chloroflexota>unclassified_Chloroflexota | GCA_014383605.1 | Chloroflexota_bacterium | H8D78_22710 | 1848..4469 | 2026724 | ATP-binding-protein-[Chloroflexi-bacterium] |
| HDQ73633.1 | AAA+ATPase+wHTH+Alpha+Beta-domain+wHTH+RRM | Type-1_BR.RM.like.systems | METHYLASE-p-><-PIN<-?\|\|?->DUF499*-> | 368 | Bacteria>Terrabacteria_group>Chloroflexota>unclassified_Chloroflexota | GCA_011046405.1 | Chloroflexota_bacterium | ENN19_16280 | 2854..3957 | 2026724 | ATP-binding-protein-[Chloroflexota-bacterium] |
| MCL5958118.1 | AAA+ATPase+wHTH+Alpha+Beta-domain+wHTH+RRM | Type-1_BR.RM.like.systems | <-HD<-DUF499*<-METHYLASE-p | 885 | Bacteria>Terrabacteria_group>Chloroflexota>unclassified_Chloroflexota | GCA_023489135.1 | Chloroflexota_bacterium | M1358_02180 | 938..3595 | 2026724 | DUF499-domain-containing-protein-[Chloroflexi-bacterium] |
| RLC91836.1 | AAA+ATPase+wHTH+Alpha+Beta-domain+wHTH+RRM | Type-1_BR.RM.like.systems | DUF499*->MNS+HTH+REase-> | 216 | Bacteria>Terrabacteria_group>Chloroflexota>unclassified_Chloroflexota | GCA_003648125.1 | Chloroflexota_bacterium | DRI77_13445 | 1..652 | 2026724 | hypothetical-protein-DRI77_13445,-partial-[Chloroflexota-bacterium] |
| KXK52491.1 | AAA+ATPase+wHTH+Alpha+Beta-domain+wHTH+RRM | Type-1_BR.RM.like.systems | <-DUF499*<-?<-?<-?<-METHYLASE-p | 1099 | Bacteria>Terrabacteria_group>Chloroflexota>unclassified_Chloroflexota | GCA_001567485.1 | Chloroflexi_bacterium_OLB13 | UZ13_01360 | 98984..102283 | 1617414 | ATPase-AAA-[Chloroflexi-bacterium-OLB13] |
| MBV7336884.1 | AAA+ATPase+wHTH+Alpha+Beta-domain+wHTH+RRM | Type-1_BR.RM.like.systems | XIS-HTH->?->?->?->TUDOR+SF1-SF2-Helicase+iREase+CTD+REase->METHYLASE->DUF499*-><-?<-DDE_Tnp_1_assoc+DDE_Tnp_1\|\|DUF433->PIN-> | 230 | Bacteria>Terrabacteria_group>Chloroflexota>unclassified_Chloroflexota | GCA_019217005.1 | Chloroflexi_bacterium_TSY | KFU94_53370 | 42959..43651 | 2834438 | DUF499-domain-containing-protein-[Chloroflexi-bacterium-TSY] |
| MBI4317402.1 | AAA+ATPase+wHTH+Alpha+Beta-domain+wHTH+RRM | Type-1_BR.RM.like.systems | <-PAIREDC-HTH<-DUF499*<-DUF499-p<-RNase<-METHYLASE<-TUDOR+SF1-SF2-Helicase+iREase+CTD+REase | 236 | Bacteria>Terrabacteria_group>Chloroflexota>unclassified_Chloroflexota | GCA_016210495.1 | Chloroflexota_bacterium | HY675_02840 | 1234..1944 | 2026724 | DUF499-domain-containing-protein-[Chloroflexota-bacterium] |
| MBI4295304.1 | AAA+ATPase+wHTH+Alpha+Beta-domain+wHTH+RRM | Type-1_BR.RM.like.systems | TUDOR+SF1-SF2-Helicase+iREase+CTD+REase->HEPN->Pkinase->METHYLASE->DUF433->PIN->DUF499*-> | 770 | Bacteria>Terrabacteria_group>Chloroflexota>unclassified_Chloroflexota | GCA_016210605.1 | Chloroflexota_bacterium | HY669_03965 | 9180..11489 | 2026724 | ATP-binding-protein-[Chloroflexi-bacterium] |
| NJD28055.1 | AAA+ATPase+wHTH+Alpha+Beta-domain+wHTH+RRM | Type-1_BR.RM.like.systems | ABC-ATPase+SbcC->?->?->?->?->TUDOR+SF1-SF2-Helicase+iREase+CTD+REase->DUF499*->?->PhdYeFM_antitox-> | 1013 | Bacteria>Terrabacteria_group>Chloroflexota>unclassified_Chloroflexota | GCA_012026785.1 | Chloroflexota_bacterium | FIA92_07135 | 27900..30941 | 2026724 | DUF499-domain-containing-protein-[Chloroflexi-bacterium] |
| MCL4370916.1 | AAA+ATPase+wHTH+Alpha+Beta-domain+wHTH+RRM | Type-1_BR.RM.like.systems | <-DUF499*<-METHYLASE<-REase<-?<-?<-Schlafen+HATPase+HTH<-TUDOR+SF1-SF2-Helicase+iREase+CTD+REase-p | 584 | Bacteria>Terrabacteria_group>Chloroflexota>unclassified_Chloroflexota | GCA_023380955.1 | Chloroflexota_bacterium | M1380_08425 | 1..1753 | 2026724 | DUF499-domain-containing-protein-[Chloroflexota-bacterium] |
| MYB84482.1 | AAA+ATPase+wHTH+Alpha+Beta-domain+wHTH+RRM | Type-1_BR.RM.like.systems | <-DUF499*<-METHYLASE<-TUDOR+SF1-SF2-Helicase+iREase+CTD+REase | 987 | Bacteria>Terrabacteria_group>Chloroflexota>unclassified_Chloroflexota | GCA_009840345.1 | Chloroflexota_bacterium | F4X54_07095 | 522..3485 | 2026724 | DUF499-domain-containing-protein-[Chloroflexi-bacterium] |
| MBI4327839.1 | AAA+ATPase+wHTH+Alpha+Beta-domain+wHTH+RRM | Type-1_BR.RM.like.systems | <-DUF499*<-TOPRIM<-OLD-ABC<-METHYLASE | 699 | Bacteria>Terrabacteria_group>Chloroflexota>unclassified_Chloroflexota | GCA_016210465.1 | Chloroflexota_bacterium | HY674_21625 | 1..2096 | 2026724 | ATP-binding-protein-[Chloroflexota-bacterium] |
| MBY0357952.1 | AAA+ATPase+wHTH+Alpha+Beta-domain+wHTH+RRM | Type-1_BR.RM.like.systems | <-SF1-SF2-Helicase+REase-p\|\|TUDOR+SF1-SF2-Helicase+iREase+CTD+REase->METHYLASE->DUF499*->REase->?->SIGMA-HTH-> | 939 | Bacteria>Terrabacteria_group>Cyanobacteriota/Melainabacteria_group>Candidatus_Melainabacteria>Candidatus_Obscuribacterales | GCA_019750335.1 | Candidatus_Obscuribacterales_bacterium | K2W82_08125 | 417056..419875 | 2364082 | DUF499-domain-containing-protein-[Candidatus-Obscuribacterales-bacterium] |

| Protein-ID | DUF499-architecture | System | Gene-Neighbourhoods | Length | Taxonomy | Assembly-ID | Strain | Locus-Tag | Genomic-coordinates | TaxID | Fasta-defline |
|---|---|---|---|---|---|---|---|---|---|---|---|
| MCS6267986.1 | AAA+ATPase+wHTH+Alpha+Beta-domain+wHTH+RRM | Type-1_BR.RM.like.systems | DOC+HTH-><-?<-DUF499*<-METHYLASE<-TUDOR+SF1-SF2-Helicase+iREase+CTD+REase\|\|DUF772+DDE_Tnp_1->TRD+TRD-> | 943 | Bacteria>Terrabacteria_group>Cyanobacteriota/Melainabacteria_group>Candidatus_Melainabacteria>Vampirovibrionales | GCA_024998755.1 | Vampirovibrio_sp. | H2174_10540 | 1332847..1335678 | 2717857 | ATP-binding-protein-[Vampirovibrio-sp.] |
| WP_245395605.1 | AAA+ATPase+wHTH+Alpha+Beta-domain+wHTH+RRM | Type-1_BR.RM.like.systems | <-DDE_Tnp_1\|\|TUDOR+SF1-SF2-Helicase+iREase+CTD+REase-><-?\|\|METHYLASE->REase->DUF499*->?->GDResolvase+HIN-HTH-><-REase+HTH | 977 | Bacteria>Terrabacteria_group>Cyanobacteriota/Melainabacteria_group>Cyanobacteriota>Cyanophyceae | GCF_018389385.1 | Anthocerotibacter_panamensis | IL331_RS05745 | 1255834..1258767 | 2857077 | DUF499-domain-containing-protein-[Anthocerotibacter-panamensis] |
| WP_242054772.1 | AAA+ATPase+wHTH+Alpha+Beta-domain+wHTH+RRM | Type-1_BR.RM.like.systems | TUDOR+SF1-SF2-Helicase+iREase+CTD+REase->Phd-YefM->PIN-><-HTH_3\|\|METHYLASE-p->DUF499*-><-TraJ-RHH\|\|PIWI->\|\|?-><-tRNA<-?<-SPOUT | 1065 | Bacteria>Terrabacteria_group>Cyanobacteriota/Melainabacteria_group>Cyanobacteriota>Cyanophyceae | GCF_014698065.1 | Nostoc_flagelliforme | H6G97_RS01005 | 252887..256084 | 1306274 | DUF499-domain-containing-protein-[Nostoc-flagelliforme] |
| MBC6477472.1 | AAA+ATPase+wHTH+Alpha+Beta-domain+wHTH+RRM | Type-1_BR.RM.like.systems | REase->REase->?->?->?->PIN-><-DUF499*<-METHYLASE<-TUDOR+SF1-SF2-Helicase+iREase+CTD+REase\|\|ABC-ATPase+SbcC->OLD-ABC->TOPRIM-> | 976 | Bacteria>Terrabacteria_group>Cyanobacteriota/Melainabacteria_group>Cyanobacteriota>Cyanophyceae | GCA_014323675.1 | Hormoscilla_sp._GM7CHS1pb | GDA56_06445 | 1108802..1111732 | 2653677 | ATP-binding-protein-[Hormoscilla-sp.-GM7CHS1pb] |
| WP_225912129.1 | AAA+ATPase+wHTH+Alpha+Beta-domain+wHTH+RRM | Type-1_BR.RM.like.systems | SF1-SF2-Helicase->?->HTH+HKD-DNase->?->SPOUT->?->tRNA-><-DUF499*\|\|HTH_3-><-TUDOR+SF1-SF2-Helicase+iREase+CTD+REase-p<-?<-DDE_Tnp_1 | 566 | Bacteria>Terrabacteria_group>Cyanobacteriota/Melainabacteria_group>Cyanobacteriota>Cyanophyceae | GCF_002813575.1 | Nostoc_flagelliforme | COO91_RS21680 | 4679917..4681617 | 1306274 | DUF499-domain-containing-protein-[Nostoc-flagelliforme] |
| WP_261362403.1 | AAA+ATPase+wHTH+Alpha+Beta-domain+wHTH+RRM | Type-1_BR.RM.like.systems | <-SIGMA-HTH<-?<-?<-DUF499*<-wHTH(SBD)+HNH<-METHYLASE-p<-HNH<-OLD-ABC<-TUDOR+SF1-SF2-Helicase+iREase+CTD+REase<-LRP-HTH | 820 | Bacteria>Terrabacteria_group>Cyanobacteriota/Melainabacteria_group>Cyanobacteriota>Cyanophyceae | - | unclassified_Cyanobium | KBZ34_RS12460 | 60686..63148 | 2627006 | MULTISPECIES:-DUF499-domain-containing-protein-[unclassified-Cyanobium] |
| WP_254953294.1 | AAA+ATPase+wHTH+Alpha+Beta-domain+wHTH+RRM | Type-1_BR.RM.like.systems | <-SIGMA-HTH<-?<-?<-DUF499*<-wHTH(SBD)+HNH<-METHYLASE-p<-HNH<-OLD-ABC<-TUDOR+SF1-SF2-Helicase+iREase+CTD+REase<-LRP-HTH | 819 | Bacteria>Terrabacteria_group>Cyanobacteriota/Melainabacteria_group>Cyanobacteriota>Cyanophyceae | GCF_024345955.1 | unclassified_Cyanobium | KBZ34_RS12405 | 60686..63145 | 2627006 | MULTISPECIES:-DUF499-domain-containing-protein-[unclassified-Cyanobium] |
| MCP9859835.1 | AAA+ATPase+wHTH+Alpha+Beta-domain+wHTH+RRM | Type-1_BR.RM.like.systems | <-SIGMA-HTH<-?<-?<-DUF499*<-wHTH(SBD)+HNH<-METHYLASE-p<-HNH<-OLD-ABC<-TUDOR+SF1-SF2-Helicase+iREase+CTD+REase<-LRP-HTH | 819 | Bacteria>Terrabacteria_group>Cyanobacteriota/Melainabacteria_group>Cyanobacteriota>Cyanophyceae | GCA_024345955.1 | Cyanobium_sp._Cruz-8H5 | KBZ34_12405 | 60686..63145 | 2823712 | ATP-binding-protein-[Cyanobium-sp.-Cruz-8H5] |
| WP_216087063.1 | AAA+ATPase+wHTH+Alpha+Beta-domain+wHTH+RRM | Type-1_BR.RM.like.systems | <-HTH_OrfB_IS605+OrfB_IS605+OrfB_Zn_ribbon\|\|METHYLASE-p->CopG/RHH->ACET-><-?\|\|HEPN->DUF499*-> | 951 | Bacteria>Terrabacteria_group>Cyanobacteriota/Melainabacteria_group>Cyanobacteriota>Cyanophyceae | GCF_000317575.1 | Stanieria_cyanosphaera | STA7437_RS18665 | 4291958..4294813 | 102116 | DUF499-domain-containing-protein-[Stanieria-cyanosphaera] |
| WP_143780302.1 | AAA+ATPase+wHTH+Alpha+Beta-domain+wHTH+RRM | Type-1_BR.RM.like.systems | <-HHH+ENDO3+NUDIX<-Peptidase_M50+PDZ\|\|?-><-METHYLASE-p<-DUF499*<-HEPN<-?<-?<-REase<-METHYLASE-p | 735 | Bacteria>Terrabacteria_group>Cyanobacteriota/Melainabacteria_group>Cyanobacteriota>Cyanophyceae | GCF_001939115.1 | Leptolyngbya_sp._'hensonii' | BST81_RS10560 | 80963..83170 | 1922337 | DUF499-domain-containing-protein-[Leptolyngbya-sp.-'hensonii'] |
| WP_236308150.1 | AAA+ATPase+wHTH+Alpha+Beta-domain+wHTH+RRM | Type-1_BR.RM.like.systems | URI-DNase+HhH-RADC-><-?<-HTH+MPTase<-HEPN<-PolB\|\|DUF499*->?->REase->?->Peptidase_C14-> | 460 | Bacteria>Terrabacteria_group>Cyanobacteriota/Melainabacteria_group>Cyanobacteriota>Cyanophyceae | GCF_021650815.1 | Cylindrospermopsis_raciborskii | L3I90_RS10045 | 2305504..2306886 | 77022 | DUF499-domain-containing-protein-[Cylindrospermopsis-raciborskii] |
| KAB8333560.1 | AAA+ATPase+wHTH+Alpha+Beta-domain+wHTH+RRM | Type-1_BR.RM.like.systems | REase->REase->SF1-SF2-Helicase+REase->METHYLASE->METHYLASE->?->DUF499*->WYL-> | 553 | Bacteria>Terrabacteria_group>Cyanobacteriota/Melainabacteria_group>Cyanobacteriota>Cyanophyceae | GCA_000828085.3 | Scytonema_tolypothrichoides_VB-61278 | SD80_012900 | 39454..41115 | 1233231 | hypothetical-protein-SD80_012900-[Scytonema-tolypothrichoides-VB-61278] |
| WP_250683155.1 | AAA+ATPase+wHTH+Alpha+Beta-domain+wHTH+RRM | Type-1_BR.RM.like.systems | <-DUF499*<-HEPN<-REase<-METHYLASE<-PemK_toxin<-AbrB<-PolB | 788 | Bacteria>Terrabacteria_group>Cyanobacteriota/Melainabacteria_group>Cyanobacteriota>Cyanophyceae | - | Thermosynechococcus_sp._HN-54 | NBE99_RS03710 | 771476..773842 | 2933959 | DUF499-domain-containing-protein-[Thermosynechococcus-sp.-HN-54] |
| WP_228036143.1 | AAA+ATPase+wHTH+Alpha+Beta-domain+wHTH+RRM | Type-1_BR.RM.like.systems | <-sGTP<-DUF29<-?<-?\|\|?->Cluster164_18clades-><-DUF499* | 856 | Bacteria>Terrabacteria_group>Cyanobacteriota/Melainabacteria_group>Cyanobacteriota>Cyanophyceae | GCF_015207925.1 | Microcystis_sp._LEGE_00066 | IQ224_RS08955 | 23232..25802 | 1828685 | DUF499-domain-containing-protein-[Microcystis-sp.-LEGE-00066] |
| KKZ10364.1 | AAA+ATPase+wHTH+Alpha+Beta-domain+wHTH+RRM | Type-1_BR.RM.like.systems | METHYLASE->HEPN->DUF499*-> | 965 | Bacteria>Terrabacteria_group>Cyanobacteriota/Melainabacteria_group>Cyanobacteriota>Cyanophyceae | GCA_001007625.1 | Candidatus_Synechococcus_spongiarum_142 | TH68_11020 | 5098..7996 | 1608213 | ATPase-AAA,-partial-[Candidatus-Synechococcus-spongiarum-142] |
| WP_227366555.1 | AAA+ATPase+wHTH+Alpha+Beta-domain+wHTH+RRM | Type-1_BR.RM.like.systems | GspA-ATPase->METHYLASE-p->REase->REase->REase->DUF499*-><-TOPRIM+SF1-SF2-Helicase\|\|Phage_int_SAM_1+Phage_integrase->?->TUDOR+SF1-SF2-Helicase+iREase+CTD->METHYLASE-p->REase-> | 1002 | Bacteria>Terrabacteria_group>Cyanobacteriota/Melainabacteria_group>Cyanobacteriota>Cyanophyceae | GCF_020594525.1 | Planktothrix_agardhii | LJE15_RS21495 | 6203..9211 | 1160 | DUF499-domain-containing-protein-[Planktothrix-agardhii] |
| WP_235753326.1 | AAA+ATPase+wHTH+Alpha+Beta-domain+wHTH+RRM | Type-1_BR.RM.like.systems | PIN->METHYLASE->REase->REase->REase->DUF499*-><-?\|\|RelB_antitoxin->HTH_OrfB_IS605+OrfB_IS605+OrfB_Zn_ribbon-><-PAIREDC-HTH | 1002 | Bacteria>Terrabacteria_group>Cyanobacteriota/Melainabacteria_group>Cyanobacteriota>Cyanophyceae | GCF_021554825.1 | Planktothrix_agardhii | L2E72_RS21865 | 12950..15958 | 1160 | DUF499-domain-containing-protein-[Planktothrix-agardhii] |
| WP_200888917.1 | AAA+ATPase+wHTH+Alpha+Beta-domain+wHTH+RRM | Type-1_BR.RM.like.systems | TUDOR+SF1-SF2-Helicase+iREase+CTD+REase->REase+HTH->?->METHYLASE-p->METHYLASE-p->DUF499*->?->Methyltransf_21->?->HTH_3->REase-><-N6_N4_MTase | 909 | Bacteria>Terrabacteria_group>Cyanobacteriota/Melainabacteria_group>Cyanobacteriota>Cyanophyceae | GCF_000789435.1 | Aphanizomenon_flos-aquae | OA07_RS10570 | 7797..10526 | 1176 | DUF499-domain-containing-protein-[Aphanizomenon-flos-aquae] |
| BAQ63189.1 | AAA+ATPase+wHTH+Alpha+Beta-domain+wHTH+RRM | Type-1_BR.RM.like.systems | TUDOR+SF1-SF2-Helicase+iREase+CTD+REase-><-ZnR\|\|METHYLASE->?->PIN->DUF499*-><-Phage_integrase<-DUF4158+DDE_Tnp_Tn3<-Phage_integrase\|\|?->N6_MTase-p-> | 1005 | Bacteria>Terrabacteria_group>Cyanobacteriota/Melainabacteria_group>Cyanobacteriota>Cyanophyceae | GCA_001548095.1 | Geminocystis_sp._NIES-3708 | GM3708_3595 | 38002..41019 | 1615909 | predicted-ATPase-[Geminocystis-sp.-NIES-3708] |
| WP_254434496.1 | AAA+ATPase+wHTH+Alpha+Beta-domain+wHTH+RRM | Type-1_BR.RM.like.systems | SF1-SF2-Helicase-p->METHYLASE-p->MetJarc-HTH->PIN->MetJarc-HTH->DUF499*->DUF499-p->?->HISKIN+MORC->Methyltransf_21->?->HTH_3-> | 386 | Bacteria>Terrabacteria_group>Cyanobacteriota/Melainabacteria_group>Cyanobacteriota>Cyanophyceae | GCF_009712065.1 | Aphanizomenon_sp._UHCC_0183 | FJR37_RS03355 | 2964..4124 | 2590028 | DUF499-domain-containing-protein-[Aphanizomenon-sp.-UHCC-0183] |
| WP_254171845.1 | AAA+ATPase+wHTH+Alpha+Beta-domain+wHTH+RRM | Type-1_BR.RM.like.systems | PIN->METHYLASE->REase->REase->REase->DUF499*-><-?\|\|RelB_antitoxin->HTH_OrfB_IS605+OrfB_IS605+OrfB_Zn_ribbon->?->RHH_1-> | 1002 | Bacteria>Terrabacteria_group>Cyanobacteriota/Melainabacteria_group>Cyanobacteriota>Cyanophyceae | GCF_904830885.1 | Planktothrix_agardhii | NMK31_RS00300 | 70820..73828 | 1160 | DUF499-domain-containing-protein-[Planktothrix-agardhii] |
| WP_189523574.1 | AAA+ATPase+wHTH+Alpha+Beta-domain+wHTH+RRM | Type-1_BR.RM.like.systems | <-ArdA\|\|MetJarc-HTH->?-><-?<-?<-DUF499*<-REase<-METHYLASE<-OLD-ABC<-TUDOR+SF1-SF2-Helicase+iREase+CTD+REase\|\|?-><-HKD-DNase+SF1-SF2-Helicase | 962 | Bacteria>Terrabacteria_group>Cyanobacteriota/Melainabacteria_group>Cyanobacteriota>Cyanophyceae | GCF_005869855.1 | Nostoc_sp._PA-18-2419 | FCR06_RS25145 | 41137..44025 | 2575443 | DUF499-domain-containing-protein-[Nostoc-sp.-PA-18-2419] |
| WP_272056151.1 | AAA+ATPase+wHTH+Alpha+Beta-domain+wHTH+RRM | Type-1_BR.RM.like.systems | <-SF1-SF2-Helicase-p<-?\|\|?->PIN->TUDOR+SF1-SF2-Helicase+iREase+CTD+REase->METHYLASE->DUF499*-><-Phage_integrase\|\|RelE-ParE->HTH_37->MORC-> | 1005 | Bacteria>Terrabacteria_group>Cyanobacteriota/Melainabacteria_group>Cyanobacteriota>Cyanophyceae | GCF_028330355.1 | Nodularia_spumigena | PN490_RS03030 | 15639..18656 | 70799 | DUF499-domain-containing-protein-[Nodularia-spumigena] |
| WP_254035908.1 | AAA+ATPase+wHTH+Alpha+Beta-domain+wHTH+RRM | Type-1_BR.RM.like.systems | PIN->PIN->METHYLASE->REase->REase->REase->DUF499*-><-?\|\|RelB_antitoxin->PemK_toxin->RHH_1-><-TOPRIM+SF1-SF2-Helicase | 999 | Bacteria>Terrabacteria_group>Cyanobacteriota/Melainabacteria_group>Cyanobacteriota>Cyanophyceae | GCF_904830935.1 | Planktothrix_agardhii | NMG88_RS22325 | 51359..54358 | 1160 | DUF499-domain-containing-protein-[Planktothrix-agardhii] |
| WP_235758599.1 | AAA+ATPase+wHTH+Alpha+Beta-domain+wHTH+RRM | Type-1_BR.RM.like.systems | TUDOR+SF1-SF2-Helicase+iREase+CTD+REase-p->PIN->METHYLASE-p->REase->REase->REase->DUF499*-><-TOPRIM+SF1-SF2-Helicase<-ZnR<-?<-?<-URI-DNase | 1002 | Bacteria>Terrabacteria_group>Cyanobacteriota/Melainabacteria_group>Cyanobacteriota>Cyanophyceae | - | Planktothrix_agardhii | L2E81_RS19965 | 6975..9983 | 1160 | DUF499-domain-containing-protein-[Planktothrix-agardhii] |
| WP_066349645.1 | AAA+ATPase+wHTH+Alpha+Beta-domain+wHTH+RRM | Type-1_BR.RM.like.systems | TUDOR+SF1-SF2-Helicase+iREase+CTD+REase-><-ZnR\|\|METHYLASE->?->PIN->DUF499*-><-DUF4158+DDE_Tnp_Tn3<-Phage_integrase\|\|?->N6_MTase-p-> | 1007 | Bacteria>Terrabacteria_group>Cyanobacteriota/Melainabacteria_group>Cyanobacteriota>Cyanophyceae | - | Geminocystis_sp._NIES-3708 | GM3708_RS17435 | 37996..41019 | 1615909 | DUF499-domain-containing-protein-[Geminocystis-sp.-NIES-3708] |
| WP_190447191.1 | AAA+ATPase+wHTH+Alpha+Beta-domain+wHTH+RRM | Type-1_BR.RM.like.systems | Prim-Pol->CHTH->TUDOR+SF1-SF2-Helicase+iREase+CTD+REase->RnaseIII+DSRBD->METHYLASE->REase->DUF499*->?->PAPS_reduct->?->ParB-><-REase | 968 | Bacteria>Terrabacteria_group>Cyanobacteriota/Melainabacteria_group>Cyanobacteriota>Cyanophyceae | GCF_014695795.1 | Phormidium_sp._FACHB-592 | H6F86_RS05945 | 30309..33215 | 2692850 | DUF499-domain-containing-protein-[Phormidium-sp.-FACHB-592] |
| MBI1240693.1 | AAA+ATPase+wHTH+Alpha+Beta-domain+wHTH+RRM | Type-1_BR.RM.like.systems | Cysteine_desulfurase->TUDOR+SF1-SF2-Helicase+iREase+CTD+REase->METHYLASE->REase->DUF499*-><-Peptidase_S51<-?<-?<-?\|\|sGTP-> | 973 | Bacteria>Terrabacteria_group>Cyanobacteriota/Melainabacteria_group>Cyanobacteriota>Cyanophyceae | GCA_016124525.1 | Nostoc_sp._RI_552 | GC195_03655 | 36340..39261 | 2654185 | DUF499-domain-containing-protein-[Nostoc-sp.-RI_552] |
| WP_227382638.1 | AAA+ATPase+wHTH+Alpha+Beta-domain+wHTH+RRM | Type-1_BR.RM.like.systems | PIN->METHYLASE->REase->REase->DUF499*-><-?\|\|RelB_antitoxin->PemK_toxin->?->RHH_1-><-TOPRIM+SF1-SF2-Helicase | 1002 | Bacteria>Terrabacteria_group>Cyanobacteriota/Melainabacteria_group>Cyanobacteriota>Cyanophyceae | GCF_020594575.1 | Planktothrix_agardhii | LJE16_RS21930 | 10143..13151 | 1160 | DUF499-domain-containing-protein-[Planktothrix-agardhii] |
| WP_227398372.1 | AAA+ATPase+wHTH+Alpha+Beta-domain+wHTH+RRM | Type-1_BR.RM.like.systems | ZnR->TOPRIM+SF1-SF2-Helicase-><-PemK_toxin<-RelB_antitoxin<-?\|\|?-><-DUF499*<-REase<-REase<-METHYLASE<-PIN | 1002 | Bacteria>Terrabacteria_group>Cyanobacteriota/Melainabacteria_group>Cyanobacteriota>Cyanophyceae | GCF_020594695.1 | Planktothrix_agardhii | L2E74_RS21495 | 55743..58751 | 1160 | DUF499-domain-containing-protein-[Planktothrix-agardhii] |
| WP_168636740.1 | AAA+ATPase+wHTH+Alpha+Beta-domain+wHTH+RRM | Type-1_BR.RM.like.systems | TUDOR+SF1-SF2-Helicase+iREase+CTD+REase->REase+HTH->?->METHYLASE->?->PIN->DUF499*->?->?->HTH_3->REase-><-N6_N4_MTase\|\|NTP_transferase-> | 911 | Bacteria>Terrabacteria_group>Cyanobacteriota/Melainabacteria_group>Cyanobacteriota>Cyanophyceae | GCF_017355725.1 | Nostocales | HEQ36_RS21575 | 43162..45897 | 1161 | DUF499-domain-containing-protein-[Dolichospermum-flos-aquae] |
| WP_053539516.1 | AAA+ATPase+wHTH+Alpha+Beta-domain+wHTH+RRM | Type-1_BR.RM.like.systems | <-NTP_transferase\|\|N6_N4_MTase-><-HTH_3<-?<-?<-DUF499*<-PIN<-?<-METHYLASE<-REase+HTH<-TUDOR+SF1-SF2-Helicase+iREase+CTD+REase<-CHTH | 911 | Bacteria>Terrabacteria_group>Cyanobacteriota/Melainabacteria_group>Cyanobacteriota>Cyanophyceae | GCF_001277295.1 | Anabaena_sp._WA102 | AA650_RS14270 | 3307169..3309904 | 1647413 | DUF499-domain-containing-protein-[Anabaena-sp.-WA102] |
| MBO1068320.1 | AAA+ATPase+wHTH+Alpha+Beta-domain+wHTH+RRM | Type-1_BR.RM.like.systems | <-NTP_transferase\|\|N6_N4_MTase-><-HTH_3<-?<-?<-DUF499*<-PIN<-?<-METHYLASE<-?<-REase+HTH<-TUDOR+SF1-SF2-Helicase+iREase+CTD+REase | 911 | Bacteria>Terrabacteria_group>Cyanobacteriota/Melainabacteria_group>Cyanobacteriota>Cyanophyceae | GCA_017355655.1 | Dolichospermum_sp._DEX189 | HEQ13_02485 | 530022..532757 | 2717870 | ATP-binding-protein-[Dolichospermum-sp.-DEX189] |
| WP_023065229.1 | AAA+ATPase+wHTH+Alpha+Beta-domain+wHTH+RRM | Type-1_BR.RM.like.systems | TUDOR+SF1-SF2-Helicase+iREase+CTD+REase->METHYLASE->REase->DUF499*-><-?\|\|?->?-><-NTP_transf_9<-TPR+TPR+TPR+TPR+TPR+TPR+TPR+TPR | 968 | Bacteria>Terrabacteria_group>Cyanobacteriota/Melainabacteria_group>Cyanobacteriota>Cyanophyceae | GCF_000478195.2 | Lyngbya_aestuarii | M595_RS06070 | 76585..79491 | 118322 | DUF499-domain-containing-protein-[Lyngbya-aestuarii] |
| WP_273769820.1 | AAA+ATPase+wHTH+Alpha+Beta-domain+wHTH+RRM | Type-1_BR.RM.like.systems | STYKinase-like->?-><-DUF499*<-METHYLASE<-?<-TUDOR+SF1-SF2-Helicase+iREase+CTD+REase<-Phage_int_SAM_4+Phage_integrase\|\|HicB-> | 968 | Bacteria>Terrabacteria_group>Cyanobacteriota/Melainabacteria_group>Cyanobacteriota>Cyanophyceae | GCF_028623165.1 | Nostoc_sp._UHCC_0926 | PQG02_RS30260 | 2075..4981 | 3025190 | DUF499-domain-containing-protein-[Nostoc-sp.-UHCC-0926] |
| WP_009787752.1 | AAA+ATPase+wHTH+Alpha+Beta-domain+wHTH+RRM | Type-1_BR.RM.like.systems | TPR+TPR+TPR+TPR+TPR+TPR+TPR+TPR->NTP_transf_9-><-?<-TPR+TPR+TPR+TPR+TPR+TPR+TPR+TPR<-?\|\|Pkinase-><-DUF499* | 968 | Bacteria>Terrabacteria_group>Cyanobacteriota/Melainabacteria_group>Cyanobacteriota>Cyanophyceae | GCF_000169095.1 | Lyngbya_sp._PCC_8106 | L8106_RS27125 | 23492..26398 | 313612 | DUF499-domain-containing-protein-[Lyngbya-sp.-PCC-8106] |
| BBD57116.1 | AAA+ATPase+wHTH+Alpha+Beta-domain+wHTH+RRM | Type-1_BR.RM.like.systems | MethyltransfD12->MethyltransfD12-><-?<-HTH_3\|\|DDE_Tnp_1->?-><-DUF499*<-REase<-REase<-REase<-METHYLASE<-PIN | 1002 | Bacteria>Terrabacteria_group>Cyanobacteriota/Melainabacteria_group>Cyanobacteriota>Cyanophyceae | GCA_003609755.1 | Planktothrix_agardhii_NIES-204 | NIES204_44520 | 39433..42441 | 282423 | AAA-family-ATPase-[Planktothrix-agardhii-NIES-204] |
| WP_096567230.1 | AAA+ATPase+wHTH+Alpha+Beta-domain+wHTH+RRM | Type-1_BR.RM.like.systems | ABC-ATPase+SbcC-><-DUF499*<-PIN<-?<-METHYLASE<-TUDOR+SF1-SF2-Helicase+iREase+CTD+REase-p<-TUDOR+SF1-SF2-Helicase+iREase+CTD+REase-p\|\|REase-> | 963 | Bacteria>Terrabacteria_group>Cyanobacteriota/Melainabacteria_group>Cyanobacteriota>Cyanophyceae | GCF_028329225.1 | Sphaerospermopsis_kisseleviana | PN497_RS06725 | 5045..7936 | 289435 | DUF499-domain-containing-protein-[Sphaerospermopsis-kisseleviana] |
| WP_204104332.1 | AAA+ATPase+wHTH+Alpha+Beta-domain+wHTH+RRM | Type-1_BR.RM.like.systems | SF1-SF2-Helicase-><-PP_kinase+HKD-DNase<-DUF499*<-METHYLASE<-?<-TUDOR+SF1-SF2-Helicase+iREase+CTD+REase<-?\|\|?-><-ABhydrolase | 968 | Bacteria>Terrabacteria_group>Cyanobacteriota/Melainabacteria_group>Cyanobacteriota>Cyanophyceae | GCF_016888125.1 | Spirulina_major | JUI58_RS13555 | 13537..16443 | 270636 | DUF499-domain-containing-protein-[Spirulina-major] |
| OIP70051.1 | AAA+ATPase+wHTH+Alpha+Beta-domain+wHTH+RRM | Type-1_BR.RM.like.systems | REase->?->TPR-><-DUF499*<-METHYLASE<-TUDOR+SF1-SF2-Helicase+iREase+CTD+REase-p<-TUDOR+SF1-SF2-Helicase+iREase+CTD+REase<-SF1-SF2-Helicase | 955 | Bacteria>Terrabacteria_group>Cyanobacteriota/Melainabacteria_group>Cyanobacteriota>Cyanophyceae | GCA_001873365.1 | Oscillatoriales_cyanobacterium_CG2_30_40_61 | AUK43_10925 | 45419..48286 | 1805291 | hypothetical-protein-AUK43_10925-[Oscillatoriales-cyanobacterium-CG2_30_40_61] |
| WP_225880403.1 | AAA+ATPase+wHTH+Alpha+Beta-domain+wHTH+RRM | Type-1_BR.RM.like.systems | <-REase<-?<-?<-DUF499<-CR-ATPase9+REase<-DUF499*<-CR-ATPase9+REase<-METHYLASE<-REase<-PIN<-MetJarc-HTH | 821 | Bacteria>Terrabacteria_group>Cyanobacteriota/Melainabacteria_group>Cyanobacteriota>Cyanophyceae | GCF_015245355.1 | Anabaenopsis_elenkinii | IM676_RS18795 | 4304033..4306498 | 156213 | DUF499-domain-containing-protein-[Anabaenopsis-elenkinii] |
| WP_141294973.1 | AAA+ATPase+wHTH+Alpha+Beta-domain+wHTH+RRM | Type-1_BR.RM.like.systems | TUDOR+SF1-SF2-Helicase+iREase+CTD+REase->REase->METHYLASE->REase->REase->DUF499*->?->?->?-><-Radical-SAM<-?<-AbrB-like | 999 | Bacteria>Terrabacteria_group>Cyanobacteriota/Melainabacteria_group>Cyanobacteriota>Cyanophyceae | GCF_005402765.1 | Planktothrix_agardhii | FLZ02_RS17820 | 16554..19553 | 1160 | DUF499-domain-containing-protein-[Planktothrix-agardhii] |
| WP_106289188.1 | AAA+ATPase+wHTH+Alpha+Beta-domain+wHTH+RRM | Type-1_BR.RM.like.systems | <-DUF499*<-REase<-METHYLASE<-OLD-ABC<-TUDOR+SF1-SF2-Helicase+iREase+CTD+REase<-CHTH\|\|OBFOLD-NUCLEASE-> | 994 | Bacteria>Terrabacteria_group>Cyanobacteriota/Melainabacteria_group>Cyanobacteriota>Cyanophyceae | GCF_003003775.1 | Merismopedia_glauca | C7B64_RS13505 | 1286..4270 | 292586 | DUF499-domain-containing-protein-[Merismopedia-glauca] |
| WP_254031531.1 | AAA+ATPase+wHTH+Alpha+Beta-domain+wHTH+RRM | Type-1_BR.RM.like.systems | <-RelB_antitoxin<-?\|\|?-><-?<-DUF499*<-REase<-REase<-REase<-METHYLASE<-DDE_Tnp_1_assoc+DDE_Tnp_1 | 999 | Bacteria>Terrabacteria_group>Cyanobacteriota/Melainabacteria_group>Cyanobacteriota>Cyanophyceae | GCF_904830945.1 | Planktothrix_agardhii | NMG75_RS16735 | 3649445..3652444 | 1160 | DUF499-domain-containing-protein-[Planktothrix-agardhii] |
| WP_190383648.1 | AAA+ATPase+wHTH+Alpha+Beta-domain+wHTH+RRM | Type-1_BR.RM.like.systems | METHYLASE->MetJarc-HTH->PIN->MetJarc-HTH->DUF499*->HISKIN+MORC->?->HTH_3->REase-><-N6_N4_MTase | 963 | Bacteria>Terrabacteria_group>Cyanobacteriota/Melainabacteria_group>Cyanobacteriota>Cyanophyceae | GCF_014696815.1 | Aphanizomenon_flos-aquae | H6F99_RS16810 | 3730..6621 | 1176 | DUF499-domain-containing-protein-[Aphanizomenon-flos-aquae] |
| WP_006634723.1 | AAA+ATPase+wHTH+Alpha+Beta-domain+wHTH+RRM | Type-1_BR.RM.like.systems | NTP_transferase->ST-Hiskin->?->TUDOR+SF1-SF2-Helicase+iREase+CTD+REase->METHYLASE->REase->DUF499*-> | 998 | Bacteria>Terrabacteria_group>Cyanobacteriota/Melainabacteria_group>Cyanobacteriota>Cyanophyceae | - | Microcoleus_vaginatus | MicvaDRAFT_4687 | 60148..63144 | 119532 | DUF499-domain-containing-protein-[Microcoleus-vaginatus] |
| EGK85305.1 | AAA+ATPase+wHTH+Alpha+Beta-domain+wHTH+RRM | Type-1_BR.RM.like.systems | NTP_transferase->ST-Hiskin->?->TUDOR+SF1-SF2-Helicase+iREase+CTD+REase->METHYLASE->REase->DUF499*-> | 998 | Bacteria>Terrabacteria_group>Cyanobacteriota/Melainabacteria_group>Cyanobacteriota>Cyanophyceae | GCA_000214075.2 | Microcoleus_vaginatus_FGP-2 | MicvaDRAFT_4687 | 60148..63144 | 756067 | AAA-family-ATPase-[Microcoleus-vaginatus-FGP-2] |
| WP_075888510.1 | AAA+ATPase+wHTH+Alpha+Beta-domain+wHTH+RRM | Type-1_BR.RM.like.systems | <-GDResolvase+HIN-HTH\|\|DUF4158+DDE_Tnp_Tn3-><-?<-?<-?<-?<-DUF499*<-Polbeta<-PolB-NTase<-PIN<-?<-METHYLASE | 965 | Bacteria>Terrabacteria_group>Cyanobacteriota/Melainabacteria_group>Cyanobacteriota>Cyanophyceae | GCF_001904615.1 | [Limnothrix_rosea]_IAM_M-220 | NIES208_RS00240 | 58146..61043 | 454133 | DUF499-domain-containing-protein-[Limnothrix-rosea] |
| QSV71060.1 | AAA+ATPase+wHTH+Alpha+Beta-domain+wHTH+RRM | Type-1_BR.RM.like.systems | REase+HTH->?->METHYLASE-p->METHYLASE-p->DUF499*->?->Methyltransf_21->?->HTH_3->REase-><-N6_N4_MTase | 911 | Bacteria>Terrabacteria_group>Cyanobacteriota/Melainabacteria_group>Cyanobacteriota>Cyanophyceae | GCA_017346875.1 | Aphanizomenon_flos-aquae_KM1D3_PB | HEQ20_10255 | 2086099..2088834 | 2723857 | ATP-binding-protein-[Aphanizomenon-flos-aquae-KM1D3_PB] |
| WP_257998857.1 | AAA+ATPase+wHTH+Alpha+Beta-domain+wHTH+RRM | Type-1_BR.RM.like.systems | ABC-ATPase+SbcC->?->?-><-?\|\|?-><-DUF499*<-REase<-METHYLASE<-OLD-ABC-p<-TUDOR+SF1-SF2-Helicase+iREase+CTD+REase | 955 | Bacteria>Terrabacteria_group>Cyanobacteriota/Melainabacteria_group>Cyanobacteriota>Cyanophyceae | GCF_002870755.1 | Fischerella_thermalis | CEN42_RS03265 | 8470..11337 | 372787 | DUF499-domain-containing-protein-[Fischerella-thermalis] |
| WP_200865503.1 | AAA+ATPase+wHTH+Alpha+Beta-domain+wHTH+RRM | Type-1_BR.RM.like.systems | TUDOR+SF1-SF2-Helicase+iREase+CTD+REase->?->PIN->METHYLASE->DUF433->PIN->DUF499*->DUF499-p->tRNA->?->?-><-?\|\|NADAR-><-MORC | 732 | Bacteria>Terrabacteria_group>Cyanobacteriota/Melainabacteria_group>Cyanobacteriota>Cyanophyceae | GCF_000505665.1 | Thermosynechococcus_sp._NK55a | NK55_RS09315 | 1926061..1928259 | 1394889 | DUF499-domain-containing-protein-[Thermosynechococcus-sp.-NK55a] |
| WP_137908144.1 | AAA+ATPase+wHTH+Alpha+Beta-domain+wHTH+RRM | Type-1_BR.RM.like.systems | <-REase+HTH<-?\|\|?-><-?<-?<-DUF499*<-PIN<-?<-MetJarc-HTH<-MetJarc-HTH<-OrfB_Zn_ribbon<-REase | 969 | Bacteria>Terrabacteria_group>Cyanobacteriota/Melainabacteria_group>Cyanobacteriota>Cyanophyceae | GCF_005402965.1 | Dolichospermum_planctonicum | FF006_RS11405 | 62234..65143 | 136072 | DUF499-domain-containing-protein-[Dolichospermum-planctonicum] |
| WP_102148297.1 | AAA+ATPase+wHTH+Alpha+Beta-domain+wHTH+RRM | Type-1_BR.RM.like.systems | ABC-ATPase+SbcC->?->?-><-?\|\|?-><-DUF499*<-REase<-METHYLASE<-OLD-ABC-p<-TUDOR+SF1-SF2-Helicase+iREase+CTD+REase | 975 | Bacteria>Terrabacteria_group>Cyanobacteriota/Melainabacteria_group>Cyanobacteriota>Cyanophyceae | - | Fischerella_thermalis | CEN42_03245 | 8470..11397 | 372787 | DUF499-domain-containing-protein-[Fischerella-thermalis] |
| PMB36920.1 | AAA+ATPase+wHTH+Alpha+Beta-domain+wHTH+RRM | Type-1_BR.RM.like.systems | ABC-ATPase+SbcC->?->?-><-?\|\|?-><-DUF499*<-REase<-METHYLASE<-OLD-ABC-p<-TUDOR+SF1-SF2-Helicase+iREase+CTD+REase | 975 | Bacteria>Terrabacteria_group>Cyanobacteriota/Melainabacteria_group>Cyanobacteriota>Cyanophyceae | GCA_002870755.1 | Fischerella_thermalis_CCMEE_5208 | CEN42_03245 | 8470..11397 | 2019669 | AAA-family-ATPase-[Fischerella-thermalis-CCMEE-5208] |
| WP_224180282.1 | AAA+ATPase+wHTH+Alpha+Beta-domain+wHTH+RRM | Type-1_BR.RM.like.systems | REase-><-?<-?<-ORC-CDC6<-ParA-Soj-PloopNTPase<-HEPN\|\|DUF499*->TUDOR+SF1-SF2-Helicase+iREase+CTD+REase-><-STYKIN | 957 | Bacteria>Terrabacteria_group>Cyanobacteriota/Melainabacteria_group>Cyanobacteriota>Cyanophyceae | - | Calothrix_sp._PCC_7716 | CAL7716_104380 | 93672..96545 | 2853227 | DUF499-domain-containing-protein-[Calothrix-sp.-PCC-7716] |
| WP_026797173.1 | AAA+ATPase+wHTH+Alpha+Beta-domain+wHTH+RRM | Type-1_BR.RM.like.systems | <-RelB_antitoxin<-?<-?<-?<-DUF499*<-REase<-REase<-REase<-METHYLASE<-REase<-TUDOR+SF1-SF2-Helicase+iREase+CTD+REase | 999 | Bacteria>Terrabacteria_group>Cyanobacteriota/Melainabacteria_group>Cyanobacteriota>Cyanophyceae | GCF_000464765.1 | Planktothrix | PLAN_RS23755 | 2806320..2809319 | 54304 | MULTISPECIES:-DUF499-domain-containing-protein-[Planktothrix] |
| CAH2571600.1 | AAA+ATPase+wHTH+Alpha+Beta-domain+wHTH+RRM | Type-1_BR.RM.like.systems | TUDOR+SF1-SF2-Helicase+iREase+CTD+REase->REase->METHYLASE->REase->REase->REase->DUF499*->?->?->RelB_antitoxin-> | 999 | Bacteria>Terrabacteria_group>Cyanobacteriota/Melainabacteria_group>Cyanobacteriota>Cyanophyceae | GCA_937024135.1 | Planktothrix_rubescens | PRNO82_00998 | 1099392..1102391 | 59512 | AAA-family-ATPase-[Planktothrix-rubescens] |
| BDA76272.1 | AAA+ATPase+wHTH+Alpha+Beta-domain+wHTH+RRM | Type-1_BR.RM.like.systems | REase-><-?<-?<-ORC-CDC6<-ParA-Soj-PloopNTPase<-HEPN\|\|DUF499*->TUDOR+SF1-SF2-Helicase+iREase+CTD+REase-><-STYKIN | 957 | Bacteria>Terrabacteria_group>Cyanobacteriota/Melainabacteria_group>Cyanobacteriota>Cyanophyceae | GCA_019977735.1 | Calothrix_sp._PCC_7716 | CAL7716_104380 | 93672..96545 | 2853227 | AAA-family-ATPase-[Calothrix-sp.-PCC-7716] |
| WP_280700572.1 | AAA+ATPase+wHTH+Alpha+Beta-domain+wHTH+RRM | Type-1_BR.RM.like.systems | Cysteine_desulfurase->METHYLASE-p->REase->DUF499*-><-Peptidase_S51<-?<-?<-?\|\|sGTP-> | 953 | Bacteria>Terrabacteria_group>Cyanobacteriota/Melainabacteria_group>Cyanobacteriota>Cyanophyceae | GCF_029891755.1 | Umezakia_ovalisporum | NWP23_RS06515 | 33643..36504 | 75695 | DUF499-domain-containing-protein-[Umezakia-ovalisporum] |
| WP_280699380.1 | AAA+ATPase+wHTH+Alpha+Beta-domain+wHTH+RRM | Type-1_BR.RM.like.systems | <-sGTP\|\|?->?->?->Peptidase_S51-><-DUF499*<-REase<-METHYLASE-p<-Cysteine_desulfurase | 953 | Bacteria>Terrabacteria_group>Cyanobacteriota/Melainabacteria_group>Cyanobacteriota>Cyanophyceae | GCF_029891685.1 | Umezakia_ovalisporum | NWP25_RS10975 | 18984..21845 | 75695 | DUF499-domain-containing-protein-[Umezakia-ovalisporum] |
| WP_280689836.1 | AAA+ATPase+wHTH+Alpha+Beta-domain+wHTH+RRM | Type-1_BR.RM.like.systems | <-sGTP\|\|?->?->?->Peptidase_S51-><-DUF499*<-REase<-METHYLASE-p<-Cysteine_desulfurase | 973 | Bacteria>Terrabacteria_group>Cyanobacteriota/Melainabacteria_group>Cyanobacteriota>Cyanophyceae | GCF_029891865.1 | Umezakia_ovalisporum | NWP15_RS06365 | 19189..22110 | 75695 | DUF499-domain-containing-protein-[Umezakia-ovalisporum] |
| WP_280687902.1 | AAA+ATPase+wHTH+Alpha+Beta-domain+wHTH+RRM | Type-1_BR.RM.like.systems | <-sGTP\|\|?->?->?->Peptidase_S51-><-DUF499*<-REase<-METHYLASE-p<-Cysteine_desulfurase | 953 | Bacteria>Terrabacteria_group>Cyanobacteriota/Melainabacteria_group>Cyanobacteriota>Cyanophyceae | GCF_029891835.1 | Umezakia_ovalisporum | NWP18_RS13190 | 14598..17459 | 75695 | DUF499-domain-containing-protein-[Umezakia-ovalisporum] |
| WP_280656504.1 | AAA+ATPase+wHTH+Alpha+Beta-domain+wHTH+RRM | Type-1_BR.RM.like.systems | <-sGTP\|\|?->?->?->Peptidase_S51-><-DUF499*<-REase<-METHYLASE-p<-Cysteine_desulfurase | 953 | Bacteria>Terrabacteria_group>Cyanobacteriota/Melainabacteria_group>Cyanobacteriota>Cyanophyceae | GCF_029891765.1 | Umezakia_ovalisporum | NWP26_RS00065 | 19111..21972 | 75695 | DUF499-domain-containing-protein-[Umezakia-ovalisporum] |
| WP_280650398.1 | AAA+ATPase+wHTH+Alpha+Beta-domain+wHTH+RRM | Type-1_BR.RM.like.systems | <-sGTP\|\|?->?->?->Peptidase_S51-><-DUF499*<-REase<-METHYLASE-p<-Cysteine_desulfurase | 953 | Bacteria>Terrabacteria_group>Cyanobacteriota/Melainabacteria_group>Cyanobacteriota>Cyanophyceae | GCF_029891655.1 | Umezakia_ovalisporum | NWP27_RS05680 | 14158..17019 | 75695 | DUF499-domain-containing-protein-[Umezakia-ovalisporum] |
| WP_046277358.1 | AAA+ATPase+wHTH+Alpha+Beta-domain+wHTH+RRM | Type-1_BR.RM.like.systems | METHYLASE-p->REase->DUF499*-><-Pkinase\|\|?->?-><-NTP_transf_9<-TPR+TPR+TPR+TPR+TPR+TPR+TPR+TPR | 968 | Bacteria>Terrabacteria_group>Cyanobacteriota/Melainabacteria_group>Cyanobacteriota>Cyanophyceae | GCF_000972705.2 | Limnoraphis_robusta | WN50_RS27095 | 1686..4592 | 1118279 | DUF499-domain-containing-protein-[Limnoraphis-robusta] |
| EHC19655.1 | AAA+ATPase+wHTH+Alpha+Beta-domain+wHTH+RRM | Type-1_BR.RM.like.systems | TUDOR+SF1-SF2-Helicase+iREase+CTD+REase->?->OLD-ABC-p->METHYLASE->REase->DUF499*->?-><-?<-?<-ABC-ATPase+SbcC | 975 | Bacteria>Terrabacteria_group>Cyanobacteriota/Melainabacteria_group>Cyanobacteriota>Cyanophyceae | GCA_000231365.2 | Fischerella_thermalis_JSC-11 | FJSC11DRAFT_0472 | 574051..576978 | 741277 | AAA-family-ATPase-[Fischerella-thermalis-JSC-11] |
| MBO1045985.1 | AAA+ATPase+wHTH+Alpha+Beta-domain+wHTH+RRM | Type-1_BR.RM.like.systems | Aldedh-><-?\|\|?->?->?-><-DUF499*<-PemK_toxin<-MetJarc-HTH<-METHYLASE<-?<-REase<-TUDOR+SF1-SF2-Helicase+iREase+CTD+REase | 965 | Bacteria>Terrabacteria_group>Cyanobacteriota/Melainabacteria_group>Cyanobacteriota>Cyanophyceae | GCA_001593825.2 | Aphanizomenon_flos-aquae_UKL13-PB | A2T80_019100 | 69616..72513 | 1803587 | DUF499-domain-containing-protein-[Aphanizomenon-flos-aquae-UKL13-PB] |
| WP_107666788.1 | AAA+ATPase+wHTH+Alpha+Beta-domain+wHTH+RRM | Type-1_BR.RM.like.systems | HicB->HicA_toxin->?->?->PIN->?->DUF499*->DDE_5-><-VirD4-TraG+VirD4-FtsK<-VirD4-TraG+VirD4-FtsK | 970 | Bacteria>Terrabacteria_group>Cyanobacteriota/Melainabacteria_group>Cyanobacteriota>Cyanophyceae | GCF_003013815.1 | Cyanothece_sp._BG0011 | C4E06_RS03245 | 1912..4824 | 2082950 | DUF499-domain-containing-protein-[Cyanothece-sp.-BG0011] |
| WP_200988278.1 | AAA+ATPase+wHTH+Alpha+Beta-domain+wHTH+RRM | Type-1_BR.RM.like.systems | <-REase<-?<-?<-DUF499*<-CR-ATPase9+REase<-DUF499<-CR-ATPase9+REase<-METHYLASE<-REase<-PIN | 950 | Bacteria>Terrabacteria_group>Cyanobacteriota/Melainabacteria_group>Cyanobacteriota>Cyanophyceae | GCF_015245355.1 | Anabaenopsis_elenkinii | IM676_RS18785 | 4299000..4301852 | 156213 | DUF499-domain-containing-protein-[Anabaenopsis-elenkinii] |
| WP_193920728.1 | AAA+ATPase+wHTH+Alpha+Beta-domain+wHTH+RRM | Type-1_BR.RM.like.systems | TUDOR+SF1-SF2-Helicase+iREase+CTD+REase->METHYLASE-><-?\|\|DUF499*-><-CHTH<-?\|\|HTH+MPTase->?-><-?<-Phage_integrase | 977 | Bacteria>Terrabacteria_group>Cyanobacteriota/Melainabacteria_group>Cyanobacteriota>Cyanophyceae | GCF_015207665.1 | Plectonema_radiosum | IQ247_RS13400 | 9838..12771 | 945768 | DUF499-domain-containing-protein-[Plectonema-radiosum] |
| WP_190667193.1 | AAA+ATPase+wHTH+Alpha+Beta-domain+wHTH+RRM | Type-1_BR.RM.like.systems | CHTH->TUDOR+SF1-SF2-Helicase+iREase+CTD+REase->METHYLASE->REase->DUF499*-><-REase<-CHTH | 1004 | Bacteria>Terrabacteria_group>Cyanobacteriota/Melainabacteria_group>Cyanobacteriota>Cyanophyceae | GCF_014695285.1 | Coleofasciculus_sp._FACHB-501 | H6F54_RS11880 | 20325..23339 | 2692786 | DUF499-domain-containing-protein-[Coleofasciculus-sp.-FACHB-501] |
| WP_069791018.1 | AAA+ATPase+wHTH+Alpha+Beta-domain+wHTH+RRM | Type-1_BR.RM.like.systems | METHYLASE->?->?->?->PIN->?->DUF499*->DUF499-p-><-GDResolvase+HIN-HTH\|\|DUF4158+DDE_Tnp_Tn3-> | 838 | Bacteria>Terrabacteria_group>Cyanobacteriota/Melainabacteria_group>Cyanobacteriota>Cyanophyceae | GCF_001747005.1 | Cyanobacterium_sp._IPPAS_B-1200 | A5482_RS09625 | 19329..21845 | 1562720 | DUF499-domain-containing-protein-[Cyanobacterium-sp.-IPPAS-B-1200] |
| MCM0594151.1 | AAA+ATPase+wHTH+Alpha+Beta-domain+wHTH+RRM | Type-1_BR.RM.like.systems | <-DUF499*<-PIN<-?<-?\|\|HTH_OrfB_IS605+OrfB_IS605+OrfB_Zn_ribbon-><-PemK_toxin<-MetJarc-HTH | 955 | Bacteria>Terrabacteria_group>Cyanobacteriota/Melainabacteria_group>Cyanobacteriota>Cyanophyceae | GCA_023657325.1 | Gloeotrichia_echinulata_DEX184 | KA716_29815 | 309224..312091 | 2824560 | DUF499-domain-containing-protein-[Gloeotrichia-echinulata-DEX184] |
| WP_181929825.1 | AAA+ATPase+wHTH+Alpha+Beta-domain+wHTH+RRM | Type-1_BR.RM.like.systems | <-TetR-HTH<-DUF499*<-METHYLASE<-METHYLASE-p<-REase<-TUDOR+SF1-SF2-Helicase+iREase+CTD+REase\|\|CHTH-> | 966 | Bacteria>Terrabacteria_group>Cyanobacteriota/Melainabacteria_group>Cyanobacteriota>Cyanophyceae | GCF_014023275.1 | Nostoc_edaphicum | HUN01_RS33840 | 7488405..7491305 | 264686 | DUF499-domain-containing-protein-[Nostoc-edaphicum] |
| WP_190397100.1 | AAA+ATPase+wHTH+Alpha+Beta-domain+wHTH+RRM | Type-1_BR.RM.like.systems | METHYLASE->REase->MetJarc-HTH->PemK_toxin->DUF499*->?->?-><-?\|\|?-><-?\|\|REase+HTH-> | 969 | Bacteria>Terrabacteria_group>Cyanobacteriota/Melainabacteria_group>Cyanobacteriota>Cyanophyceae | GCF_014696335.1 | Anabaena_sp._FACHB-1250 | H6F39_RS17030 | 5832..8741 | 2692770 | DUF499-domain-containing-protein-[Anabaena-sp.-FACHB-1250] |
| WP_214559734.1 | AAA+ATPase+wHTH+Alpha+Beta-domain+wHTH+RRM | Type-1_BR.RM.like.systems | <-RelB_antitoxin<-?<-?\|\|?-><-?<-DUF499*<-REase<-REase<-REase<-REase<-METHYLASE<-METHYLASE-p | 999 | Bacteria>Terrabacteria_group>Cyanobacteriota/Melainabacteria_group>Cyanobacteriota>Cyanophyceae | GCF_903969095.1 | Planktothrix_agardhii | KM073_RS08965 | 67038..70037 | 1160 | DUF499-domain-containing-protein-[Planktothrix-agardhii] |
| WP_193849072.1 | AAA+ATPase+wHTH+Alpha+Beta-domain+wHTH+RRM | Type-1_BR.RM.like.systems | TUDOR+SF1-SF2-Helicase+iREase+CTD+REase-><-?\|\|METHYLASE->REase->DUF499*->?-><-?\|\|?-><-?\|\|REase+HTH-> | 969 | Bacteria>Terrabacteria_group>Cyanobacteriota/Melainabacteria_group>Cyanobacteriota>Cyanophyceae | GCF_015207785.1 | Dolichospermum_flos-aquae | IQ222_RS12495 | 19378..22287 | 1166 | DUF499-domain-containing-protein-[Dolichospermum-flos-aquae] |
| WP_065711823.1 | AAA+ATPase+wHTH+Alpha+Beta-domain+wHTH+RRM | Type-1_BR.RM.like.systems | REase->TUDOR+SF1-SF2-Helicase+iREase+CTD+REase->OLD-ABC+TOPRIM->PolB->DUF499*->?->?->NACHT->?-><-HIN-HTH\|\|PIN-> | 989 | Bacteria>Terrabacteria_group>Cyanobacteriota/Melainabacteria_group>Cyanobacteriota>Cyanophyceae | GCF_001693275.1 | Picosynechococcus_sp._PCC_7117 | AWQ22_RS15050 | 32294..35263 | 195498 | DUF499-domain-containing-protein-[Synechococcus-sp.-PCC-7117] |
| WP_027249039.1 | AAA+ATPase+wHTH+Alpha+Beta-domain+wHTH+RRM | Type-1_BR.RM.like.systems | METHYLASE-p->METHYLASE->REase->REase->REase->REase->DUF499*->?-><-?\|\|?->RelB_antitoxin-> | 999 | Bacteria>Terrabacteria_group>Cyanobacteriota/Melainabacteria_group>Cyanobacteriota>Cyanophyceae | GCF_000464665.1 | Planktothrix_agardhii | N318_RS0105300 | 1257351..1260350 | 1160 | DUF499-domain-containing-protein-[Planktothrix-agardhii] |
| WP_026794456.1 | AAA+ATPase+wHTH+Alpha+Beta-domain+wHTH+RRM | Type-1_BR.RM.like.systems | <-RelB_antitoxin<-?\|\|?-><-?<-DUF499*<-REase<-REase<-REase<-REase<-METHYLASE<-METHYLASE-p | 999 | Bacteria>Terrabacteria_group>Cyanobacteriota/Melainabacteria_group>Cyanobacteriota>Cyanophyceae | GCF_000464725.1 | Planktothrix | N317_RS0110130 | 2122440..2125439 | 54304 | MULTISPECIES:-DUF499-domain-containing-protein-[Planktothrix] |
| OAB62772.1 | AAA+ATPase+wHTH+Alpha+Beta-domain+wHTH+RRM | Type-1_BR.RM.like.systems | TUDOR+SF1-SF2-Helicase+iREase+CTD+REase->PRTase+PRTase->?->METHYLASE->REase->DUF499*-> | 1013 | Bacteria>Terrabacteria_group>Cyanobacteriota/Melainabacteria_group>Cyanobacteriota>Cyanophyceae | GCA_001637395.1 | Leptolyngbya_valderiana_BDU_20041 | AY599_12440 | 19718..22759 | 322866 | AAA-family-ATPase-[Leptolyngbya-valderiana-BDU-20041] |
| CAD5939228.1 | AAA+ATPase+wHTH+Alpha+Beta-domain+wHTH+RRM | Type-1_BR.RM.like.systems | AbrB-like->?->Radical-SAM-><-?<-?<-DUF499*<-REase<-REase<-REase<-REase<-METHYLASE<-REase | 1002 | Bacteria>Terrabacteria_group>Cyanobacteriota/Melainabacteria_group>Cyanobacteriota>Cyanophyceae | GCA_904830785.1 | Planktothrix_rubescens | NO108_02187 | 2314018..2317026 | 59512 | Putative-AAA-family-ATPase-[Planktothrix-rubescens] |
| OBQ20276.1 | AAA+ATPase+wHTH+Alpha+Beta-domain+wHTH+RRM | Type-1_BR.RM.like.systems | <-DnaJ<-?<-NTP_transferase<-?<-DUF499*<-PIN<-?<-REase+HTH<-TUDOR+SF1-SF2-Helicase+iREase+CTD+REase<-CHTH | 916 | Bacteria>Terrabacteria_group>Cyanobacteriota/Melainabacteria_group>Cyanobacteriota>Cyanophyceae | GCA_001672085.1 | Anabaena_sp._AL93 | AN486_07300 | 48734..51484 | 1678133 | MAG:-ATPase-[Anabaena-sp.-AL93] |
| WP_254175247.1 | AAA+ATPase+wHTH+Alpha+Beta-domain+wHTH+RRM | Type-1_BR.RM.like.systems | <-DUF499*<-REase<-METHYLASE<-?<-SbcC<-TUDOR+SF1-SF2-Helicase+iREase+CTD+REase<-RHH_1 | 998 | Bacteria>Terrabacteria_group>Cyanobacteriota/Melainabacteria_group>Cyanobacteriota>Cyanophyceae | GCF_904830925.1 | Planktothrix_pseudagardhii | NMK36_RS28460 | 5796..8792 | 132604 | DUF499-domain-containing-protein-[Planktothrix-pseudagardhii] |
| WP_042158736.1 | AAA+ATPase+wHTH+Alpha+Beta-domain+wHTH+RRM | Type-1_BR.RM.like.systems | PIN->METHYLASE->REase->REase->REase->DUF499*-><-?\|\|RelB_antitoxin->PemK_toxin-> | 1002 | Bacteria>Terrabacteria_group>Cyanobacteriota/Melainabacteria_group>Cyanobacteriota>Cyanophyceae | GCF_000710505.1 | Planktothrix_agardhii | NMK38_RS22725 | 84961..87969 | 1160 | DUF499-domain-containing-protein-[Planktothrix-agardhii] |
| WP_168507185.1 | AAA+ATPase+wHTH+Alpha+Beta-domain+wHTH+RRM | Type-1_BR.RM.like.systems | <-DUF499*<-REase<-METHYLASE<-PIN<-TUDOR+SF1-SF2-Helicase+iREase+CTD+REase<-CHTH | 962 | Bacteria>Terrabacteria_group>Cyanobacteriota/Melainabacteria_group>Cyanobacteriota>Cyanophyceae | GCF_009712035.1 | Anabaena_sp._UHCC_0187 | FJR11_RS20795 | 2169..5057 | 2590018 | DUF499-domain-containing-protein-[Anabaena-sp.-UHCC-0187] |
| NJR56033.1 | AAA+ATPase+wHTH+Alpha+Beta-domain+wHTH+RRM | Type-1_BR.RM.like.systems | METHYLASE-p->REase->DUF499*->Phage_integrase->?->MazE_antitoxin->DOC->DUF29-> | 968 | Bacteria>Terrabacteria_group>Cyanobacteriota/Melainabacteria_group>Cyanobacteriota>Cyanophyceae | GCA_012034865.1 | Acaryochloris_sp._CRU_2_0 | HC768_16565 | 11406..14312 | 2720402 | ATP-binding-protein-[Acaryochloris-sp.-CRU_2_0] |
| WP_106370048.1 | AAA+ATPase+wHTH+Alpha+Beta-domain+wHTH+RRM | Type-1_BR.RM.like.systems | RNase-><-?\|\|?->TIR-><-HKD-DNase+SF1-SF2-Helicase<-HEPN<-DUF499*<-METHYLASE<-PIN | 1004 | Bacteria>Terrabacteria_group>Cyanobacteriota/Melainabacteria_group>Cyanobacteriota>Cyanophyceae | GCF_003003885.1 | Chlorogloea_sp._CCALA_695 | C7B70_RS16210 | 39604..42618 | 2107693 | DUF499-domain-containing-protein-[Chlorogloea-sp.-CCALA-695] |
| WP_096671658.1 | AAA+ATPase+wHTH+Alpha+Beta-domain+wHTH+RRM | Type-1_BR.RM.like.systems | <-DUF499*<-PIN<-?<-METHYLASE<-METHYLASE-p<-REase+HTH<-TUDOR+SF1-SF2-Helicase+iREase+CTD+REase | 912 | Bacteria>Terrabacteria_group>Cyanobacteriota/Melainabacteria_group>Cyanobacteriota>Cyanophyceae | GCF_002368115.1 | Dolichospermum_compactum | CA730_RS19465 | 4197332..4200070 | 136073 | DUF499-domain-containing-protein-[Dolichospermum-compactum] |
| WP_073075160.1 | AAA+ATPase+wHTH+Alpha+Beta-domain+wHTH+RRM | Type-1_BR.RM.like.systems | CR-Trypsin1->?-><-RVT+HNH<-DUF499*<-REase<-REase<-METHYLASE<-DUF4926+DUF2281 | 965 | Bacteria>Terrabacteria_group>Cyanobacteriota/Melainabacteria_group>Cyanobacteriota>Cyanophyceae | GCF_001895925.1 | Phormidesmis_priestleyi | C7B65_RS24715 | 5123..8020 | 268141 | DUF499-domain-containing-protein-[Phormidesmis-priestleyi] |
| MBV6627856.1 | AAA+ATPase+wHTH+Alpha+Beta-domain+wHTH+RRM | Type-1_BR.RM.like.systems | <-RVT+HNH<-?\|\|?-><-DUF499*<-?<-METHYLASE<-DOC+HTH<-DOC+HTH<-TUDOR+SF1-SF2-Helicase+iREase+CTD+REase | 952 | Bacteria>Terrabacteria_group>Cyanobacteriota/Melainabacteria_group>Cyanobacteriota>Cyanophyceae | GCA_019192545.1 | Rivularia_sp._MS3 | KI793_33860 | 4318..7176 | 3012223 | ATP-binding-protein-[Rivularia-sp.-(in:-Bacteria)] |
| WP_192216664.1 | AAA+ATPase+wHTH+Alpha+Beta-domain+wHTH+RRM | Type-1_BR.RM.like.systems | <-HTH\|\|?->?-><-?<-GDResolvase+Recombinase+ZnR<-?<-DUF499*<-REase<-REase<-METHYLASE-p | 969 | Bacteria>Terrabacteria_group>Cyanobacteriota/Melainabacteria_group>Cyanobacteriota>Cyanophyceae | GCF_014849525.1 | Microcoleus_sp._FACHB-SPT15 | H6F98_RS00230 | 47740..50649 | 2692830 | DUF499-domain-containing-protein-[Microcoleus-sp.-FACHB-SPT15] |
| MBW4429556.1 | AAA+ATPase+wHTH+Alpha+Beta-domain+wHTH+RRM | Type-1_BR.RM.like.systems | TUDOR+SF1-SF2-Helicase+iREase+CTD+REase->METHYLASE->REase->DUF499*->?->PIN-> | 1005 | Bacteria>Terrabacteria_group>Cyanobacteriota/Melainabacteria_group>Cyanobacteriota>Cyanophyceae | GCA_019359245.1 | Nostoc_desertorum_CM1-VF14 | KME50_35640 | 5971..8988 | 287862 | DUF499-domain-containing-protein-[Nostoc-desertorum-CM1-VF14] |
| WP_039990643.1 | AAA+ATPase+wHTH+Alpha+Beta-domain+wHTH+RRM | Type-1_BR.RM.like.systems | TUDOR+SF1-SF2-Helicase+iREase+CTD+REase->OLD-ABC-p->METHYLASE->REase->DUF499*-> | 955 | Bacteria>Terrabacteria_group>Cyanobacteriota/Melainabacteria_group>Cyanobacteriota>Cyanophyceae | GCF_000231365.1 | Fischerella_thermalis | FJSC11DRAFT_RS02260 | 574111..576978 | 372787 | DUF499-domain-containing-protein-[Fischerella-thermalis] |
| WP_180965961.1 | AAA+ATPase+wHTH+Alpha+Beta-domain+wHTH+RRM | Type-1_BR.RM.like.systems | DUF4277+DDE_Tnp_1->TUDOR+SF1-SF2-Helicase+iREase+CTD+REase->OLD-ABC-p->METHYLASE->DUF499*-> | 954 | Bacteria>Terrabacteria_group>Cyanobacteriota/Melainabacteria_group>Cyanobacteriota>Cyanophyceae | GCF_002870545.1 | Fischerella_thermalis | CBP21_RS00920 | 44648..47512 | 372787 | DUF499-domain-containing-protein-[Fischerella-thermalis] |
| MBS9772311.1 | AAA+ATPase+wHTH+Alpha+Beta-domain+wHTH+RRM | Type-1_BR.RM.like.systems | <-DUF499*<-REase<-PIN<-?<-METHYLASE<-TUDOR+SF1-SF2-Helicase+iREase+CTD+REase | 1016 | Bacteria>Terrabacteria_group>Cyanobacteriota/Melainabacteria_group>Cyanobacteriota>Cyanophyceae | GCA_018457135.1 | Trichodesmium_erythraeum_GBRTRLIN201 | J7F05_18140 | 304..3354 | 1045867 | ATP-binding-protein-[Trichodesmium-erythraeum-GBRTRLIN201] |
| WP_190347798.1 | AAA+ATPase+wHTH+Alpha+Beta-domain+wHTH+RRM | Type-1_BR.RM.like.systems | ABC-ATPase+SbcC-><-DUF499*<-PIN<-?<-METHYLASE\|\|?->?-><-TUDOR+SF1-SF2-Helicase+iREase+CTD+REase | 963 | Bacteria>Terrabacteria_group>Cyanobacteriota/Melainabacteria_group>Cyanobacteriota>Cyanophyceae | GCF_014696235.1 | Sphaerospermopsis_sp._FACHB-1194 | H6F45_RS16760 | 52263..55154 | 2692862 | DUF499-domain-containing-protein-[Sphaerospermopsis-sp.-FACHB-1194] |
| WP_168360489.1 | AAA+ATPase+wHTH+Alpha+Beta-domain+wHTH+RRM | Type-1_BR.RM.like.systems | <-REase+HTH\|\|?-><-?\|\|?-><-?<-?<-DUF499*<-?<-?<-MetJarc-HTH<-REase<-METHYLASE | 967 | Bacteria>Terrabacteria_group>Cyanobacteriota/Melainabacteria_group>Cyanobacteriota>Cyanophyceae | GCF_009712075.2 | Dolichospermum_planctonicum | FJR41_RS21600 | 23214..26117 | 136072 | DUF499-domain-containing-protein-[Dolichospermum-planctonicum] |
| WP_083617094.1 | AAA+ATPase+wHTH+Alpha+Beta-domain+wHTH+RRM | Type-1_BR.RM.like.systems | TUDOR+SF1-SF2-Helicase+iREase+CTD+REase->HTH_3->METHYLASE->DUF499*->?->?-><-FURR-HTH\|\|?-><-PIN | 955 | Bacteria>Terrabacteria_group>Cyanobacteriota/Melainabacteria_group>Cyanobacteriota>Cyanophyceae | GCF_900010725.2 | Planktothrix_serta | PL8927_RS02125 | 111808..114675 | 1678310 | DUF499-domain-containing-protein-[Planktothrix-serta] |
| WP_026788986.1 | AAA+ATPase+wHTH+Alpha+Beta-domain+wHTH+RRM | Type-1_BR.RM.like.systems | REase->METHYLASE->REase->REase->REase->REase->DUF499*->?->?->?->?-><-Radical-SAM | 999 | Bacteria>Terrabacteria_group>Cyanobacteriota/Melainabacteria_group>Cyanobacteriota>Cyanophyceae | GCF_000464785.1 | Planktothrix_rubescens | N313_RS0123225 | 1913145..1916144 | 59512 | ATP-binding-protein-[Planktothrix-rubescens] |
| WP_102148052.1 | AAA+ATPase+wHTH+Alpha+Beta-domain+wHTH+RRM | Type-1_BR.RM.like.systems | TUDOR+SF1-SF2-Helicase+iREase+CTD+REase->METHYLASE->DUF499*-><-?<-ABC-ATPase+SbcC | 974 | Bacteria>Terrabacteria_group>Cyanobacteriota/Melainabacteria_group>Cyanobacteriota>Cyanophyceae | - | Fischerella_thermalis | CBP17_11765 | 50415..53339 | 372787 | DUF499-domain-containing-protein-[Fischerella-thermalis] |
| PLZ10023.1 | AAA+ATPase+wHTH+Alpha+Beta-domain+wHTH+RRM | Type-1_BR.RM.like.systems | TUDOR+SF1-SF2-Helicase+iREase+CTD+REase->METHYLASE->DUF499*-><-?<-ABC-ATPase+SbcC | 974 | Bacteria>Terrabacteria_group>Cyanobacteriota/Melainabacteria_group>Cyanobacteriota>Cyanophyceae | GCA_002870545.1 | Fischerella_thermalis_WC114 | CBP17_11765 | 50415..53339 | 1521403 | AAA-family-ATPase-[Fischerella-thermalis-WC114] |
| WP_190397164.1 | AAA+ATPase+wHTH+Alpha+Beta-domain+wHTH+RRM | Type-1_BR.RM.like.systems | TPR+TPR-><-?\|\|?-><-DUF499*<-METHYLASE<-?<-REase+HTH<-TUDOR+SF1-SF2-Helicase+iREase+CTD+REase | 913 | Bacteria>Terrabacteria_group>Cyanobacteriota/Melainabacteria_group>Cyanobacteriota>Cyanophyceae | GCF_014696335.1 | Anabaena_sp._FACHB-1250 | H6F39_RS18020 | 6551..9292 | 2692770 | DUF499-domain-containing-protein-[Anabaena-sp.-FACHB-1250] |
| WP_190372718.1 | AAA+ATPase+wHTH+Alpha+Beta-domain+wHTH+RRM | Type-1_BR.RM.like.systems | TPR+TPR-><-?\|\|?-><-DUF499*<-METHYLASE<-?<-?<-REase+HTH<-TUDOR+SF1-SF2-Helicase+iREase+CTD+REase | 913 | Bacteria>Terrabacteria_group>Cyanobacteriota/Melainabacteria_group>Cyanobacteriota>Cyanophyceae | GCF_014696755.1 | Anabaena_sp._FACHB-1391 | H6F62_RS19000 | 6552..9293 | 2692771 | DUF499-domain-containing-protein-[Anabaena-sp.-FACHB-1391] |
| WP_137909444.1 | AAA+ATPase+wHTH+Alpha+Beta-domain+wHTH+RRM | Type-1_BR.RM.like.systems | TUDOR+SF1-SF2-Helicase+iREase+CTD+REase->REase+HTH->?->METHYLASE->DUF499*-><-?\|\|?-><-TPR+TPR | 913 | Bacteria>Terrabacteria_group>Cyanobacteriota/Melainabacteria_group>Cyanobacteriota>Cyanophyceae | GCF_005402965.1 | Dolichospermum_planctonicum | FF006_RS19070 | 11296..14037 | 136072 | DUF499-domain-containing-protein-[Dolichospermum-planctonicum] |
| WP_106371305.1 | AAA+ATPase+wHTH+Alpha+Beta-domain+wHTH+RRM | Type-1_BR.RM.like.systems | <-STYKIN\|\|?-><-SIR2+SIR2<-DUF499*<-REase<-METHYLASE<-PIN | 994 | Bacteria>Terrabacteria_group>Cyanobacteriota/Melainabacteria_group>Cyanobacteriota>Cyanophyceae | GCF_003003885.1 | Chlorogloea_sp._CCALA_695 | C7B70_RS23190 | 7463..10447 | 2107693 | DUF499-domain-containing-protein-[Chlorogloea-sp.-CCALA-695] |
| MBD1215623.1 | AAA+ATPase+wHTH+Alpha+Beta-domain+wHTH+RRM | Type-1_BR.RM.like.systems | TPR+TPR-><-?\|\|?-><-DUF499*<-METHYLASE<-?<-REase+HTH<-TUDOR+SF1-SF2-Helicase+iREase+CTD+REase | 913 | Bacteria>Terrabacteria_group>Cyanobacteriota/Melainabacteria_group>Cyanobacteriota>Cyanophyceae | GCA_014654715.1 | Dolichospermum_circinale_Clear-D4 | H9534_23500 | 3273..6014 | 2766971 | ATP-binding-protein-[Dolichospermum-circinale-Clear-D4] |
| WP_104386435.1 | AAA+ATPase+wHTH+Alpha+Beta-domain+wHTH+RRM | Type-1_BR.RM.like.systems | <-GspA-ATPase<-rve+Mu-transpos_C<-GDResolvase+HIN-HTH\|\|<-?\|\|?-><-?<-DUF499* | 709 | Bacteria>Terrabacteria_group>Cyanobacteriota/Melainabacteria_group>Cyanobacteriota>Cyanophyceae | GCF_002934005.1 | Cuspidothrix_issatschenkoi | CUN59_RS03020 | 55602..57732 | 230752 | AAA-family-ATPase,-partial-[Cuspidothrix-issatschenkoi] |
| OBQ29403.1 | AAA+ATPase+wHTH+Alpha+Beta-domain+wHTH+RRM | Type-1_BR.RM.like.systems | Aldedh-><-?\|\|?->?->?-><-DUF499*<-PemK_toxin<-MetJarc-HTH<-METHYLASE\|\|?-><-?<-REase | 965 | Bacteria>Terrabacteria_group>Cyanobacteriota/Melainabacteria_group>Cyanobacteriota>Cyanophyceae | GCA_001672095.1 | Aphanizomenon_flos-aquae_MDT14a | AN483_10570 | 7911..10808 | 1710895 | AAA-family-ATPase-[Aphanizomenon-flos-aquae-MDT14a] |
| MBW4559185.1 | AAA+ATPase+wHTH+Alpha+Beta-domain+wHTH+RRM | Type-1_BR.RM.like.systems | METHYLASE-p->REase->DUF499*-><-RVT+HNH\|\|?->?-><-?<-Phage_integrase<-REase | 990 | Bacteria>Terrabacteria_group>Cyanobacteriota/Melainabacteria_group>Cyanobacteriota>Cyanophyceae | GCA_019358915.1 | Trichormus_sp._ATA11-4-KO1 | KME59_25360 | 2404..5376 | 2835091 | DUF499-domain-containing-protein-[Trichormus-sp.-ATA11-4-KO1] |
| WP_254171832.1 | AAA+ATPase+wHTH+Alpha+Beta-domain+wHTH+RRM | Type-1_BR.RM.like.systems | TUDOR+SF1-SF2-Helicase+iREase+CTD+REase->Tox-RelE->HTH_3->METHYLASE->DUF499*-> | 957 | Bacteria>Terrabacteria_group>Cyanobacteriota/Melainabacteria_group>Cyanobacteriota>Cyanophyceae | GCF_904830885.1 | Planktothrix_agardhii | NMK31_RS00065 | 18757..21630 | 1160 | DUF499-domain-containing-protein-[Planktothrix-agardhii] |
| WP_227351566.1 | AAA+ATPase+wHTH+Alpha+Beta-domain+wHTH+RRM | Type-1_BR.RM.like.systems | TUDOR+SF1-SF2-Helicase+iREase+CTD+REase->Tox-RelE->HTH_3->METHYLASE->DUF499*-> | 957 | Bacteria>Terrabacteria_group>Cyanobacteriota/Melainabacteria_group>Cyanobacteriota>Cyanophyceae | GCF_020594555.1 | Planktothrix_agardhii | LJE14_RS21460 | 66217..69090 | 1160 | DUF499-domain-containing-protein-[Planktothrix-agardhii] |
| WP_042158676.1 | AAA+ATPase+wHTH+Alpha+Beta-domain+wHTH+RRM | Type-1_BR.RM.like.systems | TUDOR+SF1-SF2-Helicase+iREase+CTD+REase->Tox-RelE->HTH_3->METHYLASE->DUF499*-> | 957 | Bacteria>Terrabacteria_group>Cyanobacteriota/Melainabacteria_group>Cyanobacteriota>Cyanophyceae | GCF_000710505.1 | Planktothrix_agardhii | NMK38_RS22560 | 51674..54547 | 1160 | DUF499-domain-containing-protein-[Planktothrix-agardhii] |
| CEJ46775.1 | AAA+ATPase+wHTH+Alpha+Beta-domain+wHTH+RRM | Type-1_BR.RM.like.systems | <-sGTP\|\|?->?->?->Peptidase_S51-><-DUF499*<-Cysteine_desulfurase | 972 | Bacteria>Terrabacteria_group>Cyanobacteriota/Melainabacteria_group>Cyanobacteriota>Cyanophyceae | GCA_001458455.1 | Umezakia_ovalisporum | apha_02693 | 32748..35666 | 75695 | Uncharacterized-protein-apha_02693-[Chrysosporum-ovalisporum] |
| WP_193849784.1 | AAA+ATPase+wHTH+Alpha+Beta-domain+wHTH+RRM | Type-1_BR.RM.like.systems | TUDOR+SF1-SF2-Helicase+iREase+CTD+REase->REase+HTH->?->METHYLASE-><-?\|\|DUF499*-> | 913 | Bacteria>Terrabacteria_group>Cyanobacteriota/Melainabacteria_group>Cyanobacteriota>Cyanophyceae | GCF_015207785.1 | Dolichospermum_flos-aquae | IQ222_RS15370 | 28947..31688 | 1166 | DUF499-domain-containing-protein-[Dolichospermum-flos-aquae] |
| WP_190875885.1 | AAA+ATPase+wHTH+Alpha+Beta-domain+wHTH+RRM | Type-1_BR.RM.like.systems | <-TPR<-DUF499*<-METHYLASE<-TUDOR+SF1-SF2-Helicase+iREase+CTD+REase\|\|CHTH-> | 966 | Bacteria>Terrabacteria_group>Cyanobacteriota/Melainabacteria_group>Cyanobacteriota>Cyanophyceae | - | Nostoc_sp._FACHB-973 | H6G93_02510 | 170755..173655 | 2692844 | DUF499-domain-containing-protein-[Nostoc-sp.-FACHB-973] |
| MBD2513894.1 | AAA+ATPase+wHTH+Alpha+Beta-domain+wHTH+RRM | Type-1_BR.RM.like.systems | <-TPR<-DUF499*<-METHYLASE<-TUDOR+SF1-SF2-Helicase+iREase+CTD+REase\|\|CHTH-> | 966 | Bacteria>Terrabacteria_group>Cyanobacteriota/Melainabacteria_group>Cyanobacteriota>Cyanophyceae | GCA_014698115.1 | Nostoc_sp._FACHB-973 | H6G93_02510 | 170755..173655 | 2692844 | ATP-binding-protein-[Nostoc-sp.-FACHB-973] |
| WP_027255058.1 | AAA+ATPase+wHTH+Alpha+Beta-domain+wHTH+RRM | Type-1_BR.RM.like.systems | <-DUF499*<-REase<-REase<-REase<-REase<-METHYLASE<-REase | 996 | Bacteria>Terrabacteria_group>Cyanobacteriota/Melainabacteria_group>Cyanobacteriota>Cyanophyceae | GCF_000464825.1 | Planktothrix_agardhii | N316_RS0113720 | 328554..331544 | 1160 | DUF499-domain-containing-protein-[Planktothrix-agardhii] |
| MBO1062702.1 | AAA+ATPase+wHTH+Alpha+Beta-domain+wHTH+RRM | Type-1_BR.RM.like.systems | Aldedh-><-?\|\|?->?->?-><-DUF499*<-PemK_toxin<-MetJarc-HTH<-METHYLASE | 965 | Bacteria>Terrabacteria_group>Cyanobacteriota/Melainabacteria_group>Cyanobacteriota>Cyanophyceae | GCA_017355665.1 | Aphanizomenon_flos-aquae_CP01 | HEQ14_17695 | 7631..10528 | 2717881 | ATP-binding-protein-[Aphanizomenon-flos-aquae-CP01] |
| WP_219724577.1 | AAA+ATPase+wHTH+Alpha+Beta-domain+wHTH+RRM | Type-1_BR.RM.like.systems | TUDOR+SF1-SF2-Helicase+iREase+CTD+REase->OLD-ABC-p->METHYLASE-p->?->DUF499*-> | 972 | Bacteria>Terrabacteria_group>Cyanobacteriota/Melainabacteria_group>Cyanobacteriota>Cyanophyceae | GCF_002870465.1 | Fischerella_thermalis | CBP22_RS18765 | 44099..47017 | 372787 | DUF499-domain-containing-protein,-partial-[Fischerella-thermalis] |
| WP_222620775.1 | AAA+ATPase+wHTH+Alpha+Beta-domain+wHTH+RRM | Type-1_BR.RM.like.systems | <-DUF499*<-PIN<-?<-METHYLASE-p\|\|?->?-><-TUDOR+SF1-SF2-Helicase+iREase+CTD+REase | 616 | Bacteria>Terrabacteria_group>Cyanobacteriota/Melainabacteria_group>Cyanobacteriota>Cyanophyceae | GCF_014299995.1 | Sphaerospermopsis_sp._LEGE_00249 | H5968_RS22665 | 435..2285 | 1380707 | DUF499-domain-containing-protein-[Sphaerospermopsis-sp.-LEGE-00249] |
| WP_190371458.1 | AAA+ATPase+wHTH+Alpha+Beta-domain+wHTH+RRM | Type-1_BR.RM.like.systems | <-REase+HTH<-?\|\|?-><-?<-?<-DUF499*<-REase<-METHYLASE | 969 | Bacteria>Terrabacteria_group>Cyanobacteriota/Melainabacteria_group>Cyanobacteriota>Cyanophyceae | GCF_014696755.1 | Anabaena_sp._FACHB-1391 | H6F62_RS04790 | 144117..147026 | 2692771 | DUF499-domain-containing-protein-[Anabaena-sp.-FACHB-1391] |
| WP_190368028.1 | AAA+ATPase+wHTH+Alpha+Beta-domain+wHTH+RRM | Type-1_BR.RM.like.systems | <-DUF499*<-PIN<-MetJarc-HTH\|\|?->?-><-TUDOR+SF1-SF2-Helicase+iREase+CTD+REase | 977 | Bacteria>Terrabacteria_group>Cyanobacteriota/Melainabacteria_group>Cyanobacteriota>Cyanophyceae | GCA_014696225.1 | Anabaena_sp._FACHB-1237 | H6F32_RS03895 | 63906..66839 | 2692769 | ATP-binding-protein-[Anabaena-sp.-FACHB-1237] |
| MBR8828508.1 | AAA+ATPase+wHTH+Alpha+Beta-domain+wHTH+RRM | Type-1_BR.RM.like.systems | <-DUF29<-?<-?<-?\|\|TUDOR+SF1-SF2-Helicase+iREase+CTD+REase->METHYLASE->DUF499*-> | 1000 | Bacteria>Terrabacteria_group>Cyanobacteriota/Melainabacteria_group>Cyanobacteriota>Cyanophyceae | GCA_016903655.1 | Gomphosphaeria_aponina_SAG_52.96_=_DSM_107014 | DSM107014_11515 | 18878..21880 | 1521640 | hypothetical-protein-[Gomphosphaeria-aponina-SAG-52.96-=-DSM-107014] |
| WP_235656705.1 | AAA+ATPase+wHTH+Alpha+Beta-domain+wHTH+RRM | Type-1_BR.RM.like.systems | TUDOR+SF1-SF2-Helicase+iREase+CTD+REase->OLD-ABC-p->METHYLASE->DUF499*-> | 973 | Bacteria>Terrabacteria_group>Cyanobacteriota/Melainabacteria_group>Cyanobacteriota>Cyanophyceae | GCF_003346965.1 | Fischerella_thermalis | CBP14_RS00540 | 42524..45445 | 372787 | DUF499-domain-containing-protein-[Fischerella-thermalis] |

| Protein-ID | DUF499-architecture | System | Gene-Neighbourhoods | Length | Taxonomy | Assembly-ID | Strain | Locus-Tag | Genomic-coordinates | TaxID | Fasta-defline |
|---|---|---|---|---|---|---|---|---|---|---|---|
| WP_219724091.1 | AAA+ATPase+wHTH+Alpha+Beta-domain+wHTH+RRM | Type-1_BR.RM.like.systems | <-DUF499*<-METHYLASE<-OLD-ABC-p<-TUDOR+SF1-SF2-Helicase+iREase+CTD+REase | 974 | Bacteria>Terrabacteria_group>Cyanobacteriota/Melainabacteria_group>Cyanobacteriota>Cyanophyceae | GCF_002870525.1 | Fischerella_thermalis | CBP25_RS02630 | 1151..4075 | 372787 | DUF499-domain-containing-protein-[Fischerella-thermalis] |
| HAO12934.1 | AAA+ATPase+wHTH+Alpha+Beta-domain+wHTH+RRM | Type-1_BR.RM.like.systems | TUDOR+SF1-SF2-Helicase+iREase+CTD+REase->METHYLASE->DUF499*-> | 957 | Bacteria>Terrabacteria_group>Cyanobacteriota/Melainabacteria_group>Cyanobacteriota>Cyanophyceae | GCA_003456115.1 | Planktothrix_sp._UBA8407 | DCQ51_17620 | 6908..9781 | 2055758 | hypothetical-protein-[Planktothrix-sp.-UBA8407] |
| WP_190385371.1 | AAA+ATPase+wHTH+Alpha+Beta-domain+wHTH+RRM | Type-1_BR.RM.like.systems | SF1-SF2-Helicase-><-DUF499*<-METHYLASE-p | 920 | Bacteria>Terrabacteria_group>Cyanobacteriota/Melainabacteria_group>Cyanobacteriota>Cyanophyceae | GCF_014697315.1 | Aphanizomenon | H6G44_RS06925 | 119734..122496 | 1175 | MULTISPECIES:-DUF499-domain-containing-protein-[Aphanizomenon] |
| OBQ34507.1 | AAA+ATPase+wHTH+Alpha+Beta-domain+wHTH+RRM | Type-1_BR.RM.like.systems | <-DUF499*<-PIN<-MetJarc-HTH<-?<-METHYLASE-p | 966 | Bacteria>Terrabacteria_group>Cyanobacteriota/Melainabacteria_group>Cyanobacteriota>Cyanophyceae | GCA_001672075.1 | Anabaena_sp._CRKS33 | AN487_17930 | 14..2914 | 1710888 | MAG:-AAA-family-ATPase-[Anabaena-sp.-CRKS33] |
| BDA76526.1 | AAA+ATPase+wHTH+Alpha+Beta-domain+wHTH+RRM | Type-1_BR.RM.like.systems | METHYLASE-><-?\|\|DUF499*->?->?->?->?->?-><-CopG/RHH | 971 | Bacteria>Terrabacteria_group>Cyanobacteriota/Melainabacteria_group>Cyanobacteriota>Cyanophyceae | GCA_019977735.1 | Calothrix_sp._PCC_7716 | CAL7716_106920 | 5632..8547 | 2853227 | hypothetical-protein-CAL7716_106920-[Calothrix-sp.-PCC-7716] |
| HBB33200.1 | AAA+ATPase+wHTH+Alpha+Beta-domain+wHTH+RRM | Type-1_BR.RM.like.systems | TUDOR+SF1-SF2-Helicase+iREase+CTD+REase->REase->METHYLASE-p->METHYLASE->Polbeta->MqsA_antitoxin+HTH_3->DUF499*-> | 1007 | Bacteria>Terrabacteria_group>Cyanobacteriota/Melainabacteria_group>Cyanobacteriota>unclassified_Cyanobacteriota | GCA_003488275.1 | Cyanobacteria_bacterium_UBA9273 | DC064_15720 | 10674..13697 | 2055791 | AAA-family-ATPase-[Cyanobacteria-bacterium-UBA9273] |
| HBB33501.1 | AAA+ATPase+wHTH+Alpha+Beta-domain+wHTH+RRM | Type-1_BR.RM.like.systems | TUDOR+SF1-SF2-Helicase+iREase+CTD+REase-><-?<-?<-MetJarc-HTH\|\|DUF4926+DUF2281->REase->DUF499*-><-?\|\|?-><-?<-?<-?<-Pkinase | 971 | Bacteria>Terrabacteria_group>Cyanobacteriota/Melainabacteria_group>Cyanobacteriota>unclassified_Cyanobacteriota | GCA_003488275.1 | Cyanobacteria_bacterium_UBA9273 | DC064_17280 | 21890..24805 | 2055791 | hypothetical-protein-[Cyanobacteria-bacterium-UBA9273] |
| OWY64929.1 | AAA+ATPase+wHTH+Alpha+Beta-domain+wHTH+RRM | Type-1_BR.RM.like.systems | METHYLASE->REase->DUF499*-><-?\|\|HTH_3-><-?<-?<-SbcC+SidE<-METHYLASE+SF1-SF2-Helicase | 921 | Bacteria>Terrabacteria_group>Cyanobacteriota/Melainabacteria_group>Cyanobacteriota>unclassified_Cyanobacteriota | GCA_002213405.1 | cyanobacterium_TDX16 | B7486_44590 | 3155..5920 | 1503470 | AAA-family-ATPase-[cyanobacterium-TDX16] |
| MDA0267772.1 | AAA+ATPase+wHTH+Alpha+Beta-domain+wHTH+RRM | Type-1_BR.RM.like.systems | METHYLASE->?->PIN->REase->DUF499*-> | 1000 | Bacteria>Terrabacteria_group>Cyanobacteriota/Melainabacteria_group>Cyanobacteriota>unclassified_Cyanobacteriota | GCA_027592485.1 | Cyanobacteriota_bacterium | O3A14_12630 | 3513..6515 | 2847862 | MAG:-DUF499-domain-containing-protein-[Cyanobacteria-bacterium] |
| HAA26685.1 | AAA+ATPase+wHTH+Alpha+Beta-domain+wHTH+RRM | Type-1_BR.RM.like.systems | CHTH->TUDOR+SF1-SF2-Helicase+iREase+CTD+REase-p->?->SF1-SF2-Helicase->TUDOR+SF1-SF2-Helicase+iREase+CTD+REase-p->?->DUF499*->DUF499-p->DUF499-p->DUF499-p-> | 616 | Bacteria>Terrabacteria_group>Cyanobacteriota/Melainabacteria_group>Cyanobacteriota>unclassified_Cyanobacteriota | GCA_003444555.1 | Cyanobacteria_bacterium_UBA8553 | DCE56_02160 | 8645..10492 | 2055771 | hypothetical-protein-[Cyanobacteria-bacterium-UBA8553] |
| HAJ59069.1 | AAA+ATPase+wHTH+Alpha+Beta-domain+wHTH+RRM | Type-1_BR.RM.like.systems | CHTH->SMODS->?->METHYLASE->METHYLASE-p->METHYLASE-p->DUF499*->DUF499->DUF499-p-> | 258 | Bacteria>Terrabacteria_group>Cyanobacteriota/Melainabacteria_group>Cyanobacteriota>unclassified_Cyanobacteriota | GCA_003454535.1 | Cyanobacteria_bacterium_UBA8543 | DCP31_07245 | 17086..17860 | 2055766 | hypothetical-protein-[Cyanobacteria-bacterium-UBA8543] |
| WP_229782279.1 | AAA+ATPase+wHTH+Alpha+Beta-domain+wHTH+RRM | Type-1_BR.RM.like.systems | OrfB_Zn_ribbon->?->ParB-><-DUF499*<-REase<-OLD-ABC<-METHYLASE<-TUDOR+SF1-SF2-Helicase+iREase+CTD+REase | 959 | Bacteria>Terrabacteria_group>Deinococcota>Deinococci>Deinococcales | GCF_014647175.1 | Deinococcus_daejeonensis | IEY59_RS18160 | 23471..26350 | 1007098 | DUF499-domain-containing-protein-[Deinococcus-daejeonensis] |
| WP_014514547.1 | AAA+ATPase+wHTH+Alpha+Beta-domain+wHTH+RRM | Type-1_BR.RM.like.systems | 35exo+SF1-SF2-Helicase->?->?->?-><-?<-DUF499*\|\|MULE-TRANSPOSASE-><-HEPN+DUF499<-HicB_lk_antitox<-HicA_toxin<-METHYLASE<-HKD-DNase+SF1-SF2-Helicase-p | 742 | Bacteria>Terrabacteria_group>Deinococcota>Deinococci>Thermales | GCF_000236585.1 | Thermus_sp._CCB_US3_UF1 | TCCBUS3UF1_RS00625 | 142056..144284 | 1111069 | DUF499-domain-containing-protein-[Thermus-sp.-CCB_US3_UF1] |
| WP_252829645.1 | AAA+ATPase+wHTH+Alpha+Beta-domain+wHTH+RRM | Type-1_BR.RM.like.systems | <-DDE_Tnp_1<-?<-?\|\|?->?->\|\|<-HTH+DSBH\|\|<-DUF499*<-HEPN+DUF499<-HIN-HTH+AT-HTH-HYD2-2+SecB<-HTH_3<-HigB/RHH<-HicB_lk_antitox<-METHYLASE-p | 779 | Bacteria>Terrabacteria_group>Deinococcota>Deinococci>Thermales | GCF_000744155.1 | Thermus_scotoductus | N686_RS12530 | 46025..48364 | 37636 | DUF499-domain-containing-protein-[Thermus-scotoductus] |
| WP_269077414.1 | AAA+ATPase+wHTH+Alpha+Beta-domain+wHTH+RRM | Type-1_BR.RM.like.systems | HigB/RHH->HTH_3->HIN-HTH+AT-HTH-HYD2-2+SecB->?->HEPN+DUF499->DUF499->DUF499*->\|\|DDE_Tnp_1_3->RADICAL-SAM->?->?->?-><-DUF4277+DDE_Tnp_1 | 488 | Bacteria>Terrabacteria_group>Deinococcota>Deinococci>Thermales | GCF_000744175.1 | Thermus_tengchongensis | BS82_RS15530 | 77388..78854 | 1214928 | hypothetical-protein-[Thermus-tengchongensis] |
| WP_236622573.1 | AAA+ATPase+wHTH+Alpha+Beta-domain+wHTH+RRM | Type-1_BR.RM.like.systems | HicB_lk_antitox->HigB/RHH->HTH_3->HIN-HTH+AT-HTH-HYD2-2+SecB->?->HEPN+DUF499->DUF499*->DUF499->\|\|DDE_Tnp_1_3->RADICAL-SAM-> | 235 | Bacteria>Terrabacteria_group>Deinococcota>Deinococci>Thermales | GCF_000744175.1 | Thermus_tengchongensis | BS82_RS15080 | 76525..77232 | 1214928 | DUF499-domain-containing-protein-[Thermus-tengchongensis] |
| WP_259331194.1 | AAA+ATPase+wHTH+Alpha+Beta-domain+wHTH+RRM | Type-1_BR.RM.like.systems | <-ABC-ATPase+SbcC<-?<-DUF499*<-HIN-HTH+AT-HTH-HYD2-2+SecB<-HTH_3<-HicA_toxin<-METHYLASE<-DOC+HTH | 834 | Bacteria>Terrabacteria_group>Deinococcota>Deinococci>Thermales | - | Thermus_thermophilus | JQN54_RS11565 | 90341..92845 | 274 | DUF499-domain-containing-protein-[Thermus-thermophilus] |
| MCS7219263.1 | AAA+ATPase+wHTH+Alpha+Beta-domain+wHTH+RRM | Type-1_BR.RM.like.systems | METHYLASE-p->HicA_toxin->HicB_lk_antitox->DUF499->DUF499*->?-><-?<-?<-?<-?<-MarR-HTH | 683 | Bacteria>Terrabacteria_group>Deinococcota>Deinococci>Thermales | GCA_025057525.1 | Thermus_sp. | NZ846_09895 | 2521..4572 | 275 | DUF499-domain-containing-protein-[Thermus-sp.] |
| MCS7219262.1 | AAA+ATPase+wHTH+Alpha+Beta-domain+wHTH+RRM | Type-1_BR.RM.like.systems | METHYLASE-p->HicA_toxin->HicB_lk_antitox->DUF499*->DUF499-> | 273 | Bacteria>Terrabacteria_group>Deinococcota>Deinococci>Thermales | GCA_025057525.1 | Thermus_sp. | NZ846_09890 | 1812..2633 | 275 | DUF499-domain-containing-protein-[Thermus-sp.] |
| WP_150118021.1 | AAA+ATPase+wHTH+Alpha+Beta-domain+wHTH+RRM | Type-1_BR.RM.like.systems | DUF499*->ORC-CDC6+REase-> | 664 | Bacteria>Terrabacteria_group>Deinococcota>Deinococci>Thermales | GCF_001623175.1 | Meiothermus_taiwanensis | A3962_RS14295 | 1..1997 | 172827 | DUF499-domain-containing-protein,-partial-[Meiothermus-taiwanensis] |
| BDG19117.1 | AAA+ATPase+wHTH+Alpha+Beta-domain+wHTH+RRM | Type-1_BR.RM.like.systems | DDE_Tnp_1_5->?->?->?-><-HicA_toxin\|\|DDE_Tnp_1-><-DUF499*<-PolB<-METHYLASE<-Schlafen+HATPase+HTH<-TUDOR+SF1-SF2-Helicase+iREase+CTD+REase | 893 | Bacteria>Terrabacteria_group>Deinococcota>Deinococci>Thermales | GCF_022846275.1 | Thermus_thermophilus | TthSNM11_13200 | 1187360..1190041 | 274 | hypothetical-protein-TthSNM11_13200-[Thermus-thermophilus] |
| WP_244347426.1 | AAA+ATPase+wHTH+Alpha+Beta-domain+wHTH+RRM | Type-1_BR.RM.like.systems | <-HicA_toxin<-HicB\|\|DDE_Tnp_1-><-DUF499*<-?<-PolB<-METHYLASE<-Schlafen+HATPase+HTH<-TUDOR+SF1-SF2-Helicase+iREase+CTD+REase | 896 | Bacteria>Terrabacteria_group>Deinococcota>Deinococci>Thermales | GCF_022846275.1 | Thermus_thermophilus | MWM06_RS06620 | 1187360..1190050 | 274 | DUF499-domain-containing-protein-[Thermus-thermophilus] |
| WP_197525135.1 | AAA+ATPase+wHTH+Alpha+Beta-domain+wHTH+RRM | Type-1_BR.RM.like.systems | <-HicA_toxin<-HicB<-DUF499*<-?<-PolB<-METHYLASE<-Schlafen+HATPase+HTH<-MULE-TRANSPOSASE<-Schlafen | 896 | Bacteria>Terrabacteria_group>Deinococcota>Deinococci>Thermales | GCF_000091545.1 | Thermus_thermophilus | QUE39_RS05170 | 961278..963968 | 274 | DUF499-domain-containing-protein-[Thermus-thermophilus] |
| WP_259329967.1 | AAA+ATPase+wHTH+Alpha+Beta-domain+wHTH+RRM | Type-1_BR.RM.like.systems | <-HicA_toxin<-HicB<-DUF499*<-?<-PolB<-METHYLASE<-Schlafen+HATPase+HTH<-TUDOR+SF1-SF2-Helicase+iREase+CTD+REase | 896 | Bacteria>Terrabacteria_group>Deinococcota>Deinococci>Thermales | GCF_016629545.1 | Thermus_thermophilus | JMH49_RS05530 | 1035330..1038020 | 274 | ATP-binding-protein-[Thermus-thermophilus] |
| WP_201351403.1 | AAA+ATPase+wHTH+Alpha+Beta-domain+wHTH+RRM | Type-1_BR.RM.like.systems | <-HicA_toxin<-HicB<-DUF499*<-PolB<-METHYLASE<-Schlafen+HATPase+HTH<-TUDOR+SF1-SF2-Helicase+iREase+CTD+REase | 893 | Bacteria>Terrabacteria_group>Deinococcota>Deinococci>Thermales | - | Thermus_thermophilus | TthHB5018_10920 | 1035330..1038011 | 274 | DUF499-domain-containing-protein-[Thermus-thermophilus] |
| BCP66158.1 | AAA+ATPase+wHTH+Alpha+Beta-domain+wHTH+RRM | Type-1_BR.RM.like.systems | <-HicA_toxin<-HicB<-DUF499*<-PolB<-METHYLASE<-Schlafen+HATPase+HTH<-TUDOR+SF1-SF2-Helicase+iREase+CTD+REase | 893 | Bacteria>Terrabacteria_group>Deinococcota>Deinococci>Thermales | GCA_016629545.1 | Thermus_thermophilus | TthHB5018_10920 | 1035330..1038011 | 274 | hypothetical-protein-TthHB5018_10920-[Thermus-thermophilus] |
| WP_126203060.1 | AAA+ATPase+wHTH+Alpha+Beta-domain+wHTH+RRM | Type-1_BR.RM.like.systems | TUDOR+SF1-SF2-Helicase+iREase+CTD+REase->Schlafen+HATPase+HTH->METHYLASE->REase->DUF499*-><-ParB | 924 | Bacteria>Terrabacteria_group>Deinococcota>Deinococci>Thermales | GCF_003959165.1 | Thermus_scotoductus | CSW41_RS02895 | 15669..18443 | 37636 | DUF499-domain-containing-protein-[Thermus-scotoductus] |
| WP_224065205.1 | AAA+ATPase+wHTH+Alpha+Beta-domain+wHTH+RRM | Type-1_BR.RM.like.systems | <-HicA_toxin<-HicB<-DUF499*<-?<-PolB<-METHYLASE<-Schlafen+HATPase+HTH | 885 | Bacteria>Terrabacteria_group>Deinococcota>Deinococci>Thermales | GCF_019973715.1 | Thermus_thermophilus | B3N72_RS05290 | 13019..15676 | 274 | DUF499-domain-containing-protein-[Thermus-thermophilus] |
| WP_011228376.1 | AAA+ATPase+wHTH+Alpha+Beta-domain+wHTH+RRM | Type-1_BR.RM.like.systems | Schlafen+HATPase+HTH->METHYLASE->PolB->?->DUF499*->HicB->HicA_toxin-> | 893 | Bacteria>Terrabacteria_group>Deinococcota>Deinococci>Thermales | GCF_019884805.1 | Thermus | DCM56_RS07505 | 4834..7515 | 270 | DUF499-domain-containing-protein-[Thermus-thermophilus] |
| WP_126188140.1 | AAA+ATPase+wHTH+Alpha+Beta-domain+wHTH+RRM | Type-1_BR.RM.like.systems | ParB-><-DUF499*<-REase\|\|DDE_Tnp_1_assoc-> | 924 | Bacteria>Terrabacteria_group>Deinococcota>Deinococci>Thermales | GCF_003958985.1 | Thermus_scotoductus | CSW50_RS14435 | 4946..7720 | 37636 | DUF499-domain-containing-protein-[Thermus-scotoductus] |
| MBX3143487.1 | AAA+ATPase+wHTH+Alpha+Beta-domain+wHTH+RRM | Type-1_BR.RM.like.systems | XIS-HTH->HhH-RADC+JAB->TUDOR+SF1-SF2-Helicase+iREase+CTD+REase->METHYLASE->DUF499*->OLD-ABC->TOPRIM->?->McrC-NTD+REase->Classical-AAA->Peptidase-S8-> | 962 | Bacteria>Terrabacteria_group>Deinococcota>Deinococci>Trueperales | GCA_019637455.1 | Trueperaceae_bacterium | KF813_07015 | 7250..10138 | 2026798 | ATP-binding-protein-[Trueperaceae-bacterium] |
| MBO9307825.1 | AAA+ATPase+wHTH+Alpha+Beta-domain+wHTH+RRM | Type-1_BR.RM.like.systems | TUDOR+SF1-SF2-Helicase+iREase+CTD+REase-p->METHYLASE->DUF499*-> | 879 | Bacteria>Terrabacteria_group>Thermomicrobiota>Thermomicrobia>Thermomicrobiales | GCA_017743575.1 | Thermomicrobium_sp. | J7451_11395 | 2992..5631 | 1969469 | ATP-binding-protein-[Thermomicrobium-sp.] |
| MYB26426.1 | AAA+ATPase+wHTH+Alpha+Beta-domain+wHTH+RRM | Type-1_BR.RM.like.systems | <-SF1-SF2-Helicase+iREase+CTD+REase<-METHYLASE<-DUF499*\|\|DDE_Tnp_1-> | 823 | Bacteria>Thermodesulfobacteriota>Candidatus_Dadabacteria>unclassified_Candidatus_Dadabacteria | GCA_009840825.1 | Candidatus_Dadabacteria_bacterium | F4X32_02855 | 9846..12317 | 2080303 | DUF499-domain-containing-protein-[Candidatus-Dadabacteria-bacterium] |
| HEC67877.1 | AAA+ATPase+wHTH+Alpha+Beta-domain+wHTH+RRM | Type-1_BR.RM.like.systems | <-DUF499*<-METHYLASE<-?<-TOPRIM<-OLD-ABC<-HEPN<-PolB | 896 | Bacteria>Thermodesulfobacteriota>Candidatus_Desulfofervidia>Candidatus_Desulfofervidales>Candidatus_Desulfofervidaceae | GCA_011052795.1 | Candidatus_Desulfofervidus_auxilii | ENI35_03575 | 617..3307 | 1621989 | ATP-binding-protein-[Candidatus-Desulfofervidus-auxilii] |
| MDD3581155.1 | AAA+ATPase+wHTH+Alpha+Beta-domain+wHTH+RRM | Type-1_BR.RM.like.systems | <-TPR+TPR<-ORC-CDC6<-?<-DUF499*<-?<-METHYLASE<-Schlafen+HATPase+HTH<-TUDOR+SF1-SF2-Helicase+iREase+CTD+REase<-?\|\|PAIREDC-HTH-> | 911 | Bacteria>Thermodesulfobacteriota>Desulfobaccia>Desulfobaccales>Desulfobaccaceae | GCA_028695935.1 | Desulfobacca_sp. | PHW74_09060 | 4526..7261 | 2067990 | MAG:-DUF499-domain-containing-protein-[Desulfobacca-sp.] |
| MDD3580510.1 | AAA+ATPase+wHTH+Alpha+Beta-domain+wHTH+RRM | Type-1_BR.RM.like.systems | BrnT_toxin->BrnA_antitoxin->TUDOR+SF1-SF2-Helicase+iREase+CTD+REase->METHYLASE->PolB->HEPN->DUF499*-> | 920 | Bacteria>Thermodesulfobacteriota>Desulfobaccia>Desulfobaccales>Desulfobaccaceae | GCA_028695935.1 | Desulfobacca_sp. | PHW74_05770 | 25341..28103 | 2067990 | MAG:-DUF499-domain-containing-protein-[Desulfobacca-sp.] |
| KPA15639.1 | AAA+ATPase+wHTH+Alpha+Beta-domain+wHTH+RRM | Type-1_BR.RM.like.systems | TUDOR+SF1-SF2-Helicase+iREase+CTD+REase-p->HIN-HTH->TUDOR+SF1-SF2-Helicase+iREase+CTD+REase->REase+HTH->METHYLASE->DUF499*-><-REase | 941 | Bacteria>Thermodesulfobacteriota>Desulfobacteria>Desulfobacterales>Candidatus_Magnetomoraceae | GCA_001292585.1 | Candidatus_Magnetomorum_sp._HK-1 | MHK_004158 | 9986..12811 | 1509431 | ATPase-AAA-[Candidatus-Magnetomorum-sp.-HK-1] |
| WP_178366469.1 | AAA+ATPase+wHTH+Alpha+Beta-domain+wHTH+RRM | Type-1_BR.RM.like.systems | <-DUF499*<-REase<-OLD-ABC<-METHYLASE<-OLD-ABC<-TUDOR+SF1-SF2-Helicase+iREase+CTD+REase<-Radical-SAM+TRAM | 952 | Bacteria>Thermodesulfobacteriota>Desulfobacteria>Desulfobacterales>Desulfobacteraceae | GCF_013389365.1 | Desulfobacter_latus | HXW94_RS08505 | 32689..35547 | 2292 | DUF499-domain-containing-protein-[Desulfobacter-latus] |
| WP_265424070.1 | AAA+ATPase+wHTH+Alpha+Beta-domain+wHTH+RRM | Type-1_BR.RM.like.systems | <-Cysteine_desulfurase<-?<-?<-HAMP+HISKIN\|\|?-><-DUF499-p<-DUF499* | 400 | Bacteria>Thermodesulfobacteriota>Desulfobacteria>Desulfobacterales>Desulfobacteraceae | GCF_026155325.1 | Desulfobotulus_pelophilus | OOTO0_RS04340 | 141258..142460 | 2823377 | DUF499-domain-containing-protein-[Desulfobotulus-sp.-H1] |
| WP_144686771.1 | AAA+ATPase+wHTH+Alpha+Beta-domain+wHTH+RRM | Type-1_BR.RM.like.systems | <-DUF499*<-METHYLASE-p | 955 | Bacteria>Thermodesulfobacteriota>Desulfobacteria>Desulfobacterales>Desulfobacteraceae | GCF_007830435.1 | Desulfobotulus_alkaliphilus | LZ24_RS16255 | 6240..9107 | 622671 | DUF499-domain-containing-protein-[Desulfobotulus-alkaliphilus] |
| VEN75475.1 | AAA+ATPase+wHTH+Alpha+Beta-domain+wHTH+RRM | Type-1_BR.RM.like.systems | 35exo->?->Pterin_bind->?->tRNA-><-URI-DNase<-DUF499*<-METHYLASE<-HEPN<-TUDOR+SF1-SF2-Helicase+iREase+CTD+REase<-wHTH+REase<-?<-METHYLASE | 988 | Bacteria>Thermodesulfobacteriota>Desulfobacteria>Desulfobacterales>Desulfobacteraceae | GCA_900659855.1 | uncultured_Desulfobacteraceae_bacterium | EPICR_90073 | 73527..76493 | 218296 | conserved-hypothetical-protein-[uncultured-Desulfobacteraceae-bacterium] |
| OQX20475.1 | AAA+ATPase+wHTH+Alpha+Beta-domain+wHTH+RRM | Type-1_BR.RM.like.systems | METHYLASE-p->PhdYeFM_antitox->RelE-ParE-><-?\|\|MazE_antitoxin->PIN->DUF499*->?->?->?->Peptidase_M48-> | 913 | Bacteria>Thermodesulfobacteriota>Desulfobacteria>Desulfobacterales>Desulfobacteraceae | GCA_002083955.1 | Desulfobacteraceae_bacterium_IS3 | BWK80_35715 | 2623..5364 | 1934248 | hypothetical-protein-BWK80_35715-[Desulfobacteraceae-bacterium-IS3] |
| MBL7204915.1 | AAA+ATPase+wHTH+Alpha+Beta-domain+wHTH+RRM | Type-1_BR.RM.like.systems | Schlafen->METHYLASE->PolB->HEPN->DUF499*->HEPN+Mrr_N->Schlafen+HATPase+HTH-> | 945 | Bacteria>Thermodesulfobacteriota>Desulfobacteria>Desulfobacterales>Desulfobacteraceae | GCA_016783005.1 | Desulfobacteraceae_bacterium | ISS63_11385 | 7746..10583 | 2049433 | ATP-binding-protein-[Desulfobacteraceae-bacterium] |
| OQX00170.1 | AAA+ATPase+wHTH+Alpha+Beta-domain+wHTH+RRM | Type-1_BR.RM.like.systems | <-DUF499*<-HEPN<-PolB<-HEPN+REase<-?<-BirA-HTH | 918 | Bacteria>Thermodesulfobacteriota>Desulfobacteria>Desulfobacterales>Desulfobacteraceae | GCA_002083965.1 | Desulfobacteraceae_bacterium_A6 | BWK74_00360 | 10328..13084 | 1934249 | AAA-family-ATPase-[Desulfobacteraceae-bacterium-A6] |
| MBF0572396.1 | AAA+ATPase+wHTH+Alpha+Beta-domain+wHTH+RRM | Type-1_BR.RM.like.systems | <-CHASE2+ACYC<-?<-?<-DUF499*<-REase<-PIN<-?<-?<-METHYLASE-p | 936 | Bacteria>Thermodesulfobacteriota>Desulfobacteria>Desulfobacterales>Desulfobacteraceae | GCA_015233755.1 | Desulfamplus_sp. | HQK69_01355 | 13339..16149 | 2006184 | ATP-binding-protein-[Desulfamplus-sp.] |
| ETR72594.1 | AAA+ATPase+wHTH+Alpha+Beta-domain+wHTH+RRM | Type-1_BR.RM.like.systems | METHYLASE-p->METHYLASE-p->?->HicB->HicA->DUF499*-> | 983 | Bacteria>Thermodesulfobacteriota>Desulfobacteria>Desulfobacterales>Desulfobacteraceae | GCA_000516475.1 | Candidatus_Magnetoglobus_multicellularis_str._Araruama | OMM_01599 | 3345..6296 | 890399 | ATPase-AAA-[Candidatus-Magnetoglobus-multicellularis-str.-Araruama] |
| OQX25145.1 | AAA+ATPase+wHTH+Alpha+Beta-domain+wHTH+RRM | Type-1_BR.RM.like.systems | METHYLASE-p->?->?->DUF499*->?->?->?->?-><-PhdYeFM_antitox | 863 | Bacteria>Thermodesulfobacteriota>Desulfobacteria>Desulfobacterales>Desulfobacteraceae | GCA_002083955.1 | Desulfobacteraceae_bacterium_IS3 | BWK80_17135 | 4454..7045 | 1934248 | hypothetical-protein-BWK80_17135-[Desulfobacteraceae-bacterium-IS3] |
| MDA8161932.1 | AAA+ATPase+wHTH+Alpha+Beta-domain+wHTH+RRM | Type-1_BR.RM.like.systems | <-DUF499*<-HEPN<-PolB<-METHYLASE-p | 922 | Bacteria>Thermodesulfobacteriota>Desulfobacteria>Desulfobacterales>Desulfobacteraceae | GCA_028065225.1 | Desulfobacteraceae_bacterium | MO022_03365 | 1133..3901 | 2049433 | MAG:-DUF499-domain-containing-protein-[Desulfobacteraceae-bacterium] |
| MBL7205279.1 | AAA+ATPase+wHTH+Alpha+Beta-domain+wHTH+RRM | Type-1_BR.RM.like.systems | METHYLASE->DUF499*->?-><-sGTP | 943 | Bacteria>Thermodesulfobacteriota>Desulfobacteria>Desulfobacterales>Desulfobacteraceae | GCA_016783005.1 | Desulfobacteraceae_bacterium | ISS63_13275 | 2792..5623 | 2049433 | ATP-binding-protein-[Desulfobacteraceae-bacterium] |
| MBC8360097.1 | AAA+ATPase+wHTH+Alpha+Beta-domain+wHTH+RRM | Type-1_BR.RM.like.systems | TUDOR+SF1-SF2-Helicase+iREase+CTD+REase->Schlafen+HATPase+HTH->?->PHP+SbcC->METHYLASE->REase->DUF499*-> | 920 | Bacteria>Thermodesulfobacteriota>Desulfobacteria>Desulfobacterales>Desulfobacterales_incertae_sedis | GCA_014382665.1 | Candidatus_Desulfatibia_profunda | H8E23_01700 | 12931..15693 | 2841695 | ATP-binding-protein-[Desulfobacterales-bacterium] |
| MBC8432173.1 | AAA+ATPase+wHTH+Alpha+Beta-domain+wHTH+RRM | Type-1_BR.RM.like.systems | <-DUF499*<-REase<-METHYLASE<-?<-TUDOR+SF1-SF2-Helicase+iREase+CTD+REase | 979 | Bacteria>Thermodesulfobacteriota>Desulfobacteria>Desulfobacterales>Desulfobacterales_incertae_sedis | GCA_014383215.1 | Candidatus_Desulfatibia_vada | H8D96_09645 | 2855..5794 | 2841696 | DUF499-domain-containing-protein-[Desulfobacterales-bacterium] |
| SPD74420.1 | AAA+ATPase+wHTH+Alpha+Beta-domain+wHTH+RRM | Type-1_BR.RM.like.systems | TUDOR+SF1-SF2-Helicase+iREase+CTD+REase->?->?->?->URI-DNase->METHYLASE->DUF499*->MORC+HISKIN->?->DNA_methylase-><-?<-MqsA_antitoxin+HTH_3 | 970 | Bacteria>Thermodesulfobacteriota>Desulfobacteria>Desulfobacterales>Desulfobacteriaceae | GCA_900258555.1 | uncultured_Desulfobacterium_sp. | PITCH_A230106 | 637326..640238 | 201089 | ATPase-of-AAA+-superfamily-[uncultured-Desulfobacterium-sp.] |
| WP_155311025.1 | AAA+ATPase+wHTH+Alpha+Beta-domain+wHTH+RRM | Type-1_BR.RM.like.systems | XIS-HTH->TUDOR+SF1-SF2-Helicase+iREase+CTD+REase->?->MACRODOMAIN->METHYLASE->DUF499*->Classical-AAA->Peptidase_S8->HEPN+SF1-SF2-Helicase->MORC+DUF3684+HNH->REase-> | 985 | Bacteria>Thermodesulfobacteriota>Desulfobacteria>Desulfobacterales>Desulfosarcinaceae | GCF_009689005.1 | Desulfosarcina_ovata | GN112_RS15400 | 3472838..3475795 | 83564 | DUF499-domain-containing-protein-[Desulfosarcina-ovata] |
| WP_054029999.1 | AAA+ATPase+wHTH+Alpha+Beta-domain+wHTH+RRM | Type-1_BR.RM.like.systems | TraJ-RHH->?->?-><-?<-?<-DUF499*<-METHYLASE<-?<-Schlafen+HATPase+HTH<-TUDOR+SF1-SF2-Helicase+iREase+CTD+REase\|\|<-HTH<-GDResolvase+Recombinase+ZnR | 970 | Bacteria>Thermodesulfobacteriota>Desulfobacteria>Desulfobacterales>Desulfosarcinaceae | GCF_001293685.1 | Desulfatitalea_tepidiphila | DFT_RS04400 | 1009563..1012475 | 1185843 | DUF499-domain-containing-protein-[Desulfatitalea-tepidiphila] |
| WP_155314702.1 | AAA+ATPase+wHTH+Alpha+Beta-domain+wHTH+RRM | Type-1_BR.RM.like.systems | <-SF1-SF2-Helicase+REase-p<-DUF499*<-METHYLASE<-MACRODOMAIN<-?<-TUDOR+SF1-SF2-Helicase+iREase+CTD+REase<-XIS-HTH | 985 | Bacteria>Thermodesulfobacteriota>Desulfobacteria>Desulfobacterales>Desulfosarcinaceae | GCF_009688945.1 | Desulfosarcina_alkanivorans | DSCA_RS01165 | 257300..260257 | 571177 | DUF499-domain-containing-protein-[Desulfosarcina-alkanivorans] |
| PIP36530.1 | AAA+ATPase+wHTH+Alpha+Beta-domain+wHTH+RRM | Type-1_BR.RM.like.systems | <-ACET<-SAM-methylase<-?<-DUF499-p<-DUF499-p<-DUF499*<-ORC-CDC6+REase<-?<-MqsA_antitoxin+HTH_3<-MqsR_toxin | 497 | Bacteria>Thermodesulfobacteriota>Desulfobacteria>Desulfobacterales>unclassified_Desulfobacterales | GCA_002771315.1 | Desulfobacterales_bacterium_CG23_combo_of_CG06-09_8_20_14_all_52_9 | COX20_05265 | 9529..11022 | 1973984 | hypothetical-protein-COX20_05265-[Desulfobacterales-bacterium-CG23_combo_of_CG06-09_8_20_14_all_52_9] |
| MBN1931790.1 | AAA+ATPase+wHTH+Alpha+Beta-domain+wHTH+RRM | Type-1_BR.RM.like.systems | LexA-HTH+LexA-protease->?->?->Schlafen+HATPase+HTH->TUDOR+SF1-SF2-Helicase+iREase+CTD+REase->METHYLASE->DUF499*-> | 942 | Bacteria>Thermodesulfobacteriota>Desulfobacteria>Desulfobacterales>unclassified_Desulfobacterales | GCA_016931455.1 | Desulfobacterales_bacterium | JW786_09310 | 30802..33630 | 2044940 | ATP-binding-protein-[Desulfobacterales-bacterium] |
| MBF0121377.1 | AAA+ATPase+wHTH+Alpha+Beta-domain+wHTH+RRM | Type-1_BR.RM.like.systems | HNH->TUDOR+SF1-SF2-Helicase+iREase+CTD+REase->METHYLASE->?->RelE->DUF499*-> | 942 | Bacteria>Thermodesulfobacteriota>Desulfobacteria>Desulfobacterales>unclassified_Desulfobacterales | GCA_015228685.1 | Desulfobacterales_bacterium | HQK79_21310 | 12197..15025 | 2044940 | ATP-binding-protein-[Desulfobacterales-bacterium] |
| NVM22696.1 | AAA+ATPase+wHTH+Alpha+Beta-domain+wHTH+RRM | Type-1_BR.RM.like.systems | XIS-HTH->TUDOR+SF1-SF2-Helicase+iREase+CTD+REase->METHYLASE->DUF499*-> | 659 | Bacteria>Thermodesulfobacteriota>Desulfobacteria>Desulfobacterales>unclassified_Desulfobacterales | GCA_013375395.1 | Desulfobacterales_bacterium | HWN68_13065 | 6977..8955 | 2044940 | ATP-binding-protein-[Desulfobacterales-bacterium] |
| OGR23289.1 | AAA+ATPase+wHTH+Alpha+Beta-domain+wHTH+RRM | Type-1_BR.RM.like.systems | <-RelE-ParE<-MetJarc-HTH\|\|XIS-HTH->DUF499*-> | 789 | Bacteria>Thermodesulfobacteriota>Desulfobacteria>Desulfobacterales>unclassified_Desulfobacterales | GCA_001799395.1 | Desulfobacterales_bacterium_RIFOXYA12_FULL_46_15 | A2277_18960 | 4403..6770 | 1797914 | hypothetical-protein-A2277_18960,-partial-[Desulfobacterales-bacterium-RIFOXYA12_FULL_46_15] |
| MBF0121417.1 | AAA+ATPase+wHTH+Alpha+Beta-domain+wHTH+RRM | Type-1_BR.RM.like.systems | <-HNH<-DUF499*<-PIN<-?<-METHYLASE<-TUDOR+SF1-SF2-Helicase+iREase+CTD+REase | 973 | Bacteria>Thermodesulfobacteriota>Desulfobacteria>Desulfobacterales>unclassified_Desulfobacterales | GCA_015228685.1 | Desulfobacterales_bacterium | HQK79_21515 | 1603..4524 | 2044940 | ATP-binding-protein-[Desulfobacterales-bacterium] |
| MCK4391114.1 | AAA+ATPase+wHTH+Alpha+Beta-domain+wHTH+RRM | Type-1_BR.RM.like.systems | MNS+HTH-><-?\|\|METHYLASE->DUF433->PIN->HicB->DUF499*-> | 914 | Bacteria>Thermodesulfobacteriota>Desulfobacteria>Desulfobacterales>unclassified_Desulfobacterales | GCA_023129535.1 | Desulfobacterales_bacterium | KAV83_12860 | 5895..8639 | 2044940 | ATP-binding-protein-[Desulfobacterales-bacterium] |
| MCG2778998.1 | AAA+ATPase+wHTH+Alpha+Beta-domain+wHTH+RRM | Type-1_BR.RM.like.systems | <-PolB<-?<-?<-DUF499*<-REase<-METHYLASE<-VirB4-FtsK+HerA | 920 | Bacteria>Thermodesulfobacteriota>Desulfobacteria>Desulfobacterales>unclassified_Desulfobacterales | GCA_022013615.1 | Desulfobacterales_bacterium | L6406_25260 | 2158..4920 | 2044940 | DUF499-domain-containing-protein-[Desulfobacterales-bacterium] |
| MBF0398065.1 | AAA+ATPase+wHTH+Alpha+Beta-domain+wHTH+RRM | Type-1_BR.RM.like.systems | DUF499*-><-?<-?<-?<-?\|\|SF1-SF2-Helicase-><-DUF4158+DDE_Tnp_Tn3 | 712 | Bacteria>Thermodesulfobacteriota>Desulfobacteria>Desulfobacterales>unclassified_Desulfobacterales | GCA_015232515.1 | Desulfobacterales_bacterium | HQK78_14895 | 1..2141 | 2044940 | DUF499-domain-containing-protein-[Desulfobacterales-bacterium] |
| MBF0102798.1 | AAA+ATPase+wHTH+Alpha+Beta-domain+wHTH+RRM | Type-1_BR.RM.like.systems | <-DUF499-p<-DUF499*<-?<-Y1_Tnp<-METHYLASE | 601 | Bacteria>Thermodesulfobacteriota>Desulfobacteria>Desulfobacterales>unclassified_Desulfobacterales | GCA_015228675.1 | Desulfobacterales_bacterium | HQK77_17995 | 1948..3753 | 2044940 | ATP-binding-protein-[Desulfobacterales-bacterium] |
| MDD5204467.1 | AAA+ATPase+wHTH+Alpha+Beta-domain+wHTH+RRM | Type-1_BR.RM.like.systems | <-DUF499*<-METHYLASE<-ParB+HNH | 968 | Bacteria>Thermodesulfobacteriota>Desulfobacteria>Desulfobacterales>unclassified_Desulfobacterales | GCA_028713085.1 | Desulfobacterales_bacterium | PHS17_03550 | 258..3164 | 2044940 | MAG:-DUF499-domain-containing-protein-[Desulfobacterales-bacterium] |
| MBA3013967.1 | AAA+ATPase+wHTH+Alpha+Beta-domain+wHTH+RRM | Type-1_BR.RM.like.systems | DUF499*->REase->HEPN->OLD-ABC+TOPRIM->SF1-SF2-Helicase-><-METHYLASE+SF1-SF2-Helicase | 944 | Bacteria>Thermodesulfobacteriota>Desulfobulbia>Desulfobulbales>Desulfobulbaceae | GCA_013792005.1 | Desulfobulbaceae_bacterium | FP815_03320 | 13..2847 | 2053307 | ATP-binding-protein-[Desulfobulbaceae-bacterium] |
| WP_205241970.1 | AAA+ATPase+wHTH+Alpha+Beta-domain+wHTH+RRM | Type-1_BR.RM.like.systems | TUDOR+SF1-SF2-Helicase+iREase+CTD+REase->HD->DUF4065->METHYLASE->OLD-ABC->TOPRIM->DUF499*->HEPN+SF1-SF2-Helicase->MORC+DUF3684+HNH->?->?-><-?<-HEPN | 985 | Bacteria>Thermodesulfobacteriota>Desulfobulbia>Desulfobulbales>Desulfobulbaceae | GCF_016918545.1 | Desulfobulbus_alkaliphilus | JWG43_RS06300 | 125818..128775 | 869814 | DUF499-domain-containing-protein-[Desulfobulbus-alkaliphilus] |
| HHL35500.1 | AAA+ATPase+wHTH+Alpha+Beta-domain+wHTH+RRM | Type-1_BR.RM.like.systems | <-DUF499*<-DUF1828+REase | 826 | Bacteria>Thermodesulfobacteriota>Desulfobulbia>Desulfobulbales>Desulfobulbaceae | GCA_011322715.1 | Desulfobulbaceae_bacterium | ENJ30_14165 | 1..2478 | 2053307 | ATP-binding-protein-[Desulfobulbaceae-bacterium] |
| WP_205218088.1 | AAA+ATPase+wHTH+Alpha+Beta-domain+wHTH+RRM | Type-1_BR.RM.like.systems | TUDOR+SF1-SF2-Helicase+iREase+CTD+REase->METHYLASE->OLD-ABC->DUF499*->?-><-?<-?<-?\|\|?->ABhydrolase-> | 942 | Bacteria>Thermodesulfobacteriota>Desulfobulbia>Desulfobulbales>Desulfocapsaceae | GCF_016918505.1 | Desulforhopalus_vacuolatus | JWG39_RS04080 | 97099..99927 | 40414 | DUF499-domain-containing-protein-[Desulforhopalus-vacuolatus] |
| MBE0582881.1 | AAA+ATPase+wHTH+Alpha+Beta-domain+wHTH+RRM | Type-1_BR.RM.like.systems | <-DUF499*<-XIS-HTH<-?\|\|METHYLASE-p-> | 184 | Bacteria>Thermodesulfobacteriota>Desulfobulbia>Desulfobulbales>Desulfocapsaceae | GCA_014860245.1 | Desulfofustis_sp. | IH612_03870 | 1..551 | 2053546 | DUF499-domain-containing-protein-[Desulfofustis-sp.] |
| WP_027371119.1 | AAA+ATPase+wHTH+Alpha+Beta-domain+wHTH+RRM | Type-1_BR.RM.like.systems | zf-ISL3+HTH_Tnp_ISL3+DDE_Tnp_ISL3->METHYLASE->DUF499*-><-RelE-ParE<-RHH_5\|\|CHTH->?-><-?\|\|ABC-ATPase+SbcC-> | 946 | Bacteria>Thermodesulfobacteriota>Desulfovibrionia>Desulfovibrionales>Desulfohalobiaceae | GCF_000620765.1 | Desulfovermiculus_halophilus | N902_RS0111960 | 2411..5251 | 339722 | DUF499-domain-containing-protein-[Desulfovermiculus-halophilus] |
| MCF8029882.1 | AAA+ATPase+wHTH+Alpha+Beta-domain+wHTH+RRM | Type-1_BR.RM.like.systems | TUDOR+SF1-SF2-Helicase+iREase+CTD+REase->METHYLASE->DUF499*->?->REase+ParB-><-?<-?<-ZnR<-ArdCN+MPTase | 944 | Bacteria>Thermodesulfobacteriota>Desulfovibrionia>Desulfovibrionales>Desulfohalobiaceae | GCA_021737085.1 | Desulfohalobiaceae_bacterium | K9K39_03215 | 9023..11857 | 2683721 | DUF499-domain-containing-protein-[Desulfohalobiaceae-bacterium] |
| HCF05840.1 | AAA+ATPase+wHTH+Alpha+Beta-domain+wHTH+RRM | Type-1_BR.RM.like.systems | TUDOR+SF1-SF2-Helicase+iREase+CTD+REase->PIN->HEPN->METHYLASE->OLD-ABC->TOPRIM->DUF499*-> | 934 | Bacteria>Thermodesulfobacteriota>Desulfovibrionia>Desulfovibrionales>Desulfomicrobiaceae | GCA_003530935.1 | Desulfomicrobiaceae_bacterium | DEU72_06355 | 12752..15556 | 2053547 | hypothetical-protein-[Desulfomicrobiaceae-bacterium] |
| MBC7355961.1 | AAA+ATPase+wHTH+Alpha+Beta-domain+wHTH+RRM | Type-1_BR.RM.like.systems | <-DUF499*<-Schlafen+HATPase+HTH<-METHYLASE<-TOPRIM<-METHYLASE<-PemK_toxin<-PrlF_antitoxin | 934 | Bacteria>Thermodesulfobacteriota>Desulfovibrionia>Desulfovibrionales>Desulfomicrobiaceae | GCA_014359945.1 | Desulfomicrobiaceae_bacterium | H5U09_07160 | 21001..23805 | 2053547 | ATP-binding-protein-[Desulfomicrobiaceae-bacterium] |
| MBZ4685271.1 | AAA+ATPase+wHTH+Alpha+Beta-domain+wHTH+RRM | Type-1_BR.RM.like.systems | FAD-NAD-dep-oxidoreductase-><-?<-?<-?\|\|?-><-DUF499*<-TOPRIM<-OLD-ABC<-METHYLASE<-HEPN<-PIN | 934 | Bacteria>Thermodesulfobacteriota>Desulfovibrionia>Desulfovibrionales>Desulfomicrobiaceae | GCA_020063575.1 | Desulfomicrobiaceae_bacterium | JG774_1016 | 59895..62699 | 2053547 | putative-ATPase-[Desulfomicrobiaceae-bacterium] |
| WP_165177019.1 | AAA+ATPase+wHTH+Alpha+Beta-domain+wHTH+RRM | Type-1_BR.RM.like.systems | TOPRIM+SF1-SF2-Helicase+HTH->TOPRIM+SF1-SF2-Helicase+HTH->?->?->GDResolvase->GDResolvase+Recombinase+ZnR-><-DUF499*<-NERD<-OLD-ABC<-METHYLASE<-TUDOR+SF1-SF2-Helicase+iREase+CTD+REase\|\|?->DUF499-p-> | 938 | Bacteria>Thermodesulfobacteriota>Desulfovibrionia>Desulfovibrionales>Desulfovibrionaceae | GCF_011039085.1 | Desulfovibrio_sp._ZJ369 | G7Y51_RS09870 | 21109..23925 | 2709793 | DUF499-domain-containing-protein-[Desulfovibrio-sp.-ZJ369] |
| WP_179981475.1 | AAA+ATPase+wHTH+Alpha+Beta-domain+wHTH+RRM | Type-1_BR.RM.like.systems | ABC-ATPase+SbcC->?->?->GDResolvase+HIN-HTH->DUF4158+DDE_Tnp_Tn3-><-OLD-ABC<-DUF499*<-NERD<-OLD-ABC<-METHYLASE<-HEPN<-TUDOR+SF1-SF2-Helicase+iREase+CTD+REase<-SF1-SF2-Helicase-p | 944 | Bacteria>Thermodesulfobacteriota>Desulfovibrionia>Desulfovibrionales>Desulfovibrionaceae | GCF_902702915.1 | Desulfovibrio_sp._86 | QZL82_RS00255 | 53858..56692 | 2666132 | DUF499-domain-containing-protein-[Desulfovibrio-sp.-86] |
| WP_144304545.1 | AAA+ATPase+wHTH+Alpha+Beta-domain+wHTH+RRM | Type-1_BR.RM.like.systems | TUDOR+SF1-SF2-Helicase+iREase+CTD+REase-p->METHYLASE->DUF499*->?->OLD-ABC+TOPRIM->SF1-SF2-Helicase-><-Phage_integrase<-?\|\|ZnR-> | 945 | Bacteria>Thermodesulfobacteriota>Desulfovibrionia>Desulfovibrionales>Desulfovibrionaceae | GCF_007625075.1 | Oceanidesulfovibrio_indonesiensis | DPQ33_RS17545 | 4200..7037 | 54767 | DUF499-domain-containing-protein-[Oceanidesulfovibrio-indonesiensis] |
| WP_078684559.1 | AAA+ATPase+wHTH+Alpha+Beta-domain+wHTH+RRM | Type-1_BR.RM.like.systems | <-MoxR-AAA\|\|?->?-><-HIN-HTH<-DUF499*<-METHYLASE<-TUDOR+SF1-SF2-Helicase+iREase+CTD+REase\|\|JAB-><-?\|\|STYKinase-like-> | 939 | Bacteria>Thermodesulfobacteriota>Desulfovibrionia>Desulfovibrionales>Desulfovibrionaceae | GCF_900167065.1 | Desulfobaculum_bizertense | B5D23_RS06275 | 276864..279683 | 376490 | DUF499-domain-containing-protein-[Paradesulfovibrio-bizertensis] |
| WP_168935605.1 | AAA+ATPase+wHTH+Alpha+Beta-domain+wHTH+RRM | Type-1_BR.RM.like.systems | <-HTH_3\|\|?-><-?\|\|?->?-><-DUF499*<-REase<-OLD-ABC<-METHYLASE<-HNH<-OLD-ABC<-TUDOR+SF1-SF2-Helicase+iREase+CTD+REase | 933 | Bacteria>Thermodesulfobacteriota>Desulfovibrionia>Desulfovibrionales>Desulfovibrionaceae | GCF_012843875.1 | Desulfovibrio_piger | HF854_RS06725 | 17522..20323 | 901 | DUF499-domain-containing-protein-[Desulfovibrio-piger] |
| GKG92116.1 | AAA+ATPase+wHTH+Alpha+Beta-domain+wHTH+RRM | Type-1_BR.RM.like.systems | XIS-HTH->TUDOR+SF1-SF2-Helicase+iREase+CTD+REase->METHYLASE->OLD-ABC->?->DUF499*->?->Y1_Tnp-> | 941 | Bacteria>Thermodesulfobacteriota>Desulfovibrionia>Desulfovibrionales>Desulfovibrionaceae | GCA_022834915.1 | Desulfovibrionaceae_bacterium | CE91St38_01240 | 144930..147755 | 2049043 | hypothetical-protein-CE91St38_01240-[Desulfovibrionaceae-bacterium] |
| WP_144304591.1 | AAA+ATPase+wHTH+Alpha+Beta-domain+wHTH+RRM | Type-1_BR.RM.like.systems | REase->?-><-?<-?<-DUF499*<-METHYLASE<-TUDOR+SF1-SF2-Helicase+iREase+CTD+REase-p | 947 | Bacteria>Thermodesulfobacteriota>Desulfovibrionia>Desulfovibrionales>Desulfovibrionaceae | GCF_007625075.1 | Oceanidesulfovibrio_indonesiensis | DPQ33_RS17820 | 8370..11213 | 54767 | DUF499-domain-containing-protein-[Oceanidesulfovibrio-indonesiensis] |
| WP_160963929.1 | AAA+ATPase+wHTH+Alpha+Beta-domain+wHTH+RRM | Type-1_BR.RM.like.systems | TUDOR+SF1-SF2-Helicase+iREase+CTD+REase->HEPN->METHYLASE->DUF499*-> | 956 | Bacteria>Thermodesulfobacteriota>Desulfovibrionia>Desulfovibrionales>Desulfovibrionaceae | GCF_009856865.1 | Solidesulfovibrio_aerotolerans | GTA51_RS18990 | 10316..13186 | 295255 | DUF499-domain-containing-protein-[Solidesulfovibrio-aerotolerans] |
| WP_011735762.1 | AAA+ATPase+wHTH+Alpha+Beta-domain+wHTH+RRM | Type-1_BR.RM.like.systems | <-REase<-?<-SF1-SF2-Helicase<-OLD-ABC+TOPRIM<-DUF499*<-METHYLASE-p<-TUDOR+SF1-SF2-Helicase+iREase+CTD+REase | 950 | Bacteria>Thermodesulfobacteriota>Desulfuromonadia>Desulfuromonadales>Desulfuromonadaceae | GCF_000015045.1 | Pelobacter_propionicus | PPRO_RS09330 | 2017723..2020575 | 29543 | DUF499-domain-containing-protein-[Pelobacter-propionicus] |
| RII28688.1 | AAA+ATPase+wHTH+Alpha+Beta-domain+wHTH+RRM | Type-1_BR.RM.like.systems | <-REase<-RelE-ParE<-RHH_5<-Cysteine_desulfurase\|\|DUF4007->DUF499*->PAPS_reduct-><-ParB+HNH<-?<-DUF2726+ZnR+TOPRIM<-?\|\|XIS-HTH-> | 1131 | Bacteria>Thermodesulfobacteriota>Desulfuromonadia>Geobacterales>Geobacteraceae | GCA_003574885.1 | Geobacter_sp. | CXR31_02060 | 447432..450827 | 46610 | MAG:-ATP-binding-protein-[Geobacter-sp.] |
| KFN37947.1 | AAA+ATPase+wHTH+Alpha+Beta-domain+wHTH+RRM | Type-1_BR.RM.like.systems | VirB4-FtsK+HerA->METHYLASE->RNase->DUF499*-> | 922 | Bacteria>Thermodesulfobacteriota>Syntrophia>Syntrophales>Smithellaceae | GCA_000747085.1 | Smithella_sp._F21 | KN63_07070 | 8502..11270 | 1538640 | ATPase-AAA-[Smithella-sp.-F21] |
| NLA42003.1 | AAA+ATPase+wHTH+Alpha+Beta-domain+wHTH+RRM | Type-1_BR.RM.like.systems | ABC-ATPase+SbcC->?->?->?->?->?-><-DUF499*<-METHYLASE-p | 929 | Bacteria>Thermodesulfobacteriota>Syntrophia>Syntrophales>Smithellaceae | GCA_012510495.1 | Smithella_sp. | GX874_11495 | 22131..24920 | 2053614 | DUF499-domain-containing-protein-[Smithella-sp.] |
| WP_011417561.1 | AAA+ATPase+wHTH+Alpha+Beta-domain+wHTH+RRM | Type-1_BR.RM.like.systems | <-SIGMA-HTH<-?<-HerA/FtsK<-NurA<-DUF4338<-DUF499*<-METHYLASE<-URI-DNase<-SF1-SF2-Helicase | 972 | Bacteria>Thermodesulfobacteriota>Syntrophia>Syntrophales>Syntrophaceae | GCF_000013405.1 | Syntrophus_aciditrophicus | SYN_RS07905 | 1720416..1723334 | 316277 | DUF499-domain-containing-protein-[Syntrophus-aciditrophicus] |
| MBN1363357.1 | AAA+ATPase+wHTH+Alpha+Beta-domain+wHTH+RRM | Type-1_BR.RM.like.systems | Phage_int_SAM_4+Phage_integrase->?->ABhydrolase->METHYLASE->DOC+HTH->DUF499-p->DUF499*-> | 707 | Bacteria>Thermodesulfobacteriota>Syntrophia>Syntrophales>Syntrophaceae | GCA_016927935.1 | Syntrophaceae_bacterium | JW976_00995 | 9584..11707 | 2052184 | DUF499-domain-containing-protein-[Syntrophaceae-bacterium] |
| OPY16343.1 | AAA+ATPase+wHTH+Alpha+Beta-domain+wHTH+RRM | Type-1_BR.RM.like.systems | AbiEi_1->TUDOR+SF1-SF2-Helicase+iREase+CTD+REase->?->METHYLASE->DUF499*-><-?<-?<-RNase<-?<-HicB<-DHTH+WYL | 919 | Bacteria>Thermodesulfobacteriota>Syntrophia>Syntrophales>Syntrophaceae | GCA_002067825.1 | Syntrophus_sp._PtaB.Bin075 | A4E74_01896 | 11338..14097 | 1823758 | MAG:-hypothetical-protein-A4E74_01896-[Syntrophus-sp.-PtaB.Bin075] |
| OIP92318.1 | AAA+ATPase+wHTH+Alpha+Beta-domain+wHTH+RRM | Type-1_BR.RM.like.systems | OLD-ABC+TOPRIM->METHYLASE->?->RelE-ParE->DUF499*-> | 960 | Bacteria>Thermodesulfobacteriota>Syntrophia>Syntrophales>Syntrophaceae | GCA_001873745.1 | Syntrophaceae_bacterium_CG2_30_58_14 | AUK26_04980 | 8100..10982 | 1805378 | AAA-family-ATPase-[Syntrophaceae-bacterium-CG2_30_58_14] |
| OIP91485.1 | AAA+ATPase+wHTH+Alpha+Beta-domain+wHTH+RRM | Type-1_BR.RM.like.systems | METHYLASE-p->OLD-ABC->TOPRIM->?->DUF499*-> | 853 | Bacteria>Thermodesulfobacteriota>Syntrophia>Syntrophales>Syntrophaceae | GCA_001873745.1 | Syntrophaceae_bacterium_CG2_30_58_14 | AUK26_06780 | 3411..5972 | 1805378 | AAA-family-ATPase-[Syntrophaceae-bacterium-CG2_30_58_14] |
| MBN1472033.1 | AAA+ATPase+wHTH+Alpha+Beta-domain+wHTH+RRM | Type-1_BR.RM.like.systems | METHYLASE-p->PolB->HEPN->?->DUF499*-> | 791 | Bacteria>Thermodesulfobacteriota>Syntrophia>Syntrophales>Syntrophaceae | GCA_016928665.1 | Syntrophaceae_bacterium | JW925_09645 | 2090..4465 | 2052184 | ATP-binding-protein-[Syntrophaceae-bacterium] |
| OPY93422.1 | AAA+ATPase+wHTH+Alpha+Beta-domain+wHTH+RRM | Type-1_BR.RM.like.systems | <-URI-DNase<-DUF499*<-METHYLASE-p | 920 | Bacteria>Thermodesulfobacteriota>Syntrophia>Syntrophales>Syntrophaceae | GCA_002068035.1 | Syntrophaceae_bacterium_PtaU1.Bin231 | A4E73_00137 | 3688..6450 | 1811718 | hypothetical-protein-A4E73_00137-[Syntrophaceae-bacterium-PtaU1.Bin231] |
| WP_137425232.1 | AAA+ATPase+wHTH+Alpha+Beta-domain+wHTH+RRM | Type-1_BR.RM.like.systems | XIS-HTH->DUF499*->PolB->METHYLASE-p->MORC+HISKIN-> | 152 | Bacteria>Thermodesulfobacteriota>Syntrophobacteria>Syntrophobacterales>Syntrophobacteraceae | GCF_005377625.1 | Desulfoglaeba_alkanexedens | FDQ92_RS12675 | 2798739..2799197 | 361111 | hypothetical-protein-[Desulfoglaeba-alkanexedens] |
| MCU0573909.1 | AAA+ATPase+wHTH+Alpha+Beta-domain+wHTH+RRM | Type-1_BR.RM.like.systems | S1+RNase+2C-><-?\|\|?->PUA+METHYLASE-><-?\|\|tRNA->DUF499*-> | 523 | Bacteria>Thermodesulfobacteriota>Syntrophobacteria>Syntrophobacterales>Syntrophobacteraceae | GCA_025454175.1 | Syntrophobacteraceae_bacterium | MUC41_13075 | 28123..29690 | 2052185 | DUF499-domain-containing-protein-[Syntrophobacteraceae-bacterium] |
| WP_137424067.1 | AAA+ATPase+wHTH+Alpha+Beta-domain+wHTH+RRM | Type-1_BR.RM.like.systems | TUDOR+SF1-SF2-Helicase+iREase+CTD+REase->DOC+HTH->ZDNA-HTH->METHYLASE->OLD-ABC->TOPRIM->DUF499*->?-><-AbiEii\|\|HIN-HTH->?-><-AbiEii<-AbiEi_1 | 927 | Bacteria>Thermodesulfobacteriota>Syntrophobacteria>Syntrophobacterales>Syntrophobacteraceae | GCF_005377625.1 | Desulfoglaeba_alkanexedens | FDQ92_RS07910 | 1746088..1748871 | 361111 | DUF499-domain-containing-protein-[Desulfoglaeba-alkanexedens] |
| WP_258927534.1 | AAA+ATPase+wHTH+Alpha+Beta-domain+wHTH+RRM | Type-1_BR.RM.like.systems | REase+NACHT-><-Polbeta<-?<-?<-DUF499*<-REase<-METHYLASE<-?<-SF1-SF2-Helicase<-SbcC<-TUDOR+SF1-SF2-Helicase+iREase+CTD+REase | 921 | Bacteria>Thermodesulfobacteriota>Syntrophobacteria>Syntrophobacterales>Syntrophobacteraceae | GCF_022603395.1 | Desulfoferrobacter_suflitae | K1V27_RS00340 | 59530..62295 | 2865782 | DUF499-domain-containing-protein-[Desulfoferrobacter-suflitae] |
| MCK8600135.1 | AAA+ATPase+wHTH+Alpha+Beta-domain+wHTH+RRM | Type-1_BR.RM.like.systems | REase+NACHT-><-Polbeta<-?<-?<-DUF499*<-REase<-METHYLASE<-?<-SF1-SF2-Helicase<-SbcC<-TUDOR+SF1-SF2-Helicase+iREase+CTD+REase | 921 | Bacteria>Thermodesulfobacteriota>Syntrophobacteria>Syntrophobacterales>Syntrophobacteraceae | GCA_022603395.1 | Desulfoferrobacter_suflitae | K1V27_00335 | 59530..62295 | 2865782 | DUF499-domain-containing-protein-[Syntrophobacteraceae-bacterium-DRH4] |
| WP_211334870.1 | AAA+ATPase+wHTH+Alpha+Beta-domain+wHTH+RRM | Type-1_BR.RM.like.systems | TUDOR+SF1-SF2-Helicase+iREase+CTD+REase->OLD-ABC->TOPRIM->METHYLASE->OLD-ABC->TOPRIM->DUF499*->?->?-><-SF1-SF2-Helicase | 923 | Bacteria>Thermodesulfobacteriota>Syntrophobacteria>Syntrophobacterales>Syntrophobacteraceae | GCF_003751385.1 | Desulfosoma_caldarium | EDC27_RS10790 | 105380..108151 | 610254 | DUF499-domain-containing-protein-[Desulfosoma-caldarium] |
| MBC7358359.1 | AAA+ATPase+wHTH+Alpha+Beta-domain+wHTH+RRM | Type-1_BR.RM.like.systems | <-HDRnase<-RAMPs<-?<-DUF499*<-DUF4065<-HD<-TUDOR+SF1-SF2-Helicase+iREase+CTD+REase<-XIS-HTH<-?<-URI-DNase | 970 | Bacteria>Thermodesulfobacteriota>Syntrophobacteria>Syntrophobacterales>Syntrophobacteraceae | GCA_014359885.1 | Desulfacinum_sp. | H5U10_07480 | 223984..226896 | 2301220 | ATP-binding-protein-[Desulfacinum-sp.] |
| WP_137423747.1 | AAA+ATPase+wHTH+Alpha+Beta-domain+wHTH+RRM | Type-1_BR.RM.like.systems | TUDOR+SF1-SF2-Helicase+iREase+CTD+REase->OLD-ABC->TOPRIM->METHYLASE->BrnT_toxin->HTH->DUF499*-> | 932 | Bacteria>Thermodesulfobacteriota>Syntrophobacteria>Syntrophobacterales>Syntrophobacteraceae | GCF_005377625.1 | Desulfoglaeba_alkanexedens | FDQ92_RS06025 | 1288996..1291794 | 361111 | DUF499-domain-containing-protein-[Desulfoglaeba-alkanexedens] |
| WP_245795145.1 | AAA+ATPase+wHTH+Alpha+Beta-domain+wHTH+RRM | Type-1_BR.RM.like.systems | <-HDRnase<-RAMPs<-?<-DUF499*<-METHYLASE<-DUF4065<-HD<-TUDOR+SF1-SF2-Helicase+iREase+CTD+REase | 952 | Bacteria>Thermodesulfobacteriota>Syntrophobacteria>Syntrophobacterales>Syntrophobacteraceae | GCF_900129305.1 | Desulfacinum_infernum | BUB04_RS07055 | 125046..127904 | 35837 | DUF499-domain-containing-protein-[Desulfacinum-infernum] |
| WP_073037944.1 | AAA+ATPase+wHTH+Alpha+Beta-domain+wHTH+RRM | Type-1_BR.RM.like.systems | TUDOR+SF1-SF2-Helicase+iREase+CTD+REase->TIR->METHYLASE->DUF499*->PIN-> | 928 | Bacteria>Thermodesulfobacteriota>Syntrophobacteria>Syntrophobacterales>Syntrophobacteraceae | GCF_900129305.1 | Desulfacinum_infernum | BUB04_RS06060 | 31912..34698 | 35837 | DUF499-domain-containing-protein-[Desulfacinum-infernum] |
| NSW87838.1 | AAA+ATPase+wHTH+Alpha+Beta-domain+wHTH+RRM | Type-1_BR.RM.like.systems | <-DUF499*<-METHYLASE-p<-HEPN<-wHTH+REase<-REase | 698 | Bacteria>Thermodesulfobacteriota>Syntrophobacteria>Syntrophobacterales>Syntrophobacteraceae | GCA_013314815.1 | Syntrophobacteraceae_bacterium | HPY84_16135 | 1106..3202 | 2052185 | DUF499-domain-containing-protein-[Syntrophobacteraceae-bacterium] |
| WP_028895668.1 | AAA+ATPase+wHTH+Alpha+Beta-domain+wHTH+RRM | Type-1_BR.RM.like.systems | XIS-HTH->TUDOR+SF1-SF2-Helicase+iREase+CTD+REase->wHTH+REase->?->VirB4-FtsK+HerA->METHYLASE->DUF499*->Classical-AAA->?->Peptidase_S8-><-?\|\|HTH_3-> | 965 | Bacteria>Thermodesulfobacteriota>Syntrophorhabdia>Syntrophorhabdales>Syntrophorhabdaceae | GCF_000512235.1 | Syntrophorhabdus_aromaticivorans | SYNAR_RS0117510 | 3034488..3037385 | 328301 | DUF499-domain-containing-protein-[Syntrophorhabdus-aromaticivorans] |
| MDD5008580.1 | AAA+ATPase+wHTH+Alpha+Beta-domain+wHTH+RRM | Type-1_BR.RM.like.systems | <-DDE_5+DDE_Tnp_1<-?<-?\|\|METHYLASE->Mbetalac->DUF499*->HKD-DNase-> | 920 | Bacteria>Thermodesulfobacteriota>Syntrophorhabdia>Syntrophorhabdales>Syntrophorhabdaceae | GCA_028711085.1 | Syntrophorhabdaceae_bacterium | PHC68_09330 | 6929..9691 | 2660714 | MAG:-DUF499-domain-containing-protein-[Syntrophorhabdaceae-bacterium] |
| OPY63949.1 | AAA+ATPase+wHTH+Alpha+Beta-domain+wHTH+RRM | Type-1_BR.RM.like.systems | HerA/FtsK->METHYLASE->?->?->DUF499*-> | 965 | Bacteria>Thermodesulfobacteriota>Syntrophorhabdia>Syntrophorhabdales>Syntrophorhabdaceae | GCA_002067405.1 | Syntrophorhabdaceae_bacterium_PtaU1.Bin034 | A4E57_03692 | 6618..9515 | 1811715 | hypothetical-protein-A4E57_03692-[Syntrophorhabdaceae-bacterium-PtaU1.Bin034] |
| WP_038060516.1 | AAA+ATPase+wHTH+Alpha+Beta-domain+wHTH+RRM | Type-1_BR.RM.like.systems | TUDOR+SF1-SF2-Helicase+iREase+CTD+REase->METHYLASE->PolB->PolB->HEPN->METHYLASE-p->DUF499*->?->?-><-Radical-SAM<-Calcineurin<-?<-PRTase | 856 | Bacteria>Thermodesulfobacteriota>Thermodesulfobacteria>Thermodesulfobacteriales>Thermodesulfobacteriaceae | GCF_000734015.1 | Thermodesulfobacterium_commune | CC99_RS02005 | 41791..44361 | 1741 | ATP-binding-protein-[Thermodesulfobacterium-commune] |
| MBZ4682110.1 | AAA+ATPase+wHTH+Alpha+Beta-domain+wHTH+RRM | Type-1_BR.RM.like.systems | PRTase->?->Calcineurin->Radical-SAM-><-?<-?<-DUF499*<-METHYLASE-p<-HEPN<-PolB<-PolB<-METHYLASE<-TUDOR+SF1-SF2-Helicase+iREase+CTD+REase | 856 | Bacteria>Thermodesulfobacteriota>Thermodesulfobacteria>Thermodesulfobacteriales>Thermodesulfobacteriaceae | GCA_020055045.1 | Thermodesulfobacterium_sp. | JG778_1185 | 26969..29539 | 1965289 | hypothetical-protein-[Thermodesulfobacterium-sp.] |
| WP_150072659.1 | AAA+ATPase+wHTH+Alpha+Beta-domain+wHTH+RRM | Type-1_BR.RM.like.systems | TUDOR+SF1-SF2-Helicase+iREase+CTD+REase->METHYLASE->Polbeta->METHYLASE-p->DUF499*-><-?<-?\|\|?->ncRNA->tRNA->Peptidase_M48-> | 861 | Bacteria>Thermodesulfobacteriota>Thermodesulfobacteria>Thermodesulfobacteriales>Thermodesulfobacteriaceae | GCF_008630935.1 | Thermodesulfobacterium_sp._TA1 | F1847_RS08695 | 1734003..1736588 | 2234087 | DUF499-domain-containing-protein-[Thermodesulfobacterium-sp.-TA1] |
| MBZ4681882.1 | AAA+ATPase+wHTH+Alpha+Beta-domain+wHTH+RRM | Type-1_BR.RM.like.systems | <-DUF499*<-METHYLASE<-?<-TUDOR+SF1-SF2-Helicase+iREase+CTD+REase<-MULE-TRANSPOSASE | 885 | Bacteria>Thermodesulfobacteriota>Thermodesulfobacteria>Thermodesulfobacteriales>Thermodesulfobacteriaceae | GCA_020055045.1 | Thermodesulfobacterium_sp. | JG778_957 | 65921..68578 | 1965289 | ATPase-[Thermodesulfobacterium-sp.] |
| PMP69075.1 | AAA+ATPase+wHTH+Alpha+Beta-domain+wHTH+RRM | Type-1_BR.RM.like.systems | METHYLASE-p->DUF499*->?-><-?<-?<-?<-ParA-Soj-PloopNTPase\|\|PIN+PhoH-> | 840 | Bacteria>Thermodesulfobacteriota>Thermodesulfobacteria>Thermodesulfobacteriales>Thermodesulfobacteriaceae | GCA_002877985.1 | Thermodesulfobacterium_geofontis | C0190_00625 | 441..2963 | 1295609 | AAA-family-ATPase-[Thermodesulfobacterium-geofontis] |
| MCC6048503.1 | AAA+ATPase+wHTH+Alpha+Beta-domain+wHTH+RRM | Type-1_BR.RM.like.systems | <-DUF499*<-METHYLASE-p<-HEPN<-PolB<-METHYLASE-p | 454 | Bacteria>Thermodesulfobacteriota>Thermodesulfobacteria>Thermodesulfobacteriales>Thermodesulfobacteriaceae | GCA_020833455.1 | Thermodesulfobacterium_sp. | LM579_03100 | 1..1362 | 1965289 | DUF499-domain-containing-protein-[Thermodesulfobacterium-sp.] |
| WP_038060834.1 | AAA+ATPase+wHTH+Alpha+Beta-domain+wHTH+RRM | Type-1_BR.RM.like.systems | <-DUF499*<-METHYLASE<-TUDOR+SF1-SF2-Helicase+iREase+CTD+REase | 885 | Bacteria>Thermodesulfobacteriota>Thermodesulfobacteria>Thermodesulfobacteriales>Thermodesulfobacteriaceae | GCF_000734015.1 | Thermodesulfobacterium_commune | CC99_RS02810 | 2829..5486 | 1741 | DUF499-domain-containing-protein-[Thermodesulfobacterium-commune] |
| MCI4453559.1 | AAA+ATPase+wHTH+Alpha+Beta-domain+wHTH+RRM | Type-1_BR.RM.like.systems | TUDOR+SF1-SF2-Helicase+iREase+CTD+REase->METHYLASE->DUF499*-> | 377 | Bacteria>Thermodesulfobacteriota>Thermodesulfobacteria>Thermodesulfobacteriales>Thermodesulfobacteriaceae | GCA_022759245.1 | Thermodesulfobacterium_sp. | JHC25_01260 | 6131..7262 | 1965289 | ATP-binding-protein-[Thermodesulfobacterium-sp.] |
| NPA49345.1 | AAA+ATPase+wHTH+Alpha+Beta-domain+wHTH+RRM | Type-1_BR.RM.like.systems | <-VirB11+MoxR-AAA<-?<-?<-DUF499*<-HEPN<-PolB<-?<-PIN<-MazE_antitoxin<-METHYLASE | 891 | Bacteria>Thermodesulfobacteriota>unclassified_Thermodesulfobacteriota | GCA_013153725.1 | Thermodesulfobacteriota_bacterium | GXD20_05135 | 14849..17524 | 2202153 | ATP-binding-protein-[Thermodesulfobacteria-bacterium] |
| WP_121526871.1 | AAA+ATPase+wHTH+Alpha+Beta-domain+wHTH+RRM | Type-1_BR.RM.like.systems | <-HIN-HTH\|\|?->PIN-><-Schlafen+HATPase+HTH<-?<-?<-DUF499*<-DUF499-p<-METHYLASE<-METHYLASE-p<-TUDOR+SF1-SF2-Helicase+iREase+CTD+REase | 693 | Bacteria>Thermotogota>Thermotogae>Kosmotogales>Kosmotogaceae | GCF_003666285.1 | Mesotoga_sp._HF07.pep.5.2.highcov | BG32_RS11745 | 4849..6930 | 1462923 | DUF499-domain-containing-protein-[Mesotoga-sp.-HF07.pep.5.2.highcov] |
| WP_090222150.1 | AAA+ATPase+wHTH+Alpha+Beta-domain+wHTH+RRM | Type-1_BR.RM.like.systems | <-MPTase<-sGTP<-?<-?<-DUF499*<-METHYLASE<-TUDOR+SF1-SF2-Helicase+iREase+CTD+REase<-ABhydrolase<-?<-?<-CARF+HEPN+CARF | 849 | Bacteria>Thermotogota>Thermotogae>Thermotogales>Fervidobacteriaceae | GCF_004117075.1 | Fervidobacterium_changbaicum | CBS1_RS05080 | 1060470..1063019 | 310769 | DUF499-domain-containing-protein-[Fervidobacterium-changbaicum] |
| WP_231556087.1 | AAA+ATPase+wHTH+Alpha+Beta-domain+wHTH+RRM | Type-1_BR.RM.like.systems | ParB+HNH-><-TUDOR+SF1-SF2-Helicase+iREase+CTD+REase<-TUDOR+SF1-SF2-Helicase+iREase+CTD+REase-p<-Cluster35_66clades<-DUF499<-DUF499*<-METHYLASE<-?<-Peptidase_M9 | 972 | Bacteria>Thermotogota>Thermotogae>Thermotogales>Thermotogaceae | GCF_000750655.1 | Thermotoga_neapolitana | LA10_RS07745 | 292621..295539 | 2337 | ATP-binding-protein-[Thermotoga-neapolitana] |
| WP_231556086.1 | AAA+ATPase+wHTH+Alpha+Beta-domain+wHTH+RRM | Type-1_BR.RM.like.systems | ParB+HNH-><-TUDOR+SF1-SF2-Helicase+iREase+CTD+REase<-TUDOR+SF1-SF2-Helicase+iREase+CTD+REase-p<-Cluster35_66clades<-DUF499*<-DUF499<-METHYLASE<-?<-Peptidase_M9 | 150 | Bacteria>Thermotogota>Thermotogae>Thermotogales>Thermotogaceae | GCF_000750655.1 | Thermotoga_neapolitana | LA10_RS10575 | 292172..292624 | 2337 | DUF499-domain-containing-protein-[Thermotoga-neapolitana] |
| WP_255347884.1 | AAA+ATPase+wHTH+Alpha+Beta-domain+wHTH+RRM | Type-1_BR.RM.like.systems | ParB+HNH->ParB+HNH-><-REase<-TUDOR+SF1-SF2-Helicase+iREase+CTD+REase<-Cluster35_66clades<-DUF499<-DUF499*<-DUF499<-METHYLASE<-?<-Peptidase_M9 | 391 | Bacteria>Thermotogota>Thermotogae>Thermotogales>Thermotogaceae | GCF_000832145.1 | Thermotoga_sp._RQ7 | TRQ7_RS09840 | 1801298..1802473 | 126738 | DUF499-domain-containing-protein-[Thermotoga-sp.-RQ7] |
| WP_255347883.1 | AAA+ATPase+wHTH+Alpha+Beta-domain+wHTH+RRM | Type-1_BR.RM.like.systems | ParB+HNH->ParB+HNH-><-REase<-TUDOR+SF1-SF2-Helicase+iREase+CTD+REase<-Cluster35_66clades<-DUF499*<-DUF499<-DUF499<-METHYLASE<-?<-Peptidase_M9 | 78 | Bacteria>Thermotogota>Thermotogae>Thermotogales>Thermotogaceae | GCF_000832145.1 | Thermotoga_sp._RQ7 | TRQ7_RS09895 | 1801093..1801329 | 126738 | DUF499-domain-containing-protein-[Thermotoga-sp.-RQ7] |
| WP_236681711.1 | AAA+ATPase+wHTH+Alpha+Beta-domain+wHTH+RRM | Type-1_BR.RM.like.systems | ParB+HNH-><-REase<-TUDOR+SF1-SF2-Helicase+iREase+CTD+REase<-Cluster35_66clades<-DUF499<-DUF499<-DUF499*<-METHYLASE<-?<-Peptidase_M9 | 807 | Bacteria>Thermotogota>Thermotogae>Thermotogales>Thermotogaceae | GCF_000832145.1 | Thermotoga_sp._RQ7 | TRQ7_RS09855 | 1802757..1805180 | 126738 | ATP-binding-protein-[Thermotoga-sp.-RQ7] |

| Protein-ID | DUF499-architecture | System | Gene-Neighbourhoods | Length | Taxonomy | Assembly-ID | Strain | Locus-Tag | Genomic-coordinates | TaxID | Fasta-defline |
|---|---|---|---|---|---|---|---|---|---|---|---|
| MBZ4661023.1 | AAA+ATPase+wHTH+Alpha+Beta-domain+wHTH+RRM | Type-1_BR.RM.like.systems | ParB+HNH-><-TUDOR+SF1-SF2-Helicase+iREase+CTD+REase<-Cluster35_66clades<-DUF499<-DUF499*<-METHYLASE<-?<-Peptidase_M9 | 704 | Bacteria>Thermotogota>Thermotogae>Thermotogales>Thermotogaceae | GCA_020054995.1 | Thermotoga_sp. | JG779_177 | 12744..14858 | 28240 | hypothetical-protein-[Thermotoga-sp.] |
| MBZ4661022.1 | AAA+ATPase+wHTH+Alpha+Beta-domain+wHTH+RRM | Type-1_BR.RM.like.systems | ParB+HNH-><-TUDOR+SF1-SF2-Helicase+iREase+CTD+REase<-Cluster35_66clades<-DUF499<-DUF499*<-METHYLASE<-?<-Peptidase_M9 | 347 | Bacteria>Thermotogota>Thermotogae>Thermotogales>Thermotogaceae | GCA_020054995.1 | Thermotoga_sp. | JG779_176 | 11603..12646 | 28240 | hypothetical-protein-[Thermotoga-sp.] |
| WP_012311354.1 | AAA+ATPase+wHTH+Alpha+Beta-domain+wHTH+RRM | Type-1_BR.RM.like.systems | ParB+HNH-><-TUDOR+SF1-SF2-Helicase+iREase+CTD+REase<-Cluster35_66clades<-DUF499*<-METHYLASE<-?<-Peptidase_M9 | 1122 | Bacteria>Thermotogota>Thermotogae>Thermotogales>Thermotogaceae | GCF_000019625.1 | Thermotoga | TRQ2_RS09320 | 1826657..1830025 | 2335 | MULTISPECIES:-ATP-binding-protein-[Thermotoga] |
| WP_004080572.1 | AAA+ATPase+wHTH+Alpha+Beta-domain+wHTH+RRM | Type-1_BR.RM.like.systems | Peptidase_M9->?->METHYLASE->DUF499*->Cluster35_66clades->TUDOR+SF1-SF2-Helicase+iREase+CTD+REase-><-ParB+HNH | 1123 | Bacteria>Thermotogota>Thermotogae>Thermotogales>Thermotogaceae | GCF_000008545.1 | Thermotoga_maritima | TM_RS05035 | 1002898..1006269 | 2336 | ATP-binding-protein-[Thermotoga-maritima] |
| ACM23765.1 | AAA+ATPase+wHTH+Alpha+Beta-domain+wHTH+RRM | Type-1_BR.RM.like.systems | ParB+HNH-><-TUDOR+SF1-SF2-Helicase+iREase+CTD+REase<-Cluster35_66clades<-DUF499*<-METHYLASE<-?<-Peptidase_M9 | 1123 | Bacteria>Thermotogota>Thermotogae>Thermotogales>Thermotogaceae | GCA_000018945.1 | Thermotoga_neapolitana_DSM_4359 | CTN_1589 | 1557052..1560423 | 309803 | Putative-uncharacterized-protein-[Thermotoga-neapolitana-DSM-4359] |
| KUK23102.1 | AAA+ATPase+wHTH+Alpha+Beta-domain+wHTH+RRM | Type-1_BR.RM.like.systems | METHYLASE->DUF499*->Cluster35_66clades->TUDOR+SF1-SF2-Helicase+iREase+CTD+REase-><-ParB+HNH | 1124 | Bacteria>Thermotogota>Thermotogae>Thermotogales>Thermotogaceae | GCA_001508125.1 | Thermotoga_petrophila | XD57_0797 | 2049..5423 | 93929 | MAG:-Uncharacterized-protein-XD57_0797-[Thermotoga-petrophila] |
| KUK33533.1 | AAA+ATPase+wHTH+Alpha+Beta-domain+wHTH+RRM | Type-1_BR.RM.like.systems | METHYLASE-p->DUF499*->Cluster35_66clades->TUDOR+SF1-SF2-Helicase+iREase+CTD+REase-> | 1124 | Bacteria>Thermotogota>Thermotogae>Thermotogales>Thermotogaceae | GCA_001508875.1 | Thermotoga_sp._47_83 | XD64_0639 | 1472..4846 | 1635261 | MAG:-Uncharacterized-protein-XD64_0639-[Thermotoga-sp.-47_83] |
| MBC7122754.1 | AAA+ATPase+wHTH+Alpha+Beta-domain+wHTH+RRM | Type-1_BR.RM.like.systems | DUF499*->Cluster35_66clades->TUDOR+SF1-SF2-Helicase+iREase+CTD+REase-p-> | 845 | Bacteria>Thermotogota>Thermotogae>Thermotogales>Thermotogaceae | GCA_014360265.1 | Pseudothermotoga_sp. | H5T94_05935 | 1..2539 | 2033661 | ATP-binding-protein-[Pseudothermotoga-sp.] |
| WP_101513394.1 | AAA+ATPase+wHTH+Alpha+Beta-domain+wHTH+RRM | Type-1_BR.RM.like.systems | <-Phage_int_SAM_1+Phage_integrase<-?\|\|MNS+HTH+REase->HARE-HTH+REase->TUDOR+SF1-SF2-Helicase+iREase+CTD+REase->METHYLASE->DUF499*-><-HEPN<-PolB<-?\|\|?->Peptidase_M9-> | 860 | Bacteria>Thermotogota>Thermotogae>Thermotogales>Thermotogaceae | GCF_002865985.1 | Thermotoga_sp._SG1 | AS006_RS05665 | 51229..53811 | 126739 | DUF499-domain-containing-protein-[Thermotoga-sp.-SG1] |
| WP_165304882.1 | AAA+ATPase+wHTH+Alpha+Beta-domain+wHTH+RRM | Type-1_BR.RM.like.systems | Peptidase_M9->?-><-DUF499*<-METHYLASE<-TUDOR+SF1-SF2-Helicase+iREase+CTD+REase<-TUDOR+SF1-SF2-Helicase+iREase+CTD+REase-p<-ParB+HNH<-ParB+HNH | 842 | Bacteria>Thermotogota>Thermotogae>Thermotogales>Thermotogaceae | GCF_011057665.1 | Thermotoga_sp._38H-to | AS158_RS08895 | 65896..68424 | 1755812 | ATP-binding-protein-[Thermotoga-sp.-38H-to] |
| KUK23759.1 | AAA+ATPase+wHTH+Alpha+Beta-domain+wHTH+RRM | Type-1_BR.RM.like.systems | REase-><-?<-?<-?<-DUF499-p<-DUF499*<-METHYLASE-p<-HEPN<-PolB<-METHYLASE<-TUDOR+SF1-SF2-Helicase+iREase+CTD+REase | 604 | Bacteria>Thermotogota>Thermotogae>Thermotogales>Thermotogaceae | GCA_001508125.1 | Thermotoga_petrophila | XD57_0150 | 12939..14753 | 93929 | MAG:-ATPase-(AAA+-superfamily)-like-protein-[Thermotoga-petrophila] |
| WP_031503324.1 | AAA+ATPase+wHTH+Alpha+Beta-domain+wHTH+RRM | Type-1_BR.RM.like.systems | TUDOR+SF1-SF2-Helicase+iREase+CTD+REase->METHYLASE-p->PolB->HEPN->METHYLASE-p->DUF499*->?->?->?-><-REase | 847 | Bacteria>Thermotogota>Thermotogae>Thermotogales>Thermotogaceae | GCF_000504105.1 | Pseudothermotoga_hypogea | THYO1S_RS07620 | 1580734..1583277 | 57487 | ATP-binding-protein-[Pseudothermotoga-hypogea] |
| WP_211205257.1 | AAA+ATPase+wHTH+Alpha+Beta-domain+wHTH+RRM | Type-1_BR.RM.like.systems | <-HIN-HTH<-Polbeta<-?<-?<-?<-DUF499*<-METHYLASE<-TUDOR+SF1-SF2-Helicase+iREase+CTD+REase<-?\|\|?-><-Polbeta<-HEPN | 857 | Bacteria>Thermotogota>Thermotogae>Thermotogales>Thermotogaceae | GCF_000217815.1 | Pseudothermotoga_thermarum | THETH_RS07110 | 1405931..1408504 | 119394 | ATP-binding-protein-[Pseudothermotoga-thermarum] |
| AEH51463.1 | AAA+ATPase+wHTH+Alpha+Beta-domain+wHTH+RRM | Type-1_BR.RM.like.systems | <-HIN-HTH<-Polbeta<-?<-?<-?<-DUF499*<-METHYLASE<-TUDOR+SF1-SF2-Helicase+iREase+CTD+REase<-?\|\|?-><-Polbeta<-HEPN | 858 | Bacteria>Thermotogota>Thermotogae>Thermotogales>Thermotogaceae | GCA_000217815.1 | Pseudothermotoga_thermarum_DSM_5069 | Theth_1401 | 1405931..1408507 | 688269 | ATPase-(AAA+-superfamily)-like-protein-[Pseudothermotoga-thermarum-DSM-5069] |
| MBC7121832.1 | AAA+ATPase+wHTH+Alpha+Beta-domain+wHTH+RRM | Type-1_BR.RM.like.systems | REase-><-?<-?<-?<-DUF499*<-HEPN<-PolB<-METHYLASE-p | 847 | Bacteria>Thermotogota>Thermotogae>Thermotogales>Thermotogaceae | GCA_014360265.1 | Pseudothermotoga_sp. | H5T94_01065 | 9865..12408 | 2033661 | ATP-binding-protein-[Pseudothermotoga-sp.] |
| MCD6450226.1 | AAA+ATPase+wHTH+Alpha+Beta-domain+wHTH+RRM | Type-1_BR.RM.like.systems | TUDOR+SF1-SF2-Helicase+iREase+CTD+REase->METHYLASE->DUF499*-> | 848 | Bacteria>Thermotogota>Thermotogae>Thermotogales>Thermotogaceae | GCA_021161105.1 | Thermotogaceae_bacterium | J7L34_06950 | 4961..7507 | 2268206 | ATP-binding-protein-[Thermotogaceae-bacterium] |
| MBC7122504.1 | AAA+ATPase+wHTH+Alpha+Beta-domain+wHTH+RRM | Type-1_BR.RM.like.systems | TUDOR+SF1-SF2-Helicase+iREase+CTD+REase-p->METHYLASE->DUF499*-> | 845 | Bacteria>Thermotogota>Thermotogae>Thermotogales>Thermotogaceae | - | Pseudothermotoga_sp. | H5T94_04620 | 3349..5886 | 2033661 | ATP-binding-protein-[Pseudothermotoga-sp.] |
| KUK93540.1 | AAA+ATPase+wHTH+Alpha+Beta-domain+wHTH+RRM | Type-1_BR.RM.like.systems | TUDOR+SF1-SF2-Helicase+iREase+CTD+REase->METHYLASE->DUF499*-> | 787 | Bacteria>Thermotogota>Thermotogae>Thermotogales>unclassified_Thermotogales | GCA_001509135.1 | Thermotogales_bacterium_46_20 | XE05_1011 | 6831..9191 | 1635293 | ATPase-(AAA+-superfamily)-like-protein,-partial-[Thermotogales-bacterium-46_20] |
| MCD6239886.1 | AAA+ATPase+wHTH+Alpha+Beta-domain+wHTH+RRM | Type-1_BR.RM.like.systems | Peptidase_S51->?->TUDOR+SF1-SF2-Helicase+iREase+CTD+REase->METHYLASE-p->PolB->HEPN->DUF499*-> | 845 | Bacteria>Thermotogota>unclassified_Thermotogota | GCA_021158015.1 | Thermotogota_bacterium | J7K51_11330 | 17137..19674 | 2053689 | ATP-binding-protein-[Thermotogae-bacterium] |
| MCD6238356.1 | AAA+ATPase+wHTH+Alpha+Beta-domain+wHTH+RRM | Type-1_BR.RM.like.systems | <-DUF499*<-REase<-METHYLASE<-?\|\|?->?-><-PIWI | 935 | Bacteria>Thermotogota>unclassified_Thermotogota | GCA_021158015.1 | Thermotogota_bacterium | J7K51_03355 | 264..3071 | 2053689 | ATP-binding-protein-[Thermotogae-bacterium] |
| MDB4627959.1 | AAA+ATPase+wHTH+Alpha+Beta-domain+wHTH+RRM | Type-1_BR.RM.like.systems | <-ZnR+TOPRIM+D5ATPase<-?<-Phage_integrase\|\|?->REase-><-?<-DUF499*<-METHYLASE<-TUDOR+SF1-SF2-Helicase+iREase+CTD+REase\|\|?-><-?<-Phage_integrase | 954 | Bacteria>unclassified_Bacteria | GCA_028269625.1 | bacterium | OAF82_00035 | 6833..9697 | 1869227 | DUF499-domain-containing-protein-[bacterium] |
| MCY3806662.1 | AAA+ATPase+wHTH+Alpha+Beta-domain+wHTH+RRM | Type-1_BR.RM.like.systems | METHYLASE->MetJarc-HTH->PemK_toxin->HEPN+DUF499->DUF499*-><-?\|\|ABhydrolase-> | 710 | Bacteria>unclassified_Bacteria | GCA_026706585.1 | bacterium | OXG91_09245 | 5222..7356 | 1869227 | MAG:-AAA+-family-ATPase,-partial-[bacterium] |
| MBI1349440.1 | AAA+ATPase+wHTH+Alpha+Beta-domain+wHTH+RRM | Type-1_BR.RM.like.systems | HTH_3->TUDOR+SF1-SF2-Helicase+iREase+CTD+REase->METHYLASE->TIR->DUF499*-> | 704 | Bacteria>unclassified_Bacteria | GCA_016125225.1 | bacterium | GC163_24505 | 54841..56953 | 1869227 | DUF499-domain-containing-protein-[bacterium] |
| NJL57105.1 | AAA+ATPase+wHTH+Alpha+Beta-domain+wHTH+RRM | Type-1_BR.RM.like.systems | METHYLASE-p->DUF433->PIN->HEPN+DUF499->DUF499*-> | 478 | Bacteria>unclassified_Bacteria | GCA_012031555.1 | bacterium | HC928_19640 | 2241..3676 | 1869227 | ATP-binding-protein-[bacterium] |
| MBT9583937.1 | AAA+ATPase+wHTH+Alpha+Beta-domain+wHTH+RRM | Type-1_BR.RM.like.systems | <-AbiEi<-?<-DUF499-p<-DUF499*<-ZDNA-HTH<-?<-?<-?\|\|Pkinase-> | 617 | Bacteria>unclassified_Bacteria | GCA_018780265.1 | bacterium | IV102_11385 | 1322..3175 | 1869227 | ATP-binding-protein-[bacterium] |
| GBD25829.1 | AAA+ATPase+wHTH+Alpha+Beta-domain+wHTH+RRM | Type-1_BR.RM.like.systems | DUF499*->METHYLASE-p->?->?->?-><-?<-ABhydrolase | 795 | Bacteria>unclassified_Bacteria | GCA_002898795.1 | bacterium_HR30 | HRbin30_01154 | 17..2404 | 2035425 | hypothetical-protein-HRbin30_01154-[bacterium-HR30] |
| MDE0416420.1 | AAA+ATPase+wHTH+Alpha+Beta-domain+wHTH+RRM | Type-1_BR.RM.like.systems | <-ABhydrolase\|\|XIS-HTH->TUDOR+SF1-SF2-Helicase+iREase+CTD+REase->METHYLASE->DUF499*->OLD-ABC+TOPRIM->Schlafen+HATPase+HTH->?->Cysteine_desulfurase->Cysteine_desulfurase->SF1-SF2-Helicase-> | 967 | Bacteria>unclassified_Bacteria | GCA_028821455.1 | bacterium | OXI95_05715 | 17093..19996 | 1869227 | MAG:-DUF499-domain-containing-protein-[bacterium] |
| GBD43556.1 | AAA+ATPase+wHTH+Alpha+Beta-domain+wHTH+RRM | Type-1_BR.RM.like.systems | <-SF1-SF2-Helicase+DUF1998<-HEPN+Mrr_N<-METHYLASE<-TUDOR+SF1-SF2-Helicase+iREase+CTD<-DUF499*<-METHYLASE<-TUDOR+SF1-SF2-Helicase+iREase+CTD+REase<-?\|\|?->?-><-RNase | 978 | Bacteria>unclassified_Bacteria | GCA_002898275.1 | bacterium_HR40 | HRbin40_01030 | 1099894..1102830 | 2035435 | hypothetical-protein-HRbin40_01030-[bacterium-HR40] |
| GBD39155.1 | AAA+ATPase+wHTH+Alpha+Beta-domain+wHTH+RRM | Type-1_BR.RM.like.systems | <-RelE-ParE\|\|TUDOR+SF1-SF2-Helicase+iREase+CTD+REase->Schlafen->METHYLASE->PolB->HEPN->DUF499*->?-><-?\|\|?->NAD_binding_2+6PGD-> | 872 | Bacteria>unclassified_Bacteria | GCA_002898375.1 | bacterium_HR37 | HRbin37_01422 | 79215..81833 | 2035432 | hypothetical-protein-HRbin37_01422-[bacterium-HR37] |
| MCX7734094.1 | AAA+ATPase+wHTH+Alpha+Beta-domain+wHTH+RRM | Type-1_BR.RM.like.systems | Transposase_31+Yae1_N->Transposase_31+Yae1_N-><-DUF499-p<-DUF499*<-METHYLASE-p<-HKD-DNase | 416 | Bacteria>unclassified_Bacteria | GCA_026418515.1 | bacterium | N2254_04945 | 75514..76764 | 1869227 | MAG:-DUF499-domain-containing-protein-[bacterium] |
| MCK9266177.1 | AAA+ATPase+wHTH+Alpha+Beta-domain+wHTH+RRM | Type-1_BR.RM.like.systems | <-MACRODOMAIN<-REase<-?<-DUF499*<-?<-METHYLASE<-TUDOR+SF1-SF2-Helicase+iREase+CTD+REase\|\|Radical-SAM-> | 861 | Bacteria>unclassified_Bacteria | GCA_023230585.1 | bacterium | MOP98_04745 | 3522..6107 | 1869227 | DUF499-domain-containing-protein-[bacterium] |
| MBN2381437.1 | AAA+ATPase+wHTH+Alpha+Beta-domain+wHTH+RRM | Type-1_BR.RM.like.systems | GDResolvase+Recombinase+ZnR->?->TUDOR+SF1-SF2-Helicase+iREase+CTD+REase->?->PIN->METHYLASE->DUF499*-> | 970 | Bacteria>unclassified_Bacteria | GCA_016934675.1 | bacterium | JXQ70_01010 | 9963..12875 | 1869227 | ATP-binding-protein-[bacterium] |
| NUO19888.1 | AAA+ATPase+wHTH+Alpha+Beta-domain+wHTH+RRM | Type-1_BR.RM.like.systems | TUDOR+SF1-SF2-Helicase+iREase+CTD+REase->OLD-ABC+TOPRIM->METHYLASE->AbiGi->DUF499*->?->?->?->Pkinase-> | 971 | Bacteria>unclassified_Bacteria | GCA_013360195.1 | bacterium | HUU59_10600 | 10537..13452 | 1869227 | ATP-binding-protein-[bacterium] |
| MCX7795564.1 | AAA+ATPase+wHTH+Alpha+Beta-domain+wHTH+RRM | Type-1_BR.RM.like.systems | METHYLASE-p->?->DUF499*->TUDOR+SF1-SF2-Helicase+iREase+CTD+REase->?-><-?\|\|FURR-HTH->?->ABC-ATPase+SbcC-> | 825 | Bacteria>unclassified_Bacteria | GCA_026418035.1 | bacterium | N2380_03480 | 903..3380 | 1869227 | MAG:-AAA-family-ATPase-[bacterium] |
| MBN2383967.1 | AAA+ATPase+wHTH+Alpha+Beta-domain+wHTH+RRM | Type-1_BR.RM.like.systems | TUDOR+SF1-SF2-Helicase+iREase+CTD+REase->?->?->OLD-ABC+TOPRIM->METHYLASE->REase->DUF499*-><-?\|\|?-><-?\|\|Aminotran_3-> | 920 | Bacteria>unclassified_Bacteria | GCA_016934675.1 | bacterium | JXQ70_13910 | 28377..31139 | 1869227 | ATP-binding-protein-[bacterium] |
| OQA20379.1 | AAA+ATPase+wHTH+Alpha+Beta-domain+wHTH+RRM | Type-1_BR.RM.like.systems | PIN->HicB_lk_antitox->HicB_lk_antitox->?->PIN->?->DUF499*-><-NUDIX<-?<-?\|\|TPR+TPR-> | 993 | Bacteria>unclassified_Bacteria | GCA_002069435.1 | bacterium_ADurb.Bin363 | BWY64_00213 | 12863..15844 | 1866927 | hypothetical-protein-BWY64_00213-[bacterium-ADurb.Bin363] |
| MBC7327984.1 | AAA+ATPase+wHTH+Alpha+Beta-domain+wHTH+RRM | Type-1_BR.RM.like.systems | ParB+HNH->TUDOR+SF1-SF2-Helicase+iREase+CTD+REase->METHYLASE->DUF499*-><-?<-ABC-ATPase+SbcC | 922 | Bacteria>unclassified_Bacteria | GCA_014360465.1 | bacterium | H5T87_07705 | 9838..12606 | 1869227 | ATP-binding-protein-[bacterium] |
| MCP3963401.1 | AAA+ATPase+wHTH+Alpha+Beta-domain+wHTH+RRM | Type-1_BR.RM.like.systems | TUDOR+SF1-SF2-Helicase+iREase+CTD+REase-p->ParB+HNH+Schlafen->METHYLASE->DUF499*-><-?\|\|?->?-><-?<-?\|\|JAB-> | 946 | Bacteria>unclassified_Bacteria | GCA_024226315.1 | bacterium | GY719_36660 | 6844..9684 | 1869227 | ATP-binding-protein-[bacterium] |
| MCY3911431.1 | AAA+ATPase+wHTH+Alpha+Beta-domain+wHTH+RRM | Type-1_BR.RM.like.systems | <-HEPN\|\|TUDOR+SF1-SF2-Helicase+iREase+CTD+REase->METHYLASE->DUF499*->TIR-> | 933 | Bacteria>unclassified_Bacteria | GCA_026707235.1 | bacterium | OXF99_07935 | 8019..10820 | 1869227 | MAG:-DUF499-domain-containing-protein-[bacterium] |
| MCP4400157.1 | AAA+ATPase+wHTH+Alpha+Beta-domain+wHTH+RRM | Type-1_BR.RM.like.systems | <-Aminotran_1_2\|\|DUF499*->?->NTP_transf->HEPN->Peptidase-> | 1006 | Bacteria>unclassified_Bacteria | GCA_024222835.1 | bacterium | GY801_22970 | 2415..5435 | 1869227 | DUF499-domain-containing-protein-[bacterium] |
| MCI0602497.1 | AAA+ATPase+wHTH+Alpha+Beta-domain+wHTH+RRM | Type-1_BR.RM.like.systems | <-XIS-HTH\|\|?-><-?<-HNH<-DUF499*<-REase<-METHYLASE<-HEPN+REase | 922 | Bacteria>unclassified_Bacteria | GCA_022616075.1 | bacterium | LO156_05740 | 5054..7822 | 1869227 | DUF499-domain-containing-protein-[bacterium] |
| MBU2462380.1 | AAA+ATPase+wHTH+Alpha+Beta-domain+wHTH+RRM | Type-1_BR.RM.like.systems | METHYLASE-p->PolB->HEPN->OLD-ABC->TOPRIM->DUF499*-> | 441 | Bacteria>unclassified_Bacteria | GCA_018828965.1 | bacterium | KKH65_05855 | 2757..4081 | 1869227 | DUF499-domain-containing-protein-[bacterium] |
| MBU1598556.1 | AAA+ATPase+wHTH+Alpha+Beta-domain+wHTH+RRM | Type-1_BR.RM.like.systems | METHYLASE-p->PolB->HEPN->OLD-ABC->TOPRIM->DUF499*-> | 494 | Bacteria>unclassified_Bacteria | GCA_018822205.1 | bacterium | KKG61_00350 | 3347..4830 | 1869227 | DUF499-domain-containing-protein-[bacterium] |
| MBU1261806.1 | AAA+ATPase+wHTH+Alpha+Beta-domain+wHTH+RRM | Type-1_BR.RM.like.systems | <-DUF499*<-TOPRIM<-OLD-ABC<-HEPN<-PolB<-METHYLASE-p | 494 | Bacteria>unclassified_Bacteria | GCA_018819855.1 | bacterium | KJ640_02495 | 1..1484 | 1869227 | DUF499-domain-containing-protein-[bacterium] |
| MCR9296473.1 | AAA+ATPase+wHTH+Alpha+Beta-domain+wHTH+RRM | Type-1_BR.RM.like.systems | <-Phage_integrase<-?<-DUF499*<-REase | 985 | Bacteria>unclassified_Bacteria | GCA_024743355.1 | bacterium | NXI32_27500 | 1811..4768 | 1869227 | DUF499-domain-containing-protein-[bacterium] |
| GBC90889.1 | AAA+ATPase+wHTH+Alpha+Beta-domain+wHTH+RRM | Type-1_BR.RM.like.systems | <-TUDOR+SF1-SF2-Helicase+iREase+CTD+REase<-DUF499*<-?<-?<-METHYLASE-p | 879 | Bacteria>unclassified_Bacteria | - | bacterium_HR14 | HRbin14_01644 | 3380..6019 | 2035409 | hypothetical-protein-HRbin14_01644-[bacterium-HR14] |
| MBU4446673.1 | AAA+ATPase+wHTH+Alpha+Beta-domain+wHTH+RRM | Type-1_BR.RM.like.systems | TUDOR+SF1-SF2-Helicase+iREase+CTD+REase-p->METHYLASE->DUF499*-> | 582 | Bacteria>unclassified_Bacteria | GCA_018895675.1 | bacterium | KJ656_16595 | 3449..5195 | 1869227 | DUF499-domain-containing-protein-[bacterium] |
| HAW50200.1 | AAA+ATPase+wHTH+Alpha+Beta-domain+wHTH+RRM | Type-1_BR.RM.like.systems | <-DUF499*<-METHYLASE<-TUDOR+SF1-SF2-Helicase+iREase+CTD+REase-p | 858 | Bacteria>unclassified_Bacteria | GCA_003498085.1 | bacterium | DCX16_04555 | 449..3025 | 1869227 | ATPase-[bacterium] |
| MCD6327181.1 | AAA+ATPase+wHTH+Alpha+Beta-domain+wHTH+RRM | Type-1_BR.RM.like.systems | wHTH+REase->SbcC->?->METHYLASE->?->PIN->DUF499*-> | 928 | Bacteria>unclassified_Bacteria | GCA_021160245.1 | bacterium | J7M28_06475 | 15142..17928 | 1869227 | ATP-binding-protein-[bacterium] |
| MBU2461874.1 | AAA+ATPase+wHTH+Alpha+Beta-domain+wHTH+RRM | Type-1_BR.RM.like.systems | METHYLASE-p->PolB->HEPN->DUF499*-> | 513 | Bacteria>unclassified_Bacteria | GCA_018828965.1 | bacterium | KKH65_03235 | 1911..3449 | 1869227 | DUF499-domain-containing-protein-[bacterium] |
| MBU1261643.1 | AAA+ATPase+wHTH+Alpha+Beta-domain+wHTH+RRM | Type-1_BR.RM.like.systems | METHYLASE-p->PolB->HEPN->DUF499*-> | 515 | Bacteria>unclassified_Bacteria | GCA_018819855.1 | bacterium | KJ640_01640 | 1911..3456 | 1869227 | DUF499-domain-containing-protein-[bacterium] |
| MCX7759448.1 | AAA+ATPase+wHTH+Alpha+Beta-domain+wHTH+RRM | Type-1_BR.RM.like.systems | <-CR-ATPase8+REase\|\|DUF499*-> | 534 | Bacteria>unclassified_Bacteria | GCA_026418475.1 | bacterium | N2169_07590 | 877..2481 | 1869227 | MAG:-DUF499-domain-containing-protein-[bacterium] |
| MDE0231071.1 | AAA+ATPase+wHTH+Alpha+Beta-domain+wHTH+RRM | Type-1_BR.RM.like.systems | DDE_Tnp_1-><-SMODS<-?\|\|HEPN->?->CHTH-><-DUF499*<-METHYLASE<-TUDOR+SF1-SF2-Helicase+iREase+CTD+REase\|\|?-><-PAIREDC-HTH<-?\|\|Phage_integrase+Phage_integrase-> | 987 | Bacteria>unclassified_Bacteria | GCA_028820285.1 | bacterium | OXN93_02170 | 23902..26865 | 1869227 | MAG:-DUF499-domain-containing-protein-[bacterium] |
| MDE0231062.1 | AAA+ATPase+wHTH+Alpha+Beta-domain+wHTH+RRM | Type-1_BR.RM.like.systems | <-REase<-?<-SF1-SF2-Helicase<-DUF499*<-METHYLASE<-TUDOR+SF1-SF2-Helicase+iREase+CTD+REase\|\|DDE_Tnp_1-><-SMODS<-?\|\|HEPN-> | 972 | Bacteria>unclassified_Bacteria | GCA_028820285.1 | bacterium | OXN93_02125 | 4290..7208 | 1869227 | MAG:-DUF499-domain-containing-protein-[bacterium] |
| MCY4104683.1 | AAA+ATPase+wHTH+Alpha+Beta-domain+wHTH+RRM | Type-1_BR.RM.like.systems | TUDOR+SF1-SF2-Helicase+iREase+CTD+REase->DUF4527->?->?->METHYLASE->DUF499*->HEPN-><-?<-?<-35exo<-?\|\|N6_N4_MTase-> | 985 | Bacteria>unclassified_Bacteria | GCA_026708055.1 | bacterium | OXG55_15720 | 11834..14791 | 1869227 | MAG:-DUF499-domain-containing-protein-[bacterium] |
| MCY4101691.1 | AAA+ATPase+wHTH+Alpha+Beta-domain+wHTH+RRM | Type-1_BR.RM.like.systems | <-HicB\|\|TUDOR+SF1-SF2-Helicase+iREase+CTD+REase->METHYLASE->DUF499*->?-><-PIN\|\|?-><-?<-REase+HTH<-TOPRIM | 1008 | Bacteria>unclassified_Bacteria | GCA_026708055.1 | bacterium | OXG55_00260 | 15226..18252 | 1869227 | MAG:-PASTA-domain-containing-protein-[bacterium] |
| MCY4630094.1 | AAA+ATPase+wHTH+Alpha+Beta-domain+wHTH+RRM | Type-1_BR.RM.like.systems | REase->METHYLASE-p->METHYLASE-p->DUF499*-><-?\|\|CR-ATPase9+REase->?-><-?<-REase-p | 1004 | Bacteria>unclassified_Bacteria | GCA_026713745.1 | bacterium | OXE75_04305 | 3540..6554 | 1869227 | MAG:-DUF499-domain-containing-protein-[bacterium] |
| MCY4163684.1 | AAA+ATPase+wHTH+Alpha+Beta-domain+wHTH+RRM | Type-1_BR.RM.like.systems | <-DUF222+HNH\|\|TUDOR+SF1-SF2-Helicase+iREase+CTD+REase->METHYLASE->DUF499*-><-?\|\|tRNA-><-?<-?<-sGTP | 993 | Bacteria>unclassified_Bacteria | GCA_026709025.1 | bacterium | OXE93_05650 | 6917..9898 | 1869227 | MAG:-DUF499-domain-containing-protein-[bacterium] |
| MCY3952181.1 | AAA+ATPase+wHTH+Alpha+Beta-domain+wHTH+RRM | Type-1_BR.RM.like.systems | <-DUF499*<-METHYLASE-p<-METHYLASE-p<-?<-TUDOR+SF1-SF2-Helicase+iREase+CTD+REase | 197 | Bacteria>unclassified_Bacteria | GCA_026707535.1 | bacterium | OXG69_06500 | 1..590 | 1869227 | MAG:-AAA-family-ATPase,-partial-[bacterium] |
| MCY3561073.1 | AAA+ATPase+wHTH+Alpha+Beta-domain+wHTH+RRM | Type-1_BR.RM.like.systems | <-DUF499*<-METHYLASE-p<-METHYLASE-p<-?<-?<-TUDOR+SF1-SF2-Helicase+iREase+CTD+REase | 566 | Bacteria>unclassified_Bacteria | GCA_026702815.1 | bacterium | OXH20_07905 | 1..1697 | 1869227 | MAG:-DUF499-domain-containing-protein,-partial-[bacterium] |
| MCY3786398.1 | AAA+ATPase+wHTH+Alpha+Beta-domain+wHTH+RRM | Type-1_BR.RM.like.systems | <-Aminotran_1_2<-?<-?<-?<-?<-DUF499*<-?<-?<-?<-Glycos_trans_3N+Glycos_transf_3 | 214 | Bacteria>unclassified_Bacteria | GCA_026706565.1 | bacterium | OXG47_06685 | 34841..35485 | 1869227 | MAG:-DUF499-domain-containing-protein-[bacterium] |
| MCY4369700.1 | AAA+ATPase+wHTH+Alpha+Beta-domain+wHTH+RRM | Type-1_BR.RM.like.systems | ABhydrolase-><-?<-?<-DUF499*<-METHYLASE-p | 986 | Bacteria>unclassified_Bacteria | GCA_026710435.1 | bacterium | OXF41_09855 | 16238..19198 | 1869227 | MAG:-DUF499-domain-containing-protein-[bacterium] |
| MCY4369121.1 | AAA+ATPase+wHTH+Alpha+Beta-domain+wHTH+RRM | Type-1_BR.RM.like.systems | <-HEPN<-DUF499*<-METHYLASE-p | 984 | Bacteria>unclassified_Bacteria | GCA_026710435.1 | bacterium | OXF41_06785 | 3276..6230 | 1869227 | MAG:-DUF499-domain-containing-protein-[bacterium] |
| MCY2651024.1 | AAA+ATPase+wHTH+Alpha+Beta-domain+wHTH+RRM | Type-1_BR.RM.like.systems | DUF499*-><-RlpA<-?\|\|AbiEi_4->AbiEii->HIN-HTH-><-HEPN | 909 | - | - | - | OXA70_25990 | 1..2730 | - | DUF499-domain-containing-protein,-partial-[Klebsiella-pneumoniae] |
| OLS15779.1 | AAA+ATPase+wHTH+Alpha+Beta-domain+wHTH1+wHTH2+wHTH3+RRM | Type-1_BR.RM.like.systems | <-REase+HTH<-?\|\|?-><-DUF499*<-METHYLASE<-TUDOR+SF1-SF2-Helicase+iREase+CTD+REase-p | 1136 | Archaea>Asgard_group>Candidatus_Lokiarchaeota>unclassified_Lokiarchaeota | GCA_001940655.1 | Candidatus_Lokiarchaeota_archaeon_CR_4 | RBG13Loki_0616 | 17465..20875 | 1849166 | MAG:-hypothetical-protein-RBG13Loki_0616-[Candidatus-Lokiarchaeota-archaeon-CR_4] |
| WP_131007313.1 | AAA+ATPase+wHTH+Alpha+Beta-domain+wHTH1+wHTH2+wHTH3+RRM | Type-1_BR.RM.like.systems | TnsC-ATPase-><-?\|\|METHYLASE->DUF499*->?->TUDOR+SF1-SF2-Helicase+iREase+CTD+REase-><-?\|\|URI-DNase-> | 1134 | Archaea>Euryarchaeota>Methanomada_group>Methanococci>Methanococcales | GCF_004310395.1 | Methanofervidicoccus_abyssi | MHHB_RS04010 | 288294..291698 | 2082189 | DUF499-domain-containing-protein-[Methanofervidicoccus-abyssi] |
| MDA8594262.1 | AAA+ATPase+wHTH+Alpha+Beta-domain+wHTH1+wHTH2+wHTH3+RRM | Type-1_BR.RM.like.systems | ACET->?->?->?->?-><-?<-DUF499*<-METHYLASE<-SF1-SF2-Helicase+iREase+CTD+REase<-GDResolvase+Recombinase | 1089 | Archaea>Euryarchaeota>unclassified_Euryarchaeota | GCA_028097625.1 | Euryarchaeota_archaeon | N9L85_02695 | 19258..22527 | 2026739 | DUF499-domain-containing-protein-[Euryarchaeota-archaeon] |
| MCE5296808.1 | AAA+ATPase+wHTH+Alpha+Beta-domain+wHTH1+wHTH2+wHTH3+RRM | Type-1_BR.RM.like.systems | Phage_int_SAM_4+Phage_integrase->?->HIN-HTH->?-><-DUF499*<-METHYLASE | 1089 | Archaea>Euryarchaeota>unclassified_Euryarchaeota | GCA_021372315.1 | Euryarchaeota_archaeon | LLG16_06880 | 9291..12560 | 2026739 | hypothetical-protein-[Euryarchaeota-archaeon] |
| MCD6443383.1 | AAA+ATPase+wHTH+Alpha+Beta-domain+wHTH1+wHTH2+wHTH3+RRM | Type-1_BR.RM.like.systems | <-TUDOR+SF1-SF2-Helicase+iREase+CTD+REase<-Cluster35_66clades<-DUF499*<-METHYLASE<-?<-EVE\|\|HNH->REase-> | 1115 | Archaea>TACK_group>Candidatus_Bathyarchaeota>unclassified_Candidatus_Bathyarchaeota | GCA_021161255.1 | Candidatus_Bathyarchaeota_archaeon | J7L70_00020 | 4870..8217 | 2026714 | DUF499-domain-containing-protein-[Candidatus-Bathyarchaeota-archaeon] |
| OYT46229.1 | AAA+ATPase+wHTH+Alpha+Beta-domain+wHTH1+wHTH2+wHTH3+RRM | Type-1_BR.RM.like.systems | <-TUDOR+SF1-SF2-Helicase+iREase+CTD+REase<-Cluster35_66clades<-DUF499*<-METHYLASE | 1122 | Archaea>TACK_group>Candidatus_Bathyarchaeota>unclassified_Candidatus_Bathyarchaeota | GCA_002254475.1 | Candidatus_Bathyarchaeota_archaeon_ex4484_40 | B6U84_00860 | 7326..10694 | 2012513 | hypothetical-protein-B6U84_00860-[Candidatus-Bathyarchaeota-archaeon-ex4484_40] |
| MCS7366031.1 | AAA+ATPase+wHTH+Alpha+Beta-domain+wHTH1+wHTH2+wHTH3+RRM | Type-1_BR.RM.like.systems | <-TUDOR+SF1-SF2-Helicase+iREase+CTD+REase-p<-Cluster35_66clades<-DUF499*<-METHYLASE-p | 1114 | Archaea>TACK_group>Candidatus_Culexarchaeota>Candidatus_Culexarchaeia>Candidatus_Culexarchaeles | GCA_025061245.1 | Candidatus_Culexmicrobium_profundum | NDF54_11395 | 2049..5393 | 2928969 | MAG:-ATP-binding-protein-[archaeon-GB-1867-035] |
| MCI4437835.1 | AAA+ATPase+wHTH+Alpha+Beta-domain+wHTH1+wHTH2+wHTH3+RRM | Type-1_BR.RM.like.systems | <-TUDOR+SF1-SF2-Helicase+iREase+CTD+REase-p<-Cluster35_66clades<-DUF499*<-METHYLASE | 1159 | Archaea>TACK_group>Thermoproteota>Thermoprotei>Desulfurococcales | GCA_022759085.1 | Ignisphaera_sp. | JHC33_13595 | 1174..4653 | 2268142 | DUF499-domain-containing-protein-[Ignisphaera-sp.] |
| MCE4628068.1 | AAA+ATPase+wHTH+Alpha+Beta-domain+wHTH1+wHTH2+wHTH3+RRM | Type-1_BR.RM.like.systems | METHYLASE-p->DUF499*->Cluster35_66clades->TUDOR+SF1-SF2-Helicase+iREase+CTD+REase-> | 1139 | Archaea>TACK_group>Thermoproteota>Thermoprotei>Desulfurococcales | GCA_015521185.1 | Desulfurococcales_archaeon | F7C34_02830 | 1546..4965 | 2480821 | ATP-binding-protein-[Desulfurococcales-archaeon] |
| MCC6057133.1 | AAA+ATPase+wHTH+Alpha+Beta-domain+wHTH1+wHTH2+wHTH3+RRM | Type-1_BR.RM.like.systems | DUF499*->Cluster35_66clades->TUDOR+SF1-SF2-Helicase+iREase+CTD+REase-p-> | 600 | Archaea>TACK_group>Thermoproteota>Thermoprotei>Desulfurococcales | GCA_020833445.1 | Desulfurococcaceae_archaeon | LM583_10700 | 1..1805 | 2184738 | ATP-binding-protein-[Desulfurococcaceae-archaeon] |
| MCI4409925.1 | AAA+ATPase+wHTH+Alpha+Beta-domain+wHTH1+wHTH2+wHTH3+RRM | Type-1_BR.RM.like.systems | METHYLASE-p->DUF499->DUF499*->Cluster35_66clades->TUDOR+SF1-SF2-Helicase+iREase+CTD+REase-> | 671 | Archaea>TACK_group>Thermoproteota>Thermoprotei>Thermofilales | GCA_022759215.1 | Thermofilum_sp. | JHC26_12600 | 1245..3260 | 1961369 | DUF499-domain-containing-protein-[Thermofilum-sp.] |
| NAZ24253.1 | AAA+ATPase+wHTH+Alpha+Beta-domain+wHTH1+wHTH2+wHTH3+RRM | Type-1_BR.RM.like.systems | METHYLASE->DUF499*->Cluster35_66clades->TUDOR+SF1-SF2-Helicase+iREase+CTD+REase-> | 1062 | Archaea>TACK_group>Thermoproteota>Thermoprotei>Thermofilales | GCA_009903875.1 | Thermofilum_sp. | GU357_00625 | 4960..8148 | 1961369 | DUF499-domain-containing-protein-[Thermofilum-sp.] |
| HDI75066.1 | AAA+ATPase+wHTH+Alpha+Beta-domain+wHTH1+wHTH2+wHTH3+RRM | Type-1_BR.RM.like.systems | DUF499*->Cluster35_66clades->TUDOR+SF1-SF2-Helicase+iREase+CTD+REase-> | 474 | Archaea>TACK_group>Thermoproteota>Thermoprotei>unclassified_Thermoprotei | GCA_011042315.1 | Thermoprotei_archaeon | ENF55_03830 | 1..1425 | 2250277 | DUF499-domain-containing-protein-[Thermoprotei-archaeon] |
| WP_194449891.1 | AAA+ATPase+wHTH+Alpha+Beta-domain+wHTH1+wHTH2+wHTH3+RRM | Type-1_BR.RM.like.systems | <-XIS-HTH<-?<-?\|\|TUDOR+SF1-SF2-Helicase+iREase+CTD+REase->?->METHYLASE->DUF499*->HNH->?->?-><-ParB<-GDResolvase+Recombinase+ZnR<-RAMA | 1110 | Bacteria>Acidobacteriota>Terriglobia>Bryobacterales>Bryobacteraceae | GCF_015277775.1 | Paludibaculum_fermentans | IRI77_RS36750 | 9264025..9267357 | 1473598 | DUF499-domain-containing-protein-[Paludibaculum-fermentans] |
| MCC6362975.1 | AAA+ATPase+wHTH+Alpha+Beta-domain+wHTH1+wHTH2+wHTH3+RRM | Type-1_BR.RM.like.systems | PIN->?->RAMA->GDResolvase+Recombinase+ZnR->ParB-><-DUF499*<-METHYLASE<-?<-?<-?<-TUDOR+SF1-SF2-Helicase+iREase+CTD+REase | 1114 | Bacteria>Acidobacteriota>Terriglobia>Bryobacterales>unclassified_Bryobacterales | GCA_020847975.1 | Bryobacterales_bacterium | IT165_05590 | 5868..9212 | 2026791 | DUF499-domain-containing-protein-[Bryobacterales-bacterium] |
| MDE0105406.1 | AAA+ATPase+wHTH+Alpha+Beta-domain+wHTH1+wHTH2+wHTH3+RRM | Type-1_BR.RM.like.systems | <-Radical-SAM<-?<-?\|\|?-><-?<-?<-DUF499*<-METHYLASE<-TUDOR+SF1-SF2-Helicase+iREase+CTD+REase | 1101 | Bacteria>Acidobacteriota>Terriglobia>Bryobacterales>unclassified_Bryobacterales | GCA_028819235.1 | Bryobacterales_bacterium | OXN89_23770 | 12797..16102 | 2026791 | MAG:-DUF499-domain-containing-protein-[Bryobacterales-bacterium] |
| MDE0626645.1 | AAA+ATPase+wHTH+Alpha+Beta-domain+wHTH1+wHTH2+wHTH3+RRM | Type-1_BR.RM.like.systems | METHYLASE-p->?->PIN->DUF499*->REase-> | 1105 | Bacteria>Acidobacteriota>Terriglobia>Bryobacterales>unclassified_Bryobacterales | GCA_028824065.1 | Bryobacterales_bacterium | OXH99_09625 | 1430..4747 | 2026791 | MAG:-DUF499-domain-containing-protein-[Bryobacterales-bacterium] |
| MCG3190932.1 | AAA+ATPase+wHTH+Alpha+Beta-domain+wHTH1+wHTH2+wHTH3+RRM | Type-1_BR.RM.like.systems | <-DUF499*<-TOPRIM<-OLD-ABC<-METHYLASE | 1102 | Bacteria>Acidobacteriota>Thermoanaerobaculia>unclassified_Thermoanaerobaculia | GCA_022071945.1 | Thermoanaerobaculia_bacterium | DIJKHBIC_00155 | 188742..192050 | 2651171 | hypothetical-protein-[Thermoanaerobaculia-bacterium] |
| MYN68213.1 | AAA+ATPase+wHTH+Alpha+Beta-domain+wHTH1+wHTH2+wHTH3+RRM | Type-1_BR.RM.like.systems | TUDOR+SF1-SF2-Helicase+iREase+CTD+REase->Schlafen+HATPase+HTH->HEPN->METHYLASE->OLD-ABC->TOPRIM->DUF499*-><-?<-SAM-methylase<-ACET+ACET<-GspA-ATPase | 1101 | Bacteria>Acidobacteriota>unclassified_Acidobacteriota | GCA_009861545.1 | Acidobacteriota_bacterium | F4X11_24880 | 56553..59858 | 1978231 | ATP-binding-protein-[Acidobacteria-bacterium] |
| MBI4454518.1 | AAA+ATPase+wHTH+Alpha+Beta-domain+wHTH1+wHTH2+wHTH3+RRM | Type-1_BR.RM.like.systems | METHYLASE-p-><-?\|\|RADICAL-SAM->?-><-DUF499*<-?<-Schlafen+HATPase+HTH<-METHYLASE\|\|?->?-><-TUDOR+SF1-SF2-Helicase+iREase+CTD+REase | 1101 | Bacteria>Acidobacteriota>unclassified_Acidobacteriota | GCA_016211105.1 | Acidobacteriota_bacterium | HY644_01320 | 37960..41265 | 1978231 | ATP-binding-protein-[Acidobacteria-bacterium] |
| MCX6538613.1 | AAA+ATPase+wHTH+Alpha+Beta-domain+wHTH1+wHTH2+wHTH3+RRM | Type-1_BR.RM.like.systems | CHASE2+ACYC->Peptidase_M48->?->TUDOR+SF1-SF2-Helicase+iREase+CTD+REase->AbiEii->METHYLASE->DUF499*-> | 1101 | Bacteria>Acidobacteriota>unclassified_Acidobacteriota | GCA_026393675.1 | Acidobacteriota_bacterium | NT151_06745 | 30423..33728 | 1978231 | MAG:-DUF499-domain-containing-protein-[Acidobacteria-bacterium] |
| MCY3965602.1 | AAA+ATPase+wHTH+Alpha+Beta-domain+wHTH1+wHTH2+wHTH3+RRM | Type-1_BR.RM.like.systems | PIN-><-?<-?\|\|OLD-ABC->TOPRIM-><-DUF499*\|\|HEPN-><-TOPRIM<-OLD-ABC<-METHYLASE\|\|?->PAPS_reduct-> | 1095 | Bacteria>Acidobacteriota>unclassified_Acidobacteriota | GCA_026707545.1 | Acidobacteriota_bacterium | OXG83_11235 | 2822742..2826029 | 1978231 | MAG:-DUF499-domain-containing-protein-[Acidobacteria-bacterium] |
| MBX3279097.1 | AAA+ATPase+wHTH+Alpha+Beta-domain+wHTH1+wHTH2+wHTH3+RRM | Type-1_BR.RM.like.systems | HTH_3->?-><-?\|\|?-><-DUF499*<-?<-zf-ISL3+DDE_Tnp_ISL3<-GDResolvase+Recombinase+ZnR<-?\|\|?-><-HEPN | 1108 | Bacteria>Acidobacteriota>unclassified_Acidobacteriota | GCA_019639255.1 | Acidobacteriota_bacterium | KF868_13925 | 29527..32853 | 1978231 | ATP-binding-protein-[Acidobacteria-bacterium] |
| MCY3931995.1 | AAA+ATPase+wHTH+Alpha+Beta-domain+wHTH1+wHTH2+wHTH3+RRM | Type-1_BR.RM.like.systems | <-HicB_lk_antitox<-HicA_toxin\|\|METHYLASE->Phd-YefM->PIN->TPR+TPR->DUF499*-> | 1099 | Bacteria>Acidobacteriota>unclassified_Acidobacteriota | GCA_026707325.1 | Acidobacteriota_bacterium | OXH70_09765 | 12744..16043 | 1978231 | MAG:-DUF499-domain-containing-protein-[Acidobacteria-bacterium] |
| MDE2979244.1 | AAA+ATPase+wHTH+Alpha+Beta-domain+wHTH1+wHTH2+wHTH3+RRM | Type-1_BR.RM.like.systems | <-HicB_lk_antitox<-HicA_toxin\|\|METHYLASE-><-HEPN<-REase-p\|\|DUF499*-> | 1124 | Bacteria>Acidobacteriota>unclassified_Acidobacteriota | GCA_028873735.1 | Acidobacteriota_bacterium | OXU63_17235 | 11921..15295 | 1978231 | MAG:-DUF499-domain-containing-protein-[Acidobacteriota-bacterium] |
| MBI2841722.1 | AAA+ATPase+wHTH+Alpha+Beta-domain+wHTH1+wHTH2+wHTH3+RRM | Type-1_BR.RM.like.systems | TUDOR+SF1-SF2-Helicase+iREase+CTD+REase->?->AAA-ATPase+REase->METHYLASE->DUF499*-> | 1100 | Bacteria>Acidobacteriota>unclassified_Acidobacteriota | GCA_016191015.1 | Acidobacteriota_bacterium | HYX75_25690 | 46699..50001 | 1978231 | ATP-binding-protein-[Acidobacteria-bacterium] |
| MYN67290.1 | AAA+ATPase+wHTH+Alpha+Beta-domain+wHTH1+wHTH2+wHTH3+RRM | Type-1_BR.RM.like.systems | <-REase\|\|?-><-DUF499*<-TOPRIM<-OLD-ABC<-?<-TIR<-OLD-ABC<-SF1-SF2-Helicase | 1106 | Bacteria>Acidobacteriota>unclassified_Acidobacteriota | GCA_009861545.1 | Acidobacteriota_bacterium | F4X11_20045 | 66058..69378 | 1978231 | ATP-binding-protein-[Acidobacteria-bacterium] |
| MYD85169.1 | AAA+ATPase+wHTH+Alpha+Beta-domain+wHTH1+wHTH2+wHTH3+RRM | Type-1_BR.RM.like.systems | METHYLASE->METHYLASE-p->HicB_lk_antitox->DUF499*-> | 764 | Bacteria>Acidobacteriota>unclassified_Acidobacteriota | GCA_009839145.1 | Acidobacteriota_bacterium | F4Y14_02715 | 3352..5644 | 1978231 | hypothetical-protein-[Acidobacteria-bacterium] |
| OYV93187.1 | AAA+ATPase+wHTH+Alpha+Beta-domain+wHTH1+wHTH2+wHTH3+RRM | Type-1_BR.RM.like.systems | METHYLASE-p->DOC+HTH->DUF499*-> | 1072 | Bacteria>Acidobacteriota>unclassified_Acidobacteriota | GCA_002279205.1 | Acidobacteria_bacterium_21-70-11 | B7Z68_08900 | 2027..5245 | 1970485 | hypothetical-protein-B7Z68_08900-[Acidobacteria-bacterium-21-70-11] |
| MPZ20368.1 | AAA+ATPase+wHTH+Alpha+Beta-domain+wHTH1+wHTH2+wHTH3+RRM | Type-1_BR.RM.like.systems | TUDOR+SF1-SF2-Helicase+iREase+CTD+REase->PIN->?->SIR2->SIGMA-HTH->METHYLASE->DUF499*->?->HEPN-> | 1103 | Bacteria>Acidobacteriota>Vicinamibacteria>Vicinamibacterales>Vicinamibacteraceae | GCA_009377605.1 | Luteitalea_sp. | GEVO6_21000 | 17307..20618 | 2004800 | DUF499-domain-containing-protein-[Luteitalea-sp.] |
| MBN2055102.1 | AAA+ATPase+wHTH+Alpha+Beta-domain+wHTH1+wHTH2+wHTH3+RRM | Type-1_BR.RM.like.systems | <-DUF499*<-METHYLASE<-Schlafen+HATPase+HTH<-TUDOR+SF1-SF2-Helicase+iREase+CTD+REase | 1105 | Bacteria>Bacteria_incertae_sedis>Bacteria_candidate_phyla>candidate_division_CSSED10-310 | GCA_016932295.1 | candidate_division_CSSED10-310_bacterium | JW905_09275 | 5385..8702 | 2855610 | ATP-binding-protein-[bacterium] |
| RIK82262.1 | AAA+ATPase+wHTH+Alpha+Beta-domain+wHTH1+wHTH2+wHTH3+RRM | Type-1_BR.RM.like.systems | XIS-HTH->REase+SF1-SF2-Helicase->REase+N6_MTase-><-VirD4-TraG+HerA\|\|?->REase-><-DUF499*<-METHYLASE<-?<-REase+NACHT<-?<-TUDOR+SF1-SF2-Helicase+iREase+CTD+REase | 1101 | Bacteria>Bacteria_incertae_sedis>Bacteria_candidate_phyla>candidate_division_KSB1>unclassified_candidate_division_KSB1 | GCA_003576975.1 | candidate_division_KSB1_bacterium | DCC62_00525 | 28717..32022 | 2172550 | hypothetical-protein-DCC62_00525-[candidate-division-KSB1-bacterium] |
| MBZ0169160.1 | AAA+ATPase+wHTH+Alpha+Beta-domain+wHTH1+wHTH2+wHTH3+RRM | Type-1_BR.RM.like.systems | RelE-ParE-><-REase<-ORC-CDC6+REase<-BrxL<-DUF499*<-wHTH+REase<-METHYLASE<-?<-TUDOR+SF1-SF2-Helicase+iREase+CTD+REase-p<-AAA-ATPase+REase<-TUDOR+SF1-SF2-Helicase+iREase+CTD+REase | 1127 | Bacteria>Bacteria_incertae_sedis>Bacteria_candidate_phyla>candidate_division_NC10>Methylomirabilis | GCA_019912285.1 | Candidatus_Methylomirabilis_lanthanidiphila | K8F29_06945 | 3701..7084 | 2211376 | DUF499-domain-containing-protein-[Kofleriaceae-bacterium] |
| MBI4572390.1 | AAA+ATPase+wHTH+Alpha+Beta-domain+wHTH1+wHTH2+wHTH3+RRM | Type-1_BR.RM.like.systems | <-ABC-ATPase+SbcC<-?<-?\|\|Polbeta->HEPN-><-DUF499*<-AAA-ATPase+REase | 1106 | Bacteria>Bacteria_incertae_sedis>Bacteria_candidate_phyla>candidate_division_NC10>unclassified_candidate_division_NC10 | GCA_016209725.1 | candidate_division_NC10_bacterium | HY713_03775 | 5635..8955 | 2072417 | ATP-binding-protein-[candidate-division-NC10-bacterium] |
| OGB88303.1 | AAA+ATPase+wHTH+Alpha+Beta-domain+wHTH1+wHTH2+wHTH3+RRM | Type-1_BR.RM.like.systems | METHYLASE-p->AAA-ATPase+REase->DUF499*-><-?<-PIN<-?<-?<-?<-NUDIX | 1106 | Bacteria>Bacteria_incertae_sedis>Bacteria_candidate_phyla>candidate_division_NC10>unclassified_candidate_division_NC10 | GCA_001771245.1 | candidate_division_NC10_bacterium_RIFCSPLOWO2_02_FULL_66_22 | A3H39_08635 | 5174..8494 | 1801658 | hypothetical-protein-A3H39_08635-[candidate-division-NC10-bacterium-RIFCSPLOWO2_02_FULL_66_22] |
| MCG9130919.1 | AAA+ATPase+wHTH+Alpha+Beta-domain+wHTH1+wHTH2+wHTH3+RRM | Type-1_BR.RM.like.systems | TUDOR+SF1-SF2-Helicase+iREase+CTD+REase->OLD-ABC->HNH->METHYLASE->PolB->HEPN->DUF499*->HEAT_2->SAM-methylase->SIGMA-HTH->ZnR->ABC-ATPase+SbcC->ABC-ATPase+SbcC-> | 1114 | Bacteria>Bacteria_incertae_sedis>Bacteria_candidate_phyla>Candidatus_Poribacteria>unclassified_Candidatus_Poribacteria | GCA_022394775.1 | Candidatus_Poribacteria_bacterium | J5I95_04465 | 27738..31082 | 2026781 | ATP-binding-protein-[Candidatus-Poribacteria-bacterium] |
| MCY3550002.1 | AAA+ATPase+wHTH+Alpha+Beta-domain+wHTH1+wHTH2+wHTH3+RRM | Type-1_BR.RM.like.systems | <-Polbeta<-ABC-ATPase+SbcC<-ABC-ATPase+SbcC<-ZnR<-SIGMA-HTH<-HEAT_2<-DUF499*<-PolB<-METHYLASE<-?<-HNH<-OLD-ABC<-TUDOR+SF1-SF2-Helicase+iREase+CTD+REase | 1122 | Bacteria>Bacteria_incertae_sedis>Bacteria_candidate_phyla>Candidatus_Poribacteria>unclassified_Candidatus_Poribacteria | GCA_026702755.1 | Candidatus_Poribacteria_bacterium | OXH39_06030 | 100519..103887 | 2026781 | MAG:-DUF499-domain-containing-protein-[Candidatus-Poribacteria-bacterium] |
| MDE0014698.1 | AAA+ATPase+wHTH+Alpha+Beta-domain+wHTH1+wHTH2+wHTH3+RRM | Type-1_BR.RM.like.systems | <-ABC-ATPase+SbcC<-ABC-ATPase+SbcC<-ZnR<-SIGMA-HTH<-SAM-methylase<-HEAT_2<-DUF499*<-HEPN<-?<-METHYLASE<-HEPN+REase<-?<-SbcC | 1116 | Bacteria>Bacteria_incertae_sedis>Bacteria_candidate_phyla>Candidatus_Poribacteria>unclassified_Candidatus_Poribacteria | GCA_028818655.1 | Candidatus_Poribacteria_bacterium | OXU51_00820 | 15793..19143 | 2026781 | MAG:-DUF499-domain-containing-protein-[Candidatus-Poribacteria-bacterium] |
| MYC75109.1 | AAA+ATPase+wHTH+Alpha+Beta-domain+wHTH1+wHTH2+wHTH3+RRM | Type-1_BR.RM.like.systems | <-ABC-ATPase+SbcC<-ABC-ATPase+SbcC<-ZnR<-SIGMA-HTH<-HEAT_2<-DUF499*<-?<-METHYLASE<-?<-HNH<-OLD-ABC<-TUDOR+SF1-SF2-Helicase+iREase+CTD+REase | 1114 | Bacteria>Bacteria_incertae_sedis>Bacteria_candidate_phyla>Candidatus_Poribacteria>unclassified_Candidatus_Poribacteria | GCA_009841255.1 | Candidatus_Poribacteria_bacterium | F4X10_04955 | 70263..73607 | 2026781 | ATP-binding-protein-[Candidatus-Poribacteria-bacterium] |
| MDE0183895.1 | AAA+ATPase+wHTH+Alpha+Beta-domain+wHTH1+wHTH2+wHTH3+RRM | Type-1_BR.RM.like.systems | OLD-ABC->REase-><-Radical-SAM<-?<-?<-REase+35exo+Beta-Barrel+SF1-SF2-Helicase<-DUF499*<-Y1_Tnp<-TOPRIM<-OLD-ABC<-METHYLASE | 1100 | Bacteria>Bacteria_incertae_sedis>Bacteria_candidate_phyla>Candidatus_Poribacteria>unclassified_Candidatus_Poribacteria | GCA_028819845.1 | Candidatus_Poribacteria_bacterium | OXP71_00395 | 88990..92292 | 2026781 | MAG:-DUF499-domain-containing-protein-[Candidatus-Poribacteria-bacterium] |
| MYB95764.1 | AAA+ATPase+wHTH+Alpha+Beta-domain+wHTH1+wHTH2+wHTH3+RRM | Type-1_BR.RM.like.systems | TUDOR+SF1-SF2-Helicase+iREase+CTD+REase->OLD-ABC->?->METHYLASE->PolB->DUF499*->HEAT_2->SAM-methylase-> | 1114 | Bacteria>Bacteria_incertae_sedis>Bacteria_candidate_phyla>Candidatus_Poribacteria>unclassified_Candidatus_Poribacteria | GCA_009838805.1 | Candidatus_Poribacteria_bacterium | F4Y43_18100 | 10443..13787 | 2026781 | DUF499-domain-containing-protein-[Candidatus-Poribacteria-bacterium] |
| MCY4568586.1 | AAA+ATPase+wHTH+Alpha+Beta-domain+wHTH1+wHTH2+wHTH3+RRM | Type-1_BR.RM.like.systems | METHYLASE-p->DUF499*->HEAT_2->SIGMA-HTH->ZnR->?->ABC-ATPase+SbcC->ABC-ATPase+SbcC-> | 1114 | Bacteria>Bacteria_incertae_sedis>Bacteria_candidate_phyla>Candidatus_Poribacteria>unclassified_Candidatus_Poribacteria | GCA_026713385.1 | Candidatus_Poribacteria_bacterium | OXD49_09805 | 1927..5271 | 2026781 | MAG:-DUF499-domain-containing-protein-[Candidatus-Poribacteria-bacterium] |
| MDE0301131.1 | AAA+ATPase+wHTH+Alpha+Beta-domain+wHTH1+wHTH2+wHTH3+RRM | Type-1_BR.RM.like.systems | METHYLASE-p->Aminotran_3->?->?-><-DUF499*<-?<-METHYLASE<-?<-?<-?<-TUDOR+SF1-SF2-Helicase+iREase+CTD+REase | 1100 | Bacteria>Bacteria_incertae_sedis>Bacteria_candidate_phyla>Candidatus_Poribacteria>unclassified_Candidatus_Poribacteria | GCA_028820655.1 | Candidatus_Poribacteria_bacterium | OXN17_21080 | 197896..201198 | 2026781 | MAG:-DUF499-domain-containing-protein-[Candidatus-Poribacteria-bacterium] |
| MDE0324945.1 | AAA+ATPase+wHTH+Alpha+Beta-domain+wHTH1+wHTH2+wHTH3+RRM | Type-1_BR.RM.like.systems | <-HEAT_2<-DUF499*<-HEPN<-?<-METHYLASE<-?<-HNH<-OLD-ABC | 1118 | Bacteria>Bacteria_incertae_sedis>Bacteria_candidate_phyla>Candidatus_Poribacteria>unclassified_Candidatus_Poribacteria | GCA_028820875.1 | Candidatus_Poribacteria_bacterium | OXN27_13610 | 421..3777 | 2026781 | MAG:-DUF499-domain-containing-protein-[Candidatus-Poribacteria-bacterium] |
| MYB01259.1 | AAA+ATPase+wHTH+Alpha+Beta-domain+wHTH1+wHTH2+wHTH3+RRM | Type-1_BR.RM.like.systems | <-SIGMA-HTH<-SAM-methylase<-HEAT_2<-DUF499*<-PolB | 1116 | Bacteria>Bacteria_incertae_sedis>Bacteria_candidate_phyla>Candidatus_Poribacteria>unclassified_Candidatus_Poribacteria | GCA_009839665.1 | Candidatus_Poribacteria_bacterium | F4X90_16535 | 1721..5071 | 2026781 | ATP-binding-protein-[Candidatus-Poribacteria-bacterium] |
| MDE0636960.1 | AAA+ATPase+wHTH+Alpha+Beta-domain+wHTH1+wHTH2+wHTH3+RRM | Type-1_BR.RM.like.systems | TUDOR+SF1-SF2-Helicase+iREase+CTD+REase->METHYLASE->DUF499*-> | 1113 | Bacteria>Bacteria_incertae_sedis>Bacteria_candidate_phyla>Candidatus_Poribacteria>unclassified_Candidatus_Poribacteria | GCA_028824125.1 | Candidatus_Poribacteria_bacterium | OXI43_14075 | 22896..26237 | 2026781 | MAG:-DUF499-domain-containing-protein-[Candidatus-Poribacteria-bacterium] |
| MCH8291845.1 | AAA+ATPase+wHTH+Alpha+Beta-domain+wHTH1+wHTH2+wHTH3+RRM | Type-1_BR.RM.like.systems | <-REase<-HTH\|\|XIS-HTH-><-DUF499* | 1040 | Bacteria>Bacteria_incertae_sedis>Bacteria_candidate_phyla>Candidatus_Poribacteria>unclassified_Candidatus_Poribacteria | GCA_022567695.1 | Candidatus_Poribacteria_bacterium | IH992_12190 | 13707..16829 | 2026781 | DUF499-domain-containing-protein-[Candidatus-Poribacteria-bacterium] |
| MXY29380.1 | AAA+ATPase+wHTH+Alpha+Beta-domain+wHTH1+wHTH2+wHTH3+RRM | Type-1_BR.RM.like.systems | HEPN->DUF499*->REase-> | 1115 | Bacteria>Bacteria_incertae_sedis>Bacteria_candidate_phyla>Candidatus_Poribacteria>unclassified_Candidatus_Poribacteria | GCA_009838315.1 | Candidatus_Poribacteria_bacterium | F4Y59_14620 | 1265..4612 | 2026781 | ATP-binding-protein-[Candidatus-Poribacteria-bacterium] |
| MXV76534.1 | AAA+ATPase+wHTH+Alpha+Beta-domain+wHTH1+wHTH2+wHTH3+RRM | Type-1_BR.RM.like.systems | METHYLASE->PolB->DUF499*-> | 1025 | Bacteria>Bacteria_incertae_sedis>Bacteria_candidate_phyla>Candidatus_Poribacteria>unclassified_Candidatus_Poribacteria | GCA_009835515.1 | Candidatus_Poribacteria_bacterium | F4Z99_19975 | 3933..7007 | 2026781 | ATP-binding-protein-[Candidatus-Poribacteria-bacterium] |
| MBI2216638.1 | AAA+ATPase+wHTH+Alpha+Beta-domain+wHTH1+wHTH2+wHTH3+RRM | Type-1_BR.RM.like.systems | <-nSTAND3<-nSTAND3\|\|TUDOR+SF1-SF2-Helicase+iREase+CTD+REase->PHP+SbcC->Schlafen+HATPase+HTH->DUF499*->?->?-><-?<-Radical-SAM | 1102 | Bacteria>Bacteria_incertae_sedis>Bacteria_candidate_phyla>Candidatus_Rokubacteria>unclassified_Candidatus_Rokubacteria | GCA_016184125.1 | Candidatus_Rokubacteria_bacterium | HYU51_05015 | 42973..46281 | 2053607 | ATP-binding-protein-[Candidatus-Rokubacteria-bacterium] |
| OGK75197.1 | AAA+ATPase+wHTH+Alpha+Beta-domain+wHTH1+wHTH2+wHTH3+RRM | Type-1_BR.RM.like.systems | AAA-ATPase+REase->METHYLASE->DUF499*->Radical-SAM->?->?->?->?->ABC-ATPase+SbcC-> | 1096 | Bacteria>Bacteria_incertae_sedis>Bacteria_candidate_phyla>Candidatus_Rokubacteria>unclassified_Candidatus_Rokubacteria | GCA_001788395.1 | Candidatus_Rokubacteria_bacterium_GWA2_70_23 | A2X53_06550 | 6264..9554 | 1802095 | MAG:-hypothetical-protein-A2X53_06550-[Candidatus-Rokubacteria-bacterium-GWA2_70_23] |
| MCX7017989.1 | AAA+ATPase+wHTH+Alpha+Beta-domain+wHTH1+wHTH2+wHTH3+RRM | Type-1_BR.RM.like.systems | <-DUF499*<-METHYLASE<-TUDOR+SF1-SF2-Helicase+iREase+CTD+REase\|\|?-><-HEPN-p<-PolB<-HEPN | 1104 | Bacteria>Bacteria_incertae_sedis>Bacteria_candidate_phyla>Candidatus_Sumerlaeota>unclassified_Candidatus_Sumerlaeota | GCA_026398535.1 | Candidatus_Sumerlaeota_bacterium | NTY46_03120 | 388..3702 | 2107588 | MAG:-DUF499-domain-containing-protein-[Candidatus-Sumerlaeota-bacterium] |
| MCC6545645.1 | AAA+ATPase+wHTH+Alpha+Beta-domain+wHTH1+wHTH2+wHTH3+RRM | Type-1_BR.RM.like.systems | <-DUF499*<-?<-PIWI<-METHYLASE<-?<-HEPN<-METHYLASE-p | 1108 | Bacteria>Bacteria_incertae_sedis>Bacteria_candidate_phyla>Candidatus_Sumerlaeota>unclassified_Candidatus_Sumerlaeota | GCA_020848155.1 | Candidatus_Sumerlaeota_bacterium | IT570_00645 | 2114..5440 | 2107588 | ATP-binding-protein-[Candidatus-Sumerlaeota-bacterium] |
| MCP6719038.1 | AAA+ATPase+wHTH+Alpha+Beta-domain+wHTH1+wHTH2+wHTH3+RRM | Type-1_BR.RM.like.systems | Schlafen-><-?\|\|?->SF1-SF2-Helicase+iREase+CTD+REase->DUF499*->METHYLASE-> | 1106 | Bacteria>Bacteria_incertae_sedis>Bacteria_candidate_phyla>Patescibacteria_group>unclassified_Patescibacteria_group | GCA_024239015.1 | Patescibacteria_group_bacterium | KJI71_02255 | 20963..24283 | 2052139 | hypothetical-protein-[Patescibacteria-group-bacterium] |
| GAK56803.1 | AAA+ATPase+wHTH+Alpha+Beta-domain+wHTH1+wHTH2+wHTH3+RRM | Type-1_BR.RM.like.systems | RnaseIII+DSRBD->?-><-TetR-HTH<-GspA-ATPase<-DUF499*<-METHYLASE<-TUDOR+SF1-SF2-Helicase+iREase+CTD+REase | 1126 | Bacteria>Candidatus_Moduliflexota>Candidatus_Vecturitrichia>Candidatus_Vecturitrichales>Candidatus_Vecturitrichaceae | GCA_000739535.1 | Candidatus_Vecturithrix_granuli | U27_03767 | 248589..251969 | 1499967 | hypothetical-protein-Isop_3371-[Candidatus-Vecturithrix-granuli] |
| MBL0059014.1 | AAA+ATPase+wHTH+Alpha+Beta-domain+wHTH1+wHTH2+wHTH3+RRM | Type-1_BR.RM.like.systems | SMODS->?->METHYLASE->wHTH+REase->DUF499-p->DUF499*-><-rve_3<-HTH_21+rve+rve_3<-HIN-HTH | 1036 | Bacteria>Elusimicrobiota>unclassified_Elusimicrobiota | GCA_016721625.1 | Elusimicrobiota_bacterium | IPP35_07880 | 1838166..1841276 | 2030800 | ATP-binding-protein-[Elusimicrobia-bacterium] |
| MBI5244011.1 | AAA+ATPase+wHTH+Alpha+Beta-domain+wHTH1+wHTH2+wHTH3+RRM | Type-1_BR.RM.like.systems | OLD-ABC->TOPRIM->REase->SF1-SF2-Helicase+wHTH+PRTase->?->DUF499*->?-><-Methyltrans_RNA<-DnaJ | 1107 | Bacteria>Elusimicrobiota>unclassified_Elusimicrobiota | GCA_016212885.1 | Elusimicrobiota_bacterium | HY922_10100 | 5875..9198 | 2030800 | ATP-binding-protein-[Elusimicrobia-bacterium] |
| WP_089688725.1 | AAA+ATPase+wHTH+Alpha+Beta-domain+wHTH1+wHTH2+wHTH3+RRM | Type-1_BR.RM.like.systems | <-HNH\|\|TUDOR+SF1-SF2-Helicase+iREase+CTD+REase->METHYLASE->DUF499*->?->?->Phage_int_SAM_1+Phage_integrase-> | 1104 | Bacteria>FCB_group>Bacteroidota/Chlorobiota_group>Bacteroidota>Cytophagia | GCF_900100765.1 | Catalinimonas_alkaloidigena | BLR44_RS27950 | 7786..11100 | 1075417 | DUF499-domain-containing-protein-[Catalinimonas-alkaloidigena] |
| MBI4428204.1 | AAA+ATPase+wHTH+Alpha+Beta-domain+wHTH1+wHTH2+wHTH3+RRM | Type-1_BR.RM.like.systems | METHYLASE->DUF499*->?->ZnR->?->SMODS->?->MazE_antitoxin-> | 1105 | Bacteria>FCB_group>Bacteroidota/Chlorobiota_group>Ignavibacteriota>Ignavibacteria | GCA_016207415.1 | Ignavibacteriales_bacterium | HY562_03710 | 2524..5841 | 2049428 | ATP-binding-protein-[Ignavibacteriales-bacterium] |
| WP_263818943.1 | AAA+ATPase+wHTH+Alpha+Beta-domain+wHTH1+wHTH2+wHTH3+RRM | Type-1_BR.RM.like.systems | <-DUF2357+REase<-?<-HicA_toxin<-DUF499*<-METHYLASE<-TUDOR+SF1-SF2-Helicase+iREase+CTD+REase<-?<-?<-?\|\|Peptidase-> | 1098 | Bacteria>FCB_group>Bacteroidota/Chlorobiota_group>Rhodothermota>Rhodothermia | GCF_947077725.1 | Salinibacter_sp. | OJA93_RS07820 | 22881..26177 | 2065818 | DUF499-domain-containing-protein-[Salinibacter-sp.] |
| WP_259049355.1 | AAA+ATPase+wHTH+Alpha+Beta-domain+wHTH1+wHTH2+wHTH3+RRM | Type-1_BR.RM.like.systems | <-DUF2357+REase<-?<-HicA_toxin<-DUF499*<-METHYLASE<-TUDOR+SF1-SF2-Helicase+iREase+CTD+REase | 1076 | Bacteria>FCB_group>Bacteroidota/Chlorobiota_group>Rhodothermota>Rhodothermia | GCF_024808355.1 | Salinibacter_ruber | GGP97_RS06070 | 339691..342921 | 146919 | DUF499-domain-containing-protein-[Salinibacter-ruber] |

| Protein-ID | DUF499-architecture | System | Gene-Neighbourhoods | Length | Taxonomy | Assembly-ID | Strain | Locus-Tag | Genomic-coordinates | TaxID | Fasta-defline |
|---|---|---|---|---|---|---|---|---|---|---|---|
| MCC6795258.1 | AAA+ATPase+wHTH+Alpha+Beta-domain+wHTH1+wHTH2+wHTH3+RRM | Type-1_BR.RM.like.systems | TUDOR+SF1-SF2-Helicase+iREase+CTD+REase->SF1-SF2-Helicase-p->?->METHYLASE->Tox-RelE->HTH+MPTase->DUF499*->?-><-?<-TPR+TPR+TPR+TPR+TPR+TPR+TPR+TPR | 1107 | Bacteria>FCB_group>Candidatus_Hydrogenedentes>unclassified_Candidatus_Hydrogenedentes | GCA_020848835.1 | Candidatus_Hydrogenedentes_bacterium | IT366_09080 | 63574..66897 | 2030809 | ATP-binding-protein-[Candidatus-Hydrogenedentes-bacterium] |
| MBX3180306.1 | AAA+ATPase+wHTH+Alpha+Beta-domain+wHTH1+wHTH2+wHTH3+RRM | Type-1_BR.RM.like.systems | <-AbiEii<-AbiEi_4\|\|?-><-ArdCN+MPTase<-DUF499*<-PIN<-DUF433<-METHYLASE<-?<-?<-SF1-SF2-Helicase-p | 1107 | Bacteria>FCB_group>Candidatus_Hydrogenedentes>unclassified_Candidatus_Hydrogenedentes | GCA_019637335.1 | Candidatus_Hydrogenedentes_bacterium | KF886_23390 | 5509..8832 | 2030809 | ATP-binding-protein-[Candidatus-Hydrogenedentes-bacterium] |
| NLO34316.1 | AAA+ATPase+wHTH+Alpha+Beta-domain+wHTH1+wHTH2+wHTH3+RRM | Type-1_BR.RM.like.systems | <-Schlafen+HATPase+HTH<-?<-DUF499*<-METHYLASE-p | 1113 | Bacteria>FCB_group>Candidatus_Hydrogenedentes>unclassified_Candidatus_Hydrogenedentes | GCA_012523015.1 | Candidatus_Hydrogenedentes_bacterium | GX117_13355 | 3475..6816 | 2030809 | ATP-binding-protein-[Candidatus-Hydrogenedentes-bacterium] |
| MBI2505138.1 | AAA+ATPase+wHTH+Alpha+Beta-domain+wHTH1+wHTH2+wHTH3+RRM | Type-1_BR.RM.like.systems | Glycos_transf_4->?->?->?-><-DUF499*<-?<-PRTase<-SF1-SF2-Helicase-p<-wHTH+REase<-METHYLASE<-METHYLASE-p | 1105 | Bacteria>FCB_group>Candidatus_Latescibacteria>unclassified_Candidatus_Latescibacteria | GCA_016187725.1 | Candidatus_Latescibacteria_bacterium | HYW07_18115 | 13371..16688 | 2053570 | ATP-binding-protein-[Candidatus-Latescibacteria-bacterium] |
| MYC86797.1 | AAA+ATPase+wHTH+Alpha+Beta-domain+wHTH1+wHTH2+wHTH3+RRM | Type-1_BR.RM.like.systems | NTP_transferase->?->PIN->NACHT-><-?<-DUF499*<-HicB_lk_antitox<-METHYLASE<-KAP-NTPase<-MORC+REase<-DNA_methylase<-HEPN | 1101 | Bacteria>FCB_group>Gemmatimonadota>Gemmatimonadetes>Candidatus_Palauibacterales | GCA_009841015.1 | Candidatus_Palauibacter_denitrificans | F4X22_01010 | 47136..50441 | 3056734 | ATP-binding-protein-[Gemmatimonadales-bacterium] |
| MYD59776.1 | AAA+ATPase+wHTH+Alpha+Beta-domain+wHTH1+wHTH2+wHTH3+RRM | Type-1_BR.RM.like.systems | TUDOR+SF1-SF2-Helicase+iREase+CTD+REase->?->METHYLASE->HTH_3->OLD-ABC->TOPRIM->DUF499*->?->?->HigB_toxin->HTH->?->Peptidase-> | 1104 | Bacteria>FCB_group>Gemmatimonadota>unclassified_Gemmatimonadota | GCA_009841665.1 | Gemmatimonadota_bacterium | F4W91_01925 | 35019..38333 | 2026742 | ATP-binding-protein-[Gemmatimonadetes-bacterium] |
| MCY4603567.1 | AAA+ATPase+wHTH+Alpha+Beta-domain+wHTH1+wHTH2+wHTH3+RRM | Type-1_BR.RM.like.systems | CR-ATPase9+REase->PhdYeFM_antitox->METHYLASE->OLD-ABC->TOPRIM->DUF499*->Glyco_hydro_42-> | 1097 | Bacteria>FCB_group>Gemmatimonadota>unclassified_Gemmatimonadota | GCA_026713655.1 | Gemmatimonadota_bacterium | OXE49_04985 | 6802..10095 | 2026742 | MAG:-DUF499-domain-containing-protein-[Gemmatimonadetes-bacterium] |
| MXX38675.1 | AAA+ATPase+wHTH+Alpha+Beta-domain+wHTH1+wHTH2+wHTH3+RRM | Type-1_BR.RM.like.systems | OLD-ABC->TOPRIM->DUF499*->OLD-ABC->TOPRIM->Glyco_hydro_42->?-><-MqsA_antitoxin+HTH_3<-Polbeta | 1097 | Bacteria>FCB_group>Gemmatimonadota>unclassified_Gemmatimonadota | GCA_009835785.1 | Gemmatimonadota_bacterium | F4Z85_11545 | 1202..4495 | 2026742 | ATP-binding-protein-[Gemmatimonadetes-bacterium] |
| MDE2653631.1 | AAA+ATPase+wHTH+Alpha+Beta-domain+wHTH1+wHTH2+wHTH3+RRM | Type-1_BR.RM.like.systems | TUDOR+SF1-SF2-Helicase+iREase+CTD+REase->METHYLASE->METHYLASE-p->?->PIN->DUF499*-> | 1103 | Bacteria>FCB_group>Gemmatimonadota>unclassified_Gemmatimonadota | GCA_028875795.1 | Gemmatimonadota_bacterium | OXI71_07420 | 11724..15035 | 2026742 | MAG:-DUF499-domain-containing-protein-[Gemmatimonadota-bacterium] |
| MDE3256982.1 | AAA+ATPase+wHTH+Alpha+Beta-domain+wHTH1+wHTH2+wHTH3+RRM | Type-1_BR.RM.like.systems | <-SbcC<-OLD-ABC+TOPRIM<-?<-Phd-YefM<-DUF499*\|\|?-><-?<-?<-METHYLASE<-SbcC | 950 | Bacteria>FCB_group>Gemmatimonadota>unclassified_Gemmatimonadota | GCA_028877215.1 | Gemmatimonadota_bacterium | OYM47_04020 | 48493..51345 | 2026742 | MAG:-DUF499-domain-containing-protein-[Gemmatimonadota-bacterium] |
| MCY4399230.1 | AAA+ATPase+wHTH+Alpha+Beta-domain+wHTH1+wHTH2+wHTH3+RRM | Type-1_BR.RM.like.systems | METHYLASE-p-><-Peptidase\|\|?->?->?-><-DUF499* | 1023 | Bacteria>FCB_group>Gemmatimonadota>unclassified_Gemmatimonadota | GCA_026710725.1 | Gemmatimonadota_bacterium | OXE96_07830 | 57550..60621 | 2026742 | MAG:-DUF499-domain-containing-protein-[Gemmatimonadetes-bacterium] |
| WP_224365796.1 | AAA+ATPase+wHTH+Alpha+Beta-domain+wHTH1+wHTH2+wHTH3+RRM | Type-1_BR.RM.like.systems | Cas_Cmr3->?->Cas_Cmr5->?->?-><-DUF499*<-METHYLASE<-TUDOR+SF1-SF2-Helicase+iREase+CTD+REase<-CRISPR_Cas2<-Cas_Cas1 | 1126 | Bacteria>Myxococcota>Myxococcia>Myxococcales>Cystobacterineae | GCF_020103695.1 | Hyalangium_versicolor | KY501_RS24855 | 93112..96492 | 2861190 | DUF499-domain-containing-protein-[Hyalangium-sp.-H56D21] |
| NMB74978.1 | AAA+ATPase+wHTH+Alpha+Beta-domain+wHTH1+wHTH2+wHTH3+RRM | Type-1_BR.RM.like.systems | XIS-HTH->?->?-><-?<-?<-?<-DUF499*<-METHYLASE<-PIN<-?<-TUDOR+SF1-SF2-Helicase+iREase+CTD+REase<-XIS-HTH | 1126 | Bacteria>Myxococcota>Myxococcia>Myxococcales>unclassified_Myxococcales | GCA_012798435.1 | Myxococcales_bacterium | GYA21_07595 | 16528..19908 | 2026763 | ATP-binding-protein-[Myxococcales-bacterium] |
| MBN1104019.1 | AAA+ATPase+wHTH+Alpha+Beta-domain+wHTH1+wHTH2+wHTH3+RRM | Type-1_BR.RM.like.systems | <-Phage_integrase\|\|TUDOR+SF1-SF2-Helicase+iREase+CTD+REase->AAA-ATPase+REase->METHYLASE->DUF499*->MORC+DUF3684+REase-><-?<-ABC-ATPase+SbcC<-ABC-ATPase+SbcC | 1109 | Bacteria>Myxococcota>Myxococcia>unclassified_Myxococcia | GCA_016926135.1 | Deltaproteobacteria_bacterium | JXL84_11450 | 9646..12975 | 2026735 | DUF499-domain-containing-protein-[Deltaproteobacteria-bacterium] |
| MBI4514358.1 | AAA+ATPase+wHTH+Alpha+Beta-domain+wHTH1+wHTH2+wHTH3+RRM | Type-1_BR.RM.like.systems | <-sGTP<-?<-BirA-HTH<-?<-?\|\|tRNA-><-DUF499*<-REase<-METHYLASE<-AAA-ATPase+REase<-TUDOR+SF1-SF2-Helicase+iREase+CTD+REase\|\|?->\|\|tRNA->NTP_transferase+MannoseP_isomer-> | 1100 | Bacteria>Myxococcota>Myxococcia>unclassified_Myxococcia | GCA_016210005.1 | Deltaproteobacteria_bacterium | HY699_00880 | 115685..118987 | 2026735 | ATP-binding-protein-[Deltaproteobacteria-bacterium] |
| MBW2343658.1 | AAA+ATPase+wHTH+Alpha+Beta-domain+wHTH1+wHTH2+wHTH3+RRM | Type-1_BR.RM.like.systems | <-RelE-ParE<-MetJarc-HTH\|\|?->TUDOR+SF1-SF2-Helicase+iREase+CTD+REase->wHTH+REase->METHYLASE->DUF499*-> | 1106 | Bacteria>Myxococcota>Myxococcia>unclassified_Myxococcia | GCA_019310785.1 | Deltaproteobacteria_bacterium | JRF53_06530 | 24941..28261 | 2026735 | DUF499-domain-containing-protein-[Deltaproteobacteria-bacterium] |
| HBA39938.1 | AAA+ATPase+wHTH+Alpha+Beta-domain+wHTH1+wHTH2+wHTH3+RRM | Type-1_BR.RM.like.systems | TUDOR+SF1-SF2-Helicase+iREase+CTD+REase->AAA-ATPase+REase->REase-p->METHYLASE->REase->DUF499*-> | 1099 | Bacteria>Myxococcota>Myxococcia>unclassified_Myxococcia | GCA_003490035.1 | Deltaproteobacteria_bacterium | DCZ05_09410 | 9752..13051 | 2026735 | hypothetical-protein-[Deltaproteobacteria-bacterium] |
| MDD9873633.1 | AAA+ATPase+wHTH+Alpha+Beta-domain+wHTH1+wHTH2+wHTH3+RRM | Type-1_BR.RM.like.systems | TUDOR+SF1-SF2-Helicase+iREase+CTD+REase->HEPN->METHYLASE->OLD-ABC->TOPRIM->DUF499*->?->?->?->?->ABC-ATPase+SbcC-> | 1106 | Bacteria>Myxococcota>Myxococcia>unclassified_Myxococcia | GCA_028817295.1 | Deltaproteobacteria_bacterium | OXU65_09090 | 9437..12757 | 2026735 | MAG:-DUF499-domain-containing-protein-[Deltaproteobacteria-bacterium] |
| MBI4525152.1 | AAA+ATPase+wHTH+Alpha+Beta-domain+wHTH1+wHTH2+wHTH3+RRM | Type-1_BR.RM.like.systems | REase+SAVED->SMODS->UbE1+JAB->JAB->METHYLASE->REase->DUF499*->REase+nSTAND1-> | 1099 | Bacteria>Myxococcota>Myxococcia>unclassified_Myxococcia | GCA_016210115.1 | Deltaproteobacteria_bacterium | HY695_15215 | 17806..21105 | 2026735 | DUF499-domain-containing-protein-[Deltaproteobacteria-bacterium] |
| MBC8466732.1 | AAA+ATPase+wHTH+Alpha+Beta-domain+wHTH1+wHTH2+wHTH3+RRM | Type-1_BR.RM.like.systems | Phage_integrase->TUDOR+SF1-SF2-Helicase+iREase+CTD+REase->Schlafen->METHYLASE->DUF499*-> | 1095 | Bacteria>Myxococcota>Myxococcia>unclassified_Myxococcia | GCA_014384065.1 | Deltaproteobacteria_bacterium | H8D55_02725 | 8487..11774 | 2026735 | ATP-binding-protein-[Deltaproteobacteria-bacterium] |
| MBM4330413.1 | AAA+ATPase+wHTH+Alpha+Beta-domain+wHTH1+wHTH2+wHTH3+RRM | Type-1_BR.RM.like.systems | TUDOR+SF1-SF2-Helicase+iREase+CTD+REase->wHTH+REase->?->METHYLASE->DUF499*-><-?<-PIN | 1106 | Bacteria>Myxococcota>Myxococcia>unclassified_Myxococcia | GCA_016875165.1 | Deltaproteobacteria_bacterium | FJ117_04155 | 62692..66012 | 2026735 | ATP-binding-protein-[Deltaproteobacteria-bacterium] |
| MBM4396669.1 | AAA+ATPase+wHTH+Alpha+Beta-domain+wHTH1+wHTH2+wHTH3+RRM | Type-1_BR.RM.like.systems | TUDOR+SF1-SF2-Helicase+iREase+CTD+REase-p->METHYLASE->OLD-ABC->TOPRIM->DUF499*-> | 1103 | Bacteria>Myxococcota>Myxococcia>unclassified_Myxococcia | GCA_016875475.1 | Deltaproteobacteria_bacterium | FJ087_13395 | 5583..8894 | 2026735 | ATP-binding-protein-[Deltaproteobacteria-bacterium] |
| MBI3795502.1 | AAA+ATPase+wHTH+Alpha+Beta-domain+wHTH1+wHTH2+wHTH3+RRM | Type-1_BR.RM.like.systems | <-DUF499*<-TOPRIM<-OLD-ABC<-METHYLASE | 1108 | Bacteria>Myxococcota>Myxococcia>unclassified_Myxococcia | GCA_016200595.1 | Deltaproteobacteria_bacterium | HY268_00830 | 2141..5467 | 2026735 | ATP-binding-protein-[Deltaproteobacteria-bacterium] |
| MBF0565077.1 | AAA+ATPase+wHTH+Alpha+Beta-domain+wHTH1+wHTH2+wHTH3+RRM | Type-1_BR.RM.like.systems | <-DUF499*<-NTP_transf_2<-HEPN<-METHYLASE | 1116 | Bacteria>Nitrospirota>unclassified_Nitrospirota | GCA_015233615.1 | Nitrospirota_bacterium | HQK89_07535 | 20523..23873 | 2026887 | ATP-binding-protein-[Nitrospirae-bacterium] |
| MCW5716290.1 | AAA+ATPase+wHTH+Alpha+Beta-domain+wHTH1+wHTH2+wHTH3+RRM | Type-1_BR.RM.like.systems | Peptidase_M15->?-><-HEPN<-DUF499*<-METHYLASE<-SF1-SF2-Helicase<-TUDOR+SF1-SF2-Helicase+iREase+CTD+REase<-XIS-HTH<-GDResolvase+Recombinase | 1108 | Bacteria>Pseudomonadota>Alphaproteobacteria>Hyphomicrobiales>Kaistiaceae | GCA_026126085.1 | Bauldia_sp. | KIS68_00515 | 97641..100967 | 2575872 | MAG:-ATP-binding-protein-[Bauldia-sp.] |
| WP_183231813.1 | AAA+ATPase+wHTH+Alpha+Beta-domain+wHTH1+wHTH2+wHTH3+RRM | Type-1_BR.RM.like.systems | CHTH-><-?<-NUDIX\|\|?->ZnR->?-><-DUF499*<-METHYLASE<-TUDOR+SF1-SF2-Helicase+iREase+CTD+REase\|\|Calcineurin->SbcC+SbcC->SIGMA-HTH-> | 1121 | Bacteria>Pseudomonadota>Alphaproteobacteria>Hyphomicrobiales>Nitrobacteraceae | GCF_014198205.1 | Bradyrhizobium_sp._SBR1B | GGD61_RS31330 | 50239..53604 | 2663836 | DUF499-domain-containing-protein-[Bradyrhizobium-sp.-SBR1B] |
| WP_212355098.1 | AAA+ATPase+wHTH+Alpha+Beta-domain+wHTH1+wHTH2+wHTH3+RRM | Type-1_BR.RM.like.systems | <-SIGMA-HTH<-SbcC+SbcC<-Calcineurin\|\|TUDOR+SF1-SF2-Helicase+iREase+CTD+REase->METHYLASE->DUF499*-><-?<-ZnR<-?\|\|?->NUDIX-> | 1121 | Bacteria>Pseudomonadota>Alphaproteobacteria>Hyphomicrobiales>Nitrobacteraceae | GCF_018130695.1 | Bradyrhizobium_jicamae | JQ604_RS07975 | 246023..249388 | 280332 | DUF499-domain-containing-protein-[Bradyrhizobium-jicamae] |
| MCV0387876.1 | AAA+ATPase+wHTH+Alpha+Beta-domain+wHTH1+wHTH2+wHTH3+RRM | Type-1_BR.RM.like.systems | <-HIN-HTH<-?<-DnaB+SMS-KaiC\|\|?-><-XIS-HTH<-Arm-DNA-bind_3+Phage_integrase<-DUF499*<-METHYLASE-p | 1121 | Bacteria>Pseudomonadota>Alphaproteobacteria>Hyphomicrobiales>Nitrobacteraceae | GCA_025698885.1 | Nitrobacter_sp. | K5821_16025 | 7595..10960 | 29420 | MAG:-hypothetical-protein-K5821_16025-[Nitrobacter-sp.] |
| WP_247772608.1 | AAA+ATPase+wHTH+Alpha+Beta-domain+wHTH1+wHTH2+wHTH3+RRM | Type-1_BR.RM.like.systems | <-SbcC+SbcC<-Calcineurin\|\|TUDOR+SF1-SF2-Helicase+iREase+CTD+REase->METHYLASE->DUF499*-> | 1121 | Bacteria>Pseudomonadota>Alphaproteobacteria>Hyphomicrobiales>Nitrobacteraceae | GCF_023100965.1 | Bradyrhizobium_sp._192 | IVB24_RS08015 | 1720329..1723694 | 2782660 | DUF499-domain-containing-protein-[Bradyrhizobium-sp.-192] |
| WP_247341913.1 | AAA+ATPase+wHTH+Alpha+Beta-domain+wHTH1+wHTH2+wHTH3+RRM | Type-1_BR.RM.like.systems | <-DUF499*<-METHYLASE<-TUDOR+SF1-SF2-Helicase+iREase+CTD+REase\|\|Calcineurin->SbcC+SbcC-> | 1121 | Bacteria>Pseudomonadota>Alphaproteobacteria>Hyphomicrobiales>Nitrobacteraceae | GCF_023073195.1 | unclassified_Bradyrhizobium | IVB38_RS06920 | 46468..49833 | 2631580 | MULTISPECIES:-DUF499-domain-containing-protein-[unclassified-Bradyrhizobium] |
| MBY0360468.1 | AAA+ATPase+wHTH+Alpha+Beta-domain+wHTH1+wHTH2+wHTH3+RRM | Type-1_BR.RM.like.systems | <-DUF499*<-METHYLASE<-TUDOR+SF1-SF2-Helicase+iREase+CTD+REase<-HTH | 1117 | Bacteria>Pseudomonadota>Alphaproteobacteria>Hyphomicrobiales>Phreatobacteraceae | GCA_019749835.1 | Phreatobacter_sp. | K2X45_01055 | 2045..5398 | 1966341 | hypothetical-protein-[Phreatobacter-sp.] |
| MBF0339369.1 | AAA+ATPase+wHTH+Alpha+Beta-domain+wHTH1+wHTH2+wHTH3+RRM | Type-1_BR.RM.like.systems | DDE_Tnp_1_assoc+DDE_Tnp_1->TUDOR+SF1-SF2-Helicase+iREase+CTD+REase->?->NADAR->METHYLASE->DUF499*->?->OLD-ABC->REase-> | 1103 | Bacteria>Pseudomonadota>Alphaproteobacteria>Magnetococcales>unclassified_Magnetococcales | GCA_015232245.1 | Magnetococcales_bacterium | HQL95_00220 | 16379..19690 | 2026759 | ATP-binding-protein-[Magnetococcales-bacterium] |
| MBF0191611.1 | AAA+ATPase+wHTH+Alpha+Beta-domain+wHTH1+wHTH2+wHTH3+RRM | Type-1_BR.RM.like.systems | <-ABC-ATPase+SbcC\|\|?->TUDOR+SF1-SF2-Helicase+iREase+CTD+REase->?->NADAR->METHYLASE->DUF499*-> | 1104 | Bacteria>Pseudomonadota>Alphaproteobacteria>Magnetococcales>unclassified_Magnetococcales | GCA_015231375.1 | Magnetococcales_bacterium | HQL99_10815 | 25599..28913 | 2026759 | ATP-binding-protein-[Magnetococcales-bacterium] |
| MBF0296601.1 | AAA+ATPase+wHTH+Alpha+Beta-domain+wHTH1+wHTH2+wHTH3+RRM | Type-1_BR.RM.like.systems | HHH+ENDO3+NUDIX->TUDOR+SF1-SF2-Helicase+iREase+CTD+REase->?->METHYLASE->DUF499*-> | 1105 | Bacteria>Pseudomonadota>Alphaproteobacteria>Magnetococcales>unclassified_Magnetococcales | GCA_015231915.1 | Magnetococcales_bacterium | HQL96_15555 | 9273..12590 | 2026759 | ATP-binding-protein-[Magnetococcales-bacterium] |
| MBF0110513.1 | AAA+ATPase+wHTH+Alpha+Beta-domain+wHTH1+wHTH2+wHTH3+RRM | Type-1_BR.RM.like.systems | PfkB->?-><-?<-DUF499*<-METHYLASE-p | 1105 | Bacteria>Pseudomonadota>Alphaproteobacteria>Magnetococcales>unclassified_Magnetococcales | GCA_015228815.1 | Magnetococcales_bacterium | HQL76_15200 | 24782..28099 | 2026759 | ATP-binding-protein-[Magnetococcales-bacterium] |
| WP_207470981.1 | AAA+ATPase+wHTH+Alpha+Beta-domain+wHTH1+wHTH2+wHTH3+RRM | Type-1_BR.RM.like.systems | METHYLASE-p->DUF499*-><-tRNA<-?\|\|PUA+METHYLASE->PIN-> | 1119 | Bacteria>Pseudomonadota>Alphaproteobacteria>Rhodobacterales>Paracoccaceae | GCF_017356265.1 | Paracoccus_fontiphilus | JOF76_RS16065 | 1340..4699 | 1815556 | ATP-binding-protein-[Paracoccus-fontiphilus] |
| WP_244408576.1 | AAA+ATPase+wHTH+Alpha+Beta-domain+wHTH1+wHTH2+wHTH3+RRM | Type-1_BR.RM.like.systems | DEDD_Tnp_IS110+Transposase_20->?->?->TUDOR+SF1-SF2-Helicase+iREase+CTD+REase->METHYLASE->DUF499*->\|\|?->?->?->XIS-HTH-> | 1129 | Bacteria>Pseudomonadota>Alphaproteobacteria>Rhodospirillales>Acetobacteraceae | GCF_022846615.1 | Roseomonas_fluvialis | MWM08_RS21610 | 4513512..4516901 | 1750527 | hypothetical-protein-[Roseomonas-sp.-S08] |
| GIX11744.1 | AAA+ATPase+wHTH+Alpha+Beta-domain+wHTH1+wHTH2+wHTH3+RRM | Type-1_BR.RM.like.systems | <-Peptidase_S8<-Classical-AAA\|\|TUDOR+SF1-SF2-Helicase+iREase+CTD+REase->RNase->METHYLASE->DUF499*->OLD-ABC+TOPRIM-><-?<-Peptidase_M15 | 1107 | Bacteria>Pseudomonadota>Alphaproteobacteria>Rhodospirillales>Elioraeaceae | GCA_026004895.1 | Elioraea_sp. | KatS3mg116_3454 | 526065..529388 | 2185103 | MAG:-hypothetical-protein-KatS3mg116_3454-[Elioraea-sp.] |
| MDE0384393.1 | AAA+ATPase+wHTH+Alpha+Beta-domain+wHTH1+wHTH2+wHTH3+RRM | Type-1_BR.RM.like.systems | TUDOR+SF1-SF2-Helicase+iREase+CTD+REase->RNase->?->METHYLASE->OLD-ABC->TOPRIM->DUF499*-><-?<-SF1-SF2-Helicase+DUF1998<-?<-PADR-HTH | 1108 | Bacteria>Pseudomonadota>Alphaproteobacteria>Rhodospirillales>Rhodospirillaceae | GCA_028821395.1 | Defluviicoccus_sp. | OXI22_10955 | 42906..46232 | 2682137 | MAG:-DUF499-domain-containing-protein-[Defluviicoccus-sp.] |
| MBX6368568.1 | AAA+ATPase+wHTH+Alpha+Beta-domain+wHTH1+wHTH2+wHTH3+RRM | Type-1_BR.RM.like.systems | <-PIWI<-SIR2+SIR2\|\|TUDOR+SF1-SF2-Helicase+iREase+CTD+REase->METHYLASE->DUF499*-> | 1121 | Bacteria>Pseudomonadota>Alphaproteobacteria>Rhodospirillales>unclassified_Rhodospirillales | GCA_019689675.1 | Rhodospirillales_bacterium | IRZ04_11335 | 10832..14197 | 2026786 | ATP-binding-protein-[Rhodospirillales-bacterium] |
| MCY3753479.1 | AAA+ATPase+wHTH+Alpha+Beta-domain+wHTH1+wHTH2+wHTH3+RRM | Type-1_BR.RM.like.systems | <-HTH_3\|\|TUDOR+SF1-SF2-Helicase+iREase+CTD+REase->?->METHYLASE->OLD-ABC->TOPRIM->DUF499*-><-MPTase<-HTH_3\|\|SMODS->?->?-><-Arm-DNA-bind_3+Phage_int_SAM_3+Phage_integrase | 1104 | Bacteria>Pseudomonadota>Alphaproteobacteria>unclassified_Alphaproteobacteria | GCA_026706245.1 | Alphaproteobacteria_bacterium | OXG99_05240 | 13033..16347 | 1913988 | MAG:-DUF499-domain-containing-protein-[Alphaproteobacteria-bacterium] |
| NWG45252.1 | AAA+ATPase+wHTH+Alpha+Beta-domain+wHTH1+wHTH2+wHTH3+RRM | Type-1_BR.RM.like.systems | Peptidase_M15->?->?-><-?<-HEPN<-DUF499*<-METHYLASE-p | 1108 | Bacteria>Pseudomonadota>Alphaproteobacteria>unclassified_Alphaproteobacteria | GCA_013389075.1 | Alphaproteobacteria_bacterium | HXY25_01725 | 2157..5483 | 1913988 | ATP-binding-protein-[Alphaproteobacteria-bacterium] |
| WP_169070868.1 | AAA+ATPase+wHTH+Alpha+Beta-domain+wHTH1+wHTH2+wHTH3+RRM | Type-1_BR.RM.like.systems | GDResolvase+Recombinase+ZnR->HTH-><-?\|\|TUDOR+SF1-SF2-Helicase+iREase+CTD+REase->?->METHYLASE->DUF499*->?->?->LZ_Tnp_IS66+zf-IS66+DDE_Tnp_IS66+DDE_Tnp_IS66_C->?->?->\|\|DDE_Tnp_1-> | 1093 | Bacteria>Pseudomonadota>Betaproteobacteria>Betaproteobacteria_incertae_sedis>Candidatus_Accumulibacter | GCF_012940005.1 | Candidatus_Accumulibacter_contiguus | E4Q08_RS14240 | 87820..91101 | 2954381 | DUF499-domain-containing-protein-[Candidatus-Accumulibacter-contiguus] |
| QKS30678.1 | AAA+ATPase+wHTH+Alpha+Beta-domain+wHTH1+wHTH2+wHTH3+RRM | Type-1_BR.RM.like.systems | <-MerR-HTH\|\|TUDOR+SF1-SF2-Helicase+iREase+CTD+REase->?->?->METHYLASE->DUF499*->HEPN+SF1-SF2-Helicase->MORC+DUF3684+HNH->?-><-?<-TraH<-Thioredoxin | 1092 | Bacteria>Pseudomonadota>Betaproteobacteria>Betaproteobacteria_incertae_sedis>Candidatus_Accumulibacter | GCA_013347225.1 | Candidatus_Accumulibacter_similis | HT579_18190 | 4129453..4132731 | 2954388 | MAG:-ATP-binding-protein-[Candidatus-Accumulibacter-similis] |
| KFB77813.1 | AAA+ATPase+wHTH+Alpha+Beta-domain+wHTH1+wHTH2+wHTH3+RRM | Type-1_BR.RM.like.systems | Hpr-Kinase-C-><-JetD<-?<-JetB<-JetA<-DUF499*<-Schlafen+HATPase+HTH<-METHYLASE<-PIN\|\|?-><-TUDOR+SF1-SF2-Helicase+iREase+CTD+REase<-HAMP+HISKIN | 1093 | Bacteria>Pseudomonadota>Betaproteobacteria>Betaproteobacteria_incertae_sedis>Candidatus_Accumulibacter | GCA_000584975.2 | Candidatus_Accumulibacter_cognatus | AW06_000996 | 10153..13434 | 2954383 | MAG:-hypothetical-protein-AW06_000996-[Candidatus-Accumulibacter-cognatus] |
| WP_034945897.1 | AAA+ATPase+wHTH+Alpha+Beta-domain+wHTH1+wHTH2+wHTH3+RRM | Type-1_BR.RM.like.systems | Hpr-Kinase-C-><-JetD<-?<-JetB<-JetA<-DUF499*<-Schlafen+HATPase+HTH<-METHYLASE<-PIN<-TUDOR+SF1-SF2-Helicase+iREase+CTD+REase<-HAMP+HISKIN | 1093 | Bacteria>Pseudomonadota>Betaproteobacteria>Betaproteobacteria_incertae_sedis>Candidatus_Accumulibacter | GCF_000584975.1 | Candidatus_Accumulibacter_cognatus | AW06_RS03920 | 10153..13434 | 2954383 | DUF499-domain-containing-protein-[Candidatus-Accumulibacter-cognatus] |
| MCQ1548298.1 | AAA+ATPase+wHTH+Alpha+Beta-domain+wHTH1+wHTH2+wHTH3+RRM | Type-1_BR.RM.like.systems | HAMP+HISKIN->TUDOR+SF1-SF2-Helicase+iREase+CTD+REase->PIN->METHYLASE->Schlafen+HATPase+HTH->DUF499*-> | 1093 | Bacteria>Pseudomonadota>Betaproteobacteria>Betaproteobacteria_incertae_sedis>Candidatus_Accumulibacter | GCA_024380085.1 | Candidatus_Accumulibacter_phosphatis | NOF05_05620 | 18582..21863 | 327160 | DUF499-domain-containing-protein-[Candidatus-Accumulibacter-phosphatis] |
| RDE49302.1 | AAA+ATPase+wHTH+Alpha+Beta-domain+wHTH1+wHTH2+wHTH3+RRM | Type-1_BR.RM.like.systems | TUDOR+SF1-SF2-Helicase+iREase+CTD+REase->?->METHYLASE->DUF499*->HTH-> | 1093 | Bacteria>Pseudomonadota>Betaproteobacteria>Betaproteobacteria_incertae_sedis>Candidatus_Accumulibacter | GCA_003332265.1 | Candidatus_Accumulibacter_meliphilus | DVS81_17340 | 9132..12413 | 2211374 | MAG:-DUF499-domain-containing-protein-[Candidatus-Accumulibacter-meliphilus] |
| EXI65857.1 | AAA+ATPase+wHTH+Alpha+Beta-domain+wHTH1+wHTH2+wHTH3+RRM | Type-1_BR.RM.like.systems | TUDOR+SF1-SF2-Helicase+iREase+CTD+REase->PIN->?->METHYLASE->DUF499*-> | 1091 | Bacteria>Pseudomonadota>Betaproteobacteria>Betaproteobacteria_incertae_sedis>Candidatus_Accumulibacter | GCF_000585015.1 | Candidatus_Accumulibacter_adjunctus | AW08_02882 | 60181..63456 | 1454001 | MAG:-hypothetical-protein-AW08_02882-[Candidatus-Accumulibacter-adjunctus] |
| WP_105508032.1 | AAA+ATPase+wHTH+Alpha+Beta-domain+wHTH1+wHTH2+wHTH3+RRM | Type-1_BR.RM.like.systems | Phage_int_SAM_1+Phage_integrase-><-REase<-?<-HTH+MPTase\|\|?-><-?<-DUF499*<-ABhydrolase<-?\|\|HTH_3-><-Peptidase_C39<-?<-Peptidase_C39 | 837 | Bacteria>Pseudomonadota>Betaproteobacteria>Burkholderiales>Burkholderiaceae | - | Paraburkholderia_sp._BL21I4N1 | B0G83_104407 | 412905..415418 | 1938801 | sigma-54-interacting-transcriptional-regulator-[Paraburkholderia-sp.-BL21I4N1] |
| PQV52187.1 | AAA+ATPase+wHTH+Alpha+Beta-domain+wHTH1+wHTH2+wHTH3+RRM | Type-1_BR.RM.like.systems | Phage_int_SAM_1+Phage_integrase-><-REase<-?<-HTH+MPTase\|\|?-><-?<-DUF499*<-ABhydrolase<-?\|\|HTH_3-><-Peptidase_C39<-?<-Peptidase_C39 | 837 | Bacteria>Pseudomonadota>Betaproteobacteria>Burkholderiales>Burkholderiaceae | GCA_002973525.1 | Paraburkholderia_sp._BL21I4N1 | B0G83_104407 | 412905..415418 | 1938801 | sigma-54-interacting-transcriptional-regulator-[Paraburkholderia-sp.-BL21I4N1] |
| WP_145864756.1 | AAA+ATPase+wHTH+Alpha+Beta-domain+wHTH1+wHTH2+wHTH3+RRM | Type-1_BR.RM.like.systems | <-PadR-HTH\|\|TUDOR+SF1-SF2-Helicase+iREase+CTD+REase->SMODS->?->METHYLASE->DUF499*->?-><-Radical-SAM<-HTH_3<-Phage_lysozyme | 1093 | Bacteria>Pseudomonadota>Betaproteobacteria>Burkholderiales>Burkholderiaceae | GCF_007833125.1 | Pandoraea_pnomenusa | FP568_RS23045 | 5133306..5136587 | 93220 | DUF499-domain-containing-protein-[Pandoraea-pnomenusa] |
| WP_043700899.1 | AAA+ATPase+wHTH+Alpha+Beta-domain+wHTH1+wHTH2+wHTH3+RRM | Type-1_BR.RM.like.systems | <-HKD-DNase<-DUF499*<-METHYLASE<-?<-SMODS<-TUDOR+SF1-SF2-Helicase+iREase+CTD+REase\|\|PadR-HTH-> | 1093 | Bacteria>Pseudomonadota>Betaproteobacteria>Burkholderiales>Burkholderiales_genera_incertae_sedis | GCF_000807215.1 | Tepidimonas_taiwanensis | PL21_RS06510 | 396..3677 | 307486 | DUF499-domain-containing-protein-[Tepidimonas-taiwanensis] |
| WP_250960620.1 | AAA+ATPase+wHTH+Alpha+Beta-domain+wHTH1+wHTH2+wHTH3+RRM | Type-1_BR.RM.like.systems | Phage_holin->Phage_lysozyme->?->HTH_3->Radical-SAM-><-?<-DUF499*<-METHYLASE<-?<-TUDOR+SF1-SF2-Helicase+iREase+CTD+REase\|\|PadR-HTH-><-?<-HTH | 1093 | Bacteria>Pseudomonadota>Betaproteobacteria>Burkholderiales>Comamonadaceae | GCF_023702035.1 | Serpentinimonas_maccroryi | G4L37_RS07185 | 227051..230332 | 1458426 | DUF499-domain-containing-protein-[Serpentinimonas-maccroryi] |
| WP_265663937.1 | AAA+ATPase+wHTH+Alpha+Beta-domain+wHTH1+wHTH2+wHTH3+RRM | Type-1_BR.RM.like.systems | ABC-ATPase+SbcC->?->THUMP+Methylase->Radical-SAM-><-Methylase<-DUF499*<-METHYLASE<-PIN<-TUDOR+SF1-SF2-Helicase+iREase+CTD+REase | 1094 | Bacteria>Pseudomonadota>Betaproteobacteria>Burkholderiales>Comamonadaceae | GCF_026192565.1 | Verminephrobacter_aporrectodeae | D8B23_RS01230 | 11909..15193 | 1110389 | DUF499-domain-containing-protein-[Verminephrobacter-aporrectodeae] |
| WP_265283911.1 | AAA+ATPase+wHTH+Alpha+Beta-domain+wHTH1+wHTH2+wHTH3+RRM | Type-1_BR.RM.like.systems | TUDOR+SF1-SF2-Helicase+iREase+CTD+REase->PIN->METHYLASE->DUF499*->Methylase-><-Radical-SAM<-THUMP+Methylase<-?<-ABC-ATPase+SbcC | 1094 | Bacteria>Pseudomonadota>Betaproteobacteria>Burkholderiales>Comamonadaceae | GCF_026127385.1 | Verminephrobacter_aporrectodeae | D5041_RS03845 | 914052..917336 | 1110389 | DUF499-domain-containing-protein-[Verminephrobacter-aporrectodeae] |
| WP_265286858.1 | AAA+ATPase+wHTH+Alpha+Beta-domain+wHTH1+wHTH2+wHTH3+RRM | Type-1_BR.RM.like.systems | ZnR+TOPRIM+D5ATPase->TUDOR+SF1-SF2-Helicase+iREase+CTD+REase->SMODS->?->wHTH+REase->METHYLASE->DUF499*->REase->REase->?-><-GNAT | 1092 | Bacteria>Pseudomonadota>Betaproteobacteria>Burkholderiales>Comamonadaceae | GCF_026127525.1 | Verminephrobacter_eiseniae | D5047_RS05820 | 1238736..1242014 | 364317 | DUF499-domain-containing-protein-[Verminephrobacter-eiseniae] |
| WP_265653305.1 | AAA+ATPase+wHTH+Alpha+Beta-domain+wHTH1+wHTH2+wHTH3+RRM | Type-1_BR.RM.like.systems | TUDOR+SF1-SF2-Helicase+iREase+CTD+REase->PIN->METHYLASE->DUF499*->\|\|?->HerA/FtsK-><-THUMP+Methylase<-?<-ABC-ATPase+SbcC | 1094 | Bacteria>Pseudomonadota>Betaproteobacteria>Burkholderiales>Comamonadaceae | GCF_026192715.1 | Verminephrobacter_aporrectodeae | D8B27_RS01360 | 119835..123119 | 1110389 | DUF499-domain-containing-protein-[Verminephrobacter-aporrectodeae] |
| MCV0438782.1 | AAA+ATPase+wHTH+Alpha+Beta-domain+wHTH1+wHTH2+wHTH3+RRM | Type-1_BR.RM.like.systems | 35exo->tRNA->TUDOR+SF1-SF2-Helicase+iREase+CTD+REase->?->METHYLASE->DUF499*->?-><-?<-ZnR+TOPRIM+D5ATPase<-?<-XIS-HTH | 1093 | Bacteria>Pseudomonadota>Betaproteobacteria>Burkholderiales>Comamonadaceae | GCA_025699295.1 | Hydrogenophaga_sp. | K5880_09120 | 131654..134935 | 1904254 | MAG:-DUF499-domain-containing-protein-[Hydrogenophaga-sp.] |
| GIX24926.1 | AAA+ATPase+wHTH+Alpha+Beta-domain+wHTH1+wHTH2+wHTH3+RRM | Type-1_BR.RM.like.systems | Phage_lysozyme->HTH_3->Radical-SAM-><-Methylase<-?<-HKD-DNase+HKD-DNase<-DUF499*<-METHYLASE-p<-METHYLASE-p<-METHYLASE-p<-?<-?<-TUDOR+SF1-SF2-Helicase+iREase+CTD+REase-p | 1093 | Bacteria>Pseudomonadota>Betaproteobacteria>Burkholderiales>Sphaerotilaceae | GCA_026004995.1 | Caldimonas_sp. | KatS3mg122_2157 | 507671..510952 | 2838790 | MAG:-hypothetical-protein-KatS3mg122_2157-[Caldimonas-sp.] |
| MCC6533588.1 | AAA+ATPase+wHTH+Alpha+Beta-domain+wHTH1+wHTH2+wHTH3+RRM | Type-1_BR.RM.like.systems | Peptidase->?->?->?-><-?<-DUF499*<-METHYLASE<-TUDOR+SF1-SF2-Helicase+iREase+CTD+REase<-?\|\|GST_N+GST_C-> | 1088 | Bacteria>Pseudomonadota>Betaproteobacteria>Burkholderiales>unclassified_Burkholderiales | GCA_020848435.1 | Burkholderiales_bacterium | IT531_13635 | 27446..30712 | 1891238 | ATP-binding-protein-[Burkholderiales-bacterium] |
| MBE0614121.1 | AAA+ATPase+wHTH+Alpha+Beta-domain+wHTH1+wHTH2+wHTH3+RRM | Type-1_BR.RM.like.systems | <-DUF499*<-METHYLASE<-TUDOR+SF1-SF2-Helicase+iREase+CTD+REase | 1106 | Bacteria>Pseudomonadota>Betaproteobacteria>Burkholderiales>unclassified_Burkholderiales | GCA_014860445.1 | Burkholderiales_bacterium | IH604_10685 | 16325..19645 | 1891238 | ATP-binding-protein-[Burkholderiales-bacterium] |
| MCX7674148.1 | AAA+ATPase+wHTH+Alpha+Beta-domain+wHTH1+wHTH2+wHTH3+RRM | Type-1_BR.RM.like.systems | NTP_transf_2->HEPN->Mbetalac->tRNA->?-><-DUF499*<-Schlafen+HATPase+HTH<-METHYLASE<-?<-TOPRIM<-OLD-ABC<-METHYLASE | 1093 | Bacteria>Pseudomonadota>Betaproteobacteria>Nitrosomonadales>Thiobacillaceae | GCA_026419045.1 | Thiobacillaceae_bacterium | N2Z63_11245 | 10361..13642 | 2528409 | MAG:-DUF499-domain-containing-protein-[Thiobacillaceae-bacterium] |
| GJQ55613.1 | AAA+ATPase+wHTH+Alpha+Beta-domain+wHTH1+wHTH2+wHTH3+RRM | Type-1_BR.RM.like.systems | Phage_lysozyme->HTH_3->Radical-SAM-><-?<-DUF499*<-METHYLASE<-?<-SMODS<-TUDOR+SF1-SF2-Helicase+iREase+CTD+REase\|\|PadR-HTH-> | 1093 | Bacteria>Pseudomonadota>Betaproteobacteria>Rhodocyclales>Rhodocyclaceae | GCA_022181025.1 | Rhodocyclaceae_bacterium | HKUEN07_21820 | 170222..173503 | 1898103 | hypothetical-protein-HKUEN07_21820-[Rhodocyclaceae-bacterium] |
| MBI5911562.1 | AAA+ATPase+wHTH+Alpha+Beta-domain+wHTH1+wHTH2+wHTH3+RRM | Type-1_BR.RM.like.systems | <-DOC+HTH\|\|TUDOR+SF1-SF2-Helicase+iREase+CTD+REase->AAA-ATPase+REase->?->?->METHYLASE->DUF499*-><-?<-?<-Aminotran_4 | 1090 | Bacteria>Pseudomonadota>Betaproteobacteria>unclassified_Betaproteobacteria | GCA_016234705.1 | Betaproteobacteria_bacterium | HY848_16630 | 24102..27374 | 1891241 | ATP-binding-protein-[Betaproteobacteria-bacterium] |
| OGA30960.1 | AAA+ATPase+wHTH+Alpha+Beta-domain+wHTH1+wHTH2+wHTH3+RRM | Type-1_BR.RM.like.systems | TUDOR+SF1-SF2-Helicase+iREase+CTD+REase->PHP+SbcC->Schlafen+HATPase+HTH->METHYLASE->DOC+HTH->DUF499*-><-?\|\|?->?-><-?<-?\|\|SPOUT-> | 1081 | Bacteria>Pseudomonadota>Betaproteobacteria>unclassified_Betaproteobacteria | GCA_001771975.1 | Betaproteobacteria_bacterium_RIFCSPLOWO2_02_FULL_68_150 | A3I65_01690 | 23801..27046 | 1797493 | hypothetical-protein-A3I65_01690-[Betaproteobacteria-bacterium-RIFCSPLOWO2_02_FULL_68_150] |
| OGA54123.1 | AAA+ATPase+wHTH+Alpha+Beta-domain+wHTH1+wHTH2+wHTH3+RRM | Type-1_BR.RM.like.systems | <-DUF499*<-METHYLASE<-PIN<-TUDOR+SF1-SF2-Helicase+iREase+CTD+REase\|\|?->?->MqsA_antitoxin+HTH_3-> | 1091 | Bacteria>Pseudomonadota>Betaproteobacteria>unclassified_Betaproteobacteria | GCA_001772005.1 | Betaproteobacteria_bacterium_RIFCSPLOWO2_12_FULL_62_58 | A3F74_24210 | 1834..5109 | 1797497 | hypothetical-protein-A3F74_24210-[Betaproteobacteria-bacterium-RIFCSPLOWO2_12_FULL_62_58] |
| MPZ45705.1 | AAA+ATPase+wHTH+Alpha+Beta-domain+wHTH1+wHTH2+wHTH3+RRM | Type-1_BR.RM.like.systems | AAA-ATPase+REase->METHYLASE->DUF499*-> | 1088 | Bacteria>Pseudomonadota>Betaproteobacteria>unclassified_Betaproteobacteria | GCA_009377585.1 | Betaproteobacteria_bacterium | GEV05_20400 | 5826..9092 | 1891241 | DUF499-domain-containing-protein-[Betaproteobacteria-bacterium] |
| MBI3370955.1 | AAA+ATPase+wHTH+Alpha+Beta-domain+wHTH1+wHTH2+wHTH3+RRM | Type-1_BR.RM.like.systems | <-OLD-ABC+TOPRIM<-?<-DUF499*<-METHYLASE | 1089 | Bacteria>Pseudomonadota>Betaproteobacteria>unclassified_Betaproteobacteria | GCA_016195915.1 | Betaproteobacteria_bacterium | HY017_04235 | 112207..115476 | 1891241 | ATP-binding-protein-[Betaproteobacteria-bacterium] |
| GHU11234.1 | AAA+ATPase+wHTH+Alpha+Beta-domain+wHTH1+wHTH2+wHTH3+RRM | Type-1_BR.RM.like.systems | JetD-><-DUF499*<-METHYLASE-p | 1094 | Bacteria>Pseudomonadota>Betaproteobacteria>unclassified_Betaproteobacteria | GCA_026000595.1 | Betaproteobacteria_bacterium | FACS1894185_4120 | 8094..11378 | 1891241 | hypothetical-protein-FACS1894185_4120-[Betaproteobacteria-bacterium] |
| WP_096461953.1 | AAA+ATPase+wHTH+Alpha+Beta-domain+wHTH1+wHTH2+wHTH3+RRM | Type-1_BR.RM.like.systems | <-TIR<-AbiEii<-wHTH+AbiEi\|\|?-><-?<-DUF499*<-METHYLASE<-?<-SMODS<-TUDOR+SF1-SF2-Helicase+iREase+CTD+REase\|\|?->HKD-DNase-> | 1093 | Bacteria>Pseudomonadota>Gammaproteobacteria>Acidiferrobacterales>Acidiferrobacteraceae | GCF_002355415.1 | Sulfurifustis_variabilis | SVA_RS14790 | 3048682..3051963 | 1675686 | DUF499-domain-containing-protein-[Sulfurifustis-variabilis] |
| WP_157817756.1 | AAA+ATPase+wHTH+Alpha+Beta-domain+wHTH1+wHTH2+wHTH3+RRM | Type-1_BR.RM.like.systems | <-Abhydrolase_4\|\|DDE_Tnp_IS1-><-DUF29<-DUF499*<-METHYLASE<-TOPRIM<-OLD-ABC<-TUDOR+SF1-SF2-Helicase+iREase+CTD+REase | 1094 | Bacteria>Pseudomonadota>Gammaproteobacteria>Chromatiales>Chromatiaceae | GCF_002813775.1 | Candidatus_Thiodictyon_syntrophicum | THSYN_RS17695 | 4223024..4226308 | 1166950 | DUF499-domain-containing-protein-[Candidatus-Thiodictyon-syntrophicum] |
| AUB82572.1 | AAA+ATPase+wHTH+Alpha+Beta-domain+wHTH1+wHTH2+wHTH3+RRM | Type-1_BR.RM.like.systems | <-Abhydrolase_4\|\|DDE_Tnp_IS1-><-DUF29<-DUF499*<-METHYLASE<-TOPRIM<-OLD-ABC<-TUDOR+SF1-SF2-Helicase+iREase+CTD+REase | 1083 | Bacteria>Pseudomonadota>Gammaproteobacteria>Chromatiales>Chromatiaceae | GCA_002813775.1 | Candidatus_Thiodictyon_syntrophicum | THSYN_17560 | 4223024..4226275 | 1166950 | hypothetical-protein-THSYN_17560-[Candidatus-Thiodictyon-syntrophicum] |
| WP_120796693.1 | AAA+ATPase+wHTH+Alpha+Beta-domain+wHTH1+wHTH2+wHTH3+RRM | Type-1_BR.RM.like.systems | TUDOR+SF1-SF2-Helicase+iREase+CTD+REase->METHYLASE->Schlafen+HATPase+HTH->DUF499*->?->?->?->CHTH-> | 1093 | Bacteria>Pseudomonadota>Gammaproteobacteria>Chromatiales>Chromatiaceae | GCF_003634315.1 | Thiocapsa_rosea | BDD21_RS07920 | 1738895..1742176 | 69360 | DUF499-domain-containing-protein-[Thiocapsa-rosea] |
| WP_153975970.1 | AAA+ATPase+wHTH+Alpha+Beta-domain+wHTH1+wHTH2+wHTH3+RRM | Type-1_BR.RM.like.systems | ABC-ATPase->?->?-><-DUF499*<-METHYLASE<-TUDOR+SF1-SF2-Helicase+iREase+CTD+REase | 1093 | Bacteria>Pseudomonadota>Gammaproteobacteria>Chromatiales>Chromatiaceae | GCF_009664085.1 | Thermochromatium_tepidum | E6P07_RS12865 | 2782709..2785990 | 1050 | DUF499-domain-containing-protein-[Thermochromatium-tepidum] |
| MBF2761740.1 | AAA+ATPase+wHTH+Alpha+Beta-domain+wHTH1+wHTH2+wHTH3+RRM | Type-1_BR.RM.like.systems | <-TOPRIM<-OLD-ABC<-N6_N4_MTase<-DUF499*<-METHYLASE<-METHYLASE-p<-TOPRIM<-OLD-ABC<-?<-N6_N4_MTase | 1092 | Bacteria>Pseudomonadota>Gammaproteobacteria>Chromatiales>Ectothiorhodospiraceae | GCA_015278165.1 | Ectothiorhodospiraceae_bacterium_AqS1 | ISN28_16015 | 2939..6217 | 1725326 | ATP-binding-protein-[Ectothiorhodospiraceae-bacterium-AqS1] |
| TVR61882.1 | AAA+ATPase+wHTH+Alpha+Beta-domain+wHTH1+wHTH2+wHTH3+RRM | Type-1_BR.RM.like.systems | <-DUF499*<-PIN<-DUF433<-METHYLASE-p | 1105 | Bacteria>Pseudomonadota>Gammaproteobacteria>Gammaproteobacteria_incertae_sedis>Candidatus_Competibacteraceae | GCA_007695245.1 | Candidatus_Competibacteraceae_bacterium | EA420_10930 | 75..3392 | 2053538 | ATP-binding-protein-[Candidatus-Competibacteraceae-bacterium] |
| MCY4450795.1 | AAA+ATPase+wHTH+Alpha+Beta-domain+wHTH1+wHTH2+wHTH3+RRM | Type-1_BR.RM.like.systems | <-STYKinase-like<-HTH<-DUF499*<-PIN<-?<-METHYLASE\|\|HicB_lk_antitox-><-Schlafen+HATPase+HTH<-HEPN | 1074 | Bacteria>Pseudomonadota>Gammaproteobacteria>Immundisolibacterales>unclassified_Immundisolibacterales | - | Immundisolibacterales_bacterium | OXC01_02485 | 119191..122415 | 2764111 | MAG:-DUF499-domain-containing-protein-[Immundisolibacterales-bacterium] |
| MBX6421242.1 | AAA+ATPase+wHTH+Alpha+Beta-domain+wHTH1+wHTH2+wHTH3+RRM | Type-1_BR.RM.like.systems | <-HIN-HTH\|\|<-DUF499*<-METHYLASE<-?<-?<-TUDOR+SF1-SF2-Helicase+iREase+CTD+REase<-HDRnase | 1093 | Bacteria>Pseudomonadota>Gammaproteobacteria>Nevskiales>Nevskiaceae | GCA_019689635.1 | Nevskia_sp. | IRZ06_09590 | 3085..6366 | 1929292 | ATP-binding-protein-[Nevskia-sp.] |
| MCY4286579.1 | AAA+ATPase+wHTH+Alpha+Beta-domain+wHTH1+wHTH2+wHTH3+RRM | Type-1_BR.RM.like.systems | PolB->ParB+HNH->METHYLASE->?->OLD-ABC->TOPRIM->DUF499*->?->?-><-?<-?<-ABC-ATPase+SbcC | 1107 | Bacteria>Pseudomonadota>Gammaproteobacteria>Thiotrichales>unclassified_Thiotrichales | GCA_026709935.1 | Thiotrichales_bacterium | OXC65_14725 | 7747..11070 | 2026796 | MAG:-DUF499-domain-containing-protein-[Thiotrichales-bacterium] |
| MCG3203220.1 | AAA+ATPase+wHTH+Alpha+Beta-domain+wHTH1+wHTH2+wHTH3+RRM | Type-1_BR.RM.like.systems | <-NTP_transf<-MORC+DUF3684+REase<-DUF499*<-METHYLASE<-Schlafen+HATPase+HTH<-PHP+SbcC<-TUDOR+SF1-SF2-Helicase+iREase+CTD+REase\|\|MerR-HTH-> | 1092 | Bacteria>Pseudomonadota>Gammaproteobacteria>unclassified_Gammaproteobacteria | GCA_022071965.1 | Gammaproteobacteria_bacterium | NFCQHLIN_03153 | 6649..9927 | 1913989 | hypothetical-protein-[Gammaproteobacteria-bacterium] |
| MDE0258535.1 | AAA+ATPase+wHTH+Alpha+Beta-domain+wHTH1+wHTH2+wHTH3+RRM | Type-1_BR.RM.like.systems | <-TraI-VirD2-RCRE<-TraJ-RHH<-Phage_integrase<-DUF499*<-PIN<-?<-METHYLASE<-TUDOR+SF1-SF2-Helicase+iREase+CTD+REase | 1103 | Bacteria>Pseudomonadota>Gammaproteobacteria>unclassified_Gammaproteobacteria | GCA_028820325.1 | Gammaproteobacteria_bacterium | OXR82_09155 | 1794..5105 | 1913989 | MAG:-DUF499-domain-containing-protein-[Gammaproteobacteria-bacterium] |
| MDE0037092.1 | AAA+ATPase+wHTH+Alpha+Beta-domain+wHTH1+wHTH2+wHTH3+RRM | Type-1_BR.RM.like.systems | <-REase\|\|TUDOR+SF1-SF2-Helicase+iREase+CTD+REase->?->METHYLASE->METHYLASE-p->DUF499*->Radical-SAM-><-ACET | 1106 | Bacteria>Pseudomonadota>Gammaproteobacteria>unclassified_Gammaproteobacteria | GCA_028818795.1 | Gammaproteobacteria_bacterium | OXU77_05940 | 67818..71138 | 1913989 | MAG:-DUF499-domain-containing-protein-[Gammaproteobacteria-bacterium] |
| MDE0038699.1 | AAA+ATPase+wHTH+Alpha+Beta-domain+wHTH1+wHTH2+wHTH3+RRM | Type-1_BR.RM.like.systems | OLD-ABC->TOPRIM-><-?<-TIR\|\|TIR->METHYLASE->DUF499*-><-SF1-SF2-Helicase | 1089 | Bacteria>Pseudomonadota>Gammaproteobacteria>unclassified_Gammaproteobacteria | GCA_028818795.1 | Gammaproteobacteria_bacterium | OXU77_14270 | 23040..26309 | 1913989 | MAG:-hypothetical-protein-OXU77_14270-[Gammaproteobacteria-bacterium] |
| MCY4039693.1 | AAA+ATPase+wHTH+Alpha+Beta-domain+wHTH1+wHTH2+wHTH3+RRM | Type-1_BR.RM.like.systems | Aminotran_1_2-><-PIN<-?\|\|TUDOR+SF1-SF2-Helicase+iREase+CTD+REase->?->METHYLASE->DUF499*-> | 1089 | Bacteria>Pseudomonadota>Gammaproteobacteria>unclassified_Gammaproteobacteria | GCA_026708835.1 | Gammaproteobacteria_bacterium | OXF72_00370 | 11324..14593 | 1913989 | MAG:-hypothetical-protein-OXF72_00370-[Gammaproteobacteria-bacterium] |
| MCY4278451.1 | AAA+ATPase+wHTH+Alpha+Beta-domain+wHTH1+wHTH2+wHTH3+RRM | Type-1_BR.RM.like.systems | <-PIN<-?\|\|TUDOR+SF1-SF2-Helicase+iREase+CTD+REase->PIN->?->METHYLASE->DUF499*-><-?<-?<-AbiEii | 1093 | Bacteria>Pseudomonadota>Gammaproteobacteria>unclassified_Gammaproteobacteria | GCA_026709965.1 | Gammaproteobacteria_bacterium | OXE81_11575 | 12290..15571 | 1913989 | MAG:-hypothetical-protein-OXE81_11575-[Gammaproteobacteria-bacterium] |
| MCY4200247.1 | AAA+ATPase+wHTH+Alpha+Beta-domain+wHTH1+wHTH2+wHTH3+RRM | Type-1_BR.RM.like.systems | AbiEii->?->?-><-DUF499*<-METHYLASE<-?<-PIN<-TUDOR+SF1-SF2-Helicase+iREase+CTD+REase\|\|?->PIN-> | 1089 | Bacteria>Pseudomonadota>Gammaproteobacteria>unclassified_Gammaproteobacteria | GCA_026709225.1 | Gammaproteobacteria_bacterium | OXF31_10650 | 19603..22872 | 1913989 | MAG:-hypothetical-protein-OXF31_10650-[Gammaproteobacteria-bacterium] |
| MDD9817274.1 | AAA+ATPase+wHTH+Alpha+Beta-domain+wHTH1+wHTH2+wHTH3+RRM | Type-1_BR.RM.like.systems | <-DUF499*<-?<-TOPRIM<-OLD-ABC<-METHYLASE<-Schlafen+HATPase+HTH<-RelE-ParE | 1113 | Bacteria>Pseudomonadota>Gammaproteobacteria>unclassified_Gammaproteobacteria | GCA_028817495.1 | Gammaproteobacteria_bacterium | OXU61_03925 | 191..3532 | 1913989 | MAG:-hypothetical-protein-OXU61_03925-[Gammaproteobacteria-bacterium] |
| MCZ0945558.1 | AAA+ATPase+wHTH+Alpha+Beta-domain+wHTH1+wHTH2+wHTH3+RRM | Type-1_BR.RM.like.systems | TUDOR+SF1-SF2-Helicase+iREase+CTD+REase->?->BirA-HTH->?->TIR->METHYLASE->DUF499*-> | 1091 | Bacteria>Pseudomonadota>Gammaproteobacteria>unclassified_Gammaproteobacteria | GCA_026914145.1 | Gammaproteobacteria_bacterium | OXJ53_21095 | 11131..14406 | 1913989 | MAG:-hypothetical-protein-OXJ53_21095-[Gammaproteobacteria-bacterium] |
| MDD9864539.1 | AAA+ATPase+wHTH+Alpha+Beta-domain+wHTH1+wHTH2+wHTH3+RRM | Type-1_BR.RM.like.systems | METHYLASE->OLD-ABC->TOPRIM->DUF499*-><-?<-?\|\|?->?->?->ABC-ATPase+SbcC-> | 1113 | Bacteria>Pseudomonadota>Gammaproteobacteria>unclassified_Gammaproteobacteria | GCA_028817305.1 | Gammaproteobacteria_bacterium | OXU54_08205 | 4537..7878 | 1913989 | MAG:-DUF499-domain-containing-protein-[Gammaproteobacteria-bacterium] |
| MDE0441671.1 | AAA+ATPase+wHTH+Alpha+Beta-domain+wHTH1+wHTH2+wHTH3+RRM | Type-1_BR.RM.like.systems | <-TIR\|\|TIR->METHYLASE->DUF499*-><-SF1-SF2-Helicase | 1089 | Bacteria>Pseudomonadota>Gammaproteobacteria>unclassified_Gammaproteobacteria | GCA_028821355.1 | Gammaproteobacteria_bacterium | OXL38_06080 | 8041..11310 | 1913989 | MAG:-hypothetical-protein-OXL38_06080-[Gammaproteobacteria-bacterium] |
| HFD86265.1 | AAA+ATPase+wHTH+Alpha+Beta-domain+wHTH1+wHTH2+wHTH3+RRM | Type-1_BR.RM.like.systems | METHYLASE-p->DUF499*->?->UbE1+JAB-> | 1106 | Bacteria>Pseudomonadota>Gammaproteobacteria>unclassified_Gammaproteobacteria | GCA_011322685.1 | Gammaproteobacteria_bacterium | ENJ35_01145 | 1296..4616 | 1913989 | ATP-binding-protein-[Gammaproteobacteria-bacterium] |
| WP_259304308.1 | AAA+ATPase+wHTH+Alpha+Beta-domain+wHTH1+wHTH2+wHTH3+RRM | Type-1_BR.RM.like.systems | HTH_3->Radical-SAM-><-DUF772+DDE_Tnp_1<-Methylase<-HIN-HTH<-?<-DUF499*<-METHYLASE<-?<-SMODS<-TUDOR+SF1-SF2-Helicase+iREase+CTD+REase-p<-TUDOR+SF1-SF2-Helicase+iREase+CTD+REase\|\|PadR-HTH-> | 1093 | Bacteria>Pseudomonadota>Gammaproteobacteria>Xanthomonadales>Xanthomonadaceae | GCF_024704365.1 | Thermomonas_sp._S9 | LJB71_RS11670 | 2331832..2335113 | 2885203 | DUF499-domain-containing-protein-[Thermomonas-sp.-S9] |
| WP_024892241.1 | AAA+ATPase+wHTH+Alpha+Beta-domain+wHTH1+wHTH2+wHTH3+RRM | Type-1_BR.RM.like.systems | Phage_lysozyme->HTH_3->Radical-SAM-><-DUF772+DDE_Tnp_1<-Methylase<-?<-DUF499*<-METHYLASE<-?<-SMODS<-TUDOR+SF1-SF2-Helicase+iREase+CTD+REase\|\|PadR-HTH->AbiEi_4-> | 1093 | Bacteria>Pseudomonadota>Gammaproteobacteria>Xanthomonadales>Xanthomonadaceae | GCF_000559025.1 | Luteimonas_huabeiensis | Z164_RS0119095 | 27437..30718 | 1244513 | DUF499-domain-containing-protein-[Luteimonas-huabeiensis] |
| WP_218535682.1 | AAA+ATPase+wHTH+Alpha+Beta-domain+wHTH1+wHTH2+wHTH3+RRM | Type-1_BR.RM.like.systems | ZnR+TOPRIM+D5ATPase-><-PadR-HTH\|\|TUDOR+SF1-SF2-Helicase+iREase+CTD+REase->SMODS->?->METHYLASE->DUF499*->TIR->Methylase-><-Radical-SAM<-HTH_3<-Phage_lysozyme | 1093 | Bacteria>Pseudomonadota>Gammaproteobacteria>Xanthomonadales>Xanthomonadaceae | GCF_019201325.1 | Xanthomonas_euvesicatoria | KWH42_RS16780 | 68650..71931 | 456327 | DUF499-domain-containing-protein-[Xanthomonas-campestris] |
| HHW77449.1 | AAA+ATPase+wHTH+Alpha+Beta-domain+wHTH1+wHTH2+wHTH3+RRM | Type-1_BR.RM.like.systems | METHYLASE->DUF499*->DDE_Tnp_1-><-ADH_N+ADH_zinc_N<-?<-?<-?\|\|MODE-HTH+PBPII-> | 1092 | Bacteria>Pseudomonadota>Gammaproteobacteria>Xanthomonadales>Xanthomonadaceae | GCA_012840695.1 | Xanthomonadaceae_bacterium | GX399_10570 | 3262..6540 | 1926873 | ATP-binding-protein-[Xanthomonadaceae-bacterium] |
| MBU4372115.1 | AAA+ATPase+wHTH+Alpha+Beta-domain+wHTH1+wHTH2+wHTH3+RRM | Type-1_BR.RM.like.systems | Peptidase_M24->?->Aminotran_1_2->?->?->DNA_pol3_delta+DNA_pol3_gamma3-><-DUF499*<-AAA-ATPase+REase<-METHYLASE-p | 1098 | Bacteria>Pseudomonadota>unclassified_Pseudomonadota | GCA_018896355.1 | Pseudomonadota_bacterium | KKD68_09540 | 35625..38921 | 1977087 | DUF499-domain-containing-protein-[Proteobacteria-bacterium] |
| MBS0213168.1 | AAA+ATPase+wHTH+Alpha+Beta-domain+wHTH1+wHTH2+wHTH3+RRM | Type-1_BR.RM.like.systems | ZnR+TOPRIM+D5ATPase->TUDOR+SF1-SF2-Helicase+iREase+CTD+REase->SMODS->?->wHTH+REase->METHYLASE->DUF499*->REase->?->ParB+HNH-p-> | 1093 | Bacteria>Pseudomonadota>unclassified_Pseudomonadota | GCA_018242445.1 | Pseudomonadota_bacterium | JSR26_08340 | 14587..17868 | 1977087 | ATP-binding-protein-[Proteobacteria-bacterium] |
| MBU4582289.1 | AAA+ATPase+wHTH+Alpha+Beta-domain+wHTH1+wHTH2+wHTH3+RRM | Type-1_BR.RM.like.systems | AAA-ATPase+REase->DUF499*-><-DNA_pol3_delta+DNA_pol3_gamma3<-?<-?<-Aminotran_1_2<-?<-Peptidase_M24 | 1098 | Bacteria>Pseudomonadota>unclassified_Pseudomonadota | GCA_018902925.1 | Pseudomonadota_bacterium | KKH02_07770 | 1387..4683 | 1977087 | DUF499-domain-containing-protein-[Proteobacteria-bacterium] |
| MCX5809780.1 | AAA+ATPase+wHTH+Alpha+Beta-domain+wHTH1+wHTH2+wHTH3+RRM | Type-1_BR.RM.like.systems | TUDOR+SF1-SF2-Helicase+iREase+CTD+REase->METHYLASE->Schlafen+HATPase+HTH->DUF499*->ParB-> | 1105 | Bacteria>Pseudomonadota>unclassified_Pseudomonadota | GCA_026388245.1 | Pseudomonadota_bacterium | NTX36_10495 | 9204..12521 | 1977087 | MAG:-DUF499-domain-containing-protein-[Proteobacteria-bacterium] |
| MBU4121125.1 | AAA+ATPase+wHTH+Alpha+Beta-domain+wHTH1+wHTH2+wHTH3+RRM | Type-1_BR.RM.like.systems | DUF499*-><-DNA_pol3_delta+DNA_pol3_gamma3<-?<-?<-?<-Aminotran_1_2 | 871 | Bacteria>Pseudomonadota>unclassified_Pseudomonadota | GCA_018899395.1 | Pseudomonadota_bacterium | KKA48_06190 | 1..2617 | 1977087 | DUF499-domain-containing-protein-[Proteobacteria-bacterium] |
| MBU0651701.1 | AAA+ATPase+wHTH+Alpha+Beta-domain+wHTH1+wHTH2+wHTH3+RRM | Type-1_BR.RM.like.systems | DUF499*-><-DNA_pol3_delta+DNA_pol3_gamma3 | 806 | Bacteria>Pseudomonadota>unclassified_Pseudomonadota | GCA_018814185.1 | Pseudomonadota_bacterium | KKG96_02330 | 1..2421 | 1977087 | DUF499-domain-containing-protein-[Proteobacteria-bacterium] |
| MBP7275753.1 | AAA+ATPase+wHTH+Alpha+Beta-domain+wHTH1+wHTH2+wHTH3+RRM | Type-1_BR.RM.like.systems | <-DUF499*<-?<-METHYLASE<-PIN<-?<-REase | 1001 | Bacteria>PVC_group>Kiritimatiellota>Kiritimatiellia>unclassified_Kiritimatiellia | GCA_017996315.1 | Kiritimatiellia_bacterium | KBA51_06085 | 1..3004 | 2212473 | ATP-binding-protein-[Kiritimatiellae-bacterium] |
| OQC13494.1 | AAA+ATPase+wHTH+Alpha+Beta-domain+wHTH1+wHTH2+wHTH3+RRM | Type-1_BR.RM.like.systems | <-N6_N4_MTase<-HicB<-MPTase<-HTH<-?<-DUF499*<-METHYLASE<-TUDOR+SF1-SF2-Helicase+iREase+CTD+REase<-tRNA<-?<-?<-?<-S1+RNase+2C | 1109 | Bacteria>PVC_group>Lentisphaerota>unclassified_Lentisphaerota | GCA_002070945.1 | Lentisphaerae_bacterium_ADurb.Bin082 | BWX73_02299 | 9926..13255 | 1852899 | MAG:-hypothetical-protein-BWX73_02299-[Lentisphaerae-bacterium-ADurb.Bin082] |
| MBT5607699.1 | AAA+ATPase+wHTH+Alpha+Beta-domain+wHTH1+wHTH2+wHTH3+RRM | Type-1_BR.RM.like.systems | PIN->METHYLASE->DUF499*->?->DUF6361->HKD-DNase->SF1-SF2-Helicase->ORC-CDC6+HTH-> | 1109 | Bacteria>PVC_group>Lentisphaerota>unclassified_Lentisphaerota | GCA_018666605.1 | Lentisphaerota_bacterium | HOJ57_17295 | 4653..7982 | 1932692 | ATP-binding-protein-[Lentisphaerae-bacterium] |
| NUO09865.1 | AAA+ATPase+wHTH+Alpha+Beta-domain+wHTH1+wHTH2+wHTH3+RRM | Type-1_BR.RM.like.systems | <-DUF499*<-ORC-CDC6+REase<-METHYLASE<-TUDOR+SF1-SF2-Helicase+iREase+CTD+REase<-Aldedh | 1099 | Bacteria>PVC_group>Planctomycetota>Candidatus_Brocadiia>Candidatus_Brocadiales | GCA_013360945.1 | Candidatus_Brocadia_sp. | HUU08_14505 | 241..3540 | 2172549 | ATP-binding-protein-[Candidatus-Brocadia-sp.] |
| MBX3386460.1 | AAA+ATPase+wHTH+Alpha+Beta-domain+wHTH1+wHTH2+wHTH3+RRM | Type-1_BR.RM.like.systems | <-HTH+MPTase<-?\|\|TUDOR+SF1-SF2-Helicase+iREase+CTD+REase->SF1-SF2-Helicase->METHYLASE->DUF499*->?-><-Phage_integrase<-?<-?<-?\|\|MoxR-AAA-> | 1103 | Bacteria>PVC_group>Planctomycetota>Phycisphaerae>Phycisphaerales | GCA_019636535.1 | Phycisphaeraceae_bacterium | KF768_07805 | 11131..14442 | 2026777 | ATP-binding-protein-[Phycisphaeraceae-bacterium] |
| MBX3391865.1 | AAA+ATPase+wHTH+Alpha+Beta-domain+wHTH1+wHTH2+wHTH3+RRM | Type-1_BR.RM.like.systems | <-ParA-Soj-PloopNTPase\|\|?->?-><-?<-N6_N4_MTase<-DUF499*<-PIN<-METHYLASE<-?<-TUDOR+SF1-SF2-Helicase+iREase+CTD+REase | 1114 | Bacteria>PVC_group>Planctomycetota>Phycisphaerae>Phycisphaerales | GCA_019636515.1 | Phycisphaeraceae_bacterium | KF787_04385 | 1038912..1042256 | 2026777 | ATP-binding-protein-[Phycisphaeraceae-bacterium] |
| QOJ18491.1 | AAA+ATPase+wHTH+Alpha+Beta-domain+wHTH1+wHTH2+wHTH3+RRM | Type-1_BR.RM.like.systems | Methyltransf_11->?->\|\|?->TUDOR+SF1-SF2-Helicase+iREase+CTD+REase->?->DUF499*->N6_N4_MTase->?-><-?<-REase | 1114 | Bacteria>PVC_group>Planctomycetota>Phycisphaerae>Phycisphaerales | GCA_015709595.1 | Phycisphaeraceae_bacterium | HRU76_13225 | 3115373..3118717 | 2026777 | MAG:-ATP-binding-protein-[Phycisphaeraceae-bacterium] |
| MCG3177990.1 | AAA+ATPase+wHTH+Alpha+Beta-domain+wHTH1+wHTH2+wHTH3+RRM | Type-1_BR.RM.like.systems | Radical-SAM-><-DOC+HTH\|\|HTH->STYKinase-like->SF1-SF2-Helicase<-DUF499*<-METHYLASE<-HEPN<-TUDOR+SF1-SF2-Helicase+iREase+CTD+REase | 1112 | Bacteria>PVC_group>Planctomycetota>Phycisphaerae>unclassified_Phycisphaerae | GCA_022072085.1 | Phycisphaerae_bacterium | BIFFINMI_00313 | 58540..61878 | 2026778 | hypothetical-protein-[Phycisphaerae-bacterium] |
| MCK6483011.1 | AAA+ATPase+wHTH+Alpha+Beta-domain+wHTH1+wHTH2+wHTH3+RRM | Type-1_BR.RM.like.systems | N6_N4_MTase->?->?-><-ParB<-GDResolvase+Recombinase+ZnR<-RAMA<-DUF499*<-TOPRIM<-OLD-ABC<-METHYLASE<-?<-?<-TUDOR+SF1-SF2-Helicase+iREase+CTD+REase | 1108 | Bacteria>PVC_group>Planctomycetota>Phycisphaerae>unclassified_Phycisphaerae | GCA_023150595.1 | Phycisphaerae_bacterium | L6R00_02615 | 131087..134413 | 2026778 | DUF499-domain-containing-protein-[Phycisphaerae-bacterium] |
| MCG3126049.1 | AAA+ATPase+wHTH+Alpha+Beta-domain+wHTH1+wHTH2+wHTH3+RRM | Type-1_BR.RM.like.systems | HEPN+REase->METHYLASE->ParB+HNH->DUF499*->?->SF1-SF2-Helicase->?->?->CASPASE+APATPase+TPR+TPR+TPR-> | 1107 | Bacteria>PVC_group>Planctomycetota>Phycisphaerae>unclassified_Phycisphaerae | GCA_022072445.1 | Phycisphaerae_bacterium | CHACPFDD_00877 | 44646..47969 | 2026778 | hypothetical-protein-[Phycisphaerae-bacterium] |
| GJQ28342.1 | AAA+ATPase+wHTH+Alpha+Beta-domain+wHTH1+wHTH2+wHTH3+RRM | Type-1_BR.RM.like.systems | Phage_integrase-><-?<-?<-?<-?<-N6_N4_MTase<-DUF499*<-METHYLASE<-HEPN<-TUDOR+SF1-SF2-Helicase+iREase+CTD+REase | 1114 | Bacteria>PVC_group>Planctomycetota>Phycisphaerae>unclassified_Phycisphaerae | GCA_022180725.1 | Phycisphaerae_bacterium | HBSAPP03_02260 | 258751..262095 | 2026778 | hypothetical-protein-HBSAPP03_02260-[Phycisphaerae-bacterium] |
| MBN2562788.1 | AAA+ATPase+wHTH+Alpha+Beta-domain+wHTH1+wHTH2+wHTH3+RRM | Type-1_BR.RM.like.systems | <-XIS-HTH<-?<-McrC-NTD+REase<-?<-?<-DUF499*<-METHYLASE | 1108 | Bacteria>PVC_group>Planctomycetota>Phycisphaerae>unclassified_Phycisphaerae | GCA_016935885.1 | Phycisphaerae_bacterium | JXQ75_17835 | 5902..9228 | 2026778 | DUF499-domain-containing-protein-[Phycisphaerae-bacterium] |
| MBE0537368.1 | AAA+ATPase+wHTH+Alpha+Beta-domain+wHTH1+wHTH2+wHTH3+RRM | Type-1_BR.RM.like.systems | TUDOR+SF1-SF2-Helicase+iREase+CTD+REase->?->REase->DUF499*-> | 1115 | Bacteria>PVC_group>Planctomycetota>Phycisphaerae>unclassified_Phycisphaerae | GCA_014859965.1 | Phycisphaerae_bacterium | IH624_17025 | 11714..15061 | 2026778 | ATP-binding-protein-[Phycisphaerae-bacterium] |
| GIW78771.1 | AAA+ATPase+wHTH+Alpha+Beta-domain+wHTH1+wHTH2+wHTH3+RRM | Type-1_BR.RM.like.systems | <-Phage_int_SAM_4+Phage_integrase\|\|tRNA-><-?<-XIS-HTH<-?<-?<-DUF499*<-TOPRIM<-OLD-ABC<-METHYLASE<-?<-TUDOR+SF1-SF2-Helicase+iREase+CTD+REase<-SF1-SF2-Helicase-p | 1113 | Bacteria>PVC_group>Planctomycetota>Planctomycetia>Gemmatales | GCA_026004675.1 | Gemmatales_bacterium | KatS3mg105_0578 | 670726..674067 | 2838789 | MAG:-hypothetical-protein-KatS3mg105_0578-[Gemmatales-bacterium] |
| MBX3417770.1 | AAA+ATPase+wHTH+Alpha+Beta-domain+wHTH1+wHTH2+wHTH3+RRM | Type-1_BR.RM.like.systems | Pkinase-><-?<-TPR+TPR<-?\|\|SIGMA-HTH+SIGMA-HTH-><-DUF499*<-?<-DOC+HTH<-METHYLASE<-?<-DDE_Tnp_1_assoc+DDE_Tnp_1<-TUDOR+SF1-SF2-Helicase+iREase+CTD+REase | 1121 | Bacteria>PVC_group>Planctomycetota>Planctomycetia>Pirellulales | GCA_019636375.1 | Pirellulaceae_bacterium | KF851_09215 | 763861..767226 | 2699754 | ATP-binding-protein-[Pirellulaceae-bacterium] |
| MCL4190101.1 | AAA+ATPase+wHTH+Alpha+Beta-domain+wHTH1+wHTH2+wHTH3+RRM | Type-1_BR.RM.like.systems | TUDOR+SF1-SF2-Helicase+iREase+CTD+REase->?->METHYLASE->DOC+HTH->wHTH+REase->DUF499*->SbcC->?->?->LZ_Tnp_IS66+zf-IS66+DDE_Tnp_IS66+DDE_Tnp_IS66_C-> | 1115 | Bacteria>PVC_group>Planctomycetota>Planctomycetia>Pirellulales | GCA_023382995.1 | Thermoguttaceae_bacterium | KJZ87_00035 | 12107..15454 | 2691551 | DUF499-domain-containing-protein-[Thermoguttaceae-bacterium] |
| WP_146679188.1 | AAA+ATPase+wHTH+Alpha+Beta-domain+wHTH1+wHTH2+wHTH3+RRM | Type-1_BR.RM.like.systems | <-DUF772+DDE_Tnp_1+DDE_Tnp_1_6\|\|?->?->METHYLASE->DUF499->DDE_Tnp_1_4->DUF499*-><-?<-?<-?<-?\|\|?->ABC-ATPase+SbcC-> | 820 | Bacteria>PVC_group>Planctomycetota>Planctomycetia>Pirellulales | GCF_001610875.1 | Pirellula_sp._SH-Sr6A | VN12_RS23355 | 6015720..6018182 | 1632865 | hypothetical-protein-[Pirellula-sp.-SH-Sr6A] |
| MBN1912210.1 | AAA+ATPase+wHTH+Alpha+Beta-domain+wHTH1+wHTH2+wHTH3+RRM | Type-1_BR.RM.like.systems | METHYLASE->wHTH+REase->DUF499*-><-Radical-SAM | 1103 | Bacteria>PVC_group>Planctomycetota>Planctomycetia>Pirellulales | GCA_016931075.1 | Pirellulales_bacterium | JW818_20990 | 4913..8224 | 2715253 | ATP-binding-protein-[Pirellulales-bacterium] |
| NLY99303.1 | AAA+ATPase+wHTH+Alpha+Beta-domain+wHTH1+wHTH2+wHTH3+RRM | Type-1_BR.RM.like.systems | <-DUF499*<-?<-TOPRIM<-OLD-ABC<-METHYLASE-p | 1112 | Bacteria>PVC_group>Planctomycetota>Planctomycetia>Pirellulales | GCA_012744395.1 | Pirellulaceae_bacterium | GXY25_02065 | 2321..5659 | 2699754 | ATP-binding-protein-[Pirellulaceae-bacterium] |
| MDD4270717.1 | AAA+ATPase+wHTH+Alpha+Beta-domain+wHTH1+wHTH2+wHTH3+RRM | Type-1_BR.RM.like.systems | METHYLASE->OLD-ABC->TOPRIM->?->DUF499*-> | 1112 | Bacteria>PVC_group>Planctomycetota>Planctomycetia>Pirellulales | GCA_028703175.1 | Thermoguttaceae_bacterium | PHN77_21030 | 4016..7354 | 2691551 | MAG:-DUF499-domain-containing-protein-[Thermoguttaceae-bacterium] |
| GHT10305.1 | AAA+ATPase+wHTH+Alpha+Beta-domain+wHTH1+wHTH2+wHTH3+RRM | Type-1_BR.RM.like.systems | METHYLASE-p->METHYLASE-p->ZDNA-HTH->DUF499*->Peptidase->Schlafen->BirA-HTH-> | 1093 | Bacteria>PVC_group>Planctomycetota>Planctomycetia>Planctomycetales | GCA_026001785.1 | Planctomycetales_bacterium | FACS1894170_02190 | 13810..17091 | 2053591 | hypothetical-protein-FACS1894170_02190-[Planctomycetales-bacterium] |
| GHT30760.1 | AAA+ATPase+wHTH+Alpha+Beta-domain+wHTH1+wHTH2+wHTH3+RRM | Type-1_BR.RM.like.systems | TUDOR+SF1-SF2-Helicase+iREase+CTD+REase-p->SbcC->METHYLASE->ZDNA-HTH->?->DUF499*-> | 1094 | Bacteria>PVC_group>Planctomycetota>Planctomycetia>Planctomycetales | GCA_026001885.1 | Planctomycetales_bacterium | FACS1894214_1230 | 7846..11130 | 2053591 | hypothetical-protein-FACS1894214_1230-[Planctomycetales-bacterium] |
| MDA8378780.1 | AAA+ATPase+wHTH+Alpha+Beta-domain+wHTH1+wHTH2+wHTH3+RRM | Type-1_BR.RM.like.systems | TUDOR+SF1-SF2-Helicase+iREase+CTD+REase->RNase->?->METHYLASE->PIN->DUF499*-><-AAA_21 | 1112 | Bacteria>PVC_group>Planctomycetota>Planctomycetia>unclassified_Planctomycetia | GCA_028068345.1 | Planctomycetia_bacterium | MOZ50_17440 | 9415..12753 | 2052181 | MAG:-DUF499-domain-containing-protein-[Planctomycetia-bacterium] |

<table>
<tr><th>Protein-ID</th><th>DUF499-architecture</th><th>System</th><th>Gene-Neighbourhoods</th><th>Length</th><th>Taxonomy</th><th>Assembly-ID</th><th>Strain</th><th>Locus-Tag</th><th>Genomic-coordinates</th><th>TaxID</th><th>Fasta-defline</th></tr>
<tr><td>QOJ03670.1</td><td>AAA+ATPase+wHTH+Alpha+Beta-domain+wHTH1+wHTH2+wHTH3+RRM</td><td>Type-1_BR.RM.like.systems</td><td>HEPN+REase->?->METHYLASE->?->DUF499*-><-?||?->?->?->XIS-HTH-></td><td>1107</td><td>Bacteria>PVC_group>Planctomycetota>Planctomycetia>unclassified_Planctomycetia</td><td>GCA_015200345.1</td><td>Planctomycetia_bacterium</td><td>HRU71_09295</td><td>2265298..2268621</td><td>2052181</td><td>MAG:-ATP-binding-protein-[Planctomycetia-bacterium]</td></tr>
<tr><td>MBS0186570.1</td><td>AAA+ATPase+wHTH+Alpha+Beta-domain+wHTH1+wHTH2+wHTH3+RRM</td><td>Type-1_BR.RM.like.systems</td><td>TUDOR+SF1-SF2-Helicase+iREase+CTD+REase->SF1-SF2-Helicase->METHYLASE->METHYLASE->DUF499*->DUF6361->HKD-DNase->SF1-SF2-Helicase-><-REase</td><td>1104</td><td>Bacteria>PVC_group>Planctomycetota>unclassified_Planctomycetota</td><td>GCA_018242645.1</td><td>Planctomycetota_bacterium</td><td>JSS51_00780</td><td>22434..25748</td><td>2026780</td><td>ATP-binding-protein-[Planctomycetota-bacterium]</td></tr>
<tr><td>MBN8600335.1</td><td>AAA+ATPase+wHTH+Alpha+Beta-domain+wHTH1+wHTH2+wHTH3+RRM</td><td>Type-1_BR.RM.like.systems</td><td>Methyltransf_21-><-UbE1+JAB<-Mbetalac<-DUF499*<-METHYLASE<-?<-?<-TUDOR+SF1-SF2-Helicase+iREase+CTD+REase||?-><-METHYLASE</td><td>1123</td><td>Bacteria>PVC_group>Planctomycetota>unclassified_Planctomycetota</td><td>GCA_017303505.1</td><td>Planctomycetota_bacterium</td><td>JOM26_04870</td><td>9176..12547</td><td>2026780</td><td>ATP-binding-protein-[Planctomycetota-bacterium]</td></tr>
<tr><td>MBI5853022.1</td><td>AAA+ATPase+wHTH+Alpha+Beta-domain+wHTH1+wHTH2+wHTH3+RRM</td><td>Type-1_BR.RM.like.systems</td><td><-DUF499*<-METHYLASE<-TUDOR+SF1-SF2-Helicase+iREase+CTD+REase<-?||?->ABC-ATPase-></td><td>1106</td><td>Bacteria>PVC_group>Planctomycetota>unclassified_Planctomycetota</td><td>GCA_016235155.1</td><td>Planctomycetota_bacterium</td><td>HZB39_18585</td><td>15271..18591</td><td>2026780</td><td>ATP-binding-protein-[Planctomycetota-bacterium]</td></tr>
<tr><td>NQW47585.1</td><td>AAA+ATPase+wHTH+Alpha+Beta-domain+wHTH1+wHTH2+wHTH3+RRM</td><td>Type-1_BR.RM.like.systems</td><td>TUDOR+SF1-SF2-Helicase+iREase+CTD+REase-p->METHYLASE->DUF499*-></td><td>1112</td><td>Bacteria>PVC_group>Planctomycetota>unclassified_Planctomycetota</td><td>GCA_013205375.1</td><td>Planctomycetota_bacterium</td><td>HQ464_07455</td><td>5220..8558</td><td>2026780</td><td>ATP-binding-protein-[Planctomycetota-bacterium]</td></tr>
<tr><td>MBM3880845.1</td><td>AAA+ATPase+wHTH+Alpha+Beta-domain+wHTH1+wHTH2+wHTH3+RRM</td><td>Type-1_BR.RM.like.systems</td><td>TUDOR+SF1-SF2-Helicase+iREase+CTD+REase->HEPN->ParA-Soj-PloopNTPase->ORC-CDC6->REase->DUF499*-></td><td>1105</td><td>Bacteria>PVC_group>Verrucomicrobiota>unclassified_Verrucomicrobiota</td><td>GCA_016871715.1</td><td>Verrucomicrobiota_bacterium</td><td>FJ387_14200</td><td>23630..26947</td><td>2026799</td><td>DUF499-domain-containing-protein-[Verrucomicrobiota-bacterium]</td></tr>
<tr><td>NLH73611.1</td><td>AAA+ATPase+wHTH+Alpha+Beta-domain+wHTH1+wHTH2+wHTH3+RRM</td><td>Type-1_BR.RM.like.systems</td><td>TPR->METHYLASE-p->?->?-><-DUF499*<-TOPRIM<-OLD-ABC<-METHYLASE<-TUDOR+SF1-SF2-Helicase+iREase+CTD+REase</td><td>1104</td><td>Bacteria>PVC_group>Verrucomicrobiota>unclassified_Verrucomicrobiota</td><td>GCA_012517065.1</td><td>Verrucomicrobiota_bacterium</td><td>GX456_11205</td><td>15913..19227</td><td>2026799</td><td>ATP-binding-protein-[Verrucomicrobia-bacterium]</td></tr>
<tr><td>MCF7707718.1</td><td>AAA+ATPase+wHTH+Alpha+Beta-domain+wHTH1+wHTH2+wHTH3+RRM</td><td>Type-1_BR.RM.like.systems</td><td>TUDOR+SF1-SF2-Helicase+iREase+CTD+REase->OLD-ABC->TOPRIM->?->METHYLASE->Schlafen+HATPase+HTH->DUF499*-></td><td>1113</td><td>Bacteria>PVC_group>Verrucomicrobiota>unclassified_Verrucomicrobiota</td><td>GCA_021733835.1</td><td>Verrucomicrobiota_bacterium</td><td>K9N52_02315</td><td>37691..41032</td><td>2026799</td><td>DUF499-domain-containing-protein-[Verrucomicrobia-bacterium]</td></tr>
<tr><td>MBP7947571.1</td><td>AAA+ATPase+wHTH+Alpha+Beta-domain+wHTH1+wHTH2+wHTH3+RRM</td><td>Type-1_BR.RM.like.systems</td><td><-DUF499*<-TOPRIM<-OLD-ABC<-METHYLASE<-?<-TUDOR+SF1-SF2-Helicase+iREase+CTD+REase</td><td>1105</td><td>Bacteria>PVC_group>Verrucomicrobiota>unclassified_Verrucomicrobiota</td><td>GCA_018003215.1</td><td>Verrucomicrobiota_bacterium</td><td>KA191_16095</td><td>1214..4531</td><td>2026799</td><td>DUF499-domain-containing-protein-[Verrucomicrobiota-bacterium]</td></tr>
<tr><td>MBI4657986.1</td><td>AAA+ATPase+wHTH+Alpha+Beta-domain+wHTH1+wHTH2+wHTH3+RRM</td><td>Type-1_BR.RM.like.systems</td><td><-REase<-PIN<-HTH+MPTase||MetJarc-HTH->PIN-><-DUF499*</td><td>1089</td><td>Bacteria>PVC_group>Verrucomicrobiota>unclassified_Verrucomicrobiota</td><td>GCA_016209525.1</td><td>Verrucomicrobiota_bacterium</td><td>HY735_03900</td><td>7671..10940</td><td>2026799</td><td>ATP-binding-protein-[Verrucomicrobia-bacterium]</td></tr>
<tr><td>NMD22053.1</td><td>AAA+ATPase+wHTH+Alpha+Beta-domain+wHTH1+wHTH2+wHTH3+RRM</td><td>Type-1_BR.RM.like.systems</td><td>METHYLASE-p->OLD-ABC->TOPRIM->OLD-ABC->TOPRIM->DUF499*-></td><td>1068</td><td>Bacteria>PVC_group>Verrucomicrobiota>unclassified_Verrucomicrobiota</td><td>GCA_012797305.1</td><td>Verrucomicrobiota_bacterium</td><td>GYA76_17670</td><td>5686..8891</td><td>2026799</td><td>ATP-binding-protein-[Verrucomicrobia-bacterium]</td></tr>
<tr><td>MBP7947738.1</td><td>AAA+ATPase+wHTH+Alpha+Beta-domain+wHTH1+wHTH2+wHTH3+RRM</td><td>Type-1_BR.RM.like.systems</td><td><-DUF499*<-TOPRIM<-OLD-ABC<-TOPRIM<-OLD-ABC<-METHYLASE</td><td>1104</td><td>Bacteria>PVC_group>Verrucomicrobiota>unclassified_Verrucomicrobiota</td><td>GCA_018003215.1</td><td>Verrucomicrobiota_bacterium</td><td>KA191_16985</td><td>4010..7324</td><td>2026799</td><td>ATP-binding-protein-[Verrucomicrobia-bacterium]</td></tr>
<tr><td>MCI0539271.1</td><td>AAA+ATPase+wHTH+Alpha+Beta-domain+wHTH1+wHTH2+wHTH3+RRM</td><td>Type-1_BR.RM.like.systems</td><td><-SF1-SF2-Helicase<-OLD-ABC+TOPRIM||TUDOR+SF1-SF2-Helicase+iREase+CTD+REase->wHTH+REase->METHYLASE->DUF499*->PIWI->REase-></td><td>1107</td><td>Bacteria>PVC_group>Verrucomicrobiota>Verrucomicrobiae>Verrucomicrobiales</td><td>GCA_022615925.1</td><td>Verrucomicrobiales_bacterium</td><td>LOZ50_29035</td><td>12351..15674</td><td>2026801</td><td>DUF499-domain-containing-protein-[Verrucomicrobiales-bacterium]</td></tr>
<tr><td>MCF7789456.1</td><td>AAA+ATPase+wHTH+Alpha+Beta-domain+wHTH1+wHTH2+wHTH3+RRM</td><td>Type-1_BR.RM.like.systems</td><td>ABhydrolase->?-><-?<-DUF499*<-?<-METHYLASE<-TUDOR+SF1-SF2-Helicase+iREase+CTD+REase||?->?->XIS-HTH-></td><td>1099</td><td>Bacteria>PVC_group>Verrucomicrobiota>Verrucomicrobiae>Verrucomicrobiales</td><td>GCA_021733925.1</td><td>Prosthecobacter_sp.</td><td>K9N47_25260</td><td>6077..9376</td><td>1965333</td><td>DUF499-domain-containing-protein-[Prosthecobacter-sp.]</td></tr>
<tr><td>MBE7503150.1</td><td>AAA+ATPase+wHTH+Alpha+Beta-domain+wHTH1+wHTH2+wHTH3+RRM</td><td>Type-1_BR.RM.like.systems</td><td>METHYLASE->OLD-ABC->TOPRIM->DUF499->DUF499*-></td><td>747</td><td>Bacteria>PVC_group>Verrucomicrobiota>Verrucomicrobiae>Verrucomicrobiales</td><td>GCA_015075445.1</td><td>Verrucomicrobiales_bacterium</td><td>HS113_23270</td><td>56827..59070</td><td>2026801</td><td>hypothetical-protein-[Verrucomicrobiales-bacterium]</td></tr>
<tr><td>MCB1193039.1</td><td>AAA+ATPase+wHTH+Alpha+Beta-domain+wHTH1+wHTH2+wHTH3+RRM</td><td>Type-1_BR.RM.like.systems</td><td><-REase<-PIN<-?<-REase<-REase<-DUF499*<-REase<-METHYLASE<-REase<-TUDOR+SF1-SF2-Helicase+iREase+CTD+REase</td><td>1067</td><td>Bacteria>Spirochaetota>Spirochaetia>Leptospirales>Leptospiraceae</td><td>GCA_020440835.1</td><td>Leptospiraceae_bacterium</td><td>KDK90_21525</td><td>1949..5152</td><td>2026757</td><td>ATP-binding-protein-[Leptospiraceae-bacterium]</td></tr>
<tr><td>MDE0447651.1</td><td>AAA+ATPase+wHTH+Alpha+Beta-domain+wHTH1+wHTH2+wHTH3+RRM</td><td>Type-1_BR.RM.like.systems</td><td>Cluster182_14clades->TPR+TPR-><-?<-DUF499*<-TOPRIM<-OLD-ABC<-HTH_3<-METHYLASE<-SbcC<-TUDOR+SF1-SF2-Helicase+iREase+CTD+REase</td><td>1107</td><td>Bacteria>Spirochaetota>Spirochaetia>Spirochaetales>Spirochaetaceae</td><td>GCA_028821475.1</td><td>Spirochaetaceae_bacterium</td><td>OXH96_13340</td><td>24817..28140</td><td>1898206</td><td>MAG:-DUF499-domain-containing-protein-[Spirochaetaceae-bacterium]</td></tr>
<tr><td>MDD9989001.1</td><td>AAA+ATPase+wHTH+Alpha+Beta-domain+wHTH1+wHTH2+wHTH3+RRM</td><td>Type-1_BR.RM.like.systems</td><td>Cluster182_14clades-><-CR-ATPase8+REase<-?<-DUF499*<-TOPRIM<-OLD-ABC<-HTH_3<-METHYLASE<-TIR</td><td>1107</td><td>Bacteria>Spirochaetota>Spirochaetia>Spirochaetales>Spirochaetaceae</td><td>GCA_028818385.1</td><td>Spirochaetaceae_bacterium</td><td>OXQ31_22210</td><td>3073..6396</td><td>1898206</td><td>MAG:-DUF499-domain-containing-protein-[Spirochaetaceae-bacterium]</td></tr>
<tr><td>MDE0023241.1</td><td>AAA+ATPase+wHTH+Alpha+Beta-domain+wHTH1+wHTH2+wHTH3+RRM</td><td>Type-1_BR.RM.like.systems</td><td>METHYLASE-p->METHYLASE-p->Tox-RelE->HTH_3->DUF499*-><-?||?->?-><-TPR+TPR<-Cluster182_14clades</td><td>1100</td><td>Bacteria>Spirochaetota>Spirochaetia>Spirochaetales>Spirochaetaceae</td><td>GCA_028818735.1</td><td>Spirochaetaceae_bacterium</td><td>OXP69_02395</td><td>20195..23497</td><td>1898206</td><td>MAG:-DUF499-domain-containing-protein-[Spirochaetaceae-bacterium]</td></tr>
<tr><td>MCW3097907.1</td><td>AAA+ATPase+wHTH+Alpha+Beta-domain+wHTH1+wHTH2+wHTH3+RRM</td><td>Type-1_BR.RM.like.systems</td><td>TUDOR+SF1-SF2-Helicase+iREase+CTD+REase->?->METHYLASE->?->MACRODOMAIN->DUF499*->Schlafen+TUDOR+PIWI->?-><-?<-ParB-HTH</td><td>1081</td><td>Bacteria>Terrabacteria_group>Armatimonadota>Chthonomonadetes>Chthonomonadales</td><td>GCA_025963795.1</td><td>Chthonomonadaceae_bacterium</td><td>JWL77_3525</td><td>19451..22696</td><td>2739509</td><td>MAG:-hypothetical-protein-JWL77_3525-[Chthonomonadaceae-bacterium]</td></tr>
<tr><td>MBS1705314.1</td><td>AAA+ATPase+wHTH+Alpha+Beta-domain+wHTH1+wHTH2+wHTH3+RRM</td><td>Type-1_BR.RM.like.systems</td><td><-REase||TUDOR+SF1-SF2-Helicase+iREase+CTD+REase->METHYLASE->DUF499*->?-><-RelE-ParE<-GDResolvase+Recombinase+ZnR||?->MORC->GDResolvase+Recombinase+ZnR-></td><td>1095</td><td>Bacteria>Terrabacteria_group>Armatimonadota>unclassified_Armatimonadota</td><td>GCA_018268255.1</td><td>Armatimonadota_bacterium</td><td>JST40_05530</td><td>199077..202364</td><td>2033014</td><td>ATP-binding-protein-[Armatimonadetes-bacterium]</td></tr>
<tr><td>MCL6449134.1</td><td>AAA+ATPase+wHTH+Alpha+Beta-domain+wHTH1+wHTH2+wHTH3+RRM</td><td>Type-1_BR.RM.like.systems</td><td>METHYLASE->?->DUF1828+REase->DUF499*->REase->Radical-SAM-><-PIN<-PhdYeFM_antitox||?-><-PIN</td><td>1104</td><td>Bacteria>Terrabacteria_group>Armatimonadota>unclassified_Armatimonadota</td><td>GCA_023511535.1</td><td>Armatimonadota_bacterium</td><td>K6U04_13475</td><td>3761..7075</td><td>2033014</td><td>DUF499-domain-containing-protein-[Armatimonadetes-bacterium]</td></tr>
<tr><td>NCO42947.1</td><td>AAA+ATPase+wHTH+Alpha+Beta-domain+wHTH1+wHTH2+wHTH3+RRM</td><td>Type-1_BR.RM.like.systems</td><td>METHYLASE-p->DUF433->PIN->DUF499*->?->Tox-RelE->HTH_3-><-?<-HicA_toxin<-HicB_lk_antitox</td><td>1098</td><td>Bacteria>Terrabacteria_group>Armatimonadota>unclassified_Armatimonadota</td><td>GCA_009992845.1</td><td>Armatimonadota_bacterium</td><td>GW892_30120</td><td>1561..4857</td><td>2033014</td><td>ATP-binding-protein-[Armatimonadetes-bacterium]</td></tr>
<tr><td>WP_152947460.1</td><td>AAA+ATPase+wHTH+Alpha+Beta-domain+wHTH1+wHTH2+wHTH3+RRM</td><td>Type-1_BR.RM.like.systems</td><td>TUDOR+SF1-SF2-Helicase+iREase+CTD+REase->METHYLASE->?->DUF1828+REase->DUF499*->Calcineurin->SbcC+SbcC->MazG->LON-AAA->LON-AAA->sGTP-></td><td>1104</td><td>Bacteria>Terrabacteria_group>Bacillota>Clostridia>Eubacteriales</td><td>GCF_009428905.1</td><td>Desulfofundulus_thermobenzoicus</td><td>GFC01_RS12530</td><td>11768..15082</td><td>29376</td><td>DUF499-domain-containing-protein-[Desulfofundulus-thermobenzoicus]</td></tr>
<tr><td>MCR4442858.1</td><td>AAA+ATPase+wHTH+Alpha+Beta-domain+wHTH1+wHTH2+wHTH3+RRM</td><td>Type-1_BR.RM.like.systems</td><td>TUDOR+SF1-SF2-Helicase+iREase+CTD+REase->REase->SF1-SF2-Helicase->METHYLASE->OLD-ABC->DUF499*->?->?->TPR+TPR->?->PolB-></td><td>1104</td><td>Bacteria>Terrabacteria_group>Bacillota>Clostridia>Eubacteriales</td><td>GCA_024655825.1</td><td>Peptococcaceae_bacterium</td><td>NUV48_11985</td><td>21314..24628</td><td>2052179</td><td>DUF499-domain-containing-protein-[Peptococcaceae-bacterium]</td></tr>
<tr><td>WP_166349185.1</td><td>AAA+ATPase+wHTH+Alpha+Beta-domain+wHTH1+wHTH2+wHTH3+RRM</td><td>Type-1_BR.RM.like.systems</td><td>ArsR-HTH->BirA-HTH->?-><-DUF499*<-N6_N4_MTase<-?<-HEPN<-METHYLASE<-TUDOR+SF1-SF2-Helicase+iREase+CTD+REase</td><td>1104</td><td>Bacteria>Terrabacteria_group>Bacillota>Clostridia>Eubacteriales</td><td>GCF_011393055.1</td><td>Desulfofundulus_sp._TPOSR</td><td>G7K71_RS17495</td><td>347545..350859</td><td>2714340</td><td>DUF499-domain-containing-protein-[Desulfofundulus-sp.-TPOSR]</td></tr>
<tr><td>PSR20354.1</td><td>AAA+ATPase+wHTH+Alpha+Beta-domain+wHTH1+wHTH2+wHTH3+RRM</td><td>Type-1_BR.RM.like.systems</td><td>LuxR-HTH->?->?->?-><-HEPN+REase<-?<-DUF499*<-METHYLASE<-TUDOR+SF1-SF2-Helicase+iREase+CTD+REase||REase+ParB-></td><td>1101</td><td>Bacteria>Terrabacteria_group>Bacillota>Clostridia>Eubacteriales</td><td>GCA_003023745.1</td><td>Sulfobacillus_acidophilus</td><td>C7B45_15335</td><td>4761..8066</td><td>53633</td><td>hypothetical-protein-C7B45_15335-[Sulfobacillus-acidophilus]</td></tr>
<tr><td>AEG16694.1</td><td>AAA+ATPase+wHTH+Alpha+Beta-domain+wHTH1+wHTH2+wHTH3+RRM</td><td>Type-1_BR.RM.like.systems</td><td>ArsR-HTH->BirA-HTH->?-><-DUF499*<-METHYLASE<-RNase<-TUDOR+SF1-SF2-Helicase+iREase+CTD+REase<-?||?->PIN-></td><td>1104</td><td>Bacteria>Terrabacteria_group>Bacillota>Clostridia>Eubacteriales</td><td>GCA_000214705.1</td><td>Desulfofundulus_kuznetsovii_DSM_6115</td><td>Desku_3204</td><td>3212906..3216220</td><td>760568</td><td>hypothetical-protein-Desku_3204-[Desulfofundulus-kuznetsovii-DSM-6115]</td></tr>
<tr><td>MDA8204878.1</td><td>AAA+ATPase+wHTH+Alpha+Beta-domain+wHTH1+wHTH2+wHTH3+RRM</td><td>Type-1_BR.RM.like.systems</td><td>TUDOR+SF1-SF2-Helicase+iREase+CTD+REase->METHYLASE->DUF499*-></td><td>1102</td><td>Bacteria>Terrabacteria_group>Bacillota>Clostridia>Eubacteriales</td><td>GCA_028065495.1</td><td>Thermaerobacter_sp.</td><td>MOZ36_02345</td><td>5454..8762</td><td>1937527</td><td>MAG:-DUF499-domain-containing-protein-[Thermaerobacter-sp.]</td></tr>
<tr><td>WP_054937991.1</td><td>AAA+ATPase+wHTH+Alpha+Beta-domain+wHTH1+wHTH2+wHTH3+RRM</td><td>Type-1_BR.RM.like.systems</td><td>DEDD_Tnp_IS110+Transposase_20->TUDOR+SF1-SF2-Helicase+iREase+CTD+REase->?->HEPN->METHYLASE->DUF499*->ParB+HNH->?->?->?-><-?<-ArsR-HTH</td><td>1103</td><td>Bacteria>Terrabacteria_group>Bacillota>Clostridia>Moorellales</td><td>-</td><td>Moorella</td><td>MOST_RS05225</td><td>66804..70115</td><td>44260</td><td>MULTISPECIES:-DUF499-domain-containing-protein-[Moorella]</td></tr>
<tr><td>MCR4419670.1</td><td>AAA+ATPase+wHTH+Alpha+Beta-domain+wHTH1+wHTH2+wHTH3+RRM</td><td>Type-1_BR.RM.like.systems</td><td><-DUF499*<-TOPRIM<-OLD-ABC<-?<-METHYLASE<-TUDOR+SF1-SF2-Helicase+iREase+CTD+REase</td><td>1104</td><td>Bacteria>Terrabacteria_group>Bacillota>Clostridia>unclassified_Clostridia</td><td>GCA_024653205.1</td><td>Clostridia_bacterium</td><td>NUV99_05980</td><td>77810..81124</td><td>2044939</td><td>DUF499-domain-containing-protein-[Clostridia-bacterium]</td></tr>
<tr><td>MCL4516331.1</td><td>AAA+ATPase+wHTH+Alpha+Beta-domain+wHTH1+wHTH2+wHTH3+RRM</td><td>Type-1_BR.RM.like.systems</td><td><-HicB_lk_antitox<-HicA_toxin||TUDOR+SF1-SF2-Helicase+iREase+CTD+REase->SMODS->SLATT->METHYLASE->DUF499*->?-><-?||OLD-ABC-><-HTH_3||HTH-></td><td>1102</td><td>Bacteria>Terrabacteria_group>Bacillota>unclassified_Bacillota</td><td>GCA_023381695.1</td><td>Bacillota_bacterium</td><td>M1379_12200</td><td>45595..48903</td><td>1879010</td><td>DUF499-domain-containing-protein-[Firmicutes-bacterium]</td></tr>
<tr><td>MCL5054662.1</td><td>AAA+ATPase+wHTH+Alpha+Beta-domain+wHTH1+wHTH2+wHTH3+RRM</td><td>Type-1_BR.RM.like.systems</td><td>METHYLASE->DUF499*-><-HEPN<-Polbeta</td><td>1100</td><td>Bacteria>Terrabacteria_group>Bacillota>unclassified_Bacillota</td><td>GCA_023475635.1</td><td>Bacillota_bacterium</td><td>M1169_01785</td><td>1736..5038</td><td>1879010</td><td>DUF499-domain-containing-protein-[Firmicutes-bacterium]</td></tr>
<tr><td>MCR4407531.1</td><td>AAA+ATPase+wHTH+Alpha+Beta-domain+wHTH1+wHTH2+wHTH3+RRM</td><td>Type-1_BR.RM.like.systems</td><td>Phage_integrase-><-?||<-DUF499*<-TOPRIM-p<-SbcC<-METHYLASE<-PIN<-MazE_antitoxin<-TUDOR+SF1-SF2-Helicase+iREase+CTD+REase</td><td>1132</td><td>Bacteria>Terrabacteria_group>Chloroflexota>Anaerolineae>unclassified_Anaerolineae</td><td>GCA_024653325.1</td><td>Anaerolineae_bacterium</td><td>NUW24_11530</td><td>3238..6636</td><td>2052143</td><td>DUF499-domain-containing-protein-[Anaerolineae-bacterium]</td></tr>
<tr><td>MBN1139681.1</td><td>AAA+ATPase+wHTH+Alpha+Beta-domain+wHTH1+wHTH2+wHTH3+RRM</td><td>Type-1_BR.RM.like.systems</td><td><-DUF433<-DUF499*<-METHYLASE<-PIN<-CopG/RHH<-TUDOR+SF1-SF2-Helicase+iREase+CTD+REase||?-><-Methyltransf</td><td>1104</td><td>Bacteria>Terrabacteria_group>Chloroflexota>Anaerolineae>unclassified_Anaerolineae</td><td>GCA_016926595.1</td><td>Anaerolineae_bacterium</td><td>JXM73_24120</td><td>8352..11666</td><td>2052143</td><td>ATP-binding-protein-[Anaerolineae-bacterium]</td></tr>
<tr><td>MCG3212549.1</td><td>AAA+ATPase+wHTH+Alpha+Beta-domain+wHTH1+wHTH2+wHTH3+RRM</td><td>Type-1_BR.RM.like.systems</td><td>SMODS->?->METHYLASE->?->AbiEii->?->DUF499*->REase->?->?->?->?->JetA-></td><td>1128</td><td>Bacteria>Terrabacteria_group>Chloroflexota>Anaerolineae>unclassified_Anaerolineae</td><td>GCA_022071955.1</td><td>Anaerolineae_bacterium</td><td>FOGNACKC_06219</td><td>22074..25460</td><td>2052143</td><td>hypothetical-protein-[Anaerolineae-bacterium]</td></tr>
<tr><td>MDE0182657.1</td><td>AAA+ATPase+wHTH+Alpha+Beta-domain+wHTH1+wHTH2+wHTH3+RRM</td><td>Type-1_BR.RM.like.systems</td><td>PIWI->METHYLASE->OLD-ABC->TPR->?->DOC+HTH->DUF499*->?->HEPN->REase+35exo+Beta-Barrel+SF1-SF2-Helicase->?->TPR+TPR-></td><td>1109</td><td>Bacteria>Terrabacteria_group>Chloroflexota>Caldilineae>Caldilineales</td><td>GCA_028819655.1</td><td>Caldilineaceae_bacterium</td><td>OXL39_15010</td><td>68027..71356</td><td>2614250</td><td>MAG:-DUF499-domain-containing-protein-[Caldilineaceae-bacterium]</td></tr>
<tr><td>MDE0078893.1</td><td>AAA+ATPase+wHTH+Alpha+Beta-domain+wHTH1+wHTH2+wHTH3+RRM</td><td>Type-1_BR.RM.like.systems</td><td>ParB+HNH+SF1-SF2-Helicase+HDRC->?->?->HEPN-><-?<-ParB+HNH<-DUF499*<-OLD-ABC<-?<-?<-?<-HEPN<-METHYLASE</td><td>1109</td><td>Bacteria>Terrabacteria_group>Chloroflexota>Caldilineae>Caldilineales</td><td>GCA_028819025.1</td><td>Caldilineaceae_bacterium</td><td>OXO50_15340</td><td>14318..17647</td><td>2614250</td><td>MAG:-DUF499-domain-containing-protein-[Caldilineaceae-bacterium]</td></tr>
<tr><td>MDE0312394.1</td><td>AAA+ATPase+wHTH+Alpha+Beta-domain+wHTH1+wHTH2+wHTH3+RRM</td><td>Type-1_BR.RM.like.systems</td><td>TOPRIM->?->DUF499*->HTH->?-><-HEPN<-?<-?<-ParB+HNH+SF1-SF2-Helicase+HDRC</td><td>1111</td><td>Bacteria>Terrabacteria_group>Chloroflexota>Caldilineae>Caldilineales</td><td>GCA_028820765.1</td><td>Caldilineaceae_bacterium</td><td>OXI52_09030</td><td>1621..4956</td><td>2614250</td><td>MAG:-DUF499-domain-containing-protein-[Caldilineaceae-bacterium]</td></tr>
<tr><td>MDE0079138.1</td><td>AAA+ATPase+wHTH+Alpha+Beta-domain+wHTH1+wHTH2+wHTH3+RRM</td><td>Type-1_BR.RM.like.systems</td><td><-DUF499*<-METHYLASE<-PIN<-TUDOR+SF1-SF2-Helicase+iREase+CTD+REase</td><td>1096</td><td>Bacteria>Terrabacteria_group>Chloroflexota>Caldilineae>Caldilineales</td><td>GCA_028819025.1</td><td>Caldilineaceae_bacterium</td><td>OXO50_16575</td><td>525..3815</td><td>2614250</td><td>MAG:-DUF499-domain-containing-protein-[Caldilineaceae-bacterium]</td></tr>
<tr><td>MDE0077722.1</td><td>AAA+ATPase+wHTH+Alpha+Beta-domain+wHTH1+wHTH2+wHTH3+RRM</td><td>Type-1_BR.RM.like.systems</td><td>TOPRIM->?->?->?-><-Peptidase<-?||DUF499*-><-Trypsin<-?<-HTH+UbE1</td><td>1108</td><td>Bacteria>Terrabacteria_group>Chloroflexota>Caldilineae>Caldilineales</td><td>GCA_028819025.1</td><td>Caldilineaceae_bacterium</td><td>OXO50_09395</td><td>50434..53760</td><td>2614250</td><td>MAG:-DUF499-domain-containing-protein-[Caldilineaceae-bacterium]</td></tr>
<tr><td>MXX27764.1</td><td>AAA+ATPase+wHTH+Alpha+Beta-domain+wHTH1+wHTH2+wHTH3+RRM</td><td>Type-1_BR.RM.like.systems</td><td>HEPN-><-RHH<-?||?-><-?<-DUF499*<-TOPRIM<-OLD-ABC<-?<-?<-?<-PIWI</td><td>1135</td><td>Bacteria>Terrabacteria_group>Chloroflexota>Caldilineae>Caldilineales</td><td>GCA_009835815.1</td><td>Caldilineaceae_bacterium_SB0668_bin_21</td><td>F4Z82_20245</td><td>21021..24428</td><td>2605264</td><td>ATP-binding-protein-[Caldilineaceae-bacterium-SB0668_bin_21]</td></tr>
<tr><td>MDE0631425.1</td><td>AAA+ATPase+wHTH+Alpha+Beta-domain+wHTH1+wHTH2+wHTH3+RRM</td><td>Type-1_BR.RM.like.systems</td><td><-DUF499*<-TOPRIM<-OLD-ABC<-METHYLASE<-SbcC</td><td>1102</td><td>Bacteria>Terrabacteria_group>Chloroflexota>Caldilineae>Caldilineales</td><td>GCA_028824195.1</td><td>Caldilineaceae_bacterium</td><td>OXH73_07970</td><td>19208..22516</td><td>2614250</td><td>MAG:-DUF499-domain-containing-protein-[Caldilineaceae-bacterium]</td></tr>
<tr><td>WP_170293298.1</td><td>AAA+ATPase+wHTH+Alpha+Beta-domain+wHTH1+wHTH2+wHTH3+RRM</td><td>Type-1_BR.RM.like.systems</td><td><-SF1-SF2-Helicase+DUF1998<-LuxR-HTH+LuxR-HTH<-?||HTH+NUDIX+NUDIX-><-DUF499*<-METHYLASE<-TUDOR+SF1-SF2-Helicase+iREase+CTD+REase</td><td>1130</td><td>Bacteria>Terrabacteria_group>Chloroflexota>Ktedonobacteria>Thermogemmatisporales</td><td>GCF_008974285.1</td><td>Thermogemmatispora_aurantia</td><td>KTAU_RS14190</td><td>408330..411722</td><td>2045279</td><td>DUF499-domain-containing-protein-[Thermogemmatispora-aurantia]</td></tr>
<tr><td>WP_112432463.1</td><td>AAA+ATPase+wHTH+Alpha+Beta-domain+wHTH1+wHTH2+wHTH3+RRM</td><td>Type-1_BR.RM.like.systems</td><td>GNTR-HTH+UTRA->HTH+NUDIX+NUDIX->HTH_3+TPR->HTH_3->?-><-DUF499*<-METHYLASE<-TUDOR+SF1-SF2-Helicase+iREase+CTD+REase</td><td>1128</td><td>Bacteria>Terrabacteria_group>Chloroflexota>Ktedonobacteria>Thermogemmatisporales</td><td>GCF_003268475.1</td><td>Thermogemmatispora_tikiterensis</td><td>A4R35_RS19835</td><td>4947611..4950997</td><td>1825093</td><td>DUF499-domain-containing-protein-[Thermogemmatispora-tikiterensis]</td></tr>
<tr><td>MBX5458190.1</td><td>AAA+ATPase+wHTH+Alpha+Beta-domain+wHTH1+wHTH2+wHTH3+RRM</td><td>Type-1_BR.RM.like.systems</td><td>GNTR-HTH+UTRA->HTH+NUDIX+NUDIX->?->HTH_3+TPR->HTH_3->?-><-DUF499*<-METHYLASE-p</td><td>1128</td><td>Bacteria>Terrabacteria_group>Chloroflexota>Ktedonobacteria>Thermogemmatisporales</td><td>GCA_019683635.1</td><td>Thermogemmatispora_sp.</td><td>IRZ31_14945</td><td>8746..12132</td><td>1968838</td><td>ATP-binding-protein-[Thermogemmatispora-sp.]</td></tr>
<tr><td>BBH93037.1</td><td>AAA+ATPase+wHTH+Alpha+Beta-domain+wHTH1+wHTH2+wHTH3+RRM</td><td>Type-1_BR.RM.like.systems</td><td>TUDOR+SF1-SF2-Helicase+iREase+CTD+REase->METHYLASE->DUF499*-><-?||?->SF1-SF2-Helicase+DUF1998-></td><td>1130</td><td>Bacteria>Terrabacteria_group>Chloroflexota>Ktedonobacteria>Thermogemmatisporales</td><td>-</td><td>Thermogemmatispora_argillosa</td><td>KTA_12360</td><td>1574331..1577723</td><td>2045280</td><td>hypothetical-protein-KTA_12360-[Thermogemmatispora-argillosa]</td></tr>
<tr><td>WP_069805283.1</td><td>AAA+ATPase+wHTH+Alpha+Beta-domain+wHTH1+wHTH2+wHTH3+RRM</td><td>Type-1_BR.RM.like.systems</td><td>TUDOR+SF1-SF2-Helicase+iREase+CTD+REase->METHYLASE->DUF499*-></td><td>1130</td><td>Bacteria>Terrabacteria_group>Chloroflexota>Ktedonobacteria>Thermogemmatisporales</td><td>GCF_001748285.1</td><td>Thermogemmatispora_onikobensis</td><td>BGC09_RS16315</td><td>48454..51846</td><td>732234</td><td>DUF499-domain-containing-protein-[Thermogemmatispora-onikobensis]</td></tr>
<tr><td>WP_052890222.1</td><td>AAA+ATPase+wHTH+Alpha+Beta-domain+wHTH1+wHTH2+wHTH3+RRM</td><td>Type-1_BR.RM.like.systems</td><td>TUDOR+SF1-SF2-Helicase+iREase+CTD+REase->METHYLASE->DUF499*-></td><td>1129</td><td>Bacteria>Terrabacteria_group>Chloroflexota>Ktedonobacteria>Thermogemmatisporales</td><td>GCF_000702505.1</td><td>Thermogemmatispora_carboxidivorans</td><td>N688_RS16020</td><td>4079046..4082435</td><td>1382306</td><td>DUF499-domain-containing-protein-[Thermogemmatispora-carboxidivorans]</td></tr>
<tr><td>MBE7518482.1</td><td>AAA+ATPase+wHTH+Alpha+Beta-domain+wHTH1+wHTH2+wHTH3+RRM</td><td>Type-1_BR.RM.like.systems</td><td>LysM+TPR-><-?<-?<-REase+NACHT<-DUF499*<-METHYLASE<-RNase<-TUDOR+SF1-SF2-Helicase+iREase+CTD+REase<-sGTP</td><td>1100</td><td>Bacteria>Terrabacteria_group>Chloroflexota>Thermoflexia>Thermoflexales</td><td>GCA_015075555.1</td><td>Thermoflexaceae_bacterium</td><td>HS107_04470</td><td>911739..915041</td><td>2739507</td><td>ATP-binding-protein-[Thermoflexaceae-bacterium]</td></tr>
<tr><td>MXZ89877.1</td><td>AAA+ATPase+wHTH+Alpha+Beta-domain+wHTH1+wHTH2+wHTH3+RRM</td><td>Type-1_BR.RM.like.systems</td><td>AbiEii->?->TIR->TRD+TRD->?->SF1-SF2-Helicase-><-DUF499*<-TOPRIM<-OLD-ABC<-METHYLASE<-PIWI<-TIR+PNTD2<-TUDOR+SF1-SF2-Helicase+iREase+CTD+REase</td><td>1106</td><td>Bacteria>Terrabacteria_group>Chloroflexota>unclassified_Chloroflexota</td><td>GCA_009837355.1</td><td>Chloroflexota_bacterium</td><td>F4Z05_00140</td><td>18343..21663</td><td>2026724</td><td>ATP-binding-protein-[Chloroflexi-bacterium]</td></tr>
<tr><td>MBI2863489.1</td><td>AAA+ATPase+wHTH+Alpha+Beta-domain+wHTH1+wHTH2+wHTH3+RRM</td><td>Type-1_BR.RM.like.systems</td><td>TUDOR+SF1-SF2-Helicase+iREase+CTD+REase-p->HTH_37->METHYLASE->OLD-ABC->TOPRIM->DUF499*->MetJarc-HTH->PemK_toxin-><-?<-?<-?||Radical-SAM-></td><td>1111</td><td>Bacteria>Terrabacteria_group>Chloroflexota>unclassified_Chloroflexota</td><td>-</td><td>Chloroflexota_bacterium</td><td>HYX94_02885</td><td>6937..10272</td><td>2026724</td><td>ATP-binding-protein-[Chloroflexi-bacterium]</td></tr>
<tr><td>MYD49641.1</td><td>AAA+ATPase+wHTH+Alpha+Beta-domain+wHTH1+wHTH2+wHTH3+RRM</td><td>Type-1_BR.RM.like.systems</td><td>SF1-SF2-Helicase-><-DUF499*<-TOPRIM<-OLD-ABC<-METHYLASE<-PIWI<-TIR+PNTD2<-TUDOR+SF1-SF2-Helicase+iREase+CTD+REase</td><td>1106</td><td>Bacteria>Terrabacteria_group>Chloroflexota>unclassified_Chloroflexota</td><td>GCA_009841575.1</td><td>Chloroflexota_bacterium</td><td>F4W95_14345</td><td>5468..8788</td><td>2026724</td><td>ATP-binding-protein-[Chloroflexi-bacterium]</td></tr>
<tr><td>MBI2863770.1</td><td>AAA+ATPase+wHTH+Alpha+Beta-domain+wHTH1+wHTH2+wHTH3+RRM</td><td>Type-1_BR.RM.like.systems</td><td>TUDOR+SF1-SF2-Helicase+iREase+CTD+REase-p->HicB_lk_antitox->MqsA_antitoxin->?->METHYLASE->REase->DUF499*-></td><td>1112</td><td>Bacteria>Terrabacteria_group>Chloroflexota>unclassified_Chloroflexota</td><td>GCA_016190625.1</td><td>Chloroflexota_bacterium</td><td>HYX94_04330</td><td>24989..28327</td><td>2026724</td><td>ATP-binding-protein-[Chloroflexi-bacterium]</td></tr>
<tr><td>MCL5107378.1</td><td>AAA+ATPase+wHTH+Alpha+Beta-domain+wHTH1+wHTH2+wHTH3+RRM</td><td>Type-1_BR.RM.like.systems</td><td>TUDOR+SF1-SF2-Helicase+iREase+CTD+REase->?->HEPN->METHYLASE->DOC+HTH->DUF499*->Schlafen+HATPase+HTH-></td><td>1102</td><td>Bacteria>Terrabacteria_group>Chloroflexota>unclassified_Chloroflexota</td><td>GCA_023473795.1</td><td>Chloroflexota_bacterium</td><td>M1401_00595</td><td>20015..23323</td><td>2026724</td><td>DUF499-domain-containing-protein-[Chloroflexi-bacterium]</td></tr>
<tr><td>MYC29508.1</td><td>AAA+ATPase+wHTH+Alpha+Beta-domain+wHTH1+wHTH2+wHTH3+RRM</td><td>Type-1_BR.RM.like.systems</td><td><-SF1-SF2-Helicase+DUF1998<-?<-DUF499*<-TOPRIM<-OLD-ABC</td><td>1107</td><td>Bacteria>Terrabacteria_group>Chloroflexota>unclassified_Chloroflexota</td><td>GCA_009840135.1</td><td>Chloroflexota_bacterium</td><td>F4X65_05395</td><td>2421..5744</td><td>2026724</td><td>ATP-binding-protein-[Chloroflexi-bacterium]</td></tr>
<tr><td>MBI2958637.1</td><td>AAA+ATPase+wHTH+Alpha+Beta-domain+wHTH1+wHTH2+wHTH3+RRM</td><td>Type-1_BR.RM.like.systems</td><td>METHYLASE-p->DUF433->PIN->DUF499-p->DUF499*-></td><td>1006</td><td>Bacteria>Terrabacteria_group>Chloroflexota>unclassified_Chloroflexota</td><td>GCA_016189825.1</td><td>Chloroflexota_bacterium</td><td>HYY32_07340</td><td>1155..4175</td><td>2026724</td><td>DUF499-domain-containing-protein-[Chloroflexi-bacterium]</td></tr>
<tr><td>MBI4318932.1</td><td>AAA+ATPase+wHTH+Alpha+Beta-domain+wHTH1+wHTH2+wHTH3+RRM</td><td>Type-1_BR.RM.like.systems</td><td>DUF499*->?-><-?<-?||tRNA->?-><-Phage_int_SAM_4||PolB-></td><td>744</td><td>Bacteria>Terrabacteria_group>Chloroflexota>unclassified_Chloroflexota</td><td>GCA_016210495.1</td><td>Chloroflexota_bacterium</td><td>HY675_10605</td><td>237..2471</td><td>2026724</td><td>hypothetical-protein-[Chloroflexi-bacterium]</td></tr>
<tr><td>MBI2955393.1</td><td>AAA+ATPase+wHTH+Alpha+Beta-domain+wHTH1+wHTH2+wHTH3+RRM</td><td>Type-1_BR.RM.like.systems</td><td>MqsA_antitoxin->?->?->METHYLASE->?->?->DUF499*-></td><td>1103</td><td>Bacteria>Terrabacteria_group>Chloroflexota>unclassified_Chloroflexota</td><td>GCA_016189905.1</td><td>Chloroflexota_bacterium</td><td>HYY30_13850</td><td>31671..34982</td><td>2026724</td><td>ATP-binding-protein-[Chloroflexi-bacterium]</td></tr>
<tr><td>MBX9723630.1</td><td>AAA+ATPase+wHTH+Alpha+Beta-domain+wHTH1+wHTH2+wHTH3+RRM</td><td>Type-1_BR.RM.like.systems</td><td><-REase<-Prim-Pol||XIS-HTH->MORC+HisKin-><-?<-DUF499*<-METHYLASE-p</td><td>1111</td><td>Bacteria>Terrabacteria_group>Cyanobacteriota/Melainabacteria_group>Candidatus_Melainabacteria>Candidatus_Obscuribacterales</td><td>GCA_019747345.1</td><td>Candidatus_Obscuribacterales_bacterium</td><td>K2X81_19655</td><td>7987..11322</td><td>2364082</td><td>DUF499-domain-containing-protein-[Candidatus-Obscuribacterales-bacterium]</td></tr>
<tr><td>NJL38478.1</td><td>AAA+ATPase+wHTH+Alpha+Beta-domain+wHTH1+wHTH2+wHTH3+RRM</td><td>Type-1_BR.RM.like.systems</td><td>TUDOR+SF1-SF2-Helicase+iREase+CTD+REase-p->METHYLASE->DUF499-p->DUF499*-><-?<-UbE1+JAB<-?<-wHTH+REase||Phage_int_SAM_4+Phage_integrase-></td><td>1056</td><td>Bacteria>Terrabacteria_group>Cyanobacteriota/Melainabacteria_group>Cyanobacteriota>Cyanophyceae</td><td>GCA_012031415.1</td><td>Leptolyngbyaceae_cyanobacterium_SM1_4_3</td><td>HC899_18345</td><td>4580..7750</td><td>2720459</td><td>ATP-binding-protein-[Leptolyngbyaceae-cyanobacterium-SM1_4_3]</td></tr>
<tr><td>WP_096608407.1</td><td>AAA+ATPase+wHTH+Alpha+Beta-domain+wHTH1+wHTH2+wHTH3+RRM</td><td>Type-1_BR.RM.like.systems</td><td>Phage_int_SAM_4+Phage_integrase-><-?<-?<-?<-PadR-HTH||<-DUF499*<-METHYLASE<-?<-SMODS<-TUDOR+SF1-SF2-Helicase+iREase+CTD+REase<-?<-TraJ-RHH</td><td>1125</td><td>Bacteria>Terrabacteria_group>Cyanobacteriota/Melainabacteria_group>Cyanobacteriota>Cyanophyceae</td><td>-</td><td>Calothrix_sp._NIES-2100</td><td>CA734_RS39950</td><td>27837..31214</td><td>1954172</td><td>DUF499-domain-containing-protein-[Calothrix-sp.-NIES-2100]</td></tr>
<tr><td>NJO49090.1</td><td>AAA+ATPase+wHTH+Alpha+Beta-domain+wHTH1+wHTH2+wHTH3+RRM</td><td>Type-1_BR.RM.like.systems</td><td>HTH_Tnp_4+DDE_Tnp_4->TUDOR+SF1-SF2-Helicase+iREase+CTD+REase-p->METHYLASE-p->METHYLASE->DUF499*-><-?<-?<-UbE1+JAB</td><td>1120</td><td>Bacteria>Terrabacteria_group>Cyanobacteriota/Melainabacteria_group>Cyanobacteriota>Cyanophyceae</td><td>GCA_012033305.1</td><td>Leptolyngbyaceae_cyanobacterium_RM2_2_4</td><td>HC840_06015</td><td>5191..8553</td><td>2720439</td><td>ATP-binding-protein-[Leptolyngbyaceae-cyanobacterium-RM2_2_4]</td></tr>
<tr><td>HAJ59070.1</td><td>AAA+ATPase+wHTH+Alpha+Beta-domain+wHTH1+wHTH2+wHTH3+RRM</td><td>Type-1_BR.RM.like.systems</td><td>SMODS->?->METHYLASE->METHYLASE-p->METHYLASE-p->DUF499->DUF499*->DUF499-p-></td><td>730</td><td>Bacteria>Terrabacteria_group>Cyanobacteriota/Melainabacteria_group>Cyanobacteriota>unclassified_Cyanobacteriota</td><td>GCA_003454535.1</td><td>Cyanobacteria_bacterium_UBA8543</td><td>DCP31_07250</td><td>17927..20117</td><td>2055766</td><td>hypothetical-protein-[Cyanobacteria-bacterium-UBA8543]</td></tr>
<tr><td>MBC8417235.1</td><td>AAA+ATPase+wHTH+Alpha+Beta-domain+wHTH1+wHTH2+wHTH3+RRM</td><td>Type-1_BR.RM.like.systems</td><td>TUDOR+SF1-SF2-Helicase+iREase+CTD+REase->Schlafen+HATPase+HTH->METHYLASE->DUF499*-></td><td>1020</td><td>Bacteria>Thermodesulfobacteriota>Desulfobacteria>Desulfobacterales>unclassified_Desulfobacterales</td><td>GCA_014382905.1</td><td>Desulfobacterales_bacterium</td><td>H8E10_01405</td><td>14745..17804</td><td>2044940</td><td>ATP-binding-protein-[Desulfobacterales-bacterium]</td></tr>
<tr><td>MBL0712134.1</td><td>AAA+ATPase+wHTH+Alpha+Beta-domain+wHTH1+wHTH2+wHTH3+RRM</td><td>Type-1_BR.RM.like.systems</td><td><-SF1-SF2-Helicase<-?<-?<-?<-DUF499*<-METHYLASE<-TOPRIM<-OLD-ABC</td><td>1111</td><td>Bacteria>Thermodesulfobacteriota>Desulfobacteria>Desulfobacterales>unclassified_Desulfobacterales</td><td>GCA_016735415.1</td><td>Desulfobacterales_bacterium</td><td>JJV98_00405</td><td>2176..5511</td><td>2044940</td><td>ATP-binding-protein-[Desulfosarcina-sp.]</td></tr>
<tr><td>MBQ9405898.1</td><td>AAA+ATPase+wHTH+Alpha+Beta-domain+wHTH1+wHTH2+wHTH3+RRM</td><td>Type-1_BR.RM.like.systems</td><td>PfkB->?-><-DUF499*||?-><-REase<-REase<-METHYLASE-p<-METHYLASE-p<-HicB</td><td>1096</td><td>Bacteria>Thermodesulfobacteriota>Desulfovibrionia>Desulfovibrionales>Desulfovibrionaceae</td><td>GCA_017454855.1</td><td>Desulfovibrio_sp.</td><td>IJU37_04050</td><td>4592..7882</td><td>885</td><td>ATP-binding-protein-[Desulfovibrio-sp.]</td></tr>
<tr><td>MBQ7738149.1</td><td>AAA+ATPase+wHTH+Alpha+Beta-domain+wHTH1+wHTH2+wHTH3+RRM</td><td>Type-1_BR.RM.like.systems</td><td>NTP_transferase->?->?->?->?->?-><-DUF499*<-METHYLASE<-TUDOR+SF1-SF2-Helicase+iREase+CTD+REase</td><td>1112</td><td>Bacteria>Thermodesulfobacteriota>Desulfovibrionia>Desulfovibrionales>Desulfovibrionaceae</td><td>GCA_017455745.1</td><td>Desulfovibrionaceae_bacterium</td><td>IJT59_00640</td><td>29230..32568</td><td>2049043</td><td>ATP-binding-protein-[Desulfovibrionaceae-bacterium]</td></tr>
<tr><td>GFH63128.1</td><td>AAA+ATPase+wHTH+Alpha+Beta-domain+wHTH1+wHTH2+wHTH3+RRM</td><td>Type-1_BR.RM.like.systems</td><td>TUDOR+SF1-SF2-Helicase+iREase+CTD+REase->METHYLASE->DUF499*->SLATT->SMODS-></td><td>1110</td><td>Bacteria>Thermodesulfobacteriota>Desulfovibrionia>Desulfovibrionales>Desulfovibrionaceae</td><td>GCA_013374015.1</td><td>Candidatus_Desulfovibrio_kirbyi</td><td>ZNDK_0899</td><td>70330..73662</td><td>2696086</td><td>hypothetical-protein-ZNDK_0899-[Candidatus-Desulfovibrio-kirbyi]</td></tr>
<tr><td>MBS3905937.1</td><td>AAA+ATPase+wHTH+Alpha+Beta-domain+wHTH1+wHTH2+wHTH3+RRM</td><td>Type-1_BR.RM.like.systems</td><td>TOPRIM->TUDOR+SF1-SF2-Helicase+iREase+CTD+REase->Schlafen+HATPase+HTH->?->METHYLASE->AAA-ATPase+REase->DUF499*->SF1-SF2-Helicase->HTH-><-TPR</td><td>1115</td><td>Bacteria>Thermodesulfobacteriota>Syntrophia>Syntrophales>Syntrophaceae</td><td>GCA_018333715.1</td><td>Syntrophaceae_bacterium</td><td>KGZ49_02765</td><td>19111..22458</td><td>2052184</td><td>ATP-binding-protein-[Syntrophaceae-bacterium]</td></tr>
<tr><td>MCG2742124.1</td><td>AAA+ATPase+wHTH+Alpha+Beta-domain+wHTH1+wHTH2+wHTH3+RRM</td><td>Type-1_BR.RM.like.systems</td><td>Aminotran_1_2->?->?->?->DNA_pol3_delta+DNA_pol3_gamma3-><-DUF499*<-AAA-ATPase+REase<-METHYLASE<-?<-Schlafen+HATPase+HTH</td><td>1098</td><td>Bacteria>Thermodesulfobacteriota>Syntrophia>Syntrophales>Syntrophaceae</td><td>GCA_022013455.1</td><td>Syntrophaceae_bacterium</td><td>L6300_18065</td><td>15618..18914</td><td>2052184</td><td>DUF499-domain-containing-protein-[Syntrophaceae-bacterium]</td></tr>
<tr><td>KAF0159804.1</td><td>AAA+ATPase+wHTH+Alpha+Beta-domain+wHTH1+wHTH2+wHTH3+RRM</td><td>Type-1_BR.RM.like.systems</td><td><-HTH_3<-?<-DUF499*<-?<-METHYLASE<-TUDOR+SF1-SF2-Helicase+iREase+CTD+REase</td><td>1110</td><td>Bacteria>Thermodesulfobacteriota>Syntrophia>Syntrophales>Syntrophaceae</td><td>GCA_009773315.1</td><td>Syntrophaceae_bacterium</td><td>FD159_302</td><td>91265..94597</td><td>2052184</td><td>hypothetical-protein-FD159_302-[Syntrophaceae-bacterium]</td></tr>
<tr><td>OIP89123.1</td><td>AAA+ATPase+wHTH+Alpha+Beta-domain+wHTH1+wHTH2+wHTH3+RRM</td><td>Type-1_BR.RM.like.systems</td><td>METHYLASE->DUF499-p->DUF499*->HTH_31->MPTase-></td><td>882</td><td>Bacteria>Thermodesulfobacteriota>Syntrophia>Syntrophales>Syntrophaceae</td><td>GCA_001873745.1</td><td>Syntrophaceae_bacterium_CG2_30_58_14</td><td>AUK26_12340</td><td>3259..5907</td><td>1805378</td><td>hypothetical-protein-AUK26_12340-[Syntrophaceae-bacterium-CG2_30_58_14]</td></tr>
<tr><td>MBE3604313.1</td><td>AAA+ATPase+wHTH+Alpha+Beta-domain+wHTH1+wHTH2+wHTH3+RRM</td><td>Type-1_BR.RM.like.systems</td><td><-HIN-HTH||tRNA-><-DUF499*<-PIN<-DUF433||CopG_antitoxin->?-><-?<-METHYLASE</td><td>1108</td><td>Bacteria>unclassified_Bacteria</td><td>GCA_014896235.1</td><td>bacterium</td><td>IMX07_11830</td><td>1248..4574</td><td>1869227</td><td>ATP-binding-protein-[bacterium]</td></tr>
<tr><td>TRZ51818.1</td><td>AAA+ATPase+wHTH+Alpha+Beta-domain+wHTH1+wHTH2+wHTH3+RRM</td><td>Type-1_BR.RM.like.systems</td><td>METHYLASE-p->DUF499*->Calcineurin-></td><td>1104</td><td>Bacteria>unclassified_Bacteria</td><td>GCA_007279595.1</td><td>bacterium</td><td>D4S03_04460</td><td>1190..4504</td><td>1869227</td><td>ATP-binding-protein-[bacterium]</td></tr>
<tr><td>WP_283168466.1</td><td>HEPN+AAA-ATPase+Alpha+Beta-domain+wHTH+RRM</td><td>Type-1_BR.RM.like.systems</td><td>NurA->HerA/FtsK->DUF4338-><-HEPN+DUF499*<-METHYLASE-p</td><td>1093</td><td>Archaea>Euryarchaeota>Stenosarchaea_group>Methanomicrobia>Methanocellales</td><td>GCF_030019315.1</td><td>Methanocella_conradii</td><td>QMC99_RS12775</td><td>4236..7517</td><td>1175444</td><td>DUF499-domain-containing-protein-[Methanocella-conradii]</td></tr>
<tr><td>WP_167816002.1</td><td>HEPN+AAA-ATPase+Alpha+Beta-domain+wHTH+RRM</td><td>Type-1_BR.RM.like.systems</td><td>Schlafen+HATPase+HTH->?->Schlafen+HATPase+HTH->METHYLASE->OLD-ABC->TOPRIM->HEPN+DUF499*-></td><td>756</td><td>Archaea>Euryarchaeota>Stenosarchaea_group>Methanomicrobia>Methanomicrobiales</td><td>GCF_012062385.1</td><td>Methanocorpusculum_sp._GPch4</td><td>EYA77_RS04465</td><td>15957..18225</td><td>2527877</td><td>Swt1-family-HEPN-domain-containing-protein,-partial-[Methanocorpusculum-sp.-GPch4]</td></tr>
<tr><td>WP_204222940.1</td><td>HEPN+AAA-ATPase+Alpha+Beta-domain+wHTH+RRM</td><td>Type-1_BR.RM.like.systems</td><td>OLD-ABC->OLD-ABC->REase->METHYLASE->OLD-ABC->TOPRIM->HEPN+DUF499*-></td><td>1082</td><td>Archaea>Euryarchaeota>Stenosarchaea_group>Methanomicrobia>Methanomicrobiales</td><td>GCF_000013445.1</td><td>Methanospirillum_hungatei</td><td>MHUN_RS09170</td><td>1984894..1988142</td><td>2203</td><td>DUF499-domain-containing-protein-[Methanospirillum-hungatei]</td></tr>
<tr><td>ABD41477.1</td><td>HEPN+AAA-ATPase+Alpha+Beta-domain+wHTH+RRM</td><td>Type-1_BR.RM.like.systems</td><td>OLD-ABC->OLD-ABC->REase->METHYLASE->OLD-ABC->TOPRIM->HEPN+DUF499*-></td><td>1094</td><td>Archaea>Euryarchaeota>Stenosarchaea_group>Methanomicrobia>Methanomicrobiales</td><td>GCA_000013445.1</td><td>Methanospirillum_hungatei_JF-1</td><td>Mhun_1756</td><td>1984858..1988142</td><td>323259</td><td>conserved-hypothetical-protein-[Methanospirillum-hungatei-JF-1]</td></tr>
<tr><td>KUL01096.1</td><td>HEPN+AAA-ATPase+Alpha+Beta-domain+wHTH+RRM</td><td>Type-1_BR.RM.like.systems</td><td>METHYLASE-p->HEPN+DUF499*-></td><td>1106</td><td>Archaea>Euryarchaeota>Stenosarchaea_group>Methanomicrobia>Methanomicrobiales</td><td>GCA_001508695.1</td><td>Methanomicrobiales_archaeon_53_19</td><td>XE11_2241</td><td>598..3918</td><td>1641394</td><td>Uncharacterized-protein-XE11_2241-[Methanomicrobiales-archaeon-53_19]</td></tr>
<tr><td>WP_048105728.1</td><td>HEPN+AAA-ATPase+Alpha+Beta-domain+wHTH+RRM</td><td>Type-1_BR.RM.like.systems</td><td>PRTase-><-RelE-ParE||?->?->tRNA-><-HEPN+DUF499*<-METHYLASE<-TUDOR+SF1-SF2-Helicase+iREase+CTD+REase<-?<-?<-Primpol+D5N+D5ATPase</td><td>1126</td><td>Archaea>Euryarchaeota>Stenosarchaea_group>Methanomicrobia>Methanosarcinales</td><td>GCF_000970305.1</td><td>Methanosarcina_barkeri</td><td>MSBR3_RS00635</td><td>148480..151860</td><td>2208</td><td>Swt1-family-HEPN-domain-containing-protein-[Methanosarcina-barkeri]</td></tr>
<tr><td>WP_048047674.1</td><td>HEPN+AAA-ATPase+Alpha+Beta-domain+wHTH+RRM</td><td>Type-1_BR.RM.like.systems</td><td>Primpol+D5N+D5ATPase->?->?->TUDOR+SF1-SF2-Helicase+iREase+CTD+REase->METHYLASE->HEPN+DUF499*-><-?<-tRNA<-?<-?||RelE-ParE-><-PRTase</td><td>1126</td><td>Archaea>Euryarchaeota>Stenosarchaea_group>Methanomicrobia>Methanosarcinales</td><td>GCF_000979995.1</td><td>Methanosarcina_mazei</td><td>DU74_RS10215</td><td>73839..77219</td><td>2209</td><td>Swt1-family-HEPN-domain-containing-protein-[Methanosarcina-mazei]</td></tr>
<tr><td>WP_048037397.1</td><td>HEPN+AAA-ATPase+Alpha+Beta-domain+wHTH+RRM</td><td>Type-1_BR.RM.like.systems</td><td>PRTase-><-RelE-ParE||?->?->tRNA-><-HEPN+DUF499*<-METHYLASE<-TUDOR+SF1-SF2-Helicase+iREase+CTD+REase<-?<-?<-Primpol+D5N+D5ATPase</td><td>1126</td><td>Archaea>Euryarchaeota>Stenosarchaea_group>Methanomicrobia>Methanosarcinales</td><td>GCF_000978965.1</td><td>Methanosarcina_mazei</td><td>DU68_RS02805</td><td>48841..52221</td><td>2209</td><td>Swt1-family-HEPN-domain-containing-protein-[Methanosarcina-mazei]</td></tr>
<tr><td>WP_048048703.1</td><td>HEPN+AAA-ATPase+Alpha+Beta-domain+wHTH+RRM</td><td>Type-1_BR.RM.like.systems</td><td>ParA-Soj-PloopNTPase->?->?-><-HEPN+DUF499*<-METHYLASE<-TUDOR+SF1-SF2-Helicase+iREase+CTD+REase<-Peptidase_C47</td><td>1113</td><td>Archaea>Euryarchaeota>Stenosarchaea_group>Methanomicrobia>Methanosarcinales</td><td>GCF_000979015.1</td><td>Methanosarcina_mazei</td><td>DU63_RS12210</td><td>161635..164976</td><td>2209</td><td>DUF499-domain-containing-protein-[Methanosarcina-mazei]</td></tr>
<tr><td>WP_048045029.1</td><td>HEPN+AAA-ATPase+Alpha+Beta-domain+wHTH+RRM</td><td>Type-1_BR.RM.like.systems</td><td>Primpol+D5N+D5ATPase->?->?->TUDOR+SF1-SF2-Helicase+iREase+CTD+REase->METHYLASE->HEPN+DUF499*-><-?<-?<-HNH</td><td>1126</td><td>Archaea>Euryarchaeota>Stenosarchaea_group>Methanomicrobia>Methanosarcinales</td><td>GCF_000979105.1</td><td>Methanosarcina_mazei</td><td>DU65_RS07520</td><td>78194..81574</td><td>2209</td><td>Swt1-family-HEPN-domain-containing-protein-[Methanosarcina-mazei]</td></tr>
<tr><td>WP_048043068.1</td><td>HEPN+AAA-ATPase+Alpha+Beta-domain+wHTH+RRM</td><td>Type-1_BR.RM.like.systems</td><td>TUDOR+SF1-SF2-Helicase+iREase+CTD+REase->METHYLASE->HEPN+DUF499*-><-ZnR<-tRNA<-?<-?||RelE-ParE-><-PRTase</td><td>1126</td><td>Archaea>Euryarchaeota>Stenosarchaea_group>Methanomicrobia>Methanosarcinales</td><td>GCF_000970185.1</td><td>Methanosarcina_mazei</td><td>MSMAP_RS05770</td><td>1314759..1318139</td><td>2209</td><td>Swt1-family-HEPN-domain-containing-protein-[Methanosarcina-mazei]</td></tr>
<tr><td>WP_048047296.1</td><td>HEPN+AAA-ATPase+Alpha+Beta-domain+wHTH+RRM</td><td>Type-1_BR.RM.like.systems</td><td><-HEPN+DUF499*<-METHYLASE<-TUDOR+SF1-SF2-Helicase+iREase+CTD+REase<-?<-?<-Primpol+D5N+D5ATPase</td><td>513</td><td>Archaea>Euryarchaeota>Stenosarchaea_group>Methanomicrobia>Methanosarcinales</td><td>GCF_000979035.1</td><td>Methanosarcina_mazei</td><td>DU66_RS08905</td><td>1..1541</td><td>2209</td><td>Swt1-family-HEPN-domain-containing-protein,-partial-[Methanosarcina-mazei]</td></tr>
<tr><td>KXS42417.1</td><td>HEPN+AAA-ATPase+Alpha+Beta-domain+wHTH+RRM</td><td>Type-1_BR.RM.like.systems</td><td><-HEPN+DUF499*<-METHYLASE<-MORC+DUF3684<-TUDOR+SF1-SF2-Helicase+iREase+CTD+REase</td><td>1129</td><td>Archaea>Euryarchaeota>Stenosarchaea_group>Methanomicrobia>Methanosarcinales</td><td>GCA_001577565.1</td><td>Methanolobus_sp._T82-4</td><td>AWU59_1648</td><td>511..3900</td><td>1794908</td><td>MAG:-ATPase-(AAA+-superfamily)-like-protein-[Methanolobus-sp.-T82-4]</td></tr>
<tr><td>MCG7852630.1</td><td>HEPN+AAA-ATPase+Alpha+Beta-domain+wHTH+RRM</td><td>Type-1_BR.RM.like.systems</td><td>METHYLASE->HEPN+DUF499*-></td><td>1119</td><td>Archaea>Euryarchaeota>Stenosarchaea_group>Methanomicrobia>Methanosarcinales</td><td>GCA_022353215.1</td><td>Methanosarcinaceae_archaeon</td><td>MIO92_08920</td><td>2388..5747</td><td>1945595</td><td>Swt1-family-HEPN-domain-containing-protein-[Methanosarcinaceae-archaeon]</td></tr>
<tr><td>MDD4161340.1</td><td>HEPN+AAA-ATPase+Alpha+Beta-domain+wHTH+RRM</td><td>Type-1_BR.RM.like.systems</td><td>ParB+HNH+Schlafen-><-DEDD_Tnp_IS110+Transposase_20<-?||TUDOR+SF1-SF2-Helicase+iREase+CTD+REase->METHYLASE->HEPN+DUF499*->?-><-SF1-SF2-Helicase-p</td><td>1125</td><td>Archaea>Euryarchaeota>Stenosarchaea_group>Methanomicrobia>Methanotrichales</td><td>GCA_028702095.1</td><td>Methanothrix_sp.</td><td>PHW87_02550</td><td>13922..17299</td><td>90426</td><td>MAG:-Swt1-family-HEPN-domain-containing-protein-[Methanothrix-sp.]</td></tr>
<tr><td>WP_273278519.1</td><td>HEPN+AAA-ATPase+Alpha+Beta-domain+wHTH+RRM</td><td>Type-1_BR.RM.like.systems</td><td>Primpol->?->ArdCN+MPTase->?->?-><-REase<-HEPN+DUF499*<-TOPRIM<-OLD-ABC<-PolB<-METHYLASE<-HEPN<-TUDOR+SF1-SF2-Helicase+iREase+CTD+REase</td><td>1113</td><td>Archaea>Euryarchaeota>Stenosarchaea_group>Methanomicrobia>Methanotrichales</td><td>GCF_913757535.1</td><td>Methanothrix_soehngenii</td><td>NW278_RS08460</td><td>1661104..1664445</td><td>2223</td><td>DUF499-domain-containing-protein-[Methanothrix-soehngenii]</td></tr>
<tr><td>MBK7385730.1</td><td>HEPN+AAA-ATPase+Alpha+Beta-domain+wHTH+RRM</td><td>Type-1_BR.RM.like.systems</td><td>Methyltransf-><-HEPN+DUF499*<-TOPRIM<-OLD-ABC<-METHYLASE||?->?-><-TUDOR+SF1-SF2-Helicase+iREase+CTD+REase</td><td>1119</td><td>Archaea>Euryarchaeota>Stenosarchaea_group>Methanomicrobia>Methanotrichales</td><td>GCA_016706325.1</td><td>Methanothrix_sp.</td><td>IPI63_02955</td><td>568305..571664</td><td>90426</td><td>DUF499-domain-containing-protein-[Methanothrix-sp.]</td></tr>
<tr><td>MCX6676562.1</td><td>HEPN+AAA-ATPase+Alpha+Beta-domain+wHTH+RRM</td><td>Type-1_BR.RM.like.systems</td><td>TUDOR+SF1-SF2-Helicase+iREase+CTD+REase-><-?<-?<-?||METHYLASE->OLD-ABC->HEPN+DUF499*-></td><td>1111</td><td>Archaea>Euryarchaeota>Stenosarchaea_group>Methanomicrobia>Methanotrichales</td><td>GCA_026394565.1</td><td>Methanothrix_sp.</td><td>NTU95_01275</td><td>32506..35841</td><td>90426</td><td>MAG:-DUF499-domain-containing-protein-[Methanothrix-sp.]</td></tr>
<tr><td>MBC7080549.1</td><td>HEPN+AAA-ATPase+Alpha+Beta-domain+wHTH+RRM</td><td>Type-1_BR.RM.like.systems</td><td>TUDOR+SF1-SF2-Helicase+iREase+CTD+REase->METHYLASE->RNase->HEPN+DUF499*-></td><td>1102</td><td>Archaea>Euryarchaeota>Stenosarchaea_group>Methanomicrobia>Methanotrichales</td><td>GCA_014361255.1</td><td>Methanothrix_sp.</td><td>H5T42_08820</td><td>8191..11499</td><td>90426</td><td>ATP-binding-protein-[Methanothrix-sp.]</td></tr>
<tr><td>OQA55746.1</td><td>HEPN+AAA-ATPase+Alpha+Beta-domain+wHTH+RRM</td><td>Type-1_BR.RM.like.systems</td><td>METHYLASE->?->OLD-ABC->?->TOPRIM->HEPN+DUF499*-></td><td>1082</td><td>Archaea>Euryarchaeota>unclassified_Euryarchaeota</td><td>GCA_002069575.1</td><td>Euryarchaeota_archaeon_ADurb.Bin294</td><td>BWY45_02073</td><td>4487..7735</td><td>1852878</td><td>hypothetical-protein-BWY45_02073-[Euryarchaeota-archaeon-ADurb.Bin294]</td></tr>
<tr><td>MCL2643280.1</td><td>HEPN+AAA-ATPase+Alpha+Beta-domain+wHTH+RRM</td><td>Type-1_BR.RM.like.systems</td><td><-DDE_Tnp_ISAZ013<-HEPN+DUF499*<-TUDOR+SF1-SF2-Helicase+iREase+CTD+REase</td><td>1134</td><td>Archaea>TACK_group>Candidatus_Bathyarchaeota>Candidatus_Bathyarchaeia>Candidatus_Bathyarchaeia_incertae_sedis</td><td>GCA_009781395.1</td><td>Candidatus_Termiticorpusculum_sp.</td><td>FWD52_07200</td><td>1651..5055</td><td>2994959</td><td>DUF499-domain-containing-protein-[Candidatus-Bathyarchaeota-archaeon]</td></tr>
<tr><td>TSA50211.1</td><td>HEPN+AAA-ATPase+Alpha+Beta-domain+wHTH+RRM</td><td>Type-1_BR.RM.like.systems</td><td>SF1-SF2-Helicase->SF1-SF2-Helicase->METHYLASE->?->HEPN+DUF499*-></td><td>607</td><td>Archaea>unclassified_Archaea</td><td>GCA_007281275.1</td><td>archaeon</td><td>D4R47_02600</td><td>5332..7152</td><td>1906665</td><td>ATP-binding-protein,-partial-[archaeon]</td></tr>
<tr><td>MBA3712812.1</td><td>HEPN+AAA-ATPase+Alpha+Beta-domain+wHTH+RRM</td><td>Type-1_BR.RM.like.systems</td><td>Schlafen+HATPase+HTH->?->METHYLASE->DUF433->PIN->HEPN+DUF499*->TIR+PNTD2->PIWI->?->Phage_integrase-></td><td>1118</td><td>Bacteria>Acidobacteriota>Blastocatellia>Blastocatellales>Pyrinomonadaceae</td><td>GCA_013816505.1</td><td>Pyrinomonadaceae_bacterium</td><td>HOW76_10240</td><td>5538..8894</td><td>2283092</td><td>ATP-binding-protein-[Pyrinomonadaceae-bacterium]</td></tr>
<tr><td>WP_246840661.1</td><td>HEPN+AAA-ATPase+Alpha+Beta-domain+wHTH+RRM</td><td>Type-1_BR.RM.like.systems</td><td>ABhydrolase-><-?||TUDOR+SF1-SF2-Helicase+iREase+CTD+REase->METHYLASE->HEPN+DUF499*->?->Response_reg+LuxR-HTH->CarboxypepD-reg-IG->BetaPropeller+ABhydrolase-></td><td>1107</td><td>Bacteria>Acidobacteriota>Blastocatellia>Chloracidobacterium</td><td>GCF_018304765.1</td><td>Chloracidobacterium_aggregatum</td><td>J8C05_RS05820</td><td>1401266..1404589</td><td>2851959</td><td>Swt1-family-HEPN-domain-containing-protein-[Chloracidobacterium-aggregatum]</td></tr>
<tr><td>WP_246604568.1</td><td>HEPN+AAA-ATPase+Alpha+Beta-domain+wHTH+RRM</td><td>Type-1_BR.RM.like.systems</td><td>ABhydrolase-><-?||TUDOR+SF1-SF2-Helicase+iREase+CTD+REase->METHYLASE->HEPN+DUF499*->?->Response_reg+LuxR-HTH->CarboxypepD-reg-IG->BetaPropeller+ABhydrolase-></td><td>1107</td><td>Bacteria>Acidobacteriota>Blastocatellia>Chloracidobacterium</td><td>GCF_018304805.1</td><td>Chloracidobacterium_aggregatum</td><td>J8C02_RS05835</td><td>1402645..1405968</td><td>2851959</td><td>Swt1-family-HEPN-domain-containing-protein-[Chloracidobacterium-aggregatum]</td></tr>
<tr><td>WP_211431037.1</td><td>HEPN+AAA-ATPase+Alpha+Beta-domain+wHTH+RRM</td><td>Type-1_BR.RM.like.systems</td><td>ABhydrolase-><-?||TUDOR+SF1-SF2-Helicase+iREase+CTD+REase->METHYLASE->HEPN+DUF499*->?->Response_reg+LuxR-HTH->CarboxypepD-reg-IG->BetaPropeller+ABhydrolase-></td><td>1100</td><td>Bacteria>Acidobacteriota>Blastocatellia>Chloracidobacterium</td><td>GCF_018304725.1</td><td>Chloracidobacterium_aggregatum</td><td>J8C07_RS06025</td><td>1402377..1405679</td><td>2851959</td><td>Swt1-family-HEPN-domain-containing-protein-[Chloracidobacterium-aggregatum]</td></tr>
<tr><td>WP_041569618.1</td><td>HEPN+AAA-ATPase+Alpha+Beta-domain+wHTH+RRM</td><td>Type-1_BR.RM.like.systems</td><td>ABhydrolase-><-?||TUDOR+SF1-SF2-Helicase+iREase+CTD+REase->METHYLASE->HEPN+DUF499*->?->Response_reg+LuxR-HTH->CarboxypepD-reg-IG->BetaPropeller+ABhydrolase-></td><td>1110</td><td>Bacteria>Acidobacteriota>Blastocatellia>Chloracidobacterium</td><td>GCF_000226295.1</td><td>Chloracidobacterium_thermophilum</td><td>CABTHER_RS05910</td><td>1425684..1429016</td><td>458033</td><td>Swt1-family-HEPN-domain-containing-protein-[Chloracidobacterium-thermophilum]</td></tr>
<tr><td>WP_211430024.1</td><td>HEPN+AAA-ATPase+Alpha+Beta-domain+wHTH+RRM</td><td>Type-1_BR.RM.like.systems</td><td>Glyoxalase->?->?->?->TUDOR+SF1-SF2-Helicase+iREase+CTD+REase->METHYLASE->HEPN+DUF499*->?-><-?<-Glycos_trans_3N+Glycos_transf_3||HTH+WYL->PIWI-></td><td>1106</td><td>Bacteria>Acidobacteriota>Blastocatellia>Chloracidobacterium</td><td>GCF_018304825.1</td><td>Chloracidobacterium_validum</td><td>J8C06_RS13915</td><td>660771..664091</td><td>2821543</td><td>Swt1-family-HEPN-domain-containing-protein-[Chloracidobacterium-validum]</td></tr>
<tr><td>WP_058867444.1</td><td>HEPN+AAA-ATPase+Alpha+Beta-domain+wHTH+RRM</td><td>Type-1_BR.RM.like.systems</td><td>TUDOR+SF1-SF2-Helicase+iREase+CTD+REase->PrlF_antitoxin->METHYLASE->PolB->HEPN->HEPN+DUF499*->?->?->?->?->?->NUDIX-></td><td>1098</td><td>Bacteria>Acidobacteriota>Blastocatellia>Chloracidobacterium</td><td>GCF_001482755.1</td><td>Chloracidobacterium_thermophilum</td><td>AVR57_RS08855</td><td>227701..230997</td><td>458033</td><td>Swt1-family-HEPN-domain-containing-protein-[Chloracidobacterium-thermophilum]</td></tr>
<tr><td>QUW01026.1</td><td>HEPN+AAA-ATPase+Alpha+Beta-domain+wHTH+RRM</td><td>Type-1_BR.RM.like.systems</td><td>ABhydrolase-><-?||TUDOR+SF1-SF2-Helicase+iREase+CTD+REase->METHYLASE->HEPN+DUF499*->?->Response_reg+LuxR-HTH->CarboxypepD-reg-IG->BetaPropeller+ABhydrolase-></td><td>1109</td><td>Bacteria>Acidobacteriota>Blastocatellia>Chloracidobacterium>Chloracidobacterium_aggregatum</td><td>GCA_018304805.1</td><td>Chloracidobacterium_sp._MS_40/45</td><td>J8C02_05835</td><td>1402639..1405968</td><td>2821542</td><td>DUF499-domain-containing-protein-[Chloracidobacterium-sp.-MS-40/45]</td></tr>
<tr><td>QUV94961.1</td><td>HEPN+AAA-ATPase+Alpha+Beta-domain+wHTH+RRM</td><td>Type-1_BR.RM.like.systems</td><td>ABhydrolase-><-?||TUDOR+SF1-SF2-Helicase+iREase+CTD+REase->METHYLASE->HEPN+DUF499*->?->Response_reg+LuxR-HTH->CarboxypepD-reg-IG->BetaPropeller+ABhydrolase-></td><td>1109</td><td>Bacteria>Acidobacteriota>Blastocatellia>Chloracidobacterium>Chloracidobacterium_aggregatum</td><td>GCA_018304765.1</td><td>Chloracidobacterium_sp._N</td><td>J8C05_05820</td><td>1401260..1404589</td><td>2821540</td><td>DUF499-domain-containing-protein-[Chloracidobacterium-sp.-N]</td></tr>
<tr><td>AEP11956.1</td><td>HEPN+AAA-ATPase+Alpha+Beta-domain+wHTH+RRM</td><td>Type-1_BR.RM.like.systems</td><td>ABhydrolase-><-?||TUDOR+SF1-SF2-Helicase+iREase+CTD+REase->METHYLASE->HEPN+DUF499*->?->Response_reg+LuxR-HTH->CarboxypepD-reg-IG->BetaPropeller+ABhydrolase-></td><td>1108</td><td>Bacteria>Acidobacteriota>Blastocatellia>Chloracidobacterium>Chloracidobacterium_thermophilum</td><td>GCA_000226295.1</td><td>Chloracidobacterium_thermophilum_B</td><td>Cabther_A1202</td><td>1425690..1429016</td><td>981222</td><td>hypothetical-protein-Cabther_A1202-[Chloracidobacterium-thermophilum-B]</td></tr>
<tr><td>MBK9214625.1</td><td>HEPN+AAA-ATPase+Alpha+Beta-domain+wHTH+RRM</td><td>Type-1_BR.RM.like.systems</td><td>HTH->?->GDResolvase+Recombinase+ZnR-><-?<-SF1-SF2-Helicase+DUF1998+HKD-DNase+HKD-DNase<-?<-HEPN+DUF499*<-?<-METHYLASE<-ArdCN+MPTase<-TUDOR+SF1-SF2-Helicase+iREase+CTD+REase<-?||HTH-></td><td>1119</td><td>Bacteria>Acidobacteriota>Blastocatellia>Chloracidobacterium>unclassified_Chloracidobacterium</td><td>GCA_016715795.1</td><td>Chloracidobacterium_sp.</td><td>IPM59_03360</td><td>738232..741591</td><td>2099375</td><td>DUF499-domain-containing-protein-[Chloracidobacterium-sp.]</td></tr>
<tr><td>WP_211431305.1</td><td>HEPN+AAA-ATPase+Alpha+Beta-domain+wHTH+RRM</td><td>Type-1_BR.RM.like.systems</td><td>ABhydrolase-><-?||TUDOR+SF1-SF2-Helicase+iREase+CTD+REase->METHYLASE->HEPN+DUF499*->?->Response_reg+LuxR-HTH->CarboxypepD-reg-IG->BetaPropeller+ABhydrolase-></td><td>1108</td><td>Bacteria>Acidobacteriota>Blastocatellia>Chloracidobacterium>unclassified_Chloracidobacterium</td><td>GCF_018304685.1</td><td>Chloracidobacterium_sp._D</td><td>J8C01_RS05850</td><td>1398986..1402312</td><td>2821536</td><td>Swt1-family-HEPN-domain-containing-protein-[Chloracidobacterium-sp.-D]</td></tr>
<tr><td>MBK9173225.1</td><td>HEPN+AAA-ATPase+Alpha+Beta-domain+wHTH+RRM</td><td>Type-1_BR.RM.like.systems</td><td><-ZnR+TOPRIM+D5ATPase<-XIS-HTH<-?<-HEPN+DUF499*<-?<-?<-METHYLASE<-?<-?<-ParB+HNH</td><td>1118</td><td>Bacteria>Acidobacteriota>Blastocatellia>Chloracidobacterium>unclassified_Chloracidobacterium</td><td>GCA_016716305.1</td><td>Chloracidobacterium_sp.</td><td>IPM28_09505</td><td>1672158..1675514</td><td>2099375</td><td>DUF499-domain-containing-protein-[Chloracidobacterium-sp.]</td></tr>
<tr><td>MCS7081031.1</td><td>HEPN+AAA-ATPase+Alpha+Beta-domain+wHTH+RRM</td><td>Type-1_BR.RM.like.systems</td><td>HEPN+DUF499*-><-METHYLASE-p||?-><-?<-?<-?<-SAM-methylase</td><td>1113</td><td>Bacteria>Acidobacteriota>Blastocatellia>Chloracidobacterium>unclassified_Chloracidobacterium</td><td>GCA_025057975.1</td><td>Chloracidobacterium_sp.</td><td>NZ585_13400</td><td>246..3587</td><td>2099375</td><td>Swt1-family-HEPN-domain-containing-protein-[Chloracidobacterium-sp.]</td></tr>
<tr><td>MCS7080483.1</td><td>HEPN+AAA-ATPase+Alpha+Beta-domain+wHTH+RRM</td><td>Type-1_BR.RM.like.systems</td><td>METHYLASE-p->?->SF1-SF2-Helicase->?->HKD-DNase->DDE_Tnp_IS1->HEPN+DUF499*-></td><td>691</td><td>Bacteria>Acidobacteriota>Blastocatellia>Chloracidobacterium>unclassified_Chloracidobacterium</td><td>GCA_025057975.1</td><td>Chloracidobacterium_sp.</td><td>NZ585_10625</td><td>170541..172613</td><td>2099375</td><td>DUF499-domain-containing-protein-[Chloracidobacterium-sp.]</td></tr>
<tr><td>MBX7223403.1</td><td>HEPN+AAA-ATPase+Alpha+Beta-domain+wHTH+RRM</td><td>Type-1_BR.RM.like.systems</td><td><-HEPN+DUF499*<-METHYLASE<-NUDIX<-TUDOR+SF1-SF2-Helicase+iREase+CTD+REase</td><td>1127</td><td>Bacteria>Acidobacteriota>Blastocatellia>unclassified_Blastocatellia</td><td>GCA_019695185.1</td><td>Blastocatellia_bacterium</td><td>K1Y36_26050</td><td>326..3709</td><td>2052146</td><td>DUF499-domain-containing-protein-[Blastocatellia-bacterium]</td></tr>
<tr><td>MBL8148954.1</td><td>HEPN+AAA-ATPase+Alpha+Beta-domain+wHTH+RRM</td><td>Type-1_BR.RM.like.systems</td><td>VirB11+MoxR-AAA->?->Radical-SAM->Radical-SAM->?-><-HEPN+DUF499*||REase+HTH-></td><td>1094</td><td>Bacteria>Acidobacteriota>Blastocatellia>unclassified_Blastocatellia</td><td>GCA_016789295.1</td><td>Blastocatellia_bacterium</td><td>JNN15_03400</td><td>5699..8983</td><td>2052146</td><td>DUF499-domain-containing-protein-[Blastocatellia-bacterium]</td></tr>
<tr><td>MBO0720791.1</td><td>HEPN+AAA-ATPase+Alpha+Beta-domain+wHTH+RRM</td><td>Type-1_BR.RM.like.systems</td><td><-HEPN+DUF499*<-METHYLASE-p</td><td>1119</td><td>Bacteria>Acidobacteriota>Blastocatellia>unclassified_Blastocatellia</td><td>GCA_017353965.1</td><td>Blastocatellia_bacterium</td><td>J2P41_08210</td><td>1861..5220</td><td>2052146</td><td>DUF499-domain-containing-protein-[Blastocatellia-bacterium]</td></tr>
<tr><td>QQR73412.1</td><td>HEPN+AAA-ATPase+Alpha+Beta-domain+wHTH+RRM</td><td>Type-1_BR.RM.like.systems</td><td>STAS+HisKin->?->?->METHYLASE->OLD-ABC->TOPRIM->HEPN+DUF499*-><-?<-Aminotran_1_2<-?<-?||ACET-></td><td>1117</td><td>Bacteria>Acidobacteriota>Holophagae>Holophagales>unclassified_Holophagales</td><td>GCA_016699405.1</td><td>Holophagales_bacterium</td><td>IPJ17_18310</td><td>4435335..4438688</td><td>2478486</td><td>DUF499-domain-containing-protein-[Holophagales-bacterium]</td></tr>
<tr><td>MYA06552.1</td><td>HEPN+AAA-ATPase+Alpha+Beta-domain+wHTH+RRM</td><td>Type-1_BR.RM.like.systems</td><td><-Radical-SAM+TRAM<-?<-?||?->TUDOR+SF1-SF2-Helicase+iREase+CTD+REase->HEPN+DUF499*-></td><td>1134</td><td>Bacteria>Acidobacteriota>Holophagae>Holophagales>unclassified_Holophagales</td><td>GCA_009838785.1</td><td>Holophagales_bacterium</td><td>F4Y32_01005</td><td>72319..75723</td><td>2478486</td><td>ATP-binding-protein-[Holophagales-bacterium]</td></tr>
<tr><td>MBZ2178820.1</td><td>HEPN+AAA-ATPase+Alpha+Beta-domain+wHTH+RRM</td><td>Type-1_BR.RM.like.systems</td><td>TUDOR+SF1-SF2-Helicase+iREase+CTD+REase->?->METHYLASE->HEPN+DUF499*->MerR-HTH->?->KAP-NTPase->?->Cluster2725_2clades->Terminase-></td><td>1118</td><td>Bacteria>Acidobacteriota>Terriglobia>Bryobacterales>Bryobacteraceae</td><td>GCA_019931245.1</td><td>Bryobacteraceae_bacterium_CoA2_C42</td><td>K7J47_14040</td><td>14682..18038</td><td>2967959</td><td>DUF499-domain-containing-protein-[Acidobacteria-bacterium]</td></tr>
<tr><td>MBE0656246.1</td><td>HEPN+AAA-ATPase+Alpha+Beta-domain+wHTH+RRM</td><td>Type-1_BR.RM.like.systems</td><td><-HEPN+DUF499*<-METHYLASE<-sGTP<-HKD-DNase<-?<-TUDOR+SF1-SF2-Helicase+iREase+CTD+REase</td><td>1127</td><td>Bacteria>Acidobacteriota>Terriglobia>Bryobacterales>Bryobacteraceae</td><td>GCA_014860455.1</td><td>Bryobacteraceae_bacterium</td><td>IH602_01075</td><td>1306..4689</td><td>2212468</td><td>DUF499-domain-containing-protein-[Bryobacteraceae-bacterium]</td></tr>
<tr><td>MCL6508118.1</td><td>HEPN+AAA-ATPase+Alpha+Beta-domain+wHTH+RRM</td><td>Type-1_BR.RM.like.systems</td><td><-HEPN+DUF499*<-METHYLASE-p</td><td>671</td><td>Bacteria>Acidobacteriota>Terriglobia>Bryobacterales>Bryobacteraceae</td><td>GCA_023511775.1</td><td>Bryobacteraceae_bacterium</td><td>K6T59_13960</td><td>1..2014</td><td>2212468</td><td>DUF499-domain-containing-protein-[Bryobacteraceae-bacterium]</td></tr>
<tr><td>SPE35562.1</td><td>HEPN+AAA-ATPase+Alpha+Beta-domain+wHTH+RRM</td><td>Type-1_BR.RM.like.systems</td><td>METHYLASE->METHYLASE-p-><-?||ParB->PolB->HEPN->HEPN+DUF499*->DUF499-p-></td><td>906</td><td>Bacteria>Acidobacteriota>Terriglobia>Bryobacterales>Solibacteraceae</td><td>GCA_900290325.1</td><td>Candidatus_Sulfopaludibacter_sp._SbA6</td><td>SBA6_470068</td><td>62270..64990</td><td>2043166</td><td>conserved-hypothetical-protein-[Candidatus-Sulfopaludibacter-sp.-SbA6]</td></tr>
<tr><td>MDE0101184.1</td><td>HEPN+AAA-ATPase+Alpha+Beta-domain+wHTH+RRM</td><td>Type-1_BR.RM.like.systems</td><td>PadR-HTH-><-?||N6_MTase-><-HEPN+DUF499*<-HEPN<-Polbeta<-METHYLASE<-HEPN<-TUDOR+SF1-SF2-Helicase+iREase+CTD+REase||REase-></td><td>1110</td><td>Bacteria>Acidobacteriota>Terriglobia>Bryobacterales>unclassified_Bryobacterales</td><td>GCA_028819235.1</td><td>Bryobacterales_bacterium</td><td>OXN89_02285</td><td>6150..9482</td><td>2026791</td><td>MAG:-Swt1-family-HEPN-domain-containing-protein-[Bryobacterales-bacterium]</td></tr>
<tr><td>MBK9166737.1</td><td>HEPN+AAA-ATPase+Alpha+Beta-domain+wHTH+RRM</td><td>Type-1_BR.RM.like.systems</td><td>Schlafen+HATPase+HTH->METHYLASE-p-><-rve+rve_3<-HTH_Tnp_1||METHYLASE-p->ZDNA-HTH->HEPN+DUF499*->?->?->ABhydrolase-></td><td>1121</td><td>Bacteria>Acidobacteriota>Terriglobia>Bryobacterales>unclassified_Bryobacterales</td><td>GCA_016716385.1</td><td>Bryobacterales_bacterium</td><td>IPM24_04655</td><td>396638..400003</td><td>2026791</td><td>DUF499-domain-containing-protein-[Bryobacterales-bacterium]</td></tr>
<tr><td>MCL4784604.1</td><td>HEPN+AAA-ATPase+Alpha+Beta-domain+wHTH+RRM</td><td>Type-1_BR.RM.like.systems</td><td><-HTH||?->?->?-><-HEPN+DUF499*<-METHYLASE<-?<-TUDOR+SF1-SF2-Helicase+iREase+CTD+REase</td><td>1109</td><td>Bacteria>Acidobacteriota>Terriglobia>Bryobacterales>unclassified_Bryobacterales</td><td>GCA_023384735.1</td><td>Bryobacterales_bacterium</td><td>KJZ70_16335</td><td>4638..7967</td><td>2026791</td><td>DUF499-domain-containing-protein-[Bryobacterales-bacterium]</td></tr>
<tr><td>MCY4596713.1</td><td>HEPN+AAA-ATPase+Alpha+Beta-domain+wHTH+RRM</td><td>Type-1_BR.RM.like.systems</td><td>TUDOR+SF1-SF2-Helicase+iREase+CTD+REase->HEPN+DUF499*->?-><-METHYLASE-p</td><td>1110</td><td>Bacteria>Acidobacteriota>Terriglobia>Bryobacterales>unclassified_Bryobacterales</td><td>GCA_026713455.1</td><td>Bryobacterales_bacterium</td><td>OXC19_18175</td><td>3853..7185</td><td>2026791</td><td>MAG:-Swt1-family-HEPN-domain-containing-protein-[Bryobacterales-bacterium]</td></tr>
</table>

| Protein-ID | DUF499-architecture | System | Gene-Neighbourhoods | Length | Taxonomy | Assembly-ID | Strain | Locus-Tag | Genomic-coordinates | TaxID | Fasta-defline |
|---|---|---|---|---|---|---|---|---|---|---|---|
| MCU1329937.1 | HEPN+AAA-ATPase+Alpha+Beta-domain+wHTH+RRM | Type-1_BR.RM.like.systems | <-HEPN+DUF499*<-METHYLASE<-TUDOR+SF1-SF2-Helicase+iREase+CTD+REase | 1116 | Bacteria>Acidobacteriota>Terriglobia>Bryobacterales>unclassified_Bryobacterales | GCA_025462945.1 | Bryobacterales_bacterium | JWN34_5307 | 1570..4920 | 2026791 | ATPase-superfamily-[Bryobacterales-bacterium] |
| WP_038049904.1 | HEPN+AAA-ATPase+Alpha+Beta-domain+wHTH+RRM | Type-1_BR.RM.like.systems | Radical-SAM-><-?<-?<-?<-ORC-CDC6+REase\|\|?-><-HEPN+DUF499*<-METHYLASE<-MORC+DUF3684<-TUDOR+SF1-SF2-Helicase+iREase+CTD+REase\|\|ORC-CDC6+REase->?->DUF433-> | 1117 | Bacteria>Acidobacteriota>Thermoanaerobaculia>Thermoanaerobaculales>Thermoanaerobaculaceae | GCF_000687145.1 | Thermoanaerobaculum_aquaticum | EG19_RS09435 | 122506..125859 | 1312852 | Swt1-family-HEPN-domain-containing-protein-[Thermoanaerobaculum-aquaticum] |
| MBP7799276.1 | HEPN+AAA-ATPase+Alpha+Beta-domain+wHTH+RRM | Type-1_BR.RM.like.systems | <-HEPN+DUF499*<-PIN<-OLD-ABC<-METHYLASE<-TOPRIM<-OLD-ABC | 1114 | Bacteria>Acidobacteriota>Thermoanaerobaculia>Thermoanaerobaculales>Thermoanaerobaculaceae | GCA_018000955.1 | Thermoanaerobaculaceae_bacterium | KA072_11695 | 22806..26150 | 2699758 | DUF499-domain-containing-protein-[Thermoanaerobaculaceae-bacterium] |
| MCG3192277.1 | HEPN+AAA-ATPase+Alpha+Beta-domain+wHTH+RRM | Type-1_BR.RM.like.systems | TUDOR+SF1-SF2-Helicase+iREase+CTD+REase->OLD-ABC->TOPRIM->METHYLASE->PHP+SbcC->Schlafen+HATPase+HTH->HEPN+DUF499*->?->?->PAIREDC-HTH-> | 1114 | Bacteria>Acidobacteriota>Thermoanaerobaculia>unclassified_Thermoanaerobaculia | GCA_022071945.1 | Thermoanaerobaculia_bacterium | DIJKHBIC_01515 | 140917..144261 | 2651171 | hypothetical-protein-[Thermoanaerobaculia-bacterium] |
| MCK6683888.1 | HEPN+AAA-ATPase+Alpha+Beta-domain+wHTH+RRM | Type-1_BR.RM.like.systems | TUDOR+SF1-SF2-Helicase+iREase+CTD+REase->?->?->MACRODOMAIN->METHYLASE->HEPN+DUF499*->?->OLD-ABC+TOPRIM->DUF932->?->ASCH+DUF3780-> | 1093 | Bacteria>Acidobacteriota>Thermoanaerobaculia>unclassified_Thermoanaerobaculia | GCA_023151375.1 | Thermoanaerobaculia_bacterium | L6R30_15920 | 12714..15995 | 2651171 | DUF499-domain-containing-protein-[Thermoanaerobaculia-bacterium] |
| KAB2964430.1 | HEPN+AAA-ATPase+Alpha+Beta-domain+wHTH+RRM | Type-1_BR.RM.like.systems | REase->TUDOR+SF1-SF2-Helicase+iREase+CTD+REase->ParB+HNH->HEPN->METHYLASE->?->HEPN+DUF499*->HNH-> | 1109 | Bacteria>Acidobacteriota>Thermoanaerobaculia>unclassified_Thermoanaerobaculia | GCA_009026165.1 | Thermoanaerobaculia_bacterium | F9K16_04350 | 12106..15435 | 2651171 | MAG:-ATP-binding-protein-[Thermoanaerobaculia-bacterium] |
| MDE2692299.1 | HEPN+AAA-ATPase+Alpha+Beta-domain+wHTH+RRM | Type-1_BR.RM.like.systems | <-METHYLASE\|\|Phage_integrase->Phage_int_SAM_3+Phage_integrase->HEPN->METHYLASE->?-><-HEPN+DUF499*<-TUDOR+SF1-SF2-Helicase+iREase+CTD+REase<-?\|\|Arm-DNA-bind_3+Phage_integrase-> | 1104 | Bacteria>Acidobacteriota>unclassified_Acidobacteriota | GCA_028875725.1 | Acidobacteriota_bacterium | OXI49_17510 | 451971..455285 | 1978231 | MAG:-Swt1-family-HEPN-domain-containing-protein-[Acidobacteriota-bacterium] |
| MCY3930234.1 | HEPN+AAA-ATPase+Alpha+Beta-domain+wHTH+RRM | Type-1_BR.RM.like.systems | <-Phage_int_SAM_3+Phage_integrase<-Phage_int_SAM_3+Phage_integrase\|\|METHYLASE-><-HEPN+DUF499*<-TUDOR+SF1-SF2-Helicase+iREase+CTD+REase<-?<-OLD-ABC-p | 1112 | Bacteria>Acidobacteriota>unclassified_Acidobacteriota | GCA_026707325.1 | Acidobacteriota_bacterium | OXH70_00800 | 52234..55572 | 1978231 | MAG:-DUF499-domain-containing-protein-[Acidobacteria-bacterium] |
| MBI4902187.1 | HEPN+AAA-ATPase+Alpha+Beta-domain+wHTH+RRM | Type-1_BR.RM.like.systems | TUDOR+SF1-SF2-Helicase+iREase+CTD+REase->ParB+HNH->METHYLASE->OLD-ABC->TOPRIM->HEPN+DUF499*->Cluster2725_2clades->Terminase-> | 1115 | Bacteria>Acidobacteriota>unclassified_Acidobacteriota | GCA_016213775.1 | Acidobacteriota_bacterium | HY820_01040 | 17603..20950 | 1978231 | ATP-binding-protein-[Acidobacteria-bacterium] |
| ODS52398.1 | HEPN+AAA-ATPase+Alpha+Beta-domain+wHTH+RRM | Type-1_BR.RM.like.systems | <-MethyltransfD12<-ParB<-HEPN+DUF499*<-METHYLASE<-Schlafen+HATPase+HTH<-TUDOR+SF1-SF2-Helicase+iREase+CTD+REase<-?\|\|?->Peptidase_S41-> | 1114 | Bacteria>Acidobacteriota>unclassified_Acidobacteriota | GCA_001724025.1 | Acidobacteria_bacterium_SCN_69-37 | ABS36_17605 | 152750..156094 | 1660160 | AAA+-family-ATPase-[Acidobacteria-bacterium-SCN-69-37] |
| MBP7777875.1 | HEPN+AAA-ATPase+Alpha+Beta-domain+wHTH+RRM | Type-1_BR.RM.like.systems | TUDOR+SF1-SF2-Helicase+iREase+CTD+REase->DUF932->METHYLASE->HEPN+DUF499*->?->Pkinase->ParB+HNH->REase->?->HicB_lk_antitox-> | 1114 | Bacteria>Acidobacteriota>unclassified_Acidobacteriota | GCA_018000455.1 | Acidobacteriota_bacterium | KA371_12150 | 120064..123408 | 1978231 | DUF499-domain-containing-protein-[Acidobacteria-bacterium] |
| MBS1816795.1 | HEPN+AAA-ATPase+Alpha+Beta-domain+wHTH+RRM | Type-1_BR.RM.like.systems | <-XIS-HTH<-?\|\|HTH+PolB->TUDOR+SF1-SF2-Helicase+iREase+CTD+REase->METHYLASE->?->HEPN+DUF499*->?-><-OLD-ABC+TOPRIM<-?<-?<-AbiEi_4\|\|REase-> | 1111 | Bacteria>Acidobacteriota>unclassified_Acidobacteriota | GCA_018268955.1 | Acidobacteriota_bacterium | JSU08_02600 | 186783..190118 | 1978231 | DUF499-domain-containing-protein-[Acidobacteria-bacterium] |
| MCL5262739.1 | HEPN+AAA-ATPase+Alpha+Beta-domain+wHTH+RRM | Type-1_BR.RM.like.systems | <-HEPN+DUF499*<-METHYLASE<-?<-TUDOR+SF1-SF2-Helicase+iREase+CTD+REase-p<-REase<-HKD-DNase+SF1-SF2-Helicase+TRD | 1103 | Bacteria>Acidobacteriota>unclassified_Acidobacteriota | GCA_023480755.1 | Acidobacteriota_bacterium | M1568_00545 | 1..3309 | 1978231 | Swt1-family-HEPN-domain-containing-protein-[Acidobacteria-bacterium] |
| MCC6131982.1 | HEPN+AAA-ATPase+Alpha+Beta-domain+wHTH+RRM | Type-1_BR.RM.like.systems | <-HEPN+DUF499*<-Schlafen+HATPase+HTH<-PHP+SbcC<-METHYLASE<-TOPRIM<-OLD-ABC<-TUDOR+SF1-SF2-Helicase+iREase+CTD+REase | 1114 | Bacteria>Acidobacteriota>unclassified_Acidobacteriota | GCA_020845555.1 | Acidobacteriota_bacterium | IT186_18830 | 2559..5903 | 1978231 | DUF499-domain-containing-protein-[Acidobacteria-bacterium] |
| MBI1792208.1 | HEPN+AAA-ATPase+Alpha+Beta-domain+wHTH+RRM | Type-1_BR.RM.like.systems | TUDOR+SF1-SF2-Helicase+iREase+CTD+REase->Schlafen+HATPase+HTH->METHYLASE->PolB->HEPN+DUF499*->OLD-ABC+TOPRIM-> | 1093 | Bacteria>Acidobacteriota>unclassified_Acidobacteriota | GCA_016178945.1 | Acidobacteriota_bacterium | HYR60_32190 | 8610..11891 | 1978231 | ATP-binding-protein-[Acidobacteria-bacterium] |
| PYQ27742.1 | HEPN+AAA-ATPase+Alpha+Beta-domain+wHTH+RRM | Type-1_BR.RM.like.systems | ABhydrolase-><-?<-HTH\|\|?-><-?<-HEPN+DUF499*\|\|HD->DUF4065-><-METHYLASE<-TUDOR+SF1-SF2-Helicase+iREase+CTD+REase | 1105 | Bacteria>Acidobacteriota>unclassified_Acidobacteriota | GCA_003223645.1 | Acidobacteriota_bacterium | DMF56_19255 | 118621..121938 | 1978231 | AAA+-family-ATPase-[Acidobacteria-bacterium] |
| MCI0623777.1 | HEPN+AAA-ATPase+Alpha+Beta-domain+wHTH+RRM | Type-1_BR.RM.like.systems | HTH-><-HEPN+DUF499*<-METHYLASE<-PIN<-?<-STAS+HisKin<-TUDOR+SF1-SF2-Helicase+iREase+CTD+REase<-XIS-HTH | 1110 | Bacteria>Acidobacteriota>unclassified_Acidobacteriota | GCA_022615935.1 | Acidobacteriota_bacterium | L0387_19325 | 508..3840 | 1978231 | DUF499-domain-containing-protein-[Acidobacteria-bacterium] |
| TAM79789.1 | HEPN+AAA-ATPase+Alpha+Beta-domain+wHTH+RRM | Type-1_BR.RM.like.systems | PadR-HTH->?->?-><-?<-HEPN+DUF499*<-DUF433<-METHYLASE<-?<-TUDOR+SF1-SF2-Helicase+iREase+CTD+REase<-MPTase<-HTH | 1104 | Bacteria>Acidobacteriota>unclassified_Acidobacteriota | GCA_004298155.1 | Acidobacteriota_bacterium | EPN47_18525 | 398399..401713 | 1978231 | DUF499-domain-containing-protein-[Acidobacteria-bacterium] |
| MDE2851190.1 | HEPN+AAA-ATPase+Alpha+Beta-domain+wHTH+RRM | Type-1_BR.RM.like.systems | METHYLASE->?-><-HEPN+DUF499*<-TUDOR+SF1-SF2-Helicase+iREase+CTD+REase<-?<-?<-?<-?<-MqsA_antitoxin+HTH_3 | 1104 | Bacteria>Acidobacteriota>unclassified_Acidobacteriota | GCA_028874635.1 | Acidobacteriota_bacterium | OXP74_11315 | 2020..5334 | 1978231 | MAG:-Swt1-family-HEPN-domain-containing-protein-[Acidobacteriota-bacterium] |
| MDE3262363.1 | HEPN+AAA-ATPase+Alpha+Beta-domain+wHTH+RRM | Type-1_BR.RM.like.systems | <-ArdCN+MPTase<-?<-?<-XIS-HTH<-?\|\|TUDOR+SF1-SF2-Helicase+iREase+CTD+REase->HEPN+DUF499*-> | 1106 | Bacteria>Acidobacteriota>unclassified_Acidobacteriota | GCA_028877265.1 | Acidobacteriota_bacterium | OYL41_10335 | 11539..14859 | 1978231 | MAG:-Swt1-family-HEPN-domain-containing-protein-[Acidobacteriota-bacterium] |
| MYD70260.1 | HEPN+AAA-ATPase+Alpha+Beta-domain+wHTH+RRM | Type-1_BR.RM.like.systems | <-HEPN+DUF499*<-TUDOR+SF1-SF2-Helicase+iREase+CTD+REase\|\|Peptidase-> | 1104 | Bacteria>Acidobacteriota>unclassified_Acidobacteriota | GCA_009841675.1 | Acidobacteriota_bacterium | F4W89_05855 | 745..4059 | 1978231 | ATP-binding-protein-[Acidobacteria-bacterium] |
| MDD8029960.1 | HEPN+AAA-ATPase+Alpha+Beta-domain+wHTH+RRM | Type-1_BR.RM.like.systems | METHYLASE->HEPN+DUF499*-><-?<-?<-Cysteine_desulfurase\|\|Peptidase_M24-> | 1114 | Bacteria>Acidobacteriota>unclassified_Acidobacteriota | GCA_028749155.1 | Acidobacteriota_bacterium | PHE62_10480 | 3051..6395 | 1978231 | MAG:-Swt1-family-HEPN-domain-containing-protein-[Acidobacteriota-bacterium] |
| RIK10616.1 | HEPN+AAA-ATPase+Alpha+Beta-domain+wHTH+RRM | Type-1_BR.RM.like.systems | DOC->?->MPTase->METHYLASE->HEPN+DUF499*-><-?<-Schlafen<-?<-tRNA<-tRNA<-?\|\|tRNA-><-METHYLASE-p | 1113 | Bacteria>Acidobacteriota>unclassified_Acidobacteriota | GCA_003577105.1 | Acidobacteriota_bacterium | DCC49_02885 | 59433..62774 | 1978231 | AAA+-family-ATPase-[Acidobacteria-bacterium] |
| MCI0622286.1 | HEPN+AAA-ATPase+Alpha+Beta-domain+wHTH+RRM | Type-1_BR.RM.like.systems | <-REase<-HEPN+DUF499*<-?<-DUF488-l<-?<-METHYLASE<-HEPN<-METHYLASE-p | 1117 | Bacteria>Acidobacteriota>unclassified_Acidobacteriota | GCA_022615935.1 | Acidobacteriota_bacterium | L0387_11575 | 924..4277 | 1978231 | DUF499-domain-containing-protein-[Acidobacteria-bacterium] |
| MXW37283.1 | HEPN+AAA-ATPase+Alpha+Beta-domain+wHTH+RRM | Type-1_BR.RM.like.systems | TUDOR+SF1-SF2-Helicase+iREase+CTD+REase-p->HEPN+DUF499*-> | 1106 | Bacteria>Acidobacteriota>unclassified_Acidobacteriota | GCA_009836105.1 | Acidobacteriota_bacterium | F4Z65_03330 | 485..3805 | 1978231 | ATP-binding-protein-[Acidobacteria-bacterium] |
| MXW70175.1 | HEPN+AAA-ATPase+Alpha+Beta-domain+wHTH+RRM | Type-1_BR.RM.like.systems | TUDOR+SF1-SF2-Helicase+iREase+CTD+REase->HEPN+DUF499*-> | 1106 | Bacteria>Acidobacteriota>unclassified_Acidobacteriota | GCA_009835985.1 | Acidobacteriota_bacterium | F4Z74_01830 | 14208..17528 | 1978231 | ATP-binding-protein-[Acidobacteria-bacterium] |
| MBP7569108.1 | HEPN+AAA-ATPase+Alpha+Beta-domain+wHTH+RRM | Type-1_BR.RM.like.systems | Cas_Cmr3->?->Cas_Cmr5->?->CARF+HEPN->HEPN+DUF499*->Schlafen-> | 1117 | Bacteria>Acidobacteriota>unclassified_Acidobacteriota | GCA_017998655.1 | Acidobacteriota_bacterium | KBA95_03500 | 22125..25478 | 1978231 | DUF499-domain-containing-protein-[Acidobacteria-bacterium] |
| MBS1819352.1 | HEPN+AAA-ATPase+Alpha+Beta-domain+wHTH+RRM | Type-1_BR.RM.like.systems | <-SF1-SF2-Helicase<-OLD-ABC+TOPRIM<-HEPN+DUF499*<-METHYLASE-p | 1115 | Bacteria>Acidobacteriota>unclassified_Acidobacteriota | GCA_018268955.1 | Acidobacteriota_bacterium | JSU08_15565 | 76293..79640 | 1978231 | DUF499-domain-containing-protein-[Acidobacteria-bacterium] |
| MCI0388810.1 | HEPN+AAA-ATPase+Alpha+Beta-domain+wHTH+RRM | Type-1_BR.RM.like.systems | HTH+PolB->HEPN->HEPN+DUF499*-><-Peptidase | 1116 | Bacteria>Acidobacteriota>unclassified_Acidobacteriota | GCA_022614935.1 | Acidobacteriota_bacterium | MOB07_08610 | 4163..7513 | 1978231 | Swt1-family-HEPN-domain-containing-protein-[Acidobacteria-bacterium] |
| RMH20600.1 | HEPN+AAA-ATPase+Alpha+Beta-domain+wHTH+RRM | Type-1_BR.RM.like.systems | METHYLASE-p->DUF433->PIN->HEPN+DUF499*-> | 1096 | Bacteria>Acidobacteriota>unclassified_Acidobacteriota | GCA_003697015.1 | Acidobacteriota_bacterium | D6696_07825 | 1571..4858 | 1978231 | ATP-binding-protein,-partial-[Acidobacteria-bacterium] |
| HHN74473.1 | HEPN+AAA-ATPase+Alpha+Beta-domain+wHTH+RRM | Type-1_BR.RM.like.systems | <-MORC+DUF3684+REase<-HEPN+DUF499* | 1091 | Bacteria>Acidobacteriota>unclassified_Acidobacteriota | GCA_011374785.1 | Acidobacteriota_bacterium | ENK10_04515 | 4666..7941 | 1978231 | ATP-binding-protein-[Acidobacteria-bacterium] |
| NOT26039.1 | HEPN+AAA-ATPase+Alpha+Beta-domain+wHTH+RRM | Type-1_BR.RM.like.systems | <-METHYLASE-p<-?<-?<-?\|\|?->?-><-HEPN+DUF499* | 1117 | Bacteria>Acidobacteriota>unclassified_Acidobacteriota | GCA_013140775.1 | Acidobacteriota_bacterium | HOP16_08040 | 9763..13116 | 1978231 | ATP-binding-protein-[Acidobacteria-bacterium] |
| NMD09846.1 | HEPN+AAA-ATPase+Alpha+Beta-domain+wHTH+RRM | Type-1_BR.RM.like.systems | <-HEPN+DUF499*<-METHYLASE-p | 545 | Bacteria>Acidobacteriota>unclassified_Acidobacteriota | GCA_012797375.1 | Acidobacteriota_bacterium | GYA74_01585 | 1..1635 | 1978231 | ATP-binding-protein-[Acidobacteriota-bacterium] |
| NLT66835.1 | HEPN+AAA-ATPase+Alpha+Beta-domain+wHTH+RRM | Type-1_BR.RM.like.systems | METHYLASE-p->HEPN+DUF499*-> | 772 | Bacteria>Acidobacteriota>unclassified_Acidobacteriota | GCA_012719355.1 | Acidobacteriota_bacterium | GXX84_09545 | 1125..3439 | 1978231 | ATP-binding-protein-[Acidobacteria-bacterium] |
| MBM3792128.1 | HEPN+AAA-ATPase+Alpha+Beta-domain+wHTH+RRM | Type-1_BR.RM.like.systems | METHYLASE-p->HEPN+DUF499*-> | 464 | Bacteria>Acidobacteriota>unclassified_Acidobacteriota | GCA_016871075.1 | Acidobacteriota_bacterium | FJW35_17505 | 483..1875 | 1978231 | ATP-binding-protein-[Acidobacteriota-bacterium] |
| MYD85827.1 | HEPN+AAA-ATPase+Alpha+Beta-domain+wHTH+RRM | Type-1_BR.RM.like.systems | <-HEPN+DUF499*<-?<-?<-METHYLASE | 1107 | Bacteria>Acidobacteriota>unclassified_Acidobacteriota | GCA_009839145.1 | Acidobacteriota_bacterium | F4Y14_06400 | 201..3524 | 1978231 | ATP-binding-protein-[Acidobacteria-bacterium] |
| MBA4124439.1 | HEPN+AAA-ATPase+Alpha+Beta-domain+wHTH+RRM | Type-1_BR.RM.like.systems | DDE_Tnp_1_assoc+DDE_Tnp_1->CHTH->?->TUDOR+SF1-SF2-Helicase+iREase+CTD+REase->ParB+HNH->METHYLASE->HEPN+DUF499*->?->?->BirA-HTH-><-35exo<-Phage_integrase | 1091 | Bacteria>Acidobacteriota>unclassified_Acidobacteriota | GCA_013820745.1 | Acidobacteriota_bacterium | HOX72_18505 | 11663..14938 | 1978231 | DUF499-domain-containing-protein-[Acidobacteria-bacterium] |
| MBS1794111.1 | HEPN+AAA-ATPase+Alpha+Beta-domain+wHTH+RRM | Type-1_BR.RM.like.systems | <-Phage_integrase\|\|TUDOR+SF1-SF2-Helicase+iREase+CTD+REase->?->METHYLASE->?->HEPN+DUF499*->?->?->REase->?->SF1-SF2-Helicase-> | 1100 | Bacteria>Acidobacteriota>unclassified_Acidobacteriota | GCA_018269075.1 | Acidobacteriota_bacterium | JSS81_09670 | 12622..15924 | 1978231 | DUF499-domain-containing-protein-[Acidobacteria-bacterium] |
| MCL4821456.1 | HEPN+AAA-ATPase+Alpha+Beta-domain+wHTH+RRM | Type-1_BR.RM.like.systems | TUDOR+SF1-SF2-Helicase+iREase+CTD+REase-p->XIS-HTH+RVT->METHYLASE->OLD-ABC->PIN->HEPN+DUF499*->SF1-SF2-Helicase-><-?<-?\|\|Pkinase-> | 1115 | Bacteria>Acidobacteriota>Vicinamibacteria>unclassified_Vicinamibacteria | GCA_023384615.1 | Vicinamibacteria_bacterium | KJ067_20160 | 8492..11839 | 2793154 | DUF499-domain-containing-protein-[Vicinamibacteria-bacterium] |
| MCL4821144.1 | HEPN+AAA-ATPase+Alpha+Beta-domain+wHTH+RRM | Type-1_BR.RM.like.systems | STYKinase-><-HEPN+DUF499*<-?<-ORC-CDC6<-ParA-Soj-PloopNTPase<-HEPN<-METHYLASE<-PIN | 1116 | Bacteria>Acidobacteriota>Vicinamibacteria>unclassified_Vicinamibacteria | GCA_023384615.1 | Vicinamibacteria_bacterium | KJ067_18565 | 2540..5890 | 2793154 | DUF499-domain-containing-protein-[Vicinamibacteria-bacterium] |
| MBE3120494.1 | HEPN+AAA-ATPase+Alpha+Beta-domain+wHTH+RRM | Type-1_BR.RM.like.systems | <-KAP-NTPase<-HEPN+DUF499*<-METHYLASE<-Schlafen+HATPase+HTH | 1116 | Bacteria>Atribacterota>Atribacteria>unclassified_Atribacteria | GCA_014894575.1 | Candidatus_Atribacteria_bacterium | IMZ50_17295 | 4085..7435 | 2053509 | DUF499-domain-containing-protein-[Candidatus-Atribacteria-bacterium] |
| NSW76333.1 | HEPN+AAA-ATPase+Alpha+Beta-domain+wHTH+RRM | Type-1_BR.RM.like.systems | METHYLASE->HEPN+DUF499*->PolB-> | 1120 | Bacteria>Atribacterota>Atribacteria>unclassified_Atribacteria | GCA_013314795.1 | Candidatus_Atribacteria_bacterium | HPY68_06090 | 2996..6358 | 2053509 | ATP-binding-protein-[Candidatus-Atribacteria-bacterium] |
| KAA0227462.1 | HEPN+AAA-ATPase+Alpha+Beta-domain+wHTH+RRM | Type-1_BR.RM.like.systems | TUDOR+SF1-SF2-Helicase+iREase+CTD+REase->ArdCN+MPTase-><-PIN<-?\|\|HEPN+DUF499*-><-?<-STYKIN<-?<-?\|\|?->HAD-> | 1113 | Bacteria>Bacteria_incertae_sedis>Bacteria_candidate_phyla>candidate_division_KSB1>unclassified_candidate_division_KSB1 | GCA_008363195.1 | candidate_division_KSB1_bacterium | EDS67_15790 | 88921..92262 | 2172550 | ATP-binding-protein-[candidate-division-KSB1-bacterium] |
| PWB47246.1 | HEPN+AAA-ATPase+Alpha+Beta-domain+wHTH+RRM | Type-1_BR.RM.like.systems | TUDOR+SF1-SF2-Helicase+iREase+CTD+REase->Schlafen+HATPase+HTH->METHYLASE->HEPN+DUF499*->?->?-><-?<-REase<-?<-CopG_antitoxin | 1102 | Bacteria>Bacteria_incertae_sedis>Bacteria_candidate_phyla>candidate_division_NC10>unclassified_candidate_division_NC10 | GCA_003104975.1 | candidate_division_NC10_bacterium | C3F12_04520 | 184009..187317 | 2072417 | AAA+-family-ATPase-[candidate-division-NC10-bacterium] |
| MDE2058883.1 | HEPN+AAA-ATPase+Alpha+Beta-domain+wHTH+RRM | Type-1_BR.RM.like.systems | <-HEPN+DUF499*<-?<-METHYLASE<-TOPRIM<-OLD-ABC<-ORC-CDC6+REase<-TUDOR+SF1-SF2-Helicase+iREase+CTD+REase | 1122 | Bacteria>Bacteria_incertae_sedis>Bacteria_candidate_phyla>candidate_division_NC10>unclassified_candidate_division_NC10 | GCA_028864815.1 | candidate_division_NC10_bacterium | KGJ27_04040 | 4506..7874 | 2072417 | MAG:-DUF499-domain-containing-protein-[candidate-division-NC10-bacterium] |
| MCX6842110.1 | HEPN+AAA-ATPase+Alpha+Beta-domain+wHTH+RRM | Type-1_BR.RM.like.systems | NTP_transf->TUDOR+SF1-SF2-Helicase+iREase+CTD+REase->Schlafen+HATPase+HTH->?->METHYLASE-p->METHYLASE->HEPN+DUF499*-> | 1030 | Bacteria>Bacteria_incertae_sedis>Bacteria_candidate_phyla>candidate_division_WOR-3 | GCA_026397595.1 | candidate_division_WOR-3_bacterium | NTX53_07520 | 11691..14783 | 2052148 | MAG:-Swt1-family-HEPN-domain-containing-protein-[candidate-division-WOR-3-bacterium] |
| OQB57508.1 | HEPN+AAA-ATPase+Alpha+Beta-domain+wHTH+RRM | Type-1_BR.RM.like.systems | TUDOR+SF1-SF2-Helicase+iREase+CTD+REase->Schlafen+HATPase+HTH->METHYLASE->HEPN+DUF499*-><-?<-?<-Cysteine_desulfurase\|\|Peptidase_M24-> | 1114 | Bacteria>Bacteria_incertae_sedis>Bacteria_candidate_phyla>Candidatus_Aminicenantes>unclassified_Candidatus_Aminicenantes | GCA_002070465.1 | Candidatus_Aminicenantes_bacterium_ADurb.Bin147 | BWX98_01441 | 17398..20742 | 1852831 | hypothetical-protein-BWX98_01441-[Candidatus-Aminicenantes-bacterium-ADurb.Bin147] |
| WP_122031531.1 | HEPN+AAA-ATPase+Alpha+Beta-domain+wHTH+RRM | Type-1_BR.RM.like.systems | TUDOR+SF1-SF2-Helicase+iREase+CTD+REase->ORC-CDC6+REase->METHYLASE->OLD-ABC->PIN->HEPN+DUF499*->Radical-SAM+TRAM->PfkB->?->35exo+SF1-SF2-Helicase->?-><-NTP_transf_2 | 1110 | Bacteria>Bacteria_incertae_sedis>Bacteria_candidate_phyla>Candidatus_Bipolaricaulota>Bipolaricaulis | GCF_900465355.1 | Candidatus_Bipolaricaulis_anaerobius | BARAN1_RS05370 | 1116813..1120145 | 2026885 | Swt1-family-HEPN-domain-containing-protein-[Candidatus-Bipolaricaulis-anaerobius] |
| MDD5263468.1 | HEPN+AAA-ATPase+Alpha+Beta-domain+wHTH+RRM | Type-1_BR.RM.like.systems | <-HEPN+DUF499*\|\|?-><-OLD-ABC<-METHYLASE<-Schlafen+HATPase+HTH<-TUDOR+SF1-SF2-Helicase+iREase+CTD+REase-p | 1114 | Bacteria>Bacteria_incertae_sedis>Bacteria_candidate_phyla>Candidatus_Bipolaricaulota>Bipolaricaulis | GCA_028713635.1 | Candidatus_Bipolaricaulis_sp. | PHU43_01355 | 30296..33640 | 2814139 | MAG:-Swt1-family-HEPN-domain-containing-protein-[Candidatus-Bipolaricaulis-sp.] |
| QAA75938.1 | HEPN+AAA-ATPase+Alpha+Beta-domain+wHTH+RRM | Type-1_BR.RM.like.systems | <-RNaseH<-?<-?<-?<-?<-HEPN+DUF499*<-?<-METHYLASE<-TOPRIM<-OLD-ABC<-TUDOR+SF1-SF2-Helicase+iREase+CTD+REase | 1113 | Bacteria>Bacteria_incertae_sedis>Bacteria_candidate_phyla>Candidatus_Bipolaricaulota>Bipolaricaulis | GCA_004102645.1 | Candidatus_Bipolaricaulis_sibiricus | BIP78_0170 | 158925..162266 | 2501609 | hypothetical-protein-BIP78_0170-[Candidatus-Bipolaricaulis-sibiricus] |
| HDN19517.1 | HEPN+AAA-ATPase+Alpha+Beta-domain+wHTH+RRM | Type-1_BR.RM.like.systems | TUDOR+SF1-SF2-Helicase+iREase+CTD+REase->OLD-ABC+TOPRIM->METHYLASE->PolB->HEPN+DUF499*->MazE_antitoxin->PIN-> | 1111 | Bacteria>Bacteria_incertae_sedis>Bacteria_candidate_phyla>Candidatus_Bipolaricaulota>unclassified_Candidatus_Bipolaricaulota | GCA_011042375.1 | Candidatus_Acetothermia_bacterium | ENF46_00270 | 10022..13357 | 2053493 | ATP-binding-protein-[Candidatus-Acetothermia-bacterium] |
| KUK26942.1 | HEPN+AAA-ATPase+Alpha+Beta-domain+wHTH+RRM | Type-1_BR.RM.like.systems | <-DUF499-p<-HEPN+DUF499*<-PolB-NTase<-METHYLASE-p<-HEPN<-PolB<-METHYLASE-p<-PIN | 493 | Bacteria>Bacteria_incertae_sedis>Bacteria_candidate_phyla>Candidatus_Bipolaricaulota>unclassified_Candidatus_Bipolaricaulota | GCA_001508155.1 | Acetothermia_bacterium_64_32 | XD60_0831 | 38456..39937 | 1635286 | Uncharacterized-protein-XD60_0831-[Acetothermia-bacterium-64_32] |
| HAF70679.1 | HEPN+AAA-ATPase+Alpha+Beta-domain+wHTH+RRM | Type-1_BR.RM.like.systems | <-DUF499-p<-HEPN+DUF499*<-PolB-NTase<-METHYLASE-p<-HEPN<-PolB<-METHYLASE-p<-PIN | 515 | Bacteria>Bacteria_incertae_sedis>Bacteria_candidate_phyla>Candidatus_Bipolaricaulota>unclassified_Candidatus_Bipolaricaulota | GCA_003445465.1 | Candidatus_Acetothermia_bacterium | DCL37_04935 | 22341..23888 | 2053493 | hypothetical-protein-[Candidatus-Acetothermia-bacterium] |
| MBK8229368.1 | HEPN+AAA-ATPase+Alpha+Beta-domain+wHTH+RRM | Type-1_BR.RM.like.systems | <-PIN<-HTH+MPTase\|\|?->?->HTH+UbE1->TUDOR+SF1-SF2-Helicase+iREase+CTD+REase->HEPN+DUF499*->PAIREDC-HTH->DDE_3->DUF499-p->?->?->LZ_Tnp_IS66+zf-IS66+DDE_Tnp_IS66+DDE_Tnp_IS66_C-> | 799 | Bacteria>Bacteria_incertae_sedis>Bacteria_candidate_phyla>Candidatus_Eisenbacteria>unclassified_Candidatus_Eisenbacteria | GCA_016712505.1 | Candidatus_Eisenbacteria_bacterium | IPK72_02030 | 519396..521795 | 2212470 | DUF499-domain-containing-protein-[Candidatus-Eisenbacteria-bacterium] |
| MYC78293.1 | HEPN+AAA-ATPase+Alpha+Beta-domain+wHTH+RRM | Type-1_BR.RM.like.systems | <-HEPN<-URI-DNase<-?<-REase<-REase<-REase<-HEPN+DUF499*<-PIN<-HTH+MPTase<-?<-?<-?<-SF1-SF2-Helicase+REase | 1093 | Bacteria>Bacteria_incertae_sedis>Bacteria_candidate_phyla>Candidatus_Poribacteria>unclassified_Candidatus_Poribacteria | GCA_009841255.1 | Candidatus_Poribacteria_bacterium | F4X10_21230 | 47302..50583 | 2026781 | ATP-binding-protein-[Candidatus-Poribacteria-bacterium] |
| MYB95133.1 | HEPN+AAA-ATPase+Alpha+Beta-domain+wHTH+RRM | Type-1_BR.RM.like.systems | <-METHYLASE<-REase+HTH<-REase+HTH<-HEPN+DUF499*<-?<-?<-?<-SF1-SF2-Helicase+REase<-ABC-ATPase+SbcC | 1108 | Bacteria>Bacteria_incertae_sedis>Bacteria_candidate_phyla>Candidatus_Poribacteria>unclassified_Candidatus_Poribacteria | GCA_009838805.1 | Candidatus_Poribacteria_bacterium | F4Y43_14870 | 2954..6280 | 2026781 | ATP-binding-protein-[Candidatus-Poribacteria-bacterium] |
| MDE0423419.1 | HEPN+AAA-ATPase+Alpha+Beta-domain+wHTH+RRM | Type-1_BR.RM.like.systems | SF1-SF2-Helicase+REase->?->?->?->?->HEPN+DUF499*->REase+HTH->REase->METHYLASE->URI-DNase-> | 1090 | Bacteria>Bacteria_incertae_sedis>Bacteria_candidate_phyla>Candidatus_Poribacteria>unclassified_Candidatus_Poribacteria | GCA_028820845.1 | Candidatus_Poribacteria_bacterium | OXN25_00975 | 9456..12728 | 2026781 | MAG:-DUF499-domain-containing-protein-[Candidatus-Poribacteria-bacterium] |
| MDE0088638.1 | HEPN+AAA-ATPase+Alpha+Beta-domain+wHTH+RRM | Type-1_BR.RM.like.systems | ABC-ATPase+SbcC->ABC-ATPase+SbcC->?-><-URI-DNase<-METHYLASE<-HEPN+DUF499*<-Aldedh | 1101 | Bacteria>Bacteria_incertae_sedis>Bacteria_candidate_phyla>Candidatus_Poribacteria>unclassified_Candidatus_Poribacteria | GCA_028818995.1 | Candidatus_Poribacteria_bacterium | OXU23_23155 | 9286..12591 | 2026781 | MAG:-hypothetical-protein-OXU23_23155-[Candidatus-Poribacteria-bacterium] |
| MBI5115316.1 | HEPN+AAA-ATPase+Alpha+Beta-domain+wHTH+RRM | Type-1_BR.RM.like.systems | <-SF1-SF2-Helicase<-?<-HEPN+REase<-?<-PIN<-HTH+MPTase<-HEPN+DUF499*<-METHYLASE-p | 1118 | Bacteria>Bacteria_incertae_sedis>Bacteria_candidate_phyla>Candidatus_Poribacteria>unclassified_Candidatus_Poribacteria | GCA_016215705.1 | Candidatus_Poribacteria_bacterium | HZA56_02480 | 5349..8705 | 2026781 | ATP-binding-protein-[Candidatus-Poribacteria-bacterium] |
| MDE0014209.1 | HEPN+AAA-ATPase+Alpha+Beta-domain+wHTH+RRM | Type-1_BR.RM.like.systems | HTH+MPTase->PIN->HEPN+DUF499*->REase+HTH->REase->METHYLASE->URI-DNase-> | 1095 | Bacteria>Bacteria_incertae_sedis>Bacteria_candidate_phyla>Candidatus_Poribacteria>unclassified_Candidatus_Poribacteria | GCA_028818705.1 | Candidatus_Poribacteria_bacterium | OXU36_23900 | 11254..14541 | 2026781 | MAG:-DUF499-domain-containing-protein-[Candidatus-Poribacteria-bacterium] |
| MXV83700.1 | HEPN+AAA-ATPase+Alpha+Beta-domain+wHTH+RRM | Type-1_BR.RM.like.systems | <-URI-DNase<-?<-METHYLASE<-REase<-REase<-REase<-HEPN+DUF499*<-PIN | 1088 | Bacteria>Bacteria_incertae_sedis>Bacteria_candidate_phyla>Candidatus_Poribacteria>unclassified_Candidatus_Poribacteria | GCA_009835485.1 | Candidatus_Poribacteria_bacterium | F4Z98_10065 | 15224..18490 | 2026781 | ATP-binding-protein-[Candidatus-Poribacteria-bacterium] |
| MCY3724179.1 | HEPN+AAA-ATPase+Alpha+Beta-domain+wHTH+RRM | Type-1_BR.RM.like.systems | PIN->HEPN+DUF499*->REase+HTH->REase->METHYLASE->?->URI-DNase-> | 1093 | Bacteria>Bacteria_incertae_sedis>Bacteria_candidate_phyla>Candidatus_Poribacteria>unclassified_Candidatus_Poribacteria | GCA_026705955.1 | Candidatus_Poribacteria_bacterium | OXG97_18330 | 699..3980 | 2026781 | MAG:-DUF499-domain-containing-protein-[Candidatus-Poribacteria-bacterium] |
| MYK17049.1 | HEPN+AAA-ATPase+Alpha+Beta-domain+wHTH+RRM | Type-1_BR.RM.like.systems | HEPN+DUF499*->REase+HTH->REase->METHYLASE->URI-DNase-> | 1090 | Bacteria>Bacteria_incertae_sedis>Bacteria_candidate_phyla>Candidatus_Poribacteria>unclassified_Candidatus_Poribacteria | GCA_009845505.1 | Candidatus_Poribacteria_bacterium | F4055_02625 | 1958..5230 | 2026781 | ATP-binding-protein-[Candidatus-Poribacteria-bacterium] |
| MDE0681304.1 | HEPN+AAA-ATPase+Alpha+Beta-domain+wHTH+RRM | Type-1_BR.RM.like.systems | HTH+MPTase->PIN->HEPN+DUF499*->METHYLASE-> | 1094 | Bacteria>Bacteria_incertae_sedis>Bacteria_candidate_phyla>Candidatus_Poribacteria>unclassified_Candidatus_Poribacteria | GCA_028824575.1 | Candidatus_Poribacteria_bacterium | OXI63_00195 | 3088..6372 | 2026781 | MAG:-DUF499-domain-containing-protein-[Candidatus-Poribacteria-bacterium] |
| MDE0468570.1 | HEPN+AAA-ATPase+Alpha+Beta-domain+wHTH+RRM | Type-1_BR.RM.like.systems | <-URI-DNase<-METHYLASE<-?<-HEPN+DUF499* | 1092 | Bacteria>Bacteria_incertae_sedis>Bacteria_candidate_phyla>Candidatus_Poribacteria>unclassified_Candidatus_Poribacteria | GCA_028821605.1 | Candidatus_Poribacteria_bacterium | OYL97_16085 | 27558..30836 | 2026781 | MAG:-DUF499-domain-containing-protein-[Candidatus-Poribacteria-bacterium] |
| MDE0020779.1 | HEPN+AAA-ATPase+Alpha+Beta-domain+wHTH+RRM | Type-1_BR.RM.like.systems | <-METHYLASE<-HEPN+DUF499*<-?<-?<-SIGMA-HTH | 1121 | Bacteria>Bacteria_incertae_sedis>Bacteria_candidate_phyla>Candidatus_Poribacteria>unclassified_Candidatus_Poribacteria | GCA_028818695.1 | Candidatus_Poribacteria_bacterium | OXT69_05310 | 103742..107107 | 2026781 | MAG:-DUF499-domain-containing-protein-[Candidatus-Poribacteria-bacterium] |
| MBI2524457.1 | HEPN+AAA-ATPase+Alpha+Beta-domain+wHTH+RRM | Type-1_BR.RM.like.systems | REase->MORC+DUF3684->METHYLASE->?->?->HEPN+DUF499*->?->?->?->XIS-HTH->Phage_int_SAM_1+Phage_integrase-> | 1121 | Bacteria>Bacteria_incertae_sedis>Bacteria_candidate_phyla>Candidatus_Rokubacteria>unclassified_Candidatus_Rokubacteria | GCA_016188005.1 | Candidatus_Rokubacteria_bacterium | HYV93_00585 | 34529..37894 | 2053607 | ATP-binding-protein-[Candidatus-Rokubacteria-bacterium] |
| MBU0999702.1 | HEPN+AAA-ATPase+Alpha+Beta-domain+wHTH+RRM | Type-1_BR.RM.like.systems | METHYLASE-p->HEPN+DUF499*-> | 1122 | Bacteria>Bacteria_incertae_sedis>Bacteria_candidate_phyla>Patescibacteria_group>unclassified_Patescibacteria_group | GCA_018816985.1 | Patescibacteria_group_bacterium | KKG24_05405 | 385..3750 | 2052139 | DUF499-domain-containing-protein-[Patescibacteria-group-bacterium] |
| HBM97560.1 | HEPN+AAA-ATPase+Alpha+Beta-domain+wHTH+RRM | Type-1_BR.RM.like.systems | <-HEPN+DUF499*<-METHYLASE | 1146 | Bacteria>Bacteria_incertae_sedis>Bacteria_candidate_phyla>unclassified_Bacteria_candidate_phyla | GCA_003504095.1 | bacterium_UBP9_UBA11836 | DD394_08835 | 303..3743 | 2060920 | hypothetical-protein-[bacterium-UBP9_UBA11836] |
| WP_142699382.1 | HEPN+AAA-ATPase+Alpha+Beta-domain+wHTH+RRM | Type-1_BR.RM.like.systems | HTH-><-?\|\|?->TUDOR+SF1-SF2-Helicase+iREase+CTD+REase->METHYLASE->HEPN+DUF499*->REase-><-ASCH | 1099 | Bacteria>Bdellovibrionota>Bdellovibrionia>Bdellovibrionales>Pseudobdellovibrionaceae | GCF_006874645.1 | Bdellovibrio_sp._ZAP7 | DOM22_RS05315 | 1082808..1086107 | 2231053 | Swt1-family-HEPN-domain-containing-protein-[Bdellovibrio-sp.-ZAP7] |
| MBK8201825.1 | HEPN+AAA-ATPase+Alpha+Beta-domain+wHTH+RRM | Type-1_BR.RM.like.systems | <-HTH\|\|HTH->?->?->HEPN+DUF499*->DUF499->MarR-HTH-> | 351 | Bacteria>Bdellovibrionota>Bdellovibrionia>Bdellovibrionales>unclassified_Bdellovibrionales | GCA_016712575.1 | Bdellovibrionales_bacterium | IPK68_05755 | 72390..73445 | 2053517 | hypothetical-protein-[Bdellovibrionales-bacterium] |
| MCB0334043.1 | HEPN+AAA-ATPase+Alpha+Beta-domain+wHTH+RRM | Type-1_BR.RM.like.systems | METHYLASE-p->HEPN+DUF499*-> | 570 | Bacteria>Bdellovibrionota>Bdellovibrionia>Bdellovibrionales>unclassified_Bdellovibrionales | GCA_020434525.1 | Bdellovibrionales_bacterium | KDD55_11125 | 601..2311 | 2053517 | DUF499-domain-containing-protein-[Bdellovibrionales-bacterium] |
| MBN1883904.1 | HEPN+AAA-ATPase+Alpha+Beta-domain+wHTH+RRM | Type-1_BR.RM.like.systems | Peptidase_M48->TUDOR+SF1-SF2-Helicase+iREase+CTD+REase->Schlafen+HATPase+HTH->METHYLASE->OLD-ABC->TOPRIM->HEPN+DUF499*->PIN+OST-HTH-><-?<-?<-REase<-?\|\|DOC+HTH-> | 1113 | Bacteria>Candidatus_Krumholzibacteriota>unclassified_Candidatus_Krumholzibacteriota | GCA_016931295.1 | Candidatus_Krumholzibacteriota_bacterium | JW876_00100 | 28696..32037 | 2725313 | DUF499-domain-containing-protein-[Candidatus-Krumholzibacteriota-bacterium] |
| OGQ97428.1 | HEPN+AAA-ATPase+Alpha+Beta-domain+wHTH+RRM | Type-1_BR.RM.like.systems | REase->?->?->?-><-?<-SF1-SF2-Helicase<-HEPN+DUF499*<-METHYLASE<-?<-?<-ParB+HNH<-TOPRIM<-OLD-ABC | 1111 | Bacteria>delta/epsilon_subdivisions>Deltaproteobacteria>unclassified_Deltaproteobacteria | GCA_001799275.1 | Deltaproteobacteria_bacterium_RIFOXYD12_FULL_57_12 | A2521_08565 | 7962..11297 | 1797906 | AAA+-family-ATPase-[Deltaproteobacteria-bacterium-RIFOXYD12_FULL_57_12] |
| PJB39408.1 | HEPN+AAA-ATPase+Alpha+Beta-domain+wHTH+RRM | Type-1_BR.RM.like.systems | <-DUF499-p<-?<-HEPN+DUF499*<-Schlafen+HATPase+HTH<-PHP+SbcC<-METHYLASE-p | 1117 | Bacteria>delta/epsilon_subdivisions>Deltaproteobacteria>unclassified_Deltaproteobacteria | GCA_002789955.1 | Deltaproteobacteria_bacterium_CG_4_9_14_3_um_filter_63_12 | CO108_17305 | 2362..5715 | 1973967 | AAA+-family-ATPase-[Deltaproteobacteria-bacterium-CG_4_9_14_3_um_filter_63_12] |
| OQB73667.1 | HEPN+AAA-ATPase+Alpha+Beta-domain+wHTH+RRM | Type-1_BR.RM.like.systems | Classical-AAA->Peptidase_S8-><-HEPN+DUF499* | 1112 | Bacteria>delta/epsilon_subdivisions>Deltaproteobacteria>unclassified_Deltaproteobacteria | GCA_002070575.1 | Deltaproteobacteria_bacterium_ADurb.Bin135 | BWX92_03289 | 4255..7593 | 1852867 | hypothetical-protein-BWX92_03289-[Deltaproteobacteria-bacterium-ADurb.Bin135] |
| MBI4801911.1 | HEPN+AAA-ATPase+Alpha+Beta-domain+wHTH+RRM | Type-1_BR.RM.like.systems | STAS+HisKin->?->PIN->OLD-ABC->HNH->METHYLASE->HEPN+DUF499*->Glycos_trans_3N+Glycos_transf_3->Radical-SAM->Radical-SAM->?->?->RADICAL-SAM-> | 1132 | Bacteria>Elusimicrobiota>unclassified_Elusimicrobiota | GCA_016208085.1 | Elusimicrobiota_bacterium | HY796_05240 | 17514..20912 | 2030800 | ATP-binding-protein-[Elusimicrobia-bacterium] |
| MBI5243147.1 | HEPN+AAA-ATPase+Alpha+Beta-domain+wHTH+RRM | Type-1_BR.RM.like.systems | <-ORC-CDC6+REase\|\|?->PIN-><-TPR\|\|?-><-HEPN+DUF499*<-METHYLASE<-PIN<-?<-STAS+HisKin<-TUDOR+SF1-SF2-Helicase+iREase+CTD+REase | 1133 | Bacteria>Elusimicrobiota>unclassified_Elusimicrobiota | GCA_016212885.1 | Elusimicrobiota_bacterium | HY922_05595 | 9033..12434 | 2030800 | ATP-binding-protein-[Elusimicrobia-bacterium] |
| MCX5796464.1 | HEPN+AAA-ATPase+Alpha+Beta-domain+wHTH+RRM | Type-1_BR.RM.like.systems | TUDOR+SF1-SF2-Helicase+iREase+CTD+REase->STAS+HisKin->?->PIN->METHYLASE->OLD-ABC+TOPRIM->HEPN+DUF499*-> | 1133 | Bacteria>Elusimicrobiota>unclassified_Elusimicrobiota | GCA_026388095.1 | Elusimicrobiota_bacterium | NTY77_13305 | 62095..65496 | 2030800 | MAG:-Swt1-family-HEPN-domain-containing-protein-[Elusimicrobia-bacterium] |
| BAH89290.1 | HEPN+AAA-ATPase+Alpha+Beta-domain+wHTH+RRM | Type-1_BR.RM.like.systems | <-UbE1+JAB<-SMODS\|\|HTH+WYL->TUDOR+SF1-SF2-Helicase+iREase+CTD+REase->TUDOR+SF1-SF2-Helicase+iREase+CTD+REase-p->METHYLASE->HEPN+DUF499*->?-><-MODE-HTH+PBPII\|\|TetR-HTH-><-?\|\|?->MODE-HTH+PBPII-> | 1099 | Bacteria>environmental_samples | - | uncultured_bacterium | . | 18704..22003 | 77133 | conserved-hypothetical-protein-[uncultured-bacterium] |
| CAG1064741.1 | HEPN+AAA-ATPase+Alpha+Beta-domain+wHTH+RRM | Type-1_BR.RM.like.systems | <-DOC<-MazE_antitoxin<-?<-TUDOR+SF1-SF2-Helicase+iREase+CTD<-?<-HEPN+DUF499*<-METHYLASE<-ArdCN+MPTase<-Schlafen+HATPase+HTH<-ArdCN+MPTase<-TUDOR+SF1-SF2-Helicase+iREase+CTD+REase<-HTH_26 | 1109 | Bacteria>environmental_samples | GCA_905339405.1 | uncultured_bacterium | BAC1_00306 | 123828..127157 | 77133 | hypothetical-protein-BAC1_00306-[uncultured-bacterium] |
| ART37960.1 | HEPN+AAA-ATPase+Alpha+Beta-domain+wHTH+RRM | Type-1_BR.RM.like.systems | <-PemK_toxin<-?\|\|Phage_integrase-><-HEPN+DUF499* | 1098 | Bacteria>environmental_samples | - | uncultured_bacterium | . | 22115..25411 | 77133 | F28-[uncultured-bacterium] |
| MCL2726873.1 | HEPN+AAA-ATPase+Alpha+Beta-domain+wHTH+RRM | Type-1_BR.RM.like.systems | <-ABhydrolase\|\|REase->Peptidase-><-Peptidase_M24<-Peptidase<-?<-HEPN+DUF499*<-TOPRIM<-OLD-ABC<-METHYLASE<-TUDOR+SF1-SF2-Helicase+iREase+CTD+REase<-LysE | 1124 | Bacteria>FCB_group>Bacteroidota/Chlorobiota_group>Bacteroidota>Bacteroidia | GCA_009781275.1 | Bacteroidales_bacterium | FWD56_00650 | 13721..17095 | 2030927 | DUF499-domain-containing-protein-[Bacteroidales-bacterium] |
| NCB25654.1 | HEPN+AAA-ATPase+Alpha+Beta-domain+wHTH+RRM | Type-1_BR.RM.like.systems | TUDOR+SF1-SF2-Helicase+iREase+CTD+REase->METHYLASE->HEPN+DUF499*->REase->Pkinase-> | 1134 | Bacteria>FCB_group>Bacteroidota/Chlorobiota_group>Bacteroidota>Bacteroidia | GCA_009929165.1 | Bacteroidia_bacterium | EOM62_09315 | 4747..8151 | 2044936 | ATP-binding-protein-[Bacteroidia-bacterium] |
| MDD4460430.1 | HEPN+AAA-ATPase+Alpha+Beta-domain+wHTH+RRM | Type-1_BR.RM.like.systems | METHYLASE-p->HEPN+DUF499*-> | 1132 | Bacteria>FCB_group>Bacteroidota/Chlorobiota_group>Bacteroidota>Bacteroidia | GCA_028705295.1 | Proteiniphilum_sp. | PHQ24_11635 | 346..3744 | 1926877 | MAG:-DUF499-domain-containing-protein-[Proteiniphilum-sp.] |
| WP_196069848.1 | HEPN+AAA-ATPase+Alpha+Beta-domain+wHTH+RRM | Type-1_BR.RM.like.systems | <-REase+SF1-SF2-Helicase<-MazG<-METHYLASE-p<-ABhydrolase<-ACET<-DOC+HTH<-HEPN+DUF499*<-METHYLASE<-TUDOR+SF1-SF2-Helicase+iREase+CTD+REase\|\|?->HTH_3->?->Phage_int_SAM_5+Phage_integrase-> | 1099 | Bacteria>FCB_group>Bacteroidota/Chlorobiota_group>Bacteroidota>Bacteroidia | GCF_015670455.1 | Bacteroides_thetaiotaomicron | PO078_RS08820 | 41190..44489 | 818 | DUF499-domain-containing-protein-[Bacteroides-thetaiotaomicron] |
| WP_016268236.1 | HEPN+AAA-ATPase+Alpha+Beta-domain+wHTH+RRM | Type-1_BR.RM.like.systems | <-URI-DNase+SF1-SF2-Helicase<-?<-?\|\|sGTP->TUDOR+SF1-SF2-Helicase+iREase+CTD+REase->METHYLASE->HEPN+DUF499*->DOC+HTH->Glyoxalase+Glyoxalase-><-HKD-DNase<-SF1-SF2-Helicase<-SF1-SF2-Helicase | 1099 | Bacteria>FCB_group>Bacteroidota/Chlorobiota_group>Bacteroidota>Bacteroidia | GCF_000403155.2 | Bacteroides_thetaiotaomicron | C799_RS12880 | 2627963..2631262 | 818 | DUF499-domain-containing-protein-[Bacteroides-thetaiotaomicron] |
| KAB4460058.1 | HEPN+AAA-ATPase+Alpha+Beta-domain+wHTH+RRM | Type-1_BR.RM.like.systems | <-URI-DNase+SF1-SF2-Helicase<-?\|\|sGTP->TUDOR+SF1-SF2-Helicase+iREase+CTD+REase->METHYLASE->HEPN+DUF499*->DOC+HTH-><-HKD-DNase<-SF1-SF2-Helicase<-SF1-SF2-Helicase<-?<-NUDIX | 1099 | Bacteria>FCB_group>Bacteroidota/Chlorobiota_group>Bacteroidota>Bacteroidia | GCA_009021605.1 | Bacteroides_thetaiotaomicron | GAN98_17895 | 29744..33043 | 818 | ATP-binding-protein-[Bacteroides-thetaiotaomicron] |
| MCS2505372.1 | HEPN+AAA-ATPase+Alpha+Beta-domain+wHTH+RRM | Type-1_BR.RM.like.systems | TUDOR+SF1-SF2-Helicase+iREase+CTD+REase->TUDOR+SF1-SF2-Helicase+iREase+CTD+REase-p->METHYLASE-p->METHYLASE-p->METHYLASE-p->HEPN-p->HEPN+DUF499*->DUF499-><-?<-HicB<-TOPRIM<-SbcC\|\|BirA-HTH-> | 434 | Bacteria>FCB_group>Bacteroidota/Chlorobiota_group>Bacteroidota>Bacteroidia | GCA_024788865.1 | Phocaeicola_vulgatus | NXW48_05305 | 1159689..1160993 | 821 | DUF499-domain-containing-protein-[Phocaeicola-vulgatus] |
| MCS3108285.1 | HEPN+AAA-ATPase+Alpha+Beta-domain+wHTH+RRM | Type-1_BR.RM.like.systems | <-ACET<-Calcineurin<-ZDNA-HTH<-DOC+HTH<-DUF499-p<-HEPN+DUF499*<-METHYLASE<-TUDOR+SF1-SF2-Helicase+iREase+CTD+REase<-TUDOR+SF1-SF2-Helicase+iREase+CTD+REase-p<-HTH_3<-?<-TraJ-RHH | 722 | Bacteria>FCB_group>Bacteroidota/Chlorobiota_group>Bacteroidota>Bacteroidia | GCA_024792645.1 | Bacteroides_fragilis | NXV07_08905 | 2189665..2191833 | 817 | DUF499-domain-containing-protein-[Bacteroides-fragilis] |
| WP_234147452.1 | HEPN+AAA-ATPase+Alpha+Beta-domain+wHTH+RRM | Type-1_BR.RM.like.systems | sGTP->TUDOR+SF1-SF2-Helicase+iREase+CTD+REase->METHYLASE->HEPN+DUF499*->DOC+HTH->Glyoxalase+Glyoxalase-><-HKD-DNase<-SF1-SF2-Helicase<-SF1-SF2-Helicase | 1099 | Bacteria>FCB_group>Bacteroidota/Chlorobiota_group>Bacteroidota>Bacteroidia | GCF_021410605.1 | Bacteroides_thetaiotaomicron | K0G45_RS08855 | 122149..125448 | 818 | DUF499-domain-containing-protein-[Bacteroides-thetaiotaomicron] |
| WP_118211914.1 | HEPN+AAA-ATPase+Alpha+Beta-domain+wHTH+RRM | Type-1_BR.RM.like.systems | PrmC_N+Methyltransf_25->?->PRTase->?->tRNA->tRNA-><-HEPN+DUF499*<-METHYLASE<-REase+ParB<-TUDOR+SF1-SF2-Helicase+iREase+CTD+REase-p<-TUDOR+SF1-SF2-Helicase+iREase+CTD+REase-p<-HTH_3<-DnaJ | 1098 | Bacteria>FCB_group>Bacteroidota/Chlorobiota_group>Bacteroidota>Bacteroidia | GCF_027661405.1 | Phocaeicola_plebeius | DWY45_RS04060 | 51608..54904 | 310297 | DUF499-domain-containing-protein-[Phocaeicola-plebeius] |
| WP_195425521.1 | HEPN+AAA-ATPase+Alpha+Beta-domain+wHTH+RRM | Type-1_BR.RM.like.systems | STYKinase-like->SF1-SF2-Helicase-><-?<-Glyoxalase+Glyoxalase<-DOC+HTH<-HEPN+DUF499*<-METHYLASE<-TUDOR+SF1-SF2-Helicase+iREase+CTD+REase<-HTH_3<-?<-?<-TraJ-RHH | 1099 | Bacteria>FCB_group>Bacteroidota/Chlorobiota_group>Bacteroidota>Bacteroidia | GCF_015551145.1 | Bacteroides_cellulosilyticus | PQH05_RS04750 | 125017..128316 | 246787 | DUF499-domain-containing-protein-[Bacteroides-cellulosilyticus] |
| WP_258968658.1 | HEPN+AAA-ATPase+Alpha+Beta-domain+wHTH+RRM | Type-1_BR.RM.like.systems | XIS-HTH->?->?->HTH_3->TUDOR+SF1-SF2-Helicase+iREase+CTD+REase->METHYLASE->HEPN+DUF499*-><-?<-HicB<-TOPRIM<-SbcC\|\|BirA-HTH-><-Phage_int_SAM_5+Phage_integrase | 1096 | Bacteria>FCB_group>Bacteroidota/Chlorobiota_group>Bacteroidota>Bacteroidia | GCF_024790655.1 | Phocaeicola_vulgatus | NXV64_RS08565 | 2221353..2224643 | 821 | DUF499-domain-containing-protein-[Phocaeicola-vulgatus] |
| MDD7697305.1 | HEPN+AAA-ATPase+Alpha+Beta-domain+wHTH+RRM | Type-1_BR.RM.like.systems | XIS-HTH->?->?->HTH_3->TUDOR+SF1-SF2-Helicase+iREase+CTD+REase->METHYLASE->HEPN+DUF499*-><-?<-HicB<-TOPRIM<-SbcC\|\|BirA-HTH-><-Phage_int_SAM_5+Phage_integrase | 1096 | Bacteria>FCB_group>Bacteroidota/Chlorobiota_group>Bacteroidota>Bacteroidia | GCA_028727985.1 | Phocaeicola_vulgatus | PUJ58_17085 | 44225..47515 | 821 | MAG:-DUF499-domain-containing-protein-[Phocaeicola-vulgatus] |
| WP_018709397.1 | HEPN+AAA-ATPase+Alpha+Beta-domain+wHTH+RRM | Type-1_BR.RM.like.systems | REase+SF1-SF2-Helicase->?->Radical-SAM->TUDOR+SF1-SF2-Helicase+iREase+CTD+REase->REase+ParB->METHYLASE->HEPN+DUF499*->?-><-?\|\|?->Peptidase_M3-> | 1098 | Bacteria>FCB_group>Bacteroidota/Chlorobiota_group>Bacteroidota>Bacteroidia | GCF_000374585.1 | Phocaeicola_barnesiae | C510_RS0100610 | 147306..150602 | 376804 | DUF499-domain-containing-protein-[Phocaeicola-barnesiae] |
| MBP6872709.1 | HEPN+AAA-ATPase+Alpha+Beta-domain+wHTH+RRM | Type-1_BR.RM.like.systems | TUDOR+SF1-SF2-Helicase+iREase+CTD+REase->AbiEi_1->AbiEii->ParB+HNH->METHYLASE->HEPN+DUF499*->?->SbcC->?-><-?<-REase+SF1-SF2-Helicase<-MazG | 1092 | Bacteria>FCB_group>Bacteroidota/Chlorobiota_group>Bacteroidota>Bacteroidia | GCA_017991835.1 | Bacteroidales_bacterium | KBC43_11940 | 58567..61845 | 2030927 | ATP-binding-protein-[Bacteroidales-bacterium] |
| WP_229532986.1 | HEPN+AAA-ATPase+Alpha+Beta-domain+wHTH+RRM | Type-1_BR.RM.like.systems | <-SF1-SF2-Helicase<-MAD-NTD+HNH<-SF1-SF2-Helicase+DUF1998<-?<-HEPN+DUF499*<-METHYLASE<-TUDOR+SF1-SF2-Helicase+iREase+CTD+REase<-WYL+WYL-C<-HTH_3 | 1108 | Bacteria>FCB_group>Bacteroidota/Chlorobiota_group>Bacteroidota>Bacteroidia | GCF_020827855.1 | Phocaeicola_vulgatus | HZY60_RS19310 | 19399..22725 | 821 | DUF499-domain-containing-protein-[Phocaeicola-vulgatus] |
| WP_118026126.1 | HEPN+AAA-ATPase+Alpha+Beta-domain+wHTH+RRM | Type-1_BR.RM.like.systems | DnaJ->HTH_3->TUDOR+SF1-SF2-Helicase+iREase+CTD+REase->REase+ParB->METHYLASE->HEPN+DUF499*-><-tRNA<-tRNA<-?<-?<-PRTase<-PRTase<-?<-PrmC_N+Methyltransf_25 | 1098 | Bacteria>FCB_group>Bacteroidota/Chlorobiota_group>Bacteroidota>Bacteroidia | GCF_027686295.1 | Phocaeicola_plebeius | PGZ43_RS01830 | 134398..137694 | 310297 | DUF499-domain-containing-protein-[Phocaeicola-plebeius] |
| WP_071137164.1 | HEPN+AAA-ATPase+Alpha+Beta-domain+wHTH+RRM | Type-1_BR.RM.like.systems | PAIREDC-HTH-><-SF1-SF2-Helicase+DUF1998<-?<-HEPN+DUF499*<-METHYLASE<-TUDOR+SF1-SF2-Helicase+iREase+CTD+REase<-DDE_Tnp_1_3<-WYL+WYL-C<-HTH_3 | 1108 | Bacteria>FCB_group>Bacteroidota/Chlorobiota_group>Bacteroidota>Bacteroidia | GCF_900095795.1 | Petrimonas_mucosa | ING2E5A_RS09235 | 2277768..2281094 | 1642646 | DUF499-domain-containing-protein-[Petrimonas-mucosa] |
| WP_087396459.1 | HEPN+AAA-ATPase+Alpha+Beta-domain+wHTH+RRM | Type-1_BR.RM.like.systems | REase->?-><-?<-DOC+HTH<-HEPN+DUF499*<-METHYLASE<-REase+ParB<-TUDOR+SF1-SF2-Helicase+iREase+CTD+REase<-DOC<-HTH_3<-FAD-NAD-dep-oxidoreductase | 1097 | Bacteria>FCB_group>Bacteroidota/Chlorobiota_group>Bacteroidota>Bacteroidia | GCF_002161565.1 | Bacteroides_sp._An51A | B5G04_RS06890 | 141402..144695 | 1965640 | DUF499-domain-containing-protein-[Bacteroides-sp.-An51A] |
| WP_118432131.1 | HEPN+AAA-ATPase+Alpha+Beta-domain+wHTH+RRM | Type-1_BR.RM.like.systems | PrmC_N+Methyltransf_25->?->PRTase->PRTase->?->?->tRNA->tRNA-><-HEPN+DUF499*<-METHYLASE<-REase+ParB<-TUDOR+SF1-SF2-Helicase+iREase+CTD+REase<-HTH_3<-DnaJ | 1098 | Bacteria>FCB_group>Bacteroidota/Chlorobiota_group>Bacteroidota>Bacteroidia | GCF_958447975.1 | Phocaeicola_plebeius | PGY45_RS13550 | 13557..16853 | 310297 | DUF499-domain-containing-protein-[Phocaeicola-plebeius] |
| WP_074635742.1 | HEPN+AAA-ATPase+Alpha+Beta-domain+wHTH+RRM | Type-1_BR.RM.like.systems | DEDD_Tnp_IS110->HTH_3->TUDOR+SF1-SF2-Helicase+iREase+CTD+REase->METHYLASE->HEPN+DUF499*->Schlafen+HATPase+HTH->?->HTH_3->DNA_methylase->MORC-> | 1096 | Bacteria>FCB_group>Bacteroidota/Chlorobiota_group>Bacteroidota>Bacteroidia | GCF_900100465.1 | Bacteroides_ovatus | BLS71_RS01410 | 96540..99830 | 28116 | DUF499-domain-containing-protein-[Bacteroides-ovatus] |
| WP_278365558.1 | HEPN+AAA-ATPase+Alpha+Beta-domain+wHTH+RRM | Type-1_BR.RM.like.systems | <-Peptidase_M3<-?\|\|?-><-?<-HEPN+DUF499*<-METHYLASE<-REase+ParB<-TUDOR+SF1-SF2-Helicase+iREase+CTD+REase<-Radical-SAM<-?<-SF1-SF2-Helicase | 1098 | Bacteria>FCB_group>Bacteroidota/Chlorobiota_group>Bacteroidota>Bacteroidia | GCF_002359135.1 | Phocaeicola_barnesiae | B3R84_RS06870 | 398812..402108 | 376804 | DUF499-domain-containing-protein-[Phocaeicola-barnesiae] |
| CDD51558.1 | HEPN+AAA-ATPase+Alpha+Beta-domain+wHTH+RRM | Type-1_BR.RM.like.systems | <-XIS-HTH<-?<-Phage_integrase\|\|TIR->TIR-><-?<-HEPN+DUF499*<-METHYLASE<-REase+ParB<-TUDOR+SF1-SF2-Helicase+iREase+CTD+REase<-HTH_3<-DnaJ | 1098 | Bacteria>FCB_group>Bacteroidota/Chlorobiota_group>Bacteroidota>Bacteroidia | GCA_000435415.1 | Bacteroides_sp._CAG:875 | BN800_02603 | 61429..64725 | 1262752 | putative-uncharacterized-protein-[Bacteroides-sp.-CAG:875] |
| WP_235327793.1 | HEPN+AAA-ATPase+Alpha+Beta-domain+wHTH+RRM | Type-1_BR.RM.like.systems | SbcC->TOPRIM->HicB->TIR->TIR-><-?<-HEPN+DUF499*<-METHYLASE<-TUDOR+SF1-SF2-Helicase+iREase+CTD+REase<-HTH_3<-?<-?<-tRNA<-Peptidase_S41 | 1094 | Bacteria>FCB_group>Bacteroidota/Chlorobiota_group>Bacteroidota>Bacteroidia | GCF_021530995.1 | Bacteroides_caecigallinarum | I6E24_RS08290 | 4207..7491 | 1411144 | DUF499-domain-containing-protein-[Bacteroides-caecigallinarum] |
| WP_038554549.1 | HEPN+AAA-ATPase+Alpha+Beta-domain+wHTH+RRM | Type-1_BR.RM.like.systems | TUDOR+SF1-SF2-Helicase+iREase+CTD+REase->?->OLD-ABC->?->METHYLASE->HEPN+DUF499*->DUF6361->HKD-DNase->SF1-SF2-Helicase-><-?<-Primpol | 1096 | Bacteria>FCB_group>Bacteroidota/Chlorobiota_group>Bacteroidota>Bacteroidia | GCF_000626635.1 | Draconibacterium_orientale | BMW53_RS05405 | 116803..120093 | 1168034 | DUF499-domain-containing-protein-[Draconibacterium-orientale] |
| WP_284525582.1 | HEPN+AAA-ATPase+Alpha+Beta-domain+wHTH+RRM | Type-1_BR.RM.like.systems | <-DUF499-p<-DUF499-p<-HEPN+DUF499*<-METHYLASE-p<-METHYLASE<-REase+ParB<-?<-TUDOR+SF1-SF2-Helicase+iREase+CTD+REase<-HTH_3 | 518 | Bacteria>FCB_group>Bacteroidota/Chlorobiota_group>Bacteroidota>Bacteroidia | - | Phocaeicola_plebeius | DW982_RS17975 | 3045..4601 | 310297 | DUF499-domain-containing-protein-[Phocaeicola-plebeius] |
| WP_270668664.1 | HEPN+AAA-ATPase+Alpha+Beta-domain+wHTH+RRM | Type-1_BR.RM.like.systems | <-BirA-HTH\|\|SbcC->TOPRIM->HicB->?-><-?<-HEPN+DUF499*<-METHYLASE<-TUDOR+SF1-SF2-Helicase+iREase+CTD+REase<-HTH_3<-?<-?<-XIS-HTH | 1096 | Bacteria>FCB_group>Bacteroidota/Chlorobiota_group>Bacteroidota>Bacteroidia | GCF_027686505.1 | Phocaeicola_vulgatus | PG001_RS15345 | 5861..9151 | 821 | DUF499-domain-containing-protein-[Phocaeicola-vulgatus] |
| WP_151851142.1 | HEPN+AAA-ATPase+Alpha+Beta-domain+wHTH+RRM | Type-1_BR.RM.like.systems | <-BirA-HTH\|\|SbcC->TOPRIM->HicB->?->?-><-HEPN+DUF499*<-METHYLASE<-TUDOR+SF1-SF2-Helicase+iREase+CTD+REase<-HTH_3<-?<-?<-XIS-HTH | 1096 | Bacteria>FCB_group>Bacteroidota/Chlorobiota_group>Bacteroidota>Bacteroidia | GCF_009025535.1 | Phocaeicola_vulgatus | F9001_RS04885 | 42984..46274 | 821 | DUF499-domain-containing-protein-[Phocaeicola-vulgatus] |
| WP_055289369.1 | HEPN+AAA-ATPase+Alpha+Beta-domain+wHTH+RRM | Type-1_BR.RM.like.systems | REC-><-METHYLASE-p<-ABhydrolase<-ACET<-DOC+HTH<-HEPN+DUF499*<-METHYLASE<-TUDOR+SF1-SF2-Helicase+iREase+CTD+REase | 1099 | Bacteria>FCB_group>Bacteroidota/Chlorobiota_group>Bacteroidota>Bacteroidia | GCF_025914195.1 | Bacteroidales | KSY52_RS08740 | 147903..151202 | 171549 | MULTISPECIES:-DUF499-domain-containing-protein-[Bacteroidales] |
| MBE6276390.1 | HEPN+AAA-ATPase+Alpha+Beta-domain+wHTH+RRM | Type-1_BR.RM.like.systems | SbcC->TOPRIM->HicB->TIR->TIR-><-?<-HEPN+DUF499*<-METHYLASE<-TUDOR+SF1-SF2-Helicase+iREase+CTD+REase<-HTH_3<-XIS-HTH | 1095 | Bacteria>FCB_group>Bacteroidota/Chlorobiota_group>Bacteroidota>Bacteroidia | GCA_015060735.1 | Bacteroides_sp. | E7096_08645 | 4627..7914 | 29523 | ATP-binding-protein-[Bacteroides-sp.] |
| MBW6537009.1 | HEPN+AAA-ATPase+Alpha+Beta-domain+wHTH+RRM | Type-1_BR.RM.like.systems | <-HEPN+DUF499*<-METHYLASE<-AbiEii<-wHTH+AbiEi<-TUDOR+SF1-SF2-Helicase+iREase+CTD+REase<-TUDOR+SF1-SF2-Helicase+iREase+CTD+REase-p | 696 | Bacteria>FCB_group>Bacteroidota/Chlorobiota_group>Bacteroidota>Bacteroidia | GCA_019429205.1 | Mariniphaga_sp. | KOB11_18520 | 1..2089 | 1954475 | DUF499-domain-containing-protein-[Mariniphaga-sp.] |
| WP_250712821.1 | HEPN+AAA-ATPase+Alpha+Beta-domain+wHTH+RRM | Type-1_BR.RM.like.systems | SF1-SF2-Helicase->?->DEDD_Tnp_IS110->HTH_3->TUDOR+SF1-SF2-Helicase+iREase+CTD+REase->METHYLASE->HEPN+DUF499*-> | 1096 | Bacteria>FCB_group>Bacteroidota/Chlorobiota_group>Bacteroidota>Bacteroidia | GCF_023654475.1 | Bacteroides_fragilis | J8K93_RS06720 | 1584260..1587550 | 817 | DUF499-domain-containing-protein-[Bacteroides-fragilis] |
| CDD33920.1 | HEPN+AAA-ATPase+Alpha+Beta-domain+wHTH+RRM | Type-1_BR.RM.like.systems | <-Peptidase_M3<-?\|\|?-><-?<-HEPN+DUF499*<-METHYLASE<-REase+ParB<-TUDOR+SF1-SF2-Helicase+iREase+CTD+REase<-Radical-SAM | 1098 | Bacteria>FCB_group>Bacteroidota/Chlorobiota_group>Bacteroidota>Bacteroidia | GCA_000436055.1 | Bacteroides_sp._CAG:714 | BN762_01576 | 11019..14315 | 1262749 | putative-uncharacterized-protein-[Bacteroides-sp.-CAG:714] |
| WP_195204977.1 | HEPN+AAA-ATPase+Alpha+Beta-domain+wHTH+RRM | Type-1_BR.RM.like.systems | DnaJ->HTH_3->TUDOR+SF1-SF2-Helicase+iREase+CTD+REase->REase+ParB->METHYLASE-p->METHYLASE->HEPN+DUF499*-> | 1098 | Bacteria>FCB_group>Bacteroidota/Chlorobiota_group>Bacteroidota>Bacteroidia | GCF_015546785.1 | Phocaeicola_plebeius | I1Q75_RS16665 | 24617..27913 | 310297 | DUF499-domain-containing-protein-[Phocaeicola-plebeius] |
| WP_220575703.1 | HEPN+AAA-ATPase+Alpha+Beta-domain+wHTH+RRM | Type-1_BR.RM.like.systems | DEDD_Tnp_IS110->HTH_3->TUDOR+SF1-SF2-Helicase+iREase+CTD+REase->METHYLASE-p->METHYLASE->HEPN+DUF499*-> | 1096 | Bacteria>FCB_group>Bacteroidota/Chlorobiota_group>Bacteroidota>Bacteroidia | GCF_019633785.1 | Bacteroides | ISX59_RS04600 | 376325..379615 | 816 | MULTISPECIES:-DUF499-domain-containing-protein-[Bacteroides] |
| WP_032540897.1 | HEPN+AAA-ATPase+Alpha+Beta-domain+wHTH+RRM | Type-1_BR.RM.like.systems | <-HEPN+DUF499*<-METHYLASE-p<-METHYLASE<-TUDOR+SF1-SF2-Helicase+iREase+CTD+REase<-HTH_3<-DEDD_Tnp_IS110 | 1096 | Bacteria>FCB_group>Bacteroidota/Chlorobiota_group>Bacteroidota>Bacteroidia | GCF_000724815.2 | Bacteroides | IB64_RS10255 | 2443318..2446608 | 816 | DUF499-domain-containing-protein-[Bacteroides-fragilis] |
| WP_235386280.1 | HEPN+AAA-ATPase+Alpha+Beta-domain+wHTH+RRM | Type-1_BR.RM.like.systems | METHYLASE->HEPN+DUF499*->?->SF1-SF2-Helicase+DUF1998->MAD-NTD+HNH->SF1-SF2-Helicase-> | 1119 | Bacteria>FCB_group>Bacteroidota/Chlorobiota_group>Bacteroidota>Bacteroidia | GCF_021531665.1 | Prevotellamassilia_timonensis | I6E40_RS06760 | 3178..6537 | 1852370 | DUF499-domain-containing-protein-[Prevotellamassilia-timonensis] |
| MCS2502386.1 | HEPN+AAA-ATPase+Alpha+Beta-domain+wHTH+RRM | Type-1_BR.RM.like.systems | <-HEPN+DUF499*<-METHYLASE<-METHYLASE-p<-TUDOR+SF1-SF2-Helicase+iREase+CTD+REase<-DEDD_Tnp_IS110 | 1026 | Bacteria>FCB_group>Bacteroidota/Chlorobiota_group>Bacteroidota>Bacteroidia | GCA_024788905.1 | Bacteroides_ovatus | NXV80_13315 | 3514916..3517996 | 28116 | DUF499-domain-containing-protein-[Bacteroides-ovatus] |
| MCC8143016.1 | HEPN+AAA-ATPase+Alpha+Beta-domain+wHTH+RRM | Type-1_BR.RM.like.systems | <-AraC-HTH+SHS2+SHS2<-Peptidase_M28<-?<-BirA-HTH<-Schlafen<-HEPN+DUF499*<-METHYLASE | 1102 | Bacteria>FCB_group>Bacteroidota/Chlorobiota_group>Bacteroidota>Bacteroidia | GCA_020860665.1 | Tannerellaceae_bacterium | LI097_03250 | 4810..8118 | 2785930 | DUF499-domain-containing-protein-[Tannerellaceae-bacterium] |
| MBN1185441.1 | HEPN+AAA-ATPase+Alpha+Beta-domain+wHTH+RRM | Type-1_BR.RM.like.systems | METHYLASE-p->HEPN+DUF499*->TPR+TPR->?->MazG->REase+SF1-SF2-Helicase-> | 1076 | Bacteria>FCB_group>Bacteroidota/Chlorobiota_group>Bacteroidota>Bacteroidia | GCA_016926235.1 | Bacteroidales_bacterium | JXB49_24360 | 1099..4329 | 2030927 | DUF499-domain-containing-protein-[Bacteroidales-bacterium] |
| WP_235301263.1 | HEPN+AAA-ATPase+Alpha+Beta-domain+wHTH+RRM | Type-1_BR.RM.like.systems | TUDOR+SF1-SF2-Helicase+iREase+CTD+REase->REase+ParB->METHYLASE->HEPN+DUF499*->?-><-?\|\|?->Peptidase_M3-> | 1098 | Bacteria>FCB_group>Bacteroidota/Chlorobiota_group>Bacteroidota>Bacteroidia | GCF_021531035.1 | Phocaeicola_barnesiae | I6E10_RS09100 | 8385..11681 | 376804 | DUF499-domain-containing-protein-[Phocaeicola-barnesiae] |
| WP_195365726.1 | HEPN+AAA-ATPase+Alpha+Beta-domain+wHTH+RRM | Type-1_BR.RM.like.systems | <-HEPN+DUF499*<-METHYLASE<-TUDOR+SF1-SF2-Helicase+iREase+CTD+REase<-DEDD_Tnp_IS110<-?<-?\|\|HTH-> | 1096 | Bacteria>FCB_group>Bacteroidota/Chlorobiota_group>Bacteroidota>Bacteroidia | GCF_015550085.1 | Bacteroides_ovatus | PO368_RS10195 | 12026..15316 | 28116 | DUF499-domain-containing-protein-[Bacteroides-ovatus] |
| WP_101543458.1 | HEPN+AAA-ATPase+Alpha+Beta-domain+wHTH+RRM | Type-1_BR.RM.like.systems | <-HEPN+DUF499*<-METHYLASE<-TUDOR+SF1-SF2-Helicase+iREase+CTD+REase<-HTH_3<-DDE_Tnp_1<-HAD+HAD | 1096 | Bacteria>FCB_group>Bacteroidota/Chlorobiota_group>Bacteroidota>Bacteroidia | GCF_902385525.1 | Bacteroidaceae | C1A09_RS08785 | 351382..354672 | 815 | MULTISPECIES:-DUF499-domain-containing-protein-[Bacteroidaceae] |

| Protein-ID | DUF499-architecture | System | Gene-Neighbourhoods | Length | Taxonomy | Assembly-ID | Strain | Locus-Tag | Genomic-coordinates | TaxID | Fasta-defline |
|---|---|---|---|---|---|---|---|---|---|---|---|
| WP_032849356.1 | HEPN+AAA-ATPase+Alpha+Beta-domain+wHTH+RRM | Type-1_BR.RM.like.systems | <-HEPN+DUF499*<-METHYLASE<-TUDOR+SF1-SF2-Helicase+iREase+CTD+REase<-DEDD_Tnp_IS110<-?<-?\|\|HTH-> | 1096 | Bacteria>FCB_group>Bacteroidota/Chlorobiota_group>Bacteroidota>Bacteroidia | GCF_901212435.1 | Bacteroides | PO242_RS12935 | 72725..76015 | 816 | MULTISPECIES:-DUF499-domain-containing-protein-[Bacteroides] |
| WP_118338055.1 | HEPN+AAA-ATPase+Alpha+Beta-domain+wHTH+RRM | Type-1_BR.RM.like.systems | <-HEPN+DUF499*<-METHYLASE<-REase+ParB<-TUDOR+SF1-SF2-Helicase+iREase+CTD+REase<-HTH_3<-DnaJ | 1098 | Bacteria>FCB_group>Bacteroidota/Chlorobiota_group>Bacteroidota>Bacteroidia | GCF_959025615.1 | Phocaeicola_plebeius | PG845_RS15420 | 1179..4475 | 310297 | DUF499-domain-containing-protein-[Phocaeicola-plebeius] |
| WP_117747735.1 | HEPN+AAA-ATPase+Alpha+Beta-domain+wHTH+RRM | Type-1_BR.RM.like.systems | <-HEPN+DUF499*<-METHYLASE<-REase+ParB<-TUDOR+SF1-SF2-Helicase+iREase+CTD+REase<-HTH_3<-DnaJ | 1098 | Bacteria>FCB_group>Bacteroidota/Chlorobiota_group>Bacteroidota>Bacteroidia | GCF_003438135.1 | Phocaeicola_plebeius | DXC17_RS06660 | 1179..4475 | 310297 | DUF499-domain-containing-protein-[Phocaeicola-plebeius] |
| WP_032953489.1 | HEPN+AAA-ATPase+Alpha+Beta-domain+wHTH+RRM | Type-1_BR.RM.like.systems | XIS-HTH->?->?->HTH_3->TUDOR+SF1-SF2-Helicase+iREase+CTD+REase->METHYLASE->HEPN+DUF499*-><-?<-HicB | 1096 | Bacteria>FCB_group>Bacteroidota/Chlorobiota_group>Bacteroidota>Bacteroidia | GCF_001405155.1 | Bacteroidaceae | KSZ98_RS20820 | 10462..13752 | 815 | MULTISPECIES:-DUF499-domain-containing-protein-[Bacteroidaceae] |
| WP_118201547.1 | HEPN+AAA-ATPase+Alpha+Beta-domain+wHTH+RRM | Type-1_BR.RM.like.systems | <-Aminotran_1_2<-?\|\|?->?->TUDOR+SF1-SF2-Helicase+iREase+CTD+REase->METHYLASE->HEPN+DUF499*-> | 1122 | Bacteria>FCB_group>Bacteroidota/Chlorobiota_group>Bacteroidota>Bacteroidia | GCF_024330865.1 | Segatella_copri | NNC52_RS13920 | 15349..18717 | 165179 | DUF499-domain-containing-protein-[Prevotella-copri] |
| WP_118170513.1 | HEPN+AAA-ATPase+Alpha+Beta-domain+wHTH+RRM | Type-1_BR.RM.like.systems | <-HEPN+DUF499*<-METHYLASE<-TUDOR+SF1-SF2-Helicase+iREase+CTD+REase<-HTH_3<-XIS-HTH | 1096 | Bacteria>FCB_group>Bacteroidota/Chlorobiota_group>Bacteroidota>Bacteroidia | GCF_934726555.1 | Bacteroidaceae | QV389_RS11685 | 2128..5418 | 815 | MULTISPECIES:-DUF499-domain-containing-protein-[Bacteroidaceae] |
| MCR5642603.1 | HEPN+AAA-ATPase+Alpha+Beta-domain+wHTH+RRM | Type-1_BR.RM.like.systems | <-Peptidase<-DOC<-?<-HEPN+DUF499*<-METHYLASE<-TUDOR+SF1-SF2-Helicase+iREase+CTD+REase | 1106 | Bacteria>FCB_group>Bacteroidota/Chlorobiota_group>Bacteroidota>Bacteroidia | GCA_024690905.1 | Prevotella_sp. | K6G32_04600 | 31632..34952 | 59823 | DUF499-domain-containing-protein-[Prevotella-sp.] |
| KXT50696.1 | HEPN+AAA-ATPase+Alpha+Beta-domain+wHTH+RRM | Type-1_BR.RM.like.systems | DEDD_Tnp_IS110->TUDOR+SF1-SF2-Helicase+iREase+CTD+REase->METHYLASE->?->HEPN+DUF499*-> | 1098 | Bacteria>FCB_group>Bacteroidota/Chlorobiota_group>Bacteroidota>Bacteroidia | GCA_001578575.1 | Bacteroides_ovatus | HMPREF2532_00822 | 114886..118182 | 28116 | hypothetical-protein-HMPREF2532_00822-[Bacteroides-ovatus] |
| EXY75216.1 | HEPN+AAA-ATPase+Alpha+Beta-domain+wHTH+RRM | Type-1_BR.RM.like.systems | <-ACET<-Calcineurin<-ZDNA-HTH<-DOC+HTH<-HEPN+DUF499*<-METHYLASE-p | 1101 | Bacteria>FCB_group>Bacteroidota/Chlorobiota_group>Bacteroidota>Bacteroidia | GCA_000598365.1 | Bacteroides_fragilis_str._3988T(B)14 | M124_1038 | 75233..78538 | 1339315 | hypothetical-protein-M124_1038-[Bacteroides-fragilis-str.-3988T(B)14] |
| MCD8261448.1 | HEPN+AAA-ATPase+Alpha+Beta-domain+wHTH+RRM | Type-1_BR.RM.like.systems | <-STYKIN<-Schlafen<-HEPN+DUF499*<-METHYLASE-p<-METHYLASE-p | 1102 | Bacteria>FCB_group>Bacteroidota/Chlorobiota_group>Bacteroidota>Bacteroidia | GCA_021204375.1 | Bacteroides_sp. | LUD15_08080 | 1305..4613 | 29523 | DUF499-domain-containing-protein-[Bacteroides-sp.] |
| MBR1880072.1 | HEPN+AAA-ATPase+Alpha+Beta-domain+wHTH+RRM | Type-1_BR.RM.like.systems | TUDOR+SF1-SF2-Helicase+iREase+CTD+REase->METHYLASE-><-PIN<-?\|\|HEPN+DUF499*-> | 593 | Bacteria>FCB_group>Bacteroidota/Chlorobiota_group>Bacteroidota>Bacteroidia | GCA_017478995.1 | Prevotella_sp. | IJ804_04915 | 6915..8693 | 59823 | DUF499-domain-containing-protein-[Prevotella-sp.] |
| MBO5419352.1 | HEPN+AAA-ATPase+Alpha+Beta-domain+wHTH+RRM | Type-1_BR.RM.like.systems | XIS-HTH->?->?-><-?<-?\|\|ORC-CDC6+REase-><-HEPN+DUF499*<-METHYLASE-p | 1102 | Bacteria>FCB_group>Bacteroidota/Chlorobiota_group>Bacteroidota>Bacteroidia | GCA_017625725.1 | Bacteroidales_bacterium | J6A22_04620 | 7056..10364 | 2030927 | DUF499-domain-containing-protein-[Bacteroidales-bacterium] |
| WP_032587868.1 | HEPN+AAA-ATPase+Alpha+Beta-domain+wHTH+RRM | Type-1_BR.RM.like.systems | <-ACET<-Calcineurin<-ZDNA-HTH<-DOC+HTH<-HEPN+DUF499* | 1099 | Bacteria>FCB_group>Bacteroidota/Chlorobiota_group>Bacteroidota>Bacteroidia | GCF_000598365.1 | Bacteroides_fragilis | MO84_RS16910 | 33544..36843 | 817 | DUF499-domain-containing-protein-[Bacteroides-fragilis] |
| HBF05518.1 | HEPN+AAA-ATPase+Alpha+Beta-domain+wHTH+RRM | Type-1_BR.RM.like.systems | <-ORC-CDC6+REase<-HEPN+DUF499*<-METHYLASE-p | 1108 | Bacteria>FCB_group>Bacteroidota/Chlorobiota_group>Bacteroidota>Bacteroidia | GCA_003501465.1 | Candidatus_Segatella_violae | DDW28_05315 | 5202..8528 | 1947177 | hypothetical-protein-[Prevotella-sp.] |
| NLB25730.1 | HEPN+AAA-ATPase+Alpha+Beta-domain+wHTH+RRM | Type-1_BR.RM.like.systems | <-HEPN+DUF499*<-METHYLASE-p | 1096 | Bacteria>FCB_group>Bacteroidota/Chlorobiota_group>Bacteroidota>Bacteroidia | GCA_012511595.1 | Bacteroidales_bacterium | GX820_03445 | 1391..4681 | 2030927 | ATP-binding-protein-[Bacteroidales-bacterium] |
| MBQ8713170.1 | HEPN+AAA-ATPase+Alpha+Beta-domain+wHTH+RRM | Type-1_BR.RM.like.systems | METHYLASE-p->HEPN+DUF499*-> | 1120 | Bacteria>FCB_group>Bacteroidota/Chlorobiota_group>Bacteroidota>Bacteroidia | GCA_017471565.1 | Prevotella_sp. | IJ551_10195 | 948..4310 | 59823 | DUF499-domain-containing-protein-[Prevotella-sp.] |
| MBP7851039.1 | HEPN+AAA-ATPase+Alpha+Beta-domain+wHTH+RRM | Type-1_BR.RM.like.systems | METHYLASE-p->HEPN+DUF499*-> | 1097 | Bacteria>FCB_group>Bacteroidota/Chlorobiota_group>Bacteroidota>Bacteroidia | GCA_018001695.1 | Lentimicrobiaceae_bacterium | KA053_12270 | 826..4119 | 2026756 | DUF499-domain-containing-protein-[Lentimicrobiaceae-bacterium] |
| MBO6254120.1 | HEPN+AAA-ATPase+Alpha+Beta-domain+wHTH+RRM | Type-1_BR.RM.like.systems | <-HEPN+DUF499*<-METHYLASE-p | 1108 | Bacteria>FCB_group>Bacteroidota/Chlorobiota_group>Bacteroidota>Bacteroidia | GCA_017635295.1 | Bacteroidaceae_bacterium | J6O49_10775 | 248..3574 | 2212467 | DUF499-domain-containing-protein-[Bacteroidaceae-bacterium] |
| MDD4108176.1 | HEPN+AAA-ATPase+Alpha+Beta-domain+wHTH+RRM | Type-1_BR.RM.like.systems | <-HEPN+DUF499*<-?<-METHYLASE | 1002 | Bacteria>FCB_group>Bacteroidota/Chlorobiota_group>Bacteroidota>Bacteroidia | GCA_028701215.1 | Prolixibacteraceae_bacterium | PHH93_05600 | 1..3006 | 2053594 | MAG:-DUF499-domain-containing-protein,-partial-[Prolixibacteraceae-bacterium] |
| MBO7596490.1 | HEPN+AAA-ATPase+Alpha+Beta-domain+wHTH+RRM | Type-1_BR.RM.like.systems | <-HEPN+DUF499*<-METHYLASE | 1105 | Bacteria>FCB_group>Bacteroidota/Chlorobiota_group>Bacteroidota>Bacteroidia | GCA_017651345.1 | Bacteroidales_bacterium | J6T70_05520 | 1703..5020 | 2030927 | DUF499-domain-containing-protein-[Bacteroidales-bacterium] |
| MCG9899089.1 | HEPN+AAA-ATPase+Alpha+Beta-domain+wHTH+RRM | Type-1_BR.RM.like.systems | <-REase<-?\|\|?-><-?<-RVT<-?<-HEPN+DUF499*<-METHYLASE<-AbiEii<-AbiEi_1<-TUDOR+SF1-SF2-Helicase+iREase+CTD+REase | 1108 | Bacteria>FCB_group>Bacteroidota/Chlorobiota_group>Bacteroidota>Chitinophagia | GCA_022402565.1 | Hydrotalea_sp. | MH132_03730 | 9399..12725 | 2881279 | DUF499-domain-containing-protein-[Hydrotalea-sp.] |
| MBW7891560.1 | HEPN+AAA-ATPase+Alpha+Beta-domain+wHTH+RRM | Type-1_BR.RM.like.systems | TUDOR+SF1-SF2-Helicase+iREase+CTD+REase-p->AbiEi_1->METHYLASE->HEPN+DUF499*-> | 917 | Bacteria>FCB_group>Bacteroidota/Chlorobiota_group>Bacteroidota>Chitinophagia | GCA_019454915.1 | Chitinophagaceae_bacterium | H3C48_11165 | 5923..8672 | 1869212 | ATP-binding-protein-[Chitinophagaceae-bacterium] |
| MCB9047412.1 | HEPN+AAA-ATPase+Alpha+Beta-domain+wHTH+RRM | Type-1_BR.RM.like.systems | <-REase+35exo<-?<-WYL+WYL-C<-ParB+HNH<-?<-HEPN+DUF499*<-METHYLASE<-TUDOR+SF1-SF2-Helicase+iREase+CTD+REase\|\|HKD-DNase->wHTH+SAD+HNH-> | 1086 | Bacteria>FCB_group>Bacteroidota/Chlorobiota_group>Bacteroidota>Chitinophagia | GCA_020636495.1 | Chitinophagales_bacterium | H6550_14865 | 3305993..3309253 | 2448779 | DUF499-domain-containing-protein-[Chitinophagales-bacterium] |
| MBN8676639.1 | HEPN+AAA-ATPase+Alpha+Beta-domain+wHTH+RRM | Type-1_BR.RM.like.systems | REase+35exo+Beta-Barrel+SF1-SF2-Helicase->?->?->TUDOR+SF1-SF2-Helicase+iREase+CTD+REase->ParB+HNH->METHYLASE->HEPN+DUF499*->REase-> | 1087 | Bacteria>FCB_group>Bacteroidota/Chlorobiota_group>Bacteroidota>Chitinophagia | GCA_017303475.1 | Chitinophagales_bacterium | JOM29_00350 | 102288..105551 | 2448779 | DUF499-domain-containing-protein-[Chitinophagales-bacterium] |
| MBB1285247.1 | HEPN+AAA-ATPase+Alpha+Beta-domain+wHTH+RRM | Type-1_BR.RM.like.systems | <-HEPN+DUF499*<-METHYLASE<-AbiEii<-wHTH+AbiEi<-TUDOR+SF1-SF2-Helicase+iREase+CTD+REase<-?\|\|MerR-HTH-> | 1093 | Bacteria>FCB_group>Bacteroidota/Chlorobiota_group>Bacteroidota>Chitinophagia | GCA_014163555.1 | Flavisolibacter_longurius | HRH25_12755 | 53623..56904 | 2740583 | DUF499-domain-containing-protein-[Flavisolibacter-longurius] |
| WP_092673506.1 | HEPN+AAA-ATPase+Alpha+Beta-domain+wHTH+RRM | Type-1_BR.RM.like.systems | HIN-HTH->HTH_21+rve+rve_3-><-SIR2\|\|?->?-><-HEPN+DUF499*<-METHYLASE<-TUDOR+SF1-SF2-Helicase+iREase+CTD+REase<-HTH_3<-?<-URI-DNase<-URI-DNase | 1128 | Bacteria>FCB_group>Bacteroidota/Chlorobiota_group>Bacteroidota>Cytophagia | GCF_900115775.1 | Hymenobacter_arizonensis | BMZ66_RS12440 | 170708..174094 | 1227077 | DUF499-domain-containing-protein-[Siccationidurans-arizonensis] |
| WP_163384119.1 | HEPN+AAA-ATPase+Alpha+Beta-domain+wHTH+RRM | Type-1_BR.RM.like.systems | HEPN->?->?->?->Peptidase-><-HEPN+DUF499*<-METHYLASE<-TUDOR+SF1-SF2-Helicase+iREase+CTD+REase<-HTH<-?<-?<-XIS-HTH | 1103 | Bacteria>FCB_group>Bacteroidota/Chlorobiota_group>Bacteroidota>Cytophagia | GCF_030409895.1 | Cyclobacterium_jeungdonense | QWZ15_RS05330 | 512550..515861 | 708087 | Swt1-family-HEPN-domain-containing-protein-[Cyclobacterium-jeungdonense] |
| WP_013929916.1 | HEPN+AAA-ATPase+Alpha+Beta-domain+wHTH+RRM | Type-1_BR.RM.like.systems | <-DDE_Tnp_1<-BirA-HTH<-DDE_Tnp_1_assoc+DDE_Tnp_1<-MULE-TRANSPOSASE<-SF1-SF2-Helicase<-MORC+DUF3684+REase<-HEPN+DUF499*<-METHYLASE-p\|\|DDE_Tnp_1-><-METHYLASE<-REase<-TUDOR+SF1-SF2-Helicase+iREase+CTD+REase<-AbiEi_4 | 1094 | Bacteria>FCB_group>Bacteroidota/Chlorobiota_group>Bacteroidota>Cytophagia | GCF_000218895.1 | Runella_slithyformis | RUNSL_RS20985 | 4951748..4955032 | 106 | DUF499-domain-containing-protein-[Runella-slithyformis] |
| WP_027303393.1 | HEPN+AAA-ATPase+Alpha+Beta-domain+wHTH+RRM | Type-1_BR.RM.like.systems | MNS+REase->CHTH->?->?-><-?\|\|MORC+HisKin+REase-><-HEPN+DUF499*<-wHTH+REase<-METHYLASE\|\|?-><-TUDOR+SF1-SF2-Helicase+iREase+CTD+REase<-PAIREDC-HTH\|\|Phage_integrase-> | 1100 | Bacteria>FCB_group>Bacteroidota/Chlorobiota_group>Bacteroidota>Cytophagia | GCF_000383955.1 | Rudanella_lutea | RUDLU_RS0125935 | 41364..44666 | 451374 | DUF499-domain-containing-protein-[Rudanella-lutea] |
| WP_106140933.1 | HEPN+AAA-ATPase+Alpha+Beta-domain+wHTH+RRM | Type-1_BR.RM.like.systems | TUDOR+SF1-SF2-Helicase+iREase+CTD+REase->wHTH+AbiEi->AbiEii->METHYLASE->HEPN+DUF499*->?->SF1-SF2-Helicase+DUF1998+HKD-DNase+HKD-DNase-> | 1123 | Bacteria>FCB_group>Bacteroidota/Chlorobiota_group>Bacteroidota>Cytophagia | GCF_003002825.1 | Spirosoma_oryzae | CLV58_RS26865 | 12572..15943 | 1469603 | DUF499-domain-containing-protein-[Spirosoma-oryzae] |
| WP_209329595.1 | HEPN+AAA-ATPase+Alpha+Beta-domain+wHTH+RRM | Type-1_BR.RM.like.systems | <-BirA-HTH<-?<-?<-HEPN+DUF499*<-AAA-ATPase+REase<-METHYLASE<-ParB+HNH<-TUDOR+SF1-SF2-Helicase+iREase+CTD+REase | 1097 | Bacteria>FCB_group>Bacteroidota/Chlorobiota_group>Bacteroidota>Cytophagia | GCF_017811155.1 | Lunatimonas_salinarum | IAQ59_RS04505 | 160980..164273 | 1774590 | DUF499-domain-containing-protein-[Lunatimonas-salinarum] |
| WP_070725327.1 | HEPN+AAA-ATPase+Alpha+Beta-domain+wHTH+RRM | Type-1_BR.RM.like.systems | <-SF1-SF2-Helicase+REase<-?\|\|TUDOR+SF1-SF2-Helicase+iREase+CTD+REase->OLD-ABC->HNH->METHYLASE->HEPN+DUF499*-><-HTH | 1110 | Bacteria>FCB_group>Bacteroidota/Chlorobiota_group>Bacteroidota>Cytophagia | GCF_001816145.1 | Hymenobacter_lapidarius | BEN47_RS03945 | 71126..74458 | 1908237 | DUF499-domain-containing-protein-[Hymenobacter-lapidarius] |
| WP_130024109.1 | HEPN+AAA-ATPase+Alpha+Beta-domain+wHTH+RRM | Type-1_BR.RM.like.systems | <-HTH<-XIS-HTH<-?<-HEPN+DUF499*<-METHYLASE<-AbiEii<-wHTH+AbiEi<-REase<-TUDOR+SF1-SF2-Helicase+iREase+CTD+REase | 1096 | Bacteria>FCB_group>Bacteroidota/Chlorobiota_group>Bacteroidota>Cytophagia | GCF_004168285.1 | Emticicia_agri | EWM59_RS25780 | 4303..7593 | 2492393 | DUF499-domain-containing-protein-[Emticicia-agri] |
| WP_114065477.1 | HEPN+AAA-ATPase+Alpha+Beta-domain+wHTH+RRM | Type-1_BR.RM.like.systems | <-TUDOR+SF1-SF2-Helicase+iREase+CTD<-HEPN+DUF499*<-?<-OLD-ABC<-ParB+HNH<-METHYLASE<-REase<-TUDOR+SF1-SF2-Helicase+iREase+CTD+REase | 1094 | Bacteria>FCB_group>Bacteroidota/Chlorobiota_group>Bacteroidota>Cytophagia | GCF_003325355.1 | Runella_rosea | DR864_RS02565 | 579082..582366 | 2259595 | DUF499-domain-containing-protein-[Runella-rosea] |
| MBK7652070.1 | HEPN+AAA-ATPase+Alpha+Beta-domain+wHTH+RRM | Type-1_BR.RM.like.systems | HTH->REase->METHYLASE-p->HEPN+DUF499*->DUF499-p->DUF499-p-> | 498 | Bacteria>FCB_group>Bacteroidota/Chlorobiota_group>Bacteroidota>Cytophagia | GCA_016709845.1 | Flammeovirgaceae_bacterium | IPJ20_16990 | 23879..25375 | 2026740 | DUF499-domain-containing-protein-[Flammeovirgaceae-bacterium] |
| MBC7408026.1 | HEPN+AAA-ATPase+Alpha+Beta-domain+wHTH+RRM | Type-1_BR.RM.like.systems | AbiEii->METHYLASE->HEPN+DUF499*-> | 559 | Bacteria>FCB_group>Bacteroidota/Chlorobiota_group>Bacteroidota>Cytophagia | GCA_014376355.1 | Arcicella_sp. | H7339_06525 | 3538..5214 | 1872628 | DUF499-domain-containing-protein-[Arcicella-sp.] |
| MBX7052699.1 | HEPN+AAA-ATPase+Alpha+Beta-domain+wHTH+RRM | Type-1_BR.RM.like.systems | MazG->REase+SF1-SF2-Helicase->TUDOR+SF1-SF2-Helicase+iREase+CTD+REase->OLD-ABC->HNH->METHYLASE->HEPN+DUF499*-> | 1088 | Bacteria>FCB_group>Bacteroidota/Chlorobiota_group>Bacteroidota>Flavobacteriia | GCA_019694805.1 | Flavobacteriales_bacterium | K1X54_11735 | 22264..25530 | 2021391 | DUF499-domain-containing-protein-[Flavobacteriales-bacterium] |
| NEW78740.1 | HEPN+AAA-ATPase+Alpha+Beta-domain+wHTH+RRM | Type-1_BR.RM.like.systems | HKD-DNase->?-><-HEPN<-HKD-DNase<-HEPN+DUF499*<-METHYLASE<-AbiEii<-AbiEi_1<-TUDOR+SF1-SF2-Helicase+iREase+CTD+REase | 1089 | Bacteria>FCB_group>Bacteroidota/Chlorobiota_group>Bacteroidota>Flavobacteriia | GCA_010906935.1 | Gelidibacter_sp. | GZ086_04810 | 7309..10578 | 2018083 | ATP-binding-protein-[Gelidibacter-sp.] |
| HCE56000.1 | HEPN+AAA-ATPase+Alpha+Beta-domain+wHTH+RRM | Type-1_BR.RM.like.systems | <-PIN<-HEPN+DUF499*<-METHYLASE<-AbiEii<-AbiEi_1<-TUDOR+SF1-SF2-Helicase+iREase+CTD+REase<-AbiEii<-AbiEi_4 | 1089 | Bacteria>FCB_group>Bacteroidota/Chlorobiota_group>Bacteroidota>Flavobacteriia | GCA_003516285.1 | Lutibacter_sp. | DER05_13810 | 23309..26578 | 1925666 | hypothetical-protein-[Lutibacter-sp.] |
| WP_143387496.1 | HEPN+AAA-ATPase+Alpha+Beta-domain+wHTH+RRM | Type-1_BR.RM.like.systems | TUDOR+SF1-SF2-Helicase+iREase+CTD+REase->AbiEi_1->AbiEii->METHYLASE->HEPN+DUF499*->?->?->?->PIN->?-><-WYL+WYL-C | 1088 | Bacteria>FCB_group>Bacteroidota/Chlorobiota_group>Bacteroidota>Flavobacteriia | GCF_007097285.1 | Flavobacterium_gawalongense | FNW12_RS09775 | 80798..84064 | 2594432 | DUF499-domain-containing-protein-[Flavobacterium-gawalongense] |
| MAM23120.1 | HEPN+AAA-ATPase+Alpha+Beta-domain+wHTH+RRM | Type-1_BR.RM.like.systems | HTH+MPTase->PIN->HTH_3->TUDOR+SF1-SF2-Helicase+iREase+CTD+REase->METHYLASE->HEPN+DUF499*-> | 869 | Bacteria>FCB_group>Bacteroidota/Chlorobiota_group>Bacteroidota>Flavobacteriia | GCA_002695265.1 | Croceibacter_sp. | CL817_05250 | 9003..11611 | 2024829 | hypothetical-protein-[Croceibacter-sp.] |
| MCS7037545.1 | HEPN+AAA-ATPase+Alpha+Beta-domain+wHTH+RRM | Type-1_BR.RM.like.systems | TUDOR+SF1-SF2-Helicase+iREase+CTD+REase-p->METHYLASE->HEPN+DUF499*-> | 1108 | Bacteria>FCB_group>Bacteroidota/Chlorobiota_group>Bacteroidota>Saprospiria | GCA_025056735.1 | Saprospiraceae_bacterium | NZM41_12835 | 4336..7662 | 2202734 | DUF499-domain-containing-protein-[Saprospiraceae-bacterium] |
| MBK8877301.1 | HEPN+AAA-ATPase+Alpha+Beta-domain+wHTH+RRM | Type-1_BR.RM.like.systems | TUDOR+SF1-SF2-Helicase+iREase+CTD+REase->HEPN+DUF499*->?-><-?<-SMODS | 1105 | Bacteria>FCB_group>Bacteroidota/Chlorobiota_group>Bacteroidota>Saprospiria | GCA_016715005.1 | Haliscomenobacter_sp. | IPN74_01745 | 433020..436337 | 2717303 | DUF499-domain-containing-protein-[Haliscomenobacter-sp.] |
| NUQ24578.1 | HEPN+AAA-ATPase+Alpha+Beta-domain+wHTH+RRM | Type-1_BR.RM.like.systems | TUDOR+SF1-SF2-Helicase+iREase+CTD+REase->OLD-ABC->HNH->METHYLASE->AAA-ATPase+REase->HEPN+DUF499*->?->PolB->HEPN-><-ORC-CDC6+REase | 1095 | Bacteria>FCB_group>Bacteroidota/Chlorobiota_group>Bacteroidota>Saprospiria | GCA_013360405.1 | Saprospiraceae_bacterium | HUU34_11550 | 44219..47506 | 2202734 | DUF499-domain-containing-protein-[Saprospiraceae-bacterium] |
| WP_051018640.1 | HEPN+AAA-ATPase+Alpha+Beta-domain+wHTH+RRM | Type-1_BR.RM.like.systems | TUDOR+SF1-SF2-Helicase+iREase+CTD+REase->METHYLASE-p->?->DDE_Tnp_1+DDE_Tnp_1_2->METHYLASE->HEPN+DUF499*->?->?->?->?-><-?<-Peptidase | 1105 | Bacteria>FCB_group>Bacteroidota/Chlorobiota_group>Bacteroidota>Saprospiria | GCF_000275825.1 | Saprospira_grandis | SAPGRDRAFT_RS17610 | 3964671..3967988 | 1008 | DUF499-domain-containing-protein-[Saprospira-grandis] |
| MBK8920115.1 | HEPN+AAA-ATPase+Alpha+Beta-domain+wHTH+RRM | Type-1_BR.RM.like.systems | <-Glyco_hydro_1<-?<-HEPN+DUF499*<-URI-DNase<-METHYLASE-p<-METHYLASE-p<-TOPRIM<-OLD-ABC | 1088 | Bacteria>FCB_group>Bacteroidota/Chlorobiota_group>Bacteroidota>Saprospiria | GCA_016719895.1 | Saprospirales_bacterium | IPM81_01165 | 280177..283443 | 2026790 | DUF499-domain-containing-protein-[Saprospirales-bacterium] |
| MBK6932297.1 | HEPN+AAA-ATPase+Alpha+Beta-domain+wHTH+RRM | Type-1_BR.RM.like.systems | <-Glyco_hydro_1\|\|?-><-DUF499-p<-HEPN+DUF499*<-URI-DNase<-METHYLASE-p<-?<-METHYLASE-p<-HNH | 951 | Bacteria>FCB_group>Bacteroidota/Chlorobiota_group>Bacteroidota>Saprospiria | GCA_016709195.1 | Saprospirales_bacterium | IPH12_16140 | 92714..95569 | 2026790 | DUF499-domain-containing-protein-[Saprospirales-bacterium] |
| WP_270099212.1 | HEPN+AAA-ATPase+Alpha+Beta-domain+wHTH+RRM | Type-1_BR.RM.like.systems | Peptidase-><-?<-?<-?<-HEPN+DUF499*<-METHYLASE<-TUDOR+SF1-SF2-Helicase+iREase+CTD+REase\|\|?->?->Y1_Tnp-> | 1105 | Bacteria>FCB_group>Bacteroidota/Chlorobiota_group>Bacteroidota>Saprospiria | GCF_027594745.1 | Saprospira_grandis | OP864_RS16310 | 4154293..4157610 | 1008 | DUF499-domain-containing-protein-[Saprospira-grandis] |
| EJF54826.1 | HEPN+AAA-ATPase+Alpha+Beta-domain+wHTH+RRM | Type-1_BR.RM.like.systems | TUDOR+SF1-SF2-Helicase+iREase+CTD+REase->DUF4096+DDE_Tnp_1+DDE_Tnp_1_2->HEPN+DUF499*-> | 1104 | Bacteria>FCB_group>Bacteroidota/Chlorobiota_group>Bacteroidota>Saprospiria | GCA_000275825.1 | Saprospira_grandis_DSM_2844 | SapgrDRAFT_3181 | 3964674..3967988 | 694433 | putative-ATPase-(AAA+-superfamily)-[Saprospira-grandis-DSM-2844] |
| WP_222581795.1 | HEPN+AAA-ATPase+Alpha+Beta-domain+wHTH+RRM | Type-1_BR.RM.like.systems | <-HIN-HTH\|\|METHYLASE->HEPN+DUF499*-> | 1089 | Bacteria>FCB_group>Bacteroidota/Chlorobiota_group>Bacteroidota>Saprospiria | GCF_019802045.1 | Membranihabitans_marinus | KUV50_RS18995 | 2282..5551 | 1227546 | DUF499-domain-containing-protein-[Membranihabitans-marinus] |
| OWY19366.1 | HEPN+AAA-ATPase+Alpha+Beta-domain+wHTH+RRM | Type-1_BR.RM.like.systems | <-HEPN+DUF499*<-METHYLASE<-TUDOR+SF1-SF2-Helicase+iREase+CTD+REase | 1103 | Bacteria>FCB_group>Bacteroidota/Chlorobiota_group>Bacteroidota>Sphingobacteriia | GCA_002212985.1 | Sphingobacteriales_bacterium_TSM_CSS | B6N25_14540 | 46408..49719 | 1972511 | hypothetical-protein-B6N25_14540-[Sphingobacteriales-bacterium-TSM_CSS] |
| MDB5032646.1 | HEPN+AAA-ATPase+Alpha+Beta-domain+wHTH+RRM | Type-1_BR.RM.like.systems | Phage_int_SAM_5+Phage_integrase->DEDD_Tnp_IS110+Transposase_20-><-ABhydrolase<-HEPN+DUF499*<-METHYLASE<-TUDOR+SF1-SF2-Helicase+iREase+CTD+REase<-?\|\|SMODS->TIR-> | 1091 | Bacteria>FCB_group>Bacteroidota/Chlorobiota_group>Bacteroidota>Sphingobacteriia | GCA_028287845.1 | Mucilaginibacter_sp. | JWP71_3367 | 6008..9283 | 1882438 | MAG:-hypothetical-protein-JWP71_3367-[Mucilaginibacter-sp.] |
| WP_090973307.1 | HEPN+AAA-ATPase+Alpha+Beta-domain+wHTH+RRM | Type-1_BR.RM.like.systems | TUDOR+SF1-SF2-Helicase+iREase+CTD+REase->METHYLASE->?->DOC->HEPN+DUF499*->?->PIN->tRNA->?->Synaptojanin-> | 1100 | Bacteria>FCB_group>Bacteroidota/Chlorobiota_group>Bacteroidota>Sphingobacteriia | GCF_900112315.1 | Parapedobacter_composti | BM030_RS09250 | 52170..55472 | 623281 | DUF499-domain-containing-protein-[Parapedobacter-composti] |
| MCY3999495.1 | HEPN+AAA-ATPase+Alpha+Beta-domain+wHTH+RRM | Type-1_BR.RM.like.systems | REase->?->?->?->?->TUDOR+SF1-SF2-Helicase+iREase+CTD+REase->HEPN+DUF499*->Phage_int_SAM_3+Phage_integrase->Phage_int_SAM_3+Phage_integrase->METHYLASE-p-> | 1095 | Bacteria>FCB_group>Bacteroidota/Chlorobiota_group>Bacteroidota>unclassified_Bacteroidota | GCA_026708685.1 | Bacteroidota_bacterium | OXF84_01635 | 21572..24859 | 1898104 | MAG:-DUF499-domain-containing-protein-[Bacteroidetes-bacterium] |
| MCL5268276.1 | HEPN+AAA-ATPase+Alpha+Beta-domain+wHTH+RRM | Type-1_BR.RM.like.systems | HNH->?-><-?<-REase<-REase<-TIR<-HEPN+DUF499*<-METHYLASE<-DrHyd+BirA-HTH<-TUDOR+SF1-SF2-Helicase+iREase+CTD+REase<-?<-?<-METHYLASE-p | 1111 | Bacteria>FCB_group>Bacteroidota/Chlorobiota_group>Bacteroidota>unclassified_Bacteroidota | GCA_023480735.1 | Bacteroidota_bacterium | M1469_09265 | 8457..11792 | 1898104 | Swt1-family-HEPN-domain-containing-protein-[Bacteroidetes-bacterium] |
| MDE2834954.1 | HEPN+AAA-ATPase+Alpha+Beta-domain+wHTH+RRM | Type-1_BR.RM.like.systems | MNS-><-?\|\|TUDOR+SF1-SF2-Helicase+iREase+CTD+REase->HEPN+DUF499*->Phage_integrase->Phage_int_SAM_3+Phage_integrase-> | 1086 | Bacteria>FCB_group>Bacteroidota/Chlorobiota_group>Bacteroidota>unclassified_Bacteroidota | GCA_028874735.1 | Bacteroidota_bacterium | OXM02_10600 | 6360..9620 | 1898104 | MAG:-Swt1-family-HEPN-domain-containing-protein-[Bacteroidota-bacterium] |
| MCY3629603.1 | HEPN+AAA-ATPase+Alpha+Beta-domain+wHTH+RRM | Type-1_BR.RM.like.systems | <-MNS\|\|TUDOR+SF1-SF2-Helicase+iREase+CTD+REase->HEPN+DUF499*->Phage_integrase->Phage_int_SAM_3+Phage_integrase-> | 1099 | Bacteria>FCB_group>Bacteroidota/Chlorobiota_group>Bacteroidota>unclassified_Bacteroidota | GCA_026705375.1 | Bacteroidota_bacterium | OXG94_06260 | 16273..19572 | 1898104 | MAG:-DUF499-domain-containing-protein-[Bacteroidetes-bacterium] |
| MDE2731786.1 | HEPN+AAA-ATPase+Alpha+Beta-domain+wHTH+RRM | Type-1_BR.RM.like.systems | <-METHYLASE-p<-Phage_integrase\|\|?-><-HEPN+DUF499*<-TUDOR+SF1-SF2-Helicase+iREase+CTD+REase<-MNS | 1086 | Bacteria>FCB_group>Bacteroidota/Chlorobiota_group>Bacteroidota>unclassified_Bacteroidota | GCA_028875295.1 | Bacteroidota_bacterium | OXI38_10375 | 1957..5217 | 1898104 | MAG:-Swt1-family-HEPN-domain-containing-protein-[Bacteroidota-bacterium] |
| MCY4223999.1 | HEPN+AAA-ATPase+Alpha+Beta-domain+wHTH+RRM | Type-1_BR.RM.like.systems | <-Phage_int_SAM_3+Phage_integrase<-Phage_integrase<-HEPN+DUF499* | 1027 | Bacteria>FCB_group>Bacteroidota/Chlorobiota_group>Bacteroidota>unclassified_Bacteroidota | GCA_026709435.1 | Bacteroidota_bacterium | OXF06_04095 | 2186..5267 | 1898104 | MAG:-DUF499-domain-containing-protein,-partial-[Bacteroidetes-bacterium] |
| MDE2771369.1 | HEPN+AAA-ATPase+Alpha+Beta-domain+wHTH+RRM | Type-1_BR.RM.like.systems | <-DnaJ<-?<-?\|\|?->HEPN+DUF499*->Phage_integrase->Phage_integrase-> | 1094 | Bacteria>FCB_group>Bacteroidota/Chlorobiota_group>Bacteroidota>unclassified_Bacteroidota | GCA_028875125.1 | Bacteroidota_bacterium | OXI44_09415 | 24188..27472 | 1898104 | MAG:-DUF499-domain-containing-protein-[Bacteroidota-bacterium] |
| MBT3751042.1 | HEPN+AAA-ATPase+Alpha+Beta-domain+wHTH+RRM | Type-1_BR.RM.like.systems | METHYLASE-p->HEPN+DUF499*->?->?->GDResolvase+Recombinase+ZnR-> | 1103 | Bacteria>FCB_group>Bacteroidota/Chlorobiota_group>Bacteroidota>unclassified_Bacteroidota | GCA_018653945.1 | Bacteroidota_bacterium | HOG34_18835 | 220..3531 | 1898104 | ATP-binding-protein-[Bacteroidetes-bacterium] |
| RLD44971.1 | HEPN+AAA-ATPase+Alpha+Beta-domain+wHTH+RRM | Type-1_BR.RM.like.systems | <-HEPN+DUF499*<-METHYLASE-p<-METHYLASE-p | 726 | Bacteria>FCB_group>Bacteroidota/Chlorobiota_group>Bacteroidota>unclassified_Bacteroidota | GCA_003650385.1 | Bacteroidota_bacterium | DRI88_08990 | 1..2178 | 1898104 | AAA+-family-ATPase,-partial-[Bacteroidetes-bacterium] |
| MCB0540282.1 | HEPN+AAA-ATPase+Alpha+Beta-domain+wHTH+RRM | Type-1_BR.RM.like.systems | METHYLASE-p->HEPN+DUF499*-> | 769 | Bacteria>FCB_group>Bacteroidota/Chlorobiota_group>Bacteroidota>unclassified_Bacteroidota | GCA_020436205.1 | Bacteroidota_bacterium | KDE33_22385 | 287..2592 | 1898104 | DUF499-domain-containing-protein-[Bacteroidetes-bacterium] |
| WP_210512925.1 | HEPN+AAA-ATPase+Alpha+Beta-domain+wHTH+RRM | Type-1_BR.RM.like.systems | Ntox49->?->tRNA-><-XIS-HTH\|\|TUDOR+SF1-SF2-Helicase+iREase+CTD+REase->METHYLASE->HEPN+DUF499*-> | 1089 | Bacteria>FCB_group>Bacteroidota/Chlorobiota_group>Balneolota>Balneolia | GCF_017921895.1 | Natronogracilivirga_saccharolytica | NATSA_RS12455 | 139567..142836 | 2812953 | DUF499-domain-containing-protein-[Natronogracilivirga-saccharolytica] |
| GIV55345.1 | HEPN+AAA-ATPase+Alpha+Beta-domain+wHTH+RRM | Type-1_BR.RM.like.systems | <-LexA-HTH+LexA-protease\|\|?->Methyltransf_5-><-MORC+DUF3684<-HEPN+DUF499*<-METHYLASE<-TUDOR+SF1-SF2-Helicase+iREase+CTD+REase | 1140 | Bacteria>FCB_group>Bacteroidota/Chlorobiota_group>Candidatus_Kapabacteria>unclassified_Candidatus_Kapabacteria | GCA_026003495.1 | Candidatus_Kapabacteria_bacterium | KatS3mg040_0113 | 136007..139429 | 2447898 | MAG:-hypothetical-protein-KatS3mg040_0113-[Candidatus-Kapabacteria-bacterium] |
| MBL6956529.1 | HEPN+AAA-ATPase+Alpha+Beta-domain+wHTH+RRM | Type-1_BR.RM.like.systems | <-HTH+MPTase<-Tox-RelE<-?<-HEPN+DUF499*<-METHYLASE<-wHTH+REase<-TUDOR+SF1-SF2-Helicase+iREase+CTD+REase | 1115 | Bacteria>FCB_group>Bacteroidota/Chlorobiota_group>Chlorobiota>Chlorobiia | GCA_016776555.1 | Chlorobium_phaeobacteroides | ISR54_06890 | 24108..27455 | 1096 | DUF499-domain-containing-protein-[Chlorobium-phaeobacteroides] |
| WP_059138717.1 | HEPN+AAA-ATPase+Alpha+Beta-domain+wHTH+RRM | Type-1_BR.RM.like.systems | RelE-ParE->TUDOR+SF1-SF2-Helicase+iREase+CTD+REase->wHTH+REase->METHYLASE->HEPN+DUF499*->?->?->?->TraJ-RHH-> | 1115 | Bacteria>FCB_group>Bacteroidota/Chlorobiota_group>Chlorobiota>Chlorobiia | GCF_001509575.1 | Chlorobium_limicola | ASB62_RS03910 | 18289..21636 | 1092 | Swt1-family-HEPN-domain-containing-protein-[Chlorobium-limicola] |
| ARA93108.1 | HEPN+AAA-ATPase+Alpha+Beta-domain+wHTH+RRM | Type-1_BR.RM.like.systems | <-RVT<-MORC<-Methyltransf_D12<-?<-DUF4011+SF1-SF2-Helicase+REase<-HEPN+DUF499*<-METHYLASE<-MORC+DUF3684<-TUDOR+SF1-SF2-Helicase+iREase+CTD+REase<-?\|\|PIN-> | 1112 | Bacteria>FCB_group>Bacteroidota/Chlorobiota_group>Rhodothermota>Rhodothermia | GCA_001518995.2 | Rhodothermaceae_bacterium_RA | AWN76_007995 | 1928774..1932112 | 1779382 | AAA+-family-ATPase-[Rhodothermaceae-bacterium-RA] |
| MXW34066.1 | HEPN+AAA-ATPase+Alpha+Beta-domain+wHTH+RRM | Type-1_BR.RM.like.systems | <-MNS\|\|TUDOR+SF1-SF2-Helicase+iREase+CTD+REase->HEPN+DUF499*->Phage_integrase->Phage_int_SAM_3+Phage_integrase-> | 1099 | Bacteria>FCB_group>Bacteroidota/Chlorobiota_group>Rhodothermota>Rhodothermia | GCA_009836215.1 | Rhodothermaceae_bacterium | F4Z62_13320 | 16280..19579 | 2026787 | ATP-binding-protein-[Rhodothermaceae-bacterium] |
| MXW14654.1 | HEPN+AAA-ATPase+Alpha+Beta-domain+wHTH+RRM | Type-1_BR.RM.like.systems | <-MNS\|\|TUDOR+SF1-SF2-Helicase+iREase+CTD+REase->HEPN+DUF499*->Phage_integrase->Phage_int_SAM_3+Phage_integrase-> | 1099 | Bacteria>FCB_group>Bacteroidota/Chlorobiota_group>Rhodothermota>Rhodothermia | GCA_009835915.1 | Rhodothermaceae_bacterium | F4Z76_07000 | 16270..19569 | 2026787 | ATP-binding-protein-[Rhodothermaceae-bacterium] |
| WP_210373541.1 | HEPN+AAA-ATPase+Alpha+Beta-domain+wHTH+RRM | Type-1_BR.RM.like.systems | Peptidase_M28-><-?\|\|?->?->SIGMA-HTH->?-><-HEPN+DUF499*<-METHYLASE<-?<-TUDOR+SF1-SF2-Helicase+iREase+CTD+REase | 1114 | Bacteria>FCB_group>Bacteroidota/Chlorobiota_group>Rhodothermota>Rhodothermia | GCF_017908595.1 | Rhodothermus_bifroesti | J8E65_RS01005 | 262538..265882 | 2823335 | Swt1-family-HEPN-domain-containing-protein-[Rhodothermus-bifroesti] |
| MBW7874866.1 | HEPN+AAA-ATPase+Alpha+Beta-domain+wHTH+RRM | Type-1_BR.RM.like.systems | <-XIS-HTH\|\|METHYLASE->HTH+MPTase->?->Classical-AAA->HEPN+DUF499*->?->?->Polbeta->HEPN-> | 1093 | Bacteria>FCB_group>Candidatus_Cloacimonadota>unclassified_Candidatus_Cloacimonadota | GCA_019454875.1 | Candidatus_Cloacimonadota_bacterium | H3C47_02625 | 20403..23684 | 2030808 | ATP-binding-protein-[Candidatus-Cloacimonetes-bacterium] |
| MBP7719545.1 | HEPN+AAA-ATPase+Alpha+Beta-domain+wHTH+RRM | Type-1_BR.RM.like.systems | TUDOR+SF1-SF2-Helicase+iREase+CTD+REase->METHYLASE->OLD-ABC->TOPRIM->HEPN+DUF499*-> | 1111 | Bacteria>FCB_group>Candidatus_Fermentibacteria>Candidatus_Fermentibacteria_(class)>Candidatus_Fermentibacterales | GCA_017999735.1 | Candidatus_Fermentibacter_sp. | KA108_02790 | 14441..17776 | 2779163 | DUF499-domain-containing-protein-[Candidatus-Fermentibacter-sp.] |
| MCC6694832.1 | HEPN+AAA-ATPase+Alpha+Beta-domain+wHTH+RRM | Type-1_BR.RM.like.systems | TUDOR+SF1-SF2-Helicase+iREase+CTD+REase->REase->HEPN->METHYLASE-p->?->METHYLASE->HEPN+DUF499*->?->?->ArdCN+MPTase-><-Arm-DNA-bind_3+Phage_integrase | 1102 | Bacteria>FCB_group>Candidatus_Hydrogenedentes>unclassified_Candidatus_Hydrogenedentes | GCA_020849535.1 | Candidatus_Hydrogenedentes_bacterium | IT365_04280 | 13299..16607 | 2030809 | DUF499-domain-containing-protein-[Candidatus-Hydrogenedentes-bacterium] |
| MBI2502153.1 | HEPN+AAA-ATPase+Alpha+Beta-domain+wHTH+RRM | Type-1_BR.RM.like.systems | MACRODOMAIN->METHYLASE->?->DUF488-1->?->HEPN+DUF499*->HNH->?->DDE_Tnp_1_4-> | 1125 | Bacteria>FCB_group>Candidatus_Latescibacteria>unclassified_Candidatus_Latescibacteria | GCA_016187725.1 | Candidatus_Latescibacteria_bacterium | HYW07_02835 | 5709..9086 | 2053570 | ATP-binding-protein-[Candidatus-Latescibacteria-bacterium] |
| MCF7823046.1 | HEPN+AAA-ATPase+Alpha+Beta-domain+wHTH+RRM | Type-1_BR.RM.like.systems | <-MazG<-?<-?<-MORC+HISKIN<-METHYLASE<-HEPN+DUF499*<-METHYLASE<-MORC+HisKin+DUF3684<-TUDOR+SF1-SF2-Helicase+iREase+CTD+REase<-XIS-HTH<-?<-Schlafen+HATPase+HTH | 1111 | Bacteria>FCB_group>Candidatus_Marinimicrobia>unclassified_Candidatus_Marinimicrobia | GCA_021734325.1 | Candidatus_Marinimicrobia_bacterium | K9N35_02645 | 72966..76301 | 2026760 | DUF499-domain-containing-protein-[Candidatus-Marinimicrobia-bacterium] |
| MCF7837530.1 | HEPN+AAA-ATPase+Alpha+Beta-domain+wHTH+RRM | Type-1_BR.RM.like.systems | Phage_integrase-><-ZnR+TOPRIM+D5ATPase<-XIS-HTH\|\|HTH->XIS-HTH-><-HEPN+DUF499*<-OLD-ABC+TOPRIM<-METHYLASE-p | 1110 | Bacteria>FCB_group>Candidatus_Marinimicrobia>unclassified_Candidatus_Marinimicrobia | GCA_021734645.1 | Candidatus_Marinimicrobia_bacterium | K9N49_02780 | 12805..16137 | 2026760 | DUF499-domain-containing-protein-[Candidatus-Marinimicrobia-bacterium] |
| MBN1984134.1 | HEPN+AAA-ATPase+Alpha+Beta-domain+wHTH+RRM | Type-1_BR.RM.like.systems | ABC-ATPase+SbcC->?->?-><-?<-HEPN+DUF499*<-METHYLASE<-DrHyd+BirA-HTH<-TUDOR+SF1-SF2-Helicase+iREase+CTD+REase-p | 1112 | Bacteria>FCB_group>Fibrobacterota>Chitinivibrionia>Chitinivibrionales | GCA_016931815.1 | Chitinivibrionales_bacterium | JW795_21570 | 16556..19894 | 2663014 | DUF499-domain-containing-protein-[Chitinivibrionales-bacterium] |
| MCZ0935996.1 | HEPN+AAA-ATPase+Alpha+Beta-domain+wHTH+RRM | Type-1_BR.RM.like.systems | SIGMA-HTH->?->?->OLD-ABC+TOPRIM->SF1-SF2-Helicase->REase-><-HEPN+DUF499*<-METHYLASE-p\|\|Zn_Tnp_IS1595+DDE_Tnp_IS1595-><-?\|\|METHYLASE+HNH->?-><-METHYLASE | 1098 | Bacteria>FCB_group>Gemmatimonadota>Gemmatimonadetes>Candidatus_Palauibacterales | GCA_026914155.1 | Candidatus_Palauibacter_rhopaloidicola | OXJ54_12525 | 22300..25596 | 3056738 | MAG:-DUF499-domain-containing-protein-[Gemmatimonadetes-bacterium] |
| MDE2877868.1 | HEPN+AAA-ATPase+Alpha+Beta-domain+wHTH+RRM | Type-1_BR.RM.like.systems | DNA_methylase->REase->DNA_methylase->SF1-SF2-Helicase+REase->OLD-ABC->METHYLASE->HEPN+DUF499*->?->?->?->MULE-TRANSPOSASE-> | 1098 | Bacteria>FCB_group>Gemmatimonadota>Gemmatimonadetes>Candidatus_Palauibacterales | GCA_028874535.1 | Candidatus_Palauibacter_soopunensis | OXO55_03070 | 20960..24256 | 3056739 | MAG:-DUF499-domain-containing-protein-[Gemmatimonadota-bacterium] |
| MBV6522081.1 | HEPN+AAA-ATPase+Alpha+Beta-domain+wHTH+RRM | Type-1_BR.RM.like.systems | TUDOR+SF1-SF2-Helicase+iREase+CTD+REase->ORC-CDC6+REase->METHYLASE->ORC-CDC6+REase->HEPN+DUF499*-><-Pkinase | 1116 | Bacteria>FCB_group>Gemmatimonadota>Gemmatimonadetes>Gemmatimonadales | GCA_019186945.1 | Gemmatimonadaceae_bacterium | MNPFHGCM_02226 | 129597..132947 | 2026741 | hypothetical-protein-[Gemmatimonadaceae-bacterium] |
| MCC7134741.1 | HEPN+AAA-ATPase+Alpha+Beta-domain+wHTH+RRM | Type-1_BR.RM.like.systems | <-LuxR-HTH\|\|TUDOR+SF1-SF2-Helicase+iREase+CTD+REase->?->METHYLASE->HEPN+DUF499*-> | 765 | Bacteria>FCB_group>Gemmatimonadota>Gemmatimonadetes>Gemmatimonadales | GCA_020851315.1 | Gemmatimonadales_bacterium | IT352_18930 | 11948..14243 | 2448054 | DUF499-domain-containing-protein-[Gemmatimonadales-bacterium] |
| MXX78159.1 | HEPN+AAA-ATPase+Alpha+Beta-domain+wHTH+RRM | Type-1_BR.RM.like.systems | <-HEPN+DUF499*<-METHYLASE<-TUDOR+SF1-SF2-Helicase+iREase+CTD+REase<-BirA-HTH | 1138 | Bacteria>FCB_group>Gemmatimonadota>Gemmatimonadetes>Gemmatimonadales | GCA_009836785.1 | Gemmatimonadales_bacterium | F4Z33_04085 | 250..3666 | 2448054 | ATP-binding-protein-[Gemmatimonadales-bacterium] |
| MBX6330503.1 | HEPN+AAA-ATPase+Alpha+Beta-domain+wHTH+RRM | Type-1_BR.RM.like.systems | HEPN-p->Primpol+D5N+D5ATPase->HEPN+DUF499*->Glyoxalase->?->?->HTH-> | 1110 | Bacteria>FCB_group>Gemmatimonadota>Gemmatimonadetes>Gemmatimonadales | GCA_019689955.1 | Gemmatimonadaceae_bacterium | IRY91_01515 | 6551..9883 | 2026741 | DUF499-domain-containing-protein-[Gemmatimonadaceae-bacterium] |
| MCC7194161.1 | HEPN+AAA-ATPase+Alpha+Beta-domain+wHTH+RRM | Type-1_BR.RM.like.systems | METHYLASE-p->ORC-CDC6+REase->HEPN+DUF499*-><-?\|\|SIGMA-HTH-> | 1104 | Bacteria>FCB_group>Gemmatimonadota>Gemmatimonadetes>Gemmatimonadales | GCA_020851455.1 | Gemmatimonadaceae_bacterium | IT356_01265 | 1930..5244 | 2026741 | DUF499-domain-containing-protein-[Gemmatimonadaceae-bacterium] |
| MXX77944.1 | HEPN+AAA-ATPase+Alpha+Beta-domain+wHTH+RRM | Type-1_BR.RM.like.systems | OLD-ABC->TOPRIM->HEPN+DUF499*-><-RNase<-?<-REase | 1102 | Bacteria>FCB_group>Gemmatimonadota>Gemmatimonadetes>Gemmatimonadales | GCA_009836785.1 | Gemmatimonadales_bacterium | F4Z33_02945 | 1211..4519 | 2448054 | ATP-binding-protein-[Gemmatimonadales-bacterium] |
| MCR4340303.1 | HEPN+AAA-ATPase+Alpha+Beta-domain+wHTH+RRM | Type-1_BR.RM.like.systems | <-HEPN+DUF499*<-METHYLASE-p | 1109 | Bacteria>FCB_group>Gemmatimonadota>Gemmatimonadetes>Gemmatimonadales | GCA_024654005.1 | Gemmatimonadaceae_bacterium | NUWO1_10510 | 510..3839 | 2026741 | DUF499-domain-containing-protein-[Gemmatimonadaceae-bacterium] |
| ODT03617.1 | HEPN+AAA-ATPase+Alpha+Beta-domain+wHTH+RRM | Type-1_BR.RM.like.systems | TUDOR+SF1-SF2-Helicase+iREase+CTD+REase->Schlafen+HATPase+HTH->METHYLASE->HEPN+DUF499*->Primpol+D5N+D5ATPase->PAIREDC-HTH->ORC-CDC6+REase-> | 1109 | Bacteria>FCB_group>Gemmatimonadota>unclassified_Gemmatimonadota | GCA_001724275.1 | Gemmatimonadetes_bacterium_SCN_70-22 | ABS52_08400 | 10589..13918 | 1660163 | AAA+-family-ATPase-[Gemmatimonadetes-bacterium-SCN-70-22] |
| MYH20375.1 | HEPN+AAA-ATPase+Alpha+Beta-domain+wHTH+RRM | Type-1_BR.RM.like.systems | TUDOR+SF1-SF2-Helicase+iREase+CTD+REase->ParB+HNH->METHYLASE->HEPN+DUF499*->?->TPR-><-Peptidase-S8<-Classical-AAA<-Peptidase_M1 | 1108 | Bacteria>FCB_group>Gemmatimonadota>unclassified_Gemmatimonadota | GCA_009844015.1 | Gemmatimonadota_bacterium | F4132_14865 | 91560..94886 | 2026742 | ATP-binding-protein-[Gemmatimonadetes-bacterium] |
| MXZ76449.1 | HEPN+AAA-ATPase+Alpha+Beta-domain+wHTH+RRM | Type-1_BR.RM.like.systems | Synaptojanin->TUDOR+SF1-SF2-Helicase+iREase+CTD+REase->OLD-ABC->METHYLASE->HEPN+DUF499*->ParB+HNH-><-?<-MazE_antitoxin | 1113 | Bacteria>FCB_group>Gemmatimonadota>unclassified_Gemmatimonadota | GCA_009837395.1 | Gemmatimonadota_bacterium | F4Z03_15245 | 64974..68315 | 2026742 | ATP-binding-protein-[Gemmatimonadetes-bacterium] |
| MYL00076.1 | HEPN+AAA-ATPase+Alpha+Beta-domain+wHTH+RRM | Type-1_BR.RM.like.systems | TUDOR+SF1-SF2-Helicase+iREase+CTD+REase->ParB+HNH-p->METHYLASE->HEPN+DUF499*->?->TPR-><-Peptidase-S8 | 1108 | Bacteria>FCB_group>Gemmatimonadota>unclassified_Gemmatimonadota | GCA_009846305.1 | Gemmatimonadota_bacterium | F4014_15170 | 26347..29673 | 2026742 | ATP-binding-protein-[Gemmatimonadetes-bacterium] |
| MXW81679.1 | HEPN+AAA-ATPase+Alpha+Beta-domain+wHTH+RRM | Type-1_BR.RM.like.systems | <-TUDOR+SF1-SF2-Helicase+iREase+CTD+REase<-HEPN+DUF499*\|\|REase+HTH->REase+HTH-><-METHYLASE | 1112 | Bacteria>FCB_group>Gemmatimonadota>unclassified_Gemmatimonadota | GCA_009836325.1 | Gemmatimonadota_bacterium | F4Z57_22370 | 10515..13853 | 2026742 | ATP-binding-protein-[Gemmatimonadetes-bacterium] |
| MYC72435.1 | HEPN+AAA-ATPase+Alpha+Beta-domain+wHTH+RRM | Type-1_BR.RM.like.systems | TUDOR+SF1-SF2-Helicase+iREase+CTD+REase-><-?<-ParB<-DNA_methylase\|\|METHYLASE->HEPN+DUF499*-> | 1131 | Bacteria>FCB_group>Gemmatimonadota>unclassified_Gemmatimonadota | GCA_009841115.1 | Gemmatimonadota_bacterium | F4X17_17170 | 10634..14029 | 2026742 | ATP-binding-protein-[Gemmatimonadetes-bacterium] |
| MDE2875355.1 | HEPN+AAA-ATPase+Alpha+Beta-domain+wHTH+RRM | Type-1_BR.RM.like.systems | TUDOR+SF1-SF2-Helicase+iREase+CTD+REase->METHYLASE->HEPN+DUF499*-><-?\|\|?->?-><-HerA/FtsK<-?\|\|DUF429-> | 1104 | Bacteria>FCB_group>Gemmatimonadota>unclassified_Gemmatimonadota | GCA_028874445.1 | Gemmatimonadota_bacterium | OXQ93_07945 | 7211..10525 | 2026742 | MAG:-DUF499-domain-containing-protein-[Gemmatimonadota-bacterium] |
| MDE2733147.1 | HEPN+AAA-ATPase+Alpha+Beta-domain+wHTH+RRM | Type-1_BR.RM.like.systems | <-TUDOR+SF1-SF2-Helicase+iREase+CTD+REase<-HEPN+DUF499*\|\|REase->?-><-?<-METHYLASE<-REase<-OLD-ABC | 1112 | Bacteria>FCB_group>Gemmatimonadota>unclassified_Gemmatimonadota | GCA_028875335.1 | Gemmatimonadota_bacterium | OXI72_02050 | 10955..14293 | 2026742 | MAG:-Swt1-family-HEPN-domain-containing-protein-[Gemmatimonadota-bacterium] |
| MDE2984847.1 | HEPN+AAA-ATPase+Alpha+Beta-domain+wHTH+RRM | Type-1_BR.RM.like.systems | TUDOR+SF1-SF2-Helicase+iREase+CTD+REase->METHYLASE->METHYLASE-p->HEPN+DUF499*-> | 1104 | Bacteria>FCB_group>Gemmatimonadota>unclassified_Gemmatimonadota | GCA_028873845.1 | Gemmatimonadota_bacterium | OXU69_09080 | 7213..10527 | 2026742 | MAG:-DUF499-domain-containing-protein-[Gemmatimonadota-bacterium] |
| MYE70134.1 | HEPN+AAA-ATPase+Alpha+Beta-domain+wHTH+RRM | Type-1_BR.RM.like.systems | TUDOR+SF1-SF2-Helicase+iREase+CTD+REase-p->METHYLASE->HEPN+DUF499*-> | 1129 | Bacteria>FCB_group>Gemmatimonadota>unclassified_Gemmatimonadota | GCA_009841855.1 | Gemmatimonadota_bacterium | F4237_08820 | 3222..6610 | 2026742 | ATP-binding-protein-[Gemmatimonadetes-bacterium] |
| MCY4572509.1 | HEPN+AAA-ATPase+Alpha+Beta-domain+wHTH+RRM | Type-1_BR.RM.like.systems | SF1-SF2-Helicase-><-?<-HEPN+DUF499*<-OLD-ABC<-METHYLASE | 1114 | Bacteria>FCB_group>Gemmatimonadota>unclassified_Gemmatimonadota | GCA_026713345.1 | Gemmatimonadota_bacterium | OXF01_06870 | 2905..6249 | 2026742 | MAG:-Swt1-family-HEPN-domain-containing-protein-[Gemmatimonadetes-bacterium] |
| MDE2654306.1 | HEPN+AAA-ATPase+Alpha+Beta-domain+wHTH+RRM | Type-1_BR.RM.like.systems | <-REC<-HEPN+DUF499*<-TOPRIM<-OLD-ABC<-METHYLASE-p | 1116 | Bacteria>FCB_group>Gemmatimonadota>unclassified_Gemmatimonadota | GCA_028875795.1 | Gemmatimonadota_bacterium | OXI71_10900 | 83236..86586 | 2026742 | MAG:-Swt1-family-HEPN-domain-containing-protein-[Gemmatimonadota-bacterium] |
| MBU10399.1 | HEPN+AAA-ATPase+Alpha+Beta-domain+wHTH+RRM | Type-1_BR.RM.like.systems | METHYLASE->HEPN+DUF499*-><-HIN-HTH | 1090 | Bacteria>FCB_group>Gemmatimonadota>unclassified_Gemmatimonadota | GCA_002730265.1 | Gemmatimonadota_bacterium | CME13_20820 | 6712..9984 | 2026742 | AAA+-family-ATPase-[Gemmatimonadetes-bacterium] |
| MBT7594687.1 | HEPN+AAA-ATPase+Alpha+Beta-domain+wHTH+RRM | Type-1_BR.RM.like.systems | <-HEPN+DUF499*<-METHYLASE-p | 1031 | Bacteria>FCB_group>Gemmatimonadota>unclassified_Gemmatimonadota | GCA_018675915.1 | Gemmatimonadota_bacterium | HN559_07235 | 1..3094 | 2026742 | ATP-binding-protein-[Gemmatimonadetes-bacterium] |
| MCB9507837.1 | HEPN+AAA-ATPase+Alpha+Beta-domain+wHTH+RRM | Type-1_BR.RM.like.systems | REase-><-Schlafen+HATPase+HTH<-SF1-SF2-Helicase+HRDC<-?<-NERD+SF1-SF2-Helicase<-HEPN+DUF499*<-METHYLASE<-?<-TUDOR+SF1-SF2-Helicase+iREase+CTD+REase\|\|DNA-glycosylase->?->HTH-> | 1103 | Bacteria>Myxococcota>Myxococcia>Myxococcales>unclassified_Myxococcales | GCA_020633615.1 | Myxococcales_bacterium | H6697_09260 | 18345..21656 | 2026763 | ATP-binding-protein-[Myxococcales-bacterium] |
| MBI3184674.1 | HEPN+AAA-ATPase+Alpha+Beta-domain+wHTH+RRM | Type-1_BR.RM.like.systems | ParB->?->XIS-HTH-><-REase<-DNA_methylase<-MORC+REase<-HEPN+DUF499*<-METHYLASE<-?<-TUDOR+SF1-SF2-Helicase+iREase+CTD+REase<-XIS-HTH\|\|AbiEi_1->AbiEii-> | 1116 | Bacteria>Myxococcota>Myxococcia>Myxococcales>unclassified_Myxococcales | GCA_016191815.1 | Myxococcales_bacterium | HYZ28_21265 | 61053..64403 | 2026763 | ATP-binding-protein-[Myxococcales-bacterium] |
| MCC6334307.1 | HEPN+AAA-ATPase+Alpha+Beta-domain+wHTH+RRM | Type-1_BR.RM.like.systems | <-REase\|\|?-><-?<-?<-REase+CTD6<-CHTH<-HEPN+DUF499*<-METHYLASE<-TUDOR+SF1-SF2-Helicase+iREase+CTD+REase | 1100 | Bacteria>Myxococcota>Myxococcia>Myxococcales>unclassified_Myxococcales | GCA_020846875.1 | Myxococcales_bacterium | IT380_10010 | 27471..30773 | 2026763 | DUF499-domain-containing-protein-[Myxococcales-bacterium] |
| MBK7583793.1 | HEPN+AAA-ATPase+Alpha+Beta-domain+wHTH+RRM | Type-1_BR.RM.like.systems | <-HTH\|\|?->REase->TUDOR+SF1-SF2-Helicase+iREase+CTD+REase->METHYLASE->HEPN+DUF499*->REase-> | 1105 | Bacteria>Myxococcota>Myxococcia>Myxococcales>unclassified_Myxococcales | GCA_016706225.1 | Myxococcales_bacterium | IPI67_26815 | 461421..464738 | 2026763 | DUF499-domain-containing-protein-[Myxococcales-bacterium] |
| MBL9045676.1 | HEPN+AAA-ATPase+Alpha+Beta-domain+wHTH+RRM | Type-1_BR.RM.like.systems | TUDOR+SF1-SF2-Helicase+iREase+CTD+REase->METHYLASE->OLD-ABC->TOPRIM->HEPN+DUF499*-><-?<-HTH | 1115 | Bacteria>Myxococcota>Myxococcia>Myxococcales>unclassified_Myxococcales | GCA_016794465.1 | Myxococcales_bacterium | JNM83_28965 | 9575..12922 | 2026763 | DUF499-domain-containing-protein-[Myxococcales-bacterium] |
| NMB74747.1 | HEPN+AAA-ATPase+Alpha+Beta-domain+wHTH+RRM | Type-1_BR.RM.like.systems | TUDOR+SF1-SF2-Helicase+iREase+CTD+REase->METHYLASE->HEPN+DUF499*->?->?-><-?<-?<-?\|\|HAMP+ACYC-> | 1111 | Bacteria>Myxococcota>Myxococcia>Myxococcales>unclassified_Myxococcales | GCA_012798435.1 | Myxococcales_bacterium | GYA21_06410 | 6654..9989 | 2026763 | ATP-binding-protein-[Myxococcales-bacterium] |
| PXF55798.1 | HEPN+AAA-ATPase+Alpha+Beta-domain+wHTH+RRM | Type-1_BR.RM.like.systems | TUDOR+SF1-SF2-Helicase+iREase+CTD+REase->Schlafen+HATPase+HTH->?->METHYLASE-p-><-MULE-TRANSPOSASE\|\|METHYLASE-p->HEPN+DUF499*->MULE-TRANSPOSASE->?-><-LD-peptidase | 1108 | Bacteria>Myxococcota>Myxococcia>unclassified_Myxococcia | GCA_003194485.1 | Deltaproteobacteria_bacterium | C4B57_02005 | 126317..129643 | 2026735 | AAA+-family-ATPase-[Deltaproteobacteria-bacterium] |
| MBK8015115.1 | HEPN+AAA-ATPase+Alpha+Beta-domain+wHTH+RRM | Type-1_BR.RM.like.systems | DEDD_Tnp_IS110+Transposase_20->Y2_Tnp-><-REase<-?<-HEPN+DUF499*<-METHYLASE<-TUDOR+SF1-SF2-Helicase+iREase+CTD+REase<-?<-GDResolvase+Recombinase+ZnR<-HTH | 1099 | Bacteria>Myxococcota>Myxococcia>unclassified_Myxococcia | GCA_016713425.1 | Deltaproteobacteria_bacterium | IPK13_27690 | 7179186..7182485 | 2026735 | DUF499-domain-containing-protein-[Deltaproteobacteria-bacterium] |
| MBN1382687.1 | HEPN+AAA-ATPase+Alpha+Beta-domain+wHTH+RRM | Type-1_BR.RM.like.systems | <-HNH<-ZDNA-HTH<-?<-MetJarc-HTH<-?<-HEPN+DUF499*<-METHYLASE<-?<-MqsA_antitoxin+HTH_3<-MqsR_toxin<-TUDOR+SF1-SF2-Helicase+iREase+CTD+REase | 1117 | Bacteria>Myxococcota>Myxococcia>unclassified_Myxococcia | GCA_016928025.1 | Deltaproteobacteria_bacterium | JXA41_13530 | 29023..32376 | 2026735 | DUF499-domain-containing-protein-[Deltaproteobacteria-bacterium] |
| MCY4442678.1 | HEPN+AAA-ATPase+Alpha+Beta-domain+wHTH+RRM | Type-1_BR.RM.like.systems | AbiEi_1-><-HEPN+DUF499*<-METHYLASE<-MORC+DUF3684<-TUDOR+SF1-SF2-Helicase+iREase+CTD+REase<-Cysteine_desulfurase\|\|35exo-> | 1115 | Bacteria>Myxococcota>Myxococcia>unclassified_Myxococcia | GCA_026710965.1 | Deltaproteobacteria_bacterium | OXE53_21030 | 4124..7471 | 2026735 | MAG:-Swt1-family-HEPN-domain-containing-protein-[Deltaproteobacteria-bacterium] |
| MBI3073658.1 | HEPN+AAA-ATPase+Alpha+Beta-domain+wHTH+RRM | Type-1_BR.RM.like.systems | <-HTH_3\|\|TUDOR+SF1-SF2-Helicase+iREase+CTD+REase->?->METHYLASE-><-PIN<-MetJarc-HTH\|\|HEPN+DUF499*->sGTP->?->Phage_integrase-> | 1105 | Bacteria>Myxococcota>Myxococcia>unclassified_Myxococcia | GCA_016192705.1 | Deltaproteobacteria_bacterium | HYY84_16220 | 33822..37139 | 2026735 | ATP-binding-protein-[Deltaproteobacteria-bacterium] |
| MBN2299509.1 | HEPN+AAA-ATPase+Alpha+Beta-domain+wHTH+RRM | Type-1_BR.RM.like.systems | <-HEPN+DUF499*<-METHYLASE<-HKD-DNase<-Cluster1090_5clades<-DrHyd+BirA-HTH<-TUDOR+SF1-SF2-Helicase+iREase+CTD+REase | 1097 | Bacteria>Myxococcota>Myxococcia>unclassified_Myxococcia | GCA_016934135.1 | Deltaproteobacteria_bacterium | JXM72_12985 | 2480..5773 | 2026735 | DUF499-domain-containing-protein-[Deltaproteobacteria-bacterium] |
| MBW2131484.1 | HEPN+AAA-ATPase+Alpha+Beta-domain+wHTH+RRM | Type-1_BR.RM.like.systems | TUDOR+SF1-SF2-Helicase+iREase+CTD+REase->MqsR_toxin->MqsA_antitoxin+HTH_3->?->METHYLASE->HEPN+DUF499*->?->Radical-SAM-> | 1113 | Bacteria>Myxococcota>Myxococcia>unclassified_Myxococcia | GCA_019309485.1 | Deltaproteobacteria_bacterium | JRG88_02955 | 150894..154235 | 2026735 | DUF499-domain-containing-protein-[Deltaproteobacteria-bacterium] |
| MBW1956070.1 | HEPN+AAA-ATPase+Alpha+Beta-domain+wHTH+RRM | Type-1_BR.RM.like.systems | <-Radical-SAM<-?<-HEPN+DUF499*<-METHYLASE<-?<-MqsA_antitoxin+HTH_3<-MqsR_toxin<-TUDOR+SF1-SF2-Helicase+iREase+CTD+REase | 1113 | Bacteria>Myxococcota>Myxococcia>unclassified_Myxococcia | GCA_019308105.1 | Deltaproteobacteria_bacterium | JRI83_10180 | 2201..5542 | 2026735 | DUF499-domain-containing-protein-[Deltaproteobacteria-bacterium] |
| MBI5504530.1 | HEPN+AAA-ATPase+Alpha+Beta-domain+wHTH+RRM | Type-1_BR.RM.like.systems | MazE_antitoxin->DOC-><-HNH<-?<-?<-HEPN+DUF499*<-METHYLASE<-?<-ParB+HNH<-TUDOR+SF1-SF2-Helicase+iREase+CTD+REase<-?\|\|HIN-HTH-> | 1116 | Bacteria>Myxococcota>Myxococcia>unclassified_Myxococcia | GCA_016218925.1 | Deltaproteobacteria_bacterium | HY899_06990 | 27440..30790 | 2026735 | ATP-binding-protein-[Deltaproteobacteria-bacterium] |
| MBW2310290.1 | HEPN+AAA-ATPase+Alpha+Beta-domain+wHTH+RRM | Type-1_BR.RM.like.systems | TUDOR+SF1-SF2-Helicase+iREase+CTD+REase->MORC+HisKin+DUF3684->METHYLASE->URI-DNase->PolB->HEPN->HEPN+DUF499*-> | 1093 | Bacteria>Myxococcota>Myxococcia>unclassified_Myxococcia | GCA_019310505.1 | Deltaproteobacteria_bacterium | JRF35_04345 | 28703..31984 | 2026735 | DUF499-domain-containing-protein-[Deltaproteobacteria-bacterium] |
| MBW1922080.1 | HEPN+AAA-ATPase+Alpha+Beta-domain+wHTH+RRM | Type-1_BR.RM.like.systems | ORC-CDC6+REase->TUDOR+SF1-SF2-Helicase+iREase+CTD+REase-p->Schlafen+HATPase+HTH->METHYLASE->PolB->HEPN->HEPN+DUF499*-> | 1062 | Bacteria>Myxococcota>Myxococcia>unclassified_Myxococcia | GCA_019307945.1 | Deltaproteobacteria_bacterium | JRJ35_01275 | 87250..90435 | 2026735 | DUF499-domain-containing-protein-[Deltaproteobacteria-bacterium] |
| MCC7383915.1 | HEPN+AAA-ATPase+Alpha+Beta-domain+wHTH+RRM | Type-1_BR.RM.like.systems | <-SIGMA-HTH<-Peptidase_S8<-Classical-AAA<-HEPN+DUF499*<-METHYLASE<-TUDOR+SF1-SF2-Helicase+iREase+CTD+REase | 1093 | Bacteria>Myxococcota>Myxococcia>unclassified_Myxococcia | GCA_020850435.1 | Deltaproteobacteria_bacterium | IT384_18880 | 106091..109372 | 2026735 | DUF499-domain-containing-protein-[Deltaproteobacteria-bacterium] |
| MBW2077703.1 | HEPN+AAA-ATPase+Alpha+Beta-domain+wHTH+RRM | Type-1_BR.RM.like.systems | TUDOR+SF1-SF2-Helicase+iREase+CTD+REase-p->Schlafen+HATPase+HTH->METHYLASE->URI-DNase->PolB->HEPN->HEPN+DUF499*-> | 1093 | Bacteria>Myxococcota>Myxococcia>unclassified_Myxococcia | GCA_019309065.1 | Deltaproteobacteria_bacterium | JRI71_09200 | 7259..10540 | 2026735 | DUF499-domain-containing-protein-[Deltaproteobacteria-bacterium] |
| MDE0031020.1 | HEPN+AAA-ATPase+Alpha+Beta-domain+wHTH+RRM | Type-1_BR.RM.like.systems | DUF433->PIN->Trypsin-><-Trypsin\|\|Phage_integrase-><-HEPN+DUF499*<-BrnT_toxin<-BrnT_toxin<-METHYLASE-p | 1098 | Bacteria>Myxococcota>Myxococcia>unclassified_Myxococcia | GCA_028818775.1 | Deltaproteobacteria_bacterium | OXU42_16660 | 47371..50667 | 2026735 | MAG:-DUF499-domain-containing-protein-[Deltaproteobacteria-bacterium] |
| MBW1949838.1 | HEPN+AAA-ATPase+Alpha+Beta-domain+wHTH+RRM | Type-1_BR.RM.like.systems | TUDOR+SF1-SF2-Helicase+iREase+CTD+REase-p->Schlafen+HATPase+HTH->?->METHYLASE->PolB->HEPN->HEPN+DUF499*-> | 1092 | Bacteria>Myxococcota>Myxococcia>unclassified_Myxococcia | GCA_019308065.1 | Deltaproteobacteria_bacterium | JRJ18_09875 | 33770..37048 | 2026735 | DUF499-domain-containing-protein-[Deltaproteobacteria-bacterium] |
| MBI4517435.1 | HEPN+AAA-ATPase+Alpha+Beta-domain+wHTH+RRM | Type-1_BR.RM.like.systems | TUDOR+SF1-SF2-Helicase+iREase+CTD+REase-><-?\|\|?->PolB->HEPN->HEPN+DUF499*-><-?<-CHTH<-PolB<-?<-Aminotran_4 | 1154 | Bacteria>Myxococcota>Myxococcia>unclassified_Myxococcia | GCA_016210005.1 | Deltaproteobacteria_bacterium | HY699_16655 | 22012..25476 | 2026735 | ATP-binding-protein-[Deltaproteobacteria-bacterium] |
| MAG32379.1 | HEPN+AAA-ATPase+Alpha+Beta-domain+wHTH+RRM | Type-1_BR.RM.like.systems | METHYLASE-p->OLD-ABC->Schlafen+HATPase+HTH->HEPN+DUF499*->?-><-METHYLASE-p<-TOPRIM<-OLD-ABC\|\|?->METHYLASE-p-> | 1115 | Bacteria>Myxococcota>Myxococcia>unclassified_Myxococcia | GCA_002688255.1 | Deltaproteobacteria_bacterium | CL908_15970 | 4485..7832 | 2026735 | AAA+-family-ATPase-[Deltaproteobacteria-bacterium] |
| MDD3553583.1 | HEPN+AAA-ATPase+Alpha+Beta-domain+wHTH+RRM | Type-1_BR.RM.like.systems | XIS-HTH->TUDOR+SF1-SF2-Helicase+iREase+CTD+REase->METHYLASE->OLD-ABC->TOPRIM->HEPN+DUF499*-> | 1108 | Bacteria>Myxococcota>Myxococcia>unclassified_Myxococcia | GCA_028695735.1 | Deltaproteobacteria_bacterium | PHC35_03575 | 35937..39263 | 2026735 | MAG:-Swt1-family-HEPN-domain-containing-protein-[Deltaproteobacteria-bacterium] |

| Protein-ID | DUF499-architecture | System | Gene-Neighbourhoods | Length | Taxonomy | Assembly-ID | Strain | Locus-Tag | Genomic-coordinates | TaxID | Fasta-defline |
|---|---|---|---|---|---|---|---|---|---|---|---|
| MBI3768225.1 | HEPN+AAA-ATPase+Alpha+Beta-domain+wHTH+RRM | Type-1_BR.RM.like.systems | <-HNH\|\|HTH_3->TUDOR+SF1-SF2-Helicase+iREase+CTD+REase->?->?->METHYLASE->HEPN+DUF499*->?->?-><-REase | 1104 | Bacteria>Myxococcota>Myxococcia>unclassified_Myxococcia | GCA_016200545.1 | Deltaproteobacteria_bacterium | HY271_06975 | 73636..76950 | 2026735 | ATP-binding-protein-[Deltaproteobacteria-bacterium] |
| MBI4527695.1 | HEPN+AAA-ATPase+Alpha+Beta-domain+wHTH+RRM | Type-1_BR.RM.like.systems | TUDOR+SF1-SF2-Helicase+iREase+CTD+REase->PolB->HEPN->HEPN->HEPN+DUF499*->ABhydrolase-> | 1116 | Bacteria>Myxococcota>Myxococcia>unclassified_Myxococcia | GCA_016210115.1 | Deltaproteobacteria_bacterium | HY695_28195 | 8921..12271 | 2026735 | DUF499-domain-containing-protein-[Deltaproteobacteria-bacterium] |
| MCC6750891.1 | HEPN+AAA-ATPase+Alpha+Beta-domain+wHTH+RRM | Type-1_BR.RM.like.systems | ParB-><-SIR2<-HEPN+DUF499*<-METHYLASE<-TUDOR+SF1-SF2-Helicase+iREase+CTD+REase\|\|PIN-> | 1103 | Bacteria>Myxococcota>Myxococcia>unclassified_Myxococcia | GCA_020848905.1 | Deltaproteobacteria_bacterium | IT371_24750 | 4486..7797 | 2026735 | DUF499-domain-containing-protein-[Deltaproteobacteria-bacterium] |
| MDE0344143.1 | HEPN+AAA-ATPase+Alpha+Beta-domain+wHTH+RRM | Type-1_BR.RM.like.systems | <-Arm-DNA-bind_3+Phage_integrase<-?<-HNH<-HEPN+DUF499*<-?\|\|Phage_integrase->METHYLASE-p-> | 1111 | Bacteria>Myxococcota>Myxococcia>unclassified_Myxococcia | GCA_028821055.1 | Deltaproteobacteria_bacterium | OXX82_13420 | 9792..13127 | 2026735 | MAG:-Swt1-family-HEPN-domain-containing-protein-[Deltaproteobacteria-bacterium] |
| MBI3249564.1 | HEPN+AAA-ATPase+Alpha+Beta-domain+wHTH+RRM | Type-1_BR.RM.like.systems | METHYLASE->REase->HEPN+DUF499*-><-HicB_lk_antitox\|\|Phage_integrase-><-tRNA<-ABhydrolase | 1110 | Bacteria>Myxococcota>Myxococcia>unclassified_Myxococcia | GCA_016197285.1 | Deltaproteobacteria_bacterium | HYZ50_23930 | 2669..6001 | 2026735 | ATP-binding-protein-[Deltaproteobacteria-bacterium] |
| MDD9852708.1 | HEPN+AAA-ATPase+Alpha+Beta-domain+wHTH+RRM | Type-1_BR.RM.like.systems | TUDOR+SF1-SF2-Helicase+iREase+CTD+REase->?->?->METHYLASE->OLD-ABC->PIN->HEPN+DUF499*-> | 632 | Bacteria>Myxococcota>Myxococcia>unclassified_Myxococcia | GCA_028817165.1 | Deltaproteobacteria_bacterium | OXU78_02000 | 21524..23422 | 2026735 | MAG:-Swt1-family-HEPN-domain-containing-protein-[Deltaproteobacteria-bacterium] |
| MCY4488956.1 | HEPN+AAA-ATPase+Alpha+Beta-domain+wHTH+RRM | Type-1_BR.RM.like.systems | DDE_Tnp_1->BrnT_toxin->HEPN+DUF499*-><-?<-tRNA\|\|?-><-DnaB<-Aminotran_3<-Aminotran_1_2 | 1099 | Bacteria>Myxococcota>Myxococcia>unclassified_Myxococcia | GCA_026712905.1 | Deltaproteobacteria_bacterium | OXF11_17810 | 2694..5993 | 2026735 | MAG:-DUF499-domain-containing-protein-[Deltaproteobacteria-bacterium] |
| NOZ87479.1 | HEPN+AAA-ATPase+Alpha+Beta-domain+wHTH+RRM | Type-1_BR.RM.like.systems | METHYLASE->?->Peptidase_C47->HEPN+DUF499*->MethyltransfD12->HerA/FtsK-> | 1103 | Bacteria>Myxococcota>Myxococcia>unclassified_Myxococcia | GCA_013152945.1 | Deltaproteobacteria_bacterium | GXP49_14650 | 6485..9796 | 2026735 | ATP-binding-protein-[Deltaproteobacteria-bacterium] |
| MBV8135224.1 | HEPN+AAA-ATPase+Alpha+Beta-domain+wHTH+RRM | Type-1_BR.RM.like.systems | <-HEPN+DUF499*<-METHYLASE<-TUDOR+SF1-SF2-Helicase+iREase+CTD+REase-p | 478 | Bacteria>Myxococcota>Myxococcia>unclassified_Myxococcia | GCA_019237115.1 | Deltaproteobacteria_bacterium | JO121_06220 | 1..1435 | 2026735 | DUF499-domain-containing-protein-[Deltaproteobacteria-bacterium] |
| PXF60387.1 | HEPN+AAA-ATPase+Alpha+Beta-domain+wHTH+RRM | Type-1_BR.RM.like.systems | PAIREDC-HTH->Schlafen+HATPase+HTH->METHYLASE->METHYLASE-p->HEPN+DUF499*-> | 1108 | Bacteria>Myxococcota>Myxococcia>unclassified_Myxococcia | GCA_003194495.1 | Deltaproteobacteria_bacterium | C4B58_00710 | 172327..175653 | 2026735 | AAA+-family-ATPase-[Deltaproteobacteria-bacterium] |
| MBW2050448.1 | HEPN+AAA-ATPase+Alpha+Beta-domain+wHTH+RRM | Type-1_BR.RM.like.systems | <-HEPN+DUF499*<-HEPN<-PolB<-METHYLASE-p<-METHYLASE-p | 362 | Bacteria>Myxococcota>Myxococcia>unclassified_Myxococcia | GCA_019308945.1 | Deltaproteobacteria_bacterium | JRJ09_18250 | 1..1087 | 2026735 | DUF499-domain-containing-protein-[Deltaproteobacteria-bacterium] |
| MBI2357704.1 | HEPN+AAA-ATPase+Alpha+Beta-domain+wHTH+RRM | Type-1_BR.RM.like.systems | <-HEPN+DUF499*<-HEPN<-HEPN<-PolB<-Schlafen+HATPase+HTH | 747 | Bacteria>Myxococcota>Myxococcia>unclassified_Myxococcia | GCA_016183075.1 | Deltaproteobacteria_bacterium | HYV04_02095 | 1..2242 | 2026735 | ATP-binding-protein-[Deltaproteobacteria-bacterium] |
| MBI4488650.1 | HEPN+AAA-ATPase+Alpha+Beta-domain+wHTH+RRM | Type-1_BR.RM.like.systems | HEPN->NTP_transf_2->HEPN+DUF499*->?->?->DUF4527-> | 1115 | Bacteria>Myxococcota>Myxococcia>unclassified_Myxococcia | GCA_016210105.1 | Deltaproteobacteria_bacterium | HY694_06155 | 1154..4501 | 2026735 | ATP-binding-protein-[Deltaproteobacteria-bacterium] |
| TRZ75307.1 | HEPN+AAA-ATPase+Alpha+Beta-domain+wHTH+RRM | Type-1_BR.RM.like.systems | METHYLASE->OLD-ABC->TOPRIM->HEPN+DUF499*-> | 362 | Bacteria>Myxococcota>Myxococcia>unclassified_Myxococcia | GCA_007279985.1 | Deltaproteobacteria_bacterium | D4R93_05190 | 4209..5295 | 2026735 | ATP-binding-protein,-partial-[Deltaproteobacteria-bacterium] |
| RME24627.1 | HEPN+AAA-ATPase+Alpha+Beta-domain+wHTH+RRM | Type-1_BR.RM.like.systems | <-HEPN+DUF499*<-SbcC<-METHYLASE-p | 1118 | Bacteria>Myxococcota>Myxococcia>unclassified_Myxococcia | GCA_003694325.1 | Deltaproteobacteria_bacterium | D6798_10765 | 99..3455 | 2026735 | ATP-binding-protein-[Deltaproteobacteria-bacterium] |
| MBW1777804.1 | HEPN+AAA-ATPase+Alpha+Beta-domain+wHTH+RRM | Type-1_BR.RM.like.systems | METHYLASE->HEPN+DUF499*-> | 1113 | Bacteria>Myxococcota>Myxococcia>unclassified_Myxococcia | GCA_019306905.1 | Deltaproteobacteria_bacterium | JRJ54_09460 | 3651..6992 | 2026735 | DUF499-domain-containing-protein-[Deltaproteobacteria-bacterium] |
| MBM4343558.1 | HEPN+AAA-ATPase+Alpha+Beta-domain+wHTH+RRM | Type-1_BR.RM.like.systems | MPTase->?->NUDIX-><-DUF499-p<-HEPN+DUF499*<-METHYLASE\|\|?->?->\|\|CHTH-> | 454 | Bacteria>Myxococcota>Myxococcia>unclassified_Myxococcia | GCA_016875335.1 | Deltaproteobacteria_bacterium | FJ100_09290 | 14865..16229 | 2026735 | hypothetical-protein-[Deltaproteobacteria-bacterium] |
| MCC6527279.1 | HEPN+AAA-ATPase+Alpha+Beta-domain+wHTH+RRM | Type-1_BR.RM.like.systems | METHYLASE-p->HEPN+DUF499*->LZ_Tnp_IS66+zf-IS66+DDE_Tnp_IS66+DDE_Tnp_IS66_C-> | 1095 | Bacteria>Myxococcota>Polyangia>Polyangiales>Polyangiaceae | GCA_020848455.1 | Polyangiaceae_bacterium | IT373_31815 | 1148..4435 | 2268199 | DUF499-domain-containing-protein-[Polyangiaceae-bacterium] |
| MCS6898143.1 | HEPN+AAA-ATPase+Alpha+Beta-domain+wHTH+RRM | Type-1_BR.RM.like.systems | METHYLASE->HEPN+DUF499*->DUF499-p-><-?<-?<-FURR-HTH | 1001 | Bacteria>Myxococcota>Polyangia>Polyangiales>Polyangiaceae | GCA_025057345.1 | Polyangiaceae_bacterium | NZX77_00010 | 2850..5855 | 2268199 | Swt1-family-HEPN-domain-containing-protein-[Polyangiaceae-bacterium] |
| MAT27035.1 | HEPN+AAA-ATPase+Alpha+Beta-domain+wHTH+RRM | Type-1_BR.RM.like.systems | HEPN->MORC+HisKin+DUF3684-><-?\|\|TUDOR+SF1-SF2-Helicase+iREase+CTD+REase->HEPN->METHYLASE->HEPN+DUF499*->MORC+HisKin+REase->?->METHYLASE-p-> | 1093 | Bacteria>Myxococcota>Polyangia>Polyangiales>Sandaracinaceae | GCA_002699025.1 | Sandaracinus_sp. | CMN29_19140 | 67210..70491 | 2024858 | AAA+-family-ATPase-[Sandaracinus-sp.] |
| MBX3269805.1 | HEPN+AAA-ATPase+Alpha+Beta-domain+wHTH+RRM | Type-1_BR.RM.like.systems | PIN->TUDOR+SF1-SF2-Helicase+iREase+CTD+REase->?->HEPN->METHYLASE->METHYLASE->HEPN+DUF499*->?-><-HIN-HTH | 1090 | Bacteria>Myxococcota>Polyangia>Polyangiales>Sandaracinaceae | GCA_019637095.1 | Sandaracinaceae_bacterium | KF729_06070 | 123033..126305 | 2510337 | DUF499-domain-containing-protein-[Sandaracinaceae-bacterium] |
| MBP7125459.1 | HEPN+AAA-ATPase+Alpha+Beta-domain+wHTH+RRM | Type-1_BR.RM.like.systems | TUDOR+SF1-SF2-Helicase+iREase+CTD+REase->Schlafen+HATPase+HTH->METHYLASE->OLD-ABC->TOPRIM->HEPN+DUF499*-> | 1039 | Bacteria>Myxococcota>unclassified_Myxococcota | GCA_017994695.1 | Myxococcota_bacterium | KBD49_03725 | 64170..67286 | 2818507 | DUF499-domain-containing-protein-[Myxococcota-bacterium] |
| MCS6914091.1 | HEPN+AAA-ATPase+Alpha+Beta-domain+wHTH+RRM | Type-1_BR.RM.like.systems | Cas_Cmr3->?->Cas_Cmr5->?->CARF+HEPN->HEPN+DUF499*->Radical-SAM-> | 1114 | Bacteria>Myxococcota>unclassified_Myxococcota | GCA_025057315.1 | Myxococcota_bacterium | NZ890_12730 | 8654..11998 | 2818507 | DUF499-domain-containing-protein-[Myxococcota-bacterium] |
| MCG3173785.1 | HEPN+AAA-ATPase+Alpha+Beta-domain+wHTH+RRM | Type-1_BR.RM.like.systems | <-HEPN+DUF499*<-HEPN+HEPN<-RAMPs<-RAMPs<-RAMPs<-?<-METHYLASE | 1117 | Bacteria>Myxococcota>unclassified_Myxococcota | GCA_022072125.1 | Myxococcota_bacterium | GMKNLPBB_01987 | 84461..87814 | 2818507 | hypothetical-protein-[Myxococcota-bacterium] |
| MBY0401387.1 | HEPN+AAA-ATPase+Alpha+Beta-domain+wHTH+RRM | Type-1_BR.RM.like.systems | TPR->GspA-ATPase-><-Mbetalac\|\|tRNA->?->?-><-HEPN+DUF499*<-METHYLASE-p | 1114 | Bacteria>Myxococcota>unclassified_Myxococcota | GCA_019751155.1 | Myxococcota_bacterium | K2X89_13900 | 4687..8031 | 2818507 | DUF499-domain-containing-protein-[Myxococcota-bacterium] |
| MDE0208007.1 | HEPN+AAA-ATPase+Alpha+Beta-domain+wHTH+RRM | Type-1_BR.RM.like.systems | <-HEPN+DUF499*<-Schlafen+HATPase+HTH<-METHYLASE<-TUDOR+SF1-SF2-Helicase+iREase+CTD+REase<-BrnA_antitoxin | 1111 | Bacteria>Nitrospinae/Tectomicrobia_group>Candidatus_Tectomicrobia>unclassified_Candidatus_Tectomicrobia | GCA_028819935.1 | Candidatus_Tectomicrobia_bacterium | OXP66_18475 | 1406..4741 | 2528274 | MAG:-Swt1-family-HEPN-domain-containing-protein-[Candidatus-Tectomicrobia-bacterium] |
| MYA94924.1 | HEPN+AAA-ATPase+Alpha+Beta-domain+wHTH+RRM | Type-1_BR.RM.like.systems | ACET-><-?\|\|?->tRNA-><-HEPN+DUF499*<-METHYLASE<-?<-?<-?<-?\|\|Peptidase-> | 1086 | Bacteria>Nitrospinae/Tectomicrobia_group>Nitrospinota>unclassified_Nitrospinota | GCA_009839615.1 | Nitrospinota_bacterium | F4X91_00725 | 11133..14393 | 2026769 | ATP-binding-protein-[Nitrospinae-bacterium] |
| MBS0184610.1 | HEPN+AAA-ATPase+Alpha+Beta-domain+wHTH+RRM | Type-1_BR.RM.like.systems | TUDOR+SF1-SF2-Helicase+iREase+CTD+REase-p->TUDOR+SF1-SF2-Helicase+iREase+CTD+REase-p->ParB+HNH->METHYLASE->METHYLASE-p->HEPN->HEPN+DUF499*-> | 1121 | Bacteria>Nitrospirota>Nitrospiria>Nitrospirales>Nitrospiraceae | GCA_018242685.1 | Nitrospira_sp. | JSS39_19650 | 30690..34055 | 70125 | DUF499-domain-containing-protein-[Nitrospira-sp.] |
| MCS6288012.1 | HEPN+AAA-ATPase+Alpha+Beta-domain+wHTH+RRM | Type-1_BR.RM.like.systems | <-XIS-HTH\|\|TUDOR+SF1-SF2-Helicase+iREase+CTD+REase->ParB+HNH->METHYLASE->HEPN+DUF499*->RelE-ParE->HTH->SF1-SF2-Helicase-> | 1121 | Bacteria>Nitrospirota>Nitrospiria>Nitrospirales>Nitrospiraceae | GCA_024998625.1 | Nitrospira_sp. | H8K10_03430 | 260842..264207 | 70125 | ATP-binding-protein-[Nitrospira-sp.] |
| MBP6261842.1 | HEPN+AAA-ATPase+Alpha+Beta-domain+wHTH+RRM | Type-1_BR.RM.like.systems | <-XIS-HTH\|\|TUDOR+SF1-SF2-Helicase+iREase+CTD+REase->AbiGi->ParB+HNH->METHYLASE->HEPN+DUF499*->HEPN+CBS->?-><-?\|\|HIN-HTH-> | 1121 | Bacteria>Nitrospirota>Nitrospiria>Nitrospirales>Nitrospiraceae | GCA_017986555.1 | Nitrospira_sp. | KA426_01230 | 11559..14924 | 70125 | DUF499-domain-containing-protein-[Nitrospira-sp.] |
| WP_090899592.1 | HEPN+AAA-ATPase+Alpha+Beta-domain+wHTH+RRM | Type-1_BR.RM.like.systems | PIN->TUDOR+SF1-SF2-Helicase+iREase+CTD+REase->STAS+HisKin->?->PIN->METHYLASE->HEPN+DUF499*->?->KAP-NTPase->?->PAPS_reduct-> | 1117 | Bacteria>Nitrospirota>Nitrospiria>Nitrospirales>Nitrospiraceae | GCF_001458775.1 | Candidatus_Nitrospira_nitrificans | COMA2_RS14290 | 32189..35542 | 1742973 | Swt1-family-HEPN-domain-containing-protein-[Candidatus-Nitrospira-nitrificans] |
| MBZ0154906.1 | HEPN+AAA-ATPase+Alpha+Beta-domain+wHTH+RRM | Type-1_BR.RM.like.systems | TIR->PAIREDC-HTH+HIN-HTH->DDE_3-><-SF1-SF2-Helicase<-HKD-DNase<-?<-HEPN+DUF499*<-METHYLASE<-?<-?<-Schlafen+HATPase+HTH\|\|REase-> | 1108 | Bacteria>Nitrospirota>Nitrospiria>Nitrospirales>Nitrospiraceae | GCA_019912415.1 | Candidatus_Nitrobium_versatile | K8I29_01665 | 56123..59449 | 2884831 | DUF499-domain-containing-protein-[Alphaproteobacteria-bacterium] |
| MDA8090379.1 | HEPN+AAA-ATPase+Alpha+Beta-domain+wHTH+RRM | Type-1_BR.RM.like.systems | ACET-><-?<-?<-REase<-HEPN+DUF499*<-METHYLASE<-TIR+PNTD2<-TUDOR+SF1-SF2-Helicase+iREase+CTD+REase<-?\|\|Tox-RelE->HTH_3-> | 1107 | Bacteria>Nitrospirota>Nitrospiria>Nitrospirales>Nitrospiraceae | GCA_028064195.1 | Nitrospiraceae_bacterium | MOZ61_09200 | 18029..21352 | 2026770 | MAG:-Swt1-family-HEPN-domain-containing-protein-[Nitrospiraceae-bacterium] |
| NGZ08749.1 | HEPN+AAA-ATPase+Alpha+Beta-domain+wHTH+RRM | Type-1_BR.RM.like.systems | Glycos_transf_4->TUDOR+SF1-SF2-Helicase+iREase+CTD+REase->ParB+HNH->METHYLASE->HEPN+DUF499*-> | 1121 | Bacteria>Nitrospirota>Nitrospiria>Nitrospirales>Nitrospiraceae | GCA_011090425.1 | Nitrospira_sp._LK70 | CV088_05085 | 57956..61321 | 2039809 | ATP-binding-protein-[Nitrospira-sp.-LK70] |
| MBS0151038.1 | HEPN+AAA-ATPase+Alpha+Beta-domain+wHTH+RRM | Type-1_BR.RM.like.systems | TUDOR+SF1-SF2-Helicase+iREase+CTD+REase->REase->METHYLASE->?->DUF488-1->?->HEPN+DUF499*->?-><-?\|\|?-><-?<-SMODS | 1121 | Bacteria>Nitrospirota>Nitrospiria>Nitrospirales>Nitrospiraceae | GCA_018242725.1 | Nitrospira_sp. | JSR31_08860 | 34905..38270 | 70125 | DUF499-domain-containing-protein-[Nitrospira-sp.] |
| MBX3234602.1 | HEPN+AAA-ATPase+Alpha+Beta-domain+wHTH+RRM | Type-1_BR.RM.like.systems | ArdCN+MPTase->?->METHYLASE->?->DUF488-1->?->HEPN+DUF499*->?->sGTP-> | 1121 | Bacteria>Nitrospirota>Nitrospiria>Nitrospirales>Nitrospiraceae | GCA_019637075.1 | Nitrospiraceae_bacterium | KF814_00500 | 97696..101061 | 2026770 | DUF499-domain-containing-protein-[Nitrospiraceae-bacterium] |
| MBZ0154723.1 | HEPN+AAA-ATPase+Alpha+Beta-domain+wHTH+RRM | Type-1_BR.RM.like.systems | PIN->OLD-ABC->HNH->METHYLASE->?->RelE-ParE->HEPN+DUF499*-> | 1114 | Bacteria>Nitrospirota>Nitrospiria>Nitrospirales>Nitrospiraceae | - | Candidatus_Nitrobium_versatile | K8I29_00740 | 55978..59322 | 2884831 | DUF499-domain-containing-protein-[Alphaproteobacteria-bacterium] |
| KAF0183094.1 | HEPN+AAA-ATPase+Alpha+Beta-domain+wHTH+RRM | Type-1_BR.RM.like.systems | MODE-HTH+PBPII->?->?->?->?-><-HEPN+DUF499*<-METHYLASE<-?<-MqsA_antitoxin+HTH_3<-MqsR_toxin<-TUDOR+SF1-SF2-Helicase+iREase+CTD+REase | 1120 | Bacteria>Nitrospirota>unclassified_Nitrospirota | GCA_009773415.1 | Nitrospirota_bacterium | FD164_823 | 11555..14917 | 2026887 | ATPase-(AAA+-superfamily)-like-protein-[Nitrospirae-bacterium] |
| PIX82647.1 | HEPN+AAA-ATPase+Alpha+Beta-domain+wHTH+RRM | Type-1_BR.RM.like.systems | METHYLASE->OLD-ABC->TOPRIM->HEPN+DUF499*-> | 561 | Bacteria>Nitrospirota>unclassified_Nitrospirota | GCA_002783945.1 | Nitrospirae_bacterium_CG_4_10_14_3_um_filter_70_108 | CDZ33_09565 | 4155..5838 | 1974054 | AAA+-family-ATPase,-partial-[Nitrospirae-bacterium-CG_4_10_14_3_um_filter_70_108] |
| WP_035746468.1 | HEPN+AAA-ATPase+Alpha+Beta-domain+wHTH+RRM | Type-1_BR.RM.like.systems | ParG_RHH-><-TetR-HTH\|\|MODE-HTH+PBPII-><-?<-HEPN+DUF499*<-METHYLASE<-TUDOR+SF1-SF2-Helicase+iREase+CTD+REase<-HTH+WYL\|\|HicB->KAP-NTPase-> | 1099 | Bacteria>Pseudomonadota | GCF_000740775.1 | Alphaproteobacteria | CG51_RS15425 | 33987..37286 | 28211 | MULTISPECIES:-DUF499-domain-containing-protein-[Alphaproteobacteria] |
| WP_054362466.1 | HEPN+AAA-ATPase+Alpha+Beta-domain+wHTH+RRM | Type-1_BR.RM.like.systems | ParG_RHH-><-MODE-HTH+PBPII<-HEPN+DUF499*<-METHYLASE<-REase | 1098 | Bacteria>Pseudomonadota | GCF_963522535.1 | Alphaproteobacteria | ABB55_RS28500 | 24847..28143 | 28211 | MULTISPECIES:-DUF499-domain-containing-protein-[Alphaproteobacteria] |
| WP_102114747.1 | HEPN+AAA-ATPase+Alpha+Beta-domain+wHTH+RRM | Type-1_BR.RM.like.systems | METHYLASE->HEPN+DUF499*->MODE-HTH+PBPII-><-ParG_RHH | 1098 | Bacteria>Pseudomonadota | GCF_014640215.1 | Alphaproteobacteria | KF818_RS31845 | 5542..8838 | 28211 | DUF499-domain-containing-protein-[Niveispirillum-cyanobacteriorum] |
| WP_106754608.1 | HEPN+AAA-ATPase+Alpha+Beta-domain+wHTH+RRM | Type-1_BR.RM.like.systems | ParG_RHH-><-TetR-HTH\|\|MODE-HTH+PBPII-><-HEPN+DUF499*<-METHYLASE<-MORC+DUF3684<-TUDOR+SF1-SF2-Helicase+iREase+CTD+REase<-HTH+WYL\|\|SF1-SF2-Helicase+REase->SF1-SF2-Helicase+REase-> | 1098 | Bacteria>Pseudomonadota>Alphaproteobacteria | GCF_003012935.1 | Hyphomicrobiales | C8348_RS17400 | 71198..74494 | 356 | MULTISPECIES:-DUF499-domain-containing-protein-[Hyphomicrobiales] |
| WP_166015916.1 | HEPN+AAA-ATPase+Alpha+Beta-domain+wHTH+RRM | Type-1_BR.RM.like.systems | <-SF1-SF2-Helicase+REase\|\|HTH+WYL->TUDOR+SF1-SF2-Helicase+iREase+CTD+REase->YtH->MORC+DUF3684->METHYLASE->HEPN+DUF499*->?->?->GDResolvase+Recombinase+ZnR-> | 1098 | Bacteria>Pseudomonadota>Alphaproteobacteria | GCF_015326295.1 | Hyphomicrobiales | GNP00_RS08405 | 1742058..1745354 | 356 | MULTISPECIES:-DUF499-domain-containing-protein-[Hyphomicrobiales] |
| WP_066690799.1 | HEPN+AAA-ATPase+Alpha+Beta-domain+wHTH+RRM | Type-1_BR.RM.like.systems | <-HicB\|\|HTH+WYL->TUDOR+SF1-SF2-Helicase+iREase+CTD+REase->ASCH->METHYLASE->HEPN+DUF499*->MODE-HTH+PBPII-><-ParG_RHH | 1097 | Bacteria>Pseudomonadota>Alphaproteobacteria | GCF_001619955.1 | Sphingomonadales | BZF57_RS16495 | 60768..64061 | 204457 | MULTISPECIES:-DUF499-domain-containing-protein-[Sphingomonadales] |
| QLQ13550.1 | HEPN+AAA-ATPase+Alpha+Beta-domain+wHTH+RRM | Type-1_BR.RM.like.systems | VirD4-TraG+VirD4-FtsK->CopG/RHH->VirB11+VirB4-FtsK->?->?-><-MODE-HTH+PBPII<-HEPN+DUF499*<-METHYLASE<-MORC+DUF3684<-HTH+WYL\|\|SF1-SF2-Helicase+REase->SF1-SF2-Helicase+REase->SF1-SF2-helicase+RdgC+Beta-barrel+REase-> | 1096 | Bacteria>Pseudomonadota>Alphaproteobacteria>Caulobacterales>Caulobacteraceae | GCA_013693835.1 | Brevundimonas_sp. | HZY74_10140 | 2044155..2047445 | 1871086 | MAG:-ATP-binding-protein-[Brevundimonas-sp.] |
| WEK58847.1 | HEPN+AAA-ATPase+Alpha+Beta-domain+wHTH+RRM | Type-1_BR.RM.like.systems | <-SF1-SF2-Helicase+REase<-SF1-SF2-Helicase+REase\|\|HTH+WYL->TUDOR+SF1-SF2-Helicase+iREase+CTD+REase->MORC+DUF3684->METHYLASE->HEPN+DUF499*-><-?<-?<-ACYC+ACYC<-CNMP+TIR\|\|HTH+WYL-> | 1085 | Bacteria>Pseudomonadota>Alphaproteobacteria>Caulobacterales>Caulobacteraceae | GCA_029203145.1 | Brevundimonas_sp. | POY52_04750 | 987163..990420 | 1871086 | MAG:-DUF499-domain-containing-protein-[Brevundimonas-sp.] |
| WP_226745381.1 | HEPN+AAA-ATPase+Alpha+Beta-domain+wHTH+RRM | Type-1_BR.RM.like.systems | <-XIS-HTH<-SF1-SF2-helicase+RdgC+Beta-barrel+REase\|\|HTH+WYL->TUDOR+SF1-SF2-Helicase+iREase+CTD+REase->MORC+DUF3684->METHYLASE->HEPN+DUF499*->MODE-HTH+PBPII-><-ParG_RHH | 1098 | Bacteria>Pseudomonadota>Alphaproteobacteria>Caulobacterales>Caulobacteraceae | GCF_020535245.1 | Asticcacaulis_sp._AND118 | LH365_RS06640 | 1378803..1382099 | 2840468 | DUF499-domain-containing-protein-[Asticcacaulis-sp.-AND118] |
| WP_246370526.1 | HEPN+AAA-ATPase+Alpha+Beta-domain+wHTH+RRM | Type-1_BR.RM.like.systems | <-SF1-SF2-Helicase+REase<-SF1-SF2-Helicase+REase\|\|HTH+WYL->TUDOR+SF1-SF2-Helicase+iREase+CTD+REase->REase->METHYLASE->HEPN+DUF499*->MODE-HTH+PBPII-><-ParG_RHH | 1086 | Bacteria>Pseudomonadota>Alphaproteobacteria>Caulobacterales>Caulobacteraceae | GCF_014196295.1 | Phenylobacterium_haematophilum | GGQ61_RS02505 | 500790..504050 | 98513 | DUF499-domain-containing-protein-[Phenylobacterium-haematophilum] |
| WP_231576665.1 | HEPN+AAA-ATPase+Alpha+Beta-domain+wHTH+RRM | Type-1_BR.RM.like.systems | ParG_RHH-><-MODE-HTH+PBPII<-HEPN+DUF499*<-METHYLASE<-MORC+DUF3684<-YtH<-TUDOR+SF1-SF2-Helicase+iREase+CTD+REase<-HTH+WYL\|\|SF1-SF2-Helicase+REase-> | 1086 | Bacteria>Pseudomonadota>Alphaproteobacteria>Caulobacterales>Caulobacteraceae | GCF_000964075.1 | Brevundimonas_sp._KM4 | VH88_RS11620 | 37954..41214 | 1628191 | DUF499-domain-containing-protein-[Brevundimonas-sp.-KM4] |
| KJV40259.1 | HEPN+AAA-ATPase+Alpha+Beta-domain+wHTH+RRM | Type-1_BR.RM.like.systems | ParG_RHH-><-MODE-HTH+PBPII<-HEPN+DUF499*<-METHYLASE<-MORC+DUF3684<-YtH<-TUDOR+SF1-SF2-Helicase+iREase+CTD+REase<-HTH+WYL\|\|SF1-SF2-Helicase+REase-> | 1098 | Bacteria>Pseudomonadota>Alphaproteobacteria>Caulobacterales>Caulobacteraceae | GCA_000964075.1 | Brevundimonas_sp._KM4 | VH88_11615 | 37954..41250 | 1628191 | ATPase-AAA-[Brevundimonas-sp.-KM4] |
| KAF0185697.1 | HEPN+AAA-ATPase+Alpha+Beta-domain+wHTH+RRM | Type-1_BR.RM.like.systems | MazE_antitoxin->PIN->TUDOR+SF1-SF2-Helicase+iREase+CTD+REase->METHYLASE->HEPN+DUF499*-><-ParB+HNH\|\|?->?->?->Phage_AlpA-><-Arm-DNA-bind_3+Phage_integrase | 1077 | Bacteria>Pseudomonadota>Alphaproteobacteria>Caulobacterales>Caulobacteraceae | GCA_009773335.1 | Caulobacteraceae_bacterium | FD160_506 | 42091..45324 | 1978230 | ATPase--like-protein-[Caulobacteraceae-bacterium] |
| MBB3889803.1 | HEPN+AAA-ATPase+Alpha+Beta-domain+wHTH+RRM | Type-1_BR.RM.like.systems | <-SF1-SF2-Helicase+REase\|\|HTH+WYL->TUDOR+SF1-SF2-Helicase+iREase+CTD+REase->REase->METHYLASE->HEPN+DUF499*->MODE-HTH+PBPII-><-ParG_RHH | 1098 | Bacteria>Pseudomonadota>Alphaproteobacteria>Caulobacterales>Caulobacteraceae | GCA_014196295.1 | Phenylobacterium_haematophilum | GGQ61_000500 | 500754..504050 | 98513 | hypothetical-protein-[Phenylobacterium-haematophilum] |
| WP_091738393.1 | HEPN+AAA-ATPase+Alpha+Beta-domain+wHTH+RRM | Type-1_BR.RM.like.systems | <-SF1-SF2-Helicase+REase<-?\|\|HTH+WYL->TUDOR+SF1-SF2-Helicase+iREase+CTD+REase->ASCH->METHYLASE->HEPN+DUF499*->MODE-HTH+PBPII-><-ParG_RHH | 1099 | Bacteria>Pseudomonadota>Alphaproteobacteria>Caulobacterales>Caulobacteraceae | GCF_001375595.1 | Phenylobacterium_immobile | BN1313_RS07235 | 1497979..1501278 | 21 | DUF499-domain-containing-protein-[Phenylobacterium-immobile] |
| WP_039244289.1 | HEPN+AAA-ATPase+Alpha+Beta-domain+wHTH+RRM | Type-1_BR.RM.like.systems | <-SF1-SF2-Helicase+REase<-?\|\|HTH+WYL->TUDOR+SF1-SF2-Helicase+iREase+CTD+REase->?->METHYLASE->HEPN+DUF499*->MODE-HTH+PBPII-><-ParG_RHH | 1098 | Bacteria>Pseudomonadota>Alphaproteobacteria>Caulobacterales>Caulobacteraceae | GCF_000813765.1 | Brevundimonas_nasdae | RM53_RS03375 | 209717..213013 | 172043 | DUF499-domain-containing-protein-[Brevundimonas-nasdae] |
| MCA6347246.1 | HEPN+AAA-ATPase+Alpha+Beta-domain+wHTH+RRM | Type-1_BR.RM.like.systems | <-HEPN+DUF499*<-METHYLASE<-TUDOR+SF1-SF2-Helicase+iREase+CTD+REase<-XIS-HTH<-Pkinase | 1101 | Bacteria>Pseudomonadota>Alphaproteobacteria>Caulobacterales>Caulobacteraceae | GCA_020402685.1 | Phenylobacterium_sp. | IM625_01145 | 33818..37123 | 1871053 | DUF499-domain-containing-protein-[Phenylobacterium-sp.] |
| MBL8553710.1 | HEPN+AAA-ATPase+Alpha+Beta-domain+wHTH+RRM | Type-1_BR.RM.like.systems | Primpol+PriCT1-><-?<-?\|\|HTH_3->TIR+PNTD2->PIWI-><-HEPN+DUF499*<-METHYLASE-p | 1114 | Bacteria>Pseudomonadota>Alphaproteobacteria>Caulobacterales>Caulobacteraceae | GCA_016793285.1 | Phenylobacterium_sp. | JNL41_05490 | 46797..50141 | 1871053 | DUF499-domain-containing-protein-[Phenylobacterium-sp.] |
| MCA3722145.1 | HEPN+AAA-ATPase+Alpha+Beta-domain+wHTH+RRM | Type-1_BR.RM.like.systems | <-METHYLASE-p\|\|?->?-><-PIWI<-TIR+PNTD2<-HEPN+DUF499*<-METHYLASE-p | 1098 | Bacteria>Pseudomonadota>Alphaproteobacteria>Caulobacterales>Caulobacteraceae | GCA_020253645.1 | Phenylobacterium_sp. | INE98_13845 | 11536..14832 | 1871053 | DUF499-domain-containing-protein-[Phenylobacterium-sp.] |
| MDB5438829.1 | HEPN+AAA-ATPase+Alpha+Beta-domain+wHTH+RRM | Type-1_BR.RM.like.systems | METHYLASE->HEPN+DUF499*-> | 1126 | Bacteria>Pseudomonadota>Alphaproteobacteria>Caulobacterales>Caulobacteraceae | GCA_028291105.1 | Caulobacteraceae_bacterium | JWM33_1256 | 2767..6147 | 1978230 | MAG:-superfamily-helicase,-family-[Caulobacteraceae-bacterium] |
| MBN8552584.1 | HEPN+AAA-ATPase+Alpha+Beta-domain+wHTH+RRM | Type-1_BR.RM.like.systems | ParG_RHH-><-MODE-HTH+PBPII<-HEPN+DUF499*<-METHYLASE<-MORC+DUF3684<-TUDOR+SF1-SF2-Helicase+iREase+CTD+REase<-HTH+WYL\|\|SF1-SF2-Helicase+REase->SF1-SF2-Helicase+REase-> | 1098 | Bacteria>Pseudomonadota>Alphaproteobacteria>Caulobacterales>unclassified_Caulobacterales | GCA_017302105.1 | Caulobacterales_bacterium | JOL52_06785 | 4618..7914 | 2282150 | DUF499-domain-containing-protein-[Caulobacterales-bacterium] |
| MCL6609753.1 | HEPN+AAA-ATPase+Alpha+Beta-domain+wHTH+RRM | Type-1_BR.RM.like.systems | METHYLASE-p->HEPN+DUF499*-> | 1105 | Bacteria>Pseudomonadota>Alphaproteobacteria>Geminicoccales>Geminicoccaceae | GCA_023512415.1 | Geminicoccaceae_bacterium | K6T74_16850 | 1426..4743 | 2448052 | DUF499-domain-containing-protein-[Geminicoccaceae-bacterium] |
| WP_152505130.1 | HEPN+AAA-ATPase+Alpha+Beta-domain+wHTH+RRM | Type-1_BR.RM.like.systems | Arm-DNA-bind_3+Phage_integrase->CopG/RHH->ACET-><-HEPN+DUF499*<-METHYLASE<-REase<-MORC+DUF3684<-YtH<-TUDOR+SF1-SF2-Helicase+iREase+CTD+REase<-HTH+WYL | 1098 | Bacteria>Pseudomonadota>Alphaproteobacteria>Hyphomicrobiales | GCF_009363315.1 | Stappiaceae | FIV06_RS06815 | 1469673..1472969 | 2821832 | DUF499-domain-containing-protein-[Roseibium-sp.-Sym1] |
| MAS03479.1 | HEPN+AAA-ATPase+Alpha+Beta-domain+wHTH+RRM | Type-1_BR.RM.like.systems | Phage_integrase+Phage_integrase->Phage_integrase+Phage_integrase->TetR-HTH->XIS-HTH->TUDOR+SF1-SF2-Helicase+iREase+CTD+REase->METHYLASE->HEPN+DUF499*->?->?-><-GNTR-HTH\|\|?->FtsK->XIS-HTH-> | 1123 | Bacteria>Pseudomonadota>Alphaproteobacteria>Hyphomicrobiales>Ahrensiaceae | GCA_002698425.1 | Ahrensia_sp. | CL534_02085 | 25586..28957 | 1872419 | AAA-family-ATPase-[Ahrensia-sp.] |
| MAS04609.1 | HEPN+AAA-ATPase+Alpha+Beta-domain+wHTH+RRM | Type-1_BR.RM.like.systems | <-TIR<-TIR<-?<-?\|\|TUDOR+SF1-SF2-Helicase+iREase+CTD+REase-p->METHYLASE->HEPN+DUF499*->HTH_58-><-ParB+ParBC<-?<-?<-?<-HTH_36 | 1125 | Bacteria>Pseudomonadota>Alphaproteobacteria>Hyphomicrobiales>Ahrensiaceae | GCA_002698425.1 | Ahrensia_sp. | CL534_07935 | 36468..39845 | 1872419 | AAA-family-ATPase-[Ahrensia-sp.] |
| WP_202913508.1 | HEPN+AAA-ATPase+Alpha+Beta-domain+wHTH+RRM | Type-1_BR.RM.like.systems | <-ArdCN+MPTase\|\|?-><-ParB+HTH\|\|?-><-PHP+SbcC<-ParA+HTH<-HEPN+DUF499*<-METHYLASE<-?<-TUDOR+SF1-SF2-Helicase+iREase+CTD+REase<-Phage_int_SAM_1+Phage_integrase | 1115 | Bacteria>Pseudomonadota>Alphaproteobacteria>Hyphomicrobiales>Amorphaceae | GCF_003258595.1 | Acuticoccus_sediminis | DLJ53_RS33595 | 27428..30775 | 2184697 | DUF499-domain-containing-protein-[Acuticoccus-sediminis] |
| RAH96071.1 | HEPN+AAA-ATPase+Alpha+Beta-domain+wHTH+RRM | Type-1_BR.RM.like.systems | <-ArdCN+MPTase\|\|?-><-ParB+HTH<-PHP+SbcC<-ParA+HTH<-HEPN+DUF499*<-METHYLASE<-?<-TUDOR+SF1-SF2-Helicase+iREase+CTD+REase<-Phage_int_SAM_1+Phage_integrase | 1116 | Bacteria>Pseudomonadota>Alphaproteobacteria>Hyphomicrobiales>Amorphaceae | GCA_003258595.1 | Acuticoccus_sediminis | DLJ53_33525 | 27428..30778 | 2184697 | ATPase-[Acuticoccus-sediminis] |
| WP_235707141.1 | HEPN+AAA-ATPase+Alpha+Beta-domain+wHTH+RRM | Type-1_BR.RM.like.systems | HPT+HISKIN+REC->?->?->tRNA-><-HEPN+DUF499*<-METHYLASE<-?<-TUDOR+SF1-SF2-Helicase+iREase+CTD+REase<-HTH+WYL<-HTH<-GDResolvase+Recombinase+ZnR | 1098 | Bacteria>Pseudomonadota>Alphaproteobacteria>Hyphomicrobiales>Amorphaceae | GCF_021556395.1 | Acuticoccus_kalidii | L1787_RS03610 | 494431..497727 | 2910977 | DUF499-domain-containing-protein-[Acuticoccus-kalidii] |
| WP_150971178.1 | HEPN+AAA-ATPase+Alpha+Beta-domain+wHTH+RRM | Type-1_BR.RM.like.systems | TUDOR+SF1-SF2-Helicase+iREase+CTD+REase->METHYLASE->HEPN+DUF499*-><-SF1-SF2-Helicase+DUF1998+HKD-DNase+HKD-DNase<-?\|\|ParB+HNH-><-PIN<-HTH+MPTase<-PAPS_reduct | 1117 | Bacteria>Pseudomonadota>Alphaproteobacteria>Hyphomicrobiales>Aurantimonadaceae | GCF_008802405.1 | Aureimonas_leprariae | F6X38_RS15550 | 20910..24263 | 2615207 | DUF499-domain-containing-protein-[Aureimonas-leprariae] |
| WP_078709662.1 | HEPN+AAA-ATPase+Alpha+Beta-domain+wHTH+RRM | Type-1_BR.RM.like.systems | MODE-HTH+PBPII->?->ABC-ATPase+SbcC-><-METHYLASE-p<-?\|\|<-HEPN+DUF499*<-METHYLASE<-SIR2<-ParB+HNH<-ASCH<-TUDOR+SF1-SF2-Helicase+iREase+CTD+REase\|\|REase-> | 1102 | Bacteria>Pseudomonadota>Alphaproteobacteria>Hyphomicrobiales>Aurantimonadaceae | GCF_900167365.1 | Consotaella_salsifontis | B5D38_RS17130 | 40507..43815 | 1365950 | DUF499-domain-containing-protein-[Consotaella-salsifontis] |
| WP_019994997.1 | HEPN+AAA-ATPase+Alpha+Beta-domain+wHTH+RRM | Type-1_BR.RM.like.systems | REase->ArdCN+MPTase-><-HEPN+DUF499*<-METHYLASE<-?<-TUDOR+SF1-SF2-Helicase+iREase+CTD+REase<-GDResolvase<-SF1-SF2-Helicase<-REase | 1114 | Bacteria>Pseudomonadota>Alphaproteobacteria>Hyphomicrobiales>Aurantimonadaceae | GCF_000382705.1 | Aureimonas_ureilytica | B146_RS0103320 | 700135..703479 | 401562 | DUF499-domain-containing-protein-[Aureimonas-ureilytica] |
| WP_069053609.1 | HEPN+AAA-ATPase+Alpha+Beta-domain+wHTH+RRM | Type-1_BR.RM.like.systems | Arm-DNA-bind_3+Phage_int_SAM_3+Phage_integrase->TUDOR+SF1-SF2-Helicase+iREase+CTD->?->TUDOR+SF1-SF2-Helicase+iREase+CTD+REase->ASCH->METHYLASE->HEPN+DUF499*->?->XIS-HTH->Phage_int_SAM_4+Phage_integrase-><-ABC-ATPase+SbcC | 1113 | Bacteria>Pseudomonadota>Alphaproteobacteria>Hyphomicrobiales>Boseaceae | GCF_001713455.1 | Bosea_sp._RAC05 | BSY19_RS07510 | 857537..860878 | 1842539 | DUF499-domain-containing-protein-[Bosea-sp.-RAC05] |
| TAJ30270.1 | HEPN+AAA-ATPase+Alpha+Beta-domain+wHTH+RRM | Type-1_BR.RM.like.systems | ParG_RHH-><-MODE-HTH+PBPII<-HIN-HTH<-HIN-HTH<-?<-SIGMA-HTH<-HEPN+DUF499*<-METHYLASE-p | 1098 | Bacteria>Pseudomonadota>Alphaproteobacteria>Hyphomicrobiales>Boseaceae | GCA_004297045.1 | Bosea_sp._(in:_a-proteobacteria) | EPO59_12315 | 16657..19953 | 1871050 | DUF499-domain-containing-protein-[Bosea-sp.] |
| WP_069053085.1 | HEPN+AAA-ATPase+Alpha+Beta-domain+wHTH+RRM | Type-1_BR.RM.like.systems | <-Response_reg+LuxR-HTH<-?\|\|?-><-?<-?<-?<-HEPN+DUF499*<-METHYLASE-p\|\|?->?->?->LON-AAA-> | 1097 | Bacteria>Pseudomonadota>Alphaproteobacteria>Hyphomicrobiales>Boseaceae | GCF_001713455.1 | Bosea_sp._RAC05 | BSY19_RS04405 | 191351..194644 | 1842539 | DUF499-domain-containing-protein-[Bosea-sp.-RAC05] |
| WP_258125110.1 | HEPN+AAA-ATPase+Alpha+Beta-domain+wHTH+RRM | Type-1_BR.RM.like.systems | <-SF1-SF2-Helicase+REase<-SF1-SF2-Helicase+REase\|\|HTH+WYL->TUDOR+SF1-SF2-Helicase+iREase+CTD+REase->?->METHYLASE->HEPN+DUF499*->?->?->GDResolvase+Recombinase+ZnR-> | 1098 | Bacteria>Pseudomonadota>Alphaproteobacteria>Hyphomicrobiales>Brucellaceae | GCF_024720755.1 | Brucella_anthropi | NOF64_RS22210 | 68035..71331 | 529 | DUF499-domain-containing-protein-[Brucella-anthropi] |
| WP_259030292.1 | HEPN+AAA-ATPase+Alpha+Beta-domain+wHTH+RRM | Type-1_BR.RM.like.systems | <-SMODS\|\|HTH+WYL->TUDOR+SF1-SF2-Helicase+iREase+CTD+REase->ParB+HNH->?->METHYLASE->HEPN+DUF499*->GDResolvase+HIN-HTH->HEPN->?->?-><-DDE_3<-PAIREDC-HTH | 1098 | Bacteria>Pseudomonadota>Alphaproteobacteria>Hyphomicrobiales>Brucellaceae | GCF_024799785.1 | Brucella_anthropi | NW321_RS01585 | 345006..348302 | 529 | DUF499-domain-containing-protein-[Brucella-anthropi] |
| WP_210296176.1 | HEPN+AAA-ATPase+Alpha+Beta-domain+wHTH+RRM | Type-1_BR.RM.like.systems | HTH+WYL->TUDOR+SF1-SF2-Helicase+iREase+CTD+REase->ASCH->YtH->MORC+DUF3684->METHYLASE->HEPN+DUF499*->TIR+PNTD2->PIWI->HTH->STYKinase-like-> | 1086 | Bacteria>Pseudomonadota>Alphaproteobacteria>Hyphomicrobiales>Brucellaceae | GCF_014197065.1 | Brucella_pecoris | GGQ79_RS19845 | 17581..20841 | 867683 | DUF499-domain-containing-protein-[Brucella-pecoris] |
| WP_140022621.1 | HEPN+AAA-ATPase+Alpha+Beta-domain+wHTH+RRM | Type-1_BR.RM.like.systems | <-STYKinase-like<-HTH<-PIWI<-TIR+PNTD2<-HEPN+DUF499*<-METHYLASE<-MORC+DUF3684<-YtH<-ASCH<-TUDOR+SF1-SF2-Helicase+iREase+CTD+REase<-HTH+WYL | 1098 | Bacteria>Pseudomonadota>Alphaproteobacteria>Hyphomicrobiales>Brucellaceae | GCF_006376675.1 | Brucella_pecoris | FIB18_RS21290 | 91464..94760 | 867683 | DUF499-domain-containing-protein-[Brucella-pecoris] |
| MCC0038267.1 | HEPN+AAA-ATPase+Alpha+Beta-domain+wHTH+RRM | Type-1_BR.RM.like.systems | <-TUDOR+SF1-SF2-Helicase+iREase+CTD<-?<-HHH_4+SF1-SF2-Helicase+REase<-HEPN+DUF499*<-METHYLASE | 1108 | Bacteria>Pseudomonadota>Alphaproteobacteria>Hyphomicrobiales>Brucellaceae | GCA_020637695.1 | Brucellaceae_bacterium | H6893_04600 | 209797..213123 | 2021378 | ATP-binding-protein-[Brucellaceae-bacterium] |
| WP_071923269.1 | HEPN+AAA-ATPase+Alpha+Beta-domain+wHTH+RRM | Type-1_BR.RM.like.systems | GDResolvase+Recombinase+ZnR->CopG/RHH->TUDOR+SF1-SF2-Helicase+iREase+CTD+REase->METHYLASE->HEPN+DUF499*->OLD-ABC+TOPRIM->SF1-SF2-Helicase-><-HTH_17<-Phage_integrase<-HIN-HTH | 1097 | Bacteria>Pseudomonadota>Alphaproteobacteria>Hyphomicrobiales>Chelatococcaceae | GCF_001887265.1 | Chelatococcus_daeguensis | BOQ54_RS03320 | 711833..715126 | 444444 | DUF499-domain-containing-protein-[Chelatococcus-daeguensis] |
| WP_188611839.1 | HEPN+AAA-ATPase+Alpha+Beta-domain+wHTH+RRM | Type-1_BR.RM.like.systems | <-SF1-SF2-helicase+RdgC+Beta-barrel+REase\|\|HTH+WYL->TUDOR+SF1-SF2-Helicase+iREase+CTD+REase->ASCH->METHYLASE->HEPN+DUF499*-><-?\|\|?-><-?\|\|MarR-HTH-> | 1097 | Bacteria>Pseudomonadota>Alphaproteobacteria>Hyphomicrobiales>Chelatococcaceae | GCF_014640075.1 | Chelatococcus_reniformis | IEW34_RS24475 | 395088..398381 | 1494448 | DUF499-domain-containing-protein-[Chelatococcus-reniformis] |
| MBN9362843.1 | HEPN+AAA-ATPase+Alpha+Beta-domain+wHTH+RRM | Type-1_BR.RM.like.systems | <-HicB\|\|HTH+WYL->TUDOR+SF1-SF2-Helicase+iREase+CTD+REase->MORC+DUF3684->HEPN->METHYLASE->HEPN+DUF499*-><-?<-N6_MTase<-?<-Arm-DNA-bind_3+Phage_int_SAM_3+Phage_integrase<-GNTR-HTH | 1098 | Bacteria>Pseudomonadota>Alphaproteobacteria>Hyphomicrobiales>Devosiaceae | GCA_017307535.1 | Devosia_sp. | JOJ09_15200 | 310565..313861 | 1871048 | DUF499-domain-containing-protein-[Devosia-sp.] |
| WP_116654657.1 | HEPN+AAA-ATPase+Alpha+Beta-domain+wHTH+RRM | Type-1_BR.RM.like.systems | <-JAB<-UbE1+JAB<-SMODS\|\|HTH+WYL->TUDOR+SF1-SF2-Helicase+iREase+CTD+REase->METHYLASE->HEPN+DUF499*->?-><-MODE-HTH+PBPII\|\|TetR-HTH-><-ParG_RHH | 1099 | Bacteria>Pseudomonadota>Alphaproteobacteria>Hyphomicrobiales>Devosiaceae | GCF_003390885.1 | Pelagibacterium_sediminicola | DQG18_RS12765 | 56546..59845 | 2248761 | DUF499-domain-containing-protein-[Pelagibacterium-sediminicola] |
| WP_129609562.1 | HEPN+AAA-ATPase+Alpha+Beta-domain+wHTH+RRM | Type-1_BR.RM.like.systems | Phage_int_SAM_3+Phage_integrase->?->XIS-HTH->ZnR+TOPRIM->TOPRIM+SF1-SF2-Helicase+HTH-><-HEPN+DUF499*<-METHYLASE<-ASCH<-sGTP<-TUDOR+SF1-SF2-Helicase+iREase+CTD+REase | 1121 | Bacteria>Pseudomonadota>Alphaproteobacteria>Hyphomicrobiales>Hyphomicrobiaceae | GCF_900604295.1 | Rhodoplanes_serenus | RHODPL_RS14920 | 10306..13671 | 200615 | Swt1-family-HEPN-domain-containing-protein-[Rhodoplanes-piscinae] |
| MBR2536595.1 | HEPN+AAA-ATPase+Alpha+Beta-domain+wHTH+RRM | Type-1_BR.RM.like.systems | ParB+N6_N4_MTase->?->?-><-REase<-REase\|\|HTH_3-><-HEPN+DUF499*<-METHYLASE<-HEPN<-TUDOR+SF1-SF2-Helicase+iREase+CTD+REase<-HTH+WYL\|\|SMODS->SLATT-> | 1092 | Bacteria>Pseudomonadota>Alphaproteobacteria>Hyphomicrobiales>Hyphomicrobiaceae | GCA_017512425.1 | Hyphomicrobium_sp. | IKE66_11035 | 173240..176518 | 82 | DUF499-domain-containing-protein-[Hyphomicrobium-sp.] |
| MBA4172963.1 | HEPN+AAA-ATPase+Alpha+Beta-domain+wHTH+RRM | Type-1_BR.RM.like.systems | <-UbE1+JAB<-?\|\|HTH+WYL-><-HEPN+DUF499*<-METHYLASE<-TUDOR+SF1-SF2-Helicase+iREase+CTD+REase<-HTH_37 | 1110 | Bacteria>Pseudomonadota>Alphaproteobacteria>Hyphomicrobiales>Hyphomicrobiaceae | GCA_013822135.1 | Hyphomicrobium_sp. | C0511_10000 | 5338..8670 | 82 | AAA+-family-ATPase-[Hyphomicrobium-sp.] |
| UYN95212.1 | HEPN+AAA-ATPase+Alpha+Beta-domain+wHTH+RRM | Type-1_BR.RM.like.systems | TUDOR+SF1-SF2-Helicase+iREase+CTD+REase->?->ParB+HNH->METHYLASE->HEPN+DUF499*->AbiEii-><-?<-?<-SPOUT | 1104 | Bacteria>Pseudomonadota>Alphaproteobacteria>Hyphomicrobiales>Hyphomicrobiales_incertae_sedis | GCA_025808875.1 | Enhydrobacter_sp. | KIT25_24915 | 5028707..5032021 | 1894999 | MAG:-DUF499-domain-containing-protein-[Enhydrobacter-sp.] |
| WP_266282109.1 | HEPN+AAA-ATPase+Alpha+Beta-domain+wHTH+RRM | Type-1_BR.RM.like.systems | GNTR-HTH->?-><-?\|\|SIGMA-HTH-><-HEPN+DUF499*<-METHYLASE<-YtH<-MORC+DUF3684<-TUDOR+SF1-SF2-Helicase+iREase+CTD+REase<-HTH+WYL | 1098 | Bacteria>Pseudomonadota>Alphaproteobacteria>Hyphomicrobiales>Kaistiaceae | GCF_026344055.1 | Kaistia_geumhonensis | OSH08_RS14490 | 3056555..3059851 | 410839 | DUF499-domain-containing-protein-[Kaistia-geumhonensis] |
| KQQ15548.1 | HEPN+AAA-ATPase+Alpha+Beta-domain+wHTH+RRM | Type-1_BR.RM.like.systems | PAIREDC-HTH->?->TUDOR+SF1-SF2-Helicase+iREase+CTD+REase->ASCH->METHYLASE->HEPN+DUF499*->?->SF1-SF2-Helicase+DUF1998+HKD-DNase+HKD-DNase->Radical-SAM->?-><-Phage_integrase<-tRNA<-tRNA<-SIGMA-HTH+SIGMA-HTH | 1115 | Bacteria>Pseudomonadota>Alphaproteobacteria>Hyphomicrobiales>Methylobacteriaceae | GCA_001423385.1 | Methylobacterium_sp._Leaf121 | ASF59_14580 | 38300..41647 | 1736262 | ATPase-[Methylobacterium-sp.-Leaf121] |
| WP_192214889.1 | HEPN+AAA-ATPase+Alpha+Beta-domain+wHTH+RRM | Type-1_BR.RM.like.systems | URI-DNase->Phage_integrase->?->RAMA->GDResolvase+Recombinase+ZnR-><-HEPN+DUF499*<-METHYLASE<-MORC+DUF3684<-TUDOR+SF1-SF2-Helicase+iREase+CTD+REase<-HTH+WYL | 1098 | Bacteria>Pseudomonadota>Alphaproteobacteria>Hyphomicrobiales>Methylobacteriaceae | GCF_022179105.1 | Methylobacterium_bullatum | MFF03_RS22740 | 7262..10558 | 570505 | DUF499-domain-containing-protein-[Methylobacterium-bullatum] |
| OAH31268.1 | HEPN+AAA-ATPase+Alpha+Beta-domain+wHTH+RRM | Type-1_BR.RM.like.systems | <-RVT+HNH<-MODE-HTH+PBPII<-HEPN+DUF499*<-METHYLASE<-MORC+DUF3684<-YtH<-TUDOR+SF1-SF2-Helicase+iREase+CTD+REase<-HTH+WYL\|\|SF1-SF2-Helicase+REase-> | 1098 | Bacteria>Pseudomonadota>Alphaproteobacteria>Hyphomicrobiales>Methylobacteriaceae | GCA_001643095.1 | Methylorubrum_populi | AX289_21395 | 87979..91275 | 223967 | AAA-family-ATPase-[Methylorubrum-populi] |
| WP_210322592.1 | HEPN+AAA-ATPase+Alpha+Beta-domain+wHTH+RRM | Type-1_BR.RM.like.systems | TUDOR+SF1-SF2-Helicase+iREase+CTD+REase->MORC+DUF3684->METHYLASE->HEPN+DUF499*-><-GDResolvase+Recombinase+ZnR<-RAMA<-?<-Phage_integrase<-URI-DNase | 1086 | Bacteria>Pseudomonadota>Alphaproteobacteria>Hyphomicrobiales>Methylobacteriaceae | GCF_014845095.1 | Methylobacterium_bullatum | MBTS_RS00690 | 11637..14897 | 570505 | DUF499-domain-containing-protein-[Methylobacterium-bullatum] |
| MCA3671902.1 | HEPN+AAA-ATPase+Alpha+Beta-domain+wHTH+RRM | Type-1_BR.RM.like.systems | TraJ-RHH->?->?-><-HEPN<-VirD4-TraG+VirD4-FtsK\|\|GDResolvase+HIN-HTH-><-HEPN+DUF499*<-METHYLASE<-TUDOR+SF1-SF2-Helicase+iREase+CTD+REase<-?<-CHTH | 1106 | Bacteria>Pseudomonadota>Alphaproteobacteria>Hyphomicrobiales>Methylobacteriaceae | GCA_020253325.1 | Methylobacterium_sp. | INF11_08205 | 10966..14286 | 409 | DUF499-domain-containing-protein-[Methylobacterium-sp.] |
| WP_182180493.1 | HEPN+AAA-ATPase+Alpha+Beta-domain+wHTH+RRM | Type-1_BR.RM.like.systems | Arm-DNA-bind_3+Phage_int_SAM_3+Phage_integrase->TUDOR+SF1-SF2-Helicase+iREase+CTD+REase->ASCH->METHYLASE->HEPN+DUF499*-> | 1111 | Bacteria>Pseudomonadota>Alphaproteobacteria>Hyphomicrobiales>Methylobacteriaceae | GCF_903971015.1 | Methylobacterium_radiotolerans | H7B86_RS28270 | 13923..17258 | 31998 | DUF499-domain-containing-protein-[Methylobacterium-radiotolerans] |
| CAA2157197.1 | HEPN+AAA-ATPase+Alpha+Beta-domain+wHTH+RRM | Type-1_BR.RM.like.systems | SF1-SF2-Helicase-p->?-><-GDResolvase+HIN-HTH\|\|TUDOR+SF1-SF2-Helicase+iREase+CTD+REase->METHYLASE->HEPN+DUF499*-> | 1109 | Bacteria>Pseudomonadota>Alphaproteobacteria>Hyphomicrobiales>Methylobacteriaceae | - | Methylobacterium_brachiatum | MBRA_02601 | 587960..591289 | 269660 | hypothetical-protein-MBRA_02601-[Methylobacterium-brachiatum] |
| WP_207183158.1 | HEPN+AAA-ATPase+Alpha+Beta-domain+wHTH+RRM | Type-1_BR.RM.like.systems | TUDOR+SF1-SF2-Helicase+iREase+CTD+REase->ASCH->METHYLASE-p->METHYLASE-p->METHYLASE-p->HEPN+DUF499*-> | 1117 | Bacteria>Pseudomonadota>Alphaproteobacteria>Hyphomicrobiales>Methylobacteriaceae | GCF_017347565.1 | Methylobacterium_indicum | miv_RS13675 | 2915960..2919313 | 1775910 | DUF499-domain-containing-protein-[Methylobacterium-indicum] |
| WP_262268834.1 | HEPN+AAA-ATPase+Alpha+Beta-domain+wHTH+RRM | Type-1_BR.RM.like.systems | RVT-><-HEPN+DUF499*<-METHYLASE<-TUDOR+SF1-SF2-Helicase+iREase+CTD+REase\|\|?->Phage_integrase-> | 1110 | Bacteria>Pseudomonadota>Alphaproteobacteria>Hyphomicrobiales>Methylobacteriaceae | GCF_025460875.1 | Microvirga_sp._HBU65207 | NF876_RS10955 | 4117..7449 | 2953740 | DUF499-domain-containing-protein-[Microvirga-sp.-HBU65207] |
| WP_103711433.1 | HEPN+AAA-ATPase+Alpha+Beta-domain+wHTH+RRM | Type-1_BR.RM.like.systems | <-HEPN+DUF499*<-METHYLASE<-MORC+DUF3684<-TUDOR+SF1-SF2-Helicase+iREase+CTD+REase<-HTH+WYL | 1098 | Bacteria>Pseudomonadota>Alphaproteobacteria>Hyphomicrobiales>Methylobacteriaceae | GCF_002917135.1 | Methylobacterium_sp._V23 | CRT23_RS20075 | 34063..37359 | 2044878 | DUF499-domain-containing-protein-[Methylobacterium-sp.-V23] |
| WP_083920313.1 | HEPN+AAA-ATPase+Alpha+Beta-domain+wHTH+RRM | Type-1_BR.RM.like.systems | HTH+WYL-><-ParB+HNH<-SF1-SF2-Helicase+DUF1998+HKD-DNase+HKD-DNase<-?<-HEPN+DUF499* | 1053 | Bacteria>Pseudomonadota>Alphaproteobacteria>Hyphomicrobiales>Methylobacteriaceae | GCF_000364445.2 | Methylobacterium_mesophilicum | MMSR116_RS17410 | 3672852..3676013 | 39956 | DUF499-domain-containing-protein-[Methylobacterium-mesophilicum] |
| QGY06229.1 | HEPN+AAA-ATPase+Alpha+Beta-domain+wHTH+RRM | Type-1_BR.RM.like.systems | HTH+WYL-><-ParB+HNH<-ParB+HNH<-SF1-SF2-Helicase+DUF1998+HKD-DNase+HKD-DNase<-?<-HEPN+DUF499* | 1100 | Bacteria>Pseudomonadota>Alphaproteobacteria>Hyphomicrobiales>Methylobacteriaceae | GCA_000364445.2 | Methylobacterium_mesophilicum_SR1.6/6 | MMSR116_17605 | 3672852..3676154 | 908290 | ATP-binding-protein-[Methylobacterium-mesophilicum-SR1.6/6] |
| WP_064034073.1 | HEPN+AAA-ATPase+Alpha+Beta-domain+wHTH+RRM | Type-1_BR.RM.like.systems | MORC+DUF3684+REase-><-?<-SMODS\|\|?->?->MPTase-><-HEPN+DUF499*<-METHYLASE<-?<-TUDOR+SF1-SF2-Helicase+iREase+CTD+REase<-HTH+WYL | 1109 | Bacteria>Pseudomonadota>Alphaproteobacteria>Hyphomicrobiales>Methylocystaceae | GCF_001644125.1 | Methylosinus_sp._R-45379 | A1351_RS23390 | 14763..18092 | 980563 | Swt1-family-HEPN-domain-containing-protein-[Methylosinus-sp.-R-45379] |
| MBI5313198.1 | HEPN+AAA-ATPase+Alpha+Beta-domain+wHTH+RRM | Type-1_BR.RM.like.systems | TUDOR+SF1-SF2-Helicase+iREase+CTD+REase->Ntox18->Peptidase_M78->METHYLASE->OLD-ABC->PIN->HEPN+DUF499*->?-><-HTH | 1107 | Bacteria>Pseudomonadota>Alphaproteobacteria>Hyphomicrobiales>Methylocystaceae | GCA_016219665.1 | Methylocystis_sp. | HZB28_08775 | 48507..51830 | 1911079 | DUF499-domain-containing-protein-[Methylocystis-sp.] |
| WP_166143172.1 | HEPN+AAA-ATPase+Alpha+Beta-domain+wHTH+RRM | Type-1_BR.RM.like.systems | <-HEPN+DUF499*<-METHYLASE<-TUDOR+SF1-SF2-Helicase+iREase+CTD+REase<-HTH+WYL\|\|?->ParB+HTH-> | 1109 | Bacteria>Pseudomonadota>Alphaproteobacteria>Hyphomicrobiales>Methylocystaceae | GCF_011378635.1 | Methylosinus_sp._RM1 | FMZ75_RS12510 | 924..4253 | 2583817 | Swt1-family-HEPN-domain-containing-protein-[Methylosinus-sp.-RM1] |
| WP_253593718.1 | HEPN+AAA-ATPase+Alpha+Beta-domain+wHTH+RRM | Type-1_BR.RM.like.systems | LZ_Tnp_IS66+zf-IS66+DDE_Tnp_IS66+DDE_Tnp_IS66_C-><-Phage_integrase\|\|HEPN+REase->TUDOR+SF1-SF2-Helicase+iREase+CTD+REase->ASCH->METHYLASE->HEPN+DUF499*->?-><-HTH_3\|\|?-><-GDResolvase+Recombinase+ZnR | 1113 | Bacteria>Pseudomonadota>Alphaproteobacteria>Hyphomicrobiales>Nitrobacteraceae | GCF_024171365.1 | Bradyrhizobium_elkanii | M2199_RS04195 | 899505..902846 | 29448 | DUF499-domain-containing-protein-[Bradyrhizobium-elkanii] |
| TAH66859.1 | HEPN+AAA-ATPase+Alpha+Beta-domain+wHTH+RRM | Type-1_BR.RM.like.systems | <-SF1-SF2-Helicase+REase<-SF1-SF2-Helicase+REase\|\|HTH+WYL->TUDOR+SF1-SF2-Helicase+iREase+CTD+REase->MORC+DUF3684->METHYLASE->HEPN+DUF499*-><-MODE-HTH+PBPII\|\|TetR-HTH->?->DUF488-1-><-ParG_RHH | 1098 | Bacteria>Pseudomonadota>Alphaproteobacteria>Hyphomicrobiales>Nitrobacteraceae | GCA_004294885.1 | Rhodopseudomonas_palustris | EWM45_09695 | 32006..35302 | 1076 | MAG:-ATP-binding-protein-[Rhodopseudomonas-palustris] |
| WP_092298105.1 | HEPN+AAA-ATPase+Alpha+Beta-domain+wHTH+RRM | Type-1_BR.RM.like.systems | ZnR->?-><-SF1-SF2-Helicase-p<-SF1-SF2-Helicase<-Exonuc_V_gamma+REase<-HEPN+DUF499*<-METHYLASE<-MORC+DUF3684<-YtH<-TUDOR+SF1-SF2-Helicase+iREase+CTD+REase<-HTH+WYL | 1094 | Bacteria>Pseudomonadota>Alphaproteobacteria>Hyphomicrobiales>Nitrobacteraceae | GCF_900115265.1 | Bradyrhizobium_sp._Ghvi | BM374_RS24865 | 13858..17142 | 1855319 | DUF499-domain-containing-protein-[Bradyrhizobium-sp.-Ghvi] |
| WP_025032482.1 | HEPN+AAA-ATPase+Alpha+Beta-domain+wHTH+RRM | Type-1_BR.RM.like.systems | ZnR+TOPRIM->TOPRIM+SF1-SF2-Helicase+HTH->?->?->REase->REase-><-HEPN+DUF499*<-METHYLASE<-ASCH<-TUDOR+SF1-SF2-Helicase+iREase+CTD+REase<-?<-MULE-TRANSPOSASE<-tRNA<-HISKIN+MORC | 1106 | Bacteria>Pseudomonadota>Alphaproteobacteria>Hyphomicrobiales>Nitrobacteraceae | GCF_000617845.2 | Bradyrhizobium_sp._DOA9 | BDOA9_RS03050 | 657043..660363 | 1126627 | DUF499-domain-containing-protein-[Bradyrhizobium-sp.-DOA9] |
| WP_011157774.1 | HEPN+AAA-ATPase+Alpha+Beta-domain+wHTH+RRM | Type-1_BR.RM.like.systems | <-SF1-SF2-Helicase+REase<-?\|\|HTH+WYL->TUDOR+SF1-SF2-Helicase+iREase+CTD+REase->MORC+DUF3684->METHYLASE->HEPN+DUF499*->MODE-HTH+PBPII-><-ParG_RHH | 1099 | Bacteria>Pseudomonadota>Alphaproteobacteria>Hyphomicrobiales>Nitrobacteraceae | GCF_016584445.1 | Rhodopseudomonas_palustris | TX73_RS11465 | 2517133..2520432 | 1076 | DUF499-domain-containing-protein-[Rhodopseudomonas-palustris] |
| WP_247407267.1 | HEPN+AAA-ATPase+Alpha+Beta-domain+wHTH+RRM | Type-1_BR.RM.like.systems | <-AbiEii<-AbiEi_4\|\|CHTH->?-><-?\|\|REase-><-HEPN+DUF499*<-METHYLASE<-ASCH<-TUDOR+SF1-SF2-Helicase+iREase+CTD+REase<-Recombinase<-REase<-tRNA<-HISKIN+MORC | 1113 | Bacteria>Pseudomonadota>Alphaproteobacteria>Hyphomicrobiales>Nitrobacteraceae | GCF_023073575.1 | unclassified_Bradyrhizobium | IVB10_RS07520 | 491503..494844 | 2631580 | MULTISPECIES:-DUF499-domain-containing-protein-[unclassified-Bradyrhizobium] |
| WP_049832574.1 | HEPN+AAA-ATPase+Alpha+Beta-domain+wHTH+RRM | Type-1_BR.RM.like.systems | <-AbiEii<-AbiEi_4\|\|CHTH->?-><-?\|\|REase-><-HEPN+DUF499*\|\|<-TUDOR+SF1-SF2-Helicase+iREase+CTD+REase-p<-Recombinase<-REase<-tRNA<-HISKIN+MORC\|\|?->RNase-> | 1051 | Bacteria>Pseudomonadota>Alphaproteobacteria>Hyphomicrobiales>Nitrobacteraceae | GCF_011604625.1 | Bradyrhizobium_diazoefficiens | HAU57_RS27175 | 5742766..5745921 | 1355477 | DUF499-domain-containing-protein-[Bradyrhizobium-diazoefficiens] |
| WP_247438142.1 | HEPN+AAA-ATPase+Alpha+Beta-domain+wHTH+RRM | Type-1_BR.RM.like.systems | <-SF1-SF2-Helicase+REase-p\|\|?->TUDOR+SF1-SF2-Helicase+iREase+CTD+REase->?->METHYLASE->HEPN+DUF499*-><-Peptidase\|\|?->VirD4-TraG+VirD4-FtsK-> | 1113 | Bacteria>Pseudomonadota>Alphaproteobacteria>Hyphomicrobiales>Nitrobacteraceae | GCF_023074175.1 | Bradyrhizobium_sp._139 | IVA80_RS29435 | 88222..91563 | 2782616 | DUF499-domain-containing-protein-[Bradyrhizobium-sp.-139] |
| WP_079537876.1 | HEPN+AAA-ATPase+Alpha+Beta-domain+wHTH+RRM | Type-1_BR.RM.like.systems | <-ACYC\|\|?->OLD-ABC->?-><-HKD-DNase<-ParB+HNH<-HEPN+DUF499*<-METHYLASE<-ASCH<-TUDOR+SF1-SF2-Helicase+iREase+CTD+REase\|\|?-><-GDResolvase+HIN-HTH | 1108 | Bacteria>Pseudomonadota>Alphaproteobacteria>Hyphomicrobiales>Nitrobacteraceae | GCF_900141755.1 | Bradyrhizobium_lablabi | B5526_RS08940 | 1910931..1914257 | 722472 | DUF499-domain-containing-protein-[Bradyrhizobium-lablabi] |
| WP_225175154.1 | HEPN+AAA-ATPase+Alpha+Beta-domain+wHTH+RRM | Type-1_BR.RM.like.systems | REase->Recombinase->TUDOR+SF1-SF2-Helicase+iREase+CTD+REase->ASCH->METHYLASE->HEPN+DUF499*-><-REase\|\|?-><-?<-CHTH\|\|AbiEi_4->AbiEii-> | 1113 | Bacteria>Pseudomonadota>Alphaproteobacteria>Hyphomicrobiales>Nitrobacteraceae | GCF_020193975.1 | Bradyrhizobium_sp._NBAIM02 | I6F23_RS23650 | 113567..116908 | 2793817 | DUF499-domain-containing-protein-[Bradyrhizobium-sp.-NBAIM02] |
| WP_225147255.1 | HEPN+AAA-ATPase+Alpha+Beta-domain+wHTH+RRM | Type-1_BR.RM.like.systems | REase->Recombinase->TUDOR+SF1-SF2-Helicase+iREase+CTD+REase->ASCH->METHYLASE->HEPN+DUF499*-><-REase\|\|?-><-?<-CHTH\|\|AbiEi_4->AbiEii-> | 1113 | Bacteria>Pseudomonadota>Alphaproteobacteria>Hyphomicrobiales>Nitrobacteraceae | GCF_020193875.1 | Bradyrhizobium_sp._NBAIM16 | I6F29_RS15715 | 113590..116931 | 2793813 | DUF499-domain-containing-protein-[Bradyrhizobium-sp.-NBAIM16] |
| WP_028141996.1 | HEPN+AAA-ATPase+Alpha+Beta-domain+wHTH+RRM | Type-1_BR.RM.like.systems | REase-><-?<-MACRODOMAIN<-HEPN+DUF499*<-METHYLASE<-ASCH<-TUDOR+SF1-SF2-Helicase+iREase+CTD+REase<-Phage_int_SAM_1+Phage_integrase | 1103 | Bacteria>Pseudomonadota>Alphaproteobacteria>Hyphomicrobiales>Nitrobacteraceae | GCF_000472725.1 | Bradyrhizobium_ottawaense | A3AQ_RS0117025 | 231238..234549 | 931866 | DUF499-domain-containing-protein-[Bradyrhizobium-ottawaense] |
| WP_212326836.1 | HEPN+AAA-ATPase+Alpha+Beta-domain+wHTH+RRM | Type-1_BR.RM.like.systems | Phage_integrase->?->XIS-HTH->?->DnaB->HIN-HTH-><-HEPN+DUF499*<-METHYLASE<-HNH<-OLD-ABC<-TUDOR+SF1-SF2-Helicase+iREase+CTD+REase | 1108 | Bacteria>Pseudomonadota>Alphaproteobacteria>Hyphomicrobiales>Nitrobacteraceae | GCF_018131105.1 | Bradyrhizobium_manausense | JQ631_RS13545 | 2838930..2842256 | 989370 | Swt1-family-HEPN-domain-containing-protein-[Bradyrhizobium-manausense] |
| WP_225188922.1 | HEPN+AAA-ATPase+Alpha+Beta-domain+wHTH+RRM | Type-1_BR.RM.like.systems | <-REC<-REase+DUF4268\|\|?->HEPN->?->?-><-HEPN+DUF499*<-METHYLASE<-TUDOR+SF1-SF2-Helicase+iREase+CTD+REase<-HTH_3\|\|?-><-XIS-HTH | 1094 | Bacteria>Pseudomonadota>Alphaproteobacteria>Hyphomicrobiales>Nitrobacteraceae | GCF_020193585.1 | Bradyrhizobium_sp._IC3195 | I6F09_RS20850 | 200244..203528 | 2793804 | DUF499-domain-containing-protein-[Bradyrhizobium-sp.-IC3195] |
| MCV0386470.1 | HEPN+AAA-ATPase+Alpha+Beta-domain+wHTH+RRM | Type-1_BR.RM.like.systems | Phage_integrase->XIS-HTH->TUDOR+SF1-SF2-Helicase+iREase+CTD+REase->METHYLASE->HEPN+DUF499*->SF1-SF2-Helicase-p-> | 1096 | Bacteria>Pseudomonadota>Alphaproteobacteria>Hyphomicrobiales>Nitrobacteraceae | GCA_025698885.1 | Nitrobacter_sp. | K5821_08550 | 83290..86580 | 29420 | MAG:-DUF499-domain-containing-protein-[Nitrobacter-sp.] |
| WP_182868871.1 | HEPN+AAA-ATPase+Alpha+Beta-domain+wHTH+RRM | Type-1_BR.RM.like.systems | Classical-AAA->Peptidase_S8->?->?->?-><-HEPN+DUF499*<-METHYLASE<-sGTP<-TUDOR+SF1-SF2-Helicase+iREase+CTD+REase<-?<-XIS-HTH | 1117 | Bacteria>Pseudomonadota>Alphaproteobacteria>Hyphomicrobiales>Nitrobacteraceae | GCF_014163475.1 | Bradyrhizobium_diazoefficiens | H7K76_RS29690 | 6385393..6388746 | 1355477 | Swt1-family-HEPN-domain-containing-protein-[Bradyrhizobium-diazoefficiens] |
| WP_231327536.1 | HEPN+AAA-ATPase+Alpha+Beta-domain+wHTH+RRM | Type-1_BR.RM.like.systems | <-Phage_int_SAM_1+Phage_integrase\|\|GNAT->?-><-?<-?<-HEPN+DUF499*<-METHYLASE<-TUDOR+SF1-SF2-Helicase+iREase+CTD+REase | 1098 | Bacteria>Pseudomonadota>Alphaproteobacteria>Hyphomicrobiales>Nitrobacteraceae | GCF_021088345.1 | Bradyrhizobium_sp._A19 | LQG66_RS18125 | 4079105..4082401 | 2898149 | DUF499-domain-containing-protein-[Bradyrhizobium-sp.-A19] |
| OJU24986.1 | HEPN+AAA-ATPase+Alpha+Beta-domain+wHTH+RRM | Type-1_BR.RM.like.systems | PIN+OST-HTH->SF1-SF2-Helicase-p->TUDOR+SF1-SF2-Helicase+iREase+CTD+REase->HEPN+DUF499*->ArdCN+MPTase-> | 1106 | Bacteria>Pseudomonadota>Alphaproteobacteria>Hyphomicrobiales>Nitrobacteraceae | GCA_001896955.1 | Nitrobacter_sp._62-13 | BGN91_09350 | 17137..20457 | 1895797 | MAG:-ATPase-[Nitrobacter-sp.-62-13] |
| WP_172236133.1 | HEPN+AAA-ATPase+Alpha+Beta-domain+wHTH+RRM | Type-1_BR.RM.like.systems | XIS-HTH->?->?->DnaB->HIN-HTH-><-HEPN+DUF499*<-METHYLASE<-HNH<-OLD-ABC<-TUDOR+SF1-SF2-Helicase+iREase+CTD+REase | 1108 | Bacteria>Pseudomonadota>Alphaproteobacteria>Hyphomicrobiales>Nitrobacteraceae | GCF_902482845.1 | Bradyrhizobium | JQ619_RS32195 | 7694..11020 | 374 | MULTISPECIES:-Swt1-family-HEPN-domain-containing-protein-[Bradyrhizobium] |
| WP_211433350.1 | HEPN+AAA-ATPase+Alpha+Beta-domain+wHTH+RRM | Type-1_BR.RM.like.systems | <-DUF488-1<-REase\|\|TUDOR+SF1-SF2-Helicase+iREase+CTD+REase->METHYLASE-p->METHYLASE-p->HEPN+DUF499*-> | 1097 | Bacteria>Pseudomonadota>Alphaproteobacteria>Hyphomicrobiales>Nitrobacteraceae | GCF_018130635.1 | Bradyrhizobium_diazoefficiens | JQ640_RS02510 | 177861..181154 | 1355477 | DUF499-domain-containing-protein-[Bradyrhizobium-diazoefficiens] |
| WP_036012275.1 | HEPN+AAA-ATPase+Alpha+Beta-domain+wHTH+RRM | Type-1_BR.RM.like.systems | URI-DNase-><-HEPN+DUF499*<-METHYLASE<-ASCH<-TUDOR+SF1-SF2-Helicase+iREase+CTD+REase\|\|GDResolvase-> | 1104 | Bacteria>Pseudomonadota>Alphaproteobacteria>Hyphomicrobiales>Nitrobacteraceae | GCF_000261825.1 | Bradyrhizobium_yuanmingense | IC46_RS27575 | 9376..12690 | 108015 | DUF499-domain-containing-protein-[Bradyrhizobium-yuanmingense] |
| WP_247531825.1 | HEPN+AAA-ATPase+Alpha+Beta-domain+wHTH+RRM | Type-1_BR.RM.like.systems | <-GDResolvase\|\|TUDOR+SF1-SF2-Helicase+iREase+CTD+REase->ASCH->METHYLASE->HEPN+DUF499*-><-?<-PHP+SbcC | 1101 | Bacteria>Pseudomonadota>Alphaproteobacteria>Hyphomicrobiales>Nitrobacteraceae | GCF_023073775.1 | Bradyrhizobium_sp._162 | IVA99_RS22470 | 14137..17442 | 2782635 | DUF499-domain-containing-protein-[Bradyrhizobium-sp.-162] |
| WP_166354634.1 | HEPN+AAA-ATPase+Alpha+Beta-domain+wHTH+RRM | Type-1_BR.RM.like.systems | <-HEPN+DUF499*<-METHYLASE<-MORC+DUF3684<-YtH<-ASCH<-ASCH<-TUDOR+SF1-SF2-Helicase+iREase+CTD+REase | 1097 | Bacteria>Pseudomonadota>Alphaproteobacteria>Hyphomicrobiales>Nitrobacteraceae | GCF_011068405.1 | Bradyrhizobium_barranii | G6321_RS03310 | 151200..154493 | 2992140 | DUF499-domain-containing-protein-[Bradyrhizobium-barranii] |

| Protein-ID | DUF499-architecture | System | Gene-Neighbourhoods | Length | Taxonomy | Assembly-ID | Strain | Locus-Tag | Genomic-coordinates | TaxID | Fasta-defline |
|---|---|---|---|---|---|---|---|---|---|---|---|
| WP_092683743.1 | HEPN+AAA-ATPase+Alpha+Beta-domain+wHTH+RRM | Type-1_BR.RM.like.systems | TUDOR+SF1-SF2-Helicase+iREase+CTD+REase->MORC+DUF3684->METHYLASE->HEPN+DUF499*-><-?<-?<-Radical-SAM | 1116 | Bacteria>Pseudomonadota>Alphaproteobacteria>Hyphomicrobiales>Nitrobacteraceae | GCF_900110435.1 | Rhodopseudomonas_pseudopalustris | BMY71_RS11075 | 11290..14640 | 1513892 | Swt1-family-HEPN-domain-containing-protein-[Rhodopseudomonas-pseudopalustris] |
| WP_247344343.1 | HEPN+AAA-ATPase+Alpha+Beta-domain+wHTH+RRM | Type-1_BR.RM.like.systems | HTH+WYL->TUDOR+SF1-SF2-Helicase+iREase+CTD+REase->ParB+HNH->?->METHYLASE->HEPN+DUF499*->HEPN-> | 1097 | Bacteria>Pseudomonadota>Alphaproteobacteria>Hyphomicrobiales>Nitrobacteraceae | GCF_023073075.1 | Bradyrhizobium_sp._61 | IVB46_RS33345 | 520098..523391 | 2782679 | DUF499-domain-containing-protein-[Bradyrhizobium-sp.-61] |
| WP_259172384.1 | HEPN+AAA-ATPase+Alpha+Beta-domain+wHTH+RRM | Type-1_BR.RM.like.systems | <-HEPN+DUF499*<-METHYLASE<-TUDOR+SF1-SF2-Helicase+iREase+CTD+REase<-HTH_3 | 1098 | Bacteria>Pseudomonadota>Alphaproteobacteria>Hyphomicrobiales>Nitrobacteraceae | GCF_024806875.1 | Bradyrhizobium_betae | FHR88_RS05915 | 1186153..1189449 | 244734 | DUF499-domain-containing-protein-[Bradyrhizobium-betae] |
| WP_151650193.1 | HEPN+AAA-ATPase+Alpha+Beta-domain+wHTH+RRM | Type-1_BR.RM.like.systems | HTH_3->TUDOR+SF1-SF2-Helicase+iREase+CTD+REase->METHYLASE->HEPN+DUF499*-> | 1111 | Bacteria>Pseudomonadota>Alphaproteobacteria>Hyphomicrobiales>Nitrobacteraceae | GCF_008932115.1 | Bradyrhizobium_betae | F8237_RS32705 | 6799441..6802776 | 244734 | DUF499-domain-containing-protein-[Bradyrhizobium-betae] |
| WP_008960924.1 | HEPN+AAA-ATPase+Alpha+Beta-domain+wHTH+RRM | Type-1_BR.RM.like.systems | HEPN+DUF499*->?-><-Arm-DNA-bind_3+Phage_integrase | 1107 | Bacteria>Pseudomonadota>Alphaproteobacteria>Hyphomicrobiales>Nitrobacteraceae | GCF_000239795.1 | Bradyrhizobium_sp._STM_3809 | BRAS3809_RS04535 | 1765..5088 | 551936 | DUF499-domain-containing-protein-[Bradyrhizobium-sp.-STM-3809] |
| WP_145832126.1 | HEPN+AAA-ATPase+Alpha+Beta-domain+wHTH+RRM | Type-1_BR.RM.like.systems | <-HEPN+DUF499*<-METHYLASE<-DDE_3 | 1113 | Bacteria>Pseudomonadota>Alphaproteobacteria>Hyphomicrobiales>Nitrobacteraceae | GCF_007830635.1 | Bradyrhizobium_huanghuaihaiense | IQ16_RS39330 | 51033..54374 | 990078 | DUF499-domain-containing-protein-[Bradyrhizobium-huanghuaihaiense] |
| MAM94938.1 | HEPN+AAA-ATPase+Alpha+Beta-domain+wHTH+RRM | Type-1_BR.RM.like.systems | <-SF1-SF2-Helicase+REase<-SF1-SF2-Helicase+REase\|\|HTH+WYL->TUDOR+SF1-SF2-Helicase+iREase+CTD+REase->?->METHYLASE->HEPN+DUF499*-><-MODE-HTH+PBPII | 1098 | Bacteria>Pseudomonadota>Alphaproteobacteria>Hyphomicrobiales>Parvibaculaceae | GCA_002694985.1 | Parvibaculum_sp. | CMI61_09980 | 298093..301389 | 2024848 | AAA-family-ATPase-[Parvibaculum-sp.] |
| MAB12295.1 | HEPN+AAA-ATPase+Alpha+Beta-domain+wHTH+RRM | Type-1_BR.RM.like.systems | <-HEPN+DUF499*<-METHYLASE<-REase<-MORC+DUF3684<-YtH<-TUDOR+SF1-SF2-Helicase+iREase+CTD+REase<-HTH+WYL | 1098 | Bacteria>Pseudomonadota>Alphaproteobacteria>Hyphomicrobiales>Parvibaculaceae | GCA_002684405.1 | Parvibaculum_sp. | CMI59_00100 | 13399..16695 | 2024848 | AAA-family-ATPase-[Parvibaculum-sp.] |
| WP_246674663.1 | HEPN+AAA-ATPase+Alpha+Beta-domain+wHTH+RRM | Type-1_BR.RM.like.systems | <-HicB\|\|HTH+WYL->TUDOR+SF1-SF2-Helicase+iREase+CTD+REase->MORC+DUF3684->REase->METHYLASE->HEPN+DUF499*->TIR+PNTD2->PIWI-><-Arm-DNA-bind_3+Phage_int_SAM_3+Phage_integrase<-?<-?<-REase | 1086 | Bacteria>Pseudomonadota>Alphaproteobacteria>Hyphomicrobiales>Phyllobacteriaceae | GCF_006439355.1 | Mesorhizobium_sp._B4-1-3 | FJW06_RS17660 | 56881..60141 | 2589889 | DUF499-domain-containing-protein-[Mesorhizobium-sp.-B4-1-3] |
| TPI12226.1 | HEPN+AAA-ATPase+Alpha+Beta-domain+wHTH+RRM | Type-1_BR.RM.like.systems | <-HicB\|\|HTH+WYL->TUDOR+SF1-SF2-Helicase+iREase+CTD+REase->MORC+DUF3684->REase->METHYLASE->HEPN+DUF499*->TIR+PNTD2->PIWI-><-Arm-DNA-bind_3+Phage_int_SAM_3+Phage_integrase<-?<-?<-REase | 1098 | Bacteria>Pseudomonadota>Alphaproteobacteria>Hyphomicrobiales>Phyllobacteriaceae | GCA_006439355.1 | Mesorhizobium_sp._B4-1-3 | FJW06_17660 | 56845..60141 | 2589889 | ATP-binding-protein-[Mesorhizobium-sp.-B4-1-3] |
| WP_280515578.1 | HEPN+AAA-ATPase+Alpha+Beta-domain+wHTH+RRM | Type-1_BR.RM.like.systems | TUDOR+SF1-SF2-Helicase+iREase+CTD+REase-p-><-HTH_Tnp_1+HTH_21+rve+rve_3\|\|?->?->METHYLASE-p->METHYLASE->HEPN+DUF499*->DUF499->ParB+HTH->HTH_3->ParB+HNH-p->ParB+HNH-p->URI-DNase-> | 427 | Bacteria>Pseudomonadota>Alphaproteobacteria>Hyphomicrobiales>Phyllobacteriaceae | GCF_013240295.1 | Mesorhizobium_sediminum | HK436_RS32250 | 843839..845122 | 1849104 | Swt1-family-HEPN-domain-containing-protein-[Mesorhizobium-sediminum] |
| RWE54183.1 | HEPN+AAA-ATPase+Alpha+Beta-domain+wHTH+RRM | Type-1_BR.RM.like.systems | HTH+WYL->TUDOR+SF1-SF2-Helicase+iREase+CTD+REase->ASCH->YtH->METHYLASE->HEPN+DUF499*->TUDOR+SF1-SF2-Helicase+iREase+CTD->?->HHH_4+SF1-SF2-Helicase+REase->?-><-tRNA<-Peptidase_S11 | 1100 | Bacteria>Pseudomonadota>Alphaproteobacteria>Hyphomicrobiales>Phyllobacteriaceae | GCA_004018175.1 | Mesorhizobium_sp. | EOS67_26215 | 294790..298092 | 1871066 | ATP-binding-protein-[Mesorhizobium-sp.] |
| CAI2932244.1 | HEPN+AAA-ATPase+Alpha+Beta-domain+wHTH+RRM | Type-1_BR.RM.like.systems | ParG_RHH-><-MODE-HTH+PBPII<-HEPN+DUF499*<-METHYLASE<-ParB+HNH<-TUDOR+SF1-SF2-Helicase+iREase+CTD+REase<-HTH+WYL\|\|SF1-SF2-Helicase+REase->SF1-SF2-Helicase+REase-> | 1098 | Bacteria>Pseudomonadota>Alphaproteobacteria>Hyphomicrobiales>Phyllobacteriaceae | GCA_946995915.1 | Aminobacter_niigataensis | AMD1_0962 | 943205..946501 | 83265 | ATPase-AAA-[Aminobacter-niigataensis] |
| WP_284273142.1 | HEPN+AAA-ATPase+Alpha+Beta-domain+wHTH+RRM | Type-1_BR.RM.like.systems | HTH+WYL->TUDOR+SF1-SF2-Helicase+iREase+CTD+REase->ASCH->YtH->METHYLASE->HEPN+DUF499*->TUDOR+SF1-SF2-Helicase+iREase+CTD->?->HHH_4+SF1-SF2-Helicase+REase-><-?\|\|HTH-> | 1100 | Bacteria>Pseudomonadota>Alphaproteobacteria>Hyphomicrobiales>Phyllobacteriaceae | GCF_030160235.1 | Mesorhizobium_huakuii | QQK95_RS13625 | 53278..56580 | 28104 | DUF499-domain-containing-protein-[Mesorhizobium-huakuii] |
| RWL15103.1 | HEPN+AAA-ATPase+Alpha+Beta-domain+wHTH+RRM | Type-1_BR.RM.like.systems | HhH-RADC+JAB->?->\|\|Phage_integrase->?->METHYLASE->PIN-><-HEPN+DUF499*<-METHYLASE<-HEPN<-MORC+DUF3684<-TUDOR+SF1-SF2-Helicase+iREase+CTD+REase<-HTH+WYL\|\|HicB-> | 1098 | Bacteria>Pseudomonadota>Alphaproteobacteria>Hyphomicrobiales>Phyllobacteriaceae | GCA_004019595.1 | Mesorhizobium_sp. | EOR57_31095 | 10515..13811 | 1871066 | DUF499-domain-containing-protein-[Mesorhizobium-sp.] |
| GLQ79184.1 | HEPN+AAA-ATPase+Alpha+Beta-domain+wHTH+RRM | Type-1_BR.RM.like.systems | HTH+WYL->TUDOR+SF1-SF2-Helicase+iREase+CTD+REase->ASCH->YtH->METHYLASE->HEPN+DUF499*->?->HHH_4+SF1-SF2-Helicase+REase-> | 1088 | Bacteria>Pseudomonadota>Alphaproteobacteria>Hyphomicrobiales>Phyllobacteriaceae | GCA_030160235.1 | Mesorhizobium_huakuii | GCM10007881_27020 | 53314..56580 | 28104 | hypothetical-protein-GCM10007881_27020-[Mesorhizobium-huakuii] |
| WP_245247928.1 | HEPN+AAA-ATPase+Alpha+Beta-domain+wHTH+RRM | Type-1_BR.RM.like.systems | <-SF1-SF2-Helicase+REase\|\|HTH+WYL->TUDOR+SF1-SF2-Helicase+iREase+CTD+REase->ASCH->YtH->METHYLASE->HEPN+DUF499*-><-ABC-ATPase+SbcC<-ABC-ATPase+SbcC | 1084 | Bacteria>Pseudomonadota>Alphaproteobacteria>Hyphomicrobiales>Phyllobacteriaceae | GCF_017813335.1 | Tianweitania_sediminis | J5Y06_RS14970 | 36842..40096 | 1502156 | DUF499-domain-containing-protein-[Tianweitania-sediminis] |
| MBP0439959.1 | HEPN+AAA-ATPase+Alpha+Beta-domain+wHTH+RRM | Type-1_BR.RM.like.systems | <-SF1-SF2-Helicase+REase\|\|HTH+WYL->TUDOR+SF1-SF2-Helicase+iREase+CTD+REase->ASCH->YtH->METHYLASE->HEPN+DUF499*-><-ABC-ATPase+SbcC<-ABC-ATPase+SbcC | 1096 | Bacteria>Pseudomonadota>Alphaproteobacteria>Hyphomicrobiales>Phyllobacteriaceae | GCA_017813335.1 | Tianweitania_sediminis | J5Y06_14970 | 36806..40096 | 1502156 | DUF499-domain-containing-protein-[Tianweitania-sediminis] |
| WP_202379562.1 | HEPN+AAA-ATPase+Alpha+Beta-domain+wHTH+RRM | Type-1_BR.RM.like.systems | GDResolvase+Recombinase+ZnR-><-?\|\|?->?-><-?\|\|?-><-HEPN+DUF499*<-METHYLASE<-TUDOR+SF1-SF2-Helicase+iREase+CTD+REase\|\|MerR-HTH->?-><-GDResolvase+Recombinase+ZnR | 1114 | Bacteria>Pseudomonadota>Alphaproteobacteria>Hyphomicrobiales>Phyllobacteriaceae | GCF_016756555.1 | Mesorhizobium_sp._131-2-5 | JG740_RS04010 | 865607..868951 | 2744519 | DUF499-domain-containing-protein-[Mesorhizobium-sp.-131-2-5] |
| WP_095831457.1 | HEPN+AAA-ATPase+Alpha+Beta-domain+wHTH+RRM | Type-1_BR.RM.like.systems | SF1-SF2-Helicase+REase->Resolvase->REase->TUDOR+SF1-SF2-Helicase+iREase+CTD+REase->METHYLASE->HEPN+DUF499*-><-XIS-HTH\|\|?->?-><-tRNA<-?\|\|?->SAM-methylase-> | 1135 | Bacteria>Pseudomonadota>Alphaproteobacteria>Hyphomicrobiales>Phyllobacteriaceae | GCF_002294955.1 | Mesorhizobium_sp._WSM3864 | CK215_RS15785 | 236448..239855 | 2029404 | DUF499-domain-containing-protein-[Mesorhizobium-sp.-WSM3864] |
| WP_135929916.1 | HEPN+AAA-ATPase+Alpha+Beta-domain+wHTH+RRM | Type-1_BR.RM.like.systems | TUDOR+SF1-SF2-Helicase+iREase+CTD+REase->METHYLASE->OLD-ABC->PIN->HEPN+DUF499*->?-><-?\|\|Methylase-><-Radical-SAM<-Arm-DNA-bind_3+Phage_integrase | 1110 | Bacteria>Pseudomonadota>Alphaproteobacteria>Hyphomicrobiales>Phyllobacteriaceae | GCF_004791875.1 | unclassified_Mesorhizobium | EN810_RS20200 | 141593..144925 | 325217 | MULTISPECIES:-DUF499-domain-containing-protein-[unclassified-Mesorhizobium] |
| WP_263490164.1 | HEPN+AAA-ATPase+Alpha+Beta-domain+wHTH+RRM | Type-1_BR.RM.like.systems | <-BirA-HTH<-BirA-HTH<-BirA-HTH\|\|?->McrC-NTD+REase-><-REase<-HEPN+DUF499*<-?<-METHYLASE<-TUDOR+SF1-SF2-Helicase+iREase+CTD+REase\|\|?->?-><-MPTase | 1095 | Bacteria>Pseudomonadota>Alphaproteobacteria>Hyphomicrobiales>Phyllobacteriaceae | - | Mesorhizobium_sp._CA15 | LB515_RS02455 | 495752..499039 | 2876641 | DUF499-domain-containing-protein-[Mesorhizobium-sp.-CA15] |
| WP_224706639.1 | HEPN+AAA-ATPase+Alpha+Beta-domain+wHTH+RRM | Type-1_BR.RM.like.systems | <-BirA-HTH<-BirA-HTH<-BirA-HTH\|\|?->McrC-NTD+REase-><-REase<-HEPN+DUF499*<-?<-METHYLASE<-TUDOR+SF1-SF2-Helicase+iREase+CTD+REase\|\|?->?-><-MPTase | 1112 | Bacteria>Pseudomonadota>Alphaproteobacteria>Hyphomicrobiales>Phyllobacteriaceae | GCF_020165215.1 | Mesorhizobium_sp._CA15 | LB515_02450 | 495752..499090 | 2876641 | DUF499-domain-containing-protein-[Mesorhizobium-sp.-CA15] |
| WP_135867691.1 | HEPN+AAA-ATPase+Alpha+Beta-domain+wHTH+RRM | Type-1_BR.RM.like.systems | PAPS_reduct->HTH+MPTase->PIN-><-HEPN+DUF499*<-METHYLASE<-MORC+DUF3684<-TUDOR+SF1-SF2-Helicase+iREase+CTD+REase<-REase<-?<-MAD-NTD+HNH | 1097 | Bacteria>Pseudomonadota>Alphaproteobacteria>Hyphomicrobiales>Phyllobacteriaceae | GCF_004791755.1 | unclassified_Mesorhizobium | EN861_RS33190 | 64871..68164 | 325217 | MULTISPECIES:-DUF499-domain-containing-protein-[unclassified-Mesorhizobium] |
| MBZ9864227.1 | HEPN+AAA-ATPase+Alpha+Beta-domain+wHTH+RRM | Type-1_BR.RM.like.systems | <-BirA-HTH<-BirA-HTH<-BirA-HTH\|\|?->McrC-NTD+REase-><-REase<-HEPN+DUF499*<-?<-METHYLASE<-TUDOR+SF1-SF2-Helicase+iREase+CTD+REase\|\|?->?-><-MPTase | 1112 | Bacteria>Pseudomonadota>Alphaproteobacteria>Hyphomicrobiales>Phyllobacteriaceae | GCA_020165215.1 | Mesorhizobium_sp._CA15 | LB515_02450 | 495752..499090 | 2876641 | DUF499-domain-containing-protein-[Mesorhizobium-sp.-CA15] |
| WP_135861107.1 | HEPN+AAA-ATPase+Alpha+Beta-domain+wHTH+RRM | Type-1_BR.RM.like.systems | HTH+WYL->TUDOR+SF1-SF2-Helicase+iREase+CTD+REase->ASCH->YtH->MORC+DUF3684->METHYLASE->HEPN+DUF499*-><-TRPR-HTH+HTH<-?<-?<-?\|\|TraJ-RHH-> | 1082 | Bacteria>Pseudomonadota>Alphaproteobacteria>Hyphomicrobiales>Phyllobacteriaceae | GCF_004791825.1 | unclassified_Mesorhizobium | EN833_RS30760 | 107531..110779 | 325217 | MULTISPECIES:-DUF499-domain-containing-protein-[unclassified-Mesorhizobium] |
| WP_120017243.1 | HEPN+AAA-ATPase+Alpha+Beta-domain+wHTH+RRM | Type-1_BR.RM.like.systems | TRPR-HTH+HTH->?->?->?->?->?-><-HEPN+DUF499*<-METHYLASE<-MORC+DUF3684<-TUDOR+SF1-SF2-Helicase+iREase+CTD+REase<-?<-MORC+HISKIN<-METHYLASE-p | 1110 | Bacteria>Pseudomonadota>Alphaproteobacteria>Hyphomicrobiales>Phyllobacteriaceae | GCF_003601975.1 | Mesorhizobium_waimense | D3227_RS26610 | 4630..7962 | 1300307 | Swt1-family-HEPN-domain-containing-protein-[Mesorhizobium-waimense] |
| TJW08146.1 | HEPN+AAA-ATPase+Alpha+Beta-domain+wHTH+RRM | Type-1_BR.RM.like.systems | Resolvase->SIR2+STALD->?->TUDOR+SF1-SF2-Helicase+iREase+CTD+REase->METHYLASE->HEPN+DUF499*-><-XIS-HTH<-?\|\|?-><-tRNA<-?\|\|?->SAM-methylase-> | 1132 | Bacteria>Pseudomonadota>Alphaproteobacteria>Hyphomicrobiales>Phyllobacteriaceae | GCA_005047185.1 | Mesorhizobium_sp. | E5W97_02655 | 563673..567071 | 1871066 | ATP-binding-protein-[Mesorhizobium-sp.] |
| WP_224580510.1 | HEPN+AAA-ATPase+Alpha+Beta-domain+wHTH+RRM | Type-1_BR.RM.like.systems | <-HEPN+DUF499*<-METHYLASE<-Peptidase_M78<-Ntox18<-TUDOR+SF1-SF2-Helicase+iREase+CTD+REase<-Phage_integrase<-XIS-HTH | 1103 | Bacteria>Pseudomonadota>Alphaproteobacteria>Hyphomicrobiales>Phyllobacteriaceae | GCF_020165495.1 | Mesorhizobium_sp._BR1-1-16 | LB518_RS24470 | 798..4109 | 2876653 | DUF499-domain-containing-protein-[Mesorhizobium-sp.-BR1-1-16] |
| WP_125006738.1 | HEPN+AAA-ATPase+Alpha+Beta-domain+wHTH+RRM | Type-1_BR.RM.like.systems | <-SMODS\|\|HTH+WYL->?->TUDOR+SF1-SF2-Helicase+iREase+CTD+REase->MORC+DUF3684->METHYLASE->HEPN+DUF499*-><-AbiEii<-AbiEi_4 | 1110 | Bacteria>Pseudomonadota>Alphaproteobacteria>Hyphomicrobiales>Phyllobacteriaceae | GCF_003863365.1 | Mesorhizobium_tamadayense | EH240_RS33970 | 18463..21795 | 425306 | Swt1-family-HEPN-domain-containing-protein-[Mesorhizobium-tamadayense] |
| RWJ00134.1 | HEPN+AAA-ATPase+Alpha+Beta-domain+wHTH+RRM | Type-1_BR.RM.like.systems | <-ZnR\|\|?-><-?<-?<-?\|\|<-HEPN+DUF499*<-METHYLASE<-MORC+DUF3684<-TUDOR+SF1-SF2-Helicase+iREase+CTD+REase<-HTH+WYL\|\|SMODS->UbE1+JAB-> | 1110 | Bacteria>Pseudomonadota>Alphaproteobacteria>Hyphomicrobiales>Phyllobacteriaceae | GCA_004018865.1 | Mesorhizobium_sp. | EOR23_30945 | 21275..24607 | 1871066 | ATP-binding-protein-[Mesorhizobium-sp.] |
| RWD48635.1 | HEPN+AAA-ATPase+Alpha+Beta-domain+wHTH+RRM | Type-1_BR.RM.like.systems | Peptidase_S11->tRNA-><-HEPN+DUF499*<-METHYLASE<-MORC+DUF3684<-YtH<-ASCH<-TUDOR+SF1-SF2-Helicase+iREase+CTD+REase<-HTH+WYL | 1098 | Bacteria>Pseudomonadota>Alphaproteobacteria>Hyphomicrobiales>Phyllobacteriaceae | GCA_004016945.1 | Mesorhizobium_sp. | EOS59_16740 | 2840..6136 | 1871066 | ATP-binding-protein-[Mesorhizobium-sp.] |
| MBZ9939464.1 | HEPN+AAA-ATPase+Alpha+Beta-domain+wHTH+RRM | Type-1_BR.RM.like.systems | <-HEPN+DUF499*<-METHYLASE<-Peptidase_M78<-Ntox18<-TUDOR+SF1-SF2-Helicase+iREase+CTD+REase<-Phage_integrase<-XIS-HTH | 1103 | Bacteria>Pseudomonadota>Alphaproteobacteria>Hyphomicrobiales>Phyllobacteriaceae | GCA_020165495.1 | Mesorhizobium_sp._BR1-1-16 | LB518_24480 | 798..4109 | 2876653 | DUF499-domain-containing-protein-[Mesorhizobium-sp.-BR1-1-16] |
| WP_127864269.1 | HEPN+AAA-ATPase+Alpha+Beta-domain+wHTH+RRM | Type-1_BR.RM.like.systems | SF1-SF2-Helicase->TUDOR+SF1-SF2-Helicase+iREase+CTD+REase->ASCH->METHYLASE->HEPN+DUF499*-><-tRNA<-Peptidase<-?\|\|MPTase-> | 1114 | Bacteria>Pseudomonadota>Alphaproteobacteria>Hyphomicrobiales>Phyllobacteriaceae | GCF_004010405.2 | unclassified_Mesorhizobium | EN797_RS08215 | 1680820..1684164 | 325217 | MULTISPECIES:-DUF499-domain-containing-protein-[unclassified-Mesorhizobium] |
| RWQ65486.1 | HEPN+AAA-ATPase+Alpha+Beta-domain+wHTH+RRM | Type-1_BR.RM.like.systems | <-ABC-ATPase+SbcC<-?<-?\|\|MarR-HTH->?-><-?<-HEPN+DUF499*<-METHYLASE<-TUDOR+SF1-SF2-Helicase+iREase+CTD+REase<-tRNA<-Peptidase_S11 | 1103 | Bacteria>Pseudomonadota>Alphaproteobacteria>Hyphomicrobiales>Phyllobacteriaceae | GCA_004021745.1 | Mesorhizobium_sp. | EOS86_15645 | 123505..126816 | 1871066 | ATP-binding-protein-[Mesorhizobium-sp.] |
| WP_107649676.1 | HEPN+AAA-ATPase+Alpha+Beta-domain+wHTH+RRM | Type-1_BR.RM.like.systems | STYKinase-like->?-><-HEPN+DUF499*<-METHYLASE<-TUDOR+SF1-SF2-Helicase+iREase+CTD+REase<-GDResolvase+HTH\|\|?-><-?\|\|HTH-> | 1105 | Bacteria>Pseudomonadota>Alphaproteobacteria>Hyphomicrobiales>Phyllobacteriaceae | GCF_003034915.1 | Mesorhizobium_helmanticense | C9427_RS13550 | 15712..19029 | 1776423 | DUF499-domain-containing-protein-[Mesorhizobium-helmanticense] |
| WP_192357697.1 | HEPN+AAA-ATPase+Alpha+Beta-domain+wHTH+RRM | Type-1_BR.RM.like.systems | REase->?->MACRODOMAIN->STYKinase-like->?-><-HEPN+DUF499*<-METHYLASE<-TUDOR+SF1-SF2-Helicase+iREase+CTD+REase | 1105 | Bacteria>Pseudomonadota>Alphaproteobacteria>Hyphomicrobiales>Phyllobacteriaceae | GCF_014843885.1 | Mesorhizobium_mediterraneum | AK98_RS11975 | 23140..26457 | 43617 | DUF499-domain-containing-protein-[Mesorhizobium-mediterraneum] |
| RWO89814.1 | HEPN+AAA-ATPase+Alpha+Beta-domain+wHTH+RRM | Type-1_BR.RM.like.systems | MarR-HTH->?-><-?\|\|TUDOR+SF1-SF2-Helicase+iREase+CTD+REase->METHYLASE->HEPN+DUF499*-><-?\|\|REase-><-?<-?<-?<-tRNA<-Peptidase_S11 | 1105 | Bacteria>Pseudomonadota>Alphaproteobacteria>Hyphomicrobiales>Phyllobacteriaceae | GCA_004021485.1 | Mesorhizobium_sp. | EOQ95_16830 | 131837..135154 | 1871066 | ATP-binding-protein-[Mesorhizobium-sp.] |
| WP_165021935.1 | HEPN+AAA-ATPase+Alpha+Beta-domain+wHTH+RRM | Type-1_BR.RM.like.systems | TUDOR+SF1-SF2-Helicase+iREase+CTD+REase->METHYLASE->HEPN+DUF499*-><-?<-STYKinase-like<-?<-?\|\|?-><-Phage_integrase | 1105 | Bacteria>Pseudomonadota>Alphaproteobacteria>Hyphomicrobiales>Phyllobacteriaceae | GCF_011045125.1 | Mesorhizobium_camelthorni | G6N73_RS00955 | 189890..193207 | 475069 | DUF499-domain-containing-protein-[Mesorhizobium-camelthorni] |
| WP_164894099.1 | HEPN+AAA-ATPase+Alpha+Beta-domain+wHTH+RRM | Type-1_BR.RM.like.systems | <-HEPN+DUF499*<-METHYLASE<-wHTH+REase<-TUDOR+SF1-SF2-Helicase+iREase+CTD+REase<-?\|\|wHTH+SAD+HNH-><-HTH_3 | 528 | Bacteria>Pseudomonadota>Alphaproteobacteria>Hyphomicrobiales>Phyllobacteriaceae | GCF_004022915.1 | Mesorhizobium_sp._M7A.F.Ca.US.006.04.2.1 | EN925_RS00840 | 1..1586 | 2496696 | Swt1-family-HEPN-domain-containing-protein,-partial-[Mesorhizobium-sp.-M7A.F.Ca.US.006.04.2.1] |
| WP_127367955.1 | HEPN+AAA-ATPase+Alpha+Beta-domain+wHTH+RRM | Type-1_BR.RM.like.systems | HTH_3-><-wHTH+SAD+HNH\|\|?->TUDOR+SF1-SF2-Helicase+iREase+CTD+REase->wHTH+REase->METHYLASE->HEPN+DUF499*-> | 1118 | Bacteria>Pseudomonadota>Alphaproteobacteria>Hyphomicrobiales>Phyllobacteriaceae | GCF_003997975.1 | unclassified_Mesorhizobium | EN990_RS21670 | 38386..41742 | 325217 | MULTISPECIES:-DUF499-domain-containing-protein-[unclassified-Mesorhizobium] |
| RWN27570.1 | HEPN+AAA-ATPase+Alpha+Beta-domain+wHTH+RRM | Type-1_BR.RM.like.systems | <-LD-peptidase<-?<-HEPN+DUF499*<-METHYLASE<-TUDOR+SF1-SF2-Helicase+iREase+CTD+REase<-RVT<-RVT<-35exo | 1109 | Bacteria>Pseudomonadota>Alphaproteobacteria>Hyphomicrobiales>Phyllobacteriaceae | GCA_004020785.1 | Mesorhizobium_sp. | EOR95_24225 | 12019..15348 | 1871066 | ATP-binding-protein-[Mesorhizobium-sp.] |
| WP_035530026.1 | HEPN+AAA-ATPase+Alpha+Beta-domain+wHTH+RRM | Type-1_BR.RM.like.systems | Phage_integrase-><-HEPN+DUF499*<-?<-METHYLASE<-?<-SIR2<-ASCH<-TUDOR+SF1-SF2-Helicase+iREase+CTD+REase | 1103 | Bacteria>Pseudomonadota>Alphaproteobacteria>Hyphomicrobiales>Phyllobacteriaceae | GCF_000759435.1 | Hoeflea_sp._BAL378 | LL06_RS18050 | 3002..6313 | 1547437 | DUF499-domain-containing-protein-[Hoeflea-sp.-BAL378] |
| WP_023715932.1 | HEPN+AAA-ATPase+Alpha+Beta-domain+wHTH+RRM | Type-1_BR.RM.like.systems | <-ParB<-?<-REase<-HEPN+DUF499*<-PIN<-OLD-ABC<-METHYLASE<-TUDOR+SF1-SF2-Helicase+iREase+CTD+REase | 1115 | Bacteria>Pseudomonadota>Alphaproteobacteria>Hyphomicrobiales>Phyllobacteriaceae | GCF_000503075.1 | Mesorhizobium_sp._LSHC422A00 | X761_RS140575 | 209394..212741 | 1287294 | DUF499-domain-containing-protein-[Mesorhizobium-sp.-LSHC422A00] |
| WP_140871483.1 | HEPN+AAA-ATPase+Alpha+Beta-domain+wHTH+RRM | Type-1_BR.RM.like.systems | <-HEPN+DUF499*<-METHYLASE<-TUDOR+SF1-SF2-Helicase+iREase+CTD+REase<-?<-SIR2+STALD<-Resolvase | 1134 | Bacteria>Pseudomonadota>Alphaproteobacteria>Hyphomicrobiales>Phyllobacteriaceae | GCF_006440815.1 | Mesorhizobium_sp._B2-8-3 | FJ418_RS28720 | 4402..7806 | 2589905 | DUF499-domain-containing-protein-[Mesorhizobium-sp.-B2-8-3] |
| TIS54880.1 | HEPN+AAA-ATPase+Alpha+Beta-domain+wHTH+RRM | Type-1_BR.RM.like.systems | <-HEPN+DUF499*<-METHYLASE<-Peptidase_M78<-Ntox18<-TUDOR+SF1-SF2-Helicase+iREase+CTD+REase | 1115 | Bacteria>Pseudomonadota>Alphaproteobacteria>Hyphomicrobiales>Phyllobacteriaceae | GCA_004962755.1 | Mesorhizobium_sp. | E5W91_25095 | 74683..78030 | 1871066 | ATP-binding-protein-[Mesorhizobium-sp.] |
| WP_023748119.1 | HEPN+AAA-ATPase+Alpha+Beta-domain+wHTH+RRM | Type-1_BR.RM.like.systems | <-HEPN+DUF499*<-?<-METHYLASE<-?<-TUDOR+SF1-SF2-Helicase+iREase+CTD+REase<-RelE-ParE<-OLD-ABC | 1108 | Bacteria>Pseudomonadota>Alphaproteobacteria>Hyphomicrobiales>Phyllobacteriaceae | GCF_000502535.1 | Mesorhizobium_sp._LNJC391B00 | X750_RS144145 | 1284..4610 | 1287273 | Swt1-family-HEPN-domain-containing-protein-[Mesorhizobium-sp.-LNJC391B00] |
| WP_056240747.1 | HEPN+AAA-ATPase+Alpha+Beta-domain+wHTH+RRM | Type-1_BR.RM.like.systems | <-SF1-SF2-Helicase<-PIWI<-TIR+PNTD2<-HicB\|\|HTH+WYL->HEPN+DUF499*->?-><-?\|\|PADR-HTH-> | 1103 | Bacteria>Pseudomonadota>Alphaproteobacteria>Hyphomicrobiales>Phyllobacteriaceae | GCF_001427285.1 | Mesorhizobium_sp._Root552 | ASD50_RS11320 | 108386..111697 | 1736555 | DUF499-domain-containing-protein-[Mesorhizobium-sp.-Root552] |
| WP_136661243.1 | HEPN+AAA-ATPase+Alpha+Beta-domain+wHTH+RRM | Type-1_BR.RM.like.systems | TUDOR+SF1-SF2-Helicase+iREase+CTD+REase->?->METHYLASE->HEPN+DUF499*->HIN-HTH-> | 1097 | Bacteria>Pseudomonadota>Alphaproteobacteria>Hyphomicrobiales>Phyllobacteriaceae | GCF_004959855.1 | Nitratireductor_sp._XY-223 | E4M07_RS25460 | 21573..24866 | 2561926 | DUF499-domain-containing-protein-[Nitratireductor-sp.-XY-223] |
| WP_023805907.1 | HEPN+AAA-ATPase+Alpha+Beta-domain+wHTH+RRM | Type-1_BR.RM.like.systems | <-HEPN+DUF499*<-METHYLASE\|\|?->?->REase-> | 1110 | Bacteria>Pseudomonadota>Alphaproteobacteria>Hyphomicrobiales>Phyllobacteriaceae | GCF_000502355.1 | unclassified_Mesorhizobium | NLY44_RS22040 | 4450400..4453732 | 325217 | DUF499-domain-containing-protein-[Mesorhizobium-sp.-L2C089B000] |
| TIX43843.1 | HEPN+AAA-ATPase+Alpha+Beta-domain+wHTH+RRM | Type-1_BR.RM.like.systems | METHYLASE->HEPN+DUF499*-><-XIS-HTH | 1132 | Bacteria>Pseudomonadota>Alphaproteobacteria>Hyphomicrobiales>Phyllobacteriaceae | GCA_004964855.1 | Mesorhizobium_sp. | E5V40_01625 | 4247..7645 | 1871066 | ATP-binding-protein-[Mesorhizobium-sp.] |
| MBY0324746.1 | HEPN+AAA-ATPase+Alpha+Beta-domain+wHTH+RRM | Type-1_BR.RM.like.systems | <-ABC-ATPase+SbcC<-ABC-ATPase+SbcC<-ABC-ATPase+SbcC<-?\|\|?-><-?<-HEPN+DUF499*<-HEPN<-METHYLASE-p<-DDE_3<-METHYLASE<-YtH | 1111 | Bacteria>Pseudomonadota>Alphaproteobacteria>Hyphomicrobiales>Reyranellaceae | GCA_019750135.1 | Reyranella_sp. | K2X72_38955 | 16276..19611 | 1929291 | DUF499-domain-containing-protein-[Reyranella-sp.] |
| SSC67554.1 | HEPN+AAA-ATPase+Alpha+Beta-domain+wHTH+RRM | Type-1_BR.RM.like.systems | ParG_RHH-><-TetR-HTH\|\|MODE-HTH+PBPII-><-HEPN+DUF499*<-METHYLASE<-MORC+DUF3684<-TUDOR+SF1-SF2-Helicase+iREase+CTD+REase<-HTH+WYL\|\|SF1-SF2-Helicase+REase->SF1-SF2-Helicase+REase-> | 1086 | Bacteria>Pseudomonadota>Alphaproteobacteria>Hyphomicrobiales>Rhizobiaceae | GCF_900492185.1 | Ciceribacter_selenitireducens_ATCC_BAA-1503 | RHIZ70_3262 | 225722..228982 | 1336235 | unnamed-protein-product-[Ciceribacter-selenitireducens-ATCC-BAA-1503] |
| WP_174156950.1 | HEPN+AAA-ATPase+Alpha+Beta-domain+wHTH+RRM | Type-1_BR.RM.like.systems | <-XIS-HTH<-SF1-SF2-Helicase+RdgC+Beta-barrel+REase\|\|HTH+WYL->TUDOR+SF1-SF2-Helicase+iREase+CTD+REase->MORC+DUF3684->METHYLASE->HEPN+DUF499*->MODE-HTH+PBPII-><-ParG_RHH | 1098 | Bacteria>Pseudomonadota>Alphaproteobacteria>Hyphomicrobiales>Rhizobiaceae | GCF_013322055.1 | Rhizobium_lusitanum | G6L40_RS07680 | 183867..187163 | 293958 | DUF499-domain-containing-protein-[Rhizobium-lusitanum] |
| WP_127665091.1 | HEPN+AAA-ATPase+Alpha+Beta-domain+wHTH+RRM | Type-1_BR.RM.like.systems | Arm-DNA-bind_3+Phage_integrase->\|\|HTH+PolB->?-><-?<-?<-ParB+HNH<-HEPN+DUF499*<-?<-METHYLASE<-?<-TUDOR+SF1-SF2-Helicase+iREase+CTD+REase<-SF1-SF2-Helicase+DUF1998+HKD-DNase+HKD-DNase | 1104 | Bacteria>Pseudomonadota>Alphaproteobacteria>Hyphomicrobiales>Rhizobiaceae | GCF_004001475.1 | Sinorhizobium_meliloti | CN231_RS17685 | 56193..59507 | 382 | DUF499-domain-containing-protein-[Sinorhizobium-meliloti] |
| MBY3260302.1 | HEPN+AAA-ATPase+Alpha+Beta-domain+wHTH+RRM | Type-1_BR.RM.like.systems | HTH+WYL->TUDOR+SF1-SF2-Helicase+iREase+CTD+REase->YtH->MORC+DUF3684->METHYLASE->HEPN+DUF499*-><-wHTH+REase<-Arm-DNA-bind_3+Phage_integrase<-Peptidase_M24\|\|TetR-HTH-> | 1098 | Bacteria>Pseudomonadota>Alphaproteobacteria>Hyphomicrobiales>Rhizobiaceae | GCA_019779025.1 | Rhizobium_laguerreae | HFO15_01275 | 256972..260268 | 1076926 | ATP-binding-protein-[Rhizobium-laguerreae] |
| WP_149746898.1 | HEPN+AAA-ATPase+Alpha+Beta-domain+wHTH+RRM | Type-1_BR.RM.like.systems | ParG_RHH-><-MODE-HTH+PBPII<-HEPN+DUF499*<-METHYLASE<-REase<-TUDOR+SF1-SF2-Helicase+iREase+CTD+REase<-HTH+WYL\|\|SF1-SF2-helicase+RdgC+Beta-barrel+REase->XIS-HTH-> | 1098 | Bacteria>Pseudomonadota>Alphaproteobacteria>Hyphomicrobiales>Rhizobiaceae | GCF_900156005.1 | Rhizobium_sp._RU35A | F3H95_RS17305 | 253422..256718 | 1907414 | DUF499-domain-containing-protein-[Rhizobium-sp.-RU35A] |
| WP_246622248.1 | HEPN+AAA-ATPase+Alpha+Beta-domain+wHTH+RRM | Type-1_BR.RM.like.systems | HTH+WYL->TUDOR+SF1-SF2-Helicase+iREase+CTD+REase->YtH->MORC+DUF3684->METHYLASE->HEPN+DUF499*-><-wHTH+REase<-Phage_integrase<-Peptidase_M24\|\|TetR-HTH-> | 1086 | Bacteria>Pseudomonadota>Alphaproteobacteria>Hyphomicrobiales>Rhizobiaceae | - | Rhizobium_laguerreae | HFO15_RS01270 | 257008..260268 | 1076926 | DUF499-domain-containing-protein-[Rhizobium-laguerreae] |
| WP_209605924.1 | HEPN+AAA-ATPase+Alpha+Beta-domain+wHTH+RRM | Type-1_BR.RM.like.systems | <-Mbetalac<-?<-UbE1+JAB<-?\|\|HTH+WYL-><-HEPN+DUF499*<-METHYLASE<-TUDOR+SF1-SF2-Helicase+iREase+CTD+REase<-SF1-SF2-Helicase+DUF1998+HKD-DNase+HKD-DNase<-?<-?<-REase | 1109 | Bacteria>Pseudomonadota>Alphaproteobacteria>Hyphomicrobiales>Rhizobiaceae | GCF_017874595.1 | Sinorhizobium_kostiense | J2Z31_RS25385 | 30830..34159 | 76747 | DUF499-domain-containing-protein-[Sinorhizobium-kostiense] |
| WP_158499670.1 | HEPN+AAA-ATPase+Alpha+Beta-domain+wHTH+RRM | Type-1_BR.RM.like.systems | Phage_integrase->XIS-HTH->TUDOR+SF1-SF2-Helicase+iREase+CTD+REase->METHYLASE->HEPN+DUF499*->?->SF1-SF2-Helicase-><-MORC\|\|ArdCN+MPTase->ParB+HTH-> | 1096 | Bacteria>Pseudomonadota>Alphaproteobacteria>Hyphomicrobiales>Rhizobiaceae | GCF_000219415.3 | Sinorhizobium_fredii | SFGR64A_RS31000 | 11079..14369 | 380 | DUF499-domain-containing-protein-[Sinorhizobium-fredii] |
| WP_028734362.1 | HEPN+AAA-ATPase+Alpha+Beta-domain+wHTH+RRM | Type-1_BR.RM.like.systems | <-HicB\|\|HTH+WYL->TUDOR+SF1-SF2-Helicase+iREase+CTD+REase->ASCH->MORC+DUF3684->METHYLASE->HEPN+DUF499*->?-><-Arm-DNA-bind_3+Phage_integrase<-?<-?<-?<-HNH | 1098 | Bacteria>Pseudomonadota>Alphaproteobacteria>Hyphomicrobiales>Rhizobiaceae | GCF_000427705.1 | Rhizobium_leguminosarum | YSW_RS0132760 | 58681..61977 | 384 | DUF499-domain-containing-protein-[Rhizobium-leguminosarum] |
| WP_077472796.1 | HEPN+AAA-ATPase+Alpha+Beta-domain+wHTH+RRM | Type-1_BR.RM.like.systems | <-REC<-?<-DOC+HTH<-?<-GDResolvase+HIN-HTH\|\|<-HEPN+DUF499*<-METHYLASE<-ASCH<-TUDOR+SF1-SF2-Helicase+iREase+CTD+REase<-HTH+WYL<-McrC-NTD+REase | 1095 | Bacteria>Pseudomonadota>Alphaproteobacteria>Hyphomicrobiales>Rhizobiaceae | GCF_001999485.2 | Rhizobium_sp._P44RR-XXIV | BSK43_RS00425 | 6721..10008 | 1921145 | DUF499-domain-containing-protein-[Rhizobium-sp.-P44RR-XXIV] |
| WP_064332302.1 | HEPN+AAA-ATPase+Alpha+Beta-domain+wHTH+RRM | Type-1_BR.RM.like.systems | ParG_RHH-><-MODE-HTH+PBPII<-HEPN+DUF499*<-METHYLASE<-ASCH<-?<-TUDOR+SF1-SF2-Helicase+iREase+CTD+REase<-HTH+WYL\|\|SF1-SF2-Helicase+REase-> | 1098 | Bacteria>Pseudomonadota>Alphaproteobacteria>Hyphomicrobiales>Rhizobiaceae | GCF_001652565.1 | Shinella_sp._HZN7 | shn_RS22015 | 4588511..4591807 | 879274 | DUF499-domain-containing-protein-[Shinella-sp.-HZN7] |
| MCO5066228.1 | HEPN+AAA-ATPase+Alpha+Beta-domain+wHTH+RRM | Type-1_BR.RM.like.systems | RelE-ParE->ParA+HTH->?->OLD-ABC+TOPRIM->SF1-SF2-Helicase-><-TIR<-HEPN+DUF499*<-METHYLASE<-?<-TUDOR+SF1-SF2-Helicase+iREase+CTD+REase<-HTH+WYL\|\|HicB-> | 1098 | Bacteria>Pseudomonadota>Alphaproteobacteria>Hyphomicrobiales>Rhizobiaceae | GCA_023953835.1 | Rhizobiaceae_bacterium | M9924_17685 | 3745551..3748847 | 1913961 | DUF499-domain-containing-protein-[Rhizobiaceae-bacterium] |
| WP_160871739.1 | HEPN+AAA-ATPase+Alpha+Beta-domain+wHTH+RRM | Type-1_BR.RM.like.systems | Phage_integrase->?-><-ZnR\|\|GDResolvase+HIN-HTH-><-?<-HEPN+DUF499*<-METHYLASE<-TUDOR+SF1-SF2-Helicase+iREase+CTD+REase\|\|tRNA-><-?\|\|?->?->ABC-ATPase+SbcC-> | 1118 | Bacteria>Pseudomonadota>Alphaproteobacteria>Hyphomicrobiales>Rhizobiaceae | GCF_009826875.1 | Shinella_sp._AETb1-6 | GR158_RS15925 | 139338..142694 | 2692210 | DUF499-domain-containing-protein-[Shinella-sp.-AETb1-6] |
| WP_040958264.1 | HEPN+AAA-ATPase+Alpha+Beta-domain+wHTH+RRM | Type-1_BR.RM.like.systems | Tox-RelA->TUDOR+SF1-SF2-Helicase+iREase+CTD+REase->METHYLASE->HEPN+DUF499*-><-SLATT<-SMODS<-RNase<-HTH_17<-DnaB+CITB-HTH<-ZnR+TOPRIM | 1107 | Bacteria>Pseudomonadota>Alphaproteobacteria>Hyphomicrobiales>Rhizobiaceae | GCF_000219415.3 | Sinorhizobium_fredii | SFGR64A_RS01795 | 20467..23790 | 380 | DUF499-domain-containing-protein-[Sinorhizobium-fredii] |
| WP_233741236.1 | HEPN+AAA-ATPase+Alpha+Beta-domain+wHTH+RRM | Type-1_BR.RM.like.systems | <-HicB\|\|HTH+WYL->TUDOR+SF1-SF2-Helicase+iREase+CTD+REase->ASCH->MORC+DUF3684->METHYLASE->HEPN+DUF499*->?-><-?<-?<-Phage_terminase<-TetR-HTH | 1098 | Bacteria>Pseudomonadota>Alphaproteobacteria>Hyphomicrobiales>Rhizobiaceae | GCF_021387325.1 | Agrobacterium_vitis | FS799_10735 | 1060385..1063681 | 373 | DUF499-domain-containing-protein-[Agrobacterium-vitis] |
| MCE6075329.1 | HEPN+AAA-ATPase+Alpha+Beta-domain+wHTH+RRM | Type-1_BR.RM.like.systems | <-HicB\|\|HTH+WYL->TUDOR+SF1-SF2-Helicase+iREase+CTD+REase->ASCH->MORC+DUF3684->METHYLASE->HEPN+DUF499*->?-><-?<-?<-Phage_terminase<-TetR-HTH | 1098 | Bacteria>Pseudomonadota>Alphaproteobacteria>Hyphomicrobiales>Rhizobiaceae | GCA_021387325.1 | Agrobacterium_vitis | FS799_10735 | 1060385..1063681 | 373 | DUF499-domain-containing-protein-[Agrobacterium-vitis] |
| WP_142627962.1 | HEPN+AAA-ATPase+Alpha+Beta-domain+wHTH+RRM | Type-1_BR.RM.like.systems | TPR-><-RNase+SF1-SF2-Helicase<-HEPN+DUF499*<-METHYLASE<-MORC+DUF3684<-YtH<-ASCH<-TUDOR+SF1-SF2-Helicase+iREase+CTD+REase<-HTH+WYL | 1098 | Bacteria>Pseudomonadota>Alphaproteobacteria>Hyphomicrobiales>Rhizobiaceae | GCF_902153225.2 | Rhizobium_sp._P007 | FN104_RS24455 | 262558..265854 | 285908 | DUF499-domain-containing-protein-[Rhizobium-sp.-P007] |
| WP_168292930.1 | HEPN+AAA-ATPase+Alpha+Beta-domain+wHTH+RRM | Type-1_BR.RM.like.systems | <-OLD-ABC+TOPRIM\|\|?-><-YtH<-HEPN+DUF499*<-METHYLASE<-TUDOR+SF1-SF2-Helicase+iREase+CTD+REase<-?<-STYKinase-like<-HTH_3<-TetR-HTH | 1118 | Bacteria>Pseudomonadota>Alphaproteobacteria>Hyphomicrobiales>Rhizobiaceae | GCF_012275615.1 | Rhizobium_leguminosarum | GFL51_RS25220 | 66305..69661 | 384 | DUF499-domain-containing-protein-[Rhizobium-leguminosarum] |
| WP_242219593.1 | HEPN+AAA-ATPase+Alpha+Beta-domain+wHTH+RRM | Type-1_BR.RM.like.systems | ParG_RHH-><-MODE-HTH+PBPII<-HEPN+DUF499*<-METHYLASE<-?<-HIN-HTH\|\|?->Phage_AlpA-><-Arm-DNA-bind_2+Phage_int_SAM_3+Phage_integrase | 1098 | Bacteria>Pseudomonadota>Alphaproteobacteria>Hyphomicrobiales>Rhizobiaceae | GCF_022682305.1 | Shinella_zoogloeoides | MOE34_RS21255 | 4281309..4284605 | 352475 | DUF499-domain-containing-protein-[Shinella-zoogloeoides] |
| WP_221153182.1 | HEPN+AAA-ATPase+Alpha+Beta-domain+wHTH+RRM | Type-1_BR.RM.like.systems | TetR-HTH-><-tRNA<-Peptidase_M15<-?\|\|TUDOR+SF1-SF2-Helicase+iREase+CTD+REase->METHYLASE->HEPN+DUF499*-><-GDResolvase+HIN-HTH\|\|ZnR-> | 1114 | Bacteria>Pseudomonadota>Alphaproteobacteria>Hyphomicrobiales>Rhizobiaceae | - | Rhizobium_sp._NLR10a | HJB90_RS14150 | 264416..267760 | 2731105 | DUF499-domain-containing-protein-[Rhizobium-sp.-NLR10a] |
| WP_174037242.1 | HEPN+AAA-ATPase+Alpha+Beta-domain+wHTH+RRM | Type-1_BR.RM.like.systems | Phage_terminase->?-><-?<-HEPN+DUF499*<-METHYLASE<-MORC+DUF3684<-ASCH<-TUDOR+SF1-SF2-Helicase+iREase+CTD+REase<-HTH+WYL\|\|HicB-> | 1098 | Bacteria>Pseudomonadota>Alphaproteobacteria>Hyphomicrobiales>Rhizobiaceae | - | Agrobacterium_tumefaciens | G6M03_RS01055 | 214165..217461 | 358 | DUF499-domain-containing-protein-[Agrobacterium-tumefaciens] |
| WP_153320511.1 | HEPN+AAA-ATPase+Alpha+Beta-domain+wHTH+RRM | Type-1_BR.RM.like.systems | wHTH+SAD+HNH-><-HTH_3\|\|?->?->?->GDResolvase+Recombinase+ZnR-><-HEPN+DUF499*<-METHYLASE<-TUDOR+SF1-SF2-Helicase+iREase+CTD+REase | 1110 | Bacteria>Pseudomonadota>Alphaproteobacteria>Hyphomicrobiales>Rhizobiaceae | GCF_009599945.1 | Sinorhizobium_meliloti | GHJ83_RS08885 | 35044..38376 | 382 | DUF499-domain-containing-protein-[Sinorhizobium-meliloti] |
| WP_153314323.1 | HEPN+AAA-ATPase+Alpha+Beta-domain+wHTH+RRM | Type-1_BR.RM.like.systems | TUDOR+SF1-SF2-Helicase+iREase+CTD+REase->METHYLASE->HEPN+DUF499*-><-GDResolvase+Recombinase+ZnR<-?<-?\|\|HTH_3-><-wHTH+SAD+HNH | 1110 | Bacteria>Pseudomonadota>Alphaproteobacteria>Hyphomicrobiales>Rhizobiaceae | GCF_009599625.1 | Sinorhizobium_meliloti | GHK35_RS22105 | 11808..15140 | 382 | DUF499-domain-containing-protein-[Sinorhizobium-meliloti] |
| WP_132653484.1 | HEPN+AAA-ATPase+Alpha+Beta-domain+wHTH+RRM | Type-1_BR.RM.like.systems | XIS-HTH->?-><-?<-?<-?<-?<-HEPN+DUF499*<-METHYLASE<-MORC+DUF3684<-TUDOR+SF1-SF2-Helicase+iREase+CTD+REase<-?\|\|HTH_3->STYKinase-like-> | 1097 | Bacteria>Pseudomonadota>Alphaproteobacteria>Hyphomicrobiales>Rhizobiaceae | GCF_004341105.1 | Rhizobium_sp._BK418 | EV281_RS12840 | 265866..269159 | 2512120 | DUF499-domain-containing-protein-[Rhizobium-sp.-BK418] |
| WP_220337433.1 | HEPN+AAA-ATPase+Alpha+Beta-domain+wHTH+RRM | Type-1_BR.RM.like.systems | TUDOR+SF1-SF2-Helicase+iREase+CTD+REase->METHYLASE->HEPN+DUF499*->?-><-GDResolvase+HIN-HTH\|\|ZnR-><-?\|\|?-><-Phage_integrase | 1118 | Bacteria>Pseudomonadota>Alphaproteobacteria>Hyphomicrobiales>Rhizobiaceae | GCF_019511345.1 | Rhizobium_mesosinicum | JNB85_RS26630 | 6855..10211 | 335017 | DUF499-domain-containing-protein-[Rhizobium-mesosinicum] |
| CAD6420434.1 | HEPN+AAA-ATPase+Alpha+Beta-domain+wHTH+RRM | Type-1_BR.RM.like.systems | <-LuxR-HTH\|\|PIN+OST-HTH-><-SF1-SF2-Helicase<-HEPN+DUF499*<-?<-METHYLASE<-?<-TUDOR+SF1-SF2-Helicase+iREase+CTD+REase | 1104 | Bacteria>Pseudomonadota>Alphaproteobacteria>Hyphomicrobiales>Rhizobiaceae | GCA_902153285.2 | Rhizobium_sp._Q54 | REQ54_02098 | 2069964..2073278 | 1335060 | hypothetical-protein-REQ54_02098-[Rhizobium-sp.-Q54] |
| WP_045535325.1 | HEPN+AAA-ATPase+Alpha+Beta-domain+wHTH+RRM | Type-1_BR.RM.like.systems | <-MORC+HisKin+REase<-HEPN+DUF499*<-?<-METHYLASE<-SIR2<-ASCH<-TUDOR+SF1-SF2-Helicase+iREase+CTD+REase<-HTH+WYL | 1104 | Bacteria>Pseudomonadota>Alphaproteobacteria>Hyphomicrobiales>Rhizobiaceae | - | Rhizobium_sp._LMB-1 | VN14_RS23455 | 4128..7442 | 1605895 | DUF499-domain-containing-protein-[Rhizobium-sp.-LMB-1] |
| NEH32401.1 | HEPN+AAA-ATPase+Alpha+Beta-domain+wHTH+RRM | Type-1_BR.RM.like.systems | HTH+WYL->TUDOR+SF1-SF2-Helicase+iREase+CTD+REase->ASCH->YtH->MORC+DUF3684->METHYLASE->HEPN+DUF499*->HEPN-> | 1097 | Bacteria>Pseudomonadota>Alphaproteobacteria>Hyphomicrobiales>Rhizobiaceae | GCA_010668085.1 | Rhizobium_ruizarguesonis | GR203_31950 | 43158..46451 | 2081791 | DUF499-domain-containing-protein-[Rhizobium-leguminosarum] |
| WP_261323154.1 | HEPN+AAA-ATPase+Alpha+Beta-domain+wHTH+RRM | Type-1_BR.RM.like.systems | <-PIN+OST-HTH\|\|TUDOR+SF1-SF2-Helicase+iREase+CTD+REase->?->METHYLASE->?->HEPN+DUF499*-><-?<-STYKinase-like<-HTH_3 | 1091 | Bacteria>Pseudomonadota>Alphaproteobacteria>Hyphomicrobiales>Rhizobiaceae | GCF_019798485.1 | Rhizobium_leguminosarum | HFO91_RS33945 | 19091..22366 | 384 | DUF499-domain-containing-protein-[Rhizobium-leguminosarum] |
| WP_245582348.1 | HEPN+AAA-ATPase+Alpha+Beta-domain+wHTH+RRM | Type-1_BR.RM.like.systems | <-HEPN<-HEPN+DUF499*<-METHYLASE<-ASCH<-MORC+DUF3684<-YtH<-ASCH<-TUDOR+SF1-SF2-Helicase+iREase+CTD+REase | 1085 | Bacteria>Pseudomonadota>Alphaproteobacteria>Hyphomicrobiales>Rhizobiaceae | GCF_004306165.1 | Rhizobium_ruizarguesonis | ELH13_RS20435 | 4103083..4106340 | 2081791 | DUF499-domain-containing-protein-[Rhizobium-leguminosarum] |
| WP_222320084.1 | HEPN+AAA-ATPase+Alpha+Beta-domain+wHTH+RRM | Type-1_BR.RM.like.systems | <-OLD-ABC+TOPRIM<-?\|\|?-><-YtH<-HEPN+DUF499*<-METHYLASE<-METHYLASE-p<-TUDOR+SF1-SF2-Helicase+iREase+CTD+REase | 1108 | Bacteria>Pseudomonadota>Alphaproteobacteria>Hyphomicrobiales>Rhizobiaceae | GCF_019798845.1 | Rhizobium_leguminosarum | HFO61_RS22785 | 274165..277491 | 384 | DUF499-domain-containing-protein-[Rhizobium-leguminosarum] |
| WP_221096744.1 | HEPN+AAA-ATPase+Alpha+Beta-domain+wHTH+RRM | Type-1_BR.RM.like.systems | <-HicB\|\|HTH+WYL->TUDOR+SF1-SF2-Helicase+iREase+CTD+REase->ASCH->MORC+DUF3684->METHYLASE->HEPN+DUF499*-> | 1098 | Bacteria>Pseudomonadota>Alphaproteobacteria>Hyphomicrobiales>Rhizobiaceae | GCF_019684055.1 | Rhizobium_bangladeshense | HJA85_RS24965 | 76913..80209 | 1138189 | DUF499-domain-containing-protein-[Rhizobium-bangladeshense] |
| WP_173514149.1 | HEPN+AAA-ATPase+Alpha+Beta-domain+wHTH+RRM | Type-1_BR.RM.like.systems | <-UbE1+JAB<-?\|\|HTH+WYL->TUDOR+SF1-SF2-Helicase+iREase+CTD+REase->YtH->METHYLASE->HEPN+DUF499*->?-><-?\|\|?-><-?\|\|REase-> | 1101 | Bacteria>Pseudomonadota>Alphaproteobacteria>Hyphomicrobiales>Rhizobiaceae | GCF_013283645.1 | Sinorhizobium_psoraleae | ILFOPFJJ_RS29980 | 72999..76304 | 520838 | DUF499-domain-containing-protein-[Ensifer-psoraleae] |
| TBD87036.1 | HEPN+AAA-ATPase+Alpha+Beta-domain+wHTH+RRM | Type-1_BR.RM.like.systems | <-HEPN<-HEPN+DUF499*<-METHYLASE<-ASCH<-MORC+DUF3684<-YtH<-ASCH<-TUDOR+SF1-SF2-Helicase+iREase+CTD+REase | 1097 | Bacteria>Pseudomonadota>Alphaproteobacteria>Hyphomicrobiales>Rhizobiaceae | GCA_004306165.1 | Rhizobium_ruizarguesonis | ELH13_20430 | 4103083..4106376 | 2081791 | ATP-binding-protein-[Rhizobium-leguminosarum] |
| WP_064252405.1 | HEPN+AAA-ATPase+Alpha+Beta-domain+wHTH+RRM | Type-1_BR.RM.like.systems | Terminase->?-><-HTH_3\|\|?-><-?\|\|<-HEPN+DUF499*<-METHYLASE<-TUDOR+SF1-SF2-Helicase+iREase+CTD+REase<-REase\|\|35exo-> | 1109 | Bacteria>Pseudomonadota>Alphaproteobacteria>Hyphomicrobiales>Rhizobiaceae | GCF_001889105.1 | Sinorhizobium_americanum | SAMCFNEI73_RS12475 | 2595143..2598472 | 194963 | DUF499-domain-containing-protein-[Sinorhizobium-americanum] |
| WP_222374442.1 | HEPN+AAA-ATPase+Alpha+Beta-domain+wHTH+RRM | Type-1_BR.RM.like.systems | TetR-HTH->HTH_3->STYKinase-like->?->TUDOR+SF1-SF2-Helicase+iREase+CTD+REase->METHYLASE->HEPN+DUF499*-> | 1114 | Bacteria>Pseudomonadota>Alphaproteobacteria>Hyphomicrobiales>Rhizobiaceae | GCF_019799295.1 | Rhizobium_leguminosarum | HFO38_RS09705 | 496684..500028 | 384 | DUF499-domain-containing-protein-[Rhizobium-leguminosarum] |
| WP_222000518.1 | HEPN+AAA-ATPase+Alpha+Beta-domain+wHTH+RRM | Type-1_BR.RM.like.systems | TetR-HTH->HTH_3->STYKinase-like->?-><-HEPN+DUF499*<-?<-METHYLASE<-?<-TUDOR+SF1-SF2-Helicase+iREase+CTD+REase | 1103 | Bacteria>Pseudomonadota>Alphaproteobacteria>Hyphomicrobiales>Rhizobiaceae | GCF_019779395.1 | Rhizobium_laguerreae | HFN92_RS29725 | 56760..60071 | 1076926 | DUF499-domain-containing-protein-[Rhizobium-laguerreae] |
| TCS04590.1 | HEPN+AAA-ATPase+Alpha+Beta-domain+wHTH+RRM | Type-1_BR.RM.like.systems | XIS-HTH->?-><-?<-?<-?<-?<-HEPN+DUF499*<-METHYLASE<-MORC+DUF3684<-TUDOR+SF1-SF2-Helicase+iREase+CTD+REase<-?<-?\|\|HTH_3-> | 1096 | Bacteria>Pseudomonadota>Alphaproteobacteria>Hyphomicrobiales>Rhizobiaceae | GCA_004341105.1 | Rhizobium_sp._BK418 | EV281_103265 | 265866..269156 | 2512120 | hypothetical-protein-EV281_103265-[Rhizobium-sp.-BK418] |
| MBX4863812.1 | HEPN+AAA-ATPase+Alpha+Beta-domain+wHTH+RRM | Type-1_BR.RM.like.systems | <-HEPN+DUF499*<-METHYLASE<-YtH<-TUDOR+SF1-SF2-Helicase+iREase+CTD+REase<-HTH+WYL\|\|?->UbE1+JAB-> | 1098 | Bacteria>Pseudomonadota>Alphaproteobacteria>Hyphomicrobiales>Rhizobiaceae | GCA_019684005.1 | Rhizobium_bangladeshense | HJA86_28440 | 35410..38706 | 1138189 | ATP-binding-protein-[Rhizobium-bangladeshense] |
| WP_222282260.1 | HEPN+AAA-ATPase+Alpha+Beta-domain+wHTH+RRM | Type-1_BR.RM.like.systems | <-PIN+OST-HTH\|\|TUDOR+SF1-SF2-Helicase+iREase+CTD+REase->?->METHYLASE->?->HEPN+DUF499*-><-HTH_3 | 1103 | Bacteria>Pseudomonadota>Alphaproteobacteria>Hyphomicrobiales>Rhizobiaceae | - | Rhizobium_leguminosarum | HFO92_34835 | 18961..22272 | 384 | DUF499-domain-containing-protein-[Rhizobium-leguminosarum] |
| MBY5371752.1 | HEPN+AAA-ATPase+Alpha+Beta-domain+wHTH+RRM | Type-1_BR.RM.like.systems | <-PIN+OST-HTH\|\|TUDOR+SF1-SF2-Helicase+iREase+CTD+REase->?->METHYLASE->?->HEPN+DUF499*-><-HTH_3 | 1103 | Bacteria>Pseudomonadota>Alphaproteobacteria>Hyphomicrobiales>Rhizobiaceae | GCA_019798485.1 | Rhizobium_leguminosarum | HFO92_34835 | 18961..22272 | 384 | ATP-binding-protein-[Rhizobium-leguminosarum] |
| WP_166603209.1 | HEPN+AAA-ATPase+Alpha+Beta-domain+wHTH+RRM | Type-1_BR.RM.like.systems | TUDOR+SF1-SF2-Helicase+iREase+CTD+REase-><-?<-?\|\|METHYLASE->HEPN+DUF499*-> | 1119 | Bacteria>Pseudomonadota>Alphaproteobacteria>Hyphomicrobiales>Rhizobiaceae | - | Rhizobium_sp._CRIBSB | GVN24_RS23375 | 29303..32662 | 2683264 | DUF499-domain-containing-protein-[Rhizobium-sp.-CRIBSB] |
| MCB1465833.1 | HEPN+AAA-ATPase+Alpha+Beta-domain+wHTH+RRM | Type-1_BR.RM.like.systems | TUDOR+SF1-SF2-Helicase+iREase+CTD+REase-p->?->METHYLASE->HEPN+DUF499*-> | 428 | Bacteria>Pseudomonadota>Alphaproteobacteria>Hyphomicrobiales>Rhizobiaceae | GCA_020441945.1 | Rhizobiaceae_bacterium | KDK08_01465 | 5124..6408 | 1913961 | DUF499-domain-containing-protein-[Rhizobiaceae-bacterium] |
| WP_168332223.1 | HEPN+AAA-ATPase+Alpha+Beta-domain+wHTH+RRM | Type-1_BR.RM.like.systems | <-HEPN+DUF499*<-?<-METHYLASE<-?<-TUDOR+SF1-SF2-Helicase+iREase+CTD+REase | 1103 | Bacteria>Pseudomonadota>Alphaproteobacteria>Hyphomicrobiales>Rhizobiaceae | GCF_012276395.1 | Rhizobium_leguminosarum | GFL80_RS35655 | 1353..4664 | 384 | DUF499-domain-containing-protein-[Rhizobium-leguminosarum] |
| WP_097140486.1 | HEPN+AAA-ATPase+Alpha+Beta-domain+wHTH+RRM | Type-1_BR.RM.like.systems | ParG_RHH-><-MODE-HTH+PBPII<-HEPN+DUF499*<-METHYLASE | 1098 | Bacteria>Pseudomonadota>Alphaproteobacteria>Hyphomicrobiales>Rhizobiaceae | GCF_900220975.1 | Rhizobium_subbaraonis | CRO48_RS14615 | 291949..295245 | 908946 | DUF499-domain-containing-protein-[Rhizobium-subbaraonis] |
| WP_286190155.1 | HEPN+AAA-ATPase+Alpha+Beta-domain+wHTH+RRM | Type-1_BR.RM.like.systems | HTH+WYL->TUDOR+SF1-SF2-Helicase+iREase+CTD+REase->YtH->MORC+DUF3684->REase->METHYLASE->HEPN+DUF499*-><-ACET<-CopG/RHH<-Arm-DNA-bind_3+Phage_integrase<-?<-MethyltransfD12 | 1086 | Bacteria>Pseudomonadota>Alphaproteobacteria>Hyphomicrobiales>Stappiaceae | GCF_017743865.1 | Labrenzia_sp._R4_2 | J7481_RS19430 | 351068..354328 | 2821107 | DUF499-domain-containing-protein-[Labrenzia-sp.-R4_2] |
| MBO9421687.1 | HEPN+AAA-ATPase+Alpha+Beta-domain+wHTH+RRM | Type-1_BR.RM.like.systems | HTH+WYL->TUDOR+SF1-SF2-Helicase+iREase+CTD+REase->YtH->MORC+DUF3684->REase->METHYLASE->HEPN+DUF499*-><-ACET<-CopG/RHH<-Arm-DNA-bind_3+Phage_integrase<-?<-MethyltransfD12 | 1098 | Bacteria>Pseudomonadota>Alphaproteobacteria>Hyphomicrobiales>Stappiaceae | GCA_017743865.1 | Labrenzia_sp._R4_2 | J7481_19425 | 351032..354328 | 2821107 | DUF499-domain-containing-protein-[Labrenzia-sp.-R4_2] |
| WP_209006426.1 | HEPN+AAA-ATPase+Alpha+Beta-domain+wHTH+RRM | Type-1_BR.RM.like.systems | <-SF1-SF2-Helicase+REase<-SF1-SF2-Helicase+REase\|\|HTH+WYL->TUDOR+SF1-SF2-Helicase+iREase+CTD+REase->?->METHYLASE->HEPN+DUF499*-><-MODE-HTH+PBPII\|\|TetR-HTH-><-ParG_RHH | 1086 | Bacteria>Pseudomonadota>Alphaproteobacteria>Hyphomicrobiales>Stappiaceae | GCF_014252955.1 | Stappia_sp._28M-7 | H7H34_RS21795 | 99426..102686 | 2762596 | DUF499-domain-containing-protein-[Stappia-sp.-28M-7] |
| MBC2861680.1 | HEPN+AAA-ATPase+Alpha+Beta-domain+wHTH+RRM | Type-1_BR.RM.like.systems | <-SF1-SF2-Helicase+REase<-SF1-SF2-Helicase+REase\|\|HTH+WYL->TUDOR+SF1-SF2-Helicase+iREase+CTD+REase->?->METHYLASE->HEPN+DUF499*-><-MODE-HTH+PBPII\|\|TetR-HTH-><-ParG_RHH | 1098 | Bacteria>Pseudomonadota>Alphaproteobacteria>Hyphomicrobiales>Stappiaceae | GCA_014252955.1 | Stappia_sp._28M-7 | H7H34_21795 | 99390..102686 | 2762596 | ATP-binding-protein-[Stappia-sp.-28M-7] |
| WP_055658296.1 | HEPN+AAA-ATPase+Alpha+Beta-domain+wHTH+RRM | Type-1_BR.RM.like.systems | Arm-DNA-bind_3+Phage_integrase->?->Phage_AlpA->ZnR+TOPRIM->DnaB+CITB-HTH->?-><-HEPN+DUF499*<-?<-METHYLASE<-?<-ParB+HNH<-MORC+DUF3684<-TUDOR+SF1-SF2-Helicase+iREase+CTD+REase | 1112 | Bacteria>Pseudomonadota>Alphaproteobacteria>Hyphomicrobiales>Stappiaceae | GCF_001403915.1 | Roseibium_aggregatum | LAL4801_RS18620 | 1338638..1341976 | 187304 | DUF499-domain-containing-protein-[Roseibium-aggregatum] |
| WP_186398691.1 | HEPN+AAA-ATPase+Alpha+Beta-domain+wHTH+RRM | Type-1_BR.RM.like.systems | <-AbiEii<-HEPN+DUF499*<-METHYLASE<-MORC+DUF3684<-TUDOR+SF1-SF2-Helicase+iREase+CTD+REase<-HTH+WYL<-XIS-HTH | 1115 | Bacteria>Pseudomonadota>Alphaproteobacteria>Hyphomicrobiales>Stappiaceae | GCF_900185735.1 | Stappia_sp._P2PMeth1 | P2PM1_RS07495 | 1079..4426 | 2003586 | Swt1-family-HEPN-domain-containing-protein-[Stappia-sp.-P2PMeth1] |
| WP_235988002.1 | HEPN+AAA-ATPase+Alpha+Beta-domain+wHTH+RRM | Type-1_BR.RM.like.systems | ParG_RHH-><-TetR-HTH\|\|MODE-HTH+PBPII-><-HEPN+DUF499*<-METHYLASE | 1060 | Bacteria>Pseudomonadota>Alphaproteobacteria>Hyphomicrobiales>Stappiaceae | GCF_014692675.1 | Roseibium_aggregatum | HK439_RS18245 | 125935..129117 | 187304 | DUF499-domain-containing-protein-[Roseibium-aggregatum] |
| MBZ0262663.1 | HEPN+AAA-ATPase+Alpha+Beta-domain+wHTH+RRM | Type-1_BR.RM.like.systems | SF1-SF2-Helicase+REase->SF1-SF2-Helicase+REase-><-?<-ACET<-HEPN+DUF499*<-METHYLASE<-TUDOR+SF1-SF2-Helicase+iREase+CTD+REase<-HTH+WYL\|\|HicB->TIR+PNTD2->PIWI-> | 1104 | Bacteria>Pseudomonadota>Alphaproteobacteria>Hyphomicrobiales>unclassified_Hyphomicrobiales | GCA_019912605.1 | Hyphomicrobiales_bacterium | K8F90_18930 | 14586..17900 | 1909294 | DUF499-domain-containing-protein-[Hyphomicrobiales-bacterium] |
| MBN8994771.1 | HEPN+AAA-ATPase+Alpha+Beta-domain+wHTH+RRM | Type-1_BR.RM.like.systems | <-SF1-SF2-Helicase+REase\|\|HTH+WYL->TUDOR+SF1-SF2-Helicase+iREase+CTD+REase->ASCH->SIR2->METHYLASE->HEPN+DUF499*->MODE-HTH+PBPII-><-ParG_RHH | 1100 | Bacteria>Pseudomonadota>Alphaproteobacteria>Hyphomicrobiales>unclassified_Hyphomicrobiales | GCA_017305635.1 | Hyphomicrobiales_bacterium | J0H94_06075 | 362415..365717 | 1909294 | DUF499-domain-containing-protein-[Hyphomicrobiales-bacterium] |
| MBN9033564.1 | HEPN+AAA-ATPase+Alpha+Beta-domain+wHTH+RRM | Type-1_BR.RM.like.systems | ABhydrolase->REase->Peptidase_M24->HhH-RADC+JAB->Arm-DNA-bind_3+Phage_integrase->?-><-HEPN+DUF499*<-METHYLASE | 1098 | Bacteria>Pseudomonadota>Alphaproteobacteria>Hyphomicrobiales>unclassified_Hyphomicrobiales | GCA_017305935.1 | Hyphomicrobiales_bacterium | J0I23_27610 | 44902..48198 | 1909294 | DUF499-domain-containing-protein-[Hyphomicrobiales-bacterium] |
| SDR39062.1 | HEPN+AAA-ATPase+Alpha+Beta-domain+wHTH+RRM | Type-1_BR.RM.like.systems | TUDOR+SF1-SF2-Helicase+iREase+CTD+REase->YtH->METHYLASE->HEPN->HEPN+DUF499*->STAS+HisKin->?->?->?->?-><-ABhydrolase | 1116 | Bacteria>Pseudomonadota>Alphaproteobacteria>Hyphomicrobiales>unclassified_Hyphomicrobiales | GCA_900103895.1 | Rhizobiales_bacterium_GAS113 | SAMN05519103_02590 | 2751925..2755275 | 1884352 | Predicted-ATPase,-AAA+-superfamily-[Rhizobiales-bacterium-GAS113] |
| MBN9010874.1 | HEPN+AAA-ATPase+Alpha+Beta-domain+wHTH+RRM | Type-1_BR.RM.like.systems | METHYLASE-p->HEPN+DUF499*-> | 540 | Bacteria>Pseudomonadota>Alphaproteobacteria>Hyphomicrobiales>unclassified_Hyphomicrobiales | GCA_017305735.1 | Hyphomicrobiales_bacterium | J0H63_12355 | 981..2600 | 1909294 | DUF499-domain-containing-protein-[Hyphomicrobiales-bacterium] |
| MBN9001777.1 | HEPN+AAA-ATPase+Alpha+Beta-domain+wHTH+RRM | Type-1_BR.RM.like.systems | METHYLASE-p->HEPN+DUF499*-> | 379 | Bacteria>Pseudomonadota>Alphaproteobacteria>Hyphomicrobiales>unclassified_Hyphomicrobiales | GCA_017305655.1 | Hyphomicrobiales_bacterium | J0H75_06805 | 482..1619 | 1909294 | DUF499-domain-containing-protein-[Hyphomicrobiales-bacterium] |
| WP_282888615.1 | HEPN+AAA-ATPase+Alpha+Beta-domain+wHTH+RRM | Type-1_BR.RM.like.systems | <-SF1-SF2-Helicase+REase<-?\|\|HTH+WYL->TUDOR+SF1-SF2-Helicase+iREase+CTD+REase->MORC+DUF3684->METHYLASE->HEPN+DUF499*->TIR+PNTD2->PIWI->Arm-DNA-bind_3+Phage_integrase-><-XIS-HTH<-MobA_MobL+SF1-SF2-helicase\|\|TraD-> | 1098 | Bacteria>Pseudomonadota>Alphaproteobacteria>Hyphomicrobiales>Xanthobacteraceae | GCF_030015065.1 | Xanthobacter_autotrophicus | K9U40_RS02195 | 72535..75831 | 280 | DUF499-domain-containing-protein-[Xanthobacter-autotrophicus] |
| WP_159599724.1 | HEPN+AAA-ATPase+Alpha+Beta-domain+wHTH+RRM | Type-1_BR.RM.like.systems | HTH+WYL->TUDOR+SF1-SF2-Helicase+iREase+CTD+REase->YtH->MORC+DUF3684->METHYLASE->HEPN+DUF499*-><-?<-Arm-DNA-bind_3+Phage_integrase<-HhH-RADC+JAB<-Peptidase_M24<-ACET\|\|Aminotran_1_2-> | 1098 | Bacteria>Pseudomonadota>Alphaproteobacteria>Hyphomicrobiales>Xanthobacteraceae | GCF_902703545.1 | Starkeya_nomas | STARVERO_RS14985 | 3130835..3134131 | 2666134 | DUF499-domain-containing-protein-[Starkeya-sp.-HF14-78462] |
| MCZ7659369.1 | HEPN+AAA-ATPase+Alpha+Beta-domain+wHTH+RRM | Type-1_BR.RM.like.systems | DEOR-HTH->TUDOR+SF1-SF2-Helicase+iREase+CTD+REase->TUDOR+SF1-SF2-Helicase+iREase+CTD+REase-p->?->REase->METHYLASE->HEPN+DUF499*->OLD-ABC+TOPRIM->SF1-SF2-Helicase-><-REase<-?<-N6_MTase<-DEOR-HTH | 1120 | Bacteria>Pseudomonadota>Alphaproteobacteria>Hyphomicrobiales>Xanthobacteraceae | GCA_027493405.1 | Xanthobacteraceae_bacterium | M5U07_16605 | 9358..12720 | 2021379 | MAG:-Swt1-family-HEPN-domain-containing-protein-[Xanthobacteraceae-bacterium] |
| ABS65938.1 | HEPN+AAA-ATPase+Alpha+Beta-domain+wHTH+RRM | Type-1_BR.RM.like.systems | TUDOR+SF1-SF2-Helicase+iREase+CTD+REase->REase->METHYLASE-><-HTH_21+rve+rve_3<-HIN-HTH\|\|METHYLASE-p->HEPN+DUF499*-><-MODE-HTH+PBPII\|\|TetR-HTH-><-ParG_RHH | 1098 | Bacteria>Pseudomonadota>Alphaproteobacteria>Hyphomicrobiales>Xanthobacteraceae | GCA_000017645.1 | Xanthobacter_autotrophicus_Py2 | Xaut_0686 | 774507..777803 | 78245 | ATPase-(AAA+-superfamily)-like-protein-[Xanthobacter-autotrophicus-Py2] |
| MBX9650469.1 | HEPN+AAA-ATPase+Alpha+Beta-domain+wHTH+RRM | Type-1_BR.RM.like.systems | <-SF1-SF2-Helicase+REase\|\|HTH+WYL->TUDOR+SF1-SF2-Helicase+iREase+CTD+REase->MORC+DUF3684->METHYLASE->HEPN+DUF499*->MODE-HTH+PBPII-><-ParG_RHH | 1098 | Bacteria>Pseudomonadota>Alphaproteobacteria>Hyphomicrobiales>Xanthobacteraceae | GCA_019748335.1 | Xanthobacteraceae_bacterium | K2X57_25835 | 14404..17700 | 2021379 | DUF499-domain-containing-protein-[Xanthobacteraceae-bacterium] |
| ABS68223.1 | HEPN+AAA-ATPase+Alpha+Beta-domain+wHTH+RRM | Type-1_BR.RM.like.systems | <-SF1-SF2-Helicase+REase\|\|HTH+WYL->TUDOR+SF1-SF2-Helicase+iREase+CTD+REase->YtH->MORC+DUF3684->METHYLASE->HEPN+DUF499*->MODE-HTH+PBPII-> | 1098 | Bacteria>Pseudomonadota>Alphaproteobacteria>Hyphomicrobiales>Xanthobacteraceae | GCA_000017645.1 | Xanthobacter_autotrophicus_Py2 | Xaut_2992 | 3328660..3331956 | 78245 | conserved-hypothetical-protein-[Xanthobacter-autotrophicus-Py2] |
| MCW5685472.1 | HEPN+AAA-ATPase+Alpha+Beta-domain+wHTH+RRM | Type-1_BR.RM.like.systems | <-Peptidase_S11<-?\|\|tRNA-><-?<-HEPN+DUF499*<-METHYLASE<-ParB+HNH<-TUDOR+SF1-SF2-Helicase+iREase+CTD+REase<-HTH+WYL\|\|SF1-SF2-Helicase+REase-> | 1098 | Bacteria>Pseudomonadota>Alphaproteobacteria>Hyphomicrobiales>Xanthobacteraceae | GCA_026125885.1 | Pseudolabrys_sp. | KIT85_13790 | 135757..139053 | 1960880 | MAG:-DUF499-domain-containing-protein-[Pseudolabrys-sp.] |
| MBX9829560.1 | HEPN+AAA-ATPase+Alpha+Beta-domain+wHTH+RRM | Type-1_BR.RM.like.systems | TrwC+SF1-SF2-helicase->TUDOR+SF1-SF2-Helicase+iREase+CTD+REase->HEPN->METHYLASE-p->METHYLASE->?->HEPN+DUF499*->?->REase-> | 1101 | Bacteria>Pseudomonadota>Alphaproteobacteria>Hyphomicrobiales>Xanthobacteraceae | GCA_019746355.1 | Xanthobacteraceae_bacterium | K2Y27_31790 | 26343..29648 | 2021379 | DUF499-domain-containing-protein-[Xanthobacteraceae-bacterium] |
| MBX6329369.1 | HEPN+AAA-ATPase+Alpha+Beta-domain+wHTH+RRM | Type-1_BR.RM.like.systems | TUDOR+SF1-SF2-Helicase+iREase+CTD+REase-p->?->?->PolB->HEPN->HEPN+DUF499*-> | 1114 | Bacteria>Pseudomonadota>Alphaproteobacteria>Hyphomicrobiales>Xanthobacteraceae | GCA_019689975.1 | Pseudolabrys_sp. | IRY89_12420 | 7103..10447 | 1960880 | DUF499-domain-containing-protein-[Pseudolabrys-sp.] |
| MCB9972622.1 | HEPN+AAA-ATPase+Alpha+Beta-domain+wHTH+RRM | Type-1_BR.RM.like.systems | XIS-HTH->?->?-><-HEPN+DUF499*<-METHYLASE<-REase<-MORC+DUF3684<-TUDOR+SF1-SF2-Helicase+iREase+CTD+REase<-HTH+WYL\|\|VirD4-TraG+VirD4-FtsK-> | 1097 | Bacteria>Pseudomonadota>Alphaproteobacteria>Hyphomonadales>Hyphomonadaceae | GCA_020638455.1 | Hyphomonas_sp. | H6847_14020 | 101452..104745 | 87 | ATP-binding-protein-[Hyphomonas-sp.] |
| WP_188873657.1 | HEPN+AAA-ATPase+Alpha+Beta-domain+wHTH+RRM | Type-1_BR.RM.like.systems | PAPS_reduct->HTH+MPTase->PIN-><-MalT<-HEPN<-HEPN<-HEPN+DUF499*<-TIR<-METHYLASE<-?<-TUDOR+SF1-SF2-Helicase+iREase+CTD+REase<-XIS-HTH\|\|DOC+HTH-> | 1098 | Bacteria>Pseudomonadota>Alphaproteobacteria>Iodidimonadales>Iodidimonadaceae | GCF_014647255.1 | Iodidimonas_muriae | IEX96_RS06665 | 363502..366798 | 261467 | DUF499-domain-containing-protein-[Iodidimonas-muriae] |
| GER06671.1 | HEPN+AAA-ATPase+Alpha+Beta-domain+wHTH+RRM | Type-1_BR.RM.like.systems | <-DOC+HTH\|\|XIS-HTH->TUDOR+SF1-SF2-Helicase+iREase+CTD+REase->?->METHYLASE->TIR->HEPN+DUF499*->HEPN->HEPN->MalT-><-PIN<-HTH+MPTase<-PAPS_reduct | 1074 | Bacteria>Pseudomonadota>Alphaproteobacteria>Kordiimonadales>unclassified_Kordiimonadales | GCA_008579105.1 | Kordiimonadales_bacterium_JCM_17843 | JCM17843_09810 | 11822..15046 | 1236965 | hypothetical-protein-JCM17843_09810-[Kordiimonadales-bacterium-JCM-17843] |
| MBF0262096.1 | HEPN+AAA-ATPase+Alpha+Beta-domain+wHTH+RRM | Type-1_BR.RM.like.systems | XIS-HTH->TUDOR+SF1-SF2-Helicase+iREase+CTD+REase->SbcC->OLD-ABC->TOPRIM->METHYLASE->HEPN+DUF499*-> | 1048 | Bacteria>Pseudomonadota>Alphaproteobacteria>Magnetococcales>unclassified_Magnetococcales | GCA_015231775.1 | Magnetococcales_bacterium | HQL97_09720 | 11242..14384 | 2026759 | DUF499-domain-containing-protein-[Magnetococcales-bacterium] |
| MBF0108101.1 | HEPN+AAA-ATPase+Alpha+Beta-domain+wHTH+RRM | Type-1_BR.RM.like.systems | PRTase->?->?->?-><-HEPN+DUF499*<-METHYLASE | 1102 | Bacteria>Pseudomonadota>Alphaproteobacteria>Magnetococcales>unclassified_Magnetococcales | GCA_015228815.1 | Magnetococcales_bacterium | HQL76_02845 | 103029..106337 | 2026759 | DUF499-domain-containing-protein-[Magnetococcales-bacterium] |
| GGY37139.1 | HEPN+AAA-ATPase+Alpha+Beta-domain+wHTH+RRM | Type-1_BR.RM.like.systems | <-CopG/RHH<-VirD4-TraG+VirD4-FtsK\|\|HTH+WYL->TUDOR+SF1-SF2-Helicase+iREase+CTD+REase->METHYLASE->HEPN+DUF499*-><-?\|\|?-><-XIS-HTH<-?<-?<-ParB+HTH | 1097 | Bacteria>Pseudomonadota>Alphaproteobacteria>Parvularculales>Parvularculaceae | GCA_014651915.1 | Parvularcula_lutaonensis | GCM10007148_01670 | 175474..178767 | 491923 | hypothetical-protein-GCM10007148_01670-[Parvularcula-lutaonensis] |

| Protein-ID | DUF499-architecture | System | Gene-Neighbourhoods | Length | Taxonomy | Assembly-ID | Strain | Locus-Tag | Genomic-coordinates | TaxID | Fasta-defline |
|---|---|---|---|---|---|---|---|---|---|---|---|
| WP_229786002.1 | HEPN+AAA-ATPase+Alpha+Beta-domain+wHTH+RRM | Type-1_BR.RM.like.systems | <-CopG/RHH<-VirD4-TraG+VirD4-FtsK\|\|HTH+WYL->TUDOR+SF1-SF2-Helicase+iREase+CTD+REase->METHYLASE->HEPN+DUF499*-><-?<-?<-XIS-HTH<-?<-?<-ParB+HTH | 1085 | Bacteria>Pseudomonadota>Alphaproteobacteria>Parvularculales>Parvularculaceae | GCF_014651915.1 | Parvularcula_lutaonensis | IE053_RS00820 | 175510..178767 | 491923 | DUF499-domain-containing-protein-[Parvularcula-lutaonensis] |
| WP_041534905.1 | HEPN+AAA-ATPase+Alpha+Beta-domain+wHTH+RRM | Type-1_BR.RM.like.systems | <-HEPN+DUF499*<-METHYLASE<-MORC+DUF3684<-TUDOR+SF1-SF2-Helicase+iREase+CTD+REase<-HTH+WYL\|\|REase+PNTD1+PNTD2->PIWI-> | 1102 | Bacteria>Pseudomonadota>Alphaproteobacteria>Parvularculales>Parvularculaceae | GCF_000152825.2 | Parvularcula_bermudensis | PB2503_RS05500 | 1154188..1157496 | 208216 | DUF499-domain-containing-protein-[Parvularcula-bermudensis] |
| ADM09268.1 | HEPN+AAA-ATPase+Alpha+Beta-domain+wHTH+RRM | Type-1_BR.RM.like.systems | <-HEPN+DUF499*<-METHYLASE<-MORC+DUF3684<-TUDOR+SF1-SF2-Helicase+iREase+CTD+REase<-HTH+WYL\|\|REase+PNTD1+PNTD2->PIWI-> | 1090 | Bacteria>Pseudomonadota>Alphaproteobacteria>Parvularculales>Parvularculaceae | GCA_000152825.2 | Parvularcula_bermudensis_HTCC2503 | PB2503_05987 | 1154188..1157460 | 314260 | hypothetical-protein-PB2503_05987-[Parvularcula-bermudensis-HTCC2503] |
| WP_255330034.1 | HEPN+AAA-ATPase+Alpha+Beta-domain+wHTH+RRM | Type-1_BR.RM.like.systems | MODE-HTH+PBPII->\|\|?->GDResolvase+Recombinase+ZnR->\|\|DDE_Tnp_IS66->MODE-HTH+PBPII-><-HEPN+DUF499*<-METHYLASE<-ParB+HNH<-TUDOR+SF1-SF2-Helicase+iREase+CTD+REase<-HTH+WYL\|\|SF1-SF2-Helicase+REase->SF1-SF2-Helicase+REase-> | 1097 | Bacteria>Pseudomonadota>Alphaproteobacteria>Rhodobacterales>Paracoccaceae | GCF_024359525.1 | Paracoccus_albicereus | MLD63_RS11355 | 50725..54018 | 2922394 | DUF499-domain-containing-protein-[Paracoccus-sp.-TK19116] |
| MCQ0971011.1 | HEPN+AAA-ATPase+Alpha+Beta-domain+wHTH+RRM | Type-1_BR.RM.like.systems | MODE-HTH+PBPII->\|\|?->GDResolvase+Recombinase+ZnR->\|\|DDE_Tnp_IS66->MODE-HTH+PBPII-><-HEPN+DUF499*<-METHYLASE<-ParB+HNH<-TUDOR+SF1-SF2-Helicase+iREase+CTD+REase<-HTH+WYL\|\|SF1-SF2-Helicase+REase->SF1-SF2-Helicase+REase-> | 1097 | Bacteria>Pseudomonadota>Alphaproteobacteria>Rhodobacterales>Paracoccaceae | GCA_024359525.1 | Paracoccus_albicereus | MLD63_11305 | 50725..54018 | 2922394 | DUF499-domain-containing-protein-[Paracoccus-sp.-TK19116] |
| WP_035746761.1 | HEPN+AAA-ATPase+Alpha+Beta-domain+wHTH+RRM | Type-1_BR.RM.like.systems | <-DDE_Tnp_1_5+DDE_Tnp_1<-HTH_Tnp_1<-HTH_Tnp_1<-HEPN+DUF499*<-METHYLASE<-ParB+HNH<-TUDOR+SF1-SF2-Helicase+iREase+CTD+REase<-HTH+WYL\|\|SF1-SF2-Helicase+REase->SF1-SF2-Helicase+REase-> | 1098 | Bacteria>Pseudomonadota>Alphaproteobacteria>Rhodobacterales>Paracoccaceae | GCF_000740775.1 | Haematobacter_missouriensis | CG51_RS16225 | 8779..12075 | 366616 | DUF499-domain-containing-protein-[Haematobacter-missouriensis] |
| WP_105291560.1 | HEPN+AAA-ATPase+Alpha+Beta-domain+wHTH+RRM | Type-1_BR.RM.like.systems | <-SbcC+SbcC<-Calcineurin\|\|HTH+WYL->TUDOR+SF1-SF2-Helicase+iREase+CTD+REase->?->METHYLASE->HEPN+DUF499*->SbcC->?->RelE-ParE-><-?<-DUF4096+DDE_Tnp_1+DDE_Tnp<-DEDD_Tnp_IS110+Transposase_20 | 1101 | Bacteria>Pseudomonadota>Alphaproteobacteria>Rhodobacterales>Paracoccaceae | GCF_002073635.2 | Paracoccus_yeei | A6J80_RS16220 | 2818181..2821486 | 147645 | DUF499-domain-containing-protein-[Paracoccus-yeei] |
| WP_108644491.1 | HEPN+AAA-ATPase+Alpha+Beta-domain+wHTH+RRM | Type-1_BR.RM.like.systems | <-HNH<-ParB+N6_N4_MTase\|\|TUDOR+SF1-SF2-Helicase+iREase+CTD+REase->HEPN->METHYLASE->HEPN+DUF499*->?-><-LZ_Tnp_IS66+zf-IS66+DDE_Tnp_IS66+DDE_Tnp_IS66_C<-?<-HIN-HTH<-?<-MarR-HTH+REase | 1103 | Bacteria>Pseudomonadota>Alphaproteobacteria>Rhodobacterales>Paracoccaceae | GCF_003071405.1 | Fuscovulum_blasticum | B6K69_RS15915 | 3260938..3264249 | 1075 | DUF499-domain-containing-protein-[Fuscovulum-blasticum] |
| WP_233253227.1 | HEPN+AAA-ATPase+Alpha+Beta-domain+wHTH+RRM | Type-1_BR.RM.like.systems | ParG_RHH-><-TetR-HTH\|\|MODE-HTH+PBPII-><-HEPN+DUF499*<-METHYLASE<-ASCH<-TUDOR+SF1-SF2-Helicase+iREase+CTD+REase<-HTH+WYL\|\|SF1-SF2-Helicase+REase->SF1-SF2-Helicase+REase-> | 1086 | Bacteria>Pseudomonadota>Alphaproteobacteria>Rhodobacterales>Paracoccaceae | GCF_016446475.1 | Paracoccus_binzhouensis | I9P22_RS04960 | 53766..57026 | 2796149 | DUF499-domain-containing-protein-[Paracoccus-binzhouensis] |
| WP_088233907.1 | HEPN+AAA-ATPase+Alpha+Beta-domain+wHTH+RRM | Type-1_BR.RM.like.systems | <-SF1-SF2-Helicase+REase<-SF1-SF2-Helicase+REase\|\|HTH+WYL->TUDOR+SF1-SF2-Helicase+iREase+CTD+REase->ASCH->METHYLASE->HEPN+DUF499*-><-MODE-HTH+PBPII\|\|TetR-HTH-><-ParG_RHH | 1098 | Bacteria>Pseudomonadota>Alphaproteobacteria>Rhodobacterales>Paracoccaceae | GCF_943915015.1 | Haematobacter_missouriensis | CDV52_RS16275 | 17208..20504 | 366616 | DUF499-domain-containing-protein-[Haematobacter-missouriensis] |
| WP_058096461.1 | HEPN+AAA-ATPase+Alpha+Beta-domain+wHTH+RRM | Type-1_BR.RM.like.systems | <-SF1-SF2-Helicase+REase<-SF1-SF2-Helicase+REase\|\|HTH+WYL->TUDOR+SF1-SF2-Helicase+iREase+CTD+REase->MORC+DUF3684->METHYLASE->HEPN+DUF499*->MODE-HTH+PBPII-><-ParG_RHH | 1098 | Bacteria>Pseudomonadota>Alphaproteobacteria>Rhodobacterales>Paracoccaceae | GCF_001447385.1 | Paracoccus_sp._MKU1 | AQY21_RS03730 | 27830..31126 | 1745182 | DUF499-domain-containing-protein-[Paracoccus-sp.-MKU1] |
| WP_189381997.1 | HEPN+AAA-ATPase+Alpha+Beta-domain+wHTH+RRM | Type-1_BR.RM.like.systems | <-ParB+N6_N4_MTase<-ParB+N6_N4_MTase\|\|TUDOR+SF1-SF2-Helicase+iREase+CTD+REase->MACRODOMAIN->METHYLASE->?->HEPN+DUF499*-><-Enterotoxin_a<-?<-MarR-HTH+REase<-?<-?<-XIS-HTH | 1099 | Bacteria>Pseudomonadota>Alphaproteobacteria>Rhodobacterales>Paracoccaceae | GCF_014652195.1 | Gemmobacter_nanjingensis | IE126_RS20360 | 40653..43952 | 488454 | DUF499-domain-containing-protein-[Gemmobacter-nanjingensis] |
| GIX14656.1 | HEPN+AAA-ATPase+Alpha+Beta-domain+wHTH+RRM | Type-1_BR.RM.like.systems | TUDOR+SF1-SF2-Helicase+iREase+CTD+REase->TUDOR+SF1-SF2-Helicase+iREase+CTD+REase-p->METHYLASE-p->METHYLASE-p->HEPN+DUF499*-><-?<-?<-?\|\|MULE-TRANSPOSASE-> | 1104 | Bacteria>Pseudomonadota>Alphaproteobacteria>Rhodobacterales>Paracoccaceae | GCA_026004915.1 | Paracoccaceae_bacterium | KatS3mg118_2615 | 175796..179110 | 1904441 | MAG:-hypothetical-protein-KatS3mg118_2615-[Paracoccaceae-bacterium] |
| WP_149139641.1 | HEPN+AAA-ATPase+Alpha+Beta-domain+wHTH+RRM | Type-1_BR.RM.like.systems | HEPN-><-ParB+N6_N4_MTase<-ParB+N6_N4_MTase\|\|TUDOR+SF1-SF2-Helicase+iREase+CTD+REase->ASCH->METHYLASE->HEPN+DUF499*-><-Enterotoxin_a<-?<-MarR-HTH+REase<-?<-?<-XIS-HTH | 1100 | Bacteria>Pseudomonadota>Alphaproteobacteria>Rhodobacterales>Paracoccaceae | GCF_008271655.1 | Gemmobacter_caeruleus | FOA17_RS00925 | 40432..43734 | 2595004 | DUF499-domain-containing-protein-[Gemmobacter-caeruleus] |
| WP_036708698.1 | HEPN+AAA-ATPase+Alpha+Beta-domain+wHTH+RRM | Type-1_BR.RM.like.systems | XIS-HTH->?->?->MarR-HTH+REase->?->Enterotoxin_a-><-HEPN+DUF499*<-METHYLASE<-REase<-TUDOR+SF1-SF2-Helicase+iREase+CTD+REase\|\|ParB+N6_N4_MTase->ParB+N6_N4_MTase-><-HNH | 1100 | Bacteria>Pseudomonadota>Alphaproteobacteria>Rhodobacterales>Paracoccaceae | GCF_000763845.1 | Paracoccus_sanguinis | IX56_RS07055 | 10378..13680 | 1545044 | DUF499-domain-containing-protein-[Paracoccus-sanguinis] |
| KPP88834.1 | HEPN+AAA-ATPase+Alpha+Beta-domain+wHTH+RRM | Type-1_BR.RM.like.systems | VirD4-TraG+VirD4-FtsK->CopG/RHH->TUDOR+SF1-SF2-Helicase+iREase+CTD+REase->PadR-HTH->METHYLASE->HEPN+DUF499*-><-PIN<-RHH<-?<-CHTH\|\|VirB11+VirB4-FtsK->VirB2-> | 1100 | Bacteria>Pseudomonadota>Alphaproteobacteria>Rhodobacterales>Paracoccaceae | GCA_001314655.1 | Rhodobacteraceae_bacterium_HLUCCA08 | HLUCCA08_13590 | 19280..22582 | 1666913 | putative-ATPase-(AAA+-superfamily)-[Rhodobacteraceae-bacterium-HLUCCA08] |
| WP_096787622.1 | HEPN+AAA-ATPase+Alpha+Beta-domain+wHTH+RRM | Type-1_BR.RM.like.systems | METHYLASE-p-><-?\|\|?-><-SF1-SF2-Helicase+DUF1998+HKD-DNase+HKD-DNase<-?<-HEPN+DUF499*<-METHYLASE<-MACRODOMAIN<-TUDOR+SF1-SF2-Helicase+iREase+CTD+REase\|\|?->HTH+WYL | 1100 | Bacteria>Pseudomonadota>Alphaproteobacteria>Rhodobacterales>Paracoccaceae | GCF_002407205.1 | Rhodobacter_sp._CZR27 | CK951_RS18110 | 425347..428649 | 2033869 | DUF499-domain-containing-protein-[Rhodobacter-sp.-CZR27] |
| MCY4445744.1 | HEPN+AAA-ATPase+Alpha+Beta-domain+wHTH+RRM | Type-1_BR.RM.like.systems | <-Arm-DNA-bind_3+Phage_int_SAM_3+Phage_integrase<-Phage_int_SAM_3+Phage_integrase<-TUDOR+SF1-SF2-Helicase+iREase+CTD+REase<-HEPN+DUF499*\|\|METHYLASE-p-> | 1118 | Bacteria>Pseudomonadota>Alphaproteobacteria>Rhodobacterales>Paracoccaceae | GCA_026712645.1 | Paracoccaceae_bacterium | OXC02_04685 | 7553..10909 | 1904441 | MAG:-DUF499-domain-containing-protein-[Rhodobacteraceae-bacterium] |
| WP_099275178.1 | HEPN+AAA-ATPase+Alpha+Beta-domain+wHTH+RRM | Type-1_BR.RM.like.systems | rve_3+Mu-transpos_C->?-><-?<-HEPN+DUF499*<-METHYLASE<-ASCH<-TUDOR+SF1-SF2-Helicase+iREase+CTD+REase\|\|HIN-HTH->?->SF1-SF2-Helicase+DUF1998+HKD-DNase+HKD-DNase-> | 1110 | Bacteria>Pseudomonadota>Alphaproteobacteria>Rhodobacterales>Paracoccaceae | GCF_002723615.1 | Limimaricola_cinnabarinus | CJ301_RS05690 | 226587..229919 | 1125964 | DUF499-domain-containing-protein-[Limimaricola-cinnabarinus] |
| MDC0095869.1 | HEPN+AAA-ATPase+Alpha+Beta-domain+wHTH+RRM | Type-1_BR.RM.like.systems | XIS-HTH->?->TUDOR+SF1-SF2-Helicase+iREase+CTD+REase->METHYLASE->METHYLASE->HEPN+DUF499*-><-URI-DNase<-RNase35exo<-RNase+SF1-SF2-Helicase\|\|?->?->HTH-> | 1097 | Bacteria>Pseudomonadota>Alphaproteobacteria>Rhodobacterales>Paracoccaceae | GCA_028342995.1 | Amylibacter_sp. | OAI58_06255 | 84173..87466 | 1872504 | DUF499-domain-containing-protein-[Amylibacter-sp.] |
| WP_276946692.1 | HEPN+AAA-ATPase+Alpha+Beta-domain+wHTH+RRM | Type-1_BR.RM.like.systems | ParG_RHH-><-TetR-HTH\|\|MODE-HTH+PBPII-><-HEPN+DUF499*<-METHYLASE<-ASCH<-TUDOR+SF1-SF2-Helicase+iREase+CTD+REase<-HTH+WYL\|\|SMODS->UbE1+JAB-> | 1098 | Bacteria>Pseudomonadota>Alphaproteobacteria>Rhodobacterales>Paracoccaceae | GCF_946222465.1 | Haematobacter_massiliensis | P6966_RS06770 | 8482..11778 | 195105 | DUF499-domain-containing-protein-[Haematobacter-massiliensis] |
| WP_152367850.1 | HEPN+AAA-ATPase+Alpha+Beta-domain+wHTH+RRM | Type-1_BR.RM.like.systems | <-JAB<-UbE1+JAB<-SMODS\|\|HTH+WYL->TUDOR+SF1-SF2-Helicase+iREase+CTD+REase->METHYLASE->HEPN+DUF499*->?-><-MODE-HTH+PBPII\|\|TetR-HTH-><-ParG_RHH | 1099 | Bacteria>Pseudomonadota>Alphaproteobacteria>Rhodobacterales>Paracoccaceae | GCF_009301415.1 | Paracoccus_kondratievae | F8A10_RS15730 | 901669..904968 | 135740 | DUF499-domain-containing-protein-[Paracoccus-kondratievae] |
| WP_108127680.1 | HEPN+AAA-ATPase+Alpha+Beta-domain+wHTH+RRM | Type-1_BR.RM.like.systems | CopG/RHH->TUDOR+SF1-SF2-Helicase+iREase+CTD+REase->PadR-HTH->METHYLASE->HEPN+DUF499*->DUF4011+SF1-SF2-Helicase+REase-><-?\|\|?->?->NTP_transf-> | 1112 | Bacteria>Pseudomonadota>Alphaproteobacteria>Rhodobacterales>Paracoccaceae | GCF_003054195.1 | Gemmobacter_caeni | C8N34_RS03245 | 88881..92219 | 589035 | DUF499-domain-containing-protein-[Gemmobacter-caeni] |
| WP_209427763.1 | HEPN+AAA-ATPase+Alpha+Beta-domain+wHTH+RRM | Type-1_BR.RM.like.systems | <-AbiEii<-HEPN+DUF499*<-METHYLASE<-MORC+DUF3684<-TUDOR+SF1-SF2-Helicase+iREase+CTD+REase<-HTH+WYL<-Phage_int_SAM_3+Phage_integrase | 1112 | Bacteria>Pseudomonadota>Alphaproteobacteria>Rhodobacterales>Paracoccaceae | GCF_017811755.1 | Pararhodobacter_sp._SW119 | IM708_RS16910 | 1952..5290 | 2780075 | Swt1-family-HEPN-domain-containing-protein-[Pararhodobacter-sp.-SW119] |
| WP_118137057.1 | HEPN+AAA-ATPase+Alpha+Beta-domain+wHTH+RRM | Type-1_BR.RM.like.systems | <-UbE1+JAB<-?\|\|HTH+WYL->TUDOR+SF1-SF2-Helicase+iREase+CTD+REase->?->METHYLASE->HEPN+DUF499*-><-?\|\|REase->SF1-SF2-Helicase+REase->HNH->?->DOC-> | 1110 | Bacteria>Pseudomonadota>Alphaproteobacteria>Rhodobacterales>Paracoccaceae | GCF_003448965.1 | Oceanicella_sp._SM1341 | D3P35_RS19735 | 60544..63876 | 1548889 | DUF499-domain-containing-protein-[Oceanicella-sp.-SM1341] |
| MDB4017580.1 | HEPN+AAA-ATPase+Alpha+Beta-domain+wHTH+RRM | Type-1_BR.RM.like.systems | XIS-HTH->?->TUDOR+SF1-SF2-Helicase+iREase+CTD+REase->METHYLASE->METHYLASE->HEPN+DUF499*-><-URI-DNase<-RNase35exo<-Schlafen\|\|?->?->HTH-> | 1097 | Bacteria>Pseudomonadota>Alphaproteobacteria>Rhodobacterales>Paracoccaceae | GCA_028244795.1 | Amylibacter_sp. | N9493_06125 | 26402..29695 | 1872504 | DUF499-domain-containing-protein-[Amylibacter-sp.] |
| WP_181746701.1 | HEPN+AAA-ATPase+Alpha+Beta-domain+wHTH+RRM | Type-1_BR.RM.like.systems | NAD_binding_2+6PGD-><-?\|\|TUDOR+SF1-SF2-Helicase+iREase+CTD+REase->HEPN->METHYLASE->?->HEPN+DUF499*-><-Enterotoxin_a<-?<-?<-?<-Peptidase_M15 | 1099 | Bacteria>Pseudomonadota>Alphaproteobacteria>Rhodobacterales>Paracoccaceae | GCF_947502345.1 | Paracoccus_sp._S1E-3 | Q0821_RS11150 | 38877..42176 | 2756130 | DUF499-domain-containing-protein-[Paracoccus-sp.-S1E-3] |
| WP_094020052.1 | HEPN+AAA-ATPase+Alpha+Beta-domain+wHTH+RRM | Type-1_BR.RM.like.systems | XIS-HTH->HTH+WYL->ParB+HNH->TUDOR+SF1-SF2-Helicase+iREase+CTD+REase->?->METHYLASE->HEPN+DUF499*-><-RNase<-?<-?<-DUF2163+Phage_BR0599 | 1098 | Bacteria>Pseudomonadota>Alphaproteobacteria>Rhodobacterales>Paracoccaceae | GCF_900184945.1 | Maliponia_aquimaris | MAA8898_RS05760 | 37634..40930 | 1673631 | DUF499-domain-containing-protein-[Maliponia-aquimaris] |
| WP_259776874.1 | HEPN+AAA-ATPase+Alpha+Beta-domain+wHTH+RRM | Type-1_BR.RM.like.systems | REase->?->HTH+WYL->TUDOR+SF1-SF2-Helicase+iREase+CTD+REase->?->METHYLASE->HEPN+DUF499*->ArdCN+MPTase->?->?->?->?->VirD4-TraG+VirD4-FtsK-> | 1112 | Bacteria>Pseudomonadota>Alphaproteobacteria>Rhodobacterales>Paracoccaceae | GCF_025138535.1 | Aliiroseovarius_crassostreae | K3X25_RS03910 | 801884..805222 | 154981 | DUF499-domain-containing-protein-[Aliiroseovarius-crassostreae] |
| WP_188408468.1 | HEPN+AAA-ATPase+Alpha+Beta-domain+wHTH+RRM | Type-1_BR.RM.like.systems | HTH+WYL->TUDOR+SF1-SF2-Helicase+iREase+CTD+REase->ASCH->YtH->METHYLASE->HEPN+DUF499*->ParB+HTH->HTH_3->STYKinase-like-> | 1102 | Bacteria>Pseudomonadota>Alphaproteobacteria>Rhodobacterales>Paracoccaceae | GCF_014635245.1 | Agaricicola_taiwanensis | IEV58_RS04465 | 943958..947266 | 591372 | DUF499-domain-containing-protein-[Agaricicola-taiwanensis] |
| WP_136646565.1 | HEPN+AAA-ATPase+Alpha+Beta-domain+wHTH+RRM | Type-1_BR.RM.like.systems | DUF2163+Phage_BR0599->?->?->RNase-><-DUF4236+TPR<-HEPN+DUF499*<-METHYLASE<-PadR-HTH<-TUDOR+SF1-SF2-Helicase+iREase+CTD+REase | 1097 | Bacteria>Pseudomonadota>Alphaproteobacteria>Rhodobacterales>Paracoccaceae | GCF_004959785.1 | Tabrizicola_sp._YIM_78059 | EZM81_RS16965 | 12465..15758 | 2529861 | DUF499-domain-containing-protein-[Tabrizicola-sp.-YIM-78059] |
| MCC5984903.1 | HEPN+AAA-ATPase+Alpha+Beta-domain+wHTH+RRM | Type-1_BR.RM.like.systems | <-HEPN+DUF499*<-METHYLASE<-REase<-TUDOR+SF1-SF2-Helicase+iREase+CTD+REase<-CopG/RHH<-VirD4-TraG+VirD4-FtsK\|\|HNH-> | 351 | Bacteria>Pseudomonadota>Alphaproteobacteria>Rhodobacterales>Paracoccaceae | GCA_020832705.1 | Paracoccaceae_bacterium | JJU42_11110 | 1..1055 | 1904441 | AAA-family-ATPase-[Paracoccaceae-bacterium] |
| WP_104009245.1 | HEPN+AAA-ATPase+Alpha+Beta-domain+wHTH+RRM | Type-1_BR.RM.like.systems | XIS-HTH->?->Primpol->TUDOR+SF1-SF2-Helicase+iREase+CTD+REase->ASCH->METHYLASE->HEPN+DUF499*-><-?<-Peptidase_M15<-?<-RNase | 1104 | Bacteria>Pseudomonadota>Alphaproteobacteria>Rhodobacterales>Paracoccaceae | GCF_900108275.1 | Jhaorihella_thermophila | C4S70_RS16725 | 13940..17254 | 488547 | DUF499-domain-containing-protein-[Jhaorihella-thermophila] |
| WP_213544567.1 | HEPN+AAA-ATPase+Alpha+Beta-domain+wHTH+RRM | Type-1_BR.RM.like.systems | TUDOR+SF1-SF2-Helicase+iREase+CTD+REase->?->METHYLASE->HEPN+DUF499*-><-?<-?<-XIS-HTH<-?<-Arm-DNA-bind_3+Phage_integrase | 1098 | Bacteria>Pseudomonadota>Alphaproteobacteria>Rhodobacterales>Paracoccaceae | GCF_018424695.1 | Vannielia_litorea | DYI42_RS03660 | 12366..15662 | 1217970 | DUF499-domain-containing-protein-[Vannielia-litorea] |
| WP_234869545.1 | HEPN+AAA-ATPase+Alpha+Beta-domain+wHTH+RRM | Type-1_BR.RM.like.systems | <-SF1-SF2-Helicase+DUF1998<-?\|\|?-><-?<-HEPN+DUF499*<-METHYLASE<-wHTH+REase<-TUDOR+SF1-SF2-Helicase+iREase+CTD+REase | 1100 | Bacteria>Pseudomonadota>Alphaproteobacteria>Rhodobacterales>Paracoccaceae | GCF_021491355.1 | Tabrizicola_rongguiensis | L0V05_RS07760 | 379884..383186 | 2909234 | DUF499-domain-containing-protein-[Tabrizicola-rongguiensis] |
| WP_017929317.1 | HEPN+AAA-ATPase+Alpha+Beta-domain+wHTH+RRM | Type-1_BR.RM.like.systems | <-HEPN+DUF499*<-METHYLASE<-ASCH<-TUDOR+SF1-SF2-Helicase+iREase+CTD+REase<-?<-?<-SF1-SF2-Helicase-p | 1110 | Bacteria>Pseudomonadota>Alphaproteobacteria>Rhodobacterales>Paracoccaceae | GCF_000365005.1 | Limimaricola_hongkongensis | A3EY_RS0112135 | 165688..169020 | 278132 | DUF499-domain-containing-protein-[Limimaricola-hongkongensis] |
| WP_090617184.1 | HEPN+AAA-ATPase+Alpha+Beta-domain+wHTH+RRM | Type-1_BR.RM.like.systems | <-HEPN+DUF499*<-METHYLASE<-ASCH<-TUDOR+SF1-SF2-Helicase+iREase+CTD+REase<-HTH+WYL<-?<-TetR-HTH | 1110 | Bacteria>Pseudomonadota>Alphaproteobacteria>Rhodobacterales>Paracoccaceae | GCF_028553725.1 | Paracoccus_alcaliphilus | BMW58_RS19895 | 488..3820 | 34002 | DUF499-domain-containing-protein-[Paracoccus-alcaliphilus] |
| WP_072748400.1 | HEPN+AAA-ATPase+Alpha+Beta-domain+wHTH+RRM | Type-1_BR.RM.like.systems | <-HEPN+DUF499*<-METHYLASE-p<-METHYLASE<-ASCH<-TUDOR+SF1-SF2-Helicase+iREase+CTD+REase<-?<-tRNA<-HEPN | 1104 | Bacteria>Pseudomonadota>Alphaproteobacteria>Rhodobacterales>Paracoccaceae | GCF_900111545.1 | Oceanicella_actignis | BMW67_RS14465 | 1463..4777 | 1189325 | DUF499-domain-containing-protein-[Oceanicella-actignis] |
| WP_139597753.1 | HEPN+AAA-ATPase+Alpha+Beta-domain+wHTH+RRM | Type-1_BR.RM.like.systems | <-RNase\|\|TUDOR+SF1-SF2-Helicase+iREase+CTD+REase->?->METHYLASE->HEPN+DUF499*->REase-><-PIN | 1100 | Bacteria>Pseudomonadota>Alphaproteobacteria>Rhodobacterales>Paracoccaceae | GCF_006239215.1 | Paracoccus_haeundaensis | FHD67_RS02835 | 188317..191619 | 225362 | DUF499-domain-containing-protein-[Paracoccus-haeundaensis] |
| WP_189635417.1 | HEPN+AAA-ATPase+Alpha+Beta-domain+wHTH+RRM | Type-1_BR.RM.like.systems | <-HEPN+DUF499*<-?<-METHYLASE<-?<-TUDOR+SF1-SF2-Helicase+iREase+CTD+REase\|\|ParB+N6_N4_MTase-> | 1099 | Bacteria>Pseudomonadota>Alphaproteobacteria>Rhodobacterales>Paracoccaceae | GCF_014652355.1 | Gemmobacter_lanyuensis | IE101_RS18950 | 648..3947 | 1054497 | DUF499-domain-containing-protein-[Gemmobacter-lanyuensis] |
| WP_154336328.1 | HEPN+AAA-ATPase+Alpha+Beta-domain+wHTH+RRM | Type-1_BR.RM.like.systems | <-ABC-ATPase+SbcC<-?<-HEPN+DUF499*<-METHYLASE<-?<-TUDOR+SF1-SF2-Helicase+iREase+CTD+REase | 1097 | Bacteria>Pseudomonadota>Alphaproteobacteria>Rhodobacterales>Paracoccaceae | GCF_009674885.1 | Paracoccus_sp._S-4012 | GI374_RS07045 | 35979..39272 | 2665648 | DUF499-domain-containing-protein-[Paracoccus-sp.-S-4012] |
| MDE0346579.1 | HEPN+AAA-ATPase+Alpha+Beta-domain+wHTH+RRM | Type-1_BR.RM.like.systems | <-HEPN+DUF499*<-PIN<-DUF499-p<-METHYLASE<-TUDOR+SF1-SF2-Helicase+iREase+CTD+REase-p | 1102 | Bacteria>Pseudomonadota>Alphaproteobacteria>Rhodobacterales>Paracoccaceae | GCA_028821095.1 | Boseongicola_sp. | OXI66_12500 | 695..4003 | 1886600 | MAG:-DUF499-domain-containing-protein-[Boseongicola-sp.] |
| MCY4460275.1 | HEPN+AAA-ATPase+Alpha+Beta-domain+wHTH+RRM | Type-1_BR.RM.like.systems | <-HEPN+DUF499*<-Schlafen+HATPase+HTH<-METHYLASE<-TUDOR+SF1-SF2-Helicase+iREase+CTD+REase | 1099 | Bacteria>Pseudomonadota>Alphaproteobacteria>Rhodobacterales>Paracoccaceae | GCA_026712665.1 | Albidovulum_sp. | OXC26_07750 | 691..3990 | 1872424 | MAG:-DUF499-domain-containing-protein-[Albidovulum-sp.] |
| WP_119887324.1 | HEPN+AAA-ATPase+Alpha+Beta-domain+wHTH+RRM | Type-1_BR.RM.like.systems | TUDOR+SF1-SF2-Helicase+iREase+CTD+REase->?->METHYLASE->HEPN+DUF499*-> | 1100 | Bacteria>Pseudomonadota>Alphaproteobacteria>Rhodobacterales>Paracoccaceae | GCF_028553885.1 | Paracoccus_aestuarii | D3P06_RS15025 | 7971..11273 | 453842 | DUF499-domain-containing-protein-[Paracoccus-aestuarii] |
| MBC6443273.1 | HEPN+AAA-ATPase+Alpha+Beta-domain+wHTH+RRM | Type-1_BR.RM.like.systems | Glycos_transf_4->?->HEPN+DUF499*->?-><-ParB+HNH\|\|?->?->ABC-ATPase+SbcC-> | 1096 | Bacteria>Pseudomonadota>Alphaproteobacteria>Rhodobacterales>Paracoccaceae | GCA_014323795.1 | Paracoccaceae_bacterium | GDA53_09225 | 19361..22651 | 1904441 | ATP-binding-protein-[Rhodobacteraceae-bacterium] |
| MBC6437846.1 | HEPN+AAA-ATPase+Alpha+Beta-domain+wHTH+RRM | Type-1_BR.RM.like.systems | <-ABC-ATPase+SbcC\|\|?->ncRNA->?-><-HEPN+DUF499*<-METHYLASE-p | 1099 | Bacteria>Pseudomonadota>Alphaproteobacteria>Rhodobacterales>Paracoccaceae | GCA_014323785.1 | Paracoccaceae_bacterium | GDA52_06875 | 42024..45323 | 1904441 | ATP-binding-protein-[Rhodobacteraceae-bacterium] |
| MBE2259559.1 | HEPN+AAA-ATPase+Alpha+Beta-domain+wHTH+RRM | Type-1_BR.RM.like.systems | <-HEPN+DUF499*<-HEPN<-PolB<-METHYLASE-p | 485 | Bacteria>Pseudomonadota>Alphaproteobacteria>Rhodobacterales>Paracoccaceae | GCA_014879755.1 | Paracoccaceae_bacterium | IAE88_11910 | 1..1455 | 1904441 | ATP-binding-protein-[Paracoccaceae-bacterium] |
| WP_007798461.1 | HEPN+AAA-ATPase+Alpha+Beta-domain+wHTH+RRM | Type-1_BR.RM.like.systems | <-Peptidase_S11<-?\|\|tRNA->TUDOR+SF1-SF2-Helicase+iREase+CTD+REase->TUDOR+SF1-SF2-Helicase+iREase+CTD+REase-p->?->METHYLASE->HEPN+DUF499*-><-DDE_Tnp_1<-SF1-SF2-Helicase+DUF1998+HKD-DNase+HKD-DNase<-?<-?\|\|?-><-MULE-TRANSPOSASE | 1110 | Bacteria>Pseudomonadota>Alphaproteobacteria>Rhodobacterales>Roseobacteraceae | GCF_000153725.1 | Salipiger_bermudensis | R2601_RS11735 | 2461039..2464371 | 344736 | DUF499-domain-containing-protein-[Salipiger-bermudensis] |
| WP_068316335.1 | HEPN+AAA-ATPase+Alpha+Beta-domain+wHTH+RRM | Type-1_BR.RM.like.systems | <-HIN-HTH<-TPR<-HEPN+SF1-SF2-Helicase<-HIN-HTH\|\|?-><-AbiEii<-HEPN+DUF499*<-METHYLASE<-ASCH<-TUDOR+SF1-SF2-Helicase+iREase+CTD+REase<-Arm-DNA-bind_3+Phage_int_SAM_3+Phage_integrase\|\|TPR-> | 1099 | Bacteria>Pseudomonadota>Alphaproteobacteria>Rhodobacterales>Roseobacteraceae | GCF_001558155.1 | Aliiruegeria_sabulilitoris | AX756_RS20370 | 52266..55565 | 1510458 | DUF499-domain-containing-protein-[Aliiruegeria-sabulilitoris] |
| MBR2572727.1 | HEPN+AAA-ATPase+Alpha+Beta-domain+wHTH+RRM | Type-1_BR.RM.like.systems | <-ParB+N6_N4_MTase<-ParB+N6_N4_MTase\|\|TUDOR+SF1-SF2-Helicase+iREase+CTD+REase->ASCH->METHYLASE->HEPN+DUF499*-><-Enterotoxin_a<-DNA_methylase<-MarR-HTH+REase<-?<-?<-XIS-HTH | 1096 | Bacteria>Pseudomonadota>Alphaproteobacteria>Rhodobacterales>Roseobacteraceae | GCA_017542965.1 | Loktanella_sp. | IKE14_00205 | 39854..43144 | 1870901 | DUF499-domain-containing-protein-[Loktanella-sp.] |
| WP_109663884.1 | HEPN+AAA-ATPase+Alpha+Beta-domain+wHTH+RRM | Type-1_BR.RM.like.systems | VirD4-TraG+VirD4-FtsK->CopG/RHH->TUDOR+SF1-SF2-Helicase+iREase+CTD+REase->HKD-DNase->METHYLASE->HEPN+DUF499*-><-PIN<-RHH<-?<-CHTH\|\|VirB11+VirB4-FtsK->VirB2-> | 1100 | Bacteria>Pseudomonadota>Alphaproteobacteria>Rhodobacterales>Roseobacteraceae | GCF_003148775.1 | Roseicyclus_mahoneyensis | C7455_RS00150 | 34589..37891 | 164332 | DUF499-domain-containing-protein-[Roseicyclus-mahoneyensis] |
| WP_093359600.1 | HEPN+AAA-ATPase+Alpha+Beta-domain+wHTH+RRM | Type-1_BR.RM.like.systems | XIS-HTH->HTH+WYL->ParB+HNH->TUDOR+SF1-SF2-Helicase+iREase+CTD+REase->ASCH->METHYLASE->HEPN+DUF499*->RNase+SF1-SF2-Helicase-><-?<-?<-DUF2163+Phage_BR0599 | 1109 | Bacteria>Pseudomonadota>Alphaproteobacteria>Rhodobacterales>Roseobacteraceae | GCF_900112335.1 | Tropicimonas_isoalkanivorans | BM075_RS05220 | 205092..208421 | 441112 | DUF499-domain-containing-protein-[Tropicimonas-isoalkanivorans] |
| WP_189682671.1 | HEPN+AAA-ATPase+Alpha+Beta-domain+wHTH+RRM | Type-1_BR.RM.like.systems | <-HicB\|\|HTH+WYL->TUDOR+SF1-SF2-Helicase+iREase+CTD+REase->MORC+DUF3684->METHYLASE->HEPN+DUF499*-><-Phage_integrase<-?<-?\|\|?->XIS-HTH-><-Phage_integrase | 1103 | Bacteria>Pseudomonadota>Alphaproteobacteria>Rhodobacterales>Roseobacteraceae | GCF_014656415.1 | Seohaeicola_zhoushanensis | IE266_RS23965 | 15420..18731 | 1569283 | DUF499-domain-containing-protein-[Seohaeicola-zhoushanensis] |
| WP_050673791.1 | HEPN+AAA-ATPase+Alpha+Beta-domain+wHTH+RRM | Type-1_BR.RM.like.systems | Peptidase_M15->?->HTH+WYL->TUDOR+SF1-SF2-Helicase+iREase+CTD+REase->?->METHYLASE->HEPN+DUF499*->ArdCN+MPTase->?->?->?->?->VirD4-TraG+VirD4-FtsK-> | 1112 | Bacteria>Pseudomonadota>Alphaproteobacteria>Rhodobacterales>Roseobacteraceae | GCF_001251095.1 | Phaeobacter_italicus | NIT7321_RS13650 | 220898..224236 | 481446 | DUF499-domain-containing-protein-[Phaeobacter-italicus] |
| WP_026758864.1 | HEPN+AAA-ATPase+Alpha+Beta-domain+wHTH+RRM | Type-1_BR.RM.like.systems | GNTR-HTH->?->?-><-HEPN+DUF499*<-METHYLASE<-METHYLASE-p<-PadR-HTH<-TUDOR+SF1-SF2-Helicase+iREase+CTD+REase\|\|MarR-HTH->ABhydrolase-> | 1111 | Bacteria>Pseudomonadota>Alphaproteobacteria>Rhodobacterales>Roseobacteraceae | GCF_000423645.1 | Sediminimonas_qiaohouensis | G568_RS0115420 | 9768..13103 | 552061 | DUF499-domain-containing-protein-[Sediminimonas-qiaohouensis] |
| WP_170755622.1 | HEPN+AAA-ATPase+Alpha+Beta-domain+wHTH+RRM | Type-1_BR.RM.like.systems | <-REase<-DNA_methylase\|\|HNH-><-?\|\|TUDOR+SF1-SF2-Helicase+iREase+CTD+REase->METHYLASE->HEPN+DUF499*->?->?->?-><-?<-DnaB+CITB-HTH | 1101 | Bacteria>Pseudomonadota>Alphaproteobacteria>Rhodobacterales>Roseobacteraceae | GCF_013031285.1 | Ruegeria_lacuscaerulensis | GOX67_RS18140 | 83323..86628 | 55218 | DUF499-domain-containing-protein-[Ruegeria-lacuscaerulensis] |
| WP_090219105.1 | HEPN+AAA-ATPase+Alpha+Beta-domain+wHTH+RRM | Type-1_BR.RM.like.systems | <-ArdCN+MPTase\|\|TUDOR+SF1-SF2-Helicase+iREase+CTD+REase->?->METHYLASE->HEPN+DUF499*-><-?<-STYKinase-like<-HTH | 1105 | Bacteria>Pseudomonadota>Alphaproteobacteria>Rhodobacterales>Roseobacteraceae | GCF_900111945.1 | Litoreibacter_janthinus | BM352_RS16595 | 3306785..3310102 | 670154 | DUF499-domain-containing-protein-[Litoreibacter-janthinus] |
| PWJ13798.1 | HEPN+AAA-ATPase+Alpha+Beta-domain+wHTH+RRM | Type-1_BR.RM.like.systems | MODE-HTH+PBPII->tRNA->TUDOR+SF1-SF2-Helicase+iREase+CTD+REase->HEPN->METHYLASE-p->HEPN+DUF499*->?-><-?<-?<-?<-?<-XIS-HTH | 1109 | Bacteria>Pseudomonadota>Alphaproteobacteria>Rhodobacterales>Roseobacteraceae | GCA_003149265.1 | Jannaschia_seohaensis | BCF38_11329 | 37396..40725 | 475081 | hypothetical-protein-BCF38_11329-[Jannaschia-seohaensis] |
| WP_240672187.1 | HEPN+AAA-ATPase+Alpha+Beta-domain+wHTH+RRM | Type-1_BR.RM.like.systems | TUDOR+SF1-SF2-Helicase+iREase+CTD+REase->PadR-HTH->METHYLASE->HEPN+DUF499*-><-?<-?<-?<-?<-ArsR-HTH+YTCD-HTH | 1098 | Bacteria>Pseudomonadota>Alphaproteobacteria>Rhodobacterales>Roseobacteraceae | GCF_003667315.2 | Tropicibacter_alexandrii | D8Z78_RS18955 | 49995..53291 | 2267683 | DUF499-domain-containing-protein-[Tropicibacter-sp.-LMIT003] |
| WP_170364042.1 | HEPN+AAA-ATPase+Alpha+Beta-domain+wHTH+RRM | Type-1_BR.RM.like.systems | Primpol->?->?->?-><-RNase<-REase<-HEPN+DUF499*<-METHYLASE<-TUDOR+SF1-SF2-Helicase+iREase+CTD+REase | 1107 | Bacteria>Pseudomonadota>Alphaproteobacteria>Rhodobacterales>Roseobacteraceae | GCF_013030285.1 | Ruegeria_arenilitoris | GOX59_RS08670 | 247066..250389 | 1173585 | DUF499-domain-containing-protein-[Ruegeria-arenilitoris] |
| WP_245947652.1 | HEPN+AAA-ATPase+Alpha+Beta-domain+wHTH+RRM | Type-1_BR.RM.like.systems | MODE-HTH+PBPII->tRNA->TUDOR+SF1-SF2-Helicase+iREase+CTD+REase->HEPN->METHYLASE->HEPN+DUF499*-> | 1097 | Bacteria>Pseudomonadota>Alphaproteobacteria>Rhodobacterales>Roseobacteraceae | GCF_003149265.1 | Jannaschia_seohaensis | BCF38_RS18825 | 37432..40725 | 475081 | DUF499-domain-containing-protein-[Jannaschia-seohaensis] |
| HCP81323.1 | HEPN+AAA-ATPase+Alpha+Beta-domain+wHTH+RRM | Type-1_BR.RM.like.systems | TUDOR+SF1-SF2-Helicase+iREase+CTD+REase->HEPN->METHYLASE->HEPN+DUF499*-><-?<-?<-Peptidase_M15 | 1102 | Bacteria>Pseudomonadota>Alphaproteobacteria>Rhodobacterales>Roseobacteraceae | GCA_003533975.1 | Octadecabacter_sp. | DIT67_06940 | 7896..11204 | 1920174 | AAA-family-ATPase-[Octadecabacter-sp.] |
| WP_216044688.1 | HEPN+AAA-ATPase+Alpha+Beta-domain+wHTH+RRM | Type-1_BR.RM.like.systems | HTH+WYL->TUDOR+SF1-SF2-Helicase+iREase+CTD+REase->PadR-HTH->METHYLASE->HEPN+DUF499*-> | 1110 | Bacteria>Pseudomonadota>Alphaproteobacteria>Rhodobacterales>Roseobacteraceae | GCF_018860305.1 | Celeribacter_halophilus | KO492_RS09190 | 8991..12323 | 576117 | DUF499-domain-containing-protein-[Celeribacter-halophilus] |
| WP_170518600.1 | HEPN+AAA-ATPase+Alpha+Beta-domain+wHTH+RRM | Type-1_BR.RM.like.systems | <-HEPN+DUF499*<-METHYLASE<-ASCH<-TUDOR+SF1-SF2-Helicase+iREase+CTD+REase<-HEPN<-REase | 1103 | Bacteria>Pseudomonadota>Alphaproteobacteria>Rhodobacterales>Roseobacteraceae | GCF_013032825.1 | Ruegeria_atlantica | GOW75_RS18355 | 50430..53741 | 81569 | DUF499-domain-containing-protein-[Ruegeria-atlantica] |
| WP_171727684.1 | HEPN+AAA-ATPase+Alpha+Beta-domain+wHTH+RRM | Type-1_BR.RM.like.systems | HEPN-><-REase<-HEPN+DUF499*<-METHYLASE<-?<-TUDOR+SF1-SF2-Helicase+iREase+CTD+REase | 1108 | Bacteria>Pseudomonadota>Alphaproteobacteria>Rhodobacterales>Roseobacteraceae | GCF_013110755.1 | Ruegeria_sp._HKCCD7318 | GS627_RS09025 | 215066..218392 | 2683014 | DUF499-domain-containing-protein-[Ruegeria-sp.-HKCCD7318] |
| MCI5043002.1 | HEPN+AAA-ATPase+Alpha+Beta-domain+wHTH+RRM | Type-1_BR.RM.like.systems | <-HEPN+DUF499*<-METHYLASE<-?<-TUDOR+SF1-SF2-Helicase+iREase+CTD+REase<-HTH+WYL | 1111 | Bacteria>Pseudomonadota>Alphaproteobacteria>Rhodobacterales>Roseobacteraceae | GCA_022763195.1 | Donghicola_eburneus | MRY81_25480 | 833..4168 | 393278 | DUF499-domain-containing-protein-[Donghicola-eburneus] |
| MBV47914.1 | HEPN+AAA-ATPase+Alpha+Beta-domain+wHTH+RRM | Type-1_BR.RM.like.systems | METHYLASE-p->HEPN+DUF499*-><-Glyco_transf_20<-?<-Glyco_transf_20 | 1111 | Bacteria>Pseudomonadota>Alphaproteobacteria>Rhodobacterales>Roseobacteraceae | GCA_002731875.1 | Roseobacter_sp. | CMN10_05040 | 1113..4448 | 1907202 | AAA-family-ATPase-[Roseobacter-sp.] |
| MBA96874.1 | HEPN+AAA-ATPase+Alpha+Beta-domain+wHTH+RRM | Type-1_BR.RM.like.systems | METHYLASE-p->HEPN+DUF499*-><-Glyco_transf_20<-?<-Glyco_transf_20 | 1111 | Bacteria>Pseudomonadota>Alphaproteobacteria>Rhodobacterales>Roseobacteraceae | GCA_002708965.1 | Roseobacter_sp. | CMN14_05170 | 1105..4440 | 1907202 | AAA-family-ATPase-[Roseobacter-sp.] |
| WP_219436397.1 | HEPN+AAA-ATPase+Alpha+Beta-domain+wHTH+RRM | Type-1_BR.RM.like.systems | HEPN+DUF499*-><-MODE-HTH+PBPII\|\|TetR-HTH-><-ParG_RHH | 1098 | Bacteria>Pseudomonadota>Alphaproteobacteria>Rhodobacterales>Roseobacteraceae | GCF_019392295.1 | Roseovarius_mucosus | KZZ08_RS20795 | 1456..4752 | 215743 | DUF499-domain-containing-protein-[Roseovarius-mucosus] |
| MBF9033364.1 | HEPN+AAA-ATPase+Alpha+Beta-domain+wHTH+RRM | Type-1_BR.RM.like.systems | XIS-HTH->HTH+WYL->ParB+HNH->TUDOR+SF1-SF2-Helicase+iREase+CTD+REase->YtH->METHYLASE->HEPN+DUF499*->AbiEii-><-?<-Peptidase_M15 | 1098 | Bacteria>Pseudomonadota>Alphaproteobacteria>Rhodobacterales>unclassified_Rhodobacterales | GCA_015689685.1 | Rhodobacterales_bacterium_HKCCE2091 | HKCCE2091_03860 | 312307..315603 | 2792619 | DUF499-domain-containing-protein-[Rhodobacterales-bacterium-HKCCE2091] |
| QPI86373.1 | HEPN+AAA-ATPase+Alpha+Beta-domain+wHTH+RRM | Type-1_BR.RM.like.systems | TUDOR+SF1-SF2-Helicase+iREase+CTD+REase->METHYLASE->HEPN+DUF499*-><-HIN-HTH\|\|?->?->?-><-OBFOLD-NUCLEASE<-Aminotran_1_2 | 1111 | Bacteria>Pseudomonadota>Alphaproteobacteria>Rhodobacterales>unclassified_Rhodobacterales | GCA_015693905.1 | Rhodobacterales_bacterium_HKCCA1288 | I3V23_05250 | 1074611..1077946 | 2899152 | DUF499-domain-containing-protein-[Rhodobacterales-bacterium-HKCCA1288] |
| WP_153467943.1 | HEPN+AAA-ATPase+Alpha+Beta-domain+wHTH+RRM | Type-1_BR.RM.like.systems | <-GTPase-AIG<-VirB11\|\|<-VirB11<-DUF499-p<-HIN-HTH<-DUF4096+DDE_Tnp_1+DDE_Tnp_1_2<-HEPN+DUF499*<-METHYLASE<-DUF4096+DDE_Tnp_1+DDE_Tnp_1_2<-METHYLASE-p<-TUDOR+SF1-SF2-Helicase+iREase+CTD+REase<-HTH+WYL | 1047 | Bacteria>Pseudomonadota>Alphaproteobacteria>Rhodospirillales>Acetobacteraceae | GCF_009268135.1 | Komagataeibacter_medellinensis | D3W54_RS02225 | 408174..411317 | 1177712 | DUF499-domain-containing-protein-[Komagataeibacter-medellinensis] |
| WP_274966108.1 | HEPN+AAA-ATPase+Alpha+Beta-domain+wHTH+RRM | Type-1_BR.RM.like.systems | TUDOR+SF1-SF2-Helicase+iREase+CTD+REase->?->PIN->Ntox18->Peptidase_M78->METHYLASE->HEPN+DUF499*->?->HHH_5+SF1-SF2-helicase-><-RVT+HNH\|\|<-Phage_int_SAM_1+Phage_integrase<-Phage_integrase<-Phage_integrase | 1113 | Bacteria>Pseudomonadota>Alphaproteobacteria>Rhodospirillales>Acetobacteraceae | GCF_020418435.1 | Acidibrevibacterium_fodinaquatile | LGL72_RS14205 | 2856594..2859935 | 1969806 | DUF499-domain-containing-protein-[Acidibrevibacterium-fodinaquatile] |
| WP_171617018.1 | HEPN+AAA-ATPase+Alpha+Beta-domain+wHTH+RRM | Type-1_BR.RM.like.systems | Phage_integrase-><-?\|\|TUDOR+SF1-SF2-Helicase+iREase+CTD+REase->?->METHYLASE->HEPN+DUF499*->HIN-HTH->?->LZ_Tnp_IS66+zf-IS66+DDE_Tnp_IS66+DDE_Tnp_IS66_C-><-DUF4158+DDE_Tnp_Tn3\|\|GDResolvase+HIN-HTH-> | 1113 | Bacteria>Pseudomonadota>Alphaproteobacteria>Rhodospirillales>Acetobacteraceae | GCF_013112485.1 | Roseicella_sp._DB1501 | HJG45_RS27180 | 18137..21478 | 2730925 | DUF499-domain-containing-protein-[Roseicella-sp.-DB1501] |
| WP_174502310.1 | HEPN+AAA-ATPase+Alpha+Beta-domain+wHTH+RRM | Type-1_BR.RM.like.systems | NUDIX->ABhydrolase->?->Arm-DNA-bind_3+Phage_integrase-><-Phage_AlpA<-HEPN+DUF499*<-METHYLASE<-MORC+DUF3684<-TUDOR+SF1-SF2-Helicase+iREase+CTD+REase<-HTH+WYL\|\|MODE-HTH+PBPII-><-ParG_RHH | 1110 | Bacteria>Pseudomonadota>Alphaproteobacteria>Rhodospirillales>Acetobacteraceae | GCF_902712915.1 | Acidiphilium_sp._C61 | HUW10_RS03970 | 224221..227553 | 1671485 | Swt1-family-HEPN-domain-containing-protein-[Acidiphilium-sp.-C61] |
| MCA7120529.1 | HEPN+AAA-ATPase+Alpha+Beta-domain+wHTH+RRM | Type-1_BR.RM.like.systems | TUDOR+SF1-SF2-Helicase+iREase+CTD+REase->?->PIN->Ntox18->Peptidase_M78->METHYLASE->HEPN+DUF499*->?->?->HHH_5+SF1-SF2-helicase-><-RVT+HNH\|\|<-Phage_int_SAM_1+Phage_integrase<-Phage_integrase | 1113 | Bacteria>Pseudomonadota>Alphaproteobacteria>Rhodospirillales>Acetobacteraceae | GCA_020418435.1 | Acidibrevibacterium_fodinaquatile | LGL72_14265 | 2856594..2859935 | 1969806 | DUF499-domain-containing-protein-[Acidibrevibacterium-sp.] |
| WP_152832805.1 | HEPN+AAA-ATPase+Alpha+Beta-domain+wHTH+RRM | Type-1_BR.RM.like.systems | PIN->TUDOR+SF1-SF2-Helicase+iREase+CTD+REase->METHYLASE-p->METHYLASE-p->HEPN+DUF499*-><-PhdYeFM_antitox<-PIN<-Phd-YefM<-GDResolvase+Recombinase+ZnR<-XIS-HTH | 1108 | Bacteria>Pseudomonadota>Alphaproteobacteria>Rhodospirillales>Acetobacteraceae | GCF_009295745.1 | Acetobacter_senegalensis | GC248_RS02655 | 30596..33922 | 446692 | DUF499-domain-containing-protein-[Acetobacter-senegalensis] |
| WP_010666859.1 | HEPN+AAA-ATPase+Alpha+Beta-domain+wHTH+RRM | Type-1_BR.RM.like.systems | VirD4-TraG+VirD4-FtsK->CopG/RHH->?->PIN->TUDOR+SF1-SF2-Helicase+iREase+CTD+REase->METHYLASE->HEPN+DUF499*-><-?<-REase\|\|VirB11+VirB4-FtsK->VirB2->?->FtsK+VirB4-FtsK-> | 1108 | Bacteria>Pseudomonadota>Alphaproteobacteria>Rhodospirillales>Acetobacteraceae | GCF_000193495.2 | Acetobacter_aceti | EDC15_RS12660 | 41147..44473 | 435 | DUF499-domain-containing-protein-[Acetobacter-aceti] |
| WP_114912917.1 | HEPN+AAA-ATPase+Alpha+Beta-domain+wHTH+RRM | Type-1_BR.RM.like.systems | Ntox18->Peptidase_M78->METHYLASE-><-HTH_Tnp_1+HTH_21+rve+rve_3\|\|MULE-TRANSPOSASE->METHYLASE-p->HEPN+DUF499*->TPR->?->?->HHH_5+SF1-SF2-helicase->?->PAIREDC-HTH-> | 1113 | Bacteria>Pseudomonadota>Alphaproteobacteria>Rhodospirillales>Acetobacteraceae | GCF_003352165.1 | Acidibrevibacterium_fodinaquatile | DEF76_RS14375 | 3023042..3026383 | 1969806 | DUF499-domain-containing-protein-[Acidibrevibacterium-fodinaquatile] |
| OZB26691.1 | HEPN+AAA-ATPase+Alpha+Beta-domain+wHTH+RRM | Type-1_BR.RM.like.systems | TUDOR+SF1-SF2-Helicase+iREase+CTD+REase->?->PIN->Ntox18->?->METHYLASE->HEPN+DUF499*->\|FAD-NAD-dep-oxidoreductase-><-NACHT+REase\|\|?->DnaB-><-NACHT+REase | 1091 | Bacteria>Pseudomonadota>Alphaproteobacteria>Rhodospirillales>Acetobacteraceae | GCA_002282635.1 | Acidiphilium_sp._34-64-41 | B7X49_12125 | 19315..22590 | 1970297 | MAG:-ATPase-[Acidiphilium-sp.-34-64-41] |
| WP_012553516.1 | HEPN+AAA-ATPase+Alpha+Beta-domain+wHTH+RRM | Type-1_BR.RM.like.systems | <-VirB2<-VirB11+VirB4-FtsK\|\|?->REase->?-><-HEPN+DUF499*<-METHYLASE<-TUDOR+SF1-SF2-Helicase+iREase+CTD+REase<-PIN<-?<-CopG/RHH<-VirD4-TraG+VirD4-FtsK | 1108 | Bacteria>Pseudomonadota>Alphaproteobacteria>Rhodospirillales>Acetobacteraceae | GCF_000021325.1 | Gluconacetobacter_diazotrophicus | GDIA_RS05030 | 1143198..1146524 | 33996 | DUF499-domain-containing-protein-[Gluconacetobacter-diazotrophicus] |
| WP_135468376.1 | HEPN+AAA-ATPase+Alpha+Beta-domain+wHTH+RRM | Type-1_BR.RM.like.systems | <-LRP-HTH<-DnaJ<-?\|\|tRNA->TUDOR+SF1-SF2-Helicase+iREase+CTD+REase->?->METHYLASE->HEPN+DUF499*->ArdCN+MPTase-><-HTH<-GDResolvase+Recombinase+ZnR<-RAMA | 1112 | Bacteria>Pseudomonadota>Alphaproteobacteria>Rhodospirillales>Acetobacteraceae | GCF_004761865.1 | Crenalkalicoccus_roseus | ET526_RS14250 | 21738..25076 | 1485588 | DUF499-domain-containing-protein-[Crenalkalicoccus-roseus] |
| WP_211861077.1 | HEPN+AAA-ATPase+Alpha+Beta-domain+wHTH+RRM | Type-1_BR.RM.like.systems | TUDOR+SF1-SF2-Helicase+iREase+CTD+REase->HKD-DNase->?->METHYLASE->?->HEPN+DUF499*-><-HEPN+REase<-HTH<-GDResolvase+Recombinase+ZnR<-RAMA<-?<-HTH | 1110 | Bacteria>Pseudomonadota>Alphaproteobacteria>Rhodospirillales>Acetobacteraceae | GCF_018129065.1 | Neoroseomonas_soli | GXW76_RS05955 | 22990..26322 | 1081025 | DUF499-domain-containing-protein-[Roseomonas-soli] |
| MBX9595850.1 | HEPN+AAA-ATPase+Alpha+Beta-domain+wHTH+RRM | Type-1_BR.RM.like.systems | MerR-HTH->TUDOR+SF1-SF2-Helicase+iREase+CTD+REase->HKD-DNase->METHYLASE->HEPN+DUF499*->MACRODOMAIN-><-HTH<-GDResolvase+Recombinase+ZnR | 1110 | Bacteria>Pseudomonadota>Alphaproteobacteria>Rhodospirillales>Acetobacteraceae | GCA_019748015.1 | Roseomonas_sp. | K2X46_15915 | 11392..14724 | 1869190 | DUF499-domain-containing-protein-[Roseomonas-sp.] |
| WP_253545427.1 | HEPN+AAA-ATPase+Alpha+Beta-domain+wHTH+RRM | Type-1_BR.RM.like.systems | <-TraJ-RHH<-Phage_integrase\|\|?->PIN->TUDOR+SF1-SF2-Helicase+iREase+CTD+REase->METHYLASE->HEPN+DUF499*->?-><-?<-?\|\|TrwC+SF1-SF2-helicase+TOPRIM-> | 1108 | Bacteria>Pseudomonadota>Alphaproteobacteria>Rhodospirillales>Acetobacteraceae | GCF_024158445.1 | Acetobacter_lovaniensis | NKW44_RS07985 | 22199..25525 | 104100 | DUF499-domain-containing-protein-[Acetobacter-lovaniensis] |
| WP_216840234.1 | HEPN+AAA-ATPase+Alpha+Beta-domain+wHTH+RRM | Type-1_BR.RM.like.systems | TUDOR+SF1-SF2-Helicase+iREase+CTD+REase->HKD-DNase->METHYLASE->HEPN+DUF499*-><-HTH<-GDResolvase+Recombinase+ZnR<-RAMA | 1109 | Bacteria>Pseudomonadota>Alphaproteobacteria>Rhodospirillales>Acetobacteraceae | GCF_019024325.1 | Falsiroseomonas_tokyonensis | J3S20_RS28040 | 39158..42487 | 430521 | DUF499-domain-containing-protein-[Falsiroseomonas-tokyonensis] |
| WP_132287513.1 | HEPN+AAA-ATPase+Alpha+Beta-domain+wHTH+RRM | Type-1_BR.RM.like.systems | TUDOR+SF1-SF2-Helicase+iREase+CTD+REase->?->METHYLASE->HEPN+DUF499*-><-?\|\|REase-><-HTH<-GDResolvase+Recombinase+ZnR<-RAMA | 1114 | Bacteria>Pseudomonadota>Alphaproteobacteria>Rhodospirillales>Acetobacteraceae | GCF_004343615.1 | Roseicella_aquatilis | EXY23_RS09345 | 90993..94337 | 2527868 | Swt1-family-HEPN-domain-containing-protein-[Paracraurococcus-sp.-NE82] |
| MBB3175065.1 | HEPN+AAA-ATPase+Alpha+Beta-domain+wHTH+RRM | Type-1_BR.RM.like.systems | rve_3->TUDOR+SF1-SF2-Helicase+iREase+CTD+REase->METHYLASE->HEPN+DUF499*->HTH_Tnp_1->?->?-><-HIN-HTH<-HIN-HTH<-HIN-HTH | 1110 | Bacteria>Pseudomonadota>Alphaproteobacteria>Rhodospirillales>Acetobacteraceae | GCA_014192375.1 | Endobacter_medicaginis | FHR90_002912 | 7080..10412 | 1181271 | hypothetical-protein-[Endobacter-medicaginis] |
| WP_199897436.1 | HEPN+AAA-ATPase+Alpha+Beta-domain+wHTH+RRM | Type-1_BR.RM.like.systems | <-DUF499-p<-HEPN+DUF499*<-METHYLASE<-ASCH<-?<-TUDOR+SF1-SF2-Helicase+iREase+CTD+REase<-HTH+WYL<-METHYLASE-p | 1018 | Bacteria>Pseudomonadota>Alphaproteobacteria>Rhodospirillales>Acetobacteraceae | GCF_000613905.1 | Acetobacter_persici | JCM25330_RS21105 | 22243..25299 | 1076596 | DUF499-domain-containing-protein-[Acetobacter-persici] |
| MCH4022379.1 | HEPN+AAA-ATPase+Alpha+Beta-domain+wHTH+RRM | Type-1_BR.RM.like.systems | <-REase+CTD6\|\|HTH+WYL->TUDOR+SF1-SF2-Helicase+iREase+CTD+REase->?->METHYLASE->HEPN+DUF499*->?-><-?<-LexA-protease | 1097 | Bacteria>Pseudomonadota>Alphaproteobacteria>Rhodospirillales>Acetobacteraceae | GCA_022483365.1 | Acetobacter_sp. | LKE62_03305 | 718063..721356 | 440 | DUF499-domain-containing-protein-[Acetobacter-sp.] |
| WP_062273923.1 | HEPN+AAA-ATPase+Alpha+Beta-domain+wHTH+RRM | Type-1_BR.RM.like.systems | PIN->TUDOR+SF1-SF2-Helicase+iREase+CTD+REase->METHYLASE->HEPN+DUF499*-><-?<-?<-Radical-SAM<-PIN<-Phd-YefM | 1108 | Bacteria>Pseudomonadota>Alphaproteobacteria>Rhodospirillales>Acetobacteraceae | GCF_001581105.1 | Acetobacter_cerevisiae | AD954_RS12075 | 12356..15682 | 178900 | DUF499-domain-containing-protein-[Acetobacter-cerevisiae] |
| WP_018981014.1 | HEPN+AAA-ATPase+Alpha+Beta-domain+wHTH+RRM | Type-1_BR.RM.like.systems | <-SF1-SF2-Helicase<-?<-?<-ABhydrolase<-?<-HEPN+DUF499*<-METHYLASE<-TUDOR+SF1-SF2-Helicase+iREase+CTD+REase | 1099 | Bacteria>Pseudomonadota>Alphaproteobacteria>Rhodospirillales>Acetobacteraceae | GCF_025994295.1 | Saccharibacter_floricola | F457_RS0110055 | 41499..44798 | 231053 | DUF499-domain-containing-protein-[Saccharibacter-floricola] |
| WP_194301389.1 | HEPN+AAA-ATPase+Alpha+Beta-domain+wHTH+RRM | Type-1_BR.RM.like.systems | HTH+WYL->TUDOR+SF1-SF2-Helicase+iREase+CTD+REase->?->METHYLASE->HEPN+DUF499*->VirB11->GTPase-AIG-> | 1085 | Bacteria>Pseudomonadota>Alphaproteobacteria>Rhodospirillales>Acetobacteraceae | GCF_011516925.1 | Acetobacter_fabarum | GOB83_RS11695 | 8455..11712 | 483199 | DUF499-domain-containing-protein-[Acetobacter-fabarum] |
| NHO42826.1 | HEPN+AAA-ATPase+Alpha+Beta-domain+wHTH+RRM | Type-1_BR.RM.like.systems | HTH+WYL->TUDOR+SF1-SF2-Helicase+iREase+CTD+REase->?->METHYLASE->HEPN+DUF499*->VirB11->GTPase-AIG-> | 1097 | Bacteria>Pseudomonadota>Alphaproteobacteria>Rhodospirillales>Acetobacteraceae | GCA_011516925.1 | Acetobacter_fabarum | GOB83_11655 | 8419..11712 | 483199 | DUF499-domain-containing-protein-[Acetobacter-fabarum] |
| WP_165926191.1 | HEPN+AAA-ATPase+Alpha+Beta-domain+wHTH+RRM | Type-1_BR.RM.like.systems | <-GTPase-AIG<-VirB11<-HEPN+DUF499*<-METHYLASE<-?<-TUDOR+SF1-SF2-Helicase+iREase+CTD+REase-p | 1097 | Bacteria>Pseudomonadota>Alphaproteobacteria>Rhodospirillales>Acetobacteraceae | GCF_025995875.1 | Acetobacter_fabarum | AA19596_RS12715 | 13783..17076 | 483199 | DUF499-domain-containing-protein-[Acetobacter-fabarum] |
| WP_246330294.1 | HEPN+AAA-ATPase+Alpha+Beta-domain+wHTH+RRM | Type-1_BR.RM.like.systems | HIN-HTH-><-?<-HTH_Tnp_1<-HEPN+DUF499*<-METHYLASE<-TUDOR+SF1-SF2-Helicase+iREase+CTD+REase | 1093 | Bacteria>Pseudomonadota>Alphaproteobacteria>Rhodospirillales>Acetobacteraceae | GCF_014192375.1 | Endobacter_medicaginis | OR615_RS11695 | 24590..27871 | 1181271 | DUF499-domain-containing-protein-[Endobacter-medicaginis] |
| MBX6741668.1 | HEPN+AAA-ATPase+Alpha+Beta-domain+wHTH+RRM | Type-1_BR.RM.like.systems | TUDOR+SF1-SF2-Helicase+iREase+CTD+REase-p->ASCH->METHYLASE->HEPN->HEPN+DUF499*-> | 1110 | Bacteria>Pseudomonadota>Alphaproteobacteria>Rhodospirillales>Acetobacteraceae | GCA_019690105.1 | Acetobacteraceae_bacterium | IRY87_06425 | 5783..9115 | 1909293 | DUF499-domain-containing-protein-[Acetobacteraceae-bacterium] |
| WP_045542211.1 | HEPN+AAA-ATPase+Alpha+Beta-domain+wHTH+RRM | Type-1_BR.RM.like.systems | METHYLASE-p->HEPN+DUF499*-><-?<-?<-Radical-SAM<-PIN<-Phd-YefM\|\|Phage_integrase-> | 1108 | Bacteria>Pseudomonadota>Alphaproteobacteria>Rhodospirillales>Acetobacteraceae | GCF_025995375.1 | Acetobacter_tropicalis | AD944_RS02385 | 1113..4439 | 104102 | DUF499-domain-containing-protein-[Acetobacter-tropicalis] |
| MBX6741130.1 | HEPN+AAA-ATPase+Alpha+Beta-domain+wHTH+RRM | Type-1_BR.RM.like.systems | METHYLASE-p->HEPN+DUF499*->?->?->?->?-><-SF1-SF2-Helicase+REase | 1108 | Bacteria>Pseudomonadota>Alphaproteobacteria>Rhodospirillales>Acetobacteraceae | GCA_019690105.1 | Acetobacteraceae_bacterium | IRY87_03665 | 460..3786 | 1909293 | DUF499-domain-containing-protein-[Acetobacteraceae-bacterium] |
| MCA3363599.1 | HEPN+AAA-ATPase+Alpha+Beta-domain+wHTH+RRM | Type-1_BR.RM.like.systems | METHYLASE->OLD-ABC->PIN->HEPN+DUF499*-> | 483 | Bacteria>Pseudomonadota>Alphaproteobacteria>Rhodospirillales>Acetobacteraceae | GCA_020249535.1 | Roseomonas_sp. | INF74_16815 | 3977..5424 | 1869190 | DUF499-domain-containing-protein-[Roseomonas-sp.] |
| MBX6743929.1 | HEPN+AAA-ATPase+Alpha+Beta-domain+wHTH+RRM | Type-1_BR.RM.like.systems | <-HEPN+DUF499*<-HEPN<-METHYLASE-p | 1111 | Bacteria>Pseudomonadota>Alphaproteobacteria>Rhodospirillales>Acetobacteraceae | GCA_019690105.1 | Acetobacteraceae_bacterium | IRY87_17990 | 402..3737 | 1909293 | DUF499-domain-containing-protein-[Acetobacteraceae-bacterium] |
| WP_209875397.1 | HEPN+AAA-ATPase+Alpha+Beta-domain+wHTH+RRM | Type-1_BR.RM.like.systems | HIN-HTH->GDResolvase+Recombinase+ZnR->?->HIN-HTH->?->LZ_Tnp_IS66+zf-IS66+DDE_Tnp_IS66+DDE_Tnp_IS66_C-><-HEPN+DUF499*<-METHYLASE<-MORC+DUF3684<-TUDOR+SF1-SF2-Helicase+iREase+CTD+REase<-HTH+WYL | 1098 | Bacteria>Pseudomonadota>Alphaproteobacteria>Rhodospirillales>Azospirillaceae | GCF_017876165.1 | Azospirillum_soli | J2852_RS11360 | 44068..47364 | 1304799 | DUF499-domain-containing-protein-[Azospirillum-soli] |
| WP_014199726.1 | HEPN+AAA-ATPase+Alpha+Beta-domain+wHTH+RRM | Type-1_BR.RM.like.systems | MODE-HTH+PBPII-><-Arm-DNA-bind_3+Phage_int_SAM_3+Phage_integrase<-?<-HEPN+DUF499*<-HEPN<-PolB<-METHYLASE<-HNH<-OLD-ABC<-TUDOR+SF1-SF2-Helicase+iREase+CTD+REase | 1115 | Bacteria>Pseudomonadota>Alphaproteobacteria>Rhodospirillales>Azospirillaceae | GCF_000237365.1 | Azospirillum | GAY33_RS05480 | 11187..14534 | 191 | MULTISPECIES:-Swt1-family-HEPN-domain-containing-protein-[Azospirillum] |
| WP_180283596.1 | HEPN+AAA-ATPase+Alpha+Beta-domain+wHTH+RRM | Type-1_BR.RM.like.systems | <-PfkB\|\|DeoR-HTH+ISOCOT->Transposase_31+Yae1_N-><-HEPN+DUF499*<-PIN<-OLD-ABC<-METHYLASE<-TUDOR+SF1-SF2-Helicase+iREase+CTD+REase\|\|HTH->GDResolvase+PPP1-HTH-> | 1135 | Bacteria>Pseudomonadota>Alphaproteobacteria>Rhodospirillales>Azospirillaceae | GCF_013423495.1 | Azospirillum_oleiclasticum | HND93_RS19070 | 2984..6391 | 2735135 | DUF499-domain-containing-protein-[Azospirillum-oleiclasticum] |
| WP_045580668.1 | HEPN+AAA-ATPase+Alpha+Beta-domain+wHTH+RRM | Type-1_BR.RM.like.systems | <-SF1-SF2-Helicase+REase\|\|TUDOR+SF1-SF2-Helicase+iREase+CTD+REase->METHYLASE->?->HEPN+DUF499*-><-?<-?<-?<-LZ_Tnp_IS66+zf-IS66+DDE_Tnp_IS66+DDE_Tnp_IS66_C<-?<-HIN-HTH | 1118 | Bacteria>Pseudomonadota>Alphaproteobacteria>Rhodospirillales>Azospirillaceae | GCF_001305595.1 | Azospirillum_thiophilum | VY88_RS04045 | 887405..890761 | 528244 | Swt1-family-HEPN-domain-containing-protein-[Azospirillum-thiophilum] |
| TWB14201.1 | HEPN+AAA-ATPase+Alpha+Beta-domain+wHTH+RRM | Type-1_BR.RM.like.systems | <-SF1-SF2-Helicase+REase\|\|HTH+WYL->TUDOR+SF1-SF2-Helicase+iREase+CTD+REase->YtH->MORC+DUF3684->METHYLASE-p->HEPN+DUF499*->MODE-HTH+PBPII-><-ParG_RHH | 1098 | Bacteria>Pseudomonadota>Alphaproteobacteria>Rhodospirillales>Azospirillaceae | GCA_007828045.1 | Nitrospirillum_amazonense | FBZ89_11767 | 103300..106596 | 28077 | hypothetical-protein-FBZ89_11767-[Nitrospirillum-amazonense] |
| WP_246172460.1 | HEPN+AAA-ATPase+Alpha+Beta-domain+wHTH+RRM | Type-1_BR.RM.like.systems | <-SF1-SF2-Helicase+REase\|\|HTH+WYL->TUDOR+SF1-SF2-Helicase+iREase+CTD+REase->YtH->MORC+DUF3684->METHYLASE->HEPN+DUF499*->MODE-HTH+PBPII-><-ParG_RHH | 1086 | Bacteria>Pseudomonadota>Alphaproteobacteria>Rhodospirillales>Azospirillaceae | GCF_007828045.1 | Nitrospirillum_amazonense | FBZ89_RS22360 | 103336..106596 | 28077 | DUF499-domain-containing-protein-[Nitrospirillum-amazonense] |
| WP_211111227.1 | HEPN+AAA-ATPase+Alpha+Beta-domain+wHTH+RRM | Type-1_BR.RM.like.systems | <-GDResolvase+Recombinase+ZnR<-?\|\|?->?->Phage_integrase-><-HEPN+DUF499*<-METHYLASE<-TUDOR+SF1-SF2-Helicase+iREase+CTD+REase<-HTH+WYL\|\|HicB->TIR->REase-> | 1089 | Bacteria>Pseudomonadota>Alphaproteobacteria>Rhodospirillales>Azospirillaceae | GCF_016806835.2 | Skermanella_rosea | JL101_RS04580 | 983168..986437 | 1817965 | DUF499-domain-containing-protein-[Skermanella-rosea] |
| WP_238518868.1 | HEPN+AAA-ATPase+Alpha+Beta-domain+wHTH+RRM | Type-1_BR.RM.like.systems | <-SF1-SF2-Helicase+DUF1998+HKD-DNase+HKD-DNase<-?<-HEPN+DUF499*<-PIN<-OLD-ABC<-METHYLASE<-?<-Schlafen+HATPase+HTH<-TUDOR+SF1-SF2-Helicase+iREase+CTD+REase | 1154 | Bacteria>Pseudomonadota>Alphaproteobacteria>Rhodospirillales>Azospirillaceae | GCF_022214805.1 | Azospirillum_doebereinerae | MCW82_RS00020 | 10412..13876 | 92933 | DUF499-domain-containing-protein-[Azospirillum-doebereinerae] |
| WP_246149069.1 | HEPN+AAA-ATPase+Alpha+Beta-domain+wHTH+RRM | Type-1_BR.RM.like.systems | REase->TUDOR+SF1-SF2-Helicase+iREase+CTD+REase->ASCH->ASCH->METHYLASE->METHYLASE-p->HEPN+DUF499*->DUF499-p-><-?<-DDE_Tnp_1\|\|REase-> | 926 | Bacteria>Pseudomonadota>Alphaproteobacteria>Rhodospirillales>Azospirillaceae | GCF_008843145.1 | Skermanella_pratensis | DPR14_RS09205 | 2041466..2044246 | 2233999 | DUF499-domain-containing-protein-[Skermanella-pratensis] |
| WP_063633927.1 | HEPN+AAA-ATPase+Alpha+Beta-domain+wHTH+RRM | Type-1_BR.RM.like.systems | <-GDResolvase+HIN-HTH\|\|?-><-HEPN+DUF499*<-METHYLASE<-ASCH<-Cluster1090_5clades<-TUDOR+SF1-SF2-Helicase+iREase+CTD+REase<-HTH+WYL | 1115 | Bacteria>Pseudomonadota>Alphaproteobacteria>Rhodospirillales>Azospirillaceae | GCF_001639105.2 | Azospirillum_humicireducens | A6A40_RS02305 | 493921..497268 | 1226968 | Swt1-family-HEPN-domain-containing-protein-[Azospirillum-humicireducens] |
| WP_109152948.1 | HEPN+AAA-ATPase+Alpha+Beta-domain+wHTH+RRM | Type-1_BR.RM.like.systems | <-VirB4-FtsK+HerA<-?\|\|TUDOR+SF1-SF2-Helicase+iREase+CTD+REase->Cluster1090_5clades->ASCH->METHYLASE->HEPN+DUF499*->Lon_C-> | 1116 | Bacteria>Pseudomonadota>Alphaproteobacteria>Rhodospirillales>Azospirillaceae | GCF_003116065.1 | Azospirillum_sp._TS05 | TS05_RS18610 | 11435..14785 | 716760 | Swt1-family-HEPN-domain-containing-protein-[Azospirillum-sp.-TS05] |
| WP_147041135.1 | HEPN+AAA-ATPase+Alpha+Beta-domain+wHTH+RRM | Type-1_BR.RM.like.systems | TPR->?->TUDOR+SF1-SF2-Helicase+iREase+CTD+REase->METHYLASE->HEPN+DUF499*->?->OLD-ABC+TOPRIM-><-RelE-ParE<-MetJarc-HTH | 1097 | Bacteria>Pseudomonadota>Alphaproteobacteria>Rhodospirillales>Azospirillaceae | GCF_007991775.1 | Skermanella_aerolata | SAE02_RS33460 | 15952..19245 | 393310 | DUF499-domain-containing-protein-[Skermanella-aerolata] |
| WP_145610986.1 | HEPN+AAA-ATPase+Alpha+Beta-domain+wHTH+RRM | Type-1_BR.RM.like.systems | TUDOR+SF1-SF2-Helicase+iREase+CTD+REase->ParB+HNH->METHYLASE->PolB->HEPN->HEPN+DUF499*->?->PIN-> | 1115 | Bacteria>Pseudomonadota>Alphaproteobacteria>Rhodospirillales>Azospirillaceae | GCF_007828035.1 | Nitrospirillum_amazonense | FBZ87_RS14570 | 656187..659534 | 28077 | Swt1-family-HEPN-domain-containing-protein-[Nitrospirillum-amazonense] |
| WP_209991153.1 | HEPN+AAA-ATPase+Alpha+Beta-domain+wHTH+RRM | Type-1_BR.RM.like.systems | TUDOR+SF1-SF2-Helicase+iREase+CTD+REase->OLD-ABC->HNH->METHYLASE->?->HEPN+DUF499*->?->Lon_C-> | 1114 | Bacteria>Pseudomonadota>Alphaproteobacteria>Rhodospirillales>Azospirillaceae | GCF_017876115.1 | Azospirillum_picis | J2850_RS30835 | 10080..13424 | 488438 | Swt1-family-HEPN-domain-containing-protein-[Azospirillum-picis] |
| WP_218285167.1 | HEPN+AAA-ATPase+Alpha+Beta-domain+wHTH+RRM | Type-1_BR.RM.like.systems | Peptidase->Peptidase_M24->TUDOR+SF1-SF2-Helicase+iREase+CTD+REase->METHYLASE->OLD-ABC->TOPRIM->HEPN+DUF499*->?-><-PAIREDC-HTH<-DUF4096+DDE_Tnp_1+DDE_Tnp_1_2\|\|DUF4096+DDE_Tnp_1+DDE_Tnp_1_2->PAIREDC-HTH-> | 1111 | Bacteria>Pseudomonadota>Alphaproteobacteria>Rhodospirillales>Elioraeaceae | GCF_019203965.1 | Elioraea_tepida | KO353_RS13040 | 2714596..2717931 | 2843330 | Swt1-family-HEPN-domain-containing-protein-[Elioraea-tepida] |
| WP_173982139.1 | HEPN+AAA-ATPase+Alpha+Beta-domain+wHTH+RRM | Type-1_BR.RM.like.systems | HTH+WYL->TUDOR+SF1-SF2-Helicase+iREase+CTD+REase->METHYLASE->HEPN+DUF499*->TPR+TPR+PIN+PIN->TPR+TPR+PIN+PIN->REase+DUF4268-><-HTH<-GDResolvase+Recombinase+ZnR<-RAMA | 1097 | Bacteria>Pseudomonadota>Alphaproteobacteria>Rhodospirillales>Rhodospirillaceae | GCF_902729425.1 | Magnetospirillum_sp._SS-4 | MTBSS4_RS04180 | 57981..61274 | 2681465 | DUF499-domain-containing-protein-[Magnetospirillum-sp.-SS-4] |
| WP_190260570.1 | HEPN+AAA-ATPase+Alpha+Beta-domain+wHTH+RRM | Type-1_BR.RM.like.systems | <-SF1-SF2-Helicase+REase<-SF1-SF2-Helicase+REase\|\|HTH+WYL->TUDOR+SF1-SF2-Helicase+iREase+CTD+REase->YtH->METHYLASE->HEPN+DUF499*->MODE-HTH+PBPII-><-ParG_RHH | 1098 | Bacteria>Pseudomonadota>Alphaproteobacteria>Rhodospirillales>Rhodospirillaceae | GCF_014672695.1 | Defluviicoccus_vanus | HQ394_RS12700 | 2827067..2830363 | 111831 | DUF499-domain-containing-protein-[Defluviicoccus-vanus] |
| MBC8269068.1 | HEPN+AAA-ATPase+Alpha+Beta-domain+wHTH+RRM | Type-1_BR.RM.like.systems | Y1_Tnp-><-?<-?\|\|HTH+WYL->TUDOR+SF1-SF2-Helicase+iREase+CTD+REase->METHYLASE->HEPN+DUF499*-> | 565 | Bacteria>Pseudomonadota>Alphaproteobacteria>Rhodospirillales>Rhodospirillaceae | GCA_014382375.1 | Rhodospirillaceae_bacterium | H8E36_09995 | 85324..87019 | 1898112 | ATP-binding-protein-[Rhodospirillaceae-bacterium] |
| MDE0333394.1 | HEPN+AAA-ATPase+Alpha+Beta-domain+wHTH+RRM | Type-1_BR.RM.like.systems | TUDOR+SF1-SF2-Helicase+iREase+CTD+REase->?->?->?->METHYLASE->HEPN->HEPN+DUF499*->REase-> | 1096 | Bacteria>Pseudomonadota>Alphaproteobacteria>Rhodospirillales>Rhodospirillaceae | GCA_028820945.1 | Defluviicoccus_sp. | OXI64_00425 | 9805..13095 | 2682137 | MAG:-DUF499-domain-containing-protein-[Defluviicoccus-sp.] |
| MBT6311358.1 | HEPN+AAA-ATPase+Alpha+Beta-domain+wHTH+RRM | Type-1_BR.RM.like.systems | XIS-HTH->?->DnaB->?->Terminase-><-HEPN+DUF499*<-?<-?<-METHYLASE<-HEPN | 1096 | Bacteria>Pseudomonadota>Alphaproteobacteria>Rhodospirillales>Rhodospirillaceae | GCA_018701575.1 | Rhodospirillaceae_bacterium | HOJ20_16125 | 146396..149686 | 1898112 | ATP-binding-protein-[Rhodospirillaceae-bacterium] |
| MBT5356878.1 | HEPN+AAA-ATPase+Alpha+Beta-domain+wHTH+RRM | Type-1_BR.RM.like.systems | XIS-HTH->?->DnaB->?->Terminase-><-HEPN+DUF499*<-?<-?<-METHYLASE<-HEPN | 1096 | Bacteria>Pseudomonadota>Alphaproteobacteria>Rhodospirillales>Rhodospirillaceae | GCA_018663095.1 | Rhodospirillaceae_bacterium | HOL07_00895 | 173068..176358 | 1898112 | ATP-binding-protein-[Rhodospirillaceae-bacterium] |
| MDE0176181.1 | HEPN+AAA-ATPase+Alpha+Beta-domain+wHTH+RRM | Type-1_BR.RM.like.systems | METHYLASE-p->METHYLASE->?->?->HEPN+DUF499*->HEPN->?->HEPN-> | 1096 | Bacteria>Pseudomonadota>Alphaproteobacteria>Rhodospirillales>Rhodospirillaceae | GCA_028819635.1 | Defluviicoccus_sp. | OXM60_22325 | 5691..8981 | 2682137 | MAG:-DUF499-domain-containing-protein-[Defluviicoccus-sp.] |
| MCY4311819.1 | HEPN+AAA-ATPase+Alpha+Beta-domain+wHTH+RRM | Type-1_BR.RM.like.systems | <-SbcC<-HEPN+DUF499*<-?<-METHYLASE-p | 1100 | Bacteria>Pseudomonadota>Alphaproteobacteria>Rhodospirillales>Rhodospirillaceae | GCA_026710125.1 | Rhodospirillaceae_bacterium | OXC54_11025 | 2036..5338 | 1898112 | MAG:-DUF499-domain-containing-protein-[Rhodospirillaceae-bacterium] |
| MCY4239105.1 | HEPN+AAA-ATPase+Alpha+Beta-domain+wHTH+RRM | Type-1_BR.RM.like.systems | METHYLASE->?->?->HEPN+DUF499*->SbcC-> | 1100 | Bacteria>Pseudomonadota>Alphaproteobacteria>Rhodospirillales>Rhodospirillaceae | GCA_026709535.1 | Rhodospirillaceae_bacterium | OXC93_12260 | 2987..6289 | 1898112 | MAG:-DUF499-domain-containing-protein-[Rhodospirillaceae-bacterium] |

| Protein-ID | DUF499-architecture | System | Gene-Neighbourhoods | Length | Taxonomy | Assembly-ID | Strain | Locus-Tag | Genomic-coordinates | TaxID | Fasta-defline |
|---|---|---|---|---|---|---|---|---|---|---|---|
| MBK8177039.1 | HEPN+AAA-ATPase+Alpha+Beta-domain+wHTH+RRM | Type-1_BR.RM.like.systems | TrwC+SF1-SF2-helicase+TOPRIM->?->RHH->RelE-ParE-><-N6_N4_MTase<-?<-HEPN+DUF499*<-METHYLASE<-TUDOR+SF1-SF2-Helicase+iREase+CTD+REase<-?<-?<-ArdCN+MPTase\|\|DOC+HTH-> | 1106 | Bacteria>Pseudomonadota>Alphaproteobacteria>Rhodospirillales>unclassified_Rhodospirillales | GCA_016712595.1 | Rhodospirillales_bacterium | IPK66_17795 | 28556..31876 | 2026786 | DUF499-domain-containing-protein-[Rhodospirillales-bacterium] |
| TVR79626.1 | HEPN+AAA-ATPase+Alpha+Beta-domain+wHTH+RRM | Type-1_BR.RM.like.systems | HDRnase-><-HEPN+DUF499*<-?<-PIN<-ParD<-METHYLASE<-OLD-ABC+TOPRIM<-TUDOR+SF1-SF2-Helicase+iREase+CTD+REase | 1117 | Bacteria>Pseudomonadota>Alphaproteobacteria>Rhodospirillales>unclassified_Rhodospirillales | GCA_007695045.1 | Rhodospirillales_bacterium | EA405_12420 | 709..4062 | 2026786 | ATP-binding-protein-[Rhodospirillales-bacterium] |
| MBL6940543.1 | HEPN+AAA-ATPase+Alpha+Beta-domain+wHTH+RRM | Type-1_BR.RM.like.systems | Y1_Tnp-><-?<-?\|\|HTH+WYL->TUDOR+SF1-SF2-Helicase+iREase+CTD+REase->METHYLASE->HEPN+DUF499*->HAD-> | 572 | Bacteria>Pseudomonadota>Alphaproteobacteria>Rhodospirillales>unclassified_Rhodospirillales | GCA_016776585.1 | Rhodospirillales_bacterium | ISR53_00145 | 38024..39742 | 2026786 | DUF499-domain-containing-protein-[Rhodospirillales-bacterium] |
| SEP49898.1 | HEPN+AAA-ATPase+Alpha+Beta-domain+wHTH+RRM | Type-1_BR.RM.like.systems | <-HEPN+DUF499*<-PIN<-OLD-ABC<-METHYLASE<-TUDOR+SF1-SF2-Helicase+iREase+CTD+REase<-HNH | 1099 | Bacteria>Pseudomonadota>Alphaproteobacteria>Rhodospirillales>unclassified_Rhodospirillales | GCA_900110395.1 | Rhodospirillales_bacterium_URHD0017 | SAMN02990966_07447 | 5640..8939 | 1380357 | hypothetical-protein-SAMN02990966_07447-[Rhodospirillales-bacterium-URHD0017] |
| MBC6439887.1 | HEPN+AAA-ATPase+Alpha+Beta-domain+wHTH+RRM | Type-1_BR.RM.like.systems | GATase->METHYLASE->HEPN+DUF499*-><-?<-?<-?<-ABC-ATPase+SbcC | 1142 | Bacteria>Pseudomonadota>Alphaproteobacteria>Rhodospirillales>unclassified_Rhodospirillales | GCA_014323865.1 | Rhodospirillales_bacterium | GDA49_05640 | 274906..278334 | 2026786 | ATP-binding-protein-[Rhodospirillales-bacterium] |
| WP_221424189.1 | HEPN+AAA-ATPase+Alpha+Beta-domain+wHTH+RRM | Type-1_BR.RM.like.systems | CopG/RHH->VirD4-TraG+VirD4-FtsK->TrwC+SF1-SF2-helicase->wHTH+SAD+HNH-><-LexA-protease\|\|?-><-HEPN+DUF499*<-METHYLASE<-MORC+DUF3684<-TUDOR+SF1-SF2-Helicase+iREase+CTD+REase<-HTH+WYL\|\|HicB->TIR-> | 1100 | Bacteria>Pseudomonadota>Alphaproteobacteria>Sphingomonadales>Erythrobacteraceae | GCF_019711375.1 | Qipengyuania_aurantiaca | K3148_RS07230 | 1481189..1484491 | 2867233 | DUF499-domain-containing-protein-[Qipengyuania-aurantiaca] |
| WP_120325010.1 | HEPN+AAA-ATPase+Alpha+Beta-domain+wHTH+RRM | Type-1_BR.RM.like.systems | CopG/RHH->VirD4-TraG+VirD4-FtsK->TrwC+SF1-SF2-helicase->wHTH+SAD+HNH-><-LexA-protease\|\|?-><-HEPN+DUF499*<-METHYLASE<-MORC+DUF3684<-TUDOR+SF1-SF2-Helicase+iREase+CTD+REase<-HTH+WYL\|\|HicB->TIR-> | 1100 | Bacteria>Pseudomonadota>Alphaproteobacteria>Sphingomonadales>Erythrobacteraceae | GCF_003610805.1 | Altericroceibacterium_spongiae | D6851_RS11415 | 253315..256617 | 2320269 | DUF499-domain-containing-protein-[Altericroceibacterium-spongiae] |
| WP_034960305.1 | HEPN+AAA-ATPase+Alpha+Beta-domain+wHTH+RRM | Type-1_BR.RM.like.systems | <-TIR<-HicB\|\|HTH+WYL->TUDOR+SF1-SF2-Helicase+iREase+CTD+REase->MORC+DUF3684->METHYLASE->HEPN+DUF499*-><-?\|\|LexA-protease-><-wHTH+SAD+HNH<-TrwC+SF1-SF2-helicase<-VirD4-TraG+VirD4-FtsK<-CopG/RHH | 1100 | Bacteria>Pseudomonadota>Alphaproteobacteria>Sphingomonadales>Erythrobacteraceae | GCF_000715015.1 | Erythrobacter_longus | EH31_RS10535 | 101708..105010 | 1044 | DUF499-domain-containing-protein-[Erythrobacter-longus] |
| WP_017665279.1 | HEPN+AAA-ATPase+Alpha+Beta-domain+wHTH+RRM | Type-1_BR.RM.like.systems | <-HTH<-GDResolvase+Recombinase+ZnR<-RAMA<-?\|\|Peptidase_S74-><-HEPN+DUF499*<-METHYLASE<-TUDOR+SF1-SF2-Helicase+iREase+CTD+REase<-XIS-HTH | 1099 | Bacteria>Pseudomonadota>Alphaproteobacteria>Sphingomonadales>Erythrobacteraceae | GCF_000331285.1 | Porphyrobacter_sp._AAP82 | D620_RS0108005 | 32206..35505 | 1248917 | DUF499-domain-containing-protein-[Porphyrobacter-sp.-AAP82] |
| WP_073974519.1 | HEPN+AAA-ATPase+Alpha+Beta-domain+wHTH+RRM | Type-1_BR.RM.like.systems | <-HicB\|\|HTH+WYL->TUDOR+SF1-SF2-Helicase+iREase+CTD+REase->?->METHYLASE->HEPN+DUF499*->MODE-HTH+PBPII-><-ParG_RHH | 1097 | Bacteria>Pseudomonadota>Alphaproteobacteria>Sphingomonadales>Erythrobacteraceae | GCF_001556555.1 | Erythrobacter_donghaensis | AX698_RS16845 | 33517..36810 | 267135 | DUF499-domain-containing-protein-[Erythrobacter-donghaensis] |
| MBA4046061.1 | HEPN+AAA-ATPase+Alpha+Beta-domain+wHTH+RRM | Type-1_BR.RM.like.systems | TUDOR+SF1-SF2-Helicase+iREase+CTD+REase->MORC+DUF3684->METHYLASE->HEPN+DUF499*->?-><-?<-TraH<-Thioredoxin | 1099 | Bacteria>Pseudomonadota>Alphaproteobacteria>Sphingomonadales>Erythrobacteraceae | GCA_013823025.1 | Erythrobacter_sp. | C0471_16820 | 9896..13195 | 1042 | AAA-family-ATPase-[Erythrobacter-sp.] |
| WP_159791246.1 | HEPN+AAA-ATPase+Alpha+Beta-domain+wHTH+RRM | Type-1_BR.RM.like.systems | ZDNA-HTH->TUDOR+SF1-SF2-Helicase+iREase+CTD+REase->MarR-HTH->?->?->METHYLASE->HEPN+DUF499*-> | 1097 | Bacteria>Pseudomonadota>Alphaproteobacteria>Sphingomonadales>Erythrobacteraceae | GCF_009827435.1 | Croceibacterium_salegens | GRI89_RS00095 | 15268..18561 | 1737568 | DUF499-domain-containing-protein-[Croceibacterium-salegens] |
| MCV0384725.1 | HEPN+AAA-ATPase+Alpha+Beta-domain+wHTH+RRM | Type-1_BR.RM.like.systems | <-HEPN+DUF499*<-METHYLASE<-HNH<-TUDOR+SF1-SF2-Helicase+iREase+CTD+REase-p | 421 | Bacteria>Pseudomonadota>Alphaproteobacteria>Sphingomonadales>Erythrobacteraceae | GCA_025698865.1 | Erythrobacter_sp. | K5799_14985 | 1..1263 | 1042 | MAG:-DUF499-domain-containing-protein,-partial-[Erythrobacter-sp.] |
| WP_100237044.1 | HEPN+AAA-ATPase+Alpha+Beta-domain+wHTH+RRM | Type-1_BR.RM.like.systems | ParG_RHH-><-TetR-HTH\|\|MODE-HTH+PBPII-><-HEPN+DUF499*<-METHYLASE<-MORC+DUF3684<-TUDOR+SF1-SF2-Helicase+iREase+CTD+REase<-HTH+WYL\|\|SF1-SF2-Helicase+REase->SF1-SF2-Helicase+REase-> | 1098 | Bacteria>Pseudomonadota>Alphaproteobacteria>Sphingomonadales>Sphingomonadaceae | GCF_002795205.1 | Sphingobium_sp._LB126 | CAF53_RS01415 | 292890..296186 | 1983755 | DUF499-domain-containing-protein-[Sphingobium-sp.-LB126] |
| WP_025161153.1 | HEPN+AAA-ATPase+Alpha+Beta-domain+wHTH+RRM | Type-1_BR.RM.like.systems | ParG_RHH-><-TetR-HTH\|\|MODE-HTH+PBPII-><-HEPN+DUF499*<-METHYLASE<-MORC+DUF3684<-TUDOR+SF1-SF2-Helicase+iREase+CTD+REase<-HTH+WYL\|\|SF1-SF2-Helicase+REase->SF1-SF2-Helicase+REase-> | 1098 | Bacteria>Pseudomonadota>Alphaproteobacteria>Sphingomonadales>Sphingomonadaceae | GCF_000445085.1 | Sphingobium | L286_RS41375 | 45892..49188 | 165695 | MULTISPECIES:-DUF499-domain-containing-protein-[Sphingobium] |
| EPR17235.1 | HEPN+AAA-ATPase+Alpha+Beta-domain+wHTH+RRM | Type-1_BR.RM.like.systems | ParG_RHH-><-TetR-HTH\|\|MODE-HTH+PBPII-><-HEPN+DUF499*<-METHYLASE<-MORC+DUF3684<-TUDOR+SF1-SF2-Helicase+iREase+CTD+REase<-HTH+WYL\|\|SF1-SF2-Helicase+REase->SF1-SF2-Helicase+REase-> | 1086 | Bacteria>Pseudomonadota>Alphaproteobacteria>Sphingomonadales>Sphingomonadaceae | GCA_000421925.1 | Sphingobium_indicum_IP26 | M527_17965 | 110739..113999 | 1346790 | ATPase-AAA-[Sphingobium-chinhatense-IP26] |
| WP_221211510.1 | HEPN+AAA-ATPase+Alpha+Beta-domain+wHTH+RRM | Type-1_BR.RM.like.systems | <-SF1-SF2-Helicase+REase<-SF1-SF2-Helicase+REase\|\|HTH+WYL->TUDOR+SF1-SF2-Helicase+iREase+CTD+REase->REase->METHYLASE->HEPN+DUF499*->MODE-HTH+PBPII-><-ParG_RHH | 1086 | Bacteria>Pseudomonadota>Alphaproteobacteria>Sphingomonadales>Sphingomonadaceae | GCF_014196255.1 | Sphingomonas_pseudosanguinis | GGR48_RS13090 | 32803..36063 | 413712 | DUF499-domain-containing-protein-[Sphingomonas-pseudosanguinis] |
| MBB3880227.1 | HEPN+AAA-ATPase+Alpha+Beta-domain+wHTH+RRM | Type-1_BR.RM.like.systems | <-SF1-SF2-Helicase+REase<-SF1-SF2-Helicase+REase\|\|HTH+WYL->TUDOR+SF1-SF2-Helicase+iREase+CTD+REase->REase->METHYLASE->HEPN+DUF499*->MODE-HTH+PBPII-><-ParG_RHH | 1098 | Bacteria>Pseudomonadota>Alphaproteobacteria>Sphingomonadales>Sphingomonadaceae | GCA_014196255.1 | Sphingomonas_pseudosanguinis | GGR48_002670 | 32767..36063 | 413712 | hypothetical-protein-[Sphingomonas-pseudosanguinis] |
| WP_037445586.1 | HEPN+AAA-ATPase+Alpha+Beta-domain+wHTH+RRM | Type-1_BR.RM.like.systems | Thioredoxin->TraH->?-><-?<-HEPN+DUF499*<-METHYLASE<-MORC+DUF3684<-TUDOR+SF1-SF2-Helicase+iREase+CTD+REase<-HTH+WYL\|\|HicB->SF1-SF2-helicase+RdgC+Beta-barrel+REase-> | 1101 | Bacteria>Pseudomonadota>Alphaproteobacteria>Sphingomonadales>Sphingomonadaceae | GCF_000633575.2 | Sphingobium_indicum | AL00_RS20465 | 14679..17984 | 332055 | DUF499-domain-containing-protein-[Sphingobium-indicum] |
| WP_053554356.1 | HEPN+AAA-ATPase+Alpha+Beta-domain+wHTH+RRM | Type-1_BR.RM.like.systems | HIN-HTH->?-><-?\|\|\|\|HIN-HTH->?->\|\|<-DEDD_Tnp_IS110+Transposase_20<-HEPN+DUF499*<-METHYLASE<-MORC+DUF3684<-TUDOR+SF1-SF2-Helicase+iREase+CTD+REase<-HTH+WYL<-XIS-HTH | 1115 | Bacteria>Pseudomonadota>Alphaproteobacteria>Sphingomonadales>Sphingomonadaceae | GCF_001278035.1 | Sphingopyxis_sp._113P3 | LH20_RS11780 | 2402847..2406194 | 292913 | Swt1-family-HEPN-domain-containing-protein-[Sphingopyxis-sp.-113P3] |
| WP_205396700.1 | HEPN+AAA-ATPase+Alpha+Beta-domain+wHTH+RRM | Type-1_BR.RM.like.systems | ParG_RHH-><-MODE-HTH+PBPII<-HEPN+DUF499*<-METHYLASE<-?<-TUDOR+SF1-SF2-Helicase+iREase+CTD+REase<-HTH+WYL\|\|SF1-SF2-Helicase+REase->SF1-SF2-Helicase+REase-> | 1098 | Bacteria>Pseudomonadota>Alphaproteobacteria>Sphingomonadales>Sphingomonadaceae | GCF_016919545.1 | Sphingomonas_paucimobilis | JT366_RS09685 | 1990566..1993862 | 13689 | DUF499-domain-containing-protein-[Sphingomonas-paucimobilis] |
| WP_182926476.1 | HEPN+AAA-ATPase+Alpha+Beta-domain+wHTH+RRM | Type-1_BR.RM.like.systems | <-SF1-SF2-Helicase+REase<-SF1-SF2-Helicase+REase\|\|HTH+WYL->TUDOR+SF1-SF2-Helicase+iREase+CTD+REase->?->METHYLASE->HEPN+DUF499*->MODE-HTH+PBPII-><-ParG_RHH | 1098 | Bacteria>Pseudomonadota>Alphaproteobacteria>Sphingomonadales>Sphingomonadaceae | GCF_014169955.1 | Sphingomonas_paucimobilis | SPKIRA_RS16090 | 3478703..3481999 | 13689 | DUF499-domain-containing-protein-[Sphingomonas-paucimobilis] |
| WP_283938333.1 | HEPN+AAA-ATPase+Alpha+Beta-domain+wHTH+RRM | Type-1_BR.RM.like.systems | wHTH+REase->TUDOR+SF1-SF2-Helicase+iREase+CTD+REase->METHYLASE->OLD-ABC->PIN->HEPN+DUF499*-><-?<-BetaPropeller+ABhydrolase\|\|?-><-ABhydrolase<-MODE-HTH+PBPII | 1103 | Bacteria>Pseudomonadota>Alphaproteobacteria>Sphingomonadales>Sphingomonadaceae | GCF_023516505.1 | Sphingomonas_brevis | LZ518_RS11630 | 2249110..2252421 | 2908206 | DUF499-domain-containing-protein-[Sphingomonas-brevis] |
| WP_249916143.1 | HEPN+AAA-ATPase+Alpha+Beta-domain+wHTH+RRM | Type-1_BR.RM.like.systems | wHTH+REase->TUDOR+SF1-SF2-Helicase+iREase+CTD+REase->METHYLASE->OLD-ABC->PIN->HEPN+DUF499*-><-?<-BetaPropeller+ABhydrolase\|\|?-><-ABhydrolase<-MODE-HTH+PBPII | 1094 | Bacteria>Pseudomonadota>Alphaproteobacteria>Sphingomonadales>Sphingomonadaceae | - | Sphingomonas_brevis | LZ518_11630 | 2249137..2252421 | 2908206 | DUF499-domain-containing-protein-[Sphingomonas-sp.-RB56-2] |
| MCL6741777.1 | HEPN+AAA-ATPase+Alpha+Beta-domain+wHTH+RRM | Type-1_BR.RM.like.systems | wHTH+REase->TUDOR+SF1-SF2-Helicase+iREase+CTD+REase->METHYLASE->OLD-ABC->PIN->HEPN+DUF499*-><-?<-BetaPropeller+ABhydrolase\|\|?-><-ABhydrolase<-MODE-HTH+PBPII | 1094 | Bacteria>Pseudomonadota>Alphaproteobacteria>Sphingomonadales>Sphingomonadaceae | GCA_023516505.1 | Sphingomonas_brevis | LZ518_11630 | 2249137..2252421 | 2908206 | DUF499-domain-containing-protein-[Sphingomonas-sp.-RB56-2] |
| WP_079648123.1 | HEPN+AAA-ATPase+Alpha+Beta-domain+wHTH+RRM | Type-1_BR.RM.like.systems | ParG_RHH-><-MODE-HTH+PBPII<-HEPN+DUF499*<-METHYLASE<-MORC+DUF3684<-YtH<-TUDOR+SF1-SF2-Helicase+iREase+CTD+REase<-HTH+WYL\|\|SF1-SF2-Helicase+REase-> | 1099 | Bacteria>Pseudomonadota>Alphaproteobacteria>Sphingomonadales>Sphingomonadaceae | GCF_900167915.1 | Rhizorhabdus_histidinilytica | B5X82_RS10580 | 218430..221729 | 439228 | DUF499-domain-containing-protein-[Rhizorhabdus-histidinilytica] |
| WP_066778028.1 | HEPN+AAA-ATPase+Alpha+Beta-domain+wHTH+RRM | Type-1_BR.RM.like.systems | <-SF1-SF2-Helicase+REase\|\|HTH+WYL->TUDOR+SF1-SF2-Helicase+iREase+CTD+REase->YtH->MORC+DUF3684->METHYLASE->HEPN+DUF499*->MODE-HTH+PBPII-><-ParG_RHH | 1100 | Bacteria>Pseudomonadota>Alphaproteobacteria>Sphingomonadales>Sphingomonadaceae | GCF_001557395.1 | Sphingomonas_sp._CCH5-D11 | AX702_RS07275 | 80145..83447 | 1768786 | DUF499-domain-containing-protein-[Sphingomonas-sp.-CCH5-D11] |
| WP_053553316.1 | HEPN+AAA-ATPase+Alpha+Beta-domain+wHTH+RRM | Type-1_BR.RM.like.systems | <-SF1-SF2-Helicase+REase\|\|HTH+WYL->TUDOR+SF1-SF2-Helicase+iREase+CTD+REase->ASCH->SIR2->METHYLASE->HEPN+DUF499*->MODE-HTH+PBPII-><-ParG_RHH | 1100 | Bacteria>Pseudomonadota>Alphaproteobacteria>Sphingomonadales>Sphingomonadaceae | GCF_001278035.1 | Sphingopyxis_sp._113P3 | LH20_RS05260 | 1100261..1103563 | 292913 | DUF499-domain-containing-protein-[Sphingopyxis-sp.-113P3] |
| WP_226870194.1 | HEPN+AAA-ATPase+Alpha+Beta-domain+wHTH+RRM | Type-1_BR.RM.like.systems | <-SF1-SF2-helicase+RdgC+Beta-barrel+REase<-HicB\|\|HTH+WYL->TUDOR+SF1-SF2-Helicase+iREase+CTD+REase->MORC+DUF3684->METHYLASE->HEPN+DUF499*-> | 1085 | Bacteria>Pseudomonadota>Alphaproteobacteria>Sphingomonadales>Sphingomonadaceae | - | Sphingobium_sp._PNB | K7W03_RS15190 | 38977..42234 | 863934 | DUF499-domain-containing-protein-[Sphingobium-sp.-PNB] |
| MCB4860948.1 | HEPN+AAA-ATPase+Alpha+Beta-domain+wHTH+RRM | Type-1_BR.RM.like.systems | <-SF1-SF2-helicase+RdgC+Beta-barrel+REase<-HicB\|\|HTH+WYL->TUDOR+SF1-SF2-Helicase+iREase+CTD+REase->MORC+DUF3684->METHYLASE->HEPN+DUF499*-> | 1097 | Bacteria>Pseudomonadota>Alphaproteobacteria>Sphingomonadales>Sphingomonadaceae | GCA_019873065.1 | Sphingobium_sp._PNB | K7W03_15250 | 38941..42234 | 863934 | DUF499-domain-containing-protein-[Sphingobium-sp.-PNB] |
| ABQ66570.1 | HEPN+AAA-ATPase+Alpha+Beta-domain+wHTH+RRM | Type-1_BR.RM.like.systems | <-HicB\|\|HTH+WYL->TUDOR+SF1-SF2-Helicase+iREase+CTD+REase->MORC+DUF3684->METHYLASE->HEPN+DUF499*->TIR+PNTD2->PIWI-><-tRNA<-?<-HHH+ENDO3+NUDIX | 1098 | Bacteria>Pseudomonadota>Alphaproteobacteria>Sphingomonadales>Sphingomonadaceae | GCA_000016765.1 | Rhizorhabdus_wittichii_RW1 | Swit_0199 | 198092..201388 | 392499 | ATPase-(AAA+-superfamily)-like-protein-[Rhizorhabdus-wittichii-RW1] |
| WP_120216254.1 | HEPN+AAA-ATPase+Alpha+Beta-domain+wHTH+RRM | Type-1_BR.RM.like.systems | ParG_RHH-><-MODE-HTH+PBPII<-HEPN+DUF499*<-METHYLASE<-MORC+DUF3684<-YtH<-ASCH<-TUDOR+SF1-SF2-Helicase+iREase+CTD+REase<-HTH+WYL | 1098 | Bacteria>Pseudomonadota>Alphaproteobacteria>Sphingomonadales>Sphingomonadaceae | GCF_003609815.1 | Sphingopyxis_sp._EG6 | SPYCW_RS07385 | 1487775..1491071 | 1874061 | DUF499-domain-containing-protein-[Sphingopyxis-sp.-EG6] |
| WP_017183457.1 | HEPN+AAA-ATPase+Alpha+Beta-domain+wHTH+RRM | Type-1_BR.RM.like.systems | <-HicB\|\|HTH+WYL->TUDOR+SF1-SF2-Helicase+iREase+CTD+REase->ParB+HNH->ASCH->METHYLASE->HEPN+DUF499*->MODE-HTH+PBPII-><-ParG_RHH | 1097 | Bacteria>Pseudomonadota>Alphaproteobacteria>Sphingomonadales>Sphingomonadaceae | GCF_000277525.1 | Sphingobium_xenophagum | QYYP_RS0113510 | 14048..17341 | 121428 | DUF499-domain-containing-protein-[Sphingobium-xenophagum] |
| WP_066601098.1 | HEPN+AAA-ATPase+Alpha+Beta-domain+wHTH+RRM | Type-1_BR.RM.like.systems | ABhydrolase->tRNA->TUDOR+SF1-SF2-Helicase+iREase+CTD+REase->?->METHYLASE->HEPN+DUF499*-><-?<-MobA_MobL+SF1-SF2-helicase\|\|TraD->?-><-HEPN | 1100 | Bacteria>Pseudomonadota>Alphaproteobacteria>Sphingomonadales>Sphingomonadaceae | GCF_001046645.1 | Sphingobium_cupriresistens | V473_RS04750 | 974632..977934 | 1132417 | DUF499-domain-containing-protein-[Sphingobium-cupriresistens] |
| WP_156842742.1 | HEPN+AAA-ATPase+Alpha+Beta-domain+wHTH+RRM | Type-1_BR.RM.like.systems | ParG_RHH-><-MODE-HTH+PBPII<-HEPN+DUF499*<-METHYLASE<-MORC+DUF3684<-YtH<-TUDOR+SF1-SF2-Helicase+iREase+CTD+REase<-HTH+WYL | 1098 | Bacteria>Pseudomonadota>Alphaproteobacteria>Sphingomonadales>Sphingomonadaceae | GCF_009746585.1 | Novosphingobium_aquimarinum | GQ094_RS17725 | 28376..31672 | 2682494 | DUF499-domain-containing-protein-[Novosphingobium-aquimarinum] |
| WP_066967630.1 | HEPN+AAA-ATPase+Alpha+Beta-domain+wHTH+RRM | Type-1_BR.RM.like.systems | ParG_RHH-><-MODE-HTH+PBPII<-HEPN+DUF499*<-METHYLASE<-MORC+DUF3684\|\|Arm-DNA-bind_3+Phage_integrase-><-XIS-HTH<-ParG_RHH | 1098 | Bacteria>Pseudomonadota>Alphaproteobacteria>Sphingomonadales>Sphingomonadaceae | GCF_002355275.1 | Rhizorhabdus_dicambivorans | CMV14_RS12985 | 2738842..2742138 | 1850238 | DUF499-domain-containing-protein-[Rhizorhabdus-dicambivorans] |
| WP_248005921.1 | HEPN+AAA-ATPase+Alpha+Beta-domain+wHTH+RRM | Type-1_BR.RM.like.systems | Thioredoxin->TraH->?-><-?<-HEPN+DUF499*<-METHYLASE<-MORC+DUF3684<-TUDOR+SF1-SF2-Helicase+iREase+CTD+REase<-HTH+WYL\|\|HicB-> | 1085 | Bacteria>Pseudomonadota>Alphaproteobacteria>Sphingomonadales>Sphingomonadaceae | - | Novosphingobium_sp._HR1a | HWN72_RS09985 | 40208..43465 | 1395637 | DUF499-domain-containing-protein-[Novosphingobium-sp.-HR1a] |
| WP_067912291.1 | HEPN+AAA-ATPase+Alpha+Beta-domain+wHTH+RRM | Type-1_BR.RM.like.systems | <-HicB\|\|HTH+WYL->TUDOR+SF1-SF2-Helicase+iREase+CTD+REase->MORC+DUF3684->METHYLASE->HEPN+DUF499*->?-><-?<-TraH<-Thioredoxin | 1099 | Bacteria>Pseudomonadota>Alphaproteobacteria>Sphingomonadales>Sphingomonadaceae | GCF_001519075.1 | Novosphingobium_fuchskuhlense | AQZ52_RS14075 | 23149..26448 | 1117702 | DUF499-domain-containing-protein-[Novosphingobium-fuchskuhlense] |
| MBX9665921.1 | HEPN+AAA-ATPase+Alpha+Beta-domain+wHTH+RRM | Type-1_BR.RM.like.systems | <-TOPRIM+SF1-SF2-Helicase+HTH<-ZnR+TOPRIM<-?<-XIS-HTH<-?<-HEPN+DUF499*<-METHYLASE<-?<-TUDOR+SF1-SF2-Helicase+iREase+CTD+REase | 1100 | Bacteria>Pseudomonadota>Alphaproteobacteria>Sphingomonadales>Sphingomonadaceae | GCF_019748365.1 | Novosphingobium_sp. | K5Q19_21680 | 3115..6417 | 1874826 | DUF499-domain-containing-protein-[Novosphingobium-sp.] |
| MBF7011871.1 | HEPN+AAA-ATPase+Alpha+Beta-domain+wHTH+RRM | Type-1_BR.RM.like.systems | Thioredoxin->TraH->?-><-?<-HEPN+DUF499*<-METHYLASE<-MORC+DUF3684<-TUDOR+SF1-SF2-Helicase+iREase+CTD+REase<-HTH+WYL\|\|HicB-> | 1097 | Bacteria>Pseudomonadota>Alphaproteobacteria>Sphingomonadales>Sphingomonadaceae | GCA_015594425.1 | Novosphingobium_sp._HR1a | HWN72_09985 | 40208..43501 | 1395637 | ATP-binding-protein-[Novosphingobium-sp.-HR1a] |
| WP_069321300.1 | HEPN+AAA-ATPase+Alpha+Beta-domain+wHTH+RRM | Type-1_BR.RM.like.systems | ParG_RHH-><-MODE-HTH+PBPII<-HEPN+DUF499*<-METHYLASE<-MORC+DUF3684<-TUDOR+SF1-SF2-Helicase+iREase+CTD+REase<-HTH+WYL | 1099 | Bacteria>Pseudomonadota>Alphaproteobacteria>Sphingomonadales>Sphingomonadaceae | GCF_001721295.1 | Sphingomonas_turrisvirgatae | BFL28_RS14750 | 27774..31073 | 1888892 | DUF499-domain-containing-protein-[Sphingomonas-turrisvirgatae] |
| WP_201515748.1 | HEPN+AAA-ATPase+Alpha+Beta-domain+wHTH+RRM | Type-1_BR.RM.like.systems | <-TraJ-RHH<-?<-?<-?\|\|?->XIS-HTH-><-HEPN+DUF499*<-METHYLASE<-REase<-TUDOR+SF1-SF2-Helicase+iREase+CTD+REase<-HTH+WYL\|\|HicB-> | 1100 | Bacteria>Pseudomonadota>Alphaproteobacteria>Sphingomonadales>Sphingomonadaceae | GCF_904830875.1 | Sphingobium_sp._S6 | SPHS8_RS12190 | 306093..309395 | 2758386 | DUF499-domain-containing-protein-[Sphingobium-sp.-S6] |
| WP_028055914.1 | HEPN+AAA-ATPase+Alpha+Beta-domain+wHTH+RRM | Type-1_BR.RM.like.systems | <-TIR<-HicB\|\|HTH+WYL->TUDOR+SF1-SF2-Helicase+iREase+CTD+REase->MORC+DUF3684->METHYLASE->HEPN+DUF499*-><-XIS-HTH | 1100 | Bacteria>Pseudomonadota>Alphaproteobacteria>Sphingomonadales>Sphingomonadaceae | GCF_000412635.1 | Sphingobium_bisphenolivorans | L858_RS0107580 | 162523..165825 | 1335760 | DUF499-domain-containing-protein-[Sphingobium-bisphenolivorans] |
| MAF61658.1 | HEPN+AAA-ATPase+Alpha+Beta-domain+wHTH+RRM | Type-1_BR.RM.like.systems | ParG_RHH-><-MODE-HTH+PBPII<-HEPN+DUF499*<-METHYLASE<-?<-TUDOR+SF1-SF2-Helicase+iREase+CTD+REase<-HTH+WYL\|\|HicB-> | 1097 | Bacteria>Pseudomonadota>Alphaproteobacteria>Sphingomonadales>Sphingomonadaceae | GCA_002687555.1 | Blastomonas_sp. | CL680_09370 | 351225..354518 | 1909299 | DUF499-domain-containing-protein-[Blastomonas-sp.] |
| WP_066520604.1 | HEPN+AAA-ATPase+Alpha+Beta-domain+wHTH+RRM | Type-1_BR.RM.like.systems | TUDOR+SF1-SF2-Helicase+iREase+CTD+REase-p->ASCH->SIR2->METHYLASE->HEPN+DUF499*->MODE-HTH+PBPII-><-ParG_RHH | 1098 | Bacteria>Pseudomonadota>Alphaproteobacteria>Sphingomonadales>Sphingomonadaceae | GCF_002355855.1 | Sphingobium_cloacae | SC4_RS16230 | 6611..9907 | 120107 | DUF499-domain-containing-protein-[Sphingobium-cloacae] |
| WP_010188165.1 | HEPN+AAA-ATPase+Alpha+Beta-domain+wHTH+RRM | Type-1_BR.RM.like.systems | HTH+WYL->TUDOR+SF1-SF2-Helicase+iREase+CTD+REase->YtH->METHYLASE->HEPN+DUF499*->?->HIN-HTH->?->?->DOC->HTH_3-> | 1098 | Bacteria>Pseudomonadota>Alphaproteobacteria>Sphingomonadales>Sphingomonadaceae | GCF_000241485.1 | Sphingomonas_sp._PAMC_26605 | RH90_RS19210 | 226826..230122 | 1112214 | DUF499-domain-containing-protein-[Sphingomonas-sp.-PAMC-26605] |
| WP_226632678.1 | HEPN+AAA-ATPase+Alpha+Beta-domain+wHTH+RRM | Type-1_BR.RM.like.systems | HTH+WYL->TUDOR+SF1-SF2-Helicase+iREase+CTD+REase->METHYLASE->HEPN+DUF499*-><-ACET<-?\|\|<-?<-STYKinase | 1110 | Bacteria>Pseudomonadota>Alphaproteobacteria>Sphingomonadales>Sphingomonadaceae | GCF_020524485.1 | Novosphingobium_profundi | LC060_RS03305 | 760563..763895 | 1774954 | DUF499-domain-containing-protein-[Novosphingobium-profundi] |
| WP_054108969.1 | HEPN+AAA-ATPase+Alpha+Beta-domain+wHTH+RRM | Type-1_BR.RM.like.systems | HIN-HTH->?-><-HEPN+DUF499*<-METHYLASE<-TUDOR+SF1-SF2-Helicase+iREase+CTD+REase<-HTH+WYL<-?\|\|HTH_3-> | 1099 | Bacteria>Pseudomonadota>Alphaproteobacteria>Sphingomonadales>Sphingomonadaceae | GCF_001295795.1 | Novosphingobium_sp._AAP83 | IP81_RS15755 | 15292..18591 | 1523425 | DUF499-domain-containing-protein-[Novosphingobium-sp.-AAP83] |
| WP_145073130.1 | HEPN+AAA-ATPase+Alpha+Beta-domain+wHTH+RRM | Type-1_BR.RM.like.systems | TetR-HTH->XIS-HTH->TUDOR+SF1-SF2-Helicase+iREase+CTD+REase->ASCH->METHYLASE->HEPN+DUF499*-> | 1099 | Bacteria>Pseudomonadota>Alphaproteobacteria>Sphingomonadales>Sphingomonadaceae | GCF_014205735.1 | Sphingobium_wenxiniae | IQ35_RS10205 | 192114..195413 | 595605 | DUF499-domain-containing-protein-[Sphingobium-wenxiniae] |
| WP_194920702.1 | HEPN+AAA-ATPase+Alpha+Beta-domain+wHTH+RRM | Type-1_BR.RM.like.systems | METHYLASE-><-DDE_Tnp_1_4\|\|METHYLASE-p->OLD-ABC->?->HIN-HTH->HEPN+DUF499*->HIN-HTH->?-><-XIS-HTH | 1041 | Bacteria>Pseudomonadota>Alphaproteobacteria>Sphingomonadales>Sphingomonadaceae | GCF_015390225.1 | Novosphingobium_sp._NBM11 | F1640_RS18815 | 3882459..3885584 | 2596914 | DUF499-domain-containing-protein-[Novosphingobium-sp.-NBM11] |
| WP_184477099.1 | HEPN+AAA-ATPase+Alpha+Beta-domain+wHTH+RRM | Type-1_BR.RM.like.systems | N6_MTase->REase-><-?<-?<-URI-DNase<-ParB+HTH<-HEPN+DUF499*<-METHYLASE<-ASCH<-HTH+WYL | 1098 | Bacteria>Pseudomonadota>Alphaproteobacteria>Sphingomonadales>Sphingomonadaceae | GCF_014200045.1 | Rhizorhapis_suberifaciens | HNQ99_RS14965 | 12710..16006 | 13656 | DUF499-domain-containing-protein-[Rhizorhapis-suberifaciens] |
| MBP8234957.1 | HEPN+AAA-ATPase+Alpha+Beta-domain+wHTH+RRM | Type-1_BR.RM.like.systems | ParG_RHH-><-MODE-HTH+PBPII<-HEPN+DUF499*<-METHYLASE-p | 1098 | Bacteria>Pseudomonadota>Alphaproteobacteria>Sphingomonadales>Sphingomonadaceae | GCA_018006725.1 | Rhizorhabdus_sp. | KAY22_21925 | 3665..6961 | 1968843 | DUF499-domain-containing-protein-[Rhizorhabdus-sp.] |
| PZU56403.1 | HEPN+AAA-ATPase+Alpha+Beta-domain+wHTH+RRM | Type-1_BR.RM.like.systems | METHYLASE->HEPN+DUF499*-><-?<-?<-?<-LON-AAA | 1100 | Bacteria>Pseudomonadota>Alphaproteobacteria>Sphingomonadales>Sphingomonadaceae | GCA_003248935.1 | Sphingobium_sp. | DI547_16590 | 2951..6253 | 1912891 | ATPase-[Sphingobium-sp.] |
| MCX7283721.1 | HEPN+AAA-ATPase+Alpha+Beta-domain+wHTH+RRM | Type-1_BR.RM.like.systems | <-HEPN+DUF499*<-METHYLASE-p | 625 | Bacteria>Pseudomonadota>Alphaproteobacteria>Sphingomonadales>Sphingomonadaceae | GCA_026400375.1 | Novosphingobium_sp. | NTX28_06715 | 1..1874 | 1874826 | MAG:-DUF499-domain-containing-protein,-partial-[Novosphingobium-sp.] |
| WP_147041999.1 | HEPN+AAA-ATPase+Alpha+Beta-domain+wHTH+RRM | Type-1_BR.RM.like.systems | <-REase<-?\|\|HTH_3-><-?<-PIWI<-TIR+PNTD2<-HEPN+DUF499*<-METHYLASE<-?<-TUDOR+SF1-SF2-Helicase+iREase+CTD+REase<-HTH+WYL\|\|HicB-> | 1098 | Bacteria>Pseudomonadota>Alphaproteobacteria>Sphingomonadales>Sphingosinicellaceae | GCF_007995055.1 | Sphingosinicella_ginsenosidimutans | FRZ32_RS02405 | 490620..493916 | 1176539 | DUF499-domain-containing-protein-[Sphingosinicella-ginsenosidimutans] |
| WP_017667308.1 | HEPN+AAA-ATPase+Alpha+Beta-domain+wHTH+RRM | Type-1_BR.RM.like.systems | HEPN->Phage_integrase->TUDOR+SF1-SF2-Helicase+iREase+CTD+REase->METHYLASE->HEPN+DUF499*->PIN-> | 1100 | Bacteria>Pseudomonadota>Alphaproteobacteria>Sphingomonadales>Sphingosinicellaceae | GCF_000331225.1 | Sandarakinorhabdus_sp._AAP62 | D619_RS0104150 | 158738..162040 | 1248916 | DUF499-domain-containing-protein-[Sandarakinorhabdus-sp.-AAP62] |
| MBS3961271.1 | HEPN+AAA-ATPase+Alpha+Beta-domain+wHTH+RRM | Type-1_BR.RM.like.systems | <-TIR<-HEPN+DUF499*<-METHYLASE | 1098 | Bacteria>Pseudomonadota>Alphaproteobacteria>Sphingomonadales>Sphingosinicellaceae | GCA_018333935.1 | Sandarakinorhabdus_sp. | KGZ61_05485 | 446..3742 | 1916663 | DUF499-domain-containing-protein-[Sandarakinorhabdus-sp.] |
| WP_094472303.1 | HEPN+AAA-ATPase+Alpha+Beta-domain+wHTH+RRM | Type-1_BR.RM.like.systems | <-HEPN+DUF499*<-?<-METHYLASE-p | 1098 | Bacteria>Pseudomonadota>Alphaproteobacteria>Sphingomonadales>Sphingosinicellaceae | GCF_002251755.1 | Sandarakinorhabdus_cyanobacteriorum | CHU93_RS00700 | 1379..4675 | 1981098 | DUF499-domain-containing-protein-[Sandarakinorhabdus-cyanobacteriorum] |
| TXG85100.1 | HEPN+AAA-ATPase+Alpha+Beta-domain+wHTH+RRM | Type-1_BR.RM.like.systems | ParG_RHH-><-MODE-HTH+PBPII<-HEPN+DUF499*<-METHYLASE<-MORC+DUF3684<-YtH<-TUDOR+SF1-SF2-Helicase+iREase+CTD+REase<-HTH+WYL\|\|SF1-SF2-Helicase+REase-> | 1098 | Bacteria>Pseudomonadota>Alphaproteobacteria>Sphingomonadales>unclassified_Sphingomonadales | GCA_008012005.1 | Sphingomonadales_bacterium | E6R12_02055 | 22905..26201 | 1978525 | MAG:-ATP-binding-protein-[Sphingomonadales-bacterium] |
| NCP12883.1 | HEPN+AAA-ATPase+Alpha+Beta-domain+wHTH+RRM | Type-1_BR.RM.like.systems | <-TIR<-HicB\|\|HTH+WYL->TUDOR+SF1-SF2-Helicase+iREase+CTD+REase->MORC+DUF3684->METHYLASE->HEPN+DUF499*-> | 1100 | Bacteria>Pseudomonadota>Alphaproteobacteria>Sphingomonadales>unclassified_Sphingomonadales | GCA_009993565.1 | Sphingomonadales_bacterium | GW858_01815 | 17021..20323 | 1978525 | ATP-binding-protein-[Sphingomonadales-bacterium] |
| MBI1213755.1 | HEPN+AAA-ATPase+Alpha+Beta-domain+wHTH+RRM | Type-1_BR.RM.like.systems | DEOR-HTH->TUDOR+SF1-SF2-Helicase+iREase+CTD+REase->?->?->TPR->METHYLASE->HEPN+DUF499*->HEPN->DOC+HTH-><-?\|\|Phage_integrase-><-AbiEi_4\|\|AbiEi_4-> | 1117 | Bacteria>Pseudomonadota>Alphaproteobacteria>unclassified_Alphaproteobacteria | GCA_016124635.1 | Alphaproteobacteria_bacterium | GC190_20030 | 81379..84732 | 1913988 | DUF499-domain-containing-protein-[Alphaproteobacteria-bacterium] |
| MCR9130420.1 | HEPN+AAA-ATPase+Alpha+Beta-domain+wHTH+RRM | Type-1_BR.RM.like.systems | <-NTP_transferase<-?<-REase+REase<-Mbetalac<-REase<-?<-HEPN+DUF499*<-METHYLASE<-MarR-HTH<-TUDOR+SF1-SF2-Helicase+iREase+CTD+REase<-?\|\|XIS-HTH-> | 1090 | Bacteria>Pseudomonadota>Alphaproteobacteria>unclassified_Alphaproteobacteria | GCA_024742735.1 | Alphaproteobacteria_bacterium | NXI12_12945 | 97854..101126 | 1913988 | DUF499-domain-containing-protein-[Alphaproteobacteria-bacterium] |
| NBC34425.1 | HEPN+AAA-ATPase+Alpha+Beta-domain+wHTH+RRM | Type-1_BR.RM.like.systems | <-DUF29<-?\|\|?-><-HEPN+DUF499*<-METHYLASE<-TUDOR+SF1-SF2-Helicase+iREase+CTD+REase<-tRNA<-ZnR+TOPRIM+DnaB<-?\|\|?->Polbeta-> | 1148 | Bacteria>Pseudomonadota>Alphaproteobacteria>unclassified_Alphaproteobacteria | GCA_009909005.1 | Alphaproteobacteria_bacterium | GVY13_17240 | 19753..23199 | 1913988 | DUF499-domain-containing-protein-[Alphaproteobacteria-bacterium] |
| MDA8776763.1 | HEPN+AAA-ATPase+Alpha+Beta-domain+wHTH+RRM | Type-1_BR.RM.like.systems | Arm-DNA-bind_3+Phage_int_SAM_3+Phage_integrase->XIS-HTH->DnaB-><-?<-REase<-HEPN+DUF499*<-METHYLASE | 1094 | Bacteria>Pseudomonadota>Alphaproteobacteria>unclassified_Alphaproteobacteria | GCA_028101585.1 | Alphaproteobacteria_bacterium | N9N58_04660 | 21038..24322 | 1913988 | DUF499-domain-containing-protein-[Alphaproteobacteria-bacterium] |
| MCL2716067.1 | HEPN+AAA-ATPase+Alpha+Beta-domain+wHTH+RRM | Type-1_BR.RM.like.systems | TUDOR+SF1-SF2-Helicase+iREase+CTD+REase-><-?\|\|METHYLASE->HEPN+DUF499*->REase+PNTD1+PNTD2->PIWI-> | 1073 | Bacteria>Pseudomonadota>Alphaproteobacteria>unclassified_Alphaproteobacteria | GCA_009781045.1 | Alphaproteobacteria_bacterium | FWD68_16235 | 9484..12705 | 1913988 | DUF499-domain-containing-protein-[Alphaproteobacteria-bacterium] |
| MDA8675490.1 | HEPN+AAA-ATPase+Alpha+Beta-domain+wHTH+RRM | Type-1_BR.RM.like.systems | RNase->?->tRNA->Phage_integrase->?->TUDOR+SF1-SF2-Helicase+iREase+CTD+REase->METHYLASE->HEPN+DUF499*-> | 1091 | Bacteria>Pseudomonadota>Alphaproteobacteria>unclassified_Alphaproteobacteria | GCA_028099405.1 | Alphaproteobacteria_bacterium | N9M53_02250 | 53918..57193 | 1913988 | DUF499-domain-containing-protein-[Alphaproteobacteria-bacterium] |
| MBF0333240.1 | HEPN+AAA-ATPase+Alpha+Beta-domain+wHTH+RRM | Type-1_BR.RM.like.systems | TUDOR+SF1-SF2-Helicase+iREase+CTD+REase->HEPN->METHYLASE->PolB->HEPN->HEPN+DUF499*-> | 1111 | Bacteria>Pseudomonadota>Alphaproteobacteria>unclassified_Alphaproteobacteria | GCA_015232055.1 | Alphaproteobacteria_bacterium | HQL40_06270 | 8520..11855 | 1913988 | DUF499-domain-containing-protein-[Alphaproteobacteria-bacterium] |
| KUO66709.1 | HEPN+AAA-ATPase+Alpha+Beta-domain+wHTH+RRM | Type-1_BR.RM.like.systems | ZnR+TOPRIM+D5ATPase-><-?\|\|?-><-HEPN+DUF499*<-METHYLASE<-TUDOR+SF1-SF2-Helicase+iREase+CTD+REase | 1101 | Bacteria>Pseudomonadota>Alphaproteobacteria>unclassified_Alphaproteobacteria | GCA_001516065.1 | Alphaproteobacteria_bacterium_BRH_c36 | APF80_06945 | 3407..6712 | 1734406 | hypothetical-protein-APF80_06945-[Alphaproteobacteria-bacterium-BRH_c36] |
| MCB9797660.1 | HEPN+AAA-ATPase+Alpha+Beta-domain+wHTH+RRM | Type-1_BR.RM.like.systems | TUDOR+SF1-SF2-Helicase+iREase+CTD+REase->METHYLASE->OLD-ABC->TOPRIM->HEPN+DUF499*-> | 1109 | Bacteria>Pseudomonadota>Alphaproteobacteria>unclassified_Alphaproteobacteria | GCA_020632765.1 | Alphaproteobacteria_bacterium | H6741_33640 | 16618..19947 | 1913988 | ATP-binding-protein-[Alphaproteobacteria-bacterium] |
| MCB9746493.1 | HEPN+AAA-ATPase+Alpha+Beta-domain+wHTH+RRM | Type-1_BR.RM.like.systems | <-HEPN+DUF499*<-TOPRIM<-OLD-ABC<-METHYLASE<-TUDOR+SF1-SF2-Helicase+iREase+CTD+REase | 1109 | Bacteria>Pseudomonadota>Alphaproteobacteria>unclassified_Alphaproteobacteria | GCA_020632815.1 | Alphaproteobacteria_bacterium | H6740_28215 | 5203..8532 | 1913988 | ATP-binding-protein-[Alphaproteobacteria-bacterium] |
| RMH48112.1 | HEPN+AAA-ATPase+Alpha+Beta-domain+wHTH+RRM | Type-1_BR.RM.like.systems | METHYLASE-p->PolB->HEPN->HEPN+DUF499*-> | 1114 | Bacteria>Pseudomonadota>Alphaproteobacteria>unclassified_Alphaproteobacteria | GCA_003696705.1 | Alphaproteobacteria_bacterium | D6688_02640 | 1365..4709 | 1913988 | ATP-binding-protein-[Alphaproteobacteria-bacterium] |
| MBM3508286.1 | HEPN+AAA-ATPase+Alpha+Beta-domain+wHTH+RRM | Type-1_BR.RM.like.systems | <-HEPN+DUF499*<-HEPN<-PolB<-METHYLASE | 566 | Bacteria>Pseudomonadota>Alphaproteobacteria>unclassified_Alphaproteobacteria | GCA_016869235.1 | Alphaproteobacteria_bacterium | FJX64_11435 | 1..1697 | 1913988 | ATP-binding-protein-[Alphaproteobacteria-bacterium] |
| PKP67098.1 | HEPN+AAA-ATPase+Alpha+Beta-domain+wHTH+RRM | Type-1_BR.RM.like.systems | <-HEPN+DUF499*<-METHYLASE-p | 366 | Bacteria>Pseudomonadota>Alphaproteobacteria>unclassified_Alphaproteobacteria | GCA_002842795.1 | Alphaproteobacteria_bacterium_HGW-Alphaproteobacteria-8 | CVT86_02370 | 1..1098 | 2013671 | AAA-family-ATPase,-partial-[Alphaproteobacteria-bacterium-HGW-Alphaproteobacteria-8] |
| HHY48691.1 | HEPN+AAA-ATPase+Alpha+Beta-domain+wHTH+RRM | Type-1_BR.RM.like.systems | <-HEPN+DUF499*<-METHYLASE-p | 365 | Bacteria>Pseudomonadota>Alphaproteobacteria>unclassified_Alphaproteobacteria | GCA_012842825.1 | Alphaproteobacteria_bacterium | GYA10_02965 | 1..1094 | 1913988 | ATP-binding-protein-[Alphaproteobacteria-bacterium] |
| WP_169068632.1 | HEPN+AAA-ATPase+Alpha+Beta-domain+wHTH+RRM | Type-1_BR.RM.like.systems | TUDOR+SF1-SF2-Helicase+iREase+CTD+REase->MORC+DUF3684->wHTH+REase->METHYLASE->PolB->HEPN->HEPN+DUF499*->MarR-HTH->tRNA->?-><-GDResolvase+HIN-HTH\|\|MetJarc-HTH->RelE-ParE-> | 1113 | Bacteria>Pseudomonadota>Betaproteobacteria>Betaproteobacteria_incertae_sedis | GCF_907163155.1 | Candidatus_Accumulibacter | QZ648_RS13960 | 40235..43576 | 327159 | Swt1-family-HEPN-domain-containing-protein-[Candidatus-Accumulibacter-phosphatis] |
| QLH49283.1 | HEPN+AAA-ATPase+Alpha+Beta-domain+wHTH+RRM | Type-1_BR.RM.like.systems | TUDOR+SF1-SF2-Helicase+iREase+CTD+REase->OLD-ABC->TOPRIM->OLD-ABC->TOPRIM->METHYLASE->HEPN+DUF499*-><-tRNA<-?<-GNAT<-?<-?<-ABhydrolase | 1112 | Bacteria>Pseudomonadota>Betaproteobacteria>Betaproteobacteria_incertae_sedis>Candidatus_Accumulibacter | GCA_013414765.1 | Candidatus_Accumulibacter_cognatus | HWD57_05410 | 1172035..1175373 | 2954383 | MAG:-ATP-binding-protein-[Candidatus-Accumulibacter-cognatus] |
| EXI84547.1 | HEPN+AAA-ATPase+Alpha+Beta-domain+wHTH+RRM | Type-1_BR.RM.like.systems | TPR+TPR+TPR+TPR+TPR+TPR+TPR+TPR-><-HerA/FtsK<-DDE_Tnp_1<-HerA/FtsK<-?\|\|HEPN-><-HEPN+DUF499*<-METHYLASE | 1114 | Bacteria>Pseudomonadota>Betaproteobacteria>Betaproteobacteria_incertae_sedis>Candidatus_Accumulibacter | GCA_000585075.1 | Candidatus_Accumulibacter_regalis | AW11_03844 | 11867..15211 | 522306 | MAG:-hypothetical-protein-AW11_03844-[Candidatus-Accumulibacter-regalis] |
| MBK8115022.1 | HEPN+AAA-ATPase+Alpha+Beta-domain+wHTH+RRM | Type-1_BR.RM.like.systems | TRD+TRD->Schlafen+HATPase+HTH-><-?<-?<-TPR+TPR<-?<-HEPN+DUF499*<-HEPN<-PolB<-HEPN<-PolB<-METHYLASE<-TOPRIM | 1109 | Bacteria>Pseudomonadota>Betaproteobacteria>Betaproteobacteria_incertae_sedis>Candidatus_Accumulibacter | GCA_016712935.1 | Accumulibacter_sp. | IPK44_10990 | 647374..650703 | 2053492 | DUF499-domain-containing-protein-[Candidatus-Accumulibacter-sp.] |
| WP_138677185.1 | HEPN+AAA-ATPase+Alpha+Beta-domain+wHTH+RRM | Type-1_BR.RM.like.systems | ABhydrolase->?->?->GNAT->?->tRNA-><-HEPN+DUF499*<-HEPN<-PolB<-METHYLASE<-TOPRIM<-OLD-ABC<-METHYLASE-p | 1112 | Bacteria>Pseudomonadota>Betaproteobacteria>Betaproteobacteria_incertae_sedis>Candidatus_Accumulibacter | GCF_005889575.1 | Candidatus_Accumulibacter_phosphatis | ACCUM_RS00275 | 63215..66553 | 327160 | Swt1-family-HEPN-domain-containing-protein-[Candidatus-Accumulibacter-phosphatis] |
| MBN8517040.1 | HEPN+AAA-ATPase+Alpha+Beta-domain+wHTH+RRM | Type-1_BR.RM.like.systems | TOPRIM->OLD-ABC->TOPRIM->METHYLASE->PolB->HEPN->HEPN+DUF499*-><-tRNA<-?<-GNAT<-?<-?<-ABhydrolase | 1112 | Bacteria>Pseudomonadota>Betaproteobacteria>Betaproteobacteria_incertae_sedis>Candidatus_Accumulibacter | GCA_017302385.1 | Accumulibacter_sp. | JOL71_04035 | 55735..59073 | 2053492 | DUF499-domain-containing-protein-[Candidatus-Accumulibacter-sp.] |
| MBK7674932.1 | HEPN+AAA-ATPase+Alpha+Beta-domain+wHTH+RRM | Type-1_BR.RM.like.systems | PIN->PolB->HEPN->PolB->HEPN->HEPN+DUF499*->?->Response_reg+LuxR-HTH-> | 1114 | Bacteria>Pseudomonadota>Betaproteobacteria>Betaproteobacteria_incertae_sedis>Candidatus_Accumulibacter | GCA_016709675.1 | Candidatus_Accumulibacter_proximus | IPJ27_09305 | 116820..120164 | 2954385 | DUF499-domain-containing-protein-[Candidatus-Accumulibacter-sp.] |
| MCQ1549843.1 | HEPN+AAA-ATPase+Alpha+Beta-domain+wHTH+RRM | Type-1_BR.RM.like.systems | <-XIS-HTH<-?<-?<-?<-Arm-DNA-bind_3+Phage_integrase<-HEPN+DUF499*<-METHYLASE-p | 1112 | Bacteria>Pseudomonadota>Betaproteobacteria>Betaproteobacteria_incertae_sedis>Candidatus_Accumulibacter | GCA_024380085.1 | Candidatus_Accumulibacter_phosphatis | NOF05_13730 | 62282..65620 | 327160 | Swt1-family-HEPN-domain-containing-protein-[Candidatus-Accumulibacter-phosphatis] |
| MBN8437943.1 | HEPN+AAA-ATPase+Alpha+Beta-domain+wHTH+RRM | Type-1_BR.RM.like.systems | PIN->METHYLASE->PolB->HEPN->PolB->HEPN->HEPN+DUF499*->?->TPR+TPR-> | 1109 | Bacteria>Pseudomonadota>Betaproteobacteria>Betaproteobacteria_incertae_sedis>Candidatus_Accumulibacter | GCA_017302555.1 | Accumulibacter_sp. | JOL95_07850 | 90759..94088 | 2053492 | DUF499-domain-containing-protein-[Candidatus-Accumulibacter-sp.] |
| WP_169069333.1 | HEPN+AAA-ATPase+Alpha+Beta-domain+wHTH+RRM | Type-1_BR.RM.like.systems | <-HEPN+DUF499*<-HEPN<-PolB<-METHYLASE<-HNH<-OLD-ABC<-PemK_toxin | 1112 | Bacteria>Pseudomonadota>Betaproteobacteria>Betaproteobacteria_incertae_sedis>Candidatus_Accumulibacter | GCF_012940005.1 | Candidatus_Accumulibacter_contiguus | E4Q08_RS03235 | 1844..5182 | 2954381 | Swt1-family-HEPN-domain-containing-protein-[Candidatus-Accumulibacter-contiguus] |
| MCM8599234.1 | HEPN+AAA-ATPase+Alpha+Beta-domain+wHTH+RRM | Type-1_BR.RM.like.systems | MerR-HTH->wHTH+REase->METHYLASE->?->DUF488-1->?->HEPN+DUF499*-> | 512 | Bacteria>Pseudomonadota>Betaproteobacteria>Betaproteobacteria_incertae_sedis>Candidatus_Accumulibacter | GCA_023806585.1 | Accumulibacter_sp. | NFW04_11345 | 16520..18057 | 2053492 | Swt1-family-HEPN-domain-containing-protein-[Candidatus-Accumulibacter-sp.] |
| MCM8594943.1 | HEPN+AAA-ATPase+Alpha+Beta-domain+wHTH+RRM | Type-1_BR.RM.like.systems | <-NUDIX\|\|?->?-><-HEPN+DUF499*<-STAS<-HEPN<-PolB | 1108 | Bacteria>Pseudomonadota>Betaproteobacteria>Betaproteobacteria_incertae_sedis>Candidatus_Accumulibacter | GCA_023806595.1 | Accumulibacter_sp. | NFV71_06200 | 31893..35219 | 2053492 | Swt1-family-HEPN-domain-containing-protein-[Candidatus-Accumulibacter-sp.] |
| HCZ17096.1 | HEPN+AAA-ATPase+Alpha+Beta-domain+wHTH+RRM | Type-1_BR.RM.like.systems | <-HEPN+DUF499*<-HEPN<-PolB<-METHYLASE-p | 1112 | Bacteria>Pseudomonadota>Betaproteobacteria>Betaproteobacteria_incertae_sedis>Candidatus_Accumulibacter | GCA_003538495.1 | Accumulibacter_sp. | DHV85_21500 | 171..3509 | 2053492 | AAA+-family-ATPase-[Candidatus-Accumulibacter-sp.] |
| OYY21358.1 | HEPN+AAA-ATPase+Alpha+Beta-domain+wHTH+RRM | Type-1_BR.RM.like.systems | <-ParB+HNH<-REase+35exo<-HTH+WYL<-Peptidase<-HEPN+DUF499*<-METHYLASE<-TUDOR+SF1-SF2-Helicase+iREase+CTD+REase\|\|MerR-HTH->?-><-REase | 1109 | Bacteria>Pseudomonadota>Betaproteobacteria>Burkholderiales>Burkholderiaceae | GCA_002281105.1 | Polynucleobacter_sp._35-46-11 | B7Y67_02075 | 7019..10348 | 1970425 | MAG:-AAA+-family-ATPase-[Polynucleobacter-sp.-35-46-11] |
| WP_205989436.1 | HEPN+AAA-ATPase+Alpha+Beta-domain+wHTH+RRM | Type-1_BR.RM.like.systems | RelB->RelE-ParE->TUDOR+SF1-SF2-Helicase+iREase+CTD+REase->METHYLASE->HEPN+DUF499*-><-HTH+WYL\|\|?->?<-HTH+UbE1-> | 1110 | Bacteria>Pseudomonadota>Betaproteobacteria>Burkholderiales>Burkholderiaceae | GCF_017105025.1 | Paraburkholderia_sp._Ac-20336 | GXB81_RS13960 | 20393..23725 | 2703886 | Swt1-family-HEPN-domain-containing-protein-[Paraburkholderia-sp.-Ac-20336] |
| WP_035924723.1 | HEPN+AAA-ATPase+Alpha+Beta-domain+wHTH+RRM | Type-1_BR.RM.like.systems | Phage_integrase-><-HEPN+DUF499*<-METHYLASE<-TUDOR+SF1-SF2-Helicase+iREase+CTD+REase<-DUF2326<-?<-?<-HTH_37 | 1108 | Bacteria>Pseudomonadota>Betaproteobacteria>Burkholderiales>Burkholderiaceae | GCF_902833485.1 | Caballeronia_glathei | BG61_RS34410 | 1980..5306 | 60547 | Swt1-family-HEPN-domain-containing-protein-[Caballeronia-glathei] |
| WP_062175042.1 | HEPN+AAA-ATPase+Alpha+Beta-domain+wHTH+RRM | Type-1_BR.RM.like.systems | <-PIN<-SF1-SF2-Helicase<-OLD-ABC+TOPRIM<-HEPN+DUF499*<-METHYLASE<-MORC+DUF3684<-DUF2326<-?<-?<-HTH_37 | 1108 | Bacteria>Pseudomonadota>Betaproteobacteria>Burkholderiales>Burkholderiaceae | GCF_001557535.2 | Burkholderia_sp._PAMC_26561 | AXG89_RS34140 | 126894..130220 | 1795043 | Swt1-family-HEPN-domain-containing-protein-[Burkholderia-sp.-PAMC-26561] |
| MBT8573895.1 | HEPN+AAA-ATPase+Alpha+Beta-domain+wHTH+RRM | Type-1_BR.RM.like.systems | Phage_integrase-><-?\|\|?->?->?-><-HEPN+DUF499*<-METHYLASE<-TUDOR+SF1-SF2-Helicase+iREase+CTD+REase | 1100 | Bacteria>Pseudomonadota>Betaproteobacteria>Burkholderiales>Burkholderiaceae | GCA_018705385.1 | Polynucleobacter_paneuropaeus | G6701_00110 | 16919..20221 | 2527775 | ATP-binding-protein-[Polynucleobacter-paneuropaeus] |
| MBT8525405.1 | HEPN+AAA-ATPase+Alpha+Beta-domain+wHTH+RRM | Type-1_BR.RM.like.systems | Phage_integrase-><-?\|\|?->?->?-><-HEPN+DUF499*<-METHYLASE<-TUDOR+SF1-SF2-Helicase+iREase+CTD+REase | 1103 | Bacteria>Pseudomonadota>Betaproteobacteria>Burkholderiales>Burkholderiaceae | GCA_018705035.1 | Polynucleobacter_paneuropaeus | G6724_05515 | 1135363..1138674 | 2527775 | ATP-binding-protein-[Polynucleobacter-paneuropaeus] |
| MCA3183095.1 | HEPN+AAA-ATPase+Alpha+Beta-domain+wHTH+RRM | Type-1_BR.RM.like.systems | METHYLASE-p->HEPN+DUF499*-> | 993 | Bacteria>Pseudomonadota>Betaproteobacteria>Burkholderiales>Burkholderiaceae | GCA_020248745.1 | Cupriavidus_sp. | INH13_05675 | 475..3455 | 1873897 | DUF499-domain-containing-protein-[Cupriavidus-sp.] |
| MCP5281003.1 | HEPN+AAA-ATPase+Alpha+Beta-domain+wHTH+RRM | Type-1_BR.RM.like.systems | <-HEPN+DUF499*<-?<-METHYLASE<-DUF932<-ParB+HNH<-?<-TUDOR+SF1-SF2-Helicase+iREase+CTD+REase | 1107 | Bacteria>Pseudomonadota>Betaproteobacteria>Burkholderiales>Comamonadaceae | GCA_024235935.1 | Rhodoferax_sp. | H6930_05335 | 220759..224082 | 50421 | ATP-binding-protein-[Rhodoferax-sp.] |
| WP_119292278.1 | HEPN+AAA-ATPase+Alpha+Beta-domain+wHTH+RRM | Type-1_BR.RM.like.systems | <-SbcC<-?<-?<-REase<-?<-HEPN+DUF499*<-HEPN<-PolB<-SbcC<-METHYLASE<-sGTP<-TUDOR+SF1-SF2-Helicase+iREase+CTD+REase | 1114 | Bacteria>Pseudomonadota>Betaproteobacteria>Burkholderiales>Sphaerotilaceae | GCF_003574675.1 | Azohydromonas_sediminis | DRV96_RS14230 | 10149..13493 | 2259674 | Swt1-family-HEPN-domain-containing-protein-[Calidifontimicrobium-sediminis] |
| MCC6854228.1 | HEPN+AAA-ATPase+Alpha+Beta-domain+wHTH+RRM | Type-1_BR.RM.like.systems | <-HEPN+DUF499*<-HEPN<-PolB<-METHYLASE<-PemK_toxin<-PrlF_antitoxin<-TOPRIM | 1117 | Bacteria>Pseudomonadota>Betaproteobacteria>Burkholderiales>Sphaerotilaceae | GCA_020852865.1 | Rubrivivax_sp. | IT502_18195 | 74611..77964 | 50259 | DUF499-domain-containing-protein-[Rubrivivax-sp.] |
| MCC6611669.1 | HEPN+AAA-ATPase+Alpha+Beta-domain+wHTH+RRM | Type-1_BR.RM.like.systems | VirD4-TraG+VirD4-FtsK->TrwC+SF1-SF2-helicase+TOPRIM-><-PIWI<-TIR+PNTD2<-HEPN+DUF499*<-METHYLASE<-PIN<-?<-STAS+HisKin<-TUDOR+SF1-SF2-Helicase+iREase+CTD+REase<-PIN | 1116 | Bacteria>Pseudomonadota>Betaproteobacteria>Burkholderiales>unclassified_Burkholderiales | GCA_020848075.1 | Burkholderiales_bacterium | IT515_18605 | 34273..37623 | 1891238 | DUF499-domain-containing-protein-[Burkholderiales-bacterium] |
| MBE0627115.1 | HEPN+AAA-ATPase+Alpha+Beta-domain+wHTH+RRM | Type-1_BR.RM.like.systems | TUDOR+SF1-SF2-Helicase+iREase+CTD+REase->OLD-ABC->TOPRIM->HEPN->METHYLASE->HEPN+DUF499*->?->?->HEPN->REase-> | 1113 | Bacteria>Pseudomonadota>Betaproteobacteria>Burkholderiales>unclassified_Burkholderiales | GCA_014860355.1 | Burkholderiales_bacterium | IH606_20135 | 9089..12430 | 1891238 | DUF499-domain-containing-protein-[Burkholderiales-bacterium] |
| MBX9629971.1 | HEPN+AAA-ATPase+Alpha+Beta-domain+wHTH+RRM | Type-1_BR.RM.like.systems | RelE-ParE->?->TUDOR+SF1-SF2-Helicase+iREase+CTD+REase->?->?->METHYLASE->HEPN+DUF499*-><-?<-AbiEii<-AbiEi_4 | 1116 | Bacteria>Pseudomonadota>Betaproteobacteria>Burkholderiales>unclassified_Burkholderiales | GCA_019747905.1 | Burkholderiales_bacterium | K2X67_05560 | 12511..15861 | 1891238 | DUF499-domain-containing-protein-[Burkholderiales-bacterium] |
| MBE7523982.1 | HEPN+AAA-ATPase+Alpha+Beta-domain+wHTH+RRM | Type-1_BR.RM.like.systems | S1+RNase+2C->ABhydrolase->Glyoxalase-><-?<-HTH+PolB<-HEPN+DUF499*<-STAS<-?<-METHYLASE<-?<-Schlafen+HATPase+HTH | 1119 | Bacteria>Pseudomonadota>Betaproteobacteria>Burkholderiales>unclassified_Burkholderiales | GCA_015075645.1 | Burkholderiales_bacterium | HS109_16570 | 209552..212911 | 1891238 | ATP-binding-protein-[Burkholderiales-bacterium] |
| MCL4799144.1 | HEPN+AAA-ATPase+Alpha+Beta-domain+wHTH+RRM | Type-1_BR.RM.like.systems | <-HEPN+DUF499*<-HEPN<-PolB<-METHYLASE<-?<-TUDOR+SF1-SF2-Helicase+iREase+CTD+REase<-TIR | 1113 | Bacteria>Pseudomonadota>Betaproteobacteria>Burkholderiales>unclassified_Burkholderiales | GCA_023383795.1 | Burkholderiales_bacterium | KJ025_06120 | 926..4267 | 1891238 | DUF499-domain-containing-protein-[Burkholderiales-bacterium] |
| MCZ2419561.1 | HEPN+AAA-ATPase+Alpha+Beta-domain+wHTH+RRM | Type-1_BR.RM.like.systems | TPR+TPR-><-?<-HTH+UbE1<-?\|\|HTH->MPTase-><-HEPN+DUF499*<-?<-DUF488-1<-?<-METHYLASE<-TOPRIM<-OLD-ABC | 1108 | Bacteria>Pseudomonadota>Betaproteobacteria>Burkholderiales>unclassified_Burkholderiales | GCA_026931465.1 | Burkholderiales_bacterium | LC123_06960 | 80916..84242 | 1891238 | MAG:-DUF499-domain-containing-protein-[Burkholderiales-bacterium] |
| MCA3160290.1 | HEPN+AAA-ATPase+Alpha+Beta-domain+wHTH+RRM | Type-1_BR.RM.like.systems | HTH_Tnp_1+HTH_21+rve+rve_3->METHYLASE-p->METHYLASE-p->HEPN+DUF499*->?-><-SAM-methylase | 1098 | Bacteria>Pseudomonadota>Betaproteobacteria>Burkholderiales>unclassified_Burkholderiales | GCA_020248485.1 | Burkholderiales_bacterium | ING31_11990 | 9686..12982 | 1891238 | DUF499-domain-containing-protein-[Burkholderiales-bacterium] |
| MBN8477381.1 | HEPN+AAA-ATPase+Alpha+Beta-domain+wHTH+RRM | Type-1_BR.RM.like.systems | TUDOR+SF1-SF2-Helicase+iREase+CTD+REase->?->METHYLASE->?->?->HEPN+DUF499*->?->HTH->NTP_transf-> | 1117 | Bacteria>Pseudomonadota>Betaproteobacteria>Burkholderiales>unclassified_Burkholderiales | GCA_017302475.1 | Burkholderiales_bacterium | JOL91_00275 | 13639..16992 | 1891238 | DUF499-domain-containing-protein-[Burkholderiales-bacterium] |
| MCL6557122.1 | HEPN+AAA-ATPase+Alpha+Beta-domain+wHTH+RRM | Type-1_BR.RM.like.systems | <-HEPN+DUF499*<-PolB<-?<-ORC-CDC6+REase<-TUDOR+SF1-SF2-Helicase+iREase+CTD+REase-p | 1098 | Bacteria>Pseudomonadota>Betaproteobacteria>Burkholderiales>unclassified_Burkholderiales | GCA_023511995.1 | Burkholderiales_bacterium | K6T56_12270 | 1861..5157 | 1891238 | DUF499-domain-containing-protein-[Burkholderiales-bacterium] |
| GIX27884.1 | HEPN+AAA-ATPase+Alpha+Beta-domain+wHTH+RRM | Type-1_BR.RM.like.systems | METHYLASE-p-><-?\|\|METHYLASE-p->ParD->PIN->HEPN+DUF499*->DUF499-p->?->?->ORC-CDC6+REase-> | 389 | Bacteria>Pseudomonadota>Betaproteobacteria>Burkholderiales>unclassified_Burkholderiales | GCA_026005015.1 | Burkholderiales_bacterium | KatS3mg123_1765 | 1711823..1712992 | 1891238 | MAG:-hypothetical-protein-KatS3mg123_1765-[Burkholderiales-bacterium] |
| MCI4428467.1 | HEPN+AAA-ATPase+Alpha+Beta-domain+wHTH+RRM | Type-1_BR.RM.like.systems | <-HEPN+DUF499*<-HEPN<-HTH+Hiskin+STAS<-HEPN<-PolB<-METHYLASE-p | 342 | Bacteria>Pseudomonadota>Betaproteobacteria>Burkholderiales>unclassified_Burkholderiales | GCA_022758925.1 | Burkholderiales_bacterium | JHC40_04765 | 1..1025 | 1891238 | ATP-binding-protein-[Burkholderiales-bacterium] |
| MBL8529542.1 | HEPN+AAA-ATPase+Alpha+Beta-domain+wHTH+RRM | Type-1_BR.RM.like.systems | Schlafen+HATPase+HTH->?->METHYLASE-p->HTH+Hiskin+STAS->PolB->HEPN+DUF499*-> | 1108 | Bacteria>Pseudomonadota>Betaproteobacteria>Burkholderiales>unclassified_Burkholderiales | GCA_016791285.1 | Burkholderiales_bacterium | JNL68_17825 | 6694..10020 | 1891238 | DUF499-domain-containing-protein-[Burkholderiales-bacterium] |
| MCC6474792.1 | HEPN+AAA-ATPase+Alpha+Beta-domain+wHTH+RRM | Type-1_BR.RM.like.systems | <-HEPN+DUF499*<-HEPN<-PolB<-METHYLASE<-HEPN<-ParB+HNH\|\|tRNA-><-PIN | 1120 | Bacteria>Pseudomonadota>Betaproteobacteria>Burkholderiales>unclassified_Burkholderiales | GCA_020846575.1 | Burkholderiales_bacterium | IT514_13720 | 1159..4521 | 1891238 | DUF499-domain-containing-protein-[Burkholderiales-bacterium] |
| MBI5907809.1 | HEPN+AAA-ATPase+Alpha+Beta-domain+wHTH+RRM | Type-1_BR.RM.like.systems | ABC-ATPase+SbcC->ABC-ATPase+SbcC->?-><-HEPN+DUF499*<-?<-METHYLASE | 1107 | Bacteria>Pseudomonadota>Betaproteobacteria>Burkholderiales>unclassified_Burkholderiales | GCA_016234645.1 | Burkholderiales_bacterium | HY853_04415 | 2095..5418 | 1891238 | ATP-binding-protein-[Burkholderiales-bacterium] |
| RYG38846.1 | HEPN+AAA-ATPase+Alpha+Beta-domain+wHTH+RRM | Type-1_BR.RM.like.systems | METHYLASE->HEPN+DUF499*->?-><-HTH+WYL\|\|?->?->?->STYKIN-> | 1103 | Bacteria>Pseudomonadota>Betaproteobacteria>Burkholderiales>unclassified_Burkholderiales | GCA_004145455.1 | Burkholderiales_bacterium | EON93_01200 | 2210..5521 | 1891238 | ATP-binding-protein-[Burkholderiales-bacterium] |
| MCC7486352.1 | HEPN+AAA-ATPase+Alpha+Beta-domain+wHTH+RRM | Type-1_BR.RM.like.systems | ORC-CDC6+REase->HEPN+DUF499*->?-><-?<-ABhydrolase | 1107 | Bacteria>Pseudomonadota>Betaproteobacteria>Burkholderiales>unclassified_Burkholderiales | GCA_020854035.1 | Burkholderiales_bacterium | IT529_15375 | 2503..5826 | 1891238 | DUF499-domain-containing-protein-[Burkholderiales-bacterium] |
| NMA26876.1 | HEPN+AAA-ATPase+Alpha+Beta-domain+wHTH+RRM | Type-1_BR.RM.like.systems | METHYLASE-p->OLD-ABC->TOPRIM->HEPN+DUF499*-> | 448 | Bacteria>Pseudomonadota>Betaproteobacteria>Burkholderiales>unclassified_Burkholderiales | GCA_012799975.1 | Burkholderiales_bacterium | GX934_03860 | 2623..3966 | 1891238 | ATP-binding-protein-[Burkholderiales-bacterium] |
| MCA3160409.1 | HEPN+AAA-ATPase+Alpha+Beta-domain+wHTH+RRM | Type-1_BR.RM.like.systems | <-HEPN+DUF499*<-METHYLASE | 815 | Bacteria>Pseudomonadota>Betaproteobacteria>Burkholderiales>unclassified_Burkholderiales | GCA_020248485.1 | Burkholderiales_bacterium | ING31_12605 | 1..2446 | 1891238 | DUF499-domain-containing-protein-[Burkholderiales-bacterium] |
| WP_031597360.1 | HEPN+AAA-ATPase+Alpha+Beta-domain+wHTH+RRM | Type-1_BR.RM.like.systems | TUDOR+SF1-SF2-Helicase+iREase+CTD+REase->TOPRIM->METHYLASE->HEPN->ParA-Soj-PloopNTPase->ORC-CDC6->HEPN+DUF499*-><-Phage_integrase<-?<-?\|\|<-?<-HTH+UbE1 | 1109 | Bacteria>Pseudomonadota>Betaproteobacteria>Ferrovales>Ferrovaceae | GCF_000735045.1 | Ferrovum_myxofaciens | IV91_RS0107710 | 11022..14351 | 416213 | Swt1-family-HEPN-domain-containing-protein-[Ferrovum-myxofaciens] |
| NDU85854.1 | HEPN+AAA-ATPase+Alpha+Beta-domain+wHTH+RRM | Type-1_BR.RM.like.systems | <-HTH+UbE1<-?\|\|?->?->MPTase-><-HEPN+DUF499*<-METHYLASE<-TOPRIM<-OLD-ABC<-TUDOR+SF1-SF2-Helicase+iREase+CTD+REase<-SF1-SF2-Helicase<-OLD-ABC+TOPRIM | 1110 | Bacteria>Pseudomonadota>Betaproteobacteria>Ferrovales>Ferrovaceae | GCA_010436015.1 | Ferrovum_sp. | G3H99_04485 | 6034..9366 | 2609467 | ATP-binding-protein-[Ferrovum-sp.] |
| WP_273118797.1 | HEPN+AAA-ATPase+Alpha+Beta-domain+wHTH+RRM | Type-1_BR.RM.like.systems | HTH+UbE1->?->\|\|?->?->Phage_integrase-><-HEPN+DUF499*<-ORC-CDC6<-ParA-Soj-PloopNTPase<-HEPN<-METHYLASE<-TOPRIM<-OLD-ABC | 1109 | Bacteria>Pseudomonadota>Betaproteobacteria>Ferrovales>Ferrovaceae | GCF_013181415.1 | Ferrovum_myxofaciens | H0274_RS08660 | 1781795..1785124 | 416213 | Swt1-family-HEPN-domain-containing-protein-[Ferrovum-myxofaciens] |
| QKE41374.1 | HEPN+AAA-ATPase+Alpha+Beta-domain+wHTH+RRM | Type-1_BR.RM.like.systems | HTH+UbE1->?->?->?->Phage_integrase-><-HEPN+DUF499*<-ORC-CDC6<-ParA-Soj-PloopNTPase<-HEPN<-METHYLASE<-TOPRIM<-OLD-ABC | 1109 | Bacteria>Pseudomonadota>Betaproteobacteria>Ferrovales>Ferrovaceae | GCA_013181415.1 | Ferrovum_myxofaciens | H0274_08630 | 1781795..1785124 | 416213 | ATP-binding-protein-[Ferrovum-myxofaciens] |
| CAG1003423.1 | HEPN+AAA-ATPase+Alpha+Beta-domain+wHTH+RRM | Type-1_BR.RM.like.systems | TUDOR+SF1-SF2-Helicase+iREase+CTD+REase->sGTP->ParB+HNH->METHYLASE->HEPN+DUF499*-> | 1107 | Bacteria>Pseudomonadota>Betaproteobacteria>Nitrosomonadales>Methylophilaceae | GCA_905339275.1 | Methylophilaceae_bacterium | MTYP_03091 | 15685..19008 | 2030816 | hypothetical-protein-MTYP_03091-[Methylophilaceae-bacterium] |
| MBP0133287.1 | HEPN+AAA-ATPase+Alpha+Beta-domain+wHTH+RRM | Type-1_BR.RM.like.systems | XIS-HTH->?->TUDOR+SF1-SF2-Helicase+iREase+CTD+REase->Cluster1090_5clades->METHYLASE->HEPN+DUF499*-><-HTH+WYL | 1105 | Bacteria>Pseudomonadota>Betaproteobacteria>Nitrosomonadales>Nitrosomonadaceae | GCA_016106425.1 | Nitrosospira_sp. | ITD31_07875 | 10694..14011 | 38080 | DUF499-domain-containing-protein-[Nitrosospira-sp.] |
| WP_107785979.1 | HEPN+AAA-ATPase+Alpha+Beta-domain+wHTH+RRM | Type-1_BR.RM.like.systems | <-HIN-HTH\|\|TUDOR+SF1-SF2-Helicase+iREase+CTD+REase->ParB+HNH->METHYLASE->HEPN+DUF499*-><-tRNA\|\|FURR-HTH-> | 1112 | Bacteria>Pseudomonadota>Betaproteobacteria>Nitrosomonadales>Nitrosomonadaceae | GCF_003051045.1 | Nitrosomonas_ureae | C8R28_RS01190 | 8316..11654 | 44577 | Swt1-family-HEPN-domain-containing-protein-[Nitrosomonas-ureae] |
| MBK6974203.1 | HEPN+AAA-ATPase+Alpha+Beta-domain+wHTH+RRM | Type-1_BR.RM.like.systems | <-SIGMA-HTH+SIGMA-HTH\|\|DDE_Tnp_1_assoc+DDE_Tnp_1-><-35exo+SF1-SF2-Helicase<-HEPN+DUF499*<-HEPN<-PolB<-wHTH+REase<-TUDOR+SF1-SF2-Helicase+iREase+CTD+REase | 1112 | Bacteria>Pseudomonadota>Betaproteobacteria>Nitrosomonadales>Sterolibacteriaceae | GCA_016708905.1 | Candidatus_Methylophosphatis_roskildensis | IPH26_15090 | 451210..454548 | 2899263 | DUF499-domain-containing-protein-[Sterolibacteriaceae-bacterium] |
| MBK7234801.1 | HEPN+AAA-ATPase+Alpha+Beta-domain+wHTH+RRM | Type-1_BR.RM.like.systems | <-SIGMA-HTH+SIGMA-HTH\|\|DDE_Tnp_1->DDE_Tnp_1_assoc+DDE_Tnp_1->DDE_Tnp_1_assoc+DDE_Tnp_1-><-35exo+SF1-SF2-Helicase<-HEPN+DUF499*<-HEPN<-PolB<-wHTH+REase | 1112 | Bacteria>Pseudomonadota>Betaproteobacteria>Nitrosomonadales>Sterolibacteriaceae | GCA_016707455.1 | Sterolibacteriaceae_bacterium | IPI02_03880 | 820379..823717 | 2682146 | DUF499-domain-containing-protein-[Sterolibacteriaceae-bacterium] |
| MBK7665767.1 | HEPN+AAA-ATPase+Alpha+Beta-domain+wHTH+RRM | Type-1_BR.RM.like.systems | TUDOR+SF1-SF2-Helicase+iREase+CTD+REase->Schlafen+HATPase+HTH->?->PolB->HEPN->HEPN+DUF499*->35exo+SF1-SF2-Helicase->SIGMA-HTH+SIGMA-HTH-> | 1112 | Bacteria>Pseudomonadota>Betaproteobacteria>Nitrosomonadales>Sterolibacteriaceae | GCA_016709795.1 | Sterolibacteriaceae_bacterium | IPJ21_19955 | 4305507..4308845 | 2682146 | DUF499-domain-containing-protein-[Sterolibacteriaceae-bacterium] |

| Protein-ID | DUF499-architecture | System | Gene-Neighbourhoods | Length | Taxonomy | Assembly-ID | Strain | Locus-Tag | Genomic-coordinates | TaxID | Fasta-defline |
|---|---|---|---|---|---|---|---|---|---|---|---|
| ODU50157.1 | HEPN+AAA-ATPase+Alpha+Beta-domain+wHTH+RRM | Type-1_BR.RM.like.systems | <-HEPN+DUF499*<-?<-DUF488-1<-?<-METHYLASE<-sGTP<-TUDOR+SF1-SF2-Helicase+iREase+CTD+REase | 1110 | Bacteria>Pseudomonadota>Betaproteobacteria>Nitrosomonadales>Thiobacillaceae | GCA_001725135.1 | Thiobacillus_sp._SCN_63-374 | ABS92_03900 | 1302..4634 | 1660149 | MAG:-AAA+-family-ATPase-[Thiobacillus-sp.-SCN-63-374] |
| MBI1887192.1 | HEPN+AAA-ATPase+Alpha+Beta-domain+wHTH+RRM | Type-1_BR.RM.like.systems | <-Glycos_trans_3N+Glycos_transf_3<-?<-Schlafen+HATPase+HTH<-?<-HEPN+DUF499*<-METHYLASE<-HEPN<-TOPRIM<-OLD-ABC<-TUDOR+SF1-SF2-Helicase+iREase+CTD+REase<-ParB | 1112 | Bacteria>Pseudomonadota>Betaproteobacteria>Nitrosomonadales>unclassified_Nitrosomonadales | GCA_016177795.1 | Nitrosomonadales_bacterium | HYS19_02300 | 86364..89702 | 2026761 | ATP-binding-protein-[Nitrosomonadales-bacterium] |
| MBI3902214.1 | HEPN+AAA-ATPase+Alpha+Beta-domain+wHTH+RRM | Type-1_BR.RM.like.systems | <-Glycos_trans_3N+Glycos_transf_3<-?<-?<-Schlafen+HATPase+HTH<-?<-HEPN+DUF499*<-METHYLASE<-?<-?<-TOPRIM<-OLD-ABC<-TUDOR+SF1-SF2-Helicase+iREase+CTD+REase | 1112 | Bacteria>Pseudomonadota>Betaproteobacteria>Nitrosomonadales>unclassified_Nitrosomonadales | GCA_016199835.1 | Nitrosomonadales_bacterium | HY306_04630 | 118339..121677 | 2026761 | ATP-binding-protein-[Nitrosomonadales-bacterium] |
| MBI5658243.1 | HEPN+AAA-ATPase+Alpha+Beta-domain+wHTH+RRM | Type-1_BR.RM.like.systems | TUDOR+SF1-SF2-Helicase+iREase+CTD+REase->OLD-ABC->TOPRIM->METHYLASE->?->?->HEPN+DUF499*-> | 1125 | Bacteria>Pseudomonadota>Betaproteobacteria>Nitrosomonadales>unclassified_Nitrosomonadales | GCA_016223025.1 | Nitrosomonadales_bacterium | HZC43_01545 | 64771..68148 | 2026761 | ATP-binding-protein-[Nitrosomonadales-bacterium] |
| MBN8462739.1 | HEPN+AAA-ATPase+Alpha+Beta-domain+wHTH+RRM | Type-1_BR.RM.like.systems | PrlF_antitoxin->PemK_toxin->REase->METHYLASE->PolB->HEPN->HEPN+DUF499*->?-><-?<-?\|\|?->sGTP-> | 1110 | Bacteria>Pseudomonadota>Betaproteobacteria>Rhodocyclales>Azonexaceae | GCA_017302355.1 | Dechloromonas_sp. | JOM01_07975 | 12361..15693 | 1917218 | DUF499-domain-containing-protein-[Dechloromonas-sp.] |
| MCA3140850.1 | HEPN+AAA-ATPase+Alpha+Beta-domain+wHTH+RRM | Type-1_BR.RM.like.systems | HTH_3->TUDOR+SF1-SF2-Helicase+iREase+CTD+REase->Schlafen+HATPase+HTH->?->METHYLASE->HTH+Hiskin+STAS->HEPN+DUF499*->?->XIS-HTH->?->TOPRIM->MetJarc-HTH->RelE-ParE-> | 1113 | Bacteria>Pseudomonadota>Betaproteobacteria>Rhodocyclales>Rhodocyclaceae | GCA_020248385.1 | Rhodocyclaceae_bacterium | ING80_01400 | 55201..58542 | 1898103 | DUF499-domain-containing-protein-[Rhodocyclaceae-bacterium] |
| KAB2934118.1 | HEPN+AAA-ATPase+Alpha+Beta-domain+wHTH+RRM | Type-1_BR.RM.like.systems | <-PAPS_reduct<-?<-KAP-NTPase<-HEPN+DUF499*<-PIN<-OLD-ABC<-METHYLASE<-?<-TUDOR+SF1-SF2-Helicase+iREase+CTD+REase | 1109 | Bacteria>Pseudomonadota>Betaproteobacteria>Rhodocyclales>Rhodocyclaceae | GCA_008933825.1 | Rhodocyclaceae_bacterium | F9K21_11715 | 6014..9343 | 1898103 | ATP-binding-protein-[Rhodocyclaceae-bacterium] |
| WP_018989132.1 | HEPN+AAA-ATPase+Alpha+Beta-domain+wHTH+RRM | Type-1_BR.RM.like.systems | <-PIN+PhoH<-Radical-SAM+TRAM\|\|tRNA->?-><-HEPN+DUF499*<-METHYLASE<-HNH<-OLD-ABC<-PIN<-?<-STAS+HisKin | 1121 | Bacteria>Pseudomonadota>Betaproteobacteria>Rhodocyclales>Rhodocyclaceae | GCF_000378245.1 | Aromatoleum_toluclasticum | F464_RS0107275 | 366039..369404 | 92003 | Swt1-family-HEPN-domain-containing-protein-[Aromatoleum-toluclasticum] |
| KAB2933410.1 | HEPN+AAA-ATPase+Alpha+Beta-domain+wHTH+RRM | Type-1_BR.RM.like.systems | OLD-ABC->TOPRIM->?->?->METHYLASE->REase->HEPN+DUF499*->?-><-?\|\|HTH->MarR-HTH-> | 1096 | Bacteria>Pseudomonadota>Betaproteobacteria>Rhodocyclales>Rhodocyclaceae | GCA_008933825.1 | Rhodocyclaceae_bacterium | F9K21_12245 | 12685..15975 | 1898103 | ATP-binding-protein-[Rhodocyclaceae-bacterium] |
| MBP6422202.1 | HEPN+AAA-ATPase+Alpha+Beta-domain+wHTH+RRM | Type-1_BR.RM.like.systems | <-CASPASE<-TIR<-HEPN+DUF499*<-METHYLASE-p | 1110 | Bacteria>Pseudomonadota>Betaproteobacteria>Rhodocyclales>Rhodocyclaceae | GCA_017988215.1 | Propionivibrio_sp. | KA271_04855 | 1433..4765 | 2212460 | DUF499-domain-containing-protein-[Propionivibrio-sp.] |
| WP_153167206.1 | HEPN+AAA-ATPase+Alpha+Beta-domain+wHTH+RRM | Type-1_BR.RM.like.systems | MORC+DNA_gyraseB+TOPRIM-><-?<-?<-HEPN+DUF499*<-?<-METHYLASE<-?<-MORC+DUF3684<-TUDOR+SF1-SF2-Helicase+iREase+CTD+REase | 1120 | Bacteria>Pseudomonadota>Betaproteobacteria>Rhodocyclales>Zoogloeaceae | GCF_009469595.2 | Thauera_sp._2A1 | GH664_RS06300 | 29943..33305 | 2570191 | Swt1-family-HEPN-domain-containing-protein-[Thauera-sp.-2A1] |
| MBL8540233.1 | HEPN+AAA-ATPase+Alpha+Beta-domain+wHTH+RRM | Type-1_BR.RM.like.systems | TUDOR+SF1-SF2-Helicase+iREase+CTD+REase->METHYLASE->HTH+Hiskin+STAS->PolB->PolB->HEPN->HEPN+DUF499*-><-?<-?<-?<-Response_reg+LuxR-HTH<-STYKIN | 1105 | Bacteria>Pseudomonadota>Betaproteobacteria>unclassified_Betaproteobacteria | GCA_016791345.1 | Betaproteobacteria_bacterium | JNK68_07660 | 10175..13492 | 1891241 | DUF499-domain-containing-protein-[Betaproteobacteria-bacterium] |
| AGG33842.1 | HEPN+AAA-ATPase+Alpha+Beta-domain+wHTH+RRM | Type-1_BR.RM.like.systems | Peptidase_M20->PrmC_N+Methyltransf_25-><-SMS-KaiC<-?<-?\|\|tRNA-><-HEPN+DUF499*<-METHYLASE<-TUDOR+SF1-SF2-Helicase+iREase+CTD+REase\|\|MerR-HTH-> | 1110 | Bacteria>Pseudomonadota>Betaproteobacteria>unclassified_Betaproteobacteria | GCA_000342265.1 | beta_proteobacterium_CB | D521_1274 | 1257498..1260830 | 543913 | ATPase-(AAA+-superfamily)-like-protein-[beta-proteobacterium-CB] |
| MDA8257273.1 | HEPN+AAA-ATPase+Alpha+Beta-domain+wHTH+RRM | Type-1_BR.RM.like.systems | <-EVE<-HEPN+DUF499*<-HEPN<-PolB<-ORC-CDC6<-ParA-Soj-PloopNTPase<-HEPN<-METHYLASE | 1114 | Bacteria>Pseudomonadota>Betaproteobacteria>unclassified_Betaproteobacteria | GCA_028065725.1 | Betaproteobacteria_bacterium | MOZ99_16875 | 140..3484 | 1891241 | MAG:-Swt1-family-HEPN-domain-containing-protein-[Betaproteobacteria-bacterium] |
| MDE2342677.1 | HEPN+AAA-ATPase+Alpha+Beta-domain+wHTH+RRM | Type-1_BR.RM.like.systems | <-HEPN+DUF499*<-METHYLASE<-TUDOR+SF1-SF2-Helicase+iREase+CTD+REase | 1094 | Bacteria>Pseudomonadota>Betaproteobacteria>unclassified_Betaproteobacteria | GCA_028870015.1 | Betaproteobacteria_bacterium | KGL63_04675 | 257..3541 | 1891241 | MAG:-DUF499-domain-containing-protein-[Betaproteobacteria-bacterium] |
| OGA65437.1 | HEPN+AAA-ATPase+Alpha+Beta-domain+wHTH+RRM | Type-1_BR.RM.like.systems | <-HEPN+DUF499*<-PIN<-MPTase<-?<-?<-METHYLASE<-ParB+HNH | 1109 | Bacteria>Pseudomonadota>Betaproteobacteria>unclassified_Betaproteobacteria | GCA_001770585.1 | Betaproteobacteria_bacterium_RIFCSPLOWO2_12_FULL_65_14 | A3G81_12440 | 218..3547 | 1797501 | AAA+-family-ATPase-[Betaproteobacteria-bacterium-RIFCSPLOWO2_12_FULL_65_14] |
| MBI4756762.1 | HEPN+AAA-ATPase+Alpha+Beta-domain+wHTH+RRM | Type-1_BR.RM.like.systems | METHYLASE->?->?->?->PolB->HEPN->HEPN+DUF499*-> | 1123 | Bacteria>Pseudomonadota>Betaproteobacteria>unclassified_Betaproteobacteria | GCA_016208415.1 | Betaproteobacteria_bacterium | HY778_15365 | 56697..60068 | 1891241 | ATP-binding-protein-[Betaproteobacteria-bacterium] |
| WP_096360574.1 | HEPN+AAA-ATPase+Alpha+Beta-domain+wHTH+RRM | Type-1_BR.RM.like.systems | <-CHTH<-HTH_3<-RelE-ParE<-UPF0061<-?<-?<-HEPN+DUF499*<-METHYLASE<-?<-TUDOR+SF1-SF2-Helicase+iREase+CTD+REase<-?<-BirA-HTH<-HKD-DNase | 1113 | Bacteria>Pseudomonadota>Gammaproteobacteria>Acidiferrobacterales>Acidiferrobacteraceae | GCF_002355735.1 | Sulfuricaulis_limicola | SCL_RS07110 | 1437860..1441201 | 1620215 | Swt1-family-HEPN-domain-containing-protein-[Sulfuricaulis-limicola] |
| MCU0834934.1 | HEPN+AAA-ATPase+Alpha+Beta-domain+wHTH+RRM | Type-1_BR.RM.like.systems | <-DUF499-p<-HEPN+DUF499*<-HTH+Hiskin+STAS<-METHYLASE-p<-METHYLASE<-PemK_toxin<-PrlF_antitoxin<-TUDOR+SF1-SF2-Helicase+iREase+CTD+REase-p | 755 | Bacteria>Pseudomonadota>Gammaproteobacteria>Chromatiales>Chromatiaceae | GCA_025458165.1 | Chromatiaceae_bacterium | MUC77_10980 | 31032..33299 | 2049432 | Swt1-family-HEPN-domain-containing-protein-[Chromatiaceae-bacterium] |
| WP_100917910.1 | HEPN+AAA-ATPase+Alpha+Beta-domain+wHTH+RRM | Type-1_BR.RM.like.systems | TUDOR+SF1-SF2-Helicase+iREase+CTD+REase->OLD-ABC->TOPRIM->METHYLASE->PolB->HEPN+DUF499*->?-><-NUDIX<-?<-?\|\|?-><-PAIREDC-HTH | 1111 | Bacteria>Pseudomonadota>Gammaproteobacteria>Chromatiales>Chromatiaceae | GCF_002813775.1 | Candidatus_Thiodictyon_syntrophicum | THSYN_RS03450 | 792207..795542 | 1166950 | Swt1-family-HEPN-domain-containing-protein-[Candidatus-Thiodictyon-syntrophicum] |
| WP_153974954.1 | HEPN+AAA-ATPase+Alpha+Beta-domain+wHTH+RRM | Type-1_BR.RM.like.systems | <-ABhydrolase<-?<-HEPN+DUF499*<-HEPN<-PolB<-METHYLASE<-?<-TUDOR+SF1-SF2-Helicase+iREase+CTD+REase | 1103 | Bacteria>Pseudomonadota>Gammaproteobacteria>Chromatiales>Chromatiaceae | GCF_009664085.1 | Thermochromatium_tepidum | E6P07_RS07035 | 1550502..1553813 | 1050 | Swt1-family-HEPN-domain-containing-protein-[Thermochromatium-tepidum] |
| WP_020507743.1 | HEPN+AAA-ATPase+Alpha+Beta-domain+wHTH+RRM | Type-1_BR.RM.like.systems | OLD-ABC->TOPRIM->METHYLASE->PolB->PolB->HEPN->HEPN+DUF499*-><-?<-?<-?<-REC<-HPT+HISKIN+REC | 1124 | Bacteria>Pseudomonadota>Gammaproteobacteria>Chromatiales>Chromatiaceae | GCF_000379525.1 | Lamprocystis_purpurea | A390_RS0126800 | 79657..83031 | 61598 | Swt1-family-HEPN-domain-containing-protein-[Lamprocystis-purpurea] |
| WP_120796610.1 | HEPN+AAA-ATPase+Alpha+Beta-domain+wHTH+RRM | Type-1_BR.RM.like.systems | PrlF_antitoxin->RHH_6->PIN->METHYLASE-p->?->PolB->HEPN+DUF499*-><-?\|\|?->RVT+HNH->RVT+HNH-> | 1114 | Bacteria>Pseudomonadota>Gammaproteobacteria>Chromatiales>Chromatiaceae | GCF_003634315.1 | Thiocapsa_rosea | BDD21_RS07415 | 1614812..1618156 | 69360 | Swt1-family-HEPN-domain-containing-protein-[Thiocapsa-rosea] |
| QVL49217.1 | HEPN+AAA-ATPase+Alpha+Beta-domain+wHTH+RRM | Type-1_BR.RM.like.systems | OLD-ABC->TOPRIM->METHYLASE->?->PolB->HEPN->HEPN+DUF499*-><-?\|\|HicB-><-sGTP\|\|?-><-TPR+TPR | 1114 | Bacteria>Pseudomonadota>Gammaproteobacteria>Chromatiales>Chromatiaceae | GCA_018399035.1 | Thiocapsa_sp. | KFB96_01415 | 301904..305248 | 2024551 | MAG:-DUF499-domain-containing-protein-[Thiocapsa-sp.] |
| MBK8637657.1 | HEPN+AAA-ATPase+Alpha+Beta-domain+wHTH+RRM | Type-1_BR.RM.like.systems | TOPRIM->METHYLASE->PolB->HEPN->?->Y1_Tnp->HEPN+DUF499*->RADICAL-SAM-> | 1120 | Bacteria>Pseudomonadota>Gammaproteobacteria>Chromatiales>Chromatiaceae | GCA_016714645.1 | Chromatiaceae_bacterium | IPN92_04990 | 1074519..1077881 | 2049432 | DUF499-domain-containing-protein-[Chromatiaceae-bacterium] |
| WP_093189479.1 | HEPN+AAA-ATPase+Alpha+Beta-domain+wHTH+RRM | Type-1_BR.RM.like.systems | METHYLASE->PolB->HEPN->HEPN+DUF499*-><-?\|\|?->?->?-><-35exo | 1113 | Bacteria>Pseudomonadota>Gammaproteobacteria>Chromatiales>Chromatiaceae | GCF_001499735.1 | Thiocapsa_sp._KS1 | THIOKS1_RS17085 | 3720907..3724248 | 610332 | Swt1-family-HEPN-domain-containing-protein-[Thiocapsa-sp.-KS1] |
| MCC7277404.1 | HEPN+AAA-ATPase+Alpha+Beta-domain+wHTH+RRM | Type-1_BR.RM.like.systems | CHTH-><-HEPN+DUF499*<-HEPN<-PolB<-HEPN<-PolB | 1122 | Bacteria>Pseudomonadota>Gammaproteobacteria>Chromatiales>Chromatiaceae | GCA_020851955.1 | Chromatiaceae_bacterium | IT487_03685 | 4457..7825 | 2049432 | DUF499-domain-containing-protein-[Chromatiaceae-bacterium] |
| MBP6807674.1 | HEPN+AAA-ATPase+Alpha+Beta-domain+wHTH+RRM | Type-1_BR.RM.like.systems | <-HEPN+DUF499*<-?<-?<-PIN<-?<-METHYLASE | 368 | Bacteria>Pseudomonadota>Gammaproteobacteria>Chromatiales>Chromatiaceae | GCA_017991215.1 | Chromatiaceae_bacterium | KA125_05205 | 1..1104 | 2049432 | DUF499-domain-containing-protein-[Chromatiaceae-bacterium] |
| WP_091642911.1 | HEPN+AAA-ATPase+Alpha+Beta-domain+wHTH+RRM | Type-1_BR.RM.like.systems | <-AbiEi_1<-GDResolvase+HIN-HTH\|\|HIN-HTH->?->LZ_Tnp_IS66+zf-IS66+DDE_Tnp_IS66+DDE_Tnp_IS66_C->?-><-HEPN+DUF499*<-METHYLASE<-TUDOR+SF1-SF2-Helicase+iREase+CTD+REase\|\|HTH_Tnp_1+HTH_21+rve-> | 1097 | Bacteria>Pseudomonadota>Gammaproteobacteria>Chromatiales>Ectothiorhodospiraceae | GCF_900110585.1 | Aquisalimonas_asiatica | BMZ02_RS09865 | 435856..439149 | 406100 | Swt1-family-HEPN-domain-containing-protein-[Aquisalimonas-asiatica] |
| MCB1764984.1 | HEPN+AAA-ATPase+Alpha+Beta-domain+wHTH+RRM | Type-1_BR.RM.like.systems | TUDOR+SF1-SF2-Helicase+iREase+CTD+REase->?->Tox-RelE->HTH->METHYLASE->HEPN+DUF499*->?->REase->Radical-SAM-> | 1099 | Bacteria>Pseudomonadota>Gammaproteobacteria>Gammaproteobacteria_incertae_sedis>Candidatus_Competibacteraceae | GCA_020444895.1 | Candidatus_Competibacteraceae_bacterium | KDJ22_02835 | 9090..12389 | 2053538 | DUF499-domain-containing-protein-[Candidatus-Competibacteraceae-bacterium] |
| WP_048676961.1 | HEPN+AAA-ATPase+Alpha+Beta-domain+wHTH+RRM | Type-1_BR.RM.like.systems | HEPN+DUF499*->MORC-><-?<-?<-?<-Phage_integrase\|\|DUF4158+DDE_Tnp_Tn3-> | 1099 | Bacteria>Pseudomonadota>Gammaproteobacteria>Gammaproteobacteria_incertae_sedis>Candidatus_Competibacteraceae | GCF_001051235.1 | Candidatus_Competibacter_denitrificans | BN873_RS17980 | 1195..4494 | 1400862 | DUF499-domain-containing-protein-[Candidatus-Competibacter-denitrificans] |
| MBC7162924.1 | HEPN+AAA-ATPase+Alpha+Beta-domain+wHTH+RRM | Type-1_BR.RM.like.systems | PIN->HTH+Hiskin+STAS->HEPN+DUF499*-><-ZDNA-HTH<-?<-?<-?<-HTH_3<-RelE-ParE | 1109 | Bacteria>Pseudomonadota>Gammaproteobacteria>Immundisolibacterales>Immundisolibacteraceae | GCA_014359565.1 | Immundisolibacter_sp. | H5U26_12570 | 5259..8588 | 1934948 | ATP-binding-protein-[Immundisolibacter-sp.] |
| WP_169910036.1 | HEPN+AAA-ATPase+Alpha+Beta-domain+wHTH+RRM | Type-1_BR.RM.like.systems | Arm-DNA-bind_3+Phage_integrase->TOPRIM->?->Integrase_1->?->HTH+MPTase-><-HEPN+DUF499*<-METHYLASE<-TUDOR+SF1-SF2-Helicase+iREase+CTD+REase<-HTH+WYL | 1098 | Bacteria>Pseudomonadota>Gammaproteobacteria>Pseudomonadales>Pseudomonadaceae | - | Pseudomonas_proteolytica | HB002_20655 | 84255..87551 | 219574 | DUF499-domain-containing-protein-[Pseudomonas-proteolytica] |
| NMZ24831.1 | HEPN+AAA-ATPase+Alpha+Beta-domain+wHTH+RRM | Type-1_BR.RM.like.systems | Arm-DNA-bind_3+Phage_integrase->TOPRIM->?->Integrase_1->?->HTH+MPTase-><-HEPN+DUF499*<-METHYLASE<-TUDOR+SF1-SF2-Helicase+iREase+CTD+REase<-HTH+WYL | 1098 | Bacteria>Pseudomonadota>Gammaproteobacteria>Pseudomonadales>Pseudomonadaceae | GCA_012986045.1 | Pseudomonas_proteolytica | HB002_20655 | 84255..87551 | 219574 | ATP-binding-protein-[Pseudomonas-proteolytica] |
| WP_262499199.1 | HEPN+AAA-ATPase+Alpha+Beta-domain+wHTH+RRM | Type-1_BR.RM.like.systems | TOPRIM->?->Integrase_1->Phage_int_SAM->?->HTH+MPTase-><-HEPN+DUF499*<-METHYLASE<-TUDOR+SF1-SF2-Helicase+iREase+CTD+REase<-HTH+WYL | 1083 | Bacteria>Pseudomonadota>Gammaproteobacteria>Pseudomonadales>Pseudomonadaceae | GCF_012986045.1 | Pseudomonas_proteolytica | HB002_RS20625 | 84255..87506 | 219574 | DUF499-domain-containing-protein-[Pseudomonas-proteolytica] |
| MCP5148712.1 | HEPN+AAA-ATPase+Alpha+Beta-domain+wHTH+RRM | Type-1_BR.RM.like.systems | Radical-SAM-><-?<-?<-?<-HEPN+DUF499*<-METHYLASE<-?<-Cluster1090_5clades<-TUDOR+SF1-SF2-Helicase+iREase+CTD+REase\|\|GDResolvase+HIN-HTH-> | 1108 | Bacteria>Pseudomonadota>Gammaproteobacteria>Pseudomonadales>unclassified_Pseudomonadales | GCA_024234795.1 | Pseudomonadales_bacterium | H6986_11380 | 65855..69181 | 1891229 | ATP-binding-protein-[Pseudomonadales-bacterium] |
| MCP5129192.1 | HEPN+AAA-ATPase+Alpha+Beta-domain+wHTH+RRM | Type-1_BR.RM.like.systems | TraG_N-><-?<-SbcC<-?<-?<-HEPN+DUF499*<-METHYLASE<-MORC+DUF3684<-TUDOR+SF1-SF2-Helicase+iREase+CTD+REase<-XIS-HTH<-tRNA\|\|RNB+RNase->SPOUT-> | 1098 | Bacteria>Pseudomonadota>Gammaproteobacteria>Pseudomonadales>unclassified_Pseudomonadales | GCA_024234815.1 | Pseudomonadales_bacterium | H6985_06385 | 1449819..1453115 | 1891229 | ATP-binding-protein-[Pseudomonadales-bacterium] |
| MCG3168781.1 | HEPN+AAA-ATPase+Alpha+Beta-domain+wHTH+RRM | Type-1_BR.RM.like.systems | ASCH->HKD-DNase->METHYLASE->HEPN+DUF499*-> | 1110 | Bacteria>Pseudomonadota>Gammaproteobacteria>Pseudomonadales>unclassified_Pseudomonadales | GCA_022072145.1 | Pseudomonadales_bacterium | CALGDGBN_00288 | 5040..8372 | 1891229 | hypothetical-protein-[Pseudomonadales-bacterium] |
| MBI2800244.1 | HEPN+AAA-ATPase+Alpha+Beta-domain+wHTH+RRM | Type-1_BR.RM.like.systems | DUF932-><-Phage_int_SAM_2+Integrase_1\|\|TUDOR+SF1-SF2-Helicase+iREase+CTD+REase->MORC+DUF3684->METHYLASE->?->HEPN+DUF499*-><-DUF772+DDE_Tnp_1<-Arm-DNA-bind_3+Phage_integrase\|\|REase+N6_MTase-><-?\|\|SPOUT-> | 1108 | Bacteria>Pseudomonadota>Gammaproteobacteria>unclassified_Gammaproteobacteria | GCA_016191285.1 | Gammaproteobacteria_bacterium | HYX63_08145 | 89797..93123 | 1913989 | ATP-binding-protein-[Gammaproteobacteria-bacterium] |
| MBC53307.1 | HEPN+AAA-ATPase+Alpha+Beta-domain+wHTH+RRM | Type-1_BR.RM.like.systems | MerR-HTH-><-?<-?<-Calcineurin<-?<-JAB<-HEPN+DUF499*<-METHYLASE<-MORC+DUF3684<-TUDOR+SF1-SF2-Helicase+iREase+CTD+REase<-HTH+WYL\|\|SF1-SF2-helicase+RdgC+Beta-barrel+REase-> | 1098 | Bacteria>Pseudomonadota>Gammaproteobacteria>unclassified_Gammaproteobacteria | GCA_002711845.1 | Gammaproteobacteria_bacterium | CMQ34_05660 | 7518..10814 | 1913989 | AAA-family-ATPase-[Gammaproteobacteria-bacterium] |
| NKC12883.1 | HEPN+AAA-ATPase+Alpha+Beta-domain+wHTH+RRM | Type-1_BR.RM.like.systems | <-HEPN+DUF499*\|\|Phage_int_SAM_3+Phage_integrase->Phage_integrase->METHYLASE-p-><-HicA_toxin<-HicB_lk_antitox<-METHYLASE | 1107 | Bacteria>Pseudomonadota>Gammaproteobacteria>unclassified_Gammaproteobacteria | GCA_012103455.1 | Gammaproteobacteria_bacterium | GKR94_11995 | 2363179..2366502 | 1913989 | DUF499-domain-containing-protein-[Gammaproteobacteria-bacterium] |
| GIX22526.1 | HEPN+AAA-ATPase+Alpha+Beta-domain+wHTH+RRM | Type-1_BR.RM.like.systems | Response_reg+LuxR-HTH->URI-DNase+HhH-RADC->?->tRNA->tRNA->tRNA-><-HEPN+DUF499*<-METHYLASE<-REase<-TUDOR+SF1-SF2-Helicase+iREase+CTD+REase-p | 1122 | Bacteria>Pseudomonadota>Gammaproteobacteria>unclassified_Gammaproteobacteria | GCA_026004975.1 | Gammaproteobacteria_bacterium | KatS3mg121_1309 | 525909..529277 | 1913989 | MAG:-hypothetical-protein-KatS3mg121_1309-[Gammaproteobacteria-bacterium] |
| MDE0036818.1 | HEPN+AAA-ATPase+Alpha+Beta-domain+wHTH+RRM | Type-1_BR.RM.like.systems | METHYLASE-p->?-><-PIN<-OLD-ABC<-?<-HEPN+DUF499*<-METHYLASE<-OLD-ABC<-TUDOR+SF1-SF2-Helicase+iREase+CTD+REase<-?<-?\|\|SF1-SF2-Helicase-> | 1118 | Bacteria>Pseudomonadota>Gammaproteobacteria>unclassified_Gammaproteobacteria | GCA_028818795.1 | Gammaproteobacteria_bacterium | OXU77_04540 | 7629..10985 | 1913989 | MAG:-Swt1-family-HEPN-domain-containing-protein-[Gammaproteobacteria-bacterium] |
| MBU1235382.1 | HEPN+AAA-ATPase+Alpha+Beta-domain+wHTH+RRM | Type-1_BR.RM.like.systems | <-VirD4-TraG+VirD4-FtsK\|\|TUDOR+SF1-SF2-Helicase+iREase+CTD+REase->STAS+HisKin->?->PIN->METHYLASE->HEPN+DUF499*-><-?<-?<-?<-ParB | 1113 | Bacteria>Pseudomonadota>Gammaproteobacteria>unclassified_Gammaproteobacteria | GCA_018819515.1 | Gammaproteobacteria_bacterium | KJ788_01195 | 188697..192038 | 1913989 | DUF499-domain-containing-protein-[Gammaproteobacteria-bacterium] |
| MCY3939524.1 | HEPN+AAA-ATPase+Alpha+Beta-domain+wHTH+RRM | Type-1_BR.RM.like.systems | MORC+DUF3684->TUDOR+SF1-SF2-Helicase+iREase+CTD+REase->METHYLASE->HEPN+DUF499*->MazE_antitoxin->DUF433->PIN-> | 1097 | Bacteria>Pseudomonadota>Gammaproteobacteria>unclassified_Gammaproteobacteria | GCA_026707445.1 | Gammaproteobacteria_bacterium | OXG29_00400 | 16758..20051 | 1913989 | MAG:-DUF499-domain-containing-protein-[Gammaproteobacteria-bacterium] |
| MDE0510268.1 | HEPN+AAA-ATPase+Alpha+Beta-domain+wHTH+RRM | Type-1_BR.RM.like.systems | SAM-methylase->TPR+TPR-><-HEPN+DUF499*<-PemK_toxin<-MazE_antitoxin<-HEPN<-CopG_antitoxin<-BrnT_toxin\|\|HEPN-> | 1110 | Bacteria>Pseudomonadota>Gammaproteobacteria>unclassified_Gammaproteobacteria | GCA_028823865.1 | Gammaproteobacteria_bacterium | OXI88_00540 | 10766..14098 | 1913989 | MAG:-Swt1-family-HEPN-domain-containing-protein-[Gammaproteobacteria-bacterium] |
| MDE0156012.1 | HEPN+AAA-ATPase+Alpha+Beta-domain+wHTH+RRM | Type-1_BR.RM.like.systems | SAM-methylase->TPR+TPR-><-HEPN+DUF499*<-PemK_toxin<-MazE_antitoxin<-HEPN<-CopG_antitoxin<-BrnT_toxin\|\|HEPN-> | 1110 | Bacteria>Pseudomonadota>Gammaproteobacteria>unclassified_Gammaproteobacteria | GCA_028819495.1 | Gammaproteobacteria_bacterium | OXS28_10440 | 75078..78410 | 1913989 | MAG:-Swt1-family-HEPN-domain-containing-protein-[Gammaproteobacteria-bacterium] |
| MBA2592967.1 | HEPN+AAA-ATPase+Alpha+Beta-domain+wHTH+RRM | Type-1_BR.RM.like.systems | ABC-ATPase->?->TUDOR+SF1-SF2-Helicase+iREase+CTD+REase->METHYLASE->HEPN+DUF499*-><-?<-?<-ParB<-TOPRIM<-ZnR<-XIS-HTH | 1115 | Bacteria>Pseudomonadota>Gammaproteobacteria>unclassified_Gammaproteobacteria | GCA_013697045.1 | Gammaproteobacteria_bacterium | HOU97_12235 | 29064..32411 | 1913989 | ATP-binding-protein-[Gammaproteobacteria-bacterium] |
| MYE82983.1 | HEPN+AAA-ATPase+Alpha+Beta-domain+wHTH+RRM | Type-1_BR.RM.like.systems | <-Arm-DNA-bind_3+Phage_int_SAM_3+Phage_integrase<-Phage_int_SAM_3+Phage_integrase\|\|HEPN+DUF499*-> | 1107 | Bacteria>Pseudomonadota>Gammaproteobacteria>unclassified_Gammaproteobacteria | GCA_009840715.1 | Gammaproteobacteria_bacterium | F4X36_14330 | 1398..4721 | 1913989 | ATP-binding-protein-[Gammaproteobacteria-bacterium] |
| MDE0680618.1 | HEPN+AAA-ATPase+Alpha+Beta-domain+wHTH+RRM | Type-1_BR.RM.like.systems | MORC+DUF3684->TUDOR+SF1-SF2-Helicase+iREase+CTD+REase->METHYLASE->?->?->HEPN+DUF499*->MazE_antitoxin-> | 1100 | Bacteria>Pseudomonadota>Gammaproteobacteria>unclassified_Gammaproteobacteria | GCA_028824615.1 | Gammaproteobacteria_bacterium | OXI11_10470 | 19020..22322 | 1913989 | MAG:-DUF499-domain-containing-protein-[Gammaproteobacteria-bacterium] |
| TAK54308.1 | HEPN+AAA-ATPase+Alpha+Beta-domain+wHTH+RRM | Type-1_BR.RM.like.systems | TUDOR+SF1-SF2-Helicase+iREase+CTD+REase->Schlafen+HATPase+HTH->METHYLASE->OLD-ABC->TOPRIM->HEPN+DUF499*-> | 1105 | Bacteria>Pseudomonadota>Gammaproteobacteria>unclassified_Gammaproteobacteria | GCA_004297335.1 | Gammaproteobacteria_bacterium | EPO25_07530 | 13135..16452 | 1913989 | ATP-binding-protein-[Gammaproteobacteria-bacterium] |
| MCY3821975.1 | HEPN+AAA-ATPase+Alpha+Beta-domain+wHTH+RRM | Type-1_BR.RM.like.systems | <-sGTP\|\|?->?-><-?<-HEPN+DUF499*\|\|Phage_int_SAM_3+Phage_integrase->Phage_integrase->METHYLASE-p-> | 1101 | Bacteria>Pseudomonadota>Gammaproteobacteria>unclassified_Gammaproteobacteria | GCA_026706665.1 | Gammaproteobacteria_bacterium | OXH52_21920 | 46820..50125 | 1913989 | MAG:-Swt1-family-HEPN-domain-containing-protein-[Gammaproteobacteria-bacterium] |
| MDE0073871.1 | HEPN+AAA-ATPase+Alpha+Beta-domain+wHTH+RRM | Type-1_BR.RM.like.systems | METHYLASE->OLD-ABC->TOPRIM->HEPN+DUF499*-><-HicB_lk_antitox<-HicA_toxin\|\|XIS-HTH->VirB2-> | 1107 | Bacteria>Pseudomonadota>Gammaproteobacteria>unclassified_Gammaproteobacteria | GCA_028819075.1 | Gammaproteobacteria_bacterium | OXU32_07805 | 3301..6624 | 1913989 | MAG:-Swt1-family-HEPN-domain-containing-protein-[Gammaproteobacteria-bacterium] |
| NKC12747.1 | HEPN+AAA-ATPase+Alpha+Beta-domain+wHTH+RRM | Type-1_BR.RM.like.systems | <-HEPN+DUF499*<-METHYLASE<-?<-TUDOR+SF1-SF2-Helicase+iREase+CTD+REase\|\|?->?->Phage_integrase-> | 1098 | Bacteria>Pseudomonadota>Gammaproteobacteria>unclassified_Gammaproteobacteria | GCA_012103455.1 | Gammaproteobacteria_bacterium | GKR94_11270 | 2223400..2226696 | 1913989 | DUF499-domain-containing-protein-[Gammaproteobacteria-bacterium] |
| MDE0284673.1 | HEPN+AAA-ATPase+Alpha+Beta-domain+wHTH+RRM | Type-1_BR.RM.like.systems | <-HEPN\|\|BrnT_toxin->CopG_antitoxin->HEPN->MazE_antitoxin->PemK_toxin->HEPN+DUF499*-> | 949 | Bacteria>Pseudomonadota>Gammaproteobacteria>unclassified_Gammaproteobacteria | GCA_028820555.1 | Gammaproteobacteria_bacterium | OXN26_09020 | 3552..6397 | 1913989 | MAG:-Swt1-family-HEPN-domain-containing-protein,-partial-[Gammaproteobacteria-bacterium] |
| MYB35869.1 | HEPN+AAA-ATPase+Alpha+Beta-domain+wHTH+RRM | Type-1_BR.RM.like.systems | TPR+TPR+PIN+PIN->TUDOR+SF1-SF2-Helicase+iREase+CTD+REase->METHYLASE->HEPN+DUF499*-> | 1114 | Bacteria>Pseudomonadota>Gammaproteobacteria>unclassified_Gammaproteobacteria | GCA_009838905.1 | Gammaproteobacteria_bacterium | F4Y26_00435 | 10511..13855 | 1913989 | ATP-binding-protein-[Gammaproteobacteria-bacterium] |
| MDE0479832.1 | HEPN+AAA-ATPase+Alpha+Beta-domain+wHTH+RRM | Type-1_BR.RM.like.systems | <-HEPN+DUF499*<-METHYLASE<-?<-RNase<-TUDOR+SF1-SF2-Helicase+iREase+CTD+REase<-HTH+UbE1 | 1108 | Bacteria>Pseudomonadota>Gammaproteobacteria>unclassified_Gammaproteobacteria | GCA_028824825.1 | Gammaproteobacteria_bacterium | OXI13_09405 | 28778..32104 | 1913989 | MAG:-DUF499-domain-containing-protein-[Gammaproteobacteria-bacterium] |
| MCY3837525.1 | HEPN+AAA-ATPase+Alpha+Beta-domain+wHTH+RRM | Type-1_BR.RM.like.systems | METHYLASE->CR-ATPase8+REase->HEPN+DUF499*->Tox-RelE->HTH+MPTase-><-VapB_antitoxin | 1112 | Bacteria>Pseudomonadota>Gammaproteobacteria>unclassified_Gammaproteobacteria | GCA_026706425.1 | Gammaproteobacteria_bacterium | OXH09_02545 | 4546..7884 | 1913989 | MAG:-Swt1-family-HEPN-domain-containing-protein-[Gammaproteobacteria-bacterium] |
| MBM4223246.1 | HEPN+AAA-ATPase+Alpha+Beta-domain+wHTH+RRM | Type-1_BR.RM.like.systems | TUDOR+SF1-SF2-Helicase+iREase+CTD+REase->sGTP->METHYLASE->HEPN+DUF499*->REase-> | 1092 | Bacteria>Pseudomonadota>Gammaproteobacteria>unclassified_Gammaproteobacteria | GCA_016874585.1 | Gammaproteobacteria_bacterium | FJ167_00320 | 8980..12258 | 1913989 | ATP-binding-protein-[Gammaproteobacteria-bacterium] |
| MYH50428.1 | HEPN+AAA-ATPase+Alpha+Beta-domain+wHTH+RRM | Type-1_BR.RM.like.systems | METHYLASE-p->HEPN+DUF499*-><-HicB_lk_antitox<-?<-?<-LexA-protease\|\|XIS-HTH->VirB2-> | 1107 | Bacteria>Pseudomonadota>Gammaproteobacteria>unclassified_Gammaproteobacteria | GCA_009843625.1 | Gammaproteobacteria_bacterium | F4151_13115 | 723..4046 | 1913989 | ATP-binding-protein-[Gammaproteobacteria-bacterium] |
| MCY4343961.1 | HEPN+AAA-ATPase+Alpha+Beta-domain+wHTH+RRM | Type-1_BR.RM.like.systems | METHYLASE->OLD-ABC->Schlafen+HATPase+HTH->?->PIN->HEPN+DUF499*-><-MNS+ZDNA-HTH<-PRTase | 1115 | Bacteria>Pseudomonadota>Gammaproteobacteria>unclassified_Gammaproteobacteria | GCA_026710215.1 | Gammaproteobacteria_bacterium | OXE83_10365 | 11310..14657 | 1913989 | MAG:-Swt1-family-HEPN-domain-containing-protein-[Gammaproteobacteria-bacterium] |
| MCY4282353.1 | HEPN+AAA-ATPase+Alpha+Beta-domain+wHTH+RRM | Type-1_BR.RM.like.systems | SAM-methylase->TPR+TPR->DOC+DOC-><-HEPN+DUF499*\|\|?->ParB+HNH->OLD-ABC-> | 941 | Bacteria>Pseudomonadota>Gammaproteobacteria>unclassified_Gammaproteobacteria | GCA_026709915.1 | Gammaproteobacteria_bacterium | OXE03_05465 | 14307..17132 | 1913989 | MAG:-Swt1-family-HEPN-domain-containing-protein-[Gammaproteobacteria-bacterium] |
| MCG3202826.1 | HEPN+AAA-ATPase+Alpha+Beta-domain+wHTH+RRM | Type-1_BR.RM.like.systems | N6_N4_MTase-><-?<-AbiEii<-AbiEi_4<-REase<-HEPN+DUF499*<-METHYLASE-p | 1115 | Bacteria>Pseudomonadota>Gammaproteobacteria>unclassified_Gammaproteobacteria | GCA_022071965.1 | Gammaproteobacteria_bacterium | NFCOHLIN_02712 | 302948..306295 | 1913989 | hypothetical-protein-[Gammaproteobacteria-bacterium] |
| MCZ0943260.1 | HEPN+AAA-ATPase+Alpha+Beta-domain+wHTH+RRM | Type-1_BR.RM.like.systems | DUF433-><-HEPN+DUF499*<-Schlafen+HATPase+HTH<-TOPRIM<-OLD-ABC<-METHYLASE-p | 1115 | Bacteria>Pseudomonadota>Gammaproteobacteria>unclassified_Gammaproteobacteria | GCA_026914145.1 | Gammaproteobacteria_bacterium | OXJ53_09405 | 2539..5886 | 1913989 | MAG:-Swt1-family-HEPN-domain-containing-protein-[Gammaproteobacteria-bacterium] |
| MYH70086.1 | HEPN+AAA-ATPase+Alpha+Beta-domain+wHTH+RRM | Type-1_BR.RM.like.systems | METHYLASE->MazE_antitoxin->PemK_toxin->HEPN+DUF499*-><-TPR+TPR | 1110 | Bacteria>Pseudomonadota>Gammaproteobacteria>unclassified_Gammaproteobacteria | GCA_009843675.1 | Gammaproteobacteria_bacterium | F4147_08315 | 2722..6054 | 1913989 | ATP-binding-protein-[Gammaproteobacteria-bacterium] |
| MCY4265599.1 | HEPN+AAA-ATPase+Alpha+Beta-domain+wHTH+RRM | Type-1_BR.RM.like.systems | TUDOR+SF1-SF2-Helicase+iREase+CTD+REase->HEPN+DUF499*-><-MNS | 1088 | Bacteria>Pseudomonadota>Gammaproteobacteria>unclassified_Gammaproteobacteria | GCA_026709755.1 | Gammaproteobacteria_bacterium | OXE78_12210 | 4990..8256 | 1913989 | MAG:-DUF499-domain-containing-protein-[Gammaproteobacteria-bacterium] |
| MCY4211280.1 | HEPN+AAA-ATPase+Alpha+Beta-domain+wHTH+RRM | Type-1_BR.RM.like.systems | SAM-methylase->TPR+TPR->DOC+DOC-><-DUF499-p<-HEPN+DUF499* | 757 | Bacteria>Pseudomonadota>Gammaproteobacteria>unclassified_Gammaproteobacteria | GCA_026709385.1 | Gammaproteobacteria_bacterium | OXE97_09605 | 24101..26373 | 1913989 | MAG:-Swt1-family-HEPN-domain-containing-protein,-partial-[Gammaproteobacteria-bacterium] |
| MXY90447.1 | HEPN+AAA-ATPase+Alpha+Beta-domain+wHTH+RRM | Type-1_BR.RM.like.systems | <-PrmC_N+Methyltransf_25\|\|?-><-?<-?<-HEPN+DUF499*<-TOPRIM<-OLD-ABC | 1116 | Bacteria>Pseudomonadota>Gammaproteobacteria>unclassified_Gammaproteobacteria | GCA_009837675.1 | Gammaproteobacteria_bacterium | F4Y89_07905 | 34085..37435 | 1913989 | ATP-binding-protein-[Gammaproteobacteria-bacterium] |
| MCY3988741.1 | HEPN+AAA-ATPase+Alpha+Beta-domain+wHTH+RRM | Type-1_BR.RM.like.systems | METHYLASE-p->HEPN+DUF499*->MazE_antitoxin->DUF433-> | 1097 | Bacteria>Pseudomonadota>Gammaproteobacteria>unclassified_Gammaproteobacteria | GCA_026708205.1 | Gammaproteobacteria_bacterium | OXF94_09155 | 914..4207 | 1913989 | MAG:-DUF499-domain-containing-protein-[Gammaproteobacteria-bacterium] |
| MDE0678898.1 | HEPN+AAA-ATPase+Alpha+Beta-domain+wHTH+RRM | Type-1_BR.RM.like.systems | METHYLASE-p->HEPN+DUF499*->SF1-SF2-Helicase+HRDC-> | 1109 | Bacteria>Pseudomonadota>Gammaproteobacteria>unclassified_Gammaproteobacteria | GCA_028824615.1 | Gammaproteobacteria_bacterium | OXI11_01560 | 1155..4484 | 1913989 | MAG:-Swt1-family-HEPN-domain-containing-protein-[Gammaproteobacteria-bacterium] |
| MXW44722.1 | HEPN+AAA-ATPase+Alpha+Beta-domain+wHTH+RRM | Type-1_BR.RM.like.systems | MORC+DUF3684->?->?->HEPN+DUF499*->?->MazE_antitoxin-> | 959 | Bacteria>Pseudomonadota>Gammaproteobacteria>unclassified_Gammaproteobacteria | GCA_009836205.1 | Gammaproteobacteria_bacterium | F4Z63_01800 | 11560..14439 | 1913989 | ATP-binding-protein-[Gammaproteobacteria-bacterium] |
| MBT4891857.1 | HEPN+AAA-ATPase+Alpha+Beta-domain+wHTH+RRM | Type-1_BR.RM.like.systems | METHYLASE-p->HEPN+DUF499*->TIR-> | 1099 | Bacteria>Pseudomonadota>Gammaproteobacteria>unclassified_Gammaproteobacteria | GCA_018664395.1 | Gammaproteobacteria_bacterium | HON60_02830 | 645..3944 | 1913989 | ATP-binding-protein-[Gammaproteobacteria-bacterium] |
| TVS19395.1 | HEPN+AAA-ATPase+Alpha+Beta-domain+wHTH+RRM | Type-1_BR.RM.like.systems | <-HEPN+DUF499*<-METHYLASE-p | 1100 | Bacteria>Pseudomonadota>Gammaproteobacteria>unclassified_Gammaproteobacteria | GCA_007694905.1 | Gammaproteobacteria_bacterium | EA417_00670 | 96..3398 | 1913989 | ATP-binding-protein-[Gammaproteobacteria-bacterium] |
| MXY57169.1 | HEPN+AAA-ATPase+Alpha+Beta-domain+wHTH+RRM | Type-1_BR.RM.like.systems | METHYLASE-p->HEPN+DUF499*-> | 413 | Bacteria>Pseudomonadota>Gammaproteobacteria>unclassified_Gammaproteobacteria | GCA_009838585.1 | Gammaproteobacteria_bacterium | F4Y41_12365 | 1564..2802 | 1913989 | ATP-binding-protein-[Gammaproteobacteria-bacterium] |
| RMG33561.1 | HEPN+AAA-ATPase+Alpha+Beta-domain+wHTH+RRM | Type-1_BR.RM.like.systems | <-HEPN+DUF499*<-METHYLASE | 1107 | Bacteria>Pseudomonadota>Gammaproteobacteria>unclassified_Gammaproteobacteria | GCA_003696905.1 | Gammaproteobacteria_bacterium | D6720_11165 | 157..3480 | 1913989 | ATP-binding-protein-[Gammaproteobacteria-bacterium] |
| MDE0441971.1 | HEPN+AAA-ATPase+Alpha+Beta-domain+wHTH+RRM | Type-1_BR.RM.like.systems | <-HEPN+DUF499*<-METHYLASE | 570 | Bacteria>Pseudomonadota>Gammaproteobacteria>unclassified_Gammaproteobacteria | GCA_028821355.1 | Gammaproteobacteria_bacterium | OXL38_07620 | 38..1750 | 1913989 | MAG:-Swt1-family-HEPN-domain-containing-protein-[Gammaproteobacteria-bacterium] |
| MCY4181966.1 | HEPN+AAA-ATPase+Alpha+Beta-domain+wHTH+RRM | Type-1_BR.RM.like.systems | <-HEPN+DUF499*<-METHYLASE | 1105 | Bacteria>Pseudomonadota>Gammaproteobacteria>unclassified_Gammaproteobacteria | GCA_026709135.1 | Gammaproteobacteria_bacterium | OXF43_03630 | 102..3419 | 1913989 | MAG:-DUF499-domain-containing-protein-[Gammaproteobacteria-bacterium] |
| GIX32370.1 | HEPN+AAA-ATPase+Alpha+Beta-domain+wHTH+RRM | Type-1_BR.RM.like.systems | REase->METHYLASE-p->METHYLASE-p->METHYLASE->ParD->PIN->HEPN+DUF499*-><-tRNA<-SIGMA-HTH+SIGMA-HTH | 1108 | Bacteria>Pseudomonadota>Gammaproteobacteria>Xanthomonadales>unclassified_Xanthomonadales | GCA_026005055.1 | Xanthomonadales_bacterium | KatS3mg125_0326 | 361925..365251 | 2006849 | MAG:-hypothetical-protein-KatS3mg125_0326-[Xanthomonadales-bacterium] |
| RIK94108.1 | HEPN+AAA-ATPase+Alpha+Beta-domain+wHTH+RRM | Type-1_BR.RM.like.systems | DUF488->ORC-CDC6+REase->?-><-PIWI<-TIR+PNTD2<-HEPN+DUF499*<-METHYLASE<-MORC+DUF3684<-TUDOR+SF1-SF2-Helicase+iREase+CTD+REase<-HTH+WYL\|\|SF1-SF2-Helicase+REase->SF1-SF2-Helicase+REase-> | 1098 | Bacteria>Pseudomonadota>unclassified_Pseudomonadota | GCA_003577275.1 | Pseudomonadota_bacterium | DCC73_05940 | 240099..243395 | 1977087 | AAA-family-ATPase-[Proteobacteria-bacterium] |
| MCA0401434.1 | HEPN+AAA-ATPase+Alpha+Beta-domain+wHTH+RRM | Type-1_BR.RM.like.systems | ParG_RHH-><-DUF488-1<-?<-TetR-HTH\|\|MODE-HTH+PBPII-><-HEPN+DUF499*<-METHYLASE<-HEPN<-TUDOR+SF1-SF2-Helicase+iREase+CTD+REase<-HTH+WYL\|\|SF1-SF2-Helicase+REase->SF1-SF2-Helicase+REase-> | 1098 | Bacteria>Pseudomonadota>unclassified_Pseudomonadota | GCA_020161575.1 | Pseudomonadota_bacterium | LCH38_11530 | 127907..131203 | 1977087 | DUF499-domain-containing-protein-[Proteobacteria-bacterium] |
| MCA0247253.1 | HEPN+AAA-ATPase+Alpha+Beta-domain+wHTH+RRM | Type-1_BR.RM.like.systems | <-SF1-SF2-Helicase+REase<-SF1-SF2-Helicase+REase\|\|HTH+WYL->TUDOR+SF1-SF2-Helicase+iREase+CTD+REase->MORC+DUF3684->METHYLASE->HEPN+DUF499*->MODE-HTH+PBPII-><-ParG_RHH | 1098 | Bacteria>Pseudomonadota>unclassified_Pseudomonadota | GCA_020160995.1 | Pseudomonadota_bacterium | LCH93_11605 | 30891..34187 | 1977087 | DUF499-domain-containing-protein-[Proteobacteria-bacterium] |
| MBS0471294.1 | HEPN+AAA-ATPase+Alpha+Beta-domain+wHTH+RRM | Type-1_BR.RM.like.systems | <-AbiEii<-AbiEi_3_N+AbiEi_3\|\|XIS-HTH->TUDOR+SF1-SF2-Helicase+iREase+CTD+REase->MORC+DUF3684->METHYLASE->HEPN+DUF499*-><-XIS-HTH<-?<-ParG_RHH | 1118 | Bacteria>Pseudomonadota>unclassified_Pseudomonadota | GCA_018241025.1 | Pseudomonadota_bacterium | JSR60_09500 | 167872..171228 | 1977087 | DUF499-domain-containing-protein-[Proteobacteria-bacterium] |
| OQW45528.1 | HEPN+AAA-ATPase+Alpha+Beta-domain+wHTH+RRM | Type-1_BR.RM.like.systems | <-HicB\|\|HTH+WYL->TUDOR+SF1-SF2-Helicase+iREase+CTD+REase-><-?\|\|METHYLASE->HEPN+DUF499*->MODE-HTH+PBPII-><-ParG_RHH | 1097 | Bacteria>Pseudomonadota>unclassified_Pseudomonadota | GCA_002083435.1 | Proteobacteria_bacterium_SG_bin6 | A4S16_12875 | 8645..11938 | 1827383 | AAA-family-ATPase-[Proteobacteria-bacterium-SG_bin6] |
| MBU1208887.1 | HEPN+AAA-ATPase+Alpha+Beta-domain+wHTH+RRM | Type-1_BR.RM.like.systems | TUDOR+SF1-SF2-Helicase+iREase+CTD+REase-p->DrHyd+BirA-HTH->METHYLASE->HEPN+DUF499*->REase-p->?->?->Mbetalac-> | 1107 | Bacteria>Pseudomonadota>unclassified_Pseudomonadota | GCA_018819135.1 | Pseudomonadota_bacterium | KKH04_18500 | 6409..9732 | 1977087 | DUF499-domain-containing-protein-[Proteobacteria-bacterium] |
| MCX7197671.1 | HEPN+AAA-ATPase+Alpha+Beta-domain+wHTH+RRM | Type-1_BR.RM.like.systems | TUDOR+SF1-SF2-Helicase+iREase+CTD+REase-p->METHYLASE->HTH+Hiskin+STAS->HEPN+DUF499*->?->HNH->SIR2->DnaB-> | 1113 | Bacteria>Pseudomonadota>unclassified_Pseudomonadota | GCA_026399745.1 | Pseudomonadota_bacterium | NTW37_06695 | 4467..7808 | 1977087 | MAG:-Swt1-family-HEPN-domain-containing-protein-[Proteobacteria-bacterium] |
| MBU4581761.1 | HEPN+AAA-ATPase+Alpha+Beta-domain+wHTH+RRM | Type-1_BR.RM.like.systems | DDE_5+DDE_Tnp_1->?-><-HEPN+DUF499*<-METHYLASE<-?<-?<-SF1-SF2-Helicase<-DrHyd+BirA-HTH<-MORC+DUF3684 | 1124 | Bacteria>Pseudomonadota>unclassified_Pseudomonadota | GCA_018902925.1 | Pseudomonadota_bacterium | KKH02_05005 | 2369..5743 | 1977087 | DUF499-domain-containing-protein-[Proteobacteria-bacterium] |
| MBU1342385.1 | HEPN+AAA-ATPase+Alpha+Beta-domain+wHTH+RRM | Type-1_BR.RM.like.systems | MORC+DUF3684->TUDOR+SF1-SF2-Helicase+iREase+CTD+REase->HEPN->METHYLASE->HEPN+DUF499*-> | 1114 | Bacteria>Pseudomonadota>unclassified_Pseudomonadota | GCA_018820265.1 | Pseudomonadota_bacterium | KKD66_12250 | 12988..16332 | 1977087 | DUF499-domain-containing-protein-[Proteobacteria-bacterium] |
| PWT88877.1 | HEPN+AAA-ATPase+Alpha+Beta-domain+wHTH+RRM | Type-1_BR.RM.like.systems | TUDOR+SF1-SF2-Helicase+iREase+CTD+REase-p->HEPN+DUF499*-> | 1096 | Bacteria>Pseudomonadota>unclassified_Pseudomonadota | GCA_003175845.1 | Pseudomonadota_bacterium | C5B56_08055 | 3674..6960 | 1977087 | AAA+-family-ATPase,-partial-[Proteobacteria-bacterium] |
| MCA0405125.1 | HEPN+AAA-ATPase+Alpha+Beta-domain+wHTH+RRM | Type-1_BR.RM.like.systems | ASCH->METHYLASE->HEPN+DUF499*->MODE-HTH+PBPII-><-ParG_RHH | 1097 | Bacteria>Pseudomonadota>unclassified_Pseudomonadota | GCA_020161475.1 | Pseudomonadota_bacterium | LCH39_03110 | 5235..8528 | 1977087 | DUF499-domain-containing-protein-[Proteobacteria-bacterium] |
| MBS0482554.1 | HEPN+AAA-ATPase+Alpha+Beta-domain+wHTH+RRM | Type-1_BR.RM.like.systems | Thioredoxin->TraH->?-><-?<-HEPN+DUF499*<-METHYLASE-p | 1098 | Bacteria>Pseudomonadota>unclassified_Pseudomonadota | GCA_018240895.1 | Pseudomonadota_bacterium | JSR96_10460 | 15112..18408 | 1977087 | DUF499-domain-containing-protein-[Proteobacteria-bacterium] |
| MCG8554136.1 | HEPN+AAA-ATPase+Alpha+Beta-domain+wHTH+RRM | Type-1_BR.RM.like.systems | Cas_Cmr3->?->Cas_Cmr5->?->CARF+HEPN->HEPN+DUF499*-> | 1114 | Bacteria>Pseudomonadota>unclassified_Pseudomonadota | GCA_022361155.1 | Pseudomonadota_bacterium | MJD61_02430 | 12967..16311 | 1977087 | Swt1-family-HEPN-domain-containing-protein-[Proteobacteria-bacterium] |
| MBI4030489.1 | HEPN+AAA-ATPase+Alpha+Beta-domain+wHTH+RRM | Type-1_BR.RM.like.systems | Peptidase_M15_4->?->?->?->?-><-HEPN+DUF499*<-METHYLASE | 1115 | Bacteria>Pseudomonadota>unclassified_Pseudomonadota | GCA_016198545.1 | Pseudomonadota_bacterium | HY370_02330 | 8251..11598 | 1977087 | ATP-binding-protein-[Proteobacteria-bacterium] |
| NBT04715.1 | HEPN+AAA-ATPase+Alpha+Beta-domain+wHTH+RRM | Type-1_BR.RM.like.systems | <-REase<-HEPN+DUF499*<-METHYLASE-p | 1140 | Bacteria>Pseudomonadota>unclassified_Pseudomonadota | GCA_009922565.1 | Pseudomonadota_bacterium | EBS94_15100 | 2192..5614 | 1977087 | ATP-binding-protein-[Proteobacteria-bacterium] |
| MBQ9395962.1 | HEPN+AAA-ATPase+Alpha+Beta-domain+wHTH+RRM | Type-1_BR.RM.like.systems | METHYLASE-p-><-PIN<-?\|\|HEPN+DUF499*-> | 729 | Bacteria>Pseudomonadota>unclassified_Pseudomonadota | GCA_017455165.1 | Pseudomonadota_bacterium | IJU23_10695 | 3126..5311 | 1977087 | DUF499-domain-containing-protein-[Proteobacteria-bacterium] |
| MBE9569155.1 | HEPN+AAA-ATPase+Alpha+Beta-domain+wHTH+RRM | Type-1_BR.RM.like.systems | PIN->METHYLASE->HEPN+DUF499*-> | 1063 | Bacteria>Pseudomonadota>unclassified_Pseudomonadota | GCA_015222845.1 | Pseudomonadota_bacterium | IMF11_00880 | 3390..6577 | 1977087 | DUF499-domain-containing-protein-[Proteobacteria-bacterium] |
| NBQ33337.1 | HEPN+AAA-ATPase+Alpha+Beta-domain+wHTH+RRM | Type-1_BR.RM.like.systems | <-HEPN+DUF499*<-METHYLASE-p | 1140 | Bacteria>Pseudomonadota>unclassified_Pseudomonadota | GCA_009920185.1 | Pseudomonadota_bacterium | EBU21_15535 | 254..3676 | 1977087 | ATP-binding-protein-[Proteobacteria-bacterium] |
| MBU2547024.1 | HEPN+AAA-ATPase+Alpha+Beta-domain+wHTH+RRM | Type-1_BR.RM.like.systems | <-HEPN+DUF499*<-METHYLASE-p | 1093 | Bacteria>Pseudomonadota>unclassified_Pseudomonadota | GCA_018830135.1 | Pseudomonadota_bacterium | KKB20_01315 | 29847..33128 | 1977087 | DUF499-domain-containing-protein-[Proteobacteria-bacterium] |
| MBU0466040.1 | HEPN+AAA-ATPase+Alpha+Beta-domain+wHTH+RRM | Type-1_BR.RM.like.systems | <-HEPN+DUF499*<-METHYLASE-p | 1114 | Bacteria>Pseudomonadota>unclassified_Pseudomonadota | GCA_018812385.1 | Pseudomonadota_bacterium | KKE12_21020 | 197..3541 | 1977087 | DUF499-domain-containing-protein-[Proteobacteria-bacterium] |
| MDA1076682.1 | HEPN+AAA-ATPase+Alpha+Beta-domain+wHTH+RRM | Type-1_BR.RM.like.systems | <-HEPN+DUF499*<-METHYLASE | 1005 | Bacteria>Pseudomonadota>unclassified_Pseudomonadota | GCA_027622495.1 | Pseudomonadota_bacterium | O3A63_18310 | 1..3015 | 1977087 | MAG:-Swt1-family-HEPN-domain-containing-protein,-partial-[Proteobacteria-bacterium] |
| MBU2496199.1 | HEPN+AAA-ATPase+Alpha+Beta-domain+wHTH+RRM | Type-1_BR.RM.like.systems | METHYLASE->HEPN+DUF499*-> | 365 | Bacteria>PVC_group>Candidatus_Omnitrophota>unclassified_Candidatus_Omnitrophota | GCA_018829455.1 | Candidatus_Omnitrophota_bacterium | KJ935_06870 | 1920..3013 | 2035772 | DUF499-domain-containing-protein-[Candidatus-Omnitrophota-bacterium] |
| MBN1673169.1 | HEPN+AAA-ATPase+Alpha+Beta-domain+wHTH+RRM | Type-1_BR.RM.like.systems | <-DrHyd+BirA-HTH<-TUDOR+SF1-SF2-Helicase+iREase+CTD+REase-p<-?<-?\|\|?->?-><-HEPN+DUF499*<-Schlafen+HATPase+HTH<-METHYLASE-p\|\|BirA-HTH->HhH-RADC+JAB-> | 1118 | Bacteria>PVC_group>Kiritimatiellota>Kiritimatiellia>unclassified_Kiritimatiellia | GCA_016929905.1 | Kiritimatiellia_bacterium | JXR37_19145 | 44287..47643 | 2212473 | DUF499-domain-containing-protein-[Kiritimatiellae-bacterium] |
| MDD4116915.1 | HEPN+AAA-ATPase+Alpha+Beta-domain+wHTH+RRM | Type-1_BR.RM.like.systems | SAM-methylase-><-?<-?<-?\|\|tRNA-><-HEPN+DUF499*<-Peptidase_C47<-?<-METHYLASE<-ParB+HNH<-?<-TUDOR+SF1-SF2-Helicase+iREase+CTD+REase | 1113 | Bacteria>PVC_group>Kiritimatiellota>Kiritimatiellia>unclassified_Kiritimatiellia | GCA_028701395.1 | Kiritimatiellia_bacterium | PHI39_01725 | 13604..16945 | 2212473 | MAG:-Swt1-family-HEPN-domain-containing-protein-[Kiritimatiellae-bacterium] |
| MCL1920537.1 | HEPN+AAA-ATPase+Alpha+Beta-domain+wHTH+RRM | Type-1_BR.RM.like.systems | <-HEPN+DUF499*<-METHYLASE-p | 1094 | Bacteria>PVC_group>Kiritimatiellota>unclassified_Kiritimatiellota | GCA_009778475.1 | Kiritimatiellota_bacterium | FWG50_05580 | 29293..32577 | 2599938 | DUF499-domain-containing-protein-[Kiritimatiellaeota-bacterium] |
| HIV08682.1 | HEPN+AAA-ATPase+Alpha+Beta-domain+wHTH+RRM | Type-1_BR.RM.like.systems | TUDOR+SF1-SF2-Helicase+iREase+CTD+REase->HEPN+DUF499*->ParB+HNH-p-> | 1108 | Bacteria>PVC_group>Lentisphaerota>Lentisphaeria>Lentisphaerales | GCA_018713405.1 | Candidatus_Spyradenecus_faecavium | IAC79_01020 | 6980..10306 | 2840947 | DUF499-domain-containing-protein-[Candidatus-Spyradenecus-faecavium] |
| MBN2450533.1 | HEPN+AAA-ATPase+Alpha+Beta-domain+wHTH+RRM | Type-1_BR.RM.like.systems | REase->?->PIN->HEPN+DUF499*-> | 1121 | Bacteria>PVC_group>Lentisphaerota>Lentisphaeria>unclassified_Lentisphaeria | GCA_016935045.1 | Lentisphaeria_bacterium | JXR77_09085 | 1530..4895 | 2053569 | DUF499-domain-containing-protein-[Lentisphaeria-bacterium] |
| MCE9615077.1 | HEPN+AAA-ATPase+Alpha+Beta-domain+wHTH+RRM | Type-1_BR.RM.like.systems | <-XIS-HTH<-?<-?<-?<-HicB<-?<-HEPN+DUF499*<-METHYLASE<-DOC+HTH<-HTH_3<-sGTP<-TUDOR+SF1-SF2-Helicase+iREase+CTD+REase | 1128 | Bacteria>PVC_group>Lentisphaerota>unclassified_Lentisphaerota | GCA_021413965.1 | Lentisphaerota_bacterium | K8T26_12435 | 13053..16439 | 1932692 | DUF499-domain-containing-protein-[Lentisphaerae-bacterium] |
| MBC8553109.1 | HEPN+AAA-ATPase+Alpha+Beta-domain+wHTH+RRM | Type-1_BR.RM.like.systems | XIS-HTH->TUDOR+SF1-SF2-Helicase+iREase+CTD+REase->METHYLASE->HEPN+DUF499*->HNH-> | 1106 | Bacteria>PVC_group>Planctomycetota>Candidatus_Brocadiia>Candidatus_Brocadiales | GCA_014384865.1 | Candidatus_Brocadiales_bacterium | H8D23_26105 | 6653..9973 | 2760054 | ATP-binding-protein-[Candidatus-Brocadiales-bacterium] |
| MBK9119771.1 | HEPN+AAA-ATPase+Alpha+Beta-domain+wHTH+RRM | Type-1_BR.RM.like.systems | DEOR-HTH->TUDOR+SF1-SF2-Helicase+iREase+CTD+REase->Schlafen+HATPase+HTH->?->?->METHYLASE->HEPN+DUF499*->MORC->REase-><-Trypsin\|\|HEPN->HTH->MPTase-> | 1121 | Bacteria>PVC_group>Planctomycetota>Phycisphaerae>Phycisphaerales | GCA_016716475.1 | Phycisphaerales_bacterium | IPM18_09260 | 379276..382641 | 2052180 | DUF499-domain-containing-protein-[Phycisphaerales-bacterium] |
| MCL4740947.1 | HEPN+AAA-ATPase+Alpha+Beta-domain+wHTH+RRM | Type-1_BR.RM.like.systems | <-XIS-HTH<-?<-MPTase<-HTH_3\|\|?-><-HEPN+DUF499*<-METHYLASE<-sGTP<-TUDOR+SF1-SF2-Helicase+iREase+CTD+REase<-Phage_integrase<-ParB+HTH<-HTH_3 | 1114 | Bacteria>PVC_group>Planctomycetota>Phycisphaerae>Phycisphaerales | GCA_023384195.1 | Phycisphaerales_bacterium | KJZ54_01975 | 143211..146555 | 2052180 | DUF499-domain-containing-protein-[Phycisphaerales-bacterium] |
| MCC6908120.1 | HEPN+AAA-ATPase+Alpha+Beta-domain+wHTH+RRM | Type-1_BR.RM.like.systems | <-Hydrolase_3<-HEPN+DUF499*<-METHYLASE<-?<-?<-Schlafen+HATPase+HTH<-TUDOR+SF1-SF2-Helicase+iREase+CTD+REase<-DEOR-HTH | 1120 | Bacteria>PVC_group>Planctomycetota>Phycisphaerae>Phycisphaerales | GCA_020852515.1 | Phycisphaerales_bacterium | IT430_09285 | 88345..91707 | 2052180 | DUF499-domain-containing-protein-[Phycisphaerales-bacterium] |
| MCA3006076.1 | HEPN+AAA-ATPase+Alpha+Beta-domain+wHTH+RRM | Type-1_BR.RM.like.systems | METHYLASE->?->HEPN+DUF499*->?->XIS-HTH->?->?->?-><-tRNA\|\|ABC-ATPase+SbcC-> | 1114 | Bacteria>PVC_group>Planctomycetota>Phycisphaerae>Phycisphaerales | GCA_020247235.1 | Phycisphaerales_bacterium | IOD15_12050 | 3800..7144 | 2052180 | DUF499-domain-containing-protein-[Phycisphaerales-bacterium] |
| MCK6484528.1 | HEPN+AAA-ATPase+Alpha+Beta-domain+wHTH+RRM | Type-1_BR.RM.like.systems | ArdCN+MPTase->DEOR-HTH->TUDOR+SF1-SF2-Helicase+iREase+CTD+REase->wHTH+REase->REase->METHYLASE->HEPN+DUF499*->DNA_methylase->MORC+HISKIN->?->REase->CR-Kinase+ORC-CDC6-like+CARF-> | 1119 | Bacteria>PVC_group>Planctomycetota>Phycisphaerae>unclassified_Phycisphaerae | GCA_023150595.1 | Phycisphaerae_bacterium | L6R00_10395 | 56365..59724 | 2026778 | DUF499-domain-containing-protein-[Phycisphaerae-bacterium] |
| GIW77359.1 | HEPN+AAA-ATPase+Alpha+Beta-domain+wHTH+RRM | Type-1_BR.RM.like.systems | TUDOR+SF1-SF2-Helicase+iREase+CTD+REase-p->TUDOR+SF1-SF2-Helicase+iREase+CTD+REase->METHYLASE->METHYLASE-p->HEPN+DUF499*->DUF499-p->DUF499-p->?-><-tRNA\|\|?->METHYLASE-> | 529 | Bacteria>PVC_group>Planctomycetota>Phycisphaerae>unclassified_Phycisphaerae | GCA_026004655.1 | Phycisphaerae_bacterium | KatS3mg104_2422 | 7750..9339 | 2026778 | MAG:-hypothetical-protein-KatS3mg104_2422-[Phycisphaerae-bacterium] |
| MBW7904855.1 | HEPN+AAA-ATPase+Alpha+Beta-domain+wHTH+RRM | Type-1_BR.RM.like.systems | <-HTH_3\|\|DEOR-HTH->TUDOR+SF1-SF2-Helicase+iREase+CTD+REase->Schlafen+HATPase+HTH->REase->METHYLASE->HEPN+DUF499*->Hydrolase_3-> | 1120 | Bacteria>PVC_group>Planctomycetota>Phycisphaerae>unclassified_Phycisphaerae | GCA_019454965.1 | Phycisphaerae_bacterium | H3C42_04715 | 502745..506107 | 2026778 | ATP-binding-protein-[Phycisphaerae-bacterium] |
| MBN2560619.1 | HEPN+AAA-ATPase+Alpha+Beta-domain+wHTH+RRM | Type-1_BR.RM.like.systems | <-HEPN+DUF499*<-?<-METHYLASE<-wHTH+REase<-TUDOR+SF1-SF2-Helicase+iREase+CTD+REase<-?<-Cluster182_14clades | 1112 | Bacteria>PVC_group>Planctomycetota>Phycisphaerae>unclassified_Phycisphaerae | GCA_016935885.1 | Phycisphaerae_bacterium | JXQ75_06790 | 1428..4766 | 2026778 | DUF499-domain-containing-protein-[Phycisphaerae-bacterium] |
| WP_008662002.1 | HEPN+AAA-ATPase+Alpha+Beta-domain+wHTH+RRM | Type-1_BR.RM.like.systems | <-XIS-HTH\|\|URI-DNase-><-HEPN<-HEPN+DUF499*<-HEPN<-METHYLASE<-?<-TUDOR+SF1-SF2-Helicase+iREase+CTD+REase\|\|HKD-DNase+SF1-SF2-Helicase+TRD->URI-DNase-> | 1113 | Bacteria>PVC_group>Planctomycetota>Planctomycetia>Pirellulales | GCF_000338295.1 | Rhodopirellula_europaea | RE6C_RS27575 | 3450..6791 | 1263866 | DUF499-domain-containing-protein-[Rhodopirellula-europaea] |
| WP_283433169.1 | HEPN+AAA-ATPase+Alpha+Beta-domain+wHTH+RRM | Type-1_BR.RM.like.systems | <-ParB+HNH<-?<-HTH\|\|MORC+DUF3684->HKD-DNase-><-HEPN+DUF499*<-PIN<-OLD-ABC<-METHYLASE<-REase\|\|?-><-TUDOR+SF1-SF2-Helicase+iREase+CTD+REase | 1109 | Bacteria>PVC_group>Planctomycetota>Planctomycetia>Pirellulales | GCF_900182915.1 | Neorhodopirellula_lusitana | QOL80_RS14710 | 91339..94668 | 445327 | DUF499-domain-containing-protein-[Rhodopirellula-lusitana] |
| GIW99318.1 | HEPN+AAA-ATPase+Alpha+Beta-domain+wHTH+RRM | Type-1_BR.RM.like.systems | METHYLASE-p->?->MACRODOMAIN->METHYLASE->PHP+SbcC->Schlafen+HATPase+HTH->HEPN+DUF499*-><-URI-DNase<-?<-REase<-DDE_Tnp_IS66+DDE_Tnp_IS66_C | 1115 | Bacteria>PVC_group>Planctomycetota>Planctomycetia>Pirellulales | GCA_026004795.1 | Pirellulaceae_bacterium | KatS3mg111_2651 | 704885..708232 | 2699754 | MAG:-hypothetical-protein-KatS3mg111_2651-[Pirellulaceae-bacterium] |
| NLX95047.1 | HEPN+AAA-ATPase+Alpha+Beta-domain+wHTH+RRM | Type-1_BR.RM.like.systems | TPR+TPR->?->?-><-?<-PHP+SbcC<-HEPN+DUF499*<-METHYLASE<-HNH<-OLD-ABC<-TUDOR+SF1-SF2-Helicase+iREase+CTD+REase-p | 1104 | Bacteria>PVC_group>Planctomycetota>Planctomycetia>Pirellulales | GCA_012728855.1 | Rhodopirellula_sp. | GXY83_02605 | 12933..16247 | 2024855 | ATP-binding-protein-[Rhodopirellula-sp.] |
| MBL7036971.1 | HEPN+AAA-ATPase+Alpha+Beta-domain+wHTH+RRM | Type-1_BR.RM.like.systems | <-HEPN+DUF499*<-?<-METHYLASE<-?<-TUDOR+SF1-SF2-Helicase+iREase+CTD+REase\|\|?->METHYLASE-> | 498 | Bacteria>PVC_group>Planctomycetota>Planctomycetia>Pirellulales | GCA_016784735.1 | Pirellulaceae_bacterium | ISR77_00005 | 1..1496 | 2699754 | DUF499-domain-containing-protein-[Pirellulaceae-bacterium] |
| MBQ6108779.1 | HEPN+AAA-ATPase+Alpha+Beta-domain+wHTH+RRM | Type-1_BR.RM.like.systems | <-HEPN+DUF499*<-METHYLASE-p<-TUDOR+SF1-SF2-Helicase+iREase+CTD+REase-p | 927 | Bacteria>PVC_group>Planctomycetota>Planctomycetia>Pirellulales | GCA_017427905.1 | Thermoguttaceae_bacterium | IJK97_11270 | 67..2850 | 2691551 | DUF499-domain-containing-protein-[Thermoguttaceae-bacterium] |
| MBR4752420.1 | HEPN+AAA-ATPase+Alpha+Beta-domain+wHTH+RRM | Type-1_BR.RM.like.systems | TUDOR+SF1-SF2-Helicase+iREase+CTD+REase->?->METHYLASE->HEPN+DUF499*-> | 1123 | Bacteria>PVC_group>Planctomycetota>Planctomycetia>Pirellulales | GCA_017550965.1 | Thermoguttaceae_bacterium | IK077_11730 | 50107..53478 | 2691551 | DUF499-domain-containing-protein-[Thermoguttaceae-bacterium] |
| MBR2694935.1 | HEPN+AAA-ATPase+Alpha+Beta-domain+wHTH+RRM | Type-1_BR.RM.like.systems | <-HEPN+DUF499*<-METHYLASE<-TUDOR+SF1-SF2-Helicase+iREase+CTD+REase | 1133 | Bacteria>PVC_group>Planctomycetota>Planctomycetia>Pirellulales | GCA_017515305.1 | Thermoguttaceae_bacterium | IKE69_12060 | 24593..27994 | 2691551 | DUF499-domain-containing-protein-[Thermoguttaceae-bacterium] |
| MDB4650128.1 | HEPN+AAA-ATPase+Alpha+Beta-domain+wHTH+RRM | Type-1_BR.RM.like.systems | METHYLASE->HEPN+DUF499*->Synaptojanin->?-><-?<-?\|\|?->SF1-SF2-Helicase-> | 1109 | Bacteria>PVC_group>Planctomycetota>Planctomycetia>Pirellulales | GCA_028266775.1 | Pirellulaceae_bacterium | OAE37_00010 | 2073..5402 | 2699754 | DUF499-domain-containing-protein-[Pirellulaceae-bacterium] |
| MDD3586122.1 | HEPN+AAA-ATPase+Alpha+Beta-domain+wHTH+RRM | Type-1_BR.RM.like.systems | <-HEPN+DUF499*<-METHYLASE<-METHYLASE-p | 1130 | Bacteria>PVC_group>Planctomycetota>Planctomycetia>Pirellulales | GCA_028695865.1 | Thermoguttaceae_bacterium | PHQ75_02970 | 4102..7494 | 2691551 | MAG:-DUF499-domain-containing-protein-[Thermoguttaceae-bacterium] |

| Protein-ID | DUF499-architecture | System | Gene-Neighbourhoods | Length | Taxonomy | Assembly-ID | Strain | Locus-Tag | Genomic-coordinates | TaxID | Fasta-defline |
|---|---|---|---|---|---|---|---|---|---|---|---|
| MBR2005706.1 | HEPN+AAA-ATPase+Alpha+Beta-domain+wHTH+RRM | Type-1_BR.RM.like.systems | METHYLASE-p->HEPN+DUF499*->DUF499-p-> | 960 | Bacteria>PVC_group>Planctomycetota>Planctomycetia>Pirellulales | GCA_017506065.1 | Thermoguttaceae_bacterium | IJ991_16180 | 1159..4041 | 2691551 | DUF499-domain-containing-protein-[Thermoguttaceae-bacterium] |
| MBR4105130.1 | HEPN+AAA-ATPase+Alpha+Beta-domain+wHTH+RRM | Type-1_BR.RM.like.systems | <-HEPN+DUF499*<-METHYLASE-p | 819 | Bacteria>PVC_group>Planctomycetota>Planctomycetia>Pirellulales | GCA_017521545.1 | Thermoguttaceae_bacterium | IKK39_13850 | 1..2456 | 2691551 | DUF499-domain-containing-protein-[Thermoguttaceae-bacterium] |
| WP_145451931.1 | HEPN+AAA-ATPase+Alpha+Beta-domain+wHTH+RRM | Type-1_BR.RM.like.systems | HTH_3-><-?<-?<-HKD-DNase<-?<-?<-HEPN+DUF499*<-METHYLASE<-Schlafen+HATPase+HTH<-MORC+DUF3684<-TUDOR+SF1-SF2-Helicase+iREase+CTD+REase<-XIS-HTH | 1094 | Bacteria>PVC_group>Planctomycetota>Planctomycetia>Planctomycetales | GCF_007748155.1 | Gimesia_panareensis | Enr10x_RS26145 | 6953187..6956471 | 2527978 | Swt1-family-HEPN-domain-containing-protein-[Gimesia-panareensis] |
| MBV8232626.1 | HEPN+AAA-ATPase+Alpha+Beta-domain+wHTH+RRM | Type-1_BR.RM.like.systems | TUDOR+SF1-SF2-Helicase+iREase+CTD+REase-p->HicB+ZnR->METHYLASE->REase->HEPN+DUF499*->Y1_Tnp-> | 1104 | Bacteria>PVC_group>Planctomycetota>Planctomycetia>Planctomycetales | GCA_019236625.1 | Planctomycetaceae_bacterium | JO329_21805 | 5478..8792 | 2026779 | DUF499-domain-containing-protein-[Planctomycetaceae-bacterium] |
| MCA9076582.1 | HEPN+AAA-ATPase+Alpha+Beta-domain+wHTH+RRM | Type-1_BR.RM.like.systems | sGTP->METHYLASE-p->REase->METHYLASE->OLD-ABC->PIN->HEPN+DUF499*-><-PAIREDC-HTH<-?<-?<-SIGMA-HTH | 1095 | Bacteria>PVC_group>Planctomycetota>Planctomycetia>Planctomycetales | GCA_020426505.1 | Planctomycetaceae_bacterium | KDA93_16275 | 17776..21063 | 2026779 | DUF499-domain-containing-protein-[Planctomycetaceae-bacterium] |
| MBV8313321.1 | HEPN+AAA-ATPase+Alpha+Beta-domain+wHTH+RRM | Type-1_BR.RM.like.systems | <-HEPN+DUF499*<-REase<-METHYLASE<-HicB+ZnR<-TUDOR+SF1-SF2-Helicase+iREase+CTD+REase-p | 939 | Bacteria>PVC_group>Planctomycetota>Planctomycetia>Planctomycetales | GCA_019236145.1 | Planctomycetaceae_bacterium | JOZ53_00025 | 1..2817 | 2026779 | DUF499-domain-containing-protein-[Planctomycetaceae-bacterium] |
| CAG0999165.1 | HEPN+AAA-ATPase+Alpha+Beta-domain+wHTH+RRM | Type-1_BR.RM.like.systems | HKD-DNase->?->HEPN+DUF499*->NERD+SF1-SF2-Helicase-> | 1113 | Bacteria>PVC_group>Planctomycetota>Planctomycetia>Planctomycetales | GCA_905339245.1 | Planctomycetaceae_bacterium | PLCT2_02959 | 2861..6202 | 2026779 | hypothetical-protein-PLCT2_02959-[Planctomycetaceae-bacterium] |
| MBV8676907.1 | HEPN+AAA-ATPase+Alpha+Beta-domain+wHTH+RRM | Type-1_BR.RM.like.systems | METHYLASE-p->REase->HEPN+DUF499*->Y1_Tnp-> | 1104 | Bacteria>PVC_group>Planctomycetota>Planctomycetia>Planctomycetales | GCA_019232645.1 | Planctomycetaceae_bacterium | JO355_07025 | 1198..4512 | 2026779 | DUF499-domain-containing-protein-[Planctomycetaceae-bacterium] |
| MBV8269357.1 | HEPN+AAA-ATPase+Alpha+Beta-domain+wHTH+RRM | Type-1_BR.RM.like.systems | <-HEPN+DUF499*<-REase<-METHYLASE-p | 1104 | Bacteria>PVC_group>Planctomycetota>Planctomycetia>Planctomycetales | GCA_019236385.1 | Planctomycetaceae_bacterium | JO252_23790 | 243..3557 | 2026779 | DUF499-domain-containing-protein-[Planctomycetaceae-bacterium] |
| MBE7458022.1 | HEPN+AAA-ATPase+Alpha+Beta-domain+wHTH+RRM | Type-1_BR.RM.like.systems | DEOR-HTH->TUDOR+SF1-SF2-Helicase+iREase+CTD+REase->Schlafen+HATPase+HTH->?->?->METHYLASE->HEPN+DUF499*->MORC-><-?<-HTH_3<-RelE-ParE<-?<-DDE_Tnp_1_4 | 1120 | Bacteria>PVC_group>Planctomycetota>Planctomycetia>unclassified_Planctomycetia | GCA_015075705.1 | Planctomycetia_bacterium | HS102_15595 | 57764..61126 | 2052181 | ATP-binding-protein-[Planctomycetia-bacterium] |
| QDU83134.1 | HEPN+AAA-ATPase+Alpha+Beta-domain+wHTH+RRM | Type-1_BR.RM.like.systems | TUDOR+SF1-SF2-Helicase+iREase+CTD+REase->Schlafen+HATPase+HTH->?->METHYLASE->OLD-ABC->TOPRIM->HEPN+DUF499*->?-><-REase+SF1-SF2-Helicase<-DUF2357+REase<-Preader7+McrB-NTD+McrB-AAA | 1111 | Bacteria>PVC_group>Planctomycetota>unclassified_Planctomycetota | GCA_007750655.1 | Planctomycetes_bacterium_Pla163 | Pla163_02310 | 293863..297198 | 2528008 | hypothetical-protein-Pla163_02310-[Planctomycetes-bacterium-Pla163] |
| MAW61410.1 | HEPN+AAA-ATPase+Alpha+Beta-domain+wHTH+RRM | Type-1_BR.RM.like.systems | HNH->Phage_int_SAM_4+Phage_integrase-><-XIS-HTH<-Phage_int_SAM_1+Phage_integrase<-?<-?<-HEPN+DUF499*<-METHYLASE<-TUDOR+SF1-SF2-Helicase+iREase+CTD+REase | 1080 | Bacteria>PVC_group>Planctomycetota>unclassified_Planctomycetota | GCA_002705055.1 | Planctomycetota_bacterium | CMJ94_11320 | 35772..39014 | 2026780 | AAA+-family-ATPase-[Planctomycetota-bacterium] |
| TAH38052.1 | HEPN+AAA-ATPase+Alpha+Beta-domain+wHTH+RRM | Type-1_BR.RM.like.systems | <-wHTH+REase\|\|AbiEi_4->AbiEii-><-PIN<-ParB+HNH\|\|HKD-DNase-><-HEPN+DUF499*<-METHYLASE<-HNH<-OLD-ABC<-TUDOR+SF1-SF2-Helicase+iREase+CTD+REase<-HTH | 1098 | Bacteria>PVC_group>Planctomycetota>unclassified_Planctomycetota | GCA_004295085.1 | Planctomycetota_bacterium | EYC70_05360 | 66593..69889 | 2026780 | MAG:-ATP-binding-protein-[Planctomycetota-bacterium] |
| KAA3609103.1 | HEPN+AAA-ATPase+Alpha+Beta-domain+wHTH+RRM | Type-1_BR.RM.like.systems | TUDOR+SF1-SF2-Helicase+iREase+CTD+REase->Schlafen+HATPase+HTH->METHYLASE->PHP+SbcC->Schlafen+HATPase+HTH->HEPN+DUF499*->?->?-><-?<-CHTH\|\|?-><-HhH-RADC+JAB | 1120 | Bacteria>PVC_group>Planctomycetota>unclassified_Planctomycetota | GCA_008501825.1 | Planctomycetota_bacterium | DWQ01_09790 | 14904..18266 | 2026780 | MAG:-DUF499-domain-containing-protein-[Planctomycetota-bacterium] |
| PIE22549.1 | HEPN+AAA-ATPase+Alpha+Beta-domain+wHTH+RRM | Type-1_BR.RM.like.systems | SMODS->SLATT->TUDOR+SF1-SF2-Helicase+iREase+CTD+REase->METHYLASE->PHP+SbcC->Schlafen+HATPase+HTH->HEPN+DUF499*-><-tRNA\|\|?->?->?->?->SF1-SF2-Helicase-> | 1122 | Bacteria>PVC_group>Planctomycetota>unclassified_Planctomycetota | GCA_002748355.1 | Planctomycetota_bacterium | CSA62_11865 | 35983..39351 | 2026780 | MAG:-AAA+-family-ATPase-[Planctomycetota-bacterium] |
| MCZ7645664.1 | HEPN+AAA-ATPase+Alpha+Beta-domain+wHTH+RRM | Type-1_BR.RM.like.systems | HTH+MPTase->PemK_toxin->HNH+SAVED->SMODS->?-><-PIN<-HEPN+DUF499*<-METHYLASE<-?<-ParB+HNH<-TUDOR+SF1-SF2-Helicase+iREase+CTD+REase | 1122 | Bacteria>PVC_group>Planctomycetota>unclassified_Planctomycetota | GCA_027493525.1 | Planctomycetota_bacterium | M5U26_10325 | 2469927..2473295 | 2026780 | MAG:-Swt1-family-HEPN-domain-containing-protein-[Planctomycetota-bacterium] |
| MCG3135902.1 | HEPN+AAA-ATPase+Alpha+Beta-domain+wHTH+RRM | Type-1_BR.RM.like.systems | TUDOR+SF1-SF2-Helicase+iREase+CTD+REase->OLD-ABC->TOPRIM->METHYLASE->HEPN+DUF499*->KAP-NTPase->?->PAPS_reduct->?-><-TPR+TPR | 1110 | Bacteria>PVC_group>Planctomycetota>unclassified_Planctomycetota | GCA_022072415.1 | Planctomycetota_bacterium | HMLKMBBP_03678 | 15324..18656 | 2026780 | hypothetical-protein-[Planctomycetota-bacterium] |
| MBX3463236.1 | HEPN+AAA-ATPase+Alpha+Beta-domain+wHTH+RRM | Type-1_BR.RM.like.systems | <-UbE1+JAB<-?\|\|HTH_3->MPTase->TUDOR+SF1-SF2-Helicase+iREase+CTD+REase->METHYLASE->HEPN+DUF499*->HD-> | 1101 | Bacteria>PVC_group>Planctomycetota>unclassified_Planctomycetota | GCA_019636245.1 | Planctomycetota_bacterium | KF830_08695 | 93755..97060 | 2026780 | DUF499-domain-containing-protein-[Planctomycetota-bacterium] |
| MBI3462535.1 | HEPN+AAA-ATPase+Alpha+Beta-domain+wHTH+RRM | Type-1_BR.RM.like.systems | PIWI-><-ParB+HNH<-HEPN+DUF499*<-METHYLASE<-wHTH+REase<-TUDOR+SF1-SF2-Helicase+iREase+CTD+REase-p | 1111 | Bacteria>PVC_group>Planctomycetota>unclassified_Planctomycetota | GCA_016196255.1 | Planctomycetota_bacterium | HY000_05665 | 2780..6115 | 2026780 | ATP-binding-protein-[Planctomycetota-bacterium] |
| MCK6472712.1 | HEPN+AAA-ATPase+Alpha+Beta-domain+wHTH+RRM | Type-1_BR.RM.like.systems | TUDOR+SF1-SF2-Helicase+iREase+CTD+REase->?->REase-><-?\|\|?->HEPN+DUF499*->SbcC->?->?->METHYLASE-><-?<-XIS-HTH | 1124 | Bacteria>PVC_group>Planctomycetota>unclassified_Planctomycetota | GCA_023150555.1 | Planctomycetota_bacterium | L6R28_13300 | 20467..23841 | 2026780 | DUF499-domain-containing-protein-[Planctomycetota-bacterium] |
| MBK8205305.1 | HEPN+AAA-ATPase+Alpha+Beta-domain+wHTH+RRM | Type-1_BR.RM.like.systems | HEPN->METHYLASE-p->HKD-DNase->?-><-?<-?\|\|HEPN+DUF499*<-?<-?<-HTH_36<-XIS-HTH\|\|Phage_integrase-> | 1097 | Bacteria>PVC_group>Planctomycetota>unclassified_Planctomycetota | GCA_016712285.1 | Planctomycetota_bacterium | IPK87_00800 | 221312..224605 | 2026780 | DUF499-domain-containing-protein-[Planctomycetota-bacterium] |
| MBI4617608.1 | HEPN+AAA-ATPase+Alpha+Beta-domain+wHTH+RRM | Type-1_BR.RM.like.systems | Radical-SAM->?->RelE-ParE->?->?->?-><-HEPN+DUF499*<-ORC-CDC6+REase | 1115 | Bacteria>PVC_group>Planctomycetota>unclassified_Planctomycetota | GCA_016209595.1 | Planctomycetota_bacterium | HY720_28610 | 4559..7906 | 2026780 | ATP-binding-protein-[Planctomycetota-bacterium] |
| MCC7173369.1 | HEPN+AAA-ATPase+Alpha+Beta-domain+wHTH+RRM | Type-1_BR.RM.like.systems | METHYLASE-p->HEPN+DUF499*->?->?-><-PIN\|\|?->SIGMA-HTH-> | 1113 | Bacteria>PVC_group>Planctomycetota>unclassified_Planctomycetota | GCA_020851675.1 | Planctomycetota_bacterium | IT459_23190 | 1090..4431 | 2026780 | DUF499-domain-containing-protein-[Planctomycetota-bacterium] |
| MCL2001508.1 | HEPN+AAA-ATPase+Alpha+Beta-domain+wHTH+RRM | Type-1_BR.RM.like.systems | <-HEPN+DUF499*<-METHYLASE-p<-METHYLASE-p | 524 | Bacteria>PVC_group>Planctomycetota>unclassified_Planctomycetota | GCA_009778045.1 | Planctomycetota_bacterium | FWG74_08750 | 1..1573 | 2026780 | Swt1-family-HEPN-domain-containing-protein-[Planctomycetota-bacterium] |
| RMH31880.1 | HEPN+AAA-ATPase+Alpha+Beta-domain+wHTH+RRM | Type-1_BR.RM.like.systems | METHYLASE-p->HEPN+DUF499*->HIN-HTH-> | 1102 | Bacteria>PVC_group>Planctomycetota>unclassified_Planctomycetota | GCA_003696675.1 | Planctomycetota_bacterium | D6692_00155 | 1472..4780 | 2026780 | MAG:-ATP-binding-protein-[Planctomycetota-bacterium] |
| RLS88266.1 | HEPN+AAA-ATPase+Alpha+Beta-domain+wHTH+RRM | Type-1_BR.RM.like.systems | METHYLASE->HEPN+DUF499*->DUF499-p-> | 912 | Bacteria>PVC_group>Planctomycetota>unclassified_Planctomycetota | GCA_003671065.1 | Planctomycetota_bacterium | DWI10_09315 | 3242..5980 | 2026780 | MAG:-ATP-binding-protein-[Planctomycetota-bacterium] |
| RMD62695.1 | HEPN+AAA-ATPase+Alpha+Beta-domain+wHTH+RRM | Type-1_BR.RM.like.systems | <-HEPN+DUF499*<-METHYLASE-p | 397 | Bacteria>PVC_group>Planctomycetota>unclassified_Planctomycetota | GCA_003693915.1 | Planctomycetota_bacterium | D6824_06430 | 1..1192 | 2026780 | MAG:-ATP-binding-protein,-partial-[Planctomycetota-bacterium] |
| MDA1195884.1 | HEPN+AAA-ATPase+Alpha+Beta-domain+wHTH+RRM | Type-1_BR.RM.like.systems | <-HEPN+DUF499*<-?<-?<-METHYLASE | 886 | Bacteria>PVC_group>Planctomycetota>unclassified_Planctomycetota | GCA_027625315.1 | Planctomycetota_bacterium | O2894_11960 | 1..2659 | 2026780 | MAG:-Swt1-family-HEPN-domain-containing-protein,-partial-[Planctomycetota-bacterium] |
| MCK5940853.1 | HEPN+AAA-ATPase+Alpha+Beta-domain+wHTH+RRM | Type-1_BR.RM.like.systems | <-HEPN+DUF499*<-METHYLASE | 1099 | Bacteria>PVC_group>Planctomycetota>unclassified_Planctomycetota | GCA_023146405.1 | Planctomycetota_bacterium | KAI24_02690 | 255..3554 | 2026780 | DUF499-domain-containing-protein-[Planctomycetota-bacterium] |
| MBI3462766.1 | HEPN+AAA-ATPase+Alpha+Beta-domain+wHTH+RRM | Type-1_BR.RM.like.systems | <-HEPN+DUF499*<-METHYLASE | 1074 | Bacteria>PVC_group>Planctomycetota>unclassified_Planctomycetota | GCA_016196255.1 | Planctomycetota_bacterium | HY000_06860 | 1..3221 | 2026780 | ATP-binding-protein-[Planctomycetota-bacterium] |
| MCC5835099.1 | HEPN+AAA-ATPase+Alpha+Beta-domain+wHTH+RRM | Type-1_BR.RM.like.systems | RVT->METHYLASE->?->CHTH->HEPN+DUF499*-><-?\|\|wHTH+SAD+HNH-><-?\|\|REase+35exo+Beta-Barrel+SF1-SF2-Helicase-><-AbiEi_4 | 1100 | Bacteria>PVC_group>Verrucomicrobiota>Opitutae>Opitutales | GCA_020831485.1 | Opitutales_bacterium | JJU20_10230 | 15950..19252 | 2026772 | DUF499-domain-containing-protein-[Opitutales-bacterium] |
| MBW7893701.1 | HEPN+AAA-ATPase+Alpha+Beta-domain+wHTH+RRM | Type-1_BR.RM.like.systems | TUDOR+SF1-SF2-Helicase+iREase+CTD+REase->MORC+DUF3684->HKD-DNase->?->METHYLASE->HEPN+DUF499*-><-?\|\|HIN-HTH-> | 1122 | Bacteria>PVC_group>Verrucomicrobiota>Opitutae>Opitutales | GCA_019455245.1 | Opitutaceae_bacterium | H3C27_01205 | 267980..271348 | 2006848 | ATP-binding-protein-[Opitutaceae-bacterium] |
| MBX3751087.1 | HEPN+AAA-ATPase+Alpha+Beta-domain+wHTH+RRM | Type-1_BR.RM.like.systems | TUDOR+SF1-SF2-Helicase+iREase+CTD+REase->?->METHYLASE->HEPN+DUF499*-><-S1+RNase+2C | 1122 | Bacteria>PVC_group>Verrucomicrobiota>Opitutae>Opitutales | GCA_019634615.1 | Opitutaceae_bacterium | KF897_13455 | 8367..11735 | 2006848 | DUF499-domain-containing-protein-[Opitutaceae-bacterium] |
| MCC5805073.1 | HEPN+AAA-ATPase+Alpha+Beta-domain+wHTH+RRM | Type-1_BR.RM.like.systems | OLD-ABC->HNH->METHYLASE->CHTH->RHH->PIN->HEPN+DUF499*->?->PhdYeFM_antitox->PIN-> | 1107 | Bacteria>PVC_group>Verrucomicrobiota>Opitutae>Opitutales | GCA_020831525.1 | Opitutales_bacterium | JJU00_01980 | 48883..52206 | 2026772 | DUF499-domain-containing-protein-[Opitutales-bacterium] |
| WP_189517054.1 | HEPN+AAA-ATPase+Alpha+Beta-domain+wHTH+RRM | Type-1_BR.RM.like.systems | <-ParB+HNH<-HEPN+DUF499*<-?<-HTH<-?<-KAP-NTPase<-METHYLASE | 1101 | Bacteria>PVC_group>Verrucomicrobiota>Opitutae>Puniceicoccales | GCF_014651635.1 | Cerasicoccus_arenae | JIN80_RS05850 | 25614..28919 | 424488 | DUF499-domain-containing-protein-[Cerasicoccus-arenae] |
| RFC50837.1 | HEPN+AAA-ATPase+Alpha+Beta-domain+wHTH+RRM | Type-1_BR.RM.like.systems | <-UPF0061\|\|TUDOR+SF1-SF2-Helicase+iREase+CTD+REase->STAS+HisKin->?->PIN->METHYLASE->HEPN+DUF499*-><-?<-?<-?<-SIGMA-HTH | 1074 | Bacteria>PVC_group>Verrucomicrobiota>unclassified_Verrucomicrobiota | GCA_003402705.1 | Verrucomicrobiota_bacterium | DVB22_001798 | 10693..13917 | 2026799 | putative-ATPase,-AAA+-superfamily-[Verrucomicrobia-bacterium] |
| MBI5394224.1 | HEPN+AAA-ATPase+Alpha+Beta-domain+wHTH+RRM | Type-1_BR.RM.like.systems | <-PfkB<-ABC-ATPase+SbcC<-?<-HEPN+DUF499*<-HEPN<-HTH+PolB<-TUDOR+SF1-SF2-Helicase+iREase+CTD+REase | 1121 | Bacteria>PVC_group>Verrucomicrobiota>unclassified_Verrucomicrobiota | GCA_016219905.1 | Verrucomicrobiota_bacterium | HZA91_02900 | 10748..14113 | 2026799 | DUF499-domain-containing-protein-[Verrucomicrobia-bacterium] |
| MBM3880418.1 | HEPN+AAA-ATPase+Alpha+Beta-domain+wHTH+RRM | Type-1_BR.RM.like.systems | <-HEPN+DUF499*<-ParB+HNH<-METHYLASE<-TOPRIM<-OLD-ABC<-TUDOR+SF1-SF2-Helicase+iREase+CTD+REase | 1127 | Bacteria>PVC_group>Verrucomicrobiota>unclassified_Verrucomicrobiota | GCA_016871715.1 | Verrucomicrobiota_bacterium | FJ387_11990 | 332..3715 | 2026799 | ATP-binding-protein-[Verrucomicrobia-bacterium] |
| MBI3869845.1 | HEPN+AAA-ATPase+Alpha+Beta-domain+wHTH+RRM | Type-1_BR.RM.like.systems | Phage_integrase-><-?\|\|HTH-><-?<-HEPN+DUF499*<-PIWI<-TIR+PNTD2<-METHYLASE<-TIR<-SIR2+STALD<-TIR | 1119 | Bacteria>PVC_group>Verrucomicrobiota>unclassified_Verrucomicrobiota | GCA_016199985.1 | Verrucomicrobiota_bacterium | HY299_15085 | 262919..266278 | 2026799 | ATP-binding-protein-[Verrucomicrobia-bacterium] |
| MCF7668203.1 | HEPN+AAA-ATPase+Alpha+Beta-domain+wHTH+RRM | Type-1_BR.RM.like.systems | TUDOR+SF1-SF2-Helicase+iREase+CTD+REase->METHYLASE->OLD-ABC->TOPRIM->HEPN+DUF499*->?->?-><-PfkB | 1108 | Bacteria>PVC_group>Verrucomicrobiota>unclassified_Verrucomicrobiota | GCA_021733785.1 | Verrucomicrobiota_bacterium | K9N48_00295 | 29599..32925 | 2026799 | DUF499-domain-containing-protein-[Verrucomicrobia-bacterium] |
| MDE3100243.1 | HEPN+AAA-ATPase+Alpha+Beta-domain+wHTH+RRM | Type-1_BR.RM.like.systems | <-Response_reg+LuxR-HTH<-?\|\|?-><-REase<-?<-HEPN+DUF499*<-AbiEii<-AbiEi_4<-REase<-PIWI | 1114 | Bacteria>PVC_group>Verrucomicrobiota>unclassified_Verrucomicrobiota | GCA_028872915.1 | Verrucomicrobiota_bacterium | KGJ88_12300 | 8441..11785 | 2026799 | MAG:-DUF499-domain-containing-protein-[Verrucomicrobiota-bacterium] |
| MCC5848502.1 | HEPN+AAA-ATPase+Alpha+Beta-domain+wHTH+RRM | Type-1_BR.RM.like.systems | <-HEPN+DUF499*<-?<-SF1-SF2-Helicase+wHTH+PRTase<-METHYLASE<-TOPRIM<-OLD-ABC<-HTH_3 | 1113 | Bacteria>PVC_group>Verrucomicrobiota>unclassified_Verrucomicrobiota | GCA_020831635.1 | Verrucomicrobiota_bacterium | JJU29_10430 | 13119..16460 | 2026799 | DUF499-domain-containing-protein-[Verrucomicrobia-bacterium] |
| MCZ7639656.1 | HEPN+AAA-ATPase+Alpha+Beta-domain+wHTH+RRM | Type-1_BR.RM.like.systems | TIR->TIR->PIWI->AbiEi_4->AbiEii->HEPN+DUF499*->HTH->HTH->NTP_transf-> | 1127 | Bacteria>PVC_group>Verrucomicrobiota>unclassified_Verrucomicrobiota | GCA_027493615.1 | Verrucomicrobiota_bacterium | M5U12_28570 | 388044..391427 | 2026799 | MAG:-Swt1-family-HEPN-domain-containing-protein-[Verrucomicrobia-bacterium] |
| PWU21781.1 | HEPN+AAA-ATPase+Alpha+Beta-domain+wHTH+RRM | Type-1_BR.RM.like.systems | SF1-SF2-Helicase->?->HKD-DNase->?->?->?-><-HEPN+DUF499*<-?<-PIWI<-REase+PNTD1+PNTD2 | 1128 | Bacteria>PVC_group>Verrucomicrobiota>unclassified_Verrucomicrobiota | GCA_003176045.1 | Verrucomicrobiota_bacterium | C5B50_01090 | 37039..40425 | 2026799 | AAA+-family-ATPase-[Verrucomicrobia-bacterium] |
| MCP5487625.1 | HEPN+AAA-ATPase+Alpha+Beta-domain+wHTH+RRM | Type-1_BR.RM.like.systems | <-HEPN+DUF499*<-METHYLASE<-?<-SMODS<-?<-HTH_3 | 1116 | Bacteria>PVC_group>Verrucomicrobiota>unclassified_Verrucomicrobiota | GCA_024233345.1 | Verrucomicrobiota_bacterium | H7A43_03165 | 141762..145112 | 2026799 | ATP-binding-protein-[Verrucomicrobia-bacterium] |
| RME69044.1 | HEPN+AAA-ATPase+Alpha+Beta-domain+wHTH+RRM | Type-1_BR.RM.like.systems | <-HEPN+DUF499*<-METHYLASE-p | 1114 | Bacteria>PVC_group>Verrucomicrobiota>unclassified_Verrucomicrobiota | GCA_003694825.1 | Verrucomicrobiota_bacterium | D6781_09510 | 385..3729 | 2026799 | ATP-binding-protein-[Verrucomicrobia-bacterium] |
| MCD6339150.1 | HEPN+AAA-ATPase+Alpha+Beta-domain+wHTH+RRM | Type-1_BR.RM.like.systems | <-HEPN+DUF499*<-METHYLASE-p | 1113 | Bacteria>PVC_group>Verrucomicrobiota>unclassified_Verrucomicrobiota | GCA_021160185.1 | Verrucomicrobiota_bacterium | J7M29_07215 | 9794..13135 | 2026799 | DUF499-domain-containing-protein-[Verrucomicrobia-bacterium] |
| MBS0647016.1 | HEPN+AAA-ATPase+Alpha+Beta-domain+wHTH+RRM | Type-1_BR.RM.like.systems | <-HEPN+DUF499*<-METHYLASE<-SF1-SF2-Helicase-p | 1093 | Bacteria>PVC_group>Verrucomicrobiota>unclassified_Verrucomicrobiota | GCA_018239885.1 | Verrucomicrobiota_bacterium | JSR97_10575 | 874..4155 | 2026799 | DUF499-domain-containing-protein-[Verrucomicrobia-bacterium] |
| MBS0646999.1 | HEPN+AAA-ATPase+Alpha+Beta-domain+wHTH+RRM | Type-1_BR.RM.like.systems | <-HEPN+DUF499*<-METHYLASE<-ParB+HNH<-AbiEii | 1093 | Bacteria>PVC_group>Verrucomicrobiota>unclassified_Verrucomicrobiota | GCA_018239885.1 | Verrucomicrobiota_bacterium | JSR97_10490 | 874..4155 | 2026799 | DUF499-domain-containing-protein-[Verrucomicrobia-bacterium] |
| MCS1417413.1 | HEPN+AAA-ATPase+Alpha+Beta-domain+wHTH+RRM | Type-1_BR.RM.like.systems | <-HEPN+DUF499*<-?<-DUF488-l<-PIN<-?<-METHYLASE-p | 1121 | Bacteria>PVC_group>Verrucomicrobiota>Verrucomicrobiae>Limisphaerales | GCA_024761205.1 | Limisphaerales_bacterium | M2R46_05167 | 18516..21881 | 2052186 | hypothetical-protein-[Verrucomicrobia-subdivision-3-bacterium] |
| MCW5552136.1 | HEPN+AAA-ATPase+Alpha+Beta-domain+wHTH+RRM | Type-1_BR.RM.like.systems | GDResolvase+PPP1-HTH->ParB->ParB-><-MarR-HTH\|\|?->ACET-><-HEPN+DUF499*<-METHYLASE<-PIN<-?<-STAS+HisKin<-TUDOR+SF1-SF2-Helicase+iREase+CTD+REase | 1120 | Bacteria>PVC_group>Verrucomicrobiota>Verrucomicrobiae>unclassified_Verrucomicrobiae | GCA_026125285.1 | Verrucomicrobiae_bacterium | KIS67_08205 | 85951..89313 | 2499141 | MAG:-DUF499-domain-containing-protein-[Verrucomicrobiae-bacterium] |
| MBE2287190.1 | HEPN+AAA-ATPase+Alpha+Beta-domain+wHTH+RRM | Type-1_BR.RM.like.systems | TIR+PNTD2->PIWI->DUF932->TUDOR+SF1-SF2-Helicase+iREase+CTD+REase->NACHT->METHYLASE->HEPN+DUF499*->?->SPOUT-> | 1114 | Bacteria>PVC_group>Verrucomicrobiota>Verrucomicrobiae>Verrucomicrobiales | GCA_014879935.1 | Prosthecobacter_sp. | IAE77_27290 | 17760..21104 | 1965333 | ATP-binding-protein-[Prosthecobacter-sp.] |
| MBP7951461.1 | HEPN+AAA-ATPase+Alpha+Beta-domain+wHTH+RRM | Type-1_BR.RM.like.systems | ACET-><-tRNA<-?<-HEPN+DUF499*<-METHYLASE<-PIN<-?<-STAS+HisKin<-TUDOR+SF1-SF2-Helicase+iREase+CTD+REase<-DDE_Tnp_1 | 1112 | Bacteria>PVC_group>Verrucomicrobiota>Verrucomicrobiae>Verrucomicrobiales | GCA_018003315.1 | Verrucomicrobiales_bacterium | KA004_17600 | 4745..8083 | 2026801 | DUF499-domain-containing-protein-[Verrucomicrobiales-bacterium] |
| QTN34153.1 | HEPN+AAA-ATPase+Alpha+Beta-domain+wHTH+RRM | Type-1_BR.RM.like.systems | MqsA_antitoxin+HTH_3->?->METHYLASE->HEPN+DUF499*->HEPN+SF1-SF2-Helicase->HEPN+TPR+RhoN->ParB+HNH-> | 1101 | Bacteria>PVC_group>Verrucomicrobiota>Verrucomicrobiae>Verrucomicrobiales | GCA_017798145.1 | Akkermansiaceae_bacterium | HZ994_18120 | 4237936..4241241 | 2562705 | ATP-binding-protein-[Akkermansiaceae-bacterium] |
| MBL9159968.1 | HEPN+AAA-ATPase+Alpha+Beta-domain+wHTH+RRM | Type-1_BR.RM.like.systems | <-HISKIN<-ArdCN+MPTase<-HNH<-HEPN+DUF499*<-?<-Mbetalac<-?<-?<-METHYLASE | 1114 | Bacteria>PVC_group>Verrucomicrobiota>Verrucomicrobiae>Verrucomicrobiales | GCA_016795385.1 | Verrucomicrobiales_bacterium | JNJ70_20975 | 159944..163288 | 2026801 | DUF499-domain-containing-protein-[Verrucomicrobiales-bacterium] |
| HCN27197.1 | HEPN+AAA-ATPase+Alpha+Beta-domain+wHTH+RRM | Type-1_BR.RM.like.systems | METHYLASE->ParD->PIN->HEPN+DUF499*-> | 1113 | Bacteria>PVC_group>Verrucomicrobiota>Verrucomicrobiae>Verrucomicrobiales | GCA_003545395.1 | Verrucomicrobiales_bacterium | DIT64_00065 | 2852..6193 | 2026801 | AAA+-family-ATPase-[Verrucomicrobiales-bacterium] |
| AEJ18699.1 | HEPN+AAA-ATPase+Alpha+Beta-domain+wHTH+RRM | Type-1_BR.RM.like.systems | PhdYeFM_antitox->Phd-YefM->RelE-ParE->?->?->ParB+HNH->HEPN+DUF499*->ABC-ATPase+SbcC-> | 1058 | Bacteria>Spirochaetota>Spirochaetia>Spirochaetales>Breznakiellaceae | GCA_000219725.1 | Gracilinema_caldarium_DSM_7334 | Spica_0539 | 580385..583561 | 744872 | ATPase-(AAA+-superfamily)-like-protein-[Gracilinema-caldarium-DSM-7334] |
| WP_215904902.1 | HEPN+AAA-ATPase+Alpha+Beta-domain+wHTH+RRM | Type-1_BR.RM.like.systems | PIN->Phd-YefM->RelE-ParE->?->?->ParB+HNH-><-?\|\|HEPN+DUF499*->ABC-ATPase+SbcC-> | 1077 | Bacteria>Spirochaetota>Spirochaetia>Spirochaetales>Breznakiellaceae | GCF_000219725.1 | Gracilinema_caldarium | SPICA_RS02735 | 580328..583561 | 215591 | DUF499-domain-containing-protein-[Gracilinema-caldarium] |
| MCK9286308.1 | HEPN+AAA-ATPase+Alpha+Beta-domain+wHTH+RRM | Type-1_BR.RM.like.systems | <-sGTP<-SF1-SF2-Helicase<-OLD-ABC+TOPRIM<-HEPN+DUF499*<-METHYLASE<-TUDOR+SF1-SF2-Helicase+iREase+CTD+REase<-HTH_26<-SbcC+SbcC<-?<-REase+CTD6 | 1135 | Bacteria>Spirochaetota>Spirochaetia>Spirochaetales>Sphaerochaetaceae | GCA_023230445.1 | Sphaerochaetaceae_bacterium | MOP29_01920 | 121891..125298 | 2791017 | DUF499-domain-containing-protein-[Sphaerochaetaceae-bacterium] |
| MBI9094232.1 | HEPN+AAA-ATPase+Alpha+Beta-domain+wHTH+RRM | Type-1_BR.RM.like.systems | <-zf-IS66+DDE_Tnp_IS66<-?\|\|tRNA-><-?<-?<-HEPN+DUF499*<-METHYLASE<-TUDOR+SF1-SF2-Helicase+iREase+CTD+REase\|\|URI-DNase->HEPN->?-><-HTH_3 | 1134 | Bacteria>Spirochaetota>Spirochaetia>Spirochaetales>Sphaerochaetaceae | GCA_016342655.1 | Sphaerochaeta_sp. | JEY71_05065 | 29138..32542 | 1972642 | DUF499-domain-containing-protein-[Sphaerochaeta-sp.] |
| MDE0219724.1 | HEPN+AAA-ATPase+Alpha+Beta-domain+wHTH+RRM | Type-1_BR.RM.like.systems | SbcC-><-OLD-ABC\|\|TUDOR+SF1-SF2-Helicase+iREase+CTD+REase-><-?\|\|HTH+MPTase->PIN->HEPN+DUF499*->DOC+HTH->?->XIS-HTH+RVT->Glyco_hydro->DUF2961->HNH-> | 1110 | Bacteria>Spirochaetota>Spirochaetia>Spirochaetales>Spirochaetaceae | GCA_028820215.1 | Spirochaetaceae_bacterium | OXJ90_10675 | 22427..25759 | 1898206 | MAG:-Swt1-family-HEPN-domain-containing-protein-[Spirochaetaceae-bacterium] |
| WP_083048603.1 | HEPN+AAA-ATPase+Alpha+Beta-domain+wHTH+RRM | Type-1_BR.RM.like.systems | <-PfkB<-?<-?<-?<-HEPN+DUF499*<-METHYLASE<-TUDOR+SF1-SF2-Helicase+iREase+CTD+REase<-?<-ParB+HNH<-?<-SF1-SF2-Helicase+REase | 1101 | Bacteria>Spirochaetota>Spirochaetia>Spirochaetales>Spirochaetaceae | GCF_002087085.1 | Marispirochaeta_aestuarii | B4097_RS04115 | 245818..249123 | 1963862 | DUF499-domain-containing-protein-[Marispirochaeta-aestuarii] |
| MCX7774711.1 | HEPN+AAA-ATPase+Alpha+Beta-domain+wHTH+RRM | Type-1_BR.RM.like.systems | ACET->TUDOR+SF1-SF2-Helicase+iREase+CTD+REase->ORC-CDC6+REase->METHYLASE->AbiEi_4->AbiEii->HEPN+DUF499*-> | 1103 | Bacteria>Spirochaetota>Spirochaetia>Spirochaetales>Spirochaetaceae | GCA_026418115.1 | Spirochaetaceae_bacterium | N2067_01230 | 29258..32569 | 1898206 | MAG:-DUF499-domain-containing-protein-[Spirochaetaceae-bacterium] |
| HAL93442.1 | HEPN+AAA-ATPase+Alpha+Beta-domain+wHTH+RRM | Type-1_BR.RM.like.systems | SF1-SF2-Helicase+DUF1998-><-?<-PIN<-?\|\|?-><-HEPN+DUF499*<-PIN<-PhdYeFM_antitox<-?<-HEPN-p | 1142 | Bacteria>Spirochaetota>Spirochaetia>Spirochaetales>Spirochaetaceae | GCA_003452735.1 | Spirochaetaceae_bacterium | DCP56_01610 | 10292..13720 | 1898206 | AAA+-family-ATPase-[Spirochaetaceae-bacterium] |
| MDD9984792.1 | HEPN+AAA-ATPase+Alpha+Beta-domain+wHTH+RRM | Type-1_BR.RM.like.systems | <-Glyco_hydro\|\|?-><-NUDIX<-DOC+HTH\|\|?-><-?<-HEPN+DUF499*\|\|DDE_Tnp_1-> | 1110 | Bacteria>Spirochaetota>Spirochaetia>Spirochaetales>Spirochaetaceae | GCA_028818385.1 | Spirochaetaceae_bacterium | OXQ31_00820 | 20813..24145 | 1898206 | MAG:-Swt1-family-HEPN-domain-containing-protein-[Spirochaetaceae-bacterium] |
| HDQ14361.1 | HEPN+AAA-ATPase+Alpha+Beta-domain+wHTH+RRM | Type-1_BR.RM.like.systems | METHYLASE-p->METHYLASE-p->HEPN+DUF499*->?->ORC-CDC6+REase-> | 1109 | Bacteria>Spirochaetota>Spirochaetia>Spirochaetales>Spirochaetaceae | GCA_011046055.1 | Sediminispirochaeta_sp. | ENN41_06065 | 1978..5307 | 1911557 | ATP-binding-protein-[Sediminispirochaeta-sp.] |
| MCY4372401.1 | HEPN+AAA-ATPase+Alpha+Beta-domain+wHTH+RRM | Type-1_BR.RM.like.systems | MPTase->HEPN+DUF499*->?-><-?\|\|Glyco_hydro->DUF2961-> | 1110 | Bacteria>Spirochaetota>Spirochaetia>Spirochaetales>Spirochaetaceae | GCA_026710565.1 | Spirochaetaceae_bacterium | OXC31_01430 | 992..4324 | 1898206 | MAG:-Swt1-family-HEPN-domain-containing-protein-[Spirochaetaceae-bacterium] |
| OQB96942.1 | HEPN+AAA-ATPase+Alpha+Beta-domain+wHTH+RRM | Type-1_BR.RM.like.systems | SIGMA-HTH->Radical-SAM-><-?<-?\|\|?->?-><-HEPN+DUF499*<-ParB+HNH<-METHYLASE<-TOPRIM<-OLD-ABC<-HEPN<-TUDOR+SF1-SF2-Helicase+iREase+CTD+REase | 1176 | Bacteria>Spirochaetota>unclassified_Spirochaetota | GCA_002070755.1 | Spirochaetes_bacterium_ADurb.Bin110 | BWX81_01211 | 5339..8869 | 1852908 | hypothetical-protein-BWX81_01211-[Spirochaetes-bacterium-ADurb.Bin110] |
| MCL1994394.1 | HEPN+AAA-ATPase+Alpha+Beta-domain+wHTH+RRM | Type-1_BR.RM.like.systems | <-HEPN+DUF499*<-METHYLASE<-?<-OLD-ABC<-ParB+HNH<-TUDOR+SF1-SF2-Helicase+iREase+CTD+REase | 1116 | Bacteria>Spirochaetota>unclassified_Spirochaetota | GCA_009778155.1 | Spirochaetota_bacterium | FWG66_15730 | 17581..20931 | 2202144 | DUF499-domain-containing-protein-[Spirochaetes-bacterium] |
| WP_026368871.1 | HEPN+AAA-ATPase+Alpha+Beta-domain+wHTH+RRM | Type-1_BR.RM.like.systems | <-HTH_3\|\|TUDOR+SF1-SF2-Helicase+iREase+CTD+REase->METHYLASE->HEPN+DUF499*-><-ABC-ATPase+SbcC<-ABC-ATPase+SbcC\|\|<-?\|\|?->ABhydrolase-> | 1123 | Bacteria>Synergistota>Synergistia>Synergistales>Aminiphilaceae | GCF_000526375.1 | Aminiphilus_circumscriptus | K349_RS0105690 | 1079637..1083008 | 290732 | Swt1-family-HEPN-domain-containing-protein-[Aminiphilus-circumscriptus] |
| HIR18062.1 | HEPN+AAA-ATPase+Alpha+Beta-domain+wHTH+RRM | Type-1_BR.RM.like.systems | <-MNS+REase<-DOC+HTH<-HEPN+DUF499*<-METHYLASE<-TUDOR+SF1-SF2-Helicase+iREase+CTD+REase | 1115 | Bacteria>Synergistota>Synergistia>Synergistales>Synergistaceae | GCA_018712745.1 | Candidatus_Caccocola_faecigallinarum | IAD35_08445 | 169850..173197 | 2840712 | DUF499-domain-containing-protein-[Candidatus-Caccocola-faecigallinarum] |
| MBR1604261.1 | HEPN+AAA-ATPase+Alpha+Beta-domain+wHTH+RRM | Type-1_BR.RM.like.systems | TUDOR+SF1-SF2-Helicase+iREase+CTD+REase->METHYLASE->HEPN+DUF499*-><-?\|\|?->REase-> | 1094 | Bacteria>Synergistota>Synergistia>Synergistales>Synergistaceae | GCA_017470785.1 | Synergistaceae_bacterium | IJ667_12605 | 8017..11301 | 2053624 | DUF499-domain-containing-protein-[Synergistaceae-bacterium] |
| GHS93871.1 | HEPN+AAA-ATPase+Alpha+Beta-domain+wHTH+RRM | Type-1_BR.RM.like.systems | TUDOR+SF1-SF2-Helicase+iREase+CTD+REase->HEPN+REase->METHYLASE->OLD-ABC->NERD->?->HEPN+DUF499*->?->?->METHYLASE->?-><-DDE_Tnp_1 | 1116 | Bacteria>Synergistota>Synergistia>Synergistales>unclassified_Synergistales | GCA_025997955.1 | Synergistales_bacterium | AGMMS50276_05310 | 53205..56555 | 2268201 | hypothetical-protein-AGMMS50276_05310-[Synergistales-bacterium] |
| GHV35275.1 | HEPN+AAA-ATPase+Alpha+Beta-domain+wHTH+RRM | Type-1_BR.RM.like.systems | HicA_toxin->OLD-ABC->NERD->?->OLD-ABC->?->HEPN+DUF499*-> | 1117 | Bacteria>Synergistota>Synergistia>Synergistales>unclassified_Synergistales | GCA_026002715.1 | Synergistales_bacterium | FACS1894187_07640 | 12375..15728 | 2268201 | hypothetical-protein-FACS1894187_07640-[Synergistales-bacterium] |
| WP_219944160.1 | HEPN+AAA-ATPase+Alpha+Beta-domain+wHTH+RRM | Type-1_BR.RM.like.systems | TUDOR+SF1-SF2-Helicase+iREase+CTD+REase->METHYLASE->METHYLASE->?->HEPN+DUF499*->?->?-><-Phage_int_SAM_1+Phage_integrase<-?\|\|MerR-HTH-> | 1096 | Bacteria>Terrabacteria_group>Actinomycetota>Acidimicrobiia>Acidimicrobiales | GCF_019443745.1 | Iamia_sp._SCSIO_61187 | HC251_RS04740 | 976382..979672 | 2722752 | Swt1-family-HEPN-domain-containing-protein-[Iamia-sp.-SCSIO-61187] |
| MXY09521.1 | HEPN+AAA-ATPase+Alpha+Beta-domain+wHTH+RRM | Type-1_BR.RM.like.systems | TUDOR+SF1-SF2-Helicase+iREase+CTD+REase->SIR2+SIR2->PIWI->METHYLASE->MerR-HTH->PIN->HEPN+DUF499*->ASCH->HAD-><-?<-?<-ABhydrolase | 1086 | Bacteria>Terrabacteria_group>Actinomycetota>Acidimicrobiia>Acidimicrobiales | GCA_009838345.1 | Acidimicrobiaceae_bacterium | F4Y56_03045 | 76366..79626 | 2024894 | ATP-binding-protein-[Acidimicrobiaceae-bacterium] |
| MDE0699638.1 | HEPN+AAA-ATPase+Alpha+Beta-domain+wHTH+RRM | Type-1_BR.RM.like.systems | <-HTH_58\|\|?->DNA_methylase->MORC+HISKIN->?-><-wHTH+SAD+HNH\|\|HEPN+DUF499*-><-ParA+HTH<-RelE-ParE<-Creatinase_N+Peptidase_M24 | 1084 | Bacteria>Terrabacteria_group>Actinomycetota>Acidimicrobiia>Acidimicrobiales | GCA_028824665.1 | Acidimicrobiaceae_bacterium | OXH61_02820 | 85956..89210 | 2024894 | MAG:-DUF499-domain-containing-protein-[Acidimicrobiaceae-bacterium] |
| MXW76241.1 | HEPN+AAA-ATPase+Alpha+Beta-domain+wHTH+RRM | Type-1_BR.RM.like.systems | <-KAP-NTPase\|\|METHYLASE-><-HTH_58<-RNase\|\|HEPN+DUF499*-><-ParA+HTH<-RelE-ParE<-Creatinase_N+Peptidase_M24 | 1084 | Bacteria>Terrabacteria_group>Actinomycetota>Acidimicrobiia>Acidimicrobiales | GCA_009836285.1 | Acidimicrobiaceae_bacterium | F4Z58_09425 | 4280..7534 | 2024894 | ATP-binding-protein-[Acidimicrobiaceae-bacterium] |
| MCY3894210.1 | HEPN+AAA-ATPase+Alpha+Beta-domain+wHTH+RRM | Type-1_BR.RM.like.systems | <-ORC-CDC6+REase<-?<-HEPN+DUF499*<-?<-CHTH<-?<-METHYLASE<-TUDOR+SF1-SF2-Helicase+iREase+CTD+REase<-MORC+REase | 1154 | Bacteria>Terrabacteria_group>Actinomycetota>Acidimicrobiia>Acidimicrobiales | GCA_026707105.1 | Acidimicrobiaceae_bacterium | OXF65_13050 | 72169..75633 | 2024894 | MAG:-DUF499-domain-containing-protein-[Acidimicrobiaceae-bacterium] |
| MDE0498271.1 | HEPN+AAA-ATPase+Alpha+Beta-domain+wHTH+RRM | Type-1_BR.RM.like.systems | <-DOC+HTH\|\|?->?-><-?<-?\|\|?-><-HEPN+DUF499*<-?<-CHTH<-?<-METHYLASE<-TUDOR+SF1-SF2-Helicase+iREase+CTD+REase<-MORC+REase | 1152 | Bacteria>Terrabacteria_group>Actinomycetota>Acidimicrobiia>Acidimicrobiales | GCA_028823965.1 | Acidimicrobiaceae_bacterium | OXH86_13045 | 54884..58342 | 2024894 | MAG:-DUF499-domain-containing-protein-[Acidimicrobiaceae-bacterium] |
| MDE0136614.1 | HEPN+AAA-ATPase+Alpha+Beta-domain+wHTH+RRM | Type-1_BR.RM.like.systems | MORC+REase->TUDOR+SF1-SF2-Helicase+iREase+CTD+REase->METHYLASE->?->CHTH->?->HEPN+DUF499*-><-?\|\|?->?-><-?<-?\|\|DOC+HTH-> | 1152 | Bacteria>Terrabacteria_group>Actinomycetota>Acidimicrobiia>Acidimicrobiales | GCA_028819405.1 | Acidimicrobiaceae_bacterium | OXM54_17470 | 118886..122344 | 2024894 | MAG:-DUF499-domain-containing-protein-[Acidimicrobiaceae-bacterium] |
| MDE0067328.1 | HEPN+AAA-ATPase+Alpha+Beta-domain+wHTH+RRM | Type-1_BR.RM.like.systems | METHYLASE-p->?-><-PIN<-?<-HTH_58\|\|HEPN+DUF499*-><-ParA+HTH<-RelE-ParE<-Creatinase_N+Peptidase_M24 | 1084 | Bacteria>Terrabacteria_group>Actinomycetota>Acidimicrobiia>Acidimicrobiales | GCA_028819005.1 | Acidimicrobiaceae_bacterium | OXN44_10730 | 3279..6533 | 2024894 | MAG:-DUF499-domain-containing-protein-[Acidimicrobiaceae-bacterium] |
| WP_262322266.1 | HEPN+AAA-ATPase+Alpha+Beta-domain+wHTH+RRM | Type-1_BR.RM.like.systems | MPTase->?->?-><-?\|\|?-><-?<-HEPN+DUF499*<-HEPN<-PolB<-METHYLASE<-?<-TUDOR+SF1-SF2-Helicase+iREase+CTD+REase | 1118 | Bacteria>Terrabacteria_group>Actinomycetota>Acidimicrobiia>Acidimicrobiales | GCF_025499425.1 | Acidiferrimicrobium_sp._IK | K6U06_RS12505 | 271531..274887 | 2871700 | Swt1-family-HEPN-domain-containing-protein-[Acidiferrimicrobium-sp.-IK] |
| MDE0320770.1 | HEPN+AAA-ATPase+Alpha+Beta-domain+wHTH+RRM | Type-1_BR.RM.like.systems | PIN->TUDOR+SF1-SF2-Helicase+iREase+CTD+REase->METHYLASE->?->CHTH->?->HEPN+DUF499*-><-?\|\|?->?-><-?<-?\|\|DOC+HTH-> | 1152 | Bacteria>Terrabacteria_group>Actinomycetota>Acidimicrobiia>Acidimicrobiales | GCA_028820815.1 | Acidimicrobiaceae_bacterium | OXI97_12920 | 73823..77281 | 2024894 | MAG:-DUF499-domain-containing-protein-[Acidimicrobiaceae-bacterium] |
| MXW61875.1 | HEPN+AAA-ATPase+Alpha+Beta-domain+wHTH+RRM | Type-1_BR.RM.like.systems | RelE-ParE->ParA+HTH-><-HEPN+DUF499*\|\|RNase->HTH_58-><-METHYLASE\|\|KAP-NTPase->HEPN-> | 1084 | Bacteria>Terrabacteria_group>Actinomycetota>Acidimicrobiia>Acidimicrobiales | GCA_009835385.1 | Acidimicrobiaceae_bacterium | F4003_08870 | 881..4135 | 2024894 | ATP-binding-protein-[Acidimicrobiaceae-bacterium] |
| MDE0163189.1 | HEPN+AAA-ATPase+Alpha+Beta-domain+wHTH+RRM | Type-1_BR.RM.like.systems | <-HEPN+DUF499*<-?<-?<-METHYLASE<-TUDOR+SF1-SF2-Helicase+iREase+CTD+REase<-HicB | 1126 | Bacteria>Terrabacteria_group>Actinomycetota>Acidimicrobiia>Acidimicrobiales | GCA_028819505.1 | Acidimicrobiaceae_bacterium | OXL98_15820 | 25839..29219 | 2024894 | MAG:-DUF499-domain-containing-protein-[Acidimicrobiaceae-bacterium] |
| MDE0803902.1 | HEPN+AAA-ATPase+Alpha+Beta-domain+wHTH+RRM | Type-1_BR.RM.like.systems | <-HTH+SF1-SF2-Helicase+REase-p<-REase<-MORC<-HEPN+DUF499*<-METHYLASE | 1136 | Bacteria>Terrabacteria_group>Actinomycetota>Acidimicrobiia>Acidimicrobiales | GCA_028822535.1 | Acidimicrobiales_bacterium | OSA99_11315 | 4439..7849 | 2201156 | MAG:-Swt1-family-HEPN-domain-containing-protein-[Acidimicrobiales-bacterium] |
| NNO07581.1 | HEPN+AAA-ATPase+Alpha+Beta-domain+wHTH+RRM | Type-1_BR.RM.like.systems | <-HEPN+DUF499*<-METHYLASE<-TUDOR+SF1-SF2-Helicase+iREase+CTD+REase | 435 | Bacteria>Terrabacteria_group>Actinomycetota>Acidimicrobiia>Acidimicrobiales | GCA_013044265.1 | Acidimicrobiaceae_bacterium | HKL85_00090 | 1..1304 | 2024894 | ATP-binding-protein-[Acidimicrobiaceae-bacterium] |
| MYJ29885.1 | HEPN+AAA-ATPase+Alpha+Beta-domain+wHTH+RRM | Type-1_BR.RM.like.systems | ABhydrolase->?->?-><-HAD<-ASCH<-HEPN+DUF499*<-PIN<-MerR-HTH<-METHYLASE-p | 1086 | Bacteria>Terrabacteria_group>Actinomycetota>Acidimicrobiia>Acidimicrobiales | GCA_009845185.1 | Acidimicrobiaceae_bacterium | F4072_07945 | 6572..9832 | 2024894 | ATP-binding-protein-[Acidimicrobiaceae-bacterium] |
| MDE0495499.1 | HEPN+AAA-ATPase+Alpha+Beta-domain+wHTH+RRM | Type-1_BR.RM.like.systems | METHYLASE-p->MerR-HTH->PIN->HEPN+DUF499*->ASCH->HAD-><-?<-?<-ABhydrolase | 1084 | Bacteria>Terrabacteria_group>Actinomycetota>Acidimicrobiia>Acidimicrobiales | GCA_028824895.1 | Acidimicrobiaceae_bacterium | OXH54_16270 | 41170..44424 | 2024894 | MAG:-DUF499-domain-containing-protein-[Acidimicrobiaceae-bacterium] |
| MCI2976467.1 | HEPN+AAA-ATPase+Alpha+Beta-domain+wHTH+RRM | Type-1_BR.RM.like.systems | TUDOR+SF1-SF2-Helicase+iREase+CTD+REase->METHYLASE->HEPN+DUF499*-> | 1103 | Bacteria>Terrabacteria_group>Actinomycetota>Acidimicrobiia>Acidimicrobiales | GCA_022690815.1 | Ferrimicrobium_sp. | MP439_10410 | 9887..13198 | 2926050 | DUF499-domain-containing-protein-[Ferrimicrobium-sp.] |
| MXW95443.1 | HEPN+AAA-ATPase+Alpha+Beta-domain+wHTH+RRM | Type-1_BR.RM.like.systems | <-SbcC+SbcC<-HEPN+DUF499*\|\|TraJ-RHH+MarR-HTH->AbiEii-><-METHYLASE | 1096 | Bacteria>Terrabacteria_group>Actinomycetota>Acidimicrobiia>Acidimicrobiales | GCA_009836165.1 | Acidimicrobiaceae_bacterium | F4Z64_08035 | 3972..7262 | 2024894 | ATP-binding-protein-[Acidimicrobiaceae-bacterium] |
| MDE0606846.1 | HEPN+AAA-ATPase+Alpha+Beta-domain+wHTH+RRM | Type-1_BR.RM.like.systems | HEPN+DUF499*-><-ParA+HTH<-RelE-ParE<-Creatinase_N+Peptidase_M24 | 1081 | Bacteria>Terrabacteria_group>Actinomycetota>Acidimicrobiia>Acidimicrobiales | GCA_028824095.1 | Acidimicrobiaceae_bacterium | OXH78_06815 | 1..3246 | 2024894 | MAG:-DUF499-domain-containing-protein,-partial-[Acidimicrobiaceae-bacterium] |
| MCY3849326.1 | HEPN+AAA-ATPase+Alpha+Beta-domain+wHTH+RRM | Type-1_BR.RM.like.systems | <-HEPN+DUF499*<-?<-METHYLASE-p\|\|?-><-?<-?<-MORC+HISKIN<-DNA_methylase | 1088 | Bacteria>Terrabacteria_group>Actinomycetota>Acidimicrobiia>Acidimicrobiales | GCA_026706815.1 | Acidimicrobiaceae_bacterium | OXF75_00810 | 86121..89387 | 2024894 | MAG:-DUF499-domain-containing-protein-[Acidimicrobiaceae-bacterium] |
| MDE0268154.1 | HEPN+AAA-ATPase+Alpha+Beta-domain+wHTH+RRM | Type-1_BR.RM.like.systems | METHYLASE-p->HEPN+DUF499*->?-><-?\|\|?->DDE_Tnp_1->AbiEi_1-><-HTH_3 | 1111 | Bacteria>Terrabacteria_group>Actinomycetota>Acidimicrobiia>Acidimicrobiales | GCA_028820455.1 | Acidimicrobiaceae_bacterium | OXI96_03835 | 408..3743 | 2024894 | MAG:-DUF499-domain-containing-protein-[Acidimicrobiaceae-bacterium] |
| MXZ99353.1 | HEPN+AAA-ATPase+Alpha+Beta-domain+wHTH+RRM | Type-1_BR.RM.like.systems | <-AbiEii<-TraJ-RHH+MarR-HTH\|\|HEPN+DUF499*->SbcC+SbcC-> | 1096 | Bacteria>Terrabacteria_group>Actinomycetota>Acidimicrobiia>Acidimicrobiales | GCA_009836995.1 | Acidimicrobiaceae_bacterium | F4Z23_09980 | 3513..6803 | 2024894 | ATP-binding-protein-[Acidimicrobiaceae-bacterium] |
| MXW97638.1 | HEPN+AAA-ATPase+Alpha+Beta-domain+wHTH+RRM | Type-1_BR.RM.like.systems | <-HEPN+DUF499*<-?<-METHYLASE\|\|HEPN-><-PemK_toxin<-?\|\|RHH_1-> | 1085 | Bacteria>Terrabacteria_group>Actinomycetota>Acidimicrobiia>Acidimicrobiales | GCA_009835735.1 | Acidimicrobiaceae_bacterium | F4Z89_03360 | 789..4046 | 2024894 | ATP-binding-protein-[Acidimicrobiaceae-bacterium] |
| MCY3892500.1 | HEPN+AAA-ATPase+Alpha+Beta-domain+wHTH+RRM | Type-1_BR.RM.like.systems | METHYLASE-p->OLD-ABC->TOPRIM->HEPN+DUF499*->?->ABhydrolase-> | 1131 | Bacteria>Terrabacteria_group>Actinomycetota>Acidimicrobiia>Acidimicrobiales | GCA_026707105.1 | Acidimicrobiaceae_bacterium | OXF65_04295 | 3174..6569 | 2024894 | MAG:-Swt1-family-HEPN-domain-containing-protein-[Acidimicrobiaceae-bacterium] |
| NNO09184.1 | HEPN+AAA-ATPase+Alpha+Beta-domain+wHTH+RRM | Type-1_BR.RM.like.systems | METHYLASE->HEPN+DUF499*-><-?<-?<-GDResolvase<-?<-XIS-HTH | 1108 | Bacteria>Terrabacteria_group>Actinomycetota>Acidimicrobiia>Acidimicrobiales | GCA_013044265.1 | Acidimicrobiaceae_bacterium | HKL85_08335 | 1610..4936 | 2024894 | ATP-binding-protein-[Acidimicrobiaceae-bacterium] |
| MDE0497130.1 | HEPN+AAA-ATPase+Alpha+Beta-domain+wHTH+RRM | Type-1_BR.RM.like.systems | DNA_methylase->MORC+HISKIN->?->?->?->HEPN+DUF499*-> | 1089 | Bacteria>Terrabacteria_group>Actinomycetota>Acidimicrobiia>Acidimicrobiales | GCA_028823965.1 | Acidimicrobiaceae_bacterium | OXH86_07240 | 49527..52796 | 2024894 | MAG:-DUF499-domain-containing-protein-[Acidimicrobiaceae-bacterium] |
| MDE0065239.1 | HEPN+AAA-ATPase+Alpha+Beta-domain+wHTH+RRM | Type-1_BR.RM.like.systems | METHYLASE-p->?->?->HEPN+DUF499*-><-?<-?<-?<-?<-Aminotran_1_2 | 1145 | Bacteria>Terrabacteria_group>Actinomycetota>Acidimicrobiia>Acidimicrobiales | GCA_028819005.1 | Acidimicrobiaceae_bacterium | OXN44_00020 | 1449..4886 | 2024894 | MAG:-DUF499-domain-containing-protein-[Acidimicrobiaceae-bacterium] |
| MCY4517135.1 | HEPN+AAA-ATPase+Alpha+Beta-domain+wHTH+RRM | Type-1_BR.RM.like.systems | METHYLASE-p->?->?->?->HEPN+DUF499*-> | 1090 | Bacteria>Terrabacteria_group>Actinomycetota>Acidimicrobiia>Acidimicrobiales | GCA_026713105.1 | Acidimicrobiaceae_bacterium | OXB99_07860 | 4033..7305 | 2024894 | MAG:-DUF499-domain-containing-protein-[Acidimicrobiaceae-bacterium] |
| MXY89812.1 | HEPN+AAA-ATPase+Alpha+Beta-domain+wHTH+RRM | Type-1_BR.RM.like.systems | ABhydrolase-><-REase<-?<-?<-SF1-SF2-Helicase\|\|ABhydrolase-><-HEPN+DUF499*<-METHYLASE<-SF1-SF2-Helicase<-TUDOR+SF1-SF2-Helicase+iREase+CTD+REase<-?\|\|?->SF1-SF2-Helicase-> | 1083 | Bacteria>Terrabacteria_group>Actinomycetota>Acidimicrobiia>unclassified_Acidimicrobiia | GCA_009835395.1 | Acidimicrobiia_bacterium | F4004_03955 | 27589..30840 | 2080302 | ATP-binding-protein-[Acidimicrobiia-bacterium] |
| MBA2283088.1 | HEPN+AAA-ATPase+Alpha+Beta-domain+wHTH+RRM | Type-1_BR.RM.like.systems | Phage_int_SAM_1+Phage_integrase-><-NTD+SF1-SF2-Helicase\|\|TUDOR+SF1-SF2-Helicase+iREase+CTD+REase->?->METHYLASE->?->HEPN+DUF499*->REase->?-><-?<-XIS-HTH\|\|XIS-HTH-> | 1110 | Bacteria>Terrabacteria_group>Actinomycetota>Acidimicrobiia>unclassified_Acidimicrobiia | GCA_013694495.1 | Acidimicrobiia_bacterium | H0W25_17905 | 18303..21635 | 2080302 | ATP-binding-protein-[Acidimicrobiia-bacterium] |
| MXX46239.1 | HEPN+AAA-ATPase+Alpha+Beta-domain+wHTH+RRM | Type-1_BR.RM.like.systems | <-GNTR-HTH<-?\|\|ORC-CDC6+REase-><-?<-?<-RNase<-HEPN+DUF499*<-METHYLASE-p<-METHYLASE-p<-TUDOR+SF1-SF2-Helicase+iREase+CTD+REase<-CHTH | 1090 | Bacteria>Terrabacteria_group>Actinomycetota>Acidimicrobiia>unclassified_Acidimicrobiia | GCA_009836655.1 | Acidimicrobiia_bacterium | F4Z41_08550 | 57400..60672 | 2080302 | ATP-binding-protein-[Acidimicrobiia-bacterium] |
| MYB44039.1 | HEPN+AAA-ATPase+Alpha+Beta-domain+wHTH+RRM | Type-1_BR.RM.like.systems | METHYLASE-><-PIN<-?\|\|SIR2+STALD->?->PRTase+PRTase->HEPN+DUF499*-><-?\|\|PIN-><-ABC-ATPase+SbcC<-ABC-ATPase+SbcC<-Peptidase_M20 | 1097 | Bacteria>Terrabacteria_group>Actinomycetota>Acidimicrobiia>unclassified_Acidimicrobiia | GCA_009839965.1 | Acidimicrobiia_bacterium | F4X74_03760 | 19073..22366 | 2080302 | ATP-binding-protein-[Acidimicrobiia-bacterium] |
| MXZ86859.1 | HEPN+AAA-ATPase+Alpha+Beta-domain+wHTH+RRM | Type-1_BR.RM.like.systems | ABC-ATPase+SbcC->?->?->?->?-><-HEPN+DUF499*<-?\|\|?-><-METHYLASE\|\|ParB+HNH->HEPN-><-TUDOR+SF1-SF2-Helicase+iREase+CTD+REase | 1114 | Bacteria>Terrabacteria_group>Actinomycetota>Acidimicrobiia>unclassified_Acidimicrobiia | GCA_009837425.1 | Acidimicrobiia_bacterium | F4Z02_14550 | 33484..36828 | 2080302 | ATP-binding-protein-[Acidimicrobiia-bacterium] |
| MCC7078886.1 | HEPN+AAA-ATPase+Alpha+Beta-domain+wHTH+RRM | Type-1_BR.RM.like.systems | <-Schlafen+HATPase+HTH<-?<-HEPN+DUF499*<-METHYLASE<-Schlafen+HATPase+HTH<-TUDOR+SF1-SF2-Helicase+iREase+CTD+REase | 1112 | Bacteria>Terrabacteria_group>Actinomycetota>Acidimicrobiia>unclassified_Acidimicrobiia | GCA_020850945.1 | Acidimicrobiia_bacterium | IT198_17355 | 29484..32822 | 2080302 | DUF499-domain-containing-protein-[Acidimicrobiia-bacterium] |
| MCB9390232.1 | HEPN+AAA-ATPase+Alpha+Beta-domain+wHTH+RRM | Type-1_BR.RM.like.systems | <-HEPN<-XIS-HTH+RVT<-?<-DUF6361<-?<-REase<-HEPN+DUF499*<-METHYLASE\|\|?-><-?<-?<-DUF772+DDE_Tnp_1+DDE_Tnp_1_6 | 1107 | Bacteria>Terrabacteria_group>Actinomycetota>Acidimicrobiia>unclassified_Acidimicrobiia | GCA_020637365.1 | Acidimicrobiia_bacterium | H6512_11920 | 836965..840288 | 2080302 | ATP-binding-protein-[Acidimicrobiia-bacterium] |
| MYB25632.1 | HEPN+AAA-ATPase+Alpha+Beta-domain+wHTH+RRM | Type-1_BR.RM.like.systems | <-Classical-AAA\|\|ORC-CDC6+REase-><-PIN<-HTH+MPTase<-PIN\|\|METHYLASE->HEPN+DUF499*-> | 1108 | Bacteria>Terrabacteria_group>Actinomycetota>Acidimicrobiia>unclassified_Acidimicrobiia | GCA_009840735.1 | Acidimicrobiia_bacterium | F4X37_11130 | 50648..53974 | 2080302 | ATP-binding-protein-[Acidimicrobiia-bacterium] |
| MXZ78951.1 | HEPN+AAA-ATPase+Alpha+Beta-domain+wHTH+RRM | Type-1_BR.RM.like.systems | HTH_31->RNase->HTH_58-><-?<-HEPN+DUF499*\|\|?->PhdYeFM_antitox->PIN-><-METHYLASE<-?\|\|CHTH-> | 1079 | Bacteria>Terrabacteria_group>Actinomycetota>Acidimicrobiia>unclassified_Acidimicrobiia | GCA_009837345.1 | Acidimicrobiia_bacterium | F4Z06_11870 | 8705..11944 | 2080302 | ATP-binding-protein-[Acidimicrobiia-bacterium] |
| MXX00359.1 | HEPN+AAA-ATPase+Alpha+Beta-domain+wHTH+RRM | Type-1_BR.RM.like.systems | <-HEPN+DUF499*<-TUDOR+SF1-SF2-Helicase+iREase+CTD+REase<-CHTH<-?<-CHTH | 1005 | Bacteria>Terrabacteria_group>Actinomycetota>Acidimicrobiia>unclassified_Acidimicrobiia | GCA_009835875.1 | Acidimicrobiia_bacterium | F4Z79_01885 | 1..3016 | 2080302 | ATP-binding-protein-[Acidimicrobiia-bacterium] |
| MXZ30232.1 | HEPN+AAA-ATPase+Alpha+Beta-domain+wHTH+RRM | Type-1_BR.RM.like.systems | <-ParD_antitoxin<-ParD_antitoxin\|\|HEPN+DUF499*-><-ABhydrolase | 1083 | Bacteria>Terrabacteria_group>Actinomycetota>Acidimicrobiia>unclassified_Acidimicrobiia | GCA_009837005.1 | Acidimicrobiia_bacterium | F4Z22_05205 | 787..4038 | 2080302 | ATP-binding-protein-[Acidimicrobiia-bacterium] |
| MXW42977.1 | HEPN+AAA-ATPase+Alpha+Beta-domain+wHTH+RRM | Type-1_BR.RM.like.systems | <-METHYLASE<-PIN<-?<-HEPN+DUF499*<-METHYLASE<-?\|\|?-><-PIN<-DUF433 | 1079 | Bacteria>Terrabacteria_group>Actinomycetota>Acidimicrobiia>unclassified_Acidimicrobiia | GCA_009835645.1 | Acidimicrobiia_bacterium | F4Z91_07815 | 3856..7095 | 2080302 | ATP-binding-protein-[Acidimicrobiia-bacterium] |
| MYA38651.1 | HEPN+AAA-ATPase+Alpha+Beta-domain+wHTH+RRM | Type-1_BR.RM.like.systems | METHYLASE-p->HEPN+DUF499*->HTH_3->NTP_transf-><-?<-PIN | 1107 | Bacteria>Terrabacteria_group>Actinomycetota>Acidimicrobiia>unclassified_Acidimicrobiia | GCA_009838695.1 | Acidimicrobiia_bacterium | F4Y36_03440 | 1539..4862 | 2080302 | ATP-binding-protein-[Acidimicrobiia-bacterium] |
| MXY73348.1 | HEPN+AAA-ATPase+Alpha+Beta-domain+wHTH+RRM | Type-1_BR.RM.like.systems | METHYLASE-p->HEPN+DUF499*->HTH_3->NTP_transf-><-?<-PIN | 1107 | Bacteria>Terrabacteria_group>Actinomycetota>Acidimicrobiia>unclassified_Acidimicrobiia | GCA_009837815.1 | Acidimicrobiia_bacterium | F4Y83_00240 | 420..3743 | 2080302 | ATP-binding-protein-[Acidimicrobiia-bacterium] |
| MXW58366.1 | HEPN+AAA-ATPase+Alpha+Beta-domain+wHTH+RRM | Type-1_BR.RM.like.systems | ABC-ATPase+SbcC->?->?->?->?-><-HEPN+DUF499*<-?<-?<-?<-Pkinase<-METHYLASE | 1114 | Bacteria>Terrabacteria_group>Actinomycetota>Acidimicrobiia>unclassified_Acidimicrobiia | GCA_009836065.1 | Acidimicrobiia_bacterium | F4Z70_07245 | 33723..37067 | 2080302 | ATP-binding-protein-[Acidimicrobiia-bacterium] |
| MYG93975.1 | HEPN+AAA-ATPase+Alpha+Beta-domain+wHTH+RRM | Type-1_BR.RM.like.systems | METHYLASE->?->?-><-?\|\|?->HEPN+DUF499*->?->PIN->METHYLASE-> | 1079 | Bacteria>Terrabacteria_group>Actinomycetota>Acidimicrobiia>unclassified_Acidimicrobiia | GCA_009843865.1 | Acidimicrobiia_bacterium | F4138_03135 | 9114..12353 | 2080302 | ATP-binding-protein-[Acidimicrobiia-bacterium] |
| MCQ3809088.1 | HEPN+AAA-ATPase+Alpha+Beta-domain+wHTH+RRM | Type-1_BR.RM.like.systems | Radical-SAM-><-?<-?<-?<-HEPN+DUF499*\|\|?-><-?\|\|?-><-METHYLASE | 1114 | Bacteria>Terrabacteria_group>Actinomycetota>Acidimicrobiia>unclassified_Acidimicrobiia | GCA_024421205.1 | Acidimicrobiia_bacterium | KTV68_00880 | 53316..56660 | 2080302 | DUF499-domain-containing-protein-[Acidimicrobiia-bacterium] |
| MYJ15813.1 | HEPN+AAA-ATPase+Alpha+Beta-domain+wHTH+RRM | Type-1_BR.RM.like.systems | <-HTH_3<-HEPN+DUF499*<-METHYLASE-p | 1107 | Bacteria>Terrabacteria_group>Actinomycetota>Acidimicrobiia>unclassified_Acidimicrobiia | GCA_009844925.1 | Acidimicrobiia_bacterium | F4085_04615 | 343..3666 | 2080302 | ATP-binding-protein-[Acidimicrobiia-bacterium] |
| MYD40244.1 | HEPN+AAA-ATPase+Alpha+Beta-domain+wHTH+RRM | Type-1_BR.RM.like.systems | <-RNase<-HEPN+DUF499*<-METHYLASE | 1090 | Bacteria>Terrabacteria_group>Actinomycetota>Acidimicrobiia>unclassified_Acidimicrobiia | GCA_009841565.1 | Acidimicrobiia_bacterium | F4W94_00700 | 1349..4621 | 2080302 | ATP-binding-protein-[Acidimicrobiia-bacterium] |
| MXY74717.1 | HEPN+AAA-ATPase+Alpha+Beta-domain+wHTH+RRM | Type-1_BR.RM.like.systems | <-HEPN+DUF499*<-METHYLASE-p | 1044 | Bacteria>Terrabacteria_group>Actinomycetota>Acidimicrobiia>unclassified_Acidimicrobiia | GCA_009837815.1 | Acidimicrobiia_bacterium | F4Y83_07345 | 1..3131 | 2080302 | ATP-binding-protein-[Acidimicrobiia-bacterium] |
| MXW68944.1 | HEPN+AAA-ATPase+Alpha+Beta-domain+wHTH+RRM | Type-1_BR.RM.like.systems | METHYLASE-p->HEPN+DUF499*-> | 1100 | Bacteria>Terrabacteria_group>Actinomycetota>Acidimicrobiia>unclassified_Acidimicrobiia | GCA_009836245.1 | Acidimicrobiia_bacterium | F4Z61_03125 | 1254..4555 | 2080302 | ATP-binding-protein-[Acidimicrobiia-bacterium] |
| MBP8181356.1 | HEPN+AAA-ATPase+Alpha+Beta-domain+wHTH+RRM | Type-1_BR.RM.like.systems | <-HEPN+DUF499*<-METHYLASE | 1096 | Bacteria>Terrabacteria_group>Actinomycetota>Acidimicrobiia>unclassified_Acidimicrobiia | GCA_018006325.1 | Acidimicrobiia_bacterium | KAZ88_10225 | 2758..6048 | 2080302 | DUF499-domain-containing-protein-[Acidimicrobiia-bacterium] |
| WP_226436803.1 | HEPN+AAA-ATPase+Alpha+Beta-domain+wHTH+RRM | Type-1_BR.RM.like.systems | <-DUF222+HNH<-HEPN+DUF499*<-METHYLASE\|\|CHTH-><-TUDOR+SF1-SF2-Helicase+iREase+CTD+REase | 1141 | Bacteria>Terrabacteria_group>Actinomycetota>Actinomycetes | GCF_012515855.1 | Mycobacteriales | GX543_RS09675 | 1506..4931 | 85007 | Swt1-family-HEPN-domain-containing-protein-[Rhodococcus-yananensis] |

| Protein-ID | DUF499-architecture | System | Gene-Neighbourhoods | Length | Taxonomy | Assembly-ID | Strain | Locus-Tag | Genomic-coordinates | TaxID | Fasta-defline |
|---|---|---|---|---|---|---|---|---|---|---|---|
| WP_128773202.1 | HEPN+AAA-ATPase+Alpha+Beta-domain+wHTH+RRM | Type-1_BR.RM.like.systems | SF1-SF2-Helicase+REase-><-?<-TrwC+SF1-SF2-Helicase\|\|DNA_methylase-><-HEPN+REase<-REase<-HEPN+DUF499*<-wHTH+REase<-METHYLASE<-TUDOR+SF1-SF2-Helicase+iREase+CTD+REase<-XIS-HTH<-ABhydrolase | 1132 | Bacteria>Terrabacteria_group>Actinomycetota>Actinomycetes>Actinomycetales | GCF_004104015.1 | Actinomyces_oricola | EUC46_RS05555 | 19810..23208 | 206043 | DUF499-domain-containing-protein-[Actinomyces-oricola] |
| WP_114990037.1 | HEPN+AAA-ATPase+Alpha+Beta-domain+wHTH+RRM | Type-1_BR.RM.like.systems | TOPRIM->RelE-ParE->ParA+HTH->TUDOR+SF1-SF2-Helicase+iREase+CTD+REase->METHYLASE->HEPN+DUF499*-><-PIN<-DUF433\|\|RNase->TraJ-RHH->RelE-ParE->HTH_3-> | 1131 | Bacteria>Terrabacteria_group>Actinomycetota>Actinomycetes>Actinomycetales | GCF_900453275.1 | Mobiluncus_mulieris | DX043_RS05220 | 1134969..1138364 | 2052 | DUF499-domain-containing-protein-[Mobiluncus-mulieris] |
| WP_169762497.1 | HEPN+AAA-ATPase+Alpha+Beta-domain+wHTH+RRM | Type-1_BR.RM.like.systems | <-HTH_3<-RelE-ParE<-TraJ-RHH<-RNase\|\|DUF433->PIN-><-HEPN+DUF499*<-METHYLASE<-TUDOR+SF1-SF2-Helicase+iREase+CTD+REase<-ParA+HTH<-TOPRIM<-PIN | 1131 | Bacteria>Terrabacteria_group>Actinomycetota>Actinomycetes>Actinomycetales | GCF_012976645.1 | Mobiluncus_mulieris | HHJ77_RS04685 | 61231..64626 | 2052 | DUF499-domain-containing-protein-[Mobiluncus-mulieris] |
| NLI19748.1 | HEPN+AAA-ATPase+Alpha+Beta-domain+wHTH+RRM | Type-1_BR.RM.like.systems | <-HTH_31<-?\|\|UDG-conflict-><-HEPN+DUF499*<-METHYLASE<-TUDOR+SF1-SF2-Helicase+iREase+CTD+REase<-?<-TrwC+SF1-SF2-Helicase | 1114 | Bacteria>Terrabacteria_group>Actinomycetota>Actinomycetes>Actinomycetales | GCA_012517595.1 | Actinomycetales_bacterium | GX427_13235 | 2536..5880 | 1911520 | ATP-binding-protein-[Actinomycetales-bacterium] |
| WP_184451288.1 | HEPN+AAA-ATPase+Alpha+Beta-domain+wHTH+RRM | Type-1_BR.RM.like.systems | TUDOR+SF1-SF2-Helicase+iREase+CTD+REase-p-><-?\|\|TUDOR+SF1-SF2-Helicase+iREase+CTD+REase->METHYLASE->HEPN+DUF499*->HEPN->RelB->tRNA-><-Glyoxalase | 1150 | Bacteria>Terrabacteria_group>Actinomycetota>Actinomycetes>Actinomycetales | GCF_014208035.1 | Schaalia_hyovaginalis | HD592_RS00720 | 169664..173116 | 29316 | DUF499-domain-containing-protein-[Schaalia-hyovaginalis] |
| WP_103061758.1 | HEPN+AAA-ATPase+Alpha+Beta-domain+wHTH+RRM | Type-1_BR.RM.like.systems | <-REase<-HTH+SF1-SF2-Helicase+REase<-REase<-MORC<-HEPN+DUF499*<-METHYLASE\|\|DNA_methylase->MORC+HisKin->HTH_Tnp_4+DDE_Tnp_4->MORC+HEPN-> | 1138 | Bacteria>Terrabacteria_group>Actinomycetota>Actinomycetes>Actinomycetales | GCF_004102025.1 | Actinomyces_qiguomingii | CWT10_RS16110 | 3847675..3851091 | 2057800 | DUF499-domain-containing-protein-[Actinomyces-qiguomingii] |
| WP_165316371.1 | HEPN+AAA-ATPase+Alpha+Beta-domain+wHTH+RRM | Type-1_BR.RM.like.systems | ABC-ATPase+SbcC-><-RecR+TOPRIM<-?\|\|TUDOR+SF1-SF2-Helicase+iREase+CTD+REase->?->METHYLASE->HEPN+DUF499*-><-PIN\|\|?->?->DDE_Tnp_1->DDE_Tnp_1-><-RNase | 1143 | Bacteria>Terrabacteria_group>Actinomycetota>Actinomycetes>Actinomycetales | GCF_011038885.2 | Schaalia_sp._ZJ405 | G7Y41_RS09385 | 2197397..2200828 | 2709403 | DUF499-domain-containing-protein-[Schaalia-sp.-ZJ405] |
| MBE6482584.1 | HEPN+AAA-ATPase+Alpha+Beta-domain+wHTH+RRM | Type-1_BR.RM.like.systems | <-Schlafen+HATPase+HTH<-REase<-HTH+SF1-SF2-Helicase+REase<-REase<-MORC<-HEPN+DUF499*<-METHYLASE\|\|DNA_methylase->DNA_methylase->MORC+HISKIN-> | 1137 | Bacteria>Terrabacteria_group>Actinomycetota>Actinomycetes>Actinomycetales | GCA_015062785.1 | Actinomyces_ruminicola | E7Z94_09510 | 10121..13534 | 332524 | ATP-binding-protein-[Actinomyces-ruminicola] |
| WP_278972061.1 | HEPN+AAA-ATPase+Alpha+Beta-domain+wHTH+RRM | Type-1_BR.RM.like.systems | <-HTH_3<-RelE-ParE\|\|DUF433->PIN-><-HEPN+DUF499*<-METHYLASE-p<-TUDOR+SF1-SF2-Helicase+iREase+CTD+REase<-ParA+HTH<-RelE-ParE<-TOPRIM | 1131 | Bacteria>Terrabacteria_group>Actinomycetota>Actinomycetes>Actinomycetales | GCF_946891785.1 | Mobiluncus_mulieris | QDN45_RS08450 | 23006..26401 | 2052 | DUF499-domain-containing-protein-[Mobiluncus-mulieris] |
| WP_169773626.1 | HEPN+AAA-ATPase+Alpha+Beta-domain+wHTH+RRM | Type-1_BR.RM.like.systems | PIN->TOPRIM->ParA+HTH->TUDOR+SF1-SF2-Helicase+iREase+CTD+REase->METHYLASE-p->HEPN+DUF499*-><-PIN<-DUF433<-?<-HEPN\|\|RNase->TraJ-RHH-> | 1131 | Bacteria>Terrabacteria_group>Actinomycetota>Actinomycetes>Actinomycetales | GCF_012976895.1 | Mobiluncus_mulieris | HHJ79_RS06650 | 39416..42811 | 2052 | DUF499-domain-containing-protein-[Mobiluncus-mulieris] |
| WP_154475624.1 | HEPN+AAA-ATPase+Alpha+Beta-domain+wHTH+RRM | Type-1_BR.RM.like.systems | <-HEPN<-?\|\|?-><-?<-tRNA<-RelB<-HEPN<-HEPN+DUF499*<-METHYLASE<-TUDOR+SF1-SF2-Helicase+iREase+CTD+REase<-TUDOR+SF1-SF2-Helicase+iREase+CTD+REase-p | 1151 | Bacteria>Terrabacteria_group>Actinomycetota>Actinomycetes>Actinomycetales | - | Schaalia_hyovaginalis | FYJ77_RS02065 | 517238..520693 | 29316 | DUF499-domain-containing-protein-[Schaalia-hyovaginalis] |
| WP_271288667.1 | HEPN+AAA-ATPase+Alpha+Beta-domain+wHTH+RRM | Type-1_BR.RM.like.systems | ACET+ACET->HTH_3+PolB-><-?<-?<-DNA_methylase\|\|METHYLASE->HEPN+DUF499*->MORC->REase->HTH+SF1-SF2-Helicase+REase->REase-><-?\|\|METHYLASE-p-> | 1132 | Bacteria>Terrabacteria_group>Actinomycetota>Actinomycetes>Actinomycetales | GCF_027945475.1 | Actinomyces_oris | PGE45_RS00580 | 136591..139989 | 544580 | DUF499-domain-containing-protein-[Actinomyces-oris] |
| WP_075403442.1 | HEPN+AAA-ATPase+Alpha+Beta-domain+wHTH+RRM | Type-1_BR.RM.like.systems | ACET+ACET->HTH_3+PolB-><-?<-?<-DNA_methylase\|\|METHYLASE->HEPN+DUF499*->MORC->REase->HTH+SF1-SF2-Helicase+REase->REase-><-?\|\|METHYLASE-p-> | 1139 | Bacteria>Terrabacteria_group>Actinomycetota>Actinomycetes>Actinomycetales | GCF_001937505.1 | Actinomyces_oris | BKH21_RS10525 | 35903..39322 | 544580 | DUF499-domain-containing-protein-[Actinomyces-oris] |
| WP_075412471.1 | HEPN+AAA-ATPase+Alpha+Beta-domain+wHTH+RRM | Type-1_BR.RM.like.systems | <-METHYLASE-p<-REase<-HTH+SF1-SF2-Helicase+REase<-REase<-MORC<-HEPN+DUF499*<-METHYLASE\|\|DNA_methylase->?-><-HTH_3+PolB<-ACET+ACET | 1137 | Bacteria>Terrabacteria_group>Actinomycetota>Actinomycetes>Actinomycetales | GCF_001937365.1 | Actinomyces_oris | BKH31_RS11955 | 29944..33357 | 544580 | DUF499-domain-containing-protein-[Actinomyces-oris] |
| MBK8733830.1 | HEPN+AAA-ATPase+Alpha+Beta-domain+wHTH+RRM | Type-1_BR.RM.like.systems | <-DUF499-p<-?<-DUF499-p<-?<-HEPN+DUF499*<-METHYLASE<-TUDOR+SF1-SF2-Helicase+iREase+CTD+REase\|\|?->Phage_integrase+Phage_integrase->PIN-> | 347 | Bacteria>Terrabacteria_group>Actinomycetota>Actinomycetes>Actinomycetales | GCA_016710645.1 | Actinomycetales_bacterium | IPL93_13505 | 220122..221165 | 1911520 | hypothetical-protein-[Actinomycetales-bacterium] |
| WP_161924011.1 | HEPN+AAA-ATPase+Alpha+Beta-domain+wHTH+RRM | Type-1_BR.RM.like.systems | TetR-HTH->TUDOR+SF1-SF2-Helicase+iREase+CTD+REase->METHYLASE-p->METHYLASE->HEPN+DUF499*-><-HTH_Tnp_4+DDE_Tnp_4\|\|?->HAD-><-?<-HNH | 1139 | Bacteria>Terrabacteria_group>Actinomycetota>Actinomycetes>Actinomycetales | GCF_009930875.1 | Actinomyces_sp._432 | CWT12_RS05510 | 1339354..1342773 | 2057798 | DUF499-domain-containing-protein-[Actinomyces-sp.-432] |
| WP_053587850.1 | HEPN+AAA-ATPase+Alpha+Beta-domain+wHTH+RRM | Type-1_BR.RM.like.systems | <-REase<-HTH+SF1-SF2-Helicase+REase<-REase<-MORC<-HEPN+DUF499*<-METHYLASE\|\|METHYLASE->BirA-HTH->?-><-TPR<-MULE-TRANSPOSASE | 1165 | Bacteria>Terrabacteria_group>Actinomycetota>Actinomycetes>Actinomycetales | GCF_001278845.1 | Actinomyces_sp._oral_taxon_414 | AM609_RS14685 | 3673757..3677254 | 712122 | DUF499-domain-containing-protein-[Actinomyces-sp.-oral-taxon-414] |
| WP_017177796.1 | HEPN+AAA-ATPase+Alpha+Beta-domain+wHTH+RRM | Type-1_BR.RM.like.systems | Phage_holin_4_2->?-><-REase<-HTH+SF1-SF2-Helicase+REase<-REase<-MORC<-HEPN+DUF499*<-METHYLASE\|\|DNA_methylase->?->?-><-?<-HTH_3+PolB | 1137 | Bacteria>Terrabacteria_group>Actinomycetota>Actinomycetes>Actinomycetales | GCF_000295095.1 | Actinomyces_timonensis | A1QA_RS0104850 | 1087937..1091350 | 1288391 | DUF499-domain-containing-protein-[Actinomyces-timonensis] |
| WP_077240589.1 | HEPN+AAA-ATPase+Alpha+Beta-domain+wHTH+RRM | Type-1_BR.RM.like.systems | <-REase<-HTH+SF1-SF2-Helicase+REase<-REase<-MORC<-HEPN+DUF499*<-METHYLASE\|\|DNA_methylase->MORC+HEPN->\|\|?-><-?<-Peptidase_M50 | 1137 | Bacteria>Terrabacteria_group>Actinomycetota>Actinomycetes>Actinomycetales | GCF_002356195.1 | Actinomyces | DY404_RS06925 | 1656821..1660234 | 1654 | MULTISPECIES:-DUF499-domain-containing-protein-[Actinomyces] |
| WP_070662487.1 | HEPN+AAA-ATPase+Alpha+Beta-domain+wHTH+RRM | Type-1_BR.RM.like.systems | HTH_3+PolB->?-><-?<-?<-DNA_methylase\|\|METHYLASE->HEPN+DUF499*->MORC->REase->HTH+SF1-SF2-Helicase+REase->REase-><-?\|\|METHYLASE-p-> | 1137 | Bacteria>Terrabacteria_group>Actinomycetota>Actinomycetes>Actinomycetales | GCF_963519085.1 | Actinomyces | BKH20_RS04780 | 16618..20031 | 1654 | MULTISPECIES:-DUF499-domain-containing-protein-[Actinomyces] |
| WP_003788306.1 | HEPN+AAA-ATPase+Alpha+Beta-domain+wHTH+RRM | Type-1_BR.RM.like.systems | <-METHYLASE-p\|\|?-><-REase<-HTH+SF1-SF2-Helicase+REase<-REase<-MORC<-HEPN+DUF499*<-METHYLASE\|\|DNA_methylase->?->?-><-?<-HTH_3+PolB | 1139 | Bacteria>Terrabacteria_group>Actinomycetota>Actinomycetes>Actinomycetales | GCF_000175315.3 | Actinomyces_viscosus | HMPREF0059_RS11125 | 118878..122297 | 1656 | DUF499-domain-containing-protein-[Actinomyces-viscosus] |
| WP_282568806.1 | HEPN+AAA-ATPase+Alpha+Beta-domain+wHTH+RRM | Type-1_BR.RM.like.systems | ACET+ACET-><-?<-?<-DNA_methylase\|\|METHYLASE->HEPN+DUF499*->MORC->REase->HTH+SF1-SF2-Helicase+REase->REase-><-?\|\|METHYLASE-p-> | 1139 | Bacteria>Terrabacteria_group>Actinomycetota>Actinomycetes>Actinomycetales | GCF_023169925.1 | Actinomyces_oris | AoKodu_RS00635 | 147963..151382 | 544580 | DUF499-domain-containing-protein-[Actinomyces-oris] |
| BDF97829.1 | HEPN+AAA-ATPase+Alpha+Beta-domain+wHTH+RRM | Type-1_BR.RM.like.systems | ACET+ACET-><-?<-?<-DNA_methylase\|\|METHYLASE->HEPN+DUF499*->MORC->REase->HTH+SF1-SF2-Helicase+REase->REase-><-?\|\|METHYLASE-p-> | 1139 | Bacteria>Terrabacteria_group>Actinomycetota>Actinomycetes>Actinomycetales | GCA_023169925.1 | Actinomyces_oris_K20 | AoKodu_01300 | 147963..151382 | 871541 | hypothetical-protein-AoKodu_01300-[Actinomyces-oris-K20] |
| WP_268916840.1 | HEPN+AAA-ATPase+Alpha+Beta-domain+wHTH+RRM | Type-1_BR.RM.like.systems | <-REase<-HTH+SF1-SF2-Helicase+REase<-REase<-MORC<-HEPN+DUF499*<-METHYLASE\|\|DNA_methylase->MORC+HEPN->SLATT->SMODS-> | 1131 | Bacteria>Terrabacteria_group>Actinomycetota>Actinomycetes>Actinomycetales | GCF_026891875.1 | Actinomyces_israelii | OHJ16_RS04000 | 49558..52953 | 1659 | DUF499-domain-containing-protein-[Actinomyces-israelii] |
| WP_278601539.1 | HEPN+AAA-ATPase+Alpha+Beta-domain+wHTH+RRM | Type-1_BR.RM.like.systems | ACET+ACET-><-?<-?<-DNA_methylase\|\|METHYLASE->HEPN+DUF499*->MORC->REase->HTH+SF1-SF2-Helicase+REase->REase->METHYLASE-p-> | 1137 | Bacteria>Terrabacteria_group>Actinomycetota>Actinomycetes>Actinomycetales | GCF_905372775.1 | Actinomyces_naeslundii | QDH38_RS03495 | 39335..42748 | 1655 | DUF499-domain-containing-protein-[Actinomyces-naeslundii] |
| WP_108833962.1 | HEPN+AAA-ATPase+Alpha+Beta-domain+wHTH+RRM | Type-1_BR.RM.like.systems | <-METHYLASE-p<-REase<-HTH+SF1-SF2-Helicase+REase<-REase<-MORC<-HEPN+DUF499*<-METHYLASE\|\|DNA_methylase->?-><-ACET+ACET | 1138 | Bacteria>Terrabacteria_group>Actinomycetota>Actinomycetes>Actinomycetales | GCF_900323545.1 | Actinomyces_sp._Marseille-P3109 | BQ8008_RS10580 | 2440018..2443434 | 2083009 | DUF499-domain-containing-protein-[Actinomyces-sp.-Marseille-P3109] |
| WP_076255389.1 | HEPN+AAA-ATPase+Alpha+Beta-domain+wHTH+RRM | Type-1_BR.RM.like.systems | ACET+ACET-><-?<-?<-DNA_methylase\|\|METHYLASE->HEPN+DUF499*->MORC->REase->HTH+SF1-SF2-Helicase+REase->REase->METHYLASE-p-> | 1137 | Bacteria>Terrabacteria_group>Actinomycetota>Actinomycetes>Actinomycetales | GCF_001956555.1 | Actinomyces_naeslundii | BKH25_RS11260 | 118842..122255 | 1655 | DUF499-domain-containing-protein-[Actinomyces-naeslundii] |
| WP_076142650.1 | HEPN+AAA-ATPase+Alpha+Beta-domain+wHTH+RRM | Type-1_BR.RM.like.systems | <-METHYLASE-p<-REase<-HTH+SF1-SF2-Helicase+REase<-REase<-MORC<-HEPN+DUF499*<-METHYLASE\|\|DNA_methylase->?->?-><-ACET+ACET | 1139 | Bacteria>Terrabacteria_group>Actinomycetota>Actinomycetes>Actinomycetales | GCF_016127855.1 | Actinomyces_naeslundii | BKH33_RS06345 | 173705..177124 | 1655 | DUF499-domain-containing-protein-[Actinomyces-naeslundii] |
| WP_075413329.1 | HEPN+AAA-ATPase+Alpha+Beta-domain+wHTH+RRM | Type-1_BR.RM.like.systems | <-METHYLASE-p<-REase<-HTH+SF1-SF2-Helicase+REase<-REase<-MORC<-HEPN+DUF499*<-METHYLASE\|\|DNA_methylase->?->?-><-ACET+ACET | 1137 | Bacteria>Terrabacteria_group>Actinomycetota>Actinomycetes>Actinomycetales | GCF_001937475.1 | Actinomyces_naeslundii | BKH08_RS04545 | 28951..32364 | 1655 | DUF499-domain-containing-protein-[Actinomyces-naeslundii] |
| WP_136193072.1 | HEPN+AAA-ATPase+Alpha+Beta-domain+wHTH+RRM | Type-1_BR.RM.like.systems | HNH->?->ABhydrolase-><-HAD<-?<-HEPN+DUF499*<-METHYLASE<-METHYLASE-p<-TUDOR+SF1-SF2-Helicase+iREase+CTD+REase<-TetR-HTH | 1138 | Bacteria>Terrabacteria_group>Actinomycetota>Actinomycetes>Actinomycetales | GCF_004796555.1 | Actinomyces_procaprae | FAQ94_RS06890 | 215265..218681 | 2560010 | DUF499-domain-containing-protein-[Actinomyces-procaprae] |
| WP_124852157.1 | HEPN+AAA-ATPase+Alpha+Beta-domain+wHTH+RRM | Type-1_BR.RM.like.systems | <-HEPN+DUF499*<-METHYLASE<-TUDOR+SF1-SF2-Helicase+iREase+CTD+REase<-TUDOR+SF1-SF2-Helicase+iREase+CTD+REase-p | 1140 | Bacteria>Terrabacteria_group>Actinomycetota>Actinomycetes>Actinomycetales | GCF_003858495.1 | Buchananella_hordeovulneris | EII12_RS05710 | 157..3579 | 52770 | DUF499-domain-containing-protein-[Buchananella-hordeovulneris] |
| WP_073824655.1 | HEPN+AAA-ATPase+Alpha+Beta-domain+wHTH+RRM | Type-1_BR.RM.like.systems | TUDOR+SF1-SF2-Helicase+iREase+CTD+REase-p->TUDOR+SF1-SF2-Helicase+iREase+CTD+REase->METHYLASE->HEPN+DUF499*-> | 1140 | Bacteria>Terrabacteria_group>Actinomycetota>Actinomycetes>Actinomycetales | GCF_001907235.1 | Buchananella_hordeovulneris | BSZ40_RS07210 | 166478..169900 | 52770 | DUF499-domain-containing-protein-[Buchananella-hordeovulneris] |
| WP_106110118.1 | HEPN+AAA-ATPase+Alpha+Beta-domain+wHTH+RRM | Type-1_BR.RM.like.systems | <-REase\|\|GATase-><-?<-HTH_28+HTH_33+DDE_3\|\|METHYLASE->HEPN+DUF499*->?->?->Peptidase_M50->?->?->VirB11+SF1-SF2-Helicase-> | 1130 | Bacteria>Terrabacteria_group>Actinomycetota>Actinomycetes>Actinomycetales | GCF_002999235.1 | Actinomyces_sp._oral_taxon_897 | C3V41_RS09940 | 2509136..2512528 | 2081702 | DUF499-domain-containing-protein-[Actinomyces-sp.-oral-taxon-897] |
| WP_200510328.1 | HEPN+AAA-ATPase+Alpha+Beta-domain+wHTH+RRM | Type-1_BR.RM.like.systems | GATase-><-?<-Schlafen+HATPase+HTH<-?<-DNA_methylase\|\|METHYLASE->HEPN+DUF499*->?->?->Peptidase_M50->?->?->VirB11+SF1-SF2-Helicase-> | 1129 | Bacteria>Terrabacteria_group>Actinomycetota>Actinomycetes>Actinomycetales | GCF_016626125.1 | Actinomyces_sp._HMT897 | JJJ15_RS13470 | 1917676..1921065 | 2789424 | DUF499-domain-containing-protein-[Actinomyces-sp.-HMT897] |
| WP_070510861.1 | HEPN+AAA-ATPase+Alpha+Beta-domain+wHTH+RRM | Type-1_BR.RM.like.systems | <-REase<-HTH+SF1-SF2-Helicase+REase<-REase<-MORC\|\|?-><-HEPN+DUF499*<-METHYLASE\|\|DNA_methylase->?->?-><-?<-HTH_3+PolB | 1137 | Bacteria>Terrabacteria_group>Actinomycetota>Actinomycetes>Actinomycetales | GCF_001807465.1 | Actinomyces_sp._HMSC08A09 | HMPREF3165_RS03500 | 45763..49176 | 1581133 | DUF499-domain-containing-protein-[Actinomyces-sp.-HMSC08A09] |
| WP_182050741.1 | HEPN+AAA-ATPase+Alpha+Beta-domain+wHTH+RRM | Type-1_BR.RM.like.systems | SF1-SF2-Helicase->TUDOR+SF1-SF2-Helicase+iREase+CTD+REase->METHYLASE->REase->HEPN->HEPN+DUF499*->SbcC-> | 1134 | Bacteria>Terrabacteria_group>Actinomycetota>Actinomycetes>Actinomycetales | GCF_014041995.1 | Changpingibacter_yushuensis | H2017_RS05160 | 1185005..1188409 | 2758440 | Swt1-family-HEPN-domain-containing-protein-[Changpingibacter-yushuensis] |
| WP_005985559.1 | HEPN+AAA-ATPase+Alpha+Beta-domain+wHTH+RRM | Type-1_BR.RM.like.systems | <-ABC-ATPase+SbcC<-?\|\|TUDOR+SF1-SF2-Helicase+iREase+CTD+REase->METHYLASE->HEPN+DUF499*-><-?<-?<-?<-?<-ABC-ATPase+SbcC | 1128 | Bacteria>Terrabacteria_group>Actinomycetota>Actinomycetes>Actinomycetales | GCF_000239695.1 | Actinomyces_graevenitzii | HMPREF0045_RS03480 | 792644..796030 | 55565 | DUF499-domain-containing-protein-[Actinomyces-graevenitzii] |
| WP_219110028.1 | HEPN+AAA-ATPase+Alpha+Beta-domain+wHTH+RRM | Type-1_BR.RM.like.systems | HNH-><-HAD<-?<-HEPN+DUF499*<-METHYLASE<-METHYLASE-p<-TUDOR+SF1-SF2-Helicase+iREase+CTD+REase<-TetR-HTH | 1139 | Bacteria>Terrabacteria_group>Actinomycetota>Actinomycetes>Actinomycetales | GCF_019325145.1 | Actinomyces_sp._594 | GZ998_RS12335 | 131266..134685 | 2057793 | DUF499-domain-containing-protein-[Actinomyces-sp.-594] |
| WP_162678504.1 | HEPN+AAA-ATPase+Alpha+Beta-domain+wHTH+RRM | Type-1_BR.RM.like.systems | TetR-HTH->TUDOR+SF1-SF2-Helicase+iREase+CTD+REase->METHYLASE-p->METHYLASE->HEPN+DUF499*->?->HAD-><-HNH | 1139 | Bacteria>Terrabacteria_group>Actinomycetota>Actinomycetes>Actinomycetales | GCF_010365615.1 | Actinomyces_sp._565 | GZ997_RS01005 | 32665..36084 | 2057794 | DUF499-domain-containing-protein-[Actinomyces-sp.-565] |
| WP_198776534.1 | HEPN+AAA-ATPase+Alpha+Beta-domain+wHTH+RRM | Type-1_BR.RM.like.systems | sGTP-><-?<-DNA_methylase\|\|METHYLASE->HEPN+DUF499*->MORC->REase->HTH+SF1-SF2-Helicase+REase->REase-> | 1127 | Bacteria>Terrabacteria_group>Actinomycetota>Actinomycetes>Actinomycetales | GCF_016405145.1 | Schaalia_cardiffensis | JFX23_RS08485 | 21486..24869 | 181487 | DUF499-domain-containing-protein-[Schaalia-cardiffensis] |
| WP_073332337.1 | HEPN+AAA-ATPase+Alpha+Beta-domain+wHTH+RRM | Type-1_BR.RM.like.systems | <-ABC-ATPase+SbcC<-?<-?<-?<-?<-?<-HEPN+DUF499*<-METHYLASE<-TUDOR+SF1-SF2-Helicase+iREase+CTD+REase<-TetR-HTH | 1138 | Bacteria>Terrabacteria_group>Actinomycetota>Actinomycetes>Actinomycetales | GCF_900098745.1 | Actinomyces_glycerinitolerans | ACGLYG10_RS12005 | 81593..85009 | 1892869 | DUF499-domain-containing-protein-[Actinomyces-glycerinitolerans] |
| MCI6556423.1 | HEPN+AAA-ATPase+Alpha+Beta-domain+wHTH+RRM | Type-1_BR.RM.like.systems | TUDOR+SF1-SF2-Helicase+iREase+CTD+REase->METHYLASE->HEPN+DUF499*->HEPN->RelB->?->tRNA-><-?<-?<-DNA_methylase | 1149 | Bacteria>Terrabacteria_group>Actinomycetota>Actinomycetes>Actinomycetales | GCA_022780845.1 | Schaalia_hyovaginalis | MR422_02105 | 263803..267252 | 29316 | DUF499-domain-containing-protein-[Schaalia-hyovaginalis] |
| WP_251775558.1 | HEPN+AAA-ATPase+Alpha+Beta-domain+wHTH+RRM | Type-1_BR.RM.like.systems | HKD-DNase+HKD-DNase-><-?\|\|?->?-><-?\|\|?-><-HEPN+DUF499*<-METHYLASE<-TUDOR+SF1-SF2-Helicase+iREase+CTD+REase | 1129 | Bacteria>Terrabacteria_group>Actinomycetota>Actinomycetes>Actinomycetales | GCF_023716965.1 | Schaalia_meyeri | ND910_RS08155 | 14607..17996 | 52773 | DUF499-domain-containing-protein-[Schaalia-meyeri] |
| ERH33797.1 | HEPN+AAA-ATPase+Alpha+Beta-domain+wHTH+RRM | Type-1_BR.RM.like.systems | <-HEPN+DUF499*<-METHYLASE<-TUDOR+SF1-SF2-Helicase+iREase+CTD+REase<-TraJ-RHH<-?<-?\|\|TetR-HTH-> | 1139 | Bacteria>Terrabacteria_group>Actinomycetota>Actinomycetes>Actinomycetales | GCA_000466305.1 | Actinomyces_sp._oral_taxon_877_str._F0543 | HMPREF1550_00139 | 24107..27526 | 1227264 | hypothetical-protein-HMPREF1550_00139-[Actinomyces-sp.-oral-taxon-877-str.-F0543] |
| QQ076952.1 | HEPN+AAA-ATPase+Alpha+Beta-domain+wHTH+RRM | Type-1_BR.RM.like.systems | GATase-><-?<-Schlafen+HATPase+HTH<-?<-DNA_methylase\|\|METHYLASE->HEPN+DUF499*->?->?->Peptidase_M50-> | 1129 | Bacteria>Terrabacteria_group>Actinomycetota>Actinomycetes>Actinomycetales | GCF_016626125.1 | Actinomyces_sp._HMT897 | JJJ15_07540 | 1917676..1921065 | 2789424 | DUF499-domain-containing-protein-[Actinomyces-sp.-HMT897] |
| WP_204423781.1 | HEPN+AAA-ATPase+Alpha+Beta-domain+wHTH+RRM | Type-1_BR.RM.like.systems | <-HEPN+DUF499*<-METHYLASE<-TUDOR+SF1-SF2-Helicase+iREase+CTD+REase | 1130 | Bacteria>Terrabacteria_group>Actinomycetota>Actinomycetes>Actinomycetales | GCF_016904675.1 | Arcanobacterium_phocisimile | JTE88_RS06650 | 1475854..1479246 | 1302235 | DUF499-domain-containing-protein-[Arcanobacterium-phocisimile] |
| WP_124913173.1 | HEPN+AAA-ATPase+Alpha+Beta-domain+wHTH+RRM | Type-1_BR.RM.like.systems | TUDOR+SF1-SF2-Helicase+iREase+CTD+REase->METHYLASE->HEPN+DUF499*-> | 1140 | Bacteria>Terrabacteria_group>Actinomycetota>Actinomycetes>Actinomycetales | GCF_003858505.1 | Buchananella_hordeovulneris | EII13_RS09935 | 35169..38591 | 52770 | DUF499-domain-containing-protein-[Buchananella-hordeovulneris] |
| WP_034502149.1 | HEPN+AAA-ATPase+Alpha+Beta-domain+wHTH+RRM | Type-1_BR.RM.like.systems | <-HEPN+DUF499*<-METHYLASE<-TUDOR+SF1-SF2-Helicase+iREase+CTD+REase | 1128 | Bacteria>Terrabacteria_group>Actinomycetota>Actinomycetes>Actinomycetales | GCF_000758755.1 | Actinomyces_sp._S6-Spd3 | HMPREF1627_RS06325 | 1474..4860 | 1284680 | DUF499-domain-containing-protein-[Actinomyces-sp.-S6-Spd3] |
| MBF0963210.1 | HEPN+AAA-ATPase+Alpha+Beta-domain+wHTH+RRM | Type-1_BR.RM.like.systems | <-HEPN+DUF499*<-TUDOR+SF1-SF2-Helicase+iREase+CTD+REase<-TraJ-RHH | 1136 | Bacteria>Terrabacteria_group>Actinomycetota>Actinomycetes>Actinomycetales | GCA_015259385.1 | Actinomyces_sp. | HXK13_05250 | 38..3448 | 29317 | ATP-binding-protein-[Actinomyces-sp.] |
| WP_075409839.1 | HEPN+AAA-ATPase+Alpha+Beta-domain+wHTH+RRM | Type-1_BR.RM.like.systems | ACET+ACET-><-?<-?<-DNA_methylase\|\|METHYLASE->HEPN+DUF499*-> | 826 | Bacteria>Terrabacteria_group>Actinomycetota>Actinomycetes>Actinomycetales | GCF_001937735.1 | Actinomyces_naeslundii | BKH13_RS04850 | 82823..85302 | 1655 | DUF499-domain-containing-protein,-partial-[Actinomyces-naeslundii] |
| WP_003781297.1 | HEPN+AAA-ATPase+Alpha+Beta-domain+wHTH+RRM | Type-1_BR.RM.like.systems | <-HEPN+DUF499*<-METHYLASE\|\|DNA_methylase-> | 1040 | Bacteria>Terrabacteria_group>Actinomycetota>Actinomycetes>Actinomycetales | GCF_000285995.1 | Actinomyces_naeslundii | HMPREF1129_RS04410 | 1..3119 | 1655 | DUF499-domain-containing-protein,-partial-[Actinomyces-naeslundii] |
| WP_214309728.1 | HEPN+AAA-ATPase+Alpha+Beta-domain+wHTH+RRM | Type-1_BR.RM.like.systems | Phage_integrase->PadR-HTH->SF1-SF2-Helicase-><-?<-?<-HEPN+DUF499*<-METHYLASE<-TUDOR+SF1-SF2-Helicase+iREase+CTD+REase<-Cluster2498_1clades<-PIN<-XIS-HTH<-Schlafen+HATPase+HTH | 1147 | Bacteria>Terrabacteria_group>Actinomycetota>Actinomycetes>Bifidobacteriales | GCF_018555415.1 | Bifidobacterium_simiarum | JS539_RS02125 | 175290..178733 | 2045441 | Swt1-family-HEPN-domain-containing-protein-[Bifidobacterium-simiarum] |
| WP_214383368.1 | HEPN+AAA-ATPase+Alpha+Beta-domain+wHTH+RRM | Type-1_BR.RM.like.systems | ABC-ATPase+SbcC->?->Cysteine_desulfurase->?-><-Primpol+D5N+D5ATPase<-HEPN+DUF499*<-METHYLASE<-METHYLASE-p<-TUDOR+SF1-SF2-Helicase+iREase+CTD+REase<-REase-p | 1134 | Bacteria>Terrabacteria_group>Actinomycetota>Actinomycetes>Bifidobacteriales | GCF_018555535.1 | Bifidobacterium_vespertilionis | JS538_RS07550 | 34017..37421 | 2562524 | Swt1-family-HEPN-domain-containing-protein-[Bifidobacterium-vespertilionis] |
| WP_150353887.1 | HEPN+AAA-ATPase+Alpha+Beta-domain+wHTH+RRM | Type-1_BR.RM.like.systems | REase-p->TUDOR+SF1-SF2-Helicase+iREase+CTD+REase->METHYLASE-p->METHYLASE->HEPN+DUF499*->Primpol+D5N+D5ATPase-><-?<-Cysteine_desulfurase<-?<-ABC-ATPase+SbcC | 1134 | Bacteria>Terrabacteria_group>Actinomycetota>Actinomycetes>Bifidobacteriales | GCF_008698145.1 | Bifidobacterium_vespertilionis | EM090_RS02025 | 149240..152644 | 2562524 | Swt1-family-HEPN-domain-containing-protein-[Bifidobacterium-vespertilionis] |
| WP_214375421.1 | HEPN+AAA-ATPase+Alpha+Beta-domain+wHTH+RRM | Type-1_BR.RM.like.systems | Phage_integrase-><-?\|\|PadR-HTH->SF1-SF2-Helicase-><-HEPN+DUF499*<-METHYLASE<-METHYLASE-p<-TUDOR+SF1-SF2-Helicase+iREase+CTD+REase<-Cluster2498_1clades | 1147 | Bacteria>Terrabacteria_group>Actinomycetota>Actinomycetes>Bifidobacteriales | GCF_018555455.1 | Bifidobacterium_colobi | JS530_RS01265 | 287140..290583 | 2809026 | Swt1-family-HEPN-domain-containing-protein-[Bifidobacterium-colobi] |
| WP_100496559.1 | HEPN+AAA-ATPase+Alpha+Beta-domain+wHTH+RRM | Type-1_BR.RM.like.systems | Phage_integrase->?->PadR-HTH->SF1-SF2-Helicase-><-HEPN+DUF499*<-METHYLASE<-TUDOR+SF1-SF2-Helicase+iREase+CTD+REase<-Cluster2498_1clades<-?\|\|BirA-HTH-> | 1148 | Bacteria>Terrabacteria_group>Actinomycetota>Actinomycetes>Bifidobacteriales | GCF_002802865.1 | Bifidobacterium_scaligerum | CUU80_RS06820 | 234586..238032 | 2052656 | DUF499-domain-containing-protein-[Bifidobacterium-scaligerum] |
| WP_107040639.1 | HEPN+AAA-ATPase+Alpha+Beta-domain+wHTH+RRM | Type-1_BR.RM.like.systems | <-ABC-ATPase+SbcC<-?<-?<-SF1-SF2-Helicase\|\|PRTase->?-><-HEPN+DUF499*<-METHYLASE<-METHYLASE-p<-TUDOR+SF1-SF2-Helicase+iREase+CTD+REase\|\|HTH-><-TraJ-RHH | 1158 | Bacteria>Terrabacteria_group>Actinomycetota>Actinomycetes>Bifidobacteriales | - | Bifidobacterium_callitrichos | COO72_RS01265 | 282673..286149 | 762209 | Swt1-family-HEPN-domain-containing-protein-[Bifidobacterium-callitrichos] |
| WP_118262158.1 | HEPN+AAA-ATPase+Alpha+Beta-domain+wHTH+RRM | Type-1_BR.RM.like.systems | Phage_integrase-><-Schlafen+HATPase+HTH<-?\|\|?-><-?\|\|<-?<-HEPN+DUF499*<-METHYLASE<-TUDOR+SF1-SF2-Helicase+iREase+CTD+REase<-TraJ-RHH\|\|HTH_Tnp-><-?\|\|SAM-methylase-> | 1129 | Bacteria>Terrabacteria_group>Actinomycetota>Actinomycetes>Bifidobacteriales | GCF_020541925.1 | Bifidobacterium_pseudocatenulatum | PGQ74_RS03980 | 281260..284649 | 28026 | DUF499-domain-containing-protein-[Bifidobacterium-pseudocatenulatum] |
| WP_129881947.1 | HEPN+AAA-ATPase+Alpha+Beta-domain+wHTH+RRM | Type-1_BR.RM.like.systems | <-REase<-ABC-ATPase+SbcC<-tRNA\|\|ACET-><-?<-HEPN+DUF499*<-METHYLASE<-TUDOR+SF1-SF2-Helicase+iREase+CTD+REase\|\|?-><-?<-ABC-ATPase+SbcC<-GNTR-HTH-p | 1127 | Bacteria>Terrabacteria_group>Actinomycetota>Actinomycetes>Bifidobacteriales | - | Bifidobacterium_animalis | PG1528B_RS04215 | 581191..584574 | 28025 | DUF499-domain-containing-protein-[Bifidobacterium-animalis] |
| MCI2185070.1 | HEPN+AAA-ATPase+Alpha+Beta-domain+wHTH+RRM | Type-1_BR.RM.like.systems | HTH+MPTase->?->TUDOR+SF1-SF2-Helicase+iREase+CTD+REase->METHYLASE->HEPN+DUF499*->SbcC+ABC-ATPase->Methyltransf12->?->URI-DNase+RAMA-> | 1149 | Bacteria>Terrabacteria_group>Actinomycetota>Actinomycetes>Bifidobacteriales | GCA_022655455.1 | Bifidobacterium_tibiigranuli | LKK60_02355 | 563467..566916 | 2172043 | DUF499-domain-containing-protein-[Bifidobacterium-tibiigranuli] |
| WP_245834842.1 | HEPN+AAA-ATPase+Alpha+Beta-domain+wHTH+RRM | Type-1_BR.RM.like.systems | <-JetD<-HTH\|\|?->?-><-HEPN+DUF499*<-METHYLASE<-?<-TUDOR+SF1-SF2-Helicase+iREase+CTD+REase<-ABC-ATPase+SbcC<-ABC-ATPase+SbcC<-MarR-HTH | 1130 | Bacteria>Terrabacteria_group>Actinomycetota>Actinomycetes>Bifidobacteriales | GCF_002259605.1 | Pseudoscardovia_radai | PSRA_RS02315 | 23959..27351 | 987066 | DUF499-domain-containing-protein-[Pseudoscardovia-radai] |
| WP_024464663.1 | HEPN+AAA-ATPase+Alpha+Beta-domain+wHTH+RRM | Type-1_BR.RM.like.systems | Cluster2439_3clades-><-HEPN\|\|TUDOR+SF1-SF2-Helicase+iREase+CTD+REase->METHYLASE-p->METHYLASE->HEPN+DUF499*->\|\|PAIREDC-HTH-> | 1163 | Bacteria>Terrabacteria_group>Actinomycetota>Actinomycetes>Bifidobacteriales | GCF_000426405.1 | Bifidobacterium_subtile | OUO0_RS10725 | 10278..13769 | 77635 | DUF499-domain-containing-protein-[Bifidobacterium-subtile] |
| WP_118284360.1 | HEPN+AAA-ATPase+Alpha+Beta-domain+wHTH+RRM | Type-1_BR.RM.like.systems | <-Schlafen+HATPase+HTH<-?\|\|?-><-?\|\|MULE-TRANSPOSASE-><-?<-HEPN+DUF499*<-METHYLASE<-TUDOR+SF1-SF2-Helicase+iREase+CTD+REase<-TraJ-RHH\|\|HTH_Tnp-> | 1129 | Bacteria>Terrabacteria_group>Actinomycetota>Actinomycetes>Bifidobacteriales | - | Bifidobacterium_pseudocatenulatum | DW149_RS06110 | 233918..237307 | 28026 | DUF499-domain-containing-protein-[Bifidobacterium-pseudocatenulatum] |
| WP_226678304.1 | HEPN+AAA-ATPase+Alpha+Beta-domain+wHTH+RRM | Type-1_BR.RM.like.systems | <-HTH_Tnp\|\|TraJ-RHH->TUDOR+SF1-SF2-Helicase+iREase+CTD+REase->METHYLASE->HEPN+DUF499*->?->\|\|?-><-?\|\|?->Schlafen+HATPase+HTH-><-Phage_integrase | 1129 | Bacteria>Terrabacteria_group>Actinomycetota>Actinomycetes>Bifidobacteriales | GCF_020526445.1 | Bifidobacterium_pseudocatenulatum | KZD94_RS04060 | 291413..294802 | 28026 | DUF499-domain-containing-protein-[Bifidobacterium-pseudocatenulatum] |
| WP_226595328.1 | HEPN+AAA-ATPase+Alpha+Beta-domain+wHTH+RRM | Type-1_BR.RM.like.systems | <-HTH_Tnp\|\|TraJ-RHH->TUDOR+SF1-SF2-Helicase+iREase+CTD+REase->METHYLASE->HEPN+DUF499*->?->\|\|?-><-?\|\|?->Schlafen+HATPase+HTH-><-Phage_integrase | 1129 | Bacteria>Terrabacteria_group>Actinomycetota>Actinomycetes>Bifidobacteriales | GCF_020526365.1 | Bifidobacterium_pseudocatenulatum | KZP21_RS03935 | 291458..294847 | 28026 | DUF499-domain-containing-protein-[Bifidobacterium-pseudocatenulatum] |
| GDZ11402.1 | HEPN+AAA-ATPase+Alpha+Beta-domain+wHTH+RRM | Type-1_BR.RM.like.systems | MULE-TRANSPOSASE-><-?<-HEPN+DUF499*<-METHYLASE<-METHYLASE-p<-TUDOR+SF1-SF2-Helicase+iREase+CTD+REase<-TraJ-RHH\|\|HTH_Tnp-> | 1129 | Bacteria>Terrabacteria_group>Actinomycetota>Actinomycetes>Bifidobacteriales | GCA_018326745.1 | Bifidobacteriaceae_bacterium_MCC01993 | MCC01993_15150 | 22460..25849 | 2569496 | hypothetical-protein-MCC01993_15150-[Bifidobacteriaceae-bacterium-MCC01993] |
| WP_214376699.1 | HEPN+AAA-ATPase+Alpha+Beta-domain+wHTH+RRM | Type-1_BR.RM.like.systems | <-HTH_3<-?<-?\|\|Calcineurin->?-><-HEPN+DUF499*<-METHYLASE<-METHYLASE-p<-TUDOR+SF1-SF2-Helicase+iREase+CTD+REase<-FtsK\|\|DOC-> | 1148 | Bacteria>Terrabacteria_group>Actinomycetota>Actinomycetes>Bifidobacteriales | GCF_018555455.1 | Bifidobacterium_colobi | JS530_RS08230 | 76001..79447 | 2809026 | Swt1-family-HEPN-domain-containing-protein-[Bifidobacterium-colobi] |
| OZG52261.1 | HEPN+AAA-ATPase+Alpha+Beta-domain+wHTH+RRM | Type-1_BR.RM.like.systems | <-JetD<-HTH\|\|?->?->?-><-HEPN+DUF499*<-METHYLASE<-?<-TUDOR+SF1-SF2-Helicase+iREase+CTD+REase<-?<-ABC-ATPase+SbcC<-ABC-ATPase+SbcC | 1139 | Bacteria>Terrabacteria_group>Actinomycetota>Actinomycetes>Bifidobacteriales | GCA_002259605.1 | Pseudoscardovia_radai | PSRA_0450 | 23959..27378 | 987066 | ATPase-AAA-[Pseudoscardovia-radai] |
| WP_130013693.1 | HEPN+AAA-ATPase+Alpha+Beta-domain+wHTH+RRM | Type-1_BR.RM.like.systems | <-GDResolvase+HIN-HTH\|\|Mbetalac->TUDOR+SF1-SF2-Helicase+iREase+CTD+REase->METHYLASE->HEPN+DUF499*->?-><-URI-DNase | 1133 | Bacteria>Terrabacteria_group>Actinomycetota>Actinomycetes>Bifidobacteriales | GCF_004155015.1 | Bifidobacterium_pseudolongum | PG2054B_RS08290 | 18590..21991 | 1694 | DUF499-domain-containing-protein-[Bifidobacterium-pseudolongum] |
| WP_195552455.1 | HEPN+AAA-ATPase+Alpha+Beta-domain+wHTH+RRM | Type-1_BR.RM.like.systems | <-Phage_int_SAM_1+Phage_integrase\|\|?->?->?->?-><-HEPN+DUF499*<-METHYLASE<-TUDOR+SF1-SF2-Helicase+iREase+CTD+REase\|\|?->HEPN-> | 1129 | Bacteria>Terrabacteria_group>Actinomycetota>Actinomycetes>Bifidobacteriales | GCF_015559175.1 | Bifidobacterium_pseudocatenulatum | I2G66_RS01750 | 59559..62948 | 28026 | DUF499-domain-containing-protein-[Bifidobacterium-pseudocatenulatum] |
| MCI2174819.1 | HEPN+AAA-ATPase+Alpha+Beta-domain+wHTH+RRM | Type-1_BR.RM.like.systems | <-ABC-ATPase+SbcC\|\|Peptidase-><-?<-TetR-HTH<-HEPN+DUF499*<-METHYLASE<-?<-TUDOR+SF1-SF2-Helicase+iREase+CTD+REase | 1134 | Bacteria>Terrabacteria_group>Actinomycetota>Actinomycetes>Bifidobacteriales | GCA_022655115.1 | Bifidobacterium_thermacidophilum | LKK47_03040 | 24047..27451 | 246618 | DUF499-domain-containing-protein-[Bifidobacterium-thermacidophilum] |
| WP_214310646.1 | HEPN+AAA-ATPase+Alpha+Beta-domain+wHTH+RRM | Type-1_BR.RM.like.systems | <-JetB<-?<-?\|\|Phage_holin_4_2->?-><-HEPN+DUF499*<-METHYLASE<-METHYLASE-p<-?<-?<-TUDOR+SF1-SF2-Helicase+iREase+CTD+REase | 1147 | Bacteria>Terrabacteria_group>Actinomycetota>Actinomycetes>Bifidobacteriales | GCF_018555415.1 | Bifidobacterium_simiarum | JS539_RS08840 | 19593..23036 | 2045441 | Swt1-family-HEPN-domain-containing-protein-[Bifidobacterium-simiarum] |
| WP_126031245.1 | HEPN+AAA-ATPase+Alpha+Beta-domain+wHTH+RRM | Type-1_BR.RM.like.systems | <-ABC-ATPase+SbcC<-tRNA\|\|ACET-><-HEPN+DUF499*<-METHYLASE<-METHYLASE-p<-TUDOR+SF1-SF2-Helicase+iREase+CTD+REase | 1127 | Bacteria>Terrabacteria_group>Actinomycetota>Actinomycetes>Bifidobacteriales | GCF_003952025.1 | Bifidobacterium_castoris | D2E22_RS00925 | 206849..210232 | 2306972 | DUF499-domain-containing-protein-[Bifidobacterium-castoris] |
| WP_278801769.1 | HEPN+AAA-ATPase+Alpha+Beta-domain+wHTH+RRM | Type-1_BR.RM.like.systems | <-SF1-SF2-Helicase<-?<-WYL<-?<-HEPN+DUF499*<-METHYLASE<-TUDOR+SF1-SF2-Helicase+iREase+CTD+REase<-?\|\|RelB-><-tRNA<-sGTP | 1135 | Bacteria>Terrabacteria_group>Actinomycetota>Actinomycetes>Bifidobacteriales | GCF_944322225.1 | Bifidobacterium_pullorum | QDJ94_RS03760 | 68735..72142 | 78448 | DUF499-domain-containing-protein-[Bifidobacterium-pullorum] |
| WP_198229240.1 | HEPN+AAA-ATPase+Alpha+Beta-domain+wHTH+RRM | Type-1_BR.RM.like.systems | NUDIX->TetR-HTH->ABC-ATPase+SbcC->?->?-><-HEPN+DUF499*<-METHYLASE<-TUDOR+SF1-SF2-Helicase+iREase+CTD+REase | 1150 | Bacteria>Terrabacteria_group>Actinomycetota>Actinomycetes>Bifidobacteriales | GCF_016101585.1 | Bifidobacterium_polysaccharolyticum | H3T90_RS00470 | 119489..122941 | 2750967 | DUF499-domain-containing-protein-[Bifidobacterium-polysaccharolyticum] |
| WP_003839950.1 | HEPN+AAA-ATPase+Alpha+Beta-domain+wHTH+RRM | Type-1_BR.RM.like.systems | Aldedh->?->?->PRTase->tRNA->TUDOR+SF1-SF2-Helicase+iREase+CTD+REase->METHYLASE->HEPN+DUF499*->?->HNH->?->?-><-Peptidase | 1138 | Bacteria>Terrabacteria_group>Actinomycetota>Actinomycetes>Bifidobacteriales | GCF_000024445.1 | Bifidobacterium_dentium | BIFDEN_RS08405 | 2078532..2081948 | 1689 | DUF499-domain-containing-protein-[Bifidobacterium-dentium] |
| WP_214305620.1 | HEPN+AAA-ATPase+Alpha+Beta-domain+wHTH+RRM | Type-1_BR.RM.like.systems | TUDOR+SF1-SF2-Helicase+iREase+CTD+REase->METHYLASE-p->METHYLASE->HEPN+DUF499*->ABhydrolase-> | 1154 | Bacteria>Terrabacteria_group>Actinomycetota>Actinomycetes>Bifidobacteriales | GCF_018555655.1 | Bifidobacterium_callimiconis | JS532_RS00845 | 238975..242439 | 2306973 | DUF499-domain-containing-protein-[Bifidobacterium-callimiconis] |
| WP_100513170.1 | HEPN+AAA-ATPase+Alpha+Beta-domain+wHTH+RRM | Type-1_BR.RM.like.systems | TUDOR+SF1-SF2-Helicase+iREase+CTD+REase->?->METHYLASE->HEPN+DUF499*-><-?<-Phage_holin_4_2\|\|?->?->JetB-> | 1147 | Bacteria>Terrabacteria_group>Actinomycetota>Actinomycetes>Bifidobacteriales | GCF_002802905.1 | Bifidobacterium_simiarum | CSQ87_RS07025 | 60477..63920 | 2045441 | Swt1-family-HEPN-domain-containing-protein-[Bifidobacterium-simiarum] |
| WP_152234686.1 | HEPN+AAA-ATPase+Alpha+Beta-domain+wHTH+RRM | Type-1_BR.RM.like.systems | TUDOR+SF1-SF2-Helicase+iREase+CTD+REase->?->TraJ-RHH->METHYLASE-p->METHYLASE->HEPN+DUF499*-> | 1149 | Bacteria>Terrabacteria_group>Actinomycetota>Actinomycetes>Bifidobacteriales | GCF_009193355.1 | Bifidobacterium_leontopitheci | F7D09_RS07000 | 42381..45830 | 2650774 | Swt1-family-HEPN-domain-containing-protein-[Bifidobacterium-leontopitheci] |
| WP_150395254.1 | HEPN+AAA-ATPase+Alpha+Beta-domain+wHTH+RRM | Type-1_BR.RM.like.systems | TUDOR+SF1-SF2-Helicase+iREase+CTD+REase->METHYLASE->HEPN+DUF499*->ABhydrolase-> | 1156 | Bacteria>Terrabacteria_group>Actinomycetota>Actinomycetes>Bifidobacteriales | GCF_008698195.1 | Bifidobacterium_tissieri | EM849_RS05800 | 47872..51342 | 1630162 | DUF499-domain-containing-protein-[Bifidobacterium-tissieri] |
| WP_150380554.1 | HEPN+AAA-ATPase+Alpha+Beta-domain+wHTH+RRM | Type-1_BR.RM.like.systems | <-ABhydrolase<-HEPN+DUF499*<-METHYLASE<-TUDOR+SF1-SF2-Helicase+iREase+CTD+REase | 1156 | Bacteria>Terrabacteria_group>Actinomycetota>Actinomycetes>Bifidobacteriales | GCF_008698165.1 | Bifidobacterium_tissieri | EM089_RS01160 | 325418..328888 | 1630162 | DUF499-domain-containing-protein-[Bifidobacterium-tissieri] |
| MDE5641417.1 | HEPN+AAA-ATPase+Alpha+Beta-domain+wHTH+RRM | Type-1_BR.RM.like.systems | METHYLASE-p->METHYLASE-p->HEPN+DUF499*-> | 583 | Bacteria>Terrabacteria_group>Actinomycetota>Actinomycetes>Bifidobacteriales | GCA_029051405.1 | Bifidobacterium_castoris | K2I40_06975 | 1749..3499 | 2306972 | MAG:-DUF499-domain-containing-protein,-partial-[Bifidobacterium-castoris] |
| MDD6374029.1 | HEPN+AAA-ATPase+Alpha+Beta-domain+wHTH+RRM | Type-1_BR.RM.like.systems | METHYLASE->HEPN+DUF499*-> | 1120 | Bacteria>Terrabacteria_group>Actinomycetota>Actinomycetes>Bifidobacteriales | GCA_029338855.1 | Bifidobacteriaceae_bacterium | PUF97_06115 | 2208..5567 | 2291324 | MAG:-DUF499-domain-containing-protein,-partial-[Bifidobacteriaceae-bacterium] |
| MBO5627607.1 | HEPN+AAA-ATPase+Alpha+Beta-domain+wHTH+RRM | Type-1_BR.RM.like.systems | <-HEPN+DUF499*<-METHYLASE-p | 467 | Bacteria>Terrabacteria_group>Actinomycetota>Actinomycetes>Bifidobacteriales | GCA_017627815.1 | Aeriscardovia_sp. | J5965_00830 | 1..1400 | 1872397 | DUF499-domain-containing-protein-[Aeriscardovia-sp.] |
| WP_194891200.1 | HEPN+AAA-ATPase+Alpha+Beta-domain+wHTH+RRM | Type-1_BR.RM.like.systems | <-PAIREDC-HTH\|\|PAIREDC-HTH-><-?\|\|?->PAIREDC-HTH->HEPN-><-HEPN+DUF499*<-METHYLASE<-TUDOR+SF1-SF2-Helicase+iREase+CTD+REase\|\|?-><-?<-?\|\|PAIREDC-HTH-> | 1134 | Bacteria>Terrabacteria_group>Actinomycetota>Actinomycetes>Catenulisporales | GCF_015356215.1 | Catenulispora_pinisilvae | IZR00_RS01870 | 88900..92304 | 2705253 | DUF499-domain-containing-protein-[Catenulispora-pinisilvae] |
| WP_009738155.1 | HEPN+AAA-ATPase+Alpha+Beta-domain+wHTH+RRM | Type-1_BR.RM.like.systems | TUDOR+SF1-SF2-Helicase+iREase+CTD+REase->UDG-conflict->?->SF1-SF2-Helicase+REase->METHYLASE->HEPN+DUF499*-><-ParA+HTH<-RelE-ParE\|\|MULE-TRANSPOSASE->MULE-TRANSPOSASE->?-><-TPR+TPR+TPR+TPR+TPR+TPR+TPR+TPR | 1119 | Bacteria>Terrabacteria_group>Actinomycetota>Actinomycetes>Frankiales | GCF_000262465.1 | Frankia_sp._QA3 | FRAQA3DRAFT_RS05925 | 1493291..1496650 | 710111 | Swt1-family-HEPN-domain-containing-protein-[Frankia-sp.-QA3] |
| TCJ34611.1 | HEPN+AAA-ATPase+Alpha+Beta-domain+wHTH+RRM | Type-1_BR.RM.like.systems | <-ParA-Soj-PloopNTPase<-REase\|\|TUDOR+SF1-SF2-Helicase+iREase+CTD+REase->METHYLASE->HEPN+DUF499*-><-HTH+UbE1<-?<-SbcC\|\|HTH_3->MPTase-><-REase | 1116 | Bacteria>Terrabacteria_group>Actinomycetota>Actinomycetes>Frankiales | GCA_004337455.1 | Parafrankia_sp._BMG5.11 | E0504_31915 | 85407..88757 | 222540 | ATP-binding-protein-[Frankia-sp.-BMG5.11] |
| WP_203013794.1 | HEPN+AAA-ATPase+Alpha+Beta-domain+wHTH+RRM | Type-1_BR.RM.like.systems | <-NTD+SF1-SF2-Helicase\|\|TUDOR+SF1-SF2-Helicase+iREase+CTD+REase->UDG-conflict->METHYLASE->HEPN+DUF499*-><-?<-?\|\|?->?-><-?<-TetR-HTH | 1128 | Bacteria>Terrabacteria_group>Actinomycetota>Actinomycetes>Frankiales | GCF_016786345.1 | unclassified_Frankia | I6A60_RS06180 | 96819..100205 | 2632575 | MULTISPECIES:-Swt1-family-HEPN-domain-containing-protein-[unclassified-Frankia] |
| WP_248817712.1 | HEPN+AAA-ATPase+Alpha+Beta-domain+wHTH+RRM | Type-1_BR.RM.like.systems | TUDOR+SF1-SF2-Helicase+iREase+CTD+REase->UDG-conflict->METHYLASE->HEPN+DUF499*-><-ncRNA\|\|?->Pkinase-> | 1126 | Bacteria>Terrabacteria_group>Actinomycetota>Actinomycetes>Frankiales | GCF_023243045.1 | Frankia_sp._AgB32 | MXD60_RS04685 | 52343..55723 | 631119 | Swt1-family-HEPN-domain-containing-protein-[Frankia-sp.-AgB32] |
| WP_251744665.1 | HEPN+AAA-ATPase+Alpha+Beta-domain+wHTH+RRM | Type-1_BR.RM.like.systems | TUDOR+SF1-SF2-Helicase+iREase+CTD+REase->METHYLASE->HEPN+DUF499*-><-?<-ParA+HTH<-RelE-ParE | 1122 | Bacteria>Terrabacteria_group>Actinomycetota>Actinomycetes>Frankiales | GCF_023716945.1 | Frankia_sp._AiPs1 | ND748_RS06005 | 10061..13429 | 573493 | Swt1-family-HEPN-domain-containing-protein-[Frankia-sp.-AiPs1] |
| WP_261570743.1 | HEPN+AAA-ATPase+Alpha+Beta-domain+wHTH+RRM | Type-1_BR.RM.like.systems | METHYLASE->HEPN+DUF499*-><-?\|\|DDE_5+DDE_Tnp_1->?->?-><-TPR+TPR+TPR+TPR+TPR+TPR+TPR+TPR | 1120 | Bacteria>Terrabacteria_group>Actinomycetota>Actinomycetes>Frankiales | GCF_025403355.1 | Frankia_gtarii | NBZ84_RS30015 | 1969..5331 | 2950102 | Swt1-family-HEPN-domain-containing-protein-[Frankia-gtarii] |
| WP_248814552.1 | HEPN+AAA-ATPase+Alpha+Beta-domain+wHTH+RRM | Type-1_BR.RM.like.systems | METHYLASE-p->HEPN+DUF499*->?-><-TPR+TPR+TPR+TPR+TPR+TPR+TPR | 1122 | Bacteria>Terrabacteria_group>Actinomycetota>Actinomycetes>Frankiales | GCF_023243055.1 | Frankia_sp._AgPm24 | MXD61_RS17500 | 1626..4994 | 631128 | Swt1-family-HEPN-domain-containing-protein-[Frankia-sp.-AgPm24] |
| WP_113812542.1 | HEPN+AAA-ATPase+Alpha+Beta-domain+wHTH+RRM | Type-1_BR.RM.like.systems | <-ABhydrolase\|\|TUDOR+SF1-SF2-Helicase+iREase+CTD+REase->Schlafen+HATPase+HTH->METHYLASE->ParB+HNH->?->HEPN+DUF499*->NTD+SF1-SF2-Helicase->wHTH(SBD)+HNH-><-?<-?\|\|?-><-REase+35exo+Beta-Barrel+SF1-SF2-Helicase | 1114 | Bacteria>Terrabacteria_group>Actinomycetota>Actinomycetes>Geodermatophilales | GCF_003319095.1 | Blastococcus_sp._TBT05-19 | DQ244_RS16250 | 131306..134650 | 2250581 | Swt1-family-HEPN-domain-containing-protein-[Blastococcus-sp.-TBT05-19] |
| SNR82321.1 | HEPN+AAA-ATPase+Alpha+Beta-domain+wHTH+RRM | Type-1_BR.RM.like.systems | <-DNA-glycosylase\|\|?-><-METHYLASE\|\|?->TUDOR+SF1-SF2-Helicase+iREase+CTD+REase->METHYLASE->HEPN+DUF499*->tRNA-><-Arm-DNA-bind_4+Phage_int_SAM_3+Phage_integrase<-ZDNA-HTH\|\|HTH_3->?->?->DnaB-> | 1132 | Bacteria>Terrabacteria_group>Actinomycetota>Actinomycetes>Geodermatophilales | GCA_900188205.1 | Geodermatophilus_saharensis | SAMN04488107_0059 | 63726..67124 | 1137994 | Predicted-ATPase,-AAA+-superfamily-[Geodermatophilus-saharensis] |
| WP_236827009.1 | HEPN+AAA-ATPase+Alpha+Beta-domain+wHTH+RRM | Type-1_BR.RM.like.systems | HKD-DNase+SF1-SF2-Helicase+TRD->?-><-SF1-SF2-Helicase<-?\|\|TUDOR+SF1-SF2-Helicase+iREase+CTD+REase->ArdCN+MPTase->METHYLASE->HEPN+DUF499*->ParB+HNH-> | 1109 | Bacteria>Terrabacteria_group>Actinomycetota>Actinomycetes>Geodermatophilales | GCF_021698995.1 | unclassified_Blastococcus | E9565_RS02685 | 57900..61229 | 2619396 | MULTISPECIES:-Swt1-family-HEPN-domain-containing-protein-[unclassified-Blastococcus] |
| WP_176449778.1 | HEPN+AAA-ATPase+Alpha+Beta-domain+wHTH+RRM | Type-1_BR.RM.like.systems | DDE_Tnp_1_4-><-DNA-glycosylase<-METHYLASE\|\|?->TUDOR+SF1-SF2-Helicase+iREase+CTD+REase->METHYLASE->HEPN+DUF499*->tRNA->?-><-ZDNA-HTH\|\|HTH_3->?->DnaB-> | 1120 | Bacteria>Terrabacteria_group>Actinomycetota>Actinomycetes>Geodermatophilales | GCF_900188205.1 | Geodermatophilus_saharensis | CHC04_RS00285 | 63762..67124 | 1137994 | DUF499-domain-containing-protein-[Geodermatophilus-saharensis] |
| RBY97952.1 | HEPN+AAA-ATPase+Alpha+Beta-domain+wHTH+RRM | Type-1_BR.RM.like.systems | <-NUDIX\|\|?-><-ABhydrolase<-DNA_methylase\|\|REase->METHYLASE->HEPN+DUF499*->MORC->REase->HTH+SF1-SF2-Helicase+REase-> | 1128 | Bacteria>Terrabacteria_group>Actinomycetota>Actinomycetes>Geodermatophilales | GCA_003319175.1 | Blastococcus_sp._TF02-8 | DQ237_03405 | 728169..731555 | 2250574 | AAA-family-ATPase-[Blastococcus-sp.-TF02-8] |
| WP_205730045.1 | HEPN+AAA-ATPase+Alpha+Beta-domain+wHTH+RRM | Type-1_BR.RM.like.systems | <-ABhydrolase<-DNA_methylase\|\|REase->METHYLASE->HEPN+DUF499*->MORC->REase->HTH+SF1-SF2-Helicase+REase-> | 1113 | Bacteria>Terrabacteria_group>Actinomycetota>Actinomycetes>Geodermatophilales | GCF_003319175.1 | Blastococcus_sp._TF02-8 | DQ237_RS03405 | 728214..731555 | 2250574 | Swt1-family-HEPN-domain-containing-protein-[Blastococcus-sp.-TF02-8] |
| WP_284493306.1 | HEPN+AAA-ATPase+Alpha+Beta-domain+wHTH+RRM | Type-1_BR.RM.like.systems | <-HTH_19\|\|TUDOR+SF1-SF2-Helicase+iREase+CTD+REase-><-?<-?<-?\|\|?->HEPN+DUF499*->DOC+HTH-> | 1121 | Bacteria>Terrabacteria_group>Actinomycetota>Actinomycetes>Geodermatophilales | GCF_030165005.1 | Blastococcus_capsensis | QOZ86_RS17755 | 63627..66992 | 1564163 | Swt1-family-HEPN-domain-containing-protein-[Blastococcus-capsensis] |
| WP_013016869.1 | HEPN+AAA-ATPase+Alpha+Beta-domain+wHTH+RRM | Type-1_BR.RM.like.systems | MoxR-AAA->?->?->?-><-REase\|\|METHYLASE->HEPN+DUF499*-><-TUDOR+SF1-SF2-Helicase+iREase+CTD+REase | 1109 | Bacteria>Terrabacteria_group>Actinomycetota>Actinomycetes>Glycomycetales | GCF_000024545.1 | Stackebrandtia_nassauensis | SNAS_RS07855 | 1675315..1678644 | 283811 | DUF499-domain-containing-protein-[Stackebrandtia-nassauensis] |
| MBK7622561.1 | HEPN+AAA-ATPase+Alpha+Beta-domain+wHTH+RRM | Type-1_BR.RM.like.systems | Phage_int_SAM_1+Phage_integrase->?-><-HTH_Tnp_1+HTH_21+rve+rve_2<-HEPN+DUF499*<-METHYLASE\|\|TUDOR+SF1-SF2-Helicase+iREase+CTD+REase-><-UDG-conflict | 1150 | Bacteria>Terrabacteria_group>Actinomycetota>Actinomycetes>Kineosporiales | GCA_016709945.1 | Kineosporiaceae_bacterium | IPJ14_07830 | 115869..119321 | 2487580 | DUF499-domain-containing-protein-[Kineosporiaceae-bacterium] |
| MBK6873560.1 | HEPN+AAA-ATPase+Alpha+Beta-domain+wHTH+RRM | Type-1_BR.RM.like.systems | UDG-conflict-><-TUDOR+SF1-SF2-Helicase+iREase+CTD+REase\|\|METHYLASE->HEPN+DUF499*->HTH_Tnp_1+HTH_21+rve+rve_2-><-?<-?<-Phage_integrase | 1150 | Bacteria>Terrabacteria_group>Actinomycetota>Actinomycetes>Kineosporiales | GCA_016702615.1 | Kineosporiaceae_bacterium | IPG94_19890 | 267597..271049 | 2487580 | DUF499-domain-containing-protein-[Kineosporiaceae-bacterium] |
| WP_136222929.1 | HEPN+AAA-ATPase+Alpha+Beta-domain+wHTH+RRM | Type-1_BR.RM.like.systems | <-Radical-SAM<-?<-?\|\|HTH_3-><-?\|\|?-><-HEPN+DUF499*<-METHYLASE<-TUDOR+SF1-SF2-Helicase+iREase+CTD+REase\|\|?->XIS-HTH->Phage_int_SAM_1+Phage_integrase-><-DUF4011+SF1-SF2-Helicase+REase | 1135 | Bacteria>Terrabacteria_group>Actinomycetota>Actinomycetes>Kitasatosporales | GCF_004794975.1 | Streptomyces_sp._A1547 | E6W17_RS04020 | 256183..259590 | 2563105 | Swt1-family-HEPN-domain-containing-protein-[Streptomyces-sp.-A1547] |
| WP_199584383.1 | HEPN+AAA-ATPase+Alpha+Beta-domain+wHTH+RRM | Type-1_BR.RM.like.systems | DNA-glycosylase-><-HNH<-Phage_holin<-DUF6731<-HEPN+DUF499*<-METHYLASE<-TUDOR+SF1-SF2-Helicase+iREase+CTD+REase<-IclR-HTH\|\|?-><-HTH_3\|\|Phage_integrase+Phage_integrase-> | 1117 | Bacteria>Terrabacteria_group>Actinomycetota>Actinomycetes>Kitasatosporales | GCF_016482685.1 | Streptomyces_sp._DSM_110735 | IF655_RS11870 | 37375..40728 | 2775031 | DUF499-domain-containing-protein-[Streptomyces-sp.-DSM-110735] |
| WP_050490834.1 | HEPN+AAA-ATPase+Alpha+Beta-domain+wHTH+RRM | Type-1_BR.RM.like.systems | ACET-><-?\|\|<-?<-?\|\|?-><-DUF4096+DDE_Tnp_1+DDE_Tnp_1_2<-HEPN+DUF499*<-METHYLASE<-TUDOR+SF1-SF2-Helicase+iREase+CTD+REase<-ZDNA-HTH\|\|MORC->Phage_terminase-><-TRAM+Methyltransf_31 | 1115 | Bacteria>Terrabacteria_group>Actinomycetota>Actinomycetes>Kitasatosporales | GCF_000718915.1 | Streptomyces | ADK33_RS30095 | 165445..168792 | 1883 | MULTISPECIES:-DUF499-domain-containing-protein-[Streptomyces] |
| WP_093789469.1 | HEPN+AAA-ATPase+Alpha+Beta-domain+wHTH+RRM | Type-1_BR.RM.like.systems | MORC+HisKin+REase-><-Phage_holin<-DUF6731<-HEPN+DUF499*<-METHYLASE<-TUDOR+SF1-SF2-Helicase+iREase+CTD+REase\|\|VapB_antitoxin->PIN-><-DUF397<-HTH+DUF5753 | 1139 | Bacteria>Terrabacteria_group>Actinomycetota>Actinomycetes>Kitasatosporales | GCF_900114955.1 | Streptomyces_sp._cf124 | BM134_RS11480 | 95921..99340 | 1761903 | DUF499-domain-containing-protein-[Streptomyces-sp.-cf124] |
| WP_210966257.1 | HEPN+AAA-ATPase+Alpha+Beta-domain+wHTH+RRM | Type-1_BR.RM.like.systems | HTH_Tnp_4+DDE_Tnp_4-><-?<-?<-?<-Phage_holin<-DUF6731<-HEPN+DUF499*<-METHYLASE<-TUDOR+SF1-SF2-Helicase+iREase+CTD+REase<-ST-Hiskin\|\|HTH+DUF5753->DUF397-><-REase | 1129 | Bacteria>Terrabacteria_group>Actinomycetota>Actinomycetes>Kitasatosporales | GCF_018070605.1 | Streptomyces_sp._B15 | KBY19_RS19140 | 34073..37462 | 1537797 | DUF499-domain-containing-protein-[Streptomyces-sp.-B15] |
| WP_189838087.1 | HEPN+AAA-ATPase+Alpha+Beta-domain+wHTH+RRM | Type-1_BR.RM.like.systems | <-FtsK<-?\|\|HTH_3-><-NUDIX\|\|TUDOR+SF1-SF2-Helicase+iREase+CTD+REase->METHYLASE->HEPN+DUF499*->DUF6731-><-?\|\|GNTR-HTH->HTH_Tnp_4+DDE_Tnp_4->Methyltransf-> | 1150 | Bacteria>Terrabacteria_group>Actinomycetota>Actinomycetes>Kitasatosporales | GCF_014650195.1 | Streptomyces_umbrinus | IE284_RS09050 | 2097992..2101444 | 67370 | DUF499-domain-containing-protein-[Streptomyces-umbrinus] |
| WP_043673297.1 | HEPN+AAA-ATPase+Alpha+Beta-domain+wHTH+RRM | Type-1_BR.RM.like.systems | <-NTP_transferase<-?<-ABC-ATPase+SbcC\|\|tRNA->?->Phage_cap_E->UDG-conflict-><-HEPN+DUF499*<-METHYLASE<-TUDOR+SF1-SF2-Helicase+iREase+CTD+REase<-?<-REase<-SLATT | 1146 | Bacteria>Terrabacteria_group>Actinomycetota>Actinomycetes>Kitasatosporales | GCF_000716435.1 | Streptomyces_xylophagus | OO70_RS20695 | 140518..143958 | 285514 | Swt1-family-HEPN-domain-containing-protein-[Streptomyces-xylophagus] |
| WP_124289019.1 | HEPN+AAA-ATPase+Alpha+Beta-domain+wHTH+RRM | Type-1_BR.RM.like.systems | <-Pkinase<-AdoMet_Synthase<-Pkinase\|\|HTH+TPR+TPR->?-><-HEPN+DUF499*<-METHYLASE<-TUDOR+SF1-SF2-Helicase+iREase+CTD+REase<-?<-REase<-SF1-SF2-Helicase | 1148 | Bacteria>Terrabacteria_group>Actinomycetota>Actinomycetes>Kitasatosporales | GCF_003846165.1 | Streptomyces_sp._ADI91-18 | EES37_RS07515 | 1680083..1683529 | 1522755 | DUF499-domain-containing-protein-[Streptomyces-sp.-ADI91-18] |
| WP_257563884.1 | HEPN+AAA-ATPase+Alpha+Beta-domain+wHTH+RRM | Type-1_BR.RM.like.systems | REase->?->SF1-SF2-Helicase->TUDOR+SF1-SF2-Helicase+iREase+CTD+REase->METHYLASE->HEPN+DUF499*-><-DUF397<-HTH+DUF5753\|\|ST-Hiskin->?-><-PIN<-TraJ-RHH | 1137 | Bacteria>Terrabacteria_group>Actinomycetota>Actinomycetes>Kitasatosporales | GCF_024612035.1 | Streptomyces_sp._DSM_40750 | JIX55_RS15360 | 3457896..3461309 | 2801030 | Swt1-family-HEPN-domain-containing-protein-[Streptomyces-sp.-DSM-40750] |
| WP_202237096.1 | HEPN+AAA-ATPase+Alpha+Beta-domain+wHTH+RRM | Type-1_BR.RM.like.systems | <-DDE_Tnp<-?\|\|?->TUDOR+SF1-SF2-Helicase+iREase+CTD+REase->METHYLASE->HEPN+DUF499*->DUF6731->Phage_holin-><-?<-HTH+DUF5753\|\|HTH+DUF5753->METHYLASE-p-> | 1126 | Bacteria>Terrabacteria_group>Actinomycetota>Actinomycetes>Kitasatosporales | GCF_016756395.1 | Streptomyces_sp._SN-593 | RVR_RS30315 | 7009804..7013184 | 659352 | DUF499-domain-containing-protein-[Streptomyces-sp.-SN-593] |
| WP_136741406.1 | HEPN+AAA-ATPase+Alpha+Beta-domain+wHTH+RRM | Type-1_BR.RM.like.systems | HAMP+HISKIN-><-SF1-SF2-Helicase+HRDC<-tRNA<-Phage_holin<-DUF6731<-HEPN+DUF499*<-METHYLASE<-TUDOR+SF1-SF2-Helicase+iREase+CTD+REase<-?<-?<-?<-tRNA\|\|METHYLASE-p-> | 1124 | Bacteria>Terrabacteria_group>Actinomycetota>Actinomycetes>Kitasatosporales | GCF_005048155.1 | Streptomyces_piniterrae | FCH28_RS19960 | 322483..325857 | 2571125 | DUF499-domain-containing-protein-[Streptomyces-piniterrae] |
| MBX7473484.1 | HEPN+AAA-ATPase+Alpha+Beta-domain+wHTH+RRM | Type-1_BR.RM.like.systems | HTH_3->REase->?->TUDOR+SF1-SF2-Helicase+iREase+CTD+REase->METHYLASE->HEPN+DUF499*-><-UDG-conflict\|\|?->?->Phage_int_SAM_1->DDE_Tnp_1_4->DDE_Tnp_1_4-> | 1146 | Bacteria>Terrabacteria_group>Actinomycetota>Actinomycetes>Kitasatosporales | GCA_019711445.1 | Streptomyces_sp._MAG02 | K1Y80_46530 | 10650..14090 | 2866213 | DUF499-domain-containing-protein-[Streptomyces-sp.-MAG02] |
| WP_158953066.1 | HEPN+AAA-ATPase+Alpha+Beta-domain+wHTH+RRM | Type-1_BR.RM.like.systems | <-SIGMA-HTH<-ST-Hiskin<-HTH+DUF5753\|\|TUDOR+SF1-SF2-Helicase+iREase+CTD+REase->METHYLASE->HEPN+DUF499*->DUF6731-><-Phage_integrase<-XIS-HTH | 1150 | Bacteria>Terrabacteria_group>Actinomycetota>Actinomycetes>Kitasatosporales | GCF_033792825.1 | Streptomyces_scabiei | AA092_RS33625 | 12095..15547 | 1930 | DUF499-domain-containing-protein-[Streptomyces-scabiei] |
| GGS96436.1 | HEPN+AAA-ATPase+Alpha+Beta-domain+wHTH+RRM | Type-1_BR.RM.like.systems | <-ST-Hiskin\|\|HTH+DUF5753->HTH+DUF5753->DUF397-><-HEPN+DUF499*<-METHYLASE<-TUDOR+SF1-SF2-Helicase+iREase+CTD+REase<-SF1-SF2-Helicase<-?<-REase | 1137 | Bacteria>Terrabacteria_group>Actinomycetota>Actinomycetes>Kitasatosporales | GCA_014649155.1 | Streptomyces_griseoviridis | GCM10010240_32240 | 242124..245537 | 45398 | hypothetical-protein-GCM10010240_32240-[Streptomyces-griseoviridis] |
| WP_052687173.1 | HEPN+AAA-ATPase+Alpha+Beta-domain+wHTH+RRM | Type-1_BR.RM.like.systems | <-PIN<-Phage_integrase+Phage_integrase\|\|HTH_3-><-?<-DUF6731<-HEPN+DUF499*<-METHYLASE<-TUDOR+SF1-SF2-Helicase+iREase+CTD+REase<-?\|\|REase-> | 1168 | Bacteria>Terrabacteria_group>Actinomycetota>Actinomycetes>Kitasatosporales | - | Streptomyces_lydicus | T261_RS23850 | 5194255..5197761 | 47763 | DUF499-domain-containing-protein-[Streptomyces-lydicus] |
| WP_260825327.1 | HEPN+AAA-ATPase+Alpha+Beta-domain+wHTH+RRM | Type-1_BR.RM.like.systems | REase->?->SF1-SF2-Helicase->TUDOR+SF1-SF2-Helicase+iREase+CTD+REase->METHYLASE->HEPN+DUF499*-><-DUF397<-HTH+DUF5753\|\|ST-Hiskin->?->?-><-PIN | 1137 | Bacteria>Terrabacteria_group>Actinomycetota>Actinomycetes>Kitasatosporales | GCF_025290915.1 | Streptomyces_sp._15-116A | N4P33_RS06645 | 23393..26806 | 2259035 | Swt1-family-HEPN-domain-containing-protein-[Streptomyces-sp.-15-116A] |
| BBC36956.1 | HEPN+AAA-ATPase+Alpha+Beta-domain+wHTH+RRM | Type-1_BR.RM.like.systems | PIN-><-?<-?<-ST-Hiskin\|\|HTH+DUF5753->DUF397-><-HEPN+DUF499*<-METHYLASE<-TUDOR+SF1-SF2-Helicase+iREase+CTD+REase<-SF1-SF2-Helicase<-?<-REase | 1135 | Bacteria>Terrabacteria_group>Actinomycetota>Actinomycetes>Kitasatosporales | GCA_030294945.1 | Streptomyces_graminofaciens | SGFS_082500 | 9169123..9172530 | 68212 | hypothetical-protein-SGFS_082500-[Streptomyces-graminofaciens] |
| WP_224221478.1 | HEPN+AAA-ATPase+Alpha+Beta-domain+wHTH+RRM | Type-1_BR.RM.like.systems | MORC+HisKin+REase-><-Phage_holin<-DUF6731<-HEPN+DUF499*<-METHYLASE<-TUDOR+SF1-SF2-Helicase+iREase+CTD+REase<-DUF397\|\|ST-Hiskin-> | 1127 | Bacteria>Terrabacteria_group>Actinomycetota>Actinomycetes>Kitasatosporales | GCF_020092375.1 | Streptomyces_olivaceus | KVH28_RS23645 | 380730..384113 | 47716 | DUF499-domain-containing-protein-[Streptomyces-olivaceus] |
| TJZ52107.1 | HEPN+AAA-ATPase+Alpha+Beta-domain+wHTH+RRM | Type-1_BR.RM.like.systems | HAMP+HISKIN-><-Phage_holin<-DUF6731<-HEPN+DUF499*<-METHYLASE<-TUDOR+SF1-SF2-Helicase+iREase+CTD+REase<-tRNA\|\|METHYLASE-p-><-?\|\|TetR-HTH-> | 1122 | Bacteria>Terrabacteria_group>Actinomycetota>Actinomycetes>Kitasatosporales | GCA_005048155.1 | Streptomyces_piniterrae | FCH28_19955 | 322483..325851 | 2571125 | ATP-binding-protein-[Streptomyces-piniterrae] |
| WP_161236363.1 | HEPN+AAA-ATPase+Alpha+Beta-domain+wHTH+RRM | Type-1_BR.RM.like.systems | <-HEPN\|\|<-wHTH(SBD)+HNH\|\|TUDOR+SF1-SF2-Helicase+iREase+CTD+REase->METHYLASE->HEPN+DUF499*->DUF6731->Phage_holin-><-MNS+HTH+REase | 1161 | Bacteria>Terrabacteria_group>Actinomycetota>Actinomycetes>Kitasatosporales | GCF_010548075.1 | unclassified_Streptomyces | G3I23_RS06870 | 11181..14666 | 2593676 | MULTISPECIES:-DUF499-domain-containing-protein-[unclassified-Streptomyces] |
| BBB01161.1 | HEPN+AAA-ATPase+Alpha+Beta-domain+wHTH+RRM | Type-1_BR.RM.like.systems | <-DDE_Tnp<-?<-?\|\|?->TUDOR+SF1-SF2-Helicase+iREase+CTD+REase->METHYLASE->HEPN+DUF499*->DUF6731-><-?<-?<-HTH+DUF5753\|\|HTH+DUF5753->METHYLASE-p-> | 1128 | Bacteria>Terrabacteria_group>Actinomycetota>Actinomycetes>Kitasatosporales | GCA_016756395.1 | Streptomyces_sp._SN-593 | RVR_8434 | 7009798..7013184 | 659352 | hypothetical-protein-RVR_8434-[Streptomyces-sp.-SN-593] |
| WP_217467331.1 | HEPN+AAA-ATPase+Alpha+Beta-domain+wHTH+RRM | Type-1_BR.RM.like.systems | HTH+HKD-DNase->NUDIX-><-HDRnase\|\|TUDOR+SF1-SF2-Helicase+iREase+CTD+REase->METHYLASE->HEPN+DUF499*-><-?\|\|?->?-><-MerR-HTH\|\|ABhydrolase-> | 1135 | Bacteria>Terrabacteria_group>Actinomycetota>Actinomycetes>Kitasatosporales | GCF_019090125.1 | Streptomyces_sp._BV333 | KUG12_RS25645 | 32046..35453 | 2849673 | Swt1-family-HEPN-domain-containing-protein-[Streptomyces-sp.-BV333] |
| WP_064533938.1 | HEPN+AAA-ATPase+Alpha+Beta-domain+wHTH+RRM | Type-1_BR.RM.like.systems | <-SF1-SF2-Helicase<-?<-?\|\|?->tRNA->tRNA->tRNA->TUDOR+SF1-SF2-Helicase+iREase+CTD+REase->METHYLASE->HEPN+DUF499*->DUF6731->Phage_holin->tRNA->tRNA->?->?->SF1-SF2-Helicase-> | 1131 | Bacteria>Terrabacteria_group>Actinomycetota>Actinomycetes>Kitasatosporales | GCF_001654495.1 | Streptomyces_sp._SAT1 | A8713_RS14845 | 3432444..3435839 | 1849967 | DUF499-domain-containing-protein-[Streptomyces-sp.-SAT1] |

| Protein-ID | DUF499-architecture | System | Gene-Neighbourhoods | Length | Taxonomy | Assembly-ID | Strain | Locus-Tag | Genomic-coordinates | TaxID | Fasta-defline |
|---|---|---|---|---|---|---|---|---|---|---|---|
| WP_123099817.1 | HEPN+AAA-ATPase+Alpha+Beta-domain+wHTH+RRM | Type-1_BR.RM.like.systems | [illegible] | 1137 | Bacteria>Terrabacteria_group>Actinomycetota>Actinomycetes>Kitasatosporales | GCF_003721215.1 | Streptomyces_botrytidirepellens | EEJ42_RS11295 | 50582..53995 | 2486417 | Swt1-family-HEPN-domain-containing-protein-[Streptomyces-botrytidirepellens] |
| WP_075268487.1 | HEPN+AAA-ATPase+Alpha+Beta-domain+wHTH+RRM | Type-1_BR.RM.like.systems | [illegible] | 1126 | Bacteria>Terrabacteria_group>Actinomycetota>Actinomycetes>Kitasatosporales | GCF_001931635.1 | Streptomyces_sp._Tue_6075 | TK78_RS20735 | 4682915..4686295 | 1661694 | DUF499-domain-containing-protein-[Streptomyces-sp.-Tue-6075] |
| WP_275780742.1 | HEPN+AAA-ATPase+Alpha+Beta-domain+wHTH+RRM | Type-1_BR.RM.like.systems | [illegible] | 1150 | Bacteria>Terrabacteria_group>Actinomycetota>Actinomycetes>Kitasatosporales | GCF_029223485.1 | Streptomyces_coacervatus | P2Q00_RS45645 | 83477..86929 | 647381 | DUF499-domain-containing-protein-[Streptomyces-coacervatus] |
| WP_019888360.1 | HEPN+AAA-ATPase+Alpha+Beta-domain+wHTH+RRM | Type-1_BR.RM.like.systems | [illegible] | 1154 | Bacteria>Terrabacteria_group>Actinomycetota>Actinomycetes>Kitasatosporales | GCF_000384175.1 | Streptomyces_purpureus | STRPU_RS0118820 | 4066802..4070266 | 1951 | DUF499-domain-containing-protein-[Streptomyces-purpureus] |
| WP_222946975.1 | HEPN+AAA-ATPase+Alpha+Beta-domain+wHTH+RRM | Type-1_BR.RM.like.systems | [illegible] | 1153 | Bacteria>Terrabacteria_group>Actinomycetota>Actinomycetes>Kitasatosporales | GCF_008271805.1 | Streptomyces_sp._TRM49041 | FVS66_RS24245 | 260388..263849 | 2603216 | DUF499-domain-containing-protein-[Streptomyces-sp.-TRM49041] |
| KFG04509.1 | HEPN+AAA-ATPase+Alpha+Beta-domain+wHTH+RRM | Type-1_BR.RM.like.systems | [illegible] | 1123 | Bacteria>Terrabacteria_group>Actinomycetota>Actinomycetes>Kitasatosporales | GCA_000738715.1 | Streptomyces_scabiei | IQ61_35115 | 15067..18438 | 1930 | hypothetical-protein-IQ61_35115-[Streptomyces-scabiei] |
| WP_122182269.1 | HEPN+AAA-ATPase+Alpha+Beta-domain+wHTH+RRM | Type-1_BR.RM.like.systems | [illegible] | 1129 | Bacteria>Terrabacteria_group>Actinomycetota>Actinomycetes>Kitasatosporales | GCF_003696235.1 | Streptomyces_triticirhizae | EBN88_RS03395 | 53009..56398 | 2483353 | DUF499-domain-containing-protein-[Streptomyces-triticirhizae] |
| KOG77665.1 | HEPN+AAA-ATPase+Alpha+Beta-domain+wHTH+RRM | Type-1_BR.RM.like.systems | [illegible] | 1126 | Bacteria>Terrabacteria_group>Actinomycetota>Actinomycetes>Kitasatosporales | GCA_001270675.1 | Streptomyces_griseus_subsp._rhodochrous | ADK33_30195 | 165445..168825 | 285575 | hypothetical-protein-ADK33_30195-[Streptomyces-griseus-subsp.-rhodochrous] |
| WP_266631931.1 | HEPN+AAA-ATPase+Alpha+Beta-domain+wHTH+RRM | Type-1_BR.RM.like.systems | [illegible] | 1159 | Bacteria>Terrabacteria_group>Actinomycetota>Actinomycetes>Kitasatosporales | GCF_026339655.1 | unclassified_Streptomyces | OOK58_RS43255 | 9431725..9435204 | 2593676 | MULTISPECIES:-DUF499-domain-containing-protein-[unclassified-Streptomyces] |
| WP_031508838.1 | HEPN+AAA-ATPase+Alpha+Beta-domain+wHTH+RRM | Type-1_BR.RM.like.systems | [illegible] | 1145 | Bacteria>Terrabacteria_group>Actinomycetota>Actinomycetes>Kitasatosporales | GCF_000718985.1 | Streptomyces_megasporus | IF47_RS0112740 | 147736..151173 | 44060 | Swt1-family-HEPN-domain-containing-protein-[Streptomyces-megasporus] |
| WP_235049964.1 | HEPN+AAA-ATPase+Alpha+Beta-domain+wHTH+RRM | Type-1_BR.RM.like.systems | [illegible] | 1137 | Bacteria>Terrabacteria_group>Actinomycetota>Actinomycetes>Kitasatosporales | GCF_021502675.1 | Yinghuangia_soli | LZ495_RS01505 | 359747..363160 | 2908204 | Swt1-family-HEPN-domain-containing-protein-[Yinghuangia-sp.-KLBMP8922] |
| WP_208640870.1 | HEPN+AAA-ATPase+Alpha+Beta-domain+wHTH+RRM | Type-1_BR.RM.like.systems | [illegible] | 1148 | Bacteria>Terrabacteria_group>Actinomycetota>Actinomycetes>Kitasatosporales | GCF_002154505.1 | Streptomyces_carpinensis | BZY54_RS35300 | 9342..12788 | 66369 | DUF499-domain-containing-protein-[Streptomyces-carpinensis] |
| WP_122182258.1 | HEPN+AAA-ATPase+Alpha+Beta-domain+wHTH+RRM | Type-1_BR.RM.like.systems | [illegible] | 1151 | Bacteria>Terrabacteria_group>Actinomycetota>Actinomycetes>Kitasatosporales | GCF_003696235.1 | Streptomyces_triticirhizae | EBN88_RS03330 | 36535..39990 | 2483353 | Swt1-family-HEPN-domain-containing-protein-[Streptomyces-triticirhizae] |
| WP_120051255.1 | HEPN+AAA-ATPase+Alpha+Beta-domain+wHTH+RRM | Type-1_BR.RM.like.systems | [illegible] | 1131 | Bacteria>Terrabacteria_group>Actinomycetota>Actinomycetes>Kitasatosporales | GCF_003595235.1 | Streptomyces_griseorubiginosus | DWG14_RS17550 | 3948403..3951798 | 67304 | DUF499-domain-containing-protein-[Streptomyces-griseorubiginosus] |
| WP_030708645.1 | HEPN+AAA-ATPase+Alpha+Beta-domain+wHTH+RRM | Type-1_BR.RM.like.systems | [illegible] | 1136 | Bacteria>Terrabacteria_group>Actinomycetota>Actinomycetes>Kitasatosporales | GCF_000720115.1 | Streptomyces_sp._NRRL_F-2580 | IF88_RS0107850 | 389452..392862 | 1463841 | Swt1-family-HEPN-domain-containing-protein-[Streptomyces-sp.-NRRL-F-2580] |
| WP_109294977.1 | HEPN+AAA-ATPase+Alpha+Beta-domain+wHTH+RRM | Type-1_BR.RM.like.systems | [illegible] | 1137 | Bacteria>Terrabacteria_group>Actinomycetota>Actinomycetes>Kitasatosporales | GCF_003122365.1 | Streptomyces_spongiicola | DDQ41_RS15240 | 3467115..3470528 | 1690221 | Swt1-family-HEPN-domain-containing-protein-[Streptomyces-spongiicola] |
| WP_030969976.1 | HEPN+AAA-ATPase+Alpha+Beta-domain+wHTH+RRM | Type-1_BR.RM.like.systems | [illegible] | 1145 | Bacteria>Terrabacteria_group>Actinomycetota>Actinomycetes>Kitasatosporales | GCF_000719575.1 | Streptomyces_sp._NRRL_S-1824 | IG01_RS0105040 | 662425..665862 | 1463889 | Swt1-family-HEPN-domain-containing-protein-[Streptomyces-sp.-NRRL-S-1824] |
| WP_189323067.1 | HEPN+AAA-ATPase+Alpha+Beta-domain+wHTH+RRM | Type-1_BR.RM.like.systems | [illegible] | 1137 | Bacteria>Terrabacteria_group>Actinomycetota>Actinomycetes>Kitasatosporales | GCF_014647675.1 | Streptomyces_flaveus | IEZ21_RS18795 | 178873..182286 | 66370 | DUF499-domain-containing-protein-[Streptomyces-flaveus] |
| WP_250287638.1 | HEPN+AAA-ATPase+Alpha+Beta-domain+wHTH+RRM | Type-1_BR.RM.like.systems | [illegible] | 1135 | Bacteria>Terrabacteria_group>Actinomycetota>Actinomycetes>Kitasatosporales | GCF_928722305.1 | Streptomyces_atroolivaceus | NBY68_RS04240 | 83190..86597 | 66869 | Swt1-family-HEPN-domain-containing-protein-[Streptomyces-atroolivaceus] |
| WP_067071154.1 | HEPN+AAA-ATPase+Alpha+Beta-domain+wHTH+RRM | Type-1_BR.RM.like.systems | [illegible] | 1122 | Bacteria>Terrabacteria_group>Actinomycetota>Actinomycetes>Kitasatosporales | GCF_001543895.1 | Carbonactinospora_thermoautotrophica | TR74_RS19795 | 3234..6602 | 1469144 | Swt1-family-HEPN-domain-containing-protein-[Carbonactinospora-thermoautotrophica] |
| WP_073228437.1 | HEPN+AAA-ATPase+Alpha+Beta-domain+wHTH+RRM | Type-1_BR.RM.like.systems | [illegible] | 1131 | Bacteria>Terrabacteria_group>Actinomycetota>Actinomycetes>Kitasatosporales | GCF_001895105.1 | Streptomyces_sp._NBRC_110465 | BHQ08_RS31730 | 47724..51119 | 1897621 | DUF499-domain-containing-protein-[Streptomyces-sp.-NBRC-110465] |
| MCP9962608.1 | HEPN+AAA-ATPase+Alpha+Beta-domain+wHTH+RRM | Type-1_BR.RM.like.systems | [illegible] | 1115 | Bacteria>Terrabacteria_group>Actinomycetota>Actinomycetes>Kitasatosporales | GCA_024349105.1 | Streptomyces_somaliensis | LUX05_16750 | 3621706..3625053 | 78355 | DUF499-domain-containing-protein-[Streptomyces-somaliensis] |
| WP_198941888.1 | HEPN+AAA-ATPase+Alpha+Beta-domain+wHTH+RRM | Type-1_BR.RM.like.systems | [illegible] | 1115 | Bacteria>Terrabacteria_group>Actinomycetota>Actinomycetes>Kitasatosporales | GCF_001905405.1 | Streptomyces_sp._TSRI0445 | AMK14_RS33650 | 173132..176479 | 1703925 | DUF499-domain-containing-protein-[Streptomyces-sp.-TSRI0445] |
| OKI62803.1 | HEPN+AAA-ATPase+Alpha+Beta-domain+wHTH+RRM | Type-1_BR.RM.like.systems | [illegible] | 1126 | Bacteria>Terrabacteria_group>Actinomycetota>Actinomycetes>Kitasatosporales | GCA_001905405.1 | Streptomyces_sp._TSRI0445 | AMK14_33000 | 173099..176479 | 1703925 | hypothetical-protein-AMK14_33000-[Streptomyces-sp.-TSRI0445] |
| WP_103510051.1 | HEPN+AAA-ATPase+Alpha+Beta-domain+wHTH+RRM | Type-1_BR.RM.like.systems | [illegible] | 1135 | Bacteria>Terrabacteria_group>Actinomycetota>Actinomycetes>Kitasatosporales | GCF_002910875.1 | Streptomyces_sp._SM13 | B9S60_RS19800 | 69389..72796 | 1983803 | Swt1-family-HEPN-domain-containing-protein-[Streptomyces-sp.-SM13] |
| WP_225014234.1 | HEPN+AAA-ATPase+Alpha+Beta-domain+wHTH+RRM | Type-1_BR.RM.like.systems | [illegible] | 753 | Bacteria>Terrabacteria_group>Actinomycetota>Actinomycetes>Kitasatosporales | GCF_020176795.1 | Streptomyces_sp._8L | LCE31_RS12240 | 76..2337 | 2877242 | DUF499-domain-containing-protein-[Streptomyces-sp.-8L] |
| TRZ87911.1 | HEPN+AAA-ATPase+Alpha+Beta-domain+wHTH+RRM | Type-1_BR.RM.like.systems | [illegible] | 1108 | Bacteria>Terrabacteria_group>Actinomycetota>Actinomycetes>Kitasatosporales | GCA_007280125.1 | Streptomycetaceae_bacterium | D4R83_01365 | 3739..7065 | 2021369 | ATP-binding-protein-[Streptomycetaceae-bacterium] |
| WP_192910841.1 | HEPN+AAA-ATPase+Alpha+Beta-domain+wHTH+RRM | Type-1_BR.RM.like.systems | [illegible] | 1123 | Bacteria>Terrabacteria_group>Actinomycetota>Actinomycetes>Micrococcales | GCF_014883915.1 | Janibacter_indicus | IGS73_RS14740 | 2999354..3002725 | 857417 | DUF499-domain-containing-protein-[Janibacter-indicus] |
| MBK6299925.1 | HEPN+AAA-ATPase+Alpha+Beta-domain+wHTH+RRM | Type-1_BR.RM.like.systems | [illegible] | 1117 | Bacteria>Terrabacteria_group>Actinomycetota>Actinomycetes>Micrococcales | GCA_016705565.1 | Candidatus_Phosphoribacter_hodrii | IPF40_02340 | 519882..523235 | 2953743 | DUF499-domain-containing-protein-[Actinomycetales-bacterium] |
| MBK6954412.1 | HEPN+AAA-ATPase+Alpha+Beta-domain+wHTH+RRM | Type-1_BR.RM.like.systems | [illegible] | 1118 | Bacteria>Terrabacteria_group>Actinomycetota>Actinomycetes>Micrococcales | GCA_016708855.1 | Candidatus_Phosphoribacter_baldrii | IPH27_02570 | 528671..532027 | 2953744 | DUF499-domain-containing-protein-[Actinomycetales-bacterium] |
| CEA09674.1 | HEPN+AAA-ATPase+Alpha+Beta-domain+wHTH+RRM | Type-1_BR.RM.like.systems | [illegible] | 1141 | Bacteria>Terrabacteria_group>Actinomycetota>Actinomycetes>Micrococcales | GCA_900016675.1 | Arthrobacter_saudimassiliensis | BN1051_03046 | 420821..424246 | 1461584 | hypothetical-protein-BN1051_03046-[Arthrobacter-saudimassiliensis] |
| WP_137123200.1 | HEPN+AAA-ATPase+Alpha+Beta-domain+wHTH+RRM | Type-1_BR.RM.like.systems | [illegible] | 1132 | Bacteria>Terrabacteria_group>Actinomycetota>Actinomycetes>Micrococcales | GCF_005222685.1 | Ornithinicoccus_soli | FCN22_RS19310 | 12208..15606 | 2574732 | DUF499-domain-containing-protein-[Ornithinicoccus-soli] |
| WP_285140308.1 | HEPN+AAA-ATPase+Alpha+Beta-domain+wHTH+RRM | Type-1_BR.RM.like.systems | [illegible] | 1132 | Bacteria>Terrabacteria_group>Actinomycetota>Actinomycetes>Micrococcales | GCF_030226745.1 | Micrococcus_luteus | QP726_RS10470 | 9707..13105 | 1270 | Swt1-family-HEPN-domain-containing-protein-[Micrococcus-luteus] |
| WP_195692263.1 | HEPN+AAA-ATPase+Alpha+Beta-domain+wHTH+RRM | Type-1_BR.RM.like.systems | [illegible] | 1129 | Bacteria>Terrabacteria_group>Actinomycetota>Actinomycetes>Micrococcales | GCF_015565955.1 | Microbacterium | JOJ05_RS14720 | 85515..88904 | 33882 | DUF499-domain-containing-protein-[Microbacterium-schleiferi] |
| WP_169323296.1 | HEPN+AAA-ATPase+Alpha+Beta-domain+wHTH+RRM | Type-1_BR.RM.like.systems | [illegible] | 1120 | Bacteria>Terrabacteria_group>Actinomycetota>Actinomycetes>Micrococcales | GCF_012952065.1 | Cellulomonas_fimi | HIR71_RS02970 | 36548..39910 | 1708 | DUF499-domain-containing-protein-[Cellulomonas-fimi] |
| WP_055898767.1 | HEPN+AAA-ATPase+Alpha+Beta-domain+wHTH+RRM | Type-1_BR.RM.like.systems | [illegible] | 1109 | Bacteria>Terrabacteria_group>Actinomycetota>Actinomycetes>Micrococcales | GCF_001425665.1 | Yonghaparkia_sp._Root332 | ASC54_RS09975 | 479255..482584 | 1736516 | DUF499-domain-containing-protein-[Yonghaparkia-sp.-Root332] |
| WP_144831472.1 | HEPN+AAA-ATPase+Alpha+Beta-domain+wHTH+RRM | Type-1_BR.RM.like.systems | [illegible] | 1133 | Bacteria>Terrabacteria_group>Actinomycetota>Actinomycetes>Micrococcales | GCF_007677595.1 | Kocuria_rhizophila | FS577_RS10180 | 96606..100007 | 72000 | DUF499-domain-containing-protein-[Kocuria-rhizophila] |
| WP_274500272.1 | HEPN+AAA-ATPase+Alpha+Beta-domain+wHTH+RRM | Type-1_BR.RM.like.systems | [illegible] | 1131 | Bacteria>Terrabacteria_group>Actinomycetota>Actinomycetes>Micrococcales | GCF_028765365.1 | Plantibacter_flavus | PUY80_RS13440 | 281410..284805 | 150123 | DUF499-domain-containing-protein-[Plantibacter-flavus] |
| WP_255162094.1 | HEPN+AAA-ATPase+Alpha+Beta-domain+wHTH+RRM | Type-1_BR.RM.like.systems | [illegible] | 1143 | Bacteria>Terrabacteria_group>Actinomycetota>Actinomycetes>Micrococcales | GCF_024362265.1 | Microbacterium_liquefaciens | NMQ05_RS13065 | 2679727..2683158 | 33918 | DUF499-domain-containing-protein-[Microbacterium-liquefaciens] |
| WP_197156507.1 | HEPN+AAA-ATPase+Alpha+Beta-domain+wHTH+RRM | Type-1_BR.RM.like.systems | [illegible] | 1110 | Bacteria>Terrabacteria_group>Actinomycetota>Actinomycetes>Micrococcales | GCF_015864945.1 | Salinibacterium_sp._NG253 | I6E52_RS09085 | 1888777..1892109 | 2792039 | DUF499-domain-containing-protein-[Salinibacterium-sp.-NG253] |
| WP_194413436.1 | HEPN+AAA-ATPase+Alpha+Beta-domain+wHTH+RRM | Type-1_BR.RM.like.systems | [illegible] | 1119 | Bacteria>Terrabacteria_group>Actinomycetota>Actinomycetes>Micrococcales | GCF_015278355.1 | Microbacterium_yannicii | NCG93_RS19890 | 89985..93344 | 671622 | Swt1-family-HEPN-domain-containing-protein-[Microbacterium-yannicii] |
| WP_225755648.1 | HEPN+AAA-ATPase+Alpha+Beta-domain+wHTH+RRM | Type-1_BR.RM.like.systems | [illegible] | 1137 | Bacteria>Terrabacteria_group>Actinomycetota>Actinomycetes>Micrococcales | GCF_916618755.1 | Actinotalea_sp._Marseille-Q4924 | LG379_RS16320 | 268678..272091 | 2866571 | DUF499-domain-containing-protein-[Actinotalea-sp.-Marseille-Q4924] |
| WP_139142668.1 | HEPN+AAA-ATPase+Alpha+Beta-domain+wHTH+RRM | Type-1_BR.RM.like.systems | [illegible] | 1111 | Bacteria>Terrabacteria_group>Actinomycetota>Actinomycetes>Micrococcales | GCF_001758225.1 | Humibacillus_sp._DSM_29435 | BA895_RS14645 | 164736..168071 | 1869167 | Swt1-family-HEPN-domain-containing-protein-[Humibacillus-sp.-DSM-29435] |
| MCU1509501.1 | HEPN+AAA-ATPase+Alpha+Beta-domain+wHTH+RRM | Type-1_BR.RM.like.systems | [illegible] | 1121 | Bacteria>Terrabacteria_group>Actinomycetota>Actinomycetes>Micrococcales | GCA_025464095.1 | Glaciihabitans_sp. | JWQ12_1766 | 110555..113920 | 1965328 | hypothetical-protein-[Glaciihabitans-sp.] |
| MBK8435770.1 | HEPN+AAA-ATPase+Alpha+Beta-domain+wHTH+RRM | Type-1_BR.RM.like.systems | [illegible] | 1139 | Bacteria>Terrabacteria_group>Actinomycetota>Actinomycetes>Micrococcales | GCA_016711265.1 | Austwickia_sp. | IPL37_04405 | 982795..986214 | 1872664 | DUF499-domain-containing-protein-[Austwickia-sp.] |
| WP_167559767.1 | HEPN+AAA-ATPase+Alpha+Beta-domain+wHTH+RRM | Type-1_BR.RM.like.systems | [illegible] | 1132 | Bacteria>Terrabacteria_group>Actinomycetota>Actinomycetes>Micrococcales | - | Brevibacterium_sp._LS14 | EZE58_RS02710 | 16654..20052 | 2528962 | DUF499-domain-containing-protein-[Brevibacterium-sp.-LS14] |
| WP_063250773.1 | HEPN+AAA-ATPase+Alpha+Beta-domain+wHTH+RRM | Type-1_BR.RM.like.systems | [illegible] | 1132 | Bacteria>Terrabacteria_group>Actinomycetota>Actinomycetes>Micrococcales | GCF_001619685.1 | Brevibacterium_casei | AVW13_RS15020 | 17875..21273 | 33889 | DUF499-domain-containing-protein-[Brevibacterium-casei] |
| WP_246200745.1 | HEPN+AAA-ATPase+Alpha+Beta-domain+wHTH+RRM | Type-1_BR.RM.like.systems | [illegible] | 1131 | Bacteria>Terrabacteria_group>Actinomycetota>Actinomycetes>Micrococcales | GCF_009749385.1 | Agromyces_allii | GQR16_RS11060 | 282353..285748 | 393607 | Swt1-family-HEPN-domain-containing-protein-[Agromyces-allii] |
| WP_218115672.1 | HEPN+AAA-ATPase+Alpha+Beta-domain+wHTH+RRM | Type-1_BR.RM.like.systems | [illegible] | 1133 | Bacteria>Terrabacteria_group>Actinomycetota>Actinomycetes>Micrococcales | GCF_910593785.1 | Leucobacter_soli | KXF26_RS09170 | 42150..45551 | 2812850 | Swt1-family-HEPN-domain-containing-protein-[Leucobacter-soli] |
| MBN9213516.1 | HEPN+AAA-ATPase+Alpha+Beta-domain+wHTH+RRM | Type-1_BR.RM.like.systems | [illegible] | 1119 | Bacteria>Terrabacteria_group>Actinomycetota>Actinomycetes>Micrococcales | GCA_017306385.1 | Microbacterium_sp. | JOJ04_01635 | 353134..356493 | 51671 | DUF499-domain-containing-protein-[Microbacterium-sp.] |
| MCI1262624.1 | HEPN+AAA-ATPase+Alpha+Beta-domain+wHTH+RRM | Type-1_BR.RM.like.systems | [illegible] | 1116 | Bacteria>Terrabacteria_group>Actinomycetota>Actinomycetes>Micrococcales | GCA_022641475.1 | Tetrasphaera_jenkinsii | LKG20_10180 | 13344..16694 | 330834 | Swt1-family-HEPN-domain-containing-protein-[Tetrasphaera-jenkinsii] |
| WP_259878499.1 | HEPN+AAA-ATPase+Alpha+Beta-domain+wHTH+RRM | Type-1_BR.RM.like.systems | [illegible] | 1132 | Bacteria>Terrabacteria_group>Actinomycetota>Actinomycetes>Micrococcales | GCF_025148185.1 | Brachybacterium_muris | M3C58_RS02745 | 15014..18412 | 219301 | DUF499-domain-containing-protein-[Brachybacterium-muris] |
| WP_220122165.1 | HEPN+AAA-ATPase+Alpha+Beta-domain+wHTH+RRM | Type-1_BR.RM.like.systems | [illegible] | 1128 | Bacteria>Terrabacteria_group>Actinomycetota>Actinomycetes>Micrococcales | - | Ornithinimicrobium_laminariae | K0651_07785 | 644333..647719 | 2864873 | Swt1-family-HEPN-domain-containing-protein-[Ornithinimicrobium-laminariae] |
| WP_188034675.1 | HEPN+AAA-ATPase+Alpha+Beta-domain+wHTH+RRM | Type-1_BR.RM.like.systems | [illegible] | 1143 | Bacteria>Terrabacteria_group>Actinomycetota>Actinomycetes>Micrococcales | GCF_014530035.1 | unclassified_Leucobacter | KOE72_RS09585 | 56127..59558 | 2621730 | MULTISPECIES:-DUF499-domain-containing-protein-[unclassified-Leucobacter] |
| MBW8172946.1 | HEPN+AAA-ATPase+Alpha+Beta-domain+wHTH+RRM | Type-1_BR.RM.like.systems | [illegible] | 1128 | Bacteria>Terrabacteria_group>Actinomycetota>Actinomycetes>Micrococcales | GCA_019457945.1 | Ornithinimicrobium_laminariae | K0651_07785 | 644333..647719 | 2864873 | DUF499-domain-containing-protein-[Ornithinimicrobium-laminariae] |
| WP_210418449.1 | HEPN+AAA-ATPase+Alpha+Beta-domain+wHTH+RRM | Type-1_BR.RM.like.systems | [illegible] | 1116 | Bacteria>Terrabacteria_group>Actinomycetota>Actinomycetes>Micrococcales | GCF_008365295.1 | Agromyces_intestinalis | FLP10_RS00780 | 160366..163716 | 2592652 | DUF499-domain-containing-protein-[Agromyces-intestinalis] |
| WP_069599267.1 | HEPN+AAA-ATPase+Alpha+Beta-domain+wHTH+RRM | Type-1_BR.RM.like.systems | [illegible] | 1131 | Bacteria>Terrabacteria_group>Actinomycetota>Actinomycetes>Micrococcales | GCF_001729525.1 | Brevibacterium_aurantiacum | CIK60_RS17585 | 12727..16122 | 273384 | DUF499-domain-containing-protein-[Brevibacterium-aurantiacum] |
| QEO13112.1 | HEPN+AAA-ATPase+Alpha+Beta-domain+wHTH+RRM | Type-1_BR.RM.like.systems | [illegible] | 1128 | Bacteria>Terrabacteria_group>Actinomycetota>Actinomycetes>Micrococcales | GCA_008365295.1 | Agromyces_intestinalis | FLP10_00780 | 160366..163752 | 2592652 | ATP-binding-protein-[Agromyces-intestinalis] |
| WP_201671845.1 | HEPN+AAA-ATPase+Alpha+Beta-domain+wHTH+RRM | Type-1_BR.RM.like.systems | [illegible] | 1132 | Bacteria>Terrabacteria_group>Actinomycetota>Actinomycetes>Micrococcales | GCF_016726245.1 | Brevibacterium_casei | I6I57_RS03275 | 754914..758312 | 33889 | DUF499-domain-containing-protein-[Brevibacterium-casei] |
| WP_188427984.1 | HEPN+AAA-ATPase+Alpha+Beta-domain+wHTH+RRM | Type-1_BR.RM.like.systems | [illegible] | 1123 | Bacteria>Terrabacteria_group>Actinomycetota>Actinomycetes>Micrococcales | GCF_014636495.1 | Ornithinimicrobium_tianjinense | IEV65_RS02295 | 470126..473497 | 1195761 | Swt1-family-HEPN-domain-containing-protein-[Ornithinimicrobium-tianjinense] |
| WP_141842659.1 | HEPN+AAA-ATPase+Alpha+Beta-domain+wHTH+RRM | Type-1_BR.RM.like.systems | [illegible] | 1132 | Bacteria>Terrabacteria_group>Actinomycetota>Actinomycetes>Micrococcales | GCF_006704135.1 | Micrococcus_sp._R8502A1 | FLM69_RS00645 | 3870..7268 | 2583239 | Swt1-family-HEPN-domain-containing-protein-[Micrococcus-sp.-R8502A1] |
| OFE15977.1 | HEPN+AAA-ATPase+Alpha+Beta-domain+wHTH+RRM | Type-1_BR.RM.like.systems | [illegible] | 1100 | Bacteria>Terrabacteria_group>Actinomycetota>Actinomycetes>Micrococcales | GCA_001758225.1 | Humibacillus_sp._DSM_29435 | BA895_05295 | 164736..168038 | 1869167 | AAA-family-ATPase-[Humibacillus-sp.-DSM-29435] |
| MCE1178998.1 | HEPN+AAA-ATPase+Alpha+Beta-domain+wHTH+RRM | Type-1_BR.RM.like.systems | [illegible] | 1133 | Bacteria>Terrabacteria_group>Actinomycetota>Actinomycetes>Micrococcales | GCF_021287075.1 | Micrococcales_bacterium | LWW86_08210 | 123178..126579 | 2026762 | DUF499-domain-containing-protein-[Micrococcales-bacterium] |
| WP_241710057.1 | HEPN+AAA-ATPase+Alpha+Beta-domain+wHTH+RRM | Type-1_BR.RM.like.systems | [illegible] | 1123 | Bacteria>Terrabacteria_group>Actinomycetota>Actinomycetes>Micrococcales | GCF_022606295.1 | Arthrobacter_bambusae | KNN17_RS16570 | 21950..25321 | 1338426 | DUF499-domain-containing-protein-[Arthrobacter-bambusae] |
| WP_048693914.1 | HEPN+AAA-ATPase+Alpha+Beta-domain+wHTH+RRM | Type-1_BR.RM.like.systems | [illegible] | 1134 | Bacteria>Terrabacteria_group>Actinomycetota>Actinomycetes>Micrococcales | GCF_001050535.1 | Tetrasphaera_australiensis | BN11_RS02235 | 472611..476015 | 99480 | DUF499-domain-containing-protein-[Tetrasphaera-australiensis] |
| WP_034802772.1 | HEPN+AAA-ATPase+Alpha+Beta-domain+wHTH+RRM | Type-1_BR.RM.like.systems | [illegible] | 1107 | Bacteria>Terrabacteria_group>Actinomycetota>Actinomycetes>Micrococcales | GCF_000576595.1 | Intrasporangium_oryzae | N865_RS05960 | 110230..113553 | 412687 | DUF499-domain-containing-protein-[Intrasporangium-oryzae] |
| WP_137369040.1 | HEPN+AAA-ATPase+Alpha+Beta-domain+wHTH+RRM | Type-1_BR.RM.like.systems | [illegible] | 1129 | Bacteria>Terrabacteria_group>Actinomycetota>Actinomycetes>Micrococcales | GCF_005281365.1 | Arthrobacter_sp._NamB2 | FDK12_RS07770 | 280727..284116 | 2576035 | Swt1-family-HEPN-domain-containing-protein-[Arthrobacter-sp.-NamB2] |
| WP_128365158.1 | HEPN+AAA-ATPase+Alpha+Beta-domain+wHTH+RRM | Type-1_BR.RM.like.systems | [illegible] | 1135 | Bacteria>Terrabacteria_group>Actinomycetota>Actinomycetes>Micrococcales | GCF_004025965.2 | Phycicoccus_flavus | EPD83_RS11930 | 11514..14921 | 2502783 | DUF499-domain-containing-protein-[Phycicoccus-flavus] |
| WP_217316614.1 | HEPN+AAA-ATPase+Alpha+Beta-domain+wHTH+RRM | Type-1_BR.RM.like.systems | [illegible] | 1128 | Bacteria>Terrabacteria_group>Actinomycetota>Actinomycetes>Micrococcales | GCF_019056515.1 | Microbacterium_paraoxydans | IZR02_RS16970 | 42865..46251 | 199592 | DUF499-domain-containing-protein-[Microbacterium-paraoxydans] |
| WP_105034124.1 | HEPN+AAA-ATPase+Alpha+Beta-domain+wHTH+RRM | Type-1_BR.RM.like.systems | [illegible] | 1121 | Bacteria>Terrabacteria_group>Actinomycetota>Actinomycetes>Micrococcales | GCF_002954245.1 | Cryobacterium_aureum | CJ028_RS04835 | 402642..406007 | 995037 | Swt1-family-HEPN-domain-containing-protein-[Cryobacterium-aureum] |
| WP_062860586.1 | HEPN+AAA-ATPase+Alpha+Beta-domain+wHTH+RRM | Type-1_BR.RM.like.systems | [illegible] | 1132 | Bacteria>Terrabacteria_group>Actinomycetota>Actinomycetes>Micrococcales | GCF_001606005.1 | Brevibacterium_linens | A2T55_RS01955 | 443137..446535 | 1703 | DUF499-domain-containing-protein-[Brevibacterium-linens] |
| WP_112235018.1 | HEPN+AAA-ATPase+Alpha+Beta-domain+wHTH+RRM | Type-1_BR.RM.like.systems | [illegible] | 1132 | Bacteria>Terrabacteria_group>Actinomycetota>Actinomycetes>Micrococcales | GCF_003290245.1 | Kocuria_sp._BT304 | DNL27_RS06220 | 1382048..1385446 | 1702043 | DUF499-domain-containing-protein-[Kocuria-sp.-BT304] |
| WP_046529475.1 | HEPN+AAA-ATPase+Alpha+Beta-domain+wHTH+RRM | Type-1_BR.RM.like.systems | [illegible] | 1124 | Bacteria>Terrabacteria_group>Actinomycetota>Actinomycetes>Micrococcales | GCF_000986825.1 | Cellulomonas_sp._FA1 | GY42_RS11845 | 2630948..2634322 | 1346710 | Swt1-family-HEPN-domain-containing-protein-[Cellulomonas-sp.-FA1] |
| WP_156242696.1 | HEPN+AAA-ATPase+Alpha+Beta-domain+wHTH+RRM | Type-1_BR.RM.like.systems | [illegible] | 1129 | Bacteria>Terrabacteria_group>Actinomycetota>Actinomycetes>Micrococcales | GCF_009735645.1 | Microbacterium_oryzae | D7D94_RS11265 | 2403131..2406520 | 743009 | Swt1-family-HEPN-domain-containing-protein-[Microbacterium-oryzae] |
| WP_104267247.1 | HEPN+AAA-ATPase+Alpha+Beta-domain+wHTH+RRM | Type-1_BR.RM.like.systems | [illegible] | 1135 | Bacteria>Terrabacteria_group>Actinomycetota>Actinomycetes>Micrococcales | GCF_002932235.1 | Rathayibacter_sp._AY1E3 | C5C53_RS07820 | 81195..84602 | 2080551 | Swt1-family-HEPN-domain-containing-protein-[Rathayibacter-sp.-AY1E3] |
| WP_101853235.1 | HEPN+AAA-ATPase+Alpha+Beta-domain+wHTH+RRM | Type-1_BR.RM.like.systems | [illegible] | 1144 | Bacteria>Terrabacteria_group>Actinomycetota>Actinomycetes>Micrococcales | GCF_002849405.1 | Kocuria_flava | AUQ48_RS15985 | 10331..13765 | 446860 | DUF499-domain-containing-protein-[Kocuria-flava] |
| WP_191562882.1 | HEPN+AAA-ATPase+Alpha+Beta-domain+wHTH+RRM | Type-1_BR.RM.like.systems | [illegible] | 1117 | Bacteria>Terrabacteria_group>Actinomycetota>Actinomycetes>Micrococcales | GCF_904423795.1 | Janibacter_melonis | IHQ27_RS00715 | 71564..74917 | 262209 | Swt1-family-HEPN-domain-containing-protein-[Janibacter-melonis] |
| WP_147826310.1 | HEPN+AAA-ATPase+Alpha+Beta-domain+wHTH+RRM | Type-1_BR.RM.like.systems | [illegible] | 1122 | Bacteria>Terrabacteria_group>Actinomycetota>Actinomycetes>Micrococcales | GCF_008017445.1 | Microbacterium_mitrae | FVP60_RS10835 | 35691..39059 | 664640 | DUF499-domain-containing-protein-[Microbacterium-mitrae] |
| WP_138608335.1 | HEPN+AAA-ATPase+Alpha+Beta-domain+wHTH+RRM | Type-1_BR.RM.like.systems | [illegible] | 1120 | Bacteria>Terrabacteria_group>Actinomycetota>Actinomycetes>Micrococcales | GCF_005796225.1 | Pseudarthrobacter_sp._NamE5 | FDW84_RS08080 | 47724..51086 | 2576839 | DUF499-domain-containing-protein-[Pseudarthrobacter-sp.-NamE5] |
| WP_129190457.1 | HEPN+AAA-ATPase+Alpha+Beta-domain+wHTH+RRM | Type-1_BR.RM.like.systems | [illegible] | 1123 | Bacteria>Terrabacteria_group>Actinomycetota>Actinomycetes>Micrococcales | GCF_004135385.1 | Xylanimonas_protaetiae | ET471_RS17505 | 3776288..3779659 | 2509457 | Swt1-family-HEPN-domain-containing-protein-[Xylanimonas-protaetiae] |
| TLM86009.1 | HEPN+AAA-ATPase+Alpha+Beta-domain+wHTH+RRM | Type-1_BR.RM.like.systems | [illegible] | 1120 | Bacteria>Terrabacteria_group>Actinomycetota>Actinomycetes>Micrococcales | GCA_005796225.1 | Pseudarthrobacter_sp._NamE5 | FDW84_06950 | 47724..51086 | 2576839 | ATP-binding-protein-[Pseudarthrobacter-sp.-NamE5] |
| QAY71606.1 | HEPN+AAA-ATPase+Alpha+Beta-domain+wHTH+RRM | Type-1_BR.RM.like.systems | [illegible] | 1123 | Bacteria>Terrabacteria_group>Actinomycetota>Actinomycetes>Micrococcales | GCA_004135385.1 | Xylanimonas_protaetiae | ET471_17505 | 3776288..3779659 | 2509457 | ATP-binding-protein-[Xylanimonas-protaetiae] |
| WP_208008530.1 | HEPN+AAA-ATPase+Alpha+Beta-domain+wHTH+RRM | Type-1_BR.RM.like.systems | [illegible] | 1101 | Bacteria>Terrabacteria_group>Actinomycetota>Actinomycetes>Micrococcales | GCF_004135405.1 | Agromyces_protaetiae | ET445_RS03710 | 798698..802003 | 2509455 | Swt1-family-HEPN-domain-containing-protein-[Agromyces-protaetiae] |
| WP_176337038.1 | HEPN+AAA-ATPase+Alpha+Beta-domain+wHTH+RRM | Type-1_BR.RM.like.systems | [illegible] | 1149 | Bacteria>Terrabacteria_group>Actinomycetota>Actinomycetes>Micrococcales | GCF_013368965.1 | Kocuria_salina | E7Z53_RS01815 | 401527..404976 | 1929416 | DUF499-domain-containing-protein-[Kocuria-salina] |
| WP_108718589.1 | HEPN+AAA-ATPase+Alpha+Beta-domain+wHTH+RRM | Type-1_BR.RM.like.systems | [illegible] | 1134 | Bacteria>Terrabacteria_group>Actinomycetota>Actinomycetes>Micrococcales | GCF_003074035.1 | Miniimonas_sp._S16 | DCS86_RS08145 | 1885887..1889291 | 2171623 | Swt1-family-HEPN-domain-containing-protein-[Miniimonas-sp.-S16] |
| MCV7624012.1 | HEPN+AAA-ATPase+Alpha+Beta-domain+wHTH+RRM | Type-1_BR.RM.like.systems | [illegible] | 1157 | Bacteria>Terrabacteria_group>Actinomycetota>Actinomycetes>Micrococcales | GCA_025145605.2 | Micrococcus_luteus | M3B28_003235 | 32627..36100 | 1270 | Swt1-family-HEPN-domain-containing-protein-[Micrococcus-luteus] |
| WP_062529103.1 | HEPN+AAA-ATPase+Alpha+Beta-domain+wHTH+RRM | Type-1_BR.RM.like.systems | [illegible] | 1118 | Bacteria>Terrabacteria_group>Actinomycetota>Actinomycetes>Micrococcales | GCF_000971455.1 | Demequina_rhizosphaerae | AOM12_RS04465 | 350500..353856 | 1638985 | DUF499-domain-containing-protein-[Demequina-rhizosphaerae] |
| WP_043123141.1 | HEPN+AAA-ATPase+Alpha+Beta-domain+wHTH+RRM | Type-1_BR.RM.like.systems | [illegible] | 1115 | Bacteria>Terrabacteria_group>Actinomycetota>Actinomycetes>Micrococcales | GCF_000802235.1 | Sinomonas_humi | LK10_RS10225 | 159632..162979 | 1338436 | Swt1-family-HEPN-domain-containing-protein-[Sinomonas-humi] |
| WP_252619626.1 | HEPN+AAA-ATPase+Alpha+Beta-domain+wHTH+RRM | Type-1_BR.RM.like.systems | [illegible] | 1118 | Bacteria>Terrabacteria_group>Actinomycetota>Actinomycetes>Micrococcales | GCF_023923205.1 | Ornithinimicrobium_cryptoxanthini | NF557_RS11930 | 2576031..2579387 | 2934161 | DUF499-domain-containing-protein-[Ornithinimicrobium-sp.-JY.X270] |
| WP_249521258.1 | HEPN+AAA-ATPase+Alpha+Beta-domain+wHTH+RRM | Type-1_BR.RM.like.systems | [illegible] | 1131 | Bacteria>Terrabacteria_group>Actinomycetota>Actinomycetes>Micrococcales | GCF_023383675.1 | Microbacterium_sp._ER1 | BV453_RS07415 | 50443..53838 | 1932846 | DUF499-domain-containing-protein-[Microbacterium-sp.-ER1] |
| WP_187368407.1 | HEPN+AAA-ATPase+Alpha+Beta-domain+wHTH+RRM | Type-1_BR.RM.like.systems | [illegible] | 1133 | Bacteria>Terrabacteria_group>Actinomycetota>Actinomycetes>Micrococcales | GCF_008000995.1 | Ruania_zhangjianzhongii | FU260_RS03480 | 750431..753832 | 2603206 | DUF499-domain-containing-protein-[Ruania-zhangjianzhongii] |
| WP_145227579.1 | HEPN+AAA-ATPase+Alpha+Beta-domain+wHTH+RRM | Type-1_BR.RM.like.systems | [illegible] | 1121 | Bacteria>Terrabacteria_group>Actinomycetota>Actinomycetes>Micrococcales | GCF_007829035.1 | Rudaeicoccus_suwonensis | BKA23_RS09875 | 2146112..2149477 | 657409 | DUF499-domain-containing-protein-[Rudaeicoccus-suwonensis] |
| WP_134401373.1 | HEPN+AAA-ATPase+Alpha+Beta-domain+wHTH+RRM | Type-1_BR.RM.like.systems | [illegible] | 1124 | Bacteria>Terrabacteria_group>Actinomycetota>Actinomycetes>Micrococcales | GCF_004402575.1 | Cryobacterium_sp._TMT2-4 | E3054_RS08280 | 76881..80255 | 1259254 | Swt1-family-HEPN-domain-containing-protein-[Cryobacterium-sp.-TMT2-4] |
| TFC67798.1 | HEPN+AAA-ATPase+Alpha+Beta-domain+wHTH+RRM | Type-1_BR.RM.like.systems | [illegible] | 1124 | Bacteria>Terrabacteria_group>Actinomycetota>Actinomycetes>Micrococcales | GCA_004402575.1 | Cryobacterium_sp._TMT2-4 | E3054_08285 | 76881..80255 | 1259254 | ATP-binding-protein-[Cryobacterium-sp.-TMT2-4] |
| WP_074694547.1 | HEPN+AAA-ATPase+Alpha+Beta-domain+wHTH+RRM | Type-1_BR.RM.like.systems | [illegible] | 1131 | Bacteria>Terrabacteria_group>Actinomycetota>Actinomycetes>Micrococcales | GCF_900104345.1 | Microbacterium_testaceum | BLS25_RS04240 | 198218..201613 | 2033 | DUF499-domain-containing-protein-[Microbacterium-testaceum] |
| WP_144794196.1 | HEPN+AAA-ATPase+Alpha+Beta-domain+wHTH+RRM | Type-1_BR.RM.like.systems | [illegible] | 1132 | Bacteria>Terrabacteria_group>Actinomycetota>Actinomycetes>Micrococcales | GCF_007679925.1 | Microbacterium_paludicola | FS695_RS03635 | 729234..732632 | 300019 | DUF499-domain-containing-protein-[Microbacterium-paludicola] |
| WP_109686041.1 | HEPN+AAA-ATPase+Alpha+Beta-domain+wHTH+RRM | Type-1_BR.RM.like.systems | [illegible] | 1124 | Bacteria>Terrabacteria_group>Actinomycetota>Actinomycetes>Micrococcales | GCF_003149195.1 | Branchiibius_hedensis | ATK17_RS11720 | 2422832..2426206 | 672460 | Swt1-family-HEPN-domain-containing-protein-[Branchiibius-hedensis] |
| WP_106298858.1 | HEPN+AAA-ATPase+Alpha+Beta-domain+wHTH+RRM | Type-1_BR.RM.like.systems | [illegible] | 1112 | Bacteria>Terrabacteria_group>Actinomycetota>Actinomycetes>Micrococcales | GCF_003002895.1 | Knoellia_remsis | BCF74_RS15510 | 21779..25117 | 407159 | Swt1-family-HEPN-domain-containing-protein-[Knoellia-remsis] |
| AFR31400.1 | HEPN+AAA-ATPase+Alpha+Beta-domain+wHTH+RRM | Type-1_BR.RM.like.systems | [illegible] | 1120 | Bacteria>Terrabacteria_group>Actinomycetota>Actinomycetes>Micrococcales | GCA_000294695.2 | Arthrobacter_sp._Rue61a | ARUE_232p01920 | 197474..200836 | 1118963 | hypothetical-protein-ARUE_232p01920-[Arthrobacter-sp.-Rue61a] |
| WP_231831079.1 | HEPN+AAA-ATPase+Alpha+Beta-domain+wHTH+RRM | Type-1_BR.RM.like.systems | [illegible] | 1128 | Bacteria>Terrabacteria_group>Actinomycetota>Actinomycetes>Micrococcales | GCF_021168475.1 | Pseudoclavibacter_sp._13-3 | LRX77_RS01510 | 334724..338110 | 2901228 | Swt1-family-HEPN-domain-containing-protein-[Pseudoclavibacter-sp.-13-3] |
| WP_216886275.1 | HEPN+AAA-ATPase+Alpha+Beta-domain+wHTH+RRM | Type-1_BR.RM.like.systems | [illegible] | 1157 | Bacteria>Terrabacteria_group>Actinomycetota>Actinomycetes>Micrococcales | GCF_019037785.1 | Micrococcus_luteus | KM867_RS03170 | 29276..32749 | 1270 | Swt1-family-HEPN-domain-containing-protein-[Micrococcus-luteus] |
| WP_210233975.1 | HEPN+AAA-ATPase+Alpha+Beta-domain+wHTH+RRM | Type-1_BR.RM.like.systems | [illegible] | 1116 | Bacteria>Terrabacteria_group>Actinomycetota>Actinomycetes>Micrococcales | GCF_017898105.1 | Arthrobacter_jiangjiafuii | J2M53_RS14475 | 604648..607998 | 2817475 | Swt1-family-HEPN-domain-containing-protein-[Arthrobacter-jiangjiafuii] |
| WP_134416003.1 | HEPN+AAA-ATPase+Alpha+Beta-domain+wHTH+RRM | Type-1_BR.RM.like.systems | [illegible] | 1157 | Bacteria>Terrabacteria_group>Actinomycetota>Actinomycetes>Micrococcales | GCF_022664405.1 | Micrococcus_aloeverae | E2F92_RS02100 | 40430..43903 | 1391911 | Swt1-family-HEPN-domain-containing-protein-[Micrococcus-aloeverae] |
| WP_044658741.1 | HEPN+AAA-ATPase+Alpha+Beta-domain+wHTH+RRM | Type-1_BR.RM.like.systems | [illegible] | 1157 | Bacteria>Terrabacteria_group>Actinomycetota>Actinomycetes>Micrococcales | GCF_000946635.1 | Micrococcus | B8W89_RS05705 | 32626..36099 | 1269 | MULTISPECIES:-Swt1-family-HEPN-domain-containing-protein-[Micrococcus] |
| WP_101639485.1 | HEPN+AAA-ATPase+Alpha+Beta-domain+wHTH+RRM | Type-1_BR.RM.like.systems | [illegible] | 1135 | Bacteria>Terrabacteria_group>Actinomycetota>Actinomycetes>Micrococcales | GCF_900169095.1 | Brevibacterium_aurantiacum | BAUR920_RS16220 | 10995..14402 | 273384 | DUF499-domain-containing-protein-[Brevibacterium-aurantiacum] |
| WP_029263985.1 | HEPN+AAA-ATPase+Alpha+Beta-domain+wHTH+RRM | Type-1_BR.RM.like.systems | [illegible] | 1128 | Bacteria>Terrabacteria_group>Actinomycetota>Actinomycetes>Micrococcales | GCF_000708325.1 | unclassified_Microbacterium | DW303_RS18520 | 62853..66239 | 2609290 | MULTISPECIES:-DUF499-domain-containing-protein-[unclassified-Microbacterium] |
| WP_141784692.1 | HEPN+AAA-ATPase+Alpha+Beta-domain+wHTH+RRM | Type-1_BR.RM.like.systems | [illegible] | 1119 | Bacteria>Terrabacteria_group>Actinomycetota>Actinomycetes>Micrococcales | GCF_006716185.1 | Ornithinicoccus_hortensis | FB467_RS08335 | 1805419..1808778 | 82346 | DUF499-domain-containing-protein-[Ornithinicoccus-hortensis] |
| WP_054682987.1 | HEPN+AAA-ATPase+Alpha+Beta-domain+wHTH+RRM | Type-1_BR.RM.like.systems | [illegible] | 1129 | Bacteria>Terrabacteria_group>Actinomycetota>Actinomycetes>Micrococcales | GCF_001314225.1 | Microbacterium_sp._No._7 | A0A12_RS21690 | 98036..101425 | 1714373 | DUF499-domain-containing-protein-[Microbacterium-sp.-No.-7] |
| WP_251355553.1 | HEPN+AAA-ATPase+Alpha+Beta-domain+wHTH+RRM | Type-1_BR.RM.like.systems | [illegible] | 1104 | Bacteria>Terrabacteria_group>Actinomycetota>Actinomycetes>Micrococcales | - | Kocuria_rosea | M3684_RS03240 | 225275..228589 | 1275 | DUF499-domain-containing-protein-[Kocuria-rosea] |
| WP_249312439.1 | HEPN+AAA-ATPase+Alpha+Beta-domain+wHTH+RRM | Type-1_BR.RM.like.systems | [illegible] | 1140 | Bacteria>Terrabacteria_group>Actinomycetota>Actinomycetes>Micrococcales | GCF_018304855.1 | Pseudoclavibacter_sp._Marseille-Q4354 | JSQ82_RS10750 | 821875..825297 | 2810515 | DUF499-domain-containing-protein-[Pseudoclavibacter-sp.-Marseille-Q4354] |
| WP_014923538.1 | HEPN+AAA-ATPase+Alpha+Beta-domain+wHTH+RRM | Type-1_BR.RM.like.systems | [illegible] | 1120 | Bacteria>Terrabacteria_group>Actinomycetota>Actinomycetes>Micrococcales | GCF_000294695.2 | Arthrobacter_sp._Rue61a | ARUE_RS22630 | 197474..200836 | 1118963 | Swt1-family-HEPN-domain-containing-protein-[Arthrobacter-sp.-Rue61a] |
| MBS3179176.1 | HEPN+AAA-ATPase+Alpha+Beta-domain+wHTH+RRM | Type-1_BR.RM.like.systems | [illegible] | 1142 | Bacteria>Terrabacteria_group>Actinomycetota>Actinomycetes>Micrococcales | GCA_018304855.1 | Pseudoclavibacter_sp._Marseille-Q4354 | JSQ82_10750 | 821869..825297 | 2810515 | DUF499-domain-containing-protein-[Pseudoclavibacter-sp.-Marseille-Q4354] |
| WP_204409507.1 | HEPN+AAA-ATPase+Alpha+Beta-domain+wHTH+RRM | Type-1_BR.RM.like.systems | [illegible] | 1132 | Bacteria>Terrabacteria_group>Actinomycetota>Actinomycetes>Micrococcales | GCF_016907455.1 | Brachybacterium_muris | JOD52_RS08675 | 1871612..1875010 | 219301 | DUF499-domain-containing-protein-[Brachybacterium-muris] |
| WP_157002068.1 | HEPN+AAA-ATPase+Alpha+Beta-domain+wHTH+RRM | Type-1_BR.RM.like.systems | [illegible] | 1143 | Bacteria>Terrabacteria_group>Actinomycetota>Actinomycetes>Micrococcales | GCF_009749475.1 | Agromyces_laixinhei | GQQ90_RS09115 | 49199..52630 | 2585717 | DUF499-domain-containing-protein-[Agromyces-laixinhei] |
| MBD1536843.1 | HEPN+AAA-ATPase+Alpha+Beta-domain+wHTH+RRM | Type-1_BR.RM.like.systems | [illegible] | 1126 | Bacteria>Terrabacteria_group>Actinomycetota>Actinomycetes>Micrococcales | GCA_014694415.1 | Arthrobacter_sp._S13_S34 | HC749_01495 | 347639..351019 | 2720227 | ATP-binding-protein-[Arthrobacter-sp.-S13_S34] |
| WP_163540895.1 | HEPN+AAA-ATPase+Alpha+Beta-domain+wHTH+RRM | Type-1_BR.RM.like.systems | [illegible] | 1130 | Bacteria>Terrabacteria_group>Actinomycetota>Actinomycetes>Micrococcales | GCF_010994285.1 | Occultella_kanbiaonis | G7B41_RS01575 | 244678..248070 | 2675754 | Swt1-family-HEPN-domain-containing-protein-[Occultella-kanbiaonis] |
| WP_104269222.1 | HEPN+AAA-ATPase+Alpha+Beta-domain+wHTH+RRM | Type-1_BR.RM.like.systems | [illegible] | 1120 | Bacteria>Terrabacteria_group>Actinomycetota>Actinomycetes>Micrococcales | - | Rathayibacter_sp._AY1C9 | C5D09_10000 | 51639..55001 | 2080541 | Swt1-family-HEPN-domain-containing-protein-[Rathayibacter-sp.-AY1C9] |
| WP_047687850.1 | HEPN+AAA-ATPase+Alpha+Beta-domain+wHTH+RRM | Type-1_BR.RM.like.systems | [illegible] | 1133 | Bacteria>Terrabacteria_group>Actinomycetota>Actinomycetes>Micrococcales | GCF_000800955.1 | Kocuria_sp._ZOR0020 | L979_RS00020 | 3003..6404 | 1339234 | DUF499-domain-containing-protein-[Kocuria-sp.-ZOR0020] |
| PPH45732.1 | HEPN+AAA-ATPase+Alpha+Beta-domain+wHTH+RRM | Type-1_BR.RM.like.systems | [illegible] | 1120 | Bacteria>Terrabacteria_group>Actinomycetota>Actinomycetes>Micrococcales | GCA_002932315.1 | Rathayibacter_sp._AY1C9 | C5D09_10000 | 51639..55001 | 2080541 | AAA-family-ATPase-[Rathayibacter-sp.-AY1C9] |
| MBK6441773.1 | HEPN+AAA-ATPase+Alpha+Beta-domain+wHTH+RRM | Type-1_BR.RM.like.systems | [illegible] | 922 | Bacteria>Terrabacteria_group>Actinomycetota>Actinomycetes>Micrococcales | GCA_016704565.1 | Candidatus_Phosphoribacter_baldrii | IPF90_06370 | 1387205..1389971 | 2953744 | DUF499-domain-containing-protein-[Actinomycetales-bacterium] |
| WP_243591605.1 | HEPN+AAA-ATPase+Alpha+Beta-domain+wHTH+RRM | Type-1_BR.RM.like.systems | [illegible] | 1123 | Bacteria>Terrabacteria_group>Actinomycetota>Actinomycetes>Micrococcales | GCF_022815585.1 | Rathayibacter_caricis | MT349_RS09900 | 122292..125663 | 110936 | Swt1-family-HEPN-domain-containing-protein-[Rathayibacter-caricis] |
| WP_258066970.1 | HEPN+AAA-ATPase+Alpha+Beta-domain+wHTH+RRM | Type-1_BR.RM.like.systems | [illegible] | 1112 | Bacteria>Terrabacteria_group>Actinomycetota>Actinomycetes>Micrococcales | GCF_002932315.1 | Rathayibacter_sp._AY1C9 | C5D09_RS10010 | 51663..55001 | 2080541 | Swt1-family-HEPN-domain-containing-protein-[Rathayibacter-sp.-AY1C9] |
| WP_133351310.1 | HEPN+AAA-ATPase+Alpha+Beta-domain+wHTH+RRM | Type-1_BR.RM.like.systems | [illegible] | 1123 | Bacteria>Terrabacteria_group>Actinomycetota>Actinomycetes>Micrococcales | GCF_004358765.1 | Arthrobacter_nitrophenolicus | E2R57_RS17850 | 242800..246171 | 683150 | Swt1-family-HEPN-domain-containing-protein-[Arthrobacter-nitrophenolicus] |
| WP_036294304.1 | HEPN+AAA-ATPase+Alpha+Beta-domain+wHTH+RRM | Type-1_BR.RM.like.systems | [illegible] | 1131 | Bacteria>Terrabacteria_group>Actinomycetota>Actinomycetes>Micrococcales | GCF_000582705.2 | Microbacterium | MIC448_RS00375 | 9420..12815 | 33882 | MULTISPECIES:-DUF499-domain-containing-protein-[Microbacterium] |
| PZU44600.1 | HEPN+AAA-ATPase+Alpha+Beta-domain+wHTH+RRM | Type-1_BR.RM.like.systems | [illegible] | 1131 | Bacteria>Terrabacteria_group>Actinomycetota>Actinomycetes>Micrococcales | GCA_003248655.1 | Microbacterium_sp. | DI566_12010 | 17678..21073 | 51671 | AAA-family-ATPase-[Microbacterium-sp.] |

| Protein-ID | DUF499-architecture | System | Gene-Neighbourhoods | Length | Taxonomy | Assembly-ID | Strain | Locus-Tag | Genomic-coordinates | TaxID | Fasta-defline |
|---|---|---|---|---|---|---|---|---|---|---|---|
| MAT19458.1 | HEPN+AAA-ATPase+Alpha+Beta-domain+wHTH+RRM | Type-1_BR.RM.like.systems | <-ABhydrolase<-?\|\|?->?-><-SLATT+SLATT<-TIR<-HEPN+DUF499*<-METHYLASE<-TUDOR+SF1-SF2-Helicase+iREase+CTD+REase<-XIS-HTH | 1130 | Bacteria>Terrabacteria_group>Actinomycetota>Actinomycetes>Micrococcales | GCA_002702785.1 | Leifsonia_sp. | CMF56_13050 | 20496..23888 | 1870902 | AAA-family-ATPase-[Leifsonia-sp.] |
| WP_223984660.1 | HEPN+AAA-ATPase+Alpha+Beta-domain+wHTH+RRM | Type-1_BR.RM.like.systems | REase-><-?\|\|HKD-DNase+SF1-SF2-Helicase+TRD->?-><-?<-HEPN+DUF499*<-METHYLASE<-TUDOR+SF1-SF2-Helicase+iREase+CTD+REase | 1139 | Bacteria>Terrabacteria_group>Actinomycetota>Actinomycetes>Micrococcales | GCF_019977355.1 | Arthrobacter_sp._NicSoilB8 | LDO15_RS05065 | 1116619..1120038 | 2830998 | Swt1-family-HEPN-domain-containing-protein-[Arthrobacter-sp.-NicSoilB8] |
| WP_282449258.1 | HEPN+AAA-ATPase+Alpha+Beta-domain+wHTH+RRM | Type-1_BR.RM.like.systems | XIS-HTH->TUDOR+SF1-SF2-Helicase+iREase+CTD+REase->METHYLASE-p->METHYLASE->HEPN+DUF499*->HEPN-><-Glyoxalase | 1126 | Bacteria>Terrabacteria_group>Actinomycetota>Actinomycetes>Micrococcales | GCF_023155745.1 | Microbacterium_kunmingense | MA875_RS17035 | 50795..54175 | 2915939 | DUF499-domain-containing-protein-[Microbacterium-kunmingensis] |
| WP_166844731.1 | HEPN+AAA-ATPase+Alpha+Beta-domain+wHTH+RRM | Type-1_BR.RM.like.systems | <-MORC+HEPN<-HTH+SF1-SF2-Helicase+REase<-REase<-MORC<-HEPN+DUF499*<-METHYLASE\|\|DNA_methylase->?-><-?<-?<-HTH_3+PolB | 1156 | Bacteria>Terrabacteria_group>Actinomycetota>Actinomycetes>Micrococcales | GCF_011634835.1 | Isoptericola_sp._BMS4 | EYB28_RS03030 | 693115..696585 | 2527875 | Swt1-family-HEPN-domain-containing-protein-[Isoptericola-sp.-BMS4] |
| WP_229577768.1 | HEPN+AAA-ATPase+Alpha+Beta-domain+wHTH+RRM | Type-1_BR.RM.like.systems | <-HTH+SF1-SF2-Helicase+REase<-REase<-MORC<-HEPN+DUF499*<-METHYLASE-p<-METHYLASE-p<-REase\|\|DNA_methylase-> | 1141 | Bacteria>Terrabacteria_group>Actinomycetota>Actinomycetes>Micrococcales | GCF_020829765.1 | Kocuria_rhizophila | JNE43_RS06540 | 5673..9098 | 72000 | DUF499-domain-containing-protein-[Kocuria-rhizophila] |
| WP_247619184.1 | HEPN+AAA-ATPase+Alpha+Beta-domain+wHTH+RRM | Type-1_BR.RM.like.systems | <-OLD-ABC+TOPRIM<-HEPN+DUF499*<-wHTH+REase<-METHYLASE<-TUDOR+SF1-SF2-Helicase+iREase+CTD+REase<-XIS-HTH | 1135 | Bacteria>Terrabacteria_group>Actinomycetota>Actinomycetes>Micrococcales | GCF_023072605.1 | Brevibacterium_sp._R8603A2 | MTQ12_RS14500 | 1781..5188 | 2929779 | Swt1-family-HEPN-domain-containing-protein-[Brevibacterium-sp.-R8603A2] |
| WP_131132843.1 | HEPN+AAA-ATPase+Alpha+Beta-domain+wHTH+RRM | Type-1_BR.RM.like.systems | <-HTH+SF1-SF2-Helicase+REase<-REase<-MORC<-HEPN+DUF499*<-METHYLASE\|\|DNA_methylase-><-SbcC<-DUF1679<-SIR2 | 1130 | Bacteria>Terrabacteria_group>Actinomycetota>Actinomycetes>Micrococcales | GCF_004320525.1 | Arthrobacter_sp._S39 | EYS21_RS09020 | 558026..561418 | 2509720 | Swt1-family-HEPN-domain-containing-protein-[Arthrobacter-sp.-S39] |
| SNS99685.1 | HEPN+AAA-ATPase+Alpha+Beta-domain+wHTH+RRM | Type-1_BR.RM.like.systems | <-HTH_19\|\|TUDOR+SF1-SF2-Helicase+iREase+CTD+REase->Phd-YefM->RelE-ParE->METHYLASE->HEPN+DUF499*->?-><-?\|\|?->?->?->HNH-> | 1175 | Bacteria>Terrabacteria_group>Actinomycetota>Actinomycetes>Micrococcales | GCA_900188465.1 | Micrococcales_bacterium_KH10 | SAMN06309944_1702 | 505184..508711 | 1945885 | Predicted-ATPase,-AAA+-superfamily-[Micrococcales-bacterium-KH10] |
| WP_257744324.1 | HEPN+AAA-ATPase+Alpha+Beta-domain+wHTH+RRM | Type-1_BR.RM.like.systems | <-DNA_methylase\|\|REase->METHYLASE->HEPN+DUF499*->MORC->REase->HTH+SF1-SF2-Helicase+REase-><-?\|\|MarR-HTH-> | 1137 | Bacteria>Terrabacteria_group>Actinomycetota>Actinomycetes>Micrococcales | GCF_024655785.1 | Kocuria_rhizophila | NUV30_RS04805 | 31091..34504 | 72000 | DUF499-domain-containing-protein-[Kocuria-rhizophila] |
| WP_126874170.1 | HEPN+AAA-ATPase+Alpha+Beta-domain+wHTH+RRM | Type-1_BR.RM.like.systems | <-DNA_methylase\|\|REase->METHYLASE->HEPN+DUF499*->MORC->REase->HTH+SF1-SF2-Helicase+REase-><-?\|\|MarR-HTH-> | 1141 | Bacteria>Terrabacteria_group>Actinomycetota>Actinomycetes>Micrococcales | GCF_003989235.1 | unclassified_Kocuria | D8M38_RS00615 | 20820..24245 | 2649579 | MULTISPECIES:-DUF499-domain-containing-protein-[unclassified-Kocuria] |
| WP_236803031.1 | HEPN+AAA-ATPase+Alpha+Beta-domain+wHTH+RRM | Type-1_BR.RM.like.systems | <-ABC-ATPase+SbcC<-?<-HEPN+DUF499*<-METHYLASE<-TUDOR+SF1-SF2-Helicase+iREase+CTD+REase<-XIS-HTH\|\|REase-> | 1127 | Bacteria>Terrabacteria_group>Actinomycetota>Actinomycetes>Micrococcales | GCF_021801725.1 | Arthrobacter_sp._FW306-05-C | LFT44_RS20985 | 51567..54950 | 2879620 | Swt1-family-HEPN-domain-containing-protein-[Arthrobacter-sp.-FW306-05-C] |
| WP_224781058.1 | HEPN+AAA-ATPase+Alpha+Beta-domain+wHTH+RRM | Type-1_BR.RM.like.systems | <-MORC+HEPN<-DNA_methylase\|\|?->METHYLASE->HEPN+DUF499*->MORC->REase->HTH+SF1-SF2-Helicase+REase->REase-> | 1130 | Bacteria>Terrabacteria_group>Actinomycetota>Actinomycetes>Micrococcales | GCF_020096995.1 | Leucobacter_sp._Psy1 | K8P10_RS06885 | 1465839..1469231 | 2875729 | Swt1-family-HEPN-domain-containing-protein-[Leucobacter-sp.-Psy1] |
| WP_255770908.1 | HEPN+AAA-ATPase+Alpha+Beta-domain+wHTH+RRM | Type-1_BR.RM.like.systems | <-DNA_methylase\|\|METHYLASE->HEPN+DUF499*->MORC->REase->HTH+SF1-SF2-Helicase+REase->ParB+HNH-><-?<-NUDIX | 1122 | Bacteria>Terrabacteria_group>Actinomycetota>Actinomycetes>Micrococcales | GCF_024391045.1 | Pseudarthrobacter_sulfonivorans | NQJ30_RS15540 | 5007..8375 | 121292 | DUF499-domain-containing-protein-[Pseudarthrobacter-sulfonivorans] |
| WP_137320273.1 | HEPN+AAA-ATPase+Alpha+Beta-domain+wHTH+RRM | Type-1_BR.RM.like.systems | <-HEPN+DUF499*<-wHTH+REase<-METHYLASE<-TUDOR+SF1-SF2-Helicase+iREase+CTD+REase<-XIS-HTH<-ABhydrolase | 1132 | Bacteria>Terrabacteria_group>Actinomycetota>Actinomycetes>Micrococcales | GCF_005280295.1 | Brevibacterium_sp._CS2 | FDF13_RS00005 | 3239176..2834 | 2575923 | DUF499-domain-containing-protein-[Brevibacterium-sp.-CS2] |
| WP_168429935.1 | HEPN+AAA-ATPase+Alpha+Beta-domain+wHTH+RRM | Type-1_BR.RM.like.systems | <-MerR-HTH<-?\|\|MORC+DUF3684+REase-><-?<-HEPN+DUF499*<-METHYLASE<-TUDOR+SF1-SF2-Helicase+iREase+CTD+REase | 1113 | Bacteria>Terrabacteria_group>Actinomycetota>Actinomycetes>Micrococcales | GCF_012396205.1 | Cellulomonas_hominis | HGA09_RS00165 | 40962..44303 | 156981 | DUF499-domain-containing-protein-[Cellulomonas-hominis] |
| WP_104088073.1 | HEPN+AAA-ATPase+Alpha+Beta-domain+wHTH+RRM | Type-1_BR.RM.like.systems | <-HEPN+DUF499*<-METHYLASE<-TUDOR+SF1-SF2-Helicase+iREase+CTD+REase<-XIS-HTH\|\|GDResolvase+HIN-HTH-> | 1133 | Bacteria>Terrabacteria_group>Actinomycetota>Actinomycetes>Micrococcales | GCF_002929595.1 | Cryobacterium_sp._N19 | CXY78_RS03325 | 298..3699 | 2048288 | DUF499-domain-containing-protein-[Cryobacterium-sp.-N19] |
| WP_194396698.1 | HEPN+AAA-ATPase+Alpha+Beta-domain+wHTH+RRM | Type-1_BR.RM.like.systems | METHYLASE-><-HNH<-OLD-ABC+TOPRIM<-HEPN+DUF499*<-METHYLASE<-TUDOR+SF1-SF2-Helicase+iREase+CTD+REase<-XIS-HTH | 1130 | Bacteria>Terrabacteria_group>Actinomycetota>Actinomycetes>Micrococcales | GCF_015277815.1 | Microbacterium_atlanticum | IR212_RS15230 | 3329381..3332773 | 2782168 | DUF499-domain-containing-protein-[Microbacterium-atlanticum] |
| WP_253867645.1 | HEPN+AAA-ATPase+Alpha+Beta-domain+wHTH+RRM | Type-1_BR.RM.like.systems | <-HTH+SF1-SF2-Helicase+REase<-REase<-MORC<-HEPN+DUF499*<-METHYLASE\|\|HTH_3->wHTH+SAD+HNH->HTH-> | 1150 | Bacteria>Terrabacteria_group>Actinomycetota>Actinomycetes>Micrococcales | GCF_024171995.1 | Promicromonospora_umidemergens | APR04_RS02600 | 547565..551017 | 629679 | Swt1-family-HEPN-domain-containing-protein-[Promicromonospora-umidemergens] |
| WP_168888196.1 | HEPN+AAA-ATPase+Alpha+Beta-domain+wHTH+RRM | Type-1_BR.RM.like.systems | CopG_antitoxin-><-?<-HEPN+DUF499*<-wHTH+REase<-METHYLASE<-TUDOR+SF1-SF2-Helicase+iREase+CTD+REase | 1127 | Bacteria>Terrabacteria_group>Actinomycetota>Actinomycetes>Micrococcales | GCF_012641515.1 | Nesterenkonia_sedimenti | HGQ17_RS12070 | 43839..47222 | 1463632 | DUF499-domain-containing-protein-[Nesterenkonia-sedimenti] |
| WP_029212622.1 | HEPN+AAA-ATPase+Alpha+Beta-domain+wHTH+RRM | Type-1_BR.RM.like.systems | SF1-SF2-Helicase->tRNA->TUDOR+SF1-SF2-Helicase+iREase+CTD+REase->METHYLASE->ParB+HNH->HEPN+DUF499*-> | 1119 | Bacteria>Terrabacteria_group>Actinomycetota>Actinomycetes>Micrococcales | GCF_000426385.1 | Arsenicicoccus_bolidensis | K313_RS0116910 | 19529..22888 | 229480 | Swt1-family-HEPN-domain-containing-protein-[Arsenicicoccus-bolidensis] |
| MBN2177583.1 | HEPN+AAA-ATPase+Alpha+Beta-domain+wHTH+RRM | Type-1_BR.RM.like.systems | <-AbiEii<-AbiEi_4<-?<-HEPN+DUF499*<-METHYLASE\|\|?->HTH_31->?-><-TUDOR+SF1-SF2-Helicase+iREase+CTD+REase | 1113 | Bacteria>Terrabacteria_group>Actinomycetota>Actinomycetes>Micrococcales | GCA_016933155.1 | Demequinaceae_bacterium | JW722_08015 | 69321..72662 | 2683256 | DUF499-domain-containing-protein-[Demequinaceae-bacterium] |
| KQY96844.1 | HEPN+AAA-ATPase+Alpha+Beta-domain+wHTH+RRM | Type-1_BR.RM.like.systems | Phage_holin->METHYLASE->?->ACET->HEPN+DUF499*->DUF4096->DDE_Tnp_1_2-><-UDG-conflict<-?<-REase<-MobL | 1102 | Bacteria>Terrabacteria_group>Actinomycetota>Actinomycetes>Micrococcales | GCA_001426925.1 | Microbacterium_sp._Root53 | ASD19_09760 | 99084..102392 | 1736553 | AAA-family-ATPase-[Microbacterium-sp.-Root53] |
| WP_200569160.1 | HEPN+AAA-ATPase+Alpha+Beta-domain+wHTH+RRM | Type-1_BR.RM.like.systems | <-DNA_methylase\|\|REase->METHYLASE->HEPN+DUF499*->MORC->REase->HTH+SF1-SF2-Helicase+REase-> | 1144 | Bacteria>Terrabacteria_group>Actinomycetota>Actinomycetes>Micrococcales | GCF_016620465.1 | Kocuria_rhizophila | JHD97_RS04255 | 38629..42063 | 72000 | DUF499-domain-containing-protein-[Kocuria-rhizophila] |
| MCV7506294.1 | HEPN+AAA-ATPase+Alpha+Beta-domain+wHTH+RRM | Type-1_BR.RM.like.systems | <-HTH+SF1-SF2-Helicase+REase<-REase<-MORC<-HEPN+DUF499*<-METHYLASE<-DNA_methylase\|\|REase-> | 1157 | Bacteria>Terrabacteria_group>Actinomycetota>Actinomycetes>Micrococcales | GCA_025151425.2 | Micrococcus_luteus | M3D82_001030 | 202363..205836 | 1270 | Swt1-family-HEPN-domain-containing-protein-[Micrococcus-luteus] |
| WP_200941118.1 | HEPN+AAA-ATPase+Alpha+Beta-domain+wHTH+RRM | Type-1_BR.RM.like.systems | Phage_holin->METHYLASE->?->ACET->HEPN+DUF499*->DUF4096+DDE_Tnp_1+DDE_Tnp_1_2-><-?<-REase<-MobL | 1112 | Bacteria>Terrabacteria_group>Actinomycetota>Actinomycetes>Micrococcales | GCF_001426925.1 | Microbacterium_sp._Root53 | ASD19_RS13260 | 99054..102392 | 1736553 | Swt1-family-HEPN-domain-containing-protein-[Microbacterium-sp.-Root53] |
| WP_159096312.1 | HEPN+AAA-ATPase+Alpha+Beta-domain+wHTH+RRM | Type-1_BR.RM.like.systems | <-DDE_Tnp_1<-HEPN+DUF499*<-METHYLASE<-TUDOR+SF1-SF2-Helicase+iREase+CTD+REase\|\|?-><-?<-?<-REase | 1126 | Bacteria>Terrabacteria_group>Actinomycetota>Actinomycetes>Micrococcales | GCF_003074035.1 | Miniimonas_sp._S16 | DCS86_RS00720 | 144486..147866 | 2171623 | DUF499-domain-containing-protein-[Miniimonas-sp.-S16] |
| WP_260073607.1 | HEPN+AAA-ATPase+Alpha+Beta-domain+wHTH+RRM | Type-1_BR.RM.like.systems | <-HIN-HTH\|\|?->TUDOR+SF1-SF2-Helicase+iREase+CTD+REase->METHYLASE->HEPN->HEPN+DUF499*-> | 1129 | Bacteria>Terrabacteria_group>Actinomycetota>Actinomycetes>Micrococcales | GCF_025146225.1 | Nesterenkonia_massiliensis | M3B43_RS10510 | 14559..17948 | 1232429 | DUF499-domain-containing-protein-[Nesterenkonia-massiliensis] |
| WP_068492369.1 | HEPN+AAA-ATPase+Alpha+Beta-domain+wHTH+RRM | Type-1_BR.RM.like.systems | <-HTH+SF1-SF2-Helicase+REase<-REase<-MORC<-HEPN+DUF499*<-METHYLASE\|\|DNA_methylase-> | 1129 | Bacteria>Terrabacteria_group>Actinomycetota>Actinomycetes>Micrococcales | GCF_900014995.1 | Pseudoclavibacter_helvolus | BN2259_RS00510 | 5683..9072 | 255205 | Swt1-family-HEPN-domain-containing-protein-[Pseudoclavibacter-helvolus] |
| WP_285329939.1 | HEPN+AAA-ATPase+Alpha+Beta-domain+wHTH+RRM | Type-1_BR.RM.like.systems | <-RelA<-?\|\|35exo->tRNA->TUDOR+SF1-SF2-Helicase+iREase+CTD+REase->METHYLASE->HEPN+DUF499*-> | 1139 | Bacteria>Terrabacteria_group>Actinomycetota>Actinomycetes>Micrococcales | GCF_030216795.1 | Pseudoglutamicibacter_cumminsii | QP411_RS02105 | 150205..153624 | 156979 | DUF499-domain-containing-protein-[Pseudoglutamicibacter-cumminsii] |
| WP_141391091.1 | HEPN+AAA-ATPase+Alpha+Beta-domain+wHTH+RRM | Type-1_BR.RM.like.systems | <-HEPN+DUF499*<-METHYLASE<-TUDOR+SF1-SF2-Helicase+iREase+CTD+REase<-tRNA<-Aminotran_4 | 1129 | Bacteria>Terrabacteria_group>Actinomycetota>Actinomycetes>Micrococcales | GCF_006539945.1 | Cellulosimicrobium_cellulans | CCE02nite_RS18570 | 1298..4687 | 1710 | DUF499-domain-containing-protein-[Cellulosimicrobium-cellulans] |
| WP_102209830.1 | HEPN+AAA-ATPase+Alpha+Beta-domain+wHTH+RRM | Type-1_BR.RM.like.systems | <-HEPN+DUF499*<-?<-METHYLASE<-Phage_holin<-?<-?<-TUDOR+SF1-SF2-Helicase+iREase+CTD+REase | 1123 | Bacteria>Terrabacteria_group>Actinomycetota>Actinomycetes>Micrococcales | GCF_002871415.1 | Microbacterium_sp._UMB0228 | CJ226_RS16275 | 2567..5938 | 2029109 | DUF499-domain-containing-protein-[Microbacterium-sp.-UMB0228] |
| WP_247963551.1 | HEPN+AAA-ATPase+Alpha+Beta-domain+wHTH+RRM | Type-1_BR.RM.like.systems | DDE_Tnp_1->HIN-HTH->HTH_21+rve+rve_3->METHYLASE-p->HEPN+DUF499*->REase+SAVED-> | 1143 | Bacteria>Terrabacteria_group>Actinomycetota>Actinomycetes>Micrococcales | GCF_023108875.1 | Microbacterium_aerolatum | MZK47_RS16315 | 3209..6640 | 153731 | DUF499-domain-containing-protein-[Microbacterium-aerolatum] |
| WP_076404375.1 | HEPN+AAA-ATPase+Alpha+Beta-domain+wHTH+RRM | Type-1_BR.RM.like.systems | REase-><-?<-?<-?<-HEPN+DUF499*<-ParB+HNH<-METHYLASE\|\|MPTase->PIN->HTH_3->wHTH(SBD)+HNH-> | 1151 | Bacteria>Terrabacteria_group>Actinomycetota>Actinomycetes>Micrococcales | GCF_900155925.1 | Cellulosimicrobium_aquatile | BXA05_RS06490 | 508556..512011 | 1612203 | DUF499-domain-containing-protein-[Cellulosimicrobium-aquatile] |
| PZU36627.1 | HEPN+AAA-ATPase+Alpha+Beta-domain+wHTH+RRM | Type-1_BR.RM.like.systems | <-MODE-HTH+PBPII\|\|?->?->?->DUF4096+DDE_Tnp_1+DDE_Tnp_1_2-><-HEPN+DUF499*<-METHYLASE-p | 1129 | Bacteria>Terrabacteria_group>Actinomycetota>Actinomycetes>Micrococcales | GCA_003248605.1 | Microbacterium_sp. | DI573_13315 | 12880..16269 | 51671 | AAA-family-ATPase-[Microbacterium-sp.] |
| WP_247963105.1 | HEPN+AAA-ATPase+Alpha+Beta-domain+wHTH+RRM | Type-1_BR.RM.like.systems | DDE_Tnp_1->METHYLASE-p->HEPN+DUF499*->ParB+HNH->?-><-N6_MTase-p<-McrC-NTD+REase | 1132 | Bacteria>Terrabacteria_group>Actinomycetota>Actinomycetes>Micrococcales | GCF_023108875.1 | Microbacterium_aerolatum | MZK47_RS13285 | 1211..4609 | 153731 | DUF499-domain-containing-protein-[Microbacterium-aerolatum] |
| WP_232711167.1 | HEPN+AAA-ATPase+Alpha+Beta-domain+wHTH+RRM | Type-1_BR.RM.like.systems | Phage_holin->METHYLASE->?->?->HEPN+DUF499*->?->?->wHTH+REase->TrwC+SF1-SF2-Helicase-> | 1114 | Bacteria>Terrabacteria_group>Actinomycetota>Actinomycetes>Micrococcales | GCF_002812725.1 | unclassified_Microbacterium | CGU35_RS05955 | 370107..373451 | 2609290 | MULTISPECIES:-DUF499-domain-containing-protein-[unclassified-Microbacterium] |
| WP_126499814.1 | HEPN+AAA-ATPase+Alpha+Beta-domain+wHTH+RRM | Type-1_BR.RM.like.systems | Response_reg+LuxR-HTH-><-?<-TetR-HTH\|\|ABhydrolase->?->METHYLASE->HEPN+DUF499*-> | 1125 | Bacteria>Terrabacteria_group>Actinomycetota>Actinomycetes>Micrococcales | GCF_900637985.1 | Rothia_aeria | EL250_RS03685 | 841376..844753 | 172042 | DUF499-domain-containing-protein-[Rothia-aeria] |
| HBS75978.1 | HEPN+AAA-ATPase+Alpha+Beta-domain+wHTH+RRM | Type-1_BR.RM.like.systems | <-TrwC+SF1-SF2-Helicase<-wHTH+REase<-?\|\|NTD+SF1-SF2-Helicase-><-HEPN+DUF499* | 1125 | Bacteria>Terrabacteria_group>Actinomycetota>Actinomycetes>Micrococcales | GCA_003510265.1 | Microbacterium_sp. | DEB55_16590 | 13006..16383 | 51671 | AAA-family-ATPase-[Microbacterium-sp.] |
| SDF11628.1 | HEPN+AAA-ATPase+Alpha+Beta-domain+wHTH+RRM | Type-1_BR.RM.like.systems | <-wHTH(SBD)+HNH<-HTH_3<-PIN<-MPTase\|\|METHYLASE->ParB+HNH->HEPN+DUF499*-> | 1151 | Bacteria>Terrabacteria_group>Actinomycetota>Actinomycetes>Micrococcales | GCA_900102445.1 | Cellulosimicrobium_cellulans | SAMN04487781_0340 | 373492..376947 | 1710 | Predicted-ATPase,-AAA+-superfamily-[Cellulosimicrobium-cellulans] |
| PZU44169.1 | HEPN+AAA-ATPase+Alpha+Beta-domain+wHTH+RRM | Type-1_BR.RM.like.systems | TetR-HTH-><-?<-?\|\|?->\|\|HEPN+DUF499*->OLD-ABC+TOPRIM-><-?<-HIN-HTH\|\|HTH_19->Phd-YefM->PIN-> | 1129 | Bacteria>Terrabacteria_group>Actinomycetota>Actinomycetes>Micrococcales | GCA_003248655.1 | Microbacterium_sp. | DI566_12710 | 6913..10302 | 51671 | AAA-family-ATPase-[Microbacterium-sp.] |
| WP_034226753.1 | HEPN+AAA-ATPase+Alpha+Beta-domain+wHTH+RRM | Type-1_BR.RM.like.systems | <-HTH+SF1-SF2-Helicase+REase<-REase<-MORC<-HEPN+DUF499*<-METHYLASE | 1144 | Bacteria>Terrabacteria_group>Actinomycetota>Actinomycetes>Micrococcales | GCF_000603945.1 | Actinotalea_ferrariae | N866_RS12275 | 5788..9222 | 1386098 | Swt1-family-HEPN-domain-containing-protein-[Actinotalea-ferrariae] |
| NDC48896.1 | HEPN+AAA-ATPase+Alpha+Beta-domain+wHTH+RRM | Type-1_BR.RM.like.systems | TUDOR+SF1-SF2-Helicase+iREase+CTD+REase->METHYLASE->HEPN+DUF499*-> | 1108 | Bacteria>Terrabacteria_group>Actinomycetota>Actinomycetes>Micrococcales | GCA_010027415.1 | Micrococcales_bacterium | EBZ61_07440 | 7302..10628 | 2026762 | ATP-binding-protein-[Micrococcales-bacterium] |
| MCV7527924.1 | HEPN+AAA-ATPase+Alpha+Beta-domain+wHTH+RRM | Type-1_BR.RM.like.systems | <-REase\|\|DNA_methylase->METHYLASE->HEPN+DUF499*->MORC->REase-> | 1157 | Bacteria>Terrabacteria_group>Actinomycetota>Actinomycetes>Micrococcales | GCA_025149725.2 | Micrococcus_luteus | M3D91_010335 | 6032..9505 | 1270 | Swt1-family-HEPN-domain-containing-protein-[Micrococcus-luteus] |
| MCL1800684.1 | HEPN+AAA-ATPase+Alpha+Beta-domain+wHTH+RRM | Type-1_BR.RM.like.systems | REase->PhdYeFM_antitox->RelE-ParE->METHYLASE->HEPN+DUF499*-> | 635 | Bacteria>Terrabacteria_group>Actinomycetota>Actinomycetes>Micrococcales | GCA_009782595.1 | Promicromonosporaceae_bacterium | FWG25_04890 | 4477..6383 | 2304674 | DUF499-domain-containing-protein-[Promicromonosporaceae-bacterium] |
| WP_197462106.1 | HEPN+AAA-ATPase+Alpha+Beta-domain+wHTH+RRM | Type-1_BR.RM.like.systems | HEPN+DUF499*->DUF499-p->OLD-ABC+TOPRIM-p->OLD-ABC+TOPRIM-p-> | 968 | Bacteria>Terrabacteria_group>Actinomycetota>Actinomycetes>Micrococcales | GCF_001595495.1 | Microbacterium_sp._CH1 | AUV07_RS03260 | 609..3515 | 1770208 | DUF499-domain-containing-protein-[Microbacterium-sp.-CH1] |
| WP_276315051.1 | HEPN+AAA-ATPase+Alpha+Beta-domain+wHTH+RRM | Type-1_BR.RM.like.systems | REase+SF1-SF2-Helicase-><-SbcC<-HEPN+DUF499*<-METHYLASE-p | 1127 | Bacteria>Terrabacteria_group>Actinomycetota>Actinomycetes>Micrococcales | GCF_021395155.1 | Microbacterium_profundi | LZG07_RS12530 | 143258..146641 | 450380 | DUF499-domain-containing-protein-[Microbacterium-profundi] |
| WP_234001366.1 | HEPN+AAA-ATPase+Alpha+Beta-domain+wHTH+RRM | Type-1_BR.RM.like.systems | REase+SF1-SF2-Helicase-><-SbcC<-HEPN+DUF499*<-METHYLASE-p | 1108 | Bacteria>Terrabacteria_group>Actinomycetota>Actinomycetes>Micrococcales | - | Microbacterium_profundi | LZG07_12530 | 143258..146584 | 450380 | DUF499-domain-containing-protein-[Microbacterium-profundi] |
| MCE7482742.1 | HEPN+AAA-ATPase+Alpha+Beta-domain+wHTH+RRM | Type-1_BR.RM.like.systems | REase+SF1-SF2-Helicase-><-SbcC<-HEPN+DUF499*<-METHYLASE-p | 1108 | Bacteria>Terrabacteria_group>Actinomycetota>Actinomycetes>Micrococcales | GCA_021395155.1 | Microbacterium_profundi | LZG07_12530 | 143258..146584 | 450380 | DUF499-domain-containing-protein-[Microbacterium-profundi] |
| WP_009381273.1 | HEPN+AAA-ATPase+Alpha+Beta-domain+wHTH+RRM | Type-1_BR.RM.like.systems | <-HEPN+REase<-REase<-HEPN+DUF499*<-wHTH+REase<-METHYLASE | 1132 | Bacteria>Terrabacteria_group>Actinomycetota>Actinomycetes>Micrococcales | GCF_000314575.1 | Brevibacterium_casei | C272_RS16125 | 18996..22394 | 33889 | DUF499-domain-containing-protein-[Brevibacterium-casei] |
| WP_047691765.1 | HEPN+AAA-ATPase+Alpha+Beta-domain+wHTH+RRM | Type-1_BR.RM.like.systems | <-DNA_methylase\|\|REase->METHYLASE->HEPN+DUF499*-> | 452 | Bacteria>Terrabacteria_group>Actinomycetota>Actinomycetes>Micrococcales | GCF_000800955.1 | Kocuria_sp._ZOR0020 | L979_RS08865 | 18602..19957 | 1339234 | Swt1-family-HEPN-domain-containing-protein,-partial-[Kocuria-sp.-ZOR0020] |
| WP_026937806.1 | HEPN+AAA-ATPase+Alpha+Beta-domain+wHTH+RRM | Type-1_BR.RM.like.systems | <-HEPN+DUF499*<-METHYLASE\|\|DNA_methylase-> | 1131 | Bacteria>Terrabacteria_group>Actinomycetota>Actinomycetes>Micrococcales | GCF_000425685.1 | Gulosibacter_molinativorax | C7K25_RS15310 | 538..3933 | 256821 | Swt1-family-HEPN-domain-containing-protein-[Gulosibacter-molinativorax] |
| WP_282859602.1 | HEPN+AAA-ATPase+Alpha+Beta-domain+wHTH+RRM | Type-1_BR.RM.like.systems | <-SbcC<-HEPN+DUF499*<-METHYLASE<-REase | 1126 | Bacteria>Terrabacteria_group>Actinomycetota>Actinomycetes>Micrococcales | GCF_030011915.1 | Pseudoclavibacter_helvolus | QM024_RS14940 | 2044..5424 | 255205 | DUF499-domain-containing-protein-[Pseudoclavibacter-helvolus] |
| MBD3753304.1 | HEPN+AAA-ATPase+Alpha+Beta-domain+wHTH+RRM | Type-1_BR.RM.like.systems | <-HEPN+DUF499*<-METHYLASE-p | 452 | Bacteria>Terrabacteria_group>Actinomycetota>Actinomycetes>Micrococcales | GCA_014763555.1 | Micrococcales_bacterium | IE935_13015 | 1..1358 | 2026762 | ATP-binding-protein-[Micrococcales-bacterium] |
| WP_076703521.1 | HEPN+AAA-ATPase+Alpha+Beta-domain+wHTH+RRM | Type-1_BR.RM.like.systems | <-HEPN+DUF499*<-wHTH+REase | 1131 | Bacteria>Terrabacteria_group>Actinomycetota>Actinomycetes>Micrococcales | GCF_001968835.1 | Tersicoccus_phoenicis | BKD30_RS07045 | 193..3588 | 554083 | Swt1-family-HEPN-domain-containing-protein-[Tersicoccus-phoenicis] |
| WP_187693213.1 | HEPN+AAA-ATPase+Alpha+Beta-domain+wHTH+RRM | Type-1_BR.RM.like.systems | SF1-SF2-helicase+RdgC+Beta-barrel+REase->\|\|HTH_Tnp_4+DDE_Tnp_4->\|\|DUF4096+DDE_Tnp_1+DDE_Tnp_1_2->TUDOR+SF1-SF2-Helicase+iREase+CTD+REase->METHYLASE->ParB+HNH->HEPN+DUF499*-><-Phage_integrase<-?<-?<-?<-Cysteine_desulfurase | 1129 | Bacteria>Terrabacteria_group>Actinomycetota>Actinomycetes>Micromonosporales | GCF_014472405.1 | Micromonospora_chalcea | H9X95_RS12030 | 19161..22550 | 1874 | Swt1-family-HEPN-domain-containing-protein-[Micromonospora-chalcea] |
| WP_030502747.1 | HEPN+AAA-ATPase+Alpha+Beta-domain+wHTH+RRM | Type-1_BR.RM.like.systems | HTH_3->?-><-DEDD_Tnp_IS110+Transposase_20<-zf-ISL3+DDE_Tnp_ISL3\|\|MULE-TRANSPOSASE-><-HEPN+DUF499*<-ParB+HNH<-METHYLASE<-TUDOR+SF1-SF2-Helicase+iREase+CTD+REase\|\|NTD+SF1-SF2-Helicase->REase->pPIWI+pPIWI-> | 1124 | Bacteria>Terrabacteria_group>Actinomycetota>Actinomycetes>Micromonosporales | - | Micromonospora_purpureochromogenes | IH31_RS0121315 | 105944..109318 | 47872 | Swt1-family-HEPN-domain-containing-protein-[Micromonospora-purpureochromogenes] |
| WP_197494866.1 | HEPN+AAA-ATPase+Alpha+Beta-domain+wHTH+RRM | Type-1_BR.RM.like.systems | <-pPIWI+pPIWI<-REase<-NTD+SF1-SF2-Helicase\|\|TUDOR+SF1-SF2-Helicase+iREase+CTD+REase->METHYLASE->ParB+HNH->HEPN+DUF499*->?->DEDD_Tnp_IS110+Transposase_20->zf-ISL3+DDE_Tnp_ISL3-><-?<-HTH_3 | 1124 | Bacteria>Terrabacteria_group>Actinomycetota>Actinomycetes>Micromonosporales | GCF_001652995.1 | Micromonospora_sp._NBRC_110037 | A4R25_RS09725 | 2384437..2387811 | 1621261 | Swt1-family-HEPN-domain-containing-protein-[Micromonospora-sp.-NBRC-110037] |
| WP_221380246.1 | HEPN+AAA-ATPase+Alpha+Beta-domain+wHTH+RRM | Type-1_BR.RM.like.systems | sGTP->tRNA->Phage_int_SAM_4+Phage_integrase->METHYLASE-><-CARF+HEPN<-?<-ParB+HNH<-HEPN+DUF499*<-METHYLASE<-TUDOR+SF1-SF2-Helicase+iREase+CTD+REase<-SF1-SF2-Helicase<-?<-REase | 1128 | Bacteria>Terrabacteria_group>Actinomycetota>Actinomycetes>Micromonosporales | GCF_019710575.1 | Actinoplanes_polyasparticus | KXC43_RS33640 | 36558..39944 | 2856853 | Swt1-family-HEPN-domain-containing-protein-[Actinoplanes-sp.-TRM66264-DLM] |
| MBE1487514.1 | HEPN+AAA-ATPase+Alpha+Beta-domain+wHTH+RRM | Type-1_BR.RM.like.systems | HTH_17->Phage_int_SAM_1+Phage_integrase-><-?\|\|?->TUDOR+SF1-SF2-Helicase+iREase+CTD+REase->METHYLASE->HEPN+DUF499*-><-DDE_Tnp_1_2\|\|?->DNA-glycosylase-><-?<-DUF6361<-RNase | 1124 | Bacteria>Terrabacteria_group>Actinomycetota>Actinomycetes>Micromonosporales | GCA_014874095.1 | Plantactinospora_soyae | H4W31_003152 | 3518204..3521578 | 1544732 | putative-AAA+-superfamily-ATPase-[Plantactinospora-soyae] |
| WP_190252034.1 | HEPN+AAA-ATPase+Alpha+Beta-domain+wHTH+RRM | Type-1_BR.RM.like.systems | <-ABC-ATPase+SbcC\|\|tRNA->?-><-?\|\|TUDOR+SF1-SF2-Helicase+iREase+CTD+REase->METHYLASE->HEPN+DUF499*->ParB+HNH->?-><-Phage_int_SAM_1+Phage_integrase<-HTH_17\|\|?-><-BirA-HTH | 1122 | Bacteria>Terrabacteria_group>Actinomycetota>Actinomycetes>Micromonosporales | GCA_014647515.1 | Dactylosporangium_sucinum | IEZ31_RS23190 | 135414..138782 | 1424081 | Swt1-family-HEPN-domain-containing-protein-[Dactylosporangium-sucinum] |
| WP_225945547.1 | HEPN+AAA-ATPase+Alpha+Beta-domain+wHTH+RRM | Type-1_BR.RM.like.systems | HTH_17->Phage_int_SAM_1+Phage_integrase->?->TUDOR+SF1-SF2-Helicase+iREase+CTD+REase->METHYLASE->HEPN+DUF499*->\|\|?->DNA-glycosylase-><-?<-DUF6361<-RNase<-REase | 1132 | Bacteria>Terrabacteria_group>Actinomycetota>Actinomycetes>Micromonosporales | - | Plantactinospora_soyae | H4W31_RS15750 | 3518180..3521578 | 1544732 | DUF499-domain-containing-protein-[Plantactinospora-soyae] |
| WP_030271467.1 | HEPN+AAA-ATPase+Alpha+Beta-domain+wHTH+RRM | Type-1_BR.RM.like.systems | Cysteine_desulfurase->?->?->Phage_integrase-><-HEPN+DUF499*<-ParB+HNH<-METHYLASE<-TUDOR+SF1-SF2-Helicase+iREase+CTD+REase\|\|NTD+SF1-SF2-Helicase-><-?<-JetD | 1125 | Bacteria>Terrabacteria_group>Actinomycetota>Actinomycetes>Micromonosporales | GCF_000717585.1 | Micromonospora_globosa | IF06_RS0114120 | 15248..18625 | 47863 | Swt1-family-HEPN-domain-containing-protein-[Micromonospora-globosa] |
| WP_192902263.1 | HEPN+AAA-ATPase+Alpha+Beta-domain+wHTH+RRM | Type-1_BR.RM.like.systems | HTH+TPR+TPR->Pkinase-><-HEPN+DUF499*<-ParB+HNH<-METHYLASE<-TUDOR+SF1-SF2-Helicase+iREase+CTD+REase\|\|NTD+SF1-SF2-Helicase->REase->pPIWI+pPIWI-> | 1129 | Bacteria>Terrabacteria_group>Actinomycetota>Actinomycetes>Micromonosporales | GCF_003610785.1 | Micromonospora_globbae | D7I43_RS25240 | 101471..104860 | 1894969 | Swt1-family-HEPN-domain-containing-protein-[Micromonospora-globbae] |
| WP_212823736.1 | HEPN+AAA-ATPase+Alpha+Beta-domain+wHTH+RRM | Type-1_BR.RM.like.systems | Phage_int_SAM_1+Phage_integrase->?-><-HTH_21+rve+rve_3<-HIN-HTH\|\|TUDOR+SF1-SF2-Helicase+iREase+CTD+REase->METHYLASE->HEPN+DUF499*-><-?<-?\|\|?->HIN-HTH-> | 1124 | Bacteria>Terrabacteria_group>Actinomycetota>Actinomycetes>Micromonosporales | GCF_018324255.1 | Polymorphospora_rubra | Prubr_RS09020 | 1986978..1990352 | 338584 | DUF499-domain-containing-protein-[Polymorphospora-rubra] |
| WP_167666591.1 | HEPN+AAA-ATPase+Alpha+Beta-domain+wHTH+RRM | Type-1_BR.RM.like.systems | HTH+TPR+TPR->Pkinase-><-HEPN+DUF499*<-ParB+HNH<-METHYLASE<-TUDOR+SF1-SF2-Helicase+iREase+CTD+REase\|\|NTD+SF1-SF2-Helicase->\|\|HNH-> | 1124 | Bacteria>Terrabacteria_group>Actinomycetota>Actinomycetes>Micromonosporales | GCF_900089605.1 | Micromonospora_narathiwatensis | GA0070621_RS04900 | 1072112..1075486 | 299146 | Swt1-family-HEPN-domain-containing-protein-[Micromonospora-narathiwatensis] |
| WP_208576856.1 | HEPN+AAA-ATPase+Alpha+Beta-domain+wHTH+RRM | Type-1_BR.RM.like.systems | HTH+TPR+TPR->Pkinase-><-HEPN+DUF499*<-ParB+HNH<-METHYLASE<-TUDOR+SF1-SF2-Helicase+iREase+CTD+REase\|\|NTD+SF1-SF2-Helicase->HEPN-> | 1126 | Bacteria>Terrabacteria_group>Actinomycetota>Actinomycetes>Micromonosporales | GCF_017599015.1 | Micromonospora_tulbaghiae | J5U46_RS08620 | 81650..85030 | 479978 | Swt1-family-HEPN-domain-containing-protein-[Micromonospora-tulbaghiae] |
| WP_204020362.1 | HEPN+AAA-ATPase+Alpha+Beta-domain+wHTH+RRM | Type-1_BR.RM.like.systems | NUDIX->?-><-?<-REase<-ParB+HNH<-HEPN+DUF499*<-METHYLASE<-TUDOR+SF1-SF2-Helicase+iREase+CTD+REase\|\|RelE-ParE->ParA+HTH-><-PIN<-Phd-YefM | 1129 | Bacteria>Terrabacteria_group>Actinomycetota>Actinomycetes>Micromonosporales | GCF_016863595.1 | Micromonospora_andamanensis | Vwe01_RS03465 | 15617..19006 | 1287068 | Swt1-family-HEPN-domain-containing-protein-[Micromonospora-trujilloniae] |
| WP_085067314.1 | HEPN+AAA-ATPase+Alpha+Beta-domain+wHTH+RRM | Type-1_BR.RM.like.systems | <-ABhydrolase\|\|SIGMA-HTH-><-?\|\|TUDOR+SF1-SF2-Helicase+iREase+CTD+REase->METHYLASE->HEPN+DUF499*-><-?<-?<-ABhydrolase\|\|HTH+WYL+WYL-C-> | 1111 | Bacteria>Terrabacteria_group>Actinomycetota>Actinomycetes>Micromonosporales | GCF_000717135.1 | Catenuloplanes_japonicus | OP04_RS45935 | 31135..34470 | 33876 | Swt1-family-HEPN-domain-containing-protein-[Catenuloplanes-japonicus] |
| WP_203769001.1 | HEPN+AAA-ATPase+Alpha+Beta-domain+wHTH+RRM | Type-1_BR.RM.like.systems | DNA_methylase-><-REase<-HTH+SF1-SF2-Helicase+REase<-REase<-MORC<-HEPN+DUF499*<-METHYLASE\|\|\|\|?->REase-><-?\|\|REase->MarR-HTH-> | 1138 | Bacteria>Terrabacteria_group>Actinomycetota>Actinomycetes>Micromonosporales | GCF_016862275.1 | Actinoplanes_nipponensis | Ani05nite_RS15610 | 113186..116602 | 135950 | Swt1-family-HEPN-domain-containing-protein-[Actinoplanes-nipponensis] |
| WP_211192376.1 | HEPN+AAA-ATPase+Alpha+Beta-domain+wHTH+RRM | Type-1_BR.RM.like.systems | Pkinase->REase-><-?<-?<-HEPN+DUF499*<-METHYLASE<-METHYLASE-p<-TUDOR+SF1-SF2-Helicase+iREase+CTD+REase\|\|?->AAA_21-><-HEPN | 1111 | Bacteria>Terrabacteria_group>Actinomycetota>Actinomycetes>Micromonosporales | GCF_012933445.1 | Actinoplanes_sp._TBRC_11911 | HH310_RS19870 | 150452..153787 | 2729386 | Swt1-family-HEPN-domain-containing-protein-[Actinoplanes-sp.-TBRC-11911] |
| MBN1173560.1 | HEPN+AAA-ATPase+Alpha+Beta-domain+wHTH+RRM | Type-1_BR.RM.like.systems | <-Exonuc_V_gamma+REase\|\|TUDOR+SF1-SF2-Helicase+iREase+CTD+REase->METHYLASE->HEPN+DUF499*->REase-><-?\|\|?->Cas_Cmr3-> | 1155 | Bacteria>Terrabacteria_group>Actinomycetota>Actinomycetes>Micromonosporales | GCA_016926535.1 | Micromonosporaceae_bacterium | JXA67_15425 | 10595..14062 | 1873464 | DUF499-domain-containing-protein-[Micromonosporaceae-bacterium] |
| NMO53435.1 | HEPN+AAA-ATPase+Alpha+Beta-domain+wHTH+RRM | Type-1_BR.RM.like.systems | Pkinase->REase-><-?<-?<-HEPN+DUF499*<-TUDOR+SF1-SF2-Helicase+iREase+CTD+REase\|\|?->AAA_21-><-HEPN<-Primpol+D5N+D5ATPase | 1100 | Bacteria>Terrabacteria_group>Actinomycetota>Actinomycetes>Micromonosporales | GCA_012933445.1 | Actinoplanes_sp._TBRC_11911 | HH310_19870 | 150452..153754 | 2729386 | ATP-binding-protein-[Actinoplanes-sp.-TBRC-11911] |
| BCJ64829.1 | HEPN+AAA-ATPase+Alpha+Beta-domain+wHTH+RRM | Type-1_BR.RM.like.systems | <-DDE_Tnp_1<-?<-?\|\|TUDOR+SF1-SF2-Helicase+iREase+CTD+REase->METHYLASE->HEPN+DUF499*-><-?<-?\|\|?->?->?->HIN-HTH-> | 1124 | Bacteria>Terrabacteria_group>Actinomycetota>Actinomycetes>Micromonosporales | GCA_018324255.1 | Polymorphospora_rubra | Prubr_18500 | 1986978..1990352 | 338584 | hypothetical-protein-Prubr_18500-[Polymorphospora-rubra] |
| WP_184787881.1 | HEPN+AAA-ATPase+Alpha+Beta-domain+wHTH+RRM | Type-1_BR.RM.like.systems | TUDOR+SF1-SF2-Helicase+iREase+CTD+REase->METHYLASE->HEPN+DUF499*-><-TetR-HTH\|\|?-><-?\|\|ACET->?-><-HTH | 1112 | Bacteria>Terrabacteria_group>Actinomycetota>Actinomycetes>Micromonosporales | GCF_016862855.1 | Phytomonospora_endophytica | Pen01_RS22635 | 44949..48287 | 714109 | DUF499-domain-containing-protein-[Phytomonospora-endophytica] |
| WP_260034146.1 | HEPN+AAA-ATPase+Alpha+Beta-domain+wHTH+RRM | Type-1_BR.RM.like.systems | <-SF1-SF2-Helicase<-?\|\|METHYLASE->HEPN+DUF499*->MORC->REase->HTH+SF1-SF2-Helicase+REase-> | 1133 | Bacteria>Terrabacteria_group>Actinomycetota>Actinomycetes>Micromonosporales | GCF_025152325.1 | Micromonospora_chalcea | M3G91_RS14945 | 8947..12348 | 1874 | Swt1-family-HEPN-domain-containing-protein-[Micromonospora-chalcea] |
| WP_187690605.1 | HEPN+AAA-ATPase+Alpha+Beta-domain+wHTH+RRM | Type-1_BR.RM.like.systems | <-ParB+HNH+RAMA<-?<-HEPN+DUF499*<-METHYLASE<-TUDOR+SF1-SF2-Helicase+iREase+CTD+REase | 1128 | Bacteria>Terrabacteria_group>Actinomycetota>Actinomycetes>Micromonosporales | GCF_014472385.1 | Micromonospora_aurantiaca | H9X94_RS13515 | 9839..13225 | 47850 | Swt1-family-HEPN-domain-containing-protein-[Micromonospora-aurantiaca] |
| WP_107269540.1 | HEPN+AAA-ATPase+Alpha+Beta-domain+wHTH+RRM | Type-1_BR.RM.like.systems | <-ABhydrolase<-TetR-HTH\|\|?->HAMP+HISKIN->?-><-ParB+HNH<-HEPN+DUF499*\|\|?->?-><-?<-?\|\|HTH_3->Pkinase-> | 1110 | Bacteria>Terrabacteria_group>Actinomycetota>Actinomycetes>Micromonosporales | GCF_003030345.1 | Plantactinospora_sp._BC1 | C6361_RS28025 | 6382634..6385966 | 2108470 | Swt1-family-HEPN-domain-containing-protein-[Plantactinospora-sp.-BC1] |
| WP_107261307.1 | HEPN+AAA-ATPase+Alpha+Beta-domain+wHTH+RRM | Type-1_BR.RM.like.systems | <-ABhydrolase<-TetR-HTH\|\|?->HAMP+HISKIN->?-><-ParB+HNH<-HEPN+DUF499*\|\|?->?-><-?<-?<-?\|\|HTH_3-> | 1110 | Bacteria>Terrabacteria_group>Actinomycetota>Actinomycetes>Micromonosporales | GCF_003030385.1 | Plantactinospora_sp._BB1 | C6W10_RS25965 | 5945851..5949183 | 2071627 | Swt1-family-HEPN-domain-containing-protein-[Plantactinospora-sp.-BB1] |
| MBX7452224.1 | HEPN+AAA-ATPase+Alpha+Beta-domain+wHTH+RRM | Type-1_BR.RM.like.systems | <-REase\|\|DNA_methylase-><-NTD+SF1-SF2-Helicase\|\|TUDOR+SF1-SF2-Helicase+iREase+CTD+REase->METHYLASE->ParB+HNH->HEPN+DUF499*->MORC->REase->HTH+SF1-SF2-Helicase+REase->\|\|MORC+SF1-SF2-Helicase->NERD+SF1-SF2-Helicase->SF1-SF2-Helicase-> | 1122 | Bacteria>Terrabacteria_group>Actinomycetota>Actinomycetes>Mycobacteriales | GCA_019710595.1 | Mycolicibacterium_aurantiacum | GR927_29910 | 284651..288019 | 2692455 | DUF499-domain-containing-protein-[Mycolicibacterium-aurantiacum] |
| WP_094652354.1 | HEPN+AAA-ATPase+Alpha+Beta-domain+wHTH+RRM | Type-1_BR.RM.like.systems | <-ABhydrolase+ABhydrolase<-REase<-ParB+HNH+RAMA\|\|Primpol+D5N+D5ATPase->SF1-SF2-Helicase-><-HEPN+DUF499*<-ParB+HNH-p<-METHYLASE<-TUDOR+SF1-SF2-Helicase+iREase+CTD+REase\|\|?->Exonuc_V_gamma+REase->SF1-SF2-Helicase+REase-> | 1120 | Bacteria>Terrabacteria_group>Actinomycetota>Actinomycetes>Mycobacteriales | GCF_002259465.1 | Rhodococcus_fascians | CH297_RS18200 | 13414..16776 | 1828 | Swt1-family-HEPN-domain-containing-protein-[Rhodococcus-fascians] |
| WP_228232863.1 | HEPN+AAA-ATPase+Alpha+Beta-domain+wHTH+RRM | Type-1_BR.RM.like.systems | <-GDResolvase+HIN-HTH<-?\|\|ABhydrolase-><-GDResolvase+HIN-HTH\|\|zf-ISL3+DDE_Tnp_ISL3-><-?<-HEPN+DUF499*<-ParB+HNH<-METHYLASE<-UDG-conflict<-TUDOR+SF1-SF2-Helicase+iREase+CTD+REase\|\|NTD+SF1-SF2-Helicase-><-HEPN | 1121 | Bacteria>Terrabacteria_group>Actinomycetota>Actinomycetes>Mycobacteriales | GCF_020782825.1 | Rhodococcus_sp._3-2 | LLO08_RS29695 | 195028..198393 | 2890836 | Swt1-family-HEPN-domain-containing-protein-[Rhodococcus-sp.-3-2] |
| WP_214389765.1 | HEPN+AAA-ATPase+Alpha+Beta-domain+wHTH+RRM | Type-1_BR.RM.like.systems | EVE+McrB-AAA->McrC-NTD+REase-><-?<-HTH_Tnp_1+HTH_21+rve+rve_3\|\|HTH_Tnp_1+HTH_21+rve+rve_3->?-><-HEPN+DUF499*<-ParB+HNH<-METHYLASE<-TUDOR+SF1-SF2-Helicase+iREase+CTD+REase\|\|NTD+SF1-SF2-Helicase->DDE_Tnp_1_4-> | 1119 | Bacteria>Terrabacteria_group>Actinomycetota>Actinomycetes>Mycobacteriales | GCF_018524295.1 | Mycolicibacterium_goodii | KL858_RS28165 | 17594..20953 | 134601 | Swt1-family-HEPN-domain-containing-protein-[Mycolicibacterium-goodii] |
| MBU8833314.1 | HEPN+AAA-ATPase+Alpha+Beta-domain+wHTH+RRM | Type-1_BR.RM.like.systems | McrC-NTD+REase-><-?<-HTH_Tnp_1+HTH_21+rve+rve_3\|\|HTH_Tnp_1+HTH_21+rve+rve_3->?-><-REase<-HEPN+DUF499*<-ParB+HNH<-METHYLASE<-TUDOR+SF1-SF2-Helicase+iREase+CTD+REase\|\|DDE_Tnp_1_4->?->HEPN+SF1-SF2-Helicase-> | 1119 | Bacteria>Terrabacteria_group>Actinomycetota>Actinomycetes>Mycobacteriales | GCA_018524295.1 | Mycolicibacterium_goodii | KL858_28165 | 17594..20953 | 134601 | DUF499-domain-containing-protein-[Mycobacterium-goodii] |
| WP_181792003.1 | HEPN+AAA-ATPase+Alpha+Beta-domain+wHTH+RRM | Type-1_BR.RM.like.systems | <-35exo+SF1-SF2-Helicase\|\|HKD-DNase+SF1-SF2-Helicase+TRD-><-?<-?<-?<-HTH_Tnp_1+HTH_21+rve+rve_2<-HEPN+DUF499*<-ParB+HNH<-METHYLASE<-TUDOR+SF1-SF2-Helicase+iREase+CTD+REase\|\|NTD+SF1-SF2-Helicase-> | 1111 | Bacteria>Terrabacteria_group>Actinomycetota>Actinomycetes>Mycobacteriales | GCF_002086815.1 | Mycolicibacterium_parafortuitum | MPP7335_RS09255 | 1901353..1904688 | 39692 | Swt1-family-HEPN-domain-containing-protein-[Mycolicibacterium-parafortuitum] |
| RUP31532.1 | HEPN+AAA-ATPase+Alpha+Beta-domain+wHTH+RRM | Type-1_BR.RM.like.systems | <-SF1-SF2-Helicase<-TRD+TRD<-?<-SMODS<-HEPN+DUF499*<-METHYLASE<-TUDOR+SF1-SF2-Helicase+iREase+CTD+REase<-XIS-HTH<-TetR-HTH<-Phage_integrase+Phage_integrase<-Phage_integrase+Phage_integrase | 1121 | Bacteria>Terrabacteria_group>Actinomycetota>Actinomycetes>Mycobacteriales | GCA_003987785.1 | Mycolicibacterium_sp. | EKK51_14275 | 206389..209754 | 2320850 | MAG:-ATP-binding-protein-[Mycolicibacterium-sp.] |
| WP_048892803.1 | HEPN+AAA-ATPase+Alpha+Beta-domain+wHTH+RRM | Type-1_BR.RM.like.systems | DEDD_Tnp_IS110->DEDD_Tnp_IS110+Transposase_20->TUDOR+SF1-SF2-Helicase+iREase+CTD+REase->METHYLASE-><-?<-?\|\|HEPN+DUF499*-><-HKD-DNase+SF1-SF2-Helicase+TRD<-?<-PIN<-PhdYeFM_antitox<-HEPN<-Polbeta | 1117 | Bacteria>Terrabacteria_group>Actinomycetota>Actinomycetes>Mycobacteriales | GCF_016592155.1 | Mycobacterium_heckeshornense | BMW24_RS01365 | 271817..275170 | 110505 | DUF499-domain-containing-protein-[Mycobacterium-heckeshornense] |
| WP_071935222.1 | HEPN+AAA-ATPase+Alpha+Beta-domain+wHTH+RRM | Type-1_BR.RM.like.systems | <-URI-DNase<-?<-?<-?\|\|PAIREDC-HTH+HIN-HTH-><-HEPN+DUF499*\|\|HTH_Tnp_1+HTH_21+rve+rve_3-><-?\|\|DDE_Tnp_1_4-><-GDResolvase+Recombinase+ZnR<-HTH_Tnp_1+HTH_21+rve+rve_3<-HTH_Tnp_1+HTH_21+rve+rve_3 | 1131 | Bacteria>Terrabacteria_group>Actinomycetota>Actinomycetes>Mycobacteriales | GCF_001886355.1 | Rhodococcus_sp._2G | BO226_RS11040 | 2363059..2366454 | 1570939 | DUF499-domain-containing-protein-[Rhodococcus-sp.-2G] |
| WP_014358285.1 | HEPN+AAA-ATPase+Alpha+Beta-domain+wHTH+RRM | Type-1_BR.RM.like.systems | MACRODOMAIN+SIR2->SF1-SF2-Helicase-><-HTH+SF1-SF2-Helicase+REase<-REase<-MORC<-HEPN+DUF499*<-METHYLASE<-REase\|\|DNA_methylase->SAM-methylase-><-CR-Kinase+ORC-CDC6-like+CARF | 1127 | Bacteria>Terrabacteria_group>Actinomycetota>Actinomycetes>Mycobacteriales | GCF_000247715.1 | Gordonia_polyisoprenivorans | GPOL_RS00855 | 197189..200572 | 84595 | DUF499-domain-containing-protein-[Gordonia-polyisoprenivorans] |
| AHK35548.1 | HEPN+AAA-ATPase+Alpha+Beta-domain+wHTH+RRM | Type-1_BR.RM.like.systems | <-DUF499-p\|\|?-><-?<-DUF499-p<-HEPN+DUF499*<-METHYLASE<-METHYLASE-p<-TUDOR+SF1-SF2-Helicase+iREase+CTD+REase<-TUDOR+SF1-SF2-Helicase+iREase+CTD+REase-p\|\|HEPN-p->DNA-glycosylase-> | 649 | Bacteria>Terrabacteria_group>Actinomycetota>Actinomycetes>Mycobacteriales | GCA_000599545.1 | Rhodococcus_opacus_PD630 | Pd630_LPD10098 | 71803..73752 | 543736 | hypothetical-protein-Pd630_LPD10098-[Rhodococcus-opacus-PD630] |
| ORB28066.1 | HEPN+AAA-ATPase+Alpha+Beta-domain+wHTH+RRM | Type-1_BR.RM.like.systems | HKD-DNase+SF1-SF2-Helicase+TRD-><-?<-?<-?<-HTH_Tnp_1+HTH_21+rve+rve_2<-HIN-HTH<-HEPN+DUF499*<-ParB+HNH<-METHYLASE<-TUDOR+SF1-SF2-Helicase+iREase+CTD+REase\|\|NTD+SF1-SF2-Helicase-> | 1100 | Bacteria>Terrabacteria_group>Actinomycetota>Actinomycetes>Mycobacteriales | GCA_002086815.1 | Mycolicibacterium_parafortuitum | BST38_21950 | 32526..35828 | 39692 | AAA-family-ATPase-[Mycolicibacterium-parafortuitum] |
| WP_203801307.1 | HEPN+AAA-ATPase+Alpha+Beta-domain+wHTH+RRM | Type-1_BR.RM.like.systems | <-SF1-SF2-helicase+DUF1998<-?\|\|TUDOR+SF1-SF2-Helicase+iREase+CTD+REase->METHYLASE-><-?<-?\|\|HEPN+DUF499*-><-HKD-DNase+SF1-SF2-Helicase+TRD<-?<-PIN<-PhdYeFM_antitox<-HEPN<-Polbeta | 1117 | Bacteria>Terrabacteria_group>Actinomycetota>Actinomycetes>Mycobacteriales | GCF_016861545.1 | Mycobacterium_heckeshornense | JUH53_RS19960 | 4159291..4162644 | 110505 | DUF499-domain-containing-protein-[Mycobacterium-heckeshornense] |
| WP_201263267.1 | HEPN+AAA-ATPase+Alpha+Beta-domain+wHTH+RRM | Type-1_BR.RM.like.systems | <-NTD+SF1-SF2-Helicase\|\|TUDOR+SF1-SF2-Helicase+iREase+CTD+REase->METHYLASE->HEPN+DUF499*->REase->SF1-SF2-Helicase-p->HTH_Tnp_1+HTH_21+rve+rve_2->SF1-SF2-Helicase-p-> | 1109 | Bacteria>Terrabacteria_group>Actinomycetota>Actinomycetes>Mycobacteriales | GCF_002201735.1 | Mycobacterium_avium | CEV14_RS17015 | 150524..153853 | 1764 | Swt1-family-HEPN-domain-containing-protein-[Mycobacterium-avium] |
| WP_260836619.1 | HEPN+AAA-ATPase+Alpha+Beta-domain+wHTH+RRM | Type-1_BR.RM.like.systems | <-MULE-TRANSPOSASE\|\|ParA-Soj-PloopNTPase->ORC-CDC6-><-HTH+SF1-SF2-Helicase+REase<-REase<-MORC<-HEPN+DUF499*<-METHYLASE\|\|XIS-HTH+RVT-><-HNH\|\|?-><-REase\|\|DNA_methylase-> | 1138 | Bacteria>Terrabacteria_group>Actinomycetota>Actinomycetes>Mycobacteriales | GCF_025268675.1 | Gordonia_amarae | GII32_RS01225 | 287584..291000 | 36821 | DUF499-domain-containing-protein-[Gordonia-amarae] |
| WP_005189162.1 | HEPN+AAA-ATPase+Alpha+Beta-domain+wHTH+RRM | Type-1_BR.RM.like.systems | <-MULE-TRANSPOSASE\|\|ParA-Soj-PloopNTPase->ORC-CDC6-><-HTH+SF1-SF2-Helicase+REase<-REase<-MORC<-HEPN+DUF499*<-METHYLASE\|\|XIS-HTH+RVT-><-HNH\|\|?-><-REase\|\|DNA_methylase-> | 1138 | Bacteria>Terrabacteria_group>Actinomycetota>Actinomycetes>Mycobacteriales | GCF_000241345.1 | Gordonia_amarae | GII35_RS01230 | 287739..291155 | 36821 | DUF499-domain-containing-protein-[Gordonia-amarae] |
| QHN29206.1 | HEPN+AAA-ATPase+Alpha+Beta-domain+wHTH+RRM | Type-1_BR.RM.like.systems | <-MULE-TRANSPOSASE\|\|ParA-Soj-PloopNTPase->ORC-CDC6-><-HTH+SF1-SF2-Helicase+REase<-REase<-MORC<-HEPN+DUF499*<-METHYLASE\|\|XIS-HTH+RVT-><-HNH\|\|?-><-REase\|\|DNA_methylase-> | 1138 | Bacteria>Terrabacteria_group>Actinomycetota>Actinomycetes>Mycobacteriales | GCA_025268675.1 | Gordonia_amarae | GII32_01220 | 287584..291000 | 36821 | DUF499-domain-containing-protein-[Gordonia-amarae] |
| WP_193904012.1 | HEPN+AAA-ATPase+Alpha+Beta-domain+wHTH+RRM | Type-1_BR.RM.like.systems | MODE-HTH+PBPII-><-IclR-HTH\|\|?->RNase-><-HEPN+DUF499*<-METHYLASE-p<-zf-ISL3+DDE_Tnp_ISL3<-METHYLASE<-TUDOR+SF1-SF2-Helicase+iREase+CTD+REase\|\|DDE_Tnp_1_4-><-HTH+MPTase | 1142 | Bacteria>Terrabacteria_group>Actinomycetota>Actinomycetes>Mycobacteriales | GCF_015169995.1 | Rhodococcus_pyridinivorans | INP59_RS25070 | 140646..144074 | 103816 | Swt1-family-HEPN-domain-containing-protein-[Rhodococcus-pyridinivorans] |
| WP_050067339.1 | HEPN+AAA-ATPase+Alpha+Beta-domain+wHTH+RRM | Type-1_BR.RM.like.systems | MarR-HTH->tRNA-><-Arm-DNA-bind_4+Phage_int_SAM_3+Phage_integrase<-HTH\|\|XIS-HTH->?-><-HEPN+DUF499*<-METHYLASE<-TUDOR+SF1-SF2-Helicase+iREase+CTD+REase<-?<-GDResolvase+HIN-HTH | 1134 | Bacteria>Terrabacteria_group>Actinomycetota>Actinomycetes>Mycobacteriales | GCF_001040705.1 | Rhodococcus_sp._RD6.2 | RHCRD62_RS24305 | 110292..113696 | 260936 | Swt1-family-HEPN-domain-containing-protein-[Rhodococcus-sp.-RD6.2] |
| WP_259918390.1 | HEPN+AAA-ATPase+Alpha+Beta-domain+wHTH+RRM | Type-1_BR.RM.like.systems | <-ABhydrolase<-?<-Aminotran_1_2<-ncRNA\|\|tRNA->?->SF1-SF2-helicase+DUF1998-><-ORC-CDC6+REase<-HEPN+DUF499*<-METHYLASE<-TUDOR+SF1-SF2-Helicase+iREase+CTD+REase\|\|ABhydrolase-> | 1108 | Bacteria>Terrabacteria_group>Actinomycetota>Actinomycetes>Mycobacteriales | - | Corynebacterium_sanguinis | M3A76_RS07775 | 66134..69460 | 2594913 | DUF499-domain-containing-protein-[Corynebacterium-sanguinis] |
| WP_259825954.1 | HEPN+AAA-ATPase+Alpha+Beta-domain+wHTH+RRM | Type-1_BR.RM.like.systems | <-ABhydrolase<-?<-Aminotran_1_2<-ncRNA\|\|tRNA->?->SF1-SF2-helicase+DUF1998-><-ORC-CDC6+REase<-HEPN+DUF499*<-METHYLASE<-TUDOR+SF1-SF2-Helicase+iREase+CTD+REase\|\|ABhydrolase-> | 1112 | Bacteria>Terrabacteria_group>Actinomycetota>Actinomycetes>Mycobacteriales | GCF_025146125.1 | Corynebacterium_sanguinis | QON61_RS06385 | 69194..72532 | 2594913 | DUF499-domain-containing-protein-[Corynebacterium-sanguinis] |
| MCT1882934.1 | HEPN+AAA-ATPase+Alpha+Beta-domain+wHTH+RRM | Type-1_BR.RM.like.systems | <-ABhydrolase<-?<-Aminotran_1_2<-ncRNA\|\|tRNA->?->SF1-SF2-helicase+DUF1998-><-ORC-CDC6+REase<-HEPN+DUF499*<-METHYLASE<-TUDOR+SF1-SF2-Helicase+iREase+CTD+REase\|\|ABhydrolase-> | 1081 | Bacteria>Terrabacteria_group>Actinomycetota>Actinomycetes>Mycobacteriales | GCA_025144605.1 | Corynebacterium_sanguinis | M3A76_07760 | 66134..69379 | 2594913 | DUF499-domain-containing-protein-[Corynebacterium-sanguinis] |
| WP_268656219.1 | HEPN+AAA-ATPase+Alpha+Beta-domain+wHTH+RRM | Type-1_BR.RM.like.systems | DUF4158+DDE_Tnp_Tn3-><-TetR-HTH\|\|?->?->GDResolvase+HIN-HTH->\|\|RNase-><-HEPN+DUF499*<-METHYLASE<-TUDOR+SF1-SF2-Helicase+iREase+CTD+REase<-HTH+MPTase<-Tox-RelE\|\|SbcC-> | 1138 | Bacteria>Terrabacteria_group>Actinomycetota>Actinomycetes>Mycobacteriales | GCF_026802115.1 | Rhodococcus_pyridinivorans | OQN32_RS28100 | 23296..26712 | 103816 | Swt1-family-HEPN-domain-containing-protein-[Rhodococcus-pyridinivorans] |
| WP_160904533.1 | HEPN+AAA-ATPase+Alpha+Beta-domain+wHTH+RRM | Type-1_BR.RM.like.systems | <-Exonuc_V_gamma+REase<-?<-NTD+SF1-SF2-Helicase\|\|TUDOR+SF1-SF2-Helicase+iREase+CTD+REase->METHYLASE->?->HEPN+DUF499*->?->?-><-HKD-DNase+HKD-DNase<-?<-wHTH+REase<-wHTH+REase | 1121 | Bacteria>Terrabacteria_group>Actinomycetota>Actinomycetes>Mycobacteriales | GCF_024734075.1 | Gordonia_mangrovi | GIY30_RS23685 | 18846..22211 | 2665643 | Swt1-family-HEPN-domain-containing-protein-[Gordonia-mangrovi] |
| WP_106851264.1 | HEPN+AAA-ATPase+Alpha+Beta-domain+wHTH+RRM | Type-1_BR.RM.like.systems | SIGMA-HTH+SIGMA-HTH-><-wHTH(SBD)+HNH<-REase+SF1-SF2-Helicase\|\|TUDOR+SF1-SF2-Helicase+iREase+CTD+REase->METHYLASE->ParB+HNH->HEPN+DUF499*->NTD+SF1-SF2-Helicase-> | 1123 | Bacteria>Terrabacteria_group>Actinomycetota>Actinomycetes>Mycobacteriales | GCF_003013675.1 | Prescottella_equi | C7H75_RS09240 | 1958101..1961472 | 43767 | Swt1-family-HEPN-domain-containing-protein-[Prescottella-equi] |
| KUI22690.1 | HEPN+AAA-ATPase+Alpha+Beta-domain+wHTH+RRM | Type-1_BR.RM.like.systems | <-TetR-HTH\|\|DapB_N-><-?<-NTD+SF1-SF2-Helicase\|\|TUDOR+SF1-SF2-Helicase+iREase+CTD+REase->METHYLASE->HEPN+DUF499*-><-?\|\|?->?-><-Cluster182_14clades<-MACRODOMAIN<-MACRODOMAIN | 1104 | Bacteria>Terrabacteria_group>Actinomycetota>Actinomycetes>Mycobacteriales | GCA_001499905.1 | Mycobacterium_sp._GA-1285 | AU193_20905 | 131956..135270 | 1772282 | AAA-family-ATPase-[Mycobacterium-sp.-GA-1285] |
| WP_075835625.1 | HEPN+AAA-ATPase+Alpha+Beta-domain+wHTH+RRM | Type-1_BR.RM.like.systems | NUDIX->?->TUDOR+SF1-SF2-Helicase+iREase+CTD+REase->METHYLASE->HEPN+DUF499*->?-><-HEPN\|\|PAIREDC-HTH->?->GDResolvase+Recombinase+ZnR->Phage_int_SAM_4+Phage_integrase-> | 1134 | Bacteria>Terrabacteria_group>Actinomycetota>Actinomycetes>Mycobacteriales | GCF_001942025.1 | unclassified_Rhodococcus_(in:_high_G+C_Gram-positive_bacteria) | BH682_RS29985 | 71816..75220 | 192944 | MULTISPECIES:-Swt1-family-HEPN-domain-containing-protein-[unclassified-Rhodococcus] |
| OBB45420.1 | HEPN+AAA-ATPase+Alpha+Beta-domain+wHTH+RRM | Type-1_BR.RM.like.systems | ParB+HNH->?-><-HTH+SF1-SF2-Helicase+REase<-REase<-MORC<-HEPN+DUF499*<-METHYLASE\|\|DNA_methylase->OLD-ABC-><-REase<-SF1-SF2-Helicase+REase<-SF1-SF2-Helicase+REase-p | 1135 | Bacteria>Terrabacteria_group>Actinomycetota>Actinomycetes>Mycobacteriales | GCF_001665535.1 | Mycobacterium_sp._852002-51961_SCH5331710 | A5752_02020 | 17252..20659 | 1834105 | hypothetical-protein-A5752_02020-[Mycobacterium-sp.-852002-51961_SCH5331710] |
| WP_174814236.1 | HEPN+AAA-ATPase+Alpha+Beta-domain+wHTH+RRM | Type-1_BR.RM.like.systems | PAIREDC-HTH->HIN-HTH->?->TUDOR+SF1-SF2-Helicase+iREase+CTD+REase->METHYLASE->HEPN+DUF499*->?->HTH_17->GDResolvase+Recombinase->Phage_int_SAM_1+Phage_integrase-> | 1121 | Bacteria>Terrabacteria_group>Actinomycetota>Actinomycetes>Mycobacteriales | GCF_010723135.1 | Mycobacterium_kubicae | G6N40_RS26550 | 2561020..2564385 | 120959 | Swt1-family-HEPN-domain-containing-protein-[Mycobacterium-kubicae] |
| WP_208722091.1 | HEPN+AAA-ATPase+Alpha+Beta-domain+wHTH+RRM | Type-1_BR.RM.like.systems | <-TetR-HTH<-GDResolvase+HIN-HTH\|\|Lsr2->?-><-?<-HEPN+DUF499*<-ParB+HNH<-METHYLASE<-UDG-conflict<-TUDOR+SF1-SF2-Helicase+iREase+CTD+REase\|\|NTD+SF1-SF2-Helicase-> | 1122 | Bacteria>Terrabacteria_group>Actinomycetota>Actinomycetes>Mycobacteriales | GCF_006384225.1 | Rhodococcus_qingshengii | IGTS8_RS29980 | 39886..43254 | 334542 | Swt1-family-HEPN-domain-containing-protein-[Rhodococcus-qingshengii] |
| WP_153212242.1 | HEPN+AAA-ATPase+Alpha+Beta-domain+wHTH+RRM | Type-1_BR.RM.like.systems | <-NTD+SF1-SF2-Helicase\|\|TUDOR+SF1-SF2-Helicase+iREase+CTD+REase->UDG-conflict->METHYLASE->ParB+HNH->HEPN+DUF499*-><-NUDIX\|\|?->REase->?-><-HAD<-ABC-ATPase+SbcC | 1121 | Bacteria>Terrabacteria_group>Actinomycetota>Actinomycetes>Mycobacteriales | GCF_009495805.1 | Rhodococcus_erythropolis | GEO20_RS16000 | 38887..42252 | 1833 | Swt1-family-HEPN-domain-containing-protein-[Rhodococcus-erythropolis] |
| WP_085185294.1 | HEPN+AAA-ATPase+Alpha+Beta-domain+wHTH+RRM | Type-1_BR.RM.like.systems | <-PemK_toxin<-?<-?\|\|Glyoxalase-><-?<-OLD-ABC<-HEPN+DUF499*<-METHYLASE<-TUDOR+SF1-SF2-Helicase+iREase+CTD+REase\|\|NTD+SF1-SF2-Helicase-><-wHTH+SAD+HNH\|\|Synaptojanin-> | 1114 | Bacteria>Terrabacteria_group>Actinomycetota>Actinomycetes>Mycobacteriales | GCF_002101935.1 | Mycobacterium_sp._IEC1808 | AWB92_RS10440 | 126977..130321 | 1743230 | Swt1-family-HEPN-domain-containing-protein-[Mycobacterium-sp.-IEC1808] |
| WP_013470723.1 | HEPN+AAA-ATPase+Alpha+Beta-domain+wHTH+RRM | Type-1_BR.RM.like.systems | <-MORC<-HTH+SF1-SF2-Helicase+REase<-DDE_Tnp_1_4<-REase<-MORC<-HEPN+DUF499*<-ParB+HNH<-MULE-TRANSPOSASE<-TUDOR+SF1-SF2-Helicase+iREase+CTD+REase<-HIN-HTH | 1122 | Bacteria>Terrabacteria_group>Actinomycetota>Actinomycetes>Mycobacteriales | GCF_000184435.1 | Mycolicibacterium_gilvum | MSPYR1_RS04230 | 878356..881724 | 1804 | Swt1-family-HEPN-domain-containing-protein-[Mycolicibacterium-gilvum] |
| UHJ57678.1 | HEPN+AAA-ATPase+Alpha+Beta-domain+wHTH+RRM | Type-1_BR.RM.like.systems | REase->DNA-glycosylase->?-><-HTH+SF1-SF2-Helicase+REase<-REase<-MORC<-HEPN+DUF499*<-METHYLASE<-REase<-PIN\|\|DNA_methylase-><-?<-Phage_integrase+Phage_integrase | 1138 | Bacteria>Terrabacteria_group>Actinomycetota>Actinomycetes>Mycobacteriales | GCA_021249325.1 | Mycolicibacterium_fortuitum | LT337_13390 | 2620949..2624365 | 1766 | DUF499-domain-containing-protein-[Mycolicibacterium-fortuitum] |
| WP_277371905.1 | HEPN+AAA-ATPase+Alpha+Beta-domain+wHTH+RRM | Type-1_BR.RM.like.systems | MODE-HTH+PBPII-><-IclR-HTH\|\|DDE_Tnp_1_4->?->RNase-><-HEPN+DUF499*<-METHYLASE<-TUDOR+SF1-SF2-Helicase+iREase+CTD+REase\|\|SF1-SF2-Helicase-><-DNA-glycosylase | 1138 | Bacteria>Terrabacteria_group>Actinomycetota>Actinomycetes>Mycobacteriales | GCF_029457455.1 | Rhodococcus_rhodochrous | PVE95_RS25585 | 81370..84786 | 1829 | Swt1-family-HEPN-domain-containing-protein-[Rhodococcus-rhodochrous] |
| MBM4528840.1 | HEPN+AAA-ATPase+Alpha+Beta-domain+wHTH+RRM | Type-1_BR.RM.like.systems | wHTH(SBD)+HNH-><-ParB<-HEPN+DUF499*<-ParB+HNH-p<-METHYLASE<-TUDOR+SF1-SF2-Helicase+iREase+CTD+REase\|\|NTD+SF1-SF2-Helicase-><-SF1-SF2-Helicase<-HKD-DNase | 1121 | Bacteria>Terrabacteria_group>Actinomycetota>Actinomycetes>Mycobacteriales | GCA_016879355.1 | Prescottella_equi | GS563_07715 | 1641397..1644762 | 43767 | DUF499-domain-containing-protein-[Prescottella-equi] |
| WP_102999581.1 | HEPN+AAA-ATPase+Alpha+Beta-domain+wHTH+RRM | Type-1_BR.RM.like.systems | TUDOR+SF1-SF2-Helicase+iREase+CTD+REase->METHYLASE->ParB+HNH->HEPN+DUF499*->zf-ISL3+DDE_Tnp_ISL3->?-><-DNA_methylase<-HEPN<-ParB+HNH<-GDResolvase+HIN-HTH | 1121 | Bacteria>Terrabacteria_group>Actinomycetota>Actinomycetes>Mycobacteriales | GCF_002893965.1 | Rhodococcus_qingshengii | C1M55_RS31235 | 10127..13492 | 334542 | Swt1-family-HEPN-domain-containing-protein-[Rhodococcus-qingshengii] |
| WP_231782199.1 | HEPN+AAA-ATPase+Alpha+Beta-domain+wHTH+RRM | Type-1_BR.RM.like.systems | <-MULE-TRANSPOSASE<-ZnR+ZnR<-TIR\|\|TUDOR+SF1-SF2-Helicase+iREase+CTD+REase->METHYLASE->HEPN+DUF499*-><-RNase<-?\|\|GDResolvase+Recombinase+ZnR-><-?\|\|DUF4192-> | 1139 | Bacteria>Terrabacteria_group>Actinomycetota>Actinomycetes>Mycobacteriales | GCF_021165995.1 | Rhodococcus_pyridinivorans | LSF60_RS23505 | 9142..12561 | 103816 | Swt1-family-HEPN-domain-containing-protein-[Rhodococcus-pyridinivorans] |
| WP_172798208.1 | HEPN+AAA-ATPase+Alpha+Beta-domain+wHTH+RRM | Type-1_BR.RM.like.systems | DapB_N-><-?\|\|?-><-NTD+SF1-SF2-Helicase\|\|TUDOR+SF1-SF2-Helicase+iREase+CTD+REase->METHYLASE->HEPN+DUF499*-><-?\|\|?->?-><-Cluster182_14clades<-MACRODOMAIN<-MACRODOMAIN | 1115 | Bacteria>Terrabacteria_group>Actinomycetota>Actinomycetes>Mycobacteriales | GCF_001499905.1 | Mycobacterium_sp._GA-1285 | AU193_RS04065 | 131923..135270 | 1772282 | Swt1-family-HEPN-domain-containing-protein-[Mycobacterium-sp.-GA-1285] |
| TDK92004.1 | HEPN+AAA-ATPase+Alpha+Beta-domain+wHTH+RRM | Type-1_BR.RM.like.systems | <-PAIREDC-HTH<-HIN-HTH\|\|JetD-><-PIN<-VapB_antitoxin\|\|<-HEPN+DUF499*<-METHYLASE<-TUDOR+SF1-SF2-Helicase+iREase+CTD+REase\|\|HTH_31->?-><-?\|\|LIP+Peptidase_S9-> | 1099 | Bacteria>Terrabacteria_group>Actinomycetota>Actinomycetes>Mycobacteriales | GCA_004359045.1 | Mycolicibacterium_mucogenicum | EUA03_05700 | 2711..6010 | 56689 | ATP-binding-protein-[Mycolicibacterium-mucogenicum] |
| WP_205848809.1 | HEPN+AAA-ATPase+Alpha+Beta-domain+wHTH+RRM | Type-1_BR.RM.like.systems | <-PAIREDC-HTH<-HIN-HTH\|\|JetD-><-PIN<-VapB_antitoxin<-HEPN+DUF499*<-METHYLASE<-TUDOR+SF1-SF2-Helicase+iREase+CTD+REase\|\|HTH_31->?-><-?\|\|LIP+Peptidase_S9-> | 1110 | Bacteria>Terrabacteria_group>Actinomycetota>Actinomycetes>Mycobacteriales | GCF_018326145.1 | Mycolicibacterium | EUA03_RS05700 | 2711..6043 | 1866885 | MULTISPECIES:-Swt1-family-HEPN-domain-containing-protein-[Mycolicibacterium] |
| WP_011210977.1 | HEPN+AAA-ATPase+Alpha+Beta-domain+wHTH+RRM | Type-1_BR.RM.like.systems | <-HTH_21+rve+rve_3\|\|?->HIN-HTH->?->rve+rve_3-><-HEPN+DUF499*<-ParB+HNH<-METHYLASE<-TUDOR+SF1-SF2-Helicase+iREase+CTD+REase\|\|GDResolvase+Recombinase+ZnR-> | 1127 | Bacteria>Terrabacteria_group>Actinomycetota>Actinomycetes>Mycobacteriales | GCF_000009805.1 | Nocardia_farcinica | NFA_RS22030 | 4605548..4608931 | 37329 | Swt1-family-HEPN-domain-containing-protein-[Nocardia-farcinica] |
| NKZ77519.1 | HEPN+AAA-ATPase+Alpha+Beta-domain+wHTH+RRM | Type-1_BR.RM.like.systems | MarR-HTH->tRNA-><-Phage_integrase<-HTH\|\|XIS-HTH->?-><-HEPN+DUF499*<-METHYLASE<-TUDOR+SF1-SF2-Helicase+iREase+CTD+REase\|\|<-GDResolvase+HIN-HTH\|\|?-><-?\|\|ABhydrolase-> | 1133 | Bacteria>Terrabacteria_group>Actinomycetota>Actinomycetes>Mycobacteriales | GCA_012443725.1 | Prescottella_equi | GTA28_11640 | 113900..117301 | 43767 | DUF499-domain-containing-protein-[Prescottella-equi] |
| WP_229481506.1 | HEPN+AAA-ATPase+Alpha+Beta-domain+wHTH+RRM | Type-1_BR.RM.like.systems | <-NTD+SF1-SF2-Helicase\|\|TUDOR+SF1-SF2-Helicase+iREase+CTD+REase->METHYLASE->ParB+HNH->HEPN+DUF499*->?->?-><-SF1-SF2-Helicase<-OLD-ABC+TOPRIM<-?<-Primpol+D5N+D5ATPase | 1114 | Bacteria>Terrabacteria_group>Actinomycetota>Actinomycetes>Mycobacteriales | GCF_020884715.1 | Mycolicibacterium_mageritense | MTY414_RS35935 | 11397..14741 | 53462 | Swt1-family-HEPN-domain-containing-protein-[Mycolicibacterium-mageritense] |
| WP_176749585.1 | HEPN+AAA-ATPase+Alpha+Beta-domain+wHTH+RRM | Type-1_BR.RM.like.systems | <-NTD+SF1-SF2-Helicase\|\|TUDOR+SF1-SF2-Helicase+iREase+CTD+REase->METHYLASE-p->DDE_Tnp_1_4->ParB+HNH->HEPN+DUF499*->?->?->?-><-Cluster182_14clades<-?<-IclR-HTH | 1120 | Bacteria>Terrabacteria_group>Actinomycetota>Actinomycetes>Mycobacteriales | GCF_008329645.1 | Mycobacterium_grossiae | FZ046_RS21205 | 4404866..4408228 | 1552759 | Swt1-family-HEPN-domain-containing-protein-[Mycobacterium-grossiae] |
| WP_248689418.1 | HEPN+AAA-ATPase+Alpha+Beta-domain+wHTH+RRM | Type-1_BR.RM.like.systems | MULE-TRANSPOSASE->?->RNase-><-HEPN+DUF499*<-METHYLASE<-TUDOR+SF1-SF2-Helicase+iREase+CTD+REase\|\|SF1-SF2-Helicase-><-DNA-glycosylase<-DNA-glycosylase | 1138 | Bacteria>Terrabacteria_group>Actinomycetota>Actinomycetes>Mycobacteriales | GCF_023238485.1 | Rhodococcus_pyridinivorans | M1C57_RS14915 | 3265410..3268826 | 103816 | Swt1-family-HEPN-domain-containing-protein-[Rhodococcus-pyridinivorans] |
| WP_170880036.1 | HEPN+AAA-ATPase+Alpha+Beta-domain+wHTH+RRM | Type-1_BR.RM.like.systems | <-ParB+HNH<-NTD+SF1-SF2-Helicase\|\|TUDOR+SF1-SF2-Helicase+iREase+CTD+REase->METHYLASE->ParB+HNH->HEPN+DUF499*-><-Glyoxalase\|\|?-><-TetR-HTH\|\|?->?-><-Pkinase | 1111 | Bacteria>Terrabacteria_group>Actinomycetota>Actinomycetes>Mycobacteriales | GCF_001954115.1 | Mycobacterium_colombiense | A5739_RS13730 | 105112..108447 | 339268 | Swt1-family-HEPN-domain-containing-protein-[Mycobacterium-colombiense] |

| Protein-ID | DUF499-architecture | System | Gene-Neighbourhoods | Length | Taxonomy | Assembly-ID | Strain | Locus-Tag | Genomic-coordinates | TaxID | Fasta-defline |
|---|---|---|---|---|---|---|---|---|---|---|---|
| PQP40021.1 | HEPN+AAA-ATPase+Alpha+Beta-domain+wHTH+RRM | Type-1_BR.RM.like.systems | [illegible] | 1103 | Bacteria>Terrabacteria_group>Actinomycetota>Actinomycetes>Mycobacteriales | GCA_002968275.1 | Mycolicibacterium_austroafricanum | C6A88_31860 | 14193..17504 | 39687 | AAA-family-ATPase-[Mycolicibacterium-austroafricanum] |
| OMC31131.1 | HEPN+AAA-ATPase+Alpha+Beta-domain+wHTH+RRM | Type-1_BR.RM.like.systems | [illegible] | 1100 | Bacteria>Terrabacteria_group>Actinomycetota>Actinomycetes>Mycobacteriales | GCA_001954115.1 | Mycobacterium_colombiense | A5739_14025 | 105145..108447 | 339268 | AAA-family-ATPase-[Mycobacterium-colombiense] |
| WP_193376452.1 | HEPN+AAA-ATPase+Alpha+Beta-domain+wHTH+RRM | Type-1_BR.RM.like.systems | [illegible] | 1107 | Bacteria>Terrabacteria_group>Actinomycetota>Actinomycetes>Mycobacteriales | GCF_000455205.1 | Mycobacterium_sp._UM_CSW | N415_RS14335 | 81564..84887 | 1370119 | Swt1-family-HEPN-domain-containing-protein-[Mycobacterium-sp.-UM_CSW] |
| WP_116058404.1 | HEPN+AAA-ATPase+Alpha+Beta-domain+wHTH+RRM | Type-1_BR.RM.like.systems | [illegible] | 1121 | Bacteria>Terrabacteria_group>Actinomycetota>Actinomycetes>Mycobacteriales | - | Rhodococcus_erythropolis | DVG80_RS32980 | 11800..15165 | 1833 | Swt1-family-HEPN-domain-containing-protein-[Rhodococcus-erythropolis] |
| WP_094602238.1 | HEPN+AAA-ATPase+Alpha+Beta-domain+wHTH+RRM | Type-1_BR.RM.like.systems | [illegible] | 1127 | Bacteria>Terrabacteria_group>Actinomycetota>Actinomycetes>Mycobacteriales | GCF_018135605.1 | Gordonia_polyisoprenivorans | J8M97_RS16335 | 3622924..3626307 | 84595 | Swt1-family-HEPN-domain-containing-protein-[Gordonia-polyisoprenivorans] |
| KUI38022.1 | HEPN+AAA-ATPase+Alpha+Beta-domain+wHTH+RRM | Type-1_BR.RM.like.systems | [illegible] | 1104 | Bacteria>Terrabacteria_group>Actinomycetota>Actinomycetes>Mycobacteriales | GCA_001500025.1 | Mycobacterium_sp._IS-1590 | AU197_09575 | 351187..354501 | 1772286 | AAA-family-ATPase-[Mycobacterium-sp.-IS-1590] |
| WP_269614895.1 | HEPN+AAA-ATPase+Alpha+Beta-domain+wHTH+RRM | Type-1_BR.RM.like.systems | [illegible] | 1121 | Bacteria>Terrabacteria_group>Actinomycetota>Actinomycetes>Mycobacteriales | GCF_027270755.1 | Rhodococcus_qingshengii | O4271_RS30885 | 26346..29711 | 334542 | Swt1-family-HEPN-domain-containing-protein-[Rhodococcus-qingshengii] |
| WP_197506097.1 | HEPN+AAA-ATPase+Alpha+Beta-domain+wHTH+RRM | Type-1_BR.RM.like.systems | [illegible] | 1121 | Bacteria>Terrabacteria_group>Actinomycetota>Actinomycetes>Mycobacteriales | GCF_001665615.1 | Mycobacterium_sp._852014-52144_SCH5372336 | A5759_RS17450 | 92738..96103 | 1834115 | Swt1-family-HEPN-domain-containing-protein-[Mycobacterium-sp.-852014-52144_SCH5372336] |
| WP_197415592.1 | HEPN+AAA-ATPase+Alpha+Beta-domain+wHTH+RRM | Type-1_BR.RM.like.systems | [illegible] | 1115 | Bacteria>Terrabacteria_group>Actinomycetota>Actinomycetes>Mycobacteriales | GCF_001500025.1 | Mycobacterium_sp._IS-1590 | AU197_RS19315 | 351187..354534 | 1772286 | Swt1-family-HEPN-domain-containing-protein-[Mycobacterium-sp.-IS-1590] |
| WP_144823668.1 | HEPN+AAA-ATPase+Alpha+Beta-domain+wHTH+RRM | Type-1_BR.RM.like.systems | [illegible] | 1112 | Bacteria>Terrabacteria_group>Actinomycetota>Actinomycetes>Mycobacteriales | GCF_025145725.1 | Corynebacterium_sanguinis | EKI50_RS10380 | 1436..4774 | 2594913 | DUF499-domain-containing-protein-[Corynebacterium-sanguinis] |
| WP_207544692.1 | HEPN+AAA-ATPase+Alpha+Beta-domain+wHTH+RRM | Type-1_BR.RM.like.systems | [illegible] | 1114 | Bacteria>Terrabacteria_group>Actinomycetota>Actinomycetes>Mycobacteriales | GCF_001673265.1 | Mycolicibacterium_conceptionense | A5639_RS11935 | 13159..16503 | 451644 | Swt1-family-HEPN-domain-containing-protein-[Mycolicibacterium-conceptionense] |
| WP_065288965.1 | HEPN+AAA-ATPase+Alpha+Beta-domain+wHTH+RRM | Type-1_BR.RM.like.systems | [illegible] | 1141 | Bacteria>Terrabacteria_group>Actinomycetota>Actinomycetes>Mycobacteriales | GCF_001679965.1 | Mycolicibacter_kumamotonensis | ACT18_RS17285 | 28680..32105 | 354243 | DUF499-domain-containing-protein-[Mycolicibacter-kumamotonensis] |
| OBK03541.1 | HEPN+AAA-ATPase+Alpha+Beta-domain+wHTH+RRM | Type-1_BR.RM.like.systems | [illegible] | 1103 | Bacteria>Terrabacteria_group>Actinomycetota>Actinomycetes>Mycobacteriales | GCA_001673265.1 | Mycolicibacterium_conceptionense | A5639_22740 | 13192..16503 | 451644 | AAA-family-ATPase-[Mycolicibacterium-conceptionense] |
| MCB0932438.1 | HEPN+AAA-ATPase+Alpha+Beta-domain+wHTH+RRM | Type-1_BR.RM.like.systems | [illegible] | 1120 | Bacteria>Terrabacteria_group>Actinomycetota>Actinomycetes>Mycobacteriales | GCA_020438845.1 | Mycobacterium_sp. | KDB71_11130 | 13103..16465 | 1785 | DUF499-domain-containing-protein-[Mycobacterium-sp.] |
| WP_207554210.1 | HEPN+AAA-ATPase+Alpha+Beta-domain+wHTH+RRM | Type-1_BR.RM.like.systems | [illegible] | 605 | Bacteria>Terrabacteria_group>Actinomycetota>Actinomycetes>Mycobacteriales | GCF_900141575.1 | Mycobacteroides_abscessus | B5Y25_RS21325 | 33267..35081 | 36809 | Swt1-family-HEPN-domain-containing-protein,-partial-[Mycobacteroides-abscessus] |
| WP_176126766.1 | HEPN+AAA-ATPase+Alpha+Beta-domain+wHTH+RRM | Type-1_BR.RM.like.systems | [illegible] | 1120 | Bacteria>Terrabacteria_group>Actinomycetota>Actinomycetes>Mycobacteriales | - | Mycobacterium_sp._D16Q20 | B4386_RS22320 | 7597..10959 | 1855660 | Swt1-family-HEPN-domain-containing-protein-[Mycobacterium-sp.-D16Q20] |
| WP_129978122.1 | HEPN+AAA-ATPase+Alpha+Beta-domain+wHTH+RRM | Type-1_BR.RM.like.systems | [illegible] | 1138 | Bacteria>Terrabacteria_group>Actinomycetota>Actinomycetes>Mycobacteriales | GCF_004153645.1 | Rhodococcus_sp._Q1 | EQ874_RS21580 | 43740..47156 | 2508718 | Swt1-family-HEPN-domain-containing-protein-[Rhodococcus-sp.-Q1] |
| WP_094024119.1 | HEPN+AAA-ATPase+Alpha+Beta-domain+wHTH+RRM | Type-1_BR.RM.like.systems | [illegible] | 1126 | Bacteria>Terrabacteria_group>Actinomycetota>Actinomycetes>Mycobacteriales | - | Nocardia_cerradoensis | B7C42_RS01205 | 260073..263453 | 85688 | DUF499-domain-containing-protein-[Nocardia-cerradoensis] |
| WP_067488116.1 | HEPN+AAA-ATPase+Alpha+Beta-domain+wHTH+RRM | Type-1_BR.RM.like.systems | [illegible] | 1123 | Bacteria>Terrabacteria_group>Actinomycetota>Actinomycetes>Mycobacteriales | GCF_001612905.1 | Nocardia_ignorata | DFR75_RS26965 | 96694..100065 | 145285 | Swt1-family-HEPN-domain-containing-protein-[Nocardia-ignorata] |
| WP_278127257.1 | HEPN+AAA-ATPase+Alpha+Beta-domain+wHTH+RRM | Type-1_BR.RM.like.systems | [illegible] | 1124 | Bacteria>Terrabacteria_group>Actinomycetota>Actinomycetes>Mycobacteriales | GCF_029632925.1 | Gordonia_sihwensis | P5P27_RS08530 | 1762026..1765400 | 173559 | DUF499-domain-containing-protein-[Gordonia-sihwensis] |
| WP_179300635.1 | HEPN+AAA-ATPase+Alpha+Beta-domain+wHTH+RRM | Type-1_BR.RM.like.systems | [illegible] | 1107 | Bacteria>Terrabacteria_group>Actinomycetota>Actinomycetes>Mycobacteriales | GCF_002285675.1 | Mycobacterium_intracellulare | CKJ67_RS10690 | 2227205..2230528 | 1767 | Swt1-family-HEPN-domain-containing-protein-[Mycobacterium-intracellulare] |
| WP_067083703.1 | HEPN+AAA-ATPase+Alpha+Beta-domain+wHTH+RRM | Type-1_BR.RM.like.systems | [illegible] | 1135 | Bacteria>Terrabacteria_group>Actinomycetota>Actinomycetes>Mycobacteriales | GCF_001665535.1 | Mycobacterium_sp._852002-51961_SCH5331710 | A5752_RS07195 | 17252..20659 | 1834105 | Swt1-family-HEPN-domain-containing-protein-[Mycobacterium-sp.-852002-51961_SCH5331710] |
| WP_045537629.1 | HEPN+AAA-ATPase+Alpha+Beta-domain+wHTH+RRM | Type-1_BR.RM.like.systems | [illegible] | 1124 | Bacteria>Terrabacteria_group>Actinomycetota>Actinomycetes>Mycobacteriales | GCF_000960465.1 | Gordonia_sihwensis | UG54_RS06085 | 5143..8517 | 173559 | DUF499-domain-containing-protein-[Gordonia-sihwensis] |
| WP_006896364.1 | HEPN+AAA-ATPase+Alpha+Beta-domain+wHTH+RRM | Type-1_BR.RM.like.systems | [illegible] | 1124 | Bacteria>Terrabacteria_group>Actinomycetota>Actinomycetes>Mycobacteriales | GCF_000333035.1 | Gordonia | GSI01S_RS09635 | 144602..147976 | 2053 | MULTISPECIES:-DUF499-domain-containing-protein-[Gordonia] |
| TXH28009.1 | HEPN+AAA-ATPase+Alpha+Beta-domain+wHTH+RRM | Type-1_BR.RM.like.systems | [illegible] | 1098 | Bacteria>Terrabacteria_group>Actinomycetota>Actinomycetes>Mycobacteriales | GCA_008012155.1 | Mycobacterium_sp. | E6R06_02260 | 39291..42587 | 1785 | MAG:-ATP-binding-protein-[Mycobacterium-sp.] |
| WP_207546421.1 | HEPN+AAA-ATPase+Alpha+Beta-domain+wHTH+RRM | Type-1_BR.RM.like.systems | [illegible] | 1120 | Bacteria>Terrabacteria_group>Actinomycetota>Actinomycetes>Mycobacteriales | GCF_001853545.1 | Mycobacteroides_chelonae | BKG62_RS21080 | 210125..213487 | 1774 | Swt1-family-HEPN-domain-containing-protein-[Mycobacteroides-chelonae] |
| WP_205874823.1 | HEPN+AAA-ATPase+Alpha+Beta-domain+wHTH+RRM | Type-1_BR.RM.like.systems | [illegible] | 1108 | Bacteria>Terrabacteria_group>Actinomycetota>Actinomycetes>Mycobacteriales | GCF_017086405.1 | Mycobacterium_gordonae | JX552_RS26920 | 6279372..6282698 | 1778 | Swt1-family-HEPN-domain-containing-protein-[Mycobacterium-gordonae] |
| WP_207546857.1 | HEPN+AAA-ATPase+Alpha+Beta-domain+wHTH+RRM | Type-1_BR.RM.like.systems | [illegible] | 1109 | Bacteria>Terrabacteria_group>Actinomycetota>Actinomycetes>Mycobacteriales | GCF_020520165.1 | Mycolicibacterium | EKK51_RS16805 | 41858..45187 | 1866885 | MULTISPECIES:-Swt1-family-HEPN-domain-containing-protein-[Mycolicibacterium] |
| WP_204161112.1 | HEPN+AAA-ATPase+Alpha+Beta-domain+wHTH+RRM | Type-1_BR.RM.like.systems | [illegible] | 1109 | Bacteria>Terrabacteria_group>Actinomycetota>Actinomycetes>Mycobacteriales | GCF_001557045.1 | Mycolicibacterium_mucogenicum | AX749_RS05580 | 20286..23615 | 56689 | Swt1-family-HEPN-domain-containing-protein-[Mycolicibacterium-mucogenicum] |
| WP_048320042.1 | HEPN+AAA-ATPase+Alpha+Beta-domain+wHTH+RRM | Type-1_BR.RM.like.systems | [illegible] | 1137 | Bacteria>Terrabacteria_group>Actinomycetota>Actinomycetes>Mycobacteriales | GCF_001037935.1 | Rhodococcus_fascians | ACG96_RS20770 | 139814..143227 | 1828 | DUF499-domain-containing-protein-[Rhodococcus-fascians] |
| WP_041784793.1 | HEPN+AAA-ATPase+Alpha+Beta-domain+wHTH+RRM | Type-1_BR.RM.like.systems | [illegible] | 1108 | Bacteria>Terrabacteria_group>Actinomycetota>Actinomycetes>Mycobacteriales | GCF_000276825.1 | Mycobacterium_paraintracellulare | B8W68_RS04660 | 128474..131800 | 1138383 | Swt1-family-HEPN-domain-containing-protein-[Mycobacterium-paraintracellulare] |
| WP_003939259.1 | HEPN+AAA-ATPase+Alpha+Beta-domain+wHTH+RRM | Type-1_BR.RM.like.systems | [illegible] | 1138 | Bacteria>Terrabacteria_group>Actinomycetota>Actinomycetes>Mycobacteriales | GCF_000341965.1 | Rhodococcus_ruber | G352_RS25680 | 27793..31209 | 1830 | Swt1-family-HEPN-domain-containing-protein-[Rhodococcus-ruber] |
| OKH84136.1 | HEPN+AAA-ATPase+Alpha+Beta-domain+wHTH+RRM | Type-1_BR.RM.like.systems | [illegible] | 1109 | Bacteria>Terrabacteria_group>Actinomycetota>Actinomycetes>Mycobacteriales | GCA_001905655.1 | Mycobacterium_sp._ST-F2 | EB75_06540 | 184865..188194 | 1490484 | ATPase-AAA-[Mycobacterium-sp.-ST-F2] |
| WP_259916185.1 | HEPN+AAA-ATPase+Alpha+Beta-domain+wHTH+RRM | Type-1_BR.RM.like.systems | [illegible] | 1111 | Bacteria>Terrabacteria_group>Actinomycetota>Actinomycetes>Mycobacteriales | GCF_025144095.1 | Corynebacterium_sanguinis | M3G11_RS01890 | 80632..83967 | 2594913 | DUF499-domain-containing-protein-[Corynebacterium-sanguinis] |
| WP_225506778.1 | HEPN+AAA-ATPase+Alpha+Beta-domain+wHTH+RRM | Type-1_BR.RM.like.systems | [illegible] | 1108 | Bacteria>Terrabacteria_group>Actinomycetota>Actinomycetes>Mycobacteriales | GCF_020227835.1 | Mycolicibacterium_fortuitum | H8Z59_RS27040 | 5490569..5493895 | 1766 | Swt1-family-HEPN-domain-containing-protein-[Mycolicibacterium-fortuitum] |
| WP_175616187.1 | HEPN+AAA-ATPase+Alpha+Beta-domain+wHTH+RRM | Type-1_BR.RM.like.systems | [illegible] | 1114 | Bacteria>Terrabacteria_group>Actinomycetota>Actinomycetes>Mycobacteriales | GCF_001954135.1 | Mycobacterium_sp._GA-1841 | A5740_RS07670 | 288135..291479 | 1834154 | Swt1-family-HEPN-domain-containing-protein-[Mycobacterium-sp.-GA-1841] |
| WP_144317543.1 | HEPN+AAA-ATPase+Alpha+Beta-domain+wHTH+RRM | Type-1_BR.RM.like.systems | [illegible] | 1111 | Bacteria>Terrabacteria_group>Actinomycetota>Actinomycetes>Mycobacteriales | GCF_007641235.1 | Corynebacterium_sanguinis | E3227_RS03440 | 690306..693641 | 2594913 | DUF499-domain-containing-protein-[Corynebacterium-sanguinis] |
| WP_063013730.1 | HEPN+AAA-ATPase+Alpha+Beta-domain+wHTH+RRM | Type-1_BR.RM.like.systems | [illegible] | 1126 | Bacteria>Terrabacteria_group>Actinomycetota>Actinomycetes>Mycobacteriales | GCF_001613005.1 | Nocardia | NN2_RS13920 | 30731..34111 | 1817 | MULTISPECIES:-Swt1-family-HEPN-domain-containing-protein-[Nocardia] |
| OMC39072.1 | HEPN+AAA-ATPase+Alpha+Beta-domain+wHTH+RRM | Type-1_BR.RM.like.systems | [illegible] | 1103 | Bacteria>Terrabacteria_group>Actinomycetota>Actinomycetes>Mycobacteriales | GCA_001954135.1 | Mycobacterium_sp._GA-1841 | A5740_02790 | 288168..291479 | 1834154 | AAA-family-ATPase-[Mycobacterium-sp.-GA-1841] |
| OBA79911.1 | HEPN+AAA-ATPase+Alpha+Beta-domain+wHTH+RRM | Type-1_BR.RM.like.systems | [illegible] | 1098 | Bacteria>Terrabacteria_group>Actinomycetota>Actinomycetes>Mycobacteriales | GCA_001665375.1 | Mycolicibacterium_mucogenicum | A5642_03430 | 86031..89327 | 56689 | AAA-family-ATPase-[Mycolicibacterium-mucogenicum] |
| ASW98156.1 | HEPN+AAA-ATPase+Alpha+Beta-domain+wHTH+RRM | Type-1_BR.RM.like.systems | [illegible] | 1096 | Bacteria>Terrabacteria_group>Actinomycetota>Actinomycetes>Mycobacteriales | GCA_002285675.1 | Mycobacterium_intracellulare | CKJ67_10690 | 2227205..2230495 | 1767 | AAA-family-ATPase-[Mycobacterium-intracellulare] |
| WP_276761078.1 | HEPN+AAA-ATPase+Alpha+Beta-domain+wHTH+RRM | Type-1_BR.RM.like.systems | [illegible] | 1114 | Bacteria>Terrabacteria_group>Actinomycetota>Actinomycetes>Mycobacteriales | GCF_008015745.1 | Mycolicibacter_arupensis | E6Q54_RS12710 | 7339..10683 | 342002 | Swt1-family-HEPN-domain-containing-protein-[Mycolicibacter-arupensis] |
| WP_276127103.1 | HEPN+AAA-ATPase+Alpha+Beta-domain+wHTH+RRM | Type-1_BR.RM.like.systems | [illegible] | 1137 | Bacteria>Terrabacteria_group>Actinomycetota>Actinomycetes>Mycobacteriales | GCF_029270025.1 | Mycolicibacterium_septicum | P3H80_RS10735 | 384555..387968 | 98668 | DUF499-domain-containing-protein-[Mycolicibacterium-septicum] |
| WP_237211675.1 | HEPN+AAA-ATPase+Alpha+Beta-domain+wHTH+RRM | Type-1_BR.RM.like.systems | [illegible] | 1138 | Bacteria>Terrabacteria_group>Actinomycetota>Actinomycetes>Mycobacteriales | GCF_021852985.1 | Rhodococcus_ruber | K7Z00_RS28700 | 67558..70974 | 1830 | Swt1-family-HEPN-domain-containing-protein-[Rhodococcus-ruber] |
| WP_211919214.1 | HEPN+AAA-ATPase+Alpha+Beta-domain+wHTH+RRM | Type-1_BR.RM.like.systems | [illegible] | 1121 | Bacteria>Terrabacteria_group>Actinomycetota>Actinomycetes>Mycobacteriales | GCF_018283555.1 | Rhodococcus_erythropolis | AC1659_RS28475 | 46414..49779 | 1833 | Swt1-family-HEPN-domain-containing-protein-[Rhodococcus-erythropolis] |
| WP_205765478.1 | HEPN+AAA-ATPase+Alpha+Beta-domain+wHTH+RRM | Type-1_BR.RM.like.systems | [illegible] | 1114 | Bacteria>Terrabacteria_group>Actinomycetota>Actinomycetes>Mycobacteriales | GCF_002968275.1 | Mycolicibacterium_austroafricanum | C6A88_RS31870 | 14193..17537 | 39687 | Swt1-family-HEPN-domain-containing-protein-[Mycolicibacterium-austroafricanum] |
| WP_194806632.1 | HEPN+AAA-ATPase+Alpha+Beta-domain+wHTH+RRM | Type-1_BR.RM.like.systems | [illegible] | 1121 | Bacteria>Terrabacteria_group>Actinomycetota>Actinomycetes>Mycobacteriales | GCF_015354405.1 | Nocardia_sp._BSTN01 | IRT45_RS15950 | 106038..109403 | 2783665 | Swt1-family-HEPN-domain-containing-protein-[Nocardia-sp.-BSTN01] |
| WP_085075211.1 | HEPN+AAA-ATPase+Alpha+Beta-domain+wHTH+RRM | Type-1_BR.RM.like.systems | [illegible] | 1110 | Bacteria>Terrabacteria_group>Actinomycetota>Actinomycetes>Mycobacteriales | - | Mycobacterium_kubicae | H7H99_RS20905 | 74999..78331 | 120959 | Swt1-family-HEPN-domain-containing-protein-[Mycobacterium-kubicae] |
| ORV96500.1 | HEPN+AAA-ATPase+Alpha+Beta-domain+wHTH+RRM | Type-1_BR.RM.like.systems | [illegible] | 1110 | Bacteria>Terrabacteria_group>Actinomycetota>Actinomycetes>Mycobacteriales | GCA_002101745.1 | Mycobacterium_kubicae | AWC13_18590 | 6697..10029 | 120959 | AAA-family-ATPase-[Mycobacterium-kubicae] |
| MCT7366341.1 | HEPN+AAA-ATPase+Alpha+Beta-domain+wHTH+RRM | Type-1_BR.RM.like.systems | [illegible] | 1099 | Bacteria>Terrabacteria_group>Actinomycetota>Actinomycetes>Mycobacteriales | GCA_025331195.1 | Mycolicibacterium_llatzerense | A7G45_26120 | 438962..442261 | 280871 | AAA-family-ATPase-[Mycolicibacterium-llatzerense] |
| WP_204801726.1 | HEPN+AAA-ATPase+Alpha+Beta-domain+wHTH+RRM | Type-1_BR.RM.like.systems | [illegible] | 1109 | Bacteria>Terrabacteria_group>Actinomycetota>Actinomycetes>Mycobacteriales | GCF_905219495.1 | Mycobacterium_riyadhense | JX354_RS03520 | 789692..793021 | 486698 | Swt1-family-HEPN-domain-containing-protein-[Mycobacterium-riyadhense] |
| WP_085470756.1 | HEPN+AAA-ATPase+Alpha+Beta-domain+wHTH+RRM | Type-1_BR.RM.like.systems | [illegible] | 1125 | Bacteria>Terrabacteria_group>Actinomycetota>Actinomycetes>Mycobacteriales | GCF_900177695.1 | Rhodococcus_rhodochrous | M502_RS25910 | 14095..17472 | 1829 | Swt1-family-HEPN-domain-containing-protein-[Rhodococcus-rhodochrous] |
| WP_238886618.1 | HEPN+AAA-ATPase+Alpha+Beta-domain+wHTH+RRM | Type-1_BR.RM.like.systems | [illegible] | 1120 | Bacteria>Terrabacteria_group>Actinomycetota>Actinomycetes>Mycobacteriales | GCF_022317005.1 | Mycobacterium_sp._IDR2000157661 | K3G64_RS19485 | 3782160..3785522 | 2867005 | Swt1-family-HEPN-domain-containing-protein-[Mycobacterium-sp.-IDR2000157661] |
| WP_197940422.1 | HEPN+AAA-ATPase+Alpha+Beta-domain+wHTH+RRM | Type-1_BR.RM.like.systems | [illegible] | 1121 | Bacteria>Terrabacteria_group>Actinomycetota>Actinomycetes>Mycobacteriales | GCF_016019085.1 | Rhodococcus_erythropolis | I3517_RS00940 | 13889..17254 | 1833 | Swt1-family-HEPN-domain-containing-protein-[Rhodococcus-erythropolis] |
| WP_005174963.1 | HEPN+AAA-ATPase+Alpha+Beta-domain+wHTH+RRM | Type-1_BR.RM.like.systems | [illegible] | 1134 | Bacteria>Terrabacteria_group>Actinomycetota>Actinomycetes>Mycobacteriales | GCF_000332975.1 | Gordonia_aichiensis | GOACH_RS12130 | 81836..85240 | 36820 | Swt1-family-HEPN-domain-containing-protein-[Gordonia-aichiensis] |
| WP_245548345.1 | HEPN+AAA-ATPase+Alpha+Beta-domain+wHTH+RRM | Type-1_BR.RM.like.systems | [illegible] | 1116 | Bacteria>Terrabacteria_group>Actinomycetota>Actinomycetes>Mycobacteriales | GCF_000241265.1 | Gordonia_araii | GOARA_RS03580 | 82752..86102 | 263909 | Swt1-family-HEPN-domain-containing-protein-[Gordonia-araii] |
| WP_230074521.1 | HEPN+AAA-ATPase+Alpha+Beta-domain+wHTH+RRM | Type-1_BR.RM.like.systems | [illegible] | 1127 | Bacteria>Terrabacteria_group>Actinomycetota>Actinomycetes>Mycobacteriales | GCF_918697885.1 | Rhodococcus_fascians | LQV52_RS22465 | 73515..76898 | 1828 | DUF499-domain-containing-protein-[Rhodococcus-fascians] |
| WP_059384669.1 | HEPN+AAA-ATPase+Alpha+Beta-domain+wHTH+RRM | Type-1_BR.RM.like.systems | [illegible] | 1131 | Bacteria>Terrabacteria_group>Actinomycetota>Actinomycetes>Mycobacteriales | GCF_001047055.1 | Rhodococcus_rhodochrous | RR2_RS22685 | 9470..12865 | 1829 | Swt1-family-HEPN-domain-containing-protein-[Rhodococcus-rhodochrous] |
| OSC29643.1 | HEPN+AAA-ATPase+Alpha+Beta-domain+wHTH+RRM | Type-1_BR.RM.like.systems | [illegible] | 1097 | Bacteria>Terrabacteria_group>Actinomycetota>Actinomycetes>Mycobacteriales | GCA_002104735.1 | Mycobacterium_paraintracellulare | B8W68_04660 | 128474..131767 | 1138383 | AAA-family-ATPase-[Mycobacterium-paraintracellulare] |
| NNG97058.1 | HEPN+AAA-ATPase+Alpha+Beta-domain+wHTH+RRM | Type-1_BR.RM.like.systems | [illegible] | 1118 | Bacteria>Terrabacteria_group>Actinomycetota>Actinomycetes>Mycobacteriales | GCA_013004535.1 | Gordonia_araii_NBRC_100433 | HLA97_07325 | 102425..105781 | 1073574 | ATP-binding-protein-[Gordonia-araii-NBRC-100433] |
| GAB08850.1 | HEPN+AAA-ATPase+Alpha+Beta-domain+wHTH+RRM | Type-1_BR.RM.like.systems | [illegible] | 1140 | Bacteria>Terrabacteria_group>Actinomycetota>Actinomycetes>Mycobacteriales | GCA_000241265.2 | Gordonia_araii_NBRC_100433 | GOARA_021_00870 | 82752..86174 | 1073574 | hypothetical-protein-GOARA_021_00870-[Gordonia-araii-NBRC-100433] |
| WP_105942269.1 | HEPN+AAA-ATPase+Alpha+Beta-domain+wHTH+RRM | Type-1_BR.RM.like.systems | [illegible] | 1121 | Bacteria>Terrabacteria_group>Actinomycetota>Actinomycetes>Mycobacteriales | GCF_002993285.1 | Gordonia_iterans | C6V83_RS09885 | 2154114..2157479 | 1004901 | DUF499-domain-containing-protein-[Gordonia-iterans] |
| WP_185712603.1 | HEPN+AAA-ATPase+Alpha+Beta-domain+wHTH+RRM | Type-1_BR.RM.like.systems | [illegible] | 1138 | Bacteria>Terrabacteria_group>Actinomycetota>Actinomycetes>Mycobacteriales | GCF_014230025.1 | Rhodococcus_aetherivorans | H0B42_RS00215 | 49328..52744 | 191292 | Swt1-family-HEPN-domain-containing-protein-[Rhodococcus-aetherivorans] |
| WP_172936837.1 | HEPN+AAA-ATPase+Alpha+Beta-domain+wHTH+RRM | Type-1_BR.RM.like.systems | [illegible] | 1125 | Bacteria>Terrabacteria_group>Actinomycetota>Actinomycetes>Mycobacteriales | GCF_013277795.1 | Rhodococcus_sp._MS16 | FEZ60_RS20030 | 139457..142834 | 2579941 | Swt1-family-HEPN-domain-containing-protein-[Rhodococcus-sp.-MS16] |
| WP_085176531.1 | HEPN+AAA-ATPase+Alpha+Beta-domain+wHTH+RRM | Type-1_BR.RM.like.systems | [illegible] | 806 | Bacteria>Terrabacteria_group>Actinomycetota>Actinomycetes>Mycobacteriales | GCF_002101705.1 | Mycolicibacterium_iranicum | AWC12_RS19975 | 111693..114110 | 912594 | DUF499-domain-containing-protein,-partial-[Mycolicibacterium-iranicum] |
| TXI55381.1 | HEPN+AAA-ATPase+Alpha+Beta-domain+wHTH+RRM | Type-1_BR.RM.like.systems | [illegible] | 1103 | Bacteria>Terrabacteria_group>Actinomycetota>Actinomycetes>Mycobacteriales | GCA_008015745.1 | Mycolicibacter_arupensis | E6Q54_12720 | 7339..10650 | 342002 | ATP-binding-protein-[Mycolicibacter-arupensis] |
| WP_005939379.1 | HEPN+AAA-ATPase+Alpha+Beta-domain+wHTH+RRM | Type-1_BR.RM.like.systems | [illegible] | 1129 | Bacteria>Terrabacteria_group>Actinomycetota>Actinomycetes>Mycobacteriales | GCF_000420685.1 | Gordonia_hirsuta | GOHSU_RS07845 | 157749..161138 | 53427 | Swt1-family-HEPN-domain-containing-protein-[Gordonia-hirsuta] |
| WP_005139269.1 | HEPN+AAA-ATPase+Alpha+Beta-domain+wHTH+RRM | Type-1_BR.RM.like.systems | [illegible] | 1108 | Bacteria>Terrabacteria_group>Actinomycetota>Actinomycetes>Mycobacteriales | - | Mycolicibacterium_rhodesiae | MycrhDRAFT_0407 | 390606..393932 | 36814 | Swt1-family-HEPN-domain-containing-protein-[Mycolicibacterium-rhodesiae] |
| KUI43016.1 | HEPN+AAA-ATPase+Alpha+Beta-domain+wHTH+RRM | Type-1_BR.RM.like.systems | [illegible] | 1101 | Bacteria>Terrabacteria_group>Actinomycetota>Actinomycetes>Mycobacteriales | GCA_001500045.1 | Mycobacterium_sp._GA-1199 | AU198_07655 | 256994..260299 | 1772287 | AAA-family-ATPase-[Mycobacterium-sp.-GA-1199] |
| EHB57971.1 | HEPN+AAA-ATPase+Alpha+Beta-domain+wHTH+RRM | Type-1_BR.RM.like.systems | [illegible] | 1108 | Bacteria>Terrabacteria_group>Actinomycetota>Actinomycetes>Mycobacteriales | GCA_000230935.2 | Mycolicibacterium_rhodesiae_JS60 | MycrhDRAFT_0407 | 390606..393932 | 931627 | hypothetical-protein-MycrhDRAFT_0407-[Mycolicibacterium-rhodesiae-JS60] |
| WP_278720226.1 | HEPN+AAA-ATPase+Alpha+Beta-domain+wHTH+RRM | Type-1_BR.RM.like.systems | [illegible] | 1184 | Bacteria>Terrabacteria_group>Actinomycetota>Actinomycetes>Mycobacteriales | GCF_905372825.1 | Corynebacterium_matruchotii | QDH69_RS07065 | 79539..83093 | 43768 | DUF499-domain-containing-protein-[Corynebacterium-matruchotii] |
| WP_267616414.1 | HEPN+AAA-ATPase+Alpha+Beta-domain+wHTH+RRM | Type-1_BR.RM.like.systems | [illegible] | 1127 | Bacteria>Terrabacteria_group>Actinomycetota>Actinomycetes>Mycobacteriales | GCF_026616955.1 | Gordonia_bronchialis | N0B45_RS12880 | 105537..108920 | 2054 | Swt1-family-HEPN-domain-containing-protein-[Gordonia-bronchialis] |
| WP_239342823.1 | HEPN+AAA-ATPase+Alpha+Beta-domain+wHTH+RRM | Type-1_BR.RM.like.systems | [illegible] | 1109 | Bacteria>Terrabacteria_group>Actinomycetota>Actinomycetes>Mycobacteriales | GCF_022348005.1 | Mycolicibacterium_sp._OfavD-34-C | MHN82_RS23245 | 18852..22181 | 2917746 | Swt1-family-HEPN-domain-containing-protein-[Mycolicibacterium-sp.-OfavD-34-C] |
| OOK63774.1 | HEPN+AAA-ATPase+Alpha+Beta-domain+wHTH+RRM | Type-1_BR.RM.like.systems | [illegible] | 1104 | Bacteria>Terrabacteria_group>Actinomycetota>Actinomycetes>Mycobacteriales | GCA_002003585.1 | Mycobacterium_kansasii | BZL29_8433 | 3439..6753 | 1768 | hypothetical-protein-BZL29_8433-[Mycobacterium-kansasii] |
| WP_183155213.1 | HEPN+AAA-ATPase+Alpha+Beta-domain+wHTH+RRM | Type-1_BR.RM.like.systems | [illegible] | 1114 | Bacteria>Terrabacteria_group>Actinomycetota>Actinomycetes>Mycobacteriales | GCF_900565985.1 | Mycobacterium_kansasii | EET11_RS25330 | 51078..54422 | 1768 | DUF499-domain-containing-protein-[Mycobacterium-kansasii] |
| WP_174485135.1 | HEPN+AAA-ATPase+Alpha+Beta-domain+wHTH+RRM | Type-1_BR.RM.like.systems | [illegible] | 1113 | Bacteria>Terrabacteria_group>Actinomycetota>Actinomycetes>Mycobacteriales | GCF_001214605.1 | Mycobacteroides_abscessus | IUT00_RS08255 | 1657384..1660725 | 36809 | Swt1-family-HEPN-domain-containing-protein-[Mycobacteroides-abscessus] |
| PZT93668.1 | HEPN+AAA-ATPase+Alpha+Beta-domain+wHTH+RRM | Type-1_BR.RM.like.systems | [illegible] | 1133 | Bacteria>Terrabacteria_group>Actinomycetota>Actinomycetes>Mycobacteriales | GCA_003248405.1 | Gordonia_sp._(in:_high_G+C_Gram-positive_bacteria) | DI630_26685 | 9643..13044 | 84139 | AAA-family-ATPase-[Gordonia-sp.] |
| WP_179973625.1 | HEPN+AAA-ATPase+Alpha+Beta-domain+wHTH+RRM | Type-1_BR.RM.like.systems | [illegible] | 1121 | Bacteria>Terrabacteria_group>Actinomycetota>Actinomycetes>Mycobacteriales | GCF_010731815.2 | Mycolicibacterium_arabiense | H5P33_RS16145 | 691085..694450 | 1286181 | Swt1-family-HEPN-domain-containing-protein-[Mycolicibacterium-arabiense] |
| WP_149772783.1 | HEPN+AAA-ATPase+Alpha+Beta-domain+wHTH+RRM | Type-1_BR.RM.like.systems | [illegible] | 1138 | Bacteria>Terrabacteria_group>Actinomycetota>Actinomycetes>Mycobacteriales | GCF_008373105.1 | Mycolicibacter_arupensis | F0402_RS09270 | 8758..12174 | 342002 | DUF499-domain-containing-protein-[Mycolicibacter-arupensis] |
| WP_108404986.1 | HEPN+AAA-ATPase+Alpha+Beta-domain+wHTH+RRM | Type-1_BR.RM.like.systems | [illegible] | 1112 | Bacteria>Terrabacteria_group>Actinomycetota>Actinomycetes>Mycobacteriales | GCF_003070865.1 | Corynebacterium_liangguodongii | C3E79_RS00685 | 144858..148196 | 2079535 | DUF499-domain-containing-protein-[Corynebacterium-liangguodongii] |
| WP_013827697.1 | HEPN+AAA-ATPase+Alpha+Beta-domain+wHTH+RRM | Type-1_BR.RM.like.systems | [illegible] | 1138 | Bacteria>Terrabacteria_group>Actinomycetota>Actinomycetes>Mycobacteriales | GCF_000214155.1 | Mycolicibacter_sinensis | JDM601_RS03755 | 802521..805937 | 875328 | DUF499-domain-containing-protein-[Mycolicibacter-sinensis] |
| WP_228819156.1 | HEPN+AAA-ATPase+Alpha+Beta-domain+wHTH+RRM | Type-1_BR.RM.like.systems | [illegible] | 1114 | Bacteria>Terrabacteria_group>Actinomycetota>Actinomycetes>Mycobacteriales | GCF_015477815.1 | Nocardia_farcinica | QH106_RS06910 | 129402..132746 | 37329 | DUF499-domain-containing-protein-[Nocardia-farcinica] |
| WP_207554378.1 | HEPN+AAA-ATPase+Alpha+Beta-domain+wHTH+RRM | Type-1_BR.RM.like.systems | [illegible] | 1113 | Bacteria>Terrabacteria_group>Actinomycetota>Actinomycetes>Mycobacteriales | GCF_900141585.1 | Mycobacteroides_abscessus | B5Z43_RS22925 | 52451..55792 | 36809 | Swt1-family-HEPN-domain-containing-protein-[Mycobacteroides-abscessus] |
| WP_161896301.1 | HEPN+AAA-ATPase+Alpha+Beta-domain+wHTH+RRM | Type-1_BR.RM.like.systems | [illegible] | 1123 | Bacteria>Terrabacteria_group>Actinomycetota>Actinomycetes>Mycobacteriales | GCF_009932475.1 | Gordonia_spumicola | GYB83_RS15365 | 872250..875621 | 589161 | DUF499-domain-containing-protein-[Gordonia-spumicola] |
| WP_062879806.1 | HEPN+AAA-ATPase+Alpha+Beta-domain+wHTH+RRM | Type-1_BR.RM.like.systems | [illegible] | 1108 | Bacteria>Terrabacteria_group>Actinomycetota>Actinomycetes>Mycobacteriales | GCF_001606235.1 | Mycobacteroides_abscessus | B5431_RS12990 | 716837..720163 | 36809 | Swt1-family-HEPN-domain-containing-protein-[Mycobacteroides-abscessus] |
| AFC52219.1 | HEPN+AAA-ATPase+Alpha+Beta-domain+wHTH+RRM | Type-1_BR.RM.like.systems | [illegible] | 1043 | Bacteria>Terrabacteria_group>Actinomycetota>Actinomycetes>Mycobacteriales | GCA_000276825.1 | Mycobacterium_paraintracellulare | OCQ_07060 | 694865..697996 | 1138383 | hypothetical-protein-OCQ_07060-[Mycobacterium-paraintracellulare] |
| WP_141142680.1 | HEPN+AAA-ATPase+Alpha+Beta-domain+wHTH+RRM | Type-1_BR.RM.like.systems | [illegible] | 1137 | Bacteria>Terrabacteria_group>Actinomycetota>Actinomycetes>Mycobacteriales | GCF_006517415.1 | Mycolicibacterium_fortuitum | FKW78_RS12965 | 17898..21311 | 1766 | DUF499-domain-containing-protein-[Mycolicibacterium-fortuitum] |
| WP_124259269.1 | HEPN+AAA-ATPase+Alpha+Beta-domain+wHTH+RRM | Type-1_BR.RM.like.systems | [illegible] | 1120 | Bacteria>Terrabacteria_group>Actinomycetota>Actinomycetes>Mycobacteriales | GCF_003844175.1 | Rhodococcus_ruber | TN91_RS21990 | 3527..6889 | 1830 | DUF499-domain-containing-protein-[Rhodococcus-ruber] |
| WP_053777653.1 | HEPN+AAA-ATPase+Alpha+Beta-domain+wHTH+RRM | Type-1_BR.RM.like.systems | [illegible] | 1120 | Bacteria>Terrabacteria_group>Actinomycetota>Actinomycetes>Mycobacteriales | GCF_000347295.2 | Gordonia_sp._NB41Y | ISGA_RS14495 | 38931..42293 | 875808 | DUF499-domain-containing-protein-[Gordonia-sp.-NB41Y] |
| OFJ52823.1 | HEPN+AAA-ATPase+Alpha+Beta-domain+wHTH+RRM | Type-1_BR.RM.like.systems | [illegible] | 1109 | Bacteria>Terrabacteria_group>Actinomycetota>Actinomycetes>Mycobacteriales | GCA_001766635.1 | Mycobacterium_grossiae | BEL07_15685 | 42542..45871 | 1552759 | AAA-family-ATPase-[Mycobacterium-grossiae] |
| OBB72728.1 | HEPN+AAA-ATPase+Alpha+Beta-domain+wHTH+RRM | Type-1_BR.RM.like.systems | [illegible] | 1110 | Bacteria>Terrabacteria_group>Actinomycetota>Actinomycetes>Mycobacteriales | GCA_001665615.1 | Mycobacterium_sp._852014-52144_SCH5372336 | A5759_18055 | 92771..96103 | 1834115 | AAA-family-ATPase-[Mycobacterium-sp.-852014-52144_SCH5372336] |
| WP_211284609.1 | HEPN+AAA-ATPase+Alpha+Beta-domain+wHTH+RRM | Type-1_BR.RM.like.systems | [illegible] | 1113 | Bacteria>Terrabacteria_group>Actinomycetota>Actinomycetes>Mycobacteriales | GCF_025823205.1 | Mycobacterium_alsense | BST11_RS03260 | 100678..104019 | 324058 | Swt1-family-HEPN-domain-containing-protein-[Mycobacterium-alsense] |
| WP_134205957.1 | HEPN+AAA-ATPase+Alpha+Beta-domain+wHTH+RRM | Type-1_BR.RM.like.systems | [illegible] | 1137 | Bacteria>Terrabacteria_group>Actinomycetota>Actinomycetes>Mycobacteriales | GCF_004366875.1 | Mycobacteroides_salmoniphilum | CCUG62472_RS09110 | 402187..405600 | 404941 | DUF499-domain-containing-protein-[Mycobacteroides-salmoniphilum] |
| RUP02204.1 | HEPN+AAA-ATPase+Alpha+Beta-domain+wHTH+RRM | Type-1_BR.RM.like.systems | [illegible] | 1097 | Bacteria>Terrabacteria_group>Actinomycetota>Actinomycetes>Mycobacteriales | GCA_003987775.1 | Mycobacterium_sp. | EKK34_25285 | 13112..16405 | 1785 | MAG:-ATP-binding-protein-[Mycobacterium-sp.] |
| WP_260868497.1 | HEPN+AAA-ATPase+Alpha+Beta-domain+wHTH+RRM | Type-1_BR.RM.like.systems | [illegible] | 1110 | Bacteria>Terrabacteria_group>Actinomycetota>Actinomycetes>Mycobacteriales | - | Gordonia_sihwensis | ACN95_RS06460 | 9637..12969 | 173559 | DUF499-domain-containing-protein-[Gordonia-sihwensis] |
| WP_207569995.1 | HEPN+AAA-ATPase+Alpha+Beta-domain+wHTH+RRM | Type-1_BR.RM.like.systems | [illegible] | 1121 | Bacteria>Terrabacteria_group>Actinomycetota>Actinomycetes>Mycobacteriales | GCF_002086285.1 | Mycolicibacter_kumamotonensis | BST28_RS00055 | 16513..19878 | 354243 | Swt1-family-HEPN-domain-containing-protein-[Mycolicibacter-kumamotonensis] |
| WP_074345357.1 | HEPN+AAA-ATPase+Alpha+Beta-domain+wHTH+RRM | Type-1_BR.RM.like.systems | [illegible] | 1115 | Bacteria>Terrabacteria_group>Actinomycetota>Actinomycetes>Mycobacteriales | GCF_900130695.1 | Mycobacteroides_abscessus | B5Z17_RS24110 | 72774..76121 | 36809 | DUF499-domain-containing-protein-[Mycobacteroides-abscessus] |
| WP_015289521.1 | HEPN+AAA-ATPase+Alpha+Beta-domain+wHTH+RRM | Type-1_BR.RM.like.systems | [illegible] | 1115 | Bacteria>Terrabacteria_group>Actinomycetota>Actinomycetes>Mycobacteriales | GCF_000328785.1 | Mycobacterium_canettii | BN42_RS05705 | 1219765..1223112 | 78331 | Swt1-family-HEPN-domain-containing-protein-[Mycobacterium-canettii] |
| WP_009680352.1 | HEPN+AAA-ATPase+Alpha+Beta-domain+wHTH+RRM | Type-1_BR.RM.like.systems | [illegible] | 1137 | Bacteria>Terrabacteria_group>Actinomycetota>Actinomycetes>Mycobacteriales | GCF_000192435.1 | Gordonia_neofelifaecis | SCNU_RS15560 | 9488..12901 | 945692 | DUF499-domain-containing-protein-[Gordonia-neofelifaecis] |
| AYJ07722.1 | HEPN+AAA-ATPase+Alpha+Beta-domain+wHTH+RRM | Type-1_BR.RM.like.systems | [illegible] | 1097 | Bacteria>Terrabacteria_group>Actinomycetota>Actinomycetes>Mycobacteriales | GCA_003640565.1 | Mycobacterium_avium | DBO90_17285 | 3694672..3697965 | 1764 | DUF499-domain-containing-protein-[Mycobacterium-avium] |
| WP_278334519.1 | HEPN+AAA-ATPase+Alpha+Beta-domain+wHTH+RRM | Type-1_BR.RM.like.systems | [illegible] | 1138 | Bacteria>Terrabacteria_group>Actinomycetota>Actinomycetes>Mycobacteriales | GCF_029691645.1 | Rhodococcus_aetherivorans | P9K37_RS00335 | 73561..76977 | 191292 | Swt1-family-HEPN-domain-containing-protein-[Rhodococcus-aetherivorans] |
| WP_260941407.1 | HEPN+AAA-ATPase+Alpha+Beta-domain+wHTH+RRM | Type-1_BR.RM.like.systems | [illegible] | 1110 | Bacteria>Terrabacteria_group>Actinomycetota>Actinomycetes>Mycobacteriales | GCF_025331195.1 | Mycolicibacterium_llatzerense | A7G45_RS26855 | 438962..442294 | 280871 | Swt1-family-HEPN-domain-containing-protein-[Mycolicibacterium-llatzerense] |
| WP_258913985.1 | HEPN+AAA-ATPase+Alpha+Beta-domain+wHTH+RRM | Type-1_BR.RM.like.systems | [illegible] | 1138 | Bacteria>Terrabacteria_group>Actinomycetota>Actinomycetes>Mycobacteriales | GCF_024760325.1 | Rhodococcus_pyridinivorans | NXT08_RS24750 | 68689..72105 | 103816 | Swt1-family-HEPN-domain-containing-protein-[Rhodococcus-pyridinivorans] |
| WP_174714145.1 | HEPN+AAA-ATPase+Alpha+Beta-domain+wHTH+RRM | Type-1_BR.RM.like.systems | [illegible] | 1108 | Bacteria>Terrabacteria_group>Actinomycetota>Actinomycetes>Mycobacteriales | GCF_019456675.1 | Mycolicibacterium_pallens | KOO64_RS24915 | 5121618..5124944 | 370524 | MULTISPECIES:-Swt1-family-HEPN-domain-containing-protein-[Mycobacterium] |
| WP_256101683.1 | HEPN+AAA-ATPase+Alpha+Beta-domain+wHTH+RRM | Type-1_BR.RM.like.systems | [illegible] | 1118 | Bacteria>Terrabacteria_group>Actinomycetota>Actinomycetes>Mycobacteriales | GCF_024451565.1 | Mycobacterium_gordonae | KQR54_RS26625 | 18169..21525 | 1778 | Swt1-family-HEPN-domain-containing-protein-[Mycobacterium-gordonae] |
| WP_239588858.1 | HEPN+AAA-ATPase+Alpha+Beta-domain+wHTH+RRM | Type-1_BR.RM.like.systems | [illegible] | 1125 | Bacteria>Terrabacteria_group>Actinomycetota>Actinomycetes>Mycobacteriales | - | Gordonia_hydrophobica | JOC45_RS12260 | 227782..231159 | 40516 | DUF499-domain-containing-protein-[Gordonia-hydrophobica] |
| WP_167106438.1 | HEPN+AAA-ATPase+Alpha+Beta-domain+wHTH+RRM | Type-1_BR.RM.like.systems | [illegible] | 1137 | Bacteria>Terrabacteria_group>Actinomycetota>Actinomycetes>Mycobacteriales | GCF_011694515.1 | Mycobacterium_sp._DL592 | HBE64_RS20620 | 4288954..4292367 | 2675524 | DUF499-domain-containing-protein-[Mycobacterium-sp.-DL592] |
| WP_066167603.1 | HEPN+AAA-ATPase+Alpha+Beta-domain+wHTH+RRM | Type-1_BR.RM.like.systems | [illegible] | 1139 | Bacteria>Terrabacteria_group>Actinomycetota>Actinomycetes>Mycobacteriales | GCF_001592365.1 | Gordonia_hydrophobica | GH2_RS11425 | 77242..80661 | 40516 | DUF499-domain-containing-protein-[Gordonia-hydrophobica] |
| OKH62617.1 | HEPN+AAA-ATPase+Alpha+Beta-domain+wHTH+RRM | Type-1_BR.RM.like.systems | [illegible] | 1117 | Bacteria>Terrabacteria_group>Actinomycetota>Actinomycetes>Mycobacteriales | GCA_001905305.1 | Mycobacterium_sp._SWH-M5 | EB74_16030 | 8207..11560 | 1490483 | ATPase-AAA-[Mycobacterium-sp.-SWH-M5] |
| MCQ4364655.1 | HEPN+AAA-ATPase+Alpha+Beta-domain+wHTH+RRM | Type-1_BR.RM.like.systems | [illegible] | 1118 | Bacteria>Terrabacteria_group>Actinomycetota>Actinomycetes>Mycobacteriales | GCA_024451565.1 | Mycobacterium_gordonae | KQR54_26610 | 18169..21525 | 1778 | Swt1-family-HEPN-domain-containing-protein-[Mycobacterium-gordonae] |
| WP_277242420.1 | HEPN+AAA-ATPase+Alpha+Beta-domain+wHTH+RRM | Type-1_BR.RM.like.systems | [illegible] | 1119 | Bacteria>Terrabacteria_group>Actinomycetota>Actinomycetes>Mycobacteriales | GCF_029427115.1 | Gordonia_hongkongensis | L2299_RS00560 | 120321..123680 | 1701090 | Swt1-family-HEPN-domain-containing-protein-[Gordonia-hongkongensis] |
| WP_219068412.1 | HEPN+AAA-ATPase+Alpha+Beta-domain+wHTH+RRM | Type-1_BR.RM.like.systems | [illegible] | 1113 | Bacteria>Terrabacteria_group>Actinomycetota>Actinomycetes>Mycobacteriales | GCF_910591625.1 | Candidatus_Mycobacterium_methanotrophicum | M5I08_RS02450 | 450660..454001 | 2943498 | MULTISPECIES:-Swt1-family-HEPN-domain-containing-protein-[unclassified-Mycobacterium] |
| WP_174557649.1 | HEPN+AAA-ATPase+Alpha+Beta-domain+wHTH+RRM | Type-1_BR.RM.like.systems | [illegible] | 1109 | Bacteria>Terrabacteria_group>Actinomycetota>Actinomycetes>Mycobacteriales | GCF_001905655.1 | Mycobacterium_sp._ST-F2 | EB75_RS10110 | 184865..188194 | 1490484 | Swt1-family-HEPN-domain-containing-protein-[Mycobacterium-sp.-ST-F2] |
| WP_149357518.1 | HEPN+AAA-ATPase+Alpha+Beta-domain+wHTH+RRM | Type-1_BR.RM.like.systems | [illegible] | 1133 | Bacteria>Terrabacteria_group>Actinomycetota>Actinomycetes>Mycobacteriales | GCF_008306195.1 | Rhodococcus_qingshengii | D6M20_RS01760 | 367041..370442 | 334542 | DUF499-domain-containing-protein-[Rhodococcus-qingshengii] |
| SHV22416.1 | HEPN+AAA-ATPase+Alpha+Beta-domain+wHTH+RRM | Type-1_BR.RM.like.systems | [illegible] | 1122 | Bacteria>Terrabacteria_group>Actinomycetota>Actinomycetes>Mycobacteriales | GCA_900132885.1 | Mycobacteroides_abscessus_subsp._abscessus | SAMEA2259605_05819 | 13396..16764 | 1185650 | Conserved-protein-of-uncharacterised-function-(part1)-[Mycobacteroides-abscessus-subsp.-abscessus] |
| WP_193603884.1 | HEPN+AAA-ATPase+Alpha+Beta-domain+wHTH+RRM | Type-1_BR.RM.like.systems | [illegible] | 1120 | Bacteria>Terrabacteria_group>Actinomycetota>Actinomycetes>Mycobacteriales | GCF_015023815.1 | Mycobacteroides_abscessus | E3G55_RS12995 | 2677503..2680865 | 36809 | DUF499-domain-containing-protein-[Mycobacteroides-abscessus] |
| WP_037183436.1 | HEPN+AAA-ATPase+Alpha+Beta-domain+wHTH+RRM | Type-1_BR.RM.like.systems | [illegible] | 1136 | Bacteria>Terrabacteria_group>Actinomycetota>Actinomycetes>Mycobacteriales | GCF_020542785.1 | Rhodococcus_opacus | OPAG_RS41845 | 61698..65108 | 37919 | Swt1-family-HEPN-domain-containing-protein-[Rhodococcus-opacus] |
| EHI39101.1 | HEPN+AAA-ATPase+Alpha+Beta-domain+wHTH+RRM | Type-1_BR.RM.like.systems | [illegible] | 1127 | Bacteria>Terrabacteria_group>Actinomycetota>Actinomycetes>Mycobacteriales | GCA_000234335.1 | Rhodococcus_opacus_PD630 | OPAG_08353 | 61698..65081 | 543736 | hypothetical-protein-OPAG_08353-[Rhodococcus-opacus-PD630] |
| WP_083088684.1 | HEPN+AAA-ATPase+Alpha+Beta-domain+wHTH+RRM | Type-1_BR.RM.like.systems | [illegible] | 1108 | Bacteria>Terrabacteria_group>Actinomycetota>Actinomycetes>Mycobacteriales | GCF_010731635.1 | Mycobacterium_noviomagense | BST37_RS15635 | 32792..36118 | 459858 | DUF499-domain-containing-protein-[Mycobacterium-noviomagense] |
| WP_066467289.1 | HEPN+AAA-ATPase+Alpha+Beta-domain+wHTH+RRM | Type-1_BR.RM.like.systems | [illegible] | 1118 | Bacteria>Terrabacteria_group>Actinomycetota>Actinomycetes>Mycobacteriales | GCF_001552855.1 | Skermania_piniformis | SP4_RS02850 | 6456..9812 | 39122 | DUF499-domain-containing-protein-[Skermania-piniformis] |
| WP_026213067.1 | HEPN+AAA-ATPase+Alpha+Beta-domain+wHTH+RRM | Type-1_BR.RM.like.systems | [illegible] | 1116 | Bacteria>Terrabacteria_group>Actinomycetota>Actinomycetes>Mycobacteriales | GCF_000300375.1 | Mycolicibacterium_hassiacum | A3G5_RS0100810 | 158802..162152 | 46351 | Swt1-family-HEPN-domain-containing-protein-[Mycolicibacterium-hassiacum] |
| WP_014941573.1 | HEPN+AAA-ATPase+Alpha+Beta-domain+wHTH+RRM | Type-1_BR.RM.like.systems | [illegible] | 1124 | Bacteria>Terrabacteria_group>Actinomycetota>Actinomycetes>Mycobacteriales | GCF_000298095.1 | Mycobacterium_avium_complex_(MAC) | MIP_RS08165 | 1729452..1732826 | 120793 | MULTISPECIES:-DUF499-domain-containing-protein-[Mycobacterium-avium-complex-(MAC)] |
| MBF6378463.1 | HEPN+AAA-ATPase+Alpha+Beta-domain+wHTH+RRM | Type-1_BR.RM.like.systems | [illegible] | 1125 | Bacteria>Terrabacteria_group>Actinomycetota>Actinomycetes>Mycobacteriales | GCA_015477815.1 | Nocardia_farcinica | IU446_07990 | 515758..519135 | 37329 | DUF499-domain-containing-protein-[Nocardia-farcinica] |
| HJC59998.1 | HEPN+AAA-ATPase+Alpha+Beta-domain+wHTH+RRM | Type-1_BR.RM.like.systems | [illegible] | 1131 | Bacteria>Terrabacteria_group>Actinomycetota>Actinomycetes>Mycobacteriales | GCA_019119875.1 | Candidatus_Dietzia_intestinigallinarum | H9755_06660 | 12112..15507 | 2838537 | DUF499-domain-containing-protein-[Candidatus-Dietzia-intestinigallinarum] |
| EKF23780.1 | HEPN+AAA-ATPase+Alpha+Beta-domain+wHTH+RRM | Type-1_BR.RM.like.systems | [illegible] | 1105 | Bacteria>Terrabacteria_group>Actinomycetota>Actinomycetes>Mycobacteriales | GCA_000300375.1 | Mycolicibacterium_hassiacum_DSM_44199 | C731_2148 | 37750..41067 | 1122247 | hypothetical-protein-C731_2148-[Mycolicibacterium-hassiacum-DSM-44199] |

<table>
<tr><th>Protein-ID</th><th>DUF499-architecture</th><th>System</th><th>Gene-Neighbourhoods</th><th>Length</th><th>Taxonomy</th><th>Assembly-ID</th><th>Strain</th><th>Locus-Tag</th><th>Genomic-coordinates</th><th>TaxID</th><th>Fasta-defline</th></tr>
<tr><td>WP_233059597.1</td><td>HEPN+AAA-ATPase+Alpha+Beta-domain+wHTH+RRM</td><td>Type-1_BR.RM.like.systems</td><td>TIR->?->?->TUDOR+SF1-SF2-Helicase+iREase+CTD+REase-><-?||METHYLASE->HEPN+DUF499*->Schlafen+HATPase+HTH->HEPN-><-?<-NUDIX||GNTR-HTH-></td><td>1108</td><td>Bacteria>Terrabacteria_group>Actinomycetota>Actinomycetes>Mycobacteriales</td><td>GCF_021183805.1</td><td>Mycobacterium_avium</td><td>LTQ57_RS17660</td><td>3790685..3794011</td><td>1764</td><td>Swt1-family-HEPN-domain-containing-protein-[Mycobacterium-avium]</td></tr>
<tr><td>WP_228833824.1</td><td>HEPN+AAA-ATPase+Alpha+Beta-domain+wHTH+RRM</td><td>Type-1_BR.RM.like.systems</td><td>ZDNA-HTH->TUDOR+SF1-SF2-Helicase+iREase+CTD+REase->METHYLASE-p->METHYLASE->HEPN+DUF499*->?->Phage_integrase->?->STAS-></td><td>1123</td><td>Bacteria>Terrabacteria_group>Actinomycetota>Actinomycetes>Mycobacteriales</td><td>GCF_020883195.1</td><td>Nocardia</td><td>LMJ57_RS17620</td><td>3762816..3766187</td><td>1817</td><td>MULTISPECIES:-DUF499-domain-containing-protein-[Nocardia]</td></tr>
<tr><td>WP_227974732.1</td><td>HEPN+AAA-ATPase+Alpha+Beta-domain+wHTH+RRM</td><td>Type-1_BR.RM.like.systems</td><td>TIR->?->?->TUDOR+SF1-SF2-Helicase+iREase+CTD+REase-><-?||METHYLASE->HEPN+DUF499*->Schlafen+HATPase+HTH->HEPN-><-?<-NUDIX||GNTR-HTH-></td><td>1108</td><td>Bacteria>Terrabacteria_group>Actinomycetota>Actinomycetes>Mycobacteriales</td><td>GCF_020735405.1</td><td>Mycobacterium_avium</td><td>LT348_RS02565</td><td>526704..530030</td><td>1764</td><td>Swt1-family-HEPN-domain-containing-protein-[Mycobacterium-avium]</td></tr>
<tr><td>WP_009974960.1</td><td>HEPN+AAA-ATPase+Alpha+Beta-domain+wHTH+RRM</td><td>Type-1_BR.RM.like.systems</td><td>TIR->?->?->TUDOR+SF1-SF2-Helicase+iREase+CTD+REase-><-?||METHYLASE->HEPN+DUF499*->Schlafen+HATPase+HTH->HEPN-><-?<-NUDIX||GNTR-HTH-></td><td>1108</td><td>Bacteria>Terrabacteria_group>Actinomycetota>Actinomycetes>Mycobacteriales</td><td>GCF_000174035.1</td><td>Mycobacterium_avium</td><td>MAA44156_RS17120</td><td>3691227..3694553</td><td>1764</td><td>Swt1-family-HEPN-domain-containing-protein-[Mycobacterium-avium]</td></tr>
<tr><td>RUP39290.1</td><td>HEPN+AAA-ATPase+Alpha+Beta-domain+wHTH+RRM</td><td>Type-1_BR.RM.like.systems</td><td>TUDOR+SF1-SF2-Helicase+iREase+CTD+REase->METHYLASE->HEPN+DUF499*-><-HTH_3||?->TetR-HTH-><-Amidohydro_3+Amidohydro_1</td><td>1112</td><td>Bacteria>Terrabacteria_group>Actinomycetota>Actinomycetes>Mycobacteriales</td><td>GCA_003987705.1</td><td>Gordonia_sp._(in:_high_G+C_Gram-positive_bacteria)</td><td>EKK60_07255</td><td>9352..12690</td><td>84139</td><td>ATP-binding-protein-[Gordonia-sp.]</td></tr>
<tr><td>OHT48546.1</td><td>HEPN+AAA-ATPase+Alpha+Beta-domain+wHTH+RRM</td><td>Type-1_BR.RM.like.systems</td><td>PAIREDC-HTH+HIN-HTH-><-?||TUDOR+SF1-SF2-Helicase+iREase+CTD+REase->METHYLASE->ParB+HNH->HEPN+DUF499*->?->?->?-><-?<-METHYLASE-p</td><td>1109</td><td>Bacteria>Terrabacteria_group>Actinomycetota>Actinomycetes>Mycobacteriales</td><td>GCA_001853545.1</td><td>Mycobacteroides_chelonae</td><td>BKG62_21080</td><td>210158..213487</td><td>1774</td><td>AAA-family-ATPase-[Mycobacteroides-chelonae]</td></tr>
<tr><td>WP_197416164.1</td><td>HEPN+AAA-ATPase+Alpha+Beta-domain+wHTH+RRM</td><td>Type-1_BR.RM.like.systems</td><td>Cluster182_14clades-><-?<-?<-HEPN+DUF499*<-METHYLASE<-TUDOR+SF1-SF2-Helicase+iREase+CTD+REase||NTD+SF1-SF2-Helicase-></td><td>1112</td><td>Bacteria>Terrabacteria_group>Actinomycetota>Actinomycetes>Mycobacteriales</td><td>GCF_001500045.1</td><td>Mycobacterium_sp._GA-1199</td><td>AU198_RS19260</td><td>256994..260332</td><td>1772287</td><td>Swt1-family-HEPN-domain-containing-protein-[Mycobacterium-sp.-GA-1199]</td></tr>
<tr><td>TXI48376.1</td><td>HEPN+AAA-ATPase+Alpha+Beta-domain+wHTH+RRM</td><td>Type-1_BR.RM.like.systems</td><td><-HIN-HTH<-NTD+SF1-SF2-Helicase||TUDOR+SF1-SF2-Helicase+iREase+CTD+REase-><-?||METHYLASE->ParB+HNH->HEPN+DUF499*-></td><td>1102</td><td>Bacteria>Terrabacteria_group>Actinomycetota>Actinomycetes>Mycobacteriales</td><td>GCA_008015745.1</td><td>Mycolicibacter_arupensis</td><td>E6Q54_23710</td><td>12171..15479</td><td>342002</td><td>ATP-binding-protein-[Mycolicibacter-arupensis]</td></tr>
<tr><td>WP_281378415.1</td><td>HEPN+AAA-ATPase+Alpha+Beta-domain+wHTH+RRM</td><td>Type-1_BR.RM.like.systems</td><td><-ABhydrolase<-?||?-><-HEPN+DUF499*<-METHYLASE||DUF222+HNH-><-TUDOR+SF1-SF2-Helicase+iREase+CTD+REase<-?<-?<-tRNA<-tRNA<-tRNA<-tRNA<-tRNA||tRNA->DDE_Tnp_1_4-></td><td>1142</td><td>Bacteria>Terrabacteria_group>Actinomycetota>Actinomycetes>Mycobacteriales</td><td>GCF_014197435.1</td><td>Gordonia_humi</td><td>BKA16_RS13810</td><td>3005759..3009187</td><td>686429</td><td>DUF499-domain-containing-protein-[Gordonia-humi]</td></tr>
<tr><td>WP_238054205.1</td><td>HEPN+AAA-ATPase+Alpha+Beta-domain+wHTH+RRM</td><td>Type-1_BR.RM.like.systems</td><td><-REase||?->DOC->GDResolvase+HIN-HTH-><-HEPN+DUF499*<-METHYLASE<-TUDOR+SF1-SF2-Helicase+iREase+CTD+REase<-?||?-><-?<-REase</td><td>1134</td><td>Bacteria>Terrabacteria_group>Actinomycetota>Actinomycetes>Mycobacteriales</td><td>GCF_022059865.1</td><td>Rhodococcus_erythropolis</td><td>ITJ47_RS31985</td><td>3376..6780</td><td>1833</td><td>Swt1-family-HEPN-domain-containing-protein-[Rhodococcus-erythropolis]</td></tr>
<tr><td>WP_179959972.1</td><td>HEPN+AAA-ATPase+Alpha+Beta-domain+wHTH+RRM</td><td>Type-1_BR.RM.like.systems</td><td><-NTD+SF1-SF2-Helicase||TUDOR+SF1-SF2-Helicase+iREase+CTD+REase->ParB+HNH->HEPN+DUF499*->?->?->?-><-Cluster182_14clades</td><td>1122</td><td>Bacteria>Terrabacteria_group>Actinomycetota>Actinomycetes>Mycobacteriales</td><td>GCF_010728155.1</td><td>Mycolicibacterium_doricum</td><td>H7I39_RS15930</td><td>40769..44137</td><td>126673</td><td>Swt1-family-HEPN-domain-containing-protein-[Mycolicibacterium-doricum]</td></tr>
<tr><td>WP_010121755.1</td><td>HEPN+AAA-ATPase+Alpha+Beta-domain+wHTH+RRM</td><td>Type-1_BR.RM.like.systems</td><td><-HTH_3<-?||TUDOR+SF1-SF2-Helicase+iREase+CTD+REase->METHYLASE->HEPN+DUF499*->||zf-ISL3+DDE_Tnp_ISL3->HEPN+HEPN-></td><td>1182</td><td>Bacteria>Terrabacteria_group>Actinomycetota>Actinomycetes>Mycobacteriales</td><td>GCF_000213935.1</td><td>Corynebacterium_nuruki</td><td>FSW06_RS05470</td><td>1228839..1232387</td><td>1032851</td><td>DUF499-domain-containing-protein-[Corynebacterium-nuruki]</td></tr>
<tr><td>WP_161926536.1</td><td>HEPN+AAA-ATPase+Alpha+Beta-domain+wHTH+RRM</td><td>Type-1_BR.RM.like.systems</td><td><-TetR-HTH<-?||HTH_19-><-?<-HEPN+DUF499*<-DUF222+HNH<-METHYLASE<-TUDOR+SF1-SF2-Helicase+iREase+CTD+REase||DOC+HTH-></td><td>1124</td><td>Bacteria>Terrabacteria_group>Actinomycetota>Actinomycetes>Mycobacteriales</td><td>GCF_009932435.1</td><td>Gordonia_crocea</td><td>nbrcactino_RS06035</td><td>1265301..1268675</td><td>589162</td><td>DUF499-domain-containing-protein-[Gordonia-crocea]</td></tr>
<tr><td>ORA83573.1</td><td>HEPN+AAA-ATPase+Alpha+Beta-domain+wHTH+RRM</td><td>Type-1_BR.RM.like.systems</td><td>REase->?->?->?->||TUDOR+SF1-SF2-Helicase+iREase+CTD+REase->METHYLASE->HEPN+DUF499*->ParB+HNH->HEPN->?-><-SF1-SF2-Helicase</td><td>1110</td><td>Bacteria>Terrabacteria_group>Actinomycetota>Actinomycetes>Mycobacteriales</td><td>GCA_002086285.1</td><td>Mycolicibacter_kumamotonensis</td><td>BST28_00055</td><td>16546..19878</td><td>354243</td><td>AAA-family-ATPase-[Mycolicibacter-kumamotonensis]</td></tr>
<tr><td>WP_036372468.1</td><td>HEPN+AAA-ATPase+Alpha+Beta-domain+wHTH+RRM</td><td>Type-1_BR.RM.like.systems</td><td><-SF1-SF2-Helicase<-TRD+TRD<-?<-HTH_Tnp_1+HTH_21+rve+rve_3<-?<-HEPN+DUF499*<-SIR2+SIR2<-BrxL<-PglZ<-METHYLASE-PglX</td><td>1123</td><td>Bacteria>Terrabacteria_group>Actinomycetota>Actinomycetes>Mycobacteriales</td><td>GCF_000612725.1</td><td>Mycolicibacterium_austroafricanum</td><td>BN976_RS17325</td><td>44630..48001</td><td>39687</td><td>DUF499-domain-containing-protein-[Mycolicibacterium-austroafricanum]</td></tr>
<tr><td>WP_264184695.1</td><td>HEPN+AAA-ATPase+Alpha+Beta-domain+wHTH+RRM</td><td>Type-1_BR.RM.like.systems</td><td><-HTH||TUDOR+SF1-SF2-Helicase+iREase+CTD+REase-><-CHTH||METHYLASE->HEPN+DUF499*->DUF222+HNH-><-?<-?<-Peptidase_A8</td><td>1139</td><td>Bacteria>Terrabacteria_group>Actinomycetota>Actinomycetes>Mycobacteriales</td><td>-</td><td>Rhodococcus_yananensis</td><td>LDQ05_RS09980</td><td>11697..15116</td><td>2879464</td><td>Swt1-family-HEPN-domain-containing-protein-[Rhodococcus-yananensis]</td></tr>
<tr><td>WP_214515005.1</td><td>HEPN+AAA-ATPase+Alpha+Beta-domain+wHTH+RRM</td><td>Type-1_BR.RM.like.systems</td><td>ABC-ATPase+SbcC->HAD-><-?||?->TUDOR+SF1-SF2-Helicase+iREase+CTD+REase->METHYLASE->HEPN+DUF499*->?->||?-><-DDE_Tnp_IS240</td><td>1134</td><td>Bacteria>Terrabacteria_group>Actinomycetota>Actinomycetes>Mycobacteriales</td><td>GCF_018588185.1</td><td>Rhodococcus_sp._WY5</td><td>G4D79_RS33795</td><td>40006..43410</td><td>2708349</td><td>Swt1-family-HEPN-domain-containing-protein-[Rhodococcus-sp.-WY5]</td></tr>
<tr><td>WP_099384398.1</td><td>HEPN+AAA-ATPase+Alpha+Beta-domain+wHTH+RRM</td><td>Type-1_BR.RM.like.systems</td><td>TUDOR+SF1-SF2-Helicase+iREase+CTD+REase->METHYLASE->ParB+HNH->HEPN+DUF499*->NTD+SF1-SF2-Helicase-><-?<-?||REase-></td><td>1115</td><td>Bacteria>Terrabacteria_group>Actinomycetota>Actinomycetes>Mycobacteriales</td><td>GCF_002735905.1</td><td>Williamsia_marianensis</td><td>CSW57_RS21175</td><td>1638758..1642105</td><td>322609</td><td>Swt1-family-HEPN-domain-containing-protein-[Williamsia-marianensis]</td></tr>
<tr><td>WP_072804385.1</td><td>HEPN+AAA-ATPase+Alpha+Beta-domain+wHTH+RRM</td><td>Type-1_BR.RM.like.systems</td><td>TUDOR+SF1-SF2-Helicase+iREase+CTD+REase->?->AbiEii->METHYLASE-><-PIN<-MazE_antitoxin||HEPN+DUF499*->?->?-><-?<-Phd-YefM</td><td>1125</td><td>Bacteria>Terrabacteria_group>Actinomycetota>Actinomycetes>Mycobacteriales</td><td>GCF_001895005.1</td><td>Rhodococcus_yunnanensis</td><td>RY1_RS14860</td><td>83069..86446</td><td>278209</td><td>Swt1-family-HEPN-domain-containing-protein-[Rhodococcus-yunnanensis]</td></tr>
<tr><td>WP_024794486.1</td><td>HEPN+AAA-ATPase+Alpha+Beta-domain+wHTH+RRM</td><td>Type-1_BR.RM.like.systems</td><td><-TetR-HTH||?->CHTH->Glyoxalase-><-?<-HEPN+DUF499*<-METHYLASE||?->?-><-TUDOR+SF1-SF2-Helicase+iREase+CTD+REase||HTH_3-></td><td>1134</td><td>Bacteria>Terrabacteria_group>Actinomycetota>Actinomycetes>Mycobacteriales</td><td>GCF_000524475.1</td><td>Tomitella_biformata</td><td>JJ684_RS09895</td><td>229365..232769</td><td>630403</td><td>DUF499-domain-containing-protein-[Tomitella-biformata]</td></tr>
<tr><td>WP_283569686.1</td><td>HEPN+AAA-ATPase+Alpha+Beta-domain+wHTH+RRM</td><td>Type-1_BR.RM.like.systems</td><td>HEPN-><-NTD+SF1-SF2-Helicase||TUDOR+SF1-SF2-Helicase+iREase+CTD+REase->METHYLASE->ParB+HNH->HEPN+DUF499*-></td><td>1120</td><td>Bacteria>Terrabacteria_group>Actinomycetota>Actinomycetes>Mycobacteriales</td><td>GCF_030063715.1</td><td>Rhodococcus_qingshengii</td><td>QNA24_RS33665</td><td>11874..15236</td><td>334542</td><td>Swt1-family-HEPN-domain-containing-protein-[Rhodococcus-qingshengii]</td></tr>
<tr><td>WP_256083185.1</td><td>HEPN+AAA-ATPase+Alpha+Beta-domain+wHTH+RRM</td><td>Type-1_BR.RM.like.systems</td><td>HEPN-><-NTD+SF1-SF2-Helicase||TUDOR+SF1-SF2-Helicase+iREase+CTD+REase->METHYLASE->ParB+HNH->HEPN+DUF499*-></td><td>1120</td><td>Bacteria>Terrabacteria_group>Actinomycetota>Actinomycetes>Mycobacteriales</td><td>GCF_024438075.1</td><td>Rhodococcus_qingshengii</td><td>NOS52_RS32465</td><td>22838..26200</td><td>334542</td><td>Swt1-family-HEPN-domain-containing-protein-[Rhodococcus-qingshengii]</td></tr>
<tr><td>WP_119699599.1</td><td>HEPN+AAA-ATPase+Alpha+Beta-domain+wHTH+RRM</td><td>Type-1_BR.RM.like.systems</td><td><-JetA<-?<-?<-?<-?<-?<-HEPN+DUF499*<-ParB+HNH<-METHYLASE<-TUDOR+SF1-SF2-Helicase+iREase+CTD+REase||NTD+SF1-SF2-Helicase-></td><td>1118</td><td>Bacteria>Terrabacteria_group>Actinomycetota>Actinomycetes>Mycobacteriales</td><td>GCF_003586525.1</td><td>Rhodococcus_ruber</td><td>TN91_RS02590</td><td>62027..65383</td><td>1830</td><td>Swt1-family-HEPN-domain-containing-protein-[Rhodococcus-ruber]</td></tr>
<tr><td>WP_074506051.1</td><td>HEPN+AAA-ATPase+Alpha+Beta-domain+wHTH+RRM</td><td>Type-1_BR.RM.like.systems</td><td><-ABhydrolase||?-><-HEPN+DUF499*<-METHYLASE<-TUDOR+SF1-SF2-Helicase+iREase+CTD+REase||HTH_3->MODE-HTH+PBPII-></td><td>1150</td><td>Bacteria>Terrabacteria_group>Actinomycetota>Actinomycetes>Mycobacteriales</td><td>GCF_001912755.2</td><td>Corynebacterium_glutamicum</td><td>AUO95_RS02900</td><td>202510..205962</td><td>1718</td><td>DUF499-domain-containing-protein-[Corynebacterium-glutamicum]</td></tr>
<tr><td>WP_039584625.1</td><td>HEPN+AAA-ATPase+Alpha+Beta-domain+wHTH+RRM</td><td>Type-1_BR.RM.like.systems</td><td>DNA-glycosylase-><-ZnR+ZnR<-TIR||TUDOR+SF1-SF2-Helicase+iREase+CTD+REase->METHYLASE->HEPN+DUF499*-><-RNase</td><td>1138</td><td>Bacteria>Terrabacteria_group>Actinomycetota>Actinomycetes>Mycobacteriales</td><td>GCF_000801125.1</td><td>Rhodococcus_sp._Chr-9</td><td>QR64_RS06710</td><td>8521..11937</td><td>713612</td><td>Swt1-family-HEPN-domain-containing-protein-[Rhodococcus-sp.-Chr-9]</td></tr>
<tr><td>MCQ4152489.1</td><td>HEPN+AAA-ATPase+Alpha+Beta-domain+wHTH+RRM</td><td>Type-1_BR.RM.like.systems</td><td>HEPN-><-NTD+SF1-SF2-Helicase||TUDOR+SF1-SF2-Helicase+iREase+CTD+REase->METHYLASE->ParB+HNH->HEPN+DUF499*-></td><td>1120</td><td>Bacteria>Terrabacteria_group>Actinomycetota>Actinomycetes>Mycobacteriales</td><td>GCA_024438075.1</td><td>Rhodococcus_qingshengii</td><td>NOS52_32470</td><td>22838..26200</td><td>334542</td><td>Swt1-family-HEPN-domain-containing-protein-[Rhodococcus-qingshengii]</td></tr>
<tr><td>WP_207760645.1</td><td>HEPN+AAA-ATPase+Alpha+Beta-domain+wHTH+RRM</td><td>Type-1_BR.RM.like.systems</td><td>Abhydrolase_3-><-?<-?<-HEPN+DUF499*<-METHYLASE<-TUDOR+SF1-SF2-Helicase+iREase+CTD+REase||NTD+SF1-SF2-Helicase-></td><td>1120</td><td>Bacteria>Terrabacteria_group>Actinomycetota>Actinomycetes>Mycobacteriales</td><td>GCF_002250655.1</td><td>Mycolicibacterium_sphagni</td><td>CG716_RS11640</td><td>96200..99562</td><td>1786</td><td>Swt1-family-HEPN-domain-containing-protein-[Mycolicibacterium-sphagni]</td></tr>
<tr><td>WP_089250630.1</td><td>HEPN+AAA-ATPase+Alpha+Beta-domain+wHTH+RRM</td><td>Type-1_BR.RM.like.systems</td><td><-DUF433<-?<-HEPN+DUF499*||MazE_antitoxin->PIN-><-METHYLASE<-AbiEii<-?<-TUDOR+SF1-SF2-Helicase+iREase+CTD+REase</td><td>1122</td><td>Bacteria>Terrabacteria_group>Actinomycetota>Actinomycetes>Mycobacteriales</td><td>GCF_900188125.1</td><td>Rhodococcus_kyotonensis</td><td>CHB96_RS22960</td><td>65313..68681</td><td>398843</td><td>Swt1-family-HEPN-domain-containing-protein-[Rhodococcus-kyotonensis]</td></tr>
<tr><td>OYN79669.1</td><td>HEPN+AAA-ATPase+Alpha+Beta-domain+wHTH+RRM</td><td>Type-1_BR.RM.like.systems</td><td>Abhydrolase_3-><-?<-?<-HEPN+DUF499*<-METHYLASE<-TUDOR+SF1-SF2-Helicase+iREase+CTD+REase||NTD+SF1-SF2-Helicase-></td><td>1109</td><td>Bacteria>Terrabacteria_group>Actinomycetota>Actinomycetes>Mycobacteriales</td><td>GCA_002250655.1</td><td>Mycolicibacterium_sphagni</td><td>CG716_11635</td><td>96200..99529</td><td>1786</td><td>AAA-family-ATPase-[Mycolicibacterium-sphagni]</td></tr>
<tr><td>WP_284860121.1</td><td>HEPN+AAA-ATPase+Alpha+Beta-domain+wHTH+RRM</td><td>Type-1_BR.RM.like.systems</td><td>ABhydrolase->?->?->?->ABhydrolase+ABhydrolase->?-><-HEPN+DUF499*<-METHYLASE<-TUDOR+SF1-SF2-Helicase+iREase+CTD+REase</td><td>1124</td><td>Bacteria>Terrabacteria_group>Actinomycetota>Actinomycetes>Mycobacteriales</td><td>GCF_030229165.1</td><td>Corynebacterium_pseudodiphtheriticum</td><td>QP898_RS02760</td><td>58935..62309</td><td>37637</td><td>DUF499-domain-containing-protein-[Corynebacterium-pseudodiphtheriticum]</td></tr>
<tr><td>WP_284588418.1</td><td>HEPN+AAA-ATPase+Alpha+Beta-domain+wHTH+RRM</td><td>Type-1_BR.RM.like.systems</td><td>TUDOR+SF1-SF2-Helicase+iREase+CTD+REase->METHYLASE->HEPN+DUF499*-><-?<-ABhydrolase+ABhydrolase<-?<-?<-?<-ABhydrolase</td><td>1124</td><td>Bacteria>Terrabacteria_group>Actinomycetota>Actinomycetes>Mycobacteriales</td><td>GCF_030176355.1</td><td>Corynebacterium_pseudodiphtheriticum</td><td>QPX18_RS08860</td><td>15682..19056</td><td>37637</td><td>DUF499-domain-containing-protein-[Corynebacterium-pseudodiphtheriticum]</td></tr>
<tr><td>WP_249619951.1</td><td>HEPN+AAA-ATPase+Alpha+Beta-domain+wHTH+RRM</td><td>Type-1_BR.RM.like.systems</td><td>TUDOR+SF1-SF2-Helicase+iREase+CTD+REase->METHYLASE->HEPN+DUF499*-><-?<-ABhydrolase+ABhydrolase<-?<-?<-?<-ABhydrolase</td><td>1124</td><td>Bacteria>Terrabacteria_group>Actinomycetota>Actinomycetes>Mycobacteriales</td><td>GCF_023509295.1</td><td>Corynebacterium_pseudodiphtheriticum</td><td>L9H78_RS09595</td><td>2140657..2144031</td><td>37637</td><td>DUF499-domain-containing-protein-[Corynebacterium-pseudodiphtheriticum]</td></tr>
<tr><td>WP_249617808.1</td><td>HEPN+AAA-ATPase+Alpha+Beta-domain+wHTH+RRM</td><td>Type-1_BR.RM.like.systems</td><td>TUDOR+SF1-SF2-Helicase+iREase+CTD+REase->METHYLASE->HEPN+DUF499*-><-?<-ABhydrolase+ABhydrolase<-?<-?<-?<-ABhydrolase</td><td>1124</td><td>Bacteria>Terrabacteria_group>Actinomycetota>Actinomycetes>Mycobacteriales</td><td>GCF_023509275.1</td><td>Corynebacterium_pseudodiphtheriticum</td><td>L9H27_RS09685</td><td>2151202..2154576</td><td>37637</td><td>DUF499-domain-containing-protein-[Corynebacterium-pseudodiphtheriticum]</td></tr>
<tr><td>WP_182858863.1</td><td>HEPN+AAA-ATPase+Alpha+Beta-domain+wHTH+RRM</td><td>Type-1_BR.RM.like.systems</td><td>TUDOR+SF1-SF2-Helicase+iREase+CTD+REase->AbiGi->METHYLASE->HEPN+DUF499*->SbcC-><-NUDIX||GNTR-HTH->?->FtsK-></td><td>1108</td><td>Bacteria>Terrabacteria_group>Actinomycetota>Actinomycetes>Mycobacteriales</td><td>GCF_003581765.1</td><td>Mycobacteroides_abscessus</td><td>D2E28_RS06470</td><td>1322672..1325998</td><td>36809</td><td>Swt1-family-HEPN-domain-containing-protein-[Mycobacteroides-abscessus]</td></tr>
<tr><td>WP_126299909.1</td><td>HEPN+AAA-ATPase+Alpha+Beta-domain+wHTH+RRM</td><td>Type-1_BR.RM.like.systems</td><td><-HEPN||?-><-Classical-AAA<-?<-?<-Classical-AAA<-HEPN+DUF499*<-METHYLASE<-TUDOR+SF1-SF2-Helicase+iREase+CTD+REase</td><td>1141</td><td>Bacteria>Terrabacteria_group>Actinomycetota>Actinomycetes>Mycobacteriales</td><td>GCF_900638255.1</td><td>Corynebacterium_matruchotii</td><td>EL293_RS12755</td><td>2814253..2817678</td><td>43768</td><td>DUF499-domain-containing-protein-[Corynebacterium-matruchotii]</td></tr>
<tr><td>MBZ4519978.1</td><td>HEPN+AAA-ATPase+Alpha+Beta-domain+wHTH+RRM</td><td>Type-1_BR.RM.like.systems</td><td><-FtsK<-?<-GNTR-HTH||NUDIX-><-SIR2<-HEPN+DUF499*<-METHYLASE<-TUDOR+SF1-SF2-Helicase+iREase+CTD+REase<-RNase</td><td>1102</td><td>Bacteria>Terrabacteria_group>Actinomycetota>Actinomycetes>Mycobacteriales</td><td>GCA_020055125.1</td><td>Mycobacterium_avium_subsp._hominissuis</td><td>GBP86_08110</td><td>354734..358042</td><td>439334</td><td>ATP-binding-protein-[Mycobacterium-avium-subsp.-hominissuis]</td></tr>
<tr><td>WP_219849138.1</td><td>HEPN+AAA-ATPase+Alpha+Beta-domain+wHTH+RRM</td><td>Type-1_BR.RM.like.systems</td><td><-HTH+SF1-SF2-Helicase+REase<-REase<-MORC<-HEPN+DUF499*<-METHYLASE<-REase||DNA_methylase-><-METHYLASE-p</td><td>1137</td><td>Bacteria>Terrabacteria_group>Actinomycetota>Actinomycetes>Mycobacteriales</td><td>GCF_014041935.1</td><td>Gordonia_jinghuaiqii</td><td>H1R19_RS12155</td><td>2732590..2736003</td><td>2758710</td><td>Swt1-family-HEPN-domain-containing-protein-[Gordonia-jinghuaiqii]</td></tr>
<tr><td>WP_213249149.1</td><td>HEPN+AAA-ATPase+Alpha+Beta-domain+wHTH+RRM</td><td>Type-1_BR.RM.like.systems</td><td><-HTH+SF1-SF2-Helicase+REase<-REase<-MORC<-HEPN+DUF499*<-SF1-SF2-Helicase<-OLD-ABC+TOPRIM||DNA_methylase-><-?||HNH-></td><td>1138</td><td>Bacteria>Terrabacteria_group>Actinomycetota>Actinomycetes>Mycobacteriales</td><td>GCF_009914535.1</td><td>Gordonia</td><td>IDH24_RS11690</td><td>142027..145443</td><td>2053</td><td>DUF499-domain-containing-protein-[Gordonia-pseudamarae]</td></tr>
<tr><td>WP_062887256.1</td><td>HEPN+AAA-ATPase+Alpha+Beta-domain+wHTH+RRM</td><td>Type-1_BR.RM.like.systems</td><td><-FtsK<-?||NUDIX-><-HEPN<-Schlafen+HATPase+HTH<-HEPN+DUF499*<-METHYLASE||?-><-TUDOR+SF1-SF2-Helicase+iREase+CTD+REase</td><td>1107</td><td>Bacteria>Terrabacteria_group>Actinomycetota>Actinomycetes>Mycobacteriales</td><td>GCF_018524295.1</td><td>Mycobacteriaceae</td><td>KL858_RS23375</td><td>11898..15221</td><td>1762</td><td>Swt1-family-HEPN-domain-containing-protein-[Mycobacterium-avium]</td></tr>
<tr><td>MBF6311832.1</td><td>HEPN+AAA-ATPase+Alpha+Beta-domain+wHTH+RRM</td><td>Type-1_BR.RM.like.systems</td><td><-ACET<-?||?-><-STAS<-?<-HEPN+DUF499*<-METHYLASE<-METHYLASE-p<-TUDOR+SF1-SF2-Helicase+iREase+CTD+REase<-ZDNA-HTH</td><td>1125</td><td>Bacteria>Terrabacteria_group>Actinomycetota>Actinomycetes>Mycobacteriales</td><td>GCA_015477685.1</td><td>Nocardia_farcinica</td><td>IU455_12400</td><td>1148688..1152065</td><td>37329</td><td>DUF499-domain-containing-protein-[Nocardia-farcinica]</td></tr>
<tr><td>WP_276763638.1</td><td>HEPN+AAA-ATPase+Alpha+Beta-domain+wHTH+RRM</td><td>Type-1_BR.RM.like.systems</td><td><-NTD+SF1-SF2-Helicase||TUDOR+SF1-SF2-Helicase+iREase+CTD+REase-><-?||METHYLASE->ParB+HNH->HEPN+DUF499*-></td><td>1113</td><td>Bacteria>Terrabacteria_group>Actinomycetota>Actinomycetes>Mycobacteriales</td><td>GCF_008015745.1</td><td>Mycolicibacter_arupensis</td><td>E6Q54_RS23755</td><td>12138..15479</td><td>342002</td><td>Swt1-family-HEPN-domain-containing-protein-[Mycolicibacter-arupensis]</td></tr>
<tr><td>WP_205317328.1</td><td>HEPN+AAA-ATPase+Alpha+Beta-domain+wHTH+RRM</td><td>Type-1_BR.RM.like.systems</td><td>GDResolvase+HTH->?-><-?||TUDOR+SF1-SF2-Helicase+iREase+CTD+REase->METHYLASE->HEPN+DUF499*-><-?||HTH-><-tRNA<-MarR-HTH</td><td>1133</td><td>Bacteria>Terrabacteria_group>Actinomycetota>Actinomycetes>Mycobacteriales</td><td>GCF_016921075.1</td><td>Prescottella_equi</td><td>JO861_RS22230</td><td>9979..13380</td><td>43767</td><td>Swt1-family-HEPN-domain-containing-protein-[Prescottella-equi]</td></tr>
<tr><td>WP_005526420.1</td><td>HEPN+AAA-ATPase+Alpha+Beta-domain+wHTH+RRM</td><td>Type-1_BR.RM.like.systems</td><td>TUDOR+SF1-SF2-Helicase+iREase+CTD+REase->METHYLASE->HEPN+DUF499*->HEPN->Classical-AAA->?->Classical-AAA-></td><td>1181</td><td>Bacteria>Terrabacteria_group>Actinomycetota>Actinomycetes>Mycobacteriales</td><td>GCF_000175375.1</td><td>Corynebacterium_matruchotii</td><td>HBA49_RS02175</td><td>486574..490119</td><td>43768</td><td>DUF499-domain-containing-protein-[Corynebacterium-matruchotii]</td></tr>
<tr><td>WP_244804062.1</td><td>HEPN+AAA-ATPase+Alpha+Beta-domain+wHTH+RRM</td><td>Type-1_BR.RM.like.systems</td><td>ABhydrolase->?-><-?<-?||?->ORC-CDC6+REase-><-HEPN+DUF499*<-TUDOR+SF1-SF2-Helicase+iREase+CTD+REase||HTH_3->?->?->?-><-GNAT</td><td>1194</td><td>Bacteria>Terrabacteria_group>Actinomycetota>Actinomycetes>Mycobacteriales</td><td>GCF_022869005.1</td><td>Corynebacterium_kalidii</td><td>MUN33_RS06320</td><td>104934..108518</td><td>2931982</td><td>DUF499-domain-containing-protein-[Corynebacterium-sp.-LD5P10]</td></tr>
<tr><td>WP_237010003.1</td><td>HEPN+AAA-ATPase+Alpha+Beta-domain+wHTH+RRM</td><td>Type-1_BR.RM.like.systems</td><td><-ABhydrolase<-?||?-><-ACET<-HEPN+DUF499*<-METHYLASE||DUF222+HNH-><-TUDOR+SF1-SF2-Helicase+iREase+CTD+REase</td><td>1115</td><td>Bacteria>Terrabacteria_group>Actinomycetota>Actinomycetes>Mycobacteriales</td><td>GCF_021738965.1</td><td>Gordonia_zhenghanii</td><td>L5G32_05410</td><td>142683..146030</td><td>2911516</td><td>MULTISPECIES:-DUF499-domain-containing-protein-[unclassified-Gordonia]</td></tr>
<tr><td>WP_237001659.1</td><td>HEPN+AAA-ATPase+Alpha+Beta-domain+wHTH+RRM</td><td>Type-1_BR.RM.like.systems</td><td>TUDOR+SF1-SF2-Helicase+iREase+CTD+REase-><-DUF222+HNH||METHYLASE->HEPN+DUF499*->ACET-><-?||?->ABhydrolase-></td><td>1125</td><td>Bacteria>Terrabacteria_group>Actinomycetota>Actinomycetes>Mycobacteriales</td><td>GCF_021739025.1</td><td>Gordonia_liuliyuniae</td><td>L5G28_11435</td><td>131354..134731</td><td>2911517</td><td>DUF499-domain-containing-protein-[Gordonia-sp.-HY285]</td></tr>
<tr><td>WP_236997292.1</td><td>HEPN+AAA-ATPase+Alpha+Beta-domain+wHTH+RRM</td><td>Type-1_BR.RM.like.systems</td><td>TUDOR+SF1-SF2-Helicase+iREase+CTD+REase-><-DUF222+HNH||METHYLASE->HEPN+DUF499*->ACET-><-?||?->ABhydrolase-></td><td>1125</td><td>Bacteria>Terrabacteria_group>Actinomycetota>Actinomycetes>Mycobacteriales</td><td>GCF_021738805.1</td><td>Gordonia_liuliyuniae</td><td>L5G33_06210</td><td>131250..134627</td><td>2911517</td><td>DUF499-domain-containing-protein-[Gordonia-sp.-HY366]</td></tr>
<tr><td>WP_116057355.1</td><td>HEPN+AAA-ATPase+Alpha+Beta-domain+wHTH+RRM</td><td>Type-1_BR.RM.like.systems</td><td><-URI-DNase<-DNA-glycosylase||?->TUDOR+SF1-SF2-Helicase+iREase+CTD+REase->METHYLASE->HEPN+DUF499*->?-><-HNH</td><td>1122</td><td>Bacteria>Terrabacteria_group>Actinomycetota>Actinomycetes>Mycobacteriales</td><td>-</td><td>Rhodococcus_erythropolis</td><td>DVG80_RS18290</td><td>3917488..3920856</td><td>1833</td><td>Swt1-family-HEPN-domain-containing-protein-[Rhodococcus-erythropolis]</td></tr>
<tr><td>WP_047262804.1</td><td>HEPN+AAA-ATPase+Alpha+Beta-domain+wHTH+RRM</td><td>Type-1_BR.RM.like.systems</td><td>ABC-ATPase+SbcC-><-tRNA<-tRNA<-?||?-><-HEPN+DUF499*<-TUDOR+SF1-SF2-Helicase+iREase+CTD+REase||XIS-HTH->SbcC->TetR-HTH-></td><td>1140</td><td>Bacteria>Terrabacteria_group>Actinomycetota>Actinomycetes>Mycobacteriales</td><td>GCF_001020985.1</td><td>Corynebacterium_mustelae</td><td>CMUST_RS12605</td><td>2821253..2824675</td><td>571915</td><td>Swt1-family-HEPN-domain-containing-protein-[Corynebacterium-mustelae]</td></tr>
<tr><td>MCF8610758.1</td><td>HEPN+AAA-ATPase+Alpha+Beta-domain+wHTH+RRM</td><td>Type-1_BR.RM.like.systems</td><td>TUDOR+SF1-SF2-Helicase+iREase+CTD+REase-><-DUF222+HNH||METHYLASE->HEPN+DUF499*->ACET-><-?||?->ABhydrolase-></td><td>1125</td><td>Bacteria>Terrabacteria_group>Actinomycetota>Actinomycetes>Mycobacteriales</td><td>GCA_021739025.1</td><td>Gordonia_liuliyuniae</td><td>L5G28_11435</td><td>131354..134731</td><td>2911517</td><td>DUF499-domain-containing-protein-[Gordonia-liuliyuniae]</td></tr>
<tr><td>MCF8588060.1</td><td>HEPN+AAA-ATPase+Alpha+Beta-domain+wHTH+RRM</td><td>Type-1_BR.RM.like.systems</td><td>TUDOR+SF1-SF2-Helicase+iREase+CTD+REase-><-DUF222+HNH||METHYLASE->HEPN+DUF499*->ACET-><-?||?->ABhydrolase-></td><td>1125</td><td>Bacteria>Terrabacteria_group>Actinomycetota>Actinomycetes>Mycobacteriales</td><td>GCA_021738805.1</td><td>Gordonia_liuliyuniae</td><td>L5G33_06210</td><td>131250..134627</td><td>2911517</td><td>DUF499-domain-containing-protein-[Gordonia-liuliyuniae]</td></tr>
<tr><td>MCF8569704.1</td><td>HEPN+AAA-ATPase+Alpha+Beta-domain+wHTH+RRM</td><td>Type-1_BR.RM.like.systems</td><td><-ABhydrolase<-?||?-><-ACET<-HEPN+DUF499*<-METHYLASE||DUF222+HNH-><-TUDOR+SF1-SF2-Helicase+iREase+CTD+REase</td><td>1115</td><td>Bacteria>Terrabacteria_group>Actinomycetota>Actinomycetes>Mycobacteriales</td><td>GCA_021738965.1</td><td>Gordonia_zhenghanii</td><td>L5G32_05410</td><td>142683..146030</td><td>2911516</td><td>DUF499-domain-containing-protein-[Gordonia-zhenghanii]</td></tr>
<tr><td>MBB4136224.1</td><td>HEPN+AAA-ATPase+Alpha+Beta-domain+wHTH+RRM</td><td>Type-1_BR.RM.like.systems</td><td><-ABhydrolase<-?||?-><-ACET<-HEPN+DUF499*<-METHYLASE||DUF222+HNH-><-TUDOR+SF1-SF2-Helicase+iREase+CTD+REase</td><td>1130</td><td>Bacteria>Terrabacteria_group>Actinomycetota>Actinomycetes>Mycobacteriales</td><td>GCA_014197435.1</td><td>Gordonia_humi</td><td>BKA16_002776</td><td>3005759..3009151</td><td>686429</td><td>hypothetical-protein-[Gordonia-humi]</td></tr>
<tr><td>WP_207553229.1</td><td>HEPN+AAA-ATPase+Alpha+Beta-domain+wHTH+RRM</td><td>Type-1_BR.RM.like.systems</td><td><-HEPN+DUF499*<-ParB+HNH<-METHYLASE<-TUDOR+SF1-SF2-Helicase+iREase+CTD+REase||NTD+SF1-SF2-Helicase-></td><td>1114</td><td>Bacteria>Terrabacteria_group>Actinomycetota>Actinomycetes>Mycobacteriales</td><td>GCF_001954125.1</td><td>Mycolicibacterium_conceptionense</td><td>A5741_RS31960</td><td>1839..5183</td><td>451644</td><td>Swt1-family-HEPN-domain-containing-protein-[Mycolicibacterium-conceptionense]</td></tr>
<tr><td>WP_067473443.1</td><td>HEPN+AAA-ATPase+Alpha+Beta-domain+wHTH+RRM</td><td>Type-1_BR.RM.like.systems</td><td>DDE_Tnp_1-><-?||TUDOR+SF1-SF2-Helicase+iREase+CTD+REase->METHYLASE->HEPN+DUF499*-><-?<-Peptidase<-?<-?||HEPN-p-></td><td>1116</td><td>Bacteria>Terrabacteria_group>Actinomycetota>Actinomycetes>Mycobacteriales</td><td>GCF_001612745.1</td><td>Nocardia_amamiensis</td><td>NA4_RS22590</td><td>30808..34158</td><td>404578</td><td>DUF499-domain-containing-protein-[Nocardia-amamiensis]</td></tr>
<tr><td>WP_006942562.1</td><td>HEPN+AAA-ATPase+Alpha+Beta-domain+wHTH+RRM</td><td>Type-1_BR.RM.like.systems</td><td><-HEPN+DUF499*<-ParB+HNH<-METHYLASE<-TUDOR+SF1-SF2-Helicase+iREase+CTD+REase||NTD+SF1-SF2-Helicase-></td><td>1119</td><td>Bacteria>Terrabacteria_group>Actinomycetota>Actinomycetes>Mycobacteriales</td><td>GCF_000382865.1</td><td>Rhodococcus_aetherivorans</td><td>EBESD8_RS20690</td><td>433..3792</td><td>191292</td><td>Swt1-family-HEPN-domain-containing-protein-[Rhodococcus-aetherivorans]</td></tr>
<tr><td>OMB68868.1</td><td>HEPN+AAA-ATPase+Alpha+Beta-domain+wHTH+RRM</td><td>Type-1_BR.RM.like.systems</td><td><-HEPN+DUF499*<-ParB+HNH<-METHYLASE<-TUDOR+SF1-SF2-Helicase+iREase+CTD+REase||NTD+SF1-SF2-Helicase-></td><td>1103</td><td>Bacteria>Terrabacteria_group>Actinomycetota>Actinomycetes>Mycobacteriales</td><td>GCA_001954125.1</td><td>Mycolicibacterium_conceptionense</td><td>A5741_09375</td><td>1839..5150</td><td>451644</td><td>AAA-family-ATPase-[Mycolicibacterium-conceptionense]</td></tr>
<tr><td>WP_220667114.1</td><td>HEPN+AAA-ATPase+Alpha+Beta-domain+wHTH+RRM</td><td>Type-1_BR.RM.like.systems</td><td><-REase<-PRTase+PRTase<-?<-SIR2+STALD||TUDOR+SF1-SF2-Helicase+iREase+CTD+REase->HEPN+DUF499*-><-RNase</td><td>1142</td><td>Bacteria>Terrabacteria_group>Actinomycetota>Actinomycetes>Mycobacteriales</td><td>GCF_019641445.1</td><td>Rhodococcus_sp._DMU2021</td><td>K3M35_RS24125</td><td>14618..18046</td><td>2866997</td><td>Swt1-family-HEPN-domain-containing-protein-[Rhodococcus-sp.-DMU2021]</td></tr>
<tr><td>WP_044505526.1</td><td>HEPN+AAA-ATPase+Alpha+Beta-domain+wHTH+RRM</td><td>Type-1_BR.RM.like.systems</td><td>TUDOR+SF1-SF2-Helicase+iREase+CTD+REase->NERD->METHYLASE->HEPN+DUF499*->REase+TPR->?->Peptidase_M48-></td><td>1115</td><td>Bacteria>Terrabacteria_group>Actinomycetota>Actinomycetes>Mycobacteriales</td><td>GCF_000143885.2</td><td>Gordonia</td><td>KTR9_RS00485</td><td>112302..115649</td><td>2053</td><td>MULTISPECIES:-DUF499-domain-containing-protein-[Gordonia]</td></tr>
<tr><td>ADK68968.1</td><td>HEPN+AAA-ATPase+Alpha+Beta-domain+wHTH+RRM</td><td>Type-1_BR.RM.like.systems</td><td>TUDOR+SF1-SF2-Helicase+iREase+CTD+REase->NERD->?->METHYLASE->HEPN+DUF499*->REase+TPR->?->Peptidase_M48-></td><td>1120</td><td>Bacteria>Terrabacteria_group>Actinomycetota>Actinomycetes>Mycobacteriales</td><td>GCA_000143885.2</td><td>Gordonia_sp._KTR9</td><td>KTR9_4887</td><td>112287..115649</td><td>337191</td><td>Predicted-ATPase-(AAA+-superfamily)-[Gordonia-sp.-KTR9]</td></tr>
<tr><td>WP_259896744.1</td><td>HEPN+AAA-ATPase+Alpha+Beta-domain+wHTH+RRM</td><td>Type-1_BR.RM.like.systems</td><td><-ABhydrolase||TUDOR+SF1-SF2-Helicase+iREase+CTD+REase->METHYLASE->HEPN+DUF499*->ORC-CDC6+REase-></td><td>1112</td><td>Bacteria>Terrabacteria_group>Actinomycetota>Actinomycetes>Mycobacteriales</td><td>GCF_025144475.1</td><td>Corynebacterium_sanguinis</td><td>M3G45_RS07405</td><td>84077..87415</td><td>2594913</td><td>DUF499-domain-containing-protein-[Corynebacterium-sanguinis]</td></tr>
<tr><td>WP_259795206.1</td><td>HEPN+AAA-ATPase+Alpha+Beta-domain+wHTH+RRM</td><td>Type-1_BR.RM.like.systems</td><td><-ABhydrolase||TUDOR+SF1-SF2-Helicase+iREase+CTD+REase->METHYLASE->HEPN+DUF499*->ORC-CDC6+REase-></td><td>1112</td><td>Bacteria>Terrabacteria_group>Actinomycetota>Actinomycetes>Mycobacteriales</td><td>GCF_025144435.1</td><td>Corynebacterium_sanguinis</td><td>M3G32_RS10715</td><td>21189..24527</td><td>2594913</td><td>DUF499-domain-containing-protein-[Corynebacterium-sanguinis]</td></tr>
<tr><td>WP_194699134.1</td><td>HEPN+AAA-ATPase+Alpha+Beta-domain+wHTH+RRM</td><td>Type-1_BR.RM.like.systems</td><td>SF1-SF2-Helicase->||REase-><-HEPN+DUF499*<-TUDOR+SF1-SF2-Helicase+iREase+CTD+REase||PAIREDC-HTH-></td><td>1135</td><td>Bacteria>Terrabacteria_group>Actinomycetota>Actinomycetes>Mycobacteriales</td><td>-</td><td>Rhodococcus_rhodochrous</td><td>IRZ34_RS06155</td><td>243205..246612</td><td>1829</td><td>Swt1-family-HEPN-domain-containing-protein-[Rhodococcus-rhodochrous]</td></tr>
<tr><td>WP_181729591.1</td><td>HEPN+AAA-ATPase+Alpha+Beta-domain+wHTH+RRM</td><td>Type-1_BR.RM.like.systems</td><td><-ABhydrolase||TUDOR+SF1-SF2-Helicase+iREase+CTD+REase->METHYLASE->HEPN+DUF499*->ORC-CDC6+REase-></td><td>1112</td><td>Bacteria>Terrabacteria_group>Actinomycetota>Actinomycetes>Mycobacteriales</td><td>GCF_013868425.1</td><td>Corynebacterium_sanguinis</td><td>HOH28_RS04340</td><td>15252..18590</td><td>2594913</td><td>DUF499-domain-containing-protein-[Corynebacterium-sanguinis]</td></tr>
<tr><td>WP_166810353.1</td><td>HEPN+AAA-ATPase+Alpha+Beta-domain+wHTH+RRM</td><td>Type-1_BR.RM.like.systems</td><td>HEPN-><-NTD+SF1-SF2-Helicase||TUDOR+SF1-SF2-Helicase+iREase+CTD+REase->HEPN+DUF499*-><-?||?-><-?||<-GNTR-HTH</td><td>1121</td><td>Bacteria>Terrabacteria_group>Actinomycetota>Actinomycetes>Mycobacteriales</td><td>GCF_011602025.1</td><td>Rhodococcus_sp._IC4_135</td><td>G8767_RS12755</td><td>177145..180510</td><td>2715537</td><td>Swt1-family-HEPN-domain-containing-protein-[Rhodococcus-sp.-IC4_135]</td></tr>
<tr><td>WP_156666271.1</td><td>HEPN+AAA-ATPase+Alpha+Beta-domain+wHTH+RRM</td><td>Type-1_BR.RM.like.systems</td><td><-SbcC||Tox-RelE->HTH+MPTase->TUDOR+SF1-SF2-Helicase+iREase+CTD+REase->METHYLASE->HEPN+DUF499*-></td><td>536</td><td>Bacteria>Terrabacteria_group>Actinomycetota>Actinomycetes>Mycobacteriales</td><td>GCF_001651055.1</td><td>Rhodococcus_sp._HS-D2</td><td>A1E04_RS24870</td><td>10916..12523</td><td>1384636</td><td>Swt1-family-HEPN-domain-containing-protein,-partial-[Rhodococcus-sp.-HS-D2]</td></tr>
<tr><td>MCI1256747.1</td><td>HEPN+AAA-ATPase+Alpha+Beta-domain+wHTH+RRM</td><td>Type-1_BR.RM.like.systems</td><td>MPTase-><-HTH_3||TUDOR+SF1-SF2-Helicase+iREase+CTD+REase->METHYLASE->HEPN+DUF499*-><-?||HEPN+HEPN-></td><td>1166</td><td>Bacteria>Terrabacteria_group>Actinomycetota>Actinomycetes>Mycobacteriales</td><td>GCA_022641305.1</td><td>Corynebacterium_provencense</td><td>LKG15_08685</td><td>46466..49966</td><td>1737425</td><td>DUF499-domain-containing-protein-[Corynebacterium-provencense]</td></tr>
<tr><td>WP_206613802.1</td><td>HEPN+AAA-ATPase+Alpha+Beta-domain+wHTH+RRM</td><td>Type-1_BR.RM.like.systems</td><td>BirA-HTH+CRP-HTH-><-HEPN+DUF499*<-ParB+HNH<-METHYLASE<-TUDOR+SF1-SF2-Helicase+iREase+CTD+REase</td><td>1113</td><td>Bacteria>Terrabacteria_group>Actinomycetota>Actinomycetes>Mycobacteriales</td><td>GCF_004014805.1</td><td>Mycolicibacterium_elephantis</td><td>H7I95_RS08850</td><td>1305..4646</td><td>81858</td><td>Swt1-family-HEPN-domain-containing-protein-[Mycolicibacterium-elephantis]</td></tr>
<tr><td>WP_099696878.1</td><td>HEPN+AAA-ATPase+Alpha+Beta-domain+wHTH+RRM</td><td>Type-1_BR.RM.like.systems</td><td>FAD-NAD-dep-oxidoreductase->?-><-HEPN+DUF499*<-METHYLASE<-TUDOR+SF1-SF2-Helicase+iREase+CTD+REase</td><td>1134</td><td>Bacteria>Terrabacteria_group>Actinomycetota>Actinomycetes>Mycobacteriales</td><td>GCF_018588185.1</td><td>Rhodococcus</td><td>G4D79_RS05030</td><td>1105468..1108872</td><td>1827</td><td>MULTISPECIES:-Swt1-family-HEPN-domain-containing-protein-[Rhodococcus]</td></tr>
<tr><td>WP_065456397.1</td><td>HEPN+AAA-ATPase+Alpha+Beta-domain+wHTH+RRM</td><td>Type-1_BR.RM.like.systems</td><td>FAD-NAD-dep-oxidoreductase-><-HEPN+DUF499*<-METHYLASE<-TUDOR+SF1-SF2-Helicase+iREase+CTD+REase</td><td>1134</td><td>Bacteria>Terrabacteria_group>Actinomycetota>Actinomycetes>Mycobacteriales</td><td>-</td><td>Prescottella_equi</td><td>AS590_01070</td><td>160611..164015</td><td>43767</td><td>Swt1-family-HEPN-domain-containing-protein-[Prescottella-equi]</td></tr>
<tr><td>OCC17284.1</td><td>HEPN+AAA-ATPase+Alpha+Beta-domain+wHTH+RRM</td><td>Type-1_BR.RM.like.systems</td><td>FAD-NAD-dep-oxidoreductase-><-HEPN+DUF499*<-METHYLASE<-TUDOR+SF1-SF2-Helicase+iREase+CTD+REase</td><td>1134</td><td>Bacteria>Terrabacteria_group>Actinomycetota>Actinomycetes>Mycobacteriales</td><td>GCA_001687425.1</td><td>Prescottella_equi</td><td>AS590_01070</td><td>160611..164015</td><td>43767</td><td>AAA-family-ATPase-[Prescottella-equi]</td></tr>
<tr><td>WP_170194849.1</td><td>HEPN+AAA-ATPase+Alpha+Beta-domain+wHTH+RRM</td><td>Type-1_BR.RM.like.systems</td><td>HTH_3-><-?<-?||?-><-ABhydrolase+ABhydrolase<-HEPN+DUF499*||DUF222+HNH-><-AbiEii<-?||?-><-METHYLASE||HTH_19-></td><td>1113</td><td>Bacteria>Terrabacteria_group>Actinomycetota>Actinomycetes>Mycobacteriales</td><td>GCF_012933505.1</td><td>Gordonia_asplenii</td><td>HH308_RS14085</td><td>95388..98729</td><td>2725283</td><td>DUF499-domain-containing-protein-[Gordonia-asplenii]</td></tr>
<tr><td>NLG46720.1</td><td>HEPN+AAA-ATPase+Alpha+Beta-domain+wHTH+RRM</td><td>Type-1_BR.RM.like.systems</td><td><-DUF222+HNH<-HEPN+DUF499*<-METHYLASE||CHTH-><-TUDOR+SF1-SF2-Helicase+iREase+CTD+REase||HTH-></td><td>1141</td><td>Bacteria>Terrabacteria_group>Actinomycetota>Actinomycetes>Mycobacteriales</td><td>GCA_012515855.1</td><td>Gordonia_sp._(in:_high_G+C_Gram-positive_bacteria)</td><td>GX543_09665</td><td>1506..4931</td><td>84139</td><td>ATP-binding-protein-[Gordonia-sp.-(in:-high-G+C-Gram-positive-bacteria)]</td></tr>
<tr><td>WP_264721883.1</td><td>HEPN+AAA-ATPase+Alpha+Beta-domain+wHTH+RRM</td><td>Type-1_BR.RM.like.systems</td><td>TUDOR+SF1-SF2-Helicase+iREase+CTD+REase->METHYLASE->HEPN+DUF499*-><-RNase<-?||?-><-?<-Phage_integrase</td><td>1139</td><td>Bacteria>Terrabacteria_group>Actinomycetota>Actinomycetes>Mycobacteriales</td><td>GCF_025989245.1</td><td>Rhodococcus_pyridinivorans</td><td>OMK68_RS15380</td><td>34680..38099</td><td>103816</td><td>Swt1-family-HEPN-domain-containing-protein-[Rhodococcus-pyridinivorans]</td></tr>
<tr><td>WP_213184733.1</td><td>HEPN+AAA-ATPase+Alpha+Beta-domain+wHTH+RRM</td><td>Type-1_BR.RM.like.systems</td><td>TUDOR+SF1-SF2-Helicase+iREase+CTD+REase-><-?||METHYLASE->ParB+HNH->HEPN+DUF499*->ANK+ANK+RAMA-></td><td>1134</td><td>Bacteria>Terrabacteria_group>Actinomycetota>Actinomycetes>Mycobacteriales</td><td>GCF_018390595.1</td><td>Dietzia_massiliensis</td><td>KHA85_RS02755</td><td>11826..15230</td><td>2697499</td><td>Swt1-family-HEPN-domain-containing-protein-[Dietzia-massiliensis]</td></tr>
<tr><td>WP_032392811.1</td><td>HEPN+AAA-ATPase+Alpha+Beta-domain+wHTH+RRM</td><td>Type-1_BR.RM.like.systems</td><td>HIN-HTH->?-><-DNA-glycosylase||TUDOR+SF1-SF2-Helicase+iREase+CTD+REase-p->HEPN+DUF499*-><-?||?->HNH-></td><td>1123</td><td>Bacteria>Terrabacteria_group>Actinomycetota>Actinomycetes>Mycobacteriales</td><td>GCF_000760775.1</td><td>Rhodococcus_fascians</td><td>BH98_RS0123030</td><td>109500..112871</td><td>1828</td><td>Swt1-family-HEPN-domain-containing-protein-[Rhodococcus-fascians]</td></tr>
<tr><td>OAH45661.1</td><td>HEPN+AAA-ATPase+Alpha+Beta-domain+wHTH+RRM</td><td>Type-1_BR.RM.like.systems</td><td>TUDOR+SF1-SF2-Helicase+iREase+CTD+REase-><-?||METHYLASE->ParB+HNH->HEPN+DUF499*->ANK+ANK+RAMA-></td><td>1134</td><td>Bacteria>Terrabacteria_group>Actinomycetota>Actinomycetes>Mycobacteriales</td><td>GCA_001643175.1</td><td>Dietzia_cinnamea</td><td>AYJ66_16790</td><td>32668..36072</td><td>321318</td><td>AAA-family-ATPase-[Dietzia-cinnamea]</td></tr>
<tr><td>WP_145941802.1</td><td>HEPN+AAA-ATPase+Alpha+Beta-domain+wHTH+RRM</td><td>Type-1_BR.RM.like.systems</td><td>ORC-CDC6+REase-><-HEPN+DUF499*<-METHYLASE<-TUDOR+SF1-SF2-Helicase+iREase+CTD+REase||HTH_3-></td><td>1182</td><td>Bacteria>Terrabacteria_group>Actinomycetota>Actinomycetes>Mycobacteriales</td><td>GCF_007673095.1</td><td>Corynebacterium_glyciniphilum</td><td>FS507_RS05565</td><td>165529..169077</td><td>1404244</td><td>DUF499-domain-containing-protein-[Corynebacterium-glyciniphilum]</td></tr>
<tr><td>WP_079929183.1</td><td>HEPN+AAA-ATPase+Alpha+Beta-domain+wHTH+RRM</td><td>Type-1_BR.RM.like.systems</td><td>TUDOR+SF1-SF2-Helicase+iREase+CTD+REase-p-><-STYKinase-like<-HTH||METHYLASE->HEPN+DUF499*-></td><td>762</td><td>Bacteria>Terrabacteria_group>Actinomycetota>Actinomycetes>Mycobacteriales</td><td>GCF_002043085.1</td><td>Gordonia_sp._i37</td><td>B1964_RS05750</td><td>6455..8741</td><td>1961707</td><td>Swt1-family-HEPN-domain-containing-protein,-partial-[Gordonia-sp.-i37]</td></tr>
<tr><td>WP_065923600.1</td><td>HEPN+AAA-ATPase+Alpha+Beta-domain+wHTH+RRM</td><td>Type-1_BR.RM.like.systems</td><td><-RVT+HNH||?->HEPN-><-HEPN+DUF499*<-METHYLASE<-TUDOR+SF1-SF2-Helicase+iREase+CTD+REase<-?<-?<-?||HNH-></td><td>1138</td><td>Bacteria>Terrabacteria_group>Actinomycetota>Actinomycetes>Mycobacteriales</td><td>GCF_001700945.1</td><td>Rhodococcus_sp._WB1</td><td>A4U64_RS26630</td><td>5329..8745</td><td>1033922</td><td>Swt1-family-HEPN-domain-containing-protein-[Rhodococcus-sp.-WB1]</td></tr>
<tr><td>OHV00237.1</td><td>HEPN+AAA-ATPase+Alpha+Beta-domain+wHTH+RRM</td><td>Type-1_BR.RM.like.systems</td><td>TUDOR+SF1-SF2-Helicase+iREase+CTD+REase->METHYLASE->ParB+HNH->HEPN+DUF499*->SIR2-><-?||SbcC-></td><td>1095</td><td>Bacteria>Terrabacteria_group>Actinomycetota>Actinomycetes>Mycobacteriales</td><td>GCA_001854525.1</td><td>Mycobacterium_talmoniae</td><td>BKN37_18410</td><td>8707..11994</td><td>1858794</td><td>AAA-family-ATPase-[Mycobacterium-talmoniae]</td></tr>
<tr><td>WP_213934645.1</td><td>HEPN+AAA-ATPase+Alpha+Beta-domain+wHTH+RRM</td><td>Type-1_BR.RM.like.systems</td><td>McrC-NTD+REase-><-?<-?||CHTH->?-><-HEPN+DUF499*||<-TUDOR+SF1-SF2-Helicase+iREase+CTD+REase||?->?-><-ACET</td><td>1138</td><td>Bacteria>Terrabacteria_group>Actinomycetota>Actinomycetes>Mycobacteriales</td><td>GCF_013602415.1</td><td>Rhodococcus_sp._B50</td><td>GON09_RS24520</td><td>12924..16340</td><td>2682847</td><td>Swt1-family-HEPN-domain-containing-protein-[Rhodococcus-sp.-B50]</td></tr>
<tr><td>WP_207547258.1</td><td>HEPN+AAA-ATPase+Alpha+Beta-domain+wHTH+RRM</td><td>Type-1_BR.RM.like.systems</td><td>TUDOR+SF1-SF2-Helicase+iREase+CTD+REase->METHYLASE->ParB+HNH->HEPN+DUF499*->SIR2->?->SbcC-></td><td>1106</td><td>Bacteria>Terrabacteria_group>Actinomycetota>Actinomycetes>Mycobacteriales</td><td>GCF_001854525.1</td><td>Mycobacterium_talmoniae</td><td>BKN37_RS18415</td><td>8674..11994</td><td>1858794</td><td>DUF499-domain-containing-protein-[Mycobacterium-talmoniae]</td></tr>
<tr><td>WP_160903143.1</td><td>HEPN+AAA-ATPase+Alpha+Beta-domain+wHTH+RRM</td><td>Type-1_BR.RM.like.systems</td><td>RelE-ParE-><-?<-HEPN+DUF499*<-METHYLASE||?-><-TUDOR+SF1-SF2-Helicase+iREase+CTD+REase||HTH_3-></td><td>1126</td><td>Bacteria>Terrabacteria_group>Actinomycetota>Actinomycetes>Mycobacteriales</td><td>GCF_024734075.1</td><td>Gordonia_mangrovi</td><td>GIY30_RS16680</td><td>372248..375628</td><td>2665643</td><td>Swt1-family-HEPN-domain-containing-protein-[Gordonia-mangrovi]</td></tr>
<tr><td>WP_159371683.1</td><td>HEPN+AAA-ATPase+Alpha+Beta-domain+wHTH+RRM</td><td>Type-1_BR.RM.like.systems</td><td>REase+SPF-cons-><-?<-CHTH<-?<-HEPN+DUF499*<-?<-METHYLASE<-?<-TUDOR+SF1-SF2-Helicase+iREase+CTD+REase</td><td>1117</td><td>Bacteria>Terrabacteria_group>Actinomycetota>Actinomycetes>Mycobacteriales</td><td>GCF_009856645.1</td><td>Gordonia_sp._JH63</td><td>GR168_RS22070</td><td>4982408..4985761</td><td>2698900</td><td>DUF499-domain-containing-protein-[Gordonia-sp.-JH63]</td></tr>
<tr><td>MCB0937449.1</td><td>HEPN+AAA-ATPase+Alpha+Beta-domain+wHTH+RRM</td><td>Type-1_BR.RM.like.systems</td><td>ParB+ParBC-><-?<-HTH_58<-HEPN+DUF499*<-METHYLASE<-TUDOR+SF1-SF2-Helicase+iREase+CTD+REase-p</td><td>1125</td><td>Bacteria>Terrabacteria_group>Actinomycetota>Actinomycetes>Mycobacteriales</td><td>GCA_020438785.1</td><td>Mycobacterium_sp.</td><td>KDB55_15800</td><td>12247..15624</td><td>1785</td><td>DUF499-domain-containing-protein-[Mycobacterium-sp.]</td></tr>
<tr><td>WP_005520661.1</td><td>HEPN+AAA-ATPase+Alpha+Beta-domain+wHTH+RRM</td><td>Type-1_BR.RM.like.systems</td><td>TUDOR+SF1-SF2-Helicase+iREase+CTD+REase->METHYLASE->HEPN+DUF499*->HEPN-><-?<-?<-JetB<-JetA</td><td>1186</td><td>Bacteria>Terrabacteria_group>Actinomycetota>Actinomycetes>Mycobacteriales</td><td>GCF_000158635.1</td><td>Corynebacterium_matruchotii</td><td>CORMATOL_RS04405</td><td>70263..73823</td><td>43768</td><td>DUF499-domain-containing-protein-[Corynebacterium-matruchotii]</td></tr>
<tr><td>WP_247450629.1</td><td>HEPN+AAA-ATPase+Alpha+Beta-domain+wHTH+RRM</td><td>Type-1_BR.RM.like.systems</td><td>HNH-><-REase+TPR<-HEPN+DUF499*<-METHYLASE<-TUDOR+SF1-SF2-Helicase+iREase+CTD+REase</td><td>1115</td><td>Bacteria>Terrabacteria_group>Actinomycetota>Actinomycetes>Mycobacteriales</td><td>GCF_023078355.1</td><td>Gordonia_hongkongensis</td><td>MVF96_RS23000</td><td>5106874..5110221</td><td>1701090</td><td>DUF499-domain-containing-protein-[Gordonia-hongkongensis]</td></tr>
<tr><td>OCW84481.1</td><td>HEPN+AAA-ATPase+Alpha+Beta-domain+wHTH+RRM</td><td>Type-1_BR.RM.like.systems</td><td><-SbcC<-SIR2<-HEPN+DUF499*<-METHYLASE<-?<-MULE-TRANSPOSASE<-?<-HTH_28+rve+Mu-transpos_C</td><td>1113</td><td>Bacteria>Terrabacteria_group>Actinomycetota>Actinomycetes>Mycobacteriales</td><td>GCA_001704135.1</td><td>Nocardia_farcinica</td><td>A8M60_10225</td><td>5506..8847</td><td>37329</td><td>AAA-family-ATPase-[Nocardia-farcinica]</td></tr>
<tr><td>WP_006361012.1</td><td>HEPN+AAA-ATPase+Alpha+Beta-domain+wHTH+RRM</td><td>Type-1_BR.RM.like.systems</td><td><-ACYC<-ACYC<-HEPN+DUF499*<-METHYLASE||?-><-TUDOR+SF1-SF2-Helicase+iREase+CTD+REase</td><td>1113</td><td>Bacteria>Terrabacteria_group>Actinomycetota>Actinomycetes>Mycobacteriales</td><td>GCF_000225505.1</td><td>Gordonia_alkanivorans</td><td>GOALK_RS21795</td><td>2511..5852</td><td>84096</td><td>DUF499-domain-containing-protein-[Gordonia-alkanivorans]</td></tr>
<tr><td>WP_176217705.1</td><td>HEPN+AAA-ATPase+Alpha+Beta-domain+wHTH+RRM</td><td>Type-1_BR.RM.like.systems</td><td><-SIR2<-HEPN+DUF499*<-METHYLASE<-TUDOR+SF1-SF2-Helicase+iREase+CTD+REase<-RNase</td><td>1113</td><td>Bacteria>Terrabacteria_group>Actinomycetota>Actinomycetes>Mycobacteriales</td><td>GCF_002089755.1</td><td>Mycobacterium_avium</td><td>B9C05_RS24160</td><td>1185..4526</td><td>1764</td><td>Swt1-family-HEPN-domain-containing-protein-[Mycobacterium-avium]</td></tr>
<tr><td>WP_023870228.1</td><td>HEPN+AAA-ATPase+Alpha+Beta-domain+wHTH+RRM</td><td>Type-1_BR.RM.like.systems</td><td>Glyoxalase->HIN-HTH-><-DNA_methylase||REase->METHYLASE->HEPN+DUF499*-><-HIN-HTH</td><td>1138</td><td>Bacteria>Terrabacteria_group>Actinomycetota>Actinomycetes>Mycobacteriales</td><td>-</td><td>Mycobacterium_avium</td><td>BEP52_09350</td><td>1958340..1961756</td><td>1764</td><td>DUF499-domain-containing-protein-[Mycobacterium-avium]</td></tr>
<tr><td>ATO62480.2</td><td>HEPN+AAA-ATPase+Alpha+Beta-domain+wHTH+RRM</td><td>Type-1_BR.RM.like.systems</td><td>Glyoxalase->HIN-HTH-><-DNA_methylase||REase->METHYLASE->HEPN+DUF499*-><-HIN-HTH</td><td>1138</td><td>Bacteria>Terrabacteria_group>Actinomycetota>Actinomycetes>Mycobacteriales</td><td>GCA_002716905.4</td><td>Mycobacterium_avium_subsp._hominissuis</td><td>BEP52_09350</td><td>1958340..1961756</td><td>439334</td><td>DUF499-domain-containing-protein-[Mycobacterium-avium-subsp.-hominissuis]</td></tr>
<tr><td>WP_062368091.1</td><td>HEPN+AAA-ATPase+Alpha+Beta-domain+wHTH+RRM</td><td>Type-1_BR.RM.like.systems</td><td><-DUF222+HNH||METHYLASE->HEPN+DUF499*-><-?||ACET-><-Peptidase_S51<-?||?->ABhydrolase-></td><td>1124</td><td>Bacteria>Terrabacteria_group>Actinomycetota>Actinomycetes>Mycobacteriales</td><td>GCF_001578675.1</td><td>Gordonia_sp._QH-12</td><td>Y710_RS17975</td><td>9303..12677</td><td>1437876</td><td>DUF499-domain-containing-protein-[Gordonia-sp.-QH-12]</td></tr>
<tr><td>WP_202931651.1</td><td>HEPN+AAA-ATPase+Alpha+Beta-domain+wHTH+RRM</td><td>Type-1_BR.RM.like.systems</td><td><-HEPN+DUF499*<-ParB-p<-METHYLASE||?-><-TUDOR+SF1-SF2-Helicase+iREase+CTD+REase</td><td>1109</td><td>Bacteria>Terrabacteria_group>Actinomycetota>Actinomycetes>Mycobacteriales</td><td>GCF_002245615.1</td><td>Mycobacterium_neumannii</td><td>CEX88_RS25285</td><td>2442..5771</td><td>2048551</td><td>Swt1-family-HEPN-domain-containing-protein-[Mycobacterium-neumannii]</td></tr>
<tr><td>PND54237.1</td><td>HEPN+AAA-ATPase+Alpha+Beta-domain+wHTH+RRM</td><td>Type-1_BR.RM.like.systems</td><td>METHYLASE->HEPN+DUF499*-><-HIN-HTH||?-><-HTH_Tnp_1+HTH_21+rve+rve_2||DUF499-p-></td><td>654</td><td>Bacteria>Terrabacteria_group>Actinomycetota>Actinomycetes>Mycobacteriales</td><td>GCA_002887815.1</td><td>Mycobacterium_sp._ENV421</td><td>CRM90_29025</td><td>2865..4829</td><td>1213407</td><td>hypothetical-protein-CRM90_29025-[Mycobacterium-sp.-ENV421]</td></tr>
<tr><td>ORV36019.1</td><td>HEPN+AAA-ATPase+Alpha+Beta-domain+wHTH+RRM</td><td>Type-1_BR.RM.like.systems</td><td>HTH_Tnp->?->?->?->||TUDOR+SF1-SF2-Helicase+iREase+CTD+REase->ParB+HNH->HEPN+DUF499*-></td><td>1111</td><td>Bacteria>Terrabacteria_group>Actinomycetota>Actinomycetes>Mycobacteriales</td><td>GCA_002102115.1</td><td>Mycolicibacterium_doricum</td><td>AWC01_00650</td><td>40820..44155</td><td>126673</td><td>AAA-family-ATPase-[Mycobacterium-doricum]</td></tr>
<tr><td>MBZ4613710.1</td><td>HEPN+AAA-ATPase+Alpha+Beta-domain+wHTH+RRM</td><td>Type-1_BR.RM.like.systems</td><td><-HEPN+DUF499*<-ParB+HNH<-METHYLASE<-TUDOR+SF1-SF2-Helicase+iREase+CTD+REase</td><td>1103</td><td>Bacteria>Terrabacteria_group>Actinomycetota>Actinomycetes>Mycobacteriales</td><td>GCA_020055445.1</td><td>Mycobacterium_avium_subsp._hominissuis</td><td>GBP66_23845</td><td>83..3394</td><td>439334</td><td>ATP-binding-protein-[Mycobacterium-avium-subsp.-hominissuis]</td></tr>
<tr><td>WP_270484774.1</td><td>HEPN+AAA-ATPase+Alpha+Beta-domain+wHTH+RRM</td><td>Type-1_BR.RM.like.systems</td><td><-HEPN+DUF499*<-METHYLASE<-TUDOR+SF1-SF2-Helicase+iREase+CTD+REase<-?<-?<-?<-XIS-HTH</td><td>1110</td><td>Bacteria>Terrabacteria_group>Actinomycetota>Actinomycetes>Mycobacteriales</td><td>GCF_027659065.1</td><td>Gordonia_jacobaea</td><td>PGQ40_RS12000</td><td>183639..186971</td><td>122202</td><td>DUF499-domain-containing-protein-[Gordonia-jacobaea]</td></tr>
<tr><td>WP_267626547.1</td><td>HEPN+AAA-ATPase+Alpha+Beta-domain+wHTH+RRM</td><td>Type-1_BR.RM.like.systems</td><td>XIS-HTH->?->?->?->TUDOR+SF1-SF2-Helicase+iREase+CTD+REase->METHYLASE->HEPN+DUF499*-></td><td>1110</td><td>Bacteria>Terrabacteria_group>Actinomycetota>Actinomycetes>Mycobacteriales</td><td>GCF_026616935.1</td><td>Gordonia_sputi</td><td>NZL84_RS17670</td><td>45237..48569</td><td>36823</td><td>DUF499-domain-containing-protein-[Gordonia-sputi]</td></tr>
<tr><td>WP_218681519.1</td><td>HEPN+AAA-ATPase+Alpha+Beta-domain+wHTH+RRM</td><td>Type-1_BR.RM.like.systems</td><td>NUDIX->?->TUDOR+SF1-SF2-Helicase+iREase+CTD+REase->METHYLASE->HEPN+DUF499*-></td><td>1134</td><td>Bacteria>Terrabacteria_group>Actinomycetota>Actinomycetes>Mycobacteriales</td><td>GCF_019279115.1</td><td>Rhodococcus_qingshengii</td><td>KZA18_RS32415</td><td>9085..12489</td><td>334542</td><td>Swt1-family-HEPN-domain-containing-protein-[Rhodococcus-qingshengii]</td></tr>
<tr><td>WP_176213652.1</td><td>HEPN+AAA-ATPase+Alpha+Beta-domain+wHTH+RRM</td><td>Type-1_BR.RM.like.systems</td><td><-HEPN+DUF499*<-METHYLASE<-TUDOR+SF1-SF2-Helicase+iREase+CTD+REase<-RNase</td><td>1113</td><td>Bacteria>Terrabacteria_group>Actinomycetota>Actinomycetes>Mycobacteriales</td><td>GCF_002089315.1</td><td>Mycobacterium_avium_complex_(MAC)</td><td>BST46_RS22320</td><td>85..3426</td><td>120793</td><td>MULTISPECIES:-Swt1-family-HEPN-domain-containing-protein-[Mycobacterium-avium-complex-(MAC)]</td></tr>
<tr><td>MCB1297052.1</td><td>HEPN+AAA-ATPase+Alpha+Beta-domain+wHTH+RRM</td><td>Type-1_BR.RM.like.systems</td><td>Peptidase_M20-><-?||?->?->?-><-REase<-HEPN+DUF499*||MazE_antitoxin->PIN-><-METHYLASE</td><td>1119</td><td>Bacteria>Terrabacteria_group>Actinomycetota>Actinomycetes>Mycobacteriales</td><td>GCA_020438995.1</td><td>Gordonia_sp._(in:_high_G+C_Gram-positive_bacteria)</td><td>KDB67_20615</td><td>20423..23782</td><td>84139</td><td>DUF499-domain-containing-protein-[Gordonia-sp.]</td></tr>
<tr><td>WP_143907483.1</td><td>HEPN+AAA-ATPase+Alpha+Beta-domain+wHTH+RRM</td><td>Type-1_BR.RM.like.systems</td><td><-PIN<-?<-?<-HEPN+DUF499*<-Glyoxalase||ArsR-HTH->?->DNA_methylase-><-?<-METHYLASE</td><td>1132</td><td>Bacteria>Terrabacteria_group>Actinomycetota>Actinomycetes>Mycobacteriales</td><td>GCF_007559025.1</td><td>Tomitella_fengzijianii</td><td>M2W19_RS17095</td><td>32946..36344</td><td>2597660</td><td>DUF499-domain-containing-protein-[Tomitella-fengzijianii]</td></tr>
<tr><td>WP_170826146.1</td><td>HEPN+AAA-ATPase+Alpha+Beta-domain+wHTH+RRM</td><td>Type-1_BR.RM.like.systems</td><td><-HEPN+DUF499*<-METHYLASE<-TUDOR+SF1-SF2-Helicase+iREase+CTD+REase-p</td><td>1112</td><td>Bacteria>Terrabacteria_group>Actinomycetota>Actinomycetes>Mycobacteriales</td><td>-</td><td>Mycobacterium_malmoense</td><td>A9X02_RS19855</td><td>3529..6867</td><td>1780</td><td>Swt1-family-HEPN-domain-containing-protein-[Mycobacterium-malmoense]</td></tr>
<tr><td>OCB34496.1</td><td>HEPN+AAA-ATPase+Alpha+Beta-domain+wHTH+RRM</td><td>Type-1_BR.RM.like.systems</td><td><-HEPN+DUF499*<-METHYLASE<-TUDOR+SF1-SF2-Helicase+iREase+CTD+REase-p</td><td>1101</td><td>Bacteria>Terrabacteria_group>Actinomycetota>Actinomycetes>Mycobacteriales</td><td>GCA_001686725.1</td><td>Mycobacterium_malmoense</td><td>A9X02_22585</td><td>3529..6834</td><td>1780</td><td>AAA-family-ATPase-[Mycobacterium-malmoense]</td></tr>
<tr><td>WP_032373157.1</td><td>HEPN+AAA-ATPase+Alpha+Beta-domain+wHTH+RRM</td><td>Type-1_BR.RM.like.systems</td><td><-HEPN+DUF499*<-METHYLASE<-TUDOR+SF1-SF2-Helicase+iREase+CTD+REase</td><td>1122</td><td>Bacteria>Terrabacteria_group>Actinomycetota>Actinomycetes>Mycobacteriales</td><td>GCF_000760815.1</td><td>Rhodococcus_fascians</td><td>BI01_RS0106730</td><td>27469..30837</td><td>1828</td><td>Swt1-family-HEPN-domain-containing-protein-[Rhodococcus-fascians]</td></tr>
<tr><td>TXH27607.1</td><td>HEPN+AAA-ATPase+Alpha+Beta-domain+wHTH+RRM</td><td>Type-1_BR.RM.like.systems</td><td>TUDOR+SF1-SF2-Helicase+iREase+CTD+REase->METHYLASE->HEPN+DUF499*-></td><td>848</td><td>Bacteria>Terrabacteria_group>Actinomycetota>Actinomycetes>Mycobacteriales</td><td>GCA_008012155.1</td><td>Mycobacterium_sp.</td><td>E6R06_03465</td><td>7027..9570</td><td>1785</td><td>MAG:-ATP-binding-protein,-partial-[Mycobacterium-sp.]</td></tr>
<tr><td>WP_200172468.1</td><td>HEPN+AAA-ATPase+Alpha+Beta-domain+wHTH+RRM</td><td>Type-1_BR.RM.like.systems</td><td><-CHTH||METHYLASE->?->HTH+MPTase->HEPN+DUF499*->?-><-?||?->?-><-?<-ABC-ATPase+SbcC</td><td>1132</td><td>Bacteria>Terrabacteria_group>Actinomycetota>Actinomycetes>Mycobacteriales</td><td>GCF_016599145.1</td><td>Tomitella_cavernea</td><td>JJ683_RS06825</td><td>52491..55889</td><td>1387982</td><td>DUF499-domain-containing-protein-[Tomitella-cavernea]</td></tr>
<tr><td>WP_248488617.1</td><td>HEPN+AAA-ATPase+Alpha+Beta-domain+wHTH+RRM</td><td>Type-1_BR.RM.like.systems</td><td>SbcC->?-><-HTH_19<-HTH_3||<-HTH||METHYLASE->HEPN+DUF499*->?->?-><-?<-?||?-><-RVT</td><td>1120</td><td>Bacteria>Terrabacteria_group>Actinomycetota>Actinomycetes>Mycobacteriales</td><td>GCF_023168325.1</td><td>Tsukamurella_sp._PLM1</td><td>MTP03_RS12765</td><td>2471846..2475208</td><td>2929795</td><td>DUF499-domain-containing-protein-[Tsukamurella-sp.-PLM1]</td></tr>
<tr><td>WP_040525708.1</td><td>HEPN+AAA-ATPase+Alpha+Beta-domain+wHTH+RRM</td><td>Type-1_BR.RM.like.systems</td><td><-ABC-ATPase+SbcC||HTH_3-><-?<-?<-?<-HEPN+DUF499*<-METHYLASE||HTH_19-></td><td>1111</td><td>Bacteria>Terrabacteria_group>Actinomycetota>Actinomycetes>Mycobacteriales</td><td>-</td><td>Gordonia_effusa</td><td>GOEFS_RS20135</td><td>108493..111828</td><td>263908</td><td>DUF499-domain-containing-protein-[Gordonia-effusa]</td></tr>
<tr><td>GAB20465.1</td><td>HEPN+AAA-ATPase+Alpha+Beta-domain+wHTH+RRM</td><td>Type-1_BR.RM.like.systems</td><td><-ABC-ATPase+SbcC||HTH_3-><-?<-?<-?<-HEPN+DUF499*<-METHYLASE||HTH_19-></td><td>1118</td><td>Bacteria>Terrabacteria_group>Actinomycetota>Actinomycetes>Mycobacteriales</td><td>GCA_000241305.2</td><td>Gordonia_effusa_NBRC_100432</td><td>GOEFS_115_01050</td><td>108493..111849</td><td>1077974</td><td>hypothetical-protein-GOEFS_115_01050-[Gordonia-effusa-NBRC-100432]</td></tr>
<tr><td>MCB0924960.1</td><td>HEPN+AAA-ATPase+Alpha+Beta-domain+wHTH+RRM</td><td>Type-1_BR.RM.like.systems</td><td><-GDResolvase+Recombinase+ZnR<-?<-?<-HEPN+DUF499*<-METHYLASE-p</td><td>1126</td><td>Bacteria>Terrabacteria_group>Actinomycetota>Actinomycetes>Mycobacteriales</td><td>GCA_020438895.1</td><td>Mycobacterium_sp.</td><td>KDB50_10535</td><td>13047..16427</td><td>1785</td><td>DUF499-domain-containing-protein-[Mycobacterium-sp.]</td></tr>
<tr><td>GES35195.1</td><td>HEPN+AAA-ATPase+Alpha+Beta-domain+wHTH+RRM</td><td>Type-1_BR.RM.like.systems</td><td><-GDResolvase+HIN-HTH||?->RNase-><-HEPN+DUF499*<-METHYLASE</td><td>1137</td><td>Bacteria>Terrabacteria_group>Actinomycetota>Actinomycetes>Mycobacteriales</td><td>GCA_009176285.1</td><td>Rhodococcus_aetherivorans</td><td>RAJCM14343_0442</td><td>3451..6864</td><td>191292</td><td>hypothetical-protein-RAJCM14343_0442-[Rhodococcus-aetherivorans]</td></tr>
<tr><td>WP_104363764.1</td><td>HEPN+AAA-ATPase+Alpha+Beta-domain+wHTH+RRM</td><td>Type-1_BR.RM.like.systems</td><td>HIN-HTH->HTH_21+rve+rve_3-><-?<-?||ParB+HNH->HEPN+DUF499*-></td><td>1121</td><td>Bacteria>Terrabacteria_group>Actinomycetota>Actinomycetes>Mycobacteriales</td><td>GCF_002933495.1</td><td>Nocardia_nova</td><td>C5F51_RS20340</td><td>202110..205475</td><td>37330</td><td>Swt1-family-HEPN-domain-containing-protein-[Nocardia-nova]</td></tr>
<tr><td>WP_232512159.1</td><td>HEPN+AAA-ATPase+Alpha+Beta-domain+wHTH+RRM</td><td>Type-1_BR.RM.like.systems</td><td><-HEPN+DUF499*<-METHYLASE<-REase||DNA_methylase-></td><td>1128</td><td>Bacteria>Terrabacteria_group>Actinomycetota>Actinomycetes>Mycobacteriales</td><td>GCF_000829075.1</td><td>Mycobacterium_avium</td><td>B9C26_RS25090</td><td>191..3577</td><td>1764</td><td>DUF499-domain-containing-protein-[Mycobacterium-avium]</td></tr>
<tr><td>WP_210731572.1</td><td>HEPN+AAA-ATPase+Alpha+Beta-domain+wHTH+RRM</td><td>Type-1_BR.RM.like.systems</td><td>METHYLASE->ParB+HNH->HEPN+DUF499*-><-?||?->?->SbcC->SbcC-></td><td>1116</td><td>Bacteria>Terrabacteria_group>Actinomycetota>Actinomycetes>Mycobacteriales</td><td>GCF_900960235.1</td><td>Mycobacterium_intracellulare</td><td>FAX65_RS26670</td><td>4893..8243</td><td>1767</td><td>Swt1-family-HEPN-domain-containing-protein-[Mycobacterium-intracellulare]</td></tr>
<tr><td>WP_125079869.1</td><td>HEPN+AAA-ATPase+Alpha+Beta-domain+wHTH+RRM</td><td>Type-1_BR.RM.like.systems</td><td>ACET->GDResolvase+HIN-HTH-><-REase<-HEPN+DUF499*</td><td>1124</td><td>Bacteria>Terrabacteria_group>Actinomycetota>Actinomycetes>Mycobacteriales</td><td>GCF_900604305.1</td><td>Mycobacterium_sp._P7213</td><td>EJK91_RS20970</td><td>3382..6756</td><td>2478465</td><td>DUF499-domain-containing-protein-[Mycobacterium-sp.-P7213]</td></tr>
<tr><td>WP_018177518.1</td><td>HEPN+AAA-ATPase+Alpha+Beta-domain+wHTH+RRM</td><td>Type-1_BR.RM.like.systems</td><td>HTH_Tnp_1+HTH_21+rve+rve_2->HEPN+DUF499*->?-><-ACET</td><td>1113</td><td>Bacteria>Terrabacteria_group>Actinomycetota>Actinomycetes>Mycobacteriales</td><td>GCF_000380485.1</td><td>Jongsikchunia_kroppenstedtii</td><td>G341_RS0102455</td><td>2656..5997</td><td>1121721</td><td>Swt1-family-HEPN-domain-containing-protein-[Jongsikchunia-kroppenstedtii]</td></tr>
<tr><td>WP_008379160.1</td><td>HEPN+AAA-ATPase+Alpha+Beta-domain+wHTH+RRM</td><td>Type-1_BR.RM.like.systems</td><td>METHYLASE->HEPN+DUF499*->ACET-><-?||?->ABhydrolase-></td><td>1125</td><td>Bacteria>Terrabacteria_group>Actinomycetota>Actinomycetes>Mycobacteriales</td><td>GCF_000344135.1</td><td>Gordonia_malaquae</td><td>GM1_RS11930</td><td>2383..5760</td><td>410332</td><td>DUF499-domain-containing-protein-[Gordonia-malaquae]</td></tr>
<tr><td>WP_043798089.1</td><td>HEPN+AAA-ATPase+Alpha+Beta-domain+wHTH+RRM</td><td>Type-1_BR.RM.like.systems</td><td><-GDResolvase+HIN-HTH||?->RNase-><-HEPN+DUF499*</td><td>1139</td><td>Bacteria>Terrabacteria_group>Actinomycetota>Actinomycetes>Mycobacteriales</td><td>GCF_009176285.1</td><td>Rhodococcus</td><td>GDN71_RS02225</td><td>3451..6870</td><td>1827</td><td>MULTISPECIES:-Swt1-family-HEPN-domain-containing-protein-[Rhodococcus]</td></tr>
<tr><td>WP_174682959.1</td><td>HEPN+AAA-ATPase+Alpha+Beta-domain+wHTH+RRM</td><td>Type-1_BR.RM.like.systems</td><td><-HEPN+DUF499*<-ParB+HNH<-METHYLASE<-rve_3</td><td>1090</td><td>Bacteria>Terrabacteria_group>Actinomycetota>Actinomycetes>Mycobacteriales</td><td>GCF_001954235.1</td><td>Mycolicibacterium_conceptionense</td><td>A5746_RS13475</td><td>1..3270</td><td>451644</td><td>Swt1-family-HEPN-domain-containing-protein,-partial-[Mycolicibacterium-conceptionense]</td></tr>
<tr><td>WP_082921219.1</td><td>HEPN+AAA-ATPase+Alpha+Beta-domain+wHTH+RRM</td><td>Type-1_BR.RM.like.systems</td><td>METHYLASE-p->HEPN+DUF499*->MORC->REase-></td><td>1121</td><td>Bacteria>Terrabacteria_group>Actinomycetota>Actinomycetes>Mycobacteriales</td><td>GCF_001651055.1</td><td>Rhodococcus_sp._HS-D2</td><td>A1E04_RS27825</td><td>597..3962</td><td>1384636</td><td>Swt1-family-HEPN-domain-containing-protein-[Rhodococcus-sp.-HS-D2]</td></tr>
<tr><td>WP_026061790.1</td><td>HEPN+AAA-ATPase+Alpha+Beta-domain+wHTH+RRM</td><td>Type-1_BR.RM.like.systems</td><td><-DNA_methylase||REase-><-HEPN+DUF499*</td><td>1135</td><td>Bacteria>Terrabacteria_group>Actinomycetota>Actinomycetes>Mycobacteriales</td><td>GCF_000239135.1</td><td>Rhodococcus_rhodochrous</td><td>RHODODRAFT_RS0223985</td><td>1954..5361</td><td>1829</td><td>Swt1-family-HEPN-domain-containing-protein-[Rhodococcus-rhodochrous]</td></tr>
<tr><td>WP_068276445.1</td><td>HEPN+AAA-ATPase+Alpha+Beta-domain+wHTH+RRM</td><td>Type-1_BR.RM.like.systems</td><td>METHYLASE->ParB+HNH->HEPN+DUF499*-></td><td>1122</td><td>Bacteria>Terrabacteria_group>Actinomycetota>Actinomycetes>Mycobacteriales</td><td>GCF_001646865.1</td><td>Aldersonia_kunmingensis</td><td>A8G08_RS16790</td><td>4064..7432</td><td>408066</td><td>Swt1-family-HEPN-domain-containing-protein-[Aldersonia-kunmingensis]</td></tr>
</table>

| Protein-ID | DUF499-architecture | System | Gene-Neighbourhoods | Length | Taxonomy | Assembly-ID | Strain | Locus-Tag | Genomic-coordinates | TaxID | Fasta-defline |
|---|---|---|---|---|---|---|---|---|---|---|---|
| WP_021798346.1 | HEPN+AAA-ATPase+Alpha+Beta-domain+wHTH+RRM | Type-1_BR.RM.like.systems | TUDOR+SF1-SF2-Helicase+iREase+CTD+REase->METHYLASE->wHTH+REase->HEPN+DUF499*->SF1-SF2-Helicase+REase-p->SF1-SF2-Helicase+REase-p->HTH_28+HTH_33+DDE_3->SF1-SF2-Helicase-p-><-Synaptojanin | 1128 | Bacteria>Terrabacteria_group>Actinomycetota>Actinomycetes>Propionibacteriales | GCF_000478805.1 | Propionibacterium_acidifaciens | HMPREF0682_RS10330 | 20512..23898 | 556499 | Swt1-family-HEPN-domain-containing-protein-[Propionibacterium-acidifaciens] |
| WP_028701426.1 | HEPN+AAA-ATPase+Alpha+Beta-domain+wHTH+RRM | Type-1_BR.RM.like.systems | <-SF1-SF2-Helicase<-MORC+SF1-SF2-Helicase\|\|DNA_methylase-><-DNA_methylase\|\|METHYLASE->HEPN+DUF499*->MORC->REase->HTH+SF1-SF2-Helicase+REase->REase-><-ParB+HNH-p<-ParB+HNH-p | 1129 | Bacteria>Terrabacteria_group>Actinomycetota>Actinomycetes>Propionibacteriales | GCF_000427845.1 | Acidipropionibacterium_acidipropionici | ASQ49_RS03445 | 772026..775415 | 1748 | DUF499-domain-containing-protein-[Acidipropionibacterium-acidipropionici] |
| WP_207473688.1 | HEPN+AAA-ATPase+Alpha+Beta-domain+wHTH+RRM | Type-1_BR.RM.like.systems | ABhydrolase->XIS-HTH->TUDOR+SF1-SF2-Helicase+iREase+CTD+REase->METHYLASE->wHTH+REase->HEPN+DUF499*->REase->HEPN+REase-><-DNA_methylase\|\|TrwC+SF1-SF2-Helicase-> | 1131 | Bacteria>Terrabacteria_group>Actinomycetota>Actinomycetes>Propionibacteriales | GCF_017355265.1 | Tessaracoccus_sp._SD287 | IPV09_RS03380 | 69383..72778 | 2782008 | DUF499-domain-containing-protein-[Tessaracoccus-sp.-SD287] |
| WP_166139838.1 | HEPN+AAA-ATPase+Alpha+Beta-domain+wHTH+RRM | Type-1_BR.RM.like.systems | <-SLATT\|\|?-><-NTD+SF1-SF2-Helicase\|\|TUDOR+SF1-SF2-Helicase+iREase+CTD+REase->METHYLASE->ParB+HNH->HEPN+DUF499*-><-?<-HNH\|\|?->?->?->REase+35exo+Beta-Barrel+SF1-SF2-Helicase-> | 1120 | Bacteria>Terrabacteria_group>Actinomycetota>Actinomycetes>Propionibacteriales | GCF_011420305.2 | Nocardioides_ochotonae | HBO46_RS08105 | 1673933..1677295 | 2685869 | Swt1-family-HEPN-domain-containing-protein-[Nocardioides-ochotonae] |
| WP_131171419.1 | HEPN+AAA-ATPase+Alpha+Beta-domain+wHTH+RRM | Type-1_BR.RM.like.systems | <-MACRODOMAIN+SIR2<-HTH+SF1-SF2-Helicase+REase<-REase<-MORC<-HEPN+DUF499*<-METHYLASE<-METHYLASE-p\|\|DNA_methylase-><-SbcC<-SbcC\|\|TrwC+SF1-SF2-helicase+TOPRIM-> | 1146 | Bacteria>Terrabacteria_group>Actinomycetota>Actinomycetes>Propionibacteriales | GCF_900182665.1 | Propioniciclava_tarda | FN821_RS04755 | 6296..9736 | 433330 | Swt1-family-HEPN-domain-containing-protein-[Propioniciclava-tarda] |
| MBM0123458.1 | HEPN+AAA-ATPase+Alpha+Beta-domain+wHTH+RRM | Type-1_BR.RM.like.systems | <-XIS-HTH<-XIS-HTH<-TrwC+SF1-SF2-helicase\|\|HEPN+Mrr_N-><-HEPN+DUF499*<-ParB+HNH<-METHYLASE<-TUDOR+SF1-SF2-Helicase+iREase+CTD+REase\|\|NTD+SF1-SF2-Helicase-> | 1119 | Bacteria>Terrabacteria_group>Actinomycetota>Actinomycetes>Propionibacteriales | GCA_016803235.1 | Pimelobacter_simplex | IMY96_01775 | 51221..54580 | 2045 | DUF499-domain-containing-protein-[Pimelobacter-simplex] |
| WP_219081771.1 | HEPN+AAA-ATPase+Alpha+Beta-domain+wHTH+RRM | Type-1_BR.RM.like.systems | ABC-ATPase+SbcC->ABC-ATPase+SbcC-><-NTD+SF1-SF2-Helicase<-NACHT\|\|TUDOR+SF1-SF2-Helicase+iREase+CTD+REase->METHYLASE->HEPN+DUF499*->?-><-ABhydrolase<-TetR-HTH | 1121 | Bacteria>Terrabacteria_group>Actinomycetota>Actinomycetes>Propionibacteriales | GCF_019316905.1 | Tessaracoccus_palaemonis | KDB89_RS13130 | 2884932..2888297 | 2829499 | Swt1-family-HEPN-domain-containing-protein-[Tessaracoccus-sp.-J1M15] |
| MBN9103777.1 | HEPN+AAA-ATPase+Alpha+Beta-domain+wHTH+RRM | Type-1_BR.RM.like.systems | TUDOR+SF1-SF2-Helicase+iREase+CTD+REase->UDG-conflict->METHYLASE->HEPN+DUF499*->ParB+HNH->?->SF1-SF2-Helicase-><-REase+PNTD1+PNTD2\|\|ACET-><-Phage_integrase | 1117 | Bacteria>Terrabacteria_group>Actinomycetota>Actinomycetes>Propionibacteriales | GCA_017306015.1 | Propionibacteriaceae_bacterium | JOI14_02120 | 371284..374637 | 2021380 | DUF499-domain-containing-protein-[Propionibacteriaceae-bacterium] |
| WP_257750656.1 | HEPN+AAA-ATPase+Alpha+Beta-domain+wHTH+RRM | Type-1_BR.RM.like.systems | TrwC+SF1-SF2-Helicase->?-><-NTD+SF1-SF2-Helicase\|\|TUDOR+SF1-SF2-Helicase+iREase+CTD+REase->METHYLASE->HEPN+DUF499*->KAP-NTPase->?->PAPS_reduct->?->NERD-> | 1119 | Bacteria>Terrabacteria_group>Actinomycetota>Actinomycetes>Propionibacteriales | GCF_024662235.1 | Pimelobacter_simplex | MOM48_RS07490 | 1563266..1566625 | 2045 | Swt1-family-HEPN-domain-containing-protein-[Pimelobacter-simplex] |
| WP_203043111.1 | HEPN+AAA-ATPase+Alpha+Beta-domain+wHTH+RRM | Type-1_BR.RM.like.systems | <-XIS-HTH<-XIS-HTH<-TrwC+SF1-SF2-helicase<-HEPN+DUF499*<-ParB+HNH<-METHYLASE<-TUDOR+SF1-SF2-Helicase+iREase+CTD+REase\|\|NTD+SF1-SF2-Helicase-> | 1119 | Bacteria>Terrabacteria_group>Actinomycetota>Actinomycetes>Propionibacteriales | - | Pimelobacter_simplex | IMY96_RS01775 | 51221..54580 | 2045 | Swt1-family-HEPN-domain-containing-protein-[Pimelobacter-simplex] |
| WP_240236717.1 | HEPN+AAA-ATPase+Alpha+Beta-domain+wHTH+RRM | Type-1_BR.RM.like.systems | <-wHTH(SBD)+HNH<-NTD+SF1-SF2-Helicase\|\|TUDOR+SF1-SF2-Helicase+iREase+CTD+REase->METHYLASE->ParB+HNH->HEPN+DUF499*-><-Peptidase\|\|?->?->?->METHYLASE-> | 1117 | Bacteria>Terrabacteria_group>Actinomycetota>Actinomycetes>Propionibacteriales | GCF_022436705.1 | Nocardioides_sp._CFH_31398 | MJ876_RS12955 | 81599..84952 | 2919579 | Swt1-family-HEPN-domain-containing-protein-[Nocardioides-sp.-CFH-31398] |
| NNG19379.1 | HEPN+AAA-ATPase+Alpha+Beta-domain+wHTH+RRM | Type-1_BR.RM.like.systems | LexA-HTH+LexA-protease-><-?<-35exo+SF1-SF2-Helicase<-?<-HEPN+DUF499*<-METHYLASE\|\|DUF222+HNH-><-TUDOR+SF1-SF2-Helicase+iREase+CTD+REase\|\|HTH_3-> | 1110 | Bacteria>Terrabacteria_group>Actinomycetota>Actinomycetes>Propionibacteriales | GCA_013002685.1 | Naumannella_sp._ID2617S | HJ590_07255 | 1496426..1499758 | 1656893 | ATP-binding-protein-[Naumannella-sp.-ID2617S] |
| MAO81794.1 | HEPN+AAA-ATPase+Alpha+Beta-domain+wHTH+RRM | Type-1_BR.RM.like.systems | <-DUF1679<-SIR2<-NERD<-?<-?<-?<-HEPN+DUF499*<-METHYLASE<-TUDOR+SF1-SF2-Helicase+iREase+CTD+REase\|\|NTD+SF1-SF2-Helicase-><-?<-?<-TrwC+SF1-SF2-Helicase | 1117 | Bacteria>Terrabacteria_group>Actinomycetota>Actinomycetes>Propionibacteriales | GCA_002693615.1 | Nocardioides_sp. | CMH82_14275 | 37872..41225 | 35761 | AAA-family-ATPase-[Nocardioides-sp.] |
| WP_040158482.1 | HEPN+AAA-ATPase+Alpha+Beta-domain+wHTH+RRM | Type-1_BR.RM.like.systems | HTH+MPTase->?->?->HTH_Tnp_1+HTH_21+rve+rve_2-><-HEPN+DUF499*<-wHTH+REase<-METHYLASE<-TUDOR+SF1-SF2-Helicase+iREase+CTD+REase<-XIS-HTH<-tRNA<-?<-ACET | 1132 | Bacteria>Terrabacteria_group>Actinomycetota>Actinomycetes>Propionibacteriales | GCF_902375435.1 | Nigerium_massiliense | BN1187_RS02660 | 259497..262895 | 1522317 | DUF499-domain-containing-protein-[Nigerium-massiliense] |
| WP_194694759.1 | HEPN+AAA-ATPase+Alpha+Beta-domain+wHTH+RRM | Type-1_BR.RM.like.systems | <-DOC<-?<-tRNA<-DDE_Tnp_1_4<-?\|\|TUDOR+SF1-SF2-Helicase+iREase+CTD+REase->METHYLASE->HEPN+DUF499*-><-?<-?\|\|?->ParB+HNH->?->Phage_int_SAM_1+Phage_integrase-> | 1113 | Bacteria>Terrabacteria_group>Actinomycetota>Actinomycetes>Propionibacteriales | GCF_015352445.1 | Nocardioides_agariphilus | ISU10_RS02340 | 86950..90291 | 433664 | DUF499-domain-containing-protein-[Nocardioides-agariphilus] |
| WP_119715651.1 | HEPN+AAA-ATPase+Alpha+Beta-domain+wHTH+RRM | Type-1_BR.RM.like.systems | TUDOR+SF1-SF2-Helicase+iREase+CTD+REase->METHYLASE->HEPN->wHTH+REase->?->PemK_toxin->HEPN+DUF499*->?-><-?\|\|ArsR-HTH-><-ABhydrolase<-TetR-HTH | 1133 | Bacteria>Terrabacteria_group>Actinomycetota>Actinomycetes>Propionibacteriales | GCF_900289195.1 | Propionibacterium_ruminifibrarum | PROPJV5_RS07045 | 37471..40872 | 1962131 | Swt1-family-HEPN-domain-containing-protein-[Propionibacterium-ruminifibrarum] |
| MBE6477658.1 | HEPN+AAA-ATPase+Alpha+Beta-domain+wHTH+RRM | Type-1_BR.RM.like.systems | TUDOR+SF1-SF2-Helicase+iREase+CTD+REase->METHYLASE->HEPN->wHTH+REase->?->PemK_toxin->HEPN+DUF499*->?->TetR-HTH->?-><-?\|\|ArsR-HTH-><-ABhydrolase | 1088 | Bacteria>Terrabacteria_group>Actinomycetota>Actinomycetes>Propionibacteriales | GCA_015062705.1 | Propionibacteriaceae_bacterium | E7Z97_06285 | 37437..40699 | 2021380 | ATP-binding-protein-[Propionibacteriaceae-bacterium] |
| WP_277066086.1 | HEPN+AAA-ATPase+Alpha+Beta-domain+wHTH+RRM | Type-1_BR.RM.like.systems | Synaptojanin-><-SF1-SF2-Helicase+REase<-HEPN+DUF499*<-wHTH+REase<-METHYLASE<-TUDOR+SF1-SF2-Helicase+iREase+CTD+REase\|\|?-><-?\|\|Peptidase-> | 1128 | Bacteria>Terrabacteria_group>Actinomycetota>Actinomycetes>Propionibacteriales | GCF_938034675.1 | Propionibacterium_acidifaciens | P7A72_RS09150 | 66970..70356 | 556499 | Swt1-family-HEPN-domain-containing-protein-[Propionibacterium-acidifaciens] |
| WP_130874284.1 | HEPN+AAA-ATPase+Alpha+Beta-domain+wHTH+RRM | Type-1_BR.RM.like.systems | TetR-HTH-><-?\|\|?-><-ACET<-KAP-NTPase<-HEPN+DUF499*<-wHTH+REase<-METHYLASE<-TUDOR+SF1-SF2-Helicase+iREase+CTD+REase<-MORC<-METHYLASE-p | 1129 | Bacteria>Terrabacteria_group>Actinomycetota>Actinomycetes>Propionibacteriales | GCF_900607225.1 | [Pseudopropionibacterium]_massiliense | E0E39_RS08095 | 1820154..1823543 | 2220000 | DUF499-domain-containing-protein-[[Pseudopropionibacterium] |
| MAS55988.1 | HEPN+AAA-ATPase+Alpha+Beta-domain+wHTH+RRM | Type-1_BR.RM.like.systems | XIS-HTH->METHYLASE-p-><-?<-MORC+HISKIN<-HEPN+DUF499*<-ParB+HNH<-METHYLASE<-UDG-conflict<-TUDOR+SF1-SF2-Helicase+iREase+CTD+REase-p | 1116 | Bacteria>Terrabacteria_group>Actinomycetota>Actinomycetes>Propionibacteriales | GCA_002698575.1 | Pimelobacter_sp. | CMJ44_15460 | 17699..21049 | 51662 | AAA-family-ATPase-[Pimelobacter-sp.] |
| WP_119162165.1 | HEPN+AAA-ATPase+Alpha+Beta-domain+wHTH+RRM | Type-1_BR.RM.like.systems | <-TetR-HTH\|\|?-><-SF1-SF2-Helicase<-OLD-ABC+TOPRIM<-HEPN+DUF499*<-PemK_toxin<-?<-wHTH+REase<-METHYLASE<-TUDOR+SF1-SF2-Helicase+iREase+CTD+REase | 1133 | Bacteria>Terrabacteria_group>Actinomycetota>Actinomycetes>Propionibacteriales | GCF_900637915.1 | Propionibacterium_australiense | EL282_RS10960 | 2474046..2477447 | 119981 | DUF499-domain-containing-protein-[Propionibacterium-australiense] |
| WP_182784909.1 | HEPN+AAA-ATPase+Alpha+Beta-domain+wHTH+RRM | Type-1_BR.RM.like.systems | <-SF1-SF2-Helicase<-OLD-ABC+TOPRIM<-HEPN+DUF499*<-METHYLASE<-TUDOR+SF1-SF2-Helicase+iREase+CTD+REase<-SF1-SF2-Helicase-p\|\|Glyoxalase-> | 1133 | Bacteria>Terrabacteria_group>Actinomycetota>Actinomycetes>Propionibacteriales | GCF_014164725.1 | Tessaracoccus_sp._MC1627 | H5399_RS17220 | 4312..7713 | 2760312 | DUF499-domain-containing-protein-[Tessaracoccus-sp.-MC1627] |
| WP_124237129.1 | HEPN+AAA-ATPase+Alpha+Beta-domain+wHTH+RRM | Type-1_BR.RM.like.systems | <-REase\|\|NTP_transf_2->tRNA-><-NTD+SF1-SF2-Helicase\|\|TUDOR+SF1-SF2-Helicase+iREase+CTD+REase->METHYLASE->ParB+HNH->HEPN+DUF499*-> | 1124 | Bacteria>Terrabacteria_group>Actinomycetota>Actinomycetes>Propionibacteriales | GCF_003850045.1 | Aeromicrobium_camelliae | EHW97_RS10560 | 83306..86680 | 1538144 | Swt1-family-HEPN-domain-containing-protein-[Aeromicrobium-camelliae] |
| WP_154690370.1 | HEPN+AAA-ATPase+Alpha+Beta-domain+wHTH+RRM | Type-1_BR.RM.like.systems | PAIREDC-HTH-><-?<-HTH+SF1-SF2-Helicase+REase<-REase<-MORC<-HEPN+DUF499*<-METHYLASE-p<-METHYLASE<-REase<-PIN\|\|DNA_methylase-> | 1132 | Bacteria>Terrabacteria_group>Actinomycetota>Actinomycetes>Propionibacteriales | GCF_009707365.1 | Microlunatus_sp._Gsoil_973 | GJV80_RS18445 | 3917369..3920767 | 2672569 | DUF499-domain-containing-protein-[Microlunatus-sp.-Gsoil-973] |
| WP_078920357.1 | HEPN+AAA-ATPase+Alpha+Beta-domain+wHTH+RRM | Type-1_BR.RM.like.systems | <-TrwC+SF1-SF2-helicase+TOPRIM<-DNA_methylase\|\|REase->METHYLASE->HEPN+DUF499*->MORC->REase->HTH+SF1-SF2-Helicase+REase-> | 1132 | Bacteria>Terrabacteria_group>Actinomycetota>Actinomycetes>Propionibacteriales | GCF_002022225.1 | Tessaracoccus | BKM78_RS14510 | 3153779..3157177 | 72763 | MULTISPECIES:-Swt1-family-HEPN-domain-containing-protein-[Tessaracoccus] |
| WP_123224702.1 | HEPN+AAA-ATPase+Alpha+Beta-domain+wHTH+RRM | Type-1_BR.RM.like.systems | <-NERD\|\|REase+SAVED->?->?-><-TIR<-HEPN+DUF499*<-METHYLASE\|\|?-><-TUDOR+SF1-SF2-Helicase+iREase+CTD+REase\|\|NTD+SF1-SF2-Helicase-> | 1120 | Bacteria>Terrabacteria_group>Actinomycetota>Actinomycetes>Propionibacteriales | GCF_003725535.1 | Nocardioides_pocheonensis | EFL26_RS20130 | 110530..113892 | 661485 | DUF499-domain-containing-protein-[Marmoricola-pocheonensis] |
| WP_115850048.1 | HEPN+AAA-ATPase+Alpha+Beta-domain+wHTH+RRM | Type-1_BR.RM.like.systems | <-Pkinase<-NTP_transf_5\|\|Pkinase->?->?-><-SF1-SF2-Helicase<-HEPN+DUF499*<-METHYLASE<-TUDOR+SF1-SF2-Helicase+iREase+CTD+REase | 1124 | Bacteria>Terrabacteria_group>Actinomycetota>Actinomycetes>Propionibacteriales | GCF_003386795.1 | Thermasporomyces_composti | DFJ64_RS08950 | 2047152..2050526 | 696763 | Swt1-family-HEPN-domain-containing-protein-[Thermasporomyces-composti] |
| WP_255892000.1 | HEPN+AAA-ATPase+Alpha+Beta-domain+wHTH+RRM | Type-1_BR.RM.like.systems | <-NTD+SF1-SF2-Helicase\|\|TUDOR+SF1-SF2-Helicase+iREase+CTD+REase->METHYLASE->HEPN+DUF499*->MORC+HISKIN->?-><-METHYLASE-p | 1104 | Bacteria>Terrabacteria_group>Actinomycetota>Actinomycetes>Propionibacteriales | GCF_024436375.1 | Nocardioides_astragali | J3P44_RS18825 | 368600..371914 | 1776736 | Swt1-family-HEPN-domain-containing-protein-[Nocardioides-astragali] |
| WP_061786876.1 | HEPN+AAA-ATPase+Alpha+Beta-domain+wHTH+RRM | Type-1_BR.RM.like.systems | REase->TUDOR+SF1-SF2-Helicase+iREase+CTD+REase->METHYLASE->wHTH+REase->HEPN+DUF499*->KAP-NTPase->ACET-><-?\|\|?-><-TetR-HTH | 1129 | Bacteria>Terrabacteria_group>Actinomycetota>Actinomycetes>Propionibacteriales | GCF_001592325.1 | Arachnia_propionica | PP13_RS01005 | 243859..247248 | 1750 | DUF499-domain-containing-protein-[Arachnia-propionica] |
| MBK8462833.1 | HEPN+AAA-ATPase+Alpha+Beta-domain+wHTH+RRM | Type-1_BR.RM.like.systems | <-REase<-?<-HEPN\|\|TUDOR+SF1-SF2-Helicase+iREase+CTD+REase->METHYLASE->ParB+HNH->HEPN+DUF499*->?-><-Peptidase_M48<-?<-?\|\|TPR-> | 1117 | Bacteria>Terrabacteria_group>Actinomycetota>Actinomycetes>Propionibacteriales | GCA_016711325.1 | Nigerium_sp. | IPL36_07335 | 626569..629922 | 2042655 | DUF499-domain-containing-protein-[Nigerium-sp.] |
| WP_126378241.1 | HEPN+AAA-ATPase+Alpha+Beta-domain+wHTH+RRM | Type-1_BR.RM.like.systems | TetR-HTH-><-?\|\|?-><-ACET<-?<-HEPN+DUF499*<-wHTH+REase<-METHYLASE-p<-METHYLASE-p<-TUDOR+SF1-SF2-Helicase+iREase+CTD+REase | 1129 | Bacteria>Terrabacteria_group>Actinomycetota>Actinomycetes>Propionibacteriales | GCF_900638645.1 | Arachnia_propionica | EL396_RS08525 | 1937379..1940768 | 1750 | DUF499-domain-containing-protein-[Arachnia-propionica] |
| WP_094363616.1 | HEPN+AAA-ATPase+Alpha+Beta-domain+wHTH+RRM | Type-1_BR.RM.like.systems | <-rve_3<-HIN-HTH\|\|?-><-?\|\|TUDOR+SF1-SF2-Helicase+iREase+CTD+REase->METHYLASE->HEPN+DUF499*->SbcC-><-?<-?\|\|?->HEPN->DOC+HTH-> | 1125 | Bacteria>Terrabacteria_group>Actinomycetota>Actinomycetes>Propionibacteriales | GCF_002250625.1 | Enemella_dayhoffiae | CGZ93_RS07965 | 9213..12590 | 2016507 | DUF499-domain-containing-protein-[Enemella-dayhoffiae] |
| WP_246322323.1 | HEPN+AAA-ATPase+Alpha+Beta-domain+wHTH+RRM | Type-1_BR.RM.like.systems | <-REase<-?\|\|?->SIGMA-HTH-><-HEPN+DUF499*<-ParB+HNH<-METHYLASE<-TUDOR+SF1-SF2-Helicase+iREase+CTD+REase\|\|REase-> | 1096 | Bacteria>Terrabacteria_group>Actinomycetota>Actinomycetes>Propionibacteriales | GCF_013407955.1 | Microlunatus_parietis | BKA15_RS11400 | 2395766..2399056 | 682979 | DUF499-domain-containing-protein-[Microlunatus-parietis] |
| WP_232416936.1 | HEPN+AAA-ATPase+Alpha+Beta-domain+wHTH+RRM | Type-1_BR.RM.like.systems | <-HTH+SF1-SF2-Helicase+REase<-REase<-MORC<-HEPN+DUF499*<-METHYLASE<-REase\|\|DNA_methylase-><-?\|\|DUF222+HNH-> | 1136 | Bacteria>Terrabacteria_group>Actinomycetota>Actinomycetes>Propionibacteriales | GCF_024508295.1 | Aeromicrobium_duanguangcaii | NP095_RS05665 | 1110875..1114285 | 2968086 | MULTISPECIES:-Swt1-family-HEPN-domain-containing-protein-[unclassified-Aeromicrobium] |
| WP_014846821.1 | HEPN+AAA-ATPase+Alpha+Beta-domain+wHTH+RRM | Type-1_BR.RM.like.systems | REase->TUDOR+SF1-SF2-Helicase+iREase+CTD+REase->METHYLASE->wHTH+REase->HEPN+DUF499*->?->ACET-><-?\|\|?-><-TetR-HTH | 1129 | Bacteria>Terrabacteria_group>Actinomycetota>Actinomycetes>Propionibacteriales | GCF_000277715.1 | Arachnia_propionica | J5A53_RS06805 | 1506504..1509893 | 1750 | DUF499-domain-containing-protein-[Arachnia-propionica] |
| WP_187768939.1 | HEPN+AAA-ATPase+Alpha+Beta-domain+wHTH+RRM | Type-1_BR.RM.like.systems | <-PIN+PhoH<-?\|\|?-><-?<-?<-HEPN+DUF499*<-METHYLASE<-TUDOR+SF1-SF2-Helicase+iREase+CTD+REase\|\|NTD+SF1-SF2-Helicase-> | 1119 | Bacteria>Terrabacteria_group>Actinomycetota>Actinomycetes>Propionibacteriales | GCF_014490085.1 | Aeromicrobium | GL325_RS06400 | 1283810..1287169 | 2040 | MULTISPECIES:-Swt1-family-HEPN-domain-containing-protein-[unclassified-Aeromicrobium] |
| WP_020385147.1 | HEPN+AAA-ATPase+Alpha+Beta-domain+wHTH+RRM | Type-1_BR.RM.like.systems | NUDIX->tRNA-><-?<-?\|\|TUDOR+SF1-SF2-Helicase+iREase+CTD+REase->METHYLASE->ParB+HNH->HEPN+DUF499*-><-?<-?<-?<-METHYLASE-p | 1116 | Bacteria>Terrabacteria_group>Actinomycetota>Actinomycetes>Propionibacteriales | GCF_000372465.1 | Kribbella_catacumbae | A3ES_RS0102270 | 28038..31388 | 460086 | Swt1-family-HEPN-domain-containing-protein-[Kribbella-catacumbae] |
| WP_167165953.1 | HEPN+AAA-ATPase+Alpha+Beta-domain+wHTH+RRM | Type-1_BR.RM.like.systems | <-Pkinase<-?<-?\|\|TUDOR+SF1-SF2-Helicase+iREase+CTD+REase->METHYLASE->HEPN+DUF499*->?->PIN->?->?-><-Aminotran_1_2 | 1131 | Bacteria>Terrabacteria_group>Actinomycetota>Actinomycetes>Propionibacteriales | GCF_011758645.1 | Brooklawnia_cerclae | FB473_RS07080 | 1539391..1542786 | 349934 | Swt1-family-HEPN-domain-containing-protein-[Brooklawnia-cerclae] |
| WP_125568181.1 | HEPN+AAA-ATPase+Alpha+Beta-domain+wHTH+RRM | Type-1_BR.RM.like.systems | HTH_3->?->?-><-HEPN+DUF499*<-METHYLASE<-TUDOR+SF1-SF2-Helicase+iREase+CTD+REase\|\|NTD+SF1-SF2-Helicase-> | 1121 | Bacteria>Terrabacteria_group>Actinomycetota>Actinomycetes>Propionibacteriales | GCF_003945325.1 | Nocardioides_baekrokdamisoli | KCTC_RS07355 | 1515846..1519211 | 1804624 | Swt1-family-HEPN-domain-containing-protein-[Nocardioides-baekrokdamisoli] |
| WP_211798072.1 | HEPN+AAA-ATPase+Alpha+Beta-domain+wHTH+RRM | Type-1_BR.RM.like.systems | TetR-HTH-><-?\|\|?-><-ACET<-?<-HEPN+DUF499*<-wHTH+REase<-METHYLASE<-TUDOR+SF1-SF2-Helicase+iREase+CTD+REase | 1129 | Bacteria>Terrabacteria_group>Actinomycetota>Actinomycetes>Propionibacteriales | GCF_018128345.1 | Arachnia_propionica | J5A61_RS08580 | 1914777..1918166 | 1750 | DUF499-domain-containing-protein-[Arachnia-propionica] |
| WP_227735039.1 | HEPN+AAA-ATPase+Alpha+Beta-domain+wHTH+RRM | Type-1_BR.RM.like.systems | TUDOR+SF1-SF2-Helicase+iREase+CTD+REase-><-PIN<-DUF433\|\|METHYLASE-><-DOC+HTH\|\|HEPN+DUF499*-> | 1104 | Bacteria>Terrabacteria_group>Actinomycetota>Actinomycetes>Propionibacteriales | GCF_020687685.1 | Tessaracoccus_sp._OS52 | LKO27_RS00980 | 227595..230909 | 2886691 | DUF499-domain-containing-protein-[Tessaracoccus-sp.-OS52] |
| WP_278693137.1 | HEPN+AAA-ATPase+Alpha+Beta-domain+wHTH+RRM | Type-1_BR.RM.like.systems | REase->TUDOR+SF1-SF2-Helicase+iREase+CTD+REase->METHYLASE->wHTH+REase->HEPN+DUF499*-> | 1129 | Bacteria>Terrabacteria_group>Actinomycetota>Actinomycetes>Propionibacteriales | GCF_915063725.1 | Arachnia_propionica | QDI34_RS08815 | 15722..19111 | 1750 | DUF499-domain-containing-protein-[Arachnia-propionica] |
| WP_182766932.1 | HEPN+AAA-ATPase+Alpha+Beta-domain+wHTH+RRM | Type-1_BR.RM.like.systems | <-HTH+SF1-SF2-Helicase+REase<-REase<-HEPN+DUF499*<-METHYLASE<-REase\|\|DNA_methylase-> | 1129 | Bacteria>Terrabacteria_group>Actinomycetota>Actinomycetes>Propionibacteriales | GCF_014164735.1 | Tessaracoccus_sp._MC1756 | H5393_RS12480 | 209092..212481 | 2760311 | Swt1-family-HEPN-domain-containing-protein-[Tessaracoccus-sp.-MC1756] |
| WP_013863116.1 | HEPN+AAA-ATPase+Alpha+Beta-domain+wHTH+RRM | Type-1_BR.RM.like.systems | PglZ-><-?\|\|?-><-DUF222+HNH\|\|?-><-HEPN+DUF499*<-ParB+HNH<-?<-METHYLASE\|\|wHTH(SBD)+HNH->HTH_19-> | 1117 | Bacteria>Terrabacteria_group>Actinomycetota>Actinomycetes>Propionibacteriales | GCF_000270245.1 | Microlunatus_phosphovorus | MLP_RS10775 | 2379873..2383226 | 29405 | Swt1-family-HEPN-domain-containing-protein-[Microlunatus-phosphovorus] |
| WP_094451589.1 | HEPN+AAA-ATPase+Alpha+Beta-domain+wHTH+RRM | Type-1_BR.RM.like.systems | <-SbcC<-HEPN+DUF499*<-METHYLASE<-TUDOR+SF1-SF2-Helicase+iREase+CTD+REase<-?\|\|HTH-> | 1135 | Bacteria>Terrabacteria_group>Actinomycetota>Actinomycetes>Propionibacteriales | GCF_002250475.1 | Parenemella_sanctibonifatiensis | CGZ92_RS11810 | 2653..6060 | 2016505 | DUF499-domain-containing-protein-[Parenemella-sanctibonifatiensis] |
| MCL2489469.1 | HEPN+AAA-ATPase+Alpha+Beta-domain+wHTH+RRM | Type-1_BR.RM.like.systems | <-TrwC+SF1-SF2-Helicase<-?<-PAPS_reduct<-?<-KAP-NTPase<-HEPN+DUF499*<-HEPN<-METHYLASE | 1120 | Bacteria>Terrabacteria_group>Actinomycetota>Actinomycetes>Propionibacteriales | GCA_009778875.1 | Propionibacteriaceae_bacterium | FWF36_01890 | 13226..16588 | 2021380 | DUF499-domain-containing-protein-[Propionibacteriaceae-bacterium] |
| WP_146174093.1 | HEPN+AAA-ATPase+Alpha+Beta-domain+wHTH+RRM | Type-1_BR.RM.like.systems | HTH+DUF5753->ZnR->?->TPR+TPR+TPR-><-?<-HEPN+DUF499*<-METHYLASE<-TUDOR+SF1-SF2-Helicase+iREase+CTD+REase\|\|XIS-HTH->Phage_int_SAM_1+Phage_integrase-> | 1099 | Bacteria>Terrabacteria_group>Actinomycetota>Actinomycetes>Pseudonocardiales | - | Saccharothrix_carnea | BOI31_12144 | 38628..41927 | 1280637 | DUF499-domain-containing-protein-[Saccharothrix-carnea] |
| WP_106619913.1 | HEPN+AAA-ATPase+Alpha+Beta-domain+wHTH+RRM | Type-1_BR.RM.like.systems | HTH+DUF5753->ZnR->?->TPR+TPR+TPR-><-?<-HEPN+DUF499*<-METHYLASE<-TUDOR+SF1-SF2-Helicase+iREase+CTD+REase\|\|XIS-HTH->Phage_int_SAM_1+Phage_integrase-> | 1114 | Bacteria>Terrabacteria_group>Actinomycetota>Actinomycetes>Pseudonocardiales | GCF_003014735.1 | Saccharothrix_carnea | BOI31_RS36295 | 38628..41972 | 1280637 | DUF499-domain-containing-protein-[Saccharothrix-carnea] |
| PSL51415.1 | HEPN+AAA-ATPase+Alpha+Beta-domain+wHTH+RRM | Type-1_BR.RM.like.systems | HTH+DUF5753->ZnR->?->TPR+TPR+TPR-><-?<-HEPN+DUF499*<-METHYLASE<-TUDOR+SF1-SF2-Helicase+iREase+CTD+REase\|\|XIS-HTH->Phage_int_SAM_1+Phage_integrase-> | 1099 | Bacteria>Terrabacteria_group>Actinomycetota>Actinomycetes>Pseudonocardiales | GCA_003014735.1 | Saccharothrix_carnea | BOI31_12144 | 38628..41927 | 1280637 | putative-AAA+-superfamily-ATPase-[Saccharothrix-carnea] |
| WP_139094500.1 | HEPN+AAA-ATPase+Alpha+Beta-domain+wHTH+RRM | Type-1_BR.RM.like.systems | <-HIN-HTH+HIN-HTH\|\|?->?->?->?->?-><-HEPN+DUF499*<-METHYLASE<-TUDOR+SF1-SF2-Helicase+iREase+CTD+REase<-?<-HIN-HTH+HIN-HTH\|\|ABC-ATPase+SbcC->IclR-HTH-> | 1111 | Bacteria>Terrabacteria_group>Actinomycetota>Actinomycetes>Pseudonocardiales | GCF_006152065.1 | Amycolatopsis_alkalitolerans | FG385_RS00135 | 35621..38956 | 2547244 | DUF499-domain-containing-protein-[Amycolatopsis-alkalitolerans] |
| WP_218590226.1 | HEPN+AAA-ATPase+Alpha+Beta-domain+wHTH+RRM | Type-1_BR.RM.like.systems | TUDOR+SF1-SF2-Helicase+iREase+CTD+REase->METHYLASE->ParB+HNH->HEPN+DUF499*->?-><-TrwC+SF1-SF2-helicase+TOPRIM<-?<-VirD4-TraG+VirD4-FtsK | 1126 | Bacteria>Terrabacteria_group>Actinomycetota>Actinomycetes>Pseudonocardiales | GCF_019263625.1 | Pseudonocardia_oceani | I4I79_RS18830 | 8746..12126 | 2792013 | Swt1-family-HEPN-domain-containing-protein-[Pseudonocardia-oceani] |
| WP_184862258.1 | HEPN+AAA-ATPase+Alpha+Beta-domain+wHTH+RRM | Type-1_BR.RM.like.systems | HIN-HTH+HIN-HTH->?->TUDOR+SF1-SF2-Helicase+iREase+CTD+REase->UDG-conflict->METHYLASE->HEPN+DUF499*-><-?<-?<-PAIREDC-HTH<-?<-HTH<-SIGMA-HTH | 1111 | Bacteria>Terrabacteria_group>Actinomycetota>Actinomycetes>Pseudonocardiales | GCF_014203355.1 | Kutzneria_kofuensis | BJ998_RS15140 | 3332051..3335386 | 103725 | DUF499-domain-containing-protein-[Kutzneria-kofuensis] |
| WP_085916424.1 | HEPN+AAA-ATPase+Alpha+Beta-domain+wHTH+RRM | Type-1_BR.RM.like.systems | DNA-glycosylase->TUDOR+SF1-SF2-Helicase+iREase+CTD+REase->METHYLASE->ParB+HNH->HEPN+DUF499*->?-><-TrwC+SF1-SF2-helicase+TOPRIM | 1135 | Bacteria>Terrabacteria_group>Actinomycetota>Actinomycetes>Pseudonocardiales | GCF_003945385.1 | Pseudonocardia | PSA01_RS26720 | 8547..11954 | 1847 | MULTISPECIES:-Swt1-family-HEPN-domain-containing-protein-[Pseudonocardia] |
| WP_061989441.1 | HEPN+AAA-ATPase+Alpha+Beta-domain+wHTH+RRM | Type-1_BR.RM.like.systems | HIN-HTH+HIN-HTH->?->TUDOR+SF1-SF2-Helicase+iREase+CTD+REase->UDG-conflict->METHYLASE->HEPN+DUF499*-><-?\|\|Phage_integrase->?->?-><-tRNA<-?<-ACET | 1112 | Bacteria>Terrabacteria_group>Actinomycetota>Actinomycetes>Pseudonocardiales | GCF_900113515.1 | Amycolatopsis_regifaucium | AVL48_RS05675 | 397234..400572 | 546365 | DUF499-domain-containing-protein-[Amycolatopsis-regifaucium] |
| MBV9142989.1 | HEPN+AAA-ATPase+Alpha+Beta-domain+wHTH+RRM | Type-1_BR.RM.like.systems | <-HEPN+DUF499*<-METHYLASE-p<-TUDOR+SF1-SF2-Helicase+iREase+CTD+REase<-Schlafen+HATPase+HTH<-Methyltransf\|\|METHYLASE-p-><-PIN | 1073 | Bacteria>Terrabacteria_group>Actinomycetota>Actinomycetes>Pseudonocardiales | GCA_019247305.1 | Pseudonocardiales_bacterium | JO115_19100 | 30..3251 | 2201155 | DUF499-domain-containing-protein-[Pseudonocardiales-bacterium] |
| WP_211261518.1 | HEPN+AAA-ATPase+Alpha+Beta-domain+wHTH+RRM | Type-1_BR.RM.like.systems | <-ArsR-HTH<-?<-?\|\|?->TUDOR+SF1-SF2-Helicase+iREase+CTD+REase->METHYLASE->HEPN+DUF499*->?-><-?<-HTH+DUF5753<-TetR-HTH | 1100 | Bacteria>Terrabacteria_group>Actinomycetota>Actinomycetes>Pseudonocardiales | GCF_000620785.1 | Pseudonocardia_acaciae | N912_RS0136125 | 176490..179792 | 551276 | DUF499-domain-containing-protein-[Pseudonocardia-acaciae] |
| RAS61288.1 | HEPN+AAA-ATPase+Alpha+Beta-domain+wHTH+RRM | Type-1_BR.RM.like.systems | TUDOR+SF1-SF2-Helicase+iREase+CTD+REase->METHYLASE-p->METHYLASE-p->HEPN+DUF499*-><-?<-HTH+DUF5753\|\|?->?->?->HTH-> | 1111 | Bacteria>Terrabacteria_group>Actinomycetota>Actinomycetes>Pseudonocardiales | GCA_003269295.1 | Lentzea_atacamensis | C8D87_110236 | 241893..245228 | 531938 | putative-AAA+-superfamily-ATPase-[Lentzea-atacamensis] |
| WP_215732591.1 | HEPN+AAA-ATPase+Alpha+Beta-domain+wHTH+RRM | Type-1_BR.RM.like.systems | TUDOR+SF1-SF2-Helicase+iREase+CTD+REase->METHYLASE-p->METHYLASE-p->HEPN+DUF499*->?-><-?<-HTH+DUF5753 | 1100 | Bacteria>Terrabacteria_group>Actinomycetota>Actinomycetes>Pseudonocardiales | GCF_003269295.1 | Lentzea_atacamensis | C8D87_RS36395 | 241926..245228 | 531938 | DUF499-domain-containing-protein-[Lentzea-atacamensis] |
| WP_019873098.1 | HEPN+AAA-ATPase+Alpha+Beta-domain+wHTH+RRM | Type-1_BR.RM.like.systems | <-ArdCN+MPTase<-?<-HEPN+DUF499*<-METHYLASE<-TUDOR+SF1-SF2-Helicase+iREase+CTD+REase-p<-DnaB+DnaB<-VirB4-FtsK<-?<-LysM+TPR | 1113 | Bacteria>Terrabacteria_group>Actinomycetota>Actinomycetes>Sporichthyales | GCF_000384115.1 | Sporichthya_polymorpha | SPOPO_RS0101950 | 384127..387468 | 35751 | DUF499-domain-containing-protein-[Sporichthya-polymorpha] |
| WP_131964568.1 | HEPN+AAA-ATPase+Alpha+Beta-domain+wHTH+RRM | Type-1_BR.RM.like.systems | <-Schlafen+HATPase+HTH<-Schlafen+HATPase+HTH<-ParB+HNH\|\|TUDOR+SF1-SF2-Helicase+iREase+CTD+REase->METHYLASE->HEPN+DUF499*->Phage_int_SAM_1+Phage_integrase->Phage_integrase-><-NUDIX<-SAM-methylase | 1135 | Bacteria>Terrabacteria_group>Actinomycetota>Actinomycetes>Streptosporangiales | GCF_004348735.1 | Actinomadura_sp._KC06 | E1287_RS09895 | 20684..24091 | 2530369 | DUF499-domain-containing-protein-[Actinomadura-sp.-KC06] |
| WP_076440368.1 | HEPN+AAA-ATPase+Alpha+Beta-domain+wHTH+RRM | Type-1_BR.RM.like.systems | <-Primpol+D5N+D5ATPase<-DUF4011+SF1-SF2-Helicase+REase<-SF1-SF2-helicase+DUF1998<-?<-?<-HEPN+DUF499*<-METHYLASE<-TUDOR+SF1-SF2-Helicase+iREase+CTD+REase<-?<-REase+N6_MTase<-Pkinase | 1125 | Bacteria>Terrabacteria_group>Actinomycetota>Actinomycetes>Streptosporangiales | GCF_900156315.1 | Microbispora_rosea | BXA11_RS31445 | 30499..33876 | 58117 | Swt1-family-HEPN-domain-containing-protein-[Microbispora-rosea] |
| GIH50617.1 | HEPN+AAA-ATPase+Alpha+Beta-domain+wHTH+RRM | Type-1_BR.RM.like.systems | <-Primpol+D5N+D5ATPase<-DUF4011+SF1-SF2-Helicase+REase<-SF1-SF2-helicase+DUF1998<-?<-?<-HEPN+DUF499*<-METHYLASE<-TUDOR+SF1-SF2-Helicase+iREase+CTD+REase<-?<-REase+N6_MTase<-Pkinase | 1125 | Bacteria>Terrabacteria_group>Actinomycetota>Actinomycetes>Streptosporangiales | GCA_016863055.1 | Microbispora_rosea_subsp._rosea | Mro03_57960 | 50675..54052 | 58118 | hypothetical-protein-Mro03_57960-[Microbispora-rosea-subsp.-rosea] |
| WP_184983644.1 | HEPN+AAA-ATPase+Alpha+Beta-domain+wHTH+RRM | Type-1_BR.RM.like.systems | 35exo-><-wHTH(SBD)+HNH<-?<-SF1-SF2-helicase+DUF1998<-?<-ParB+HNH<-HEPN+DUF499*<-METHYLASE<-TUDOR+SF1-SF2-Helicase+iREase+CTD+REase<-?<-REase+SF1-SF2-Helicase<-HTH_3<-ABhydrolase | 1124 | Bacteria>Terrabacteria_group>Actinomycetota>Actinomycetes>Streptosporangiales | GCF_014207705.1 | Sphaerisporangium_rubeum | BJ992_RS21150 | 4926161..4929535 | 321317 | Swt1-family-HEPN-domain-containing-protein-[Sphaerisporangium-rubeum] |
| WP_132634803.1 | HEPN+AAA-ATPase+Alpha+Beta-domain+wHTH+RRM | Type-1_BR.RM.like.systems | <-REase+N6_MTase<-TRD+TRD<-REase+SF1-SF2-Helicase<-REase+SF1-SF2-Helicase\|\|TUDOR+SF1-SF2-Helicase+iREase+CTD+REase->METHYLASE->HEPN+DUF499*-><-DndB | 1126 | Bacteria>Terrabacteria_group>Actinomycetota>Actinomycetes>Streptosporangiales | GCF_004352805.1 | Nonomuraea_mesophila | E1295_RS29355 | 30406..33786 | 2530382 | Swt1-family-HEPN-domain-containing-protein-[Nonomuraea-mesophila] |
| WP_030913330.1 | HEPN+AAA-ATPase+Alpha+Beta-domain+wHTH+RRM | Type-1_BR.RM.like.systems | <-REase+SF1-SF2-Helicase<-MazG\|\|UDG-conflict->?->TUDOR+SF1-SF2-Helicase+iREase+CTD+REase->METHYLASE->HEPN+DUF499*->?->?->?-><-DUF397<-HTH+DUF5753\|\|ST-Hiskin-> | 1133 | Bacteria>Terrabacteria_group>Actinomycetota>Actinomycetes>Streptosporangiales | GCF_000721075.1 | Streptosporangium_amethystogenes | IG98_RS0118225 | 190587..193988 | 2002 | Swt1-family-HEPN-domain-containing-protein-[Streptosporangium-amethystogenes] |
| WP_185051091.1 | HEPN+AAA-ATPase+Alpha+Beta-domain+wHTH+RRM | Type-1_BR.RM.like.systems | <-HDRnase<-GNTR-HTH<-?\|\|HTH+HKD-DNase->TUDOR+SF1-SF2-Helicase+iREase+CTD+REase->METHYLASE->HEPN+DUF499*->tRNA-><-?\|\|MODE-HTH+PBPII->ABhydrolase-> | 1126 | Bacteria>Terrabacteria_group>Actinomycetota>Actinomycetes>Streptosporangiales | GCF_014202315.1 | Thermocatellispora_tengchongensis | HNP84_RS19220 | 438143..441523 | 1073253 | DUF499-domain-containing-protein-[Thermocatellispora-tengchongensis] |
| WP_203946296.1 | HEPN+AAA-ATPase+Alpha+Beta-domain+wHTH+RRM | Type-1_BR.RM.like.systems | REase+N6_MTase->?->TUDOR+SF1-SF2-Helicase+iREase+CTD+REase->METHYLASE->HEPN+DUF499*->?->ParB+HNH->?->SF1-SF2-helicase+DUF1998-><-?\|\|OLD-ABC+TOPRIM-> | 1129 | Bacteria>Terrabacteria_group>Actinomycetota>Actinomycetes>Streptosporangiales | GCF_016863335.1 | Planotetraspora_thailandica | Pth03_RS22295 | 43771..47160 | 487172 | Swt1-family-HEPN-domain-containing-protein-[Planotetraspora-thailandica] |
| WP_179824532.1 | HEPN+AAA-ATPase+Alpha+Beta-domain+wHTH+RRM | Type-1_BR.RM.like.systems | <-ST-Hiskin\|\|HTH+DUF5753->DUF397-><-?<-ParB+HNH<-HEPN+DUF499*<-METHYLASE<-TUDOR+SF1-SF2-Helicase+iREase+CTD+REase<-?<-UDG-conflict | 1132 | Bacteria>Terrabacteria_group>Actinomycetota>Actinomycetes>Streptosporangiales | GCF_013409935.1 | Streptosporangium_sandarakinum | HDA43_RS21565 | 651273..654671 | 1260955 | Swt1-family-HEPN-domain-containing-protein-[Streptosporangium-sandarakinum] |
| GHJ54460.1 | HEPN+AAA-ATPase+Alpha+Beta-domain+wHTH+RRM | Type-1_BR.RM.like.systems | <-REase<-?<-NTD+SF1-SF2-Helicase\|\|TUDOR+SF1-SF2-Helicase+iREase+CTD+REase->METHYLASE->ParB+HNH->HEPN+DUF499*->?->?->?-><-?<-SbcC\|\|HIN-HTH-> | 1122 | Bacteria>Terrabacteria_group>Actinomycetota>Actinomycetes>Streptosporangiales | GCA_018327725.1 | Nonomuraea_sp._TT08I-71 | Nm8I071_37670 | 3194026..3197394 | 2733864 | hypothetical-protein-Nm8I071_37670-[Nonomuraea-sp.-TT08I-71] |
| WP_203980270.1 | HEPN+AAA-ATPase+Alpha+Beta-domain+wHTH+RRM | Type-1_BR.RM.like.systems | <-SF1-SF2-helicase+DUF1998<-?<-ParB+HNH<-?<-HEPN+DUF499*<-METHYLASE<-TUDOR+SF1-SF2-Helicase+iREase+CTD+REase<-?<-REase+N6_MTase | 1129 | Bacteria>Terrabacteria_group>Actinomycetota>Actinomycetes>Streptosporangiales | GCF_016863315.1 | Planotetraspora_silvatica | Psi02_RS35595 | 24804..28193 | 234614 | Swt1-family-HEPN-domain-containing-protein-[Planotetraspora-silvatica] |
| WP_237104850.1 | HEPN+AAA-ATPase+Alpha+Beta-domain+wHTH+RRM | Type-1_BR.RM.like.systems | GNTR-HTH-p->NUDIX->TUDOR+SF1-SF2-Helicase+iREase+CTD+REase->METHYLASE->HEPN+DUF499*->ParB+HNH->?->REase-><-?<-?\|\|SIGMA-HTH-> | 1137 | Bacteria>Terrabacteria_group>Actinomycetota>Actinomycetes>Streptosporangiales | GCF_021699195.1 | Nonomuraea_sp._MG754425 | FAF44_RS20630 | 17644..21057 | 2570319 | DUF499-domain-containing-protein-[Nonomuraea-sp.-MG754425] |
| MCF6470725.1 | HEPN+AAA-ATPase+Alpha+Beta-domain+wHTH+RRM | Type-1_BR.RM.like.systems | GNTR-HTH-p->NUDIX->TUDOR+SF1-SF2-Helicase+iREase+CTD+REase->METHYLASE->HEPN+DUF499*->ParB+HNH->?->REase-><-?\|\|SIGMA-HTH-> | 1139 | Bacteria>Terrabacteria_group>Actinomycetota>Actinomycetes>Streptosporangiales | GCA_021699195.1 | Nonomuraea_sp._MG754425 | FAF44_20335 | 17638..21057 | 2570319 | ATP-binding-protein-[Nonomuraea-sp.-MG754425] |
| WP_061082585.1 | HEPN+AAA-ATPase+Alpha+Beta-domain+wHTH+RRM | Type-1_BR.RM.like.systems | REase->?->McrC-NTD+REase->TUDOR+SF1-SF2-Helicase+iREase+CTD+REase->METHYLASE->HEPN+DUF499*-><-?<-ST-Hiskin\|\|HTH+DUF5753-> | 1135 | Bacteria>Terrabacteria_group>Actinomycetota>Actinomycetes>Streptosporangiales | GCF_001552695.1 | Nocardiopsis_alborubida | ND1_RS24360 | 24019..27426 | 146802 | Swt1-family-HEPN-domain-containing-protein-[Nocardiopsis-alborubida] |
| WP_017595710.1 | HEPN+AAA-ATPase+Alpha+Beta-domain+wHTH+RRM | Type-1_BR.RM.like.systems | <-JetB<-JetA<-HEPN+DUF499*<-METHYLASE<-TUDOR+SF1-SF2-Helicase+iREase+CTD+REase<-PIN<-XIS-HTH<-tRNA<-SIGMA-HTH | 1136 | Bacteria>Terrabacteria_group>Actinomycetota>Actinomycetes>Streptosporangiales | GCF_000341105.1 | Nocardiopsis_potens | D459_RS0124835 | 11224..14634 | 1246458 | Swt1-family-HEPN-domain-containing-protein-[Nocardiopsis-potens] |
| WP_191055573.1 | HEPN+AAA-ATPase+Alpha+Beta-domain+wHTH+RRM | Type-1_BR.RM.like.systems | HTH+HKD-DNase->?->?->TUDOR+SF1-SF2-Helicase+iREase+CTD+REase->METHYLASE->HEPN+DUF499*->ParB+HNH->HEPN-> | 1138 | Bacteria>Terrabacteria_group>Actinomycetota>Actinomycetes>Streptosporangiales | GCF_014712745.1 | Microbispora_camponoti | IEQ31_RS35675 | 16717..20133 | 1677852 | DUF499-domain-containing-protein-[Microbispora-camponoti] |
| WP_204029818.1 | HEPN+AAA-ATPase+Alpha+Beta-domain+wHTH+RRM | Type-1_BR.RM.like.systems | <-GNTR-HTH-p<-?<-?<-?<-HEPN<-ParB+HNH<-HEPN+DUF499*<-METHYLASE<-TUDOR+SF1-SF2-Helicase+iREase+CTD+REase | 1127 | Bacteria>Terrabacteria_group>Actinomycetota>Actinomycetes>Streptosporangiales | GCF_016863435.1 | Sinosporangium_siamense | Ssi02_RS24940 | 167948..171331 | 1367973 | DUF499-domain-containing-protein-[Sinosporangium-siamense] |
| GHJ56382.1 | HEPN+AAA-ATPase+Alpha+Beta-domain+wHTH+RRM | Type-1_BR.RM.like.systems | <-HEPN+DUF499*<-METHYLASE<-TUDOR+SF1-SF2-Helicase+iREase+CTD+REase<-?\|\|SAD+HNH-><-Peptidase | 1124 | Bacteria>Terrabacteria_group>Actinomycetota>Actinomycetes>Streptosporangiales | GCA_018327725.1 | Nonomuraea_sp._TT08I-71 | Nm8I071_56890 | 5264667..5268041 | 2733864 | hypothetical-protein-Nm8I071_56890-[Nonomuraea-sp.-TT08I-71] |
| WP_155999755.1 | HEPN+AAA-ATPase+Alpha+Beta-domain+wHTH+RRM | Type-1_BR.RM.like.systems | <-HTH+DUF5753\|\|?-><-?<-?<-METHYLASE<-?\|\|HEPN+DUF499*->ParB+HNH-><-?<-Glycos_trans_3N+Glycos_transf_3 | 1141 | Bacteria>Terrabacteria_group>Actinomycetota>Actinomycetes>Streptosporangiales | GCF_009728995.1 | Streptomonospora_sp._PA3 | FZ103_RS07070 | 271290..274715 | 2607326 | Swt1-family-HEPN-domain-containing-protein-[Streptomonospora-sp.-PA3] |
| WP_189237444.1 | HEPN+AAA-ATPase+Alpha+Beta-domain+wHTH+RRM | Type-1_BR.RM.like.systems | METHYLASE-p->HEPN+DUF499*-><-DUF397<-HTH+DUF5753\|\|ST-Hiskin->SIGMA-HTH->ParB+HNH-> | 1133 | Bacteria>Terrabacteria_group>Actinomycetota>Actinomycetes>Streptosporangiales | GCF_014647955.1 | Planomonospora_parontospora | Ppa05_RS15805 | 840..4241 | 58119 | Swt1-family-HEPN-domain-containing-protein-[Planomonospora-parontospora] |
| GII94898.1 | HEPN+AAA-ATPase+Alpha+Beta-domain+wHTH+RRM | Type-1_BR.RM.like.systems | <-HEPN<-ParB+HNH<-HEPN+DUF499*<-METHYLASE<-TUDOR+SF1-SF2-Helicase+iREase+CTD+REase | 1137 | Bacteria>Terrabacteria_group>Actinomycetota>Actinomycetes>Streptosporangiales | GCA_016863435.1 | Sinosporangium_siamense | Ssi02_51290 | 167948..171361 | 1367973 | hypothetical-protein-Ssi02_51290-[Sinosporangium-siamense] |
| EAR26285.1 | HEPN+AAA-ATPase+Alpha+Beta-domain+wHTH+RRM | Type-1_BR.RM.like.systems | REase-><-REase<-DNA_methylase<-MORC+SIGMA-HTH\|\|METHYLASE->HEPN+DUF499*->MORC->REase->HTH+SF1-SF2-Helicase+REase->?-><-?<-?<-ABC-ATPase+SbcC | 1127 | Bacteria>Terrabacteria_group>Actinomycetota>Actinomycetes>unclassified_Actinomycetes | GCA_000153145.1 | marine_actinobacterium_PHSC20C1 | A20C1_10399 | 2131040..2134423 | 312284 | hypothetical-protein-A20C1_10399-[marine-actinobacterium-PHSC20C1] |
| MCH9729512.1 | HEPN+AAA-ATPase+Alpha+Beta-domain+wHTH+RRM | Type-1_BR.RM.like.systems | PIN->TUDOR+SF1-SF2-Helicase+iREase+CTD+REase->METHYLASE->ParB+HNH->HEPN+DUF499*->SIR2-><-DndB\|\|?->Recombinase-><-NUDIX\|\|GNTR-HTH-> | 1114 | Bacteria>Terrabacteria_group>Actinomycetota>Actinomycetes>unclassified_Actinomycetes | GCA_022600605.1 | Actinomycetes_bacterium | KOU84_07525 | 8492..11836 | 1883427 | DUF499-domain-containing-protein-[Actinomycetia-bacterium] |
| KLR61150.1 | HEPN+AAA-ATPase+Alpha+Beta-domain+wHTH+RRM | Type-1_BR.RM.like.systems | <-ParB+Schlafen\|\|TUDOR+SF1-SF2-Helicase+iREase+CTD+REase->METHYLASE-p->METHYLASE-p->HEPN+DUF499*->tRNA-><-?\|\|RelE-ParE->HTH-> | 1095 | Bacteria>Terrabacteria_group>Actinomycetota>Actinomycetes>unclassified_Actinomycetes | GCA_001025035.1 | Actinobacteria_bacterium_IMCC26207 | IMCC26207_109668 | 707438..710725 | 1641811 | putative-ATPase-(AAA+-superfamily)-[Actinobacteria-bacterium-IMCC26207] |
| MCL2317098.1 | HEPN+AAA-ATPase+Alpha+Beta-domain+wHTH+RRM | Type-1_BR.RM.like.systems | <-HEPN+DUF499*<-PIN<-PhdYeFM_antitox<-METHYLASE-p | 623 | Bacteria>Terrabacteria_group>Actinomycetota>Actinomycetes>unclassified_Actinomycetes | GCA_009785885.1 | Actinomycetes_bacterium | FWC46_08465 | 1..1871 | 1883427 | Swt1-family-HEPN-domain-containing-protein-[Actinomycetia-bacterium] |
| PKQ27862.1 | HEPN+AAA-ATPase+Alpha+Beta-domain+wHTH+RRM | Type-1_BR.RM.like.systems | REase->TUDOR+SF1-SF2-Helicase+iREase+CTD+REase->Schlafen+HATPase+HTH->METHYLASE->HEPN+DUF499*->?->TIR-> | 1105 | Bacteria>Terrabacteria_group>Actinomycetota>Candidatus_Geothermincolia>Candidatus_Geothermincolales | GCA_002843005.1 | Candidatus_Anoxymicrobium_japonicum | CVT63_05765 | 8955..12272 | 2013648 | AAA+-family-ATPase-[Candidatus-Anoxymicrobium-japonicum] |
| WP_267192478.1 | HEPN+AAA-ATPase+Alpha+Beta-domain+wHTH+RRM | Type-1_BR.RM.like.systems | <-Peptidase_C39_2<-?<-?\|\|PspA+PIN->?-><-HEPN+DUF499*<-?<-MqsA_antitoxin+HTH_3<-?<-METHYLASE<-ParB+HNH<-TUDOR+SF1-SF2-Helicase+iREase+CTD+REase | 1106 | Bacteria>Terrabacteria_group>Actinomycetota>Coriobacteriia>Anaerosomatales | - | Parvivirga_hydrogeniphila | MX659_RS00500 | 98516..101836 | 2939460 | Swt1-family-HEPN-domain-containing-protein-[Parvivirga-hydrogeniphila] |
| MCL4078096.1 | HEPN+AAA-ATPase+Alpha+Beta-domain+wHTH+RRM | Type-1_BR.RM.like.systems | <-Peptidase_C39_2<-?<-?\|\|PspA+PIN->?-><-HEPN+DUF499*<-?<-MqsA_antitoxin+HTH_3<-?<-METHYLASE<-ParB+HNH<-TUDOR+SF1-SF2-Helicase+iREase+CTD+REase | 1106 | Bacteria>Terrabacteria_group>Actinomycetota>Coriobacteriia>Anaerosomatales | GCA_023371205.1 | Parvivirga_hydrogeniphila | MX659_00500 | 98516..101836 | 2939460 | Swt1-family-HEPN-domain-containing-protein-[Coriobacteriia-bacterium-Es71-Z0120] |
| WP_273395403.1 | HEPN+AAA-ATPase+Alpha+Beta-domain+wHTH+RRM | Type-1_BR.RM.like.systems | <-ABC-ATPase+SbcC<-MarR-HTH<-?\|\|ORC-CDC6+REase-><-ORC-CDC6+REase<-?<-HEPN+DUF499*<-METHYLASE<-TUDOR+SF1-SF2-Helicase+iREase+CTD+REase\|\|HTH_3->HAD-> | 1133 | Bacteria>Terrabacteria_group>Actinomycetota>Coriobacteriia>Coriobacteriales | GCF_944322105.1 | Thermophilibacter_mediterraneus | PUQ24_RS02970 | 23710..27111 | 1871031 | DUF499-domain-containing-protein-[Thermophilibacter-mediterraneus] |
| WP_072371784.1 | HEPN+AAA-ATPase+Alpha+Beta-domain+wHTH+RRM | Type-1_BR.RM.like.systems | <-ABC-ATPase+SbcC<-MarR-HTH<-?\|\|ORC-CDC6+REase-><-ORC-CDC6+REase<-?<-HEPN+DUF499*<-METHYLASE<-TUDOR+SF1-SF2-Helicase+iREase+CTD+REase\|\|HTH_3->HAD-> | 1133 | Bacteria>Terrabacteria_group>Actinomycetota>Coriobacteriia>Coriobacteriales | GCF_900119385.1 | Thermophilibacter_mediterraneus | BQ7106_RS04355 | 522048..525449 | 1871031 | DUF499-domain-containing-protein-[Thermophilibacter-mediterraneus] |
| TGY60542.1 | HEPN+AAA-ATPase+Alpha+Beta-domain+wHTH+RRM | Type-1_BR.RM.like.systems | MazG->URI-DNase+SF1-SF2-Helicase->HEPN->HEPN->TUDOR+SF1-SF2-Helicase+iREase+CTD+REase->METHYLASE->HEPN+DUF499*->ORC-CDC6+REase->?-><-?\|\|?->?->REase-> | 1126 | Bacteria>Terrabacteria_group>Actinomycetota>Coriobacteriia>Coriobacteriales | GCA_004793665.1 | Coriobacteriaceae_bacterium | E5335_00400 | 90399..93779 | 2011094 | ATP-binding-protein-[Coriobacteriaceae-bacterium] |
| WP_265590834.1 | HEPN+AAA-ATPase+Alpha+Beta-domain+wHTH+RRM | Type-1_BR.RM.like.systems | <-REase<-?<-?\|\|?-><-?<-ORC-CDC6+REase<-HEPN+DUF499*<-METHYLASE<-TUDOR+SF1-SF2-Helicase+iREase+CTD+REase<-HEPN<-URI-DNase+SF1-SF2-Helicase<-MazG | 1126 | Bacteria>Terrabacteria_group>Actinomycetota>Coriobacteriia>Coriobacteriales | GCF_022834715.1 | Granulimonas_faecalis | OR600_RS06205 | 1358904..1362284 | 2894155 | DUF499-domain-containing-protein-[Granulimonas-faecalis] |
| GJM55560.1 | HEPN+AAA-ATPase+Alpha+Beta-domain+wHTH+RRM | Type-1_BR.RM.like.systems | <-REase<-?<-?\|\|?-><-?<-ORC-CDC6+REase<-HEPN+DUF499*<-METHYLASE<-TUDOR+SF1-SF2-Helicase+iREase+CTD+REase<-HEPN<-URI-DNase+SF1-SF2-Helicase<-MazG | 1126 | Bacteria>Terrabacteria_group>Actinomycetota>Coriobacteriia>Coriobacteriales | GCA_022834715.1 | Granulimonas_faecalis | ATOP_12150 | 1358904..1362284 | 2894155 | hypothetical-protein-ATOP_12150-[Granulimonas-faecalis] |
| WP_251158236.1 | HEPN+AAA-ATPase+Alpha+Beta-domain+wHTH+RRM | Type-1_BR.RM.like.systems | <-ORC-CDC6+REase<-HEPN+DUF499*<-METHYLASE<-TUDOR+SF1-SF2-Helicase+iREase+CTD+REase<-HEPN<-URI-DNase+SF1-SF2-Helicase<-MazG | 1126 | Bacteria>Terrabacteria_group>Actinomycetota>Coriobacteriia>Coriobacteriales | GCF_943193035.1 | Caniella_muris | NFR49_RS03270 | 716481..719861 | 2941502 | DUF499-domain-containing-protein-[Caniella-sp.-DSM-110323] |
| WP_218957899.1 | HEPN+AAA-ATPase+Alpha+Beta-domain+wHTH+RRM | Type-1_BR.RM.like.systems | <-Peptidase<-?<-?<-?<-?<-DUF499-p<-HEPN+DUF499*<-METHYLASE<-TUDOR+SF1-SF2-Helicase+iREase+CTD+REase\|\|OLD-ABC+TOPRIM->SF1-SF2-Helicase-> | 864 | Bacteria>Terrabacteria_group>Actinomycetota>Coriobacteriia>Coriobacteriales | GCF_007655325.1 | Enorma_shizhengliae | GJE22_RS07705 | 20738..23332 | 2606615 | DUF499-domain-containing-protein-[Enorma-shizhengliae] |
| WP_204556581.1 | HEPN+AAA-ATPase+Alpha+Beta-domain+wHTH+RRM | Type-1_BR.RM.like.systems | HTH_3->?->?-><-?\|\|TUDOR+SF1-SF2-Helicase+iREase+CTD+REase->METHYLASE->HEPN+DUF499*->?->XIS-HTH-><-SF1-SF2-Helicase\|\|?->?->MNS+HTH-> | 1118 | Bacteria>Terrabacteria_group>Actinomycetota>Coriobacteriia>Coriobacteriales | GCF_016901585.1 | Collinsella_intestinalis | H7U33_RS06515 | 78334..81690 | 147207 | DUF499-domain-containing-protein-[Collinsella-intestinalis] |
| WP_270472295.1 | HEPN+AAA-ATPase+Alpha+Beta-domain+wHTH+RRM | Type-1_BR.RM.like.systems | HISKIN-><-?<-ORC-CDC6+REase\|\|TUDOR+SF1-SF2-Helicase+iREase+CTD+REase->METHYLASE->HEPN+DUF499*->?->?->?->?->ABC-ATPase+SbcC-> | 1129 | Bacteria>Terrabacteria_group>Actinomycetota>Coriobacteriia>Coriobacteriales | GCF_027683425.1 | Collinsella_intestinalis | PGX73_RS07325 | 61994..65383 | 147207 | DUF499-domain-containing-protein-[Collinsella-intestinalis] |
| WP_154324032.1 | HEPN+AAA-ATPase+Alpha+Beta-domain+wHTH+RRM | Type-1_BR.RM.like.systems | Classical-AAA->Peptidase_S8-><-HEPN+DUF499*<-METHYLASE<-TUDOR+SF1-SF2-Helicase+iREase+CTD+REase<-?<-Phage-portal | 1146 | Bacteria>Terrabacteria_group>Actinomycetota>Coriobacteriia>Coriobacteriales | GCF_009676995.1 | Enorma_shizhengliae | GJE22_RS00805 | 201758..205198 | 2606615 | DUF499-domain-containing-protein-[Enorma-shizhengliae] |
| WP_117794170.1 | HEPN+AAA-ATPase+Alpha+Beta-domain+wHTH+RRM | Type-1_BR.RM.like.systems | <-MNS+REase\|\|?-><-?<-HEPN+DUF499*<-METHYLASE<-TUDOR+SF1-SF2-Helicase+iREase+CTD+REase\|\|?-><-HEPN<-REase-p<-REase-p | 1137 | Bacteria>Terrabacteria_group>Actinomycetota>Coriobacteriia>Coriobacteriales | GCF_003466445.1 | Collinsella_sp._TM05-38 | DXD49_RS03005 | 54584..57997 | 2292341 | DUF499-domain-containing-protein-[Collinsella-sp.-TM05-38] |
| MBS6858281.1 | HEPN+AAA-ATPase+Alpha+Beta-domain+wHTH+RRM | Type-1_BR.RM.like.systems | MNS+HTH+REase->35exo->TUDOR+SF1-SF2-Helicase+iREase+CTD+REase->METHYLASE->HEPN+DUF499*-><-?<-?<-?\|\|?->Peptidase_M20-> | 1137 | Bacteria>Terrabacteria_group>Actinomycetota>Coriobacteriia>Coriobacteriales | GCA_018380015.1 | Collinsella_sp. | KH187_03945 | 112102..115515 | 1965294 | DUF499-domain-containing-protein-[Collinsella-sp.] |
| WP_195644120.1 | HEPN+AAA-ATPase+Alpha+Beta-domain+wHTH+RRM | Type-1_BR.RM.like.systems | <-MNS+REase\|\|?-><-?<-HEPN+DUF499*<-METHYLASE<-TUDOR+SF1-SF2-Helicase+iREase+CTD+REase<-?<-35exo<-MNS+HTH+REase | 1137 | Bacteria>Terrabacteria_group>Actinomycetota>Coriobacteriia>Coriobacteriales | GCF_015561195.1 | Collinsella_aerofaciens | PMX13_RS01910 | 103698..107111 | 74426 | DUF499-domain-containing-protein-[Collinsella-aerofaciens] |
| WP_059053651.1 | HEPN+AAA-ATPase+Alpha+Beta-domain+wHTH+RRM | Type-1_BR.RM.like.systems | <-HEPN+DUF499*<-METHYLASE<-TUDOR+SF1-SF2-Helicase+iREase+CTD+REase\|\|REase->?-><-Phage-portal | 1133 | Bacteria>Terrabacteria_group>Actinomycetota>Coriobacteriia>Coriobacteriales | GCF_001494635.1 | Tractidigestivibacter_scatoligenes | AUL39_RS03515 | 793561..796962 | 1299998 | DUF499-domain-containing-protein-[Tractidigestivibacter-scatoligenes] |
| MCI1646461.1 | HEPN+AAA-ATPase+Alpha+Beta-domain+wHTH+RRM | Type-1_BR.RM.like.systems | Phage-portal-><-?<-REase<-?\|\|TUDOR+SF1-SF2-Helicase+iREase+CTD+REase->METHYLASE->HEPN+DUF499*-> | 1135 | Bacteria>Terrabacteria_group>Actinomycetota>Coriobacteriia>Coriobacteriales | GCA_022648325.1 | Olsenella_sp. | LKI40_09700 | 33450..36857 | 2042683 | Swt1-family-HEPN-domain-containing-protein-[Olsenella-sp.] |
| MBR3315601.1 | HEPN+AAA-ATPase+Alpha+Beta-domain+wHTH+RRM | Type-1_BR.RM.like.systems | ASCH-><-PIN+ACET+EVE<-HEPN+DUF499*<-METHYLASE<-TUDOR+SF1-SF2-Helicase+iREase+CTD+REase | 1137 | Bacteria>Terrabacteria_group>Actinomycetota>Coriobacteriia>Coriobacteriales | GCA_017510745.1 | Atopobiaceae_bacterium | IKG18_15850 | 6540..9953 | 2485926 | DUF499-domain-containing-protein-[Atopobiaceae-bacterium] |
| MBQ6523654.1 | HEPN+AAA-ATPase+Alpha+Beta-domain+wHTH+RRM | Type-1_BR.RM.like.systems | <-HEPN+DUF499*<-METHYLASE-p | 1129 | Bacteria>Terrabacteria_group>Actinomycetota>Coriobacteriia>Coriobacteriales | GCA_017433555.1 | Atopobiaceae_bacterium | IJI12_03205 | 1663..5052 | 2485926 | DUF499-domain-containing-protein-[Atopobiaceae-bacterium] |
| MCD8317268.1 | HEPN+AAA-ATPase+Alpha+Beta-domain+wHTH+RRM | Type-1_BR.RM.like.systems | TUDOR+SF1-SF2-Helicase+iREase+CTD+REase->METHYLASE-p-><-Phage_int_SAM_1+Phage_integrase<-Phage_integrase<-Phage_integrase\|\|METHYLASE-p->HEPN+DUF499*-> | 1113 | Bacteria>Terrabacteria_group>Actinomycetota>Coriobacteriia>Eggerthellales | GCA_021204985.1 | Eggerthellaceae_bacterium | LUB61_07720 | 22349..25690 | 1972561 | DUF499-domain-containing-protein-[Eggerthellaceae-bacterium] |
| WP_206214419.1 | HEPN+AAA-ATPase+Alpha+Beta-domain+wHTH+RRM | Type-1_BR.RM.like.systems | <-wHTH+REase<-HTH_3<-Tox-RelE\|\|Classical-AAA->Peptidase_S8-><-HEPN+DUF499*<-METHYLASE<-TUDOR+SF1-SF2-Helicase+iREase+CTD+REase<-?\|\|?-><-?<-ABC-ATPase | 1145 | Bacteria>Terrabacteria_group>Actinomycetota>Coriobacteriia>Eggerthellales | GCF_011038965.1 | Adlercreutzia_sp._ZJ141 | G7Y44_RS06490 | 70626..74063 | 2709406 | DUF499-domain-containing-protein-[Adlercreutzia-sp.-ZJ141] |
| MBQ9000666.1 | HEPN+AAA-ATPase+Alpha+Beta-domain+wHTH+RRM | Type-1_BR.RM.like.systems | TUDOR+SF1-SF2-Helicase+iREase+CTD+REase->METHYLASE->HEPN+DUF499*->ParB-> | 1133 | Bacteria>Terrabacteria_group>Actinomycetota>Coriobacteriia>Eggerthellales | GCA_017460365.1 | Eggerthellaceae_bacterium | IJ087_02285 | 5494..8895 | 1972561 | DUF499-domain-containing-protein-[Eggerthellaceae-bacterium] |
| WP_165173327.1 | HEPN+AAA-ATPase+Alpha+Beta-domain+wHTH+RRM | Type-1_BR.RM.like.systems | TUDOR+SF1-SF2-Helicase+iREase+CTD+REase->METHYLASE->HEPN+DUF499*->?->UDG-> | 1132 | Bacteria>Terrabacteria_group>Actinomycetota>Coriobacteriia>Eggerthellales | GCF_011038995.1 | Adlercreutzia_sp._ZJ242 | G7Y49_RS09770 | 7632..11030 | 2709409 | DUF499-domain-containing-protein-[Adlercreutzia-sp.-ZJ242] |
| MDD5805721.1 | HEPN+AAA-ATPase+Alpha+Beta-domain+wHTH+RRM | Type-1_BR.RM.like.systems | METHYLASE->HEPN+DUF499*-> | 557 | Bacteria>Terrabacteria_group>Actinomycetota>Coriobacteriia>Eggerthellales | GCA_029101185.1 | Eggerthellales_bacterium | PUD02_00035 | 2228..3899 | 2137879 | MAG:-Swt1-family-HEPN-domain-containing-protein,-partial-[Eggerthellales-bacterium] |
| MBC7265381.1 | HEPN+AAA-ATPase+Alpha+Beta-domain+wHTH+RRM | Type-1_BR.RM.like.systems | <-Peptidase_C39_2<-?\|\|PspA+PIN->?-><-HEPN+DUF499*<-?<-MqsA_antitoxin+HTH_3<-?<-METHYLASE<-ParB+HNH<-TUDOR+SF1-SF2-Helicase+iREase+CTD+REase | 1106 | Bacteria>Terrabacteria_group>Actinomycetota>Coriobacteriia>unclassified_Coriobacteriia | GCA_014360605.1 | Coriobacteriia_bacterium | H5T75_00145 | 31010..34330 | 2052159 | ATP-binding-protein-[Coriobacteriia-bacterium] |
| MBE0418219.1 | HEPN+AAA-ATPase+Alpha+Beta-domain+wHTH+RRM | Type-1_BR.RM.like.systems | METHYLASE-p->HEPN+DUF499*-> | 630 | Bacteria>Terrabacteria_group>Actinomycetota>Coriobacteriia>unclassified_Coriobacteriia | GCA_014859265.1 | Coriobacteriia_bacterium | IBX63_10685 | 617..2507 | 2052159 | ATP-binding-protein-[Coriobacteriia-bacterium] |
| GIU94958.1 | HEPN+AAA-ATPase+Alpha+Beta-domain+wHTH+RRM | Type-1_BR.RM.like.systems | Schlafen+HATPase+HTH->?->OLD-ABC->TOPRIM->METHYLASE->METHYLASE-p->HEPN+DUF499*->?->TPR->DUF222+HNH->REase+35exo+Beta-Barrel+SF1-SF2-Helicase-> | 1082 | Bacteria>Terrabacteria_group>Actinomycetota>Rubrobacteria>Gaiellales | GCA_026002995.1 | Gaiellaceae_bacterium | KatS3mg012_1415 | 1404220..1407468 | 2838783 | MAG:-hypothetical-protein-KatS3mg012_1415-[Gaiellaceae-bacterium] |
| MCP9484920.1 | HEPN+AAA-ATPase+Alpha+Beta-domain+wHTH+RRM | Type-1_BR.RM.like.systems | METHYLASE-p->?->METHYLASE-p->SbcC->PIN->?->HEPN+DUF499*->HIN-HTH->?->\|\|?-><-Peptidase_M20 | 1087 | Bacteria>Terrabacteria_group>Actinomycetota>Rubrobacteria>Gaiellales | GCA_024330215.1 | Candidatus_Gaiellasilicea_maunaloa | MSC30_03590 | 46535..49798 | 2928748 | DUF499-domain-containing-protein-[Candidatus-Gaiellasilicea-maunaloa] |
| WP_219975492.1 | HEPN+AAA-ATPase+Alpha+Beta-domain+wHTH+RRM | Type-1_BR.RM.like.systems | <-SIGMA-HTH<-?<-MULE-TRANSPOSASE\|\|?->?-><-REase<-HEPN+DUF499*<-PIN<-MazE_antitoxin<-METHYLASE<-PIN<-TUDOR+SF1-SF2-Helicase+iREase+CTD+REase\|\|DUF4096-> | 1106 | Bacteria>Terrabacteria_group>Actinomycetota>Rubrobacteria>Rubrobacterales | GCF_019448335.1 | Rubrobacter_xylanophilus | Rxycam_RS04975 | 969402..972722 | 49319 | Swt1-family-HEPN-domain-containing-protein-[Rubrobacter-xylanophilus] |
| MBX6764706.1 | HEPN+AAA-ATPase+Alpha+Beta-domain+wHTH+RRM | Type-1_BR.RM.like.systems | METHYLASE-p->MazE_antitoxin->PIN->HEPN+DUF499*->REase-> | 1106 | Bacteria>Terrabacteria_group>Actinomycetota>Rubrobacteria>Rubrobacterales | GCA_019690035.1 | Rubrobacteraceae_bacterium | IRY88_13660 | 1711..5031 | 2740537 | DUF499-domain-containing-protein-[Rubrobacteraceae-bacterium] |
| WP_217923107.1 | HEPN+AAA-ATPase+Alpha+Beta-domain+wHTH+RRM | Type-1_BR.RM.like.systems | <-HEPN+DUF499*<-METHYLASE<-TUDOR+SF1-SF2-Helicase+iREase+CTD+REase | 1108 | Bacteria>Terrabacteria_group>Actinomycetota>Thermoleophilia>Miltoncostaeales | GCF_018141545.1 | Miltoncostaea_oceani | IU369_RS03095 | 622243..625569 | 2843216 | Swt1-family-HEPN-domain-containing-protein-[Miltoncostaea-oceani] |
| MBI2692601.1 | HEPN+AAA-ATPase+Alpha+Beta-domain+wHTH+RRM | Type-1_BR.RM.like.systems | <-HTH+UbE1<-?\|\|HTH_31->MPTase->SF1-SF2-Helicase+REase-p->?-><-HEPN+DUF499*<-ParB+HNH<-METHYLASE<-TUDOR+SF1-SF2-Helicase+iREase+CTD+REase<-HTH_3\|\|?-><-HTH | 1109 | Bacteria>Terrabacteria_group>Actinomycetota>Thermoleophilia>Solirubrobacterales | GCA_016185345.1 | Solirubrobacterales_bacterium | HYX29_11735 | 159016..162345 | 1909295 | ATP-binding-protein-[Solirubrobacterales-bacterium] |
| MCW3047079.1 | HEPN+AAA-ATPase+Alpha+Beta-domain+wHTH+RRM | Type-1_BR.RM.like.systems | TUDOR+SF1-SF2-Helicase+iREase+CTD+REase->METHYLASE->HEPN+DUF499*->?->?-><-SF1-SF2-helicase<-SF1-SF2-Helicase+REase | 1109 | Bacteria>Terrabacteria_group>Actinomycetota>Thermoleophilia>Solirubrobacterales | GCA_025963905.1 | Solirubrobacterales_bacterium | JW074_1363 | 11829..15158 | 1909295 | MAG:-hypothetical-protein-JW074_1363-[Solirubrobacterales-bacterium] |
| MCZ7663785.1 | HEPN+AAA-ATPase+Alpha+Beta-domain+wHTH+RRM | Type-1_BR.RM.like.systems | <-NTP_transf<-HEPN<-PolB<-?<-TPR<-HEPN+DUF499*<-RelE-ParE<-?<-PIN<-?<-PIN | 1114 | Bacteria>Terrabacteria_group>Actinomycetota>Thermoleophilia>unclassified_Thermoleophilia | GCA_027493445.1 | Thermoleophilia_bacterium | M5U22_13030 | 2549788..2553132 | 2026888 | MAG:-Swt1-family-HEPN-domain-containing-protein-[Thermoleophilia-bacterium] |
| MSW27210.1 | HEPN+AAA-ATPase+Alpha+Beta-domain+wHTH+RRM | Type-1_BR.RM.like.systems | Phage_int_SAM_1+Phage_integrase->BirA-HTH-><-REase+DUF4268\|\|TUDOR+SF1-SF2-Helicase+iREase+CTD+REase->METHYLASE->METHYLASE->HEPN+DUF499*->RNase+SF1-SF2-Helicase->HEPN+SF1-SF2-Helicase->?->REase-> | 1092 | Bacteria>Terrabacteria_group>Actinomycetota>unclassified_Actinomycetota | GCA_009698865.1 | Actinomycetota_bacterium | F2849_00700 | 14320..17598 | 2900548 | DUF499-domain-containing-protein-[Actinomycetia-bacterium] |
| MCA0180524.1 | HEPN+AAA-ATPase+Alpha+Beta-domain+wHTH+RRM | Type-1_BR.RM.like.systems | <-SMODS<-?<-TrwC+SF1-SF2-Helicase\|\|DNA_methylase-><-SF1-SF2-Helicase<-OLD-ABC+TOPRIM<-HEPN+DUF499*<-wHTH+REase<-METHYLASE<-TUDOR+SF1-SF2-Helicase+iREase+CTD+REase<-XIS-HTH<-?\|\|GDResolvase+Recombinase+ZnR-> | 1132 | Bacteria>Terrabacteria_group>Actinomycetota>unclassified_Actinomycetota | GCA_020160805.1 | Actinomycetota_bacterium | LCH77_13240 | 54204..57602 | 2900548 | DUF499-domain-containing-protein-[Actinomycetia-bacterium] |
| MCA0294159.1 | HEPN+AAA-ATPase+Alpha+Beta-domain+wHTH+RRM | Type-1_BR.RM.like.systems | <-NTD+SF1-SF2-Helicase\|\|TUDOR+SF1-SF2-Helicase+iREase+CTD+REase->METHYLASE->HEPN+DUF499*->ParB+HNH->?->SF1-SF2-Helicase-><-REase+PNTD1+PNTD2\|\|MULE-TRANSPOSASE->ACET-> | 1117 | Bacteria>Terrabacteria_group>Actinomycetota>unclassified_Actinomycetota | GCA_020161755.1 | Actinomycetota_bacterium | LCH96_02385 | 200193..203546 | 2900548 | DUF499-domain-containing-protein-[Actinomycetia-bacterium] |
| TAK69428.1 | HEPN+AAA-ATPase+Alpha+Beta-domain+wHTH+RRM | Type-1_BR.RM.like.systems | ParB+HNH->HEPN-><-TUDOR+SF1-SF2-Helicase+iREase+CTD+REase\|\|?->?->HEPN+DUF499*->SF1-SF2-Helicase->?-><-?<-REase+35exo+Beta-Barrel+SF1-SF2-Helicase | 1114 | Bacteria>Terrabacteria_group>Actinomycetota>unclassified_Actinomycetota | GCA_004297305.1 | Actinomycetota_bacterium | EPO13_05945 | 61030..64374 | 2900548 | ATP-binding-protein-[Actinobacteria-bacterium] |
| TAK68533.1 | HEPN+AAA-ATPase+Alpha+Beta-domain+wHTH+RRM | Type-1_BR.RM.like.systems | ABC-ATPase+SbcC->?-><-ACET+ACET\|\|?->?-><-HEPN+DUF499*<-METHYLASE<-TUDOR+SF1-SF2-Helicase+iREase+CTD+REase<-?\|\|SIR2+SIR2->PIWI-><-SF1-SF2-Helicase | 1113 | Bacteria>Terrabacteria_group>Actinomycetota>unclassified_Actinomycetota | GCA_004297305.1 | Actinomycetota_bacterium | EPO13_10510 | 141712..145053 | 2900548 | ATP-binding-protein-[Actinobacteria-bacterium] |
| QQS24220.1 | HEPN+AAA-ATPase+Alpha+Beta-domain+wHTH+RRM | Type-1_BR.RM.like.systems | TUDOR+SF1-SF2-Helicase+iREase+CTD+REase->METHYLASE->HEPN+DUF499*->NTD+SF1-SF2-Helicase->REase-><-ACET\|\|METHYLASE-p-><-PIN<-Phd-YefM | 1101 | Bacteria>Terrabacteria_group>Actinomycetota>unclassified_Actinomycetota | GCA_016700055.1 | Actinomycetota_bacterium | IPM43_12505 | 2586473..2589778 | 2900548 | DUF499-domain-containing-protein-[Actinobacteria-bacterium] |

| Protein-ID | DUF499-architecture | System | Gene-Neighbourhoods | Length | Taxonomy | Assembly-ID | Strain | Locus-Tag | Genomic-coordinates | TaxID | Fasta-defline |
|---|---|---|---|---|---|---|---|---|---|---|---|
| MCA0180887.1 | HEPN+AAA-ATPase+Alpha+Beta-domain+wHTH+RRM | Type-1_BR.RM.like.systems | <-VirD4-TraG+VirD4-FtsK<-?\|\|XIS-HTH->TUDOR+SF1-SF2-Helicase+iREase+CTD+REase->METHYLASE->wHTH+REase->HEPN+DUF499*->REase->HEPN+REase-> | 1132 | Bacteria>Terrabacteria_group>Actinomycetota>unclassified_Actinomycetota | GCA_020160805.1 | Actinomycetota_bacterium | LCH77_15145 | 46365..49763 | 2900548 | DUF499-domain-containing-protein-[Actinomycetia-bacterium] |
| MBS1906106.1 | HEPN+AAA-ATPase+Alpha+Beta-domain+wHTH+RRM | Type-1_BR.RM.like.systems | N6_MTase-p->?->HTH+MPTase-><-SF1-SF2-Helicase<-REase<-HEPN+DUF499*<-METHYLASE<-TUDOR+SF1-SF2-Helicase+iREase+CTD+REase<-XIS-HTH<-?<-?<-HTH_3 | 1133 | Bacteria>Terrabacteria_group>Actinomycetota>unclassified_Actinomycetota | GCA_018268515.1 | Actinomycetota_bacterium | JST33_05955 | 17244..20645 | 2900548 | DUF499-domain-containing-protein-[Actinomycetia-bacterium] |
| MDA8358565.1 | HEPN+AAA-ATPase+Alpha+Beta-domain+wHTH+RRM | Type-1_BR.RM.like.systems | TUDOR+SF1-SF2-Helicase+iREase+CTD+REase->METHYLASE->HEPN+DUF499*->DNA-glycosylase-><-PIWI<-SIR2+SIR2\|\|SbcC->SF1-SF2-Helicase-> | 1122 | Bacteria>Terrabacteria_group>Actinomycetota>unclassified_Actinomycetota | GCA_028067995.1 | Actinomycetota_bacterium | M0Z95_20250 | 17381..20749 | 2900548 | MAG:-DUF499-domain-containing-protein-[Actinomycetota-bacterium] |
| NCD16052.1 | HEPN+AAA-ATPase+Alpha+Beta-domain+wHTH+RRM | Type-1_BR.RM.like.systems | <-MORC+REase<-HTH+SF1-SF2-Helicase+REase<-REase<-MORC<-HEPN+DUF499*<-METHYLASE<-REase\|\|DNA_methylase-><-?<-?<-TrwC+SF1-SF2-Helicase | 1131 | Bacteria>Terrabacteria_group>Actinomycetota>unclassified_Actinomycetota | GCA_009930385.1 | Actinomycetota_bacterium | EOL91_01830 | 7837..11232 | 2900548 | ATP-binding-protein-[Actinomycetia-bacterium] |
| MBU2074163.1 | HEPN+AAA-ATPase+Alpha+Beta-domain+wHTH+RRM | Type-1_BR.RM.like.systems | <-HTH+SF1-SF2-Helicase+REase<-REase<-MORC<-HEPN+DUF499*<-METHYLASE<-REase\|\|DNA_methylase->MORC+HEPN-><-?<-TrwC+SF1-SF2-Helicase | 1135 | Bacteria>Terrabacteria_group>Actinomycetota>unclassified_Actinomycetota | GCA_018826065.1 | Actinomycetota_bacterium | KJ938_06990 | 8952..12359 | 2900548 | DUF499-domain-containing-protein-[Actinomycetia-bacterium] |
| MCL4554262.1 | HEPN+AAA-ATPase+Alpha+Beta-domain+wHTH+RRM | Type-1_BR.RM.like.systems | TUDOR+SF1-SF2-Helicase+iREase+CTD+REase-p->OLD-ABC+TOPRIM->METHYLASE->HEPN->ParA-Soj-PloopNTPase->ORC-CDC6->HEPN+DUF499*-> | 468 | Bacteria>Terrabacteria_group>Actinomycetota>unclassified_Actinomycetota | GCA_023382015.1 | Actinomycetota_bacterium | M1565_03890 | 9031..10434 | 2900548 | Swt1-family-HEPN-domain-containing-protein-[Actinomycetota-bacterium] |
| MSW31301.1 | HEPN+AAA-ATPase+Alpha+Beta-domain+wHTH+RRM | Type-1_BR.RM.like.systems | PadR-HTH->?->?-><-?<-RNase<-REase+DUF4268<-HEPN+DUF499*<-METHYLASE<-TUDOR+SF1-SF2-Helicase+iREase+CTD+REase<-?\|\|?->?->Phage_int_SAM_1-> | 1098 | Bacteria>Terrabacteria_group>Actinomycetota>unclassified_Actinomycetota | GCA_009698915.1 | Actinomycetota_bacterium | F2847_00635 | 27684..30980 | 2900548 | DUF499-domain-containing-protein-[Actinomycetia-bacterium] |
| MCL5734830.1 | HEPN+AAA-ATPase+Alpha+Beta-domain+wHTH+RRM | Type-1_BR.RM.like.systems | TUDOR+SF1-SF2-Helicase+iREase+CTD+REase-p->MORC+DUF3684->METHYLASE->REase->SF1-SF2-Helicase->HEPN+DUF499*-><-HTH+DSBH | 1116 | Bacteria>Terrabacteria_group>Actinomycetota>unclassified_Actinomycetota | GCA_023483915.1 | Actinomycetota_bacterium | M1274_04450 | 22427..25777 | 2900548 | Swt1-family-HEPN-domain-containing-protein-[Actinobacteria-bacterium] |
| MBC7252723.1 | HEPN+AAA-ATPase+Alpha+Beta-domain+wHTH+RRM | Type-1_BR.RM.like.systems | HIN-HTH->?-><-?<-Schlafen+HATPase+HTH<-ParB+HNH+Schlafen<-HEPN+DUF499*<-METHYLASE<-PIN<-TUDOR+SF1-SF2-Helicase+iREase+CTD+REase | 1115 | Bacteria>Terrabacteria_group>Actinomycetota>unclassified_Actinomycetota | GCA_014360685.1 | Actinomycetota_bacterium | H5T72_01965 | 75820..79167 | 2900548 | ATP-binding-protein-[Actinomycetia-bacterium] |
| NLE73128.1 | HEPN+AAA-ATPase+Alpha+Beta-domain+wHTH+RRM | Type-1_BR.RM.like.systems | AbiEi_1->AbiEii->HIN-HTH-><-?<-Schlafen<-HEPN+DUF499*<-DOC+DOC<-HTH_3<-Tox-RelE<-RelE-ParE<-?<-METHYLASE-p | 1130 | Bacteria>Terrabacteria_group>Actinomycetota>unclassified_Actinomycetota | GCA_012514635.1 | Actinomycetota_bacterium | GX604_00625 | 12665..16057 | 2900548 | ATP-binding-protein-[Actinomycetia-bacterium] |
| MDA8316555.1 | HEPN+AAA-ATPase+Alpha+Beta-domain+wHTH+RRM | Type-1_BR.RM.like.systems | TUDOR+SF1-SF2-Helicase+iREase+CTD+REase->METHYLASE->PolB->HEPN->PolB->HEPN+DUF499*-><-RNase<-XIS-HTH | 1117 | Bacteria>Terrabacteria_group>Actinomycetota>unclassified_Actinomycetota | GCA_028067765.1 | Actinomycetota_bacterium | M0010_15505 | 14614..17967 | 2900548 | MAG:-Swt1-family-HEPN-domain-containing-protein-[Actinomycetota-bacterium] |
| MBU2601803.1 | HEPN+AAA-ATPase+Alpha+Beta-domain+wHTH+RRM | Type-1_BR.RM.like.systems | <-Mbetalac<-?<-?<-Radical-SAM<-HEPN+DUF499*<-METHYLASE<-?<-?<-TUDOR+SF1-SF2-Helicase+iREase+CTD+REase<-AbiEii | 1115 | Bacteria>Terrabacteria_group>Actinomycetota>unclassified_Actinomycetota | GCA_018830645.1 | Actinomycetota_bacterium | KKA32_06500 | 57657..61004 | 2900548 | DUF499-domain-containing-protein-[Actinomycetia-bacterium] |
| MDA8312150.1 | HEPN+AAA-ATPase+Alpha+Beta-domain+wHTH+RRM | Type-1_BR.RM.like.systems | Phage_integrase-><-HEPN+DUF499*<-METHYLASE<-TUDOR+SF1-SF2-Helicase+iREase+CTD+REase<-SIR2 | 1097 | Bacteria>Terrabacteria_group>Actinomycetota>unclassified_Actinomycetota | GCA_028067795.1 | Actinomycetota_bacterium | M0Z46_16395 | 628..3921 | 2900548 | MAG:-Swt1-family-HEPN-domain-containing-protein-[Actinomycetota-bacterium] |
| MCL5447091.1 | HEPN+AAA-ATPase+Alpha+Beta-domain+wHTH+RRM | Type-1_BR.RM.like.systems | <-HEPN+DUF499*<-METHYLASE<-?<-TUDOR+SF1-SF2-Helicase+iREase+CTD+REase<-REase<-?<-rve+rve_3 | 1178 | Bacteria>Terrabacteria_group>Actinomycetota>unclassified_Actinomycetota | GCA_023486005.1 | Actinomycetota_bacterium | M1350_06795 | 8812..12348 | 2900548 | DUF499-domain-containing-protein-[Actinobacteria-bacterium] |
| MDA8317108.1 | HEPN+AAA-ATPase+Alpha+Beta-domain+wHTH+RRM | Type-1_BR.RM.like.systems | <-HEPN+DUF499*<-METHYLASE\|\|?-><-TUDOR+SF1-SF2-Helicase+iREase+CTD+REase<-HTH+MPTase | 493 | Bacteria>Terrabacteria_group>Actinomycetota>unclassified_Actinomycetota | GCA_028067765.1 | Actinomycetota_bacterium | M0010_18355 | 6..1487 | 2900548 | MAG:-Swt1-family-HEPN-domain-containing-protein-[Actinomycetota-bacterium] |
| MSW80815.1 | HEPN+AAA-ATPase+Alpha+Beta-domain+wHTH+RRM | Type-1_BR.RM.like.systems | <-REase<-?\|\|TUDOR+SF1-SF2-Helicase+iREase+CTD+REase->METHYLASE->HEPN+DUF499*-> | 1104 | Bacteria>Terrabacteria_group>Actinomycetota>unclassified_Actinomycetota | GCA_009699015.1 | Actinomycetota_bacterium | F2840_06650 | 17897..21211 | 2900548 | DUF499-domain-containing-protein-[Actinomycetia-bacterium] |
| MDA8117679.1 | HEPN+AAA-ATPase+Alpha+Beta-domain+wHTH+RRM | Type-1_BR.RM.like.systems | <-MerR-HTH<-?<-HEPN+DUF499*<-TOPRIM<-OLD-ABC<-METHYLASE<-HEPN<-ParB+HNH<-HicB+ZnR | 1122 | Bacteria>Terrabacteria_group>Actinomycetota>unclassified_Actinomycetota | GCA_028064745.1 | Actinomycetota_bacterium | M0000_09960 | 1809..5177 | 2900548 | MAG:-Swt1-family-HEPN-domain-containing-protein-[Actinomycetota-bacterium] |
| TXH45529.1 | HEPN+AAA-ATPase+Alpha+Beta-domain+wHTH+RRM | Type-1_BR.RM.like.systems | Mbetalac-><-?<-?\|\|Peptidase-><-HEPN+DUF499*\|\|Phd-YefM->PIN-><-?<-METHYLASE\|\|?->PIN-> | 1130 | Bacteria>Terrabacteria_group>Actinomycetota>unclassified_Actinomycetota | GCA_008012465.1 | Actinomycetota_bacterium | E6Q90_00220 | 20268..23660 | 2900548 | ATP-binding-protein-[Actinobacteria-bacterium] |
| MDA8316124.1 | HEPN+AAA-ATPase+Alpha+Beta-domain+wHTH+RRM | Type-1_BR.RM.like.systems | <-HEPN+DUF499*<-METHYLASE<-TUDOR+SF1-SF2-Helicase+iREase+CTD+REase-p | 1027 | Bacteria>Terrabacteria_group>Actinomycetota>unclassified_Actinomycetota | GCA_028067765.1 | Actinomycetota_bacterium | M0010_13295 | 109..3192 | 2900548 | MAG:-Swt1-family-HEPN-domain-containing-protein-[Actinomycetota-bacterium] |
| NCD16598.1 | HEPN+AAA-ATPase+Alpha+Beta-domain+wHTH+RRM | Type-1_BR.RM.like.systems | <-HEPN+REase<-REase<-HEPN+DUF499*<-wHTH+REase<-METHYLASE | 1133 | Bacteria>Terrabacteria_group>Actinomycetota>unclassified_Actinomycetota | GCA_009930385.1 | Actinomycetota_bacterium | EOL91_04665 | 3104..6505 | 2900548 | ATP-binding-protein-[Actinomycetia-bacterium] |
| MBU2602478.1 | HEPN+AAA-ATPase+Alpha+Beta-domain+wHTH+RRM | Type-1_BR.RM.like.systems | <-MNS+HTH\|\|?->HerA/FtsK-><-?<-HEPN+DUF499*<-?<-NTP_transf<-HTH | 1112 | Bacteria>Terrabacteria_group>Actinomycetota>unclassified_Actinomycetota | GCA_018830645.1 | Actinomycetota_bacterium | KKA32_09965 | 120264..123602 | 2900548 | DUF499-domain-containing-protein-[Actinomycetia-bacterium] |
| MDA8063428.1 | HEPN+AAA-ATPase+Alpha+Beta-domain+wHTH+RRM | Type-1_BR.RM.like.systems | <-HEPN+DUF499*<-HEPN<-PolB<-METHYLASE-p | 641 | Bacteria>Terrabacteria_group>Actinomycetota>unclassified_Actinomycetota | GCA_028064035.1 | Actinomycetota_bacterium | M0T80_13560 | 1..1922 | 2900548 | MAG:-Swt1-family-HEPN-domain-containing-protein,-partial-[Actinomycetota-bacterium] |
| MBW3642892.1 | HEPN+AAA-ATPase+Alpha+Beta-domain+wHTH+RRM | Type-1_BR.RM.like.systems | <-HEPN+DUF499*<-?<-PIN<-SbcC<-METHYLASE | 1106 | Bacteria>Terrabacteria_group>Actinomycetota>unclassified_Actinomycetota | GCA_019348235.1 | Actinomycetota_bacterium | KY447_08275 | 1669..4989 | 2900548 | DUF499-domain-containing-protein-[Actinomycetia-bacterium] |
| NLO26762.1 | HEPN+AAA-ATPase+Alpha+Beta-domain+wHTH+RRM | Type-1_BR.RM.like.systems | PIN->METHYLASE->PIN->?->?->?->HEPN+DUF499*-> | 401 | Bacteria>Terrabacteria_group>Actinomycetota>unclassified_Actinomycetota | GCA_012523095.1 | Actinomycetota_bacterium | GX113_01060 | 16345..17548 | 2900548 | ATP-binding-protein-[Actinomycetia-bacterium] |
| MCX6403861.1 | HEPN+AAA-ATPase+Alpha+Beta-domain+wHTH+RRM | Type-1_BR.RM.like.systems | REase->METHYLASE->HEPN+DUF499*-> | 1088 | Bacteria>Terrabacteria_group>Actinomycetota>unclassified_Actinomycetota | GCA_026392535.1 | Actinomycetota_bacterium | NT032_08225 | 3485..6751 | 2900548 | MAG:-DUF499-domain-containing-protein-[Actinobacteria-bacterium] |
| MDA8210090.1 | HEPN+AAA-ATPase+Alpha+Beta-domain+wHTH+RRM | Type-1_BR.RM.like.systems | METHYLASE->HEPN+DUF499*-> | 900 | Bacteria>Terrabacteria_group>Actinomycetota>unclassified_Actinomycetota | GCA_028065575.1 | Actinomycetota_bacterium | M0Z92_13975 | 2687..5388 | 2900548 | MAG:-Swt1-family-HEPN-domain-containing-protein,-partial-[Actinomycetota-bacterium] |
| NLH98010.1 | HEPN+AAA-ATPase+Alpha+Beta-domain+wHTH+RRM | Type-1_BR.RM.like.systems | TUDOR+SF1-SF2-Helicase+iREase+CTD+REase->?->METHYLASE->OLD-ABC->TOPRIM->HEPN+DUF499*-> | 1111 | Bacteria>Terrabacteria_group>Armatimonadota>Chthonomonadetes>Chthonomonadales | GCA_012517255.1 | Chthonomonadales_bacterium | GX446_00800 | 20611..23946 | 2282151 | ATP-binding-protein-[Chthonomonadales-bacterium] |
| MBL8066214.1 | HEPN+AAA-ATPase+Alpha+Beta-domain+wHTH+RRM | Type-1_BR.RM.like.systems | <-HEPN+DUF499*<-HEPN<-METHYLASE-p | 1119 | Bacteria>Terrabacteria_group>Armatimonadota>Chthonomonadetes>Chthonomonadales | GCA_016788345.1 | Chthonomonadaceae_bacterium | JNM34_10200 | 1590..4949 | 2739509 | DUF499-domain-containing-protein-[Chthonomonadaceae-bacterium] |
| MBX7131086.1 | HEPN+AAA-ATPase+Alpha+Beta-domain+wHTH+RRM | Type-1_BR.RM.like.systems | Phage_int_SAM_1+Phage_integrase-><-Peptidase<-HEPN+DUF499*<-?<-HTH<-HKD-DNase<-METHYLASE<-ParB+HNH<-TUDOR+SF1-SF2-Helicase+iREase+CTD+REase | 1104 | Bacteria>Terrabacteria_group>Armatimonadota>Fimbriimonadia>Fimbriimonadales | GCA_019694835.1 | Fimbriimonadaceae_bacterium | K1X67_00255 | 55329..58643 | 2682144 | DUF499-domain-containing-protein-[Fimbriimonadaceae-bacterium] |
| MCC7230625.1 | HEPN+AAA-ATPase+Alpha+Beta-domain+wHTH+RRM | Type-1_BR.RM.like.systems | TIR->?->?->TUDOR+SF1-SF2-Helicase+iREase+CTD+REase->Schlafen+HATPase+HTH->METHYLASE->HEPN+DUF499*-><-JAB\|\|?->ArdCN+MPTase->?->N6_N4_MTase-><-PIN | 1116 | Bacteria>Terrabacteria_group>Armatimonadota>Fimbriimonadia>Fimbriimonadales | GCA_020851885.1 | Fimbriimonadaceae_bacterium | IT203_09555 | 16596..19946 | 2682144 | DUF499-domain-containing-protein-[Fimbriimonadaceae-bacterium] |
| MCX6342680.1 | HEPN+AAA-ATPase+Alpha+Beta-domain+wHTH+RRM | Type-1_BR.RM.like.systems | <-URI-DNase\|\|MazG->URI-DNase+SF1-SF2-Helicase-><-?<-REase<-HEPN+DUF499*<-METHYLASE<-TUDOR+SF1-SF2-Helicase+iREase+CTD+REase | 1119 | Bacteria>Terrabacteria_group>Armatimonadota>Fimbriimonadia>Fimbriimonadales | GCA_026392145.1 | Fimbriimonadales_bacterium | NTU72_09890 | 5893..9252 | 2682145 | MAG:-Swt1-family-HEPN-domain-containing-protein-[Fimbriimonadales-bacterium] |
| MBW7926628.1 | HEPN+AAA-ATPase+Alpha+Beta-domain+wHTH+RRM | Type-1_BR.RM.like.systems | TUDOR+SF1-SF2-Helicase+iREase+CTD+REase->?->SMODS->METHYLASE->HEPN+DUF499*->?->?->JAB-><-D5N+viral-AAA | 1119 | Bacteria>Terrabacteria_group>Armatimonadota>Fimbriimonadia>Fimbriimonadales | GCA_019454665.1 | Fimbriimonadaceae_bacterium | H3C58_00785 | 10553..13912 | 2682144 | ATP-binding-protein-[Fimbriimonadaceae-bacterium] |
| GIV04239.1 | HEPN+AAA-ATPase+Alpha+Beta-domain+wHTH+RRM | Type-1_BR.RM.like.systems | METHYLASE-p->?->RAMPs->RAMPs->RAMPs->HEPN+HEPN->HEPN+DUF499*->MORC+DUF3684-> | 891 | Bacteria>Terrabacteria_group>Armatimonadota>Fimbriimonadia>Fimbriimonadales | GCA_026003055.1 | Fimbriimonadales_bacterium | KatS3mg015_3069 | 13495..16170 | 2682145 | MAG:-hypothetical-protein-KatS3mg015_3069-[Fimbriimonadales-bacterium] |
| MBS1724660.1 | HEPN+AAA-ATPase+Alpha+Beta-domain+wHTH+RRM | Type-1_BR.RM.like.systems | TUDOR+SF1-SF2-Helicase+iREase+CTD+REase->METHYLASE->HEPN+DUF499*->MORC-><-?\|\|N6_N4_MTase-><-Phage_integrase<-MAD-NTD+HNH | 1107 | Bacteria>Terrabacteria_group>Armatimonadota>unclassified_Armatimonadota | GCA_018268375.1 | Armatimonadota_bacterium | JSS66_17085 | 550844..554167 | 2033014 | DUF499-domain-containing-protein-[Armatimonadetes-bacterium] |
| MBI3912159.1 | HEPN+AAA-ATPase+Alpha+Beta-domain+wHTH+RRM | Type-1_BR.RM.like.systems | METHYLASE-p->DUF433->PIN->HEPN+DUF499*->?-><-NTP_transf<-Peptidase<-?<-Radical-SAM | 1099 | Bacteria>Terrabacteria_group>Armatimonadota>unclassified_Armatimonadota | GCA_016199525.1 | Armatimonadota_bacterium | HY320_14665 | 1429..4728 | 2033014 | ATP-binding-protein-[Armatimonadetes-bacterium] |
| HCD99868.1 | HEPN+AAA-ATPase+Alpha+Beta-domain+wHTH+RRM | Type-1_BR.RM.like.systems | METHYLASE-p->HEPN+DUF499*->TUDOR+SF1-SF2-Helicase+iREase+CTD+REase-p-> | 1116 | Bacteria>Terrabacteria_group>Armatimonadota>unclassified_Armatimonadota | GCA_003516955.1 | Armatimonadota_bacterium | DER07_02370 | 679..4029 | 2033014 | AAA+-family-ATPase-[Armatimonadetes-bacterium] |
| MCX6379947.1 | HEPN+AAA-ATPase+Alpha+Beta-domain+wHTH+RRM | Type-1_BR.RM.like.systems | REase->HEPN+DUF499*-><-?<-?<-?<-?<-?\|\|Phage_int_SAM_1+Phage_integrase-> | 1119 | Bacteria>Terrabacteria_group>Armatimonadota>unclassified_Armatimonadota | GCA_026392135.1 | Armatimonadota_bacterium | NT023_10805 | 3514..6873 | 2033014 | MAG:-Swt1-family-HEPN-domain-containing-protein-[Armatimonadetes-bacterium] |
| MBM3457180.1 | HEPN+AAA-ATPase+Alpha+Beta-domain+wHTH+RRM | Type-1_BR.RM.like.systems | METHYLASE-p->HEPN+DUF499*-> | 1128 | Bacteria>Terrabacteria_group>Armatimonadota>unclassified_Armatimonadota | GCA_016868995.1 | Armatimonadota_bacterium | FJX77_01400 | 1230..4616 | 2033014 | DUF499-domain-containing-protein-[Armatimonadetes-bacterium] |
| AKM18157.1 | HEPN+AAA-ATPase+Alpha+Beta-domain+wHTH+RRM | Type-1_BR.RM.like.systems | <-GspA-ATPase<-HTH_Tnp<-ZnR\|\|?->TUDOR+SF1-SF2-Helicase+iREase+CTD+REase->METHYLASE->HEPN+DUF499*-><-?<-DDE_5+DDE_Tnp_2\|\|REase-><-Arm-DNA-bind_4+Phage_int_SAM_3+Phage_integrase<-DDE_5+DDE_Tnp_2 | 1126 | Bacteria>Terrabacteria_group>Bacillota>Bacilli>Bacillales | GCA_001028085.1 | Geobacillus_sp._12AMOR1 | GARCT_00860 | 838569..841949 | 1629723 | hypothetical-protein-GARCT_00860-[Geobacillus-sp.-12AMOR1] |
| WP_220678456.1 | HEPN+AAA-ATPase+Alpha+Beta-domain+wHTH+RRM | Type-1_BR.RM.like.systems | TUDOR+SF1-SF2-Helicase+iREase+CTD+REase->METHYLASE-p-><-MULE-TRANSPOSASE\|\|METHYLASE->HEPN+DUF499*->?->DUF772+DDE_Tnp_1+DDE_Tnp_1_6->?->zf-ISL3+DDE_Tnp_ISL3->?->Glycos_trans_3N+Glycos_transf_3-> | 1124 | Bacteria>Terrabacteria_group>Bacillota>Bacilli>Bacillales | GCF_019645935.1 | Hydrogenibacillus_sp._N12 | K2M58_RS04470 | 965696..969070 | 2866627 | Swt1-family-HEPN-domain-containing-protein-[Hydrogenibacillus-sp.-N12] |
| STO36143.1 | HEPN+AAA-ATPase+Alpha+Beta-domain+wHTH+RRM | Type-1_BR.RM.like.systems | <-REase\|\|DDE_5+DDE_Tnp_2->HIN-HTH->?->?-><-HEPN+DUF499*<-METHYLASE<-DUF4277+DDE_Tnp_1<-METHYLASE-p<-TUDOR+SF1-SF2-Helicase+iREase+CTD+REase<-?\|\|ZnR-> | 1026 | Bacteria>Terrabacteria_group>Bacillota>Bacilli>Bacillales | GCA_900450595.1 | [Flavobacterium]_thermophilum | NCTC11526_03101 | 455797..458877 | 1414643 | Protein-of-uncharacterised-function-(DUF499)-[[Flavobacterium] |
| WP_129929317.1 | HEPN+AAA-ATPase+Alpha+Beta-domain+wHTH+RRM | Type-1_BR.RM.like.systems | REase+N6_MTase->?->TUDOR+SF1-SF2-Helicase+iREase+CTD+REase->METHYLASE->HEPN+DUF499*->ParB+HNH-><-DDE_Tnp_1<-?<-?\|\|DDE_Tnp_1-> | 1123 | Bacteria>Terrabacteria_group>Bacillota>Bacilli>Bacillales | - | Sporolactobacillus_sp._THM19-2 | EWH91_06945 | 33831..37202 | 2511171 | DUF499-domain-containing-protein-[Sporolactobacillus-sp.-THM19-2] |
| RYL92593.1 | HEPN+AAA-ATPase+Alpha+Beta-domain+wHTH+RRM | Type-1_BR.RM.like.systems | REase+N6_MTase->?->TUDOR+SF1-SF2-Helicase+iREase+CTD+REase->METHYLASE->HEPN+DUF499*->ParB+HNH-><-DDE_Tnp_1<-?<-?\|\|DDE_Tnp_1-> | 1123 | Bacteria>Terrabacteria_group>Bacillota>Bacilli>Bacillales | GCA_004153195.1 | Sporolactobacillus_sp._THM19-2 | EWH91_06945 | 33831..37202 | 2511171 | ATP-binding-protein-[Sporolactobacillus-sp.-THM19-2] |
| WP_054669274.1 | HEPN+AAA-ATPase+Alpha+Beta-domain+wHTH+RRM | Type-1_BR.RM.like.systems | <-DUF499-p<-HEPN+DUF499*<-METHYLASE-p<-METHYLASE-p<-METHYLASE-p<-TUDOR+SF1-SF2-Helicase+iREase+CTD+REase-p | 833 | Bacteria>Terrabacteria_group>Bacillota>Bacilli>Bacillales | GCF_001311905.1 | Calditerricola_satsumensis | JCM14719_RS01550 | 8456..10957 | 373054 | Swt1-family-HEPN-domain-containing-protein-[Calditerricola-satsumensis] |
| WP_124727907.1 | HEPN+AAA-ATPase+Alpha+Beta-domain+wHTH+RRM | Type-1_BR.RM.like.systems | Mbetalac->?-><-?<-?<-ABhydrolase<-HEPN+DUF499*<-METHYLASE<-TUDOR+SF1-SF2-Helicase+iREase+CTD+REase\|\|?->ACET-><-CHTH | 1107 | Bacteria>Terrabacteria_group>Bacillota>Bacilli>Bacillales | GCF_003856495.1 | Staphylospora_marina | EG886_RS09450 | 1890646..1893969 | 2490858 | Swt1-family-HEPN-domain-containing-protein-[Staphylospora-marina] |
| WP_188816795.1 | HEPN+AAA-ATPase+Alpha+Beta-domain+wHTH+RRM | Type-1_BR.RM.like.systems | <-Radical-SAM+TRAM<-?<-?<-?<-?<-?<-HEPN+DUF499*<-METHYLASE<-TUDOR+SF1-SF2-Helicase+iREase+CTD+REase<-Peptidase | 1124 | Bacteria>Terrabacteria_group>Bacillota>Bacilli>Bacillales | GCF_014646935.1 | Calditerricola_satsumensis | IEX61_RS03410 | 40382..43756 | 373054 | DUF499-domain-containing-protein-[Calditerricola-satsumensis] |
| PTQ56106.1 | HEPN+AAA-ATPase+Alpha+Beta-domain+wHTH+RRM | Type-1_BR.RM.like.systems | TUDOR+SF1-SF2-Helicase+iREase+CTD+REase->METHYLASE->HEPN+DUF499*-><-?\|\|?-><-?<-?<-Primpol+D5N+D5ATPase | 1116 | Bacteria>Terrabacteria_group>Bacillota>Bacilli>Bacillales | GCA_003050645.1 | Candidatus_Carbobacillus_altaicus | BSOLF_0824 | 6920..10270 | 2163959 | hypothetical-protein-BSOLF_0824-[Candidatus-Carbobacillus-altaicus] |
| WP_244410622.1 | HEPN+AAA-ATPase+Alpha+Beta-domain+wHTH+RRM | Type-1_BR.RM.like.systems | <-HEPN+DUF499*<-METHYLASE<-TUDOR+SF1-SF2-Helicase+iREase+CTD+REase<-?\|\|?-><-MULE-TRANSPOSASE | 1126 | Bacteria>Terrabacteria_group>Bacillota>Bacilli>Bacillales | GCF_022846475.1 | Parageobacillus_thermoglucosidasius | MWM14_RS16385 | 3230827..3234207 | 1426 | Swt1-family-HEPN-domain-containing-protein-[Parageobacillus-thermoglucosidasius] |
| WP_236408338.1 | HEPN+AAA-ATPase+Alpha+Beta-domain+wHTH+RRM | Type-1_BR.RM.like.systems | XIS-HTH->TUDOR+SF1-SF2-Helicase+iREase+CTD+REase->METHYLASE->HEPN+DUF499*-> | 1134 | Bacteria>Terrabacteria_group>Bacillota>Bacilli>Bacillales | GCF_021608135.1 | Microaerobacter_geothermalis | L1765_RS15200 | 6759..10163 | 674972 | Swt1-family-HEPN-domain-containing-protein-[Microaerobacter-geothermalis] |
| WP_258111975.1 | HEPN+AAA-ATPase+Alpha+Beta-domain+wHTH+RRM | Type-1_BR.RM.like.systems | <-HicB<-HicA_toxin<-?<-PIN<-?\|\|METHYLASE-p->HEPN+DUF499*->RVT-> | 1128 | Bacteria>Terrabacteria_group>Bacillota>Bacilli>Bacillales | GCF_024706985.1 | Alicyclobacillus_sp._SP_1 | M4I36_RS11945 | 3650..7036 | 2942475 | DUF499-domain-containing-protein-[Alicyclobacillus-sp.-SP_1] |
| MBO2534075.1 | HEPN+AAA-ATPase+Alpha+Beta-domain+wHTH+RRM | Type-1_BR.RM.like.systems | <-MqsA_antitoxin+HTH_3<-?<-HEPN+DUF499*<-METHYLASE-p | 1117 | Bacteria>Terrabacteria_group>Bacillota>Bacilli>Bacillales | GCA_017577425.1 | Thermoactinomycetaceae_bacterium | CW342_14595 | 2341..5694 | 2056641 | AAA+-family-ATPase-[Thermoactinomycetaceae-bacterium] |
| MCG0314785.1 | HEPN+AAA-ATPase+Alpha+Beta-domain+wHTH+RRM | Type-1_BR.RM.like.systems | <-HEPN+DUF499*<-METHYLASE-p | 616 | Bacteria>Terrabacteria_group>Bacillota>Bacilli>Bacillales | GCA_021890815.1 | Calditerricola_sp. | L5657_09065 | 1..1847 | 2885161 | DUF499-domain-containing-protein-[Calditerricola-sp.] |
| WP_181339998.1 | HEPN+AAA-ATPase+Alpha+Beta-domain+wHTH+RRM | Type-1_BR.RM.like.systems | DeoR-HTH+ISOCOT->?->?->?->?-><-HEPN+DUF499*<-METHYLASE<-TUDOR+SF1-SF2-Helicase+iREase+CTD+REase\|\|OLD-ABC+TOPRIM-> | 1104 | Bacteria>Terrabacteria_group>Bacillota>Bacillota_sensu_stricto_incertae_sedis>Capillibacterium | GCF_013664685.1 | Capillibacterium_thermochitinicola | G5B42_RS08290 | 130685..133999 | 2699427 | Swt1-family-HEPN-domain-containing-protein-[Capillibacterium-thermochitinicola] |
| WP_087282257.1 | HEPN+AAA-ATPase+Alpha+Beta-domain+wHTH+RRM | Type-1_BR.RM.like.systems | XIS-HTH->TUDOR+SF1-SF2-Helicase+iREase+CTD+REase->METHYLASE->METHYLASE->HEPN+DUF499*->?->SbcC->SF1-SF2-Helicase->WYL->?->LexA-HTH+LexA-protease-> | 1134 | Bacteria>Terrabacteria_group>Bacillota>Clostridia | GCF_016902345.1 | Eubacteriales | H6B11_RS01180 | 245745..249149 | 186802 | MULTISPECIES:-DUF499-domain-containing-protein-[Eubacteriales] |
| HIR07141.1 | HEPN+AAA-ATPase+Alpha+Beta-domain+wHTH+RRM | Type-1_BR.RM.like.systems | TUDOR+SF1-SF2-Helicase+iREase+CTD+REase->METHYLASE->HEPN+DUF499*-> | 987 | Bacteria>Terrabacteria_group>Bacillota>Clostridia>Clostridia_incertae_sedis | GCA_018712885.1 | Candidatus_Pullichristensenella_stercoripullorum | IAA71_02960 | 11277..14237 | 2840910 | DUF499-domain-containing-protein-[Candidatus-Pullichristensenella-stercoripullorum] |
| WP_156058230.1 | HEPN+AAA-ATPase+Alpha+Beta-domain+wHTH+RRM | Type-1_BR.RM.like.systems | AbiGi->GDResolvase+Recombinase+ZnR->TUDOR+SF1-SF2-Helicase+iREase+CTD+REase->TUDOR+SF1-SF2-Helicase+iREase+CTD+REase-p->MNS->METHYLASE->HEPN+DUF499*->?->ParB->CRP-HTH+GNTR-HTH->?->CHTH->GDResolvase+Recombinase+ZnR-> | 1130 | Bacteria>Terrabacteria_group>Bacillota>Clostridia>Eubacteriales | GCF_019969385.1 | Faecalibacterium_prausnitzii | GO822_RS02960 | 605262..608654 | 853 | Swt1-family-HEPN-domain-containing-protein-[Faecalibacterium-prausnitzii] |
| WP_015942906.1 | HEPN+AAA-ATPase+Alpha+Beta-domain+wHTH+RRM | Type-1_BR.RM.like.systems | SF1-SF2-Helicase+HKD-DNase->MORC+SF1-SF2-Helicase->?->HTH_26->TUDOR+SF1-SF2-Helicase+iREase+CTD+REase->METHYLASE->HEPN+DUF499*->OLD-ABC+TOPRIM->SF1-SF2-Helicase->METHYLASE-p->ParB+HNH->HEPN->pPIWI+pPIWI-> | 1137 | Bacteria>Terrabacteria_group>Bacillota>Clostridia>Eubacteriales | GCF_000021925.1 | Desulfitobacterium_hafniense | DPCES_RS04010 | 854740..858153 | 49338 | DUF499-domain-containing-protein-[Desulfitobacterium-hafniense] |
| WP_018214270.1 | HEPN+AAA-ATPase+Alpha+Beta-domain+wHTH+RRM | Type-1_BR.RM.like.systems | <-pPIWI+pPIWI<-HEPN<-ParB+HNH<-METHYLASE-p<-Schlafen<-MORC<-HEPN+DUF499*<-METHYLASE<-TUDOR+SF1-SF2-Helicase+iREase+CTD+REase<-HTH_26<-?<-MORC+SF1-SF2-Helicase<-SF1-SF2-Helicase+HKD-DNase | 1137 | Bacteria>Terrabacteria_group>Bacillota>Clostridia>Eubacteriales | GCF_000378805.1 | Desulfitobacterium_hafniense | DESHAF_RS0122600 | 1962937..1966350 | 49338 | DUF499-domain-containing-protein-[Desulfitobacterium-hafniense] |
| WP_207684444.1 | HEPN+AAA-ATPase+Alpha+Beta-domain+wHTH+RRM | Type-1_BR.RM.like.systems | GDResolvase->AbiGi->GDResolvase+Recombinase+ZnR->TUDOR+SF1-SF2-Helicase+iREase+CTD+REase->?->METHYLASE->HEPN+DUF499*->ParB->CRP-HTH+GNTR-HTH->?->?->ZDNA-HTH->GDResolvase+Recombinase+ZnR-> | 1129 | Bacteria>Terrabacteria_group>Bacillota>Clostridia>Eubacteriales | GCF_017377535.1 | Faecalibacterium_sp._Marseille-Q4137 | J3353_RS06385 | 45904..49293 | 2817021 | DUF499-domain-containing-protein-[Faecalibacterium-sp.-Marseille-Q4137] |
| MBS4922099.1 | HEPN+AAA-ATPase+Alpha+Beta-domain+wHTH+RRM | Type-1_BR.RM.like.systems | AbiGi->GDResolvase+Recombinase+ZnR->TUDOR+SF1-SF2-Helicase+iREase+CTD+REase->MNS->METHYLASE->HEPN+DUF499*->ParB->CRP-HTH+GNTR-HTH->?->ZDNA-HTH->GDResolvase+Recombinase+ZnR-> | 1129 | Bacteria>Terrabacteria_group>Bacillota>Clostridia>Eubacteriales | GCA_018364195.1 | Faecalibacterium_prausnitzii | KHZ84_09455 | 17444..20833 | 853 | DUF499-domain-containing-protein-[Faecalibacterium-prausnitzii] |
| UYJ02009.1 | HEPN+AAA-ATPase+Alpha+Beta-domain+wHTH+RRM | Type-1_BR.RM.like.systems | TRAM+Methyltransf_31->AbiGi->TUDOR+SF1-SF2-Helicase+iREase+CTD+REase->MNS->METHYLASE->HEPN+DUF499*->ParB->CRP-HTH+GNTR-HTH->CRP-HTH+GNTR-HTH->?->?->ZDNA-HTH-> | 1130 | Bacteria>Terrabacteria_group>Bacillota>Clostridia>Eubacteriales | GCA_025757765.1 | Faecalibacterium_prausnitzii | OGM65_02940 | 601933..605325 | 853 | MAG:-Swt1-family-HEPN-domain-containing-protein-[Faecalibacterium-prausnitzii] |
| WP_166512732.1 | HEPN+AAA-ATPase+Alpha+Beta-domain+wHTH+RRM | Type-1_BR.RM.like.systems | <-SF1-SF2-Helicase+DUF1998<-HEPN+Mrr_N<-HDRnase<-?<-Primpol+D5N+D5ATPase<-?<-HEPN+DUF499*<-METHYLASE<-TUDOR+SF1-SF2-Helicase+iREase+CTD+REase<-Peptidase_M15 | 1110 | Bacteria>Terrabacteria_group>Bacillota>Clostridia>Eubacteriales | GCF_008124625.1 | Desulfallas_thermosapovorans | LX24_RS13825 | 12605..15937 | 58137 | Swt1-family-HEPN-domain-containing-protein-[Desulfallas-thermosapovorans] |
| WP_076780282.1 | HEPN+AAA-ATPase+Alpha+Beta-domain+wHTH+RRM | Type-1_BR.RM.like.systems | HDRnase->TRAM+Methyltransf_31-><-?<-?<-HEPN+DUF499*<-METHYLASE<-METHYLASE<-TUDOR+SF1-SF2-Helicase+iREase+CTD+REase<-HTH_26<-SIGMA-HTH+SIGMA-HTH<-HTH | 1134 | Bacteria>Terrabacteria_group>Bacillota>Clostridia>Eubacteriales | GCF_900120345.1 | Lachnoclostridium_phocaeense | BQ7370_RS15745 | 3233710..3237114 | 1871021 | DUF499-domain-containing-protein-[Lachnoclostridium-phocaeense] |
| MBU5249610.1 | HEPN+AAA-ATPase+Alpha+Beta-domain+wHTH+RRM | Type-1_BR.RM.like.systems | DUF772+DDE_Tnp_1_6->Schlafen+HATPase+HTH->TUDOR+SF1-SF2-Helicase+iREase+CTD+REase->METHYLASE->METHYLASE->HEPN+DUF499*->?->CRP-HTH+GNTR-HTH->?->?->?->REase-> | 1128 | Bacteria>Terrabacteria_group>Bacillota>Clostridia>Eubacteriales | GCA_018916865.1 | Coprococcus_comes | KQH93_07505 | 29424..32810 | 410072 | DUF499-domain-containing-protein-[Coprococcus-comes] |
| WP_033117941.1 | HEPN+AAA-ATPase+Alpha+Beta-domain+wHTH+RRM | Type-1_BR.RM.like.systems | <-HTH\|\|?-><-?<-Response_reg+LuxR-HTH<-HISKIN+MORC<-?<-HEPN+DUF499*<-METHYLASE<-TUDOR+SF1-SF2-Helicase+iREase+CTD+REase<-XIS-HTH<-?\|\|?->GDResolvase-> | 1116 | Bacteria>Terrabacteria_group>Bacillota>Clostridia>Eubacteriales | GCF_000765135.1 | Intestinimonas | E0E41_RS12870 | 2605213..2608563 | 1392389 | MULTISPECIES:-DUF499-domain-containing-protein-[Intestinimonas] |
| MBO6231158.1 | HEPN+AAA-ATPase+Alpha+Beta-domain+wHTH+RRM | Type-1_BR.RM.like.systems | <-HNH+HNH\|\|?-><-HTH_3\|\|TUDOR+SF1-SF2-Helicase+iREase+CTD+REase->METHYLASE->HEPN+DUF499*->?->Classical-AAA->REase->SMODS->HTH_3-><-Phd-YefM | 1114 | Bacteria>Terrabacteria_group>Bacillota>Clostridia>Eubacteriales | GCA_017635095.1 | Ruminiclostridium_sp. | J6050_11400 | 23257..26601 | 2053608 | DUF499-domain-containing-protein-[Ruminiclostridium-sp.] |
| WP_204889591.1 | HEPN+AAA-ATPase+Alpha+Beta-domain+wHTH+RRM | Type-1_BR.RM.like.systems | HTH_26->TUDOR+SF1-SF2-Helicase+iREase+CTD+REase->METHYLASE->Schlafen+HATPase+HTH->HEPN+DUF499*-><-?<-HIN-HTH\|\|DUF772+DDE_Tnp_1+DDE_Tnp_1_6-> | 1123 | Bacteria>Terrabacteria_group>Bacillota>Clostridia>Eubacteriales | GCF_016902175.1 | Pseudoflavonifractor_phocaeensis | H6B27_RS11850 | 9172..12543 | 1870988 | Swt1-family-HEPN-domain-containing-protein-[Pseudoflavonifractor-phocaeensis] |
| MCD7717395.1 | HEPN+AAA-ATPase+Alpha+Beta-domain+wHTH+RRM | Type-1_BR.RM.like.systems | REase->?->?-><-NACHT\|\|?-><-REase+ParB<-HEPN+DUF499*<-METHYLASE<-Schlafen+HATPase+HTH<-METHYLASE<-TUDOR+SF1-SF2-Helicase+iREase+CTD+REase<-XIS-HTH | 1132 | Bacteria>Terrabacteria_group>Bacillota>Clostridia>Eubacteriales | GCA_021199235.1 | Lachnospiraceae_bacterium | LUI39_13305 | 18966..22364 | 1898203 | DUF499-domain-containing-protein-[Lachnospiraceae-bacterium] |
| MBD9257204.1 | HEPN+AAA-ATPase+Alpha+Beta-domain+wHTH+RRM | Type-1_BR.RM.like.systems | TRAM+Methyltransf_31->AbiGi->GDResolvase+Recombinase+ZnR->TUDOR+SF1-SF2-Helicase+iREase+CTD+REase->?->METHYLASE->HEPN+DUF499*-> | 1129 | Bacteria>Terrabacteria_group>Bacillota>Clostridia>Eubacteriales | GCA_014848725.1 | Faecalibacterium_sp. | EGR99_01350 | 18059..21448 | 1971605 | ATP-binding-protein-[Faecalibacterium-sp.] |
| WP_195975511.1 | HEPN+AAA-ATPase+Alpha+Beta-domain+wHTH+RRM | Type-1_BR.RM.like.systems | HTH_26->TUDOR+SF1-SF2-Helicase+iREase+CTD+REase->METHYLASE->METHYLASE->HEPN+DUF499*-><-?<-GDResolvase+Recombinase+ZnR<-CHTH | 1134 | Bacteria>Terrabacteria_group>Bacillota>Clostridia>Eubacteriales | GCF_015670165.1 | Hydrogeniiclostridium_mannosilyticum | I4V57_RS07180 | 144125..147529 | 2764322 | DUF499-domain-containing-protein-[Hydrogeniiclostridium-mannosilyticum] |
| HHW47352.1 | HEPN+AAA-ATPase+Alpha+Beta-domain+wHTH+RRM | Type-1_BR.RM.like.systems | XIS-HTH->TUDOR+SF1-SF2-Helicase+iREase+CTD+REase->METHYLASE->HEPN+DUF499*->?->ParB+HNH-><-Aminotran_1_2\|\|?->ABC-ATPase+SbcC-> | 1135 | Bacteria>Terrabacteria_group>Bacillota>Clostridia>Eubacteriales | GCA_012840535.1 | Clostridiaceae_bacterium | GXX14_01895 | 11508..14915 | 1898204 | ATP-binding-protein-[Clostridiaceae-bacterium] |
| WP_211802648.1 | HEPN+AAA-ATPase+Alpha+Beta-domain+wHTH+RRM | Type-1_BR.RM.like.systems | REase+CTD6->?->SbcC+SbcC->HTH_26->TUDOR+SF1-SF2-Helicase+iREase+CTD+REase->METHYLASE->HEPN+DUF499*->SF1-SF2-Helicase-p-> | 1134 | Bacteria>Terrabacteria_group>Bacillota>Clostridia>Eubacteriales | GCF_018128025.1 | Proteiniclasticum_sediminis | KCG48_RS13075 | 34929..38333 | 2804028 | DUF499-domain-containing-protein-[Proteiniclasticum-sediminis] |
| MCH3999398.1 | HEPN+AAA-ATPase+Alpha+Beta-domain+wHTH+RRM | Type-1_BR.RM.like.systems | SbcC->HTH_26->TUDOR+SF1-SF2-Helicase+iREase+CTD+REase->METHYLASE->METHYLASE->HEPN+DUF499*->?->REase->HNH->?-><-?\|\|MarR-HTH-> | 1138 | Bacteria>Terrabacteria_group>Bacillota>Clostridia>Eubacteriales | GCA_022483165.1 | Lachnospiraceae_bacterium | LKE66_07755 | 1772421..1775837 | 1898203 | DUF499-domain-containing-protein-[Lachnospiraceae-bacterium] |
| MCI7107135.1 | HEPN+AAA-ATPase+Alpha+Beta-domain+wHTH+RRM | Type-1_BR.RM.like.systems | REase->?->?->?-><-ParB+HNH<-HEPN+DUF499*<-METHYLASE<-METHYLASE<-TUDOR+SF1-SF2-Helicase+iREase+CTD+REase<-HTH_26<-SbcC+SbcC | 1129 | Bacteria>Terrabacteria_group>Bacillota>Clostridia>Eubacteriales | GCA_022775045.1 | Agathobacter_sp. | MR945_04560 | 37097..40486 | 2021311 | DUF499-domain-containing-protein-[Agathobacter-sp.] |
| MBZ4669808.1 | HEPN+AAA-ATPase+Alpha+Beta-domain+wHTH+RRM | Type-1_BR.RM.like.systems | ABhydrolase+ABhydrolase->?->?->TUDOR+SF1-SF2-Helicase+iREase+CTD+REase->METHYLASE->HEPN+DUF499*->?-><-PIN<-PhdYeFM_antitox | 1133 | Bacteria>Terrabacteria_group>Bacillota>Clostridia>Eubacteriales | GCA_020054675.1 | Oscillospiraceae_bacterium | JG769_112 | 107395..110796 | 2485925 | hypothetical-protein-[Oscillospiraceae-bacterium] |
| WP_073238382.1 | HEPN+AAA-ATPase+Alpha+Beta-domain+wHTH+RRM | Type-1_BR.RM.like.systems | <-TPR<-?<-Radical-SAM<-?<-tRNA<-?<-HEPN+DUF499*<-METHYLASE<-TUDOR+SF1-SF2-Helicase+iREase+CTD+REase\|\|tRNA->RNaseH->ZnR->PAIREDC-HTH-> | 1123 | Bacteria>Terrabacteria_group>Bacillota>Clostridia>Eubacteriales | - | Desulforamulus_putei | BUB67_RS07930 | 94636..98007 | 74701 | Swt1-family-HEPN-domain-containing-protein-[Desulforamulus-putei] |
| WP_204872058.1 | HEPN+AAA-ATPase+Alpha+Beta-domain+wHTH+RRM | Type-1_BR.RM.like.systems | SIGMA-HTH->HTH_26->TUDOR+SF1-SF2-Helicase+iREase+CTD+REase->Schlafen+HATPase+HTH->METHYLASE->METHYLASE->HEPN+DUF499*-> | 1132 | Bacteria>Terrabacteria_group>Bacillota>Clostridia>Eubacteriales | GCF_016899615.1 | Pseudoflavonifractor_capillosus | H9X87_RS10190 | 14789..18187 | 106588 | DUF499-domain-containing-protein-[Pseudoflavonifractor-capillosus] |
| WP_168124741.1 | HEPN+AAA-ATPase+Alpha+Beta-domain+wHTH+RRM | Type-1_BR.RM.like.systems | <-HEPN+DUF499*<-METHYLASE<-METHYLASE<-Schlafen+HATPase+HTH<-TUDOR+SF1-SF2-Helicase+iREase+CTD+REase<-HTH_26<-SIGMA-HTH | 1132 | Bacteria>Terrabacteria_group>Bacillota>Clostridia>Eubacteriales | GCF_012027765.1 | Pseudoflavonifractor_sp._SW1122 | E0L15_RS04955 | 37276..40674 | 2530044 | DUF499-domain-containing-protein-[Pseudoflavonifractor-sp.-SW1122] |
| MBQ3841838.1 | HEPN+AAA-ATPase+Alpha+Beta-domain+wHTH+RRM | Type-1_BR.RM.like.systems | <-XIS-HTH<-HNH+HNH\|\|?-><-HTH_3\|\|TUDOR+SF1-SF2-Helicase+iREase+CTD+REase->METHYLASE->HEPN+DUF499*->?->?->HTH_3-><-?<-HTH_3 | 1114 | Bacteria>Terrabacteria_group>Bacillota>Clostridia>Eubacteriales | GCA_017395555.1 | Ruminiclostridium_sp. | II820_04000 | 63719..67063 | 2053608 | DUF499-domain-containing-protein-[Ruminiclostridium-sp.] |
| KLU38984.1 | HEPN+AAA-ATPase+Alpha+Beta-domain+wHTH+RRM | Type-1_BR.RM.like.systems | TUDOR+SF1-SF2-Helicase+iREase+CTD+REase->METHYLASE->METHYLASE->METHYLASE->HEPN+DUF499*->?->?->METHYLASE-p->?-><-?\|\|CHTH-> | 1130 | Bacteria>Terrabacteria_group>Bacillota>Clostridia>Eubacteriales | GCA_001028815.1 | Peptococcaceae_bacterium_1109 | AA931_11700 | 75012..78404 | 1655638 | hypothetical-protein-AA931_11700-[Peptococcaceae-bacterium-1109] |
| WP_087179373.1 | HEPN+AAA-ATPase+Alpha+Beta-domain+wHTH+RRM | Type-1_BR.RM.like.systems | <-NACHT\|\|PemK_toxin->?-><-HEPN+DUF499*<-METHYLASE<-METHYLASE<-TUDOR+SF1-SF2-Helicase+iREase+CTD+REase<-XIS-HTH | 1128 | Bacteria>Terrabacteria_group>Bacillota>Clostridia>Eubacteriales | GCF_002161355.1 | Lachnoclostridium_sp._An76 | B5G27_RS13410 | 49376..52762 | 1965654 | DUF499-domain-containing-protein-[Lachnoclostridium-sp.-An76] |
| NQS74830.1 | HEPN+AAA-ATPase+Alpha+Beta-domain+wHTH+RRM | Type-1_BR.RM.like.systems | Peptidase_M24->?->ZnR->?->TUDOR+SF1-SF2-Helicase+iREase+CTD+REase->METHYLASE->HEPN+DUF499*->?->XIS-HTH-><-HTH+MPTase | 1135 | Bacteria>Terrabacteria_group>Bacillota>Clostridia>Eubacteriales | GCA_013201395.1 | Candidatus_Syntrophopropionicum_ammoniitolerans | HQP61_00110 | 30439..33846 | 2807340 | ATP-binding-protein-[Peptococcaceae-bacterium] |
| MBC7075381.1 | HEPN+AAA-ATPase+Alpha+Beta-domain+wHTH+RRM | Type-1_BR.RM.like.systems | MazE_antitoxin->RelE-ParE->\|\|?->?->TUDOR+SF1-SF2-Helicase+iREase+CTD+REase->METHYLASE->HEPN+DUF499*->ParB+HNH-> | 1122 | Bacteria>Terrabacteria_group>Bacillota>Clostridia>Eubacteriales | GCA_014360155.1 | Syntrophomonadaceae_bacterium | H5T98_04805 | 11477..14845 | 2093811 | ATP-binding-protein-[Syntrophomonadaceae-bacterium] |
| MCI7280004.1 | HEPN+AAA-ATPase+Alpha+Beta-domain+wHTH+RRM | Type-1_BR.RM.like.systems | METHYLASE->HEPN+DUF499*-><-?\|\|ZDNA-HTH->GDResolvase+Recombinase+ZnR-><-tRNA<-Peptidase_U32+Peptidase_U32_C<-?<-GNAT | 1129 | Bacteria>Terrabacteria_group>Bacillota>Clostridia>Eubacteriales | GCA_022783405.1 | Faecalibacterium_prausnitzii | MR538_06465 | 2347..5736 | 853 | DUF499-domain-containing-protein-[Faecalibacterium-prausnitzii] |
| MBR3354625.1 | HEPN+AAA-ATPase+Alpha+Beta-domain+wHTH+RRM | Type-1_BR.RM.like.systems | <-HTH\|\|HTH_26->TUDOR+SF1-SF2-Helicase+iREase+CTD+REase->METHYLASE->METHYLASE->METHYLASE->HEPN+DUF499*-> | 1125 | Bacteria>Terrabacteria_group>Bacillota>Clostridia>Eubacteriales | GCA_017510465.1 | Oscillospiraceae_bacterium | IKG47_04625 | 20932..24309 | 2485925 | DUF499-domain-containing-protein-[Oscillospiraceae-bacterium] |
| WP_228724972.1 | HEPN+AAA-ATPase+Alpha+Beta-domain+wHTH+RRM | Type-1_BR.RM.like.systems | HTH_36-><-?<-?<-?<-JetA<-HEPN+DUF499*<-METHYLASE<-METHYLASE-p<-TUDOR+SF1-SF2-Helicase+iREase+CTD+REase<-CHTH | 1117 | Bacteria>Terrabacteria_group>Bacillota>Clostridia>Eubacteriales | GCF_014303955.1 | Caproicibacter_fermentans | HCR03_RS14035 | 2844426..2847779 | 2576756 | DUF499-domain-containing-protein-[Caproicibacter-fermentans] |
| WP_008413292.1 | HEPN+AAA-ATPase+Alpha+Beta-domain+wHTH+RRM | Type-1_BR.RM.like.systems | <-HIN-HTH<-?<-TraJ-RHH<-HEPN+DUF499*<-METHYLASE<-TUDOR+SF1-SF2-Helicase+iREase+CTD+REase<-XIS-HTH<-?<-?<-HTH | 1135 | Bacteria>Terrabacteria_group>Bacillota>Clostridia>Eubacteriales | GCF_000315365.1 | Desulforamulus_hydrothermalis | BUA68_RS05960 | 1191..4598 | 412895 | Swt1-family-HEPN-domain-containing-protein-[Desulforamulus-hydrothermalis] |
| MCD8011085.1 | HEPN+AAA-ATPase+Alpha+Beta-domain+wHTH+RRM | Type-1_BR.RM.like.systems | <-HEPN+DUF499*<-RelE-ParE<-RelB<-METHYLASE-p<-METHYLASE-p<-GDResolvase<-GDResolvase+Recombinase+ZnR | 819 | Bacteria>Terrabacteria_group>Bacillota>Clostridia>Eubacteriales | GCA_021202235.1 | Lachnospiraceae_bacterium | LUF34_10055 | 34..2493 | 1898203 | DUF499-domain-containing-protein-[Lachnospiraceae-bacterium] |
| WP_092487583.1 | HEPN+AAA-ATPase+Alpha+Beta-domain+wHTH+RRM | Type-1_BR.RM.like.systems | HIN-HTH-><-DDE_Tnp_1<-HEPN+DUF499*<-PIN<-SbcC<-METHYLASE<-TUDOR+SF1-SF2-Helicase+iREase+CTD+REase | 888 | Bacteria>Terrabacteria_group>Bacillota>Clostridia>Eubacteriales | GCF_900115975.1 | Desulfoscipio_geothermicus | BM299_RS17670 | 2600..5266 | 39060 | Swt1-family-HEPN-domain-containing-protein-[Desulfoscipio-geothermicus] |
| MCI8809285.1 | HEPN+AAA-ATPase+Alpha+Beta-domain+wHTH+RRM | Type-1_BR.RM.like.systems | <-ABC-ATPase+SbcC<-?<-?\|\|TUDOR+SF1-SF2-Helicase+iREase+CTD+REase->METHYLASE->HEPN+DUF499*->Resolvase-> | 1137 | Bacteria>Terrabacteria_group>Bacillota>Clostridia>Eubacteriales | GCA_022795445.1 | Oscillibacter_sp. | HFF84_04010 | 14987..18400 | 1945593 | ATP-binding-protein-[Oscillibacter-sp.] |
| WP_135225812.1 | HEPN+AAA-ATPase+Alpha+Beta-domain+wHTH+RRM | Type-1_BR.RM.like.systems | <-HEPN+DUF499*<-METHYLASE<-TUDOR+SF1-SF2-Helicase+iREase+CTD+REase\|\|MarR-HTH->ABC-ATPase+SbcC-> | 1129 | Bacteria>Terrabacteria_group>Bacillota>Clostridia>Eubacteriales | GCF_004634385.1 | Thermaerobacter_sp._FW80 | E1B22_RS11885 | 2873077..2876466 | 2546351 | Swt1-family-HEPN-domain-containing-protein-[Thermaerobacter-sp.-FW80] |
| MBR3240816.1 | HEPN+AAA-ATPase+Alpha+Beta-domain+wHTH+RRM | Type-1_BR.RM.like.systems | <-MORC<-HEPN+DUF499*<-METHYLASE<-TUDOR+SF1-SF2-Helicase+iREase+CTD+REase<-?\|\|HTH->HTH_31->MPTase-> | 1133 | Bacteria>Terrabacteria_group>Bacillota>Clostridia>Eubacteriales | GCA_017511105.1 | Oscillospiraceae_bacterium | IKF99_20575 | 19684..23085 | 2485925 | DUF499-domain-containing-protein-[Oscillospiraceae-bacterium] |
| NLW38082.1 | HEPN+AAA-ATPase+Alpha+Beta-domain+wHTH+RRM | Type-1_BR.RM.like.systems | <-MethyltransfD12\|\|?->PIN-><-HEPN+DUF499*<-METHYLASE<-TPR+TPR<-MetJarc-HTH<-SF1-SF2-Helicase<-EVE | 1120 | Bacteria>Terrabacteria_group>Bacillota>Clostridia>Eubacteriales | GCA_012728715.1 | Peptococcaceae_bacterium | GXY90_05855 | 3552..6914 | 2052179 | ATP-binding-protein-[Peptococcaceae-bacterium] |
| MCL1790896.1 | HEPN+AAA-ATPase+Alpha+Beta-domain+wHTH+RRM | Type-1_BR.RM.like.systems | <-Peptidase\|\|TUDOR+SF1-SF2-Helicase+iREase+CTD+REase->METHYLASE->Schlafen+HATPase+HTH->HEPN+DUF499*-> | 1122 | Bacteria>Terrabacteria_group>Bacillota>Clostridia>Eubacteriales | GCA_009778595.1 | Peptococcaceae_bacterium | FWG40_05990 | 9595..12963 | 2052179 | Swt1-family-HEPN-domain-containing-protein-[Peptococcaceae-bacterium] |
| MBS3975902.1 | HEPN+AAA-ATPase+Alpha+Beta-domain+wHTH+RRM | Type-1_BR.RM.like.systems | <-Phage_terminase<-NACHT<-HEPN+DUF499*<-METHYLASE<-TUDOR+SF1-SF2-Helicase+iREase+CTD+REase-p | 1133 | Bacteria>Terrabacteria_group>Bacillota>Clostridia>Eubacteriales | GCA_018333865.1 | Syntrophomonadaceae_bacterium | KGZ75_04135 | 3808..7209 | 2093811 | DUF499-domain-containing-protein-[Syntrophomonadaceae-bacterium] |
| WP_166510781.1 | HEPN+AAA-ATPase+Alpha+Beta-domain+wHTH+RRM | Type-1_BR.RM.like.systems | <-HISKIN<-?<-?<-MPTase<-HTH<-?<-HEPN+DUF499*<-METHYLASE-p<-TUDOR+SF1-SF2-Helicase+iREase+CTD+REase-p | 1122 | Bacteria>Terrabacteria_group>Bacillota>Clostridia>Eubacteriales | GCF_008124625.1 | Desulfallas_thermosapovorans | LX24_RS03620 | 91122..94490 | 58137 | Swt1-family-HEPN-domain-containing-protein-[Desulfallas-thermosapovorans] |
| WP_066668116.1 | HEPN+AAA-ATPase+Alpha+Beta-domain+wHTH+RRM | Type-1_BR.RM.like.systems | <-HD<-?\|\|?->?->?-><-HEPN+DUF499*<-METHYLASE<-PIN<-MetJarc-HTH<-TUDOR+SF1-SF2-Helicase+iREase+CTD+REase | 1130 | Bacteria>Terrabacteria_group>Bacillota>Clostridia>Eubacteriales | GCF_001655685.1 | Desulfotomaculum_copahuensis | A6M21_RS09370 | 52458..55850 | 1838280 | Swt1-family-HEPN-domain-containing-protein-[Desulfotomaculum-copahuensis] |
| MBR3354573.1 | HEPN+AAA-ATPase+Alpha+Beta-domain+wHTH+RRM | Type-1_BR.RM.like.systems | HTH->tRNA->tRNA->tRNA->tRNA->tRNA->tRNA->tRNA->tRNA->?->TUDOR+SF1-SF2-Helicase+iREase+CTD+REase->METHYLASE->METHYLASE->HEPN+DUF499*->REase-> | 1111 | Bacteria>Terrabacteria_group>Bacillota>Clostridia>Eubacteriales | GCA_017510465.1 | Oscillospiraceae_bacterium | IKG47_04355 | 36923..40258 | 2485925 | DUF499-domain-containing-protein-[Oscillospiraceae-bacterium] |
| NLI53236.1 | HEPN+AAA-ATPase+Alpha+Beta-domain+wHTH+RRM | Type-1_BR.RM.like.systems | WYL->?->TUDOR+SF1-SF2-Helicase+iREase+CTD+REase->METHYLASE->HEPN+DUF499*->OLD-ABC+TOPRIM->?->?-><-tRNA<-SMS-KaiC | 1124 | Bacteria>Terrabacteria_group>Bacillota>Clostridia>Eubacteriales | GCA_012517905.1 | Clostridiales_bacterium | GX417_02810 | 9381..12755 | 1898207 | ATP-binding-protein-[Clostridiales-bacterium] |
| WP_161846233.1 | HEPN+AAA-ATPase+Alpha+Beta-domain+wHTH+RRM | Type-1_BR.RM.like.systems | TUDOR+SF1-SF2-Helicase+iREase+CTD+REase->?->METHYLASE->HEPN+DUF499*->REase-><-GDResolvase | 1134 | Bacteria>Terrabacteria_group>Bacillota>Clostridia>Eubacteriales | GCF_009935845.1 | Pseudoflavonifractor_sp._524-17 | D1159_RS13725 | 7029..10433 | 2304577 | DUF499-domain-containing-protein-[Pseudoflavonifractor-sp.-524-17] |
| MCI6268195.1 | HEPN+AAA-ATPase+Alpha+Beta-domain+wHTH+RRM | Type-1_BR.RM.like.systems | <-MerR-HTH<-?\|\|?->tRNA->HTH_26->TUDOR+SF1-SF2-Helicase+iREase+CTD+REase->METHYLASE->HEPN+DUF499*-> | 1130 | Bacteria>Terrabacteria_group>Bacillota>Clostridia>Eubacteriales | GCA_022772795.1 | Clostridiales_bacterium | MR637_06175 | 165246..168638 | 1898207 | DUF499-domain-containing-protein-[Clostridiales-bacterium] |
| NLV15838.1 | HEPN+AAA-ATPase+Alpha+Beta-domain+wHTH+RRM | Type-1_BR.RM.like.systems | PRTase-><-?\|\|TUDOR+SF1-SF2-Helicase+iREase+CTD+REase->METHYLASE->HEPN+DUF499*->REase-> | 1154 | Bacteria>Terrabacteria_group>Bacillota>Clostridia>Eubacteriales | GCA_012729505.1 | Syntrophomonadaceae_bacterium | GXY50_01325 | 75931..79395 | 2093811 | ATP-binding-protein-[Syntrophomonadaceae-bacterium] |
| MBQ9662099.1 | HEPN+AAA-ATPase+Alpha+Beta-domain+wHTH+RRM | Type-1_BR.RM.like.systems | TUDOR+SF1-SF2-Helicase+iREase+CTD+REase->METHYLASE->HEPN+DUF499*-><-?<-?<-?<-Glycos_transf_4 | 1119 | Bacteria>Terrabacteria_group>Bacillota>Clostridia>Eubacteriales | GCA_017458065.1 | Oscillospiraceae_bacterium | IJV40_02965 | 18565..21924 | 2485925 | DUF499-domain-containing-protein-[Oscillospiraceae-bacterium] |
| MCD8117856.1 | HEPN+AAA-ATPase+Alpha+Beta-domain+wHTH+RRM | Type-1_BR.RM.like.systems | <-HEPN+DUF499*<-ORC-CDC6+REase<-METHYLASE<-?<-?<-TUDOR+SF1-SF2-Helicase+iREase+CTD+REase | 1126 | Bacteria>Terrabacteria_group>Bacillota>Clostridia>Eubacteriales | GCA_021203045.1 | Oscillospiraceae_bacterium | LUE21_12275 | 415..3795 | 2485925 | DUF499-domain-containing-protein-[Oscillospiraceae-bacterium] |
| MCD7750767.1 | HEPN+AAA-ATPase+Alpha+Beta-domain+wHTH+RRM | Type-1_BR.RM.like.systems | <-NERD<-?<-?<-HEPN+DUF499*<-METHYLASE<-TUDOR+SF1-SF2-Helicase+iREase+CTD+REase<-XIS-HTH | 1127 | Bacteria>Terrabacteria_group>Bacillota>Clostridia>Eubacteriales | GCA_021199495.1 | Lachnospiraceae_bacterium | LUI10_03350 | 11139..14522 | 1898203 | DUF499-domain-containing-protein-[Lachnospiraceae-bacterium] |
| QNK42634.1 | HEPN+AAA-ATPase+Alpha+Beta-domain+wHTH+RRM | Type-1_BR.RM.like.systems | HTH_36-><-?<-?<-?<-JetA<-HEPN+DUF499*<-TUDOR+SF1-SF2-Helicase+iREase+CTD+REase<-CHTH | 1113 | Bacteria>Terrabacteria_group>Bacillota>Clostridia>Eubacteriales | GCA_014303955.1 | Caproicibacter_fermentans | HCR03_14035 | 2844426..2847767 | 2576756 | ATP-binding-protein-[Caproicibacter-fermentans] |
| MCL1794475.1 | HEPN+AAA-ATPase+Alpha+Beta-domain+wHTH+RRM | Type-1_BR.RM.like.systems | <-RelE-ParE<-MazE_antitoxin<-HEPN+DUF499*\|\|?->GDResolvase->?-><-DUF772+DDE_Tnp_1 | 1135 | Bacteria>Terrabacteria_group>Bacillota>Clostridia>Eubacteriales | GCA_009782405.1 | Oscillospiraceae_bacterium | FWG34_11475 | 15838..19245 | 2485925 | DUF499-domain-containing-protein-[Oscillospiraceae-bacterium] |
| MCR5263712.1 | HEPN+AAA-ATPase+Alpha+Beta-domain+wHTH+RRM | Type-1_BR.RM.like.systems | <-HTH\|\|?->?->TUDOR+SF1-SF2-Helicase+iREase+CTD+REase->METHYLASE->HEPN+DUF499*-> | 1118 | Bacteria>Terrabacteria_group>Bacillota>Clostridia>Eubacteriales | GCA_024695945.1 | Clostridiales_bacterium | K6D94_07550 | 8144..11500 | 1898207 | DUF499-domain-containing-protein-[Clostridiales-bacterium] |
| MCI9350231.1 | HEPN+AAA-ATPase+Alpha+Beta-domain+wHTH+RRM | Type-1_BR.RM.like.systems | TUDOR+SF1-SF2-Helicase+iREase+CTD+REase-p->?->METHYLASE->HEPN+DUF499*-> | 1134 | Bacteria>Terrabacteria_group>Bacillota>Clostridia>Eubacteriales | GCA_022799735.1 | Oscillibacter_sp. | HFF83_13605 | 3993..7397 | 1945593 | ATP-binding-protein-[Oscillibacter-sp.] |
| WP_006903458.1 | HEPN+AAA-ATPase+Alpha+Beta-domain+wHTH+RRM | Type-1_BR.RM.like.systems | TUDOR+SF1-SF2-Helicase+iREase+CTD+REase->METHYLASE->HEPN+DUF499*-> | 1128 | Bacteria>Terrabacteria_group>Bacillota>Clostridia>Eubacteriales | GCF_000183545.2 | Thermaerobacter_subterraneus | THESUDRAFT_RS04080 | 969973..973359 | 175696 | Swt1-family-HEPN-domain-containing-protein-[Thermaerobacter-subterraneus] |
| MCD7762788.1 | HEPN+AAA-ATPase+Alpha+Beta-domain+wHTH+RRM | Type-1_BR.RM.like.systems | <-NACHT\|\|?-><-MORC<-HEPN+DUF499*<-METHYLASE<-Schlafen+HATPase+HTH<-METHYLASE | 1132 | Bacteria>Terrabacteria_group>Bacillota>Clostridia>Eubacteriales | GCA_021199655.1 | Lachnospiraceae_bacterium | LUI14_06240 | 6036..9434 | 1898203 | DUF499-domain-containing-protein-[Lachnospiraceae-bacterium] |
| MDA8201143.1 | HEPN+AAA-ATPase+Alpha+Beta-domain+wHTH+RRM | Type-1_BR.RM.like.systems | <-RelE-ParE<-PhdYeFM_antitox\|\|DOC+HTH-><-?<-HEPN+DUF499*<-METHYLASE | 1093 | Bacteria>Terrabacteria_group>Bacillota>Clostridia>Eubacteriales | GCA_028065455.1 | Thermaerobacter_sp. | M0Z54_17205 | 4106..7387 | 1937527 | MAG:-Swt1-family-HEPN-domain-containing-protein-[Thermaerobacter-sp.] |
| HJB61498.1 | HEPN+AAA-ATPase+Alpha+Beta-domain+wHTH+RRM | Type-1_BR.RM.like.systems | TRAM+Methyltransf_31-><-HEPN<-HEPN+DUF499*<-METHYLASE-p | 1134 | Bacteria>Terrabacteria_group>Bacillota>Clostridia>Eubacteriales | GCA_019119145.1 | Candidatus_Ruminococcus_gallistercoris | IAA38_07560 | 7302..10706 | 2838749 | DUF499-domain-containing-protein-[Candidatus-Ruminococcus-gallistercoris] |
| MCC8151343.1 | HEPN+AAA-ATPase+Alpha+Beta-domain+wHTH+RRM | Type-1_BR.RM.like.systems | METHYLASE->HEPN+DUF499*->?->?->HTH+MPTase->PIN->SIGMA-HTH-> | 1132 | Bacteria>Terrabacteria_group>Bacillota>Clostridia>Eubacteriales | GCA_020860745.1 | Lachnospiraceae_bacterium | LIO96_07800 | 2635..6033 | 1898203 | DUF499-domain-containing-protein-[Lachnospiraceae-bacterium] |
| HJB96184.1 | HEPN+AAA-ATPase+Alpha+Beta-domain+wHTH+RRM | Type-1_BR.RM.like.systems | METHYLASE-p->HEPN+DUF499*-><-?<-PemK_toxin\|\|NACHT-> | 1128 | Bacteria>Terrabacteria_group>Bacillota>Clostridia>Eubacteriales | GCA_019119265.1 | Candidatus_Mediterraneibacter_intestinigallinarum | H9765_11315 | 720..4106 | 2838674 | DUF499-domain-containing-protein-[Candidatus-Mediterraneibacter-intestinigallinarum] |
| HCW03906.1 | HEPN+AAA-ATPase+Alpha+Beta-domain+wHTH+RRM | Type-1_BR.RM.like.systems | <-HEPN+DUF499*<-?<-?<-?<-ABC-ATPase+SbcC<-ABC-ATPase+SbcC | 1132 | Bacteria>Terrabacteria_group>Bacillota>Clostridia>Eubacteriales | GCA_003541575.1 | Clostridium_sp. | DGK91_04815 | 19..3417 | 1506 | hypothetical-protein-[Clostridium-sp.] |
| NLJ76878.1 | HEPN+AAA-ATPase+Alpha+Beta-domain+wHTH+RRM | Type-1_BR.RM.like.systems | <-ParB+HNH<-TPR<-?<-HEPN+DUF499*<-METHYLASE-p | 1133 | Bacteria>Terrabacteria_group>Bacillota>Clostridia>Eubacteriales | GCA_012518875.1 | Peptococcaceae_bacterium | GX325_06435 | 3363..6764 | 2052179 | ATP-binding-protein-[Peptococcaceae-bacterium] |
| MCD7843035.1 | HEPN+AAA-ATPase+Alpha+Beta-domain+wHTH+RRM | Type-1_BR.RM.like.systems | METHYLASE->RelB->RelE-ParE->HEPN+DUF499*-> | 1126 | Bacteria>Terrabacteria_group>Bacillota>Clostridia>Eubacteriales | GCA_021200795.1 | Clostridiales_bacterium | LUG17_00205 | 3743..7123 | 1898207 | DUF499-domain-containing-protein-[Clostridiales-bacterium] |
| MCR5807676.1 | HEPN+AAA-ATPase+Alpha+Beta-domain+wHTH+RRM | Type-1_BR.RM.like.systems | BirA-HTH->METHYLASE->HEPN+DUF499*-> | 732 | Bacteria>Terrabacteria_group>Bacillota>Clostridia>Eubacteriales | GCA_024698785.1 | Oscillospiraceae_bacterium | K6G68_11685 | 3761..5956 | 2485925 | DUF499-domain-containing-protein-[Oscillospiraceae-bacterium] |
| NLI59701.1 | HEPN+AAA-ATPase+Alpha+Beta-domain+wHTH+RRM | Type-1_BR.RM.like.systems | METHYLASE-p->HEPN+DUF499*-> | 985 | Bacteria>Terrabacteria_group>Bacillota>Clostridia>Eubacteriales | GCA_012517995.1 | Clostridium_sp. | GX387_14560 | 352..3307 | 1506 | ATP-binding-protein-[Clostridium-sp.] |
| MCR5122540.1 | HEPN+AAA-ATPase+Alpha+Beta-domain+wHTH+RRM | Type-1_BR.RM.like.systems | METHYLASE-p->HEPN+DUF499*-> | 458 | Bacteria>Terrabacteria_group>Bacillota>Clostridia>Eubacteriales | GCA_024697705.1 | Ruminococcus_sp. | K6B74_09000 | 1756..3129 | 41978 | DUF499-domain-containing-protein-[Ruminococcus-sp.] |
| MCM1579586.1 | HEPN+AAA-ATPase+Alpha+Beta-domain+wHTH+RRM | Type-1_BR.RM.like.systems | METHYLASE-p->HEPN+DUF499*-> | 410 | Bacteria>Terrabacteria_group>Bacillota>Clostridia>Eubacteriales | GCA_023667745.1 | Ruminococcus_sp. | NC078_12395 | 2153..3383 | 41978 | Swt1-family-HEPN-domain-containing-protein-[Ruminococcus-sp.] |
| MCD8379771.1 | HEPN+AAA-ATPase+Alpha+Beta-domain+wHTH+RRM | Type-1_BR.RM.like.systems | <-HEPN+DUF499*<-METHYLASE-p | 474 | Bacteria>Terrabacteria_group>Bacillota>Clostridia>Eubacteriales | GCA_021205565.1 | Lachnospiraceae_bacterium | LUC95_05545 | 1..1422 | 1898203 | DUF499-domain-containing-protein-[Lachnospiraceae-bacterium] |
| MCR5825960.1 | HEPN+AAA-ATPase+Alpha+Beta-domain+wHTH+RRM | Type-1_BR.RM.like.systems | <-HEPN+DUF499*<-METHYLASE | 853 | Bacteria>Terrabacteria_group>Bacillota>Clostridia>Eubacteriales | GCA_024698225.1 | Oscillospiraceae_bacterium | K6G54_05305 | 1..2559 | 2485925 | DUF499-domain-containing-protein-[Oscillospiraceae-bacterium] |
| MCL2699418.1 | HEPN+AAA-ATPase+Alpha+Beta-domain+wHTH+RRM | Type-1_BR.RM.like.systems | <-HEPN+DUF499*<-METHYLASE | 645 | Bacteria>Terrabacteria_group>Bacillota>Clostridia>Eubacteriales | GCA_009783765.1 | Defluviitaleaceae_bacterium | FWE68_03815 | 1..1937 | 2660712 | Swt1-family-HEPN-domain-containing-protein-[Defluviitaleaceae-bacterium] |

| Protein-ID | DUF499-architecture | System | Gene-Neighbourhoods | Length | Taxonomy | Assembly-ID | Strain | Locus-Tag | Genomic-coordinates | TaxID | Fasta-defline |
|---|---|---|---|---|---|---|---|---|---|---|---|
| MCI9425971.1 | HEPN+AAA-ATPase+Alpha+Beta-domain+wHTH+RRM | Type-1_BR.RM.like.systems | METHYLASE->HEPN+DUF499*-> | 444 | Bacteria>Terrabacteria_group>Bacillota>Clostridia>Eubacteriales | GCA_022800375.1 | Flavonifractor_sp. | HFF30_10520 | 3535..4866 | 2049025 | ATP-binding-protein-[Flavonifractor-sp.] |
| WP_156275504.1 | HEPN+AAA-ATPase+Alpha+Beta-domain+wHTH+RRM | Type-1_BR.RM.like.systems | <-CopG_antitoxin\|\|NTP_transf_2-><-?<-?<-Schlafen<-HEPN+DUF499*<-METHYLASE<-TUDOR+SF1-SF2-Helicase+iREase+CTD+REase\|\|?-><-?<-Phage_terminase | 1122 | Bacteria>Terrabacteria_group>Bacillota>Clostridia>Moorellales | GCF_009735625.1 | Moorella_glycerini | MGLY_RS15905 | 3178613..3181981 | 55779 | DUF499-domain-containing-protein-[Moorella-glycerini] |
| WP_071522349.1 | HEPN+AAA-ATPase+Alpha+Beta-domain+wHTH+RRM | Type-1_BR.RM.like.systems | PolB->HEPN->\|\|?->?->TUDOR+SF1-SF2-Helicase+iREase+CTD+REase->METHYLASE->HEPN+DUF499*->NERD-> | 1122 | Bacteria>Terrabacteria_group>Bacillota>Clostridia>Moorellales | GCF_001874085.1 | Moorella_thermoacetica | MOOTH_RS08230 | 71675..75043 | 1525 | Swt1-family-HEPN-domain-containing-protein-[Moorella-thermoacetica] |
| WP_013782413.1 | HEPN+AAA-ATPase+Alpha+Beta-domain+wHTH+RRM | Type-1_BR.RM.like.systems | TUDOR+SF1-SF2-Helicase+iREase+CTD+REase->METHYLASE->HEPN+DUF499*->?->XIS-HTH->BirA-HTH->?->Primpol+PriCT1->ZnR+TOPRIM+D5ATPase-> | 1118 | Bacteria>Terrabacteria_group>Bacillota>Clostridia>Thermoanaerobacterales | GCF_000213255.1 | Mahella_australiensis | MAHAU_RS14260 | 3047947..3051303 | 252966 | Swt1-family-HEPN-domain-containing-protein-[Mahella-australiensis] |
| WP_221039750.1 | HEPN+AAA-ATPase+Alpha+Beta-domain+wHTH+RRM | Type-1_BR.RM.like.systems | <-REase<-HEPN+DUF499*<-METHYLASE<-TUDOR+SF1-SF2-Helicase+iREase+CTD+REase<-McrC-NTD+REase | 1133 | Bacteria>Terrabacteria_group>Bacillota>Clostridia>Thermoanaerobacterales | GCF_019668485.1 | Gelria_sp._Kuro-4 | K5554_RS03460 | 654715..658116 | 2796927 | Swt1-family-HEPN-domain-containing-protein-[Gelria-sp.-Kuro-4] |
| MBQ6961345.1 | HEPN+AAA-ATPase+Alpha+Beta-domain+wHTH+RRM | Type-1_BR.RM.like.systems | <-SF1-SF2-Helicase-p<-OLD-ABC+TOPRIM<-HEPN+DUF499*<-HEPN<-PolB<-METHYLASE<-MNS<-TUDOR+SF1-SF2-Helicase+iREase+CTD+REase<-HTH | 1126 | Bacteria>Terrabacteria_group>Bacillota>Clostridia>unclassified_Clostridia | GCA_017448975.1 | Clostridia_bacterium | IJP78_10255 | 13529..16909 | 2044939 | DUF499-domain-containing-protein-[Clostridia-bacterium] |
| GHU66463.1 | HEPN+AAA-ATPase+Alpha+Beta-domain+wHTH+RRM | Type-1_BR.RM.like.systems | ABhydrolase->TUDOR+SF1-SF2-Helicase+iREase+CTD+REase->NTP_transf->HEPN->METHYLASE->HEPN+DUF499*->?->?->?-><-PADR-HTH | 1148 | Bacteria>Terrabacteria_group>Bacillota>Clostridia>unclassified_Clostridia | GCA_026000995.1 | Clostridia_bacterium | FACS1894184_04510 | 37803..41249 | 2044939 | hypothetical-protein-FACS1894184_04510-[Clostridia-bacterium] |
| HHY05947.1 | HEPN+AAA-ATPase+Alpha+Beta-domain+wHTH+RRM | Type-1_BR.RM.like.systems | <-TRAM+Methyltransf_31<-?<-HEPN+DUF499*<-METHYLASE<-TUDOR+SF1-SF2-Helicase+iREase+CTD+REase | 1125 | Bacteria>Terrabacteria_group>Bacillota>Clostridia>unclassified_Clostridia | GCA_012842035.1 | Clostridia_bacterium | GX532_03075 | 2394..5771 | 2044939 | ATP-binding-protein-[Clostridia-bacterium] |
| MBS3947521.1 | HEPN+AAA-ATPase+Alpha+Beta-domain+wHTH+RRM | Type-1_BR.RM.like.systems | <-HEPN+DUF499*<-HEPN<-Polbeta<-METHYLASE-p | 1118 | Bacteria>Terrabacteria_group>Bacillota>Dethiobacteria>Dethiobacterales | GCA_018333395.1 | Dethiobacter_sp. | KGZ57_04310 | 478..3834 | 2093370 | DUF499-domain-containing-protein-[Dethiobacter-sp.] |
| WP_285944649.1 | HEPN+AAA-ATPase+Alpha+Beta-domain+wHTH+RRM | Type-1_BR.RM.like.systems | <-HEPN+DUF499*<-METHYLASE | 1106 | Bacteria>Terrabacteria_group>Bacillota>Erysipelotrichia>Erysipelotrichales | GCF_949383055.1 | Faecalibaculum_rodentium | QTM32_RS09735 | 250..3570 | 1702221 | DUF499-domain-containing-protein-[Faecalibaculum-rodentium] |
| WP_281790372.1 | HEPN+AAA-ATPase+Alpha+Beta-domain+wHTH+RRM | Type-1_BR.RM.like.systems | <-HEPN+DUF499*<-METHYLASE | 1106 | Bacteria>Terrabacteria_group>Bacillota>Erysipelotrichia>Erysipelotrichales | GCF_937916425.1 | Faecalibaculum_rodentium | QU443_RS10445 | 250..3570 | 1702221 | DUF499-domain-containing-protein-[Faecalibaculum-rodentium] |
| MBE3584147.1 | HEPN+AAA-ATPase+Alpha+Beta-domain+wHTH+RRM | Type-1_BR.RM.like.systems | <-METHYLASE-p\|\|?->TUDOR+SF1-SF2-Helicase+iREase+CTD+REase->OLD-ABC+TOPRIM->METHYLASE->HEPN+DUF499*-><-?\|\|ABC-ATPase+SbcC->ABC-ATPase+SbcC-><-MazE_antitoxin | 1113 | Bacteria>Terrabacteria_group>Bacillota>Limnochordia>Limnochordales | GCA_014896305.1 | Limnochordaceae_bacterium | IMX01_08555 | 27896..31237 | 2773236 | DUF499-domain-containing-protein-[Limnochordaceae-bacterium] |
| MCK9524538.1 | HEPN+AAA-ATPase+Alpha+Beta-domain+wHTH+RRM | Type-1_BR.RM.like.systems | ArdCN+MPTase->?-><-?<-REase<-?<-REase<-HEPN+DUF499*<-METHYLASE<-TUDOR+SF1-SF2-Helicase+iREase+CTD+REase-p | 1113 | Bacteria>Terrabacteria_group>Bacillota>Limnochordia>unclassified_Limnochordia | GCA_023228555.1 | Limnochordia_bacterium | MOR49_01235 | 6495..9836 | 2497621 | Swt1-family-HEPN-domain-containing-protein-[Limnochordia-bacterium] |
| WP_232709101.1 | HEPN+AAA-ATPase+Alpha+Beta-domain+wHTH+RRM | Type-1_BR.RM.like.systems | NUDIX->TUDOR+SF1-SF2-Helicase+iREase+CTD+REase->METHYLASE->METHYLASE-p->HIN-HTH->METHYLASE->HEPN+DUF499*->HTH_3->?->SIGMA-HTH-> | 1130 | Bacteria>Terrabacteria_group>Bacillota>Negativicutes>Acidaminococcales | GCF_902376135.1 | Acidaminococcus_massiliensis | BQ2657_RS09005 | 1875868..1879260 | 1852375 | DUF499-domain-containing-protein-[Acidaminococcus-massiliensis] |
| PWM79769.1 | HEPN+AAA-ATPase+Alpha+Beta-domain+wHTH+RRM | Type-1_BR.RM.like.systems | Peptidase_S8->?-><-?<-?<-STYKinase<-HEPN+DUF499*<-METHYLASE<-Schlafen+HATPase+HTH<-TUDOR+SF1-SF2-Helicase+iREase+CTD+REase<-HTH_26 | 1130 | Bacteria>Terrabacteria_group>Bacillota>Negativicutes>Acidaminococcales | GCA_003150755.1 | Phascolarctobacterium_sp. | DBY32_02635 | 546389..549781 | 2049039 | MAG:-hypothetical-protein-DBY32_02635-[Phascolarctobacterium-sp.] |
| MBR6861287.1 | HEPN+AAA-ATPase+Alpha+Beta-domain+wHTH+RRM | Type-1_BR.RM.like.systems | RADICAL-SAM-><-HEPN+DUF499*<-METHYLASE<-METHYLASE<-TUDOR+SF1-SF2-Helicase+iREase+CTD+REase<-?<-?<-REase+35exo | 1123 | Bacteria>Terrabacteria_group>Bacillota>Negativicutes>Acidaminococcales | GCA_017531245.1 | Acidaminococcaceae_bacterium | IKM73_08200 | 27143..30514 | 2049052 | DUF499-domain-containing-protein-[Acidaminococcaceae-bacterium] |
| WP_279002420.1 | HEPN+AAA-ATPase+Alpha+Beta-domain+wHTH+RRM | Type-1_BR.RM.like.systems | NUDIX->?->TUDOR+SF1-SF2-Helicase+iREase+CTD+REase->METHYLASE->METHYLASE->HEPN+DUF499*->HTH_3-> | 1145 | Bacteria>Terrabacteria_group>Bacillota>Negativicutes>Acidaminococcales | GCF_945915305.1 | Acidaminococcus_fermentans | QDM86_RS09090 | 15622..19059 | 905 | DUF499-domain-containing-protein-[Acidaminococcus-fermentans] |
| WP_270837292.1 | HEPN+AAA-ATPase+Alpha+Beta-domain+wHTH+RRM | Type-1_BR.RM.like.systems | <-MULE-TRANSPOSASE\|\|HTH_OrfB_IS605+OrfB_IS605+OrfB_Zn_ribbon->?->TUDOR+SF1-SF2-Helicase+iREase+CTD+REase->METHYLASE->METHYLASE->HEPN+DUF499*-><-DDE_Tnp_IS66+DDE_Tnp_IS66_C<-?<-HIN-HTH | 1134 | Bacteria>Terrabacteria_group>Bacillota>Negativicutes>Selenomonadales | GCF_027670005.1 | Mitsuokella_multacida | PHA28_RS02250 | 15218..18622 | 52226 | DUF499-domain-containing-protein-[Mitsuokella-multacida] |
| WP_278503476.1 | HEPN+AAA-ATPase+Alpha+Beta-domain+wHTH+RRM | Type-1_BR.RM.like.systems | Pkinase-><-?<-HEPN+DUF499*<-METHYLASE<-METHYLASE<-TUDOR+SF1-SF2-Helicase+iREase+CTD+REase<-?<-HTH_OrfB_IS605+OrfB_IS605+OrfB_Zn_ribbon<-PhdYeFM_antitox | 1134 | Bacteria>Terrabacteria_group>Bacillota>Negativicutes>Selenomonadales | GCF_905193285.1 | Mitsuokella_multacida | QMF84_RS08000 | 50365..53769 | 52226 | DUF499-domain-containing-protein-[Mitsuokella-multacida] |
| WP_264918710.1 | HEPN+AAA-ATPase+Alpha+Beta-domain+wHTH+RRM | Type-1_BR.RM.like.systems | <-Mbetalac\|\|GDResolvase+HIN-HTH->TUDOR+SF1-SF2-Helicase+iREase+CTD+REase->METHYLASE->HEPN+DUF499*->?->Peptidase->?->sGTP->?->Aldedh-> | 1127 | Bacteria>Terrabacteria_group>Bacillota>Negativicutes>Selenomonadales | GCF_026013845.1 | Selenomonas_sputigena | OL240_RS12405 | 2695256..2698639 | 69823 | DUF499-domain-containing-protein-[Selenomonas-sputigena] |
| MBR7025847.1 | HEPN+AAA-ATPase+Alpha+Beta-domain+wHTH+RRM | Type-1_BR.RM.like.systems | ABC-ATPase+SbcC-><-?\|\|TUDOR+SF1-SF2-Helicase+iREase+CTD+REase->METHYLASE->HEPN+DUF499*-><-ABC-ATPase+SbcC<-ABC-ATPase+SbcC | 1077 | Bacteria>Terrabacteria_group>Bacillota>Negativicutes>Selenomonadales | GCA_017518905.1 | Selenomonadaceae_bacterium | IKI08_07585 | 26267..29500 | 2652289 | DUF499-domain-containing-protein-[Selenomonadaceae-bacterium] |
| WP_009350157.1 | HEPN+AAA-ATPase+Alpha+Beta-domain+wHTH+RRM | Type-1_BR.RM.like.systems | XIS-HTH->HIN-HTH->?-><-CHTH<-HEPN+DUF499*<-METHYLASE<-TUDOR+SF1-SF2-Helicase+iREase+CTD+REase<-?\|\|DOC+HTH-><-MACRODOMAIN | 1142 | Bacteria>Terrabacteria_group>Bacillota>Negativicutes>Selenomonadales | GCF_000187125.1 | Selenomonas_artemidis | HMPREF9555_RS07020 | 6182..9610 | 671224 | DUF499-domain-containing-protein-[Selenomonas-artemidis] |
| WP_061945835.1 | HEPN+AAA-ATPase+Alpha+Beta-domain+wHTH+RRM | Type-1_BR.RM.like.systems | METHYLASE-p->?->METHYLASE-p->METHYLASE-p->METHYLASE-p->HEPN+DUF499*->tRNA-><-REase<-Phage_integrase\|\|?-><-?\|\|DUF772+DDE_Tnp_1_6-> | 377 | Bacteria>Terrabacteria_group>Bacillota>Negativicutes>Selenomonadales | GCF_001554015.1 | Selenomonas_sp._oral_taxon_136 | AXE86_RS06860 | 1418306..1419439 | 713030 | Swt1-family-HEPN-domain-containing-protein-[Selenomonas-sp.-oral-taxon-136] |
| MBF1662768.1 | HEPN+AAA-ATPase+Alpha+Beta-domain+wHTH+RRM | Type-1_BR.RM.like.systems | Classical-AAA->Peptidase_S8-><-?<-?<-?<-HEPN+DUF499*<-METHYLASE<-TUDOR+SF1-SF2-Helicase+iREase+CTD+REase<-?<-PfkB | 1145 | Bacteria>Terrabacteria_group>Bacillota>Negativicutes>Selenomonadales | GCA_015259755.1 | Selenomonas_noxia | HX076_05010 | 11846..15283 | 135083 | ATP-binding-protein-[Selenomonas-noxia] |
| WP_007288290.1 | HEPN+AAA-ATPase+Alpha+Beta-domain+wHTH+RRM | Type-1_BR.RM.like.systems | Calcineurin->SbcC+SbcC-><-?<-?<-HEPN+DUF499*<-METHYLASE\|\|DDE_Tnp-><-?<-HTH_21+rve+rve_3<-HIN-HTH<-RNase | 1126 | Bacteria>Terrabacteria_group>Bacillota>Negativicutes>Selenomonadales | GCF_000169155.1 | Thermosinus_carboxydivorans | TCARDRAFT_RS01740 | 31494..34874 | 261685 | Swt1-family-HEPN-domain-containing-protein-[Thermosinus-carboxydivorans] |
| MBF1693417.1 | HEPN+AAA-ATPase+Alpha+Beta-domain+wHTH+RRM | Type-1_BR.RM.like.systems | TUDOR+SF1-SF2-Helicase+iREase+CTD+REase-p->HEPN+DUF499*-><-ASCH<-PIN+ACET+EVE<-?<-Classical-AAA | 1142 | Bacteria>Terrabacteria_group>Bacillota>Negativicutes>Selenomonadales | GCA_015265235.1 | Selenomonas_sp. | HX082_03045 | 3480..6908 | 2053611 | ATP-binding-protein-[Selenomonas-sp.] |
| WP_270528574.1 | HEPN+AAA-ATPase+Alpha+Beta-domain+wHTH+RRM | Type-1_BR.RM.like.systems | <-35exo\|\|?-><-?\|\|?->?-><-HEPN+DUF499*<-METHYLASE<-TUDOR+SF1-SF2-Helicase+iREase+CTD+REase<-?<-?<-?<-Methylase | 1161 | Bacteria>Terrabacteria_group>Bacillota>Negativicutes>Selenomonadales | GCF_027681165.1 | Mitsuokella_jalaludinii | PGX61_RS07200 | 41318..44803 | 187979 | DUF499-domain-containing-protein-[Mitsuokella-jalaludinii] |
| MCI6231067.1 | HEPN+AAA-ATPase+Alpha+Beta-domain+wHTH+RRM | Type-1_BR.RM.like.systems | TUDOR+SF1-SF2-Helicase+iREase+CTD+REase->Transposase_31->METHYLASE->HEPN+DUF499*-> | 1147 | Bacteria>Terrabacteria_group>Bacillota>Negativicutes>Selenomonadales | GCA_022773585.1 | Selenomonas_sp. | MR630_00380 | 8523..11966 | 2053611 | DUF499-domain-containing-protein-[Selenomonas-sp.] |
| MCI6099016.1 | HEPN+AAA-ATPase+Alpha+Beta-domain+wHTH+RRM | Type-1_BR.RM.like.systems | TUDOR+SF1-SF2-Helicase+iREase+CTD+REase->Transposase_31->METHYLASE->HEPN+DUF499*-> | 1147 | Bacteria>Terrabacteria_group>Bacillota>Negativicutes>Selenomonadales | GCA_022770905.1 | Selenomonas_sp. | MR698_00025 | 8556..11999 | 2053611 | DUF499-domain-containing-protein-[Selenomonas-sp.] |
| MBR0290429.1 | HEPN+AAA-ATPase+Alpha+Beta-domain+wHTH+RRM | Type-1_BR.RM.like.systems | TUDOR+SF1-SF2-Helicase+iREase+CTD+REase->METHYLASE->HEPN+DUF499*->DUF499-p-> | 334 | Bacteria>Terrabacteria_group>Bacillota>Negativicutes>Selenomonadales | GCA_017446965.1 | Selenomonadaceae_bacterium | IJQ82_15765 | 5468..6472 | 2652289 | hypothetical-protein-[Selenomonadaceae-bacterium] |
| MBR0260489.1 | HEPN+AAA-ATPase+Alpha+Beta-domain+wHTH+RRM | Type-1_BR.RM.like.systems | METHYLASE->HEPN+DUF499*->sGTP->SIGMA-HTH+SIGMA-HTH->SIGMA-HTH+SIGMA-HTH-> | 1077 | Bacteria>Terrabacteria_group>Bacillota>Negativicutes>Selenomonadales | GCA_017446845.1 | Selenomonadaceae_bacterium | IJQ85_01715 | 15691..18924 | 2652289 | DUF499-domain-containing-protein-[Selenomonadaceae-bacterium] |
| MBF1688626.1 | HEPN+AAA-ATPase+Alpha+Beta-domain+wHTH+RRM | Type-1_BR.RM.like.systems | TUDOR+SF1-SF2-Helicase+iREase+CTD+REase-p->METHYLASE->HEPN+DUF499*-> | 887 | Bacteria>Terrabacteria_group>Bacillota>Negativicutes>Selenomonadales | GCA_015265195.1 | Selenomonas_sp. | HX083_10375 | 4201..6861 | 2053611 | ATP-binding-protein-[Selenomonas-sp.] |
| WP_031587665.1 | HEPN+AAA-ATPase+Alpha+Beta-domain+wHTH+RRM | Type-1_BR.RM.like.systems | <-HEPN+DUF499*<-METHYLASE<-?<-TUDOR+SF1-SF2-Helicase+iREase+CTD+REase | 1132 | Bacteria>Terrabacteria_group>Bacillota>Negativicutes>Selenomonadales | GCF_000701885.1 | Selenomonas_bovis | T485_RS0110635 | 1077..4475 | 416586 | DUF499-domain-containing-protein-[Selenomonas-bovis] |
| MBQ6005594.1 | HEPN+AAA-ATPase+Alpha+Beta-domain+wHTH+RRM | Type-1_BR.RM.like.systems | TUDOR+SF1-SF2-Helicase+iREase+CTD+REase->METHYLASE->HEPN+DUF499*-> | 1081 | Bacteria>Terrabacteria_group>Bacillota>Negativicutes>Selenomonadales | GCA_017427545.1 | Selenomonadaceae_bacterium | IJL14_05025 | 8141..11386 | 2652289 | DUF499-domain-containing-protein-[Selenomonadaceae-bacterium] |
| MBR4642051.1 | HEPN+AAA-ATPase+Alpha+Beta-domain+wHTH+RRM | Type-1_BR.RM.like.systems | SF1-SF2-Helicase-p->REase->?->?->METHYLASE-p->METHYLASE->HEPN+DUF499*-> | 1118 | Bacteria>Terrabacteria_group>Bacillota>Negativicutes>Selenomonadales | GCA_017535925.1 | Selenomonadaceae_bacterium | IKO74_04930 | 6896..10252 | 2652289 | hypothetical-protein-[Selenomonadaceae-bacterium] |
| MBQ7497714.1 | HEPN+AAA-ATPase+Alpha+Beta-domain+wHTH+RRM | Type-1_BR.RM.like.systems | <-DOC+HTH\|\|DeoR-HTH+ISOCOT-><-HEPN+DUF499*<-METHYLASE | 1125 | Bacteria>Terrabacteria_group>Bacillota>Negativicutes>Selenomonadales | GCA_017455465.1 | Selenomonas_sp. | IJU00_07780 | 3332..6709 | 2053611 | DUF499-domain-containing-protein-[Selenomonas-sp.] |
| MBR1580270.1 | HEPN+AAA-ATPase+Alpha+Beta-domain+wHTH+RRM | Type-1_BR.RM.like.systems | <-Peptidase_M30<-HEPN+DUF499*<-?<-NTP_transf<-METHYLASE | 1098 | Bacteria>Terrabacteria_group>Bacillota>Negativicutes>Selenomonadales | GCA_017477125.1 | Selenomonadaceae_bacterium | IJ668_07210 | 12971..16267 | 2652289 | hypothetical-protein-[Selenomonadaceae-bacterium] |
| MBR3051424.1 | HEPN+AAA-ATPase+Alpha+Beta-domain+wHTH+RRM | Type-1_BR.RM.like.systems | ABC-ATPase+SbcC-><-HEPN+DUF499*<-METHYLASE-p | 1077 | Bacteria>Terrabacteria_group>Bacillota>Negativicutes>Selenomonadales | GCA_017510185.1 | Selenomonadaceae_bacterium | IKG61_08245 | 624..3857 | 2652289 | DUF499-domain-containing-protein-[Selenomonadaceae-bacterium] |
| MBR3623059.1 | HEPN+AAA-ATPase+Alpha+Beta-domain+wHTH+RRM | Type-1_BR.RM.like.systems | <-HEPN+DUF499*<-NERD<-OLD-ABC<-METHYLASE-p | 509 | Bacteria>Terrabacteria_group>Bacillota>Negativicutes>Selenomonadales | GCA_017529085.1 | Selenomonadaceae_bacterium | IKN43_06910 | 1..1529 | 2652289 | DUF499-domain-containing-protein-[Selenomonadaceae-bacterium] |
| MBQ9480498.1 | HEPN+AAA-ATPase+Alpha+Beta-domain+wHTH+RRM | Type-1_BR.RM.like.systems | <-HEPN+DUF499*<-?<-BrnT_toxin<-METHYLASE-p | 465 | Bacteria>Terrabacteria_group>Bacillota>Negativicutes>Selenomonadales | GCA_017454185.1 | Selenomonadaceae_bacterium | IJU71_13220 | 1..1395 | 2652289 | DUF499-domain-containing-protein-[Selenomonadaceae-bacterium] |
| MBR1807330.1 | HEPN+AAA-ATPase+Alpha+Beta-domain+wHTH+RRM | Type-1_BR.RM.like.systems | <-HEPN+DUF499*<-METHYLASE<-METHYLASE-p | 1085 | Bacteria>Terrabacteria_group>Bacillota>Negativicutes>Selenomonadales | GCA_017479925.1 | Selenomonadaceae_bacterium | IJ774_13210 | 2314..5571 | 2652289 | DUF499-domain-containing-protein-[Selenomonadaceae-bacterium] |
| MBR3723747.1 | HEPN+AAA-ATPase+Alpha+Beta-domain+wHTH+RRM | Type-1_BR.RM.like.systems | <-HEPN+DUF499*<-NERD<-OLD-ABC<-METHYLASE | 503 | Bacteria>Terrabacteria_group>Bacillota>Negativicutes>Selenomonadales | GCA_017530025.1 | Selenomonadaceae_bacterium | IKN12_13455 | 1..1509 | 2652289 | DUF499-domain-containing-protein-[Selenomonadaceae-bacterium] |
| MBR3746746.1 | HEPN+AAA-ATPase+Alpha+Beta-domain+wHTH+RRM | Type-1_BR.RM.like.systems | <-HEPN+DUF499*<-METHYLASE-p | 1077 | Bacteria>Terrabacteria_group>Bacillota>Negativicutes>Selenomonadales | GCA_017529545.1 | Selenomonadaceae_bacterium | IKN27_07285 | 344..3577 | 2652289 | DUF499-domain-containing-protein-[Selenomonadaceae-bacterium] |
| MBR0060910.1 | HEPN+AAA-ATPase+Alpha+Beta-domain+wHTH+RRM | Type-1_BR.RM.like.systems | METHYLASE->HEPN+DUF499*-> | 1084 | Bacteria>Terrabacteria_group>Bacillota>Negativicutes>Selenomonadales | GCA_017449105.1 | Selenomonadaceae_bacterium | IJP68_05465 | 1758..5012 | 2652289 | DUF499-domain-containing-protein-[Selenomonadaceae-bacterium] |
| WP_008860390.1 | HEPN+AAA-ATPase+Alpha+Beta-domain+wHTH+RRM | Type-1_BR.RM.like.systems | Phage_integrase->Phage_integrase->?->TUDOR+SF1-SF2-Helicase+iREase+CTD+REase->METHYLASE->METHYLASE->HEPN+DUF499*->?-><-REase+SF1-SF2-Helicase<-REase+SF1-SF2-Helicase\|\|MazG-> | 1135 | Bacteria>Terrabacteria_group>Bacillota>Negativicutes>Veillonellales | GCF_902374635.1 | Dialister_succinatiphilus | HMPREF9453_RS09730 | 179520..182927 | 487173 | DUF499-domain-containing-protein-[Dialister-succinatiphilus] |
| WP_204967821.1 | HEPN+AAA-ATPase+Alpha+Beta-domain+wHTH+RRM | Type-1_BR.RM.like.systems | TRD+TRD->GDResolvase+Recombinase+ZnR->HTH_26->TUDOR+SF1-SF2-Helicase+iREase+CTD+REase->METHYLASE->METHYLASE->HEPN+DUF499*->?->?->?->TPR+TPR-> | 1132 | Bacteria>Terrabacteria_group>Bacillota>Negativicutes>Veillonellales | GCF_016900675.1 | Megasphaera_stantonii | H6G85_RS06205 | 65633..69031 | 2144175 | DUF499-domain-containing-protein-[Megasphaera-stantonii] |
| HBL50377.1 | HEPN+AAA-ATPase+Alpha+Beta-domain+wHTH+RRM | Type-1_BR.RM.like.systems | <-SF1-SF2-Helicase<-OLD-ABC+TOPRIM<-HEPN+DUF499*<-METHYLASE<-TUDOR+SF1-SF2-Helicase+iREase+CTD+REase<-?<-MORC+DUF3684+HNH<-SF1-SF2-Helicase-p<-HEPN+SF1-SF2-Helicase | 1129 | Bacteria>Terrabacteria_group>Bacillota>unclassified_Bacillota | GCA_003504715.1 | Bacillota_bacterium | DDZ65_09075 | 4304..7693 | 1879010 | hypothetical-protein-[Firmicutes-bacterium] |
| NPV89870.1 | HEPN+AAA-ATPase+Alpha+Beta-domain+wHTH+RRM | Type-1_BR.RM.like.systems | GDResolvase->?->GDResolvase->TUDOR+SF1-SF2-Helicase+iREase+CTD+REase->AbiGi->METHYLASE->HEPN+DUF499*->OLD-ABC+TOPRIM->SF1-SF2-Helicase->?->Peptidase_S8-><-HTH_3 | 1137 | Bacteria>Terrabacteria_group>Bacillota>unclassified_Bacillota | GCA_013178275.1 | Bacillota_bacterium | HPY50_03730 | 179885..183298 | 1879010 | ATP-binding-protein-[Firmicutes-bacterium] |
| MCL6590158.1 | HEPN+AAA-ATPase+Alpha+Beta-domain+wHTH+RRM | Type-1_BR.RM.like.systems | HicB_lk_antitox->TUDOR+SF1-SF2-Helicase+iREase+CTD+REase->?->MACRODOMAIN->METHYLASE->Y1_Tnp->HEPN+DUF499*->REase->SF1-SF2-Helicase->JetD-> | 1116 | Bacteria>Terrabacteria_group>Bacillota>unclassified_Bacillota | GCA_023512195.1 | Bacillota_bacterium | K6U80_09420 | 20552..23902 | 1879010 | DUF499-domain-containing-protein-[Firmicutes-bacterium] |
| NPV80543.1 | HEPN+AAA-ATPase+Alpha+Beta-domain+wHTH+RRM | Type-1_BR.RM.like.systems | REase->?-><-DOC\|\|TUDOR+SF1-SF2-Helicase+iREase+CTD+REase->Schlafen+HATPase+HTH->METHYLASE->HEPN+DUF499*->?-><-HEPN-p | 1115 | Bacteria>Terrabacteria_group>Bacillota>unclassified_Bacillota | GCA_013178415.1 | Bacillota_bacterium | HPY52_09760 | 58381..61728 | 1879010 | ATP-binding-protein-[Firmicutes-bacterium] |
| NLG83533.1 | HEPN+AAA-ATPase+Alpha+Beta-domain+wHTH+RRM | Type-1_BR.RM.like.systems | REase-><-?<-HEPN+DUF499*<-METHYLASE<-Schlafen+HATPase+HTH<-TUDOR+SF1-SF2-Helicase+iREase+CTD+REase | 1127 | Bacteria>Terrabacteria_group>Bacillota>unclassified_Bacillota | GCA_012516115.1 | Bacillota_bacterium | GX493_02745 | 877..4260 | 1879010 | ATP-binding-protein-[Firmicutes-bacterium] |
| OQB47592.1 | HEPN+AAA-ATPase+Alpha+Beta-domain+wHTH+RRM | Type-1_BR.RM.like.systems | DSBH-><-?\|\|tRNA->REase-><-HEPN+DUF499*<-METHYLASE-p<-TUDOR+SF1-SF2-Helicase+iREase+CTD+REase | 1103 | Bacteria>Terrabacteria_group>Bacillota>unclassified_Bacillota | GCA_002070415.1 | Firmicutes_bacterium_ADurb.Bin153 | BWY00_01229 | 26495..29806 | 1852883 | hypothetical-protein-BWY00_01229-[Firmicutes-bacterium-ADurb.Bin153] |
| HHW13082.1 | HEPN+AAA-ATPase+Alpha+Beta-domain+wHTH+RRM | Type-1_BR.RM.like.systems | TUDOR+SF1-SF2-Helicase+iREase+CTD+REase->METHYLASE->HEPN+DUF499*-><-?<-?<-?<-DeoR-HTH+ISOCOT | 1104 | Bacteria>Terrabacteria_group>Bacillota>unclassified_Bacillota | GCA_012840155.1 | Bacillota_bacterium | GXX33_08800 | 5884..9198 | 1879010 | ATP-binding-protein-[Firmicutes-bacterium] |
| NLN07229.1 | HEPN+AAA-ATPase+Alpha+Beta-domain+wHTH+RRM | Type-1_BR.RM.like.systems | <-HEPN+DUF499*<-METHYLASE<-TUDOR+SF1-SF2-Helicase+iREase+CTD+REase<-REase | 1130 | Bacteria>Terrabacteria_group>Bacillota>unclassified_Bacillota | GCA_012522075.1 | Bacillota_bacterium | GX167_06390 | 552..3944 | 1879010 | ATP-binding-protein-[Firmicutes-bacterium] |
| NLG85967.1 | HEPN+AAA-ATPase+Alpha+Beta-domain+wHTH+RRM | Type-1_BR.RM.like.systems | <-HHH_5+SF1-SF2-helicase<-?<-?<-?<-Cysteine_desulfurase<-HEPN+DUF499*<-METHYLASE-p | 1132 | Bacteria>Terrabacteria_group>Bacillota>unclassified_Bacillota | GCA_012516165.1 | Bacillota_bacterium | GX489_01905 | 9843..13241 | 1879010 | ATP-binding-protein-[Firmicutes-bacterium] |
| HHV07782.1 | HEPN+AAA-ATPase+Alpha+Beta-domain+wHTH+RRM | Type-1_BR.RM.like.systems | TUDOR+SF1-SF2-Helicase+iREase+CTD+REase-p->?->METHYLASE->HEPN+DUF499*-> | 441 | Bacteria>Terrabacteria_group>Bacillota>unclassified_Bacillota | GCA_012839445.1 | Bacillota_bacterium | GXX69_06175 | 4070..5392 | 1879010 | ATP-binding-protein-[Bacillota-bacterium] |
| HHV56346.1 | HEPN+AAA-ATPase+Alpha+Beta-domain+wHTH+RRM | Type-1_BR.RM.like.systems | <-DUF772+DDE_Tnp_1+DDE_Tnp_1_6\|\|METHYLASE-p->HEPN+DUF499*->REase-> | 1131 | Bacteria>Terrabacteria_group>Bacillota>unclassified_Bacillota | GCA_012839755.1 | Bacillota_bacterium | GXX50_01080 | 3217..6612 | 1879010 | ATP-binding-protein-[Firmicutes-bacterium] |
| MBO2495132.1 | HEPN+AAA-ATPase+Alpha+Beta-domain+wHTH+RRM | Type-1_BR.RM.like.systems | METHYLASE-p->HEPN+DUF499*->?->MqsA_antitoxin+HTH_3->ABhydrolase-> | 1117 | Bacteria>Terrabacteria_group>Bacillota>unclassified_Bacillota | GCA_017577845.1 | Bacillota_bacterium | COP67_00010 | 2134..5487 | 1879010 | AAA+-family-ATPase-[Firmicutes-bacterium] |
| MTI83942.1 | HEPN+AAA-ATPase+Alpha+Beta-domain+wHTH+RRM | Type-1_BR.RM.like.systems | <-SF1-SF2-Helicase+REase<-PIWI<-HEPN+DUF499*<-METHYLASE-p | 1111 | Bacteria>Terrabacteria_group>Bacillota>unclassified_Bacillota | GCA_009711705.1 | Bacillota_bacterium | FH756_08540 | 83034..86369 | 1879010 | ATP-binding-protein-[Firmicutes-bacterium] |
| MCL5982191.1 | HEPN+AAA-ATPase+Alpha+Beta-domain+wHTH+RRM | Type-1_BR.RM.like.systems | <-HEPN+DUF499*<-METHYLASE-p | 1131 | Bacteria>Terrabacteria_group>Bacillota>unclassified_Bacillota | GCA_023488665.1 | Bacillota_bacterium | M1571_06745 | 19839..23234 | 1879010 | DUF499-domain-containing-protein-[Firmicutes-bacterium] |
| HHU75981.1 | HEPN+AAA-ATPase+Alpha+Beta-domain+wHTH+RRM | Type-1_BR.RM.like.systems | <-HEPN+DUF499*<-METHYLASE-p | 1124 | Bacteria>Terrabacteria_group>Bacillota>unclassified_Bacillota | GCA_012839215.1 | Bacillota_bacterium | GXZ24_03690 | 12668..16042 | 1879010 | ATP-binding-protein-[Firmicutes-bacterium] |
| HHU30916.1 | HEPN+AAA-ATPase+Alpha+Beta-domain+wHTH+RRM | Type-1_BR.RM.like.systems | METHYLASE->HEPN+DUF499*-> | 1077 | Bacteria>Terrabacteria_group>Bacillota>unclassified_Bacillota | GCA_012838605.1 | Bacillota_bacterium | GXZ53_09580 | 2024..5256 | 1879010 | ATP-binding-protein-[Firmicutes-bacterium] |
| MBJ7608107.1 | HEPN+AAA-ATPase+Alpha+Beta-domain+wHTH+RRM | Type-1_BR.RM.like.systems | Phd-YefM->PIN->?->?->?-><-HEPN+DUF499*<-METHYLASE<-TUDOR+SF1-SF2-Helicase+iREase+CTD+REase<-MPTase<-HTH\|\|?->SMODS-> | 1110 | Bacteria>Terrabacteria_group>Candidatus_Dormibacteraeota>unclassified_Candidatus_Dormibacteraeota | GCA_016464795.1 | Candidatus_Dormibacteraeota_bacterium | JF887_01575 | 11366..14698 | 2052315 | DUF499-domain-containing-protein-[Candidatus-Dormibacteraeota-bacterium] |
| MDA8332521.1 | HEPN+AAA-ATPase+Alpha+Beta-domain+wHTH+RRM | Type-1_BR.RM.like.systems | <-HEPN+DUF499*<-METHYLASE<-?<-TUDOR+SF1-SF2-Helicase+iREase+CTD+REase<-TIR | 485 | Bacteria>Terrabacteria_group>Candidatus_Dormibacteraeota>unclassified_Candidatus_Dormibacteraeota | GCA_028067975.1 | Candidatus_Dormibacteraeota_bacterium | M0027_15220 | 1..1455 | 2052315 | MAG:-Swt1-family-HEPN-domain-containing-protein,-partial-[Candidatus-Dormibacteraeota-bacterium] |
| MDA8331677.1 | HEPN+AAA-ATPase+Alpha+Beta-domain+wHTH+RRM | Type-1_BR.RM.like.systems | TUDOR+SF1-SF2-Helicase+iREase+CTD+REase->?->METHYLASE->HEPN+DUF499*-> | 1075 | Bacteria>Terrabacteria_group>Candidatus_Dormibacteraeota>unclassified_Candidatus_Dormibacteraeota | GCA_028067975.1 | Candidatus_Dormibacteraeota_bacterium | M0027_10845 | 5705..8930 | 2052315 | MAG:-DUF499-domain-containing-protein,-partial-[Candidatus-Dormibacteraeota-bacterium] |
| MCB9452414.1 | HEPN+AAA-ATPase+Alpha+Beta-domain+wHTH+RRM | Type-1_BR.RM.like.systems | <-MetJarc-HTH\|\|?->TPR+TPR+TPR+TPR+TPR+TPR+TPR+TPR-><-HEPN+DUF499*<-ParB+HNH<-METHYLASE<-Schlafen+HATPase+HTH<-PIN<-?<-TUDOR+SF1-SF2-Helicase+iREase+CTD+REase | 1106 | Bacteria>Terrabacteria_group>Chloroflexota>Anaerolineae>Anaerolineales | GCA_020634115.1 | Anaerolineaceae_bacterium | H6672_13335 | 112307..115627 | 2024896 | ATP-binding-protein-[Anaerolineaceae-bacterium] |
| MBE7511263.1 | HEPN+AAA-ATPase+Alpha+Beta-domain+wHTH+RRM | Type-1_BR.RM.like.systems | Peptidase_M1->?->TUDOR+SF1-SF2-Helicase+iREase+CTD+REase->?->PIN->METHYLASE-p->HEPN+DUF499*-><-tRNA<-?<-ABhydrolase\|\|TPR+TPR->ACET+ACET->ACET-> | 1098 | Bacteria>Terrabacteria_group>Chloroflexota>Anaerolineae>Anaerolineales | GCA_015075625.1 | Anaerolineales_bacterium | HS103_00415 | 94825..98121 | 2073117 | ATP-binding-protein-[Anaerolineales-bacterium] |
| MCZ7550961.1 | HEPN+AAA-ATPase+Alpha+Beta-domain+wHTH+RRM | Type-1_BR.RM.like.systems | <-NACHT\|\|<-?<-?<-ParB+HNH<-?<-HEPN+DUF499*<-METHYLASE<-MORC+DUF3684<-MORC+DUF3684<-TUDOR+SF1-SF2-Helicase+iREase+CTD+REase-p | 1112 | Bacteria>Terrabacteria_group>Chloroflexota>Anaerolineae>Anaerolineales | GCA_027494025.1 | Anaerolineales_bacterium | M5U11_17685 | 3656371..3659709 | 2073117 | MAG:-Swt1-family-HEPN-domain-containing-protein-[Anaerolineales-bacterium] |
| MCB0010245.1 | HEPN+AAA-ATPase+Alpha+Beta-domain+wHTH+RRM | Type-1_BR.RM.like.systems | METHYLASE-p->TIR->TPR->?->?-><-?\|\|HEPN+DUF499*->?->HKD-DNase+HKD-DNase-><-nSTAND1<-?<-SIR2+CASPASE<-SIR2+nSTAND1+FGS | 1088 | Bacteria>Terrabacteria_group>Chloroflexota>Anaerolineae>Anaerolineales | GCA_020433005.1 | Anaerolineales_bacterium | KDE34_00040 | 5582..8848 | 2073117 | DUF499-domain-containing-protein-[Anaerolineales-bacterium] |
| QYK51438.1 | HEPN+AAA-ATPase+Alpha+Beta-domain+wHTH+RRM | Type-1_BR.RM.like.systems | DOC+HTH-><-HEPN+DUF499*<-METHYLASE<-SIR2<-TUDOR+SF1-SF2-Helicase+iREase+CTD+REase<-?<-TIR\|\|PRTase-> | 1117 | Bacteria>Terrabacteria_group>Chloroflexota>Anaerolineae>Anaerolineales | GCA_019454165.1 | Anaerolineales_bacterium | KF701_02830 | 580231..583584 | 2073117 | MAG:-DUF499-domain-containing-protein-[Anaerolineales-bacterium] |
| MCW5886897.1 | HEPN+AAA-ATPase+Alpha+Beta-domain+wHTH+RRM | Type-1_BR.RM.like.systems | <-PRTase\|\|TIR->?->TUDOR+SF1-SF2-Helicase+iREase+CTD+REase->METHYLASE->HEPN+DUF499*-><-DOC+HTH | 1117 | Bacteria>Terrabacteria_group>Chloroflexota>Anaerolineae>Anaerolineales | GCA_026129325.1 | Anaerolineales_bacterium | KIT07_02110 | 8706..12059 | 2073117 | MAG:-DUF499-domain-containing-protein-[Anaerolineales-bacterium] |
| NLF02984.1 | HEPN+AAA-ATPase+Alpha+Beta-domain+wHTH+RRM | Type-1_BR.RM.like.systems | <-HEPN+DUF499*<-METHYLASE<-?<-MORC<-TUDOR+SF1-SF2-Helicase+iREase+CTD+REase | 1091 | Bacteria>Terrabacteria_group>Chloroflexota>Anaerolineae>Anaerolineales | GCA_012514695.1 | Anaerolineales_bacterium | GX601_18630 | 3316..6591 | 2073117 | ATP-binding-protein-[Anaerolineales-bacterium] |
| MAS36305.1 | HEPN+AAA-ATPase+Alpha+Beta-domain+wHTH+RRM | Type-1_BR.RM.like.systems | Glyoxalase->?-><-PIN<-?<-HEPN+DUF499*<-REase<-?<-METHYLASE<-Schlafen+HATPase+HTH | 1094 | Bacteria>Terrabacteria_group>Chloroflexota>Anaerolineae>Anaerolineales | GCA_002702065.1 | Anaerolineaceae_bacterium | CL610_20035 | 35853..39137 | 2024896 | AAA+-family-ATPase-[Anaerolineaceae-bacterium] |
| MCB9078004.1 | HEPN+AAA-ATPase+Alpha+Beta-domain+wHTH+RRM | Type-1_BR.RM.like.systems | <-HEPN+DUF499*<-REase<-?<-METHYLASE-p<-METHYLASE<-PIN<-DUF433 | 385 | Bacteria>Terrabacteria_group>Chloroflexota>Anaerolineae>Anaerolineales | GCA_020634955.1 | Anaerolineaceae_bacterium | H6631_10465 | 1..1157 | 2024896 | ATP-binding-protein-[Anaerolineaceae-bacterium] |
| MBQ6503986.1 | HEPN+AAA-ATPase+Alpha+Beta-domain+wHTH+RRM | Type-1_BR.RM.like.systems | METHYLASE-p->METHYLASE->HEPN+DUF499*->KAP-NTPase-> | 1120 | Bacteria>Terrabacteria_group>Chloroflexota>Anaerolineae>Anaerolineales | GCA_017432685.1 | Flexilinea_sp. | IJI57_08765 | 4499..7861 | 2017390 | DUF499-domain-containing-protein-[Flexilinea-sp.] |
| MCC6958025.1 | HEPN+AAA-ATPase+Alpha+Beta-domain+wHTH+RRM | Type-1_BR.RM.like.systems | <-MarR-HTH<-HEPN+DUF499*<-?<-METHYLASE<-?<-PIN<-REase | 1114 | Bacteria>Terrabacteria_group>Chloroflexota>Anaerolineae>Anaerolineales | GCA_020853835.1 | Anaerolineales_bacterium | IT316_14590 | 1878..5222 | 2073117 | DUF499-domain-containing-protein-[Anaerolineales-bacterium] |
| MBL6983853.1 | HEPN+AAA-ATPase+Alpha+Beta-domain+wHTH+RRM | Type-1_BR.RM.like.systems | <-REase<-HEPN+DUF499*<-SF1-SF2-Helicase-p | 1095 | Bacteria>Terrabacteria_group>Chloroflexota>Anaerolineae>Anaerolineales | GCA_016785015.1 | Anaerolineales_bacterium | ISR58_21930 | 2169..5456 | 2073117 | DUF499-domain-containing-protein-[Anaerolineales-bacterium] |
| MBE2198717.1 | HEPN+AAA-ATPase+Alpha+Beta-domain+wHTH+RRM | Type-1_BR.RM.like.systems | METHYLASE-p->?->HEPN+DUF499*-> | 1099 | Bacteria>Terrabacteria_group>Chloroflexota>Anaerolineae>Anaerolineales | GCA_014879875.1 | Anaerolinea_sp. | IAE79_08915 | 3861..7160 | 1872519 | ATP-binding-protein-[Anaerolinea-sp.] |
| MBK9093182.1 | HEPN+AAA-ATPase+Alpha+Beta-domain+wHTH+RRM | Type-1_BR.RM.like.systems | <-ABhydrolase+ABhydrolase<-?<-ABhydrolase<-?<-?<-HEPN+DUF499*<-Schlafen+HATPase+HTH<-METHYLASE-p<-Schlafen+HATPase+HTH<-TUDOR+SF1-SF2-Helicase+iREase+CTD+REase<-TOPRIM<-OLD-ABC | 1133 | Bacteria>Terrabacteria_group>Chloroflexota>Anaerolineae>unclassified_Anaerolineae | GCA_016719785.1 | Anaerolineae_bacterium | IPM84_10425 | 127540..130941 | 2052143 | DUF499-domain-containing-protein-[Anaerolineae-bacterium] |
| MCL4871680.1 | HEPN+AAA-ATPase+Alpha+Beta-domain+wHTH+RRM | Type-1_BR.RM.like.systems | <-AbiEii<-AbiEii<-AbiEi_4\|\|TUDOR+SF1-SF2-Helicase+iREase+CTD+REase->REase->METHYLASE->HEPN+DUF499*-><-?<-?<-SIGMA-HTH<-RNB+RNase\|\|?->PadR-HTH-> | 1115 | Bacteria>Terrabacteria_group>Chloroflexota>Anaerolineae>unclassified_Anaerolineae | GCA_023384415.1 | Anaerolineae_bacterium | KJ063_22200 | 16187..19534 | 2052143 | DUF499-domain-containing-protein-[Anaerolineae-bacterium] |
| MCL4237388.1 | HEPN+AAA-ATPase+Alpha+Beta-domain+wHTH+RRM | Type-1_BR.RM.like.systems | MULE-TRANSPOSASE->REase->SF1-SF2-Helicase-p->MULE-TRANSPOSASE-><-HEPN+DUF499*<-METHYLASE<-PIN<-?<-TUDOR+SF1-SF2-Helicase+iREase+CTD+REase | 1111 | Bacteria>Terrabacteria_group>Chloroflexota>Anaerolineae>unclassified_Anaerolineae | GCA_023382715.1 | Anaerolineae_bacterium | KJ047_03985 | 66352..69687 | 2052143 | DUF499-domain-containing-protein-[Anaerolineae-bacterium] |
| MBK6429951.1 | HEPN+AAA-ATPase+Alpha+Beta-domain+wHTH+RRM | Type-1_BR.RM.like.systems | OLD-ABC->TOPRIM->Schlafen+HATPase+HTH->METHYLASE-p->Schlafen+HATPase+HTH->REase->HEPN+DUF499*->?->?->PemK_toxin->ABhydrolase->?->ABhydrolase+ABhydrolase-> | 1113 | Bacteria>Terrabacteria_group>Chloroflexota>Anaerolineae>unclassified_Anaerolineae | GCA_016704685.1 | Anaerolineae_bacterium | IPF85_04125 | 226121..229462 | 2052143 | DUF499-domain-containing-protein-[Anaerolineae-bacterium] |
| MCZ2114537.1 | HEPN+AAA-ATPase+Alpha+Beta-domain+wHTH+RRM | Type-1_BR.RM.like.systems | <-HTH+UbE1<-?\|\|HTH->MPTase-><-HEPN+DUF499*<-PIN<-OLD-ABC<-METHYLASE<-MORC+DUF3684<-TUDOR+SF1-SF2-Helicase+iREase+CTD+REase | 1108 | Bacteria>Terrabacteria_group>Chloroflexota>Anaerolineae>unclassified_Anaerolineae | GCA_026954395.1 | Anaerolineae_bacterium | LC131_12000 | 4338..7664 | 2052143 | MAG:-DUF499-domain-containing-protein-[Anaerolineae-bacterium] |
| MCK6580687.1 | HEPN+AAA-ATPase+Alpha+Beta-domain+wHTH+RRM | Type-1_BR.RM.like.systems | <-GDResolvase<-?<-PIN<-HEPN+DUF499*<-PIN<-DUF433<-?<-METHYLASE<-?<-TUDOR+SF1-SF2-Helicase+iREase+CTD+REase | 1104 | Bacteria>Terrabacteria_group>Chloroflexota>Anaerolineae>unclassified_Anaerolineae | GCA_023151025.1 | Anaerolineae_bacterium | L6Q98_21555 | 21655..24969 | 2052143 | DUF499-domain-containing-protein-[Anaerolineae-bacterium] |
| MBN2002049.1 | HEPN+AAA-ATPase+Alpha+Beta-domain+wHTH+RRM | Type-1_BR.RM.like.systems | <-HEPN+DUF499*<-Y1_Tnp<-METHYLASE<-MACRODOMAIN<-?<-?<-TUDOR+SF1-SF2-Helicase+iREase+CTD+REase | 1119 | Bacteria>Terrabacteria_group>Chloroflexota>Anaerolineae>unclassified_Anaerolineae | GCA_016931835.1 | Anaerolineae_bacterium | JXA21_01725 | 49309..52668 | 2052143 | DUF499-domain-containing-protein-[Anaerolineae-bacterium] |
| MCL4805199.1 | HEPN+AAA-ATPase+Alpha+Beta-domain+wHTH+RRM | Type-1_BR.RM.like.systems | <-HEPN+DUF499*<-METHYLASE<-?<-TUDOR+SF1-SF2-Helicase+iREase+CTD+REase<-McrC-NTD+REase | 1098 | Bacteria>Terrabacteria_group>Chloroflexota>Anaerolineae>unclassified_Anaerolineae | GCA_023384355.1 | Anaerolineae_bacterium | KJ046_12935 | 9107..12403 | 2052143 | DUF499-domain-containing-protein-[Anaerolineae-bacterium] |
| NPV86424.1 | HEPN+AAA-ATPase+Alpha+Beta-domain+wHTH+RRM | Type-1_BR.RM.like.systems | <-REC<-tRNA\|\|TUDOR+SF1-SF2-Helicase+iREase+CTD+REase->PIN->METHYLASE->HEPN+DUF499*-> | 1115 | Bacteria>Terrabacteria_group>Chloroflexota>Anaerolineae>unclassified_Anaerolineae | GCA_013178295.1 | Anaerolineae_bacterium | HPY45_10505 | 184598..187945 | 2052143 | ATP-binding-protein-[Anaerolineae-bacterium] |
| MDD3828881.1 | HEPN+AAA-ATPase+Alpha+Beta-domain+wHTH+RRM | Type-1_BR.RM.like.systems | TUDOR+SF1-SF2-Helicase+iREase+CTD+REase->METHYLASE->HEPN+DUF499*-> | 350 | Bacteria>Terrabacteria_group>Chloroflexota>Anaerolineae>unclassified_Anaerolineae | GCA_028698515.1 | Anaerolineae_bacterium | PHY79_23180 | 6005..7056 | 2052143 | MAG:-Swt1-family-HEPN-domain-containing-protein,-partial-[Anaerolineae-bacterium] |
| MCZ2096700.1 | HEPN+AAA-ATPase+Alpha+Beta-domain+wHTH+RRM | Type-1_BR.RM.like.systems | SF1-SF2-Helicase->wHTH+SAD+HNH->ORC-CDC6+REase->HEPN+DUF499*-> | 1112 | Bacteria>Terrabacteria_group>Chloroflexota>Anaerolineae>unclassified_Anaerolineae | GCA_026954235.1 | Anaerolineae_bacterium | LC121_10630 | 39414..42752 | 2052143 | MAG:-DUF499-domain-containing-protein-[Anaerolineae-bacterium] |
| MCL4299450.1 | HEPN+AAA-ATPase+Alpha+Beta-domain+wHTH+RRM | Type-1_BR.RM.like.systems | TPR->?->?->METHYLASE->URI-DNase->HEPN+DUF499*->DUF2726+ZnR+TOPRIM-> | 1096 | Bacteria>Terrabacteria_group>Chloroflexota>Anaerolineae>unclassified_Anaerolineae | GCA_023382535.1 | Anaerolineae_bacterium | KJ077_27185 | 26645..29935 | 2052143 | DUF499-domain-containing-protein-[Anaerolineae-bacterium] |
| MCC6801844.1 | HEPN+AAA-ATPase+Alpha+Beta-domain+wHTH+RRM | Type-1_BR.RM.like.systems | <-HNH<-CopG/RHH<-DUF2726+ZnR+TOPRIM<-HEPN+DUF499*<-?<-METHYLASE | 1095 | Bacteria>Terrabacteria_group>Chloroflexota>Anaerolineae>unclassified_Anaerolineae | GCA_020848855.1 | Anaerolineae_bacterium | IT319_03080 | 2285..5572 | 2052143 | DUF499-domain-containing-protein-[Anaerolineae-bacterium] |
| MCU0500065.1 | HEPN+AAA-ATPase+Alpha+Beta-domain+wHTH+RRM | Type-1_BR.RM.like.systems | <-Glyco_transf_28<-TetR-HTH\|\|?-><-HEPN+DUF499*<-METHYLASE-p | 1097 | Bacteria>Terrabacteria_group>Chloroflexota>Anaerolineae>unclassified_Anaerolineae | GCA_025454525.1 | Anaerolineae_bacterium | MUF87_22175 | 4596..7889 | 2052143 | DUF499-domain-containing-protein-[Anaerolineae-bacterium] |
| MCW5847519.1 | HEPN+AAA-ATPase+Alpha+Beta-domain+wHTH+RRM | Type-1_BR.RM.like.systems | METHYLASE-p->REase->HEPN+DUF499*-><-?<-?<-?<-?<-ABC-ATPase+SbcC | 1097 | Bacteria>Terrabacteria_group>Chloroflexota>Anaerolineae>unclassified_Anaerolineae | GCA_026129205.1 | Anaerolineae_bacterium | KIS95_09830 | 2201..5494 | 2052143 | MAG:-DUF499-domain-containing-protein-[Anaerolineae-bacterium] |
| MCC6973771.1 | HEPN+AAA-ATPase+Alpha+Beta-domain+wHTH+RRM | Type-1_BR.RM.like.systems | PIN->?->METHYLASE->?->HEPN+DUF499*->HEPN-> | 1096 | Bacteria>Terrabacteria_group>Chloroflexota>Anaerolineae>unclassified_Anaerolineae | GCA_020853695.1 | Anaerolineae_bacterium | IT322_07165 | 5474..8764 | 2052143 | DUF499-domain-containing-protein-[Anaerolineae-bacterium] |
| MCB0250740.1 | HEPN+AAA-ATPase+Alpha+Beta-domain+wHTH+RRM | Type-1_BR.RM.like.systems | <-HEPN+DUF499*<-Y1_Tnp<-METHYLASE-p | 1124 | Bacteria>Terrabacteria_group>Chloroflexota>Anaerolineae>unclassified_Anaerolineae | GCA_020434015.1 | Anaerolineae_bacterium | KDI07_19355 | 427..3801 | 2052143 | DUF499-domain-containing-protein-[Anaerolineae-bacterium] |
| MCU0476247.1 | HEPN+AAA-ATPase+Alpha+Beta-domain+wHTH+RRM | Type-1_BR.RM.like.systems | <-HEPN+DUF499*<-METHYLASE-p | 1076 | Bacteria>Terrabacteria_group>Chloroflexota>Anaerolineae>unclassified_Anaerolineae | GCA_025454715.1 | Anaerolineae_bacterium | MUC99_09085 | 257..3487 | 2052143 | DUF499-domain-containing-protein-[Anaerolineae-bacterium] |
| MBN1429664.1 | HEPN+AAA-ATPase+Alpha+Beta-domain+wHTH+RRM | Type-1_BR.RM.like.systems | <-HEPN+DUF499*<-?<-?<-?<-?<-METHYLASE | 352 | Bacteria>Terrabacteria_group>Chloroflexota>Anaerolineae>unclassified_Anaerolineae | GCA_016928235.1 | Anaerolineae_bacterium | JXB07_14940 | 1..1058 | 2052143 | AAA-family-ATPase-[Anaerolineae-bacterium] |
| MCL4302340.1 | HEPN+AAA-ATPase+Alpha+Beta-domain+wHTH+RRM | Type-1_BR.RM.like.systems | <-TraJ-RHH\|\|TUDOR+SF1-SF2-Helicase+iREase+CTD+REase->?->?->METHYLASE->REase->HEPN+DUF499*-><-?\|\|?->GDResolvase+Recombinase+ZnR-> | 1117 | Bacteria>Terrabacteria_group>Chloroflexota>Anaerolineae>unclassified_Anaerolineae | GCA_023382535.1 | Anaerolineae_bacterium | KJ077_42015 | 34544..37897 | 2052143 | DUF499-domain-containing-protein-[Anaerolineae-bacterium] |
| MCS6835397.1 | HEPN+AAA-ATPase+Alpha+Beta-domain+wHTH+RRM | Type-1_BR.RM.like.systems | METHYLASE-p->OLD-ABC+TOPRIM-p->HEPN+DUF499*-> | 1089 | Bacteria>Terrabacteria_group>Chloroflexota>Anaerolineae>unclassified_Anaerolineae | GCA_025055175.1 | Anaerolineae_bacterium | NZ750_05190 | 1306..4575 | 2052143 | DUF499-domain-containing-protein-[Anaerolineae-bacterium] |
| MBN1136865.1 | HEPN+AAA-ATPase+Alpha+Beta-domain+wHTH+RRM | Type-1_BR.RM.like.systems | APATPase+iSTAND3->METHYLASE->?->?->HEPN+DUF499*-> | 684 | Bacteria>Terrabacteria_group>Chloroflexota>Anaerolineae>unclassified_Anaerolineae | GCA_016926595.1 | Anaerolineae_bacterium | JXM73_09775 | 6509..8561 | 2052143 | DUF499-domain-containing-protein-[Anaerolineae-bacterium] |
| MCZ7570815.1 | HEPN+AAA-ATPase+Alpha+Beta-domain+wHTH+RRM | Type-1_BR.RM.like.systems | DEDD_Tnp_IS110+Transposase_20->?->TUDOR+SF1-SF2-Helicase+iREase+CTD+REase->METHYLASE->REase->HEPN+DUF499*-><-PIN<-PhdYeFM_antitox<-?<-HAD | 1117 | Bacteria>Terrabacteria_group>Chloroflexota>Ardenticatenia>Ardenticatenales | GCA_027493995.1 | Ardenticatenaceae_bacterium | M5U01_19825 | 4779576..4782929 | 2614247 | MAG:-Swt1-family-HEPN-domain-containing-protein-[Ardenticatenaceae-bacterium] |
| MCB9423196.1 | HEPN+AAA-ATPase+Alpha+Beta-domain+wHTH+RRM | Type-1_BR.RM.like.systems | TUDOR+SF1-SF2-Helicase+iREase+CTD+REase->?->SF1-SF2-Helicase+HRDC->?->METHYLASE->TIR->HEPN+DUF499*->XIS-HTH-> | 1096 | Bacteria>Terrabacteria_group>Chloroflexota>Ardenticatenia>Ardenticatenales | GCA_020634245.1 | Ardenticatenaceae_bacterium | H6667_25585 | 35052..38342 | 2614247 | ATP-binding-protein-[Ardenticatenaceae-bacterium] |
| MCB8944346.1 | HEPN+AAA-ATPase+Alpha+Beta-domain+wHTH+RRM | Type-1_BR.RM.like.systems | GDResolvase+HIN-HTH-><-?\|\|?-><-?\|\|<-?<-?<-HEPN+DUF499*<-?<-METHYLASE<-ParB+HNH<-TUDOR+SF1-SF2-Helicase+iREase+CTD+REase | 1098 | Bacteria>Terrabacteria_group>Chloroflexota>Ardenticatenia>Ardenticatenales | GCA_020634415.1 | Ardenticatenaceae_bacterium | H6658_11405 | 5286..8582 | 2614247 | ATP-binding-protein-[Ardenticatenaceae-bacterium] |
| MBP7961438.1 | HEPN+AAA-ATPase+Alpha+Beta-domain+wHTH+RRM | Type-1_BR.RM.like.systems | TUDOR+SF1-SF2-Helicase+iREase+CTD+REase->?->?->METHYLASE->OLD-ABC->PIN->HEPN+DUF499*->?->SF1-SF2-Helicase->SF1-SF2-Helicase+DUF1998-> | 1102 | Bacteria>Terrabacteria_group>Chloroflexota>Caldilineae>Caldilineales | GCA_018003515.1 | Caldilineaceae_bacterium | KA003_05670 | 15367..18675 | 2614250 | DUF499-domain-containing-protein-[Caldilineaceae-bacterium] |
| MDE0338616.1 | HEPN+AAA-ATPase+Alpha+Beta-domain+wHTH+RRM | Type-1_BR.RM.like.systems | <-HAD+HAD\|\|?-><-?\|\|TUDOR+SF1-SF2-Helicase+iREase+CTD+REase->METHYLASE->PolB->HEPN+DUF499*->?->?->MazE_antitoxin-><-DSBH | 1121 | Bacteria>Terrabacteria_group>Chloroflexota>Caldilineae>Caldilineales | GCA_028820895.1 | Caldilineaceae_bacterium | OXI80_13170 | 100490..103855 | 2614250 | MAG:-Swt1-family-HEPN-domain-containing-protein-[Caldilineaceae-bacterium] |
| MCY4117637.1 | HEPN+AAA-ATPase+Alpha+Beta-domain+wHTH+RRM | Type-1_BR.RM.like.systems | TUDOR+SF1-SF2-Helicase+iREase+CTD+REase->METHYLASE->PolB->HEPN->OLD-ABC->TOPRIM->HEPN+DUF499*->?->?->?-><-DSBH | 1124 | Bacteria>Terrabacteria_group>Chloroflexota>Caldilineae>Caldilineales | GCA_026708105.1 | Caldilineaceae_bacterium | OXF55_12190 | 34492..37866 | 2614250 | MAG:-Swt1-family-HEPN-domain-containing-protein-[Caldilineaceae-bacterium] |
| MCY3902338.1 | HEPN+AAA-ATPase+Alpha+Beta-domain+wHTH+RRM | Type-1_BR.RM.like.systems | TIR->TUDOR+SF1-SF2-Helicase+iREase+CTD+REase-p->PIN->METHYLASE->PolB->HEPN->HEPN+DUF499*->?->?-><-?<-DSBH | 1125 | Bacteria>Terrabacteria_group>Chloroflexota>Caldilineae>Caldilineales | GCA_026707125.1 | Caldilineaceae_bacterium | OXF76_04160 | 61061..64438 | 2614250 | MAG:-Swt1-family-HEPN-domain-containing-protein-[Caldilineaceae-bacterium] |
| WP_014433685.1 | HEPN+AAA-ATPase+Alpha+Beta-domain+wHTH+RRM | Type-1_BR.RM.like.systems | <-Peptidase\|\|TUDOR+SF1-SF2-Helicase+iREase+CTD+REase->METHYLASE->METHYLASE->HEPN+DUF499*-> | 1108 | Bacteria>Terrabacteria_group>Chloroflexota>Caldilineae>Caldilineales | GCF_000281175.1 | Caldilinea | QVE71_RS01780 | 444218..447544 | 233191 | Swt1-family-HEPN-domain-containing-protein-[Caldilinea-aerophila] |
| MCY4079348.1 | HEPN+AAA-ATPase+Alpha+Beta-domain+wHTH+RRM | Type-1_BR.RM.like.systems | METHYLASE->PolB->?->?->OLD-ABC->TOPRIM->HEPN+DUF499*->?->?-><-?\|\|MazE_antitoxin-><-DSBH | 1121 | Bacteria>Terrabacteria_group>Chloroflexota>Caldilineae>Caldilineales | GCA_026707715.1 | Caldilineaceae_bacterium | OXF54_03840 | 16452..19817 | 2614250 | MAG:-Swt1-family-HEPN-domain-containing-protein-[Caldilineaceae-bacterium] |
| MYA05496.1 | HEPN+AAA-ATPase+Alpha+Beta-domain+wHTH+RRM | Type-1_BR.RM.like.systems | <-METHYLASE<-TUDOR+SF1-SF2-Helicase+iREase+CTD+REase<-HEPN+DUF499* | 1079 | Bacteria>Terrabacteria_group>Chloroflexota>Caldilineae>Caldilineales | GCA_009838685.1 | Caldilineaceae_bacterium_SB0664_bin_22 | F4Y37_12945 | 5308..8547 | 2605259 | ATP-binding-protein-[Caldilineaceae-bacterium-SB0664_bin_22] |
| MCE7987213.1 | HEPN+AAA-ATPase+Alpha+Beta-domain+wHTH+RRM | Type-1_BR.RM.like.systems | <-HAMP+HISKIN<-?<-HEPN+DUF499*<-Y1_Tnp<-METHYLASE<-Schlafen+HATPase+HTH | 1167 | Bacteria>Terrabacteria_group>Chloroflexota>Caldilineae>Caldilineales | GCA_021405325.1 | Caldilinea_sp._CFX5 | DYG89_39080 | 104570..108073 | 2293554 | ATP-binding-protein-[Caldilinea-sp.-CFX5] |
| MDE0431259.1 | HEPN+AAA-ATPase+Alpha+Beta-domain+wHTH+RRM | Type-1_BR.RM.like.systems | DSBH->?-><-?<-?<-HEPN+DUF499*<-TOPRIM<-OLD-ABC<-?<-HEPN<-PolB<-METHYLASE | 1123 | Bacteria>Terrabacteria_group>Chloroflexota>Caldilineae>Caldilineales | GCA_028821555.1 | Caldilineaceae_bacterium | OXH98_15965 | 76934..80305 | 2614250 | MAG:-Swt1-family-HEPN-domain-containing-protein-[Caldilineaceae-bacterium] |
| MCZ0940408.1 | HEPN+AAA-ATPase+Alpha+Beta-domain+wHTH+RRM | Type-1_BR.RM.like.systems | REase+NACHT-><-?\|\|?->?-><-MAD-NTD+HNH<-HEPN+DUF499*<-METHYLASE-p | 1114 | Bacteria>Terrabacteria_group>Chloroflexota>Caldilineae>Caldilineales | GCA_026914085.1 | Caldilineaceae_bacterium | OXJ55_17325 | 15852..19196 | 2614250 | MAG:-DUF499-domain-containing-protein-[Caldilineaceae-bacterium] |
| MXX28115.1 | HEPN+AAA-ATPase+Alpha+Beta-domain+wHTH+RRM | Type-1_BR.RM.like.systems | <-HEPN+DUF499*<-TOPRIM<-OLD-ABC<-HEPN<-PolB<-METHYLASE | 1121 | Bacteria>Terrabacteria_group>Chloroflexota>Caldilineae>Caldilineales | GCA_009835815.1 | Caldilineaceae_bacterium_SB0668_bin_21 | F4Z82_22075 | 67..3432 | 2605264 | ATP-binding-protein-[Caldilineaceae-bacterium-SB0668_bin_21] |
| MDE0200338.1 | HEPN+AAA-ATPase+Alpha+Beta-domain+wHTH+RRM | Type-1_BR.RM.like.systems | METHYLASE->PolB->HEPN->OLD-ABC->TOPRIM->HEPN+DUF499*-> | 1112 | Bacteria>Terrabacteria_group>Chloroflexota>Caldilineae>Caldilineales | GCA_028819675.1 | Caldilineaceae_bacterium | OXK78_19185 | 5066..8404 | 2614250 | MAG:-Swt1-family-HEPN-domain-containing-protein-[Caldilineaceae-bacterium] |
| MDE0069281.1 | HEPN+AAA-ATPase+Alpha+Beta-domain+wHTH+RRM | Type-1_BR.RM.like.systems | DSBH->?-><-?<-?<-HEPN+DUF499*<-?<-?<-METHYLASE-p | 1125 | Bacteria>Terrabacteria_group>Chloroflexota>Caldilineae>Caldilineales | GCA_028819015.1 | Caldilineaceae_bacterium | OXO48_06170 | 52227..55604 | 2614250 | MAG:-Swt1-family-HEPN-domain-containing-protein-[Caldilineaceae-bacterium] |
| MDE0183518.1 | HEPN+AAA-ATPase+Alpha+Beta-domain+wHTH+RRM | Type-1_BR.RM.like.systems | METHYLASE->?->?->HEPN+DUF499*->?->?-><-?<-DSBH | 1122 | Bacteria>Terrabacteria_group>Chloroflexota>Caldilineae>Caldilineales | GCA_028819655.1 | Caldilineaceae_bacterium | OXL39_19355 | 3803..7171 | 2614250 | MAG:-Swt1-family-HEPN-domain-containing-protein-[Caldilineaceae-bacterium] |
| MCC6169408.1 | HEPN+AAA-ATPase+Alpha+Beta-domain+wHTH+RRM | Type-1_BR.RM.like.systems | PIN->METHYLASE->HEPN+DUF499*-> | 1104 | Bacteria>Terrabacteria_group>Chloroflexota>Caldilineae>Caldilineales | GCA_028845455.1 | Caldilineaceae_bacterium | IT329_19460 | 3543..6857 | 2614250 | DUF499-domain-containing-protein-[Caldilineaceae-bacterium] |
| WP_097642123.1 | HEPN+AAA-ATPase+Alpha+Beta-domain+wHTH+RRM | Type-1_BR.RM.like.systems | <-DUF488-l<-?<-?<-Phage_integrase<-HKD-DNase+HKD-DNase<-?<-HEPN+DUF499*<-Polbeta<-PolB-NTase<-METHYLASE<-HEPN<-?<-TUDOR+SF1-SF2-Helicase+iREase+CTD+REase | 1103 | Bacteria>Terrabacteria_group>Chloroflexota>Chloroflexia>Chloroflexales | GCF_002532535.1 | Candidatus_Viridilinea_mediisalina | CJ255_RS00480 | 82670..85981 | 2024553 | Swt1-family-HEPN-domain-containing-protein-[Candidatus-Viridilinea-mediisalina] |
| WP_097643072.1 | HEPN+AAA-ATPase+Alpha+Beta-domain+wHTH+RRM | Type-1_BR.RM.like.systems | <-PIN\|\|TUDOR+SF1-SF2-Helicase+iREase+CTD+REase-><-?\|\|?->METHYLASE->DUF433->HEPN+DUF499*->?-><-MarR-HTH\|\|?-><-?<-ST-Hiskin<-NTP_transferase | 1128 | Bacteria>Terrabacteria_group>Chloroflexota>Chloroflexia>Chloroflexales | GCF_002532535.1 | Candidatus_Viridilinea_mediisalina | CJ255_RS05405 | 56277..59663 | 2024553 | Swt1-family-HEPN-domain-containing-protein-[Candidatus-Viridilinea-mediisalina] |
| MBK9709775.1 | HEPN+AAA-ATPase+Alpha+Beta-domain+wHTH+RRM | Type-1_BR.RM.like.systems | TUDOR+SF1-SF2-Helicase+iREase+CTD+REase->REase->METHYLASE->URI-DNase->HEPN+DUF499*->?->?-><-tRNA<-?<-?<-RnaseIII+DSRBD | 1115 | Bacteria>Terrabacteria_group>Chloroflexota>Chloroflexia>Chloroflexales | GCA_016717335.1 | Kouleothrix_sp. | IPO81_00330 | 71152..74499 | 2779161 | DUF499-domain-containing-protein-[Kouleothrix-sp.] |
| WP_066790944.1 | HEPN+AAA-ATPase+Alpha+Beta-domain+wHTH+RRM | Type-1_BR.RM.like.systems | <-HEPN+DUF499*<-METHYLASE<-METHYLASE<-REase<-TUDOR+SF1-SF2-Helicase+iREase+CTD+REase | 1104 | Bacteria>Terrabacteria_group>Chloroflexota>Chloroflexia>Chloroflexales | GCF_001650695.1 | Chloroflexus_islandicus | A6A03_RS17620 | 125816..129130 | 1707952 | Swt1-family-HEPN-domain-containing-protein-[Chloroflexus-islandicus] |
| WP_011958262.1 | HEPN+AAA-ATPase+Alpha+Beta-domain+wHTH+RRM | Type-1_BR.RM.like.systems | HEPN->?->?->DDE_Tnp_1->DDE_Tnp_1->METHYLASE-p->HEPN+DUF499*->ABC-ATPase+SbcC-> | 1122 | Bacteria>Terrabacteria_group>Chloroflexota>Chloroflexia>Chloroflexales | GCF_000016665.1 | Roseiflexus_sp._RS-1 | ROSERS_RS18370 | 4458924..4462292 | 357808 | Swt1-family-HEPN-domain-containing-protein-[Roseiflexus-sp.-RS-1] |
| WP_031460600.1 | HEPN+AAA-ATPase+Alpha+Beta-domain+wHTH+RRM | Type-1_BR.RM.like.systems | <-STAS\|\|?-><-?<-?<-?\|\|HEPN+DUF499*->TUDOR+SF1-SF2-Helicase+iREase+CTD+REase->METHYLASE-> | 1112 | Bacteria>Terrabacteria_group>Chloroflexota>Chloroflexia>Chloroflexales | GCF_000735195.1 | Chloroflexus_sp._MS-G | JE47_RS0115345 | 6861..10199 | 1521187 | Swt1-family-HEPN-domain-containing-protein-[Chloroflexus-sp.-MS-G] |
| MBO9390433.1 | HEPN+AAA-ATPase+Alpha+Beta-domain+wHTH+RRM | Type-1_BR.RM.like.systems | METHYLASE-p->?->RAMPs->RAMPs->RAMPs->HEPN+HEPN->HEPN+DUF499*-> | 1118 | Bacteria>Terrabacteria_group>Chloroflexota>Chloroflexia>Chloroflexales | GCA_017743475.1 | Roseiflexus_sp. | J7482_16995 | 17677..21033 | 2562120 | DUF499-domain-containing-protein-[Roseiflexus-sp.] |

| Protein-ID | DUF499-architecture | System | Gene-Neighbourhoods | Length | Taxonomy | Assembly-ID | Strain | Locus-Tag | Genomic-coordinates | TaxID | Fasta-defline |
|---|---|---|---|---|---|---|---|---|---|---|---|
| MCS6883598.1 | HEPN+AAA-ATPase+Alpha+Beta-domain+wHTH+RRM | Type-1_BR.RM.like.systems | PolB->HEPN->?->METHYLASE->METHYLASE-p->REase->HEPN+DUF499*-> | 1130 | Bacteria>Terrabacteria_group>Chloroflexota>Chloroflexia>Chloroflexales | GCA_025057155.1 | Chloroflexaceae_bacterium | NZU74_19900 | 13790..17182 | 2562117 | Swt1-family-HEPN-domain-containing-protein-[Chloroflexaceae-bacterium] |
| MBO9374240.1 | HEPN+AAA-ATPase+Alpha+Beta-domain+wHTH+RRM | Type-1_BR.RM.like.systems | <-TUDOR+SF1-SF2-Helicase+iREase+CTD+REase<-HEPN+DUF499* | 1112 | Bacteria>Terrabacteria_group>Chloroflexota>Chloroflexia>Chloroflexales | GCA_017743265.1 | Chloroflexus_sp. | J7459_12885 | 2503..5841 | 1904827 | DUF499-domain-containing-protein-[Chloroflexus-sp.] |
| MBO9321562.1 | HEPN+AAA-ATPase+Alpha+Beta-domain+wHTH+RRM | Type-1_BR.RM.like.systems | <-TUDOR+SF1-SF2-Helicase+iREase+CTD+REase<-HEPN+DUF499* | 1112 | Bacteria>Terrabacteria_group>Chloroflexota>Chloroflexia>Chloroflexales | GCA_017743635.1 | Roseiflexus_sp. | J7457_05465 | 3598..6936 | 2562120 | DUF499-domain-containing-protein-[Roseiflexus-sp.] |
| WP_255346773.1 | HEPN+AAA-ATPase+Alpha+Beta-domain+wHTH+RRM | Type-1_BR.RM.like.systems | HEPN+DUF499*-><-?<-MODE-HTH+PBPII\|\|?->?->?->ABC-ATPase+SbcC-> | 501 | Bacteria>Terrabacteria_group>Chloroflexota>Chloroflexia>Chloroflexales | GCF_000516515.1 | Chloroflexus_sp._Y-396-1 | CHY396_RS19730 | 597760..599265 | 867845 | Swt1-family-HEPN-domain-containing-protein-[Chloroflexus-sp.-Y-396-1] |
| WP_205703984.1 | HEPN+AAA-ATPase+Alpha+Beta-domain+wHTH+RRM | Type-1_BR.RM.like.systems | RAMPs->?->?->?->HEPN+DUF499*->?->?->Cas_Cas1->CRISPR_Cas2-> | 1169 | Bacteria>Terrabacteria_group>Chloroflexota>Chloroflexia>Chloroflexales | GCF_004138165.1 | Candidatus_Chloroploca_sp._Khr17 | EKD05_RS19000 | 28531..32040 | 2496869 | Swt1-family-HEPN-domain-containing-protein-[Candidatus-Chloroploca-sp.-Khr17] |
| MCX7860988.1 | HEPN+AAA-ATPase+Alpha+Beta-domain+wHTH+RRM | Type-1_BR.RM.like.systems | <-HEPN+DUF499*<-METHYLASE-p | 1115 | Bacteria>Terrabacteria_group>Chloroflexota>Chloroflexia>Chloroflexales | GCA_026417255.1 | Chloroflexus_sp. | N2385_12910 | 223..3570 | 1904827 | MAG:-Swt1-family-HEPN-domain-containing-protein-[Chloroflexus-sp.] |
| MCS6937885.1 | HEPN+AAA-ATPase+Alpha+Beta-domain+wHTH+RRM | Type-1_BR.RM.like.systems | METHYLASE-p->HEPN+DUF499*-> | 531 | Bacteria>Terrabacteria_group>Chloroflexota>Chloroflexia>Chloroflexales | GCA_025055265.1 | Roseiflexus_sp. | NZM94_01360 | 1244..2838 | 2562120 | Swt1-family-HEPN-domain-containing-protein-[Roseiflexus-sp.] |
| MCS6889332.1 | HEPN+AAA-ATPase+Alpha+Beta-domain+wHTH+RRM | Type-1_BR.RM.like.systems | <-HEPN+DUF499*<-METHYLASE-p | 1115 | Bacteria>Terrabacteria_group>Chloroflexota>Chloroflexia>Chloroflexales | GCA_025057175.1 | Chloroflexus_sp. | NZQ09_14085 | 968..4315 | 1904827 | Swt1-family-HEPN-domain-containing-protein-[Chloroflexus-sp.] |
| WP_026369943.1 | HEPN+AAA-ATPase+Alpha+Beta-domain+wHTH+RRM | Type-1_BR.RM.like.systems | ABC-ATPase+SbcC->ABC-ATPase+SbcC->?->tRNA-><-HEPN+DUF499*\|\|PAIREDC-HTH->?->?->?->?-><-METHYLASE | 1108 | Bacteria>Terrabacteria_group>Chloroflexota>Chloroflexia>Kallotenuales | GCF_000526415.1 | Kallotenue_papyrolyticum | K361_RS0107050 | 1033569..1036895 | 1325125 | Swt1-family-HEPN-domain-containing-protein-[Kallotenue-papyrolyticum] |
| KAB8142895.1 | HEPN+AAA-ATPase+Alpha+Beta-domain+wHTH+RRM | Type-1_BR.RM.like.systems | PAPS_reduct->?->TUDOR+SF1-SF2-Helicase+iREase+CTD+REase->?->METHYLASE->HEPN+DUF499*-><-?<-HNH\|\|?->Phage_int_SAM_4+Phage_integrase-> | 1122 | Bacteria>Terrabacteria_group>Chloroflexota>Chloroflexia>unclassified_Chloroflexia | GCA_009268125.1 | Chloroflexia_bacterium_SDU3-3 | F8S13_11620 | 110122..113490 | 2651839 | ATP-binding-protein-[Chloroflexia-bacterium-SDU3-3] |
| KAB8139622.1 | HEPN+AAA-ATPase+Alpha+Beta-domain+wHTH+RRM | Type-1_BR.RM.like.systems | <-HEPN+DUF499*<-METHYLASE<-TUDOR+SF1-SF2-Helicase+iREase+CTD+REase | 1122 | Bacteria>Terrabacteria_group>Chloroflexota>Chloroflexia>unclassified_Chloroflexia | GCA_009268125.1 | Chloroflexia_bacterium_SDU3-3 | F8S13_27385 | 758..4126 | 2651839 | ATP-binding-protein-[Chloroflexia-bacterium-SDU3-3] |
| WP_012876078.1 | HEPN+AAA-ATPase+Alpha+Beta-domain+wHTH+RRM | Type-1_BR.RM.like.systems | <-GNTR-HTH\|\|?->TUDOR+SF1-SF2-Helicase+iREase+CTD+REase->METHYLASE->HEPN+DUF499*-> | 1099 | Bacteria>Terrabacteria_group>Chloroflexota>Chloroflexota_incertae_sedis>Thermobaculum | GCF_000025005.1 | Thermobaculum_terrenum | TTER_RS10895 | 291668..294967 | 166501 | Swt1-family-HEPN-domain-containing-protein-[Thermobaculum-terrenum] |
| WP_072555801.1 | HEPN+AAA-ATPase+Alpha+Beta-domain+wHTH+RRM | Type-1_BR.RM.like.systems | <-XIS-HTH<-HEPN+DUF499*<-METHYLASE<-MORC+DUF3684<-TUDOR+SF1-SF2-Helicase+iREase+CTD+REase<-HTH<-REase+SF1-SF2-Helicase<-HTH+WYL | 1114 | Bacteria>Terrabacteria_group>Chloroflexota>Dehalococcoidia>Dehalococcoidales | GCF_001889305.1 | Dehalococcoides_mccartyi | ASJ33_RS07850 | 1416330..1419674 | 61435 | Swt1-family-HEPN-domain-containing-protein-[Dehalococcoides-mccartyi] |
| WP_046961399.1 | HEPN+AAA-ATPase+Alpha+Beta-domain+wHTH+RRM | Type-1_BR.RM.like.systems | <-XIS-HTH<-HEPN+DUF499*<-METHYLASE<-MORC+DUF3684<-TUDOR+SF1-SF2-Helicase+iREase+CTD+REase<-HTH<-REase+SF1-SF2-Helicase<-HTH+WYL | 1114 | Bacteria>Terrabacteria_group>Chloroflexota>Dehalococcoidia>Dehalococcoidales | GCF_001010485.1 | Dehalococcoides | CWV2_RS06675 | 1251842..1255186 | 61434 | MULTISPECIES:-Swt1-family-HEPN-domain-containing-protein-[Dehalococcoides] |
| MBN4064100.1 | HEPN+AAA-ATPase+Alpha+Beta-domain+wHTH+RRM | Type-1_BR.RM.like.systems | METHYLASE-p->HEPN+DUF499*-><-REase\|\|XIS-HTH-> | 1109 | Bacteria>Terrabacteria_group>Chloroflexota>Dehalococcoidia>Dehalococcoidales | GCA_017114025.1 | Dehalococcoides_mccartyi | JYU04_00010 | 654..3983 | 61435 | DUF499-domain-containing-protein-[Dehalococcoides-mccartyi] |
| MCL4239733.1 | HEPN+AAA-ATPase+Alpha+Beta-domain+wHTH+RRM | Type-1_BR.RM.like.systems | <-SF1-SF2-Helicase+REase<-?<-SIGMA-HTH<-?<-?<-HEPN+DUF499*<-METHYLASE<-?<-MORC+DUF3684<-TUDOR+SF1-SF2-Helicase+iREase+CTD+REase\|\|HEPN-><-LysM+TPR | 1122 | Bacteria>Terrabacteria_group>Chloroflexota>Dehalococcoidia>unclassified_Dehalococcoidia | GCA_023382755.1 | Dehalococcoidia_bacterium | KJ048_00120 | 28707..32075 | 2026734 | DUF499-domain-containing-protein-[Dehalococcoidia-bacterium] |
| MXX18045.1 | HEPN+AAA-ATPase+Alpha+Beta-domain+wHTH+RRM | Type-1_BR.RM.like.systems | TUDOR+SF1-SF2-Helicase+iREase+CTD+REase->Trypsin->METHYLASE->PolB->HEPN->HEPN+DUF499*->?-><-ABhydrolase\|\|STYKinase-> | 1125 | Bacteria>Terrabacteria_group>Chloroflexota>Dehalococcoidia>unclassified_Dehalococcoidia | GCA_009836455.1 | Dehalococcoidia_bacterium | F4Z48_00945 | 10112..13489 | 2026734 | ATP-binding-protein-[Dehalococcoidia-bacterium] |
| PWB45748.1 | HEPN+AAA-ATPase+Alpha+Beta-domain+wHTH+RRM | Type-1_BR.RM.like.systems | LysM+TPR-><-HEPN\|\|TUDOR+SF1-SF2-Helicase+iREase+CTD+REase->MORC+DUF3684->HEPN+DUF499*-><-?<-HISKIN<-SbcC<-sGTP | 1122 | Bacteria>Terrabacteria_group>Chloroflexota>Dehalococcoidia>unclassified_Dehalococcoidia | GCA_003104995.1 | Dehalococcoidia_bacterium | C3F10_05490 | 126089..129457 | 2026734 | AAA+-family-ATPase-[Dehalococcoidia-bacterium] |
| MCL4230333.1 | HEPN+AAA-ATPase+Alpha+Beta-domain+wHTH+RRM | Type-1_BR.RM.like.systems | <-HTH_17\|\|HTH->Peptidase-><-HEPN+DUF499*<-METHYLASE<-HEPN<-TUDOR+SF1-SF2-Helicase+iREase+CTD+REase | 1139 | Bacteria>Terrabacteria_group>Chloroflexota>Dehalococcoidia>unclassified_Dehalococcoidia | GCA_023382735.1 | Dehalococcoidia_bacterium | KJ053_02015 | 402494..405913 | 2026734 | DUF499-domain-containing-protein-[Dehalococcoidia-bacterium] |
| MXY35173.1 | HEPN+AAA-ATPase+Alpha+Beta-domain+wHTH+RRM | Type-1_BR.RM.like.systems | <-HEPN+DUF499*<-HEPN<-PolB<-METHYLASE<-METHYLASE<-HEPN<-TUDOR+SF1-SF2-Helicase+iREase+CTD+REase | 1125 | Bacteria>Terrabacteria_group>Chloroflexota>Dehalococcoidia>unclassified_Dehalococcoidia | GCA_009838385.1 | Dehalococcoidia_bacterium | F4Y54_00090 | 1779..5156 | 2026734 | ATP-binding-protein-[Dehalococcoidia-bacterium] |
| MXW25933.1 | HEPN+AAA-ATPase+Alpha+Beta-domain+wHTH+RRM | Type-1_BR.RM.like.systems | ABhydrolase-><-?<-URI-DNase\|\|tRNA->\|\|?-><-PadR-HTH\|\|?-><-HEPN+DUF499*<-HEPN<-PolB<-METHYLASE-p | 1120 | Bacteria>Terrabacteria_group>Chloroflexota>Dehalococcoidia>unclassified_Dehalococcoidia | GCA_009835935.1 | Dehalococcoidia_bacterium | F4Z77_06550 | 10102..13464 | 2026734 | ATP-binding-protein-[Dehalococcoidia-bacterium] |
| MCL4240066.1 | HEPN+AAA-ATPase+Alpha+Beta-domain+wHTH+RRM | Type-1_BR.RM.like.systems | <-HEPN+DUF499*<-?<-HTH<-?<-METHYLASE<-ParB+HNH | 1138 | Bacteria>Terrabacteria_group>Chloroflexota>Dehalococcoidia>unclassified_Dehalococcoidia | GCA_023382755.1 | Dehalococcoidia_bacterium | KJ048_01835 | 17815..21231 | 2026734 | DUF499-domain-containing-protein-[Dehalococcoidia-bacterium] |
| MCS6803437.1 | HEPN+AAA-ATPase+Alpha+Beta-domain+wHTH+RRM | Type-1_BR.RM.like.systems | METHYLASE-><-?\|\|HEPN+DUF499*-> | 1109 | Bacteria>Terrabacteria_group>Chloroflexota>Dehalococcoidia>unclassified_Dehalococcoidia | GCA_025054935.1 | Dehalococcoidia_bacterium | NZ773_16035 | 2535..5864 | 2026734 | Swt1-family-HEPN-domain-containing-protein-[Dehalococcoidia-bacterium] |
| MPZ22055.1 | HEPN+AAA-ATPase+Alpha+Beta-domain+wHTH+RRM | Type-1_BR.RM.like.systems | <-CR-Kinase<-?<-HEPN+DUF499*\|\|<-TUDOR+SF1-SF2-Helicase+iREase+CTD+REase\|\|Peptidase_M20->?-><-DNA-glycosylase<-HAD | 1112 | Bacteria>Terrabacteria_group>Chloroflexota>Dehalococcoidia>unclassified_Dehalococcoidia | GCA_009377665.1 | Dehalococcoidia_bacterium | GEU28_00570 | 3174..6512 | 2026734 | DUF499-domain-containing-protein-[Dehalococcoidia-bacterium] |
| MCA9831553.1 | HEPN+AAA-ATPase+Alpha+Beta-domain+wHTH+RRM | Type-1_BR.RM.like.systems | TUDOR+SF1-SF2-Helicase+iREase+CTD+REase->REase->?->?->?->METHYLASE->HEPN+DUF499*->?->HerA/FtsK->PIN-> | 1134 | Bacteria>Terrabacteria_group>Chloroflexota>Dehalococcoidia>unclassified_Dehalococcoidia | GCA_020432045.1 | Dehalococcoidia_bacterium | KC495_13860 | 27821..31225 | 2026734 | ATP-binding-protein-[Dehalococcoidia-bacterium] |
| WP_126627899.1 | HEPN+AAA-ATPase+Alpha+Beta-domain+wHTH+RRM | Type-1_BR.RM.like.systems | ParB->TUDOR+SF1-SF2-Helicase+iREase+CTD+REase->METHYLASE-p->DDE_Tnp_1->DDE_Tnp_1+DDE_Tnp_1_2->METHYLASE->HEPN+DUF499*->?->?-><-?<-?<-NERD+Pkinase+Pkinase+nSTAND3 | 1124 | Bacteria>Terrabacteria_group>Chloroflexota>Ktedonobacteria>Ktedonobacterales | GCF_003967575.1 | Dictyobacter_alpinus | EL633_RS14700 | 3690225..3693599 | 2014873 | DUF499-domain-containing-protein-[Dictyobacter-alpinus] |
| WP_201387653.1 | HEPN+AAA-ATPase+Alpha+Beta-domain+wHTH+RRM | Type-1_BR.RM.like.systems | TUDOR+SF1-SF2-Helicase+iREase+CTD+REase->METHYLASE-p-><-DDE_3<-PAIREDC-HTH<-PAIREDC-HTH\|\|METHYLASE->HEPN+DUF499*->ParB+HNH-><-?<-?<-?\|\|HISKIN->REC-> | 1128 | Bacteria>Terrabacteria_group>Chloroflexota>Ktedonobacteria>Ktedonobacterales | GCF_016587395.1 | Ktedonobacter_sp._SOSP1-52 | JMJ16_RS30300 | 1126374..1129760 | 2778366 | DUF499-domain-containing-protein-[Ktedonobacter-sp.-SOSP1-52] |
| WP_129891798.1 | HEPN+AAA-ATPase+Alpha+Beta-domain+wHTH+RRM | Type-1_BR.RM.like.systems | MPTase-><-METHYLASE-p<-SF1-SF2-helicase+DUF1998<-?\|\|TUDOR+SF1-SF2-Helicase+iREase+CTD+REase->METHYLASE->HEPN+DUF499*-><-?<-?<-?\|\|?->ABhydrolase-> | 1138 | Bacteria>Terrabacteria_group>Chloroflexota>Ktedonobacteria>Ktedonobacterales | GCF_004208415.1 | Ktedonosporobacter_rubrisoli | EPA93_RS34115 | 7851540..7854956 | 2509675 | Swt1-family-HEPN-domain-containing-protein-[Ktedonosporobacter-rubrisoli] |
| WP_201359963.1 | HEPN+AAA-ATPase+Alpha+Beta-domain+wHTH+RRM | Type-1_BR.RM.like.systems | NERD+Pkinase+Pkinase+nSTAND3->?->?->APATPase+iSTAND3-><-HEPN+DUF499*<-METHYLASE<-TUDOR+SF1-SF2-Helicase+iREase+CTD+REase\|\|?->?->?->STYKIN-> | 1127 | Bacteria>Terrabacteria_group>Chloroflexota>Ktedonobacteria>Ktedonobacterales | GCF_016587435.1 | Dictyobacter_formicarum | JMI88_RS01395 | 354947..358330 | 2778368 | DUF499-domain-containing-protein-[Dictyobacter-formicarum] |
| BBH87966.1 | HEPN+AAA-ATPase+Alpha+Beta-domain+wHTH+RRM | Type-1_BR.RM.like.systems | ACET->TUDOR+SF1-SF2-Helicase+iREase+CTD+REase->METHYLASE->HEPN+DUF499*-><-PADR-HTH | 1126 | Bacteria>Terrabacteria_group>Chloroflexota>Ktedonobacteria>Ktedonobacterales | - | Thermosporothrix_sp._COM3 | KTC_27170 | 3109376..3112756 | 2490863 | hypothetical-protein-KTC_27170-[Thermosporothrix-sp.-COM3] |
| WP_201365551.1 | HEPN+AAA-ATPase+Alpha+Beta-domain+wHTH+RRM | Type-1_BR.RM.like.systems | <-NUDIX<-?<-?<-HEPN+DUF499*<-METHYLASE<-TUDOR+SF1-SF2-Helicase+iREase+CTD+REase | 1131 | Bacteria>Terrabacteria_group>Chloroflexota>Ktedonobacteria>Ktedonobacterales | GCF_016587435.1 | Dictyobacter_formicarum | JMI88_RS29565 | 30848..34243 | 2778368 | Swt1-family-HEPN-domain-containing-protein-[Dictyobacter-formicarum] |
| WP_111324027.1 | HEPN+AAA-ATPase+Alpha+Beta-domain+wHTH+RRM | Type-1_BR.RM.like.systems | TUDOR+SF1-SF2-Helicase+iREase+CTD+REase->METHYLASE->HEPN+DUF499*-> | 1126 | Bacteria>Terrabacteria_group>Chloroflexota>Ktedonobacteria>Ktedonobacterales | GCF_005402645.1 | Thermosporothrix_hazakensis | EI42_RS18025 | 56767..60147 | 644383 | Swt1-family-HEPN-domain-containing-protein-[Thermosporothrix-hazakensis] |
| WP_069804546.1 | HEPN+AAA-ATPase+Alpha+Beta-domain+wHTH+RRM | Type-1_BR.RM.like.systems | TUDOR+SF1-SF2-Helicase+iREase+CTD+REase->METHYLASE->HEPN+DUF499*-><-?<-LuxR-HTH+LuxR-HTH<-?\|\|XIS-HTH->?->HTH+TPR+TPR-> | 1138 | Bacteria>Terrabacteria_group>Chloroflexota>Ktedonobacteria>Thermogemmatisporales | GCF_001748285.1 | Thermogemmatispora_onikobensis | BGC09_RS13650 | 65405..68821 | 732234 | Swt1-family-HEPN-domain-containing-protein-[Thermogemmatispora-onikobensis] |
| WP_151729028.1 | HEPN+AAA-ATPase+Alpha+Beta-domain+wHTH+RRM | Type-1_BR.RM.like.systems | <-HTH+TPR+TPR\|\|?->LuxR-HTH+LuxR-HTH->?-><-HEPN+DUF499*<-METHYLASE<-TUDOR+SF1-SF2-Helicase+iREase+CTD+REase | 1139 | Bacteria>Terrabacteria_group>Chloroflexota>Ktedonobacteria>Thermogemmatisporales | GCF_008974285.1 | Thermogemmatispora_aurantia | KTAU_RS14380 | 459463..462882 | 2045279 | Swt1-family-HEPN-domain-containing-protein-[Thermogemmatispora-aurantia] |
| BBH92994.1 | HEPN+AAA-ATPase+Alpha+Beta-domain+wHTH+RRM | Type-1_BR.RM.like.systems | TUDOR+SF1-SF2-Helicase+iREase+CTD+REase->METHYLASE->HEPN+DUF499*-><-?<-LuxR-HTH+LuxR-HTH<-?\|\|HTH+TPR+TPR-> | 1130 | Bacteria>Terrabacteria_group>Chloroflexota>Ktedonobacteria>Thermogemmatisporales | - | Thermogemmatispora_argillosa | KTA_11930 | 1512698..1516090 | 2045280 | hypothetical-protein-KTA_11930-[Thermogemmatispora-argillosa] |
| MBX5450396.1 | HEPN+AAA-ATPase+Alpha+Beta-domain+wHTH+RRM | Type-1_BR.RM.like.systems | TUDOR+SF1-SF2-Helicase+iREase+CTD+REase->METHYLASE->HEPN+DUF499*-><-?<-LuxR-HTH+LuxR-HTH | 1139 | Bacteria>Terrabacteria_group>Chloroflexota>Ktedonobacteria>Thermogemmatisporales | GCA_019683755.1 | Thermogemmatispora_sp. | IRZ24_10040 | 7541..10960 | 1968838 | DUF499-domain-containing-protein-[Thermogemmatispora-sp.] |
| MBX5459182.1 | HEPN+AAA-ATPase+Alpha+Beta-domain+wHTH+RRM | Type-1_BR.RM.like.systems | LuxR-HTH+LuxR-HTH->?-><-HEPN+DUF499*<-METHYLASE-p | 1136 | Bacteria>Terrabacteria_group>Chloroflexota>Ktedonobacteria>Thermogemmatisporales | GCA_019683635.1 | Thermogemmatispora_sp. | IRZ31_20000 | 1031..4441 | 1968838 | DUF499-domain-containing-protein-[Thermogemmatispora-sp.] |
| WP_158067647.1 | HEPN+AAA-ATPase+Alpha+Beta-domain+wHTH+RRM | Type-1_BR.RM.like.systems | RNase+SF1-SF2-Helicase->CRISPR_Cas2->TUDOR+SF1-SF2-Helicase+iREase+CTD+REase->?->METHYLASE-p->METHYLASE->HEPN+DUF499*-><-?<-?\|\|?-><-?\|\|?-><-ZnR | 1109 | Bacteria>Terrabacteria_group>Chloroflexota>Tepidiformia>Tepidiformales | GCF_008838325.1 | Tepidiforma_bonchosmolovskayae | Tbon_RS10440 | 2086716..2090045 | 2601677 | Swt1-family-HEPN-domain-containing-protein-[Tepidiforma-bonchosmolovskayae] |
| WP_098503018.1 | HEPN+AAA-ATPase+Alpha+Beta-domain+wHTH+RRM | Type-1_BR.RM.like.systems | PolB->HEPN->?->TUDOR+SF1-SF2-Helicase+iREase+CTD+REase->OLD-ABC+TOPRIM->METHYLASE->HEPN+DUF499*->?-><-HAD | 1111 | Bacteria>Terrabacteria_group>Chloroflexota>Tepidiformia>Tepidiformales | GCF_002563855.1 | Tepidiforma_thermophila | A9A59_RS03825 | 812331..815666 | 2761530 | Swt1-family-HEPN-domain-containing-protein-[Tepidiforma-thermophila] |
| PFG73567.1 | HEPN+AAA-ATPase+Alpha+Beta-domain+wHTH+RRM | Type-1_BR.RM.like.systems | PolB->HEPN->?->TUDOR+SF1-SF2-Helicase+iREase+CTD+REase->OLD-ABC+TOPRIM->METHYLASE->HEPN+DUF499*->?-><-HAD | 1111 | Bacteria>Terrabacteria_group>Chloroflexota>Tepidiformia>Tepidiformales | GCF_002563855.1 | Tepidiforma_thermophila | A9A59_0767 | 812331..815666 | 2761530 | putative-AAA+-superfamily-ATPase-[Chloroflexi-bacterium-G233] |
| QWK10277.1 | HEPN+AAA-ATPase+Alpha+Beta-domain+wHTH+RRM | Type-1_BR.RM.like.systems | <-OrfB_IS605+OrfB_Zn_ribbon<-?\|\|ABC-ATPase+SbcC-><-?<-?<-?<-HEPN+DUF499*<-HEPN<-PolB<-METHYLASE<-REase<-OLD-ABC+TOPRIM<-TUDOR+SF1-SF2-Helicase+iREase+CTD+REase | 1119 | Bacteria>Terrabacteria_group>Chloroflexota>Thermoflexia>Thermoflexales | GCA_018771565.1 | Thermoflexus_hugenholtzii | KNN16_13035 | 2858619..2861978 | 1495650 | MAG:-DUF499-domain-containing-protein-[Thermoflexus-hugenholtzii] |
| MBO9362157.1 | HEPN+AAA-ATPase+Alpha+Beta-domain+wHTH+RRM | Type-1_BR.RM.like.systems | <-HEPN+DUF499*<-HEPN<-PolB<-METHYLASE | 1119 | Bacteria>Terrabacteria_group>Chloroflexota>Thermoflexia>Thermoflexales | GCA_017743605.1 | Thermoflexus_sp. | J7452_08145 | 638..3997 | 1969742 | DUF499-domain-containing-protein-[Thermoflexus-sp.] |
| MBE7520719.1 | HEPN+AAA-ATPase+Alpha+Beta-domain+wHTH+RRM | Type-1_BR.RM.like.systems | sGTP-><-PAIREDC-HTH<-?<-PAPS_reduct<-?<-KAP-NTPase<-HEPN+DUF499*<-METHYLASE<-?<-?<-TUDOR+SF1-SF2-Helicase+iREase+CTD+REase<-LysM+TPR | 1132 | Bacteria>Terrabacteria_group>Chloroflexota>Thermoflexia>Thermoflexales | GCA_015075555.1 | Thermoflexaceae_bacterium | HS107_15925 | 106294..109692 | 2739507 | DUF499-domain-containing-protein-[Thermoflexaceae-bacterium] |
| MCQ3935712.1 | HEPN+AAA-ATPase+Alpha+Beta-domain+wHTH+RRM | Type-1_BR.RM.like.systems | SF1-SF2-Helicase-><-PIWI<-REase+PNTD1+PNTD2<-HEPN+DUF499*<-METHYLASE<-SF1-SF2-Helicase<-?<-TUDOR+SF1-SF2-Helicase+iREase+CTD+REase\|\|OLD-ABC+TOPRIM->SF1-SF2-Helicase-> | 1108 | Bacteria>Terrabacteria_group>Chloroflexota>unclassified_Chloroflexota | GCA_024434165.1 | Chloroflexota_bacterium | DPW18_01570 | 11480..14806 | 2026724 | AAA+-family-ATPase-[Chloroflexi-bacterium] |
| MXZ46491.1 | HEPN+AAA-ATPase+Alpha+Beta-domain+wHTH+RRM | Type-1_BR.RM.like.systems | <-GDResolvase+Recombinase+ZnR<-?\|\|?->?-><-BirA-HTH+CRP-HTH\|\|?-><-HEPN+DUF499*<-HEPN<-PolB<-METHYLASE<-TUDOR+SF1-SF2-Helicase+iREase+CTD+REase\|\|PIN+ACET+EVE->ASCH-> | 1125 | Bacteria>Terrabacteria_group>Chloroflexota>unclassified_Chloroflexota | GCA_009837285.1 | Chloroflexota_bacterium | F4Z08_05790 | 8633..12010 | 2026724 | ATP-binding-protein-[Chloroflexi-bacterium] |
| MCC6178177.1 | HEPN+AAA-ATPase+Alpha+Beta-domain+wHTH+RRM | Type-1_BR.RM.like.systems | <-HTH_3<-Tox-RelE<-SF1-SF2-Helicase<-OLD-ABC+TOPRIM<-REase<-HEPN+DUF499*<-METHYLASE<-wHTH+REase<-TUDOR+SF1-SF2-Helicase+iREase+CTD+REase\|\|?->Peptidase_S41-> | 1117 | Bacteria>Terrabacteria_group>Chloroflexota>unclassified_Chloroflexota | GCA_020845495.1 | Chloroflexota_bacterium | IT305_22975 | 6439..9792 | 2026724 | DUF499-domain-containing-protein-[Chloroflexi-bacterium] |
| MXX47637.1 | HEPN+AAA-ATPase+Alpha+Beta-domain+wHTH+RRM | Type-1_BR.RM.like.systems | VirD4-TraG+HerA->PadR-HTH->RADICAL-SAM->?->?-><-HEPN+DUF499*<-HEPN<-Polbeta<-METHYLASE<-Trypsin<-TUDOR+SF1-SF2-Helicase+iREase+CTD+REase\|\|REase-> | 1112 | Bacteria>Terrabacteria_group>Chloroflexota>unclassified_Chloroflexota | GCA_009836695.1 | Chloroflexota_bacterium | F4Z38_04900 | 11069..14407 | 2026724 | ATP-binding-protein-[Chloroflexi-bacterium] |
| MDE2892396.1 | HEPN+AAA-ATPase+Alpha+Beta-domain+wHTH+RRM | Type-1_BR.RM.like.systems | <-REase\|\|TUDOR+SF1-SF2-Helicase+iREase+CTD+REase->Trypsin->METHYLASE->Polbeta->HEPN->HEPN+DUF499*-><-?<-?<-McrC-NTD+REase<-?<-wHTH+REase<-PadR-HTH | 1110 | Bacteria>Terrabacteria_group>Chloroflexota>unclassified_Chloroflexota | GCA_028874365.1 | Chloroflexota_bacterium | OXN86_07855 | 154158..157490 | 2026724 | MAG:-Swt1-family-HEPN-domain-containing-protein-[Chloroflexota-bacterium] |
| MCY3570526.1 | HEPN+AAA-ATPase+Alpha+Beta-domain+wHTH+RRM | Type-1_BR.RM.like.systems | TUDOR+SF1-SF2-Helicase+iREase+CTD+REase-p->HEPN->METHYLASE->Polbeta->HEPN->HEPN+DUF499*-><-?<-METHYLASE-p<-?<-ParB+HNH<-wHTH+REase<-PadR-HTH | 1110 | Bacteria>Terrabacteria_group>Chloroflexota>unclassified_Chloroflexota | GCA_026705025.1 | Chloroflexota_bacterium | OXH19_04200 | 6390..9722 | 2026724 | MAG:-Swt1-family-HEPN-domain-containing-protein-[Chloroflexi-bacterium] |
| MBP8000880.1 | HEPN+AAA-ATPase+Alpha+Beta-domain+wHTH+RRM | Type-1_BR.RM.like.systems | <-XIS-HTH<-?\|\|SF1-SF2-helicase+RdgC+Beta-barrel+REase-><-HEPN+DUF499*<-?<-?<-METHYLASE<-ParB+HNH<-TUDOR+SF1-SF2-Helicase+iREase+CTD+REase | 1097 | Bacteria>Terrabacteria_group>Chloroflexota>unclassified_Chloroflexota | GCA_018004095.1 | Chloroflexota_bacterium | KA338_16630 | 13590..16883 | 2026724 | DUF499-domain-containing-protein-[Chloroflexi-bacterium] |
| MCE7858272.1 | HEPN+AAA-ATPase+Alpha+Beta-domain+wHTH+RRM | Type-1_BR.RM.like.systems | HTH+UbE1-><-Schlafen<-?<-NACHT<-ParB+HNH<-HEPN+DUF499*<-METHYLASE<-MORC+DUF3684<-TUDOR+SF1-SF2-Helicase+iREase+CTD+REase<-?<-?<-HTH+WYL | 1112 | Bacteria>Terrabacteria_group>Chloroflexota>unclassified_Chloroflexota | GCA_021404705.1 | Chloroflexi_bacterium_CFX2 | DYG86_00620 | 111078..114416 | 2293708 | ATP-binding-protein-[Chloroflexi-bacterium-CFX2] |
| NJD66806.1 | HEPN+AAA-ATPase+Alpha+Beta-domain+wHTH+RRM | Type-1_BR.RM.like.systems | sGTP-><-SF1-SF2-Helicase<-OLD-ABC+TOPRIM<-HEPN+DUF499*<-METHYLASE<-DUF4326<-?<-DUF488-l<-TUDOR+SF1-SF2-Helicase+iREase+CTD+REase<-LysM+TPR | 1122 | Bacteria>Terrabacteria_group>Chloroflexota>unclassified_Chloroflexota | GCA_012027015.1 | Chloroflexota_bacterium | FIB00_16460 | 5569..8937 | 2026724 | ATP-binding-protein-[Chloroflexi-bacterium] |
| MCY3559222.1 | HEPN+AAA-ATPase+Alpha+Beta-domain+wHTH+RRM | Type-1_BR.RM.like.systems | TUDOR+SF1-SF2-Helicase+iREase+CTD+REase-p->Trypsin->METHYLASE->Polbeta->HEPN->HEPN+DUF499*->MORC+REase-><-REase<-PadR-HTH | 1111 | Bacteria>Terrabacteria_group>Chloroflexota>unclassified_Chloroflexota | GCA_026702785.1 | Chloroflexota_bacterium | OXH13_11370 | 6173..9508 | 2026724 | MAG:-Swt1-family-HEPN-domain-containing-protein-[Chloroflexi-bacterium] |
| MCA0453853.1 | HEPN+AAA-ATPase+Alpha+Beta-domain+wHTH+RRM | Type-1_BR.RM.like.systems | <-GDResolvase<-?<-zf-ISL3+DDE_Tnp_ISL3<-?<-?<-?<-HEPN+DUF499*<-PIN<-DUF433<-METHYLASE<-PIN<-?<-TUDOR+SF1-SF2-Helicase+iREase+CTD+REase | 1103 | Bacteria>Terrabacteria_group>Chloroflexota>unclassified_Chloroflexota | GCA_020161935.1 | Chloroflexota_bacterium | LCI00_07765 | 53613..56924 | 2026724 | DUF499-domain-containing-protein-[Chloroflexi-bacterium] |
| MDE2935729.1 | HEPN+AAA-ATPase+Alpha+Beta-domain+wHTH+RRM | Type-1_BR.RM.like.systems | <-HNH<-?<-OrfB_IS605+OrfB_Zn_ribbon\|\|?-><-?<-HEPN+DUF499*<-METHYLASE-p<-METHYLASE-p<-TUDOR+SF1-SF2-Helicase+iREase+CTD+REase | 1054 | Bacteria>Terrabacteria_group>Chloroflexota>unclassified_Chloroflexota | GCA_028874095.1 | Chloroflexota_bacterium | OXP37_02700 | 49007..52171 | 2026724 | MAG:-DUF499-domain-containing-protein-[Chloroflexota-bacterium] |
| MCY4393097.1 | HEPN+AAA-ATPase+Alpha+Beta-domain+wHTH+RRM | Type-1_BR.RM.like.systems | SF1-SF2-Helicase-p-><-REase<-N6_MTase<-HEPN+DUF499*<-HEPN<-?<-METHYLASE<-HEPN<-TUDOR+SF1-SF2-Helicase+iREase+CTD+REase | 1125 | Bacteria>Terrabacteria_group>Chloroflexota>unclassified_Chloroflexota | GCA_026710655.1 | Chloroflexota_bacterium | OXE43_13775 | 6581..9958 | 2026724 | MAG:-Swt1-family-HEPN-domain-containing-protein-[Chloroflexi-bacterium] |
| MBK8988277.1 | HEPN+AAA-ATPase+Alpha+Beta-domain+wHTH+RRM | Type-1_BR.RM.like.systems | TUDOR+SF1-SF2-Helicase+iREase+CTD+REase->?->PIN->CopG/RHH->PIN->METHYLASE-p->HEPN+DUF499*-><-AbiEii<-?\|\|?->?->?->ABC-ATPase+SbcC-> | 1097 | Bacteria>Terrabacteria_group>Chloroflexota>unclassified_Chloroflexota | GCA_016716065.1 | Chloroflexota_bacterium | IPM39_19790 | 148627..151920 | 2026724 | DUF499-domain-containing-protein-[Chloroflexi-bacterium] |
| MCQ3930198.1 | HEPN+AAA-ATPase+Alpha+Beta-domain+wHTH+RRM | Type-1_BR.RM.like.systems | ParB->?-><-?<-?<-STYKinase<-TIR<-HEPN+DUF499*<-HEPN<-PolB<-METHYLASE<-Pkinase<-TUDOR+SF1-SF2-Helicase+iREase+CTD+REase | 1098 | Bacteria>Terrabacteria_group>Chloroflexota>unclassified_Chloroflexota | GCA_024434115.1 | Chloroflexota_bacterium | DPW16_07035 | 13947..17243 | 2026724 | AAA+-family-ATPase-[Chloroflexi-bacterium] |
| MBI4332978.1 | HEPN+AAA-ATPase+Alpha+Beta-domain+wHTH+RRM | Type-1_BR.RM.like.systems | <-PIN<-MazE_antitoxin<-HicA_toxin<-HEPN+DUF499*<-METHYLASE<-Schlafen+HATPase+HTH<-TUDOR+SF1-SF2-Helicase+iREase+CTD+REase | 1127 | Bacteria>Terrabacteria_group>Chloroflexota>unclassified_Chloroflexota | GCA_016210565.1 | Chloroflexota_bacterium | HY673_17055 | 9786..13169 | 2026724 | ATP-binding-protein-[Chloroflexi-bacterium] |
| MBF6606506.1 | HEPN+AAA-ATPase+Alpha+Beta-domain+wHTH+RRM | Type-1_BR.RM.like.systems | TUDOR+SF1-SF2-Helicase+iREase+CTD+REase->SIGMA-HTH+CR-ATPase9+Schlafen->METHYLASE->HEPN+DUF499*->?->?-><-Schlafen+HATPase+HTH | 1085 | Bacteria>Terrabacteria_group>Chloroflexota>unclassified_Chloroflexota | GCA_015478725.1 | Chloroflexota_bacterium | IVW53_13110 | 14708..17965 | 2026724 | DUF499-domain-containing-protein-[Chloroflexi-bacterium] |
| HAF11542.1 | HEPN+AAA-ATPase+Alpha+Beta-domain+wHTH+RRM | Type-1_BR.RM.like.systems | <-GDResolvase+Recombinase+ZnR<-?<-HEPN+DUF499*<-METHYLASE<-TUDOR+SF1-SF2-Helicase+iREase+CTD+REase<-Schlafen+HATPase+HTH | 1109 | Bacteria>Terrabacteria_group>Chloroflexota>unclassified_Chloroflexota | GCA_003446795.1 | Chloroflexota_bacterium | DCK98_15880 | 2135..5464 | 2026724 | AAA+-family-ATPase-[Chloroflexi-bacterium] |
| NWJ47051.1 | HEPN+AAA-ATPase+Alpha+Beta-domain+wHTH+RRM | Type-1_BR.RM.like.systems | <-Peptidase<-?<-SIGMA-HTH<-?<-?<-?<-HEPN+DUF499*<-Y1_Tnp<-?<-TUDOR+SF1-SF2-Helicase+iREase+CTD+REase<-?<-CRISPR_Cas2<-Cas_Cas1 | 1122 | Bacteria>Terrabacteria_group>Chloroflexota>unclassified_Chloroflexota | GCA_013390565.1 | Chloroflexota_bacterium | HXX08_14415 | 53237..56605 | 2026724 | ATP-binding-protein-[Chloroflexi-bacterium] |
| MCY3923477.1 | HEPN+AAA-ATPase+Alpha+Beta-domain+wHTH+RRM | Type-1_BR.RM.like.systems | SF1-SF2-Helicase+REase-><-HEPN+DUF499*<-HEPN<-PolB<-METHYLASE<-HEPN<-TUDOR+SF1-SF2-Helicase+iREase+CTD+REase | 1114 | Bacteria>Terrabacteria_group>Chloroflexota>unclassified_Chloroflexota | GCA_026707265.1 | Chloroflexota_bacterium | OXG27_13945 | 4231..7575 | 2026724 | MAG:-Swt1-family-HEPN-domain-containing-protein-[Chloroflexi-bacterium] |
| MCC6176203.1 | HEPN+AAA-ATPase+Alpha+Beta-domain+wHTH+RRM | Type-1_BR.RM.like.systems | TUDOR+SF1-SF2-Helicase+iREase+CTD+REase->Schlafen+HATPase+HTH->Cluster1090_5clades->METHYLASE->HEPN+DUF499*-><-REase | 1129 | Bacteria>Terrabacteria_group>Chloroflexota>unclassified_Chloroflexota | GCA_020845495.1 | Chloroflexota_bacterium | IT305_12930 | 12450..15839 | 2026724 | DUF499-domain-containing-protein-[Chloroflexi-bacterium] |
| NJD65682.1 | HEPN+AAA-ATPase+Alpha+Beta-domain+wHTH+RRM | Type-1_BR.RM.like.systems | sGTP->SbcC->HISKIN->?-><-HEPN+DUF499*<-MORC+DUF3684<-TUDOR+SF1-SF2-Helicase+iREase+CTD+REase\|\|HEPN-><-LysM+TPR | 1122 | Bacteria>Terrabacteria_group>Chloroflexota>unclassified_Chloroflexota | GCA_012027015.1 | Chloroflexota_bacterium | FIB00_10630 | 11912..15280 | 2026724 | ATP-binding-protein-[Chloroflexi-bacterium] |
| MCC6174343.1 | HEPN+AAA-ATPase+Alpha+Beta-domain+wHTH+RRM | Type-1_BR.RM.like.systems | TUDOR+SF1-SF2-Helicase+iREase+CTD+REase->Schlafen->BirA-HTH->METHYLASE->ParD->PIN->HEPN+DUF499*-><-?\|\|?->Aldedh-> | 1160 | Bacteria>Terrabacteria_group>Chloroflexota>unclassified_Chloroflexota | GCA_020845495.1 | Chloroflexota_bacterium | IT305_03475 | 199239..202721 | 2026724 | DUF499-domain-containing-protein-[Chloroflexi-bacterium] |
| MXY00663.1 | HEPN+AAA-ATPase+Alpha+Beta-domain+wHTH+RRM | Type-1_BR.RM.like.systems | <-REase\|\|TUDOR+SF1-SF2-Helicase+iREase+CTD+REase->METHYLASE->METHYLASE-p->HEPN+DUF499*-><-?\|\|?-><-ABhydrolase | 1058 | Bacteria>Terrabacteria_group>Chloroflexota>unclassified_Chloroflexota | GCA_009838085.1 | Chloroflexota_bacterium | F4Y67_07570 | 133498..136674 | 2026724 | ATP-binding-protein-[Chloroflexi-bacterium] |
| MBX3030766.1 | HEPN+AAA-ATPase+Alpha+Beta-domain+wHTH+RRM | Type-1_BR.RM.like.systems | <-NUDIX<-?<-HEPN+DUF499*<-?\|\|HTH_31-><-TUDOR+SF1-SF2-Helicase+iREase+CTD+REase<-?<-DOC+HTH\|\|ABC-ATPase+SbcC-> | 1095 | Bacteria>Terrabacteria_group>Chloroflexota>unclassified_Chloroflexota | GCA_019637815.1 | Chloroflexota_bacterium | KF809_11435 | 95688..98975 | 2026724 | DUF499-domain-containing-protein-[Chloroflexi-bacterium] |
| MBK7919048.1 | HEPN+AAA-ATPase+Alpha+Beta-domain+wHTH+RRM | Type-1_BR.RM.like.systems | DDE_5-><-DDE_Tnp_IS1\|\|METHYLASE-p->?->?->HEPN-p->HEPN+DUF499*->DUF499-><-DDE_5<-DDE_5\|\|?-><-DDE_5+DDE_Tnp_2 | 582 | Bacteria>Terrabacteria_group>Chloroflexota>unclassified_Chloroflexota | GCA_016713795.1 | Chloroflexota_bacterium | IPJ94_22860 | 29678..31426 | 2026724 | DUF499-domain-containing-protein-[Chloroflexota-bacterium] |
| NLG97290.1 | HEPN+AAA-ATPase+Alpha+Beta-domain+wHTH+RRM | Type-1_BR.RM.like.systems | METHYLASE->Peptidase->HEPN+DUF499*-><-MULE-TRANSPOSASE<-MULE-TRANSPOSASE<-?<-tRNA<-REase<-Abi<-Glyoxalase | 1131 | Bacteria>Terrabacteria_group>Chloroflexota>unclassified_Chloroflexota | GCA_012516155.1 | Chloroflexota_bacterium | GX491_08015 | 33163..36558 | 2026724 | ATP-binding-protein-[Chloroflexi-bacterium] |
| MXX31451.1 | HEPN+AAA-ATPase+Alpha+Beta-domain+wHTH+RRM | Type-1_BR.RM.like.systems | <-REase\|\|TUDOR+SF1-SF2-Helicase+iREase+CTD+REase->HEPN->METHYLASE->Polbeta->HEPN->HEPN+DUF499*-> | 1112 | Bacteria>Terrabacteria_group>Chloroflexota>unclassified_Chloroflexota | GCA_009836445.1 | Chloroflexota_bacterium | F4Z51_04295 | 8660..11998 | 2026724 | ATP-binding-protein-[Chloroflexi-bacterium] |
| MYD93547.1 | HEPN+AAA-ATPase+Alpha+Beta-domain+wHTH+RRM | Type-1_BR.RM.like.systems | DUF429->DOC-><-HEPN+DUF499*<-METHYLASE<-TUDOR+SF1-SF2-Helicase+iREase+CTD+REase\|\|MORC+DUF3684-> | 1098 | Bacteria>Terrabacteria_group>Chloroflexota>unclassified_Chloroflexota | GCA_009839425.1 | Chloroflexota_bacterium | F4Y02_07625 | 2005..5301 | 2026724 | ATP-binding-protein-[Chloroflexi-bacterium] |
| MCY4557669.1 | HEPN+AAA-ATPase+Alpha+Beta-domain+wHTH+RRM | Type-1_BR.RM.like.systems | <-NADAR<-?<-HIN-HTH<-HEPN+DUF499*<-METHYLASE<-OLD-ABC<-TUDOR+SF1-SF2-Helicase+iREase+CTD+REase<-?<-CHTH | 1117 | Bacteria>Terrabacteria_group>Chloroflexota>unclassified_Chloroflexota | GCA_026713265.1 | Chloroflexota_bacterium | OXF79_15050 | 3333..6686 | 2026724 | MAG:-Swt1-family-HEPN-domain-containing-protein-[Chloroflexi-bacterium] |
| MBN9390798.1 | HEPN+AAA-ATPase+Alpha+Beta-domain+wHTH+RRM | Type-1_BR.RM.like.systems | <-HEPN+DUF499*<-METHYLASE<-YtH<-?<-TUDOR+SF1-SF2-Helicase+iREase+CTD+REase<-HTH_17<-ArdCN+MPTase | 1111 | Bacteria>Terrabacteria_group>Chloroflexota>unclassified_Chloroflexota | GCA_017307685.1 | Chloroflexota_bacterium | JOI20_22395 | 2393..5728 | 2026724 | DUF499-domain-containing-protein-[Chloroflexi-bacterium] |
| MBI4673984.1 | HEPN+AAA-ATPase+Alpha+Beta-domain+wHTH+RRM | Type-1_BR.RM.like.systems | TPR+TPR+TPR+TPR-><-HEPN+DUF499*<-NTP_transf<-?<-DOC+HTH<-TUDOR+SF1-SF2-Helicase+iREase+CTD+REase | 1130 | Bacteria>Terrabacteria_group>Chloroflexota>unclassified_Chloroflexota | GCA_016209205.1 | Chloroflexota_bacterium | HY741_20245 | 1845..5237 | 2026724 | ATP-binding-protein-[Chloroflexi-bacterium] |
| MCC6567532.1 | HEPN+AAA-ATPase+Alpha+Beta-domain+wHTH+RRM | Type-1_BR.RM.like.systems | <-SF1-SF2-Helicase<-?<-?<-HEPN+DUF499*<-REase<-?<-METHYLASE<-?<-?<-TUDOR+SF1-SF2-Helicase+iREase+CTD+REase | 1096 | Bacteria>Terrabacteria_group>Chloroflexota>unclassified_Chloroflexota | GCA_020848335.1 | Chloroflexota_bacterium | IT298_17380 | 16185..19475 | 2026724 | DUF499-domain-containing-protein-[Chloroflexi-bacterium] |
| NOK58906.1 | HEPN+AAA-ATPase+Alpha+Beta-domain+wHTH+RRM | Type-1_BR.RM.like.systems | STAS->SAM-methylase->?-><-HEPN+DUF499*<-?<-METHYLASE<-TUDOR+SF1-SF2-Helicase+iREase+CTD+REase | 1106 | Bacteria>Terrabacteria_group>Chloroflexota>unclassified_Chloroflexota | GCA_013122435.1 | Chloroflexi_bacterium_AL-N1 | GFH23_1086616n269 | 267583..270903 | 2686367 | ATP-binding-protein-[Chloroflexi-bacterium-AL-N1] |
| MBI5949977.1 | HEPN+AAA-ATPase+Alpha+Beta-domain+wHTH+RRM | Type-1_BR.RM.like.systems | SIGMA-HTH-><-Pkinase<-?<-Phage_integrase<-Phage_integrase<-HEPN+DUF499*<-?<-METHYLASE<-?<-?<-?<-YtH | 1133 | Bacteria>Terrabacteria_group>Chloroflexota>unclassified_Chloroflexota | GCA_016234445.1 | Chloroflexota_bacterium | HY875_17715 | 143137..146538 | 2026724 | ATP-binding-protein-[Chloroflexi-bacterium] |
| NWF79207.1 | HEPN+AAA-ATPase+Alpha+Beta-domain+wHTH+RRM | Type-1_BR.RM.like.systems | TUDOR+SF1-SF2-Helicase+iREase+CTD+REase-><-?\|\|METHYLASE-><-PIN<-DUF433\|\|HEPN+DUF499*-> | 825 | Bacteria>Terrabacteria_group>Chloroflexota>unclassified_Chloroflexota | GCA_013388805.1 | Chloroflexota_bacterium | HXY37_04055 | 14161..16634 | 2026724 | ATP-binding-protein-[Chloroflexi-bacterium] |
| MCC6175191.1 | HEPN+AAA-ATPase+Alpha+Beta-domain+wHTH+RRM | Type-1_BR.RM.like.systems | <-METHYLASE-p<-HEPN+DUF499*<-METHYLASE<-TUDOR+SF1-SF2-Helicase+iREase+CTD+REase | 1129 | Bacteria>Terrabacteria_group>Chloroflexota>unclassified_Chloroflexota | GCA_020845495.1 | Chloroflexota_bacterium | IT305_07815 | 409..3798 | 2026724 | DUF499-domain-containing-protein-[Chloroflexi-bacterium] |
| MBV9281297.1 | HEPN+AAA-ATPase+Alpha+Beta-domain+wHTH+RRM | Type-1_BR.RM.like.systems | METHYLASE-p->Glyoxalase->HEPN+DUF499*->REase-><-SIGMA-HTH+SIGMA-HTH<-?<-Glyoxalase | 1102 | Bacteria>Terrabacteria_group>Chloroflexota>unclassified_Chloroflexota | GCA_019245855.1 | Chloroflexota_bacterium | JOZ41_14550 | 2324..5632 | 2026724 | DUF499-domain-containing-protein-[Chloroflexi-bacterium] |
| MBM2811008.1 | HEPN+AAA-ATPase+Alpha+Beta-domain+wHTH+RRM | Type-1_BR.RM.like.systems | METHYLASE-p->METHYLASE-p-><-?\|\|?->HicB_lk_antitox->HicA_toxin->HEPN+DUF499*->DUF499-> | 384 | Bacteria>Terrabacteria_group>Chloroflexota>unclassified_Chloroflexota | GCA_016844585.1 | Chloroflexota_bacterium | HW416_1767 | 3649..4803 | 2026724 | ATPase-of-the-superfamily-[Chloroflexota-bacterium] |
| MCA0453442.1 | HEPN+AAA-ATPase+Alpha+Beta-domain+wHTH+RRM | Type-1_BR.RM.like.systems | HTH-><-?<-?<-?<-HEPN+DUF499*<-METHYLASE<-HEPN<-TUDOR+SF1-SF2-Helicase+iREase+CTD+REase | 1099 | Bacteria>Terrabacteria_group>Chloroflexota>unclassified_Chloroflexota | GCA_020161935.1 | Chloroflexota_bacterium | LCI00_05665 | 1284524..1287823 | 2026724 | DUF499-domain-containing-protein-[Chloroflexi-bacterium] |
| MBU1746748.1 | HEPN+AAA-ATPase+Alpha+Beta-domain+wHTH+RRM | Type-1_BR.RM.like.systems | <-HEPN+DUF499*<-METHYLASE<-?<-ParB+HNH<-TUDOR+SF1-SF2-Helicase+iREase+CTD+REase | 551 | Bacteria>Terrabacteria_group>Chloroflexota>unclassified_Chloroflexota | GCA_018823415.1 | Chloroflexota_bacterium | KKA73_03595 | 1..1652 | 2026724 | DUF499-domain-containing-protein-[Chloroflexota-bacterium] |
| MCY3938336.1 | HEPN+AAA-ATPase+Alpha+Beta-domain+wHTH+RRM | Type-1_BR.RM.like.systems | HEPN+DUF499*->METHYLASE->URI-DNase-><-?\|\|METHYLASE-p-><-ABhydrolase\|\|NUDIX-> | 1121 | Bacteria>Terrabacteria_group>Chloroflexota>unclassified_Chloroflexota | GCA_026707405.1 | Chloroflexota_bacterium | OXG22_08290 | 17127..20492 | 2026724 | MAG:-DUF499-domain-containing-protein-[Chloroflexi-bacterium] |
| MCY3801423.1 | HEPN+AAA-ATPase+Alpha+Beta-domain+wHTH+RRM | Type-1_BR.RM.like.systems | HEPN+DUF499*->METHYLASE->URI-DNase-><-?\|\|METHYLASE-p-><-ABhydrolase\|\|NUDIX-> | 1121 | Bacteria>Terrabacteria_group>Chloroflexota>unclassified_Chloroflexota | GCA_026706785.1 | Chloroflexota_bacterium | OXG46_07570 | 255383..258748 | 2026724 | MAG:-DUF499-domain-containing-protein-[Chloroflexi-bacterium] |
| MBG54827.1 | HEPN+AAA-ATPase+Alpha+Beta-domain+wHTH+RRM | Type-1_BR.RM.like.systems | TUDOR+SF1-SF2-Helicase+iREase+CTD+REase->METHYLASE->HEPN+DUF499*-><-tRNA<-?<-?<-?<-?<-?<-RNase | 1093 | Bacteria>Terrabacteria_group>Chloroflexota>unclassified_Chloroflexota | GCA_002715705.1 | Chloroflexota_bacterium | CL795_03415 | 7874..11155 | 2026724 | AAA+-family-ATPase-[Chloroflexi-bacterium] |
| MXW23202.1 | HEPN+AAA-ATPase+Alpha+Beta-domain+wHTH+RRM | Type-1_BR.RM.like.systems | TUDOR+SF1-SF2-Helicase+iREase+CTD+REase-p->PolB->HEPN->HEPN+DUF499*-> | 1125 | Bacteria>Terrabacteria_group>Chloroflexota>unclassified_Chloroflexota | GCA_009835505.1 | Chloroflexota_bacterium | F4Z96_00705 | 5471..8848 | 2026724 | ATP-binding-protein-[Chloroflexi-bacterium] |
| NLE83803.1 | HEPN+AAA-ATPase+Alpha+Beta-domain+wHTH+RRM | Type-1_BR.RM.like.systems | TUDOR+SF1-SF2-Helicase+iREase+CTD+REase-p->METHYLASE->HEPN+DUF499*-> | 944 | Bacteria>Terrabacteria_group>Chloroflexota>unclassified_Chloroflexota | GCA_012514645.1 | Chloroflexota_bacterium | GX603_04735 | 3343..6175 | 2026724 | ATP-binding-protein-[Chloroflexi-bacterium] |
| MCY3747797.1 | HEPN+AAA-ATPase+Alpha+Beta-domain+wHTH+RRM | Type-1_BR.RM.like.systems | PolB->METHYLASE->PolB->?->?->HEPN+DUF499*->?->PADR-HTH->?-><-REase<-?<-HTH+UbE1 | 1113 | Bacteria>Terrabacteria_group>Chloroflexota>unclassified_Chloroflexota | GCA_026706265.1 | Chloroflexota_bacterium | OXG64_00755 | 49727..53068 | 2026724 | MAG:-Swt1-family-HEPN-domain-containing-protein-[Chloroflexi-bacterium] |
| MYD10935.1 | HEPN+AAA-ATPase+Alpha+Beta-domain+wHTH+RRM | Type-1_BR.RM.like.systems | HEPN+DUF499*-><-DUF2961<-ABC-ATPase+SbcC<-ABC-ATPase+SbcC | 1106 | Bacteria>Terrabacteria_group>Chloroflexota>unclassified_Chloroflexota | GCA_009841405.1 | Chloroflexota_bacterium | F4X02_12950 | 92..3412 | 2026724 | ATP-binding-protein-[Chloroflexi-bacterium] |
| MCX6066946.1 | HEPN+AAA-ATPase+Alpha+Beta-domain+wHTH+RRM | Type-1_BR.RM.like.systems | <-REase<-?<-?<-?<-DUF499-p<-HEPN+DUF499*<-PIN<-MazE_antitoxin | 972 | Bacteria>Terrabacteria_group>Chloroflexota>unclassified_Chloroflexota | GCA_026389995.1 | Chloroflexota_bacterium | NT121_14520 | 4682..7598 | 2026724 | MAG:-Swt1-family-HEPN-domain-containing-protein,-partial-[Chloroflexi-bacterium] |
| MCE7937169.1 | HEPN+AAA-ATPase+Alpha+Beta-domain+wHTH+RRM | Type-1_BR.RM.like.systems | <-HEPN+DUF499*<-SF1-SF2-Helicase+HNH<-?<-?<-ORC-CDC6+REase | 1145 | Bacteria>Terrabacteria_group>Chloroflexota>unclassified_Chloroflexota | - | Chloroflexi_bacterium_CFX6 | DYG90_00940 | 4..3441 | 2293710 | ATP-binding-protein-[Chloroflexi-bacterium-CFX6] |
| MDC0253131.1 | HEPN+AAA-ATPase+Alpha+Beta-domain+wHTH+RRM | Type-1_BR.RM.like.systems | REase->METHYLASE->HEPN+DUF499*-><-Mbetalac\|\|?-><-?<-?<-RNase | 1088 | Bacteria>Terrabacteria_group>Chloroflexota>unclassified_Chloroflexota | GCA_028346115.1 | Chloroflexota_bacterium | OAK52_03100 | 3358..6624 | 2026724 | DUF499-domain-containing-protein-[Chloroflexi-bacterium] |
| MBI3979239.1 | HEPN+AAA-ATPase+Alpha+Beta-domain+wHTH+RRM | Type-1_BR.RM.like.systems | <-PemK_toxin<-?\|\|Peptidase_M20-><-?<-?<-HEPN+DUF499*<-REase | 1136 | Bacteria>Terrabacteria_group>Chloroflexota>unclassified_Chloroflexota | GCA_016199325.1 | Chloroflexota_bacterium | HY331_13735 | 11922..15332 | 2026724 | ATP-binding-protein-[Chloroflexi-bacterium] |
| MYJ01661.1 | HEPN+AAA-ATPase+Alpha+Beta-domain+wHTH+RRM | Type-1_BR.RM.like.systems | METHYLASE-p->Polbeta->HEPN->HEPN+DUF499*-><-PIN | 1112 | Bacteria>Terrabacteria_group>Chloroflexota>unclassified_Chloroflexota | GCA_009844585.1 | Chloroflexota_bacterium | F4102_05150 | 3394..6732 | 2026724 | ATP-binding-protein-[Chloroflexi-bacterium] |
| MBI1882242.1 | HEPN+AAA-ATPase+Alpha+Beta-domain+wHTH+RRM | Type-1_BR.RM.like.systems | <-HEPN+DUF499*<-REase<-METHYLASE-p<-METHYLASE-p | 401 | Bacteria>Terrabacteria_group>Chloroflexota>unclassified_Chloroflexota | GCA_016178315.1 | Chloroflexota_bacterium | HYR94_29060 | 1..1204 | 2026724 | ATP-binding-protein-[Chloroflexota-bacterium] |
| MCY4583299.1 | HEPN+AAA-ATPase+Alpha+Beta-domain+wHTH+RRM | Type-1_BR.RM.like.systems | METHYLASE->HEPN+DUF499*->?->?-><-ABC-ATPase+SbcC | 1128 | Bacteria>Terrabacteria_group>Chloroflexota>unclassified_Chloroflexota | GCA_026713505.1 | Chloroflexota_bacterium | OXE50_10970 | 2507..5893 | 2026724 | MAG:-Swt1-family-HEPN-domain-containing-protein-[Chloroflexi-bacterium] |
| MCY4474618.1 | HEPN+AAA-ATPase+Alpha+Beta-domain+wHTH+RRM | Type-1_BR.RM.like.systems | <-METHYLASE<-PIN<-?<-?<-HEPN<-HEPN<-HEPN+DUF499* | 1144 | Bacteria>Terrabacteria_group>Chloroflexota>unclassified_Chloroflexota | GCA_026712855.1 | Chloroflexota_bacterium | OXC83_04195 | 178388..181822 | 2026724 | MAG:-DUF499-domain-containing-protein-[Chloroflexi-bacterium] |
| MBC8453263.1 | HEPN+AAA-ATPase+Alpha+Beta-domain+wHTH+RRM | Type-1_BR.RM.like.systems | METHYLASE-p->METHYLASE-p->HEPN+DUF499*-> | 404 | Bacteria>Terrabacteria_group>Chloroflexota>unclassified_Chloroflexota | GCA_014383785.1 | Chloroflexota_bacterium | H8D69_02180 | 1481..2692 | 2026724 | ATP-binding-protein-[Chloroflexota-bacterium] |
| MCX6028868.1 | HEPN+AAA-ATPase+Alpha+Beta-domain+wHTH+RRM | Type-1_BR.RM.like.systems | TPR+TPR->?->HEPN-p->RNaseH->?->HEPN+DUF499*-> | 1119 | Bacteria>Terrabacteria_group>Chloroflexota>unclassified_Chloroflexota | GCA_026389845.1 | Chloroflexota_bacterium | NT169_06150 | 82672..86031 | 2026724 | MAG:-Swt1-family-HEPN-domain-containing-protein-[Chloroflexi-bacterium] |
| MBI3961042.1 | HEPN+AAA-ATPase+Alpha+Beta-domain+wHTH+RRM | Type-1_BR.RM.like.systems | METHYLASE-p->OLD-ABC->PIN->HEPN+DUF499*-> | 1102 | Bacteria>Terrabacteria_group>Chloroflexota>unclassified_Chloroflexota | GCA_016199425.1 | Chloroflexota_bacterium | HY328_19695 | 3483..6791 | 2026724 | ATP-binding-protein-[Chloroflexi-bacterium] |
| MDE2824105.1 | HEPN+AAA-ATPase+Alpha+Beta-domain+wHTH+RRM | Type-1_BR.RM.like.systems | METHYLASE->HEPN+DUF499*->SIR2-> | 1105 | Bacteria>Terrabacteria_group>Chloroflexota>unclassified_Chloroflexota | GCA_028874805.1 | Chloroflexota_bacterium | OXK79_11455 | 2958..6275 | 2026724 | MAG:-DUF499-domain-containing-protein-[Chloroflexota-bacterium] |
| NLS78478.1 | HEPN+AAA-ATPase+Alpha+Beta-domain+wHTH+RRM | Type-1_BR.RM.like.systems | <-HEPN+DUF499*<-?<-METHYLASE-p | 1075 | Bacteria>Terrabacteria_group>Chloroflexota>unclassified_Chloroflexota | GCA_012689325.1 | Chloroflexota_bacterium | GXY76_14575 | 1..3224 | 2026724 | ATP-binding-protein-[Chloroflexi-bacterium] |
| NLH06920.1 | HEPN+AAA-ATPase+Alpha+Beta-domain+wHTH+RRM | Type-1_BR.RM.like.systems | METHYLASE-p->?->HEPN+DUF499*-> | 445 | Bacteria>Terrabacteria_group>Chloroflexota>unclassified_Chloroflexota | GCA_012516245.1 | Chloroflexota_bacterium | GX484_04875 | 1862..3195 | 2026724 | ATP-binding-protein-[Chloroflexota-bacterium] |
| MXY13704.1 | HEPN+AAA-ATPase+Alpha+Beta-domain+wHTH+RRM | Type-1_BR.RM.like.systems | METHYLASE-p->HEPN+DUF499*-> | 1057 | Bacteria>Terrabacteria_group>Chloroflexota>unclassified_Chloroflexota | GCA_009838555.1 | Chloroflexota_bacterium | F4Y46_06845 | 861..4034 | 2026724 | ATP-binding-protein-[Chloroflexi-bacterium] |
| MCC6627902.1 | HEPN+AAA-ATPase+Alpha+Beta-domain+wHTH+RRM | Type-1_BR.RM.like.systems | METHYLASE->?->?->HEPN+DUF499*-> | 524 | Bacteria>Terrabacteria_group>Chloroflexota>unclassified_Chloroflexota | GCA_020849725.1 | Chloroflexota_bacterium | IT340_10930 | 2805..4377 | 2026724 | DUF499-domain-containing-protein-[Chloroflexota-bacterium] |
| MBX6773122.1 | HEPN+AAA-ATPase+Alpha+Beta-domain+wHTH+RRM | Type-1_BR.RM.like.systems | <-HEPN+DUF499*<-METHYLASE | 328 | Bacteria>Terrabacteria_group>Chloroflexota>unclassified_Chloroflexota | GCA_019690095.1 | Chloroflexota_bacterium | IRY83_15455 | 1..985 | 2026724 | AAA+-family-ATPase-[Chloroflexota-bacterium] |
| MBU6150404.1 | HEPN+AAA-ATPase+Alpha+Beta-domain+wHTH+RRM | Type-1_BR.RM.like.systems | <-HEPN+DUF499*<-METHYLASE | 1130 | Bacteria>Terrabacteria_group>Chloroflexota>unclassified_Chloroflexota | GCA_018969665.1 | Chloroflexota_bacterium | KGR25_03080 | 672..4064 | 2026724 | DUF499-domain-containing-protein-[Chloroflexi-bacterium] |
| MDE2668025.1 | HEPN+AAA-ATPase+Alpha+Beta-domain+wHTH+RRM | Type-1_BR.RM.like.systems | <-PIN<-AbrB<-HEPN+DUF499*<-METHYLASE<-TUDOR+SF1-SF2-Helicase+iREase+CTD+REase<-REase<-MORC+REase<-DNA_methylase\|\|AbiEi_4-> | 1121 | Bacteria>Terrabacteria_group>Chloroflexota>unclassified_Chloroflexota | GCA_028875515.1 | Chloroflexota_bacterium | OXI51_00025 | 2709..6074 | 2026724 | MAG:-DUF499-domain-containing-protein-[Chloroflexota-bacterium] |
| MCY4618411.1 | HEPN+AAA-ATPase+Alpha+Beta-domain+wHTH+RRM | Type-1_BR.RM.like.systems | <-SF1-SF2-Helicase-p<-SF1-SF2-Helicase<-OLD-ABC+TOPRIM<-?<-?<-?<-HEPN+DUF499*<-METHYLASE<-TUDOR+SF1-SF2-Helicase+iREase+CTD+REase<-?<-CHTH | 1114 | Bacteria>Terrabacteria_group>Chloroflexota>unclassified_Chloroflexota | GCA_026713765.1 | Chloroflexota_bacterium | OXD50_07650 | 54962..58306 | 2026724 | MAG:-DUF499-domain-containing-protein-[Chloroflexi-bacterium] |
| MCY3957107.1 | HEPN+AAA-ATPase+Alpha+Beta-domain+wHTH+RRM | Type-1_BR.RM.like.systems | TUDOR+SF1-SF2-Helicase+iREase+CTD+REase->METHYLASE->HEPN+DUF499*-> | 1113 | Bacteria>Terrabacteria_group>Chloroflexota>unclassified_Chloroflexota | GCA_026707585.1 | Chloroflexota_bacterium | OXG65_02325 | 8124..11465 | 2026724 | MAG:-DUF499-domain-containing-protein-[Chloroflexi-bacterium] |
| MBI3978251.1 | HEPN+AAA-ATPase+Alpha+Beta-domain+wHTH+RRM | Type-1_BR.RM.like.systems | <-RNase<-?\|\|?-><-?<-HEPN+DUF499*<-METHYLASE | 1129 | Bacteria>Terrabacteria_group>Chloroflexota>unclassified_Chloroflexota | GCA_016199325.1 | Chloroflexota_bacterium | HY331_08705 | 13275..16664 | 2026724 | ATP-binding-protein-[Chloroflexi-bacterium] |
| WP_115538087.1 | HEPN+AAA-ATPase+Alpha+Beta-domain+wHTH+RRM | Type-1_BR.RM.like.systems | <-Peptidase_C14<-?<-REase<-DUF499-p\|\|DDE_Tnp_IS1-><-HEPN+DUF499*<-METHYLASE<-HTH_OrfB_IS605+OrfB_IS605+OrfB_Zn_ribbon\|\|<-TUDOR+SF1-SF2-Helicase+iREase+CTD+REase\|\|HTH+MPTase-><-?<-TUDOR+SF1-SF2-Helicase+iREase+CTD+REase-p | 1013 | Bacteria>Terrabacteria_group>Cyanobacteriota/Melainabacteria_group>Cyanobacteriota>Cyanophyceae | GCF_003367075.2 | Cylindrospermopsis_raciborskii | C6N34_RS14575 | 3182311..3185352 | 77022 | Swt1-family-HEPN-domain-containing-protein-[Cylindrospermopsis-raciborskii] |
| BAZ89419.1 | HEPN+AAA-ATPase+Alpha+Beta-domain+wHTH+RRM | Type-1_BR.RM.like.systems | <-PolB\|\|TUDOR+SF1-SF2-Helicase+iREase+CTD+REase-p->?-><-HTH+MPTase<-Tox-RelE\|\|TUDOR+SF1-SF2-Helicase+iREase+CTD+REase->HEPN+DUF499*-><-CR-ATPase9+REase\|\|ABhydrolase->?->?->Cysteine_desulfurase-> | 1015 | Bacteria>Terrabacteria_group>Cyanobacteriota/Melainabacteria_group>Cyanobacteriota>Cyanophyceae | GCA_002368135.1 | Raphidiopsis_curvata_NIES-932 | NIES932_08970 | 961455..964502 | 1973482 | hypothetical-protein-NIES932_08970-[Raphidiopsis-curvata-NIES-932] |
| WP_190676125.1 | HEPN+AAA-ATPase+Alpha+Beta-domain+wHTH+RRM | Type-1_BR.RM.like.systems | <-pPIWI+pPIWI\|\|ParB-><-DDE_3<-PAIREDC-HTH\|\|ParB-><-TUDOR+SF1-SF2-Helicase+iREase+CTD+REase<-HEPN+DUF499*<-PIN<-OLD-ABC<-METHYLASE<-REase\|\|?-><-TUDOR+SF1-SF2-Helicase+iREase+CTD+REase | 1112 | Bacteria>Terrabacteria_group>Cyanobacteriota/Melainabacteria_group>Cyanobacteriota>Cyanophyceae | GCF_014695435.1 | Coleofasciculus_sp._FACHB-129 | H6F60_RS14220 | 70669..74007 | 2692785 | Swt1-family-HEPN-domain-containing-protein-[Coleofasciculus-sp.-FACHB-129] |
| WP_087068561.1 | HEPN+AAA-ATPase+Alpha+Beta-domain+wHTH+RRM | Type-1_BR.RM.like.systems | TUDOR+SF1-SF2-Helicase+iREase+CTD+REase->?->PIN->METHYLASE-p->DUF772+DDE_Tnp_1_2->METHYLASE->HEPN+DUF499*->SF1-SF2-Helicase+REase->SF1-SF2-Helicase-p-><-?<-DUF772+DDE_Tnp_1_2 | 1101 | Bacteria>Terrabacteria_group>Cyanobacteriota/Melainabacteria_group>Cyanobacteriota>Cyanophyceae | GCF_900088535.1 | Cyanobium_sp._NIES-981 | CBM981_RS11760 | 2335245..2338550 | 1851505 | DUF499-domain-containing-protein-[Cyanobium-sp.-NIES-981] |
| WP_045875055.1 | HEPN+AAA-ATPase+Alpha+Beta-domain+wHTH+RRM | Type-1_BR.RM.like.systems | <-HTH_37<-GDResolvase+Recombinase<-TraJ-RHH\|\|?->Phage_int_SAM_1+Phage_integrase-><-?<-HEPN+DUF499*<-REase<-?<-REase<-METHYLASE<-HicB+ZnR<-TUDOR+SF1-SF2-Helicase+iREase+CTD+REase | 1112 | Bacteria>Terrabacteria_group>Cyanobacteriota/Melainabacteria_group>Cyanobacteriota>Cyanophyceae | GCF_025860405.1 | unclassified_Tolypothrix | FDUTEX481_RS39715 | 28585..31923 | 2649714 | MULTISPECIES:-Swt1-family-HEPN-domain-containing-protein-[unclassified-Tolypothrix] |
| WP_163928210.1 | HEPN+AAA-ATPase+Alpha+Beta-domain+wHTH+RRM | Type-1_BR.RM.like.systems | <-TPR+TPR\|\|TUDOR+SF1-SF2-Helicase+iREase+CTD+REase->Phd-YefM->PIN-><-HTH_3\|\|METHYLASE->HEPN+DUF499*-><-TraJ-RHH\|\|PIWI-><-?<-?\|\|?->HTH_OrfB_IS605+OrfB_IS605+OrfB_Zn_ribbon-> | 1108 | Bacteria>Terrabacteria_group>Cyanobacteriota/Melainabacteria_group>Cyanobacteriota>Cyanophyceae | GCF_010747425.1 | Nostoc_sp._UIC_10630 | G1D98_RS07815 | 12096..15422 | 2100146 | Swt1-family-HEPN-domain-containing-protein-[Nostoc-sp.-UIC-10630] |
| MBW4455562.1 | HEPN+AAA-ATPase+Alpha+Beta-domain+wHTH+RRM | Type-1_BR.RM.like.systems | HEPN-><-?\|\|TUDOR+SF1-SF2-Helicase+iREase+CTD+REase->HicB+ZnR->METHYLASE-p->REase->HEPN+DUF499*-><-Phage_int_SAM_1+Phage_integrase\|\|?->?-><-Response_reg+LuxR-HTH<-HerA/FtsK | 1112 | Bacteria>Terrabacteria_group>Cyanobacteriota/Melainabacteria_group>Cyanobacteriota>Cyanophyceae | GCA_019359215.1 | Nostoc_indistinguendum_CM1-VF10 | KME55_24210 | 27093..30431 | 287863 | DUF499-domain-containing-protein-[Nostoc-indistinguendum-CM1-VF10] |
| WP_011318979.1 | HEPN+AAA-ATPase+Alpha+Beta-domain+wHTH+RRM | Type-1_BR.RM.like.systems | HTH_28+rve+Mu-transpos_C->?->?->TUDOR+SF1-SF2-Helicase+iREase+CTD+REase->METHYLASE->METHYLASE-p->HEPN+DUF499*-><-TraJ-RHH\|\|PIWI-><-HTH_3\|\|?-><-tRNA\|\|?->LexA-HTH+LexA-protease-> | 1109 | Bacteria>Terrabacteria_group>Cyanobacteriota/Melainabacteria_group>Cyanobacteriota>Cyanophyceae | GCF_000204075.1 | Nostocaceae | GSQ19_RS16260 | 3995693..3999022 | 1162 | MULTISPECIES:-Swt1-family-HEPN-domain-containing-protein-[Nostocaceae] |
| ARV61678.1 | HEPN+AAA-ATPase+Alpha+Beta-domain+wHTH+RRM | Type-1_BR.RM.like.systems | <-Trypsin\|\|?-><-?\|\|DDE_Tnp_1_3->TUDOR+SF1-SF2-Helicase+iREase+CTD+REase->METHYLASE->HEPN+DUF499*-><-tRNA\|\|?->LexA-HTH+LexA-protease->SF1-SF2-Helicase->SbcC->REase-> | 1115 | Bacteria>Terrabacteria_group>Cyanobacteriota/Melainabacteria_group>Cyanobacteriota>Cyanophyceae | GCA_002163975.1 | Nostocales_cyanobacterium_HT-58-2 | BZZ01_26385 | 6380631..6383978 | 1940762 | AAA+-family-ATPase-[Nostocales-cyanobacterium-HT-58-2] |
| WP_190513827.1 | HEPN+AAA-ATPase+Alpha+Beta-domain+wHTH+RRM | Type-1_BR.RM.like.systems | CHTH->TUDOR+SF1-SF2-Helicase+iREase+CTD+REase->?->METHYLASE->RHH_6->PIN->HEPN+DUF499*->URI-DNase-><-RelE-ParE<-?\|\|pPIWI+pPIWI->pPIWI->pPIWI+MID+RNaseH-> | 1117 | Bacteria>Terrabacteria_group>Cyanobacteriota/Melainabacteria_group>Cyanobacteriota>Cyanophyceae | GCF_014697545.1 | Oscillatoria_sp._FACHB-1407 | H6G89_RS30695 | 49800..53153 | 2692847 | Swt1-family-HEPN-domain-containing-protein-[Oscillatoria-sp.-FACHB-1407] |
| WP_244349767.1 | HEPN+AAA-ATPase+Alpha+Beta-domain+wHTH+RRM | Type-1_BR.RM.like.systems | TUDOR+SF1-SF2-Helicase+iREase+CTD+REase->METHYLASE->?->PemK_toxin->REase->REase->HEPN+DUF499*->?->ABC-ATPase+SbcC->?->?-><-ABC-ATPase+SbcC<-ABC-ATPase+SbcC | 1107 | Bacteria>Terrabacteria_group>Cyanobacteriota/Melainabacteria_group>Cyanobacteriota>Cyanophyceae | GCF_022848905.1 | Thermostichus_vulcanus | JX360_RS06145 | 28078..31401 | 32053 | Swt1-family-HEPN-domain-containing-protein-[Thermostichus-vulcanus] |
| WP_196514030.1 | HEPN+AAA-ATPase+Alpha+Beta-domain+wHTH+RRM | Type-1_BR.RM.like.systems | HTH+HKD-DNase->SPOUT->?->tRNA-><-?\|\|<-PIWI\|\|TraJ-RHH-><-HEPN+DUF499*<-METHYLASE\|\|HTH_3-><-TUDOR+SF1-SF2-Helicase+iREase+CTD+REase<-?<-?<-HTH_28+rve+Mu-transpos_C | 1107 | Bacteria>Terrabacteria_group>Cyanobacteriota/Melainabacteria_group>Cyanobacteriota>Cyanophyceae | GCF_015714705.1 | Nostoc_sp._NZL | F8R90_RS31675 | 50056..53379 | 2650612 | Swt1-family-HEPN-domain-containing-protein-[Nostoc-sp.-NZL] |

| Protein-ID | DUF499-architecture | System | Gene-Neighbourhoods | Length | Taxonomy | Assembly-ID | Strain | Locus-Tag | Genomic-coordinates | TaxID | Fasta-defline |
|---|---|---|---|---|---|---|---|---|---|---|---|
| MCD8490337.1 | HEPN+AAA-ATPase+Alpha+Beta-domain+wHTH+RRM | Type-1_BR.RM.like.systems | pPIWI+pPIWI->pPIWI->pPIWI+MID+RNaseH->?->?->?-><-HEPN+DUF499*<-METHYLASE<-PIN<-AbrB<-TUDOR+SF1-SF2-Helicase+iREase+CTD+REase<-TOPRIM+SF1-SF2-Helicase | 1118 | Bacteria>Terrabacteria_group>Cyanobacteriota/Melainabacteria_group>Cyanobacteriota>Cyanophyceae | GCA_021208125.1 | Desertifilum_sp. | LRZ84_27325 | 10272..13628 | 2497424 | DUF499-domain-containing-protein-[Desertifilum-sp.] |
| WP_231828504.1 | HEPN+AAA-ATPase+Alpha+Beta-domain+wHTH+RRM | Type-1_BR.RM.like.systems | TUDOR+SF1-SF2-Helicase+iREase+CTD+REase->REase->METHYLASE->PAIREDC-HTH->PIN->HEPN+DUF499*-><-Cluster164_18clades<-?\|\|?->?->DUF29-><-Y1_Tnp | 1129 | Bacteria>Terrabacteria_group>Cyanobacteriota/Melainabacteria_group>Cyanobacteriota>Cyanophyceae | GCF_002095975.1 | Microcystis_aeruginosa | BH695_RS02870 | 541185..544574 | 1126 | Swt1-family-HEPN-domain-containing-protein-[Microcystis-aeruginosa] |
| WP_190359815.1 | HEPN+AAA-ATPase+Alpha+Beta-domain+wHTH+RRM | Type-1_BR.RM.like.systems | REase-><-DUF29<-?<-?\|\|?->Cluster164_18clades-><-HEPN+DUF499*<-PIN<-RHH_6<-METHYLASE<-REase<-TUDOR+SF1-SF2-Helicase+iREase+CTD+REase<-ABhydrolase | 1117 | Bacteria>Terrabacteria_group>Cyanobacteriota/Melainabacteria_group>Cyanobacteriota>Cyanophyceae | GCF_026222535.1 | Microcystis | H6F92_RS24355 | 16318..19671 | 1125 | Swt1-family-HEPN-domain-containing-protein-[Microcystis-wesenbergii] |
| WP_255090875.1 | HEPN+AAA-ATPase+Alpha+Beta-domain+wHTH+RRM | Type-1_BR.RM.like.systems | <-ACET<-CopG/RHH<-WYL\|\|LRP-HTH->TUDOR+SF1-SF2-Helicase+iREase+CTD+REase->METHYLASE->HEPN+DUF499*->Peptidase_M48-><-HNH\|\|ArdCN+MPTase-> | 1122 | Bacteria>Terrabacteria_group>Cyanobacteriota/Melainabacteria_group>Cyanobacteriota>Cyanophyceae | GCF_024346175.1 | Vulcanococcus_limneticus | KBZ20_RS12915 | 25528..28896 | 2170428 | DUF499-domain-containing-protein-[Vulcanococcus-limneticus] |
| WP_211941813.1 | HEPN+AAA-ATPase+Alpha+Beta-domain+wHTH+RRM | Type-1_BR.RM.like.systems | HKD-DNase+SF1-SF2-Helicase-p-><-HTH+MPTase<-?<-?\|\|TUDOR+SF1-SF2-Helicase+iREase+CTD+REase->METHYLASE->HEPN+DUF499*->REase->?->Peptidase_C14-> | 1103 | Bacteria>Terrabacteria_group>Cyanobacteriota/Melainabacteria_group>Cyanobacteriota>Cyanophyceae | GCF_018139025.1 | Cylindrospermopsis_raciborskii | J9B92_RS12735 | 2844714..2848025 | 77022 | Swt1-family-HEPN-domain-containing-protein-[Cylindrospermopsis-raciborskii] |
| WP_057177165.1 | HEPN+AAA-ATPase+Alpha+Beta-domain+wHTH+RRM | Type-1_BR.RM.like.systems | HKD-DNase+SF1-SF2-Helicase-p-><-HTH+MPTase<-?<-?\|\|TUDOR+SF1-SF2-Helicase+iREase+CTD+REase->METHYLASE->HEPN+DUF499*->REase->?->Peptidase_C14-> | 1103 | Bacteria>Terrabacteria_group>Cyanobacteriota/Melainabacteria_group>Cyanobacteriota>Cyanophyceae | GCF_001432185.1 | Cylindrospermopsis_sp._CR12 | ASL19_RS02560 | 94062..97373 | 1747196 | Swt1-family-HEPN-domain-containing-protein-[Cylindrospermopsis-sp.-CR12] |
| WP_094555433.1 | HEPN+AAA-ATPase+Alpha+Beta-domain+wHTH+RRM | Type-1_BR.RM.like.systems | GDResolvase-><-D5N+D5ATPase\|\|?->ArdCN+MPTase-><-?<-HEPN+DUF499*<-METHYLASE<-?<-PIN<-RHH<-TUDOR+SF1-SF2-Helicase+iREase+CTD+REase\|\|DUF4236-> | 1103 | Bacteria>Terrabacteria_group>Cyanobacteriota/Melainabacteria_group>Cyanobacteriota>Cyanophyceae | GCF_002252625.1 | Synechococcus_sp._1G10 | CJZ82_RS08935 | 55076..58387 | 2025605 | DUF499-domain-containing-protein-[Synechococcus-sp.-1G10] |
| WP_230840692.1 | HEPN+AAA-ATPase+Alpha+Beta-domain+wHTH+RRM | Type-1_BR.RM.like.systems | TUDOR+SF1-SF2-Helicase+iREase+CTD+REase->REase->METHYLASE->DDE_Tnp_1->METHYLASE-p->?->HEPN+DUF499*-><-?<-Peptidase_U32+Peptidase_U32_C | 1101 | Bacteria>Terrabacteria_group>Cyanobacteriota/Melainabacteria_group>Cyanobacteriota>Cyanophyceae | GCF_021018745.1 | Gloeobacter_morelensis | ISF26_RS17965 | 3744120..3747425 | 2907343 | Swt1-family-HEPN-domain-containing-protein-[Gloeobacter-morelensis] |
| ARI79893.1 | HEPN+AAA-ATPase+Alpha+Beta-domain+wHTH+RRM | Type-1_BR.RM.like.systems | TUDOR+SF1-SF2-Helicase+iREase+CTD+REase->REase->REase->METHYLASE->PAIREDC-HTH->PIN->HEPN+DUF499*-><-Cluster164_18clades<-?\|\|?->?->DUF29-> | 1155 | Bacteria>Terrabacteria_group>Cyanobacteriota/Melainabacteria_group>Cyanobacteriota>Cyanophyceae | GCA_002095975.1 | Microcystis_aeruginosa_PCC_7806SL | BH695_0612 | 541107..544574 | 1903187 | hypothetical-protein-BH695_0612-[Microcystis-aeruginosa-PCC-7806SL] |
| WP_159816100.1 | HEPN+AAA-ATPase+Alpha+Beta-domain+wHTH+RRM | Type-1_BR.RM.like.systems | <-DUF772+DDE_Tnp_1<-?<-HIN-HTH<-MORC+DUF3684<-HEPN+DUF499*<-METHYLASE-p<-DUF772+DDE_Tnp_1+DDE_Tnp_1_6<-METHYLASE<-METHYLASE-p<-PIN | 1104 | Bacteria>Terrabacteria_group>Cyanobacteriota/Melainabacteria_group>Cyanobacteriota>Cyanophyceae | GCF_009834675.1 | Cyanobium_sp._Copco_Reservoir_LC18 | L107_RS01345 | 7355..10669 | 1328305 | DUF499-domain-containing-protein-[Cyanobium-sp.-Copco_Reservoir_LC18] |
| TRU24939.1 | HEPN+AAA-ATPase+Alpha+Beta-domain+wHTH+RRM | Type-1_BR.RM.like.systems | ABhydrolase->TUDOR+SF1-SF2-Helicase+iREase+CTD+REase->REase->METHYLASE->?->PIN->HEPN+DUF499*-><-Cluster164_18clades<-?\|\|?->?->DUF29->PolB-> | 1130 | Bacteria>Terrabacteria_group>Cyanobacteriota/Melainabacteria_group>Cyanobacteriota>Cyanophyceae | GCA_007095795.1 | Microcystis_aeruginosa_Ma_MB_S_20031200_S102D | EWV79_09040 | 24486..27878 | 2486255 | ATP-binding-protein-[Microcystis-aeruginosa-Ma_MB_S_20031200_S102D] |
| QJB25544.1 | HEPN+AAA-ATPase+Alpha+Beta-domain+wHTH+RRM | Type-1_BR.RM.like.systems | SF1-SF2-Helicase-><-THUMP+Methylase<-ABhydrolase\|\|TUDOR+SF1-SF2-Helicase+iREase+CTD+REase->METHYLASE-><-?\|\|HEPN+DUF499*-><-?\|\|?->?->?->?->SPOUT-> | 1123 | Bacteria>Terrabacteria_group>Cyanobacteriota/Melainabacteria_group>Cyanobacteriota>Cyanophyceae | GCA_012516315.1 | Limnospira_fusiformis_SAG_85.79 | HFV01_06765 | 1374121..1377492 | 1521533 | ATP-binding-protein-[Limnospira-fusiformis-SAG-85.79] |
| WP_216916461.1 | HEPN+AAA-ATPase+Alpha+Beta-domain+wHTH+RRM | Type-1_BR.RM.like.systems | <-ArdCN+MPTase\|\|HNH-><-Peptidase_M48<-?<-?<-?<-HEPN+DUF499*<-METHYLASE<-TUDOR+SF1-SF2-Helicase+iREase+CTD+REase\|\|DUF4236->?->?-><-ABhydrolase | 1099 | Bacteria>Terrabacteria_group>Cyanobacteriota/Melainabacteria_group>Cyanobacteriota>Cyanophyceae | GCF_019038475.1 | unclassified_Synechococcus | KUG39_RS15775 | 99576..102875 | 2626047 | MULTISPECIES:-DUF499-domain-containing-protein-[unclassified-Synechococcus] |
| WP_059001129.1 | HEPN+AAA-ATPase+Alpha+Beta-domain+wHTH+RRM | Type-1_BR.RM.like.systems | PIN->?-><-HEPN+DUF499*<-METHYLASE\|\|REase+HTH-><-TUDOR+SF1-SF2-Helicase+iREase+CTD+REase<-TrwC+SF1-SF2-helicase+TOPRIM\|\|?->ParB+HNH-> | 1124 | Bacteria>Terrabacteria_group>Cyanobacteriota/Melainabacteria_group>Cyanobacteriota>Cyanophyceae | GCF_001485215.1 | Leptolyngbya_sp._NIES-2104 | NIES2104_RS25560 | 5320050..5323424 | 1552121 | Swt1-family-HEPN-domain-containing-protein-[Leptolyngbya-sp.-NIES-2104] |
| NJO72425.1 | HEPN+AAA-ATPase+Alpha+Beta-domain+wHTH+RRM | Type-1_BR.RM.like.systems | CHTH->TUDOR+SF1-SF2-Helicase+iREase+CTD+REase->REase->METHYLASE->?->PIN->HEPN+DUF499*->?-><-?<-?<-REase\|\|Phage_int_SAM_4+Phage_integrase-> | 1125 | Bacteria>Terrabacteria_group>Cyanobacteriota/Melainabacteria_group>Cyanobacteriota>Cyanophyceae | GCA_012033355.1 | Leptolyngbyaceae_cyanobacterium_RM1_406_9 | HC833_00760 | 33344..36721 | 2720434 | ATP-binding-protein-[Leptolyngbyaceae-cyanobacterium-RM1_406_9] |
| WP_187757730.1 | HEPN+AAA-ATPase+Alpha+Beta-domain+wHTH+RRM | Type-1_BR.RM.like.systems | SF1-SF2-Helicase-><-THUMP+Methylase<-ABhydrolase\|\|TUDOR+SF1-SF2-Helicase+iREase+CTD+REase->METHYLASE->HEPN+DUF499*-><-?\|\|?->?->?->?->SPOUT-> | 1123 | Bacteria>Terrabacteria_group>Cyanobacteriota/Melainabacteria_group>Cyanobacteriota>Cyanophyceae | GCF_014489865.2 | Limnospira | H4N54_RS05785 | 1241805..1245176 | 2596745 | Swt1-family-HEPN-domain-containing-protein-[Limnospira-fusiformis] |
| WP_008052582.1 | HEPN+AAA-ATPase+Alpha+Beta-domain+wHTH+RRM | Type-1_BR.RM.like.systems | <-SPOUT<-?<-?<-?<-?\|\|?-><-HEPN+DUF499*<-METHYLASE<-TUDOR+SF1-SF2-Helicase+iREase+CTD+REase\|\|ABhydrolase->THUMP+Methylase-><-SF1-SF2-Helicase | 1120 | Bacteria>Terrabacteria_group>Cyanobacteriota/Melainabacteria_group>Cyanobacteriota>Cyanophyceae | GCF_000973065.1 | Limnospira_indica | XZ46_RS18610 | 4259944..4263306 | 147322 | Swt1-family-HEPN-domain-containing-protein-[Limnospira-indica] |
| REJ55686.1 | HEPN+AAA-ATPase+Alpha+Beta-domain+wHTH+RRM | Type-1_BR.RM.like.systems | TUDOR+SF1-SF2-Helicase+iREase+CTD+REase->\|\|AbrB->PIN->METHYLASE->RHH_6->PIN->HEPN+DUF499*->?-><-Cluster164_18clades<-?\|\|?->?->DUF29-> | 1111 | Bacteria>Terrabacteria_group>Cyanobacteriota/Melainabacteria_group>Cyanobacteriota>Cyanophyceae | GCA_003388675.1 | Microcystis_aeruginosa_DA14 | DWQ56_15555 | 244128..247463 | 1987506 | DUF499-domain-containing-protein-[Microcystis-aeruginosa-DA14] |
| WP_152590042.1 | HEPN+AAA-ATPase+Alpha+Beta-domain+wHTH+RRM | Type-1_BR.RM.like.systems | TUDOR+SF1-SF2-Helicase+iREase+CTD+REase-><-HTH_3<-RelE-ParE\|\|METHYLASE->HEPN+DUF499*-><-TraJ-RHH\|\|PIWI-><-HTH_3\|\|?-><-tRNA<-?<-SPOUT | 1108 | Bacteria>Terrabacteria_group>Cyanobacteriota/Melainabacteria_group>Cyanobacteriota>Cyanophyceae | GCF_009372195.1 | Nostoc_sphaeroides | GXM_RS22730 | 5029149..5032475 | 446679 | Swt1-family-HEPN-domain-containing-protein-[Nostoc-sphaeroides] |
| WP_015155777.1 | HEPN+AAA-ATPase+Alpha+Beta-domain+wHTH+RRM | Type-1_BR.RM.like.systems | Phage_integrase-><-Phage_integrase\|\|?-><-HEPN+DUF499*<-?<-METHYLASE<-REase<-PIN<-AbrB<-TUDOR+SF1-SF2-Helicase+iREase+CTD+REase | 1113 | Bacteria>Terrabacteria_group>Cyanobacteriota/Melainabacteria_group>Cyanobacteriota>Cyanophyceae | GCF_000317125.1 | Chroococcidiopsis_thermalis | CHRO_RS18600 | 4302197..4305538 | 54299 | Swt1-family-HEPN-domain-containing-protein-[Chroococcidiopsis-thermalis] |
| WP_190449861.1 | HEPN+AAA-ATPase+Alpha+Beta-domain+wHTH+RRM | Type-1_BR.RM.like.systems | CHTH->TUDOR+SF1-SF2-Helicase+iREase+CTD+REase->HEPN->METHYLASE->OLD-ABC->PIN->HEPN+DUF499*->ParB+HNH->?->CopG/RHH->HNH-><-RHH | 1121 | Bacteria>Terrabacteria_group>Cyanobacteriota/Melainabacteria_group>Cyanobacteriota>Cyanophyceae | GCF_014695795.1 | Phormidium_sp._FACHB-592 | H6F86_RS16680 | 58473..61838 | 2692850 | Swt1-family-HEPN-domain-containing-protein-[Phormidium-sp.-FACHB-592] |
| WP_006624320.1 | HEPN+AAA-ATPase+Alpha+Beta-domain+wHTH+RRM | Type-1_BR.RM.like.systems | <-ABhydrolase\|\|TUDOR+SF1-SF2-Helicase+iREase+CTD+REase->METHYLASE-p->METHYLASE->METHYLASE-p-><-?\|\|HEPN+DUF499*-><-?\|\|?->?->?->SPOUT-> | 1123 | Bacteria>Terrabacteria_group>Cyanobacteriota/Melainabacteria_group>Cyanobacteriota>Cyanophyceae | - | Arthrospira_platensis | APLC1_RS09865 | 2007550..2010921 | 118562 | Swt1-family-HEPN-domain-containing-protein-[Arthrospira-platensis] |
| WP_229465572.1 | HEPN+AAA-ATPase+Alpha+Beta-domain+wHTH+RRM | Type-1_BR.RM.like.systems | <-SIGMA-HTH\|\|?-><-?<-PIWI\|\|TraJ-RHH-><-HEPN+DUF499*<-METHYLASE-p<-METHYLASE-p<-DOC<-?<-TUDOR+SF1-SF2-Helicase+iREase+CTD+REase | 1108 | Bacteria>Terrabacteria_group>Cyanobacteriota/Melainabacteria_group>Cyanobacteriota>Cyanophyceae | - | Nostoc_sp._CHAB_5824 | LC607_29645 | 82754..86080 | 2780402 | Swt1-family-HEPN-domain-containing-protein-[Nostoc-sp.-CHAB-5824] |
| MCC5647010.1 | HEPN+AAA-ATPase+Alpha+Beta-domain+wHTH+RRM | Type-1_BR.RM.like.systems | <-SIGMA-HTH\|\|?-><-?<-PIWI\|\|TraJ-RHH-><-HEPN+DUF499*<-METHYLASE-p<-METHYLASE-p<-DOC<-?<-TUDOR+SF1-SF2-Helicase+iREase+CTD+REase | 1108 | Bacteria>Terrabacteria_group>Cyanobacteriota/Melainabacteria_group>Cyanobacteriota>Cyanophyceae | GCA_020829665.1 | Nostoc_sp._CHAB_5824 | LC607_29645 | 82754..86080 | 2780402 | DUF499-domain-containing-protein-[Nostoc-sp.-CHAB-5824] |
| WP_006668432.1 | HEPN+AAA-ATPase+Alpha+Beta-domain+wHTH+RRM | Type-1_BR.RM.like.systems | SF1-SF2-Helicase-><-THUMP+Methylase\|\|TUDOR+SF1-SF2-Helicase+iREase+CTD+REase-><-?\|\|METHYLASE-><-?\|\|HEPN+DUF499*-><-?\|\|?->?->?->?->SPOUT-> | 1123 | Bacteria>Terrabacteria_group>Cyanobacteriota/Melainabacteria_group>Cyanobacteriota>Cyanophyceae | GCF_000173555.1 | Limnospira_maxima | AMAXDRAFT_RS04610 | 145363..148734 | 129910 | Swt1-family-HEPN-domain-containing-protein-[Limnospira-maxima] |
| WP_223278246.1 | HEPN+AAA-ATPase+Alpha+Beta-domain+wHTH+RRM | Type-1_BR.RM.like.systems | TUDOR+SF1-SF2-Helicase+iREase+CTD+REase->Phd-YefM->PIN-><-HTH_3\|\|METHYLASE->HEPN+DUF499*-><-TraJ-RHH\|\|PIWI-><-HTH_3 | 1023 | Bacteria>Terrabacteria_group>Cyanobacteriota/Melainabacteria_group>Cyanobacteriota>Cyanophyceae | GCF_002245985.1 | Nostoc_sp._'Peltigera_membranacea_cyanobiont'_232 | CDG79_RS21465 | 10493..13564 | 2014531 | DUF499-domain-containing-protein-[Nostoc-sp.-'Peltigera-membranacea-cyanobiont'-232] |
| WP_066030118.1 | HEPN+AAA-ATPase+Alpha+Beta-domain+wHTH+RRM | Type-1_BR.RM.like.systems | PP_kinase+HKD-DNase-><-?<-?\|\|TUDOR+SF1-SF2-Helicase+iREase+CTD+REase->REase->METHYLASE->HEPN+DUF499*-><-?<-?\|\|?->?-><-THUMP+Methylase | 1130 | Bacteria>Terrabacteria_group>Cyanobacteriota/Melainabacteria_group>Cyanobacteriota>Cyanophyceae | GCF_001704955.2 | Microcystis_aeruginosa | amyaer_RS18255 | 3708666..3712058 | 1126 | Swt1-family-HEPN-domain-containing-protein-[Microcystis-aeruginosa] |
| WP_046663117.1 | HEPN+AAA-ATPase+Alpha+Beta-domain+wHTH+RRM | Type-1_BR.RM.like.systems | PP_kinase+HKD-DNase-><-?<-?\|\|TUDOR+SF1-SF2-Helicase+iREase+CTD+REase->REase->METHYLASE->HEPN+DUF499*-><-?<-?\|\|?->?-><-THUMP+Methylase | 1130 | Bacteria>Terrabacteria_group>Cyanobacteriota/Melainabacteria_group>Cyanobacteriota>Cyanophyceae | GCF_000981785.2 | Microcystis_aeruginosa | myaer_RS18245 | 3707749..3711141 | 1126 | Swt1-family-HEPN-domain-containing-protein-[Microcystis-aeruginosa] |
| WP_149977735.1 | HEPN+AAA-ATPase+Alpha+Beta-domain+wHTH+RRM | Type-1_BR.RM.like.systems | Cluster164_18clades-><-?<-?<-HEPN+DUF499*<-ACET<-?<-METHYLASE<-REase<-TUDOR+SF1-SF2-Helicase+iREase+CTD+REase<-ABhydrolase | 1130 | Bacteria>Terrabacteria_group>Cyanobacteriota/Melainabacteria_group>Cyanobacteriota>Cyanophyceae | GCF_008579685.1 | Microcystis_aeruginosa | MiTa_RS10465 | 30953..34345 | 1126 | Swt1-family-HEPN-domain-containing-protein-[Microcystis-aeruginosa] |
| WP_110545657.1 | HEPN+AAA-ATPase+Alpha+Beta-domain+wHTH+RRM | Type-1_BR.RM.like.systems | TUDOR+SF1-SF2-Helicase+iREase+CTD+REase->REase->METHYLASE->METHYLASE-p->?->ACET->HEPN+DUF499*-><-Cluster164_18clades | 1111 | Bacteria>Terrabacteria_group>Cyanobacteriota/Melainabacteria_group>Cyanobacteriota>Cyanophyceae | GCF_003206625.1 | Microcystis_aeruginosa | MTo_RS18515 | 13505..16840 | 1126 | Swt1-family-HEPN-domain-containing-protein-[Microcystis-aeruginosa] |
| NCR40139.1 | HEPN+AAA-ATPase+Alpha+Beta-domain+wHTH+RRM | Type-1_BR.RM.like.systems | ABhydrolase->TUDOR+SF1-SF2-Helicase+iREase+CTD+REase->REase->METHYLASE->?->PIN->HEPN+DUF499*-><-Cluster164_18clades | 1130 | Bacteria>Terrabacteria_group>Cyanobacteriota/Melainabacteria_group>Cyanobacteriota>Cyanophyceae | GCA_009995705.1 | Microcystis_aeruginosa_W13-11 | GPJ21_09840 | 22820..26212 | 2685991 | ATP-binding-protein-[Microcystis-aeruginosa-W13-11] |
| WP_104397640.1 | HEPN+AAA-ATPase+Alpha+Beta-domain+wHTH+RRM | Type-1_BR.RM.like.systems | REase->AbrB->PIN->METHYLASE->RHH_6->PIN->HEPN+DUF499*-><-Cluster164_18clades<-?\|\|?->?->?-><-TPR+TPR+TPR+TPR+TPR+TPR+TPR+TPR | 1111 | Bacteria>Terrabacteria_group>Cyanobacteriota/Melainabacteria_group>Cyanobacteriota>Cyanophyceae | - | Microcystis_aeruginosa | myaer87_32740 | 119164..122499 | 1126 | Swt1-family-HEPN-domain-containing-protein-[Microcystis-aeruginosa] |
| GBE76047.1 | HEPN+AAA-ATPase+Alpha+Beta-domain+wHTH+RRM | Type-1_BR.RM.like.systems | REase->AbrB->PIN->METHYLASE->RHH_6->PIN->HEPN+DUF499*-><-Cluster164_18clades<-?\|\|?->?->?-><-TPR+TPR+TPR+TPR+TPR+TPR+TPR+TPR | 1111 | Bacteria>Terrabacteria_group>Cyanobacteriota/Melainabacteria_group>Cyanobacteriota>Cyanophyceae | GCA_002933835.1 | Microcystis_aeruginosa_NIES-87 | myaer87_32740 | 119164..122499 | 449440 | hypothetical-protein-myaer87_32740-[Microcystis-aeruginosa-NIES-87] |
| WP_200988810.1 | HEPN+AAA-ATPase+Alpha+Beta-domain+wHTH+RRM | Type-1_BR.RM.like.systems | PRTase-><-Y1_Tnp\|\|?->?->?-><-HEPN+DUF499*<-METHYLASE\|\|REase+HTH-><-TUDOR+SF1-SF2-Helicase+iREase+CTD+REase | 1108 | Bacteria>Terrabacteria_group>Cyanobacteriota/Melainabacteria_group>Cyanobacteriota>Cyanophyceae | GCF_015245355.1 | Anabaenopsis_elenkinii | IM676_RS02385 | 515316..518642 | 156213 | Swt1-family-HEPN-domain-containing-protein-[Anabaenopsis-elenkinii] |
| WP_273247370.1 | HEPN+AAA-ATPase+Alpha+Beta-domain+wHTH+RRM | Type-1_BR.RM.like.systems | <-SPOUT<-?<-?<-?<-?\|\|?-><-HEPN+DUF499*<-?<-METHYLASE<-TUDOR+SF1-SF2-Helicase+iREase+CTD+REase<-?\|\|ABhydrolase->THUMP+Methylase-> | 1056 | Bacteria>Terrabacteria_group>Cyanobacteriota/Melainabacteria_group>Cyanobacteriota>Cyanophyceae | GCF_014250495.1 | Limnospira_indica | H2674_RS19805 | 4259944..4263114 | 147322 | DUF499-domain-containing-protein,-partial-[Limnospira-indica] |
| WP_251956292.1 | HEPN+AAA-ATPase+Alpha+Beta-domain+wHTH+RRM | Type-1_BR.RM.like.systems | <-HEPN+DUF499*<-METHYLASE-p<-METHYLASE-p\|\|HTH_3-><-PIN<-Phd-YefM<-TUDOR+SF1-SF2-Helicase+iREase+CTD+REase-p | 1108 | Bacteria>Terrabacteria_group>Cyanobacteriota/Melainabacteria_group>Cyanobacteriota>Cyanophyceae | GCF_023734775.1 | Nostoc_commune | ANSO36C_RS22370 | 4938684..4942010 | 1178 | Swt1-family-HEPN-domain-containing-protein-[Nostoc-commune] |
| KAB8314079.1 | HEPN+AAA-ATPase+Alpha+Beta-domain+wHTH+RRM | Type-1_BR.RM.like.systems | TUDOR+SF1-SF2-Helicase+iREase+CTD+REase-p->REase-><-?\|\|METHYLASE->HEPN+DUF499*-><-?<-?\|\|FAD-NAD-dep-oxidoreductase-> | 1113 | Bacteria>Terrabacteria_group>Cyanobacteriota/Melainabacteria_group>Cyanobacteriota>Cyanophyceae | GCA_000828075.3 | Tolypothrix_campylonemoides_VB511288 | SD81_039415 | 98698..102039 | 1245935 | ATP-binding-protein-[Tolypothrix-campylonemoides-VB511288] |
| EKD07858.1 | HEPN+AAA-ATPase+Alpha+Beta-domain+wHTH+RRM | Type-1_BR.RM.like.systems | <-ABhydrolase<-ABhydrolase\|\|TUDOR+SF1-SF2-Helicase+iREase+CTD+REase-><-?\|\|METHYLASE-><-?\|\|HEPN+DUF499*-><-?\|\|?->?->?->?->SPOUT-> | 1123 | Bacteria>Terrabacteria_group>Cyanobacteriota/Melainabacteria_group>Cyanobacteriota>Cyanophyceae | GCA_000307915.1 | Arthrospira_platensis_C1 | SPLC1_S361100 | 3984646..3988017 | 459495 | hypothetical-protein-SPLC1_S361100-[Arthrospira-platensis-C1] |
| MCA2716396.1 | HEPN+AAA-ATPase+Alpha+Beta-domain+wHTH+RRM | Type-1_BR.RM.like.systems | REase->AbrB->PIN->METHYLASE->RHH_6->HEPN+DUF499*-><-Cluster164_18clades<-?\|\|?->?->?-><-TPR+TPR+TPR+TPR+TPR+TPR+TPR+TPR | 1130 | Bacteria>Terrabacteria_group>Cyanobacteriota/Melainabacteria_group>Cyanobacteriota>Cyanophyceae | GCA_020245865.1 | Microcystis_sp._M169S2 | IM485_01040 | 5502..8894 | 2771157 | DUF499-domain-containing-protein-[Microcystis-sp.-M169S2] |
| WP_159297746.1 | HEPN+AAA-ATPase+Alpha+Beta-domain+wHTH+RRM | Type-1_BR.RM.like.systems | TUDOR+SF1-SF2-Helicase+iREase+CTD+REase->REase->METHYLASE->OLD-ABC->PIN->HEPN+DUF499*-><-?<-?\|\|?->?-><-THUMP+Methylase | 1130 | Bacteria>Terrabacteria_group>Cyanobacteriota/Melainabacteria_group>Cyanobacteriota>Cyanophyceae | GCF_009811875.1 | Microcystis_aeruginosa | GVO69_RS15755 | 16338..19730 | 1126 | Swt1-family-HEPN-domain-containing-protein-[Microcystis-aeruginosa] |
| WP_159253859.1 | HEPN+AAA-ATPase+Alpha+Beta-domain+wHTH+RRM | Type-1_BR.RM.like.systems | TUDOR+SF1-SF2-Helicase+iREase+CTD+REase->REase->METHYLASE->OLD-ABC->PIN->HEPN+DUF499*-><-?<-?\|\|?->?-><-THUMP+Methylase | 1130 | Bacteria>Terrabacteria_group>Cyanobacteriota/Melainabacteria_group>Cyanobacteriota>Cyanophyceae | GCF_009811855.1 | Microcystis_aeruginosa | GVO78_RS16540 | 16290..19682 | 1126 | Swt1-family-HEPN-domain-containing-protein-[Microcystis-aeruginosa] |
| WP_149989607.1 | HEPN+AAA-ATPase+Alpha+Beta-domain+wHTH+RRM | Type-1_BR.RM.like.systems | Cluster164_18clades-><-?<-HEPN+DUF499*<-ACET<-?<-METHYLASE<-REase\|\|?-><-TUDOR+SF1-SF2-Helicase+iREase+CTD+REase | 1117 | Bacteria>Terrabacteria_group>Cyanobacteriota/Melainabacteria_group>Cyanobacteriota>Cyanophyceae | GCF_008579565.1 | Microcystis_aeruginosa | MiHa_RS15955 | 12499..15852 | 1126 | Swt1-family-HEPN-domain-containing-protein-[Microcystis-aeruginosa] |
| WP_149979248.1 | HEPN+AAA-ATPase+Alpha+Beta-domain+wHTH+RRM | Type-1_BR.RM.like.systems | Cluster164_18clades-><-HEPN+DUF499*<-ACET<-?<-METHYLASE<-REase\|\|?-><-TUDOR+SF1-SF2-Helicase+iREase+CTD+REase | 1130 | Bacteria>Terrabacteria_group>Cyanobacteriota/Melainabacteria_group>Cyanobacteriota>Cyanophyceae | GCF_008579225.1 | Microcystis_aeruginosa | MiYa_RS12425 | 10420..13812 | 1126 | Swt1-family-HEPN-domain-containing-protein-[Microcystis-aeruginosa] |
| WP_106908219.1 | HEPN+AAA-ATPase+Alpha+Beta-domain+wHTH+RRM | Type-1_BR.RM.like.systems | Cluster164_18clades-><-?<-HEPN+DUF499*<-ACET<-?<-METHYLASE<-REase\|\|?-><-TUDOR+SF1-SF2-Helicase+iREase+CTD+REase | 1130 | Bacteria>Terrabacteria_group>Cyanobacteriota/Melainabacteria_group>Cyanobacteriota>Cyanophyceae | GCF_003019735.1 | Microcystis_sp._MC19 | B5D77_RS02535 | 488054..491446 | 1967666 | Swt1-family-HEPN-domain-containing-protein-[Microcystis-sp.-MC19] |
| WP_106502064.1 | HEPN+AAA-ATPase+Alpha+Beta-domain+wHTH+RRM | Type-1_BR.RM.like.systems | <-WYL\|\|LRP-HTH->TUDOR+SF1-SF2-Helicase+iREase+CTD+REase->METHYLASE->HEPN+DUF499*->?->?->?->?->Peptidase_M48-><-HNH | 1113 | Bacteria>Terrabacteria_group>Cyanobacteriota/Melainabacteria_group>Cyanobacteriota>Cyanophyceae | GCF_024345435.1 | Cyanobium | C7K55_RS03685 | 97796..101137 | 167375 | MULTISPECIES:-DUF499-domain-containing-protein-[Cyanobium] |
| KPQ40173.1 | HEPN+AAA-ATPase+Alpha+Beta-domain+wHTH+RRM | Type-1_BR.RM.like.systems | <-CARF+HEPN+CARF<-REase<-HEPN+DUF499*<-?<-?<-?<-METHYLASE\|\|METHYLASE-><-TUDOR+SF1-SF2-Helicase+iREase+CTD+REase | 1133 | Bacteria>Terrabacteria_group>Cyanobacteriota/Melainabacteria_group>Cyanobacteriota>Cyanophyceae | GCA_001314905.1 | Phormidium_sp._OSCR | HLUCCO16_04190 | 45328..48729 | 1666905 | putative-ATPase-(AAA+-superfamily)-[Phormidium-sp.-OSCR] |
| WP_190576377.1 | HEPN+AAA-ATPase+Alpha+Beta-domain+wHTH+RRM | Type-1_BR.RM.like.systems | RelE-ParE-><-?<-URI-DNase<-?<-?<-HEPN+DUF499*<-METHYLASE<-REase<-AbrB<-TUDOR+SF1-SF2-Helicase+iREase+CTD+REase | 1117 | Bacteria>Terrabacteria_group>Cyanobacteriota/Melainabacteria_group>Cyanobacteriota>Cyanophyceae | GCF_014697025.1 | Alkalinema_sp._FACHB-956 | H6G21_RS22810 | 35052..38405 | 2692768 | Swt1-family-HEPN-domain-containing-protein-[Alkalinema-sp.-FACHB-956] |
| WP_254935737.1 | HEPN+AAA-ATPase+Alpha+Beta-domain+wHTH+RRM | Type-1_BR.RM.like.systems | <-HEPN+DUF499*<-METHYLASE<-SF1-SF2-Helicase<-TUDOR+SF1-SF2-Helicase+iREase+CTD+REase<-LRP-HTH\|\|WYL-> | 1105 | Bacteria>Terrabacteria_group>Cyanobacteriota/Melainabacteria_group>Cyanobacteriota>Cyanophyceae | GCF_024345395.1 | unclassified_Cyanobium | KBZ17_RS00995 | 40704..44021 | 2627006 | MULTISPECIES:-DUF499-domain-containing-protein-[unclassified-Cyanobium] |
| MCA6503991.1 | HEPN+AAA-ATPase+Alpha+Beta-domain+wHTH+RRM | Type-1_BR.RM.like.systems | METHYLASE-p->?->PIN->REase->REase->REase->HEPN+DUF499*-><-Thioredoxin\|\|tRNA-><-PIN<-Phd-YefM\|\|?->?->ABhydrolase-> | 1118 | Bacteria>Terrabacteria_group>Cyanobacteriota/Melainabacteria_group>Cyanobacteriota>Cyanophyceae | GCA_020403835.1 | Pseudanabaena_sp._M090S1SP2A07QC | IM580_15730 | 4170..7526 | 2771254 | DUF499-domain-containing-protein-[Pseudanabaena-sp.-M090S1SP2A07QC] |
| MBV6627478.1 | HEPN+AAA-ATPase+Alpha+Beta-domain+wHTH+RRM | Type-1_BR.RM.like.systems | <-HEPN+DUF499*<-REase<-METHYLASE-p\|\|wHTH+REase-><-TUDOR+SF1-SF2-Helicase+iREase+CTD+REase-p<-REase | 1116 | Bacteria>Terrabacteria_group>Cyanobacteriota/Melainabacteria_group>Cyanobacteriota>Cyanophyceae | GCA_019192545.1 | Rivularia_sp._MS3 | KI793_31915 | 3971..7321 | 3012223 | DUF499-domain-containing-protein-[Rivularia-sp.-(in:-Bacteria)] |
| AMW31767.1 | HEPN+AAA-ATPase+Alpha+Beta-domain+wHTH+RRM | Type-1_BR.RM.like.systems | <-THUMP+Methylase<-ABhydrolase<-?\|\|TUDOR+SF1-SF2-Helicase+iREase+CTD+REase->METHYLASE->HEPN+DUF499*-> | 1119 | Bacteria>Terrabacteria_group>Cyanobacteriota/Melainabacteria_group>Cyanobacteriota>Cyanophyceae | GCA_001611905.1 | Arthrospira_platensis_YZ | AP285_21895 | 4862859..4866218 | 1738638 | AAA+-family-ATPase-[Arthrospira-platensis-YZ] |
| WP_272078194.1 | HEPN+AAA-ATPase+Alpha+Beta-domain+wHTH+RRM | Type-1_BR.RM.like.systems | DDE_3-><-HEPN+DUF499*<-PIN<-PIN<-RHH_6<-METHYLASE<-REase<-TUDOR+SF1-SF2-Helicase+iREase+CTD+REase | 1111 | Bacteria>Terrabacteria_group>Cyanobacteriota/Melainabacteria_group>Cyanobacteriota>Cyanophyceae | GCF_028331105.1 | Microcystis_aeruginosa | PN487_RS19700 | 7822..11157 | 1126 | Swt1-family-HEPN-domain-containing-protein-[Microcystis-aeruginosa] |
| WP_104908185.1 | HEPN+AAA-ATPase+Alpha+Beta-domain+wHTH+RRM | Type-1_BR.RM.like.systems | TUDOR+SF1-SF2-Helicase+iREase+CTD+REase->Phd-YefM->PIN-><-HTH_3\|\|METHYLASE->REase->HEPN+DUF499*-> | 1109 | Bacteria>Terrabacteria_group>Cyanobacteriota/Melainabacteria_group>Cyanobacteriota>Cyanophyceae | GCF_002949795.1 | Nostoc_sp._'Lobaria_pulmonaria_(5183)_cyanobiont' | NLP_RS21725 | 4933532..4936861 | 1618022 | Swt1-family-HEPN-domain-containing-protein-[Nostoc-sp.-'Lobaria-pulmonaria-(5183)-cyanobiont'] |
| MBW4532455.1 | HEPN+AAA-ATPase+Alpha+Beta-domain+wHTH+RRM | Type-1_BR.RM.like.systems | <-HTH_OrfB_IS605+OrfB_IS605+OrfB_Zn_ribbon\|\|<-?<-?<-?<-HEPN+DUF499*<-CopG/RHH\|\|DDE_Tnp_1-><-PIN<-?<-?<-METHYLASE | 1104 | Bacteria>Terrabacteria_group>Cyanobacteriota/Melainabacteria_group>Cyanobacteriota>Cyanophyceae | GCA_019359095.1 | Pleurocapsa_minor_HA4230-MV1 | KME09_00805 | 42655..45969 | 2839664 | DUF499-domain-containing-protein-[Pleurocapsa-minor-HA4230-MV1] |
| WP_254940710.1 | HEPN+AAA-ATPase+Alpha+Beta-domain+wHTH+RRM | Type-1_BR.RM.like.systems | <-DUF4236\|\|TUDOR+SF1-SF2-Helicase+iREase+CTD+REase->?->PIN->METHYLASE->HEPN+DUF499*->?->?-><-ArdCN+MPTase | 1103 | Bacteria>Terrabacteria_group>Cyanobacteriota/Melainabacteria_group>Cyanobacteriota>Cyanophyceae | GCF_024345575.1 | Cyanobium_sp._Morenito_9A2 | KBY88_RS14795 | 41582..44893 | 2823718 | DUF499-domain-containing-protein-[Cyanobium-sp.-Morenito-9A2] |
| WP_149107756.1 | HEPN+AAA-ATPase+Alpha+Beta-domain+wHTH+RRM | Type-1_BR.RM.like.systems | <-sGTP<-DUF29<-?\|\|?->Cluster164_18clades-><-HEPN+DUF499*<-PIN<-RHH_6<-METHYLASE<-PemK_toxin<-AbrB<-PIN | 1130 | Bacteria>Terrabacteria_group>Cyanobacteriota/Melainabacteria_group>Cyanobacteriota>Cyanophyceae | GCF_008257445.1 | Microcystis_aeruginosa | FXO09_RS20205 | 21434..24826 | 1126 | Swt1-family-HEPN-domain-containing-protein-[Microcystis-aeruginosa] |
| WP_103113079.1 | HEPN+AAA-ATPase+Alpha+Beta-domain+wHTH+RRM | Type-1_BR.RM.like.systems | PemK_toxin->METHYLASE->RHH_6->PIN->OLD-ABC->PIN->HEPN+DUF499*-><-Cluster164_18clades<-?\|\|?->?->DUF29->sGTP-> | 1111 | Bacteria>Terrabacteria_group>Cyanobacteriota/Melainabacteria_group>Cyanobacteriota>Cyanophyceae | GCF_002897275.1 | Microcystis_aeruginosa | D3800_RS23535 | 4968568..4971903 | 1126 | Swt1-family-HEPN-domain-containing-protein-[Microcystis-aeruginosa] |
| WP_026733571.1 | HEPN+AAA-ATPase+Alpha+Beta-domain+wHTH+RRM | Type-1_BR.RM.like.systems | TUDOR+SF1-SF2-Helicase+iREase+CTD+REase->AbrB->PIN->METHYLASE->REase->HEPN+DUF499*->?->TraJ-RHH-> | 1122 | Bacteria>Terrabacteria_group>Cyanobacteriota/Melainabacteria_group>Cyanobacteriota>Cyanophyceae | GCF_000517105.1 | Fischerella_sp._PCC_9605 | FIS9605_RS0116420 | 337165..340533 | 1173024 | Swt1-family-HEPN-domain-containing-protein-[Fischerella-sp.-PCC-9605] |
| WP_194014089.1 | HEPN+AAA-ATPase+Alpha+Beta-domain+wHTH+RRM | Type-1_BR.RM.like.systems | AbrB->PIN->AbrB->METHYLASE->?->ACET->HEPN+DUF499*-><-?\|\|?->?->?-><-TPR+TPR+TPR+TPR+TPR+TPR+TPR+TPR<-ORC-CDC6 | 1130 | Bacteria>Terrabacteria_group>Cyanobacteriota/Melainabacteria_group>Cyanobacteriota>Cyanophyceae | GCF_015207115.1 | Microcystis_aeruginosa | IQ232_RS04140 | 10022..13414 | 1126 | Swt1-family-HEPN-domain-containing-protein-[Microcystis-aeruginosa] |
| WP_130756579.1 | HEPN+AAA-ATPase+Alpha+Beta-domain+wHTH+RRM | Type-1_BR.RM.like.systems | REase-p->AbrB->PIN->METHYLASE->RHH_6->PIN->HEPN+DUF499*-><-Cluster164_18clades<-?\|\|?->?->DUF29->sGTP-> | 1111 | Bacteria>Terrabacteria_group>Cyanobacteriota/Melainabacteria_group>Cyanobacteriota>Cyanophyceae | GCF_004305995.1 | Microcystis_aeruginosa | MiAbB_RS04285 | 58496..61831 | 1126 | Swt1-family-HEPN-domain-containing-protein-[Microcystis-aeruginosa] |
| WJI23189.1 | HEPN+AAA-ATPase+Alpha+Beta-domain+wHTH+RRM | Type-1_BR.RM.like.systems | <-HEPN+DUF499*<-REase<-METHYLASE<-TUDOR+SF1-SF2-Helicase+iREase+CTD+REase\|\|REase-><-DDE_Tnp_1 | 1098 | Bacteria>Terrabacteria_group>Cyanobacteriota/Melainabacteria_group>Cyanobacteriota>Cyanophyceae | GCA_030322965.1 | Thermosynechococcus_sp._B0 | MZ909_08100 | 1635834..1639130 | 2937284 | Swt1-family-HEPN-domain-containing-protein-[Thermosynechococcus-sp.-B0] |
| WP_250012765.1 | HEPN+AAA-ATPase+Alpha+Beta-domain+wHTH+RRM | Type-1_BR.RM.like.systems | AbrB->PIN->REase->METHYLASE-p->METHYLASE->?->HEPN+DUF499*->?-><-?\|\|Phage_integrase-><-Phage_integrase | 1113 | Bacteria>Terrabacteria_group>Cyanobacteriota/Melainabacteria_group>Cyanobacteriota>Cyanophyceae | GCF_023546805.1 | Chroococcidiopsis_sp._CCNUC1 | M5J74_RS19090 | 4105692..4109033 | 2653189 | Swt1-family-HEPN-domain-containing-protein-[Chroococcidiopsis-sp.-CCNUC1] |
| HBH74016.1 | HEPN+AAA-ATPase+Alpha+Beta-domain+wHTH+RRM | Type-1_BR.RM.like.systems | TUDOR+SF1-SF2-Helicase+iREase+CTD+REase-><-PIN<-MazE_antitoxin\|\|?->wHTH(SBD)+HNH->HEPN+DUF499*-> | 1110 | Bacteria>Terrabacteria_group>Cyanobacteriota/Melainabacteria_group>Cyanobacteriota>Cyanophyceae | GCA_003508015.1 | Synechococcales_bacterium_UBA10510 | DDY43_11405 | 28893..32225 | 2055795 | AAA+-family-ATPase-[Synechococcales-bacterium-UBA10510] |
| WP_240867511.1 | HEPN+AAA-ATPase+Alpha+Beta-domain+wHTH+RRM | Type-1_BR.RM.like.systems | <-PolB<-?<-DDE_Tnp_1<-?<-?\|\|TUDOR+SF1-SF2-Helicase+iREase+CTD+REase->HEPN+DUF499*->?->?->?-><-?\|\|Peptidase_C14-> | 1102 | Bacteria>Terrabacteria_group>Cyanobacteriota/Melainabacteria_group>Cyanobacteriota>Cyanophyceae | GCF_022496815.1 | Cylindrospermopsis_raciborskii | GSN00_RS07900 | 80955..84263 | 77022 | Swt1-family-HEPN-domain-containing-protein-[Cylindrospermopsis-raciborskii] |
| WP_002754187.1 | HEPN+AAA-ATPase+Alpha+Beta-domain+wHTH+RRM | Type-1_BR.RM.like.systems | AbrB->PemK_toxin->METHYLASE->RHH_6->PIN->HEPN+DUF499*-><-Cluster164_18clades<-?\|\|?->?->DUF29->sGTP-> | 1130 | Bacteria>Terrabacteria_group>Cyanobacteriota/Melainabacteria_group>Cyanobacteriota>Cyanophyceae | GCF_000307995.1 | Microcystis_aeruginosa | MICCA_RS15375 | 7336..10728 | 1126 | Swt1-family-HEPN-domain-containing-protein-[Microcystis-aeruginosa] |
| WP_271935711.1 | HEPN+AAA-ATPase+Alpha+Beta-domain+wHTH+RRM | Type-1_BR.RM.like.systems | AbrB->METHYLASE->RHH_6->PIN->OLD-ABC->PIN->HEPN+DUF499*-><-Cluster164_18clades<-?\|\|?->?->DUF29->sGTP-> | 1111 | Bacteria>Terrabacteria_group>Cyanobacteriota/Melainabacteria_group>Cyanobacteriota>Cyanophyceae | GCF_028329095.1 | Microcystis | PN470_RS21190 | 25157..28492 | 1125 | MULTISPECIES:-Swt1-family-HEPN-domain-containing-protein-[Microcystis] |
| WP_096720868.1 | HEPN+AAA-ATPase+Alpha+Beta-domain+wHTH+RRM | Type-1_BR.RM.like.systems | DUF29-><-?<-?<-?\|\|?-><-HEPN+DUF499*<-PIN<-?<-METHYLASE\|\|REase-><-TUDOR+SF1-SF2-Helicase+iREase+CTD+REase | 1125 | Bacteria>Terrabacteria_group>Cyanobacteriota/Melainabacteria_group>Cyanobacteriota>Cyanophyceae | - | Chondrocystis_sp._NIES-4102 | CDC35_RS11850 | 2748341..2751718 | 2005460 | Swt1-family-HEPN-domain-containing-protein-[Chondrocystis-sp.-NIES-4102] |
| WP_235278697.1 | HEPN+AAA-ATPase+Alpha+Beta-domain+wHTH+RRM | Type-1_BR.RM.like.systems | METHYLASE->?->PemK_toxin->REase->REase->CopG/RHH->HEPN+DUF499*->?->?->MORC+DNA_gyraseB+TOPRIM-> | 1109 | Bacteria>Terrabacteria_group>Cyanobacteriota/Melainabacteria_group>Cyanobacteriota>Cyanophyceae | GCF_021533835.1 | Synechococcus_sp._Nb3U1 | L1047_RS09935 | 2086840..2090169 | 1914529 | Swt1-family-HEPN-domain-containing-protein-[Synechococcus-sp.-Nb3U1] |
| WP_073632965.1 | HEPN+AAA-ATPase+Alpha+Beta-domain+wHTH+RRM | Type-1_BR.RM.like.systems | TUDOR+SF1-SF2-Helicase+iREase+CTD+REase->METHYLASE->REase->?->HEPN+DUF499*-><-HTH\|\|PIWI-> | 1113 | Bacteria>Terrabacteria_group>Cyanobacteriota/Melainabacteria_group>Cyanobacteriota>Cyanophyceae | GCF_001904675.1 | Scytonema_sp._HK-05 | NIES2130_RS19405 | 80454..83795 | 1137095 | Swt1-family-HEPN-domain-containing-protein-[Scytonema-sp.-HK-05] |
| WP_247214939.1 | HEPN+AAA-ATPase+Alpha+Beta-domain+wHTH+RRM | Type-1_BR.RM.like.systems | DDE_Tnp_1-><-Peptidase_M24\|\|?->tRNA-><-HEPN+DUF499*<-REase<-REase<-REase<-METHYLASE<-PIN<-AbrB | 1107 | Bacteria>Terrabacteria_group>Cyanobacteriota/Melainabacteria_group>Cyanobacteriota>Cyanophyceae | GCF_022984075.1 | Synechococcus_sp._C9 | MLD66_RS00195 | 39889..43212 | 102119 | Swt1-family-HEPN-domain-containing-protein-[Synechococcus-sp.-C9] |
| WP_075601490.1 | HEPN+AAA-ATPase+Alpha+Beta-domain+wHTH+RRM | Type-1_BR.RM.like.systems | <-HEPN+DUF499*<-METHYLASE<-?<-SbcC<-TUDOR+SF1-SF2-Helicase+iREase+CTD+REase<-MPTase<-HTH | 834 | Bacteria>Terrabacteria_group>Cyanobacteriota/Melainabacteria_group>Cyanobacteriota>Cyanophyceae | GCF_001939115.1 | Leptolyngbya_sp._'hensonii' | BST81_RS26530 | 1..2502 | 1922337 | Swt1-family-HEPN-domain-containing-protein,-partial-[Leptolyngbya-sp.-'hensonii'] |
| WP_017744970.1 | HEPN+AAA-ATPase+Alpha+Beta-domain+wHTH+RRM | Type-1_BR.RM.like.systems | OrfB_IS605+OrfB_Zn_ribbon-><-HEPN+DUF499*\|\|?-><-MqsA_antitoxin+HTH_3<-Polbeta<-?<-METHYLASE | 1113 | Bacteria>Terrabacteria_group>Cyanobacteriota/Melainabacteria_group>Cyanobacteriota>Cyanophyceae | GCF_000346485.2 | Scytonema_hofmannii | WA1_RS46765 | 11110546..11113887 | 34078 | Swt1-family-HEPN-domain-containing-protein-[Scytonema-hofmannii] |
| WP_002777134.1 | HEPN+AAA-ATPase+Alpha+Beta-domain+wHTH+RRM | Type-1_BR.RM.like.systems | METHYLASE->RHH_6->PIN->OLD-ABC->PIN->HEPN+DUF499*-><-Cluster164_18clades<-?\|\|?->?->DUF29->sGTP-> | 1130 | Bacteria>Terrabacteria_group>Cyanobacteriota/Melainabacteria_group>Cyanobacteriota>Cyanophyceae | GCF_000312205.1 | Microcystis_aeruginosa | MICAD_RS14460 | 204713..208105 | 1126 | Swt1-family-HEPN-domain-containing-protein-[Microcystis-aeruginosa] |
| PSB44266.1 | HEPN+AAA-ATPase+Alpha+Beta-domain+wHTH+RRM | Type-1_BR.RM.like.systems | <-HEPN+DUF499*<-?<-METHYLASE<-REase<-PIN<-AbrB<-TUDOR+SF1-SF2-Helicase+iREase+CTD+REase | 1113 | Bacteria>Terrabacteria_group>Cyanobacteriota/Melainabacteria_group>Cyanobacteriota>Cyanophyceae | GCA_003004015.1 | Cyanosarcina_cf._burmensis_CCALA_770 | C7B80_21165 | 280..3621 | 2107697 | AAA+-family-ATPase-[Cyanosarcina-cf.-burmensis-CCALA-770] |
| WP_265264586.1 | HEPN+AAA-ATPase+Alpha+Beta-domain+wHTH+RRM | Type-1_BR.RM.like.systems | NTP_transferase-><-HEPN+DUF499*<-PIN<-AbrB<-TUDOR+SF1-SF2-Helicase+iREase+CTD+REase | 1112 | Bacteria>Terrabacteria_group>Cyanobacteriota/Melainabacteria_group>Cyanobacteriota>Cyanophyceae | GCF_026130165.1 | Spirulina_subsalsa | K4A83_RS11005 | 13314..16652 | 54311 | Swt1-family-HEPN-domain-containing-protein-[Spirulina-subsalsa] |
| WP_100904446.1 | HEPN+AAA-ATPase+Alpha+Beta-domain+wHTH+RRM | Type-1_BR.RM.like.systems | <-DDE_Tnp_1_3<-HEPN+DUF499*<-REase<-TUDOR+SF1-SF2-Helicase+iREase+CTD+REase<-?<-?\|\|DndB-> | 1112 | Bacteria>Terrabacteria_group>Cyanobacteriota/Melainabacteria_group>Cyanobacteriota>Cyanophyceae | GCF_002813575.1 | Nostoc_flagelliforme | CO091_RS47960 | 745879..749217 | 1306274 | Swt1-family-HEPN-domain-containing-protein-[Nostoc-flagelliforme] |
| WP_006616387.1 | HEPN+AAA-ATPase+Alpha+Beta-domain+wHTH+RRM | Type-1_BR.RM.like.systems | <-ABhydrolase<-?\|\|TUDOR+SF1-SF2-Helicase+iREase+CTD+REase->METHYLASE->?-><-?\|\|HEPN+DUF499*-> | 1135 | Bacteria>Terrabacteria_group>Cyanobacteriota/Melainabacteria_group>Cyanobacteriota>Cyanophyceae | - | Arthrospira_platensis | AP285_RS21915 | 4862811..4866218 | 118562 | Swt1-family-HEPN-domain-containing-protein-[Arthrospira-platensis] |
| TGH19526.1 | HEPN+AAA-ATPase+Alpha+Beta-domain+wHTH+RRM | Type-1_BR.RM.like.systems | TUDOR+SF1-SF2-Helicase+iREase+CTD+REase-p->METHYLASE-p->METHYLASE-p->HEPN+DUF499*-> | 1094 | Bacteria>Terrabacteria_group>Cyanobacteriota/Melainabacteria_group>Cyanobacteriota>Cyanophyceae | GCA_004768385.2 | Aphanocapsa_feldmannii_277cI | ERJ68_08375 | 4147..7431 | 2507554 | ATP-binding-protein-[Aphanocapsa-feldmannii-277cI] |
| MCA6600378.1 | HEPN+AAA-ATPase+Alpha+Beta-domain+wHTH+RRM | Type-1_BR.RM.like.systems | Phd-YefM->PIN-><-tRNA\|\|Thioredoxin-><-HEPN+DUF499*<-REase<-REase<-REase<-PIN<-?<-METHYLASE-p | 1118 | Bacteria>Terrabacteria_group>Cyanobacteriota/Melainabacteria_group>Cyanobacteriota>Cyanophyceae | GCA_020404965.1 | Pseudanabaena_sp._M57BS1SP1A06MG | IM537_09260 | 3869..7225 | 2771237 | DUF499-domain-containing-protein-[Pseudanabaena-sp.-M57BS1SP1A06MG] |
| EKQ69986.1 | HEPN+AAA-ATPase+Alpha+Beta-domain+wHTH+RRM | Type-1_BR.RM.like.systems | TUDOR+SF1-SF2-Helicase+iREase+CTD+REase->REase->PolB->HEPN-><-?\|\|REase->HEPN+DUF499*-> | 1113 | Bacteria>Terrabacteria_group>Cyanobacteriota/Melainabacteria_group>Cyanobacteriota>Cyanophyceae | GCA_000309945.1 | Leptolyngbyaceae_cyanobacterium_JSC-12 | OsccyDRAFT_0242 | 268586..271927 | 864702 | putative-ATPase-(AAA+-superfamily)-[Leptolyngbyaceae-cyanobacterium-JSC-12] |
| RFP57752.1 | HEPN+AAA-ATPase+Alpha+Beta-domain+wHTH+RRM | Type-1_BR.RM.like.systems | HISKIN+MORC->?-><-?\|\|TUDOR+SF1-SF2-Helicase+iREase+CTD+REase->METHYLASE->HEPN+DUF499*-> | 1119 | Bacteria>Terrabacteria_group>Cyanobacteriota/Melainabacteria_group>Cyanobacteriota>Cyanophyceae | GCA_001913845.2 | Limnothrix_sp._CACIAM_69d | BJG00_013040 | 63508..66867 | 1890733 | MAG:-ATP-binding-protein-[Limnothrix-sp.-CACIAM-69d] |
| WP_255548761.1 | HEPN+AAA-ATPase+Alpha+Beta-domain+wHTH+RRM | Type-1_BR.RM.like.systems | REase-><-?<-CR-Trypsin1<-?\|\|REase-><-?<-HEPN+DUF499*<-?<-METHYLASE<-PIN<-AbrB<-REase<-REase | 1078 | Bacteria>Terrabacteria_group>Cyanobacteriota/Melainabacteria_group>Cyanobacteriota>Cyanophyceae | - | Thermoleptolyngbya_sp._PKUAC-SCTB121 | JTM80_RS13275 | 3238289..3241525 | 2811482 | DUF499-domain-containing-protein-[Thermoleptolyngbya-sp.-PKUAC-SCTB121] |
| WP_172355891.1 | HEPN+AAA-ATPase+Alpha+Beta-domain+wHTH+RRM | Type-1_BR.RM.like.systems | REase->REase->AbrB->PIN->METHYLASE->?->HEPN+DUF499*->?-><-REase\|\|?->CR-Trypsin1->?-><-REase | 1105 | Bacteria>Terrabacteria_group>Cyanobacteriota/Melainabacteria_group>Cyanobacteriota>Cyanophyceae | GCF_013177315.1 | Thermoleptolyngbya_sichuanensis | HPC62_RS11805 | 2814598..2817915 | 2885951 | Swt1-family-HEPN-domain-containing-protein-[Thermoleptolyngbya-sichuanensis] |
| MCA6597113.1 | HEPN+AAA-ATPase+Alpha+Beta-domain+wHTH+RRM | Type-1_BR.RM.like.systems | Phd-YefM->PIN-><-tRNA\|\|Thioredoxin-><-HEPN+DUF499*<-REase<-REase<-?<-REase<-REase | 1118 | Bacteria>Terrabacteria_group>Cyanobacteriota/Melainabacteria_group>Cyanobacteriota>Cyanophyceae | GCA_020405005.1 | Pseudanabaena_sp._M046S1SP1A06QC | IM539_12615 | 33014..36370 | 2771236 | DUF499-domain-containing-protein-[Pseudanabaena-sp.-M046S1SP1A06QC] |
| TRU11470.1 | HEPN+AAA-ATPase+Alpha+Beta-domain+wHTH+RRM | Type-1_BR.RM.like.systems | <-DUF29<-?<-?\|\|?->Cluster164_18clades-><-?<-HEPN+DUF499*<-ACET<-?<-METHYLASE<-AbrB<-PIN<-AbrB | 1117 | Bacteria>Terrabacteria_group>Cyanobacteriota/Melainabacteria_group>Cyanobacteriota>Cyanophyceae | GCA_007095705.1 | Microcystis_sp._Msp_OC_L_20101000_S702 | EWV60_07295 | 28722..32075 | 2486218 | MAG:-ATP-binding-protein-[Microcystis-sp.-Msp_OC_L_20101000_S702] |
| WP_170190015.1 | HEPN+AAA-ATPase+Alpha+Beta-domain+wHTH+RRM | Type-1_BR.RM.like.systems | TUDOR+SF1-SF2-Helicase+iREase+CTD+REase->SbcC->?->METHYLASE->?->PIN->HEPN+DUF499*-> | 1130 | Bacteria>Terrabacteria_group>Cyanobacteriota/Melainabacteria_group>Cyanobacteriota>Cyanophyceae | GCF_012911965.1 | Geitlerinema_sp._P-1104 | E1H12_RS13025 | 9854..13246 | 2546230 | Swt1-family-HEPN-domain-containing-protein-[Geitlerinema-sp.-P-1104] |
| WP_015227928.1 | HEPN+AAA-ATPase+Alpha+Beta-domain+wHTH+RRM | Type-1_BR.RM.like.systems | <-HEPN+DUF499*<-HEPN<-PolB<-METHYLASE<-?<-HicB_lk_antitox\|\|BrnA_antitoxin-> | 1113 | Bacteria>Terrabacteria_group>Cyanobacteriota/Melainabacteria_group>Cyanobacteriota>Cyanophyceae | GCF_000317615.1 | Dactylococcopsis_salina | DACSA_RS00655 | 98261..101602 | 292566 | Swt1-family-HEPN-domain-containing-protein-[Dactylococcopsis-salina] |
| NMG59376.1 | HEPN+AAA-ATPase+Alpha+Beta-domain+wHTH+RRM | Type-1_BR.RM.like.systems | TUDOR+SF1-SF2-Helicase+iREase+CTD+REase->SbcC->?->METHYLASE->?->PIN->HEPN+DUF499*-> | 1130 | Bacteria>Terrabacteria_group>Cyanobacteriota/Melainabacteria_group>Cyanobacteriota>Cyanophyceae | GCA_012911965.1 | Geitlerinema_sp._P-1104 | E1H12_12830 | 9854..13246 | 2546230 | ATP-binding-protein-[Geitlerinema-sp.-P-1104] |
| WP_002801405.1 | HEPN+AAA-ATPase+Alpha+Beta-domain+wHTH+RRM | Type-1_BR.RM.like.systems | REase->METHYLASE->RHH_6->PIN->OLD-ABC->PIN->HEPN+DUF499*-><-Cluster164_18clades | 1117 | Bacteria>Terrabacteria_group>Cyanobacteriota/Melainabacteria_group>Cyanobacteriota>Cyanophyceae | GCF_000312285.1 | Microcystis_aeruginosa | MICAK_RS07420 | 6579..9932 | 1126 | Swt1-family-HEPN-domain-containing-protein-[Microcystis-aeruginosa] |
| MCE2669063.1 | HEPN+AAA-ATPase+Alpha+Beta-domain+wHTH+RRM | Type-1_BR.RM.like.systems | Cluster164_18clades-><-HEPN+DUF499*<-PIN<-OLD-ABC<-PIN<-RHH_6<-METHYLASE<-REase | 1130 | Bacteria>Terrabacteria_group>Cyanobacteriota/Melainabacteria_group>Cyanobacteriota>Cyanophyceae | GCA_021300285.1 | Microcystis_sp._49638_E5 | LW728_07560 | 10523..13915 | 2904986 | DUF499-domain-containing-protein-[Microcystis-sp.-49638_E5] |
| MCA2627579.1 | HEPN+AAA-ATPase+Alpha+Beta-domain+wHTH+RRM | Type-1_BR.RM.like.systems | REase->METHYLASE->RHH_6->PIN->OLD-ABC->PIN->HEPN+DUF499*-><-Cluster164_18clades | 1130 | Bacteria>Terrabacteria_group>Cyanobacteriota/Melainabacteria_group>Cyanobacteriota>Cyanophyceae | GCA_020245485.1 | Microcystis_sp._M091S2 | IM505_05380 | 45571..48963 | 2771177 | DUF499-domain-containing-protein-[Microcystis-sp.-M091S2] |
| MBR8832157.1 | HEPN+AAA-ATPase+Alpha+Beta-domain+wHTH+RRM | Type-1_BR.RM.like.systems | <-PIN<-RHH<-HEPN+DUF499*<-METHYLASE<-?<-?<-?<-?<-TUDOR+SF1-SF2-Helicase+iREase+CTD+REase | 1101 | Bacteria>Terrabacteria_group>Cyanobacteriota/Melainabacteria_group>Cyanobacteriota>Cyanophyceae | GCA_016903695.1 | Chlorogloea_purpurea_SAG_13.99 | Cpurp_13245 | 34415..37720 | 1521637 | DUF499-domain-containing-protein-[Chlorogloea-purpurea-SAG-13.99] |
| WP_272014891.1 | HEPN+AAA-ATPase+Alpha+Beta-domain+wHTH+RRM | Type-1_BR.RM.like.systems | METHYLASE-p->RHH_6->PIN->HEPN+DUF499*-><-Cluster164_18clades<-?\|\|?->?->DUF29->sGTP-> | 1111 | Bacteria>Terrabacteria_group>Cyanobacteriota/Melainabacteria_group>Cyanobacteriota>Cyanophyceae | GCF_028330745.1 | Microcystis_aeruginosa | PN482_RS11750 | 1666..5001 | 1126 | Swt1-family-HEPN-domain-containing-protein-[Microcystis-aeruginosa] |
| WP_016515479.1 | HEPN+AAA-ATPase+Alpha+Beta-domain+wHTH+RRM | Type-1_BR.RM.like.systems | Cluster164_18clades-><-?<-HEPN+DUF499*<-PIN<-OLD-ABC<-PIN<-RHH_6<-METHYLASE<-AbrB | 1130 | Bacteria>Terrabacteria_group>Cyanobacteriota/Melainabacteria_group>Cyanobacteriota>Cyanophyceae | GCF_000412595.1 | Microcystis_aeruginosa | MAESPC_RS19915 | 50545..53937 | 1126 | Swt1-family-HEPN-domain-containing-protein-[Microcystis-aeruginosa] |
| MCA2653775.1 | HEPN+AAA-ATPase+Alpha+Beta-domain+wHTH+RRM | Type-1_BR.RM.like.systems | Cluster164_18clades-><-HEPN+DUF499*<-PIN<-RHH_6<-METHYLASE<-AbrB<-PIN<-AbrB | 1130 | Bacteria>Terrabacteria_group>Cyanobacteriota/Melainabacteria_group>Cyanobacteriota>Cyanophyceae | GCA_020245605.1 | Microcystis_sp._M061S2 | IM475_05875 | 1597..4989 | 2771171 | DUF499-domain-containing-protein-[Microcystis-sp.-M061S2] |
| WP_149976793.1 | HEPN+AAA-ATPase+Alpha+Beta-domain+wHTH+RRM | Type-1_BR.RM.like.systems | <-HEPN+DUF499*<-METHYLASE<-REase<-TUDOR+SF1-SF2-Helicase+iREase+CTD+REase | 1130 | Bacteria>Terrabacteria_group>Cyanobacteriota/Melainabacteria_group>Cyanobacteriota>Cyanophyceae | GCF_008579445.1 | Microcystis_aeruginosa | MiTs_RS18945 | 1418..4810 | 1126 | Swt1-family-HEPN-domain-containing-protein-[Microcystis-aeruginosa] |
| WP_130128840.1 | HEPN+AAA-ATPase+Alpha+Beta-domain+wHTH+RRM | Type-1_BR.RM.like.systems | <-MORC+DUF3684<-HEPN+DUF499*<-METHYLASE<-METHYLASE<-PIN<-RHH\|\|?->RelE-ParE-> | 1113 | Bacteria>Terrabacteria_group>Cyanobacteriota/Melainabacteria_group>Cyanobacteriota>Cyanophyceae | GCF_004209775.1 | Synechococcus_sp._WH_8101 | SynWH8101_RS05090 | 1007085..1010426 | 59932 | DUF499-domain-containing-protein-[Synechococcus-sp.-WH-8101] |
| MCE2977896.1 | HEPN+AAA-ATPase+Alpha+Beta-domain+wHTH+RRM | Type-1_BR.RM.like.systems | METHYLASE-p->?->PIN->REase->REase->REase->HEPN+DUF499*-><-Thioredoxin\|\|tRNA-><-PIN | 1118 | Bacteria>Terrabacteria_group>Cyanobacteriota/Melainabacteria_group>Cyanobacteriota>Cyanophyceae | GCA_021298775.1 | Pseudanabaena_sp._CoA8_M7 | LW851_16625 | 3735..7091 | 2904992 | DUF499-domain-containing-protein-[Pseudanabaena-sp.-CoA8_M7] |
| WP_233748345.1 | HEPN+AAA-ATPase+Alpha+Beta-domain+wHTH+RRM | Type-1_BR.RM.like.systems | <-Peptidase<-HEPN+DUF499*<-CopG/RHH<-PemK_toxin<-?<-REase<-REase<-METHYLASE | 1125 | Bacteria>Terrabacteria_group>Cyanobacteriota/Melainabacteria_group>Cyanobacteriota>Cyanophyceae | GCF_021379005.1 | Leptodesmis_sichuanensis | KIK02_RS09445 | 2037020..2040397 | 2906798 | Swt1-family-HEPN-domain-containing-protein-[Leptodesmis-sichuanensis] |
| TRT82535.1 | HEPN+AAA-ATPase+Alpha+Beta-domain+wHTH+RRM | Type-1_BR.RM.like.systems | Cluster164_18clades-><-?<-HEPN+DUF499*<-ACET<-?<-METHYLASE<-PIN<-AbrB<-REase | 1130 | Bacteria>Terrabacteria_group>Cyanobacteriota/Melainabacteria_group>Cyanobacteriota>Cyanophyceae | GCA_007095595.1 | Microcystis_aeruginosa_Ma_AC_P_19900807_S299 | EWV82_11170 | 1786..5178 | 2486264 | ATP-binding-protein-[Microcystis-aeruginosa-Ma_AC_P_19900807_S299] |
| WP_285965028.1 | HEPN+AAA-ATPase+Alpha+Beta-domain+wHTH+RRM | Type-1_BR.RM.like.systems | XIS-HTH-><-?<-Phage_integrase\|\|tRNA-><-?\|\|REase-><-DUF499-p<-HEPN+DUF499*<-AbiEii | 1005 | Bacteria>Terrabacteria_group>Cyanobacteriota/Melainabacteria_group>Cyanobacteriota>Cyanophyceae | GCF_030285465.1 | Oscillatoria_laete-virens | QQ056_RS17150 | 57885..60902 | 1565316 | Swt1-family-HEPN-domain-containing-protein-[Oscillatoria-laete-virens] |
| WP_193965432.1 | HEPN+AAA-ATPase+Alpha+Beta-domain+wHTH+RRM | Type-1_BR.RM.like.systems | Phage_integrase-><-?\|\|?->?-><-HTH\|\|?-><-HEPN+DUF499*\|\|?-><-ParB+HNH<-?<-PIN<-?<-METHYLASE | 1114 | Bacteria>Terrabacteria_group>Cyanobacteriota/Melainabacteria_group>Cyanobacteriota>Cyanophyceae | GCF_015207395.1 | Nodosilinea_sp._LEGE_07088 | IQ254_RS06885 | 5859..9203 | 2777968 | Swt1-family-HEPN-domain-containing-protein-[Nodosilinea-sp.-LEGE-07088] |
| MBF2075191.1 | HEPN+AAA-ATPase+Alpha+Beta-domain+wHTH+RRM | Type-1_BR.RM.like.systems | AbrB->HicB_lk_antitox->REase->REase->REase->REase->HEPN+DUF499*-> | 1114 | Bacteria>Terrabacteria_group>Cyanobacteriota/Melainabacteria_group>Cyanobacteriota>Cyanophyceae | GCA_015272325.1 | Synechococcales_cyanobacterium_C42_A2020_086 | IGS50_15730 | 92250..95594 | 2774344 | DUF499-domain-containing-protein-[Synechococcales-cyanobacterium-C42_A2020_086] |
| WP_019503187.1 | HEPN+AAA-ATPase+Alpha+Beta-domain+wHTH+RRM | Type-1_BR.RM.like.systems | <-DDE_3<-PAIREDC-HTH\|\|?->\|\|<-?\|\|PIWI-><-HEPN+DUF499*\|\|?-><-PIN<-?<-METHYLASE | 1127 | Bacteria>Terrabacteria_group>Cyanobacteriota/Melainabacteria_group>Cyanobacteriota>Cyanophyceae | GCF_000332195.1 | Pleurocapsa_sp._PCC_7319 | PLEUR7319_RS0100190 | 37513..40896 | 118161 | Swt1-family-HEPN-domain-containing-protein-[Pleurocapsa-sp.-PCC-7319] |
| WP_150976007.1 | HEPN+AAA-ATPase+Alpha+Beta-domain+wHTH+RRM | Type-1_BR.RM.like.systems | REase->AbrB->PIN->METHYLASE->RHH_6->PIN->HEPN+DUF499*-><-DDE_3 | 1111 | Bacteria>Terrabacteria_group>Cyanobacteriota/Melainabacteria_group>Cyanobacteriota>Cyanophyceae | GCF_008757435.1 | Microcystis_aeruginosa | EZJ55_RS08120 | 1414057..1417392 | 1126 | Swt1-family-HEPN-domain-containing-protein-[Microcystis-aeruginosa] |
| WP_002748938.1 | HEPN+AAA-ATPase+Alpha+Beta-domain+wHTH+RRM | Type-1_BR.RM.like.systems | PAIREDC-HTH->PIN->HEPN+DUF499*-><-Cluster164_18clades<-?\|\|?->?->DUF29-> | 1111 | Bacteria>Terrabacteria_group>Cyanobacteriota/Melainabacteria_group>Cyanobacteriota>Cyanophyceae | GCF_000332585.1 | Microcystis_aeruginosa | C789_RS21375 | 1161..4496 | 1126 | Swt1-family-HEPN-domain-containing-protein-[Microcystis-aeruginosa] |
| WP_015211462.1 | HEPN+AAA-ATPase+Alpha+Beta-domain+wHTH+RRM | Type-1_BR.RM.like.systems | DUF29->AbrB->OLD-ABC->?->METHYLASE->?->HEPN+DUF499*-><-?<-MobL+ZnR+TOPRIM | 1117 | Bacteria>Terrabacteria_group>Cyanobacteriota/Melainabacteria_group>Cyanobacteriota>Cyanophyceae | GCF_000317555.1 | Gloeocapsa_sp._PCC_7428 | GLO7428_RS25580 | 12645..15998 | 1173026 | Swt1-family-HEPN-domain-containing-protein-[Gloeocapsa-sp.-PCC-7428] |
| WP_265900465.1 | HEPN+AAA-ATPase+Alpha+Beta-domain+wHTH+RRM | Type-1_BR.RM.like.systems | REase->REase->?->?-><-sGTP<-HEPN+DUF499*<-PIN<-?<-REase<-REase<-REase | 1128 | Bacteria>Terrabacteria_group>Cyanobacteriota/Melainabacteria_group>Cyanobacteriota>Cyanophyceae | GCF_026240635.1 | Trichothermofontia_sichuanensis | OOK60_RS10505 | 2477245..2480631 | 3045816 | DUF499-domain-containing-protein-[Leptolyngbya-sp.-PKUAC-SCTB231] |
| WP_008200990.1 | HEPN+AAA-ATPase+Alpha+Beta-domain+wHTH+RRM | Type-1_BR.RM.like.systems | REase->METHYLASE->?->ACET->HEPN+DUF499*->?-><-Cluster164_18clades | 1130 | Bacteria>Terrabacteria_group>Cyanobacteriota/Melainabacteria_group>Cyanobacteriota>Cyanophyceae | GCF_000297435.1 | Microcystis_sp._T1-4 | MICAI_RS09635 | 4446..7838 | 1160279 | Swt1-family-HEPN-domain-containing-protein-[Microcystis-sp.-T1-4] |
| WP_272094882.1 | HEPN+AAA-ATPase+Alpha+Beta-domain+wHTH+RRM | Type-1_BR.RM.like.systems | <-HEPN+DUF499*<-PIN<-OLD-ABC<-PIN<-RHH_6<-METHYLASE<-REase | 1065 | Bacteria>Terrabacteria_group>Cyanobacteriota/Melainabacteria_group>Cyanobacteriota>Cyanophyceae | GCF_028330705.1 | Microcystis_aeruginosa | PN479_RS02395 | 1..3194 | 1126 | Swt1-family-HEPN-domain-containing-protein,-partial-[Microcystis-aeruginosa] |
| WP_024971076.1 | HEPN+AAA-ATPase+Alpha+Beta-domain+wHTH+RRM | Type-1_BR.RM.like.systems | REase->METHYLASE->RHH_6->PIN->OLD-ABC->PIN->HEPN+DUF499*-> | 1130 | Bacteria>Terrabacteria_group>Cyanobacteriota/Melainabacteria_group>Cyanobacteriota>Cyanophyceae | GCF_000599945.1 | Microcystis_aeruginosa | MIC7005_RS0124495 | 6350..9742 | 1126 | Swt1-family-HEPN-domain-containing-protein-[Microcystis-aeruginosa] |
| TGG91622.1 | HEPN+AAA-ATPase+Alpha+Beta-domain+wHTH+RRM | Type-1_BR.RM.like.systems | REase+NACHT-><-?<-?<-HEPN+DUF499*<-METHYLASE<-REase<-?<-?<-ParB+HNH | 1109 | Bacteria>Terrabacteria_group>Cyanobacteriota/Melainabacteria_group>Cyanobacteriota>Cyanophyceae | GCA_004768415.1 | Aphanocapsa_feldmannii_277cV | ERJ67_07705 | 17520..20849 | 2507553 | ATP-binding-protein-[Aphanocapsa-feldmannii-277cV] |
| WP_190546091.1 | HEPN+AAA-ATPase+Alpha+Beta-domain+wHTH+RRM | Type-1_BR.RM.like.systems | LRP-HTH->METHYLASE->HEPN+DUF499*->?->?->?->?->Peptidase_M48-><-HNH | 1126 | Bacteria>Terrabacteria_group>Cyanobacteriota/Melainabacteria_group>Cyanobacteriota>Cyanophyceae | - | Microcystis_elabens | H6G65_10800 | 7559..10939 | 44824 | DUF499-domain-containing-protein-[Microcystis-elabens] |
| MBD2550060.1 | HEPN+AAA-ATPase+Alpha+Beta-domain+wHTH+RRM | Type-1_BR.RM.like.systems | LRP-HTH->METHYLASE->HEPN+DUF499*->?->?->?->?->Peptidase_M48-><-HNH | 1126 | Bacteria>Terrabacteria_group>Cyanobacteriota/Melainabacteria_group>Cyanobacteriota>Cyanophyceae | GCA_014698445.1 | Microcystis_elabens_FACHB-917 | H6G65_10800 | 7559..10939 | 2692898 | ATP-binding-protein-[Microcystis-elabens-FACHB-917] |
| MCA2660027.1 | HEPN+AAA-ATPase+Alpha+Beta-domain+wHTH+RRM | Type-1_BR.RM.like.systems | Cluster164_18clades-><-HEPN+DUF499*<-ACET<-?<-METHYLASE | 1130 | Bacteria>Terrabacteria_group>Cyanobacteriota/Melainabacteria_group>Cyanobacteriota>Cyanophyceae | GCA_020245625.1 | Microcystis_sp._M049S2 | IM496_16500 | 8003..11395 | 2771169 | DUF499-domain-containing-protein-[Microcystis-sp.-M049S2] |
| WP_281006617.1 | HEPN+AAA-ATPase+Alpha+Beta-domain+wHTH+RRM | Type-1_BR.RM.like.systems | METHYLASE->?->PIN->REase->REase->REase->HEPN+DUF499*-> | 1118 | Bacteria>Terrabacteria_group>Cyanobacteriota/Melainabacteria_group>Cyanobacteriota>Cyanophyceae | GCF_029910235.1 | Pseudanabaena_galeata | OA858_RS18405 | 4059827..4063183 | 1112103 | Swt1-family-HEPN-domain-containing-protein-[Pseudanabaena-galeata] |
| WP_151696808.1 | HEPN+AAA-ATPase+Alpha+Beta-domain+wHTH+RRM | Type-1_BR.RM.like.systems | <-PRTase<-sGTP<-?\|\|?->?->?-><-HEPN+DUF499*<-PIN<-?<-METHYLASE-p | 1130 | Bacteria>Terrabacteria_group>Cyanobacteriota/Melainabacteria_group>Cyanobacteriota>Cyanophyceae | GCF_008974145.1 | Microcystis_aeruginosa | MiAbW_RS15325 | 39137..42529 | 1126 | Swt1-family-HEPN-domain-containing-protein-[Microcystis-aeruginosa] |
| WP_006527613.1 | HEPN+AAA-ATPase+Alpha+Beta-domain+wHTH+RRM | Type-1_BR.RM.like.systems | <-METHYLASE-p<-HEPN+DUF499*<-?<-ABhydrolase\|\|sGTP-> | 1093 | Bacteria>Terrabacteria_group>Cyanobacteriota/Melainabacteria_group>Cyanobacteriota>Cyanophyceae | GCF_000332035.1 | Gloeocapsa_sp._PCC_73106 | GLO73106_RS03445 | 648..3929 | 102232 | Swt1-family-HEPN-domain-containing-protein-[Gloeocapsa-sp.-PCC-73106] |
| MXW41265.1 | HEPN+AAA-ATPase+Alpha+Beta-domain+wHTH+RRM | Type-1_BR.RM.like.systems | HEPN->METHYLASE->HEPN+DUF499*-><-?<-N6_N4_MTase | 1110 | Bacteria>Terrabacteria_group>Cyanobacteriota/Melainabacteria_group>Cyanobacteriota>Cyanophyceae | GCA_009835965.1 | Synechococcus_sp._SB0668_bin_15 | F4Z75_09040 | 35350..38682 | 2604860 | ATP-binding-protein-[Synechococcus-sp.-SB0668_bin_15] |
| WP_271953564.1 | HEPN+AAA-ATPase+Alpha+Beta-domain+wHTH+RRM | Type-1_BR.RM.like.systems | PIN->HEPN+DUF499*-><-Cluster164_18clades | 1130 | Bacteria>Terrabacteria_group>Cyanobacteriota/Melainabacteria_group>Cyanobacteriota>Cyanophyceae | GCF_028331205.1 | Microcystis_aeruginosa | PN437_RS04350 | 1001..4393 | 1126 | Swt1-family-HEPN-domain-containing-protein-[Microcystis-aeruginosa] |
| MCA6605401.1 | HEPN+AAA-ATPase+Alpha+Beta-domain+wHTH+RRM | Type-1_BR.RM.like.systems | PIN->REase->REase->REase->HEPN+DUF499*-> | 895 | Bacteria>Terrabacteria_group>Cyanobacteriota/Melainabacteria_group>Cyanobacteriota>Cyanophyceae | GCA_020405035.1 | Pseudanabaena_sp._M007S1SP1A06QC | IM533_13245 | 2810..5496 | 2771232 | DUF499-domain-containing-protein-[Pseudanabaena-sp.-M007S1SP1A06QC] |
| MCA6581759.1 | HEPN+AAA-ATPase+Alpha+Beta-domain+wHTH+RRM | Type-1_BR.RM.like.systems | <-HEPN+DUF499*<-REase<-REase<-REase<-PIN | 1118 | Bacteria>Terrabacteria_group>Cyanobacteriota/Melainabacteria_group>Cyanobacteriota>Cyanophyceae | GCA_020404925.1 | Pseudanabaena_sp._M34BS1SP1A06MG | IM536_06930 | 164..3520 | 2771240 | DUF499-domain-containing-protein-[Pseudanabaena-sp.-M34BS1SP1A06MG] |
| NJO76740.1 | HEPN+AAA-ATPase+Alpha+Beta-domain+wHTH+RRM | Type-1_BR.RM.like.systems | METHYLASE-p->REase->HEPN+DUF499*-> | 1125 | Bacteria>Terrabacteria_group>Cyanobacteriota/Melainabacteria_group>Cyanobacteriota>Cyanophyceae | GCA_012033355.1 | Leptolyngbyaceae_cyanobacterium_RM1_406_9 | HC833_25190 | 1227..4604 | 2720434 | ATP-binding-protein-[Leptolyngbyaceae-cyanobacterium-RM1_406_9] |
| WP_280653420.1 | HEPN+AAA-ATPase+Alpha+Beta-domain+wHTH+RRM | Type-1_BR.RM.like.systems | DNA_methylase->?->?-><-?<-HEPN+DUF499* | 1111 | Bacteria>Terrabacteria_group>Cyanobacteriota/Melainabacteria_group>Cyanobacteriota>Cyanophyceae | GCF_029891735.1 | Chrysosporum_bergii | NWP17_RS02960 | 23805..27140 | 105352 | Swt1-family-HEPN-domain-containing-protein-[Chrysosporum-bergii] |
| WP_137669391.1 | HEPN+AAA-ATPase+Alpha+Beta-domain+wHTH+RRM | Type-1_BR.RM.like.systems | <-METHYLASE-p\|\|HEPN+DUF499*-> | 965 | Bacteria>Terrabacteria_group>Cyanobacteriota/Melainabacteria_group>Cyanobacteriota>Cyanophyceae | GCF_005402885.1 | Sphaerospermopsis_reniformis | FFR48_RS26320 | 561..3458 | 531300 | Swt1-family-HEPN-domain-containing-protein-[Sphaerospermopsis-reniformis] |

| Protein-ID | DUF499-architecture | System | Gene-Neighbourhoods | Length | Taxonomy | Assembly-ID | Strain | Locus-Tag | Genomic-coordinates | TaxID | Fasta-define |
|---|---|---|---|---|---|---|---|---|---|---|---|
| GCL39965.1 | HEPN+AAA-ATPase+Alpha+Beta-domain+wHTH+RRM | Type-1_BR.RM.like.systems | <-METHYLASE-p\|\|HEPN+DUF499*-> | 931 | Bacteria>Terrabacteria_group>Cyanobacteriota/Melainabacteria_group>Cyanobacteriota>Cyanophyceae | GCA_005402885.1 | Sphaerospermopsis_reniformis | SR1949_50960 | 663..3458 | 531300 | hypothetical-protein-SR1949_50960-[Sphaerospermopsis-reniformis] |
| MBF2000219.1 | HEPN+AAA-ATPase+Alpha+Beta-domain+wHTH+RRM | Type-1_BR.RM.like.systems | <-HEPN+DUF499*<-REase<-REase | 1113 | Bacteria>Terrabacteria_group>Cyanobacteriota/Melainabacteria_group>Cyanobacteriota>Cyanophyceae | GCA_015272295.1 | Synechococcales_cyanobacterium_M58_A2018_015 | IGS38_05830 | 38711..42052 | 2774348 | DUF499-domain-containing-protein-[Synechococcales-cyanobacterium-M58_A2018_015] |
| TVR05251.1 | HEPN+AAA-ATPase+Alpha+Beta-domain+wHTH+RRM | Type-1_BR.RM.like.systems | METHYLASE-p->?->?->HEPN+DUF499*-> | 1026 | Bacteria>Terrabacteria_group>Cyanobacteriota/Melainabacteria_group>Cyanobacteriota>Cyanophyceae | GCA_007693465.1 | Phormidium_sp._GEM2.Bin31 | EA395_15935 | 1874..4951 | 2480216 | ATP-binding-protein,-partial-[Phormidium-sp.-GEM2.Bin31] |
| TAE60284.1 | HEPN+AAA-ATPase+Alpha+Beta-domain+wHTH+RRM | Type-1_BR.RM.like.systems | <-HEPN+DUF499*\|\|<-METHYLASE | 1113 | Bacteria>Terrabacteria_group>Cyanobacteriota/Melainabacteria_group>Cyanobacteriota>Cyanophyceae | GCA_004292695.1 | Nostocales_cyanobacterium | EAZ87_06610 | 35276..38617 | 2447902 | ATP-binding-protein-[Nostocales-cyanobacterium] |
| TAD76280.1 | HEPN+AAA-ATPase+Alpha+Beta-domain+wHTH+RRM | Type-1_BR.RM.like.systems | METHYLASE-p->HEPN+DUF499*-> | 1114 | Bacteria>Terrabacteria_group>Cyanobacteriota/Melainabacteria_group>Cyanobacteriota>Cyanophyceae | GCA_004292515.1 | Oscillatoriales_cyanobacterium | EA001_12365 | 1816..5160 | 1151 | ATP-binding-protein-[Oscillatoriales-cyanobacterium] |
| MCL6436540.1 | HEPN+AAA-ATPase+Alpha+Beta-domain+wHTH+RRM | Type-1_BR.RM.like.systems | <-HEPN+DUF499*<-METHYLASE-p | 1113 | Bacteria>Terrabacteria_group>Cyanobacteriota/Melainabacteria_group>Cyanobacteriota>Cyanophyceae | GCA_023511615.1 | Leptolyngbyaceae_cyanobacterium_HOT.MB2.61 | K6T90_20505 | 755..4096 | 2855467 | DUF499-domain-containing-protein-[Leptolyngbyaceae-cyanobacterium-HOT.MB2.61] |
| WP_094561768.1 | HEPN+AAA-ATPase+Alpha+Beta-domain+wHTH+RRM | Type-1_BR.RM.like.systems | METHYLASE->HEPN+DUF499*-> | 902 | Bacteria>Terrabacteria_group>Cyanobacteriota/Melainabacteria_group>Cyanobacteriota>Cyanophyceae | GCF_002252665.1 | Synechococcus_sp._8F6 | CJZ78_RS13255 | 2049..4755 | 2025606 | DUF499-domain-containing-protein,-partial-[Synechococcus-sp.-8F6] |
| WP_014276704.1 | HEPN+AAA-ATPase+Alpha+Beta-domain+wHTH+RRM | Type-1_BR.RM.like.systems | METHYLASE->HEPN+DUF499*-> | 1119 | Bacteria>Terrabacteria_group>Cyanobacteriota/Melainabacteria_group>Cyanobacteriota>Cyanophyceae | GCF_009176225.1 | Arthrospira_platensis | H6G70_RS20095 | 3941..7300 | 118562 | Swt1-family-HEPN-domain-containing-protein-[Arthrospira-platensis] |
| NJM67133.1 | HEPN+AAA-ATPase+Alpha+Beta-domain+wHTH+RRM | Type-1_BR.RM.like.systems | METHYLASE->HEPN+DUF499*-> | 1106 | Bacteria>Terrabacteria_group>Cyanobacteriota/Melainabacteria_group>Cyanobacteriota>Cyanophyceae | GCA_012032145.1 | Acaryochloris_sp._RU_4_1 | HC851_16415 | 2448..5768 | 2720442 | ATP-binding-protein-[Acaryochloris-sp.-RU_4_1] |
| HBP53069.1 | HEPN+AAA-ATPase+Alpha+Beta-domain+wHTH+RRM | Type-1_BR.RM.like.systems | METHYLASE->HEPN+DUF499*-> | 1057 | Bacteria>Terrabacteria_group>Cyanobacteriota/Melainabacteria_group>Cyanobacteriota>Cyanophyceae | GCA_003523965.1 | Synechococcus_sp._UBA8638 | DD643_01360 | 4940..8110 | 2055762 | AAA-family-ATPase-[Synechococcus-sp.-UBA8638] |
| WP_250682698.1 | HEPN+AAA-ATPase+Alpha+Beta-domain+wHTH+RRM | Type-1_BR.RM.like.systems | <-HEPN+DUF499*<-METHYLASE<-?<-GATase<-?\|\|?->RNB+RNase-> | 1053 | Bacteria>Terrabacteria_group>Cyanobacteriota/Melainabacteria_group>Cyanobacteriota>Cyanophyceae | GCF_023650955.1 | Thermosynechococcus_sp._HN-54 | NBE99_RS01185 | 246244..249405 | 2933959 | DUF499-domain-containing-protein-[Thermosynechococcus-sp.-HN-54] |
| OWY65325.1 | HEPN+AAA-ATPase+Alpha+Beta-domain+wHTH+RRM | Type-1_BR.RM.like.systems | Phage_integrase-><-Phage_integrase<-HEPN+DUF499*<-DUF29<-REase<-PIN<-AbrB<-TUDOR+SF1-SF2-Helicase+iREase+CTD+REase\|\|Classical-AAA-> | 1119 | Bacteria>Terrabacteria_group>Cyanobacteriota/Melainabacteria_group>Cyanobacteriota>unclassified_Cyanobacteriota | GCA_002213405.1 | cyanobacterium_TDX16 | B7486_42500 | 6012..9371 | 1503470 | AAA+-family-ATPase-[cyanobacterium-TDX16] |
| MBJ7900527.1 | HEPN+AAA-ATPase+Alpha+Beta-domain+wHTH+RRM | Type-1_BR.RM.like.systems | <-Glyco_hydro_57+DUF3536<-?\|\|?->HAD-><-HEPN+DUF499*<-REase<-PIN<-OLD-ABC<-CopG/RHH<-?<-TUDOR+SF1-SF2-Helicase+iREase+CTD+REase | 1111 | Bacteria>Terrabacteria_group>Cyanobacteriota/Melainabacteria_group>Cyanobacteriota>unclassified_Cyanobacteriota | GCA_016471375.1 | Cyanobacteria_bacterium_RI_101 | GC158_11540 | 10083..13418 | 2775578 | DUF499-domain-containing-protein-[Cyanobacteria-bacterium-RI_101] |
| OWY70620.1 | HEPN+AAA-ATPase+Alpha+Beta-domain+wHTH+RRM | Type-1_BR.RM.like.systems | TUDOR+SF1-SF2-Helicase+iREase+CTD+REase->?->METHYLASE->?->DUF488-1->?->HEPN+DUF499*->?-><-?\|\|ZnR+TOPRIM+D5ATPase-> | 1113 | Bacteria>Terrabacteria_group>Cyanobacteriota/Melainabacteria_group>Cyanobacteriota>unclassified_Cyanobacteriota | GCA_002213405.1 | cyanobacterium_TDX16 | B7486_13460 | 127054..130395 | 1503470 | AAA+-family-ATPase-[cyanobacterium-TDX16] |
| MCY4173696.1 | HEPN+AAA-ATPase+Alpha+Beta-domain+wHTH+RRM | Type-1_BR.RM.like.systems | ABC-ATPase+SbcC-><-HEPN+DUF499*<-METHYLASE<-SIR2<-?\|\|?->?-><-TUDOR+SF1-SF2-Helicase+iREase+CTD+REase | 1112 | Bacteria>Terrabacteria_group>Cyanobacteriota/Melainabacteria_group>Cyanobacteriota>unclassified_Cyanobacteriota | GCA_026709085.1 | Cyanobacteria_bacterium_MAG_CAR3_bin_5 | OXF25_06475 | 22273..25611 | 2775591 | MAG:-DUF499-domain-containing-protein-[Cyanobacteria-bacterium-MAG-CAR3_bin_5] |
| PSO73463.1 | HEPN+AAA-ATPase+Alpha+Beta-domain+wHTH+RRM | Type-1_BR.RM.like.systems | <-HEPN+DUF499*<-TUDOR+SF1-SF2-Helicase+iREase+CTD+REase<-?\|\|?-><-BrnA_antitoxin\|\|?-><-DDE_Tnp_ISL3 | 1112 | Bacteria>Terrabacteria_group>Cyanobacteriota/Melainabacteria_group>Cyanobacteriota>unclassified_Cyanobacteriota | GCA_003021535.1 | Cyanobacteria_bacterium_QH_3_48_40 | BRC37_09455 | 1052..4390 | 1919122 | AAA+-family-ATPase-[Cyanobacteria-bacterium-QH_3_48_40] |
| PSO64590.1 | HEPN+AAA-ATPase+Alpha+Beta-domain+wHTH+RRM | Type-1_BR.RM.like.systems | <-HEPN+DUF499*<-TUDOR+SF1-SF2-Helicase+iREase+CTD+REase<-?\|\|?-><-BrnA_antitoxin\|\|?-><-DDE_Tnp_ISL3 | 1112 | Bacteria>Terrabacteria_group>Cyanobacteriota/Melainabacteria_group>Cyanobacteriota>unclassified_Cyanobacteriota | GCA_003022265.1 | Cyanobacteria_bacterium_QH_6_48_35 | BRC38_11110 | 999..4337 | 1919123 | AAA+-family-ATPase-[Cyanobacteria-bacterium-QH_6_48_35] |
| MCY4332863.1 | HEPN+AAA-ATPase+Alpha+Beta-domain+wHTH+RRM | Type-1_BR.RM.like.systems | ABC-ATPase+SbcC-><-HEPN+DUF499*<-METHYLASE<-wHTH+REase | 1113 | Bacteria>Terrabacteria_group>Cyanobacteriota/Melainabacteria_group>Cyanobacteriota>unclassified_Cyanobacteriota | GCA_026710375.1 | Cyanobacteria_bacterium_MAG_CAR1_bin_15 | OXC96_10300 | 26378..29719 | 2775589 | MAG:-DUF499-domain-containing-protein-[Cyanobacteria-bacterium-MAG-CAR1_bin_15] |
| MCY4055150.1 | HEPN+AAA-ATPase+Alpha+Beta-domain+wHTH+RRM | Type-1_BR.RM.like.systems | ABC-ATPase+SbcC-><-HEPN+DUF499*<-METHYLASE<-SIR2 | 1113 | Bacteria>Terrabacteria_group>Cyanobacteriota/Melainabacteria_group>Cyanobacteriota>unclassified_Cyanobacteriota | GCA_026708625.1 | Cyanobacteria_bacterium_MAG_CAR4_bin_6 | OXF67_04645 | 13192..16533 | 2775592 | MAG:-DUF499-domain-containing-protein-[Cyanobacteria-bacterium-MAG-CAR4_bin_6] |
| MBU6346401.1 | HEPN+AAA-ATPase+Alpha+Beta-domain+wHTH+RRM | Type-1_BR.RM.like.systems | <-Peptidase_C14<-?<-REase<-HEPN+DUF499*<-METHYLASE | 1103 | Bacteria>Terrabacteria_group>Cyanobacteriota/Melainabacteria_group>Cyanobacteriota>unclassified_Cyanobacteriota | GCA_018969285.1 | Cyanobacteria_bacterium_REEB494 | KGR70_14045 | 15811..19122 | 2800107 | DUF499-domain-containing-protein-[Cyanobacteria-bacterium-REEB494] |
| PSP10430.1 | HEPN+AAA-ATPase+Alpha+Beta-domain+wHTH+RRM | Type-1_BR.RM.like.systems | DDE_Tnp_ISL3->BrnA_antitoxin-><-?\|\|?->HEPN+DUF499*-> | 1112 | Bacteria>Terrabacteria_group>Cyanobacteriota/Melainabacteria_group>Cyanobacteriota>unclassified_Cyanobacteriota | GCA_003022175.1 | Cyanobacteria_bacterium_SW_10_48_33 | BRC49_10060 | 15348..18686 | 1919136 | AAA+-family-ATPase-[Cyanobacteria-bacterium-SW_10_48_33] |
| MBM5817894.1 | HEPN+AAA-ATPase+Alpha+Beta-domain+wHTH+RRM | Type-1_BR.RM.like.systems | HicB_lk_antitox->HicA_toxin->HEPN+DUF499*-> | 1124 | Bacteria>Terrabacteria_group>Cyanobacteriota/Melainabacteria_group>Cyanobacteriota>unclassified_Cyanobacteriota | GCA_016882305.1 | Cyanobacteria_bacterium_K_Offshore_surface_m2_239 | FJO83_15335 | 1681..5055 | 2775555 | ATP-binding-protein-[Cyanobacteria-bacterium-K_Offshore_surface_m2_239] |
| MCY4235602.1 | HEPN+AAA-ATPase+Alpha+Beta-domain+wHTH+RRM | Type-1_BR.RM.like.systems | SIR2->METHYLASE->HEPN+DUF499*-> | 348 | Bacteria>Terrabacteria_group>Cyanobacteriota/Melainabacteria_group>Cyanobacteriota>unclassified_Cyanobacteriota | GCA_026709585.1 | Cyanobacteria_bacterium_MAG_CAR2_bin_4 | OXE74_04750 | 4969..6012 | 2775590 | MAG:-Swt1-family-HEPN-domain-containing-protein,-partial-[Cyanobacteria-bacterium-MAG-CAR2_bin_4] |
| MCY4234708.1 | HEPN+AAA-ATPase+Alpha+Beta-domain+wHTH+RRM | Type-1_BR.RM.like.systems | HEPN->METHYLASE->HEPN+DUF499*-> | 347 | Bacteria>Terrabacteria_group>Cyanobacteriota/Melainabacteria_group>Cyanobacteriota>unclassified_Cyanobacteriota | GCA_026709585.1 | Cyanobacteria_bacterium_MAG_CAR2_bin_4 | OXE74_00055 | 3576..4616 | 2775590 | MAG:-Swt1-family-HEPN-domain-containing-protein,-partial-[Cyanobacteria-bacterium-MAG-CAR2_bin_4] |
| MBY0405416.1 | HEPN+AAA-ATPase+Alpha+Beta-domain+wHTH+RRM | Type-1_BR.RM.like.systems | METHYLASE-p->HEPN+DUF499*-> | 1058 | Bacteria>Terrabacteria_group>Cyanobacteriota/Melainabacteria_group>Cyanobacteriota>unclassified_Cyanobacteriota | GCA_019751245.1 | Cyanobacteriota_bacterium | K2X66_16060 | 763..3936 | 2847862 | DUF499-domain-containing-protein-[Cyanobacteria-bacterium] |
| PSO86698.1 | HEPN+AAA-ATPase+Alpha+Beta-domain+wHTH+RRM | Type-1_BR.RM.like.systems | REase->?-><-?\|\|HEPN+DUF499*-> | 1112 | Bacteria>Terrabacteria_group>Cyanobacteriota/Melainabacteria_group>Cyanobacteriota>unclassified_Cyanobacteriota | GCA_003021925.1 | Cyanobacteria_bacterium_QH_9_48_43 | BRC41_06380 | 3561..6899 | 1919126 | AAA+-family-ATPase-[Cyanobacteria-bacterium-QH_9_48_43] |
| WP_184025303.1 | HEPN+AAA-ATPase+Alpha+Beta-domain+wHTH+RRM | Type-1_BR.RM.like.systems | Phage_int_SAM_1+Phage_integrase->TUDOR+SF1-SF2-Helicase+iREase+CTD+REase->DUF433->PIN->METHYLASE->?->HEPN+DUF499*-><-?<-TOPRIM+primpol<-OrfB_Zn_ribbon<-XIS-HTH\|\|HTH->Phage_integrase-> | 1097 | Bacteria>Terrabacteria_group>Deinococcota>Deinococci>Deinococcales | GCF_014201885.1 | Deinococcus_budaensis | HNQ09_RS02950 | 29745..33038 | 1665626 | DUF499-domain-containing-protein-[Deinococcus-budaensis] |
| WP_103130569.1 | HEPN+AAA-ATPase+Alpha+Beta-domain+wHTH+RRM | Type-1_BR.RM.like.systems | ABhydrolase->NTP_transferase->?->IclR-HTH-><-TraJ-RHH<-HEPN+DUF499*\|\|?-><-METHYLASE\|\|?->?-><-TUDOR+SF1-SF2-Helicase+iREase+CTD+REase | 1099 | Bacteria>Terrabacteria_group>Deinococcota>Deinococci>Deinococcales | GCF_002897375.1 | Deinococcus_aerius | DAERI_RS16670 | 55118..58417 | 200253 | DUF499-domain-containing-protein-[Deinococcus-aerius] |
| WP_075830034.1 | HEPN+AAA-ATPase+Alpha+Beta-domain+wHTH+RRM | Type-1_BR.RM.like.systems | TUDOR+SF1-SF2-Helicase+iREase+CTD+REase-><-?<-?\|\|METHYLASE->?->HEPN+DUF499*->?->?-><-NERD+PspA | 1100 | Bacteria>Terrabacteria_group>Deinococcota>Deinococci>Deinococcales | GCF_001949125.1 | Deinococcus_marmoris | BOO71_RS00480 | 38514..41816 | 249408 | DUF499-domain-containing-protein-[Deinococcus-marmoris] |
| OLV20266.1 | HEPN+AAA-ATPase+Alpha+Beta-domain+wHTH+RRM | Type-1_BR.RM.like.systems | TUDOR+SF1-SF2-Helicase+iREase+CTD+REase-><-?<-?\|\|METHYLASE->?->HEPN+DUF499*->?-><-NERD+PspA | 1092 | Bacteria>Terrabacteria_group>Deinococcota>Deinococci>Deinococcales | GCA_001949125.1 | Deinococcus_marmoris | BOO71_0000262 | 38538..41816 | 249408 | hypothetical-protein-BOO71_0000262-[Deinococcus-marmoris] |
| WP_189066307.1 | HEPN+AAA-ATPase+Alpha+Beta-domain+wHTH+RRM | Type-1_BR.RM.like.systems | Phage_int_SAM_4+Phage_integrase->?-><-HIN-HTH<-?<-HEPN+DUF499*<-METHYLASE-p<-METHYLASE<-?<-Schlafen+HATPase+HTH<-HKD-DNase<-TUDOR+SF1-SF2-Helicase+iREase+CTD+REase | 1082 | Bacteria>Terrabacteria_group>Deinococcota>Deinococci>Deinococcales | GCF_014648115.1 | Deinococcus_seoulensis | IEY70_RS17415 | 18426..21674 | 1837379 | DUF499-domain-containing-protein-[Deinococcus-seoulensis] |
| WP_239048765.1 | HEPN+AAA-ATPase+Alpha+Beta-domain+wHTH+RRM | Type-1_BR.RM.like.systems | <-HIN-HTH<-?<-?<-McrC-NTD+REase<-DUF499-p<-HEPN+DUF499*<-HEPN-p<-METHYLASE\|\|?-><-Schlafen+HATPase+HTH<-HKD-DNase<-TUDOR+SF1-SF2-Helicase+iREase+CTD+REase | 559 | Bacteria>Terrabacteria_group>Deinococcota>Deinococci>Deinococcales | GCF_022344005.1 | Deinococcus_sp._KNUC1210 | MF271_RS10550 | 1554668..1556347 | 2917691 | DUF499-domain-containing-protein-[Deinococcus-sp.-KNUC1210] |
| WP_107138393.1 | HEPN+AAA-ATPase+Alpha+Beta-domain+wHTH+RRM | Type-1_BR.RM.like.systems | TUDOR+SF1-SF2-Helicase+iREase+CTD+REase->HKD-DNase->Schlafen+HATPase+HTH->?->METHYLASE->HEPN+DUF499*->?->HIN-HTH-><-Phage_int_SAM_4+Phage_integrase | 1079 | Bacteria>Terrabacteria_group>Deinococcota>Deinococci>Deinococcales | GCF_003028415.1 | Deinococcus_arcticus | C8263_RS12115 | 79347..82586 | 2136176 | DUF499-domain-containing-protein-[Deinococcus-arcticus] |
| WP_135230540.1 | HEPN+AAA-ATPase+Alpha+Beta-domain+wHTH+RRM | Type-1_BR.RM.like.systems | Phage_integrase-><-?<-GDResolvase+HIN-HTH<-HEPN+DUF499*<-METHYLASE\|\|?-><-Schlafen+HATPase+HTH<-HKD-DNase<-Phage_integrase | 1098 | Bacteria>Terrabacteria_group>Deinococcota>Deinococci>Deinococcales | GCF_004634215.1 | Deinococcus_fonticola | E5Z01_RS17500 | 5706..9002 | 2528713 | DUF499-domain-containing-protein-[Deinococcus-fonticola] |
| WP_253581497.1 | HEPN+AAA-ATPase+Alpha+Beta-domain+wHTH+RRM | Type-1_BR.RM.like.systems | TUDOR+SF1-SF2-Helicase+iREase+CTD+REase->HKD-DNase->Schlafen+HATPase+HTH-><-?\|\|METHYLASE->HEPN+DUF499*-> | 1096 | Bacteria>Terrabacteria_group>Deinococcota>Deinococci>Deinococcales | GCF_024171495.1 | Deinococcus_sp._HSC-46F16 | L1280_RS07655 | 450556..453846 | 2910968 | DUF499-domain-containing-protein-[Deinococcus-sp.-HSC-46F16] |
| WP_129117970.1 | HEPN+AAA-ATPase+Alpha+Beta-domain+wHTH+RRM | Type-1_BR.RM.like.systems | <-HEPN+DUF499*<-METHYLASE-p<-METHYLASE-p<-?<-Schlafen+HATPase+HTH<-?<-HKD-DNase | 1076 | Bacteria>Terrabacteria_group>Deinococcota>Deinococci>Deinococcales | GCF_030161515.1 | Deinococcus_metallilatus | QQK08_RS18410 | 2112..5342 | 1211322 | DUF499-domain-containing-protein-[Deinococcus-metallilatus] |
| WP_275974710.1 | HEPN+AAA-ATPase+Alpha+Beta-domain+wHTH+RRM | Type-1_BR.RM.like.systems | <-TUDOR+SF1-SF2-Helicase+iREase+CTD+REase\|\|?->METHYLASE->HicA_toxin->HTH_3->HIN-HTH+AT-HTH-HYD2-2+SecB->HEPN+DUF499*-><-DDE_Tnp_1_3\|\|HTH_3->HIN-HTH+AT-HTH-HYD2-2+SecB->?->ABC-ATPase+SbcC-> | 1084 | Bacteria>Terrabacteria_group>Deinococcota>Deinococci>Thermales | GCF_022846355.1 | Thermus_thermophilus | MWM12_RS12790 | 105744..108998 | 274 | Swt1-family-HEPN-domain-containing-protein-[Thermus-thermophilus] |
| WP_038055761.1 | HEPN+AAA-ATPase+Alpha+Beta-domain+wHTH+RRM | Type-1_BR.RM.like.systems | <-Radical-SAM<-tRNA<-rRNA<-rRNA\|\|DDE_Tnp_1_5->tRNA-><-HEPN+DUF499*<-HIN-HTH+SecB<-HTH_3<-HigB/RHH<-METHYLASE<-TUDOR+SF1-SF2-Helicase+iREase+CTD+REase-p<-TUDOR+SF1-SF2-Helicase+iREase+CTD+REase | 1086 | Bacteria>Terrabacteria_group>Deinococcota>Deinococci>Thermales | GCF_000744885.1 | Thermus_amyloliquefaciens | BS74_RS12920 | 400983..404243 | 1449080 | Swt1-family-HEPN-domain-containing-protein-[Thermus-amyloliquefaciens] |
| BCZ90733.1 | HEPN+AAA-ATPase+Alpha+Beta-domain+wHTH+RRM | Type-1_BR.RM.like.systems | <-MULE-TRANSPOSASE\|\|HIN-HTH->HTH_21+rve+rve_3-><-?\|\|PAIREDC-HTH+HIN-HTH->DDE_3-><-HEPN+DUF499*<-HIN-HTH+AT-HTH-HYD2-2+SecB<-HTH_3<-HicB_lk_antitox<-HicA_toxin<-METHYLASE<-DOC+HTH | 1084 | Bacteria>Terrabacteria_group>Deinococcota>Deinococci>Thermales | GCA_019974355.1 | Thermus_thermophilus | TthAA22_25380 | 104954..108208 | 274 | hypothetical-protein-TthAA22_25380-[Thermus-thermophilus] |
| WP_224065568.1 | HEPN+AAA-ATPase+Alpha+Beta-domain+wHTH+RRM | Type-1_BR.RM.like.systems | <-TUDOR+SF1-SF2-Helicase+iREase+CTD+REase\|\|DOC+HTH->METHYLASE->HicA_toxin->HTH_3->HIN-HTH+AT-HTH-HYD2-2+SecB->HEPN+DUF499*->DDE_Tnp_1+DDE_Tnp_1_2->?-><-HIN-HTH\|\|?-><-?\|\|DDE_Tnp_1_3-> | 1084 | Bacteria>Terrabacteria_group>Deinococcota>Deinococci>Thermales | GCF_019974375.1 | Thermus_thermophilus | LDD80_RS12530 | 90011..93265 | 274 | Swt1-family-HEPN-domain-containing-protein-[Thermus-thermophilus] |
| WP_275974790.1 | HEPN+AAA-ATPase+Alpha+Beta-domain+wHTH+RRM | Type-1_BR.RM.like.systems | DUF4096+DDE_Tnp_1+DDE_Tnp->PAIREDC-HTH+HIN-HTH->DDE_3-><-HicA_toxin<-HicB<-DDE_Tnp_1<-HEPN+DUF499*<-HIN-HTH+AT-HTH-HYD2-2+SecB<-HTH_3<-HigB/RHH<-HicB_lk_antitox<-METHYLASE | 1084 | Bacteria>Terrabacteria_group>Deinococcota>Deinococci>Thermales | GCF_022846315.1 | Thermus_thermophilus | SNM_RS12085 | 50871..54125 | 274 | Swt1-family-HEPN-domain-containing-protein-[Thermus-thermophilus] |
| BDG27741.1 | HEPN+AAA-ATPase+Alpha+Beta-domain+wHTH+RRM | Type-1_BR.RM.like.systems | DDE_Tnp_1+DDE_Tnp_1_2->PAIREDC-HTH+HIN-HTH->DDE_3-><-HicA_toxin<-HicB<-DDE_Tnp_1<-HEPN+DUF499*<-HIN-HTH+AT-HTH-HYD2-2+SecB<-HTH_3<-HigB/RHH<-HicB_lk_antitox<-METHYLASE | 1084 | Bacteria>Terrabacteria_group>Deinococcota>Deinococci>Thermales | GCA_022846315.1 | Thermus_thermophilus | TthSNM66_23770 | 50871..54125 | 274 | hypothetical-protein-TthSNM66_23770-[Thermus-thermophilus] |
| WP_267461447.1 | HEPN+AAA-ATPase+Alpha+Beta-domain+wHTH+RRM | Type-1_BR.RM.like.systems | HIN-HTH-><-?\|\|PAIREDC-HTH+HIN-HTH->DDE_3-><-HEPN+DUF499*<-HIN-HTH+AT-HTH-HYD2-2+SecB<-HTH_3<-HicA_toxin<-METHYLASE<-DOC+HTH\|\|TUDOR+SF1-SF2-Helicase+iREase+CTD+REase-> | 1084 | Bacteria>Terrabacteria_group>Deinococcota>Deinococci>Thermales | GCF_019974355.1 | Thermus_thermophilus | LDD10_RS13400 | 104954..108208 | 274 | Swt1-family-HEPN-domain-containing-protein-[Thermus-thermophilus] |
| BDG22724.1 | HEPN+AAA-ATPase+Alpha+Beta-domain+wHTH+RRM | Type-1_BR.RM.like.systems | METHYLASE->HicA_toxin->HicB_lk_antitox->HTH_3->HIN-HTH+AT-HTH-HYD2-2+SecB->HEPN+DUF499*-><-DDE_Tnp_1_3\|\|HTH_3->HIN-HTH+AT-HTH-HYD2-2+SecB->?->ABC-ATPase+SbcC-> | 1084 | Bacteria>Terrabacteria_group>Deinococcota>Deinococci>Thermales | GCA_022846355.1 | Thermus_thermophilus | TthSNM17_23860 | 105744..108998 | 274 | hypothetical-protein-TthSNM17_23860-[Thermus-thermophilus] |
| WP_275973822.1 | HEPN+AAA-ATPase+Alpha+Beta-domain+wHTH+RRM | Type-1_BR.RM.like.systems | <-HTH_3<-?\|\|DDE_Tnp_1_3->DDE_Tnp_1->DDE_Tnp_1_3->DDE_Tnp_1-><-HEPN+DUF499*<-HIN-HTH+AT-HTH-HYD2-2+SecB<-HTH_3<-HigB/RHH<-HicB_lk_antitox<-METHYLASE | 1084 | Bacteria>Terrabacteria_group>Deinococcota>Deinococci>Thermales | GCF_022846275.1 | Thermus_thermophilus | MWM06_RS12915 | 20817..24071 | 274 | Swt1-family-HEPN-domain-containing-protein-[Thermus-thermophilus] |
| WP_014632301.1 | HEPN+AAA-ATPase+Alpha+Beta-domain+wHTH+RRM | Type-1_BR.RM.like.systems | <-SF1-SF2-Helicase+REase<-Exonuc_V_gamma+REase<-?<-?<-?<-?<-HEPN+DUF499*<-HIN-HTH+AT-HTH-HYD2-2+SecB<-HTH_3<-HigB/RHH<-HicB_lk_antitox<-METHYLASE<-DOC+HTH | 1084 | Bacteria>Terrabacteria_group>Deinococcota>Deinococci>Thermales | GCF_000258245.1 | Thermus_thermophilus | TTJL18_RS12200 | 95863..99117 | 274 | Swt1-family-HEPN-domain-containing-protein-[Thermus-thermophilus] |
| WP_224065603.1 | HEPN+AAA-ATPase+Alpha+Beta-domain+wHTH+RRM | Type-1_BR.RM.like.systems | PP_kinase+HKD-DNase-><-HEPN+DUF499*<-HIN-HTH+AT-HTH-HYD2-2+SecB<-HTH_3<-HicB_lk_antitox<-METHYLASE<-?\|\|TUDOR+SF1-SF2-Helicase+iREase+CTD+REase-p-> | 1086 | Bacteria>Terrabacteria_group>Deinococcota>Deinococci>Thermales | GCF_019974375.1 | Thermus_thermophilus | LDD80_RS12010 | 2426..5686 | 274 | Swt1-family-HEPN-domain-containing-protein-[Thermus-thermophilus] |
| WP_014511084.1 | HEPN+AAA-ATPase+Alpha+Beta-domain+wHTH+RRM | Type-1_BR.RM.like.systems | METHYLASE->HicB_lk_antitox->?->HigB/RHH->HTH_3->HIN-HTH+AT-HTH-HYD2-2+SecB->HEPN+DUF499*->?->?->?->?->Exonuc_V_gamma+REase->SF1-SF2-Helicase+REase-> | 1084 | Bacteria>Terrabacteria_group>Deinococcota>Deinococci>Thermales | GCF_000214845.1 | Thermus_thermophilus | THTHE16_RS13550 | 92847..96101 | 274 | Swt1-family-HEPN-domain-containing-protein-[Thermus-thermophilus] |
| BDG20195.1 | HEPN+AAA-ATPase+Alpha+Beta-domain+wHTH+RRM | Type-1_BR.RM.like.systems | DDE_Tnp_1_3->DDE_Tnp_1->?->DDE_Tnp_1_3->DDE_Tnp_1-><-HEPN+DUF499*<-HIN-HTH+AT-HTH-HYD2-2+SecB<-HTH_3<-HigB/RHH<-HicB_lk_antitox<-METHYLASE | 1084 | Bacteria>Terrabacteria_group>Deinococcota>Deinococci>Thermales | GCA_022846275.1 | Thermus_thermophilus | TthSNM11_23980 | 20817..24071 | 274 | hypothetical-protein-TthSNM11_23980-[Thermus-thermophilus] |
| AEG34458.1 | HEPN+AAA-ATPase+Alpha+Beta-domain+wHTH+RRM | Type-1_BR.RM.like.systems | METHYLASE->HicB_lk_antitox->?->HigB/RHH->HTH_3->HIN-HTH+AT-HTH-HYD2-2+SecB->HEPN+DUF499*->?->?->?->?->Exonuc_V_gamma+REase->SF1-SF2-Helicase+REase-> | 1084 | Bacteria>Terrabacteria_group>Deinococcota>Deinococci>Thermales | GCA_000214845.1 | Thermus_thermophilus_SG0.5JP17-16 | Tthe16_2072 | 92847..96101 | 762633 | ATPase-(AAA+-superfamily)-like-protein-[Thermus-thermophilus-SG0.5JP17-16] |
| WP_269077413.1 | HEPN+AAA-ATPase+Alpha+Beta-domain+wHTH+RRM | Type-1_BR.RM.like.systems | METHYLASE-p->HicB_lk_antitox->HigB/RHH->HTH_3->HIN-HTH+AT-HTH-HYD2-2+SecB->?->HEPN+DUF499*->DUF499->DUF499->\|\|DDE_Tnp_1_3->RADICAL-SAM-> | 335 | Bacteria>Terrabacteria_group>Deinococcota>Deinococci>Thermales | GCF_000744175.1 | Thermus_tengchongensis | BS82_RS15525 | 75605..76612 | 1214928 | Swt1-family-HEPN-domain-containing-protein-[Thermus-tengchongensis] |
| MBF6594441.1 | HEPN+AAA-ATPase+Alpha+Beta-domain+wHTH+RRM | Type-1_BR.RM.like.systems | TUDOR+SF1-SF2-Helicase+iREase+CTD+REase->Schlafen+HATPase+HTH->METHYLASE-><-PIN<-MazE_antitoxin\|\|HEPN+DUF499*-><-HTH_Tnp_4\|\|<-?<-?\|\|METHYLASE-p-> | 1117 | Bacteria>Terrabacteria_group>Deinococcota>Deinococci>Thermales | GCA_015478585.1 | Thermaceae_bacterium | IVW51_08335 | 21188..24541 | 2614257 | DUF499-domain-containing-protein-[Thermaceae-bacterium] |
| WP_202819508.1 | HEPN+AAA-ATPase+Alpha+Beta-domain+wHTH+RRM | Type-1_BR.RM.like.systems | <-ABC-ATPase+SbcC<-?\|\|?-><-DDE_Tnp_1<-HEPN+DUF499*<-HIN-HTH+AT-HTH-HYD2-2+SecB<-HTH_3<-HicB_lk_antitox<-HicA_toxin<-METHYLASE<-DOC+HTH | 1083 | Bacteria>Terrabacteria_group>Deinococcota>Deinococci>Thermales | GCF_016865275.1 | Thermus_thermophilus | TthHB5002_b22180 | 17958..21209 | 274 | Swt1-family-HEPN-domain-containing-protein-[Thermus-thermophilus] |
| BCP99115.1 | HEPN+AAA-ATPase+Alpha+Beta-domain+wHTH+RRM | Type-1_BR.RM.like.systems | <-ABC-ATPase+SbcC<-?\|\|?-><-DDE_Tnp_1<-HEPN+DUF499*<-HIN-HTH+AT-HTH-HYD2-2+SecB<-HTH_3<-HicB_lk_antitox<-HicA_toxin<-METHYLASE<-DOC+HTH | 1083 | Bacteria>Terrabacteria_group>Deinococcota>Deinococci>Thermales | GCA_016865275.1 | Thermus_thermophilus | TthHB5002_b22180 | 17958..21209 | 274 | hypothetical-protein-TthHB5002_b22180-[Thermus-thermophilus] |
| WP_027888723.1 | HEPN+AAA-ATPase+Alpha+Beta-domain+wHTH+RRM | Type-1_BR.RM.like.systems | <-PAIREDC-HTH<-BrxL<-?<-?<-?<-ORC-CDC6+REase<-HEPN+DUF499*<-METHYLASE<-TUDOR+SF1-SF2-Helicase+iREase+CTD+REase\|\|DDE_Tnp_IS1-><-?\|\|?->TPR+TPR-> | 1105 | Bacteria>Terrabacteria_group>Deinococcota>Deinococci>Thermales | GCF_003568585.1 | Meiothermus_taiwanensis | MTAI_RS14270 | 92295..95612 | 172827 | Swt1-family-HEPN-domain-containing-protein-[Meiothermus-taiwanensis] |
| WP_252829575.1 | HEPN+AAA-ATPase+Alpha+Beta-domain+wHTH+RRM | Type-1_BR.RM.like.systems | <-HTH+DSBH\|\|<-DUF499<-HEPN+DUF499*<-HIN-HTH+AT-HTH-HYD2-2+SecB<-HTH_3<-HigB/RHH<-HicB_lk_antitox<-METHYLASE-p<-METHYLASE | 335 | Bacteria>Terrabacteria_group>Deinococcota>Deinococci>Thermales | GCF_000744155.1 | Thermus_scotoductus | N686_RS14355 | 48271..49278 | 37636 | Swt1-family-HEPN-domain-containing-protein-[Thermus-scotoductus] |
| WP_243092374.1 | HEPN+AAA-ATPase+Alpha+Beta-domain+wHTH+RRM | Type-1_BR.RM.like.systems | Hpr-Kinase-C-><-HEPN+DUF499*<-HIN-HTH+SecB<-HTH_3<-HigB/RHH<-METHYLASE<-DOC+HTH<-TUDOR+SF1-SF2-Helicase+iREase+CTD+REase | 1092 | Bacteria>Terrabacteria_group>Deinococcota>Deinococci>Thermales | GCF_022760925.1 | Thermus_hydrothermalis | L0C60_RS00500 | 88186..91464 | 2908148 | DUF499-domain-containing-protein-[Thermus-hydrothermalis] |
| WP_071678256.1 | HEPN+AAA-ATPase+Alpha+Beta-domain+wHTH+RRM | Type-1_BR.RM.like.systems | TUDOR+SF1-SF2-Helicase+iREase+CTD+REase->DOC+HTH->METHYLASE->HigB/RHH->HTH_3->HIN-HTH+SecB->HEPN+DUF499*-><-Hpr-Kinase-C | 1092 | Bacteria>Terrabacteria_group>Deinococcota>Deinococci>Thermales | GCF_022846375.1 | Thermus_brockianus | MWM05_RS11465 | 93356..96634 | 56956 | DUF499-domain-containing-protein-[Thermus-brockianus] |
| WP_155983233.1 | HEPN+AAA-ATPase+Alpha+Beta-domain+wHTH+RRM | Type-1_BR.RM.like.systems | <-DUF499\|\|MULE-TRANSPOSASE-><-HEPN+DUF499*<-HicB_lk_antitox<-HicA_toxin<-METHYLASE<-HKD-DNase+SF1-SF2-Helicase-p\|\|?->PIN-> | 341 | Bacteria>Terrabacteria_group>Deinococcota>Deinococci>Thermales | GCF_000236585.1 | Thermus_sp._CCB_US3_UF1 | TCCBUS3UF1_RS00635 | 146394..147419 | 1111069 | Swt1-family-HEPN-domain-containing-protein-[Thermus-sp.-CCB_US3_UF1] |
| GIW33264.1 | HEPN+AAA-ATPase+Alpha+Beta-domain+wHTH+RRM | Type-1_BR.RM.like.systems | Pkinase->DUF4388-><-?<-?\|\|?-><-HEPN+DUF499*<-HEPN<-PolB<-METHYLASE<-ORC-CDC6+REase<-TUDOR+SF1-SF2-Helicase+iREase+CTD+REase | 1104 | Bacteria>Terrabacteria_group>Deinococcota>Deinococci>Thermales | GCA_026004115.1 | Meiothermus_sp. | KatS3mg072_0597 | 655659..658973 | 1955249 | MAG:-hypothetical-protein-KatS3mg072_0597-[Meiothermus-sp.] |
| WP_244348266.1 | HEPN+AAA-ATPase+Alpha+Beta-domain+wHTH+RRM | Type-1_BR.RM.like.systems | DOC+HTH->METHYLASE->HicB_lk_antitox->HigB/RHH->HTH_3->HIN-HTH+AT-HTH-HYD2-2+SecB->HEPN+DUF499*->HNH->Peptidase_M48-> | 1084 | Bacteria>Terrabacteria_group>Deinococcota>Deinococci>Thermales | GCF_022846295.1 | Thermus_thermophilus | MWM13_RS12110 | 24920..28174 | 274 | Swt1-family-HEPN-domain-containing-protein-[Thermus-thermophilus] |
| WP_243028145.1 | HEPN+AAA-ATPase+Alpha+Beta-domain+wHTH+RRM | Type-1_BR.RM.like.systems | RNB+RNase-><-?\|\|?->?->tRNA-><-HEPN+DUF499*<-HicB_lk_antitox<-METHYLASE<-TUDOR+SF1-SF2-Helicase+iREase+CTD+REase\|\|Radical-SAM+TRAM-> | 1076 | Bacteria>Terrabacteria_group>Deinococcota>Deinococci>Thermales | GCF_022760855.1 | Thermus_albus | L0D18_RS06900 | 53799..57029 | 2908146 | DUF499-domain-containing-protein-[Thermus-albus] |
| WP_015716390.1 | HEPN+AAA-ATPase+Alpha+Beta-domain+wHTH+RRM | Type-1_BR.RM.like.systems | <-Phage_integrase\|\|?->?-><-?\|\|OLD-ABC+TOPRIM-><-HEPN+DUF499*<-METHYLASE<-MACRODOMAIN<-?<-TUDOR+SF1-SF2-Helicase+iREase+CTD+REase<-?<-ArsR-HTH | 1116 | Bacteria>Terrabacteria_group>Deinococcota>Deinococci>Thermales | GCF_000187005.1 | Thermus_scotoductus | TSC_RS02330 | 419073..422423 | 37636 | Swt1-family-HEPN-domain-containing-protein-[Thermus-scotoductus] |
| WP_013458840.1 | HEPN+AAA-ATPase+Alpha+Beta-domain+wHTH+RRM | Type-1_BR.RM.like.systems | TUDOR+SF1-SF2-Helicase+iREase+CTD+REase->HicA_toxin->HicB_lk_antitox->PIN->METHYLASE->HEPN+DUF499*->?-><-?<-?<-?\|\|?->METHYLASE-p-> | 1120 | Bacteria>Terrabacteria_group>Deinococcota>Deinococci>Thermales | GCF_000183745.1 | Oceanithermus_profundus | OCEPR_RS11190 | 2275579..2278941 | 187137 | DUF499-domain-containing-protein-[Oceanithermus-profundus] |
| AEV15179.1 | HEPN+AAA-ATPase+Alpha+Beta-domain+wHTH+RRM | Type-1_BR.RM.like.systems | <-DUF499-p\|\|MULE-TRANSPOSASE-><-HEPN+DUF499*<-HicB_lk_antitox<-HicA_toxin<-METHYLASE<-HKD-DNase+SF1-SF2-Helicase-p | 355 | Bacteria>Terrabacteria_group>Deinococcota>Deinococci>Thermales | GCA_000236585.1 | Thermus_sp._CCB_US3_UF1 | TCCBUS3UF1_1290 | 146394..147461 | 1111069 | hypothetical-protein-TCCBUS3UF1_1290-[Thermus-sp.-CCB_US3_UF1] |
| WP_126209517.1 | HEPN+AAA-ATPase+Alpha+Beta-domain+wHTH+RRM | Type-1_BR.RM.like.systems | METHYLASE->HicA_toxin->HicB_lk_antitox->HIN-HTH+AT-HTH-HYD2-2+SecB->HEPN+DUF499*->DDE_Tnp_1_3-><-MULE-TRANSPOSASE | 1085 | Bacteria>Terrabacteria_group>Deinococcota>Deinococci>Thermales | GCF_003959225.1 | Thermus_scotoductus | CSW37_RS13630 | 3217..6474 | 37636 | DUF499-domain-containing-protein-[Thermus-scotoductus] |
| WP_018110668.1 | HEPN+AAA-ATPase+Alpha+Beta-domain+wHTH+RRM | Type-1_BR.RM.like.systems | <-MULE-TRANSPOSASE\|\|METHYLASE->HigB/RHH->HTH_3->HIN-HTH+SecB->HEPN+DUF499*->?-><-?<-XIS-HTH\|\|Primpol->TOPRIM+primpol-> | 1095 | Bacteria>Terrabacteria_group>Deinococcota>Deinococci>Thermales | GCF_000376265.1 | Thermus_igniterrae | B128_RS0102275 | 11501..14788 | 88189 | Swt1-family-HEPN-domain-containing-protein-[Thermus-igniterrae] |
| WP_018461866.1 | HEPN+AAA-ATPase+Alpha+Beta-domain+wHTH+RRM | Type-1_BR.RM.like.systems | ParB->TUDOR+SF1-SF2-Helicase+iREase+CTD+REase->PIN->METHYLASE->HEPN+DUF499*->SF1-SF2-Helicase->?-><-?<-Peptidase_C26 | 1097 | Bacteria>Terrabacteria_group>Deinococcota>Deinococci>Thermales | GCF_000373145.1 | Thermus_oshimai | B043_RS0109800 | 87375..90668 | 56957 | Swt1-family-HEPN-domain-containing-protein-[Thermus-oshimai] |
| WP_130842394.1 | HEPN+AAA-ATPase+Alpha+Beta-domain+wHTH+RRM | Type-1_BR.RM.like.systems | TUDOR+SF1-SF2-Helicase+iREase+CTD+REase->DOC+HTH->METHYLASE->HigB/RHH->HTH_3->HIN-HTH+SecB->HEPN+DUF499*-> | 1088 | Bacteria>Terrabacteria_group>Deinococcota>Deinococci>Thermales | GCF_004307015.1 | Thermus_thermamylovorans | ETP66_RS09490 | 48126..51392 | 2509362 | Swt1-family-HEPN-domain-containing-protein-[Thermus-thermamylovorans] |
| WP_015065389.1 | HEPN+AAA-ATPase+Alpha+Beta-domain+wHTH+RRM | Type-1_BR.RM.like.systems | ParB->TUDOR+SF1-SF2-Helicase+iREase+CTD+REase->PIN->DDE_Tnp_1->HEPN+DUF499*->SF1-SF2-Helicase->IclR-HTH-> | 1099 | Bacteria>Terrabacteria_group>Deinococcota>Deinococci>Thermales | GCF_000309885.1 | Thermus_oshimai | THEOS_RS11985 | 210648..213947 | 56957 | Swt1-family-HEPN-domain-containing-protein-[Thermus-oshimai] |
| WP_119356451.1 | HEPN+AAA-ATPase+Alpha+Beta-domain+wHTH+RRM | Type-1_BR.RM.like.systems | PIN->TUDOR+SF1-SF2-Helicase+iREase+CTD+REase->XIS-HTH->METHYLASE-><-PIN<-MazE_antitoxin\|\|HEPN+DUF499*-> | 1110 | Bacteria>Terrabacteria_group>Deinococcota>Deinococci>Thermales | GCF_007990995.1 | Meiothermus_granaticius | MGR01S_RS15620 | 8907..12239 | 863370 | Swt1-family-HEPN-domain-containing-protein-[Meiothermus-granaticius] |
| WP_119341253.1 | HEPN+AAA-ATPase+Alpha+Beta-domain+wHTH+RRM | Type-1_BR.RM.like.systems | Pkinase->DUF4388-><-?<-?\|\|?-><-RHH<-HEPN+DUF499*<-METHYLASE<-TUDOR+SF1-SF2-Helicase+iREase+CTD+REase\|\|?-><-?<-PfkB | 1099 | Bacteria>Terrabacteria_group>Deinococcota>Deinococci>Thermales | GCF_007990975.1 | Meiothermus_hypogaeus | MHY01S_RS03040 | 15508..18807 | 884155 | Swt1-family-HEPN-domain-containing-protein-[Meiothermus-hypogaeus] |
| WP_278359523.1 | HEPN+AAA-ATPase+Alpha+Beta-domain+wHTH+RRM | Type-1_BR.RM.like.systems | <-HEPN+DUF499*<-HIN-HTH+AT-HTH-HYD2-2+SecB<-HTH_3<-HicB_lk_antitox<-HicA_toxin<-METHYLASE<-DOC+HTH | 601 | Bacteria>Terrabacteria_group>Deinococcota>Deinococci>Thermales | GCF_002378965.1 | Thermus_thermophilus | B3N72_RS06335 | 2095..3898 | 274 | Swt1-family-HEPN-domain-containing-protein,-partial-[Thermus-thermophilus] |
| WP_014677588.1 | HEPN+AAA-ATPase+Alpha+Beta-domain+wHTH+RRM | Type-1_BR.RM.like.systems | <-HEPN+DUF499*<-HIN-HTH+AT-HTH-HYD2-2+SecB<-HTH_3<-HicB_lk_antitox<-HicA_toxin<-METHYLASE<-DOC+HTH | 1084 | Bacteria>Terrabacteria_group>Deinococcota>Deinococci>Thermales | - | Thermus | DCM56_RS11460 | 979..4233 | 270 | Swt1-family-HEPN-domain-containing-protein-[Thermus-thermophilus] |
| WP_267262111.1 | HEPN+AAA-ATPase+Alpha+Beta-domain+wHTH+RRM | Type-1_BR.RM.like.systems | <-HEPN<-DDE_3<-?<-?<-HEPN+DUF499*<-METHYLASE<-OLD-ABC+TOPRIM<-TUDOR+SF1-SF2-Helicase+iREase+CTD+REase<-?<-ArsR-HTH | 1116 | Bacteria>Terrabacteria_group>Deinococcota>Deinococci>Thermales | GCF_026427635.1 | Thermus_sp._PS18 | KQ693_RS05000 | 868393..871743 | 2849039 | Swt1-family-HEPN-domain-containing-protein-[Thermus-sp.-PS18] |
| WP_119360281.1 | HEPN+AAA-ATPase+Alpha+Beta-domain+wHTH+RRM | Type-1_BR.RM.like.systems | ACET->?-><-?<-ABhydrolase<-HEPN+DUF499*<-?<-HTH_3<-METHYLASE<-?<-TUDOR+SF1-SF2-Helicase+iREase+CTD+REase | 1091 | Bacteria>Terrabacteria_group>Deinococcota>Deinococci>Thermales | GCF_003574085.1 | Meiothermus_luteus | Mlute_RS08485 | 5782..9057 | 2026184 | Swt1-family-HEPN-domain-containing-protein-[Meiothermus-luteus] |
| WP_119313343.1 | HEPN+AAA-ATPase+Alpha+Beta-domain+wHTH+RRM | Type-1_BR.RM.like.systems | TUDOR+SF1-SF2-Helicase+iREase+CTD+REase->METHYLASE->HEPN+DUF499*->ZnR+TOPRIM+primpol+HTH-> | 1117 | Bacteria>Terrabacteria_group>Deinococcota>Deinococci>Thermales | GCF_003574345.1 | Calidithermus_terrae | Mterra_RS00325 | 80239..83592 | 1408545 | Swt1-family-HEPN-domain-containing-protein-[Calidithermus-terrae] |
| WP_119361752.1 | HEPN+AAA-ATPase+Alpha+Beta-domain+wHTH+RRM | Type-1_BR.RM.like.systems | TUDOR+SF1-SF2-Helicase+iREase+CTD+REase->METHYLASE->HEPN+DUF499*->ORC-CDC6+REase-> | 1108 | Bacteria>Terrabacteria_group>Deinococcota>Deinococci>Thermales | GCF_003574045.1 | Meiothermus_taiwanensis | Mcate_RS11630 | 6951..10277 | 172827 | Swt1-family-HEPN-domain-containing-protein-[Meiothermus-taiwanensis] |
| WP_234507999.1 | HEPN+AAA-ATPase+Alpha+Beta-domain+wHTH+RRM | Type-1_BR.RM.like.systems | DOC+HTH->METHYLASE->HigB/RHH->HTH_3->HIN-HTH+SecB->HEPN+DUF499*-><-Hpr-Kinase-C | 1090 | Bacteria>Terrabacteria_group>Deinococcota>Deinococci>Thermales | GCF_021462505.1 | Thermus_brockianus | LXN00_RS10430 | 5041..8313 | 56956 | DUF499-domain-containing-protein-[Thermus-brockianus] |
| WP_135343460.1 | HEPN+AAA-ATPase+Alpha+Beta-domain+wHTH+RRM | Type-1_BR.RM.like.systems | <-HEPN+DUF499*<-METHYLASE<-PIN<-TUDOR+SF1-SF2-Helicase+iREase+CTD+REase<-ParB | 1099 | Bacteria>Terrabacteria_group>Deinococcota>Deinococci>Thermales | GCF_004685085.1 | Thermus_tengchongensis | E0489_RS07460 | 252..3551 | 1214928 | Swt1-family-HEPN-domain-containing-protein-[Thermus-tengchongensis] |
| GIW32506.1 | HEPN+AAA-ATPase+Alpha+Beta-domain+wHTH+RRM | Type-1_BR.RM.like.systems | MerR-HTH->?->TUDOR+SF1-SF2-Helicase+iREase+CTD+REase->METHYLASE->HEPN+DUF499*-> | 1105 | Bacteria>Terrabacteria_group>Deinococcota>Deinococci>Thermales | GCA_026004095.1 | Meiothermus_sp. | KatS3mg071_2680 | 19805..23122 | 1955249 | MAG:-hypothetical-protein-KatS3mg071_2680-[Meiothermus-sp.] |
| AEG34698.1 | HEPN+AAA-ATPase+Alpha+Beta-domain+wHTH+RRM | Type-1_BR.RM.like.systems | DDE_Tnp_1->HTH_3->HIN-HTH+SecB->HEPN+DUF499*->HTH+DSBH->?->?->?->?->ABC-ATPase+SbcC-> | 1084 | Bacteria>Terrabacteria_group>Deinococcota>Deinococci>Thermales | GCA_000214845.1 | Thermus_thermophilus_SG0.5JP17-16 | Tthe16_2339 | 352760..356014 | 762633 | ATPase-(AAA+-superfamily)-like-protein-[Thermus-thermophilus-SG0.5JP17-16] |
| WP_223299558.1 | HEPN+AAA-ATPase+Alpha+Beta-domain+wHTH+RRM | Type-1_BR.RM.like.systems | <-HEPN+DUF499*<-METHYLASE<-TUDOR+SF1-SF2-Helicase+iREase+CTD+REase-p | 451 | Bacteria>Terrabacteria_group>Deinococcota>Deinococci>Thermales | GCF_001623175.1 | Meiothermus_taiwanensis | A3962_RS01225 | 1..1353 | 172827 | Swt1-family-HEPN-domain-containing-protein,-partial-[Meiothermus-taiwanensis] |
| WP_014511304.1 | HEPN+AAA-ATPase+Alpha+Beta-domain+wHTH+RRM | Type-1_BR.RM.like.systems | DDE_Tnp_1->HTH_3->HIN-HTH+SecB->HEPN+DUF499*->\|\|HTH+DSBH->DSBH-> | 1084 | Bacteria>Terrabacteria_group>Deinococcota>Deinococci>Thermales | GCF_000214845.1 | Thermus_thermophilus | THTHE16_RS11840 | 352760..356014 | 274 | Swt1-family-HEPN-domain-containing-protein-[Thermus-thermophilus] |
| WP_253730420.1 | HEPN+AAA-ATPase+Alpha+Beta-domain+wHTH+RRM | Type-1_BR.RM.like.systems | METHYLASE-p->HigB/RHH->HTH_3->HIN-HTH+SecB->HEPN+DUF499*-> | 1004 | Bacteria>Terrabacteria_group>Deinococcota>Deinococci>Thermales | GCF_003959405.1 | Thermus_scotoductus | CSW25_RS02145 | 3152..6166 | 37636 | Swt1-family-HEPN-domain-containing-protein-[Thermus-scotoductus] |
| WP_253724412.1 | HEPN+AAA-ATPase+Alpha+Beta-domain+wHTH+RRM | Type-1_BR.RM.like.systems | <-HEPN+DUF499*<-HIN-HTH+SecB<-HTH_3<-HigB/RHH<-METHYLASE | 424 | Bacteria>Terrabacteria_group>Deinococcota>Deinococci>Thermales | GCF_003959115.1 | Thermus_scotoductus | CSW43_RS02865 | 82..1356 | 37636 | Swt1-family-HEPN-domain-containing-protein-[Thermus-scotoductus] |
| WP_253665784.1 | HEPN+AAA-ATPase+Alpha+Beta-domain+wHTH+RRM | Type-1_BR.RM.like.systems | <-HEPN+DUF499*<-HIN-HTH+SecB<-HTH_3<-HigB/RHH<-METHYLASE | 429 | Bacteria>Terrabacteria_group>Deinococcota>Deinococci>Thermales | GCF_003959565.1 | Thermus_scotoductus | CSW39_RS03125 | 1..1288 | 37636 | Swt1-family-HEPN-domain-containing-protein,-partial-[Thermus-scotoductus] |
| WP_153185240.1 | HEPN+AAA-ATPase+Alpha+Beta-domain+wHTH+RRM | Type-1_BR.RM.like.systems | <-HEPN+DUF499*<-HIN-HTH+SecB<-HTH_3<-HigB/RHH<-METHYLASE | 1011 | Bacteria>Terrabacteria_group>Deinococcota>Deinococci>Thermales | GCF_003959445.1 | Thermus_scotoductus | CSW20_RS00300 | 1..3032 | 37636 | Swt1-family-HEPN-domain-containing-protein,-partial-[Thermus-scotoductus] |
| WP_126212240.1 | HEPN+AAA-ATPase+Alpha+Beta-domain+wHTH+RRM | Type-1_BR.RM.like.systems | METHYLASE->HigB/RHH->HTH_3->HIN-HTH+SecB->HEPN+DUF499*-> | 418 | Bacteria>Terrabacteria_group>Deinococcota>Deinococci>Thermales | GCF_003959525.1 | Thermus_scotoductus | CSW17_RS01375 | 4891..6146 | 37636 | Swt1-family-HEPN-domain-containing-protein,-partial-[Thermus-scotoductus] |
| WP_126193807.1 | HEPN+AAA-ATPase+Alpha+Beta-domain+wHTH+RRM | Type-1_BR.RM.like.systems | METHYLASE->HigB/RHH->HTH_3->HIN-HTH+SecB->HEPN+DUF499*-> | 416 | Bacteria>Terrabacteria_group>Deinococcota>Deinococci>Thermales | GCF_003959075.1 | Thermus_scotoductus | CSW46_RS01260 | 4967..6215 | 37636 | Swt1-family-HEPN-domain-containing-protein,-partial-[Thermus-scotoductus] |
| WP_126164480.1 | HEPN+AAA-ATPase+Alpha+Beta-domain+wHTH+RRM | Type-1_BR.RM.like.systems | METHYLASE->HigB/RHH->HTH_3->HIN-HTH+SecB->HEPN+DUF499*-> | 412 | Bacteria>Terrabacteria_group>Deinococcota>Deinococci>Thermales | GCF_003958975.1 | Thermus_scotoductus | CSW48_RS02070 | 4415..5652 | 37636 | Swt1-family-HEPN-domain-containing-protein,-partial-[Thermus-scotoductus] |
| WP_253724062.1 | HEPN+AAA-ATPase+Alpha+Beta-domain+wHTH+RRM | Type-1_BR.RM.like.systems | <-HEPN+DUF499*<-HIN-HTH+SecB<-HTH_3<-HigB/RHH | 446 | Bacteria>Terrabacteria_group>Deinococcota>Deinococci>Thermales | GCF_003959305.1 | Thermus_scotoductus | CSW32_RS03250 | 10..1350 | 37636 | Swt1-family-HEPN-domain-containing-protein-[Thermus-scotoductus] |
| WP_253665106.1 | HEPN+AAA-ATPase+Alpha+Beta-domain+wHTH+RRM | Type-1_BR.RM.like.systems | HigB/RHH->HTH_3->HIN-HTH+SecB->HEPN+DUF499*-> | 1022 | Bacteria>Terrabacteria_group>Deinococcota>Deinococci>Thermales | GCF_003959545.1 | Thermus_scotoductus | CSW16_RS01280 | 2718..5783 | 37636 | Swt1-family-HEPN-domain-containing-protein,-partial-[Thermus-scotoductus] |
| WP_126252475.1 | HEPN+AAA-ATPase+Alpha+Beta-domain+wHTH+RRM | Type-1_BR.RM.like.systems | HigB/RHH->HTH_3->HIN-HTH+SecB->HEPN+DUF499*-> | 415 | Bacteria>Terrabacteria_group>Deinococcota>Deinococci>Thermales | GCF_003959675.1 | Thermus_scotoductus | CSW15_RS03250 | 2694..3937 | 37636 | Swt1-family-HEPN-domain-containing-protein,-partial-[Thermus-scotoductus] |
| WP_126216336.1 | HEPN+AAA-ATPase+Alpha+Beta-domain+wHTH+RRM | Type-1_BR.RM.like.systems | <-HEPN+DUF499*<-HIN-HTH+SecB<-HTH_3<-HigB/RHH | 413 | Bacteria>Terrabacteria_group>Deinococcota>Deinococci>Thermales | GCF_003959575.1 | Thermus_scotoductus | CSW36_RS03285 | 1..1238 | 37636 | Swt1-family-HEPN-domain-containing-protein,-partial-[Thermus-scotoductus] |
| WP_126212451.1 | HEPN+AAA-ATPase+Alpha+Beta-domain+wHTH+RRM | Type-1_BR.RM.like.systems | HigB/RHH->HTH_3->HIN-HTH+SecB->HEPN+DUF499*-> | 419 | Bacteria>Terrabacteria_group>Deinococcota>Deinococci>Thermales | GCF_003959265.1 | Thermus_scotoductus | CSW34_RS03430 | 2738..3995 | 37636 | Swt1-family-HEPN-domain-containing-protein,-partial-[Thermus-scotoductus] |
| WP_126211477.1 | HEPN+AAA-ATPase+Alpha+Beta-domain+wHTH+RRM | Type-1_BR.RM.like.systems | <-HEPN+DUF499*<-HIN-HTH+SecB<-HTH_3<-HigB/RHH | 421 | Bacteria>Terrabacteria_group>Deinococcota>Deinococci>Thermales | - | Thermus_scotoductus | CSW19_RS03320 | 1..1262 | 37636 | Swt1-family-HEPN-domain-containing-protein,-partial-[Thermus-scotoductus] |
| WP_126210662.1 | HEPN+AAA-ATPase+Alpha+Beta-domain+wHTH+RRM | Type-1_BR.RM.like.systems | <-HEPN+DUF499*<-HIN-HTH+SecB<-HTH_3<-HigB/RHH | 424 | Bacteria>Terrabacteria_group>Deinococcota>Deinococci>Thermales | GCF_003959395.1 | Thermus_scotoductus | CSW26_RS03555 | 1..1273 | 37636 | Swt1-family-HEPN-domain-containing-protein,-partial-[Thermus-scotoductus] |
| WP_126199710.1 | HEPN+AAA-ATPase+Alpha+Beta-domain+wHTH+RRM | Type-1_BR.RM.like.systems | HigB/RHH->HTH_3->HIN-HTH+SecB->HEPN+DUF499*-> | 425 | Bacteria>Terrabacteria_group>Deinococcota>Deinococci>Thermales | GCF_003959105.1 | Thermus_scotoductus | CSW44_RS03290 | 2683..3958 | 37636 | Swt1-family-HEPN-domain-containing-protein,-partial-[Thermus-scotoductus] |
| WP_028494585.1 | HEPN+AAA-ATPase+Alpha+Beta-domain+wHTH+RRM | Type-1_BR.RM.like.systems | <-HEPN+DUF499*<-HIN-HTH+SecB<-HTH_3<-HigB/RHH | 1095 | Bacteria>Terrabacteria_group>Deinococcota>Deinococci>Thermales | GCF_000423905.1 | Thermus | CSW24_RS01285 | 852..4139 | 270 | MULTISPECIES:-Swt1-family-HEPN-domain-containing-protein-[Thermus] |
| WP_219760305.1 | HEPN+AAA-ATPase+Alpha+Beta-domain+wHTH+RRM | Type-1_BR.RM.like.systems | SLATT->METHYLASE->HEPN+DUF499*->?->?->DDE_3-> | 1116 | Bacteria>Terrabacteria_group>Deinococcota>Deinococci>Thermales | GCF_019430925.1 | Thermus_brevis | KZX47_RS11605 | 3607..6957 | 2862456 | Swt1-family-HEPN-domain-containing-protein-[Thermus-brevis] |
| GIW28837.1 | HEPN+AAA-ATPase+Alpha+Beta-domain+wHTH+RRM | Type-1_BR.RM.like.systems | TPR-><-ABhydrolase<-HDRnase<-?\|\|HEPN+DUF499*->TUDOR+SF1-SF2-Helicase+iREase+CTD+REase->SF1-SF2-Helicase+wHTH+PRTase->?->Schlafen+HATPase+HTH->METHYLASE-><-TPR | 1063 | Bacteria>Terrabacteria_group>Deinococcota>Deinococci>Thermales | GCA_026004075.1 | Meiothermus_sp. | KatS3mg070_2200 | 2304208..2307399 | 1955249 | MAG:-hypothetical-protein-KatS3mg070_2200-[Meiothermus-sp.] |
| WP_240637265.1 | HEPN+AAA-ATPase+Alpha+Beta-domain+wHTH+RRM | Type-1_BR.RM.like.systems | DDE_Tnp_1->HEPN+DUF499*->TUDOR+SF1-SF2-Helicase+iREase+CTD+REase->SF1-SF2-Helicase+wHTH+PRTase->?->Schlafen+HATPase+HTH->METHYLASE-> | 1066 | Bacteria>Terrabacteria_group>Deinococcota>Deinococci>Thermales | GCF_007990975.1 | Meiothermus_hypogaeus | MHY01S_RS11115 | 3652..6852 | 884155 | DUF499-domain-containing-protein-[Meiothermus-hypogaeus] |
| WP_083922075.1 | HEPN+AAA-ATPase+Alpha+Beta-domain+wHTH+RRM | Type-1_BR.RM.like.systems | Hpr-Kinase-C->?->?->?->?->HEPN+DUF499*->TUDOR+SF1-SF2-Helicase+iREase+CTD+REase->METHYLASE-><-?\|\|?->TraJ-RHH->PIN-> | 1058 | Bacteria>Terrabacteria_group>Deinococcota>Deinococci>Thermales | GCF_000376265.1 | Thermus_igniterrae | B128_RS0103820 | 7701..10877 | 88189 | DUF499-domain-containing-protein-[Thermus-igniterrae] |
| WP_245587643.1 | HEPN+AAA-ATPase+Alpha+Beta-domain+wHTH+RRM | Type-1_BR.RM.like.systems | <-REase<-?\|\|HNH-><-?\|\|HEPN+DUF499*->TUDOR+SF1-SF2-Helicase+iREase+CTD+REase->?->MACRODOMAIN->METHYLASE-> | 1065 | Bacteria>Terrabacteria_group>Deinococcota>Deinococci>Thermales | GCF_000430045.1 | Calidithermus_chliarophilus | MEICH_RS0111510 | 2368714..2371911 | 52023 | DUF499-domain-containing-protein-[Calidithermus-chliarophilus] |
| WP_114799026.1 | HEPN+AAA-ATPase+Alpha+Beta-domain+wHTH+RRM | Type-1_BR.RM.like.systems | ABC-ATPase+SbcC-><-?\|\|?->?->?-><-?\|\|tRNA->tRNA->HEPN+DUF499*->TUDOR+SF1-SF2-Helicase+iREase+CTD+REase->METHYLASE-><-TPR | 1057 | Bacteria>Terrabacteria_group>Deinococcota>Deinococci>Thermales | GCF_003351145.1 | Meiothermus_sp._QL-1 | DV704_RS07840 | 44133..47306 | 2058095 | DUF499-domain-containing-protein-[Meiothermus-sp.-QL-1] |
| WP_036270380.1 | HEPN+AAA-ATPase+Alpha+Beta-domain+wHTH+RRM | Type-1_BR.RM.like.systems | <-METHYLASE<-MACRODOMAIN<-?<-TUDOR+SF1-SF2-Helicase+iREase+CTD+REase<-HEPN+DUF499*<-?<-BirA-HTH | 1062 | Bacteria>Terrabacteria_group>Deinococcota>Deinococci>Thermales | GCF_000423425.1 | Meiothermus_rufus | G552_RS0106115 | 16057..19245 | 604332 | DUF499-domain-containing-protein-[Meiothermus-rufus] |
| MCY4354897.1 | HEPN+AAA-ATPase+Alpha+Beta-domain+wHTH+RRM | Type-1_BR.RM.like.systems | HicA_toxin->HicB->TUDOR+SF1-SF2-Helicase+iREase+CTD+REase->METHYLASE->PHP+SbcC->Schlafen+HATPase+HTH->HEPN+DUF499*->?->PIN->?->?->PrmC_N+Methyltransf_25-><-Peptidase_A8 | 1107 | Bacteria>Terrabacteria_group>Deinococcota>Deinococci>Trueperales | GCA_026710445.1 | Truepera_sp. | OXC09_08995 | 25870..29193 | 2478917 | MAG:-Swt1-family-HEPN-domain-containing-protein-[Truepera-sp.] |
| WP_013177687.1 | HEPN+AAA-ATPase+Alpha+Beta-domain+wHTH+RRM | Type-1_BR.RM.like.systems | PUA+METHYLASE->?->?->?-><-sGTP<-Aminotran_3<-HEPN+DUF499*<-METHYLASE-p\|\|Y1_Tnp-><-METHYLASE<-PIN | 1108 | Bacteria>Terrabacteria_group>Deinococcota>Deinococci>Trueperales | GCF_000092425.1 | Truepera_radiovictrix | TRAD_RS05920 | 1278601..1281927 | 332249 | Swt1-family-HEPN-domain-containing-protein-[Truepera-radiovictrix] |
| WP_013179180.1 | HEPN+AAA-ATPase+Alpha+Beta-domain+wHTH+RRM | Type-1_BR.RM.like.systems | TUDOR+SF1-SF2-Helicase+iREase+CTD+REase->Phd-YefM->PIN->METHYLASE->?->PIN->HEPN+DUF499*-><-Peptidase_M20 | 1069 | Bacteria>Terrabacteria_group>Deinococcota>Deinococci>Trueperales | GCF_000092425.1 | Truepera_radiovictrix | TRAD_RS13560 | 2957587..2960796 | 332249 | DUF499-domain-containing-protein-[Truepera-radiovictrix] |
| RMH56280.1 | HEPN+AAA-ATPase+Alpha+Beta-domain+wHTH+RRM | Type-1_BR.RM.like.systems | <-HAD<-?<-?\|\|TUDOR+SF1-SF2-Helicase+iREase+CTD+REase->METHYLASE->HTH_3->HEPN+DUF499*->ABhydrolase->?-><-?<-ACET | 1091 | Bacteria>Terrabacteria_group>Deinococcota>unclassified_Deinococcota | GCA_003695925.1 | Deinococcota_bacterium | D6684_05775 | 12315..15590 | 1913990 | ATP-binding-protein-[Deinococcus-Thermus-bacterium] |
| MCA9833507.1 | HEPN+AAA-ATPase+Alpha+Beta-domain+wHTH+RRM | Type-1_BR.RM.like.systems | Peptidase-><-?<-?\|\|TUDOR+SF1-SF2-Helicase+iREase+CTD+REase->HEPN->METHYLASE->HEPN+DUF499*->DDE_3->?-><-tRNA<-?\|\|?-><-?<-NACHT | 1122 | Bacteria>Terrabacteria_group>Thermomicrobiota>Thermomicrobia>Thermomicrobiales | GCA_020432165.1 | Thermomicrobiales_bacterium | KC435_06155 | 74622..77990 | 2482790 | DUF499-domain-containing-protein-[Thermomicrobiales-bacterium] |
| MCC6945870.1 | HEPN+AAA-ATPase+Alpha+Beta-domain+wHTH+RRM | Type-1_BR.RM.like.systems | METHYLASE-p->HEPN+DUF499*-> | 1058 | Bacteria>Terrabacteria_group>Thermomicrobiota>Thermomicrobia>Thermomicrobiales | GCA_020853875.1 | Thermomicrobiales_bacterium | IT335_14920 | 865..4040 | 2482790 | DUF499-domain-containing-protein-[Thermomicrobiales-bacterium] |
| MCC6945442.1 | HEPN+AAA-ATPase+Alpha+Beta-domain+wHTH+RRM | Type-1_BR.RM.like.systems | METHYLASE-p->HEPN+DUF499*-> | 1060 | Bacteria>Terrabacteria_group>Thermomicrobiota>Thermomicrobia>Thermomicrobiales | GCA_020853875.1 | Thermomicrobiales_bacterium | IT335_12740 | 291..3470 | 2482790 | DUF499-domain-containing-protein-[Thermomicrobiales-bacterium] |
| MBI5524652.1 | HEPN+AAA-ATPase+Alpha+Beta-domain+wHTH+RRM | Type-1_BR.RM.like.systems | <-Peptidase\|\|tRNA->TUDOR+SF1-SF2-Helicase+iREase+CTD+REase->DOC+HTH->METHYLASE->HEPN+DUF499*-> | 1110 | Bacteria>Thermodesulfobacteriota>Desulfarculia>Desulfarculales>Desulfarculaceae | GCA_016218705.1 | Desulfarculus_sp. | HY910_18660 | 18095..21427 | 2093367 | ATP-binding-protein-[Desulfarculus-sp.] |
| VBB42850.1 | HEPN+AAA-ATPase+Alpha+Beta-domain+wHTH+RRM | Type-1_BR.RM.like.systems | <-Phage_integrase<-?\|\|TUDOR+SF1-SF2-Helicase+iREase+CTD+REase->ParB+HNH->METHYLASE->HEPN+DUF499*->?->?->?-><-DDE_Tnp_1\|\|HNH-> | 1117 | Bacteria>Thermodesulfobacteriota>Desulfobacteria>Desulfatiglandales>Desulfatiglandaceae | GCA_900498135.1 | uncultured_Desulfatiglans_sp. | TRIP_B220098 | 96105..99458 | 1748965 | conserved-hypothetical-protein-[uncultured-Desulfatiglans-sp.] |
| WP_028322621.1 | HEPN+AAA-ATPase+Alpha+Beta-domain+wHTH+RRM | Type-1_BR.RM.like.systems | METHYLASE->HEPN+DUF499*->?->?->Primpol+D5N+D5ATPase-> | 1114 | Bacteria>Thermodesulfobacteriota>Desulfobacteria>Desulfatiglandales>Desulfatiglandaceae | GCF_000422285.1 | Desulfatiglans_anilini | H567_RS0119135 | 2085..5429 | 90728 | Swt1-family-HEPN-domain-containing-protein-[Desulfatiglans-anilini] |
| MBA3017256.1 | HEPN+AAA-ATPase+Alpha+Beta-domain+wHTH+RRM | Type-1_BR.RM.like.systems | <-N6_N4_MTase<-Peptidase_C47<-?\|\|TUDOR+SF1-SF2-Helicase+iREase+CTD+REase->MORC+HisKin+DUF3684->HEPN+DUF499*->RNase->ParB+HNH->HEPN-> | 1092 | Bacteria>Thermodesulfobacteriota>Desulfobacteria>Desulfobacterales>Desulfobacteraceae | GCA_013792055.1 | Desulfobacteraceae_bacterium | FP811_03735 | 27065..30343 | 2049433 | ATP-binding-protein-[Desulfobacteraceae-bacterium] |
| MBL7176677.1 | HEPN+AAA-ATPase+Alpha+Beta-domain+wHTH+RRM | Type-1_BR.RM.like.systems | TUDOR+SF1-SF2-Helicase+iREase+CTD+REase-p->OLD-ABC->PIN->?->HEPN+DUF499*->?->OLD-ABC->SF1-SF2-Helicase->?->MNS+HTH-> | 1094 | Bacteria>Thermodesulfobacteriota>Desulfobacteria>Desulfobacterales>Desulfobacteraceae | GCA_016782925.1 | Desulfobacteraceae_bacterium | ISS66_12705 | 13240..16524 | 2049433 | DUF499-domain-containing-protein-[Desulfobacteraceae-bacterium] |
| WP_004073680.1 | HEPN+AAA-ATPase+Alpha+Beta-domain+wHTH+RRM | Type-1_BR.RM.like.systems | SPOUT->PRTase->TUDOR+SF1-SF2-Helicase+iREase+CTD+REase->HEPN->METHYLASE->HEPN+DUF499*-> | 1113 | Bacteria>Thermodesulfobacteriota>Desulfobacteria>Desulfobacterales>Desulfobacteraceae | GCF_000233695.2 | Desulfobacter_postgatei | DESPODRAFT_RS11525 | 2528550..2531891 | 2293 | Swt1-family-HEPN-domain-containing-protein-[Desulfobacter-postgatei] |
| HHO48401.1 | HEPN+AAA-ATPase+Alpha+Beta-domain+wHTH+RRM | Type-1_BR.RM.like.systems | <-HEPN+DUF499*<-METHYLASE<-TUDOR+SF1-SF2-Helicase+iREase+CTD+REase | 1112 | Bacteria>Thermodesulfobacteriota>Desulfobacteria>Desulfobacterales>Desulfobacteraceae | GCA_011372855.1 | Desulfobacteraceae_bacterium | ENN06_08115 | 325..3663 | 2049433 | ATP-binding-protein-[Desulfobacteraceae-bacterium] |
| OPX33848.1 | HEPN+AAA-ATPase+Alpha+Beta-domain+wHTH+RRM | Type-1_BR.RM.like.systems | <-HicB_lk_antitox<-?<-HEPN+DUF499*<-METHYLASE-p | 957 | Bacteria>Thermodesulfobacteriota>Desulfobacteria>Desulfobacterales>Desulfobacteraceae | GCA_002049905.1 | Desulfobacteraceae_bacterium_4484_190.1 | B1H11_11870 | 1782..4655 | 1956167 | hypothetical-protein-B1H11_11870-[Desulfobacteraceae-bacterium-4484_190.1] |
| CBX30683.1 | HEPN+AAA-ATPase+Alpha+Beta-domain+wHTH+RRM | Type-1_BR.RM.like.systems | <-SbcC\|\|TUDOR+SF1-SF2-Helicase+iREase+CTD+REase->MqsR_toxin->MqsA_antitoxin+HTH_3->?->METHYLASE->HEPN+DUF499*-> | 1111 | Bacteria>Thermodesulfobacteriota>Desulfobacteria>Desulfobacterales>Desulfobacteriaceae | - | uncultured_Desulfobacterium_sp. | N47_E41950 | 104821..108156 | 201089 | hypothetical-protein-N47_E41950-[uncultured-Desulfobacterium-sp.] |
| MBC2710211.1 | HEPN+AAA-ATPase+Alpha+Beta-domain+wHTH+RRM | Type-1_BR.RM.like.systems | REase-><-Phage_integrase<-?\|\|?->?->?-><-HEPN+DUF499*<-METHYLASE<-TOPRIM<-OLD-ABC<-MORC+DUF3684<-TUDOR+SF1-SF2-Helicase+iREase+CTD+REase\|\|DDE_Tnp_1-> | 1102 | Bacteria>Thermodesulfobacteriota>Desulfobacteria>Desulfobacterales>Desulfosarcinaceae | GCA_014237405.1 | Desulfosarcina_sp. | HGJ94_04170 | 9166..12474 | 2027861 | ATP-binding-protein-[Desulfosarcina-sp.] |
| MBC2711122.1 | HEPN+AAA-ATPase+Alpha+Beta-domain+wHTH+RRM | Type-1_BR.RM.like.systems | TUDOR+SF1-SF2-Helicase+iREase+CTD+REase->?->OLD-ABC->HNH->METHYLASE->HEPN+DUF499*->?->GNTR-HTH->?->?->ABC-ATPase+SbcC->ABC-ATPase+SbcC-> | 1122 | Bacteria>Thermodesulfobacteriota>Desulfobacteria>Desulfobacterales>Desulfosarcinaceae | GCA_014237405.1 | Desulfosarcina_sp. | HGJ94_09035 | 43660..47028 | 2027861 | ATP-binding-protein-[Desulfosarcina-sp.] |

| Protein-ID | DUF499-architecture | System | Gene-Neighbourhoods | Length | Taxonomy | Assembly-ID | Strain | Locus-Tag | Genomic-coordinates | TaxID | Fasta-defline |
|---|---|---|---|---|---|---|---|---|---|---|---|
| PIU50309.1 | HEPN+AAA-ATPase+Alpha+Beta-domain+wHTH+RRM | Type-1_BR.RM.like.systems | METHYLASE->?->HEPN+DUF499*->?-><-?\|\|SF1-SF2-Helicase-> | 1109 | Bacteria>Thermodesulfobacteriota>Desulfobacteria>Desulfobacterales>unclassified_Desulfobacterales | GCA_002779465.1 | Desulfobacterales_bacterium_CG07_land_8_20_14_0_80_52_14 | COS92_01840 | 3954..7283 | 1973982 | AAA+-family-ATPase-[Desulfobacterales-bacterium-CG07_land_8_20_14_0_80_52_14] |
| MCU0558984.1 | HEPN+AAA-ATPase+Alpha+Beta-domain+wHTH+RRM | Type-1_BR.RM.like.systems | <-HEPN+DUF499*<-TOPRIM<-OLD-ABC<-METHYLASE-p | 1112 | Bacteria>Thermodesulfobacteriota>Desulfobacteria>Desulfobacterales>unclassified_Desulfobacterales | GCA_025454255.1 | Desulfobacterales_bacterium | MUD16_02165 | 66226..69564 | 2044940 | Swt1-family-HEPN-domain-containing-protein-[Desulfobacterales-bacterium] |
| WP_028319484.1 | HEPN+AAA-ATPase+Alpha+Beta-domain+wHTH+RRM | Type-1_BR.RM.like.systems | DDE_Tnp_1_2-><-HEPN+DUF499*<-METHYLASE<-TOPRIM<-OLD-ABC<-MORC+DUF3684<-TUDOR+SF1-SF2-Helicase+iREase+CTD+REase<-XIS-HTH | 1112 | Bacteria>Thermodesulfobacteriota>Desulfobulbia>Desulfobulbales>Desulfobulbaceae | GCF_000621145.1 | Desulfobulbus_elongatus | Q362_RS0116010 | 475..3813 | 53332 | Swt1-family-HEPN-domain-containing-protein-[Desulfobulbus-elongatus] |
| MDD6180986.1 | HEPN+AAA-ATPase+Alpha+Beta-domain+wHTH+RRM | Type-1_BR.RM.like.systems | CR-ATPase8+REase-><-HEPN+DUF499*<-METHYLASE<-TUDOR+SF1-SF2-Helicase+iREase+CTD+REase<-GDResolvase+Recombinase+ZnR<-Recombinase+ZnR<-GDResolvase+Recombinase+ZnR<-RAMA | 1147 | Bacteria>Thermodesulfobacteriota>Desulfovibrionia>Desulfovibrionales>Desulfovibrionaceae | GCA_029008115.1 | Desulfovibrionaceae_bacterium | PUB01_02780 | 4125..7568 | 2049043 | MAG:-Swt1-family-HEPN-domain-containing-protein-[Desulfovibrionaceae-bacterium] |
| WP_165067758.1 | HEPN+AAA-ATPase+Alpha+Beta-domain+wHTH+RRM | Type-1_BR.RM.like.systems | GDResolvase+Recombinase+ZnR->TUDOR+SF1-SF2-Helicase+iREase+CTD+REase-><-?\|\|METHYLASE->HEPN+DUF499*-><-ACET<-?<-?<-?<-?\|\|Arm-DNA-bind_3+Phage_integrase-> | 1129 | Bacteria>Thermodesulfobacteriota>Desulfovibrionia>Desulfovibrionales>Desulfovibrionaceae | GCF_011039095.1 | Desulfovibrio_sp._ZJ200 | G7Y52_RS05935 | 42436..45825 | 2709792 | Swt1-family-HEPN-domain-containing-protein-[Desulfovibrio-sp.-ZJ200] |
| WP_273524160.1 | HEPN+AAA-ATPase+Alpha+Beta-domain+wHTH+RRM | Type-1_BR.RM.like.systems | DOC+HTH->REase+SAVED->?->?-><-?<-HEPN+DUF499*<-METHYLASE<-TUDOR+SF1-SF2-Helicase+iREase+CTD+REase<-GDResolvase+Recombinase+ZnR<-?<-?<-Phage_AlpA | 1120 | Bacteria>Thermodesulfobacteriota>Desulfovibrionia>Desulfovibrionales>Desulfovibrionaceae | GCF_944324605.1 | Mailhella_massiliensis | PUQ52_RS06600 | 138638..142000 | 1903261 | Swt1-family-HEPN-domain-containing-protein-[Mailhella-massiliensis] |
| WP_077072828.1 | HEPN+AAA-ATPase+Alpha+Beta-domain+wHTH+RRM | Type-1_BR.RM.like.systems | DOC+HTH->REase+SAVED->?->?-><-?<-HEPN+DUF499*<-METHYLASE<-TUDOR+SF1-SF2-Helicase+iREase+CTD+REase<-GDResolvase+Recombinase+ZnR<-?<-?<-Phage_AlpA | 1120 | Bacteria>Thermodesulfobacteriota>Desulfovibrionia>Desulfovibrionales>Desulfovibrionaceae | GCF_944324615.1 | Mailhella_massiliensis | Q0K07_RS04875 | 39206..42568 | 1903261 | Swt1-family-HEPN-domain-containing-protein-[Mailhella-massiliensis] |
| WP_278732740.1 | HEPN+AAA-ATPase+Alpha+Beta-domain+wHTH+RRM | Type-1_BR.RM.like.systems | TOPRIM+SF1-SF2-Helicase+HTH->?->?->GDResolvase+Recombinase+ZnR->TUDOR+SF1-SF2-Helicase+iREase+CTD+REase->METHYLASE->HEPN+DUF499*-> | 865 | Bacteria>Thermodesulfobacteriota>Desulfovibrionia>Desulfovibrionales>Desulfovibrionaceae | GCF_020743955.1 | Bilophila_wadsworthia | K8U76_RS10810 | 19353..21949 | 35833 | Swt1-family-HEPN-domain-containing-protein,-partial-[Bilophila-wadsworthia] |
| HJH15746.1 | HEPN+AAA-ATPase+Alpha+Beta-domain+wHTH+RRM | Type-1_BR.RM.like.systems | TOPRIM+SF1-SF2-Helicase+HTH->?->?->GDResolvase+Recombinase+ZnR->TUDOR+SF1-SF2-Helicase+iREase+CTD+REase->METHYLASE->HEPN+DUF499*-> | 865 | Bacteria>Thermodesulfobacteriota>Desulfovibrionia>Desulfovibrionales>Desulfovibrionaceae | GCA_020743955.1 | Bilophila_wadsworthia | K8U76_10820 | 19353..21949 | 35833 | DUF499-domain-containing-protein-[Bilophila-wadsworthia] |
| WP_168936425.1 | HEPN+AAA-ATPase+Alpha+Beta-domain+wHTH+RRM | Type-1_BR.RM.like.systems | <-HEPN+DUF499*<-METHYLASE<-TUDOR+SF1-SF2-Helicase+iREase+CTD+REase-p<-TUDOR+SF1-SF2-Helicase+iREase+CTD+REase | 1121 | Bacteria>Thermodesulfobacteriota>Desulfovibrionia>Desulfovibrionales>Desulfovibrionaceae | GCF_012843875.1 | Desulfovibrio_piger | HF854_RS11535 | 110..3475 | 901 | Swt1-family-HEPN-domain-containing-protein-[Desulfovibrio-piger] |
| WP_279060950.1 | HEPN+AAA-ATPase+Alpha+Beta-domain+wHTH+RRM | Type-1_BR.RM.like.systems | GDResolvase+Recombinase+ZnR->TUDOR+SF1-SF2-Helicase+iREase+CTD+REase->METHYLASE->METHYLASE-p->?->HEPN+DUF499*-> | 1135 | Bacteria>Thermodesulfobacteriota>Desulfovibrionia>Desulfovibrionales>Desulfovibrionaceae | GCF_025543515.1 | Bilophila_wadsworthia | QDO56_RS08660 | 91751..95158 | 35833 | DUF499-domain-containing-protein-[Bilophila-wadsworthia] |
| MBD5607241.1 | HEPN+AAA-ATPase+Alpha+Beta-domain+wHTH+RRM | Type-1_BR.RM.like.systems | TUDOR+SF1-SF2-Helicase+iREase+CTD+REase->METHYLASE->METHYLASE->HEPN+DUF499*->?->?->SF1-SF2-Helicase+REase-p-> | 1123 | Bacteria>Thermodesulfobacteriota>Desulfovibrionia>Desulfovibrionales>Desulfovibrionaceae | GCA_014804865.1 | Desulfovibrio_sp. | HDQ93_00095 | 9665..13036 | 885 | ATP-binding-protein-[Desulfovibrio-sp.] |
| HJB66433.1 | HEPN+AAA-ATPase+Alpha+Beta-domain+wHTH+RRM | Type-1_BR.RM.like.systems | TraJ-RHH->TraI-VirD2-RCRE->?->?->?-><-HEPN<-HEPN+DUF499*<-METHYLASE<-TOPRIM | 1112 | Bacteria>Thermodesulfobacteriota>Desulfovibrionia>Desulfovibrionales>Desulfovibrionaceae | GCA_019119045.1 | Candidatus_Mailhella_merdavium | H9768_09155 | 39301..42639 | 2838657 | DUF499-domain-containing-protein-[Candidatus-Mailhella-merdavium] |
| MBR3880305.1 | HEPN+AAA-ATPase+Alpha+Beta-domain+wHTH+RRM | Type-1_BR.RM.like.systems | METHYLASE->HEPN+DUF499*-> | 1086 | Bacteria>Thermodesulfobacteriota>Desulfovibrionia>Desulfovibrionales>Desulfovibrionaceae | GCA_017522745.1 | Mailhella_sp. | IKJ34_01740 | 2844..6102 | 1981029 | DUF499-domain-containing-protein-[Mailhella-sp.] |
| TVR02140.1 | HEPN+AAA-ATPase+Alpha+Beta-domain+wHTH+RRM | Type-1_BR.RM.like.systems | <-HEPN+DUF499*<-?<-METHYLASE<-PemK_toxin<-?<-Y1_Tnp<-TUDOR+SF1-SF2-Helicase+iREase+CTD+REase | 1114 | Bacteria>Thermodesulfobacteriota>Desulfovibrionia>Desulfovibrionales>unclassified_Desulfovibrionales | GCA_007693505.1 | Desulfovibrionales_bacterium | EA399_00610 | 1087..4431 | 2448050 | ATP-binding-protein-[Desulfovibrionales-bacterium] |
| WP_067619295.1 | HEPN+AAA-ATPase+Alpha+Beta-domain+wHTH+RRM | Type-1_BR.RM.like.systems | LexA-HTH+LexA-protease->TUDOR+SF1-SF2-Helicase+iREase+CTD+REase->Schlafen+HATPase+HTH->?->METHYLASE->DOC+HTH->HEPN+DUF499*->?->Peptidase_C47->?->PIN-> | 1099 | Bacteria>Thermodesulfobacteriota>Dissulfuribacteria>Dissulfuribacterales>Dissulfuribacteraceae | GCF_001687335.1 | Dissulfuribacter_thermophilus | DBT_RS08725 | 80186..83485 | 1156395 | Swt1-family-HEPN-domain-containing-protein-[Dissulfuribacter-thermophilus] |
| NLD79955.1 | HEPN+AAA-ATPase+Alpha+Beta-domain+wHTH+RRM | Type-1_BR.RM.like.systems | <-HEPN+DUF499*<-METHYLASE-p | 1123 | Bacteria>Thermodesulfobacteriota>Syntrophia>Syntrophales>Smithellaceae | GCA_012513865.1 | Smithella_sp. | GX642_02135 | 555..3926 | 2053614 | ATP-binding-protein-[Smithella-sp.] |
| NLA40358.1 | HEPN+AAA-ATPase+Alpha+Beta-domain+wHTH+RRM | Type-1_BR.RM.like.systems | <-HEPN+DUF499*<-METHYLASE | 1123 | Bacteria>Thermodesulfobacteriota>Syntrophia>Syntrophales>Smithellaceae | GCA_012510495.1 | Smithella_sp. | GX874_02955 | 108..3479 | 2053614 | ATP-binding-protein-[Smithella-sp.] |
| MCG2740517.1 | HEPN+AAA-ATPase+Alpha+Beta-domain+wHTH+RRM | Type-1_BR.RM.like.systems | MerR-HTH->METHYLASE->PIN->?-><-HEPN+DUF499*<-METHYLASE<-MerR-HTH<-TUDOR+SF1-SF2-Helicase+iREase+CTD+REase | 1109 | Bacteria>Thermodesulfobacteriota>Syntrophia>Syntrophales>Syntrophaceae | GCA_022013455.1 | Syntrophaceae_bacterium | L6300_09825 | 5770..9099 | 2052184 | DUF499-domain-containing-protein-[Syntrophaceae-bacterium] |
| ABC75909.1 | HEPN+AAA-ATPase+Alpha+Beta-domain+wHTH+RRM | Type-1_BR.RM.like.systems | <-sGTP\|\|?-><-DrHyd+BirA-HTH<-DUF499-p<-HEPN+DUF499*<-TOPRIM<-OLD-ABC<-METHYLASE-p<-METHYLASE-p<-HKD-DNase | 700 | Bacteria>Thermodesulfobacteriota>Syntrophia>Syntrophales>Syntrophaceae | GCA_000013405.1 | Syntrophus_aciditrophicus_SB | SYN_00008 | 32188..34290 | 56780 | ATPase-of-the-AAA+-superfamily-[Syntrophus-aciditrophicus-SB] |
| WP_011415944.1 | HEPN+AAA-ATPase+Alpha+Beta-domain+wHTH+RRM | Type-1_BR.RM.like.systems | <-sGTP<-DrHyd+BirA-HTH<-DUF499-p<-HEPN+DUF499*<-TOPRIM<-OLD-ABC<-METHYLASE-p<-METHYLASE-p<-HKD-DNase | 700 | Bacteria>Thermodesulfobacteriota>Syntrophia>Syntrophales>Syntrophaceae | GCF_000013405.1 | Syntrophus_aciditrophicus | SYN_RS16675 | 32188..34290 | 316277 | Swt1-family-HEPN-domain-containing-protein-[Syntrophus-aciditrophicus] |
| HBL52555.1 | HEPN+AAA-ATPase+Alpha+Beta-domain+wHTH+RRM | Type-1_BR.RM.like.systems | TUDOR+SF1-SF2-Helicase+iREase+CTD+REase-p->ORC-CDC6+REase->MqsA_antitoxin+HTH_3->HEPN+DUF499*-> | 1127 | Bacteria>Thermodesulfobacteriota>Syntrophia>Syntrophales>Syntrophaceae | GCA_003506745.1 | Syntrophaceae_bacterium | DDZ34_00420 | 4849..8232 | 2052184 | AAA+-family-ATPase-[Syntrophaceae-bacterium] |
| MCG2738715.1 | HEPN+AAA-ATPase+Alpha+Beta-domain+wHTH+RRM | Type-1_BR.RM.like.systems | MORC+DUF3684->DrHyd+BirA-HTH->SF1-SF2-Helicase->?->?->METHYLASE->HEPN+DUF499*-> | 1124 | Bacteria>Thermodesulfobacteriota>Syntrophia>Syntrophales>Syntrophaceae | GCA_022013455.1 | Syntrophaceae_bacterium | L6300_00555 | 19371..22745 | 2052184 | DUF499-domain-containing-protein-[Syntrophaceae-bacterium] |
| MBN1365905.1 | HEPN+AAA-ATPase+Alpha+Beta-domain+wHTH+RRM | Type-1_BR.RM.like.systems | HKD-DNase->METHYLASE->HEPN+DUF499*->?->?->REase-><-?\|\|HNH-><-REase | 1119 | Bacteria>Thermodesulfobacteriota>Syntrophia>Syntrophales>Syntrophaceae | GCA_016927935.1 | Syntrophaceae_bacterium | JW976_13940 | 4360..7719 | 2052184 | DUF499-domain-containing-protein-[Syntrophaceae-bacterium] |
| MCK9528171.1 | HEPN+AAA-ATPase+Alpha+Beta-domain+wHTH+RRM | Type-1_BR.RM.like.systems | TUDOR+SF1-SF2-Helicase+iREase+CTD+REase->PHP+SbcC->Schlafen+HATPase+HTH->METHYLASE->OLD-ABC->TOPRIM->HEPN+DUF499*->ABhydrolase-> | 1108 | Bacteria>Thermodesulfobacteriota>Syntrophia>Syntrophales>unclassified_Syntrophales | GCA_023228425.1 | Syntrophales_bacterium | MOQ23_05915 | 66900..70226 | 2928017 | Swt1-family-HEPN-domain-containing-protein-[Syntrophales-bacterium] |
| MCK9230422.1 | HEPN+AAA-ATPase+Alpha+Beta-domain+wHTH+RRM | Type-1_BR.RM.like.systems | <-HEPN+DUF499*<-TOPRIM<-OLD-ABC<-METHYLASE<-Schlafen+HATPase+HTH<-PHP+SbcC<-TUDOR+SF1-SF2-Helicase+iREase+CTD+REase | 638 | Bacteria>Thermodesulfobacteriota>Syntrophia>Syntrophales>unclassified_Syntrophales | GCA_023230845.1 | Syntrophales_bacterium | MOP18_07070 | 1..1913 | 2928017 | Swt1-family-HEPN-domain-containing-protein-[Syntrophales-bacterium] |
| SPF49808.1 | HEPN+AAA-ATPase+Alpha+Beta-domain+wHTH+RRM | Type-1_BR.RM.like.systems | ParB+HNH->DrHyd+BirA-HTH->STAS+HisKin->METHYLASE->HEPN+DUF499*-> | 1116 | Bacteria>Thermodesulfobacteriota>Syntrophobacteria>Syntrophobacterales>Syntrophobacteraceae | GCA_900290365.1 | Syntrophobacter_sp._SbD1 | SBDP1_830032 | 43636..46986 | 2043167 | conserved-hypothetical-protein-[Syntrophobacter-sp.-SbD1] |
| WP_028893190.1 | HEPN+AAA-ATPase+Alpha+Beta-domain+wHTH+RRM | Type-1_BR.RM.like.systems | TUDOR+SF1-SF2-Helicase+iREase+CTD+REase->MORC+DUF3684->SbcC->OLD-ABC->TOPRIM->METHYLASE->HEPN+DUF499*->DUF4338->NurA->HerA/FtsK-><-?<-AbiEii<-AbiEi_1 | 1108 | Bacteria>Thermodesulfobacteriota>Syntrophorhabdia>Syntrophorhabdales>Syntrophorhabdaceae | GCF_000512235.1 | Syntrophorhabdus_aromaticivorans | SYNAR_RS0100650 | 131724..135050 | 328301 | Swt1-family-HEPN-domain-containing-protein-[Syntrophorhabdus-aromaticivorans] |
| MDD3846877.1 | HEPN+AAA-ATPase+Alpha+Beta-domain+wHTH+RRM | Type-1_BR.RM.like.systems | <-HEPN<-ParB+HNH<-HEPN+DUF499*<-METHYLASE<-MORC+DUF3684<-TUDOR+SF1-SF2-Helicase+iREase+CTD+REase<-DUF932 | 1114 | Bacteria>Thermodesulfobacteriota>Syntrophorhabdia>Syntrophorhabdales>Syntrophorhabdaceae | GCA_028698615.1 | Syntrophorhabdaceae_bacterium | PHC90_11025 | 3963..7307 | 2660714 | MAG:-Swt1-family-HEPN-domain-containing-protein-[Syntrophorhabdaceae-bacterium] |
| OPY85236.1 | HEPN+AAA-ATPase+Alpha+Beta-domain+wHTH+RRM | Type-1_BR.RM.like.systems | METHYLASE->HEPN+DUF499*-> | 1175 | Bacteria>Thermodesulfobacteriota>Syntrophorhabdia>Syntrophorhabdales>Syntrophorhabdaceae | GCA_002067475.1 | Syntrophorhabdus_sp._PtaU1.Bin153 | A4E65_00163 | 2092..5619 | 1811705 | MAG:-hypothetical-protein-A4E65_00163-[Syntrophorhabdus-sp.-PtaU1.Bin153] |
| MDE2480824.1 | HEPN+AAA-ATPase+Alpha+Beta-domain+wHTH+RRM | Type-1_BR.RM.like.systems | <-SF1-SF2-Helicase<-OLD-ABC+TOPRIM<-HEPN+DUF499*<-METHYLASE<-TUDOR+SF1-SF2-Helicase+iREase+CTD+REase\|\|PIN->AbiEi_3_N+AbiEi_3-><-SF1-SF2-helicase | 1110 | Bacteria>unclassified_Bacteria | GCA_028869385.1 | bacterium | KGN02_01370 | 63070..66402 | 1869227 | MAG:-DUF499-domain-containing-protein-[bacterium] |
| MCG3151370.1 | HEPN+AAA-ATPase+Alpha+Beta-domain+wHTH+RRM | Type-1_BR.RM.like.systems | TUDOR+SF1-SF2-Helicase+iREase+CTD+REase->SIR2->METHYLASE->HEPN+DUF499*->HTH->MPTase->?->?->Primpol+D5N+D5ATPase->HIN-HTH-> | 1134 | Bacteria>unclassified_Bacteria | GCA_022072165.1 | bacterium | GEEBNDBF_00641 | 748382..751786 | 1869227 | hypothetical-protein-[bacterium] |
| MCY4258519.1 | HEPN+AAA-ATPase+Alpha+Beta-domain+wHTH+RRM | Type-1_BR.RM.like.systems | <-DUF222+HNH<-PIN<-DUF433\|\|?-><-HEPN+DUF499*<-?\|\|?-><-METHYLASE<-SbcC<-SbcC<-TUDOR+SF1-SF2-Helicase+iREase+CTD+REase | 1091 | Bacteria>unclassified_Bacteria | GCA_026709785.1 | bacterium | OXE04_09620 | 60604..63879 | 1869227 | MAG:-DUF499-domain-containing-protein-[bacterium] |
| OQA20075.1 | HEPN+AAA-ATPase+Alpha+Beta-domain+wHTH+RRM | Type-1_BR.RM.like.systems | DUF4236+TPR-><-Peptidase_M20<-?<-HEPN+DUF499*<-?<-DUF433<-METHYLASE<-TUDOR+SF1-SF2-Helicase+iREase+CTD+REase | 1128 | Bacteria>unclassified_Bacteria | GCA_002069435.1 | bacterium_ADurb.Bin363 | BWY64_00439 | 13366..16752 | 1866927 | hypothetical-protein-BWY64_00439-[bacterium-ADurb.Bin363] |
| KAF0250487.1 | HEPN+AAA-ATPase+Alpha+Beta-domain+wHTH+RRM | Type-1_BR.RM.like.systems | Phd-YefM->PIN->METHYLASE->REase->OLD-ABC->TOPRIM->HEPN+DUF499*->?-><-Phage_integrase\|\|?->MqsA_antitoxin+HTH_3-> | 1130 | Bacteria>unclassified_Bacteria | GCA_009773835.1 | bacterium | FD167_113 | 11053..14445 | 1869227 | hypothetical-protein-FD167_113-[bacterium] |
| MCG3156467.1 | HEPN+AAA-ATPase+Alpha+Beta-domain+wHTH+RRM | Type-1_BR.RM.like.systems | TUDOR+SF1-SF2-Helicase+iREase+CTD+REase->Schlafen+HATPase+HTH->DOC+HTH->METHYLASE-><-PIN<-?\|\|HEPN+DUF499*->TIR-> | 1113 | Bacteria>unclassified_Bacteria | GCA_022072305.1 | bacterium | DKINENOH_03091 | 110830..114171 | 1869227 | hypothetical-protein-[bacterium] |
| MDE0368588.1 | HEPN+AAA-ATPase+Alpha+Beta-domain+wHTH+RRM | Type-1_BR.RM.like.systems | <-FAD-NAD-dep-oxidoreductase<-?<-?<-?\|\|?-><-HEPN+DUF499*<-METHYLASE\|\|?-><-MORC+HisKin+REase<-MNS+HTH<-REase\|\|HEPN-> | 1095 | Bacteria>unclassified_Bacteria | GCA_028821265.1 | bacterium | OXI84_00435 | 13176..16463 | 1869227 | MAG:-DUF499-domain-containing-protein-[bacterium] |
| MBK8128933.1 | HEPN+AAA-ATPase+Alpha+Beta-domain+wHTH+RRM | Type-1_BR.RM.like.systems | DDE_3->?-><-?\|\|?-><-?<-?<-HEPN+DUF499*<-METHYLASE-p<-PIN<-CopG/RHH<-PIN<-?<-TUDOR+SF1-SF2-Helicase+iREase+CTD+REase-p | 1097 | Bacteria>unclassified_Bacteria | GCA_016712825.1 | bacterium | IPK53_08290 | 27062..30355 | 1869227 | DUF499-domain-containing-protein-[bacterium] |
| MDD3925571.1 | HEPN+AAA-ATPase+Alpha+Beta-domain+wHTH+RRM | Type-1_BR.RM.like.systems | <-SF1-SF2-Helicase<-HEPN+DUF499*<-METHYLASE<-Schlafen+HATPase+HTH<-TUDOR+SF1-SF2-Helicase+iREase+CTD+REase | 1106 | Bacteria>unclassified_Bacteria | GCA_028699365.1 | bacterium | PHT33_02815 | 1591..4911 | 1869227 | MAG:-Swt1-family-HEPN-domain-containing-protein-[bacterium] |
| MCY4102170.1 | HEPN+AAA-ATPase+Alpha+Beta-domain+wHTH+RRM | Type-1_BR.RM.like.systems | CopG/RHH->PemK_toxin->REase-p-><-PIN<-CopG/RHH\|\|METHYLASE->HEPN+DUF499*->?->?->NUDIX-><-?<-?<-ABC-ATPase+SbcC | 1108 | Bacteria>unclassified_Bacteria | GCA_026708055.1 | bacterium | OXG55_02720 | 179181..182507 | 1869227 | MAG:-Swt1-family-HEPN-domain-containing-protein-[bacterium] |
| MCY3924850.1 | HEPN+AAA-ATPase+Alpha+Beta-domain+wHTH+RRM | Type-1_BR.RM.like.systems | PIN->DUF433->PIN->METHYLASE->MetJarc-HTH->PemK_toxin->HEPN+DUF499*-><-?\|\|ABhydrolase->?-><-?<-Glyoxalase | 1099 | Bacteria>unclassified_Bacteria | GCA_026707305.1 | bacterium | OXG52_04985 | 27802..31101 | 1869227 | MAG:-DUF499-domain-containing-protein-[bacterium] |
| PZN89122.1 | HEPN+AAA-ATPase+Alpha+Beta-domain+wHTH+RRM | Type-1_BR.RM.like.systems | SF1-SF2-Helicase+REase-p->SF1-SF2-Helicase+REase-><-?\|\|?-><-?<-?<-HEPN+DUF499*<-RNaseH<-METHYLASE-p | 1114 | Bacteria>unclassified_Bacteria | GCA_003242735.1 | bacterium | DIU52_15115 | 17855..21199 | 1869227 | AAA+-family-ATPase-[bacterium] |
| MCY4103460.1 | HEPN+AAA-ATPase+Alpha+Beta-domain+wHTH+RRM | Type-1_BR.RM.like.systems | DUF433->PIN->METHYLASE->MetJarc-HTH->PemK_toxin->HEPN+DUF499*-><-?\|\|ABhydrolase-><-?\|\|?-><-?\|\|REase-> | 1099 | Bacteria>unclassified_Bacteria | GCA_026708055.1 | bacterium | OXG55_09400 | 103234..106533 | 1869227 | MAG:-DUF499-domain-containing-protein-[bacterium] |
| MBQ7502581.1 | HEPN+AAA-ATPase+Alpha+Beta-domain+wHTH+RRM | Type-1_BR.RM.like.systems | <-HEPN+DUF499*\|\|REase->?->?-><-METHYLASE<-TUDOR+SF1-SF2-Helicase+iREase+CTD+REase\|\|REase-> | 1127 | Bacteria>unclassified_Bacteria | GCA_017458675.1 | bacterium | IJT93_07695 | 1961..5344 | 1869227 | DUF499-domain-containing-protein-[bacterium] |
| MCY3951956.1 | HEPN+AAA-ATPase+Alpha+Beta-domain+wHTH+RRM | Type-1_BR.RM.like.systems | <-NUDIX<-?<-?<-HEPN+DUF499*<-METHYLASE\|\|CopG/RHH->PIN-><-REase-p<-PemK_toxin<-CopG/RHH | 1108 | Bacteria>unclassified_Bacteria | GCA_026707535.1 | bacterium | OXG69_05350 | 3578..6904 | 1869227 | MAG:-Swt1-family-HEPN-domain-containing-protein-[bacterium] |
| MCY4163400.1 | HEPN+AAA-ATPase+Alpha+Beta-domain+wHTH+RRM | Type-1_BR.RM.like.systems | DUF433->PIN->METHYLASE->METHYLASE-p-><-?\|\|?->HEPN+DUF499*-><-?\|\|?->?->?->ABC-ATPase+SbcC->PfkB-> | 1091 | Bacteria>unclassified_Bacteria | GCA_026709025.1 | bacterium | OXE93_04200 | 10413..13688 | 1869227 | MAG:-DUF499-domain-containing-protein-[bacterium] |
| MCY3953257.1 | HEPN+AAA-ATPase+Alpha+Beta-domain+wHTH+RRM | Type-1_BR.RM.like.systems | DUF433->PIN->METHYLASE->MetJarc-HTH->PemK_toxin->HEPN+DUF499*-><-?\|\|ABhydrolase-> | 1099 | Bacteria>unclassified_Bacteria | GCA_026707535.1 | bacterium | OXG69_12060 | 23481..26780 | 1869227 | MAG:-DUF499-domain-containing-protein-[bacterium] |
| MCY3806661.1 | HEPN+AAA-ATPase+Alpha+Beta-domain+wHTH+RRM | Type-1_BR.RM.like.systems | METHYLASE->MetJarc-HTH->PemK_toxin->HEPN+DUF499*->DUF499-><-?\|\|ABhydrolase-> | 355 | Bacteria>unclassified_Bacteria | GCA_026706585.1 | bacterium | OXG91_09240 | 4057..5121 | 1869227 | MAG:-Swt1-family-HEPN-domain-containing-protein,-partial-[bacterium] |
| MCY3889489.1 | HEPN+AAA-ATPase+Alpha+Beta-domain+wHTH+RRM | Type-1_BR.RM.like.systems | <-PIN<-PhdYeFM_antitox<-?\|\|DUF222+HNH->?->PIN->HEPN+DUF499*-><-?<-DOC+DOC<-HTH_31 | 1079 | Bacteria>unclassified_Bacteria | GCA_026707085.1 | bacterium | OXG50_06285 | 19658..22897 | 1869227 | MAG:-DUF499-domain-containing-protein-[bacterium] |
| MCY3862385.1 | HEPN+AAA-ATPase+Alpha+Beta-domain+wHTH+RRM | Type-1_BR.RM.like.systems | <-PIN<-PhdYeFM_antitox<-?\|\|DUF222+HNH->?->PIN->HEPN+DUF499*-><-?<-DOC+DOC<-HTH_31 | 1079 | Bacteria>unclassified_Bacteria | GCA_026706925.1 | bacterium | OXG67_10535 | 19896..23135 | 1869227 | MAG:-DUF499-domain-containing-protein-[bacterium] |
| MCY3562206.1 | HEPN+AAA-ATPase+Alpha+Beta-domain+wHTH+RRM | Type-1_BR.RM.like.systems | <-ABhydrolase\|\|?->?-><-HEPN+DUF499*\|\|ParD_antitoxin-><-?<-METHYLASE<-PIN<-DUF433 | 1084 | Bacteria>unclassified_Bacteria | GCA_026702815.1 | bacterium | OXH20_13745 | 3210..6464 | 1869227 | MAG:-DUF499-domain-containing-protein-[bacterium] |
| MCY4258449.1 | HEPN+AAA-ATPase+Alpha+Beta-domain+wHTH+RRM | Type-1_BR.RM.like.systems | <-HEPN+DUF499*<-?<-METHYLASE<-?<-TUDOR+SF1-SF2-Helicase+iREase+CTD+REase | 1116 | Bacteria>unclassified_Bacteria | GCA_026709785.1 | bacterium | OXE04_09265 | 10514..13864 | 1869227 | MAG:-Swt1-family-HEPN-domain-containing-protein-[bacterium] |
| MDE0668308.1 | HEPN+AAA-ATPase+Alpha+Beta-domain+wHTH+RRM | Type-1_BR.RM.like.systems | CopG/RHH-><-PIN<-MPTase<-PIN<-CopG/RHH\|\|METHYLASE->HEPN+DUF499*-> | 1108 | Bacteria>unclassified_Bacteria | GCA_028824385.1 | bacterium | OXI48_04695 | 5107..8433 | 1869227 | MAG:-Swt1-family-HEPN-domain-containing-protein-[bacterium] |
| MCY3963126.1 | HEPN+AAA-ATPase+Alpha+Beta-domain+wHTH+RRM | Type-1_BR.RM.like.systems | <-PIN<-PhdYeFM_antitox<-?\|\|DUF222+HNH->?->PIN->HEPN+DUF499*-><-?<-HTH_31 | 1079 | Bacteria>unclassified_Bacteria | GCA_026707505.1 | bacterium | OXG34_15910 | 28430..31669 | 1869227 | MAG:-DUF499-domain-containing-protein-[bacterium] |
| MCP3995426.1 | HEPN+AAA-ATPase+Alpha+Beta-domain+wHTH+RRM | Type-1_BR.RM.like.systems | zf-ISL3+HTH_Tnp_ISL3+DDE_Tnp_ISL3->METHYLASE-p->HEPN+DUF499*-> | 479 | Bacteria>unclassified_Bacteria | GCA_024226295.1 | bacterium | GY722_10220 | 2321..3757 | 1869227 | ATP-binding-protein-[bacterium] |
| MDE0115147.1 | HEPN+AAA-ATPase+Alpha+Beta-domain+wHTH+RRM | Type-1_BR.RM.like.systems | <-HEPN+DUF499*<-DUF222+HNH\|\|?->PhdYeFM_antitox->PIN->?-><-METHYLASE | 1079 | Bacteria>unclassified_Bacteria | GCA_028819295.1 | bacterium | OXT07_00780 | 18092..21331 | 1869227 | MAG:-DUF499-domain-containing-protein-[bacterium] |
| MCY4370194.1 | HEPN+AAA-ATPase+Alpha+Beta-domain+wHTH+RRM | Type-1_BR.RM.like.systems | <-HEPN+DUF499*<-METHYLASE\|\|HEPN->DDE_Tnp_1-><-?<-HerA/FtsK | 1095 | Bacteria>unclassified_Bacteria | GCA_026710435.1 | bacterium | OXF41_12460 | 874..4161 | 1869227 | MAG:-DUF499-domain-containing-protein-[bacterium] |
| MCY3785681.1 | HEPN+AAA-ATPase+Alpha+Beta-domain+wHTH+RRM | Type-1_BR.RM.like.systems | <-CHTH<-?\|\|BirA-HTH->CopG/RHH->METHYLASE->HEPN+DUF499*-> | 1126 | Bacteria>unclassified_Bacteria | GCA_026706565.1 | bacterium | OXG47_03025 | 15510..18890 | 1869227 | MAG:-Swt1-family-HEPN-domain-containing-protein-[bacterium] |
| MCY3562591.1 | HEPN+AAA-ATPase+Alpha+Beta-domain+wHTH+RRM | Type-1_BR.RM.like.systems | METHYLASE->HEPN+DUF499*->HTH_3->NTP_transf-><-?<-PIN | 1107 | Bacteria>unclassified_Bacteria | GCA_026702805.1 | bacterium | OXH26_01515 | 1911..5234 | 1869227 | MAG:-DUF499-domain-containing-protein-[bacterium] |
| NJL57104.1 | HEPN+AAA-ATPase+Alpha+Beta-domain+wHTH+RRM | Type-1_BR.RM.like.systems | METHYLASE-p->DUF433->PIN->HEPN+DUF499*->DUF499-> | 411 | Bacteria>unclassified_Bacteria | GCA_012031555.1 | bacterium | HC928_19635 | 992..2227 | 1869227 | ATP-binding-protein-[bacterium] |
| MCY3923936.1 | HEPN+AAA-ATPase+Alpha+Beta-domain+wHTH+RRM | Type-1_BR.RM.like.systems | <-CHTH\|\|DOC->CopG/RHH->METHYLASE->HEPN+DUF499*-> | 1131 | Bacteria>unclassified_Bacteria | GCA_026707305.1 | bacterium | OXG52_00270 | 5305..8700 | 1869227 | MAG:-Swt1-family-HEPN-domain-containing-protein-[bacterium] |
| MCY4135752.1 | HEPN+AAA-ATPase+Alpha+Beta-domain+wHTH+RRM | Type-1_BR.RM.like.systems | <-HEPN+DUF499*\|\|?-><-METHYLASE\|\|DOC-><-HEPN+REase | 1079 | Bacteria>unclassified_Bacteria | GCA_026708015.1 | bacterium | OXG30_12705 | 27919..31158 | 1869227 | MAG:-DUF499-domain-containing-protein-[bacterium] |
| MDE0170777.1 | HEPN+AAA-ATPase+Alpha+Beta-domain+wHTH+RRM | Type-1_BR.RM.like.systems | <-HEPN+DUF499*<-METHYLASE<-XIS-HTH+RVT<-?<-?<-?<-REase | 1096 | Bacteria>unclassified_Bacteria | GCA_028819595.1 | bacterium | OXS29_14930 | 550..3840 | 1869227 | MAG:-DUF499-domain-containing-protein-[bacterium] |
| MCY3637183.1 | HEPN+AAA-ATPase+Alpha+Beta-domain+wHTH+RRM | Type-1_BR.RM.like.systems | <-HEPN+DUF499*<-METHYLASE<-OLD-ABC+TOPRIM<-?<-TOPRIM<-OLD-ABC | 1079 | Bacteria>unclassified_Bacteria | GCA_026705625.1 | bacterium | OXH23_16400 | 13524..16763 | 1869227 | MAG:-DUF499-domain-containing-protein-[bacterium] |
| MDE0377403.1 | HEPN+AAA-ATPase+Alpha+Beta-domain+wHTH+RRM | Type-1_BR.RM.like.systems | <-Glyoxalase<-?\|\|?->?-><-?<-HEPN+DUF499*<-METHYLASE | 1095 | Bacteria>unclassified_Bacteria | GCA_028821235.1 | bacterium | OXK16_15785 | 33048..36335 | 1869227 | MAG:-DUF499-domain-containing-protein-[bacterium] |
| MDE0578824.1 | HEPN+AAA-ATPase+Alpha+Beta-domain+wHTH+RRM | Type-1_BR.RM.like.systems | <-HEPN+DUF499*\|\|?-><-METHYLASE\|\|HTH_58-> | 1079 | Bacteria>unclassified_Bacteria | GCA_028824815.1 | bacterium | OXI29_03025 | 42794..46033 | 1869227 | MAG:-DUF499-domain-containing-protein-[bacterium] |
| MCR5845017.1 | HEPN+AAA-ATPase+Alpha+Beta-domain+wHTH+RRM | Type-1_BR.RM.like.systems | REase-><-?\|\|?-><-?<-HEPN+DUF499*<-METHYLASE | 1130 | Bacteria>unclassified_Bacteria | GCA_024698685.1 | bacterium | K6G78_00395 | 19668..23060 | 1869227 | DUF499-domain-containing-protein-[bacterium] |
| MDE0193912.1 | HEPN+AAA-ATPase+Alpha+Beta-domain+wHTH+RRM | Type-1_BR.RM.like.systems | METHYLASE->HEPN+DUF499*->?->?->NUDIX-> | 1108 | Bacteria>unclassified_Bacteria | GCA_028819945.1 | bacterium | OXP08_00010 | 2300..5626 | 1869227 | MAG:-Swt1-family-HEPN-domain-containing-protein-[bacterium] |
| MDC0992039.1 | HEPN+AAA-ATPase+Alpha+Beta-domain+wHTH+RRM | Type-1_BR.RM.like.systems | <-OLD-ABC<-HEPN+DUF499*<-METHYLASE | 1135 | Bacteria>unclassified_Bacteria | GCA_028375025.1 | bacterium | OAR33_00540 | 2470..5877 | 1869227 | Swt1-family-HEPN-domain-containing-protein-[bacterium] |
| MCI0414205.1 | HEPN+AAA-ATPase+Alpha+Beta-domain+wHTH+RRM | Type-1_BR.RM.like.systems | <-HEPN<-HEPN+DUF499*<-METHYLASE | 1107 | Bacteria>unclassified_Bacteria | GCA_022615055.1 | bacterium | L0222_15590 | 1218..4541 | 1869227 | DUF499-domain-containing-protein-[bacterium] |
| TRZ47091.1 | HEPN+AAA-ATPase+Alpha+Beta-domain+wHTH+RRM | Type-1_BR.RM.like.systems | <-HEPN+DUF499*<-?<-METHYLASE-p | 1109 | Bacteria>unclassified_Bacteria | GCA_007279595.1 | bacterium | D4S03_11515 | 927..4256 | 1869227 | ATP-binding-protein-[bacterium] |
| MDD2328800.1 | HEPN+AAA-ATPase+Alpha+Beta-domain+wHTH+RRM | Type-1_BR.RM.like.systems | METHYLASE-p->HEPN+DUF499*-> | 1132 | Bacteria>unclassified_Bacteria | GCA_028683535.1 | bacterium | PHZ13_10910 | 283..3681 | 1869227 | MAG:-DUF499-domain-containing-protein-[bacterium] |
| MCY3563828.1 | HEPN+AAA-ATPase+Alpha+Beta-domain+wHTH+RRM | Type-1_BR.RM.like.systems | <-HEPN+DUF499*<-METHYLASE-p | 754 | Bacteria>unclassified_Bacteria | GCA_026702805.1 | bacterium | OXH26_07915 | 1..2262 | 1869227 | MAG:-DUF499-domain-containing-protein,-partial-[bacterium] |
| MCY3560578.1 | HEPN+AAA-ATPase+Alpha+Beta-domain+wHTH+RRM | Type-1_BR.RM.like.systems | METHYLASE-p->HEPN+DUF499*-> | 1108 | Bacteria>unclassified_Bacteria | GCA_026702815.1 | bacterium | OXH20_05350 | 698..4024 | 1869227 | MAG:-Swt1-family-HEPN-domain-containing-protein-[bacterium] |
| MCP4656470.1 | HEPN+AAA-ATPase+Alpha+Beta-domain+wHTH+RRM | Type-1_BR.RM.like.systems | <-HEPN+DUF499*<-?<-METHYLASE-p | 1058 | Bacteria>unclassified_Bacteria | GCA_024231315.1 | bacterium | GY856_13735 | 1..3174 | 1869227 | ATP-binding-protein-[bacterium] |
| MCY4632748.1 | HEPN+AAA-ATPase+Alpha+Beta-domain+wHTH+RRM | Type-1_BR.RM.like.systems | <-HEPN+DUF499*<-METHYLASE | 487 | Bacteria>unclassified_Bacteria | GCA_026713745.1 | bacterium | OXE75_17955 | 1..1460 | 1869227 | MAG:-Swt1-family-HEPN-domain-containing-protein,-partial-[bacterium] |
| MCY4630164.1 | HEPN+AAA-ATPase+Alpha+Beta-domain+wHTH+RRM | Type-1_BR.RM.like.systems | <-HEPN+DUF499*<-METHYLASE | 332 | Bacteria>unclassified_Bacteria | GCA_026713745.1 | bacterium | OXE75_04665 | 1..996 | 1869227 | MAG:-Swt1-family-HEPN-domain-containing-protein,-partial-[bacterium] |
| MBU2648361.1 | HEPN+AAA-ATPase+Alpha+Beta-domain+wHTH+RRM | Type-1_BR.RM.like.systems | TUDOR+SF1-SF2-Helicase+iREase+CTD+REase->wHTH+REase->METHYLASE->HEPN+DUF499*->?->Phage_integrase-><-Arm-DNA-bind_3+Phage_int_SAM_3+Phage_integrase | 1091 | Bacteria>unclassified_Bacteria | GCA_018830885.1 | bacterium | KKI24_26885 | 8390..11665 | 1869227 | DUF499-domain-containing-protein-[bacterium] |
| MCP4152612.1 | HEPN+AAA-ATPase+Alpha+Beta-domain+wHTH+RRM | Type-1_BR.RM.like.systems | Radical-SAM->?-><-HEPN+DUF499*<-METHYLASE-p | 1101 | Bacteria>unclassified_Bacteria | GCA_024224895.1 | bacterium | GY757_33055 | 6237..9542 | 1869227 | ATP-binding-protein-[bacterium] |
| WP_248539435.1 | HEPN+AAA-ATPase+Alpha+Beta-domain+wHTH+RRM | Type-1_BR.RM.like.systems | MerR-HTH-><-?\|\|N6_MTase-p-><-OLD-ABC+TOPRIM<-HEPN+DUF499*<-METHYLASE<-TUDOR+SF1-SF2-Helicase+iREase+CTD+REase<-XIS-HTH<-?<-HTH_Tnp_1+HTH_21+rve+rve_2 | 1132 | - | GCF_023195655.1 | Not | M0721_RS02800 | 592569..595967 | 2851643 | DUF499-domain-containing-protein-[Microbacterium-sp.-SSW1-7] |

| Protein-ID | DUF499-architecture | System | Gene-Neighbourhoods | Length | Taxonomy | Assembly-ID | Strain | Locus-Tag | Genomic-coordinates | TaxID | Fasta-defline |
|---|---|---|---|---|---|---|---|---|---|---|---|
| -- | | | | | | | | | | | |
| --BR-2.RM.like.systems | | | | | | | | | | | |
| -- | | | | | | | | | | | |
| PMP75322.1 | AAA+ATPase+wHTH+Alpha+Beta-domain+wHTH+RRM | Type-2_BR.RM.like.systems | <-SF1-SF2-Helicase+iREase+CTD+REase<-METHYLASE<-DUF3780<-DUF499* | 883 | Archaea>Candidatus_Thermoplasmatota>DHVE2_group>Aciduliprofundum>unclassified_Aciduliprofundum | GCA_002878135.1 | Aciduliprofundum_sp. | C0180_01660 | 6420..9071 | 2060325 | MAG:-hypothetical-protein-C0180_01660-[Aciduliprofundum-sp.] |
| OPY34213.1 | AAA+ATPase+wHTH+Alpha+Beta-domain+wHTH+RRM | Type-2_BR.RM.like.systems | Phage_int_SAM_4+Phage_integrase-><-ASCH<-PIN+ACET+EVE\|\|DUF499*->DUF3780->METHYLASE->TUDOR+SF1-SF2-Helicase+iREase+CTD-><-ZetaToxin<-?<-OLD-ABC+TOPRIM | 981 | Archaea>Candidatus_Thermoplasmatota>Thermoplasmata>Methanomassiliicoccales>unclassified_Methanomassiliicoccales | GCA_002067865.1 | Methanomassiliicoccales_archaeon_PtaU1.Bin124 | A4E32_00485 | 156463..159408 | 1811730 | MAG:-hypothetical-protein-A4E32_00485-[Methanomassiliicoccales-archaeon-PtaU1.Bin124] |
| MBD6956366.1 | AAA+ATPase+wHTH+Alpha+Beta-domain+wHTH+RRM | Type-2_BR.RM.like.systems | <-SF1-SF2-Helicase+iREase+CTD+REase<-METHYLASE<-DUF3780<-DUF499* | 890 | Archaea>Candidatus_Thermoplasmatota>Thermoplasmata>Thermoplasmatales>unclassified_Thermoplasmatales | GCA_014836265.1 | Thermoplasmatales_archaeon | F9Y92_07015 | 7804..10476 | 2268204 | DUF499-domain-containing-protein-[Thermoplasmatales-archaeon] |
| PKL69081.1 | AAA+ATPase+wHTH+Alpha+Beta-domain+wHTH+RRM | Type-2_BR.RM.like.systems | <-REase<-?<-EVE<-SF1-SF2-Helicase+iREase+CTD+REase<-METHYLASE<-DUF3780<-DUF499*\|\|REase->?->?-><-REase\|\|?->ACET-> | 1018 | Archaea>Euryarchaeota>Methanomada_group>Methanobacteria>Methanobacteriales | GCA_002839705.1 | Methanobacteriales_archaeon_HGW-Methanobacteriales-1 | CVV28_02915 | 553077..556133 | 2013815 | hypothetical-protein-CVV28_02915-[Methanobacteriales-archaeon-HGW-Methanobacteriales-1] |
| WP_211533304.1 | AAA+ATPase+wHTH+Alpha+Beta-domain+wHTH+RRM | Type-2_BR.RM.like.systems | <-REase<-?<-EVE<-SF1-SF2-Helicase+iREase+CTD+REase<-METHYLASE<-DUF3780<-DUF499*\|\|REase->?->?->?-><-?<-TPR+TPR | 965 | Archaea>Euryarchaeota>Methanomada_group>Methanobacteria>Methanobacteriales | GCF_018141185.1 | Methanobacterium_alkalithermotolerans | HYG87_RS00570 | 112143..115040 | 2731220 | DUF499-domain-containing-protein-[Methanobacterium-alkalithermotolerans] |
| WP_004594465.1 | AAA+ATPase+wHTH+Alpha+Beta-domain+wHTH+RRM | Type-2_BR.RM.like.systems | PIWI-><-Cluster197_21clades<-HKD-DNase+SF1-SF2-Helicase+iREase+CTD<-?<-METHYLASE<-DUF3780<-DUF499* | 774 | Archaea>Euryarchaeota>Stenosarchaea_group>Halobacteria>Halobacteriales | GCF_000336635.1 | Haloarcula_japonica | C444_RS17915 | 26155..28480 | 29282 | hypothetical-protein,-partial-[Haloarcula-japonica] |
| WP_007743304.1 | AAA+ATPase+wHTH+Alpha+Beta-domain+wHTH+RRM | Type-2_BR.RM.like.systems | <-HKD-DNase+SF1-SF2-Helicase+iREase+CTD<-METHYLASE<-DUF499*<-AbrB-HTH<-PadR-HTH | 1099 | Archaea>Euryarchaeota>Stenosarchaea_group>Halobacteria>Halobacteriales | GCF_000336715.1 | Halococcus_thailandensis | C451_RS19280 | 34174..37473 | 335952 | DUF499-domain-containing-protein-[Halococcus-thailandensis] |
| WP_008524485.1 | AAA+ATPase+wHTH+Alpha+Beta-domain+wHTH+RRM | Type-2_BR.RM.like.systems | <-HKD-DNase+SF1-SF2-Helicase+iREase+CTD<-METHYLASE<-DUF3780<-DUF499*<-?\|\|Cluster197_21clades-><-SbcC+SbcC<-?\|\|?->ZetaToxin-> | 1097 | Archaea>Euryarchaeota>Stenosarchaea_group>Halobacteria>Halobacteriales | GCF_000470655.1 | Halorhabdus_tiamatea | HTIA_RS08715 | 1770607..1773900 | 430914 | DUF499-domain-containing-protein-[Halorhabdus-tiamatea] |
| WP_015408759.1 | AAA+ATPase+wHTH+Alpha+Beta-domain+wHTH+RRM | Type-2_BR.RM.like.systems | <-HKD-DNase+SF1-SF2-Helicase+iREase+CTD<-METHYLASE<-zf-IS66+DDE_Tnp_IS66<-METHYLASE-p<-DUF3780<-DUF499*<-Phage_integrase | 1124 | Archaea>Euryarchaeota>Stenosarchaea_group>Halobacteria>Halobacteriales | GCF_000591055.1 | Natronomonas_moolapensis | NMLP_RS03550 | 712690..716064 | 416273 | DUF499-domain-containing-protein-[Natronomonas-moolapensis] |
| WP_228841862.1 | AAA+ATPase+wHTH+Alpha+Beta-domain+wHTH+RRM | Type-2_BR.RM.like.systems | <-HKD-DNase+SF1-SF2-Helicase+iREase+CTD<-HEPN<-METHYLASE<-DUF3780<-DUF499*<-AbrB-HTH | 1108 | Archaea>Euryarchaeota>Stenosarchaea_group>Halobacteria>Halobacteriales | GCF_001485575.1 | Haloarcula_sp._CBA1127 | AV059_RS20935 | 42218..45544 | 1765055 | DUF499-domain-containing-protein-[Haloarcula-sp.-CBA1127] |
| WP_232820588.1 | AAA+ATPase+wHTH+Alpha+Beta-domain+wHTH+RRM | Type-2_BR.RM.like.systems | <-HKD-DNase+SF1-SF2-Helicase+iREase+CTD<-?<-?<-METHYLASE-p<-DUF3780<-DUF499*<-DUF4011+SF1-SF2-Helicase+REase\|\|REase->SF1-SF2-Helicase+REase->Phage_integrase-><-ParB+HNH\|\|ZDNA-HTH-> | 1046 | Archaea>Euryarchaeota>Stenosarchaea_group>Halobacteria>Halobacteriales | GCF_003382685.1 | Halorussus_litoreus | DPQ64_RS14205 | 17558..20698 | 1710536 | DUF499-domain-containing-protein-[Halorussus-litoreus] |
| WP_123538893.1 | AAA+ATPase+wHTH+Alpha+Beta-domain+wHTH+RRM | Type-2_BR.RM.like.systems | <-HKD-DNase+SF1-SF2-Helicase+iREase+CTD<-METHYLASE<-DUF3780<-DUF499*<-?\|\|Cluster197_21clades-><-PIWI<-REase+PNTD1+PNTD2\|\|?->DDE_Tnp_1-> | 1097 | Archaea>Euryarchaeota>Stenosarchaea_group>Halobacteria>Halobacteriales | GCF_003730195.1 | Halosimplex_salinum | D1409_RS20665 | 23577..26870 | 1710538 | DUF499-domain-containing-protein-[Halosimplex-salinum] |
| WP_208023500.1 | AAA+ATPase+wHTH+Alpha+Beta-domain+wHTH+RRM | Type-2_BR.RM.like.systems | Aminotran_1_2->?->?->?->CHTH-><-DUF772+DDE_Tnp_1\|\|DUF499*->METHYLASE->HKD-DNase+SF1-SF2-Helicase+iREase+CTD-><-Phage_int_SAM_4+Phage_integrase<-TraJ-RHH | 1108 | Archaea>Euryarchaeota>Stenosarchaea_group>Halobacteria>Halobacteriales | GCF_004087835.1 | Halorussus_pelagicus | EP007_RS13895 | 2742258..2745584 | 2505977 | DUF499-domain-containing-protein-[Halorussus-pelagicus] |
| WP_209309559.1 | AAA+ATPase+wHTH+Alpha+Beta-domain+wHTH+RRM | Type-2_BR.RM.like.systems | <-PadR-HTH\|\|?->?->?->AbrB-HTH->DUF499*->DUF3780->METHYLASE->HKD-DNase+SF1-SF2-Helicase+iREase+CTD->Cluster197_21clades-><-PIWI<-REase+PNTD1+PNTD2 | 1099 | Archaea>Euryarchaeota>Stenosarchaea_group>Halobacteria>Halobacteriales | GCF_004681185.1 | Halomicroarcula_amylolytica | E4S61_RS18885 | 132255..135554 | 2562280 | DUF499-domain-containing-protein-[Halomicroarcula-amylolytica] |
| WP_135536459.1 | AAA+ATPase+wHTH+Alpha+Beta-domain+wHTH+RRM | Type-2_BR.RM.like.systems | <-Cluster197_21clades\|\|DUF499*->DUF3780->METHYLASE->HKD-DNase+SF1-SF2-Helicase+iREase+CTD->SMODS-> | 1098 | Archaea>Euryarchaeota>Stenosarchaea_group>Halobacteria>Halobacteriales | GCF_004762015.1 | Halostella_pelagica | EYW42_RS16390 | 56792..60088 | 2583824 | DUF499-domain-containing-protein-[Halostella-pelagica] |
| WP_136689357.1 | AAA+ATPase+wHTH+Alpha+Beta-domain+wHTH+RRM | Type-2_BR.RM.like.systems | REase+PNTD1+PNTD2->PIWI-><-Cluster197_21clades\|\|?->DUF499*->DUF3780->METHYLASE->HKD-DNase+SF1-SF2-Helicase+iREase+CTD->ORC/CDC6+HTH-> | 1096 | Archaea>Euryarchaeota>Stenosarchaea_group>Halobacteria>Halobacteriales | GCF_004944975.1 | Halorhabdus_amylolytica | E3374_RS12585 | 68541..71831 | 2559573 | DUF499-domain-containing-protein-[Halorhabdus-amylolytica] |
| WP_152156194.1 | AAA+ATPase+wHTH+Alpha+Beta-domain+wHTH+RRM | Type-2_BR.RM.like.systems | RHH+RAMA-><-REase\|\|?-><-HKD-DNase+SF1-SF2-Helicase+iREase+CTD<-METHYLASE<-DUF3780<-DUF499* | 1087 | Archaea>Euryarchaeota>Stenosarchaea_group>Halobacteria>Halobacteriales | GCF_009184555.1 | Halosegnis_rubeus | DP108_RS06505 | 166762..170025 | 2212850 | DUF499-domain-containing-protein-[Halosegnis-rubeus] |
| WP_157511186.1 | AAA+ATPase+wHTH+Alpha+Beta-domain+wHTH+RRM | Type-2_BR.RM.like.systems | <-HKD-DNase+SF1-SF2-Helicase+iREase+CTD-p<-HKD-DNase+SF1-SF2-Helicase+iREase+CTD-p<-?<-METHYLASE<-METHYLASE-p<-DUF3780<-DUF499*<-REase+HTH<-?\|\|REase->?->ORC/CDC6+HTH-> | 1097 | Archaea>Euryarchaeota>Stenosarchaea_group>Halobacteria>Halobacteriales | GCF_009741825.1 | Halapricum_sp._CBA1109 | GJ629_RS00110 | 29598..32891 | 2668068 | DUF499-domain-containing-protein-[Halapricum-sp.-CBA1109] |
| WP_179908254.1 | AAA+ATPase+wHTH+Alpha+Beta-domain+wHTH+RRM | Type-2_BR.RM.like.systems | <-SMODS<-ORC/CDC6+HTH<-?<-?<-?\|\|AbrB-HTH->DUF499*->DUF3780->METHYLASE->HKD-DNase+SF1-SF2-Helicase+iREase+CTD->Cluster197_21clades-> | 1099 | Archaea>Euryarchaeota>Stenosarchaea_group>Halobacteria>Halobacteriales | GCF_013415885.1 | Halosimplex_rubrum | HZS55_RS14055 | 2806959..2810258 | 869889 | DUF499-domain-containing-protein-[Halosimplex-rubrum] |
| WP_179922873.1 | AAA+ATPase+wHTH+Alpha+Beta-domain+wHTH+RRM | Type-2_BR.RM.like.systems | Mbetalac-><-ORC/CDC6+HTH\|\|ORC/CDC6+HTH-><-HKD-DNase+SF1-SF2-Helicase+iREase+CTD<-METHYLASE<-DUF3780<-DUF499*<-AbrB-HTH<-?\|\|Phage_int_SAM_5+Phage_integrase-><-DUF772+DDE_Tnp_1\|\|?->OrfB_IS605+OrfB_Zn_ribbon-> | 1099 | Archaea>Euryarchaeota>Stenosarchaea_group>Halobacteria>Halobacteriales | GCF_013415905.1 | Halosimplex_pelagicum | HZS54_RS12595 | 2543503..2546802 | 869886 | DUF499-domain-containing-protein-[Halosimplex-pelagicum] |
| WP_229772914.1 | AAA+ATPase+wHTH+Alpha+Beta-domain+wHTH+RRM | Type-2_BR.RM.like.systems | <-ASCH<-PIN+ACET+EVE\|\|AbrB-HTH->DUF499*->DUF3780->METHYLASE->SF1-SF2-Helicase-><-ZnR+REase\|\|HKD-DNase+SF1-SF2-Helicase+iREase+CTD->MORC+DUF3684+REase-> | 1109 | Archaea>Euryarchaeota>Stenosarchaea_group>Halobacteria>Halobacteriales | GCF_014647155.1 | Halarchaeum_nitratireducens | IEX91_RS14675 | 40696..44025 | 489913 | DUF499-domain-containing-protein-[Halarchaeum-nitratireducens] |
| WP_229871353.1 | AAA+ATPase+wHTH+Alpha+Beta-domain+wHTH+RRM | Type-2_BR.RM.like.systems | DDE_Tnp_IS240->\|\|?->RADICAL-SAM->?->AbrB-HTH->DUF499*->DUF3780->METHYLASE->SF1-SF2-Helicase-><-?<-SWACOS\|\|HKD-DNase+SF1-SF2-Helicase+iREase+CTD-> | 1108 | Archaea>Euryarchaeota>Stenosarchaea_group>Halobacteria>Halobacteriales | GCF_014647455.2 | Halarchaeum_grantii | IEY12_RS13405 | 16136..19462 | 1193105 | DUF499-domain-containing-protein-[Halarchaeum-grantii] |
| WP_220589967.1 | AAA+ATPase+wHTH+Alpha+Beta-domain+wHTH+RRM | Type-2_BR.RM.like.systems | SWACOS-><-?<-?<-HNH\|\|REase->DUF499*->DUF3780->METHYLASE->HKD-DNase+SF1-SF2-Helicase+iREase+CTD->Resolvase-><-MarR-HTH | 1098 | Archaea>Euryarchaeota>Stenosarchaea_group>Halobacteria>Halobacteriales | GCF_019599405.1 | Halomicroarcula_salinisoli | EGD98_RS19205 | 187933..191229 | 2487746 | DUF499-domain-containing-protein-[Halomicroarcula-salinisoli] |
| WP_220600311.1 | AAA+ATPase+wHTH+Alpha+Beta-domain+wHTH+RRM | Type-2_BR.RM.like.systems | <-HKD-DNase+SF1-SF2-Helicase+iREase+CTD<-?<-?<-METHYLASE<-DUF3780<-DUF499*\|\|?->BirA-HTH->?->?-><-CoiA | 1109 | Archaea>Euryarchaeota>Stenosarchaea_group>Halobacteria>Halobacteriales | GCF_019599485.1 | Halomicroarcula_salinisoli | EGH22_RS20450 | 35568..38897 | 2487746 | DUF499-domain-containing-protein-[Halomicroarcula-salinisoli] |
| WP_220581564.1 | AAA+ATPase+wHTH+Alpha+Beta-domain+wHTH+RRM | Type-2_BR.RM.like.systems | <-PadR-HTH\|\|HTH->?->?->AbrB-HTH->DUF499*->DUF3780->METHYLASE->HKD-DNase+SF1-SF2-Helicase+iREase+CTD->CoiA-> | 1099 | Archaea>Euryarchaeota>Stenosarchaea_group>Halobacteria>Halobacteriales | GCF_019599505.1 | Halomicroarcula_nitratireducens | EGH23_RS19010 | 75081..78380 | 2487749 | DUF499-domain-containing-protein-[Halomicroarcula-nitratireducens] |
| WP_226011842.1 | AAA+ATPase+wHTH+Alpha+Beta-domain+wHTH+RRM | Type-2_BR.RM.like.systems | GDResolvase+Recombinase->?->DUF499*->DUF3780->METHYLASE-><-?\|\|ParB+HNH->HKD-DNase+SF1-SF2-Helicase+iREase+CTD-> | 1101 | Archaea>Euryarchaeota>Stenosarchaea_group>Halobacteria>Halobacteriales | GCF_020405185.1 | Halomicrobium_salinisoli | LE162_RS01580 | 297593..300898 | 2878391 | DUF499-domain-containing-protein-[Halomicrobium-salinisoli] |
| WP_232590526.1 | AAA+ATPase+wHTH+Alpha+Beta-domain+wHTH+RRM | Type-2_BR.RM.like.systems | <-ORC/CDC6+HTH<-SbcC+SbcC<-?\|\|AbrB-HTH->DUF499*->DUF3780->METHYLASE->HKD-DNase+SF1-SF2-Helicase+iREase+CTD->?->?->RADICAL-SAM-> | 1099 | Archaea>Euryarchaeota>Stenosarchaea_group>Halobacteria>Halobacteriales | GCF_021233435.1 | Halobacterium_noricense | LT974_RS17375 | 346912..350211 | 223182 | DUF499-domain-containing-protein-[Halobacterium-noricense] |
| WP_232688937.1 | AAA+ATPase+wHTH+Alpha+Beta-domain+wHTH+RRM | Type-2_BR.RM.like.systems | <-ORC/CDC6+HTH<-?<-?<-?\|\|AbrB-HTH->DUF499*->DUF3780->METHYLASE->HKD-DNase+SF1-SF2-Helicase+iREase+CTD->Cluster197_21clades-> | 1099 | Archaea>Euryarchaeota>Stenosarchaea_group>Halobacteria>Halobacteriales | GCF_021249405.1 | Halobacterium_zhouii | LT970_RS14545 | 104100..107399 | 2902624 | DUF499-domain-containing-protein-[Halobacterium-sp.-XZYJT26] |
| WP_248653039.1 | AAA+ATPase+wHTH+Alpha+Beta-domain+wHTH+RRM | Type-2_BR.RM.like.systems | <-ParB+HNH\|\|?-><-ORC/CDC6+HTH<-HKD-DNase+SF1-SF2-Helicase+iREase+CTD<-METHYLASE<-DUF3780<-DUF499*<-?\|\|Cluster197_21clades-><-PIWI<-REase+PNTD1+PNTD2 | 1096 | Archaea>Euryarchaeota>Stenosarchaea_group>Halobacteria>Halobacteriales | GCF_023238205.1 | Halorussus_limi | MOR89_RS22505 | 156540..159830 | 2938695 | DUF499-domain-containing-protein-[Halorussus-sp.-XZYJT49] |
| WP_255150364.1 | AAA+ATPase+wHTH+Alpha+Beta-domain+wHTH+RRM | Type-2_BR.RM.like.systems | Resolvase->?-><-?\|\|REase+REase->?->DUF499*->DUF3780->METHYLASE->HKD-DNase+SF1-SF2-Helicase+iREase+CTD-><-REase<-?<-ACYC | 1084 | Archaea>Euryarchaeota>Stenosarchaea_group>Halobacteria>Halobacteriales | GCF_024298865.1 | Halorarius_halobius | NL112_RS18705 | 3543298..3546552 | 2962671 | DUF499-domain-containing-protein-[Halosegnis-sp.-ZY10] |
| WP_254824893.1 | AAA+ATPase+wHTH+Alpha+Beta-domain+wHTH+RRM | Type-2_BR.RM.like.systems | ZnR+REase-><-HKD-DNase+SF1-SF2-Helicase+iREase+CTD\|\|ParB+HNH-><-METHYLASE<-DUF3780<-DUF499*\|\|ZnR+REase-><-ParA-Soj-PloopNTPase | 1107 | Archaea>Euryarchaeota>Stenosarchaea_group>Halobacteria>Halobacteriales | GCF_024298885.1 | Haloglomus_halophilum | NL114_RS21530 | 126665..129988 | 2962672 | DUF499-domain-containing-protein-[Haloglomus-sp.-ZY41] |
| WP_254860685.1 | AAA+ATPase+wHTH+Alpha+Beta-domain+wHTH+RRM | Type-2_BR.RM.like.systems | NERD+SF1-SF2-Helicase-><-BirA-HTH\|\|?-><-SF1-SF2-Helicase+REase<-REase\|\|DUF499*->DUF3780-><-?\|\|METHYLASE->HKD-DNase+SF1-SF2-Helicase+iREase+CTD-><-?<-ParB+HNH+Schlafen | 1123 | Archaea>Euryarchaeota>Stenosarchaea_group>Halobacteria>Halobacteriales | GCF_024300825.1 | Natronomonas_gomsonensis | NMP98_RS06305 | 1133633..1137004 | 1046043 | DUF499-domain-containing-protein-[Natronomonas-gomsonensis] |
| WP_260644222.1 | AAA+ATPase+wHTH+Alpha+Beta-domain+wHTH+RRM | Type-2_BR.RM.like.systems | <-ORC/CDC6+HTH\|\|Exonuc_V_gamma+REase->SF1-SF2-Helicase+REase-><-HKD-DNase+SF1-SF2-Helicase+iREase+CTD<-METHYLASE<-DUF3780<-DUF499*<-AbrB-HTH<-ORC/CDC6+REase\|\|PIWI-> | 925 | Archaea>Euryarchaeota>Stenosarchaea_group>Halobacteria>Halobacteriales | GCF_025231485.1 | Salinirubellus_salinus | NOB31_RS21715 | 52598..55375 | 1364945 | ATP-binding-protein-[Salinirubellus-salinus] |
| WP_285255572.1 | AAA+ATPase+wHTH+Alpha+Beta-domain+wHTH+RRM | Type-2_BR.RM.like.systems | PIN-><-?<-?<-Phage_integrase\|\|DUF499*->?->METHYLASE->HKD-DNase+SF1-SF2-Helicase+iREase+CTD-> | 1093 | Archaea>Euryarchaeota>Stenosarchaea_group>Halobacteria>Halobacteriales | GCF_030250325.1 | Halobacterium_salinarum | PNP83_RS11130 | 2979..6260 | 2242 | DUF499-domain-containing-protein-[Halobacterium-salinarum] |
| WP_135665718.1 | AAA+ATPase+wHTH+Alpha+Beta-domain+wHTH+RRM | Type-2_BR.RM.like.systems | <-SF1-SF2-Helicase+REase<-Exonuc_V_gamma+REase\|\|?->SbcC+SbcC-><-Cluster197_21clades\|\|?->DUF499*->DUF3780->METHYLASE->HKD-DNase+SF1-SF2-Helicase+iREase+CTD->ORC/CDC6+HTH-><-HEPN<-METHYLASE | 1096 | Archaea>Euryarchaeota>Stenosarchaea_group>Halobacteria>Halobacteriales | GCF_900880625.1 | Halorhabdus_rudnickae | BN2694_RS11750 | 66442..69732 | 1775544 | DUF499-domain-containing-protein-[Halorhabdus-rudnickae] |
| WP_210408989.1 | AAA+ATPase+wHTH+Alpha+Beta-domain+wHTH+RRM | Type-2_BR.RM.like.systems | <-DUF772+DDE_Tnp_1\|\|?-><-?<-?<-OrfB_Zn_ribbon\|\|AbrB-HTH->DUF499*->DUF3780->METHYLASE->HKD-DNase+SF1-SF2-Helicase+iREase+CTD-> | 1098 | Archaea>Euryarchaeota>Stenosarchaea_group>Halobacteria>Halobacteriales | GCF_900880625.1 | Halorhabdus_rudnickae | BN2694_RS14135 | 149108..152404 | 1775544 | DUF499-domain-containing-protein-[Halorhabdus-rudnickae] |
| WP_176705305.1 | AAA+ATPase+wHTH+Alpha+Beta-domain+wHTH+RRM | Type-2_BR.RM.like.systems | <-CoiA\|\|?-><-?\|\|HNH-><-Cluster197_21clades\|\|?->DUF499*->DUF3780->METHYLASE->HKD-DNase+SF1-SF2-Helicase+iREase+CTD->ORC/CDC6+HTH-> | 1097 | Archaea>Euryarchaeota>Stenosarchaea_group>Halobacteria>Halobacteriales | - | Halobacterium_sp._GN101 | HXZ12_RS01050 | 222081..225374 | 88773 | DUF499-domain-containing-protein-[Halobacterium-sp.-GN101] |
| WP_070365960.1 | AAA+ATPase+wHTH+Alpha+Beta-domain+wHTH+RRM | Type-2_BR.RM.like.systems | <-REase\|\|ParB+HNH-p->?-><-METHYLASE<-DUF3780<-DUF499* | 933 | Archaea>Euryarchaeota>Stenosarchaea_group>Halobacteria>Halobacteriales | GCF_001767315.1 | Halodesulfurarchaeum_formicicum | HTSR_RS08490 | 1660006..1662807 | 1873524 | DUF499-domain-containing-protein-[Halodesulfurarchaeum-formicicum] |
| WP_211087444.1 | AAA+ATPase+wHTH+Alpha+Beta-domain+wHTH+RRM | Type-2_BR.RM.like.systems | <-METHYLASE<-DUF3780<-?<-?\|\|?->?-><-DUF499*<-?\|\|?-><-?<-PIWI<-REase+PNTD1+PNTD2 | 985 | Archaea>Euryarchaeota>Stenosarchaea_group>Halobacteria>Halobacteriales | GCF_017874695.1 | Halarchaeum_solikamskense | J2754_RS10890 | 121277..124234 | 1073081 | DUF499-domain-containing-protein-[Halarchaeum-solikamskense] |
| AOW80904.1 | AAA+ATPase+wHTH+Alpha+Beta-domain+wHTH+RRM | Type-2_BR.RM.like.systems | <-REase\|\|ParB+HNH-p->?-><-METHYLASE<-DUF3780<-DUF499* | 1107 | Archaea>Euryarchaeota>Stenosarchaea_group>Halobacteria>Halobacteriales | GCA_001767315.1 | Halodesulfurarchaeum_formicicum | HTSR_1734 | 1660006..1663329 | 1873524 | putative-ATPase-(AAA+-superfamily)-[Halodesulfurarchaeum-formicicum] |
| WP_011222498.1 | AAA+ATPase+wHTH+Alpha+Beta-domain+wHTH+RRM | Type-2_BR.RM.like.systems | <-METHYLASE<-DUF3780<-DUF499*<-?\|\|Cluster197_21clades-><-PIWI<-REase+PNTD1+PNTD2 | 1097 | Archaea>Euryarchaeota>Stenosarchaea_group>Halobacteria>Halobacteriales | GCF_000011085.1 | Haloarcula | E6P14_RS00430 | 3013..6306 | 2237 | MULTISPECIES:-DUF499-domain-containing-protein-[Haloarcula] |
| AEN07321.1 | AAA+ATPase+wHTH+Alpha+Beta-domain+wHTH+RRM | Type-2_BR.RM.like.systems | Schlafen->?->?-><-ParB+HNH\|\|ParB+HNH+Schlafen-><-?\|\|DUF499*->DUF3780->METHYLASE->HKD-DNase+SF1-SF2-Helicase+iREase+CTD->?->Resolvase->WYL-> | 1103 | Archaea>Euryarchaeota>Stenosarchaea_group>Halobacteria>Haloferacales | GCA_000224475.1 | halophilic_archaeon_DL31 | Halar_0046 | 17393..20704 | 756883 | hypothetical-protein-Halar_0046-[halophilic-archaeon-DL31] |
| ELZ45915.1 | AAA+ATPase+wHTH+Alpha+Beta-domain+wHTH+RRM | Type-2_BR.RM.like.systems | REase-><-HKD-DNase+SF1-SF2-Helicase+iREase+CTD<-METHYLASE-p<-DUF3780<-DUF499*\|\|BirA-HTH-><-Phage_integrase\|\|HTH-><-?\|\|ParB+HNH-> | 1085 | Archaea>Euryarchaeota>Stenosarchaea_group>Halobacteria>Haloferacales | GCA_000336875.1 | Halorubrum_californiense_DSM_19288 | C463_05775 | 141068..144325 | 1227465 | hypothetical-protein-C463_05775-[Halorubrum-californiensis-DSM-19288] |
| ELZ44610.1 | AAA+ATPase+wHTH+Alpha+Beta-domain+wHTH+RRM | Type-2_BR.RM.like.systems | <-HKD-DNase+SF1-SF2-Helicase+iREase+CTD<-METHYLASE-p<-DUF3780<-DUF499*\|\|?-><-ORC/CDC6+HTH\|\|REase+DUF4268->ParB+HNH-> | 934 | Archaea>Euryarchaeota>Stenosarchaea_group>Halobacteria>Haloferacales | GCA_000337055.1 | Halorubrum_distributum_JCM_9100 | C465_15856 | 15249..18053 | 1227467 | hypothetical-protein-C465_15856-[Halorubrum-distributum-JCM-9100] |
| ELZ56264.1 | AAA+ATPase+wHTH+Alpha+Beta-domain+wHTH+RRM | Type-2_BR.RM.like.systems | DDE_Tnp_1-><-HKD-DNase+SF1-SF2-Helicase+iREase+CTD<-HEPN<-METHYLASE-p<-DUF3780<-DUF499*<-?\|\|Resolvase->?->PadR-HTH->?->BirA-HTH-> | 935 | Archaea>Euryarchaeota>Stenosarchaea_group>Halobacteria>Haloferacales | GCA_000337075.1 | Halorubrum_hochstenium_ATCC_700873 | C467_08795 | 110652..113459 | 1227481 | hypothetical-protein-C467_08795-[Halorubrum-hochstenium-ATCC-700873] |
| POG53947.1 | AAA+ATPase+wHTH+Alpha+Beta-domain+wHTH+RRM | Type-2_BR.RM.like.systems | <-HTH\|\|?->?->Phage_int_SAM_1+Phage_integrase-><-ParB+HNH\|\|DUF499*->DUF3780->METHYLASE->HKD-DNase+SF1-SF2-Helicase+iREase+CTD-><-Phage_int_SAM_1+Phage_integrase\|\|?-><-Phage_integrase | 909 | Archaea>Euryarchaeota>Stenosarchaea_group>Halobacteria>Haloferacales | GCA_001469875.2 | Haloferax_marisrubri | AUR65_018735 | 249618..252347 | 1544719 | DUF499-domain-containing-protein-[Haloferax-marisrubri] |
| OYR84711.1 | AAA+ATPase+wHTH+Alpha+Beta-domain+wHTH+RRM | Type-2_BR.RM.like.systems | <-REase\|\|DUF499*->DUF3780->METHYLASE->HKD-DNase+SF1-SF2-Helicase+iREase+CTD-> | 1098 | Archaea>Euryarchaeota>Stenosarchaea_group>Halobacteria>Haloferacales | GCA_002252955.1 | Halorubrum_sp._E3 | DJ71_08490 | 3071..6367 | 1483400 | hypothetical-protein-DJ71_08490-[Halorubrum-sp.-E3] |
| OYR54791.1 | AAA+ATPase+wHTH+Alpha+Beta-domain+wHTH+RRM | Type-2_BR.RM.like.systems | <-HKD-DNase+SF1-SF2-Helicase+iREase+CTD<-METHYLASE<-DUF3780<-DUF499*<-?<-BirA-HTH\|\|<-ORC/CDC6+HTH\|\|?-><-REase\|\|URI-DNase-> | 1110 | Archaea>Euryarchaeota>Stenosarchaea_group>Halobacteria>Haloferacales | GCA_002252985.1 | Halorubrum_halodurans | DJ70_13120 | 25437..28769 | 1383851 | hypothetical-protein-DJ70_13120-[Halorubrum-halodurans] |
| RLM76840.1 | AAA+ATPase+wHTH+Alpha+Beta-domain+wHTH+RRM | Type-2_BR.RM.like.systems | REase+PNTD1+PNTD2->PIWI-><-Cluster197_21clades<-HKD-DNase+SF1-SF2-Helicase+iREase+CTD<-METHYLASE<-DUF3780<-DUF499*<-AbrB-HTH\|\|?->?->ORC/CDC6+HTH-><-?\|\|HTH+IclR-HTH-> | 926 | Archaea>Euryarchaeota>Stenosarchaea_group>Halobacteria>Haloferacales | GCA_003666025.1 | Halorubrum_sp._Atlit-8R | DVK05_15580 | 67088..69868 | 2282126 | DUF499-domain-containing-protein-[Halorubrum-sp.-Atlit-8R] |
| QCC49701.1 | AAA+ATPase+wHTH+Alpha+Beta-domain+wHTH+RRM | Type-2_BR.RM.like.systems | <-ORC/CDC6+HTH<-ORC/CDC6+HTH<-?<-?<-?\|\|AbrB-HTH->DUF499*->DUF3780->METHYLASE->HKD-DNase+SF1-SF2-Helicase+iREase+CTD-><-?<-?\|\|CoiA-> | 926 | Archaea>Euryarchaeota>Stenosarchaea_group>Halobacteria>Haloferacales | GCA_004799685.1 | Halobellus_limi | DV707_18320 | 167715..170495 | 699433 | DUF499-domain-containing-protein-[Halobellus-limi] |
| QGX93495.1 | AAA+ATPase+wHTH+Alpha+Beta-domain+wHTH+RRM | Type-2_BR.RM.like.systems | <-ORC/CDC6+HTH<-?<-?<-?\|\|AbrB-HTH->DUF499*->DUF3780->METHYLASE->HKD-DNase+SF1-SF2-Helicase+iREase+CTD->Cluster197_21clades-><-PIWI<-REase+PNTD1+PNTD2 | 926 | Archaea>Euryarchaeota>Stenosarchaea_group>Halobacteria>Haloferacales | GCA_009762275.1 | Haloplanus_rallus | EI982_00350 | 69646..72426 | 1816183 | DUF499-domain-containing-protein-[Haloplanus-rallus] |
| MBB6645927.1 | AAA+ATPase+wHTH+Alpha+Beta-domain+wHTH+RRM | Type-2_BR.RM.like.systems | Phage_integrase-><-HKD-DNase+SF1-SF2-Helicase+iREase+CTD<-HEPN<-METHYLASE<-DUF3780<-DUF499* | 935 | Archaea>Euryarchaeota>Stenosarchaea_group>Halobacteria>Haloferacales | GCA_014212355.1 | Halobellus_ruber | H5V44_06430 | 284116..286923 | 2761102 | DUF499-domain-containing-protein-[Halobellus-ruber] |
| MBC9987873.1 | AAA+ATPase+wHTH+Alpha+Beta-domain+wHTH+RRM | Type-2_BR.RM.like.systems | <-Peptidase_M48<-?<-SF1-SF2-Helicase+REase<-Exonuc_V_gamma+REase<-CBS+HEPN\|\|?->DUF499*->DUF3780->METHYLASE-><-ParB+HNH\|\|HKD-DNase+SF1-SF2-Helicase+iREase+CTD-><-ZnR+REase | 933 | Archaea>Euryarchaeota>Stenosarchaea_group>Halobacteria>Haloferacales | GCA_014530665.1 | Haloferax_sp._AS1 | E4P24_16065 | 306929..309730 | 2562277 | DUF499-domain-containing-protein-[Haloferax-sp.-AS1] |
| SFR56296.1 | AAA+ATPase+wHTH+Alpha+Beta-domain+wHTH+RRM | Type-2_BR.RM.like.systems | <-ABC-ATPase+SbcC<-?<-?<-?\|\|?-><-?\|\|tRNA->DUF499*->DUF3780->METHYLASE->HKD-DNase+SF1-SF2-Helicase+iREase+CTD-><-REase\|\|?->SbcC+SbcC-> | 1121 | Archaea>Euryarchaeota>Stenosarchaea_group>Halobacteria>Haloferacales | GCA_900111935.1 | Halorubrum_sodomense | SAMN04487937_2790 | 92085..95450 | 35743 | Protein-of-unknown-function-[Halorubrum-sodomense] |
| SNR42929.1 | AAA+ATPase+wHTH+Alpha+Beta-domain+wHTH+RRM | Type-2_BR.RM.like.systems | <-HKD-DNase+SF1-SF2-Helicase+iREase+CTD<-METHYLASE<-DUF3780<-DUF499*\|\|?-><-nSTAND3\|\|?->GDResolvase+Recombinase+ZnR-> | 1099 | Archaea>Euryarchaeota>Stenosarchaea_group>Halobacteria>Haloferacales | GCA_900188065.1 | Halorubrum_vacuolatum | SAMN06264855_10645 | 53783..57082 | 63740 | Protein-of-unknown-function-[Halorubrum-vacuolatum] |
| WP_014555848.1 | AAA+ATPase+wHTH+Alpha+Beta-domain+wHTH+RRM | Type-2_BR.RM.like.systems | <-Zn_Tnp_IS1595+DDE_Tnp_IS1595<-?<-?<-?<-SF1-SF2-Helicase\|\|DUF499*->DUF3780->METHYLASE->HKD-DNase+SF1-SF2-Helicase+iREase+CTD-><-?<-SF1-SF2-Helicase | 1150 | Archaea>Euryarchaeota>Stenosarchaea_group>Halobacteria>Haloferacales | GCF_000237865.1 | Haloquadratum_walsbyi | HQRW_RS05760 | 1307615..1311067 | 293091 | DUF499-domain-containing-protein-[Haloquadratum-walsbyi] |
| WP_008095328.1 | AAA+ATPase+wHTH+Alpha+Beta-domain+wHTH+RRM | Type-2_BR.RM.like.systems | <-HKD-DNase+SF1-SF2-Helicase+iREase+CTD<-METHYLASE<-DUF3780<-DUF499*\|\|REase-><-?<-?<-?\|\|PADR-HTH-> | 1115 | Archaea>Euryarchaeota>Stenosarchaea_group>Halobacteria>Haloferacales | GCF_000336815.1 | Haloferax_prahovense | C457_RS13190 | 268523..271870 | 381852 | DUF499-domain-containing-protein-[Haloferax-prahovense] |
| WP_239638853.1 | AAA+ATPase+wHTH+Alpha+Beta-domain+wHTH+RRM | Type-2_BR.RM.like.systems | <-SF1-SF2-Helicase<-?\|\|REase-><-HKD-DNase+SF1-SF2-Helicase+iREase+CTD<-METHYLASE<-DUF3780<-DUF499*\|\|?->BirA-HTH-><-Phage_integrase<-?\|\|HNH->HTH-> | 1121 | Archaea>Euryarchaeota>Stenosarchaea_group>Halobacteria>Haloferacales | GCF_000336875.1 | Halorubrum_californiense | C463_RS05650 | 141068..144433 | 416585 | DUF499-domain-containing-protein-[Halorubrum-californiense] |
| WP_241431564.1 | AAA+ATPase+wHTH+Alpha+Beta-domain+wHTH+RRM | Type-2_BR.RM.like.systems | <-HKD-DNase+SF1-SF2-Helicase+iREase+CTD\|\|?-><-METHYLASE-p<-DUF3780<-DUF499*<-ORC/CDC6+HTH\|\|REase+DUF4268->?->ParB+HNH-> | 1054 | Archaea>Euryarchaeota>Stenosarchaea_group>Halobacteria>Haloferacales | GCF_000337055.1 | Halorubrum_distributum | C465_RS15555 | 15249..18413 | 29283 | DUF499-domain-containing-protein-[Halorubrum-distributum] |
| WP_239638715.1 | AAA+ATPase+wHTH+Alpha+Beta-domain+wHTH+RRM | Type-2_BR.RM.like.systems | <-PadR-HTH\|\|?->?-><-Resolvase\|\|?->AbrB-HTH->DUF499*->DUF3780->METHYLASE->HEPN->HKD-DNase+SF1-SF2-Helicase+iREase+CTD->REase-><-DDE_Tnp_1 | 1054 | Archaea>Euryarchaeota>Stenosarchaea_group>Halobacteria>Haloferacales | GCF_000337075.1 | Halorubrum | NAF06_RS08930 | 1779031..1782195 | 56688 | MULTISPECIES:-DUF499-domain-containing-protein-[Halorubrum] |
| WP_080506006.1 | AAA+ATPase+wHTH+Alpha+Beta-domain+wHTH+RRM | Type-2_BR.RM.like.systems | <-ParB+HNH<-?<-REase+DUF4268\|\|ORC/CDC6+HTH->DUF499*->DUF3780->METHYLASE-p-><-?\|\|HKD-DNase+SF1-SF2-Helicase+iREase+CTD-> | 1109 | Archaea>Euryarchaeota>Stenosarchaea_group>Halobacteria>Haloferacales | GCF_000337335.1 | Halorubrum_distributum | C466_RS13155 | 9015..12344 | 29283 | DUF499-domain-containing-protein-[Halorubrum-distributum] |
| WP_245575955.1 | AAA+ATPase+wHTH+Alpha+Beta-domain+wHTH+RRM | Type-2_BR.RM.like.systems | SF1-SF2-Helicase+REase->?-><-HKD-DNase+SF1-SF2-Helicase+iREase+CTD<-?<-METHYLASE<-DUF3780<-DUF499*<-AbrB-HTH<-ORC/CDC6+REase\|\|PIWI-> | 1102 | Archaea>Euryarchaeota>Stenosarchaea_group>Halobacteria>Haloferacales | GCF_000427685.1 | Haloplanus_natans | HALNA_RS00240 | 57388..60696 | 376171 | DUF499-domain-containing-protein-[Haloplanus-natans] |
| WP_195155664.1 | AAA+ATPase+wHTH+Alpha+Beta-domain+wHTH+RRM | Type-2_BR.RM.like.systems | Zn_Tnp_IS1595+DDE_Tnp_IS1595->HEPN->HEPN-><-?\|\|?->AbrB-HTH->DUF499*->DUF3780->METHYLASE->HKD-DNase+SF1-SF2-Helicase+iREase+CTD->REase-p->HEPN-> | 1099 | Archaea>Euryarchaeota>Stenosarchaea_group>Halobacteria>Haloferacales | GCF_000513435.1 | Halorubrum_sp._AJ67 | BN903_RS00695 | 145466..148765 | 1173487 | DUF499-domain-containing-protein-[Halorubrum-sp.-AJ67] |
| WP_082222573.1 | AAA+ATPase+wHTH+Alpha+Beta-domain+wHTH+RRM | Type-2_BR.RM.like.systems | <-REase<-REase\|\|HicA_toxin->?->?->AbrB-HTH->DUF499*->DUF3780->METHYLASE->HKD-DNase+SF1-SF2-Helicase+iREase+CTD->GDResolvase+Recombinase->?->DOC-> | 1099 | Archaea>Euryarchaeota>Stenosarchaea_group>Halobacteria>Haloferacales | GCF_000739595.1 | Halorubrum_halophilum | ACP96_RS14010 | 6716..10015 | 413816 | DUF499-domain-containing-protein-[Halorubrum-halophilum] |
| WP_050460473.1 | AAA+ATPase+wHTH+Alpha+Beta-domain+wHTH+RRM | Type-2_BR.RM.like.systems | <-MarR-HTH<-HKD-DNase+SF1-SF2-Helicase+iREase+CTD<-METHYLASE<-DUF3780<-DUF499*<-Schlafen+HATPase+HTH<-SWACOS\|\|?->RADICAL-SAM->HTH+IclR-HTH->NTP_transf+wHTH-> | 1097 | Archaea>Euryarchaeota>Stenosarchaea_group>Halobacteria>Haloferacales | GCF_001190965.1 | Haloferax_gibbonsii | ABY42_RS18700 | 86264..89557 | 35746 | DUF499-domain-containing-protein-[Haloferax-gibbonsii] |
| WP_233194430.1 | AAA+ATPase+wHTH+Alpha+Beta-domain+wHTH+RRM | Type-2_BR.RM.like.systems | REase->?->Phage_int_SAM_1+Phage_integrase-><-ParB+HNH\|\|?->DUF499*->DUF3780->METHYLASE->HKD-DNase+SF1-SF2-Helicase+iREase+CTD-><-Phage_int_SAM_1+Phage_integrase<-?<-Phage_integrase | 1080 | Archaea>Euryarchaeota>Stenosarchaea_group>Halobacteria>Haloferacales | GCF_001469875.2 | Haloferax_marisrubri | AUR65_RS18765 | 249105..252347 | 1544719 | DUF499-domain-containing-protein-[Haloferax-marisrubri] |
| WP_082230522.1 | AAA+ATPase+wHTH+Alpha+Beta-domain+wHTH+RRM | Type-2_BR.RM.like.systems | MORC+DUF3684+REase-><-?\|\|DUF499*->DUF3780->METHYLASE->?->?->HKD-DNase+SF1-SF2-Helicase+iREase+CTD-> | 1102 | Archaea>Euryarchaeota>Stenosarchaea_group>Halobacteria>Haloferacales | GCF_001469955.1 | Haloprofundus_marisrubri | AUR64_RS01320 | 118093..121401 | 1514971 | DUF499-domain-containing-protein-[Haloprofundus-marisrubri] |
| WP_232745144.1 | AAA+ATPase+wHTH+Alpha+Beta-domain+wHTH+RRM | Type-2_BR.RM.like.systems | ORC/CDC6+HTH->AbrB-HTH->DUF499*->DUF3780->METHYLASE->HKD-DNase+SF1-SF2-Helicase+iREase+CTD-> | 1099 | Archaea>Euryarchaeota>Stenosarchaea_group>Halobacteria>Haloferacales | GCF_001542905.1 | Halorubrum_aethiopicum | AXA68_RS15345 | 3320..6619 | 1758255 | DUF499-domain-containing-protein-[Halorubrum-aethiopicum] |
| WP_079233154.1 | AAA+ATPase+wHTH+Alpha+Beta-domain+wHTH+RRM | Type-2_BR.RM.like.systems | BirA-HTH-><-?<-HKD-DNase+SF1-SF2-Helicase+iREase+CTD<-METHYLASE<-DUF3780<-DUF499*\|\|?->?-><-?<-?\|\|RHH+RAMA-> | 1095 | Archaea>Euryarchaeota>Stenosarchaea_group>Halobacteria>Haloferacales | GCF_002025255.2 | Halolamina_sp._CBA1230 | B4589_RS04535 | 891070..894357 | 1853690 | DUF499-domain-containing-protein-[Halolamina-sp.-CBA1230] |
| WP_094580341.1 | AAA+ATPase+wHTH+Alpha+Beta-domain+wHTH+RRM | Type-2_BR.RM.like.systems | <-REase\|\|?->DUF499*->DUF3780->METHYLASE->HKD-DNase+SF1-SF2-Helicase+iREase+CTD-><-?<-MarR-HTH | 1097 | Archaea>Euryarchaeota>Stenosarchaea_group>Halobacteria>Haloferacales | GCF_002252725.1 | Halorubrum_ezzemoulense | DJ83_RS13005 | 1301..4594 | 337243 | DUF499-domain-containing-protein-[Halorubrum-ezzemoulense] |
| WP_094522756.1 | AAA+ATPase+wHTH+Alpha+Beta-domain+wHTH+RRM | Type-2_BR.RM.like.systems | <-SF1-SF2-Helicase<-?\|\|REase-><-HKD-DNase+SF1-SF2-Helicase+iREase+CTD<-METHYLASE<-DUF3780<-DUF499*<-?<-?\|\|BirA-HTH->HTH-><-Phage_integrase | 1102 | Archaea>Euryarchaeota>Stenosarchaea_group>Halobacteria>Haloferacales | GCF_002252895.1 | Halorubrum_sp._Eb13 | DJ75_RS03965 | 27103..30411 | 1383843 | DUF499-domain-containing-protein-[Halorubrum-sp.-Eb13] |
| WP_245810850.1 | AAA+ATPase+wHTH+Alpha+Beta-domain+wHTH+RRM | Type-2_BR.RM.like.systems | <-HKD-DNase+SF1-SF2-Helicase+iREase+CTD\|\|?-><-METHYLASE<-DUF3780<-DUF499*<-?<-BirA-HTH\|\|?-><-ORC/CDC6+HTH\|\|?-><-REase | 1055 | Archaea>Euryarchaeota>Stenosarchaea_group>Halobacteria>Haloferacales | GCF_002252985.1 | Halorubrum_halodurans | DJ70_RS13160 | 25437..28604 | 1383851 | DUF499-domain-containing-protein-[Halorubrum-halodurans] |
| WP_120102742.1 | AAA+ATPase+wHTH+Alpha+Beta-domain+wHTH+RRM | Type-2_BR.RM.like.systems | BirA-HTH->?->REase-><-ORC-CDC6+HTH<-HKD-DNase+SF1-SF2-Helicase+iREase+CTD<-METHYLASE<-DUF3780<-DUF499*<-?<-?\|\|?->ZnR+REase-> | 1101 | Archaea>Euryarchaeota>Stenosarchaea_group>Halobacteria>Haloferacales | GCF_003605575.1 | Halonotius_aquaticus | DM826_RS07280 | 18673..21978 | 2216978 | DUF499-domain-containing-protein-[Halonotius-aquaticus] |
| WP_233735731.1 | AAA+ATPase+wHTH+Alpha+Beta-domain+wHTH+RRM | Type-2_BR.RM.like.systems | SbcC+SbcC-><-Cluster197_21clades<-HKD-DNase+SF1-SF2-Helicase+iREase+CTD<-METHYLASE<-DUF3780<-DUF499*<-AbrB-HTH\|\|?->?->?->ORC/CDC6+HTH-><-ORC/CDC6+REase | 1104 | Archaea>Euryarchaeota>Stenosarchaea_group>Halobacteria>Haloferacales | GCF_003605575.1 | Halonotius_aquaticus | DM826_RS08055 | 57803..61117 | 2216978 | DUF499-domain-containing-protein-[Halonotius-aquaticus] |
| WP_121571189.1 | AAA+ATPase+wHTH+Alpha+Beta-domain+wHTH+RRM | Type-2_BR.RM.like.systems | CARF-><-ZnR\|\|?->DUF499*->DUF3780->METHYLASE->HKD-DNase+SF1-SF2-Helicase+iREase+CTD-><-Radical-SAM\|\|Radical-SAM-> | 1101 | Archaea>Euryarchaeota>Stenosarchaea_group>Halobacteria>Haloferacales | GCF_003665925.1 | Halobellus_sp._Atlit-38R | D3D02_RS17200 | 8638..11943 | 2282131 | DUF499-domain-containing-protein-[Halobellus-sp.-Atlit-38R] |
| WP_233562229.1 | AAA+ATPase+wHTH+Alpha+Beta-domain+wHTH+RRM | Type-2_BR.RM.like.systems | REase+PNTD1+PNTD2->PIWI-><-Cluster197_21clades<-HKD-DNase+SF1-SF2-Helicase+iREase+CTD<-METHYLASE<-DUF3780<-DUF499*<-AbrB-HTH\|\|?->?->ORC/CDC6+HTH-><-?\|\|HTH+IclR-HTH-> | 1099 | Archaea>Euryarchaeota>Stenosarchaea_group>Halobacteria>Haloferacales | GCF_003666025.1 | Halorubrum_sp._Atlit-8R | DVK05_RS15570 | 67088..70387 | 2282126 | DUF499-domain-containing-protein-[Halorubrum-sp.-Atlit-8R] |
| WP_129452722.1 | AAA+ATPase+wHTH+Alpha+Beta-domain+wHTH+RRM | Type-2_BR.RM.like.systems | <-Resolvase<-Resolvase<-?<-HKD-DNase+SF1-SF2-Helicase+iREase+CTD<-METHYLASE<-DUF3780<-DUF499*<-?<-ZetaToxin<-?\|\|REase- | 1097 | Archaea>Euryarchaeota>Stenosarchaea_group>Halobacteria>Haloferacales | GCF_004126515.1 | Halorubrum_ezzemoulense | EO776_RS18665 | 45317..48610 | 337243 | DUF499-domain-containing-protein-[Halorubrum-ezzemoulense] |
| WP_205742829.1 | AAA+ATPase+wHTH+Alpha+Beta-domain+wHTH+RRM | Type-2_BR.RM.like.systems | <-ORC/CDC6+HTH<-ORC/CDC6+HTH<-?<-?<-?\|\|AbrB-HTH->DUF499*->DUF3780->METHYLASE->HKD-DNase+SF1-SF2-Helicase+iREase+CTD->REase-> | 1099 | Archaea>Euryarchaeota>Stenosarchaea_group>Halobacteria>Haloferacales | GCF_004799685.1 | Halobellus_limi | DV707_RS18320 | 167196..170495 | 699433 | DUF499-domain-containing-protein-[Halobellus-limi] |
| WP_137695329.1 | AAA+ATPase+wHTH+Alpha+Beta-domain+wHTH+RRM | Type-2_BR.RM.like.systems | <-REase\|\|?->ZetaToxin->?->DUF499*->DUF3780->METHYLASE->HKD-DNase+SF1-SF2-Helicase+iREase+CTD->?->?->MPTase-> | 1097 | Archaea>Euryarchaeota>Stenosarchaea_group>Halobacteria>Haloferacales | GCF_005435175.1 | Halorubrum_sp._GN11GM_10-3_MGM | EXE40_RS03790 | 28191..31484 | 2518111 | DUF499-domain-containing-protein-[Halorubrum-sp.-GN11GM_10-3_MGM] |
| WP_137705915.1 | AAA+ATPase+wHTH+Alpha+Beta-domain+wHTH+RRM | Type-2_BR.RM.like.systems | PRTase->?->?-><-HKD-DNase+SF1-SF2-Helicase+iREase+CTD<-METHYLASE<-DUF3780<-DUF499*\|\|?->BirA-HTH-><-Phage_integrase<-?<-?<-Schlafen+HATPase+HTH | 1102 | Archaea>Euryarchaeota>Stenosarchaea_group>Halobacteria>Haloferacales | GCF_005435195.1 | Halorubrum_sp._SP3 | EXE42_RS03640 | 182542..185850 | 1537265 | DUF499-domain-containing-protein-[Halorubrum-sp.-SP3] |
| WP_199243581.1 | AAA+ATPase+wHTH+Alpha+Beta-domain+wHTH+RRM | Type-2_BR.RM.like.systems | <-ORC/CDC6+HTH<-?<-?<-?\|\|AbrB-HTH->DUF499*->DUF3780->METHYLASE->HKD-DNase+SF1-SF2-Helicase+iREase+CTD->Cluster197_21clades-><-PIWI<-REase+PNTD1+PNTD2 | 1099 | Archaea>Euryarchaeota>Stenosarchaea_group>Halobacteria>Haloferacales | GCF_009762275.1 | Haloplanus_rallus | EI982_RS00330 | 69127..72426 | 1816183 | DUF499-domain-containing-protein-[Haloplanus-rallus] |
| WP_238992092.1 | AAA+ATPase+wHTH+Alpha+Beta-domain+wHTH+RRM | Type-2_BR.RM.like.systems | <-HKD-DNase+SF1-SF2-Helicase+iREase+CTD<-?<-METHYLASE<-DUF3780<-DUF499* | 1110 | Archaea>Euryarchaeota>Stenosarchaea_group>Halobacteria>Haloferacales | GCF_009856455.1 | Halorubrum_terrestre | GLW30_RS04820 | 197870..201202 | 267368 | DUF499-domain-containing-protein-[Halorubrum-terrestre] |
| WP_246403805.1 | AAA+ATPase+wHTH+Alpha+Beta-domain+wHTH+RRM | Type-2_BR.RM.like.systems | Phage_integrase-><-HKD-DNase+SF1-SF2-Helicase+iREase+CTD<-HEPN<-METHYLASE<-DUF3780<-DUF499* | 1109 | Archaea>Euryarchaeota>Stenosarchaea_group>Halobacteria>Haloferacales | GCF_014212355.1 | Halobellus_ruber | H5V44_RS06435 | 284116..287445 | 2761102 | DUF499-domain-containing-protein-[Halobellus-ruber] |
| WP_188148783.1 | AAA+ATPase+wHTH+Alpha+Beta-domain+wHTH+RRM | Type-2_BR.RM.like.systems | <-Peptidase_M48<-?<-SF1-SF2-Helicase+REase<-Exonuc_V_gamma+REase<-CBS+HEPN\|\|?->DUF499*->DUF3780->METHYLASE-><-ParB+HNH\|\|HKD-DNase+SF1-SF2-Helicase+iREase+CTD-><-ZnR+REase | 1107 | Archaea>Euryarchaeota>Stenosarchaea_group>Halobacteria>Haloferacales | GCF_014530665.1 | Haloferax_sp._AS1 | E4P24_RS16200 | 306407..309730 | 2562277 | DUF499-domain-containing-protein-[Haloferax-sp.-AS1] |
| WP_234699407.1 | AAA+ATPase+wHTH+Alpha+Beta-domain+wHTH+RRM | Type-2_BR.RM.like.systems | AbrB-HTH->DUF499*->DUF3780->METHYLASE->HKD-DNase+SF1-SF2-Helicase+iREase+CTD->REase-><-?<-BirA-HTH | 1099 | Archaea>Euryarchaeota>Stenosarchaea_group>Halobacteria>Haloferacales | GCF_016635065.1 | Halorubrum_sp._LN27 | JIN95_RS16120 | 50683..53982 | 2801032 | DUF499-domain-containing-protein-[Halorubrum-sp.-LN27] |
| WP_251344662.1 | AAA+ATPase+wHTH+Alpha+Beta-domain+wHTH+RRM | Type-2_BR.RM.like.systems | <-CoiA\|\|HNH-><-Cluster197_21clades<-?<-HKD-DNase+SF1-SF2-Helicase+iREase+CTD<-METHYLASE<-DUF499*<-AbrB-HTH\|\|?->?->?->ORC/CDC6+HTH->SMODS-> | 1099 | Archaea>Euryarchaeota>Stenosarchaea_group>Halobacteria>Haloferacales | GCF_023703775.1 | Haloplanus_sp._GDY1 | NBT67_RS17100 | 71897..75196 | 2949993 | DUF499-domain-containing-protein-[Haloplanus-sp.-GDY1] |
| WP_256397181.1 | AAA+ATPase+wHTH+Alpha+Beta-domain+wHTH+RRM | Type-2_BR.RM.like.systems | <-DDE_Tnp_IS240\|\|AbrB-HTH->DUF499*->DUF3780->METHYLASE->HKD-DNase+SF1-SF2-Helicase+iREase+CTD->HEPN-><-?\|\|ORC/CDC6+HTH-> | 1045 | Archaea>Euryarchaeota>Stenosarchaea_group>Halobacteria>Haloferacales | GCF_024494645.1 | Halorubraceae | NO887_RS09745 | 233066..236203 | 1963271 | MULTISPECIES:-DUF499-domain-containing-protein-[Halorubraceae] |
| WP_256406460.1 | AAA+ATPase+wHTH+Alpha+Beta-domain+wHTH+RRM | Type-2_BR.RM.like.systems | HEPN-><-?\|\|?->AbrB-HTH->DUF499*->DUF3780->METHYLASE->HKD-DNase+SF1-SF2-Helicase+iREase+CTD->\|\|?->OrfB_Zn_ribbon-> | 1099 | Archaea>Euryarchaeota>Stenosarchaea_group>Halobacteria>Haloferacales | GCF_024494725.1 | Haloplanus_ruber | NO878_RS13275 | 61244..64543 | 869892 | DUF499-domain-containing-protein-[Haloplanus-ruber] |
| WP_267623061.1 | AAA+ATPase+wHTH+Alpha+Beta-domain+wHTH+RRM | Type-2_BR.RM.like.systems | <-REase<-ORC/CDC6+HTH\|\|CoiA-><-DOC\|\|AbrB-HTH->DUF499*->DUF3780->METHYLASE->HEPN->HKD-DNase+SF1-SF2-Helicase+iREase+CTD->Cluster197_21clades-> | 1103 | Archaea>Euryarchaeota>Stenosarchaea_group>Halobacteria>Haloferacales | GCF_026586625.1 | Halobium_salinum | N5095_RS11700 | 30435..33746 | 1364940 | DUF499-domain-containing-protein-[Halobium-salinum] |
| WP_267624427.1 | AAA+ATPase+wHTH+Alpha+Beta-domain+wHTH+RRM | Type-2_BR.RM.like.systems | <-Phage_integrase<-?<-HKD-DNase+SF1-SF2-Helicase+iREase+CTD<-METHYLASE<-DUF3780<-DUF499*<-?<-?<-?\|\|?->REase+DUF4268-> | 1101 | Archaea>Euryarchaeota>Stenosarchaea_group>Halobacteria>Haloferacales | GCF_026586625.1 | Halobium_salinum | N5095_RS17535 | 1056167..1059472 | 1364940 | DUF499-domain-containing-protein-[Halobium-salinum] |
| WP_267646551.1 | AAA+ATPase+wHTH+Alpha+Beta-domain+wHTH+RRM | Type-2_BR.RM.like.systems | BirA-HTH->REase-><-HKD-DNase+SF1-SF2-Helicase+iREase+CTD<-METHYLASE<-DUF3780<-DUF499*<-?\|\|?->HTH->?->RelE-ParE-> | 1108 | Archaea>Euryarchaeota>Stenosarchaea_group>Halobacteria>Haloferacales | GCF_026616205.1 | Halolamina_litorea | NO998_RS07830 | 1504199..1507525 | 1515593 | DUF499-domain-containing-protein-[Halolamina-litorea] |
| WP_271946610.1 | AAA+ATPase+wHTH+Alpha+Beta-domain+wHTH+RRM | Type-2_BR.RM.like.systems | <-HKD-DNase+SF1-SF2-Helicase+iREase+CTD<-METHYLASE<-DUF3780<-DUF499*<-?\|\|REase->?->?->REase-><-HEPN | 1097 | Archaea>Euryarchaeota>Stenosarchaea_group>Halobacteria>Haloferacales | GCF_028211275.1 | Halorubrum_ezzemoulense | PM036_RS16555 | 16669..19962 | 337243 | DUF499-domain-containing-protein-[Halorubrum-ezzemoulense] |
| WP_271963542.1 | AAA+ATPase+wHTH+Alpha+Beta-domain+wHTH+RRM | Type-2_BR.RM.like.systems | <-Schlafen<-HKD-DNase+SF1-SF2-Helicase+iREase+CTD\|\|ParB+HNH-><-METHYLASE<-DUF3780<-DUF499*\|\|?->?->?-><-Phage_integrase\|\|PadR-HTH->PadR-HTH-> | 1107 | Archaea>Euryarchaeota>Stenosarchaea_group>Halobacteria>Haloferacales | GCF_028211285.1 | Halorubrum_ezzemoulense | PM025_RS15440 | 14437..17760 | 337243 | DUF499-domain-containing-protein-[Halorubrum-ezzemoulense] |
| WP_092923569.1 | AAA+ATPase+wHTH+Alpha+Beta-domain+wHTH+RRM | Type-2_BR.RM.like.systems | DUF499*->DUF3780->METHYLASE->HKD-DNase+SF1-SF2-Helicase+iREase+CTD-><-REase\|\|?->SbcC+SbcC-> | 1124 | Archaea>Euryarchaeota>Stenosarchaea_group>Halobacteria>Haloferacales | GCF_900111935.1 | Halorubrum_sodomense | BM286_RS13660 | 92076..95450 | 35743 | DUF499-domain-containing-protein-[Halorubrum-sodomense] |
| WP_245809927.1 | AAA+ATPase+wHTH+Alpha+Beta-domain+wHTH+RRM | Type-2_BR.RM.like.systems | <-HKD-DNase+SF1-SF2-Helicase+iREase+CTD<-METHYLASE<-DUF499*\|\|LysR-HTH->?-><-nSTAND3\|\|?->GDResolvase+Recombinase+ZnR-> | 1104 | Archaea>Euryarchaeota>Stenosarchaea_group>Halobacteria>Haloferacales | GCF_900188065.1 | Halorubrum_vacuolatum | CHB65_RS07630 | 53783..57097 | 63740 | DUF499-domain-containing-protein-[Halorubrum-vacuolatum] |
| WP_114606940.1 | AAA+ATPase+wHTH+Alpha+Beta-domain+wHTH+RRM | Type-2_BR.RM.like.systems | <-BirA-HTH\|\|?->AbrB-HTH->DUF499*->DUF3780->METHYLASE->?->HKD-DNase+SF1-SF2-Helicase+iREase+CTD->DDE_Tnp_IS240-><-MarR-HTH | 928 | Archaea>Euryarchaeota>Stenosarchaea_group>Halobacteria>Haloferacales | - | Haloplanus_rubicundus | DU484_RS18925 | 82703..85489 | 1547898 | DUF499-domain-containing-protein-[Haloplanus-rubicundus] |
| WP_158293466.1 | AAA+ATPase+wHTH+Alpha+Beta-domain+wHTH+RRM | Type-2_BR.RM.like.systems | DUF499*->DUF3780->METHYLASE->HKD-DNase+SF1-SF2-Helicase+iREase+CTD->HKD-DNase+SF1-SF2-Helicase+iREase+CTD-p-> | 633 | Archaea>Euryarchaeota>Stenosarchaea_group>Halobacteria>Haloferacales | - | Halorubrum_sp._SS7 | EXE44_RS15475 | 1..1902 | 2518119 | DUF499-domain-containing-protein,-partial-[Halorubrum-sp.-SS7] |
| WP_233139259.1 | AAA+ATPase+wHTH+Alpha+Beta-domain+wHTH+RRM | Type-2_BR.RM.like.systems | <-ORC/CDC6+HTH<-REase<-?\|\|DUF4011+SF1-SF2-Helicase+REase-><-?\|\|DUF499*->DUF3780->METHYLASE-p-><-DDE_Tnp_IS66 | 1055 | Archaea>Euryarchaeota>Stenosarchaea_group>Halobacteria>Haloferacales | GCF_002286985.1 | Halorubrum_salipaludis | CK500_RS16405 | 17917..21084 | 2032630 | DUF499-domain-containing-protein-[Halorubrum-salipaludis] |
| KTG11418.1 | AAA+ATPase+wHTH+Alpha+Beta-domain+wHTH+RRM | Type-2_BR.RM.like.systems | HNH-><-?<-?\|\|MORC+DUF3684+REase-><-?\|\|AbrB-HTH->DUF499*->DUF3780-><-?<-?\|\|METHYLASE-p-> | 929 | Archaea>Euryarchaeota>Stenosarchaea_group>Halobacteria>Haloferacales | GCA_001469955.1 | Haloprofundus_marisrubri | AUR64_03955 | 118612..121401 | 1514971 | hypothetical-protein-AUR64_03955-[Haloprofundus-marisrubri] |
| PAU78613.1 | AAA+ATPase+wHTH+Alpha+Beta-domain+wHTH+RRM | Type-2_BR.RM.like.systems | <-ORC/CDC6+HTH<-REase<-?\|\|DUF4011+SF1-SF2-Helicase+REase-><-?\|\|DUF499*->DUF3780->METHYLASE-p-><-DDE_Tnp_IS66 | 1110 | Archaea>Euryarchaeota>Stenosarchaea_group>Halobacteria>Haloferacales | GCA_002286985.1 | Halorubrum_salipaludis | CK500_16385 | 17752..21084 | 2032630 | hypothetical-protein-CK500_16385-[Halorubrum-salipaludis] |
| OAQ51496.1 | AAA+ATPase+wHTH+Alpha+Beta-domain+wHTH+RRM | Type-2_BR.RM.like.systems | ZetaToxin-><-HEPN-p\|\|?->?->AbrB-HTH->DUF499*->DUF3780->METHYLASE-p->HKD-DNase+SF1-SF2-Helicase+iREase+CTD->REase-><-?\|\|CoiA-> | 1099 | Archaea>Euryarchaeota>Stenosarchaea_group>Halobacteria>Natrialbales | GCA_000690595.2 | Natrinema_mahii | HTG_19000 | 43326..46625 | 1416969 | ATPase-AAA-[Haloterrigena-mahii] |
| WP_008458790.1 | AAA+ATPase+wHTH+Alpha+Beta-domain+wHTH+RRM | Type-2_BR.RM.like.systems | <-ORC/CDC6+HTH<-HKD-DNase+SF1-SF2-Helicase+iREase+CTD<-METHYLASE<-DUF3780<-DUF499*<-?\|\|Cluster197_21clades-><-PIWI | 1097 | Archaea>Euryarchaeota>Stenosarchaea_group>Halobacteria>Natrialbales | GCF_000337175.1 | Natrinema_gari | C486_RS18175 | 86707..90000 | 419186 | DUF499-domain-containing-protein-[Natrinema-gari] |
| WP_006826449.1 | AAA+ATPase+wHTH+Alpha+Beta-domain+wHTH+RRM | Type-2_BR.RM.like.systems | NTP_transf+wHTH-><-?\|\|?->?->Phage_integrase->AbrB-HTH->DUF499*->DUF3780->METHYLASE-><-?<-?\|\|HKD-DNase+SF1-SF2-Helicase+iREase+CTD-><-CoiA | 1080 | Archaea>Euryarchaeota>Stenosarchaea_group>Halobacteria>Natrialbales | GCF_000337595.1 | Natrialba_taiwanensis | C484_RS13375 | 74471..77713 | 160846 | DUF499-domain-containing-protein-[Natrialba-taiwanensis] |
| WP_138247001.1 | AAA+ATPase+wHTH+Alpha+Beta-domain+wHTH+RRM | Type-2_BR.RM.like.systems | <-ORC/CDC6+HTH<-HKD-DNase+SF1-SF2-Helicase+iREase+CTD<-METHYLASE<-DUF3780<-DUF499*\|\|Cluster197_21clades-><-PIWI<-REase+PNTD1+PNTD2 | 1097 | Archaea>Euryarchaeota>Stenosarchaea_group>Halobacteria>Natrialbales | GCF_005576615.1 | Natrinema_versiforme | FEJ81_RS19880 | 258413..261706 | 88724 | DUF499-domain-containing-protein-[Natrinema-versiforme] |
| WP_160063634.1 | AAA+ATPase+wHTH+Alpha+Beta-domain+wHTH+RRM | Type-2_BR.RM.like.systems | <-REase<-HKD-DNase+SF1-SF2-Helicase+iREase+CTD<-METHYLASE<-DUF3780<-DUF499*<-AbrB-HTH<-?<-?\|\|?->HNH-> | 926 | Archaea>Euryarchaeota>Stenosarchaea_group>Halobacteria>Natrialbales | GCF_009834785.1 | Natronorubrum_halalkaliphilum | GS429_RS05865 | 14261..17041 | 2691917 | DUF499-domain-containing-protein-[Natronorubrum-halalkaliphilum] |
| WP_226778690.1 | AAA+ATPase+wHTH+Alpha+Beta-domain+wHTH+RRM | Type-2_BR.RM.like.systems | PadR-HTH->?->DUF499*->DUF3780->METHYLASE->SF1-SF2-Helicase->HKD-DNase+SF1-SF2-Helicase+iREase+CTD-><-?<-SF1-SF2-Helicase | 909 | Archaea>Euryarchaeota>Stenosarchaea_group>Halobacteria>Natrialbales | GCF_013456555.2 | Natrinema_zhouii | HYG81_RS42930 | 110101..112830 | 1710539 | DUF499-domain-containing-protein-[Natrinema-sp.-YPL30] |
| WP_204749924.1 | AAA+ATPase+wHTH+Alpha+Beta-domain+wHTH+RRM | Type-2_BR.RM.like.systems | REase+PNTD1+PNTD2->PIWI-><-Cluster197_21clades<-HKD-DNase+SF1-SF2-Helicase+iREase+CTD<-METHYLASE<-DUF3780<-DUF499*<-AbrB-HTH\|\|?->?->?->ORC/CDC6+HTH-> | 1099 | Archaea>Euryarchaeota>Stenosarchaea_group>Halobacteria>Natrialbales | GCF_016906025.1 | Haloterrigena_salifodinae | JMJ58_RS24560 | 74729..78028 | 2675099 | DUF499-domain-containing-protein-[Haloterrigena-salifodinae] |
| WP_265109158.1 | AAA+ATPase+wHTH+Alpha+Beta-domain+wHTH+RRM | Type-2_BR.RM.like.systems | Glyoxalase-><-?<-?<-?\|\|BirA-HTH->?->DUF499*->DUF3780->METHYLASE->HKD-DNase+SF1-SF2-Helicase+iREase+CTD-> | 1102 | Archaea>Euryarchaeota>Stenosarchaea_group>Halobacteria>Natrialbales | GCF_022869805.1 | Halosolutus_halophilus | MUG98_RS19875 | 4018249..4021557 | 1552990 | DUF499-domain-containing-protein-[Halosolutus-halophilus] |
| WP_254862272.1 | AAA+ATPase+wHTH+Alpha+Beta-domain+wHTH+RRM | Type-2_BR.RM.like.systems | <-HNH\|\|HEPN-><-?<-?\|\|REase-><-REase\|\|DUF499*->DUF3780->METHYLASE-><-PHP+SbcC+SidE+CR-ATPase9\|\|HKD-DNase+SF1-SF2-Helicase+iREase+CTD-> | 1084 | Archaea>Euryarchaeota>Stenosarchaea_group>Halobacteria>Natrialbales | GCF_024300625.1 | Halovivax_gelatinilyticus | NKH31_RS13270 | 2785012..2788266 | 2961597 | DUF499-domain-containing-protein-[Halovivax-sp.-CGA30] |
| WP_256532744.1 | AAA+ATPase+wHTH+Alpha+Beta-domain+wHTH+RRM | Type-2_BR.RM.like.systems | REase->SF1-SF2-Helicase+REase->DUF499*->DUF3780->METHYLASE->SF1-SF2-Helicase->?->HKD-DNase+SF1-SF2-Helicase+iREase+CTD->iSTAND-> | 986 | Archaea>Euryarchaeota>Stenosarchaea_group>Halobacteria>Natrialbales | GCF_024498195.1 | Halovivax_cerinus | NO366_RS02535 | 531191..534151 | 1487865 | DUF499-domain-containing-protein-[Halovivax-cerinus] |
| WP_049990607.1 | AAA+ATPase+wHTH+Alpha+Beta-domain+wHTH+RRM | Type-2_BR.RM.like.systems | <-CITB-HTH<-?<-HKD-DNase+SF1-SF2-Helicase+iREase+CTD<-METHYLASE<-DUF3780<-DUF499* | 1094 | Archaea>Euryarchaeota>Stenosarchaea_group>Halobacteria>Natrialbales | GCF_900110455.1 | Natrinema_salifodinae | BMY29_RS05885 | 83396..86680 | 1202768 | DUF499-domain-containing-protein-[Halopiger-salifodinae] |
| WP_216819686.1 | AAA+ATPase+wHTH+Alpha+Beta-domain+wHTH+RRM | Type-2_BR.RM.like.systems | <-HKD-DNase+SF1-SF2-Helicase+iREase+CTD<-SF1-SF2-Helicase<-METHYLASE<-DUF3780<-DUF499*<-CARF\|\|?-><-?<-BirA-HTH | 1076 | Archaea>Euryarchaeota>Stenosarchaea_group>Halobacteria>Natrialbales | GCF_900156475.1 | Natronorubrum_thiooxidans | BW940_RS19225 | 17133..20363 | 308853 | DUF499-domain-containing-protein-[Natronorubrum-thiooxidans] |
| NLA99643.1 | AAA+ATPase+wHTH+Alpha+Beta-domain+wHTH+RRM | Type-2_BR.RM.like.systems | Schlafen+HATPase+HTH->?-><-DUF3780<-DUF499*\|\|METHYLASE-><-?<-HKD-DNase+SF1-SF2-Helicase+iREase+CTD<-?<-ORC/CDC6+HTH | 950 | Archaea>Euryarchaeota>Stenosarchaea_group>Methanomicrobia>Methanomicrobiales | GCA_012511245.1 | Methanomicrobiales_archaeon | GX837_01550 | 2570..5422 | 2184052 | DUF499-domain-containing-protein-[Methanomicrobiales-archaeon] |
| CAG0963808.1 | AAA+ATPase+wHTH+Alpha+Beta-domain+wHTH+RRM | Type-2_BR.RM.like.systems | HKD-DNase+SF1-SF2-Helicase+iREase+CTD->?-><-?<-METHYLASE\|\|DUF499*->DUF3780-> | 994 | Archaea>Euryarchaeota>Stenosarchaea_group>Methanomicrobia>Methanosarcinales | GCA_905339305.1 | Methanosarcinales_archaeon | METP3_01010 | 17343..20327 | 2250255 | hypothetical-protein-METP3_01010-[Methanosarcinales-archaeon] |
| WAH99918.1 | AAA+ATPase+wHTH+Alpha+Beta-domain+wHTH+RRM | Type-2_BR.RM.like.systems | <-SF1-SF2-Helicase+iREase+CTD+REase<-METHYLASE<-?<-DUF3780<-DUF499* | 978 | Archaea>Euryarchaeota>Stenosarchaea_group>Methanomicrobia>Methanosarcinales | - | Candidatus_Methanoperedens_sp. | OI716_00725 | 153031..155967 | 2032688 | MAG:-DUF499-domain-containing-protein-[Candidatus-Methanoperedens-sp.] |
| MCZ7400325.1 | AAA+ATPase+wHTH+Alpha+Beta-domain+wHTH+RRM | Type-2_BR.RM.like.systems | RelE-ParE->?->?->DUF499*->DUF3780->PADR-HTH->HD->METHYLASE->NUDIX->SF1-SF2-Helicase+iREase+CTD+REase-> | 980 | Archaea>Euryarchaeota>Stenosarchaea_group>Methanomicrobia>Methanosarcinales | GCA_027460605.1 | Candidatus_Methanoperedens_sp. | O8C62_11750 | 3097..6039 | 2032688 | MAG:-DUF499-domain-containing-protein-[Candidatus-Methanoperedens-sp.] |
| CAG0952062.1 | AAA+ATPase+wHTH+Alpha+Beta-domain+wHTH+RRM | Type-2_BR.RM.like.systems | <-Primpol+D5N+D5ATPase\|\|DUF499*->DUF3780->PADR-HTH->HD->METHYLASE->SF1-SF2-Helicase+iREase+CTD+REase->ParB+HNH-> | 980 | Archaea>Euryarchaeota>Stenosarchaea_group>Methanomicrobia>Methanosarcinales | GCA_905339305.1 | Methanosarcinales_archaeon | METP3_00272 | 13443..16385 | 2250255 | hypothetical-protein-METP3_00272-[Methanosarcinales-archaeon] |
| WP_048043536.1 | AAA+ATPase+wHTH+Alpha+Beta-domain+wHTH+RRM | Type-2_BR.RM.like.systems | <-SF1-SF2-Helicase+iREase+CTD+REase<-METHYLASE<-DUF3780<-DUF499*<-?<-RelE-ParE | 983 | Archaea>Euryarchaeota>Stenosarchaea_group>Methanomicrobia>Methanosarcinales | GCF_000970185.1 | Methanosarcina_mazei | MSMAP_RS10730 | 2555385..2558336 | 2209 | DUF499-domain-containing-protein-[Methanosarcina-mazei] |
| WP_221068977.1 | AAA+ATPase+wHTH+Alpha+Beta-domain+wHTH+RRM | Type-2_BR.RM.like.systems | <-HTH\|\|?->?-><-?<-?\|\|Phage_integrase+ZnR->DUF499*->DUF3780->METHYLASE->SF1-SF2-Helicase+iREase+CTD+REase-> | 983 | Archaea>Euryarchaeota>Stenosarchaea_group>Methanomicrobia>Methanosarcinales | GCF_019669945.1 | Methanosarcina_mazei | MmazTMA_RS08615 | 2012144..2015095 | 2209 | DUF499-domain-containing-protein-[Methanosarcina-mazei] |
| MBP7066715.1 | AAA+ATPase+wHTH+Alpha+Beta-domain+wHTH+RRM | Type-2_BR.RM.like.systems | TPR+TPR->?->?-><-DUF3780<-DUF499*\|\|METHYLASE-><-?<-?<-HKD-DNase+SF1-SF2-Helicase+iREase+CTD<-ParB+HNH<-ArdCN+MPTase | 1011 | Archaea>Euryarchaeota>Stenosarchaea_group>Methanomicrobia>Methanotrichales | GCA_017994095.1 | Methanothrix_sp. | KBB04_00320 | 64883..67918 | 90426 | ATP-binding-protein-[Methanothrix-sp.] |
| MBA3046419.1 | AAA+ATPase+wHTH+Alpha+Beta-domain+wHTH+RRM | Type-2_BR.RM.like.systems | DUF499*->DUF3780->METHYLASE->TUDOR+SF1-SF2-Helicase+iREase+CTD->GDResolvase->OLD-ABC->TOPRIM-> | 1035 | Archaea>Euryarchaeota>unclassified_Euryarchaeota | GCA_013791785.1 | Euryarchaeota_archaeon | FP824_09440 | 19887..22984 | 2026739 | ATP-binding-protein-[Euryarchaeota-archaeon] |
| MBN1683203.1 | AAA+ATPase+wHTH+Alpha+Beta-domain+wHTH+RRM | Type-2_BR.RM.like.systems | DUF499*->DUF3780->METHYLASE->SF1-SF2-Helicase+iREase+CTD+REase-> | 678 | Archaea>TACK_group>Candidatus_Bathyarchaeota>unclassified_Candidatus_Bathyarchaeota | GCA_016930055.1 | Candidatus_Bathyarchaeota_archaeon | JW865_06595 | 1..2037 | 2026714 | hypothetical-protein-[Candidatus-Bathyarchaeota-archaeon] |

| Protein-ID | DUF499-architecture | System | Gene-Neighbourhoods | Length | Taxonomy | Assembly-ID | Strain | Locus-Tag | Genomic-coordinates | TaxID | Fasta-defline |
|---|---|---|---|---|---|---|---|---|---|---|---|
| MCW3988812.1 | AAA+ATPase+wHTH+Alpha+Beta-domain+wHTH+RRM | Type-2_BR.RM.like.systems | <-URI-DNase\|\|?->35exo->?-><-SF1-SF2-Helicase<-?\|\|DUF499*->DUF3780->METHYLASE->SF1-SF2-Helicase+iREase+CTD+REase->?->?->HIN-HTH-> | 885 | Archaea>TACK_group>Candidatus_Bathyarchaeota>unclassified_Candidatus_Bathyarchaeota | GCA_026014875.1 | Candidatus_Bathyarchaeota_archaeon | NWE88_01905 | 126678..129335 | 2026714 | MAG:-DUF499-domain-containing-protein-[Candidatus-Bathyarchaeota-archaeon] |
| RWX74207.1 | AAA+ATPase+wHTH+Alpha+Beta-domain+wHTH+RRM | Type-2_BR.RM.like.systems | PIN->?-><-?\|\|DUF499*->DUF3780->METHYLASE->SF1-SF2-Helicase+iREase+CTD+REase->?->?-><-PIN | 892 | Archaea>TACK_group>Candidatus_Verstraetearchaeota>Candidatus_Methanomethylicia>Candidatus_Methanomethylicales | GCA_004028775.1 | Candidatus_Methanosuratincola_subterraneus | Metus_0232 | 214945..217623 | 2593994 | hypothetical-protein-Metus_0232-[Candidatus-Methanosuratincola-subterraneus] |
| WP_103287092.1 | AAA+ATPase+wHTH+Alpha+Beta-domain+wHTH+RRM | Type-2_BR.RM.like.systems | VirD4-TraG+HerA->?->HEPN->EVE-><-?<-DUF3780<-DUF499*\|\|METHYLASE-><-?<-HKD-DNase+SF1-SF2-Helicase+iREase+CTD\|\|OLD-ABC+TOPRIM-> | 996 | Archaea>TACK_group>Nitrososphaerota>Nitrososphaeria>Candidatus_Nitrosocaldales | GCF_002906215.1 | Candidatus_Nitrosocaldus | CRG89_RS04275 | 822422..825412 | 498374 | MULTISPECIES:-DUF499-domain-containing-protein-[Candidatus-Nitrosocaldus] |
| NHV98109.1 | AAA+ATPase+wHTH+Alpha+Beta-domain+wHTH+RRM | Type-2_BR.RM.like.systems | <-Radical-SAM\|\|Radical-SAM-><-EVE\|\|ParB+HNH-><-DUF3780<-DUF499*\|\|METHYLASE-><-?<-HKD-DNase+SF1-SF2-Helicase+iREase+CTD\|\|?-><-DNA-glycosylase | 982 | Archaea>TACK_group>Nitrososphaerota>unclassified_Nitrososphaerota | GCA_011605725.1 | Nitrososphaerota_archaeon | HA496_00520 | 58939..61887 | 2026795 | ATP-binding-protein-[Thaumarchaeota-archaeon] |
| WP_054837388.1 | AAA+ATPase+wHTH+Alpha+Beta-domain+wHTH+RRM | Type-2_BR.RM.like.systems | DUF499*->DUF3780->METHYLASE-p->METHYLASE-p->SF1-SF2-Helicase+iREase+CTD+REase-p->REase-> | 877 | Archaea>TACK_group>Thermoproteota>Thermoprotei>Sulfolobales | GCF_001315825.1 | Metallosphaera_hakonensis | JCM8857_RS10980 | 481..3114 | 79601 | hypothetical-protein-[Metallosphaera-hakonensis] |
| MCL4850298.1 | AAA+ATPase+wHTH+Alpha+Beta-domain+wHTH+RRM | Type-2_BR.RM.like.systems | ParB->ParB->DUF499*->DUF3780->METHYLASE->HKD-DNase+SF1-SF2-Helicase+iREase+CTD-> | 1078 | Bacteria>Acidobacteriota>Terriglobia>Bryobacterales>Bryobacteraceae | GCA_023384715.1 | Bryobacteraceae_bacterium | KJZ78_02835 | 2179..5415 | 2212468 | DUF499-domain-containing-protein-[Bryobacteraceae-bacterium] |
| GIU73087.1 | AAA+ATPase+wHTH+Alpha+Beta-domain+wHTH+RRM | Type-2_BR.RM.like.systems | <-XIS-HTH<-GDResolvase+HIN-HTH<-?<-HKD-DNase+SF1-SF2-Helicase+iREase+CTD<-METHYLASE<-DUF3780<-DUF499*\|\|?->?-><-HTH_Tnp_1+HTH_21+rve<-ACET | 1076 | Bacteria>Acidobacteriota>Terriglobia>Bryobacterales>Bryobacteraceae | GCA_026002855.1 | Bryobacteraceae_bacterium | KatS3mg004_0174 | 202425..205655 | 2212468 | MAG:-hypothetical-protein-KatS3mg004_0174-[Bryobacteraceae-bacterium] |
| GIU73302.1 | AAA+ATPase+wHTH+Alpha+Beta-domain+wHTH+RRM | Type-2_BR.RM.like.systems | <-XIS-HTH<-GDResolvase+HIN-HTH<-HKD-DNase+SF1-SF2-Helicase+iREase+CTD<-METHYLASE<-DUF3780<-DUF499* | 1076 | Bacteria>Acidobacteriota>Terriglobia>Bryobacterales>Bryobacteraceae | GCA_026002855.1 | Bryobacteraceae_bacterium | KatS3mg004_0389 | 448361..451591 | 2212468 | MAG:-hypothetical-protein-KatS3mg004_0389-[Bryobacteraceae-bacterium] |
| GIU78629.1 | AAA+ATPase+wHTH+Alpha+Beta-domain+wHTH+RRM | Type-2_BR.RM.like.systems | Arm-DNA-bind_3+Phage_integrase->DUF772+DDE_Tnp_1+DDE_Tnp_1_6-><-?\|\|DUF499*->DUF3780->HTH+MPTase->PIN->METHYLASE->HKD-DNase+SF1-SF2-Helicase+iREase+CTD-> | 1076 | Bacteria>Acidobacteriota>Terriglobia>Bryobacterales>Bryobacteraceae | GCA_026002875.1 | Bryobacteraceae_bacterium | KatS3mg005_1867 | 2212823..2216053 | 2212468 | MAG:-hypothetical-protein-KatS3mg005_1867-[Bryobacteraceae-bacterium] |
| MCS6954437.1 | AAA+ATPase+wHTH+Alpha+Beta-domain+wHTH+RRM | Type-2_BR.RM.like.systems | <-HicA_toxin<-HicB_lk_antitox\|\|?->?->DUF499*->DUF3780->MPTase->PIN->METHYLASE-p-> | 1077 | Bacteria>Acidobacteriota>Terriglobia>Bryobacterales>Bryobacteraceae | GCA_025055455.1 | Bryobacteraceae_bacterium | NZM33_16450 | 10420..13653 | 2212468 | DUF499-domain-containing-protein-[Bryobacteraceae-bacterium] |
| SPE38928.1 | AAA+ATPase+wHTH+Alpha+Beta-domain+wHTH+RRM | Type-2_BR.RM.like.systems | <-PIN<-?<-?<-Ntox49<-METHYLASE<-DUF3780<-DUF499*<-?<-?<-DSBH | 1092 | Bacteria>Acidobacteriota>Terriglobia>Bryobacterales>Solibacteraceae | GCA_900290325.1 | Candidatus_Sulfopaludibacter_sp._SbA6 | SBA6_970023 | 16637..19915 | 2043166 | hypothetical-protein-SBA6_970023-[Candidatus-Sulfopaludibacter-sp.-SbA6] |
| MBV9397053.1 | AAA+ATPase+wHTH+Alpha+Beta-domain+wHTH+RRM | Type-2_BR.RM.like.systems | <-Cluster2725_2clades\|\|Terminase->PolB->HEPN->DUF499*->DUF3780->METHYLASE->HKD-DNase+SF1-SF2-Helicase+iREase+CTD-> | 1087 | Bacteria>Acidobacteriota>Terriglobia>Bryobacterales>unclassified_Bryobacterales | GCA_019261985.1 | Bryobacterales_bacterium | JO062_03680 | 122094..125357 | 2026791 | ATP-binding-protein-[Bryobacterales-bacterium] |
| MCY4597199.1 | AAA+ATPase+wHTH+Alpha+Beta-domain+wHTH+RRM | Type-2_BR.RM.like.systems | <-HKD-DNase+SF1-SF2-Helicase+iREase+CTD<-PIN<-?<-METHYLASE<-TOPRIM<-OLD-ABC\|\|DUF499*->DUF3780-> | 1006 | Bacteria>Acidobacteriota>Terriglobia>Bryobacterales>unclassified_Bryobacterales | GCA_026713455.1 | Bryobacterales_bacterium | OXC19_20655 | 14043..17063 | 2026791 | MAG:-DUF499-domain-containing-protein-[Bryobacterales-bacterium] |
| MCY4585111.1 | AAA+ATPase+wHTH+Alpha+Beta-domain+wHTH+RRM | Type-2_BR.RM.like.systems | <-HKD-DNase+SF1-SF2-Helicase+iREase+CTD<-METHYLASE<-?<-OLD-ABC<-ParB+HNH<-DUF3780<-DUF499*\|\|Phage_integrase-> | 1018 | Bacteria>Acidobacteriota>Terriglobia>Bryobacterales>unclassified_Bryobacterales | GCA_026713545.1 | Bryobacterales_bacterium | OXB98_03645 | 8005..11061 | 2026791 | MAG:-DUF499-domain-containing-protein-[Bryobacterales-bacterium] |
| MXY68843.1 | AAA+ATPase+wHTH+Alpha+Beta-domain+wHTH+RRM | Type-2_BR.RM.like.systems | <-Methyltransf<-?\|\|?-><-ArdCN+MPTase<-DUF3780<-DUF499*\|\|OLD-ABC->TOPRIM->METHYLASE->HKD-DNase+SF1-SF2-Helicase+iREase+CTD-> | 995 | Bacteria>Acidobacteriota>Terriglobia>unclassified_Terriglobia | GCA_009838515.1 | Terriglobia_bacterium | F4Y47_09825 | 221199..224186 | 2081523 | DUF499-domain-containing-protein-[Acidobacteriia-bacterium] |
| ODS53422.1 | AAA+ATPase+wHTH+Alpha+Beta-domain+wHTH+RRM | Type-2_BR.RM.like.systems | RVT-><-HEPN<-HKD-DNase+SF1-SF2-Helicase+iREase+CTD<-METHYLASE<-DOC+HTH<-DUF3780<-DUF499* | 997 | Bacteria>Acidobacteriota>unclassified_Acidobacteriota | GCA_001724025.1 | Acidobacteria_bacterium_SCN_69-37 | ABS36_12275 | 12058..15051 | 1660160 | hypothetical-protein-ABS36_12275-[Acidobacteria-bacterium-SCN-69-37] |
| MXX85702.1 | AAA+ATPase+wHTH+Alpha+Beta-domain+wHTH+RRM | Type-2_BR.RM.like.systems | <-TraI-VirD2-RCRE<-TraJ-RHH<-HicA_toxin<-HicB_lk_antitox\|\|DUF499*->DUF3780->METHYLASE-p->METHYLASE->HKD-DNase+SF1-SF2-Helicase+iREase+CTD-> | 1043 | Bacteria>Acidobacteriota>unclassified_Acidobacteriota | GCA_009838025.1 | Acidobacteriota_bacterium | F4Y71_04530 | 2244..5375 | 1978231 | ATP-binding-protein-[Acidobacteria-bacterium] |
| MCY4637129.1 | AAA+ATPase+wHTH+Alpha+Beta-domain+wHTH+RRM | Type-2_BR.RM.like.systems | <-DUF3780<-DUF499*\|\|<-?\|\|METHYLASE->HKD-DNase+SF1-SF2-Helicase+iREase+CTD->?-><-HEPN<-TniQ | 1037 | Bacteria>Acidobacteriota>unclassified_Acidobacteriota | GCA_026713685.1 | Acidobacteriota_bacterium | OXG04_21975 | 2884..5997 | 1978231 | MAG:-DUF499-domain-containing-protein-[Acidobacteria-bacterium] |
| MDE3261994.1 | AAA+ATPase+wHTH+Alpha+Beta-domain+wHTH+RRM | Type-2_BR.RM.like.systems | REase->DUF499*->DUF3780->OLD-ABC->HNH->METHYLASE->HKD-DNase+SF1-SF2-Helicase+iREase+CTD-> | 1006 | Bacteria>Acidobacteriota>unclassified_Acidobacteriota | GCA_028877265.1 | Acidobacteriota_bacterium | OYL41_08435 | 3377..6397 | 1978231 | MAG:-DUF499-domain-containing-protein-[Acidobacteriota-bacterium] |
| MCY4634540.1 | AAA+ATPase+wHTH+Alpha+Beta-domain+wHTH+RRM | Type-2_BR.RM.like.systems | <-REase+NACHT\|\|?-><-DUF3780<-DUF499*\|\|METHYLASE-> | 963 | Bacteria>Acidobacteriota>unclassified_Acidobacteriota | GCA_026713685.1 | Acidobacteriota_bacterium | OXG04_08580 | 8124..11015 | 1978231 | MAG:-DUF499-domain-containing-protein-[Acidobacteria-bacterium] |
| MCY3933638.1 | AAA+ATPase+wHTH+Alpha+Beta-domain+wHTH+RRM | Type-2_BR.RM.like.systems | <-HNH<-OLD-ABC<-?<-OLD-ABC<-ParB+HNH<-DUF3780<-DUF499*<-?<-?<-SMODS\|\|HTH+MPTase-><-?<-Phd-YefM | 1021 | Bacteria>Acidobacteriota>unclassified_Acidobacteriota | GCA_026707325.1 | Acidobacteriota_bacterium | OXH70_18150 | 6925..9990 | 1978231 | MAG:-DUF499-domain-containing-protein-[Acidobacteria-bacterium] |
| MDE2976517.1 | AAA+ATPase+wHTH+Alpha+Beta-domain+wHTH+RRM | Type-2_BR.RM.like.systems | MetJarc-HTH->?->?->DUF499*->DUF3780->ParB+HNH->OLD-ABC->?->OLD-ABC->HNH-> | 1021 | Bacteria>Acidobacteriota>unclassified_Acidobacteriota | GCA_028873735.1 | Acidobacteriota_bacterium | OXU63_03230 | 1513..4578 | 1978231 | MAG:-DUF499-domain-containing-protein-[Acidobacteriota-bacterium] |
| MCY3776603.1 | AAA+ATPase+wHTH+Alpha+Beta-domain+wHTH+RRM | Type-2_BR.RM.like.systems | <-OLD-ABC<-DUF3780<-DUF499*<-CHTH<-MazE_antitoxin<-?<-?\|\|?->PIN-> | 1004 | Bacteria>Bacteria_incertae_sedis>Bacteria_candidate_phyla>Candidatus_Aminicenantes>unclassified_Candidatus_Aminicenantes | GCA_026706325.1 | Candidatus_Aminicenantes_bacterium | OXH11_11535 | 1231..4245 | 2052149 | MAG:-DUF499-domain-containing-protein-[Candidatus-Aminicenantes-bacterium] |
| NCA71196.1 | AAA+ATPase+wHTH+Alpha+Beta-domain+wHTH+RRM | Type-2_BR.RM.like.systems | MODE-HTH+PBPII-><-DUF29\|\|DUF499*->DUF3780->Schlafen+HATPase+HTH->METHYLASE-> | 1037 | Bacteria>FCB_group>Bacteroidota/Chlorobiota_group>Bacteroidota>Sphingobacteriia | GCA_009928635.1 | Sphingobacteriia_bacterium | EOM91_14035 | 5029..8142 | 2035813 | ATP-binding-protein-[Sphingobacteriia-bacterium] |
| NLV45647.1 | AAA+ATPase+wHTH+Alpha+Beta-domain+wHTH+RRM | Type-2_BR.RM.like.systems | PolB->HEPN->?-><-?<-?\|\|DUF499*->?->DUF3780->?->HD->METHYLASE-> | 1012 | Bacteria>FCB_group>Candidatus_Hydrogenedentes>unclassified_Candidatus_Hydrogenedentes | GCA_012730105.1 | Candidatus_Hydrogenedentes_bacterium | GXY07_14260 | 8243..11281 | 2030809 | ATP-binding-protein-[Candidatus-Hydrogenedentes-bacterium] |
| MCY3601004.1 | AAA+ATPase+wHTH+Alpha+Beta-domain+wHTH+RRM | Type-2_BR.RM.like.systems | <-HKD-DNase+SF1-SF2-Helicase+iREase+CTD<-METHYLASE<-DUF3780<-DUF499*\|\|HicB_lk_antitox->HicA_toxin-> | 1043 | Bacteria>FCB_group>Gemmatimonadota>Gemmatimonadetes>Candidatus_Palauibacterales | GCA_026705295.1 | Candidatus_Palauibacter_australiensis | OXN85_13645 | 7648..10779 | 3056737 | MAG:-DUF499-domain-containing-protein-[Gemmatimonadetes-bacterium] |
| MBY0489937.1 | AAA+ATPase+wHTH+Alpha+Beta-domain+wHTH+RRM | Type-2_BR.RM.like.systems | <-REase<-HKD-DNase+SF1-SF2-Helicase+iREase+CTD<-METHYLASE<-DUF3780<-DUF499*<-?\|\|?->?-><-MazF | 974 | Bacteria>FCB_group>Gemmatimonadota>Gemmatimonadetes>Gemmatimonadales | GCA_019752115.1 | Gemmatimonadaceae_bacterium | K2R93_08880 | 95795..98719 | 2026741 | DUF499-domain-containing-protein-[Gemmatimonadaceae-bacterium] |
| MCC6769769.1 | AAA+ATPase+wHTH+Alpha+Beta-domain+wHTH+RRM | Type-2_BR.RM.like.systems | <-DUF772+DDE_Tnp_1+DDE_Tnp_1_6<-?<-?\|\|DUF499*->DUF3780->METHYLASE->Schlafen+HATPase+HTH->HKD-DNase+SF1-SF2-Helicase+iREase+CTD-><-CHTH | 1013 | Bacteria>FCB_group>Gemmatimonadota>Gemmatimonadetes>Gemmatimonadales | GCA_020848865.1 | Gemmatimonadaceae_bacterium | IT360_01040 | 3136..6177 | 2026741 | ATP-binding-protein-[Gemmatimonadaceae-bacterium] |
| GIW50768.1 | AAA+ATPase+wHTH+Alpha+Beta-domain+wHTH+RRM | Type-2_BR.RM.like.systems | DUF499*->DUF3780->?->METHYLASE-p->METHYLASE-p->METHYLASE-p->HKD-DNase+SF1-SF2-Helicase+iREase+CTD-> | 998 | Bacteria>FCB_group>Gemmatimonadota>Gemmatimonadetes>Gemmatimonadales | GCA_026004295.1 | Gemmatimonadales_bacterium | KatS3mg081_0123 | 136420..139416 | 2448054 | MAG:-hypothetical-protein-KatS3mg081_0123-[Gemmatimonadales-bacterium] |
| MCZ8206518.1 | AAA+ATPase+wHTH+Alpha+Beta-domain+wHTH+RRM | Type-2_BR.RM.like.systems | <-REase+NACHT\|\|DUF499*->DUF3780->METHYLASE->Schlafen+HATPase+HTH->HKD-DNase+SF1-SF2-Helicase+iREase+CTD-> | 1013 | Bacteria>FCB_group>Gemmatimonadota>Gemmatimonadetes>Gemmatimonadales | GCA_027531815.1 | Gemmatimonas_sp. | O9271_18135 | 3551..6592 | 1962908 | MAG:-DUF499-domain-containing-protein-[Gemmatimonas-sp.] |
| MBK6307121.1 | AAA+ATPase+wHTH+Alpha+Beta-domain+wHTH+RRM | Type-2_BR.RM.like.systems | <-GDResolvase<-?\|\|?-><-HKD-DNase+SF1-SF2-Helicase+iREase+CTD<-?<-METHYLASE<-DUF499*<-Schlafen+HATPase+HTH | 957 | Bacteria>FCB_group>Gemmatimonadota>unclassified_Gemmatimonadota | GCA_016705425.1 | Gemmatimonadota_bacterium | IPF47_15915 | 119468..122341 | 2026742 | ATP-binding-protein-[Gemmatimonadetes-bacterium] |
| MBK8059521.1 | AAA+ATPase+wHTH+Alpha+Beta-domain+wHTH+RRM | Type-2_BR.RM.like.systems | GDResolvase->GDResolvase-><-HKD-DNase+SF1-SF2-Helicase+iREase+CTD<-ParB+HNH<-METHYLASE<-DUF3780<-DUF499*\|\|HTH_3-><-?<-PAPS_reduct<-?<-KAP-NTPase<-HTH | 1026 | Bacteria>FCB_group>Gemmatimonadota>unclassified_Gemmatimonadota | GCA_016713015.1 | Gemmatimonadota_bacterium | IPK33_17185 | 755230..758310 | 2026742 | ATP-binding-protein-[Gemmatimonadetes-bacterium] |
| MCE2900724.1 | AAA+ATPase+wHTH+Alpha+Beta-domain+wHTH+RRM | Type-2_BR.RM.like.systems | DUF499*->DUF3780->METHYLASE-p->DUF772+DDE_Tnp_1->METHYLASE-p->Schlafen+HATPase+HTH->HKD-DNase+SF1-SF2-Helicase+iREase+CTD-> | 1013 | Bacteria>FCB_group>Gemmatimonadota>unclassified_Gemmatimonadota | GCA_021299375.1 | Gemmatimonadota_bacterium | LXA09_05735 | 68806..71847 | 2026742 | DUF499-domain-containing-protein-[Gemmatimonadetes-bacterium] |
| MCY3677712.1 | AAA+ATPase+wHTH+Alpha+Beta-domain+wHTH+RRM | Type-2_BR.RM.like.systems | <-HKD-DNase+SF1-SF2-Helicase+iREase+CTD<-METHYLASE<-DUF3780<-DUF499*\|\|HicB_lk_antitox->HicA_toxin-> | 1044 | Bacteria>FCB_group>Gemmatimonadota>unclassified_Gemmatimonadota | GCA_026705785.1 | Gemmatimonadota_bacterium | OXH66_08930 | 26252..29386 | 2026742 | MAG:-DUF499-domain-containing-protein-[Gemmatimonadetes-bacterium] |
| MCY4606377.1 | AAA+ATPase+wHTH+Alpha+Beta-domain+wHTH+RRM | Type-2_BR.RM.like.systems | OLD-ABC->HNH->DUF499*->DUF3780->?->?->?->METHYLASE->HKD-DNase+SF1-SF2-Helicase+iREase+CTD-> | 995 | Bacteria>FCB_group>Gemmatimonadota>unclassified_Gemmatimonadota | GCA_026713655.1 | Gemmatimonadota_bacterium | OXE49_19245 | 3266..6253 | 2026742 | MAG:-DUF499-domain-containing-protein-[Gemmatimonadetes-bacterium] |
| MDE2982966.1 | AAA+ATPase+wHTH+Alpha+Beta-domain+wHTH+RRM | Type-2_BR.RM.like.systems | DUF499*->DUF3780->METHYLASE->HKD-DNase+SF1-SF2-Helicase+iREase+CTD-> | 1016 | Bacteria>FCB_group>Gemmatimonadota>unclassified_Gemmatimonadota | GCA_028873815.1 | Gemmatimonadota_bacterium | OXU74_17385 | 23059..26109 | 2026742 | MAG:-DUF499-domain-containing-protein-[Gemmatimonadota-bacterium] |
| MDE2870422.1 | AAA+ATPase+wHTH+Alpha+Beta-domain+wHTH+RRM | Type-2_BR.RM.like.systems | <-HKD-DNase+SF1-SF2-Helicase+iREase+CTD<-METHYLASE<-?<-?<-?<-DUF3780<-DUF499*\|\|N6_MTase-> | 1002 | Bacteria>FCB_group>Gemmatimonadota>unclassified_Gemmatimonadota | GCA_028874435.1 | Gemmatimonadota_bacterium | OXQ94_01840 | 10684..13692 | 2026742 | MAG:-DUF499-domain-containing-protein-[Gemmatimonadota-bacterium] |
| MDE3259718.1 | AAA+ATPase+wHTH+Alpha+Beta-domain+wHTH+RRM | Type-2_BR.RM.like.systems | DUF499*->DUF3780->?->OLD-ABC->HNH->ParB+HNH->OLD-ABC-> | 955 | Bacteria>FCB_group>Gemmatimonadota>unclassified_Gemmatimonadota | GCA_028877215.1 | Gemmatimonadota_bacterium | OYM47_17950 | 187414..190281 | 2026742 | MAG:-DUF499-domain-containing-protein-[Gemmatimonadota-bacterium] |
| TAK25717.1 | AAA+ATPase+wHTH+Alpha+Beta-domain+wHTH+RRM | Type-2_BR.RM.like.systems | <-HKD-DNase+SF1-SF2-Helicase+iREase+CTD<-METHYLASE<-DUF3780<-DUF499*<-?<-?\|\|Synaptojanin->?-><-HTH_3\|\|TPR+TPR-> | 1011 | Bacteria>Myxococcota>Myxococcia>Myxococcales>Cystobacterineae | GCA_004297725.1 | Myxococcaceae_bacterium | EPO40_21965 | 13686..16721 | 2484252 | DUF499-domain-containing-protein-[Myxococcaceae-bacterium] |
| WP_164002142.1 | AAA+ATPase+wHTH+Alpha+Beta-domain+wHTH+RRM | Type-2_BR.RM.like.systems | <-HKD-DNase+SF1-SF2-Helicase+iREase+CTD<-METHYLASE<-DUF3780<-DUF499*<-?<-?\|\|DDE_3-><-HKD-DNase<-Pkinase+SF1-SF2-Helicase | 1272 | Bacteria>Myxococcota>Myxococcia>Myxococcales>Cystobacterineae | GCF_010894405.1 | Pyxidicoccus_caerfyrddinensis | G4D86_RS47295 | 8527..12345 | 2709663 | DUF499-domain-containing-protein-[Pyxidicoccus-caerfyrddinensis] |
| MCY4441764.1 | AAA+ATPase+wHTH+Alpha+Beta-domain+wHTH+RRM | Type-2_BR.RM.like.systems | DUF499*->DUF3780-><-HTH_3<-Tox-RelE\|\|?->METHYLASE->HKD-DNase+SF1-SF2-Helicase+iREase+CTD-> | 1032 | Bacteria>Myxococcota>Myxococcia>unclassified_Myxococcia | GCA_026710965.1 | Deltaproteobacteria_bacterium | OXE53_16355 | 908..4006 | 2026735 | MAG:-DUF499-domain-containing-protein-[Deltaproteobacteria-bacterium] |
| MDE0030313.1 | AAA+ATPase+wHTH+Alpha+Beta-domain+wHTH+RRM | Type-2_BR.RM.like.systems | <-HKD-DNase+SF1-SF2-Helicase+iREase+CTD<-METHYLASE<-?<-OLD-ABC<-ParB+HNH<-DUF3780<-DUF499*\|\|?-><-HEPN\|\|HEPN-><-DUF772+DDE_Tnp_1+DDE_Tnp_1_6 | 1005 | Bacteria>Myxococcota>Myxococcia>unclassified_Myxococcia | GCA_028818775.1 | Deltaproteobacteria_bacterium | OXU42_13040 | 19997..23014 | 2026735 | MAG:-DUF499-domain-containing-protein-[Deltaproteobacteria-bacterium] |
| MCK6511429.1 | AAA+ATPase+wHTH+Alpha+Beta-domain+wHTH+RRM | Type-2_BR.RM.like.systems | <-HKD-DNase+SF1-SF2-Helicase+iREase+CTD<-METHYLASE<-?<-?<-DUF3780<-DUF499*\|\|EVE-><-tRNA<-tRNA<-S1+RNase+2C\|\|?-><-SMS-KaiC | 1029 | Bacteria>Myxococcota>unclassified_Myxococcota | GCA_023150835.1 | Myxococcota_bacterium | L6R29_15840 | 12120..15209 | 2818507 | DUF499-domain-containing-protein-[Myxococcota-bacterium] |
| MDE0206328.1 | AAA+ATPase+wHTH+Alpha+Beta-domain+wHTH+RRM | Type-2_BR.RM.like.systems | <-PAS+LuxR-HTH<-PAS+LuxR-HTH\|\|?->DUF499*->DUF3780->OLD-ABC->?->REase+NACHT->Ntox18->Peptidase_M78-> | 1012 | Bacteria>Nitrospinae/Tectomicrobia_group>Candidatus_Tectomicrobia>unclassified_Candidatus_Tectomicrobia | GCA_028819935.1 | Candidatus_Tectomicrobia_bacterium | OXP66_09920 | 9081..12119 | 2528274 | MAG:-DUF499-domain-containing-protein-[Candidatus-Tectomicrobia-bacterium] |
| MCY4382893.1 | AAA+ATPase+wHTH+Alpha+Beta-domain+wHTH+RRM | Type-2_BR.RM.like.systems | PIN->?->?->?->DUF499*->DUF3780->METHYLASE->HKD-DNase+SF1-SF2-Helicase+iREase+CTD-><-TraJ-RHH | 998 | Bacteria>Nitrospinae/Tectomicrobia_group>Nitrospinota>unclassified_Nitrospinota | GCA_026710535.1 | Nitrospinota_bacterium | OXE44_07060 | 3249..6245 | 2026769 | MAG:-DUF499-domain-containing-protein-[Nitrospinae-bacterium] |
| GIW55576.1 | AAA+ATPase+wHTH+Alpha+Beta-domain+wHTH+RRM | Type-2_BR.RM.like.systems | XIS-HTH->DUF499-p->DUF499*->DUF3780->HD->PADR-HTH->METHYLASE->HKD-DNase+SF1-SF2-Helicase+iREase+CTD-> | 899 | Bacteria>Nitrospirota>Nitrospiria>Nitrospirales>Nitrospiraceae | GCA_026004315.1 | Nitrospiraceae_bacterium | KatS3mg082_1980 | 35349..38048 | 2026770 | MAG:-hypothetical-protein-KatS3mg082_1980-[Nitrospiraceae-bacterium] |
| WP_246765470.1 | AAA+ATPase+wHTH+Alpha+Beta-domain+wHTH+RRM | Type-2_BR.RM.like.systems | BirA-HTH->?->?->HisKin->?->DUF499*->DUF3780->METHYLASE->HKD-DNase+SF1-SF2-Helicase+iREase+CTD-><-OBFOLD-NUCLEASE | 1007 | Bacteria>Pseudomonadota | GCF_014843375.1 | Alphaproteobacteria | N7E70_RS30355 | 86433..89456 | 28211 | MULTISPECIES:-DUF499-domain-containing-protein-[Alphaproteobacteria] |
| MCB1884749.1 | AAA+ATPase+wHTH+Alpha+Beta-domain+wHTH+RRM | Type-2_BR.RM.like.systems | XIS-HTH->?->?->?->?->DUF499*->DUF3780->REase->METHYLASE->HKD-DNase+SF1-SF2-Helicase+iREase+CTD-><-Arm-DNA-bind_2+Phage_int_SAM_3+Phage_integrase | 1009 | Bacteria>Pseudomonadota>Alphaproteobacteria>Geminicoccales>Geminicoccaceae | GCA_020445645.1 | Geminicoccaceae_bacterium | KDG89_12270 | 5302..8331 | 2448052 | ATP-binding-protein-[Geminicoccaceae-bacterium] |
| WP_090598154.1 | AAA+ATPase+wHTH+Alpha+Beta-domain+wHTH+RRM | Type-2_BR.RM.like.systems | BirA-HTH->?->?->HisKin-><-?\|\|?->DUF499*->DUF3780->METHYLASE->HKD-DNase+SF1-SF2-Helicase+iREase+CTD-><-OBFOLD-NUCLEASE | 1021 | Bacteria>Pseudomonadota>Alphaproteobacteria>Hyphomicrobiales>Devosiaceae | GCF_900100665.1 | Pelagibacterium_luteolum | BLR31_RS17080 | 50153..53218 | 440168 | DUF499-domain-containing-protein-[Pelagibacterium-luteolum] |
| WP_203443311.1 | AAA+ATPase+wHTH+Alpha+Beta-domain+wHTH+RRM | Type-2_BR.RM.like.systems | <-GDResolvase+HIN-HTH<-HEPN<-HKD-DNase+SF1-SF2-Helicase+iREase+CTD-p<-DUF932<-METHYLASE<-DUF3780<-DUF499*\|\|?->?->?->?-><-?\|\|MarR-HTH-> | 1021 | Bacteria>Pseudomonadota>Alphaproteobacteria>Hyphomicrobiales>Methylobacteriaceae | GCF_016864375.1 | Microvirga_sp._VF16 | JO965_RS08880 | 1887910..1890975 | 2807101 | DUF499-domain-containing-protein-[Microvirga-sp.-VF16] |
| WP_230534162.1 | AAA+ATPase+wHTH+Alpha+Beta-domain+wHTH+RRM | Type-2_BR.RM.like.systems | <-HEPN<-HKD-DNase+SF1-SF2-Helicase+iREase+CTD<-DUF932<-METHYLASE<-DUF3780<-DUF499* | 1021 | Bacteria>Pseudomonadota>Alphaproteobacteria>Hyphomicrobiales>Methylobacteriaceae | GCF_020866965.1 | Microvirga_roseola | LG661_RS22040 | 8580..11645 | 2883126 | DUF499-domain-containing-protein-[Microvirga-roseola] |
| WP_003612121.1 | AAA+ATPase+wHTH+Alpha+Beta-domain+wHTH+RRM | Type-2_BR.RM.like.systems | <-HKD-DNase+SF1-SF2-Helicase+iREase+CTD<-METHYLASE<-DUF3780<-DUF499*\|\|?->?->?->LON-AAA-> | 1016 | Bacteria>Pseudomonadota>Alphaproteobacteria>Hyphomicrobiales>Methylocystaceae | GCF_000178815.2 | Methylosinus | P3C36_RS01035 | 199571..202621 | 425 | MULTISPECIES:-DUF499-domain-containing-protein-[Methylosinus] |
| WP_196404653.1 | AAA+ATPase+wHTH+Alpha+Beta-domain+wHTH+RRM | Type-2_BR.RM.like.systems | Phage_int_SAM_1+Phage_integrase->GDResolvase+Recombinase+ZnR->DUF499*->DUF3780->METHYLASE->HKD-DNase+SF1-SF2-Helicase+iREase+CTD-><-Arm-DNA-bind_3+Phage_integrase | 1017 | Bacteria>Pseudomonadota>Alphaproteobacteria>Hyphomicrobiales>Methylocystaceae | GCF_015709515.1 | Methylocystis_sp._H62 | IYY11_RS15350 | 2722..5775 | 2785789 | DUF499-domain-containing-protein-[Methylocystis-sp.-H62] |
| WP_245162922.1 | AAA+ATPase+wHTH+Alpha+Beta-domain+wHTH+RRM | Type-2_BR.RM.like.systems | <-HTH+UbE1<-?<-SF1-SF2-Helicase+REase<-?<-SIGMA-HTH\|\|CopG/RHH->DUF499*->DUF3780->METHYLASE->HKD-DNase+SF1-SF2-Helicase+iREase+CTD->wHTH+SAD+HNH->ParB+HNH-p->ParB+HNH-> | 1014 | Bacteria>Pseudomonadota>Alphaproteobacteria>Hyphomicrobiales>Nitrobacteraceae | - | Bradyrhizobium_vignae | JWS04_RS17155 | 6362..9406 | 1549949 | DUF499-domain-containing-protein-[Bradyrhizobium-vignae] |
| MBP0112785.1 | AAA+ATPase+wHTH+Alpha+Beta-domain+wHTH+RRM | Type-2_BR.RM.like.systems | <-HTH+UbE1<-?<-SF1-SF2-Helicase+REase<-?<-SIGMA-HTH\|\|CopG/RHH->DUF499*->DUF3780->METHYLASE->HKD-DNase+SF1-SF2-Helicase+iREase+CTD->wHTH(SBD)+HNH-> | 1029 | Bacteria>Pseudomonadota>Alphaproteobacteria>Hyphomicrobiales>Nitrobacteraceae | GCA_017788805.1 | Bradyrhizobium_vignae | JWS04_17170 | 6317..9406 | 1549949 | ATP-binding-protein-[Bradyrhizobium-vignae] |
| WP_076863827.1 | AAA+ATPase+wHTH+Alpha+Beta-domain+wHTH+RRM | Type-2_BR.RM.like.systems | <-HKD-DNase+SF1-SF2-Helicase+iREase+CTD<-METHYLASE<-DUF3780<-DUF499*<-?<-?<-?<-?<-?<-Phage_integrase | 1023 | Bacteria>Pseudomonadota>Alphaproteobacteria>Hyphomicrobiales>Nitrobacteraceae | GCF_001982635.1 | Bradyrhizobium_mercantei | BI327_RS24760 | 444106..447177 | 1904807 | DUF499-domain-containing-protein-[Bradyrhizobium-mercantei] |
| WP_178372757.1 | AAA+ATPase+wHTH+Alpha+Beta-domain+wHTH+RRM | Type-2_BR.RM.like.systems | <-NUDIX<-?<-?<-?\|\|DUF499*->DUF3780->METHYLASE->HKD-DNase+SF1-SF2-Helicase+iREase+CTD->REase+PNTD1+PNTD2->PIWI->HEPN-> | 1014 | Bacteria>Pseudomonadota>Alphaproteobacteria>Hyphomicrobiales>Nitrobacteraceae | GCF_013390305.1 | Bradyrhizobium_diazoefficiens | HUW42_RS33505 | 7080156..7083200 | 1355477 | DUF499-domain-containing-protein-[Bradyrhizobium-diazoefficiens] |
| WP_212504742.1 | AAA+ATPase+wHTH+Alpha+Beta-domain+wHTH+RRM | Type-2_BR.RM.like.systems | <-HKD-DNase<-Schlafen\|\|Phage_integrase->?->?->?->DUF499*->DUF3780->METHYLASE->HKD-DNase+SF1-SF2-Helicase+iREase+CTD-> | 1023 | Bacteria>Pseudomonadota>Alphaproteobacteria>Hyphomicrobiales>Nitrobacteraceae | GCF_018130735.1 | Bradyrhizobium_japonicum | JQ613_RS14815 | 103564..106635 | 375 | DUF499-domain-containing-protein-[Bradyrhizobium-japonicum] |
| WP_259243044.1 | AAA+ATPase+wHTH+Alpha+Beta-domain+wHTH+RRM | Type-2_BR.RM.like.systems | <-HIN-HTH\|\|?-><-HKD-DNase+SF1-SF2-Helicase+iREase+CTD<-METHYLASE<-DUF3780<-DUF499*<-CopG/RHH<-?<-PIN | 1012 | Bacteria>Pseudomonadota>Alphaproteobacteria>Hyphomicrobiales>Nitrobacteraceae | GCF_024808955.1 | Bradyrhizobium_elkanii | M2219_RS32005 | 6850109..6853147 | 29448 | DUF499-domain-containing-protein-[Bradyrhizobium-elkanii] |
| WP_271672235.1 | AAA+ATPase+wHTH+Alpha+Beta-domain+wHTH+RRM | Type-2_BR.RM.like.systems | HisKin-><-HKD-DNase+SF1-SF2-Helicase+iREase+CTD<-METHYLASE<-DUF3780<-DUF499*<-CopG/RHH\|\|SIGMA-HTH->?->SF1-SF2-Helicase+REase->?->HTH+UbE1-> | 1014 | Bacteria>Pseudomonadota>Alphaproteobacteria>Hyphomicrobiales>Nitrobacteraceae | GCF_028128765.1 | Bradyrhizobium_sp._CCBAU_51627 | XH88_RS32200 | 149327..152371 | 1325088 | DUF499-domain-containing-protein-[Bradyrhizobium-sp.-CCBAU-51627] |
| RWB32387.1 | AAA+ATPase+wHTH+Alpha+Beta-domain+wHTH+RRM | Type-2_BR.RM.like.systems | Peptidase_S11-><-HKD-DNase+SF1-SF2-Helicase+iREase+CTD<-METHYLASE<-DUF3780<-DUF499*\|\|tRNA->ParB+HNH->?-><-?<-?<-ACYC<-TetR-HTH | 1018 | Bacteria>Pseudomonadota>Alphaproteobacteria>Hyphomicrobiales>Phyllobacteriaceae | GCA_004016585.1 | Mesorhizobium_sp. | EOQ43_07600 | 7908..10964 | 1871066 | ATP-binding-protein-[Mesorhizobium-sp.] |
| RWA80837.1 | AAA+ATPase+wHTH+Alpha+Beta-domain+wHTH+RRM | Type-2_BR.RM.like.systems | XIS-HTH->?->ZnR+TOPRIM-><-HKD-DNase+SF1-SF2-Helicase+iREase+CTD<-METHYLASE<-DUF3780<-DUF499*\|\|?-><-?<-?<-XIS-HTH<-?\|\|Resolvase-> | 1015 | Bacteria>Pseudomonadota>Alphaproteobacteria>Hyphomicrobiales>Phyllobacteriaceae | GCA_004016695.1 | Mesorhizobium_sp. | EOQ30_21525 | 141970..145017 | 1871066 | ATP-binding-protein-[Mesorhizobium-sp.] |
| RWG07594.1 | AAA+ATPase+wHTH+Alpha+Beta-domain+wHTH+RRM | Type-2_BR.RM.like.systems | Phage_integrase-><-HKD-DNase+SF1-SF2-Helicase+iREase+CTD<-METHYLASE<-DUF3780<-DUF499*<-XIS-HTH\|\|GDResolvase+HTH-> | 1015 | Bacteria>Pseudomonadota>Alphaproteobacteria>Hyphomicrobiales>Phyllobacteriaceae | GCA_004018525.1 | Mesorhizobium_sp. | EOQ54_04025 | 85569..88616 | 1871066 | DUF499-domain-containing-protein-[Mesorhizobium-sp.] |
| WP_063171915.1 | AAA+ATPase+wHTH+Alpha+Beta-domain+wHTH+RRM | Type-2_BR.RM.like.systems | HTH->?-><-REase<-?\|\|SIR2->Phage_terminase->DUF499*->DUF3780->METHYLASE->HKD-DNase+SF1-SF2-Helicase+iREase+CTD->?->HNH->HNH-> | 1017 | Bacteria>Pseudomonadota>Alphaproteobacteria>Hyphomicrobiales>Phyllobacteriaceae | GCF_001618845.1 | Mesorhizobium_ciceri | A4R29_RS11120 | 2446069..2449122 | 39645 | DUF499-domain-containing-protein-[Mesorhizobium-ciceri] |
| WP_127299913.1 | AAA+ATPase+wHTH+Alpha+Beta-domain+wHTH+RRM | Type-2_BR.RM.like.systems | <-HKD-DNase+SF1-SF2-Helicase+iREase+CTD<-METHYLASE<-DUF3780<-DUF499*<-?<-HEPN | 1017 | Bacteria>Pseudomonadota>Alphaproteobacteria>Hyphomicrobiales>Phyllobacteriaceae | GCF_003997355.1 | Mesorhizobium_sp._M1E.F.Ca.ET.041.01.1.1 | EOA38_RS18715 | 4307..7360 | 2496759 | DUF499-domain-containing-protein-[Mesorhizobium-sp.-M1E.F.Ca.ET.041.01.1.1] |
| WP_154386575.1 | AAA+ATPase+wHTH+Alpha+Beta-domain+wHTH+RRM | Type-2_BR.RM.like.systems | BirA-HTH->?->?->HisKin->?->DUF499*->DUF3780->METHYLASE->HKD-DNase+SF1-SF2-Helicase+iREase+CTD-><-OBFOLD-NUCLEASE | 1022 | Bacteria>Pseudomonadota>Alphaproteobacteria>Hyphomicrobiales>Phyllobacteriaceae | GCF_014250155.1 | Aminobacter_sp._MDW-2 | GJR93_RS23015 | 44156..47224 | 2666139 | MULTISPECIES:-DUF499-domain-containing-protein-[unclassified-Aminobacter] |
| TBA20192.1 | AAA+ATPase+wHTH+Alpha+Beta-domain+wHTH+RRM | Type-2_BR.RM.like.systems | <-SF1-SF2-Helicase\|\|DUF499*->DUF3780->METHYLASE->HKD-DNase+SF1-SF2-Helicase+iREase+CTD-><-?<-TetR-HTH | 1020 | Bacteria>Pseudomonadota>Alphaproteobacteria>Hyphomicrobiales>Rhizobiaceae | GCA_004304965.1 | Rhizobium_ruizarguesonis | ELH66_03755 | 758245..761307 | 2081791 | ATP-binding-protein-[Rhizobium-leguminosarum] |
| TBZ80384.1 | AAA+ATPase+wHTH+Alpha+Beta-domain+wHTH+RRM | Type-2_BR.RM.like.systems | Glyco_transf_8->Glyco_trans_1_2->?-><-?\|\|?->Phage_integrase->DUF499*->DUF3780->METHYLASE->HKD-DNase+SF1-SF2-Helicase+iREase+CTD-><-?\|\|?->HIN-HTH-> | 1020 | Bacteria>Pseudomonadota>Alphaproteobacteria>Hyphomicrobiales>Rhizobiaceae | GCA_004330515.1 | Rhizobium_leguminosarum_bv._viciae | EOH61_17530 | 191921..194983 | 387 | ATP-binding-protein-[Rhizobium-leguminosarum-bv.-viciae] |
| MBO1965284.1 | AAA+ATPase+wHTH+Alpha+Beta-domain+wHTH+RRM | Type-2_BR.RM.like.systems | <-Glyoxalase_8<-PIN+OST-HTH\|\|REase-><-?\|\|<-HIN-HTH\|\|DUF499*->DUF3780->METHYLASE->HKD-DNase+SF1-SF2-Helicase+iREase+CTD-> | 1020 | Bacteria>Pseudomonadota>Alphaproteobacteria>Hyphomicrobiales>Rhizobiaceae | GCA_017571595.1 | Sinorhizobium_medicae | J4T85_34045 | 70691..73753 | 110321 | ATP-binding-protein-[Sinorhizobium-medicae] |
| WP_245488900.1 | AAA+ATPase+wHTH+Alpha+Beta-domain+wHTH+RRM | Type-2_BR.RM.like.systems | <-SF1-SF2-Helicase\|\|tRNA->DUF499*->DUF3780->METHYLASE->HKD-DNase+SF1-SF2-Helicase+iREase+CTD-><-?<-TetR-HTH | 1018 | Bacteria>Pseudomonadota>Alphaproteobacteria>Hyphomicrobiales>Rhizobiaceae | GCF_004304965.1 | Rhizobium_ruizarguesonis | ELH66_RS03755 | 758251..761307 | 2081791 | DUF499-domain-containing-protein-[Rhizobium-leguminosarum] |
| WP_245515890.1 | AAA+ATPase+wHTH+Alpha+Beta-domain+wHTH+RRM | Type-2_BR.RM.like.systems | Glyco_transf_8->Glyco_trans_1_2->?-><-?\|\|?->Phage_integrase->DUF499*->DUF3780->METHYLASE->HKD-DNase+SF1-SF2-Helicase+iREase+CTD-><-?\|\|?->HIN-HTH-> | 942 | Bacteria>Pseudomonadota>Alphaproteobacteria>Hyphomicrobiales>Rhizobiaceae | GCF_004330515.1 | Rhizobium_leguminosarum | EOH61_RS17530 | 192155..194983 | 384 | DUF499-domain-containing-protein-[Rhizobium-leguminosarum] |
| WP_157214161.1 | AAA+ATPase+wHTH+Alpha+Beta-domain+wHTH+RRM | Type-2_BR.RM.like.systems | Phage_integrase->DUF499*->DUF3780->METHYLASE->HKD-DNase+SF1-SF2-Helicase+iREase+CTD-> | 1020 | Bacteria>Pseudomonadota>Alphaproteobacteria>Hyphomicrobiales>Rhizobiaceae | GCF_009755765.1 | Rhizobium_leguminosarum | FML87_RS35050 | 2445..5507 | 384 | DUF499-domain-containing-protein-[Rhizobium-leguminosarum] |
| WP_221130431.1 | AAA+ATPase+wHTH+Alpha+Beta-domain+wHTH+RRM | Type-2_BR.RM.like.systems | TetR-HTH->?->REase->Tox-RelE->HTH+MPTase-><-XIS-HTH+RVT<-tRNA\|\|DUF499*->DUF3780->METHYLASE->HKD-DNase+SF1-SF2-Helicase+iREase+CTD->?-><-URI-DNase\|\|<-GDResolvase+HIN-HTH | 1018 | Bacteria>Pseudomonadota>Alphaproteobacteria>Hyphomicrobiales>Rhizobiaceae | GCF_019684795.1 | Rhizobium_lentis | HJB59_RS06875 | 255609..258665 | 1138194 | DUF499-domain-containing-protein-[Rhizobium-lentis] |
| WP_221160088.1 | AAA+ATPase+wHTH+Alpha+Beta-domain+wHTH+RRM | Type-2_BR.RM.like.systems | ABC-ATPase+SbcC-><-?<-?<-?<-?\|\|TetR-HTH-><-tRNA\|\|DUF499*->DUF3780->METHYLASE->HKD-DNase+SF1-SF2-Helicase+iREase+CTD->tRNA-><-GDResolvase+HIN-HTH\|\|ZnR-> | 1021 | Bacteria>Pseudomonadota>Alphaproteobacteria>Hyphomicrobiales>Rhizobiaceae | GCF_019685215.1 | Rhizobium_sp._NLR4b | HJC03_RS05950 | 256658..259723 | 2731118 | DUF499-domain-containing-protein-[Rhizobium-sp.-NLR4b] |
| WP_234942434.1 | AAA+ATPase+wHTH+Alpha+Beta-domain+wHTH+RRM | Type-2_BR.RM.like.systems | <-Glyoxalase_8<-PIN+OST-HTH\|\|REase-><-?\|\|<-HIN-HTH\|\|DUF499*->DUF3780->METHYLASE->HKD-DNase+SF1-SF2-Helicase+iREase+CTD-> | 973 | Bacteria>Pseudomonadota>Alphaproteobacteria>Hyphomicrobiales>Rhizobiaceae | GCF_034629665.1 | Sinorhizobium_medicae | J4T85_RS34045 | 70832..73753 | 110321 | DUF499-domain-containing-protein-[Sinorhizobium-medicae] |
| WP_149252800.1 | AAA+ATPase+wHTH+Alpha+Beta-domain+wHTH+RRM | Type-2_BR.RM.like.systems | Glyoxalase->tRNA->Arm-DNA-bind_3+Phage_integrase->DUF499*->DUF3780->METHYLASE->HKD-DNase+SF1-SF2-Helicase+iREase+CTD->?->XIS-HTH->HTH_36-> | 1022 | Bacteria>Pseudomonadota>Alphaproteobacteria>Hyphomicrobiales>Xanthobacteraceae | GCF_008330945.1 | Labrys_sp._KNU-23 | FZC33_RS20755 | 2431934..2435002 | 2789216 | DUF499-domain-containing-protein-[Labrys-sp.-KNU-23] |
| WP_119001174.1 | AAA+ATPase+wHTH+Alpha+Beta-domain+wHTH+RRM | Type-2_BR.RM.like.systems | <-HKD-DNase+SF1-SF2-Helicase+iREase+CTD<-METHYLASE<-DUF3780<-DUF499*<-MODE-HTH+PBPII\|\|TetR-HTH-><-ParG_RHH | 1019 | Bacteria>Pseudomonadota>Alphaproteobacteria>Rhodobacterales>Paracoccaceae | GCF_003547135.1 | Cereibacter_sphaeroides | D1114_RS19755 | 8373..11432 | 1063 | DUF499-domain-containing-protein-[Cereibacter-sphaeroides] |
| WP_161763477.1 | AAA+ATPase+wHTH+Alpha+Beta-domain+wHTH+RRM | Type-2_BR.RM.like.systems | Peptidase_M15->?->?->DUF499*->DUF3780->METHYLASE->HKD-DNase+SF1-SF2-Helicase+iREase+CTD-> | 1019 | Bacteria>Pseudomonadota>Alphaproteobacteria>Rhodobacterales>Paracoccaceae | GCF_009908265.2 | Rhodobacter_amnigenus | KTV87_RS16080 | 175141..178200 | 2852097 | DUF499-domain-containing-protein-[Rhodobacter-amnigenus] |
| WP_217900503.1 | AAA+ATPase+wHTH+Alpha+Beta-domain+wHTH+RRM | Type-2_BR.RM.like.systems | RAMA->GDResolvase+Recombinase+ZnR->HTH->?->DUF499*->DUF3780->METHYLASE->HKD-DNase+SF1-SF2-Helicase+iREase+CTD-> | 1045 | Bacteria>Pseudomonadota>Alphaproteobacteria>Rhodobacterales>Paracoccaceae | GCF_900184945.1 | Maliponia_aquimaris | MAA8898_RS15965 | 28146..31283 | 1673631 | DUF499-domain-containing-protein-[Maliponia-aquimaris] |
| MBQ5516158.1 | AAA+ATPase+wHTH+Alpha+Beta-domain+wHTH+RRM | Type-2_BR.RM.like.systems | HKD-DNase+HKD-DNase-><-Synaptojanin\|\|?->DUF499*->DUF3780->METHYLASE-> | 1034 | Bacteria>Pseudomonadota>Alphaproteobacteria>Rhodospirillales>Acetobacteraceae | GCA_017385515.1 | Acetobacter_sp. | IIT88_03630 | 7546..10650 | 440 | ATP-binding-protein-[Acetobacter-sp.] |
| MBQ5479180.1 | AAA+ATPase+wHTH+Alpha+Beta-domain+wHTH+RRM | Type-2_BR.RM.like.systems | HKD-DNase+HKD-DNase-><-Synaptojanin\|\|?->DUF499*->DUF3780-> | 1034 | Bacteria>Pseudomonadota>Alphaproteobacteria>Rhodospirillales>Acetobacteraceae | GCA_017386005.1 | Acetobacter_sp. | IIT66_03080 | 12505..15609 | 440 | ATP-binding-protein-[Acetobacter-sp.] |
| MBQ5469892.1 | AAA+ATPase+wHTH+Alpha+Beta-domain+wHTH+RRM | Type-2_BR.RM.like.systems | <-DUF3780<-DUF499*<-?\|\|Synaptojanin-><-HKD-DNase+HKD-DNase | 1031 | Bacteria>Pseudomonadota>Alphaproteobacteria>Rhodospirillales>Acetobacteraceae | GCA_017386225.1 | Acetobacter_sp. | IIT54_04720 | 323..3418 | 440 | ATP-binding-protein-[Acetobacter-sp.] |
| MBR2123973.1 | AAA+ATPase+wHTH+Alpha+Beta-domain+wHTH+RRM | Type-2_BR.RM.like.systems | <-METHYLASE-p<-DUF3780<-DUF499*<-?\|\|Synaptojanin-><-HKD-DNase+HKD-DNase | 1047 | Bacteria>Pseudomonadota>Alphaproteobacteria>Rhodospirillales>Acetobacteraceae | GCA_017474285.1 | Acetobacter_sp. | IJ934_02160 | 1080..4223 | 440 | ATP-binding-protein-[Acetobacter-sp.] |
| MBO6035783.1 | AAA+ATPase+wHTH+Alpha+Beta-domain+wHTH+RRM | Type-2_BR.RM.like.systems | <-METHYLASE-p<-DUF3780<-DUF499*<-?\|\|Synaptojanin-><-HKD-DNase+HKD-DNase | 1032 | Bacteria>Pseudomonadota>Alphaproteobacteria>Rhodospirillales>Acetobacteraceae | GCA_017632995.1 | Acetobacter_sp. | J6P38_01670 | 2020..5118 | 440 | ATP-binding-protein-[Acetobacter-sp.] |
| MBO6091752.1 | AAA+ATPase+wHTH+Alpha+Beta-domain+wHTH+RRM | Type-2_BR.RM.like.systems | <-DUF499*<-?\|\|Synaptojanin-><-HKD-DNase+HKD-DNase | 1001 | Bacteria>Pseudomonadota>Alphaproteobacteria>Rhodospirillales>Acetobacteraceae | GCA_017633685.1 | Acetobacter_sp. | J6P00_06015 | 1..3003 | 440 | ATP-binding-protein-[Acetobacter-sp.] |
| MCA3396953.1 | AAA+ATPase+wHTH+Alpha+Beta-domain+wHTH+RRM | Type-2_BR.RM.like.systems | <-HIN-HTH<-DUF499*\|\|TIR->METHYLASE->HKD-DNase+SF1-SF2-Helicase+iREase+CTD-> | 1031 | Bacteria>Pseudomonadota>Alphaproteobacteria>Rhodospirillales>Acetobacteraceae | GCA_020249805.1 | Roseomonas_sp. | INF85_15065 | 1743..4838 | 1869190 | ATP-binding-protein-[Roseomonas-sp.] |
| MCZ8277167.1 | AAA+ATPase+wHTH+Alpha+Beta-domain+wHTH+RRM | Type-2_BR.RM.like.systems | HTH-><-HKD-DNase+SF1-SF2-Helicase+iREase+CTD<-METHYLASE<-?<-?<-DUF3780<-DUF499* | 940 | Bacteria>Pseudomonadota>Alphaproteobacteria>Rhodospirillales>Acetobacteraceae | GCA_027532445.1 | Acetobacteraceae_bacterium | O9313_02395 | 10685..13507 | 1909293 | MAG:-DUF499-domain-containing-protein-[Acetobacteraceae-bacterium] |
| WP_198375108.1 | AAA+ATPase+wHTH+Alpha+Beta-domain+wHTH+RRM | Type-2_BR.RM.like.systems | <-HKD-DNase+SF1-SF2-Helicase+iREase+CTD<-DUF932<-METHYLASE<-HNH<-OLD-ABC<-DUF3780<-DUF499*<-DnaB<-HTH_17<-?<-Arm-DNA-bind_3+Phage_integrase\|\|Methyltransf_21->Methyltransf-> | 1033 | Bacteria>Pseudomonadota>Alphaproteobacteria>Rhodospirillales>Acetobacteraceae | GCF_016106015.1 | Roseomonas_rubea | HW577_RS05730 | 12708..15809 | 2748666 | DUF499-domain-containing-protein-[Roseomonas-rubea] |
| WP_211844700.1 | AAA+ATPase+wHTH+Alpha+Beta-domain+wHTH+RRM | Type-2_BR.RM.like.systems | GDResolvase+Recombinase+ZnR->HTH->?->HTH-><-HKD-DNase+SF1-SF2-Helicase+iREase+CTD<-METHYLASE\|\|DUF499*->DUF3780-><-tRNA<-?<-sGTP | 1030 | Bacteria>Pseudomonadota>Alphaproteobacteria>Rhodospirillales>Acetobacteraceae | GCF_018129005.1 | Neoroseomonas_eburnea | GXW74_RS02465 | 27272..30364 | 1346889 | DUF499-domain-containing-protein-[Roseomonas-eburnea] |
| MCI0431032.1 | AAA+ATPase+wHTH+Alpha+Beta-domain+wHTH+RRM | Type-2_BR.RM.like.systems | DUF499*->DUF3780->METHYLASE->HKD-DNase+SF1-SF2-Helicase+iREase+CTD-> | 983 | Bacteria>Pseudomonadota>Alphaproteobacteria>Rhodospirillales>unclassified_Rhodospirillales | GCA_022614775.1 | Rhodospirillales_bacterium | L0210_10875 | 3961..6912 | 2026786 | DUF499-domain-containing-protein-[Rhodospirillales-bacterium] |
| MDD9991563.1 | AAA+ATPase+wHTH+Alpha+Beta-domain+wHTH+RRM | Type-2_BR.RM.like.systems | <-SF1-SF2-Helicase<-OLD-ABC+TOPRIM<-?<-?\|\|DUF499*->DUF3780->METHYLASE->HKD-DNase+SF1-SF2-Helicase+iREase+CTD->DUF222+HNH-> | 1041 | Bacteria>Pseudomonadota>Alphaproteobacteria>Rhodospirillales>unclassified_Rhodospirillales | GCA_028818565.1 | Rhodospirillales_bacterium | OXP75_07170 | 4041..7166 | 2026786 | MAG:-DUF499-domain-containing-protein-[Rhodospirillales-bacterium] |
| MDA8048919.1 | AAA+ATPase+wHTH+Alpha+Beta-domain+wHTH+RRM | Type-2_BR.RM.like.systems | <-SF1-SF2-Helicase<-?<-TRD+TRD<-REase+N6_MTase<-PIN\|\|DUF499*->DUF3780->HD->DUF4065-><-?<-REase\|\|ORC-CDC6+REase-> | 1006 | Bacteria>Pseudomonadota>Alphaproteobacteria>Rhodospirillales>unclassified_Rhodospirillales | GCA_028063655.1 | Rhodospirillales_bacterium | M0002_02755 | 5882..8902 | 2026786 | MAG:-DUF499-domain-containing-protein-[Rhodospirillales-bacterium] |
| WP_160766410.1 | AAA+ATPase+wHTH+Alpha+Beta-domain+wHTH+RRM | Type-2_BR.RM.like.systems | <-Zn_Tnp_IS1595+DDE_Tnp_IS1595\|\|?->DUF499*->DUF3780->METHYLASE->?->HKD-DNase+SF1-SF2-Helicase+iREase+CTD->TPR->RNase+SF1-SF2-Helicase-> | 1022 | Bacteria>Pseudomonadota>Alphaproteobacteria>Sphingomonadales>Erythrobacteraceae | GCF_030815185.1 | Qipengyuania_citrea | GRI55_RS05380 | 1121606..1124674 | 225971 | DUF499-domain-containing-protein-[Qipengyuania-citrea] |
| AOF98012.1 | AAA+ATPase+wHTH+Alpha+Beta-domain+wHTH+RRM | Type-2_BR.RM.like.systems | <-HIN-HTH<-HKD-DNase+SF1-SF2-Helicase+iREase+CTD<-METHYLASE-p<-DUF3780<-DUF499*<-REase-p<-?<-?<-OLD-ABC<-ParB+HNH | 1026 | Bacteria>Pseudomonadota>Alphaproteobacteria>Sphingomonadales>Sphingomonadaceae | GCA_001713415.1 | Sphingobium_sp._RAC03 | BSY17_1605 | 1643040..1646120 | 1843368 | hypothetical-protein-BSY17_1605-[Sphingobium-sp.-RAC03] |
| OYY89817.1 | AAA+ATPase+wHTH+Alpha+Beta-domain+wHTH+RRM | Type-2_BR.RM.like.systems | <-SF1-SF2-Helicase+REase<-?<-SMODS<-?\|\|CHTH->MPTase->DUF499*->DUF3780->METHYLASE->HKD-DNase+SF1-SF2-Helicase+iREase+CTD-><-?<-HTH | 1004 | Bacteria>Pseudomonadota>Alphaproteobacteria>Sphingomonadales>Sphingomonadaceae | GCA_002281485.1 | Sphingomonas_sp._28-66-16 | B7Y45_10350 | 25240..28254 | 1970435 | hypothetical-protein-B7Y45_10350-[Sphingomonas-sp.-28-66-16] |
| RSV33259.1 | AAA+ATPase+wHTH+Alpha+Beta-domain+wHTH+RRM | Type-2_BR.RM.like.systems | Phage_integrase->ZnR+TOPRIM->?->?-><-ParB+HNH\|\|?->DUF499*->DUF3780->METHYLASE-><-?\|\|HKD-DNase+SF1-SF2-Helicase+iREase+CTD-><-Schlafen<-PIWI | 1005 | Bacteria>Pseudomonadota>Alphaproteobacteria>Sphingomonadales>Sphingomonadaceae | GCA_003951715.1 | Sphingomonas_sp._ABOLH | CA237_01110 | 31273..34290 | 1985881 | DUF499-domain-containing-protein-[Sphingomonas-sp.-ABOLH] |
| MBX9859748.1 | AAA+ATPase+wHTH+Alpha+Beta-domain+wHTH+RRM | Type-2_BR.RM.like.systems | <-SMODS<-?<-HKD-DNase+SF1-SF2-Helicase-p<-HKD-DNase+SF1-SF2-Helicase+iREase+CTD<-METHYLASE<-DUF3780<-DUF499*<-?<-?\|\|HTH->HEPN-><-ArsR-HTH+YTCD-HTH | 1024 | Bacteria>Pseudomonadota>Alphaproteobacteria>Sphingomonadales>Sphingomonadaceae | GCA_019744885.1 | Sphingomonas_sp. | K2Y20_09195 | 11117..14191 | 28214 | DUF499-domain-containing-protein-[Sphingomonas-sp.] |
| WP_209445045.1 | AAA+ATPase+wHTH+Alpha+Beta-domain+wHTH+RRM | Type-2_BR.RM.like.systems | PIWI->Schlafen-><-HKD-DNase+SF1-SF2-Helicase+iREase+CTD<-?<-METHYLASE<-DUF3780<-DUF499*<-?\|\|ParB+HNH-><-?<-?<-ZnR+TOPRIM<-Phage_integrase | 1025 | Bacteria>Pseudomonadota>Alphaproteobacteria>Sphingomonadales>Sphingomonadaceae | GCF_001477495.1 | Sphingomonas_yabuuchiae | NS355_RS08100 | 26418..29495 | 172044 | DUF499-domain-containing-protein-[Sphingomonas-yabuuchiae] |
| WP_061939617.1 | AAA+ATPase+wHTH+Alpha+Beta-domain+wHTH+RRM | Type-2_BR.RM.like.systems | XIS-HTH-><-?<-DUF488-1\|\|DUF499*->DUF3780->METHYLASE->HKD-DNase+SF1-SF2-Helicase+iREase+CTD->?-><-Arm-DNA-bind_3+Phage_integrase | 1025 | Bacteria>Pseudomonadota>Alphaproteobacteria>Sphingomonadales>Sphingomonadaceae | GCF_001550165.1 | unclassified_Sphingobium | AXW74_RS19420 | 19251..22328 | 2611147 | MULTISPECIES:-DUF499-domain-containing-protein-[unclassified-Sphingobium] |
| WP_065846240.1 | AAA+ATPase+wHTH+Alpha+Beta-domain+wHTH+RRM | Type-2_BR.RM.like.systems | Arm-DNA-bind_3+Phage_integrase-><-?<-HKD-DNase+SF1-SF2-Helicase+iREase+CTD<-METHYLASE<-DUF3780<-DUF499*\|\|DUF488-1->?-><-XIS-HTH | 1025 | Bacteria>Pseudomonadota>Alphaproteobacteria>Sphingomonadales>Sphingomonadaceae | GCF_001699185.1 | Sphingobium_sp._Ndbn-10 | BFF74_RS09030 | 21448..24525 | 1667223 | DUF499-domain-containing-protein-[Sphingobium-sp.-Ndbn-10] |
| WP_237236295.1 | AAA+ATPase+wHTH+Alpha+Beta-domain+wHTH+RRM | Type-2_BR.RM.like.systems | <-HIN-HTH<-XIS-HTH<-?<-HKD-DNase+SF1-SF2-Helicase+iREase+CTD<-METHYLASE<-DUF3780<-DUF499*<-REase-p<-?<-?<-OLD-ABC<-ParB+HNH | 1006 | Bacteria>Pseudomonadota>Alphaproteobacteria>Sphingomonadales>Sphingomonadaceae | GCF_001713415.1 | Sphingobium_sp._RAC03 | BSY17_RS12535 | 1643040..1646060 | 1843368 | DUF499-domain-containing-protein-[Sphingobium-sp.-RAC03] |
| WP_206432931.1 | AAA+ATPase+wHTH+Alpha+Beta-domain+wHTH+RRM | Type-2_BR.RM.like.systems | XIS-HTH->Phage_integrase->ZnR+TOPRIM->?->?-><-ParB+HNH\|\|DUF499*->DUF3780->METHYLASE->?->HKD-DNase+SF1-SF2-Helicase+iREase+CTD-><-Schlafen<-PIWI | 1025 | Bacteria>Pseudomonadota>Alphaproteobacteria>Sphingomonadales>Sphingomonadaceae | GCF_003951715.1 | Sphingomonas_sp._ABOLH | CA237_RS01115 | 31213..34290 | 1985881 | DUF499-domain-containing-protein-[Sphingomonas-sp.-ABOLH] |
| WP_167952885.1 | AAA+ATPase+wHTH+Alpha+Beta-domain+wHTH+RRM | Type-2_BR.RM.like.systems | PIWI-><-?<-HKD-DNase+SF1-SF2-Helicase+iREase+CTD<-?<-METHYLASE<-DUF3780<-DUF499*<-?\|\|?->?->?-><-TniQ | 1029 | Bacteria>Pseudomonadota>Alphaproteobacteria>Sphingomonadales>Sphingomonadaceae | GCF_011927695.1 | Sphingomonas_jejuensis | GGR88_RS03065 | 607476..610565 | 904715 | DUF499-domain-containing-protein-[Sphingomonas-jejuensis] |
| WP_168134876.1 | AAA+ATPase+wHTH+Alpha+Beta-domain+wHTH+RRM | Type-2_BR.RM.like.systems | PIWI-><-?<-HKD-DNase+SF1-SF2-Helicase+iREase+CTD<-?<-METHYLASE<-DUF3780<-DUF499*<-?<-HEPN\|\|AAA_21->?-><-TniQ | 1031 | Bacteria>Pseudomonadota>Alphaproteobacteria>Sphingomonadales>Sphingomonadaceae | GCF_012035195.1 | Sphingomonas_corticis | HBH26_RS12185 | 283937..287032 | 2722791 | DUF499-domain-containing-protein-[Sphingomonas-corticis] |
| WP_168603135.1 | AAA+ATPase+wHTH+Alpha+Beta-domain+wHTH+RRM | Type-2_BR.RM.like.systems | <-REase<-?\|\|?-><-HKD-DNase+SF1-SF2-Helicase+iREase+CTD<-METHYLASE<-DUF3780<-DUF499*<-?\|\|SMODS->?-><-tRNA\|\|?->SAM-methylase-> | 1023 | Bacteria>Pseudomonadota>Alphaproteobacteria>Sphingomonadales>Sphingomonadaceae | GCF_012275515.1 | Novosphingobium_sp._SG707 | FHW92_RS10095 | 286102..289173 | 2586996 | DUF499-domain-containing-protein-[Novosphingobium-sp.-SG707] |
| WP_175310430.1 | AAA+ATPase+wHTH+Alpha+Beta-domain+wHTH+RRM | Type-2_BR.RM.like.systems | <-SbcC<-?<-SIR2<-HKD-DNase+SF1-SF2-Helicase+iREase+CTD<-METHYLASE<-DUF3780<-DUF499*<-?<-?<-?<-REase+35exo+Beta-Barrel+SF1-SF2-Helicase\|\|HTH+WYL-> | 1028 | Bacteria>Pseudomonadota>Alphaproteobacteria>Sphingomonadales>Sphingomonadaceae | GCF_013359795.1 | Sphingomonas_zeae | HP438_RS01360 | 31429..34515 | 1646122 | DUF499-domain-containing-protein-[Sphingomonas-zeae] |
| WP_088441309.1 | AAA+ATPase+wHTH+Alpha+Beta-domain+wHTH+RRM | Type-2_BR.RM.like.systems | <-HKD-DNase+SF1-SF2-Helicase+iREase+CTD<-METHYLASE<-DUF3780<-DUF499*<-HHH_4+SF1-SF2-Helicase+REase<-REase+35exo+Beta-Barrel+SF1-SF2-Helicase<-Ntox49\|\|HTH+WYL-> | 1022 | Bacteria>Pseudomonadota>Alphaproteobacteria>Sphingomonadales>Sphingomonadaceae | GCF_014645635.1 | Sphingopyxis_bauzanensis | CDQ92_RS10605 | 849090..852158 | 651663 | DUF499-domain-containing-protein-[Sphingopyxis-bauzanensis] |
| WP_223010001.1 | AAA+ATPase+wHTH+Alpha+Beta-domain+wHTH+RRM | Type-2_BR.RM.like.systems | <-SbcC<-?<-SIR2<-HKD-DNase+SF1-SF2-Helicase+iREase+CTD<-METHYLASE<-DUF3780<-DUF499*\|\|?-><-HKD-DNase | 1023 | Bacteria>Pseudomonadota>Alphaproteobacteria>Sphingomonadales>Sphingomonadaceae | GCF_019898765.1 | Sphingomonas_yunnanensis | K7957_RS00115 | 31463..34534 | 310400 | DUF499-domain-containing-protein-[Sphingomonas-yunnanensis] |
| MBN9423714.1 | AAA+ATPase+wHTH+Alpha+Beta-domain+wHTH+RRM | Type-2_BR.RM.like.systems | <-HKD-DNase<-HD<-?<-NUDIX\|\|HTH+WYL->DUF499*->DUF3780->METHYLASE->HKD-DNase+SF1-SF2-Helicase+iREase+CTD-><-HTH_Tnp_1+HTH_21+rve+rve_3\|\|ParB+HNH-> | 1262 | Bacteria>Pseudomonadota>Betaproteobacteria>Betaproteobacteria_incertae_sedis>Candidatus_Accumulibacter | GCA_017307795.1 | Accumulibacter_sp. | JOI91_14740 | 18980..22768 | 2053492 | ATP-binding-protein-[Candidatus-Accumulibacter-sp.] |
| MCM8580151.1 | AAA+ATPase+wHTH+Alpha+Beta-domain+wHTH+RRM | Type-2_BR.RM.like.systems | <-METHYLASE<-Schlafen+HATPase+HTH<-DUF3780<-DUF499*<-SF1-SF2-Helicase+DUF1998+HKD-DNase+HKD-DNase<-SF1-SF2-Helicase+DUF1998+HKD-DNase+HKD-DNase | 1010 | Bacteria>Pseudomonadota>Betaproteobacteria>Betaproteobacteria_incertae_sedis>Candidatus_Accumulibacter | GCA_023806705.1 | Accumulibacter_sp. | NFV56_11820 | 5127..8159 | 2053492 | DUF499-domain-containing-protein-[Candidatus-Accumulibacter-sp.] |
| MCQ1547870.1 | AAA+ATPase+wHTH+Alpha+Beta-domain+wHTH+RRM | Type-2_BR.RM.like.systems | <-Classical-AAA\|\|MetJarc-HTH-><-HKD-DNase+SF1-SF2-Helicase+iREase+CTD<-METHYLASE<-Schlafen+HATPase+HTH<-DUF3780<-DUF499*\|\|DUF6361->HKD-DNase->SF1-SF2-Helicase-><-?<-?<-RelE-ParE | 1010 | Bacteria>Pseudomonadota>Betaproteobacteria>Betaproteobacteria_incertae_sedis>Candidatus_Accumulibacter | GCA_024380085.1 | Candidatus_Accumulibacter_phosphatis | NOF05_03365 | 13347..16379 | 327160 | DUF499-domain-containing-protein-[Candidatus-Accumulibacter-phosphatis] |
| WP_138677516.1 | AAA+ATPase+wHTH+Alpha+Beta-domain+wHTH+RRM | Type-2_BR.RM.like.systems | RelE-ParE->?->?-><-SF1-SF2-Helicase<-HKD-DNase<-DUF6361\|\|DUF499*->DUF3780->Schlafen+HATPase+HTH->METHYLASE->HKD-DNase+SF1-SF2-Helicase+iREase+CTD-><-MetJarc-HTH\|\|Classical-AAA-> | 1010 | Bacteria>Pseudomonadota>Betaproteobacteria>Betaproteobacteria_incertae_sedis>Candidatus_Accumulibacter | GCF_005889575.1 | Candidatus_Accumulibacter_phosphatis | ACCUM_RS04320 | 32152..35184 | 327160 | DUF499-domain-containing-protein-[Candidatus-Accumulibacter-phosphatis] |
| WP_138678982.1 | AAA+ATPase+wHTH+Alpha+Beta-domain+wHTH+RRM | Type-2_BR.RM.like.systems | <-HKD-DNase+SF1-SF2-Helicase+iREase+CTD<-METHYLASE<-Schlafen+HATPase+HTH<-DUF3780<-DUF499* | 944 | Bacteria>Pseudomonadota>Betaproteobacteria>Betaproteobacteria_incertae_sedis>Candidatus_Accumulibacter | GCF_005889575.1 | Candidatus_Accumulibacter_phosphatis | ACCUM_RS19260 | 7387..10221 | 327160 | DUF499-domain-containing-protein,-partial-[Candidatus-Accumulibacter-phosphatis] |
| WP_248595601.1 | AAA+ATPase+wHTH+Alpha+Beta-domain+wHTH+RRM | Type-2_BR.RM.like.systems | HKD-DNase->SIR2+SIR2->DUF499-p->DUF499*->DUF3780->Schlafen+HATPase+HTH->METHYLASE->METHYLASE-p->HKD-DNase+SF1-SF2-Helicase+iREase+CTD-><-RVT | 845 | Bacteria>Pseudomonadota>Betaproteobacteria>Betaproteobacteria_incertae_sedis>Candidatus_Accumulibacter | GCF_012939955.1 | Candidatus_Accumulibacter_phosphatis | E4Q23_RS19540 | 3573..6110 | 327160 | DUF499-domain-containing-protein-[Candidatus-Accumulibacter-phosphatis] |
| MCC2867811.1 | AAA+ATPase+wHTH+Alpha+Beta-domain+wHTH+RRM | Type-2_BR.RM.like.systems | <-DUF3780<-DUF499*<-?<-NERD<-ZDNA-HTH<-ZnR+TOPRIM<-REase+SF1-SF2-Helicase | 1010 | Bacteria>Pseudomonadota>Betaproteobacteria>Betaproteobacteria_incertae_sedis>Candidatus_Accumulibacter | GCA_020709745.1 | Candidatus_Accumulibacter_phosphatis | LK562_08495 | 219..3251 | 327160 | DUF499-domain-containing-protein-[Candidatus-Accumulibacter-phosphatis] |
| TMQ74963.1 | AAA+ATPase+wHTH+Alpha+Beta-domain+wHTH+RRM | Type-2_BR.RM.like.systems | <-METHYLASE\|\|?-><-?<-Schlafen+HATPase+HTH<-DUF3780<-DUF499* | 911 | Bacteria>Pseudomonadota>Betaproteobacteria>Betaproteobacteria_incertae_sedis>Candidatus_Accumulibacter | GCA_005889575.1 | Candidatus_Accumulibacter_phosphatis | ACCUM_2270 | 7387..10122 | 327160 | putative-ATPase-(AAA+-superfamily)-[Candidatus-Accumulibacter-phosphatis] |
| WP_212161746.1 | AAA+ATPase+wHTH+Alpha+Beta-domain+wHTH+RRM | Type-2_BR.RM.like.systems | <-REase\|\|?-><-HKD-DNase+SF1-SF2-Helicase+iREase+CTD<-METHYLASE<-Schlafen+HATPase+HTH<-DUF3780<-DUF499*<-Methylase\|\|?-><-Radical-SAM<-MORC+DNA_gyraseB+TOPRIM | 1012 | Bacteria>Pseudomonadota>Betaproteobacteria>Burkholderiales>Burkholderiaceae | - | Burkholderia_vietnamiensis | KDW46_26565 | 46825..49863 | 60552 | DUF499-domain-containing-protein-[Burkholderia-vietnamiensis] |
| MBR8145923.1 | AAA+ATPase+wHTH+Alpha+Beta-domain+wHTH+RRM | Type-2_BR.RM.like.systems | <-REase\|\|?-><-HKD-DNase+SF1-SF2-Helicase+iREase+CTD<-METHYLASE<-Schlafen+HATPase+HTH<-DUF3780<-DUF499*<-Methylase\|\|?-><-Radical-SAM<-MORC+DNA_gyraseB+TOPRIM | 1012 | Bacteria>Pseudomonadota>Betaproteobacteria>Burkholderiales>Burkholderiaceae | GCA_018147135.1 | Burkholderia_vietnamiensis | KDW46_26565 | 46825..49863 | 60552 | ATP-binding-protein-[Burkholderia-vietnamiensis] |
| MCA8374011.1 | AAA+ATPase+wHTH+Alpha+Beta-domain+wHTH+RRM | Type-2_BR.RM.like.systems | HIN-HTH-><-HKD-DNase+SF1-SF2-Helicase+iREase+CTD<-METHYLASE<-Schlafen+HATPase+HTH<-DUF3780<-DUF499*<-?\|\|?->?->Pkinase-> | 1011 | Bacteria>Pseudomonadota>Betaproteobacteria>Burkholderiales>Burkholderiaceae | GCA_020420025.1 | Burkholderia_multivorans | LGM80_11600 | 411823..414858 | 87883 | DUF499-domain-containing-protein-[Burkholderia-multivorans] |
| MCO5106862.1 | AAA+ATPase+wHTH+Alpha+Beta-domain+wHTH+RRM | Type-2_BR.RM.like.systems | <-LON-AAA<-?<-?\|\|?->DUF499*->DUF3780->Schlafen+HATPase+HTH->?->METHYLASE->HKD-DNase+SF1-SF2-Helicase+iREase+CTD-> | 1008 | Bacteria>Pseudomonadota>Betaproteobacteria>Burkholderiales>Burkholderiaceae | GCA_023954045.1 | Burkholderiaceae_bacterium | M9907_07245 | 32418..35444 | 2030806 | DUF499-domain-containing-protein-[Burkholderiaceae-bacterium] |
| WP_027788794.1 | AAA+ATPase+wHTH+Alpha+Beta-domain+wHTH+RRM | Type-2_BR.RM.like.systems | <-REase\|\|?-><-HKD-DNase+SF1-SF2-Helicase+iREase+CTD<-METHYLASE<-Schlafen+HATPase+HTH<-DUF3780<-DUF499* | 1012 | Bacteria>Pseudomonadota>Betaproteobacteria>Burkholderiales>Burkholderiaceae | GCF_001528845.1 | Burkholderia_cepacia | BCE01_RS37115 | 24596..27634 | 292 | DUF499-domain-containing-protein-[Burkholderia-cepacia] |
| WP_088931578.1 | AAA+ATPase+wHTH+Alpha+Beta-domain+wHTH+RRM | Type-2_BR.RM.like.systems | DUF499*->DUF3780->Schlafen+HATPase+HTH->METHYLASE->HKD-DNase+SF1-SF2-Helicase+iREase+CTD-> | 1013 | Bacteria>Pseudomonadota>Betaproteobacteria>Burkholderiales>Burkholderiaceae | GCF_003812585.1 | Burkholderia_multivorans | UA17_RS04225 | 849369..852410 | 87883 | DUF499-domain-containing-protein-[Burkholderia-multivorans] |
| WP_124581834.1 | AAA+ATPase+wHTH+Alpha+Beta-domain+wHTH+RRM | Type-2_BR.RM.like.systems | <-HKD-DNase+SF1-SF2-Helicase+iREase+CTD<-METHYLASE<-Schlafen+HATPase+HTH<-DUF3780<-DUF499*<-Methylase\|\|?-><-Radical-SAM<-MORC+DNA_gyraseB+TOPRIM | 1012 | Bacteria>Pseudomonadota>Betaproteobacteria>Burkholderiales>Burkholderiaceae | GCF_003853875.1 | Burkholderia_contaminans | DF051_RS23135 | 9301..12339 | 488447 | DUF499-domain-containing-protein-[Burkholderia-contaminans] |
| WP_207876957.1 | AAA+ATPase+wHTH+Alpha+Beta-domain+wHTH+RRM | Type-2_BR.RM.like.systems | <-HKD-DNase+SF1-SF2-Helicase+iREase+CTD<-METHYLASE<-Schlafen+HATPase+HTH<-DUF3780<-DUF499*<-?\|\|?->?->DEDD_Tnp_IS110+Transposase_20->SIR2-> | 1011 | Bacteria>Pseudomonadota>Betaproteobacteria>Burkholderiales>Burkholderiaceae | GCF_017378435.1 | Burkholderia_pseudomallei | J3L14_RS17850 | 4003230..4006265 | 28450 | DUF499-domain-containing-protein-[Burkholderia-pseudomallei] |
| WP_259032552.1 | AAA+ATPase+wHTH+Alpha+Beta-domain+wHTH+RRM | Type-2_BR.RM.like.systems | <-HKD-DNase+SF1-SF2-Helicase+iREase+CTD<-METHYLASE<-Schlafen+HATPase+HTH<-DUF3780<-DUF499*<-Methylase\|\|?-><-Radical-SAM<-HIN-HTH<-?<-Phage_AlpA | 1012 | Bacteria>Pseudomonadota>Betaproteobacteria>Burkholderiales>Burkholderiaceae | GCF_024793955.1 | Burkholderia_thailandensis | NX841_RS02865 | 24753..27791 | 57975 | DUF499-domain-containing-protein-[Burkholderia-thailandensis] |

| Protein-ID | DUF499-architecture | System | Gene-Neighbourhoods | Length | Taxonomy | Assembly-ID | Strain | Locus-Tag | Genomic-coordinates | TaxID | Fasta-define |
|---|---|---|---|---|---|---|---|---|---|---|---|
| WP_122877385.1 | AAA+ATPase+wHTH+Alpha+Beta-domain+wHTH+RRM | Type-2_BR.RM.like.systems | MORC+DNA_gyraseB+TOPRIM->Radical-SAM-><-?\|\|Methylase->DUF499*->DUF3780->Schlafen+HATPase+HTH->METHYLASE->HKD-DNase+SF1-SF2-Helicase+iREase+CTD-><-?\|\|REase-> | 1012 | Bacteria>Pseudomonadota>Betaproteobacteria>Burkholderiales>Burkholderiaceae | GCF_900591725.1 | Burkholderia_pseudomallei | EFU85_RS04200 | 244549..247587 | 28450 | DUF499-domain-containing-protein-[Burkholderia-pseudomallei] |
| WP_173125224.1 | AAA+ATPase+wHTH+Alpha+Beta-domain+wHTH+RRM | Type-2_BR.RM.like.systems | <-HKD-DNase+SF1-SF2-Helicase+iREase+CTD<-?<-METHYLASE<-Schlafen+HATPase+HTH<-DUF3780<-DUF499*<-?<-?<-?<-XIS-HTH<-?\|\|HTH-> | 1009 | Bacteria>Pseudomonadota>Betaproteobacteria>Burkholderiales>Burkholderiales_genera_incertae_sedis | GCF_013266755.1 | Aquabacterium_terrae | HLB44_RS18520 | 560708..563737 | 2732868 | DUF499-domain-containing-protein-[Aquabacterium-terrae] |
| MCW5609142.1 | AAA+ATPase+wHTH+Alpha+Beta-domain+wHTH+RRM | Type-2_BR.RM.like.systems | <-ParB+HNH<-?<-OLD-ABC<-METHYLASE<-Schlafen+HATPase+HTH<-DUF3780<-DUF499*<-Arm-DNA-bind_3+Phage_integrase\|\|Phage_int_SAM_2+Integrase_1-><-HTH_3\|\|?-><-HTH+MPTase | 1009 | Bacteria>Pseudomonadota>Betaproteobacteria>Burkholderiales>Sphaerotilaceae | GCA_026125605.1 | Rubrivivax_sp. | KIS83_00415 | 81577..84606 | 50259 | MAG:-ATP-binding-protein-[Rubrivivax-sp.] |
| MBK7768857.1 | AAA+ATPase+wHTH+Alpha+Beta-domain+wHTH+RRM | Type-2_BR.RM.like.systems | <-METHYLASE-p<-Schlafen+HATPase+HTH<-DUF3780<-DUF499*\|\|?->?-><-NTP_transf_3<-ZnR | 1005 | Bacteria>Pseudomonadota>Betaproteobacteria>Nitrosomonadales>Sterolibacteriaceae | GCA_016706645.1 | Sulfuritalea_sp. | IPI44_24195 | 5665..8682 | 2480090 | ATP-binding-protein-[Sulfuritalea-sp.] |
| MBL8482095.1 | AAA+ATPase+wHTH+Alpha+Beta-domain+wHTH+RRM | Type-2_BR.RM.like.systems | <-ZnR<-?\|\|?->Arm-DNA-bind_3+Phage_integrase->DUF499*->DUF3780->Schlafen+HATPase+HTH->METHYLASE->HKD-DNase+SF1-SF2-Helicase+iREase+CTD->REase+PNTD1+PNTD2->PIWI-> | 1008 | Bacteria>Pseudomonadota>Betaproteobacteria>Rhodocyclales>Rhodocyclaceae | GCA_016791085.1 | Rhodocyclaceae_bacterium | JNJ60_07865 | 13203..16229 | 1898103 | ATP-binding-protein-[Rhodocyclaceae-bacterium] |
| OHC61993.1 | AAA+ATPase+wHTH+Alpha+Beta-domain+wHTH+RRM | Type-2_BR.RM.like.systems | <-HKD-DNase+SF1-SF2-Helicase+iREase+CTD<-METHYLASE<-Schlafen+HATPase+HTH<-DUF3780<-DUF499* | 1005 | Bacteria>Pseudomonadota>Betaproteobacteria>Rhodocyclales>unclassified_Rhodocyclales | GCA_001828915.1 | Rhodocyclales_bacterium_GWA2_65_19 | A2040_16480 | 43406..46423 | 1802016 | MAG:-hypothetical-protein-A2040_16480-[Rhodocyclales-bacterium-GWA2_65_19] |
| KIN88580.1 | AAA+ATPase+wHTH+Alpha+Beta-domain+wHTH+RRM | Type-2_BR.RM.like.systems | <-MORC+HisKin<-sGTP\|\|HTH+WYL->?->DUF6079-p->DUF499*->DUF3780->METHYLASE->HKD-DNase+SF1-SF2-Helicase+iREase+CTD-> | 1244 | Bacteria>Pseudomonadota>Betaproteobacteria>Rhodocyclales>Zoogloeaceae | GCA_000831325.1 | Thauera_sp._SWB20 | P078_251 | 260874..264608 | 1572758 | hypothetical-protein-P078_251-[Thauera-sp.-SWB20] |
| WP_043745923.1 | AAA+ATPase+wHTH+Alpha+Beta-domain+wHTH+RRM | Type-2_BR.RM.like.systems | <-MORC+HisKin<-sGTP\|\|HTH+WYL->?->DUF6079-p->DUF499*->DUF3780->METHYLASE->HKD-DNase+SF1-SF2-Helicase+iREase+CTD-> | 1259 | Bacteria>Pseudomonadota>Betaproteobacteria>Rhodocyclales>Zoogloeaceae | GCF_000831325.1 | Thauera_sp._SWB20 | P078_RS18625 | 260829..264608 | 1572758 | DUF499-domain-containing-protein-[Thauera-sp.-SWB20] |
| WP_159690410.1 | AAA+ATPase+wHTH+Alpha+Beta-domain+wHTH+RRM | Type-2_BR.RM.like.systems | <-HKD-DNase+SF1-SF2-Helicase+iREase+CTD<-METHYLASE<-DUF3780<-DUF499*<-DUF6079-p<-?<-HTH+WYL\|\|sGTP->MORC+HisKin-> | 1244 | Bacteria>Pseudomonadota>Betaproteobacteria>Rhodocyclales>Zoogloeaceae | GCF_009800965.1 | Cognatazoarcus_halotolerans | GQB03_RS13810 | 94058..97792 | 2686016 | DUF499-domain-containing-protein-[Cognatazoarcus-halotolerans] |
| WP_007193607.1 | AAA+ATPase+wHTH+Alpha+Beta-domain+wHTH+RRM | Type-2_BR.RM.like.systems | <-HKD-DNase+SF1-SF2-Helicase+iREase+CTD<-ParB+HNH<-METHYLASE-p<-METHYLASE<-Schlafen+HATPase+HTH<-DUF3780<-DUF499*\|\|?->?->?->CHTH-> | 1041 | Bacteria>Pseudomonadota>Gammaproteobacteria>Chromatiales>Chromatiaceae | GCF_000223985.1 | Thiocapsa_marina | THIMADRAFT_RS13475 | 269670..272795 | 244573 | DUF499-domain-containing-protein-[Thiocapsa-marina] |
| QVL47522.1 | AAA+ATPase+wHTH+Alpha+Beta-domain+wHTH+RRM | Type-2_BR.RM.like.systems | <-CHTH<-?<-?<-?\|\|DUF499*->DUF3780->?->PIN->?-><-DDE_5+DDE_Tnp_1<-DDE_Tnp_IS66+DDE_Tnp_IS66_C | 828 | Bacteria>Pseudomonadota>Gammaproteobacteria>Chromatiales>Chromatiaceae | GCA_018399035.1 | Thiocapsa_sp. | KFB96_17705 | 3898302..3900788 | 2024551 | MAG:-DUF499-domain-containing-protein-[Thiocapsa-sp.] |
| WP_082673870.1 | AAA+ATPase+wHTH+Alpha+Beta-domain+wHTH+RRM | Type-2_BR.RM.like.systems | <-ABC-ATPase+SbcC<-?\|\|FtsK+HerA+A32->DUF499*->?->REase->DUF3780-><-?<-HIN-HTH | 1004 | Bacteria>Pseudomonadota>Gammaproteobacteria>Chromatiales>Chromatiaceae | GCF_001469165.1 | Thiohalocapsa_sp._ML1 | ML01_RS02170 | 202750..205764 | 1431688 | DUF499-domain-containing-protein-[Thiohalocapsa-sp.-ML1] |
| PZN82194.1 | AAA+ATPase+wHTH+Alpha+Beta-domain+wHTH+RRM | Type-2_BR.RM.like.systems | <-METHYLASE-p<-?<-HNH<-OLD-ABC<-DUF3780<-DUF499* | 1015 | Bacteria>Pseudomonadota>Gammaproteobacteria>Methylococcales>Methylococcales_incertae_sedis | GCA_003242955.1 | Candidatus_Methylumidiphilus_alinenensis | DM484_06870 | 3894..6941 | 2202197 | hypothetical-protein-DM484_06870-[Candidatus-Methylumidiphilus-alinenensis] |
| MCG3169754.1 | AAA+ATPase+wHTH+Alpha+Beta-domain+wHTH+RRM | Type-2_BR.RM.like.systems | <-sGTP\|\|?->?->?->?->DUF772+DDE_Tnp_1+DDE_Tnp_1_6->DUF499*->DUF3780->Schlafen+HATPase+HTH->METHYLASE->HKD-DNase+SF1-SF2-Helicase+iREase+CTD-> | 1010 | Bacteria>Pseudomonadota>Gammaproteobacteria>Pseudomonadales>unclassified_Pseudomonadales | GCA_022072145.1 | Pseudomonadales_bacterium | CALGDGBN_01277 | 15067..18099 | 1891229 | hypothetical-protein-[Pseudomonadales-bacterium] |
| MXX05925.1 | AAA+ATPase+wHTH+Alpha+Beta-domain+wHTH+RRM | Type-2_BR.RM.like.systems | <-HEPN<-?\|\|MazE_antitoxin->PIN->DUF499*->DUF3780->?->?->?->METHYLASE->HKD-DNase+SF1-SF2-Helicase+iREase+CTD-> | 1000 | Bacteria>Pseudomonadota>Gammaproteobacteria>unclassified_Gammaproteobacteria | GCA_009836015.1 | Gammaproteobacteria_bacterium | F4Z71_04120 | 93593..96595 | 1913989 | DUF499-domain-containing-protein-[Gammaproteobacteria-bacterium] |
| MXY53195.1 | AAA+ATPase+wHTH+Alpha+Beta-domain+wHTH+RRM | Type-2_BR.RM.like.systems | <-TPR<-?\|\|?->?->?->DUF499*->DUF3780->OLD-ABC->HNH->METHYLASE->HKD-DNase+SF1-SF2-Helicase+iREase+CTD->OLD-ABC-> | 1009 | Bacteria>Pseudomonadota>Gammaproteobacteria>unclassified_Gammaproteobacteria | GCA_009837725.1 | Gammaproteobacteria_bacterium | F4Y86_11820 | 5670..8699 | 1913989 | ATP-binding-protein-[Gammaproteobacteria-bacterium] |
| MYB38346.1 | AAA+ATPase+wHTH+Alpha+Beta-domain+wHTH+RRM | Type-2_BR.RM.like.systems | <-OLD-ABC<-HKD-DNase+SF1-SF2-Helicase+iREase+CTD<-METHYLASE<-HNH<-OLD-ABC<-DUF3780<-DUF499* | 1009 | Bacteria>Pseudomonadota>Gammaproteobacteria>unclassified_Gammaproteobacteria | GCA_009838905.1 | Gammaproteobacteria_bacterium | F4Y26_13400 | 18076..21105 | 1913989 | DUF499-domain-containing-protein-[Gammaproteobacteria-bacterium] |
| MCY4011318.1 | AAA+ATPase+wHTH+Alpha+Beta-domain+wHTH+RRM | Type-2_BR.RM.like.systems | <-HKD-DNase+SF1-SF2-Helicase+iREase+CTD<-METHYLASE-p<-METHYLASE<-DUF3780<-DUF499*<-REase\|\|Arm-DNA-bind_3+Phage_integrase-> | 1000 | Bacteria>Pseudomonadota>Gammaproteobacteria>unclassified_Gammaproteobacteria | GCA_026708185.1 | Gammaproteobacteria_bacterium | OXG82_01240 | 7722..10724 | 1913989 | MAG:-DUF499-domain-containing-protein-[Gammaproteobacteria-bacterium] |
| MCY4165205.1 | AAA+ATPase+wHTH+Alpha+Beta-domain+wHTH+RRM | Type-2_BR.RM.like.systems | <-HKD-DNase+SF1-SF2-Helicase+iREase+CTD<-METHYLASE<-?<-?<-?<-DUF3780<-DUF499* | 907 | Bacteria>Pseudomonadota>Gammaproteobacteria>unclassified_Gammaproteobacteria | GCA_026709045.1 | Gammaproteobacteria_bacterium | OXF03_03550 | 9312..12035 | 1913989 | MAG:-DUF499-domain-containing-protein-[Gammaproteobacteria-bacterium] |
| MCY4646294.1 | AAA+ATPase+wHTH+Alpha+Beta-domain+wHTH+RRM | Type-2_BR.RM.like.systems | <-HKD-DNase+SF1-SF2-Helicase+iREase+CTD<-METHYLASE<-?<-?<-?<-DUF3780<-DUF499*<-LexA-protease\|\|XIS-HTH->VirB2->?->FtsK+VirB4-FtsK-> | 1002 | Bacteria>Pseudomonadota>Gammaproteobacteria>unclassified_Gammaproteobacteria | GCA_026713895.1 | Gammaproteobacteria_bacterium | OXE73_05415 | 129919..132927 | 1913989 | MAG:-DUF499-domain-containing-protein-[Gammaproteobacteria-bacterium] |
| MDE0055389.1 | AAA+ATPase+wHTH+Alpha+Beta-domain+wHTH+RRM | Type-2_BR.RM.like.systems | DUF499*->DUF3780->METHYLASE->HKD-DNase+SF1-SF2-Helicase+iREase+CTD->?->SF1-SF2-Helicase+DUF1998->REase-> | 947 | Bacteria>Pseudomonadota>Gammaproteobacteria>unclassified_Gammaproteobacteria | GCA_028818875.1 | Gammaproteobacteria_bacterium | OXT64_14225 | 5722..8565 | 1913989 | MAG:-DUF499-domain-containing-protein-[Gammaproteobacteria-bacterium] |
| MDE0357070.1 | AAA+ATPase+wHTH+Alpha+Beta-domain+wHTH+RRM | Type-2_BR.RM.like.systems | <-HKD-DNase+SF1-SF2-Helicase+iREase+CTD<-METHYLASE<-DUF3780<-DUF499*\|\|HicB_lk_antitox->HicA_toxin-> | 1042 | Bacteria>Pseudomonadota>Gammaproteobacteria>unclassified_Gammaproteobacteria | GCA_028821195.1 | Gammaproteobacteria_bacterium | OXN92_04925 | 28607..31735 | 1913989 | MAG:-DUF499-domain-containing-protein-[Gammaproteobacteria-bacterium] |
| MDE0441395.1 | AAA+ATPase+wHTH+Alpha+Beta-domain+wHTH+RRM | Type-2_BR.RM.like.systems | <-SF1-SF2-Helicase+DUF1998<-?<-HKD-DNase+SF1-SF2-Helicase+iREase+CTD<-METHYLASE<-DUF3780<-DUF499* | 998 | Bacteria>Pseudomonadota>Gammaproteobacteria>unclassified_Gammaproteobacteria | GCA_028821355.1 | Gammaproteobacteria_bacterium | OXL38_04655 | 10994..13990 | 1913989 | MAG:-DUF499-domain-containing-protein-[Gammaproteobacteria-bacterium] |
| MDE0444961.1 | AAA+ATPase+wHTH+Alpha+Beta-domain+wHTH+RRM | Type-2_BR.RM.like.systems | <-HKD-DNase+SF1-SF2-Helicase+iREase+CTD<-METHYLASE<-PIN<-MPTase<-DUF3780<-DUF499*\|\|?-><-HicA_toxin<-SF1-SF2-Helicase+REase<-SF1-SF2-Helicase+REase<-?<-PIN | 1015 | Bacteria>Pseudomonadota>Gammaproteobacteria>unclassified_Gammaproteobacteria | GCA_028821355.1 | Gammaproteobacteria_bacterium | OXL38_23000 | 9764..12811 | 1913989 | MAG:-DUF499-domain-containing-protein-[Gammaproteobacteria-bacterium] |
| MDE0038229.1 | AAA+ATPase+wHTH+Alpha+Beta-domain+wHTH+RRM | Type-2_BR.RM.like.systems | <-METHYLASE<-PIN<-MPTase<-DUF3780<-DUF499*\|\|?-><-HicA_toxin<-SF1-SF2-Helicase+REase<-SF1-SF2-Helicase+REase<-?<-PIN | 1015 | Bacteria>Pseudomonadota>Gammaproteobacteria>unclassified_Gammaproteobacteria | GCA_028818795.1 | Gammaproteobacteria_bacterium | OXU77_11860 | 4307..7354 | 1913989 | MAG:-DUF499-domain-containing-protein-[Gammaproteobacteria-bacterium] |
| MDE0478449.1 | AAA+ATPase+wHTH+Alpha+Beta-domain+wHTH+RRM | Type-2_BR.RM.like.systems | <-METHYLASE<-?<-?<-?<-DUF3780<-DUF499*<-PIN<-MazE_antitoxin\|\|?->HEPN-> | 1000 | Bacteria>Pseudomonadota>Gammaproteobacteria>unclassified_Gammaproteobacteria | GCA_028824825.1 | Gammaproteobacteria_bacterium | OXI13_02365 | 5014..8016 | 1913989 | MAG:-DUF499-domain-containing-protein-[Gammaproteobacteria-bacterium] |
| HAI59970.1 | AAA+ATPase+wHTH+Alpha+Beta-domain+wHTH+RRM | Type-2_BR.RM.like.systems | SF1-SF2-Helicase-><-Pkinase\|\|?-><-Synaptojanin\|\|DUF499*->DUF3780->Schlafen+HATPase+HTH->?->METHYLASE->HKD-DNase+SF1-SF2-Helicase+iREase+CTD-> | 1013 | Bacteria>Pseudomonadota>Gammaproteobacteria>Xanthomonadales>Xanthomonadaceae | GCA_003449515.1 | Xanthomonadaceae_bacterium | DCM32_08860 | 4597..7638 | 1926873 | hypothetical-protein-[Xanthomonadaceae-bacterium] |
| MCZ8164568.1 | AAA+ATPase+wHTH+Alpha+Beta-domain+wHTH+RRM | Type-2_BR.RM.like.systems | XIS-HTH->HTH_36->?->?-><-?<-Phage_int_SAM_2+Integrase_1\|\|DUF499*->DUF3780->Schlafen+HATPase+HTH->?->METHYLASE->HKD-DNase+SF1-SF2-Helicase+iREase+CTD-> | 1013 | Bacteria>Pseudomonadota>Gammaproteobacteria>Xanthomonadales>Xanthomonadaceae | GCA_027531545.1 | Silanimonas_sp. | O9280_02335 | 527918..530959 | 1929290 | MAG:-DUF499-domain-containing-protein-[Silanimonas-sp.] |
| WP_146472112.1 | AAA+ATPase+wHTH+Alpha+Beta-domain+wHTH+RRM | Type-2_BR.RM.like.systems | <-ABC-ATPase+SbcC\|\|?->ABhydrolase->ABC-ATPase+SbcC->?-><-?\|\|DUF499*->DUF3780->Schlafen+HATPase+HTH->?->?->METHYLASE->HKD-DNase+SF1-SF2-Helicase+iREase+CTD-> | 1012 | Bacteria>Pseudomonadota>Gammaproteobacteria>Xanthomonadales>Xanthomonadaceae | GCF_014656335.1 | Vulcaniibacterium_thermophilum | IE210_RS02820 | 81268..84306 | 1169913 | DUF499-domain-containing-protein-[Vulcaniibacterium-thermophilum] |
| NCC04604.1 | AAA+ATPase+wHTH+Alpha+Beta-domain+wHTH+RRM | Type-2_BR.RM.like.systems | <-HKD-DNase+SF1-SF2-Helicase+iREase+CTD<-METHYLASE<-PIN<-DUF3780<-DUF499* | 1015 | Bacteria>Pseudomonadota>unclassified_Pseudomonadota | GCA_009929575.1 | Pseudomonadota_bacterium | EOM37_11320 | 14987..18034 | 1977087 | ATP-binding-protein-[Proteobacteria-bacterium] |
| MBS0382004.1 | AAA+ATPase+wHTH+Alpha+Beta-domain+wHTH+RRM | Type-2_BR.RM.like.systems | DUF499*->DUF3780->Schlafen+HATPase+HTH->?->METHYLASE-p-> | 889 | Bacteria>Pseudomonadota>unclassified_Pseudomonadota | GCA_018241365.1 | Pseudomonadota_bacterium | JSR56_06185 | 1..2671 | 1977087 | DUF499-domain-containing-protein-[Proteobacteria-bacterium] |
| MBS0338394.1 | AAA+ATPase+wHTH+Alpha+Beta-domain+wHTH+RRM | Type-2_BR.RM.like.systems | <-Schlafen+HATPase+HTH<-Schlafen+HATPase+HTH<-DUF3780<-DUF499* | 1016 | Bacteria>Pseudomonadota>unclassified_Pseudomonadota | GCA_018241645.1 | Pseudomonadota_bacterium | JSS40_16645 | 2322..5372 | 1977087 | ATP-binding-protein-[Proteobacteria-bacterium] |
| MCE9614797.1 | AAA+ATPase+wHTH+Alpha+Beta-domain+wHTH+RRM | Type-2_BR.RM.like.systems | <-HTH_3\|\|?->?->?->?->HTH->DUF499*->DUF3780->REase->?->DNA_methylase->METHYLASE->HKD-DNase+SF1-SF2-Helicase+iREase+CTD-> | 986 | Bacteria>PVC_group>Lentisphaerota>unclassified_Lentisphaerota | GCA_021413965.1 | Lentisphaerota_bacterium | K8T26_11010 | 329228..332188 | 1932692 | DUF499-domain-containing-protein-[Lentisphaerae-bacterium] |
| NLX59557.1 | AAA+ATPase+wHTH+Alpha+Beta-domain+wHTH+RRM | Type-2_BR.RM.like.systems | <-SbcC\|\|?->HTH->RVT->CHTH->DUF499*->DUF3780->METHYLASE->HKD-DNase+SF1-SF2-Helicase+iREase+CTD-> | 1011 | Bacteria>PVC_group>Planctomycetota>Phycisphaerae>unclassified_Phycisphaerae | GCA_012729015.1 | Phycisphaerae_bacterium | GXY74_10770 | 18997..22032 | 2026778 | ATP-binding-protein-[Phycisphaerae-bacterium] |
| MBY0527363.1 | AAA+ATPase+wHTH+Alpha+Beta-domain+wHTH+RRM | Type-2_BR.RM.like.systems | DUF499*->DUF3780->OLD-ABC->TOPRIM->METHYLASE->HKD-DNase+SF1-SF2-Helicase+iREase+CTD-> | 1065 | Bacteria>PVC_group>Planctomycetota>Planctomycetia>Gemmatales | GCA_019752375.1 | Gemmataceae_bacterium | K2R98_28465 | 27968..31165 | 2052164 | DUF499-domain-containing-protein-[Gemmataceae-bacterium] |
| WP_194535999.1 | AAA+ATPase+wHTH+Alpha+Beta-domain+wHTH+RRM | Type-2_BR.RM.like.systems | <-HKD-DNase+SF1-SF2-Helicase+iREase+CTD<-PIN<-?<-?<-METHYLASE<-DUF3780<-DUF499*<-?\|\|?->?->PAIREDC-HTH-> | 1013 | Bacteria>PVC_group>Planctomycetota>Planctomycetia>Gemmatales | GCF_013694095.1 | Thermogemmata_fonticola | H0921_RS00060 | 23590..26631 | 2755323 | DUF499-domain-containing-protein-[Thermogemmata-fonticola] |
| WP_169978696.1 | AAA+ATPase+wHTH+Alpha+Beta-domain+wHTH+RRM | Type-2_BR.RM.like.systems | Peptidase->PAIREDC-HTH+HIN-HTH-><-HKD-DNase+SF1-SF2-Helicase+iREase+CTD<-METHYLASE<-DUF3780<-DUF499*\|\|?->ZnR+TOPRIM+D5ATPase->?-><-Phage_integrase | 1272 | Bacteria>PVC_group>Planctomycetota>Planctomycetia>Isosphaerales | GCF_012958305.1 | Tautonia_rosea | H1B33_RS19750 | 70038..73856 | 2728037 | DUF499-domain-containing-protein-[Tautonia-rosea] |
| MBX3412588.1 | AAA+ATPase+wHTH+Alpha+Beta-domain+wHTH+RRM | Type-2_BR.RM.like.systems | ArdCN+MPTase->?->?->NTD+SF1-SF2-Helicase->Phage_int_SAM_1+Phage_integrase->XIS-HTH->DUF499*->DUF3780->REase->?->METHYLASE->?->HKD-DNase+SF1-SF2-Helicase+iREase+CTD-> | 1023 | Bacteria>PVC_group>Planctomycetota>Planctomycetia>Pirellulales | GCA_019636335.1 | Pirellulales_bacterium | KF708_07865 | 38769..41840 | 2715253 | ATP-binding-protein-[Pirellulales-bacterium] |
| NLF73182.1 | AAA+ATPase+wHTH+Alpha+Beta-domain+wHTH+RRM | Type-2_BR.RM.like.systems | TUDOR+SF1-SF2-Helicase+iREase+CTD+REase->SF1-SF2-Helicase->METHYLASE->?->DUF499*->?->ASCH+DUF3780-><-?\|\|?->XIS-HTH-> | 975 | Bacteria>PVC_group>Planctomycetota>Planctomycetia>Pirellulales | GCA_012515235.1 | Candidatus_Anammoximicrobium_sp. | GX575_29450 | 8853..11780 | 2699742 | ATP-binding-protein-[Candidatus-Anammoximicrobium-sp.] |
| OJW08007.1 | AAA+ATPase+wHTH+Alpha+Beta-domain+wHTH+RRM | Type-2_BR.RM.like.systems | DUF499*->DUF3780->OLD-ABC->TOPRIM->METHYLASE-p->HKD-DNase+SF1-SF2-Helicase+iREase+CTD->OLD-ABC+TOPRIM-> | 1008 | Bacteria>PVC_group>Planctomycetota>Planctomycetia>Planctomycetales | GCA_001898385.1 | Planctomycetales_bacterium_71-10 | BG049_20675 | 18469..21495 | 1895807 | MAG:-hypothetical-protein-BG049_20675-[Planctomycetales-bacterium-71-10] |
| RLT16925.1 | AAA+ATPase+wHTH+Alpha+Beta-domain+wHTH+RRM | Type-2_BR.RM.like.systems | DUF499*->DUF3780->METHYLASE->HKD-DNase+SF1-SF2-Helicase+iREase+CTD-> | 731 | Bacteria>PVC_group>Planctomycetota>unclassified_Planctomycetota | GCA_003672035.1 | Planctomycetota_bacterium | DWI28_06925 | 1..2198 | 2026780 | MAG:-DUF499-domain-containing-protein,-partial-[Planctomycetota-bacterium] |
| MCE9554575.1 | AAA+ATPase+wHTH+Alpha+Beta-domain+wHTH+RRM | Type-2_BR.RM.like.systems | DUF499*->DUF3780->ParB+HNH->METHYLASE->?->HKD-DNase+SF1-SF2-Helicase+iREase+CTD-> | 885 | Bacteria>PVC_group>Planctomycetota>unclassified_Planctomycetota | GCA_021413705.1 | Planctomycetota_bacterium | K8T91_14550 | 155..2812 | 2026780 | ATP-binding-protein-[Planctomycetota-bacterium] |
| MCK6447147.1 | AAA+ATPase+wHTH+Alpha+Beta-domain+wHTH+RRM | Type-2_BR.RM.like.systems | <-ABhydrolase\|\|Phage_integrase->?->?->XIS-HTH->DUF499*->DUF3780->METHYLASE->?->?->PIN->HKD-DNase+SF1-SF2-Helicase+iREase+CTD-> | 1029 | Bacteria>PVC_group>Planctomycetota>unclassified_Planctomycetota | GCA_023150475.1 | Planctomycetota_bacterium | L6Q99_12220 | 25859..28948 | 2026780 | ATP-binding-protein-[Planctomycetota-bacterium] |
| MCB9932998.1 | AAA+ATPase+wHTH+Alpha+Beta-domain+wHTH+RRM | Type-2_BR.RM.like.systems | SF1-SF2-Helicase-p->?->DUF499*->DUF3780->?->REase+HTH->NTP_transf->XIS-HTH->PIN-> | 1008 | Bacteria>PVC_group>Planctomycetota>unclassified_Planctomycetota | GCA_020638605.1 | Planctomycetota_bacterium | H6841_06205 | 57026..60052 | 2026780 | ATP-binding-protein-[Planctomycetota-bacterium] |
| MCY2987606.1 | AAA+ATPase+wHTH+Alpha+Beta-domain+wHTH+RRM | Type-2_BR.RM.like.systems | <-DOC+HTH<-HNH<-DUF3780<-DUF499*<-XIS-HTH | 1025 | Bacteria>PVC_group>Planctomycetota>unclassified_Planctomycetota | GCA_026692885.1 | Planctomycetota_bacterium | NTY19_07045 | 3408..6485 | 2026780 | MAG:-DUF499-domain-containing-protein-[Planctomycetota-bacterium] |
| MBM3879657.1 | AAA+ATPase+wHTH+Alpha+Beta-domain+wHTH+RRM | Type-2_BR.RM.like.systems | <-ABC-ATPase+SbcC<-?<-?<-?<-?<-ABC-ATPase+SbcC\|\|DUF499*->DUF3780->Schlafen+HATPase+HTH->?->Schlafen+HATPase+HTH->METHYLASE->HEPN-> | 1019 | Bacteria>PVC_group>Verrucomicrobiota>unclassified_Verrucomicrobiota | GCA_016871715.1 | Verrucomicrobiota_bacterium | FJ387_08035 | 7884..10943 | 2026799 | ATP-binding-protein-[Verrucomicrobia-bacterium] |
| MCP5524364.1 | AAA+ATPase+wHTH+Alpha+Beta-domain+wHTH+RRM | Type-2_BR.RM.like.systems | <-HKD-DNase+SF1-SF2-Helicase+iREase+CTD<-?<-?<-?<-?<-DUF3780<-DUF499* | 1019 | Bacteria>PVC_group>Verrucomicrobiota>Verrucomicrobiae>Verrucomicrobiales | GCA_024233215.1 | Verrucomicrobiales_bacterium | H7A46_22760 | 69953..73012 | 2026801 | ATP-binding-protein-[Verrucomicrobiales-bacterium] |
| MBP6739048.1 | AAA+ATPase+wHTH+Alpha+Beta-domain+wHTH+RRM | Type-2_BR.RM.like.systems | DUF499*->DUF3780->ParB+HNH->OLD-ABC->?->Y1_Tnp->METHYLASE-> | 1072 | Bacteria>Spirochaetota>Spirochaetia>Leptospirales>Leptospiraceae | GCA_017990775.1 | Leptospiraceae_bacterium | KA146_03605 | 102..3320 | 2026757 | ATP-binding-protein-[Leptospiraceae-bacterium] |
| MDE0024272.1 | AAA+ATPase+wHTH+Alpha+Beta-domain+wHTH+RRM | Type-2_BR.RM.like.systems | <-HKD-DNase+SF1-SF2-Helicase+iREase+CTD<-METHYLASE<-?<-?<-?<-DUF3780<-DUF499*<-Glyoxalase<-HicB_lk_antitox | 1012 | Bacteria>Spirochaetota>Spirochaetia>Spirochaetales>Spirochaetaceae | GCA_028818735.1 | Spirochaetaceae_bacterium | OXP69_07645 | 37630..40668 | 1898206 | MAG:-DUF499-domain-containing-protein-[Spirochaetaceae-bacterium] |
| MCY4377157.1 | AAA+ATPase+wHTH+Alpha+Beta-domain+wHTH+RRM | Type-2_BR.RM.like.systems | <-METHYLASE<-SF1-SF2-Helicase+REase-p\|\|NACHT->SbcC-><-?<-DUF3780<-DUF499* | 1005 | Bacteria>Spirochaetota>Spirochaetia>Spirochaetales>Spirochaetaceae | GCA_026710565.1 | Spirochaetaceae_bacterium | OXC31_25645 | 10086..13103 | 1898206 | MAG:-DUF499-domain-containing-protein-[Spirochaetaceae-bacterium] |
| MDD9986492.1 | AAA+ATPase+wHTH+Alpha+Beta-domain+wHTH+RRM | Type-2_BR.RM.like.systems | <-HEPN<-PolB<-?\|\|?->?->?->DUF499*->DUF3780->HEPN->?->?->?->METHYLASE-p-> | 1007 | Bacteria>Spirochaetota>Spirochaetia>Spirochaetales>Spirochaetaceae | GCA_028818385.1 | Spirochaetaceae_bacterium | OXQ31_09490 | 10176..13199 | 1898206 | MAG:-DUF499-domain-containing-protein-[Spirochaetaceae-bacterium] |
| MXW62749.1 | AAA+ATPase+wHTH+Alpha+Beta-domain+wHTH+RRM | Type-2_BR.RM.like.systems | <-sGTP<-sGTP<-?<-?<-?<-?\|\|DUF499*->DUF3780->METHYLASE->HKD-DNase+SF1-SF2-Helicase+iREase+CTD->HKD-DNase+SF1-SF2-Helicase+iREase+CTD->TIR-><-PIWI | 1087 | Bacteria>Terrabacteria_group>Actinomycetota>Acidimicrobiia>Acidimicrobiales | GCA_009835385.1 | Acidimicrobiaceae_bacterium | F4003_13330 | 50978..54241 | 2024894 | ATP-binding-protein-[Acidimicrobiaceae-bacterium] |
| MXW96686.1 | AAA+ATPase+wHTH+Alpha+Beta-domain+wHTH+RRM | Type-2_BR.RM.like.systems | <-RnasePH+RnasePH+KH+S1<-?<-HKD-DNase+SF1-SF2-Helicase+iREase+CTD<-METHYLASE<-METHYLASE-p<-DUF3780<-DUF499* | 986 | Bacteria>Terrabacteria_group>Actinomycetota>Acidimicrobiia>Acidimicrobiales | GCA_009836165.1 | Acidimicrobiaceae_bacterium | F4Z64_14470 | 45717..48678 | 2024894 | ATP-binding-protein-[Acidimicrobiaceae-bacterium] |
| MXW76679.1 | AAA+ATPase+wHTH+Alpha+Beta-domain+wHTH+RRM | Type-2_BR.RM.like.systems | PIWI-><-TIR<-HKD-DNase+SF1-SF2-Helicase+iREase+CTD-p<-HKD-DNase+SF1-SF2-Helicase+iREase+CTD<-METHYLASE<-DUF3780<-DUF499*\|\|?->?->?->?->sGTP->sGTP-> | 1087 | Bacteria>Terrabacteria_group>Actinomycetota>Acidimicrobiia>Acidimicrobiales | GCA_009836285.1 | Acidimicrobiaceae_bacterium | F4Z58_11720 | 147353..150616 | 2024894 | ATP-binding-protein-[Acidimicrobiaceae-bacterium] |
| MXX42195.1 | AAA+ATPase+wHTH+Alpha+Beta-domain+wHTH+RRM | Type-2_BR.RM.like.systems | MODE-HTH+PBPII-><-PIN<-?<-HKD-DNase+SF1-SF2-Helicase+iREase+CTD<-METHYLASE<-DUF3780<-DUF499*\|\|REase-> | 1088 | Bacteria>Terrabacteria_group>Actinomycetota>Acidimicrobiia>Acidimicrobiales | GCA_009836375.1 | Acidimicrobiales_bacterium | F4Z53_04000 | 47327..50593 | 2201156 | ATP-binding-protein-[Acidimicrobiales-bacterium] |
| MXZ98360.1 | AAA+ATPase+wHTH+Alpha+Beta-domain+wHTH+RRM | Type-2_BR.RM.like.systems | DUF499*->DUF3780->METHYLASE->HKD-DNase+SF1-SF2-Helicase+iREase+CTD-> | 987 | Bacteria>Terrabacteria_group>Actinomycetota>Acidimicrobiia>Acidimicrobiales | GCA_009836995.1 | Acidimicrobiaceae_bacterium | F4Z23_04905 | 1..2965 | 2024894 | DUF499-domain-containing-protein-[Acidimicrobiaceae-bacterium] |
| MYE76343.1 | AAA+ATPase+wHTH+Alpha+Beta-domain+wHTH+RRM | Type-2_BR.RM.like.systems | <-HKD-DNase+SF1-SF2-Helicase+iREase+CTD<-METHYLASE<-DUF3780<-DUF499* | 1053 | Bacteria>Terrabacteria_group>Actinomycetota>Acidimicrobiia>Acidimicrobiales | GCA_009841965.1 | Acidimicrobiaceae_bacterium | F4232_08040 | 7307..10468 | 2024894 | ATP-binding-protein-[Acidimicrobiaceae-bacterium] |
| MYH42527.1 | AAA+ATPase+wHTH+Alpha+Beta-domain+wHTH+RRM | Type-2_BR.RM.like.systems | <-RnasePH+RnasePH+KH+S1<-?<-HKD-DNase+SF1-SF2-Helicase+iREase+CTD<-METHYLASE<-DUF3780<-DUF499*<-DUF499-p\|\|?->?-><-PAIREDC-HTH+HIN-HTH\|\|N6_N4_MTase-> | 1033 | Bacteria>Terrabacteria_group>Actinomycetota>Acidimicrobiia>Acidimicrobiales | GCA_009843995.1 | Acidimicrobiaceae_bacterium | F4131_02330 | 14500..17601 | 2024894 | ATP-binding-protein-[Acidimicrobiaceae-bacterium] |
| MYI55289.1 | AAA+ATPase+wHTH+Alpha+Beta-domain+wHTH+RRM | Type-2_BR.RM.like.systems | <-RnasePH+RnasePH+KH+S1<-?<-HKD-DNase+SF1-SF2-Helicase+iREase+CTD<-METHYLASE<-METHYLASE-p<-DUF3780<-DUF499* | 1022 | Bacteria>Terrabacteria_group>Actinomycetota>Acidimicrobiia>Acidimicrobiales | GCA_009844405.1 | Acidimicrobiaceae_bacterium | F4110_15145 | 14637..17705 | 2024894 | ATP-binding-protein-[Acidimicrobiaceae-bacterium] |
| MYJ42091.1 | AAA+ATPase+wHTH+Alpha+Beta-domain+wHTH+RRM | Type-2_BR.RM.like.systems | <-HKD-DNase+SF1-SF2-Helicase+iREase+CTD<-METHYLASE<-DUF3780<-DUF499* | 940 | Bacteria>Terrabacteria_group>Actinomycetota>Acidimicrobiia>Acidimicrobiales | GCA_009845165.1 | Acidimicrobiaceae_bacterium | F4076_06550 | 7397..10219 | 2024894 | DUF499-domain-containing-protein-[Acidimicrobiaceae-bacterium] |
| MYJ81972.1 | AAA+ATPase+wHTH+Alpha+Beta-domain+wHTH+RRM | Type-2_BR.RM.like.systems | <-HKD-DNase+SF1-SF2-Helicase+iREase+CTD<-METHYLASE<-DUF3780<-DUF499* | 988 | Bacteria>Terrabacteria_group>Actinomycetota>Acidimicrobiia>Acidimicrobiales | GCA_009845665.1 | Acidimicrobiaceae_bacterium | F4046_09180 | 7300..10266 | 2024894 | ATP-binding-protein-[Acidimicrobiaceae-bacterium] |
| MYK73531.1 | AAA+ATPase+wHTH+Alpha+Beta-domain+wHTH+RRM | Type-2_BR.RM.like.systems | DUF499*->DUF3780->METHYLASE->HKD-DNase+SF1-SF2-Helicase+iREase+CTD->?->RnasePH+RnasePH+KH+S1-> | 997 | Bacteria>Terrabacteria_group>Actinomycetota>Acidimicrobiia>Acidimicrobiales | GCA_009846275.1 | Acidimicrobiaceae_bacterium | F4017_02885 | 1..2994 | 2024894 | ATP-binding-protein-[Acidimicrobiaceae-bacterium] |
| MCY3619217.1 | AAA+ATPase+wHTH+Alpha+Beta-domain+wHTH+RRM | Type-2_BR.RM.like.systems | <-HKD-DNase+SF1-SF2-Helicase+iREase+CTD<-METHYLASE<-DUF3780<-DUF499*<-DUF499-p | 904 | Bacteria>Terrabacteria_group>Actinomycetota>Acidimicrobiia>Acidimicrobiales | GCA_026705265.1 | Acidimicrobiaceae_bacterium | OXG66_16190 | 5597..8311 | 2024894 | MAG:-DUF499-domain-containing-protein-[Acidimicrobiaceae-bacterium] |
| MCY3892071.1 | AAA+ATPase+wHTH+Alpha+Beta-domain+wHTH+RRM | Type-2_BR.RM.like.systems | <-Mbetalac\|\|?-><-REase\|\|DUF499*->DUF3780->METHYLASE->HKD-DNase+SF1-SF2-Helicase+iREase+CTD-><-FtsK | 1088 | Bacteria>Terrabacteria_group>Actinomycetota>Acidimicrobiia>Acidimicrobiales | GCA_026707105.1 | Acidimicrobiaceae_bacterium | OXF65_02125 | 22259..25525 | 2024894 | MAG:-DUF499-domain-containing-protein-[Acidimicrobiaceae-bacterium] |
| MCY4068111.1 | AAA+ATPase+wHTH+Alpha+Beta-domain+wHTH+RRM | Type-2_BR.RM.like.systems | DUF499-p->DUF499*->METHYLASE->HKD-DNase+SF1-SF2-Helicase+iREase+CTD-><-?<-?\|\|?->RnasePH+RnasePH+KH+S1-> | 889 | Bacteria>Terrabacteria_group>Actinomycetota>Acidimicrobiia>Acidimicrobiales | GCA_026708645.1 | Acidimicrobiaceae_bacterium | OXE79_03230 | 135527..138196 | 2024894 | MAG:-DUF499-domain-containing-protein-[Acidimicrobiaceae-bacterium] |
| MCY4174992.1 | AAA+ATPase+wHTH+Alpha+Beta-domain+wHTH+RRM | Type-2_BR.RM.like.systems | DUF499-p->DUF499*->METHYLASE->HKD-DNase+SF1-SF2-Helicase+iREase+CTD-><-?<-?\|\|?->RnasePH+RnasePH+KH+S1-> | 890 | Bacteria>Terrabacteria_group>Actinomycetota>Acidimicrobiia>Acidimicrobiales | GCA_026709125.1 | Acidimicrobiaceae_bacterium | OXD37_01465 | 24384..27056 | 2024894 | MAG:-DUF499-domain-containing-protein-[Acidimicrobiaceae-bacterium] |
| MCY4280945.1 | AAA+ATPase+wHTH+Alpha+Beta-domain+wHTH+RRM | Type-2_BR.RM.like.systems | DUF499*->METHYLASE->HKD-DNase+SF1-SF2-Helicase+iREase+CTD-> | 957 | Bacteria>Terrabacteria_group>Actinomycetota>Acidimicrobiia>Acidimicrobiales | GCA_026709905.1 | Acidimicrobiaceae_bacterium | OXC59_09605 | 1..2874 | 2024894 | MAG:-DUF499-domain-containing-protein,-partial-[Acidimicrobiaceae-bacterium] |
| MCY4293669.1 | AAA+ATPase+wHTH+Alpha+Beta-domain+wHTH+RRM | Type-2_BR.RM.like.systems | DUF499-p->DUF499*->METHYLASE->HKD-DNase+SF1-SF2-Helicase+iREase+CTD-><-?<-?\|\|?->RnasePH+RnasePH+KH+S1-> | 889 | Bacteria>Terrabacteria_group>Actinomycetota>Acidimicrobiia>Acidimicrobiales | GCA_026710015.1 | Acidimicrobiaceae_bacterium | OXC58_02335 | 25005..27674 | 2024894 | MAG:-DUF499-domain-containing-protein-[Acidimicrobiaceae-bacterium] |
| MDE0135358.1 | AAA+ATPase+wHTH+Alpha+Beta-domain+wHTH+RRM | Type-2_BR.RM.like.systems | <-HKD-DNase+SF1-SF2-Helicase+iREase+CTD<-METHYLASE<-DUF3780<-DUF499*\|\|REase-> | 1088 | Bacteria>Terrabacteria_group>Actinomycetota>Acidimicrobiia>Acidimicrobiales | GCA_028819405.1 | Acidimicrobiaceae_bacterium | OXM54_11015 | 23317..26583 | 2024894 | MAG:-DUF499-domain-containing-protein-[Acidimicrobiaceae-bacterium] |
| MDE0496286.1 | AAA+ATPase+wHTH+Alpha+Beta-domain+wHTH+RRM | Type-2_BR.RM.like.systems | <-REase\|\|DUF499*->DUF3780->METHYLASE->HKD-DNase+SF1-SF2-Helicase+iREase+CTD-> | 1088 | Bacteria>Terrabacteria_group>Actinomycetota>Acidimicrobiia>Acidimicrobiales | GCA_028823965.1 | Acidimicrobiaceae_bacterium | OXH86_02940 | 41213..44479 | 2024894 | MAG:-DUF499-domain-containing-protein-[Acidimicrobiaceae-bacterium] |
| MYD03946.1 | AAA+ATPase+wHTH+Alpha+Beta-domain+wHTH+RRM | Type-2_BR.RM.like.systems | <-HKD-DNase+SF1-SF2-Helicase+iREase+CTD<-METHYLASE<-DUF3780<-DUF499* | 986 | Bacteria>Terrabacteria_group>Actinomycetota>Acidimicrobiia>unclassified_Acidimicrobiia | GCA_009841075.1 | Acidimicrobiia_bacterium | F4X21_03265 | 7005..9965 | 2080302 | ATP-binding-protein-[Acidimicrobiia-bacterium] |
| KJE19503.1 | AAA+ATPase+wHTH+Alpha+Beta-domain+wHTH+RRM | Type-2_BR.RM.like.systems | DUF4096->DDE_Tnp_1_2-><-?\|\|DDE_Tnp_1_4->DUF499*->DUF3780->METHYLASE->HKD-DNase+SF1-SF2-Helicase+iREase+CTD-><-DDE_Tnp_1_2<-DUF4096 | 1073 | Bacteria>Terrabacteria_group>Actinomycetota>Actinomycetes>Frankiales | GCA_000948395.1 | Frankia_torreyi | FF36_06214 | 3173..6394 | 1856 | Protein-of-unknown-function-(DUF499)-[Frankia-torreyi] |
| KQM01919.1 | AAA+ATPase+wHTH+Alpha+Beta-domain+wHTH+RRM | Type-2_BR.RM.like.systems | DDE_Tnp_1+DDE_Tnp_1_2-><-HKD-DNase+SF1-SF2-Helicase+iREase+CTD<-METHYLASE<-DUF3780<-DUF499*<-DDE_Tnp_1_4\|\|?-><-DDE_Tnp_1_2 | 1073 | Bacteria>Terrabacteria_group>Actinomycetota>Actinomycetes>Frankiales | GCA_001421075.1 | Frankia_sp._CpI1-P | FF86_10935 | 7159..10380 | 1502734 | Protein-of-unknown-function-(DUF499)-[Frankia-sp.-CpI1-P] |
| WP_138732028.1 | AAA+ATPase+wHTH+Alpha+Beta-domain+wHTH+RRM | Type-2_BR.RM.like.systems | ABC-ATPase+SbcC->?->?->?-><-?\|\|DUF499*->DUF3780->METHYLASE->HKD-DNase+SF1-SF2-Helicase+iREase+CTD-><-SF1-SF2-Helicase+REase<-Exonuc_V_gamma+REase\|\|ABhydrolase+ABhydrolase-> | 994 | Bacteria>Terrabacteria_group>Actinomycetota>Actinomycetes>Geodermatophilales | GCF_005930495.1 | Modestobacter_excelsi | FHL80_RS03265 | 33106..36090 | 2213161 | DUF499-domain-containing-protein-[Modestobacter-excelsi] |
| QES21410.1 | AAA+ATPase+wHTH+Alpha+Beta-domain+wHTH+RRM | Type-2_BR.RM.like.systems | Methyltransf_11->ST-Hiskin->DUF499*-><-?\|\|METHYLASE->HKD-DNase+SF1-SF2-Helicase+iREase+CTD->?->LuxR-HTH-> | 1070 | Bacteria>Terrabacteria_group>Actinomycetota>Actinomycetes>Kitasatosporales | GCA_008642375.1 | Streptomyces_venezuelae | DEJ46_21795 | 4741407..4744619 | 54571 | hypothetical-protein-DEJ46_21795-[Streptomyces-venezuelae] |
| MTE20048.1 | AAA+ATPase+wHTH+Alpha+Beta-domain+wHTH+RRM | Type-2_BR.RM.like.systems | Pkinase-><-HTH_3\|\|ST-Hiskin->?->?->?->DUF499*->DUF3780->METHYLASE->HKD-DNase+SF1-SF2-Helicase+iREase+CTD-><-GNTR-HTH\|\|?->FtsK-> | 1022 | Bacteria>Terrabacteria_group>Actinomycetota>Actinomycetes>Kitasatosporales | GCA_009709575.1 | Streptomyces_taklimakanensis | FOL17_13155 | 3018420..3021488 | 2569853 | DUF499-domain-containing-protein-[Streptomyces-taklimakanensis] |
| MBE4799174.1 | AAA+ATPase+wHTH+Alpha+Beta-domain+wHTH+RRM | Type-2_BR.RM.like.systems | <-HKD-DNase+SF1-SF2-Helicase+iREase+CTD<-METHYLASE<-DUF3780<-DUF499*\|\|REase+DUF5655-><-N6_N4_MTase | 1002 | Bacteria>Terrabacteria_group>Actinomycetota>Actinomycetes>Kitasatosporales | GCA_014930365.1 | Streptomyces_caniscabiei | HW837_43475 | 7242..10250 | 2746961 | ATP-binding-protein-[Streptomyces-caniscabiei] |
| MBM7088758.1 | AAA+ATPase+wHTH+Alpha+Beta-domain+wHTH+RRM | Type-2_BR.RM.like.systems | XIS-HTH->Phage_integrase-><-HTH_Tnp_4+DDE_Tnp_4\|\|DUF499*->DUF3780->METHYLASE->HKD-DNase+SF1-SF2-Helicase+iREase+CTD-><-?\|\|GNTR-HTH->ST-Hiskin-> | 1001 | Bacteria>Terrabacteria_group>Actinomycetota>Actinomycetes>Kitasatosporales | GCA_016900935.1 | Streptomyces_sp._S12 | JTP67_09410 | 34483..37488 | 1399828 | ATP-binding-protein-[Streptomyces-sp.-S12] |
| WP_067071388.1 | AAA+ATPase+wHTH+Alpha+Beta-domain+wHTH+RRM | Type-2_BR.RM.like.systems | <-HKD-DNase+SF1-SF2-Helicase+iREase+CTD<-METHYLASE<-DUF3780<-DUF499*\|\|?->?->?->?-><-?<-HKD-DNase | 1001 | Bacteria>Terrabacteria_group>Actinomycetota>Actinomycetes>Kitasatosporales | GCF_001543895.1 | Carbonactinospora_thermoautotrophica | TH66_RS19600 | 537285..540290 | 1469144 | DUF499-domain-containing-protein-[Carbonactinospora-thermoautotrophica] |
| WP_158013629.1 | AAA+ATPase+wHTH+Alpha+Beta-domain+wHTH+RRM | Type-2_BR.RM.like.systems | DUF499*->DUF3780->METHYLASE->HKD-DNase+SF1-SF2-Helicase+iREase+CTD-> | 700 | Bacteria>Terrabacteria_group>Actinomycetota>Actinomycetes>Kitasatosporales | GCF_001543915.1 | Carbonactinospora_thermoautotrophica | TR74_RS22760 | 1..2104 | 1469144 | DUF499-domain-containing-protein,-partial-[Carbonactinospora-thermoautotrophica] |
| WP_066883461.1 | AAA+ATPase+wHTH+Alpha+Beta-domain+wHTH+RRM | Type-2_BR.RM.like.systems | HKD-DNase->?-><-?<-?<-?<-?\|\|DUF499*->DUF3780->METHYLASE->HKD-DNase+SF1-SF2-Helicase+iREase+CTD-> | 1001 | Bacteria>Terrabacteria_group>Actinomycetota>Actinomycetes>Kitasatosporales | GCF_001543925.1 | Carbonactinospora_thermoautotrophica | LI90_RS01235 | 264009..267014 | 1469144 | DUF499-domain-containing-protein-[Carbonactinospora-thermoautotrophica] |
| WP_094211291.1 | AAA+ATPase+wHTH+Alpha+Beta-domain+wHTH+RRM | Type-2_BR.RM.like.systems | <-Methyltransf_25<-?<-?\|\|HTH_3->NUDIX->DUF499*->DUF3780->METHYLASE->HKD-DNase+SF1-SF2-Helicase+iREase+CTD-> | 1002 | Bacteria>Terrabacteria_group>Actinomycetota>Actinomycetes>Kitasatosporales | GCF_002242735.1 | Streptomyces_sp._2R | BEH93_RS05710 | 6682..9690 | 1883452 | DUF499-domain-containing-protein-[Streptomyces-sp.-2R] |
| WP_190622806.1 | AAA+ATPase+wHTH+Alpha+Beta-domain+wHTH+RRM | Type-2_BR.RM.like.systems | Methyltransf_11->ST-Hiskin->DUF499*-><-?\|\|METHYLASE->HKD-DNase+SF1-SF2-Helicase+iREase+CTD->?->LuxR-HTH-> | 1002 | Bacteria>Terrabacteria_group>Actinomycetota>Actinomycetes>Kitasatosporales | GCF_008642375.1 | Streptomyces_venezuelae | DEJ46_RS21805 | 4741611..4744619 | 54571 | DUF499-domain-containing-protein-[Streptomyces-venezuelae] |
| WP_162466148.1 | AAA+ATPase+wHTH+Alpha+Beta-domain+wHTH+RRM | Type-2_BR.RM.like.systems | Pkinase-><-HTH_3\|\|ST-Hiskin->?->?->?->DUF499*->DUF3780->METHYLASE->HKD-DNase+SF1-SF2-Helicase+iREase+CTD-><-GNTR-HTH\|\|?->FtsK-> | 999 | Bacteria>Terrabacteria_group>Actinomycetota>Actinomycetes>Kitasatosporales | GCF_009709575.1 | Streptomyces_taklimakanensis | FOL17_RS13180 | 3018489..3021488 | 2569853 | DUF499-domain-containing-protein-[Streptomyces-taklimakanensis] |
| WP_189301311.1 | AAA+ATPase+wHTH+Alpha+Beta-domain+wHTH+RRM | Type-2_BR.RM.like.systems | XIS-HTH-><-HKD-DNase+SF1-SF2-Helicase+iREase+CTD<-METHYLASE<-DUF3780<-DUF499*<-ABhydrolase\|\|?->?->?->ParA-Soj-PloopNTPase-> | 1093 | Bacteria>Terrabacteria_group>Actinomycetota>Actinomycetes>Kitasatosporales | GCF_014648695.1 | Streptomyces_albospinus | IEZ17_RS17930 | 11197..14478 | 285515 | DUF499-domain-containing-protein-[Streptomyces-albospinus] |
| WP_237270650.1 | AAA+ATPase+wHTH+Alpha+Beta-domain+wHTH+RRM | Type-2_BR.RM.like.systems | N6_N4_MTase-><-REase+DUF5655\|\|DUF499*->DUF3780->METHYLASE->HKD-DNase+SF1-SF2-Helicase+iREase+CTD-> | 999 | Bacteria>Terrabacteria_group>Actinomycetota>Actinomycetes>Kitasatosporales | GCF_014930395.1 | Streptomyces_caniscabiei | HW845_RS42820 | 91168..94167 | 2746961 | DUF499-domain-containing-protein-[Streptomyces-caniscabiei] |
| WP_093894449.1 | AAA+ATPase+wHTH+Alpha+Beta-domain+wHTH+RRM | Type-2_BR.RM.like.systems | <-DUF397<-HTH+DUF5753\|\|ST-Hiskin->DUF499*->DUF3780->METHYLASE->HKD-DNase+SF1-SF2-Helicase+iREase+CTD-> | 1093 | Bacteria>Terrabacteria_group>Actinomycetota>Actinomycetes>Kitasatosporales | GCF_900091955.1 | Streptomyces_sp._Ncost-T10-10d | GA0115254_RS07180 | 39686..42967 | 1839774 | DUF499-domain-containing-protein-[Streptomyces-sp.-Ncost-T10-10d] |
| WP_210775023.1 | AAA+ATPase+wHTH+Alpha+Beta-domain+wHTH+RRM | Type-2_BR.RM.like.systems | HTH_17->Phage_int_SAM_1+Phage_integrase->?-><-HKD-DNase+SF1-SF2-Helicase+iREase+CTD-p<-METHYLASE<-DUF3780<-DUF499*\|\|<-Phage_integrase\|\|?->?-><-?\|\|MULE-TRANSPOSASE-> | 1075 | Bacteria>Terrabacteria_group>Actinomycetota>Actinomycetes>Micromonosporales | GCF_018070095.1 | Micromonospora_sp._U56 | KBX37_RS06580 | 75892..79119 | 2824900 | DUF499-domain-containing-protein-[Micromonospora-sp.-U56] |
| WP_265004266.1 | AAA+ATPase+wHTH+Alpha+Beta-domain+wHTH+RRM | Type-2_BR.RM.like.systems | <-HKD-DNase+SF1-SF2-Helicase+iREase+CTD<-METHYLASE<-DUF3780<-DUF499* | 1001 | Bacteria>Terrabacteria_group>Actinomycetota>Actinomycetes>Micromonosporales | GCF_026013475.1 | Micromonospora_yasonensis | ONA70_RS00435 | 78334..81339 | 1128667 | DUF499-domain-containing-protein-[Micromonospora-yasonensis] |
| WP_091105663.1 | AAA+ATPase+wHTH+Alpha+Beta-domain+wHTH+RRM | Type-2_BR.RM.like.systems | <-URI-DNase\|\|?-><-?\|\|?->DUF499*->DUF3780->METHYLASE->HKD-DNase+SF1-SF2-Helicase+iREase+CTD-><-HEPN<-XIS-HTH+RVT | 1075 | Bacteria>Terrabacteria_group>Actinomycetota>Actinomycetes>Micromonosporales | GCF_900090315.1 | Micromonospora_citrea | GA0070606_RS26660 | 5834350..5837577 | 47855 | DUF499-domain-containing-protein-[Micromonospora-citrea] |
| WP_172896457.1 | AAA+ATPase+wHTH+Alpha+Beta-domain+wHTH+RRM | Type-2_BR.RM.like.systems | DUF499*->DUF3780->METHYLASE->HKD-DNase+SF1-SF2-Helicase+iREase+CTD->REase-> | 989 | Bacteria>Terrabacteria_group>Actinomycetota>Actinomycetes>Micromonosporales | GCF_900091445.1 | Micromonospora_echinofusca | GA0070610_RS08520 | 1858371..1861340 | 47858 | DUF499-domain-containing-protein-[Micromonospora-echinofusca] |
| AHD24334.1 | AAA+ATPase+wHTH+Alpha+Beta-domain+wHTH+RRM | Type-2_BR.RM.like.systems | DUF499*->DUF3780->METHYLASE-p->HKD-DNase+SF1-SF2-Helicase+iREase+CTD-><-?\|\|PIN-> | 1067 | Bacteria>Terrabacteria_group>Actinomycetota>Actinomycetes>Mycobacteriales | GCA_000511305.1 | Rhodococcus_pyridinivorans_SB3094 | Y013_26415 | 338597..341800 | 1435356 | hypothetical-protein-Y013_26415-[Rhodococcus-pyridinivorans-SB3094] |
| MCT7364800.1 | AAA+ATPase+wHTH+Alpha+Beta-domain+wHTH+RRM | Type-2_BR.RM.like.systems | <-HTH_21+rve+rve_3<-HIN-HTH<-HKD-DNase+SF1-SF2-Helicase+iREase+CTD<-METHYLASE<-DUF3780<-DUF499*<-XIS-HTH<-TetR-HTH<-Phage_integrase+Phage_integrase<-Phage_integrase+Phage_integrase<-Phage_integrase<-DNA-glycosylase | 981 | Bacteria>Terrabacteria_group>Actinomycetota>Actinomycetes>Mycobacteriales | GCA_025331195.1 | Mycolicibacterium_llatzerense | A7G45_18170 | 801835..804780 | 280871 | hypothetical-protein-[Mycolicibacterium-llatzerense] |
| MCV7153468.1 | AAA+ATPase+wHTH+Alpha+Beta-domain+wHTH+RRM | Type-2_BR.RM.like.systems | <-TetR-HTH\|\|?-><-?\|\|Phage_integrase+Phage_integrase->Phage_integrase+Phage_integrase->TetR-HTH->DUF499*->DUF3780->METHYLASE->HKD-DNase+SF1-SF2-Helicase+iREase+CTD->\|\|HTH_Tnp_1+HTH_21+rve+rve_3->\|\|DDE_Tnp_1-> | 993 | Bacteria>Terrabacteria_group>Actinomycetota>Actinomycetes>Mycobacteriales | GCA_025821625.1 | Mycolicibacterium_pyrenivorans | H7I57_20005 | 26013..28994 | 187102 | DUF499-domain-containing-protein-[Mycobacterium-pyrenivorans] |
| MCW0191653.1 | AAA+ATPase+wHTH+Alpha+Beta-domain+wHTH+RRM | Type-2_BR.RM.like.systems | HTH_Tnp->DUF499*->DUF3780->METHYLASE->HKD-DNase+SF1-SF2-Helicase+iREase+CTD-> | 1089 | Bacteria>Terrabacteria_group>Actinomycetota>Actinomycetes>Mycobacteriales | GCA_025869315.1 | Rhodococcus_sp._(in:_high_G+C_Gram-positive_bacteria) | OJJ55_10175 | 803..4072 | 1831 | MAG:-ATP-binding-protein-[Rhodococcus-sp.] |
| WP_083847075.1 | AAA+ATPase+wHTH+Alpha+Beta-domain+wHTH+RRM | Type-2_BR.RM.like.systems | <-Phage_integrase<-?<-?<-HKD-DNase+SF1-SF2-Helicase+iREase+CTD<-METHYLASE<-DUF3780<-DUF499*\|\|HicA_toxin-><-HNH<-REase | 989 | Bacteria>Terrabacteria_group>Actinomycetota>Actinomycetes>Mycobacteriales | GCF_000256505.1 | Rhodococcus_sp._P14 | OEY_RS30125 | 9685..12654 | 450821 | DUF499-domain-containing-protein-[Rhodococcus-sp.-P14] |
| WP_158455960.1 | AAA+ATPase+wHTH+Alpha+Beta-domain+wHTH+RRM | Type-2_BR.RM.like.systems | DUF499*->DUF3780->METHYLASE->HKD-DNase+SF1-SF2-Helicase+iREase+CTD-><-?\|\|?->PIN-> | 1080 | Bacteria>Terrabacteria_group>Actinomycetota>Actinomycetes>Mycobacteriales | GCF_000511305.1 | Rhodococcus_pyridinivorans | Y013_RS25450 | 338558..341800 | 103816 | DUF499-domain-containing-protein-[Rhodococcus-pyridinivorans] |
| WP_138874062.1 | AAA+ATPase+wHTH+Alpha+Beta-domain+wHTH+RRM | Type-2_BR.RM.like.systems | Peptidase->?-><-REase\|\|?-><-?\|\|HTH_Tnp->DUF499*->DUF3780->METHYLASE->HKD-DNase+SF1-SF2-Helicase+iREase+CTD-> | 1089 | Bacteria>Terrabacteria_group>Actinomycetota>Actinomycetes>Mycobacteriales | GCF_005938745.2 | Rhodococcus_sp._KBS0724 | FFI94_RS33030 | 462712..465981 | 1179674 | DUF499-domain-containing-protein-[Rhodococcus-sp.-KBS0724] |
| WP_145239328.1 | AAA+ATPase+wHTH+Alpha+Beta-domain+wHTH+RRM | Type-2_BR.RM.like.systems | wHTH(SBD)+HNH->DUF499*->DUF3780->METHYLASE->HKD-DNase+SF1-SF2-Helicase+iREase+CTD->?->?->OLD-ABC+TOPRIM-> | 1089 | Bacteria>Terrabacteria_group>Actinomycetota>Actinomycetes>Mycobacteriales | GCF_007672705.1 | Prescottella_equi | FS672_RS22145 | 117636..120905 | 43767 | DUF499-domain-containing-protein-[Prescottella-equi] |
| WP_168707290.1 | AAA+ATPase+wHTH+Alpha+Beta-domain+wHTH+RRM | Type-2_BR.RM.like.systems | <-SF1-SF2-Helicase<-Primpol+D5N+D5ATPase<-HKD-DNase+SF1-SF2-Helicase+iREase+CTD<-METHYLASE<-DUF3780<-DUF499*<-?<-?<-JetA\|\|ABhydrolase-> | 1088 | Bacteria>Terrabacteria_group>Actinomycetota>Actinomycetes>Mycobacteriales | GCF_012498585.1 | Gordonia_paraffinivorans | HH035_RS01955 | 91655..94921 | 175628 | DUF499-domain-containing-protein-[Gordonia-paraffinivorans] |
| WP_209907589.1 | AAA+ATPase+wHTH+Alpha+Beta-domain+wHTH+RRM | Type-2_BR.RM.like.systems | <-HKD-DNase+SF1-SF2-Helicase+iREase+CTD<-METHYLASE<-DUF3780<-DUF499*<-HTH_Tnp\|\|HIN-HTH-><-?\|\|URI-DNase-> | 1090 | Bacteria>Terrabacteria_group>Actinomycetota>Actinomycetes>Mycobacteriales | GCF_017876985.1 | Rhodococcus_sp._PvP104 | J3A74_RS00025 | 8376..11648 | 2817911 | DUF499-domain-containing-protein-[Rhodococcus-sp.-PvP104] |
| WP_157892521.1 | AAA+ATPase+wHTH+Alpha+Beta-domain+wHTH+RRM | Type-2_BR.RM.like.systems | ABC-ATPase+SbcC->?-><-?\|\|?->DUF499*->DUF3780->METHYLASE->HKD-DNase+SF1-SF2-Helicase+iREase+CTD-> | 1088 | Bacteria>Terrabacteria_group>Actinomycetota>Actinomycetes>Mycobacteriales | GCF_019930745.1 | Mycolicibacterium_smegmatis | PXH78_RS30725 | 15102..18368 | 1772 | DUF499-domain-containing-protein-[Mycolicibacterium-smegmatis] |
| WP_275760079.1 | AAA+ATPase+wHTH+Alpha+Beta-domain+wHTH+RRM | Type-2_BR.RM.like.systems | ABC-ATPase+SbcC->?-><-?\|\|?->DUF499*->DUF3780->METHYLASE->HKD-DNase+SF1-SF2-Helicase+iREase+CTD-> | 1067 | Bacteria>Terrabacteria_group>Actinomycetota>Actinomycetes>Mycobacteriales | GCF_029221935.1 | Mycolicibacterium_smegmatis | PXH80_RS30425 | 15165..18368 | 1772 | DUF499-domain-containing-protein-[Mycolicibacterium-smegmatis] |
| WP_182263237.1 | AAA+ATPase+wHTH+Alpha+Beta-domain+wHTH+RRM | Type-2_BR.RM.like.systems | HTH_Tnp->DUF499*->DUF3780->METHYLASE->HKD-DNase+SF1-SF2-Helicase+iREase+CTD-><-?<-?\|\|GDResolvase+HIN-HTH-> | 1089 | Bacteria>Terrabacteria_group>Actinomycetota>Actinomycetes>Mycobacteriales | GCF_903819475.1 | Rhodococcus_sp._UFZ-B548 | H7C86_RS34245 | 1125..4394 | 2742212 | DUF499-domain-containing-protein-[Rhodococcus-sp.-UFZ-B548] |
| WP_276555405.1 | AAA+ATPase+wHTH+Alpha+Beta-domain+wHTH+RRM | Type-2_BR.RM.like.systems | <-HIN-HTH<-HKD-DNase+SF1-SF2-Helicase+iREase+CTD-p<-METHYLASE<-DUF3780<-DUF499*<-XIS-HTH<-TetR-HTH<-Phage_integrase+Phage_integrase<-Phage_integrase+Phage_integrase<-Phage_integrase<-DNA-glycosylase | 992 | Bacteria>Terrabacteria_group>Actinomycetota>Actinomycetes>Mycobacteriales | - | Mycolicibacterium_llatzerense | A7G45_RS18760 | 801835..804813 | 280871 | DUF499-domain-containing-protein-[Mycolicibacterium-llatzerense] |
| WP_275081497.1 | AAA+ATPase+wHTH+Alpha+Beta-domain+wHTH+RRM | Type-2_BR.RM.like.systems | CHTH->Phage_integrase+Phage_integrase->Phage_integrase+Phage_integrase->TetR-HTH->DUF499*->DUF3780->METHYLASE->HKD-DNase+SF1-SF2-Helicase+iREase+CTD-><-METHYLASE-p\|\|HTH_Tnp_1+HTH_21+rve+rve_3-> | 883 | Bacteria>Terrabacteria_group>Actinomycetota>Actinomycetes>Mycobacteriales | - | Mycolicibacterium_pyrenivorans | H7I57_RS20155 | 26343..28994 | 187102 | DUF499-domain-containing-protein-[Mycolicibacterium-pyrenivorans] |
| WP_012476995.1 | AAA+ATPase+wHTH+Alpha+Beta-domain+wHTH+RRM | Type-2_BR.RM.like.systems | DUF499*->DUF3780->METHYLASE->HKD-DNase+SF1-SF2-Helicase+iREase+CTD-> | 1088 | Bacteria>Terrabacteria_group>Actinomycetota>Actinomycetes>Mycobacteriales | - | Rhodococcus_sp._NS1 | HS530_RS00365 | 66277..69543 | 402236 | DUF499-domain-containing-protein-[Rhodococcus-sp.-NS1] |
| WP_274196404.1 | AAA+ATPase+wHTH+Alpha+Beta-domain+wHTH+RRM | Type-2_BR.RM.like.systems | REase->Exonuc_V_gamma+REase->SF1-SF2-Helicase+REase-><-HKD-DNase+SF1-SF2-Helicase+iREase+CTD<-METHYLASE<-DUF3780<-DUF499* | 1019 | Bacteria>Terrabacteria_group>Actinomycetota>Actinomycetes>Pseudonocardiales | GCF_028737265.1 | Actinomycetospora_callitridis | PH191_RS20135 | 253278..256337 | 913944 | DUF499-domain-containing-protein-[Actinomycetospora-callitridis] |
| WP_142649215.1 | AAA+ATPase+wHTH+Alpha+Beta-domain+wHTH+RRM | Type-2_BR.RM.like.systems | NUDIX->?-><-HDRnase<-GNTR-HTH<-?\|\|HTH+HKD-DNase->DUF499*->DUF3780->METHYLASE->HKD-DNase+SF1-SF2-Helicase+iREase+CTD->XIS-HTH-> | 1098 | Bacteria>Terrabacteria_group>Actinomycetota>Actinomycetes>Streptosporangiales | GCF_006874455.1 | Microbispora_sp._SCL1-1 | FLW53_RS21915 | 132280..135576 | 2592812 | DUF499-domain-containing-protein-[Microbispora-sp.-SCL1-1] |
| WP_240971672.1 | AAA+ATPase+wHTH+Alpha+Beta-domain+wHTH+RRM | Type-2_BR.RM.like.systems | NUDIX->?-><-HDRnase<-GNTR-HTH<-?\|\|HTH+HKD-DNase->DUF499*->DUF3780->METHYLASE->HKD-DNase+SF1-SF2-Helicase+iREase+CTD->XIS-HTH-> | 1091 | Bacteria>Terrabacteria_group>Actinomycetota>Actinomycetes>Streptosporangiales | GCF_012034195.1 | Microbispora_sp._CL1-1 | HCK04_RS21825 | 132301..135576 | 2720026 | DUF499-domain-containing-protein-[Microbispora-sp.-CL1-1] |
| WP_165634759.1 | AAA+ATPase+wHTH+Alpha+Beta-domain+wHTH+RRM | Type-2_BR.RM.like.systems | <-GNTR-HTH<-?<-?\|\|HTH_3-><-NUDIX\|\|DUF499*->DUF3780->METHYLASE->HKD-DNase+SF1-SF2-Helicase+iREase+CTD-><-?\|\|?-><-PIN | 991 | Bacteria>Terrabacteria_group>Actinomycetota>Actinomycetes>Streptosporangiales | GCF_900099985.1 | Thermostaphylospora_chromogena | BLS31_RS10180 | 2220186..2223161 | 35622 | DUF499-domain-containing-protein-[Thermostaphylospora-chromogena] |
| MCB0857325.1 | AAA+ATPase+wHTH+Alpha+Beta-domain+wHTH+RRM | Type-2_BR.RM.like.systems | <-TUDOR+SF1-SF2-Helicase+iREase+CTD<-?<-?\|\|ParB+HNH->DUF499*->DUF3780->METHYLASE->HKD-DNase+SF1-SF2-Helicase+iREase+CTD->?-><-REase | 976 | Bacteria>Terrabacteria_group>Actinomycetota>Thermoleophilia>Solirubrobacterales | GCA_020438355.1 | Solirubrobacterales_bacterium | KDB57_04290 | 63088..66018 | 1909295 | DUF499-domain-containing-protein-[Solirubrobacterales-bacterium] |
| MDA8309413.1 | AAA+ATPase+wHTH+Alpha+Beta-domain+wHTH+RRM | Type-2_BR.RM.like.systems | <-sGTP-MglA+MORC+REase\|\|?->DDE_Tnp_ISAZ013-><-?\|\|DUF499*->DUF3780->METHYLASE->HKD-DNase+SF1-SF2-Helicase+iREase+CTD-><-MODE-HTH+PBPII | 978 | Bacteria>Terrabacteria_group>Actinomycetota>unclassified_Actinomycetota | GCA_028067795.1 | Actinomycetota_bacterium | MOZ46_02150 | 8496..11432 | 2900548 | MAG:-DUF499-domain-containing-protein-[Actinomycetota-bacterium] |
| MDA8372907.1 | AAA+ATPase+wHTH+Alpha+Beta-domain+wHTH+RRM | Type-2_BR.RM.like.systems | <-Phage_int_SAM_1+Phage_integrase\|\|DUF499*->?->METHYLASE->HKD-DNase+SF1-SF2-Helicase+iREase+CTD-><-?<-?<-DUF4011+SF1-SF2-Helicase+REase | 978 | Bacteria>Terrabacteria_group>Actinomycetota>unclassified_Actinomycetota | GCA_028068365.1 | Actinomycetota_bacterium | MOZ91_01415 | 1480..4416 | 2900548 | MAG:-DUF499-domain-containing-protein-[Actinomycetota-bacterium] |
| MBE3519594.1 | AAA+ATPase+wHTH+Alpha+Beta-domain+wHTH+RRM | Type-2_BR.RM.like.systems | <-ABC-ATPase+SbcC<-PAIREDC-HTH\|\|tRNA-><-Phage_integrase\|\|MazE_antitoxin->PIN->DUF499*->DUF3780->METHYLASE->HKD-DNase+SF1-SF2-Helicase+iREase+CTD-> | 1008 | Bacteria>Terrabacteria_group>Bacillota>Candidatus_Fermentithermobacillia>Candidatus_Fermentithermobacillales | GCA_014896535.1 | Candidatus_Fermentithermobacillaceae_bacterium | IMW97_04750 | 107753..110779 | 3085339 | ATP-binding-protein-[Firmicutes-bacterium] |
| WP_073234041.1 | AAA+ATPase+wHTH+Alpha+Beta-domain+wHTH+RRM | Type-2_BR.RM.like.systems | <-HTH_3\|\|?->NERD->?->?->DUF499*->DUF3780->METHYLASE->HKD-DNase+SF1-SF2-Helicase+iREase+CTD-> | 1013 | Bacteria>Terrabacteria_group>Bacillota>Clostridia>Eubacteriales | GCF_900129195.1 | Desulforamulus_putei | BUB67_RS00525 | 118587..121628 | 74701 | DUF499-domain-containing-protein-[Desulforamulus-putei] |
| WP_028962636.1 | AAA+ATPase+wHTH+Alpha+Beta-domain+wHTH+RRM | Type-2_BR.RM.like.systems | Peptidase_M20->rve_3->HTH->DUF499*->DUF3780->METHYLASE->TUDOR+SF1-SF2-Helicase+iREase+CTD->?->MethyltransfD12-> | 992 | Bacteria>Terrabacteria_group>Bacillota>Clostridia>Eubacteriales | GCF_000497695.1 | Sulfobacillus_thermosulfidooxidans | L801_RS0106580 | 9020..11998 | 28034 | DUF499-domain-containing-protein-[Sulfobacillus-thermosulfidooxidans] |
| AEJ38296.1 | AAA+ATPase+wHTH+Alpha+Beta-domain+wHTH+RRM | Type-2_BR.RM.like.systems | REase->DNA_methylase->SF1-SF2-Helicase+REase->?->DUF499*->DUF3780->METHYLASE->TUDOR+SF1-SF2-Helicase+iREase+CTD-> | 999 | Bacteria>Terrabacteria_group>Bacillota>Clostridia>Eubacteriales | GCA_000219855.1 | Sulfobacillus_acidophilus_TPY | TPY_0069 | 67681..70680 | 1051632 | ATPase-(AAA+-superfamily)-like-protein-[Sulfobacillus-acidophilus-TPY] |
| MBE3577570.1 | AAA+ATPase+wHTH+Alpha+Beta-domain+wHTH+RRM | Type-2_BR.RM.like.systems | <-HKD-DNase+SF1-SF2-Helicase+iREase+CTD<-METHYLASE<-DUF3780<-DUF499* | 1008 | Bacteria>Terrabacteria_group>Bacillota>Limnochordia>Limnochordales | GCA_014896315.1 | Limnochordales_bacterium | IMX00_07740 | 22208..25234 | 2602711 | ATP-binding-protein-[Limnochordales-bacterium] |
| HHV54946.1 | AAA+ATPase+wHTH+Alpha+Beta-domain+wHTH+RRM | Type-2_BR.RM.like.systems | DUF499*->DUF3780->METHYLASE->HKD-DNase+SF1-SF2-Helicase+iREase+CTD-> | 1011 | Bacteria>Terrabacteria_group>Bacillota>unclassified_Bacillota | GCA_012839805.1 | Bacillota_bacterium | GXX55_05835 | 3376..6411 | 1879010 | ATP-binding-protein-[Firmicutes-bacterium] |
| NLJ36704.1 | AAA+ATPase+wHTH+Alpha+Beta-domain+wHTH+RRM | Type-2_BR.RM.like.systems | DUF499*->DUF3780->METHYLASE->HKD-DNase+SF1-SF2-Helicase+iREase+CTD-><-?\|\|Phage_integrase-> | 1049 | Bacteria>Terrabacteria_group>candidate_division_WS1>unclassified_candidate_division_WS1 | GCA_012518355.1 | candidate_division_WS1_bacterium | GX358_10820 | 2331..5480 | 2099669 | ATP-binding-protein-[candidate-division-WS1-bacterium] |
| MDA8395862.1 | AAA+ATPase+wHTH+Alpha+Beta-domain+wHTH+RRM | Type-2_BR.RM.like.systems | Phage_integrase-><-?<-?<-HKD-DNase+SF1-SF2-Helicase+iREase+CTD<-METHYLASE<-DUF3780<-DUF499*<-MetJarc-HTH\|\|ParA-Soj-PloopNTPase->?->?-><-RelE-ParE<-PhdYeFM_antitox | 995 | Bacteria>Terrabacteria_group>Candidatus_Dormibacteraeota>unclassified_Candidatus_Dormibacteraeota | GCA_028068535.1 | Candidatus_Dormibacteraeota_bacterium | MOT72_11630 | 9348..12335 | 2052315 | MAG:-DUF499-domain-containing-protein-[Candidatus-Dormibacteraeota-bacterium] |
| NPV68429.1 | AAA+ATPase+wHTH+Alpha+Beta-domain+wHTH+RRM | Type-2_BR.RM.like.systems | <-ABC-ATPase+SbcC<-?<-?\|\|?->Phage_int_SAM_1+Phage_integrase->DUF499*->ASCH+DUF3780->MazE_antitoxin->PIN->METHYLASE->HKD-DNase+SF1-SF2-Helicase+iREase+CTD-> | 1008 | Bacteria>Terrabacteria_group>Chloroflexota>Anaerolineae>unclassified_Anaerolineae | GCA_013178165.1 | Anaerolineae_bacterium | HPY64_14900 | 23774..26800 | 2052143 | ATP-binding-protein-[Anaerolineae-bacterium] |
| MBN1262662.1 | AAA+ATPase+wHTH+Alpha+Beta-domain+wHTH+RRM | Type-2_BR.RM.like.systems | <-HKD-DNase+SF1-SF2-Helicase+iREase+CTD<-METHYLASE<-ASCH+DUF3780<-DUF499* | 1016 | Bacteria>Terrabacteria_group>Chloroflexota>Anaerolineae>unclassified_Anaerolineae | GCA_016926915.1 | Anaerolineae_bacterium | JXB35_18415 | 7964..11014 | 2052143 | ATP-binding-protein-[Anaerolineae-bacterium] |
| MCL4868352.1 | AAA+ATPase+wHTH+Alpha+Beta-domain+wHTH+RRM | Type-2_BR.RM.like.systems | <-METHYLASE<-TOPRIM<-OLD-ABC<-ASCH+DUF3780<-PIN<-PhdYeFM_antitox<-DUF499* | 1019 | Bacteria>Terrabacteria_group>Chloroflexota>Anaerolineae>unclassified_Anaerolineae | GCA_023384415.1 | Anaerolineae_bacterium | KJ063_05240 | 198944..202003 | 2052143 | DUF499-domain-containing-protein-[Anaerolineae-bacterium] |

<table>
<tr><th>Protein-ID</th><th>DUF499-architecture</th><th>System</th><th>Gene-Neighbourhoods</th><th>Length</th><th>Taxonomy</th><th>Assembly-ID</th><th>Strain</th><th>Locus-Tag</th><th>Genomic-coordinates</th><th>TaxID</th><th>Fasta-defline</th></tr>
<tr><td>MCX7854723.1</td><td>AAA+ATPase+wHTH+Alpha+Beta-domain+wHTH+RRM</td><td>Type-2_BR.RM.like.systems</td><td><-MazE_antitoxin<-Peptidase_C14||DUF499*->ASCH+DUF3780->OLD-ABC->TOPRIM->METHYLASE-p-></td><td>1008</td><td>Bacteria>Terrabacteria_group>Chloroflexota>Anaerolineae>unclassified_Anaerolineae</td><td>GCA_026417235.1</td><td>Anaerolineae_bacterium</td><td>N2556_01880</td><td>1669..4695</td><td>2052143</td><td>MAG:-DUF499-domain-containing-protein-[Anaerolineae-bacterium]</td></tr>
<tr><td>MCZ7568046.1</td><td>AAA+ATPase+wHTH+Alpha+Beta-domain+wHTH+RRM</td><td>Type-2_BR.RM.like.systems</td><td>NTP_transf_5->Peptidase_M24->ACET->?-><-?||DUF499*->DUF3780->?->OLD-ABC->TOPRIM->METHYLASE->METHYLASE-p-></td><td>1024</td><td>Bacteria>Terrabacteria_group>Chloroflexota>Ardenticatenia>Ardenticatenales</td><td>GCA_027493995.1</td><td>Ardenticatenaceae_bacterium</td><td>M5U01_05505</td><td>1311111..1314185</td><td>2614247</td><td>MAG:-ATP-binding-protein-[Ardenticatenaceae-bacterium]</td></tr>
<tr><td>MYC64339.1</td><td>AAA+ATPase+wHTH+Alpha+Beta-domain+wHTH+RRM</td><td>Type-2_BR.RM.like.systems</td><td><-NERD<-?<-DUF772+DDE_Tnp_1+DDE_Tnp_1_6||DUF499*->DUF3780->METHYLASE->HKD-DNase+SF1-SF2-Helicase+iREase+CTD-></td><td>985</td><td>Bacteria>Terrabacteria_group>Chloroflexota>Caldilineae>Caldilineales</td><td>GCA_009841145.1</td><td>Caldilineaceae_bacterium_SB0661_bin_34</td><td>F4X16_16165</td><td>38045..41002</td><td>2605256</td><td>ATP-binding-protein-[Caldilineaceae-bacterium-SB0661_bin_34]</td></tr>
<tr><td>MCY4522925.1</td><td>AAA+ATPase+wHTH+Alpha+Beta-domain+wHTH+RRM</td><td>Type-2_BR.RM.like.systems</td><td>DUF499*->DUF3780->METHYLASE->HKD-DNase+SF1-SF2-Helicase+iREase+CTD->?->?->DDE_3-></td><td>764</td><td>Bacteria>Terrabacteria_group>Chloroflexota>Caldilineae>Caldilineales</td><td>GCA_026713025.1</td><td>Caldilineaceae_bacterium</td><td>OXC13_19395</td><td>1..2295</td><td>2614250</td><td>MAG:-DUF499-domain-containing-protein,-partial-[Caldilineaceae-bacterium]</td></tr>
<tr><td>MCB9124684.1</td><td>AAA+ATPase+wHTH+Alpha+Beta-domain+wHTH+RRM</td><td>Type-2_BR.RM.like.systems</td><td><-METHYLASE-p<-HTH_Tnp_1+HTH_21+rve<-TOPRIM<-OLD-ABC<-DUF4276<-ASCH+DUF3780<-DUF499*||?->?-><-?<-NTP_transf_2</td><td>1027</td><td>Bacteria>Terrabacteria_group>Chloroflexota>Caldilineae>Caldilineales</td><td>GCA_020634715.1</td><td>Caldilineaceae_bacterium</td><td>H6643_09260</td><td>39017..42100</td><td>2614250</td><td>ATP-binding-protein-[Caldilineaceae-bacterium]</td></tr>
<tr><td>MCB9118705.1</td><td>AAA+ATPase+wHTH+Alpha+Beta-domain+wHTH+RRM</td><td>Type-2_BR.RM.like.systems</td><td>NTP_transf_2->?-><-?<-?||DUF499*->ASCH+DUF3780->DUF4276->OLD-ABC->TOPRIM->HTH_Tnp_1+HTH_21+rve->METHYLASE-p-></td><td>1027</td><td>Bacteria>Terrabacteria_group>Chloroflexota>Caldilineae>Caldilineales</td><td>GCA_020634765.1</td><td>Caldilineaceae_bacterium</td><td>H6640_03130</td><td>11971..15054</td><td>2614250</td><td>ATP-binding-protein-[Caldilineaceae-bacterium]</td></tr>
<tr><td>MCB0025417.1</td><td>AAA+ATPase+wHTH+Alpha+Beta-domain+wHTH+RRM</td><td>Type-2_BR.RM.like.systems</td><td><-METHYLASE-p<-TOPRIM<-OLD-ABC<-DUF4276<-ASCH+DUF3780<-DUF499*</td><td>1027</td><td>Bacteria>Terrabacteria_group>Chloroflexota>Caldilineae>Caldilineales</td><td>GCA_020433165.1</td><td>Caldilinea_sp.</td><td>KDD91_20435</td><td>3611..6694</td><td>2293560</td><td>ATP-binding-protein-[Caldilinea-sp.]</td></tr>
<tr><td>MBN1856440.1</td><td>AAA+ATPase+wHTH+Alpha+Beta-domain+wHTH+RRM</td><td>Type-2_BR.RM.like.systems</td><td><-HKD-DNase+SF1-SF2-Helicase+iREase+CTD<-METHYLASE<-DUF3780<-DUF499*<-?<-ZnR</td><td>1006</td><td>Bacteria>Terrabacteria_group>Chloroflexota>Dehalococcoidia>unclassified_Dehalococcoidia</td><td>GCA_016930935.1</td><td>Dehalococcoidia_bacterium</td><td>JW846_05735</td><td>9362..12382</td><td>2026734</td><td>ATP-binding-protein-[Dehalococcoidia-bacterium]</td></tr>
<tr><td>MCY4653293.1</td><td>AAA+ATPase+wHTH+Alpha+Beta-domain+wHTH+RRM</td><td>Type-2_BR.RM.like.systems</td><td><-Phage_integrase||DUF499*->DUF3780->METHYLASE->HKD-DNase+SF1-SF2-Helicase+iREase+CTD-></td><td>1005</td><td>Bacteria>Terrabacteria_group>Chloroflexota>Dehalococcoidia>unclassified_Dehalococcoidia</td><td>GCA_026713885.1</td><td>Dehalococcoidia_bacterium</td><td>OXC95_09035</td><td>1209..4226</td><td>2026734</td><td>MAG:-DUF499-domain-containing-protein-[Dehalococcoidia-bacterium]</td></tr>
<tr><td>HEY77166.1</td><td>AAA+ATPase+wHTH+Alpha+Beta-domain+wHTH+RRM</td><td>Type-2_BR.RM.like.systems</td><td>PhdYeFM_antitox-><-HTH<-XIS-HTH<-GDResolvase+Recombinase+ZnR<-GDResolvase||?->DUF499*->ASCH+DUF3780->Schlafen+HATPase+HTH->HicB_lk_antitox->METHYLASE->SF1-SF2-Helicase->HKD-DNase+SF1-SF2-Helicase+iREase+CTD-></td><td>1010</td><td>Bacteria>Terrabacteria_group>Chloroflexota>Thermoflexia>unclassified_Thermoflexia</td><td>GCA_011192125.1</td><td>Thermoflexia_bacterium</td><td>G4O00_13500</td><td>6629..9661</td><td>2420334</td><td>ATP-binding-protein-[Thermoflexia-bacterium]</td></tr>
<tr><td>OQB28032.1</td><td>AAA+ATPase+wHTH+Alpha+Beta-domain+wHTH+RRM</td><td>Type-2_BR.RM.like.systems</td><td><-PolB||?-><-HEPN+Mrr_N<-HKD-DNase+SF1-SF2-Helicase+iREase+CTD<-METHYLASE<-DUF3780<-DUF499*<-?<-?<-REase<-?<-HHH_4+SF1-SF2-Helicase+REase</td><td>1020</td><td>Bacteria>Terrabacteria_group>Chloroflexota>unclassified_Chloroflexota</td><td>GCA_002070215.1</td><td>Chloroflexi_bacterium_ADurb.Bin180</td><td>BWY10_00836</td><td>30087..33149</td><td>1852857</td><td>hypothetical-protein-BWY10_00836-[Chloroflexi-bacterium-ADurb.Bin180]</td></tr>
<tr><td>MBI3972516.1</td><td>AAA+ATPase+wHTH+Alpha+Beta-domain+wHTH+RRM</td><td>Type-2_BR.RM.like.systems</td><td>Phage_integrase-><-HKD-DNase+SF1-SF2-Helicase+iREase+CTD<-METHYLASE<-DUF3780<-DUF499*<-zf-IS66+DDE_Tnp_IS66||Phd-YefM->PIN-><-ABhydrolase<-DDE_Tnp_1</td><td>1046</td><td>Bacteria>Terrabacteria_group>Chloroflexota>unclassified_Chloroflexota</td><td>GCA_016199265.1</td><td>Chloroflexota_bacterium</td><td>HY332_14650</td><td>10576..13716</td><td>2026724</td><td>ATP-binding-protein-[Chloroflexi-bacterium]</td></tr>
<tr><td>MBX6770938.1</td><td>AAA+ATPase+wHTH+Alpha+Beta-domain+wHTH+RRM</td><td>Type-2_BR.RM.like.systems</td><td><-HKD-DNase+SF1-SF2-Helicase+iREase+CTD<-METHYLASE<-SbcC<-DUF3780<-DUF499*</td><td>1016</td><td>Bacteria>Terrabacteria_group>Chloroflexota>unclassified_Chloroflexota</td><td>GCA_019690095.1</td><td>Chloroflexota_bacterium</td><td>IRY83_04380</td><td>23664..26714</td><td>2026724</td><td>ATP-binding-protein-[Chloroflexi-bacterium]</td></tr>
<tr><td>MCY4559676.1</td><td>AAA+ATPase+wHTH+Alpha+Beta-domain+wHTH+RRM</td><td>Type-2_BR.RM.like.systems</td><td><-REase<-REase<-?||SF1-SF2-Helicase->DUF932->XIS-HTH->DUF499*->DUF3780->OLD-ABC->HNH->METHYLASE->HKD-DNase+SF1-SF2-Helicase+iREase+CTD-></td><td>1013</td><td>Bacteria>Terrabacteria_group>Chloroflexota>unclassified_Chloroflexota</td><td>GCA_026713265.1</td><td>Chloroflexota_bacterium</td><td>OXF79_25580</td><td>10138..13179</td><td>2026724</td><td>MAG:-DUF499-domain-containing-protein-[Chloroflexi-bacterium]</td></tr>
<tr><td>MBC8448144.1</td><td>AAA+ATPase+wHTH+Alpha+Beta-domain+wHTH+RRM</td><td>Type-2_BR.RM.like.systems</td><td>PhdYeFM_antitox->RelE-ParE->DUF499*->ASCH+DUF3780->DUF433->PIN-></td><td>1019</td><td>Bacteria>Terrabacteria_group>Chloroflexota>unclassified_Chloroflexota</td><td>GCA_014383605.1</td><td>Chloroflexota_bacterium</td><td>H8D78_10365</td><td>9726..12785</td><td>2026724</td><td>ATP-binding-protein-[Chloroflexi-bacterium]</td></tr>
<tr><td>OQA44952.1</td><td>AAA+ATPase+wHTH+Alpha+Beta-domain+wHTH+RRM</td><td>Type-2_BR.RM.like.systems</td><td><-METHYLASE<-HEPN<-PolB<-PhdYeFM_antitox<-RelE-ParE<-ASCH+DUF3780<-DUF499*</td><td>1029</td><td>Bacteria>Terrabacteria_group>Chloroflexota>unclassified_Chloroflexota</td><td>GCA_002068935.1</td><td>Chloroflexi_bacterium_ADurb.Bin325</td><td>BWY52_01458</td><td>8239..11328</td><td>1852859</td><td>hypothetical-protein-BWY52_01458-[Chloroflexi-bacterium-ADurb.Bin325]</td></tr>
<tr><td>WP_234991458.1</td><td>AAA+ATPase+wHTH+Alpha+Beta-domain+wHTH+RRM</td><td>Type-2_BR.RM.like.systems</td><td><-METHYLASE-p<-METHYLASE-p<-REase<-PIN<-?<-DUF3780<-DUF499*<-?<-GDResolvase+HIN-HTH</td><td>1043</td><td>Bacteria>Terrabacteria_group>Cyanobacteriota/Melainabacteria_group>Cyanobacteriota</td><td>GCF_000019485.1</td><td>Cyanophyceae</td><td>CCK85_RS15790</td><td>115639..118770</td><td>3028117</td><td>MULTISPECIES:-DUF499-domain-containing-protein-[unclassified-Synechococcus]</td></tr>
<tr><td>WP_190626291.1</td><td>AAA+ATPase+wHTH+Alpha+Beta-domain+wHTH+RRM</td><td>Type-2_BR.RM.like.systems</td><td><-NERD+SF1-SF2-Helicase<-METHYLASE||?-><-?<-?<-DUF3780<-DUF499*<-REase<-?<-?||HTH_26-></td><td>1072</td><td>Bacteria>Terrabacteria_group>Cyanobacteriota/Melainabacteria_group>Cyanobacteriota</td><td>GCF_014695595.1</td><td>Cyanophyceae</td><td>H6F55_RS16010</td><td>31589..34807</td><td>3028117</td><td>MULTISPECIES:-DUF499-domain-containing-protein-[Cyanobacteria]</td></tr>
<tr><td>WP_224174984.1</td><td>AAA+ATPase+wHTH+Alpha+Beta-domain+wHTH+RRM</td><td>Type-2_BR.RM.like.systems</td><td><-DDE_Tnp_1<-HKD-DNase+SF1-SF2-Helicase+iREase+CTD<-ParB+HNH<-METHYLASE<-DUF3780<-DUF499*<-CHTH||STYKIN-></td><td>1013</td><td>Bacteria>Terrabacteria_group>Cyanobacteriota/Melainabacteria_group>Cyanobacteriota>Cyanophyceae</td><td>-</td><td>Calothrix_sp._PCC_7716</td><td>CAL7716_066380</td><td>7620880..7623921</td><td>2853227</td><td>DUF499-domain-containing-protein-[Calothrix-sp.-PCC-7716]</td></tr>
<tr><td>CEJ45544.1</td><td>AAA+ATPase+wHTH+Alpha+Beta-domain+wHTH+RRM</td><td>Type-2_BR.RM.like.systems</td><td>URI-DNase-><-HKD-DNase+SF1-SF2-Helicase+iREase+CTD<-?<-METHYLASE<-DUF499*||?->HAD-><-NUDIX<-?||<-?<-Methyltransf_5</td><td>1088</td><td>Bacteria>Terrabacteria_group>Cyanobacteriota/Melainabacteria_group>Cyanobacteriota>Cyanophyceae</td><td>GCA_001458455.1</td><td>Umezakia_ovalisporum</td><td>apha_01996</td><td>54460..57726</td><td>75695</td><td>Putative-ATPase-(AAA+-superfamily)-[Chrysosporum-ovalisporum]</td></tr>
<tr><td>OSO90289.1</td><td>AAA+ATPase+wHTH+Alpha+Beta-domain+wHTH+RRM</td><td>Type-2_BR.RM.like.systems</td><td><-HEPN<-PolB||METHYLASE->HKD-DNase+SF1-SF2-Helicase+iREase+CTD-><-?<-?||DUF499*->DUF3780-></td><td>1075</td><td>Bacteria>Terrabacteria_group>Cyanobacteriota/Melainabacteria_group>Cyanobacteriota>Cyanophyceae</td><td>GCA_002114155.1</td><td>Cylindrospermopsis_raciborskii_CENA303</td><td>B7O87_09645</td><td>23367..26594</td><td>1170769</td><td>AAA-family-ATPase-[Cylindrospermopsis-raciborskii-CENA303]</td></tr>
<tr><td>PSB37932.1</td><td>AAA+ATPase+wHTH+Alpha+Beta-domain+wHTH+RRM</td><td>Type-2_BR.RM.like.systems</td><td>Phage_integrase->DUF499*->DUF3780->METHYLASE->HKD-DNase+SF1-SF2-Helicase+iREase+CTD-></td><td>1086</td><td>Bacteria>Terrabacteria_group>Cyanobacteriota/Melainabacteria_group>Cyanobacteriota>Cyanophyceae</td><td>GCA_003003925.1</td><td>Aphanothece_cf._minutissima_CCALA_015</td><td>C7B81_07485</td><td>152093..155353</td><td>2107695</td><td>hypothetical-protein-C7B81_07485-[Aphanothece-cf.-minutissima-CCALA-015]</td></tr>
<tr><td>PZO60790.1</td><td>AAA+ATPase+wHTH+Alpha+Beta-domain+wHTH+RRM</td><td>Type-2_BR.RM.like.systems</td><td><-RNB+RNase<-DDE_Tnp_1_5+DDE_Tnp_1<-OLD-ABC+TOPRIM||?->DUF499*->DUF3780-><-MACRODOMAIN||METHYLASE->?->PIN->HKD-DNase+SF1-SF2-Helicase+iREase+CTD-></td><td>1079</td><td>Bacteria>Terrabacteria_group>Cyanobacteriota/Melainabacteria_group>Cyanobacteriota>Cyanophyceae</td><td>GCA_003242115.1</td><td>Phormidesmis_priestleyi</td><td>DCF15_01370</td><td>14594..17833</td><td>268141</td><td>AAA-family-ATPase-[Phormidesmis-priestleyi]</td></tr>
<tr><td>PZV04176.1</td><td>AAA+ATPase+wHTH+Alpha+Beta-domain+wHTH+RRM</td><td>Type-2_BR.RM.like.systems</td><td><-HEPN<-HEPN||?->DUF499*->DUF3780->?->METHYLASE->HKD-DNase+SF1-SF2-Helicase+iREase+CTD->RNaseH+RNaseH-></td><td>1009</td><td>Bacteria>Terrabacteria_group>Cyanobacteriota/Melainabacteria_group>Cyanobacteriota>Cyanophyceae</td><td>GCA_003248995.1</td><td>Cyanobium_sp.</td><td>DCF23_07060</td><td>23229..26258</td><td>2164130</td><td>hypothetical-protein-DCF23_07060-[Cyanobium-sp.]</td></tr>
<tr><td>TVQ60424.1</td><td>AAA+ATPase+wHTH+Alpha+Beta-domain+wHTH+RRM</td><td>Type-2_BR.RM.like.systems</td><td>DUF499*->DUF3780->HKD-DNase+SF1-SF2-Helicase+iREase+CTD-></td><td>1080</td><td>Bacteria>Terrabacteria_group>Cyanobacteriota/Melainabacteria_group>Cyanobacteriota>Cyanophyceae</td><td>GCA_007693875.1</td><td>Spirulina_sp._DLM2.Bin59</td><td>EA366_03660</td><td>2326..5568</td><td>2480215</td><td>ATP-binding-protein-[Spirulina-sp.-DLM2.Bin59]</td></tr>
<tr><td>NJN12005.1</td><td>AAA+ATPase+wHTH+Alpha+Beta-domain+wHTH+RRM</td><td>Type-2_BR.RM.like.systems</td><td>MazE_antitoxin->Phd-YefM->MazE_antitoxin->PIN-><-?||DUF499*->DUF3780->?->?-><-REase||METHYLASE->HKD-DNase+SF1-SF2-Helicase+iREase+CTD-></td><td>1079</td><td>Bacteria>Terrabacteria_group>Cyanobacteriota/Melainabacteria_group>Cyanobacteriota>Cyanophyceae</td><td>GCA_012032385.1</td><td>Richelia_sp._RM1_1_1</td><td>HC815_30215</td><td>3465..6704</td><td>2720426</td><td>ATP-binding-protein-[Richelia-sp.-RM1_1_1]</td></tr>
<tr><td>MBD1193924.1</td><td>AAA+ATPase+wHTH+Alpha+Beta-domain+wHTH+RRM</td><td>Type-2_BR.RM.like.systems</td><td>DUF499*->DUF3780->?->METHYLASE->HKD-DNase+SF1-SF2-Helicase+iREase+CTD-></td><td>1033</td><td>Bacteria>Terrabacteria_group>Cyanobacteriota/Melainabacteria_group>Cyanobacteriota>Cyanophyceae</td><td>GCA_014654725.1</td><td>Vulcanococcus_sp._Clear-D1</td><td>H9532_07300</td><td>91187..94288</td><td>2766970</td><td>ATP-binding-protein-[Vulcanococcus-sp.-Clear-D1]</td></tr>
<tr><td>MBD1812659.1</td><td>AAA+ATPase+wHTH+Alpha+Beta-domain+wHTH+RRM</td><td>Type-2_BR.RM.like.systems</td><td><-RelE-ParE<-PhdYeFM_antitox<-?<-HKD-DNase+SF1-SF2-Helicase+iREase+CTD<-METHYLASE<-DUF3780<-DUF499*</td><td>1075</td><td>Bacteria>Terrabacteria_group>Cyanobacteriota/Melainabacteria_group>Cyanobacteriota>Cyanophyceae</td><td>GCA_014695175.1</td><td>Microcoleus_sp._FACHB-DQ6</td><td>H6F65_13345</td><td>15046..18273</td><td>2692829</td><td>ATP-binding-protein-[Microcoleus-sp.-FACHB-DQ6]</td></tr>
<tr><td>MBD2438833.1</td><td>AAA+ATPase+wHTH+Alpha+Beta-domain+wHTH+RRM</td><td>Type-2_BR.RM.like.systems</td><td><-Phage_int_SAM_1+Phage_integrase||?->?->?->DUF499*->DUF3780->METHYLASE->ParB+HNH->HKD-DNase+SF1-SF2-Helicase+iREase+CTD-></td><td>1019</td><td>Bacteria>Terrabacteria_group>Cyanobacteriota/Melainabacteria_group>Cyanobacteriota>Cyanophyceae</td><td>GCA_014697425.1</td><td>Nostoc_sp._FACHB-110</td><td>H6G69_18770</td><td>61408..64467</td><td>2692834</td><td>ATP-binding-protein-[Nostoc-sp.-FACHB-110]</td></tr>
<tr><td>MBD2467864.1</td><td>AAA+ATPase+wHTH+Alpha+Beta-domain+wHTH+RRM</td><td>Type-2_BR.RM.like.systems</td><td><-CHTH<-TPR||<-?||?->?->||DUF499*->DUF3780->METHYLASE->ParB+HNH->HKD-DNase+SF1-SF2-Helicase+iREase+CTD-></td><td>1019</td><td>Bacteria>Terrabacteria_group>Cyanobacteriota/Melainabacteria_group>Cyanobacteriota>Cyanophyceae</td><td>GCA_014697455.1</td><td>Nostoc_sp._FACHB-145</td><td>H6G90_09455</td><td>23220..26279</td><td>2692836</td><td>ATP-binding-protein-[Nostoc-sp.-FACHB-145]</td></tr>
<tr><td>MBE9018013.1</td><td>AAA+ATPase+wHTH+Alpha+Beta-domain+wHTH+RRM</td><td>Type-2_BR.RM.like.systems</td><td><-HKD-DNase+SF1-SF2-Helicase+iREase+CTD<-ParB+HNH<-METHYLASE<-DUF3780<-DUF499*</td><td>1026</td><td>Bacteria>Terrabacteria_group>Cyanobacteriota/Melainabacteria_group>Cyanobacteriota>Cyanophyceae</td><td>GCA_015206905.1</td><td>Chroococcidiopsidales_cyanobacterium_LEGE_13417</td><td>IQ272_18060</td><td>6249..9329</td><td>1828829</td><td>ATP-binding-protein-[Chroococcidiopsidales-cyanobacterium-LEGE-13417]</td></tr>
<tr><td>MBE9044407.1</td><td>AAA+ATPase+wHTH+Alpha+Beta-domain+wHTH+RRM</td><td>Type-2_BR.RM.like.systems</td><td>METHYLASE->?-><-REase||HKD-DNase+SF1-SF2-Helicase+iREase+CTD->?-><-DUF3780<-DUF499*</td><td>1077</td><td>Bacteria>Terrabacteria_group>Cyanobacteriota/Melainabacteria_group>Cyanobacteriota>Cyanophyceae</td><td>GCA_015207195.1</td><td>Pleurocapsales_cyanobacterium_LEGE_10410</td><td>IQ255_08325</td><td>15194..18427</td><td>1828828</td><td>ATP-binding-protein-[Pleurocapsales-cyanobacterium-LEGE-10410]</td></tr>
<tr><td>MBN3957901.1</td><td>AAA+ATPase+wHTH+Alpha+Beta-domain+wHTH+RRM</td><td>Type-2_BR.RM.like.systems</td><td><-HKD-DNase+SF1-SF2-Helicase+iREase+CTD<-PIN<-?<-?<-METHYLASE<-DUF3780<-DUF499*<-SF1-SF2-Helicase||ParB+HNH->Pkinase+SF1-SF2-Helicase->HTH+HKD-DNase->||?-><-PAPS_reduct</td><td>1078</td><td>Bacteria>Terrabacteria_group>Cyanobacteriota/Melainabacteria_group>Cyanobacteriota>Cyanophyceae</td><td>GCA_017115045.1</td><td>Nostoc_sp._NMS8</td><td>HWQ39_03310</td><td>19363..22599</td><td>2815392</td><td>ATP-binding-protein-[Nostoc-sp.-NMS8]</td></tr>
<tr><td>BDA72472.1</td><td>AAA+ATPase+wHTH+Alpha+Beta-domain+wHTH+RRM</td><td>Type-2_BR.RM.like.systems</td><td><-DDE_Tnp_1<-HKD-DNase+SF1-SF2-Helicase+iREase+CTD<-ParB+HNH<-METHYLASE<-DUF3780<-DUF499*<-CHTH||STYKIN-></td><td>1013</td><td>Bacteria>Terrabacteria_group>Cyanobacteriota/Melainabacteria_group>Cyanobacteriota>Cyanophyceae</td><td>GCA_019977735.1</td><td>Calothrix_sp._PCC_7716</td><td>CAL7716_066380</td><td>7620880..7623921</td><td>2853227</td><td>hypothetical-protein-CAL7716_066380-[Calothrix-sp.-PCC-7716]</td></tr>
<tr><td>MCC5898057.1</td><td>AAA+ATPase+wHTH+Alpha+Beta-domain+wHTH+RRM</td><td>Type-2_BR.RM.like.systems</td><td><-HKD-DNase+SF1-SF2-Helicase+iREase+CTD<-METHYLASE<-DUF3780<-DUF499*<-REase<-REase</td><td>1079</td><td>Bacteria>Terrabacteria_group>Cyanobacteriota/Melainabacteria_group>Cyanobacteriota>Cyanophyceae</td><td>GCA_020832215.1</td><td>Phormidium_sp._BM_Day4_Bin.17</td><td>JJU32_09090</td><td>28948..32187</td><td>2813767</td><td>ATP-binding-protein-[Phormidium-sp.-BM_Day4_Bin.17]</td></tr>
<tr><td>MCD8487854.1</td><td>AAA+ATPase+wHTH+Alpha+Beta-domain+wHTH+RRM</td><td>Type-2_BR.RM.like.systems</td><td><-HKD-DNase+SF1-SF2-Helicase+iREase+CTD<-PIN<-?<-METHYLASE<-DUF3780<-REase<-DUF499*</td><td>1077</td><td>Bacteria>Terrabacteria_group>Cyanobacteriota/Melainabacteria_group>Cyanobacteriota>Cyanophyceae</td><td>GCA_021208125.1</td><td>Desertifilum_sp.</td><td>LRZ84_14250</td><td>9885..13118</td><td>2497424</td><td>DUF499-domain-containing-protein-[Desertifilum-sp.]</td></tr>
<tr><td>UNU27023.1</td><td>AAA+ATPase+wHTH+Alpha+Beta-domain+wHTH+RRM</td><td>Type-2_BR.RM.like.systems</td><td><-AbrB-like<-?<-?<-?||?->DUF499*->DUF3780-><-?||?->METHYLASE->HKD-DNase+SF1-SF2-Helicase+iREase+CTD-></td><td>1075</td><td>Bacteria>Terrabacteria_group>Cyanobacteriota/Melainabacteria_group>Cyanobacteriota>Cyanophyceae</td><td>GCA_022701255.1</td><td>Microcoleus_vaginatus_HSN003</td><td>DOA37_27340</td><td>6455094..6458321</td><td>2303329</td><td>ATP-binding-protein-[Microcoleus-vaginatus-HSN003]</td></tr>
<tr><td>WP_009789601.1</td><td>AAA+ATPase+wHTH+Alpha+Beta-domain+wHTH+RRM</td><td>Type-2_BR.RM.like.systems</td><td><-HKD-DNase+SF1-SF2-Helicase+iREase+CTD<-METHYLASE<-DUF3780<-DUF499*<-REase+35exo+Beta-Barrel+SF1-SF2-Helicase||?-><-HD</td><td>1027</td><td>Bacteria>Terrabacteria_group>Cyanobacteriota/Melainabacteria_group>Cyanobacteriota>Cyanophyceae</td><td>GCF_000153805.1</td><td>Synechococcus_sp._BL107</td><td>BL107_RS07010</td><td>1349787..1352870</td><td>313625</td><td>DUF499-domain-containing-protein-[Synechococcus-sp.-BL107]</td></tr>
<tr><td>WP_015178384.1</td><td>AAA+ATPase+wHTH+Alpha+Beta-domain+wHTH+RRM</td><td>Type-2_BR.RM.like.systems</td><td><-HKD-DNase+SF1-SF2-Helicase+iREase+CTD<-?<-METHYLASE||?-><-DUF3780<-DUF499*</td><td>1080</td><td>Bacteria>Terrabacteria_group>Cyanobacteriota/Melainabacteria_group>Cyanobacteriota>Cyanophyceae</td><td>GCF_000317475.1</td><td>Oscillatoria_nigro-viridis</td><td>OSC7112_RS24310</td><td>5835081..5838323</td><td>482564</td><td>DUF499-domain-containing-protein-[Oscillatoria-nigro-viridis]</td></tr>
<tr><td>WP_052378119.1</td><td>AAA+ATPase+wHTH+Alpha+Beta-domain+wHTH+RRM</td><td>Type-2_BR.RM.like.systems</td><td><-MerR-HTH||?->?-><-HKD-DNase+SF1-SF2-Helicase+iREase+CTD<-METHYLASE<-DUF3780<-DUF499*<-REase+35exo+Beta-Barrel+SF1-SF2-Helicase<-?<-?<-?<-35exo</td><td>1015</td><td>Bacteria>Terrabacteria_group>Cyanobacteriota/Melainabacteria_group>Cyanobacteriota>Cyanophyceae</td><td>GCF_000737575.1</td><td>Synechococcus_sp._KORDI-49</td><td>KR49_RS14345</td><td>181511..184558</td><td>585423</td><td>DUF499-domain-containing-protein-[Synechococcus-sp.-KORDI-49]</td></tr>
<tr><td>WP_054466655.1</td><td>AAA+ATPase+wHTH+Alpha+Beta-domain+wHTH+RRM</td><td>Type-2_BR.RM.like.systems</td><td><-HKD-DNase+SF1-SF2-Helicase+iREase+CTD<-METHYLASE<-DUF3780<-DUF499*<-?||?->?-><-?||Calcineurin->SbcC-></td><td>1110</td><td>Bacteria>Terrabacteria_group>Cyanobacteriota/Melainabacteria_group>Cyanobacteriota>Cyanophyceae</td><td>GCF_001276715.1</td><td>Planktothricoides_sp._SR001</td><td>AM228_RS13635</td><td>37590..40922</td><td>1705388</td><td>DUF499-domain-containing-protein-[Planktothricoides-sp.-SR001]</td></tr>
<tr><td>WP_057177419.1</td><td>AAA+ATPase+wHTH+Alpha+Beta-domain+wHTH+RRM</td><td>Type-2_BR.RM.like.systems</td><td>HKD-DNase+SF1-SF2-Helicase+iREase+CTD->DUF499*-></td><td>719</td><td>Bacteria>Terrabacteria_group>Cyanobacteriota/Melainabacteria_group>Cyanobacteriota>Cyanophyceae</td><td>GCF_001432185.1</td><td>Cylindrospermopsis_sp._CR12</td><td>ASL19_RS04260</td><td>5486..7644</td><td>1747196</td><td>DUF499-domain-containing-protein,-partial-[Cylindrospermopsis-sp.-CR12]</td></tr>
<tr><td>WP_179147480.1</td><td>AAA+ATPase+wHTH+Alpha+Beta-domain+wHTH+RRM</td><td>Type-2_BR.RM.like.systems</td><td>PIN-><-?<-HEPN<-PolB||METHYLASE->HKD-DNase+SF1-SF2-Helicase+iREase+CTD->DUF499*->DUF3780-></td><td>1090</td><td>Bacteria>Terrabacteria_group>Cyanobacteriota/Melainabacteria_group>Cyanobacteriota>Cyanophyceae</td><td>GCF_002114155.1</td><td>Cylindrospermopsis_raciborskii</td><td>B7O87_RS09640</td><td>23322..26594</td><td>77022</td><td>DUF499-domain-containing-protein-[Cylindrospermopsis-raciborskii]</td></tr>
<tr><td>WP_181280075.1</td><td>AAA+ATPase+wHTH+Alpha+Beta-domain+wHTH+RRM</td><td>Type-2_BR.RM.like.systems</td><td>Phage_integrase-><-?||DUF499*->DUF3780->METHYLASE->HKD-DNase+SF1-SF2-Helicase+iREase+CTD->?->?->Pkinase-></td><td>1032</td><td>Bacteria>Terrabacteria_group>Cyanobacteriota/Melainabacteria_group>Cyanobacteriota>Cyanophyceae</td><td>GCF_003003925.1</td><td>Aphanothece_minutissima</td><td>C7B81_RS07500</td><td>152255..155353</td><td>543815</td><td>DUF499-domain-containing-protein-[Aphanothece-minutissima]</td></tr>
<tr><td>WP_236107436.1</td><td>AAA+ATPase+wHTH+Alpha+Beta-domain+wHTH+RRM</td><td>Type-2_BR.RM.like.systems</td><td><-DDE_Tnp_IS1||METHYLASE->HKD-DNase+SF1-SF2-Helicase+iREase+CTD->PAIREDC-HTH->DDE_3->DUF499*->DUF3780-><-?||MORC-></td><td>1108</td><td>Bacteria>Terrabacteria_group>Cyanobacteriota/Melainabacteria_group>Cyanobacteriota>Cyanophyceae</td><td>GCF_003367075.2</td><td>Cylindrospermopsis_raciborskii</td><td>C6N34_RS05820</td><td>1276239..1279565</td><td>77022</td><td>DUF499-domain-containing-protein-[Cylindrospermopsis-raciborskii]</td></tr>
<tr><td>WP_181495442.1</td><td>AAA+ATPase+wHTH+Alpha+Beta-domain+wHTH+RRM</td><td>Type-2_BR.RM.like.systems</td><td><-REase<-?||?->?-><-Peptidase<-?||DUF499*->DUF3780-><-REase||?->METHYLASE->HKD-DNase+SF1-SF2-Helicase+iREase+CTD-></td><td>1012</td><td>Bacteria>Terrabacteria_group>Cyanobacteriota/Melainabacteria_group>Cyanobacteriota>Cyanophyceae</td><td>GCF_003555505.1</td><td>Thermosynechococcus_vestitus</td><td>D3A95_RS00385</td><td>85196..88234</td><td>146786</td><td>DUF499-domain-containing-protein-[Thermosynechococcus-vestitus]</td></tr>
<tr><td>WP_258316377.1</td><td>AAA+ATPase+wHTH+Alpha+Beta-domain+wHTH+RRM</td><td>Type-2_BR.RM.like.systems</td><td><-MORC||?-><-DUF3780<-?<-?<-DUF499*<-DUF499<-HKD-DNase+SF1-SF2-Helicase-p<-?<-?<-?<-SF1-SF2-Helicase-p</td><td>609</td><td>Bacteria>Terrabacteria_group>Cyanobacteriota/Melainabacteria_group>Cyanobacteriota>Cyanophyceae</td><td>GCF_006523545.1</td><td>Cylindrospermopsis_raciborskii</td><td>FIV49_RS17545</td><td>245967..247796</td><td>77022</td><td>DUF499-domain-containing-protein-[Cylindrospermopsis-raciborskii]</td></tr>
<tr><td>WP_155084576.1</td><td>AAA+ATPase+wHTH+Alpha+Beta-domain+wHTH+RRM</td><td>Type-2_BR.RM.like.systems</td><td><-Primpol||?->Phage_int_SAM_4+Phage_integrase->DUF499*->REase+NACHT-><-METHYLASE<-DUF3780||DUF433->PIN->HKD-DNase+SF1-SF2-Helicase+iREase+CTD-></td><td>1100</td><td>Bacteria>Terrabacteria_group>Cyanobacteriota/Melainabacteria_group>Cyanobacteriota>Cyanophyceae</td><td>GCF_009710015.1</td><td>Cyanobacterium_aponinum</td><td>GGC33_RS15815</td><td>17362..20664</td><td>379064</td><td>DUF499-domain-containing-protein-[Cyanobacterium-aponinum]</td></tr>
<tr><td>WP_168466096.1</td><td>AAA+ATPase+wHTH+Alpha+Beta-domain+wHTH+RRM</td><td>Type-2_BR.RM.like.systems</td><td>DUF29->DUF3780->?->DUF499*->DUF3780->?->REase->METHYLASE->?->HKD-DNase+SF1-SF2-Helicase+iREase+CTD-></td><td>1088</td><td>Bacteria>Terrabacteria_group>Cyanobacteriota/Melainabacteria_group>Cyanobacteriota>Cyanophyceae</td><td>GCF_009712065.1</td><td>Aphanizomenon_sp._UHCC_0183</td><td>FJR37_RS11780</td><td>50771..54037</td><td>2590028</td><td>DUF499-domain-containing-protein-[Aphanizomenon-sp.-UHCC-0183]</td></tr>
<tr><td>WP_168359778.1</td><td>AAA+ATPase+wHTH+Alpha+Beta-domain+wHTH+RRM</td><td>Type-2_BR.RM.like.systems</td><td><-Pterin_bind<-?<-NUDIX<-Glyoxalase||DUF499*->DUF3780->HNH->?->METHYLASE-p->METHYLASE-p->HKD-DNase+SF1-SF2-Helicase+iREase+CTD-></td><td>1093</td><td>Bacteria>Terrabacteria_group>Cyanobacteriota/Melainabacteria_group>Cyanobacteriota>Cyanophyceae</td><td>GCF_009712075.2</td><td>Dolichospermum_planctonicum</td><td>FJR41_RS17240</td><td>43574..46855</td><td>136072</td><td>DUF499-domain-containing-protein-[Dolichospermum-planctonicum]</td></tr>
<tr><td>WP_179052056.1</td><td>AAA+ATPase+wHTH+Alpha+Beta-domain+wHTH+RRM</td><td>Type-2_BR.RM.like.systems</td><td>PIN-><-HKD-DNase+SF1-SF2-Helicase+iREase+CTD<-PIN<-?<-METHYLASE<-DUF3780<-DUF499*<-TPR<-?<-sGTP<-MetJarc-HTH||ArdA-></td><td>1017</td><td>Bacteria>Terrabacteria_group>Cyanobacteriota/Melainabacteria_group>Cyanobacteriota>Cyanophyceae</td><td>GCF_013393945.1</td><td>Nostoc_sp._TCL26-01</td><td>FD725_RS31200</td><td>89491..92544</td><td>2576904</td><td>DUF499-domain-containing-protein-[Nostoc-sp.-TCL26-01]</td></tr>
<tr><td>WP_187705251.1</td><td>AAA+ATPase+wHTH+Alpha+Beta-domain+wHTH+RRM</td><td>Type-2_BR.RM.like.systems</td><td><-CRP-HTH||?->?->METHYLASE->HKD-DNase+SF1-SF2-Helicase+iREase+CTD->DUF499*->DUF3780-><-?||MORC-></td><td>1092</td><td>Bacteria>Terrabacteria_group>Cyanobacteriota/Melainabacteria_group>Cyanobacteriota>Cyanophyceae</td><td>GCF_014489415.1</td><td>Cylindrospermopsis_curvispora</td><td>IAR63_RS10645</td><td>2396129..2399407</td><td>747548</td><td>DUF499-domain-containing-protein-[Cylindrospermopsis-curvispora]</td></tr>
<tr><td>WP_187705597.1</td><td>AAA+ATPase+wHTH+Alpha+Beta-domain+wHTH+RRM</td><td>Type-2_BR.RM.like.systems</td><td><-REase<-?<-?<-?<-DUF3780<-DUF499*<-HKD-DNase+SF1-SF2-Helicase+iREase+CTD<-METHYLASE</td><td>1092</td><td>Bacteria>Terrabacteria_group>Cyanobacteriota/Melainabacteria_group>Cyanobacteriota>Cyanophyceae</td><td>GCF_014489415.1</td><td>Cylindrospermopsis_curvispora</td><td>IAR63_RS13335</td><td>2992458..2995736</td><td>747548</td><td>DUF499-domain-containing-protein-[Cylindrospermopsis-curvispora]</td></tr>
<tr><td>WP_199338599.1</td><td>AAA+ATPase+wHTH+Alpha+Beta-domain+wHTH+RRM</td><td>Type-2_BR.RM.like.systems</td><td><-CHTH<-TPR||<-?||?->?->||DUF499*->DUF3780->METHYLASE->ParB+HNH->HKD-DNase+SF1-SF2-Helicase+iREase+CTD-></td><td>1045</td><td>Bacteria>Terrabacteria_group>Cyanobacteriota/Melainabacteria_group>Cyanobacteriota>Cyanophyceae</td><td>GCF_014697455.1</td><td>Nostoc_sp._FACHB-145</td><td>H6G90_RS09455</td><td>23142..26279</td><td>2692836</td><td>DUF499-domain-containing-protein-[Nostoc-sp.-FACHB-145]</td></tr>
<tr><td>WP_193877251.1</td><td>AAA+ATPase+wHTH+Alpha+Beta-domain+wHTH+RRM</td><td>Type-2_BR.RM.like.systems</td><td>DUF499*->DUF3780->METHYLASE->HKD-DNase+SF1-SF2-Helicase+iREase+CTD-><-RHH_1||Tn7_Tnp_TnsA_N+Tn7_Tnp_TnsA_C-></td><td>1024</td><td>Bacteria>Terrabacteria_group>Cyanobacteriota/Melainabacteria_group>Cyanobacteriota>Cyanophyceae</td><td>GCF_015207605.1</td><td>Oculatella_sp._LEGE_06141</td><td>IQ268_RS07955</td><td>1375..4449</td><td>1828648</td><td>DUF499-domain-containing-protein-[Oculatella-sp.-LEGE-06141]</td></tr>
<tr><td>WP_193799565.1</td><td>AAA+ATPase+wHTH+Alpha+Beta-domain+wHTH+RRM</td><td>Type-2_BR.RM.like.systems</td><td>SF1-SF2-Helicase-><-SF1-SF2-Helicase||ABC-ATPase+SbcC->?->DUF499*-><-Y1_Tnp||?-><-METHYLASE||?-><-DUF3780||HKD-DNase+SF1-SF2-Helicase+iREase+CTD-></td><td>1091</td><td>Bacteria>Terrabacteria_group>Cyanobacteriota/Melainabacteria_group>Cyanobacteriota>Cyanophyceae</td><td>GCF_015207825.1</td><td>Cyanobacterium_stanieri</td><td>IQ215_RS01555</td><td>126055..129330</td><td>102235</td><td>DUF499-domain-containing-protein-[Cyanobacterium-stanieri]</td></tr>
<tr><td>WP_211941997.1</td><td>AAA+ATPase+wHTH+Alpha+Beta-domain+wHTH+RRM</td><td>Type-2_BR.RM.like.systems</td><td>SbcC-><-CRP-HTH||?->?->METHYLASE->HKD-DNase+SF1-SF2-Helicase+iREase+CTD->DUF499*->DUF3780-><-?||MORC-></td><td>1092</td><td>Bacteria>Terrabacteria_group>Cyanobacteriota/Melainabacteria_group>Cyanobacteriota>Cyanophyceae</td><td>GCF_018139025.1</td><td>Cylindrospermopsis_raciborskii</td><td>J9B92_RS16930</td><td>3783988..3787266</td><td>77022</td><td>DUF499-domain-containing-protein-[Cylindrospermopsis-raciborskii]</td></tr>
<tr><td>WP_236308314.1</td><td>AAA+ATPase+wHTH+Alpha+Beta-domain+wHTH+RRM</td><td>Type-2_BR.RM.like.systems</td><td>SbcC-><-CRP-HTH||?->?->METHYLASE->HKD-DNase+SF1-SF2-Helicase+iREase+CTD->DUF499*->DUF3780-><-?||MORC-></td><td>1092</td><td>Bacteria>Terrabacteria_group>Cyanobacteriota/Melainabacteria_group>Cyanobacteriota>Cyanophyceae</td><td>GCF_021650815.1</td><td>Cylindrospermopsis_raciborskii</td><td>L3I90_RS14150</td><td>3213455..3216733</td><td>77022</td><td>DUF499-domain-containing-protein-[Cylindrospermopsis-raciborskii]</td></tr>
<tr><td>WP_240866046.1</td><td>AAA+ATPase+wHTH+Alpha+Beta-domain+wHTH+RRM</td><td>Type-2_BR.RM.like.systems</td><td>METHYLASE-p->METHYLASE-p->HKD-DNase+SF1-SF2-Helicase+iREase+CTD-p->DUF499*->DUF3780-><-?||MORC-></td><td>1092</td><td>Bacteria>Terrabacteria_group>Cyanobacteriota/Melainabacteria_group>Cyanobacteriota>Cyanophyceae</td><td>GCF_022496815.1</td><td>Cylindrospermopsis_raciborskii</td><td>GSN00_RS02230</td><td>6039..9317</td><td>77022</td><td>DUF499-domain-containing-protein-[Cylindrospermopsis-raciborskii]</td></tr>
<tr><td>WP_252662978.1</td><td>AAA+ATPase+wHTH+Alpha+Beta-domain+wHTH+RRM</td><td>Type-2_BR.RM.like.systems</td><td><-HKD-DNase+SF1-SF2-Helicase+iREase+CTD<-METHYLASE<-?<-?<-DUF3780<-DUF499*<-?<-PIN<-DUF433<-MetJarc-HTH<-?<-HEPN</td><td>1079</td><td>Bacteria>Terrabacteria_group>Cyanobacteriota/Melainabacteria_group>Cyanobacteriota>Cyanophyceae</td><td>GCF_023983615.1</td><td>Phormidium_yuhuli</td><td>NEA10_RS19360</td><td>4490827..4494066</td><td>2974039</td><td>DUF499-domain-containing-protein-[Phormidium-yuhuli]</td></tr>
<tr><td>WP_259702440.1</td><td>AAA+ATPase+wHTH+Alpha+Beta-domain+wHTH+RRM</td><td>Type-2_BR.RM.like.systems</td><td><-RNaseH+RNaseH<-HKD-DNase+SF1-SF2-Helicase+iREase+CTD<-METHYLASE<-?<-DUF3780<-DUF499*<-?||HEPN->HEPN-></td><td>1009</td><td>Bacteria>Terrabacteria_group>Cyanobacteriota/Melainabacteria_group>Cyanobacteriota>Cyanophyceae</td><td>GCF_025128155.1</td><td>Synechococcus_sp._CS-1324</td><td>KQ306_RS08900</td><td>95996..99025</td><td>2847980</td><td>DUF499-domain-containing-protein-[Synechococcus-sp.-CS-1324]</td></tr>
<tr><td>WP_272033246.1</td><td>AAA+ATPase+wHTH+Alpha+Beta-domain+wHTH+RRM</td><td>Type-2_BR.RM.like.systems</td><td><-HKD-DNase+SF1-SF2-Helicase+iREase+CTD<-REase<-METHYLASE<-?<-DUF3780<-REase<-DUF499*||?-><-ACET<-?<-RHH<-RHH</td><td>1079</td><td>Bacteria>Terrabacteria_group>Cyanobacteriota/Melainabacteria_group>Cyanobacteriota>Cyanophyceae</td><td>GCF_028330895.1</td><td>Kamptonema_animale</td><td>PN499_RS16795</td><td>10473..13712</td><td>92934</td><td>DUF499-domain-containing-protein-[Kamptonema-animale]</td></tr>
<tr><td>WP_280701012.1</td><td>AAA+ATPase+wHTH+Alpha+Beta-domain+wHTH+RRM</td><td>Type-2_BR.RM.like.systems</td><td>URI-DNase->?-><-HKD-DNase+SF1-SF2-Helicase+iREase+CTD<-REase<-METHYLASE<-HicA_toxin<-DUF499*||?->HAD-><-NUDIX</td><td>1088</td><td>Bacteria>Terrabacteria_group>Cyanobacteriota/Melainabacteria_group>Cyanobacteriota>Cyanophyceae</td><td>GCF_029891755.1</td><td>Umezakia_ovalisporum</td><td>NWP23_RS16815</td><td>9300..12566</td><td>75695</td><td>DUF499-domain-containing-protein-[Umezakia-ovalisporum]</td></tr>
<tr><td>WP_280686613.1</td><td>AAA+ATPase+wHTH+Alpha+Beta-domain+wHTH+RRM</td><td>Type-2_BR.RM.like.systems</td><td>URI-DNase-><-HKD-DNase+SF1-SF2-Helicase+iREase+CTD<-REase<-METHYLASE<-?<-HicA_toxin<-DUF499*||?->HAD-><-NUDIX</td><td>1088</td><td>Bacteria>Terrabacteria_group>Cyanobacteriota/Melainabacteria_group>Cyanobacteriota>Cyanophyceae</td><td>GCF_029891765.1</td><td>Umezakia_ovalisporum</td><td>NWP30_RS07830</td><td>9432..12698</td><td>75695</td><td>DUF499-domain-containing-protein-[Umezakia-ovalisporum]</td></tr>
<tr><td>WP_280657425.1</td><td>AAA+ATPase+wHTH+Alpha+Beta-domain+wHTH+RRM</td><td>Type-2_BR.RM.like.systems</td><td>URI-DNase-><-HKD-DNase+SF1-SF2-Helicase+iREase+CTD<-REase<-METHYLASE<-?<-HicA_toxin<-DUF499*||?->HAD-><-NUDIX</td><td>1088</td><td>Bacteria>Terrabacteria_group>Cyanobacteriota/Melainabacteria_group>Cyanobacteriota>Cyanophyceae</td><td>GCF_029891785.1</td><td>Umezakia_ovalisporum</td><td>NWP24_RS13330</td><td>9520..12786</td><td>75695</td><td>DUF499-domain-containing-protein-[Umezakia-ovalisporum]</td></tr>
<tr><td>WP_280688181.1</td><td>AAA+ATPase+wHTH+Alpha+Beta-domain+wHTH+RRM</td><td>Type-2_BR.RM.like.systems</td><td>NUDIX-><-HAD<-?||DUF499*->HicA_toxin->METHYLASE->REase->HKD-DNase+SF1-SF2-Helicase+iREase+CTD-><-URI-DNase</td><td>1102</td><td>Bacteria>Terrabacteria_group>Cyanobacteriota/Melainabacteria_group>Cyanobacteriota>Cyanophyceae</td><td>GCF_029891835.1</td><td>Umezakia_ovalisporum</td><td>NWP16_RS11415</td><td>21389..24697</td><td>75695</td><td>DUF499-domain-containing-protein-[Umezakia-ovalisporum]</td></tr>
<tr><td>WP_280692849.1</td><td>AAA+ATPase+wHTH+Alpha+Beta-domain+wHTH+RRM</td><td>Type-2_BR.RM.like.systems</td><td>URI-DNase-><-HKD-DNase+SF1-SF2-Helicase+iREase+CTD<-REase<-METHYLASE<-HicA_toxin<-DUF499*||?->HAD-><-NUDIX</td><td>1088</td><td>Bacteria>Terrabacteria_group>Cyanobacteriota/Melainabacteria_group>Cyanobacteriota>Cyanophyceae</td><td>GCF_029891915.1</td><td>Umezakia_ovalisporum</td><td>NWP18_RS15355</td><td>9410..12676</td><td>75695</td><td>DUF499-domain-containing-protein-[Umezakia-ovalisporum]</td></tr>
<tr><td>WP_280702141.1</td><td>AAA+ATPase+wHTH+Alpha+Beta-domain+wHTH+RRM</td><td>Type-2_BR.RM.like.systems</td><td>URI-DNase-><-HKD-DNase+SF1-SF2-Helicase+iREase+CTD<-REase<-METHYLASE<-?<-HicA_toxin<-DUF499*||?->HAD-><-NUDIX</td><td>1088</td><td>Bacteria>Terrabacteria_group>Cyanobacteriota/Melainabacteria_group>Cyanobacteriota>Cyanophyceae</td><td>GCF_029891925.1</td><td>Umezakia_ovalisporum</td><td>NWP26_RS18515</td><td>9506..12772</td><td>75695</td><td>DUF499-domain-containing-protein-[Umezakia-ovalisporum]</td></tr>
<tr><td>WP_281009999.1</td><td>AAA+ATPase+wHTH+Alpha+Beta-domain+wHTH+RRM</td><td>Type-2_BR.RM.like.systems</td><td><-HKD-DNase+SF1-SF2-Helicase+iREase+CTD<-REase<-METHYLASE||MetJarc-HTH->PemK_toxin-><-DUF3780<-DUF499*||?->?-><-CopG/RHH</td><td>1074</td><td>Bacteria>Terrabacteria_group>Cyanobacteriota/Melainabacteria_group>Cyanobacteriota>Cyanophyceae</td><td>GCF_029910235.1</td><td>Pseudanabaena_galeata</td><td>OA858_RS25990</td><td>135028..138252</td><td>1112103</td><td>DUF499-domain-containing-protein-[Pseudanabaena-galeata]</td></tr>
<tr><td>WP_114988660.1</td><td>AAA+ATPase+wHTH+Alpha+Beta-domain+wHTH+RRM</td><td>Type-2_BR.RM.like.systems</td><td>REase-><-HTH_17<-HKD-DNase+SF1-SF2-Helicase+iREase+CTD<-METHYLASE<-DUF3780<-DUF499*<-?<-?<-?<-?||THUMP+Methylase-></td><td>1027</td><td>Bacteria>Terrabacteria_group>Cyanobacteriota/Melainabacteria_group>Cyanobacteriota>Cyanophyceae</td><td>GCF_900474045.1</td><td>Synechococcus_sp._N19</td><td>DXY02_RS06950</td><td>10966..14049</td><td>2575512</td><td>DUF499-domain-containing-protein-[Synechococcus-sp.-N19]</td></tr>
<tr><td>WP_254033282.1</td><td>AAA+ATPase+wHTH+Alpha+Beta-domain+wHTH+RRM</td><td>Type-2_BR.RM.like.systems</td><td>SAM-methylase->?->?->?->DUF499*->DUF3780->?->PIN->METHYLASE->METHYLASE-p->HKD-DNase+SF1-SF2-Helicase+iREase+CTD-></td><td>1076</td><td>Bacteria>Terrabacteria_group>Cyanobacteriota/Melainabacteria_group>Cyanobacteriota>Cyanophyceae</td><td>GCF_904830845.1</td><td>Planktothrix_agardhii</td><td>NMG87_RS03740</td><td>860760..863990</td><td>1160</td><td>DUF499-domain-containing-protein-[Planktothrix-agardhii]</td></tr>
<tr><td>WP_254034264.1</td><td>AAA+ATPase+wHTH+Alpha+Beta-domain+wHTH+RRM</td><td>Type-2_BR.RM.like.systems</td><td><-NTP_transferase||?->SAM-methylase->?->?->?->DUF499*->DUF3780->?->PIN->METHYLASE->METHYLASE-p->HKD-DNase+SF1-SF2-Helicase+iREase+CTD-></td><td>1076</td><td>Bacteria>Terrabacteria_group>Cyanobacteriota/Melainabacteria_group>Cyanobacteriota>Cyanophyceae</td><td>GCF_904830915.1</td><td>Planktothrix_agardhii</td><td>NMG85_RS01800</td><td>410077..413307</td><td>1160</td><td>DUF499-domain-containing-protein-[Planktothrix-agardhii]</td></tr>
<tr><td>WP_242719410.1</td><td>AAA+ATPase+wHTH+Alpha+Beta-domain+wHTH+RRM</td><td>Type-2_BR.RM.like.systems</td><td><-AbrB-like<-?<-?<-?||?->DUF499*->DUF3780-><-?||?->METHYLASE->HKD-DNase+SF1-SF2-Helicase+iREase+CTD-></td><td>1075</td><td>Bacteria>Terrabacteria_group>Cyanobacteriota/Melainabacteria_group>Cyanobacteriota>Cyanophyceae</td><td>-</td><td>Microcoleus_vaginatus</td><td>DOA37_27340</td><td>6455094..6458321</td><td>119532</td><td>DUF499-domain-containing-protein-[Microcoleus-vaginatus]</td></tr>
<tr><td>WP_096542894.1</td><td>AAA+ATPase+wHTH+Alpha+Beta-domain+wHTH+RRM</td><td>Type-2_BR.RM.like.systems</td><td><-HTH+MPTase||ZnR->?->?->HTH+UbE1-><-DUF3780<-DUF499*||TPR+TPR-><-?||?->ParB+HTH->Pkinase+SF1-SF2-Helicase->HTH+HKD-DNase-></td><td>1077</td><td>Bacteria>Terrabacteria_group>Cyanobacteriota/Melainabacteria_group>Cyanobacteriota>Cyanophyceae</td><td>-</td><td>Nostoc_linckia</td><td>CA726_RS34720</td><td>13848..17081</td><td>92942</td><td>DUF499-domain-containing-protein-[Nostoc-linckia]</td></tr>
<tr><td>WP_242072110.1</td><td>AAA+ATPase+wHTH+Alpha+Beta-domain+wHTH+RRM</td><td>Type-2_BR.RM.like.systems</td><td><-Phage_int_SAM_1+Phage_integrase||?->?->?->DUF499*->DUF3780->METHYLASE->ParB+HNH->HKD-DNase+SF1-SF2-Helicase+iREase+CTD-></td><td>920</td><td>Bacteria>Terrabacteria_group>Cyanobacteriota/Melainabacteria_group>Cyanobacteriota>Cyanophyceae</td><td>-</td><td>Nostoc_sp._FACHB-110</td><td>H6G69_RS18775</td><td>61705..64467</td><td>2692834</td><td>DUF499-domain-containing-protein-[Nostoc-sp.-FACHB-110]</td></tr>
<tr><td>WP_235759069.1</td><td>AAA+ATPase+wHTH+Alpha+Beta-domain+wHTH+RRM</td><td>Type-2_BR.RM.like.systems</td><td><-HKD-DNase+SF1-SF2-Helicase+iREase+CTD<-METHYLASE-p<-PIN<-?<-DUF3780<-DUF499*</td><td>1076</td><td>Bacteria>Terrabacteria_group>Cyanobacteriota/Melainabacteria_group>Cyanobacteriota>Cyanophyceae</td><td>-</td><td>Planktothrix_agardhii</td><td>L2E81_RS23100</td><td>5349..8579</td><td>1160</td><td>DUF499-domain-containing-protein-[Planktothrix-agardhii]</td></tr>
<tr><td>WP_251960876.1</td><td>AAA+ATPase+wHTH+Alpha+Beta-domain+wHTH+RRM</td><td>Type-2_BR.RM.like.systems</td><td><-PIN<-MPTase<-HTH_3||ZnR->HTH+UbE1-><-DUF3780<-DUF499*<-?||Phage_int_SAM_4+Phage_integrase->Phage_integrase->Phage_integrase-></td><td>1085</td><td>Bacteria>Terrabacteria_group>Cyanobacteriota/Melainabacteria_group>Cyanobacteriota>Cyanophyceae</td><td>GCF_023734775.1</td><td>Nostoc_commune</td><td>ANS036C_RS32215</td><td>25490..28747</td><td>1178</td><td>DUF499-domain-containing-protein-[Nostoc-commune]</td></tr>
<tr><td>WP_088889980.1</td><td>AAA+ATPase+wHTH+Alpha+Beta-domain+wHTH+RRM</td><td>Type-2_BR.RM.like.systems</td><td>DEDD_Tnp_IS110+Transposase_20-><-REase<-PIN<-?<-DUF3780<-DUF499*||wHTH+REase-><-?<-URI-DNase<-Prim-Pol</td><td>1072</td><td>Bacteria>Terrabacteria_group>Cyanobacteriota/Melainabacteria_group>Cyanobacteriota>Cyanophyceae</td><td>GCF_002215035.1</td><td>Leptolyngbya_ohadii</td><td>CDV24_RS06815</td><td>190985..194203</td><td>1962290</td><td>DUF499-domain-containing-protein-[Leptolyngbya-ohadii]</td></tr>
<tr><td>WP_017747474.1</td><td>AAA+ATPase+wHTH+Alpha+Beta-domain+wHTH+RRM</td><td>Type-2_BR.RM.like.systems</td><td><-PIN<-?<-METHYLASE<-Schlafen+TUDOR+PIWI<-DUF3780<-DUF499*<-?<-ABhydrolase||?->MetJarc-HTH->RelE-ParE-><-REase</td><td>1072</td><td>Bacteria>Terrabacteria_group>Cyanobacteriota/Melainabacteria_group>Cyanobacteriota>Cyanophyceae</td><td>GCF_000346485.2</td><td>Scytonema_hofmannii</td><td>WA1_RS47720</td><td>11345103..11348321</td><td>34078</td><td>DUF499-domain-containing-protein-[Scytonema-hofmannii]</td></tr>
<tr><td>MBW4588299.1</td><td>AAA+ATPase+wHTH+Alpha+Beta-domain+wHTH+RRM</td><td>Type-2_BR.RM.like.systems</td><td><-REase<-METHYLASE<-PIN<-?<-?<-DUF3780<-DUF499*<-ABC-ATPase+SbcC||||OrfB_IS605+OrfB_Zn_ribbon->?-><-ACET<-?<-VapB_antitoxin</td><td>1077</td><td>Bacteria>Terrabacteria_group>Cyanobacteriota/Melainabacteria_group>Cyanobacteriota>Cyanophyceae</td><td>GCA_019359795.1</td><td>Aetokthonos_hydrillicola_CCALA_1050</td><td>KME33_24335</td><td>51118..54351</td><td>2710484</td><td>DUF499-domain-containing-protein-[Aetokthonos-hydrillicola-CCALA-1050]</td></tr>
<tr><td>WP_109013093.1</td><td>AAA+ATPase+wHTH+Alpha+Beta-domain+wHTH+RRM</td><td>Type-2_BR.RM.like.systems</td><td><-Phage_int_SAM_4+Phage_integrase||?-><-PIN<-?<-DUF3780<-DUF499*<-HNH<-?||?-><-PAIREDC-HTH||?->Prim-Pol-></td><td>1102</td><td>Bacteria>Terrabacteria_group>Cyanobacteriota/Melainabacteria_group>Cyanobacteriota>Cyanophyceae</td><td>GCF_003113895.1</td><td>Nostoc_commune</td><td>CDC33_RS34475</td><td>462428..465736</td><td>1178</td><td>DUF499-domain-containing-protein-[Nostoc-commune]</td></tr>
<tr><td>WP_012308559.1</td><td>AAA+ATPase+wHTH+Alpha+Beta-domain+wHTH+RRM</td><td>Type-2_BR.RM.like.systems</td><td><-HicA_toxin<-HicB<-?<-?||GDResolvase+HIN-HTH->?->DUF499*->DUF3780->?->PIN->REase->METHYLASE->HigB_toxin-></td><td>1066</td><td>Bacteria>Terrabacteria_group>Cyanobacteriota/Melainabacteria_group>Cyanobacteriota>Cyanophyceae</td><td>-</td><td>Picosynechococcus_sp._PCC_7002</td><td>SYNPCC7002_RS14755</td><td>16276..19476</td><td>32049</td><td>DUF499-domain-containing-protein-[Synechococcus-sp.-PCC-7002]</td></tr>
<tr><td>WP_235751988.1</td><td>AAA+ATPase+wHTH+Alpha+Beta-domain+wHTH+RRM</td><td>Type-2_BR.RM.like.systems</td><td>SAM-methylase->?->?->?->?->DUF499*->DUF3780->?->PIN-><-rRNA<-?<-Peptidase_M16+Peptidase_M16_C</td><td>1076</td><td>Bacteria>Terrabacteria_group>Cyanobacteriota/Melainabacteria_group>Cyanobacteriota>Cyanophyceae</td><td>GCF_021554875.1</td><td>Planktothrix_agardhii</td><td>L2E65_RS17630</td><td>3802221..3805451</td><td>1160</td><td>DUF499-domain-containing-protein-[Planktothrix-agardhii]</td></tr>
<tr><td>MBS9391445.1</td><td>AAA+ATPase+wHTH+Alpha+Beta-domain+wHTH+RRM</td><td>Type-2_BR.RM.like.systems</td><td>METHYLASE-p->?->ABC-ATPase+SbcC->?-><-DDE_Tnp_1||?->||DUF499*->DUF3780->?->REase->?->?->PIN-></td><td>1083</td><td>Bacteria>Terrabacteria_group>Cyanobacteriota/Melainabacteria_group>Cyanobacteriota>Cyanophyceae</td><td>GCA_018447775.1</td><td>Dolichospermum_sp._WA123</td><td>HEQ33_22150</td><td>4813071..4816322</td><td>2717878</td><td>ATP-binding-protein-[Dolichospermum-sp.-WA123]</td></tr>
<tr><td>WP_094345566.1</td><td>AAA+ATPase+wHTH+Alpha+Beta-domain+wHTH+RRM</td><td>Type-2_BR.RM.like.systems</td><td><-ParB||PAIREDC-HTH->Phage_int_SAM_1+Phage_integrase->?->HNH->DUF499*->DUF3780->?->PIN->METHYLASE-></td><td>1076</td><td>Bacteria>Terrabacteria_group>Cyanobacteriota/Melainabacteria_group>Cyanobacteriota>Cyanophyceae</td><td>GCF_002245985.1</td><td>Nostoc_sp._'Peltigera_membranacea_cyanobiont'_232</td><td>CDG79_RS31090</td><td>5976..9206</td><td>2014531</td><td>DUF499-domain-containing-protein-[Nostoc-sp.-'Peltigera-membranacea-cyanobiont'-232]</td></tr>
<tr><td>WP_017302292.1</td><td>AAA+ATPase+wHTH+Alpha+Beta-domain+wHTH+RRM</td><td>Type-2_BR.RM.like.systems</td><td>NUDIX->DUF499*->DUF3780->?->PIN-><-PAIREDC-HTH||PIN-></td><td>1075</td><td>Bacteria>Terrabacteria_group>Cyanobacteriota/Melainabacteria_group>Cyanobacteriota>Cyanophyceae</td><td>GCF_000309385.1</td><td>Nodosilinea_nodulosa</td><td>LEPTO7104_RS0130115</td><td>429641..432868</td><td>416001</td><td>DUF499-domain-containing-protein-[Nodosilinea-nodulosa]</td></tr>
<tr><td>WP_015130212.1</td><td>AAA+ATPase+wHTH+Alpha+Beta-domain+wHTH+RRM</td><td>Type-2_BR.RM.like.systems</td><td><-METHYLASE-p<-PIN<-?<-DUF3780<-DUF499*<-?<-?<-Thioredoxin</td><td>1078</td><td>Bacteria>Terrabacteria_group>Cyanobacteriota/Melainabacteria_group>Cyanobacteriota>Cyanophyceae</td><td>GCF_000316575.1</td><td>Calothrix_sp._PCC_7507</td><td>CAL7507_RS19505</td><td>4557851..4561087</td><td>99598</td><td>DUF499-domain-containing-protein-[Calothrix-sp.-PCC-7507]</td></tr>
<tr><td>WP_019504690.1</td><td>AAA+ATPase+wHTH+Alpha+Beta-domain+wHTH+RRM</td><td>Type-2_BR.RM.like.systems</td><td>DUF499*->DUF3780-><-PIN<-?<-PemK_toxin<-MetJarc-HTH||METHYLASE-></td><td>1082</td><td>Bacteria>Terrabacteria_group>Cyanobacteriota/Melainabacteria_group>Cyanobacteriota>Cyanophyceae</td><td>GCF_000332195.1</td><td>Pleurocapsa_sp._PCC_7319</td><td>PLEUR7319_RS0107970</td><td>842087..845335</td><td>118161</td><td>DUF499-domain-containing-protein-[Pleurocapsa-sp.-PCC-7319]</td></tr>
<tr><td>WP_216915326.1</td><td>AAA+ATPase+wHTH+Alpha+Beta-domain+wHTH+RRM</td><td>Type-2_BR.RM.like.systems</td><td>PIN-><-?<-?<-?<-DUF4236||DUF499*->DUF3780->RelE-ParE->HTH_3->?->?->METHYLASE-></td><td>1028</td><td>Bacteria>Terrabacteria_group>Cyanobacteriota/Melainabacteria_group>Cyanobacteriota>Cyanophyceae</td><td>GCF_019038475.1</td><td>unclassified_Synechococcus</td><td>KUG39_RS12575</td><td>13470..16556</td><td>2626047</td><td>MULTISPECIES:-DUF499-domain-containing-protein-[unclassified-Synechococcus]</td></tr>
<tr><td>WP_106236198.1</td><td>AAA+ATPase+wHTH+Alpha+Beta-domain+wHTH+RRM</td><td>Type-2_BR.RM.like.systems</td><td>MetJarc-HTH->PemK_toxin->?->PIN-><-DUF3780||?-><-DUF499*</td><td>1094</td><td>Bacteria>Terrabacteria_group>Cyanobacteriota/Melainabacteria_group>Cyanobacteriota>Cyanophyceae</td><td>GCF_003003995.1</td><td>Pleurocapsa_sp._CCALA_161</td><td>C7B62_RS11765</td><td>10633..13917</td><td>2107688</td><td>DUF499-domain-containing-protein-[Pleurocapsa-sp.-CCALA-161]</td></tr>
<tr><td>WP_268611164.1</td><td>AAA+ATPase+wHTH+Alpha+Beta-domain+wHTH+RRM</td><td>Type-2_BR.RM.like.systems</td><td>DUF499*->DUF3780-><-?<-ZnR||HTH+MPTase->PIN->METHYLASE-p-></td><td>1072</td><td>Bacteria>Terrabacteria_group>Cyanobacteriota/Melainabacteria_group>Cyanobacteriota>Cyanophyceae</td><td>GCF_026802175.1</td><td>Thermocoleostomius_sinensis</td><td>OXH18_RS04190</td><td>963011..966229</td><td>3065396</td><td>DUF499-domain-containing-protein-[Leptolyngbya-sp.-PKUAC-SCTA174]</td></tr>
<tr><td>WP_190566691.1</td><td>AAA+ATPase+wHTH+Alpha+Beta-domain+wHTH+RRM</td><td>Type-2_BR.RM.like.systems</td><td>DUF499*->REase->DUF3780->METHYLASE-p->METHYLASE->?->PIN-></td><td>1077</td><td>Bacteria>Terrabacteria_group>Cyanobacteriota/Melainabacteria_group>Cyanobacteriota>Cyanophyceae</td><td>GCF_014696935.1</td><td>unclassified_Desertifilum</td><td>H6G23_RS20070</td><td>60642..63875</td><td>2621682</td><td>MULTISPECIES:-DUF499-domain-containing-protein-[unclassified-Desertifilum]</td></tr>
<tr><td>WP_068394464.1</td><td>AAA+ATPase+wHTH+Alpha+Beta-domain+wHTH+RRM</td><td>Type-2_BR.RM.like.systems</td><td>Peptidase-S8->DUF433->PIN-><-HNH||?-><-DUF3780<-DUF499*</td><td>1071</td><td>Bacteria>Terrabacteria_group>Cyanobacteriota/Melainabacteria_group>Cyanobacteriota>Cyanophyceae</td><td>-</td><td>Leptolyngbya_sp._NIES-3755</td><td>LEP3755_RS32315</td><td>59687..62902</td><td>1752064</td><td>DUF499-domain-containing-protein-[Leptolyngbya-sp.-NIES-3755]</td></tr>
<tr><td>WP_015186345.1</td><td>AAA+ATPase+wHTH+Alpha+Beta-domain+wHTH+RRM</td><td>Type-2_BR.RM.like.systems</td><td>D5N+viral-AAA-><-?<-Phage_integrase||?->?->CHTH->DUF499*->DUF3780->?->PIN->METHYLASE-></td><td>1078</td><td>Bacteria>Terrabacteria_group>Cyanobacteriota/Melainabacteria_group>Cyanobacteriota>Cyanophyceae</td><td>GCF_000317515.1</td><td>Allocoleopsis_franciscana</td><td>MIC7113_RS32605</td><td>20770..24006</td><td>2886352</td><td>DUF499-domain-containing-protein-[Allocoleopsis-franciscana]</td></tr>
<tr><td>WP_271941641.1</td><td>AAA+ATPase+wHTH+Alpha+Beta-domain+wHTH+RRM</td><td>Type-2_BR.RM.like.systems</td><td><-ParB<-?||DUF499*->DUF3780->?->PIN->METHYLASE->REase-></td><td>1128</td><td>Bacteria>Terrabacteria_group>Cyanobacteriota/Melainabacteria_group>Cyanobacteriota>Cyanophyceae</td><td>GCF_028329345.1</td><td>Nodularia_spumigena</td><td>PN454_RS18900</td><td>6974..10360</td><td>70799</td><td>DUF499-domain-containing-protein-[Nodularia-spumigena]</td></tr>
<tr><td>WP_198808387.1</td><td>AAA+ATPase+wHTH+Alpha+Beta-domain+wHTH+RRM</td><td>Type-2_BR.RM.like.systems</td><td>NACHT->?->?->Peptidase->?->EVE->DUF499*->?->DUF3780->?->PIN->?-><-PIN</td><td>1073</td><td>Bacteria>Terrabacteria_group>Cyanobacteriota/Melainabacteria_group>Cyanobacteriota>Cyanophyceae</td><td>GCF_016403105.1</td><td>Leptolyngbya_sp._BL0902</td><td>GFS31_RS19740</td><td>138456..141677</td><td>1115757</td><td>DUF499-domain-containing-protein-[Leptolyngbya-sp.-BL0902]</td></tr>
<tr><td>NUN66948.1</td><td>AAA+ATPase+wHTH+Alpha+Beta-domain+wHTH+RRM</td><td>Type-2_BR.RM.like.systems</td><td><-RVT+HNH<-LPD36+METHYLASE+SF1-SF2-Helicase||?->DUF499*->DUF3780-><-?||HTH+MPTase->PIN->METHYLASE->HigB_toxin-></td><td>1074</td><td>Bacteria>Terrabacteria_group>Cyanobacteriota/Melainabacteria_group>Cyanobacteriota>Cyanophyceae</td><td>GCA_013361095.1</td><td>Pseudanabaena_biceps</td><td>HCU40_19855</td><td>93752..96976</td><td>927669</td><td>ATP-binding-protein-[Pseudanabaena-biceps]</td></tr>
<tr><td>NEP85621.1</td><td>AAA+ATPase+wHTH+Alpha+Beta-domain+wHTH+RRM</td><td>Type-2_BR.RM.like.systems</td><td>RHH->PolB->HEPN->SF1-SF2-Helicase->CHTH->DUF499*->DUF3780->HTH_Tnp->PIN->AbrB-like-></td><td>1085</td><td>Bacteria>Terrabacteria_group>Cyanobacteriota/Melainabacteria_group>Cyanobacteriota>Cyanophyceae</td><td>GCA_010692625.1</td><td>Okeania_sp._SIO2C2</td><td>F6K18_01580</td><td>185519..188776</td><td>2607787</td><td>ATP-binding-protein-[Okeania-sp.-SIO2C2]</td></tr>
<tr><td>MBS3030295.1</td><td>AAA+ATPase+wHTH+Alpha+Beta-domain+wHTH+RRM</td><td>Type-2_BR.RM.like.systems</td><td><-TPR<-?<-CR-ATPase9+REase||?->?->DUF499*->?->DUF3780->REase->METHYLASE->?->PIN-></td><td>1088</td><td>Bacteria>Terrabacteria_group>Cyanobacteriota/Melainabacteria_group>Cyanobacteriota>Cyanophyceae</td><td>GCA_018295985.1</td><td>Dolichospermum_sp._DET66</td><td>HCA26_25880</td><td>5395527..5398793</td><td>2704415</td><td>ATP-binding-protein-[Dolichospermum-sp.-DET66]</td></tr>
<tr><td>MBW4466337.1</td><td>AAA+ATPase+wHTH+Alpha+Beta-domain+wHTH+RRM</td><td>Type-2_BR.RM.like.systems</td><td>DUF499*->DUF3780->?->PIN-><-Peptidase-S8<-Classical-AAA||METHYLASE-></td><td>1072</td><td>Bacteria>Terrabacteria_group>Cyanobacteriota/Melainabacteria_group>Cyanobacteriota>Cyanophyceae</td><td>GCA_019359155.1</td><td>Pegethrix_bostrychoides_GSE-TBD4-15B</td><td>KME07_13005</td><td>91225..94443</td><td>2839662</td><td>DUF499-domain-containing-protein-[Pegethrix-bostrychoides-GSE-TBD4-15B]</td></tr>
<tr><td>WP_190499965.1</td><td>AAA+ATPase+wHTH+Alpha+Beta-domain+wHTH+RRM</td><td>Type-2_BR.RM.like.systems</td><td>MACRODOMAIN-><-?<-?<-PIN<-?<-DUF3780<-DUF499*<-EVE<-?||RHH-></td><td>1073</td><td>Bacteria>Terrabacteria_group>Cyanobacteriota/Melainabacteria_group>Cyanobacteriota>Cyanophyceae</td><td>GCF_014697375.1</td><td>Phormidium_sp._FACHB-1136</td><td>H6G62_RS24145</td><td>5974..9195</td><td>2692848</td><td>DUF499-domain-containing-protein-[Phormidium-sp.-FACHB-1136]</td></tr>
<tr><td>MBW4537488.1</td><td>AAA+ATPase+wHTH+Alpha+Beta-domain+wHTH+RRM</td><td>Type-2_BR.RM.like.systems</td><td><-sGTP||?->?->?-><-?||DUF499*-><-?||DUF3780-><-PIN<-?<-PemK_toxin<-MetJarc-HTH</td><td>1097</td><td>Bacteria>Terrabacteria_group>Cyanobacteriota/Melainabacteria_group>Cyanobacteriota>Cyanophyceae</td><td>GCA_019359095.1</td><td>Pleurocapsa_minor_HA4230-MV1</td><td>KME09_26475</td><td>587119..590412</td><td>2839664</td><td>DUF499-domain-containing-protein-[Pleurocapsa-minor-HA4230-MV1]</td></tr>
<tr><td>WP_185582347.1</td><td>AAA+ATPase+wHTH+Alpha+Beta-domain+wHTH+RRM</td><td>Type-2_BR.RM.like.systems</td><td><-CopG/RHH||Peptidase-><-METHYLASE<-PIN<-?<-DUF3780<-DUF499*</td><td>1077</td><td>Bacteria>Terrabacteria_group>Cyanobacteriota/Melainabacteria_group>Cyanobacteriota>Cyanophyceae</td><td>GCF_014222255.1</td><td>unclassified_Nostoc</td><td>GNF10_RS25055</td><td>11550..14783</td><td>2593658</td><td>MULTISPECIES:-DUF499-domain-containing-protein-[unclassified-Nostoc]</td></tr>
<tr><td>WP_193963259.1</td><td>AAA+ATPase+wHTH+Alpha+Beta-domain+wHTH+RRM</td><td>Type-2_BR.RM.like.systems</td><td>MACRODOMAIN-><-MACRODOMAIN<-?<-PIN<-?<-DUF3780<-DUF499*<-?||ParD_antitoxin->RelE-ParE-></td><td>1075</td><td>Bacteria>Terrabacteria_group>Cyanobacteriota/Melainabacteria_group>Cyanobacteriota>Cyanophyceae</td><td>GCF_015207395.1</td><td>Nodosilinea_sp._LEGE_07088</td><td>IQ254_RS00715</td><td>151941..155168</td><td>2777968</td><td>DUF499-domain-containing-protein-[Nodosilinea-sp.-LEGE-07088]</td></tr>
<tr><td>NJO66191.1</td><td>AAA+ATPase+wHTH+Alpha+Beta-domain+wHTH+RRM</td><td>Type-2_BR.RM.like.systems</td><td><-REase<-REase<-PIN<-?<-DUF3780<-?<-DUF499*||?->REase+ParB-></td><td>1069</td><td>Bacteria>Terrabacteria_group>Cyanobacteriota/Melainabacteria_group>Cyanobacteriota>Cyanophyceae</td><td>GCA_012033405.1</td><td>Leptolyngbyaceae_cyanobacterium_RM1_405_57</td><td>HC832_00965</td><td>7160..10369</td><td>2720433</td><td>ATP-binding-protein-[Leptolyngbyaceae-cyanobacterium-RM1_405_57]</td></tr>
<tr><td>NJK39172.1</td><td>AAA+ATPase+wHTH+Alpha+Beta-domain+wHTH+RRM</td><td>Type-2_BR.RM.like.systems</td><td><-REase<-?<-?<-?<-?||DUF499*->DUF3780->HTH_Tnp->?->PIN->REase-></td><td>1079</td><td>Bacteria>Terrabacteria_group>Cyanobacteriota/Melainabacteria_group>Cyanobacteriota>Cyanophyceae</td><td>GCA_012030745.1</td><td>Oscillatoriales_cyanobacterium_SM2_3_0</td><td>HC920_16470</td><td>6917..10156</td><td>2720468</td><td>DUF499-domain-containing-protein-[Oscillatoriales-cyanobacterium-SM2_3_0]</td></tr>
<tr><td>MBV5261741.1</td><td>AAA+ATPase+wHTH+Alpha+Beta-domain+wHTH+RRM</td><td>Type-2_BR.RM.like.systems</td><td><-HTH<-HigB_toxin<-METHYLASE<-PIN<-AbrB<-DUF3780<-DUF499*||MazE_antitoxin->?-><-?<-REase</td><td>1080</td><td>Bacteria>Terrabacteria_group>Cyanobacteriota/Melainabacteria_group>Cyanobacteriota>Cyanophyceae</td><td>GCA_019155215.1</td><td>Synechococcus_moorigangaii_CMS01</td><td>FLX56_25345</td><td>11889..15131</td><td>2025807</td><td>ATP-binding-protein-[Synechococcus-moorigangaii-CMS01]</td></tr>
<tr><td>WP_019497040.1</td><td>AAA+ATPase+wHTH+Alpha+Beta-domain+wHTH+RRM</td><td>Type-2_BR.RM.like.systems</td><td><-PIN<-?<-?<-SMODS<-BirA-HTH<-HISKIN||DUF499*->DUF3780-><-?<-?<-?<-?||METHYLASE-></td><td>1076</td><td>Bacteria>Terrabacteria_group>Cyanobacteriota/Melainabacteria_group>Cyanobacteriota>Cyanophyceae</td><td>-</td><td>Nostocales</td><td>RIVM261_048610</td><td>654121..657351</td><td>1161</td><td>MULTISPECIES:-DUF499-domain-containing-protein-[Nostocales]</td></tr>
<tr><td>MCE2718757.1</td><td>AAA+ATPase+wHTH+Alpha+Beta-domain+wHTH+RRM</td><td>Type-2_BR.RM.like.systems</td><td><-METHYLASE-p<-PIN<-?<-DUF3780<-DUF499*||Glyoxalase->NUDIX->?->Pterin_bind-></td><td>1083</td><td>Bacteria>Terrabacteria_group>Cyanobacteriota/Melainabacteria_group>Cyanobacteriota>Cyanophyceae</td><td>GCA_021300315.1</td><td>Anabaena_sp._49628_E55</td><td>LW708_08020</td><td>3060..6311</td><td>2904988</td><td>DUF499-domain-containing-protein-[Anabaena-sp.-49628_E55]</td></tr>
<tr><td>GJD19905.1</td><td>AAA+ATPase+wHTH+Alpha+Beta-domain+wHTH+RRM</td><td>Type-2_BR.RM.like.systems</td><td><-PIN<-?<-?<-SMODS<-BirA-HTH<-HISKIN||DUF499*->DUF3780-><-?<-?<-?<-?||METHYLASE-></td><td>1076</td><td>Bacteria>Terrabacteria_group>Cyanobacteriota/Melainabacteria_group>Cyanobacteriota>Cyanophyceae</td><td>GCA_019973055.1</td><td>Rivularia_sp._IAM_M-261</td><td>RIVM261_048610</td><td>654121..657351</td><td>448371</td><td>hypothetical-protein-RIVM261_048610-[Rivularia-sp.-IAM-M-261]</td></tr>
<tr><td>WP_278615280.1</td><td>AAA+ATPase+wHTH+Alpha+Beta-domain+wHTH+RRM</td><td>Type-2_BR.RM.like.systems</td><td><-PIN<-?<-DUF3780<-DUF499*<-?<-?<-?<-?<-SAM-methylase</td><td>1076</td><td>Bacteria>Terrabacteria_group>Cyanobacteriota/Melainabacteria_group>Cyanobacteriota>Cyanophyceae</td><td>GCF_015708255.1</td><td>Planktothrix_agardhii</td><td>I4E98_RS06875</td><td>1485..4715</td><td>1160</td><td>DUF499-domain-containing-protein-[Planktothrix-agardhii]</td></tr>
<tr><td>WP_235766086.1</td><td>AAA+ATPase+wHTH+Alpha+Beta-domain+wHTH+RRM</td><td>Type-2_BR.RM.like.systems</td><td>SAM-methylase->?->?->?->?->DUF499*->DUF3780->?->PIN-></td><td>1076</td><td>Bacteria>Terrabacteria_group>Cyanobacteriota/Melainabacteria_group>Cyanobacteriota>Cyanophyceae</td><td>GCF_021554705.1</td><td>Planktothrix_agardhii</td><td>L2E80_RS22970</td><td>30217..33447</td><td>1160</td><td>DUF499-domain-containing-protein-[Planktothrix-agardhii]</td></tr>
<tr><td>WP_202895460.1</td><td>AAA+ATPase+wHTH+Alpha+Beta-domain+wHTH+RRM</td><td>Type-2_BR.RM.like.systems</td><td><-METHYLASE-p<-DUF3780<-DUF499*||?->PIN-></td><td>1018</td><td>Bacteria>Terrabacteria_group>Cyanobacteriota/Melainabacteria_group>Cyanobacteriota>Cyanophyceae</td><td>GCF_014698965.1</td><td>Iningainema_tapete</td><td>ICL16_RS08685</td><td>918..3974</td><td>2806730</td><td>DUF499-domain-containing-protein-[Iningainema-tapete]</td></tr>
<tr><td>MBG0746295.1</td><td>AAA+ATPase+wHTH+Alpha+Beta-domain+wHTH+RRM</td><td>Type-2_BR.RM.like.systems</td><td><-PIN<-?<-DUF3780<-DUF499*<-?<-?<-?<-?<-SAM-methylase</td><td>1076</td><td>Bacteria>Terrabacteria_group>Cyanobacteriota/Melainabacteria_group>Cyanobacteriota>Cyanophyceae</td><td>GCA_015708255.1</td><td>Planktothrix_agardhii_KL2</td><td>I4E98_06850</td><td>1485..4715</td><td>2789715</td><td>ATP-binding-protein-[Planktothrix-agardhii-KL2]</td></tr>
<tr><td>MCG9890253.1</td><td>AAA+ATPase+wHTH+Alpha+Beta-domain+wHTH+RRM</td><td>Type-2_BR.RM.like.systems</td><td>CHTH->DUF499*->DUF3780->CHTH->?->METHYLASE->?->PIN-></td><td>1071</td><td>Bacteria>Terrabacteria_group>Cyanobacteriota/Melainabacteria_group>Cyanobacteriota>Cyanophyceae</td><td>GCA_022402525.1</td><td>Thermosynechococcaceae_cyanobacterium_MS004</td><td>MH252_04185</td><td>862..4077</td><td>2881380</td><td>DUF499-domain-containing-protein-[Thermosynechococcaceae-cyanobacterium-MS004]</td></tr>
<tr><td>WP_223209979.1</td><td>AAA+ATPase+wHTH+Alpha+Beta-domain+wHTH+RRM</td><td>Type-2_BR.RM.like.systems</td><td><-REase<-DUF433<-?<-PIN<-?<-DUF3780<-DUF499*</td><td>1044</td><td>Bacteria>Terrabacteria_group>Cyanobacteriota/Melainabacteria_group>Cyanobacteriota>Cyanophyceae</td><td>GCF_000715475.1</td><td>Picosynechococcus_sp._NKBG042902</td><td>BG2_RS15810</td><td>13294..16428</td><td>490193</td><td>DUF499-domain-containing-protein-[Synechococcus-sp.-NKBG042902]</td></tr>
<tr><td>NJM00138.1</td><td>AAA+ATPase+wHTH+Alpha+Beta-domain+wHTH+RRM</td><td>Type-2_BR.RM.like.systems</td><td>DUF499*->DUF3780-><-?<-AbrB-HTH||?->PIN->RNase-></td><td>844</td><td>Bacteria>Terrabacteria_group>Cyanobacteriota/Melainabacteria_group>Cyanobacteriota>Cyanophyceae</td><td>GCA_012031565.1</td><td>Synechococcaceae_cyanobacterium_SM2_3_2</td><td>HC924_15665</td><td>1..2537</td><td>2720471</td><td>DUF499-domain-containing-protein-[Synechococcaceae-cyanobacterium-SM2_3_2]</td></tr>
<tr><td>MBF2090291.1</td><td>AAA+ATPase+wHTH+Alpha+Beta-domain+wHTH+RRM</td><td>Type-2_BR.RM.like.systems</td><td>TOPRIM+SF1-SF2-Helicase-><-TPR+TPR+TPR+TPR+TPR+TPR+TPR+TPR<-?<-RHH_1<-TPR+TPR+TPR+TPR+TPR+TPR+TPR+TPR||DUF499*->DUF3780->REase+PNTD1+PNTD2->PIWI-><-MACRODOMAIN<-PemK_toxin<-MetJarc-HTH</td><td>1082</td><td>Bacteria>Terrabacteria_group>Cyanobacteriota/Melainabacteria_group>Cyanobacteriota>Cyanophyceae</td><td>GCA_015272375.1</td><td>Synechococcales_cyanobacterium_K32_A2020_035</td><td>IGR90_15410</td><td>40350..43598</td><td>2774345</td><td>ATP-binding-protein-[Synechococcales-cyanobacterium-K32_A2020_035]</td></tr>
<tr><td>MBW4483623.1</td><td>AAA+ATPase+wHTH+Alpha+Beta-domain+wHTH+RRM</td><td>Type-2_BR.RM.like.systems</td><td>Arm-DNA-bind_2+Phage_int_SAM_3+Phage_integrase-><-LexA-protease||HTH->?->?-><-ParD_antitoxin||DUF499*->DUF3780->?->?->?->REase+PNTD1+PNTD2->PIWI-></td><td>1075</td><td>Bacteria>Terrabacteria_group>Cyanobacteriota/Melainabacteria_group>Cyanobacteriota>Cyanophyceae</td><td>GCA_019358835.1</td><td>Tildeniella_torsiva_UHER_1998/13D</td><td>KME14_13865</td><td>174768..177995</td><td>337900</td><td>DUF499-domain-containing-protein-[Tildeniella-torsiva-UHER-1998/13D]</td></tr>
<tr><td>WP_065715514.1</td><td>AAA+ATPase+wHTH+Alpha+Beta-domain+wHTH+RRM</td><td>Type-2_BR.RM.like.systems</td><td><-PemK_toxin<-MazE_antitoxin<-?||Phage_int_SAM_4+Phage_integrase->DUF499*->DUF3780->?->DUF433->REase->METHYLASE-></td><td>1075</td><td>Bacteria>Terrabacteria_group>Cyanobacteriota/Melainabacteria_group>Cyanobacteriota>Cyanophyceae</td><td>GCF_001693255.1</td><td>Picosynechococcus_sp._PCC_7003</td><td>AWQ21_RS15070</td><td>95173..98400</td><td>374981</td><td>DUF499-domain-containing-protein-[Synechococcus-sp.-PCC-7003]</td></tr>
<tr><td>WP_190748058.1</td><td>AAA+ATPase+wHTH+Alpha+Beta-domain+wHTH+RRM</td><td>Type-2_BR.RM.like.systems</td><td><-METHYLASE||?-><-?<-Zn_Tnp_IS1595+DDE_Tnp_IS1595<-?<-DUF3780<-DUF499*<-REase<-?||?->HTH_26->?->Phage_integrase-></td><td>1074</td><td>Bacteria>Terrabacteria_group>Cyanobacteriota/Melainabacteria_group>Cyanobacteriota>Cyanophyceae</td><td>GCF_014695395.1</td><td>Nodosilinea_sp._FACHB-131</td><td>H6F75_RS00215</td><td>20757..23981</td><td>2692832</td><td>DUF499-domain-containing-protein-[Nodosilinea-sp.-FACHB-131]</td></tr>
<tr><td>WP_015328425.1</td><td>AAA+ATPase+wHTH+Alpha+Beta-domain+wHTH+RRM</td><td>Type-2_BR.RM.like.systems</td><td><-CopG/RHH<-REase||?->Peptidase-><-METHYLASE<-DUF3780<-DUF499*<-Prim-Pol<-?<-Phage_int_SAM_4+Phage_integrase</td><td>1110</td><td>Bacteria>Terrabacteria_group>Cyanobacteriota/Melainabacteria_group>Cyanobacteriota>Cyanophyceae</td><td>GCF_000317535.1</td><td>Cylindrospermum_stagnale</td><td>CYLST_RS30675</td><td>267763..271095</td><td>142864</td><td>DUF499-domain-containing-protein-[Cylindrospermum-stagnale]</td></tr>
<tr><td>WP_026798357.1</td><td>AAA+ATPase+wHTH+Alpha+Beta-domain+wHTH+RRM</td><td>Type-2_BR.RM.like.systems</td><td><-METHYLASE-p<-?<-METHYLASE<-HTH_Tnp<-DUF3780<-DUF499*<-?||?->?-><-REase<-Phage_int_SAM_1+Phage_integrase<-REase</td><td>1076</td><td>Bacteria>Terrabacteria_group>Cyanobacteriota/Melainabacteria_group>Cyanobacteriota>Cyanophyceae</td><td>GCF_900009275.2</td><td>Planktothrix</td><td>PLAN_RS00560</td><td>89888..93118</td><td>54304</td><td>MULTISPECIES:-DUF499-domain-containing-protein-[Planktothrix]</td></tr>
<tr><td>WP_026789283.1</td><td>AAA+ATPase+wHTH+Alpha+Beta-domain+wHTH+RRM</td><td>Type-2_BR.RM.like.systems</td><td><-METHYLASE-p<-?<-METHYLASE<-HTH_Tnp<-DUF3780<-DUF499*<-?<-REase<-Phage_int_SAM_1+Phage_integrase<-REase</td><td>1076</td><td>Bacteria>Terrabacteria_group>Cyanobacteriota/Melainabacteria_group>Cyanobacteriota>Cyanophyceae</td><td>GCF_000464785.1</td><td>Planktothrix_rubescens</td><td>N313_RS0125125</td><td>92611..95841</td><td>59512</td><td>DUF499-domain-containing-protein-[Planktothrix-rubescens]</td></tr>
<tr><td>WP_242016564.1</td><td>AAA+ATPase+wHTH+Alpha+Beta-domain+wHTH+RRM</td><td>Type-2_BR.RM.like.systems</td><td><-REase<-?<-METHYLASE||Classical-AAA->Peptidase-S8-><-DUF3780<-DUF499*||?->?->HPT+HISKIN+REC-></td><td>1132</td><td>Bacteria>Terrabacteria_group>Cyanobacteriota/Melainabacteria_group>Cyanobacteriota>Cyanophyceae</td><td>GCF_014696345.1</td><td>Pseudanabaena_cinerea</td><td>H6F44_RS13260</td><td>24620..28018</td><td>2661616</td><td>DUF499-domain-containing-protein-[Pseudanabaena-sp.-FACHB-1277]</td></tr>
</table>

| Protein-ID | DUF499-architecture | System | Gene-Neighbourhoods | Length | Taxonomy | Assembly-ID | Strain | Locus-Tag | Genomic-coordinates | TaxID | Fasta-defline |
|---|---|---|---|---|---|---|---|---|---|---|---|
| WP_199304646.1 | AAA+ATPase+wHTH+Alpha+Beta-domain+wHTH+RRM | Type-2_BR.RM.like.systems | <-REase<-METHYLASE\|\|Classical-AAA->Peptidase-S8-><-REase<-DUF3780<-DUF499* | 1067 | Bacteria>Terrabacteria_group>Cyanobacteriota/Melainabacteria_group>Cyanobacteriota>Cyanophyceae | - | Microcoleus_sp._FACHB-1515 | H6F67_RS25815 | 77444..80647 | 2692821 | DUF499-domain-containing-protein-[Microcoleus-sp.-FACHB-1515] |
| WP_027400718.1 | AAA+ATPase+wHTH+Alpha+Beta-domain+wHTH+RRM | Type-2_BR.RM.like.systems | ParA-Soj-PloopNTPase->?->?->?->DUF499*-><-?\|\|?->DUF3780->?->REase-> | 1116 | Bacteria>Terrabacteria_group>Cyanobacteriota/Melainabacteria_group>Cyanobacteriota>Cyanophyceae | GCF_000521175.1 | Aphanizomenon_flos-aquae | Y931_RS0101135 | 108477..111827 | 1176 | DUF499-domain-containing-protein-[Aphanizomenon-flos-aquae] |
| WP_103670645.1 | AAA+ATPase+wHTH+Alpha+Beta-domain+wHTH+RRM | Type-2_BR.RM.like.systems | <-METHYLASE\|\|MetJarc-HTH->PemK_toxin-><-REase<-?<-DUF3780<-DUF499*\|\|?->?->?->ParB-> | 1074 | Bacteria>Terrabacteria_group>Cyanobacteriota/Melainabacteria_group>Cyanobacteriota>Cyanophyceae | GCF_002914585.1 | Pseudanabaena_sp._BC1403 | CQ839_RS23050 | 18348..21572 | 2043171 | DUF499-domain-containing-protein-[Pseudanabaena-sp.-BC1403] |
| CAH2575563.1 | AAA+ATPase+wHTH+Alpha+Beta-domain+wHTH+RRM | Type-2_BR.RM.like.systems | REase->Phage_int_SAM_1+Phage_integrase->REase-><-?<-?\|\|?->DUF499*->DUF3780->HTH_Tnp->?->METHYLASE->?->METHYLASE-p-> | 1076 | Bacteria>Terrabacteria_group>Cyanobacteriota/Melainabacteria_group>Cyanobacteriota>Cyanophyceae | GCA_937024135.1 | Planktothrix_rubescens | PRNO82_05038 | 19838..23068 | 59512 | hypothetical-protein-PRNO82_05038-[Planktothrix-rubescens] |
| CAD5985010.1 | AAA+ATPase+wHTH+Alpha+Beta-domain+wHTH+RRM | Type-2_BR.RM.like.systems | REase->Phage_int_SAM_1+Phage_integrase->REase-><-?<-?\|\|?->DUF499*->DUF3780->HTH_Tnp->METHYLASE->?->METHYLASE-p-> | 1076 | Bacteria>Terrabacteria_group>Cyanobacteriota/Melainabacteria_group>Cyanobacteriota>Cyanophyceae | GCA_904830785.1 | Planktothrix_rubescens | NO108_04990 | 18013..21243 | 59512 | hypothetical-protein-NO108_04990-[Planktothrix-rubescens] |
| WP_190577353.1 | AAA+ATPase+wHTH+Alpha+Beta-domain+wHTH+RRM | Type-2_BR.RM.like.systems | Prim-Pol->?->?->CopG/RHH-><-?<-?\|\|DUF499*->DUF3780-><-MORC<-?\|\|METHYLASE->REase->HTH-> | 1074 | Bacteria>Terrabacteria_group>Cyanobacteriota/Melainabacteria_group>Cyanobacteriota>Cyanophyceae | GCF_014696925.1 | Phormidium_tenue | H6G05_RS06495 | 52293..55517 | 126344 | DUF499-domain-containing-protein-[Pseudanabaena-sp.-FACHB-1050] |
| WP_242049455.1 | AAA+ATPase+wHTH+Alpha+Beta-domain+wHTH+RRM | Type-2_BR.RM.like.systems | Classical-AAA->Peptidase-S8-><-?<-?\|\|?-><-DUF3780<-DUF499*<-wHTH+REase | 1070 | Bacteria>Terrabacteria_group>Cyanobacteriota/Melainabacteria_group>Cyanobacteriota>Cyanophyceae | GCF_014697245.1 | unclassified_Leptolyngbya | H6G61_RS21985 | 21330..24542 | 2650499 | MULTISPECIES:-DUF499-domain-containing-protein-[unclassified-Leptolyngbya] |
| WP_026149087.1 | AAA+ATPase+wHTH+Alpha+Beta-domain+wHTH+RRM | Type-2_BR.RM.like.systems | Classical-AAA->Peptidase-S8-><-?<-?\|\|?-><-DUF3780<-DUF499*<-wHTH+REase | 1071 | Bacteria>Terrabacteria_group>Cyanobacteriota/Melainabacteria_group>Cyanobacteriota>Cyanophyceae | GCF_002142475.1 | Leptolyngbya_boryana | MCP04_RS33160 | 388752..391967 | 1184 | DUF499-domain-containing-protein-[Leptolyngbya-boryana] |
| WP_017291799.1 | AAA+ATPase+wHTH+Alpha+Beta-domain+wHTH+RRM | Type-2_BR.RM.like.systems | wHTH+REase->DUF499*->DUF3780-><-?\|\|?->?-><-Peptidase-S8<-Classical-AAA | 1073 | Bacteria>Terrabacteria_group>Cyanobacteriota/Melainabacteria_group>Cyanobacteriota>Cyanophyceae | GCF_014697225.1 | Leptolyngbya | H6G55_RS21500 | 113566..116787 | 47251 | MULTISPECIES:-DUF499-domain-containing-protein-[Leptolyngbya] |
| WP_227788789.1 | AAA+ATPase+wHTH+Alpha+Beta-domain+wHTH+RRM | Type-2_BR.RM.like.systems | <-REase<-METHYLASE<-?<-METHYLASE-p<-DUF3780<-DUF499*\|\|?->ParB-> | 1077 | Bacteria>Terrabacteria_group>Cyanobacteriota/Melainabacteria_group>Cyanobacteriota>Cyanophyceae | GCF_020702635.1 | Nodularia_sp._LEGE_04288 | IQ240_RS03265 | 103901..107134 | 1828639 | DUF499-domain-containing-protein-[Nodularia-sp.-LEGE-04288] |
| MBD2151080.1 | AAA+ATPase+wHTH+Alpha+Beta-domain+wHTH+RRM | Type-2_BR.RM.like.systems | <-REase<-?<-METHYLASE\|\|Classical-AAA->Peptidase-S8-><-DUF3780<-DUF499*\|\|?->?->HPT+HISKIN+REC-> | 1085 | Bacteria>Terrabacteria_group>Cyanobacteriota/Melainabacteria_group>Cyanobacteriota>Cyanophyceae | GCA_014696345.1 | Pseudanabaena_cinerea_FACHB-1277 | H6F44_13265 | 24620..27877 | 2949581 | ATP-binding-protein-[Pseudanabaena-sp.-FACHB-1277] |
| MBW4617375.1 | AAA+ATPase+wHTH+Alpha+Beta-domain+wHTH+RRM | Type-2_BR.RM.like.systems | <-URI-DNase<-?\|\|?->DUF499*->DUF3780->?->METHYLASE-p-><-?\|\|REase->CopG/RHH-> | 1078 | Bacteria>Terrabacteria_group>Cyanobacteriota/Melainabacteria_group>Cyanobacteriota>Cyanophyceae | GCA_019359515.1 | Desmonostoc_vinosum_HA7617-LM4 | KME21_29995 | 8094..11330 | 1382956 | DUF499-domain-containing-protein-[Desmonostoc-vinosum-HA7617-LM4] |
| MBD2093275.1 | AAA+ATPase+wHTH+Alpha+Beta-domain+wHTH+RRM | Type-2_BR.RM.like.systems | <-METHYLASE\|\|Classical-AAA->Peptidase-S8-><-REase<-MACRODOMAIN<-DUF3780<-DUF499* | 1068 | Bacteria>Terrabacteria_group>Cyanobacteriota/Melainabacteria_group>Cyanobacteriota>Cyanophyceae | GCA_014696105.1 | Microcoleus_sp._FACHB-1515 | H6F67_25860 | 77444..80650 | 2692821 | ATP-binding-protein-[Microcoleus-sp.-FACHB-1515] |
| BAZ33968.1 | AAA+ATPase+wHTH+Alpha+Beta-domain+wHTH+RRM | Type-2_BR.RM.like.systems | <-CopG/RHH<-REase\|\|?->?-><-METHYLASE<-DUF3780<-DUF499*<-?<-Prim-Pol<-?<-ParB\|\|PAIREDC-HTH->Phage_int_SAM_1+Phage_integrase-> | 1131 | Bacteria>Terrabacteria_group>Cyanobacteriota/Melainabacteria_group>Cyanobacteriota>Cyanophyceae | GCA_003994795.1 | Cylindrospermum_sp._NIES-4074 | NIES4074_64820 | 86312..89707 | 2005457 | AAA-family-ATPase-[Cylindrospermum-sp.-NIES-4074] |
| WP_257121428.1 | AAA+ATPase+wHTH+Alpha+Beta-domain+wHTH+RRM | Type-2_BR.RM.like.systems | <-REase<-?<-DUF3780<-?\|\|?-><-DUF499*<-DUF29 | 1137 | Bacteria>Terrabacteria_group>Cyanobacteriota/Melainabacteria_group>Cyanobacteriota>Cyanophyceae | GCF_024584745.1 | Dolichospermum_heterosporum | NG743_RS02235 | 513982..517395 | 747522 | DUF499-domain-containing-protein-[Dolichospermum-heterosporum] |
| WP_017721096.1 | AAA+ATPase+wHTH+Alpha+Beta-domain+wHTH+RRM | Type-2_BR.RM.like.systems | DUF499*->DUF3780->REase->?->?->METHYLASE-> | 1078 | Bacteria>Terrabacteria_group>Cyanobacteriota/Melainabacteria_group>Cyanobacteriota>Cyanophyceae | GCF_000332335.1 | Oscillatoria_sp._PCC_10802 | OSC10802_RS0133315 | 641156..644392 | 1173028 | DUF499-domain-containing-protein-[Oscillatoria-sp.-PCC-10802] |
| WP_284081457.1 | AAA+ATPase+wHTH+Alpha+Beta-domain+wHTH+RRM | Type-2_BR.RM.like.systems | <-REase<-?<-DUF3780<-?\|\|?-><-DUF499*<-DUF29 | 1104 | Bacteria>Terrabacteria_group>Cyanobacteriota/Melainabacteria_group>Cyanobacteriota>Cyanophyceae | - | Aphanizomenon_sp._PH219 | QHH03_03540 | 3211..6525 | 2027336 | DUF499-domain-containing-protein-[Aphanizomenon-sp.-PH219] |
| WP_190384561.1 | AAA+ATPase+wHTH+Alpha+Beta-domain+wHTH+RRM | Type-2_BR.RM.like.systems | ParA-Soj-PloopNTPase->?->?->?->?->DUF499*->DUF3780->REase->?->REase-> | 1088 | Bacteria>Terrabacteria_group>Cyanobacteriota/Melainabacteria_group>Cyanobacteriota>Cyanophyceae | GCF_014696815.1 | Aphanizomenon_flos-aquae | H6F99_RS24975 | 5640..8906 | 1176 | DUF499-domain-containing-protein-[Aphanizomenon-flos-aquae] |
| WP_190398670.1 | AAA+ATPase+wHTH+Alpha+Beta-domain+wHTH+RRM | Type-2_BR.RM.like.systems | <-REase<-?<-METHYLASE\|\|Classical-AAA->Peptidase-S8-><-DUF3780<-DUF499* | 1074 | Bacteria>Terrabacteria_group>Cyanobacteriota/Melainabacteria_group>Cyanobacteriota>Cyanophyceae | GCF_014696355.1 | Pseudanabaena_sp._FACHB-1998 | H6F42_RS07140 | 12844..16068 | 2692858 | DUF499-domain-containing-protein-[Pseudanabaena-sp.-FACHB-1998] |
| WP_254567384.1 | AAA+ATPase+wHTH+Alpha+Beta-domain+wHTH+RRM | Type-2_BR.RM.like.systems | <-METHYLASE<-?<-DUF3780<-DUF499*\|\|REase-><-?<-?<-?\|\|Peptidase_M50-> | 1075 | Bacteria>Terrabacteria_group>Cyanobacteriota/Melainabacteria_group>Cyanobacteriota>Cyanophyceae | GCF_024220515.1 | Oscillatoria_sp._HE19RPO | NG833_RS21820 | 4599..7826 | 2954806 | DUF499-domain-containing-protein-[Oscillatoria-sp.-HE19RPO] |
| MCA6573123.1 | AAA+ATPase+wHTH+Alpha+Beta-domain+wHTH+RRM | Type-2_BR.RM.like.systems | <-METHYLASE<-?<-?<-?<-DUF3780<-DUF499*\|\|REase->?->?-><-SF1-SF2-Helicase+DUF1998 | 1074 | Bacteria>Terrabacteria_group>Cyanobacteriota/Melainabacteria_group>Cyanobacteriota>Cyanophyceae | GCA_020404105.1 | Pseudanabaena_sp._M53BS1SP1A06MG | IM549_07735 | 10744..13968 | 2771239 | ATP-binding-protein-[Pseudanabaena-sp.-M53BS1SP1A06MG] |
| MXW11415.1 | AAA+ATPase+wHTH+Alpha+Beta-domain+wHTH+RRM | Type-2_BR.RM.like.systems | <-AbrB-like\|\|?->?->DUF499*->DUF3780->REase->?->OLD-ABC->METHYLASE-> | 1020 | Bacteria>Terrabacteria_group>Cyanobacteriota/Melainabacteria_group>Cyanobacteriota>Cyanophyceae | GCA_009836025.1 | Synechococcus_sp._SB0668_bin_13 | F4Z73_00765 | 8681..11743 | 2604859 | ATP-binding-protein-[Synechococcus-sp.-SB0668_bin_13] |
| WP_039200355.1 | AAA+ATPase+wHTH+Alpha+Beta-domain+wHTH+RRM | Type-2_BR.RM.like.systems | <-METHYLASE-p<-REase<-?<-REase<-DUF3780\|\|?-><-DUF499* | 1088 | Bacteria>Terrabacteria_group>Cyanobacteriota/Melainabacteria_group>Cyanobacteriota>Cyanophyceae | GCF_000789435.1 | Aphanizomenon_flos-aquae | OA07_RS04085 | 6997..10263 | 1176 | DUF499-domain-containing-protein-[Aphanizomenon-flos-aquae] |
| OOV26578.1 | AAA+ATPase+wHTH+Alpha+Beta-domain+wHTH+RRM | Type-2_BR.RM.like.systems | DUF499*->DUF3780->REase->?->ParB+HNH-> | 1040 | Bacteria>Terrabacteria_group>Cyanobacteriota/Melainabacteria_group>Cyanobacteriota>Cyanophyceae | GCA_002018045.1 | Candidatus_Synechococcus_spongiarum_LMB_bulk15M | BV61_06570 | 12460..15582 | 1943582 | hypothetical-protein-BV61_06570-[Candidatus-Synechococcus-spongiarum-LMB-bulk15M] |
| PZV10243.1 | AAA+ATPase+wHTH+Alpha+Beta-domain+wHTH+RRM | Type-2_BR.RM.like.systems | DUF499*->DUF3780->REase+PNTD1+PNTD2->PIWI-><-MACRODOMAIN<-?\|\|METHYLASE-> | 1076 | Bacteria>Terrabacteria_group>Cyanobacteriota/Melainabacteria_group>Cyanobacteriota>Cyanophyceae | GCA_003249105.1 | Leptolyngbya_sp. | DCF32_00020 | 1234..4464 | 47254 | AAA-family-ATPase-[Leptolyngbya-sp.] |
| MYG64556.1 | AAA+ATPase+wHTH+Alpha+Beta-domain+wHTH+RRM | Type-2_BR.RM.like.systems | <-REase<-DUF3780<-DUF499*<-?<-?\|\|AbrB-like-> | 1025 | Bacteria>Terrabacteria_group>Cyanobacteriota/Melainabacteria_group>Cyanobacteriota>Cyanophyceae | GCA_009843455.1 | Synechococcus_sp._SB0675_bin_7 | F4158_08385 | 1344..4421 | 2604868 | ATP-binding-protein-[Synechococcus-sp.-SB0675_bin_7] |
| MXY61915.1 | AAA+ATPase+wHTH+Alpha+Beta-domain+wHTH+RRM | Type-2_BR.RM.like.systems | DUF499*->DUF3780->REase->?->?->ParB+HNH->OLD-ABC-> | 1035 | Bacteria>Terrabacteria_group>Cyanobacteriota/Melainabacteria_group>Cyanobacteriota>Cyanophyceae | GCA_009837735.1 | Synechococcus_sp._SB0665_bin_28 | F4Y87_00410 | 1971..5078 | 2604856 | ATP-binding-protein-[Synechococcus-sp.-SB0665_bin_28] |
| WP_277867662.1 | AAA+ATPase+wHTH+Alpha+Beta-domain+wHTH+RRM | Type-2_BR.RM.like.systems | <-DDE_Tnp_1\|\|DDE_Tnp_1-><-REase<-DUF3780<-DUF499* | 1012 | Bacteria>Terrabacteria_group>Cyanobacteriota/Melainabacteria_group>Cyanobacteriota>Cyanophyceae | GCF_029582805.1 | Candidatus_Synechococcus_calcipolaris | L3556_RS12450 | 3668..6706 | 1522304 | DUF499-domain-containing-protein-[Candidatus-Synechococcus-calcipolaris] |
| KKZ13991.1 | AAA+ATPase+wHTH+Alpha+Beta-domain+wHTH+RRM | Type-2_BR.RM.like.systems | HEPN->DUF499*->DUF3780->REase->?->ParB+HNH->OLD-ABC-> | 1040 | Bacteria>Terrabacteria_group>Cyanobacteriota/Melainabacteria_group>Cyanobacteriota>Cyanophyceae | GCA_001007635.1 | Candidatus_Synechococcus_spongiarum_15L | TQ37_02560 | 1477..4599 | 1608419 | hypothetical-protein-TQ37_02560-[Candidatus-Synechococcus-spongiarum-15L] |
| MYI71486.1 | AAA+ATPase+wHTH+Alpha+Beta-domain+wHTH+RRM | Type-2_BR.RM.like.systems | DUF499*->DUF3780->REase->?->?->OLD-ABC-> | 1020 | Bacteria>Terrabacteria_group>Cyanobacteriota/Melainabacteria_group>Cyanobacteriota>Cyanophyceae | GCA_009844645.1 | Synechococcus_sp._SB0673_bin_10 | F4099_03010 | 1152..4214 | 2604866 | ATP-binding-protein-[Synechococcus-sp.-SB0673_bin_10] |
| KKZ14064.1 | AAA+ATPase+wHTH+Alpha+Beta-domain+wHTH+RRM | Type-2_BR.RM.like.systems | <-REase<-?<-DUF3780<-DUF499* | 1040 | Bacteria>Terrabacteria_group>Cyanobacteriota/Melainabacteria_group>Cyanobacteriota>Cyanophyceae | GCA_001007625.1 | Candidatus_Synechococcus_spongiarum_142 | TH68_05775 | 1342..4464 | 1608213 | hypothetical-protein-TH68_05775-[Candidatus-Synechococcus-spongiarum-142] |
| WP_271835223.1 | AAA+ATPase+wHTH+Alpha+Beta-domain+wHTH+RRM | Type-2_BR.RM.like.systems | DUF499*->DUF3780->REase-> | 1083 | Bacteria>Terrabacteria_group>Cyanobacteriota/Melainabacteria_group>Cyanobacteriota>Cyanophyceae | GCF_028330815.1 | Dolichospermum_lemmermannii | PN450_RS01660 | 685..3936 | 54295 | DUF499-domain-containing-protein-[Dolichospermum-lemmermannii] |
| WP_255094986.1 | AAA+ATPase+wHTH+Alpha+Beta-domain+wHTH+RRM | Type-2_BR.RM.like.systems | AbrB-like->?->HD-><-?<-DUF4236\|\|DUF499*->DUF3780->ParB+HNH->METHYLASE-p-> | 1037 | Bacteria>Terrabacteria_group>Cyanobacteriota/Melainabacteria_group>Cyanobacteriota>Cyanophyceae | GCF_024345795.1 | Synechococcus_sp._J7-Johnson | KBY93_RS13695 | 20943..24056 | 2823737 | DUF499-domain-containing-protein-[Synechococcus-sp.-J7-Johnson] |
| MCA6595420.1 | AAA+ATPase+wHTH+Alpha+Beta-domain+wHTH+RRM | Type-2_BR.RM.like.systems | <-METHYLASE\|\|Classical-AAA->Peptidase-S8-><-DUF3780<-DUF499*<-?<-?<-HTH+HEPN<-SF1-SF2-Helicase<-STYKIN | 1074 | Bacteria>Terrabacteria_group>Cyanobacteriota/Melainabacteria_group>Cyanobacteriota>Cyanophyceae | GCA_020405005.1 | Pseudanabaena_sp._M046S1SP1A06QC | IM539_03680 | 19554..22778 | 2771236 | ATP-binding-protein-[Pseudanabaena-sp.-M046S1SP1A06QC] |
| MCA6522844.1 | AAA+ATPase+wHTH+Alpha+Beta-domain+wHTH+RRM | Type-2_BR.RM.like.systems | STYKIN->SF1-SF2-Helicase->HTH+HEPN->?->?->DUF499*->DUF3780->?->?->?->METHYLASE-p-> | 1074 | Bacteria>Terrabacteria_group>Cyanobacteriota/Melainabacteria_group>Cyanobacteriota>Cyanophyceae | GCA_020403865.1 | Pseudanabaena_sp._M051S1SP2A07QC | IM596_13630 | 16100..19324 | 2771253 | ATP-binding-protein-[Pseudanabaena-sp.-M051S1SP2A07QC] |
| WP_017654864.1 | AAA+ATPase+wHTH+Alpha+Beta-domain+wHTH+RRM | Type-2_BR.RM.like.systems | <-HNH<-?<-?<-VirD4-TraG+VirD4-FtsK\|\|<-DDE_Tnp_1\|\|SF1-SF2-Helicase->DUF499*->DUF3780-><-HTH+UbE1<-?<-?<-ZnR\|\|HTH+MPTase-> | 1077 | Bacteria>Terrabacteria_group>Cyanobacteriota/Melainabacteria_group>Cyanobacteriota>Cyanophyceae | GCF_000332295.1 | Fortiea_contorta | MIC7126_RS0119575 | 4220906..4224139 | 1892405 | DUF499-domain-containing-protein-[Fortiea-contorta] |
| WP_190371129.1 | AAA+ATPase+wHTH+Alpha+Beta-domain+wHTH+RRM | Type-2_BR.RM.like.systems | <-Pterin_bind<-?<-NUDIX<-Glyoxalase\|\|DUF499*->DUF3780->HNH->?->METHYLASE-p->?->METHYLASE-> | 1093 | Bacteria>Terrabacteria_group>Cyanobacteriota/Melainabacteria_group>Cyanobacteriota>Cyanophyceae | GCF_014696755.1 | Anabaena_sp._FACHB-1391 | H6F62_RS01330 | 59822..63103 | 2692771 | DUF499-domain-containing-protein-[Anabaena-sp.-FACHB-1391] |
| WP_146297647.1 | AAA+ATPase+wHTH+Alpha+Beta-domain+wHTH+RRM | Type-2_BR.RM.like.systems | <-BirA-HTH<-?<-TOPRIM+SF1-SF2-Helicase<-?<-BrnT_toxin<-DUF499*\|\|?->?->CopG/RHH-> | 1074 | Bacteria>Terrabacteria_group>Cyanobacteriota/Melainabacteria_group>Cyanobacteriota>Cyanophyceae | GCF_007904085.1 | Euhalothece_natronophila | FRE64_RS17115 | 29221..32445 | 577489 | DUF499-domain-containing-protein-[Euhalothece-natronophila] |
| WP_027255649.1 | AAA+ATPase+wHTH+Alpha+Beta-domain+wHTH+RRM | Type-2_BR.RM.like.systems | RVT+HNH->?->DUF499*->DUF3780->HTH_Tnp->METHYLASE->?->METHYLASE-p-> | 1076 | Bacteria>Terrabacteria_group>Cyanobacteriota/Melainabacteria_group>Cyanobacteriota>Cyanophyceae | GCF_000464825.1 | Planktothrix_agardhii | N316_RS0124610 | 93302..96532 | 1160 | DUF499-domain-containing-protein-[Planktothrix-agardhii] |
| WP_073634985.1 | AAA+ATPase+wHTH+Alpha+Beta-domain+wHTH+RRM | Type-2_BR.RM.like.systems | HNH->Mbetalac->?->DUF499*->DUF3780-><-?\|\|?->METHYLASE-p->METHYLASE-p->METHYLASE-> | 1092 | Bacteria>Terrabacteria_group>Cyanobacteriota/Melainabacteria_group>Cyanobacteriota>Cyanophyceae | GCF_001904675.1 | Scytonema_sp._HK-05 | NIES2130_RS30570 | 7897..11175 | 1137095 | DUF499-domain-containing-protein-[Scytonema-sp.-HK-05] |
| NJO63285.1 | AAA+ATPase+wHTH+Alpha+Beta-domain+wHTH+RRM | Type-2_BR.RM.like.systems | RVT+HNH->RVT+HNH->HNH->?-><-?\|\|DUF499*->DUF3780-> | 1079 | Bacteria>Terrabacteria_group>Cyanobacteriota/Melainabacteria_group>Cyanobacteriota>Cyanophyceae | GCA_012033295.1 | Richelia_sp._RM2_1_2 | HC836_35240 | 9019..12258 | 2720436 | ATP-binding-protein-[Richelia-sp.-RM2_1_2] |
| PSB50413.1 | AAA+ATPase+wHTH+Alpha+Beta-domain+wHTH+RRM | Type-2_BR.RM.like.systems | <-HKD-DNase+SF1-SF2-Helicase+iREase+CTD<-?<-METHYLASE<-DUF3780<-DUF499*\|\|?->ABC-ATPase+SbcC-> | 1076 | Bacteria>Terrabacteria_group>Cyanobacteriota/Melainabacteria_group>Cyanobacteriota>unclassified_Cyanobacteriota | GCA_003003725.1 | filamentous_cyanobacterium_Phorm_6 | C7B67_14560 | 4819..8049 | 2107706 | AAA-family-ATPase-[filamentous-cyanobacterium-Phorm-6] |
| MBU6346613.1 | AAA+ATPase+wHTH+Alpha+Beta-domain+wHTH+RRM | Type-2_BR.RM.like.systems | METHYLASE->HKD-DNase+SF1-SF2-Helicase+iREase+CTD->DUF499*->DUF3780-> | 1093 | Bacteria>Terrabacteria_group>Cyanobacteriota/Melainabacteria_group>Cyanobacteriota>unclassified_Cyanobacteriota | GCA_018969285.1 | Cyanobacteria_bacterium_REEB494 | KGR70_15160 | 7181..10462 | 2800107 | DUF499-domain-containing-protein-[Cyanobacteria-bacterium-REEB494] |
| MCY4173473.1 | AAA+ATPase+wHTH+Alpha+Beta-domain+wHTH+RRM | Type-2_BR.RM.like.systems | <-HTH_39\|\|<-?\|\|DUF499*->DUF3780->?->METHYLASE-><-PIN\|\|HKD-DNase+SF1-SF2-Helicase+iREase+CTD-> | 1020 | Bacteria>Terrabacteria_group>Cyanobacteriota/Melainabacteria_group>Cyanobacteriota>unclassified_Cyanobacteriota | GCA_026709085.1 | Cyanobacteria_bacterium_MAG_CAR3_bin_5 | OXF25_05325 | 614..3676 | 2775591 | MAG:-DUF499-domain-containing-protein-[Cyanobacteria-bacterium-MAG-CAR3_bin_5] |
| MCY4234851.1 | AAA+ATPase+wHTH+Alpha+Beta-domain+wHTH+RRM | Type-2_BR.RM.like.systems | DUF499*->DUF3780->?->METHYLASE-><-HEPN<-?\|\|HKD-DNase+SF1-SF2-Helicase+iREase+CTD-> | 1020 | Bacteria>Terrabacteria_group>Cyanobacteriota/Melainabacteria_group>Cyanobacteriota>unclassified_Cyanobacteriota | GCA_026709585.1 | Cyanobacteria_bacterium_MAG_CAR2_bin_4 | OXE74_00800 | 134..3196 | 2775590 | MAG:-DUF499-domain-containing-protein-[Cyanobacteria-bacterium-MAG-CAR2_bin_4] |
| HBB35643.1 | AAA+ATPase+wHTH+Alpha+Beta-domain+wHTH+RRM | Type-2_BR.RM.like.systems | FAD-NAD-dep-oxidoreductase->?->?->DUF499*->DUF3780-><-ZnR\|\|HTH+MPTase->PIN->METHYLASE-> | 1077 | Bacteria>Terrabacteria_group>Cyanobacteriota/Melainabacteria_group>Cyanobacteriota>unclassified_Cyanobacteriota | GCA_003488275.1 | Cyanobacteria_bacterium_UBA9273 | DC064_28650 | 3837..7070 | 2055791 | AAA-family-ATPase-[Cyanobacteria-bacterium-UBA9273] |
| MBM5817556.1 | AAA+ATPase+wHTH+Alpha+Beta-domain+wHTH+RRM | Type-2_BR.RM.like.systems | <-METHYLASE-p<-PIN<-RHH<-PIN<-TraJ-RHH<-DUF3780<-DUF499*<-Y2_Tnp<-?<-Y2_Tnp | 1031 | Bacteria>Terrabacteria_group>Cyanobacteriota/Melainabacteria_group>Cyanobacteriota>unclassified_Cyanobacteriota | GCA_016882305.1 | Cyanobacteria_bacterium_K_Offshore_surface_m2_239 | FJ083_13530 | 2227..5322 | 2775555 | ATP-binding-protein-[Cyanobacteria-bacterium-K_Offshore_surface_m2_239] |
| MCX5965989.1 | AAA+ATPase+wHTH+Alpha+Beta-domain+wHTH+RRM | Type-2_BR.RM.like.systems | DUF499*->DUF3780->Phage_integrase->METHYLASE->?->PIN-> | 1079 | Bacteria>Terrabacteria_group>Cyanobacteriota/Melainabacteria_group>Cyanobacteriota>unclassified_Cyanobacteriota | GCA_026389365.1 | Cyanobacteriota_bacterium | NT070_23555 | 754..3993 | 2847862 | MAG:-DUF499-domain-containing-protein-[Cyanobacteria-bacterium] |
| MCY4331636.1 | AAA+ATPase+wHTH+Alpha+Beta-domain+wHTH+RRM | Type-2_BR.RM.like.systems | <-OLD-ABC<-ParB+HNH<-?<-REase<-DUF3780<-DUF499*\|\|?->\|\|HTH_39-><-MethyltransfD12\|\|?->NUDIX-> | 1020 | Bacteria>Terrabacteria_group>Cyanobacteriota/Melainabacteria_group>Cyanobacteriota>unclassified_Cyanobacteriota | GCA_026710375.1 | Cyanobacteria_bacterium_MAG_CAR1_bin_15 | OXC96_03845 | 58664..61726 | 2775589 | MAG:-DUF499-domain-containing-protein-[Cyanobacteria-bacterium-MAG-CAR1_bin_15] |
| MDE0648516.1 | AAA+ATPase+wHTH+Alpha+Beta-domain+wHTH+RRM | Type-2_BR.RM.like.systems | <-AbrB-like\|\|?->METHYLASE-p->?->DUF499*->DUF3780->REase->?->OLD-ABC->METHYLASE-> | 1025 | Bacteria>Terrabacteria_group>Cyanobacteriota/Melainabacteria_group>Cyanobacteriota>unclassified_Cyanobacteriota | GCA_028824215.1 | Cyanobacteria_bacterium_MAG_IRC4_bin_6 | OXI08_10885 | 8218..11295 | 2775598 | MAG:-DUF499-domain-containing-protein-[Cyanobacteria-bacterium-MAG-IRC4_bin_6] |
| MCX5946177.1 | AAA+ATPase+wHTH+Alpha+Beta-domain+wHTH+RRM | Type-2_BR.RM.like.systems | REase->?->?->?->SIGMA-HTH->DUF499*->DUF3780->?->METHYLASE-p-> | 1059 | Bacteria>Terrabacteria_group>Cyanobacteriota/Melainabacteria_group>Cyanobacteriota>unclassified_Cyanobacteriota | GCA_026389235.1 | Cyanobacteriota_bacterium | NTZ53_12960 | 51039..54218 | 2847862 | MAG:-DUF499-domain-containing-protein-[Cyanobacteria-bacterium] |
| MCY4360135.1 | AAA+ATPase+wHTH+Alpha+Beta-domain+wHTH+RRM | Type-2_BR.RM.like.systems | <-REase<-DUF3780<-DUF499* | 1040 | Bacteria>Terrabacteria_group>Cyanobacteriota/Melainabacteria_group>Cyanobacteriota>unclassified_Cyanobacteriota | GCA_026710605.1 | Cyanobacteria_bacterium_MAG_APA_bin_95 | OXC47_05670 | 648..3770 | 2775588 | MAG:-DUF499-domain-containing-protein-[Cyanobacteria-bacterium-MAG-APA_bin_95] |
| MCY3653787.1 | AAA+ATPase+wHTH+Alpha+Beta-domain+wHTH+RRM | Type-2_BR.RM.like.systems | DUF499*->DUF3780->REase-> | 877 | Bacteria>Terrabacteria_group>Cyanobacteriota/Melainabacteria_group>Cyanobacteriota>unclassified_Cyanobacteriota | GCA_026705665.1 | Cyanobacteria_bacterium_MAG_IRC1_bin_28 | OXG70_01955 | 1..2635 | 2775596 | MAG:-DUF499-domain-containing-protein,-partial-[Cyanobacteria-bacterium-MAG-IRC1_bin_28] |
| WP_012873276.1 | AAA+ATPase+wHTH+Alpha+Beta-domain+wHTH+RRM | Type-2_BR.RM.like.systems | HAD+HAD->?->?-><-DUF6361\|\|DDE_Tnp_1_3->DUF499*->DUF3780->METHYLASE->HKD-DNase+SF1-SF2-Helicase+iREase+CTD->tRNA-><-Phage_int_SAM_1+Phage_integrase<-?\|\|XIS-HTH-> | 1020 | Bacteria>Terrabacteria_group>Thermomicrobiota>Thermomicrobia>Sphaerobacterales | GCF_000024985.1 | Sphaerobacter_thermophilus | STHE_RS14160 | 440471..443533 | 2057 | DUF499-domain-containing-protein-[Sphaerobacter-thermophilus] |
| MBX6754321.1 | AAA+ATPase+wHTH+Alpha+Beta-domain+wHTH+RRM | Type-2_BR.RM.like.systems | DUF499*->DUF3780->SF1-SF2-Helicase+DUF1998->METHYLASE->HKD-DNase+SF1-SF2-Helicase+iREase+CTD-> | 1014 | Bacteria>Terrabacteria_group>Thermomicrobiota>Thermomicrobia>Thermomicrobia_incertae_sedis | GCA_019690065.1 | Thermorudis_peleae | IRY86_08820 | 311..3355 | 1382356 | ATP-binding-protein-[Thermorudis-peleae] |
| MCX7623923.1 | AAA+ATPase+wHTH+Alpha+Beta-domain+wHTH+RRM | Type-2_BR.RM.like.systems | DUF499*->DUF3780->?->MazE_antitoxin->METHYLASE->HKD-DNase+SF1-SF2-Helicase+iREase+CTD-> | 1015 | Bacteria>Terrabacteria_group>Thermomicrobiota>Thermomicrobia>Thermomicrobiales | GCA_026419575.1 | Thermomicrobium_sp. | N2Z82_09275 | 210..3257 | 1969469 | MAG:-DUF499-domain-containing-protein-[Thermomicrobium-sp.] |
| WP_252129089.1 | AAA+ATPase+wHTH+Alpha+Beta-domain+wHTH+RRM | Type-2_BR.RM.like.systems | <-HKD-DNase+SF1-SF2-Helicase+iREase+CTD<-METHYLASE<-?<-?<-DUF3780<-DUF499*<-?<-?\|\|ABC-ATPase+SbcC-> | 1004 | Bacteria>Terrabacteria_group>Thermomicrobiota>Thermomicrobia>Thermomicrobiales | GCF_023806465.1 | Thermomicrobium_sp._CFH_73360 | NETO3_RS11605 | 91122..94136 | 2951987 | DUF499-domain-containing-protein-[Thermomicrobium-sp.-CFH-73360] |
| WP_284055791.1 | AAA+ATPase+wHTH+Alpha+Beta-domain+wHTH+RRM | Type-2_BR.RM.like.systems | <-Phage_integrase\|\|DUF499*->DUF3780->HEPN+Mrr_N->?->?->REase->MazE_antitoxin-> | 1016 | Bacteria>Terrabacteria_group>Thermomicrobiota>Thermomicrobia>Thermomicrobiales | GCF_023806485.1 | Thermalbibacter_longus | NETO2_RS02500 | 568134..571184 | 2951981 | DUF499-domain-containing-protein-[Thermalbibacter-longus] |
| MCM8748012.1 | AAA+ATPase+wHTH+Alpha+Beta-domain+wHTH+RRM | Type-2_BR.RM.like.systems | <-Phage_integrase\|\|DUF499*->DUF3780->HEPN+Mrr_N->?->?->REase->MazE_antitoxin-> | 1016 | Bacteria>Terrabacteria_group>Thermomicrobiota>Thermomicrobia>Thermomicrobiales | GCA_023806485.1 | Thermalbibacter_longus | NETO2_02505 | 568134..571184 | 2951981 | DUF499-domain-containing-protein-[Thermomicrobiaceae-bacterium-CFH-74404] |
| WP_051375312.1 | AAA+ATPase+wHTH+Alpha+Beta-domain+wHTH+RRM | Type-2_BR.RM.like.systems | <-DDE_Tnp_1\|\|DUF499*->DUF499-p->DUF3780->METHYLASE->Schlafen+HATPase+HTH->HKD-DNase+SF1-SF2-Helicase->TIR-> | 790 | Bacteria>Thermodesulfobacteriota>Desulfobacteria>Desulfobacterales>Desulfosarcinaceae | GCF_028752835.1 | Desulfosarcina_sp._BuS5 | A39W_RS22855 | 2496..4868 | 933262 | DUF499-domain-containing-protein-[Desulfosarcina-sp.-BuS5] |
| WP_027369815.1 | AAA+ATPase+wHTH+Alpha+Beta-domain+wHTH+RRM | Type-2_BR.RM.like.systems | <-DUF433\|\|Schlafen+HATPase+HTH->REase-><-?<-DUF433\|\|DUF499*->DUF3780->DOC+HTH->METHYLASE->HicA_toxin->HicB_lk_antitox->HKD-DNase+SF1-SF2-Helicase+iREase+CTD-> | 1022 | Bacteria>Thermodesulfobacteriota>Desulfovibrionia>Desulfovibrionales>Desulfohalobiaceae | GCF_000620765.1 | Desulfovermiculus_halophilus | N902_RS0103555 | 234313..237381 | 339722 | DUF499-domain-containing-protein-[Desulfovermiculus-halophilus] |
| MBD5553419.1 | AAA+ATPase+wHTH+Alpha+Beta-domain+wHTH+RRM | Type-2_BR.RM.like.systems | <-HKD-DNase+SF1-SF2-Helicase+iREase+CTD<-METHYLASE<-DUF3780<-DUF499* | 1049 | Bacteria>Thermodesulfobacteriota>Desulfovibrionia>Desulfovibrionales>Desulfovibrionaceae | GCA_014804885.1 | Desulfovibrio_sp. | HDQ44_03680 | 24575..27724 | 885 | ATP-binding-protein-[Desulfovibrio-sp.] |
| MBQ7457436.1 | AAA+ATPase+wHTH+Alpha+Beta-domain+wHTH+RRM | Type-2_BR.RM.like.systems | <-HKD-DNase+SF1-SF2-Helicase+iREase+CTD<-METHYLASE<-DUF3780<-DUF499*<-CR-ATPase8+REase<-?\|\|HDRnase-> | 1008 | Bacteria>Thermodesulfobacteriota>Desulfovibrionia>Desulfovibrionales>Desulfovibrionaceae | GCA_017450275.1 | Desulfovibrio_sp. | IJS54_07515 | 6253..9279 | 885 | ATP-binding-protein-[Desulfovibrio-sp.] |
| MCR5814188.1 | AAA+ATPase+wHTH+Alpha+Beta-domain+wHTH+RRM | Type-2_BR.RM.like.systems | DUF499*->DUF3780->RelE-ParE->HTH->METHYLASE-p->METHYLASE->HKD-DNase+SF1-SF2-Helicase+iREase+CTD-> | 1014 | Bacteria>Thermodesulfobacteriota>Desulfovibrionia>Desulfovibrionales>Desulfovibrionaceae | GCA_024699605.1 | Desulfovibrio_sp. | K6G15_06825 | 3599..6643 | 885 | DUF499-domain-containing-protein-[Desulfovibrio-sp.] |
| SBW09990.1 | AAA+ATPase+wHTH+Alpha+Beta-domain+wHTH+RRM | Type-2_BR.RM.like.systems | DUF499*->DUF3780->HTH+MPTase->PIN->METHYLASE->HKD-DNase+SF1-SF2-Helicase+iREase+CTD-> | 1024 | Bacteria>Thermodesulfobacteriota>Desulfovibrionia>Desulfovibrionales>Desulfovibrionaceae | GCA_900079715.1 | uncultured_Desulfovibrio_sp. | KM92DES2_12846 | 2964314..2967388 | 167968 | conserved-hypothetical-protein-[uncultured-Desulfovibrio-sp.] |
| WP_206214249.1 | AAA+ATPase+wHTH+Alpha+Beta-domain+wHTH+RRM | Type-2_BR.RM.like.systems | <-HIN-HTH\|\|TraJ-RHH->?->RVT->?->DUF499*->DUF3780->HTH+MPTase->PIN->METHYLASE->HKD-DNase+SF1-SF2-Helicase+iREase+CTD-> | 1012 | Bacteria>Thermodesulfobacteriota>Desulfovibrionia>Desulfovibrionales>Desulfovibrionaceae | GCF_011039085.1 | Desulfovibrio_sp._ZJ369 | G7Y51_RS04960 | 82456..85494 | 2709793 | DUF499-domain-containing-protein-[Desulfovibrio-sp.-ZJ369] |
| WP_163302422.1 | AAA+ATPase+wHTH+Alpha+Beta-domain+wHTH+RRM | Type-2_BR.RM.like.systems | Phage_integrase-><-DUF932<-HKD-DNase+SF1-SF2-Helicase+iREase+CTD<-METHYLASE<-Schlafen+HATPase+HTH<-DUF3780<-DUF499*<-?\|\|?-><-PAIREDC-HTH\|\|?->ZnR+TOPRIM+primpol+HTH-><-DUF772+DDE_Tnp_1 | 1014 | Bacteria>Thermodesulfobacteriota>Desulfovibrionia>Desulfovibrionales>Desulfovibrionaceae | GCF_013376455.1 | Desulfolutivibrio_sulfodismutans | GD606_RS12045 | 2614561..2617605 | 63561 | DUF499-domain-containing-protein-[Desulfovibrio-sulfodismutans] |
| GBD21619.1 | AAA+ATPase+wHTH+Alpha+Beta-domain+wHTH+RRM | Type-2_BR.RM.like.systems | <-ABC-ATPase+SbcC\|\|?->?->DUF499*->DUF3780->METHYLASE->?->HKD-DNase+SF1-SF2-Helicase+iREase+CTD-> | 1018 | Bacteria>unclassified_Bacteria | GCA_002898255.1 | bacterium_HR28 | HRbin28_02070 | 193704..196760 | 2035423 | hypothetical-protein-HRbin28_02070-[bacterium-HR28] |
| GBD14658.1 | AAA+ATPase+wHTH+Alpha+Beta-domain+wHTH+RRM | Type-2_BR.RM.like.systems | DUF499*->DUF3780->METHYLASE->HKD-DNase+SF1-SF2-Helicase+iREase+CTD-> | 1011 | Bacteria>unclassified_Bacteria | GCA_002898615.1 | bacterium_HR25 | HRbin25_00537 | 680..3715 | 2035420 | hypothetical-protein-HRbin25_00537-[bacterium-HR25] |
| MCY4162822.1 | AAA+ATPase+wHTH+Alpha+Beta-domain+wHTH+RRM | Type-2_BR.RM.like.systems | <-HKD-DNase+SF1-SF2-Helicase+iREase+CTD<-METHYLASE<-DUF3780<-DUF499* | 1110 | Bacteria>unclassified_Bacteria | GCA_026709025.1 | bacterium | OXE93_01185 | 15478..18810 | 1869227 | MAG:-DUF499-domain-containing-protein-[bacterium] |
| MCY4257860.1 | AAA+ATPase+wHTH+Alpha+Beta-domain+wHTH+RRM | Type-2_BR.RM.like.systems | DUF499*->DUF499-p->DUF3780->METHYLASE->HKD-DNase+SF1-SF2-Helicase+iREase+CTD-> | 934 | Bacteria>unclassified_Bacteria | GCA_026709785.1 | bacterium | OXE04_06160 | 17115..19918 | 1869227 | MAG:-DUF499-domain-containing-protein,-partial-[bacterium] |
| MDE0417756.1 | AAA+ATPase+wHTH+Alpha+Beta-domain+wHTH+RRM | Type-2_BR.RM.like.systems | <-Peptidase_M20\|\|HAD->TraJ-RHH->?->TraJ-RHH-><-TrwC+SF1-SF2-Helicase+TOPRIM\|\|DUF499*->DUF3780->?->?->?->METHYLASE->HKD-DNase+SF1-SF2-Helicase+iREase+CTD-> | 1009 | Bacteria>unclassified_Bacteria | GCA_028821455.1 | bacterium | OXI95_12600 | 159017..162046 | 1869227 | MAG:-DUF499-domain-containing-protein-[bacterium] |
| MDE0603886.1 | AAA+ATPase+wHTH+Alpha+Beta-domain+wHTH+RRM | Type-2_BR.RM.like.systems | <-HKD-DNase+SF1-SF2-Helicase+iREase+CTD<-METHYLASE<-DUF3780<-DUF499* | 984 | Bacteria>unclassified_Bacteria | GCA_028824745.1 | bacterium | OXI18_05720 | 7136..10090 | 1869227 | MAG:-DUF499-domain-containing-protein-[bacterium] |
| GBE30120.1 | AAA+ATPase+wHTH+Alpha+Beta-domain+wHTH+RRM | Type-2_BR.RM.like.systems | <-PIN<-?<-DUF3780<-DUF499*\|\|?-><-REase\|\|?->DUF932-> | 1005 | Bacteria>unclassified_Bacteria | GCA_002898195.1 | bacterium_BMS3Bbin04 | BMS3Bbin04_01146 | 2034..5051 | 2005727 | hypothetical-protein-BMS3Bbin04_01146-[bacterium-BMS3Bbin04] |
| MTJ27308.1 | AAA+ATPase+wHTH+Alpha+Beta-domain+wHTH+RRM | Type-2_BR.RM.like.systems | <-Pterin_bind<-?<-NUDIX<-Glyoxalase\|\|DUF499*->DUF3780->HNH->?->METHYLASE-p->METHYLASE-p->HKD-DNase+SF1-SF2-Helicase+iREase+CTD-> | 1093 | - | - | - | FJR41_RS17240 | 43574..46855 | - | ATP-binding-protein-[Dolichospermum-planctonicum-UHCC-0167] |
| WP_235225417.1 | AAA+ATPase+wHTH+Alpha+Beta-domain+wHTH+RRM | Type-2_BR.RM.like.systems | <-MAD-NTD+HNH\|\|<-?\|\|zf-ISL3+DDE_Tnp_ISL3->DUF499*->DUF499-p->DUF3780->METHYLASE->HKD-DNase+SF1-SF2-Helicase-><-NACHT+SF1-SF2-Helicase+HKD-DNase<-RNaseH | 910 | Bacteria>Pseudomonadota>Gammaproteobacteria>Enterobacterales>Enterobacteriaceae | GCF_021498025.2 | Escherichia_coli | HPY99_RS03315 | 704972..707704 | 562 | DUF499-domain-containing-protein-[Escherichia-coli] |
| HCK2681827.1 | AAA+ATPase+wHTH+Alpha+Beta-domain+wHTH+RRM | Type-2_BR.RM.like.systems | <-DUF499*\|\|?->MAD-NTD+HNH-><-?<-?\|\|Phage_integrase-> | 801 | Bacteria>Pseudomonadota>Gammaproteobacteria>Enterobacterales>Enterobacteriaceae | GCA_024776365.1 | Escherichia_coli | NY534_005063 | 1..2402 | 562 | ATP-binding-protein-[Escherichia-coli] |
| MBU4356577.1 | AAA+ATPase+wHTH+Alpha+Beta-domain+wHTH+RRM | Type-2_BR.RM.like.systems | <-Phage_int_SAM_4+Phage_integrase<-XIS-HTH<-?<-TOPRIM<-HTH<-?\|\|DUF499*-> | 873 | Bacteria>Pseudomonadota>unclassified_Pseudomonadota | GCA_018896435.1 | Pseudomonadota_bacterium | KJ822_14745 | 11884..14502 | 1977087 | DUF499-domain-containing-protein-[Proteobacteria-bacterium] |
| MCZ7647980.1 | AAA+ATPase+wHTH+Alpha+Beta-domain+wHTH+RRM | Type-2_BR.RM.like.systems | <-XIS-HTH<-?<-?<-?<-HTH_3\|\|?->DUF499*->DUF3780->METHYLASE->HKD-DNase+SF1-SF2-Helicase-><-Trypsin<-?<-PIWI | 941 | Bacteria>PVC_group>Planctomycetota>unclassified_Planctomycetota | GCA_027493525.1 | Planctomycetota_bacterium | M5U26_22435 | 8207..11032 | 2026780 | MAG:-DUF499-domain-containing-protein-[Planctomycetota-bacterium] |
| HJI24373.1 | AAA+ATPase+wHTH+Alpha+Beta-domain+wHTH+RRM | Type-2_BR.RM.like.systems | <-Peptidase_M28<-?<-?<-?<-Phage_integrase\|\|DUF499*->DUF499-p->DUF3780->METHYLASE->HKD-DNase+SF1-SF2-Helicase-> | 923 | Bacteria>PVC_group>Verrucomicrobiota>Verrucomicrobiae>Verrucomicrobiales | GCA_025993435.1 | Akkermansia_muciniphila | OIL89_05300 | 1236852..1239623 | 239935 | MAG-TPA:-DUF499-domain-containing-protein-[Akkermansia-muciniphila] |
| EJU29934.1 | AAA+ATPase+wHTH+Alpha+Beta-domain+wHTH+RRM | Type-2_BR.RM.like.systems | Peptidase_S8-><-?<-HKD-DNase+SF1-SF2-Helicase<-METHYLASE-p<-METHYLASE-p<-METHYLASE-p<-DUF499* | 964 | Bacteria>Terrabacteria_group>Bacillota>Negativicutes>Selenomonadales | GCA_000292955.1 | Selenomonas_sp._CM52 | HMPREF1153_1707 | 45364..48258 | 936381 | chitobiase/beta-hexosaminidase-C-terminal-domain-protein-[Selenomonas-sp.-CM52] |
| WP_198003306.1 | AAA+ATPase+wHTH+Alpha+Beta-domain+wHTH+RRM | Type-2_BR.RM.like.systems | Peptidase_S8-><-?<-HKD-DNase+SF1-SF2-Helicase<-METHYLASE-p<-METHYLASE-p<-METHYLASE-p<-DUF499* | 958 | Bacteria>Terrabacteria_group>Bacillota>Negativicutes>Selenomonadales | GCF_000292955.1 | Selenomonas_sp._CM52 | HMPREF1153_RS02155 | 45364..48240 | 936381 | DUF499-domain-containing-protein-[Selenomonas-sp.-CM52] |
| MCX7789960.1 | AAA+ATPase+wHTH+Alpha+Beta-domain+wHTH+RRM | Type-2_BR.RM.like.systems | ZnR->?->PolB->DUF499*->DUF499-p->DUF3780->PolB->HEPN-> | 910 | Bacteria>Terrabacteria_group>Chloroflexota>Chloroflexia>Chloroflexales | GCA_026417815.1 | Chloroflexaceae_bacterium | N2378_04905 | 2392..5124 | 2562117 | MAG:-DUF499-domain-containing-protein-[Chloroflexaceae-bacterium] |
| WP_260644223.1 | AAA+ATPase+wHTH+Alpha+Beta-domain+wHTH+RRM | Type-2_BR.RM.like.systems | REase-><-HKD-DNase+SF1-SF2-Helicase+iREase+CTD<-METHYLASE<-DUF3780<-?<-?<-DUF499* | 601 | Archaea>Euryarchaeota>Stenosarchaea_group>Halobacteria>Halobacteriales | GCF_025231485.1 | Salinirubellus_salinus | N0B31_RS22540 | 247172..248977 | 1364945 | DUF499-domain-containing-protein-[Salinirubellus-salinus] |
| WP_209019655.1 | AAA+ATPase+wHTH+Alpha+Beta-domain+wHTH+RRM | Type-2_BR.RM.like.systems | HicA_toxin-><-?\|\|<-DDE_Tnp_1<-ZnR\|\|DUF772+DDE_Tnp_1-><-DUF499*\|\|?->MORC+DUF3684+REase->ORC/CDC6+HTH-><-REase | 638 | Archaea>Euryarchaeota>Stenosarchaea_group>Halobacteria>Halobacteriales | GCF_004521895.1 | Halorussus_marinus | EPL04_RS16715 | 110620..112536 | 2505976 | DUF499-domain-containing-protein-[Halorussus-marinus] |
| KPN31954.1 | AAA+ATPase+wHTH+Alpha+Beta-domain+wHTH+RRM | Type-2_BR.RM.like.systems | <-HKD-DNase+SF1-SF2-Helicase+iREase+CTD<-METHYLASE-p<-METHYLASE<-DUF3780<-?<-DUF499-p<-DUF499*\|\|?-><-?\|\|?->?->HTH-> | 486 | Archaea>Euryarchaeota>Stenosarchaea_group>Halobacteria>Haloferacales | GCA_001307315.1 | Halolamina_pelagica | SY89_02711 | 2262976..2264436 | 699431 | hypothetical-protein-SY89_02711-[Halolamina-pelagica] |
| WP_239685613.1 | AAA+ATPase+wHTH+Alpha+Beta-domain+wHTH+RRM | Type-2_BR.RM.like.systems | <-HKD-DNase+SF1-SF2-Helicase+iREase+CTD<-METHYLASE-p<-METHYLASE<-DUF3780<-?<-DUF499*<-DUF499\|\|?-><-?\|\|?->?->HTH-> | 448 | Archaea>Euryarchaeota>Stenosarchaea_group>Halobacteria>Haloferacales | GCF_001307315.1 | Halolamina_pelagica | SY89_RS18610 | 2261633..2262979 | 699431 | DUF499-domain-containing-protein-[Halolamina-pelagica] |
| WP_239685614.1 | AAA+ATPase+wHTH+Alpha+Beta-domain+wHTH+RRM | Type-2_BR.RM.like.systems | <-HKD-DNase+SF1-SF2-Helicase+iREase+CTD<-METHYLASE-p<-METHYLASE<-DUF3780<-?<-DUF499<-DUF499*\|\|?-><-?\|\|?->?->HTH-> | 469 | Archaea>Euryarchaeota>Stenosarchaea_group>Halobacteria>Haloferacales | GCF_001307315.1 | Halolamina_pelagica | SY89_RS18615 | 2262976..2264385 | 699431 | DUF499-domain-containing-protein-[Halolamina-pelagica] |
| WP_006169677.1 | AAA+ATPase+wHTH+Alpha+Beta-domain+wHTH+RRM | Type-2_BR.RM.like.systems | REase+PNTD1+PNTD2->PIWI-><-Cluster197_21clades\|\|?->DUF499*-> | 827 | Archaea>Euryarchaeota>Stenosarchaea_group>Halobacteria>Natrialbales | GCF_000337135.1 | Natrialba_chahannaoensis | C482_RS20060 | 11469..13949 | 68911 | DUF499-domain-containing-protein,-partial-[Natrialba-chahannaoensis] |
| MCE8429039.1 | AAA+ATPase+wHTH+Alpha+Beta-domain+wHTH+RRM | Type-2_BR.RM.like.systems | DUF499*->DUF499-p->DUF3780->METHYLASE->HKD-DNase+SF1-SF2-Helicase-><-EVE | 399 | Archaea>Euryarchaeota>Stenosarchaea_group>Methanomicrobia>Methanosarcinales | GCA_021409285.1 | Candidatus_Methanoperedens_sp. | J5U19_11695 | 1..1202 | 2032688 | DUF499-domain-containing-protein-[Candidatus-Methanoperedens-sp.] |
| GIU72777.1 | AAA+ATPase+wHTH+Alpha+Beta-domain+wHTH+RRM | Type-2_BR.RM.like.systems | DUF488->DUF499*->DUF3780-><-PIN<-?<-?\|\|METHYLASE->SF1-SF2-Helicase+iREase+CTD+REase-> | 894 | Archaea>TACK_group>Nitrososphaerota>Nitrososphaeria>Candidatus_Nitrosocaldales | GCA_026002835.1 | Candidatus_Nitrosocaldaceae_archaeon | KatS3mg003_2256 | 14146..16830 | 2838776 | MAG:-hypothetical-protein-KatS3mg003_2256-[Candidatus-Nitrosocaldaceae-archaeon] |
| MDE0525348.1 | AAA+ATPase+wHTH+Alpha+Beta-domain+wHTH+RRM | Type-2_BR.RM.like.systems | <-MarR-HTH\|\|DUF499*->DUF3780->METHYLASE->SF1-SF2-Helicase+iREase+CTD+REase-> | 885 | Archaea>TACK_group>Nitrososphaerota>unclassified_Nitrososphaerota | GCA_028823795.1 | Nitrososphaerota_archaeon | OXI27_01935 | 36993..39650 | 2026795 | MAG:-DUF499-domain-containing-protein-[Thaumarchaeota-archaeon] |
| MYK11747.1 | AAA+ATPase+wHTH+Alpha+Beta-domain+wHTH+RRM | Type-2_BR.RM.like.systems | <-DUF499*\|\|OLD-ABC->TOPRIM->METHYLASE->HKD-DNase+SF1-SF2-Helicase+iREase+CTD-> | 815 | Bacteria>Acidobacteriota>Terriglobia>unclassified_Terriglobia | GCA_009845805.1 | Terriglobia_bacterium | F4041_19170 | 1..2445 | 2081523 | ATP-binding-protein-[Acidobacteriia-bacterium] |
| MCI0393005.1 | AAA+ATPase+wHTH+Alpha+Beta-domain+wHTH+RRM | Type-2_BR.RM.like.systems | DUF499*->DUF499-p->DUF3780->METHYLASE->HKD-DNase+SF1-SF2-Helicase-> | 798 | Bacteria>Acidobacteriota>unclassified_Acidobacteriota | GCA_022614935.1 | Acidobacteriota_bacterium | MOB07_30090 | 151..2547 | 1978231 | DUF499-domain-containing-protein-[Acidobacteria-bacterium] |
| MCG2812992.1 | AAA+ATPase+wHTH+Alpha+Beta-domain+wHTH+RRM | Type-2_BR.RM.like.systems | <-DUF499*<-TOPRIM<-OLD-ABC<-METHYLASE<-ASCH+DUF3780<-PAPS_reduct | 916 | Bacteria>Bacteria_incertae_sedis>Bacteria_candidate_phyla>Candidatus_Aminicenantes>unclassified_Candidatus_Aminicenantes | GCA_022013915.1 | Candidatus_Aminicenantes_bacterium | L6428_16300 | 153..2903 | 2052149 | DUF499-domain-containing-protein-[Candidatus-Aminicenantes-bacterium] |
| QQS19533.1 | AAA+ATPase+wHTH+Alpha+Beta-domain+wHTH+RRM | Type-2_BR.RM.like.systems | DDE_Tnp_1->Peptidase_S58-><-?\|\|DEDD_Tnp_IS110+Transposase_20->FtsK->ORC-CDC6+REase->DUF499*-><-?\|\|?->?-><-?\|\|ABhydrolase-> | 766 | Bacteria>Bacteria_incertae_sedis>Bacteria_candidate_phyla>Candidatus_Saccharibacteria>unclassified_Candidatus_Saccharibacteria | GCA_016699955.1 | Candidatus_Saccharibacteria_bacterium | IPL85_04630 | 901666..903966 | 2026720 | ATP-binding-protein-[Candidatus-Saccharibacteria-bacterium] |
| MBI2571108.1 | AAA+ATPase+wHTH+Alpha+Beta-domain+wHTH+RRM | Type-2_BR.RM.like.systems | <-DUF499*\|\|REase->AbiEi_1->AbiEi_1-><-?<-REase | 526 | Bacteria>Bacteria_incertae_sedis>Bacteria_candidate_phyla>Candidatus_Schekmanbacteria>unclassified_Candidatus_Schekmanbacteria | GCA_016188575.1 | Candidatus_Schekmanbacteria_bacterium | HYV63_29230 | 1..1578 | 2419984 | ATP-binding-protein-[Candidatus-Schekmanbacteria-bacterium] |
| MBX2928866.1 | AAA+ATPase+wHTH+Alpha+Beta-domain+wHTH+RRM | Type-2_BR.RM.like.systems | DUF499*->RVT+HNH->DUF499-p->DUF3780->?->METHYLASE-> | 507 | Bacteria>FCB_group>Bacteroidota/Chlorobiota_group>Bacteroidota>Saprospiria | GCA_019638385.1 | Saprospiraceae_bacterium | KF852_13605 | 113678..115201 | 2202734 | ATP-binding-protein-[Saprospiraceae-bacterium] |
| MBV5303481.1 | AAA+ATPase+wHTH+Alpha+Beta-domain+wHTH+RRM | Type-2_BR.RM.like.systems | <-HTH\|\|ORC-CDC6+REase->MazE_antitoxin->PemK_toxin-><-?\|\|DUF499*-> | 622 | Bacteria>FCB_group>Bacteroidota/Chlorobiota_group>Chlorobiota>Chlorobiia | GCA_019163115.1 | Chlorobium_sp. | JZU70_04665 | 96654..98520 | 1095 | ATP-binding-protein-[Chlorobium-sp.] |
| MCD4708414.1 | AAA+ATPase+wHTH+Alpha+Beta-domain+wHTH+RRM | Type-2_BR.RM.like.systems | MqsA_antitoxin+HTH_3->HTH+MPTase->PIN->?->DUF499*->DUF499-p->DUF3780->METHYLASE->RADICAL-SAM-><-METHYLASE | 647 | Bacteria>FCB_group>Candidatus_Fermentibacteria>Sabulitectum>unclassified_Candidatus_Sabulitectum | GCA_021107435.1 | Candidatus_Sabulitectum_sp. | K8S62_11845 | 5005..6948 | 2822438 | DUF499-domain-containing-protein-[Candidatus-Sabulitectum-sp.] |
| MBK9409374.1 | AAA+ATPase+wHTH+Alpha+Beta-domain+wHTH+RRM | Type-2_BR.RM.like.systems | <-HKD-DNase+SF1-SF2-Helicase+iREase+CTD-p<-HKD-DNase+SF1-SF2-Helicase+iREase+CTD-p<-METHYLASE<-DUF499-p<-DUF499*<-HD | 399 | Bacteria>FCB_group>Gemmatimonadota>unclassified_Gemmatimonadota | GCA_016719105.1 | Gemmatimonadota_bacterium | IPN47_15260 | 139828..141027 | 2026742 | ATP-binding-protein-[Gemmatimonadota-bacterium] |
| MBA4156892.1 | AAA+ATPase+wHTH+Alpha+Beta-domain+wHTH+RRM | Type-2_BR.RM.like.systems | <-HTH+MPTase<-?<-TIR\|\|DUF499*->DUF3780->METHYLASE->?->?->HicB_lk_antitox->REase-> | 312 | Bacteria>FCB_group>Gemmatimonadota>unclassified_Gemmatimonadota | GCA_013820915.1 | Gemmatimonadota_bacterium | HOX65_05385 | 4079..5017 | 2026742 | ATP-binding-protein-[Gemmatimonadota-bacterium] |
| MBW2046233.1 | AAA+ATPase+wHTH+Alpha+Beta-domain+wHTH+RRM | Type-2_BR.RM.like.systems | DUF499-p->DUF499*->DUF499-p->DUF3780-><-DDE_5+DDE_Tnp_1\|\|RHH->PIN->HEPN-p-> | 455 | Bacteria>Myxococcota>Myxococcia>unclassified_Myxococcia | GCA_019309025.1 | Deltaproteobacteria_bacterium | JRI96_15365 | 321..1688 | 2026735 | ATP-binding-protein-[Deltaproteobacteria-bacterium] |
| RLB16925.1 | AAA+ATPase+wHTH+Alpha+Beta-domain+wHTH+RRM | Type-2_BR.RM.like.systems | <-DUF499*<-ORC-CDC6+REase<-?<-ABhydrolase | 348 | Bacteria>Myxococcota>Myxococcia>unclassified_Myxococcia | GCA_003646815.1 | Deltaproteobacteria_bacterium | DRG82_07820 | 1..1043 | 2026735 | glycosyl-transferase,-partial-[Deltaproteobacteria-bacterium] |
| ETX04990.1 | AAA+ATPase+wHTH+Alpha+Beta-domain+wHTH+RRM | Type-2_BR.RM.like.systems | <-HKD-DNase+SF1-SF2-Helicase<-PIN<-?<-METHYLASE<-DUF3780<-DUF499-p<-DUF499* | 495 | Bacteria>Nitrospinae/Tectomicrobia_group>Candidatus_Tectomicrobia>Entotheonella | GCA_000522445.1 | Candidatus_Entotheonella_gemina | ETSY2_25620 | 8747..10231 | 1429439 | hypothetical-protein-ETSY2_25620,-partial-[Candidatus-Entotheonella-gemina] |
| GIX46075.1 | AAA+ATPase+wHTH+Alpha+Beta-domain+wHTH+RRM | Type-2_BR.RM.like.systems | <-PIN<-?<-MULE-TRANSPOSASE<-?\|\|DUF499-p->DUF499*->?->?->PHP+SbcC+PspA+SbcC->PHP+SbcC+PspA+SbcC->Schlafen+HATPase+HTH-> | 476 | Bacteria>Nitrospinae/Tectomicrobia_group>Candidatus_Tectomicrobia>unclassified_Candidatus_Tectomicrobia | GCA_026005175.1 | Candidatus_Tectomicrobia_bacterium | KatS3mg131_0286 | 296359..297789 | 2528274 | MAG:-hypothetical-protein-KatS3mg131_0286-[Candidatus-Tectomicrobia-bacterium] |
| MCS6305125.1 | AAA+ATPase+wHTH+Alpha+Beta-domain+wHTH+RRM | Type-2_BR.RM.like.systems | MetJarc-HTH->?->?->?->PIN->CopG/RHH->DUF499*-> | 297 | Bacteria>Nitrospirota>Nitrospiria>Nitrospirales>Nitrospiraceae | GCA_024998685.1 | Nitrospira_sp. | H8K07_15900 | 51882..52775 | 70125 | DUF499-domain-containing-protein-[Nitrospira-sp.] |
| RWC26997.1 | AAA+ATPase+wHTH+Alpha+Beta-domain+wHTH+RRM | Type-2_BR.RM.like.systems | PIN+ACET+EVE->?-><-HNH\|\|SIR2-><-wHTH+SAD+HNH\|\|?->DUF499*->DUF499->DUF3780->METHYLASE->HKD-DNase+SF1-SF2-Helicase-><-SF1-SF2-Helicase+DUF1998+HKD-DNase+HKD-DNase | 230 | Bacteria>Pseudomonadota>Alphaproteobacteria>Hyphomicrobiales>Phyllobacteriaceae | GCA_004017365.1 | Mesorhizobium_sp. | EOS70_30590 | 20112..20804 | 1871066 | DUF499-domain-containing-protein-[Mesorhizobium-sp.] |
| WP_233943241.1 | AAA+ATPase+wHTH+Alpha+Beta-domain+wHTH+RRM | Type-2_BR.RM.like.systems | <-HKD-DNase+SF1-SF2-Helicase<-METHYLASE<-DUF499-p<-DUF499*\|\|wHTH+REase->\|\|?-><-?<-PIN+OST-HTH<-?<-Arm-DNA-bind_3+Phage_integrase | 526 | Bacteria>Pseudomonadota>Alphaproteobacteria>Hyphomicrobiales>Rhizobiaceae | GCF_021391595.1 | Rhizobium_leguminosarum | LZK78_RS32110 | 712264..713844 | 384 | DUF499-domain-containing-protein-[Rhizobium-leguminosarum] |
| MBO7350942.1 | AAA+ATPase+wHTH+Alpha+Beta-domain+wHTH+RRM | Type-2_BR.RM.like.systems | <-DUF499*<-?\|\|Synaptojanin-><-HKD-DNase+HKD-DNase | 770 | Bacteria>Pseudomonadota>Alphaproteobacteria>Rhodospirillales>Acetobacteraceae | GCA_017648865.1 | Acetobacter_sp. | J6U18_05075 | 1..2310 | 440 | ATP-binding-protein-[Acetobacter-sp.] |
| MBV8524768.1 | AAA+ATPase+wHTH+Alpha+Beta-domain+wHTH+RRM | Type-2_BR.RM.like.systems | <-DUF499<-DUF499*\|\|REase->?-><-?<-ABC-ATPase+SbcC | 433 | Bacteria>Pseudomonadota>Alphaproteobacteria>Rhodospirillales>Acetobacteraceae | GCA_019234085.1 | Acetobacteraceae_bacterium | JOY71_22065 | 913..2214 | 1909293 | ATP-binding-protein-[Acetobacteraceae-bacterium] |
| MBV8524767.1 | AAA+ATPase+wHTH+Alpha+Beta-domain+wHTH+RRM | Type-2_BR.RM.like.systems | <-DUF499*<-DUF499\|\|REase->?-><-?<-ABC-ATPase+SbcC | 324 | Bacteria>Pseudomonadota>Alphaproteobacteria>Rhodospirillales>Acetobacteraceae | GCA_019234085.1 | Acetobacteraceae_bacterium | JOY71_22060 | 1..973 | 1909293 | hypothetical-protein-[Acetobacteraceae-bacterium] |
| MBV8590611.1 | AAA+ATPase+wHTH+Alpha+Beta-domain+wHTH+RRM | Type-2_BR.RM.like.systems | <-DUF499*\|\|REase->?-><-?<-ABC-ATPase+SbcC | 433 | Bacteria>Pseudomonadota>Alphaproteobacteria>Rhodospirillales>Acetobacteraceae | GCA_019233525.1 | Acetobacteraceae_bacterium | JO212_11260 | 226..1527 | 1909293 | ATP-binding-protein-[Acetobacteraceae-bacterium] |
| MBK9010719.1 | AAA+ATPase+wHTH+Alpha+Beta-domain+wHTH+RRM | Type-2_BR.RM.like.systems | <-Calcineurin<-HIN-HTH<-Calcineurin<-HKD-DNase+SF1-SF2-Helicase+iREase+CTD<-DUF3780<-DUF499-p<-DUF499*<-RNase+35exo<-RVT<-RVT<-ZnR | 837 | Bacteria>Pseudomonadota>Alphaproteobacteria>Sphingomonadales>Sphingomonadaceae | GCA_016719755.1 | Novosphingobium_sp. | IPM87_08350 | 118615..121128 | 1874826 | ATP-binding-protein-[Novosphingobium-sp.] |
| WP_272798655.1 | AAA+ATPase+wHTH+Alpha+Beta-domain+wHTH+RRM | Type-2_BR.RM.like.systems | <-HKD-DNase+SF1-SF2-Helicase<-METHYLASE<-DUF3780<-DUF499-p<-?<-HIN-HTH<-DUF499*<-?\|\|REase+PNTD1+PNTD2->PIWI->DDE_Tnp_1_4->?->HIN-HTH-> | 663 | Bacteria>Pseudomonadota>Alphaproteobacteria>Sphingomonadales>Sphingomonadaceae | GCF_028538045.1 | Sphingobium_sp._AntQ-1 | sphantq_RS12565 | 2511854..2513845 | 2930091 | DUF499-domain-containing-protein-[Sphingobium-sp.-AntQ-1] |
| MCM8596898.1 | AAA+ATPase+wHTH+Alpha+Beta-domain+wHTH+RRM | Type-2_BR.RM.like.systems | <-XIS-HTH<-?\|\|DUF499*->DUF499-p->DUF3780->METHYLASE->HKD-DNase+SF1-SF2-Helicase-> | 747 | Bacteria>Pseudomonadota>Betaproteobacteria>Betaproteobacteria_incertae_sedis>Candidatus_Accumulibacter | GCA_023806595.1 | Accumulibacter_sp. | NFV71_16345 | 9561..11800 | 2053492 | DUF499-domain-containing-protein-[Candidatus-Accumulibacter-sp.] |
| WP_258231469.1 | AAA+ATPase+wHTH+Alpha+Beta-domain+wHTH+RRM | Type-2_BR.RM.like.systems | MarR-HTH->Phage_int_SAM_2+Integrase_1->RHH_1->Phage_int_SAM_2+Integrase_1->RHH_1->DUF499*->DUF499->DUF3780->?->MACRODOMAIN->METHYLASE->HKD-DNase+SF1-SF2-Helicase-> | 359 | Bacteria>Pseudomonadota>Betaproteobacteria>Burkholderiales>Alcaligenaceae | GCF_002885955.2 | Achromobacter_pulmonis | C1I89_RS17900 | 576242..577318 | 1389932 | ATP-binding-protein,-partial-[Achromobacter-pulmonis] |
| WP_090140695.1 | AAA+ATPase+wHTH+Alpha+Beta-domain+wHTH+RRM | Type-2_BR.RM.like.systems | <-HKD-DNase+SF1-SF2-Helicase<-URI-DNase+RAMA<-METHYLASE<-DUF3780<-DUF499-p<-DUF499*\|\|RelE-ParE->HTH_3->?-><-PIN<-THUMP+Methylase | 375 | Bacteria>Pseudomonadota>Betaproteobacteria>Burkholderiales>Comamonadaceae | GCF_001269385.1 | Limnohabitans_sp._DM1 | AMM33_RS09415 | 63880..65007 | 1597955 | DUF499-domain-containing-protein-[Limnohabitans-sp.-DM1] |
| MBK7421883.1 | AAA+ATPase+wHTH+Alpha+Beta-domain+wHTH+RRM | Type-2_BR.RM.like.systems | Phage_int_SAM_2+Integrase_1-><-?\|\|DUF499-p->DUF499*->DUF499-p->DUF499-p->DUF3780->Schlafen+HATPase+HTH->Pkinase->TPR-> | 487 | Bacteria>Pseudomonadota>Betaproteobacteria>Rhodocyclales>Rhodocyclaceae | GCA_016709335.1 | Candidatus_Propionivibrio_dominans | IPJ48_01605 | 217179..218642 | 2954373 | DUF499-domain-containing-protein-[Propionivibrio-sp.] |
| WP_256684210.1 | AAA+ATPase+wHTH+Alpha+Beta-domain+wHTH+RRM | Type-2_BR.RM.like.systems | RHH->DUF499-p->DUF499*->DUF499-p->DUF3780+RAMA->METHYLASE->HKD-DNase+SF1-SF2-Helicase-> | 356 | Bacteria>Pseudomonadota>Gammaproteobacteria>Aeromonadales>Aeromonadaceae | GCF_024508255.1 | Aeromonas_salmonicida | NP805_RS14160 | 3078633..3079703 | 645 | DUF499-domain-containing-protein-[Aeromonas-salmonicida] |
| WP_216593132.1 | AAA+ATPase+wHTH+Alpha+Beta-domain+wHTH+RRM | Type-2_BR.RM.like.systems | <-Phage_integrase<-JAB\|\|XIS-HTH->RHH_1->DUF499*->DUF499-p->RAMA->METHYLASE->METHYLASE-p->HKD-DNase+SF1-SF2-Helicase->HKD-DNase+SF1-SF2-Helicase-p-> | 715 | Bacteria>Pseudomonadota>Gammaproteobacteria>Alteromonadales>Alteromonadaceae | - | Alteromonas_australica | EP12_RS19375 | 4393285..4395432 | 589873 | DUF499-domain-containing-protein-[Alteromonas-australica] |
| WP_246237869.1 | AAA+ATPase+wHTH+Alpha+Beta-domain+wHTH+RRM | Type-2_BR.RM.like.systems | <-SF1-SF2-Helicase-p<-HKD-DNase+SF1-SF2-Helicase<-HKD-DNase+SF1-SF2-Helicase-p<-URI-DNase<-DUF499-p<-?<-DUF499*<-DUF499\|\|?->ParB+N6_N4_MTase-><-RelE-ParE<-ParD_like | 263 | Bacteria>Pseudomonadota>Gammaproteobacteria>Chromatiales>Chromatiaceae | GCF_011290485.1 | Caldichromatium_japonicum | GWK36_RS14650 | 150616..151407 | 2699430 | DUF499-domain-containing-protein-[Caldichromatium-japonicum] |
| WP_210756809.1 | AAA+ATPase+wHTH+Alpha+Beta-domain+wHTH+RRM | Type-2_BR.RM.like.systems | <-HKD-DNase+SF1-SF2-Helicase<-HKD-DNase+SF1-SF2-Helicase-p<-URI-DNase<-DUF499-p<-?<-DUF499<-DUF499*\|\|?->ParB+N6_N4_MTase-><-RelE-ParE<-ParD_like | 109 | Bacteria>Pseudomonadota>Gammaproteobacteria>Chromatiales>Chromatiaceae | GCF_011290485.1 | Caldichromatium_japonicum | GWK36_RS14655 | 151446..151775 | 2699430 | hypothetical-protein-[Caldichromatium-japonicum] |
| MBV5274819.1 | AAA+ATPase+wHTH+Alpha+Beta-domain+wHTH+RRM | Type-2_BR.RM.like.systems | <-SF1-SF2-Helicase\|\|?->NTP_transferase->DUF499*->?->DUF3780-><-PIN<-?\|\|METHYLASE->OLD-ABC-p-> | 898 | Bacteria>Pseudomonadota>Gammaproteobacteria>Chromatiales>Chromatiaceae | GCA_019163055.1 | Lamprocystis_purpurea | JZU52_14660 | 4546..7239 | 61598 | ATP-binding-protein-[Lamprocystis-purpurea] |
| WP_242523202.1 | AAA+ATPase+wHTH+Alpha+Beta-domain+wHTH+RRM | Type-2_BR.RM.like.systems | <-DUF499*<-IclR-HTH<-?<-?<-REase<-?\|\|TP_methylase-> | 508 | Bacteria>Pseudomonadota>Gammaproteobacteria>Chromatiales>Chromatiaceae | GCF_016653465.1 | Thiocystis_minor | CCR95_RS11640 | 78..1604 | 61597 | DUF499-domain-containing-protein-[Thiocystis-minor] |

<table>
<tr><th>Protein-ID</th><th>DUF499-architecture</th><th>System</th><th>Gene-Neighbourhoods</th><th>Length</th><th>Taxonomy</th><th>Assembly-ID</th><th>Strain</th><th>Locus-Tag</th><th>Genomic-coordinates</th><th>TaxID</th><th>Fasta-define</th></tr>
<tr><td>UZF63336.1</td><td>AAA+ATPase+wHTH+Alpha+Beta-domain+wHTH+RRM</td><td>Type-2_BR.RM.like.systems</td><td><-HIN-HTH<-wHTH+REase<-HKD-DNase+SF1-SF2-Helicase<-METHYLASE<-DUF3780+RAMA<-DUF499-p<-DUF499*<-DUF499-p<-RHH_5<-?<-DEDD_Tnp_IS110+Transposase_20<-HTH_Tnp_1+HTH_21+rve+rve_3||<-Arm-DNA-bind_3+Phage_integrase</td><td>355</td><td>Bacteria>Pseudomonadota>Gammaproteobacteria>Enterobacterales>Enterobacteriaceae</td><td>GCA_026016605.1</td><td>Escherichia_coli</td><td>OM434_11360</td><td>2373962..2375029</td><td>562</td><td>ATP-binding-protein-[Escherichia-coli]</td></tr>
<tr><td>VTN25635.1</td><td>AAA+ATPase+wHTH+Alpha+Beta-domain+wHTH+RRM</td><td>Type-2_BR.RM.like.systems</td><td>HTH+MPTase-><-wHTH+REase<-HKD-DNase+SF1-SF2-Helicase<-METHYLASE<-DUF3780+RAMA<-DUF499-p<-DUF499*<-RHH<-Arm-DNA-bind_3+Phage_integrase<-tRNA||?->?->ADH_N+ADH_zinc_N-></td><td>247</td><td>Bacteria>Pseudomonadota>Gammaproteobacteria>Enterobacterales>Enterobacteriaceae</td><td>GCA_901420815.1</td><td>Klebsiella_pneumoniae</td><td>NCTC9129_01967</td><td>1513305..1514048</td><td>573</td><td>Uncharacterised-protein-[Klebsiella-pneumoniae]</td></tr>
<tr><td>WP_250216358.1</td><td>AAA+ATPase+wHTH+Alpha+Beta-domain+wHTH+RRM</td><td>Type-2_BR.RM.like.systems</td><td><-HKD-DNase+SF1-SF2-Helicase<-METHYLASE<-DUF3780<-DUF499||HIN-HTH->?-><-DUF499*<-zf-ISL3+DDE_Tnp_ISL3||?->||MAD-NTD+HNH-></td><td>340</td><td>Bacteria>Pseudomonadota>Gammaproteobacteria>Enterobacterales>Enterobacteriaceae</td><td>GCF_005381525.1</td><td>Escherichia_coli</td><td>FE858_RS22010</td><td>11270..12292</td><td>562</td><td>DUF499-domain-containing-protein-[Escherichia-coli]</td></tr>
<tr><td>CNI93772.1</td><td>AAA+ATPase+wHTH+Alpha+Beta-domain+wHTH+RRM</td><td>Type-2_BR.RM.like.systems</td><td><-Peptidase||?->?->tRNA->Arm-DNA-bind_3+Phage_integrase->RHH->DUF499*->METHYLASE-p->HKD-DNase+SF1-SF2-Helicase-p-><-OLD-ABC+TOPRIM<-HTH+MPTase<-?||DDE_3-></td><td>758</td><td>Bacteria>Pseudomonadota>Gammaproteobacteria>Enterobacterales>Yersiniaceae</td><td>GCA_001319685.1</td><td>Yersinia_pseudotuberculosis</td><td>ERS008465_02107</td><td>77957..80233</td><td>633</td><td>Protein-of-uncharacterised-function-(DUF499)-[Yersinia-pseudotuberculosis]</td></tr>
<tr><td>WP_226974938.1</td><td>AAA+ATPase+wHTH+Alpha+Beta-domain+wHTH+RRM</td><td>Type-2_BR.RM.like.systems</td><td><-Peptidase||?->?->tRNA->Arm-DNA-bind_3+Phage_integrase->RHH->DUF499*->METHYLASE-p->HKD-DNase+SF1-SF2-Helicase->wHTH+REase-><-OLD-ABC+TOPRIM<-SbcC<-HTH+MPTase</td><td>777</td><td>Bacteria>Pseudomonadota>Gammaproteobacteria>Enterobacterales>Yersiniaceae</td><td>GCF_001319705.1</td><td>Yersinia_pseudotuberculosis</td><td>EGX52_RS11275</td><td>2325348..2327681</td><td>633</td><td>DUF499-domain-containing-protein-[Yersinia-pseudotuberculosis]</td></tr>
<tr><td>WP_235045586.1</td><td>AAA+ATPase+wHTH+Alpha+Beta-domain+wHTH+RRM</td><td>Type-2_BR.RM.like.systems</td><td>Phage_integrase->Phage_integrase->?->?->LON-AAA->RHH->DUF499*->DUF499-p->DUF3780->METHYLASE-p->METHYLASE->HKD-DNase+SF1-SF2-Helicase->wHTH+REase-></td><td>776</td><td>Bacteria>Pseudomonadota>Gammaproteobacteria>Thiotrichales>Piscirickettsiaceae</td><td>GCF_000349205.1</td><td>Methylophaga_lonarensis</td><td>MPL1_RS12025</td><td>17805..20135</td><td>999151</td><td>DUF499-domain-containing-protein-[Methylophaga-lonarensis]</td></tr>
<tr><td>MDE0094449.1</td><td>AAA+ATPase+wHTH+Alpha+Beta-domain+wHTH+RRM</td><td>Type-2_BR.RM.like.systems</td><td>Schlafen+HATPase+HTH->DUF499*->DUF499-p->DUF3780->METHYLASE->HKD-DNase+SF1-SF2-Helicase+iREase+CTD-></td><td>782</td><td>Bacteria>Pseudomonadota>Gammaproteobacteria>unclassified_Gammaproteobacteria</td><td>GCA_028819195.1</td><td>Gammaproteobacteria_bacterium</td><td>OXO49_08130</td><td>171410..173758</td><td>1913989</td><td>MAG:-DUF499-domain-containing-protein-[Gammaproteobacteria-bacterium]</td></tr>
<tr><td>MDE0179377.1</td><td>AAA+ATPase+wHTH+Alpha+Beta-domain+wHTH+RRM</td><td>Type-2_BR.RM.like.systems</td><td><-HKD-DNase+SF1-SF2-Helicase+iREase+CTD<-?<-?<-DUF3780<-DUF499-p<-DUF499*<-Glyoxalase<-HicB_lk_antitox<-HicA_toxin</td><td>208</td><td>Bacteria>Pseudomonadota>Gammaproteobacteria>unclassified_Gammaproteobacteria</td><td>GCA_028819775.1</td><td>Gammaproteobacteria_bacterium</td><td>OXP36_12380</td><td>3318..3944</td><td>1913989</td><td>MAG:-DUF499-domain-containing-protein-[Gammaproteobacteria-bacterium]</td></tr>
<tr><td>MDE0649696.1</td><td>AAA+ATPase+wHTH+Alpha+Beta-domain+wHTH+RRM</td><td>Type-2_BR.RM.like.systems</td><td><-HKD-DNase+SF1-SF2-Helicase+iREase+CTD<-METHYLASE<-DUF3780<-DUF499-p<-DUF499*||HicB_lk_antitox->HicA_toxin-></td><td>453</td><td>Bacteria>Pseudomonadota>Gammaproteobacteria>unclassified_Gammaproteobacteria</td><td>GCA_028824315.1</td><td>Gammaproteobacteria_bacterium</td><td>OXI12_05060</td><td>9541..10901</td><td>1913989</td><td>MAG:-DUF499-domain-containing-protein,-partial-[Gammaproteobacteria-bacterium]</td></tr>
<tr><td>HBJ34236.1</td><td>AAA+ATPase+wHTH+Alpha+Beta-domain+wHTH+RRM</td><td>Type-2_BR.RM.like.systems</td><td><-REase<-HTH||DUF499*-><-?||?->DUF3780->METHYLASE-></td><td>516</td><td>Bacteria>PVC_group>Planctomycetota>Planctomycetia>Planctomycetales</td><td>GCA_003506095.1</td><td>Planctomycetaceae_bacterium</td><td>DDZ51_05610</td><td>2380..3930</td><td>2026779</td><td>hypothetical-protein-[Planctomycetaceae-bacterium]</td></tr>
<tr><td>MCX5673909.1</td><td>AAA+ATPase+wHTH+Alpha+Beta-domain+wHTH+RRM</td><td>Type-2_BR.RM.like.systems</td><td><-REase<-REase+NACHT<-REase+NACHT||DUF499*->DUF3780->Schlafen+HATPase+HTH->METHYLASE->HKD-DNase+SF1-SF2-Helicase+iREase+CTD->?->HKD-DNase-></td><td>978</td><td>Bacteria>PVC_group>Planctomycetota>unclassified_Planctomycetota</td><td>GCA_026387015.1</td><td>Planctomycetota_bacterium</td><td>NTX87_02775</td><td>8015..10951</td><td>2026780</td><td>MAG:-DUF499-domain-containing-protein-[Planctomycetota-bacterium]</td></tr>
<tr><td>MCA9314335.1</td><td>AAA+ATPase+wHTH+Alpha+Beta-domain+wHTH+RRM</td><td>Type-2_BR.RM.like.systems</td><td><-McrC-NTD+REase<-?<-?<-DUF488-l<-?<-DUF499*<-XIS-HTH<-HNH<-?<-?<-rve+rve_3<-HTH_Tnp_1</td><td>500</td><td>Bacteria>PVC_group>Planctomycetota>unclassified_Planctomycetota</td><td>GCA_020427815.1</td><td>Planctomycetota_bacterium</td><td>KDB73_02465</td><td>8962..10464</td><td>2026780</td><td>ATP-binding-protein-[Planctomycetota-bacterium]</td></tr>
<tr><td>WP_182916264.1</td><td>AAA+ATPase+wHTH+Alpha+Beta-domain+wHTH+RRM</td><td>Type-2_BR.RM.like.systems</td><td><-HKD-DNase+SF1-SF2-Helicase<-METHYLASE-p<-METHYLASE<-DUF3780<-DUF499-p<-DUF499*||Phage_integrase->?->?->?->Peptidase_M28-></td><td>684</td><td>Bacteria>PVC_group>Verrucomicrobiota>Verrucomicrobiae>Verrucomicrobiales</td><td>GCF_014170075.1</td><td>Akkermansia_muciniphila</td><td>HXS70_RS11830</td><td>2746914..2748968</td><td>239935</td><td>DUF499-domain-containing-protein-[Akkermansia-muciniphila]</td></tr>
<tr><td>WP_257229194.1</td><td>AAA+ATPase+wHTH+Alpha+Beta-domain+wHTH+RRM</td><td>Type-2_BR.RM.like.systems</td><td><-HKD-DNase+SF1-SF2-Helicase<-DUF3780<-DUF499-p<-DUF499*<-DUF499-p||Phage_integrase->?->?->?->Peptidase_M28-></td><td>371</td><td>Bacteria>PVC_group>Verrucomicrobiota>Verrucomicrobiae>Verrucomicrobiales</td><td>GCF_018847275.1</td><td>Akkermansia_muciniphila</td><td>J5W48_RS12770</td><td>23240..24355</td><td>239935</td><td>ATP-binding-protein-[Akkermansia-muciniphila]</td></tr>
<tr><td>WP_257229149.1</td><td>AAA+ATPase+wHTH+Alpha+Beta-domain+wHTH+RRM</td><td>Type-2_BR.RM.like.systems</td><td><-HKD-DNase+SF1-SF2-Helicase<-HKD-DNase+SF1-SF2-Helicase-p<-DUF3780<-DUF499-p<-DUF499*<-DUF499-p||Phage_integrase->?->?->?->Peptidase_M28-></td><td>371</td><td>Bacteria>PVC_group>Verrucomicrobiota>Verrucomicrobiae>Verrucomicrobiales</td><td>GCF_018847535.1</td><td>Akkermansia_muciniphila</td><td>J5W77_RS12760</td><td>23250..24365</td><td>239935</td><td>ATP-binding-protein-[Akkermansia-muciniphila]</td></tr>
<tr><td>MBK9497792.1</td><td>AAA+ATPase+wHTH+Alpha+Beta-domain+wHTH+RRM</td><td>Type-2_BR.RM.like.systems</td><td><-OLD-ABC<-?<-OLD-ABC<-ParB+HNH<-DUF3780<-DUF499<-DUF499*||?->?->?->Peptidase-></td><td>294</td><td>Bacteria>Spirochaetota>Spirochaetia>Leptospirales>Leptospiraceae</td><td>GCA_016718635.1</td><td>Leptospiraceae_bacterium</td><td>IPO06_00160</td><td>44986..45870</td><td>2026757</td><td>DUF499-domain-containing-protein-[Leptospiraceae-bacterium]</td></tr>
<tr><td>MXZ29854.1</td><td>AAA+ATPase+wHTH+Alpha+Beta-domain+wHTH+RRM</td><td>Type-2_BR.RM.like.systems</td><td><-HKD-DNase+SF1-SF2-Helicase+iREase+CTD<-METHYLASE<-DUF3780<-DUF499*</td><td>482</td><td>Bacteria>Terrabacteria_group>Actinomycetota>Acidimicrobiia>unclassified_Acidimicrobiia</td><td>GCA_009837005.1</td><td>Acidimicrobiia_bacterium</td><td>F4Z22_03260</td><td>6357..7807</td><td>2080302</td><td>DUF499-domain-containing-protein-[Acidimicrobiia-bacterium]</td></tr>
<tr><td>MYB24502.1</td><td>AAA+ATPase+wHTH+Alpha+Beta-domain+wHTH+RRM</td><td>Type-2_BR.RM.like.systems</td><td>ORC-CDC6+REase->?->MACRODOMAIN->DUF499*->DUF3780->METHYLASE->HKD-DNase+SF1-SF2-Helicase+iREase+CTD-><-?<-?<-DUF4527</td><td>703</td><td>Bacteria>Terrabacteria_group>Actinomycetota>Acidimicrobiia>unclassified_Acidimicrobiia</td><td>GCA_009840735.1</td><td>Acidimicrobiia_bacterium</td><td>F4X37_05290</td><td>24179..26290</td><td>2080302</td><td>ATP-binding-protein-[Acidimicrobiia-bacterium]</td></tr>
<tr><td>WP_036280506.1</td><td>AAA+ATPase+wHTH+Alpha+Beta-domain+wHTH+RRM</td><td>Type-2_BR.RM.like.systems</td><td><-HTH_3||?->?->XIS-HTH->DUF499*->?->DUF3780->METHYLASE->METHYLASE-p->HKD-DNase+SF1-SF2-Helicase+iREase+CTD-><-N6_MTase-p</td><td>818</td><td>Bacteria>Terrabacteria_group>Actinomycetota>Actinomycetes>Micrococcales</td><td>GCF_000685355.1</td><td>Microbacterium_sp._CH12i</td><td>DC31_RS12080</td><td>19282..21738</td><td>1479651</td><td>DUF499-domain-containing-protein-[Microbacterium-sp.-CH12i]</td></tr>
<tr><td>ETI92890.1</td><td>AAA+ATPase+wHTH+Alpha+Beta-domain+wHTH+RRM</td><td>Type-2_BR.RM.like.systems</td><td>DUF499*->DUF499-p->DUF3780->METHYLASE->HKD-DNase+SF1-SF2-Helicase-></td><td>550</td><td>Bacteria>Terrabacteria_group>Bacillota>Clostridia>Eubacteriales</td><td>GCA_000508445.1</td><td>Intestinibacter_bartlettii_DORA_8_9</td><td>Q606_CBAC00367G0002</td><td>198..1850</td><td>1403940</td><td>hypothetical-protein-Q606_CBAC00367G0002-[Intestinibacter-bartlettii-DORA_8_9]</td></tr>
<tr><td>HCS72475.1</td><td>AAA+ATPase+wHTH+Alpha+Beta-domain+wHTH+RRM</td><td>Type-2_BR.RM.like.systems</td><td><-HKD-DNase+SF1-SF2-Helicase<-HKD-DNase+SF1-SF2-Helicase-p<-METHYLASE<-DUF3780<-DUF499-p<-DUF499*<-HTH_26<-?<-MPTase<-CHTH</td><td>715</td><td>Bacteria>Terrabacteria_group>Bacillota>Clostridia>Eubacteriales</td><td>GCA_003524825.1</td><td>Clostridiales_bacterium</td><td>DIW17_01185</td><td>14149..16292</td><td>1898207</td><td>glycosyl-transferase-[Clostridiales-bacterium]</td></tr>
<tr><td>WP_252891859.1</td><td>AAA+ATPase+wHTH+Alpha+Beta-domain+wHTH+RRM</td><td>Type-2_BR.RM.like.systems</td><td>DUF499-p->DUF499*->DUF499-p->DUF499-p->METHYLASE->REase->HKD-DNase+SF1-SF2-Helicase-p->HKD-DNase+SF1-SF2-Helicase-></td><td>195</td><td>Bacteria>Terrabacteria_group>Bacillota>Clostridia>Eubacteriales</td><td>GCF_001310995.1</td><td>Thermoclostridium_stercorarium</td><td>JCM9324_RS16190</td><td>584..1171</td><td>1510</td><td>DUF499-domain-containing-protein-[Thermoclostridium-stercorarium]</td></tr>
<tr><td>WP_099082940.1</td><td>AAA+ATPase+wHTH+Alpha+Beta-domain+wHTH+RRM</td><td>Type-2_BR.RM.like.systems</td><td><-HKD-DNase+SF1-SF2-Helicase+iREase+CTD-p<-HKD-DNase+SF1-SF2-Helicase+iREase+CTD<-METHYLASE<-METHYLASE-p<-DUF3780<-DUF499-p<-DUF499*<-?||?->SIGMA-HTH->ZnR-></td><td>461</td><td>Bacteria>Terrabacteria_group>Bacillota>Clostridia>Eubacteriales</td><td>GCF_002607855.1</td><td>Desulforamulus_profundi</td><td>P378_RS08455</td><td>21622..23007</td><td>1383067</td><td>DUF499-domain-containing-protein-[Desulforamulus-profundi]</td></tr>
<tr><td>HCC32356.1</td><td>AAA+ATPase+wHTH+Alpha+Beta-domain+wHTH+RRM</td><td>Type-2_BR.RM.like.systems</td><td><-DUF499*<-?||?-><-PIN<-HTH+MPTase||ParA-Soj-PloopNTPase->ORC-CDC6-></td><td>810</td><td>Bacteria>Terrabacteria_group>Bacillota>Clostridia>Eubacteriales</td><td>GCA_003517845.1</td><td>Clostridiales_bacterium</td><td>DEQ28_00385</td><td>1..2429</td><td>1898207</td><td>AAA-family-ATPase-[Clostridiales-bacterium]</td></tr>
<tr><td>HHV96419.1</td><td>AAA+ATPase+wHTH+Alpha+Beta-domain+wHTH+RRM</td><td>Type-2_BR.RM.like.systems</td><td>METHYLASE->?->REase->?->SF1-SF2-Helicase->DUF499*-></td><td>401</td><td>Bacteria>Terrabacteria_group>Bacillota>Clostridia>Eubacteriales</td><td>GCA_012840075.1</td><td>Clostridiaceae_bacterium</td><td>GXX37_08125</td><td>11739..12942</td><td>1898204</td><td>ATP-binding-protein-[Clostridiaceae-bacterium]</td></tr>
<tr><td>MBE7435441.1</td><td>AAA+ATPase+wHTH+Alpha+Beta-domain+wHTH+RRM</td><td>Type-2_BR.RM.like.systems</td><td><-URI-DNase<-ParB+HNH<-DUF499-p<-ASCH+DUF3780<-?<-?<-DUF499*<-?<-METHYLASE<-TUDOR+SF1-SF2-Helicase+iREase+CTD+REase</td><td>1032</td><td>Bacteria>Terrabacteria_group>Chloroflexota>Anaerolineae>Anaerolineales</td><td>GCA_015075725.1</td><td>Anaerolineales_bacterium</td><td>HS100_16115</td><td>3331332..3334430</td><td>2073117</td><td>DUF499-domain-containing-protein-[Anaerolineales-bacterium]</td></tr>
<tr><td>WP_258316378.1</td><td>AAA+ATPase+wHTH+Alpha+Beta-domain+wHTH+RRM</td><td>Type-2_BR.RM.like.systems</td><td><-MORC||?-><-DUF3780<-?<-?<-DUF499<-DUF499*<-HKD-DNase+SF1-SF2-Helicase-p<-?<-?<-SF1-SF2-Helicase-p<-HKD-DNase+SF1-SF2-Helicase+iREase+CTD</td><td>311</td><td>Bacteria>Terrabacteria_group>Cyanobacteriota/Melainabacteria_group>Cyanobacteriota>Cyanophyceae</td><td>GCF_006523545.1</td><td>Cylindrospermopsis_raciborskii</td><td>FIV49_RS17550</td><td>247769..248704</td><td>77022</td><td>DUF499-domain-containing-protein-[Cylindrospermopsis-raciborskii]</td></tr>
<tr><td>WP_230967088.1</td><td>AAA+ATPase+wHTH+Alpha+Beta-domain+wHTH+RRM</td><td>Type-2_BR.RM.like.systems</td><td><-HKD-DNase+SF1-SF2-Helicase+iREase+CTD-p<-HKD-DNase+SF1-SF2-Helicase+iREase+CTD<-RelE-ParE<-CopG/RHH<-DUF3780<-DUF499*</td><td>640</td><td>Bacteria>Terrabacteria_group>Cyanobacteriota/Melainabacteria_group>Cyanobacteriota>Cyanophyceae</td><td>GCF_015714765.1</td><td>Nostoc_commune</td><td>F8S20_RS18355</td><td>6312..8234</td><td>1178</td><td>DUF499-domain-containing-protein-[Nostoc-commune]</td></tr>
<tr><td>WP_163936881.1</td><td>AAA+ATPase+wHTH+Alpha+Beta-domain+wHTH+RRM</td><td>Type-2_BR.RM.like.systems</td><td><-FAD-NAD-dep-oxidoreductase<-REase||HTH_28+rve+Mu-transpos_C->?->?->DUF499*-></td><td>596</td><td>Bacteria>Terrabacteria_group>Cyanobacteriota/Melainabacteria_group>Cyanobacteriota>Cyanophyceae</td><td>GCF_010747425.1</td><td>Nostoc_sp._UIC_10630</td><td>G1O98_RS31925</td><td>10178..11968</td><td>2100146</td><td>DUF499-domain-containing-protein-[Nostoc-sp.-UIC-10630]</td></tr>
<tr><td>MBC8359968.1</td><td>AAA+ATPase+wHTH+Alpha+Beta-domain+wHTH+RRM</td><td>Type-2_BR.RM.like.systems</td><td>Schlafen+HATPase+HTH->BirA-HTH->?->PolB->HEPN->DUF499*-></td><td>637</td><td>Bacteria>Thermodesulfobacteriota>Desulfobacteria>Desulfobacterales>Desulfobacterales_incertae_sedis</td><td>GCA_014382665.1</td><td>Candidatus_Desulfatibia_profunda</td><td>H8E23_01035</td><td>3975..5887</td><td>2841695</td><td>ATP-binding-protein-[Desulfobacterales-bacterium]</td></tr>
<tr><td>MCY4104732.1</td><td>AAA+ATPase+wHTH+Alpha+Beta-domain+wHTH+RRM</td><td>Type-2_BR.RM.like.systems</td><td>DUF499*->DUF499-p->DUF3780->METHYLASE->HKD-DNase+SF1-SF2-Helicase+iREase+CTD->HKD-DNase+SF1-SF2-Helicase+iREase+CTD-p-><-PIN</td><td>678</td><td>Bacteria>unclassified_Bacteria</td><td>GCA_026708055.1</td><td>bacterium</td><td>OXG55_15965</td><td>64138..66174</td><td>1869227</td><td>MAG:-DUF499-domain-containing-protein-[bacterium]</td></tr>
<tr><td>MCY4631878.1</td><td>AAA+ATPase+wHTH+Alpha+Beta-domain+wHTH+RRM</td><td>Type-2_BR.RM.like.systems</td><td>MACRODOMAIN->DUF499*->DUF3780->METHYLASE->HKD-DNase+SF1-SF2-Helicase+iREase+CTD->?->CHTH-></td><td>756</td><td>Bacteria>unclassified_Bacteria</td><td>GCA_026713745.1</td><td>bacterium</td><td>OXE75_13500</td><td>3556..5826</td><td>1869227</td><td>MAG:-DUF499-domain-containing-protein-[bacterium]</td></tr>
<tr><td>MDE0667717.1</td><td>AAA+ATPase+wHTH+Alpha+Beta-domain+wHTH+RRM</td><td>Type-2_BR.RM.like.systems</td><td><-DUF499*<-MACRODOMAIN<-?<-ORC-CDC6+REase</td><td>733</td><td>Bacteria>unclassified_Bacteria</td><td>GCA_028824385.1</td><td>bacterium</td><td>OXI48_01630</td><td>1..2198</td><td>1869227</td><td>MAG:-DUF499-domain-containing-protein,-partial-[bacterium]</td></tr>
<tr><td>MCY3561261.1</td><td>AAA+ATPase+wHTH+Alpha+Beta-domain+wHTH+RRM</td><td>Type-2_BR.RM.like.systems</td><td><-DUF499*<-MACRODOMAIN<-?<-ORC-CDC6+REase</td><td>695</td><td>Bacteria>unclassified_Bacteria</td><td>GCA_026702815.1</td><td>bacterium</td><td>OXH20_08865</td><td>1..2084</td><td>1869227</td><td>MAG:-DUF499-domain-containing-protein,-partial-[bacterium]</td></tr>
<tr><td>MBI4750004.1</td><td>AAA+ATPase+wHTH+Alpha+Beta-domain+wHTH+RRM</td><td>Type-2_BR.RM.like.systems</td><td>TUDOR+SF1-SF2-Helicase+iREase+CTD+REase->HicB_lk_antitox->REase->METHYLASE->HicB_lk_antitox->HicA_toxin->DUF499*->ABhydrolase-><-?<-?<-?<-BirA-HTH</td><td>939</td><td>Bacteria>Acidobacteriota>unclassified_Acidobacteriota</td><td>GCA_016208495.1</td><td>Acidobacteriota_bacterium</td><td>HY774_16075</td><td>69401..72220</td><td>1978231</td><td>ATP-binding-protein-[Acidobacteria-bacterium]</td></tr>
<tr><td>WP_202318894.1</td><td>AAA+ATPase+wHTH+Alpha+Beta-domain+wHTH+wHTH+RRM</td><td>Type-2_BR.RM.like.systems</td><td><-METHYLASE-p<-PIN<-?<-MazE_antitoxin||?-><-DUF3780<-DUF499*<-?<-tRNA||?->?->Cysteine_desulfurase-></td><td>773</td><td>Archaea>Euryarchaeota>Archaeoglobi>Archaeoglobales>Archaeoglobaceae</td><td>GCF_016757965.1</td><td>Archaeoglobus_neptunius</td><td>JFQ59_RS02795</td><td>87809..90130</td><td>2798580</td><td>ATP-binding-protein-[Archaeoglobus-neptunius]</td></tr>
<tr><td>WP_013749028.1</td><td>AAA+ATPase+wHTH+Alpha+Beta-domain+wHTH+wHTH+RRM</td><td>Type-2_BR.RM.like.systems</td><td><-SF1-SF2-Helicase+iREase+CTD+REase<-METHYLASE<-DUF3780<-DUF499*<-ORC-CDC6+REase<-?<-?<-?<-LRP-HTH<-Peptidase</td><td>1059</td><td>Archaea>Euryarchaeota>Thermococci>Thermococcales>Thermococcaceae</td><td>GCF_000211475.1</td><td>Pyrococcus_sp._NA2</td><td>PNA2_RS07895</td><td>1443844..1447023</td><td>342949</td><td>ATP-binding-protein-[Pyrococcus-sp.-NA2]</td></tr>
<tr><td>NQW03349.1</td><td>AAA+ATPase+wHTH+Alpha+Beta-domain+wHTH+wHTH+RRM</td><td>Type-2_BR.RM.like.systems</td><td>DUF499*->DUF3780->METHYLASE->HKD-DNase+SF1-SF2-Helicase+iREase+CTD-></td><td>1084</td><td>Bacteria>Acidobacteriota>unclassified_Acidobacteriota</td><td>GCA_013203475.1</td><td>Acidobacteriota_bacterium</td><td>HQ485_04920</td><td>424..3678</td><td>1978231</td><td>ATP-binding-protein-[Acidobacteria-bacterium]</td></tr>
<tr><td>MBI1799725.1</td><td>AAA+ATPase+wHTH+Alpha+Beta-domain+wHTH+wHTH+RRM</td><td>Type-2_BR.RM.like.systems</td><td><-ParA-Soj-PloopNTPase||Radical-SAM->?-><-HKD-DNase+SF1-SF2-Helicase+iREase+CTD<-METHYLASE<-PIN<-DUF499*<-?<-ABC-ATPase+SbcC<-?<-?<-PRTase</td><td>972</td><td>Bacteria>Bacteria_incertae_sedis>Bacteria_candidate_phyla>Candidatus_Eisenbacteria>unclassified_Candidatus_Eisenbacteria</td><td>GCA_016178685.1</td><td>Candidatus_Eisenbacteria_bacterium</td><td>HYR73_08560</td><td>14195..17113</td><td>2212470</td><td>ATP-binding-protein-[Candidatus-Eisenbacteria-bacterium]</td></tr>
<tr><td>MCB9367519.1</td><td>AAA+ATPase+wHTH+Alpha+Beta-domain+wHTH+wHTH+RRM</td><td>Type-2_BR.RM.like.systems</td><td>DUF499*->DUF3780->METHYLASE->HKD-DNase+SF1-SF2-Helicase+iREase+CTD->ParB+HNH->OLD-ABC-></td><td>970</td><td>Bacteria>Calditrichota>unclassified_Calditrichota</td><td>GCA_020637445.1</td><td>Calditrichota_bacterium</td><td>H6507_00185</td><td>43427..46339</td><td>2212469</td><td>ATP-binding-protein-[Calditrichaeota-bacterium]</td></tr>
<tr><td>NCA71589.1</td><td>AAA+ATPase+wHTH+Alpha+Beta-domain+wHTH+wHTH+RRM</td><td>Type-2_BR.RM.like.systems</td><td>DUF499*->DUF3780->?->?->?->METHYLASE->PIN-></td><td>942</td><td>Bacteria>FCB_group>Bacteroidota/Chlorobiota_group>Bacteroidota>Sphingobacteriia</td><td>GCA_009928635.1</td><td>Sphingobacteriia_bacterium</td><td>EOM91_16190</td><td>1..2830</td><td>2035813</td><td>DUF499-domain-containing-protein-[Sphingobacteriia-bacterium]</td></tr>
<tr><td>MBM4287682.1</td><td>AAA+ATPase+wHTH+Alpha+Beta-domain+wHTH+wHTH+RRM</td><td>Type-2_BR.RM.like.systems</td><td><-HKD-DNase+SF1-SF2-Helicase+iREase+CTD<-METHYLASE<-RNaseH<-DUF3780<-DUF499*<-?||?-><-HTH+PolB</td><td>1019</td><td>Bacteria>Myxococcota>Myxococcia>unclassified_Myxococcia</td><td>GCA_016874895.1</td><td>Deltaproteobacteria_bacterium</td><td>FJ135_05975</td><td>8432..11491</td><td>2026735</td><td>ATP-binding-protein-[Deltaproteobacteria-bacterium]</td></tr>
<tr><td>MBM4273482.1</td><td>AAA+ATPase+wHTH+Alpha+Beta-domain+wHTH+wHTH+RRM</td><td>Type-2_BR.RM.like.systems</td><td>TPR+TPR+TPR+TPR+TPR+TPR+TPR+TPR->?->?->DUF499*->DUF3780->RNaseH->METHYLASE->HKD-DNase+SF1-SF2-Helicase+iREase+CTD-></td><td>973</td><td>Bacteria>Myxococcota>Myxococcia>unclassified_Myxococcia</td><td>GCA_016874915.1</td><td>Deltaproteobacteria_bacterium</td><td>FJ134_03345</td><td>24713..27634</td><td>2026735</td><td>ATP-binding-protein-[Deltaproteobacteria-bacterium]</td></tr>
<tr><td>MBW1961245.1</td><td>AAA+ATPase+wHTH+Alpha+Beta-domain+wHTH+wHTH+RRM</td><td>Type-2_BR.RM.like.systems</td><td><-HKD-DNase+SF1-SF2-Helicase+iREase+CTD<-METHYLASE<-DUF3780<-DUF499*||?->?-><-?<-?||?->AbiEii-></td><td>1082</td><td>Bacteria>Myxococcota>Myxococcia>unclassified_Myxococcia</td><td>GCA_019308005.1</td><td>Deltaproteobacteria_bacterium</td><td>JRJ04_07300</td><td>8076..11324</td><td>2026735</td><td>DUF499-domain-containing-protein-[Deltaproteobacteria-bacterium]</td></tr>
<tr><td>MCD6261593.1</td><td>AAA+ATPase+wHTH+Alpha+Beta-domain+wHTH+wHTH+RRM</td><td>Type-2_BR.RM.like.systems</td><td>Calcineurin->?->DUF499*->DUF3780->METHYLASE->HKD-DNase+SF1-SF2-Helicase+iREase+CTD-><-?<-?||sGTP-></td><td>1081</td><td>Bacteria>Myxococcota>Myxococcia>unclassified_Myxococcia</td><td>GCA_021163815.1</td><td>Deltaproteobacteria_bacterium</td><td>J7J52_01035</td><td>3419..6664</td><td>2026735</td><td>ATP-binding-protein-[Deltaproteobacteria-bacterium]</td></tr>
<tr><td>GIW54142.1</td><td>AAA+ATPase+wHTH+Alpha+Beta-domain+wHTH+wHTH+RRM</td><td>Type-2_BR.RM.like.systems</td><td>REase+HTH->NTP_transf-><-?<-?||AbiEii->?->DUF499*->DUF3780->PIN->METHYLASE-></td><td>972</td><td>Bacteria>Nitrospirota>Nitrospiria>Nitrospirales>Nitrospiraceae</td><td>GCA_026004315.1</td><td>Nitrospiraceae_bacterium</td><td>KatS3mg082_0546</td><td>99014..101932</td><td>2026770</td><td>MAG:-hypothetical-protein-KatS3mg082_0546-[Nitrospiraceae-bacterium]</td></tr>
<tr><td>MTW18316.1</td><td>AAA+ATPase+wHTH+Alpha+Beta-domain+wHTH+wHTH+RRM</td><td>Type-2_BR.RM.like.systems</td><td>TnsC-ATPase-><-NTP_transf_3||tRNA-><-HKD-DNase+SF1-SF2-Helicase+iREase+CTD<-METHYLASE<-DUF3780<-DUF499*<-HTH<-GDResolvase+Recombinase+ZnR<-RAMA||?->?->SIGMA-HTH-></td><td>969</td><td>Bacteria>Pseudomonadota>Alphaproteobacteria>Hyphomicrobiales>Hyphomicrobiaceae</td><td>GCA_009720755.1</td><td>Rhodoplanes_serenus</td><td>GJ689_19120</td><td>50553..53462</td><td>200615</td><td>DUF499-domain-containing-protein-[Rhodoplanes-serenus]</td></tr>
<tr><td>WP_155480776.1</td><td>AAA+ATPase+wHTH+Alpha+Beta-domain+wHTH+wHTH+RRM</td><td>Type-2_BR.RM.like.systems</td><td>TnsC-ATPase-><-NTP_transf_3||tRNA-><-HKD-DNase+SF1-SF2-Helicase+iREase+CTD<-METHYLASE<-DUF3780<-DUF499*<-HTH<-GDResolvase+Recombinase+ZnR<-RAMA</td><td>969</td><td>Bacteria>Pseudomonadota>Alphaproteobacteria>Hyphomicrobiales>Hyphomicrobiaceae</td><td>GCF_009720755.1</td><td>Rhodoplanes_serenus</td><td>GJ689_RS19115</td><td>50553..53462</td><td>200615</td><td>DUF499-domain-containing-protein-[Rhodoplanes-serenus]</td></tr>
<tr><td>WP_247297741.1</td><td>AAA+ATPase+wHTH+Alpha+Beta-domain+wHTH+wHTH+RRM</td><td>Type-2_BR.RM.like.systems</td><td>Glyco_transf_20->Glyco_transf_20-><-?||tRNA->DUF499*->METHYLASE->HKD-DNase+SF1-SF2-Helicase+iREase+CTD-></td><td>969</td><td>Bacteria>Pseudomonadota>Alphaproteobacteria>Hyphomicrobiales>Nitrobacteraceae</td><td>GCF_023073535.1</td><td>Bradyrhizobium_sp._179</td><td>IVB12_RS08740</td><td>17828..20737</td><td>2782648</td><td>DUF499-domain-containing-protein-[Bradyrhizobium-sp.-179]</td></tr>
<tr><td>TIL33927.1</td><td>AAA+ATPase+wHTH+Alpha+Beta-domain+wHTH+wHTH+RRM</td><td>Type-2_BR.RM.like.systems</td><td><-Peptidase<-?||?->Methyltransf_25-><-REase||DUF499*->DUF3780->HKD-DNase+SF1-SF2-Helicase+iREase+CTD-><-SF1-SF2-Helicase+DUF1998</td><td>1038</td><td>Bacteria>Pseudomonadota>Alphaproteobacteria>Hyphomicrobiales>Phyllobacteriaceae</td><td>GCA_004961305.1</td><td>Mesorhizobium_sp.</td><td>E5Y85_12100</td><td>117884..121000</td><td>1871066</td><td>ATP-binding-protein-[Mesorhizobium-sp.]</td></tr>
<tr><td>WP_183367784.1</td><td>AAA+ATPase+wHTH+Alpha+Beta-domain+wHTH+wHTH+RRM</td><td>Type-2_BR.RM.like.systems</td><td><-HNH+SAVED<-HKD-DNase+SF1-SF2-Helicase+iREase+CTD<-METHYLASE<-DUF3780<-DUF499*<-HTH+WYL<-XIS-HTH||Phage_integrase-><-?||HNH-></td><td>1101</td><td>Bacteria>Pseudomonadota>Alphaproteobacteria>Hyphomicrobiales>Rhizobiaceae</td><td>GCF_014197145.1</td><td>Gellertiella_hungarica</td><td>GGR23_RS18445</td><td>165891..169196</td><td>1572859</td><td>DUF499-domain-containing-protein-[Gellertiella-hungarica]</td></tr>
<tr><td>WP_135313433.1</td><td>AAA+ATPase+wHTH+Alpha+Beta-domain+wHTH+wHTH+RRM</td><td>Type-2_BR.RM.like.systems</td><td>XIS-HTH->?->RAMA->GDResolvase+Recombinase+ZnR->HTH->?->DUF499*->DUF3780->METHYLASE->?->HKD-DNase+SF1-SF2-Helicase+iREase+CTD->Arm-DNA-bind_3+Phage_integrase-></td><td>969</td><td>Bacteria>Pseudomonadota>Alphaproteobacteria>Rhodobacterales>Paracoccaceae</td><td>GCF_004683865.2</td><td>Paracoccus_liaowanqingii</td><td>E4191_RS10950</td><td>2265090..2267999</td><td>2560053</td><td>DUF499-domain-containing-protein-[Paracoccus-liaowanqingii]</td></tr>
<tr><td>WP_036720035.1</td><td>AAA+ATPase+wHTH+Alpha+Beta-domain+wHTH+wHTH+RRM</td><td>Type-2_BR.RM.like.systems</td><td>XIS-HTH->HTH+WYL->DUF499*->DUF3780->METHYLASE->HKD-DNase+SF1-SF2-Helicase+iREase+CTD-></td><td>1086</td><td>Bacteria>Pseudomonadota>Alphaproteobacteria>Rhodobacterales>Paracoccaceae</td><td>GCF_000763805.1</td><td>Paracoccus_sphaerophysae</td><td>IC63_RS10560</td><td>2078..5338</td><td>690417</td><td>DUF499-domain-containing-protein-[Paracoccus-sphaerophysae]</td></tr>
<tr><td>WP_096434278.1</td><td>AAA+ATPase+wHTH+Alpha+Beta-domain+wHTH+wHTH+RRM</td><td>Type-2_BR.RM.like.systems</td><td><-XIS-HTH<-?||LexA-protease-><-ParD_antitoxin||HTH+WYL->DUF499*->DUF3780->METHYLASE->HKD-DNase+SF1-SF2-Helicase+iREase+CTD-><-RNase</td><td>1086</td><td>Bacteria>Pseudomonadota>Alphaproteobacteria>Rhodobacterales>Paracoccaceae</td><td>GCF_002358085.1</td><td>Pseudothioclava_arenosa</td><td>CLN94_RS12435</td><td>22675..25935</td><td>1795308</td><td>DUF499-domain-containing-protein-[Pseudothioclava-arenosa]</td></tr>
<tr><td>WP_252930533.1</td><td>AAA+ATPase+wHTH+Alpha+Beta-domain+wHTH+wHTH+RRM</td><td>Type-2_BR.RM.like.systems</td><td><-DDE_Tnp_1_4||CopG/RHH->DUF499*->DUF3780->METHYLASE->HKD-DNase+SF1-SF2-Helicase+iREase+CTD-><-GDResolvase+HIN-HTH<-Phage_integrase<-Phage_int_SAM_1+Phage_integrase</td><td>1086</td><td>Bacteria>Pseudomonadota>Alphaproteobacteria>Rhodobacterales>Paracoccaceae</td><td>GCF_024105395.1</td><td>Paracoccus_sp._08</td><td>GB037_RS19525</td><td>2113..5370</td><td>2606624</td><td>DUF499-domain-containing-protein-[Paracoccus-sp.-08]</td></tr>
<tr><td>WP_261193162.1</td><td>AAA+ATPase+wHTH+Alpha+Beta-domain+wHTH+wHTH+RRM</td><td>Type-2_BR.RM.like.systems</td><td>RNase-><-HKD-DNase+SF1-SF2-Helicase+iREase+CTD<-METHYLASE<-DUF3780<-DUF499*<-HTH+WYL||ParD_antitoxin-><-XIS-HTH||?-><-MODE-HTH+PBPII</td><td>1086</td><td>Bacteria>Pseudomonadota>Alphaproteobacteria>Rhodobacterales>Roseobacteraceae</td><td>GCF_025376885.1</td><td>Pseudoruegeria_sp._SHC-113</td><td>KVX96_RS04850</td><td>968995..972255</td><td>2855439</td><td>DUF499-domain-containing-protein-[Pseudoruegeria-sp.-SHC-113]</td></tr>
<tr><td>QDJ08642.1</td><td>AAA+ATPase+wHTH+Alpha+Beta-domain+wHTH+wHTH+RRM</td><td>Type-2_BR.RM.like.systems</td><td><-N6_N4_MTase<-MPTase<-HTH_3||RAMA->GDResolvase+Recombinase+ZnR->HTH->DUF499*->DUF3780->METHYLASE->?->HKD-DNase+SF1-SF2-Helicase+iREase+CTD-></td><td>969</td><td>Bacteria>Pseudomonadota>Alphaproteobacteria>Rhodospirillales>Acetobacteraceae</td><td>GCF_006771025.1</td><td>Roseomonas_mucosa</td><td>HVPorG_04654</td><td>1098172..1101081</td><td>207340</td><td>Hypothetical-protein-HVPorG_04654-[Roseomonas-mucosa]</td></tr>
<tr><td>MCZ8140259.1</td><td>AAA+ATPase+wHTH+Alpha+Beta-domain+wHTH+wHTH+RRM</td><td>Type-2_BR.RM.like.systems</td><td><-HKD-DNase<-HKD-DNase+SF1-SF2-Helicase+iREase+CTD<-?<-METHYLASE<-DUF3780<-DUF499*<-?<-?<-HTH<-GDResolvase+Recombinase+ZnR<-RAMA<-LexA-protease</td><td>969</td><td>Bacteria>Pseudomonadota>Alphaproteobacteria>Rhodospirillales>Acetobacteraceae</td><td>GCA_027531275.1</td><td>Acetobacteraceae_bacterium</td><td>O9273_01165</td><td>231571..234480</td><td>1909293</td><td>MAG:-DUF499-domain-containing-protein-[Acetobacteraceae-bacterium]</td></tr>
<tr><td>WP_237719425.1</td><td>AAA+ATPase+wHTH+Alpha+Beta-domain+wHTH+wHTH+RRM</td><td>Type-2_BR.RM.like.systems</td><td>MazE_antitoxin->PIN-><-HKD-DNase+SF1-SF2-Helicase+iREase+CTD<-?<-METHYLASE<-DUF3780<-DUF499*</td><td>411</td><td>Bacteria>Pseudomonadota>Alphaproteobacteria>Rhodospirillales>Acetobacteraceae</td><td>GCF_000292225.1</td><td>Roseomonas_sp._B5</td><td>B555_RS0120340</td><td>9698..10935</td><td>1217720</td><td>ATP-binding-protein,-partial-[Roseomonas-sp.-B5]</td></tr>
<tr><td>WP_146422921.1</td><td>AAA+ATPase+wHTH+Alpha+Beta-domain+wHTH+wHTH+RRM</td><td>Type-2_BR.RM.like.systems</td><td><-XIS-HTH<-PAPS_reduct<-?||RAMA->GDResolvase+Recombinase+ZnR->HTH->DUF499*->DUF3780->METHYLASE->?->HKD-DNase+SF1-SF2-Helicase+iREase+CTD-></td><td>969</td><td>Bacteria>Pseudomonadota>Alphaproteobacteria>Rhodospirillales>Acetobacteraceae</td><td>GCF_003253705.1</td><td>Humitalea_rosea</td><td>C8P66_RS16710</td><td>26262..29171</td><td>990373</td><td>DUF499-domain-containing-protein-[Humitalea-rosea]</td></tr>
<tr><td>WP_019463153.1</td><td>AAA+ATPase+wHTH+Alpha+Beta-domain+wHTH+wHTH+RRM</td><td>Type-2_BR.RM.like.systems</td><td>DUF499*->DUF3780->METHYLASE->?->HKD-DNase+SF1-SF2-Helicase+iREase+CTD-><-PIN<-MazE_antitoxin</td><td>399</td><td>Bacteria>Pseudomonadota>Alphaproteobacteria>Rhodospirillales>Acetobacteraceae</td><td>GCF_006384895.1</td><td>Roseomonas_mucosa</td><td>BT98_RS0119345</td><td>343..1542</td><td>207340</td><td>hypothetical-protein-[Roseomonas-mucosa]</td></tr>
<tr><td>WP_265547898.1</td><td>AAA+ATPase+wHTH+Alpha+Beta-domain+wHTH+wHTH+RRM</td><td>Type-2_BR.RM.like.systems</td><td><-N6_N4_MTase<-MPTase<-HTH_3||RAMA->GDResolvase+Recombinase+ZnR->HTH->DUF499*->DUF3780->METHYLASE->?->HKD-DNase+SF1-SF2-Helicase+iREase+CTD-></td><td>969</td><td>Bacteria>Pseudomonadota>Alphaproteobacteria>Rhodospirillales>Acetobacteraceae</td><td>GCF_006771025.1</td><td>Roseomonas_mucosa</td><td>HVPorG_RS04810</td><td>1098172..1101081</td><td>207340</td><td>DUF499-domain-containing-protein-[Roseomonas-mucosa]</td></tr>
<tr><td>WP_184481416.1</td><td>AAA+ATPase+wHTH+Alpha+Beta-domain+wHTH+wHTH+RRM</td><td>Type-2_BR.RM.like.systems</td><td><-N6_N4_MTase<-MPTase<-HTH_3||RAMA->GDResolvase+Recombinase+ZnR->HTH->DUF499*->DUF3780->METHYLASE->HKD-DNase+SF1-SF2-Helicase+iREase+CTD-><-tRNA<-ABhydrolase</td><td>990</td><td>Bacteria>Pseudomonadota>Alphaproteobacteria>Rhodospirillales>Acetobacteraceae</td><td>GCF_014199195.1</td><td>Neoroseomonas_alkaliterrae</td><td>FHS88_RS03560</td><td>32349..35321</td><td>1452450</td><td>DUF499-domain-containing-protein-[Roseomonas-alkaliterrae]</td></tr>
<tr><td>WP_081055878.1</td><td>AAA+ATPase+wHTH+Alpha+Beta-domain+wHTH+wHTH+RRM</td><td>Type-2_BR.RM.like.systems</td><td><-SymE_toxin||?-><-HKD-DNase+SF1-SF2-Helicase+iREase+CTD<-METHYLASE<-DUF3780<-DUF499*<-HTH+WYL||HEPN-p-></td><td>1073</td><td>Bacteria>Pseudomonadota>Betaproteobacteria>Burkholderiales>Burkholderiaceae</td><td>GCF_001524505.1</td><td>Burkholderia_pseudomultivorans</td><td>WS80_RS30005</td><td>9016..12237</td><td>1207504</td><td>DUF499-domain-containing-protein-[Burkholderia-pseudomultivorans]</td></tr>
<tr><td>WP_239294343.1</td><td>AAA+ATPase+wHTH+Alpha+Beta-domain+wHTH+wHTH+RRM</td><td>Type-2_BR.RM.like.systems</td><td><-HKD-DNase+SF1-SF2-Helicase+iREase+CTD<-METHYLASE<-DUF3780<-DUF499*<-RHH||?-><-Phage_AlpA<-Arm-DNA-bind_3+Phage_int_SAM_3+Phage_integrase<-Arm-DNA-bind_3+Phage_integrase</td><td>1073</td><td>Bacteria>Pseudomonadota>Betaproteobacteria>Burkholderiales>Burkholderiaceae</td><td>GCF_022346825.1</td><td>Caballeronia_zhejiangensis</td><td>MHM92_RS02640</td><td>565275..568496</td><td>871203</td><td>DUF499-domain-containing-protein-[Caballeronia-zhejiangensis]</td></tr>
<tr><td>TAK83982.1</td><td>AAA+ATPase+wHTH+Alpha+Beta-domain+wHTH+wHTH+RRM</td><td>Type-2_BR.RM.like.systems</td><td><-MethyltransfD12<-ParB<-HKD-DNase+SF1-SF2-Helicase+iREase+CTD<-METHYLASE<-DUF3780<-DUF499*||?->?-><-TetR-HTH||?->TIR-></td><td>966</td><td>Bacteria>Pseudomonadota>Betaproteobacteria>unclassified_Betaproteobacteria</td><td>GCA_004297625.1</td><td>Betaproteobacteria_bacterium</td><td>EPO20_16110</td><td>30839..33739</td><td>1891241</td><td>ATP-binding-protein-[Betaproteobacteria-bacterium]</td></tr>
<tr><td>WP_094037979.1</td><td>AAA+ATPase+wHTH+Alpha+Beta-domain+wHTH+wHTH+RRM</td><td>Type-2_BR.RM.like.systems</td><td><-MNS+HTH||OLD-ABC->RlpA-><-HKD-DNase+SF1-SF2-Helicase+iREase+CTD<-METHYLASE<-DUF3780<-DUF499*||?->?-><-?||?-><-HTH+WYL</td><td>1077</td><td>Bacteria>Pseudomonadota>Gammaproteobacteria>Aeromonadales>Aeromonadaceae</td><td>GCF_002237365.1</td><td>Zobellella_denitrificans</td><td>CGX12_RS00725</td><td>23792..27025</td><td>347534</td><td>DUF499-domain-containing-protein-[Zobellella-denitrificans]</td></tr>
<tr><td>TMO42407.1</td><td>AAA+ATPase+wHTH+Alpha+Beta-domain+wHTH+wHTH+RRM</td><td>Type-2_BR.RM.like.systems</td><td><-MNS+REase||HTH->STYKinase-like->XIS-HTH->ZnR+TOPRIM+primpol+HTH->HTH+WYL->DUF499*->DUF3780->METHYLASE->ParB+HNH->HKD-DNase+SF1-SF2-Helicase+iREase+CTD->?->MqsA_antitoxin+HTH_3-></td><td>1081</td><td>Bacteria>Pseudomonadota>Gammaproteobacteria>Alteromonadales>Pseudoalteromonadaceae</td><td>GCA_005886815.1</td><td>Pseudoalteromonas_sp._S4389</td><td>CWC25_15720</td><td>9911..13156</td><td>579556</td><td>AAA-family-ATPase-[Pseudoalteromonas-sp.-S4389]</td></tr>
<tr><td>WP_158678953.1</td><td>AAA+ATPase+wHTH+Alpha+Beta-domain+wHTH+wHTH+RRM</td><td>Type-2_BR.RM.like.systems</td><td><-Phage_integrase<-JAB||XIS-HTH->PIN+OST-HTH->DUF499*->DUF3780->METHYLASE->HKD-DNase+SF1-SF2-Helicase+iREase+CTD-><-Arm-DNA-bind_3+Phage_integrase<-35exo</td><td>1076</td><td>Bacteria>Pseudomonadota>Gammaproteobacteria>Alteromonadales>Pseudoalteromonadaceae</td><td>GCF_002964705.1</td><td>Pseudoalteromonas_sp._T1lg75</td><td>C2M36_RS13740</td><td>243061..246291</td><td>2077102</td><td>DUF499-domain-containing-protein-[Pseudoalteromonas-sp.-T1lg75]</td></tr>
<tr><td>WP_171040448.1</td><td>AAA+ATPase+wHTH+Alpha+Beta-domain+wHTH+wHTH+RRM</td><td>Type-2_BR.RM.like.systems</td><td><-MNS+REase||HTH->STYKinase-like->XIS-HTH->ZnR+TOPRIM+primpol+HTH->HTH+WYL->DUF499*->DUF3780->METHYLASE->ParB+HNH->HKD-DNase+SF1-SF2-Helicase+iREase+CTD->?->MqsA_antitoxin+HTH_3-></td><td>1077</td><td>Bacteria>Pseudomonadota>Gammaproteobacteria>Alteromonadales>Pseudoalteromonadaceae</td><td>GCF_005886815.1</td><td>Pseudoalteromonas_sp._S4389</td><td>CWC25_RS15645</td><td>9923..13156</td><td>579556</td><td>DUF499-domain-containing-protein-[Pseudoalteromonas-sp.-S4389]</td></tr>
<tr><td>WP_199525347.1</td><td>AAA+ATPase+wHTH+Alpha+Beta-domain+wHTH+wHTH+RRM</td><td>Type-2_BR.RM.like.systems</td><td><-MqsA_antitoxin+HTH_3<-?<-HKD-DNase+SF1-SF2-Helicase+iREase+CTD<-ParB+HNH<-METHYLASE<-DUF3780<-DUF499*<-HTH+WYL<-ZnR+TOPRIM+primpol+HTH<-XIS-HTH<-STYKinase-like<-HTH||MNS+REase-></td><td>1077</td><td>Bacteria>Pseudomonadota>Gammaproteobacteria>Alteromonadales>Pseudoalteromonadaceae</td><td>GCF_016464045.1</td><td>Pseudoalteromonas_sp._bablab_jr011</td><td>JIG41_RS20360</td><td>15866..19099</td><td>2755062</td><td>DUF499-domain-containing-protein-[Pseudoalteromonas-sp.-bablab_jr011]</td></tr>
<tr><td>WP_076922918.1</td><td>AAA+ATPase+wHTH+Alpha+Beta-domain+wHTH+wHTH+RRM</td><td>Type-2_BR.RM.like.systems</td><td><-MqsA_antitoxin+HTH_3<-?<-HKD-DNase+SF1-SF2-Helicase+iREase+CTD<-ParB+HNH<-METHYLASE<-DUF3780<-DUF499*<-HTH+WYL<-ZnR+TOPRIM+primpol+HTH<-XIS-HTH<-STYKinase-like<-HTH||MNS+REase-></td><td>1077</td><td>Bacteria>Pseudomonadota>Gammaproteobacteria>Alteromonadales>Pseudoalteromonadaceae</td><td>GCF_030407565.1</td><td>unclassified_Pseudoalteromonas</td><td>BW140_RS15975</td><td>19057..22290</td><td>194690</td><td>DUF499-domain-containing-protein-[Pseudoalteromonas-sp.-SK20]</td></tr>
<tr><td>MAL94418.1</td><td>AAA+ATPase+wHTH+Alpha+Beta-domain+wHTH+wHTH+RRM</td><td>Type-2_BR.RM.like.systems</td><td><-HKD-DNase+SF1-SF2-Helicase+iREase+CTD<-?<-PIN<-OLD-ABC<-METHYLASE<-DUF3780<-DUF499*||SF1-SF2-Helicase-></td><td>1088</td><td>Bacteria>Pseudomonadota>Gammaproteobacteria>Cellvibrionales>Halieaceae</td><td>GCA_002692965.1</td><td>Haliea_sp.</td><td>CME40_05000</td><td>41677..44943</td><td>1932666</td><td>hypothetical-protein-[Haliea-sp.]</td></tr>
<tr><td>ESQ16532.1</td><td>AAA+ATPase+wHTH+Alpha+Beta-domain+wHTH+wHTH+RRM</td><td>Type-2_BR.RM.like.systems</td><td><-HKD-DNase+SF1-SF2-Helicase+iREase+CTD<-HEPN<-PolB<-METHYLASE<-DUF3780<-REase<-DUF499*</td><td>1087</td><td>Bacteria>Pseudomonadota>Gammaproteobacteria>Chromatiales>Chromatiaceae</td><td>GCA_000495615.1</td><td>Thiohalocapsa_sp._PB-PSB1</td><td>N838_05685</td><td>9740..13003</td><td>1385625</td><td>MAG:-hypothetical-protein-N838_05685-[Thiohalocapsa-sp.-PB-PSB1]</td></tr>
<tr><td>QQO57135.1</td><td>AAA+ATPase+wHTH+Alpha+Beta-domain+wHTH+wHTH+RRM</td><td>Type-2_BR.RM.like.systems</td><td>DDE_Tnp_1->DUF499*->REase->DUF3780->METHYLASE->PolB->HEPN->HKD-DNase+SF1-SF2-Helicase+iREase+CTD-></td><td>1085</td><td>Bacteria>Pseudomonadota>Gammaproteobacteria>Chromatiales>Chromatiaceae</td><td>GCA_016745215.1</td><td>Thiohalocapsa_sp._PB-PSB1</td><td>N838_31100</td><td>7373531..7376788</td><td>1385625</td><td>MAG:-ATP-binding-protein-[Thiohalocapsa-sp.-PB-PSB1]</td></tr>
<tr><td>WP_020506762.1</td><td>AAA+ATPase+wHTH+Alpha+Beta-domain+wHTH+wHTH+RRM</td><td>Type-2_BR.RM.like.systems</td><td><-HKD-DNase+SF1-SF2-Helicase+iREase+CTD<-Zn_Tnp_IS1595+DDE_Tnp_IS1595<-OLD-ABC<-METHYLASE<-HKD-DNase<-DUF3780<-DUF499*</td><td>1089</td><td>Bacteria>Pseudomonadota>Gammaproteobacteria>Chromatiales>Chromatiaceae</td><td>GCF_000379525.1</td><td>Lamprocystis_purpurea</td><td>A390_RS0121645</td><td>37320..40589</td><td>61598</td><td>DUF499-domain-containing-protein-[Lamprocystis-purpurea]</td></tr>
<tr><td>WP_100917768.1</td><td>AAA+ATPase+wHTH+Alpha+Beta-domain+wHTH+wHTH+RRM</td><td>Type-2_BR.RM.like.systems</td><td>Phage_int_SAM_1+Phage_integrase-><-HKD-DNase+SF1-SF2-Helicase+iREase+CTD<-HNH<-OLD-ABC<-METHYLASE<-DUF3780<-DUF499*||?->?-><-?<-?<-Aminotran_4</td><td>1083</td><td>Bacteria>Pseudomonadota>Gammaproteobacteria>Chromatiales>Chromatiaceae</td><td>GCF_002813775.1</td><td>Candidatus_Thiodictyon_syntrophicum</td><td>THSYN_RS02615</td><td>607511..610762</td><td>1166950</td><td>DUF499-domain-containing-protein-[Candidatus-Thiodictyon-syntrophicum]</td></tr>
<tr><td>WP_100919617.1</td><td>AAA+ATPase+wHTH+Alpha+Beta-domain+wHTH+wHTH+RRM</td><td>Type-2_BR.RM.like.systems</td><td><-HKD-DNase+SF1-SF2-Helicase+iREase+CTD<-?<-OLD-ABC<-METHYLASE<-?<-DUF3780<-DUF499*||?->?->?->?->?->SMS-KaiC-></td><td>1084</td><td>Bacteria>Pseudomonadota>Gammaproteobacteria>Chromatiales>Chromatiaceae</td><td>GCF_002813775.1</td><td>Candidatus_Thiodictyon_syntrophicum</td><td>THSYN_RS13530</td><td>3196797..3200051</td><td>1166950</td><td>DUF499-domain-containing-protein-[Candidatus-Thiodictyon-syntrophicum]</td></tr>
<tr><td>WP_093031934.1</td><td>AAA+ATPase+wHTH+Alpha+Beta-domain+wHTH+wHTH+RRM</td><td>Type-2_BR.RM.like.systems</td><td>DnaB->?->DUF499*->DUF3780->?->?->?->METHYLASE->HKD-DNase+SF1-SF2-Helicase+iREase+CTD-></td><td>1084</td><td>Bacteria>Pseudomonadota>Gammaproteobacteria>Chromatiales>Chromatiaceae</td><td>GCF_900106925.1</td><td>Thiocapsa_roseopersicina</td><td>BLV04_RS11740</td><td>2639..5893</td><td>1058</td><td>DUF499-domain-containing-protein-[Thiocapsa-roseopersicina]</td></tr>
<tr><td>WP_020503208.1</td><td>AAA+ATPase+wHTH+Alpha+Beta-domain+wHTH+wHTH+RRM</td><td>Type-2_BR.RM.like.systems</td><td><-NUDIX||?-><-SF1-SF2-Helicase||?->?->NTP_transferase->DUF499*->DUF3780-><-PIN<-?||METHYLASE->OLD-ABC-p->OLD-ABC-></td><td>1081</td><td>Bacteria>Pseudomonadota>Gammaproteobacteria>Chromatiales>Chromatiaceae</td><td>GCF_000379525.1</td><td>Lamprocystis_purpurea</td><td>A390_RS0102830</td><td>110926..114171</td><td>61598</td><td>DUF499-domain-containing-protein-[Lamprocystis-purpurea]</td></tr>
<tr><td>WP_132976384.1</td><td>AAA+ATPase+wHTH+Alpha+Beta-domain+wHTH+wHTH+RRM</td><td>Type-2_BR.RM.like.systems</td><td><-HAD<-?<-METHYLASE<-PIN<-MazE_antitoxin<-DUF3780<-DUF499*||?-><-?<-?<-HISKIN<-REC</td><td>1086</td><td>Bacteria>Pseudomonadota>Gammaproteobacteria>Chromatiales>Chromatiaceae</td><td>GCF_004342175.1</td><td>Thiobaca_trueperi</td><td>EDC35_RS06210</td><td>66133..69393</td><td>127458</td><td>DUF499-domain-containing-protein-[Thiobaca-trueperi]</td></tr>
<tr><td>WP_201244800.1</td><td>AAA+ATPase+wHTH+Alpha+Beta-domain+wHTH+wHTH+RRM</td><td>Type-2_BR.RM.like.systems</td><td>PrmC_N+Methyltransf_25->?->?-><-?||DUF499*->DUF3780->HicA_toxin->?->PolB->HEPN->METHYLASE-></td><td>1086</td><td>Bacteria>Pseudomonadota>Gammaproteobacteria>Chromatiales>Chromatiaceae</td><td>GCF_016653295.1</td><td>Halochromatium_salexigens</td><td>CCR82_RS07690</td><td>81879..85139</td><td>49447</td><td>DUF499-domain-containing-protein-[Halochromatium-salexigens]</td></tr>
<tr><td>MCO6441615.1</td><td>AAA+ATPase+wHTH+Alpha+Beta-domain+wHTH+wHTH+RRM</td><td>Type-2_BR.RM.like.systems</td><td><-HKD-DNase+SF1-SF2-Helicase+iREase+CTD<-HicB_lk_antitox<-METHYLASE<-DUF3780<-DUF499*||ABC-ATPase+SbcC-></td><td>1087</td><td>Bacteria>Pseudomonadota>Gammaproteobacteria>Chromatiales>Ectothiorhodospiraceae</td><td>GCA_024102725.1</td><td>Nitrococcus_mobilis</td><td>J5I81_11110</td><td>16313..19576</td><td>35797</td><td>ATP-binding-protein-[Nitrococcus-mobilis]</td></tr>
<tr><td>WP_005369522.1</td><td>AAA+ATPase+wHTH+Alpha+Beta-domain+wHTH+wHTH+RRM</td><td>Type-2_BR.RM.like.systems</td><td>SbcC->?->MetJarc-HTH->RelE-ParE->DUF499*->DUF3780->METHYLASE->ParB+HNH->HKD-DNase+SF1-SF2-Helicase+iREase+CTD->OLD-ABC+TOPRIM->SF1-SF2-Helicase-></td><td>1077</td><td>Bacteria>Pseudomonadota>Gammaproteobacteria>Methylococcales>Methylococcaceae</td><td>GCF_000214275.2</td><td>Methylomicrobium_album</td><td>METAL_RS03085</td><td>732822..736055</td><td>39775</td><td>DUF499-domain-containing-protein-[Methylomicrobium-album]</td></tr>
<tr><td>WP_249692546.1</td><td>AAA+ATPase+wHTH+Alpha+Beta-domain+wHTH+wHTH+RRM</td><td>Type-2_BR.RM.like.systems</td><td>HTH+WYL->DUF499*->DUF3780->METHYLASE->ParB+HNH->HKD-DNase+SF1-SF2-Helicase+iREase+CTD-><-Phage_lysozyme</td><td>1077</td><td>Bacteria>Pseudomonadota>Gammaproteobacteria>Oceanospirillales>Aestuariirhabdaceae</td><td>GCF_023509475.1</td><td>Aestuariirhabdus_haliotis</td><td>MIB40_RS07390</td><td>56644..59877</td><td>2918751</td><td>DUF499-domain-containing-protein-[Aestuariirhabdus-haliotis]</td></tr>
<tr><td>WP_086622422.1</td><td>AAA+ATPase+wHTH+Alpha+Beta-domain+wHTH+wHTH+RRM</td><td>Type-2_BR.RM.like.systems</td><td>DUF932->?->?->?->DUF499*->DUF3780->METHYLASE->OLD-ABC->PIN->?->HKD-DNase+SF1-SF2-Helicase+iREase+CTD-></td><td>1088</td><td>Bacteria>Pseudomonadota>Gammaproteobacteria>Oceanospirillales>Halomonadaceae</td><td>GCF_002155145.1</td><td>Kushneria_konosiri</td><td>B9G99_RS12330</td><td>2656346..2659612</td><td>698828</td><td>DUF499-domain-containing-protein-[Kushneria-konosiri]</td></tr>
<tr><td>WP_153843409.1</td><td>AAA+ATPase+wHTH+Alpha+Beta-domain+wHTH+wHTH+RRM</td><td>Type-2_BR.RM.like.systems</td><td>NERD+SF1-SF2-Helicase->DUF499*->DUF3780->METHYLASE->ParB+HNH->REase+ParB->HKD-DNase+SF1-SF2-Helicase+iREase+CTD-></td><td>1106</td><td>Bacteria>Pseudomonadota>Gammaproteobacteria>Oceanospirillales>Halomonadaceae</td><td>GCF_012427705.1</td><td>Halomonas_piezotolerans</td><td>GYM47_RS02295</td><td>484423..487743</td><td>2609667</td><td>DUF499-domain-containing-protein-[Halomonas-piezotolerans]</td></tr>
<tr><td>WP_213115175.1</td><td>AAA+ATPase+wHTH+Alpha+Beta-domain+wHTH+wHTH+RRM</td><td>Type-2_BR.RM.like.systems</td><td>MODE-HTH+PBPII-><-?||DUF499*->DUF3780->METHYLASE->?->HKD-DNase+SF1-SF2-Helicase+iREase+CTD-></td><td>1088</td><td>Bacteria>Pseudomonadota>Gammaproteobacteria>Oceanospirillales>Halomonadaceae</td><td>GCF_018340035.1</td><td>Cobetia_sp._MC34</td><td>IZU87_RS15975</td><td>7021..10287</td><td>2785080</td><td>DUF499-domain-containing-protein-[Cobetia-sp.-MC34]</td></tr>
<tr><td>WP_247619860.1</td><td>AAA+ATPase+wHTH+Alpha+Beta-domain+wHTH+wHTH+RRM</td><td>Type-2_BR.RM.like.systems</td><td>DUF499*->DUF3780->METHYLASE->ParB+HNH->?->HKD-DNase+SF1-SF2-Helicase+iREase+CTD-></td><td>1088</td><td>Bacteria>Pseudomonadota>Gammaproteobacteria>Oceanospirillales>Halomonadaceae</td><td>GCF_025393375.1</td><td>Chromohalobacter</td><td>KZO79_RS04210</td><td>936140..939406</td><td>42054</td><td>MULTISPECIES:-DUF499-domain-containing-protein-[Chromohalobacter]</td></tr>
<tr><td>WP_092844114.1</td><td>AAA+ATPase+wHTH+Alpha+Beta-domain+wHTH+wHTH+RRM</td><td>Type-2_BR.RM.like.systems</td><td>CHTH-><-?||?->XIS-HTH->DUF499*->DUF3780->METHYLASE->PolB->HEPN->HKD-DNase+SF1-SF2-Helicase+iREase+CTD->ParB+HNH-></td><td>1088</td><td>Bacteria>Pseudomonadota>Gammaproteobacteria>Oceanospirillales>Halomonadaceae</td><td>GCF_900113605.1</td><td>Halomonas_xianhensis</td><td>BM194_RS04450</td><td>210680..213946</td><td>442341</td><td>DUF499-domain-containing-protein-[Halomonas-xianhensis]</td></tr>
<tr><td>WP_136067364.1</td><td>AAA+ATPase+wHTH+Alpha+Beta-domain+wHTH+wHTH+RRM</td><td>Type-2_BR.RM.like.systems</td><td><-HKD-DNase+SF1-SF2-Helicase+iREase+CTD<-?<-?<-METHYLASE<-DUF3780<-DUF499*<-JAB<-?<-?<-?<-DUF932</td><td>1088</td><td>Bacteria>Pseudomonadota>Gammaproteobacteria>Oceanospirillales>Halomonadaceae</td><td>GCF_900961225.1</td><td>Halomonas_sp._EAR18</td><td>HR9077_RS12645</td><td>16845..20111</td><td>2518972</td><td>DUF499-domain-containing-protein-[Halomonas-sp.-EAR18]</td></tr>
<tr><td>WP_133483952.1</td><td>AAA+ATPase+wHTH+Alpha+Beta-domain+wHTH+wHTH+RRM</td><td>Type-2_BR.RM.like.systems</td><td>TraH->?->TraG_N-><-?<-CHTH||DUF4236+TPR->DUF499*->DUF3780->METHYLASE->OLD-ABC->PIN->?->ParB+HNH-></td><td>1088</td><td>Bacteria>Pseudomonadota>Gammaproteobacteria>Oceanospirillales>Halomonadaceae</td><td>GCF_024137965.1</td><td>Halomonas</td><td>NHL60_RS01495</td><td>294251..297517</td><td>2745</td><td>MULTISPECIES:-DUF499-domain-containing-protein-[Halomonas]</td></tr>
<tr><td>WP_110069957.1</td><td>AAA+ATPase+wHTH+Alpha+Beta-domain+wHTH+wHTH+RRM</td><td>Type-2_BR.RM.like.systems</td><td>TraH->?->TraG_N-><-?||DUF4236+TPR->DUF499*->DUF3780->METHYLASE->OLD-ABC->PIN->?->ParB+HNH-></td><td>1088</td><td>Bacteria>Pseudomonadota>Gammaproteobacteria>Oceanospirillales>Halomonadaceae</td><td>GCF_003182195.1</td><td>Halomonas_sp._A11-A</td><td>DER72_RS10560</td><td>14381..17647</td><td>2183985</td><td>DUF499-domain-containing-protein-[Halomonas-sp.-A11-A]</td></tr>
<tr><td>WP_147183730.1</td><td>AAA+ATPase+wHTH+Alpha+Beta-domain+wHTH+wHTH+RRM</td><td>Type-2_BR.RM.like.systems</td><td><-PIN<-DUF433||DUF4236+TPR->DUF499*->DUF3780->METHYLASE->OLD-ABC->PIN->?->ParB+HNH-></td><td>1088</td><td>Bacteria>Pseudomonadota>Gammaproteobacteria>Oceanospirillales>Halomonadaceae</td><td>GCF_007954585.1</td><td>Pistricoccus_aurantiacus</td><td>FGL86_RS05990</td><td>1230922..1234188</td><td>1883414</td><td>DUF499-domain-containing-protein-[Pistricoccus-aurantiacus]</td></tr>
<tr><td>WP_163558801.1</td><td>AAA+ATPase+wHTH+Alpha+Beta-domain+wHTH+wHTH+RRM</td><td>Type-2_BR.RM.like.systems</td><td><-ParB+HNH<-?<-PIN<-OLD-ABC<-METHYLASE<-DUF3780<-DUF499*<-SH3||?-><-CHTH<-?||?->DUF932-></td><td>1088</td><td>Bacteria>Pseudomonadota>Gammaproteobacteria>Oceanospirillales>Halomonadaceae</td><td>GCF_010994375.1</td><td>Halomonas_sp._NO4</td><td>FEF15_RS08845</td><td>14465..17731</td><td>2484813</td><td>DUF499-domain-containing-protein-[Halomonas-sp.-NO4]</td></tr>
<tr><td>WP_234259726.1</td><td>AAA+ATPase+wHTH+Alpha+Beta-domain+wHTH+wHTH+RRM</td><td>Type-2_BR.RM.like.systems</td><td><-ParB+HNH<-?<-PIN<-OLD-ABC<-METHYLASE<-DUF3780<-DUF499*<-SH3</td><td>1088</td><td>Bacteria>Pseudomonadota>Gammaproteobacteria>Oceanospirillales>Halomonadaceae</td><td>GCF_021404355.1</td><td>Halomonas_sp._MCCC_1A11062</td><td>HOP58_RS04945</td><td>176236..179502</td><td>2733485</td><td>DUF499-domain-containing-protein-[Halomonas-sp.-MCCC-1A11062]</td></tr>
<tr><td>WP_168444437.1</td><td>AAA+ATPase+wHTH+Alpha+Beta-domain+wHTH+wHTH+RRM</td><td>Type-2_BR.RM.like.systems</td><td><-DUF932||?->?-><-?||SH3->DUF499*->DUF3780->METHYLASE->OLD-ABC->PIN->?->ParB+HNH-></td><td>1088</td><td>Bacteria>Pseudomonadota>Gammaproteobacteria>Oceanospirillales>Halomonadaceae</td><td>GCF_012427705.1</td><td>Halomonas_piezotolerans</td><td>GYM47_RS05600</td><td>1204301..1207567</td><td>2609667</td><td>DUF499-domain-containing-protein-[Halomonas-piezotolerans]</td></tr>
<tr><td>WP_223136851.1</td><td>AAA+ATPase+wHTH+Alpha+Beta-domain+wHTH+wHTH+RRM</td><td>Type-2_BR.RM.like.systems</td><td>CHTH->?-><-?||SH3->DUF499*->DUF3780->METHYLASE->OLD-ABC->PIN->?->ParB+HNH-></td><td>1088</td><td>Bacteria>Pseudomonadota>Gammaproteobacteria>Oceanospirillales>Halomonadaceae</td><td>GCF_019915905.1</td><td>Halomonas_sp._ANAO-440</td><td>KZO25_RS10115</td><td>168620..171886</td><td>2861360</td><td>DUF499-domain-containing-protein-[Halomonas-sp.-ANAO-440]</td></tr>
<tr><td>KAE8440079.1</td><td>AAA+ATPase+wHTH+Alpha+Beta-domain+wHTH+wHTH+RRM</td><td>Type-2_BR.RM.like.systems</td><td><-DUF932<-?||<-?||SH3->DUF499*->DUF3780->METHYLASE->OLD-ABC->PIN->?->ParB+HNH-></td><td>1113</td><td>Bacteria>Pseudomonadota>Gammaproteobacteria>Oceanospirillales>Halomonadaceae</td><td>GCA_009660035.1</td><td>Halomonas_piezotolerans</td><td>F1978_02220</td><td>465970..469311</td><td>2609667</td><td>ATP-binding-protein-[Halomonas-piezotolerans]</td></tr>
<tr><td>WP_068642258.1</td><td>AAA+ATPase+wHTH+Alpha+Beta-domain+wHTH+wHTH+RRM</td><td>Type-2_BR.RM.like.systems</td><td><-SbcC<-JetB<-JetA||HTH+WYL->DUF499*->DUF3780->METHYLASE->HKD-DNase+SF1-SF2-Helicase+iREase+CTD-></td><td>1077</td><td>Bacteria>Pseudomonadota>Gammaproteobacteria>Oceanospirillales>Oleiphilaceae</td><td>GCF_001635845.1</td><td>Oleiphilus_sp._HI0132</td><td>A3766_RS11865</td><td>4418..7651</td><td>1822270</td><td>DUF499-domain-containing-protein-[Oleiphilus-sp.-HI0132]</td></tr>
<tr><td>WP_200371316.1</td><td>AAA+ATPase+wHTH+Alpha+Beta-domain+wHTH+wHTH+RRM</td><td>Type-2_BR.RM.like.systems</td><td><-DUF4236+TPR<-?||?->?->?->?->DUF499*->DUF3780->METHYLASE->ParB+HNH->OLD-ABC->PIN-></td><td>1081</td><td>Bacteria>Pseudomonadota>Gammaproteobacteria>Pseudomonadales>Marinobacteraceae</td><td>GCF_016597815.1</td><td>Marinobacter_sp._DY40_1A1</td><td>FFI61_RS14505</td><td>345780..349025</td><td>2583229</td><td>DUF499-domain-containing-protein-[Marinobacter-sp.-DY40_1A1]</td></tr>
<tr><td>WP_090215787.1</td><td>AAA+ATPase+wHTH+Alpha+Beta-domain+wHTH+wHTH+RRM</td><td>Type-2_BR.RM.like.systems</td><td>RlpA-><-HKD-DNase+SF1-SF2-Helicase+iREase+CTD<-METHYLASE<-DUF3780<-DUF499*<-?<-?<-?<-?<-?<-XIS-HTH</td><td>1082</td><td>Bacteria>Pseudomonadota>Gammaproteobacteria>Pseudomonadales>Pseudomonadaceae</td><td>GCF_900105885.1</td><td>Pseudomonas_guangdongensis</td><td>BLU22_RS14145</td><td>3036479..3039727</td><td>1245526</td><td>DUF499-domain-containing-protein-[Pseudomonas-guangdongensis]</td></tr>
<tr><td>WP_046619147.1</td><td>AAA+ATPase+wHTH+Alpha+Beta-domain+wHTH+wHTH+RRM</td><td>Type-2_BR.RM.like.systems</td><td>MarR-HTH->XIS-HTH->?->DUF499*->DUF3780->METHYLASE->HKD-DNase+SF1-SF2-Helicase+iREase+CTD-><-RlpA<-?<-JAB</td><td>1082</td><td>Bacteria>Pseudomonadota>Gammaproteobacteria>Pseudomonadales>Pseudomonadaceae</td><td>-</td><td>Stutzerimonas_stutzeri</td><td>PK34_RS09970</td><td>34155..37403</td><td>316</td><td>DUF499-domain-containing-protein-[Stutzerimonas-stutzeri]</td></tr>
<tr><td>NCA89397.1</td><td>AAA+ATPase+wHTH+Alpha+Beta-domain+wHTH+wHTH+RRM</td><td>Type-2_BR.RM.like.systems</td><td><-HTH||RHH->DUF499*->DUF3780->METHYLASE->?->HKD-DNase+SF1-SF2-Helicase+iREase+CTD-></td><td>1084</td><td>Bacteria>Pseudomonadota>Gammaproteobacteria>unclassified_Gammaproteobacteria</td><td>GCA_009928645.1</td><td>Gammaproteobacteria_bacterium</td><td>EOM92_10935</td><td>816..4070</td><td>1913989</td><td>ATP-binding-protein-[Gammaproteobacteria-bacterium]</td></tr>
<tr><td>HCG7747321.1</td><td>AAA+ATPase+wHTH+Alpha+Beta-domain+wHTH+wHTH+RRM</td><td>Type-2_BR.RM.like.systems</td><td><-HKD-DNase+SF1-SF2-Helicase+iREase+CTD<-METHYLASE<-DUF3780<-DUF499*<-XIS-HTH||HEPN-><-?<-HNH||JAB-></td><td>1075</td><td>Bacteria>Pseudomonadota>Gammaproteobacteria>Vibrionales>Vibrionaceae</td><td>GCA_024045415.1</td><td>Vibrio_parahaemolyticus</td><td>NJ442_001228</td><td>553706..556933</td><td>670</td><td>ATP-binding-protein-[Vibrio-parahaemolyticus]</td></tr>
<tr><td>HCK0611955.1</td><td>AAA+ATPase+wHTH+Alpha+Beta-domain+wHTH+wHTH+RRM</td><td>Type-2_BR.RM.like.systems</td><td><-MORC+DUF3684+REase||RlpA-><-HKD-DNase+SF1-SF2-Helicase+iREase+CTD<-METHYLASE<-DUF3780<-DUF499*||HEPN-><-?<-HNH<-?<-McrC-NTD+REase</td><td>1076</td><td>Bacteria>Pseudomonadota>Gammaproteobacteria>Vibrionales>Vibrionaceae</td><td>GCA_024739345.1</td><td>Vibrio_parahaemolyticus</td><td>NGT52_002945</td><td>240200..243430</td><td>670</td><td>ATP-binding-protein-[Vibrio-parahaemolyticus]</td></tr>
<tr><td>WP_064353772.1</td><td>AAA+ATPase+wHTH+Alpha+Beta-domain+wHTH+wHTH+RRM</td><td>Type-2_BR.RM.like.systems</td><td><-HEPN||XIS-HTH->DUF499*->DUF3780->METHYLASE->HKD-DNase+SF1-SF2-Helicase+iREase+CTD-><-RlpA</td><td>1075</td><td>Bacteria>Pseudomonadota>Gammaproteobacteria>Vibrionales>Vibrionaceae</td><td>GCF_001617475.1</td><td>Vibrio_alginolyticus</td><td>XM68_RS05165</td><td>20692..23919</td><td>663</td><td>DUF499-domain-containing-protein-[Vibrio-alginolyticus]</td></tr>
<tr><td>WP_069494520.1</td><td>AAA+ATPase+wHTH+Alpha+Beta-domain+wHTH+wHTH+RRM</td><td>Type-2_BR.RM.like.systems</td><td><-JAB||HNH->?-><-HEPN||XIS-HTH->DUF499*->DUF3780->METHYLASE->HKD-DNase+SF1-SF2-Helicase+iREase+CTD-><-RlpA</td><td>1075</td><td>Bacteria>Pseudomonadota>Gammaproteobacteria>Vibrionales>Vibrionaceae</td><td>GCF_001726325.1</td><td>Vibrio_parahaemolyticus</td><td>BBL88_RS06400</td><td>350486..353713</td><td>670</td><td>DUF499-domain-containing-protein-[Vibrio-parahaemolyticus]</td></tr>
<tr><td>WP_033905940.1</td><td>AAA+ATPase+wHTH+Alpha+Beta-domain+wHTH+wHTH+RRM</td><td>Type-2_BR.RM.like.systems</td><td><-JAB||RNase->XIS-HTH->HTH+WYL->DUF499*->DUF3780->METHYLASE->HKD-DNase+SF1-SF2-Helicase+iREase+CTD->?-><-?<-Arm-DNA-bind_3+Phage_integrase</td><td>1077</td><td>Bacteria>Pseudomonadota>Gammaproteobacteria>Vibrionales>Vibrionaceae</td><td>GCF_017313445.1</td><td>Vibrio</td><td>J4H45_RS11285</td><td>94869..98102</td><td>662</td><td>MULTISPECIES:-DUF499-domain-containing-protein-[Vibrio]</td></tr>
<tr><td>WP_228067030.1</td><td>AAA+ATPase+wHTH+Alpha+Beta-domain+wHTH+wHTH+RRM</td><td>Type-2_BR.RM.like.systems</td><td><-JAB||HNH->?-><-HEPN||?->DUF499*->DUF3780->METHYLASE->HKD-DNase+SF1-SF2-Helicase+iREase+CTD-><-RlpA||MORC+DUF3684+REase-></td><td>1076</td><td>Bacteria>Pseudomonadota>Gammaproteobacteria>Vibrionales>Vibrionaceae</td><td>GCF_020740875.1</td><td>Vibrio_parahaemolyticus</td><td>IB288_RS04445</td><td>157878..161108</td><td>670</td><td>DUF499-domain-containing-protein-[Vibrio-parahaemolyticus]</td></tr>
<tr><td>WP_285401339.1</td><td>AAA+ATPase+wHTH+Alpha+Beta-domain+wHTH+wHTH+RRM</td><td>Type-2_BR.RM.like.systems</td><td><-JAB||XIS-HTH->?->XIS-HTH->DUF499*->DUF3780->METHYLASE->ParB+HNH->HKD-DNase+SF1-SF2-Helicase+iREase+CTD-><-RlpA<-Arm-DNA-bind_3+Phage_integrase</td><td>1076</td><td>Bacteria>Pseudomonadota>Gammaproteobacteria>Vibrionales>Vibrionaceae</td><td>GCF_030248535.1</td><td>unclassified_Vibrio</td><td>KJB41_RS10780</td><td>100358..103588</td><td>2614977</td><td>MULTISPECIES:-DUF499-domain-containing-protein-[unclassified-Vibrio]</td></tr>
<tr><td>MCA9273427.1</td><td>AAA+ATPase+wHTH+Alpha+Beta-domain+wHTH+wHTH+RRM</td><td>Type-2_BR.RM.like.systems</td><td>XIS-HTH->DUF499*->DUF3780->METHYLASE->HKD-DNase+SF1-SF2-Helicase+iREase+CTD-></td><td>1087</td><td>Bacteria>PVC_group>Planctomycetota>Phycisphaerae>Phycisphaerales</td><td>GCA_020427465.1</td><td>Phycisphaerales_bacterium</td><td>KDA31_10305</td><td>493..3756</td><td>2052180</td><td>ATP-binding-protein-[Phycisphaerales-bacterium]</td></tr>
<tr><td>MCB9841661.1</td><td>AAA+ATPase+wHTH+Alpha+Beta-domain+wHTH+wHTH+RRM</td><td>Type-2_BR.RM.like.systems</td><td><-XIS-HTH||SIR2->DUF1679->SbcC->XIS-HTH->DUF499*->DUF3780->METHYLASE->HKD-DNase+SF1-SF2-Helicase+iREase+CTD-></td><td>1086</td><td>Bacteria>PVC_group>Planctomycetota>Phycisphaerae>Phycisphaerales</td><td>GCA_020639235.1</td><td>Phycisphaeraceae_bacterium</td><td>H6812_00215</td><td>68785..72045</td><td>2026777</td><td>ATP-binding-protein-[Phycisphaeraceae-bacterium]</td></tr>
<tr><td>QDV88977.1</td><td>AAA+ATPase+wHTH+Alpha+Beta-domain+wHTH+wHTH+RRM</td><td>Type-2_BR.RM.like.systems</td><td><-HKD-DNase+SF1-SF2-Helicase+iREase+CTD<-METHYLASE<-SIR2<-ArdCN+MPTase<-MORC<-DUF3780<-DUF499*<-ArdCN+MPTase<-?<-?<-?<-?<-PadR-HTH</td><td>982</td><td>Bacteria>PVC_group>Planctomycetota>Phycisphaerae>unclassified_Phycisphaerae</td><td>GCA_007753915.1</td><td>Phycisphaerae_bacterium_RAS2</td><td>RAS2_00360</td><td>48471..51419</td><td>2528035</td><td>hypothetical-protein-RAS2_00360-[Phycisphaerae-bacterium-RAS2]</td></tr>
<tr><td>MBI3099182.1</td><td>AAA+ATPase+wHTH+Alpha+Beta-domain+wHTH+wHTH+RRM</td><td>Type-2_BR.RM.like.systems</td><td><-HKD-DNase+SF1-SF2-Helicase+iREase+CTD<-METHYLASE<-DUF3780<-DUF499*<-HEPN<-?<-Phage_integrase||tRNA->TPR-></td><td>977</td><td>Bacteria>PVC_group>Planctomycetota>unclassified_Planctomycetota</td><td>GCA_016192525.1</td><td>Planctomycetota_bacterium</td><td>HYY93_13215</td><td>4536..7469</td><td>2026780</td><td>ATP-binding-protein-[Planctomycetota-bacterium]</td></tr>
<tr><td>MCY2953678.1</td><td>AAA+ATPase+wHTH+Alpha+Beta-domain+wHTH+wHTH+RRM</td><td>Type-2_BR.RM.like.systems</td><td>DUF499*->DUF3780->PIN->METHYLASE-></td><td>977</td><td>Bacteria>PVC_group>Planctomycetota>unclassified_Planctomycetota</td><td>GCA_026693085.1</td><td>Planctomycetota_bacterium</td><td>NTU53_17115</td><td>20539..23472</td><td>2026780</td><td>MAG:-DUF499-domain-containing-protein-[Planctomycetota-bacterium]</td></tr>
<tr><td>AKC83089.1</td><td>AAA+ATPase+wHTH+Alpha+Beta-domain+wHTH+wHTH+RRM</td><td>Type-2_BR.RM.like.systems</td><td>SbcC+SbcC-><-?||DUF499*->DUF3780->SMODS->SLATT->METHYLASE->HKD-DNase+SF1-SF2-Helicase+iREase+CTD->PIN-></td><td>988</td><td>Bacteria>PVC_group>Verrucomicrobiota>unclassified_Verrucomicrobiota</td><td>GCA_000972765.1</td><td>Verrucomicrobia_bacterium_IMCC26134</td><td>IMCC26134_10435</td><td>2769967..2772933</td><td>1637999</td><td>hypothetical-protein-IMCC26134_10435-[Verrucomicrobia-bacterium-IMCC26134]</td></tr>
<tr><td>MCF7668420.1</td><td>AAA+ATPase+wHTH+Alpha+Beta-domain+wHTH+wHTH+RRM</td><td>Type-2_BR.RM.like.systems</td><td><-HKD-DNase+SF1-SF2-Helicase+iREase+CTD<-HTH+MPTase<-Schlafen+HATPase+HTH<-SbcC+SidE<-METHYLASE<-DUF3780<-DUF499*</td><td>1083</td><td>Bacteria>PVC_group>Verrucomicrobiota>unclassified_Verrucomicrobiota</td><td>GCA_021733785.1</td><td>Verrucomicrobiota_bacterium</td><td>K9N48_01435</td><td>17156..20407</td><td>2026799</td><td>DUF499-domain-containing-protein-[Verrucomicrobia-bacterium]</td></tr>
<tr><td>MBL9158412.1</td><td>AAA+ATPase+wHTH+Alpha+Beta-domain+wHTH+wHTH+RRM</td><td>Type-2_BR.RM.like.systems</td><td><-HKD-DNase+SF1-SF2-Helicase+iREase+CTD<-?<-CHTH<-METHYLASE<-?<-DUF3780<-DUF499*</td><td>1081</td><td>Bacteria>PVC_group>Verrucomicrobiota>Verrucomicrobiae>Verrucomicrobiales</td><td>GCA_016795385.1</td><td>Verrucomicrobiales_bacterium</td><td>JNJ70_13120</td><td>81865..85110</td><td>2026801</td><td>ATP-binding-protein-[Verrucomicrobiales-bacterium]</td></tr>
<tr><td>WP_029165421.1</td><td>AAA+ATPase+wHTH+Alpha+Beta-domain+wHTH+wHTH+RRM</td><td>Type-2_BR.RM.like.systems</td><td><-HKD-DNase+SF1-SF2-Helicase+iREase+CTD<-METHYLASE<-DUF3780<-DUF499*||?-><-?<-ACET</td><td>1076</td><td>Bacteria>Synergistota>Synergistia>Synergistales>Aminiphilaceae</td><td>GCF_000526375.1</td><td>Aminiphilus_circumscriptus</td><td>K349_RS0108485</td><td>287707..290937</td><td>290732</td><td>DUF499-domain-containing-protein-[Aminiphilus-circumscriptus]</td></tr>
<tr><td>WP_085543648.1</td><td>AAA+ATPase+wHTH+Alpha+Beta-domain+wHTH+wHTH+RRM</td><td>Type-2_BR.RM.like.systems</td><td>OLD-ABC->?-><-?<-HKD-DNase+SF1-SF2-Helicase+iREase+CTD<-METHYLASE<-DUF3780<-DUF499*<-?<-?||?-><-Peptidase_M20</td><td>1083</td><td>Bacteria>Synergistota>Synergistia>Synergistales>Dethiosulfovibrionaceae</td><td>GCF_900177735.1</td><td>Dethiosulfovibrio_salsuginis</td><td>B9Y55_RS01800</td><td>28225..31476</td><td>561720</td><td>DUF499-domain-containing-protein-[Dethiosulfovibrio-salsuginis]</td></tr>
<tr><td>MDD3916530.1</td><td>AAA+ATPase+wHTH+Alpha+Beta-domain+wHTH+wHTH+RRM</td><td>Type-2_BR.RM.like.systems</td><td>DUF932->DUF499*->DUF3780->METHYLASE->HKD-DNase+SF1-SF2-Helicase+iREase+CTD->HKD-DNase+SF1-SF2-Helicase+iREase+CTD->VapB_antitoxin->PIN-></td><td>1079</td><td>Bacteria>Synergistota>Synergistia>Synergistales>Synergistaceae</td><td>GCA_028699385.1</td><td>Synergistaceae_bacterium</td><td>PHX00_05210</td><td>3629..6868</td><td>2053624</td><td>MAG:-DUF499-domain-containing-protein-[Synergistaceae-bacterium]</td></tr>
<tr><td>WP_273491645.1</td><td>AAA+ATPase+wHTH+Alpha+Beta-domain+wHTH+wHTH+RRM</td><td>Type-2_BR.RM.like.systems</td><td><-REase||MarR-HTH->?->?->?->XIS-HTH->DUF499*->DUF3780->METHYLASE->HKD-DNase+SF1-SF2-Helicase+iREase+CTD-><-DnaB+SMS-KaiC</td><td>1092</td><td>Bacteria>Terrabacteria_group>Actinomycetota</td><td>GCF_936918145.1</td><td>Actinomycetes</td><td>Q0172_RS03030</td><td>30769..34047</td><td>1760</td><td>MULTISPECIES:-DUF499-domain-containing-protein-[Actinomycetes]</td></tr>
<tr><td>MBF6555079.1</td><td>AAA+ATPase+wHTH+Alpha+Beta-domain+wHTH+wHTH+RRM</td><td>Type-2_BR.RM.like.systems</td><td>REase+35exo+Beta-Barrel+SF1-SF2-Helicase->?-><-HKD-DNase+SF1-SF2-Helicase+iREase+CTD<-METHYLASE<-DUF3780<-DUF499*<-?<-?||?->?->ParA-Soj-PloopNTPase-></td><td>1077</td><td>Bacteria>Terrabacteria_group>Actinomycetota>Acidimicrobiia>Acidimicrobiales</td><td>GCA_015478655.1</td><td>Acidimicrobiales_bacterium</td><td>IVW52_02790</td><td>45007..48240</td><td>2201156</td><td>ATP-binding-protein-[Acidimicrobiales-bacterium]</td></tr>
<tr><td>MCZ7532618.1</td><td>AAA+ATPase+wHTH+Alpha+Beta-domain+wHTH+wHTH+RRM</td><td>Type-2_BR.RM.like.systems</td><td>TrwC+SF1-SF2-helicase+TOPRIM-><-?||?-><-?<-HTH+MPTase<-Tox-RelE||DUF499*->DUF3780->METHYLASE->HKD-DNase+SF1-SF2-Helicase+iREase+CTD->OLD-ABC->?-><-SIR2</td><td>1074</td><td>Bacteria>Terrabacteria_group>Actinomycetota>Acidimicrobiia>unclassified_Acidimicrobiia</td><td>GCA_027494075.1</td><td>Acidimicrobiia_bacterium</td><td>M5U23_04305</td><td>852296..855520</td><td>2080302</td><td>MAG:-DUF499-domain-containing-protein-[Acidimicrobiia-bacterium]</td></tr>
</table>

| Protein-ID | DUF499-architecture | System | Gene-Neighbourhoods | Length | Taxonomy | Assembly-ID | Strain | Locus-Tag | Genomic-coordinates | TaxID | Fasta-defline |
|---|---|---|---|---|---|---|---|---|---|---|---|
| KUL69962.1 | AAA+ATPase+wHTH+Alpha+Beta-domain+wHTH+wHTH+RRM | Type-2_BR.RM.like.systems | <-HTH+DUF5753<-DUF397\|\|ST-Hiskin->?->?->DUF499*->DUF3780->METHYLASE->HKD-DNase+SF1-SF2-Helicase+iREase+CTD->?-><-REase | 1070 | Bacteria>Terrabacteria_group>Actinomycetota>Actinomycetes>Kitasatosporales | GCA_001509795.1 | Streptomyces_sp._NRRL_WC-3605 | ADL34_30365 | 368586..371798 | 1609103 | hypothetical-protein-ADL34_30365-[Streptomyces-sp.-NRRL-WC-3605] |
| TRZ84781.1 | AAA+ATPase+wHTH+Alpha+Beta-domain+wHTH+wHTH+RRM | Type-2_BR.RM.like.systems | <-HKD-DNase+SF1-SF2-Helicase+iREase+CTD<-METHYLASE<-DUF3780<-DUF499* | 1084 | Bacteria>Terrabacteria_group>Actinomycetota>Actinomycetes>Kitasatosporales | GCA_007280125.1 | Streptomycetaceae_bacterium | D4R83_07970 | 6018..9272 | 2021369 | DUF499-domain-containing-protein-[Streptomycetaceae-bacterium] |
| MBP0459504.1 | AAA+ATPase+wHTH+Alpha+Beta-domain+wHTH+wHTH+RRM | Type-2_BR.RM.like.systems | HTH+DUF5753->DUF397-><-HTH_Tnp_4+DDE_Tnp_4<-HKD-DNase+SF1-SF2-Helicase+iREase+CTD<-METHYLASE<-DUF3780<-DUF499*<-tRNA<-?\|\|?->ZnR-> | 1091 | Bacteria>Terrabacteria_group>Actinomycetota>Actinomycetes>Kitasatosporales | GCA_017813245.1 | Streptomyces_montanisoli | JFN87_18620 | 20884..24159 | 2798581 | ATP-binding-protein-[Streptomyces-montanisoli] |
| MCZ1002042.1 | AAA+ATPase+wHTH+Alpha+Beta-domain+wHTH+wHTH+RRM | Type-2_BR.RM.like.systems | XIS-HTH->?->Phage_integrase-><-?\|\|DUF499*->DUF3780->METHYLASE->HKD-DNase+SF1-SF2-Helicase+iREase+CTD-> | 1091 | Bacteria>Terrabacteria_group>Actinomycetota>Actinomycetes>Kitasatosporales | GCA_026913145.1 | Streptomyces_mirabilis | O1M63_34880 | 1588072..1591347 | 68239 | DUF499-domain-containing-protein-[Streptomyces-mirabilis] |
| WP_078656197.1 | AAA+ATPase+wHTH+Alpha+Beta-domain+wHTH+wHTH+RRM | Type-2_BR.RM.like.systems | <-HKD-DNase+SF1-SF2-Helicase+iREase+CTD<-METHYLASE<-DUF3780<-DUF499*\|\|?-><-ABhydrolase\|\|?-><-Response_reg+LuxR-HTH | 1091 | Bacteria>Terrabacteria_group>Actinomycetota>Actinomycetes>Kitasatosporales | GCF_000716885.1 | Kitasatospora_aureofaciens | OP17_RS00890 | 204481..207756 | 1894 | DUF499-domain-containing-protein-[Kitasatospora-aureofaciens] |
| WP_079034155.1 | AAA+ATPase+wHTH+Alpha+Beta-domain+wHTH+wHTH+RRM | Type-2_BR.RM.like.systems | <-HTH+DUF5753<-DUF397\|\|ST-Hiskin->?->?->?->DUF499*->DUF3780->METHYLASE->HKD-DNase+SF1-SF2-Helicase+iREase+CTD-><-?\|\|?-><-REase | 1093 | Bacteria>Terrabacteria_group>Actinomycetota>Actinomycetes>Kitasatosporales | GCF_001509785.1 | unclassified_Streptomyces | ADL34_RS30385 | 368517..371798 | 2593676 | MULTISPECIES:-DUF499-domain-containing-protein-[unclassified-Streptomyces] |
| WP_159398890.1 | AAA+ATPase+wHTH+Alpha+Beta-domain+wHTH+wHTH+RRM | Type-2_BR.RM.like.systems | REase->?->SF1-SF2-Helicase->DUF499*->DUF3780->METHYLASE->HKD-DNase+SF1-SF2-Helicase+iREase+CTD-><-ST-Hiskin\|\|HTH+DUF5753-> | 1091 | Bacteria>Terrabacteria_group>Actinomycetota>Actinomycetes>Kitasatosporales | GCF_002812405.1 | Streptomyces_sp._M56 | CFP59_RS16970 | 4223222..4226497 | 1495638 | DUF499-domain-containing-protein-[Streptomyces-sp.-M56] |
| WP_237508703.1 | AAA+ATPase+wHTH+Alpha+Beta-domain+wHTH+wHTH+RRM | Type-2_BR.RM.like.systems | <-HTH+DUF5753\|\|ST-Hiskin-><-HKD-DNase+SF1-SF2-Helicase+iREase+CTD<-METHYLASE<-DUF3780<-DUF499*<-SF1-SF2-Helicase<-?<-REase | 1088 | Bacteria>Terrabacteria_group>Actinomycetota>Actinomycetes>Kitasatosporales | GCF_009864805.1 | Streptomyces_sp._SID8382 | GTZ89_RS09435 | 22128..25394 | 2690362 | DUF499-domain-containing-protein-[Streptomyces-sp.-SID8382] |
| WP_262057850.1 | AAA+ATPase+wHTH+Alpha+Beta-domain+wHTH+wHTH+RRM | Type-2_BR.RM.like.systems | <-ST-Hiskin\|\|HTH+DUF5753->DUF397-><-HKD-DNase+SF1-SF2-Helicase+iREase+CTD<-METHYLASE<-DUF3780<-DUF499*<-SF1-SF2-Helicase<-?<-REase<-?\|\|REase->pPIWI+pPIWI-> | 1093 | Bacteria>Terrabacteria_group>Actinomycetota>Actinomycetes>Kitasatosporales | GCF_024519455.1 | Streptomyces_sp._STR69 | OC490_RS11915 | 311195..314476 | 1796942 | DUF499-domain-containing-protein-[Streptomyces-sp.-STR69] |
| WP_266371749.1 | AAA+ATPase+wHTH+Alpha+Beta-domain+wHTH+wHTH+RRM | Type-2_BR.RM.like.systems | <-N6_N4_MTase<-?\|\|?->DUF499*->DUF3780->METHYLASE->HKD-DNase+SF1-SF2-Helicase+iREase+CTD->?->RVT->HTH+DUF5753-> | 1093 | Bacteria>Terrabacteria_group>Actinomycetota>Actinomycetes>Kitasatosporales | GCF_026342305.1 | Streptomyces_antibioticus | OG616_RS26230 | 5685167..5688448 | 1890 | DUF499-domain-containing-protein-[Streptomyces-antibioticus] |
| WP_275817440.1 | AAA+ATPase+wHTH+Alpha+Beta-domain+wHTH+wHTH+RRM | Type-2_BR.RM.like.systems | <-Radical-SAM\|\|HTH_20->?-><-HKD-DNase+SF1-SF2-Helicase+iREase+CTD<-METHYLASE<-DUF3780<-DUF499*\|\|?-><-OLD-ABC+TOPRIM<-?\|\|?->HTH_Tnp_4+DDE_Tnp_4-> | 1093 | Bacteria>Terrabacteria_group>Actinomycetota>Actinomycetes>Kitasatosporales | GCF_029223525.1 | Streptantibioticus_ferralitis | P2L57_RS22720 | 141423..144704 | 236510 | DUF499-domain-containing-protein-[Streptomyces-ferralitis] |
| WP_284505300.1 | AAA+ATPase+wHTH+Alpha+Beta-domain+wHTH+wHTH+RRM | Type-2_BR.RM.like.systems | Glyoxalase->?->?->?->\|\|<-?\|\|DDE_Tnp_1->\|\|DUF499*->METHYLASE->HKD-DNase+SF1-SF2-Helicase+iREase+CTD->\|\|Phage_integrase-> | 1070 | Bacteria>Terrabacteria_group>Actinomycetota>Actinomycetes>Kitasatosporales | - | Streptomyces_cellostaticus | MXD57_RS08755 | 66377..69589 | 67285 | DUF499-domain-containing-protein-[Streptomyces-cellostaticus] |
| WP_245180283.1 | AAA+ATPase+wHTH+Alpha+Beta-domain+wHTH+wHTH+RRM | Type-2_BR.RM.like.systems | HTH+DUF5753->DUF397-><-HTH_Tnp_4+DDE_Tnp_4<-HKD-DNase+SF1-SF2-Helicase+iREase+CTD<-METHYLASE<-DUF3780<-DUF499*<-tRNA<-?\|\|?->ZnR-> | 1088 | Bacteria>Terrabacteria_group>Actinomycetota>Actinomycetes>Kitasatosporales | - | Streptomyces_montanisoli | JFN87_RS18620 | 20884..24150 | 2798581 | DUF499-domain-containing-protein-[Streptomyces-montanisoli] |
| EKU49302.1 | AAA+ATPase+wHTH+Alpha+Beta-domain+wHTH+wHTH+RRM | Type-2_BR.RM.like.systems | DUF499*->DUF3780->METHYLASE->HKD-DNase+SF1-SF2-Helicase+iREase+CTD->GDResolvase+Recombinase->ArdA-> | 1007 | Bacteria>Terrabacteria_group>Actinomycetota>Actinomycetes>Micrococcales | GCA_000314575.1 | Brevibacterium_casei_S18 | C272_01160 | 266972..269995 | 1229781 | putative-protein-kinase,-ATPase-[Brevibacterium-casei-S18] |
| OMH22920.1 | AAA+ATPase+wHTH+Alpha+Beta-domain+wHTH+wHTH+RRM | Type-2_BR.RM.like.systems | <-DDE_Tnp_1\|\|DDE_Tnp_1_2-><-HKD-DNase+SF1-SF2-Helicase+iREase+CTD<-METHYLASE<-DUF3780<-DUF499*<-XIS-HTH<-ABhydrolase<-?<-MPTase<-HTH\|\|ArdCN+MPTase-> | 1074 | Bacteria>Terrabacteria_group>Actinomycetota>Actinomycetes>Micrococcales | GCA_001968835.1 | Tersicoccus_phoenicis | BKD30_15225 | 8674..11898 | 554083 | hypothetical-protein-BKD30_15225-[Tersicoccus-phoenicis] |
| AXH34464.1 | AAA+ATPase+wHTH+Alpha+Beta-domain+wHTH+wHTH+RRM | Type-2_BR.RM.like.systems | DnaB+SMS-KaiC-><-HKD-DNase+SF1-SF2-Helicase+iREase+CTD<-METHYLASE<-DUF3780<-DUF499*<-XIS-HTH<-?<-?<-?<-?<-MarR-HTH | 1092 | Bacteria>Terrabacteria_group>Actinomycetota>Actinomycetes>Micrococcales | GCA_003347035.1 | Humibacter_sp._BT305 | DVJ78_02650 | 553287..556565 | 2282656 | DUF499-domain-containing-protein-[Humibacter-sp.-BT305] |
| MBI5161479.1 | AAA+ATPase+wHTH+Alpha+Beta-domain+wHTH+wHTH+RRM | Type-2_BR.RM.like.systems | TrwC+SF1-SF2-helicase-><-MORC+HisKin+REase\|\|XIS-HTH->DUF499*->DUF3780->METHYLASE->HKD-DNase+SF1-SF2-Helicase+iREase+CTD->PolB->HEPN->TraJ-RHH-> | 1092 | Bacteria>Terrabacteria_group>Actinomycetota>Actinomycetes>Micrococcales | GCA_016212425.1 | Micrococcales_bacterium | HY996_08740 | 41373..44651 | 2026762 | ATP-binding-protein-[Micrococcales-bacterium] |
| WP_081590932.1 | AAA+ATPase+wHTH+Alpha+Beta-domain+wHTH+wHTH+RRM | Type-2_BR.RM.like.systems | DUF499*->DUF3780->METHYLASE->HKD-DNase+SF1-SF2-Helicase+iREase+CTD->GDResolvase+Recombinase->ArdA-> | 1090 | Bacteria>Terrabacteria_group>Actinomycetota>Actinomycetes>Micrococcales | GCF_000314575.1 | Brevibacterium_casei | C272_RS01165 | 266723..269995 | 33889 | DUF499-domain-containing-protein-[Brevibacterium-casei] |
| WP_082067576.1 | AAA+ATPase+wHTH+Alpha+Beta-domain+wHTH+wHTH+RRM | Type-2_BR.RM.like.systems | PAIREDC-HTH-><-?\|\|<-HKD-DNase+SF1-SF2-Helicase+iREase+CTD<-METHYLASE<-DUF3780<-DUF499*<-?\|\|<-?\|\|ABC-ATPase+SbcC->?->?->Response_reg+LuxR-HTH-> | 1090 | Bacteria>Terrabacteria_group>Actinomycetota>Actinomycetes>Micrococcales | GCF_000946635.1 | Micrococcus | CIT32_RS01920 | 441439..444711 | 1269 | MULTISPECIES:-DUF499-domain-containing-protein-[Micrococcus] |
| WP_084343725.1 | AAA+ATPase+wHTH+Alpha+Beta-domain+wHTH+wHTH+RRM | Type-2_BR.RM.like.systems | DUF499*->DUF3780->METHYLASE->HKD-DNase+SF1-SF2-Helicase+iREase+CTD-> | 1090 | Bacteria>Terrabacteria_group>Actinomycetota>Actinomycetes>Micrococcales | GCF_001515525.1 | Piscicoccus_intestinalis | PINTE_RS17100 | 2367..5639 | 746033 | DUF499-domain-containing-protein-[Piscicoccus-intestinalis] |
| WP_083956610.1 | AAA+ATPase+wHTH+Alpha+Beta-domain+wHTH+wHTH+RRM | Type-2_BR.RM.like.systems | <-DDE_Tnp_1\|\|<-HKD-DNase+SF1-SF2-Helicase+iREase+CTD<-METHYLASE<-DUF3780<-DUF499*<-XIS-HTH<-ABhydrolase<-?<-MPTase<-HTH\|\|ArdCN+MPTase-> | 1092 | Bacteria>Terrabacteria_group>Actinomycetota>Actinomycetes>Micrococcales | GCF_001968835.1 | Tersicoccus_phoenicis | BKD30_RS15250 | 8674..11952 | 554083 | DUF499-domain-containing-protein-[Tersicoccus-phoenicis] |
| WP_104257506.1 | AAA+ATPase+wHTH+Alpha+Beta-domain+wHTH+wHTH+RRM | Type-2_BR.RM.like.systems | <-HKD-DNase+SF1-SF2-Helicase+iREase+CTD<-METHYLASE<-DUF3780<-DUF499*<-XIS-HTH | 1092 | Bacteria>Terrabacteria_group>Actinomycetota>Actinomycetes>Micrococcales | GCF_002932355.1 | unclassified_Rathayibacter | C5C67_RS05330 | 42447..45725 | 2609250 | MULTISPECIES:-DUF499-domain-containing-protein-[unclassified-Rathayibacter] |
| WP_104326310.1 | AAA+ATPase+wHTH+Alpha+Beta-domain+wHTH+wHTH+RRM | Type-2_BR.RM.like.systems | <-ArdA\|\|NERD->NERD-><-HKD-DNase+SF1-SF2-Helicase+iREase+CTD<-METHYLASE<-DUF3780<-DUF499*<-XIS-HTH<-tRNA<-ABhydrolase\|\|SMS-KaiC-> | 1092 | Bacteria>Terrabacteria_group>Actinomycetota>Actinomycetes>Micrococcales | GCF_002932775.1 | Rathayibacter_sp._AY1B8 | C5C63_RS14380 | 39547..42825 | 2080533 | DUF499-domain-containing-protein-[Rathayibacter-sp.-AY1B8] |
| WP_138926566.1 | AAA+ATPase+wHTH+Alpha+Beta-domain+wHTH+wHTH+RRM | Type-2_BR.RM.like.systems | <-PhdYeFM_antitox<-ArdA<-GDResolvase+Recombinase<-HKD-DNase+SF1-SF2-Helicase+iREase+CTD<-METHYLASE<-DUF3780<-DUF499*<-?<-?\|\|?->?-><-?<-Aminotran_1_2 | 1092 | Bacteria>Terrabacteria_group>Actinomycetota>Actinomycetes>Micrococcales | GCF_006094275.1 | Glutamicibacter_creatinolyticus | GcLGCM259_RS10305 | 2255876..2259154 | 162496 | DUF499-domain-containing-protein-[Glutamicibacter-creatinolyticus] |
| WP_144831113.1 | AAA+ATPase+wHTH+Alpha+Beta-domain+wHTH+wHTH+RRM | Type-2_BR.RM.like.systems | DnaB+SMS-KaiC-><-HKD-DNase+SF1-SF2-Helicase+iREase+CTD<-METHYLASE<-DUF3780<-DUF499*<-XIS-HTH<-?\|\|REase->?-><-URI-DNase<-ABhydrolase+ABhydrolase | 1092 | Bacteria>Terrabacteria_group>Actinomycetota>Actinomycetes>Micrococcales | GCF_007679045.1 | Microbacterium_sp._BH-3-3-3 | FS769_RS06770 | 1449456..1452734 | 1906742 | DUF499-domain-containing-protein-[Microbacterium-sp.-BH-3-3-3] |
| WP_248642106.1 | AAA+ATPase+wHTH+Alpha+Beta-domain+wHTH+wHTH+RRM | Type-2_BR.RM.like.systems | <-PAIREDC-HTH<-REase+SF1-SF2-Helicase\|\|MazG-><-HKD-DNase+SF1-SF2-Helicase+iREase+CTD<-METHYLASE<-DUF3780<-DUF499*\|\|ParB+HNH-><-REase\|\|DUF429->NERD-> | 1092 | Bacteria>Terrabacteria_group>Actinomycetota>Actinomycetes>Micrococcales | GCF_020447105.2 | unclassified_Demequina | LGT36_RS12815 | 2648905..2652183 | 2620311 | MULTISPECIES:-DUF499-domain-containing-protein-[unclassified-Demequina] |
| WP_248257239.1 | AAA+ATPase+wHTH+Alpha+Beta-domain+wHTH+wHTH+RRM | Type-2_BR.RM.like.systems | Phage_integrase-><-HKD-DNase+SF1-SF2-Helicase+iREase+CTD<-METHYLASE<-DUF3780<-DUF499*\|\|HTH->?-><-?<-?<-KAP-NTPase | 1084 | Bacteria>Terrabacteria_group>Actinomycetota>Actinomycetes>Micrococcales | GCF_023148685.1 | Georgenia_sp._EYE_87 | KZX45_RS08185 | 46684..49938 | 2853448 | DUF499-domain-containing-protein-[Georgenia-sp.-EYE_87] |
| WP_253034129.1 | AAA+ATPase+wHTH+Alpha+Beta-domain+wHTH+wHTH+RRM | Type-2_BR.RM.like.systems | Cluster2439_3clades->DUF499*->DUF3780->HKD-DNase+SF1-SF2-Helicase+iREase+CTD->HIN-HTH+HIN-HTH-><-HEPN<-?<-PAPS_reduct | 1092 | Bacteria>Terrabacteria_group>Actinomycetota>Actinomycetes>Micrococcales | GCF_024124465.1 | Microbacterium_sp._CnD16-F | NH287_RS01095 | 84562..87840 | 2954493 | DUF499-domain-containing-protein-[Microbacterium-sp.-CnD16-F] |
| WP_259917334.1 | AAA+ATPase+wHTH+Alpha+Beta-domain+wHTH+wHTH+RRM | Type-2_BR.RM.like.systems | <-HKD-DNase+SF1-SF2-Helicase+iREase+CTD<-METHYLASE<-DUF3780<-DUF499*<-?\|\|HTH_21+rve+rve_3-><-PAIREDC-HTH | 1074 | Bacteria>Terrabacteria_group>Actinomycetota>Actinomycetes>Micrococcales | GCF_025145075.1 | Kocuria_carniphila | M3B51_RS13295 | 9499..12723 | 262208 | DUF499-domain-containing-protein-[Kocuria-carniphila] |
| WP_186368756.1 | AAA+ATPase+wHTH+Alpha+Beta-domain+wHTH+wHTH+RRM | Type-2_BR.RM.like.systems | <-GNAT\|\|<-HKD-DNase+SF1-SF2-Helicase+iREase+CTD<-METHYLASE-p<-METHYLASE<-DUF3780<-DUF499*<-?\|\|ABC-ATPase+SbcC->?->?->Response_reg+LuxR-HTH-> | 1090 | Bacteria>Terrabacteria_group>Actinomycetota>Actinomycetes>Micrococcales | GCF_025149565.1 | Kocuria | FS750_RS00535 | 109526..112798 | 57493 | DUF499-domain-containing-protein-[Kocuria-indica] |
| WP_259894632.1 | AAA+ATPase+wHTH+Alpha+Beta-domain+wHTH+wHTH+RRM | Type-2_BR.RM.like.systems | DUF499*->DUF3780->METHYLASE->HKD-DNase+SF1-SF2-Helicase+iREase+CTD-><-RelE-ParE<-Phd-YefM | 1090 | Bacteria>Terrabacteria_group>Actinomycetota>Actinomycetes>Micrococcales | GCF_025151625.1 | unclassified_Kocuria | M3E18_RS01545 | 2496..5768 | 2649579 | MULTISPECIES:-DUF499-domain-containing-protein-[unclassified-Kocuria] |
| WP_282837860.1 | AAA+ATPase+wHTH+Alpha+Beta-domain+wHTH+wHTH+RRM | Type-2_BR.RM.like.systems | <-HKD-DNase+SF1-SF2-Helicase+iREase+CTD<-METHYLASE<-DUF3780<-DUF499*<-XIS-HTH<-?<-?<-DDE_5 | 1092 | Bacteria>Terrabacteria_group>Actinomycetota>Actinomycetes>Micrococcales | GCF_030011835.1 | Microbacterium_flavum | QM026_RS12475 | 6925..10203 | 415216 | DUF499-domain-containing-protein-[Microbacterium-flavum] |
| WP_125106250.1 | AAA+ATPase+wHTH+Alpha+Beta-domain+wHTH+wHTH+RRM | Type-2_BR.RM.like.systems | <-ArdA<-GDResolvase+Recombinase<-HKD-DNase+SF1-SF2-Helicase+iREase+CTD<-METHYLASE<-DUF3780<-DUF499*<-?\|\|35exo-> | 1090 | Bacteria>Terrabacteria_group>Actinomycetota>Actinomycetes>Micrococcales | GCF_900626105.1 | Gulosibacter_massiliensis | EJL01_RS03165 | 640383..643655 | 2479839 | DUF499-domain-containing-protein-[Gulosibacter-massiliensis] |
| NED58711.1 | AAA+ATPase+wHTH+Alpha+Beta-domain+wHTH+wHTH+RRM | Type-2_BR.RM.like.systems | <-HKD-DNase+SF1-SF2-Helicase+iREase+CTD<-METHYLASE<-DUF3780<-DUF499*\|\|GDResolvase+Recombinase+ZnR-> | 1093 | Bacteria>Terrabacteria_group>Actinomycetota>Actinomycetes>Micromonosporales | GCA_010550585.1 | Micromonospora_aurantiaca | G3I24_48480 | 12699..15980 | 47850 | ATP-binding-protein-[Micromonospora-aurantiaca] |
| WP_063007180.1 | AAA+ATPase+wHTH+Alpha+Beta-domain+wHTH+wHTH+RRM | Type-2_BR.RM.like.systems | <-HTH<-?<-?<-HKD-DNase+SF1-SF2-Helicase+iREase+CTD<-METHYLASE<-DUF3780<-DUF499*<-NUDIX<-?<-?<-SF1-SF2-Helicase | 1093 | Bacteria>Terrabacteria_group>Actinomycetota>Actinomycetes>Mycobacteriales | GCF_001612965.1 | Nocardia_kruczakiae | NK1_RS03875 | 189470..192751 | 261477 | DUF499-domain-containing-protein-[Nocardia-kruczakiae] |
| WP_064862403.1 | AAA+ATPase+wHTH+Alpha+Beta-domain+wHTH+wHTH+RRM | Type-2_BR.RM.like.systems | <-ArdCN+MPTase\|\|HTH->MPTase->?->ABhydrolase->XIS-HTH->DUF499*->DUF3780->METHYLASE->HKD-DNase+SF1-SF2-Helicase+iREase+CTD-><-?<-DNA_methylase\|\|TrwC+SF1-SF2-Helicase-> | 1092 | Bacteria>Terrabacteria_group>Actinomycetota>Actinomycetes>Mycobacteriales | GCF_001665715.1 | Gordonia_sp._852002-10350_SCH5691597 | A5777_RS08980 | 4876..8154 | 1834085 | DUF499-domain-containing-protein-[Gordonia-sp.-852002-10350_SCH5691597] |
| WP_096907611.1 | AAA+ATPase+wHTH+Alpha+Beta-domain+wHTH+wHTH+RRM | Type-2_BR.RM.like.systems | HTH->?-><-?<-HKD-DNase+SF1-SF2-Helicase+iREase+CTD<-METHYLASE<-DUF3780<-DUF499*<-HTH_Tnp\|\|GDResolvase+HIN-HTH-><-?<-MODE-HTH+PBPII | 1089 | Bacteria>Terrabacteria_group>Actinomycetota>Actinomycetes>Mycobacteriales | GCF_002441635.1 | Dietzia_sp._WMMA184 | CK206_RS18720 | 83413..86682 | 2039808 | DUF499-domain-containing-protein-[Dietzia-sp.-WMMA184] |
| WP_144689647.1 | AAA+ATPase+wHTH+Alpha+Beta-domain+wHTH+wHTH+RRM | Type-2_BR.RM.like.systems | <-RelE-ParE<-Phd-YefM<-HKD-DNase+SF1-SF2-Helicase+iREase+CTD-p\|\|?-><-METHYLASE<-DUF3780<-DUF499* | 1090 | Bacteria>Terrabacteria_group>Actinomycetota>Actinomycetes>Mycobacteriales | GCF_025144605.1 | Corynebacterium_sanguinis | M3A76_RS07250 | 53141..56413 | 2594913 | MULTISPECIES:-DUF499-domain-containing-protein-[Corynebacterium] |
| WP_136651048.1 | AAA+ATPase+wHTH+Alpha+Beta-domain+wHTH+wHTH+RRM | Type-2_BR.RM.like.systems | DUF499*->DUF3780->METHYLASE->HKD-DNase+SF1-SF2-Helicase+iREase+CTD->Phd-YefM->RelE-ParE-> | 1090 | Bacteria>Terrabacteria_group>Actinomycetota>Actinomycetes>Mycobacteriales | GCF_025144815.1 | Corynebacterium_sanguinis | M3B20_RS07455 | 35434..38706 | 2594913 | DUF499-domain-containing-protein-[Corynebacterium-sanguinis] |
| WP_279938925.1 | AAA+ATPase+wHTH+Alpha+Beta-domain+wHTH+wHTH+RRM | Type-2_BR.RM.like.systems | DUF499*->DUF3780->METHYLASE->HKD-DNase+SF1-SF2-Helicase+iREase+CTD->GDResolvase+Recombinase-><-?<-TraJ-RHH | 1090 | Bacteria>Terrabacteria_group>Actinomycetota>Actinomycetes>Mycobacteriales | GCF_029853935.1 | Gordonia_alkanivorans | QEN38_RS07200 | 33675..36947 | 84096 | DUF499-domain-containing-protein-[Gordonia-alkanivorans] |
| WP_259787954.1 | AAA+ATPase+wHTH+Alpha+Beta-domain+wHTH+wHTH+RRM | Type-2_BR.RM.like.systems | DUF499*->DUF3780->METHYLASE->HKD-DNase+SF1-SF2-Helicase+iREase+CTD->Phd-YefM->RelE-ParE-> | 1090 | Bacteria>Terrabacteria_group>Actinomycetota>Actinomycetes>Mycobacteriales | GCF_943914675.1 | Corynebacterium | QV646_RS01785 | 35219..38491 | 1716 | DUF499-domain-containing-protein-[Corynebacterium-sanguinis] |
| WP_243580595.1 | AAA+ATPase+wHTH+Alpha+Beta-domain+wHTH+wHTH+RRM | Type-2_BR.RM.like.systems | HNH-><-HKD-DNase+SF1-SF2-Helicase+iREase+CTD<-METHYLASE<-DUF3780<-DUF499*<-HTH_Tnp<-ACET<-?\|\|SIR2-> | 1089 | Bacteria>Terrabacteria_group>Actinomycetota>Actinomycetes>Mycobacteriales | GCF_022810365.1 | unclassified_Rhodococcus_(in:_high_G+C_Gram-positive_bacteria) | MTX38_RS19005 | 53191..56460 | 192944 | MULTISPECIES:-DUF499-domain-containing-protein-[unclassified-Rhodococcus] |
| WP_094623222.1 | AAA+ATPase+wHTH+Alpha+Beta-domain+wHTH+wHTH+RRM | Type-2_BR.RM.like.systems | <-REase<-SF1-SF2-Helicase+REase<-Exonuc_V_gamma+REase\|\|DUF499*->DUF3780->METHYLASE->HKD-DNase+SF1-SF2-Helicase+iREase+CTD->PAIREDC-HTH->?-><-REase | 1090 | Bacteria>Terrabacteria_group>Actinomycetota>Actinomycetes>Mycobacteriales | - | unclassified_Rhodococcus_(in:_high_G+C_Gram-positive_bacteria) | CH269_RS10940 | 142592..145864 | 192944 | MULTISPECIES:-DUF499-domain-containing-protein-[unclassified-Rhodococcus] |
| TCO33382.1 | AAA+ATPase+wHTH+Alpha+Beta-domain+wHTH+wHTH+RRM | Type-2_BR.RM.like.systems | CHTH->?->Primpol+PriCT1-><-?<-HEPN\|\|SIR2->DUF499*->DUF3780->METHYLASE->HKD-DNase+SF1-SF2-Helicase+iREase+CTD-><-HKD-DNase+HKD-DNase<-?<-HTH+HKD-DNase | 1103 | Bacteria>Terrabacteria_group>Actinomycetota>Actinomycetes>Propionibacteriales | GCA_004342025.1 | Kribbella_steppae | EV652_103383 | 388366..391677 | 2512223 | uncharacterized-protein-DUF499-[Kribbella-sp.-VKM-Ac-2572] |
| SCQ62922.1 | AAA+ATPase+wHTH+Alpha+Beta-domain+wHTH+wHTH+RRM | Type-2_BR.RM.like.systems | <-ArdCN+MPTase\|\|HTH->MPTase->?->ABhydrolase->XIS-HTH->DUF499*-><-?\|\|?->METHYLASE->HKD-DNase+SF1-SF2-Helicase+iREase+CTD-><-DNA_methylase\|\|TrwC+SF1-SF2-Helicase-> | 1092 | Bacteria>Terrabacteria_group>Actinomycetota>Actinomycetes>Propionibacteriales | GCA_900097265.1 | Propionibacterium_freudenreichii | PFR_JS9-1_1587 | 1777278..1780556 | 1744 | AAA-family-ATPase-[Propionibacterium-freudenreichii] |
| WP_158441140.1 | AAA+ATPase+wHTH+Alpha+Beta-domain+wHTH+wHTH+RRM | Type-2_BR.RM.like.systems | CHTH->?->Primpol+PriCT1-><-?<-HEPN\|\|SIR2->DUF499*->METHYLASE->HKD-DNase+SF1-SF2-Helicase+iREase+CTD-><-?<-HTH+HKD-DNase<-Pkinase+SF1-SF2-Helicase | 1084 | Bacteria>Terrabacteria_group>Actinomycetota>Actinomycetes>Propionibacteriales | GCF_004342025.1 | Kribbella_steppae | EV652_RS10505 | 388423..391677 | 2512223 | DUF499-domain-containing-protein-[Kribbella-sp.-VKM-Ac-2572] |
| WP_260289258.1 | AAA+ATPase+wHTH+Alpha+Beta-domain+wHTH+wHTH+RRM | Type-2_BR.RM.like.systems | <-ParB<-TrwC+SF1-SF2-Helicase<-REase<-HKD-DNase+SF1-SF2-Helicase+iREase+CTD<-METHYLASE<-DUF3780<-DUF499*<-XIS-HTH\|\|?->ArdCN+MPTase->Primpol+PriCT1-> | 1092 | Bacteria>Terrabacteria_group>Actinomycetota>Actinomycetes>Propionibacteriales | GCF_025187725.1 | Propionibacterium_freudenreichii | EFN01_RS07350 | 14264..17542 | 1744 | DUF499-domain-containing-protein-[Propionibacterium-freudenreichii] |
| WP_081083807.1 | AAA+ATPase+wHTH+Alpha+Beta-domain+wHTH+wHTH+RRM | Type-2_BR.RM.like.systems | XIS-HTH->DUF499*->DUF3780->METHYLASE->HKD-DNase+SF1-SF2-Helicase+iREase+CTD->REase->TrwC+SF1-SF2-Helicase->ParB-> | 1092 | Bacteria>Terrabacteria_group>Actinomycetota>Actinomycetes>Propionibacteriales | GCF_900000055.1 | Propionibacterium_freudenreichii | FAM23868_RS05705 | 570..3848 | 1744 | DUF499-domain-containing-protein-[Propionibacterium-freudenreichii] |
| MBB6545858.1 | AAA+ATPase+wHTH+Alpha+Beta-domain+wHTH+wHTH+RRM | Type-2_BR.RM.like.systems | NUDIX->?-><-HKD-DNase+SF1-SF2-Helicase+iREase+CTD<-METHYLASE<-DUF3780<-DUF499*\|\|?-><-GNTR-HTH-p\|\|?-><-?<-?<-GNTR-HTH-p | 1091 | Bacteria>Terrabacteria_group>Actinomycetota>Actinomycetes>Streptosporangiales | GCA_014207985.1 | Nonomuraea_rubra | HD593_000653 | 715510..718785 | 46180 | hypothetical-protein-[Nonomuraea-rubra] |
| GII79103.1 | AAA+ATPase+wHTH+Alpha+Beta-domain+wHTH+wHTH+RRM | Type-2_BR.RM.like.systems | <-HEPN<-HTH_Tnp_4+DDE_Tnp_4<-MULE-TRANSPOSASE\|\|DDE_Tnp_ISAZ013-><-MULE-TRANSPOSASE<-MULE-TRANSPOSASE\|\|DUF499*->DUF3780->METHYLASE->HKD-DNase+SF1-SF2-Helicase+iREase+CTD-><-?<-RelE-ParE | 1096 | Bacteria>Terrabacteria_group>Actinomycetota>Actinomycetes>Streptosporangiales | GCA_016863395.1 | Sphaerisporangium_rufum | Sru01_40850 | 146394..149684 | 1381558 | hypothetical-protein-Sru01_40850-[Sphaerisporangium-rufum] |
| MBX6386284.1 | AAA+ATPase+wHTH+Alpha+Beta-domain+wHTH+wHTH+RRM | Type-2_BR.RM.like.systems | DUF499*->?->?->HKD-DNase+SF1-SF2-Helicase+iREase+CTD-> | 582 | Bacteria>Terrabacteria_group>Actinomycetota>Actinomycetes>Streptosporangiales | GCA_019689655.1 | Microbispora_sp. | IRZ07_25480 | 114..1862 | 1898321 | hypothetical-protein-[Microbispora-sp.] |
| WP_203988837.1 | AAA+ATPase+wHTH+Alpha+Beta-domain+wHTH+wHTH+RRM | Type-2_BR.RM.like.systems | HEPN->REase->HAD-><-HEPN<-HTH_Tnp_4+DDE_Tnp_4\|\|DDE_Tnp_ISAZ013->\|\|DUF499*->DUF3780->METHYLASE->HKD-DNase+SF1-SF2-Helicase+iREase+CTD-><-?<-RelE-ParE | 1070 | Bacteria>Terrabacteria_group>Actinomycetota>Actinomycetes>Streptosporangiales | GCF_016863395.1 | Sphaerisporangium_rufum | Sru01_RS20350 | 146472..149684 | 1381558 | DUF499-domain-containing-protein-[Sphaerisporangium-rufum] |
| WP_246547501.1 | AAA+ATPase+wHTH+Alpha+Beta-domain+wHTH+wHTH+RRM | Type-2_BR.RM.like.systems | NUDIX->?-><-HKD-DNase+SF1-SF2-Helicase+iREase+CTD<-METHYLASE<-DUF3780<-DUF499*\|\|?->\|\|?-><-?<-GNTR-HTH-p | 1088 | Bacteria>Terrabacteria_group>Actinomycetota>Actinomycetes>Streptosporangiales | - | Nonomuraea_rubra | HD593_RS03230 | 715510..718776 | 46180 | DUF499-domain-containing-protein-[Nonomuraea-rubra] |
| MCC5947763.1 | AAA+ATPase+wHTH+Alpha+Beta-domain+wHTH+wHTH+RRM | Type-2_BR.RM.like.systems | <-HTH<-?\|\|tRNA-><-?<-TUDOR+SF1-SF2-Helicase+iREase+CTD<-METHYLASE<-DUF3780<-DUF499*\|\|?->?->Phage_integrase+Phage_integrase->Phage_integrase+Phage_integrase->?->PIN-> | 1068 | Bacteria>Terrabacteria_group>Actinomycetota>Nitriliruptoria>Nitriliruptorales | GCA_020832455.1 | Nitriliruptoraceae_bacterium | JJT89_04835 | 21790..24996 | 2093808 | ATP-binding-protein-[Nitriliruptoraceae-bacterium] |
| TXH43022.1 | AAA+ATPase+wHTH+Alpha+Beta-domain+wHTH+wHTH+RRM | Type-2_BR.RM.like.systems | <-Primpol+PriCT1<-ArdCN+MPTase\|\|HTH->MPTase->ABhydrolase->XIS-HTH->DUF499*->DUF3780->METHYLASE->HKD-DNase+SF1-SF2-Helicase+iREase+CTD-> | 1092 | Bacteria>Terrabacteria_group>Actinomycetota>unclassified_Actinomycetota | GCA_008012465.1 | Actinomycetota_bacterium | E6Q90_07820 | 6765..10043 | 2900548 | ATP-binding-protein-[Actinobacteria-bacterium] |
| MBU1800851.1 | AAA+ATPase+wHTH+Alpha+Beta-domain+wHTH+wHTH+RRM | Type-2_BR.RM.like.systems | DUF499*->DUF3780->METHYLASE->HKD-DNase+SF1-SF2-Helicase+iREase+CTD-p-><-?<-ArdA<-GDResolvase+HIN-HTH | 861 | Bacteria>Terrabacteria_group>Actinomycetota>unclassified_Actinomycetota | GCA_018823685.1 | Actinomycetota_bacterium | KKA97_01120 | 1..2587 | 2900548 | ATP-binding-protein-[Actinomycetia-bacterium] |
| PJB73371.1 | AAA+ATPase+wHTH+Alpha+Beta-domain+wHTH+wHTH+RRM | Type-2_BR.RM.like.systems | <-Radical-SAM\|\|?->?-><-?<-?<-?\|\|DUF499*->DUF3780->OLD-ABC->TOPRIM-> | 974 | Bacteria>Terrabacteria_group>Armatimonadota>unclassified_Armatimonadota | GCA_002789735.1 | Armatimonadetes_bacterium_CG_4_9_14_3_um_filter_66_14 | CO096_06105 | 32491..35415 | 1973914 | hypothetical-protein-CO096_06105-[Armatimonadetes-bacterium-CG_4_9_14_3_um_filter_66_14] |
| PIZ50111.1 | AAA+ATPase+wHTH+Alpha+Beta-domain+wHTH+wHTH+RRM | Type-2_BR.RM.like.systems | <-METHYLASE-p<-TOPRIM<-OLD-ABC<-DUF3780<-DUF499* | 974 | Bacteria>Terrabacteria_group>Armatimonadota>unclassified_Armatimonadota | GCA_002791775.1 | Armatimonadetes_bacterium_CG_4_10_14_0_8_um_filter_66_14 | COY42_02315 | 3146..6070 | 1973908 | hypothetical-protein-COY42_02315-[Armatimonadetes-bacterium-CG_4_10_14_0_8_um_filter_66_14] |
| OIO95800.1 | AAA+ATPase+wHTH+Alpha+Beta-domain+wHTH+wHTH+RRM | Type-2_BR.RM.like.systems | DUF499*->DUF3780->OLD-ABC->TOPRIM-> | 974 | Bacteria>Terrabacteria_group>Armatimonadota>unclassified_Armatimonadota | GCA_001871675.1 | Armatimonadetes_bacterium_CG2_30_66_41 | AUJ96_25895 | 5778..8702 | 1805028 | hypothetical-protein-AUJ96_25895-[Armatimonadetes-bacterium-CG2_30_66_41] |
| PIU89760.1 | AAA+ATPase+wHTH+Alpha+Beta-domain+wHTH+wHTH+RRM | Type-2_BR.RM.like.systems | <-METHYLASE-p<-TOPRIM<-OLD-ABC<-DUF3780<-DUF499* | 974 | Bacteria>Terrabacteria_group>Armatimonadota>unclassified_Armatimonadota | GCA_002779825.1 | Armatimonadetes_bacterium_CG06_land_8_20_14_3_00_66_21 | COS65_27530 | 3126..6050 | 1973915 | hypothetical-protein-COS65_27530-[Armatimonadetes-bacterium-CG06_land_8_20_14_3_00_66_21] |
| NDK16278.1 | AAA+ATPase+wHTH+Alpha+Beta-domain+wHTH+wHTH+RRM | Type-2_BR.RM.like.systems | <-TOPRIM<-OLD-ABC<-DUF3780<-DUF499* | 974 | Bacteria>Terrabacteria_group>Armatimonadota>unclassified_Armatimonadota | GCA_010119115.1 | Armatimonadota_bacterium | GW911_29980 | 2382..5306 | 2033014 | ATP-binding-protein-[Armatimonadetes-bacterium] |
| NCO91848.1 | AAA+ATPase+wHTH+Alpha+Beta-domain+wHTH+wHTH+RRM | Type-2_BR.RM.like.systems | <-TOPRIM<-OLD-ABC<-DUF3780<-DUF499* | 974 | Bacteria>Terrabacteria_group>Armatimonadota>unclassified_Armatimonadota | GCA_009993145.1 | Armatimonadota_bacterium | GW880_11055 | 2472..5396 | 2033014 | ATP-binding-protein-[Armatimonadetes-bacterium] |
| KPC75471.1 | AAA+ATPase+wHTH+Alpha+Beta-domain+wHTH+wHTH+RRM | Type-2_BR.RM.like.systems | <-HKD-DNase+SF1-SF2-Helicase+iREase+CTD<-METHYLASE<-DUF3780<-DUF499* | 1076 | Bacteria>Terrabacteria_group>Bacillota>Bacilli>Bacillales | GCA_001294365.1 | Thermoactinomyces_vulgaris | ADL26_07600 | 8634..11864 | 2026 | ATPase-AAA-[Thermoactinomyces-vulgaris] |
| OUM88560.1 | AAA+ATPase+wHTH+Alpha+Beta-domain+wHTH+wHTH+RRM | Type-2_BR.RM.like.systems | <-HKD-DNase+SF1-SF2-Helicase+iREase+CTD<-METHYLASE<-DUF3780<-DUF499*\|\|pPIWI+pPIWI->pPIWI->pPIWI+MID+RNaseH->MULE-TRANSPOSASE-> | 1074 | Bacteria>Terrabacteria_group>Bacillota>Bacilli>Bacillales | GCA_002159075.1 | Bacillus_thermozeamaize | BAA01_05365 | 48594..51818 | 230954 | AAA-family-ATPase-[Bacillus-thermozeamaize] |
| PDO10504.1 | AAA+ATPase+wHTH+Alpha+Beta-domain+wHTH+wHTH+RRM | Type-2_BR.RM.like.systems | <-HKD-DNase+SF1-SF2-Helicase+iREase+CTD<-DUF3780<-DUF499*<-?<-Polbeta\|\|HIN-HTH-> | 1083 | Bacteria>Terrabacteria_group>Bacillota>Bacilli>Bacillales | GCA_002507565.1 | Candidatus_Reconcilbacillus_cellulovorans | BLM47_07210 | 108136..111387 | 1906605 | AAA-family-ATPase-[Candidatus-Reconcilbacillus-cellulovorans] |
| PRZ12344.1 | AAA+ATPase+wHTH+Alpha+Beta-domain+wHTH+wHTH+RRM | Type-2_BR.RM.like.systems | rve_3-><-?<-HKD-DNase+SF1-SF2-Helicase+iREase+CTD<-METHYLASE<-DUF3780<-DUF499*<-HD\|\|?-><-?<-Methyltransf_25 | 1076 | Bacteria>Terrabacteria_group>Bacillota>Bacilli>Bacillales | GCA_003003125.1 | Laceyella_sediminis | CLV36_1147 | 10508..13738 | 573074 | uncharacterized-protein-DUF499-[Laceyella-sediminis] |
| MCL6597625.1 | AAA+ATPase+wHTH+Alpha+Beta-domain+wHTH+wHTH+RRM | Type-2_BR.RM.like.systems | <-HKD-DNase+SF1-SF2-Helicase+iREase+CTD<-METHYLASE<-DUF3780<-DUF499*<-REase+REase<-REase<-?<-?\|\|XIS-HTH->Arm-DNA-bind_4+Phage_int_SAM_3+Phage_integrase-> | 1071 | Bacteria>Terrabacteria_group>Bacillota>Bacilli>Bacillales | GCA_023512275.1 | Alicyclobacillus_macrosporangiidus | K6T81_02655 | 38493..41708 | 392015 | DUF499-domain-containing-protein-[Alicyclobacillus-macrosporangiidus] |
| WP_013076776.1 | AAA+ATPase+wHTH+Alpha+Beta-domain+wHTH+wHTH+RRM | Type-2_BR.RM.like.systems | <-HKD-DNase+SF1-SF2-Helicase+iREase+CTD<-METHYLASE<-DUF3780<-DUF499*\|\|ABhydrolase->ABhydrolase->?->?-><-ABC-ATPase+SbcC | 1074 | Bacteria>Terrabacteria_group>Bacillota>Bacilli>Bacillales | GCF_000092905.1 | Kyrpidia_tusciae | BTUS_RS14310 | 2935912..2939136 | 33943 | DUF499-domain-containing-protein-[Kyrpidia-tusciae] |
| WP_067849130.1 | AAA+ATPase+wHTH+Alpha+Beta-domain+wHTH+wHTH+RRM | Type-2_BR.RM.like.systems | DUF499*->DUF3780->METHYLASE->HKD-DNase+SF1-SF2-Helicase+iREase+CTD-> | 1111 | Bacteria>Terrabacteria_group>Bacillota>Bacilli>Bacillales | GCF_001570745.1 | Alicyclobacillus_mali_(ex_Roth_et_al._2021) | AMA01_RS06480 | 1550..4885 | 1123961 | DUF499-domain-containing-protein-[Alicyclobacillus-mali] |
| WP_245888378.1 | AAA+ATPase+wHTH+Alpha+Beta-domain+wHTH+wHTH+RRM | Type-2_BR.RM.like.systems | rve_3-><-?<-HKD-DNase+SF1-SF2-Helicase+iREase+CTD<-METHYLASE<-DUF3780<-DUF499*<-HD\|\|?-><-?<-Methyltransf_25 | 1062 | Bacteria>Terrabacteria_group>Bacillota>Bacilli>Bacillales | GCF_003003125.1 | Laceyella_sediminis | CLV36_RS14160 | 10508..13696 | 573074 | DUF499-domain-containing-protein-[Laceyella-sediminis] |
| WP_181754673.1 | AAA+ATPase+wHTH+Alpha+Beta-domain+wHTH+wHTH+RRM | Type-2_BR.RM.like.systems | <-HKD-DNase+SF1-SF2-Helicase+iREase+CTD<-METHYLASE<-DUF3780<-DUF499*\|\|?->DDE_Tnp_1-> | 1076 | Bacteria>Terrabacteria_group>Bacillota>Bacilli>Bacillales | GCF_013867725.1 | Paenactinomyces_guangxiensis | I8U27_RS05455 | 222049..225279 | 1490290 | DUF499-domain-containing-protein-[Paenactinomyces-guangxiensis] |
| WP_033099138.1 | AAA+ATPase+wHTH+Alpha+Beta-domain+wHTH+wHTH+RRM | Type-2_BR.RM.like.systems | HD->DUF499*->DUF3780->METHYLASE->HKD-DNase+SF1-SF2-Helicase+iREase+CTD-> | 1076 | Bacteria>Terrabacteria_group>Bacillota>Bacilli>Bacillales | GCF_013867735.1 | Thermoactinomyces_daqus | JG50_RS0100370 | 4293..7523 | 1329516 | DUF499-domain-containing-protein-[Thermoactinomyces-daqus] |
| WP_198056113.1 | AAA+ATPase+wHTH+Alpha+Beta-domain+wHTH+wHTH+RRM | Type-2_BR.RM.like.systems | PAIREDC-HTH-><-?<-?<-HKD-DNase+SF1-SF2-Helicase+iREase+CTD<-METHYLASE<-DUF3780<-DUF499*<-HD | 1076 | Bacteria>Terrabacteria_group>Bacillota>Bacilli>Bacillales | GCF_016055105.1 | Thermoactinomyces_sp._CICC_10523 | I8U19_RS13150 | 12194..15424 | 2767428 | DUF499-domain-containing-protein-[Thermoactinomyces-sp.-CICC-10523] |
| WP_242549801.1 | AAA+ATPase+wHTH+Alpha+Beta-domain+wHTH+wHTH+RRM | Type-2_BR.RM.like.systems | SbcC->tRNA-><-?<-?\|\|DUF499*->DUF3780->METHYLASE->HKD-DNase+SF1-SF2-Helicase+iREase+CTD-> | 984 | Bacteria>Terrabacteria_group>Bacillota>Bacilli>Bacillales | GCF_017329565.1 | Alicyclobacillus_mali_(ex_Roth_et_al._2021) | J3479_RS12595 | 9539..12493 | 1123961 | DUF499-domain-containing-protein-[Alicyclobacillus-mali] |
| WP_242549237.1 | AAA+ATPase+wHTH+Alpha+Beta-domain+wHTH+wHTH+RRM | Type-2_BR.RM.like.systems | <-HKD-DNase+SF1-SF2-Helicase+iREase+CTD<-METHYLASE<-DUF3780<-DUF499* | 984 | Bacteria>Terrabacteria_group>Bacillota>Bacilli>Bacillales | GCF_017329625.1 | Alicyclobacillus_mali_(ex_Roth_et_al._2021) | J3484_RS08440 | 65687..68641 | 1123961 | DUF499-domain-containing-protein-[Alicyclobacillus-mali] |
| WP_236011684.1 | AAA+ATPase+wHTH+Alpha+Beta-domain+wHTH+wHTH+RRM | Type-2_BR.RM.like.systems | <-HKD-DNase+SF1-SF2-Helicase+iREase+CTD<-METHYLASE<-DUF3780<-DUF499* | 1035 | Bacteria>Terrabacteria_group>Bacillota>Bacilli>Bacillales | GCF_017329635.1 | Alicyclobacillus_fructus | J3481_RS13160 | 17542..20649 | 2816082 | DUF499-domain-containing-protein-[Alicyclobacillus-fructus] |
| WP_170099813.1 | AAA+ATPase+wHTH+Alpha+Beta-domain+wHTH+wHTH+RRM | Type-2_BR.RM.like.systems | <-HKD-DNase+SF1-SF2-Helicase+iREase+CTD<-METHYLASE<-DUF3780<-DUF499*\|\|ABhydrolase->?->?-><-ABC-ATPase+SbcC | 1074 | Bacteria>Terrabacteria_group>Bacillota>Bacilli>Bacillales | GCF_018802225.1 | Kyrpidia_spormannii | GWK40_RS14140 | 2871659..2874883 | 2055160 | DUF499-domain-containing-protein-[Kyrpidia-spormannii] |
| WP_170086420.1 | AAA+ATPase+wHTH+Alpha+Beta-domain+wHTH+wHTH+RRM | Type-2_BR.RM.like.systems | <-HKD-DNase+SF1-SF2-Helicase+iREase+CTD<-?<-METHYLASE<-DUF3780<-DUF499*\|\|ABhydrolase-> | 1074 | Bacteria>Terrabacteria_group>Bacillota>Bacilli>Bacillales | GCF_018802245.1 | Kyrpidia_spormannii | GWK42_RS14095 | 2869994..2873218 | 2055160 | DUF499-domain-containing-protein-[Kyrpidia-spormannii] |
| WP_259436284.1 | AAA+ATPase+wHTH+Alpha+Beta-domain+wHTH+wHTH+RRM | Type-2_BR.RM.like.systems | DUF499*->DUF3780->METHYLASE->HKD-DNase+SF1-SF2-Helicase+iREase+CTD-> | 1076 | Bacteria>Terrabacteria_group>Bacillota>Bacilli>Bacillales | GCF_024927925.1 | Laceyella_sacchari | NYR52_RS03490 | 660153..663383 | 37482 | DUF499-domain-containing-protein-[Laceyella-sacchari] |
| WP_102991502.1 | AAA+ATPase+wHTH+Alpha+Beta-domain+wHTH+wHTH+RRM | Type-2_BR.RM.like.systems | Methyltransf_25->?-><-?\|\|HD->DUF499*->DUF3780->METHYLASE->HKD-DNase+SF1-SF2-Helicase+iREase+CTD->?-><-rve_3 | 1076 | Bacteria>Terrabacteria_group>Bacillota>Bacilli>Bacillales | GCF_900182735.1 | Laceyella_tengchongensis | QQZ12_RS00295 | 54408..57638 | 574699 | DUF499-domain-containing-protein-[Laceyella-sacchari] |
| WP_243276991.1 | AAA+ATPase+wHTH+Alpha+Beta-domain+wHTH+wHTH+RRM | Type-2_BR.RM.like.systems | REase-><-?<-?\|\|DUF499*->DUF3780->METHYLASE->HKD-DNase+SF1-SF2-Helicase+iREase+CTD->PolB->?->PemK_toxin-> | 1067 | Bacteria>Terrabacteria_group>Bacillota>Clostridia>Eubacteriales | - | Desulfallas_sp._Bu1-1 | IT084_RS14240 | 47390..50593 | 2787620 | DUF499-domain-containing-protein-[Desulfallas-sp.-Bu1-1] |
| WP_207712783.1 | AAA+ATPase+wHTH+Alpha+Beta-domain+wHTH+wHTH+RRM | Type-2_BR.RM.like.systems | PolB-NTase->Polbeta-><-Peptidase_S51<-HKD-DNase+SF1-SF2-Helicase+iREase+CTD<-METHYLASE<-DUF3780<-DUF499*<-PIN<-MetJarc-HTH<-REase | 1069 | Bacteria>Terrabacteria_group>Bacillota>Clostridia>Eubacteriales | - | Desulfofundulus_sp._TPOSR | G7K71_RS06320 | 1300314..1303523 | 2714340 | DUF499-domain-containing-protein-[Desulfofundulus-sp.-TPOSR] |
| AEG13741.1 | AAA+ATPase+wHTH+Alpha+Beta-domain+wHTH+wHTH+RRM | Type-2_BR.RM.like.systems | REase->?->MetJarc-HTH->PIN->DUF499*->DUF3780->METHYLASE->HKD-DNase+SF1-SF2-Helicase+iREase+CTD->Peptidase_S51-><-Polbeta<-PolB-NTase | 1073 | Bacteria>Terrabacteria_group>Bacillota>Clostridia>Eubacteriales | GCA_000214705.1 | Desulfofundulus_kuznetsovii_DSM_6115 | Desku_0093 | 89841..93062 | 760568 | ATPase-(AAA+-superfamily)-like-protein-[Desulfofundulus-kuznetsovii-DSM-6115] |
| KKM12713.1 | AAA+ATPase+wHTH+Alpha+Beta-domain+wHTH+wHTH+RRM | Type-2_BR.RM.like.systems | ParA-Soj-PloopNTPase->?-><-RNaseH<-HKD-DNase+SF1-SF2-Helicase+iREase+CTD<-METHYLASE<-DUF3780<-DUF499*<-PrlF_antitoxin<-NTP_transf_2<-HEPN<-tRNA<-tRNA<-tRNA<-SIGMA-HTH+SIGMA-HTH<-PIN<-SPOUT | 1072 | Bacteria>Terrabacteria_group>Bacillota>Clostridia>Eubacteriales | GCA_000987045.1 | Clostridiales_bacterium_PH28_bin88 | SY88_02270 | 81238..84456 | 1605376 | ATPase-AAA-[Clostridiales-bacterium-PH28_bin88] |
| NHM26610.1 | AAA+ATPase+wHTH+Alpha+Beta-domain+wHTH+wHTH+RRM | Type-2_BR.RM.like.systems | PolB-NTase->Polbeta-><-Peptidase_S51<-HKD-DNase+SF1-SF2-Helicase+iREase+CTD<-METHYLASE<-DUF3780<-DUF499*<-PIN<-MetJarc-HTH<-REase | 1073 | Bacteria>Terrabacteria_group>Bacillota>Clostridia>Eubacteriales | GCA_011393055.1 | Desulfofundulus_sp._TPOSR | G7K71_06325 | 1300314..1303535 | 2714340 | ATP-binding-protein-[Desulfofundulus-sp.-TPOSR] |
| NLW91010.1 | AAA+ATPase+wHTH+Alpha+Beta-domain+wHTH+wHTH+RRM | Type-2_BR.RM.like.systems | HTH+WYL->HDRnase-><-REase\|\|DUF499*->DUF3780->METHYLASE->HKD-DNase+SF1-SF2-Helicase+iREase+CTD->?->MqsA_antitoxin+HTH_3-><-DUF932 | 1076 | Bacteria>Terrabacteria_group>Bacillota>Clostridia>Eubacteriales | GCA_012729835.1 | Syntrophomonadaceae_bacterium | GXY34_05360 | 7265..10495 | 2093811 | ATP-binding-protein-[Syntrophomonadaceae-bacterium] |
| MBF7084128.1 | AAA+ATPase+wHTH+Alpha+Beta-domain+wHTH+wHTH+RRM | Type-2_BR.RM.like.systems | REase-><-?<-?\|\|DUF499*->DUF3780->METHYLASE->HKD-DNase+SF1-SF2-Helicase+iREase+CTD->PolB->?->PemK_toxin-> | 1081 | Bacteria>Terrabacteria_group>Bacillota>Clostridia>Eubacteriales | GCA_015594625.1 | Desulfallas_sp._Bu1-1 | IT084_14245 | 47348..50593 | 2787620 | ATP-binding-protein-[Desulfallas-sp.-Bu1-1] |
| MCF8011662.1 | AAA+ATPase+wHTH+Alpha+Beta-domain+wHTH+wHTH+RRM | Type-2_BR.RM.like.systems | <-TPR<-?<-ParB+HNH<-HKD-DNase+SF1-SF2-Helicase+iREase+CTD<-METHYLASE<-DUF3780<-DUF499*<-PrlF_antitoxin | 1080 | Bacteria>Terrabacteria_group>Bacillota>Clostridia>Eubacteriales | GCA_021736925.1 | Clostridiales_bacterium | K9L56_10290 | 19311..22553 | 1898207 | DUF499-domain-containing-protein-[Clostridiales-bacterium] |
| WP_027363592.1 | AAA+ATPase+wHTH+Alpha+Beta-domain+wHTH+wHTH+RRM | Type-2_BR.RM.like.systems | <-HKD-DNase+SF1-SF2-Helicase+iREase+CTD<-METHYLASE<-DUF3780<-DUF499*<-?<-?<-?<-?<-Peptidase_M24 | 1075 | Bacteria>Terrabacteria_group>Bacillota>Clostridia>Eubacteriales | GCF_000430885.1 | Desulfotruncus_alcoholivorax | H569_RS0101950 | 125864..129091 | 265477 | DUF499-domain-containing-protein-[Desulfotruncus-alcoholivorax] |
| WP_066640200.1 | AAA+ATPase+wHTH+Alpha+Beta-domain+wHTH+wHTH+RRM | Type-2_BR.RM.like.systems | <-HKD-DNase+SF1-SF2-Helicase+iREase+CTD<-METHYLASE<-DUF3780<-DUF499*<-PrlF_antitoxin\|\|HIN-HTH-> | 1079 | Bacteria>Terrabacteria_group>Bacillota>Clostridia>Eubacteriales | GCF_001592105.1 | Desulfolucanica_intricata | DIN01_RS13810 | 34611..37850 | 1285191 | DUF499-domain-containing-protein-[Desulfolucianica-intricata] |
| WP_078664716.1 | AAA+ATPase+wHTH+Alpha+Beta-domain+wHTH+wHTH+RRM | Type-2_BR.RM.like.systems | <-REase<-?<-?<-HKD-DNase+SF1-SF2-Helicase+iREase+CTD<-METHYLASE<-DUF3780<-DUF499*<-DndB\|\|HTH_3-> | 1075 | Bacteria>Terrabacteria_group>Bacillota>Clostridia>Eubacteriales | GCF_003047205.1 | Carboxydocella | CFE_RS10845 | 2156909..2160136 | 178898 | MULTISPECIES:-DUF499-domain-containing-protein-[Carboxydocella] |
| WP_134215527.1 | AAA+ATPase+wHTH+Alpha+Beta-domain+wHTH+wHTH+RRM | Type-2_BR.RM.like.systems | <-SMODS<-HKD-DNase+SF1-SF2-Helicase+iREase+CTD<-METHYLASE<-DUF3780<-DUF499*<-PrlF_antitoxin<-SF1-SF2-Helicase<-DUF429<-DnaB | 1074 | Bacteria>Terrabacteria_group>Bacillota>Clostridia>Eubacteriales | GCF_004369225.1 | Pelotomaculum_propionicicum | Pmgp_RS17020 | 9683..12907 | 258475 | DUF499-domain-containing-protein-[Pelotomaculum-propionicicum] |
| WP_161821648.1 | AAA+ATPase+wHTH+Alpha+Beta-domain+wHTH+wHTH+RRM | Type-2_BR.RM.like.systems | <-wHTH+SAD+HNH<-REase<-HEPN<-HKD-DNase+SF1-SF2-Helicase+iREase+CTD<-METHYLASE<-DUF3780<-DUF499*\|\|?-><-?<-?<-Glycos_trans_3N+Glycos_transf_3\|\|?->HAMP+HISKIN-> | 1082 | Bacteria>Terrabacteria_group>Bacillota>Clostridia>Eubacteriales | GCF_009932395.1 | Sporotomaculum_syntrophicum | SPSYN_RS06335 | 287290..290538 | 182264 | DUF499-domain-containing-protein-[Sporotomaculum-syntrophicum] |
| WP_161844019.1 | AAA+ATPase+wHTH+Alpha+Beta-domain+wHTH+wHTH+RRM | Type-2_BR.RM.like.systems | TetR-HTH-><-HKD-DNase+SF1-SF2-Helicase+iREase+CTD<-METHYLASE<-DUF3780<-DUF499*\|\|?->ABhydrolase-><-?<-HHH+ENDO3+NUDIX | 1078 | Bacteria>Terrabacteria_group>Bacillota>Clostridia>Eubacteriales | GCF_009935845.1 | Pseudoflavonifractor_sp._524-17 | D1159_RS02025 | 177996..181232 | 2304577 | DUF499-domain-containing-protein-[Pseudoflavonifractor-sp.-524-17] |
| WP_073162464.1 | AAA+ATPase+wHTH+Alpha+Beta-domain+wHTH+wHTH+RRM | Type-2_BR.RM.like.systems | <-Peptidase_S51<-HKD-DNase+SF1-SF2-Helicase+iREase+CTD<-METHYLASE<-DUF3780<-DUF499*<-PIN<-MetJarc-HTH\|\|DEDD_Tnp_IS110+Transposase_20-> | 1072 | Bacteria>Terrabacteria_group>Bacillota>Clostridia>Eubacteriales | GCF_900129285.1 | Desulfofundulus_australicus | BUB44_RS00895 | 187151..190369 | 1566 | DUF499-domain-containing-protein-[Desulfofundulus-australicus] |
| WP_072867394.1 | AAA+ATPase+wHTH+Alpha+Beta-domain+wHTH+wHTH+RRM | Type-2_BR.RM.like.systems | OrfB_IS605+OrfB_Zn_ribbon->REase->?->Radical-SAM->MetJarc-HTH->PIN->DUF499*->DUF3780->METHYLASE->HKD-DNase+SF1-SF2-Helicase+iREase+CTD->Peptidase_S51-> | 1073 | Bacteria>Terrabacteria_group>Bacillota>Clostridia>Eubacteriales | GCF_900142025.1 | Desulfofundulus_thermosubterraneus | BUA65_RS03535 | 7382..10603 | 348840 | DUF499-domain-containing-protein-[Desulfofundulus-thermosubterraneus] |
| WP_141263005.1 | AAA+ATPase+wHTH+Alpha+Beta-domain+wHTH+wHTH+RRM | Type-2_BR.RM.like.systems | <-HKD-DNase+SF1-SF2-Helicase+iREase+CTD<-METHYLASE<-DUF3780<-DUF499*\|\|?-><-?\|\|Phd-YefM->PIN->REase->Polbeta-> | 1084 | Bacteria>Terrabacteria_group>Bacillota>Clostridia>Moorellales | GCF_006538365.1 | Moorella_sp._E308F | E308F_RS01830 | 387704..390958 | 2572682 | DUF499-domain-containing-protein-[Moorella-sp.-E308F] |
| WP_156271943.1 | AAA+ATPase+wHTH+Alpha+Beta-domain+wHTH+wHTH+RRM | Type-2_BR.RM.like.systems | DSBH-><-HKD-DNase+SF1-SF2-Helicase+iREase+CTD<-METHYLASE<-DUF3780<-DUF499*<-MNS+HTH+REase<-RVT<-?<-?\|\|OLD-ABC-p->OLD-ABC-> | 1074 | Bacteria>Terrabacteria_group>Bacillota>Clostridia>Moorellales | GCF_009735625.1 | Moorella_glycerini | MGLY_RS03970 | 823454..826678 | 55779 | DUF499-domain-containing-protein-[Moorella-glycerini] |
| WP_075515826.1 | AAA+ATPase+wHTH+Alpha+Beta-domain+wHTH+wHTH+RRM | Type-2_BR.RM.like.systems | DUF499*->DUF3780->METHYLASE->HKD-DNase+SF1-SF2-Helicase+iREase+CTD-> | 1074 | Bacteria>Terrabacteria_group>Bacillota>Clostridia>Moorellales | - | Moorella_thermoacetica | MOTE_RS01400 | 88708..91932 | 1525 | DUF499-domain-containing-protein-[Moorella-thermoacetica] |
| NLL17843.1 | AAA+ATPase+wHTH+Alpha+Beta-domain+wHTH+wHTH+RRM | Type-2_BR.RM.like.systems | TOPRIM->RelE-ParE->?-><-SF1-SF2-Helicase\|\|DUF499*->DUF3780->METHYLASE-> | 1076 | Bacteria>Terrabacteria_group>Bacillota>Clostridia>unclassified_Clostridia | GCA_012520135.1 | Clostridia_bacterium | GX262_02325 | 5472..8702 | 2044939 | ATP-binding-protein-[Clostridia-bacterium] |
| MBS5425877.1 | AAA+ATPase+wHTH+Alpha+Beta-domain+wHTH+wHTH+RRM | Type-2_BR.RM.like.systems | HTH_3->tRNA-><-HTH_3\|\|XIS-HTH->Phage_int_SAM_1+Phage_integrase->?->GDResolvase+Recombinase+ZnR->DUF499*->DUF3780->METHYLASE->HKD-DNase+SF1-SF2-Helicase+iREase+CTD-> | 1089 | Bacteria>Terrabacteria_group>Bacillota>Negativicutes>Acidaminococcales | GCA_018368575.1 | Phascolarctobacterium_succinatutens | KHX39_02005 | 174252..177521 | 626940 | ATP-binding-protein-[Phascolarctobacterium-succinatutens] |
| MCI7613152.1 | AAA+ATPase+wHTH+Alpha+Beta-domain+wHTH+wHTH+RRM | Type-2_BR.RM.like.systems | DUF499*->DUF3780->METHYLASE->HKD-DNase+SF1-SF2-Helicase+iREase+CTD-> | 1079 | Bacteria>Terrabacteria_group>Bacillota>Negativicutes>Selenomonadales | GCA_022786545.1 | Selenomonadaceae_bacterium | MSS66_09270 | 1698..4937 | 2652289 | DUF499-domain-containing-protein-[Selenomonadaceae-bacterium] |
| WP_080657027.1 | AAA+ATPase+wHTH+Alpha+Beta-domain+wHTH+wHTH+RRM | Type-2_BR.RM.like.systems | <-HKD-DNase+SF1-SF2-Helicase+iREase+CTD<-HEPN<-METHYLASE<-DUF3780<-DUF499*<-RelE-ParE<-Phd-YefM<-?<-?<-?<-ABC-ATPase+SbcC | 1126 | Bacteria>Terrabacteria_group>Bacillota>Negativicutes>Selenomonadales | GCF_000424065.1 | Selenomonas_ruminantium | G598_RS0111655 | 42037..45417 | 971 | DUF499-domain-containing-protein-[Selenomonas-ruminantium] |
| WP_074570871.1 | AAA+ATPase+wHTH+Alpha+Beta-domain+wHTH+wHTH+RRM | Type-2_BR.RM.like.systems | <-HIN-HTH\|\|?->DUF499*->DUF3780->METHYLASE->HEPN->HKD-DNase+SF1-SF2-Helicase+iREase+CTD-><-DOC+HTH | 1085 | Bacteria>Terrabacteria_group>Bacillota>Negativicutes>Selenomonadales | GCF_900104055.1 | Selenomonas_ruminantium | BLQ59_RS00930 | 972..4229 | 971 | DUF499-domain-containing-protein-[Selenomonas-ruminantium] |
| WP_278520348.1 | AAA+ATPase+wHTH+Alpha+Beta-domain+wHTH+wHTH+RRM | Type-2_BR.RM.like.systems | <-HTH_OrfB_IS605+OrfB_IS605+OrfB_Zn_ribbon<-?<-?<-HKD-DNase+SF1-SF2-Helicase+iREase+CTD<-METHYLASE<-DUF3780<-DUF499* | 1085 | Bacteria>Terrabacteria_group>Bacillota>Negativicutes>Selenomonadales | GCF_905194285.1 | Mitsuokella_jalaludinii | QWB65_RS08355 | 93245..96502 | 187979 | DUF499-domain-containing-protein-[Mitsuokella-jalaludinii] |
| WP_107196386.1 | AAA+ATPase+wHTH+Alpha+Beta-domain+wHTH+wHTH+RRM | Type-2_BR.RM.like.systems | SPOUT_MTase->?->?-><-Phage_integrase<-?\|\|DUF499*->DUF3780->METHYLASE->HKD-DNase+SF1-SF2-Helicase+iREase+CTD->Schlafen->BirA-HTH->DOC+HTH-> | 1081 | Bacteria>Terrabacteria_group>Bacillota>Negativicutes>Veillonellales | GCF_003367905.1 | Megasphaera_stantonii | DKB62_RS10425 | 2193335..2196580 | 2144175 | DUF499-domain-containing-protein-[Megasphaera-stantonii] |
| MCY3660412.1 | AAA+ATPase+wHTH+Alpha+Beta-domain+wHTH+wHTH+RRM | Type-2_BR.RM.like.systems | <-DUF499*<-CR-ATPase8+REase | 943 | Bacteria>Terrabacteria_group>Chloroflexota>Caldilineae>Caldilineales | GCA_026705565.1 | Caldilineaceae_bacterium | OXG36_15520 | 517..3348 | 2614250 | MAG:-DUF499-domain-containing-protein-[Caldilineaceae-bacterium] |
| MBC7234301.1 | AAA+ATPase+wHTH+Alpha+Beta-domain+wHTH+wHTH+RRM | Type-2_BR.RM.like.systems | Peptidase_M20-><-?\|\|?->DUF499*->DUF3780->METHYLASE->HKD-DNase+SF1-SF2-Helicase+iREase+CTD-> | 1024 | Bacteria>Terrabacteria_group>Chloroflexota>unclassified_Chloroflexota | GCA_014360745.1 | Chloroflexota_bacterium | H5T69_00550 | 12856..15930 | 2026724 | ATP-binding-protein-[Chloroflexi-bacterium] |
| WP_013706311.1 | AAA+ATPase+wHTH+Alpha+Beta-domain+wHTH+wHTH+RRM | Type-2_BR.RM.like.systems | DUF499*->DUF3780->?->METHYLASE->?->HKD-DNase+SF1-SF2-Helicase+iREase+CTD-><-HKD-DNase | 1084 | Bacteria>Thermodesulfobacteriota>Desulfobaccia>Desulfobaccales>Desulfobaccaceae | GCF_000195295.1 | Desulfobacca_acetoxidans | DESAC_RS06700 | 1547695..1550949 | 60893 | DUF499-domain-containing-protein-[Desulfobacca-acetoxidans] |
| WP_028324165.1 | AAA+ATPase+wHTH+Alpha+Beta-domain+wHTH+wHTH+RRM | Type-2_BR.RM.like.systems | <-METHYLASE-p<-METHYLASE<-TOPRIM<-OLD-ABC<-DUF3780<-?<-DUF499*\|\|?-><-?<-ORC-CDC6+REase\|\|ACET+ACET-> | 1083 | Bacteria>Thermodesulfobacteriota>Desulfobacteria>Desulfobacterales>Desulfatirhabdiaceae | GCF_000429925.1 | Desulfatirhabdium_butyrativorans | G492_RS0107615 | 74798..78049 | 340467 | DUF499-domain-containing-protein-[Desulfatirhabdium-butyrativorans] |
| MCF8137378.1 | AAA+ATPase+wHTH+Alpha+Beta-domain+wHTH+wHTH+RRM | Type-2_BR.RM.like.systems | <-HKD-DNase+SF1-SF2-Helicase+iREase+CTD<-METHYLASE<-DUF3780<-DUF499*<-AbiEii<-AbiEi_3_N+AbiEi_3<-URI-DNase\|\|ParD_like->RelE-ParE-> | 1080 | Bacteria>Thermodesulfobacteriota>Desulfobacteria>Desulfobacterales>Desulfobacteraceae | GCA_021737695.1 | Desulfotignum_sp. | K9K63_08725 | 19501..22743 | 2049434 | DUF499-domain-containing-protein-[Desulfotignum-sp.] |
| WP_111958333.1 | AAA+ATPase+wHTH+Alpha+Beta-domain+wHTH+wHTH+RRM | Type-2_BR.RM.like.systems | <-HEPN\|\|NACHT-><-HKD-DNase+SF1-SF2-Helicase+iREase+CTD<-METHYLASE<-DUF3780<-DUF499*<-?\|\|Phage_int_SAM_4+Phage_integrase->?->?->?->SMODS-> | 1076 | Bacteria>Thermodesulfobacteriota>Desulfobacteria>Desulfobacterales>Desulfobacteraceae | GCF_004319545.1 | Desulfobacter_hydrogenophilus | G3N28_RS13820 | 47377..50607 | 2291 | DUF499-domain-containing-protein-[Desulfobacter-hydrogenophilus] |
| WP_178368050.1 | AAA+ATPase+wHTH+Alpha+Beta-domain+wHTH+wHTH+RRM | Type-2_BR.RM.like.systems | DUF499*->DUF3780->METHYLASE->HKD-DNase+SF1-SF2-Helicase+iREase+CTD-><-SF1-SF2-Helicase | 1079 | Bacteria>Thermodesulfobacteriota>Desulfobacteria>Desulfobacterales>Desulfobacteraceae | GCF_013389365.1 | Desulfobacter_latus | HXW94_RS16750 | 239..3478 | 2292 | DUF499-domain-containing-protein-[Desulfobacter-latus] |
| WP_205241274.1 | AAA+ATPase+wHTH+Alpha+Beta-domain+wHTH+wHTH+RRM | Type-2_BR.RM.like.systems | <-HKD-DNase+SF1-SF2-Helicase+iREase+CTD<-METHYLASE<-DUF3780<-DUF499*<-ParB+HNH-p<-?<-DOC+DOC<-HTH_31 | 1077 | Bacteria>Thermodesulfobacteriota>Desulfobulbia>Desulfobulbales>Desulfobulbaceae | GCF_016918545.1 | Desulfobulbus_alkaliphilus | JWG43_RS02730 | 133485..136718 | 869814 | DUF499-domain-containing-protein-[Desulfobulbus-alkaliphilus] |
| WP_045216075.1 | AAA+ATPase+wHTH+Alpha+Beta-domain+wHTH+wHTH+RRM | Type-2_BR.RM.like.systems | TUDOR+SF1-SF2-Helicase+iREase+CTD+REase->?->PIN->?->?->DUF499*->DUF3780->METHYLASE->PolB->HEPN-> | 1078 | Bacteria>Thermodesulfobacteriota>Desulfovibrionia>Desulfovibrionales>Desulfonatronovibrionaceae | GCF_000934755.1 | Desulfonatronovibrio_magnus | LZ23_RS16895 | 146926..150162 | 698827 | DUF499-domain-containing-protein-[Desulfonatronovibrio-magnus] |
| MCY4434133.1 | AAA+ATPase+wHTH+Alpha+Beta-domain+wHTH+wHTH+RRM | Type-2_BR.RM.like.systems | HTH+UbE1->?-><-?<-?\|\|DUF499-p->DUF499*->DUF3780->METHYLASE->HKD-DNase+SF1-SF2-Helicase+iREase+CTD->?-><-SF1-SF2-Helicase | 768 | Bacteria>unclassified_Bacteria | GCA_026710925.1 | bacterium | OXE45_10725 | 18441..20747 | 1869227 | MAG:-hypothetical-protein-OXE45_10725-[bacterium] |
| MCB2391437.1 | AAA+ATPase+wHTH+Alpha+Beta-domain+wHTH+wHTH+RRM | Type-2_BR.RM.like.systems | <-PAIREDC-HTH<-REase+SF1-SF2-Helicase\|\|MazG-><-HKD-DNase+SF1-SF2-Helicase+iREase+CTD<-METHYLASE<-DUF3780<-DUF499*\|\|ParB+HNH-><-REase\|\|DUF429->NERD-> | 1092 | - | - | - | LGT36_RS12815 | 2648905..2652183 | - | DUF499-domain-containing-protein-[Demequina-sp.-TMPB413] |
| WP_229109768.1 | AAA+ATPase+wHTH+Alpha+Beta-domain+wHTH+Fn-III+Fn-III+RRM | Type-2_BR.RM.like.systems | Phage_integrase->DUF499*->DUF3780->METHYLASE->HKD-DNase+SF1-SF2-Helicase+iREase+CTD->?->REase-><-CoiA | 1115 | Archaea>Euryarchaeota>Stenosarchaea_group>Halobacteria>Halobacteriales | GCF_017094525.1 | Halapricum_desulfuricans | HSR122_RS10640 | 2060643..2063990 | 2841257 | DUF499-domain-containing-protein-[Halapricum-desulfuricans] |
| WP_285253509.1 | AAA+ATPase+wHTH+Alpha+Beta-domain+wHTH+Fn-III+Fn-III+RRM | Type-2_BR.RM.like.systems | <-Phage_integrase\|\|?-><-HKD-DNase+SF1-SF2-Helicase+iREase+CTD<-METHYLASE<-DUF3780<-DUF499*<-?<-?<-?<-?\|\|Phage_integrase-> | 1030 | Archaea>Euryarchaeota>Stenosarchaea_group>Halobacteria>Halobacteriales | GCF_030250185.1 | Halobacterium_salinarum | PNP84_RS07925 | 14471..17563 | 2242 | DUF499-domain-containing-protein-[Halobacterium-salinarum] |
| ELY67372.1 | AAA+ATPase+wHTH+Alpha+Beta-domain+wHTH+Fn-III+Fn-III+RRM | Type-2_BR.RM.like.systems | DUF499*->DUF3780-><-?<-?\|\|METHYLASE-p->HKD-DNase+SF1-SF2-Helicase+iREase+CTD-> | 963 | Archaea>Euryarchaeota>Stenosarchaea_group>Halobacteria>Natrialbales | GCA_000337695.1 | Natronococcus_jeotgali_DSM_18795 | C492_00120 | 208..3099 | 1227498 | hypothetical-protein-C492_00120-[Natronococcus-jeotgali-DSM-18795] |
| WP_204365588.1 | AAA+ATPase+wHTH+Alpha+Beta-domain+wHTH+Fn-III+Fn-III+RRM | Type-2_BR.RM.like.systems | DUF499*->DUF3780->METHYLASE->HKD-DNase+SF1-SF2-Helicase+iREase+CTD-> | 989 | Archaea>Euryarchaeota>Stenosarchaea_group>Halobacteria>Natrialbales | GCF_000337695.1 | Natronococcus_jeotgali | C492_RS00100 | 130..3099 | 413812 | DUF499-domain-containing-protein-[Natronococcus-jeotgali] |
| WP_119720856.1 | AAA+ATPase+wHTH+Alpha+Beta-domain+wHTH+Fn-III+Fn-III+RRM | Type-2_BR.RM.like.systems | <-Methyltransf<-?<-EVE<-HKD-DNase+SF1-SF2-Helicase<-METHYLASE<-DUF3780<-DUF499*<-CARF+HTH\|\|MORC-><-SF1-SF2-Helicase+REase<-REase\|\|?->McrC-NTD+REase-> | 1047 | Archaea>Euryarchaeota>Methanomada_group>Methanococci>Methanococcales | GCF_000828555.1 | Methanococcus_maripaludis | B5K38_RS04185 | 779454..782597 | 39152 | DUF499-domain-containing-protein-[Methanococcus-maripaludis] |
| KKG11630.1 | AAA+ATPase+wHTH+Alpha+Beta-domain+wHTH+Fn-III+Fn-III+RRM | Type-2_BR.RM.like.systems | <-ParA-Soj-PloopNTPase\|\|?->HD-><-?\|\|DUF499*->DUF3780->METHYLASE->HKD-DNase+SF1-SF2-Helicase-> | 1033 | Archaea>Euryarchaeota>Stenosarchaea_group>Methanomicrobia>Methanosarcinales | GCA_000979415.1 | Methanosarcina_sp._2.H.T.1A.15 | EO97_16340 | 24902..28003 | 1483596 | hypothetical-protein-EO97_16340-[Methanosarcina-sp.-2.H.T.1A.15] |
| OPY16698.1 | AAA+ATPase+wHTH+Alpha+Beta-domain+wHTH+Fn-III+Fn-III+RRM | Type-2_BR.RM.like.systems | DUF499*->DUF3780->METHYLASE->HKD-DNase+SF1-SF2-Helicase-><-SF1-SF2-Helicase<-?<-SF1-SF2-Helicase-p | 1020 | Archaea>Euryarchaeota>Stenosarchaea_group>Methanomicrobia>Methanosarcinales | GCA_002067265.1 | Methanomethylovorans_sp._PtaU1.Bin073 | A4E23_01684 | 10719..13781 | 1811678 | MAG:-hypothetical-protein-A4E23_01684-[Methanomethylovorans-sp.-PtaU1.Bin073] |
| MBP1909178.1 | AAA+ATPase+wHTH+Alpha+Beta-domain+wHTH+Fn-III+Fn-III+RRM | Type-2_BR.RM.like.systems | <-CARF+HTH\|\|SF1-SF2-Helicase+REase->?->?->DUF499*->DUF3780->METHYLASE->HKD-DNase+SF1-SF2-Helicase-> | 1020 | Archaea>Euryarchaeota>Stenosarchaea_group>Methanomicrobia>Methanosarcinales | GCA_017873415.1 | Methanolobus_bombayensis | J2746_001390 | 249854..252916 | 38023 | hypothetical-protein-[Methanolobus-bombayensis] |

| Protein-ID | DUF499-architecture | System | Gene-Neighbourhoods | Length | Taxonomy | Assembly-ID | Strain | Locus-Tag | Genomic-coordinates | TaxID | Fasta-define |
|---|---|---|---|---|---|---|---|---|---|---|---|
| WP_156161107.1 | AAA+ATPase+wHTH+Alpha+Beta-domain+wHTH+Fn-III+Fn-III+RRM | Type-2_BR.RM.like.systems | <-ParA-Soj-PloopNTPase\|\|?->HD-><-?\|\|ParB+HNH->?->DUF499*->DUF3780->METHYLASE->HKD-DNase+SF1-SF2-Helicase-> | 1034 | Archaea>Euryarchaeota>Stenosarchaea_group>Methanomicrobia>Methanosarcinales | GCF_000979415.1 | unclassified_Methanosarcina | EO97_RS16030 | 24899..28003 | 2644672 | MULTISPECIES:-DUF499-domain-containing-protein-[unclassified-Methanosarcina] |
| WP_245250036.1 | AAA+ATPase+wHTH+Alpha+Beta-domain+wHTH+Fn-III+Fn-III+RRM | Type-2_BR.RM.like.systems | <-CARF+HTH\|\|SF1-SF2-Helicase+REase->?->?->DUF499*->METHYLASE->HKD-DNase+SF1-SF2-Helicase->?->?->?->SF1-SF2-Helicase-> | 921 | Archaea>Euryarchaeota>Stenosarchaea_group>Methanomicrobia>Methanosarcinales | GCF_017873415.1 | Methanolobus_bombayensis | J2746_RS06815 | 250151..252916 | 38023 | DUF499-domain-containing-protein-[Methanolobus-bombayensis] |
| WP_091690449.1 | AAA+ATPase+wHTH+Alpha+Beta-domain+wHTH+Fn-III+Fn-III+RRM | Type-2_BR.RM.like.systems | Phage_integrase->?->SF1-SF2-Helicase->URI-DNase+RAMA->DUF499*->DUF3780->METHYLASE->?->HKD-DNase+SF1-SF2-Helicase-><-?<-DUF499-p | 1021 | Archaea>Euryarchaeota>Stenosarchaea_group>Methanomicrobia>Methanosarcinales | GCF_900773955.1 | Methanococcoides_vulcani | BMX02_RS09485 | 150182..153247 | 1353158 | DUF499-domain-containing-protein-[Methanococcoides-vulcani] |
| QNO47110.1 | AAA+ATPase+wHTH+Alpha+Beta-domain+wHTH+Fn-III+Fn-III+RRM | Type-2_BR.RM.like.systems | <-METHYLASE<-DUF3780<-DUF499*<-METHYLASE-p<-?<-?<-Aminotran_1_2\|\|REase-> | 1021 | Archaea>Euryarchaeota>Stenosarchaea_group>Methanomicrobia>Methanosarcinales | - | Methanosarcinales_archaeon_ANME-2c_ERB4 | PLINODHC_00004 | 4889..7954 | 2759911 | hypothetical-protein-PLINODHC_00004-[Methanosarcinales-archaeon-ANME-2c-ERB4] |
| MCK9404871.1 | AAA+ATPase+wHTH+Alpha+Beta-domain+wHTH+Fn-III+Fn-III+RRM | Type-2_BR.RM.like.systems | DUF499*->DUF3780->METHYLASE->HKD-DNase+SF1-SF2-Helicase-><-SF1-SF2-Helicase<-REase\|\|PIN-> | 1027 | Archaea>Euryarchaeota>Stenosarchaea_group>Methanomicrobia>Methanotrichales | GCA_023229265.1 | Methanothrix_sp. | M0Q47_00445 | 3470..6553 | 90426 | DUF499-domain-containing-protein-[Methanothrix-sp.] |
| WP_211424620.1 | AAA+ATPase+wHTH+Alpha+Beta-domain+wHTH+Fn-III+Fn-III+RRM | Type-2_BR.RM.like.systems | <-HEPN<-PemK_toxin<-OrfB_IS605+OrfB_Zn_ribbon<-?<-?<-DUF3780<-DUF499*<-?<-?<-?<-SIGMA-HTH | 1041 | Bacteria>Acidobacteriota>Blastocatellia>Chloracidobacterium | GCF_018304705.1 | Chloracidobacterium_aggregatum | J8C07_RS14540 | 703515..706640 | 2851959 | DUF499-domain-containing-protein-[Chloracidobacterium-aggregatum] |
| WP_211427121.1 | AAA+ATPase+wHTH+Alpha+Beta-domain+wHTH+Fn-III+Fn-III+RRM | Type-2_BR.RM.like.systems | SIGMA-HTH->?->?->DUF499*->DUF3780->?->?->PemK_toxin->METHYLASE->HEPN-> | 1041 | Bacteria>Acidobacteriota>Blastocatellia>Chloracidobacterium | GCF_018304805.1 | Chloracidobacterium_aggregatum | J8C02_RS13600 | 632551..635676 | 2851959 | DUF499-domain-containing-protein-[Chloracidobacterium-aggregatum] |
| MCA2968203.1 | AAA+ATPase+wHTH+Alpha+Beta-domain+wHTH+Fn-III+Fn-III+RRM | Type-2_BR.RM.like.systems | XIS-HTH->DUF499*->DUF3780->METHYLASE->HKD-DNase+SF1-SF2-Helicase->?-><-?<-TraH | 1042 | Bacteria>Acidobacteriota>Terriglobia>Terriglobales>Acidobacteriaceae | GCA_020247055.1 | Acidobacteriaceae_bacterium | INH43_06790 | 1502..4630 | 2052142 | ATP-binding-protein-[Acidobacteriaceae-bacterium] |
| MBM63538.1 | AAA+ATPase+wHTH+Alpha+Beta-domain+wHTH+Fn-III+Fn-III+RRM | Type-2_BR.RM.like.systems | DUF499*->DUF3780->METHYLASE->HKD-DNase+SF1-SF2-Helicase-> | 1033 | Bacteria>Acidobacteriota>unclassified_Acidobacteriota | GCA_002722875.1 | Acidobacteriota_bacterium | CL484_11380 | 512..3613 | 1978231 | hypothetical-protein-[Acidobacteria-bacterium] |
| MBN1569786.1 | AAA+ATPase+wHTH+Alpha+Beta-domain+wHTH+Fn-III+Fn-III+RRM | Type-2_BR.RM.like.systems | <-wHTH+SAD+HNH<-?<-HKD-DNase+SF1-SF2-Helicase<-Schlafen+HATPase+HTH<-METHYLASE<-DUF3780<-DUF499*<-?<-HTH+WYL | 1021 | Bacteria>Acidobacteriota>unclassified_Acidobacteriota | GCA_016929335.1 | Acidobacteriota_bacterium | JXA73_18215 | 77162..80227 | 1978231 | DUF499-domain-containing-protein-[Acidobacteria-bacterium] |
| MDE2058553.1 | AAA+ATPase+wHTH+Alpha+Beta-domain+wHTH+Fn-III+Fn-III+RRM | Type-2_BR.RM.like.systems | HTH_31->Peptidase-><-HKD-DNase+SF1-SF2-Helicase<-?<-METHYLASE<-DUF3780<-DUF499* | 1027 | Bacteria>Bacteria_incertae_sedis>Bacteria_candidate_phyla>candidate_division_NC10>unclassified_candidate_division_NC10 | GCA_028864815.1 | candidate_division_NC10_bacterium | KGJ27_02330 | 29205..32288 | 2072417 | MAG:-DUF499-domain-containing-protein-[candidate-division-NC10-bacterium] |
| MBF0543186.1 | AAA+ATPase+wHTH+Alpha+Beta-domain+wHTH+Fn-III+Fn-III+RRM | Type-2_BR.RM.like.systems | ABhydrolase->HAMP+ACYC->?-><-TPR+TPR+TPR+TPR<-?\|\|?->DUF499*->DUF3780->METHYLASE->HKD-DNase+SF1-SF2-Helicase-><-ACET<-?<-TPR+TPR+TPR+TPR | 1022 | Bacteria>Bacteria_incertae_sedis>Bacteria_candidate_phyla>Candidatus_Riflebacteria>unclassified_Candidatus_Riflebacteria | GCA_015233565.1 | Candidatus_Riflebacteria_bacterium | HQM08_02070 | 217810..220878 | 2053586 | ATP-binding-protein-[Candidatus-Riflebacteria-bacterium] |
| NLT83945.1 | AAA+ATPase+wHTH+Alpha+Beta-domain+wHTH+Fn-III+Fn-III+RRM | Type-2_BR.RM.like.systems | <-REase+DUF5655\|\|DUF499*->DUF3780->METHYLASE->HKD-DNase+SF1-SF2-Helicase-> | 1103 | Bacteria>Coprothermobacterota>Coprothermobacteria>Coprothermobacterales>Coprothermobacteraceae | GCA_012719895.1 | Coprothermobacter_proteolyticus | GXW92_04120 | 966..4277 | 35786 | ATP-binding-protein-[Coprothermobacter-proteolyticus] |
| WP_018962942.1 | AAA+ATPase+wHTH+Alpha+Beta-domain+wHTH+Fn-III+Fn-III+RRM | Type-2_BR.RM.like.systems | ZnR-><-?<-HKD-DNase+SF1-SF2-Helicase<-METHYLASE<-DUF3780<-DUF499*\|\|?->?->?->?->?->DnaB+SMS-KaiC-> | 1104 | Bacteria>Coprothermobacterota>Coprothermobacteria>Coprothermobacterales>Coprothermobacteraceae | GCF_000378005.1 | Coprothermobacter_platensis | D890_RS0102405 | 440158..443472 | 108819 | DUF499-domain-containing-protein-[Coprothermobacter-platensis] |
| MBL4824170.1 | AAA+ATPase+wHTH+Alpha+Beta-domain+wHTH+Fn-III+Fn-III+RRM | Type-2_BR.RM.like.systems | <-RelE-ParE<-?<-?<-HKD-DNase+SF1-SF2-Helicase<-DUF3780<-DUF499*<-?<-?<-?<-ABC-ATPase+SbcC<-?<-Aminotran_1_2 | 1039 | Bacteria>delta/epsilon_subdivisions>Deltaproteobacteria>Deltaproteobacteria_incertae_sedis>SAR324_cluster | GCA_016764995.1 | SAR324_cluster_bacterium | JKY07_02795 | 10847..13966 | 2024889 | DUF499-domain-containing-protein-[SAR324-cluster-bacterium] |
| NLT02438.1 | AAA+ATPase+wHTH+Alpha+Beta-domain+wHTH+Fn-III+Fn-III+RRM | Type-2_BR.RM.like.systems | <-ParB+HNH<-?<-HKD-DNase+SF1-SF2-Helicase<-Schlafen+HATPase+HTH<-METHYLASE<-DUF3780<-DUF499*<-tRNA\|\|?->?->NADAR->?->Aminotran_1_2-> | 1038 | Bacteria>FCB_group>Bacteroidota/Chlorobiota_group>Bacteroidota>Bacteroidia | GCA_012718945.1 | Bacteroidales_bacterium | GXY09_01085 | 15955..19071 | 2030927 | DUF499-domain-containing-protein-[Bacteroidales-bacterium] |
| MDD3040647.1 | AAA+ATPase+wHTH+Alpha+Beta-domain+wHTH+Fn-III+Fn-III+RRM | Type-2_BR.RM.like.systems | <-HKD-DNase+SF1-SF2-Helicase<-NERD<-SbcC<-METHYLASE<-DUF3780<-DUF499*<-?<-?<-?<-JetA | 1037 | Bacteria>FCB_group>Bacteroidota/Chlorobiota_group>Bacteroidota>Bacteroidia | GCA_028690125.1 | Bacteroides_sp. | PHO71_22645 | 8586..11699 | 29523 | MAG:-DUF499-domain-containing-protein-[Bacteroides-sp.] |
| AEW00270.1 | AAA+ATPase+wHTH+Alpha+Beta-domain+wHTH+Fn-III+Fn-III+RRM | Type-2_BR.RM.like.systems | <-HTH_19\|\|HEPN->?-><-HEPN<-?<-?\|\|DUF499*->DUF3780->METHYLASE->HKD-DNase+SF1-SF2-Helicase-> | 1036 | Bacteria>FCB_group>Bacteroidota/Chlorobiota_group>Bacteroidota>Chitinophagia | GCA_000246855.1 | Niastella_koreensis_GR20-10 | Niako_3988 | 5013582..5016692 | 700598 | hypothetical-protein-Niako_3988-[Niastella-koreensis-GR20-10] |
| WP_207622366.1 | AAA+ATPase+wHTH+Alpha+Beta-domain+wHTH+Fn-III+Fn-III+RRM | Type-2_BR.RM.like.systems | <-HTH_19\|\|HEPN->?-><-HEPN<-?<-?\|\|DUF499*->DUF3780->METHYLASE->HKD-DNase+SF1-SF2-Helicase-> | 992 | Bacteria>FCB_group>Bacteroidota/Chlorobiota_group>Bacteroidota>Chitinophagia | GCF_000246855.1 | Niastella_koreensis | NIAKO_RS19575 | 5013714..5016692 | 354356 | DUF499-domain-containing-protein-[Niastella-koreensis] |
| WP_085411740.1 | AAA+ATPase+wHTH+Alpha+Beta-domain+wHTH+Fn-III+Fn-III+RRM | Type-2_BR.RM.like.systems | <-HKD-DNase+SF1-SF2-Helicase<-?<-HTH<-?<-METHYLASE<-DUF3780<-DUF499*<-tRNA<-Peptidase_S74\|\|?-><-?<-?<-Hpr-Kinase-N | 1029 | Bacteria>FCB_group>Bacteroidota/Chlorobiota_group>Bacteroidota>Cytophagia | - | Fibrella_sp._ES10-3-2-2 | A6C57_RS08645 | 2083632..2086721 | 1834519 | DUF499-domain-containing-protein-[Fibrella-sp.-ES10-3-2-2] |
| WP_135464405.1 | AAA+ATPase+wHTH+Alpha+Beta-domain+wHTH+Fn-III+Fn-III+RRM | Type-2_BR.RM.like.systems | <-URI-DNase<-HKD-DNase<-AbiEii<-AbiEii<-?\|\|DUF499*->DUF3780->METHYLASE->HKD-DNase+SF1-SF2-Helicase-><-tRNA\|\|?->?->ACET-> | 1028 | Bacteria>FCB_group>Bacteroidota/Chlorobiota_group>Bacteroidota>Cytophagia | GCF_004765605.1 | Hymenobacter_aquaticus | E5K00_RS16470 | 902079..905165 | 1867101 | DUF499-domain-containing-protein-[Hymenobacter-aquaticus] |
| MBF8982611.1 | AAA+ATPase+wHTH+Alpha+Beta-domain+wHTH+Fn-III+Fn-III+RRM | Type-2_BR.RM.like.systems | HIN-HTH->RVT+HNH->?->HIN-HTH->?->?->DUF499*->DUF3780->METHYLASE->HKD-DNase+SF1-SF2-Helicase->?->ABhydrolase-> | 1031 | Bacteria>FCB_group>Bacteroidota/Chlorobiota_group>Bacteroidota>Flavobacteriia | GCA_015686845.1 | Lutibacter_sp._B2 | IZY60_03570 | 21818..24913 | 2789328 | DUF499-domain-containing-protein-[Lutibacter-sp.-B2] |
| MBP7407818.1 | AAA+ATPase+wHTH+Alpha+Beta-domain+wHTH+Fn-III+Fn-III+RRM | Type-2_BR.RM.like.systems | <-HKD-DNase+SF1-SF2-Helicase<-?<-?<-METHYLASE<-DUF3780<-DUF499* | 1000 | Bacteria>FCB_group>Bacteroidota/Chlorobiota_group>Bacteroidota>Flavobacteriia | GCA_017997375.1 | Flavobacteriales_bacterium | KA941_03585 | 17100..20104 | 2021391 | DUF499-domain-containing-protein-[Flavobacteriales-bacterium] |
| MCB0784563.1 | AAA+ATPase+wHTH+Alpha+Beta-domain+wHTH+Fn-III+Fn-III+RRM | Type-2_BR.RM.like.systems | DUF499*->DUF3780->METHYLASE->?->HTH->?->HKD-DNase+SF1-SF2-Helicase-> | 1024 | Bacteria>FCB_group>Bacteroidota/Chlorobiota_group>Bacteroidota>Flavobacteriia | GCA_020437915.1 | Flavobacteriales_bacterium | KDC02_10170 | 977..4051 | 2021391 | DUF499-domain-containing-protein-[Flavobacteriales-bacterium] |
| TAK42618.1 | AAA+ATPase+wHTH+Alpha+Beta-domain+wHTH+Fn-III+Fn-III+RRM | Type-2_BR.RM.like.systems | BirA-HTH->?->wHTH+REase->METHYLASE-p->DUF499*->DUF3780->ParB+HNH->OLD-ABC->RelE-ParE->METHYLASE-p-> | 1024 | Bacteria>FCB_group>Bacteroidota/Chlorobiota_group>Bacteroidota>Saprospiria | GCA_004297645.1 | Saprospiraceae_bacterium | EPO28_07065 | 17499..20573 | 2202734 | DUF499-domain-containing-protein-[Saprospiraceae-bacterium] |
| NUQ24407.1 | AAA+ATPase+wHTH+Alpha+Beta-domain+wHTH+Fn-III+Fn-III+RRM | Type-2_BR.RM.like.systems | <-ABhydrolase\|\|?->?-><-?<-?\|\|?->DUF499*->DUF3780->ParB+HNH->OLD-ABC->RelE-ParE->METHYLASE->OLD-ABC-> | 1024 | Bacteria>FCB_group>Bacteroidota/Chlorobiota_group>Bacteroidota>Saprospiria | GCA_013360405.1 | Saprospiraceae_bacterium | HUU34_10665 | 13446..16520 | 2202734 | DUF499-domain-containing-protein-[Saprospiraceae-bacterium] |
| NBB84709.1 | AAA+ATPase+wHTH+Alpha+Beta-domain+wHTH+Fn-III+Fn-III+RRM | Type-2_BR.RM.like.systems | <-HKD-DNase+SF1-SF2-Helicase<-?\|\|?-><-HTH+MPTase<-DUF3780<-DUF499* | 1021 | Bacteria>FCB_group>Bacteroidota/Chlorobiota_group>Bacteroidota>unclassified_Bacteroidota | GCA_009908985.1 | Bacteroidota_bacterium | GVY12_00600 | 31223..34288 | 1898104 | DUF499-domain-containing-protein-[Bacteroidetes-bacterium] |
| MCX6251131.1 | AAA+ATPase+wHTH+Alpha+Beta-domain+wHTH+Fn-III+Fn-III+RRM | Type-2_BR.RM.like.systems | DUF499*->DUF3780->METHYLASE->HKD-DNase+SF1-SF2-Helicase-> | 1027 | Bacteria>FCB_group>Bacteroidota/Chlorobiota_group>Bacteroidota>unclassified_Bacteroidota | GCA_026391355.1 | Bacteroidota_bacterium | NTX61_10310 | 145885..148968 | 1898104 | MAG:-DUF499-domain-containing-protein-[Bacteroidetes-bacterium] |
| WP_109646585.1 | AAA+ATPase+wHTH+Alpha+Beta-domain+wHTH+Fn-III+Fn-III+RRM | Type-2_BR.RM.like.systems | ParB+HNH->DUF499*->DUF3780->METHYLASE->HKD-DNase+SF1-SF2-Helicase->HEPN+CBS-> | 1023 | Bacteria>FCB_group>Bacteroidota/Chlorobiota_group>Balneolota>Balneolia | GCF_003150675.1 | Rhodohalobacter_mucosus | DDZ15_RS08075 | 64787..67858 | 2079485 | DUF499-domain-containing-protein-[Rhodohalobacter-mucosus] |
| WP_012507105.1 | AAA+ATPase+wHTH+Alpha+Beta-domain+wHTH+Fn-III+Fn-III+RRM | Type-2_BR.RM.like.systems | NUDIX-><-?<-TIR\|\|DUF499*->DUF3780->METHYLASE-p->DDE_Tnp_1->METHYLASE->HKD-DNase+SF1-SF2-Helicase-><-RelE-ParE | 1037 | Bacteria>FCB_group>Bacteroidota/Chlorobiota_group>Chlorobiota>Chlorobiia | GCF_000020645.1 | Pelodictyon_phaeoclathratiforme | PPHA_RS01390 | 269156..272269 | 34090 | DUF499-domain-containing-protein-[Pelodictyon-phaeoclathratiforme] |
| MBM2814383.1 | AAA+ATPase+wHTH+Alpha+Beta-domain+wHTH+Fn-III+Fn-III+RRM | Type-2_BR.RM.like.systems | <-ParB+HNH<-?<-HKD-DNase+SF1-SF2-Helicase<-METHYLASE<-DUF3780<-DUF499*<-Peptidase<-Peptidase | 1037 | Bacteria>FCB_group>Bacteroidota/Chlorobiota_group>Ignavibacteriota>Ignavibacteria | GCA_016844745.1 | Ignavibacteria_bacterium | HW421_1145 | 16386..19499 | 2053306 | hypothetical-protein-[Ignavibacteria-bacterium] |
| WP_183957933.1 | AAA+ATPase+wHTH+Alpha+Beta-domain+wHTH+Fn-III+Fn-III+RRM | Type-2_BR.RM.like.systems | <-REase<-URI-DNase\|\|RNase->DUF499*->DUF3780->METHYLASE->HKD-DNase+SF1-SF2-Helicase->URI-DNase+RAMA-> | 1021 | Bacteria>FCB_group>Bacteroidota/Chlorobiota_group>Rhodothermota>Rhodothermia | GCF_014197195.1 | Salinibacter_ruber | GGQ18_RS15525 | 40084..43149 | 146919 | DUF499-domain-containing-protein-[Salinibacter-ruber] |
| WP_259060935.1 | AAA+ATPase+wHTH+Alpha+Beta-domain+wHTH+Fn-III+Fn-III+RRM | Type-2_BR.RM.like.systems | <-URI-DNase+RAMA<-HKD-DNase+SF1-SF2-Helicase<-METHYLASE<-PIN<-HTH+MPTase<-DUF3780<-DUF499*<-RNase\|\|URI-DNase->REase-> | 1021 | Bacteria>FCB_group>Bacteroidota/Chlorobiota_group>Rhodothermota>Rhodothermia | GCF_024806505.1 | Salinibacter_ruber | GGQ15_RS12220 | 47661..50726 | 146919 | DUF499-domain-containing-protein-[Salinibacter-ruber] |
| WP_259082418.1 | AAA+ATPase+wHTH+Alpha+Beta-domain+wHTH+Fn-III+Fn-III+RRM | Type-2_BR.RM.like.systems | <-HKD-DNase+SF1-SF2-Helicase-p<-METHYLASE<-METHYLASE<-PIN<-HTH+MPTase<-DUF3780<-DUF499*<-RNase<-?\|\|REase-> | 1021 | Bacteria>FCB_group>Bacteroidota/Chlorobiota_group>Rhodothermota>Rhodothermia | GCF_024806735.1 | Salinibacter_ruber | GGP83_RS14580 | 9847..12912 | 146919 | DUF499-domain-containing-protein-[Salinibacter-ruber] |
| WP_259170624.1 | AAA+ATPase+wHTH+Alpha+Beta-domain+wHTH+Fn-III+Fn-III+RRM | Type-2_BR.RM.like.systems | <-HKD-DNase+SF1-SF2-Helicase<-METHYLASE<-METHYLASE<-PIN<-HTH+MPTase<-DUF3780<-DUF499*<-REase+35exo<-?<-?\|\|DEDD_Tnp_IS110+Transposase_20->?->DUF4096+DDE_Tnp_1+DDE_Tnp-> | 1021 | Bacteria>FCB_group>Bacteroidota/Chlorobiota_group>Rhodothermota>Rhodothermia | GCF_024807095.1 | Salinibacter_ruber | GGP42_RS07650 | 508801..511866 | 146919 | DUF499-domain-containing-protein-[Salinibacter-ruber] |
| WP_259058979.1 | AAA+ATPase+wHTH+Alpha+Beta-domain+wHTH+Fn-III+Fn-III+RRM | Type-2_BR.RM.like.systems | <-PAIREDC-HTH<-?<-ZnR<-sGTP<-?<-?\|\|tRNA->tRNA->DUF499*->DUF3780->METHYLASE->HKD-DNase+SF1-SF2-Helicase-><-?<-?<-HD | 1021 | Bacteria>FCB_group>Bacteroidota/Chlorobiota_group>Rhodothermota>Rhodothermia | GCF_024809535.1 | Salinibacter_ruber | GGQ20_RS08045 | 268747..271812 | 146919 | DUF499-domain-containing-protein-[Salinibacter-ruber] |
| WP_259066518.1 | AAA+ATPase+wHTH+Alpha+Beta-domain+wHTH+Fn-III+Fn-III+RRM | Type-2_BR.RM.like.systems | <-HKD-DNase+SF1-SF2-Helicase<-METHYLASE<-PIN<-HTH+MPTase<-DUF3780<-DUF499*<-?\|\|URI-DNase->REase-> | 1021 | Bacteria>FCB_group>Bacteroidota/Chlorobiota_group>Rhodothermota>Rhodothermia | GCF_024809675.1 | Salinibacter_ruber | GGP46_RS15095 | 82348..85413 | 146919 | DUF499-domain-containing-protein-[Salinibacter-ruber] |
| WP_251930233.1 | AAA+ATPase+wHTH+Alpha+Beta-domain+wHTH+Fn-III+Fn-III+RRM | Type-2_BR.RM.like.systems | <-REase<-URI-DNase\|\|?->RNase->DUF499*->DUF3780->HTH+MPTase->PIN->METHYLASE->HKD-DNase+SF1-SF2-Helicase-> | 1021 | Bacteria>FCB_group>Bacteroidota/Chlorobiota_group>Rhodothermota>Rhodothermia | GCF_943908145.1 | Salinibacter_ruber | NGL32_RS11815 | 75461..78526 | 146919 | DUF499-domain-containing-protein-[Salinibacter-ruber] |
| NLD91450.1 | AAA+ATPase+wHTH+Alpha+Beta-domain+wHTH+Fn-III+Fn-III+RRM | Type-2_BR.RM.like.systems | <-METHYLASE<-Glycos_transf_4\|\|?->SF1-SF2-Helicase->Peptidase_M48->?->DUF499*->DUF3780->?->METHYLASE->OLD-ABC->TOPRIM->HKD-DNase+SF1-SF2-Helicase-> | 1024 | Bacteria>FCB_group>Fibrobacterota>Fibrobacteria>Fibrobacterales | GCA_012513915.1 | Fibrobacter_sp. | GX639_02155 | 52641..55715 | 35828 | DUF499-domain-containing-protein-[Fibrobacter-sp.] |
| AKU92642.1 | AAA+ATPase+wHTH+Alpha+Beta-domain+wHTH+Fn-III+Fn-III+RRM | Type-2_BR.RM.like.systems | ParB+HNH-><-?<-?<-?\|\|?->DUF499*->DUF3780->METHYLASE-p->METHYLASE->HKD-DNase+SF1-SF2-Helicase-> | 1066 | Bacteria>Myxococcota>Myxococcia>Myxococcales>Cystobacterineae | GCA_001263175.1 | Vulgatibacter_incomptus | AKJ08_3029 | 3557369..3560569 | 1391653 | hypothetical-protein-AKJ08_3029-[Vulgatibacter-incomptus] |
| WP_050726781.1 | AAA+ATPase+wHTH+Alpha+Beta-domain+wHTH+Fn-III+Fn-III+RRM | Type-2_BR.RM.like.systems | ParB+HNH-><-?<-?\|\|?->DUF499*->DUF3780->METHYLASE-p-><-?<-?\|\|METHYLASE-p->HKD-DNase+SF1-SF2-Helicase-> | 1032 | Bacteria>Myxococcota>Myxococcia>Myxococcales>Cystobacterineae | GCF_001263175.1 | Vulgatibacter_incomptus | AKJ08_RS14855 | 3557471..3560569 | 1391653 | DUF499-domain-containing-protein-[Vulgatibacter-incomptus] |
| MBF0106878.1 | AAA+ATPase+wHTH+Alpha+Beta-domain+wHTH+Fn-III+Fn-III+RRM | Type-2_BR.RM.like.systems | DUF499*->DUF3780->METHYLASE->HKD-DNase+SF1-SF2-Helicase->TIR-> | 1043 | Bacteria>Myxococcota>Myxococcia>unclassified_Myxococcia | GCA_015228875.1 | Deltaproteobacteria_bacterium | HQM16_16320 | 2366..5497 | 2026735 | DUF499-domain-containing-protein-[Deltaproteobacteria-bacterium] |
| MBM4287907.1 | AAA+ATPase+wHTH+Alpha+Beta-domain+wHTH+Fn-III+Fn-III+RRM | Type-2_BR.RM.like.systems | DUF499*->DUF3780->METHYLASE->HKD-DNase+SF1-SF2-Helicase-> | 1023 | Bacteria>Myxococcota>Myxococcia>unclassified_Myxococcia | GCA_016874895.1 | Deltaproteobacteria_bacterium | FJ135_07120 | 2551..5622 | 2026735 | DUF499-domain-containing-protein-[Deltaproteobacteria-bacterium] |
| MBN1335625.1 | AAA+ATPase+wHTH+Alpha+Beta-domain+wHTH+Fn-III+Fn-III+RRM | Type-2_BR.RM.like.systems | REase->SF1-SF2-Helicase-><-ParB+HNH<-HKD-DNase+SF1-SF2-Helicase<-METHYLASE<-DUF3780<-DUF499*<-XIS-HTH<-?<-?<-HEPN | 1020 | Bacteria>Myxococcota>Myxococcia>unclassified_Myxococcia | GCA_016927815.1 | Deltaproteobacteria_bacterium | JXB39_06665 | 17124..20186 | 2026735 | DUF499-domain-containing-protein-[Deltaproteobacteria-bacterium] |
| MBW2095694.1 | AAA+ATPase+wHTH+Alpha+Beta-domain+wHTH+Fn-III+Fn-III+RRM | Type-2_BR.RM.like.systems | <-DUF932\|\|?->?->DUF499*->DUF3780->?->?->?->METHYLASE->HKD-DNase+SF1-SF2-Helicase-p-> | 1031 | Bacteria>Myxococcota>Myxococcia>unclassified_Myxococcia | GCA_019309145.1 | Deltaproteobacteria_bacterium | JRI80_12480 | 2126..5221 | 2026735 | DUF499-domain-containing-protein-[Deltaproteobacteria-bacterium] |
| MCY4488568.1 | AAA+ATPase+wHTH+Alpha+Beta-domain+wHTH+Fn-III+Fn-III+RRM | Type-2_BR.RM.like.systems | LD-peptidase-><-?\|\|DUF499*->DUF3780->METHYLASE->N6_MTase->REase->HKD-DNase+SF1-SF2-Helicase-> | 1042 | Bacteria>Myxococcota>Myxococcia>unclassified_Myxococcia | GCA_026712905.1 | Deltaproteobacteria_bacterium | OXF11_15860 | 11359..14487 | 2026735 | MAG:-DUF499-domain-containing-protein-[Deltaproteobacteria-bacterium] |
| MBI5442127.1 | AAA+ATPase+wHTH+Alpha+Beta-domain+wHTH+Fn-III+Fn-III+RRM | Type-2_BR.RM.like.systems | <-METHYLASE-p<-?<-PIN<-METHYLASE<-DUF3780<-DUF499* | 842 | Bacteria>Myxococcota>Myxococcia>unclassified_Myxococcia | GCA_016218905.1 | Deltaproteobacteria_bacterium | HY900_13060 | 8281..10809 | 2026735 | DUF499-domain-containing-protein-[Deltaproteobacteria-bacterium] |
| MBW2065551.1 | AAA+ATPase+wHTH+Alpha+Beta-domain+wHTH+Fn-III+Fn-III+RRM | Type-2_BR.RM.like.systems | <-HEPN<-PolB<-METHYLASE<-DUF3780<-DUF499*<-tRNA\|\|?->TUDOR+SF1-SF2-Helicase+iREase+CTD->METHYLASE-> | 1025 | Bacteria>Myxococcota>Myxococcia>unclassified_Myxococcia | GCA_019308985.1 | Deltaproteobacteria_bacterium | JRJ03_11535 | 4380..7457 | 2026735 | DUF499-domain-containing-protein-[Deltaproteobacteria-bacterium] |
| MDD3554381.1 | AAA+ATPase+wHTH+Alpha+Beta-domain+wHTH+Fn-III+Fn-III+RRM | Type-2_BR.RM.like.systems | <-ParA-Soj-PloopNTPase<-ParA-Soj-PloopNTPase<-?\|\|DUF499*->DUF3780->?->MACRODOMAIN->METHYLASE->HEPN->ParA-Soj-PloopNTPase-> | 1040 | Bacteria>Myxococcota>Myxococcia>unclassified_Myxococcia | GCA_028695735.1 | Deltaproteobacteria_bacterium | PHC35_07690 | 76771..79893 | 2026735 | MAG:-DUF499-domain-containing-protein-[Deltaproteobacteria-bacterium] |
| MCD6274302.1 | AAA+ATPase+wHTH+Alpha+Beta-domain+wHTH+Fn-III+Fn-III+RRM | Type-2_BR.RM.like.systems | DUF499*->DUF3780->METHYLASE->Schlafen-> | 1032 | Bacteria>Myxococcota>Myxococcia>unclassified_Myxococcia | GCA_021163685.1 | Deltaproteobacteria_bacterium | J7K30_16050 | 3210..6308 | 2026735 | DUF499-domain-containing-protein-[Deltaproteobacteria-bacterium] |
| MCC6665145.1 | AAA+ATPase+wHTH+Alpha+Beta-domain+wHTH+Fn-III+Fn-III+RRM | Type-2_BR.RM.like.systems | <-REase+SAVED<-HKD-DNase+SF1-SF2-Helicase<-METHYLASE<-DUF3780<-DUF499*<-XIS-HTH<-ParB | 1027 | Bacteria>Myxococcota>Polyangia>Polyangiales>Polyangiaceae | GCA_020849115.1 | Polyangiaceae_bacterium | IT375_15450 | 209262..212345 | 2268199 | DUF499-domain-containing-protein-[Polyangiaceae-bacterium] |
| MCK6571620.1 | AAA+ATPase+wHTH+Alpha+Beta-domain+wHTH+Fn-III+Fn-III+RRM | Type-2_BR.RM.like.systems | TPR+TPR->?->?->HTH_3->?->?->DUF499*->DUF3780->METHYLASE->HKD-DNase+SF1-SF2-Helicase-><-HHH+ENDO3+NUDIX | 1034 | Bacteria>Myxococcota>unclassified_Myxococcota | GCA_023150895.1 | Myxococcota_bacterium | L6V77_11080 | 5114..8218 | 2818507 | DUF499-domain-containing-protein-[Myxococcota-bacterium] |
| MBT3366889.1 | AAA+ATPase+wHTH+Alpha+Beta-domain+wHTH+Fn-III+Fn-III+RRM | Type-2_BR.RM.like.systems | ParB+HNH+Schlafen->DUF499*->DUF3780->METHYLASE->HicB_lk_antitox->HicA_toxin->HKD-DNase+SF1-SF2-Helicase->HNH-> | 1035 | Bacteria>Nitrospinae/Tectomicrobia_group>Nitrospinota>Nitrospinia>Nitrospinales | GCA_018648125.1 | Nitrospina_sp. | HN416_07030 | 2974..6081 | 2024844 | DUF499-domain-containing-protein-[Nitrospina-sp.] |
| CBK39915.1 | AAA+ATPase+wHTH+Alpha+Beta-domain+wHTH+Fn-III+Fn-III+RRM | Type-2_BR.RM.like.systems | <-HKD-DNase+SF1-SF2-Helicase<-METHYLASE<-DUF3780<-DUF499*<-RHH\|\|PhdYeFM_antitox->RelE-ParE->?->REase+SF1-SF2-Helicase->TRD+TRD-> | 1040 | Bacteria>Nitrospirota>Nitrospiria>Nitrospirales>Nitrospiraceae | GCA_000196815.1 | Nitrospira_defluvii | NIDE0131 | 105559..108681 | 330214 | conserved-protein-of-unknown-function-[Nitrospira-defluvii] |
| HBP88391.1 | AAA+ATPase+wHTH+Alpha+Beta-domain+wHTH+Fn-III+Fn-III+RRM | Type-2_BR.RM.like.systems | DUF499*->DUF3780->METHYLASE->HKD-DNase+SF1-SF2-Helicase->ParB+HNH+RAMA->HEPN-> | 1027 | Bacteria>Nitrospirota>Nitrospiria>Nitrospirales>Nitrospiraceae | GCA_003523945.1 | Nitrospiraceae_bacterium | DD706_11920 | 779..3862 | 2026770 | hypothetical-protein-[Nitrospiraceae-bacterium] |
| MCI0428429.1 | AAA+ATPase+wHTH+Alpha+Beta-domain+wHTH+Fn-III+Fn-III+RRM | Type-2_BR.RM.like.systems | <-HKD-DNase+SF1-SF2-Helicase<-METHYLASE<-DUF3780<-DUF499* | 1031 | Bacteria>Nitrospirota>Nitrospiria>Nitrospirales>Nitrospiraceae | GCA_022614835.1 | Nitrospiraceae_bacterium | LOZ46_10480 | 8376..11471 | 2026770 | DUF499-domain-containing-protein-[Nitrospiraceae-bacterium] |
| MCS6304978.1 | AAA+ATPase+wHTH+Alpha+Beta-domain+wHTH+Fn-III+Fn-III+RRM | Type-2_BR.RM.like.systems | HicA_toxin->?->PIN->?-><-HKD-DNase+SF1-SF2-Helicase<-DUF3780<-DUF499* | 759 | Bacteria>Nitrospirota>Nitrospiria>Nitrospirales>Nitrospiraceae | GCA_024998685.1 | Nitrospira_sp. | H8K07_15150 | 53882..56161 | 70125 | DUF499-domain-containing-protein-[Nitrospira-sp.] |
| MDA2911232.1 | AAA+ATPase+wHTH+Alpha+Beta-domain+wHTH+Fn-III+Fn-III+RRM | Type-2_BR.RM.like.systems | DUF499*->DUF3780->METHYLASE->HKD-DNase+SF1-SF2-Helicase->HKD-DNase+SF1-SF2-Helicase-p->?->MethyltransfD12-> | 1031 | Bacteria>Nitrospirota>Nitrospiria>Nitrospirales>Nitrospiraceae | GCA_027717545.1 | Nitrospiraceae_bacterium_AH_259_D15_M11_P09 | MYX04_09910 | 1638..4733 | 2935764 | DUF499-domain-containing-protein-[Nitrospiraceae-bacterium-AH_259_D15_M11_P09] |
| MCS6286386.1 | AAA+ATPase+wHTH+Alpha+Beta-domain+wHTH+Fn-III+Fn-III+RRM | Type-2_BR.RM.like.systems | <-METHYLASE-p<-DUF3780<-DUF499*<-REase+35exo+Beta-Barrel+SF1-SF2-Helicase | 1044 | Bacteria>Nitrospirota>Nitrospiria>Nitrospirales>Nitrospiraceae | GCA_024998665.1 | Nitrospira_sp. | H8K08_13255 | 985..4119 | 70125 | ATP-binding-protein-[Nitrospira-sp.] |
| WP_108717267.1 | AAA+ATPase+wHTH+Alpha+Beta-domain+wHTH+Fn-III+Fn-III+RRM | Type-2_BR.RM.like.systems | <-VirD4-TraG+VirB4-FtsK<-HDRNase<-wHTH+REase<-HKD-DNase+SF1-SF2-Helicase<-METHYLASE<-DUF3780+RAMA<-DUF499*<-RHH_1<-35exo<-DDE_Tnp_1 | 1041 | Bacteria>Pseudomonadota | GCF_003073935.1 | Gammaproteobacteria | QQJ65_RS05090 | 258052..261177 | 1236 | MULTISPECIES:-DUF499-domain-containing-protein-[Gammaproteobacteria] |
| WP_222132889.1 | AAA+ATPase+wHTH+Alpha+Beta-domain+wHTH+Fn-III+Fn-III+RRM | Type-2_BR.RM.like.systems | DDE_Tnp_1-><-Phage_integrase<-?\|\|35exo->XIS-HTH->DUF499*->DUF3780->METHYLASE->HKD-DNase+SF1-SF2-Helicase->wHTH+REase->HDRNase->VirD4-TraG+VirB4-FtsK-> | 1022 | Bacteria>Pseudomonadota | GCF_030388465.1 | Gammaproteobacteria | FG482_RS02520 | 511750..514818 | 1236 | DUF499-domain-containing-protein-[Vibrio-parahaemolyticus] |
| WP_215851339.1 | AAA+ATPase+wHTH+Alpha+Beta-domain+wHTH+Fn-III+Fn-III+RRM | Type-2_BR.RM.like.systems | Phage_int_SAM_4+Phage_integrase->RHH->DUF499*->DUF3780->?->?->PIN->METHYLASE->REase+ParB-> | 1041 | Bacteria>Pseudomonadota>Acidithiobacillia>Acidithiobacillales>Acidithiobacillaceae | GCF_031200535.1 | Acidithiobacillus_caldus | HFV02_RS10450 | 3946..7071 | 33059 | DUF499-domain-containing-protein-[Acidithiobacillus-caldus] |
| MDD5374995.1 | AAA+ATPase+wHTH+Alpha+Beta-domain+wHTH+Fn-III+Fn-III+RRM | Type-2_BR.RM.like.systems | <-ParA-Soj-PloopNTPase<-HEPN<-METHYLASE<-MACRODOMAIN<-?<-DUF3780<-DUF499*<-RHH_5 | 1041 | Bacteria>Pseudomonadota>Acidithiobacillia>Acidithiobacillales>Acidithiobacillaceae | GCA_028714835.1 | Acidithiobacillus_sp. | PHO38_05125 | 22024..25149 | 1872118 | MAG:-DUF499-domain-containing-protein-[Acidithiobacillus-sp.] |
| ESQ73727.1 | AAA+ATPase+wHTH+Alpha+Beta-domain+wHTH+Fn-III+Fn-III+RRM | Type-2_BR.RM.like.systems | HTH->Cluster182_14clades->?->DUF499*->DUF3780->METHYLASE->HKD-DNase+SF1-SF2-Helicase-><-?<-Arm-DNA-bind_3+Phage_integrase<-tRNA<-ANK+ANK | 1044 | Bacteria>Pseudomonadota>Alphaproteobacteria>Caulobacterales>Caulobacteraceae | GCA_000495835.1 | Asticcacaulis_sp._AC402 | ABAC402_17840 | 26712..29846 | 1282361 | hypothetical-protein-ABAC402_17840-[Asticcacaulis-sp.-AC402] |
| MCZ8170923.1 | AAA+ATPase+wHTH+Alpha+Beta-domain+wHTH+Fn-III+Fn-III+RRM | Type-2_BR.RM.like.systems | <-SMODS<-OBFOLD-NUCLEASE<-HEPN+REase<-HKD-DNase+SF1-SF2-Helicase<-METHYLASE<-DUF3780<-DUF499*<-CopG/RHH\|\|?-><-?<-HTH_3 | 1034 | Bacteria>Pseudomonadota>Alphaproteobacteria>Caulobacterales>Caulobacteraceae | GCA_027531485.1 | Brevundimonas_sp. | O9272_04190 | 53052..56156 | 1871086 | MAG:-DUF499-domain-containing-protein-[Brevundimonas-sp.] |
| WP_162148546.1 | AAA+ATPase+wHTH+Alpha+Beta-domain+wHTH+Fn-III+Fn-III+RRM | Type-2_BR.RM.like.systems | HTH->Cluster182_14clades->?->DUF499*->DUF3780->METHYLASE->HKD-DNase+SF1-SF2-Helicase-><-?<-Arm-DNA-bind_3+Phage_integrase<-tRNA<-ANK+ANK | 1020 | Bacteria>Pseudomonadota>Alphaproteobacteria>Caulobacterales>Caulobacteraceae | GCF_000495835.1 | Asticcacaulis_sp._AC402 | ABAC402_RS37545 | 26784..29846 | 1282361 | DUF499-domain-containing-protein-[Asticcacaulis-sp.-AC402] |
| WP_264595813.1 | AAA+ATPase+wHTH+Alpha+Beta-domain+wHTH+Fn-III+Fn-III+RRM | Type-2_BR.RM.like.systems | ABhydrolase-><-?<-HKD-DNase+SF1-SF2-Helicase<-METHYLASE<-DUF3780<-DUF499*\|\|?->?->?->REase->rve+Mu-transpos_C-> | 1043 | Bacteria>Pseudomonadota>Alphaproteobacteria>Hyphomicrobiales>Beijerinckiaceae | GCF_025961575.1 | Rhodoblastus_acidophilus | M2323_RS01700 | 101217..104348 | 1074 | DUF499-domain-containing-protein-[Rhodoblastus-acidophilus] |
| TWH01684.1 | AAA+ATPase+wHTH+Alpha+Beta-domain+wHTH+Fn-III+Fn-III+RRM | Type-2_BR.RM.like.systems | Peptidase_M15->?-><-URI-DNase+RAMA<-HKD-DNase+SF1-SF2-Helicase<-METHYLASE<-DUF3780<-DUF499*<-RHH<-XIS-HTH\|\|?-><-GST_N+GST_C | 1040 | Bacteria>Pseudomonadota>Alphaproteobacteria>Hyphomicrobiales>Brucellaceae | GCA_007829595.1 | Ochrobactrum_sp._J50 | L614_002100000140 | 13220..16342 | 936132 | putative-AAA+-superfamily-ATPase-[Ochrobactrum-sp.-J50] |
| WP_144897301.1 | AAA+ATPase+wHTH+Alpha+Beta-domain+wHTH+Fn-III+Fn-III+RRM | Type-2_BR.RM.like.systems | Peptidase_M15->?-><-URI-DNase+RAMA<-HKD-DNase+SF1-SF2-Helicase<-METHYLASE<-DUF3780<-DUF499*<-RHH<-XIS-HTH\|\|?-><-GST_N+GST_C | 1040 | Bacteria>Pseudomonadota>Alphaproteobacteria>Hyphomicrobiales>Brucellaceae | GCF_007829595.1 | Ochrobactrum_sp._J50 | L614_RS09275 | 13220..16342 | 936132 | DUF499-domain-containing-protein-[Ochrobactrum-sp.-J50] |
| WP_210189959.1 | AAA+ATPase+wHTH+Alpha+Beta-domain+wHTH+Fn-III+Fn-III+RRM | Type-2_BR.RM.like.systems | HKD-DNase-><-ParB+HNH+RAMA\|\|?->DUF499*->DUF3780->METHYLASE->HKD-DNase+SF1-SF2-Helicase-> | 1039 | Bacteria>Pseudomonadota>Alphaproteobacteria>Hyphomicrobiales>Devosiaceae | GCF_900177655.1 | Devosia_lucknowensis | CCK88_RS15845 | 727170..730289 | 1096929 | DUF499-domain-containing-protein-[Devosia-lucknowensis] |
| MCB1501366.1 | AAA+ATPase+wHTH+Alpha+Beta-domain+wHTH+Fn-III+Fn-III+RRM | Type-2_BR.RM.like.systems | RNase->?->?->?->?->DUF499*->DUF3780->METHYLASE->HKD-DNase+SF1-SF2-Helicase-><-Arm-DNA-bind_3+Phage_int_SAM_3+Phage_integrase<-ParB<-ParB | 1040 | Bacteria>Pseudomonadota>Alphaproteobacteria>Hyphomicrobiales>Kaistiaceae | GCA_020442065.1 | Bauldia_sp. | KDK07_16535 | 19619..22741 | 2575872 | DUF499-domain-containing-protein-[Bauldia-sp.] |
| MCG5246879.1 | AAA+ATPase+wHTH+Alpha+Beta-domain+wHTH+Fn-III+Fn-III+RRM | Type-2_BR.RM.like.systems | <-DDE_Tnp_IS66\|\|CopG/RHH->DUF499*->DUF3780->METHYLASE->HKD-DNase+SF1-SF2-Helicase-> | 1037 | Bacteria>Pseudomonadota>Alphaproteobacteria>Hyphomicrobiales>Methylobacteriaceae | GCA_022214735.1 | Methylorubrum_extorquens | MCX33_13055 | 24088..27201 | 408 | DUF499-domain-containing-protein-[Methylorubrum-extorquens] |
| WP_019904786.1 | AAA+ATPase+wHTH+Alpha+Beta-domain+wHTH+Fn-III+Fn-III+RRM | Type-2_BR.RM.like.systems | HTH_3->MPTase-><-?<-HKD-DNase+SF1-SF2-Helicase<-METHYLASE<-DUF3780<-DUF499*<-CopG/RHH\|\|?->Calcineurin-><-MODE-HTH+PBPII\|\|?-><-ABhydrolase | 1037 | Bacteria>Pseudomonadota>Alphaproteobacteria>Hyphomicrobiales>Methylobacteriaceae | GCF_000372825.1 | Methylobacterium_sp._77 | A3OK_RS0110320 | 2170970..2174083 | 1101192 | DUF499-domain-containing-protein-[Methylobacterium-sp.-77] |
| WP_060847076.1 | AAA+ATPase+wHTH+Alpha+Beta-domain+wHTH+Fn-III+Fn-III+RRM | Type-2_BR.RM.like.systems | <-HKD-DNase+SF1-SF2-Helicase<-METHYLASE<-DUF3780<-DUF499*<-CopG/RHH<-?<-SAM-methylase | 1034 | Bacteria>Pseudomonadota>Alphaproteobacteria>Hyphomicrobiales>Methylobacteriaceae | GCF_001548015.1 | Methylobacterium_aquaticum | Maq22A_RS11360 | 2477154..2480258 | 270351 | DUF499-domain-containing-protein-[Methylobacterium-aquaticum] |
| WP_231343733.1 | AAA+ATPase+wHTH+Alpha+Beta-domain+wHTH+Fn-III+Fn-III+RRM | Type-2_BR.RM.like.systems | <-HKD-DNase+SF1-SF2-Helicase<-METHYLASE<-DUF3780<-DUF499*<-CopG/RHH\|\|DDE_Tnp_IS66-> | 1037 | Bacteria>Pseudomonadota>Alphaproteobacteria>Hyphomicrobiales>Methylobacteriaceae | GCF_021117295.1 | Methylorubrum_sp._B1-46 | LPC10_RS17665 | 3807873..3810986 | 2897334 | DUF499-domain-containing-protein-[Methylorubrum-sp.-B1-46] |
| WP_274382743.1 | AAA+ATPase+wHTH+Alpha+Beta-domain+wHTH+Fn-III+Fn-III+RRM | Type-2_BR.RM.like.systems | <-DDE_Tnp_IS66\|\|CopG/RHH->DUF499*->DUF3780->METHYLASE->HKD-DNase+SF1-SF2-Helicase-> | 961 | Bacteria>Pseudomonadota>Alphaproteobacteria>Hyphomicrobiales>Methylobacteriaceae | GCF_022214735.1 | Methylorubrum_extorquens | MCX33_RS13055 | 24316..27201 | 408 | DUF499-domain-containing-protein-[Methylorubrum-extorquens] |
| WP_243984965.1 | AAA+ATPase+wHTH+Alpha+Beta-domain+wHTH+Fn-III+Fn-III+RRM | Type-2_BR.RM.like.systems | CopG/RHH->DUF499*->DUF3780->METHYLASE->HKD-DNase+SF1-SF2-Helicase->SIGMA-HTH->?->SF1-SF2-Helicase+REase-> | 1035 | Bacteria>Pseudomonadota>Alphaproteobacteria>Hyphomicrobiales>Methylobacteriaceae | GCF_022829325.1 | Methylobacterium_sp._J-092 | MKK60_RS08115 | 733..3840 | 2836667 | DUF499-domain-containing-protein-[Methylobacterium-sp.-J-092] |
| WP_243957552.1 | AAA+ATPase+wHTH+Alpha+Beta-domain+wHTH+Fn-III+Fn-III+RRM | Type-2_BR.RM.like.systems | LexA-HTH+LexA-protease->\|\|GDResolvase+Recombinase+ZnR-><-HKD-DNase+SF1-SF2-Helicase<-METHYLASE<-DUF3780<-DUF499*<-CopG/RHH<-URI-DNase<-?<-SF1-SF2-Helicase | 1034 | Bacteria>Pseudomonadota>Alphaproteobacteria>Hyphomicrobiales>Methylobacteriaceae | GCF_022829565.1 | Methylobacterium_sp._J-001 | MKK65_RS10575 | 98896..102000 | 2836609 | DUF499-domain-containing-protein-[Methylobacterium-sp.-J-001] |
| WP_244032334.1 | AAA+ATPase+wHTH+Alpha+Beta-domain+wHTH+Fn-III+Fn-III+RRM | Type-2_BR.RM.like.systems | <-KAP-NTPase<-RnaseHII\|\|?->?->MPTase->CopG/RHH->DUF499*->DUF3780->METHYLASE->HKD-DNase+SF1-SF2-Helicase-> | 1045 | Bacteria>Pseudomonadota>Alphaproteobacteria>Hyphomicrobiales>Methylobacteriaceae | GCF_022829745.1 | Methylobacterium_sp._E-016 | MKK68_RS18845 | 76366..79503 | 2836556 | DUF499-domain-containing-protein-[Methylobacterium-sp.-E-016] |
| WP_238507602.1 | AAA+ATPase+wHTH+Alpha+Beta-domain+wHTH+Fn-III+Fn-III+RRM | Type-2_BR.RM.like.systems | <-DDE_Tnp_IS66\|\|CopG/RHH->DUF499*->DUF3780->METHYLASE->HKD-DNase+SF1-SF2-Helicase-> | 1037 | Bacteria>Pseudomonadota>Alphaproteobacteria>Hyphomicrobiales>Methylobacteriaceae | GCF_031851355.1 | Methylorubrum_extorquens | MCX33_13055 | 24088..27201 | 408 | DUF499-domain-containing-protein-[Methylorubrum-extorquens] |
| WP_091970185.1 | AAA+ATPase+wHTH+Alpha+Beta-domain+wHTH+Fn-III+Fn-III+RRM | Type-2_BR.RM.like.systems | PIWI->REase-><-HKD-DNase+SF1-SF2-Helicase<-METHYLASE<-DUF3780<-DUF499*<-CopG/RHH<-Phage_integrase | 1035 | Bacteria>Pseudomonadota>Alphaproteobacteria>Hyphomicrobiales>Methylobacteriaceae | GCF_900113485.1 | Methylobacterium_gossipiicola | BM249_RS06630 | 32386..35493 | 582675 | DUF499-domain-containing-protein-[Methylobacterium-gossipiicola] |
| WP_136498122.1 | AAA+ATPase+wHTH+Alpha+Beta-domain+wHTH+Fn-III+Fn-III+RRM | Type-2_BR.RM.like.systems | Phd-YefM->PIN->Phage_integrase->CopG/RHH->DUF499*->DUF3780->METHYLASE->HKD-DNase+SF1-SF2-Helicase->ParB+HNH-p->ParB+HNH-p-> | 1045 | Bacteria>Pseudomonadota>Alphaproteobacteria>Hyphomicrobiales>Methylocystaceae | GCF_004802635.2 | Methylocystis_heyeri | H2LOC_RS20845 | 59150..62287 | 391905 | DUF499-domain-containing-protein-[Methylocystis-heyeri] |
| WP_283816843.1 | AAA+ATPase+wHTH+Alpha+Beta-domain+wHTH+Fn-III+Fn-III+RRM | Type-2_BR.RM.like.systems | Phage_integrase-><-?\|\|GDResolvase+Recombinase+ZnR-><-?<-?<-HTH_36\|\|DUF499*->DUF3780->METHYLASE->HKD-DNase+SF1-SF2-Helicase-><-JAB<-UbE1+JAB | 956 | Bacteria>Pseudomonadota>Alphaproteobacteria>Hyphomicrobiales>Nitrobacteraceae | - | Bradyrhizobium_liaoningense | JQ584_RS26270 | 27003..29873 | 43992 | DUF499-domain-containing-protein-[Bradyrhizobium-liaoningense] |
| ABE37255.1 | AAA+ATPase+wHTH+Alpha+Beta-domain+wHTH+Fn-III+Fn-III+RRM | Type-2_BR.RM.like.systems | ZnR->?-><-?<-HKD-DNase+SF1-SF2-Helicase<-METHYLASE<-DUF3780<-DUF499*<-CopG/RHH\|\|?->Calcineurin->REase+wHTH-> | 1050 | Bacteria>Pseudomonadota>Alphaproteobacteria>Hyphomicrobiales>Nitrobacteraceae | GCA_000013685.1 | Rhodopseudomonas_palustris_BisB5 | RPD_0015 | 19336..22488 | 316057 | conserved-hypothetical-protein-[Rhodopseudomonas-palustris-BisB5] |
| ABE62179.1 | AAA+ATPase+wHTH+Alpha+Beta-domain+wHTH+Fn-III+Fn-III+RRM | Type-2_BR.RM.like.systems | <-HTH+MPTase<-?<-HKD-DNase+SF1-SF2-Helicase<-METHYLASE<-DUF3780<-DUF499*<-CopG/RHH\|\|?->?->Calcineurin->REase+wHTH-> | 1062 | Bacteria>Pseudomonadota>Alphaproteobacteria>Hyphomicrobiales>Nitrobacteraceae | GCA_000013885.1 | Nitrobacter_hamburgensis_X14 | Nham_1355 | 1536266..1539454 | 323097 | conserved-hypothetical-protein-[Nitrobacter-hamburgensis-X14] |
| EHR03038.1 | AAA+ATPase+wHTH+Alpha+Beta-domain+wHTH+Fn-III+Fn-III+RRM | Type-2_BR.RM.like.systems | <-REase+wHTH<-Calcineurin<-?\|\|CopG/RHH->DUF499*->DUF3780->METHYLASE->HKD-DNase+SF1-SF2-Helicase-><-JAB<-UbE1+JAB<-SMODS | 1045 | Bacteria>Pseudomonadota>Alphaproteobacteria>Hyphomicrobiales>Nitrobacteraceae | GCA_000244915.1 | Bradyrhizobium_sp._WSM471 | Bra471DRAFT_03804 | 4063635..4066772 | 319017 | putative-ATPase-(AAA+-superfamily)-[Bradyrhizobium-sp.-WSM471] |
| BAR58173.1 | AAA+ATPase+wHTH+Alpha+Beta-domain+wHTH+Fn-III+Fn-III+RRM | Type-2_BR.RM.like.systems | GDResolvase+PPP1-HTH-><-35exo<-?<-DEDD_Tnp_IS110+Transposase_20<-ZnR\|\|DUF499*->DUF3780->HKD-DNase+SF1-SF2-Helicase->?->UbE1+JAB->DDE_Tnp_1_4-> | 714 | Bacteria>Pseudomonadota>Alphaproteobacteria>Hyphomicrobiales>Nitrobacteraceae | GCA_001549695.1 | Bradyrhizobium_diazoefficiens | NK6_5014 | 4915194..4917338 | 1355477 | hypothetical-protein-NK6_5014-[Bradyrhizobium-diazoefficiens] |
| RQH12683.1 | AAA+ATPase+wHTH+Alpha+Beta-domain+wHTH+Fn-III+Fn-III+RRM | Type-2_BR.RM.like.systems | CopG/RHH->DUF499*->DUF3780->METHYLASE->HKD-DNase+SF1-SF2-Helicase-><-?<-MPTase | 1064 | Bacteria>Pseudomonadota>Alphaproteobacteria>Hyphomicrobiales>Nitrobacteraceae | GCA_003837805.1 | Bradyrhizobium_sp._RP6 | EHH60_14420 | 35819..39013 | 2489596 | ATP-binding-protein-[Bradyrhizobium-sp.-RP6] |
| MBR1030295.1 | AAA+ATPase+wHTH+Alpha+Beta-domain+wHTH+Fn-III+Fn-III+RRM | Type-2_BR.RM.like.systems | Phage_integrase-><-?\|\|GDResolvase+Recombinase+ZnR-><-?<-?<-HTH_36\|\|DUF499*->DUF3780->METHYLASE->HKD-DNase+SF1-SF2-Helicase-><-JAB<-UbE1+JAB | 1038 | Bacteria>Pseudomonadota>Alphaproteobacteria>Hyphomicrobiales>Nitrobacteraceae | GCA_018130275.1 | Bradyrhizobium_liaoningense | JQ584_26260 | 26757..29873 | 43992 | ATP-binding-protein-[Bradyrhizobium-liaoningense] |
| MCW5702705.1 | AAA+ATPase+wHTH+Alpha+Beta-domain+wHTH+Fn-III+Fn-III+RRM | Type-2_BR.RM.like.systems | <-HKD-DNase+SF1-SF2-Helicase<-METHYLASE<-DUF3780<-DUF499*\|\|NADAR-><-?<-Hydrolase_3 | 991 | Bacteria>Pseudomonadota>Alphaproteobacteria>Hyphomicrobiales>Nitrobacteraceae | GCA_026125985.1 | Bradyrhizobium_sp. | KIT82_08985 | 6309..9284 | 376 | MAG:-ATP-binding-protein-[Bradyrhizobium-sp.] |
| GLR99270.1 | AAA+ATPase+wHTH+Alpha+Beta-domain+wHTH+Fn-III+Fn-III+RRM | Type-2_BR.RM.like.systems | GDResolvase+Recombinase+ZnR-><-?<-?<-?<-HTH_36<-MetJarc-HTH\|\|DUF499*->DUF3780->METHYLASE->HKD-DNase+SF1-SF2-Helicase-><-MPTase | 1047 | Bacteria>Pseudomonadota>Alphaproteobacteria>Hyphomicrobiales>Nitrobacteraceae | GCA_030160735.1 | Bradyrhizobium_liaoningense | GCM10007858_69130 | 27511..30654 | 43992 | hypothetical-protein-GCM10007858_69130-[Bradyrhizobium-liaoningense] |
| WP_041357804.1 | AAA+ATPase+wHTH+Alpha+Beta-domain+wHTH+Fn-III+Fn-III+RRM | Type-2_BR.RM.like.systems | <-HTH+MPTase<-?<-HKD-DNase+SF1-SF2-Helicase<-METHYLASE<-DUF3780<-DUF499*<-CopG/RHH\|\|?->?->?->Calcineurin->REase+wHTH-> | 1045 | Bacteria>Pseudomonadota>Alphaproteobacteria>Hyphomicrobiales>Nitrobacteraceae | GCF_000013885.1 | Nitrobacter_hamburgensis | NHAM_RS06920 | 1536266..1539403 | 912 | DUF499-domain-containing-protein-[Nitrobacter-hamburgensis] |
| WP_231170937.1 | AAA+ATPase+wHTH+Alpha+Beta-domain+wHTH+Fn-III+Fn-III+RRM | Type-2_BR.RM.like.systems | <-REase+wHTH<-Calcineurin\|\|DUF499*->DUF3780->METHYLASE->HKD-DNase+SF1-SF2-Helicase-><-JAB<-UbE1+JAB<-SMODS | 1150 | Bacteria>Pseudomonadota>Alphaproteobacteria>Hyphomicrobiales>Nitrobacteraceae | GCF_000244915.1 | Bradyrhizobium | BcanWSM471_RS18650 | 4063659..4067111 | 374 | MULTISPECIES:-DUF499-domain-containing-protein-[Bradyrhizobium] |
| WP_063198232.1 | AAA+ATPase+wHTH+Alpha+Beta-domain+wHTH+Fn-III+Fn-III+RRM | Type-2_BR.RM.like.systems | HTH_3-><-REase<-?\|\|REase->?->DUF499*->DUF3780->METHYLASE->HKD-DNase+SF1-SF2-Helicase->?-><-tRNA\|\|?-><-Glyco_transf_20 | 1047 | Bacteria>Pseudomonadota>Alphaproteobacteria>Hyphomicrobiales>Nitrobacteraceae | GCF_001590795.1 | Bradyrhizobium_sp._AT1 | SE92_RS21720 | 4672667..4675810 | 574934 | DUF499-domain-containing-protein-[Bradyrhizobium-sp.-AT1] |
| WP_085395240.1 | AAA+ATPase+wHTH+Alpha+Beta-domain+wHTH+Fn-III+Fn-III+RRM | Type-2_BR.RM.like.systems | HTH_31->MPTase-><-HKD-DNase+SF1-SF2-Helicase<-METHYLASE<-DUF3780<-DUF499*<-CopG/RHH\|\|?->?->REase+wHTH-> | 1047 | Bacteria>Pseudomonadota>Alphaproteobacteria>Hyphomicrobiales>Nitrobacteraceae | GCF_002108825.1 | Bradyrhizobium_canariense | BSZ21_RS21955 | 70717..73860 | 255045 | DUF499-domain-containing-protein-[Bradyrhizobium-canariense] |
| WP_245454495.1 | AAA+ATPase+wHTH+Alpha+Beta-domain+wHTH+Fn-III+Fn-III+RRM | Type-2_BR.RM.like.systems | CopG/RHH->DUF499*->DUF3780->METHYLASE->HKD-DNase+SF1-SF2-Helicase-><-?<-MPTase | 1043 | Bacteria>Pseudomonadota>Alphaproteobacteria>Hyphomicrobiales>Nitrobacteraceae | GCF_003837805.1 | Bradyrhizobium_sp._RP6 | EHH60_RS14420 | 35882..39013 | 2489596 | DUF499-domain-containing-protein-[Bradyrhizobium-sp.-RP6] |
| WP_211433491.1 | AAA+ATPase+wHTH+Alpha+Beta-domain+wHTH+Fn-III+Fn-III+RRM | Type-2_BR.RM.like.systems | GDResolvase+HIN-HTH-><-?<-?<-HTH_36\|\|DUF499*->METHYLASE->HKD-DNase+SF1-SF2-Helicase-><-MPTase<-?<-?\|\|ABhydrolase-> | 1047 | Bacteria>Pseudomonadota>Alphaproteobacteria>Hyphomicrobiales>Nitrobacteraceae | GCF_018130635.1 | Bradyrhizobium_diazoefficiens | JQ640_RS08145 | 54329..57472 | 1355477 | DUF499-domain-containing-protein-[Bradyrhizobium-diazoefficiens] |
| WP_283823355.1 | AAA+ATPase+wHTH+Alpha+Beta-domain+wHTH+Fn-III+Fn-III+RRM | Type-2_BR.RM.like.systems | <-ZnR\|\|DUF499*->DUF3780->HKD-DNase+SF1-SF2-Helicase->?->UbE1+JAB-> | 825 | Bacteria>Pseudomonadota>Alphaproteobacteria>Hyphomicrobiales>Nitrobacteraceae | GCF_018130635.1 | Bradyrhizobium_diazoefficiens | JQ640_RS39765 | 1315..3792 | 1355477 | DUF499-domain-containing-protein-[Bradyrhizobium-diazoefficiens] |
| WP_247467598.1 | AAA+ATPase+wHTH+Alpha+Beta-domain+wHTH+Fn-III+Fn-III+RRM | Type-2_BR.RM.like.systems | MPTase-><-HKD-DNase+SF1-SF2-Helicase<-METHYLASE<-DUF3780<-DUF499*\|\|MetJarc-HTH->HTH_36-> | 1047 | Bacteria>Pseudomonadota>Alphaproteobacteria>Hyphomicrobiales>Nitrobacteraceae | GCF_023074095.1 | Bradyrhizobium_sp._137 | IVA78_RS21415 | 74673..77816 | 2782614 | DUF499-domain-containing-protein-[Bradyrhizobium-sp.-137] |
| WP_247821020.1 | AAA+ATPase+wHTH+Alpha+Beta-domain+wHTH+Fn-III+Fn-III+RRM | Type-2_BR.RM.like.systems | MPTase-><-HKD-DNase+SF1-SF2-Helicase<-METHYLASE<-DUF3780<-DUF499*\|\|MetJarc-HTH-> | 1047 | Bacteria>Pseudomonadota>Alphaproteobacteria>Hyphomicrobiales>Nitrobacteraceae | GCF_023101005.1 | Bradyrhizobium_sp._187 | IVB19_RS02895 | 596600..599743 | 2782655 | DUF499-domain-containing-protein-[Bradyrhizobium-sp.-187] |
| WP_253579688.1 | AAA+ATPase+wHTH+Alpha+Beta-domain+wHTH+Fn-III+Fn-III+RRM | Type-2_BR.RM.like.systems | GDResolvase+Recombinase+ZnR-><-?<-?<-?<-?\|\|DUF499*->DUF3780->METHYLASE->HKD-DNase+SF1-SF2-Helicase-><-?\|\|KAP-NTPase-> | 1043 | Bacteria>Pseudomonadota>Alphaproteobacteria>Hyphomicrobiales>Nitrobacteraceae | GCF_024170955.1 | Bradyrhizobium_japonicum | J2R91_RS00790 | 164869..168000 | 375 | DUF499-domain-containing-protein-[Bradyrhizobium-japonicum] |
| WP_271613183.1 | AAA+ATPase+wHTH+Alpha+Beta-domain+wHTH+Fn-III+Fn-III+RRM | Type-2_BR.RM.like.systems | MPTase-><-HKD-DNase+SF1-SF2-Helicase<-METHYLASE<-DUF3780<-DUF499*\|\|HTH_36->?->?->?->?-><-GDResolvase+Recombinase+ZnR | 1047 | Bacteria>Pseudomonadota>Alphaproteobacteria>Hyphomicrobiales>Nitrobacteraceae | GCF_028129015.1 | Bradyrhizobium_sp._CCBAU_21362 | ACM41_RS41390 | 456252..459395 | 1325082 | DUF499-domain-containing-protein-[Bradyrhizobium-sp.-CCBAU-21362] |
| WP_284282971.1 | AAA+ATPase+wHTH+Alpha+Beta-domain+wHTH+Fn-III+Fn-III+RRM | Type-2_BR.RM.like.systems | GDResolvase+Recombinase+ZnR-><-?<-?<-HTH_36\|\|?->DUF499*->DUF3780->METHYLASE->HKD-DNase+SF1-SF2-Helicase-><-MPTase | 1047 | Bacteria>Pseudomonadota>Alphaproteobacteria>Hyphomicrobiales>Nitrobacteraceae | GCF_030160735.1 | Bradyrhizobium_liaoningense | QQL14_RS35055 | 27511..30654 | 43992 | DUF499-domain-containing-protein-[Bradyrhizobium-liaoningense] |
| PAQ03744.1 | AAA+ATPase+wHTH+Alpha+Beta-domain+wHTH+Fn-III+Fn-III+RRM | Type-2_BR.RM.like.systems | HISKIN+MORC-><-?<-Resolvase<-?\|\|?->DUF499*->DUF3780->METHYLASE->HKD-DNase+SF1-SF2-Helicase-> | 1034 | Bacteria>Pseudomonadota>Alphaproteobacteria>Hyphomicrobiales>Phyllobacteriaceae | GCA_002284565.1 | Mesorhizobium_mediterraneum | CIT25_02795 | 3278..6382 | 43617 | hypothetical-protein-CIT25_02795-[Mesorhizobium-mediterraneum] |
| RWB54343.1 | AAA+ATPase+wHTH+Alpha+Beta-domain+wHTH+Fn-III+Fn-III+RRM | Type-2_BR.RM.like.systems | <-DUF433<-?\|\|<-HTH_Tnp\|\|CopG/RHH->DUF499*->DUF3780->METHYLASE->HKD-DNase+SF1-SF2-Helicase->URI-DNase->sGTP-MglA+MORC+REase-> | 1034 | Bacteria>Pseudomonadota>Alphaproteobacteria>Hyphomicrobiales>Phyllobacteriaceae | GCA_004016745.1 | Mesorhizobium_sp. | EOQ48_31970 | 14136..17240 | 1871066 | ATP-binding-protein-[Mesorhizobium-sp.] |
| RWA88262.1 | AAA+ATPase+wHTH+Alpha+Beta-domain+wHTH+Fn-III+Fn-III+RRM | Type-2_BR.RM.like.systems | <-DEDD_Tnp_IS110+Transposase_20<-MODE-HTH+PBPII<-DUF433<-?\|\|<-HTH_Tnp\|\|CopG/RHH->DUF499*->DUF3780->METHYLASE->HKD-DNase+SF1-SF2-Helicase->URI-DNase->sGTP-MglA+MORC+REase-> | 1034 | Bacteria>Pseudomonadota>Alphaproteobacteria>Hyphomicrobiales>Phyllobacteriaceae | GCA_004016895.1 | Mesorhizobium_sp. | EOQ31_22360 | 210553..213657 | 1871066 | ATP-binding-protein-[Mesorhizobium-sp.] |
| RWD32092.1 | AAA+ATPase+wHTH+Alpha+Beta-domain+wHTH+Fn-III+Fn-III+RRM | Type-2_BR.RM.like.systems | <-DUF433<-?<-DDE_Tnp_1\|\|CopG/RHH->DUF499*->DUF3780->METHYLASE->HKD-DNase+SF1-SF2-Helicase->URI-DNase->DDE_3-> | 1034 | Bacteria>Pseudomonadota>Alphaproteobacteria>Hyphomicrobiales>Phyllobacteriaceae | GCA_004017065.1 | Mesorhizobium_sp. | EOS22_03615 | 83872..86976 | 1871066 | ATP-binding-protein-[Mesorhizobium-sp.] |
| RWC26998.1 | AAA+ATPase+wHTH+Alpha+Beta-domain+wHTH+Fn-III+Fn-III+RRM | Type-2_BR.RM.like.systems | <-HNH\|\|SIR2-><-wHTH+SAD+HNH\|\|?->DUF499->DUF499*->DUF3780->METHYLASE->HKD-DNase+SF1-SF2-Helicase-><-SF1-SF2-Helicase+DUF1998+HKD-DNase+HKD-DNase | 819 | Bacteria>Pseudomonadota>Alphaproteobacteria>Hyphomicrobiales>Phyllobacteriaceae | GCA_004017365.1 | Mesorhizobium_sp. | EOS70_30595 | 20753..23212 | 1871066 | DUF499-domain-containing-protein-[Mesorhizobium-sp.] |
| RWG75421.1 | AAA+ATPase+wHTH+Alpha+Beta-domain+wHTH+Fn-III+Fn-III+RRM | Type-2_BR.RM.like.systems | UbE1+JAB->?-><-HEPN<-HKD-DNase+SF1-SF2-Helicase<-METHYLASE<-DUF3780<-DUF499*<-CopG/RHH<-HEPN | 830 | Bacteria>Pseudomonadota>Alphaproteobacteria>Hyphomicrobiales>Phyllobacteriaceae | GCA_004018415.1 | Mesorhizobium_sp. | EOQ66_00050 | 16547..19039 | 1871066 | DUF499-domain-containing-protein-[Mesorhizobium-sp.] |
| RWK94990.1 | AAA+ATPase+wHTH+Alpha+Beta-domain+wHTH+Fn-III+Fn-III+RRM | Type-2_BR.RM.like.systems | MULE-TRANSPOSASE-><-Resolvase<-?<-HKD-DNase+SF1-SF2-Helicase<-METHYLASE<-DUF3780<-DUF499*\|\|HEPN->?->HTH+MPTase-><-?<-TRPR-HTH+HTH | 1036 | Bacteria>Pseudomonadota>Alphaproteobacteria>Hyphomicrobiales>Phyllobacteriaceae | GCA_004020105.1 | Mesorhizobium_sp. | EOR53_16295 | 117341..120451 | 1871066 | ATP-binding-protein-[Mesorhizobium-sp.] |
| RWI67698.1 | AAA+ATPase+wHTH+Alpha+Beta-domain+wHTH+Fn-III+Fn-III+RRM | Type-2_BR.RM.like.systems | <-HKD-DNase+SF1-SF2-Helicase<-METHYLASE<-DUF3780<-DUF499*<-?<-ParB+HNH+RAMA\|\|?->Resolvase-> | 1033 | Bacteria>Pseudomonadota>Alphaproteobacteria>Hyphomicrobiales>Phyllobacteriaceae | GCA_004020235.1 | Mesorhizobium_sp. | EOR19_30450 | 12024..15125 | 1871066 | ATP-binding-protein-[Mesorhizobium-sp.] |
| RWL80664.1 | AAA+ATPase+wHTH+Alpha+Beta-domain+wHTH+Fn-III+Fn-III+RRM | Type-2_BR.RM.like.systems | <-HKD-DNase+SF1-SF2-Helicase<-METHYLASE<-DUF3780<-DUF499*<-CopG/RHH<-HEPN\|\|<-SF1-SF2-Helicase+HRDC | 1042 | Bacteria>Pseudomonadota>Alphaproteobacteria>Hyphomicrobiales>Phyllobacteriaceae | GCA_004020855.1 | Mesorhizobium_sp. | EOR67_30945 | 18644..21772 | 1871066 | ATP-binding-protein-[Mesorhizobium-sp.] |
| RWQ62425.1 | AAA+ATPase+wHTH+Alpha+Beta-domain+wHTH+Fn-III+Fn-III+RRM | Type-2_BR.RM.like.systems | <-NTP_transf<-REase+HTH<-URI-DNase<-HKD-DNase+SF1-SF2-Helicase<-METHYLASE<-DUF3780<-DUF499*<-CopG/RHH\|\|HIN-HTH-> | 1034 | Bacteria>Pseudomonadota>Alphaproteobacteria>Hyphomicrobiales>Phyllobacteriaceae | GCA_004021745.1 | Mesorhizobium_sp. | EOS86_30155 | 35225..38329 | 1871066 | ATP-binding-protein-[Mesorhizobium-sp.] |
| TIW61818.1 | AAA+ATPase+wHTH+Alpha+Beta-domain+wHTH+Fn-III+Fn-III+RRM | Type-2_BR.RM.like.systems | ABhydrolase-><-?\|\|GDResolvase+HIN-HTH-><-?\|\|REase->CopG/RHH->DUF499*->DUF3780->METHYLASE->HKD-DNase+SF1-SF2-Helicase->URI-DNase-> | 1042 | Bacteria>Pseudomonadota>Alphaproteobacteria>Hyphomicrobiales>Phyllobacteriaceae | GCA_004964595.1 | Mesorhizobium_sp. | E5V48_07640 | 44498..47626 | 1871066 | ATP-binding-protein-[Mesorhizobium-sp.] |
| TIU01605.1 | AAA+ATPase+wHTH+Alpha+Beta-domain+wHTH+Fn-III+Fn-III+RRM | Type-2_BR.RM.like.systems | <-HKD-DNase+SF1-SF2-Helicase<-METHYLASE<-DUF3780<-DUF499*<-CopG/RHH\|\|HTH_Tnp->\|\|?->DUF433-> | 1034 | Bacteria>Pseudomonadota>Alphaproteobacteria>Hyphomicrobiales>Phyllobacteriaceae | GCA_004965025.1 | Mesorhizobium_sp. | E5W43_04420 | 5162..8266 | 1871066 | ATP-binding-protein-[Mesorhizobium-sp.] |
| WP_158302499.1 | AAA+ATPase+wHTH+Alpha+Beta-domain+wHTH+Fn-III+Fn-III+RRM | Type-2_BR.RM.like.systems | HISKIN+MORC-><-?<-Resolvase<-?\|\|DUF499*->DUF3780->METHYLASE->HKD-DNase+SF1-SF2-Helicase-><-?<-?<-TRPR-HTH+HTH | 1033 | Bacteria>Pseudomonadota>Alphaproteobacteria>Hyphomicrobiales>Phyllobacteriaceae | GCF_002284565.1 | Mesorhizobium_mediterraneum | CIT25_RS02815 | 3281..6382 | 43617 | DUF499-domain-containing-protein-[Mesorhizobium-mediterraneum] |
| WP_126092843.1 | AAA+ATPase+wHTH+Alpha+Beta-domain+wHTH+Fn-III+Fn-III+RRM | Type-2_BR.RM.like.systems | <-DDE_3<-URI-DNase<-HKD-DNase+SF1-SF2-Helicase<-METHYLASE<-DUF3780<-DUF499*<-CopG/RHH<-TRPR-HTH+HTH\|\|HTH_Tnp->MODE-HTH+PBPII-> | 1034 | Bacteria>Pseudomonadota>Alphaproteobacteria>Hyphomicrobiales>Phyllobacteriaceae | GCF_003952485.1 | Mesorhizobium_sp._M1E.F.Ca.ET.045.02.1.1 | EJ070_RS19855 | 4114898..4118002 | 2493672 | DUF499-domain-containing-protein-[Mesorhizobium-sp.-M1E.F.Ca.ET.045.02.1.1] |
| WP_128294455.1 | AAA+ATPase+wHTH+Alpha+Beta-domain+wHTH+Fn-III+Fn-III+RRM | Type-2_BR.RM.like.systems | HEPN->CopG/RHH->DUF499*->DUF3780->METHYLASE->HKD-DNase+SF1-SF2-Helicase-p->HEPN-> | 1042 | Bacteria>Pseudomonadota>Alphaproteobacteria>Hyphomicrobiales>Phyllobacteriaceae | GCF_004023225.1 | unclassified_Mesorhizobium | EOQ59_RS13085 | 2651..5779 | 325217 | DUF499-domain-containing-protein-[Mesorhizobium-sp.-M1A.F.Ca.IN.020.30.1.1] |
| WP_267322120.1 | AAA+ATPase+wHTH+Alpha+Beta-domain+wHTH+Fn-III+Fn-III+RRM | Type-2_BR.RM.like.systems | <-GDResolvase+HIN-HTH<-?<-?<-HTH_28+rve+Mu-transpos_C\|\|DUF499*->DUF3780->METHYLASE->HKD-DNase+SF1-SF2-Helicase-> | 1034 | Bacteria>Pseudomonadota>Alphaproteobacteria>Hyphomicrobiales>Phyllobacteriaceae | GCF_026521175.1 | Aminobacter_sp._MET-1 | ND048_RS29605 | 5575..8679 | 2951085 | DUF499-domain-containing-protein-[Aminobacter-sp.-MET-1] |
| WP_095483057.1 | AAA+ATPase+wHTH+Alpha+Beta-domain+wHTH+Fn-III+Fn-III+RRM | Type-2_BR.RM.like.systems | METHYLASE->?->HISKIN+MORC-><-?<-Resolvase<-?\|\|DUF499*->DUF3780->METHYLASE->HKD-DNase+SF1-SF2-Helicase-><-?<-?<-TRPR-HTH+HTH | 1034 | Bacteria>Pseudomonadota>Alphaproteobacteria>Hyphomicrobiales>Phyllobacteriaceae | GCF_030272285.1 | Mesorhizobium_mediterraneum | LRP31_RS35030 | 32637..35741 | 43617 | DUF499-domain-containing-protein-[Mesorhizobium-mediterraneum] |
| RWP46882.1 | AAA+ATPase+wHTH+Alpha+Beta-domain+wHTH+Fn-III+Fn-III+RRM | Type-2_BR.RM.like.systems | <-NTP_transf<-REase+HTH<-URI-DNase<-HKD-DNase+SF1-SF2-Helicase<-METHYLASE<-DUF3780<-DUF499*<-CopG/RHH<-REase<-?<-?\|\|?->Phage_integrase-> | 1034 | Bacteria>Pseudomonadota>Alphaproteobacteria>Hyphomicrobiales>Phyllobacteriaceae | - | Mesorhizobium_sp. | EOR05_19895 | 25334..28438 | 1871066 | ATP-binding-protein-[Mesorhizobium-sp.] |
| WP_127405482.1 | AAA+ATPase+wHTH+Alpha+Beta-domain+wHTH+Fn-III+Fn-III+RRM | Type-2_BR.RM.like.systems | <-METHYLASE<-DUF3780<-DUF499*<-?\|\|wHTH+SAD+HNH-> | 1033 | Bacteria>Pseudomonadota>Alphaproteobacteria>Hyphomicrobiales>Phyllobacteriaceae | GCF_003997725.1 | Mesorhizobium_sp._M4A.F.Ca.ET.050.02.1.1 | EOA33_RS09415 | 2546..5647 | 2496754 | DUF499-domain-containing-protein-[Mesorhizobium-sp.-M4A.F.Ca.ET.050.02.1.1] |
| PDS57769.1 | AAA+ATPase+wHTH+Alpha+Beta-domain+wHTH+Fn-III+Fn-III+RRM | Type-2_BR.RM.like.systems | METHYLASE-p-><-?<-Phage_integrase+Phage_integrase<-Phage_integrase+Phage_integrase<-Phage_integrase\|\|DUF499*->DUF3780->METHYLASE->HKD-DNase+SF1-SF2-Helicase->Pkinase+SF1-SF2-Helicase->HTH+HKD-DNase-> | 1041 | Bacteria>Pseudomonadota>Alphaproteobacteria>Hyphomicrobiales>Rhizobiaceae | GCA_002531635.1 | Rhizobium_anhuiense | CO663_16960 | 123338..126463 | 1184720 | AAA-family-ATPase-[Rhizobium-anhuiense] |
| MBY5896032.1 | AAA+ATPase+wHTH+Alpha+Beta-domain+wHTH+Fn-III+Fn-III+RRM | Type-2_BR.RM.like.systems | DUF499*->DUF3780->METHYLASE->HKD-DNase+SF1-SF2-Helicase-><-?<-TetR-HTH | 1041 | Bacteria>Pseudomonadota>Alphaproteobacteria>Hyphomicrobiales>Rhizobiaceae | GCA_019799885.1 | Rhizobium_leguminosarum | HFN45_21935 | 190896..194021 | 384 | DUF499-domain-containing-protein-[Rhizobium-leguminosarum] |
| WP_037456478.1 | AAA+ATPase+wHTH+Alpha+Beta-domain+wHTH+Fn-III+Fn-III+RRM | Type-2_BR.RM.like.systems | <-HKD-DNase+SF1-SF2-Helicase-p<-HKD-DNase+SF1-SF2-Helicase<-METHYLASE<-DUF3780<-DUF499* | 1033 | Bacteria>Pseudomonadota>Alphaproteobacteria>Hyphomicrobiales>Rhizobiaceae | GCF_000261885.1 | Sinorhizobium_fredii | AB395_RS33470 | 113083..116184 | 380 | DUF499-domain-containing-protein-[Sinorhizobium-fredii] |

| Protein-ID | DUF499-architecture | System | Gene-Neighbourhoods | Length | Taxonomy | Assembly-ID | Strain | Locus-Tag | Genomic-coordinates | TaxID | Fasta-defline |
|---|---|---|---|---|---|---|---|---|---|---|---|
| WP_088199411.1 | AAA+ATPase+wHTH+Alpha+Beta-domain+wHTH+Fn-III+Fn-III+RRM | Type-2_BR.RM.like.systems | DNA_methylase->?->?-><-HKD-DNase+SF1-SF2-Helicase<-METHYLASE<-DUF3780<-DUF499*<-?\|\|?-><-ParB+HNH\|\|?-><-REase | 1039 | Bacteria>Pseudomonadota>Alphaproteobacteria>Hyphomicrobiales>Rhizobiaceae | GCF_002197465.1 | Sinorhizobium_meliloti | GHK31_RS12210 | 25646..28765 | 382 | DUF499-domain-containing-protein-[Sinorhizobium-meliloti] |
| WP_245445203.1 | AAA+ATPase+wHTH+Alpha+Beta-domain+wHTH+Fn-III+Fn-III+RRM | Type-2_BR.RM.like.systems | METHYLASE-p-><-?<-Phage_integrase+Phage_integrase<-Phage_integrase+Phage_integrase<-Phage_integrase+Phage_integrase<-Phage_integrase\|\|DUF499*->DUF3780->METHYLASE->HKD-DNase+SF1-SF2-Helicase->Pkinase+SF1-SF2-Helicase->HTH+HKD-DNase-> | 989 | Bacteria>Pseudomonadota>Alphaproteobacteria>Hyphomicrobiales>Rhizobiaceae | GCF_002531635.1 | Rhizobium_anhuiense | CO663_RS16955 | 123494..126463 | 1184720 | DUF499-domain-containing-protein-[Rhizobium-anhuiense] |
| WP_127660879.1 | AAA+ATPase+wHTH+Alpha+Beta-domain+wHTH+Fn-III+Fn-III+RRM | Type-2_BR.RM.like.systems | <-HIN-HTH<-DDE_Tnp_1\|\|?-><-HKD-DNase+SF1-SF2-Helicase<-METHYLASE<-DUF3780<-DUF499*\|\|?->?-><-?\|\|?->PIN->Phage_integrase-> | 1042 | Bacteria>Pseudomonadota>Alphaproteobacteria>Hyphomicrobiales>Rhizobiaceae | GCF_004002365.1 | Sinorhizobium_medicae | CN168_RS28120 | 8678..11806 | 110321 | DUF499-domain-containing-protein-[Sinorhizobium-medicae] |
| WP_245497535.1 | AAA+ATPase+wHTH+Alpha+Beta-domain+wHTH+Fn-III+Fn-III+RRM | Type-2_BR.RM.like.systems | DUF499*->DUF3780->METHYLASE->HKD-DNase+SF1-SF2-Helicase-><-?<-TetR-HTH | 1026 | Bacteria>Pseudomonadota>Alphaproteobacteria>Hyphomicrobiales>Rhizobiaceae | GCF_004306385.1 | Rhizobium_ruizarguesonis | ELH20_RS03710 | 755502..758582 | 2081791 | DUF499-domain-containing-protein-[Rhizobium-leguminosarum] |
| WP_131704163.1 | AAA+ATPase+wHTH+Alpha+Beta-domain+wHTH+Fn-III+Fn-III+RRM | Type-2_BR.RM.like.systems | <-REase\|\|RHH->DUF499*->DUF3780->METHYLASE->HKD-DNase+SF1-SF2-Helicase->ParB->REase-> | 1040 | Bacteria>Pseudomonadota>Alphaproteobacteria>Hyphomicrobiales>Rhizobiaceae | GCF_004330715.1 | Rhizobium_leguminosarum | E0H72_RS11610 | 3872..6994 | 384 | DUF499-domain-containing-protein-[Rhizobium-leguminosarum] |
| WP_153518275.1 | AAA+ATPase+wHTH+Alpha+Beta-domain+wHTH+Fn-III+Fn-III+RRM | Type-2_BR.RM.like.systems | <-PIN<-PhdYeFM_antitox<-HKD-DNase+SF1-SF2-Helicase<-METHYLASE<-DUF3780<-DUF499*<-Methyltransf_25<-ParB+HNH<-HTH+WYL\|\|SF1-SF2-Helicase+REase->SF1-SF2-Helicase+REase->HKD-DNase-> | 1040 | Bacteria>Pseudomonadota>Alphaproteobacteria>Hyphomicrobiales>Rhizobiaceae | GCF_009599805.1 | Sinorhizobium_meliloti | GHK65_RS07990 | 24886..28008 | 382 | DUF499-domain-containing-protein-[Sinorhizobium-meliloti] |
| WP_153520038.1 | AAA+ATPase+wHTH+Alpha+Beta-domain+wHTH+Fn-III+Fn-III+RRM | Type-2_BR.RM.like.systems | URI-DNase-><-?\|\|DDE_Tnp_1-><-HKD-DNase+SF1-SF2-Helicase<-METHYLASE<-DUF3780<-DUF499*\|\|?-><-PemK_toxin<-?\|\|?-><-PIN+OST-HTH | 1042 | Bacteria>Pseudomonadota>Alphaproteobacteria>Hyphomicrobiales>Rhizobiaceae | GCF_009600005.1 | Sinorhizobium_meliloti | GHL01_RS04135 | 66008..69136 | 382 | DUF499-domain-containing-protein-[Sinorhizobium-meliloti] |
| WP_170263459.1 | AAA+ATPase+wHTH+Alpha+Beta-domain+wHTH+Fn-III+Fn-III+RRM | Type-2_BR.RM.like.systems | <-DOC+HTH\|\|?-><-?<-HKD-DNase+SF1-SF2-Helicase<-METHYLASE<-DUF3780<-DUF499* | 1041 | Bacteria>Pseudomonadota>Alphaproteobacteria>Hyphomicrobiales>Rhizobiaceae | GCF_013004175.1 | Rhizobium_laguerreae | HLI18_RS32230 | 10035..13160 | 1076926 | DUF499-domain-containing-protein-[Rhizobium-laguerreae] |
| WP_222407811.1 | AAA+ATPase+wHTH+Alpha+Beta-domain+wHTH+Fn-III+Fn-III+RRM | Type-2_BR.RM.like.systems | <-REase<-ParB<-HKD-DNase+SF1-SF2-Helicase<-METHYLASE<-DUF3780<-DUF499*<-RHH\|\|REase->?->?-><-?<-TUDOR+SF1-SF2-Helicase+iREase+CTD | 1040 | Bacteria>Pseudomonadota>Alphaproteobacteria>Hyphomicrobiales>Rhizobiaceae | GCF_019798725.1 | Rhizobium_leguminosarum | HF082_RS21825 | 332651..335773 | 384 | DUF499-domain-containing-protein-[Rhizobium-leguminosarum] |
| WP_221975247.1 | AAA+ATPase+wHTH+Alpha+Beta-domain+wHTH+Fn-III+Fn-III+RRM | Type-2_BR.RM.like.systems | DUF499*->DUF3780->METHYLASE->HKD-DNase+SF1-SF2-Helicase-><-?<-TetR-HTH | 1041 | Bacteria>Pseudomonadota>Alphaproteobacteria>Hyphomicrobiales>Rhizobiaceae | - | Rhizobium_leguminosarum | HF204_RS31495 | 121770..124895 | 384 | DUF499-domain-containing-protein-[Rhizobium-leguminosarum] |
| WP_068408954.1 | AAA+ATPase+wHTH+Alpha+Beta-domain+wHTH+Fn-III+Fn-III+RRM | Type-2_BR.RM.like.systems | <-Phage_integrase\|\|HTH-><-STYKinase-like\|\|ParB+HNH+RAMA->SF1-SF2-Helicase->DUF499*->DUF3780->METHYLASE->HKD-DNase+SF1-SF2-Helicase-><-TraI-VirD2-RCRE<-TraJ-RHH<-HTH_36 | 1040 | Bacteria>Pseudomonadota>Alphaproteobacteria>Hyphomicrobiales>Stappiaceae | GCF_001624695.1 | Labrenzia_sp._OB1 | OA90_RS07080 | 119206..122328 | 1561204 | DUF499-domain-containing-protein-[Labrenzia-sp.-OB1] |
| MBN8978630.1 | AAA+ATPase+wHTH+Alpha+Beta-domain+wHTH+Fn-III+Fn-III+RRM | Type-2_BR.RM.like.systems | <-METHYLASE<-DUF3780<-DUF499*<-CopG/RHH\|\|?->?->Calcineurin->REase+wHTH-> | 1046 | Bacteria>Pseudomonadota>Alphaproteobacteria>Hyphomicrobiales>unclassified_Hyphomicrobiales | GCA_017305555.1 | Hyphomicrobiales_bacterium | JOI08_19330 | 3242..6382 | 1909294 | DUF499-domain-containing-protein-[Hyphomicrobiales-bacterium] |
| MAW82209.1 | AAA+ATPase+wHTH+Alpha+Beta-domain+wHTH+Fn-III+Fn-III+RRM | Type-2_BR.RM.like.systems | <-HKD-DNase+SF1-SF2-Helicase<-METHYLASE<-DUF3780<-DUF499*<-RHH_1\|\|?->ArdA->?->RNase->ParB+HTH-> | 1039 | Bacteria>Pseudomonadota>Alphaproteobacteria>Parvularculales>Parvularculaceae | GCA_002704915.1 | Parvularcula_sp. | CMI63_18385 | 35385..38504 | 1979207 | AAA-family-ATPase-[Parvularcula-sp.] |
| MAY89501.1 | AAA+ATPase+wHTH+Alpha+Beta-domain+wHTH+Fn-III+Fn-III+RRM | Type-2_BR.RM.like.systems | Peptidase_M15->?->?-><-URI-DNase+RAMA<-HKD-DNase+SF1-SF2-Helicase<-DUF499*<-RHH<-ParB | 973 | Bacteria>Pseudomonadota>Alphaproteobacteria>Rhodobacterales>Paracoccaceae | GCA_002706905.1 | Pseudooceanicola_sp. | CML02_22640 | 66522..69443 | 1914328 | AAA-family-ATPase-[Pseudooceanicola-sp.] |
| RDD72166.1 | AAA+ATPase+wHTH+Alpha+Beta-domain+wHTH+Fn-III+Fn-III+RRM | Type-2_BR.RM.like.systems | RNase-><-URI-DNase+RAMA<-HKD-DNase+SF1-SF2-Helicase<-METHYLASE<-DUF3780<-DUF499*<-RHH<-Arm-DNA-bind_3+Phage_int_SAM_3+Phage_integrase\|\|TPR-><-GDResolvase+Recombinase+ZnR | 1020 | Bacteria>Pseudomonadota>Alphaproteobacteria>Rhodobacterales>Paracoccaceae | GCA_003344805.1 | Paracoccus_versutus | DVR11_07365 | 55704..58766 | 34007 | DUF499-domain-containing-protein-[Paracoccus-versutus] |
| QYK40247.1 | AAA+ATPase+wHTH+Alpha+Beta-domain+wHTH+Fn-III+Fn-III+RRM | Type-2_BR.RM.like.systems | <-ACET<-?<-HEPN+REase<-HKD-DNase+SF1-SF2-Helicase<-METHYLASE<-DUF3780<-DUF499*<-CopG/RHH\|\|?-><-tRNA\|\|?->?-><-tRNA\|\|?->Peptidase_M48-> | 1034 | Bacteria>Pseudomonadota>Alphaproteobacteria>Rhodobacterales>Paracoccaceae | GCA_019454225.1 | Paracoccaceae_bacterium | KF887_12470 | 2639699..2642803 | 1904441 | MAG:-DUF499-domain-containing-protein-[Rhodobacteraceae-bacterium] |
| WP_036718023.1 | AAA+ATPase+wHTH+Alpha+Beta-domain+wHTH+Fn-III+Fn-III+RRM | Type-2_BR.RM.like.systems | RNase-><-URI-DNase+RAMA<-HKD-DNase+SF1-SF2-Helicase<-METHYLASE<-DUF3780<-DUF499*<-RHH<-Arm-DNA-bind_3+Phage_int_SAM_3+Phage_integrase\|\|TPR->?-><-MPTase | 1039 | Bacteria>Pseudomonadota>Alphaproteobacteria>Rhodobacterales>Paracoccaceae | GCF_000763805.1 | Paracoccus_sphaerophysae | IC63_RS06155 | 51780..54899 | 690417 | DUF499-domain-containing-protein-[Paracoccus-sphaerophysae] |
| WP_107325843.1 | AAA+ATPase+wHTH+Alpha+Beta-domain+wHTH+Fn-III+Fn-III+RRM | Type-2_BR.RM.like.systems | REase->?->?-><-?\|\|CopG/RHH->DUF499*->DUF3780->METHYLASE->HKD-DNase+SF1-SF2-Helicase-> | 1034 | Bacteria>Pseudomonadota>Alphaproteobacteria>Rhodobacterales>Paracoccaceae | GCF_003034995.1 | Phaeovulum_veldkampii | EV658_RS14470 | 14082..17186 | 33049 | DUF499-domain-containing-protein-[Phaeovulum-veldkampii] |
| WP_111405614.1 | AAA+ATPase+wHTH+Alpha+Beta-domain+wHTH+Fn-III+Fn-III+RRM | Type-2_BR.RM.like.systems | RNase-><-URI-DNase+RAMA<-HKD-DNase+SF1-SF2-Helicase<-METHYLASE<-DUF3780<-DUF499*<-RHH<-XIS-HTH | 1039 | Bacteria>Pseudomonadota>Alphaproteobacteria>Rhodobacterales>Paracoccaceae | GCF_003254465.1 | Phycocomes_zhengii | DDE13_RS17795 | 16902..20021 | 2056810 | DUF499-domain-containing-protein-[Phycocomes-zhengii] |
| WP_220180866.1 | AAA+ATPase+wHTH+Alpha+Beta-domain+wHTH+Fn-III+Fn-III+RRM | Type-2_BR.RM.like.systems | RNase-><-URI-DNase+RAMA<-HKD-DNase+SF1-SF2-Helicase<-METHYLASE<-DUF3780<-DUF499*<-RHH<-Arm-DNA-bind_3+Phage_int_SAM_3+Phage_integrase\|\|TPR-><-GDResolvase+Recombinase+ZnR | 1039 | Bacteria>Pseudomonadota>Alphaproteobacteria>Rhodobacterales>Paracoccaceae | GCF_003344805.1 | Paracoccus_versutus | DVR11_RS07370 | 55704..58823 | 34007 | DUF499-domain-containing-protein-[Paracoccus-versutus] |
| WP_152828752.1 | AAA+ATPase+wHTH+Alpha+Beta-domain+wHTH+Fn-III+Fn-III+RRM | Type-2_BR.RM.like.systems | Peptidase_S11-><-HKD-DNase+SF1-SF2-Helicase<-METHYLASE<-DUF3780<-DUF499*<-PIN<-RHH\|\|RNase-> | 1044 | Bacteria>Pseudomonadota>Alphaproteobacteria>Rhodobacterales>Paracoccaceae | GCF_009380135.2 | Fertoeibacter_niger | GEU84_RS20380 | 38813..41947 | 2656921 | DUF499-domain-containing-protein-[Fertoeibacter-niger] |
| RLJ36255.1 | AAA+ATPase+wHTH+Alpha+Beta-domain+wHTH+Fn-III+Fn-III+RRM | Type-2_BR.RM.like.systems | <-ParB+HTH<-?<-ZDNA-HTH<-?<-?\|\|CopG/RHH->DUF499*->DUF3780->METHYLASE->HKD-DNase+SF1-SF2-Helicase-><-ABhydrolase | 1034 | Bacteria>Pseudomonadota>Alphaproteobacteria>Rhodobacterales>Roseobacteraceae | GCA_003663885.1 | Litoreibacter_meonggei | BCF46_3837 | 30311..33415 | 1049199 | uncharacterized-protein-DUF499-[Litoreibacter-meonggei] |
| WP_211332850.1 | AAA+ATPase+wHTH+Alpha+Beta-domain+wHTH+Fn-III+Fn-III+RRM | Type-2_BR.RM.like.systems | <-ParB+HTH<-?<-?<-ZDNA-HTH<-?<-?\|\|DUF499*->DUF3780->METHYLASE->HKD-DNase+SF1-SF2-Helicase-><-ABhydrolase | 979 | Bacteria>Pseudomonadota>Alphaproteobacteria>Rhodobacterales>Roseobacteraceae | GCF_003663885.1 | Litoreibacter_meonggei | BCF46_RS18880 | 30476..33415 | 1049199 | DUF499-domain-containing-protein-[Litoreibacter-meonggei] |
| WP_217644466.1 | AAA+ATPase+wHTH+Alpha+Beta-domain+wHTH+Fn-III+Fn-III+RRM | Type-2_BR.RM.like.systems | DDE_Tnp_IS1595->DUF499*->DUF3780->METHYLASE->HKD-DNase+SF1-SF2-Helicase-><-URI-DNase+RAMA\|\|ArdCN+MPTase-> | 925 | Bacteria>Pseudomonadota>Alphaproteobacteria>Rhodobacterales>Roseobacteraceae | GCF_900109855.1 | Roseovarius_tolerans | BMX04_RS11590 | 280..3057 | 74031 | DUF499-domain-containing-protein-[Roseovarius-tolerans] |
| SEN02116.1 | AAA+ATPase+wHTH+Alpha+Beta-domain+wHTH+Fn-III+Fn-III+RRM | Type-2_BR.RM.like.systems | DDE_Tnp_IS1595->DUF499*->DUF3780->METHYLASE->HKD-DNase+SF1-SF2-Helicase-><-URI-DNase+RAMA\|\|ArdCN+MPTase-> | 871 | Bacteria>Pseudomonadota>Alphaproteobacteria>Rhodobacterales>Roseobacteraceae | - | Roseovarius_tolerans | SAMN04488077_1112 | 442..3057 | 74031 | Protein-of-unknown-function-[Roseovarius-tolerans] |
| TPG55477.1 | AAA+ATPase+wHTH+Alpha+Beta-domain+wHTH+Fn-III+Fn-III+RRM | Type-2_BR.RM.like.systems | <-HKD-DNase+SF1-SF2-Helicase<-MACRODOMAIN<-ParB+HNH<-METHYLASE<-DUF3780<-DUF499*<-35exo<-?<-?\|\|HEPN+REase-> | 1086 | Bacteria>Pseudomonadota>Alphaproteobacteria>Rhodospirillales>Acetobacteraceae | GCA_006438825.1 | Roseomonas_nepalensis | EAH89_14595 | 46471..49731 | 1854500 | DUF499-domain-containing-protein-[Roseomonas-nepalensis] |
| MBV1800582.1 | AAA+ATPase+wHTH+Alpha+Beta-domain+wHTH+Fn-III+Fn-III+RRM | Type-2_BR.RM.like.systems | DUF499*->DUF3780->METHYLASE->HKD-DNase+SF1-SF2-Helicase-p->?->HKD-DNase+SF1-SF2-Helicase-p-><-HicB | 858 | Bacteria>Pseudomonadota>Alphaproteobacteria>Rhodospirillales>Acetobacteraceae | GCA_019084075.1 | Siccirubricoccus_sp._G192 | KSF81_27995 | 969..3545 | 2849651 | DUF499-domain-containing-protein-[Siccirubricoccus-sp.-G192] |
| MCA3288087.1 | AAA+ATPase+wHTH+Alpha+Beta-domain+wHTH+Fn-III+Fn-III+RRM | Type-2_BR.RM.like.systems | <-sGTP<-?<-?\|\|tRNA->DUF499*->DUF3780->METHYLASE->HKD-DNase+SF1-SF2-Helicase-><-ParB+HNH<-HTH<-GDResolvase+Recombinase+ZnR | 1045 | Bacteria>Pseudomonadota>Alphaproteobacteria>Rhodospirillales>Acetobacteraceae | GCA_020249245.1 | Roseomonas_sp. | ING09_16215 | 121046..124183 | 1869190 | ATP-binding-protein-[Roseomonas-sp.] |
| MCA4920336.1 | AAA+ATPase+wHTH+Alpha+Beta-domain+wHTH+Fn-III+Fn-III+RRM | Type-2_BR.RM.like.systems | DUF499*->DUF3780->METHYLASE->HKD-DNase+SF1-SF2-Helicase-><-ParB+HNH<-HTH<-GDResolvase+Recombinase+ZnR | 1045 | Bacteria>Pseudomonadota>Alphaproteobacteria>Rhodospirillales>Acetobacteraceae | GCA_020285315.1 | Roseomonas_sp. | ING82_12990 | 1209..4346 | 1869190 | ATP-binding-protein-[Roseomonas-sp.] |
| WP_010510756.1 | AAA+ATPase+wHTH+Alpha+Beta-domain+wHTH+Fn-III+Fn-III+RRM | Type-2_BR.RM.like.systems | ParB+HNH->ParB+HNH-><-?<-HKD-DNase+SF1-SF2-Helicase<-METHYLASE<-DUF3780<-DUF499*<-CopG/RHH<-PAIREDC-HTH+HIN-HTH<-DUF772+DDE_Tnp_1+DDE_Tnp_1_6 | 1032 | Bacteria>Pseudomonadota>Alphaproteobacteria>Rhodospirillales>Acetobacteraceae | GCF_000227545.1 | Komagataeibacter_europaeus | GEU02_RS13955 | 17786..20884 | 33995 | DUF499-domain-containing-protein-[Komagataeibacter-europaeus] |
| WP_075822782.1 | AAA+ATPase+wHTH+Alpha+Beta-domain+wHTH+Fn-III+Fn-III+RRM | Type-2_BR.RM.like.systems | <-DDE_Tnp_1<-?<-?<-?<-?\|\|DUF499*->DUF3780->METHYLASE->ParB+HNH->MACRODOMAIN->HKD-DNase+SF1-SF2-Helicase-> | 1066 | Bacteria>Pseudomonadota>Alphaproteobacteria>Rhodospirillales>Acetobacteraceae | GCF_001950415.1 | Roseomonas | MHL39_RS17975 | 5795..8995 | 125216 | MULTISPECIES:-DUF499-domain-containing-protein-[Roseomonas] |
| WP_161993601.1 | AAA+ATPase+wHTH+Alpha+Beta-domain+wHTH+Fn-III+Fn-III+RRM | Type-2_BR.RM.like.systems | <-HKD-DNase+SF1-SF2-Helicase<-MACRODOMAIN<-ParB+HNH<-METHYLASE<-DUF3780<-DUF499*<-35exo<-?<-?\|\|HEPN+REase-> | 1022 | Bacteria>Pseudomonadota>Alphaproteobacteria>Rhodospirillales>Acetobacteraceae | GCF_006438825.1 | Roseomonas_nepalensis | EAH89_RS14655 | 46471..49539 | 1854500 | DUF499-domain-containing-protein-[Roseomonas-nepalensis] |
| WP_149536631.1 | AAA+ATPase+wHTH+Alpha+Beta-domain+wHTH+Fn-III+Fn-III+RRM | Type-2_BR.RM.like.systems | <-MetJarc-HTH<-SAM-methylase<-HKD-DNase+SF1-SF2-Helicase<-?<-METHYLASE<-DUF3780<-DUF499*<-?\|\|DUF4096+DDE_Tnp_1+DDE_Tnp_1_2-> | 1059 | Bacteria>Pseudomonadota>Alphaproteobacteria>Rhodospirillales>Acetobacteraceae | GCF_008360935.1 | Siccirubricoccus_phaeus | E5N76_RS07285 | 142618..145797 | 2595053 | DUF499-domain-containing-protein-[Siccirubricoccus-phaeus] |
| WP_171833827.1 | AAA+ATPase+wHTH+Alpha+Beta-domain+wHTH+Fn-III+Fn-III+RRM | Type-2_BR.RM.like.systems | ParB+HNH+RAMA-><-GDResolvase<-HKD-DNase+SF1-SF2-Helicase<-METHYLASE<-DUF3780<-DUF499*<-?<-REase<-SF1-SF2-Helicase<-MobL+ZnR+TOPRIM\|\|REase-><-PIN | 1040 | Bacteria>Pseudomonadota>Alphaproteobacteria>Rhodospirillales>Acetobacteraceae | GCF_013201075.1 | Lichenicola_cladoniae | HN018_RS23720 | 375896..379018 | 1484109 | DUF499-domain-containing-protein-[Lichenicola-cladoniae] |
| WP_203075822.1 | AAA+ATPase+wHTH+Alpha+Beta-domain+wHTH+Fn-III+Fn-III+RRM | Type-2_BR.RM.like.systems | RAMA->GDResolvase+Recombinase+ZnR->HTH->?->DUF499*->DUF3780->METHYLASE->HKD-DNase+SF1-SF2-Helicase->HEPN-> | 1045 | Bacteria>Pseudomonadota>Alphaproteobacteria>Rhodospirillales>Acetobacteraceae | GCF_016805305.1 | Roseomonas_ponticola | JJI54_RS23740 | 272470..275607 | 2786951 | DUF499-domain-containing-protein-[Roseomonas-ponticola] |
| WP_213612192.1 | AAA+ATPase+wHTH+Alpha+Beta-domain+wHTH+Fn-III+Fn-III+RRM | Type-2_BR.RM.like.systems | HTH->?-><-MalT\|\|DUF499*->DUF3780->METHYLASE->HKD-DNase+SF1-SF2-Helicase-> | 1045 | Bacteria>Pseudomonadota>Alphaproteobacteria>Rhodospirillales>Acetobacteraceae | GCF_018413585.1 | Roseococcus_sp._SDR | KTR66_RS08730 | 93597..96734 | 2835532 | DUF499-domain-containing-protein-[Roseococcus-sp.-SDR] |
| WP_256444398.1 | AAA+ATPase+wHTH+Alpha+Beta-domain+wHTH+Fn-III+Fn-III+RRM | Type-2_BR.RM.like.systems | DUF499*->DUF3780->METHYLASE->ParB+HNH->?->?->HKD-DNase+SF1-SF2-Helicase-p-> | 869 | Bacteria>Pseudomonadota>Alphaproteobacteria>Rhodospirillales>Acetobacteraceae | GCF_019084075.1 | Siccirubricoccus_sp._G192 | KSF81_RS27995 | 936..3545 | 2849651 | DUF499-domain-containing-protein-[Siccirubricoccus-sp.-G192] |
| WP_217382201.1 | AAA+ATPase+wHTH+Alpha+Beta-domain+wHTH+Fn-III+Fn-III+RRM | Type-2_BR.RM.like.systems | Phage_integrase-><-?<-?<-ParA-Soj-PloopNTPase\|\|SF1-SF2-Helicase->?->\|\|DUF499*->DUF3780->METHYLASE-p->METHYLASE->ParB+HNH->HKD-DNase+SF1-SF2-Helicase-><-HicB | 1072 | Bacteria>Pseudomonadota>Alphaproteobacteria>Rhodospirillales>Acetobacteraceae | GCF_019084075.1 | Siccirubricoccus_sp._G192 | KSF81_RS28660 | 165691..168909 | 2849651 | DUF499-domain-containing-protein-[Siccirubricoccus-sp.-G192] |
| WP_090663037.1 | AAA+ATPase+wHTH+Alpha+Beta-domain+wHTH+Fn-III+Fn-III+RRM | Type-2_BR.RM.like.systems | GDResolvase+Recombinase+ZnR->HTH->REase-><-HKD-DNase+SF1-SF2-Helicase<-METHYLASE<-DUF3780<-DUF499*<-tRNA<-sGTP | 1059 | Bacteria>Pseudomonadota>Alphaproteobacteria>Rhodospirillales>Acetobacteraceae | GCF_900101615.1 | Belnapia_rosea | BLR02_RS10830 | 87446..90625 | 938405 | DUF499-domain-containing-protein-[Belnapia-rosea] |
| WP_217382140.1 | AAA+ATPase+wHTH+Alpha+Beta-domain+wHTH+Fn-III+Fn-III+RRM | Type-2_BR.RM.like.systems | DUF499*->DUF3780->METHYLASE->HKD-DNase+SF1-SF2-Helicase-p->?->HKD-DNase+SF1-SF2-Helicase-p-><-HicB | 858 | Bacteria>Pseudomonadota>Alphaproteobacteria>Rhodospirillales>Acetobacteraceae | - | Siccirubricoccus_sp._G192 | KSF81_27995 | 969..3545 | 2849651 | DUF499-domain-containing-protein-[Siccirubricoccus-sp.-G192] |
| MBV8096531.1 | AAA+ATPase+wHTH+Alpha+Beta-domain+wHTH+Fn-III+Fn-III+RRM | Type-2_BR.RM.like.systems | <-DUF3780<-DUF499*\|\|REase->?-><-?<-ABC-ATPase+SbcC | 1063 | Bacteria>Pseudomonadota>Alphaproteobacteria>Rhodospirillales>Acetobacteraceae | GCA_019237355.1 | Acetobacteraceae_bacterium | JO110_25480 | 497..3688 | 1909293 | DUF499-domain-containing-protein-[Acetobacteraceae-bacterium] |
| MBE7157121.1 | AAA+ATPase+wHTH+Alpha+Beta-domain+wHTH+Fn-III+Fn-III+RRM | Type-2_BR.RM.like.systems | HTH->ParB+HNH->ParB+HNH->DUF499*->DUF3780-> | 1031 | Bacteria>Pseudomonadota>Alphaproteobacteria>Rhodospirillales>Acetobacteraceae | GCA_015069705.1 | Acetobacter_sp. | INR62_01575 | 6375..9470 | 440 | DUF499-domain-containing-protein-[Rhodospirillales-bacterium] |
| WP_019647020.1 | AAA+ATPase+wHTH+Alpha+Beta-domain+wHTH+Fn-III+Fn-III+RRM | Type-2_BR.RM.like.systems | <-HKD-DNase+SF1-SF2-Helicase<-METHYLASE<-DUF3780<-DUF499*<-CopG/RHH\|\|XIS-HTH-> | 1032 | Bacteria>Pseudomonadota>Alphaproteobacteria>Rhodospirillales>Novispirillaceae | GCF_000381985.1 | Novispirillum_itersonii | G365_RS0118445 | 37771..40869 | 189 | DUF499-domain-containing-protein-[Novispirillum-itersonii] |
| MDD9997662.1 | AAA+ATPase+wHTH+Alpha+Beta-domain+wHTH+Fn-III+Fn-III+RRM | Type-2_BR.RM.like.systems | <-ParB+HNH\|\|?->DUF499*->DUF3780->METHYLASE->HKD-DNase+SF1-SF2-Helicase-><-?\|\|?-><-PIWI | 1022 | Bacteria>Pseudomonadota>Alphaproteobacteria>Rhodospirillales>Rhodospirillaceae | GCA_028818525.1 | Rhodospirillaceae_bacterium | OXQ89_07950 | 12303..15371 | 1898112 | MAG:-DUF499-domain-containing-protein-[Rhodospirillaceae-bacterium] |
| SUS04809.1 | AAA+ATPase+wHTH+Alpha+Beta-domain+wHTH+Fn-III+Fn-III+RRM | Type-2_BR.RM.like.systems | ParB+HNH->DUF499*->DUF3780->RelE-ParE->ParA+HTH->METHYLASE->HKD-DNase+SF1-SF2-Helicase-> | 1024 | Bacteria>Pseudomonadota>Alphaproteobacteria>Rhodospirillales>Rhodospirillaceae | GCA_900498315.1 | uncultured_Defluviicoccus_sp. | DF3PB_150002 | 1531..4605 | 543062 | conserved-hypothetical-protein-[uncultured-Defluviicoccus-sp.] |
| MBX3517862.1 | AAA+ATPase+wHTH+Alpha+Beta-domain+wHTH+Fn-III+Fn-III+RRM | Type-2_BR.RM.like.systems | <-HKD-DNase+SF1-SF2-Helicase<-METHYLASE<-DUF3780<-DUF499*\|\|35exo->?->?->?-><-D5N+D5ATPase | 1055 | Bacteria>Pseudomonadota>Alphaproteobacteria>Rhodospirillales>unclassified_Rhodospirillales | GCA_019635555.1 | Rhodospirillales_bacterium | KF815_13070 | 97074..100241 | 2026786 | DUF499-domain-containing-protein-[Rhodospirillales-bacterium] |
| MAW91001.1 | AAA+ATPase+wHTH+Alpha+Beta-domain+wHTH+Fn-III+Fn-III+RRM | Type-2_BR.RM.like.systems | REase->HTH_Tnp->?->TniQ-><-?\|\|?->DUF499*->DUF3780->METHYLASE->HKD-DNase+SF1-SF2-Helicase->ACYC->CNMP+TIR->ACYC+ACYC-> | 1033 | Bacteria>Pseudomonadota>Alphaproteobacteria>Sphingomonadales>Erythrobacteraceae | GCA_002710955.1 | Altererythrobacter_sp. | CL574_07895 | 51084..54185 | 1872480 | hypothetical-protein-[Altererythrobacter-sp.] |
| MBK63127.1 | AAA+ATPase+wHTH+Alpha+Beta-domain+wHTH+Fn-III+Fn-III+RRM | Type-2_BR.RM.like.systems | <-ACYC+ACYC<-CNMP+TIR<-ACYC<-HKD-DNase+SF1-SF2-Helicase<-METHYLASE<-DUF3780<-DUF499*<-?\|\|?-><-TniQ<-?<-HTH_Tnp<-REase | 1033 | Bacteria>Pseudomonadota>Alphaproteobacteria>Sphingomonadales>Erythrobacteraceae | GCA_002719495.1 | Altererythrobacter_sp. | CL575_09345 | 28296..31397 | 1872480 | hypothetical-protein-[Altererythrobacter-sp.] |
| WP_119081271.1 | AAA+ATPase+wHTH+Alpha+Beta-domain+wHTH+Fn-III+Fn-III+RRM | Type-2_BR.RM.like.systems | <-ABhydrolase<-HKD-DNase+SF1-SF2-Helicase<-METHYLASE<-DUF3780<-DUF499*\|\|?->DDE_Tnp_1_4->?->?-><-TniQ | 1033 | Bacteria>Pseudomonadota>Alphaproteobacteria>Sphingomonadales>Erythrobacteraceae | GCF_003569745.1 | Altererythrobacter_sp._B11 | AEB_RS00150 | 30987..34088 | 2060312 | DUF499-domain-containing-protein-[Altererythrobacter-sp.-B11] |
| WP_144094495.1 | AAA+ATPase+wHTH+Alpha+Beta-domain+wHTH+Fn-III+Fn-III+RRM | Type-2_BR.RM.like.systems | <-LexA-protease\|\|HTH+WYL->DUF499*->DUF3780->METHYLASE->HEPN->HKD-DNase+SF1-SF2-Helicase->?->TUDOR+SF1-SF2-Helicase+iREase+CTD-> | 1034 | Bacteria>Pseudomonadota>Alphaproteobacteria>Sphingomonadales>Erythrobacteraceae | GCF_007570835.1 | Croceicoccus_sediminis | FAX90_RS00355 | 90290..93394 | 2571150 | DUF499-domain-containing-protein-[Croceicoccus-sediminis] |
| EQB10242.1 | AAA+ATPase+wHTH+Alpha+Beta-domain+wHTH+Fn-III+Fn-III+RRM | Type-2_BR.RM.like.systems | ZnR+TOPRIM->METHYLASE+SF1-SF2-Helicase->ParB+HTH->ArdCN+MPTase->?->?->DUF499*->DUF3780->METHYLASE->HKD-DNase+SF1-SF2-Helicase-> | 981 | Bacteria>Pseudomonadota>Alphaproteobacteria>Sphingomonadales>Sphingomonadaceae | GCA_000445125.1 | Novosphingobium_lindaniclasticum_LE124 | L284_17630 | 11560..14505 | 1096930 | hypothetical-protein-L284_17630-[Novosphingobium-lindaniclasticum-LE124] |
| MAO03351.1 | AAA+ATPase+wHTH+Alpha+Beta-domain+wHTH+Fn-III+Fn-III+RRM | Type-2_BR.RM.like.systems | <-ACYC+ACYC<-CNMP+TIR<-ACYC<-HKD-DNase+SF1-SF2-Helicase<-METHYLASE<-DUF3780<-DUF499*<-?\|\|?-><-TniQ<-?<-HTH_Tnp<-REase | 1033 | Bacteria>Pseudomonadota>Alphaproteobacteria>Sphingomonadales>Sphingomonadaceae | GCA_002693455.1 | Citromicrobium_sp. | CL804_02295 | 38836..41937 | 2024827 | hypothetical-protein-[Citromicrobium-sp.] |
| MCZ8320051.1 | AAA+ATPase+wHTH+Alpha+Beta-domain+wHTH+Fn-III+Fn-III+RRM | Type-2_BR.RM.like.systems | Thioredoxin->TraH->?-><-HKD-DNase+SF1-SF2-Helicase<-METHYLASE<-DUF3780<-DUF499*<-XIS-HTH\|\|?->?->SF1-SF2-Helicase-> | 1043 | Bacteria>Pseudomonadota>Alphaproteobacteria>Sphingomonadales>Sphingomonadaceae | GCA_027533085.1 | Novosphingobium_sp. | O9296_00505 | 39480..42611 | 1874826 | MAG:-DUF499-domain-containing-protein-[Novosphingobium-sp.] |
| WP_011445823.1 | AAA+ATPase+wHTH+Alpha+Beta-domain+wHTH+Fn-III+Fn-III+RRM | Type-2_BR.RM.like.systems | TraH->?-><-?<-HKD-DNase+SF1-SF2-Helicase<-METHYLASE<-DUF3780<-DUF499*<-HTH+WYL\|\|LexA-protease-> | 1034 | Bacteria>Pseudomonadota>Alphaproteobacteria>Sphingomonadales>Sphingomonadaceae | GCF_000013325.1 | Novosphingobium_aromaticivorans | SARO_RS10955 | 2317040..2320144 | 48935 | DUF499-domain-containing-protein-[Novosphingobium-aromaticivorans] |
| WP_160172144.1 | AAA+ATPase+wHTH+Alpha+Beta-domain+wHTH+Fn-III+Fn-III+RRM | Type-2_BR.RM.like.systems | ZnR+TOPRIM->METHYLASE+SF1-SF2-Helicase->ParB+HTH->ArdCN+MPTase->?-><-?\|\|DUF499*->DUF3780->METHYLASE->HKD-DNase+SF1-SF2-Helicase-> | 897 | Bacteria>Pseudomonadota>Alphaproteobacteria>Sphingomonadales>Sphingomonadaceae | GCF_000445125.1 | Novosphingobium_lindaniclasticum | L284_RS41460 | 11812..14505 | 1329895 | DUF499-domain-containing-protein-[Novosphingobium-lindaniclasticum] |
| WP_114686204.1 | AAA+ATPase+wHTH+Alpha+Beta-domain+wHTH+Fn-III+Fn-III+RRM | Type-2_BR.RM.like.systems | ZnR+TOPRIM->?->?-><-?\|\|DNA_methylase-><-MORC\|\|DUF499*->DUF3780->METHYLASE->HKD-DNase+SF1-SF2-Helicase->TPR->REase->RNase+SF1-SF2-Helicase-> | 1034 | Bacteria>Pseudomonadota>Alphaproteobacteria>Sphingomonadales>Sphingomonadaceae | GCF_003345355.1 | Sphingomonas_aracearum | DVW87_RS02650 | 537271..540375 | 2283317 | DUF499-domain-containing-protein-[Sphingomonas-aracearum] |
| WP_150424836.1 | AAA+ATPase+wHTH+Alpha+Beta-domain+wHTH+Fn-III+Fn-III+RRM | Type-2_BR.RM.like.systems | ZnR->?->?->DUF499*->DUF3780->METHYLASE->HKD-DNase+SF1-SF2-Helicase->HIN-HTH->?->LZ_Tnp_IS66+zf-IS66+DDE_Tnp_IS66+DDE_Tnp_IS66_C-> | 1032 | Bacteria>Pseudomonadota>Alphaproteobacteria>Sphingomonadales>Sphingomonadaceae | GCF_008710165.1 | Sphingobium_limneticum | F4U95_RS05095 | 85246..88344 | 1007511 | DUF499-domain-containing-protein-[Sphingobium-limneticum] |
| WP_144901136.1 | AAA+ATPase+wHTH+Alpha+Beta-domain+wHTH+Fn-III+Fn-III+RRM | Type-2_BR.RM.like.systems | <-LexA-protease\|\|HTH+WYL->DUF499*->DUF3780->METHYLASE->HKD-DNase+SF1-SF2-Helicase->?-><-?<-TraH | 1034 | Bacteria>Pseudomonadota>Alphaproteobacteria>Sphingomonadales>Sphingomonadaceae | GCF_014199635.1 | Novosphingobium_taihuense | GGR37_RS04765 | 980502..983606 | 260085 | DUF499-domain-containing-protein-[Novosphingobium-taihuense] |
| WP_242095370.1 | AAA+ATPase+wHTH+Alpha+Beta-domain+wHTH+Fn-III+Fn-III+RRM | Type-2_BR.RM.like.systems | <-HKD-DNase+SF1-SF2-Helicase<-METHYLASE<-DUF3780<-DUF499* | 1033 | Bacteria>Pseudomonadota>Alphaproteobacteria>Sphingomonadales>Sphingomonadaceae | GCF_022662335.1 | Sphingomonas_sp._CROZ-RG-20F-R02-07 | L8H00_RS01065 | 237791..240892 | 2914832 | DUF499-domain-containing-protein-[Sphingomonas-sp.-CROZ-RG-20F-R02-07] |
| WP_094472367.1 | AAA+ATPase+wHTH+Alpha+Beta-domain+wHTH+Fn-III+Fn-III+RRM | Type-2_BR.RM.like.systems | <-Arm-DNA-bind_3\|\|HTH+WYL->DUF499*->DUF3780->METHYLASE->HKD-DNase+SF1-SF2-Helicase->URI-DNase+RAMA-> | 1034 | Bacteria>Pseudomonadota>Alphaproteobacteria>Sphingomonadales>Sphingosinicellaceae | GCF_002251755.1 | Sandarakinorhabdus_cyanobacteriorum | CHU93_RS01065 | 1818..4922 | 1981098 | DUF499-domain-containing-protein-[Sandarakinorhabdus-cyanobacteriorum] |
| MBK6720173.1 | AAA+ATPase+wHTH+Alpha+Beta-domain+wHTH+Fn-III+Fn-III+RRM | Type-2_BR.RM.like.systems | HTH->?->?->?->?->PhdYeFM_antitox->DUF499*->DUF3780->HKD-DNase+SF1-SF2-Helicase-><-Peptidase_S74<-Peptidase_S74 | 1044 | Bacteria>Pseudomonadota>Alphaproteobacteria>Sphingomonadales>unclassified_Sphingomonadales | GCA_016703235.1 | Sphingomonadales_bacterium | IPG62_09975 | 520657..523791 | 1978525 | DUF499-domain-containing-protein-[Sphingomonadales-bacterium] |
| MBK6492958.1 | AAA+ATPase+wHTH+Alpha+Beta-domain+wHTH+Fn-III+Fn-III+RRM | Type-2_BR.RM.like.systems | Peptidase_S74->Peptidase_S74-><-HKD-DNase+SF1-SF2-Helicase<-METHYLASE<-DUF3780<-DUF499*<-PhdYeFM_antitox<-?<-?<-?<-HTH<-GDResolvase+Recombinase+ZnR | 1044 | Bacteria>Pseudomonadota>Alphaproteobacteria>Sphingomonadales>unclassified_Sphingomonadales | GCA_016704425.1 | Sphingomonadales_bacterium | IPF97_15015 | 762153..765287 | 1978525 | DUF499-domain-containing-protein-[Sphingomonadales-bacterium] |
| EXI78520.1 | AAA+ATPase+wHTH+Alpha+Beta-domain+wHTH+Fn-III+Fn-III+RRM | Type-2_BR.RM.like.systems | DUF499*->DUF3780->METHYLASE->HKD-DNase+SF1-SF2-Helicase-> | 1045 | Bacteria>Pseudomonadota>Betaproteobacteria>Betaproteobacteria_incertae_sedis>Candidatus_Accumulibacter | GCA_000585055.1 | Candidatus_Accumulibacter_appositus | AW10_03004 | 2832..5969 | 1454003 | MAG:-hypothetical-protein-AW10_03004-[Candidatus-Accumulibacter-appositus] |
| RDE52523.1 | AAA+ATPase+wHTH+Alpha+Beta-domain+wHTH+Fn-III+Fn-III+RRM | Type-2_BR.RM.like.systems | RAMA->GDResolvase+Recombinase+ZnR->HTH->TIR->DUF499*->DUF3780->METHYLASE->HKD-DNase+SF1-SF2-Helicase-><-tRNA<-?\|\|?-><-SbcC+SbcC | 1050 | Bacteria>Pseudomonadota>Betaproteobacteria>Betaproteobacteria_incertae_sedis>Candidatus_Accumulibacter | GCA_003332265.1 | Candidatus_Accumulibacter_meliphilus | DVS81_01880 | 389951..393103 | 2211374 | MAG:-DUF499-domain-containing-protein-[Candidatus-Accumulibacter-meliphilus] |
| TLD44573.1 | AAA+ATPase+wHTH+Alpha+Beta-domain+wHTH+Fn-III+Fn-III+RRM | Type-2_BR.RM.like.systems | ZnR+TOPRIM+D5ATPase-><-HKD-DNase+SF1-SF2-Helicase<-METHYLASE<-DUF3780<-DUF499* | 1044 | Bacteria>Pseudomonadota>Betaproteobacteria>Betaproteobacteria_incertae_sedis>Candidatus_Accumulibacter | GCA_005524045.1 | Accumulibacter_sp. | FAZ92_03175 | 9813..12947 | 2053492 | MAG:-hypothetical-protein-FAZ92_03175-[Candidatus-Accumulibacter-sp.] |
| MBK8387660.1 | AAA+ATPase+wHTH+Alpha+Beta-domain+wHTH+Fn-III+Fn-III+RRM | Type-2_BR.RM.like.systems | <-HKD-DNase+SF1-SF2-Helicase-p<-TOPRIM<-METHYLASE<-PIN<-MazE_antitoxin<-DUF3780<-DUF499*<-DSBH\|\|?->?-><-?<-Methyltransf_25 | 1044 | Bacteria>Pseudomonadota>Betaproteobacteria>Betaproteobacteria_incertae_sedis>Candidatus_Accumulibacter | GCA_016711725.1 | Accumulibacter_sp. | IPL11_19685 | 6141..9275 | 2053492 | DUF499-domain-containing-protein-[Candidatus-Accumulibacter-sp.] |
| MBK8117091.1 | AAA+ATPase+wHTH+Alpha+Beta-domain+wHTH+Fn-III+Fn-III+RRM | Type-2_BR.RM.like.systems | <-HKD-DNase+SF1-SF2-Helicase<-?<-METHYLASE<-PIN<-MazE_antitoxin<-DUF3780<-DUF499*<-DSBH\|\|?->?-><-?<-Methyltransf_25 | 1044 | Bacteria>Pseudomonadota>Betaproteobacteria>Betaproteobacteria_incertae_sedis>Candidatus_Accumulibacter | GCA_016712935.1 | Accumulibacter_sp. | IPK44_22565 | 276336..279470 | 2053492 | DUF499-domain-containing-protein-[Candidatus-Accumulibacter-sp.] |
| MBN8450450.1 | AAA+ATPase+wHTH+Alpha+Beta-domain+wHTH+Fn-III+Fn-III+RRM | Type-2_BR.RM.like.systems | <-HKD-DNase+SF1-SF2-Helicase<-PIN<-?<-METHYLASE<-DUF3780<-DUF499* | 1034 | Bacteria>Pseudomonadota>Betaproteobacteria>Betaproteobacteria_incertae_sedis>Candidatus_Accumulibacter | GCA_017302435.1 | Candidatus_Accumulibacter_necessarius | JOM13_15880 | 43840..46944 | 2954386 | ATP-binding-protein-[Candidatus-Accumulibacter-sp.] |
| MCM8612468.1 | AAA+ATPase+wHTH+Alpha+Beta-domain+wHTH+Fn-III+Fn-III+RRM | Type-2_BR.RM.like.systems | ZnR+TOPRIM+D5ATPase-><-?<-HKD-DNase+SF1-SF2-Helicase<-METHYLASE<-DUF3780<-DUF499* | 930 | Bacteria>Pseudomonadota>Betaproteobacteria>Betaproteobacteria_incertae_sedis>Candidatus_Accumulibacter | GCA_023806505.1 | Accumulibacter_sp. | NFW17_10295 | 117648..120440 | 2053492 | DUF499-domain-containing-protein-[Candidatus-Accumulibacter-sp.] |
| MCM8636865.1 | AAA+ATPase+wHTH+Alpha+Beta-domain+wHTH+Fn-III+Fn-III+RRM | Type-2_BR.RM.like.systems | ZnR+TOPRIM+D5ATPase-><-?<-HKD-DNase+SF1-SF2-Helicase<-METHYLASE<-DUF3780<-DUF499* | 935 | Bacteria>Pseudomonadota>Betaproteobacteria>Betaproteobacteria_incertae_sedis>Candidatus_Accumulibacter | GCA_023806545.1 | Accumulibacter_sp. | NFW15_13160 | 75797..78604 | 2053492 | DUF499-domain-containing-protein-[Candidatus-Accumulibacter-sp.] |
| MCM8594871.1 | AAA+ATPase+wHTH+Alpha+Beta-domain+wHTH+Fn-III+Fn-III+RRM | Type-2_BR.RM.like.systems | VirB11+MoxR-AAA->?->tRNA-><-?<-?<-?\|\|DUF499*->DUF3780->METHYLASE->HKD-DNase+SF1-SF2-Helicase-p-> | 1048 | Bacteria>Pseudomonadota>Betaproteobacteria>Betaproteobacteria_incertae_sedis>Candidatus_Accumulibacter | GCA_023806595.1 | Accumulibacter_sp. | NFV71_05830 | 29949..33095 | 2053492 | DUF499-domain-containing-protein-[Candidatus-Accumulibacter-sp.] |
| MCM8624397.1 | AAA+ATPase+wHTH+Alpha+Beta-domain+wHTH+Fn-III+Fn-III+RRM | Type-2_BR.RM.like.systems | <-XIS-HTH<-?\|\|DUF499*->DUF3780->METHYLASE->HKD-DNase+SF1-SF2-Helicase->?-><-?<-Phage_integrase | 1027 | Bacteria>Pseudomonadota>Betaproteobacteria>Betaproteobacteria_incertae_sedis>Candidatus_Accumulibacter | GCA_023806625.1 | Accumulibacter_sp. | NFW05_00285 | 9161..12244 | 2053492 | DUF499-domain-containing-protein-[Candidatus-Accumulibacter-sp.] |
| MCM8626847.1 | AAA+ATPase+wHTH+Alpha+Beta-domain+wHTH+Fn-III+Fn-III+RRM | Type-2_BR.RM.like.systems | VirB11+MoxR-AAA->?->tRNA-><-?<-?<-?\|\|DUF499*->DUF3780->METHYLASE->HKD-DNase+SF1-SF2-Helicase-p-> | 1028 | Bacteria>Pseudomonadota>Betaproteobacteria>Betaproteobacteria_incertae_sedis>Candidatus_Accumulibacter | GCA_023806625.1 | Accumulibacter_sp. | NFW05_12950 | 6812..9898 | 2053492 | DUF499-domain-containing-protein-[Candidatus-Accumulibacter-sp.] |
| MCP5230244.1 | AAA+ATPase+wHTH+Alpha+Beta-domain+wHTH+Fn-III+Fn-III+RRM | Type-2_BR.RM.like.systems | Phage_integrase->?-><-?\|\|GDResolvase+HIN-HTH-><-URI-DNase\|\|PhdYeFM_antitox->DUF499*->DUF3780->METHYLASE->HKD-DNase+SF1-SF2-Helicase-><-Schlafen<-?\|\|ArdCN+MPTase-> | 1037 | Bacteria>Pseudomonadota>Betaproteobacteria>Betaproteobacteria_incertae_sedis>Candidatus_Accumulibacter | GCA_024235715.1 | Accumulibacter_sp. | H6941_16790 | 59686..62799 | 2053492 | ATP-binding-protein-[Candidatus-Accumulibacter-sp.] |
| WP_186409049.1 | AAA+ATPase+wHTH+Alpha+Beta-domain+wHTH+Fn-III+Fn-III+RRM | Type-2_BR.RM.like.systems | DnaB->?->?->?->DUF499*->DUF3780->MazE_antitoxin->PIN->METHYLASE->?->PIN-> | 1044 | Bacteria>Pseudomonadota>Betaproteobacteria>Betaproteobacteria_incertae_sedis>Candidatus_Accumulibacter | GCF_900089955.1 | Candidatus_Accumulibacter_aalborgensis | ACCAA_RS20210 | 12255..15389 | 1860102 | DUF499-domain-containing-protein-[Candidatus-Accumulibacter-aalborgensis] |
| MBN8436880.1 | AAA+ATPase+wHTH+Alpha+Beta-domain+wHTH+Fn-III+Fn-III+RRM | Type-2_BR.RM.like.systems | Methyltransf_25->?-><-?<-?\|\|?->DSBH->DUF499*->DUF3780->MazE_antitoxin->PIN->METHYLASE->?->PIN-> | 1044 | Bacteria>Pseudomonadota>Betaproteobacteria>Betaproteobacteria_incertae_sedis>Candidatus_Accumulibacter | GCA_017302555.1 | Accumulibacter_sp. | JOL95_02360 | 92935..96069 | 2053492 | DUF499-domain-containing-protein-[Candidatus-Accumulibacter-sp.] |
| MBK8576832.1 | AAA+ATPase+wHTH+Alpha+Beta-domain+wHTH+Fn-III+Fn-III+RRM | Type-2_BR.RM.like.systems | <-HicB<-PIN<-METHYLASE<-PIN<-MazE_antitoxin<-DUF3780<-DUF499*<-DSBH\|\|?->?-><-?<-Methyltransf_25 | 1044 | Bacteria>Pseudomonadota>Betaproteobacteria>Betaproteobacteria_incertae_sedis>Candidatus_Accumulibacter | GCA_016714935.1 | Candidatus_Accumulibacter_propinquus | IPN78_00515 | 111101..114235 | 2954380 | DUF499-domain-containing-protein-[Candidatus-Accumulibacter-sp.] |
| EXI76478.1 | AAA+ATPase+wHTH+Alpha+Beta-domain+wHTH+Fn-III+Fn-III+RRM | Type-2_BR.RM.like.systems | <-PIN<-?<-METHYLASE<-PIN<-MazE_antitoxin<-DUF3780<-DUF499*<-?<-?\|\|?->?-><-?<-Peptidase | 1044 | Bacteria>Pseudomonadota>Betaproteobacteria>Betaproteobacteria_incertae_sedis>Candidatus_Accumulibacter | GCA_000584995.1 | Candidatus_Accumulibacter_sp._SK-11 | AW07_00426 | 125131..128265 | 1454000 | MAG:-hypothetical-protein-AW07_00426-[Candidatus-Accumulibacter-sp.-SK-11] |
| MCM8613692.1 | AAA+ATPase+wHTH+Alpha+Beta-domain+wHTH+Fn-III+Fn-III+RRM | Type-2_BR.RM.like.systems | <-METHYLASE<-PIN<-MazE_antitoxin<-DUF3780<-DUF499*\|\|?->?-><-?<-?<-Radical-SAM | 1044 | Bacteria>Pseudomonadota>Betaproteobacteria>Betaproteobacteria_incertae_sedis>Candidatus_Accumulibacter | GCA_023806505.1 | Accumulibacter_sp. | NFW17_16745 | 3715..6849 | 2053492 | DUF499-domain-containing-protein-[Candidatus-Accumulibacter-sp.] |
| MBN8455770.1 | AAA+ATPase+wHTH+Alpha+Beta-domain+wHTH+Fn-III+Fn-III+RRM | Type-2_BR.RM.like.systems | <-Schlafen+HATPase+HTH<-DUF3780<-DUF499* | 1039 | Bacteria>Pseudomonadota>Betaproteobacteria>Betaproteobacteria_incertae_sedis>Candidatus_Accumulibacter | GCA_017302415.1 | Accumulibacter_sp. | JOM31_22260 | 5561..8680 | 2053492 | ATP-binding-protein-[Candidatus-Accumulibacter-sp.] |
| TAL91807.1 | AAA+ATPase+wHTH+Alpha+Beta-domain+wHTH+Fn-III+Fn-III+RRM | Type-2_BR.RM.like.systems | <-PhdYeFM_antitox<-HKD-DNase+SF1-SF2-Helicase<-METHYLASE<-MACRODOMAIN<-?<-DUF3780<-DUF499*<-RHH_5\|\|?-><-Arm-DNA-bind_3+Phage_integrase | 1041 | Bacteria>Pseudomonadota>Betaproteobacteria>Burkholderiales>Alcaligenaceae | GCA_004322185.1 | Candidimonas_sp. | EPN77_00110 | 8692..11817 | 1912597 | MAG:-DUF499-domain-containing-protein-[Candidimonas-sp.] |
| NGM85844.1 | AAA+ATPase+wHTH+Alpha+Beta-domain+wHTH+Fn-III+Fn-III+RRM | Type-2_BR.RM.like.systems | <-SbcC<-JAB\|\|RHH_5->DUF499*->DUF3780->?->MACRODOMAIN->METHYLASE->HKD-DNase+SF1-SF2-Helicase->HTH_19-> | 1041 | Bacteria>Pseudomonadota>Betaproteobacteria>Burkholderiales>Alcaligenaceae | GCA_011043535.1 | Parapusillimonas_sp._SGNA-6 | G5B35_00845 | 169912..173037 | 2710884 | DUF499-domain-containing-protein-[Parapusillimonas-sp.-SGNA-6] |
| WP_008167870.1 | AAA+ATPase+wHTH+Alpha+Beta-domain+wHTH+Fn-III+Fn-III+RRM | Type-2_BR.RM.like.systems | <-HTH_3<-?\|\|?->tRNA-><-HKD-DNase+SF1-SF2-Helicase<-METHYLASE<-DUF3780<-DUF499* | 1043 | Bacteria>Pseudomonadota>Betaproteobacteria>Burkholderiales>Alcaligenaceae | GCF_000236785.1 | Achromobacter_arsenitoxydans | KYC_RS25370 | 56027..59158 | 1147684 | DUF499-domain-containing-protein-[Achromobacter-arsenitoxydans] |
| WP_054473216.1 | AAA+ATPase+wHTH+Alpha+Beta-domain+wHTH+Fn-III+Fn-III+RRM | Type-2_BR.RM.like.systems | RHH->DUF499*->DUF3780->?->MACRODOMAIN->METHYLASE->HKD-DNase+SF1-SF2-Helicase->MORC+REase-> | 1041 | Bacteria>Pseudomonadota>Betaproteobacteria>Burkholderiales>Alcaligenaceae | GCF_001298895.1 | Achromobacter_xylosoxidans | N7677_RS16680 | 47740..50865 | 85698 | DUF499-domain-containing-protein-[Achromobacter-xylosoxidans] |
| WP_066667991.1 | AAA+ATPase+wHTH+Alpha+Beta-domain+wHTH+Fn-III+Fn-III+RRM | Type-2_BR.RM.like.systems | Arm-DNA-bind_3+Phage_int_SAM_3+Phage_integrase-><-HKD-DNase+SF1-SF2-Helicase<-METHYLASE<-MACRODOMAIN<-?<-DUF3780<-DUF499*<-RHH_1<-Integrase_1<-MarR-HTH\|\|ABhydrolase->DOC+HTH-><-MODE-HTH+PBPII | 1041 | Bacteria>Pseudomonadota>Betaproteobacteria>Burkholderiales>Alcaligenaceae | GCF_001676745.1 | Bordetella_bronchialis | BAU08_RS02720 | 616565..619690 | 463025 | DUF499-domain-containing-protein-[Bordetella-bronchialis] |
| WP_076468429.1 | AAA+ATPase+wHTH+Alpha+Beta-domain+wHTH+Fn-III+Fn-III+RRM | Type-2_BR.RM.like.systems | <-HAD\|\|?-><-?\|\|RHH->DUF499*->DUF3780->?->MACRODOMAIN->METHYLASE->HKD-DNase+SF1-SF2-Helicase->MORC+REase-> | 1041 | Bacteria>Pseudomonadota>Betaproteobacteria>Burkholderiales>Alcaligenaceae | GCF_001957045.1 | Achromobacter_xylosoxidans | BI147_RS15725 | 74082..77207 | 85698 | DUF499-domain-containing-protein-[Achromobacter-xylosoxidans] |
| WP_102773984.1 | AAA+ATPase+wHTH+Alpha+Beta-domain+wHTH+Fn-III+Fn-III+RRM | Type-2_BR.RM.like.systems | MarR-HTH->Phage_int_SAM_2+Integrase_1->RHH_1->Phage_int_SAM_2+Integrase_1->RHH_1->DUF499->DUF499*->DUF3780->?->MACRODOMAIN->METHYLASE->HKD-DNase+SF1-SF2-Helicase-><-Arm-DNA-bind_3+Phage_int_SAM_3+Phage_integrase | 904 | Bacteria>Pseudomonadota>Betaproteobacteria>Burkholderiales>Alcaligenaceae | GCF_002885955.2 | Achromobacter_pulmonis | C1I89_RS17905 | 578102..580818 | 1389932 | DUF499-domain-containing-protein,-partial-[Achromobacter-pulmonis] |
| WP_130302965.1 | AAA+ATPase+wHTH+Alpha+Beta-domain+wHTH+Fn-III+Fn-III+RRM | Type-2_BR.RM.like.systems | <-RelB\|\|?->XIS-HTH->DUF499*->DUF3780->METHYLASE->HKD-DNase+SF1-SF2-Helicase-><-ParB+HNH<-MORC+REase\|\|HEPN-> | 1022 | Bacteria>Pseudomonadota>Betaproteobacteria>Burkholderiales>Alcaligenaceae | GCF_004217095.1 | Advenella_incenata | EV681_RS00025 | 2520..5588 | 267800 | DUF499-domain-containing-protein-[Advenella-incenata] |
| WP_153229102.1 | AAA+ATPase+wHTH+Alpha+Beta-domain+wHTH+Fn-III+Fn-III+RRM | Type-2_BR.RM.like.systems | Arm-DNA-bind_3+Phage_int_SAM_3+Phage_integrase-><-HKD-DNase+SF1-SF2-Helicase<-METHYLASE<-MACRODOMAIN<-?<-DUF3780<-DUF499*<-RHH_1<-Phage_int_SAM_2+Integrase_1\|\|?-><-MarR-HTH\|\|RelE->CHTH-> | 1041 | Bacteria>Pseudomonadota>Betaproteobacteria>Burkholderiales>Alcaligenaceae | GCF_009497775.1 | Alcaligenes_faecalis | DUD43_RS03025 | 665460..668585 | 511 | DUF499-domain-containing-protein-[Alcaligenes-faecalis] |
| WP_185780770.1 | AAA+ATPase+wHTH+Alpha+Beta-domain+wHTH+Fn-III+Fn-III+RRM | Type-2_BR.RM.like.systems | <-HKD-DNase+SF1-SF2-Helicase<-METHYLASE<-MACRODOMAIN<-?<-DUF3780<-DUF499*<-RHH_5\|\|JAB-> | 1041 | Bacteria>Pseudomonadota>Betaproteobacteria>Burkholderiales>Alcaligenaceae | GCF_014237865.1 | Pusillimonas_minor | GTU67_RS14785 | 10023..13148 | 2697024 | DUF499-domain-containing-protein-[Pusillimonas-minor] |
| WP_269498091.1 | AAA+ATPase+wHTH+Alpha+Beta-domain+wHTH+Fn-III+Fn-III+RRM | Type-2_BR.RM.like.systems | RVT->?-><-HKD-DNase+SF1-SF2-Helicase<-METHYLASE<-Schlafen+HATPase+HTH<-DUF3780<-DUF499*<-RHH_5<-CHTH\|\|XIS-HTH->HTH_17-><-Arm-DNA-bind_3+Phage_integrase | 1041 | Bacteria>Pseudomonadota>Betaproteobacteria>Burkholderiales>Alcaligenaceae | GCF_027257175.1 | Castellaniella_sp._S9 | OOZ48_RS10600 | 328374..331499 | 2993652 | DUF499-domain-containing-protein-[Castellaniella-sp.-S9] |
| WP_102776215.1 | AAA+ATPase+wHTH+Alpha+Beta-domain+wHTH+Fn-III+Fn-III+RRM | Type-2_BR.RM.like.systems | TrwC+SF1-SF2-helicase+TOPRIM-><-?<-?<-SbcC<-JAB\|\|RHH_5->DUF499*->DUF3780->?->MACRODOMAIN->METHYLASE-p->DUF772+DDE_Tnp_1->METHYLASE-> | 1041 | Bacteria>Pseudomonadota>Betaproteobacteria>Burkholderiales>Alcaligenaceae | GCF_002885955.2 | Achromobacter_pulmonis | C1I89_RS32060 | 50774..53899 | 1389932 | DUF499-domain-containing-protein-[Achromobacter-pulmonis] |
| KOE22418.1 | AAA+ATPase+wHTH+Alpha+Beta-domain+wHTH+Fn-III+Fn-III+RRM | Type-2_BR.RM.like.systems | <-ACET<-TraJ-RHH\|\|RHH->DUF499*->DUF3780->METHYLASE->HKD-DNase+SF1-SF2-Helicase-><-XIS-HTH | 1040 | Bacteria>Pseudomonadota>Betaproteobacteria>Burkholderiales>Burkholderiaceae | GCA_001267755.1 | Burkholderia_multivorans_R-20526 | AI46_30005 | 2327..5449 | 1449334 | ATPase-AAA-[Burkholderia-multivorans-R-20526] |
| ODS98135.1 | AAA+ATPase+wHTH+Alpha+Beta-domain+wHTH+Fn-III+Fn-III+RRM | Type-2_BR.RM.like.systems | <-MORC+DUF3684+REase<-HKD-DNase+SF1-SF2-Helicase<-?<-METHYLASE<-DUF3780<-DUF499*<-HTH+WYL\|\|DNA-glycosylase-><-?\|\|Phage_int_SAM_2+Integrase_1-><-?<-MerR-HTH | 1027 | Bacteria>Pseudomonadota>Betaproteobacteria>Burkholderiales>Burkholderiaceae | GCA_001724855.1 | Lautropia_sp._SCN_69-89 | ABS56_05865 | 14072..17155 | 1660104 | MAG:-hypothetical-protein-ABS56_05865-[Lautropia-sp.-SCN-69-89] |
| MBU65595.1 | AAA+ATPase+wHTH+Alpha+Beta-domain+wHTH+Fn-III+Fn-III+RRM | Type-2_BR.RM.like.systems | <-Methylase<-HKD-DNase+SF1-SF2-Helicase<-METHYLASE<-MACRODOMAIN<-?<-DUF3780<-DUF499*<-RHH_5<-ZnR+TOPRIM+D5ATPase<-ParB<-ParB<-GDResolvase+PPP1-HTH | 1041 | Bacteria>Pseudomonadota>Betaproteobacteria>Burkholderiales>Burkholderiaceae | GCA_002730895.1 | Cupriavidus_sp. | CL858_09105 | 332114..335239 | 1873897 | AAA-family-ATPase-[Cupriavidus-sp.] |
| TXH35855.1 | AAA+ATPase+wHTH+Alpha+Beta-domain+wHTH+Fn-III+Fn-III+RRM | Type-2_BR.RM.like.systems | <-HKD-DNase+SF1-SF2-Helicase<-METHYLASE<-MACRODOMAIN<-?<-DUF3780<-DUF499*<-RHH_5 | 1041 | Bacteria>Pseudomonadota>Betaproteobacteria>Burkholderiales>Burkholderiaceae | GCA_008012415.1 | Burkholderiaceae_bacterium | E6Q92_11760 | 8695..11820 | 2030806 | DUF499-domain-containing-protein-[Burkholderiaceae-bacterium] |
| MBP6645096.1 | AAA+ATPase+wHTH+Alpha+Beta-domain+wHTH+Fn-III+Fn-III+RRM | Type-2_BR.RM.like.systems | <-XIS-HTH<-?\|\|DUF499*->DUF3780->METHYLASE->HKD-DNase+SF1-SF2-Helicase->?->SF1-SF2-Helicase->HEPN+TPR+RhoN-> | 1053 | Bacteria>Pseudomonadota>Betaproteobacteria>Burkholderiales>Burkholderiaceae | GCA_017989995.1 | Burkholderiaceae_bacterium | KA207_04515 | 1913..5074 | 2030806 | DUF499-domain-containing-protein-[Burkholderiaceae-bacterium] |
| MBW0450648.1 | AAA+ATPase+wHTH+Alpha+Beta-domain+wHTH+Fn-III+Fn-III+RRM | Type-2_BR.RM.like.systems | CHTH-><-HTH+HKD-DNase<-Pkinase+SF1-SF2-Helicase\|\|RHH_5->DUF499*->DUF3780->METHYLASE->HKD-DNase+SF1-SF2-Helicase->?-><-DUF488-l | 1041 | Bacteria>Pseudomonadota>Betaproteobacteria>Burkholderiales>Burkholderiaceae | GCA_019278015.1 | Paraburkholderia_phenoliruptrix | JRV94_28610 | 13392..16517 | 252970 | DUF499-domain-containing-protein-[Paraburkholderia-phenoliruptrix] |
| MCC7226161.1 | AAA+ATPase+wHTH+Alpha+Beta-domain+wHTH+Fn-III+Fn-III+RRM | Type-2_BR.RM.like.systems | XIS-HTH->ZnR+TOPRIM+primpol+HTH->DUF499*->DUF3780->METHYLASE->HKD-DNase+SF1-SF2-Helicase->?->?-><-ParB | 1044 | Bacteria>Pseudomonadota>Betaproteobacteria>Burkholderiales>Burkholderiaceae | GCA_020851995.1 | Burkholderiaceae_bacterium | IT507_05645 | 24088..27222 | 2030806 | DUF499-domain-containing-protein-[Burkholderiaceae-bacterium] |
| WP_053299181.1 | AAA+ATPase+wHTH+Alpha+Beta-domain+wHTH+Fn-III+Fn-III+RRM | Type-2_BR.RM.like.systems | <-ACET<-TraJ-RHH\|\|RHH->DUF499*->DUF3780->METHYLASE->HKD-DNase+SF1-SF2-Helicase-><-XIS-HTH | 1040 | Bacteria>Pseudomonadota>Betaproteobacteria>Burkholderiales>Burkholderiaceae | GCF_001267755.1 | Burkholderia_multivorans | AI46_30005 | 2327..5449 | 87883 | DUF499-domain-containing-protein-[Burkholderia-multivorans] |
| WP_059825747.1 | AAA+ATPase+wHTH+Alpha+Beta-domain+wHTH+Fn-III+Fn-III+RRM | Type-2_BR.RM.like.systems | <-HKD-DNase+SF1-SF2-Helicase<-METHYLASE<-MACRODOMAIN<-DUF3780<-DUF499*<-RHH_5 | 1041 | Bacteria>Pseudomonadota>Betaproteobacteria>Burkholderiales>Burkholderiaceae | GCF_001525845.1 | Burkholderia_ubonensis | WI72_RS11080 | 7508..10633 | 101571 | DUF499-domain-containing-protein-[Burkholderia-ubonensis] |
| WP_076934373.1 | AAA+ATPase+wHTH+Alpha+Beta-domain+wHTH+Fn-III+Fn-III+RRM | Type-2_BR.RM.like.systems | GDResolvase+PPP1-HTH->ParB->ParB->RHH_5->DUF499*->DUF3780->?->MACRODOMAIN->METHYLASE->HKD-DNase+SF1-SF2-Helicase-> | 1041 | Bacteria>Pseudomonadota>Betaproteobacteria>Burkholderiales>Burkholderiaceae | GCF_001977385.1 | Burkholderia_pseudomallei | EFM92_RS18385 | 41595..44720 | 28450 | DUF499-domain-containing-protein-[Burkholderia-pseudomallei] |
| WP_137330769.1 | AAA+ATPase+wHTH+Alpha+Beta-domain+wHTH+Fn-III+Fn-III+RRM | Type-2_BR.RM.like.systems | TUDOR+SF1-SF2-Helicase+iREase+CTD->?->HHH_4+SF1-SF2-Helicase+REase-><-?\|\|RHH_5->DUF499*->DUF3780->MACRODOMAIN->METHYLASE->HKD-DNase+SF1-SF2-Helicase-> | 1041 | Bacteria>Pseudomonadota>Betaproteobacteria>Burkholderiales>Burkholderiaceae | GCF_005280735.1 | Trinickia_violacea | FAZ95_RS01270 | 296654..299779 | 2571746 | DUF499-domain-containing-protein-[Burkholderia-sp.-DHOD12] |
| WP_216254672.1 | AAA+ATPase+wHTH+Alpha+Beta-domain+wHTH+Fn-III+Fn-III+RRM | Type-2_BR.RM.like.systems | <-HTH+WYL<-?\|\|?->?-><-?<-MerR-HTH\|\|DUF499*->DUF3780->METHYLASE->HKD-DNase+SF1-SF2-Helicase->ParB+HNH+Schlafen-><-PIN | 1038 | Bacteria>Pseudomonadota>Betaproteobacteria>Burkholderiales>Burkholderiaceae | GCF_018882365.1 | Polynucleobacter_sp._71A-WALBACH | ICN42_RS10945 | 39602..42718 | 2689097 | DUF499-domain-containing-protein-[Polynucleobacter-sp.-71A-WALBACH] |
| WP_217114508.1 | AAA+ATPase+wHTH+Alpha+Beta-domain+wHTH+Fn-III+Fn-III+RRM | Type-2_BR.RM.like.systems | RHH->DUF499*->DUF3780->METHYLASE->HKD-DNase+SF1-SF2-Helicase-><-XIS-HTH | 1040 | Bacteria>Pseudomonadota>Betaproteobacteria>Burkholderiales>Burkholderiaceae | GCF_019042455.1 | Burkholderia_multivorans | KTE70_RS07490 | 172913..176035 | 87883 | DUF499-domain-containing-protein-[Burkholderia-multivorans] |
| WP_265206756.1 | AAA+ATPase+wHTH+Alpha+Beta-domain+wHTH+Fn-III+Fn-III+RRM | Type-2_BR.RM.like.systems | <-HNH\|\|\|\|CopG_antitoxin->DUF499*->DUF3780->METHYLASE->HKD-DNase+SF1-SF2-Helicase-><-?<-MORC+DNA_gyraseB+TOPRIM | 1040 | Bacteria>Pseudomonadota>Betaproteobacteria>Burkholderiales>Burkholderiaceae | GCF_019266405.1 | Burkholderia_cenocepacia | B7L14_RS29735 | 5744..8866 | 95486 | DUF499-domain-containing-protein-[Burkholderia-cenocepacia] |
| WP_222858495.1 | AAA+ATPase+wHTH+Alpha+Beta-domain+wHTH+Fn-III+Fn-III+RRM | Type-2_BR.RM.like.systems | DUF488-l-><-?<-HKD-DNase+SF1-SF2-Helicase<-METHYLASE<-?<-DUF3780<-DUF499*<-RHH_5\|\|Pkinase+SF1-SF2-Helicase->HTH+HKD-DNase-><-CHTH | 1022 | Bacteria>Pseudomonadota>Betaproteobacteria>Burkholderiales>Burkholderiaceae | GCF_019511465.1 | Paraburkholderia_phenoliruptrix | JNC06_RS12015 | 480243..483311 | 252970 | DUF499-domain-containing-protein-[Paraburkholderia-phenoliruptrix] |
| WP_227893206.1 | AAA+ATPase+wHTH+Alpha+Beta-domain+wHTH+Fn-III+Fn-III+RRM | Type-2_BR.RM.like.systems | <-URI-DNase+RAMA<-HKD-DNase+SF1-SF2-Helicase<-METHYLASE<-MACRODOMAIN<-?<-DUF3780<-DUF499*<-RHH_5\|\|?-><-?<-?\|\|HIN-HTH-> | 1041 | Bacteria>Pseudomonadota>Betaproteobacteria>Burkholderiales>Burkholderiaceae | GCF_020736425.1 | Burkholderia_vietnamiensis | LK462_RS15635 | 2189797..2192922 | 60552 | DUF499-domain-containing-protein-[Burkholderia-vietnamiensis] |
| WP_234744912.1 | AAA+ATPase+wHTH+Alpha+Beta-domain+wHTH+Fn-III+Fn-III+RRM | Type-2_BR.RM.like.systems | <-HKD-DNase+SF1-SF2-Helicase<-DUF3780<-DUF499*<-RHH_5<-ParB<-ParB<-GDResolvase+PPP1-HTH<-?\|\|MACRODOMAIN-> | 1041 | Bacteria>Pseudomonadota>Betaproteobacteria>Burkholderiales>Burkholderiaceae | GCF_021483755.1 | Burkholderia_sp._WTPI3 | J7024_RS06765 | 56093..59218 | 2822167 | DUF499-domain-containing-protein-[Burkholderia-sp.-WTPI3] |
| WP_250481419.1 | AAA+ATPase+wHTH+Alpha+Beta-domain+wHTH+Fn-III+Fn-III+RRM | Type-2_BR.RM.like.systems | HNH->?->?->DUF499*->DUF3780->METHYLASE->HKD-DNase+SF1-SF2-Helicase->Glyoxalase-> | 1047 | Bacteria>Pseudomonadota>Betaproteobacteria>Burkholderiales>Burkholderiaceae | GCF_023631185.1 | Caballeronia_sp._NCTM5 | NCS64_RS01050 | 209772..212915 | 2921755 | DUF499-domain-containing-protein-[Caballeronia-sp.-NCTM5] |
| WP_250459453.1 | AAA+ATPase+wHTH+Alpha+Beta-domain+wHTH+Fn-III+Fn-III+RRM | Type-2_BR.RM.like.systems | DUF499*->DUF3780->METHYLASE->HKD-DNase+SF1-SF2-Helicase->Glyoxalase-> | 1047 | Bacteria>Pseudomonadota>Betaproteobacteria>Burkholderiales>Burkholderiaceae | GCF_023631285.1 | unclassified_Caballeronia | NCS62_RS27085 | 1232..4375 | 2646786 | DUF499-domain-containing-protein-[Caballeronia-sp.-INML5] |
| WP_160111850.1 | AAA+ATPase+wHTH+Alpha+Beta-domain+wHTH+Fn-III+Fn-III+RRM | Type-2_BR.RM.like.systems | <-Phage_integrase\|\|tRNA->Arm-DNA-bind_3+Phage_integrase->DUF499*->DUF3780->METHYLASE->HKD-DNase+SF1-SF2-Helicase-><-HTH+MPTase<-Tox-RelE | 1034 | Bacteria>Pseudomonadota>Betaproteobacteria>Burkholderiales>Burkholderiaceae | GCF_900007165.1 | Paraburkholderia_piptadeniae | BN2476_RS40865 | 50131..53235 | 1701573 | DUF499-domain-containing-protein-[Paraburkholderia-piptadeniae] |
| WP_084286022.1 | AAA+ATPase+wHTH+Alpha+Beta-domain+wHTH+Fn-III+Fn-III+RRM | Type-2_BR.RM.like.systems | <-HIN-HTH<-?<-?\|\|DUF499*->DUF3780->METHYLASE->HKD-DNase+SF1-SF2-Helicase-> | 1043 | Bacteria>Pseudomonadota>Betaproteobacteria>Burkholderiales>Burkholderiaceae | GCF_900176405.1 | Polynucleobacter_kasalickyi | B8983_RS13500 | 3154..6285 | 1938817 | DUF499-domain-containing-protein-[Polynucleobacter-kasalickyi] |
| TXH35005.1 | AAA+ATPase+wHTH+Alpha+Beta-domain+wHTH+Fn-III+Fn-III+RRM | Type-2_BR.RM.like.systems | <-ParA-Soj-PloopNTPase<-HEPN<-METHYLASE<-MACRODOMAIN<-?<-DUF3780<-DUF499* | 828 | Bacteria>Pseudomonadota>Betaproteobacteria>Burkholderiales>Burkholderiaceae | GCA_008012415.1 | Burkholderiaceae_bacterium | E6Q92_12395 | 11663..14151 | 2030806 | DUF499-domain-containing-protein,-partial-[Burkholderiaceae-bacterium] |
| ODS67619.1 | AAA+ATPase+wHTH+Alpha+Beta-domain+wHTH+Fn-III+Fn-III+RRM | Type-2_BR.RM.like.systems | <-HKD-DNase+SF1-SF2-Helicase<-METHYLASE<-PIN<-MazE_antitoxin<-DUF3780<-DUF499*\|\|OLD-ABC+TOPRIM->SF1-SF2-Helicase-><-HTH_21+rve+rve_3<-HIN-HTH | 1045 | Bacteria>Pseudomonadota>Betaproteobacteria>Burkholderiales>Comamonadaceae | GCA_001724795.1 | Acidovorax_sp._SCN_65-28 | ABS39_19765 | 13942..17079 | 1660086 | MAG:-hypothetical-protein-ABS39_19765-[Acidovorax-sp.-SCN-65-28] |
| MBA3056686.1 | AAA+ATPase+wHTH+Alpha+Beta-domain+wHTH+Fn-III+Fn-III+RRM | Type-2_BR.RM.like.systems | <-HKD-DNase+SF1-SF2-Helicase<-METHYLASE<-DUF3780<-DUF499*<-OLD-ABC+TOPRIM<-HTH_3\|\|?->?->RNase-> | 1049 | Bacteria>Pseudomonadota>Betaproteobacteria>Burkholderiales>Comamonadaceae | GCA_013791765.1 | Rhodoferax_sp. | FP823_02085 | 51974..55123 | 50421 | DUF499-domain-containing-protein-[Rhodoferax-sp.] |
| MBC7619868.1 | AAA+ATPase+wHTH+Alpha+Beta-domain+wHTH+Fn-III+Fn-III+RRM | Type-2_BR.RM.like.systems | <-XIS-HTH<-?\|\|DUF499*->DUF3780->METHYLASE->?->HKD-DNase+SF1-SF2-Helicase-p-> | 1057 | Bacteria>Pseudomonadota>Betaproteobacteria>Burkholderiales>Comamonadaceae | GCA_014377745.1 | Rhodoferax_sp. | H7293_12945 | 1752..4925 | 50421 | DUF499-domain-containing-protein-[Candidatus-Saccharibacteria-bacterium] |
| MBN9628764.1 | AAA+ATPase+wHTH+Alpha+Beta-domain+wHTH+Fn-III+Fn-III+RRM | Type-2_BR.RM.like.systems | HTH_3-><-?<-?\|\|HTH_Tnp_1+HTH_21+rve+rve_2-><-SF1-SF2-Helicase<-OLD-ABC+TOPRIM\|\|DUF499*->DUF3780->MazE_antitoxin->PIN->METHYLASE->HKD-DNase+SF1-SF2-Helicase-> | 1024 | Bacteria>Pseudomonadota>Betaproteobacteria>Burkholderiales>Comamonadaceae | GCA_017307415.1 | Acidovorax_sp. | JOG38_21240 | 6814..9888 | 1872122 | DUF499-domain-containing-protein-[Acidovorax-sp.] |
| MCF8209762.1 | AAA+ATPase+wHTH+Alpha+Beta-domain+wHTH+Fn-III+Fn-III+RRM | Type-2_BR.RM.like.systems | DUF499*->?->HKD-DNase+SF1-SF2-Helicase-p->?->SF1-SF2-Helicase-p-><-?<-ACET | 991 | Bacteria>Pseudomonadota>Betaproteobacteria>Burkholderiales>Comamonadaceae | GCA_021738345.1 | Rhodoferax_sp. | K9K38_10205 | 597..3572 | 50421 | DUF499-domain-containing-protein-[Rhodoferax-sp.] |
| WP_169741491.1 | AAA+ATPase+wHTH+Alpha+Beta-domain+wHTH+Fn-III+Fn-III+RRM | Type-2_BR.RM.like.systems | <-REase<-?\|\|?->DUF499*->DUF3780->MazE_antitoxin->PIN->METHYLASE->HKD-DNase+SF1-SF2-Helicase-> | 1024 | Bacteria>Pseudomonadota>Betaproteobacteria>Burkholderiales>Comamonadaceae | GCF_000739875.1 | Comamonas_aquatica | CA2_RS16400 | 2260..5334 | 225991 | DUF499-domain-containing-protein-[Comamonas-aquatica] |
| WP_042427321.1 | AAA+ATPase+wHTH+Alpha+Beta-domain+wHTH+Fn-III+Fn-III+RRM | Type-2_BR.RM.like.systems | <-Methylase<-HKD-DNase+SF1-SF2-Helicase<-METHYLASE<-MACRODOMAIN<-?<-DUF3780<-DUF499*<-RHH_5<-ZnR+TOPRIM+D5ATPase<-ParB<-ParB<-GDResolvase+PPP1-HTH | 1041 | Bacteria>Pseudomonadota>Betaproteobacteria>Burkholderiales>Comamonadaceae | GCF_000739995.1 | Comamonas_granuli | CGR01S_RS12125 | 152499..155624 | 290309 | DUF499-domain-containing-protein-[Comamonas-granuli] |
| WP_090142652.1 | AAA+ATPase+wHTH+Alpha+Beta-domain+wHTH+Fn-III+Fn-III+RRM | Type-2_BR.RM.like.systems | HTH+MPTase-><-HKD-DNase+SF1-SF2-Helicase<-METHYLASE<-DUF3780<-DUF499*<-RHH_1\|\|HTH-><-?\|\|HIN-HTH->?->RVT-> | 1033 | Bacteria>Pseudomonadota>Betaproteobacteria>Burkholderiales>Comamonadaceae | GCF_001269385.1 | Limnohabitans_sp._DM1 | AMM33_RS17975 | 10059..13160 | 1597955 | DUF499-domain-containing-protein-[Limnohabitans-sp.-DM1] |
| WP_009518701.1 | AAA+ATPase+wHTH+Alpha+Beta-domain+wHTH+Fn-III+Fn-III+RRM | Type-2_BR.RM.like.systems | ZnR+TOPRIM+D5ATPase-><-?<-?<-HTH+MPTase<-?\|\|RHH_5->DUF499*->DUF3780->?->MACRODOMAIN->METHYLASE->HKD-DNase+SF1-SF2-Helicase->REase-> | 1041 | Bacteria>Pseudomonadota>Betaproteobacteria>Burkholderiales>Comamonadaceae | GCF_001571205.1 | Hydrogenophaga | FGJ01_RS07155 | 285549..288674 | 47420 | MULTISPECIES:-DUF499-domain-containing-protein-[Hydrogenophaga] |
| WP_114969907.1 | AAA+ATPase+wHTH+Alpha+Beta-domain+wHTH+Fn-III+Fn-III+RRM | Type-2_BR.RM.like.systems | HTH_17->?-><-?\|\|?->DUF499*->DUF3780->METHYLASE->HKD-DNase+SF1-SF2-Helicase->sGTP-> | 1030 | Bacteria>Pseudomonadota>Betaproteobacteria>Burkholderiales>Comamonadaceae | GCF_003347515.1 | Rhodoferax_ferrireducens | C5P05_RS11095 | 35271..38363 | 192843 | DUF499-domain-containing-protein-[Rhodoferax-ferrireducens] |
| WP_182324144.1 | AAA+ATPase+wHTH+Alpha+Beta-domain+wHTH+Fn-III+Fn-III+RRM | Type-2_BR.RM.like.systems | <-HKD-DNase+SF1-SF2-Helicase<-METHYLASE\|\|?-><-DUF3780<-DUF499*<-?<-?<-TOPRIM+SF1-SF2-Helicase+HTH<-?<-XIS-HTH | 1013 | Bacteria>Pseudomonadota>Betaproteobacteria>Burkholderiales>Comamonadaceae | GCF_014109725.1 | Comamonas_piscis | HS961_RS17370 | 3854444..3857485 | 1562974 | DUF499-domain-containing-protein-[Comamonas-piscis] |

| Protein-ID | DUF499-architecture | System | Gene-Neighbourhoods | Length | Taxonomy | Assembly-ID | Strain | Locus-Tag | Genomic-coordinates | TaxID | Fasta-defline |
|---|---|---|---|---|---|---|---|---|---|---|---|
| WP_210931298.1 | AAA+ATPase+wHTH+Alpha+Beta-domain+wHTH+Fn-III+Fn-III+RRM | Type-2_BR.RM.like.systems | XIS-HTH->?->?->?->DUF499*->DUF3780->MazE_antitoxin->PIN->METHYLASE->HKD-DNase+SF1-SF2-Helicase-><-STYKinase-like | 1024 | Bacteria>Pseudomonadota>Betaproteobacteria>Burkholderiales>Comamonadaceae | GCF_018064205.1 | Comamonas_aquatica | KAQ61_RS08175 | 1732991..1736065 | 225991 | DUF499-domain-containing-protein-[Comamonas-aquatica] |
| WP_284213693.1 | AAA+ATPase+wHTH+Alpha+Beta-domain+wHTH+Fn-III+Fn-III+RRM | Type-2_BR.RM.like.systems | <-HNH+SAVED<-JAB<-UbE1+JAB<-SMODS<-GDResolvase+HIN-HTH\|\|RHH_1->DUF499*->DUF3780->METHYLASE->URI-DNase+RAMA->HKD-DNase+SF1-SF2-Helicase-><-RVT | 1031 | Bacteria>Pseudomonadota>Betaproteobacteria>Burkholderiales>Comamonadaceae | GCF_030162955.1 | Comamonas_jiangduensis | QEP12_RS06200 | 174293..177388 | 1194168 | DUF499-domain-containing-protein-[Comamonas-jiangduensis] |
| WP_022771796.1 | AAA+ATPase+wHTH+Alpha+Beta-domain+wHTH+Fn-III+Fn-III+RRM | Type-2_BR.RM.like.systems | <-PIN<-?<-METHYLASE<-PIN<-MazE_antitoxin<-DUF3780<-DUF499*<-RHH<-tRNA<-?<-RecR+TOPRIM | 1061 | Bacteria>Pseudomonadota>Betaproteobacteria>Burkholderiales>Comamonadaceae | GCF_000477435.1 | Candidatus_Symbiobacter_mobilis | CENROD_RS03870 | 937551..940736 | 1436290 | DUF499-domain-containing-protein-[Candidatus-Symbiobacter-mobilis] |
| MBC7621638.1 | AAA+ATPase+wHTH+Alpha+Beta-domain+wHTH+Fn-III+Fn-III+RRM | Type-2_BR.RM.like.systems | <-METHYLASE-p<-PIN<-?<-DUF3780<-DUF499* | 1048 | Bacteria>Pseudomonadota>Betaproteobacteria>Burkholderiales>Comamonadaceae | GCA_014377745.1 | Rhodoferax_sp. | H7293_22115 | 2441..5587 | 50421 | DUF499-domain-containing-protein-[Candidatus-Saccharibacteria-bacterium] |
| MBN9355826.1 | AAA+ATPase+wHTH+Alpha+Beta-domain+wHTH+Fn-III+Fn-III+RRM | Type-2_BR.RM.like.systems | <-STAS<-ABC-ATPase+SbcC\|\|?->RHH_5->DUF499*->DUF3780->?->MACRODOMAIN->METHYLASE->HKD-DNase+SF1-SF2-Helicase->REase-> | 1041 | Bacteria>Pseudomonadota>Betaproteobacteria>Burkholderiales>Oxalobacteraceae | GCA_017307195.1 | Herbaspirillum_huttiense | J0I15_05180 | 142983..146108 | 863372 | DUF499-domain-containing-protein-[Herbaspirillum-huttiense] |
| WP_186922122.1 | AAA+ATPase+wHTH+Alpha+Beta-domain+wHTH+Fn-III+Fn-III+RRM | Type-2_BR.RM.like.systems | <-URI-DNase+RAMA<-HKD-DNase+SF1-SF2-Helicase<-METHYLASE<-MACRODOMAIN<-?<-DUF3780<-DUF499*<-RHH_5<-HTH+WYL\|\|NERD+SF1-SF2-Helicase->SF1-SF2-Helicase-p-> | 1041 | Bacteria>Pseudomonadota>Betaproteobacteria>Burkholderiales>Oxalobacteraceae | GCF_014284305.1 | Undibacterium_seohonense | H8K52_RS06745 | 169046..172171 | 1344950 | DUF499-domain-containing-protein-[Undibacterium-seohonense] |
| WP_276838598.1 | AAA+ATPase+wHTH+Alpha+Beta-domain+wHTH+Fn-III+Fn-III+RRM | Type-2_BR.RM.like.systems | <-STAS<-ABC-ATPase+SbcC\|\|?->RHH_5->DUF499*->DUF3780->?->MACRODOMAIN->METHYLASE->HKD-DNase+SF1-SF2-Helicase->REase-> | 1041 | Bacteria>Pseudomonadota>Betaproteobacteria>Burkholderiales>Oxalobacteraceae | GCF_017307195.1 | Herbaspirillum_huttiense | J0I15_RS05200 | 142983..146108 | 863372 | DUF499-domain-containing-protein-[Herbaspirillum-huttiense] |
| RVT52743.1 | AAA+ATPase+wHTH+Alpha+Beta-domain+wHTH+Fn-III+Fn-III+RRM | Type-2_BR.RM.like.systems | sGTP->Arm-DNA-bind_3+Phage_integrase->?->MqsA_antitoxin+HTH_3->DUF499*->DUF3780->METHYLASE-p-><-MULE-TRANSPOSASE\|\|METHYLASE-p->HKD-DNase+SF1-SF2-Helicase-> | 1045 | Bacteria>Pseudomonadota>Betaproteobacteria>Burkholderiales>Sphaerotilaceae | GCA_004016515.1 | Rubrivivax_albus | ENE75_10045 | 837812..840949 | 2499835 | DUF499-domain-containing-protein-[Rubrivivax-albus] |
| NYG35432.1 | AAA+ATPase+wHTH+Alpha+Beta-domain+wHTH+Fn-III+Fn-III+RRM | Type-2_BR.RM.like.systems | <-AbiEii<-AbiEi_4<-?\|\|?-><-35exo+SF1-SF2-Helicase\|\|DUF499*->DUF3780->METHYLASE->HKD-DNase+SF1-SF2-Helicase-> | 1045 | Bacteria>Pseudomonadota>Betaproteobacteria>Burkholderiales>Sphaerotilaceae | GCA_013410775.1 | Sphaerotilus_montanus | BDD16_004494 | 55356..58493 | 522889 | hypothetical-protein-[Sphaerotilus-montanus] |
| MBL0730627.1 | AAA+ATPase+wHTH+Alpha+Beta-domain+wHTH+Fn-III+Fn-III+RRM | Type-2_BR.RM.like.systems | <-REase<-?\|\|RAMA->?->?->RHH_1->DUF499*->DUF3780->METHYLASE->HKD-DNase+SF1-SF2-Helicase-><-?<-TetR-HTH | 1040 | Bacteria>Pseudomonadota>Betaproteobacteria>Burkholderiales>Sphaerotilaceae | GCA_016735515.1 | Piscinibacter_sp._HJYY11 | JI745_23165 | 4948200..4951322 | 2801333 | DUF499-domain-containing-protein-[Piscinibacter-sp.-HJYY11] |
| MBL8328914.1 | AAA+ATPase+wHTH+Alpha+Beta-domain+wHTH+Fn-III+Fn-III+RRM | Type-2_BR.RM.like.systems | <-HKD-DNase+SF1-SF2-Helicase<-METHYLASE<-Schlafen+HATPase+HTH<-DUF3780<-DUF499*<-RHH_1\|\|HNH->?->CopG_antitoxin-> | 1040 | Bacteria>Pseudomonadota>Betaproteobacteria>Burkholderiales>Sphaerotilaceae | GCA_016790225.1 | Rubrivivax_sp. | JNJ71_08675 | 122219..125341 | 50259 | DUF499-domain-containing-protein-[Rubrivivax-sp.] |
| MBX3641768.1 | AAA+ATPase+wHTH+Alpha+Beta-domain+wHTH+Fn-III+Fn-III+RRM | Type-2_BR.RM.like.systems | <-HIN-HTH\|\|PhdYeFM_antitox->RelE-ParE->DUF499*->DUF3780->METHYLASE->HKD-DNase+SF1-SF2-Helicase-> | 1013 | Bacteria>Pseudomonadota>Betaproteobacteria>Burkholderiales>Sphaerotilaceae | GCA_019635285.1 | Rubrivivax_sp. | KF720_01710 | 134268..137309 | 50259 | DUF499-domain-containing-protein-[Rubrivivax-sp.] |
| MCE2689652.1 | AAA+ATPase+wHTH+Alpha+Beta-domain+wHTH+Fn-III+Fn-III+RRM | Type-2_BR.RM.like.systems | <-HKD-DNase+SF1-SF2-Helicase<-?<-METHYLASE<-DUF3780<-DUF499*\|\|?-><-?\|\|RelE->CHTH-> | 1027 | Bacteria>Pseudomonadota>Betaproteobacteria>Burkholderiales>Sphaerotilaceae | GCA_021300575.1 | Rubrivivax_sp. | LW862_07365 | 8331..11414 | 50259 | DUF499-domain-containing-protein-[Rubrivivax-sp.] |
| WP_104302216.1 | AAA+ATPase+wHTH+Alpha+Beta-domain+wHTH+Fn-III+Fn-III+RRM | Type-2_BR.RM.like.systems | <-HKD-DNase+SF1-SF2-Helicase<-METHYLASE<-MACRODOMAIN<-?<-DUF3780<-DUF499*<-RHH\|\|?-><-MerR-HTH\|\|<-HAMP+HISKIN | 1041 | Bacteria>Pseudomonadota>Betaproteobacteria>Burkholderiales>Sphaerotilaceae | GCF_002930615.1 | Caldimonas_caldifontis | C1704_RS08060 | 51594..54719 | 1452508 | DUF499-domain-containing-protein-[Zhizhongheella-caldifontis] |
| WP_119157530.1 | AAA+ATPase+wHTH+Alpha+Beta-domain+wHTH+Fn-III+Fn-III+RRM | Type-2_BR.RM.like.systems | RHH->DUF499*->DUF3780->METHYLASE->HKD-DNase+SF1-SF2-Helicase-><-HIN-HTH<-HTH+WYL | 1034 | Bacteria>Pseudomonadota>Betaproteobacteria>Burkholderiales>Sphaerotilaceae | GCF_003569765.1 | Caldimonas_tepidiphila | D3A59_RS22310 | 1272..4376 | 2315841 | DUF499-domain-containing-protein-[Aquabacterium-tepidiphilum] |
| WP_158288907.1 | AAA+ATPase+wHTH+Alpha+Beta-domain+wHTH+Fn-III+Fn-III+RRM | Type-2_BR.RM.like.systems | sGTP->Arm-DNA-bind_3+Phage_integrase->?->MqsA_antitoxin+HTH_3->DUF499*->DUF3780->METHYLASE-p-><-MULE-TRANSPOSASE\|\|METHYLASE-p->HKD-DNase+SF1-SF2-Helicase-> | 963 | Bacteria>Pseudomonadota>Betaproteobacteria>Burkholderiales>Sphaerotilaceae | GCF_004016515.1 | Rubrivivax_albus | ENE75_RS10045 | 838058..840949 | 2499835 | DUF499-domain-containing-protein-[Rubrivivax-albus] |
| WP_132762965.1 | AAA+ATPase+wHTH+Alpha+Beta-domain+wHTH+Fn-III+Fn-III+RRM | Type-2_BR.RM.like.systems | <-HKD-DNase+SF1-SF2-Helicase<-METHYLASE<-MACRODOMAIN<-?<-DUF3780<-DUF499*<-RHH | 1041 | Bacteria>Pseudomonadota>Betaproteobacteria>Burkholderiales>Sphaerotilaceae | GCF_026057715.1 | Caldimonas_thermodepolymerans | ONS87_RS07545 | 1526756..1529881 | 215580 | DUF499-domain-containing-protein-[Schlegelella-thermodepolymerans] |
| WP_247652921.1 | AAA+ATPase+wHTH+Alpha+Beta-domain+wHTH+Fn-III+Fn-III+RRM | Type-2_BR.RM.like.systems | CopG/RHH->RHH_1->DUF499*->DUF3780->Schlafen+HATPase+HTH->METHYLASE->HKD-DNase+SF1-SF2-Helicase-> | 1021 | Bacteria>Pseudomonadota>Betaproteobacteria>Burkholderiales>Sphaerotilaceae | - | Ideonella | KAK07_RS23525 | 27586..30651 | 36862 | MULTISPECIES:-DUF499-domain-containing-protein-[Ideonella] |
| WP_236675415.1 | AAA+ATPase+wHTH+Alpha+Beta-domain+wHTH+Fn-III+Fn-III+RRM | Type-2_BR.RM.like.systems | <-REase<-?\|\|RAMA->?->?->RHH_1->DUF499*->DUF3780->METHYLASE->HKD-DNase+SF1-SF2-Helicase-><-?<-TetR-HTH | 1021 | Bacteria>Pseudomonadota>Betaproteobacteria>Burkholderiales>Sphaerotilaceae | - | Piscinibacter_sp._HJYY11 | JI745_RS23165 | 4948257..4951322 | 2801333 | DUF499-domain-containing-protein-[Piscinibacter-sp.-HJYY11] |
| WP_246332866.1 | AAA+ATPase+wHTH+Alpha+Beta-domain+wHTH+Fn-III+Fn-III+RRM | Type-2_BR.RM.like.systems | <-AbiEii<-AbiEi_4<-?\|\|?-><-35exo+SF1-SF2-Helicase\|\|DUF499*->DUF3780->METHYLASE->HKD-DNase+SF1-SF2-Helicase-> | 963 | Bacteria>Pseudomonadota>Betaproteobacteria>Burkholderiales>Sphaerotilaceae | - | Sphaerotilus_montanus | BDD16_RS22275 | 55602..58493 | 522889 | DUF499-domain-containing-protein-[Sphaerotilus-montanus] |
| WP_138938924.1 | AAA+ATPase+wHTH+Alpha+Beta-domain+wHTH+Fn-III+Fn-III+RRM | Type-2_BR.RM.like.systems | <-SIGMA-HTH<-Transposase_mut\|\|HTH_3->RHH_1->DUF499*->DUF3780->Schlafen+HATPase+HTH->METHYLASE->HEPN->ParA-Soj-PloopNTPase->ORC-CDC6-> | 1022 | Bacteria>Pseudomonadota>Betaproteobacteria>Burkholderiales>Sphaerotilaceae | GCF_011365705.1 | Rubrivivax_benzoatilyticus | G7087_RS13130 | 24930..27998 | 316997 | DUF499-domain-containing-protein-[Rubrivivax-benzoatilyticus] |
| WP_009858743.1 | AAA+ATPase+wHTH+Alpha+Beta-domain+wHTH+Fn-III+Fn-III+RRM | Type-2_BR.RM.like.systems | <-HIN-HTH<-METHYLASE<-Schlafen+HATPase+HTH<-DUF3780<-DUF499*<-RHH_1<-HTH_3\|\|Transposase_mut->SIGMA-HTH-> | 1041 | Bacteria>Pseudomonadota>Betaproteobacteria>Burkholderiales>Sphaerotilaceae | GCF_000190375.1 | Rubrivivax_benzoatilyticus | F799_RS0117815 | 6811..9936 | 316997 | DUF499-domain-containing-protein-[Rubrivivax-benzoatilyticus] |
| MCC7099553.1 | AAA+ATPase+wHTH+Alpha+Beta-domain+wHTH+Fn-III+Fn-III+RRM | Type-2_BR.RM.like.systems | <-Schlafen<-DUF3780<-DUF499*<-RHH_1 | 1040 | Bacteria>Pseudomonadota>Betaproteobacteria>Burkholderiales>Sphaerotilaceae | GCA_020850905.1 | Rubrivivax_sp. | IT500_08190 | 1682..4804 | 50259 | DUF499-domain-containing-protein-[Rubrivivax-sp.] |
| MBE0548353.1 | AAA+ATPase+wHTH+Alpha+Beta-domain+wHTH+Fn-III+Fn-III+RRM | Type-2_BR.RM.like.systems | <-DUF3780<-DUF499*<-RHH_1\|\|?->TOPRIM-> | 1040 | Bacteria>Pseudomonadota>Betaproteobacteria>Burkholderiales>Sphaerotilaceae | GCA_014859905.1 | Rubrivivax_sp. | IH627_12045 | 2233..5355 | 50259 | DUF499-domain-containing-protein-[Rubrivivax-sp.] |
| OGA80233.1 | AAA+ATPase+wHTH+Alpha+Beta-domain+wHTH+Fn-III+Fn-III+RRM | Type-2_BR.RM.like.systems | MPTase-><-HKD-DNase+SF1-SF2-Helicase<-METHYLASE<-MACRODOMAIN<-?<-DUF3780<-DUF499* | 1041 | Bacteria>Pseudomonadota>Betaproteobacteria>Burkholderiales>unclassified_Burkholderiales | GCA_001770725.1 | Burkholderiales_bacterium_RIFCSPHIGHO2_01_FULL_63_240 | A2711_11415 | 41165..44290 | 1797552 | AAA-family-ATPase-[Burkholderiales-bacterium-RIFCSPHIGHO2_01_FULL_63_240] |
| NNM63254.1 | AAA+ATPase+wHTH+Alpha+Beta-domain+wHTH+Fn-III+Fn-III+RRM | Type-2_BR.RM.like.systems | <-HKD-DNase+SF1-SF2-Helicase<-URI-DNase+RAMA<-METHYLASE<-DUF3780<-DUF499*<-RHH\|\|HTH+WYL-><-?\|\|?->?->Zn_Tnp_IS1595+DDE_Tnp_IS1595-> | 1031 | Bacteria>Pseudomonadota>Betaproteobacteria>Burkholderiales>unclassified_Burkholderiales | GCA_013044035.1 | Burkholderiales_bacterium | HKL99_01325 | 194447..197542 | 1891238 | DUF499-domain-containing-protein-[Burkholderiales-bacterium] |
| MBI3155186.1 | AAA+ATPase+wHTH+Alpha+Beta-domain+wHTH+Fn-III+Fn-III+RRM | Type-2_BR.RM.like.systems | <-HKD-DNase+SF1-SF2-Helicase<-METHYLASE<-Schlafen+HATPase+HTH<-DUF3780<-DUF499*<-RHH_1<-ABC-ATPase+SbcC<-PrmC_N+Methyltransf_25<-Peptidase_M20 | 1040 | Bacteria>Pseudomonadota>Betaproteobacteria>Burkholderiales>unclassified_Burkholderiales | GCA_016191995.1 | Burkholderiales_bacterium | HYZ20_07235 | 13809..16931 | 1891238 | DUF499-domain-containing-protein-[Burkholderiales-bacterium] |
| MCA3175693.1 | AAA+ATPase+wHTH+Alpha+Beta-domain+wHTH+Fn-III+Fn-III+RRM | Type-2_BR.RM.like.systems | DOC+HTH-><-HKD-DNase+SF1-SF2-Helicase<-METHYLASE<-MACRODOMAIN<-?<-DUF3780<-DUF499*<-RHH_5 | 1041 | Bacteria>Pseudomonadota>Betaproteobacteria>Burkholderiales>unclassified_Burkholderiales | GCA_020248585.1 | Burkholderiales_bacterium | ING36_09175 | 8666..11791 | 1891238 | DUF499-domain-containing-protein-[Burkholderiales-bacterium] |
| MCA3223201.1 | AAA+ATPase+wHTH+Alpha+Beta-domain+wHTH+Fn-III+Fn-III+RRM | Type-2_BR.RM.like.systems | sGTP->tRNA->?->DUF499*->DUF3780->METHYLASE->HKD-DNase+SF1-SF2-Helicase->HEPN->?->URI-DNase+RAMA-> | 1027 | Bacteria>Pseudomonadota>Betaproteobacteria>Burkholderiales>unclassified_Burkholderiales | GCA_020248865.1 | Burkholderiales_bacterium | ING41_12560 | 90260..93343 | 1891238 | DUF499-domain-containing-protein-[Burkholderiales-bacterium] |
| MDE2076666.1 | AAA+ATPase+wHTH+Alpha+Beta-domain+wHTH+Fn-III+Fn-III+RRM | Type-2_BR.RM.like.systems | <-HTH+WYL\|\|?->?->DUF499*->DUF3780->?->PIN->METHYLASE->HKD-DNase+SF1-SF2-Helicase-p-> | 1056 | Bacteria>Pseudomonadota>Betaproteobacteria>Burkholderiales>unclassified_Burkholderiales | GCA_028864885.1 | Burkholderiales_bacterium | KGI91_06275 | 36073..39243 | 1891238 | MAG:-DUF499-domain-containing-protein-[Burkholderiales-bacterium] |
| MBK7616707.1 | AAA+ATPase+wHTH+Alpha+Beta-domain+wHTH+Fn-III+Fn-III+RRM | Type-2_BR.RM.like.systems | <-HTH+WYL\|\|HNH-><-?\|\|Arm-DNA-bind_3+Phage_integrase-><-ParB+HNH<-DUF3780<-DUF499*<-SF1-SF2-Helicase+DUF1998<-?<-?<-?\|\|HNH-> | 1045 | Bacteria>Pseudomonadota>Betaproteobacteria>Burkholderiales>unclassified_Burkholderiales | GCA_016710065.1 | Burkholderiales_bacterium | IPJ08_20475 | 317100..320237 | 1891238 | ATP-binding-protein-[Burkholderiales-bacterium] |
| MDE2080727.1 | AAA+ATPase+wHTH+Alpha+Beta-domain+wHTH+Fn-III+Fn-III+RRM | Type-2_BR.RM.like.systems | ABC-ATPase->HAD-><-?\|\|?-><-Schlafen+HATPase+HTH<-DUF3780<-DUF499*<-RHH_1<-?<-MORC\|\|?->?->Glycos_trans_3N+Glycos_transf_3-> | 1040 | Bacteria>Pseudomonadota>Betaproteobacteria>Burkholderiales>unclassified_Burkholderiales | GCA_028872015.1 | Burkholderiales_bacterium | KGI90_05185 | 52053..55175 | 1891238 | MAG:-DUF499-domain-containing-protein-[Burkholderiales-bacterium] |
| WP_239905585.1 | AAA+ATPase+wHTH+Alpha+Beta-domain+wHTH+Fn-III+Fn-III+RRM | Type-2_BR.RM.like.systems | HTH_3->?->?->RHH_5->DUF499*->DUF3780->?->MACRODOMAIN->METHYLASE->HKD-DNase+SF1-SF2-Helicase-> | 1041 | Bacteria>Pseudomonadota>Betaproteobacteria>Neisseriales>Aquaspirillaceae | GCF_022394635.1 | Laribacter_hongkongensis | LH433_RS03065 | 103644..106769 | 168471 | DUF499-domain-containing-protein-[Laribacter-hongkongensis] |
| WP_128581437.1 | AAA+ATPase+wHTH+Alpha+Beta-domain+wHTH+Fn-III+Fn-III+RRM | Type-2_BR.RM.like.systems | Aminotran_1_2->?->?-><-HKD-DNase+SF1-SF2-Helicase<-METHYLASE<-DUF3780<-DUF499*<-RHH | 1040 | Bacteria>Pseudomonadota>Betaproteobacteria>Neisseriales>Neisseriaceae | GCF_001618065.1 | Neisseria_flavescens | TW89_RS10890 | 66415..69537 | 484 | DUF499-domain-containing-protein-[Neisseria-flavescens] |
| WP_080614041.1 | AAA+ATPase+wHTH+Alpha+Beta-domain+wHTH+Fn-III+Fn-III+RRM | Type-2_BR.RM.like.systems | <-HKD-DNase+SF1-SF2-Helicase<-METHYLASE<-MACRODOMAIN<-?<-DUF3780<-DUF499*<-RHH\|\|?->?-><-?\|\|ABhydrolase-> | 1041 | Bacteria>Pseudomonadota>Betaproteobacteria>Neisseriales>Neisseriaceae | GCF_002073715.2 | Neisseria_mucosa | A6J88_RS22875 | 1744324..1747449 | 488 | DUF499-domain-containing-protein-[Neisseria-mucosa] |
| WP_074747016.1 | AAA+ATPase+wHTH+Alpha+Beta-domain+wHTH+Fn-III+Fn-III+RRM | Type-2_BR.RM.like.systems | <-Phage_integrase<-?\|\|?->DUF499*->DUF3780->METHYLASE->HKD-DNase+SF1-SF2-Helicase-><-McrC-NTD+REase | 1043 | Bacteria>Pseudomonadota>Betaproteobacteria>Nitrosomonadales>Nitrosomonadaceae | GCF_900110185.1 | Nitrosospira_multiformis | BMY01_RS10795 | 146430..149561 | 1231 | DUF499-domain-containing-protein-[Nitrosospira-multiformis] |
| MBW8328694.1 | AAA+ATPase+wHTH+Alpha+Beta-domain+wHTH+Fn-III+Fn-III+RRM | Type-2_BR.RM.like.systems | Calcineurin->MORC+DUF3684+REase->HNH-><-?\|\|DUF499*->DUF3780->METHYLASE->HKD-DNase+SF1-SF2-Helicase->HicA_toxin->HicB_lk_antitox-> | 1034 | Bacteria>Pseudomonadota>Betaproteobacteria>Nitrosomonadales>Thiobacillaceae | GCA_019459705.1 | Thiobacillus_sp. | K0M48_05945 | 15328..18432 | 924 | DUF499-domain-containing-protein-[Thiobacillus-sp.] |
| MBL0353835.1 | AAA+ATPase+wHTH+Alpha+Beta-domain+wHTH+Fn-III+Fn-III+RRM | Type-2_BR.RM.like.systems | NUDIX-><-?<-REase<-ParB+HNH<-?\|\|?->DUF499*->DUF3780->METHYLASE->HKD-DNase+SF1-SF2-Helicase-><-OLD-ABC<-DDE_Tnp_1 | 1049 | Bacteria>Pseudomonadota>Betaproteobacteria>Rhodocyclales>Azonexaceae | GCA_016722705.1 | Candidatus_Dechloromonas_phosphoritropha | IPP03_14705 | 1844732..1847881 | 1946401 | DUF499-domain-containing-protein-[Dechloromonas-sp.] |
| WP_274740690.1 | AAA+ATPase+wHTH+Alpha+Beta-domain+wHTH+Fn-III+Fn-III+RRM | Type-2_BR.RM.like.systems | GDResolvase+PPP1-HTH->ParB->ParB->RHH_5->DUF499*->DUF3780->?->MACRODOMAIN->METHYLASE->HKD-DNase+SF1-SF2-Helicase->URI-DNase+RAMA-> | 1041 | Bacteria>Pseudomonadota>Betaproteobacteria>Rhodocyclales>Azonexaceae | GCF_028864255.1 | Dechloromonas_agitata | PTW32_RS19040 | 3225..6350 | 73030 | DUF499-domain-containing-protein-[Dechloromonas-agitata] |
| TXH01181.1 | AAA+ATPase+wHTH+Alpha+Beta-domain+wHTH+Fn-III+Fn-III+RRM | Type-2_BR.RM.like.systems | <-HKD-DNase+SF1-SF2-Helicase<-METHYLASE<-DUF3780<-DUF499* | 1033 | Bacteria>Pseudomonadota>Betaproteobacteria>Rhodocyclales>Rhodocyclaceae | GCA_008012085.1 | Rhodocyclaceae_bacterium | E6R09_07140 | 11061..14162 | 1898103 | DUF499-domain-containing-protein-[Rhodocyclaceae-bacterium] |
| MBY0340665.1 | AAA+ATPase+wHTH+Alpha+Beta-domain+wHTH+Fn-III+Fn-III+RRM | Type-2_BR.RM.like.systems | <-URI-DNase+RAMA<-HKD-DNase+SF1-SF2-Helicase<-METHYLASE<-MACRODOMAIN<-?<-DUF3780<-DUF499*<-RHH_5\|\|?->?->STYKIN-> | 1022 | Bacteria>Pseudomonadota>Betaproteobacteria>Rhodocyclales>Rhodocyclaceae | GCA_019750575.1 | Rhodocyclaceae_bacterium | K2Q19_05825 | 10584..13652 | 1898103 | DUF499-domain-containing-protein-[Rhodocyclaceae-bacterium] |
| MDD2741615.1 | AAA+ATPase+wHTH+Alpha+Beta-domain+wHTH+Fn-III+Fn-III+RRM | Type-2_BR.RM.like.systems | HHH_4+SF1-SF2-helicase-><-HKD-DNase+SF1-SF2-Helicase<-METHYLASE<-DUF3780<-DUF499* | 1034 | Bacteria>Pseudomonadota>Betaproteobacteria>Rhodocyclales>Rhodocyclaceae | GCA_028687645.1 | Rhodocyclaceae_bacterium | PHV02_05040 | 30899..34003 | 1898103 | MAG:-DUF499-domain-containing-protein-[Rhodocyclaceae-bacterium] |
| TMW74654.1 | AAA+ATPase+wHTH+Alpha+Beta-domain+wHTH+Fn-III+Fn-III+RRM | Type-2_BR.RM.like.systems | <-CHTH\|\|?->?->?->tRNA-><-?\|\|DUF499*->DUF3780->METHYLASE->HKD-DNase+SF1-SF2-Helicase-> | 1049 | Bacteria>Pseudomonadota>Betaproteobacteria>Rhodocyclales>Zoogloeaceae | GCA_006007845.1 | Thauera_sp._UPWRP | FG147_06645 | 11672..14821 | 2488794 | DUF499-domain-containing-protein-[Thauera-sp.-UPWRP] |
| MBP7790325.1 | AAA+ATPase+wHTH+Alpha+Beta-domain+wHTH+Fn-III+Fn-III+RRM | Type-2_BR.RM.like.systems | DUF499*->DUF3780->METHYLASE->HKD-DNase+SF1-SF2-Helicase-><-REase+nSTAND3<-SbcC+SbcC<-Phage_int_SAM_1+Phage_integrase | 1044 | Bacteria>Pseudomonadota>Betaproteobacteria>Rhodocyclales>Zoogloeaceae | GCA_018000855.1 | Zoogloea_sp. | KA024_00675 | 1365..4499 | 49181 | DUF499-domain-containing-protein-[Zoogloea-sp.] |
| OGA48883.1 | AAA+ATPase+wHTH+Alpha+Beta-domain+wHTH+Fn-III+Fn-III+RRM | Type-2_BR.RM.like.systems | DUF499*->DUF3780->METHYLASE->?->HKD-DNase+SF1-SF2-Helicase-><-HTH+UbE1 | 1027 | Bacteria>Pseudomonadota>Betaproteobacteria>unclassified_Betaproteobacteria | GCA_001772005.1 | Betaproteobacteria_bacterium_RIFCSPLOWO2_12_FULL_62_58 | A3F74_27660 | 2223..5306 | 1797497 | hypothetical-protein-A3F74_27660-[Betaproteobacteria-bacterium-RIFCSPLOWO2_12_FULL_62_58] |
| MPZ42202.1 | AAA+ATPase+wHTH+Alpha+Beta-domain+wHTH+Fn-III+Fn-III+RRM | Type-2_BR.RM.like.systems | <-HKD-DNase+SF1-SF2-Helicase<-?<-Ntox49<-DUF3780<-PIN<-?<-DUF499* | 1021 | Bacteria>Pseudomonadota>Betaproteobacteria>unclassified_Betaproteobacteria | GCA_009377585.1 | Betaproteobacteria_bacterium | GEV05_02130 | 145657..148722 | 1891241 | DUF499-domain-containing-protein-[Betaproteobacteria-bacterium] |
| MBI3148522.1 | AAA+ATPase+wHTH+Alpha+Beta-domain+wHTH+Fn-III+Fn-III+RRM | Type-2_BR.RM.like.systems | <-Integrase_1+Phage_integrase\|\|?->?->RHH_5->DUF499*->DUF3780->?->MACRODOMAIN->METHYLASE->HKD-DNase+SF1-SF2-Helicase->DOC+HTH-> | 1041 | Bacteria>Pseudomonadota>Betaproteobacteria>unclassified_Betaproteobacteria | GCA_016192065.1 | Betaproteobacteria_bacterium | HYZ17_08415 | 3794..6919 | 1891241 | DUF499-domain-containing-protein-[Betaproteobacteria-bacterium] |
| MBI3370534.1 | AAA+ATPase+wHTH+Alpha+Beta-domain+wHTH+Fn-III+Fn-III+RRM | Type-2_BR.RM.like.systems | <-MqsA_antitoxin+HTH_3\|\|?->HEPN->?->DUF499*->DUF3780->METHYLASE->?->?->HKD-DNase+SF1-SF2-Helicase-> | 1026 | Bacteria>Pseudomonadota>Betaproteobacteria>unclassified_Betaproteobacteria | GCA_016195915.1 | Betaproteobacteria_bacterium | HY017_02090 | 33561..36641 | 1891241 | DUF499-domain-containing-protein-[Betaproteobacteria-bacterium] |
| MBK7647592.1 | AAA+ATPase+wHTH+Alpha+Beta-domain+wHTH+Fn-III+Fn-III+RRM | Type-2_BR.RM.like.systems | <-Phage_int_SAM_2+Integrase_1\|\|?->?->RHH_5->DUF499*->DUF3780->?->MACRODOMAIN->METHYLASE->HKD-DNase+SF1-SF2-Helicase-> | 1041 | Bacteria>Pseudomonadota>Betaproteobacteria>unclassified_Betaproteobacteria | GCA_016709965.1 | Betaproteobacteria_bacterium | IPJ12_10575 | 2144439..2147564 | 1891241 | DUF499-domain-containing-protein-[Betaproteobacteria-bacterium] |
| GHU10669.1 | AAA+ATPase+wHTH+Alpha+Beta-domain+wHTH+Fn-III+Fn-III+RRM | Type-2_BR.RM.like.systems | <-ParA-Soj-PloopNTPase<-HEPN<-METHYLASE<-MACRODOMAIN<-?<-DUF3780<-DUF499*<-RHH_5 | 1041 | Bacteria>Pseudomonadota>Betaproteobacteria>unclassified_Betaproteobacteria | GCA_025997015.1 | Betaproteobacteria_bacterium | AGMMS50225_14630 | 22175..25300 | 1891241 | hypothetical-protein-AGMMS50225_14630-[Betaproteobacteria-bacterium] |
| WP_024250705.1 | AAA+ATPase+wHTH+Alpha+Beta-domain+wHTH+Fn-III+Fn-III+RRM | Type-2_BR.RM.like.systems | <-HIN-HTH<-wHTH+REase<-HKD-DNase+SF1-SF2-Helicase<-METHYLASE<-DUF3780+RAMA<-DUF499*<-RHH_5<-?<-DEDD_Tnp_IS110+Transposase_20<-HTH_Tnp_1+HTH_21+rve+rve_3\|\|<-Arm-DNA-bind_3+Phage_integrase | 1041 | Bacteria>Pseudomonadota>Gammaproteobacteria | GCF_000212715.2 | Enterobacterales | KT011_RS04530 | 922314..925439 | 91347 | MULTISPECIES:-DUF499-domain-containing-protein-[Enterobacterales] |
| NJI20146.1 | AAA+ATPase+wHTH+Alpha+Beta-domain+wHTH+Fn-III+Fn-III+RRM | Type-2_BR.RM.like.systems | <-GNTR-HTH<-?<-HKD-DNase+SF1-SF2-Helicase<-METHYLASE<-DUF3780+RAMA<-DUF499*<-RHH<-?<-?<-?<-?<-CbeA_antitoxin | 1041 | Bacteria>Pseudomonadota>Gammaproteobacteria>Aeromonadales>Aeromonadaceae | GCA_012029575.1 | Aeromonas_veronii | HBJ00_15870 | 37049..40174 | 654 | DUF499-domain-containing-protein-[Aeromonas-veronii] |
| HAT2492740.1 | AAA+ATPase+wHTH+Alpha+Beta-domain+wHTH+Fn-III+Fn-III+RRM | Type-2_BR.RM.like.systems | RHH->DUF499*->DUF3780+RAMA->METHYLASE->HKD-DNase+SF1-SF2-Helicase-p-> | 1041 | Bacteria>Pseudomonadota>Gammaproteobacteria>Aeromonadales>Aeromonadaceae | GCA_015897745.1 | Aeromonas_hydrophila | I8210_004528 | 330..3455 | 644 | DUF499-domain-containing-protein-[Aeromonas-hydrophila] |
| EIM1709203.1 | AAA+ATPase+wHTH+Alpha+Beta-domain+wHTH+Fn-III+Fn-III+RRM | Type-2_BR.RM.like.systems | RHH->DUF499*->DUF3780+RAMA->METHYLASE->HKD-DNase+SF1-SF2-Helicase-><-tRNA<-?\|\|TetR-HTH-> | 1041 | Bacteria>Pseudomonadota>Gammaproteobacteria>Aeromonadales>Aeromonadaceae | GCA_020765715.1 | Aeromonas_dhakensis | JFQ88_003073 | 114609..117734 | 196024 | DUF499-domain-containing-protein-[Aeromonas-dhakensis] |
| WP_016349971.1 | AAA+ATPase+wHTH+Alpha+Beta-domain+wHTH+Fn-III+Fn-III+RRM | Type-2_BR.RM.like.systems | <-LD-peptidase\|\|tRNA-><-HKD-DNase+SF1-SF2-Helicase<-METHYLASE<-DUF3780+RAMA<-DUF499*<-RHH | 1041 | Bacteria>Pseudomonadota>Gammaproteobacteria>Aeromonadales>Aeromonadaceae | GCF_000401555.1 | Aeromonas | V428_RS06965 | 1532835..1535960 | 642 | MULTISPECIES:-DUF499-domain-containing-protein-[Aeromonas] |
| WP_127008751.1 | AAA+ATPase+wHTH+Alpha+Beta-domain+wHTH+Fn-III+Fn-III+RRM | Type-2_BR.RM.like.systems | <-Phage_integrase<-HKD-DNase+SF1-SF2-Helicase<-METHYLASE<-DUF3780+RAMA<-DUF499*<-RHH | 1041 | Bacteria>Pseudomonadota>Gammaproteobacteria>Aeromonadales>Aeromonadaceae | GCF_003990305.1 | Aeromonas_veronii | ELS78_RS22195 | 13228..16353 | 654 | DUF499-domain-containing-protein-[Aeromonas-veronii] |
| WP_136501404.1 | AAA+ATPase+wHTH+Alpha+Beta-domain+wHTH+Fn-III+Fn-III+RRM | Type-2_BR.RM.like.systems | <-TetR-HTH\|\|?->tRNA-><-HKD-DNase+SF1-SF2-Helicase<-METHYLASE<-DUF3780+RAMA<-DUF499*<-RHH | 1041 | Bacteria>Pseudomonadota>Gammaproteobacteria>Aeromonadales>Aeromonadaceae | GCF_004803495.1 | Aeromonas_veronii | E8Q35_RS05065 | 517343..520468 | 654 | DUF499-domain-containing-protein-[Aeromonas-veronii] |
| WP_167567093.1 | AAA+ATPase+wHTH+Alpha+Beta-domain+wHTH+Fn-III+Fn-III+RRM | Type-2_BR.RM.like.systems | <-GNTR-HTH<-?<-HKD-DNase+SF1-SF2-Helicase<-METHYLASE<-DUF3780+RAMA<-DUF499*<-RHH<-?<-?<-?<-?<-CbeA_antitoxin | 1041 | Bacteria>Pseudomonadota>Gammaproteobacteria>Aeromonadales>Aeromonadaceae | GCF_012029575.1 | Aeromonas_veronii | HBJ00_15870 | 37049..40174 | 654 | DUF499-domain-containing-protein-[Aeromonas-veronii] |
| WP_182962959.1 | AAA+ATPase+wHTH+Alpha+Beta-domain+wHTH+Fn-III+Fn-III+RRM | Type-2_BR.RM.like.systems | HIN-HTH-><-SF1-SF2-Helicase-p\|\|PAIREDC-HTH-><-HKD-DNase+SF1-SF2-Helicase<-METHYLASE<-DUF3780<-DUF499*<-XIS-HTH\|\|<-HIN-HTH<-MazE_antitoxin | 1022 | Bacteria>Pseudomonadota>Gammaproteobacteria>Aeromonadales>Aeromonadaceae | GCF_014169835.1 | Aeromonas_veronii | H7R74_RS15935 | 3511914..3514982 | 654 | DUF499-domain-containing-protein-[Aeromonas-veronii] |
| WP_223953095.1 | AAA+ATPase+wHTH+Alpha+Beta-domain+wHTH+Fn-III+Fn-III+RRM | Type-2_BR.RM.like.systems | <-HKD-DNase+SF1-SF2-Helicase<-METHYLASE<-DUF3780<-DUF499*<-XIS-HTH | 1022 | Bacteria>Pseudomonadota>Gammaproteobacteria>Aeromonadales>Aeromonadaceae | GCF_019971735.1 | Aeromonas | MWA36_RS19625 | 6312..9380 | 642 | DUF499-domain-containing-protein-[Aeromonas-caviae] |
| WP_256687664.1 | AAA+ATPase+wHTH+Alpha+Beta-domain+wHTH+Fn-III+Fn-III+RRM | Type-2_BR.RM.like.systems | <-HKD-DNase+SF1-SF2-Helicase<-METHYLASE<-DUF3780+RAMA<-DUF499*<-DUF499-p<-RHH | 847 | Bacteria>Pseudomonadota>Gammaproteobacteria>Aeromonadales>Aeromonadaceae | GCF_024508255.1 | Aeromonas_salmonicida | NP805_RS05835 | 1211766..1214309 | 645 | DUF499-domain-containing-protein-[Aeromonas-salmonicida] |
| WP_257726076.1 | AAA+ATPase+wHTH+Alpha+Beta-domain+wHTH+Fn-III+Fn-III+RRM | Type-2_BR.RM.like.systems | <-HKD-DNase+SF1-SF2-Helicase<-METHYLASE<-DUF3780+RAMA<-DUF499*<-RHH<-?<-?<-?<-?<-CbeA_antitoxin | 1041 | Bacteria>Pseudomonadota>Gammaproteobacteria>Aeromonadales>Aeromonadaceae | GCF_024655985.1 | Aeromonas_veronii | NS965_RS21105 | 7370..10495 | 654 | DUF499-domain-containing-protein-[Aeromonas-veronii] |
| WP_265431750.1 | AAA+ATPase+wHTH+Alpha+Beta-domain+wHTH+Fn-III+Fn-III+RRM | Type-2_BR.RM.like.systems | CbeA_antitoxin->?->?->?->?->RHH->DUF499*->DUF3780+RAMA->METHYLASE->HKD-DNase+SF1-SF2-Helicase-> | 1041 | Bacteria>Pseudomonadota>Gammaproteobacteria>Aeromonadales>Aeromonadaceae | GCF_026173065.1 | Aeromonas_salmonicida | ORM39_RS16575 | 7047..10172 | 645 | DUF499-domain-containing-protein-[Aeromonas-salmonicida] |
| WP_265763441.1 | AAA+ATPase+wHTH+Alpha+Beta-domain+wHTH+Fn-III+Fn-III+RRM | Type-2_BR.RM.like.systems | RHH->DUF499*->DUF3780+RAMA->METHYLASE->HKD-DNase+SF1-SF2-Helicase-><-tRNA<-?\|\|TetR-HTH-> | 1041 | Bacteria>Pseudomonadota>Gammaproteobacteria>Aeromonadales>Aeromonadaceae | GCF_026230465.1 | Aeromonas_veronii | OD806_RS18940 | 27776..30901 | 654 | DUF499-domain-containing-protein-[Aeromonas-veronii] |
| WP_277096146.1 | AAA+ATPase+wHTH+Alpha+Beta-domain+wHTH+Fn-III+Fn-III+RRM | Type-2_BR.RM.like.systems | RHH->DUF499-p->DUF499*->DUF3780+RAMA->METHYLASE->HKD-DNase+SF1-SF2-Helicase-> | 834 | Bacteria>Pseudomonadota>Gammaproteobacteria>Aeromonadales>Aeromonadaceae | GCF_029436615.1 | Aeromonas_salmonicida | P5G63_RS24895 | 4870375..4872879 | 645 | DUF499-domain-containing-protein-[Aeromonas-salmonicida] |
| AFT74739.1 | AAA+ATPase+wHTH+Alpha+Beta-domain+wHTH+Fn-III+Fn-III+RRM | Type-2_BR.RM.like.systems | HEPN-><-Phage_integrase<-JAB<-HKD-DNase+SF1-SF2-Helicase<-METHYLASE<-DUF3780+RAMA<-DUF499*<-Phage_integrase<-Arm-DNA-bind_3<-tRNA<-tRNA<-tRNA<-?<-URI-DNase+HhH-RADC<-Response_reg+LuxR-HTH | 1022 | Bacteria>Pseudomonadota>Gammaproteobacteria>Alteromonadales>Alteromonadaceae | GCA_000299955.1 | Alteromonas_macleodii_str._'English_Channel_673' | AMEC673_10225 | 2405443..2408511 | 1004788 | hypothetical-protein-AMEC673_10225-[Alteromonas-macleodii-str.-'English-Channel-673'] |
| RPH13128.1 | AAA+ATPase+wHTH+Alpha+Beta-domain+wHTH+Fn-III+Fn-III+RRM | Type-2_BR.RM.like.systems | <-OLD-ABC+TOPRIM-p<-HKD-DNase+SF1-SF2-Helicase<-METHYLASE<-DUF3780+RAMA<-DUF499*<-RHH_1<-XIS-HTH | 1041 | Bacteria>Pseudomonadota>Gammaproteobacteria>Alteromonadales>Alteromonadaceae | GCA_002173075.2 | Alteromonadaceae_bacterium_TMED7 | CBB67_021520 | 7937..11062 | 1986650 | DUF499-domain-containing-protein-[Alteromonadaceae-bacterium-TMED7] |
| WP_044583752.1 | AAA+ATPase+wHTH+Alpha+Beta-domain+wHTH+Fn-III+Fn-III+RRM | Type-2_BR.RM.like.systems | HEPN-><-Phage_integrase<-JAB<-HKD-DNase+SF1-SF2-Helicase<-METHYLASE<-DUF3780+RAMA<-DUF499*<-Phage_integrase<-Arm-DNA-bind_3<-tRNA<-tRNA<-tRNA<-?<-URI-DNase+HhH-RADC<-Response_reg+LuxR-HTH | 1041 | Bacteria>Pseudomonadota>Gammaproteobacteria>Alteromonadales>Alteromonadaceae | GCF_000299955.1 | Alteromonas_macleodii | AMEC673_RS10395 | 2405443..2408568 | 28108 | DUF499-domain-containing-protein-[Alteromonas-macleodii] |
| MBY6226422.1 | AAA+ATPase+wHTH+Alpha+Beta-domain+wHTH+Fn-III+Fn-III+RRM | Type-2_BR.RM.like.systems | <-HEPN<-Tox-RelA\|\|tRNA-><-HKD-DNase+SF1-SF2-Helicase<-METHYLASE<-DUF3780+RAMA<-DUF499*<-RHH<-XIS-HTH\|\|SAM-methylase-><-?\|\|Cysteine_desulfurase-> | 1022 | Bacteria>Pseudomonadota>Gammaproteobacteria>Alteromonadales>Ferrimonadaceae | GCA_019800765.1 | Ferrimonas_balearica | KUW13_18305 | 80575..83643 | 44012 | DUF499-domain-containing-protein-[Ferrimonas-balearica] |
| WP_246814860.1 | AAA+ATPase+wHTH+Alpha+Beta-domain+wHTH+Fn-III+Fn-III+RRM | Type-2_BR.RM.like.systems | <-HEPN<-Tox-RelA\|\|tRNA-><-HKD-DNase+SF1-SF2-Helicase<-METHYLASE<-DUF3780+RAMA<-DUF499*<-RHH<-XIS-HTH\|\|SAM-methylase-><-?\|\|Cysteine_desulfurase-> | 1041 | Bacteria>Pseudomonadota>Gammaproteobacteria>Alteromonadales>Ferrimonadaceae | GCF_019800765.1 | Ferrimonas_balearica | KUW13_RS18310 | 80575..83700 | 44012 | DUF499-domain-containing-protein-[Ferrimonas-balearica] |
| QTL37180.1 | AAA+ATPase+wHTH+Alpha+Beta-domain+wHTH+Fn-III+Fn-III+RRM | Type-2_BR.RM.like.systems | <-VirD4-TraG+VirB4-FtsK<-HDRNase<-wHTH+REase<-HKD-DNase+SF1-SF2-Helicase<-METHYLASE<-DUF3780+RAMA<-DUF499*<-RHH_1<-35exo\|\|?->?->Classical-AAA->Peptidase_S8-> | 1022 | Bacteria>Pseudomonadota>Gammaproteobacteria>Alteromonadales>Pseudoalteromonadaceae | GCA_017742995.1 | Pseudoalteromonas_viridis | J5X90_01860 | 434218..437286 | 339617 | DUF499-domain-containing-protein-[Pseudoalteromonas-viridis] |
| WP_165744672.1 | AAA+ATPase+wHTH+Alpha+Beta-domain+wHTH+Fn-III+Fn-III+RRM | Type-2_BR.RM.like.systems | <-JAB\|\|RNase->XIS-HTH->RHH->DUF499*->DUF3780+RAMA->METHYLASE->HKD-DNase+SF1-SF2-Helicase->?->MACRODOMAIN-><-Arm-DNA-bind_3+Phage_integrase | 1041 | Bacteria>Pseudomonadota>Gammaproteobacteria>Alteromonadales>Pseudoalteromonadaceae | GCF_011378805.1 | Pseudoalteromonas_sp._Z9A6 | GQR53_RS17200 | 6833..9958 | 2686352 | DUF499-domain-containing-protein-[Pseudoalteromonas-sp.-Z9A6] |
| WP_248443748.1 | AAA+ATPase+wHTH+Alpha+Beta-domain+wHTH+Fn-III+Fn-III+RRM | Type-2_BR.RM.like.systems | <-MACRODOMAIN<-?<-?<-HKD-DNase+SF1-SF2-Helicase<-METHYLASE<-DUF3780+RAMA<-DUF499*<-XIS-HTH\|\|JAB->?->Phage_integrase-> | 1041 | Bacteria>Pseudomonadota>Gammaproteobacteria>Alteromonadales>Pseudoalteromonadaceae | GCF_023161805.1 | Pseudoalteromonas_sp._1CM17D | MSC25_RS13675 | 176224..179349 | 2929162 | DUF499-domain-containing-protein-[Pseudoalteromonas-sp.-1CM17D] |
| WP_099295148.1 | AAA+ATPase+wHTH+Alpha+Beta-domain+wHTH+Fn-III+Fn-III+RRM | Type-2_BR.RM.like.systems | <-Phage_integrase<-?<-JAB\|\|XIS-HTH->DUF499*->DUF3780+RAMA->METHYLASE->HKD-DNase+SF1-SF2-Helicase-> | 1041 | Bacteria>Pseudomonadota>Gammaproteobacteria>Alteromonadales>Pseudoalteromonadaceae | GCF_030689005.1 | unclassified_Pseudoalteromonas | CSC79_RS07135 | 94445..97570 | 194690 | DUF499-domain-containing-protein-[Pseudoalteromonas-sp.-3D05] |
| WP_036957217.1 | AAA+ATPase+wHTH+Alpha+Beta-domain+wHTH+Fn-III+Fn-III+RRM | Type-2_BR.RM.like.systems | <-HNH<-ABC-ATPase<-wHTH+REase<-HKD-DNase+SF1-SF2-Helicase<-METHYLASE<-DUF3780+RAMA<-DUF499*<-RHH<-?<-XIS-HTH<-?<-XIS-HTH\|\|JAB-> | 1041 | Bacteria>Pseudomonadota>Gammaproteobacteria>Alteromonadales>Pseudoalteromonadaceae | GCF_913044265.1 | Pseudoalteromonas | QL994_RS18060 | 15258..18383 | 53246 | DUF499-domain-containing-protein-[Pseudoalteromonas-sp.-A2] |
| QYK03187.1 | AAA+ATPase+wHTH+Alpha+Beta-domain+wHTH+Fn-III+Fn-III+RRM | Type-2_BR.RM.like.systems | <-XIS-HTH<-Phage_integrase<-?<-JAB\|\|XIS-HTH->DUF499*->DUF3780+RAMA->METHYLASE->HKD-DNase+SF1-SF2-Helicase->SF1-SF2-Helicase->?-><-Arm-DNA-bind_3+Phage_integrase | 1022 | Bacteria>Pseudomonadota>Gammaproteobacteria>Alteromonadales>Shewanellaceae | GCA_019457595.1 | Shewanella_psychrotolerans | K0I62_01745 | 370192..373260 | 2864206 | DUF499-domain-containing-protein-[Shewanella-sp.-FJAT-53749] |
| WP_218997779.1 | AAA+ATPase+wHTH+Alpha+Beta-domain+wHTH+Fn-III+Fn-III+RRM | Type-2_BR.RM.like.systems | METHYLASE->REase->rve+rve_3+Mu-transpos_C->?->TetR-HTH->XIS-HTH->DUF499*->DUF3780->METHYLASE->HKD-DNase+SF1-SF2-Helicase->SbcC->\|\|?->SF1-SF2-Helicase-> | 1022 | Bacteria>Pseudomonadota>Gammaproteobacteria>Alteromonadales>Shewanellaceae | GCF_019303335.1 | Shewanella_algae | K0E68_RS03920 | 10181..13249 | 38313 | DUF499-domain-containing-protein-[Shewanella-algae] |
| WP_258405063.1 | AAA+ATPase+wHTH+Alpha+Beta-domain+wHTH+Fn-III+Fn-III+RRM | Type-2_BR.RM.like.systems | <-XIS-HTH<-Phage_integrase<-?<-JAB\|\|XIS-HTH->DUF499*->DUF3780+RAMA->METHYLASE->HKD-DNase+SF1-SF2-Helicase->SF1-SF2-Helicase->?-><-Arm-DNA-bind_3+Phage_integrase | 1041 | Bacteria>Pseudomonadota>Gammaproteobacteria>Alteromonadales>Shewanellaceae | GCF_019457595.1 | Shewanella_psychrotolerans | K0I62_RS01745 | 370135..373260 | 2864206 | DUF499-domain-containing-protein-[Shewanella-sp.-FJAT-53749] |
| WP_259215849.1 | AAA+ATPase+wHTH+Alpha+Beta-domain+wHTH+Fn-III+Fn-III+RRM | Type-2_BR.RM.like.systems | MerR-HTH->?->?->?->?->\|\|XIS-HTH->DUF499*->DUF3780+RAMA->METHYLASE->HKD-DNase+SF1-SF2-Helicase->?-><-HTH_3 | 1041 | Bacteria>Pseudomonadota>Gammaproteobacteria>Alteromonadales>Shewanellaceae | GCF_024803605.1 | Shewanella_baltica | G3492_RS15830 | 133246..136371 | 62322 | DUF499-domain-containing-protein-[Shewanella-baltica] |
| WP_126492065.1 | AAA+ATPase+wHTH+Alpha+Beta-domain+wHTH+Fn-III+Fn-III+RRM | Type-2_BR.RM.like.systems | HTH_3-><-?<-HKD-DNase+SF1-SF2-Helicase<-METHYLASE<-DUF3780+RAMA<-DUF499*<-XIS-HTH\|\|JAB->?->CbeA_antitoxin-> | 1041 | Bacteria>Pseudomonadota>Gammaproteobacteria>Alteromonadales>Shewanellaceae | GCF_900636855.1 | Shewanella_baltica | EL185_RS11920 | 2770503..2773628 | 62322 | DUF499-domain-containing-protein-[Shewanella-baltica] |
| WP_220071247.1 | AAA+ATPase+wHTH+Alpha+Beta-domain+wHTH+Fn-III+Fn-III+RRM | Type-2_BR.RM.like.systems | <-XIS-HTH<-Phage_integrase<-?<-JAB\|\|XIS-HTH->DUF499*->DUF3780+RAMA->METHYLASE->HKD-DNase+SF1-SF2-Helicase->SF1-SF2-Helicase->?-><-Arm-DNA-bind_3+Phage_integrase | 1022 | Bacteria>Pseudomonadota>Gammaproteobacteria>Alteromonadales>Shewanellaceae | - | Shewanella_psychrotolerans | K0I62_01745 | 370192..373260 | 2864206 | DUF499-domain-containing-protein-[Shewanella-sp.-FJAT-53749] |
| AUB85626.1 | AAA+ATPase+wHTH+Alpha+Beta-domain+wHTH+Fn-III+Fn-III+RRM | Type-2_BR.RM.like.systems | <-HKD-DNase+SF1-SF2-Helicase<-PIN<-?<-?<-METHYLASE<-DUF3780<-DUF499* | 1026 | Bacteria>Pseudomonadota>Gammaproteobacteria>Chromatiales>Chromatiaceae | GCA_002813775.1 | Candidatus_Thiodictyon_syntrophicum | THSYN_32535 | 358287..361367 | 1166950 | AAA-family-ATPase-[Candidatus-Thiodictyon-syntrophicum] |
| MCF7983262.1 | AAA+ATPase+wHTH+Alpha+Beta-domain+wHTH+Fn-III+Fn-III+RRM | Type-2_BR.RM.like.systems | <-HKD-DNase+SF1-SF2-Helicase-p<-?<-STYKinase-like<-METHYLASE<-DUF3780<-DUF499*\|\|?->?->MazE_antitoxin-><-Peptidase<-ABC-ATPase+SbcC\|\|ABC-ATPase+SbcC-> | 1021 | Bacteria>Pseudomonadota>Gammaproteobacteria>Chromatiales>Chromatiaceae | GCA_021736665.1 | Thiohalocapsa_sp. | K9L70_02570 | 9369..12434 | 2497641 | DUF499-domain-containing-protein-[Thiohalocapsa-sp.] |
| MCF7997122.1 | AAA+ATPase+wHTH+Alpha+Beta-domain+wHTH+Fn-III+Fn-III+RRM | Type-2_BR.RM.like.systems | DUF499*->?->HKD-DNase+SF1-SF2-Helicase-> | 1047 | Bacteria>Pseudomonadota>Gammaproteobacteria>Chromatiales>Chromatiaceae | GCA_021736825.1 | Chromatiaceae_bacterium | K9L88_20170 | 286..3429 | 2049432 | DUF499-domain-containing-protein-[Chromatiaceae-bacterium] |
| MCF8003666.1 | AAA+ATPase+wHTH+Alpha+Beta-domain+wHTH+Fn-III+Fn-III+RRM | Type-2_BR.RM.like.systems | <-DUF29<-HKD-DNase+SF1-SF2-Helicase<-PIN<-?<-METHYLASE<-DUF3780<-DUF499*<-CR-ATPase9+REase | 1047 | Bacteria>Pseudomonadota>Gammaproteobacteria>Chromatiales>Chromatiaceae | GCA_021736845.1 | Chromatiaceae_bacterium | K9L32_05580 | 9009..12152 | 2049432 | DUF499-domain-containing-protein-[Chromatiaceae-bacterium] |
| MCF8004660.1 | AAA+ATPase+wHTH+Alpha+Beta-domain+wHTH+Fn-III+Fn-III+RRM | Type-2_BR.RM.like.systems | <-HKD-DNase+SF1-SF2-Helicase<-?<-DUF499* | 1047 | Bacteria>Pseudomonadota>Gammaproteobacteria>Chromatiales>Chromatiaceae | GCA_021736845.1 | Chromatiaceae_bacterium | K9L32_10745 | 4477..7620 | 2049432 | DUF499-domain-containing-protein-[Chromatiaceae-bacterium] |
| MCP5417446.1 | AAA+ATPase+wHTH+Alpha+Beta-domain+wHTH+Fn-III+Fn-III+RRM | Type-2_BR.RM.like.systems | TraH->?->TraG_N-><-?\|\|XIS-HTH->ParB+HNH->DUF499*->DUF3780->METHYLASE->HKD-DNase+SF1-SF2-Helicase->?->Hydrolase_3-> | 1043 | Bacteria>Pseudomonadota>Gammaproteobacteria>Chromatiales>Chromatiaceae | GCA_024235215.1 | Chromatiaceae_bacterium | H6965_10165 | 390461..393592 | 2049432 | DUF499-domain-containing-protein-[Chromatiaceae-bacterium] |
| MCU0833529.1 | AAA+ATPase+wHTH+Alpha+Beta-domain+wHTH+Fn-III+Fn-III+RRM | Type-2_BR.RM.like.systems | DUF2026-><-HKD-DNase+SF1-SF2-Helicase<-?<-STYKinase-like<-METHYLASE<-DUF3780<-DUF499*\|\|Thioredoxin-> | 1022 | Bacteria>Pseudomonadota>Gammaproteobacteria>Chromatiales>Chromatiaceae | GCA_025458165.1 | Chromatiaceae_bacterium | MUC77_03690 | 57036..60104 | 2049432 | DUF499-domain-containing-protein-[Chromatiaceae-bacterium] |
| WP_236849074.1 | AAA+ATPase+wHTH+Alpha+Beta-domain+wHTH+Fn-III+Fn-III+RRM | Type-2_BR.RM.like.systems | <-HKD-DNase+SF1-SF2-Helicase<-PIN<-?<-?<-METHYLASE<-DUF3780<-DUF499* | 1039 | Bacteria>Pseudomonadota>Gammaproteobacteria>Chromatiales>Chromatiaceae | GCF_002813775.1 | Candidatus_Thiodictyon_syntrophicum | THSYN_RS32810 | 358287..361406 | 1166950 | DUF499-domain-containing-protein-[Candidatus-Thiodictyon-syntrophicum] |
| WP_134356845.1 | AAA+ATPase+wHTH+Alpha+Beta-domain+wHTH+Fn-III+Fn-III+RRM | Type-2_BR.RM.like.systems | <-HNH\|\|DUF499*->DUF3780->METHYLASE->HKD-DNase+SF1-SF2-Helicase-> | 1032 | Bacteria>Pseudomonadota>Gammaproteobacteria>Chromatiales>Chromatiaceae | GCF_004421105.1 | Nitrosococcus_wardiae | E3U44_RS04390 | 903911..907009 | 1814290 | DUF499-domain-containing-protein-[Nitrosococcus-wardiae] |
| WP_153975105.1 | AAA+ATPase+wHTH+Alpha+Beta-domain+wHTH+Fn-III+Fn-III+RRM | Type-2_BR.RM.like.systems | <-HKD-DNase+SF1-SF2-Helicase<-METHYLASE<-PIN<-?<-MACRODOMAIN<-DUF3780<-DUF499* | 1040 | Bacteria>Pseudomonadota>Gammaproteobacteria>Chromatiales>Chromatiaceae | GCF_009664085.1 | Thermochromatium_tepidum | E6P07_RS07905 | 1743171..1746293 | 1050 | DUF499-domain-containing-protein-[Thermochromatium-tepidum] |
| WP_201219751.1 | AAA+ATPase+wHTH+Alpha+Beta-domain+wHTH+Fn-III+Fn-III+RRM | Type-2_BR.RM.like.systems | CR-ATPase9+REase->DUF499*->DUF3780->?->PIN->HKD-DNase+SF1-SF2-Helicase->DUF29-> | 1047 | Bacteria>Pseudomonadota>Gammaproteobacteria>Chromatiales>Chromatiaceae | GCF_016653315.1 | Halochromatium_roseum | CCR96_RS08785 | 123675..126818 | 391920 | DUF499-domain-containing-protein-[Halochromatium-roseum] |
| WP_026199823.1 | AAA+ATPase+wHTH+Alpha+Beta-domain+wHTH+Fn-III+Fn-III+RRM | Type-2_BR.RM.like.systems | REase->DUF499-p-><-HicB_lk_antitox\|\|Phd-YefM->PIN->DUF499*->DUF3780-><-HicB_lk_antitox<-HicA_toxin\|\|METHYLASE-p->METHYLASE->REase-> | 1035 | Bacteria>Pseudomonadota>Gammaproteobacteria>Chromatiales>Chromatiaceae | GCF_000379525.1 | Lamprocystis_purpurea | A390_RS0128810 | 112256..115363 | 61598 | DUF499-domain-containing-protein-[Lamprocystis-purpurea] |
| WP_232410096.1 | AAA+ATPase+wHTH+Alpha+Beta-domain+wHTH+Fn-III+Fn-III+RRM | Type-2_BR.RM.like.systems | <-PIN\|\|CopG/RHH->ACET-><-DUF3780<-HTH_29+rve+rve_3<-DUF499* | 1036 | Bacteria>Pseudomonadota>Gammaproteobacteria>Chromatiales>Chromatiaceae | GCF_021228795.1 | Thiocapsa_bogorovii | LT988_RS10530 | 2321159..2324269 | 521689 | DUF499-domain-containing-protein-[Thiocapsa-bogorovii] |
| WP_245969509.1 | AAA+ATPase+wHTH+Alpha+Beta-domain+wHTH+Fn-III+Fn-III+RRM | Type-2_BR.RM.like.systems | <-ABhydrolase<-?\|\|ABhydrolase-><-?\|\|?->DUF499*->REase->DUF3780->?->PIN->METHYLASE->METHYLASE-p-> | 1016 | Bacteria>Pseudomonadota>Gammaproteobacteria>Chromatiales>Chromatiaceae | GCF_003634315.1 | Thiocapsa_rosea | BDD21_RS09910 | 2229310..2232360 | 69360 | DUF499-domain-containing-protein-[Thiocapsa-rosea] |
| WP_242518638.1 | AAA+ATPase+wHTH+Alpha+Beta-domain+wHTH+Fn-III+Fn-III+RRM | Type-2_BR.RM.like.systems | DUF499*->DUF3780->PIN->METHYLASE->DDE_Tnp_1-><-GDResolvase+Recombinase+ZnR | 1021 | Bacteria>Pseudomonadota>Gammaproteobacteria>Chromatiales>Chromatiaceae | GCF_016653315.1 | Halochromatium_roseum | CCR96_RS05805 | 17129..20194 | 391920 | DUF499-domain-containing-protein-[Halochromatium-roseum] |
| MBK5938754.1 | AAA+ATPase+wHTH+Alpha+Beta-domain+wHTH+Fn-III+Fn-III+RRM | Type-2_BR.RM.like.systems | DUF499*->DUF3780->PIN->DDE_Tnp_1-><-GDResolvase+Recombinase+ZnR<-?\|\|<-TraJ-RHH | 1040 | Bacteria>Pseudomonadota>Gammaproteobacteria>Chromatiales>Chromatiaceae | GCA_016653315.1 | Halochromatium_roseum | CCR96_05695 | 17072..20194 | 391920 | AAA-family-ATPase-[Halochromatium-roseum] |
| WP_245970045.1 | AAA+ATPase+wHTH+Alpha+Beta-domain+wHTH+Fn-III+Fn-III+RRM | Type-2_BR.RM.like.systems | <-PIN<-?<-METHYLASE<-PIN<-?<-DUF3780<-DUF499*<-?<-REase+NACHT<-GDResolvase+HIN-HTH | 1018 | Bacteria>Pseudomonadota>Gammaproteobacteria>Chromatiales>Chromatiaceae | GCF_003634315.1 | Thiocapsa_rosea | BDD21_RS27140 | 182504..185560 | 69360 | DUF499-domain-containing-protein-[Thiocapsa-rosea] |
| WP_200375560.1 | AAA+ATPase+wHTH+Alpha+Beta-domain+wHTH+Fn-III+Fn-III+RRM | Type-2_BR.RM.like.systems | <-MerR-HTH<-?\|\|?->?->?->SMS-KaiC->DUF499*->DUF3780->?->PIN->OLD-ABC->TOPRIM->METHYLASE-> | 1027 | Bacteria>Pseudomonadota>Gammaproteobacteria>Chromatiales>Chromatiaceae | GCF_016583575.1 | Thiocystis_violacea | CKO27_RS14205 | 21670..24753 | 13725 | DUF499-domain-containing-protein-[Thiocystis-violacea] |
| MCU0836365.1 | AAA+ATPase+wHTH+Alpha+Beta-domain+wHTH+Fn-III+Fn-III+RRM | Type-2_BR.RM.like.systems | <-METHYLASE-p<-HEPN<-PolB<-PIN<-?<-DUF3780<-DUF499*<-RHH\|\|REase-><-Cluster182_14clades\|\|?->?->ZnR+TOPRIM-> | 1037 | Bacteria>Pseudomonadota>Gammaproteobacteria>Chromatiales>Chromatiaceae | GCA_025458165.1 | Chromatiaceae_bacterium | MUC77_18335 | 8879..11992 | 2049432 | DUF499-domain-containing-protein-[Chromatiaceae-bacterium] |
| WP_207148756.1 | AAA+ATPase+wHTH+Alpha+Beta-domain+wHTH+Fn-III+Fn-III+RRM | Type-2_BR.RM.like.systems | <-METHYLASE<-PIN<-?<-DUF3780<-PIN<-MazE_antitoxin<-DUF499* | 1020 | Bacteria>Pseudomonadota>Gammaproteobacteria>Chromatiales>Chromatiaceae | GCF_016583905.1 | Lamprobacter_modestohalophilus | CKO42_RS16530 | 30403..33465 | 1064514 | DUF499-domain-containing-protein-[Lamprobacter-modestohalophilus] |
| WP_009148233.1 | AAA+ATPase+wHTH+Alpha+Beta-domain+wHTH+Fn-III+Fn-III+RRM | Type-2_BR.RM.like.systems | <-ACET<-?\|\|DUF499*->DUF3780->RHH_1->?->PIN->ParB+HNH->METHYLASE-> | 1040 | Bacteria>Pseudomonadota>Gammaproteobacteria>Chromatiales>Chromatiaceae | GCF_033954835.1 | Thiorhodovibrio_frisius | THI970DRAFT_RS08785 | 535191..538313 | 631362 | DUF499-domain-containing-protein-[Thiorhodovibrio-sp.-970] |
| WP_093189452.1 | AAA+ATPase+wHTH+Alpha+Beta-domain+wHTH+Fn-III+Fn-III+RRM | Type-2_BR.RM.like.systems | <-Aminotran_3<-?\|\|DUF499*->?->DUF3780->?->PIN->?->PIN-> | 1025 | Bacteria>Pseudomonadota>Gammaproteobacteria>Chromatiales>Chromatiaceae | GCF_001499735.1 | Thiocapsa_sp._KS1 | THIOKS1_RS16995 | 3698794..3701871 | 610332 | DUF499-domain-containing-protein-[Thiocapsa-sp.-KS1] |
| MBK1620020.1 | AAA+ATPase+wHTH+Alpha+Beta-domain+wHTH+Fn-III+Fn-III+RRM | Type-2_BR.RM.like.systems | <-METHYLASE<-PIN<-?<-DUF3780<-PIN<-MazE_antitoxin<-DUF499* | 1039 | Bacteria>Pseudomonadota>Gammaproteobacteria>Chromatiales>Chromatiaceae | GCA_016583905.1 | Lamprobacter_modestohalophilus | CKO42_16545 | 30403..33522 | 1064514 | AAA-family-ATPase-[Lamprobacter-modestohalophilus] |
| TVQ88665.1 | AAA+ATPase+wHTH+Alpha+Beta-domain+wHTH+Fn-III+Fn-III+RRM | Type-2_BR.RM.like.systems | ABhydrolase-><-?<-?\|\|?->DUF499*->MazE_antitoxin->PIN->DUF3780->?->PIN->PolB-> | 1030 | Bacteria>Pseudomonadota>Gammaproteobacteria>Chromatiales>Chromatiaceae | GCA_007693535.1 | Chromatiaceae_bacterium | EA400_08510 | 10879..13971 | 2049432 | ATP-binding-protein-[Chromatiaceae-bacterium] |
| QQO52729.1 | AAA+ATPase+wHTH+Alpha+Beta-domain+wHTH+Fn-III+Fn-III+RRM | Type-2_BR.RM.like.systems | 35exo-><-?<-?<-?\|\|?->DUF499*->DUF3780-><-?\|\|?->?->PIN->METHYLASE-> | 1040 | Bacteria>Pseudomonadota>Gammaproteobacteria>Chromatiales>Chromatiaceae | GCA_016745215.1 | Thiohalocapsa_sp._PB-PSB1 | N838_04400 | 1035281..1038403 | 1385625 | MAG:-DUF499-domain-containing-protein-[Thiohalocapsa-sp.-PB-PSB1] |
| NEX15887.1 | AAA+ATPase+wHTH+Alpha+Beta-domain+wHTH+Fn-III+Fn-III+RRM | Type-2_BR.RM.like.systems | <-PolB<-ParB+HNH<-?<-?<-RHH_1<-DUF3780<-DUF499*<-?<-PIN | 1040 | Bacteria>Pseudomonadota>Gammaproteobacteria>Chromatiales>Chromatiaceae | GCA_010912705.1 | Halochromatium_sp. | C1943_04405 | 213833..216955 | 2049430 | AAA-family-ATPase-[Halochromatium-sp.] |
| MCF8005387.1 | AAA+ATPase+wHTH+Alpha+Beta-domain+wHTH+Fn-III+Fn-III+RRM | Type-2_BR.RM.like.systems | <-DUF3780<-PIN<-MazE_antitoxin<-DUF499* | 1026 | Bacteria>Pseudomonadota>Gammaproteobacteria>Chromatiales>Chromatiaceae | GCA_021736845.1 | Chromatiaceae_bacterium | K9L32_14520 | 1092..4172 | 2049432 | DUF499-domain-containing-protein-[Chromatiaceae-bacterium] |
| QVL47709.1 | AAA+ATPase+wHTH+Alpha+Beta-domain+wHTH+Fn-III+Fn-III+RRM | Type-2_BR.RM.like.systems | <-METHYLASE<-METHYLASE-p<-DUF3780<-?<-HTH_OrfB_IS605+OrfB_IS605+OrfB_Zn_ribbon<-MerR-HTH+GDResolvase<-DUF499*\|\|REase-> | 1025 | Bacteria>Pseudomonadota>Gammaproteobacteria>Chromatiales>Chromatiaceae | GCA_018399035.1 | Thiocapsa_sp. | KFB96_18760 | 4106445..4109522 | 2024551 | MAG:-DUF499-domain-containing-protein-[Thiocapsa-sp.] |
| WP_211211735.1 | AAA+ATPase+wHTH+Alpha+Beta-domain+wHTH+Fn-III+Fn-III+RRM | Type-2_BR.RM.like.systems | DUF499*->REase->DUF3780->?->METHYLASE->STYKinase-like-> | 1034 | Bacteria>Pseudomonadota>Gammaproteobacteria>Chromatiales>Chromatiaceae | GCF_000379525.1 | Lamprocystis_purpurea | A390_RS0125930 | 13980..17084 | 61598 | DUF499-domain-containing-protein-[Lamprocystis-purpurea] |
| MBV5273604.1 | AAA+ATPase+wHTH+Alpha+Beta-domain+wHTH+Fn-III+Fn-III+RRM | Type-2_BR.RM.like.systems | DUF499*->REase->DUF3780->?->?->METHYLASE->STYKinase-like-> | 804 | Bacteria>Pseudomonadota>Gammaproteobacteria>Chromatiales>Chromatiaceae | GCA_019163055.1 | Lamprocystis_purpurea | JZU52_08160 | 1..2416 | 61598 | DUF499-domain-containing-protein-[Lamprocystis-purpurea] |
| QVL47943.1 | AAA+ATPase+wHTH+Alpha+Beta-domain+wHTH+Fn-III+Fn-III+RRM | Type-2_BR.RM.like.systems | <-ABhydrolase<-?\|\|ABhydrolase-><-?\|\|?->DUF499*->OLD-ABC->TOPRIM->DUF3780->?->METHYLASE-> | 1028 | Bacteria>Pseudomonadota>Gammaproteobacteria>Chromatiales>Chromatiaceae | GCA_018399035.1 | Thiocapsa_sp. | KFB96_20135 | 4418562..4421648 | 2024551 | MAG:-DUF499-domain-containing-protein-[Thiocapsa-sp.] |
| MBL1275688.1 | AAA+ATPase+wHTH+Alpha+Beta-domain+wHTH+Fn-III+Fn-III+RRM | Type-2_BR.RM.like.systems | <-HKD-DNase+SF1-SF2-Helicase<-Schlafen+HATPase+HTH<-METHYLASE<-DUF3780<-DUF499*<-RHH_5<-REase\|\|SF1-SF2-Helicase->?-><-?\|\|PADR-HTH-> | 1041 | Bacteria>Pseudomonadota>Gammaproteobacteria>Chromatiales>Ectothiorhodospiraceae | GCA_002401775.2 | Ectothiorhodospiraceae_bacterium | COB30_006350 | 9914..13039 | 2026737 | DUF499-domain-containing-protein-[Ectothiorhodospiraceae-bacterium] |
| PYG00798.1 | AAA+ATPase+wHTH+Alpha+Beta-domain+wHTH+Fn-III+Fn-III+RRM | Type-2_BR.RM.like.systems | rve_3->rve_3-><-HKD-DNase+SF1-SF2-Helicase<-METHYLASE<-DUF3780<-DUF499*<-?<-Arm-DNA-bind_3+Phage_integrase | 1073 | Bacteria>Pseudomonadota>Gammaproteobacteria>Chromatiales>Ectothiorhodospiraceae | GCA_003208255.1 | Thioalkalivibrio_sp._ALE21 | D893_02198 | 7289..10510 | 1158175 | putative-AAA+-superfamily-ATPase-[Thioalkalivibrio-sp.-ALE21] |
| TQE98850.1 | AAA+ATPase+wHTH+Alpha+Beta-domain+wHTH+Fn-III+Fn-III+RRM | Type-2_BR.RM.like.systems | <-HKD-DNase+SF1-SF2-Helicase-p<-METHYLASE<-DUF3780<-DUF499* | 1040 | Bacteria>Pseudomonadota>Gammaproteobacteria>Chromatiales>Ectothiorhodospiraceae | GCA_006569195.1 | Spiribacter_salinus | FKY71_11670 | 4270..7392 | 1335746 | DUF499-domain-containing-protein-[Spiribacter-salinus] |
| WP_018995154.1 | AAA+ATPase+wHTH+Alpha+Beta-domain+wHTH+Fn-III+Fn-III+RRM | Type-2_BR.RM.like.systems | <-ArsR-HTH\|\|Arm-DNA-bind_3+Phage_integrase->DUF499*->DUF3780->METHYLASE->HKD-DNase+SF1-SF2-Helicase->?->XIS-HTH-> | 1039 | Bacteria>Pseudomonadota>Gammaproteobacteria>Chromatiales>Ectothiorhodospiraceae | GCF_000378325.1 | Thioalkalivibrio_sp._ALJ2 | F468_RS0110855 | 71234..74353 | 1261622 | DUF499-domain-containing-protein-[Thioalkalivibrio-sp.-ALJ2] |
| WP_019568718.1 | AAA+ATPase+wHTH+Alpha+Beta-domain+wHTH+Fn-III+Fn-III+RRM | Type-2_BR.RM.like.systems | <-XIS-HTH<-?<-HKD-DNase+SF1-SF2-Helicase<-METHYLASE<-DUF3780<-DUF499*<-Arm-DNA-bind_3+Phage_integrase<-ABC-ATPase+SbcC<-ABC-ATPase+SbcC | 1039 | Bacteria>Pseudomonadota>Gammaproteobacteria>Chromatiales>Ectothiorhodospiraceae | GCF_000381185.1 | Thioalkalivibrio_sp._ALMg13-2 | F618_RS0112140 | 8832..11951 | 1158167 | DUF499-domain-containing-protein-[Thioalkalivibrio-sp.-ALMg13-2] |
| WP_110619825.1 | AAA+ATPase+wHTH+Alpha+Beta-domain+wHTH+Fn-III+Fn-III+RRM | Type-2_BR.RM.like.systems | rve_3->rve_3-><-HKD-DNase+SF1-SF2-Helicase<-METHYLASE<-DUF3780<-DUF499*<-Arm-DNA-bind_3+Phage_integrase | 1039 | Bacteria>Pseudomonadota>Gammaproteobacteria>Chromatiales>Ectothiorhodospiraceae | GCF_003208255.1 | Thioalkalivibrio_sp._ALE21 | D893_RS11030 | 7289..10408 | 1158175 | DUF499-domain-containing-protein-[Thioalkalivibrio-sp.-ALE21] |
| OZB36169.1 | AAA+ATPase+wHTH+Alpha+Beta-domain+wHTH+Fn-III+Fn-III+RRM | Type-2_BR.RM.like.systems | <-ParA-Soj-PloopNTPase<-HEPN<-METHYLASE<-MACRODOMAIN<-?<-DUF3780<-DUF499*<-RHH_5<-?<-?\|\|XIS-HTH->DnaB-><-HAMP+HISKIN | 1041 | Bacteria>Pseudomonadota>Gammaproteobacteria>Chromatiales>Halothiobacillaceae | GCA_002282715.1 | Halothiobacillus_sp._15-55-196 | B7X44_07150 | 24460..27585 | 1970382 | MAG:-AAA-family-ATPase-[Halothiobacillus-sp.-15-55-196] |
| WP_040132773.1 | AAA+ATPase+wHTH+Alpha+Beta-domain+wHTH+Fn-III+Fn-III+RRM | Type-2_BR.RM.like.systems | 35exo->RHH_1->DUF499*->DUF3780+RAMA->METHYLASE->HKD-DNase+SF1-SF2-Helicase->wHTH+REase->HDRNase->VirD4-TraG+VirB4-FtsK-> | 1041 | Bacteria>Pseudomonadota>Gammaproteobacteria>Enterobacterales | GCF_000754345.1 | Morganellaceae | I6H92_RS05395 | 1203958..1207083 | 1903414 | MULTISPECIES:-DUF499-domain-containing-protein-[Morganellaceae] |
| WP_032437326.1 | AAA+ATPase+wHTH+Alpha+Beta-domain+wHTH+Fn-III+Fn-III+RRM | Type-2_BR.RM.like.systems | SF1-SF2-Helicase->?-><-wHTH+REase<-HKD-DNase+SF1-SF2-Helicase<-METHYLASE<-DUF3780+RAMA<-DUF499*<-RHH<-Arm-DNA-bind_3+Phage_integrase<-tRNA\|\|?->?->ADH_N+ADH_zinc_N-> | 1041 | Bacteria>Pseudomonadota>Gammaproteobacteria>Enterobacterales | GCF_009387945.1 | Enterobacteriaceae | OMD39_RS22365 | 4570598..4573723 | 543 | MULTISPECIES:-DUF499-domain-containing-protein-[Enterobacteriaceae] |
| WP_000860009.1 | AAA+ATPase+wHTH+Alpha+Beta-domain+wHTH+Fn-III+Fn-III+RRM | Type-2_BR.RM.like.systems | <-HKD-DNase+SF1-SF2-Helicase<-METHYLASE<-DUF3780<-DUF499*<-zf-ISL3+DDE_Tnp_ISL3\|\|?->\|\|MAD-NTD+HNH-> | 1022 | Bacteria>Pseudomonadota>Gammaproteobacteria>Enterobacterales | GCF_014168595.1 | Enterobacteriaceae | KUM08_RS17055 | 7220..10288 | 543 | MULTISPECIES:-DUF499-domain-containing-protein-[Enterobacteriaceae] |
| WP_272536813.1 | AAA+ATPase+wHTH+Alpha+Beta-domain+wHTH+Fn-III+Fn-III+RRM | Type-2_BR.RM.like.systems | XIS-HTH->Inovirus_Gp2-><-?\|\|ZnR->XIS-HTH->RHH->DUF499*->DUF3780+RAMA->METHYLASE->HKD-DNase+SF1-SF2-Helicase->Zn_Tnp_IS1595+DDE_Tnp_IS1595-> | 1041 | Bacteria>Pseudomonadota>Gammaproteobacteria>Enterobacterales | GCF_028480325.1 | Morganellaceae | PVW56_RS12320 | 55801..58926 | 1903414 | MULTISPECIES:-DUF499-domain-containing-protein-[Morganellaceae] |
| WP_230608232.1 | AAA+ATPase+wHTH+Alpha+Beta-domain+wHTH+Fn-III+Fn-III+RRM | Type-2_BR.RM.like.systems | 35exo->RHH_1->DUF499*->DUF3780+RAMA->METHYLASE->HKD-DNase+SF1-SF2-Helicase->wHTH+REase->HDRNase->VirD4-TraG+VirB4-FtsK-> | 1041 | Bacteria>Pseudomonadota>Gammaproteobacteria>Enterobacterales>Budviciaceae | GCF_021010575.1 | Limnobaculum_eriocheiris | LPW36_RS04285 | 61896..65021 | 2897391 | DUF499-domain-containing-protein-[Jinshanibacter-sp.-LJY008] |
| WP_200893021.1 | AAA+ATPase+wHTH+Alpha+Beta-domain+wHTH+Fn-III+Fn-III+RRM | Type-2_BR.RM.like.systems | Arm-DNA-bind_3+Phage_integrase->RHH->DUF499*->DUF3780+RAMA->METHYLASE->HKD-DNase+SF1-SF2-Helicase->wHTH+REase-><-?\|\|HTH_Tnp_1+HTH_21+rve+rve_2-> | 1022 | Bacteria>Pseudomonadota>Gammaproteobacteria>Enterobacterales>Enterobacteriaceae | - | Enterobacter_hormaechei | I4A92_000735 | 150340..153408 | 158836 | DUF499-domain-containing-protein-[Enterobacter-hormaechei] |

| Protein-ID | DUF499-architecture | System | Gene-Neighbourhoods | Length | Taxonomy | Assembly-ID | Strain | Locus-Tag | Genomic-coordinates | TaxID | Fasta-defline |
|---|---|---|---|---|---|---|---|---|---|---|---|
| WP_218968419.1 | AAA+ATPase+wHTH+Alpha+Beta-domain+wHTH+Fn-III+Fn-III+RRM | Type-2_BR.RM.like.systems | <-wHTH+REase<-HKD-DNase+SF1-SF2-Helicase<-METHYLASE<-DUF3780+RAMA<-DUF499*<-RHH<-Arm-DNA-bind_3+Phage_integrase | 1022 | Bacteria>Pseudomonadota>Gammaproteobacteria>Enterobacterales>Enterobacteriaceae | - | Enterobacter_hormaechei | SAMEA2053862_03698 | 27017..30085 | 158836 | DUF499-domain-containing-protein-[Enterobacter-hormaechei] |
| WP_219810711.1 | AAA+ATPase+wHTH+Alpha+Beta-domain+wHTH+Fn-III+Fn-III+RRM | Type-2_BR.RM.like.systems | HIN-HTH->HIN-HTH-><-REase<-HKD-DNase+SF1-SF2-Helicase<-METHYLASE<-DUF3780+RAMA<-DUF499*<-RHH<-Arm-DNA-bind_3+Phage_integrase | 1022 | Bacteria>Pseudomonadota>Gammaproteobacteria>Enterobacterales>Enterobacteriaceae | - | Enterobacter_sp._BIDMC_29 | L465_03720 | 733767..736835 | 1329841 | DUF499-domain-containing-protein-[Enterobacter-sp.-BIDMC-29] |
| EUM08187.1 | AAA+ATPase+wHTH+Alpha+Beta-domain+wHTH+Fn-III+Fn-III+RRM | Type-2_BR.RM.like.systems | HIN-HTH->HIN-HTH-><-REase<-HKD-DNase+SF1-SF2-Helicase<-METHYLASE<-DUF3780+RAMA<-DUF499*<-RHH<-Arm-DNA-bind_3+Phage_integrase | 1022 | Bacteria>Pseudomonadota>Gammaproteobacteria>Enterobacterales>Enterobacteriaceae | GCA_000534375.1 | Enterobacter_sp._BIDMC_29 | L465_03720 | 733767..736835 | 1329841 | hypothetical-protein-L465_03720-[Enterobacter-sp.-BIDMC-29] |
| KDL80228.1 | AAA+ATPase+wHTH+Alpha+Beta-domain+wHTH+Fn-III+Fn-III+RRM | Type-2_BR.RM.like.systems | <-ADH_N+ADH_zinc_N<-?<-?\|\|tRNA->Arm-DNA-bind_3+Phage_integrase->RHH->DUF499*->DUF3780+RAMA->METHYLASE->HKD-DNase+SF1-SF2-Helicase->wHTH+REase-> | 1022 | Bacteria>Pseudomonadota>Gammaproteobacteria>Enterobacterales>Enterobacteriaceae | GCA_000694675.1 | Klebsiella_pneumoniae_MGH_72 | AD98_04564 | 185426..188494 | 1438795 | hypothetical-protein-AD98_04564-[Klebsiella-pneumoniae-MGH-72] |
| ARA42377.1 | AAA+ATPase+wHTH+Alpha+Beta-domain+wHTH+Fn-III+Fn-III+RRM | Type-2_BR.RM.like.systems | SF1-SF2-Helicase->?-><-wHTH+REase<-HKD-DNase+SF1-SF2-Helicase<-METHYLASE<-DUF3780+RAMA<-DUF499*<-DDE_Tnp_1<-DUF499-p<-RHH<-Arm-DNA-bind_3+Phage_integrase | 984 | Bacteria>Pseudomonadota>Gammaproteobacteria>Enterobacterales>Enterobacteriaceae | GCA_002055765.1 | Klebsiella_pneumoniae | AM364_19165 | 3781580..3784534 | 573 | AAA-family-ATPase-[Klebsiella-pneumoniae] |
| PMD76486.1 | AAA+ATPase+wHTH+Alpha+Beta-domain+wHTH+Fn-III+Fn-III+RRM | Type-2_BR.RM.like.systems | <-ADH_N+ADH_zinc_N<-?<-?\|\|tRNA->Arm-DNA-bind_3+Phage_integrase->RHH->DUF499*->DUF3780+RAMA->METHYLASE->HKD-DNase+SF1-SF2-Helicase->wHTH+REase-><-?\|\|<-XIS-HTH | 1041 | Bacteria>Pseudomonadota>Gammaproteobacteria>Enterobacterales>Enterobacteriaceae | GCA_002872205.1 | Klebsiella_pneumoniae | A8A10_21890 | 132525..135650 | 573 | AAA-family-ATPase-[Klebsiella-pneumoniae] |
| ECA3794685.1 | AAA+ATPase+wHTH+Alpha+Beta-domain+wHTH+Fn-III+Fn-III+RRM | Type-2_BR.RM.like.systems | <-DnaB+SMS-KaiC<-?<-ParA-Soj-PloopNTPase\|\|Arm-DNA-bind_3+Phage_integrase->RHH->DUF499*->DUF3780+RAMA->METHYLASE->HKD-DNase+SF1-SF2-Helicase->wHTH+REase-> | 1041 | Bacteria>Pseudomonadota>Gammaproteobacteria>Enterobacterales>Enterobacteriaceae | GCA_007378915.1 | Salmonella_enterica_subsp._enterica_serovar_Aqua | EKG95_23275 | 13505..16630 | 1302615 | DUF499-domain-containing-protein-[Salmonella-enterica-subsp.-enterica-serovar-Aqua] |
| EES4626560.1 | AAA+ATPase+wHTH+Alpha+Beta-domain+wHTH+Fn-III+Fn-III+RRM | Type-2_BR.RM.like.systems | <-Phage_integrase\|\|?->?->?-><-HNH<-?\|\|DUF499*->DUF3780->METHYLASE->HKD-DNase+SF1-SF2-Helicase-><-REase<-NACHT+SF1-SF2-Helicase+HKD-DNase<-RNaseH | 1022 | Bacteria>Pseudomonadota>Gammaproteobacteria>Enterobacterales>Enterobacteriaceae | GCA_011897625.1 | Escherichia_coli | EZR16_001805 | 93004..96072 | 562 | DUF499-domain-containing-protein-[Escherichia-coli] |
| EFA6413129.1 | AAA+ATPase+wHTH+Alpha+Beta-domain+wHTH+Fn-III+Fn-III+RRM | Type-2_BR.RM.like.systems | <-HIN-HTH<-wHTH+REase<-HKD-DNase+SF1-SF2-Helicase<-METHYLASE<-DUF3780+RAMA<-DUF499*<-RHH_5\|\|<-?<-Aminotran_1_2\|\|<-Arm-DNA-bind_3+Phage_integrase<-tRNA\|\|ADH_N+ADH_zinc_N->FtsK+HerA+A32-> | 1041 | Bacteria>Pseudomonadota>Gammaproteobacteria>Enterobacterales>Enterobacteriaceae | GCA_012256865.1 | Escherichia_coli | CIW06_000361 | 54690..57815 | 562 | DUF499-domain-containing-protein-[Escherichia-coli] |
| MBK1516290.1 | AAA+ATPase+wHTH+Alpha+Beta-domain+wHTH+Fn-III+Fn-III+RRM | Type-2_BR.RM.like.systems | <-ADH_N+ADH_zinc_N<-?<-?\|\|tRNA->Arm-DNA-bind_3+Phage_integrase->RHH->DUF499*->DUF3780+RAMA->METHYLASE->HKD-DNase+SF1-SF2-Helicase->wHTH+REase-><-HTH+MPTase | 1022 | Bacteria>Pseudomonadota>Gammaproteobacteria>Enterobacterales>Enterobacteriaceae | GCA_014235385.1 | Klebsiella_variicola | H8S48_22830 | 314255..317323 | 244366 | DUF499-domain-containing-protein-[Klebsiella-variicola] |
| HAM9542477.1 | AAA+ATPase+wHTH+Alpha+Beta-domain+wHTH+Fn-III+Fn-III+RRM | Type-2_BR.RM.like.systems | <-Phage_integrase\|\|?->?->?-><-MAD-NTD+HNH<-?\|\|DUF499*->DUF3780->METHYLASE->HKD-DNase+SF1-SF2-Helicase-><-REase<-NACHT+SF1-SF2-Helicase+HKD-DNase<-RNaseH | 1022 | Bacteria>Pseudomonadota>Gammaproteobacteria>Enterobacterales>Enterobacteriaceae | GCA_014583135.1 | Escherichia_coli | IDY85_002513 | 90940..94008 | 562 | DUF499-domain-containing-protein-[Escherichia-coli] |
| HAT2182700.1 | AAA+ATPase+wHTH+Alpha+Beta-domain+wHTH+Fn-III+Fn-III+RRM | Type-2_BR.RM.like.systems | OLD-ABC+TOPRIM->SF1-SF2-Helicase-><-HTH_Tnp_1+HTH_21+rve+rve_2<-HKD-DNase+SF1-SF2-Helicase<-METHYLASE<-DUF3780+RAMA<-DUF499*<-RHH<-Arm-DNA-bind_3+Phage_integrase<-tRNA\|\|ADH_N+ADH_zinc_N->FtsK+HerA+A32-> | 1041 | Bacteria>Pseudomonadota>Gammaproteobacteria>Enterobacterales>Enterobacteriaceae | GCA_015897445.1 | Citrobacter_freundii | I8143_001064 | 210810..213935 | 546 | DUF499-domain-containing-protein-[Citrobacter-freundii] |
| HAV2294932.1 | AAA+ATPase+wHTH+Alpha+Beta-domain+wHTH+Fn-III+Fn-III+RRM | Type-2_BR.RM.like.systems | HTH+MPTase-><-wHTH+REase<-HKD-DNase+SF1-SF2-Helicase<-METHYLASE<-DUF3780+RAMA<-DUF499*<-RHH<-Arm-DNA-bind_3+Phage_integrase<-tRNA\|\|ADH_N+ADH_zinc_N->FtsK+HerA+A32-> | 1041 | Bacteria>Pseudomonadota>Gammaproteobacteria>Enterobacterales>Enterobacteriaceae | GCA_016445565.1 | Citrobacter_koseri | JHK52_001290 | 177195..180320 | 545 | DUF499-domain-containing-protein-[Citrobacter-koseri] |
| MBJ8391833.1 | AAA+ATPase+wHTH+Alpha+Beta-domain+wHTH+Fn-III+Fn-III+RRM | Type-2_BR.RM.like.systems | <-HIN-HTH<-wHTH+REase<-HKD-DNase+SF1-SF2-Helicase<-METHYLASE<-DUF3780+RAMA<-DUF499*<-RHH<-Arm-DNA-bind_3+Phage_integrase<-tRNA\|\|ADH_N+ADH_zinc_N-> | 1022 | Bacteria>Pseudomonadota>Gammaproteobacteria>Enterobacterales>Enterobacteriaceae | GCA_016501115.1 | Citrobacter_cronae | I6M80_16495 | 16784..19852 | 1748967 | DUF499-domain-containing-protein-[Citrobacter-cronae] |
| MBK2656985.1 | AAA+ATPase+wHTH+Alpha+Beta-domain+wHTH+Fn-III+Fn-III+RRM | Type-2_BR.RM.like.systems | HTH+MPTase-><-wHTH+REase<-HKD-DNase+SF1-SF2-Helicase<-METHYLASE<-DUF3780+RAMA<-DUF499*<-RHH<-Arm-DNA-bind_3+Phage_integrase<-tRNA\|\|?->?->ADH_N+ADH_zinc_N-> | 1039 | Bacteria>Pseudomonadota>Gammaproteobacteria>Enterobacterales>Enterobacteriaceae | GCA_016614755.1 | Klebsiella_pneumoniae | FKL12_18780 | 12401..15520 | 573 | DUF499-domain-containing-protein-[Klebsiella-pneumoniae] |
| MBK5711249.1 | AAA+ATPase+wHTH+Alpha+Beta-domain+wHTH+Fn-III+Fn-III+RRM | Type-2_BR.RM.like.systems | HTH+MPTase-><-wHTH+REase<-HKD-DNase+SF1-SF2-Helicase<-METHYLASE<-DUF3780+RAMA<-DUF499*<-RHH<-Arm-DNA-bind_3+Phage_integrase<-tRNA\|\|?->?->ADH_N+ADH_zinc_N-> | 1022 | Bacteria>Pseudomonadota>Gammaproteobacteria>Enterobacterales>Enterobacteriaceae | GCA_016651545.1 | Klebsiella_pneumoniae | JKA42_15120 | 66258..69326 | 573 | DUF499-domain-containing-protein-[Klebsiella-pneumoniae] |
| EHF0011153.1 | AAA+ATPase+wHTH+Alpha+Beta-domain+wHTH+Fn-III+Fn-III+RRM | Type-2_BR.RM.like.systems | DUF499*->DUF3780->METHYLASE->HKD-DNase+SF1-SF2-Helicase-> | 826 | Bacteria>Pseudomonadota>Gammaproteobacteria>Enterobacterales>Enterobacteriaceae | GCA_017278235.1 | Escherichia_coli | IFX60_004912 | 1..2483 | 562 | DUF499-domain-containing-protein-[Escherichia-coli] |
| HBB8152959.1 | AAA+ATPase+wHTH+Alpha+Beta-domain+wHTH+Fn-III+Fn-III+RRM | Type-2_BR.RM.like.systems | DEDD_Tnp_IS110+Transposase_20->?->RHH_1->DUF499-p->DUF499*->DUF3780+RAMA->METHYLASE->HKD-DNase+SF1-SF2-Helicase->wHTH+REase->HIN-HTH-> | 958 | Bacteria>Pseudomonadota>Gammaproteobacteria>Enterobacterales>Enterobacteriaceae | GCA_018015815.1 | Escherichia_coli | J8710_004813 | 1587..4463 | 562 | DUF499-domain-containing-protein-[Escherichia-coli] |
| HBC5231502.1 | AAA+ATPase+wHTH+Alpha+Beta-domain+wHTH+Fn-III+Fn-III+RRM | Type-2_BR.RM.like.systems | SF1-SF2-Helicase->?-><-wHTH+REase<-HKD-DNase+SF1-SF2-Helicase<-METHYLASE<-DUF3780+RAMA<-DUF499*<-DUF499-p<-RHH<-Arm-DNA-bind_3+Phage_integrase<-tRNA\|\|ADH_N+ADH_zinc_N-> | 959 | Bacteria>Pseudomonadota>Gammaproteobacteria>Enterobacterales>Enterobacteriaceae | GCA_018155305.1 | Klebsiella_oxytoca | KEU23_000991 | 23793..26672 | 571 | DUF499-domain-containing-protein-[Klebsiella-oxytoca] |
| MBS9010073.1 | AAA+ATPase+wHTH+Alpha+Beta-domain+wHTH+Fn-III+Fn-III+RRM | Type-2_BR.RM.like.systems | <-Phage_integrase\|\|?->?->?-><-MAD-NTD+HNH<-?\|\|DUF499*->DUF3780->METHYLASE->HKD-DNase+SF1-SF2-Helicase-><-REase<-NACHT+SF1-SF2-Helicase+HKD-DNase<-RNaseH | 1022 | Bacteria>Pseudomonadota>Gammaproteobacteria>Enterobacterales>Enterobacteriaceae | GCA_018428565.1 | Escherichia_coli | KEP54_04330 | 349575..352643 | 562 | DUF499-domain-containing-protein-[Escherichia-coli] |
| EHN8715928.1 | AAA+ATPase+wHTH+Alpha+Beta-domain+wHTH+Fn-III+Fn-III+RRM | Type-2_BR.RM.like.systems | Arm-DNA-bind_3+Phage_integrase->RHH->DUF499*->DUF3780+RAMA->METHYLASE->HKD-DNase+SF1-SF2-Helicase->wHTH+REase-><-?\|\|HTH_Tnp_1+HTH_21+rve+rve_2-> | 1022 | Bacteria>Pseudomonadota>Gammaproteobacteria>Enterobacterales>Enterobacteriaceae | GCA_018454885.1 | Enterobacter_hormaechei | I4A92_000735 | 150340..153408 | 158836 | DUF499-domain-containing-protein-[Enterobacter-hormaechei] |
| HBE2428354.1 | AAA+ATPase+wHTH+Alpha+Beta-domain+wHTH+Fn-III+Fn-III+RRM | Type-2_BR.RM.like.systems | RHH_1->DUF499*->DUF3780+RAMA->METHYLASE->HKD-DNase+SF1-SF2-Helicase->wHTH+REase-> | 1022 | Bacteria>Pseudomonadota>Gammaproteobacteria>Enterobacterales>Enterobacteriaceae | GCA_018545885.1 | Escherichia_coli | KL339_004657 | 1620..4688 | 562 | DUF499-domain-containing-protein-[Escherichia-coli] |
| MBT1774403.1 | AAA+ATPase+wHTH+Alpha+Beta-domain+wHTH+Fn-III+Fn-III+RRM | Type-2_BR.RM.like.systems | Arm-DNA-bind_3+Phage_integrase->RHH->DUF499*->DUF3780+RAMA->METHYLASE->HKD-DNase+SF1-SF2-Helicase-> | 1022 | Bacteria>Pseudomonadota>Gammaproteobacteria>Enterobacterales>Enterobacteriaceae | GCA_018596995.1 | Enterobacter_hormaechei_subsp._xiangfangensis | KK000_19420 | 150119..153187 | 1296536 | DUF499-domain-containing-protein-[Enterobacter-hormaechei-subsp.-xiangfangensis] |
| EHR1235094.1 | AAA+ATPase+wHTH+Alpha+Beta-domain+wHTH+Fn-III+Fn-III+RRM | Type-2_BR.RM.like.systems | DUF499*->DUF3780->METHYLASE->HKD-DNase+SF1-SF2-Helicase-><-REase<-SF1-SF2-Helicase+HKD-DNase | 788 | Bacteria>Pseudomonadota>Gammaproteobacteria>Enterobacterales>Enterobacteriaceae | GCA_019000485.1 | Escherichia_coli | KS907_004964 | 1..2368 | 562 | DUF499-domain-containing-protein-[Escherichia-coli] |
| MBV4586514.1 | AAA+ATPase+wHTH+Alpha+Beta-domain+wHTH+Fn-III+Fn-III+RRM | Type-2_BR.RM.like.systems | ZnR->ZnR->HTH_Tnp_1+HTH_21+rve+rve_3->?->RHH_1->DUF499*->DUF3780+RAMA->METHYLASE->HKD-DNase+SF1-SF2-Helicase->wHTH+REase->HIN-HTH-> | 1022 | Bacteria>Pseudomonadota>Gammaproteobacteria>Enterobacterales>Enterobacteriaceae | GCA_019145775.1 | Escherichia_coli | I2922_14300 | 166455..169523 | 562 | DUF499-domain-containing-protein-[Escherichia-coli] |
| EHS9957778.1 | AAA+ATPase+wHTH+Alpha+Beta-domain+wHTH+Fn-III+Fn-III+RRM | Type-2_BR.RM.like.systems | <-Phage_integrase\|\|?->?->?-><-MAD-NTD+HNH<-?\|\|DUF499*->DUF3780->METHYLASE->HKD-DNase+SF1-SF2-Helicase-><-REase | 1022 | Bacteria>Pseudomonadota>Gammaproteobacteria>Enterobacterales>Enterobacteriaceae | GCA_019181685.1 | Escherichia_coli | KV153_004362 | 9266..12334 | 562 | DUF499-domain-containing-protein-[Escherichia-coli] |
| EHT5191361.1 | AAA+ATPase+wHTH+Alpha+Beta-domain+wHTH+Fn-III+Fn-III+RRM | Type-2_BR.RM.like.systems | <-Phage_integrase\|\|?->?->?-><-MAD-NTD+HNH<-?\|\|DUF499*->DUF3780->METHYLASE->HKD-DNase+SF1-SF2-Helicase-><-REase | 1022 | Bacteria>Pseudomonadota>Gammaproteobacteria>Enterobacterales>Enterobacteriaceae | GCA_019222045.1 | Escherichia_coli | KFA52_004662 | 10352..13420 | 562 | DUF499-domain-containing-protein-[Escherichia-coli] |
| HBK6101728.1 | AAA+ATPase+wHTH+Alpha+Beta-domain+wHTH+Fn-III+Fn-III+RRM | Type-2_BR.RM.like.systems | OLD-ABC+TOPRIM->SF1-SF2-Helicase-><-HTH_Tnp_1+HTH_21+rve+rve_2<-HKD-DNase+SF1-SF2-Helicase<-METHYLASE<-DUF3780+RAMA<-DUF499*<-RHH<-Arm-DNA-bind_3+Phage_integrase<-tRNA\|\|ADH_N+ADH_zinc_N->FtsK+HerA+A32-> | 1022 | Bacteria>Pseudomonadota>Gammaproteobacteria>Enterobacterales>Enterobacteriaceae | GCA_020776975.1 | Citrobacter_freundii | LMY18_000638 | 210810..213878 | 546 | DUF499-domain-containing-protein-[Citrobacter-freundii] |
| EIQ9407871.1 | AAA+ATPase+wHTH+Alpha+Beta-domain+wHTH+Fn-III+Fn-III+RRM | Type-2_BR.RM.like.systems | DUF499*->DUF3780->METHYLASE->HKD-DNase+SF1-SF2-Helicase-><-REase | 986 | Bacteria>Pseudomonadota>Gammaproteobacteria>Enterobacterales>Enterobacteriaceae | GCA_021226995.1 | Escherichia_coli | LV737_004815 | 1..2962 | 562 | DUF499-domain-containing-protein-[Escherichia-coli] |
| HBM8437234.1 | AAA+ATPase+wHTH+Alpha+Beta-domain+wHTH+Fn-III+Fn-III+RRM | Type-2_BR.RM.like.systems | <-HIN-HTH<-wHTH+REase<-HKD-DNase+SF1-SF2-Helicase<-METHYLASE<-DUF3780+RAMA<-DUF499*<-RHH_5\|\|<-?<-Aminotran_1_2 | 1022 | Bacteria>Pseudomonadota>Gammaproteobacteria>Enterobacterales>Enterobacteriaceae | GCA_021425605.1 | Escherichia_coli | LOV60_003801 | 18779..21847 | 562 | DUF499-domain-containing-protein-[Escherichia-coli] |
| HBQ4762183.1 | AAA+ATPase+wHTH+Alpha+Beta-domain+wHTH+Fn-III+Fn-III+RRM | Type-2_BR.RM.like.systems | DUF499*->DUF3780->METHYLASE->HKD-DNase+SF1-SF2-Helicase-><-REase<-NACHT+SF1-SF2-Helicase+HKD-DNase<-RNaseH | 1019 | Bacteria>Pseudomonadota>Gammaproteobacteria>Enterobacterales>Enterobacteriaceae | GCA_021860145.1 | Escherichia_coli | L7317_004528 | 1..3060 | 562 | DUF499-domain-containing-protein-[Escherichia-coli] |
| HBQ8501082.1 | AAA+ATPase+wHTH+Alpha+Beta-domain+wHTH+Fn-III+Fn-III+RRM | Type-2_BR.RM.like.systems | SF1-SF2-Helicase->?-><-wHTH+REase<-HKD-DNase+SF1-SF2-Helicase<-METHYLASE<-DUF3780+RAMA<-DUF499*<-RHH<-Arm-DNA-bind_3+Phage_integrase | 1022 | Bacteria>Pseudomonadota>Gammaproteobacteria>Enterobacterales>Enterobacteriaceae | GCA_021895715.1 | Klebsiella_pneumoniae | L9A44_004853 | 54041..57109 | 573 | DUF499-domain-containing-protein-[Klebsiella-pneumoniae] |
| HBX9967586.1 | AAA+ATPase+wHTH+Alpha+Beta-domain+wHTH+Fn-III+Fn-III+RRM | Type-2_BR.RM.like.systems | HTH+MPTase-><-wHTH+REase<-HKD-DNase+SF1-SF2-Helicase<-METHYLASE<-DUF3780+RAMA<-DUF499*<-RHH<-Arm-DNA-bind_3+Phage_integrase<-tRNA\|\|?->?->ADH_N+ADH_zinc_N-> | 1041 | Bacteria>Pseudomonadota>Gammaproteobacteria>Enterobacterales>Enterobacteriaceae | GCA_022266445.1 | Klebsiella_variicola | MIM73_23195 | 35789..38914 | 244366 | DUF499-domain-containing-protein-[Klebsiella-variicola] |
| HBY8019970.1 | AAA+ATPase+wHTH+Alpha+Beta-domain+wHTH+Fn-III+Fn-III+RRM | Type-2_BR.RM.like.systems | SF1-SF2-Helicase->?-><-wHTH+REase<-HKD-DNase+SF1-SF2-Helicase<-METHYLASE<-DUF3780+RAMA<-DUF499* | 982 | Bacteria>Pseudomonadota>Gammaproteobacteria>Enterobacterales>Enterobacteriaceae | GCA_022301155.1 | Klebsiella_pneumoniae | MI822_22070 | 54206..57154 | 573 | DUF499-domain-containing-protein-[Klebsiella-pneumoniae] |
| EIY1898012.1 | AAA+ATPase+wHTH+Alpha+Beta-domain+wHTH+Fn-III+Fn-III+RRM | Type-2_BR.RM.like.systems | SF1-SF2-Helicase->?-><-wHTH+REase<-HKD-DNase+SF1-SF2-Helicase<-METHYLASE<-DUF3780+RAMA<-DUF499*<-RHH<-Arm-DNA-bind_3+Phage_integrase<-tRNA\|\|?->?->ADH_N+ADH_zinc_N-> | 1022 | Bacteria>Pseudomonadota>Gammaproteobacteria>Enterobacterales>Enterobacteriaceae | GCA_022464035.1 | Klebsiella_pneumoniae | MMH51_002463 | 54491..57559 | 573 | DUF499-domain-containing-protein-[Klebsiella-pneumoniae] |
| HBZ7479225.1 | AAA+ATPase+wHTH+Alpha+Beta-domain+wHTH+Fn-III+Fn-III+RRM | Type-2_BR.RM.like.systems | <-ADH_N+ADH_zinc_N<-?<-?\|\|tRNA->Arm-DNA-bind_3+Phage_integrase->RHH->DUF499*->DUF3780+RAMA->METHYLASE->HKD-DNase+SF1-SF2-Helicase->wHTH+REase-><-HTH+MPTase | 1022 | Bacteria>Pseudomonadota>Gammaproteobacteria>Enterobacterales>Enterobacteriaceae | GCA_022497945.1 | Klebsiella_pneumoniae | MM066_002839 | 198069..201137 | 573 | DUF499-domain-containing-protein-[Klebsiella-pneumoniae] |
| EIY6173593.1 | AAA+ATPase+wHTH+Alpha+Beta-domain+wHTH+Fn-III+Fn-III+RRM | Type-2_BR.RM.like.systems | <-Phage_integrase\|\|?->?->?-><-MAD-NTD+HNH<-?\|\|DUF499*->DUF3780->METHYLASE->HKD-DNase+SF1-SF2-Helicase-><-REase | 1022 | Bacteria>Pseudomonadota>Gammaproteobacteria>Enterobacterales>Enterobacteriaceae | GCA_022521285.1 | Escherichia_coli | MND03_004216 | 10313..13381 | 562 | DUF499-domain-containing-protein-[Escherichia-coli] |
| MCM7420480.1 | AAA+ATPase+wHTH+Alpha+Beta-domain+wHTH+Fn-III+Fn-III+RRM | Type-2_BR.RM.like.systems | SF1-SF2-Helicase->?-><-wHTH+REase<-HKD-DNase+SF1-SF2-Helicase<-METHYLASE<-DUF3780+RAMA<-DUF499*<-RHH<-Arm-DNA-bind_3+Phage_integrase | 1041 | Bacteria>Pseudomonadota>Gammaproteobacteria>Enterobacterales>Enterobacteriaceae | GCA_023751025.1 | Enterobacter_hormaechei | M8R61_11745 | 350884..354009 | 158836 | DUF499-domain-containing-protein-[Enterobacter-hormaechei] |
| HCD9026459.1 | AAA+ATPase+wHTH+Alpha+Beta-domain+wHTH+Fn-III+Fn-III+RRM | Type-2_BR.RM.like.systems | HTH+MPTase-><-HNH+SAVED<-wHTH+REase<-HKD-DNase+SF1-SF2-Helicase<-METHYLASE<-DUF3780+RAMA<-DUF499*<-RHH<-Arm-DNA-bind_3+Phage_integrase<-tRNA\|\|wHTH+HNH->SAM-methylase-> | 1022 | Bacteria>Pseudomonadota>Gammaproteobacteria>Enterobacterales>Enterobacteriaceae | GCA_023760145.1 | Klebsiella_oxytoca | NEF91_003870 | 43874..46942 | 571 | DUF499-domain-containing-protein-[Klebsiella-oxytoca] |
| HCJ8634260.1 | AAA+ATPase+wHTH+Alpha+Beta-domain+wHTH+Fn-III+Fn-III+RRM | Type-2_BR.RM.like.systems | <-ADH_N+ADH_zinc_N<-?<-?\|\|tRNA->RHH->DUF499*->DUF3780+RAMA->METHYLASE->HKD-DNase+SF1-SF2-Helicase->wHTH+REase-><-?<-SF1-SF2-Helicase | 1022 | Bacteria>Pseudomonadota>Gammaproteobacteria>Enterobacterales>Enterobacteriaceae | GCA_024674695.1 | Klebsiella_pneumoniae | NV748_003018 | 174251..177319 | 573 | DUF499-domain-containing-protein-[Klebsiella-pneumoniae] |
| HCM9136403.1 | AAA+ATPase+wHTH+Alpha+Beta-domain+wHTH+Fn-III+Fn-III+RRM | Type-2_BR.RM.like.systems | ZnR->ZnR-><-wHTH+REase<-HKD-DNase+SF1-SF2-Helicase<-METHYLASE<-DUF3780+RAMA<-DUF499*<-RHH<-Arm-DNA-bind_3+Phage_integrase | 1022 | Bacteria>Pseudomonadota>Gammaproteobacteria>Enterobacterales>Enterobacteriaceae | GCA_025209595.1 | Enterobacter_cloacae_subsp._cloacae | N5L02_000891 | 375093..378161 | 336306 | DUF499-domain-containing-protein-[Enterobacter-cloacae-subsp.-cloacae] |
| HCN6859046.1 | AAA+ATPase+wHTH+Alpha+Beta-domain+wHTH+Fn-III+Fn-III+RRM | Type-2_BR.RM.like.systems | DUF499*->DUF3780->METHYLASE->HKD-DNase+SF1-SF2-Helicase-><-REase<-NACHT+SF1-SF2-Helicase+HKD-DNase<-RNaseH | 746 | Bacteria>Pseudomonadota>Gammaproteobacteria>Enterobacterales>Enterobacteriaceae | GCA_025285375.1 | Escherichia_coli | N6101_002250 | 1..2242 | 562 | DUF499-domain-containing-protein-[Escherichia-coli] |
| HCT3226645.1 | AAA+ATPase+wHTH+Alpha+Beta-domain+wHTH+Fn-III+Fn-III+RRM | Type-2_BR.RM.like.systems | <-SymE_toxin\|\|wHTH(SBD)+HNH-><-HKD-DNase+SF1-SF2-Helicase<-METHYLASE<-DUF3780+RAMA<-DUF499*<-RHH<-wHTH(SBD)+HNH\|\|?->XIS-HTH->ParB+BAM-> | 1022 | Bacteria>Pseudomonadota>Gammaproteobacteria>Enterobacterales>Enterobacteriaceae | GCA_026310715.1 | Enterobacter_hormaechei | OTS66_001787 | 86000..89068 | 158836 | TPA:-DUF499-domain-containing-protein-[Enterobacter-hormaechei] |
| HCW2948669.1 | AAA+ATPase+wHTH+Alpha+Beta-domain+wHTH+Fn-III+Fn-III+RRM | Type-2_BR.RM.like.systems | <-HIN-HTH<-wHTH+REase<-HKD-DNase+SF1-SF2-Helicase<-METHYLASE<-DUF3780+RAMA<-DUF499*<-RHH_1 | 1022 | Bacteria>Pseudomonadota>Gammaproteobacteria>Enterobacterales>Enterobacteriaceae | GCA_026548625.1 | Escherichia_coli | OXB36_004615 | 17018..20086 | 562 | TPA:-DUF499-domain-containing-protein-[Escherichia-coli] |
| HCW3139350.1 | AAA+ATPase+wHTH+Alpha+Beta-domain+wHTH+Fn-III+Fn-III+RRM | Type-2_BR.RM.like.systems | <-FtsK+HerA+A32<-ADH_N+ADH_zinc_N\|\|tRNA->Arm-DNA-bind_3+Phage_integrase->RHH->DUF499*->DUF3780+RAMA->METHYLASE->HKD-DNase+SF1-SF2-Helicase->wHTH+REase-><-HTH+MPTase | 1022 | Bacteria>Pseudomonadota>Gammaproteobacteria>Enterobacterales>Enterobacteriaceae | GCA_026549635.1 | Citrobacter_freundii | OXC88_001869 | 152209..155277 | 546 | TPA:-DUF499-domain-containing-protein-[Citrobacter-freundii] |
| EKK5919799.1 | AAA+ATPase+wHTH+Alpha+Beta-domain+wHTH+Fn-III+Fn-III+RRM | Type-2_BR.RM.like.systems | Arm-DNA-bind_3+Phage_integrase->RHH->DUF499*->DUF3780+RAMA->METHYLASE->HKD-DNase+SF1-SF2-Helicase->wHTH+REase-><-?<-SF1-SF2-Helicase | 1022 | Bacteria>Pseudomonadota>Gammaproteobacteria>Enterobacterales>Enterobacteriaceae | GCA_028182685.1 | Enterobacter_hormaechei | POC94_000676 | 161243..164311 | 158836 | DUF499-domain-containing-protein-[Enterobacter-hormaechei] |
| HDK9936257.1 | AAA+ATPase+wHTH+Alpha+Beta-domain+wHTH+Fn-III+Fn-III+RRM | Type-2_BR.RM.like.systems | DUF499*->DUF3780->METHYLASE->HKD-DNase+SF1-SF2-Helicase-><-REase<-NACHT+SF1-SF2-Helicase+HKD-DNase<-RNaseH | 794 | Bacteria>Pseudomonadota>Gammaproteobacteria>Enterobacterales>Enterobacteriaceae | GCA_028671755.1 | Escherichia_coli | PVT51_004404 | 1..2385 | 562 | TPA:-DUF499-domain-containing-protein,-partial-[Escherichia-coli] |
| WIJ56059.1 | AAA+ATPase+wHTH+Alpha+Beta-domain+wHTH+Fn-III+Fn-III+RRM | Type-2_BR.RM.like.systems | Arm-DNA-bind_3+Phage_integrase->RHH->DUF499-p->DUF499*->DUF3780+RAMA->METHYLASE->HKD-DNase+SF1-SF2-Helicase->wHTH+REase-><-?\|\|HTH_Tnp_1+HTH_21+rve+rve_2-> | 894 | Bacteria>Pseudomonadota>Gammaproteobacteria>Enterobacterales>Enterobacteriaceae | GCA_030179615.1 | Enterobacter_hormaechei | OI979_06745 | 1401626..1404310 | 158836 | DUF499-domain-containing-protein-[Enterobacter-hormaechei] |
| STR56259.1 | AAA+ATPase+wHTH+Alpha+Beta-domain+wHTH+Fn-III+Fn-III+RRM | Type-2_BR.RM.like.systems | SF1-SF2-Helicase->?-><-wHTH+REase<-HKD-DNase+SF1-SF2-Helicase<-METHYLASE<-DUF3780+RAMA<-DUF499*<-DUF499-p<-RHH<-Arm-DNA-bind_3+Phage_integrase<-tRNA\|\|ADH_N+ADH_zinc_N-> | 984 | Bacteria>Pseudomonadota>Gammaproteobacteria>Enterobacterales>Enterobacteriaceae | GCA_900451255.1 | Klebsiella_oxytoca | NCTC11696_05224 | 5162019..5164973 | 571 | Protein-of-uncharacterised-function-(DUF499)-[Klebsiella-oxytoca] |
| VAE22474.1 | AAA+ATPase+wHTH+Alpha+Beta-domain+wHTH+Fn-III+Fn-III+RRM | Type-2_BR.RM.like.systems | <-wHTH+REase<-HKD-DNase+SF1-SF2-Helicase<-METHYLASE<-DUF3780+RAMA<-DUF499*<-RHH<-Arm-DNA-bind_3+Phage_integrase | 1022 | Bacteria>Pseudomonadota>Gammaproteobacteria>Enterobacterales>Enterobacteriaceae | GCA_900558075.1 | Enterobacter_hormaechei | SAMEA2053862_03698 | 27017..30085 | 158836 | Protein-of-uncharacterised-function-(DUF499)-[Enterobacter-hormaechei] |
| WP_000860008.1 | AAA+ATPase+wHTH+Alpha+Beta-domain+wHTH+Fn-III+Fn-III+RRM | Type-2_BR.RM.like.systems | <-MAD-NTD+HNH\|\|<-?\|\|zf-ISL3+DDE_Tnp_ISL3->DUF499*->DUF3780->METHYLASE->HKD-DNase+SF1-SF2-Helicase-> | 1022 | Bacteria>Pseudomonadota>Gammaproteobacteria>Enterobacterales>Enterobacteriaceae | GCF_000259385.1 | Escherichia_coli | WA3_RS0115475 | 167802..170870 | 562 | DUF499-domain-containing-protein-[Escherichia-coli] |
| WP_024008845.1 | AAA+ATPase+wHTH+Alpha+Beta-domain+wHTH+Fn-III+Fn-III+RRM | Type-2_BR.RM.like.systems | <-VirD4-TraG+VirB4-FtsK<-HDRNase<-wHTH+REase<-HKD-DNase+SF1-SF2-Helicase<-METHYLASE<-DUF3780+RAMA<-DUF499*<-RHH_1<-35exo\|\|?->Phage_integrase-> | 1041 | Bacteria>Pseudomonadota>Gammaproteobacteria>Enterobacterales>Enterobacteriaceae | GCF_000507645.1 | Escherichia_coli | G831_RS00450 | 65264..68389 | 562 | DUF499-domain-containing-protein-[Escherichia-coli] |
| WP_047743274.1 | AAA+ATPase+wHTH+Alpha+Beta-domain+wHTH+Fn-III+Fn-III+RRM | Type-2_BR.RM.like.systems | wHTH+SAD+HNH->NACHT+SF1-SF2-Helicase+HKD-DNase->REase-><-HKD-DNase+SF1-SF2-Helicase<-METHYLASE<-DUF3780<-DUF499*<-?<-wHTH+SAD+HNH\|\|SymE_toxin-><-Methyltransf_25 | 1022 | Bacteria>Pseudomonadota>Gammaproteobacteria>Enterobacterales>Enterobacteriaceae | GCF_001022445.2 | Enterobacter_ludwigii | ABR30_RS08135 | 180323..183391 | 299767 | DUF499-domain-containing-protein-[Enterobacter-ludwigii] |
| WP_059294450.1 | AAA+ATPase+wHTH+Alpha+Beta-domain+wHTH+Fn-III+Fn-III+RRM | Type-2_BR.RM.like.systems | Arm-DNA-bind_3+Phage_integrase->RHH->DUF499*->DUF3780+RAMA->METHYLASE->HKD-DNase+SF1-SF2-Helicase->REase-> | 1041 | Bacteria>Pseudomonadota>Gammaproteobacteria>Enterobacterales>Enterobacteriaceae | GCF_001518515.1 | Enterobacter_hormaechei | AWI06_RS12030 | 36830..39955 | 158836 | DUF499-domain-containing-protein-[Enterobacter-hormaechei] |
| WP_249533074.1 | AAA+ATPase+wHTH+Alpha+Beta-domain+wHTH+Fn-III+Fn-III+RRM | Type-2_BR.RM.like.systems | <-HKD-DNase+SF1-SF2-Helicase<-METHYLASE<-DUF3780<-DUF499* | 822 | Bacteria>Pseudomonadota>Gammaproteobacteria>Enterobacterales>Enterobacteriaceae | GCF_002016495.1 | Escherichia_coli | BT088_RS26550 | 5064..7532 | 562 | DUF499-domain-containing-protein-[Escherichia-coli] |
| WP_227518579.1 | AAA+ATPase+wHTH+Alpha+Beta-domain+wHTH+Fn-III+Fn-III+RRM | Type-2_BR.RM.like.systems | SF1-SF2-Helicase->?-><-wHTH+REase<-HKD-DNase+SF1-SF2-Helicase<-METHYLASE<-DUF3780+RAMA<-DUF499*<-DDE_Tnp_1<-DUF499-p<-RHH<-Arm-DNA-bind_3+Phage_integrase | 973 | Bacteria>Pseudomonadota>Gammaproteobacteria>Enterobacterales>Enterobacteriaceae | GCF_002055765.1 | Klebsiella_pneumoniae | AM364_RS18410 | 3781580..3784501 | 573 | DUF499-domain-containing-protein-[Klebsiella-pneumoniae] |
| WP_094163055.1 | AAA+ATPase+wHTH+Alpha+Beta-domain+wHTH+Fn-III+Fn-III+RRM | Type-2_BR.RM.like.systems | <-MAD-NTD+HNH\|\|<-?\|\|zf-ISL3+DDE_Tnp_ISL3->DUF499*->DUF3780->METHYLASE->HKD-DNase+SF1-SF2-Helicase-><-REase<-NACHT+SF1-SF2-Helicase+HKD-DNase<-RNaseH | 1022 | Bacteria>Pseudomonadota>Gammaproteobacteria>Enterobacterales>Enterobacteriaceae | GCF_002240935.1 | Escherichia_coli | CDL55_RS21695 | 10424..13492 | 562 | DUF499-domain-containing-protein-[Escherichia-coli] |
| WP_101744256.1 | AAA+ATPase+wHTH+Alpha+Beta-domain+wHTH+Fn-III+Fn-III+RRM | Type-2_BR.RM.like.systems | zf-ISL3+DDE_Tnp_ISL3->DUF499*->DUF3780->METHYLASE->HKD-DNase+SF1-SF2-Helicase-><-REase<-NACHT+SF1-SF2-Helicase+HKD-DNase<-wHTH+SAD+HNH | 1022 | Bacteria>Pseudomonadota>Gammaproteobacteria>Enterobacterales>Enterobacteriaceae | GCF_002863965.1 | Citrobacter_koseri | C1H48_RS13720 | 303487..306555 | 545 | DUF499-domain-containing-protein-[Citrobacter-koseri] |
| WP_102217321.1 | AAA+ATPase+wHTH+Alpha+Beta-domain+wHTH+Fn-III+Fn-III+RRM | Type-2_BR.RM.like.systems | <-ADH_N+ADH_zinc_N<-?<-?\|\|tRNA->Arm-DNA-bind_3+Phage_integrase->RHH->DUF499*->DUF3780+RAMA->METHYLASE->HKD-DNase+SF1-SF2-Helicase->wHTH+REase-><-HTH+MPTase | 1041 | Bacteria>Pseudomonadota>Gammaproteobacteria>Enterobacterales>Enterobacteriaceae | GCF_002872205.1 | Klebsiella_pneumoniae | A8A10_RS01535 | 132525..135650 | 573 | DUF499-domain-containing-protein-[Klebsiella-pneumoniae] |
| WP_105476848.1 | AAA+ATPase+wHTH+Alpha+Beta-domain+wHTH+Fn-III+Fn-III+RRM | Type-2_BR.RM.like.systems | RNaseH->NACHT+SF1-SF2-Helicase+HKD-DNase->REase-><-HKD-DNase+SF1-SF2-Helicase<-METHYLASE<-DUF3780<-DUF499*\|\|<-zf-ISL3+DDE_Tnp_ISL3 | 1022 | Bacteria>Pseudomonadota>Gammaproteobacteria>Enterobacterales>Enterobacteriaceae | GCF_002972735.1 | Escherichia_coli | VH93_RS04625 | 62417..65485 | 562 | DUF499-domain-containing-protein-[Escherichia-coli] |
| WP_120159929.1 | AAA+ATPase+wHTH+Alpha+Beta-domain+wHTH+Fn-III+Fn-III+RRM | Type-2_BR.RM.like.systems | RHH_5->DUF499*->DUF3780+RAMA->METHYLASE->HKD-DNase+SF1-SF2-Helicase->wHTH+REase->HIN-HTH-> | 1041 | Bacteria>Pseudomonadota>Gammaproteobacteria>Enterobacterales>Enterobacteriaceae | GCF_003605695.1 | Escherichia_coli | D3Y67_RS09395 | 1782..4907 | 562 | DUF499-domain-containing-protein-[Escherichia-coli] |
| WP_131009058.1 | AAA+ATPase+wHTH+Alpha+Beta-domain+wHTH+Fn-III+Fn-III+RRM | Type-2_BR.RM.like.systems | HTH+MPTase-><-wHTH+REase<-HKD-DNase+SF1-SF2-Helicase<-METHYLASE<-DUF3780+RAMA<-DUF499*<-RHH<-Arm-DNA-bind_3+Phage_integrase<-tRNA\|\|?->?->ADH_N+ADH_zinc_N-> | 1041 | Bacteria>Pseudomonadota>Gammaproteobacteria>Enterobacterales>Enterobacteriaceae | GCF_004312385.1 | Klebsiella_variicola | HZ923_RS12940 | 35774..38899 | 244366 | DUF499-domain-containing-protein-[Klebsiella-variicola] |
| WP_250216357.1 | AAA+ATPase+wHTH+Alpha+Beta-domain+wHTH+Fn-III+Fn-III+RRM | Type-2_BR.RM.like.systems | <-HKD-DNase+SF1-SF2-Helicase<-METHYLASE<-DUF3780<-DUF499*\|\|HIN-HTH->?-><-DUF499<-zf-ISL3+DDE_Tnp_ISL3\|\|?->\|\|MAD-NTD+HNH-> | 711 | Bacteria>Pseudomonadota>Gammaproteobacteria>Enterobacterales>Enterobacteriaceae | GCF_005381525.1 | Escherichia_coli | FE858_RS21990 | 7265..9400 | 562 | chitobiase/beta-hexosaminidase-C-terminal-domain-containing-protein-[Escherichia-coli] |
| WP_000077518.1 | AAA+ATPase+wHTH+Alpha+Beta-domain+wHTH+Fn-III+Fn-III+RRM | Type-2_BR.RM.like.systems | DEDD_Tnp_IS110+Transposase_20->?->RHH_5->DUF499*->DUF3780+RAMA->METHYLASE->HKD-DNase+SF1-SF2-Helicase->wHTH+REase->HIN-HTH-> | 1022 | Bacteria>Pseudomonadota>Gammaproteobacteria>Enterobacterales>Enterobacteriaceae | GCF_005390065.1 | Escherichia_coli | EXY46_RS20920 | 4095..7163 | 562 | DUF499-domain-containing-protein-[Escherichia-coli] |
| WP_153674499.1 | AAA+ATPase+wHTH+Alpha+Beta-domain+wHTH+Fn-III+Fn-III+RRM | Type-2_BR.RM.like.systems | rve+rve_3-><-HIN-HTH<-wHTH+REase<-HKD-DNase+SF1-SF2-Helicase<-METHYLASE<-DUF3780+RAMA<-DUF499*<-RHH_5<-?\|\|<-?<-HTH_Tnp_1+HTH_21+rve+rve_3<-ZnR | 1041 | Bacteria>Pseudomonadota>Gammaproteobacteria>Enterobacterales>Enterobacteriaceae | GCF_009648255.1 | Escherichia_coli | GIJ00_RS05185 | 9599..12724 | 562 | DUF499-domain-containing-protein-[Escherichia-coli] |
| WP_185774377.1 | AAA+ATPase+wHTH+Alpha+Beta-domain+wHTH+Fn-III+Fn-III+RRM | Type-2_BR.RM.like.systems | <-ADH_N+ADH_zinc_N<-?\|\|tRNA->Arm-DNA-bind_3+Phage_integrase->RHH->DUF499*->DUF3780+RAMA->METHYLASE->HKD-DNase+SF1-SF2-Helicase->wHTH+REase-><-HTH+MPTase | 1041 | Bacteria>Pseudomonadota>Gammaproteobacteria>Enterobacterales>Enterobacteriaceae | GCF_014235385.1 | Klebsiella_variicola | H8S48_RS22840 | 314198..317323 | 244366 | DUF499-domain-containing-protein-[Klebsiella-variicola] |
| WP_041911426.1 | AAA+ATPase+wHTH+Alpha+Beta-domain+wHTH+Fn-III+Fn-III+RRM | Type-2_BR.RM.like.systems | HIN-HTH-><-REase<-HKD-DNase+SF1-SF2-Helicase<-METHYLASE<-DUF3780+RAMA<-DUF499*<-RHH<-Arm-DNA-bind_3+Phage_integrase | 1041 | Bacteria>Pseudomonadota>Gammaproteobacteria>Enterobacterales>Enterobacteriaceae | GCF_014596715.1 | Enterobacter | L465_RS18340 | 3972931..3976056 | 547 | MULTISPECIES:-DUF499-domain-containing-protein-[Enterobacter] |
| WP_154813114.1 | AAA+ATPase+wHTH+Alpha+Beta-domain+wHTH+Fn-III+Fn-III+RRM | Type-2_BR.RM.like.systems | <-MAD-NTD+HNH\|\|<-?\|\|zf-ISL3+DDE_Tnp_ISL3->DUF499*->DUF3780->METHYLASE->HKD-DNase+SF1-SF2-Helicase-> | 1022 | Bacteria>Pseudomonadota>Gammaproteobacteria>Enterobacterales>Enterobacteriaceae | GCF_015234845.1 | Shigella_flexneri | E4T96_RS13560 | 111927..114995 | 623 | DUF499-domain-containing-protein-[Shigella-flexneri] |
| WP_227628590.1 | AAA+ATPase+wHTH+Alpha+Beta-domain+wHTH+Fn-III+Fn-III+RRM | Type-2_BR.RM.like.systems | <-ADH_N+ADH_zinc_N\|\|tRNA->Arm-DNA-bind_3+Phage_integrase->RHH->DUF499*->DUF3780+RAMA->METHYLASE->HKD-DNase+SF1-SF2-Helicase->wHTH+REase-> | 1041 | Bacteria>Pseudomonadota>Gammaproteobacteria>Enterobacterales>Enterobacteriaceae | GCF_015721555.1 | Klebsiella_oxytoca | QP770_RS13995 | 142941..146066 | 571 | DUF499-domain-containing-protein-[Klebsiella-oxytoca] |
| WP_233593808.1 | AAA+ATPase+wHTH+Alpha+Beta-domain+wHTH+Fn-III+Fn-III+RRM | Type-2_BR.RM.like.systems | <-wHTH+REase<-HKD-DNase+SF1-SF2-Helicase<-METHYLASE<-DUF3780+RAMA<-DUF499*<-RHH<-Arm-DNA-bind_3+Phage_integrase<-tRNA\|\|ADH_N+ADH_zinc_N-> | 1041 | Bacteria>Pseudomonadota>Gammaproteobacteria>Enterobacterales>Enterobacteriaceae | GCF_016501115.1 | Citrobacter | OGW14_RS13170 | 9006..12131 | 544 | DUF499-domain-containing-protein-[Citrobacter-cronae] |
| WP_227639547.1 | AAA+ATPase+wHTH+Alpha+Beta-domain+wHTH+Fn-III+Fn-III+RRM | Type-2_BR.RM.like.systems | HTH+MPTase-><-wHTH+REase<-HKD-DNase+SF1-SF2-Helicase<-METHYLASE<-DUF3780+RAMA<-DUF499*<-RHH<-Arm-DNA-bind_3+Phage_integrase<-tRNA\|\|?->?->ADH_N+ADH_zinc_N-> | 1041 | Bacteria>Pseudomonadota>Gammaproteobacteria>Enterobacterales>Enterobacteriaceae | GCF_016651545.1 | Klebsiella_pneumoniae | JKA42_RS15120 | 66258..69383 | 573 | DUF499-domain-containing-protein-[Klebsiella-pneumoniae] |
| WP_213752381.1 | AAA+ATPase+wHTH+Alpha+Beta-domain+wHTH+Fn-III+Fn-III+RRM | Type-2_BR.RM.like.systems | wHTH(SBD)+HNH->?->DUF499*->DUF3780->METHYLASE->HKD-DNase+SF1-SF2-Helicase-><-REase\|\|HIN-HTH-><-Arm-DNA-bind_3+Phage_integrase | 1022 | Bacteria>Pseudomonadota>Gammaproteobacteria>Enterobacterales>Enterobacteriaceae | GCF_018446995.1 | Citrobacter | OGY13_RS10400 | 358156..361224 | 544 | DUF499-domain-containing-protein-[Citrobacter-portucalensis] |
| WP_089553231.1 | AAA+ATPase+wHTH+Alpha+Beta-domain+wHTH+Fn-III+Fn-III+RRM | Type-2_BR.RM.like.systems | <-HIN-HTH<-wHTH+REase<-HKD-DNase+SF1-SF2-Helicase<-METHYLASE<-DUF3780+RAMA<-DUF499*<-RHH_5\|\|<-?<-Aminotran_1_2\|\|<-Arm-DNA-bind_3+Phage_integrase<-tRNA\|\|ADH_N+ADH_zinc_N->FtsK+HerA+A32-> | 1041 | Bacteria>Pseudomonadota>Gammaproteobacteria>Enterobacterales>Enterobacteriaceae | GCF_019957165.1 | Escherichia_coli | BEC12_RS18680 | 17132..20257 | 562 | DUF499-domain-containing-protein-[Escherichia-coli] |
| WP_196557776.1 | AAA+ATPase+wHTH+Alpha+Beta-domain+wHTH+Fn-III+Fn-III+RRM | Type-2_BR.RM.like.systems | Arm-DNA-bind_3+Phage_integrase->RHH->DUF499*->DUF3780+RAMA->METHYLASE->HKD-DNase+SF1-SF2-Helicase->wHTH+REase-><-?<-SF1-SF2-Helicase | 1022 | Bacteria>Pseudomonadota>Gammaproteobacteria>Enterobacterales>Enterobacteriaceae | GCF_020116575.1 | Klebsiella_oxytoca | FML32_RS25495 | 2110..5178 | 571 | DUF499-domain-containing-protein-[Klebsiella-oxytoca] |
| WP_232904627.1 | AAA+ATPase+wHTH+Alpha+Beta-domain+wHTH+Fn-III+Fn-III+RRM | Type-2_BR.RM.like.systems | PIN->?-><-wHTH+REase<-HKD-DNase+SF1-SF2-Helicase<-METHYLASE<-DUF3780+RAMA<-DUF499*<-RHH<-Arm-DNA-bind_3+Phage_integrase | 1041 | Bacteria>Pseudomonadota>Gammaproteobacteria>Enterobacterales>Enterobacteriaceae | GCF_021282435.1 | Buttiauxella_sp._W03-F01 | LVQ77_RS12925 | 38707..41832 | 2904524 | DUF499-domain-containing-protein-[Buttiauxella-sp.-W03-F01] |
| WP_233395631.1 | AAA+ATPase+wHTH+Alpha+Beta-domain+wHTH+Fn-III+Fn-III+RRM | Type-2_BR.RM.like.systems | <-MAD-NTD+HNH\|\|<-?\|\|zf-ISL3+DDE_Tnp_ISL3->DUF499*->DUF3780->METHYLASE->HKD-DNase+SF1-SF2-Helicase-> | 1022 | Bacteria>Pseudomonadota>Gammaproteobacteria>Enterobacterales>Enterobacteriaceae | GCF_021354515.1 | Escherichia_coli | I5143_RS13495 | 121544..124612 | 562 | DUF499-domain-containing-protein-[Escherichia-coli] |
| WP_201792481.1 | AAA+ATPase+wHTH+Alpha+Beta-domain+wHTH+Fn-III+Fn-III+RRM | Type-2_BR.RM.like.systems | RHH_5->DUF499*->DUF3780+RAMA->METHYLASE->HKD-DNase+SF1-SF2-Helicase->wHTH+REase->HIN-HTH-> | 1022 | Bacteria>Pseudomonadota>Gammaproteobacteria>Enterobacterales>Enterobacteriaceae | GCF_023516095.1 | Escherichia_coli | IXW39_RS10870 | 2714..5782 | 562 | DUF499-domain-containing-protein-[Escherichia-coli] |
| WP_252076363.1 | AAA+ATPase+wHTH+Alpha+Beta-domain+wHTH+Fn-III+Fn-III+RRM | Type-2_BR.RM.like.systems | Arm-DNA-bind_3+Phage_integrase->RHH->DUF499*->DUF3780+RAMA->METHYLASE->HKD-DNase+SF1-SF2-Helicase->wHTH+REase-><-HTH+MPTase | 1041 | Bacteria>Pseudomonadota>Gammaproteobacteria>Enterobacterales>Enterobacteriaceae | GCF_023751425.1 | Enterobacter_kobei | M8R77_RS12255 | 159585..162710 | 208224 | DUF499-domain-containing-protein-[Enterobacter-kobei] |
| WP_258301444.1 | AAA+ATPase+wHTH+Alpha+Beta-domain+wHTH+Fn-III+Fn-III+RRM | Type-2_BR.RM.like.systems | <-HIN-HTH<-wHTH+REase<-HKD-DNase+SF1-SF2-Helicase<-METHYLASE<-DUF3780+RAMA<-DUF499*<-RHH_5\|\|<-?<-Aminotran_1_2\|\|<-Arm-DNA-bind_3+Phage_integrase<-tRNA\|\|ADH_N+ADH_zinc_N->FtsK+HerA+A32-> | 1041 | Bacteria>Pseudomonadota>Gammaproteobacteria>Enterobacterales>Enterobacteriaceae | GCF_024733045.1 | Escherichia_coli | NX108_RS16810 | 9461..12586 | 562 | DUF499-domain-containing-protein-[Escherichia-coli] |
| WP_258442601.1 | AAA+ATPase+wHTH+Alpha+Beta-domain+wHTH+Fn-III+Fn-III+RRM | Type-2_BR.RM.like.systems | <-ADH_N+ADH_zinc_N\|\|tRNA->Arm-DNA-bind_3+Phage_integrase->RHH->DUF499*->DUF3780+RAMA->METHYLASE->HKD-DNase+SF1-SF2-Helicase->wHTH+REase-><-?<-SF1-SF2-Helicase | 1041 | Bacteria>Pseudomonadota>Gammaproteobacteria>Enterobacterales>Enterobacteriaceae | GCF_024741195.1 | Citrobacter_amalonaticus | NXA99_RS21100 | 70776..73901 | 35703 | DUF499-domain-containing-protein-[Citrobacter-amalonaticus] |
| WP_273924537.1 | AAA+ATPase+wHTH+Alpha+Beta-domain+wHTH+Fn-III+Fn-III+RRM | Type-2_BR.RM.like.systems | <-MAD-NTD+HNH\|\|<-?\|\|zf-ISL3+DDE_Tnp_ISL3->DUF499-p-><-DDE_Tnp_1\|\|DUF499*->DUF3780->METHYLASE->HKD-DNase+SF1-SF2-Helicase-><-REase<-NACHT+SF1-SF2-Helicase+HKD-DNase<-RNaseH | 790 | Bacteria>Pseudomonadota>Gammaproteobacteria>Enterobacterales>Enterobacteriaceae | GCF_028623065.1 | Escherichia_coli | PTA69_RS21070 | 4298151..4300523 | 562 | DUF499-domain-containing-protein-[Escherichia-coli] |
| WP_101974273.1 | AAA+ATPase+wHTH+Alpha+Beta-domain+wHTH+Fn-III+Fn-III+RRM | Type-2_BR.RM.like.systems | HTH+MPTase-><-wHTH+REase<-HKD-DNase+SF1-SF2-Helicase<-METHYLASE<-DUF3780+RAMA<-DUF499*<-RHH<-Arm-DNA-bind_3+Phage_integrase<-tRNA\|\|?->?->ADH_N+ADH_zinc_N-> | 1041 | Bacteria>Pseudomonadota>Gammaproteobacteria>Enterobacterales>Enterobacteriaceae | GCF_028894585.1 | Klebsiella_pneumoniae | PXW64_RS18870 | 24255..27380 | 573 | DUF499-domain-containing-protein-[Klebsiella-pneumoniae] |
| WP_275399056.1 | AAA+ATPase+wHTH+Alpha+Beta-domain+wHTH+Fn-III+Fn-III+RRM | Type-2_BR.RM.like.systems | <-FtsK+HerA+A32<-ADH_N+ADH_zinc_N\|\|tRNA->Arm-DNA-bind_3+Phage_integrase->RHH->DUF499*->DUF3780+RAMA->METHYLASE->HKD-DNase+SF1-SF2-Helicase->wHTH+REase-><-SF1-SF2-Helicase<-OLD-ABC+TOPRIM | 1041 | Bacteria>Pseudomonadota>Gammaproteobacteria>Enterobacterales>Enterobacteriaceae | GCF_029104325.1 | Citrobacter_portucalensis | L2173_RS05340 | 153527..156652 | 1639133 | DUF499-domain-containing-protein-[Citrobacter-portucalensis] |
| WP_045337753.1 | AAA+ATPase+wHTH+Alpha+Beta-domain+wHTH+Fn-III+Fn-III+RRM | Type-2_BR.RM.like.systems | Arm-DNA-bind_3+Phage_integrase->RHH->DUF499*->DUF3780+RAMA->METHYLASE->HKD-DNase+SF1-SF2-Helicase->wHTH+REase-><-?\|\|HTH_Tnp_1+HTH_21+rve+rve_2-> | 1041 | Bacteria>Pseudomonadota>Gammaproteobacteria>Enterobacterales>Enterobacteriaceae | GCF_029207635.1 | Enterobacter_hormaechei | H0762_RS10605 | 150283..153408 | 158836 | DUF499-domain-containing-protein-[Enterobacter-hormaechei] |
| WP_284106250.1 | AAA+ATPase+wHTH+Alpha+Beta-domain+wHTH+Fn-III+Fn-III+RRM | Type-2_BR.RM.like.systems | HTH+MPTase-><-HNH+SAVED<-wHTH+REase<-HKD-DNase+SF1-SF2-Helicase<-METHYLASE<-DUF3780+RAMA<-DUF499*<-RHH<-Arm-DNA-bind_3+Phage_integrase<-tRNA\|\|ADH_N+ADH_zinc_N->FtsK+HerA+A32-> | 1041 | Bacteria>Pseudomonadota>Gammaproteobacteria>Enterobacterales>Enterobacteriaceae | GCF_030132145.1 | Citrobacter_portucalensis | QL289_RS16600 | 141668..144793 | 1639133 | DUF499-domain-containing-protein-[Citrobacter-portucalensis] |
| WP_285003151.1 | AAA+ATPase+wHTH+Alpha+Beta-domain+wHTH+Fn-III+Fn-III+RRM | Type-2_BR.RM.like.systems | DUF499*->DUF3780->METHYLASE->HKD-DNase+SF1-SF2-Helicase-><-REase<-NACHT+SF1-SF2-Helicase+HKD-DNase<-RNaseH | 801 | Bacteria>Pseudomonadota>Gammaproteobacteria>Enterobacterales>Enterobacteriaceae | GCF_030208715.1 | Escherichia_coli | QRN35_RS23695 | 1..2407 | 562 | DUF499-domain-containing-protein,-partial-[Escherichia-coli] |
| WP_285145128.1 | AAA+ATPase+wHTH+Alpha+Beta-domain+wHTH+Fn-III+Fn-III+RRM | Type-2_BR.RM.like.systems | XIS-HTH->Phage_ASH->?->?->TOPRIM+DNAG+D5N+D5ATPase+P4PRIMASE-HTH->RHH_5->DUF499*->DUF3780+RAMA->METHYLASE->HKD-DNase+SF1-SF2-Helicase->wHTH+REase-><-ParB+HNH<-SymE_toxin | 1041 | Bacteria>Pseudomonadota>Gammaproteobacteria>Enterobacterales>Enterobacteriaceae | GCF_030234145.1 | unclassified_Lelliottia | QQF24_RS22360 | 130751..133876 | 2642424 | MULTISPECIES:-DUF499-domain-containing-protein-[unclassified-Lelliottia] |
| WP_180245885.1 | AAA+ATPase+wHTH+Alpha+Beta-domain+wHTH+Fn-III+Fn-III+RRM | Type-2_BR.RM.like.systems | SF1-SF2-Helicase->?-><-wHTH+REase<-HKD-DNase+SF1-SF2-Helicase<-METHYLASE<-DUF3780+RAMA<-DUF499*<-RHH<-Arm-DNA-bind_3+Phage_integrase | 1041 | Bacteria>Pseudomonadota>Gammaproteobacteria>Enterobacterales>Enterobacteriaceae | GCF_900558075.1 | Enterobacter_hormaechei | H0785_RS18375 | 27017..30142 | 158836 | DUF499-domain-containing-protein-[Enterobacter-hormaechei] |
| WP_204372837.1 | AAA+ATPase+wHTH+Alpha+Beta-domain+wHTH+Fn-III+Fn-III+RRM | Type-2_BR.RM.like.systems | SF1-SF2-Helicase->?-><-wHTH+REase<-HKD-DNase+SF1-SF2-Helicase<-METHYLASE<-DUF3780+RAMA<-DUF499*<-RHH<-Arm-DNA-bind_3+Phage_integrase<-tRNA\|\|?->?->ADH_N+ADH_zinc_N-> | 1022 | Bacteria>Pseudomonadota>Gammaproteobacteria>Enterobacterales>Enterobacteriaceae | - | Klebsiella_pneumoniae | MMH51_002463 | 54491..57559 | 573 | DUF499-domain-containing-protein-[Klebsiella-pneumoniae] |
| WP_198740177.1 | AAA+ATPase+wHTH+Alpha+Beta-domain+wHTH+Fn-III+Fn-III+RRM | Type-2_BR.RM.like.systems | Arm-DNA-bind_3+Phage_integrase->RHH->DUF499*->DUF3780+RAMA->METHYLASE->HKD-DNase+SF1-SF2-Helicase->wHTH+REase-><-?<-SF1-SF2-Helicase | 1022 | Bacteria>Pseudomonadota>Gammaproteobacteria>Enterobacterales>Enterobacteriaceae | - | Klebsiella_pneumoniae | POC94_000676 | 161243..164311 | 573 | MULTISPECIES:-DUF499-domain-containing-protein-[Enterobacteriaceae] |
| WP_048997689.1 | AAA+ATPase+wHTH+Alpha+Beta-domain+wHTH+Fn-III+Fn-III+RRM | Type-2_BR.RM.like.systems | <-ADH_N+ADH_zinc_N<-?<-?\|\|tRNA->Arm-DNA-bind_3+Phage_integrase->RHH->DUF499*->DUF3780+RAMA->METHYLASE->HKD-DNase+SF1-SF2-Helicase->wHTH+REase-><-?<-SF1-SF2-Helicase | 1041 | Bacteria>Pseudomonadota>Gammaproteobacteria>Enterobacterales>Enterobacteriaceae | - | Klebsiella_pneumoniae | QVO21_RS15045 | 200942..204067 | 573 | DUF499-domain-containing-protein-[Klebsiella-pneumoniae] |
| WP_116197723.1 | AAA+ATPase+wHTH+Alpha+Beta-domain+wHTH+Fn-III+Fn-III+RRM | Type-2_BR.RM.like.systems | <-NACHT+SF1-SF2-Helicase+HKD-DNase\|\|ParB+HNH-><-wHTH+REase<-HKD-DNase+SF1-SF2-Helicase<-METHYLASE<-DUF3780+RAMA<-DUF499*<-RHH | 1041 | Bacteria>Pseudomonadota>Gammaproteobacteria>Enterobacterales>Erwiniaceae | GCF_017920215.1 | Pantoea_ananatis | HG975_RS10020 | 359371..362496 | 553 | DUF499-domain-containing-protein-[Pantoea-ananatis] |
| WP_266093834.1 | AAA+ATPase+wHTH+Alpha+Beta-domain+wHTH+Fn-III+Fn-III+RRM | Type-2_BR.RM.like.systems | DUF1176->tRNA->RHH->DUF499*->DUF3780+RAMA->METHYLASE->HKD-DNase+SF1-SF2-Helicase->wHTH+REase-> | 1041 | Bacteria>Pseudomonadota>Gammaproteobacteria>Enterobacterales>Erwiniaceae | GCF_026241965.1 | [Curtobacterium]_plantarum | ORG41_RS07860 | 286061..289186 | 221276 | DUF499-domain-containing-protein-[[Curtobacterium] |
| WP_035596320.1 | AAA+ATPase+wHTH+Alpha+Beta-domain+wHTH+Fn-III+Fn-III+RRM | Type-2_BR.RM.like.systems | <-XIS-HTH\|\|DDE_Tnp_1_5+DDE_Tnp_1->\|\|DDE_Tnp_1_5+DDE_Tnp_1-><-HKD-DNase+SF1-SF2-Helicase<-METHYLASE<-DUF3780<-DUF499* | 1022 | Bacteria>Pseudomonadota>Gammaproteobacteria>Enterobacterales>Hafniaceae | GCF_000264805.1 | Edwardsiella_tarda | ETATCC_RS01040 | 59957..63025 | 636 | DUF499-domain-containing-protein-[Edwardsiella-tarda] |
| MBK1419820.1 | AAA+ATPase+wHTH+Alpha+Beta-domain+wHTH+Fn-III+Fn-III+RRM | Type-2_BR.RM.like.systems | 35exo->RHH_1->DUF499*->DUF3780+RAMA->METHYLASE->HKD-DNase+SF1-SF2-Helicase->wHTH+REase->HDRNase->VirD4-TraG+VirB4-FtsK-> | 1022 | Bacteria>Pseudomonadota>Gammaproteobacteria>Enterobacterales>Morganellaceae | GCA_016623625.1 | Providencia_stuartii | I6H92_05395 | 1204015..1207083 | 588 | DUF499-domain-containing-protein-[Providencia-stuartii] |
| MCL8535002.1 | AAA+ATPase+wHTH+Alpha+Beta-domain+wHTH+Fn-III+Fn-III+RRM | Type-2_BR.RM.like.systems | DUF499*->DUF3780->METHYLASE->HKD-DNase+SF1-SF2-Helicase->wHTH+REase->HDRNase->VirD4-TraG+VirB4-FtsK-> | 850 | Bacteria>Pseudomonadota>Gammaproteobacteria>Enterobacterales>Morganellaceae | GCA_023604365.1 | Proteus_mirabilis | NAE57_17165 | 121..2673 | 584 | ATP-binding-protein-[Proteus-mirabilis] |
| EKK5375336.1 | AAA+ATPase+wHTH+Alpha+Beta-domain+wHTH+Fn-III+Fn-III+RRM | Type-2_BR.RM.like.systems | <-Zn_Tnp_IS1595+DDE_Tnp_IS1595<-HKD-DNase+SF1-SF2-Helicase<-METHYLASE<-DUF3780+RAMA<-DUF499*<-RHH<-XIS-HTH<-ZnR\|\|?-><-Inovirus_Gp2<-XIS-HTH | 1022 | Bacteria>Pseudomonadota>Gammaproteobacteria>Enterobacterales>Morganellaceae | GCA_028172445.1 | Morganella_morganii | PN738_000385 | 133802..136870 | 582 | DUF499-domain-containing-protein-[Morganella-morganii] |
| WP_004234628.1 | AAA+ATPase+wHTH+Alpha+Beta-domain+wHTH+Fn-III+Fn-III+RRM | Type-2_BR.RM.like.systems | <-ArdB<-nSTAND1+sGTP<-?\|\|?->XIS-HTH->XIS-HTH->DUF499*->DUF3780->METHYLASE->HKD-DNase+SF1-SF2-Helicase-><-?<-Arm-DNA-bind_3+Phage_integrase<-tRNA\|\|Beta-lactamase-> | 1022 | Bacteria>Pseudomonadota>Gammaproteobacteria>Enterobacterales>Morganellaceae | GCF_000286435.2 | Morganella_morganii | MU9_RS00630 | 138231..141299 | 582 | DUF499-domain-containing-protein-[Morganella-morganii] |
| WP_038193804.1 | AAA+ATPase+wHTH+Alpha+Beta-domain+wHTH+Fn-III+Fn-III+RRM | Type-2_BR.RM.like.systems | <-wHTH+REase<-HKD-DNase+SF1-SF2-Helicase<-METHYLASE<-DUF3780+RAMA<-DUF499*<-RHH<-tRNA\|\|Beta-lactamase-><-?<-?\|\|Peptidase-> | 1041 | Bacteria>Pseudomonadota>Gammaproteobacteria>Enterobacterales>Morganellaceae | GCF_000736695.1 | Xenorhabdus_bovienii | XBKB1_RS06355 | 8159..11284 | 40576 | DUF499-domain-containing-protein-[Xenorhabdus-bovienii] |
| WP_282555203.1 | AAA+ATPase+wHTH+Alpha+Beta-domain+wHTH+Fn-III+Fn-III+RRM | Type-2_BR.RM.like.systems | <-ParB+HNH<-N6_MTase-p<-SbcC<-HKD-DNase+SF1-SF2-Helicase<-METHYLASE<-DUF3780<-DUF499*<-XIS-HTH<-NACHT\|\|DDE_Tnp_IS240-><-?\|\|Zn_Tnp_IS1+HTH_Tnp_IS1+DDE_Tnp_IS1-> | 921 | Bacteria>Pseudomonadota>Gammaproteobacteria>Enterobacterales>Morganellaceae | GCF_002947315.1 | Providencia_stuartii | AM353_RS02810 | 621695..624460 | 588 | DUF499-domain-containing-protein-[Providencia-stuartii] |
| WP_107337129.1 | AAA+ATPase+wHTH+Alpha+Beta-domain+wHTH+Fn-III+Fn-III+RRM | Type-2_BR.RM.like.systems | <-ArdB<-nSTAND1+sGTP<-CopG/RHH\|\|?->XIS-HTH->XIS-HTH->DUF499*->DUF3780->METHYLASE->HKD-DNase+SF1-SF2-Helicase-><-?<-Arm-DNA-bind_3+Phage_integrase<-tRNA\|\|Beta-lactamase-> | 1022 | Bacteria>Pseudomonadota>Gammaproteobacteria>Enterobacterales>Morganellaceae | GCF_003030945.1 | Proteus | O9355_RS16120 | 9799..12867 | 583 | MULTISPECIES:-DUF499-domain-containing-protein-[Proteus] |
| WP_152110353.1 | AAA+ATPase+wHTH+Alpha+Beta-domain+wHTH+Fn-III+Fn-III+RRM | Type-2_BR.RM.like.systems | <-Peptidase_S8<-?\|\|35exo->RHH_1->DUF499*->DUF3780+RAMA->METHYLASE->HKD-DNase+SF1-SF2-Helicase->wHTH+REase->HDRNase->VirD4-TraG+VirB4-FtsK-> | 1041 | Bacteria>Pseudomonadota>Gammaproteobacteria>Enterobacterales>Morganellaceae | GCF_009183685.1 | Proteus_mirabilis | GBN12_RS03240 | 63661..66786 | 584 | DUF499-domain-containing-protein-[Proteus-mirabilis] |
| WP_104872941.1 | AAA+ATPase+wHTH+Alpha+Beta-domain+wHTH+Fn-III+Fn-III+RRM | Type-2_BR.RM.like.systems | <-DDE_Tnp_IS240\|\|NACHT->TetR-HTH->XIS-HTH->DUF499*->DUF3780->METHYLASE->HKD-DNase+SF1-SF2-Helicase->SbcC->N6_MTase-p->ParB+HNH-> | 1022 | Bacteria>Pseudomonadota>Gammaproteobacteria>Enterobacterales>Morganellaceae | GCF_009706655.1 | Providencia_stuartii | GKR72_RS00025 | 4280..7348 | 588 | DUF499-domain-containing-protein-[Providencia-stuartii] |
| WP_159262627.1 | AAA+ATPase+wHTH+Alpha+Beta-domain+wHTH+Fn-III+Fn-III+RRM | Type-2_BR.RM.like.systems | <-VirD4-TraG+VirB4-FtsK<-HDRNase<-wHTH+REase<-HKD-DNase+SF1-SF2-Helicase<-METHYLASE<-DUF3780<-DUF499*<-XIS-HTH<-35exo | 1022 | Bacteria>Pseudomonadota>Gammaproteobacteria>Enterobacterales>Morganellaceae | GCF_010692865.1 | Proteus_mirabilis | QN307_RS16755 | 34681..37749 | 584 | DUF499-domain-containing-protein-[Proteus-mirabilis] |
| WP_064483386.1 | AAA+ATPase+wHTH+Alpha+Beta-domain+wHTH+Fn-III+Fn-III+RRM | Type-2_BR.RM.like.systems | <-CopG/RHH\|\|?->XIS-HTH->XIS-HTH->DUF499*->DUF3780->METHYLASE->HKD-DNase+SF1-SF2-Helicase-><-Arm-DNA-bind_3+Phage_integrase<-tRNA\|\|Beta-lactamase-> | 1022 | Bacteria>Pseudomonadota>Gammaproteobacteria>Enterobacterales>Morganellaceae | GCF_020790175.1 | Morganella_morganii | AYO06_RS06545 | 4112..7180 | 582 | DUF499-domain-containing-protein-[Morganella-morganii] |
| WP_275358359.1 | AAA+ATPase+wHTH+Alpha+Beta-domain+wHTH+Fn-III+Fn-III+RRM | Type-2_BR.RM.like.systems | <-Peptidase\|\|?->?-><-Beta-lactamase\|\|tRNA->RHH->DUF499*->DUF3780+RAMA->METHYLASE->HKD-DNase+SF1-SF2-Helicase->wHTH+REase-> | 1041 | Bacteria>Pseudomonadota>Gammaproteobacteria>Enterobacterales>Morganellaceae | GCF_029104585.1 | Xenorhabdus_bovienii | KKI95_RS09990 | 130588..133713 | 40576 | DUF499-domain-containing-protein-[Xenorhabdus-bovienii] |
| WP_196578769.1 | AAA+ATPase+wHTH+Alpha+Beta-domain+wHTH+Fn-III+Fn-III+RRM | Type-2_BR.RM.like.systems | 35exo->RHH_1->DUF499*->DUF3780+RAMA->METHYLASE->HKD-DNase+SF1-SF2-Helicase->wHTH+REase->HDRNase->VirD4-TraG+VirB4-FtsK-> | 1022 | Bacteria>Pseudomonadota>Gammaproteobacteria>Enterobacterales>Morganellaceae | - | Proteus_mirabilis | I6H92_05395 | 1204015..1207083 | 584 | DUF499-domain-containing-protein-[Proteus-mirabilis] |
| WP_250269692.1 | AAA+ATPase+wHTH+Alpha+Beta-domain+wHTH+Fn-III+Fn-III+RRM | Type-2_BR.RM.like.systems | DUF499*->DUF3780->METHYLASE->HKD-DNase+SF1-SF2-Helicase->wHTH+REase->HDRNase->VirD4-TraG+VirB4-FtsK-> | 850 | Bacteria>Pseudomonadota>Gammaproteobacteria>Enterobacterales>Morganellaceae | - | Proteus_mirabilis | NAE57_17165 | 121..2673 | 584 | DUF499-domain-containing-protein-[Proteus-mirabilis] |
| WP_095924507.1 | AAA+ATPase+wHTH+Alpha+Beta-domain+wHTH+Fn-III+Fn-III+RRM | Type-2_BR.RM.like.systems | <-Peptidase\|\|?->?-><-Beta-lactamase\|\|tRNA->Arm-DNA-bind_3+Phage_integrase->RHH->DUF499*->DUF3780+RAMA->METHYLASE->HKD-DNase+SF1-SF2-Helicase->wHTH+REase-><-HIN-HTH | 1041 | Bacteria>Pseudomonadota>Gammaproteobacteria>Enterobacterales>Yersiniaceae | GCF_002303785.1 | Rahnella_victoriana | CO635_RS19350 | 4088068..4091193 | 1510570 | DUF499-domain-containing-protein-[Rahnella-victoriana] |
| RUQ39484.1 | AAA+ATPase+wHTH+Alpha+Beta-domain+wHTH+Fn-III+Fn-III+RRM | Type-2_BR.RM.like.systems | <-FURR-HTH\|\|?-><-?<-?\|\|?->RHH_5->DUF499*->DUF3780->?->PIN->METHYLASE-p-> | 1041 | Bacteria>Pseudomonadota>Gammaproteobacteria>Gammaproteobacteria_incertae_sedis>Candidatus_Competibacteraceae | GCA_003989075.1 | Candidatus_Competibacteraceae_bacterium | EKK69_09085 | 13765..16890 | 2053538 | DUF499-domain-containing-protein-[Candidatus-Competibacteraceae-bacterium] |
| HAU1604805.1 | AAA+ATPase+wHTH+Alpha+Beta-domain+wHTH+Fn-III+Fn-III+RRM | Type-2_BR.RM.like.systems | <-RelE-ParE<-ParD_like<-?<-Y1_Tnp\|\|RHH->DUF499*->DUF3780->METHYLASE->HKD-DNase+SF1-SF2-Helicase->HTH_3->RVT->REase-> | 1041 | Bacteria>Pseudomonadota>Gammaproteobacteria>Legionellales>Legionellaceae | GCA_015989465.1 | Legionella_pneumophila | JBJ19_08155 | 5208..8333 | 446 | DUF499-domain-containing-protein-[Legionella-pneumophila] |
| TXK95834.1 | AAA+ATPase+wHTH+Alpha+Beta-domain+wHTH+Fn-III+Fn-III+RRM | Type-2_BR.RM.like.systems | <-ParA-Soj-PloopNTPase<-?\|\|DUF499*->DUF3780->METHYLASE->HKD-DNase+SF1-SF2-Helicase-><-Phage_integrase+Phage_integrase<-Phage_integrase<-Phage_integrase | 1021 | Bacteria>Pseudomonadota>Gammaproteobacteria>Methylococcales>Methylococcaceae | GCA_008030125.1 | Methylococcaceae_bacterium_CS5 | BMR11_12830 | 5291..8356 | 1916070 | hypothetical-protein-BMR11_12830-[Methylococcaceae-bacterium-CS5] |
| TXL03596.1 | AAA+ATPase+wHTH+Alpha+Beta-domain+wHTH+Fn-III+Fn-III+RRM | Type-2_BR.RM.like.systems | DUF499*->DUF3780->HKD-DNase+SF1-SF2-Helicase-><-Phage_integrase+Phage_integrase | 716 | Bacteria>Pseudomonadota>Gammaproteobacteria>Methylococcales>Methylococcaceae | GCA_008030145.1 | Methylococcaceae_bacterium_CS3 | BMR09_14555 | 35..2185 | 1916068 | hypothetical-protein-BMR09_14555-[Methylococcaceae-bacterium-CS3] |
| TXL04644.1 | AAA+ATPase+wHTH+Alpha+Beta-domain+wHTH+Fn-III+Fn-III+RRM | Type-2_BR.RM.like.systems | DUF499*->DUF3780->METHYLASE->HKD-DNase+SF1-SF2-Helicase-><-Phage_integrase+Phage_integrase<-Phage_integrase<-Phage_integrase | 1021 | Bacteria>Pseudomonadota>Gammaproteobacteria>Methylococcales>Methylococcaceae | GCA_008030215.1 | Methylococcaceae_bacterium_CS1 | BMR07_11835 | 1562..4627 | 1916066 | hypothetical-protein-BMR07_11835-[Methylococcaceae-bacterium-CS1] |
| CAI8823424.1 | AAA+ATPase+wHTH+Alpha+Beta-domain+wHTH+Fn-III+Fn-III+RRM | Type-2_BR.RM.like.systems | <-HKD-DNase+SF1-SF2-Helicase<-METHYLASE<-DUF3780<-DUF499*<-?<-?\|\|?-><-tRNA\|\|Phage_integrase-> | 1039 | Bacteria>Pseudomonadota>Gammaproteobacteria>Methylococcales>Methylococcaceae | GCA_949769195.1 | Methylocaldum_szegediense | MSZNOR_2007 | 1923893..1927012 | 73780 | AAA-family-ATPase-[Methylocaldum-szegediense] |
| WP_159657692.1 | AAA+ATPase+wHTH+Alpha+Beta-domain+wHTH+Fn-III+Fn-III+RRM | Type-2_BR.RM.like.systems | <-HNH<-ParB+HTH\|\|MNS+HTH->DUF499*->DUF3780->METHYLASE->HKD-DNase+SF1-SF2-Helicase->RVT-> | 1022 | Bacteria>Pseudomonadota>Gammaproteobacteria>Methylococcales>Methylococcaceae | GCF_009828925.2 | Methylicorpusculum_oleiharenae | GO003_RS24720 | 281775..284843 | 1338687 | DUF499-domain-containing-protein-[Methylicorpusculum-oleiharenae] |
| WP_169601231.1 | AAA+ATPase+wHTH+Alpha+Beta-domain+wHTH+Fn-III+Fn-III+RRM | Type-2_BR.RM.like.systems | <-HKD-DNase+SF1-SF2-Helicase<-METHYLASE<-MACRODOMAIN<-?<-DUF3780<-DUF499*<-RHH_1\|\|HTH_Tnp_1+HTH_21+rve+rve_3->?->ORC-CDC6+REase->JetA->JetB-> | 1041 | Bacteria>Pseudomonadota>Gammaproteobacteria>Methylococcales>Methylococcaceae | GCF_012769535.1 | Methylococcus_geothermalis | GNH96_RS00310 | 69324..72449 | 2681310 | DUF499-domain-containing-protein-[Methylococcus-geothermalis] |
| WP_005373819.1 | AAA+ATPase+wHTH+Alpha+Beta-domain+wHTH+Fn-III+Fn-III+RRM | Type-2_BR.RM.like.systems | SPOUT->?->Cysteine_desulfurase->?->?->DUF499*->DUF3780->?->RelE-ParE->?->PIN->URI-DNase-> | 1039 | Bacteria>Pseudomonadota>Gammaproteobacteria>Methylococcales>Methylococcaceae | GCF_000214275.2 | Methylomicrobium | CC94_RS0103015 | 650450..653569 | 39773 | MULTISPECIES:-DUF499-domain-containing-protein-[Methylomicrobium] |
| WP_017842534.1 | AAA+ATPase+wHTH+Alpha+Beta-domain+wHTH+Fn-III+Fn-III+RRM | Type-2_BR.RM.like.systems | HNH-><-?\|\|?->?->MetJarc-HTH->RelE-ParE->DUF499*->DUF3780->?->PIN->?->PIN->METHYLASE-> | 1021 | Bacteria>Pseudomonadota>Gammaproteobacteria>Methylococcales>Methylococcaceae | GCF_005931095.1 | Methylotuvimicrobium_buryatense | EQU24_RS16455 | 3692788..3695853 | 95641 | DUF499-domain-containing-protein-[Methylotuvimicrobium-buryatense] |
| WP_026223744.1 | AAA+ATPase+wHTH+Alpha+Beta-domain+wHTH+Fn-III+Fn-III+RRM | Type-2_BR.RM.like.systems | <-METHYLASE<-PIN<-VapB_antitoxin<-PIN<-?<-DUF3780<-DUF499*\|\|?->?->?-><-?<-?<-LON-AAA | 1042 | Bacteria>Pseudomonadota>Gammaproteobacteria>Methylococcales>Methylococcaceae | GCF_000372865.1 | Methylosarcina_fibrata | A3OW_RS0119500 | 4143557..4146685 | 105972 | DUF499-domain-containing-protein-[Methylosarcina-fibrata] |
| MBS3952485.1 | AAA+ATPase+wHTH+Alpha+Beta-domain+wHTH+Fn-III+Fn-III+RRM | Type-2_BR.RM.like.systems | HDRnase->?->DUF499*->DUF3780->?->RelE-ParE-><-?\|\|?->PIN-> | 1039 | Bacteria>Pseudomonadota>Gammaproteobacteria>Methylococcales>Methylococcaceae | GCA_018333785.1 | Methylomicrobium_sp. | KGZ88_06015 | 4130..7249 | 2303549 | DUF499-domain-containing-protein-[Methylomicrobium-sp.] |
| WP_198323836.1 | AAA+ATPase+wHTH+Alpha+Beta-domain+wHTH+Fn-III+Fn-III+RRM | Type-2_BR.RM.like.systems | ADH_N+ADH_zinc_N-><-SF1-SF2-Helicase\|\|?->?->RHH->DUF499*->DUF3780->MACRODOMAIN->METHYLASE->HEPN->ParA-Soj-PloopNTPase->ORC-CDC6-> | 1041 | Bacteria>Pseudomonadota>Gammaproteobacteria>Methylococcales>Methylococcaceae | GCF_016106025.1 | Methylococcus | ELQ36_RS13470 | 36069..39194 | 413 | DUF499-domain-containing-protein-[Methylococcus-sp.-BF19-07] |
| WP_218806438.1 | AAA+ATPase+wHTH+Alpha+Beta-domain+wHTH+Fn-III+Fn-III+RRM | Type-2_BR.RM.like.systems | ADH_N+ADH_zinc_N-><-SF1-SF2-Helicase\|\|?->?->CopG_antitoxin->DUF499*->DUF3780->METHYLASE->Schlafen+HATPase+HTH->?->METHYLASE->HEPN-> | 1041 | Bacteria>Pseudomonadota>Gammaproteobacteria>Methylococcales>Methylococcaceae | GCF_019285515.1 | Methylococcus_sp._Mc7 | KW115_RS14805 | 3051261..3054386 | 2860258 | DUF499-domain-containing-protein-[Methylococcus-sp.-Mc7] |
| MSP27870.1 | AAA+ATPase+wHTH+Alpha+Beta-domain+wHTH+Fn-III+Fn-III+RRM | Type-2_BR.RM.like.systems | HIN-HTH->DUF499*->DUF3780->?->PIN->METHYLASE->HKD-DNase+SF1-SF2-Helicase-> | 1021 | Bacteria>Pseudomonadota>Gammaproteobacteria>Methylococcales>unclassified_Methylococcales | GCA_009692395.1 | Methylococcales_bacterium | EXR80_05400 | 1625..4690 | 2304002 | DUF499-domain-containing-protein-[Methylococcales-bacterium] |
| MDD5268641.1 | AAA+ATPase+wHTH+Alpha+Beta-domain+wHTH+Fn-III+Fn-III+RRM | Type-2_BR.RM.like.systems | <-HKD-DNase+SF1-SF2-Helicase<-PIN<-VapB_antitoxin<-METHYLASE<-DUF3780<-DUF499*<-DOC+HTH | 1043 | Bacteria>Pseudomonadota>Gammaproteobacteria>Methylococcales>unclassified_Methylococcales | GCA_028712795.1 | Methylococcales_bacterium | PHO08_16185 | 15697..18828 | 2304002 | MAG:-DUF499-domain-containing-protein-[Methylococcales-bacterium] |

| Protein-ID | DUF499-architecture | System | Gene-Neighbourhoods | Length | Taxonomy | Assembly-ID | Strain | Locus-Tag | Genomic-coordinates | TaxID | Fasta-defline |
|---|---|---|---|---|---|---|---|---|---|---|---|
| MDD5321594.1 | AAA+ATPase+wHTH+Alpha+Beta-domain+wHTH+Fn-III+Fn-III+RRM | Type-2_BR.RM.like.systems | <-HKD-DNase+SF1-SF2-Helicase<-?<-PIN<-?<-METHYLASE<-DUF3780<-DUF499*<-tRNA<-MerR-HTH | 1047 | Bacteria>Pseudomonadota>Gammaproteobacteria>Methylococcales>unclassified_Methylococcales | GCA_028714175.1 | Methylococcales_bacterium | PHD43_13460 | 8074..11217 | 2304002 | MAG:-DUF499-domain-containing-protein-[Methylococcales-bacterium] |
| MDD5320442.1 | AAA+ATPase+wHTH+Alpha+Beta-domain+wHTH+Fn-III+Fn-III+RRM | Type-2_BR.RM.like.systems | DUF499*->DUF3780->REase->?->PIN->REase->METHYLASE-> | 1041 | Bacteria>Pseudomonadota>Gammaproteobacteria>Methylococcales>unclassified_Methylococcales | GCA_028714175.1 | Methylococcales_bacterium | PHD43_07485 | 17817..20942 | 2304002 | MAG:-DUF499-domain-containing-protein-[Methylococcales-bacterium] |
| WP_214288673.1 | AAA+ATPase+wHTH+Alpha+Beta-domain+wHTH+Fn-III+Fn-III+RRM | Type-2_BR.RM.like.systems | rve+Mu-transpos_C->TnsC-ATPase->?->?->Cluster1976_1clades->HTH+MPTase->DUF499*->DUF3780->METHYLASE->HKD-DNase+SF1-SF2-Helicase-> | 1013 | Bacteria>Pseudomonadota>Gammaproteobacteria>Moraxellales>Moraxellaceae | - | Acinetobacter_pittii | JD510_RS16135 | 28076..31117 | 48296 | DUF499-domain-containing-protein-[Acinetobacter-pittii] |
| RUP42373.1 | AAA+ATPase+wHTH+Alpha+Beta-domain+wHTH+Fn-III+Fn-III+RRM | Type-2_BR.RM.like.systems | LRP-HTH->?-><-HKD-DNase+SF1-SF2-Helicase<-METHYLASE<-DUF3780<-DUF499*<-HTH+MPTase<-Cluster1976_1clades<-?<-?<-TnsC-ATPase<-rve+Mu-transpos_C | 1022 | Bacteria>Pseudomonadota>Gammaproteobacteria>Moraxellales>Moraxellaceae | GCA_003987695.1 | Acinetobacter_sp. | EKK63_01560 | 43281..46349 | 472 | DUF499-domain-containing-protein-[Acinetobacter-sp.] |
| WP_019767350.1 | AAA+ATPase+wHTH+Alpha+Beta-domain+wHTH+Fn-III+Fn-III+RRM | Type-2_BR.RM.like.systems | rve+Mu-transpos_C->TnsC-ATPase->?->?->Cluster1976_1clades->HTH+MPTase->DUF499*->DUF3780->METHYLASE->HKD-DNase+SF1-SF2-Helicase-> | 1022 | Bacteria>Pseudomonadota>Gammaproteobacteria>Moraxellales>Moraxellaceae | GCF_000230465.1 | Acinetobacter_pittii | QY040_RS16165 | 28008..31076 | 48296 | MULTISPECIES:-DUF499-domain-containing-protein-[Acinetobacter-calcoaceticus/baumannii-complex] |
| WP_004965096.1 | AAA+ATPase+wHTH+Alpha+Beta-domain+wHTH+Fn-III+Fn-III+RRM | Type-2_BR.RM.like.systems | LRP-HTH->?-><-HKD-DNase+SF1-SF2-Helicase<-METHYLASE<-DUF3780<-DUF499*<-HTH+MPTase<-Cluster1976_1clades<-?<-?<-TnsC-ATPase<-rve+Mu-transpos_C | 1022 | Bacteria>Pseudomonadota>Gammaproteobacteria>Moraxellales>Moraxellaceae | GCF_000302355.1 | Acinetobacter | AJU02_RS05515 | 41078..44146 | 469 | MULTISPECIES:-DUF499-domain-containing-protein-[Acinetobacter] |
| WP_033854088.1 | AAA+ATPase+wHTH+Alpha+Beta-domain+wHTH+Fn-III+Fn-III+RRM | Type-2_BR.RM.like.systems | rve+Mu-transpos_C->TnsC-ATPase->?->?->Cluster1976_1clades->HTH+MPTase->DUF499*->DUF3780->METHYLASE->HKD-DNase+SF1-SF2-Helicase-> | 1022 | Bacteria>Pseudomonadota>Gammaproteobacteria>Moraxellales>Moraxellaceae | GCF_008979865.1 | Acinetobacter | GBV81_RS05730 | 228883..231951 | 469 | MULTISPECIES:-DUF499-domain-containing-protein-[Acinetobacter] |
| WP_151738990.1 | AAA+ATPase+wHTH+Alpha+Beta-domain+wHTH+Fn-III+Fn-III+RRM | Type-2_BR.RM.like.systems | DUF499*->DUF3780->METHYLASE->HKD-DNase+SF1-SF2-Helicase-> | 1022 | Bacteria>Pseudomonadota>Gammaproteobacteria>Moraxellales>Moraxellaceae | GCF_008982085.1 | Acinetobacter_sp._TUM15099 | GBJ83_RS07185 | 19..3087 | 2609136 | DUF499-domain-containing-protein-[Acinetobacter-sp.-TUM15099] |
| WP_228276565.1 | AAA+ATPase+wHTH+Alpha+Beta-domain+wHTH+Fn-III+Fn-III+RRM | Type-2_BR.RM.like.systems | <-HKD-DNase+SF1-SF2-Helicase<-METHYLASE<-DUF3780<-DUF499* | 883 | Bacteria>Pseudomonadota>Gammaproteobacteria>Moraxellales>Moraxellaceae | GCF_008982165.1 | Acinetobacter_sp._TUM15103 | GBJ92_RS15120 | 7611..10262 | 2609137 | chitobiase/beta-hexosaminidase-C-terminal-domain-containing-protein-[Acinetobacter-sp.-TUM15103] |
| WP_236765703.1 | AAA+ATPase+wHTH+Alpha+Beta-domain+wHTH+Fn-III+Fn-III+RRM | Type-2_BR.RM.like.systems | LRP-HTH->?-><-HKD-DNase+SF1-SF2-Helicase<-METHYLASE<-DUF3780<-DUF499*<-HTH+MPTase<-Cluster1976_1clades<-?<-?<-TnsC-ATPase<-rve+Mu-transpos_C | 1013 | Bacteria>Pseudomonadota>Gammaproteobacteria>Moraxellales>Moraxellaceae | GCF_009004115.1 | Acinetobacter_junii | GBR74_RS00175 | 36032..39073 | 40215 | DUF499-domain-containing-protein-[Acinetobacter-junii] |
| WP_168416643.1 | AAA+ATPase+wHTH+Alpha+Beta-domain+wHTH+Fn-III+Fn-III+RRM | Type-2_BR.RM.like.systems | ZnR+ZnR->DUF932->?->ParB->XIS-HTH->RHH_1->DUF499*->DUF3780->METHYLASE->HKD-DNase+SF1-SF2-Helicase->sGTP-> | 1041 | Bacteria>Pseudomonadota>Gammaproteobacteria>Moraxellales>Moraxellaceae | GCF_012371205.1 | Acinetobacter_indicus | HHC89_RS06080 | 207809..210934 | 756892 | DUF499-domain-containing-protein-[Acinetobacter-indicus] |
| WP_180127358.1 | AAA+ATPase+wHTH+Alpha+Beta-domain+wHTH+Fn-III+Fn-III+RRM | Type-2_BR.RM.like.systems | ParB->XIS-HTH->SAM-methylase->wHTH(SBD)+HNH->RHH_1->DUF499*->DUF3780->METHYLASE->HKD-DNase+SF1-SF2-Helicase-p->?-><-Arm-DNA-bind_3+Phage_int_SAM_3+Phage_integrase | 1040 | Bacteria>Pseudomonadota>Gammaproteobacteria>Moraxellales>Moraxellaceae | GCF_013419465.1 | unclassified_Acinetobacter | FTU25_RS07965 | 188161..191283 | 196816 | MULTISPECIES:-DUF499-domain-containing-protein-[unclassified-Acinetobacter] |
| WP_200150193.1 | AAA+ATPase+wHTH+Alpha+Beta-domain+wHTH+Fn-III+Fn-III+RRM | Type-2_BR.RM.like.systems | <-HKD-DNase+SF1-SF2-Helicase<-METHYLASE<-DUF3780<-DUF499*<-XIS-HTH\|\|?-><-DUF932 | 1022 | Bacteria>Pseudomonadota>Gammaproteobacteria>Moraxellales>Moraxellaceae | GCF_016576965.1 | Acinetobacter_bereziniae | P7L54_RS16685 | 49736..52804 | 106648 | DUF499-domain-containing-protein-[Acinetobacter-bereziniae] |
| WP_156188412.1 | AAA+ATPase+wHTH+Alpha+Beta-domain+wHTH+Fn-III+Fn-III+RRM | Type-2_BR.RM.like.systems | <-HKD-DNase+SF1-SF2-Helicase<-METHYLASE<-DUF3780<-DUF499*<-HTH+MPTase<-Cluster1976_1clades<-?<-?<-TnsC-ATPase<-rve+Mu-transpos_C | 1013 | Bacteria>Pseudomonadota>Gammaproteobacteria>Moraxellales>Moraxellaceae | GCF_017837665.1 | Acinetobacter_nosocomialis | JRZ80_RS04920 | 78315..81356 | 106654 | MULTISPECIES:-DUF499-domain-containing-protein-[Acinetobacter-calcoaceticus/baumannii-complex] |
| WP_227062135.1 | AAA+ATPase+wHTH+Alpha+Beta-domain+wHTH+Fn-III+Fn-III+RRM | Type-2_BR.RM.like.systems | <-Peptidase_S51\|\|?-><-HKD-DNase+SF1-SF2-Helicase<-METHYLASE<-DUF3780<-DUF499*<-HTH+MPTase<-Cluster1976_1clades | 1022 | Bacteria>Pseudomonadota>Gammaproteobacteria>Moraxellales>Moraxellaceae | GCF_020544425.1 | Acinetobacter_pittii | JC624_RS16535 | 3433264..3436332 | 48296 | DUF499-domain-containing-protein-[Acinetobacter-pittii] |
| MDB5972090.1 | AAA+ATPase+wHTH+Alpha+Beta-domain+wHTH+Fn-III+Fn-III+RRM | Type-2_BR.RM.like.systems | <-MACRODOMAIN<-?<-HKD-DNase+SF1-SF2-Helicase<-METHYLASE<-DUF3780<-DUF499*<-Phage_int_SAM_2+Integrase_1<-?<-HHH+ENDO3+NUDIX | 1027 | Bacteria>Pseudomonadota>Gammaproteobacteria>Nevskiales>Nevskiaceae | GCA_028291725.1 | Hydrocarboniphaga_sp. | JWQ90_4540 | 26375..29458 | 2033016 | MAG:-hypothetical-protein-JWQ90_4540-[Hydrocarboniphaga-sp.] |
| WP_262597092.1 | AAA+ATPase+wHTH+Alpha+Beta-domain+wHTH+Fn-III+Fn-III+RRM | Type-2_BR.RM.like.systems | XIS-HTH->DUF499*->DUF3780->METHYLASE-><-?\|\|HKD-DNase+SF1-SF2-Helicase->?->RVT-> | 1040 | Bacteria>Pseudomonadota>Gammaproteobacteria>Oceanospirillales>Endozoicomonadaceae | GCF_025562755.1 | Endozoicomonas_euniceicola | NX720_RS20380 | 5029732..5032854 | 1234143 | DUF499-domain-containing-protein-[Endozoicomonas-euniceicola] |
| WP_257264226.1 | AAA+ATPase+wHTH+Alpha+Beta-domain+wHTH+Fn-III+Fn-III+RRM | Type-2_BR.RM.like.systems | DUF499*->DUF3780->?->METHYLASE-p->METHYLASE->PIN->XIS-HTH-> | 740 | Bacteria>Pseudomonadota>Gammaproteobacteria>Oceanospirillales>Endozoicomonadaceae | GCF_024606365.1 | Endozoicomonas_sp._ONNA2 | KDK14_RS06570 | 43..2265 | 2828741 | FN3-associated-domain-containing-protein-[Endozoicomonas-sp.-ONNA2] |
| MBP5978753.1 | AAA+ATPase+wHTH+Alpha+Beta-domain+wHTH+Fn-III+Fn-III+RRM | Type-2_BR.RM.like.systems | <-MODE-HTH+PBPII\|\|?->ABC-ATPase+SbcC-><-HKD-DNase+SF1-SF2-Helicase<-METHYLASE<-DUF3780<-DUF499*<-RHH | 1041 | Bacteria>Pseudomonadota>Gammaproteobacteria>Oceanospirillales>Halomonadaceae | GCA_017983445.1 | Halomonas_sp. | KA748_00935 | 220755..223880 | 1486246 | DUF499-domain-containing-protein-[Halomonas-sp.] |
| WP_035565412.1 | AAA+ATPase+wHTH+Alpha+Beta-domain+wHTH+Fn-III+Fn-III+RRM | Type-2_BR.RM.like.systems | <-HKD-DNase+SF1-SF2-Helicase<-METHYLASE<-DUF3780<-DUF499*<-?<-?<-?<-?<-PAIREDC-HTH | 1034 | Bacteria>Pseudomonadota>Gammaproteobacteria>Oceanospirillales>Halomonadaceae | GCF_000734975.2 | Halomonas_sp._KO116 | KO116_RS23500 | 103563..106667 | 1504981 | DUF499-domain-containing-protein-[Halomonas-sp.-KO116] |
| WP_125749599.1 | AAA+ATPase+wHTH+Alpha+Beta-domain+wHTH+Fn-III+Fn-III+RRM | Type-2_BR.RM.like.systems | DUF499*->DUF3780->METHYLASE->HKD-DNase+SF1-SF2-Helicase-><-?<-HTH_OrfB_IS605+OrfB_IS605+OrfB_Zn_ribbon | 1034 | Bacteria>Pseudomonadota>Gammaproteobacteria>Oceanospirillales>Halomonadaceae | GCF_003950215.1 | Halomonas_venusta | DDR56_RS06095 | 352926..356030 | 44935 | DUF499-domain-containing-protein-[Halomonas-venusta] |
| WP_193092566.1 | AAA+ATPase+wHTH+Alpha+Beta-domain+wHTH+Fn-III+Fn-III+RRM | Type-2_BR.RM.like.systems | <-REase<-?<-?<-HKD-DNase+SF1-SF2-Helicase<-METHYLASE<-DUF3780<-DUF499*<-?<-?\|\|?->?->CR-ATPase8+REase-> | 1034 | Bacteria>Pseudomonadota>Gammaproteobacteria>Oceanospirillales>Halomonadaceae | GCF_014897985.1 | Halomonas_sp._FME66 | INT33_RS17400 | 16589..19693 | 2742615 | DUF499-domain-containing-protein-[Halomonas-sp.-FME66] |
| WP_110715559.1 | AAA+ATPase+wHTH+Alpha+Beta-domain+wHTH+Fn-III+Fn-III+RRM | Type-2_BR.RM.like.systems | Classical-AAA->Peptidase_S8-><-HKD-DNase+SF1-SF2-Helicase<-METHYLASE<-DUF3780<-DUF499*<-RHH_5<-?\|\|?->?-><-?\|\|FURR-HTH-> | 1041 | Bacteria>Pseudomonadota>Gammaproteobacteria>Oceanospirillales>Halomonadaceae | GCF_029870975.1 | Salinicola_acroporae | CUR86_RS04115 | 820457..823582 | 1541440 | DUF499-domain-containing-protein-[Salinicola-acroporae] |
| WP_090726653.1 | AAA+ATPase+wHTH+Alpha+Beta-domain+wHTH+Fn-III+Fn-III+RRM | Type-2_BR.RM.like.systems | <-SF1-SF2-Helicase<-HKD-DNase+SF1-SF2-Helicase<-METHYLASE<-DUF3780<-DUF499*<-RHH_1\|\|JAB->Phage_integrase-><-?<-?<-ACET | 1041 | Bacteria>Pseudomonadota>Gammaproteobacteria>Oceanospirillales>Oceanospirillaceae | GCF_016082375.1 | Neptunomonas_qingdaonensis | Nqin_RS15525 | 312285..315410 | 1045558 | DUF499-domain-containing-protein-[Neptunomonas-qingdaonensis] |
| WP_218150140.1 | AAA+ATPase+wHTH+Alpha+Beta-domain+wHTH+Fn-III+Fn-III+RRM | Type-2_BR.RM.like.systems | <-SF1-SF2-Helicase<-HKD-DNase+SF1-SF2-Helicase<-METHYLASE<-DUF3780<-DUF499*<-RHH_1\|\|JAB->Phage_integrase-><-?<-?<-ACET | 1022 | Bacteria>Pseudomonadota>Gammaproteobacteria>Oceanospirillales>Oceanospirillaceae | GCF_900113275.1 | Neptunomonas_qingdaonensis | BM099_RS08335 | 312372..315440 | 1045558 | DUF499-domain-containing-protein-[Neptunomonas-qingdaonensis] |
| WP_043884792.1 | AAA+ATPase+wHTH+Alpha+Beta-domain+wHTH+Fn-III+Fn-III+RRM | Type-2_BR.RM.like.systems | MazG->REase+SF1-SF2-Helicase->Arm-DNA-bind_3+Phage_integrase-><-HKD-DNase+SF1-SF2-Helicase<-METHYLASE<-DUF3780<-DUF499* | 1038 | Bacteria>Pseudomonadota>Gammaproteobacteria>Pseudomonadales | GCF_003836565.1 | Pseudomonadaceae | PA2G_RS09470 | 2116753..2119869 | 135621 | MULTISPECIES:-DUF499-domain-containing-protein-[Pseudomonadaceae] |
| MAB53981.1 | AAA+ATPase+wHTH+Alpha+Beta-domain+wHTH+Fn-III+Fn-III+RRM | Type-2_BR.RM.like.systems | DUF499*->DUF3780->METHYLASE->HKD-DNase+SF1-SF2-Helicase->?->AbiEi->CHTH-> | 1034 | Bacteria>Pseudomonadota>Gammaproteobacteria>Pseudomonadales>Marinobacteraceae | GCA_002683435.1 | Marinobacter_sp. | CMG80_19940 | 487..3591 | 50741 | AAA-family-ATPase-[Marinobacter-sp.] |
| MAK49681.1 | AAA+ATPase+wHTH+Alpha+Beta-domain+wHTH+Fn-III+Fn-III+RRM | Type-2_BR.RM.like.systems | <-CHTH<-AbiEi<-?<-HKD-DNase+SF1-SF2-Helicase<-METHYLASE<-DUF3780<-DUF499*<-GDResolvase+Recombinase+ZnR\|\|ParB->REase+SAVED-> | 1034 | Bacteria>Pseudomonadota>Gammaproteobacteria>Pseudomonadales>Marinobacteraceae | GCA_002693995.1 | Marinobacter_sp. | CMG85_10440 | 142423..145527 | 50741 | AAA-family-ATPase-[Marinobacter-sp.] |
| MBE93922.1 | AAA+ATPase+wHTH+Alpha+Beta-domain+wHTH+Fn-III+Fn-III+RRM | Type-2_BR.RM.like.systems | ArsR-HTH+YTCD-HTH-><-?<-?<-?\|\|MerR-HTH->GDResolvase+Recombinase+ZnR->DUF499*->DUF3780->METHYLASE->HKD-DNase+SF1-SF2-Helicase-> | 1034 | Bacteria>Pseudomonadota>Gammaproteobacteria>Pseudomonadales>Marinobacteraceae | GCA_002714765.1 | Marinobacter_sp. | CMG89_02520 | 5039..8143 | 50741 | AAA-family-ATPase-[Marinobacter-sp.] |
| WP_076724732.1 | AAA+ATPase+wHTH+Alpha+Beta-domain+wHTH+Fn-III+Fn-III+RRM | Type-2_BR.RM.like.systems | RHH_5->DUF499*->DUF3780->METHYLASE->HKD-DNase+SF1-SF2-Helicase-><-Peptidase_S8<-Classical-AAA<-Phage_lysozyme | 1041 | Bacteria>Pseudomonadota>Gammaproteobacteria>Pseudomonadales>Marinobacteraceae | GCF_001981305.1 | Marinobacter_lutaoensis | BT032_RS11255 | 364358..367483 | 135739 | DUF499-domain-containing-protein-[Marinobacter-lutaoensis] |
| WP_273070524.1 | AAA+ATPase+wHTH+Alpha+Beta-domain+wHTH+Fn-III+Fn-III+RRM | Type-2_BR.RM.like.systems | <-AbiEi<-?<-HKD-DNase+SF1-SF2-Helicase<-METHYLASE<-DUF3780<-DUF499*<-GDResolvase+Recombinase+ZnR\|\|ParB->REase+SAVED-> | 1034 | Bacteria>Pseudomonadota>Gammaproteobacteria>Pseudomonadales>Marinobacteraceae | GCF_002341575.1 | unclassified_Marinobacter | CMG85_RS10390 | 142423..145527 | 83889 | DUF499-domain-containing-protein-[Marinobacter-sp.] |
| EAZ58601.1 | AAA+ATPase+wHTH+Alpha+Beta-domain+wHTH+Fn-III+Fn-III+RRM | Type-2_BR.RM.like.systems | MazG->REase+SF1-SF2-Helicase->Arm-DNA-bind_3+Phage_integrase-><-HKD-DNase+SF1-SF2-Helicase<-METHYLASE<-DUF499*<-?<-?<-?<-?\|\|HTH+WYL-> | 1072 | Bacteria>Pseudomonadota>Gammaproteobacteria>Pseudomonadales>Pseudomonadaceae | GCA_000152545.1 | Pseudomonas_aeruginosa_2192 | PA2G_01849 | 2116753..2119971 | 350703 | hypothetical-protein-PA2G_01849-[Pseudomonas-aeruginosa-2192] |
| KJS67130.1 | AAA+ATPase+wHTH+Alpha+Beta-domain+wHTH+Fn-III+Fn-III+RRM | Type-2_BR.RM.like.systems | MerR-HTH->MerR-HTH->DUF4158+DDE_Tnp_Tn3-><-HKD-DNase+SF1-SF2-Helicase<-METHYLASE<-DUF3780<-DUF499*<-?<-Phage_integrase\|\|Peptidase_M15-> | 1038 | Bacteria>Pseudomonadota>Gammaproteobacteria>Pseudomonadales>Pseudomonadaceae | GCA_000802035.2 | [Pseudomonas]_sp._BICA1-14 | JL55_36155 | 12259..15375 | 1619235 | MAG:-ATPase-AAA-[Pseudomonas-sp.-BICA1-14] |
| RUB64276.1 | AAA+ATPase+wHTH+Alpha+Beta-domain+wHTH+Fn-III+Fn-III+RRM | Type-2_BR.RM.like.systems | MerR-HTH->tRNA->\|\|TUDOR+SF1-SF2-Helicase+iREase+CTD->?->DUF499*->DUF3780->METHYLASE->HKD-DNase+SF1-SF2-Helicase-><-?<-ZNR+Pkinase+SF1-SF2-Helicase+REase | 1038 | Bacteria>Pseudomonadota>Gammaproteobacteria>Pseudomonadales>Pseudomonadaceae | GCA_003973995.1 | Pseudomonas_aeruginosa | IPC1413_22670 | 96006..99122 | 287 | DUF499-domain-containing-protein-[Pseudomonas-aeruginosa] |
| MBG6817109.1 | AAA+ATPase+wHTH+Alpha+Beta-domain+wHTH+Fn-III+Fn-III+RRM | Type-2_BR.RM.like.systems | RHH_1->DUF499*->DUF3780->METHYLASE->Schlafen+HATPase+HTH->METHYLASE->HKD-DNase+SF1-SF2-Helicase->MORC+REase-> | 1041 | Bacteria>Pseudomonadota>Gammaproteobacteria>Pseudomonadales>Pseudomonadaceae | GCA_015764555.1 | Pseudomonas_aeruginosa | I5J00_12285 | 138318..141443 | 287 | DUF499-domain-containing-protein-[Pseudomonas-aeruginosa] |
| MBH9417281.1 | AAA+ATPase+wHTH+Alpha+Beta-domain+wHTH+Fn-III+Fn-III+RRM | Type-2_BR.RM.like.systems | HIN-HTH->Peptidase_M15->?->TUDOR+SF1-SF2-Helicase+iREase+CTD->?->?->DUF499*->DUF3780->METHYLASE->HKD-DNase+SF1-SF2-Helicase-><-Arm-DNA-bind_3+Phage_integrase<-REase+SF1-SF2-Helicase<-MazG | 1038 | Bacteria>Pseudomonadota>Gammaproteobacteria>Pseudomonadales>Pseudomonadaceae | GCA_016092775.1 | Pseudomonas_aeruginosa | GHU28_23410 | 8846..11962 | 287 | DUF499-domain-containing-protein-[Pseudomonas-aeruginosa] |
| MBS7414067.1 | AAA+ATPase+wHTH+Alpha+Beta-domain+wHTH+Fn-III+Fn-III+RRM | Type-2_BR.RM.like.systems | HTH+MPTase->OLD-ABC+TOPRIM->SF1-SF2-Helicase-><-HKD-DNase+SF1-SF2-Helicase<-METHYLASE<-DUF3780<-DUF499*<-?\|\|?-><-?\|\|HTH_3-><-DUF932 | 1072 | Bacteria>Pseudomonadota>Gammaproteobacteria>Pseudomonadales>Pseudomonadaceae | GCA_018383755.1 | Pseudomonas_syringae | KID89_13640 | 3099163..3102381 | 317 | DUF499-domain-containing-protein-[Pseudomonas-syringae] |
| MBX5883026.1 | AAA+ATPase+wHTH+Alpha+Beta-domain+wHTH+Fn-III+Fn-III+RRM | Type-2_BR.RM.like.systems | <-URI-DNase+RAMA<-HKD-DNase+SF1-SF2-Helicase<-METHYLASE<-MACRODOMAIN<-?<-DUF3780<-DUF499*<-RHH_5\|\|?->TOPRIM+SF1-SF2-Helicase+HTH-> | 1041 | Bacteria>Pseudomonadota>Gammaproteobacteria>Pseudomonadales>Pseudomonadaceae | GCA_019688335.1 | Pseudomonas_aeruginosa | ISE51_20275 | 196634..199759 | 287 | DUF499-domain-containing-protein-[Pseudomonas-aeruginosa] |
| HJH19646.1 | AAA+ATPase+wHTH+Alpha+Beta-domain+wHTH+Fn-III+Fn-III+RRM | Type-2_BR.RM.like.systems | ZnR+ZnR-><-HTH_3\|\|?->?->?->DUF499*->DUF3780->METHYLASE->HKD-DNase+SF1-SF2-Helicase-><-SF1-SF2-Helicase<-OLD-ABC+TOPRIM | 1038 | Bacteria>Pseudomonadota>Gammaproteobacteria>Pseudomonadales>Pseudomonadaceae | GCA_020744095.1 | Pseudomonas_lactis | K8W20_13130 | 6705..9821 | 1615674 | DUF499-domain-containing-protein-[Pseudomonas-lactis] |
| EIU2715573.1 | AAA+ATPase+wHTH+Alpha+Beta-domain+wHTH+Fn-III+Fn-III+RRM | Type-2_BR.RM.like.systems | SF1-SF2-Helicase->HIN-HTH->?->TUDOR+SF1-SF2-Helicase+iREase+CTD->?->?->DUF499*->DUF3780->METHYLASE->HKD-DNase+SF1-SF2-Helicase-><-Arm-DNA-bind_3+Phage_integrase<-REase+SF1-SF2-Helicase<-MazG | 1072 | Bacteria>Pseudomonadota>Gammaproteobacteria>Pseudomonadales>Pseudomonadaceae | GCA_021614535.1 | Pseudomonas_aeruginosa | L4O33_002031 | 46678..49896 | 287 | DUF499-domain-containing-protein-[Pseudomonas-aeruginosa] |
| HBP0460596.1 | AAA+ATPase+wHTH+Alpha+Beta-domain+wHTH+Fn-III+Fn-III+RRM | Type-2_BR.RM.like.systems | ParB->JAB-><-HEPN<-HKD-DNase+SF1-SF2-Helicase<-METHYLASE<-DUF3780<-DUF499*<-RHH\|\|?->Phage_integrase->Phage_integrase-> | 1040 | Bacteria>Pseudomonadota>Gammaproteobacteria>Pseudomonadales>Pseudomonadaceae | GCA_021685755.1 | Pseudomonas_aeruginosa | L5Q34_003641 | 43212..46334 | 287 | DUF499-domain-containing-protein-[Pseudomonas-aeruginosa] |
| HBP5533303.1 | AAA+ATPase+wHTH+Alpha+Beta-domain+wHTH+Fn-III+Fn-III+RRM | Type-2_BR.RM.like.systems | ParB->JAB-><-HKD-DNase+SF1-SF2-Helicase<-METHYLASE<-DUF3780<-DUF499*<-RHH\|\|<-?<-XIS-HTH\|\|CopG/RHH-><-tRNA<-MerR-HTH | 1040 | Bacteria>Pseudomonadota>Gammaproteobacteria>Pseudomonadales>Pseudomonadaceae | GCA_021712375.1 | Pseudomonas_aeruginosa | L6B68_08675 | 333439..336561 | 287 | DUF499-domain-containing-protein-[Pseudomonas-aeruginosa] |
| MCF6783903.1 | AAA+ATPase+wHTH+Alpha+Beta-domain+wHTH+Fn-III+Fn-III+RRM | Type-2_BR.RM.like.systems | DUF499*->DUF3780->METHYLASE->HKD-DNase+SF1-SF2-Helicase->\|\|SF1-SF2-Helicase-><-CHTH<-SF1-SF2-Helicase | 1072 | Bacteria>Pseudomonadota>Gammaproteobacteria>Pseudomonadales>Pseudomonadaceae | GCA_021713665.1 | Stutzerimonas_stutzeri | L3X17_19300 | 544..3762 | 316 | DUF499-domain-containing-protein-[Stutzerimonas-stutzeri] |
| HCE5959030.1 | AAA+ATPase+wHTH+Alpha+Beta-domain+wHTH+Fn-III+Fn-III+RRM | Type-2_BR.RM.like.systems | CNMP+TIR->ACYC+ACYC->?->DUF499*->DUF3780->METHYLASE->HKD-DNase+SF1-SF2-Helicase-><-SF1-SF2-Helicase<-OLD-ABC+TOPRIM<-CHTH | 1072 | Bacteria>Pseudomonadota>Gammaproteobacteria>Pseudomonadales>Pseudomonadaceae | GCA_023904655.1 | Pseudomonas_aeruginosa | NHK89_004125 | 34242..37460 | 287 | DUF499-domain-containing-protein-[Pseudomonas-aeruginosa] |
| HCF5250768.1 | AAA+ATPase+wHTH+Alpha+Beta-domain+wHTH+Fn-III+Fn-III+RRM | Type-2_BR.RM.like.systems | Peptidase_M15->?->TUDOR+SF1-SF2-Helicase+iREase+CTD->?->?->DUF499*->DUF3780->METHYLASE->HKD-DNase+SF1-SF2-Helicase-><-Arm-DNA-bind_3+Phage_integrase | 1072 | Bacteria>Pseudomonadota>Gammaproteobacteria>Pseudomonadales>Pseudomonadaceae | GCA_023953455.1 | Pseudomonas_aeruginosa | NIG42_002592 | 72601..75819 | 287 | DUF499-domain-containing-protein-[Pseudomonas-aeruginosa] |
| HCG0874251.1 | AAA+ATPase+wHTH+Alpha+Beta-domain+wHTH+Fn-III+Fn-III+RRM | Type-2_BR.RM.like.systems | ZNR+Pkinase+SF1-SF2-Helicase+REase->?-><-HKD-DNase+SF1-SF2-Helicase<-METHYLASE<-DUF3780<-DUF499*<-?<-?<-TUDOR+SF1-SF2-Helicase+iREase+CTD | 1072 | Bacteria>Pseudomonadota>Gammaproteobacteria>Pseudomonadales>Pseudomonadaceae | GCA_024004315.1 | Pseudomonas_aeruginosa | NJF72_002791 | 346926..350144 | 287 | DUF499-domain-containing-protein-[Pseudomonas-aeruginosa] |
| MCQ4281551.1 | AAA+ATPase+wHTH+Alpha+Beta-domain+wHTH+Fn-III+Fn-III+RRM | Type-2_BR.RM.like.systems | Glyoxalase-><-?\|\|?->CNMP+TIR->ACYC+ACYC->?->DUF499*->DUF3780->METHYLASE->HKD-DNase+SF1-SF2-Helicase-><-CHTH<-?<-SF1-SF2-Helicase | 1038 | Bacteria>Pseudomonadota>Gammaproteobacteria>Pseudomonadales>Pseudomonadaceae | GCA_024448855.1 | Stutzerimonas_stutzeri | NA633_00395 | 79686..82802 | 316 | DUF499-domain-containing-protein-[Pseudomonas-stutzeri] |
| HCR1362767.1 | AAA+ATPase+wHTH+Alpha+Beta-domain+wHTH+Fn-III+Fn-III+RRM | Type-2_BR.RM.like.systems | <-HIN-HTH\|\|TUDOR+SF1-SF2-Helicase+iREase+CTD->?->?->DUF499*->DUF3780->METHYLASE->HKD-DNase+SF1-SF2-Helicase-><-?<-ZNR+Pkinase+SF1-SF2-Helicase+REase | 1072 | Bacteria>Pseudomonadota>Gammaproteobacteria>Pseudomonadales>Pseudomonadaceae | GCA_025921445.1 | Pseudomonas_aeruginosa | ONA48_004000 | 6159..9377 | 287 | TPA:-DUF499-domain-containing-protein-[Pseudomonas-aeruginosa] |
| WP_003280170.1 | AAA+ATPase+wHTH+Alpha+Beta-domain+wHTH+Fn-III+Fn-III+RRM | Type-2_BR.RM.like.systems | DUF499*->DUF3780->METHYLASE->HKD-DNase+SF1-SF2-Helicase-> | 1038 | Bacteria>Pseudomonadota>Gammaproteobacteria>Pseudomonadales>Pseudomonadaceae | GCF_000237885.1 | Stutzerimonas_stutzeri | PSTZOBELL_RS01535 | 2264..5380 | 316 | DUF499-domain-containing-protein-[Stutzerimonas-stutzeri] |
| WP_028615088.1 | AAA+ATPase+wHTH+Alpha+Beta-domain+wHTH+Fn-III+Fn-III+RRM | Type-2_BR.RM.like.systems | <-HKD-DNase+SF1-SF2-Helicase<-METHYLASE<-DUF3780<-DUF499*<-RHH<-JAB<-ParB<-?<-DUF932 | 1041 | Bacteria>Pseudomonadota>Gammaproteobacteria>Pseudomonadales>Pseudomonadaceae | GCF_000410875.1 | Halopseudomonas_pelagia | K787_RS0118500 | 63461..66586 | 553151 | DUF499-domain-containing-protein-[Halopseudomonas-pelagia] |
| WP_023443648.1 | AAA+ATPase+wHTH+Alpha+Beta-domain+wHTH+Fn-III+Fn-III+RRM | Type-2_BR.RM.like.systems | <-Phage_integrase<-?\|\|RHH->DUF499*->DUF3780->METHYLASE->HKD-DNase+SF1-SF2-Helicase->HEPN-><-?<-JAB | 1040 | Bacteria>Pseudomonadota>Gammaproteobacteria>Pseudomonadales>Pseudomonadaceae | GCF_000495915.1 | Stutzerimonas_stutzeri_group | F753_RS25405 | 1974..5096 | 136846 | MULTISPECIES:-DUF499-domain-containing-protein-[Stutzerimonas-stutzeri-group] |
| WP_028697645.1 | AAA+ATPase+wHTH+Alpha+Beta-domain+wHTH+Fn-III+Fn-III+RRM | Type-2_BR.RM.like.systems | <-RHH_6\|\|GDResolvase+Recombinase+ZnR->CHTH-><-HKD-DNase+SF1-SF2-Helicase<-METHYLASE<-DUF3780<-DUF499*<-?<-ACYC+ACYC<-CNMP+TIR<-?\|\|?->PIN-> | 1038 | Bacteria>Pseudomonadota>Gammaproteobacteria>Pseudomonadales>Pseudomonadaceae | GCF_000633915.1 | Pseudomonas_putida_group | AOQ85_RS15325 | 12239..15355 | 136845 | MULTISPECIES:-DUF499-domain-containing-protein-[Pseudomonas-putida-group] |
| WP_047595839.1 | AAA+ATPase+wHTH+Alpha+Beta-domain+wHTH+Fn-III+Fn-III+RRM | Type-2_BR.RM.like.systems | DUF932->?->ParB-><-HKD-DNase+SF1-SF2-Helicase<-METHYLASE<-DUF3780<-DUF499*<-?<-?<-XIS-HTH\|\|CopG/RHH-><-?\|\|MODE-HTH+PBPII-> | 1038 | Bacteria>Pseudomonadota>Gammaproteobacteria>Pseudomonadales>Pseudomonadaceae | GCF_000764405.1 | Pseudomonas_plecoglossicida | GT93_RS22520 | 4924039..4927155 | 70775 | DUF499-domain-containing-protein-[Pseudomonas-plecoglossicida] |
| WP_034022384.1 | AAA+ATPase+wHTH+Alpha+Beta-domain+wHTH+Fn-III+Fn-III+RRM | Type-2_BR.RM.like.systems | CNMP+TIR->ACYC+ACYC->?->DUF499*->DUF3780->METHYLASE->HKD-DNase+SF1-SF2-Helicase-><-?\|\|MerR-HTH-> | 1038 | Bacteria>Pseudomonadota>Gammaproteobacteria>Pseudomonadales>Pseudomonadaceae | GCF_000792265.1 | Pseudomonas_aeruginosa | NT44_RS12690 | 35095..38211 | 287 | DUF499-domain-containing-protein-[Pseudomonas-aeruginosa] |
| WP_034126986.1 | AAA+ATPase+wHTH+Alpha+Beta-domain+wHTH+Fn-III+Fn-III+RRM | Type-2_BR.RM.like.systems | HTH+MPTase-><-REase<-?<-HKD-DNase+SF1-SF2-Helicase<-METHYLASE<-DUF3780<-DUF499*<-?\|\|TOPRIM->?->Integrase_1->Phage_int_SAM-><-Arm-DNA-bind_3+Phage_integrase | 1038 | Bacteria>Pseudomonadota>Gammaproteobacteria>Pseudomonadales>Pseudomonadaceae | GCF_000801795.1 | Pseudomonas_fluorescens | MZ32_RS11545 | 750418..753534 | 294 | DUF499-domain-containing-protein-[Pseudomonas-fluorescens] |
| WP_058147188.1 | AAA+ATPase+wHTH+Alpha+Beta-domain+wHTH+Fn-III+Fn-III+RRM | Type-2_BR.RM.like.systems | <-MPTase<-?\|\|?-><-?<-Arm-DNA-bind_3+Phage_integrase\|\|RHH->DUF499*->DUF3780->METHYLASE->HKD-DNase+SF1-SF2-Helicase->HEPN-><-?<-Phage_lysozyme | 1040 | Bacteria>Pseudomonadota>Gammaproteobacteria>Pseudomonadales>Pseudomonadaceae | GCF_001449155.1 | Pseudomonas_aeruginosa | AO881_RS09155 | 11696..14818 | 287 | DUF499-domain-containing-protein-[Pseudomonas-aeruginosa] |
| WP_058152388.1 | AAA+ATPase+wHTH+Alpha+Beta-domain+wHTH+Fn-III+Fn-III+RRM | Type-2_BR.RM.like.systems | <-CopG/RHH\|\|XIS-HTH->?->RHH->DUF499*->DUF3780->?->MACRODOMAIN->METHYLASE->HKD-DNase+SF1-SF2-Helicase-> | 1041 | Bacteria>Pseudomonadota>Gammaproteobacteria>Pseudomonadales>Pseudomonadaceae | GCF_001449485.1 | Pseudomonas_aeruginosa | AO904_RS13505 | 165327..168452 | 287 | DUF499-domain-containing-protein-[Pseudomonas-aeruginosa] |
| WP_058181793.1 | AAA+ATPase+wHTH+Alpha+Beta-domain+wHTH+Fn-III+Fn-III+RRM | Type-2_BR.RM.like.systems | <-SF1-SF2-Helicase<-STYKIN<-?<-?<-?\|\|RHH_5->DUF499*->DUF3780->?->MACRODOMAIN->METHYLASE->HKD-DNase+SF1-SF2-Helicase->ParB+HNH-> | 1041 | Bacteria>Pseudomonadota>Gammaproteobacteria>Pseudomonadales>Pseudomonadaceae | GCF_001453595.1 | Pseudomonas_aeruginosa | APA84_RS22695 | 17823..20948 | 287 | DUF499-domain-containing-protein-[Pseudomonas-aeruginosa] |
| WP_059317285.1 | AAA+ATPase+wHTH+Alpha+Beta-domain+wHTH+Fn-III+Fn-III+RRM | Type-2_BR.RM.like.systems | MerR-HTH->tRNA->\|\|TUDOR+SF1-SF2-Helicase+iREase+CTD->?->?->DUF499*->DUF3780->METHYLASE-p->METHYLASE->HKD-DNase+SF1-SF2-Helicase-><-?<-ZNR+Pkinase+SF1-SF2-Helicase+REase | 1038 | Bacteria>Pseudomonadota>Gammaproteobacteria>Pseudomonadales>Pseudomonadaceae | GCF_001516005.1 | Pseudomonas_aeruginosa | HW00_RS22675 | 4884088..4887204 | 287 | DUF499-domain-containing-protein-[Pseudomonas-aeruginosa] |
| WP_043096885.1 | AAA+ATPase+wHTH+Alpha+Beta-domain+wHTH+Fn-III+Fn-III+RRM | Type-2_BR.RM.like.systems | CNMP+TIR->ACYC+ACYC->?->DUF499*->DUF3780->METHYLASE->HKD-DNase+SF1-SF2-Helicase-><-SF1-SF2-Helicase<-OLD-ABC+TOPRIM<-CHTH | 1038 | Bacteria>Pseudomonadota>Gammaproteobacteria>Pseudomonadales>Pseudomonadaceae | GCF_001516245.2 | Pseudomonas_aeruginosa | B8B65_RS21600 | 126942..130058 | 287 | DUF499-domain-containing-protein-[Pseudomonas-aeruginosa] |
| WP_085633597.1 | AAA+ATPase+wHTH+Alpha+Beta-domain+wHTH+Fn-III+Fn-III+RRM | Type-2_BR.RM.like.systems | <-HKD-DNase+SF1-SF2-Helicase<-METHYLASE<-MACRODOMAIN<-?<-DUF3780<-DUF499*<-RHH<-?<-XIS-HTH\|\|CopG/RHH-><-REase | 1041 | Bacteria>Pseudomonadota>Gammaproteobacteria>Pseudomonadales>Pseudomonadaceae | GCF_002113495.1 | Pseudomonas_sp._R16(2017) | CAF93_RS24955 | 18818..21943 | 1981704 | DUF499-domain-containing-protein-[Pseudomonas-sp.-R16(2017)] |
| WP_095039130.1 | AAA+ATPase+wHTH+Alpha+Beta-domain+wHTH+Fn-III+Fn-III+RRM | Type-2_BR.RM.like.systems | XIS-HTH->ZnR+TOPRIM->?->?-><-?\|\|DUF499*->DUF3780->METHYLASE->HKD-DNase+SF1-SF2-Helicase-> | 1049 | Bacteria>Pseudomonadota>Gammaproteobacteria>Pseudomonadales>Pseudomonadaceae | GCF_002269595.1 | Pseudomonas_fragi | CJU72_RS15135 | 99455..102604 | 296 | DUF499-domain-containing-protein-[Pseudomonas-fragi] |
| WP_095598174.1 | AAA+ATPase+wHTH+Alpha+Beta-domain+wHTH+Fn-III+Fn-III+RRM | Type-2_BR.RM.like.systems | <-MORC+REase<-HKD-DNase+SF1-SF2-Helicase<-METHYLASE<-Schlafen+HATPase+HTH<-METHYLASE<-DUF3780<-DUF499*<-RHH | 1041 | Bacteria>Pseudomonadota>Gammaproteobacteria>Pseudomonadales>Pseudomonadaceae | GCF_002286785.1 | Pseudomonas_sp._PIC25 | BZL41_RS04445 | 60362..63487 | 1958773 | DUF499-domain-containing-protein-[Pseudomonas-sp.-PIC25] |
| WP_110748134.1 | AAA+ATPase+wHTH+Alpha+Beta-domain+wHTH+Fn-III+Fn-III+RRM | Type-2_BR.RM.like.systems | <-HTH+MPTase<-?<-?<-HKD-DNase+SF1-SF2-Helicase<-METHYLASE<-DUF3780<-DUF499*<-?<-Phage_int_SAM<-Integrase_1<-?<-TOPRIM<-Arm-DNA-bind_3+Phage_integrase | 1038 | Bacteria>Pseudomonadota>Gammaproteobacteria>Pseudomonadales>Pseudomonadaceae | GCF_003205205.1 | Pseudomonas_sp._LB-090624 | DMX11_RS24825 | 9199..12315 | 2213079 | DUF499-domain-containing-protein-[Pseudomonas-sp.-LB-090624] |
| WP_121373084.1 | AAA+ATPase+wHTH+Alpha+Beta-domain+wHTH+Fn-III+Fn-III+RRM | Type-2_BR.RM.like.systems | REase+SF1-SF2-Helicase->Arm-DNA-bind_3+Phage_integrase-><-HKD-DNase+SF1-SF2-Helicase<-METHYLASE<-DUF3780<-DUF499* | 737 | Bacteria>Pseudomonadota>Gammaproteobacteria>Pseudomonadales>Pseudomonadaceae | GCF_003630125.1 | Pseudomonas_aeruginosa | DQI18_RS19110 | 9389..11604 | 287 | DUF499-domain-containing-protein,-partial-[Pseudomonas-aeruginosa] |
| WP_126517622.1 | AAA+ATPase+wHTH+Alpha+Beta-domain+wHTH+Fn-III+Fn-III+RRM | Type-2_BR.RM.like.systems | ZNR+Pkinase+SF1-SF2-Helicase+REase->?->SymE_toxin->\|\|<-HKD-DNase+SF1-SF2-Helicase<-METHYLASE<-DUF3780<-DUF499*<-RHH_5\|\|?-><-tRNA<-MerR-HTH | 1041 | Bacteria>Pseudomonadota>Gammaproteobacteria>Pseudomonadales>Pseudomonadaceae | GCF_003969255.1 | Pseudomonas_aeruginosa | DY989_RS01785 | 49004..52129 | 287 | DUF499-domain-containing-protein-[Pseudomonas-aeruginosa] |
| WP_128663668.1 | AAA+ATPase+wHTH+Alpha+Beta-domain+wHTH+Fn-III+Fn-III+RRM | Type-2_BR.RM.like.systems | MerR-HTH->tRNA->\|\|TUDOR+SF1-SF2-Helicase+iREase+CTD->?->?->DUF499*->DUF3780->METHYLASE->HKD-DNase+SF1-SF2-Helicase-><-?<-ZNR+Pkinase+SF1-SF2-Helicase+REase | 1038 | Bacteria>Pseudomonadota>Gammaproteobacteria>Pseudomonadales>Pseudomonadaceae | GCF_003973995.1 | Pseudomonas_aeruginosa | IPC1413_RS22670 | 96006..99122 | 287 | DUF499-domain-containing-protein-[Pseudomonas-aeruginosa] |
| WP_134269529.1 | AAA+ATPase+wHTH+Alpha+Beta-domain+wHTH+Fn-III+Fn-III+RRM | Type-2_BR.RM.like.systems | MazG->REase+SF1-SF2-Helicase->Arm-DNA-bind_3+Phage_integrase-><-HKD-DNase+SF1-SF2-Helicase<-METHYLASE<-DUF3780<-DUF499*<-?<-?<-TUDOR+SF1-SF2-Helicase+iREase+CTD<-?<-Peptidase_M15<-HIN-HTH | 1038 | Bacteria>Pseudomonadota>Gammaproteobacteria>Pseudomonadales>Pseudomonadaceae | GCF_004373135.1 | Pseudomonas_aeruginosa | IPC162_RS08525 | 440785..443901 | 287 | DUF499-domain-containing-protein-[Pseudomonas-aeruginosa] |
| WP_134252683.1 | AAA+ATPase+wHTH+Alpha+Beta-domain+wHTH+Fn-III+Fn-III+RRM | Type-2_BR.RM.like.systems | <-HIN-HTH\|\|TUDOR+SF1-SF2-Helicase+iREase+CTD->?->?->DUF499*->DUF3780->METHYLASE->HKD-DNase+SF1-SF2-Helicase->HIN-HTH-> | 1038 | Bacteria>Pseudomonadota>Gammaproteobacteria>Pseudomonadales>Pseudomonadaceae | GCF_004375175.1 | Pseudomonas_aeruginosa | IPC48_RS30590 | 6361..9477 | 287 | DUF499-domain-containing-protein-[Pseudomonas-aeruginosa] |
| WP_153986190.1 | AAA+ATPase+wHTH+Alpha+Beta-domain+wHTH+Fn-III+Fn-III+RRM | Type-2_BR.RM.like.systems | Peptidase_M15->?->Ntox49->TUDOR+SF1-SF2-Helicase+iREase+CTD->?->?->DUF499*->DUF3780->METHYLASE->HKD-DNase+SF1-SF2-Helicase-><-Arm-DNA-bind_3+Phage_integrase<-REase+SF1-SF2-Helicase<-MazG | 1038 | Bacteria>Pseudomonadota>Gammaproteobacteria>Pseudomonadales>Pseudomonadaceae | GCF_009664165.1 | Pseudomonas_aeruginosa | FOZ66_RS12385 | 2584610..2587726 | 287 | DUF499-domain-containing-protein-[Pseudomonas-aeruginosa] |
| WP_158190235.1 | AAA+ATPase+wHTH+Alpha+Beta-domain+wHTH+Fn-III+Fn-III+RRM | Type-2_BR.RM.like.systems | SF1-SF2-Helicase->?->CHTH-><-HKD-DNase+SF1-SF2-Helicase<-METHYLASE<-DUF3780<-DUF499*<-?<-ACYC+ACYC<-CNMP+TIR | 1038 | Bacteria>Pseudomonadota>Gammaproteobacteria>Pseudomonadales>Pseudomonadaceae | GCF_009789555.1 | Stutzerimonas_stutzeri | GQA94_RS22905 | 195258..198374 | 316 | DUF499-domain-containing-protein-[Stutzerimonas-stutzeri] |
| WP_237047019.1 | AAA+ATPase+wHTH+Alpha+Beta-domain+wHTH+Fn-III+Fn-III+RRM | Type-2_BR.RM.like.systems | MazG->REase+SF1-SF2-Helicase->Arm-DNA-bind_3+Phage_integrase-><-HKD-DNase+SF1-SF2-Helicase<-METHYLASE<-DUF3780<-DUF499* | 908 | Bacteria>Pseudomonadota>Gammaproteobacteria>Pseudomonadales>Pseudomonadaceae | GCF_009791355.1 | Pseudomonas_aeruginosa | GQF67_RS10665 | 323805..326532 | 287 | DUF499-domain-containing-protein,-partial-[Pseudomonas-aeruginosa] |
| WP_169989884.1 | AAA+ATPase+wHTH+Alpha+Beta-domain+wHTH+Fn-III+Fn-III+RRM | Type-2_BR.RM.like.systems | REase->?->HTH+MPTase-><-HKD-DNase+SF1-SF2-Helicase<-METHYLASE<-DUF3780<-DUF499*<-?\|\|?->TOPRIM->?->Integrase_1->Phage_int_SAM-> | 1038 | Bacteria>Pseudomonadota>Gammaproteobacteria>Pseudomonadales>Pseudomonadaceae | GCF_012985575.1 | Pseudomonas_sp._WS_5413 | HBO21_RS13365 | 120078..123194 | 2717488 | DUF499-domain-containing-protein-[Pseudomonas-sp.-WS-5413] |
| WP_123186567.1 | AAA+ATPase+wHTH+Alpha+Beta-domain+wHTH+Fn-III+Fn-III+RRM | Type-2_BR.RM.like.systems | Ntox49->TUDOR+SF1-SF2-Helicase+iREase+CTD->?->DUF499*->DUF3780->METHYLASE->HKD-DNase+SF1-SF2-Helicase-><-Arm-DNA-bind_3+Phage_integrase<-REase+SF1-SF2-Helicase<-MazG | 1038 | Bacteria>Pseudomonadota>Gammaproteobacteria>Pseudomonadales>Pseudomonadaceae | GCF_013255565.1 | Pseudomonas | NLZ23_RS29065 | 7522..10638 | 286 | DUF499-domain-containing-protein-[Pseudomonas-aeruginosa] |
| WP_178952339.1 | AAA+ATPase+wHTH+Alpha+Beta-domain+wHTH+Fn-III+Fn-III+RRM | Type-2_BR.RM.like.systems | ParA-Soj-PloopNTPase->?->?->?->DUF499*->DUF3780->METHYLASE->HKD-DNase+SF1-SF2-Helicase-> | 1038 | Bacteria>Pseudomonadota>Gammaproteobacteria>Pseudomonadales>Pseudomonadaceae | GCF_013386335.1 | Pseudomonas_reactans | HX891_RS10935 | 21467..24583 | 117680 | DUF499-domain-containing-protein-[Pseudomonas-reactans] |
| WP_031634902.1 | AAA+ATPase+wHTH+Alpha+Beta-domain+wHTH+Fn-III+Fn-III+RRM | Type-2_BR.RM.like.systems | HIN-HTH->Peptidase_M15->?->TUDOR+SF1-SF2-Helicase+iREase+CTD->?->?->DUF499*->DUF3780->METHYLASE->HKD-DNase+SF1-SF2-Helicase-><-Arm-DNA-bind_3+Phage_integrase<-MazG<-SymE_toxin | 1038 | Bacteria>Pseudomonadota>Gammaproteobacteria>Pseudomonadales>Pseudomonadaceae | GCF_016105665.1 | Pseudomonas | CJU36_RS21560 | 8845..11961 | 286 | MULTISPECIES:-DUF499-domain-containing-protein-[Pseudomonas] |
| WP_210799995.1 | AAA+ATPase+wHTH+Alpha+Beta-domain+wHTH+Fn-III+Fn-III+RRM | Type-2_BR.RM.like.systems | RHH_1->DUF499*->DUF3780->METHYLASE->Schlafen+HATPase+HTH->METHYLASE->HKD-DNase+SF1-SF2-Helicase->MORC+REase-> | 1041 | Bacteria>Pseudomonadota>Gammaproteobacteria>Pseudomonadales>Pseudomonadaceae | GCF_018026755.1 | Pseudomonas_aeruginosa | J5F08_RS20235 | 53337..56462 | 287 | DUF499-domain-containing-protein-[Pseudomonas-aeruginosa] |
| WP_249721383.1 | AAA+ATPase+wHTH+Alpha+Beta-domain+wHTH+Fn-III+Fn-III+RRM | Type-2_BR.RM.like.systems | HTH+MPTase->OLD-ABC+TOPRIM->SF1-SF2-Helicase-><-HKD-DNase+SF1-SF2-Helicase<-METHYLASE<-DUF3780<-DUF499*<-?\|\|?-><-?\|\|HTH_3-><-DUF932 | 1038 | Bacteria>Pseudomonadota>Gammaproteobacteria>Pseudomonadales>Pseudomonadaceae | GCF_018383755.1 | Pseudomonas_syringae | KID89_RS13640 | 3099163..3102279 | 317 | DUF499-domain-containing-protein-[Pseudomonas-syringae] |
| WP_278917457.1 | AAA+ATPase+wHTH+Alpha+Beta-domain+wHTH+Fn-III+Fn-III+RRM | Type-2_BR.RM.like.systems | ZnR+ZnR-><-HTH_3\|\|?->?->?->DUF499*->DUF3780->METHYLASE->HKD-DNase+SF1-SF2-Helicase-><-SF1-SF2-Helicase<-OLD-ABC+TOPRIM | 1038 | Bacteria>Pseudomonadota>Gammaproteobacteria>Pseudomonadales>Pseudomonadaceae | GCF_020744095.1 | Pseudomonas_lactis | K8W20_RS13160 | 6705..9821 | 1615674 | DUF499-domain-containing-protein-[Pseudomonas-lactis] |
| WP_232893124.1 | AAA+ATPase+wHTH+Alpha+Beta-domain+wHTH+Fn-III+Fn-III+RRM | Type-2_BR.RM.like.systems | ParB+HNH-p-><-HKD-DNase+SF1-SF2-Helicase<-METHYLASE<-DUF3780<-DUF499*<-HTH_21+rve+rve_3<-HIN-HTH<-?<-?<-?\|\|MarR-HTH-> | 967 | Bacteria>Pseudomonadota>Gammaproteobacteria>Pseudomonadales>Pseudomonadaceae | GCF_021282385.2 | Pseudomonas_juntendi | LU686_RS25220 | 1011606..1014509 | 2666183 | DUF499-domain-containing-protein-[Pseudomonas-juntendi] |
| WP_236167518.1 | AAA+ATPase+wHTH+Alpha+Beta-domain+wHTH+Fn-III+Fn-III+RRM | Type-2_BR.RM.like.systems | <-ParB+HNH<-ParB+HNH<-HKD-DNase+SF1-SF2-Helicase<-METHYLASE<-DUF3780<-DUF499*<-?<-Phage_int_SAM<-Integrase_1<-?<-TOPRIM<-Arm-DNA-bind_3+Phage_integrase | 1038 | Bacteria>Pseudomonadota>Gammaproteobacteria>Pseudomonadales>Pseudomonadaceae | GCF_021602545.1 | Pseudomonas_fulva | L6D33_RS20200 | 11642..14758 | 47880 | DUF499-domain-containing-protein-[Pseudomonas-fulva] |
| WP_045424232.1 | AAA+ATPase+wHTH+Alpha+Beta-domain+wHTH+Fn-III+Fn-III+RRM | Type-2_BR.RM.like.systems | Glyoxalase-><-?\|\|?->CNMP+TIR->ACYC+ACYC->?->DUF499*->DUF3780->METHYLASE->HKD-DNase+SF1-SF2-Helicase-><-CHTH<-?<-SF1-SF2-Helicase | 1038 | Bacteria>Pseudomonadota>Gammaproteobacteria>Pseudomonadales>Pseudomonadaceae | GCF_024448855.1 | Stutzerimonas_stutzeri | NA633_RS00395 | 79686..82802 | 316 | DUF499-domain-containing-protein-[Stutzerimonas-stutzeri] |
| WP_257375001.1 | AAA+ATPase+wHTH+Alpha+Beta-domain+wHTH+Fn-III+Fn-III+RRM | Type-2_BR.RM.like.systems | ZnR+ZnR-><-HTH_3\|\|?->?->?->DUF499*->DUF3780->METHYLASE->HKD-DNase+SF1-SF2-Helicase-><-SF1-SF2-Helicase<-OLD-ABC+TOPRIM | 1038 | Bacteria>Pseudomonadota>Gammaproteobacteria>Pseudomonadales>Pseudomonadaceae | GCF_024610635.1 | Pseudomonas_sp._T8 | NRG23_RS15195 | 3369528..3372644 | 645292 | DUF499-domain-containing-protein-[Pseudomonas-sp.-T8] |
| WP_262155685.1 | AAA+ATPase+wHTH+Alpha+Beta-domain+wHTH+Fn-III+Fn-III+RRM | Type-2_BR.RM.like.systems | Phage_integrase->?->DUF499*->DUF3780->METHYLASE->HKD-DNase+SF1-SF2-Helicase->HTH_Tnp_1+HTH_21+rve+rve_2-> | 1022 | Bacteria>Pseudomonadota>Gammaproteobacteria>Pseudomonadales>Pseudomonadaceae | GCF_025466035.1 | unclassified_Pseudomonas | NTD88_RS17485 | 21326..24394 | 196821 | MULTISPECIES:-DUF499-domain-containing-protein-[unclassified-Pseudomonas] |
| WP_263227050.1 | AAA+ATPase+wHTH+Alpha+Beta-domain+wHTH+Fn-III+Fn-III+RRM | Type-2_BR.RM.like.systems | DUF932->?->ParB-><-HKD-DNase+SF1-SF2-Helicase<-METHYLASE<-DUF3780<-DUF499*<-?<-?<-XIS-HTH\|\|CopG/RHH-> | 1038 | Bacteria>Pseudomonadota>Gammaproteobacteria>Pseudomonadales>Pseudomonadaceae | GCF_025642795.1 | Pseudomonas_entomophila | OGL91_RS16850 | 37362..40478 | 312306 | DUF499-domain-containing-protein-[Pseudomonas-entomophila] |
| WP_273622718.1 | AAA+ATPase+wHTH+Alpha+Beta-domain+wHTH+Fn-III+Fn-III+RRM | Type-2_BR.RM.like.systems | DUF499*->DUF3780->METHYLASE->HKD-DNase+SF1-SF2-Helicase-><-Arm-DNA-bind_3+Phage_integrase | 1038 | Bacteria>Pseudomonadota>Gammaproteobacteria>Pseudomonadales>Pseudomonadaceae | GCF_028595385.1 | Pseudomonas_aeruginosa | PUL38_RS31525 | 996..4112 | 287 | DUF499-domain-containing-protein-[Pseudomonas-aeruginosa] |
| WP_172693438.1 | AAA+ATPase+wHTH+Alpha+Beta-domain+wHTH+Fn-III+Fn-III+RRM | Type-2_BR.RM.like.systems | RelE-ParE->HTH_3->SbcC-><-HKD-DNase+SF1-SF2-Helicase<-METHYLASE<-DUF3780<-DUF499* | 1038 | Bacteria>Pseudomonadota>Gammaproteobacteria>Pseudomonadales>Pseudomonadaceae | GCF_035019465.1 | Pseudomonas | HTS36_RS00745 | 135940..139056 | 286 | DUF499-domain-containing-protein-[Pseudomonas-fluorescens] |
| WP_069673157.1 | AAA+ATPase+wHTH+Alpha+Beta-domain+wHTH+Fn-III+Fn-III+RRM | Type-2_BR.RM.like.systems | <-Phage_int_SAM<-Integrase_1<-?<-TOPRIM<-?\|\|?->DUF499*->DUF3780->METHYLASE->HKD-DNase+SF1-SF2-Helicase-><-HTH+MPTase<-?<-REase | 1038 | Bacteria>Pseudomonadota>Gammaproteobacteria>Pseudomonadales>Pseudomonadaceae | GCF_900111895.1 | Pseudomonas_simiae | BMZ57_RS20825 | 83316..86432 | 321846 | DUF499-domain-containing-protein-[Pseudomonas-simiae] |
| WP_109421334.1 | AAA+ATPase+wHTH+Alpha+Beta-domain+wHTH+Fn-III+Fn-III+RRM | Type-2_BR.RM.like.systems | <-HNH+SAVED\|\|?->DUF499*->DUF3780->METHYLASE->HKD-DNase+SF1-SF2-Helicase-><-SF1-SF2-Helicase<-OLD-ABC+TOPRIM<-CHTH | 1038 | Bacteria>Pseudomonadota>Gammaproteobacteria>Pseudomonadales>Pseudomonadaceae | GCF_900406905.1 | Pseudomonas_aeruginosa | DK331_RS20265 | 35149..38265 | 287 | DUF499-domain-containing-protein-[Pseudomonas-aeruginosa] |
| WP_202808302.1 | AAA+ATPase+wHTH+Alpha+Beta-domain+wHTH+Fn-III+Fn-III+RRM | Type-2_BR.RM.like.systems | CNMP+TIR->ACYC+ACYC->?->DUF499*->DUF3780->METHYLASE->HKD-DNase+SF1-SF2-Helicase-><-SF1-SF2-Helicase<-OLD-ABC+TOPRIM<-CHTH | 1072 | Bacteria>Pseudomonadota>Gammaproteobacteria>Pseudomonadales>Pseudomonadaceae | - | Pseudomonas_aeruginosa | NHK89_004125 | 34242..37460 | 287 | DUF499-domain-containing-protein-[Pseudomonas-aeruginosa] |
| WP_213155768.1 | AAA+ATPase+wHTH+Alpha+Beta-domain+wHTH+Fn-III+Fn-III+RRM | Type-2_BR.RM.like.systems | ZNR+Pkinase+SF1-SF2-Helicase+REase->?-><-HKD-DNase+SF1-SF2-Helicase<-METHYLASE<-DUF3780<-DUF499*<-?<-?<-TUDOR+SF1-SF2-Helicase+iREase+CTD | 1072 | Bacteria>Pseudomonadota>Gammaproteobacteria>Pseudomonadales>Pseudomonadaceae | - | Pseudomonas_aeruginosa | NJF72_002791 | 346926..350144 | 287 | DUF499-domain-containing-protein-[Pseudomonas-aeruginosa] |
| WP_221768843.1 | AAA+ATPase+wHTH+Alpha+Beta-domain+wHTH+Fn-III+Fn-III+RRM | Type-2_BR.RM.like.systems | <-HIN-HTH\|\|TUDOR+SF1-SF2-Helicase+iREase+CTD->?->?->DUF499*->DUF3780->METHYLASE->HKD-DNase+SF1-SF2-Helicase-><-?<-ZNR+Pkinase+SF1-SF2-Helicase+REase | 1072 | Bacteria>Pseudomonadota>Gammaproteobacteria>Pseudomonadales>Pseudomonadaceae | - | Pseudomonas_aeruginosa | ONA48_004000 | 6159..9377 | 287 | DUF499-domain-containing-protein-[Pseudomonas-aeruginosa] |
| WP_206437318.1 | AAA+ATPase+wHTH+Alpha+Beta-domain+wHTH+Fn-III+Fn-III+RRM | Type-2_BR.RM.like.systems | Peptidase_M15->?->Ntox49->TUDOR+SF1-SF2-Helicase+iREase+CTD->?->?->DUF499*->DUF3780->METHYLASE->HKD-DNase+SF1-SF2-Helicase-><-Arm-DNA-bind_3+Phage_integrase<-REase+SF1-SF2-Helicase<-MazG | 1072 | Bacteria>Pseudomonadota>Gammaproteobacteria>Pseudomonadales>Pseudomonadaceae | - | Pseudomonas_aeruginosa | PQC20_002029 | 43524..46742 | 287 | DUF499-domain-containing-protein-[Pseudomonas-aeruginosa] |
| WP_254873384.1 | AAA+ATPase+wHTH+Alpha+Beta-domain+wHTH+Fn-III+Fn-III+RRM | Type-2_BR.RM.like.systems | <-ParA-Soj-PloopNTPase<-HEPN<-METHYLASE<-MACRODOMAIN<-?<-DUF3780<-DUF499*<-RHH | 1036 | Bacteria>Pseudomonadota>Gammaproteobacteria>Pseudomonadales>Pseudomonadaceae | GCF_003411965.2 | Pseudomonas_aeruginosa | DZ913_RS12395 | 105397..108507 | 287 | DUF499-domain-containing-protein-[Pseudomonas-aeruginosa] |
| WP_073667316.1 | AAA+ATPase+wHTH+Alpha+Beta-domain+wHTH+Fn-III+Fn-III+RRM | Type-2_BR.RM.like.systems | RHH_5->DUF499*->DUF3780->?->MACRODOMAIN->METHYLASE->HEPN->ParA-Soj-PloopNTPase-> | 1041 | Bacteria>Pseudomonadota>Gammaproteobacteria>Pseudomonadales>Pseudomonadaceae | GCF_900708645.1 | Pseudomonas_aeruginosa | CHH44_RS18170 | 263427..266552 | 287 | DUF499-domain-containing-protein-[Pseudomonas-aeruginosa] |
| MBV1952380.1 | AAA+ATPase+wHTH+Alpha+Beta-domain+wHTH+Fn-III+Fn-III+RRM | Type-2_BR.RM.like.systems | RHH_1->DUF499*->DUF3780->METHYLASE->HKD-DNase+SF1-SF2-Helicase->SF1-SF2-Helicase->REase-> | 1043 | Bacteria>Pseudomonadota>Gammaproteobacteria>Thiotrichales>Piscirickettsiaceae | GCA_019090295.1 | Cycloclasticus_sp. | KUG64_09355 | 786..3917 | 2024830 | DUF499-domain-containing-protein-[Cycloclasticus-sp.] |
| WP_277642465.1 | AAA+ATPase+wHTH+Alpha+Beta-domain+wHTH+Fn-III+Fn-III+RRM | Type-2_BR.RM.like.systems | <-HKD-DNase+SF1-SF2-Helicase<-METHYLASE<-DUF3780<-DUF499*\|\|?-><-?<-XIS-HTH<-?<-Arm-DNA-bind_3+Phage_integrase | 962 | Bacteria>Pseudomonadota>Gammaproteobacteria>Thiotrichales>Piscirickettsiaceae | GCF_029212605.1 | Cycloclasticus_pugetii | P1U51_RS07825 | 6293..9181 | 34068 | DUF499-domain-containing-protein,-partial-[Cycloclasticus-pugetii] |
| GAW87350.1 | AAA+ATPase+wHTH+Alpha+Beta-domain+wHTH+Fn-III+Fn-III+RRM | Type-2_BR.RM.like.systems | DUF499*->DUF3780->METHYLASE->HKD-DNase+SF1-SF2-Helicase-> | 703 | Bacteria>Pseudomonadota>Gammaproteobacteria>unclassified_Gammaproteobacteria | GCA_002189065.1 | Bathymodiolus_platifrons_methanotrophic_gill_symbiont | bplSymb_SCF07702P001 | 1761..3872 | 113268 | hypothetical-protein-bplSymb_SCF07702P001-[Bathymodiolus-platifrons-methanotrophic-gill-symbiont] |
| MBK80531.1 | AAA+ATPase+wHTH+Alpha+Beta-domain+wHTH+Fn-III+Fn-III+RRM | Type-2_BR.RM.like.systems | TraH->?->TraG_N-><-HKD-DNase+SF1-SF2-Helicase<-METHYLASE<-DUF3780<-DUF499*<-RHH_5\|\|?->?->HerA/FtsK->?-><-Arm-DNA-bind_3+Phage_integrase | 1040 | Bacteria>Pseudomonadota>Gammaproteobacteria>unclassified_Gammaproteobacteria | GCA_002720645.1 | Gammaproteobacteria_bacterium | CMQ43_06390 | 60391..63513 | 1913989 | AAA-family-ATPase-[Gammaproteobacteria-bacterium] |
| PUB90401.1 | AAA+ATPase+wHTH+Alpha+Beta-domain+wHTH+Fn-III+Fn-III+RRM | Type-2_BR.RM.like.systems | DUF499*->DUF3780->METHYLASE->HKD-DNase+SF1-SF2-Helicase->wHTH+REase->HTH-> | 1012 | Bacteria>Pseudomonadota>Gammaproteobacteria>unclassified_Gammaproteobacteria | GCA_003058535.1 | gamma_proteobacterium_symbiont_of_Ctena_orbiculata | DBP01_07750 | 1..3039 | 1968598 | AAA-family-ATPase,-partial-[gamma-proteobacterium-symbiont-of-Ctena-orbiculata] |
| QFY90794.1 | AAA+ATPase+wHTH+Alpha+Beta-domain+wHTH+Fn-III+Fn-III+RRM | Type-2_BR.RM.like.systems | XIS-HTH->DUF499*->DUF3780->METHYLASE->HEPN->ParA-Soj-PloopNTPase->ORC-CDC6->HKD-DNase+SF1-SF2-Helicase-> | 1040 | Bacteria>Pseudomonadota>Gammaproteobacteria>unclassified_Gammaproteobacteria | GCA_009497875.2 | gamma_proteobacterium_SS-5 | D5125_15735 | 2296694..2299816 | 947516 | DUF499-domain-containing-protein-[gamma-proteobacterium-SS-5] |
| MXW50605.1 | AAA+ATPase+wHTH+Alpha+Beta-domain+wHTH+Fn-III+Fn-III+RRM | Type-2_BR.RM.like.systems | <-HicB\|\|DUF499*->DUF3780->METHYLASE->PHP+SbcC->Schlafen+HATPase+HTH->HKD-DNase+SF1-SF2-Helicase-> | 1019 | Bacteria>Pseudomonadota>Gammaproteobacteria>unclassified_Gammaproteobacteria | GCA_009835805.1 | Gammaproteobacteria_bacterium | F4Z84_10190 | 3046..6105 | 1913989 | DUF499-domain-containing-protein-[Gammaproteobacteria-bacterium] |
| MBU0657159.1 | AAA+ATPase+wHTH+Alpha+Beta-domain+wHTH+Fn-III+Fn-III+RRM | Type-2_BR.RM.like.systems | Phage_integrase-><-SF1-SF2-Helicase<-?<-HKD-DNase+SF1-SF2-Helicase<-METHYLASE<-DUF3780<-DUF499*<-ParB+HNH+Schlafen\|\|N6_N4_MTase->?->Mbetalac->?->XIS-HTH-> | 1043 | Bacteria>Pseudomonadota>Gammaproteobacteria>unclassified_Gammaproteobacteria | GCA_018813855.1 | Gammaproteobacteria_bacterium | KJ914_18710 | 11621..14752 | 1913989 | DUF499-domain-containing-protein-[Gammaproteobacteria-bacterium] |
| MCB1859932.1 | AAA+ATPase+wHTH+Alpha+Beta-domain+wHTH+Fn-III+Fn-III+RRM | Type-2_BR.RM.like.systems | <-HKD-DNase+SF1-SF2-Helicase<-METHYLASE<-DUF3780<-DUF499*<-ParB+HNH<-XIS-HTH\|\|TIR-><-TraG_N | 1043 | Bacteria>Pseudomonadota>Gammaproteobacteria>unclassified_Gammaproteobacteria | GCA_020445415.1 | Gammaproteobacteria_bacterium | KDI63_16790 | 5935..9066 | 1913989 | DUF499-domain-containing-protein-[Gammaproteobacteria-bacterium] |
| MCY4014266.1 | AAA+ATPase+wHTH+Alpha+Beta-domain+wHTH+Fn-III+Fn-III+RRM | Type-2_BR.RM.like.systems | <-HKD-DNase+SF1-SF2-Helicase<-Schlafen+HATPase+HTH<-PHP+SbcC<-METHYLASE<-DUF3780<-DUF499* | 1023 | Bacteria>Pseudomonadota>Gammaproteobacteria>unclassified_Gammaproteobacteria | GCA_026708185.1 | Gammaproteobacteria_bacterium | OXG82_16305 | 63884..66955 | 1913989 | MAG:-DUF499-domain-containing-protein-[Gammaproteobacteria-bacterium] |
| MDE0192036.1 | AAA+ATPase+wHTH+Alpha+Beta-domain+wHTH+Fn-III+Fn-III+RRM | Type-2_BR.RM.like.systems | <-HKD-DNase+SF1-SF2-Helicase<-REase<-N6_MTase<-METHYLASE<-DUF3780<-DUF499*<-RelE-ParE<-OLD-ABC<-ParB+HNH | 1021 | Bacteria>Pseudomonadota>Gammaproteobacteria>unclassified_Gammaproteobacteria | GCA_028819695.1 | Gammaproteobacteria_bacterium | OXQ90_11825 | 72946..76011 | 1913989 | MAG:-DUF499-domain-containing-protein-[Gammaproteobacteria-bacterium] |
| WP_087868382.1 | AAA+ATPase+wHTH+Alpha+Beta-domain+wHTH+Fn-III+Fn-III+RRM | Type-2_BR.RM.like.systems | <-wHTH+REase<-HKD-DNase+SF1-SF2-Helicase<-METHYLASE<-?<-DUF3780<-DUF499*<-RHH<-XIS-HTH | 1041 | Bacteria>Pseudomonadota>Gammaproteobacteria>unclassified_Gammaproteobacteria | GCF_002189065.1 | Bathymodiolus_platifrons_methanotrophic_gill_symbiont | CDZ78_RS01565 | 51590..54715 | 113268 | DUF499-domain-containing-protein-[Bathymodiolus-platifrons-methanotrophic-gill-symbiont] |
| MDD9983184.1 | AAA+ATPase+wHTH+Alpha+Beta-domain+wHTH+Fn-III+Fn-III+RRM | Type-2_BR.RM.like.systems | ABhydrolase->DUF499*->DUF3780->METHYLASE-p->?->METHYLASE-p->ParB+HNH->HEPN-> | 1021 | Bacteria>Pseudomonadota>Gammaproteobacteria>unclassified_Gammaproteobacteria | GCA_028818395.1 | Gammaproteobacteria_bacterium | OXU81_17805 | 2420..5485 | 1913989 | MAG:-DUF499-domain-containing-protein-[Gammaproteobacteria-bacterium] |
| MCC5517983.1 | AAA+ATPase+wHTH+Alpha+Beta-domain+wHTH+Fn-III+Fn-III+RRM | Type-2_BR.RM.like.systems | <-JAB\|\|RNase->XIS-HTH->RHH->DUF499*->DUF3780+RAMA->METHYLASE->HKD-DNase+SF1-SF2-Helicase-><-Arm-DNA-bind_3+Phage_int_SAM_3+Phage_integrase | 1041 | Bacteria>Pseudomonadota>Gammaproteobacteria>Vibrionales>Vibrionaceae | GCA_002876945.2 | Vibrio_splendidus | BCT65_013080 | 411206..414331 | 29497 | DUF499-domain-containing-protein-[Vibrio-splendidus] |
| QLK44250.1 | AAA+ATPase+wHTH+Alpha+Beta-domain+wHTH+Fn-III+Fn-III+RRM | Type-2_BR.RM.like.systems | HTH+WYL-><-HKD-DNase+SF1-SF2-Helicase<-METHYLASE<-DUF3780+RAMA<-DUF499*<-RHH\|\|HEPN-><-HNH<-XIS-HTH\|\|JAB-> | 1041 | Bacteria>Pseudomonadota>Gammaproteobacteria>Vibrionales>Vibrionaceae | GCA_013459835.1 | Vibrio_owensii | DR996_02410 | 549792..552917 | 696485 | AAA-family-ATPase-[Vibrio-owensii] |
| MBE4228752.1 | AAA+ATPase+wHTH+Alpha+Beta-domain+wHTH+Fn-III+Fn-III+RRM | Type-2_BR.RM.like.systems | <-VirD4-TraG+VirB4-FtsK<-HDRNase<-wHTH+REase<-HKD-DNase+SF1-SF2-Helicase<-DUF3780+RAMA<-DUF499*<-RHH_1<-35exo | 1041 | Bacteria>Pseudomonadota>Gammaproteobacteria>Vibrionales>Vibrionaceae | GCA_014920775.1 | Vibrio_parahaemolyticus | HJ082_06195 | 253692..256817 | 670 | DUF499-domain-containing-protein-[Vibrio-parahaemolyticus] |
| EGQ8040269.1 | AAA+ATPase+wHTH+Alpha+Beta-domain+wHTH+Fn-III+Fn-III+RRM | Type-2_BR.RM.like.systems | Arm-DNA-bind_3+Phage_integrase-><-MACRODOMAIN<-?<-HKD-DNase+SF1-SF2-Helicase<-METHYLASE<-DUF3780+RAMA<-DUF499*<-RHH<-XIS-HTH<-RNase\|\|JAB-> | 1041 | Bacteria>Pseudomonadota>Gammaproteobacteria>Vibrionales>Vibrionaceae | GCA_015955145.1 | Vibrio_alginolyticus | HJA73_001778 | 380130..383255 | 663 | DUF499-domain-containing-protein-[Vibrio-alginolyticus] |
| MBS9942123.1 | AAA+ATPase+wHTH+Alpha+Beta-domain+wHTH+Fn-III+Fn-III+RRM | Type-2_BR.RM.like.systems | Arm-DNA-bind_3+Phage_integrase-><-MACRODOMAIN<-?<-HKD-DNase+SF1-SF2-Helicase<-METHYLASE<-DUF3780+RAMA<-DUF499*<-RHH<-XIS-HTH<-RNase\|\|JAB-> | 1041 | Bacteria>Pseudomonadota>Gammaproteobacteria>Vibrionales>Vibrionaceae | GCA_018458105.1 | Vibrio_alginolyticus | J4H27_09040 | 373054..376179 | 663 | DUF499-domain-containing-protein-[Vibrio-alginolyticus] |
| MBY7816961.1 | AAA+ATPase+wHTH+Alpha+Beta-domain+wHTH+Fn-III+Fn-III+RRM | Type-2_BR.RM.like.systems | 35exo->XIS-HTH->DUF499*->DUF3780->METHYLASE->HKD-DNase+SF1-SF2-Helicase->wHTH+REase->HDRNase->VirD4-TraG+VirB4-FtsK-> | 1022 | Bacteria>Pseudomonadota>Gammaproteobacteria>Vibrionales>Vibrionaceae | GCA_019852895.1 | Vibrio_fluvialis | KW416_13045 | 310292..313360 | 676 | DUF499-domain-containing-protein-[Vibrio-fluvialis] |
| MCE9842953.1 | AAA+ATPase+wHTH+Alpha+Beta-domain+wHTH+Fn-III+Fn-III+RRM | Type-2_BR.RM.like.systems | <-JAB\|\|?-><-DDE_Tnp_1_5+DDE_Tnp_1\|\|XIS-HTH->RHH->DUF499*->DUF3780+RAMA->METHYLASE->HKD-DNase+SF1-SF2-Helicase->?->MACRODOMAIN-><-Arm-DNA-bind_3+Phage_integrase | 1022 | Bacteria>Pseudomonadota>Gammaproteobacteria>Vibrionales>Vibrionaceae | GCA_021440305.1 | Vibrio_antiquarius | LZ666_05745 | 166857..169925 | 150340 | DUF499-domain-containing-protein-[Vibrio-antiquarius] |
| MCR9497773.1 | AAA+ATPase+wHTH+Alpha+Beta-domain+wHTH+Fn-III+Fn-III+RRM | Type-2_BR.RM.like.systems | Arm-DNA-bind_3+Phage_integrase-><-MACRODOMAIN<-?<-HKD-DNase+SF1-SF2-Helicase<-METHYLASE<-DUF3780+RAMA<-DUF499*<-RHH<-XIS-HTH<-RNase\|\|JAB-> | 1022 | Bacteria>Pseudomonadota>Gammaproteobacteria>Vibrionales>Vibrionaceae | GCA_024744525.1 | Vibrio_alginolyticus | NB493_20505 | 383209..386277 | 663 | DUF499-domain-containing-protein-[Vibrio-alginolyticus] |
| EKO3646393.1 | AAA+ATPase+wHTH+Alpha+Beta-domain+wHTH+Fn-III+Fn-III+RRM | Type-2_BR.RM.like.systems | <-VirD4-TraG+VirB4-FtsK<-HDRNase<-wHTH+REase<-HKD-DNase+SF1-SF2-Helicase<-METHYLASE<-DUF3780+RAMA<-DUF499*<-RHH_1<-35exo | 1041 | Bacteria>Pseudomonadota>Gammaproteobacteria>Vibrionales>Vibrionaceae | GCA_028989625.1 | Vibrio_metschnikovii | M3925_003006 | 63497..66622 | 28172 | DUF499-domain-containing-protein-[Vibrio-metschnikovii] |
| EKO3692967.1 | AAA+ATPase+wHTH+Alpha+Beta-domain+wHTH+Fn-III+Fn-III+RRM | Type-2_BR.RM.like.systems | 35exo->XIS-HTH->DUF499*->DUF3780->METHYLASE->HKD-DNase+SF1-SF2-Helicase->wHTH+REase->HDRNase->VirD4-TraG+VirB4-FtsK-> | 1022 | Bacteria>Pseudomonadota>Gammaproteobacteria>Vibrionales>Vibrionaceae | GCA_028990005.1 | Vibrio_metschnikovii | POF02_001010 | 192750..195818 | 28172 | ATP-binding-protein-[Vibrio-metschnikovii] |
| WP_017062798.1 | AAA+ATPase+wHTH+Alpha+Beta-domain+wHTH+Fn-III+Fn-III+RRM | Type-2_BR.RM.like.systems | Arm-DNA-bind_3+Phage_integrase-><-HKD-DNase+SF1-SF2-Helicase<-DUF3780+RAMA<-DUF499*<-RHH<-XIS-HTH<-RNase\|\|JAB-> | 1041 | Bacteria>Pseudomonadota>Gammaproteobacteria>Vibrionales>Vibrionaceae | GCF_000272045.2 | Vibrio_crassostreae | A136_RS10915 | 27154..30279 | 246167 | DUF499-domain-containing-protein-[Vibrio-crassostreae] |
| WP_036795924.1 | AAA+ATPase+wHTH+Alpha+Beta-domain+wHTH+Fn-III+Fn-III+RRM | Type-2_BR.RM.like.systems | Arm-DNA-bind_3+Phage_int_SAM_3+Phage_integrase->MULE-TRANSPOSASE-><-HKD-DNase+SF1-SF2-Helicase<-METHYLASE<-DUF3780+RAMA<-DUF499*<-RHH_1\|\|JAB->?->?-><-XIS-HTH | 1041 | Bacteria>Pseudomonadota>Gammaproteobacteria>Vibrionales>Vibrionaceae | GCF_000613045.3 | Photobacterium_kishitanii | PPBDW_RS12780 | 2735530..2738655 | 318456 | DUF499-domain-containing-protein-[Photobacterium-kishitanii] |
| WP_203342017.1 | AAA+ATPase+wHTH+Alpha+Beta-domain+wHTH+Fn-III+Fn-III+RRM | Type-2_BR.RM.like.systems | <-VirD4-TraG+VirB4-FtsK<-HDRNase<-wHTH+REase<-HKD-DNase+SF1-SF2-Helicase<-METHYLASE<-DUF3780+RAMA<-DUF499*<-?<-DUF499-p<-RHH_1<-35exo | 782 | Bacteria>Pseudomonadota>Gammaproteobacteria>Vibrionales>Vibrionaceae | GCF_014525255.1 | Vibrio_metschnikovii | GO180_RS04985 | 107142..109490 | 28172 | DUF499-domain-containing-protein-[Vibrio-metschnikovii] |
| WP_250156807.1 | AAA+ATPase+wHTH+Alpha+Beta-domain+wHTH+Fn-III+Fn-III+RRM | Type-2_BR.RM.like.systems | Arm-DNA-bind_3+Phage_integrase-><-MACRODOMAIN<-?<-HKD-DNase+SF1-SF2-Helicase<-METHYLASE<-DUF3780+RAMA<-DUF499*<-RHH<-XIS-HTH<-RNase\|\|JAB-> | 1022 | Bacteria>Pseudomonadota>Gammaproteobacteria>Vibrionales>Vibrionaceae | GCF_018458105.1 | Vibrio_alginolyticus | J4H27_RS09040 | 373054..376122 | 663 | DUF499-domain-containing-protein-[Vibrio-alginolyticus] |
| WP_086597666.1 | AAA+ATPase+wHTH+Alpha+Beta-domain+wHTH+Fn-III+Fn-III+RRM | Type-2_BR.RM.like.systems | <-JAB\|\|RNase->XIS-HTH->RHH->DUF499*->DUF3780+RAMA->METHYLASE->HKD-DNase+SF1-SF2-Helicase->?->MACRODOMAIN-><-Arm-DNA-bind_3+Phage_integrase | 1041 | Bacteria>Pseudomonadota>Gammaproteobacteria>Vibrionales>Vibrionaceae | GCF_019846795.1 | Vibrio_parahaemolyticus | D5E77_RS03755 | 157307..160432 | 670 | DUF499-domain-containing-protein-[Vibrio-parahaemolyticus] |
| WP_258489585.1 | AAA+ATPase+wHTH+Alpha+Beta-domain+wHTH+Fn-III+Fn-III+RRM | Type-2_BR.RM.like.systems | <-JAB\|\|?-><-DDE_Tnp_1_5+DDE_Tnp_1\|\|XIS-HTH->RHH->DUF499*->DUF3780+RAMA->METHYLASE->HKD-DNase+SF1-SF2-Helicase->?->MACRODOMAIN-><-Arm-DNA-bind_3+Phage_integrase | 1022 | Bacteria>Pseudomonadota>Gammaproteobacteria>Vibrionales>Vibrionaceae | GCF_021440305.1 | Vibrio_harveyi_group | LZ666_RS05760 | 166857..169925 | 717610 | DUF499-domain-containing-protein-[Vibrio-alginolyticus] |
| WP_238130455.1 | AAA+ATPase+wHTH+Alpha+Beta-domain+wHTH+Fn-III+Fn-III+RRM | Type-2_BR.RM.like.systems | <-VirD4-TraG+VirB4-FtsK<-HDRNase<-wHTH+REase<-HKD-DNase+SF1-SF2-Helicase<-METHYLASE<-DUF3780+RAMA<-DUF499*<-RHH_1<-35exo | 1041 | Bacteria>Pseudomonadota>Gammaproteobacteria>Vibrionales>Vibrionaceae | GCF_022114045.1 | Vibrio_cincinnatiensis | EXA22_RS10090 | 12757..15882 | 675 | DUF499-domain-containing-protein-[Vibrio-cincinnatiensis] |

| Protein-ID | DUF499-architecture | System | Gene-Neighbourhoods | Length | Taxonomy | Assembly-ID | Strain | Locus-Tag | Genomic-coordinates | TaxID | Fasta-define |
|---|---|---|---|---|---|---|---|---|---|---|---|
| WP_123015107.1 | AAA+ATPase+wHTH+Alpha+Beta-domain+wHTH+Fn-III+Fn-III+RRM | Type-2_BR.RM.like.systems | <-JAB\|\|RNase->XIS-HTH->RHH->DUF499*->DUF3780+RAMA->METHYLASE->HKD-DNase+SF1-SF2-Helicase->?->MACRODOMAIN-><-Arm-DNA-bind_3+Phage_integrase | 1041 | Bacteria>Pseudomonadota>Gammaproteobacteria>Vibrionales>Vibrionaceae | GCF_024347155.1 | Vibrio_zhugei | EAE30_RS05630 | 117440..120565 | 2479546 | DUF499-domain-containing-protein-[Vibrio-zhugei] |
| WP_269977406.1 | AAA+ATPase+wHTH+Alpha+Beta-domain+wHTH+Fn-III+Fn-III+RRM | Type-2_BR.RM.like.systems | <-JAB\|\|RNase->XIS-HTH->RHH->DUF499*->DUF3780+RAMA->METHYLASE->HKD-DNase+SF1-SF2-Helicase-><-Arm-DNA-bind_3+Phage_integrase | 1041 | Bacteria>Pseudomonadota>Gammaproteobacteria>Vibrionales>Vibrionaceae | GCF_027587055.1 | Vibrio_sp._MarTm2 | OH458_RS04465 | 5022..8147 | 2998831 | DUF499-domain-containing-protein-[Vibrio-sp.-MarTm2] |
| WP_274682768.1 | AAA+ATPase+wHTH+Alpha+Beta-domain+wHTH+Fn-III+Fn-III+RRM | Type-2_BR.RM.like.systems | <-JAB\|\|RHH_1->DUF499*->DUF3780+RAMA->METHYLASE->HKD-DNase+SF1-SF2-Helicase->?-><-?<-Arm-DNA-bind_3+Phage_integrase | 1041 | Bacteria>Pseudomonadota>Gammaproteobacteria>Vibrionales>Vibrionaceae | GCF_028858065.1 | Vibrio_aestuarianus | L9W94_RS04800 | 100542..103667 | 28171 | DUF499-domain-containing-protein-[Vibrio-aestuarianus] |
| WP_228607499.1 | AAA+ATPase+wHTH+Alpha+Beta-domain+wHTH+Fn-III+Fn-III+RRM | Type-2_BR.RM.like.systems | <-JAB\|\|RNase->XIS-HTH->RHH->DUF499*->DUF3780+RAMA->METHYLASE->HKD-DNase+SF1-SF2-Helicase-><-Arm-DNA-bind_3+Phage_int_SAM_3+Phage_integrase | 1022 | Bacteria>Pseudomonadota>Gammaproteobacteria>Vibrionales>Vibrionaceae | - | Vibrio_splendidus | BCT65_RS13195 | 411263..414331 | 29497 | DUF499-domain-containing-protein-[Vibrio-splendidus] |
| MBK8067117.1 | AAA+ATPase+wHTH+Alpha+Beta-domain+wHTH+Fn-III+Fn-III+RRM | Type-2_BR.RM.like.systems | <-REase+35exo<-REase+35exo\|\|?->HTH+WYL->XIS-HTH->DUF499*->DUF3780->METHYLASE->HKD-DNase+SF1-SF2-Helicase->ParB+HNH-> | 1046 | Bacteria>Pseudomonadota>Gammaproteobacteria>Xanthomonadales>Rhodanobacteraceae | GCA_016713135.1 | Rhodanobacteraceae_bacterium | IPK27_05685 | 321286..324426 | 2052182 | ATP-binding-protein-[Rhodanobacteraceae-bacterium] |
| MBL8298151.1 | AAA+ATPase+wHTH+Alpha+Beta-domain+wHTH+Fn-III+Fn-III+RRM | Type-2_BR.RM.like.systems | <-HKD-DNase+SF1-SF2-Helicase<-METHYLASE<-MACRODOMAIN<-?<-DUF3780<-DUF499*<-RHH_1<-?<-TPR+TPR | 1040 | Bacteria>Pseudomonadota>Gammaproteobacteria>Xanthomonadales>Rhodanobacteraceae | GCA_016790075.1 | Rhodanobacteraceae_bacterium | JNN30_07385 | 50639..53761 | 2052182 | DUF499-domain-containing-protein-[Rhodanobacteraceae-bacterium] |
| MCW0201519.1 | AAA+ATPase+wHTH+Alpha+Beta-domain+wHTH+Fn-III+Fn-III+RRM | Type-2_BR.RM.like.systems | <-HKD-DNase+SF1-SF2-Helicase<-METHYLASE<-DUF3780<-DUF499*<-?<-HTH\|\|HTH->STYKinase-like->?->HTH+WYL-> | 1043 | Bacteria>Pseudomonadota>Gammaproteobacteria>Xanthomonadales>Rhodanobacteraceae | GCA_025869415.1 | Rhodanobacter_thiooxydans | OJI74_06590 | 59621..62752 | 416169 | MAG:-DUF499-domain-containing-protein-[Rhodanobacter-thiooxydans] |
| WP_039006157.1 | AAA+ATPase+wHTH+Alpha+Beta-domain+wHTH+Fn-III+Fn-III+RRM | Type-2_BR.RM.like.systems | <-HKD-DNase+SF1-SF2-Helicase<-METHYLASE<-MACRODOMAIN<-?<-DUF3780<-DUF499*<-RHH_5\|\|?->ABhydrolase->?-><-?<-TetR-HTH | 1041 | Bacteria>Pseudomonadota>Gammaproteobacteria>Xanthomonadales>Xanthomonadaceae | GCF_000807145.1 | Xanthomonas_translucens | KHA79_RS02625 | 616013..619138 | 343 | DUF499-domain-containing-protein-[Xanthomonas-translucens] |
| WP_076739482.1 | AAA+ATPase+wHTH+Alpha+Beta-domain+wHTH+Fn-III+Fn-III+RRM | Type-2_BR.RM.like.systems | Methyltransf_11->RNase->35exo-><-?\|\|tRNA->RHH_5->DUF499*->DUF3780->?->MACRODOMAIN->METHYLASE->HKD-DNase+SF1-SF2-Helicase->DUF932-> | 1041 | Bacteria>Pseudomonadota>Gammaproteobacteria>Xanthomonadales>Xanthomonadaceae | GCF_001974685.1 | Stenotrophomonas_sp._MB339 | BU225_RS12360 | 15100..18225 | 1663558 | DUF499-domain-containing-protein-[Stenotrophomonas-sp.-MB339] |
| WP_072756491.1 | AAA+ATPase+wHTH+Alpha+Beta-domain+wHTH+Fn-III+Fn-III+RRM | Type-2_BR.RM.like.systems | DDE_Tnp_1->RHH->DUF499*->DUF3780->?->MACRODOMAIN->METHYLASE->HKD-DNase+SF1-SF2-Helicase->Endonuclease_5-> | 1041 | Bacteria>Pseudomonadota>Gammaproteobacteria>Xanthomonadales>Xanthomonadaceae | GCF_900129205.1 | Thermomonas_hydrothermalis | BUA29_RS10400 | 869..3994 | 213588 | DUF499-domain-containing-protein-[Thermomonas-hydrothermalis] |
| WP_261997736.1 | AAA+ATPase+wHTH+Alpha+Beta-domain+wHTH+Fn-III+Fn-III+RRM | Type-2_BR.RM.like.systems | Methyltransf_11->RNase->35exo-><-?\|\|tRNA->RHH_5->DUF499*->DUF3780->?->MACRODOMAIN->METHYLASE->HEPN->ParA-Soj-PloopNTPase-> | 1041 | Bacteria>Pseudomonadota>Gammaproteobacteria>Xanthomonadales>Xanthomonadaceae | GCF_024519495.1 | Stenotrophomonas_sp._Ste96 | OC373_RS11855 | 536568..539693 | 2926029 | DUF499-domain-containing-protein-[Stenotrophomonas-sp.-Ste96] |
| WP_053462285.1 | AAA+ATPase+wHTH+Alpha+Beta-domain+wHTH+Fn-III+Fn-III+RRM | Type-2_BR.RM.like.systems | Methyltransf_11->RNase->35exo-><-?\|\|tRNA->RHH_5->DUF499*->DUF3780->?->MACRODOMAIN->METHYLASE->HEPN->ParA-Soj-PloopNTPase-> | 1041 | Bacteria>Pseudomonadota>Gammaproteobacteria>Xanthomonadales>Xanthomonadaceae | GCF_021554495.1 | Stenotrophomonas | N5C25_RS09560 | 28481..31606 | 40323 | DUF499-domain-containing-protein-[Stenotrophomonas-maltophilia] |
| WP_032129329.1 | AAA+ATPase+wHTH+Alpha+Beta-domain+wHTH+Fn-III+Fn-III+RRM | Type-2_BR.RM.like.systems | Methyltransf_11->RNase->35exo-><-?\|\|tRNA->RHH_5->DUF499*->DUF3780->?->METHYLASE->HEPN->ParA-Soj-PloopNTPase->ORC-CDC6-> | 1044 | Bacteria>Pseudomonadota>Gammaproteobacteria>Xanthomonadales>Xanthomonadaceae | GCF_000758465.1 | Stenotrophomonas_maltophilia | LI87_RS0113700 | 18740..21874 | 40324 | DUF499-domain-containing-protein-[Stenotrophomonas-maltophilia] |
| WP_099589177.1 | AAA+ATPase+wHTH+Alpha+Beta-domain+wHTH+Fn-III+Fn-III+RRM | Type-2_BR.RM.like.systems | RHH_5->DUF499*->DUF3780->?->MACRODOMAIN->METHYLASE->HEPN->ParA-Soj-PloopNTPase-> | 1041 | Bacteria>Pseudomonadota>Gammaproteobacteria>Xanthomonadales>Xanthomonadaceae | GCF_002738905.1 | Stenotrophomonas_maltophilia | BRB41_RS09090 | 797..3922 | 40324 | DUF499-domain-containing-protein-[Stenotrophomonas-maltophilia] |
| WP_055423214.1 | AAA+ATPase+wHTH+Alpha+Beta-domain+wHTH+Fn-III+Fn-III+RRM | Type-2_BR.RM.like.systems | <-HKD-DNase+SF1-SF2-Helicase<-REase+ParB<-METHYLASE<-MACRODOMAIN<-?<-DUF3780<-DUF499*<-RHH\|\|?->\|\|NAD_binding_2+6PGD-> | 1041 | Bacteria>Pseudomonadota>Hydrogenophilia>Hydrogenophilales>Hydrogenophilaceae | GCF_001517365.1 | Tepidiphilus_thermophilus | ASC37_RS04380 | 10728..13853 | 876478 | DUF499-domain-containing-protein-[Tepidiphilus-thermophilus] |
| NCC04614.1 | AAA+ATPase+wHTH+Alpha+Beta-domain+wHTH+Fn-III+Fn-III+RRM | Type-2_BR.RM.like.systems | <-HKD-DNase+SF1-SF2-Helicase<-METHYLASE<-DUF3780<-DUF499* | 1043 | Bacteria>Pseudomonadota>unclassified_Pseudomonadota | GCA_009929575.1 | Pseudomonadota_bacterium | EOM37_11375 | 11977..15108 | 1977087 | DUF499-domain-containing-protein-[Proteobacteria-bacterium] |
| MBS0593923.1 | AAA+ATPase+wHTH+Alpha+Beta-domain+wHTH+Fn-III+Fn-III+RRM | Type-2_BR.RM.like.systems | <-HKD-DNase+SF1-SF2-Helicase<-HicB<-PIN<-METHYLASE<-DUF3780<-DUF499*<-?\|\|?->ABhydrolase-><-MerR-HTH | 1044 | Bacteria>Pseudomonadota>unclassified_Pseudomonadota | GCA_018240325.1 | Pseudomonadota_bacterium | JSR84_10695 | 72059..75193 | 1977087 | DUF499-domain-containing-protein-[Proteobacteria-bacterium] |
| MBU1052860.1 | AAA+ATPase+wHTH+Alpha+Beta-domain+wHTH+Fn-III+Fn-III+RRM | Type-2_BR.RM.like.systems | <-HKD-DNase+SF1-SF2-Helicase<-MACRODOMAIN<-?<-METHYLASE<-DUF3780<-DUF499* | 1027 | Bacteria>Pseudomonadota>unclassified_Pseudomonadota | GCA_018817425.1 | Pseudomonadota_bacterium | KKC46_03390 | 75459..78542 | 1977087 | DUF499-domain-containing-protein-[Proteobacteria-bacterium] |
| MBQ6473948.1 | AAA+ATPase+wHTH+Alpha+Beta-domain+wHTH+Fn-III+Fn-III+RRM | Type-2_BR.RM.like.systems | DUF499*->DUF3780->METHYLASE->HKD-DNase+SF1-SF2-Helicase-> | 994 | Bacteria>PVC_group>Lentisphaerota>Lentisphaeria>Victivallales | GCA_017431185.1 | Victivallales_bacterium | IJJ33_18310 | 728..3712 | 2212475 | DUF499-domain-containing-protein-[Victivallales-bacterium] |
| OGV20214.1 | AAA+ATPase+wHTH+Alpha+Beta-domain+wHTH+Fn-III+Fn-III+RRM | Type-2_BR.RM.like.systems | <-HKD-DNase+SF1-SF2-Helicase<-Schlafen+HATPase+HTH<-METHYLASE<-DUF3780<-DUF499*\|\|?-><-?<-?\|\|?->STAS->ABC-ATPase+SbcC-> | 1036 | Bacteria>PVC_group>Lentisphaerota>unclassified_Lentisphaerota | GCA_001803205.1 | Lentisphaerae_bacterium_GWF2_38_69 | A2X47_06010 | 8891..12001 | 1798570 | MAG:-glycosyl-transferase-[Lentisphaerae-bacterium-GWF2_38_69] |
| WP_169704427.1 | AAA+ATPase+wHTH+Alpha+Beta-domain+wHTH+Fn-III+Fn-III+RRM | Type-2_BR.RM.like.systems | <-HKD-DNase+SF1-SF2-Helicase<-METHYLASE<-DUF3780<-DUF499*\|\|?->?->?->?->OLD-ABC+TOPRIM->SF1-SF2-Helicase-> | 1036 | Bacteria>PVC_group>Planctomycetota>Candidatus_Brocadiia>Candidatus_Brocadiales | GCF_900232175.1 | Candidatus_Kuenenia_stuttgartiensis | HL046_RS36405 | 60109..63219 | 174633 | DUF499-domain-containing-protein-[Candidatus-Kuenenia-stuttgartiensis] |
| MCC5786917.1 | AAA+ATPase+wHTH+Alpha+Beta-domain+wHTH+Fn-III+Fn-III+RRM | Type-2_BR.RM.like.systems | <-N6_N4_MTase<-MPTase<-HTH\|\|DUF499*->DUF3780->METHYLASE->HKD-DNase+SF1-SF2-Helicase->URI-DNase+RAMA-><-ParB<-GDResolvase+Recombinase+ZnR | 1033 | Bacteria>PVC_group>Planctomycetota>Phycisphaerae>Phycisphaerales | GCA_020831775.1 | Phycisphaerales_bacterium | JJU33_09470 | 30542..33643 | 2052180 | DUF499-domain-containing-protein-[Phycisphaerales-bacterium] |
| MAO19196.1 | AAA+ATPase+wHTH+Alpha+Beta-domain+wHTH+Fn-III+Fn-III+RRM | Type-2_BR.RM.like.systems | <-XIS-HTH<-?\|\|?-><-HKD-DNase+SF1-SF2-Helicase<-METHYLASE<-DUF3780<-DUF499*<-XIS-HTH | 1033 | Bacteria>PVC_group>Planctomycetota>Phycisphaerae>unclassified_Phycisphaerae | GCA_002696165.1 | Phycisphaerae_bacterium | CMJ25_00465 | 51602..54703 | 2026778 | hypothetical-protein-[Phycisphaerae-bacterium] |
| WP_011117919.1 | AAA+ATPase+wHTH+Alpha+Beta-domain+wHTH+Fn-III+Fn-III+RRM | Type-2_BR.RM.like.systems | <-Schlafen<-HKD-DNase+SF1-SF2-Helicase<-TOPRIM<-OLD-ABC<-METHYLASE-p<-DUF3780<-DUF499*<-XIS-HTH | 1031 | Bacteria>PVC_group>Planctomycetota>Planctomycetia>Pirellulales | GCF_000196115.1 | Rhodopirellula_baltica | DDX19_RS10420 | 136482..139577 | 265606 | DUF499-domain-containing-protein-[Rhodopirellula-baltica] |
| WP_146459050.1 | AAA+ATPase+wHTH+Alpha+Beta-domain+wHTH+Fn-III+Fn-III+RRM | Type-2_BR.RM.like.systems | <-HKD-DNase<-?<-Schlafen<-HKD-DNase+SF1-SF2-Helicase<-METHYLASE<-DUF3780<-DUF499*<-XIS-HTH<-?<-SF1-SF2-Helicase<-?<-NERD+SF1-SF2-Helicase<-DUF2726+ZnR+TOPRIM | 1031 | Bacteria>PVC_group>Planctomycetota>Planctomycetia>Pirellulales | GCF_007860125.1 | Rubripirellula_tenax | Poly51_RS17440 | 744140..747235 | 2528015 | DUF499-domain-containing-protein-[Rubripirellula-tenax] |
| WP_253156603.1 | AAA+ATPase+wHTH+Alpha+Beta-domain+wHTH+Fn-III+Fn-III+RRM | Type-2_BR.RM.like.systems | <-HEPN<-?<-HKD-DNase+SF1-SF2-Helicase<-METHYLASE<-DUF3780<-DUF499*\|\|HTH-><-REase<-?\|\|GDResolvase->ParB+HTH->ParB-> | 1050 | Bacteria>PVC_group>Planctomycetota>Planctomycetia>Pirellulales | GCF_024129865.1 | Stieleria_tagensis | NHH03_RS12675 | 165548..168700 | 2956795 | DUF499-domain-containing-protein-[Stieleria-sp.-TO1_6] |
| MDB5387810.1 | AAA+ATPase+wHTH+Alpha+Beta-domain+wHTH+Fn-III+Fn-III+RRM | Type-2_BR.RM.like.systems | HNH->?->MetJarc-HTH->?->DUF499*->DUF3780->METHYLASE-> | 1027 | Bacteria>PVC_group>Planctomycetota>Planctomycetia>Planctomycetales | GCA_028291565.1 | Planctomycetaceae_bacterium | JWM11_3456 | 3579..6662 | 2026779 | MAG:-hypothetical-protein-JWM11_3456-[Planctomycetaceae-bacterium] |
| QKK08601.1 | AAA+ATPase+wHTH+Alpha+Beta-domain+wHTH+Fn-III+Fn-III+RRM | Type-2_BR.RM.like.systems | <-REase+SF1-SF2-Helicase<-?\|\|<-tRNA\|\|?->?->?->XIS-HTH->DUF499*->DUF3780->METHYLASE->HKD-DNase+SF1-SF2-Helicase-> | 1033 | Bacteria>PVC_group>Planctomycetota>unclassified_Planctomycetota | GCA_013285505.1 | Planctomycetota_bacterium | HND58_10800 | 2323544..2326645 | 2026780 | MAG:-DUF499-domain-containing-protein-[Planctomycetota-bacterium] |
| MCY2993648.1 | AAA+ATPase+wHTH+Alpha+Beta-domain+wHTH+Fn-III+Fn-III+RRM | Type-2_BR.RM.like.systems | <-ParA-Soj-PloopNTPase<-HKD-DNase+SF1-SF2-Helicase<-METHYLASE-p\|\|?-><-DUF3780<-DUF499*<-HDRnase<-SIGMA-HTH+SIGMA-HTH | 999 | Bacteria>PVC_group>Planctomycetota>unclassified_Planctomycetota | GCA_026692885.1 | Planctomycetota_bacterium | NTY19_38085 | 6695..9694 | 2026780 | MAG:-DUF499-domain-containing-protein-[Planctomycetota-bacterium] |
| MBL9192080.1 | AAA+ATPase+wHTH+Alpha+Beta-domain+wHTH+Fn-III+Fn-III+RRM | Type-2_BR.RM.like.systems | METHYLASE-p-><-?\|\|?-><-?\|\|?->DUF499*->DUF3780->METHYLASE->HKD-DNase+SF1-SF2-Helicase->HTH-> | 1049 | Bacteria>PVC_group>Verrucomicrobiota>Opitutae>Opitutales | GCA_016794625.1 | Opitutaceae_bacterium | JNJ82_06960 | 6491..9640 | 2006848 | DUF499-domain-containing-protein-[Opitutaceae-bacterium] |
| MDD2765364.1 | AAA+ATPase+wHTH+Alpha+Beta-domain+wHTH+Fn-III+Fn-III+RRM | Type-2_BR.RM.like.systems | <-HKD-DNase+SF1-SF2-Helicase<-METHYLASE<-DUF3780<-DUF499*<-?<-?\|\|?->?-><-HTH_17 | 1063 | Bacteria>PVC_group>Verrucomicrobiota>Opitutae>Opitutales | GCA_028688225.1 | Opitutaceae_bacterium | PHE83_15480 | 23391..26582 | 2006848 | MAG:-DUF499-domain-containing-protein-[Opitutaceae-bacterium] |
| MCH8539497.1 | AAA+ATPase+wHTH+Alpha+Beta-domain+wHTH+Fn-III+Fn-III+RRM | Type-2_BR.RM.like.systems | Radical-SAM->?->?->DUF499*->DUF3780->PIN->OLD-ABC+TOPRIM->METHYLASE->OLD-ABC->TOPRIM-> | 1021 | Bacteria>PVC_group>Verrucomicrobiota>Opitutae>Opitutales | GCA_022565675.1 | Opitutales_bacterium | LAT58_01865 | 10372..13437 | 2026772 | DUF499-domain-containing-protein-[Opitutales-bacterium] |
| MCC5838834.1 | AAA+ATPase+wHTH+Alpha+Beta-domain+wHTH+Fn-III+Fn-III+RRM | Type-2_BR.RM.like.systems | <-Radical-SAM<-Classical-AAA<-PIN<-?<-DUF3780<-DUF499*<-?<-PIN<-RHH\|\|?-><-?<-XIS-HTH+RVT | 1043 | Bacteria>PVC_group>Verrucomicrobiota>Opitutae>Opitutales | GCA_020831585.1 | Opitutales_bacterium | JJT96_01820 | 59223..62354 | 2026772 | DUF499-domain-containing-protein-[Opitutales-bacterium] |
| MBC8243979.1 | AAA+ATPase+wHTH+Alpha+Beta-domain+wHTH+Fn-III+Fn-III+RRM | Type-2_BR.RM.like.systems | DUF499*->DUF3780->METHYLASE->HKD-DNase+SF1-SF2-Helicase->?->RelE-ParE-> | 933 | Bacteria>PVC_group>Verrucomicrobiota>unclassified_Verrucomicrobiota | GCA_014382705.1 | Verrucomicrobiota_bacterium | H8E20_06270 | 3711..6512 | 2026799 | DUF499-domain-containing-protein-[Verrucomicrobiota-bacterium] |
| MBI4028162.1 | AAA+ATPase+wHTH+Alpha+Beta-domain+wHTH+Fn-III+Fn-III+RRM | Type-2_BR.RM.like.systems | <-HKD-DNase+SF1-SF2-Helicase<-METHYLASE<-DUF3780<-DUF499*\|\|?->PIN-> | 1032 | Bacteria>PVC_group>Verrucomicrobiota>unclassified_Verrucomicrobiota | GCA_016198745.1 | Verrucomicrobiota_bacterium | HY360_24470 | 6469..9567 | 2026799 | DUF499-domain-containing-protein-[Verrucomicrobia-bacterium] |
| MBS0652531.1 | AAA+ATPase+wHTH+Alpha+Beta-domain+wHTH+Fn-III+Fn-III+RRM | Type-2_BR.RM.like.systems | ABhydrolase-><-HEPN+CBS\|\|?-><-HKD-DNase+SF1-SF2-Helicase<-METHYLASE<-DUF3780<-DUF499* | 1029 | Bacteria>PVC_group>Verrucomicrobiota>unclassified_Verrucomicrobiota | GCA_018239665.1 | Verrucomicrobiota_bacterium | JSR39_03280 | 41454..44543 | 2026799 | ATP-binding-protein-[Verrucomicrobia-bacterium] |
| MCF7762652.1 | AAA+ATPase+wHTH+Alpha+Beta-domain+wHTH+Fn-III+Fn-III+RRM | Type-2_BR.RM.like.systems | <-HKD-DNase+SF1-SF2-Helicase<-Schlafen+HATPase+HTH<-METHYLASE<-wHTH+REase<-DUF3780<-DUF499*<-XIS-HTH\|\|SF1-SF2-Helicase-><-Glyoxalase<-?<-REC | 1027 | Bacteria>PVC_group>Verrucomicrobiota>unclassified_Verrucomicrobiota | GCA_021733945.1 | Verrucomicrobiota_bacterium | K9N62_03175 | 25626..28709 | 2026799 | DUF499-domain-containing-protein-[Verrucomicrobia-bacterium] |
| MCC5848622.1 | AAA+ATPase+wHTH+Alpha+Beta-domain+wHTH+Fn-III+Fn-III+RRM | Type-2_BR.RM.like.systems | <-Schlafen+HATPase+HTH<-DUF3780<-DUF499*<-?<-?\|\|?->?->?-><-tRNA<-ABhydrolase | 1021 | Bacteria>PVC_group>Verrucomicrobiota>unclassified_Verrucomicrobiota | GCA_020831635.1 | Verrucomicrobiota_bacterium | JJU29_11045 | 2236..5301 | 2026799 | DUF499-domain-containing-protein-[Verrucomicrobia-bacterium] |
| NCC60914.1 | AAA+ATPase+wHTH+Alpha+Beta-domain+wHTH+Fn-III+Fn-III+RRM | Type-2_BR.RM.like.systems | DUF499*->DUF3780->METHYLASE->HKD-DNase+SF1-SF2-Helicase-> | 1022 | Bacteria>PVC_group>Verrucomicrobiota>Verrucomicrobiae>unclassified_Verrucomicrobiae | GCA_009930015.1 | Verrucomicrobiae_bacterium | EOM12_08220 | 1214..4282 | 2499141 | DUF499-domain-containing-protein-[Verrucomicrobiae-bacterium] |
| WP_217730384.1 | AAA+ATPase+wHTH+Alpha+Beta-domain+wHTH+Fn-III+Fn-III+RRM | Type-2_BR.RM.like.systems | <-HKD-DNase+SF1-SF2-Helicase<-METHYLASE<-DUF3780<-DUF499*\|\|Phage_integrase->?->?->?->Peptidase_M28-> | 1031 | Bacteria>PVC_group>Verrucomicrobiota>Verrucomicrobiae>Verrucomicrobiales | - | Akkermansia_muciniphila | NE654_RS08875 | 77960..81055 | 239935 | DUF499-domain-containing-protein-[Akkermansia-muciniphila] |
| WP_253822299.1 | AAA+ATPase+wHTH+Alpha+Beta-domain+wHTH+Fn-III+Fn-III+RRM | Type-2_BR.RM.like.systems | <-Peptidase_M28<-?<-?<-?<-Phage_integrase\|\|DUF499*->DUF3780->METHYLASE->HKD-DNase+SF1-SF2-Helicase-> | 1031 | Bacteria>PVC_group>Verrucomicrobiota>Verrucomicrobiae>Verrucomicrobiales | - | Akkermansia_muciniphila | NLA09_07035 | 42923..46018 | 239935 | DUF499-domain-containing-protein-[Akkermansia-muciniphila] |
| OLA90112.1 | AAA+ATPase+wHTH+Alpha+Beta-domain+wHTH+Fn-III+Fn-III+RRM | Type-2_BR.RM.like.systems | <-Peptidase_M28<-?<-?<-?<-Phage_integrase\|\|DUF499*->DUF3780->METHYLASE->HKD-DNase+SF1-SF2-Helicase-> | 1031 | Bacteria>PVC_group>Verrucomicrobiota>Verrucomicrobiae>Verrucomicrobiales | GCA_001917295.1 | Akkermansia_sp._54_46 | BHW66_03800 | 27081..30176 | 1896967 | hypothetical-protein-BHW66_03800-[Akkermansia-sp.-54_46] |
| PNC55947.1 | AAA+ATPase+wHTH+Alpha+Beta-domain+wHTH+Fn-III+Fn-III+RRM | Type-2_BR.RM.like.systems | <-Peptidase_M28<-?<-?<-?<-Phage_integrase\|\|DUF499*->DUF3780->METHYLASE->HKD-DNase+SF1-SF2-Helicase-> | 1022 | Bacteria>PVC_group>Verrucomicrobiota>Verrucomicrobiae>Verrucomicrobiales | GCA_002885155.1 | Akkermansia_muciniphila | CXU09_07630 | 32573..35641 | 239935 | hypothetical-protein-CXU09_07630-[Akkermansia-muciniphila] |
| PNC89921.1 | AAA+ATPase+wHTH+Alpha+Beta-domain+wHTH+Fn-III+Fn-III+RRM | Type-2_BR.RM.like.systems | <-Peptidase_M28<-?<-?<-?<-Phage_integrase\|\|DUF499*->DUF3780->METHYLASE->HKD-DNase+SF1-SF2-Helicase-> | 1022 | Bacteria>PVC_group>Verrucomicrobiota>Verrucomicrobiae>Verrucomicrobiales | GCA_002885375.1 | Akkermansia_muciniphila | CXT91_10285 | 189606..192674 | 239935 | hypothetical-protein-CXT91_10285-[Akkermansia-muciniphila] |
| PNC60733.1 | AAA+ATPase+wHTH+Alpha+Beta-domain+wHTH+Fn-III+Fn-III+RRM | Type-2_BR.RM.like.systems | <-HKD-DNase+SF1-SF2-Helicase<-METHYLASE<-DUF3780<-DUF499*\|\|Phage_integrase->?->?->?->Peptidase_M28-> | 1022 | Bacteria>PVC_group>Verrucomicrobiota>Verrucomicrobiae>Verrucomicrobiales | GCA_002885575.1 | Akkermansia_muciniphila | CXU07_10505 | 76886..79954 | 239935 | hypothetical-protein-CXU07_10505-[Akkermansia-muciniphila] |
| MCL6686250.1 | AAA+ATPase+wHTH+Alpha+Beta-domain+wHTH+Fn-III+Fn-III+RRM | Type-2_BR.RM.like.systems | <-Peptidase_M28<-?<-?<-?<-Phage_integrase\|\|DUF499*->DUF3780->HKD-DNase+SF1-SF2-Helicase-> | 1031 | Bacteria>PVC_group>Verrucomicrobiota>Verrucomicrobiae>Verrucomicrobiales | GCA_023516675.1 | Akkermansia_muciniphila | M8N40_07390 | 46448..49543 | 239935 | DUF499-domain-containing-protein-[Akkermansia-muciniphila] |
| MCX8496484.1 | AAA+ATPase+wHTH+Alpha+Beta-domain+wHTH+Fn-III+Fn-III+RRM | Type-2_BR.RM.like.systems | <-REase<-?\|\|HTH->MPTase->?->?->DUF499*->DUF3780->METHYLASE->METHYLASE->HKD-DNase+SF1-SF2-Helicase->REase->HEPN-> | 1044 | Bacteria>PVC_group>Verrucomicrobiota>Verrucomicrobiae>Verrucomicrobiales | GCA_026461385.1 | Akkermansiaceae_bacterium | ORN51_09920 | 5504..8638 | 2562705 | MAG:-DUF499-domain-containing-protein-[Akkermansiaceae-bacterium] |
| MDA7889828.1 | AAA+ATPase+wHTH+Alpha+Beta-domain+wHTH+Fn-III+Fn-III+RRM | Type-2_BR.RM.like.systems | <-HKD-DNase+SF1-SF2-Helicase<-METHYLASE<-DUF3780<-DUF499*<-?\|\|?-><-PAIREDC-HTH | 1025 | Bacteria>PVC_group>Verrucomicrobiota>Verrucomicrobiae>Verrucomicrobiales | GCA_028046845.1 | Akkermansiaceae_bacterium | N9A78_01310 | 7498..10575 | 2562705 | DUF499-domain-containing-protein-[Akkermansiaceae-bacterium] |
| WP_102739763.1 | AAA+ATPase+wHTH+Alpha+Beta-domain+wHTH+Fn-III+Fn-III+RRM | Type-2_BR.RM.like.systems | <-HKD-DNase+SF1-SF2-Helicase<-METHYLASE<-DUF3780<-DUF499*\|\|Phage_integrase->?->?->?->Peptidase_M28-> | 1031 | Bacteria>PVC_group>Verrucomicrobiota>Verrucomicrobiae>Verrucomicrobiales | GCF_002885235.1 | Akkermansia_muciniphila | CXU03_RS10880 | 164258..167353 | 239935 | DUF499-domain-containing-protein-[Akkermansia-muciniphila] |
| WP_146020897.1 | AAA+ATPase+wHTH+Alpha+Beta-domain+wHTH+Fn-III+Fn-III+RRM | Type-2_BR.RM.like.systems | <-Peptidase_M28<-?<-?<-?<-Phage_integrase\|\|DUF499*->DUF3780->METHYLASE->HKD-DNase+SF1-SF2-Helicase-> | 1031 | Bacteria>PVC_group>Verrucomicrobiota>Verrucomicrobiae>Verrucomicrobiales | GCF_002885375.1 | Akkermansia_muciniphila | CXT91_RS10270 | 189579..192674 | 239935 | DUF499-domain-containing-protein-[Akkermansia-muciniphila] |
| WP_146018571.1 | AAA+ATPase+wHTH+Alpha+Beta-domain+wHTH+Fn-III+Fn-III+RRM | Type-2_BR.RM.like.systems | <-HKD-DNase+SF1-SF2-Helicase<-METHYLASE<-DUF3780<-DUF499*\|\|Phage_integrase->?->?->?->Peptidase_M28-> | 1031 | Bacteria>PVC_group>Verrucomicrobiota>Verrucomicrobiae>Verrucomicrobiales | GCF_002885575.1 | Akkermansia_muciniphila | CXU07_RS10500 | 76886..79981 | 239935 | DUF499-domain-containing-protein-[Akkermansia-muciniphila] |
| WP_123052235.1 | AAA+ATPase+wHTH+Alpha+Beta-domain+wHTH+Fn-III+Fn-III+RRM | Type-2_BR.RM.like.systems | <-Peptidase_M28<-?<-?<-?<-Phage_integrase\|\|DUF499*->DUF3780->METHYLASE->HKD-DNase+SF1-SF2-Helicase-> | 1031 | Bacteria>PVC_group>Verrucomicrobiota>Verrucomicrobiae>Verrucomicrobiales | GCF_003716915.1 | Akkermansia_muciniphila | CUB89_RS06705 | 1494864..1497959 | 239935 | DUF499-domain-containing-protein-[Akkermansia-muciniphila] |
| WP_155844469.1 | AAA+ATPase+wHTH+Alpha+Beta-domain+wHTH+Fn-III+Fn-III+RRM | Type-2_BR.RM.like.systems | <-HKD-DNase+SF1-SF2-Helicase<-METHYLASE<-DUF3780<-DUF499*\|\|Phage_integrase->?->?->?->Peptidase_M28-> | 1031 | Bacteria>PVC_group>Verrucomicrobiota>Verrucomicrobiae>Verrucomicrobiales | GCF_009731575.1 | Akkermansia_muciniphila | GOZ73_RS00075 | 21492..24587 | 239935 | DUF499-domain-containing-protein-[Akkermansia-muciniphila] |
| WP_251841163.1 | AAA+ATPase+wHTH+Alpha+Beta-domain+wHTH+Fn-III+Fn-III+RRM | Type-2_BR.RM.like.systems | <-HKD-DNase+SF1-SF2-Helicase<-METHYLASE<-DUF3780<-DUF499*\|\|Phage_integrase->?->?->?->Peptidase_M28-> | 933 | Bacteria>PVC_group>Verrucomicrobiota>Verrucomicrobiae>Verrucomicrobiales | GCF_018709665.1 | Akkermansia_muciniphila | HHJ08_RS08625 | 63705..66506 | 239935 | DUF499-domain-containing-protein-[Akkermansia-muciniphila] |
| WP_146018440.1 | AAA+ATPase+wHTH+Alpha+Beta-domain+wHTH+Fn-III+Fn-III+RRM | Type-2_BR.RM.like.systems | <-Peptidase_M28<-?<-?<-?<-Phage_integrase\|\|DUF499*->DUF3780->METHYLASE->HKD-DNase+SF1-SF2-Helicase-> | 1031 | Bacteria>PVC_group>Verrucomicrobiota>Verrucomicrobiae>Verrucomicrobiales | GCF_018847215.1 | Akkermansia_muciniphila | CXU09_RS07625 | 32546..35641 | 239935 | DUF499-domain-containing-protein-[Akkermansia-muciniphila] |
| WP_215490137.1 | AAA+ATPase+wHTH+Alpha+Beta-domain+wHTH+Fn-III+Fn-III+RRM | Type-2_BR.RM.like.systems | <-HKD-DNase+SF1-SF2-Helicase<-DUF3780<-DUF499*\|\|Phage_integrase->?->?->?->Peptidase_M28-> | 1031 | Bacteria>PVC_group>Verrucomicrobiota>Verrucomicrobiae>Verrucomicrobiales | GCF_018847255.1 | Akkermansia_muciniphila | J5W54_RS00075 | 21500..24595 | 239935 | DUF499-domain-containing-protein-[Akkermansia-muciniphila] |
| WP_215833905.1 | AAA+ATPase+wHTH+Alpha+Beta-domain+wHTH+Fn-III+Fn-III+RRM | Type-2_BR.RM.like.systems | <-HKD-DNase+SF1-SF2-Helicase<-DUF3780<-DUF499*\|\|Phage_integrase->?->?->?->Peptidase_M28-> | 1031 | Bacteria>PVC_group>Verrucomicrobiota>Verrucomicrobiae>Verrucomicrobiales | GCF_018847355.1 | Akkermansia_muciniphila | J5W80_RS00075 | 21512..24607 | 239935 | DUF499-domain-containing-protein-[Akkermansia-muciniphila] |
| WP_273701145.1 | AAA+ATPase+wHTH+Alpha+Beta-domain+wHTH+Fn-III+Fn-III+RRM | Type-2_BR.RM.like.systems | <-HKD-DNase+SF1-SF2-Helicase<-METHYLASE<-DUF3780<-DUF499*\|\|Phage_integrase->?->?->?->Peptidase_M28-> | 933 | Bacteria>PVC_group>Verrucomicrobiota>Verrucomicrobiae>Verrucomicrobiales | GCF_019132835.1 | Akkermansia_muciniphila | LON27_RS02830 | 77960..80761 | 239935 | DUF499-domain-containing-protein-[Akkermansia-muciniphila] |
| WP_254005885.1 | AAA+ATPase+wHTH+Alpha+Beta-domain+wHTH+Fn-III+Fn-III+RRM | Type-2_BR.RM.like.systems | <-Peptidase_M28<-?<-?<-?<-Phage_integrase\|\|DUF499*->DUF3780->METHYLASE->HKD-DNase+SF1-SF2-Helicase-> | 933 | Bacteria>PVC_group>Verrucomicrobiota>Verrucomicrobiae>Verrucomicrobiales | GCF_023516675.1 | Akkermansia_muciniphila | M8N40_RS07405 | 46742..49543 | 239935 | DUF499-domain-containing-protein-[Akkermansia-muciniphila] |
| WP_285605586.1 | AAA+ATPase+wHTH+Alpha+Beta-domain+wHTH+Fn-III+Fn-III+RRM | Type-2_BR.RM.like.systems | <-HKD-DNase+SF1-SF2-Helicase<-METHYLASE<-DUF3780<-DUF499*\|\|Phage_integrase->?->?->?->Peptidase_M28-> | 1031 | Bacteria>PVC_group>Verrucomicrobiota>Verrucomicrobiae>Verrucomicrobiales | GCF_030268525.1 | Akkermansia_muciniphila | QSJ39_RS10215 | 50121..53216 | 239935 | DUF499-domain-containing-protein-[Akkermansia-muciniphila] |
| AFM14204.1 | AAA+ATPase+wHTH+Alpha+Beta-domain+wHTH+Fn-III+Fn-III+RRM | Type-2_BR.RM.like.systems | <-TIR<-TIR<-TIR_2<-HKD-DNase+SF1-SF2-Helicase<-METHYLASE<-DUF3780<-DUF499*\|\|HIN-HTH->DDE_Tnp_IS1595->DDE_Tnp_IS1595->?->?->HNH-> | 1050 | Bacteria>Spirochaetota>Spirochaetia>Leptospirales>Leptospiraceae | GCA_000266885.1 | Turneriella_parva_DSM_21527 | Turpa_3569 | 3707986..3711138 | 869212 | hypothetical-protein-Turpa_3569-[Turneriella-parva-DSM-21527] |
| TGM42334.1 | AAA+ATPase+wHTH+Alpha+Beta-domain+wHTH+Fn-III+Fn-III+RRM | Type-2_BR.RM.like.systems | DUF499*->DUF3780->MPTase->PIN->METHYLASE->HKD-DNase+SF1-SF2-Helicase-> | 1030 | Bacteria>Spirochaetota>Spirochaetia>Leptospirales>Leptospiraceae | GCA_004770805.1 | Leptospira_biflexa | EHQ92_16185 | 338..3430 | 172 | DUF499-domain-containing-protein-[Leptospira-biflexa] |
| MCB1144576.1 | AAA+ATPase+wHTH+Alpha+Beta-domain+wHTH+Fn-III+Fn-III+RRM | Type-2_BR.RM.like.systems | <-HKD-DNase+SF1-SF2-Helicase<-METHYLASE<-?<-PIN<-MPTase<-DUF3780<-DUF499*<-?<-PIN<-ParB+HNH<-rve+rve_3 | 1030 | Bacteria>Spirochaetota>Spirochaetia>Leptospirales>Leptospiraceae | GCA_020440865.1 | Leptospiraceae_bacterium | KDK54_20140 | 8602..11694 | 2026757 | ATP-binding-protein-[Leptospiraceae-bacterium] |
| WP_041948680.1 | AAA+ATPase+wHTH+Alpha+Beta-domain+wHTH+Fn-III+Fn-III+RRM | Type-2_BR.RM.like.systems | <-TIR<-TIR<-TIR_2<-HKD-DNase+SF1-SF2-Helicase<-METHYLASE<-DUF3780<-DUF499*\|\|ParB+HNH->HTH_29+rve+rve_3->DDE_Tnp_IS1595->DDE_Tnp_IS1595->?->HNH-> | 1046 | Bacteria>Spirochaetota>Spirochaetia>Leptospirales>Leptospiraceae | GCF_000266885.1 | Turneriella_parva | TURPA_RS17950 | 3707986..3711126 | 29510 | DUF499-domain-containing-protein-[Turneriella-parva] |
| WP_014804717.1 | AAA+ATPase+wHTH+Alpha+Beta-domain+wHTH+Fn-III+Fn-III+RRM | Type-2_BR.RM.like.systems | <-TIR<-TIR<-TIR_2<-HKD-DNase+SF1-SF2-Helicase<-METHYLASE<-DUF3780<-DUF499*\|\|REase->DDE_Tnp_IS1595-> | 1050 | Bacteria>Spirochaetota>Spirochaetia>Leptospirales>Leptospiraceae | GCF_000266885.1 | Turneriella_parva | TURPA_RS18115 | 3751659..3754811 | 29510 | DUF499-domain-containing-protein-[Turneriella-parva] |
| WP_135769228.1 | AAA+ATPase+wHTH+Alpha+Beta-domain+wHTH+Fn-III+Fn-III+RRM | Type-2_BR.RM.like.systems | <-HKD-DNase+SF1-SF2-Helicase<-METHYLASE<-DOC+HTH<-DOC+HTH<-DUF3780<-DUF499* | 1034 | Bacteria>Spirochaetota>Spirochaetia>Leptospirales>Leptospiraceae | GCF_004769195.1 | Leptospira_fletcheri | EHO60_RS16115 | 293955..297059 | 2484981 | DUF499-domain-containing-protein-[Leptospira-fletcheri] |
| WP_167881406.1 | AAA+ATPase+wHTH+Alpha+Beta-domain+wHTH+Fn-III+Fn-III+RRM | Type-2_BR.RM.like.systems | DUF499*->DUF3780->MPTase->PIN->METHYLASE->HKD-DNase+SF1-SF2-Helicase-> | 998 | Bacteria>Spirochaetota>Spirochaetia>Leptospirales>Leptospiraceae | GCF_004770325.1 | Leptospira_biflexa | EHQ92_RS16185 | 434..3430 | 172 | DUF499-domain-containing-protein-[Leptospira-biflexa] |
| WP_135740495.1 | AAA+ATPase+wHTH+Alpha+Beta-domain+wHTH+Fn-III+Fn-III+RRM | Type-2_BR.RM.like.systems | DUF499*->DUF3780->MPTase->PIN->METHYLASE->HKD-DNase+SF1-SF2-Helicase-> | 1030 | Bacteria>Spirochaetota>Spirochaetia>Leptospirales>Leptospiraceae | GCF_004770445.1 | Leptospira_bouyouniensis | EHQ99_RS04855 | 387..3479 | 2484911 | DUF499-domain-containing-protein-[Leptospira-bouyouniensis] |
| MBK9497791.1 | AAA+ATPase+wHTH+Alpha+Beta-domain+wHTH+Fn-III+Fn-III+RRM | Type-2_BR.RM.like.systems | <-HNH<-OLD-ABC<-?<-OLD-ABC<-ParB+HNH<-DUF3780<-DUF499*<-DUF499\|\|?->?->?->Peptidase-> | 736 | Bacteria>Spirochaetota>Spirochaetia>Leptospirales>Leptospiraceae | GCA_016718635.1 | Leptospiraceae_bacterium | IPO06_00155 | 42788..44998 | 2026757 | DUF499-domain-containing-protein-[Leptospiraceae-bacterium] |
| MBX7059766.1 | AAA+ATPase+wHTH+Alpha+Beta-domain+wHTH+Fn-III+Fn-III+RRM | Type-2_BR.RM.like.systems | RelE->CHTH->?->?->?->RelE-ParE->DUF499*->DUF3780->DOC+HTH->METHYLASE->HKD-DNase+SF1-SF2-Helicase-><-DOC+HTH | 1027 | Bacteria>Spirochaetota>Spirochaetia>Leptospirales>unclassified_Leptospirales | GCA_019694655.1 | Leptospirales_bacterium | K1X75_17010 | 16945..20028 | 2604146 | DUF499-domain-containing-protein-[Leptospirales-bacterium] |
| AEJ20205.1 | AAA+ATPase+wHTH+Alpha+Beta-domain+wHTH+Fn-III+Fn-III+RRM | Type-2_BR.RM.like.systems | <-HTH+UbE1\|\|DUF499*->DUF3780->OLD-ABC+TOPRIM->DUF772+DDE_Tnp_1_2->HKD-DNase+SF1-SF2-Helicase->HTH+MPTase->PIN-> | 1024 | Bacteria>Spirochaetota>Spirochaetia>Spirochaetales>Breznakiellaceae | GCA_000219725.1 | Gracilinema_caldarium_DSM_7334 | Spica_2078 | 2335967..2339041 | 744872 | hypothetical-protein-Spica_2078-[Gracilinema-caldarium-DSM-7334] |
| SLM14732.1 | AAA+ATPase+wHTH+Alpha+Beta-domain+wHTH+Fn-III+Fn-III+RRM | Type-2_BR.RM.like.systems | <-HKD-DNase+SF1-SF2-Helicase<-Schlafen+HATPase+HTH<-METHYLASE<-OLD-ABC+TOPRIM<-DUF3780<-DUF499*\|\|wHTH+SAD+HNH-><-?\|\|HEPN->PolB-> | 1036 | Bacteria>Spirochaetota>Spirochaetia>Spirochaetales>environmental_samples | GCA_900163985.1 | uncultured_spirochete | SPIROBIBN47_350053 | 62416..65526 | 156406 | conserved-hypothetical-protein-[uncultured-spirochete] |
| ORC33988.1 | AAA+ATPase+wHTH+Alpha+Beta-domain+wHTH+Fn-III+Fn-III+RRM | Type-2_BR.RM.like.systems | <-HKD-DNase+SF1-SF2-Helicase<-METHYLASE<-ParB+HNH<-DUF3780<-DUF499*<-?<-Phage_integrase<-tRNA\|\|?-><-NTP_transf_3 | 1091 | Bacteria>Spirochaetota>Spirochaetia>Spirochaetales>Spirochaetaceae | GCA_002087085.1 | Marispirochaeta_aestuarii | B4O97_13955 | 8718..11993 | 1963862 | hypothetical-protein-B4O97_13955-[Marispirochaeta-aestuarii] |
| WP_158084306.1 | AAA+ATPase+wHTH+Alpha+Beta-domain+wHTH+Fn-III+Fn-III+RRM | Type-2_BR.RM.like.systems | <-HKD-DNase+SF1-SF2-Helicase<-METHYLASE<-ParB+HNH<-DUF3780<-DUF499*<-?<-Phage_integrase<-tRNA\|\|?-><-NTP_transf_3 | 1039 | Bacteria>Spirochaetota>Spirochaetia>Spirochaetales>Spirochaetaceae | GCF_002087085.1 | Marispirochaeta_aestuarii | B4O97_RS13955 | 8718..11837 | 1963862 | DUF499-domain-containing-protein-[Marispirochaeta-aestuarii] |
| HCX97050.1 | AAA+ATPase+wHTH+Alpha+Beta-domain+wHTH+Fn-III+Fn-III+RRM | Type-2_BR.RM.like.systems | <-PolB<-HEPN\|\|?->\|\|DUF499*->DUF3780-> | 1036 | Bacteria>Spirochaetota>Spirochaetia>Spirochaetales>Spirochaetaceae | GCA_003539135.1 | Spirochaetaceae_bacterium | DHU26_08820 | 5566..8676 | 1898206 | glycosyl-transferase-[Spirochaetaceae-bacterium] |
| WP_132781615.1 | AAA+ATPase+wHTH+Alpha+Beta-domain+wHTH+Fn-III+Fn-III+RRM | Type-2_BR.RM.like.systems | <-N6_N4_MTase<-Aminotran_1_2\|\|TIR->DUF499*->DUF3780->OLD-ABC+TOPRIM->METHYLASE->Schlafen+HATPase+HTH->HKD-DNase+SF1-SF2-Helicase-> | 1038 | Bacteria>Spirochaetota>Spirochaetia>Spirochaetales>Treponemataceae | GCF_004343725.1 | Treponema_sp._J25 | C5O22_RS10535 | 15156..18272 | 2094121 | DUF499-domain-containing-protein-[Treponema-sp.-J25] |
| MCA1950497.1 | AAA+ATPase+wHTH+Alpha+Beta-domain+wHTH+Fn-III+Fn-III+RRM | Type-2_BR.RM.like.systems | REase->?-><-?\|\|HTH+WYL->?->DUF499*->DUF3780->OLD-ABC+TOPRIM->METHYLASE-p-> | 1038 | Bacteria>Spirochaetota>Spirochaetia>Spirochaetales>Treponemataceae | GCA_020216655.1 | Treponema_sp. | LDL24_07980 | 9395..12511 | 166 | DUF499-domain-containing-protein-[Treponema-sp.] |
| NCN05110.1 | AAA+ATPase+wHTH+Alpha+Beta-domain+wHTH+Fn-III+Fn-III+RRM | Type-2_BR.RM.like.systems | REase-><-MORC<-?<-HKD-DNase+SF1-SF2-Helicase<-METHYLASE<-DUF3780<-DUF499*<-XIS-HTH\|\|?->AbiEi_4->OLD-ABC-><-TPR+TPR\|\|AbiEi_1-> | 1027 | Bacteria>Spirochaetota>Spirochaetia>Spirochaetales>unclassified_Spirochaetales | GCA_009991955.1 | Spirochaetales_bacterium | GW949_05720 | 40118..43201 | 2026792 | ATP-binding-protein-[Spirochaetales-bacterium] |
| MBN1524672.1 | AAA+ATPase+wHTH+Alpha+Beta-domain+wHTH+Fn-III+Fn-III+RRM | Type-2_BR.RM.like.systems | RADICAL-SAM->?-><-?<-?<-?\|\|?->DUF499*->DUF3780->METHYLASE->BirA-HTH->AbiEii->HKD-DNase+SF1-SF2-Helicase-> | 1039 | Bacteria>Spirochaetota>Spirochaetia>Spirochaetales>unclassified_Spirochaetales | GCA_016929065.1 | Spirochaetales_bacterium | JW904_09325 | 65076..68195 | 2026792 | ATP-binding-protein-[Spirochaetales-bacterium] |
| MBN2352085.1 | AAA+ATPase+wHTH+Alpha+Beta-domain+wHTH+Fn-III+Fn-III+RRM | Type-2_BR.RM.like.systems | <-HKD-DNase+SF1-SF2-Helicase<-AbiEii<-BirA-HTH<-HEPN<-METHYLASE<-DUF3780<-DUF499*\|\|?->?-><-?<-?<-HNH | 1039 | Bacteria>Spirochaetota>Spirochaetia>Spirochaetales>unclassified_Spirochaetales | GCA_016934415.1 | Spirochaetales_bacterium | JXD23_05895 | 38302..41421 | 2026792 | ATP-binding-protein-[Spirochaetales-bacterium] |
| MDA3832123.1 | AAA+ATPase+wHTH+Alpha+Beta-domain+wHTH+Fn-III+Fn-III+RRM | Type-2_BR.RM.like.systems | DUF499*->DUF3780->Schlafen+HATPase+HTH->METHYLASE-p-> | 1006 | Bacteria>Spirochaetota>Spirochaetia>Spirochaetales>unclassified_Spirochaetales | GCA_027858175.1 | Spirochaetales_bacterium | PF495_01880 | 1..3021 | 2026792 | MAG:-DUF499-domain-containing-protein,-partial-[Spirochaetales-bacterium] |
| PKL23537.1 | AAA+ATPase+wHTH+Alpha+Beta-domain+wHTH+Fn-III+Fn-III+RRM | Type-2_BR.RM.like.systems | <-HKD-DNase+SF1-SF2-Helicase<-DOC+HTH<-MORC<-METHYLASE<-?<-DUF3780<-DUF499*\|\|?-><-ParB\|\|JAB-> | 1032 | Bacteria>Spirochaetota>unclassified_Spirochaetota | GCA_002839315.1 | Spirochaetae_bacterium_HGW-Spirochaetae-3 | CVV47_15000 | 172085..175183 | 2013835 | glycosyl-transferase-[Spirochaetae-bacterium-HGW-Spirochaetae-3] |
| MCX7023764.1 | AAA+ATPase+wHTH+Alpha+Beta-domain+wHTH+Fn-III+Fn-III+RRM | Type-2_BR.RM.like.systems | SF1-SF2-Helicase-><-?<-?<-?<-?<-ACET\|\|DUF499*->DUF3780->BirA-HTH->METHYLASE->DOC+HTH->HKD-DNase+SF1-SF2-Helicase-> | 1032 | Bacteria>Spirochaetota>unclassified_Spirochaetota | GCA_026398615.1 | Spirochaetota_bacterium | NT080_04000 | 27702..30800 | 2202144 | MAG:-DUF499-domain-containing-protein-[Spirochaetes-bacterium] |
| WP_276978228.1 | AAA+ATPase+wHTH+Alpha+Beta-domain+wHTH+Fn-III+Fn-III+RRM | Type-2_BR.RM.like.systems | SF1-SF2-Helicase-><-LON-AAA<-LON-AAA\|\|?->?->?->DUF499*->DUF3780->METHYLASE->HKD-DNase+SF1-SF2-Helicase-><-GDResolvase+HIN-HTH | 1046 | Bacteria>Terrabacteria_group>Actinomycetota>Acidimicrobiia>Acidimicrobiales | GCF_027412955.1 | Ferrimicrobium_acidiphilum | M7P62_RS13375 | 9535..12675 | 121039 | DUF499-domain-containing-protein-[Ferrimicrobium-acidiphilum] |
| WP_201732542.1 | AAA+ATPase+wHTH+Alpha+Beta-domain+wHTH+Fn-III+Fn-III+RRM | Type-2_BR.RM.like.systems | PIN->?-><-HKD-DNase+SF1-SF2-Helicase<-URI-DNase<-METHYLASE<-DUF3780<-DUF499*\|\|?->?->MazE_antitoxin->PIN-><-wHTH+SAD+HNH\|\|SIGMA-HTH-> | 1046 | Bacteria>Terrabacteria_group>Actinomycetota>Acidimicrobiia>Acidimicrobiales | GCF_904848545.2 | Acidithrix_sp._C25 | ACIDITHRIX25_RS38085 | 815227..818367 | 1671482 | DUF499-domain-containing-protein-[Acidithrix-sp.-C25] |
| KOX56111.1 | AAA+ATPase+wHTH+Alpha+Beta-domain+wHTH+Fn-III+Fn-III+RRM | Type-2_BR.RM.like.systems | <-ABC-ATPase+SbcC\|\|?->?->DUF499*->METHYLASE->ParB+HNH->HKD-DNase+SF1-SF2-Helicase-> | 1045 | Bacteria>Terrabacteria_group>Actinomycetota>Actinomycetes>Kitasatosporales | GCA_001280155.1 | Streptomyces_purpurogeneiscleroticus | ADL19_10425 | 17672..20809 | 68259 | hypothetical-protein-ADL19_10425-[Streptomyces-purpurogeneiscleroticus] |
| MBX3108928.1 | AAA+ATPase+wHTH+Alpha+Beta-domain+wHTH+Fn-III+Fn-III+RRM | Type-2_BR.RM.like.systems | <-HTH_3\|\|?->?->GDResolvase+HIN-HTH-><-HerA/FtsK<-?\|\|DUF499*->DUF3780->METHYLASE->Schlafen+HATPase+HTH->HKD-DNase+SF1-SF2-Helicase-><-?<-DNA_methylase | 1037 | Bacteria>Terrabacteria_group>Armatimonadota>Fimbriimonadia>Fimbriimonadales | GCA_019638755.1 | Fimbriimonadaceae_bacterium | KF743_07090 | 1426545..1429658 | 2682144 | DUF499-domain-containing-protein-[Fimbriimonadaceae-bacterium] |
| WP_098986579.1 | AAA+ATPase+wHTH+Alpha+Beta-domain+wHTH+Fn-III+Fn-III+RRM | Type-2_BR.RM.like.systems | <-NERD+SF1-SF2-Helicase\|\|Preader7+McrB-NTD+McrB-AAA->McrC-NTD+REase->?-><-HEPN\|\|?->DUF499*->DUF3780->METHYLASE->HKD-DNase+SF1-SF2-Helicase->?->?->MerR-HTH-> | 1022 | Bacteria>Terrabacteria_group>Bacillota>Bacilli>Bacillales | - | Bacillus_cereus | COD90_RS22430 | 19622..22690 | 1396 | DUF499-domain-containing-protein-[Bacillus-cereus] |
| PGW11809.1 | AAA+ATPase+wHTH+Alpha+Beta-domain+wHTH+Fn-III+Fn-III+RRM | Type-2_BR.RM.like.systems | <-NERD+SF1-SF2-Helicase\|\|Preader7+McrB-NTD+McrB-AAA->McrC-NTD+REase->?-><-HEPN\|\|?->DUF499*->DUF3780->METHYLASE->HKD-DNase+SF1-SF2-Helicase->?->?->MerR-HTH-> | 1022 | Bacteria>Terrabacteria_group>Bacillota>Bacilli>Bacillales | GCA_002585495.1 | Bacillus_cereus | COD90_22430 | 19622..22690 | 1396 | hypothetical-protein-COD90_22430-[Bacillus-cereus] |
| MBB2482470.1 | AAA+ATPase+wHTH+Alpha+Beta-domain+wHTH+Fn-III+Fn-III+RRM | Type-2_BR.RM.like.systems | REase+DUF4268->?->DUF499*->DUF3780->METHYLASE->HKD-DNase+SF1-SF2-Helicase->HEPN-> | 1031 | Bacteria>Terrabacteria_group>Bacillota>Bacilli>Bacillales | GCA_014174455.1 | Bacillus_sp._APMAM | H5P36_20065 | 1869..4964 | 2761103 | DUF499-domain-containing-protein-[Bacillus-sp.-APMAM] |
| MBN8209372.1 | AAA+ATPase+wHTH+Alpha+Beta-domain+wHTH+Fn-III+Fn-III+RRM | Type-2_BR.RM.like.systems | DUF499*->DUF3780->METHYLASE->HKD-DNase+SF1-SF2-Helicase-> | 1028 | Bacteria>Terrabacteria_group>Bacillota>Bacilli>Bacillales | GCA_017303315.1 | Bacillus_sp._NTK071 | JI666_11505 | 25787..28873 | 2802175 | DUF499-domain-containing-protein-[Bacillus-sp.-NTK071] |
| MBP3040009.1 | AAA+ATPase+wHTH+Alpha+Beta-domain+wHTH+Fn-III+Fn-III+RRM | Type-2_BR.RM.like.systems | CHTH->tRNA->?->DDE_Tnp->XIS-HTH->DUF499*->DUF3780->METHYLASE->HKD-DNase+SF1-SF2-Helicase-><-ZnR\|\|?->REase-> | 1022 | Bacteria>Terrabacteria_group>Bacillota>Bacilli>Bacillales | GCA_017898095.1 | Bacillaceae_bacterium_Marseille-Q3522 | J9303_10980 | 4040..7108 | 2759695 | DUF499-domain-containing-protein-[Bacillaceae-bacterium-Marseille-Q3522] |
| MBT2694631.1 | AAA+ATPase+wHTH+Alpha+Beta-domain+wHTH+Fn-III+Fn-III+RRM | Type-2_BR.RM.like.systems | <-ZnR+HTH\|\|?->?-><-HKD-DNase+SF1-SF2-Helicase<-METHYLASE<-DUF3780<-DUF499*<-?<-REase+DUF4268<-HTH_31<-REase | 1032 | Bacteria>Terrabacteria_group>Bacillota>Bacilli>Bacillales | GCA_018613155.1 | Bacillus_sp._ISL-55 | J7E32_16715 | 33533..36631 | 2819134 | DUF499-domain-containing-protein-[Bacillus-sp.-ISL-55] |
| MBT2667307.1 | AAA+ATPase+wHTH+Alpha+Beta-domain+wHTH+Fn-III+Fn-III+RRM | Type-2_BR.RM.like.systems | <-Phage_int_SAM_5<-URI-DNase\|\|?-><-HKD-DNase+SF1-SF2-Helicase<-METHYLASE<-DUF3780<-DUF499*<-?<-?<-GDResolvase+Recombinase+ZnR | 1033 | Bacteria>Terrabacteria_group>Bacillota>Bacilli>Bacillales | GCA_018613275.1 | Bacillus_sp._ISL-4 | J7J00_17650 | 67951..71052 | 2819125 | DUF499-domain-containing-protein-[Bacillus-sp.-ISL-4] |
| WP_027725741.1 | AAA+ATPase+wHTH+Alpha+Beta-domain+wHTH+Fn-III+Fn-III+RRM | Type-2_BR.RM.like.systems | HerA/FtsK->?->?->DUF499*->DUF3780->METHYLASE->HKD-DNase+SF1-SF2-Helicase->HIN-HTH->\|\|Peptidase-> | 1035 | Bacteria>Terrabacteria_group>Bacillota>Bacilli>Bacillales | GCF_000430685.1 | Tuberibacillus_calidus | H532_RS0112350 | 4411..7518 | 340097 | DUF499-domain-containing-protein-[Tuberibacillus-calidus] |
| WP_025333200.1 | AAA+ATPase+wHTH+Alpha+Beta-domain+wHTH+Fn-III+Fn-III+RRM | Type-2_BR.RM.like.systems | HTH+WYL->?->?->?->Exonuc_V_gamma+REase->SF1-SF2-Helicase+REase->DUF499*->DUF3780->METHYLASE->HKD-DNase+SF1-SF2-Helicase->?->REase->pPIWI+pPIWI-> | 1022 | Bacteria>Terrabacteria_group>Bacillota>Bacilli>Bacillales | GCF_000612505.1 | Paenibacillus_sabinae | PSAB_RS03435 | 744096..747164 | 365617 | DUF499-domain-containing-protein-[Paenibacillus-sabinae] |
| WP_042210529.1 | AAA+ATPase+wHTH+Alpha+Beta-domain+wHTH+Fn-III+Fn-III+RRM | Type-2_BR.RM.like.systems | TPR->?->?->HTH+WYL->Exonuc_V_gamma+REase->SF1-SF2-Helicase+REase->DUF499*->DUF3780->METHYLASE->HKD-DNase+SF1-SF2-Helicase->Calcineurin->SbcC+SbcC-> | 1022 | Bacteria>Terrabacteria_group>Bacillota>Bacilli>Bacillales | GCF_000758665.1 | Paenibacillus_borealis | PBOR_RS03715 | 849881..852949 | 160799 | DUF499-domain-containing-protein-[Paenibacillus-borealis] |
| WP_040209692.1 | AAA+ATPase+wHTH+Alpha+Beta-domain+wHTH+Fn-III+Fn-III+RRM | Type-2_BR.RM.like.systems | <-GspA-ATPase<-Mu-transpos_C<-HKD-DNase+SF1-SF2-Helicase<-METHYLASE<-DUF3780<-DUF499*<-HhH-RADC+JAB<-?<-?<-?<-?<-GDResolvase+Recombinase+ZnR | 1029 | Bacteria>Terrabacteria_group>Bacillota>Bacilli>Bacillales | GCF_000820865.2 | Neobacillus_jeddahensis | TA93_RS24225 | 1723854..1726943 | 1461580 | DUF499-domain-containing-protein-[Bacillus-sp.-JCA] |
| WP_047154476.1 | AAA+ATPase+wHTH+Alpha+Beta-domain+wHTH+Fn-III+Fn-III+RRM | Type-2_BR.RM.like.systems | <-HKD-DNase+SF1-SF2-Helicase<-METHYLASE<-DUF3780<-DUF499*\|\|<-METHYLASE-p<-?\|\|GDResolvase-><-?<-35exo | 1022 | Bacteria>Terrabacteria_group>Bacillota>Bacilli>Bacillales | GCF_001015055.1 | Aneurinibacillus_tyrosinisolvens | MBEAL2_RS22455 | 13443..16511 | 1443435 | DUF499-domain-containing-protein-[Aneurinibacillus-tyrosinisolvens] |
| WP_075310335.1 | AAA+ATPase+wHTH+Alpha+Beta-domain+wHTH+Fn-III+Fn-III+RRM | Type-2_BR.RM.like.systems | Preader7+McrB-NTD+McrB-AAA->McrC-NTD+REase->REase->HNH-><-HEPN<-HEPN-p\|\|DUF499*->DUF3780->METHYLASE->HKD-DNase+SF1-SF2-Helicase->?->?->MerR-HTH-> | 1022 | Bacteria>Terrabacteria_group>Bacillota>Bacilli>Bacillales | GCF_001932005.1 | Bacillus_wiedmannii | BUC33_RS14890 | 28645..31713 | 1890302 | DUF499-domain-containing-protein-[Bacillus-wiedmannii] |
| WP_098896857.1 | AAA+ATPase+wHTH+Alpha+Beta-domain+wHTH+Fn-III+Fn-III+RRM | Type-2_BR.RM.like.systems | <-FtsK<-?<-?\|\|?-><-URI-DNase+RAMA\|\|DUF499*->DUF3780->METHYLASE->HKD-DNase+SF1-SF2-Helicase-> | 1022 | Bacteria>Terrabacteria_group>Bacillota>Bacilli>Bacillales | GCF_002552565.1 | Priestia_megaterium | CON20_RS27020 | 16730..19798 | 1404 | DUF499-domain-containing-protein-[Priestia-megaterium] |
| WP_098128686.1 | AAA+ATPase+wHTH+Alpha+Beta-domain+wHTH+Fn-III+Fn-III+RRM | Type-2_BR.RM.like.systems | HTH+WYL->?->Preader7+McrB-NTD+McrB-AAA->McrC-NTD+REase->REase->?->DUF499*->DUF3780->METHYLASE->HKD-DNase+SF1-SF2-Helicase-> | 1033 | Bacteria>Terrabacteria_group>Bacillota>Bacilli>Bacillales | GCF_002556745.1 | Bacillus_wiedmannii | CN576_RS24165 | 43423..46524 | 1890302 | DUF499-domain-containing-protein-[Bacillus-wiedmannii] |
| WP_098807661.1 | AAA+ATPase+wHTH+Alpha+Beta-domain+wHTH+Fn-III+Fn-III+RRM | Type-2_BR.RM.like.systems | HEPN-p-><-HKD-DNase\|\|?->?->DUF499*->DUF3780->METHYLASE->HKD-DNase+SF1-SF2-Helicase->?->?->HNH-> | 1022 | Bacteria>Terrabacteria_group>Bacillota>Bacilli>Bacillales | GCF_002579615.1 | Bacillus_toyonensis | COF68_RS16765 | 26144..29212 | 155322 | DUF499-domain-containing-protein-[Bacillus-toyonensis] |
| WP_098836958.1 | AAA+ATPase+wHTH+Alpha+Beta-domain+wHTH+Fn-III+Fn-III+RRM | Type-2_BR.RM.like.systems | <-HNH<-?<-?<-HKD-DNase+SF1-SF2-Helicase<-METHYLASE<-DUF3780<-DUF499*<-?\|\|HKD-DNase-><-HEPN-p | 1022 | Bacteria>Terrabacteria_group>Bacillota>Bacilli>Bacillales | GCF_002580305.1 | Bacillus_toyonensis | COF04_RS03765 | 130813..133881 | 155322 | DUF499-domain-containing-protein-[Bacillus-toyonensis] |
| WP_119923933.1 | AAA+ATPase+wHTH+Alpha+Beta-domain+wHTH+Fn-III+Fn-III+RRM | Type-2_BR.RM.like.systems | <-HKD-DNase+SF1-SF2-Helicase<-METHYLASE<-DUF3780<-DUF499*<-HKD-DNase+SF1-SF2-Helicase+TRD<-?\|\|?->?-><-?\|\|Peptidase-> | 1022 | Bacteria>Terrabacteria_group>Bacillota>Bacilli>Bacillales | GCF_003598705.1 | Exiguobacterium_sp._RIT452 | D3D03_RS14170 | 138411..141479 | 2315552 | DUF499-domain-containing-protein-[Exiguobacterium-sp.-RIT452] |
| WP_122925303.1 | AAA+ATPase+wHTH+Alpha+Beta-domain+wHTH+Fn-III+Fn-III+RRM | Type-2_BR.RM.like.systems | SF1-SF2-Helicase->?->DUF2791->?->?->SF1-SF2-Helicase+DUF1998->DUF499*->DUF3780->METHYLASE->HKD-DNase+SF1-SF2-Helicase->?->JetD-> | 1022 | Bacteria>Terrabacteria_group>Bacillota>Bacilli>Bacillales | GCF_003710965.1 | Brevibacillus_nitrificans | EDM59_RS20400 | 101341..104409 | 651560 | DUF499-domain-containing-protein-[Brevibacillus-nitrificans] |
| WP_131490690.1 | AAA+ATPase+wHTH+Alpha+Beta-domain+wHTH+Fn-III+Fn-III+RRM | Type-2_BR.RM.like.systems | DUF499*->DUF3780->METHYLASE->HKD-DNase+SF1-SF2-Helicase->HIN-HTH->PIWI-> | 1022 | Bacteria>Terrabacteria_group>Bacillota>Bacilli>Bacillales | GCF_004337185.1 | Exiguobacterium_sp._SHOS2 | EVJ21_RS13505 | 28496..31564 | 2510950 | DUF499-domain-containing-protein-[Exiguobacterium-sp.-SHOS2] |
| WP_151627816.1 | AAA+ATPase+wHTH+Alpha+Beta-domain+wHTH+Fn-III+Fn-III+RRM | Type-2_BR.RM.like.systems | HNH-><-HKD-DNase+SF1-SF2-Helicase<-METHYLASE<-DUF3780<-DUF499*<-?\|\|HEPN-><-HNH<-REase<-McrC-NTD+REase<-Preader7+McrB-NTD+McrB-AAA | 1022 | Bacteria>Terrabacteria_group>Bacillota>Bacilli>Bacillales | GCF_008923665.1 | Bacillus_luti | F8163_RS24235 | 197732..200800 | 2026191 | DUF499-domain-containing-protein-[Bacillus-luti] |
| WP_153736957.1 | AAA+ATPase+wHTH+Alpha+Beta-domain+wHTH+Fn-III+Fn-III+RRM | Type-2_BR.RM.like.systems | <-REase\|\|SbcC->ParB+HNH->?->?->DUF499*->DUF3780->METHYLASE->HKD-DNase+SF1-SF2-Helicase-><-?<-?<-HTH | 1022 | Bacteria>Terrabacteria_group>Bacillota>Bacilli>Bacillales | GCF_009649745.1 | Aquibacillus_halophilus | GH741_RS11625 | 116271..119339 | 930132 | DUF499-domain-containing-protein-[Aquibacillus-halophilus] |
| WP_179034754.1 | AAA+ATPase+wHTH+Alpha+Beta-domain+wHTH+Fn-III+Fn-III+RRM | Type-2_BR.RM.like.systems | HTH+WYL->Exonuc_V_gamma+REase->SF1-SF2-Helicase+REase->?->?->DUF499*->DUF3780->METHYLASE->HKD-DNase+SF1-SF2-Helicase->?->Calcineurin->SbcC+SbcC-> | 1022 | Bacteria>Terrabacteria_group>Bacillota>Bacilli>Bacillales | GCF_013393385.1 | Paenibacillus_sp._URB8-2 | PUR_RS07815 | 1668256..1671324 | 2741301 | DUF499-domain-containing-protein-[Paenibacillus-sp.-URB8-2] |
| WP_188856936.1 | AAA+ATPase+wHTH+Alpha+Beta-domain+wHTH+Fn-III+Fn-III+RRM | Type-2_BR.RM.like.systems | <-HKD-DNase+SF1-SF2-Helicase<-METHYLASE<-DUF3780<-DUF499*<-?<-SPOUT_MTase<-TnsC-ATPase | 1032 | Bacteria>Terrabacteria_group>Bacillota>Bacilli>Bacillales | GCF_014647195.1 | Oceanobacillus_indicireducens | IEX94_RS09185 | 11196..14294 | 1004261 | DUF499-domain-containing-protein-[Oceanobacillus-indicireducens] |
| WP_190867336.1 | AAA+ATPase+wHTH+Alpha+Beta-domain+wHTH+Fn-III+Fn-III+RRM | Type-2_BR.RM.like.systems | <-DUF2357+REase<-?<-?<-HKD-DNase+SF1-SF2-Helicase<-METHYLASE<-DUF3780<-DUF499*<-?<-?\|\|35exo-><-REase+REase<-SF1-SF2-Helicase+REase<-Exonuc_V_gamma+REase | 1022 | Bacteria>Terrabacteria_group>Bacillota>Bacilli>Bacillales | GCF_014705655.1 | Paenibacillus_arenilitoris | IDH41_RS28595 | 20504..23572 | 2772299 | DUF499-domain-containing-protein-[Paenibacillus-arenilitoris] |
| WP_191705089.1 | AAA+ATPase+wHTH+Alpha+Beta-domain+wHTH+Fn-III+Fn-III+RRM | Type-2_BR.RM.like.systems | <-HKD-DNase+SF1-SF2-Helicase<-METHYLASE<-DUF3780<-DUF499*<-?<-REase+DUF4268<-HTH_31<-?<-SF1-SF2-Helicase+HNH | 1031 | Bacteria>Terrabacteria_group>Bacillota>Bacilli>Bacillales | GCF_014836935.1 | Solibacillus_merdavium | H9632_RS16075 | 8152..11247 | 2762218 | DUF499-domain-containing-protein-[Solibacillus-merdavium] |
| WP_192951079.1 | AAA+ATPase+wHTH+Alpha+Beta-domain+wHTH+Fn-III+Fn-III+RRM | Type-2_BR.RM.like.systems | TRAM+Methyltransf_31->?->REase->MULE-TRANSPOSASE->DUF499*->DUF3780->METHYLASE->HKD-DNase+SF1-SF2-Helicase-><-HTH+MPTase | 1034 | Bacteria>Terrabacteria_group>Bacillota>Bacilli>Bacillales | GCF_014890455.1 | Anoxybacillus_flavithermus | HPK01_RS05390 | 42811..45915 | 33934 | DUF499-domain-containing-protein-[Anoxybacillus-flavithermus] |
| WP_206891906.1 | AAA+ATPase+wHTH+Alpha+Beta-domain+wHTH+Fn-III+Fn-III+RRM | Type-2_BR.RM.like.systems | Peptidase->?->DUF499*->DUF3780->METHYLASE->HKD-DNase+SF1-SF2-Helicase->?->?->HD-> | 1028 | Bacteria>Terrabacteria_group>Bacillota>Bacilli>Bacillales | GCF_017303315.1 | Bacillus_sp._NTK071 | JI666_RS11505 | 25787..28873 | 2802175 | DUF499-domain-containing-protein-[Bacillus-sp.-NTK071] |
| WP_209852303.1 | AAA+ATPase+wHTH+Alpha+Beta-domain+wHTH+Fn-III+Fn-III+RRM | Type-2_BR.RM.like.systems | <-HKD-DNase+SF1-SF2-Helicase<-METHYLASE<-DUF3780<-DUF499*<-?<-?<-HerA/FtsK<-?<-SPOUT_MTase | 1035 | Bacteria>Terrabacteria_group>Bacillota>Bacilli>Bacillales | GCF_017873765.1 | Paenibacillus_sediminis | J2Z20_RS15375 | 6485..9592 | 664909 | DUF499-domain-containing-protein-[Paenibacillus-sediminis] |
| WP_209976485.1 | AAA+ATPase+wHTH+Alpha+Beta-domain+wHTH+Fn-III+Fn-III+RRM | Type-2_BR.RM.like.systems | SPOUT_MTase-><-?<-?\|\|Recombinase+ZnR->?->DUF499*->DUF3780->METHYLASE->HKD-DNase+SF1-SF2-Helicase->?->?->SMODS-> | 1022 | Bacteria>Terrabacteria_group>Bacillota>Bacilli>Bacillales | GCF_017874215.1 | Paenibacillus_eucommiae | J2Z66_RS30220 | 56106..59174 | 1355755 | DUF499-domain-containing-protein-[Paenibacillus-eucommiae] |
| WP_213147727.1 | AAA+ATPase+wHTH+Alpha+Beta-domain+wHTH+Fn-III+Fn-III+RRM | Type-2_BR.RM.like.systems | HNH->?->REase->HTH_31->?->?->DUF499*->DUF3780->METHYLASE->HKD-DNase+SF1-SF2-Helicase-><-?<-?\|\|ZnR+HTH-> | 1033 | Bacteria>Terrabacteria_group>Bacillota>Bacilli>Bacillales | GCF_018343545.2 | Neobacillus_citreus | KHB02_RS06060 | 38173..41274 | 2833578 | DUF499-domain-containing-protein-[Neobacillus-citreus] |
| WP_214909476.1 | AAA+ATPase+wHTH+Alpha+Beta-domain+wHTH+Fn-III+Fn-III+RRM | Type-2_BR.RM.like.systems | <-ZnR+HTH\|\|?->?-><-HKD-DNase+SF1-SF2-Helicase<-METHYLASE<-DUF3780<-DUF499*<-?<-REase+DUF4268<-HTH_31<-REase | 1032 | Bacteria>Terrabacteria_group>Bacillota>Bacilli>Bacillales | GCF_018613155.1 | Bacillus_sp._ISL-55 | J7E32_RS16635 | 33533..36631 | 2819134 | DUF499-domain-containing-protein-[Bacillus-sp.-ISL-55] |
| WP_214792875.1 | AAA+ATPase+wHTH+Alpha+Beta-domain+wHTH+Fn-III+Fn-III+RRM | Type-2_BR.RM.like.systems | <-Phage_int_SAM_5<-URI-DNase\|\|?-><-HKD-DNase+SF1-SF2-Helicase<-METHYLASE<-DUF3780<-DUF499*<-?<-?<-GDResolvase+Recombinase+ZnR | 1033 | Bacteria>Terrabacteria_group>Bacillota>Bacilli>Bacillales | GCF_018613275.1 | Bacillus_sp._ISL-4 | J7J00_RS17675 | 67951..71052 | 2819125 | DUF499-domain-containing-protein-[Bacillus-sp.-ISL-4] |
| WP_220775373.1 | AAA+ATPase+wHTH+Alpha+Beta-domain+wHTH+Fn-III+Fn-III+RRM | Type-2_BR.RM.like.systems | <-GDResolvase<-GDResolvase+Recombinase+ZnR<-?<-HKD-DNase+SF1-SF2-Helicase<-METHYLASE<-DUF3780<-DUF499*<-?<-LuxR-HTH<-?<-?\|\|?-><-HTH+DSBH | 1032 | Bacteria>Terrabacteria_group>Bacillota>Bacilli>Bacillales | GCF_021655155.1 | Bacillus | L6486_RS22210 | 4362036..4365134 | 1386 | MULTISPECIES:-DUF499-domain-containing-protein-[Bacillus] |
| WP_236805686.1 | AAA+ATPase+wHTH+Alpha+Beta-domain+wHTH+Fn-III+Fn-III+RRM | Type-2_BR.RM.like.systems | XIS-HTH->Arm-DNA-bind_4+Phage_int_SAM_3+Phage_integrase->DUF499*->DUF3780->METHYLASE->HKD-DNase+SF1-SF2-Helicase-><-tRNA<-HTH_19\|\|CHTH-> | 1022 | Bacteria>Terrabacteria_group>Bacillota>Bacilli>Bacillales | GCF_021655315.1 | Bacillus | M4D74_RS24360 | 2857..5925 | 1386 | MULTISPECIES:-DUF499-domain-containing-protein-[Bacillus] |
| WP_241475128.1 | AAA+ATPase+wHTH+Alpha+Beta-domain+wHTH+Fn-III+Fn-III+RRM | Type-2_BR.RM.like.systems | XIS-HTH->?->HTH+MPTase-><-HKD-DNase+SF1-SF2-Helicase<-METHYLASE<-DUF3780<-DUF499*\|\|ParB+HNH->?-><-SPOUT_MTase | 1023 | Bacteria>Terrabacteria_group>Bacillota>Bacilli>Bacillales | GCF_022559225.1 | Priestia_flexa | H1W68_RS20670 | 3864077..3867148 | 86664 | DUF499-domain-containing-protein-[Priestia-flexa] |
| WP_097858041.1 | AAA+ATPase+wHTH+Alpha+Beta-domain+wHTH+Fn-III+Fn-III+RRM | Type-2_BR.RM.like.systems | Preader7+McrB-NTD+McrB-AAA->McrC-NTD+REase->REase->HNH-><-HEPN\|\|?->DUF499*->DUF3780->METHYLASE->HKD-DNase+SF1-SF2-Helicase->?->?->MerR-HTH-> | 1022 | Bacteria>Terrabacteria_group>Bacillota>Bacilli>Bacillales | GCF_025516925.1 | Bacillus_wiedmannii | CON82_RS17990 | 124407..127475 | 1890302 | DUF499-domain-containing-protein-[Bacillus-wiedmannii] |
| WP_270407228.1 | AAA+ATPase+wHTH+Alpha+Beta-domain+wHTH+Fn-III+Fn-III+RRM | Type-2_BR.RM.like.systems | MULE-TRANSPOSASE->DUF772+DDE_Tnp_1_2->?->?->DUF499*->DUF3780->METHYLASE->HKD-DNase+SF1-SF2-Helicase-> | 1035 | Bacteria>Terrabacteria_group>Bacillota>Bacilli>Bacillales | GCF_027665165.1 | Paenibacillus_timonensis | PGS16_RS22725 | 6488..9595 | 225915 | DUF499-domain-containing-protein-[Paenibacillus-timonensis] |

| Protein-ID | DUF499-architecture | System | Gene-Neighbourhoods | Length | Taxonomy | Assembly-ID | Strain | Locus-Tag | Genomic-coordinates | TaxID | Fasta-define |
|---|---|---|---|---|---|---|---|---|---|---|---|
| WP_274435612.1 | AAA+ATPase+wHTH+Alpha+Beta-domain+wHTH+Fn-III+Fn-III+RRM | Type-2_BR.RM.like.systems | <-RVT+HNH<-DUF4277+DDE_Tnp_1<-RVT<-HKD-DNase+SF1-SF2-Helicase<-METHYLASE<-DUF3780<-DUF499*<-TUDOR+SF1-SF2-Helicase+iREase+CTD+REase-p | 1058 | Bacteria>Terrabacteria_group>Bacillota>Bacilli>Bacillales | GCF_028751195.1 | Alicyclobacillus_sp._ALC3 | JC200_RS07360 | 1554106..1557282 | 2796143 | DUF499-domain-containing-protein-[Alicyclobacillus-sp.-ALC3] |
| WP_230127616.1 | AAA+ATPase+wHTH+Alpha+Beta-domain+wHTH+Fn-III+Fn-III+RRM | Type-2_BR.RM.like.systems | SPOUT_MTase-><-HTH+MPTase<-?\|\|DUF499*->DUF3780->METHYLASE->HKD-DNase+SF1-SF2-Helicase->HEPN-> | 1022 | Bacteria>Terrabacteria_group>Bacillota>Bacilli>Bacillales | GCF_918741275.1 | Bacillus_sp._CECT_9360 | LQ216_RS05715 | 45654..48722 | 2845821 | DUF499-domain-containing-protein-[Bacillus-sp.-CECT-9360] |
| NLJ18207.1 | AAA+ATPase+wHTH+Alpha+Beta-domain+wHTH+Fn-III+Fn-III+RRM | Type-2_BR.RM.like.systems | <-HKD-DNase+SF1-SF2-Helicase<-METHYLASE<-DUF3780<-DUF499* | 1031 | Bacteria>Terrabacteria_group>Bacillota>Bacilli>Lactobacillales | GCA_012518335.1 | Globicatella_sulfidifaciens | GX355_05025 | 6421..9516 | 136093 | DUF499-domain-containing-protein-[Globicatella-sulfidifaciens] |
| WP_276647681.1 | AAA+ATPase+wHTH+Alpha+Beta-domain+wHTH+Fn-III+Fn-III+RRM | Type-2_BR.RM.like.systems | <-HKD-DNase+SF1-SF2-Helicase<-METHYLASE<-DUF3780<-DUF499* | 1031 | Bacteria>Terrabacteria_group>Bacillota>Bacilli>Lactobacillales | GCF_012518335.1 | Globicatella_sulfidifaciens | GX355_RS05005 | 6421..9516 | 136093 | DUF499-domain-containing-protein-[Globicatella-sulfidifaciens] |
| WP_015908040.1 | AAA+ATPase+wHTH+Alpha+Beta-domain+wHTH+Fn-III+Fn-III+RRM | Type-2_BR.RM.like.systems | <-OLD-ABC+TOPRIM<-ParB+HNH<-HKD-DNase+SF1-SF2-Helicase<-METHYLASE<-DUF3780<-DUF499*<-SF1-SF2-Helicase<-OLD-ABC+TOPRIM<-NACHT<-?<-NERD | 1080 | Bacteria>Terrabacteria_group>Bacillota>Clostridia>Caldicellulosiruptorales | GCF_000022325.1 | Caldicellulosiruptor_bescii | ATHE_RS08025 | 1698593..1701835 | 31899 | DUF499-domain-containing-protein-[Caldicellulosiruptor-bescii] |
| WP_013403049.1 | AAA+ATPase+wHTH+Alpha+Beta-domain+wHTH+Fn-III+Fn-III+RRM | Type-2_BR.RM.like.systems | NERD->?->NACHT->OLD-ABC+TOPRIM->SF1-SF2-Helicase->DUF499*->DUF3780->METHYLASE->HKD-DNase+SF1-SF2-Helicase->ParB+HNH-> | 1080 | Bacteria>Terrabacteria_group>Bacillota>Clostridia>Caldicellulosiruptorales | GCF_000166355.1 | Caldicellulosiruptor_hydrothermalis | CALHY_RS05800 | 1166413..1169655 | 413888 | DUF499-domain-containing-protein-[Caldicellulosiruptor-hydrothermalis] |
| WP_013432875.1 | AAA+ATPase+wHTH+Alpha+Beta-domain+wHTH+Fn-III+Fn-III+RRM | Type-2_BR.RM.like.systems | <-REase<-MethyltransfD12<-SbcC<-HKD-DNase+SF1-SF2-Helicase<-METHYLASE<-DUF3780<-DUF499*<-SbcC<-SF1-SF2-Helicase<-NERD<-?<-?<-Peptidase_M56+Peptidase_M48 | 1080 | Bacteria>Terrabacteria_group>Bacillota>Clostridia>Caldicellulosiruptorales | GCF_000166695.1 | Caldicellulosiruptor_acetigenus | CALKR_RS08085 | 1684950..1688192 | 301953 | DUF499-domain-containing-protein-[Caldicellulosiruptor-acetigenus] |
| WP_013430101.1 | AAA+ATPase+wHTH+Alpha+Beta-domain+wHTH+Fn-III+Fn-III+RRM | Type-2_BR.RM.like.systems | MerR-HTH+GDResolvase->NERD->SF1-SF2-Helicase->?->?->?->DUF499*->DUF3780->METHYLASE->HKD-DNase+SF1-SF2-Helicase->OLD-ABC+TOPRIM-> | 1080 | Bacteria>Terrabacteria_group>Bacillota>Clostridia>Caldicellulosiruptorales | GCF_000166775.1 | Caldicellulosiruptor_kronotskyensis | CALKRO_RS05655 | 1282748..1285990 | 413889 | DUF499-domain-containing-protein-[Caldicellulosiruptor-kronotskyensis] |
| WP_014042422.1 | AAA+ATPase+wHTH+Alpha+Beta-domain+wHTH+Fn-III+Fn-III+RRM | Type-2_BR.RM.like.systems | <-MethyltransfD12<-?<-ParB+HNH<-HKD-DNase+SF1-SF2-Helicase<-METHYLASE<-DUF3780<-DUF499*<-SbcC<-SF1-SF2-Helicase<-NERD<-?<-?<-Peptidase_M56+Peptidase_M48 | 1080 | Bacteria>Terrabacteria_group>Bacillota>Clostridia>Caldicellulosiruptorales | GCF_000193435.2 | Caldicellulosiruptor_acetigenus | CALLA_RS05120 | 1088104..1091346 | 301953 | DUF499-domain-containing-protein-[Caldicellulosiruptor-acetigenus] |
| WP_029228888.1 | AAA+ATPase+wHTH+Alpha+Beta-domain+wHTH+Fn-III+Fn-III+RRM | Type-2_BR.RM.like.systems | Peptidase_M56+Peptidase_M48-><-?\|\|?->?->OrfB_IS605+OrfB_Zn_ribbon->?->DUF499*->DUF3780->METHYLASE->HKD-DNase+SF1-SF2-Helicase->SbcC->MethyltransfD12->REase-> | 1080 | Bacteria>Terrabacteria_group>Bacillota>Clostridia>Caldicellulosiruptorales | GCF_000421725.1 | Caldicellulosiruptor_acetigenus | OTK01_RS05715 | 1153589..1156831 | 301953 | DUF499-domain-containing-protein-[Caldicellulosiruptor-acetigenus] |
| WP_207181944.1 | AAA+ATPase+wHTH+Alpha+Beta-domain+wHTH+Fn-III+Fn-III+RRM | Type-2_BR.RM.like.systems | SF1-SF2-Helicase->SbcC->DUF499*->DUF3780->METHYLASE-p-><-MULE-TRANSPOSASE\|\|METHYLASE->?->HKD-DNase+SF1-SF2-Helicase-> | 1072 | Bacteria>Terrabacteria_group>Bacillota>Clostridia>Caldicellulosiruptorales | GCF_017347585.1 | Caldicellulosiruptor_diazotrophicus | CaldiYA01_RS04625 | 973296..976514 | 2806205 | DUF499-domain-containing-protein-[Caldicellulosiruptor-diazotrophicus] |
| WP_045169275.1 | AAA+ATPase+wHTH+Alpha+Beta-domain+wHTH+Fn-III+Fn-III+RRM | Type-2_BR.RM.like.systems | LON+AAA->?->?->tRNA->tRNA->tRNA->tRNA->tRNA->\|\|tRNA->?->?->DUF499*->DUF3780->METHYLASE->HKD-DNase+SF1-SF2-Helicase-><-?\|\|?->SIGMA-HTH-> | 1081 | Bacteria>Terrabacteria_group>Bacillota>Clostridia>Caldicellulosiruptorales | GCF_026810225.1 | Caldicellulosiruptor_morganii | OTK00_RS04800 | 989349..992594 | 1387555 | DUF499-domain-containing-protein-[Caldicellulosiruptor-morganii] |
| WP_045165444.1 | AAA+ATPase+wHTH+Alpha+Beta-domain+wHTH+Fn-III+Fn-III+RRM | Type-2_BR.RM.like.systems | <-REase<-MethyltransfD12<-?<-HKD-DNase+SF1-SF2-Helicase<-METHYLASE<-DUF3780<-DUF499*<-HEPN<-PolB | 1080 | Bacteria>Terrabacteria_group>Bacillota>Clostridia>Caldicellulosiruptorales | GCF_026914285.1 | Caldicellulosiruptor_naganoensis | N907_RS06310 | 1198788..1202030 | 29324 | DUF499-domain-containing-protein-[Caldicellulosiruptor-naganoensis] |
| WP_045175072.1 | AAA+ATPase+wHTH+Alpha+Beta-domain+wHTH+Fn-III+Fn-III+RRM | Type-2_BR.RM.like.systems | <-PIN<-MPTase<-SbcC<-HKD-DNase+SF1-SF2-Helicase<-METHYLASE<-DUF3780<-DUF499*<-?<-HEPN<-PolB | 1081 | Bacteria>Terrabacteria_group>Bacillota>Clostridia>Caldicellulosiruptorales | GCF_034343125.1 | Caldicellulosiruptor_danielii | N913_RS08680 | 1859027..1862272 | 1387557 | DUF499-domain-containing-protein-[Caldicellulosiruptor-danielii] |
| WP_236914906.1 | AAA+ATPase+wHTH+Alpha+Beta-domain+wHTH+Fn-III+Fn-III+RRM | Type-2_BR.RM.like.systems | HKD-DNase->?->SF1-SF2-Helicase+DUF1998->?->?->DUF499*->DUF3780->METHYLASE->HKD-DNase+SF1-SF2-Helicase-> | 1038 | Bacteria>Terrabacteria_group>Bacillota>Clostridia>Eubacteriales | - | Clostridium_sp._Cult2 | C3E90_06760 | 35293..38409 | 2079003 | DUF499-domain-containing-protein-[Clostridium-sp.-Cult2] |
| AGC68793.1 | AAA+ATPase+wHTH+Alpha+Beta-domain+wHTH+Fn-III+Fn-III+RRM | Type-2_BR.RM.like.systems | MULE-TRANSPOSASE->MULE-TRANSPOSASE-><-HKD-DNase+SF1-SF2-Helicase<-METHYLASE<-DUF3780<-DUF499*\|\|HIN-HTH->MULE-TRANSPOSASE-><-?<-?<-?\|\|MULE-TRANSPOSASE-> | 1100 | Bacteria>Terrabacteria_group>Bacillota>Clostridia>Eubacteriales | GCA_000331995.1 | Thermoclostridium_stercorarium_subsp._stercorarium_DSM_8532 | Cst_c18150 | 1976764..1980066 | 1121335 | hypothetical-protein-Cst_c18150-[Thermoclostridium-stercorarium-subsp.-stercorarium-DSM-8532] |
| KUK65379.1 | AAA+ATPase+wHTH+Alpha+Beta-domain+wHTH+Fn-III+Fn-III+RRM | Type-2_BR.RM.like.systems | <-Radical-SAM<-HKD-DNase+SF1-SF2-Helicase<-METHYLASE<-DUF3780<-DUF499*\|\|DUF772+DDE_Tnp_1_6-><-?\|\|?->?-><-?<-TOPRIM | 1020 | Bacteria>Terrabacteria_group>Bacillota>Clostridia>Eubacteriales | GCA_001508995.1 | Desulfotomaculum_sp._46_80 | XD84_0583 | 17278..20340 | 1641375 | MAG:-Uncharacterized-protein-XD84_0583-[Desulfotomaculum-sp.-46_80] |
| ODM26534.1 | AAA+ATPase+wHTH+Alpha+Beta-domain+wHTH+Fn-III+Fn-III+RRM | Type-2_BR.RM.like.systems | <-HIN-HTH<-HKD-DNase+SF1-SF2-Helicase<-METHYLASE-p<-DDE_5<-METHYLASE<-DUF3780<-DUF499*<-HTH_26<-Peptidase_S8<-Classical-AAA<-METHYLASE-p<-TRAM+Methyltransf_31<-ABC-ATPase+SbcC | 1033 | Bacteria>Terrabacteria_group>Bacillota>Clostridia>Eubacteriales | GCA_001717745.1 | Clostridium_sp._Bc-iso-3 | A7W90_10065 | 2361220..2364321 | 1848158 | glycosyl-transferase-[Clostridium-sp.-Bc-iso-3] |
| PHS35622.1 | AAA+ATPase+wHTH+Alpha+Beta-domain+wHTH+Fn-III+Fn-III+RRM | Type-2_BR.RM.like.systems | <-Pkinase<-REase<-HKD-DNase+SF1-SF2-Helicase<-METHYLASE<-DUF3780<-DUF499*<-?<-?<-DOC+HTH<-BirA-HTH | 1030 | Bacteria>Terrabacteria_group>Bacillota>Clostridia>Eubacteriales | GCA_002733545.1 | Alkaliphilus_sp. | COA82_04085 | 77629..80721 | 1872455 | MAG:-glycosyl-transferase-[Alkaliphilus-sp.] |
| AVX31525.1 | AAA+ATPase+wHTH+Alpha+Beta-domain+wHTH+Fn-III+Fn-III+RRM | Type-2_BR.RM.like.systems | <-REase<-PIN<-REase+NACHT<-HKD-DNase+SF1-SF2-Helicase<-METHYLASE<-DUF3780<-DUF499*<-Phage_int_SAM_4+Phage_integrase<-DEDD_Tnp_IS110+Transposase_20 | 1026 | Bacteria>Terrabacteria_group>Bacillota>Clostridia>Eubacteriales | GCA_003054495.1 | Carboxydocella_thermautotrophica | CTH_1954 | 1949811..1952891 | 178899 | putative-ATPase,-AAA+-superfamily-[Carboxydocella-thermautotrophica] |
| HAG11399.1 | AAA+ATPase+wHTH+Alpha+Beta-domain+wHTH+Fn-III+Fn-III+RRM | Type-2_BR.RM.like.systems | <-Mbetalac<-?<-Radical-SAM<-HKD-DNase+SF1-SF2-Helicase<-METHYLASE<-DUF3780<-DUF499*\|\|DUF772+DDE_Tnp_1_6-><-?\|\|?->?-><-?<-TOPRIM | 1025 | Bacteria>Terrabacteria_group>Bacillota>Clostridia>Eubacteriales | GCA_003445665.1 | Desulfotomaculum_sp. | DCK76_08485 | 61319..64396 | 41211 | glycosyl-transferase-[Desulfotomaculum-sp.] |
| HAS91822.1 | AAA+ATPase+wHTH+Alpha+Beta-domain+wHTH+Fn-III+Fn-III+RRM | Type-2_BR.RM.like.systems | <-wHTH+SAD+HNH<-SF1-SF2-Helicase+REase<-?<-HKD-DNase+SF1-SF2-Helicase<-METHYLASE<-DUF3780<-DUF499*<-?<-Peptidase_S8<-Classical-AAA | 1034 | Bacteria>Terrabacteria_group>Bacillota>Clostridia>Eubacteriales | GCA_003483345.1 | Clostridiales_bacterium | DCS12_06155 | 17914..21018 | 1898207 | glycosyl-transferase-[Clostridiales-bacterium] |
| HBQ28796.1 | AAA+ATPase+wHTH+Alpha+Beta-domain+wHTH+Fn-III+Fn-III+RRM | Type-2_BR.RM.like.systems | <-HKD-DNase+SF1-SF2-Helicase<-METHYLASE<-DUF3780<-DUF499*<-?\|\|DNA-glycosylase-> | 1045 | Bacteria>Terrabacteria_group>Bacillota>Clostridia>Eubacteriales | GCA_003535365.1 | Desulfotomaculum_sp. | DD719_05275 | 9557..12694 | 41211 | glycosyl-transferase-[Desulfotomaculum-sp.] |
| NFR87735.1 | AAA+ATPase+wHTH+Alpha+Beta-domain+wHTH+Fn-III+Fn-III+RRM | Type-2_BR.RM.like.systems | SIGMA-HTH-><-HKD-DNase+SF1-SF2-Helicase<-METHYLASE<-DUF3780<-DUF499* | 1032 | Bacteria>Terrabacteria_group>Bacillota>Clostridia>Eubacteriales | GCA_011018715.1 | Clostridium_botulinum | FDE85_13385 | 8947..12045 | 1491 | DUF499-domain-containing-protein-[Clostridium-botulinum] |
| NJD03475.1 | AAA+ATPase+wHTH+Alpha+Beta-domain+wHTH+Fn-III+Fn-III+RRM | Type-2_BR.RM.like.systems | <-NUDIX<-?<-?<-HKD-DNase+SF1-SF2-Helicase<-METHYLASE<-DUF3780<-DUF499* | 1029 | Bacteria>Terrabacteria_group>Bacillota>Clostridia>Eubacteriales | GCA_012027035.1 | Ruminiclostridium_sp. | FIA99_12975 | 13665..16754 | 2053608 | DUF499-domain-containing-protein-[Ruminiclostridium-sp.] |
| HHY76914.1 | AAA+ATPase+wHTH+Alpha+Beta-domain+wHTH+Fn-III+Fn-III+RRM | Type-2_BR.RM.like.systems | <-URI-DNase<-ParB+HNH<-SbcC<-HKD-DNase+SF1-SF2-Helicase<-METHYLASE<-DUF3780<-DUF499*<-TraJ-RHH | 1098 | Bacteria>Terrabacteria_group>Bacillota>Clostridia>Eubacteriales | GCA_012842725.1 | Clostridiales_bacterium | GX498_00205 | 10822..14118 | 1898207 | DUF499-domain-containing-protein-[Clostridiales-bacterium] |
| MBC7074494.1 | AAA+ATPase+wHTH+Alpha+Beta-domain+wHTH+Fn-III+Fn-III+RRM | Type-2_BR.RM.like.systems | HHH_5+SF1-SF2-helicase->LON-AAA->?-><-?<-ZnR\|\|?->DUF499*->DUF3780->METHYLASE->HKD-DNase+SF1-SF2-Helicase->?->REase+RAMA->DUF6361-> | 1041 | Bacteria>Terrabacteria_group>Bacillota>Clostridia>Eubacteriales | GCA_014360155.1 | Syntrophomonadaceae_bacterium | H5T98_00170 | 31047..34172 | 2093811 | DUF499-domain-containing-protein-[Syntrophomonadaceae-bacterium] |
| MBD5170374.1 | AAA+ATPase+wHTH+Alpha+Beta-domain+wHTH+Fn-III+Fn-III+RRM | Type-2_BR.RM.like.systems | ACET->NTP_transf_6->?->NTP_transf_2->?->?->DUF499*->DUF3780->METHYLASE->HKD-DNase+SF1-SF2-Helicase-> | 1039 | Bacteria>Terrabacteria_group>Bacillota>Clostridia>Eubacteriales | GCA_014799795.1 | Oscillibacter_sp. | HDT20_09715 | 9924..13043 | 1945593 | DUF499-domain-containing-protein-[Oscillibacter-sp.] |
| MBD5152197.1 | AAA+ATPase+wHTH+Alpha+Beta-domain+wHTH+Fn-III+Fn-III+RRM | Type-2_BR.RM.like.systems | NTP_transf_6->?->NTP_transf_2->?->?->DUF499*->DUF3780->METHYLASE->HKD-DNase+SF1-SF2-Helicase-> | 1039 | Bacteria>Terrabacteria_group>Bacillota>Clostridia>Eubacteriales | GCA_014799885.1 | Oscillibacter_sp. | HDT16_06850 | 18982..22101 | 1945593 | DUF499-domain-containing-protein-[Oscillibacter-sp.] |
| MBD5154273.1 | AAA+ATPase+wHTH+Alpha+Beta-domain+wHTH+Fn-III+Fn-III+RRM | Type-2_BR.RM.like.systems | ACET->NTP_transf_6->?->NTP_transf_2->Peptidase_S51->?->DUF499*->DUF3780->METHYLASE->HKD-DNase+SF1-SF2-Helicase-> | 1039 | Bacteria>Terrabacteria_group>Bacillota>Clostridia>Eubacteriales | GCA_014799915.1 | Oscillibacter_sp. | HDT15_04160 | 19111..22230 | 1945593 | DUF499-domain-containing-protein-[Oscillibacter-sp.] |
| MBD8933315.1 | AAA+ATPase+wHTH+Alpha+Beta-domain+wHTH+Fn-III+Fn-III+RRM | Type-2_BR.RM.like.systems | DUF499*->DUF3780->METHYLASE->HKD-DNase+SF1-SF2-Helicase->SLATT->SMODS-> | 1039 | Bacteria>Terrabacteria_group>Bacillota>Clostridia>Eubacteriales | GCA_014858285.1 | Lachnospiraceae_bacterium | EGR91_05070 | 108..3227 | 1898203 | DUF499-domain-containing-protein-[Lachnospiraceae-bacterium] |
| MBE6059264.1 | AAA+ATPase+wHTH+Alpha+Beta-domain+wHTH+Fn-III+Fn-III+RRM | Type-2_BR.RM.like.systems | <-Radical-SAM<-?<-TetR-HTH\|\|TRAM+Methyltransf_31->?->DUF499*->DUF3780->METHYLASE->HKD-DNase+SF1-SF2-Helicase->REase+RAMA->?-><-HTH_3 | 1031 | Bacteria>Terrabacteria_group>Bacillota>Clostridia>Eubacteriales | GCA_015058365.1 | Clostridium_sulfidigenes | E7215_03695 | 31892..34987 | 318464 | DUF499-domain-containing-protein-[Clostridium-sulfidigenes] |
| MBM7867397.1 | AAA+ATPase+wHTH+Alpha+Beta-domain+wHTH+Fn-III+Fn-III+RRM | Type-2_BR.RM.like.systems | REase->DUF499*->DUF3780->METHYLASE->HKD-DNase+SF1-SF2-Helicase->ParB+HNH-p->?->Classical-AAA-> | 1023 | Bacteria>Terrabacteria_group>Bacillota>Clostridia>Eubacteriales | GCA_016908025.1 | Heliomicrobium_gestii | JOC69_002303 | 75173..78244 | 2699 | hypothetical-protein-[Heliomicrobium-gestii] |
| MBR2257284.1 | AAA+ATPase+wHTH+Alpha+Beta-domain+wHTH+Fn-III+Fn-III+RRM | Type-2_BR.RM.like.systems | HTH_26->DUF499*->DUF3780->METHYLASE->HKD-DNase+SF1-SF2-Helicase-> | 1043 | Bacteria>Terrabacteria_group>Bacillota>Clostridia>Eubacteriales | GCA_017475035.1 | Blautia_sp. | IJ899_08090 | 51779..54910 | 1955243 | DUF499-domain-containing-protein-[Blautia-sp.] |
| MBP3806185.1 | AAA+ATPase+wHTH+Alpha+Beta-domain+wHTH+Fn-III+Fn-III+RRM | Type-2_BR.RM.like.systems | <-GNAT<-SbcC\|\|35exo->HTH_26->DUF499*->DUF3780->METHYLASE->HKD-DNase+SF1-SF2-Helicase->SMODS->?->URI-DNase-> | 1042 | Bacteria>Terrabacteria_group>Bacillota>Clostridia>Eubacteriales | GCA_017940955.1 | Oribacterium_sp. | J6I76_20230 | 3385..6513 | 1969407 | DUF499-domain-containing-protein-[Oribacterium-sp.] |
| MCC0639043.1 | AAA+ATPase+wHTH+Alpha+Beta-domain+wHTH+Fn-III+Fn-III+RRM | Type-2_BR.RM.like.systems | <-HKD-DNase+SF1-SF2-Helicase<-METHYLASE<-DUF3780<-DUF499*\|\|<-?<-?<-?<-PRTase | 1031 | Bacteria>Terrabacteria_group>Bacillota>Clostridia>Eubacteriales | GCA_020658435.1 | Clostridioides_sp._ES-S-0049-03 | IC212_02425 | 567103..570198 | 2770779 | DUF499-domain-containing-protein-[Clostridioides-sp.-ES-S-0049-03] |
| MCF6465586.1 | AAA+ATPase+wHTH+Alpha+Beta-domain+wHTH+Fn-III+Fn-III+RRM | Type-2_BR.RM.like.systems | HKD-DNase->?->SF1-SF2-Helicase+DUF1998->?->?->DUF499*->DUF3780->METHYLASE->HKD-DNase+SF1-SF2-Helicase-> | 1038 | Bacteria>Terrabacteria_group>Bacillota>Clostridia>Eubacteriales | GCA_021655075.1 | Clostridium_sp._Cult2 | C3E90_06760 | 35293..38409 | 2079003 | glycosyl-transferase-[Clostridium-sp.-Cult2] |
| MDA8227626.1 | AAA+ATPase+wHTH+Alpha+Beta-domain+wHTH+Fn-III+Fn-III+RRM | Type-2_BR.RM.like.systems | DUF499*->DUF3780->METHYLASE->HKD-DNase+SF1-SF2-Helicase-p-> | 1038 | Bacteria>Terrabacteria_group>Bacillota>Clostridia>Eubacteriales | GCA_028065655.1 | Desulfitobacterium_hafniense | MOT74_07965 | 111..3227 | 49338 | MAG:-DUF499-domain-containing-protein-[Desulfitobacterium-hafniense] |
| MDA8346579.1 | AAA+ATPase+wHTH+Alpha+Beta-domain+wHTH+Fn-III+Fn-III+RRM | Type-2_BR.RM.like.systems | SIGMA-HTH->?->PolB->HEPN->?->DUF499*->DUF3780->METHYLASE->HKD-DNase+SF1-SF2-Helicase-> | 1028 | Bacteria>Terrabacteria_group>Bacillota>Clostridia>Eubacteriales | GCA_028068135.1 | Thermaerobacter_sp. | MOZ66_14030 | 66749..69835 | 1937527 | MAG:-DUF499-domain-containing-protein-[Thermaerobacter-sp.] |
| MDC7291477.1 | AAA+ATPase+wHTH+Alpha+Beta-domain+wHTH+Fn-III+Fn-III+RRM | Type-2_BR.RM.like.systems | <-ParB+HNH<-?<-HKD-DNase+SF1-SF2-Helicase<-METHYLASE<-DUF3780<-DUF499*<-HTH_26<-DDE_Tnp_1<-?<-GDResolvase<-GDResolvase<-GDResolvase | 1039 | Bacteria>Terrabacteria_group>Bacillota>Clostridia>Eubacteriales | GCA_028539195.1 | Blautia_schinkii | NXH76_27225 | 12059..15178 | 180164 | DUF499-domain-containing-protein-[Blautia-schinkii] |
| WP_003361845.1 | AAA+ATPase+wHTH+Alpha+Beta-domain+wHTH+Fn-III+Fn-III+RRM | Type-2_BR.RM.like.systems | <-PADR-HTH\|\|wHTH(SBD)+HNH->?->DUF499*->DUF3780->METHYLASE->HKD-DNase+SF1-SF2-Helicase->REase+RAMA->?->Radical-SAM-> | 1031 | Bacteria>Terrabacteria_group>Bacillota>Clostridia>Eubacteriales | GCF_000020345.1 | Clostridium_botulinum | AGE29_RS02755 | 251614..254709 | 1491 | DUF499-domain-containing-protein-[Clostridium-botulinum] |
| WP_015260687.1 | AAA+ATPase+wHTH+Alpha+Beta-domain+wHTH+Fn-III+Fn-III+RRM | Type-2_BR.RM.like.systems | HTH_26->DUF499*->DUF3780->METHYLASE->HKD-DNase+SF1-SF2-Helicase->?->OLD-ABC+TOPRIM->SF1-SF2-Helicase-> | 1038 | Bacteria>Terrabacteria_group>Bacillota>Clostridia>Eubacteriales | GCF_000243135.2 | Desulfitobacterium_dichloroeliminans | DESDI_RS00585 | 115746..118862 | 233055 | DUF499-domain-containing-protein-[Desulfitobacterium-dichloroeliminans] |
| WP_008908701.1 | AAA+ATPase+wHTH+Alpha+Beta-domain+wHTH+Fn-III+Fn-III+RRM | Type-2_BR.RM.like.systems | DUF499*->METHYLASE->HKD-DNase+SF1-SF2-Helicase->?->?->?-><-ZnR | 1035 | Bacteria>Terrabacteria_group>Bacillota>Clostridia>Eubacteriales | GCF_000297115.1 | Caloramator_australicus | CAAU_RS06845 | 1386..4493 | 515264 | DUF499-domain-containing-protein-[Caloramator-australicus] |
| WP_242823541.1 | AAA+ATPase+wHTH+Alpha+Beta-domain+wHTH+Fn-III+Fn-III+RRM | Type-2_BR.RM.like.systems | <-HKD-DNase+SF1-SF2-Helicase<-METHYLASE<-DUF3780<-DUF499*\|\|MULE-TRANSPOSASE-> | 1085 | Bacteria>Terrabacteria_group>Bacillota>Clostridia>Eubacteriales | GCF_000331995.1 | Thermoclostridium_stercorarium | CLST_RS08500 | 1974405..1977662 | 1510 | DUF499-domain-containing-protein-[Thermoclostridium-stercorarium] |
| WP_018086521.1 | AAA+ATPase+wHTH+Alpha+Beta-domain+wHTH+Fn-III+Fn-III+RRM | Type-2_BR.RM.like.systems | Polbeta->?-><-HKD-DNase+SF1-SF2-Helicase<-METHYLASE<-DUF3780<-DUF499* | 1035 | Bacteria>Terrabacteria_group>Bacillota>Clostridia>Eubacteriales | GCF_000376385.1 | Desulfurispora_thermophila | B064_RS0111620 | 166713..169820 | 265470 | DUF499-domain-containing-protein-[Desulfurispora-thermophila] |
| WP_029232559.1 | AAA+ATPase+wHTH+Alpha+Beta-domain+wHTH+Fn-III+Fn-III+RRM | Type-2_BR.RM.like.systems | <-ParB+HNH<-ParB+HNH<-HKD-DNase+SF1-SF2-Helicase<-METHYLASE<-DUF3780<-DUF499*<-XIS-HTH<-?<-?<-ZnR | 1039 | Bacteria>Terrabacteria_group>Bacillota>Clostridia>Eubacteriales | GCF_000420945.1 | Butyrivibrio_sp._VCB2006 | G613_RS0110145 | 46570..49689 | 1280679 | DUF499-domain-containing-protein-[Butyrivibrio-sp.-VCB2006] |
| WP_018666427.1 | AAA+ATPase+wHTH+Alpha+Beta-domain+wHTH+Fn-III+Fn-III+RRM | Type-2_BR.RM.like.systems | <-METHYLASE<-METHYLASE-p<-HKD-DNase+SF1-SF2-Helicase<-METHYLASE-p<-DUF3780<-DUF499* | 1035 | Bacteria>Terrabacteria_group>Bacillota>Clostridia>Eubacteriales | GCF_000430995.1 | Thermobrachium_celere | TCEL_RS11875 | 61747..64854 | 53422 | DUF499-domain-containing-protein-[Thermobrachium-celere] |
| WP_023162377.1 | AAA+ATPase+wHTH+Alpha+Beta-domain+wHTH+Fn-III+Fn-III+RRM | Type-2_BR.RM.like.systems | SIGMA-HTH->?->?->?->?->?->DUF499*->DUF3780->METHYLASE->HKD-DNase+SF1-SF2-Helicase->Schlafen+HATPase+HTH-> | 1033 | Bacteria>Terrabacteria_group>Bacillota>Clostridia>Eubacteriales | GCF_000484505.2 | Clostridium_autoethanogenum | WX72_RS00855 | 181813..184914 | 84023 | DUF499-domain-containing-protein-[Clostridium-autoethanogenum] |
| WP_039231429.1 | AAA+ATPase+wHTH+Alpha+Beta-domain+wHTH+Fn-III+Fn-III+RRM | Type-2_BR.RM.like.systems | OrfB_IS605+OrfB_Zn_ribbon->SAM-methylase->?->tRNA->tRNA->tRNA->tRNA->tRNA->?->?->DUF499*->DUF3780->METHYLASE->HKD-DNase+SF1-SF2-Helicase->REase+RAMA-> | 1031 | Bacteria>Terrabacteria_group>Bacillota>Clostridia>Eubacteriales | GCF_000724265.2 | Clostridium_novyi | Z958_RS01340 | 4777..7872 | 1542 | DUF499-domain-containing-protein-[Clostridium-novyi] |
| WP_036935476.1 | AAA+ATPase+wHTH+Alpha+Beta-domain+wHTH+Fn-III+Fn-III+RRM | Type-2_BR.RM.like.systems | CHTH->MPTase-><-HKD-DNase+SF1-SF2-Helicase<-METHYLASE<-DUF3780<-DUF499*<-HTH_36<-?<-?<-Peptidase | 1036 | Bacteria>Terrabacteria_group>Bacillota>Clostridia>Eubacteriales | GCF_000745845.1 | Pseudobacteroides_cellulosolvens | Bccel_RS23015 | 5550747..5553857 | 35825 | DUF499-domain-containing-protein-[Pseudobacteroides-cellulosolvens] |
| WP_039680700.1 | AAA+ATPase+wHTH+Alpha+Beta-domain+wHTH+Fn-III+Fn-III+RRM | Type-2_BR.RM.like.systems | <-HKD-DNase+SF1-SF2-Helicase<-METHYLASE<-DUF3780<-DUF499* | 1032 | Bacteria>Terrabacteria_group>Bacillota>Clostridia>Eubacteriales | GCF_000808015.1 | Terrisporobacter_othiniensis | QX51_RS14940 | 7169..10267 | 1577792 | DUF499-domain-containing-protein-[Terrisporobacter-othiniensis] |
| WP_039774266.1 | AAA+ATPase+wHTH+Alpha+Beta-domain+wHTH+Fn-III+Fn-III+RRM | Type-2_BR.RM.like.systems | <-PADR-HTH\|\|wHTH(SBD)+HNH->?->DUF499*->DUF3780->METHYLASE->HKD-DNase+SF1-SF2-Helicase->REase+RAMA->?->Radical-SAM-> | 1031 | Bacteria>Terrabacteria_group>Bacillota>Clostridia>Eubacteriales | GCF_000817935.1 | Clostridium_botulinum | T259_RS11600 | 2303984..2307079 | 1491 | DUF499-domain-containing-protein-[Clostridium-botulinum] |
| WP_061317398.1 | AAA+ATPase+wHTH+Alpha+Beta-domain+wHTH+Fn-III+Fn-III+RRM | Type-2_BR.RM.like.systems | <-Radical-SAM<-REase+RAMA<-HKD-DNase+SF1-SF2-Helicase<-METHYLASE-p<-METHYLASE<-DUF3780<-DUF499*<-?<-wHTH(SBD)+HNH\|\|PADR-HTH-> | 1031 | Bacteria>Terrabacteria_group>Bacillota>Clostridia>Eubacteriales | GCF_001573255.1 | Clostridium_botulinum | AZI51_RS06445 | 49483..52578 | 1491 | DUF499-domain-containing-protein-[Clostridium-botulinum] |
| WP_061320311.1 | AAA+ATPase+wHTH+Alpha+Beta-domain+wHTH+Fn-III+Fn-III+RRM | Type-2_BR.RM.like.systems | wHTH(SBD)+HNH->?->DUF499*->DUF3780->METHYLASE->HKD-DNase+SF1-SF2-Helicase->REase+RAMA->?->Radical-SAM-> | 1031 | Bacteria>Terrabacteria_group>Bacillota>Clostridia>Eubacteriales | GCF_001573935.1 | Clostridium_botulinum | AZI72_RS02380 | 8952..12047 | 1491 | DUF499-domain-containing-protein-[Clostridium-botulinum] |
| WP_065821436.1 | AAA+ATPase+wHTH+Alpha+Beta-domain+wHTH+Fn-III+Fn-III+RRM | Type-2_BR.RM.like.systems | MULE-TRANSPOSASE-><-HD<-HKD-DNase+SF1-SF2-Helicase<-REase<-METHYLASE<-DUF3780<-DUF499*\|\|MULE-TRANSPOSASE-><-?<-?\|\|MULE-TRANSPOSASE-> | 1036 | Bacteria>Terrabacteria_group>Bacillota>Clostridia>Eubacteriales | GCF_001697245.1 | Thermoclostridium_stercorarium | CSTERTH_RS08760 | 2032374..2035484 | 1510 | DUF499-domain-containing-protein-[Thermoclostridium-stercorarium] |
| WP_034844279.1 | AAA+ATPase+wHTH+Alpha+Beta-domain+wHTH+Fn-III+Fn-III+RRM | Type-2_BR.RM.like.systems | HIN-HTH-><-?<-HIN-HTH<-HIN-HTH<-ZnR\|\|TraJ-RHH->DUF499*->DUF3780->METHYLASE->REase->HKD-DNase+SF1-SF2-Helicase->?->URI-DNase-> | 1036 | Bacteria>Terrabacteria_group>Bacillota>Clostridia>Eubacteriales | GCF_001697285.1 | Thermoclostridium_stercorarium | CSTERLE_RS00935 | 220886..223996 | 1510 | DUF499-domain-containing-protein-[Thermoclostridium-stercorarium] |
| WP_070957121.1 | AAA+ATPase+wHTH+Alpha+Beta-domain+wHTH+Fn-III+Fn-III+RRM | Type-2_BR.RM.like.systems | <-Peptidase-S8+Subtilisin<-Peptidase-S8+Subtilisin<-HTH_3<-HKD-DNase+SF1-SF2-Helicase<-METHYLASE<-DUF3780<-DUF499*\|\|GDResolvase+Recombinase+ZnR-><-?\|\|?->GDResolvase-><-TRAM+Methyltransf_31 | 1031 | Bacteria>Terrabacteria_group>Bacillota>Clostridia>Eubacteriales | GCF_001854085.1 | Clostridium_perfringens | FXZ97_RS12165 | 2722294..2725389 | 1502 | DUF499-domain-containing-protein-[Clostridium-perfringens] |
| WP_101515073.1 | AAA+ATPase+wHTH+Alpha+Beta-domain+wHTH+Fn-III+Fn-III+RRM | Type-2_BR.RM.like.systems | <-Radical-SAM<-?<-REase+RAMA<-HKD-DNase+SF1-SF2-Helicase<-METHYLASE<-DUF3780<-DUF499*<-?<-wHTH(SBD)+HNH | 1031 | Bacteria>Terrabacteria_group>Bacillota>Clostridia>Eubacteriales | GCF_002866225.1 | Clostridium_botulinum | RSJ21_RS11400 | 2254845..2257940 | 1491 | DUF499-domain-containing-protein-[Clostridium-botulinum] |
| WP_106024681.1 | AAA+ATPase+wHTH+Alpha+Beta-domain+wHTH+Fn-III+Fn-III+RRM | Type-2_BR.RM.like.systems | TnsC-ATPase->?->?->RNase->?->?->DUF499*->DUF3780->METHYLASE->HKD-DNase+SF1-SF2-Helicase->?->SMODS->UbE1+JAB-> | 1036 | Bacteria>Terrabacteria_group>Bacillota>Clostridia>Eubacteriales | GCF_002995795.1 | Clostridium_thermopalmarium | LX19_RS07985 | 59154..62264 | 29373 | DUF499-domain-containing-protein-[Clostridium-thermopalmarium] |
| WP_106574724.1 | AAA+ATPase+wHTH+Alpha+Beta-domain+wHTH+Fn-III+Fn-III+RRM | Type-2_BR.RM.like.systems | <-Radical-SAM<-REase+RAMA<-HKD-DNase+SF1-SF2-Helicase<-METHYLASE<-DUF3780<-DUF499*<-?<-?\|\|?->?-><-sGTP | 1031 | Bacteria>Terrabacteria_group>Bacillota>Clostridia>Eubacteriales | GCF_003014955.1 | Clostridium_botulinum | C6C12_RS10395 | 111152..114247 | 1491 | DUF499-domain-containing-protein-[Clostridium-botulinum] |
| WP_110463402.1 | AAA+ATPase+wHTH+Alpha+Beta-domain+wHTH+Fn-III+Fn-III+RRM | Type-2_BR.RM.like.systems | McrC-NTD+REase-><-HKD-DNase+SF1-SF2-Helicase<-METHYLASE<-DUF3780<-DUF499*<-?<-?<-?<-GDResolvase<-TOPRIM<-OLD-ABC | 1032 | Bacteria>Terrabacteria_group>Bacillota>Clostridia>Eubacteriales | GCF_003208175.1 | Ruminiclostridium_sufflavum | LY28_RS17135 | 12744..15842 | 396504 | DUF499-domain-containing-protein-[Ruminiclostridium-sufflavum] |
| WP_117567862.1 | AAA+ATPase+wHTH+Alpha+Beta-domain+wHTH+Fn-III+Fn-III+RRM | Type-2_BR.RM.like.systems | HTH_26->DUF499*->DUF3780->METHYLASE->HKD-DNase+SF1-SF2-Helicase->SLATT->SMODS-> | 1039 | Bacteria>Terrabacteria_group>Bacillota>Clostridia>Eubacteriales | GCF_003435125.1 | Clostridium_sp._AM25-23AC | DW677_RS16735 | 1639..4758 | 2305240 | DUF499-domain-containing-protein-[Clostridium-sp.-AM25-23AC] |
| WP_138979355.1 | AAA+ATPase+wHTH+Alpha+Beta-domain+wHTH+Fn-III+Fn-III+RRM | Type-2_BR.RM.like.systems | <-FAD-NAD-dep-oxidoreductase<-?<-?<-HKD-DNase+SF1-SF2-Helicase<-METHYLASE<-DUF3780<-DUF499*<-?<-?<-SIGMA-HTH<-?<-?<-MACRODOMAIN | 1035 | Bacteria>Terrabacteria_group>Bacillota>Clostridia>Eubacteriales | GCF_006016075.1 | Caloramator_sp._E03 | FDN13_RS05925 | 1176985..1180092 | 2576307 | DUF499-domain-containing-protein-[Caloramator-sp.-E03] |
| WP_161840113.1 | AAA+ATPase+wHTH+Alpha+Beta-domain+wHTH+Fn-III+Fn-III+RRM | Type-2_BR.RM.like.systems | <-HKD-DNase+SF1-SF2-Helicase<-METHYLASE<-DUF3780<-DUF499*<-HNH<-wHTH(SBD)+HNH<-wHTH(SBD)+HNH<-SF1-SF2-Helicase+REase<-SF1-SF2-Helicase+REase<-HAMP+HISKIN | 1027 | Bacteria>Terrabacteria_group>Bacillota>Clostridia>Eubacteriales | GCF_009931695.1 | Anaerocolumna_sedimenticola | Ana3638_RS22990 | 5160581..5163664 | 2696063 | DUF499-domain-containing-protein-[Anaerocolumna-sedimenticola] |
| WP_164948359.1 | AAA+ATPase+wHTH+Alpha+Beta-domain+wHTH+Fn-III+Fn-III+RRM | Type-2_BR.RM.like.systems | zf-IS66+DDE_Tnp_IS66->DUF499*->DUF3780->METHYLASE->OLD-ABC->NERD->HKD-DNase+SF1-SF2-Helicase-> | 1031 | Bacteria>Terrabacteria_group>Bacillota>Clostridia>Eubacteriales | GCF_011065955.2 | Clostridium_sp._FP2 | G9F71_RS10335 | 2159683..2162778 | 2724481 | DUF499-domain-containing-protein-[Clostridium-sp.-FP2] |
| WP_235920429.1 | AAA+ATPase+wHTH+Alpha+Beta-domain+wHTH+Fn-III+Fn-III+RRM | Type-2_BR.RM.like.systems | REase->DUF499*->DUF3780->METHYLASE->HKD-DNase+SF1-SF2-Helicase->ParB+HNH-p->?->Classical-AAA-> | 942 | Bacteria>Terrabacteria_group>Bacillota>Clostridia>Eubacteriales | GCF_016908025.1 | Heliomicrobium_gestii | JOC69_RS11260 | 75416..78244 | 2699 | DUF499-domain-containing-protein-[Heliomicrobium-gestii] |
| WP_206812928.1 | AAA+ATPase+wHTH+Alpha+Beta-domain+wHTH+Fn-III+Fn-III+RRM | Type-2_BR.RM.like.systems | <-SIR2<-HKD-DNase+SF1-SF2-Helicase<-METHYLASE<-DUF3780<-DUF499*<-CHTH | 1039 | Bacteria>Terrabacteria_group>Bacillota>Clostridia>Eubacteriales | GCF_017310645.1 | Paradesulfitobacterium_ferrireducens | JOX21_RS14505 | 49501..52620 | 2816476 | DUF499-domain-containing-protein-[Paradesulfitobacterium-ferrireducens] |
| WP_211802356.1 | AAA+ATPase+wHTH+Alpha+Beta-domain+wHTH+Fn-III+Fn-III+RRM | Type-2_BR.RM.like.systems | <-PAPS_reduct<-?<-KAP-NTPase<-HKD-DNase+SF1-SF2-Helicase<-METHYLASE<-DUF3780<-DUF499*<-HTH_26<-MPTase<-CHTH\|\|?->?->SIGMA-HTH-> | 1038 | Bacteria>Terrabacteria_group>Bacillota>Clostridia>Eubacteriales | GCF_018128025.1 | Proteiniclasticum_sediminis | KCG48_RS11535 | 23162..26278 | 2804028 | DUF499-domain-containing-protein-[Proteiniclasticum-sediminis] |
| WP_216398042.1 | AAA+ATPase+wHTH+Alpha+Beta-domain+wHTH+Fn-III+Fn-III+RRM | Type-2_BR.RM.like.systems | HTH_26->DUF499*->DUF3780->METHYLASE->HKD-DNase+SF1-SF2-Helicase-> | 1033 | Bacteria>Terrabacteria_group>Bacillota>Clostridia>Eubacteriales | GCF_018917405.1 | Anaerocolumna_aminovalerica | KQI61_RS20640 | 728..3829 | 1527 | DUF499-domain-containing-protein-[Anaerocolumna-aminovalerica] |
| WP_272879084.1 | AAA+ATPase+wHTH+Alpha+Beta-domain+wHTH+Fn-III+Fn-III+RRM | Type-2_BR.RM.like.systems | HKD-DNase->?->SF1-SF2-Helicase+DUF1998->?->DUF499*->DUF3780->METHYLASE->HKD-DNase+SF1-SF2-Helicase-> | 956 | Bacteria>Terrabacteria_group>Bacillota>Clostridia>Eubacteriales | GCF_021655075.1 | Clostridium_sp._Cult2 | C3E90_RS06775 | 35539..38409 | 2079003 | DUF499-domain-containing-protein-[Clostridium-sp.-Cult2] |
| WP_245124324.1 | AAA+ATPase+wHTH+Alpha+Beta-domain+wHTH+Fn-III+Fn-III+RRM | Type-2_BR.RM.like.systems | sGTP-><-?<-?\|\|?->wHTH(SBD)+HNH->?->DUF499*->DUF3780->METHYLASE->HKD-DNase+SF1-SF2-Helicase->REase+RAMA->Radical-SAM-> | 1031 | Bacteria>Terrabacteria_group>Bacillota>Clostridia>Eubacteriales | GCF_022936445.1 | Clostridium_botulinum | LO064_RS12750 | 357814..360909 | 1491 | DUF499-domain-containing-protein-[Clostridium-botulinum] |
| WP_255228679.1 | AAA+ATPase+wHTH+Alpha+Beta-domain+wHTH+Fn-III+Fn-III+RRM | Type-2_BR.RM.like.systems | JetD->?->?->?->OLD-ABC->TOPRIM->DUF499*->DUF3780->METHYLASE->HKD-DNase+SF1-SF2-Helicase->GNTR-HTH->Aminotran_1_2-> | 1039 | Bacteria>Terrabacteria_group>Bacillota>Clostridia>Eubacteriales | GCF_024363565.1 | Lutispora_saccharofermentans | LJD61_RS16670 | 81318..84437 | 3024236 | DUF499-domain-containing-protein-[Lutispora-thermophila] |
| WP_163278136.1 | AAA+ATPase+wHTH+Alpha+Beta-domain+wHTH+Fn-III+Fn-III+RRM | Type-2_BR.RM.like.systems | <-HIN-HTH<-HKD-DNase+SF1-SF2-Helicase<-METHYLASE<-DUF3780<-DUF499*<-HTH_26<-Peptidase_S8<-Classical-AAA<-?<-TRAM+Methyltransf_31 | 1033 | Bacteria>Terrabacteria_group>Bacillota>Clostridia>Eubacteriales | GCF_026130265.1 | Clostridium | FCV37_RS07265 | 127477..130578 | 1485 | DUF499-domain-containing-protein-[Clostridium-sporogenes] |
| WP_180703528.1 | AAA+ATPase+wHTH+Alpha+Beta-domain+wHTH+Fn-III+Fn-III+RRM | Type-2_BR.RM.like.systems | <-Phage_int_SAM_1+Phage_integrase\|\|Methyltransf_4-><-?\|\|?->DUF499*->DUF3780->METHYLASE->HKD-DNase+SF1-SF2-Helicase->REase->Phage_integrase->LexA-HTH+LexA-protease-> | 1029 | Bacteria>Terrabacteria_group>Bacillota>Clostridia>Eubacteriales | GCF_900015215.1 | Romboutsia_ilealis | CRIB_RS05530 | 1166039..1169128 | 1115758 | DUF499-domain-containing-protein-[Romboutsia-ilealis] |
| WP_092330897.1 | AAA+ATPase+wHTH+Alpha+Beta-domain+wHTH+Fn-III+Fn-III+RRM | Type-2_BR.RM.like.systems | <-wHTH(SBD)+HNH<-?<-HKD-DNase+SF1-SF2-Helicase<-METHYLASE<-DUF3780<-DUF499*<-JetD<-?<-HTH\|\|REC-> | 1022 | Bacteria>Terrabacteria_group>Bacillota>Clostridia>Eubacteriales | GCF_900100785.1 | Desulfosporosinus_hippei | BLS16_RS05460 | 173725..176793 | 569859 | DUF499-domain-containing-protein-[Desulfosporosinus-hippei] |
| WP_092282215.1 | AAA+ATPase+wHTH+Alpha+Beta-domain+wHTH+Fn-III+Fn-III+RRM | Type-2_BR.RM.like.systems | <-SF1-SF2-Helicase<-SIGMA-HTH+SIGMA-HTH<-?<-HKD-DNase+SF1-SF2-Helicase<-METHYLASE<-DUF3780<-DUF499*<-?<-RNB+RNase<-HD | 1035 | Bacteria>Terrabacteria_group>Bacillota>Clostridia>Eubacteriales | GCF_900115765.1 | Caldicoprobacter_faecalis | BM241_RS05315 | 48180..51287 | 937334 | DUF499-domain-containing-protein-[Caldicoprobacter-faecalis] |
| WP_226926778.1 | AAA+ATPase+wHTH+Alpha+Beta-domain+wHTH+Fn-III+Fn-III+RRM | Type-2_BR.RM.like.systems | <-HKD-DNase+SF1-SF2-Helicase<-METHYLASE<-DUF3780<-DUF499* | 1032 | Bacteria>Terrabacteria_group>Bacillota>Clostridia>Eubacteriales | GCF_937997615.1 | Intestinibacter_bartlettii | LIQ82_RS01500 | 333183..336281 | 261299 | DUF499-domain-containing-protein-[Intestinibacter-bartlettii] |
| WP_265923632.1 | AAA+ATPase+wHTH+Alpha+Beta-domain+wHTH+Fn-III+Fn-III+RRM | Type-2_BR.RM.like.systems | <-Peptidase-S8+Subtilisin<-Peptidase-S8+Subtilisin<-HTH_3<-HKD-DNase+SF1-SF2-Helicase<-METHYLASE<-DUF3780<-DUF499*\|\|GDResolvase+Recombinase+ZnR-><-?\|\|?->GDResolvase-><-TRAM+Methyltransf_31 | 1031 | Bacteria>Terrabacteria_group>Bacillota>Clostridia>Eubacteriales | GCF_951336995.1 | Clostridium_perfringens | QDX72_RS05140 | 1151785..1154880 | 1502 | DUF499-domain-containing-protein-[Clostridium-perfringens] |
| NJP39667.1 | AAA+ATPase+wHTH+Alpha+Beta-domain+wHTH+Fn-III+Fn-III+RRM | Type-2_BR.RM.like.systems | MULE-TRANSPOSASE-><-SF1-SF2-Helicase<-OLD-ABC+TOPRIM<-HKD-DNase+SF1-SF2-Helicase<-METHYLASE<-DUF3780<-DUF499*<-XIS-HTH<-?\|\|HTH_3-><-MPTase<-CHTH | 1039 | Bacteria>Terrabacteria_group>Bacillota>Clostridia>Eubacteriales | - | Oscillospiraceae_bacterium_HV4-5-C5C | HCH52_01160 | 266770..269889 | 2721777 | DUF499-domain-containing-protein-[Oscillospiraceae-bacterium-HV4-5-C5C] |
| WP_227830710.1 | AAA+ATPase+wHTH+Alpha+Beta-domain+wHTH+Fn-III+Fn-III+RRM | Type-2_BR.RM.like.systems | <-HKD-DNase+SF1-SF2-Helicase<-METHYLASE<-DUF3780+RAMA<-DUF499*\|\|<-?<-?<-?<-PRTase | 1031 | Bacteria>Terrabacteria_group>Bacillota>Clostridia>Eubacteriales | - | unclassified_Clostridioides | IC219_RS04610 | 1030948..1034043 | 2635829 | MULTISPECIES:-DUF499-domain-containing-protein-[unclassified-Clostridioides] |
| WP_005488051.1 | AAA+ATPase+wHTH+Alpha+Beta-domain+wHTH+Fn-III+Fn-III+RRM | Type-2_BR.RM.like.systems | SF1-SF2-Helicase->OLD-ABC+TOPRIM->SF1-SF2-Helicase->DUF499*->DUF3780->METHYLASE->HKD-DNase+SF1-SF2-Helicase->MORC->URI-DNase+RAMA-><-MazF_antitoxin | 1032 | Bacteria>Terrabacteria_group>Bacillota>Clostridia>Halanaerobiales | GCF_000350165.1 | Halanaerobium_saccharolyticum | HSACCH_RS04045 | 795760..798858 | 43595 | DUF499-domain-containing-protein-[Halanaerobium-saccharolyticum] |
| WP_108141012.1 | AAA+ATPase+wHTH+Alpha+Beta-domain+wHTH+Fn-III+Fn-III+RRM | Type-2_BR.RM.like.systems | MazF_antitoxin-><-RAMA<-?<-HKD-DNase+SF1-SF2-Helicase<-METHYLASE<-DUF3780<-DUF499*<-SF1-SF2-Helicase<-OLD-ABC+TOPRIM<-SF1-SF2-Helicase | 1030 | Bacteria>Terrabacteria_group>Bacillota>Clostridia>Halanaerobiales | GCF_003053565.1 | Halanaerobium_saccharolyticum | C8C76_RS09355 | 24676..27768 | 43595 | DUF499-domain-containing-protein-[Halanaerobium-saccharolyticum] |
| WP_133505831.1 | AAA+ATPase+wHTH+Alpha+Beta-domain+wHTH+Fn-III+Fn-III+RRM | Type-2_BR.RM.like.systems | SF1-SF2-Helicase->OLD-ABC+TOPRIM->SF1-SF2-Helicase->DUF499*->DUF3780->METHYLASE->HKD-DNase+SF1-SF2-Helicase->RVT->RAMA->RAMA-> | 1030 | Bacteria>Terrabacteria_group>Bacillota>Clostridia>Halanaerobiales | GCF_004362065.1 | Halanaerobium_congolense | C8C79_RS10220 | 14213..17305 | 54121 | DUF499-domain-containing-protein-[Halanaerobium-congolense] |
| WP_229346644.1 | AAA+ATPase+wHTH+Alpha+Beta-domain+wHTH+Fn-III+Fn-III+RRM | Type-2_BR.RM.like.systems | <-URI-DNase+RAMA<-HEPN+CBS<-?<-HKD-DNase+SF1-SF2-Helicase<-METHYLASE<-DUF3780<-DUF499*<-SF1-SF2-Helicase<-OLD-ABC+TOPRIM-p | 1030 | Bacteria>Terrabacteria_group>Bacillota>Clostridia>Halanaerobiales | GCF_020711195.1 | Halanaerobium_polyolivorans | LJ207_RS11500 | 8208..11300 | 2886943 | DUF499-domain-containing-protein-[Halanaerobium-sp.-Z-7514] |
| WP_089655503.1 | AAA+ATPase+wHTH+Alpha+Beta-domain+wHTH+Fn-III+Fn-III+RRM | Type-2_BR.RM.like.systems | MazF_antitoxin-><-RAMA<-?<-HKD-DNase+SF1-SF2-Helicase<-METHYLASE<-DUF3780<-DUF499*<-ParB+HNH<-SF1-SF2-Helicase | 1030 | Bacteria>Terrabacteria_group>Bacillota>Clostridia>Halanaerobiales | GCF_900100575.1 | Halanaerobium_congolense | BLS10_RS04565 | 105480..108572 | 54121 | DUF499-domain-containing-protein-[Halanaerobium-congolense] |
| WP_089859264.1 | AAA+ATPase+wHTH+Alpha+Beta-domain+wHTH+Fn-III+Fn-III+RRM | Type-2_BR.RM.like.systems | DUF499*->DUF3780->METHYLASE->HKD-DNase+SF1-SF2-Helicase->Phage_int_SAM_4+Phage_integrase-> | 1030 | Bacteria>Terrabacteria_group>Bacillota>Clostridia>Halanaerobiales | GCF_900114545.1 | Halanaerobium_salsuginis | BMZ74_RS02640 | 52151..55243 | 29563 | DUF499-domain-containing-protein-[Halanaerobium-salsuginis] |
| WP_097019088.1 | AAA+ATPase+wHTH+Alpha+Beta-domain+wHTH+Fn-III+Fn-III+RRM | Type-2_BR.RM.like.systems | <-HEPN+CBS<-HKD-DNase+SF1-SF2-Helicase<-METHYLASE<-DUF3780<-DUF499*<-REase<-?<-?<-?<-Methyltransf<-METHYLASE | 1029 | Bacteria>Terrabacteria_group>Bacillota>Clostridia>Halanaerobiales | GCF_900215265.1 | Orenia_metallireducens | CRN99_RS15855 | 34717..37806 | 1413210 | DUF499-domain-containing-protein-[Orenia-metallireducens] |
| WP_071541194.1 | AAA+ATPase+wHTH+Alpha+Beta-domain+wHTH+Fn-III+Fn-III+RRM | Type-2_BR.RM.like.systems | RNase->?->tRNA->tRNA->tRNA->tRNA->tRNA->tRNA->Phage_int_SAM_1+Phage_integrase->?-><-DDE_Tnp\|\|DUF499*->DUF3780->METHYLASE->HKD-DNase+SF1-SF2-Helicase->?->Phage_terminase-> | 1036 | Bacteria>Terrabacteria_group>Bacillota>Clostridia>Moorellales | GCF_001874605.1 | Moorella_thermoacetica | MTJW_RS02200 | 446381..449491 | 1525 | DUF499-domain-containing-protein-[Moorella-thermoacetica] |
| WP_227764536.1 | AAA+ATPase+wHTH+Alpha+Beta-domain+wHTH+Fn-III+Fn-III+RRM | Type-2_BR.RM.like.systems | DUF499*->DUF3780->METHYLASE->HKD-DNase+SF1-SF2-Helicase->?->?->HIN-HTH-> | 1036 | Bacteria>Terrabacteria_group>Bacillota>Clostridia>Moorellales | GCF_004353525.1 | Zhaonella_formicivorans | EYS13_RS15595 | 3197652..3200762 | 2528593 | DUF499-domain-containing-protein-[Zhaonella-formicivorans] |
| WP_075517732.1 | AAA+ATPase+wHTH+Alpha+Beta-domain+wHTH+Fn-III+Fn-III+RRM | Type-2_BR.RM.like.systems | <-MULE-TRANSPOSASE<-?\|\|DUF499*->DUF3780->METHYLASE->HKD-DNase+SF1-SF2-Helicase->REase+NACHT->Mbetalac-> | 1026 | Bacteria>Terrabacteria_group>Bacillota>Clostridia>Moorellales | - | Moorella_thermoacetica | MOTE_RS11890 | 1060..4140 | 1525 | DUF499-domain-containing-protein-[Moorella-thermoacetica] |
| HHV75152.1 | AAA+ATPase+wHTH+Alpha+Beta-domain+wHTH+Fn-III+Fn-III+RRM | Type-2_BR.RM.like.systems | <-McrC-NTD+REase<-?<-HKD-DNase+SF1-SF2-Helicase<-METHYLASE<-DUF3780<-DUF499*<-SF1-SF2-Helicase<-SIGMA-HTH+SIGMA-HTH<-?<-ACET | 1037 | Bacteria>Terrabacteria_group>Bacillota>Clostridia>Thermoanaerobacterales | GCA_012839995.1 | Thermoanaerobacterium_sp. | GXX41_11045 | 45158..48271 | 40549 | DUF499-domain-containing-protein-[Thermoanaerobacterium-sp.] |
| HHW03029.1 | AAA+ATPase+wHTH+Alpha+Beta-domain+wHTH+Fn-III+Fn-III+RRM | Type-2_BR.RM.like.systems | HTH_3-><-?<-?<-HKD-DNase+SF1-SF2-Helicase<-METHYLASE<-DUF3780<-DUF499*<-?<-SF1-SF2-helicase+DUF1998<-?\|\|HDRnase-> | 1035 | Bacteria>Terrabacteria_group>Bacillota>Clostridia>Thermoanaerobacterales | GCA_012840135.1 | Thermoanaerobacterales_bacterium | GXX35_09500 | 366720..369827 | 2304039 | DUF499-domain-containing-protein-[Thermoanaerobacterales-bacterium] |
| WP_014063447.1 | AAA+ATPase+wHTH+Alpha+Beta-domain+wHTH+Fn-III+Fn-III+RRM | Type-2_BR.RM.like.systems | <-Primpol+D5N+D5ATPase<-ParB+HNH<-HKD-DNase+SF1-SF2-Helicase<-METHYLASE<-DUF3780<-DUF499*<-TraJ-RHH\|\|DDE_Tnp_1-> | 1037 | Bacteria>Terrabacteria_group>Bacillota>Clostridia>Thermoanaerobacterales | GCF_000147695.2 | Thermoanaerobacter_wiegelii | THEWI_RS10775 | 2213989..2217102 | 46354 | DUF499-domain-containing-protein-[Thermoanaerobacter-wiegelii] |
| WP_235837234.1 | AAA+ATPase+wHTH+Alpha+Beta-domain+wHTH+Fn-III+Fn-III+RRM | Type-2_BR.RM.like.systems | DUF499*->DUF3780->METHYLASE->HKD-DNase+SF1-SF2-Helicase-><-wHTH(SBD)+HNH<-SF1-SF2-Helicase+DUF1998 | 1038 | Bacteria>Terrabacteria_group>Bacillota>Clostridia>Thermoanaerobacterales | GCF_001950325.1 | Carboxydothermus_islandicus | ciss_RS03990 | 2222..5338 | 661089 | DUF499-domain-containing-protein-[Carboxydothermus-islandicus] |
| NLU09544.1 | AAA+ATPase+wHTH+Alpha+Beta-domain+wHTH+Fn-III+Fn-III+RRM | Type-2_BR.RM.like.systems | <-HKD-DNase+SF1-SF2-Helicase<-METHYLASE<-DUF3780<-DUF499*<-?<-?<-?<-?<-?<-MORC | 1036 | Bacteria>Terrabacteria_group>Bacillota>Clostridia>Thermosediminibacterales | GCA_012719935.1 | Tepidanaerobacter_acetatoxydans | GXW90_01115 | 23411..26521 | 499229 | DUF499-domain-containing-protein-[Tepidanaerobacter-acetatoxydans] |
| NCB42796.1 | AAA+ATPase+wHTH+Alpha+Beta-domain+wHTH+Fn-III+Fn-III+RRM | Type-2_BR.RM.like.systems | SF1-SF2-Helicase+DUF1998->DUF499*->DUF3780->METHYLASE->HKD-DNase+SF1-SF2-Helicase->?->SF1-SF2-Helicase+REase->SF1-SF2-Helicase+REase-> | 1038 | Bacteria>Terrabacteria_group>Bacillota>Clostridia>unclassified_Clostridia | GCA_009929155.1 | Clostridia_bacterium | EOM59_09270 | 22400..25516 | 2044939 | DUF499-domain-containing-protein-[Clostridia-bacterium] |
| NCB71009.1 | AAA+ATPase+wHTH+Alpha+Beta-domain+wHTH+Fn-III+Fn-III+RRM | Type-2_BR.RM.like.systems | <-HTH_31<-HKD-DNase+SF1-SF2-Helicase<-METHYLASE<-DUF3780<-DUF499* | 1047 | Bacteria>Terrabacteria_group>Bacillota>Clostridia>unclassified_Clostridia | GCA_009929415.1 | Clostridia_bacterium | EOM45_04370 | 7042..10185 | 2044939 | DUF499-domain-containing-protein-[Clostridia-bacterium] |
| MBO2494343.1 | AAA+ATPase+wHTH+Alpha+Beta-domain+wHTH+Fn-III+Fn-III+RRM | Type-2_BR.RM.like.systems | HD->RNB+RNase->?->Phd-YefM->PIN->\|\|DUF499*->DUF3780->METHYLASE->HKD-DNase+SF1-SF2-Helicase-> | 1035 | Bacteria>Terrabacteria_group>Bacillota>Clostridia>unclassified_Clostridia | GCA_017577715.1 | Clostridia_bacterium | COP72_06005 | 40310..43417 | 2044939 | glycosyl-transferase-[Clostridia-bacterium] |
| MDD2401874.1 | AAA+ATPase+wHTH+Alpha+Beta-domain+wHTH+Fn-III+Fn-III+RRM | Type-2_BR.RM.like.systems | DUF499*->DUF3780->METHYLASE->HKD-DNase+SF1-SF2-Helicase->?->AbiEii-> | 964 | Bacteria>Terrabacteria_group>Bacillota>Clostridia>unclassified_Clostridia | GCA_028684195.1 | Clostridia_bacterium | PHD60_06710 | 1..2897 | 2044939 | MAG:-DUF499-domain-containing-protein,-partial-[Clostridia-bacterium] |
| WP_281737584.1 | AAA+ATPase+wHTH+Alpha+Beta-domain+wHTH+Fn-III+Fn-III+RRM | Type-2_BR.RM.like.systems | <-SIGMA-HTH<-?<-HKD-DNase+SF1-SF2-Helicase<-METHYLASE<-DUF3780<-DUF499*<-?<-?<-?<-?<-?<-ACET | 1046 | Bacteria>Terrabacteria_group>Bacillota>Negativicutes>Acidaminococcales | GCF_938046715.1 | Phascolarctobacterium_succinatutens | QMF16_RS00400 | 30323..33463 | 626940 | DUF499-domain-containing-protein-[Phascolarctobacterium-succinatutens] |
| WP_277249689.1 | AAA+ATPase+wHTH+Alpha+Beta-domain+wHTH+Fn-III+Fn-III+RRM | Type-2_BR.RM.like.systems | <-HKD-DNase+SF1-SF2-Helicase<-METHYLASE<-DUF3780<-DUF499*<-GDResolvase+Recombinase+ZnR<-?\|\|?-><-?<-tRNA<-HTH_3 | 1056 | Bacteria>Terrabacteria_group>Bacillota>Negativicutes>Acidaminococcales | GCF_945908285.1 | Phascolarctobacterium_succinatutens | P6850_RS02330 | 6337..9507 | 626940 | DUF499-domain-containing-protein-[Phascolarctobacterium-succinatutens] |
| WP_277247933.1 | AAA+ATPase+wHTH+Alpha+Beta-domain+wHTH+Fn-III+Fn-III+RRM | Type-2_BR.RM.like.systems | <-HKD-DNase+SF1-SF2-Helicase<-METHYLASE<-DUF3780<-DUF499*<-GDResolvase+Recombinase+ZnR<-?\|\|?-><-?<-tRNA<-HTH_3 | 1056 | Bacteria>Terrabacteria_group>Bacillota>Negativicutes>Acidaminococcales | GCF_946012055.1 | Phascolarctobacterium_succinatutens | P6838_RS01020 | 12800..15970 | 626940 | DUF499-domain-containing-protein-[Phascolarctobacterium-succinatutens] |
| MCI6483250.1 | AAA+ATPase+wHTH+Alpha+Beta-domain+wHTH+Fn-III+Fn-III+RRM | Type-2_BR.RM.like.systems | DUF499*->DUF3780->METHYLASE->HKD-DNase+SF1-SF2-Helicase-> | 970 | Bacteria>Terrabacteria_group>Bacillota>Negativicutes>Selenomonadales | GCA_022774045.1 | Selenomonadaceae_bacterium | MSA24_02620 | 23575..26487 | 2652289 | DUF499-domain-containing-protein-[Selenomonadaceae-bacterium] |
| WP_154407767.1 | AAA+ATPase+wHTH+Alpha+Beta-domain+wHTH+Fn-III+Fn-III+RRM | Type-2_BR.RM.like.systems | DUF499*->DUF3780->METHYLASE->HKD-DNase+SF1-SF2-Helicase-><-REase<-Glyoxalase\|\|ACET-> | 1040 | Bacteria>Terrabacteria_group>Bacillota>Negativicutes>Selenomonadales | GCF_009695585.1 | Anaerovibrio_slackiae | FYJ84_RS11575 | 24516..27638 | 2652309 | DUF499-domain-containing-protein-[Anaerovibrio-slackiae] |
| NLY45482.1 | AAA+ATPase+wHTH+Alpha+Beta-domain+wHTH+Fn-III+Fn-III+RRM | Type-2_BR.RM.like.systems | <-MORC<-PIN<-HKD-DNase+SF1-SF2-Helicase<-METHYLASE<-DUF3780<-DUF499*<-HTH_26 | 1033 | Bacteria>Terrabacteria_group>Bacillota>Tissierellia>Tissierellales | GCA_012735585.1 | Tissierella_sp. | GX053_05775 | 9074..12175 | 41274 | DUF499-domain-containing-protein-[Tissierella-sp.] |
| MCK9444924.1 | AAA+ATPase+wHTH+Alpha+Beta-domain+wHTH+Fn-III+Fn-III+RRM | Type-2_BR.RM.like.systems | <-DEDD_Tnp_IS110+Transposase_20<-HEPN<-?<-HKD-DNase+SF1-SF2-Helicase<-METHYLASE<-DUF3780<-DUF499*<-HTH_26 | 1033 | Bacteria>Terrabacteria_group>Bacillota>Tissierellia>Tissierellales | GCA_023229175.1 | Tissierellaceae_bacterium | MOQ14_10415 | 14002..17103 | 2562714 | DUF499-domain-containing-protein-[Tissierellaceae-bacterium] |
| WP_216384534.1 | AAA+ATPase+wHTH+Alpha+Beta-domain+wHTH+Fn-III+Fn-III+RRM | Type-2_BR.RM.like.systems | HTH_26->DUF499*->DUF3780->METHYLASE->HKD-DNase+SF1-SF2-Helicase->ParB+HNH->?->Cysteine_desulfurase-> | 1038 | Bacteria>Terrabacteria_group>Bacillota>Tissierellia>Tissierellales | GCF_018917195.1 | Anaerosalibacter_bizertensis | KQH90_RS07250 | 53730..56846 | 932217 | DUF499-domain-containing-protein-[Anaerosalibacter-bizertensis] |
| NMB40363.1 | AAA+ATPase+wHTH+Alpha+Beta-domain+wHTH+Fn-III+Fn-III+RRM | Type-2_BR.RM.like.systems | <-HKD-DNase+SF1-SF2-Helicase<-METHYLASE<-DUF3780<-DUF499*<-METHYLASE-p<-HTH_3<-Tox-RelE | 1037 | Bacteria>Terrabacteria_group>Bacillota>unclassified_Bacillota | GCA_012798755.1 | Bacillota_bacterium | GX996_00275 | 6889..10002 | 1879010 | DUF499-domain-containing-protein-[Firmicutes-bacterium] |
| MCK9217630.1 | AAA+ATPase+wHTH+Alpha+Beta-domain+wHTH+Fn-III+Fn-III+RRM | Type-2_BR.RM.like.systems | DUF499*->DUF3780->METHYLASE->HKD-DNase+SF1-SF2-Helicase-> | 1041 | Bacteria>Terrabacteria_group>Bacillota>unclassified_Bacillota | GCA_023230985.1 | Bacillota_bacterium | MOP77_06905 | 1..3128 | 1879010 | DUF499-domain-containing-protein-[Firmicutes-bacterium] |
| MBI9046792.1 | AAA+ATPase+wHTH+Alpha+Beta-domain+wHTH+Fn-III+Fn-III+RRM | Type-2_BR.RM.like.systems | <-HKD-DNase+SF1-SF2-Helicase<-METHYLASE<-DUF3780<-DUF499* | 1023 | Bacteria>Terrabacteria_group>Chloroflexota>Anaerolineae>Anaerolineales | GCA_016342475.1 | Anaerolineaceae_bacterium | JEZ06_20065 | 6553..9624 | 2024896 | DUF499-domain-containing-protein-[Anaerolineaceae-bacterium] |
| MCA9900782.1 | AAA+ATPase+wHTH+Alpha+Beta-domain+wHTH+Fn-III+Fn-III+RRM | Type-2_BR.RM.like.systems | DUF499*->DUF3780->METHYLASE->METHYLASE-p->HKD-DNase+SF1-SF2-Helicase->ParB+HNH-p-> | 1027 | Bacteria>Terrabacteria_group>Chloroflexota>Anaerolineae>Anaerolineales | GCA_020432515.1 | Anaerolineales_bacterium | KC433_21490 | 560..3643 | 2073117 | DUF499-domain-containing-protein-[Anaerolineales-bacterium] |
| GIV64890.1 | AAA+ATPase+wHTH+Alpha+Beta-domain+wHTH+Fn-III+Fn-III+RRM | Type-2_BR.RM.like.systems | <-Peptidase<-HKD-DNase+SF1-SF2-Helicase<-METHYLASE<-DUF3780<-DUF499*<-REase+DUF5655\|\|Phage_int_SAM_4+Phage_integrase-> | 1029 | Bacteria>Terrabacteria_group>Chloroflexota>Anaerolineae>Anaerolineales | GCA_026003595.1 | Bellilinea_sp. | KatS3mg046_150 | 153425..156514 | 2838785 | MAG:-hypothetical-protein-KatS3mg046_150-[Bellilinea-sp.] |
| MBQ6482232.1 | AAA+ATPase+wHTH+Alpha+Beta-domain+wHTH+Fn-III+Fn-III+RRM | Type-2_BR.RM.like.systems | PAIREDC-HTH->DDE_3-><-METHYLASE-p<-ORC-CDC6+REase<-DUF3780<-DUF499*<-?\|\|REase+SPF-cons-> | 1041 | Bacteria>Terrabacteria_group>Chloroflexota>Anaerolineae>Anaerolineales | GCA_017432945.1 | Anaerolineaceae_bacterium | IJI45_14030 | 4377..7502 | 2024896 | DUF499-domain-containing-protein-[Anaerolineaceae-bacterium] |
| NPV42307.1 | AAA+ATPase+wHTH+Alpha+Beta-domain+wHTH+Fn-III+Fn-III+RRM | Type-2_BR.RM.like.systems | <-HKD-DNase+SF1-SF2-Helicase<-HNH<-METHYLASE<-DUF3780<-DUF499*<-XIS-HTH<-DOC+HTH\|\|XIS-HTH->?->Phage_integrase-> | 1030 | Bacteria>Terrabacteria_group>Chloroflexota>Anaerolineae>unclassified_Anaerolineae | GCA_013178095.1 | Anaerolineae_bacterium | HPY72_13330 | 39338..42430 | 2052143 | DUF499-domain-containing-protein-[Anaerolineae-bacterium] |
| MBN1991939.1 | AAA+ATPase+wHTH+Alpha+Beta-domain+wHTH+Fn-III+Fn-III+RRM | Type-2_BR.RM.like.systems | LON-AAA-><-GDResolvase<-?<-?<-?<-XIS-HTH\|\|DUF499*->DUF3780->METHYLASE->HKD-DNase+SF1-SF2-Helicase->OLD-ABC+TOPRIM->SF1-SF2-Helicase-> | 1027 | Bacteria>Terrabacteria_group>Chloroflexota>Anaerolineae>unclassified_Anaerolineae | GCA_016931895.1 | Anaerolineae_bacterium | JW953_04500 | 49654..52737 | 2052143 | DUF499-domain-containing-protein-[Anaerolineae-bacterium] |
| MCB0192590.1 | AAA+ATPase+wHTH+Alpha+Beta-domain+wHTH+Fn-III+Fn-III+RRM | Type-2_BR.RM.like.systems | <-DDE_Tnp_IS1<-ABC-ATPase+SbcC<-tRNA<-Phage_int_SAM_1+Phage_integrase\|\|PIWI->DUF499*->DUF3780->METHYLASE->HKD-DNase+SF1-SF2-Helicase-><-METHYLASE<-?<-TnsC-ATPase | 1028 | Bacteria>Terrabacteria_group>Chloroflexota>Anaerolineae>unclassified_Anaerolineae | GCA_020433805.1 | Anaerolineae_bacterium | KDJ65_11660 | 8073..11159 | 2052143 | DUF499-domain-containing-protein-[Anaerolineae-bacterium] |
| MCB8978659.1 | AAA+ATPase+wHTH+Alpha+Beta-domain+wHTH+Fn-III+Fn-III+RRM | Type-2_BR.RM.like.systems | XIS-HTH-><-?<-?\|\|TPR+TPR->?->DUF499*->DUF3780->METHYLASE->HKD-DNase+SF1-SF2-Helicase-><-?<-DDE_3<-PAIREDC-HTH | 1020 | Bacteria>Terrabacteria_group>Chloroflexota>Ardenticatenia>Ardenticatenales | GCA_020634435.1 | Ardenticatenaceae_bacterium | H6657_14660 | 499064..502126 | 2614247 | DUF499-domain-containing-protein-[Ardenticatenaceae-bacterium] |
| MCE7987123.1 | AAA+ATPase+wHTH+Alpha+Beta-domain+wHTH+Fn-III+Fn-III+RRM | Type-2_BR.RM.like.systems | RNaseH-><-HKD-DNase+SF1-SF2-Helicase<-wHTH+REase<-METHYLASE<-DUF3780<-DUF499* | 1028 | Bacteria>Terrabacteria_group>Chloroflexota>Caldilineae>Caldilineales | GCA_021405325.1 | Caldilinea_sp._CFX5 | DYG89_38625 | 47315..50401 | 2293554 | DUF499-domain-containing-protein-[Caldilinea-sp.-CFX5] |
| WP_211358664.1 | AAA+ATPase+wHTH+Alpha+Beta-domain+wHTH+Fn-III+Fn-III+RRM | Type-2_BR.RM.like.systems | <-DHTH+WYL<-?<-?\|\|DUF499*->DUF3780->METHYLASE->HKD-DNase+SF1-SF2-Helicase-><-McrC-NTD+REase<-?\|\|Pkinase-> | 1026 | Bacteria>Terrabacteria_group>Chloroflexota>Caldilineae>Caldilineales | GCF_006569185.2 | Litorilinea_aerophila | FKZ61_RS20575 | 18600..21680 | 1204385 | DUF499-domain-containing-protein-[Litorilinea-aerophila] |
| MCC6456909.1 | AAA+ATPase+wHTH+Alpha+Beta-domain+wHTH+Fn-III+Fn-III+RRM | Type-2_BR.RM.like.systems | <-HIN-HTH<-HIN-HTH<-?\|\|MazE_antitoxin->DUF499*->MAD-NTD+HNH->DUF3780->DUF3780->ParB+HNH->METHYLASE->Calcineurin-> | 1021 | Bacteria>Terrabacteria_group>Chloroflexota>Caldilineae>Caldilineales | GCA_020846635.1 | Caldilineaceae_bacterium | IT328_18275 | 1966..5031 | 2614250 | DUF499-domain-containing-protein-[Caldilineaceae-bacterium] |
| NTV99826.1 | AAA+ATPase+wHTH+Alpha+Beta-domain+wHTH+Fn-III+Fn-III+RRM | Type-2_BR.RM.like.systems | DUF499*->DUF3780->METHYLASE->HKD-DNase+SF1-SF2-Helicase-><-REase\|\|DDE_Tnp_1_assoc+DDE_Tnp_1->DDE_Tnp_1-> | 816 | Bacteria>Terrabacteria_group>Chloroflexota>Chloroflexia>Chloroflexales | GCA_013336555.1 | Oscillochloris_sp. | HGA19_00810 | 1512..3962 | 2547424 | DUF499-domain-containing-protein-[Oscillochloris-sp.] |
| WP_097655028.1 | AAA+ATPase+wHTH+Alpha+Beta-domain+wHTH+Fn-III+Fn-III+RRM | Type-2_BR.RM.like.systems | DUF4096+DDE_Tnp_1+DDE_Tnp->Trypsin->?->?->?->?->DUF499*->DUF3780->METHYLASE->HKD-DNase+SF1-SF2-Helicase-> | 1027 | Bacteria>Terrabacteria_group>Chloroflexota>Chloroflexia>Chloroflexales | GCF_002532075.1 | Candidatus_Chloroploca_asiatica | A9Q02_RS20960 | 9491..12574 | 1506545 | DUF499-domain-containing-protein-[Candidatus-Chloroploca-asiatica] |
| WP_129673421.1 | AAA+ATPase+wHTH+Alpha+Beta-domain+wHTH+Fn-III+Fn-III+RRM | Type-2_BR.RM.like.systems | DUF499*->DUF3780->METHYLASE->OLD-ABC->TOPRIM-p->HKD-DNase+SF1-SF2-Helicase->NAD_binding_2+6PGD-> | 1028 | Bacteria>Terrabacteria_group>Chloroflexota>Chloroflexia>Chloroflexales | GCF_004138165.1 | Candidatus_Chloroploca_sp._Khr17 | EKD05_RS13120 | 232..3318 | 2496869 | DUF499-domain-containing-protein-[Candidatus-Chloroploca-sp.-Khr17] |
| PFG75312.1 | AAA+ATPase+wHTH+Alpha+Beta-domain+wHTH+Fn-III+Fn-III+RRM | Type-2_BR.RM.like.systems | Terminase_4-><-?\|\|XIS-HTH->XIS-HTH->Primpol+PriCT1->?->DUF499*->DUF3780->METHYLASE->HKD-DNase+SF1-SF2-Helicase->Terminase_4->Phage_int_SAM_1+Phage_integrase-> | 1033 | Bacteria>Terrabacteria_group>Chloroflexota>Tepidiformia>Tepidiformales | GCA_002563855.1 | Tepidiforma_thermophila | A9A59_2580 | 2632628..2635729 | 2761530 | putative-AAA+-superfamily-ATPase-[Chloroflexi-bacterium-G233] |
| WP_278286911.1 | AAA+ATPase+wHTH+Alpha+Beta-domain+wHTH+Fn-III+Fn-III+RRM | Type-2_BR.RM.like.systems | Terminase_4-><-?\|\|XIS-HTH->XIS-HTH->Primpol+PriCT1->?->DUF499*->DUF3780->METHYLASE->HKD-DNase+SF1-SF2-Helicase->Terminase_4->Phage_int_SAM_1+Phage_integrase-> | 1025 | Bacteria>Terrabacteria_group>Chloroflexota>Tepidiformia>Tepidiformales | GCF_002563855.1 | Tepidiforma_thermophila | A9A59_RS12785 | 2632652..2635729 | 2761530 | DUF499-domain-containing-protein-[Tepidiforma-thermophila] |

| Protein-ID | DUF499-architecture | System | Gene-Neighbourhoods | Length | Taxonomy | Assembly-ID | Strain | Locus-Tag | Genomic-coordinates | TaxID | Fasta-defline |
|---|---|---|---|---|---|---|---|---|---|---|---|
| HCG29367.1 | AAA+ATPase+wHTH+Alpha+Beta-domain+wHTH+Fn-III+Fn-III+RRM | Type-2_BR.RM.like.systems | REase->DUF499*->DUF3780->METHYLASE->HKD-DNase+SF1-SF2-Helicase->HicA_toxin->HicB_lk_antitox-> | 1024 | Bacteria>Terrabacteria_group>Chloroflexota>unclassified_Chloroflexota | GCA_003534145.1 | Chloroflexota_bacterium | DEU95_06405 | 3683..6757 | 2026724 | hypothetical-protein-[Chloroflexi-bacterium] |
| MCI0728575.1 | AAA+ATPase+wHTH+Alpha+Beta-domain+wHTH+Fn-III+Fn-III+RRM | Type-2_BR.RM.like.systems | ArdCN+MPTase->?->?->?->DUF499*->DUF3780->METHYLASE->HKD-DNase+SF1-SF2-Helicase-> | 1026 | Bacteria>Terrabacteria_group>Chloroflexota>unclassified_Chloroflexota | GCA_022616635.1 | Chloroflexota_bacterium | L0332_17900 | 5300..8380 | 2026724 | DUF499-domain-containing-protein-[Chloroflexi-bacterium] |
| MCL4561934.1 | AAA+ATPase+wHTH+Alpha+Beta-domain+wHTH+Fn-III+Fn-III+RRM | Type-2_BR.RM.like.systems | <-D5N+D5ATPase<-?<-XIS-HTH\|\|HIN-HTH-><-?<-?\|\|DUF499*->ZnR->DUF3780->?->METHYLASE->?->HKD-DNase+SF1-SF2-Helicase-> | 1035 | Bacteria>Terrabacteria_group>Chloroflexota>unclassified_Chloroflexota | GCA_023381565.1 | Chloroflexota_bacterium | M1281_15165 | 7329..10436 | 2026724 | DUF499-domain-containing-protein-[Chloroflexi-bacterium] |
| MCX5991391.1 | AAA+ATPase+wHTH+Alpha+Beta-domain+wHTH+Fn-III+Fn-III+RRM | Type-2_BR.RM.like.systems | <-HTH+PolB<-?<-ParB+HNH<-HKD-DNase+SF1-SF2-Helicase<-METHYLASE<-DUF3780<-DUF499*\|\|HD->PADR-HTH-><-PIN<-HTH+MPTase | 1031 | Bacteria>Terrabacteria_group>Chloroflexota>unclassified_Chloroflexota | GCA_026389585.1 | Chloroflexota_bacterium | NTZ04_03550 | 87097..90192 | 2026724 | MAG:-DUF499-domain-containing-protein-[Chloroflexi-bacterium] |
| MCA0313165.1 | AAA+ATPase+wHTH+Alpha+Beta-domain+wHTH+Fn-III+Fn-III+RRM | Type-2_BR.RM.like.systems | DDE_Tnp_1-><-HKD-DNase+SF1-SF2-Helicase<-METHYLASE<-DUF3780<-DUF499* | 1027 | Bacteria>Terrabacteria_group>Cyanobacteriota/Melainabacteria_group>Candidatus_Melainabacteria>unclassified_Candidatus_Melainabacteria | GCA_020161235.1 | Candidatus_Melainabacteria_bacterium | LCH63_04905 | 6694..9777 | 2052166 | DUF499-domain-containing-protein-[Candidatus-Melainabacteria-bacterium] |
| MBD2718328.1 | AAA+ATPase+wHTH+Alpha+Beta-domain+wHTH+Fn-III+Fn-III+RRM | Type-2_BR.RM.like.systems | <-ParA-Soj-PloopNTPase\|\|?->?->?->DUF499*->DUF3780->METHYLASE->HKD-DNase+SF1-SF2-Helicase->?-><-DOC+HTH | 1047 | Bacteria>Terrabacteria_group>Cyanobacteriota/Melainabacteria_group>Cyanobacteriota>Cyanophyceae | GCA_014698945.1 | Synechococcus_sp._FACHB-909 | H6G99_05105 | 362265..365408 | 2692863 | DUF499-domain-containing-protein-[Synechococcus-sp.-FACHB-909] |
| MBW4531835.1 | AAA+ATPase+wHTH+Alpha+Beta-domain+wHTH+Fn-III+Fn-III+RRM | Type-2_BR.RM.like.systems | PIN->?-><-?<-?<-?\|\|DUF499*->DUF3780->METHYLASE->HKD-DNase+SF1-SF2-Helicase->?-><-REase | 1026 | Bacteria>Terrabacteria_group>Cyanobacteriota/Melainabacteria_group>Cyanobacteriota>Cyanophyceae | GCA_019359745.1 | Aphanothece_saxicola_GSE-SYN-MK-01-06B | KME02_14215 | 37139..40219 | 2835092 | DUF499-domain-containing-protein-[Aphanothece-saxicola-GSE-SYN-MK-01-06B] |
| BCX11788.1 | AAA+ATPase+wHTH+Alpha+Beta-domain+wHTH+Fn-III+Fn-III+RRM | Type-2_BR.RM.like.systems | SF1-SF2-Helicase-><-REase\|\|TUDOR+SF1-SF2-Helicase+iREase+CTD+REase-p->DUF499*->DUF3780->METHYLASE->HKD-DNase+SF1-SF2-Helicase->HEPN+Mrr_N->HEPN+REase-> | 1027 | Bacteria>Terrabacteria_group>Cyanobacteriota/Melainabacteria_group>Cyanobacteriota>Cyanophyceae | GCA_025999095.1 | Thermosynechococcus_sp. | KatS3mg067_0726 | 755896..758979 | 2814275 | MAG:-hypothetical-protein-KatS3mg067_0726-[Thermosynechococcus-sp.] |
| WP_075596948.1 | AAA+ATPase+wHTH+Alpha+Beta-domain+wHTH+Fn-III+Fn-III+RRM | Type-2_BR.RM.like.systems | <-HKD-DNase+SF1-SF2-Helicase<-PIN<-?<-METHYLASE<-REase<-DUF3780<-DUF499*\|\|?->Phage_integrase->?->ABC-ATPase+SbcC->ABC-ATPase+SbcC-> | 1026 | Bacteria>Terrabacteria_group>Cyanobacteriota/Melainabacteria_group>Cyanobacteriota>Cyanophyceae | GCF_001939115.1 | Leptolyngbya_sp._'hensonii' | BST81_RS02385 | 92870..95950 | 1922337 | DUF499-domain-containing-protein-[Leptolyngbya-sp.-'hensonii'] |
| WP_106500153.1 | AAA+ATPase+wHTH+Alpha+Beta-domain+wHTH+Fn-III+Fn-III+RRM | Type-2_BR.RM.like.systems | SF1-SF2-Helicase+DUF1998-><-HKD-DNase+SF1-SF2-Helicase<-METHYLASE<-DUF3780<-DUF499* | 1047 | Bacteria>Terrabacteria_group>Cyanobacteriota/Melainabacteria_group>Cyanobacteriota>Cyanophyceae | GCF_003011125.1 | Synechococcus_lacustris | C7K08_RS08195 | 18616..21759 | 2116544 | DUF499-domain-containing-protein-[Synechococcus-lacustris] |
| WP_181494812.1 | AAA+ATPase+wHTH+Alpha+Beta-domain+wHTH+Fn-III+Fn-III+RRM | Type-2_BR.RM.like.systems | <-HKD-DNase+SF1-SF2-Helicase<-METHYLASE<-?\|\|DUF499*->DUF3780-> | 1031 | Bacteria>Terrabacteria_group>Cyanobacteriota/Melainabacteria_group>Cyanobacteriota>Cyanophyceae | GCF_003555505.1 | Thermosynechococcus_vestitus | D3A95_RS09535 | 1951205..1954300 | 146786 | DUF499-domain-containing-protein-[Thermosynechococcus-vestitus] |
| WP_168652560.1 | AAA+ATPase+wHTH+Alpha+Beta-domain+wHTH+Fn-III+Fn-III+RRM | Type-2_BR.RM.like.systems | HNH-><-HKD-DNase+SF1-SF2-Helicase<-METHYLASE<-DUF499*<-Phage_int_SAM_1+Phage_integrase<-?\|\|?->PIN->REC-> | 828 | Bacteria>Terrabacteria_group>Cyanobacteriota/Melainabacteria_group>Cyanobacteriota>Cyanophyceae | GCF_009711965.1 | unclassified_Dolichospermum | FJR09_RS23710 | 6633..9119 | 2622029 | MULTISPECIES:-DUF499-domain-containing-protein-[unclassified-Dolichospermum] |
| WP_192161916.1 | AAA+ATPase+wHTH+Alpha+Beta-domain+wHTH+Fn-III+Fn-III+RRM | Type-2_BR.RM.like.systems | <-HKD-DNase+SF1-SF2-Helicase-p<-HKD-DNase+SF1-SF2-Helicase<-?<-METHYLASE-p<-DUF3780<-DUF499*\|\|HKD-DNase+SF1-SF2-Helicase-p-><-?<-AbrB-HTH | 1039 | Bacteria>Terrabacteria_group>Cyanobacteriota/Melainabacteria_group>Cyanobacteriota>Cyanophyceae | GCF_014696895.1 | Chroococcidiopsis_sp._[FACHB-1243] | H6G17_RS28475 | 10763..13882 | 2692781 | DUF499-domain-containing-protein-[Chroococcidiopsis-sp.-FACHB-1243] |
| WP_190610329.1 | AAA+ATPase+wHTH+Alpha+Beta-domain+wHTH+Fn-III+Fn-III+RRM | Type-2_BR.RM.like.systems | CHASE2+ACYC-><-?\|\|?-><-ParA-Soj-PloopNTPase\|\|?->?->DUF499*->DUF3780->METHYLASE->HKD-DNase+SF1-SF2-Helicase->?-><-DOC+HTH | 1102 | Bacteria>Terrabacteria_group>Cyanobacteriota/Melainabacteria_group>Cyanobacteriota>Cyanophyceae | GCF_014698945.1 | Synechococcus_sp._FACHB-909 | H6G99_RS05105 | 362100..365408 | 2692863 | DUF499-domain-containing-protein-[Synechococcus-sp.-FACHB-909] |
| WP_201323620.1 | AAA+ATPase+wHTH+Alpha+Beta-domain+wHTH+Fn-III+Fn-III+RRM | Type-2_BR.RM.like.systems | <-HKD-DNase+SF1-SF2-Helicase<-ParB+HNH<-METHYLASE<-REase<-DUF3780<-DUF499* | 1027 | Bacteria>Terrabacteria_group>Cyanobacteriota/Melainabacteria_group>Cyanobacteriota>Cyanophyceae | GCF_016584665.1 | Pseudanabaena_sp._lw0831 | APA_RS17915 | 65254..68337 | 1357935 | DUF499-domain-containing-protein-[Pseudanabaena-sp.-lw0831] |
| WP_255125046.1 | AAA+ATPase+wHTH+Alpha+Beta-domain+wHTH+Fn-III+Fn-III+RRM | Type-2_BR.RM.like.systems | DUF499*->DUF3780->METHYLASE->HKD-DNase+SF1-SF2-Helicase-><-SF1-SF2-Helicase+DUF1998 | 1047 | Bacteria>Terrabacteria_group>Cyanobacteriota/Melainabacteria_group>Cyanobacteriota>Cyanophyceae | GCF_024346215.1 | Synechococcus_lacustris | KBY79_RS10925 | 397..3540 | 2116544 | DUF499-domain-containing-protein-[Synechococcus-lacustris] |
| WP_255001865.1 | AAA+ATPase+wHTH+Alpha+Beta-domain+wHTH+Fn-III+Fn-III+RRM | Type-2_BR.RM.like.systems | DUF499*->DUF3780->METHYLASE->METHYLASE-p->HKD-DNase+SF1-SF2-Helicase-><-ParB+HNH | 1047 | Bacteria>Terrabacteria_group>Cyanobacteriota/Melainabacteria_group>Cyanobacteriota>Cyanophyceae | GCF_024346435.1 | Cyanobium_sp._Alchichica_3B3-8F6 | KBY65_RS13005 | 1719..4862 | 2823696 | DUF499-domain-containing-protein-[Cyanobium-sp.-Alchichica-3B3-8F6] |
| WP_268243228.1 | AAA+ATPase+wHTH+Alpha+Beta-domain+wHTH+Fn-III+Fn-III+RRM | Type-2_BR.RM.like.systems | <-HKD-DNase+SF1-SF2-Helicase<-ParB+HNH<-METHYLASE<-DUF3780<-DUF499*\|\|?->SF1-SF2-Helicase-><-Phage_int_SAM_4+Phage_integrase | 1031 | Bacteria>Terrabacteria_group>Cyanobacteriota/Melainabacteria_group>Cyanobacteriota>Cyanophyceae | GCF_026735145.1 | Leptolyngbya_sp._GGD | OU814_RS30730 | 9328..12423 | 2997907 | DUF499-domain-containing-protein-[Leptolyngbya-sp.-GGD] |
| MCL1491775.1 | AAA+ATPase+wHTH+Alpha+Beta-domain+wHTH+Fn-III+Fn-III+RRM | Type-2_BR.RM.like.systems | <-RNaseH\|\|PspA+ParA-Soj-PloopNTPase-><-?\|\|?->DDE_3->D5N+D5ATPase->DUF499*->DUF3780->STAS+HisKin->?->PIN->MPTase->PIN-> | 1027 | Bacteria>Terrabacteria_group>Cyanobacteriota/Melainabacteria_group>Cyanobacteriota>Cyanophyceae | GCA_023336205.1 | Pseudanabaena_sp._Salubria-1 | M1G31_13655 | 139342..142425 | 2938918 | DUF499-domain-containing-protein-[Pseudanabaena-sp.-Salubria-1] |
| WP_026099180.1 | AAA+ATPase+wHTH+Alpha+Beta-domain+wHTH+Fn-III+Fn-III+RRM | Type-2_BR.RM.like.systems | <-METHYLASE-p\|\|DDE_Tnp_1-><-METHYLASE-p<-PIN<-DUF433<-DUF3780<-DUF499*\|\|?->REase-> | 1030 | Bacteria>Terrabacteria_group>Cyanobacteriota/Melainabacteria_group>Cyanobacteriota>Cyanophyceae | GCF_000332315.1 | Prochlorothrix_hollandica | PRO9006_RS0100215 | 51714..54806 | 1223 | DUF499-domain-containing-protein-[Prochlorothrix-hollandica] |
| WP_011433321.1 | AAA+ATPase+wHTH+Alpha+Beta-domain+wHTH+Fn-III+Fn-III+RRM | Type-2_BR.RM.like.systems | TPR->tRNA->?->DUF499*->DUF3780->?-><-PIN<-OrfB_Zn_ribbon<-DUF433\|\|METHYLASE-> | 1026 | Bacteria>Terrabacteria_group>Cyanobacteriota/Melainabacteria_group>Cyanobacteriota>Cyanophyceae | GCF_000013225.1 | Synechococcus_sp._JA-2-3B'a(2-13) | CYB_RS08180 | 1798790..1801870 | 321332 | DUF499-domain-containing-protein-[Synechococcus-sp.-JA-2-3B'a(2-13)] |
| WP_190598812.1 | AAA+ATPase+wHTH+Alpha+Beta-domain+wHTH+Fn-III+Fn-III+RRM | Type-2_BR.RM.like.systems | AbrB-HTH->DUF433->PIN-><-DUF29<-DUF3780<-DUF499*\|\|?->Radical-SAM-><-REase | 1031 | Bacteria>Terrabacteria_group>Cyanobacteriota/Melainabacteria_group>Cyanobacteriota>Cyanophyceae | GCF_014698605.1 | Synechocystis_sp._FACHB-383 | H6G45_RS12825 | 4289..7384 | 2692864 | DUF499-domain-containing-protein-[Synechocystis-sp.-FACHB-383] |
| WP_009624988.1 | AAA+ATPase+wHTH+Alpha+Beta-domain+wHTH+Fn-III+Fn-III+RRM | Type-2_BR.RM.like.systems | <-PIN<-MPTase<-PIN<-?<-STAS+HisKin<-DUF3780<-DUF499*\|\|?->?->DDE_Tnp_1->DDE_5-> | 1027 | Bacteria>Terrabacteria_group>Cyanobacteriota/Melainabacteria_group>Cyanobacteriota>Cyanophyceae | GCF_000332215.1 | Pseudanabaena | FEV09_RS00155 | 30199..33282 | 1152 | DUF499-domain-containing-protein-[Pseudanabaena-biceps] |
| WP_239112532.1 | AAA+ATPase+wHTH+Alpha+Beta-domain+wHTH+Fn-III+Fn-III+RRM | Type-2_BR.RM.like.systems | <-DUF4236+TPR<-?<-?<-?\|\|DUF499*->DUF3780->DUF433->PIN->DUF433->PIN->METHYLASE-> | 1031 | Bacteria>Terrabacteria_group>Cyanobacteriota/Melainabacteria_group>Cyanobacteriota>Cyanophyceae | GCF_016888165.1 | Halomicronema_sp._CCY15110 | JUK32_RS10475 | 44838..47933 | 2767773 | DUF499-domain-containing-protein-[Halomicronema-sp.-CCY15110] |
| WP_212717131.1 | AAA+ATPase+wHTH+Alpha+Beta-domain+wHTH+Fn-III+Fn-III+RRM | Type-2_BR.RM.like.systems | AbrB-HTH->?->DUF433->PIN-><-?<-DUF3780<-DUF499*\|\|?->Radical-SAM->?-><-REase | 1032 | Bacteria>Terrabacteria_group>Cyanobacteriota/Melainabacteria_group>Cyanobacteriota>Cyanophyceae | GCF_018282115.1 | unclassified_Synechocystis | HTZ78_RS15005 | 3211021..3214119 | 2640012 | MULTISPECIES:-DUF499-domain-containing-protein-[unclassified-Synechocystis] |
| WP_249369271.1 | AAA+ATPase+wHTH+Alpha+Beta-domain+wHTH+Fn-III+Fn-III+RRM | Type-2_BR.RM.like.systems | <-RHH_1\|\|?->DUF499*->DUF3780->DUF433->DUF433->PIN->METHYLASE-p-> | 1033 | Bacteria>Terrabacteria_group>Cyanobacteriota/Melainabacteria_group>Cyanobacteriota>Cyanophyceae | GCF_018336915.1 | Acaryochloris_marina | I1H34_RS15275 | 3282074..3285175 | 155978 | DUF499-domain-containing-protein-[Acaryochloris-marina] |
| PZV21714.1 | AAA+ATPase+wHTH+Alpha+Beta-domain+wHTH+Fn-III+Fn-III+RRM | Type-2_BR.RM.like.systems | <-PIN<-?<-?\|\|Y1_Tnp-><-METHYLASE<-DUF3780<-DUF499*\|\|?-><-ParB+HTH<-?\|\|?-><-MULE-TRANSPOSASE\|\|DDE_Tnp_ISAZ013-> | 1035 | Bacteria>Terrabacteria_group>Cyanobacteriota/Melainabacteria_group>Cyanobacteriota>Cyanophyceae | GCA_003249155.1 | Leptolyngbya_sp. | DCF21_01735 | 19345..22452 | 47254 | AAA-family-ATPase-[Leptolyngbya-sp.] |
| PZO60752.1 | AAA+ATPase+wHTH+Alpha+Beta-domain+wHTH+Fn-III+Fn-III+RRM | Type-2_BR.RM.like.systems | <-METHYLASE-p\|\|?-><-?<-?\|\|?->?->tRNA->DUF499*->DUF3780->?->PIN->METHYLASE-><-PIN<-MPTase | 1029 | Bacteria>Terrabacteria_group>Cyanobacteriota/Melainabacteria_group>Cyanobacteriota>Cyanophyceae | GCA_003242115.1 | Phormidesmis_priestleyi | DCF15_01580 | 22660..25749 | 268141 | AAA-family-ATPase-[Phormidesmis-priestleyi] |
| NJL98833.1 | AAA+ATPase+wHTH+Alpha+Beta-domain+wHTH+Fn-III+Fn-III+RRM | Type-2_BR.RM.like.systems | <-PIN<-DUF433<-DUF433<-DUF3780<-DUF499* | 1031 | Bacteria>Terrabacteria_group>Cyanobacteriota/Melainabacteria_group>Cyanobacteriota>Cyanophyceae | GCA_012031565.1 | Synechococcaceae_cyanobacterium_SM2_3_2 | HC924_08500 | 13502..16597 | 2720471 | DUF499-domain-containing-protein-[Synechococcaceae-cyanobacterium-SM2_3_2] |
| KPQ33158.1 | AAA+ATPase+wHTH+Alpha+Beta-domain+wHTH+Fn-III+Fn-III+RRM | Type-2_BR.RM.like.systems | DUF499*->DUF3780->DUF433->?->PIN->METHYLASE->OLD-ABC-> | 1030 | Bacteria>Terrabacteria_group>Cyanobacteriota/Melainabacteria_group>Cyanobacteriota>Cyanophyceae | GCA_001314865.1 | Phormidesmis_priestleyi_Ana | HLUCCA11_19330 | 2367..5459 | 1666911 | putative-ATPase-(AAA+-superfamily)-[Phormidesmis-priestleyi-Ana] |
| WP_190499833.1 | AAA+ATPase+wHTH+Alpha+Beta-domain+wHTH+Fn-III+Fn-III+RRM | Type-2_BR.RM.like.systems | <-ACET<-?\|\|DUF499*->DUF3780->REase-> | 1025 | Bacteria>Terrabacteria_group>Cyanobacteriota/Melainabacteria_group>Cyanobacteriota>Cyanophyceae | GCF_014697375.1 | Phormidium_sp._FACHB-1136 | H6G62_RS23765 | 9557..12634 | 2692848 | DUF499-domain-containing-protein-[Phormidium-sp.-FACHB-1136] |
| MBW4553105.1 | AAA+ATPase+wHTH+Alpha+Beta-domain+wHTH+Fn-III+Fn-III+RRM | Type-2_BR.RM.like.systems | <-DUF499*<-DUF499-p<-?<-?<-REase<-ABhydrolase | 774 | Bacteria>Terrabacteria_group>Cyanobacteriota/Melainabacteria_group>Cyanobacteriota>Cyanophyceae | GCA_019359715.1 | Aphanocapsa_sp._GSE-SYN-MK-11-07L | KME35_18635 | 88..2412 | 2835086 | DUF499-domain-containing-protein-[Aphanocapsa-sp.-GSE-SYN-MK-11-07L] |
| MBM5816147.1 | AAA+ATPase+wHTH+Alpha+Beta-domain+wHTH+Fn-III+Fn-III+RRM | Type-2_BR.RM.like.systems | METHYLASE->OLD-ABC->PIN->?-><-?\|\|<-?\|\|DUF499*->DUF3780->METHYLASE->HKD-DNase+SF1-SF2-Helicase-><-ArdCN+MPTase\|\|SIGMA-HTH-> | 1011 | Bacteria>Terrabacteria_group>Cyanobacteriota/Melainabacteria_group>Cyanobacteriota>unclassified_Cyanobacteriota | GCA_016882305.1 | Cyanobacteria_bacterium_K_Offshore_surface_m2_239 | FJ083_06030 | 12810..15845 | 2775555 | DUF499-domain-containing-protein-[Cyanobacteria-bacterium-K_Offshore_surface_m2_239] |
| MCY4056140.1 | AAA+ATPase+wHTH+Alpha+Beta-domain+wHTH+Fn-III+Fn-III+RRM | Type-2_BR.RM.like.systems | <-HKD-DNase+SF1-SF2-Helicase<-METHYLASE<-DUF3780<-DUF499*<-Peptidase_M22<-?\|\|Zn_Tnp_IS1595+DDE_Tnp_IS1595->?->METHYLASE+HNH-> | 1050 | Bacteria>Terrabacteria_group>Cyanobacteriota/Melainabacteria_group>Cyanobacteriota>unclassified_Cyanobacteriota | GCA_026708625.1 | Cyanobacteria_bacterium_MAG_CAR4_bin_6 | OXF67_09885 | 8757..11909 | 2775592 | MAG:-DUF499-domain-containing-protein-[Cyanobacteria-bacterium-MAG-CAR4_bin_6] |
| MCY4173285.1 | AAA+ATPase+wHTH+Alpha+Beta-domain+wHTH+Fn-III+Fn-III+RRM | Type-2_BR.RM.like.systems | Peptidase_M22->DUF499*->DUF3780->METHYLASE->HKD-DNase+SF1-SF2-Helicase-> | 1050 | Bacteria>Terrabacteria_group>Cyanobacteriota/Melainabacteria_group>Cyanobacteriota>unclassified_Cyanobacteriota | GCA_026709085.1 | Cyanobacteria_bacterium_MAG_CAR3_bin_5 | OXF25_04340 | 2970..6122 | 2775591 | MAG:-DUF499-domain-containing-protein-[Cyanobacteria-bacterium-MAG-CAR3_bin_5] |
| HAN46918.1 | AAA+ATPase+wHTH+Alpha+Beta-domain+wHTH+Fn-III+Fn-III+RRM | Type-2_BR.RM.like.systems | <-ABhydrolase<-TOPRIM+ZnR<-?\|\|?->METHYLASE-><-?\|\|DUF499*->DUF3780->?->RelE-ParE-> | 1026 | Bacteria>Terrabacteria_group>Cyanobacteriota/Melainabacteria_group>Cyanobacteriota>unclassified_Cyanobacteriota | GCA_003456125.1 | Cyanobacteria_bacterium_UBA8156 | DCQ32_10280 | 6756..9836 | 2055767 | hypothetical-protein-[Cyanobacteria-bacterium-UBA8156] |
| WP_012692622.1 | AAA+ATPase+wHTH+Alpha+Beta-domain+wHTH+Fn-III+Fn-III+RRM | Type-2_BR.RM.like.systems | <-HKD-DNase+SF1-SF2-Helicase<-METHYLASE<-METHYLASE<-DUF3780<-DUF499*\|\|35exo+SF1-SF2-Helicase->?-><-SF1-SF2-Helicase\|\|SIR2->DDE_Tnp_1-> | 1024 | Bacteria>Terrabacteria_group>Deinococcota>Deinococci>Deinococcales | GCF_000020685.1 | Deinococcus_deserti | DEIDE_RS03790 | 776943..780017 | 310783 | DUF499-domain-containing-protein-[Deinococcus-deserti] |
| WP_017869320.1 | AAA+ATPase+wHTH+Alpha+Beta-domain+wHTH+Fn-III+Fn-III+RRM | Type-2_BR.RM.like.systems | <-HKD-DNase+SF1-SF2-Helicase<-METHYLASE<-METHYLASE<-DUF3780<-DUF499*<-MetJarc-HTH\|\|HTH+HNH->35exo+SF1-SF2-Helicase->?-><-SF1-SF2-Helicase | 1024 | Bacteria>Terrabacteria_group>Deinococcota>Deinococci>Deinococcales | GCF_000348665.1 | Deinococcus_wulumuqiensis | G6R31_RS03660 | 752437..755511 | 980427 | DUF499-domain-containing-protein-[Deinococcus-wulumuqiensis] |
| WP_119673927.1 | AAA+ATPase+wHTH+Alpha+Beta-domain+wHTH+Fn-III+Fn-III+RRM | Type-2_BR.RM.like.systems | <-PIN<-?\|\|SF1-SF2-Helicase-><-?<-35exo+SF1-SF2-Helicase\|\|MetJarc-HTH->DUF499*->DUF3780->METHYLASE->METHYLASE->HKD-DNase+SF1-SF2-Helicase-> | 1023 | Bacteria>Terrabacteria_group>Deinococcota>Deinococci>Deinococcales | GCF_003583865.1 | Deinococcus_sp._RM | D3W47_RS12435 | 13530..16601 | 2316359 | DUF499-domain-containing-protein-[Deinococcus-sp.-RM] |
| WP_119762732.1 | AAA+ATPase+wHTH+Alpha+Beta-domain+wHTH+Fn-III+Fn-III+RRM | Type-2_BR.RM.like.systems | <-Phage_integrase<-XIS-HTH<-?\|\|?->?->DUF499*->DUF3780->METHYLASE->METHYLASE->HKD-DNase+SF1-SF2-Helicase-> | 1024 | Bacteria>Terrabacteria_group>Deinococcota>Deinococci>Deinococcales | GCF_003590815.1 | Deinococcus_cavernae | D3875_RS07880 | 676200..679274 | 2320857 | DUF499-domain-containing-protein-[Deinococcus-cavernae] |
| WP_184024653.1 | AAA+ATPase+wHTH+Alpha+Beta-domain+wHTH+Fn-III+Fn-III+RRM | Type-2_BR.RM.like.systems | HTH-><-?<-HNH+SAVED<-JAB<-UbE1+JAB<-SMODS\|\|DUF499*->DUF3780->METHYLASE->HKD-DNase+SF1-SF2-Helicase-><-?<-?<-MarR-HTH | 1024 | Bacteria>Terrabacteria_group>Deinococcota>Deinococci>Deinococcales | GCF_014201885.1 | Deinococcus_budaensis | HNQ09_RS01910 | 398460..401534 | 1665626 | DUF499-domain-containing-protein-[Deinococcus-budaensis] |
| WP_229659273.1 | AAA+ATPase+wHTH+Alpha+Beta-domain+wHTH+Fn-III+Fn-III+RRM | Type-2_BR.RM.like.systems | SF1-SF2-Helicase-><-?<-35exo+SF1-SF2-Helicase<-HTH+HNH\|\|MetJarc-HTH-><-?\|\|DUF499*->DUF3780->METHYLASE->METHYLASE->HKD-DNase+SF1-SF2-Helicase-> | 945 | Bacteria>Terrabacteria_group>Deinococcota>Deinococci>Deinococcales | GCF_014645575.1 | Deinococcus_wulumuqiensis | IEX15_RS07550 | 43264..46101 | 980427 | DUF499-domain-containing-protein-[Deinococcus-wulumuqiensis] |
| WP_189055329.1 | AAA+ATPase+wHTH+Alpha+Beta-domain+wHTH+Fn-III+Fn-III+RRM | Type-2_BR.RM.like.systems | XIS-HTH-><-35exo+SF1-SF2-Helicase<-HKD-DNase+SF1-SF2-Helicase<-METHYLASE<-METHYLASE<-DUF3780<-DUF499*<-?\|\|35exo+SF1-SF2-Helicase-> | 1024 | Bacteria>Terrabacteria_group>Deinococcota>Deinococci>Deinococcales | GCF_014647175.1 | Deinococcus_daejeonensis | IEY59_RS06775 | 64413..67487 | 1007098 | DUF499-domain-containing-protein-[Deinococcus-daejeonensis] |
| WP_234009276.1 | AAA+ATPase+wHTH+Alpha+Beta-domain+wHTH+Fn-III+Fn-III+RRM | Type-2_BR.RM.like.systems | <-DDE_Tnp_1<-PAIREDC-HTH\|\|DDE_5+DDE_Tnp_1-><-?\|\|DUF499*->DUF3780->?->PIN->DDE_Tnp_1-><-HIN-HTH\|\|DDE_5+DDE_Tnp_1-> | 1025 | Bacteria>Terrabacteria_group>Deinococcota>Deinococci>Deinococcales | GCF_002953415.1 | Deinococcus_sp._NW-56 | C3K08_RS17170 | 342216..345293 | 2080419 | DUF499-domain-containing-protein-[Deinococcus-sp.-NW-56] |
| WP_124875512.1 | AAA+ATPase+wHTH+Alpha+Beta-domain+wHTH+Fn-III+Fn-III+RRM | Type-2_BR.RM.like.systems | <-METHYLASE<-PIN<-?<-DUF3780<-DUF499*<-HTH+WYL+WYL-C<-?<-SF1-SF2-Helicase+REase<-Exonuc_V_gamma+REase | 1031 | Bacteria>Terrabacteria_group>Deinococcota>Deinococci>Deinococcales | GCF_003860465.1 | Deinococcus_psychrotolerans | EHF33_RS19840 | 23522..26617 | 2489213 | DUF499-domain-containing-protein-[Deinococcus-psychrotolerans] |
| NLH45761.1 | AAA+ATPase+wHTH+Alpha+Beta-domain+wHTH+Fn-III+Fn-III+RRM | Type-2_BR.RM.like.systems | DUF499*->DUF3780->METHYLASE->HKD-DNase+SF1-SF2-Helicase-> | 1038 | Bacteria>Terrabacteria_group>Mycoplasmatota>Mollicutes>Acholeplasmatales | GCA_012517145.1 | Acholeplasmataceae_bacterium | GX451_06515 | 3138..6254 | 1898209 | DUF499-domain-containing-protein-[Acholeplasmataceae-bacterium] |
| WP_038039187.1 | AAA+ATPase+wHTH+Alpha+Beta-domain+wHTH+Fn-III+Fn-III+RRM | Type-2_BR.RM.like.systems | PfkB->?->?->DeoR-HTH+ISOCOT->DUF499*->DUF3780->METHYLASE->HKD-DNase+SF1-SF2-Helicase-> | 1024 | Bacteria>Terrabacteria_group>Thermomicrobiota>Thermomicrobia>Thermomicrobia_incertae_sedis | GCF_000744775.1 | Thermorudis_peleae | N675_RS09860 | 1494378..1497452 | 1382356 | DUF499-domain-containing-protein-[Thermorudis-peleae] |
| WP_012642488.1 | AAA+ATPase+wHTH+Alpha+Beta-domain+wHTH+Fn-III+Fn-III+RRM | Type-2_BR.RM.like.systems | <-RNaseH<-?\|\|DUF499*->DUF3780->METHYLASE->HKD-DNase+SF1-SF2-Helicase-> | 1024 | Bacteria>Terrabacteria_group>Thermomicrobiota>Thermomicrobia>Thermomicrobiales | GCF_000021685.1 | Thermomicrobium_roseum | TRD_RS10465 | 261946..265020 | 500 | DUF499-domain-containing-protein-[Thermomicrobium-roseum] |
| KPA18631.1 | AAA+ATPase+wHTH+Alpha+Beta-domain+wHTH+Fn-III+Fn-III+RRM | Type-2_BR.RM.like.systems | <-HKD-DNase+SF1-SF2-Helicase<-wHTH+REase<-METHYLASE<-DUF3780<-DUF499* | 1032 | Bacteria>Thermodesulfobacteriota>Desulfobacteria>Desulfobacterales>Candidatus_Magnetomoraceae | GCA_001292585.1 | Candidatus_Magnetomorum_sp._HK-1 | MHK_001159 | 10914..14012 | 1509431 | ATPase-AAA-[Candidatus-Magnetomorum-sp.-HK-1] |
| KPA12292.1 | AAA+ATPase+wHTH+Alpha+Beta-domain+wHTH+Fn-III+Fn-III+RRM | Type-2_BR.RM.like.systems | <-HTH<-HKD-DNase+SF1-SF2-Helicase<-?<-?<-METHYLASE<-DUF3780<-DUF499*<-ZDNA-HTH | 876 | Bacteria>Thermodesulfobacteriota>Desulfobacteria>Desulfobacterales>Candidatus_Magnetomoraceae | GCA_001292585.1 | Candidatus_Magnetomorum_sp._HK-1 | MHK_007537 | 35476..38106 | 1509431 | glycosyl-transferase-[Candidatus-Magnetomorum-sp.-HK-1] |
| KPA09724.1 | AAA+ATPase+wHTH+Alpha+Beta-domain+wHTH+Fn-III+Fn-III+RRM | Type-2_BR.RM.like.systems | DUF499*->DUF3780->Schlafen+HATPase+HTH->METHYLASE->ParB+HNH->OLD-ABC-> | 1021 | Bacteria>Thermodesulfobacteriota>Desulfobacteria>Desulfobacterales>Candidatus_Magnetomoraceae | GCA_001292585.1 | Candidatus_Magnetomorum_sp._HK-1 | MHK_010064 | 367..3432 | 1509431 | ATPase-AAA-[Candidatus-Magnetomorum-sp.-HK-1] |
| RZB36988.1 | AAA+ATPase+wHTH+Alpha+Beta-domain+wHTH+Fn-III+Fn-III+RRM | Type-2_BR.RM.like.systems | <-HKD-DNase+SF1-SF2-Helicase<-?<-RNaseH<-METHYLASE<-DUF3780<-?<-DUF499*<-?<-?<-N6_MTase<-?<-HTH_3 | 1032 | Bacteria>Thermodesulfobacteriota>Desulfobacteria>Desulfobacterales>Desulfobacteraceae | GCA_004193595.1 | Desulfobacteraceae_bacterium_Eth-SRB2 | SRB2_01492 | 19212..22310 | 2528438 | hypothetical-protein-SRB2_01492-[Desulfobacteraceae-bacterium-Eth-SRB2] |
| MBA3018061.1 | AAA+ATPase+wHTH+Alpha+Beta-domain+wHTH+Fn-III+Fn-III+RRM | Type-2_BR.RM.like.systems | <-FtsK<-?<-?<-?\|\|DUF499*->DUF3780->METHYLASE->Schlafen+HATPase+HTH->HKD-DNase+SF1-SF2-Helicase-> | 1032 | Bacteria>Thermodesulfobacteriota>Desulfobacteria>Desulfobacterales>Desulfobacteraceae | GCA_013792055.1 | Desulfobacteraceae_bacterium | FP811_07930 | 7617..10715 | 2049433 | DUF499-domain-containing-protein-[Desulfobacteraceae-bacterium] |
| MBC2696453.1 | AAA+ATPase+wHTH+Alpha+Beta-domain+wHTH+Fn-III+Fn-III+RRM | Type-2_BR.RM.like.systems | <-MNS+REase\|\|DUF499*->DUF3780->METHYLASE->HKD-DNase+SF1-SF2-Helicase-> | 1032 | Bacteria>Thermodesulfobacteriota>Desulfobacteria>Desulfobacterales>Desulfobacteraceae | GCA_014237195.1 | Desulfobacteraceae_bacterium | HF982_14490 | 2508..5606 | 2049433 | DUF499-domain-containing-protein-[Desulfobacteraceae-bacterium] |
| MCF8025627.1 | AAA+ATPase+wHTH+Alpha+Beta-domain+wHTH+Fn-III+Fn-III+RRM | Type-2_BR.RM.like.systems | DUF499*->DUF3780->METHYLASE->HKD-DNase+SF1-SF2-Helicase->ParB+HNH-> | 1031 | Bacteria>Thermodesulfobacteriota>Desulfobacteria>Desulfobacterales>Desulfobacteraceae | GCA_021737095.1 | Desulfobacteraceae_bacterium | K9K82_09080 | 113..3208 | 2049433 | DUF499-domain-containing-protein-[Desulfobacteraceae-bacterium] |
| MCF8096019.1 | AAA+ATPase+wHTH+Alpha+Beta-domain+wHTH+Fn-III+Fn-III+RRM | Type-2_BR.RM.like.systems | DUF499*->DUF3780->METHYLASE->HKD-DNase+SF1-SF2-Helicase-p-> | 707 | Bacteria>Thermodesulfobacteriota>Desulfobacteria>Desulfobacterales>Desulfobacteraceae | GCA_021737545.1 | Desulfobacteraceae_bacterium | K9J79_11740 | 1..2126 | 2049433 | chitobiase/beta-hexosaminidase-C-terminal-domain-containing-protein-[Desulfobacteraceae-bacterium] |
| WP_178365842.1 | AAA+ATPase+wHTH+Alpha+Beta-domain+wHTH+Fn-III+Fn-III+RRM | Type-2_BR.RM.like.systems | <-TPR+TPR\|\|?->?-><-HKD-DNase+SF1-SF2-Helicase<-METHYLASE-p<-DUF3780<-DUF499* | 1021 | Bacteria>Thermodesulfobacteriota>Desulfobacteria>Desulfobacterales>Desulfobacteraceae | GCF_013389365.1 | Desulfobacter_latus | HXW94_RS05225 | 23224..26289 | 2292 | DUF499-domain-containing-protein-[Desulfobacter-latus] |
| OQY49645.1 | AAA+ATPase+wHTH+Alpha+Beta-domain+wHTH+Fn-III+Fn-III+RRM | Type-2_BR.RM.like.systems | <-OLD-ABC<-ParB+HNH<-DUF3780<-DUF499*<-HTH+WYL\|\|SMODS-> | 1021 | Bacteria>Thermodesulfobacteriota>Desulfobacteria>Desulfobacterales>Desulfobacteraceae | GCA_002085465.1 | Desulfobacteraceae_bacterium_4572_89 | B6230_07770 | 5853..8918 | 1972445 | hypothetical-protein-B6230_07770-[Desulfobacteraceae-bacterium-4572_89] |
| OIP44020.1 | AAA+ATPase+wHTH+Alpha+Beta-domain+wHTH+Fn-III+Fn-III+RRM | Type-2_BR.RM.like.systems | HTH-><-HKD-DNase+SF1-SF2-Helicase<-MACRODOMAIN<-?<-METHYLASE<-DUF3780<-DUF499*<-XIS-HTH | 1027 | Bacteria>Thermodesulfobacteriota>Desulfobacteria>Desulfobacterales>unclassified_Desulfobacterales | GCA_001873115.1 | Desulfobacterales_bacterium_CG2_30_60_27 | AUK28_11250 | 9145..12228 | 1805129 | hypothetical-protein-AUK28_11250-[Desulfobacterales-bacterium-CG2_30_60_27] |
| MCP4350268.1 | AAA+ATPase+wHTH+Alpha+Beta-domain+wHTH+Fn-III+Fn-III+RRM | Type-2_BR.RM.like.systems | <-DUF3780<-DUF499*\|\|REase-> | 1057 | Bacteria>Thermodesulfobacteriota>Desulfobacteria>Desulfobacterales>unclassified_Desulfobacterales | GCA_024244725.1 | Desulfobacterales_bacterium | GY795_32745 | 4306..7479 | 2044940 | DUF499-domain-containing-protein-[Desulfobacterales-bacterium] |
| MBF0120192.1 | AAA+ATPase+wHTH+Alpha+Beta-domain+wHTH+Fn-III+Fn-III+RRM | Type-2_BR.RM.like.systems | <-NERD<-OLD-ABC<-METHYLASE<-HEPN<-PolB<-DUF3780<-DUF499*<-?<-?<-ABC-ATPase+SbcC | 1037 | Bacteria>Thermodesulfobacteriota>Desulfobacteria>Desulfobacterales>unclassified_Desulfobacterales | GCA_015228685.1 | Desulfobacterales_bacterium | HQK79_15260 | 23007..26120 | 2044940 | DUF499-domain-containing-protein-[Desulfobacterales-bacterium] |
| MCK9293532.1 | AAA+ATPase+wHTH+Alpha+Beta-domain+wHTH+Fn-III+Fn-III+RRM | Type-2_BR.RM.like.systems | JAB->?-><-HKD-DNase+SF1-SF2-Helicase<-Schlafen+HATPase+HTH<-METHYLASE<-DUF3780<-DUF499*<-XIS-HTH | 1027 | Bacteria>Thermodesulfobacteriota>Desulfobulbia>Desulfobulbales>Desulfobulbaceae | GCA_023230525.1 | Desulfobulbaceae_bacterium | MOP70_00480 | 103201..106284 | 2053307 | DUF499-domain-containing-protein-[Desulfobulbaceae-bacterium] |
| MCK9175498.1 | AAA+ATPase+wHTH+Alpha+Beta-domain+wHTH+Fn-III+Fn-III+RRM | Type-2_BR.RM.like.systems | DUF499*->DUF3780->Schlafen+HATPase+HTH->METHYLASE->HKD-DNase+SF1-SF2-Helicase->URI-DNase+RAMA-> | 1021 | Bacteria>Thermodesulfobacteriota>Desulfobulbia>Desulfobulbales>Desulfocapsaceae | GCA_023231425.1 | Desulforhopalus_sp. | MOO96_09510 | 4806..7871 | 2099376 | DUF499-domain-containing-protein-[Desulforhopalus-sp.] |
| WP_136815950.1 | AAA+ATPase+wHTH+Alpha+Beta-domain+wHTH+Fn-III+Fn-III+RRM | Type-2_BR.RM.like.systems | MarR-HTH-><-HKD-DNase+SF1-SF2-Helicase<-METHYLASE<-DUF3780<-DUF499*<-?<-?<-?<-?<-TraJ-RHH | 1033 | Bacteria>Thermodesulfobacteriota>Desulfobulbia>Desulfobulbales>Desulfocapsaceae | GCF_005116645.1 | Desulfopila_sp._IMCC35006 | FCL47_RS22410 | 56979..60080 | 2569542 | DUF499-domain-containing-protein-[Desulfopila-sp.-IMCC35006] |
| WP_205219501.1 | AAA+ATPase+wHTH+Alpha+Beta-domain+wHTH+Fn-III+Fn-III+RRM | Type-2_BR.RM.like.systems | DUF499*->DUF3780->METHYLASE->HKD-DNase+SF1-SF2-Helicase->?->AbiEi_4-> | 1021 | Bacteria>Thermodesulfobacteriota>Desulfobulbia>Desulfobulbales>Desulfocapsaceae | GCF_016918505.1 | Desulforhopalus_vacuolatus | JWG39_RS11235 | 7446..10511 | 40414 | DUF499-domain-containing-protein-[Desulforhopalus-vacuolatus] |
| WP_045217121.1 | AAA+ATPase+wHTH+Alpha+Beta-domain+wHTH+Fn-III+Fn-III+RRM | Type-2_BR.RM.like.systems | <-HKD-DNase+SF1-SF2-Helicase<-CHTH<-?<-MetJarc-HTH<-METHYLASE<-DUF3780<-DUF499*<-Transposase_31+DUF4351<-?<-?<-ORC-CDC6+REase<-ORC-CDC6+REase | 1032 | Bacteria>Thermodesulfobacteriota>Desulfovibrionia>Desulfovibrionales>Desulfonatronovibrionaceae | GCF_000934755.1 | Desulfonatronovibrio_magnus | LZ23_RS20095 | 32185..35283 | 698827 | DUF499-domain-containing-protein-[Desulfonatronovibrio-magnus] |
| WP_243366428.1 | AAA+ATPase+wHTH+Alpha+Beta-domain+wHTH+Fn-III+Fn-III+RRM | Type-2_BR.RM.like.systems | <-METHYLASE<-HKD-DNase+SF1-SF2-Helicase<-METHYLASE<-DUF3780<-DUF499*<-ParB+HNH<-?<-?<-REase | 1022 | Bacteria>Thermodesulfobacteriota>Desulfovibrionia>Desulfovibrionales>Desulfovibrionaceae | GCF_022808695.1 | Fundidesulfovibrio_soli | MLE18_RS00810 | 167395..170463 | 2922716 | DUF499-domain-containing-protein-[Fundidesulfovibrio-sp.-SG60] |
| TVQ96846.1 | AAA+ATPase+wHTH+Alpha+Beta-domain+wHTH+Fn-III+Fn-III+RRM | Type-2_BR.RM.like.systems | DUF4236+TPR->DUF499*->DUF3780-><-HEPN<-PolB<-?\|\|METHYLASE-p->REase+N6_MTase-> | 438 | Bacteria>Thermodesulfobacteriota>Desulfovibrionia>Desulfovibrionales>unclassified_Desulfovibrionales | GCA_007693505.1 | Desulfovibrionales_bacterium | EA399_14900 | 820..2136 | 2448050 | hypothetical-protein-EA399_14900-[Desulfovibrionales-bacterium] |
| HIJ86203.1 | AAA+ATPase+wHTH+Alpha+Beta-domain+wHTH+Fn-III+Fn-III+RRM | Type-2_BR.RM.like.systems | <-HKD-DNase+SF1-SF2-Helicase\|\|?->?-><-METHYLASE<-DUF3780<-DUF499*\|\|?->MNS+HTH->?-><-Radical-SAM\|\|MetJarc-HTH->PemK_toxin-> | 1043 | Bacteria>Thermodesulfobacteriota>Desulfuromonadia>Desulfuromonadales>unclassified_Desulfuromonadales | GCA_013349405.1 | Desulfuromonadales_bacterium | HPP97_00805 | 167050..170181 | 2099678 | DUF499-domain-containing-protein-[Desulfuromonadales-bacterium] |
| MBK7550235.1 | AAA+ATPase+wHTH+Alpha+Beta-domain+wHTH+Fn-III+Fn-III+RRM | Type-2_BR.RM.like.systems | <-HKD-DNase+SF1-SF2-Helicase<-DUF3780<-DUF499*<-wHTH+REase<-PIN<-?<-CHTH | 1026 | Bacteria>Thermodesulfobacteriota>Syntrophia>Syntrophales>Syntrophaceae | - | Syntrophaceae_bacterium | IPI61_00440 | 88025..91105 | 2052184 | DUF499-domain-containing-protein-[Syntrophaceae-bacterium] |
| MDD2900810.1 | AAA+ATPase+wHTH+Alpha+Beta-domain+wHTH+Fn-III+Fn-III+RRM | Type-2_BR.RM.like.systems | OLD-ABC->HNH->?->?->RADICAL-SAM->DUF499*->DUF3780->METHYLASE->HKD-DNase+SF1-SF2-Helicase->?->Peptidase-> | 1027 | Bacteria>Thermodesulfobacteriota>Syntrophia>Syntrophales>unclassified_Syntrophales | GCA_028688935.1 | Syntrophales_bacterium | PHU44_00075 | 18845..21928 | 2928017 | MAG:-DUF499-domain-containing-protein-[Syntrophales-bacterium] |
| MCK9390838.1 | AAA+ATPase+wHTH+Alpha+Beta-domain+wHTH+Fn-III+Fn-III+RRM | Type-2_BR.RM.like.systems | <-MORC+HISKIN<-MORC<-REase<-DNA_methylase<-METHYLASE<-DUF3780<-DUF499* | 1026 | Bacteria>Thermodesulfobacteriota>Syntrophia>Syntrophales>unclassified_Syntrophales | GCA_023229565.1 | Syntrophales_bacterium | MOQ01_04645 | 15666..18746 | 2928017 | DUF499-domain-containing-protein-[Syntrophales-bacterium] |
| PIP07168.1 | AAA+ATPase+wHTH+Alpha+Beta-domain+wHTH+Fn-III+Fn-III+RRM | Type-2_BR.RM.like.systems | <-HKD-DNase+SF1-SF2-Helicase<-DUF3780<-DUF499* | 1026 | Bacteria>Thermodesulfobacteriota>Syntrophobacteria>Syntrophobacterales>unclassified_Syntrophobacterales | GCA_002771815.1 | Syntrophobacterales_bacterium_CG23_combo_of_CG06-09_8_20_14_all_48_27 | COX52_04725 | 9017..12097 | 1974101 | hypothetical-protein-COX52_04725-[Syntrophobacterales-bacterium-CG23_combo_of_CG06-09_8_20_14_all_48_27] |
| OPX97686.1 | AAA+ATPase+wHTH+Alpha+Beta-domain+wHTH+Fn-III+Fn-III+RRM | Type-2_BR.RM.like.systems | <-REase<-?\|\|?->HEPN+Mrr_N->?->DUF499-p->DUF499*->DUF3780->METHYLASE->HKD-DNase+SF1-SF2-Helicase->ParB+HNH-> | 861 | Bacteria>Thermodesulfobacteriota>Syntrophorhabdia>Syntrophorhabdales>Syntrophorhabdaceae | GCA_002067585.1 | Syntrophorhabdus_sp._PtaB.Bin027 | A4E59_00667 | 16354..18939 | 1811699 | MAG:-hypothetical-protein-A4E59_00667-[Syntrophorhabdus-sp.-PtaB.Bin027] |
| WP_033191784.1 | AAA+ATPase+wHTH+Alpha+Beta-domain+wHTH+Fn-III+Fn-III+RRM | Type-2_BR.RM.like.systems | DUF499*->DUF3780->METHYLASE->HKD-DNase+SF1-SF2-Helicase->?->?->Peptidase_M22-> | 1034 | Bacteria>Thermotogota>Thermotogae>Thermotogales>Fervidobacteriaceae | GCF_000767275.3 | Fervidobacterium | CBS1_RS02775 | 604999..608103 | 2422 | MULTISPECIES:-DUF499-domain-containing-protein-[Fervidobacterium] |
| MBI1298957.1 | AAA+ATPase+wHTH+Alpha+Beta-domain+wHTH+Fn-III+Fn-III+RRM | Type-2_BR.RM.like.systems | <-Trypsin<-?<-?\|\|DUF499*->DUF3780->METHYLASE->?->PIN->?->HKD-DNase+SF1-SF2-Helicase-> | 1026 | Bacteria>unclassified_Bacteria | GCA_016124945.1 | bacterium | GC175_28840 | 7719..10799 | 1869227 | DUF499-domain-containing-protein-[bacterium] |
| MBW6503451.1 | AAA+ATPase+wHTH+Alpha+Beta-domain+wHTH+Fn-III+Fn-III+RRM | Type-2_BR.RM.like.systems | DUF499*->DUF3780->METHYLASE->HKD-DNase+SF1-SF2-Helicase->?->?->Peptidase_M16+Peptidase_M16_C-> | 996 | Bacteria>unclassified_Bacteria | GCA_019429245.1 | bacterium | KOB90_04135 | 121..3111 | 1869227 | DUF499-domain-containing-protein-[bacterium] |
| MBX3024939.1 | AAA+ATPase+wHTH+Alpha+Beta-domain+wHTH+Fn-III+Fn-III+RRM | Type-2_BR.RM.like.systems | <-HKD-DNase+SF1-SF2-Helicase<-METHYLASE<-?<-PIN<-DUF3780<-DUF499*<-?<-?\|\|DUF2163+Phage_BR0599-> | 1021 | Bacteria>unclassified_Bacteria | GCA_019637795.1 | bacterium | KF840_08515 | 743696..746761 | 1869227 | DUF499-domain-containing-protein-[bacterium] |
| MCE5223354.1 | AAA+ATPase+wHTH+Alpha+Beta-domain+wHTH+Fn-III+Fn-III+RRM | Type-2_BR.RM.like.systems | <-ParB+HNH<-RNase<-?<-HKD-DNase+SF1-SF2-Helicase<-METHYLASE<-DUF3780<-DUF499* | 1030 | Bacteria>unclassified_Bacteria | GCA_021372585.1 | bacterium | LLG10_04080 | 11946..15038 | 1869227 | DUF499-domain-containing-protein-[bacterium] |
| EIP1037433.1 | AAA+ATPase+wHTH+Alpha+Beta-domain+wHTH+Fn-III+Fn-III+RRM | Type-2_BR.RM.like.systems | <-Phage_integrase\|\|?->?->?-><-MAD-NTD+HNH<-?\|\|DUF499*->DUF3780->METHYLASE->HKD-DNase+SF1-SF2-Helicase-><-REase | 1022 | - | - | - | LRR03_002276 | 218063..221131 | - | DUF499-domain-containing-protein-[Escherichia-coli] |
| EKJ6866575.1 | AAA+ATPase+wHTH+Alpha+Beta-domain+wHTH+Fn-III+Fn-III+RRM | Type-2_BR.RM.like.systems | Peptidase_M15->?->Ntox49->TUDOR+SF1-SF2-Helicase+iREase+CTD->?->?->DUF499*->DUF3780->METHYLASE->HKD-DNase+SF1-SF2-Helicase-><-Arm-DNA-bind_3+Phage_integrase<-REase+SF1-SF2-Helicase<-MazG | 1072 | - | - | - | O8Q49_001735 | 84420..87638 | - | DUF499-domain-containing-protein-[Pseudomonas-aeruginosa] |
| EJE9402911.1 | AAA+ATPase+wHTH+Alpha+Beta-domain+wHTH+Fn-III+Fn-III+RRM | Type-2_BR.RM.like.systems | Peptidase_M15->?->Ntox49->TUDOR+SF1-SF2-Helicase+iREase+CTD->?->?->DUF499*->DUF3780->METHYLASE->HKD-DNase+SF1-SF2-Helicase-><-Arm-DNA-bind_3+Phage_integrase<-REase+SF1-SF2-Helicase<-MazG | 1072 | - | - | - | PQC20_002029 | 43524..46742 | - | DUF499-domain-containing-protein-[Pseudomonas-aeruginosa] |

<table>
<tr><th>Protein-ID</th><th>DUF499-architecture</th><th>System</th><th>Gene-Neighbourhoods</th><th>Length</th><th>Taxonomy</th><th>Assembly-ID</th><th>Strain</th><th>Locus-Tag</th><th>Genomic-coordinates</th><th>TaxID</th><th>Fasta-defline</th></tr>
<tr><td>--</td><td></td><td></td><td></td><td></td><td></td><td></td><td></td><td></td><td></td><td></td><td></td></tr>
<tr><td>--BR-3.PglZ.associated.systems</td><td></td><td></td><td></td><td></td><td></td><td></td><td></td><td></td><td></td><td></td><td></td></tr>
<tr><td>--</td><td></td><td></td><td></td><td></td><td></td><td></td><td></td><td></td><td></td><td></td><td></td></tr>
<tr><td>RLJ04319.1</td><td>AAA+ATPase+wHTH+Alpha+Beta-domain+wHTH+RRM</td><td>Type-3_BR.PglZ.associated.systems</td><td><-METHYLASE-p<-PglZ<-DUF499*<-Phage_integrase</td><td>809</td><td>Archaea>DPANN_group>Candidatus_Aenigmarchaeota>unclassified_Candidatus_Aenigmarchaeota</td><td>GCA_003663285.1</td><td>Candidatus_Aenigmarchaeota_archaeon</td><td>DRP08_02010</td><td>1892..4321</td><td>2093792</td><td>hypothetical-protein-DRP08_02010-[Candidatus-Aenigmarchaeota-archaeon]</td></tr>
<tr><td>MCL6500835.1</td><td>AAA+ATPase+wHTH+Alpha+Beta-domain+wHTH+RRM</td><td>Type-3_BR.PglZ.associated.systems</td><td><-PolB<-PglZ<-DUF499*<-XIS-HTH</td><td>807</td><td>Archaea>DPANN_group>Candidatus_Pacearchaeota>unclassified_Candidatus_Pacearchaeota</td><td>GCA_023511755.1</td><td>Candidatus_Pacearchaeota_archaeon</td><td>K6T16_02265</td><td>1045..3468</td><td>2026773</td><td>hypothetical-protein-[Candidatus-Pacearchaeota-archaeon]</td></tr>
<tr><td>KXA95606.1</td><td>AAA+ATPase+wHTH+Alpha+Beta-domain+wHTH+RRM</td><td>Type-3_BR.PglZ.associated.systems</td><td>DNA-glycosylase-><-SF1-SF2-Helicase<-iSTAND<-METHYLASE<-PglZ<-DUF499*||ORC/CDC6+HTH-><-?<-REase<-?||HTH->PADR-HTH-></td><td>802</td><td>Archaea>Euryarchaeota>Euryarchaeota_incertae_sedis>candidate_division_MSBL1</td><td>GCA_001548965.1</td><td>candidate_division_MSBL1_archaeon_SCGC-AAA259I07</td><td>AKJ36_00090</td><td>15715..18123</td><td>1698266</td><td>hypothetical-protein-AKJ36_00090-[candidate-division-MSBL1-archaeon-SCGC-AAA259I07]</td></tr>
<tr><td>WP_246986818.1</td><td>AAA+ATPase+wHTH+Alpha+Beta-domain+wHTH+RRM</td><td>Type-3_BR.PglZ.associated.systems</td><td>ZnR->URI-DNase->McrB-AAA->McrC-NTD+REase-><-iSTAND<-DUF499*||METHYLASE-><-?||SF1-SF2-Helicase->?-><-?<-SWACOS</td><td>1086</td><td>Archaea>Euryarchaeota>Stenosarchaea_group>Halobacteria>Halobacteriales</td><td>GCF_023028345.1</td><td>Halorientalis_marina</td><td>MUG96_RS04275</td><td>804107..807367</td><td>2931976</td><td>DUF499-domain-containing-protein-[Halorientalis-marina]</td></tr>
<tr><td>WP_080508865.1</td><td>AAA+ATPase+wHTH+Alpha+Beta-domain+wHTH+RRM</td><td>Type-3_BR.PglZ.associated.systems</td><td>SF1-SF2-Helicase->?-><-?||DUF499*->PglZ->METHYLASE->iSTAND-></td><td>800</td><td>Archaea>Euryarchaeota>Stenosarchaea_group>Halobacteria>Halobacteriales</td><td>GCF_001280425.1</td><td>Haloarcula_rubripromontorii</td><td>AMS69_RS17865</td><td>34834..37236</td><td>1705562</td><td>DUF499-domain-containing-protein-[Haloarcula-rubripromontorii]</td></tr>
<tr><td>KOX91592.1</td><td>AAA+ATPase+wHTH+Alpha+Beta-domain+wHTH+RRM</td><td>Type-3_BR.PglZ.associated.systems</td><td>SF1-SF2-Helicase->?->?->DUF499*->PglZ->METHYLASE->iSTAND-></td><td>843</td><td>Archaea>Euryarchaeota>Stenosarchaea_group>Halobacteria>Halobacteriales</td><td>GCA_001280425.1</td><td>Haloarcula_rubripromontorii</td><td>AMS69_17885</td><td>34705..37236</td><td>1705562</td><td>AAA-family-ATPase-[Haloarcula-rubripromontorii]</td></tr>
<tr><td>WP_152120851.1</td><td>AAA+ATPase+wHTH+Alpha+Beta-domain+wHTH+RRM</td><td>Type-3_BR.PglZ.associated.systems</td><td>SbcC-><-?<-SF1-SF2-Helicase+REase<-Exonuc_V_gamma+REase<-?||METHYLASE->DUF499*->METHYLASE->NACHT->iSTAND-><-McrC-NTD+REase<-McrB-AAA||ORC/CDC6+REase-></td><td>1071</td><td>Archaea>Euryarchaeota>Stenosarchaea_group>Halobacteria>Halobacteriales</td><td>GCF_009184545.1</td><td>Halosegnis_rubeus</td><td>DMP03_RS12320</td><td>55278..58493</td><td>2212850</td><td>DUF499-domain-containing-protein-[Halosegnis-rubeus]</td></tr>
<tr><td>WP_274323786.1</td><td>AAA+ATPase+wHTH+Alpha+Beta-domain+wHTH+RRM</td><td>Type-3_BR.PglZ.associated.systems</td><td>REase+PNTD1+PNTD2->PIWI-><-iSTAND<-METHYLASE<-PglZ<-DUF499*<-?||ORC/CDC6+HTH->REase-><-?||DUF772+DDE_Tnp_1-></td><td>934</td><td>Archaea>Euryarchaeota>Stenosarchaea_group>Halobacteria>Halobacteriales</td><td>GCF_028747785.1</td><td>Halosimplex_sp._XZYJT29</td><td>PSC72_RS23170</td><td>4705748..4708552</td><td>3026162</td><td>DUF499-domain-containing-protein-[Halosimplex-sp.-XZYJT29]</td></tr>
<tr><td>WP_260594793.1</td><td>AAA+ATPase+wHTH+Alpha+Beta-domain+wHTH+RRM</td><td>Type-3_BR.PglZ.associated.systems</td><td>REase+PNTD1+PNTD2->PIWI-><-35exo||?-><-ORC/CDC6+HTH||DUF499*->PglZ->METHYLASE->?-><-?||DUF772+DDE_Tnp_1-></td><td>909</td><td>Archaea>Euryarchaeota>Stenosarchaea_group>Halobacteria>Halobacteriales</td><td>GCF_025231485.1</td><td>Salinirubellus_salinus</td><td>NOB31_RS05225</td><td>942571..945300</td><td>1364945</td><td>ATP-binding-protein-[Salinirubellus-salinus]</td></tr>
<tr><td>WP_007188154.1</td><td>AAA+ATPase+wHTH+Alpha+Beta-domain+wHTH+RRM</td><td>Type-3_BR.PglZ.associated.systems</td><td>RADICAL-SAM-><-?||HEPN->METHYLASE-><-HD<-?||DUF499*->METHYLASE->iSTAND-><-McrC-NTD+REase<-McrB-AAA||REase+DUF5655->SF1-SF2-Helicase-></td><td>1075</td><td>Archaea>Euryarchaeota>Stenosarchaea_group>Halobacteria>Halobacteriales</td><td>GCF_000337755.1</td><td>Haloarcula_californiae</td><td>C435_RS03615</td><td>18358..21585</td><td>244363</td><td>DUF499-domain-containing-protein-[Haloarcula-californiae]</td></tr>
<tr><td>WP_256392866.1</td><td>AAA+ATPase+wHTH+Alpha+Beta-domain+wHTH+RRM</td><td>Type-3_BR.PglZ.associated.systems</td><td>RADICAL-SAM->||McrB-AAA->McrC-NTD+REase-><-iSTAND<-DUF499*||?->PIN->?->ParB+HNH-><-REase||METHYLASE-></td><td>1089</td><td>Archaea>Euryarchaeota>Stenosarchaea_group>Halobacteria>Halobacteriales</td><td>GCF_024494695.1</td><td>Natronoarchaeum_rubrum</td><td>NO931_RS13565</td><td>560055..563324</td><td>755311</td><td>DUF499-domain-containing-protein-[Natronoarchaeum-rubrum]</td></tr>
<tr><td>WP_235158035.1</td><td>AAA+ATPase+wHTH+Alpha+Beta-domain+wHTH+RRM</td><td>Type-3_BR.PglZ.associated.systems</td><td>Phage_integrase->HEPN->DUF499*->PglZ->METHYLASE-p-></td><td>833</td><td>Archaea>Euryarchaeota>Stenosarchaea_group>Halobacteria>Halobacteriales</td><td>GCF_021504045.1</td><td>Halobacterium_salinarum</td><td>KV539_RS04330</td><td>69355..71856</td><td>2242</td><td>hypothetical-protein-[Halobacterium-salinarum]</td></tr>
<tr><td>WP_248655467.1</td><td>AAA+ATPase+wHTH+Alpha+Beta-domain+wHTH+RRM</td><td>Type-3_BR.PglZ.associated.systems</td><td>Phage_integrase-><-?<-?||BirA-HTH-><-?<-SF1-SF2-Helicase||DUF499*-><-?<-ParB+HNH||?->METHYLASE->?->iSTAND-></td><td>1068</td><td>Archaea>Euryarchaeota>Stenosarchaea_group>Halobacteria>Halobacteriales</td><td>GCF_023238445.1</td><td>Halorussus_gelatinilyticus</td><td>MOR88_RS02920</td><td>556415..559621</td><td>2937524</td><td>DUF499-domain-containing-protein-[Halorussus-sp.-XZYJT40]</td></tr>
<tr><td>WP_153952805.1</td><td>AAA+ATPase+wHTH+Alpha+Beta-domain+wHTH+RRM</td><td>Type-3_BR.PglZ.associated.systems</td><td>Phage_int_SAM_5+Phage_integrase-><-?<-?<-MTaX1+HNH<-iSTAND<-DUF499*<-METHYLASE||SF1-SF2-Helicase-></td><td>1075</td><td>Archaea>Euryarchaeota>Stenosarchaea_group>Halobacteria>Halobacteriales</td><td>GCF_009663395.1</td><td>Halosegnis_longus</td><td>DM818_RS13820</td><td>128498..131725</td><td>2216012</td><td>DUF499-domain-containing-protein-[Halosegnis-longus]</td></tr>
<tr><td>WP_225334475.1</td><td>AAA+ATPase+wHTH+Alpha+Beta-domain+wHTH+RRM</td><td>Type-3_BR.PglZ.associated.systems</td><td>PadR-HTH->?-><-?<-?<-?<-tRNA||DUF499*->PglZ->METHYLASE->iSTAND-><-BirA-HTH</td><td>943</td><td>Archaea>Euryarchaeota>Stenosarchaea_group>Halobacteria>Halobacteriales</td><td>GCF_020217425.1</td><td>Halomicrobium_urmianum</td><td>LCY71_RS00840</td><td>162225..165056</td><td>1586233</td><td>hypothetical-protein-[Halosiccatus-urmianus]</td></tr>
<tr><td>EMA25693.1</td><td>AAA+ATPase+wHTH+Alpha+Beta-domain+wHTH+RRM</td><td>Type-3_BR.PglZ.associated.systems</td><td>ORC/CDC6+HTH-><-?<-SF1-SF2-Helicase||DUF499*->?->RNase->iSTAND->?-><-McrC-NTD+REase<-McrB-AAA</td><td>1068</td><td>Archaea>Euryarchaeota>Stenosarchaea_group>Halobacteria>Halobacteriales</td><td>GCA_000336895.1</td><td>Haloarcula_argentinensis_DSM_12282</td><td>C443_02749</td><td>44456..47662</td><td>1230451</td><td>ATPase-(AAA+-superfamily)-like-protein-[Haloarcula-argentinensis-DSM-12282]</td></tr>
<tr><td>WP_274326254.1</td><td>AAA+ATPase+wHTH+Alpha+Beta-domain+wHTH+RRM</td><td>Type-3_BR.PglZ.associated.systems</td><td>MORC-><-zf-IS66+DDE_Tnp_IS66||MORC+DUF3684+REase->METHYLASE-><-?||DUF499*->METHYLASE->iSTAND-><-McrC-NTD+REase<-McrB-AAA||ORC/CDC6+REase-></td><td>984</td><td>Archaea>Euryarchaeota>Stenosarchaea_group>Halobacteria>Halobacteriales</td><td>GCF_028747785.1</td><td>Halosimplex_sp._XZYJT29</td><td>PSC72_RS24190</td><td>111156..114110</td><td>3026162</td><td>DUF499-domain-containing-protein-[Halosimplex-sp.-XZYJT29]</td></tr>
<tr><td>WP_254824803.1</td><td>AAA+ATPase+wHTH+Alpha+Beta-domain+wHTH+RRM</td><td>Type-3_BR.PglZ.associated.systems</td><td>McrC-NTD+REase->ZnR+REase-><-iSTAND<-ParB+HNH<-METHYLASE<-MTaX1+HNH+ColE7<-DUF499*||SF1-SF2-Helicase->?-><-SF1-SF2-Helicase+REase<-REase||SMODS-></td><td>1071</td><td>Archaea>Euryarchaeota>Stenosarchaea_group>Halobacteria>Halobacteriales</td><td>GCF_024298885.1</td><td>Haloglomus_halophilum</td><td>NL114_RS20950</td><td>108014..111229</td><td>2962672</td><td>DUF499-domain-containing-protein-[Haloglomus-sp.-ZY41]</td></tr>
<tr><td>WP_135823131.1</td><td>AAA+ATPase+wHTH+Alpha+Beta-domain+wHTH+RRM</td><td>Type-3_BR.PglZ.associated.systems</td><td>McrC-NTD+REase->ZnR+REase-><-iSTAND<-ParB+HNH<-METHYLASE||?-><-DUF499*||SF1-SF2-Helicase->?-><-SF1-SF2-Helicase+REase<-REase</td><td>1071</td><td>Archaea>Euryarchaeota>Stenosarchaea_group>Halobacteria>Halobacteriales</td><td>GCF_004785955.1</td><td>Halostella_litorea</td><td>EYW40_RS18725</td><td>43618..46833</td><td>2528831</td><td>DUF499-domain-containing-protein-[Halostella-litorea]</td></tr>
<tr><td>WP_229123016.1</td><td>AAA+ATPase+wHTH+Alpha+Beta-domain+wHTH+RRM</td><td>Type-3_BR.PglZ.associated.systems</td><td>McrC-NTD+REase-><-iSTAND<-RNase<-METHYLASE||PHP+SbcC+SidE+CR-ATPase9->ParB+HNH-><-DUF499*||SF1-SF2-Helicase->?->?-><-SF1-SF2-Helicase</td><td>1069</td><td>Archaea>Euryarchaeota>Stenosarchaea_group>Halobacteria>Halobacteriales</td><td>GCF_017094465.1</td><td>Halapricum_desulfuricans</td><td>HSEST_RS14120</td><td>98603..101812</td><td>2841257</td><td>DUF499-domain-containing-protein-[Halapricum-desulfuricans]</td></tr>
<tr><td>WP_170084520.1</td><td>AAA+ATPase+wHTH+Alpha+Beta-domain+wHTH+RRM</td><td>Type-3_BR.PglZ.associated.systems</td><td>McrC-NTD+REase-><-?<-iSTAND<-RNase<-METHYLASE||REase-><-DUF499*||SF1-SF2-Helicase->?-><-ORC/CDC6+HTH||?->?->ZDNA-HTH-></td><td>1070</td><td>Archaea>Euryarchaeota>Stenosarchaea_group>Halobacteria>Halobacteriales</td><td>GCF_012726155.1</td><td>Haloarcula_rubripromontorii</td><td>GOC83_RS19580</td><td>48744..51956</td><td>1705562</td><td>DUF499-domain-containing-protein-[Haloarcula-rubripromontorii]</td></tr>
<tr><td>WP_247010670.1</td><td>AAA+ATPase+wHTH+Alpha+Beta-domain+wHTH+RRM</td><td>Type-3_BR.PglZ.associated.systems</td><td>McrC-NTD+REase-><-?<-?||METHYLASE->?->DUF499*->METHYLASE->iSTAND-><-McrC-NTD+REase<-McrB-AAA||ORC/CDC6+REase->METHYLASE-></td><td>1078</td><td>Archaea>Euryarchaeota>Stenosarchaea_group>Halobacteria>Halobacteriales</td><td>GCF_023028225.1</td><td>Halorientalis_litorea</td><td>MUG95_RS16045</td><td>63979..67215</td><td>2931977</td><td>DUF499-domain-containing-protein-[Halorientalis-litorea]</td></tr>
<tr><td>WP_135829423.1</td><td>AAA+ATPase+wHTH+Alpha+Beta-domain+wHTH+RRM</td><td>Type-3_BR.PglZ.associated.systems</td><td>McrC-NTD+REase->?-><-iSTAND<-METHYLASE<-?||?-><-DUF499*||SF1-SF2-Helicase->?->SF1-SF2-Helicase-><-DUF772+DDE_Tnp_1_6</td><td>1068</td><td>Archaea>Euryarchaeota>Stenosarchaea_group>Halobacteria>Halobacteriales</td><td>GCF_004765805.1</td><td>Halorussus_halobius</td><td>EPL03_RS11400</td><td>932771..935977</td><td>1710537</td><td>DUF499-domain-containing-protein-[Halorussus-halobius]</td></tr>
<tr><td>WP_285253701.1</td><td>AAA+ATPase+wHTH+Alpha+Beta-domain+wHTH+RRM</td><td>Type-3_BR.PglZ.associated.systems</td><td>McrC-NTD+REase->?-><-iSTAND<-?<-METHYLASE<-?<-DUF499*</td><td>1062</td><td>Archaea>Euryarchaeota>Stenosarchaea_group>Halobacteria>Halobacteriales</td><td>GCF_030250185.1</td><td>Halobacterium_salinarum</td><td>PNP84_RS10970</td><td>9207..12395</td><td>2242</td><td>DUF499-domain-containing-protein-[Halobacterium-salinarum]</td></tr>
<tr><td>WP_015409830.1</td><td>AAA+ATPase+wHTH+Alpha+Beta-domain+wHTH+RRM</td><td>Type-3_BR.PglZ.associated.systems</td><td>McrB-AAA->McrC-NTD+REase-><-iSTAND<-METHYLASE<-NTP_transf+wHTH<-HTH+IclR-HTH<-DUF499*||SF1-SF2-Helicase->?->Phage_integrase->?-><-PadR-HTH||<-DEDD_Tnp_IS110+Transposase_20</td><td>1068</td><td>Archaea>Euryarchaeota>Stenosarchaea_group>Halobacteria>Halobacteriales</td><td>GCF_000591055.1</td><td>Natronomonas_moolapensis</td><td>NMLP_RS09150</td><td>1887636..1890842</td><td>416273</td><td>DUF499-domain-containing-protein-[Natronomonas-moolapensis]</td></tr>
<tr><td>WP_020446091.1</td><td>AAA+ATPase+wHTH+Alpha+Beta-domain+wHTH+RRM</td><td>Type-3_BR.PglZ.associated.systems</td><td>McrB-AAA->McrC-NTD+REase-><-iSTAND<-DUF499*<-ParB+HNH<-?||METHYLASE->SbcC-></td><td>1086</td><td>Archaea>Euryarchaeota>Stenosarchaea_group>Halobacteria>Halobacteriales</td><td>GCF_000403645.1</td><td>Salinarchaeum_sp._Harcht-Bsk1</td><td>L593_RS06220</td><td>1342919..1346179</td><td>1333523</td><td>DUF499-domain-containing-protein-[Salinarchaeum-sp.-Harcht-Bsk1]</td></tr>
<tr><td>WP_053969685.1</td><td>AAA+ATPase+wHTH+Alpha+Beta-domain+wHTH+RRM</td><td>Type-3_BR.PglZ.associated.systems</td><td>McrB-AAA->McrC-NTD+REase-><-?<-iSTAND<-RNase<-METHYLASE<-DUF499*||SF1-SF2-Helicase->?-><-ORC/CDC6+HTH</td><td>1068</td><td>Archaea>Euryarchaeota>Stenosarchaea_group>Halobacteria>Halobacteriales</td><td>GCF_001280425.1</td><td>Haloarcula_rubripromontorii</td><td>AMS69_RS19455</td><td>14199..17405</td><td>1705562</td><td>DUF499-domain-containing-protein-[Haloarcula-rubripromontorii]</td></tr>
<tr><td>WP_004516383.1</td><td>AAA+ATPase+wHTH+Alpha+Beta-domain+wHTH+RRM</td><td>Type-3_BR.PglZ.associated.systems</td><td>McrB-AAA->McrC-NTD+REase-><-?||CARF-><-iSTAND<-METHYLASE<-DUF499*||?->HD-><-METHYLASE<-Phage_integrase<-METHYLASE</td><td>1075</td><td>Archaea>Euryarchaeota>Stenosarchaea_group>Halobacteria>Halobacteriales</td><td>GCF_000337775.1</td><td>Haloarcula_vallismortis</td><td>BLV10_RS17025</td><td>26097..29324</td><td>28442</td><td>DUF499-domain-containing-protein-[Haloarcula-vallismortis]</td></tr>
<tr><td>WP_227355755.1</td><td>AAA+ATPase+wHTH+Alpha+Beta-domain+wHTH+RRM</td><td>Type-3_BR.PglZ.associated.systems</td><td>MarR-HTH-><-ParB+HNH<-NACHT<-McrC-NTD+REase<-EVE+SF1-SF2-Helicase+McrB||REase+DUF4268->DUF499*->PglZ->METHYLASE->iSTAND-><-SF1-SF2-Helicase-p||?->Radical-SAM-></td><td>938</td><td>Archaea>Euryarchaeota>Stenosarchaea_group>Halobacteria>Halobacteriales</td><td>GCF_020614375.1</td><td>Haladaptatus_salinisoli</td><td>LIX00_RS17100</td><td>281449..284265</td><td>2884876</td><td>DUF499-domain-containing-protein-[Haladaptatus-salinisoli]</td></tr>
<tr><td>WP_135853937.1</td><td>AAA+ATPase+wHTH+Alpha+Beta-domain+wHTH+RRM</td><td>Type-3_BR.PglZ.associated.systems</td><td>MarR-HTH->?->McrB-AAA->McrC-NTD+REase-><-iSTAND<-METHYLASE<-DUF499*<-METHYLASE||SF1-SF2-Helicase->?->METHYLASE-><-McrC-NTD+REase</td><td>1069</td><td>Archaea>Euryarchaeota>Stenosarchaea_group>Halobacteria>Halobacteriales</td><td>GCF_004765815.2</td><td>Halorussus_salinus</td><td>EPL00_RS19250</td><td>484788..487997</td><td>1364935</td><td>DUF499-domain-containing-protein-[Halorussus-salinus]</td></tr>
<tr><td>WP_076432494.1</td><td>AAA+ATPase+wHTH+Alpha+Beta-domain+wHTH+RRM</td><td>Type-3_BR.PglZ.associated.systems</td><td>LysR-HTH->McrB-AAA->McrC-NTD+REase-><-iSTAND<-METHYLASE||ParB+HNH-><-DUF499*||SF1-SF2-Helicase->?-><-?||?->EVE+SF1-SF2-Helicase+McrB->McrC-NTD+REase-></td><td>1069</td><td>Archaea>Euryarchaeota>Stenosarchaea_group>Halobacteria>Halobacteriales</td><td>GCF_900156425.1</td><td>Haladaptatus_litoreus</td><td>BW974_RS20500</td><td>117928..121137</td><td>553468</td><td>DUF499-domain-containing-protein-[Haladaptatus-litoreus]</td></tr>
<tr><td>WP_135807376.1</td><td>AAA+ATPase+wHTH+Alpha+Beta-domain+wHTH+RRM</td><td>Type-3_BR.PglZ.associated.systems</td><td>DUF499*->PglZ->METHYLASE->iSTAND-><-?||Phage_integrase-></td><td>931</td><td>Archaea>Euryarchaeota>Stenosarchaea_group>Halobacteria>Halobacteriales</td><td>GCF_004765795.1</td><td>Halorussus_marinus</td><td>EPY72_RS18550</td><td>3253..6048</td><td>2505976</td><td>DUF499-domain-containing-protein-[Halorussus-marinus]</td></tr>
<tr><td>WP_285255286.1</td><td>AAA+ATPase+wHTH+Alpha+Beta-domain+wHTH+RRM</td><td>Type-3_BR.PglZ.associated.systems</td><td>DUF499*->PglZ->METHYLASE->iSTAND->?->Resolvase-></td><td>930</td><td>Archaea>Euryarchaeota>Stenosarchaea_group>Halobacteria>Halobacteriales</td><td>GCF_030250325.1</td><td>Halobacterium_salinarum</td><td>PNP83_RS03665</td><td>67..2859</td><td>2242</td><td>DUF499-domain-containing-protein-[Halobacterium-salinarum]</td></tr>
<tr><td>WP_170845500.1</td><td>AAA+ATPase+wHTH+Alpha+Beta-domain+wHTH+RRM</td><td>Type-3_BR.PglZ.associated.systems</td><td>DDE_Tnp_1->?->McrC-NTD+REase-><-?||METHYLASE->DUF499*->METHYLASE->iSTAND-><-McrC-NTD+REase<-McrB-AAA||ORC/CDC6+REase->METHYLASE-></td><td>1078</td><td>Archaea>Euryarchaeota>Stenosarchaea_group>Halobacteria>Halobacteriales</td><td>GCF_900110215.1</td><td>Halorientalis_persicus</td><td>BMX17_RS20575</td><td>8615..11851</td><td>1367881</td><td>DUF499-domain-containing-protein-[Halorientalis-persicus]</td></tr>
<tr><td>WP_237561076.1</td><td>AAA+ATPase+wHTH+Alpha+Beta-domain+wHTH+RRM</td><td>Type-3_BR.PglZ.associated.systems</td><td>ABhydrolase-><-?||REase-><-?<-SF1-SF2-Helicase||DUF499*->METHYLASE->iSTAND-><-McrC-NTD+REase<-McrB-AAA<-REase</td><td>935</td><td>Archaea>Euryarchaeota>Stenosarchaea_group>Halobacteria>Halobacteriales</td><td>GCF_000755245.1</td><td>Halococcus_sediminicola</td><td>ACP97_RS02845</td><td>208503..211310</td><td>1264579</td><td>ATP-binding-protein-[Halococcus-sediminicola]</td></tr>
<tr><td>WP_004591998.1</td><td>AAA+ATPase+wHTH+Alpha+Beta-domain+wHTH+RRM</td><td>Type-3_BR.PglZ.associated.systems</td><td><-Zn_Tnp_IS1595+DDE_Tnp_IS1595||?->?->||?->?->DUF499*->PglZ->METHYLASE->iSTAND-><-?||SF1-SF2-Helicase-p-></td><td>931</td><td>Archaea>Euryarchaeota>Stenosarchaea_group>Halobacteria>Halobacteriales</td><td>GCF_000336635.1</td><td>Haloarcula_japonica</td><td>C444_RS07455</td><td>21015..23810</td><td>29282</td><td>DUF499-domain-containing-protein-[Haloarcula-japonica]</td></tr>
<tr><td>WP_158056534.1</td><td>AAA+ATPase+wHTH+Alpha+Beta-domain+wHTH+RRM</td><td>Type-3_BR.PglZ.associated.systems</td><td><-SF1-SF2-Helicase<-REase<-METHYLASE<-?||DUF499*->iSTAND-><-McrC-NTD+REase<-McrB-AAA||REase-><-HEPN</td><td>1090</td><td>Archaea>Euryarchaeota>Stenosarchaea_group>Halobacteria>Halobacteriales</td><td>GCF_008831545.1</td><td>Halorussus_halophilus</td><td>F7R90_RS06970</td><td>1419992..1423264</td><td>2650975</td><td>DUF499-domain-containing-protein-[Halorussus-halophilus]</td></tr>
<tr><td>WP_050038544.1</td><td>AAA+ATPase+wHTH+Alpha+Beta-domain+wHTH+RRM</td><td>Type-3_BR.PglZ.associated.systems</td><td><-SF1-SF2-Helicase<-ORC/CDC6+REase||McrB-AAA->McrC-NTD+REase-><-iSTAND<-METHYLASE<-DUF499*<-?<-METHYLASE<-MORC+DUF3684+REase||?-><-CoiA||ORC/CDC6+HTH-></td><td>1078</td><td>Archaea>Euryarchaeota>Stenosarchaea_group>Halobacteria>Halobacteriales</td><td>GCF_001485555.1</td><td>unclassified_Haloarcula</td><td>AVZ98_RS18980</td><td>49445..52681</td><td>2624677</td><td>MULTISPECIES:-DUF499-domain-containing-protein-[unclassified-Haloarcula]</td></tr>
<tr><td>WP_129069634.1</td><td>AAA+ATPase+wHTH+Alpha+Beta-domain+wHTH+RRM</td><td>Type-3_BR.PglZ.associated.systems</td><td><-SF1-SF2-Helicase+REase<-REase||McrB-AAA->McrC-NTD+REase->METHYLASE-><-iSTAND<-DUF499*<-NTP_transf_5<-?||?->PIN-><-HEPN||METHYLASE-></td><td>1086</td><td>Archaea>Euryarchaeota>Stenosarchaea_group>Halobacteria>Halobacteriales</td><td>-</td><td>Halorientalis_pallida</td><td>EAF64_RS14100</td><td>110036..113296</td><td>2479928</td><td>DUF499-domain-containing-protein-[Halorientalis-pallida]</td></tr>
<tr><td>WP_137275290.1</td><td>AAA+ATPase+wHTH+Alpha+Beta-domain+wHTH+RRM</td><td>Type-3_BR.PglZ.associated.systems</td><td><-SF1-SF2-Helicase||McrB-AAA->McrC-NTD+REase-><-iSTAND<-METHYLASE<-DUF499*<-METHYLASE||?->ParB+HNH-></td><td>1076</td><td>Archaea>Euryarchaeota>Stenosarchaea_group>Halobacteria>Halobacteriales</td><td>GCF_005239135.1</td><td>Natronomonas_salsuginis</td><td>DM868_RS02605</td><td>463806..467036</td><td>2217661</td><td>DUF499-domain-containing-protein-[Natronomonas-salsuginis]</td></tr>
<tr><td>WP_245998557.1</td><td>AAA+ATPase+wHTH+Alpha+Beta-domain+wHTH+RRM</td><td>Type-3_BR.PglZ.associated.systems</td><td><-SF1-SF2-Helicase||DUF499*->MTaX1+HNH+ColE7->METHYLASE->iSTAND->?->ZetaToxin-><-McrC-NTD+REase</td><td>1057</td><td>Archaea>Euryarchaeota>Stenosarchaea_group>Halobacteria>Halobacteriales</td><td>GCF_003697815.1</td><td>Halalkalicoccus_subterraneus</td><td>EAO80_RS10110</td><td>56658..59831</td><td>2675002</td><td>DUF499-domain-containing-protein-[Halalkalicoccus-subterraneus]</td></tr>
<tr><td>WP_257299380.1</td><td>AAA+ATPase+wHTH+Alpha+Beta-domain+wHTH+RRM</td><td>Type-3_BR.PglZ.associated.systems</td><td><-SF1-SF2-Helicase||DUF499*-><-?||METHYLASE->CITB-HTH->?->iSTAND-><-McrC-NTD+REase</td><td>1068</td><td>Archaea>Euryarchaeota>Stenosarchaea_group>Halobacteria>Halobacteriales</td><td>-</td><td>Haloarchaeobius_sp._FL176</td><td>NO978_RS09655</td><td>221264..224470</td><td>2967129</td><td>DUF499-domain-containing-protein-[Haloarchaeobius-sp.-FL176]</td></tr>
<tr><td>WP_188980552.1</td><td>AAA+ATPase+wHTH+Alpha+Beta-domain+wHTH+RRM</td><td>Type-3_BR.PglZ.associated.systems</td><td><-SF1-SF2-Helicase||DUF499*-><-?||METHYLASE->?->?->PIN->iSTAND-></td><td>1070</td><td>Archaea>Euryarchaeota>Stenosarchaea_group>Halobacteria>Halobacteriales</td><td>GCF_014647415.1</td><td>Haloarcula_sebkhae</td><td>IEY38_RS19870</td><td>57205..60417</td><td>932660</td><td>DUF499-domain-containing-protein-[Haloarcula-sebkhae]</td></tr>
<tr><td>WP_233553836.1</td><td>AAA+ATPase+wHTH+Alpha+Beta-domain+wHTH+RRM</td><td>Type-3_BR.PglZ.associated.systems</td><td><-SF1-SF2-Helicase||?-><-METHYLASE<-?||DUF499*->iSTAND-><-McrC-NTD+REase<-McrB-AAA<-?<-?<-PadR-HTH</td><td>843</td><td>Archaea>Euryarchaeota>Stenosarchaea_group>Halobacteria>Halobacteriales</td><td>GCF_003602035.1</td><td>Halococcus_sp._IIIV-5B</td><td>D3261_RS02505</td><td>109531..112062</td><td>2321230</td><td>DUF499-domain-containing-protein-[Halococcus-sp.-IIIV-5B]</td></tr>
<tr><td>SDG40939.1</td><td>AAA+ATPase+wHTH+Alpha+Beta-domain+wHTH+RRM</td><td>Type-3_BR.PglZ.associated.systems</td><td><-Resolvase||McrB-AAA->McrC-NTD+REase-><-iSTAND<-DUF499*</td><td>242</td><td>Archaea>Euryarchaeota>Stenosarchaea_group>Halobacteria>Halobacteriales</td><td>GCA_900102305.1</td><td>Halorientalis_regularis</td><td>SAMN05216218_13410</td><td>8735..9465</td><td>660518</td><td>hypothetical-protein-SAMN05216218_13410,-partial-[Halorientalis-regularis]</td></tr>
<tr><td>WP_285254229.1</td><td>AAA+ATPase+wHTH+Alpha+Beta-domain+wHTH+RRM</td><td>Type-3_BR.PglZ.associated.systems</td><td><-Phage_integrase||?->?->?->?->DUF499*->PglZ->METHYLASE->iSTAND-></td><td>931</td><td>Archaea>Euryarchaeota>Stenosarchaea_group>Halobacteria>Halobacteriales</td><td>GCF_030250225.1</td><td>Halobacterium_salinarum</td><td>PNP03_RS08435</td><td>4236..7031</td><td>2242</td><td>DUF499-domain-containing-protein-[Halobacterium-salinarum]</td></tr>
<tr><td>WP_227130964.1</td><td>AAA+ATPase+wHTH+Alpha+Beta-domain+wHTH+RRM</td><td>Type-3_BR.PglZ.associated.systems</td><td><-ParB+METHYLASE<-ParB+METHYLASE||McrB-AAA->McrC-NTD+REase-><-iSTAND<-DUF499*<-NTP_transf_5<-?||?->ParB+HNH-><-REase||METHYLASE-></td><td>1089</td><td>Archaea>Euryarchaeota>Stenosarchaea_group>Halobacteria>Halobacteriales</td><td>GCF_020567525.1</td><td>Halorubellus_salinus</td><td>LI336_RS02105</td><td>417083..420352</td><td>755309</td><td>DUF499-domain-containing-protein-[Halorubellus-salinus]</td></tr>
<tr><td>WP_285254565.1</td><td>AAA+ATPase+wHTH+Alpha+Beta-domain+wHTH+RRM</td><td>Type-3_BR.PglZ.associated.systems</td><td><-ORC/CDC6+REase||McrB-AAA->McrC-NTD+REase-><-iSTAND<-METHYLASE<-DUF499*<-?||?-><-BirA-HTH</td><td>1079</td><td>Archaea>Euryarchaeota>Stenosarchaea_group>Halobacteria>Halobacteriales</td><td>GCF_030250245.1</td><td>Halobacterium_salinarum</td><td>PNP85_RS03280</td><td>8159..11398</td><td>2242</td><td>DUF499-domain-containing-protein-[Halobacterium-salinarum]</td></tr>
<tr><td>WP_158854013.1</td><td>AAA+ATPase+wHTH+Alpha+Beta-domain+wHTH+RRM</td><td>Type-3_BR.PglZ.associated.systems</td><td><-METHYLASE<-?<-PIN<-?<-Zn_Tnp_IS1595+DDE_Tnp_IS1595||DUF499*-><-?||METHYLASE->iSTAND-><-McrC-NTD+REase<-McrB-AAA</td><td>1086</td><td>Archaea>Euryarchaeota>Stenosarchaea_group>Halobacteria>Halobacteriales</td><td>GCF_009797925.1</td><td>Halorhabdus_sp._CUG00001</td><td>FR699_RS03320</td><td>667983..671243</td><td>2600297</td><td>DUF499-domain-containing-protein-[Halorhabdus-sp.-CUG00001]</td></tr>
<tr><td>MCD2204530.1</td><td>AAA+ATPase+wHTH+Alpha+Beta-domain+wHTH+RRM</td><td>Type-3_BR.PglZ.associated.systems</td><td><-MarR-HTH||?->McrB-AAA->McrC-NTD+REase-><-iSTAND<-METHYLASE<-DUF499*||Phage_integrase->METHYLASE-p->METHYLASE-p->SF1-SF2-Helicase->?-><-McrC-NTD+REase</td><td>1033</td><td>Archaea>Euryarchaeota>Stenosarchaea_group>Halobacteria>Halobacteriales</td><td>GCA_021029015.1</td><td>Halobacterium_sp._KA-6</td><td>LPA46_14445</td><td>35375..38476</td><td>2896368</td><td>DUF499-domain-containing-protein-[Halobacterium-sp.-KA-6]</td></tr>
<tr><td>WP_269073710.1</td><td>AAA+ATPase+wHTH+Alpha+Beta-domain+wHTH+RRM</td><td>Type-3_BR.PglZ.associated.systems</td><td><-MarR-HTH||?->McrB-AAA->McrC-NTD+REase-><-iSTAND<-METHYLASE<-DUF499*||Phage_integrase->METHYLASE-p->METHYLASE-p->SF1-SF2-Helicase->?-><-McrC-NTD+REase</td><td>1068</td><td>Archaea>Euryarchaeota>Stenosarchaea_group>Halobacteria>Halobacteriales</td><td>GCF_021029015.1</td><td>Halobacterium_sp._KA-6</td><td>LPA46_RS14450</td><td>35375..38581</td><td>2896368</td><td>DUF499-domain-containing-protein-[Halobacterium-sp.-KA-6]</td></tr>
<tr><td>WP_230893090.1</td><td>AAA+ATPase+wHTH+Alpha+Beta-domain+wHTH+RRM</td><td>Type-3_BR.PglZ.associated.systems</td><td><-MarR-HTH||?->McrB-AAA->McrC-NTD+REase-><-iSTAND<-METHYLASE<-DUF499*||Phage_integrase->METHYLASE-p->METHYLASE-p->SF1-SF2-Helicase->?-><-McrC-NTD+REase</td><td>1033</td><td>Archaea>Euryarchaeota>Stenosarchaea_group>Halobacteria>Halobacteriales</td><td>-</td><td>Halobacterium_sp._KA-6</td><td>LPA46_14445</td><td>35375..38476</td><td>2896368</td><td>DUF499-domain-containing-protein-[Halobacterium-sp.-KA-6]</td></tr>
<tr><td>WP_285254334.1</td><td>AAA+ATPase+wHTH+Alpha+Beta-domain+wHTH+RRM</td><td>Type-3_BR.PglZ.associated.systems</td><td><-iSTAND<-METHYLASE<-PglZ<-DUF499*<-HEPN<-Phage_integrase</td><td>833</td><td>Archaea>Euryarchaeota>Stenosarchaea_group>Halobacteria>Halobacteriales</td><td>GCF_030250225.1</td><td>Halobacterium_salinarum</td><td>PNP03_RS10250</td><td>14588..17089</td><td>2242</td><td>hypothetical-protein-[Halobacterium-salinarum]</td></tr>
<tr><td>WP_229109461.1</td><td>AAA+ATPase+wHTH+Alpha+Beta-domain+wHTH+RRM</td><td>Type-3_BR.PglZ.associated.systems</td><td><-iSTAND<-METHYLASE<-PglZ<-DUF499*<-?||?-><-CHTH<-OrfB_Zn_ribbon</td><td>841</td><td>Archaea>Euryarchaeota>Stenosarchaea_group>Halobacteria>Halobacteriales</td><td>GCF_017094525.1</td><td>Halapricum_desulfuricans</td><td>HSR122_RS09455</td><td>1830239..1832764</td><td>2841257</td><td>DUF499-domain-containing-protein-[Halapricum-desulfuricans]</td></tr>
<tr><td>WP_064287641.1</td><td>AAA+ATPase+wHTH+Alpha+Beta-domain+wHTH+RRM</td><td>Type-3_BR.PglZ.associated.systems</td><td><-iSTAND<-METHYLASE<-PglZ<-DUF499*||?-><-Phage_integrase</td><td>837</td><td>Archaea>Euryarchaeota>Stenosarchaea_group>Halobacteria>Halobacteriales</td><td>GCF_001647155.1</td><td>Haloarcula_sp._K1</td><td>AV929_RS05370</td><td>105139..107652</td><td>1622207</td><td>DUF499-domain-containing-protein-[Haloarcula-sp.-K1]</td></tr>
<tr><td>WP_121816683.1</td><td>AAA+ATPase+wHTH+Alpha+Beta-domain+wHTH+RRM</td><td>Type-3_BR.PglZ.associated.systems</td><td><-iSTAND<-METHYLASE<-PglZ<-DUF499*||?->?->?-><-Phage_integrase</td><td>836</td><td>Archaea>Euryarchaeota>Stenosarchaea_group>Halobacteria>Halobacteriales</td><td>GCF_011440285.1</td><td>Halostella</td><td>G9464_RS02095</td><td>431416..433926</td><td>1843185</td><td>MULTISPECIES:-DUF499-domain-containing-protein-[Halostella]</td></tr>
<tr><td>WP_135806761.1</td><td>AAA+ATPase+wHTH+Alpha+Beta-domain+wHTH+RRM</td><td>Type-3_BR.PglZ.associated.systems</td><td><-iSTAND<-?<-METHYLASE<-?||?-><-DUF499*||SF1-SF2-Helicase->?->SF1-SF2-Helicase-><-?<-?||MarR-HTH-></td><td>1068</td><td>Archaea>Euryarchaeota>Stenosarchaea_group>Halobacteria>Halobacteriales</td><td>GCF_004765795.1</td><td>Halorussus_marinus</td><td>EPY72_RS13530</td><td>232726..235932</td><td>2505976</td><td>DUF499-domain-containing-protein-[Halorussus-marinus]</td></tr>
<tr><td>WP_135823667.1</td><td>AAA+ATPase+wHTH+Alpha+Beta-domain+wHTH+RRM</td><td>Type-3_BR.PglZ.associated.systems</td><td><-HTH||McrB-AAA->McrC-NTD+REase-><-iSTAND<-METHYLASE<-DUF499*<-METHYLASE||SF1-SF2-Helicase->?->METHYLASE-><-McrC-NTD+REase<-EVE+SF1-SF2-Helicase+McrB</td><td>1071</td><td>Archaea>Euryarchaeota>Stenosarchaea_group>Halobacteria>Halobacteriales</td><td>GCF_004765785.1</td><td>Halorussus_ruber</td><td>EPL01_RS02355</td><td>27346..30561</td><td>1126238</td><td>DUF499-domain-containing-protein-[Halorussus-ruber]</td></tr>
<tr><td>WP_179918061.1</td><td>AAA+ATPase+wHTH+Alpha+Beta-domain+wHTH+RRM</td><td>Type-3_BR.PglZ.associated.systems</td><td><-HEPN<-?<-SF1-SF2-Helicase||DUF499*->METHYLASE->?->iSTAND->?-><-McrC-NTD+REase<-McrB-AAA</td><td>1068</td><td>Archaea>Euryarchaeota>Stenosarchaea_group>Halobacteria>Halobacteriales</td><td>GCF_013415905.1</td><td>Halosimplex_pelagicum</td><td>HZS54_RS15820</td><td>3167949..3171155</td><td>869886</td><td>DUF499-domain-containing-protein-[Halosimplex-pelagicum]</td></tr>
<tr><td>AHG02823.1</td><td>AAA+ATPase+wHTH+Alpha+Beta-domain+wHTH+RRM</td><td>Type-3_BR.PglZ.associated.systems</td><td><-DUF772+DDE_Tnp_1||RHH_6+RHH_6-><-?<-SF1-SF2-Helicase<-?||DUF499*-><-?||?->METHYLASE->?->CITB-HTH->iSTAND-></td><td>1068</td><td>Archaea>Euryarchaeota>Stenosarchaea_group>Halobacteria>Halobacteriales</td><td>GCA_000230955.3</td><td>Halobacterium_sp._DL1</td><td>HALDL1_03655</td><td>404623..407829</td><td>751944</td><td>ATPase-AAA-[Halobacterium-sp.-DL1]</td></tr>
<tr><td>AKH97510.1</td><td>AAA+ATPase+wHTH+Alpha+Beta-domain+wHTH+RRM</td><td>Type-3_BR.PglZ.associated.systems</td><td><-DUF772+DDE_Tnp_1||HTH_5->?->PAPS_reduct-><-ORC/CDC6+HTH||DUF499*->PglZ->METHYLASE->iSTAND->REase-></td><td>936</td><td>Archaea>Euryarchaeota>Stenosarchaea_group>Halobacteria>Halobacteriales</td><td>GCA_001011115.1</td><td>Halanaeroarchaeum_sulfurireducens</td><td>HLASF_1021</td><td>991096..993906</td><td>1604004</td><td>ATPase-AAA-[Halanaeroarchaeum-sulfurireducens]</td></tr>
<tr><td>ALG81906.1</td><td>AAA+ATPase+wHTH+Alpha+Beta-domain+wHTH+RRM</td><td>Type-3_BR.PglZ.associated.systems</td><td><-DUF772+DDE_Tnp_1||HTH_5->?->PAPS_reduct-><-ORC/CDC6+HTH||DUF499*->PglZ->METHYLASE->iSTAND->REase-></td><td>936</td><td>Archaea>Euryarchaeota>Stenosarchaea_group>Halobacteria>Halobacteriales</td><td>GCA_001305655.1</td><td>Halanaeroarchaeum_sulfurireducens</td><td>HLASA_1010</td><td>987895..990705</td><td>1604004</td><td>ATPase-AAA-[Halanaeroarchaeum-sulfurireducens]</td></tr>
<tr><td>WP_050048259.1</td><td>AAA+ATPase+wHTH+Alpha+Beta-domain+wHTH+RRM</td><td>Type-3_BR.PglZ.associated.systems</td><td><-DUF772+DDE_Tnp_1||HTH_5->?->?->PAPS_reduct-><-ORC/CDC6+HTH||DUF499*->PglZ->METHYLASE->iSTAND->REase-></td><td>930</td><td>Archaea>Euryarchaeota>Stenosarchaea_group>Halobacteria>Halobacteriales</td><td>GCF_001011115.1</td><td>Halanaeroarchaeum_sulfurireducens</td><td>HLASF_RS04940</td><td>991114..993906</td><td>1604004</td><td>DUF499-domain-containing-protein-[Halanaeroarchaeum-sulfurireducens]</td></tr>
<tr><td>WP_054519645.1</td><td>AAA+ATPase+wHTH+Alpha+Beta-domain+wHTH+RRM</td><td>Type-3_BR.PglZ.associated.systems</td><td><-DUF772+DDE_Tnp_1||HTH_5->?->?->PAPS_reduct-><-ORC/CDC6+HTH||DUF499*->PglZ->METHYLASE->iSTAND->REase-></td><td>930</td><td>Archaea>Euryarchaeota>Stenosarchaea_group>Halobacteria>Halobacteriales</td><td>GCF_001305655.1</td><td>Halanaeroarchaeum_sulfurireducens</td><td>HLASA_RS04935</td><td>987913..990705</td><td>1604004</td><td>DUF499-domain-containing-protein-[Halanaeroarchaeum-sulfurireducens]</td></tr>
<tr><td>WP_230889378.1</td><td>AAA+ATPase+wHTH+Alpha+Beta-domain+wHTH+RRM</td><td>Type-3_BR.PglZ.associated.systems</td><td><-Calcineurin<-Phage_integrase<-?||REase->SF1-SF2-Helicase->DUF499*->PglZ->METHYLASE->iSTAND-><-PadR-HTH<-DndE<-SbcC</td><td>929</td><td>Archaea>Euryarchaeota>Stenosarchaea_group>Halobacteria>Halobacteriales</td><td>GCF_021028995.1</td><td>Halobacterium_sp._KA-4</td><td>LPA44_RS09645</td><td>32260..35049</td><td>2896367</td><td>DUF499-domain-containing-protein-[Halobacterium-sp.-KA-4]</td></tr>
<tr><td>WP_137275743.1</td><td>AAA+ATPase+wHTH+Alpha+Beta-domain+wHTH+RRM</td><td>Type-3_BR.PglZ.associated.systems</td><td><-BirA-HTH||ZnR->SF1-SF2-Helicase-><-?<-METHYLASE<-PglZ<-DUF499*||?->?->?->?->?-><-Phage_integrase</td><td>942</td><td>Archaea>Euryarchaeota>Stenosarchaea_group>Halobacteria>Halobacteriales</td><td>GCF_005239135.1</td><td>Natronomonas_salsuginis</td><td>DM868_RS04875</td><td>29184..32012</td><td>2217661</td><td>hypothetical-protein-[Natronomonas-salsuginis]</td></tr>
<tr><td>WP_310901786.1</td><td>AAA+ATPase+wHTH+Alpha+Beta-domain+wHTH+RRM</td><td>Type-3_BR.PglZ.associated.systems</td><td><-35exo||DUF499+RRM*->PglZ->METHYLASE-></td><td>930</td><td>Archaea>Euryarchaeota>Stenosarchaea_group>Halobacteria>Halobacteriales>Haloarculaceae</td><td>GCF_031624605.1</td><td>Halomicroarcula_onubensis</td><td>NDI86_RS18205</td><td>39949..42741</td><td>2950539</td><td>DUF499-domain-containing-protein-[Halomicroarcula-sp.-S3CR25-11]</td></tr>
<tr><td>WWX76766.1</td><td>AAA+ATPase+wHTH+Alpha+Beta-domain+wHTH+RRM</td><td>Type-3_BR.PglZ.associated.systems</td><td><-GDResolvase||DUF499+RRM*->PglZ->METHYLASE->||SF1-SF2-Helicase+REase-></td><td>942</td><td>Archaea>Euryarchaeota>Stenosarchaea_group>Halobacteria>Halobacteriales>Halobacteriaceae</td><td>GCA_037094595.1</td><td>Halobacterium_salinarum</td><td>V9T20_12280</td><td>7348..10176</td><td>2242</td><td>DUF499-domain-containing-protein-[Halobacterium-salinarum]</td></tr>
<tr><td>WP_193493617.1</td><td>AAA+ATPase+wHTH+Alpha+Beta-domain+wHTH+RRM</td><td>Type-3_BR.PglZ.associated.systems</td><td>ZnR+REase-><-iSTAND<-METHYLASE+CBS+HEPN<-METHYLASE||ParB+HNH-p->ParB+HNH-><-DUF499*||SF1-SF2-Helicase->?-><-?<-ORC/CDC6+HTH||?-><-ZnR</td><td>1069</td><td>Archaea>Euryarchaeota>Stenosarchaea_group>Halobacteria>Haloferacales</td><td>GCF_014969745.1</td><td>Haloferax_gibbonsii</td><td>HfgLR_RS17320</td><td>401617..404826</td><td>35746</td><td>DUF499-domain-containing-protein-[Haloferax-gibbonsii]</td></tr>
<tr><td>WP_257627366.1</td><td>AAA+ATPase+wHTH+Alpha+Beta-domain+wHTH+RRM</td><td>Type-3_BR.PglZ.associated.systems</td><td>SF1-SF2-Helicase->DUF499*->PglZ->METHYLASE-><-?<-?||DDE_Tnp_IS240-></td><td>949</td><td>Archaea>Euryarchaeota>Stenosarchaea_group>Halobacteria>Haloferacales</td><td>GCF_024498175.1</td><td>Haloplanus_salinarum</td><td>NO364_RS09470</td><td>1827336..1830185</td><td>1912324</td><td>hypothetical-protein-[Haloplanus-salinarum]</td></tr>
<tr><td>WP_123622887.1</td><td>AAA+ATPase+wHTH+Alpha+Beta-domain+wHTH+RRM</td><td>Type-3_BR.PglZ.associated.systems</td><td>REase->SF1-SF2-Helicase+REase-><-?<-SF1-SF2-Helicase||DUF499*->MTaX1+McrB->REase->METHYLASE->ParB+HNH->iSTAND-><-ZnR+REase</td><td>1071</td><td>Archaea>Euryarchaeota>Stenosarchaea_group>Halobacteria>Haloferacales</td><td>GCF_003781945.1</td><td>Halorubrum_sp._CSM-61</td><td>EDM04_RS13285</td><td>102150..105365</td><td>2485838</td><td>DUF499-domain-containing-protein-[Halorubrum-sp.-CSM-61]</td></tr>
<tr><td>WP_170830671.1</td><td>AAA+ATPase+wHTH+Alpha+Beta-domain+wHTH+RRM</td><td>Type-3_BR.PglZ.associated.systems</td><td>REase->SF1-SF2-Helicase+REase-><-?||METHYLASE->DUF499*->METHYLASE->iSTAND-><-McrC-NTD+REase<-McrB-AAA||ORC/CDC6+REase->SF1-SF2-Helicase-></td><td>1077</td><td>Archaea>Euryarchaeota>Stenosarchaea_group>Halobacteria>Haloferacales</td><td>GCF_900103715.1</td><td>Halogranum_gelatinilyticum</td><td>BLR57_RS15895</td><td>52884..56117</td><td>660521</td><td>DUF499-domain-containing-protein-[Halogranum-gelatinilyticum]</td></tr>
<tr><td>WP_241897866.1</td><td>AAA+ATPase+wHTH+Alpha+Beta-domain+wHTH+RRM</td><td>Type-3_BR.PglZ.associated.systems</td><td>REase-><-TIR+PNTD2<-?<-SF1-SF2-Helicase<-?||DUF499*-><-?||METHYLASE->HNH->?->iSTAND-><-McrC-NTD+REase</td><td>1038</td><td>Archaea>Euryarchaeota>Stenosarchaea_group>Halobacteria>Haloferacales</td><td>-</td><td>Halorubrum_sp._C191</td><td>Z052_RS00925</td><td>21444..24560</td><td>1383842</td><td>DUF499-domain-containing-protein-[Halorubrum-sp.-C191]</td></tr>
<tr><td>PHQ44026.1</td><td>AAA+ATPase+wHTH+Alpha+Beta-domain+wHTH+RRM</td><td>Type-3_BR.PglZ.associated.systems</td><td>REase-><-TIR+PNTD2<-?<-SF1-SF2-Helicase||?->DUF499*-><-?||METHYLASE->HNH->?->iSTAND-><-McrC-NTD+REase</td><td>1068</td><td>Archaea>Euryarchaeota>Stenosarchaea_group>Halobacteria>Haloferacales</td><td>GCA_002727095.1</td><td>Halorubrum_sp._C191</td><td>Z052_00910</td><td>21354..24560</td><td>1383842</td><td>DUF499-domain-containing-protein-[Halorubrum-sp.-C191]</td></tr>
<tr><td>WP_089699572.1</td><td>AAA+ATPase+wHTH+Alpha+Beta-domain+wHTH+RRM</td><td>Type-3_BR.PglZ.associated.systems</td><td>REase-><-PIN||?-><-?<-METHYLASE||REase->DUF499*->iSTAND-><-McrC-NTD+REase<-McrB-AAA</td><td>1086</td><td>Archaea>Euryarchaeota>Stenosarchaea_group>Halobacteria>Haloferacales</td><td>GCF_900103715.1</td><td>Halogranum_gelatinilyticum</td><td>BLR57_RS16980</td><td>80220..83480</td><td>660521</td><td>DUF499-domain-containing-protein-[Halogranum-gelatinilyticum]</td></tr>
<tr><td>WP_012660078.1</td><td>AAA+ATPase+wHTH+Alpha+Beta-domain+wHTH+RRM</td><td>Type-3_BR.PglZ.associated.systems</td><td>PIWI->DUF499*->PglZ->METHYLASE->iSTAND->MethyltransfD12->||DUF772+DDE_Tnp_1-><-MTaX1+McrB</td><td>834</td><td>Archaea>Euryarchaeota>Stenosarchaea_group>Halobacteria>Haloferacales</td><td>GCF_000022205.1</td><td>Halorubrum_lacusprofundi</td><td>HLAC_RS16270</td><td>97421..99925</td><td>2247</td><td>DUF499-domain-containing-protein-[Halorubrum-lacusprofundi]</td></tr>
<tr><td>WP_245957329.1</td><td>AAA+ATPase+wHTH+Alpha+Beta-domain+wHTH+RRM</td><td>Type-3_BR.PglZ.associated.systems</td><td>PIWI-><-Peptidase_M48<-?<-SF1-SF2-Helicase+REase<-Exonuc_V_gamma+REase||DUF499*->PglZ->METHYLASE->iSTAND->MethyltransfD12-><-McrC-NTD+REase<-MTaX1+McrB</td><td>834</td><td>Archaea>Euryarchaeota>Stenosarchaea_group>Halobacteria>Haloferacales</td><td>GCF_003342695.1</td><td>Haloplanus_rubicundus</td><td>DU500_RS17090</td><td>20722..23226</td><td>1547898</td><td>DUF499-domain-containing-protein-[Haloplanus-rubicundus]</td></tr>
<tr><td>AXG08207.1</td><td>AAA+ATPase+wHTH+Alpha+Beta-domain+wHTH+RRM</td><td>Type-3_BR.PglZ.associated.systems</td><td>PIWI-><-?<-?<-SF1-SF2-Helicase+REase<-Exonuc_V_gamma+REase||DUF499*->PglZ->METHYLASE->iSTAND->MethyltransfD12-><-McrC-NTD+REase<-MTaX1+McrB</td><td>791</td><td>Archaea>Euryarchaeota>Stenosarchaea_group>Halobacteria>Haloferacales</td><td>GCA_003342695.1</td><td>Haloplanus_rubicundus</td><td>DU500_17090</td><td>20851..23226</td><td>1547898</td><td>DUF499-domain-containing-protein-[Haloplanus-rubicundus]</td></tr>
<tr><td>WP_271967209.1</td><td>AAA+ATPase+wHTH+Alpha+Beta-domain+wHTH+RRM</td><td>Type-3_BR.PglZ.associated.systems</td><td>PIN-><-?<-SF1-SF2-Helicase+REase<-Exonuc_V_gamma+REase<-?||METHYLASE->DUF499*->METHYLASE->iSTAND-><-McrC-NTD+REase<-McrB-AAA||ORC/CDC6+REase->METHYLASE-></td><td>1071</td><td>Archaea>Euryarchaeota>Stenosarchaea_group>Halobacteria>Haloferacales</td><td>GCF_028211085.1</td><td>Halorubrum_ezzemoulense</td><td>PM032_RS14240</td><td>26916..30131</td><td>337243</td><td>DUF499-domain-containing-protein-[Halorubrum-ezzemoulense]</td></tr>
<tr><td>WP_245902853.1</td><td>AAA+ATPase+wHTH+Alpha+Beta-domain+wHTH+RRM</td><td>Type-3_BR.PglZ.associated.systems</td><td>Phage_integrase-><-iSTAND||HKD-DNase-><-METHYLASE||HKD-DNase-><-PglZ<-DUF499*<-DUF499<-?<-tRNA||<-?<-sGTP</td><td>549</td><td>Archaea>Euryarchaeota>Stenosarchaea_group>Halobacteria>Haloferacales</td><td>GCF_002906575.1</td><td>Salinigranum_rubrum</td><td>C2R22_RS01985</td><td>409942..411591</td><td>755307</td><td>hypothetical-protein-[Salinigranum-rubrum]</td></tr>
<tr><td>WP_185194288.1</td><td>AAA+ATPase+wHTH+Alpha+Beta-domain+wHTH+RRM</td><td>Type-3_BR.PglZ.associated.systems</td><td>Phage_integrase->?-><-Phage_integrase+ZnR||McrB-AAA->McrC-NTD+REase-><-iSTAND<-DUF499*||HEPN->?->METHYLASE-><-?||SF1-SF2-Helicase-></td><td>1086</td><td>Archaea>Euryarchaeota>Stenosarchaea_group>Halobacteria>Haloferacales</td><td>GCF_014212355.1</td><td>Halobellus_ruber</td><td>H5V44_RS16775</td><td>47153..50413</td><td>2761102</td><td>DUF499-domain-containing-protein-[Halobellus-ruber]</td></tr>
<tr><td>WP_121597923.1</td><td>AAA+ATPase+wHTH+Alpha+Beta-domain+wHTH+RRM</td><td>Type-3_BR.PglZ.associated.systems</td><td>Phage_integrase->?-><-?||DUF499*->PglZ->METHYLASE->?-><-?<-BirA-HTH</td><td>946</td><td>Archaea>Euryarchaeota>Stenosarchaea_group>Halobacteria>Haloferacales</td><td>GCF_003666015.1</td><td>Halorubrum_sp._Atlit-26R</td><td>DVK07_RS15665</td><td>60497..63337</td><td>2282128</td><td>hypothetical-protein-[Halorubrum-sp.-Atlit-26R]</td></tr>
<tr><td>WP_234298121.1</td><td>AAA+ATPase+wHTH+Alpha+Beta-domain+wHTH+RRM</td><td>Type-3_BR.PglZ.associated.systems</td><td>Phage_integrase+ZnR->?-><-Phage_integrase+ZnR||McrB-AAA->McrC-NTD+REase-><-iSTAND<-DUF499*||HEPN->?->METHYLASE->Schlafen->SF1-SF2-Helicase-></td><td>1086</td><td>Archaea>Euryarchaeota>Stenosarchaea_group>Halobacteria>Haloferacales</td><td>GCF_021431925.1</td><td>Halobaculum_sp._CBA1158</td><td>Hbl1158_RS00445</td><td>95193..98453</td><td>2904243</td><td>DUF499-domain-containing-protein-[Halobaculum-sp.-CBA1158]</td></tr>
<tr><td>WP_159483888.1</td><td>AAA+ATPase+wHTH+Alpha+Beta-domain+wHTH+RRM</td><td>Type-3_BR.PglZ.associated.systems</td><td>Phage_int_SAM_4+Phage_integrase->?-><-?||?->?->DUF499*->PglZ->METHYLASE->iSTAND-><-Resolvase</td><td>930</td><td>Archaea>Euryarchaeota>Stenosarchaea_group>Halobacteria>Haloferacales</td><td>GCF_009806985.1</td><td>Halorubrum_sp._JWXQ-INN_858</td><td>GRS48_RS04310</td><td>197977..200769</td><td>2690782</td><td>DUF499-domain-containing-protein-[Halorubrum-sp.-JWXQ-INN-858]</td></tr>
<tr><td>WP_089768776.1</td><td>AAA+ATPase+wHTH+Alpha+Beta-domain+wHTH+RRM</td><td>Type-3_BR.PglZ.associated.systems</td><td>N6_MTase-><-?||Phage_integrase->?->?->?->DUF499*->PglZ->METHYLASE->iSTAND-><-?<-tRNA<-?<-sGTP</td><td>942</td><td>Archaea>Euryarchaeota>Stenosarchaea_group>Halobacteria>Haloferacales</td><td>GCF_900107195.1</td><td>Halobellus_clavatus</td><td>BLU74_RS14980</td><td>71837..74665</td><td>660517</td><td>hypothetical-protein-[Halobellus-clavatus]</td></tr>
<tr><td>ELZ84490.1</td><td>AAA+ATPase+wHTH+Alpha+Beta-domain+wHTH+RRM</td><td>Type-3_BR.PglZ.associated.systems</td><td>MTaX1+McrB->McrC-NTD+REase-><-MethyltransfD12<-METHYLASE<-PglZ<-DUF499*<-ORC/CDC6+REase||SF1-SF2-Helicase->REase->SF1-SF2-Helicase+REase-><-RNase</td><td>791</td><td>Archaea>Euryarchaeota>Stenosarchaea_group>Halobacteria>Haloferacales</td><td>GCA_000336955.1</td><td>Haloferax_larsenii_JCM_13917</td><td>C455_00117</td><td>28316..30691</td><td>1227460</td><td>AAA-family-ATPase-[Haloferax-larsenii-JCM-13917]</td></tr>
<tr><td>WP_049918509.1</td><td>AAA+ATPase+wHTH+Alpha+Beta-domain+wHTH+RRM</td><td>Type-3_BR.PglZ.associated.systems</td><td>MTaX1+McrB->McrC-NTD+REase-><-MethyltransfD12<-iSTAND<-METHYLASE<-PglZ<-DUF499*<-ORC/CDC6+REase||SF1-SF2-Helicase->REase->SF1-SF2-Helicase+REase-><-RNase</td><td>834</td><td>Archaea>Euryarchaeota>Stenosarchaea_group>Halobacteria>Haloferacales</td><td>GCF_000336955.1</td><td>Haloferax_larsenii</td><td>C455_RS00110</td><td>28316..30820</td><td>302484</td><td>DUF499-domain-containing-protein-[Haloferax-larsenii]</td></tr>
<tr><td>WP_240137805.1</td><td>AAA+ATPase+wHTH+Alpha+Beta-domain+wHTH+RRM</td><td>Type-3_BR.PglZ.associated.systems</td><td>MTaX1+McrB->McrC-NTD+REase-><-MethyltransfD12<-iSTAND<-METHYLASE<-PglZ<-DUF499*<-?<-?<-?<-ORC/CDC6+HTH||ORC/CDC6+HTH-></td><td>832</td><td>Archaea>Euryarchaeota>Stenosarchaea_group>Halobacteria>Haloferacales</td><td>GCF_009176545.1</td><td>Salinigranum_salinum</td><td>FZP69_RS21685</td><td>58691..61189</td><td>1364937</td><td>DUF499-domain-containing-protein-[Salinigranum-salinum]</td></tr>
<tr><td>WP_137692123.1</td><td>AAA+ATPase+wHTH+Alpha+Beta-domain+wHTH+RRM</td><td>Type-3_BR.PglZ.associated.systems</td><td>METHYLASE->DUF499*->METHYLASE->iSTAND-><-McrC-NTD+REase<-McrB-AAA<-METHYLASE||SF1-SF2-Helicase-></td><td>1076</td><td>Archaea>Euryarchaeota>Stenosarchaea_group>Halobacteria>Haloferacales</td><td>GCF_005435095.1</td><td>Halorubrum_sp._CGM5_25_10-8B</td><td>EXE51_RS05025</td><td>124677..127907</td><td>2518115</td><td>DUF499-domain-containing-protein-[Halorubrum-sp.-CGM5_25_10-8B]</td></tr>
<tr><td>WP_233509326.1</td><td>AAA+ATPase+wHTH+Alpha+Beta-domain+wHTH+RRM</td><td>Type-3_BR.PglZ.associated.systems</td><td>McrC-NTD+REase-><-iSTAND<-RNase||?-><-METHYLASE<-?<-DUF499*||SF1-SF2-Helicase->?-><-?<-?<-UbE1+JAB<-SMODS</td><td>1015</td><td>Archaea>Euryarchaeota>Stenosarchaea_group>Halobacteria>Haloferacales</td><td>GCF_003383015.1</td><td>Haloferax_sp._Atlit-48N</td><td>DEQ67_RS14625</td><td>37469..40516</td><td>2077198</td><td>DUF499-domain-containing-protein-[Haloferax-sp.-Atlit-48N]</td></tr>
<tr><td>AUX08315.1</td><td>AAA+ATPase+wHTH+Alpha+Beta-domain+wHTH+RRM</td><td>Type-3_BR.PglZ.associated.systems</td><td>McrC-NTD+REase-><-iSTAND<-DOC<-METHYLASE||REase-><-HTH+IclR-HTH<-DUF499*||?->Resolvase-><-?||?->RelE-ParE->ACYC-></td><td>1068</td><td>Archaea>Euryarchaeota>Stenosarchaea_group>Halobacteria>Haloferacales</td><td>GCA_002952775.1</td><td>Halalkaliarchaeum_desulfuricum</td><td>AArcSl_0665</td><td>701854..705060</td><td>2055893</td><td>ATPase-(AAA+-superfamily)-like-protein-[Halalkaliarchaeum-desulfuricum]</td></tr>
<tr><td>WP_122104307.1</td><td>AAA+ATPase+wHTH+Alpha+Beta-domain+wHTH+RRM</td><td>Type-3_BR.PglZ.associated.systems</td><td>McrC-NTD+REase-><-iSTAND<-?<-DOC<-METHYLASE<-?<-DUF499*||SF1-SF2-Helicase->?-><-HHH+ENDO3+NUDIX<-Aminotran_3<-PadR-HTH</td><td>1071</td><td>Archaea>Euryarchaeota>Stenosarchaea_group>Halobacteria>Haloferacales</td><td>GCF_003697845.1</td><td>Halobellus_captivus</td><td>EA078_RS02610</td><td>112806..116021</td><td>2592614</td><td>DUF499-domain-containing-protein-[Halobellus-captivus]</td></tr>
<tr><td>WP_271969326.1</td><td>AAA+ATPase+wHTH+Alpha+Beta-domain+wHTH+RRM</td><td>Type-3_BR.PglZ.associated.systems</td><td>McrB-AAA->McrC-NTD+REase->PIN+ACET+EVE->ASCH-><-iSTAND<-METHYLASE<-DUF499*||SF1-SF2-Helicase->?->SF1-SF2-Helicase-><-?||BirA-HTH-></td><td>1068</td><td>Archaea>Euryarchaeota>Stenosarchaea_group>Halobacteria>Haloferacales</td><td>GCF_028211405.1</td><td>Halorubrum_ezzemoulense</td><td>PM038_RS04335</td><td>798473..801679</td><td>337243</td><td>DUF499-domain-containing-protein-[Halorubrum-ezzemoulense]</td></tr>
<tr><td>WP_193308745.1</td><td>AAA+ATPase+wHTH+Alpha+Beta-domain+wHTH+RRM</td><td>Type-3_BR.PglZ.associated.systems</td><td>McrB-AAA->McrC-NTD+REase-><-iSTAND<-PIN<-METHYLASE<-MTaX1+HNH+ColE7<-DUF499*||SF1-SF2-Helicase-></td><td>1071</td><td>Archaea>Euryarchaeota>Stenosarchaea_group>Halobacteria>Haloferacales</td><td>GCF_008806895.1</td><td>Halorubrum_halophilum</td><td>HrrH1147_RS02390</td><td>471375..474590</td><td>413816</td><td>DUF499-domain-containing-protein-[Halorubrum-halophilum]</td></tr>
<tr><td>WP_017342169.1</td><td>AAA+ATPase+wHTH+Alpha+Beta-domain+wHTH+RRM</td><td>Type-3_BR.PglZ.associated.systems</td><td>McrB-AAA->McrC-NTD+REase-><-iSTAND<-METHYLASE||?-><-DUF499*||METHYLASE-><-REase<-REase||SF1-SF2-Helicase-></td><td>1086</td><td>Archaea>Euryarchaeota>Stenosarchaea_group>Halobacteria>Haloferacales</td><td>GCF_000296615.1</td><td>Halorubrum_sp._T3</td><td>B378_RS0102055</td><td>69263..72523</td><td>1194088</td><td>DUF499-domain-containing-protein-[Halorubrum-sp.-T3]</td></tr>
<tr><td>WP_115805855.1</td><td>AAA+ATPase+wHTH+Alpha+Beta-domain+wHTH+RRM</td><td>Type-3_BR.PglZ.associated.systems</td><td>McrB-AAA->McrC-NTD+REase-><-iSTAND<-DUF499*<-?<-?<-?<-METHYLASE||?->REase+HTH-></td><td>1073</td><td>Archaea>Euryarchaeota>Stenosarchaea_group>Halobacteria>Haloferacales</td><td>GCF_003383025.1</td><td>Haloferax_sp._Atlit-4N</td><td>C5C07_RS19165</td><td>27711..30932</td><td>2077206</td><td>DUF499-domain-containing-protein-[Haloferax-sp.-Atlit-4N]</td></tr>
<tr><td>ELZ41075.1</td><td>AAA+ATPase+wHTH+Alpha+Beta-domain+wHTH+RRM</td><td>Type-3_BR.PglZ.associated.systems</td><td>McrB-AAA->McrC-NTD+REase-><-iSTAND<-?<-METHYLASE<-?<-DUF499*||Phage_integrase->?->SF1-SF2-Helicase->?->?-><-CITB-HTH</td><td>1033</td><td>Archaea>Euryarchaeota>Stenosarchaea_group>Halobacteria>Haloferacales</td><td>GCA_000337915.1</td><td>Halorubrum_saccharovorum_DSM_1137</td><td>C471_06248</td><td>15260..18361</td><td>1227484</td><td>ATPase-[Halorubrum-saccharovorum-DSM-1137]</td></tr>
<tr><td>WP_142980433.1</td><td>AAA+ATPase+wHTH+Alpha+Beta-domain+wHTH+RRM</td><td>Type-3_BR.PglZ.associated.systems</td><td>McrB-AAA->McrC-NTD+REase-><-iSTAND||?->SbcC+SbcC-><-METHYLASE<-DUF499*||SF1-SF2-Helicase->?-><-SF1-SF2-Helicase+REase<-REase<-SbcC+SbcC</td><td>1071</td><td>Archaea>Euryarchaeota>Stenosarchaea_group>Halobacteria>Haloferacales</td><td>GCF_006861665.1</td><td>Halonotius_terrestris</td><td>EGH24_RS12285</td><td>61033..64248</td><td>2487750</td><td>DUF499-domain-containing-protein-[Halonotius-terrestris]</td></tr>
<tr><td>WP_120083768.1</td><td>AAA+ATPase+wHTH+Alpha+Beta-domain+wHTH+RRM</td><td>Type-3_BR.PglZ.associated.systems</td><td>McrB-AAA->McrC-NTD+REase->?-><-iSTAND<-METHYLASE<-DUF499*||SF1-SF2-Helicase->?->?->?-><-?<-HNH</td><td>1071</td><td>Archaea>Euryarchaeota>Stenosarchaea_group>Halobacteria>Haloferacales</td><td>GCF_003605635.1</td><td>Halonotius_pteroides</td><td>DP106_RS04765</td><td>53600..56815</td><td>268735</td><td>DUF499-domain-containing-protein-[Halonotius-pteroides]</td></tr>
<tr><td>WP_137703666.1</td><td>AAA+ATPase+wHTH+Alpha+Beta-domain+wHTH+RRM</td><td>Type-3_BR.PglZ.associated.systems</td><td>HTH_OrfB_IS605+OrfB_IS605+OrfB_Zn_ribbon-><-?||<-DDE_Tnp_1||?->REase-><-?||DUF499*->METHYLASE->iSTAND-><-McrC-NTD+REase<-McrB-AAA<-METHYLASE||SF1-SF2-Helicase-></td><td>1074</td><td>Archaea>Euryarchaeota>Stenosarchaea_group>Halobacteria>Haloferacales</td><td>GCF_005435025.1</td><td>Halorubrum_sp._CGM4_25_10-8A</td><td>EXE52_RS08795</td><td>123511..126735</td><td>2518116</td><td>DUF499-domain-containing-protein-[Halorubrum-sp.-CGM4_25_10-8A]</td></tr>
<tr><td>PAU80010.1</td><td>AAA+ATPase+wHTH+Alpha+Beta-domain+wHTH+RRM</td><td>Type-3_BR.PglZ.associated.systems</td><td>DUF499*->PglZ->METHYLASE->iSTAND->?->||?->REase-></td><td>788</td><td>Archaea>Euryarchaeota>Stenosarchaea_group>Halobacteria>Haloferacales</td><td>GCA_002286985.1</td><td>Halorubrum_salipaludis</td><td>CK500_15950</td><td>32163..34529</td><td>2032630</td><td>DUF499-domain-containing-protein-[Halorubrum-salipaludis]</td></tr>
<tr><td>WP_233139255.1</td><td>AAA+ATPase+wHTH+Alpha+Beta-domain+wHTH+RRM</td><td>Type-3_BR.PglZ.associated.systems</td><td>DUF499*->PglZ->METHYLASE->iSTAND->?->||?->REase-></td><td>832</td><td>Archaea>Euryarchaeota>Stenosarchaea_group>Halobacteria>Haloferacales</td><td>GCF_002286985.1</td><td>Halorubrum_salipaludis</td><td>CK500_RS15970</td><td>32031..34529</td><td>2032630</td><td>DUF499-domain-containing-protein-[Halorubrum-salipaludis]</td></tr>
<tr><td>WP_138007004.1</td><td>AAA+ATPase+wHTH+Alpha+Beta-domain+wHTH+RRM</td><td>Type-3_BR.PglZ.associated.systems</td><td>BirA-HTH-><-?<-SF1-SF2-Helicase||?-><-Phage_integrase||tRNA->DUF499*->METHYLASE->iSTAND-><-McrC-NTD+REase<-McrB-AAA||REase-></td><td>1068</td><td>Archaea>Euryarchaeota>Stenosarchaea_group>Halobacteria>Haloferacales</td><td>GCF_005543295.1</td><td>Halalkalirubrum_salinum</td><td>E5K03_RS11590</td><td>23030..26236</td><td>2563889</td><td>DUF499-domain-containing-protein-[Halalkalirubrum-salinum]</td></tr>
<tr><td>WP_123619536.1</td><td>AAA+ATPase+wHTH+Alpha+Beta-domain+wHTH+RRM</td><td>Type-3_BR.PglZ.associated.systems</td><td>BirA-HTH-><-?<-?<-METHYLASE<-PglZ<-DUF499*<-?||?-><-ZnR+REase||?-><-?<-Schlafen+HATPase+HTH</td><td>934</td><td>Archaea>Euryarchaeota>Stenosarchaea_group>Halobacteria>Haloferacales</td><td>GCF_003781945.1</td><td>Halorubrum_sp._CSM-61</td><td>EDM04_RS02225</td><td>428083..430887</td><td>2485838</td><td>DUF499-domain-containing-protein-[Halorubrum-sp.-CSM-61]</td></tr>
<tr><td>TKX86475.1</td><td>AAA+ATPase+wHTH+Alpha+Beta-domain+wHTH+RRM</td><td>Type-3_BR.PglZ.associated.systems</td><td>BirA-HTH->?->?-><-?||?-><-Phage_integrase||DUF499*->PglZ->METHYLASE-></td><td>946</td><td>Archaea>Euryarchaeota>Stenosarchaea_group>Halobacteria>Haloferacales</td><td>GCA_005435045.1</td><td>Halorubrum_sp._SS5</td><td>EXE43_08255</td><td>9934..12774</td><td>2518118</td><td>hypothetical-protein-EXE43_08255-[Halorubrum-sp.-SS5]</td></tr>
<tr><td>WP_156588024.1</td><td>AAA+ATPase+wHTH+Alpha+Beta-domain+wHTH+RRM</td><td>Type-3_BR.PglZ.associated.systems</td><td><-SF1-SF2-Helicase<-SF1-SF2-Helicase||DUF499*->MTaX1+HNH-><-MethyltransfD12||METHYLASE->iSTAND-></td><td>1068</td><td>Archaea>Euryarchaeota>Stenosarchaea_group>Halobacteria>Haloferacales</td><td>GCF_009742825.1</td><td>Halorubrum_sp._CBA1125</td><td>GJ633_RS06035</td><td>9367..12573</td><td>2668072</td><td>DUF499-domain-containing-protein-[Halorubrum-sp.-CBA1125]</td></tr>
<tr><td>WP_007344265.1</td><td>AAA+ATPase+wHTH+Alpha+Beta-domain+wHTH+RRM</td><td>Type-3_BR.PglZ.associated.systems</td><td><-SF1-SF2-Helicase<-METHYLASE||McrB-AAA->McrC-NTD+REase-><-iSTAND<-METHYLASE<-DUF499*||?-><-?<-DDE_Tnp_1</td><td>1076</td><td>Archaea>Euryarchaeota>Stenosarchaea_group>Halobacteria>Haloferacales</td><td>GCF_000337435.1</td><td>Halorubrum_terrestre</td><td>C473_RS01625</td><td>14122..17352</td><td>267368</td><td>DUF499-domain-containing-protein-[Halorubrum-terrestre]</td></tr>
<tr><td>ELZ46378.1</td><td>AAA+ATPase+wHTH+Alpha+Beta-domain+wHTH+RRM</td><td>Type-3_BR.PglZ.associated.systems</td><td><-SF1-SF2-Helicase<-HTH+ACET+HNH<-?||DUF499*->?-><-DUF772+DDE_Tnp_1<-?<-BirA-HTH||?->iSTAND-></td><td>937</td><td>Archaea>Euryarchaeota>Stenosarchaea_group>Halobacteria>Haloferacales</td><td>GCA_000336875.1</td><td>Halorubrum_californiense_DSM_19288</td><td>C463_04179</td><td>16388..19201</td><td>1227465</td><td>ATPase-[Halorubrum-californiensis-DSM-19288]</td></tr>
<tr><td>WP_271970501.1</td><td>AAA+ATPase+wHTH+Alpha+Beta-domain+wHTH+RRM</td><td>Type-3_BR.PglZ.associated.systems</td><td><-SF1-SF2-Helicase||METHYLASE->McrB-AAA->McrC-NTD+REase-><-iSTAND<-METHYLASE<-DUF499*||?->?-><-?<-?||REase-><-HicA_toxin</td><td>1074</td><td>Archaea>Euryarchaeota>Stenosarchaea_group>Halobacteria>Haloferacales</td><td>GCF_028210945.1</td><td>Halorubrum_ezzemoulense</td><td>PM085_RS15895</td><td>13658..16882</td><td>337243</td><td>DUF499-domain-containing-protein-[Halorubrum-ezzemoulense]</td></tr>
<tr><td>WP_251330422.1</td><td>AAA+ATPase+wHTH+Alpha+Beta-domain+wHTH+RRM</td><td>Type-3_BR.PglZ.associated.systems</td><td><-REase<-?||?-><-SF1-SF2-Helicase<-?||DUF499*->PglZ->METHYLASE->iSTAND->REase-><-?<-Phage_integrase</td><td>832</td><td>Archaea>Euryarchaeota>Stenosarchaea_group>Halobacteria>Haloferacales</td><td>GCF_023703795.1</td><td>Haloplanus_sp._HW8-1</td><td>NBT82_RS04725</td><td>877082..879580</td><td>2949995</td><td>DUF499-domain-containing-protein-[Haloplanus-sp.-HW8-1]</td></tr>
<tr><td>WP_008322212.1</td><td>AAA+ATPase+wHTH+Alpha+Beta-domain+wHTH+RRM</td><td>Type-3_BR.PglZ.associated.systems</td><td><-PIN<-?||REase+35exo+Beta-Barrel+SF1-SF2-Helicase-><-?<-?<-SF1-SF2-Helicase||DUF499*->?->METHYLASE-><-?||iSTAND->?->ZetaToxin-></td><td>1069</td><td>Archaea>Euryarchaeota>Stenosarchaea_group>Halobacteria>Haloferacales</td><td>GCF_000336755.1</td><td>Haloferax_elongans</td><td>C453_RS01405</td><td>21218..24427</td><td>403191</td><td>DUF499-domain-containing-protein-[Haloferax-elongans]</td></tr>
<tr><td>WP_157687718.1</td><td>AAA+ATPase+wHTH+Alpha+Beta-domain+wHTH+RRM</td><td>Type-3_BR.PglZ.associated.systems</td><td><-ORC/CDC6+HTH<-?<-?<-?<-?<-ORC/CDC6+HTH||DUF499*->PglZ->METHYLASE->iSTAND->MethyltransfD12-><-REase<-MTaX1+McrB</td><td>834</td><td>Archaea>Euryarchaeota>Stenosarchaea_group>Halobacteria>Haloferacales</td><td>GCF_009762275.1</td><td>Haloplanus_rallus</td><td>EI982_RS00925</td><td>188623..191127</td><td>1816183</td><td>DUF499-domain-containing-protein-[Haloplanus-rallus]</td></tr>
<tr><td>WP_159369583.1</td><td>AAA+ATPase+wHTH+Alpha+Beta-domain+wHTH+RRM</td><td>Type-3_BR.PglZ.associated.systems</td><td><-N6_N4_MTase<-MORC<-HEPN||DUF499*->PglZ->METHYLASE->?-><-BirA-HTH||Glyoxalase-></td><td>944</td><td>Archaea>Euryarchaeota>Stenosarchaea_group>Halobacteria>Haloferacales</td><td>GCF_009856205.1</td><td>Halorubrum_terrestre</td><td>GLW36_RS15285</td><td>11268..14102</td><td>267368</td><td>hypothetical-protein-[Halorubrum-terrestre]</td></tr>
<tr><td>WP_114586574.1</td><td>AAA+ATPase+wHTH+Alpha+Beta-domain+wHTH+RRM</td><td>Type-3_BR.PglZ.associated.systems</td><td><-METHYLASE<-PglZ<-DUF499*||?-><-?<-?<-RNase</td><td>949</td><td>Archaea>Euryarchaeota>Stenosarchaea_group>Halobacteria>Haloferacales</td><td>GCF_003342695.1</td><td>Haloplanus_rubicundus</td><td>DU500_RS14000</td><td>2749829..2752678</td><td>1547898</td><td>hypothetical-protein-[Haloplanus-rubicundus]</td></tr>
<tr><td>WP_271946307.1</td><td>AAA+ATPase+wHTH+Alpha+Beta-domain+wHTH+RRM</td><td>Type-3_BR.PglZ.associated.systems</td><td><-METHYLASE<-ORC/CDC6+REase||McrB-AAA->McrC-NTD+REase-><-iSTAND<-METHYLASE<-DUF499*<-METHYLASE||?->Exonuc_V_gamma+REase->SF1-SF2-Helicase+REase->?-><-SbcC</td><td>1071</td><td>Archaea>Euryarchaeota>Stenosarchaea_group>Halobacteria>Haloferacales</td><td>GCF_028211295.1</td><td>Halorubrum_ezzemoulense</td><td>PM026_RS15325</td><td>38110..41325</td><td>337243</td><td>DUF499-domain-containing-protein-[Halorubrum-ezzemoulense]</td></tr>
<tr><td>RDZ30299.1</td><td>AAA+ATPase+wHTH+Alpha+Beta-domain+wHTH+RRM</td><td>Type-3_BR.PglZ.associated.systems</td><td><-iSTAND<-RNase||?-><-METHYLASE<-REase<-?<-DUF499*||SF1-SF2-Helicase->?-><-?<-UbE1+JAB<-SMODS</td><td>1069</td><td>Archaea>Euryarchaeota>Stenosarchaea_group>Halobacteria>Haloferacales</td><td>GCA_003383015.1</td><td>Haloferax_sp._Atlit-48N</td><td>DEQ67_14625</td><td>37469..40678</td><td>2077198</td><td>DUF499-domain-containing-protein-[Haloferax-sp.-Atlit-48N]</td></tr>
<tr><td>WP_121919202.1</td><td>AAA+ATPase+wHTH+Alpha+Beta-domain+wHTH+RRM</td><td>Type-3_BR.PglZ.associated.systems</td><td><-iSTAND<-METHYLASE<-PglZ<-DUF499*||?->Primpol-><-NACHT||?-><-Phage_integrase</td><td>833</td><td>Archaea>Euryarchaeota>Stenosarchaea_group>Halobacteria>Haloferacales</td><td>GCF_003856835.1</td><td>Haloplanus_aerogenes</td><td>DU502_RS09875</td><td>1940385..1942886</td><td>660522</td><td>DUF499-domain-containing-protein-[Haloplanus-aerogenes]</td></tr>
<tr><td>WP_256417376.1</td><td>AAA+ATPase+wHTH+Alpha+Beta-domain+wHTH+RRM</td><td>Type-3_BR.PglZ.associated.systems</td><td><-iSTAND<-METHYLASE<-PglZ<-DUF499*||?->HEPN-></td><td>930</td><td>Archaea>Euryarchaeota>Stenosarchaea_group>Halobacteria>Haloferacales</td><td>GCF_024494655.1</td><td>Halorubrum_laminariae</td><td>NO887_RS05075</td><td>406335..409127</td><td>1433523</td><td>DUF499-domain-containing-protein-[Halorubrum-laminariae]</td></tr>
<tr><td>WP_092817109.1</td><td>AAA+ATPase+wHTH+Alpha+Beta-domain+wHTH+RRM</td><td>Type-3_BR.PglZ.associated.systems</td><td><-iSTAND<-METHYLASE||?-><-NTP_transf+wHTH<-HTH+IclR-HTH<-DUF499*||SF1-SF2-Helicase->?-><-?<-?||REase+DUF4268-><-HEPN</td><td>1068</td><td>Archaea>Euryarchaeota>Stenosarchaea_group>Halobacteria>Haloferacales</td><td>GCF_900108505.1</td><td>Halopenitus_malekzadehii</td><td>BLW38_RS09985</td><td>83792..86998</td><td>1267564</td><td>DUF499-domain-containing-protein-[Halopenitus-malekzadehii]</td></tr>
<tr><td>WP_049982438.1</td><td>AAA+ATPase+wHTH+Alpha+Beta-domain+wHTH+RRM</td><td>Type-3_BR.PglZ.associated.systems</td><td><-iSTAND<-?<-SbcC<-?<-?<-METHYLASE<-DUF499*||SF1-SF2-Helicase->?-><-?<-?||?-><-Phage_integrase</td><td>1069</td><td>Archaea>Euryarchaeota>Stenosarchaea_group>Halobacteria>Haloferacales</td><td>GCF_000746205.1</td><td>Halorubrum_sp._BV1</td><td>EP28_RS02610</td><td>519644..522853</td><td>1498500</td><td>DUF499-domain-containing-protein-[Halorubrum-sp.-BV1]</td></tr>
<tr><td>WP_049909041.1</td><td>AAA+ATPase+wHTH+Alpha+Beta-domain+wHTH+RRM</td><td>Type-3_BR.PglZ.associated.systems</td><td><-iSTAND<-?<-HNH<-METHYLASE<-REase<-MTaX1+McrB<-DUF499*||Phage_integrase->METHYLASE->SF1-SF2-Helicase-></td><td>1068</td><td>Archaea>Euryarchaeota>Stenosarchaea_group>Halobacteria>Haloferacales</td><td>GCF_000337915.1</td><td>Halorubrum_saccharovorum</td><td>C471_RS06270</td><td>15260..18466</td><td>2248</td><td>DUF499-domain-containing-protein-[Halorubrum-saccharovorum]</td></tr>
<tr><td>WP_245883345.1</td><td>AAA+ATPase+wHTH+Alpha+Beta-domain+wHTH+RRM</td><td>Type-3_BR.PglZ.associated.systems</td><td><-iSTAND<-?<-DOC<-METHYLASE||REase-><-HTH+IclR-HTH<-DUF499*<-?<-?||?->Resolvase->?->RelE-ParE-></td><td>1033</td><td>Archaea>Euryarchaeota>Stenosarchaea_group>Halobacteria>Haloferacales</td><td>-</td><td>Halalkaliarchaeum_desulfuricum</td><td>AArcSl_RS03355</td><td>701854..704955</td><td>2055893</td><td>DUF499-domain-containing-protein-[Halalkaliarchaeum-desulfuricum]</td></tr>
</table>

<table>
<thead>
<tr><th>Protein-ID</th><th>DUF499-architecture</th><th>System</th><th>Gene-Neighbourhoods</th><th>Length</th><th>Taxonomy</th><th>Assembly-ID</th><th>Strain</th><th>Locus-Tag</th><th>Genomic-coordinates</th><th>TaxID</th><th>Fasta-defline</th></tr>
</thead>
<tbody>
<tr><td>WP_245902854.1</td><td>AAA+ATPase+wHTH+Alpha+Beta-domain+wHTH+RRM</td><td>Type-3_BR.PglZ.associated.systems</td><td><-iSTAND||HKD-DNase-><-METHYLASE||HKD-DNase-><-PglZ<-DUF499<-DUF499*<-?<-tRNA||<-?<-sGTP</td><td>323</td><td>Archaea>Euryarchaeota>Stenosarchaea_group>Halobacteria>Haloferacales</td><td>GCF_002906575.1</td><td>Salinigranum_rubrum</td><td>C2R22_RS26070</td><td>411495..412466</td><td>755307</td><td>hypothetical-protein-[Salinigranum-rubrum]</td></tr>
<tr><td>WP_181861797.1</td><td>AAA+ATPase+wHTH+Alpha+Beta-domain+wHTH+RRM</td><td>Type-3_BR.PglZ.associated.systems</td><td><-DDE_Tnp_1<-ORC/CDC6+REase||McrB-AAA->McrC-NTD+REase-><-iSTAND<-METHYLASE<-DUF499*<-METHYLASE||?-><-McrC-NTD+REase<-?||MORC+DUF3684+REase-><-MarR-HTH</td><td>1078</td><td>Archaea>Euryarchaeota>Stenosarchaea_group>Halobacteria>Haloferacales</td><td>GCF_003336245.1</td><td>Haloplanus_salinus</td><td>DU504_RS17490</td><td>28986..32222</td><td>1126245</td><td>DUF499-domain-containing-protein-[Haloplanus-salinus]</td></tr>
<tr><td>RCU43688.1</td><td>AAA+ATPase+wHTH+Alpha+Beta-domain+wHTH+RRM</td><td>Type-3_BR.PglZ.associated.systems</td><td><-DDE_Tnp_1<-ORC/CDC6+REase||McrB-AAA->McrC-NTD+REase-><-iSTAND<-METHYLASE<-DUF499*||?-><-METHYLASE||?-><-McrC-NTD+REase</td><td>984</td><td>Archaea>Euryarchaeota>Stenosarchaea_group>Halobacteria>Haloferacales</td><td>GCA_003336245.1</td><td>Haloplanus_salinus</td><td>DU504_17495</td><td>28986..31940</td><td>1126245</td><td>DUF499-domain-containing-protein-[Haloplanus-salinus]</td></tr>
<tr><td>PHQ40460.1</td><td>AAA+ATPase+wHTH+Alpha+Beta-domain+wHTH+RRM</td><td>Type-3_BR.PglZ.associated.systems</td><td><-DDE_Tnp_1<-iSTAND<-METHYLASE<-?<-NTP_transf+wHTH<-HTH+IclR-HTH<-DUF499*<-?||SF1-SF2-Helicase-></td><td>1069</td><td>Archaea>Euryarchaeota>Stenosarchaea_group>Halobacteria>Haloferacales</td><td>GCA_002727125.1</td><td>Halorubrum_persicum</td><td>DJ69_01230</td><td>85507..88716</td><td>1383844</td><td>DUF499-domain-containing-protein-[Halorubrum-persicum]</td></tr>
<tr><td>WP_245851434.1</td><td>AAA+ATPase+wHTH+Alpha+Beta-domain+wHTH+RRM</td><td>Type-3_BR.PglZ.associated.systems</td><td><-DDE_Tnp_1<-iSTAND<-METHYLASE<-?<-NTP_transf+wHTH<-HTH+IclR-HTH<-DUF499*||SF1-SF2-Helicase-></td><td>1081</td><td>Archaea>Euryarchaeota>Stenosarchaea_group>Halobacteria>Haloferacales</td><td>-</td><td>Halorubrum_persicum</td><td>DJ69_RS01250</td><td>85507..88752</td><td>1383844</td><td>DUF499-domain-containing-protein-[Halorubrum-persicum]</td></tr>
<tr><td>WP_050034071.1</td><td>AAA+ATPase+wHTH+Alpha+Beta-domain+wHTH+RRM</td><td>Type-3_BR.PglZ.associated.systems</td><td><-Calcineurin<-?<-SF1-SF2-Helicase||DUF499*->METHYLASE->iSTAND->?-><-HEPN</td><td>1068</td><td>Archaea>Euryarchaeota>Stenosarchaea_group>Halobacteria>Haloferacales</td><td>GCF_000739595.1</td><td>Halorubrum_halophilum</td><td>ACP96_RS15860</td><td>8001..11207</td><td>413816</td><td>DUF499-domain-containing-protein-[Halorubrum-halophilum]</td></tr>
<tr><td>WP_121564026.1</td><td>AAA+ATPase+wHTH+Alpha+Beta-domain+wHTH+RRM</td><td>Type-3_BR.PglZ.associated.systems</td><td><-BirA-HTH<-iSTAND<-METHYLASE<-METHYLASE-p<-PglZ<-DUF499*||?->Phage_integrase-></td><td>936</td><td>Archaea>Euryarchaeota>Stenosarchaea_group>Halobacteria>Haloferacales</td><td>GCF_003666005.1</td><td>unclassified_Halorubrum</td><td>DVK08_RS15025</td><td>123925..126735</td><td>2642239</td><td>MULTISPECIES:-DUF499-domain-containing-protein-[unclassified-Halorubrum]</td></tr>
<tr><td>WP_272044658.1</td><td>AAA+ATPase+wHTH+Alpha+Beta-domain+wHTH+RRM</td><td>Type-3_BR.PglZ.associated.systems</td><td><-BirA-HTH||?-><-SF1-SF2-Helicase<-?<-SF1-SF2-Helicase||DUF499*->?->?->METHYLASE->iSTAND-><-ASCH<-PIN+ACET+EVE</td><td>1068</td><td>Archaea>Euryarchaeota>Stenosarchaea_group>Halobacteria>Haloferacales</td><td>GCF_028329875.1</td><td>Halorubrum_ezzemoulense</td><td>PN409_RS07550</td><td>110586..113792</td><td>337243</td><td>DUF499-domain-containing-protein-[Halorubrum-ezzemoulense]</td></tr>
<tr><td>WP_310922358.1</td><td>AAA+ATPase+wHTH+Alpha+Beta-domain+wHTH+RRM</td><td>Type-3_BR.PglZ.associated.systems</td><td>GDResolvase-><-METHYLASE<-PglZ<-DUF499+RRM*||<-PolB</td><td>935</td><td>Archaea>Euryarchaeota>Stenosarchaea_group>Halobacteria>Haloferacales>Haloferacaceae</td><td>GCF_031624495.1</td><td>Halogeometricum_sp._S1BR25-6</td><td>NDI76_RS02150</td><td>417326..420133</td><td>2950536</td><td>DUF499-domain-containing-protein-[Halogeometricum-sp.-S1BR25-6]</td></tr>
<tr><td>WP_310930530.1</td><td>AAA+ATPase+wHTH+Alpha+Beta-domain+wHTH+RRM</td><td>Type-3_BR.PglZ.associated.systems</td><td>PolB->||DUF499+RRM*->PglZ->METHYLASE-><-GDResolvase</td><td>937</td><td>Archaea>Euryarchaeota>Stenosarchaea_group>Halobacteria>Haloferacales>Haloferacaceae</td><td>GCF_031624635.1</td><td>Halogeometricum_sp._S3BR5-2</td><td>NDI79_RS20400</td><td>309465..312278</td><td>2950537</td><td>DUF499-domain-containing-protein-[Halogeometricum-sp.-S3BR5-2]</td></tr>
<tr><td>GAA0541242.1</td><td>AAA+ATPase+wHTH+Alpha+Beta-domain+wHTH+RRM</td><td>Type-3_BR.PglZ.associated.systems</td><td><-SF1-SF2-Helicase+REase||<-METHYLASE<-PglZ<-DUF499+RRM*||GDResolvase-></td><td>942</td><td>Archaea>Euryarchaeota>Stenosarchaea_group>Halobacteria>Haloferacales>Halorubraceae</td><td>GCA_039522565.1</td><td>Halorubrum_ejinorense</td><td>GCM10008994_15410</td><td>94881..97709</td><td>425309</td><td>hypothetical-protein-GCM10008994_15410-[Halorubrum-ejinorense]</td></tr>
<tr><td>WP_280585078.1</td><td>AAA+ATPase+wHTH+Alpha+Beta-domain+wHTH+RRM</td><td>Type-3_BR.PglZ.associated.systems</td><td><-METHYLASE<-PglZ<-DUF499+RRM*<-URI-DNase||<-Phage_integrase</td><td>827</td><td>Archaea>Euryarchaeota>Stenosarchaea_group>Halobacteria>Haloferacales>Halorubraceae</td><td>GCF_029889805.1</td><td>Halorubrum_sp._Boch-26</td><td>OQF71_RS00510</td><td>23624..26107</td><td>2994426</td><td>DUF499-domain-containing-protein-[Halorubrum-sp.-Boch-26]</td></tr>
<tr><td>WP_207290710.1</td><td>AAA+ATPase+wHTH+Alpha+Beta-domain+wHTH+RRM</td><td>Type-3_BR.PglZ.associated.systems</td><td>REase->?->?->tRNA-><-?<-SF1-SF2-Helicase||DUF499*-><-?||?-><-ParB+HNH<-?||METHYLASE->iSTAND-></td><td>1064</td><td>Archaea>Euryarchaeota>Stenosarchaea_group>Halobacteria>Natrialbales</td><td>GCF_017352155.2</td><td>Haloterrigena_alkaliphila</td><td>JOX25_RS27460</td><td>1775468..1778662</td><td>2816475</td><td>DUF499-domain-containing-protein-[Haloterrigena-alkaliphila]</td></tr>
<tr><td>WP_006065907.1</td><td>AAA+ATPase+wHTH+Alpha+Beta-domain+wHTH+RRM</td><td>Type-3_BR.PglZ.associated.systems</td><td>METHYLASE-><-ORC/CDC6+REase||McrB-AAA->McrC-NTD+REase-><-iSTAND<-METHYLASE<-DUF499*||?->?-><-METHYLASE-p||zf-IS66+DDE_Tnp_IS66-><-MORC+DUF3684+REase</td><td>1077</td><td>Archaea>Euryarchaeota>Stenosarchaea_group>Halobacteria>Natrialbales</td><td>GCF_000337715.1</td><td>Natronorubrum_bangense</td><td>DV706_RS20705</td><td>103313..106546</td><td>61858</td><td>DUF499-domain-containing-protein-[Natronorubrum-bangense]</td></tr>
<tr><td>QCC57013.1</td><td>AAA+ATPase+wHTH+Alpha+Beta-domain+wHTH+RRM</td><td>Type-3_BR.PglZ.associated.systems</td><td>METHYLASE-><-ORC/CDC6+REase||McrB-AAA->McrC-NTD+REase-><-iSTAND<-METHYLASE<-DUF499*||?->?->?->||zf-IS66+DDE_Tnp_IS66-><-MORC+DUF3684+REase</td><td>983</td><td>Archaea>Euryarchaeota>Stenosarchaea_group>Halobacteria>Natrialbales</td><td>GCA_004799645.1</td><td>Natronorubrum_bangense</td><td>DV706_20890</td><td>103313..106264</td><td>61858</td><td>DUF499-domain-containing-protein-[Natronorubrum-bangense]</td></tr>
<tr><td>WP_117362386.1</td><td>AAA+ATPase+wHTH+Alpha+Beta-domain+wHTH+RRM</td><td>Type-3_BR.PglZ.associated.systems</td><td>McrC-NTD+REase-><-MethyltransfD12<-iSTAND<-METHYLASE<-PglZ<-DUF499*||?-><-?<-?<-?||?->HEPN-></td><td>834</td><td>Archaea>Euryarchaeota>Stenosarchaea_group>Halobacteria>Natrialbales</td><td>GCF_003430825.1</td><td>Natrarchaeobaculum_sulfurireducens</td><td>AArc1_RS00285</td><td>75312..77816</td><td>2044521</td><td>DUF499-domain-containing-protein-[Natrarchaeobaculum-sulfurireducens]</td></tr>
<tr><td>ELY46666.1</td><td>AAA+ATPase+wHTH+Alpha+Beta-domain+wHTH+RRM</td><td>Type-3_BR.PglZ.associated.systems</td><td>McrB-AAA->McrC-NTD+REase-><-iSTAND<-SF1-SF2-Helicase<-METHYLASE||ParB+HNH-><-DUF499*||SF1-SF2-Helicase->?->?->RNase-></td><td>1068</td><td>Archaea>Euryarchaeota>Stenosarchaea_group>Halobacteria>Natrialbales</td><td>GCA_000337735.1</td><td>Natronorubrum_sulfidifaciens_JCM_14089</td><td>C495_06148</td><td>92654..95860</td><td>1230460</td><td>ATPase-[Natronorubrum-sulfidifaciens-JCM-14089]</td></tr>
<tr><td>WP_004213898.1</td><td>AAA+ATPase+wHTH+Alpha+Beta-domain+wHTH+RRM</td><td>Type-3_BR.PglZ.associated.systems</td><td>McrB-AAA->McrC-NTD+REase-><-iSTAND||METHYLASE-><-?<-?<-DUF499*||SF1-SF2-Helicase-></td><td>1068</td><td>Archaea>Euryarchaeota>Stenosarchaea_group>Halobacteria>Natrialbales</td><td>GCF_000025625.1</td><td>Natrialba_magadii</td><td>C500_RS02195</td><td>143164..146370</td><td>13769</td><td>DUF499-domain-containing-protein-[Natrialba-magadii]</td></tr>
<tr><td>TYL37429.1</td><td>AAA+ATPase+wHTH+Alpha+Beta-domain+wHTH+RRM</td><td>Type-3_BR.PglZ.associated.systems</td><td>McrB-AAA->McrC-NTD+REase->?-><-iSTAND<-METHYLASE<-DUF499*||SF1-SF2-Helicase-></td><td>1068</td><td>Archaea>Euryarchaeota>Stenosarchaea_group>Halobacteria>Natrialbales</td><td>GCA_008122205.1</td><td>Natronococcus_pandeyae</td><td>CV102_17630</td><td>157597..160803</td><td>2055836</td><td>DUF499-domain-containing-protein-[Natronococcus-pandeyae]</td></tr>
<tr><td>WP_233710727.1</td><td>AAA+ATPase+wHTH+Alpha+Beta-domain+wHTH+RRM</td><td>Type-3_BR.PglZ.associated.systems</td><td>McrB-AAA->McrC-NTD+REase->?-><-iSTAND<-METHYLASE<-DUF499*||SF1-SF2-Helicase-></td><td>976</td><td>Archaea>Euryarchaeota>Stenosarchaea_group>Halobacteria>Natrialbales</td><td>GCF_008122205.1</td><td>Natronococcus_pandeyae</td><td>CV102_RS17540</td><td>157597..160527</td><td>2055836</td><td>DUF499-domain-containing-protein-[Natronococcus-pandeyae]</td></tr>
<tr><td>WP_252490024.1</td><td>AAA+ATPase+wHTH+Alpha+Beta-domain+wHTH+RRM</td><td>Type-3_BR.PglZ.associated.systems</td><td><-SF1-SF2-Helicase||DUF499*->HEPN+REase->METHYLASE->DUF429->ParB+HNH->iSTAND-><-McrC-NTD+REase</td><td>1070</td><td>Archaea>Euryarchaeota>Stenosarchaea_group>Halobacteria>Natrialbales</td><td>GCF_023913515.1</td><td>Natronococcus_sp._CG52</td><td>NED97_RS07150</td><td>1394012..1397224</td><td>2951804</td><td>DUF499-domain-containing-protein-[Natronococcus-sp.-CG52]</td></tr>
<tr><td>SFC23913.1</td><td>AAA+ATPase+wHTH+Alpha+Beta-domain+wHTH+RRM</td><td>Type-3_BR.PglZ.associated.systems</td><td><-SF1-SF2-Helicase||?->?-><-METHYLASE<-PglZ<-DUF499*<-?<-HEPN||?-><-?<-?||Resolvase-></td><td>955</td><td>Archaea>Euryarchaeota>Stenosarchaea_group>Halobacteria>Natrialbales</td><td>GCA_900112205.1</td><td>Halobiforma_haloterrestris</td><td>SAMN05444422_10610</td><td>10310..13177</td><td>148448</td><td>hypothetical-protein-SAMN05444422_10610-[Halobiforma-haloterrestris]</td></tr>
<tr><td>WP_143095830.1</td><td>AAA+ATPase+wHTH+Alpha+Beta-domain+wHTH+RRM</td><td>Type-3_BR.PglZ.associated.systems</td><td><-SF1-SF2-Helicase||?->?-><-iSTAND<-METHYLASE<-PglZ<-DUF499*<-HEPN||?-><-?||Resolvase-></td><td>952</td><td>Archaea>Euryarchaeota>Stenosarchaea_group>Halobacteria>Natrialbales</td><td>GCF_900112205.1</td><td>Halobiforma_haloterrestris</td><td>BM113_RS12255</td><td>10310..13168</td><td>148448</td><td>hypothetical-protein-[Halobiforma-haloterrestris]</td></tr>
<tr><td>WP_029601777.1</td><td>AAA+ATPase+wHTH+Alpha+Beta-domain+wHTH+RRM</td><td>Type-3_BR.PglZ.associated.systems</td><td><-REase<-ORC/CDC6+REase||McrB-AAA->McrC-NTD+REase-><-iSTAND<-METHYLASE<-DUF499*||?->?-><-METHYLASE</td><td>1077</td><td>Archaea>Euryarchaeota>Stenosarchaea_group>Halobacteria>Natrialbales</td><td>GCF_000336655.1</td><td>Halobiforma_lacisalsi</td><td>C445_RS16630</td><td>18229..21462</td><td>229731</td><td>DUF499-domain-containing-protein-[Halobiforma-lacisalsi]</td></tr>
<tr><td>EMA30005.1</td><td>AAA+ATPase+wHTH+Alpha+Beta-domain+wHTH+RRM</td><td>Type-3_BR.PglZ.associated.systems</td><td><-ORC/CDC6+REase||McrB-AAA->McrC-NTD+REase-><-iSTAND<-METHYLASE<-DUF499*</td><td>983</td><td>Archaea>Euryarchaeota>Stenosarchaea_group>Halobacteria>Natrialbales</td><td>GCA_000336655.1</td><td>Halobiforma_lacisalsi_AJ5</td><td>C445_16904</td><td>18229..21180</td><td>358396</td><td>ATPase-(AAA+-superfamily)-like-protein-[Halobiforma-lacisalsi-AJ5]</td></tr>
<tr><td>ELY39843.1</td><td>AAA+ATPase+wHTH+Alpha+Beta-domain+wHTH+RRM</td><td>Type-3_BR.PglZ.associated.systems</td><td><-ORC/CDC6+REase||McrB-AAA-><-?||McrC-NTD+REase-><-iSTAND<-METHYLASE<-DUF499*||?-><-METHYLASE||?->ParB+HNH-></td><td>1074</td><td>Archaea>Euryarchaeota>Stenosarchaea_group>Halobacteria>Natrialbales</td><td>GCA_000337235.1</td><td>Natronorubrum_tibetense_GA33</td><td>C496_14236</td><td>262588..265812</td><td>1114856</td><td>ATPase-[Natronorubrum-tibetense-GA33]</td></tr>
<tr><td>WP_027119197.1</td><td>AAA+ATPase+wHTH+Alpha+Beta-domain+wHTH+RRM</td><td>Type-3_BR.PglZ.associated.systems</td><td><-METHYLASE<-ORC/CDC6+REase||McrB-AAA->McrC-NTD+REase-><-iSTAND<-METHYLASE<-DUF499*||?->?-><-METHYLASE||?->ParB+HNH-></td><td>1076</td><td>Archaea>Euryarchaeota>Stenosarchaea_group>Halobacteria>Natrialbales</td><td>GCF_000337235.1</td><td>Natronorubrum_tibetense</td><td>NATTI_RS0117070</td><td>3305760..3308990</td><td>63128</td><td>ATPase-[Natronorubrum-tibetense]</td></tr>
<tr><td>WP_130170137.1</td><td>AAA+ATPase+wHTH+Alpha+Beta-domain+wHTH+RRM</td><td>Type-3_BR.PglZ.associated.systems</td><td><-iSTAND<-METHYLASE<-PglZ<-DUF499*<-Resolvase||<-?<-Phage_integrase||rRNA->tRNA->rRNA->rRNA->rRNA-><-?<-PadR-HTH</td><td>950</td><td>Archaea>Euryarchaeota>Stenosarchaea_group>Halobacteria>Natrialbales</td><td>GCF_004209855.1</td><td>Natrinema_altunense</td><td>ELS17_RS07420</td><td>1518703..1521555</td><td>222984</td><td>hypothetical-protein-[Natrinema-altunense]</td></tr>
<tr><td>WP_076610738.1</td><td>AAA+ATPase+wHTH+Alpha+Beta-domain+wHTH+RRM</td><td>Type-3_BR.PglZ.associated.systems</td><td><-iSTAND<-METHYLASE<-PglZ<-DUF499*</td><td>843</td><td>Archaea>Euryarchaeota>Stenosarchaea_group>Halobacteria>Natrialbales</td><td>GCF_900156475.1</td><td>Natronorubrum_thiooxidans</td><td>BW940_RS18885</td><td>31270..33801</td><td>308853</td><td>DUF499-domain-containing-protein-[Natronorubrum-thiooxidans]</td></tr>
<tr><td>WP_241430589.1</td><td>AAA+ATPase+wHTH+Alpha+Beta-domain+wHTH+RRM</td><td>Type-3_BR.PglZ.associated.systems</td><td><-iSTAND<-?<-SF1-SF2-Helicase<-METHYLASE||ParB+HNH->SIGMA-HTH-><-DUF499*||SF1-SF2-Helicase->?-><-?||RNase-></td><td>1033</td><td>Archaea>Euryarchaeota>Stenosarchaea_group>Halobacteria>Natrialbales</td><td>GCF_000337735.1</td><td>Natronorubrum_sulfidifaciens</td><td>C495_RS06085</td><td>92654..95755</td><td>388259</td><td>DUF499-domain-containing-protein-[Natronorubrum-sulfidifaciens]</td></tr>
<tr><td>WP_263020920.1</td><td>AAA+ATPase+wHTH+Alpha+Beta-domain+wHTH+RRM</td><td>Type-3_BR.PglZ.associated.systems</td><td><-ABC-ATPase+SbcC<-?||?->?->DUF499*->PglZ->METHYLASE->iSTAND-></td><td>832</td><td>Archaea>Euryarchaeota>Stenosarchaea_group>Halobacteria>Natrialbales</td><td>GCF_025629245.1</td><td>Natronobiforma_cellulositropha</td><td>NOB15_RS17085</td><td>46291..48789</td><td>1679076</td><td>DUF499-domain-containing-protein-[Natronobiforma-cellulositropha]</td></tr>
<tr><td>WP_312910163.1</td><td>AAA+ATPase+wHTH+Alpha+Beta-domain+wHTH+RRM</td><td>Type-3_BR.PglZ.associated.systems</td><td><-METHYLASE<-PglZ<-DUF499+RRM*</td><td>929</td><td>Archaea>Euryarchaeota>Stenosarchaea_group>Halobacteria>Natrialbales>Natrialbaceae</td><td>GCF_017357105.1</td><td>Natronosalvus_caseinilyticus</td><td>J1N60_RS01630</td><td>320944..323733</td><td>2953747</td><td>hypothetical-protein-[Natronosalvus-caseinilyticus]</td></tr>
<tr><td>HJH26897.1</td><td>AAA+ATPase+wHTH+Alpha+Beta-domain+wHTH+RRM</td><td>Type-3_BR.PglZ.associated.systems</td><td><-METHYLASE-p<-REase<-?<-PglZ<-DUF499*</td><td>832</td><td>Archaea>Euryarchaeota>Stenosarchaea_group>Methanomicrobia>Methanophagales</td><td>GCA_020793595.1</td><td>Methanophagales_archaeon</td><td>C5S37_09040</td><td>4404..6902</td><td>2056316</td><td>hypothetical-protein-[Methanophagales-archaeon]</td></tr>
<tr><td>MCW3138460.1</td><td>AAA+ATPase+wHTH+Alpha+Beta-domain+wHTH+RRM</td><td>Type-3_BR.PglZ.associated.systems</td><td><-METHYLASE-p<-?<-?<-DUF499*<-?<-?<-?<-iSTAND<-METHYLASE<-PglZ</td><td>244</td><td>Archaea>Euryarchaeota>Stenosarchaea_group>Methanomicrobia>Methanophagales</td><td>GCA_025966495.1</td><td>Methanophagales_archaeon</td><td>N2V76_08630</td><td>1224..1958</td><td>2056316</td><td>MAG:-hypothetical-protein-N2V76_08630-[Methanophagales-archaeon]</td></tr>
<tr><td>NMX21982.1</td><td>AAA+ATPase+wHTH+Alpha+Beta-domain+wHTH+RRM</td><td>Type-3_BR.PglZ.associated.systems</td><td><-iSTAND<-METHYLASE<-PglZ<-DUF499*<-JAB</td><td>829</td><td>Archaea>Euryarchaeota>Stenosarchaea_group>Methanomicrobia>Methanophagales</td><td>GCA_012979255.1</td><td>ANME-1_cluster_archaeon_GoMg4</td><td>C5S30_06035</td><td>4462..6951</td><td>2081357</td><td>hypothetical-protein-[ANME-1-cluster-archaeon-GoMg4]</td></tr>
<tr><td>MCW3138467.1</td><td>AAA+ATPase+wHTH+Alpha+Beta-domain+wHTH+RRM</td><td>Type-3_BR.PglZ.associated.systems</td><td><-iSTAND<-METHYLASE<-PglZ<-DUF499*<-HNH</td><td>999</td><td>Archaea>Euryarchaeota>Stenosarchaea_group>Methanomicrobia>Methanophagales</td><td>GCA_025966495.1</td><td>Methanophagales_archaeon</td><td>N2V76_08665</td><td>7672..10671</td><td>2056316</td><td>MAG:-DUF499-domain-containing-protein-[Methanophagales-archaeon]</td></tr>
<tr><td>OFV67936.1</td><td>AAA+ATPase+wHTH+Alpha+Beta-domain+wHTH+RRM</td><td>Type-3_BR.PglZ.associated.systems</td><td>REase->?->?->DUF499*->PglZ->METHYLASE->iSTAND-></td><td>1023</td><td>Archaea>Euryarchaeota>Stenosarchaea_group>Methanomicrobia>Methanosarcinales</td><td>GCA_001766815.1</td><td>Candidatus_Syntrophoarchaeum_caldarius</td><td>SCAL_000576</td><td>27693..30764</td><td>1838285</td><td>AAA-family-ATPase-[Candidatus-Syntrophoarchaeum-caldarius]</td></tr>
<tr><td>WP_270095351.1</td><td>AAA+ATPase+wHTH+Alpha+Beta-domain+wHTH+RRM</td><td>Type-3_BR.PglZ.associated.systems</td><td>MarR-HTH->?->tRNA-><-TUDOR+SF1-SF2-Helicase+iREase+CTD<-iSTAND<-METHYLASE<-PglZ<-DUF499*||?-><-GDResolvase+Recombinase+ZnR</td><td>928</td><td>Archaea>Euryarchaeota>Stenosarchaea_group>Methanomicrobia>Methanosarcinales</td><td>GCF_027594145.1</td><td>Methanococcoides_alaskense</td><td>PBV53_RS01130</td><td>228076..230862</td><td>325778</td><td>hypothetical-protein-[Methanococcoides-alaskense]</td></tr>
<tr><td>PXF60464.1</td><td>AAA+ATPase+wHTH+Alpha+Beta-domain+wHTH+RRM</td><td>Type-3_BR.PglZ.associated.systems</td><td>DUF433->HicB_lk_antitox->HhH-RADC+JAB->DUF499*->PglZ->?->REase->METHYLASE->iSTAND-></td><td>869</td><td>Archaea>Euryarchaeota>Stenosarchaea_group>Methanomicrobia>Methanosarcinales</td><td>GCA_003194445.1</td><td>ANME-2_cluster_archaeon</td><td>C4B59_09325</td><td>28064..30673</td><td>2056317</td><td>hypothetical-protein-C4B59_09325-[ANME-2-cluster-archaeon]</td></tr>
<tr><td>RLG35041.1</td><td>AAA+ATPase+wHTH+Alpha+Beta-domain+wHTH+RRM</td><td>Type-3_BR.PglZ.associated.systems</td><td><-METHYLASE<-PglZ<-DUF499*<-HhH-RADC+JAB<-?<-tRNA<-DNA-glycosylase</td><td>829</td><td>Archaea>Euryarchaeota>Stenosarchaea_group>Methanomicrobia>Methanosarcinales</td><td>GCA_003661125.1</td><td>Methanosarcinales_archaeon</td><td>DRN97_00825</td><td>2857..5346</td><td>2250255</td><td>hypothetical-protein-DRN97_00825-[Methanosarcinales-archaeon]</td></tr>
<tr><td>CAD6492704.1</td><td>AAA+ATPase+wHTH+Alpha+Beta-domain+wHTH+RRM</td><td>Type-3_BR.PglZ.associated.systems</td><td><-iSTAND<-METHYLASE<-REase<-?<-PglZ<-DUF499*<-HhH-RADC+JAB</td><td>870</td><td>Archaea>Euryarchaeota>Stenosarchaea_group>Methanomicrobia>Methanosarcinales</td><td>GCA_905067565.1</td><td>Candidatus_Argoarchaeum_ethanivorans</td><td>CHKLHMKO_00332</td><td>6578..9190</td><td>2608793</td><td>hypothetical-protein-CHKLHMKO_00332-[Candidatus-Argoarchaeum-ethanivorans]</td></tr>
<tr><td>MDR6223666.1</td><td>AAA+ATPase+wHTH+Alpha+Beta-domain+wHTH+RRM</td><td>Type-3_BR.PglZ.associated.systems</td><td>GDResolvase+HIN-HTH->||DUF499+RRM*->PglZ->METHYLASE->iSTAND->TUDOR+SF1-SF2-Helicase+iREase+CTD-></td><td>935</td><td>Archaea>Euryarchaeota>Stenosarchaea_group>Methanomicrobia>Methanosarcinales>Methanosarcinaceae</td><td>GCA_031454185.1</td><td>Methanococcoides_alaskense</td><td>J2750_002139</td><td>41269..44076</td><td>325778</td><td>hypothetical-protein-[Methanococcoides-alaskense]</td></tr>
<tr><td>RLG24730.1</td><td>AAA+ATPase+wHTH+Alpha+Beta-domain+wHTH+RRM</td><td>Type-3_BR.PglZ.associated.systems</td><td><-PglZ<-OB-Fold+AAA-ATPase+LonP-BrxL<-DUF499+RRM*</td><td>1014</td><td>Archaea>Euryarchaeota>Stenosarchaea_group>Methanomicrobia>Methanosarcinales>unclassified_Methanosarcinales</td><td>GCA_003661045.1</td><td>Methanosarcinales_archaeon</td><td>DRN85_07235</td><td>3188..6232</td><td>2250255</td><td>MAG:-hypothetical-protein-DRN85_07235,-partial-[Methanosarcinales-archaeon]</td></tr>
<tr><td>MBU7037419.1</td><td>AAA+ATPase+wHTH+Alpha+Beta-domain+wHTH+RRM</td><td>Type-3_BR.PglZ.associated.systems</td><td><-CR-Kinase<-?<-iSTAND<-METHYLASE<-PglZ<-DUF499*</td><td>999</td><td>Archaea>Euryarchaeota>Theionarchaea>unclassified_Theionarchaea</td><td>GCA_018992685.1</td><td>Theionarchaea_archaeon</td><td>HXS52_05780</td><td>146119..149118</td><td>2747605</td><td>DUF499-domain-containing-protein-[Theionarchaea-archaeon]</td></tr>
<tr><td>WP_167886533.1</td><td>AAA+ATPase+wHTH+Alpha+Beta-domain+wHTH+RRM</td><td>Type-3_BR.PglZ.associated.systems</td><td>SF1-SF2-Helicase->?->?->DndE-><-?<-PglZ<-DUF499*||METHYLASE->?->?->REase-><-EVE||EVE-></td><td>1105</td><td>Archaea>Euryarchaeota>Thermococci>Thermococcales>Thermococcaceae</td><td>GCF_012027325.1</td><td>Thermococcus_sp._M39</td><td>E3E31_RS08155</td><td>9394..12711</td><td>1638262</td><td>DUF499-domain-containing-protein-[Thermococcus-sp.-M39]</td></tr>
<tr><td>WP_297420748.1</td><td>AAA+ATPase+wHTH+Alpha+Beta-domain+wHTH+RRM</td><td>Type-3_BR.PglZ.associated.systems</td><td>SbcC->SbcC-><-PglZ<-DUF499+RRM*||METHYLASE-></td><td>1105</td><td>Archaea>Euryarchaeota>Thermococci>Thermococcales>Thermococcaceae>Thermococcus</td><td>GCF_027019385.1</td><td>Thermococcus_sp.</td><td>MV421_RS00875</td><td>76122..79439</td><td>35749</td><td>DUF499-domain-containing-protein-[Thermococcus-sp.]</td></tr>
<tr><td>RLF82338.1</td><td>AAA+ATPase+wHTH+Alpha+Beta-domain+wHTH+RRM</td><td>Type-3_BR.PglZ.associated.systems</td><td>SF1-SF2-Helicase->?->?->DndE-><-?<-PglZ<-DUF499*||METHYLASE->?->?-><-EVE||EVE-></td><td>1106</td><td>Archaea>Euryarchaeota>Thermococci>unclassified_Thermococci</td><td>GCA_003650065.1</td><td>Thermococci_archaeon</td><td>DRN38_00300</td><td>6104..9424</td><td>2250254</td><td>hypothetical-protein-DRN38_00300-[Thermococci-archaeon]</td></tr>
<tr><td>WP_125672294.1</td><td>AAA+ATPase+wHTH+Alpha+Beta-domain+wHTH+RRM</td><td>Type-3_BR.PglZ.associated.systems</td><td><-METHYLASE||DUF499*->BrxL->PglZ-></td><td>1122</td><td>Archaea>TACK_group>Candidatus_Korarchaeota>Candidatus_Methanodesulfokores</td><td>GCF_003947435.1</td><td>Candidatus_Methanodesulfokores_washburnensis</td><td>D6D85_RS12065</td><td>3774..7142</td><td>2478471</td><td>DUF499-domain-containing-protein-[Candidatus-Methanodesulfokores-washburnensis]</td></tr>
<tr><td>WP_125670094.1</td><td>AAA+ATPase+wHTH+Alpha+Beta-domain+wHTH+RRM</td><td>Type-3_BR.PglZ.associated.systems</td><td><-iSTAND<-METHYLASE<-PglZ<-DUF499*||?->DndB->?->?->SbcC-></td><td>847</td><td>Archaea>TACK_group>Candidatus_Korarchaeota>Candidatus_Methanodesulfokores</td><td>GCF_003947435.1</td><td>Candidatus_Methanodesulfokores_washburnensis</td><td>D6D85_RS00580</td><td>4292..6835</td><td>2478471</td><td>hypothetical-protein-[Candidatus-Methanodesulfokores-washburnensis]</td></tr>
<tr><td>MCC6023981.1</td><td>AAA+ATPase+wHTH+Alpha+Beta-domain+wHTH+RRM</td><td>Type-3_BR.PglZ.associated.systems</td><td>METHYLASE-p->DUF499+RRM*->PglZ->Cluster81_1clades-></td><td>1075</td><td>Archaea>TACK_group>Nitrososphaerota>unclassified_Nitrososphaerota</td><td>GCA_020833125.1</td><td>Nitrososphaerota_archaeon</td><td>LM600_04705</td><td>1748..4975</td><td>2026795</td><td>hypothetical-protein-[Nitrososphaerota-archaeon]</td></tr>
<tr><td>WP_012608229.1</td><td>AAA+ATPase+wHTH+Alpha+Beta-domain+wHTH+RRM</td><td>Type-3_BR.PglZ.associated.systems</td><td>XIS-HTH-><-PglZ<-DUF499+RRM*<-METHYLASE</td><td>1070</td><td>Archaea>TACK_group>Thermoproteota>Thermoprotei>Desulfurococcales>Desulfurococcaceae</td><td>GCF_000020905.1</td><td>Desulfurococcus_amylolyticus</td><td>DKAM_RS02675</td><td>523466..526678</td><td>94694</td><td>DUF499-domain-containing-protein-[Desulfurococcus-amylolyticus]</td></tr>
<tr><td>MEB3780842.1</td><td>AAA+ATPase+wHTH+Alpha+Beta-domain+wHTH+RRM</td><td>Type-3_BR.PglZ.associated.systems</td><td><-PglZ<-OB-Fold+AAA-ATPase+LonP-BrxL<-DUF499+RRM*<-METHYLASE||AAA-ATPase+REase-></td><td>1100</td><td>Archaea>TACK_group>Thermoproteota>Thermoprotei>Desulfurococcales>unclassified_Desulfurococcales</td><td>GCA_035615675.1</td><td>Desulfurococcales_archaeon</td><td>GSR85_11550</td><td>165012..168314</td><td>2480821</td><td>hypothetical-protein-[Desulfurococcales-archaeon]</td></tr>
<tr><td>MEB3759607.1</td><td>AAA+ATPase+wHTH+Alpha+Beta-domain+wHTH+RRM</td><td>Type-3_BR.PglZ.associated.systems</td><td><-PglZ<-OB-Fold+AAA-ATPase+LonP-BrxL<-DUF499+RRM*<-METHYLASE||<-HEPN</td><td>1100</td><td>Archaea>TACK_group>Thermoproteota>Thermoprotei>Desulfurococcales>unclassified_Desulfurococcales</td><td>GCA_035615755.1</td><td>Desulfurococcales_archaeon</td><td>GSR81_02015</td><td>170120..173422</td><td>2480821</td><td>hypothetical-protein-[Desulfurococcales-archaeon]</td></tr>
<tr><td>MEB3788750.1</td><td>AAA+ATPase+wHTH+Alpha+Beta-domain+wHTH+RRM</td><td>Type-3_BR.PglZ.associated.systems</td><td><-PglZ<-OB-Fold+AAA-ATPase+LonP-BrxL<-DUF499+RRM*<-METHYLASE||<-HEPN</td><td>1100</td><td>Archaea>TACK_group>Thermoproteota>Thermoprotei>Desulfurococcales>unclassified_Desulfurococcales</td><td>GCA_035615915.1</td><td>Desulfurococcales_archaeon</td><td>GSR72_02520</td><td>3073..6375</td><td>2480821</td><td>hypothetical-protein-[Desulfurococcales-archaeon]</td></tr>
<tr><td>MCG2864876.1</td><td>AAA+ATPase+wHTH+Alpha+Beta-domain+wHTH+RRM</td><td>Type-3_BR.PglZ.associated.systems</td><td>METHYLASE->DUF499+RRM*->PglZ->Cluster81_1clades-></td><td>1080</td><td>Archaea>TACK_group>Thermoproteota>Thermoprotei>Thermoproteales>Thermoproteaceae</td><td>GCA_022014895.1</td><td>Vulcanisaeta_sp.</td><td>L7H04_03325</td><td>3346..6588</td><td>2020871</td><td>hypothetical-protein-[Vulcanisaeta-sp.]</td></tr>
<tr><td>RLF12260.1</td><td>AAA+ATPase+wHTH+Alpha+Beta-domain+wHTH+RRM</td><td>Type-3_BR.PglZ.associated.systems</td><td><-Phage_integrase||?->DUF499*->PglZ->METHYLASE->iSTAND-></td><td>832</td><td>Archaea>TACK_group>Thermoproteota>Thermoprotei>unclassified_Thermoprotei</td><td>GCA_003649605.1</td><td>Thermoprotei_archaeon</td><td>DRJ69_01100</td><td>1173..3671</td><td>2250277</td><td>MAG:-hypothetical-protein-DRJ69_01100-[Thermoprotei-archaeon]</td></tr>
<tr><td>MDM7274691.1</td><td>AAA+ATPase+wHTH+Alpha+Beta-domain+wHTH+RRM</td><td>Type-3_BR.PglZ.associated.systems</td><td>METHYLASE->DUF499+RRM*->OB-Fold+AAA-ATPase+LonP-BrxL->PglZ-><-HEPN</td><td>1081</td><td>Archaea>TACK_group>Thermoproteota>Thermoprotei>unclassified_Thermoprotei</td><td>GCA_029335875.2</td><td>Thermoprotei_archaeon</td><td>P3X22_001000</td><td>3466..6711</td><td>2250277</td><td>hypothetical-protein-[Thermoprotei-archaeon]</td></tr>
<tr><td>MCR8454632.1</td><td>AAA+ATPase+wHTH+Alpha+Beta-domain+wHTH+RRM</td><td>Type-3_BR.PglZ.associated.systems</td><td>DUF499*->PglZ->METHYLASE->iSTAND-></td><td>845</td><td>Archaea>TACK_group>Thermoproteota>unclassified_Thermoproteota</td><td>GCA_024720955.1</td><td>Thermoproteota_archaeon</td><td>NDP13_06570</td><td>1996..4533</td><td>2056631</td><td>hypothetical-protein-[Thermoproteota-archaeon]</td></tr>
<tr><td>MCR8433831.1</td><td>AAA+ATPase+wHTH+Alpha+Beta-domain+wHTH+RRM</td><td>Type-3_BR.PglZ.associated.systems</td><td><-PglZ<-OB-Fold+AAA-ATPase+LonP-BrxL<-DUF499+RRM*<-METHYLASE-p</td><td>1079</td><td>Archaea>TACK_group>Thermoproteota>unclassified_Thermoproteota</td><td>GCA_024720975.1</td><td>Thermoproteota_archaeon</td><td>NDP24_06350</td><td>3366..6605</td><td>2056631</td><td>hypothetical-protein-[Thermoproteota-archaeon]</td></tr>
<tr><td>WP_279735282.1</td><td>AAA+ATPase+wHTH+Alpha+Beta-domain+wHTH+RRM</td><td>Type-3_BR.PglZ.associated.systems</td><td><-TUDOR+SF1-SF2-Helicase+iREase+CTD<-iSTAND<-METHYLASE<-Peptidase_S8<-Classical-AAA<-PglZ<-DUF499*<-XIS-HTH||MODE-HTH+PBPII->?->VirD4-TraG+VirD4-FtsK->CopG/RHH->VirB11+VirB4-FtsK-></td><td>922</td><td>Bacteria</td><td>GCF_030845255.1</td><td>Pseudomonadota</td><td>Q2343_RS07055</td><td>402338..405106</td><td>1224</td><td>DUF499-domain-containing-protein-[Stenotrophomonas-maltophilia]</td></tr>
<tr><td>MBA2528240.1</td><td>AAA+ATPase+wHTH+Alpha+Beta-domain+wHTH+RRM</td><td>Type-3_BR.PglZ.associated.systems</td><td><-REase+35exo+Beta-Barrel+SF1-SF2-Helicase<-TUDOR+SF1-SF2-Helicase+iREase+CTD<-iSTAND<-URI-DNase<-METHYLASE<-PglZ<-DUF499*</td><td>901</td><td>Bacteria>Acidobacteriota>Blastocatellia>Blastocatellales>Pyrinomonadaceae</td><td>GCA_013696625.1</td><td>Pyrinomonadaceae_bacterium</td><td>HOV18_21020</td><td>15268..17973</td><td>2283092</td><td>DUF499-domain-containing-protein-[Pyrinomonadaceae-bacterium]</td></tr>
<tr><td>PYP82692.1</td><td>AAA+ATPase+wHTH+Alpha+Beta-domain+wHTH+RRM</td><td>Type-3_BR.PglZ.associated.systems</td><td><-XIS-HTH||Phage_int_SAM_1+Phage_integrase-><-TUDOR+SF1-SF2-Helicase+iREase+CTD<-iSTAND<-METHYLASE<-PglZ<-DUF499*</td><td>346</td><td>Bacteria>Acidobacteriota>Blastocatellia>unclassified_Blastocatellia</td><td>GCA_003222245.1</td><td>Blastocatellia_bacterium_AA13</td><td>DMF61_25860</td><td>53723..54763</td><td>2169413</td><td>hypothetical-protein-DMF61_25860-[Blastocatellia-bacterium-AA13]</td></tr>
<tr><td>HEX8998547.1</td><td>AAA+ATPase+wHTH+Alpha+Beta-domain+wHTH+RRM</td><td>Type-3_BR.PglZ.associated.systems</td><td><-METHYLASE-p<-PglZ<-DUF499+RRM*</td><td>903</td><td>Bacteria>Acidobacteriota>Blastocatellia>unclassified_Blastocatellia</td><td>GCA_036392235.1</td><td>Blastocatellia_bacterium</td><td>VGB07_01535</td><td>3765..6476</td><td>2052146</td><td>DUF499-domain-containing-protein-[Blastocatellia-bacterium]</td></tr>
<tr><td>MDR2561287.1</td><td>AAA+ATPase+wHTH+Alpha+Beta-domain+wHTH+RRM</td><td>Type-3_BR.PglZ.associated.systems</td><td><-PglZ<-DUF499+RRM*</td><td>902</td><td>Bacteria>Acidobacteriota>Holophagae>Holophagales>unclassified_Holophagales</td><td>GCA_031281385.1</td><td>Holophagales_bacterium</td><td>LBC63_05920</td><td>651..3359</td><td>2478486</td><td>DUF499-domain-containing-protein-[Holophagales-bacterium]</td></tr>
<tr><td>WP_263354240.1</td><td>AAA+ATPase+wHTH+Alpha+Beta-domain+wHTH+RRM</td><td>Type-3_BR.PglZ.associated.systems</td><td>DEDD_Tnp_IS110+Transposase_20->ParB+HTH-><-TUDOR+SF1-SF2-Helicase+iREase+CTD<-iSTAND<-METHYLASE<-PglZ<-DUF499*||HTH->STYKinase-like->?->?->?-><-MODE-HTH+PBPII</td><td>917</td><td>Bacteria>Acidobacteriota>Terriglobia>Terriglobales>Acidobacteriaceae</td><td>GCF_025685625.1</td><td>Acidicapsa_acidisoli</td><td>OHL23_RS22435</td><td>96896..99649</td><td>1615681</td><td>DUF499-domain-containing-protein-[Acidicapsa-acidisoli]</td></tr>
<tr><td>MCC6128740.1</td><td>AAA+ATPase+wHTH+Alpha+Beta-domain+wHTH+RRM</td><td>Type-3_BR.PglZ.associated.systems</td><td>HTH+WYL->DUF499*->PglZ->METHYLASE-p-></td><td>898</td><td>Bacteria>Acidobacteriota>unclassified_Acidobacteriota</td><td>GCA_020845555.1</td><td>Acidobacteriota_bacterium</td><td>IT186_02330</td><td>824..3520</td><td>1978231</td><td>hypothetical-protein-[Acidobacteria-bacterium]</td></tr>
<tr><td>MBI4455133.1</td><td>AAA+ATPase+wHTH+Alpha+Beta-domain+wHTH+RRM</td><td>Type-3_BR.PglZ.associated.systems</td><td><-PIN<-DUF433||?->?-><-?||DUF499*->PglZ->CopG/RHH->PIN->METHYLASE-p->METHYLASE->iSTAND-></td><td>872</td><td>Bacteria>Acidobacteriota>unclassified_Acidobacteriota</td><td>GCA_016211105.1</td><td>Acidobacteriota_bacterium</td><td>HY644_04455</td><td>210412..213030</td><td>1978231</td><td>hypothetical-protein-[Acidobacteria-bacterium]</td></tr>
<tr><td>KAA6461889.1</td><td>AAA+ATPase+wHTH+Alpha+Beta-domain+wHTH+RRM</td><td>Type-3_BR.PglZ.associated.systems</td><td><-CHTH<-TUDOR+SF1-SF2-Helicase+iREase+CTD<-iSTAND<-METHYLASE<-PglZ<-DUF499*||ArdCN+MPTase->ParB->Phage_int_SAM_1+Phage_integrase->Phage_integrase-></td><td>916</td><td>Bacteria>Acidobacteriota>unclassified_Acidobacteriota</td><td>GCA_008682415.1</td><td>Acidobacteria_bacterium_AB60</td><td>DYQ86_09580</td><td>197055..199805</td><td>1230846</td><td>DUF499-domain-containing-protein-[Acidobacteria-bacterium-AB60]</td></tr>
<tr><td>MEA2005693.1</td><td>AAA+ATPase+wHTH+Alpha+Beta-domain+wHTH+RRM</td><td>Type-3_BR.PglZ.associated.systems</td><td><-PglZ<-DUF499+RRM*<-XIS-HTH<-MerR-HTH</td><td>808</td><td>Bacteria>Acidobacteriota>unclassified_Acidobacteriota</td><td>GCA_034660295.1</td><td>Acidobacteriota_bacterium</td><td>U9O50_05480</td><td>2926..5352</td><td>1978231</td><td>hypothetical-protein-[Acidobacteriota-bacterium]</td></tr>
<tr><td>MBP6702582.1</td><td>AAA+ATPase+wHTH+Alpha+Beta-domain+wHTH+RRM</td><td>Type-3_BR.PglZ.associated.systems</td><td><-MORC+HisKin<-sGTP<-TUDOR+SF1-SF2-Helicase+iREase+CTD<-iSTAND<-METHYLASE<-PglZ<-DUF499*<-HTH+WYL||OLD-ABC+TOPRIM->SF1-SF2-Helicase-></td><td>904</td><td>Bacteria>Acidobacteriota>Vicinamibacteria>unclassified_Vicinamibacteria</td><td>GCA_017990495.1</td><td>Vicinamibacteria_bacterium</td><td>KA385_03705</td><td>18565..21279</td><td>2793154</td><td>DUF499-domain-containing-protein-[Vicinamibacteria-bacterium]</td></tr>
<tr><td>NLY06143.1</td><td>AAA+ATPase+wHTH+Alpha+Beta-domain+wHTH+RRM</td><td>Type-3_BR.PglZ.associated.systems</td><td>XIS-HTH->DUF499*->PglZ->PolB->HEPN->METHYLASE->iSTAND->SF1-SF2-Helicase-></td><td>812</td><td>Bacteria>Atribacterota>Atribacteria>unclassified_Atribacteria</td><td>GCA_012728195.1</td><td>Candidatus_Atribacteria_bacterium</td><td>GXZ17_08485</td><td>474..2912</td><td>2053509</td><td>hypothetical-protein-[Candidatus-Atribacteria-bacterium]</td></tr>
<tr><td>MDY0134309.1</td><td>AAA+ATPase+wHTH+Alpha+Beta-domain+wHTH+RRM</td><td>Type-3_BR.PglZ.associated.systems</td><td><-METHYLASE<-PemK_toxin<-PglZ<-DUF499+RRM*<-XIS-HTH</td><td>917</td><td>Bacteria>Atribacterota>unclassified_Atribacterota</td><td>GCA_034005935.1</td><td>Atribacterota_bacterium</td><td>RBS14_01300</td><td>9425..12178</td><td>2952890</td><td>DUF499-domain-containing-protein-[Atribacterota-bacterium]</td></tr>
<tr><td>MDZ7363404.1</td><td>AAA+ATPase+wHTH+Alpha+Beta-domain+wHTH+RRM</td><td>Type-3_BR.PglZ.associated.systems</td><td><-PIN<-CopG-RHH<-PglZ<-DUF499+RRM*||REase-></td><td>866</td><td>Bacteria>Bacteria_incertae_sedis>Bacteria_candidate_phyla>candidate_division_KSB1>unclassified_candidate_division_KSB1</td><td>GCA_034506175.1</td><td>candidate_division_KSB1_bacterium</td><td>ONB46_22195</td><td>42829..45429</td><td>2172550</td><td>ATP-binding-protein-[candidate-division-KSB1-bacterium]</td></tr>
<tr><td>WP_337288695.1</td><td>AAA+ATPase+wHTH+Alpha+Beta-domain+wHTH+RRM</td><td>Type-3_BR.PglZ.associated.systems</td><td><-METHYLASE<-PIN<-CopG-RHH<-PglZ<-DUF499+RRM*</td><td>815</td><td>Bacteria>Bacteria_incertae_sedis>Bacteria_candidate_phyla>candidate_division_NC10>Candidatus_Methylomirabilis>unclassified_Candidatus_Methylomirabilis</td><td>GCF_002634395.1</td><td>Candidatus_Methylomirabilis_sp.</td><td>CI804_RS12515</td><td>8396..10843</td><td>2032687</td><td>hypothetical-protein-[Candidatus-Methylomirabilis-sp.]</td></tr>
<tr><td>MCK4308254.1</td><td>AAA+ATPase+wHTH+Alpha+Beta-domain+wHTH+RRM</td><td>Type-3_BR.PglZ.associated.systems</td><td>DUF499*->PglZ->METHYLASE-p-></td><td>750</td><td>Bacteria>Bacteria_incertae_sedis>Bacteria_candidate_phyla>candidate_division_WOR-3</td><td>GCA_023129235.1</td><td>candidate_division_WOR-3_bacterium</td><td>KAW50_08525</td><td>1..2255</td><td>2052148</td><td>hypothetical-protein-[candidate-division-WOR-3-bacterium]</td></tr>
<tr><td>MBE0478554.1</td><td>AAA+ATPase+wHTH+Alpha+Beta-domain+wHTH+RRM</td><td>Type-3_BR.PglZ.associated.systems</td><td><-Radical-SAM||DUF499*->PglZ->METHYLASE->Polbeta-></td><td>809</td><td>Bacteria>Bacteria_incertae_sedis>Bacteria_candidate_phyla>Candidatus_Aerophobetes>unclassified_Candidatus_Aerophobetes</td><td>GCA_014859225.1</td><td>Candidatus_Aerophobetes_bacterium</td><td>IBX65_05470</td><td>1499..3928</td><td>2030807</td><td>hypothetical-protein-[Candidatus-Aerophobetes-bacterium]</td></tr>
<tr><td>MCD6232554.1</td><td>AAA+ATPase+wHTH+Alpha+Beta-domain+wHTH+RRM</td><td>Type-3_BR.PglZ.associated.systems</td><td><-HD<-HEPN<-PolB<-PglZ<-PglZ<-DUF499*<-DUF499-p<-XIS-HTH</td><td>586</td><td>Bacteria>Bacteria_incertae_sedis>Bacteria_candidate_phyla>Candidatus_Aerophobetes>unclassified_Candidatus_Aerophobetes</td><td>GCA_021158005.1</td><td>Candidatus_Aerophobetes_bacterium</td><td>J7K28_08055</td><td>2638..4398</td><td>2030807</td><td>hypothetical-protein-[Candidatus-Aerophobetes-bacterium]</td></tr>
<tr><td>WP_304820872.1</td><td>AAA+ATPase+wHTH+Alpha+Beta-domain+wHTH+RRM</td><td>Type-3_BR.PglZ.associated.systems</td><td><-TUDOR+SF1-SF2-Helicase+iREase+CTD||DUF499+RRM*->PglZ-></td><td>892</td><td>Bacteria>Bacteria_incertae_sedis>Bacteria_candidate_phyla>Candidatus_Binatota>Candidatus_Binatia>Candidatus_Binatales</td><td>GCF_030646925.1</td><td>Candidatus_Binatus_sp.</td><td>Q7S58_RS03535</td><td>10994..13672</td><td>2811406</td><td>DUF499-domain-containing-protein-[Candidatus-Binatus-sp.]</td></tr>
<tr><td>HXK27027.1</td><td>AAA+ATPase+wHTH+Alpha+Beta-domain+wHTH+RRM</td><td>Type-3_BR.PglZ.associated.systems</td><td><-DUF499+RRM*<-METHYLASE<-OLD-ABC+TOPRIM<-TUDOR+SF1-SF2-Helicase+iREase+CTD<-PglZ<-Schlafen+HATPase+wHTH</td><td>949</td><td>Bacteria>Bacteria_incertae_sedis>Bacteria_candidate_phyla>Candidatus_Binatota>Candidatus_Binatia>unclassified_Candidatus_Binatia</td><td>GCA_035583555.1</td><td>Candidatus_Binatia_bacterium</td><td>VJ646_02090</td><td>262..3111</td><td>2838779</td><td>DUF499-domain-containing-protein-[Candidatus-Binatia-bacterium]</td></tr>
<tr><td>HWO41905.1</td><td>AAA+ATPase+wHTH+Alpha+Beta-domain+wHTH+RRM</td><td>Type-3_BR.PglZ.associated.systems</td><td><-PglZ<-DUF499+RRM*</td><td>795</td><td>Bacteria>Bacteria_incertae_sedis>Bacteria_candidate_phyla>Candidatus_Eisenbacteria>unclassified_Candidatus_Eisenbacteria</td><td>GCA_035559295.1</td><td>Candidatus_Eisenbacteria_bacterium</td><td>VNO43_08895</td><td>883..3270</td><td>2212470</td><td>hypothetical-protein-[Candidatus-Eisenbacteria-bacterium]</td></tr>
<tr><td>WP_259101344.1</td><td>AAA+ATPase+wHTH+Alpha+Beta-domain+wHTH+RRM</td><td>Type-3_BR.PglZ.associated.systems</td><td>METHYLASE-p->METHYLASE-p->?->OLD-ABC->?->?->DUF499*->PglZ-><-?<-?<-DDE_Tnp_1_3<-Peptidase_M50+PDZ</td><td>1011</td><td>Bacteria>Bacteria_incertae_sedis>Bacteria_candidate_phyla>Candidatus_Fervidibacteria>Candidatus_Fervidibacter</td><td>GCF_030520105.1</td><td>Candidatus_Fervidibacter_sacchari</td><td>M2350_RS16405</td><td>12362..15397</td><td>1448929</td><td>hypothetical-protein-[Candidatus-Fervidibacter-sacchari]</td></tr>
<tr><td>MCD6507458.1</td><td>AAA+ATPase+wHTH+Alpha+Beta-domain+wHTH+RRM</td><td>Type-3_BR.PglZ.associated.systems</td><td>METHYLASE->?->?->DUF499*->PglZ->?-><-?<-sGTP</td><td>1015</td><td>Bacteria>Bacteria_incertae_sedis>Bacteria_candidate_phyla>Candidatus_Poribacteria>unclassified_Candidatus_Poribacteria</td><td>GCA_021162805.1</td><td>Candidatus_Poribacteria_bacterium</td><td>J7M22_12660</td><td>4844..7891</td><td>2026781</td><td>DUF499-domain-containing-protein-[Candidatus-Poribacteria-bacterium]</td></tr>
<tr><td>HIE29204.1</td><td>AAA+ATPase+wHTH+Alpha+Beta-domain+wHTH+RRM</td><td>Type-3_BR.PglZ.associated.systems</td><td>DUF499*->PglZ->METHYLASE-></td><td>848</td><td>Bacteria>Bacteria_incertae_sedis>Bacteria_candidate_phyla>Candidatus_Poribacteria>unclassified_Candidatus_Poribacteria</td><td>GCA_012961655.1</td><td>Candidatus_Poribacteria_bacterium</td><td>EYP66_18180</td><td>406..2952</td><td>2026781</td><td>hypothetical-protein-[Candidatus-Poribacteria-bacterium]</td></tr>
<tr><td>HZM03188.1</td><td>AAA+ATPase+wHTH+Alpha+Beta-domain+wHTH+RRM</td><td>Type-3_BR.PglZ.associated.systems</td><td><-METHYLASE<-REase<-PglZ<-DUF499+RRM*</td><td>927</td><td>Bacteria>Bacteria_incertae_sedis>Bacteria_candidate_phyla>Candidatus_Saccharibacteria>Candidatus_Saccharimonadia>Candidatus_Saccharimonadales</td><td>GCA_035648615.1</td><td>Candidatus_Saccharimonadales_bacterium</td><td>VFC44_09205</td><td>7434..10217</td><td>2973664</td><td>hypothetical-protein-[Candidatus-Saccharimonadales-bacterium]</td></tr>
<tr><td>MBI2571916.1</td><td>AAA+ATPase+wHTH+Alpha+Beta-domain+wHTH+RRM</td><td>Type-3_BR.PglZ.associated.systems</td><td><-TUDOR+SF1-SF2-Helicase+iREase+CTD<-iSTAND<-METHYLASE<-PglZ<-DUF499*<-XIS-HTH</td><td>914</td><td>Bacteria>Bacteria_incertae_sedis>Bacteria_candidate_phyla>Candidatus_Schekmanbacteria>unclassified_Candidatus_Schekmanbacteria</td><td>GCA_016188575.1</td><td>Candidatus_Schekmanbacteria_bacterium</td><td>HYV63_33310</td><td>16963..19707</td><td>2419984</td><td>DUF499-domain-containing-protein-[Candidatus-Schekmanbacteria-bacterium]</td></tr>
<tr><td>MBI2569499.1</td><td>AAA+ATPase+wHTH+Alpha+Beta-domain+wHTH+RRM</td><td>Type-3_BR.PglZ.associated.systems</td><td><-OLD-ABC<-HEPN<-PolB<-PglZ<-DUF499*<-Peptidase_M24<-?<-ABC-ATPase+SbcC<-ABC-ATPase+SbcC</td><td>909</td><td>Bacteria>Bacteria_incertae_sedis>Bacteria_candidate_phyla>Candidatus_Schekmanbacteria>unclassified_Candidatus_Schekmanbacteria</td><td>GCA_016188575.1</td><td>Candidatus_Schekmanbacteria_bacterium</td><td>HYV63_21030</td><td>5126..7855</td><td>2419984</td><td>DUF499-domain-containing-protein-[Candidatus-Schekmanbacteria-bacterium]</td></tr>
<tr><td>HNY28787.1</td><td>AAA+ATPase+wHTH+Alpha+Beta-domain+wHTH+RRM</td><td>Type-3_BR.PglZ.associated.systems</td><td><-PglZ<-DUF499+RRM*</td><td>929</td><td>Bacteria>Bacteria_incertae_sedis>Bacteria_candidate_phyla>Candidatus_Sumerlaeota>unclassified_Candidatus_Sumerlaeota</td><td>GCA_035353095.1</td><td>Candidatus_Sumerlaeota_bacterium</td><td>PKH31_15550</td><td>294..3083</td><td>2107588</td><td>DUF499-domain-containing-protein-[Candidatus-Sumerlaeota-bacterium]</td></tr>
<tr><td>HRZ47307.1</td><td>AAA+ATPase+wHTH+Alpha+Beta-domain+wHTH+RRM</td><td>Type-3_BR.PglZ.associated.systems</td><td><-XIS-HTH<-HTH_3||DUF499+RRM*->PglZ->PIN->METHYLASE-p-></td><td>869</td><td>Bacteria>Bacteria_incertae_sedis>Bacteria_candidate_phyla>Patescibacteria_group>Microgenomates_group>Candidatus_Paceibacterota</td><td>GCA_035453485.1</td><td>Candidatus_Paceibacterota_bacterium</td><td>P5555_19175</td><td>13269..15878</td><td>2053589</td><td>hypothetical-protein-[Candidatus-Paceibacterota-bacterium]</td></tr>
<tr><td>MBC7429623.1</td><td>AAA+ATPase+wHTH+Alpha+Beta-domain+wHTH+RRM</td><td>Type-3_BR.PglZ.associated.systems</td><td>DUF499+RRM*->PglZ->METHYLASE->iSTAND->TUDOR+SF1-SF2-Helicase+iREase+CTD-></td><td>890</td><td>Bacteria>Bdellovibrionota>Bacteriovoracia>Bacteriovoracales>Bacteriovoracaceae>Bacteriovorax</td><td>GCA_014376445.1</td><td>Bacteriovorax_sp.</td><td>H7336_13480</td><td>202..2874</td><td>2608087</td><td>hypothetical-protein-[Bacteriovorax-sp.]</td></tr>
<tr><td>OGQ80565.1</td><td>AAA+ATPase+wHTH+Alpha+Beta-domain+wHTH+RRM</td><td>Type-3_BR.PglZ.associated.systems</td><td>DUF499*->PglZ->CopG/RHH->PIN->TUDOR+SF1-SF2-Helicase+iREase+CTD-></td><td>333</td><td>Bacteria>delta/epsilon_subdivisions>Deltaproteobacteria>unclassified_Deltaproteobacteria</td><td>GCA_001798595.1</td><td>Deltaproteobacteria_bacterium_RIFCSPLOWO2_12_FULL_60_19</td><td>A3F90_01965</td><td>176..1177</td><td>1797894</td><td>hypothetical-protein-A3F90_01965-[Deltaproteobacteria-bacterium-RIFCSPLOWO2_12_FULL_60_19]</td></tr>
<tr><td>MBI5243200.1</td><td>AAA+ATPase+wHTH+Alpha+Beta-domain+wHTH+RRM</td><td>Type-3_BR.PglZ.associated.systems</td><td>DUF499*->PglZ->CopG/RHH->PIN->RNaseH->?->METHYLASE-p-></td><td>846</td><td>Bacteria>Elusimicrobiota>unclassified_Elusimicrobiota</td><td>GCA_016212885.1</td><td>Elusimicrobiota_bacterium</td><td>HY922_05860</td><td>504..3044</td><td>2030800</td><td>hypothetical-protein-[Elusimicrobia-bacterium]</td></tr>
<tr><td>MCZ2342044.1</td><td>AAA+ATPase+wHTH+Alpha+Beta-domain+wHTH+RRM</td><td>Type-3_BR.PglZ.associated.systems</td><td>CRP-HTH->?->AbiEi_1->AbiEii->HTH_3->DUF499*->PglZ->SIR2->VirB4-FtsK+HerA-></td><td>908</td><td>Bacteria>FCB_group>Bacteroidota/Chlorobiota_group>Bacteroidota>Bacteroidia</td><td>GCA_026931725.1</td><td>Bacteroidales_bacterium</td><td>LC104_09655</td><td>11640..14366</td><td>2030927</td><td>MAG:-DUF499-domain-containing-protein-[Bacteroidales-bacterium]</td></tr>
<tr><td>WP_012508809.1</td><td>AAA+ATPase+wHTH+Alpha+Beta-domain+wHTH+RRM</td><td>Type-3_BR.PglZ.associated.systems</td><td>REase->REase-><-BirA-HTH||?->HTH_3->DUF499*->PglZ->?->?->PemK_toxin->METHYLASE->iSTAND-></td><td>911</td><td>Bacteria>FCB_group>Bacteroidota/Chlorobiota_group>Chlorobiota>Chlorobiia</td><td>GCF_000020645.1</td><td>Pelodictyon_phaeoclathratiforme</td><td>PPHA_RS10545</td><td>2206349..2209084</td><td>34090</td><td>DUF499-domain-containing-protein-[Pelodictyon-phaeoclathratiforme]</td></tr>
<tr><td>MBL7986431.1</td><td>AAA+ATPase+wHTH+Alpha+Beta-domain+wHTH+RRM</td><td>Type-3_BR.PglZ.associated.systems</td><td><-METHYLASE<-PglZ<-DUF499*<-?||TPR+TPR-></td><td>911</td><td>Bacteria>FCB_group>Bacteroidota/Chlorobiota_group>Chlorobiota>unclassified_Chlorobiota</td><td>GCA_016788565.1</td><td>Chlorobiota_bacterium</td><td>JNJ94_00015</td><td>3158..5893</td><td>2268192</td><td>DUF499-domain-containing-protein-[Chlorobi-bacterium]</td></tr>
<tr><td>MCF8267317.1</td><td>AAA+ATPase+wHTH+Alpha+Beta-domain+wHTH+RRM</td><td>Type-3_BR.PglZ.associated.systems</td><td>DUF499*->PglZ->CopG/RHH->PIN->METHYLASE->iSTAND->TUDOR+SF1-SF2-Helicase+iREase+CTD-></td><td>875</td><td>Bacteria>FCB_group>Bacteroidota/Chlorobiota_group>Ignavibacteriota>Ignavibacteria</td><td>GCA_021738695.1</td><td>Ignavibacteriales_bacterium</td><td>K9I69_04480</td><td>425..3052</td><td>2049428</td><td>hypothetical-protein-[Ignavibacteriales-bacterium]</td></tr>
<tr><td>MBK6914962.1</td><td>AAA+ATPase+wHTH+Alpha+Beta-domain+wHTH+RRM</td><td>Type-3_BR.PglZ.associated.systems</td><td>ACET->?->?->?->?->||DUF499*->PglZ->SMODS->SLATT->CopG/RHH->PIN->METHYLASE-p-></td><td>878</td><td>Bacteria>FCB_group>Bacteroidota/Chlorobiota_group>Ignavibacteriota>Ignavibacteria</td><td>GCA_016709155.1</td><td>Ignavibacteriales_bacterium</td><td>IPH11_15375</td><td>469821..472457</td><td>2049428</td><td>hypothetical-protein-[Ignavibacteriales-bacterium]</td></tr>
<tr><td>OIO18559.1</td><td>AAA+ATPase+wHTH+Alpha+Beta-domain+wHTH+RRM</td><td>Type-3_BR.PglZ.associated.systems</td><td><-iSTAND<-METHYLASE<-PIN<-CopG/RHH<-PglZ<-DUF499*||?->?->?->?->?->DUF932-></td><td>875</td><td>Bacteria>FCB_group>Bacteroidota/Chlorobiota_group>Ignavibacteriota>Ignavibacteria</td><td>GCA_001871325.1</td><td>Ignavibacteria_bacterium_CG1_02_37_35</td><td>AUJ54_07725</td><td>5254..7881</td><td>1805220</td><td>hypothetical-protein-AUJ54_07725-[Ignavibacteria-bacterium-CG1_02_37_35]</td></tr>
<tr><td>MDP2302477.1</td><td>AAA+ATPase+wHTH+Alpha+Beta-domain+wHTH+RRM</td><td>Type-3_BR.PglZ.associated.systems</td><td>DUF499+RRM*->PglZ->CopG-RHH->PIN->METHYLASE-></td><td>877</td><td>Bacteria>FCB_group>Bacteroidota/Chlorobiota_group>Ignavibacteriota>Ignavibacteria>unclassified_Ignavibacteria</td><td>GCA_030684555.1</td><td>Ignavibacteria_bacterium</td><td>Q8N03_08640</td><td>475..3108</td><td>2053306</td><td>hypothetical-protein-[Ignavibacteria-bacterium]</td></tr>
<tr><td>TET76899.1</td><td>AAA+ATPase+wHTH+Alpha+Beta-domain+wHTH+RRM</td><td>Type-3_BR.PglZ.associated.systems</td><td><-TUDOR+SF1-SF2-Helicase+iREase+CTD<-iSTAND<-METHYLASE<-PglZ<-DUF499*</td><td>538</td><td>Bacteria>FCB_group>Candidatus_Cloacimonadota>unclassified_Candidatus_Cloacimonadota</td><td>GCA_004375965.1</td><td>Candidatus_Cloacimonadota_bacterium</td><td>E3J41_08420</td><td>4840..6456</td><td>2030808</td><td>MAG:-hypothetical-protein-E3J41_08420-[Candidatus-Cloacimonadota-bacterium]</td></tr>
<tr><td>HOF41518.1</td><td>AAA+ATPase+wHTH+Alpha+Beta-domain+wHTH+RRM</td><td>Type-3_BR.PglZ.associated.systems</td><td><-METHYLASE-p<-PglZ<-DUF499+RRM*<-XIS-HTH</td><td>921</td><td>Bacteria>FCB_group>Candidatus_Hydrogenedentota>unclassified_Candidatus_Hydrogenedentota</td><td>GCA_035367355.1</td><td>Candidatus_Hydrogenedentota_bacterium</td><td>PLD73_15720</td><td>1326..4091</td><td>2030809</td><td>DUF499-domain-containing-protein-[Candidatus-Hydrogenedentota-bacterium]</td></tr>
<tr><td>RKY62550.1</td><td>AAA+ATPase+wHTH+Alpha+Beta-domain+wHTH+RRM</td><td>Type-3_BR.PglZ.associated.systems</td><td>METHYLASE-p->?->Radical-SAM->iSTAND->DUF499*-></td><td>462</td><td>Bacteria>FCB_group>Candidatus_Latescibacteria>unclassified_Candidatus_Latescibacteria</td><td>GCA_003644635.1</td><td>Candidatus_Latescibacteria_bacterium</td><td>DRP95_01360</td><td>4105..5489</td><td>2053570</td><td>hypothetical-protein-DRP95_01360,-partial-[Candidatus-Latescibacteria-bacterium]</td></tr>
<tr><td>MBI2504357.1</td><td>AAA+ATPase+wHTH+Alpha+Beta-domain+wHTH+RRM</td><td>Type-3_BR.PglZ.associated.systems</td><td><-Peptidase_M55<-?||DUF499*->PglZ->PolB->METHYLASE-p-></td><td>706</td><td>Bacteria>FCB_group>Candidatus_Latescibacteria>unclassified_Candidatus_Latescibacteria</td><td>GCA_016187725.1</td><td>Candidatus_Latescibacteria_bacterium</td><td>HYW07_14125</td><td>11765..13885</td><td>2053570</td><td>hypothetical-protein-[Candidatus-Latescibacteria-bacterium]</td></tr>
<tr><td>HDN59916.1</td><td>AAA+ATPase+wHTH+Alpha+Beta-domain+wHTH+RRM</td><td>Type-3_BR.PglZ.associated.systems</td><td>DUF499*->PglZ->METHYLASE-p-></td><td>999</td><td>Bacteria>FCB_group>Candidatus_Marinimicrobia>unclassified_Candidatus_Marinimicrobia</td><td>GCA_011043495.1</td><td>Candidatus_Marinimicrobia_bacterium</td><td>ENF20_08155</td><td>340..3339</td><td>2026760</td><td>DUF499-domain-containing-protein-[Candidatus-Marinimicrobia-bacterium]</td></tr>
<tr><td>MCD6099024.1</td><td>AAA+ATPase+wHTH+Alpha+Beta-domain+wHTH+RRM</td><td>Type-3_BR.PglZ.associated.systems</td><td><-PglZ<-DUF499*<-XIS-HTH</td><td>809</td><td>Bacteria>FCB_group>Candidatus_Marinimicrobia>unclassified_Candidatus_Marinimicrobia</td><td>GCA_021158485.1</td><td>Candidatus_Marinimicrobia_bacterium</td><td>J7K33_00415</td><td>1042..3471</td><td>2026760</td><td>hypothetical-protein-[Candidatus-Marinimicrobia-bacterium]</td></tr>
<tr><td>HLV33113.1</td><td>AAA+ATPase+wHTH+Alpha+Beta-domain+wHTH+RRM</td><td>Type-3_BR.PglZ.associated.systems</td><td><-SbcC<-PglZ<-DUF499+RRM*</td><td>930</td><td>Bacteria>FCB_group>Fibrobacterota>Chitinispirillia>Chitinispirillales>Chitinispirillaceae</td><td>GCA_035306725.1</td><td>Chitinispirillaceae_bacterium</td><td>VKY57_16215</td><td>1459..4251</td><td>2813024</td><td>hypothetical-protein-[Chitinispirillaceae-bacterium]</td></tr>
<tr><td>HLP43276.1</td><td>AAA+ATPase+wHTH+Alpha+Beta-domain+wHTH+RRM</td><td>Type-3_BR.PglZ.associated.systems</td><td>DUF499+RRM*->PglZ-></td><td>781</td><td>Bacteria>FCB_group>Fibrobacterota>Fibrobacteria>unclassified_Fibrobacteria</td><td>GCA_035299185.1</td><td>Fibrobacteria_bacterium</td><td>VK465_17360</td><td>3220..5565</td><td>2725311</td><td>DUF499-domain-containing-protein-[Fibrobacteria-bacterium]</td></tr>
<tr><td>HEX8359295.1</td><td>AAA+ATPase+wHTH+Alpha+Beta-domain+wHTH+RRM</td><td>Type-3_BR.PglZ.associated.systems</td><td>DUF499+RRM*->PglZ->METHYLASE->iSTAND->||TUDOR+SF1-SF2-Helicase+iREase+CTD-></td><td>907</td><td>Bacteria>FCB_group>Gemmatimonadota>Longimicrobiia>Longimicrobiales>Longimicrobiaceae</td><td>GCA_036388335.1</td><td>Longimicrobium_sp.</td><td>VF613_04280</td><td>493..3216</td><td>2029185</td><td>hypothetical-protein-[Longimicrobium-sp.]</td></tr>
<tr><td>WP_309896349.1</td><td>AAA+ATPase+wHTH+Alpha+Beta-domain+wHTH+RRM</td><td>Type-3_BR.PglZ.associated.systems</td><td><-TUDOR+SF1-SF2-Helicase+iREase+CTD<-iSTAND<-METHYLASE<-PglZ<-DUF499+RRM*</td><td>902</td><td>Bacteria>Myxococcota>Myxococcia>Myxococcales>Cystobacterineae>Archangiaceae</td><td>GCF_031412795.1</td><td>Archangium_sp.</td><td>RMO48_RS40080</td><td>50428..53136</td><td>1872627</td><td>DUF499-domain-containing-protein-[Archangium-sp.]</td></tr>
<tr><td>MBK8217908.1</td><td>AAA+ATPase+wHTH+Alpha+Beta-domain+wHTH+RRM</td><td>Type-3_BR.PglZ.associated.systems</td><td>GDResolvase+Recombinase+ZnR->ParB->?->?->XIS-HTH->DUF499*->PglZ->METHYLASE->iSTAND->TUDOR+SF1-SF2-Helicase+iREase+CTD->HHH_4+SF1-SF2-Helicase+REase->REase+SAVED-></td><td>912</td><td>Bacteria>Myxococcota>Myxococcia>Myxococcales>unclassified_Myxococcales</td><td>GCA_016712525.1</td><td>Myxococcales_bacterium</td><td>IPK71_29630</td><td>19752..22490</td><td>2026763</td><td>DUF499-domain-containing-protein-[Myxococcales-bacterium]</td></tr>
<tr><td>MBK8999613.1</td><td>AAA+ATPase+wHTH+Alpha+Beta-domain+wHTH+RRM</td><td>Type-3_BR.PglZ.associated.systems</td><td><-TUDOR+SF1-SF2-Helicase+iREase+CTD<-?<-iSTAND<-METHYLASE<-PglZ<-DUF499*<-?<-HTH||TPR->?->?-><-Phage_int_SAM_1+Phage_integrase</td><td>929</td><td>Bacteria>Myxococcota>Myxococcia>Myxococcales>unclassified_Myxococcales</td><td>GCA_016716175.1</td><td>Myxococcales_bacterium</td><td>IPM35_28170</td><td>15207..17996</td><td>2026763</td><td>DUF499-domain-containing-protein-[Myxococcales-bacterium]</td></tr>
<tr><td>MBK7078211.1</td><td>AAA+ATPase+wHTH+Alpha+Beta-domain+wHTH+RRM</td><td>Type-3_BR.PglZ.associated.systems</td><td><-Tox-RelA||DUF499*->PglZ->METHYLASE->iSTAND->TUDOR+SF1-SF2-Helicase+iREase+CTD->MORC+SF1-SF2-Helicase->HHH_4+SF1-SF2-Helicase+REase-></td><td>1397</td><td>Bacteria>Myxococcota>Myxococcia>Myxococcales>unclassified_Myxococcales</td><td>GCA_016708565.1</td><td>Myxococcales_bacterium</td><td>IPH44_38690</td><td>590888..595081</td><td>2026763</td><td>AAA-family-ATPase-[Myxococcales-bacterium]</td></tr>
<tr><td>MCA9629693.1</td><td>AAA+ATPase+wHTH+Alpha+Beta-domain+wHTH+RRM</td><td>Type-3_BR.PglZ.associated.systems</td><td><-METHYLASE<-PglZ<-DUF499*<-XIS-HTH</td><td>918</td><td>Bacteria>Myxococcota>Myxococcia>Myxococcales>unclassified_Myxococcales</td><td>GCA_020430865.1</td><td>Myxococcales_bacterium</td><td>KC766_18590</td><td>3656..6412</td><td>2026763</td><td>hypothetical-protein-[Myxococcales-bacterium]</td></tr>
<tr><td>MBI5511445.1</td><td>AAA+ATPase+wHTH+Alpha+Beta-domain+wHTH+RRM</td><td>Type-3_BR.PglZ.associated.systems</td><td>XIS-HTH->Phage_int_SAM_1+Phage_integrase-><-?<-Phage_int_SAM_1+Phage_integrase||DUF499*->PglZ->METHYLASE->iSTAND->TUDOR+SF1-SF2-Helicase+iREase+CTD-><-SF1-SF2-Helicase+DUF1998<-REase</td><td>896</td><td>Bacteria>Myxococcota>Myxococcia>unclassified_Myxococcia</td><td>GCA_016218825.1</td><td>Deltaproteobacteria_bacterium</td><td>HY903_22020</td><td>61793..64483</td><td>2026735</td><td>DUF499-domain-containing-protein-[Deltaproteobacteria-bacterium]</td></tr>
<tr><td>MBW1935297.1</td><td>AAA+ATPase+wHTH+Alpha+Beta-domain+wHTH+RRM</td><td>Type-3_BR.PglZ.associated.systems</td><td>XIS-HTH->DUF499*->?->PglZ->Schlafen+HATPase+HTH->METHYLASE->iSTAND->TUDOR+SF1-SF2-Helicase+iREase+CTD-></td><td>780</td><td>Bacteria>Myxococcota>Myxococcia>unclassified_Myxococcia</td><td>GCA_019307915.1</td><td>Deltaproteobacteria_bacterium</td><td>JRI84_07075</td><td>1782..4124</td><td>2026735</td><td>hypothetical-protein-[Deltaproteobacteria-bacterium]</td></tr>
<tr><td>MBW2329469.1</td><td>AAA+ATPase+wHTH+Alpha+Beta-domain+wHTH+RRM</td><td>Type-3_BR.PglZ.associated.systems</td><td>PolB->HEPN->ACET->?->RelB->TIR->DUF499*->PglZ->METHYLASE->?->?->REase+PNTD1+PNTD2->DUF433-></td><td>882</td><td>Bacteria>Myxococcota>Myxococcia>unclassified_Myxococcia</td><td>GCA_019310695.1</td><td>Deltaproteobacteria_bacterium</td><td>JRF30_00695</td><td>6296..8944</td><td>2026735</td><td>AAA-family-ATPase-[Deltaproteobacteria-bacterium]</td></tr>
<tr><td>MBW1717201.1</td><td>AAA+ATPase+wHTH+Alpha+Beta-domain+wHTH+RRM</td><td>Type-3_BR.PglZ.associated.systems</td><td>DUF499*->Schlafen+HATPase+HTH->METHYLASE-p-></td><td>457</td><td>Bacteria>Myxococcota>Myxococcia>unclassified_Myxococcia</td><td>GCA_019306255.1</td><td>Deltaproteobacteria_bacterium</td><td>JRJ77_15495</td><td>1..1374</td><td>2026735</td><td>hypothetical-protein-[Deltaproteobacteria-bacterium]</td></tr>
<tr><td>MBM4322257.1</td><td>AAA+ATPase+wHTH+Alpha+Beta-domain+wHTH+RRM</td><td>Type-3_BR.PglZ.associated.systems</td><td>DUF499*->PglZ->PIN->METHYLASE->iSTAND->TUDOR+SF1-SF2-Helicase+iREase+CTD-></td><td>825</td><td>Bacteria>Myxococcota>Myxococcia>unclassified_Myxococcia</td><td>GCA_016875155.1</td><td>Deltaproteobacteria_bacterium</td><td>FJ115_01740</td><td>605..3082</td><td>2026735</td><td>hypothetical-protein-[Deltaproteobacteria-bacterium]</td></tr>
<tr><td>MBW1708134.1</td><td>AAA+ATPase+wHTH+Alpha+Beta-domain+wHTH+RRM</td><td>Type-3_BR.PglZ.associated.systems</td><td>DUF499*->PglZ->METHYLASE-p-></td><td>498</td><td>Bacteria>Myxococcota>Myxococcia>unclassified_Myxococcia</td><td>GCA_019306165.1</td><td>Deltaproteobacteria_bacterium</td><td>JRJ86_23765</td><td>1..1499</td><td>2026735</td><td>hypothetical-protein-[Deltaproteobacteria-bacterium]</td></tr>
<tr><td>MBW1716465.1</td><td>AAA+ATPase+wHTH+Alpha+Beta-domain+wHTH+RRM</td><td>Type-3_BR.PglZ.associated.systems</td><td>DUF499*->PglZ->METHYLASE-><-DDE_Tnp_1<-Peptidase</td><td>882</td><td>Bacteria>Myxococcota>Myxococcia>unclassified_Myxococcia</td><td>GCA_019306255.1</td><td>Deltaproteobacteria_bacterium</td><td>JRJ77_11700</td><td>173..2821</td><td>2026735</td><td>AAA-family-ATPase-[Deltaproteobacteria-bacterium]</td></tr>
<tr><td>MBW1895891.1</td><td>AAA+ATPase+wHTH+Alpha+Beta-domain+wHTH+RRM</td><td>Type-3_BR.PglZ.associated.systems</td><td>DUF499*->PglZ->CopG/RHH->PIN->SF1-SF2-Helicase-></td><td>620</td><td>Bacteria>Myxococcota>Myxococcia>unclassified_Myxococcia</td><td>GCA_019307715.1</td><td>Deltaproteobacteria_bacterium</td><td>JRI47_02425</td><td>1..1865</td><td>2026735</td><td>hypothetical-protein-[Deltaproteobacteria-bacterium]</td></tr>
<tr><td>MBW1706453.1</td><td>AAA+ATPase+wHTH+Alpha+Beta-domain+wHTH+RRM</td><td>Type-3_BR.PglZ.associated.systems</td><td>DUF499*->PglZ->CopG/RHH->PIN->RelB->RelE->PolB-></td><td>844</td><td>Bacteria>Myxococcota>Myxococcia>unclassified_Myxococcia</td><td>GCA_019306165.1</td><td>Deltaproteobacteria_bacterium</td><td>JRJ86_14995</td><td>652..3186</td><td>2026735</td><td>hypothetical-protein-[Deltaproteobacteria-bacterium]</td></tr>
<tr><td>MBW2117192.1</td><td>AAA+ATPase+wHTH+Alpha+Beta-domain+wHTH+RRM</td><td>Type-3_BR.PglZ.associated.systems</td><td>DUF499*->PglZ->CopG/RHH->PIN->RelB->RelE->PolB-></td><td>844</td><td>Bacteria>Myxococcota>Myxococcia>unclassified_Myxococcia</td><td>GCA_019309305.1</td><td>Deltaproteobacteria_bacterium</td><td>JRH09_04635</td><td>652..3186</td><td>2026735</td><td>hypothetical-protein-[Deltaproteobacteria-bacterium]</td></tr>
<tr><td>MBL8614386.1</td><td>AAA+ATPase+wHTH+Alpha+Beta-domain+wHTH+RRM</td><td>Type-3_BR.PglZ.associated.systems</td><td>DUF499*->PglZ->?->SF1-SF2-Helicase+DUF1998-><-NACHT</td><td>1347</td><td>Bacteria>Myxococcota>Myxococcia>unclassified_Myxococcia</td><td>GCA_016792625.1</td><td>Deltaproteobacteria_bacterium</td><td>JNM72_02140</td><td>283..4326</td><td>2026735</td><td>hypothetical-protein-[Deltaproteobacteria-bacterium]</td></tr>
<tr><td>MBW1697962.1</td><td>AAA+ATPase+wHTH+Alpha+Beta-domain+wHTH+RRM</td><td>Type-3_BR.PglZ.associated.systems</td><td>ABC-ATPase+SbcC->?->?->?->DUF499*->PglZ->METHYLASE-></td><td>462</td><td>Bacteria>Myxococcota>Myxococcia>unclassified_Myxococcia</td><td>GCA_019306465.1</td><td>Deltaproteobacteria_bacterium</td><td>JRK26_14315</td><td>44501..45889</td><td>2026735</td><td>hypothetical-protein-[Deltaproteobacteria-bacterium]</td></tr>
<tr><td>MBM4329656.1</td><td>AAA+ATPase+wHTH+Alpha+Beta-domain+wHTH+RRM</td><td>Type-3_BR.PglZ.associated.systems</td><td><-TUDOR+SF1-SF2-Helicase+iREase+CTD<-METHYLASE-p<-?<-METHYLASE<-DOC+HTH<-PglZ<-DUF499*</td><td>883</td><td>Bacteria>Myxococcota>Myxococcia>unclassified_Myxococcia</td><td>GCA_016875165.1</td><td>Deltaproteobacteria_bacterium</td><td>FJ117_00285</td><td>60971..63622</td><td>2026735</td><td>hypothetical-protein-[Deltaproteobacteria-bacterium]</td></tr>
</tbody>
</table>

| Protein-ID | DUF499-architecture | System | Gene-Neighbourhoods | Length | Taxonomy | Assembly-ID | Strain | Locus-Tag | Genomic-coordinates | TaxID | Fasta-defline |
|---|---|---|---|---|---|---|---|---|---|---|---|
| MCX5862619.1 | AAA+ATPase+wHTH+Alpha+Beta-domain+wHTH+RRM | Type-3_BR.PglZ.associated.systems | <-TUDOR+SF1-SF2-Helicase+iREase+CTD<-iSTAND<-METHYLASE<-REase<-OLD-ABC<-PglZ<-DUF499* | 922 | Bacteria>Myxococcota>Myxococcia>unclassified_Myxococcia | GCA_026388625.1 | Deltaproteobacteria_bacterium | NTW27_10950 | 92147..94915 | 2026735 | MAG:-DUF499-domain-containing-protein-[Deltaproteobacteria-bacterium] |
| MBI4488888.1 | AAA+ATPase+wHTH+Alpha+Beta-domain+wHTH+RRM | Type-3_BR.PglZ.associated.systems | <-TUDOR+SF1-SF2-Helicase+iREase+CTD<-iSTAND<-METHYLASE<-PIN<-CopG/RHH<-PglZ<-DUF499*<-URI-DNase+HhH-RADC\|\|?->ABhydrolase-> | 867 | Bacteria>Myxococcota>Myxococcia>unclassified_Myxococcia | GCA_016210105.1 | Deltaproteobacteria_bacterium | HY694_07360 | 23521..26124 | 2026735 | hypothetical-protein-[Deltaproteobacteria-bacterium] |
| MBI4523256.1 | AAA+ATPase+wHTH+Alpha+Beta-domain+wHTH+RRM | Type-3_BR.PglZ.associated.systems | <-REase<-TUDOR+SF1-SF2-Helicase+iREase+CTD<-METHYLASE-p<-PIN<-CopG/RHH<-PglZ<-DUF499*\|\|?-><-PIN | 793 | Bacteria>Myxococcota>Myxococcia>unclassified_Myxococcia | GCA_016210115.1 | Deltaproteobacteria_bacterium | HY695_05515 | 11284..13665 | 2026735 | hypothetical-protein-[Deltaproteobacteria-bacterium] |
| MBW2342104.1 | AAA+ATPase+wHTH+Alpha+Beta-domain+wHTH+RRM | Type-3_BR.PglZ.associated.systems | <-METHYLASE-p<-Schlafen+HATPase+HTH<-DUF499* | 616 | Bacteria>Myxococcota>Myxococcia>unclassified_Myxococcia | GCA_019310725.1 | Deltaproteobacteria_bacterium | JRF50_17480 | 2066..3918 | 2026735 | hypothetical-protein-[Deltaproteobacteria-bacterium] |
| MBW2737991.1 | AAA+ATPase+wHTH+Alpha+Beta-domain+wHTH+RRM | Type-3_BR.PglZ.associated.systems | <-METHYLASE<-PglZ<-DUF499* | 535 | Bacteria>Myxococcota>Myxococcia>unclassified_Myxococcia | GCA_019314485.1 | Deltaproteobacteria_bacterium | JRE64_03890 | 3447..5054 | 2026735 | hypothetical-protein-[Deltaproteobacteria-bacterium] |
| MBP1716773.1 | AAA+ATPase+wHTH+Alpha+Beta-domain+wHTH+RRM | Type-3_BR.PglZ.associated.systems | <-iSTAND<-?<-METHYLASE<-PIN<-CopG/RHH<-PglZ<-DUF499* | 807 | Bacteria>Myxococcota>Myxococcia>unclassified_Myxococcia | GCA_017860815.1 | Deltaproteobacteria_bacterium | H6Q43_211 | 7959..10382 | 2026735 | ATPase-superfamily-like-protein-[Deltaproteobacteria-bacterium] |
| MBI1949370.1 | AAA+ATPase+wHTH+Alpha+Beta-domain+wHTH+RRM | Type-3_BR.PglZ.associated.systems | <-HHH_4+SF1-SF2-Helicase+REase<-TUDOR+SF1-SF2-Helicase+iREase+CTD<-iSTAND<-METHYLASE<-PglZ<-DUF499*<-XIS-HTH<-?<-DUF488-1<-XIS-HTH<-ParB<-GDResolvase+Recombinase+ZnR | 912 | Bacteria>Myxococcota>Myxococcia>unclassified_Myxococcia | GCA_016177705.1 | Deltaproteobacteria_bacterium | HYS27_27030 | 302678..305416 | 2026735 | DUF499-domain-containing-protein-[Deltaproteobacteria-bacterium] |
| MBN1842856.1 | AAA+ATPase+wHTH+Alpha+Beta-domain+wHTH+RRM | Type-3_BR.PglZ.associated.systems | <-HEPN<-PolB<-PIN<-CopG/RHH<-PglZ<-DUF499*<-?\|\|?->?-><-?<-?\|\|DSBH-> | 872 | Bacteria>Myxococcota>Myxococcia>unclassified_Myxococcia | GCA_016930875.1 | Deltaproteobacteria_bacterium | JW883_11325 | 21739..24357 | 2026735 | hypothetical-protein-[Deltaproteobacteria-bacterium] |
| MBN2034137.1 | AAA+ATPase+wHTH+Alpha+Beta-domain+wHTH+RRM | Type-3_BR.PglZ.associated.systems | <-PglZ<-DUF499+RRM*\|\|<-PIN | 859 | Bacteria>Myxococcota>Myxococcia>unclassified_Myxococcia | GCA_016932115.1 | Deltaproteobacteria_bacterium | JW836_12740 | 1133..3712 | 2026735 | hypothetical-protein-[Deltaproteobacteria-bacterium] |
| MDO8527161.1 | AAA+ATPase+wHTH+Alpha+Beta-domain+wHTH+RRM | Type-3_BR.PglZ.associated.systems | <-TUDOR+SF1-SF2-Helicase+iREase+CTD<-iSTAND<-METHYLASE<-PglZ<-DUF499+RRM*\|\|<-RHH | 896 | Bacteria>Myxococcota>Myxococcia>unclassified_Myxococcia | GCA_030648035.1 | Deltaproteobacteria_bacterium | Q7T03_05675 | 23313..26003 | 2026735 | DUF499-domain-containing-protein-[Deltaproteobacteria-bacterium] |
| MDR1607839.1 | AAA+ATPase+wHTH+Alpha+Beta-domain+wHTH+RRM | Type-3_BR.PglZ.associated.systems | DUF499+RRM*->PglZ->METHYLASE->iSTAND->TUDOR+SF1-SF2-Helicase+iREase+CTD-> | 923 | Bacteria>Myxococcota>Myxococcia>unclassified_Myxococcia | GCA_031270705.1 | Deltaproteobacteria_bacterium | LBT38_05460 | 37700..40471 | 2026735 | hypothetical-protein-[Deltaproteobacteria-bacterium] |
| MBX3129295.1 | AAA+ATPase+wHTH+Alpha+Beta-domain+wHTH+RRM | Type-3_BR.PglZ.associated.systems | XIS-HTH->TUDOR+SF1-SF2-Helicase+iREase+CTD+REase->METHYLASE->BirA-HTH->METHYLASE->REase->DUF499*-><-?<-ACET<-DNA-glycosylase<-?<-PglZ<-SF1-SF2-Helicase+iREase+CTD+REase-p | 936 | Bacteria>Myxococcota>Polyangia>Polyangiales>Polyangiaceae | GCA_019637295.1 | Polyangiaceae_bacterium | KF718_21435 | 16411..19221 | 2268199 | ATP-binding-protein-[Polyangiaceae-bacterium] |
| KYF71594.1 | AAA+ATPase+wHTH+Alpha+Beta-domain+wHTH+RRM | Type-3_BR.PglZ.associated.systems | ABC-ATPase+SbcC->N6_N4_MTase->DUF499*->PglZ->METHYLASE->iSTAND->TUDOR+SF1-SF2-Helicase+iREase+CTD-> | 907 | Bacteria>Myxococcota>Polyangia>Polyangiales>Polyangiaceae | GCA_001589185.1 | Sorangium_cellulosum | BE17_05970 | 3194..5917 | 56 | hypothetical-protein-BE17_05970-[Sorangium-cellulosum] |
| WP_104982305.1 | AAA+ATPase+wHTH+Alpha+Beta-domain+wHTH+RRM | Type-3_BR.PglZ.associated.systems | <-SF1-SF2-Helicase+DUF1998<-?<-PglZ<-DUF499*<-METHYLASE-PglX | 1270 | Bacteria>Myxococcota>Polyangia>Polyangiales>Polyangiaceae | GCF_002950945.1 | Sorangium_cellulosum | SOCE26_RS25435 | 7076861..7080673 | 56 | hypothetical-protein-[Sorangium-cellulosum] |
| HTN83164.1 | AAA+ATPase+wHTH+Alpha+Beta-domain+wHTH+RRM | Type-3_BR.PglZ.associated.systems | METHYLASE-p->DUF499+RRM*->PglZ-> | 908 | Bacteria>Myxococcota>Polyangia>Polyangiales>Polyangiaceae>Sorangium | GCA_035497415.1 | Sorangium_sp. | VL242_05750 | 932..3658 | 2093377 | hypothetical-protein-[Sorangium-sp.] |
| MBK8173470.1 | AAA+ATPase+wHTH+Alpha+Beta-domain+wHTH+RRM | Type-3_BR.PglZ.associated.systems | DUF499*->PglZ->iSTAND->TUDOR+SF1-SF2-Helicase+iREase+CTD-><-REase+35exo | 909 | Bacteria>Myxococcota>Polyangia>Polyangiales>Sandaracinaceae | GCA_016712745.1 | Sandaracinaceae_bacterium | IPK60_24475 | 716247..718976 | 2510337 | DUF499-domain-containing-protein-[Sandaracinaceae-bacterium] |
| OGW17481.1 | AAA+ATPase+wHTH+Alpha+Beta-domain+wHTH+RRM | Type-3_BR.PglZ.associated.systems | <-Schlafen+HATPase+HTH<-PglZ<-DUF499*<-XIS-HTH | 872 | Bacteria>Nitrospinae/Tectomicrobia_group>Nitrospinota>unclassified_Nitrospinota | GCA_001803565.1 | Nitrospinae_bacterium_RIFCSPLOWO2_12_FULL_45_22 | A3G93_06415 | 4599..7217 | 1801687 | hypothetical-protein-A3G93_06415-[Nitrospinae-bacterium-RIFCSPLOWO2_12_FULL_45_22] |
| MBX9659010.1 | AAA+ATPase+wHTH+Alpha+Beta-domain+wHTH+RRM | Type-3_BR.PglZ.associated.systems | SbcC+SbcC->HTH+WYL->DUF499*->PglZ->METHYLASE->iSTAND->TUDOR+SF1-SF2-Helicase+iREase+CTD-> | 914 | Bacteria>Nitrospirota>Nitrospiria>Nitrospirales>Nitrospiraceae | GCA_019748255.1 | Nitrospiraceae_bacterium | K2X00_10620 | 2595..5339 | 2026770 | DUF499-domain-containing-protein-[Nitrospiraceae-bacterium] |
| MBP8199110.1 | AAA+ATPase+wHTH+Alpha+Beta-domain+wHTH+RRM | Type-3_BR.PglZ.associated.systems | HTH+WYL->DUF499*->PglZ->METHYLASE->iSTAND->TUDOR+SF1-SF2-Helicase+iREase+CTD->MORC+HisKin-> | 897 | Bacteria>Nitrospirota>Nitrospiria>Nitrospirales>Nitrospiraceae | GCA_018006455.1 | Nitrospira_sp. | KAX98_02165 | 16781..19474 | 70125 | hypothetical-protein-[Nitrospira-sp.] |
| KJR40413.1 | AAA+ATPase+wHTH+Alpha+Beta-domain+wHTH+RRM | Type-3_BR.PglZ.associated.systems | <-Transposase_31+Yae1_N<-Transposase_31+Yae1_N<-TUDOR+SF1-SF2-Helicase+iREase+CTD<-iSTAND<-METHYLASE<-PglZ<-DUF499* | 877 | Bacteria>Nitrospirota>Nitrospiria>Nitrospirales>Nitrospiraceae | GCA_000960725.1 | Candidatus_Magnetoovum_chiemensis | MCHI_003705 | 16876..19509 | 1609970 | ATPase-[Candidatus-Magnetoovum-chiemensis] |
| MBA5868094.1 | AAA+ATPase+wHTH+Alpha+Beta-domain+wHTH+RRM | Type-3_BR.PglZ.associated.systems | <-REase\|\|?->?->?->DUF499*->PglZ->METHYLASE->iSTAND->TUDOR+SF1-SF2-Helicase+iREase+CTD-><-SbcC+SbcC | 930 | Bacteria>Nitrospirota>Nitrospiria>Nitrospirales>Nitrospiraceae | GCA_014055525.1 | Nitrospira_sp._CR1.3 | GDA67_15485 | 24238..27030 | 2656758 | DUF499-domain-containing-protein-[Nitrospira-sp.-CR1.3] |
| MDR4485159.1 | AAA+ATPase+wHTH+Alpha+Beta-domain+wHTH+RRM | Type-3_BR.PglZ.associated.systems | DUF499+RRM*->PglZ->METHYLASE->iSTAND->TUDOR+SF1-SF2-Helicase+iREase+CTD-> | 894 | Bacteria>Nitrospirota>Nitrospiria>Nitrospirales>unclassified_Nitrospirales | GCA_031315875.1 | Nitrospirales_bacterium | MRJ94_18240 | 874..3558 | 2358460 | hypothetical-protein-[Nitrospirales-bacterium] |
| MDR4496269.1 | AAA+ATPase+wHTH+Alpha+Beta-domain+wHTH+RRM | Type-3_BR.PglZ.associated.systems | TUDOR+SF1-SF2-Helicase+iREase+CTD->\|\|DUF499+RRM*->PglZ->CopG-RHH->PIN->Schlafen+HATPase+wHTH-> | 821 | Bacteria>Nitrospirota>Nitrospiria>Nitrospirales>unclassified_Nitrospirales | GCA_031315915.1 | Nitrospirales_bacterium | MRJ99_17830 | 3988..6453 | 2358460 | hypothetical-protein-[Nitrospirales-bacterium] |
| OHE58167.1 | AAA+ATPase+wHTH+Alpha+Beta-domain+wHTH+RRM | Type-3_BR.PglZ.associated.systems | <-TUDOR+SF1-SF2-Helicase+iREase+CTD<-iSTAND<-?<-METHYLASE<-PglZ<-DUF499<-DUF499*\|\|METHYLASE->?->?->ParB+HNH-p-> | 214 | Bacteria>Nitrospirota>Thermodesulfovibrionia>Thermodesulfovibrionales>Thermodesulfovibrionaceae | GCA_001830985.1 | Thermodesulfovibrio_sp._RBG_19FT_COMBO_42_12 | A2Z47_05835 | 11195..11839 | 1802373 | MAG:-hypothetical-protein-A2Z47_05835-[Thermodesulfovibrio-sp.-RBG_19FT_COMBO_42_12] |
| OHE58166.1 | AAA+ATPase+wHTH+Alpha+Beta-domain+wHTH+RRM | Type-3_BR.PglZ.associated.systems | <-TUDOR+SF1-SF2-Helicase+iREase+CTD<-iSTAND<-?<-METHYLASE<-PglZ<-DUF499*<-DUF499\|\|METHYLASE->?->?->ParB+HNH-p-> | 621 | Bacteria>Nitrospirota>Thermodesulfovibrionia>Thermodesulfovibrionales>Thermodesulfovibrionaceae | GCA_001830985.1 | Thermodesulfovibrio_sp._RBG_19FT_COMBO_42_12 | A2Z47_05830 | 9175..11040 | 1802373 | MAG:-hypothetical-protein-A2Z47_05830-[Thermodesulfovibrio-sp.-RBG_19FT_COMBO_42_12] |
| MCC7203047.1 | AAA+ATPase+wHTH+Alpha+Beta-domain+wHTH+RRM | Type-3_BR.PglZ.associated.systems | <-TUDOR+SF1-SF2-Helicase+iREase+CTD<-?<-RelE-ParE<-?<-METHYLASE<-PglZ<-DUF499*\|\|?->ABC-ATPase+SbcC-> | 889 | Bacteria>Nitrospirota>unclassified_Nitrospirota | GCA_020851375.1 | Nitrospirota_bacterium | IT393_10370 | 196208..198877 | 2026887 | hypothetical-protein-[Nitrospirae-bacterium] |
| MCL5421046.1 | AAA+ATPase+wHTH+Alpha+Beta-domain+wHTH+RRM | Type-3_BR.PglZ.associated.systems | <-METHYLASE<-HEPN<-HEPN<-PIN<-CopG/RHH<-PglZ<-DUF499* | 877 | Bacteria>Nitrospirota>unclassified_Nitrospirota | GCA_023485285.1 | Nitrospirota_bacterium | M1461_01045 | 9759..12392 | 2026887 | hypothetical-protein-[Nitrospirae-bacterium] |
| MDH4162607.1 | AAA+ATPase+wHTH+Alpha+Beta-domain+wHTH+RRM | Type-3_BR.PglZ.associated.systems | <-PIN<-CopG-RHH<-PglZ<-DUF499+RRM*<-AAA-ATPase+REase | 882 | Bacteria>Nitrospirota>unclassified_Nitrospirota | GCA_029858565.1 | Nitrospirota_bacterium | OEW15_07945 | 49708..52356 | 2026887 | DUF499-domain-containing-protein-[Nitrospirota-bacterium] |
| HUJ16438.1 | AAA+ATPase+wHTH+Alpha+Beta-domain+wHTH+RRM | Type-3_BR.PglZ.associated.systems | Arm-DNA-bind_3+Phage_integrase->DUF499+RRM*->PglZ-> | 807 | Bacteria>Nitrospirota>unclassified_Nitrospirota | GCA_035516965.1 | Nitrospirota_bacterium | VL197_00415 | 17959..20382 | 2026887 | DUF499-domain-containing-protein-[Nitrospirota-bacterium] |
| HZX48546.1 | AAA+ATPase+wHTH+Alpha+Beta-domain+wHTH+RRM | Type-3_BR.PglZ.associated.systems | <-ParB+HNH<-METHYLASE-p\|\|DUF499+RRM*->PglZ->METHYLASE->TUDOR+SF1-SF2-Helicase+iREase+CTD-> | 665 | Bacteria>Nitrospirota>unclassified_Nitrospirota | GCA_035655335.1 | Nitrospirota_bacterium | VFF47_04945 | 7229..9226 | 2026887 | hypothetical-protein-[Nitrospirota-bacterium] |
| WP_040227130.1 | AAA+ATPase+wHTH+Alpha+Beta-domain+wHTH+RRM | Type-3_BR.PglZ.associated.systems | REase->Mu-transpos_C->TnsC-ATPase->?->?->XIS-HTH->DUF499*->PglZ->SIR2->VirB4-FtsK+HerA->METHYLASE->iSTAND->TUDOR+SF1-SF2-Helicase+iREase+CTD-> | 921 | Bacteria>Pseudomonadota | GCF_003115575.1 | Gammaproteobacteria | L1D06_RS23305 | 38708..41473 | 1236 | MULTISPECIES:-DUF499-domain-containing-protein-[Enterobacteriaceae] |
| WP_204157998.1 | AAA+ATPase+wHTH+Alpha+Beta-domain+wHTH+RRM | Type-3_BR.PglZ.associated.systems | DUF772+DDE_Tnp_1-><-TUDOR+SF1-SF2-Helicase+iREase+CTD<-iSTAND<-METHYLASE<-PglZ<-DUF499*<-HTH+WYL<-McrC-NTD+REase<-?\|\|ArdCN+MPTase->?->ParB+HTH-> | 917 | Bacteria>Pseudomonadota | GCF_016887905.1 | Alphaproteobacteria | L6Q21_RS14595 | 26503..29256 | 28211 | MULTISPECIES:-DUF499-domain-containing-protein-[Alphaproteobacteria] |
| WP_047289057.1 | AAA+ATPase+wHTH+Alpha+Beta-domain+wHTH+RRM | Type-3_BR.PglZ.associated.systems | <-VirB11+VirB4-FtsK<-CopG/RHH<-VirD4-TraG+VirD4-FtsK<-?<-MODE-HTH+PBPII\|\|XIS-HTH->DUF499*->PglZ->Classical-AAA->Peptidase_S8->METHYLASE->iSTAND->TUDOR+SF1-SF2-Helicase+iREase+CTD-> | 941 | Bacteria>Pseudomonadota | GCF_029837305.1 | Gammaproteobacteria | N7609_RS06520 | 27132..29957 | 1236 | MULTISPECIES:-DUF499-domain-containing-protein-[Gammaproteobacteria] |
| WP_180883211.1 | AAA+ATPase+wHTH+Alpha+Beta-domain+wHTH+RRM | Type-3_BR.PglZ.associated.systems | <-TUDOR+SF1-SF2-Helicase+iREase+CTD<-iSTAND<-METHYLASE<-VirB4-FtsK+HerA<-SIR2<-PglZ<-DUF499*<-XIS-HTH\|\|MODE-HTH+PBPII->?->VirD4-TraG+VirD4-FtsK->CopG/RHH->VirB11+VirB4-FtsK-> | 922 | Bacteria>Pseudomonadota | GCF_030236855.1 | Gammaproteobacteria | QRO95_RS19435 | 15633..18401 | 1236 | DUF499-domain-containing-protein-[Stenotrophomonas-maltophilia] |
| WP_257539443.1 | AAA+ATPase+wHTH+Alpha+Beta-domain+wHTH+RRM | Type-3_BR.PglZ.associated.systems | <-TUDOR+SF1-SF2-Helicase+iREase+CTD<-iSTAND<-METHYLASE<-PglZ<-DUF499*<-HTH+WYL<-McrC-NTD+REase<-?\|\|ArdCN+MPTase->?->ParB+HTH-> | 917 | Bacteria>Pseudomonadota | GCF_024611115.1 | Alphaproteobacteria | M2352_RS17095 | 502515..505268 | 28211 | MULTISPECIES:-ATP-binding-protein-[Alphaproteobacteria] |
| WP_279703502.1 | AAA+ATPase+wHTH+Alpha+Beta-domain+wHTH+RRM | Type-3_BR.PglZ.associated.systems | <-TUDOR+SF1-SF2-Helicase+iREase+CTD<-iSTAND<-METHYLASE<-Peptidase_S8<-Classical-AAA<-PglZ<-DUF499*<-XIS-HTH\|\|MODE-HTH+PBPII->?->VirD4-TraG+VirD4-FtsK->CopG/RHH->VirB11+VirB4-FtsK-> | 922 | Bacteria>Pseudomonadota | GCF_032231025.1 | Gammaproteobacteria | N5C47_RS04580 | 13342..16110 | 1236 | DUF499-domain-containing-protein-[Stenotrophomonas-maltophilia] |
| WP_046802276.1 | AAA+ATPase+wHTH+Alpha+Beta-domain+wHTH+RRM | Type-3_BR.PglZ.associated.systems | DUF4158+DDE_Tnp_Tn3-><-?<-TUDOR+SF1-SF2-Helicase+iREase+CTD<-iSTAND<-METHYLASE<-PglZ<-DUF499*<-HTH_3\|\|?->?->Phage_int_SAM_1+Phage_integrase->?-><-HTH+MPTase | 914 | Bacteria>Pseudomonadota>Alphaproteobacteria | GCF_014843375.1 | Hyphomicrobiales | YA62_RS01425 | 293757..296501 | 356 | MULTISPECIES:-DUF499-domain-containing-protein-[Hyphomicrobiales] |
| WP_242915516.1 | AAA+ATPase+wHTH+Alpha+Beta-domain+wHTH+RRM | Type-3_BR.PglZ.associated.systems | Peptidase_M20-><-DDE_Tnp_IS240<-TUDOR+SF1-SF2-Helicase+iREase+CTD<-iSTAND<-METHYLASE<-PglZ<-DUF499*<-HTH+WYL<-DDE_Tnp_IS240<-?\|\|DEDD_Tnp_IS110+Transposase_20-> | 917 | Bacteria>Pseudomonadota>Alphaproteobacteria>Caulobacterales>Caulobacteraceae | GCF_022750635.1 | Brevundimonas_pishanensis | LOX95_RS13580 | 11838..14591 | 2896315 | DUF499-domain-containing-protein-[Brevundimonas-pishanensis] |
| MDZ4061052.1 | AAA+ATPase+wHTH+Alpha+Beta-domain+wHTH+RRM | Type-3_BR.PglZ.associated.systems | DUF499+RRM*->PglZ->METHYLASE->iSTAND->TUDOR+SF1-SF2-Helicase+iREase+CTD-><-HTH_Tnp_1+HTH_21+rve+rve_3 | 911 | Bacteria>Pseudomonadota>Alphaproteobacteria>Caulobacterales>Caulobacteraceae>Brevundimonas | GCA_034366565.1 | Brevundimonas_sp. | U1C75_07700 | 37415..40150 | 1871086 | DUF499-domain-containing-protein-[Brevundimonas-sp.] |
| WP_298128243.1 | AAA+ATPase+wHTH+Alpha+Beta-domain+wHTH+RRM | Type-3_BR.PglZ.associated.systems | DUF499+RRM*->PglZ->METHYLASE->iSTAND->TUDOR+SF1-SF2-Helicase+iREase+CTD-> | 914 | Bacteria>Pseudomonadota>Alphaproteobacteria>Caulobacterales>Caulobacteraceae>Brevundimonas | GCF_027490075.1 | Brevundimonas_sp. | Q0045_RS14405 | 23621..26365 | 1871086 | DUF499-domain-containing-protein-[Brevundimonas-sp.] |
| MBN8607693.1 | AAA+ATPase+wHTH+Alpha+Beta-domain+wHTH+RRM | Type-3_BR.PglZ.associated.systems | HTH+WYL->REase+35exo+Beta-Barrel+SF1-SF2-Helicase-><-TUDOR+SF1-SF2-Helicase+iREase+CTD<-iSTAND<-METHYLASE<-PglZ<-DUF499*<-PIN<-MazE_antitoxin<-?<-ArdCN+MPTase\|\|HIN-HTH-> | 915 | Bacteria>Pseudomonadota>Alphaproteobacteria>Caulobacterales>unclassified_Caulobacterales | GCA_017303355.1 | Caulobacterales_bacterium | JOL81_12320 | 130564..133311 | 2282150 | DUF499-domain-containing-protein-[Caulobacterales-bacterium] |
| WP_206455647.1 | AAA+ATPase+wHTH+Alpha+Beta-domain+wHTH+RRM | Type-3_BR.PglZ.associated.systems | OLD-ABC+TOPRIM->SF1-SF2-Helicase-><-TUDOR+SF1-SF2-Helicase+iREase+CTD<-iSTAND<-METHYLASE<-PglZ<-DUF499*<-HTH+WYL\|\|SF1-SF2-Helicase+REase->SF1-SF2-Helicase+REase->SF1-SF2-helicase+RdgC+Beta-barrel+REase->DUF6361->HKD-DNase-> | 915 | Bacteria>Pseudomonadota>Alphaproteobacteria>Hyphomicrobiales>Aurantimonadaceae | GCF_017183135.1 | Aurantimonas_marina | ING15_RS14985 | 13646..16393 | 2780508 | DUF499-domain-containing-protein-[Aurantimonas-marina] |
| WP_286167649.1 | AAA+ATPase+wHTH+Alpha+Beta-domain+wHTH+RRM | Type-3_BR.PglZ.associated.systems | ABhydrolase->HISKIN+MORC->HTH+WYL->DUF499*->PglZ->METHYLASE->iSTAND->TUDOR+SF1-SF2-Helicase+iREase+CTD->?->HTH+MPTase-> | 949 | Bacteria>Pseudomonadota>Alphaproteobacteria>Hyphomicrobiales>Brucellaceae | GCF_003332115.1 | Ochrobactrum_sp._3-3 | A4U69_RS21825 | 57437..60286 | 1830124 | DUF499-domain-containing-protein-[Ochrobactrum-sp.-3-3] |
| RRY02343.1 | AAA+ATPase+wHTH+Alpha+Beta-domain+wHTH+RRM | Type-3_BR.PglZ.associated.systems | <-PIN+OST-HTH\|\|MobA_MobL+SF1-SF2-helicase-><-TUDOR+SF1-SF2-Helicase+iREase+CTD<-iSTAND<-METHYLASE<-PglZ<-DUF499*<-HTH_3\|\|HEPN->?->?->Phage_int_SAM_1+Phage_integrase-> | 913 | Bacteria>Pseudomonadota>Alphaproteobacteria>Hyphomicrobiales>Brucellaceae | GCA_003937425.1 | Brucella_anthropi | EGJ58_24130 | 19251..21992 | 529 | DUF499-domain-containing-protein-[Brucella-anthropi] |
| WP_235824343.1 | AAA+ATPase+wHTH+Alpha+Beta-domain+wHTH+RRM | Type-3_BR.PglZ.associated.systems | <-PIN+OST-HTH\|\|MobA_MobL+SF1-SF2-helicase-><-TUDOR+SF1-SF2-Helicase+iREase+CTD<-iSTAND<-METHYLASE<-PglZ<-DUF499*<-HTH_3\|\|HEPN->?->?->Phage_int_SAM_1+Phage_integrase-> | 813 | Bacteria>Pseudomonadota>Alphaproteobacteria>Hyphomicrobiales>Brucellaceae | GCF_003937425.1 | Brucella_anthropi | EGJ58_RS23965 | 19251..21692 | 529 | DUF499-domain-containing-protein-[Brucella-anthropi] |
| WP_321442581.1 | AAA+ATPase+wHTH+Alpha+Beta-domain+wHTH+RRM | Type-3_BR.PglZ.associated.systems | DUF499+RRM*->PglZ->METHYLASE->iSTAND->TUDOR+SF1-SF2-Helicase+iREase+CTD->GDResolvase+HIN-HTH-> | 914 | Bacteria>Pseudomonadota>Alphaproteobacteria>Hyphomicrobiales>Cohaesibacteraceae>Cohaesibacter | GCF_963677725.1 | uncultured_Cohaesibacter_sp. | U2957_RS10480 | 2457317..2460061 | 1002546 | DUF499-domain-containing-protein-[uncultured-Cohaesibacter-sp.] |
| WP_201657149.1 | AAA+ATPase+wHTH+Alpha+Beta-domain+wHTH+RRM | Type-3_BR.PglZ.associated.systems | HHH_4+SF1-SF2-Helicase+REase-><-TUDOR+SF1-SF2-Helicase+iREase+CTD<-iSTAND<-METHYLASE<-PglZ<-DUF499*<-?<-TUDOR+SF1-SF2-Helicase+iREase+CTD\|\|Recombinase+ZnR-><-tRNA<-HISKIN+MORC | 907 | Bacteria>Pseudomonadota>Alphaproteobacteria>Hyphomicrobiales>Devosiaceae | GCF_016698645.1 | Devosia_oryziradicis | JI749_RS00420 | 95377..98100 | 2801335 | DUF499-domain-containing-protein-[Devosia-oryziradicis] |
| WP_172122492.1 | AAA+ATPase+wHTH+Alpha+Beta-domain+wHTH+RRM | Type-3_BR.PglZ.associated.systems | ABhydrolase->?-><-?\|\|CopG/RHH->DUF499*->PglZ->METHYLASE->TUDOR+SF1-SF2-Helicase+iREase+CTD->\|\|?->?-><-PIN+OST-HTH | 907 | Bacteria>Pseudomonadota>Alphaproteobacteria>Hyphomicrobiales>Devosiaceae | GCF_013185965.1 | Devosia_sp._919 | HRQ64_RS05340 | 1105701..1108424 | 2726065 | DUF499-domain-containing-protein-[Devosia-sp.-919] |
| MCA3679746.1 | AAA+ATPase+wHTH+Alpha+Beta-domain+wHTH+RRM | Type-3_BR.PglZ.associated.systems | XIS-HTH->DUF499*->PglZ->METHYLASE->iSTAND->TUDOR+SF1-SF2-Helicase+iREase+CTD-><-?<-Peptidase_M15 | 903 | Bacteria>Pseudomonadota>Alphaproteobacteria>Hyphomicrobiales>Methylobacteriaceae | GCA_020253465.1 | Methylobacterium_sp. | IDB85_02240 | 1160..3871 | 409 | DUF499-domain-containing-protein-[Methylobacterium-sp.] |
| WP_012454975.1 | AAA+ATPase+wHTH+Alpha+Beta-domain+wHTH+RRM | Type-3_BR.PglZ.associated.systems | ParG_RHH-><-MODE-HTH+PBPII<-TUDOR+SF1-SF2-Helicase+iREase+CTD<-iSTAND<-METHYLASE<-PglZ<-DUF499*<-HTH+WYL<-?\|\|SF1-SF2-Helicase+REase->DUF6361-><-GDResolvase+PPP1-HTH | 914 | Bacteria>Pseudomonadota>Alphaproteobacteria>Hyphomicrobiales>Methylobacteriaceae | GCF_000019945.1 | Methylorubrum_populi | MPOP_RS15440 | 3329238..3331982 | 223967 | DUF499-domain-containing-protein-[Methylorubrum-populi] |
| WP_271520342.1 | AAA+ATPase+wHTH+Alpha+Beta-domain+wHTH+RRM | Type-3_BR.PglZ.associated.systems | HTH_3-><-?<-GDResolvase<-?\|\|DUF499*->PglZ->METHYLASE->iSTAND->TUDOR+SF1-SF2-Helicase+iREase+CTD-><-HIN-HTH | 909 | Bacteria>Pseudomonadota>Alphaproteobacteria>Hyphomicrobiales>Nitrobacteraceae | GCF_028128755.1 | Bradyrhizobium_sp._CCBAU_53380 | XH97_RS05860 | 41970..44699 | 1325117 | DUF499-domain-containing-protein-[Bradyrhizobium-sp.-CCBAU-53380] |
| WP_260345495.1 | AAA+ATPase+wHTH+Alpha+Beta-domain+wHTH+RRM | Type-3_BR.PglZ.associated.systems | Glyco_transf_20->Glyco_transf_20-><-?\|\|tRNA-><-?\|\|HTH_3-><-?\|\|DUF499*->PglZ->METHYLASE->iSTAND->TUDOR+SF1-SF2-Helicase+iREase+CTD-> | 910 | Bacteria>Pseudomonadota>Alphaproteobacteria>Hyphomicrobiales>Nitrobacteraceae | GCF_025200905.1 | Bradyrhizobium_sp._CB1024 | N2605_RS00120 | 26209..28941 | 2976823 | DUF499-domain-containing-protein-[Bradyrhizobium-sp.-CB1024] |
| WP_212428804.1 | AAA+ATPase+wHTH+Alpha+Beta-domain+wHTH+RRM | Type-3_BR.PglZ.associated.systems | DUF499*->PglZ->METHYLASE->iSTAND->SF1-SF2-Helicase+iREase+CTD->?->HTH_36-> | 909 | Bacteria>Pseudomonadota>Alphaproteobacteria>Hyphomicrobiales>Nitrobacteraceae | GCF_018130405.1 | Bradyrhizobium_ottawaense | JQ556_RS08060 | 406554..409283 | 931866 | DUF499-domain-containing-protein-[Bradyrhizobium-ottawaense] |
| WP_212491123.1 | AAA+ATPase+wHTH+Alpha+Beta-domain+wHTH+RRM | Type-3_BR.PglZ.associated.systems | <-SF1-SF2-Helicase+iREase+CTD<-iSTAND<-METHYLASE<-PglZ<-DUF499*\|\|?-><-HTH_3\|\|?-><-?\|\|?-><-XIS-HTH | 910 | Bacteria>Pseudomonadota>Alphaproteobacteria>Hyphomicrobiales>Nitrobacteraceae | GCF_018130445.1 | Bradyrhizobium | J2R95_RS40410 | 8398545..8401277 | 374 | MULTISPECIES:-DUF499-domain-containing-protein-[Bradyrhizobium] |
| QIP02981.1 | AAA+ATPase+wHTH+Alpha+Beta-domain+wHTH+RRM | Type-3_BR.PglZ.associated.systems | <-RHH_1<-?<-TUDOR+SF1-SF2-Helicase+iREase+CTD<-iSTAND<-METHYLASE<-PglZ<-DUF499* | 910 | Bacteria>Pseudomonadota>Alphaproteobacteria>Hyphomicrobiales>Nitrobacteraceae | GCA_011604645.1 | Bradyrhizobium_symbiodeficiens | HAU86_25750 | 5451333..5454065 | 1404367 | DUF499-domain-containing-protein-[Bradyrhizobium-symbiodeficiens] |
| WP_283812243.1 | AAA+ATPase+wHTH+Alpha+Beta-domain+wHTH+RRM | Type-3_BR.PglZ.associated.systems | <-RHH_1<-?<-TUDOR+SF1-SF2-Helicase+iREase+CTD<-iSTAND<-METHYLASE<-PglZ<-DUF499* | 889 | Bacteria>Pseudomonadota>Alphaproteobacteria>Hyphomicrobiales>Nitrobacteraceae | GCF_011604645.1 | Bradyrhizobium_symbiodeficiens | HAU86_RS33435 | 5451333..5454002 | 1404367 | DUF499-domain-containing-protein-[Bradyrhizobium-symbiodeficiens] |
| WP_283022678.1 | AAA+ATPase+wHTH+Alpha+Beta-domain+wHTH+RRM | Type-3_BR.PglZ.associated.systems | DUF499+RRM*->PglZ->METHYLASE->iSTAND->TUDOR+SF1-SF2-Helicase+iREase+CTD-> | 897 | Bacteria>Pseudomonadota>Alphaproteobacteria>Hyphomicrobiales>Nitrobacteraceae>Bradyrhizobium | GCF_029714325.1 | Bradyrhizobium_sp._CB1717 | QA649_RS01650 | 350857..353550 | 3039154 | DUF499-domain-containing-protein-[Bradyrhizobium-sp.-CB1717] |
| WP_304499825.1 | AAA+ATPase+wHTH+Alpha+Beta-domain+wHTH+RRM | Type-3_BR.PglZ.associated.systems | <-SF1-SF2-Helicase<-SF1-SF2-Helicase+REase<-REase\|\|WYL->DUF499+RRM*->PglZ->METHYLASE->iSTAND->TUDOR+SF1-SF2-Helicase+iREase+CTD-> | 914 | Bacteria>Pseudomonadota>Alphaproteobacteria>Hyphomicrobiales>Nitrobacteraceae>Bradyrhizobium | GCF_030584725.1 | Bradyrhizobium_elkanii | QIH83_RS18795 | 4112604..4115348 | 29448 | DUF499-domain-containing-protein-[Bradyrhizobium-elkanii] |
| WP_315722376.1 | AAA+ATPase+wHTH+Alpha+Beta-domain+wHTH+RRM | Type-3_BR.PglZ.associated.systems | <-HTH_3\|\|DUF499+RRM*->PglZ->METHYLASE->iSTAND->TUDOR+SF1-SF2-Helicase+iREase+CTD->RHH-> | 909 | Bacteria>Pseudomonadota>Alphaproteobacteria>Hyphomicrobiales>Nitrobacteraceae>Bradyrhizobium | GCF_032484555.1 | unclassified_Bradyrhizobium | QXZ82_RS12810 | 2865047..2867776 | 2631580 | MULTISPECIES:-DUF499-domain-containing-protein-[unclassified-Bradyrhizobium] |
| WP_066996950.1 | AAA+ATPase+wHTH+Alpha+Beta-domain+wHTH+RRM | Type-3_BR.PglZ.associated.systems | REase-><-TUDOR+SF1-SF2-Helicase+iREase+CTD<-iSTAND<-METHYLASE<-PglZ<-DUF499*<-REase<-ParB+HNH<-ParB+HNH\|\|REase+35exo-><-HTH+WYL | 910 | Bacteria>Pseudomonadota>Alphaproteobacteria>Hyphomicrobiales>Phyllobacteriaceae | GCF_001672445.1 | Mesorhizobium_sp._WSM3873 | A9K71_RS15155 | 40507..43239 | 1854056 | DUF499-domain-containing-protein-[Mesorhizobium-sp.-WSM3873] |
| CDX58767.1 | AAA+ATPase+wHTH+Alpha+Beta-domain+wHTH+RRM | Type-3_BR.PglZ.associated.systems | <-REase<-TUDOR+SF1-SF2-Helicase+iREase+CTD<-iSTAND<-METHYLASE<-PglZ<-DUF499*<-?<-REase\|\|?-><-ParB+HNH<-ParB+HNH\|\|REase+35exo-> | 910 | Bacteria>Pseudomonadota>Alphaproteobacteria>Hyphomicrobiales>Phyllobacteriaceae | GCA_000824765.1 | Mesorhizobium_plurifarium | MPL3365_30298 | 299651..302383 | 69974 | conserved-hypothetical-protein-[Mesorhizobium-plurifarium] |
| WP_245517160.1 | AAA+ATPase+wHTH+Alpha+Beta-domain+wHTH+RRM | Type-3_BR.PglZ.associated.systems | <-N6_MTase<-?<-TUDOR+SF1-SF2-Helicase+iREase+CTD<-iSTAND<-METHYLASE<-PglZ<-DUF499*<-?<-URI-DNase<-?<-PemK_toxin<-?\|\|Phage_integrase-> | 917 | Bacteria>Pseudomonadota>Alphaproteobacteria>Hyphomicrobiales>Phyllobacteriaceae | GCF_004022625.2 | unclassified_Mesorhizobium | EN838_RS23485 | 18883..21636 | 325217 | MULTISPECIES:-DUF499-domain-containing-protein-[unclassified-Mesorhizobium] |
| TGQ51611.1 | AAA+ATPase+wHTH+Alpha+Beta-domain+wHTH+RRM | Type-3_BR.PglZ.associated.systems | <-N6_MTase<-?<-TUDOR+SF1-SF2-Helicase+iREase+CTD<-iSTAND<-METHYLASE<-PglZ<-DUF499*<-?<-URI-DNase<-?<-PemK_toxin\|\|Phage_integrase-> | 914 | Bacteria>Pseudomonadota>Alphaproteobacteria>Hyphomicrobiales>Phyllobacteriaceae | GCA_004791585.1 | Mesorhizobium_sp._M1C.F.Ca.ET.210.01.1.1 | EN853_23480 | 18883..21627 | 2563930 | DUF499-domain-containing-protein-[Mesorhizobium-sp.-M1C.F.Ca.ET.210.01.1.1] |
| WP_252913873.1 | AAA+ATPase+wHTH+Alpha+Beta-domain+wHTH+RRM | Type-3_BR.PglZ.associated.systems | <-HIN-HTH<-TUDOR+SF1-SF2-Helicase+iREase+CTD<-iSTAND<-METHYLASE<-PglZ<-DUF499*<-HTH+WYL<-McrC-NTD+REase<-?\|\|ArdCN+MPTase->?->ParB+HTH-> | 917 | Bacteria>Pseudomonadota>Alphaproteobacteria>Hyphomicrobiales>Phyllobacteriaceae | GCF_024105665.1 | Aliihoeflea_aestuarii | GTW25_RS19045 | 13819..16572 | 453840 | DUF499-domain-containing-protein-[Aliihoeflea-aestuarii] |
| WP_227345053.1 | AAA+ATPase+wHTH+Alpha+Beta-domain+wHTH+RRM | Type-3_BR.PglZ.associated.systems | <-GDResolvase+HIN-HTH<-?<-?\|\|?->?->DUF499*->PglZ->METHYLASE->iSTAND->TUDOR+SF1-SF2-Helicase+iREase+CTD-><-tRNA<-Peptidase_M20\|\|Peptidase_S11-> | 891 | Bacteria>Pseudomonadota>Alphaproteobacteria>Hyphomicrobiales>Phyllobacteriaceae | GCF_020616515.1 | Mesorhizobium_sp._PAMC28654 | LGH82_RS23595 | 4797492..4800167 | 2880934 | DUF499-domain-containing-protein-[Mesorhizobium-sp.-PAMC28654] |
| MCF1450408.1 | AAA+ATPase+wHTH+Alpha+Beta-domain+wHTH+RRM | Type-3_BR.PglZ.associated.systems | RVT-><-?<-TUDOR+SF1-SF2-Helicase+iREase+CTD<-iSTAND<-METHYLASE<-PglZ<-DUF499*<-HTH_3\|\|HEPN->?->?->Phage_int_SAM_1+Phage_integrase-> | 914 | Bacteria>Pseudomonadota>Alphaproteobacteria>Hyphomicrobiales>Rhizobiaceae | GCA_021462735.1 | Agrobacterium_vitis | FS815_26875 | 27279..30023 | 373 | DUF499-domain-containing-protein-[Agrobacterium-vitis] |
| WP_234689564.1 | AAA+ATPase+wHTH+Alpha+Beta-domain+wHTH+RRM | Type-3_BR.PglZ.associated.systems | RVT-><-?<-SF1-SF2-Helicase+iREase+CTD<-iSTAND<-METHYLASE<-PglZ<-DUF499*<-HTH_3\|\|HEPN->?->?->Phage_int_SAM_1+Phage_integrase-> | 838 | Bacteria>Pseudomonadota>Alphaproteobacteria>Hyphomicrobiales>Rhizobiaceae | GCF_021462735.1 | Allorhizobium_ampelinum | FS815_RS27030 | 27279..29795 | 3025782 | DUF499-domain-containing-protein-[Agrobacterium-vitis] |
| WP_260297856.1 | AAA+ATPase+wHTH+Alpha+Beta-domain+wHTH+RRM | Type-3_BR.PglZ.associated.systems | Phage_integrase->?->DUF499*->PglZ->METHYLASE->iSTAND->TUDOR+SF1-SF2-Helicase+iREase+CTD-> | 911 | Bacteria>Pseudomonadota>Alphaproteobacteria>Hyphomicrobiales>Rhizobiaceae | GCF_025200775.1 | Rhizobium_leguminosarum | N2600_RS02420 | 499502..502237 | 384 | DUF499-domain-containing-protein-[Rhizobium-leguminosarum] |
| WP_168336271.1 | AAA+ATPase+wHTH+Alpha+Beta-domain+wHTH+RRM | Type-3_BR.PglZ.associated.systems | Peptidase->?->CHTH-><-?<-?\|\|HTH_3->DUF499*->PglZ->METHYLASE->iSTAND->SF1-SF2-Helicase+iREase+CTD-> | 914 | Bacteria>Pseudomonadota>Alphaproteobacteria>Hyphomicrobiales>Rhizobiaceae | GCF_012275785.1 | Rhizobium_laguerreae | GFM01_RS22355 | 99011..101755 | 1076926 | DUF499-domain-containing-protein-[Rhizobium-laguerreae] |
| WP_142846515.1 | AAA+ATPase+wHTH+Alpha+Beta-domain+wHTH+RRM | Type-3_BR.PglZ.associated.systems | MobA_MobL+SF1-SF2-helicase->?-><-TUDOR+SF1-SF2-Helicase+iREase+CTD<-iSTAND<-METHYLASE<-PglZ<-DUF499*<-HTH_3\|\|?->?->Phage_int_SAM_1+Phage_integrase->HTH_Tnp_1+HTH_21+rve-> | 913 | Bacteria>Pseudomonadota>Alphaproteobacteria>Hyphomicrobiales>Rhizobiaceae | GCF_027921025.1 | Agrobacterium | EXN32_RS24485 | 16537..19278 | 357 | DUF499-domain-containing-protein-[Agrobacterium-tumefaciens] |
| MCB1467157.1 | AAA+ATPase+wHTH+Alpha+Beta-domain+wHTH+RRM | Type-3_BR.PglZ.associated.systems | HTH+WYL->DUF499*->PglZ->METHYLASE->iSTAND->TUDOR+SF1-SF2-Helicase+iREase+CTD-> | 913 | Bacteria>Pseudomonadota>Alphaproteobacteria>Hyphomicrobiales>Rhizobiaceae | GCA_020441945.1 | Rhizobiaceae_bacterium | KDK08_08450 | 1309..4050 | 1913961 | DUF499-domain-containing-protein-[Rhizobiaceae-bacterium] |
| WP_153473550.1 | AAA+ATPase+wHTH+Alpha+Beta-domain+wHTH+RRM | Type-3_BR.PglZ.associated.systems | DUF499*->PglZ->METHYLASE->iSTAND->SF1-SF2-Helicase+iREase+CTD-> | 916 | Bacteria>Pseudomonadota>Alphaproteobacteria>Hyphomicrobiales>Rhizobiaceae | GCF_009599615.1 | Sinorhizobium_meliloti | GHK14_RS10815 | 1237..3987 | 382 | DUF499-domain-containing-protein-[Sinorhizobium-meliloti] |
| WP_064697201.1 | AAA+ATPase+wHTH+Alpha+Beta-domain+wHTH+RRM | Type-3_BR.PglZ.associated.systems | ABhydrolase-><-HEPN<-TUDOR+SF1-SF2-Helicase+iREase+CTD<-iSTAND<-METHYLASE<-PglZ<-DUF499*<-HTH+WYL\|\|Radical-SAM->?->?->NADAR-><-ACET | 915 | Bacteria>Pseudomonadota>Alphaproteobacteria>Hyphomicrobiales>Rhizobiaceae | GCF_001662105.1 | Rhizobium_aegyptiacum | A7X79_RS30920 | 89703..92450 | 1764550 | DUF499-domain-containing-protein-[Rhizobium-aegyptiacum] |
| WP_132665734.1 | AAA+ATPase+wHTH+Alpha+Beta-domain+wHTH+RRM | Type-3_BR.PglZ.associated.systems | <-TUDOR+SF1-SF2-Helicase+iREase+CTD<-iSTAND<-METHYLASE<-PglZ<-DUF499*<-HTH_3<-?<-HISKIN<-SbcC<-GDResolvase+HTH | 913 | Bacteria>Pseudomonadota>Alphaproteobacteria>Hyphomicrobiales>Rhizobiaceae | - | Rhizobium_sp._PP-CC-3G-465 | C8J33_RS24990 | 8087..10828 | 2135648 | DUF499-domain-containing-protein-[Rhizobium-sp.-PP-CC-3G-465] |
| WP_234936258.1 | AAA+ATPase+wHTH+Alpha+Beta-domain+wHTH+RRM | Type-3_BR.PglZ.associated.systems | <-TUDOR+SF1-SF2-Helicase+iREase+CTD<-iSTAND<-METHYLASE<-PglZ<-DUF499*<-HTH_3\|\|?->?->Phage_int_SAM_1+Phage_integrase->?->MetJarc-HTH-> | 838 | Bacteria>Pseudomonadota>Alphaproteobacteria>Hyphomicrobiales>Rhizobiaceae | GCF_000739935.1 | Agrobacterium_rubi | RRU01S_RS24275 | 4977..7493 | 28099 | DUF499-domain-containing-protein-[Agrobacterium-rubi] |
| WP_222277391.1 | AAA+ATPase+wHTH+Alpha+Beta-domain+wHTH+RRM | Type-3_BR.PglZ.associated.systems | <-TUDOR+SF1-SF2-Helicase+iREase+CTD<-iSTAND<-METHYLASE<-PglZ<-DUF499*<-?<-Phage_integrase | 913 | Bacteria>Pseudomonadota>Alphaproteobacteria>Hyphomicrobiales>Rhizobiaceae | GCF_019798225.1 | Rhizobium_leguminosarum | HFN10_RS00615 | 134552..137293 | 384 | DUF499-domain-containing-protein-[Rhizobium-leguminosarum] |
| WP_203529913.1 | AAA+ATPase+wHTH+Alpha+Beta-domain+wHTH+RRM | Type-3_BR.PglZ.associated.systems | <-TUDOR+SF1-SF2-Helicase+iREase+CTD<-iSTAND<-METHYLASE<-PglZ<-DUF499*\|\|RHH->PIN-><-?<-MODE-HTH+PBPII | 914 | Bacteria>Pseudomonadota>Alphaproteobacteria>Hyphomicrobiales>Rhizobiaceae | GCF_017488845.2 | Ensifer_canadensis | J3R84_RS30765 | 229977..232721 | 555315 | DUF499-domain-containing-protein-[Ensifer-sp.-T173] |
| WP_097143356.1 | AAA+ATPase+wHTH+Alpha+Beta-domain+wHTH+RRM | Type-3_BR.PglZ.associated.systems | <-Phage_int_SAM_1+Phage_integrase<-?<-?\|\|HTH_3->DUF499*->PglZ->METHYLASE->iSTAND->TUDOR+SF1-SF2-Helicase+iREase+CTD-> | 913 | Bacteria>Pseudomonadota>Alphaproteobacteria>Hyphomicrobiales>Rhizobiaceae | GCF_900220975.1 | Rhizobium_subbaraonis | CRO48_RS30165 | 5920..8661 | 908946 | DUF499-domain-containing-protein-[Rhizobium-subbaraonis] |
| GAK73401.1 | AAA+ATPase+wHTH+Alpha+Beta-domain+wHTH+RRM | Type-3_BR.PglZ.associated.systems | <-iSTAND<-METHYLASE<-PglZ<-DUF499*\|\|?->?->Phage_int_SAM_1+Phage_integrase->?->MetJarc-HTH-> | 914 | Bacteria>Pseudomonadota>Alphaproteobacteria>Hyphomicrobiales>Rhizobiaceae | GCA_000739935.1 | Agrobacterium_rubi_TR3_=_NBRC_13261 | RRU01S_38_00050 | 4977..7721 | 1368415 | hypothetical-protein-RRU01S_38_00050-[Agrobacterium-rubi-TR3-=-NBRC-13261] |
| WP_183613692.1 | AAA+ATPase+wHTH+Alpha+Beta-domain+wHTH+RRM | Type-3_BR.PglZ.associated.systems | <-HEPN<-PAIREDC-HTH+HIN-HTH<-TUDOR+SF1-SF2-Helicase+iREase+CTD<-iSTAND<-METHYLASE<-PglZ<-DUF499*<-HTH+WYL\|\|Radical-SAM->?->?->NADAR-> | 913 | Bacteria>Pseudomonadota>Alphaproteobacteria>Hyphomicrobiales>Rhizobiaceae | GCF_014198655.1 | Rhizobium_leguminosarum | GGE50_RS31045 | 10727..13468 | 384 | DUF499-domain-containing-protein-[Rhizobium-leguminosarum] |
| WP_130765314.1 | AAA+ATPase+wHTH+Alpha+Beta-domain+wHTH+RRM | Type-3_BR.PglZ.associated.systems | <-Calcineurin\|\|HTH+WYL-><-TUDOR+SF1-SF2-Helicase+iREase+CTD<-iSTAND<-METHYLASE<-PglZ<-DUF499*<-?<-?<-?<-?<-REase<-ParB | 913 | Bacteria>Pseudomonadota>Alphaproteobacteria>Hyphomicrobiales>Rhizobiaceae | GCF_004304965.1 | Rhizobium_ruizarguesonis | ELH66_RS18325 | 3654017..3656758 | 2081791 | DUF499-domain-containing-protein-[Rhizobium-leguminosarum] |
| WP_131112880.1 | AAA+ATPase+wHTH+Alpha+Beta-domain+wHTH+RRM | Type-3_BR.PglZ.associated.systems | <-Calcineurin\|\|HTH+WYL-><-TUDOR+SF1-SF2-Helicase+iREase+CTD<-iSTAND<-METHYLASE<-PglZ<-DUF499* | 913 | Bacteria>Pseudomonadota>Alphaproteobacteria>Hyphomicrobiales>Rhizobiaceae | GCF_004322935.1 | Rhizobium_ruizarguesonis | ELH18_RS20215 | 4083448..4086189 | 2081791 | DUF499-domain-containing-protein-[Rhizobium-leguminosarum] |
| WP_203192678.1 | AAA+ATPase+wHTH+Alpha+Beta-domain+wHTH+RRM | Type-3_BR.PglZ.associated.systems | <-HKD-DNase<-DUF6361<-SF1-SF2-helicase+RdgC+Beta-barrel+REase<-SF1-SF2-Helicase+REase<-SF1-SF2-Helicase+REase\|\|HTH+WYL->DUF499*->PglZ->METHYLASE->iSTAND->TUDOR+SF1-SF2-Helicase+iREase+CTD-> | 915 | Bacteria>Pseudomonadota>Alphaproteobacteria>Hyphomicrobiales>Xanthobacteraceae | GCF_016807805.1 | Xanthobacter_dioxanivorans | EZH22_RS22705 | 4862984..4865731 | 2528964 | DUF499-domain-containing-protein-[Xanthobacter-dioxanivorans] |
| MBF0460762.1 | AAA+ATPase+wHTH+Alpha+Beta-domain+wHTH+RRM | Type-3_BR.PglZ.associated.systems | DDE_Tnp_IS1595-><-?<-?<-?<-PglZ<-DUF499*<-?<-?<-Peptidase | 909 | Bacteria>Pseudomonadota>Alphaproteobacteria>Magnetococcales>unclassified_Magnetococcales | GCA_015233035.1 | Magnetococcales_bacterium | HQL87_05140 | 3919..6648 | 2026759 | DUF499-domain-containing-protein-[Magnetococcales-bacterium] |
| MBF0628342.1 | AAA+ATPase+wHTH+Alpha+Beta-domain+wHTH+RRM | Type-3_BR.PglZ.associated.systems | <-REase+NACHT\|\|HTH+WYL->DUF499*->PglZ->?->PemK_toxin->METHYLASE->iSTAND->OLD-ABC-> | 914 | Bacteria>Pseudomonadota>Alphaproteobacteria>Magnetococcales>unclassified_Magnetococcales | GCA_015234115.1 | Magnetococcales_bacterium | HQL91_08975 | 6934..9678 | 2026759 | DUF499-domain-containing-protein-[Magnetococcales-bacterium] |
| MBF0422814.1 | AAA+ATPase+wHTH+Alpha+Beta-domain+wHTH+RRM | Type-3_BR.PglZ.associated.systems | <-Peptidase_S8<-Classical-AAA<-PglZ<-DUF499*<-HTH+WYL<-PAIREDC-HTH | 915 | Bacteria>Pseudomonadota>Alphaproteobacteria>Magnetococcales>unclassified_Magnetococcales | GCA_015232645.1 | Magnetococcales_bacterium | HQL73_07465 | 5425..8172 | 2026759 | DUF499-domain-containing-protein-[Magnetococcales-bacterium] |
| GLS88419.1 | AAA+ATPase+wHTH+Alpha+Beta-domain+wHTH+RRM | Type-3_BR.PglZ.associated.systems | RNase-><-TUDOR+SF1-SF2-Helicase+iREase+CTD<-iSTAND<-METHYLASE<-PglZ<-DUF499*<-HTH+WYL<-ParB<-ParB<-GDResolvase+PPP1-HTH\|\|?->HD-> | 914 | Bacteria>Pseudomonadota>Alphaproteobacteria>Rhodobacterales>Paracoccaceae | GCA_030161095.1 | Cypionkella_aquatica | GCM10010873_33930 | 355568..358312 | 1756042 | hypothetical-protein-GCM10010873_33930-[Cypionkella-aquatica] |
| WP_284326581.1 | AAA+ATPase+wHTH+Alpha+Beta-domain+wHTH+RRM | Type-3_BR.PglZ.associated.systems | RNase-><-TUDOR+SF1-SF2-Helicase+iREase+CTD<-iSTAND<-METHYLASE<-PglZ<-DUF499*<-HTH+WYL<-ParB<-ParB<-GDResolvase+PPP1-HTH\|\|?->HD-> | 914 | Bacteria>Pseudomonadota>Alphaproteobacteria>Rhodobacterales>Paracoccaceae | GCF_030161095.1 | Cypionkella_aquatica | QQJ98_RS17135 | 355568..358312 | 1756042 | DUF499-domain-containing-protein-[Cypionkella-aquatica] |
| MCR9157071.1 | AAA+ATPase+wHTH+Alpha+Beta-domain+wHTH+RRM | Type-3_BR.PglZ.associated.systems | RADICAL-SAM->?-><-TUDOR+SF1-SF2-Helicase+iREase+CTD<-iSTAND<-METHYLASE<-PglZ<-DUF499*<-HTH+WYL<-HEPN\|\|REase+35exo+Beta-Barrel+SF1-SF2-Helicase-> | 914 | Bacteria>Pseudomonadota>Alphaproteobacteria>Rhodobacterales>Paracoccaceae | GCA_024742575.1 | Paracoccaceae_bacterium | NXH80_07485 | 25301..28045 | 1904441 | DUF499-domain-containing-protein-[Paracoccaceae-bacterium] |
| WP_173488294.1 | AAA+ATPase+wHTH+Alpha+Beta-domain+wHTH+RRM | Type-3_BR.PglZ.associated.systems | HTH-><-?<-?<-?\|\|TetR-HTH->?->DUF499*->PglZ->METHYLASE->iSTAND->TUDOR+SF1-SF2-Helicase+iREase+CTD-> | 914 | Bacteria>Pseudomonadota>Alphaproteobacteria>Rhodobacterales>Paracoccaceae | GCF_013283585.1 | unclassified_Aliiroseovarius | XMD517_RS04540 | 962406..965150 | 2623558 | MULTISPECIES:-DUF499-domain-containing-protein-[unclassified-Aliiroseovarius] |
| PZQ97082.1 | AAA+ATPase+wHTH+Alpha+Beta-domain+wHTH+RRM | Type-3_BR.PglZ.associated.systems | GDResolvase+PPP1-HTH->ParB->ParB->HTH+WYL->DUF499*->PglZ->METHYLASE->iSTAND->TUDOR+SF1-SF2-Helicase+iREase+CTD-> | 914 | Bacteria>Pseudomonadota>Alphaproteobacteria>Rhodobacterales>Paracoccaceae | GCA_003243455.1 | Cereibacter_sphaeroides | DI533_16205 | 628782..631526 | 1063 | DUF499-domain-containing-protein-[Cereibacter-sphaeroides] |
| WP_206771951.1 | AAA+ATPase+wHTH+Alpha+Beta-domain+wHTH+RRM | Type-3_BR.PglZ.associated.systems | GDResolvase+HIN-HTH-><-TUDOR+SF1-SF2-Helicase+iREase+CTD<-iSTAND<-METHYLASE<-PglZ<-DUF499*<-HTH+WYL<-?<-HTH_17<-Phage_integrase | 917 | Bacteria>Pseudomonadota>Alphaproteobacteria>Rhodobacterales>Paracoccaceae | GCF_000192745.1 | Polymorphum_gilvum | SL003B_RS02635 | 542544..545297 | 991904 | DUF499-domain-containing-protein-[Polymorphum-gilvum] |
| ADZ68952.1 | AAA+ATPase+wHTH+Alpha+Beta-domain+wHTH+RRM | Type-3_BR.PglZ.associated.systems | <-VirD4-TraG+VirD4-FtsK\|\|?->?-><-TUDOR+SF1-SF2-Helicase+iREase+CTD<-METHYLASE<-PglZ<-DUF499*<-HTH+WYL<-HTH_17<-Phage_integrase\|\|?->?-><-HTH_21+rve+rve_3 | 766 | Bacteria>Pseudomonadota>Alphaproteobacteria>Rhodobacterales>Paracoccaceae | GCA_000192745.1 | Polymorphum_gilvum_SL003B-26A1 | SL003B_0519 | 542544..544844 | 991905 | hypothetical-protein-SL003B_0519-[Polymorphum-gilvum-SL003B-26A1] |
| WP_189381906.1 | AAA+ATPase+wHTH+Alpha+Beta-domain+wHTH+RRM | Type-3_BR.PglZ.associated.systems | <-TUDOR+SF1-SF2-Helicase+iREase+CTD<-iSTAND<-METHYLASE<-PglZ<-DUF499*<-HTH+WYL\|\|MerR-HTH->?->?->?->PhdYeFM_antitox-> | 914 | Bacteria>Pseudomonadota>Alphaproteobacteria>Rhodobacterales>Paracoccaceae | GCF_014652195.1 | Gemmobacter_nanjingensis | IE126_RS19310 | 35968..38712 | 488454 | DUF499-domain-containing-protein-[Gemmobacter-nanjingensis] |
| WRH63789.1 | AAA+ATPase+wHTH+Alpha+Beta-domain+wHTH+RRM | Type-3_BR.PglZ.associated.systems | <-MerR-HTH\|\|WYL->DUF499+RRM*->PglZ->METHYLASE->iSTAND->TUDOR+SF1-SF2-Helicase+iREase+CTD-> | 914 | Bacteria>Pseudomonadota>Alphaproteobacteria>Rhodobacterales>Paracoccaceae>Fuscovulum | GCA_035192965.1 | Fuscovulum_sp. | RSE12_05475 | 1109757..1112501 | 2818471 | MAG:-DUF499-domain-containing-protein-[Fuscovulum-sp.] |
| WP_170773962.1 | AAA+ATPase+wHTH+Alpha+Beta-domain+wHTH+RRM | Type-3_BR.PglZ.associated.systems | REase-><-SF1-SF2-Helicase<-TUDOR+SF1-SF2-Helicase+iREase+CTD<-iSTAND<-METHYLASE<-PglZ<-DUF499*\|\|?-><-tRNA<-ABhydrolase | 914 | Bacteria>Pseudomonadota>Alphaproteobacteria>Rhodobacterales>Roseobacteraceae | GCF_013030225.1 | Ruegeria_lacuscaerulensis | GOX68_RS16715 | 238882..241626 | 55218 | DUF499-domain-containing-protein-[Ruegeria-lacuscaerulensis] |
| WP_200879667.1 | AAA+ATPase+wHTH+Alpha+Beta-domain+wHTH+RRM | Type-3_BR.PglZ.associated.systems | Peptidase_M15->?-><-TUDOR+SF1-SF2-Helicase+iREase+CTD<-iSTAND<-METHYLASE<-PglZ<-DUF499*<-HTH+WYL<-McrC-NTD+REase<-?<-ParB<-ParB<-GDResolvase+PPP1-HTH | 924 | Bacteria>Pseudomonadota>Alphaproteobacteria>Rhodobacterales>Roseobacteraceae | GCF_000743715.1 | Palleronia_rufa | KW83_RS07530 | 22758..25532 | 1530186 | DUF499-domain-containing-protein-[Palleronia-rufa] |
| WP_171230676.1 | AAA+ATPase+wHTH+Alpha+Beta-domain+wHTH+RRM | Type-3_BR.PglZ.associated.systems | McrC-NTD+REase->HTH-><-RVT<-?<-RNase+35exo\|\|DUF499*->PglZ->METHYLASE->iSTAND->TUDOR+SF1-SF2-Helicase+iREase+CTD-> | 907 | Bacteria>Pseudomonadota>Alphaproteobacteria>Rhodobacterales>Roseobacteraceae | GCF_013033125.1 | Ruegeria_sp._HKCCA4008 | GGW60_RS20085 | 13608..16331 | 2682999 | DUF499-domain-containing-protein-[Ruegeria-sp.-HKCCA4008] |
| WP_034927475.1 | AAA+ATPase+wHTH+Alpha+Beta-domain+wHTH+RRM | Type-3_BR.PglZ.associated.systems | <-HTH<-ArsR-HTH+YTCD-HTH<-?\|\|?->GDResolvase+HIN-HTH->DUF499*->PglZ->METHYLASE-p-> | 912 | Bacteria>Pseudomonadota>Alphaproteobacteria>Rhodospirillales | GCF_000700985.1 | Acetobacteraceae | BIV10_RS03765 | 7225..9963 | 433 | MULTISPECIES:-DUF499-domain-containing-protein-[Acetobacteraceae] |
| MBR2124997.1 | AAA+ATPase+wHTH+Alpha+Beta-domain+wHTH+RRM | Type-3_BR.PglZ.associated.systems | <-SF1-SF2-Helicase+iREase+CTD<-TUDOR+SF1-SF2-Helicase+iREase+CTD<-iSTAND<-METHYLASE<-PglZ<-DUF499*<-?<-?\|\|?->Aminotran_1_2-> | 931 | Bacteria>Pseudomonadota>Alphaproteobacteria>Rhodospirillales>Acetobacteraceae | GCA_017474285.1 | Acetobacter_sp. | IJ934_07580 | 13175..15970 | 440 | DUF499-domain-containing-protein-[Acetobacter-sp.] |
| WP_176643704.1 | AAA+ATPase+wHTH+Alpha+Beta-domain+wHTH+RRM | Type-3_BR.PglZ.associated.systems | <-SF1-SF2-Helicase+iREase+CTD<-iSTAND<-METHYLASE<-Peptidase_S8<-Classical-AAA<-PglZ<-DUF499*<-GDResolvase+HIN-HTH\|\|?-><-TetR-HTH | 909 | Bacteria>Pseudomonadota>Alphaproteobacteria>Rhodospirillales>Acetobacteraceae | GCF_013376245.1 | Komagataeibacter_swingsii | HUK81_RS13370 | 11766..14495 | 215220 | DUF499-domain-containing-protein-[Komagataeibacter-swingsii] |
| WP_149537232.1 | AAA+ATPase+wHTH+Alpha+Beta-domain+wHTH+RRM | Type-3_BR.PglZ.associated.systems | <-MPTase<-HTH_3<-?\|\|RAMA->GDResolvase+Recombinase+ZnR->HTH->DUF499*->PglZ->METHYLASE->iSTAND->TUDOR+SF1-SF2-Helicase+iREase+CTD-><-ASCH | 944 | Bacteria>Pseudomonadota>Alphaproteobacteria>Rhodospirillales>Acetobacteraceae | GCF_008360935.1 | Siccirubricoccus_phaeus | E5N76_RS10395 | 67681..70515 | 2595053 | DUF499-domain-containing-protein-[Siccirubricoccus-phaeus] |
| MBQ5470382.1 | AAA+ATPase+wHTH+Alpha+Beta-domain+wHTH+RRM | Type-3_BR.PglZ.associated.systems | <-METHYLASE-p<-PglZ<-DUF499<-DUF499* | 427 | Bacteria>Pseudomonadota>Alphaproteobacteria>Rhodospirillales>Acetobacteraceae | GCA_017386225.1 | Acetobacter_sp. | IIT54_07235 | 3403..4686 | 440 | AAA-family-ATPase-[Acetobacter-sp.] |
| MBQ5516117.1 | AAA+ATPase+wHTH+Alpha+Beta-domain+wHTH+RRM | Type-3_BR.PglZ.associated.systems | <-METHYLASE-p<-PglZ<-DUF499*<-DUF499-p | 523 | Bacteria>Pseudomonadota>Alphaproteobacteria>Rhodospirillales>Acetobacteraceae | GCA_017385515.1 | Acetobacter_sp. | IIT88_03420 | 2106..3677 | 440 | hypothetical-protein-[Acetobacter-sp.] |
| MBQ5470381.1 | AAA+ATPase+wHTH+Alpha+Beta-domain+wHTH+RRM | Type-3_BR.PglZ.associated.systems | <-METHYLASE-p<-PglZ<-DUF499*<-DUF499 | 523 | Bacteria>Pseudomonadota>Alphaproteobacteria>Rhodospirillales>Acetobacteraceae | GCA_017386225.1 | Acetobacter_sp. | IIT54_07230 | 1902..3473 | 440 | hypothetical-protein-[Acetobacter-sp.] |
| WP_064275617.1 | AAA+ATPase+wHTH+Alpha+Beta-domain+wHTH+RRM | Type-3_BR.PglZ.associated.systems | <-GDResolvase+Recombinase+ZnR<-?\|\|GDResolvase+HIN-HTH->DUF499*->PglZ->METHYLASE->iSTAND->TUDOR+SF1-SF2-Helicase+iREase+CTD->?->N6_N4_MTase-> | 905 | Bacteria>Pseudomonadota>Alphaproteobacteria>Rhodospirillales>Acetobacteraceae | GCF_001645165.1 | Gluconobacter_cerinus | A0123_RS16035 | 3609..6326 | 38307 | DUF499-domain-containing-protein-[Gluconobacter-cerinus] |
| WP_288005656.1 | AAA+ATPase+wHTH+Alpha+Beta-domain+wHTH+RRM | Type-3_BR.PglZ.associated.systems | <-SF1-SF2-Helicase+REase\|\|Arm-DNA-bind_3+Phage_integrase->\|\|WYL->DUF499+RRM*->PglZ->METHYLASE->iSTAND->TUDOR+SF1-SF2-Helicase+iREase+CTD-> | 916 | Bacteria>Pseudomonadota>Alphaproteobacteria>Rhodospirillales>Acetobacteraceae | GCF_027308825.1 | Acidiphilium | M3W77_RS08825 | 11167..13917 | 522 | MULTISPECIES:-DUF499-domain-containing-protein-[Acidiphilium] |
| WP_294509802.1 | AAA+ATPase+wHTH+Alpha+Beta-domain+wHTH+RRM | Type-3_BR.PglZ.associated.systems | <-TUDOR+SF1-SF2-Helicase+iREase+CTD<-iSTAND<-METHYLASE<-PglZ<-DUF499+RRM*\|\|ParB+HNH-><-SF1-SF2-Helicase | 768 | Bacteria>Pseudomonadota>Alphaproteobacteria>Rhodospirillales>Acetobacteraceae>Rhodopila | GCF_903841135.1 | uncultured_Rhodopila_sp. | QZY05_RS24400 | 7644..9950 | 310091 | DUF499-domain-containing-protein-[uncultured-Rhodopila-sp.] |
| WP_318650451.1 | AAA+ATPase+wHTH+Alpha+Beta-domain+wHTH+RRM | Type-3_BR.PglZ.associated.systems | GDResolvase->Cluster83_1clades->DUF499+RRM*->PglZ->METHYLASE->iSTAND->TUDOR+SF1-SF2-Helicase+iREase+CTD-> | 884 | Bacteria>Pseudomonadota>Alphaproteobacteria>Rhodospirillales>Acetobacteraceae>Sediminicoccus | GCF_033547095.1 | Sediminicoccus_rosea | R9Z33_RS06275 | 1310521..1313175 | 1225128 | hypothetical-protein-[Sediminicoccus-rosea] |
| WP_109076132.1 | AAA+ATPase+wHTH+Alpha+Beta-domain+wHTH+RRM | Type-3_BR.PglZ.associated.systems | HNH-><-?\|\|?->?->Peptidase-><-?\|\|DUF499*->PglZ->METHYLASE->iSTAND->TUDOR+SF1-SF2-Helicase+iREase+CTD->Phage_integrase->SF1-SF2-Helicase-p-> | 916 | Bacteria>Pseudomonadota>Alphaproteobacteria>Rhodospirillales>Azospirillaceae | GCF_003115945.1 | Azospirillum_sp._TSH7 | TSH7_RS25855 | 9619..12369 | 652751 | DUF499-domain-containing-protein-[Azospirillum-sp.-TSH7] |
| WP_199232438.1 | AAA+ATPase+wHTH+Alpha+Beta-domain+wHTH+RRM | Type-3_BR.PglZ.associated.systems | <-SF1-SF2-Helicase-p<-Phage_integrase<-TUDOR+SF1-SF2-Helicase+iREase+CTD<-iSTAND<-METHYLASE<-PglZ<-DUF499*\|\|?-><-Peptidase<-?<-?\|\|?-><-HNH | 902 | Bacteria>Pseudomonadota>Alphaproteobacteria>Rhodospirillales>Azospirillaceae | GCF_003115935.1 | Azospirillum_sp._TSH20 | TSH20_RS18720 | 15042..17750 | 652754 | DUF499-domain-containing-protein-[Azospirillum-sp.-TSH20] |
| PZR07944.1 | AAA+ATPase+wHTH+Alpha+Beta-domain+wHTH+RRM | Type-3_BR.PglZ.associated.systems | <-Phage_integrase<-TUDOR+SF1-SF2-Helicase+iREase+CTD<-iSTAND<-METHYLASE<-PglZ<-DUF499*<-REase<-N6_MTase | 909 | Bacteria>Pseudomonadota>Alphaproteobacteria>Rhodospirillales>Azospirillaceae | GCA_003243305.1 | Azospirillum_brasilense | DI532_22855 | 8674..11403 | 192 | DUF499-domain-containing-protein-[Azospirillum-brasilense] |
| WP_322332028.1 | AAA+ATPase+wHTH+Alpha+Beta-domain+wHTH+RRM | Type-3_BR.PglZ.associated.systems | GDResolvase+HIN-HTH->Phage_integrase->DUF499+RRM*->PglZ->METHYLASE->iSTAND->TUDOR+SF1-SF2-Helicase+iREase+CTD-><-GDResolvase+HIN-HTH | 913 | Bacteria>Pseudomonadota>Alphaproteobacteria>Rhodospirillales>Thalassobaculaceae>Thalassobaculum | GCF_034377285.1 | Thalassobaculum_sp._OXR-137 | T8K17_RS22795 | 4921987..4924728 | 3100173 | DUF499-domain-containing-protein-[Thalassobaculum-sp.-OXR-137] |
| WP_267619446.1 | AAA+ATPase+wHTH+Alpha+Beta-domain+wHTH+RRM | Type-3_BR.PglZ.associated.systems | HKD-DNase+HKD-DNase-><-REase+35exo+Beta-Barrel+SF1-SF2-Helicase<-?<-TUDOR+SF1-SF2-Helicase+iREase+CTD\|\|XIS-HTH->DUF499*->PglZ->METHYLASE->iSTAND->TUDOR+SF1-SF2-Helicase+iREase+CTD-> | 914 | Bacteria>Pseudomonadota>Alphaproteobacteria>Sphingomonadales>Erythrobacteraceae | GCF_026586795.1 | Erythrobacter_sp._WG | OPU67_RS10295 | 68588..71332 | 2985510 | DUF499-domain-containing-protein-[Erythrobacter-sp.-WG] |
| MBX7482086.1 | AAA+ATPase+wHTH+Alpha+Beta-domain+wHTH+RRM | Type-3_BR.PglZ.associated.systems | <-REase+35exo<-?<-TUDOR+SF1-SF2-Helicase+iREase+CTD<-Ntox49\|\|HTH+WYL->XIS-HTH->DUF499*->PglZ->METHYLASE-p->iSTAND->SF1-SF2-Helicase+iREase+CTD->LexA-protease-><-wHTH+SAD+HNH | 764 | Bacteria>Pseudomonadota>Alphaproteobacteria>Sphingomonadales>Erythrobacteraceae | GCA_019711515.1 | Qipengyuania_qiaonensis | K3174_06055 | 80764..83058 | 2867240 | ATP-binding-protein-[Qipengyuania-qiaonensis] |
| WP_247711953.1 | AAA+ATPase+wHTH+Alpha+Beta-domain+wHTH+RRM | Type-3_BR.PglZ.associated.systems | <-REase+35exo<-?<-TUDOR+SF1-SF2-Helicase+iREase+CTD<-Ntox49\|\|HTH+WYL->XIS-HTH->DUF499*->PglZ->METHYLASE->iSTAND->SF1-SF2-Helicase+iREase+CTD->LexA-protease-><-wHTH+SAD+HNH | 914 | Bacteria>Pseudomonadota>Alphaproteobacteria>Sphingomonadales>Erythrobacteraceae | GCF_019711515.1 | Qipengyuania_qiaonensis | K3174_RS06055 | 80314..83058 | 2867240 | DUF499-domain-containing-protein-[Qipengyuania-qiaonensis] |
| WP_246386365.1 | AAA+ATPase+wHTH+Alpha+Beta-domain+wHTH+RRM | Type-3_BR.PglZ.associated.systems | XIS-HTH->DUF499*->PglZ->METHYLASE->iSTAND->TUDOR+SF1-SF2-Helicase+iREase+CTD->HIN-HTH-> | 838 | Bacteria>Pseudomonadota>Alphaproteobacteria>Sphingomonadales>Sphingomonadaceae | GCF_014196055.1 | Novosphingobium_hassiacum | GGQ88_RS19045 | 885..3401 | 173676 | DUF499-domain-containing-protein-[Novosphingobium-hassiacum] |
| WP_093314938.1 | AAA+ATPase+wHTH+Alpha+Beta-domain+wHTH+RRM | Type-3_BR.PglZ.associated.systems | HTH-><-TUDOR+SF1-SF2-Helicase+iREase+CTD<-iSTAND<-METHYLASE<-PglZ<-DUF499*<-GDResolvase+HIN-HTH<-?<-?<-?<-MODE-HTH+PBPII | 913 | Bacteria>Pseudomonadota>Alphaproteobacteria>Sphingomonadales>Sphingomonadaceae | GCF_900113315.1 | Sphingomonas_jatrophae | BM360_RS12030 | 577357..580098 | 1166337 | nucleoporin-[Sphingomonas-jatrophae] |
| WP_037491255.1 | AAA+ATPase+wHTH+Alpha+Beta-domain+wHTH+RRM | Type-3_BR.PglZ.associated.systems | DUF499*->PglZ->METHYLASE-> | 717 | Bacteria>Pseudomonadota>Alphaproteobacteria>Sphingomonadales>Sphingomonadaceae | GCF_000382885.1 | Sphingobium_indicum | EBBID32_RS16170 | 1..2156 | 332055 | hypothetical-protein,-partial-[Sphingobium-indicum] |
| WP_230188972.1 | AAA+ATPase+wHTH+Alpha+Beta-domain+wHTH+RRM | Type-3_BR.PglZ.associated.systems | AbiEii<-Phage_integrase<-TUDOR+SF1-SF2-Helicase+iREase+CTD<-iSTAND<-METHYLASE<-PglZ<-DUF499*<-HTH_3\|\|?->METHYLASE->?-><-?<-Calcineurin | 911 | Bacteria>Pseudomonadota>Alphaproteobacteria>Sphingomonadales>Sphingomonadaceae | GCF_918741285.1 | Sphingobium_sp._CECT_9361 | LQ220_RS14860 | 69359..72094 | 2845384 | DUF499-domain-containing-protein-[Sphingobium-sp.-CECT-9361] |
| WP_056450793.1 | AAA+ATPase+wHTH+Alpha+Beta-domain+wHTH+RRM | Type-3_BR.PglZ.associated.systems | <-RNase<-?<-?\|\|<-HIN-HTH\|\|GDResolvase+HTH->DUF499*->PglZ->METHYLASE->iSTAND->TUDOR+SF1-SF2-Helicase+iREase+CTD-> | 914 | Bacteria>Pseudomonadota>Alphaproteobacteria>Sphingomonadales>Sphingomonadaceae | GCF_001421245.1 | Sphingomonas_sp._Leaf10 | ASE59_RS00525 | 106933..109677 | 1735676 | DUF499-domain-containing-protein-[Sphingomonas-sp.-Leaf10] |
| WP_070156238.1 | AAA+ATPase+wHTH+Alpha+Beta-domain+wHTH+RRM | Type-3_BR.PglZ.associated.systems | <-REase+35exo<-?<-TUDOR+SF1-SF2-Helicase+iREase+CTD<-Ntox49\|\|HTH+WYL->XIS-HTH->DUF499*->PglZ->METHYLASE->iSTAND->TUDOR+SF1-SF2-Helicase+iREase+CTD->LexA-protease-><-wHTH+SAD+HNH | 914 | Bacteria>Pseudomonadota>Alphaproteobacteria>Sphingomonadales>Sphingomonadaceae | GCF_001757355.1 | Sphingobium_phenoxybenzoativorans | BHQ40_RS12645 | 29293..32037 | 1592790 | DUF499-domain-containing-protein-[Sphingobium-phenoxybenzoativorans] |
| WP_231471110.1 | AAA+ATPase+wHTH+Alpha+Beta-domain+wHTH+RRM | Type-3_BR.PglZ.associated.systems | <-MODE-HTH+PBPII\|\|PIN+OST-HTH-><-TUDOR+SF1-SF2-Helicase+iREase+CTD<-iSTAND<-METHYLASE<-PglZ<-DUF499*<-HTH+WYL<-SbcC+SbcC<-Calcineurin\|\|ArdCN+MPTase->ParB+HTH-> | 914 | Bacteria>Pseudomonadota>Alphaproteobacteria>Sphingomonadales>Sphingomonadaceae | GCF_920987055.1 | Novosphingobium_sp._CECT_9465 | LUA85_RS15260 | 3120008..3122752 | 2829794 | DUF499-domain-containing-protein-[Novosphingobium-sp.-CECT-9465] |
| WP_169574502.1 | AAA+ATPase+wHTH+Alpha+Beta-domain+wHTH+RRM | Type-3_BR.PglZ.associated.systems | <-MODE-HTH+PBPII\|\|PIN+OST-HTH-><-TUDOR+SF1-SF2-Helicase+iREase+CTD<-iSTAND<-METHYLASE<-PglZ<-DUF499*<-HTH+WYL<-SbcC+SbcC<-Calcineurin\|\|ArdCN+MPTase->?->ParB+HTH-> | 914 | Bacteria>Pseudomonadota>Alphaproteobacteria>Sphingomonadales>Sphingomonadaceae | GCF_012927105.1 | Sphingobium_psychrophilum | HHL08_RS18425 | 42120..44864 | 2728834 | DUF499-domain-containing-protein-[Sphingobium-psychrophilum] |
| WP_298166666.1 | AAA+ATPase+wHTH+Alpha+Beta-domain+wHTH+RRM | Type-3_BR.PglZ.associated.systems | <-TUDOR+SF1-SF2-Helicase+iREase+CTD\|\|WYL+wHTH->XIS-HTH->DUF499+RRM*->PglZ->METHYLASE->iSTAND->TUDOR+SF1-SF2-Helicase+iREase+CTD-> | 914 | Bacteria>Pseudomonadota>Alphaproteobacteria>Sphingomonadales>Sphingomonadaceae>Novosphingobium | GCF_027486195.1 | Novosphingobium_sp. | Q0071_RS06905 | 47304..50048 | 1874826 | DUF499-domain-containing-protein-[Novosphingobium-sp.] |

| Protein-ID | DUF499-architecture | System | Gene-Neighbourhoods | Length | Taxonomy | Assembly-ID | Strain | Locus-Tag | Genomic-coordinates | TaxID | Fasta-defline |
|---|---|---|---|---|---|---|---|---|---|---|---|
| MEC6699672.1 | AAA+ATPase+wHTH+Alpha+Beta-domain+wHTH+RRM | Type-3_BR.PglZ.associated.systems | XIS-HTH-><-TUDOR+SF1-SF2-Helicase+iREase+CTD<-iSTAND<-METHYLASE<-PglZ<-DUF499+RRM*<-XIS-HTH<-Arm-DNA-bind_3+Phage_integrase | 914 | Bacteria>Pseudomonadota>Alphaproteobacteria>Sphingomonadales>Sphingomonadaceae>Sphingobium | - | Sphingobium_sp._SJ10-10 | VVT58_RS11490 | 2369358..2372102 | 3114999 | DUF499-domain-containing-protein-[Sphingobium-sp.-SJ10-10] |
| WP_317514299.1 | AAA+ATPase+wHTH+Alpha+Beta-domain+wHTH+RRM | Type-3_BR.PglZ.associated.systems | XIS-HTH->DUF499+RRM*->PglZ->METHYLASE->iSTAND->TUDOR+SF1-SF2-Helicase+iREase+CTD-> | 914 | Bacteria>Pseudomonadota>Alphaproteobacteria>Sphingomonadales>Sphingomonadaceae>Sphingomonas | GCF_033025475.1 | Sphingomonas_sp._C3-2 | QYC26_RS05005 | 1053743..1056487 | 3062169 | DUF499-domain-containing-protein-[Sphingomonas-sp.-C3-2] |
| MBK6492143.1 | AAA+ATPase+wHTH+Alpha+Beta-domain+wHTH+RRM | Type-3_BR.PglZ.associated.systems | <-DDE_Tnp_1_4\|\|GDResolvase+HIN-HTH->DDE_Tnp_1->XIS-HTH->DUF499*->PglZ->METHYLASE->iSTAND->TUDOR+SF1-SF2-Helicase+iREase+CTD-><-SF1-SF2-Helicase | 914 | Bacteria>Pseudomonadota>Alphaproteobacteria>Sphingomonadales>unclassified_Sphingomonadales | GCA_016704425.1 | Sphingomonadales_bacterium | IPF97_10750 | 490462..493206 | 1978525 | DUF499-domain-containing-protein-[Sphingomonadales-bacterium] |
| NCC23132.1 | AAA+ATPase+wHTH+Alpha+Beta-domain+wHTH+RRM | Type-3_BR.PglZ.associated.systems | Phage_integrase-><-TUDOR+SF1-SF2-Helicase+iREase+CTD<-iSTAND<-METHYLASE<-PglZ<-DUF499* | 906 | Bacteria>Pseudomonadota>Alphaproteobacteria>unclassified_Alphaproteobacteria | GCA_009929775.1 | Alphaproteobacteria_bacterium | EOM26_11845 | 7748..10468 | 1913988 | DUF499-domain-containing-protein-[Alphaproteobacteria-bacterium] |
| TLD46490.1 | AAA+ATPase+wHTH+Alpha+Beta-domain+wHTH+RRM | Type-3_BR.PglZ.associated.systems | XIS-HTH->DUF499*->PglZ->ParB+HNH->METHYLASE->iSTAND->OLD-ABC->TOPRIM-> | 909 | Bacteria>Pseudomonadota>Betaproteobacteria>Betaproteobacteria_incertae_sedis>Candidatus_Accumulibacter | GCA_005524045.1 | Accumulibacter_sp. | FAZ92_01227 | 1323..4052 | 2053492 | MAG:-hypothetical-protein-FAZ92_01227-[Candidatus-Accumulibacter-sp.] |
| EXI88111.1 | AAA+ATPase+wHTH+Alpha+Beta-domain+wHTH+RRM | Type-3_BR.PglZ.associated.systems | <-VirB11+VirB4-FtsK<-CopG/RHH<-VirD4-TraG+VirD4-FtsK<-?<-MODE-HTH+PBPII\|\|XIS-HTH->DUF499*->PglZ->Classical-AAA->Peptidase_S8->METHYLASE->iSTAND->TUDOR+SF1-SF2-Helicase+iREase+CTD-> | 922 | Bacteria>Pseudomonadota>Betaproteobacteria>Betaproteobacteria_incertae_sedis>Candidatus_Accumulibacter | GCA_000585075.1 | Candidatus_Accumulibacter_regalis | AW11_02234 | 10498..13266 | 522306 | MAG:-hypothetical-protein-AW11_02234-[Candidatus-Accumulibacter-regalis] |
| QLH49694.1 | AAA+ATPase+wHTH+Alpha+Beta-domain+wHTH+RRM | Type-3_BR.PglZ.associated.systems | <-OLD-ABC<-iSTAND<-METHYLASE<-PIN<-?<-PglZ<-DUF499*<-XIS-HTH<-?<-RVT<-?<-TOPRIM | 909 | Bacteria>Pseudomonadota>Betaproteobacteria>Betaproteobacteria_incertae_sedis>Candidatus_Accumulibacter | GCA_013414765.1 | Candidatus_Accumulibacter_cognatus | HWD57_07800 | 1710495..1713224 | 2954383 | MAG:-DUF499-domain-containing-protein-[Candidatus-Accumulibacter-cognatus] |
| MDR3298567.1 | AAA+ATPase+wHTH+Alpha+Beta-domain+wHTH+RRM | Type-3_BR.PglZ.associated.systems | DUF499+RRM*->PglZ->Classical-AAA->METHYLASE->iSTAND-> | 908 | Bacteria>Pseudomonadota>Betaproteobacteria>Betaproteobacteria_incertae_sedis>Candidatus_Accumulibacter>unclassified_Candidatus_Accumulibacter | GCA_031290495.1 | Accumulibacter_sp. | LBU43_00825 | 11976..14702 | 2053492 | DUF499-domain-containing-protein-[Accumulibacter-sp.] |
| HPT50269.1 | AAA+ATPase+wHTH+Alpha+Beta-domain+wHTH+RRM | Type-3_BR.PglZ.associated.systems | <-DUF4276<-SbcC+SbcC<-iSTAND<-METHYLASE<-PglZ<-DUF499+RRM*<-XIS-HTH | 912 | Bacteria>Pseudomonadota>Betaproteobacteria>Betaproteobacteria_incertae_sedis>Candidatus_Accumulibacter>unclassified_Candidatus_Accumulibacter | GCA_035402605.1 | Accumulibacter_sp. | PKZ22_08725 | 20536..23274 | 2053492 | DUF499-domain-containing-protein-[Accumulibacter-sp.] |
| HRI93224.1 | AAA+ATPase+wHTH+Alpha+Beta-domain+wHTH+RRM | Type-3_BR.PglZ.associated.systems | <-METHYLASE-p<-PglZ<-DUF499+RRM* | 915 | Bacteria>Pseudomonadota>Betaproteobacteria>Betaproteobacteria_incertae_sedis>Candidatus_Accumulibacter>unclassified_Candidatus_Accumulibacter | GCA_035437285.1 | Accumulibacter_sp. | PLS93_16455 | 1800..4547 | 2053492 | DUF499-domain-containing-protein-[Accumulibacter-sp.] |
| WP_287461342.1 | AAA+ATPase+wHTH+Alpha+Beta-domain+wHTH+RRM | Type-3_BR.PglZ.associated.systems | XIS-HTH->DUF499+RRM*->PglZ->ParB+HNH->METHYLASE->iSTAND->SbcC+SbcC-> | 909 | Bacteria>Pseudomonadota>Betaproteobacteria>Betaproteobacteria_incertae_sedis>Candidatus_Accumulibacter>unclassified_Candidatus_Accumulibacter | GCF_005524045.1 | Accumulibacter_sp. | FAZ92_RS06335 | 1323..4052 | 2053492 | DUF499-domain-containing-protein-[Accumulibacter-sp.] |
| WP_043566948.1 | AAA+ATPase+wHTH+Alpha+Beta-domain+wHTH+RRM | Type-3_BR.PglZ.associated.systems | <-iSTAND<-METHYLASE<-Classical-AAA<-PglZ<-DUF499+RRM*<-XIS-HTH\|\|MODE-HTH+PBPII->RHH-> | 922 | Bacteria>Pseudomonadota>Betaproteobacteria>Betaproteobacteria_incertae_sedis>Candidatus_Accumulibacter>unclassified_Candidatus_Accumulibacter | GCF_035440265.1 | Accumulibacter_sp. | PLC14_RS15270 | 47480..50248 | 2053492 | DUF499-domain-containing-protein-[Accumulibacter-sp.] |
| WP_291996720.1 | AAA+ATPase+wHTH+Alpha+Beta-domain+wHTH+RRM | Type-3_BR.PglZ.associated.systems | <-RelE-ParE\|\|XIS-HTH->DUF499+RRM*->PglZ->METHYLASE->iSTAND->TUDOR+SF1-SF2-Helicase+iREase+CTD-> | 908 | Bacteria>Pseudomonadota>Betaproteobacteria>Betaproteobacteria_incertae_sedis>Candidatus_Accumulibacter>unclassified_Candidatus_Accumulibacter | GCF_907163155.1 | Candidatus_Accumulibacter_sp._ACC012 | QZ648_RS08530 | 35661..38387 | 2823332 | DUF499-domain-containing-protein-[Candidatus-Accumulibacter-sp.-ACC012] |
| WP_024973516.1 | AAA+ATPase+wHTH+Alpha+Beta-domain+wHTH+RRM | Type-3_BR.PglZ.associated.systems | XIS-HTH->DUF499*->PglZ->METHYLASE->iSTAND->TUDOR+SF1-SF2-Helicase+iREase+CTD-><-MORC+HisKin\|\|TraJ-RHH-> | 907 | Bacteria>Pseudomonadota>Betaproteobacteria>Burkholderiales | GCF_000974605.1 | Burkholderiaceae | BEK68_RS19805 | 1502..4225 | 119060 | MULTISPECIES:-DUF499-domain-containing-protein-[Burkholderiaceae] |
| WP_099722418.1 | AAA+ATPase+wHTH+Alpha+Beta-domain+wHTH+RRM | Type-3_BR.PglZ.associated.systems | TetR-HTH->?-><-?\|\|MarR-HTH->Phage_int_SAM_2+Integrase_1->XIS-HTH->DUF499*->PglZ->SIR2->VirB4-FtsK+HerA->?->METHYLASE->iSTAND-> | 909 | Bacteria>Pseudomonadota>Betaproteobacteria>Burkholderiales>Alcaligenaceae | GCF_002750775.1 | Bordetella_bronchiseptica | CJ995_RS20390 | 15086..17815 | 518 | DUF499-domain-containing-protein-[Bordetella-bronchiseptica] |
| WP_099722243.1 | AAA+ATPase+wHTH+Alpha+Beta-domain+wHTH+RRM | Type-3_BR.PglZ.associated.systems | TetR-HTH->?-><-?\|\|MarR-HTH->Phage_int_SAM_2+Integrase_1->XIS-HTH->DUF499*->PglZ->SIR2->VirB4-FtsK+HerA->?->METHYLASE->iSTAND-> | 909 | Bacteria>Pseudomonadota>Betaproteobacteria>Burkholderiales>Alcaligenaceae | GCF_002751245.1 | Bordetella_bronchiseptica | CJJ25_RS21415 | 14968..17697 | 518 | DUF499-domain-containing-protein-[Bordetella-bronchiseptica] |
| WP_144630727.1 | AAA+ATPase+wHTH+Alpha+Beta-domain+wHTH+RRM | Type-3_BR.PglZ.associated.systems | Arm-DNA-bind_3+Phage_int_SAM_3+Phage_integrase-><-TUDOR+SF1-SF2-Helicase+iREase+CTD<-Schlafen+TPR<-iSTAND<-METHYLASE<-PglZ<-DUF499*<-XIS-HTH<-Integrase_1<-Phage_int_SAM_2+Integrase_1<-MarR-HTH | 910 | Bacteria>Pseudomonadota>Betaproteobacteria>Burkholderiales>Alcaligenaceae | GCF_007679965.1 | Bordetella_genomosp._13 | FS845_RS03495 | 757641..760373 | 463040 | DUF499-domain-containing-protein-[Bordetella-genomosp.-13] |
| WP_033451751.1 | AAA+ATPase+wHTH+Alpha+Beta-domain+wHTH+RRM | Type-3_BR.PglZ.associated.systems | <-VirB4-FtsK+HerA<-SIR2<-PglZ<-DUF499*<-XIS-HTH<-Phage_int_SAM_2+Integrase_1<-MarR-HTH\|\|?-><-?<-TetR-HTH | 909 | Bacteria>Pseudomonadota>Betaproteobacteria>Burkholderiales>Alcaligenaceae | GCF_003443855.2 | Bordetella_bronchiseptica | KM22_RS04625 | 3023..5752 | 518 | DUF499-domain-containing-protein-[Bordetella-bronchiseptica] |
| WP_015064471.1 | AAA+ATPase+wHTH+Alpha+Beta-domain+wHTH+RRM | Type-3_BR.PglZ.associated.systems | <-iSTAND<-METHYLASE<-?<-VirB4-FtsK+HerA<-SIR2<-PglZ<-DUF499*<-XIS-HTH<-Phage_int_SAM_2+Integrase_1<-MarR-HTH\|\|?-><-?<-TetR-HTH | 909 | Bacteria>Pseudomonadota>Betaproteobacteria>Burkholderiales>Alcaligenaceae | GCF_000318015.1 | Bordetella_bronchiseptica | BB1_RS13020 | 39685..42414 | 518 | DUF499-domain-containing-protein-[Bordetella-bronchiseptica] |
| WP_033474477.1 | AAA+ATPase+wHTH+Alpha+Beta-domain+wHTH+RRM | Type-3_BR.PglZ.associated.systems | <-iSTAND<-METHYLASE<-?<-VirB4-FtsK+HerA<-SIR2<-PglZ<-DUF499*<-XIS-HTH<-Phage_int_SAM_2+Integrase_1<-MarR-HTH\|\|?-><-?<-TetR-HTH | 909 | Bacteria>Pseudomonadota>Betaproteobacteria>Burkholderiales>Alcaligenaceae | GCF_000662095.1 | Bordetella_bronchiseptica | L490_RS08805 | 21773..24502 | 518 | DUF499-domain-containing-protein-[Bordetella-bronchiseptica] |
| WP_110139123.1 | AAA+ATPase+wHTH+Alpha+Beta-domain+wHTH+RRM | Type-3_BR.PglZ.associated.systems | <-iSTAND<-METHYLASE<-?<-VirB4-FtsK+HerA<-SIR2<-PglZ<-DUF499*<-XIS-HTH<-Phage_int_SAM_2+Integrase_1<-MarR-HTH\|\|?-><-?<-TetR-HTH | 909 | Bacteria>Pseudomonadota>Betaproteobacteria>Burkholderiales>Alcaligenaceae | GCF_003186685.1 | Bordetella_bronchiseptica | B7P00_RS04520 | 964851..967580 | 518 | DUF499-domain-containing-protein-[Bordetella-bronchiseptica] |
| WP_185780658.1 | AAA+ATPase+wHTH+Alpha+Beta-domain+wHTH+RRM | Type-3_BR.PglZ.associated.systems | <-HIN-HTH<-?\|\|?->?->XIS-HTH->DUF499*->PglZ->SIR2->VirB4-FtsK+HerA->METHYLASE->iSTAND->OLD-ABC-> | 912 | Bacteria>Pseudomonadota>Betaproteobacteria>Burkholderiales>Alcaligenaceae | GCF_014237865.1 | Pusillimonas_minor | GTU67_RS13985 | 22665..25403 | 2697024 | DUF499-domain-containing-protein-[Pusillimonas-minor] |
| WP_186025513.1 | AAA+ATPase+wHTH+Alpha+Beta-domain+wHTH+RRM | Type-3_BR.PglZ.associated.systems | TOPRIM+ZnR-><-MODE-HTH+PBPII\|\|?-><-?<-?\|\|tRNA->DUF499*->PglZ->METHYLASE->iSTAND->SF1-SF2-Helicase+iREase+CTD->HTH_Tnp_1+HTH_21+rve+rve_2-> | 909 | Bacteria>Pseudomonadota>Betaproteobacteria>Burkholderiales>Burkholderiaceae | GCF_902830185.1 | Burkholderia_gladioli | KTE24_RS33185 | 25102..27831 | 28095 | DUF499-domain-containing-protein-[Burkholderia-gladioli] |
| WP_091926080.1 | AAA+ATPase+wHTH+Alpha+Beta-domain+wHTH+RRM | Type-3_BR.PglZ.associated.systems | rve+rve_3-><-PRTase\|\|?->?->DUF499*->PglZ->METHYLASE->iSTAND->TUDOR+SF1-SF2-Helicase+iREase+CTD-> | 909 | Bacteria>Pseudomonadota>Betaproteobacteria>Burkholderiales>Burkholderiaceae | GCF_900111715.1 | Burkholderia_cepacia | BMW81_RS40255 | 2792..5521 | 292 | DUF499-domain-containing-protein-[Burkholderia-cepacia] |
| WP_200622122.1 | AAA+ATPase+wHTH+Alpha+Beta-domain+wHTH+RRM | Type-3_BR.PglZ.associated.systems | RelE-ParE->DUF499*->PglZ->METHYLASE->iSTAND-> | 922 | Bacteria>Pseudomonadota>Betaproteobacteria>Burkholderiales>Burkholderiaceae | GCF_905220685.1 | Paraburkholderia_aspalathi | KHO25_RS35560 | 11281..14049 | 1324617 | DUF499-domain-containing-protein-[Paraburkholderia-aspalathi] |
| AET95554.1 | AAA+ATPase+wHTH+Alpha+Beta-domain+wHTH+RRM | Type-3_BR.PglZ.associated.systems | Phage_integrase->XIS-HTH->DUF499*->PglZ->METHYLASE->iSTAND->TUDOR+SF1-SF2-Helicase+iREase+CTD->?-><-HEPN | 915 | Bacteria>Pseudomonadota>Betaproteobacteria>Burkholderiales>Burkholderiaceae | GCA_000236065.1 | Burkholderia_sp._YI23 | BYI23_F000030 | 1550..4297 | 1097668 | hypothetical-protein-BYI23_F000030-[Burkholderia-sp.-YI23] |
| WP_176328878.1 | AAA+ATPase+wHTH+Alpha+Beta-domain+wHTH+RRM | Type-3_BR.PglZ.associated.systems | ParB->?->?->XIS-HTH->DUF499*->PglZ->Classical-AAA->Peptidase_S8->METHYLASE->iSTAND->TUDOR+SF1-SF2-Helicase+iREase+CTD-> | 910 | Bacteria>Pseudomonadota>Betaproteobacteria>Burkholderiales>Burkholderiaceae | GCF_902833435.1 | Burkholderia_vietnamiensis | HW535_RS33775 | 184037..186769 | 60552 | DUF499-domain-containing-protein-[Burkholderia-vietnamiensis] |
| RZF25887.1 | AAA+ATPase+wHTH+Alpha+Beta-domain+wHTH+RRM | Type-3_BR.PglZ.associated.systems | MPTase->?->?-><-?\|\|?->Phage_int_SAM_1+Phage_integrase->DUF499*->PglZ->METHYLASE->iSTAND->TUDOR+SF1-SF2-Helicase+iREase+CTD->STYKinase-like-> | 911 | Bacteria>Pseudomonadota>Betaproteobacteria>Burkholderiales>Burkholderiaceae | GCA_004197395.1 | Paraburkholderia_sp._UYCP14C | EVC45_31230 | 48884..51619 | 2511130 | DUF499-domain-containing-protein-[Paraburkholderia-sp.-UYCP14C] |
| WP_129962032.1 | AAA+ATPase+wHTH+Alpha+Beta-domain+wHTH+RRM | Type-3_BR.PglZ.associated.systems | MPTase->?->?-><-?\|\|?->Phage_int_SAM_1+Phage_integrase->DUF499*->PglZ->METHYLASE->iSTAND->TUDOR+SF1-SF2-Helicase+iREase+CTD->STYKinase-like-> | 911 | Bacteria>Pseudomonadota>Betaproteobacteria>Burkholderiales>Burkholderiaceae | GCF_004197395.1 | Paraburkholderia_sp._UYCP14C | EVC45_31230 | 48884..51619 | 2511130 | DUF499-domain-containing-protein-[Paraburkholderia-sp.-UYCP14C] |
| WP_229061130.1 | AAA+ATPase+wHTH+Alpha+Beta-domain+wHTH+RRM | Type-3_BR.PglZ.associated.systems | MORC+DNA_gyraseB+TOPRIM->DUF499->DUF499*->PglZ->METHYLASE->iSTAND->TUDOR+SF1-SF2-Helicase+iREase+CTD->SF1-SF2-Helicase+iREase+CTD->SymE_toxin-> | 250 | Bacteria>Pseudomonadota>Betaproteobacteria>Burkholderiales>Burkholderiaceae | GCF_020843905.1 | Burkholderia_ambifaria | LL999_RS00030 | 8240..8992 | 152480 | hypothetical-protein-[Burkholderia-ambifaria] |
| WP_229061128.1 | AAA+ATPase+wHTH+Alpha+Beta-domain+wHTH+RRM | Type-3_BR.PglZ.associated.systems | MORC+DNA_gyraseB+TOPRIM->DUF499*->DUF499->PglZ->METHYLASE->iSTAND->TUDOR+SF1-SF2-Helicase+iREase+CTD->SF1-SF2-Helicase+iREase+CTD-> | 661 | Bacteria>Pseudomonadota>Betaproteobacteria>Burkholderiales>Burkholderiaceae | GCF_020843905.1 | Burkholderia_ambifaria | LL999_RS00025 | 6262..8247 | 152480 | DUF499-domain-containing-protein-[Burkholderia-ambifaria] |
| WP_059855476.1 | AAA+ATPase+wHTH+Alpha+Beta-domain+wHTH+RRM | Type-3_BR.PglZ.associated.systems | MORC+DNA_gyraseB+TOPRIM-><-TUDOR+SF1-SF2-Helicase+iREase+CTD<-iSTAND<-METHYLASE<-PglZ<-DUF499*<-XIS-HTH<-?<-STYKinase-like\|\|HTH_31->REase->rve+rve_3+Mu-transpos_C-> | 907 | Bacteria>Pseudomonadota>Betaproteobacteria>Burkholderiales>Burkholderiaceae | GCF_001526755.1 | Burkholderia_cepacia | WK03_RS11770 | 46492..49215 | 292 | DUF499-domain-containing-protein-[Burkholderia-cepacia] |
| WP_124520597.1 | AAA+ATPase+wHTH+Alpha+Beta-domain+wHTH+RRM | Type-3_BR.PglZ.associated.systems | MORC+DNA_gyraseB+TOPRIM->\|\|?-><-TUDOR+SF1-SF2-Helicase+iREase+CTD<-iSTAND<-METHYLASE<-PglZ<-DUF499*<-XIS-HTH<-?<-STYKinase-like\|\|HTH_31->REase->rve+rve_3+Mu-transpos_C-> | 915 | Bacteria>Pseudomonadota>Betaproteobacteria>Burkholderiales>Burkholderiaceae | GCF_003853695.1 | unclassified_Burkholderia | DIE00_RS27530 | 72749..75496 | 2613784 | MULTISPECIES:-DUF499-domain-containing-protein-[unclassified-Burkholderia] |
| WP_060222787.1 | AAA+ATPase+wHTH+Alpha+Beta-domain+wHTH+RRM | Type-3_BR.PglZ.associated.systems | MazG->HNH-><-SymE_toxin<-HTH_3\|\|XIS-HTH->DUF499*->PglZ->METHYLASE->iSTAND->TUDOR+SF1-SF2-Helicase+iREase+CTD-> | 915 | Bacteria>Pseudomonadota>Betaproteobacteria>Burkholderiales>Burkholderiaceae | GCF_001531715.1 | Burkholderia_ubonensis | WL44_RS01920 | 9515..12262 | 101571 | DUF499-domain-containing-protein-[Burkholderia-ubonensis] |
| KJJ99199.1 | AAA+ATPase+wHTH+Alpha+Beta-domain+wHTH+RRM | Type-3_BR.PglZ.associated.systems | CHTH-><-?<-?<-Phage_int_SAM_2+Integrase_1<-?\|\|?->DUF499*->PglZ->?->METHYLASE->iSTAND->TUDOR+SF1-SF2-Helicase+iREase+CTD-> | 910 | Bacteria>Pseudomonadota>Betaproteobacteria>Burkholderiales>Burkholderiaceae | GCA_000955795.1 | Burkholderiaceae_bacterium_26 | UB44_13745 | 405465..408197 | 1619951 | hypothetical-protein-UB44_13745-[Burkholderiaceae-bacterium-26] |
| WP_230938041.1 | AAA+ATPase+wHTH+Alpha+Beta-domain+wHTH+RRM | Type-3_BR.PglZ.associated.systems | <-TUDOR+SF1-SF2-Helicase+iREase+CTD<-iSTAND<-METHYLASE<-VirB4-FtsK+HerA<-SIR2<-PglZ<-DUF499* | 586 | Bacteria>Pseudomonadota>Betaproteobacteria>Burkholderiales>Burkholderiaceae | GCF_001528605.1 | Burkholderia_multivorans | WK33_RS10475 | 225111..226871 | 87883 | hypothetical-protein-[Burkholderia-multivorans] |
| WP_124982684.1 | AAA+ATPase+wHTH+Alpha+Beta-domain+wHTH+RRM | Type-3_BR.PglZ.associated.systems | <-TUDOR+SF1-SF2-Helicase+iREase+CTD<-iSTAND<-METHYLASE<-PglZ<-DUF499*<-XIS-HTH<-ZnR+TOPRIM+D5ATPase<-ParB<-ParB<-GDResolvase+PPP1-HTH | 910 | Bacteria>Pseudomonadota>Betaproteobacteria>Burkholderiales>Burkholderiaceae | GCF_003860725.1 | Ralstonia_solanacearum | EIH14_RS14875 | 3259002..3261734 | 305 | DUF499-domain-containing-protein-[Ralstonia-solanacearum] |
| WP_059636912.1 | AAA+ATPase+wHTH+Alpha+Beta-domain+wHTH+RRM | Type-3_BR.PglZ.associated.systems | <-TUDOR+SF1-SF2-Helicase+iREase+CTD<-iSTAND<-METHYLASE<-PglZ<-DUF499*<-XIS-HTH\|\|HTH_3->SymE_toxin-><-HNH<-MazG | 915 | Bacteria>Pseudomonadota>Betaproteobacteria>Burkholderiales>Burkholderiaceae | GCF_001718355.1 | Burkholderia_ubonensis | WI31_RS04605 | 15203..17950 | 101571 | DUF499-domain-containing-protein-[Burkholderia-ubonensis] |
| WP_186124915.1 | AAA+ATPase+wHTH+Alpha+Beta-domain+wHTH+RRM | Type-3_BR.PglZ.associated.systems | <-TUDOR+SF1-SF2-Helicase+iREase+CTD<-iSTAND<-METHYLASE<-PglZ<-DUF499*<-tRNA\|\|?->?-><-?\|\|MODE-HTH+PBPII-><-TOPRIM+ZnR | 909 | Bacteria>Pseudomonadota>Betaproteobacteria>Burkholderiales>Burkholderiaceae | GCF_902831955.1 | Burkholderia_gladioli | H9E10_RS01495 | 330557..333286 | 28095 | DUF499-domain-containing-protein-[Burkholderia-gladioli] |
| WP_175976581.1 | AAA+ATPase+wHTH+Alpha+Beta-domain+wHTH+RRM | Type-3_BR.PglZ.associated.systems | <-TUDOR+SF1-SF2-Helicase+iREase+CTD<-iSTAND<-METHYLASE<-PglZ<-DUF499*<-?<-?\|\|PRTase-> | 910 | Bacteria>Pseudomonadota>Betaproteobacteria>Burkholderiales>Burkholderiaceae | GCF_902833155.1 | Burkholderia_sp._BCC1047 | HWX55_RS17145 | 46438..49170 | 2676299 | DUF499-domain-containing-protein-[Burkholderia-sp.-BCC1047] |
| WP_212202038.1 | AAA+ATPase+wHTH+Alpha+Beta-domain+wHTH+RRM | Type-3_BR.PglZ.associated.systems | <-TUDOR+SF1-SF2-Helicase+iREase+CTD<-iSTAND<-METHYLASE<-PglZ<-DUF499*\|\|<-MORC+DNA_gyraseB+TOPRIM | 909 | Bacteria>Pseudomonadota>Betaproteobacteria>Burkholderiales>Burkholderiaceae | GCF_018148135.1 | Burkholderia_vietnamiensis | KDW10_RS29905 | 10367..13096 | 60552 | DUF499-domain-containing-protein-[Burkholderia-vietnamiensis] |
| WP_267600860.1 | AAA+ATPase+wHTH+Alpha+Beta-domain+wHTH+RRM | Type-3_BR.PglZ.associated.systems | <-TUDOR+SF1-SF2-Helicase+iREase+CTD<-iSTAND<-?<-METHYLASE<-METHYLASE<-PglZ<-DUF499*<-HTH+WYL\|\|?->?-><-?<-HicB | 908 | Bacteria>Pseudomonadota>Betaproteobacteria>Burkholderiales>Burkholderiaceae | GCF_026627925.1 | Cupriavidus_sp._D39 | OMK73_RS03905 | 81354..84080 | 2997877 | DUF499-domain-containing-protein-[Cupriavidus-sp.-D39] |
| WP_105857077.1 | AAA+ATPase+wHTH+Alpha+Beta-domain+wHTH+RRM | Type-3_BR.PglZ.associated.systems | <-TraJ-RHH\|\|MORC+HisKin-><-TUDOR+SF1-SF2-Helicase+iREase+CTD<-iSTAND<-METHYLASE<-PglZ<-DUF499*<-XIS-HTH\|\|?-><-?\|\|PAIREDC-HTH-> | 907 | Bacteria>Pseudomonadota>Betaproteobacteria>Burkholderiales>Burkholderiaceae | GCF_002981575.1 | Burkholderia_vietnamiensis | C6T65_RS31540 | 70531..73254 | 60552 | DUF499-domain-containing-protein-[Burkholderia-vietnamiensis] |
| WP_254170749.1 | AAA+ATPase+wHTH+Alpha+Beta-domain+wHTH+RRM | Type-3_BR.PglZ.associated.systems | <-TraJ-RHH\|\|MORC+HisKin-><-SF1-SF2-Helicase+iREase+CTD<-iSTAND<-METHYLASE<-PglZ<-DUF499*\|\|?-><-XIS-HTH\|\|?-><-?\|\|?->ParB-> | 778 | Bacteria>Pseudomonadota>Betaproteobacteria>Burkholderiales>Burkholderiaceae | GCF_018726805.1 | Ralstonia_sp._B265 | KOL72_RS03995 | 817362..819698 | 2836825 | DUF499-domain-containing-protein-[Ralstonia-sp.-B265] |
| WP_060275749.1 | AAA+ATPase+wHTH+Alpha+Beta-domain+wHTH+RRM | Type-3_BR.PglZ.associated.systems | <-SymE_toxin<-TUDOR+SF1-SF2-Helicase+iREase+CTD<-iSTAND<-METHYLASE<-PglZ<-DUF499*\|\|<-MORC+DNA_gyraseB+TOPRIM<-?<-?\|\|?->RNaseP-> | 909 | Bacteria>Pseudomonadota>Betaproteobacteria>Burkholderiales>Burkholderiaceae | GCF_001532125.1 | Burkholderia_cepacia | WL99_RS07440 | 8342..11071 | 292 | nucleoporin-[Burkholderia-cepacia] |
| WP_212184034.1 | AAA+ATPase+wHTH+Alpha+Beta-domain+wHTH+RRM | Type-3_BR.PglZ.associated.systems | <-SF1-SF2-Helicase+iREase+CTD<-iSTAND<-METHYLASE<-PglZ<-DUF499*\|\|Phage_integrase-><-MORC+DNA_gyraseB+TOPRIM<-?<-?\|\|?->RNaseP-> | 909 | Bacteria>Pseudomonadota>Betaproteobacteria>Burkholderiales>Burkholderiaceae | GCF_018147225.1 | Burkholderia_vietnamiensis | KDW84_RS23745 | 8248..10977 | 60552 | DUF499-domain-containing-protein-[Burkholderia-vietnamiensis] |
| WP_261533397.1 | AAA+ATPase+wHTH+Alpha+Beta-domain+wHTH+RRM | Type-3_BR.PglZ.associated.systems | <-rve+rve_3+Mu-transpos_C<-HTH_31\|\|XIS-HTH->DUF499*->PglZ->METHYLASE->iSTAND->TUDOR+SF1-SF2-Helicase+iREase+CTD-> | 914 | Bacteria>Pseudomonadota>Betaproteobacteria>Burkholderiales>Burkholderiaceae | GCF_905332655.1 | Burkholderia_multivorans | OCJ25_RS11095 | 7266..10010 | 87883 | DUF499-domain-containing-protein-[Burkholderia-multivorans] |
| WP_160473034.1 | AAA+ATPase+wHTH+Alpha+Beta-domain+wHTH+RRM | Type-3_BR.PglZ.associated.systems | <-REase<-TUDOR+SF1-SF2-Helicase+iREase+CTD<-iSTAND<-METHYLASE<-PglZ<-DUF499*<-PIWI<-TIR+PNTD2<-HTH+WYL\|\|?->DUF932-> | 903 | Bacteria>Pseudomonadota>Betaproteobacteria>Burkholderiales>Burkholderiaceae | GCF_009793345.1 | Cupriavidus_sp._SW-Y-13 | GA566_RS14170 | 3175309..3178020 | 2653854 | DUF499-domain-containing-protein-[Cupriavidus-sp.-SW-Y-13] |
| KGX52783.1 | AAA+ATPase+wHTH+Alpha+Beta-domain+wHTH+RRM | Type-3_BR.PglZ.associated.systems | <-PRTase<-?<-?\|\|?->DUF499*->PglZ->METHYLASE->iSTAND->TUDOR+SF1-SF2-Helicase+iREase+CTD-> | 862 | Bacteria>Pseudomonadota>Betaproteobacteria>Burkholderiales>Burkholderiaceae | GCA_000775115.1 | Burkholderia_pseudomallei_TSV44 | Y024_5460 | 20528..23116 | 1439858 | hypothetical-protein-Y024_5460-[Burkholderia-pseudomallei-TSV44] |
| WP_043304690.1 | AAA+ATPase+wHTH+Alpha+Beta-domain+wHTH+RRM | Type-3_BR.PglZ.associated.systems | <-PRTase\|\|?->DUF499*->PglZ->METHYLASE->iSTAND->TUDOR+SF1-SF2-Helicase+iREase+CTD-> | 909 | Bacteria>Pseudomonadota>Betaproteobacteria>Burkholderiales>Burkholderiaceae | GCF_000775115.1 | Burkholderia_pseudomallei | Y024_RS25770 | 20387..23116 | 28450 | DUF499-domain-containing-protein-[Burkholderia-pseudomallei] |
| WP_133098465.1 | AAA+ATPase+wHTH+Alpha+Beta-domain+wHTH+RRM | Type-3_BR.PglZ.associated.systems | <-PglZ<-DUF499*<-XIS-HTH | 913 | Bacteria>Pseudomonadota>Betaproteobacteria>Burkholderiales>Burkholderiaceae | GCF_004353535.1 | Cupriavidus_sp._L7L | E1J61_RS35410 | 980..3721 | 2546443 | DUF499-domain-containing-protein-[Cupriavidus-sp.-L7L] |
| WP_165590571.1 | AAA+ATPase+wHTH+Alpha+Beta-domain+wHTH+RRM | Type-3_BR.PglZ.associated.systems | <-ParB<-?<-DUF932<-?\|\|XIS-HTH->DUF499*->PglZ->SIR2->VirB4-FtsK+HerA->METHYLASE->iSTAND->TUDOR+SF1-SF2-Helicase+iREase+CTD-> | 914 | Bacteria>Pseudomonadota>Betaproteobacteria>Burkholderiales>Burkholderiaceae | - | Ralstonia_solanacearum | G7947_RS02400 | 517859..520603 | 305 | nucleoporin-[Ralstonia-solanacearum] |
| WP_193027477.1 | AAA+ATPase+wHTH+Alpha+Beta-domain+wHTH+RRM | Type-3_BR.PglZ.associated.systems | <-ParB<-?<-DUF932<-?\|\|XIS-HTH->DUF499*->PglZ->METHYLASE->iSTAND->TUDOR+SF1-SF2-Helicase+iREase+CTD->HEPN-><-Arm-DNA-bind_3+Phage_integrase+Phage_integr_3 | 915 | Bacteria>Pseudomonadota>Betaproteobacteria>Burkholderiales>Burkholderiaceae | GCF_014884705.1 | Ralstonia_pseudosolanacearum | HF908_RS02775 | 577051..579798 | 1310165 | DUF499-domain-containing-protein-[Ralstonia-solanacearum] |
| NKJ47212.1 | AAA+ATPase+wHTH+Alpha+Beta-domain+wHTH+RRM | Type-3_BR.PglZ.associated.systems | <-ParB<-?<-DUF932<-?\|\|HTH+WYL->TIR+PNTD2->DUF499*->PglZ->METHYLASE->iSTAND->TUDOR+SF1-SF2-Helicase+iREase+CTD->PHP+SbcC->Schlafen+HATPase+HTH-> | 908 | Bacteria>Pseudomonadota>Betaproteobacteria>Burkholderiales>Burkholderiaceae | GCA_012275575.1 | Paraburkholderia_sp._SG-MS1 | CIC12_10750 | 68504..71230 | 2023741 | hypothetical-protein-[Burkholderia-sp.-SG-MS1] |
| WP_168664091.1 | AAA+ATPase+wHTH+Alpha+Beta-domain+wHTH+RRM | Type-3_BR.PglZ.associated.systems | <-ParB<-?<-DUF932<-?\|\|HTH+WYL->TIR+PNTD2->DUF499*->PglZ->METHYLASE->iSTAND->TUDOR+SF1-SF2-Helicase+iREase+CTD->PHP+SbcC->Schlafen+HATPase+HTH-> | 908 | Bacteria>Pseudomonadota>Betaproteobacteria>Burkholderiales>Burkholderiaceae | GCF_012275575.1 | Paraburkholderia_sp._SG-MS1 | CIC12_10750 | 68504..71230 | 2023741 | DUF499-domain-containing-protein-[Burkholderia-sp.-SG-MS1] |
| MCA8456511.1 | AAA+ATPase+wHTH+Alpha+Beta-domain+wHTH+RRM | Type-3_BR.PglZ.associated.systems | <-LuxR-HTH\|\|Phage_AlpA->?->TraJ-RHH->RHH->XIS-HTH->DUF499*->PglZ->SIR2->VirB4-FtsK+HerA->METHYLASE->iSTAND->TUDOR+SF1-SF2-Helicase+iREase+CTD-> | 909 | Bacteria>Pseudomonadota>Betaproteobacteria>Burkholderiales>Burkholderiaceae | GCA_020421375.1 | Burkholderia_multivorans | LGN04_21610 | 112280..115009 | 87883 | DUF499-domain-containing-protein-[Burkholderia-multivorans] |
| WP_176314319.1 | AAA+ATPase+wHTH+Alpha+Beta-domain+wHTH+RRM | Type-3_BR.PglZ.associated.systems | <-HIN-HTH<-STYKinase-like<-TUDOR+SF1-SF2-Helicase+iREase+CTD<-iSTAND<-METHYLASE<-PglZ<-DUF499*<-RelE-ParE<-?<-?<-?\|\|HEPN-> | 922 | Bacteria>Pseudomonadota>Betaproteobacteria>Burkholderiales>Burkholderiaceae | GCF_902833535.1 | Burkholderia_vietnamiensis | HW467_RS06840 | 18032..20800 | 60552 | DUF499-domain-containing-protein-[Burkholderia-vietnamiensis] |
| WP_239687442.1 | AAA+ATPase+wHTH+Alpha+Beta-domain+wHTH+RRM | Type-3_BR.PglZ.associated.systems | <-HerA/FtsK<-?<-ParB<-?\|\|?-><-?\|\|DUF499*->PglZ-> | 780 | Bacteria>Pseudomonadota>Betaproteobacteria>Burkholderiales>Burkholderiaceae | - | Burkholderia_cenocepacia | ADM12_RS28385 | 16194..18533 | 95486 | DUF499-domain-containing-protein,-partial-[Burkholderia-cenocepacia] |
| WP_183716828.1 | AAA+ATPase+wHTH+Alpha+Beta-domain+wHTH+RRM | Type-3_BR.PglZ.associated.systems | <-DUF932<-?\|\|HTH+WYL->TIR+PNTD2->PIWI->DUF499*->PglZ->METHYLASE->iSTAND->TUDOR+SF1-SF2-Helicase+iREase+CTD->PHP+SbcC->Schlafen+HATPase+HTH-> | 908 | Bacteria>Pseudomonadota>Betaproteobacteria>Burkholderiales>Burkholderiaceae | GCF_014191085.1 | Paraburkholderia_tropica | FHX57_RS30195 | 108403..111129 | 92647 | DUF499-domain-containing-protein-[Paraburkholderia-tropica] |
| WP_036033575.1 | AAA+ATPase+wHTH+Alpha+Beta-domain+wHTH+RRM | Type-3_BR.PglZ.associated.systems | <-DUF932<-?\|\|HTH+WYL->TIR+PNTD2->PIWI->DUF499*->PglZ->METHYLASE->iSTAND->TUDOR+SF1-SF2-Helicase+iREase+CTD-> | 908 | Bacteria>Pseudomonadota>Betaproteobacteria>Burkholderiales>Burkholderiaceae | GCF_000757585.1 | Burkholderia_gladioli | LA03_RS13700 | 112083..114809 | 28095 | hypothetical-protein-[Burkholderia-gladioli] |
| HDR9280232.1 | AAA+ATPase+wHTH+Alpha+Beta-domain+wHTH+RRM | Type-3_BR.PglZ.associated.systems | DUF499+RRM*->PglZ->METHYLASE->iSTAND-> | 909 | Bacteria>Pseudomonadota>Betaproteobacteria>Burkholderiales>Burkholderiaceae>Burkholderia | GCA_029727675.1 | Burkholderia_vietnamiensis | QDB45_001755 | 129454..132183 | 60552 | DUF499-domain-containing-protein-[Burkholderia-vietnamiensis] |
| CAJ5018355.1 | AAA+ATPase+wHTH+Alpha+Beta-domain+wHTH+RRM | Type-3_BR.PglZ.associated.systems | <-SbcC<-TUDOR+SF1-SF2-Helicase+iREase+CTD<-iSTAND<-METHYLASE-p<-PglZ<-DUF499+RRM*<-WYL\|\|DUF932-> | 908 | Bacteria>Pseudomonadota>Betaproteobacteria>Burkholderiales>Burkholderiaceae>Burkholderia | GCA_963563145.1 | Burkholderia_pseudomallei | SAMEA4874637_04387 | 10651..13377 | 28450 | Uncharacterised-protein-[Burkholderia-pseudomallei] |
| WP_317922488.1 | AAA+ATPase+wHTH+Alpha+Beta-domain+wHTH+RRM | Type-3_BR.PglZ.associated.systems | DUF499+RRM*->PglZ->METHYLASE->iSTAND->TUDOR+SF1-SF2-Helicase+iREase+CTD-> | 917 | Bacteria>Pseudomonadota>Betaproteobacteria>Burkholderiales>Burkholderiaceae>Cupriavidus | GCF_033238665.1 | Cupriavidus_sp._TA19 | Tamer99_RS17225 | 3741..6494 | 701108 | DUF499-domain-containing-protein-[Cupriavidus-sp.-TA19] |
| CAJ0808540.1 | AAA+ATPase+wHTH+Alpha+Beta-domain+wHTH+RRM | Type-3_BR.PglZ.associated.systems | <-Phage_int_SAM_2+Integrase_1\|\|DUF499+RRM*->PglZ->METHYLASE->iSTAND->TUDOR+SF1-SF2-Helicase+iREase+CTD-> | 910 | Bacteria>Pseudomonadota>Betaproteobacteria>Burkholderiales>Burkholderiaceae>Ralstonia | GCA_958415835.1 | Ralstonia_wenshanensis | LMG19087_00247 | 262051..264783 | 2842456 | hypothetical-protein-LMG19087_00247-[Ralstonia-wenshanensis] |
| WP_316877888.1 | AAA+ATPase+wHTH+Alpha+Beta-domain+wHTH+RRM | Type-3_BR.PglZ.associated.systems | HEPN-><-Phage_int_SAM_2+Integrase_1\|\|DUF499+RRM*->PglZ->METHYLASE->iSTAND->TUDOR+SF1-SF2-Helicase+iREase+CTD-> | 889 | Bacteria>Pseudomonadota>Betaproteobacteria>Burkholderiales>Burkholderiaceae>Ralstonia | GCF_958415835.1 | Ralstonia_wenshanensis | R2N09_RS01235 | 262114..264783 | 2842456 | DUF499-domain-containing-protein-[Ralstonia-wenshanensis] |
| WP_316867537.1 | AAA+ATPase+wHTH+Alpha+Beta-domain+wHTH+RRM | Type-3_BR.PglZ.associated.systems | GDResolvase->Cluster83_1clades->\|\|XIS-HTH->DUF499+RRM*->PglZ->METHYLASE->iSTAND->TUDOR+SF1-SF2-Helicase+iREase+CTD-> | 903 | Bacteria>Pseudomonadota>Betaproteobacteria>Burkholderiales>Burkholderiaceae>Ralstonia | GCF_958437825.1 | Ralstonia_mannitolilytica | R2663_RS14955 | 46989..49700 | 105219 | DUF499-domain-containing-protein-[Ralstonia-mannitolilytica] |
| WP_311528734.1 | AAA+ATPase+wHTH+Alpha+Beta-domain+wHTH+RRM | Type-3_BR.PglZ.associated.systems | <-TUDOR+SF1-SF2-Helicase+iREase+CTD<-iSTAND<-METHYLASE<-PglZ<-DUF499+RRM*\|\|<-XIS-HTH<-ParB<-ParB<-GDResolvase+Recombinase | 812 | Bacteria>Pseudomonadota>Betaproteobacteria>Burkholderiales>Burkholderiaceae>Ralstonia | GCF_963512445.1 | uncultured_Ralstonia_sp. | RRB00_RS12225 | 60430..62868 | 114715 | DUF499-domain-containing-protein-[uncultured-Ralstonia-sp.] |
| HJW53974.1 | AAA+ATPase+wHTH+Alpha+Beta-domain+wHTH+RRM | Type-3_BR.PglZ.associated.systems | <-TUDOR+SF1-SF2-Helicase+iREase+CTD<-iSTAND<-METHYLASE<-PglZ<-DUF499+RRM*<-XIS-HTH<-PIN | 909 | Bacteria>Pseudomonadota>Betaproteobacteria>Burkholderiales>Burkholderiaceae>unclassified_Burkholderiaceae | GCA_035255485.1 | Burkholderiaceae_bacterium | VJ577_01750 | 10931..13660 | 2030806 | DUF499-domain-containing-protein-[Burkholderiaceae-bacterium] |
| MBI3381001.1 | AAA+ATPase+wHTH+Alpha+Beta-domain+wHTH+RRM | Type-3_BR.PglZ.associated.systems | HTH+WYL->?->?->XIS-HTH->DUF499*->PglZ->METHYLASE->iSTAND->TUDOR+SF1-SF2-Helicase+iREase+CTD-><-?<-TUDOR+SF1-SF2-Helicase+iREase+CTD | 924 | Bacteria>Pseudomonadota>Betaproteobacteria>Burkholderiales>Burkholderiales_genera_incertae_sedis | GCA_016195855.1 | Aquabacterium_sp. | HYO19_03245 | 359496..362270 | 1872578 | DUF499-domain-containing-protein-[Aquabacterium-sp.] |
| WP_290688310.1 | AAA+ATPase+wHTH+Alpha+Beta-domain+wHTH+RRM | Type-3_BR.PglZ.associated.systems | WYL+wHTH->XIS-HTH->DUF499+RRM*->PglZ->METHYLASE->iSTAND->TUDOR+SF1-SF2-Helicase+iREase+CTD-> | 924 | Bacteria>Pseudomonadota>Betaproteobacteria>Burkholderiales>Burkholderiales_genera_incertae_sedis>Aquabacterium | GCF_016195855.1 | Aquabacterium_sp. | HYO19_RS03255 | 359496..362270 | 1872578 | DUF499-domain-containing-protein-[Aquabacterium-sp.] |
| WP_288127761.1 | AAA+ATPase+wHTH+Alpha+Beta-domain+wHTH+RRM | Type-3_BR.PglZ.associated.systems | XIS-HTH->DUF499+RRM*->PglZ->METHYLASE->iSTAND->TUDOR+SF1-SF2-Helicase+iREase+CTD-> | 911 | Bacteria>Pseudomonadota>Betaproteobacteria>Burkholderiales>Burkholderiales_genera_incertae_sedis>Thiomonas | GCF_027318795.1 | Thiomonas_sp. | M6N45_RS07505 | 1558..4293 | 2047785 | DUF499-domain-containing-protein-[Thiomonas-sp.] |
| WP_265300095.1 | AAA+ATPase+wHTH+Alpha+Beta-domain+wHTH+RRM | Type-3_BR.PglZ.associated.systems | ZnR+TOPRIM+DnaB->XIS-HTH->DUF499*->PglZ->?->METHYLASE-p->iSTAND->Schlafen+TPR->TUDOR+SF1-SF2-Helicase+iREase+CTD-> | 910 | Bacteria>Pseudomonadota>Betaproteobacteria>Burkholderiales>Comamonadaceae | GCF_026127465.1 | Verminephrobacter_eiseniae | D5045_RS13495 | 2764640..2767372 | 364317 | DUF499-domain-containing-protein-[Verminephrobacter-eiseniae] |
| WP_265285930.1 | AAA+ATPase+wHTH+Alpha+Beta-domain+wHTH+RRM | Type-3_BR.PglZ.associated.systems | ZnR+TOPRIM+DnaB->XIS-HTH->DUF499*->PglZ->?->?->PemK_toxin->METHYLASE-p->METHYLASE-> | 913 | Bacteria>Pseudomonadota>Betaproteobacteria>Burkholderiales>Comamonadaceae | GCF_026127525.1 | Verminephrobacter_eiseniae | D5047_RS00495 | 110351..113092 | 364317 | DUF499-domain-containing-protein-[Verminephrobacter-eiseniae] |
| WP_265283655.1 | AAA+ATPase+wHTH+Alpha+Beta-domain+wHTH+RRM | Type-3_BR.PglZ.associated.systems | XIS-HTH->DUF499*->PglZ->Classical-AAA->Peptidase_S8->METHYLASE->iSTAND->OLD-ABC-> | 913 | Bacteria>Pseudomonadota>Betaproteobacteria>Burkholderiales>Comamonadaceae | GCF_026127385.1 | Verminephrobacter_aporrectodeae | D5041_RS01105 | 337752..340493 | 1110389 | DUF499-domain-containing-protein-[Verminephrobacter-aporrectodeae] |
| WP_232081151.1 | AAA+ATPase+wHTH+Alpha+Beta-domain+wHTH+RRM | Type-3_BR.PglZ.associated.systems | Phage_integrase-><-?\|\|HTH_3->DUF499->DUF499*->PglZ->METHYLASE-> | 692 | Bacteria>Pseudomonadota>Betaproteobacteria>Burkholderiales>Comamonadaceae | GCF_001541225.1 | Variovorax_sp._WDL1 | APY03_RS11680 | 2406..4484 | 207745 | DUF499-domain-containing-protein-[Variovorax-sp.-WDL1] |
| WP_145894925.1 | AAA+ATPase+wHTH+Alpha+Beta-domain+wHTH+RRM | Type-3_BR.PglZ.associated.systems | MORC+WLM->?->Phage_integrase-><-REase\|\|XIS-HTH->DUF499*->PglZ->METHYLASE->iSTAND->TUDOR+SF1-SF2-Helicase+iREase+CTD-> | 909 | Bacteria>Pseudomonadota>Betaproteobacteria>Burkholderiales>Comamonadaceae | GCF_007833165.1 | Caenimonas_sedimenti | FN976_RS20490 | 131533..134262 | 2596921 | DUF499-domain-containing-protein-[Caenimonas-sp.-HX-9-20] |
| WP_241520345.1 | AAA+ATPase+wHTH+Alpha+Beta-domain+wHTH+RRM | Type-3_BR.PglZ.associated.systems | LZ_Tnp_IS66+zf-IS66+DDE_Tnp_IS66+DDE_Tnp_IS66_C->Phage_integrase-><-?\|\|HTH_3->DUF499->DUF499*->PglZ->METHYLASE-><-DUF772+DDE_Tnp_1\|\|METHYLASE-p->iSTAND->TUDOR+SF1-SF2-Helicase+iREase+CTD-> | 694 | Bacteria>Pseudomonadota>Betaproteobacteria>Burkholderiales>Comamonadaceae | GCF_002891725.1 | unclassified_Variovorax | CHC06_RS23100 | 371277..373361 | 663243 | MULTISPECIES:-DUF499-domain-containing-protein-[unclassified-Variovorax] |
| WP_232081150.1 | AAA+ATPase+wHTH+Alpha+Beta-domain+wHTH+RRM | Type-3_BR.PglZ.associated.systems | HIN-HTH->?->LZ_Tnp_IS66+zf-IS66+DDE_Tnp_IS66+DDE_Tnp_IS66_C->Phage_integrase-><-?\|\|HTH_3->DUF499*->DUF499->PglZ->METHYLASE-><-DUF772+DDE_Tnp_1\|\|METHYLASE-p->iSTAND-> | 180 | Bacteria>Pseudomonadota>Betaproteobacteria>Burkholderiales>Comamonadaceae | GCF_001541225.1 | unclassified_Variovorax | APY03_RS43635 | 1749..2291 | 663243 | MULTISPECIES:-hypothetical-protein-[unclassified-Variovorax] |
| MCW5655148.1 | AAA+ATPase+wHTH+Alpha+Beta-domain+wHTH+RRM | Type-3_BR.PglZ.associated.systems | GDResolvase+PPP1-HTH->ParB->ParB->XIS-HTH->DUF499*->PglZ->SIR2->VirB4-FtsK+HerA->METHYLASE->iSTAND->TUDOR+SF1-SF2-Helicase+iREase+CTD-> | 914 | Bacteria>Pseudomonadota>Betaproteobacteria>Burkholderiales>Comamonadaceae | GCA_026125785.1 | Hydrogenophaga_sp. | KIT53_15520 | 6474..9218 | 1904254 | MAG:-DUF499-domain-containing-protein-[Hydrogenophaga-sp.] |
| WP_003058386.1 | AAA+ATPase+wHTH+Alpha+Beta-domain+wHTH+RRM | Type-3_BR.PglZ.associated.systems | <-TUDOR+SF1-SF2-Helicase+iREase+CTD<-iSTAND<-METHYLASE<-Peptidase_S8<-Classical-AAA<-PglZ<-DUF499*<-XIS-HTH<-?<-?\|\|?-><-tRNA<-?<-GDResolvase+Recombinase+ZnR | 920 | Bacteria>Pseudomonadota>Betaproteobacteria>Burkholderiales>Comamonadaceae | GCF_000168855.1 | Comamonas_testosteroni | CTESDRAFT_RS21085 | 4612982..4615744 | 285 | DUF499-domain-containing-protein-[Comamonas-testosteroni] |
| WP_061287703.1 | AAA+ATPase+wHTH+Alpha+Beta-domain+wHTH+RRM | Type-3_BR.PglZ.associated.systems | <-TUDOR+SF1-SF2-Helicase+iREase+CTD<-iSTAND<-METHYLASE<-Peptidase_S8<-Classical-AAA<-PglZ<-DUF499*<-XIS-HTH<-?\|\|?->DUF932->?->ParB-> | 920 | Bacteria>Pseudomonadota>Betaproteobacteria>Burkholderiales>Comamonadaceae | GCF_022846195.1 | Delftia | DTS01S_RS10800 | 83630..86392 | 80865 | MULTISPECIES:-DUF499-domain-containing-protein-[Delftia] |
| WP_243854304.1 | AAA+ATPase+wHTH+Alpha+Beta-domain+wHTH+RRM | Type-3_BR.PglZ.associated.systems | <-TUDOR+SF1-SF2-Helicase+iREase+CTD<-iSTAND<-METHYLASE<-?<-SIR2<-PglZ<-DUF499*<-GDResolvase+HIN-HTH<-tRNA<-?<-TPR+TPR<-?<-HAD | 943 | Bacteria>Pseudomonadota>Betaproteobacteria>Burkholderiales>Comamonadaceae | GCF_004364775.1 | Comamonas_sp._JUb58 | EDF71_RS13840 | 170163..172994 | 2485114 | DUF499-domain-containing-protein-[Comamonas-sp.-JUb58] |
| MBY4597996.1 | AAA+ATPase+wHTH+Alpha+Beta-domain+wHTH+RRM | Type-3_BR.PglZ.associated.systems | <-TUDOR+SF1-SF2-Helicase+iREase+CTD<-iSTAND<-?<-Schlafen+HATPase+HTH<-METHYLASE<-PglZ<-DUF499*<-HTH_26\|\|?->?->HDRnase-> | 911 | Bacteria>Pseudomonadota>Betaproteobacteria>Burkholderiales>Comamonadaceae | GCA_019793415.1 | Ottowia_caeni | K3217_20965 | 100483..103218 | 2870339 | DUF499-domain-containing-protein-[Ottowia-caeni] |
| WP_265306803.1 | AAA+ATPase+wHTH+Alpha+Beta-domain+wHTH+RRM | Type-3_BR.PglZ.associated.systems | <-Schlafen+TPR<-iSTAND<-METHYLASE<-VirB4-FtsK+HerA<-SIR2<-PglZ<-DUF499*<-XIS-HTH<-ZnR+TOPRIM+DnaB<-?\|\|?->METHYLASE-p-> | 910 | Bacteria>Pseudomonadota>Betaproteobacteria>Burkholderiales>Comamonadaceae | GCF_026127485.1 | Verminephrobacter_eiseniae | D8B31_RS14985 | 27555..30287 | 364317 | DUF499-domain-containing-protein-[Verminephrobacter-eiseniae] |
| KAB7584466.1 | AAA+ATPase+wHTH+Alpha+Beta-domain+wHTH+RRM | Type-3_BR.PglZ.associated.systems | <-PglZ<-DUF499*<-XIS-HTH<-ZnR+TOPRIM+DnaB | 910 | Bacteria>Pseudomonadota>Betaproteobacteria>Burkholderiales>Comamonadaceae | GCA_004134765.2 | Verminephrobacter_sp._Larva24 | ET532_016930 | 278..3010 | 2509434 | DUF499-domain-containing-protein-[Verminephrobacter-sp.-Larva24] |
| WP_207375510.1 | AAA+ATPase+wHTH+Alpha+Beta-domain+wHTH+RRM | Type-3_BR.PglZ.associated.systems | <-PAIREDC-HTH<-?<-PglZ<-DUF499*<-XIS-HTH<-Phage_integrase<-?<-?<-ABhydrolase | 917 | Bacteria>Pseudomonadota>Betaproteobacteria>Burkholderiales>Comamonadaceae | GCF_017353205.1 | Acidovorax_temperans | J1N44_RS20445 | 1935..4688 | 80878 | DUF499-domain-containing-protein-[Acidovorax-temperans] |
| WP_265258634.1 | AAA+ATPase+wHTH+Alpha+Beta-domain+wHTH+RRM | Type-3_BR.PglZ.associated.systems | <-OLD-ABC<-iSTAND<-METHYLASE<-Peptidase_S8<-Classical-AAA<-PglZ<-DUF499*<-XIS-HTH | 914 | Bacteria>Pseudomonadota>Betaproteobacteria>Burkholderiales>Comamonadaceae | GCF_026127315.1 | Verminephrobacter_aporrectodeae | D5038_RS14895 | 1638771..1641515 | 1110389 | DUF499-domain-containing-protein-[Verminephrobacter-aporrectodeae] |
| WP_011807907.1 | AAA+ATPase+wHTH+Alpha+Beta-domain+wHTH+RRM | Type-3_BR.PglZ.associated.systems | <-METHYLASE-p<-?\|\|?->ZnR+TOPRIM+DnaB->XIS-HTH->DUF499*->PglZ->Classical-AAA->Peptidase_S8->METHYLASE->iSTAND->Schlafen+TPR-> | 910 | Bacteria>Pseudomonadota>Betaproteobacteria>Burkholderiales>Comamonadaceae | GCF_000015565.1 | Verminephrobacter_eiseniae | VEIS_RS00490 | 112310..115042 | 364317 | DUF499-domain-containing-protein-[Verminephrobacter-eiseniae] |
| WP_279849361.1 | AAA+ATPase+wHTH+Alpha+Beta-domain+wHTH+RRM | Type-3_BR.PglZ.associated.systems | <-METHYLASE-p\|\|DUF772+DDE_Tnp_1-><-METHYLASE<-Peptidase_S8<-Classical-AAA<-PglZ<-DUF499*<-XIS-HTH<-ParB<-ParB<-GDResolvase+PPP1-HTH<-?<-AbiEii | 915 | Bacteria>Pseudomonadota>Betaproteobacteria>Burkholderiales>Comamonadaceae | GCF_029841605.1 | Comamonas_aquatica | N5D37_RS08145 | 1644651..1647398 | 225991 | DUF499-domain-containing-protein-[Comamonas-aquatica] |
| QIL80928.1 | AAA+ATPase+wHTH+Alpha+Beta-domain+wHTH+RRM | Type-3_BR.PglZ.associated.systems | <-HTH_3\|\|?->ABhydrolase->?->XIS-HTH->DUF499*->PglZ->METHYLASE->iSTAND->OLD-ABC->TOPRIM->TUDOR+SF1-SF2-Helicase+iREase+CTD-> | 918 | Bacteria>Pseudomonadota>Betaproteobacteria>Burkholderiales>Comamonadaceae | GCA_011305995.1 | Diaphorobacter_sp._HDW4A | G7047_14235 | 3121074..3123830 | 2714924 | DUF499-domain-containing-protein-[Diaphorobacter-sp.-HDW4A] |
| WP_240939489.1 | AAA+ATPase+wHTH+Alpha+Beta-domain+wHTH+RRM | Type-3_BR.PglZ.associated.systems | <-HTH_3\|\|?->ABhydrolase->?->XIS-HTH->DUF499*->PglZ->METHYLASE->iSTAND->OLD-ABC->TOPRIM->TUDOR+SF1-SF2-Helicase+iREase+CTD-> | 855 | Bacteria>Pseudomonadota>Betaproteobacteria>Burkholderiales>Comamonadaceae | GCA_011305995.1 | Diaphorobacter_sp._HDW4A | G7047_RS14230 | 3121263..3123830 | 2714924 | DUF499-domain-containing-protein-[Diaphorobacter-sp.-HDW4A] |
| WP_265289617.1 | AAA+ATPase+wHTH+Alpha+Beta-domain+wHTH+RRM | Type-3_BR.PglZ.associated.systems | <-DOC+HTH\|\|XIS-HTH->DUF499*->PglZ->Classical-AAA->Peptidase_S8->METHYLASE->iSTAND->Schlafen+TPR-> | 879 | Bacteria>Pseudomonadota>Betaproteobacteria>Burkholderiales>Comamonadaceae | GCF_026127525.1 | Verminephrobacter_eiseniae | D5047_RS22040 | 4723398..4726037 | 364317 | DUF499-domain-containing-protein-[Verminephrobacter-eiseniae] |
| WP_287803793.1 | AAA+ATPase+wHTH+Alpha+Beta-domain+wHTH+RRM | Type-3_BR.PglZ.associated.systems | XIS-HTH->DUF499+RRM*->PglZ->METHYLASE->iSTAND->TUDOR+SF1-SF2-Helicase+iREase+CTD-> | 917 | Bacteria>Pseudomonadota>Betaproteobacteria>Burkholderiales>Comamonadaceae>Diaphorobacter | GCF_022729045.1 | Diaphorobacter_sp. | NW886_RS05350 | 99441..102194 | 1934310 | DUF499-domain-containing-protein-[Diaphorobacter-sp.] |
| WP_297754296.1 | AAA+ATPase+wHTH+Alpha+Beta-domain+wHTH+RRM | Type-3_BR.PglZ.associated.systems | GDResolvase+Recombinase->ParB->ParB->XIS-HTH->DUF499+RRM*->PglZ->SIR2->VirB4-FtsK+HerA->METHYLASE->iSTAND-> | 914 | Bacteria>Pseudomonadota>Betaproteobacteria>Burkholderiales>Comamonadaceae>Hydrogenophaga | GCF_026125785.1 | Hydrogenophaga_sp. | KIT53_RS15595 | 6474..9218 | 1904254 | DUF499-domain-containing-protein-[Hydrogenophaga-sp.] |
| HOE40136.1 | AAA+ATPase+wHTH+Alpha+Beta-domain+wHTH+RRM | Type-3_BR.PglZ.associated.systems | <-TUDOR+SF1-SF2-Helicase+iREase+CTD<-iSTAND<-METHYLASE<-PglZ<-DUF499+RRM*<-XIS-HTH | 914 | Bacteria>Pseudomonadota>Betaproteobacteria>Burkholderiales>Comamonadaceae>Rhodoferax | GCA_035368905.1 | Rhodoferax_sp. | PLB25_00645 | 16244..18988 | 50421 | DUF499-domain-containing-protein-[Rhodoferax-sp.] |
| WP_307642257.1 | AAA+ATPase+wHTH+Alpha+Beta-domain+wHTH+RRM | Type-3_BR.PglZ.associated.systems | <-TUDOR+SF1-SF2-Helicase+iREase+CTD<-iSTAND<-METHYLASE<-PglZ<-DUF499+RRM*<-CHTH<-HTH_3\|\|METHYLASE-p-> | 911 | Bacteria>Pseudomonadota>Betaproteobacteria>Burkholderiales>Comamonadaceae>Variovorax | GCF_030811995.1 | Variovorax_paradoxus | J2X90_RS28335 | 20502..23237 | 34073 | DUF499-domain-containing-protein-[Variovorax-paradoxus] |
| WP_241453801.1 | AAA+ATPase+wHTH+Alpha+Beta-domain+wHTH+RRM | Type-3_BR.PglZ.associated.systems | MORC+DNA_gyraseB+TOPRIM-><-ParB+HNH-p<-TUDOR+SF1-SF2-Helicase+iREase+CTD<-iSTAND<-METHYLASE<-PglZ<-DUF499*<-HTH+WYL\|\|STYKinase-like->?-><-DUF2026<-HEPN<-HTH_3 | 905 | Bacteria>Pseudomonadota>Betaproteobacteria>Burkholderiales>Oxalobacteraceae | GCF_022568815.1 | Undibacterium_sp._TS12 | LZ463_RS03270 | 735782..738499 | 2908202 | DUF499-domain-containing-protein-[Undibacterium-sp.-TS12] |
| WP_092435923.1 | AAA+ATPase+wHTH+Alpha+Beta-domain+wHTH+RRM | Type-3_BR.PglZ.associated.systems | <-URI-DNase\|\|?->GDResolvase+PPP1-HTH->ParB->ParB->XIS-HTH->DUF499*->PglZ->METHYLASE->iSTAND->TUDOR+SF1-SF2-Helicase+iREase+CTD-> | 941 | Bacteria>Pseudomonadota>Betaproteobacteria>Burkholderiales>Oxalobacteraceae | GCF_900111995.1 | Collimonas_sp._OK607 | BM029_RS10985 | 20800..23625 | 1798194 | DUF499-domain-containing-protein-[Collimonas-sp.-OK607] |
| RYF23847.1 | AAA+ATPase+wHTH+Alpha+Beta-domain+wHTH+RRM | Type-3_BR.PglZ.associated.systems | <-TUDOR+SF1-SF2-Helicase+iREase+CTD<-iSTAND<-METHYLASE<-PglZ<-DUF499* | 451 | Bacteria>Pseudomonadota>Betaproteobacteria>Burkholderiales>Oxalobacteraceae | GCA_004144765.1 | Oxalobacteraceae_bacterium | EOO77_01345 | 7709..9066 | 1871070 | DUF499-domain-containing-protein,-partial-[Oxalobacteraceae-bacterium] |
| WP_262835447.1 | AAA+ATPase+wHTH+Alpha+Beta-domain+wHTH+RRM | Type-3_BR.PglZ.associated.systems | <-SIGMA-HTH\|\|?->?->XIS-HTH->DUF499*->PglZ->METHYLASE->iSTAND->TUDOR+SF1-SF2-Helicase+iREase+CTD->?-><-DUF488-l | 915 | Bacteria>Pseudomonadota>Betaproteobacteria>Burkholderiales>Oxalobacteraceae | GCF_025548535.1 | Undibacterium_sp._Jales_W-56 | LPB67_RS02300 | 520019..522766 | 2897325 | ATP-binding-protein-[Undibacterium-sp.-Jales-W-56] |
| WP_228894839.1 | AAA+ATPase+wHTH+Alpha+Beta-domain+wHTH+RRM | Type-3_BR.PglZ.associated.systems | <-RADICAL-SAM<-?<-?\|\|HTH+WYL->DUF499*->PglZ->METHYLASE->iSTAND->?->TUDOR+SF1-SF2-Helicase+iREase+CTD-> | 917 | Bacteria>Pseudomonadota>Betaproteobacteria>Burkholderiales>Oxalobacteraceae | GCF_020865585.1 | Pseudoduganella_aquatica | KT882_RS05635 | 1286945..1289698 | 2660641 | DUF499-domain-containing-protein-[Massilia-sp.-HC52] |
| WP_322248650.1 | AAA+ATPase+wHTH+Alpha+Beta-domain+wHTH+RRM | Type-3_BR.PglZ.associated.systems | <-RHH\|\|WYL->DUF499+RRM*->PglZ->METHYLASE->iSTAND->TUDOR+SF1-SF2-Helicase+iREase+CTD-> | 904 | Bacteria>Pseudomonadota>Betaproteobacteria>Burkholderiales>Oxalobacteraceae>Undibacterium | GCF_034347425.1 | unclassified_Undibacterium | QN364_RS06135 | 35909..38623 | 2630295 | MULTISPECIES:-DUF499-domain-containing-protein-[unclassified-Undibacterium] |
| ODV10321.1 | AAA+ATPase+wHTH+Alpha+Beta-domain+wHTH+RRM | Type-3_BR.PglZ.associated.systems | <-PglZ<-DUF499*<-XIS-HTH<-?<-?<-?<-ParB | 909 | Bacteria>Pseudomonadota>Betaproteobacteria>Burkholderiales>Sphaerotilaceae | GCA_001725505.1 | Rubrivivax_sp._SCN_70-15 | ABT20_08395 | 1159..3888 | 1660139 | MAG:-DUF499-domain-containing-protein-[Rubrivivax-sp.-SCN-70-15] |
| WP_322586885.1 | AAA+ATPase+wHTH+Alpha+Beta-domain+wHTH+RRM | Type-3_BR.PglZ.associated.systems | XIS-HTH->DUF499+RRM*->PglZ->METHYLASE-p->iSTAND-> | 922 | Bacteria>Pseudomonadota>Betaproteobacteria>Burkholderiales>Sphaerotilaceae>Sphaerotilus | GCF_034519685.1 | Sphaerotilus_sp. | U5044_RS02520 | 443079..445847 | 2093942 | nucleoporin-[Sphaerotilus-sp.] |
| MBK6594595.1 | AAA+ATPase+wHTH+Alpha+Beta-domain+wHTH+RRM | Type-3_BR.PglZ.associated.systems | PIN-><-TUDOR+SF1-SF2-Helicase+iREase+CTD<-iSTAND<-HEPN<-METHYLASE<-PglZ<-DUF499*\|\|?-><-Phage_int_SAM_1+Phage_integrase | 917 | Bacteria>Pseudomonadota>Betaproteobacteria>Burkholderiales>unclassified_Burkholderiales | GCA_016703945.1 | Burkholderiales_bacterium | IPG23_18880 | 28989..31742 | 1891238 | DUF499-domain-containing-protein-[Burkholderiales-bacterium] |
| WP_193583345.1 | AAA+ATPase+wHTH+Alpha+Beta-domain+wHTH+RRM | Type-3_BR.PglZ.associated.systems | Arm-DNA-bind_3+Phage_integrase-><-CopG/RHH\|\|XIS-HTH->?->DOC+DOC->\|\|XIS-HTH->DUF499*->PglZ->Classical-AAA->Peptidase_S8->METHYLASE->iSTAND->OLD-ABC-> | 912 | Bacteria>Pseudomonadota>Betaproteobacteria>Neisseriales>Aquaspirillaceae | GCF_022394265.1 | Laribacter_hongkongensis | LH451_RS13700 | 37041..39779 | 168471 | DUF499-domain-containing-protein-[Laribacter-hongkongensis] |
| WP_199067553.1 | AAA+ATPase+wHTH+Alpha+Beta-domain+wHTH+RRM | Type-3_BR.PglZ.associated.systems | <-HIN-HTH\|\|XIS-HTH->DUF499*->PglZ->Classical-AAA->Peptidase_S8->METHYLASE->iSTAND->SF1-SF2-Helicase+iREase+CTD-> | 915 | Bacteria>Pseudomonadota>Betaproteobacteria>Neisseriales>Chromobacteriaceae | GCF_016429325.1 | Chromobacterium_sp._ASV5 | JHD66_RS14020 | 717..3464 | 2795109 | DUF499-domain-containing-protein-[Chromobacterium-sp.-ASV5] |
| WP_058355577.1 | AAA+ATPase+wHTH+Alpha+Beta-domain+wHTH+RRM | Type-3_BR.PglZ.associated.systems | <-REase+N6_MTase<-TUDOR+SF1-SF2-Helicase+iREase+CTD<-iSTAND<-METHYLASE<-PglZ<-DUF499*<-XIS-HTH<-HTH_Tnp\|\|HTH_3-><-TetR-HTH<-NACHT<-HTH_Tnp_1+HTH_21+rve+rve_2 | 893 | Bacteria>Pseudomonadota>Betaproteobacteria>Neisseriales>Neisseriaceae | GCF_902375805.1 | Vitreoscilla_massiliensis | LVJ82_RS01920 | 368282..370963 | 1689272 | hypothetical-protein-[Vitreoscilla-massiliensis] |
| MBY0576970.1 | AAA+ATPase+wHTH+Alpha+Beta-domain+wHTH+RRM | Type-3_BR.PglZ.associated.systems | AbiEi_3_N+AbiEi_3->AbiEii->DUF499*->PglZ->METHYLASE->iSTAND->TUDOR+SF1-SF2-Helicase+iREase+CTD->REase-> | 908 | Bacteria>Pseudomonadota>Betaproteobacteria>Nitrosomonadales>Gallionellaceae | GCA_019753125.1 | Gallionellaceae_bacterium | K2P67_10320 | 1411..4137 | 2052162 | DUF499-domain-containing-protein-[Gallionellaceae-bacterium] |
| WP_074776128.1 | AAA+ATPase+wHTH+Alpha+Beta-domain+wHTH+RRM | Type-3_BR.PglZ.associated.systems | ParA+HTH-><-?<-TUDOR+SF1-SF2-Helicase+iREase+CTD<-iSTAND<-METHYLASE<-PglZ<-DUF499*<-XIS-HTH<-GDResolvase+HIN-HTH<-ParB+HNH\|\|DDE_Tnp_1-> | 910 | Bacteria>Pseudomonadota>Betaproteobacteria>Nitrosomonadales>Nitrosomonadaceae | GCF_900107715.1 | Nitrosospira_multiformis | BLV78_RS14485 | 8979..11711 | 1231 | DUF499-domain-containing-protein-[Nitrosospira-multiformis] |
| WP_074456054.1 | AAA+ATPase+wHTH+Alpha+Beta-domain+wHTH+RRM | Type-3_BR.PglZ.associated.systems | <-TUDOR+SF1-SF2-Helicase+iREase+CTD<-iSTAND<-METHYLASE<-PglZ<-DUF499*<-XIS-HTH | 911 | Bacteria>Pseudomonadota>Betaproteobacteria>Nitrosomonadales>Nitrosomonadaceae | GCF_900100815.1 | Nitrosomonas_eutropha | BLP42_RS07295 | 27222..29957 | 916 | DUF499-domain-containing-protein-[Nitrosomonas-eutropha] |
| MBK8336964.1 | AAA+ATPase+wHTH+Alpha+Beta-domain+wHTH+RRM | Type-3_BR.PglZ.associated.systems | XIS-HTH->DUF499*->PglZ->METHYLASE->iSTAND->TUDOR+SF1-SF2-Helicase+iREase+CTD->HEPN-> | 914 | Bacteria>Pseudomonadota>Betaproteobacteria>Nitrosomonadales>Sterolibacteriaceae | GCA_016711925.1 | Candidatus_Methylophosphatis_haderslevensis | IPL03_10375 | 329..3073 | 2899264 | DUF499-domain-containing-protein-[Sterolibacteriaceae-bacterium] |
| MDP2795245.1 | AAA+ATPase+wHTH+Alpha+Beta-domain+wHTH+RRM | Type-3_BR.PglZ.associated.systems | <-DUF4276<-SbcC+SbcC<-iSTAND<-METHYLASE<-PglZ<-DUF499+RRM*<-XIS-HTH\|\|HTH_3-><-RelE-ParE | 912 | Bacteria>Pseudomonadota>Betaproteobacteria>Nitrosomonadales>Sterolibacteriaceae>Sulfurisoma | GCA_030691075.1 | Sulfurisoma_sp. | Q8O25_14410 | 15256..17994 | 1965311 | DUF499-domain-containing-protein-[Sulfurisoma-sp.] |
| MBN8761840.1 | AAA+ATPase+wHTH+Alpha+Beta-domain+wHTH+RRM | Type-3_BR.PglZ.associated.systems | SF1-SF2-Helicase-><-SbcC+SbcC<-Calcineurin\|\|XIS-HTH->DUF499*->PglZ->SIR2->VirB4-FtsK+HerA->METHYLASE->iSTAND->TUDOR+SF1-SF2-Helicase+iREase+CTD-> | 939 | Bacteria>Pseudomonadota>Betaproteobacteria>Nitrosomonadales>Thiobacillaceae | GCA_017309235.1 | Thiobacillus_sp. | JOI63_00595 | 15907..18726 | 924 | DUF499-domain-containing-protein-[Thiobacillus-sp.] |
| MBN8780443.1 | AAA+ATPase+wHTH+Alpha+Beta-domain+wHTH+RRM | Type-3_BR.PglZ.associated.systems | <-TPR+TPR+TPR+TPR<-?<-?<-NERD+SF1-SF2-Helicase<-REase\|\|XIS-HTH->DUF499*->PglZ->METHYLASE->iSTAND->TUDOR+SF1-SF2-Helicase+iREase+CTD->ParB+HNH->Methylase-> | 912 | Bacteria>Pseudomonadota>Betaproteobacteria>Nitrosomonadales>Thiobacillaceae | GCA_017304205.1 | Thiobacillus_sp. | JOJ07_12925 | 541953..544691 | 924 | DUF499-domain-containing-protein-[Thiobacillus-sp.] |
| WP_296652459.1 | AAA+ATPase+wHTH+Alpha+Beta-domain+wHTH+RRM | Type-3_BR.PglZ.associated.systems | <-iSTAND<-METHYLASE<-VirB4-FtsK+HerA<-SIR2<-PglZ<-DUF499+RRM*<-XIS-HTH\|\|SbcC-><-TUDOR+SF1-SF2-Helicase+iREase+CTD | 939 | Bacteria>Pseudomonadota>Betaproteobacteria>Nitrosomonadales>Thiobacillaceae>Thiobacillus | GCF_001899765.1 | Thiobacillus_sp._63-78 | BGP20_RS13050 | 14820..17639 | 1895859 | DUF499-domain-containing-protein-[Thiobacillus-sp.-63-78] |
| WP_296844157.1 | AAA+ATPase+wHTH+Alpha+Beta-domain+wHTH+RRM | Type-3_BR.PglZ.associated.systems | <-AAA-ATPase+REase<-AAA-ATPase+REase\|\|XIS-HTH->DUF499+RRM*->PglZ->METHYLASE->iSTAND->TUDOR+SF1-SF2-Helicase+iREase+CTD->ParB+HNH-> | 912 | Bacteria>Pseudomonadota>Betaproteobacteria>Nitrosomonadales>Thiobacillaceae>Thiobacillus | GCF_017304205.1 | Thiobacillus_sp. | JOJ07_RS12940 | 541953..544691 | 924 | DUF499-domain-containing-protein-[Thiobacillus-sp.] |
| MBI5006212.1 | AAA+ATPase+wHTH+Alpha+Beta-domain+wHTH+RRM | Type-3_BR.PglZ.associated.systems | <-TUDOR+SF1-SF2-Helicase+iREase+CTD<-iSTAND<-METHYLASE<-?<-PglZ<-DUF499*\|\|HD->?-><-?<-HEPN<-?<-CHTH | 904 | Bacteria>Pseudomonadota>Betaproteobacteria>Nitrosomonadales>unclassified_Nitrosomonadales | GCA_016214695.1 | Nitrosomonadales_bacterium | HZB95_03705 | 753813..756527 | 2026761 | DUF499-domain-containing-protein-[Nitrosomonadales-bacterium] |
| WP_211164044.1 | AAA+ATPase+wHTH+Alpha+Beta-domain+wHTH+RRM | Type-3_BR.PglZ.associated.systems | <-METHYLASE<-Peptidase_S8<-Classical-AAA<-PglZ<-DUF499-p<-DUF499*<-XIS-HTH | 320 | Bacteria>Pseudomonadota>Betaproteobacteria>Rhodocyclales | GCF_029982735.1 | Zoogloeaceae | GO613_RS23385 | 10194..11156 | 2008794 | hypothetical-protein-[Parazoarcus-communis] |
| WP_274740636.1 | AAA+ATPase+wHTH+Alpha+Beta-domain+wHTH+RRM | Type-3_BR.PglZ.associated.systems | GDResolvase+PPP1-HTH->ParB->ParB->XIS-HTH->DUF499*->PglZ->METHYLASE->METHYLASE->iSTAND->TUDOR+SF1-SF2-Helicase+iREase+CTD-> | 912 | Bacteria>Pseudomonadota>Betaproteobacteria>Rhodocyclales>Azonexaceae | GCF_028864255.1 | Dechloromonas_agitata | PTW32_RS18765 | 3097..5835 | 73030 | DUF499-domain-containing-protein-[Dechloromonas-agitata] |
| MBX9847895.1 | AAA+ATPase+wHTH+Alpha+Beta-domain+wHTH+RRM | Type-3_BR.PglZ.associated.systems | <-METHYLASE-p<-Peptidase_S8<-Classical-AAA<-PglZ<-DUF499* | 881 | Bacteria>Pseudomonadota>Betaproteobacteria>Rhodocyclales>Rhodocyclaceae | GCA_019746535.1 | Rhodocyclaceae_bacterium | K2X64_01275 | 5860..8505 | 1898103 | DUF499-domain-containing-protein-[Rhodocyclaceae-bacterium] |
| MDY0012930.1 | AAA+ATPase+wHTH+Alpha+Beta-domain+wHTH+RRM | Type-3_BR.PglZ.associated.systems | Arm-DNA-bind_3+Phage_integrase-><-Phage_int_SAM_2+Integrase_1\|\|XIS-HTH->DUF499+RRM*->PglZ->SIR2->VirB4-FtsK+HerA->METHYLASE->iSTAND-> | 914 | Bacteria>Pseudomonadota>Betaproteobacteria>Rhodocyclales>Rhodocyclaceae>unclassified_Rhodocyclaceae | GCA_034004685.1 | Rhodocyclaceae_bacterium | RBS40_08585 | 41209..43953 | 1898103 | DUF499-domain-containing-protein-[Rhodocyclaceae-bacterium] |
| HEY3431396.1 | AAA+ATPase+wHTH+Alpha+Beta-domain+wHTH+RRM | Type-3_BR.PglZ.associated.systems | <-TUDOR+SF1-SF2-Helicase+iREase+CTD<-iSTAND<-METHYLASE<-PglZ<-DUF499+RRM*<-XIS-HTH | 915 | Bacteria>Pseudomonadota>Betaproteobacteria>Rhodocyclales>Rhodocyclaceae>unclassified_Rhodocyclaceae | GCA_036504515.1 | Rhodocyclaceae_bacterium | VGK09_02500 | 539128..541875 | 1898103 | DUF499-domain-containing-protein-[Rhodocyclaceae-bacterium] |
| MBI5901627.1 | AAA+ATPase+wHTH+Alpha+Beta-domain+wHTH+RRM | Type-3_BR.PglZ.associated.systems | <-TUDOR+SF1-SF2-Helicase+iREase+CTD<-iSTAND<-METHYLASE<-PIN<-CopG/RHH<-PglZ<-DUF499* | 798 | Bacteria>Pseudomonadota>Betaproteobacteria>Rhodocyclales>unclassified_Rhodocyclales | GCA_016235165.1 | Rhodocyclales_bacterium | HZB40_20700 | 26495..28891 | 2072421 | hypothetical-protein-[Rhodocyclales-bacterium] |
| WP_211164043.1 | AAA+ATPase+wHTH+Alpha+Beta-domain+wHTH+RRM | Type-3_BR.PglZ.associated.systems | <-METHYLASE<-Peptidase_S8<-Classical-AAA<-PglZ<-DUF499*<-DUF499<-XIS-HTH | 599 | Bacteria>Pseudomonadota>Betaproteobacteria>Rhodocyclales>Zoogloeaceae | GCF_012910825.1 | Parazoarcus_communis | GO613_RS14065 | 8425..10224 | 41977 | hypothetical-protein-[Parazoarcus-communis] |

| Protein-ID | DUF499-architecture | System | Gene-Neighbourhoods | Length | Taxonomy | Assembly-ID | Strain | Locus-Tag | Genomic-coordinates | TaxID | Fasta-define |
|---|---|---|---|---|---|---|---|---|---|---|---|
| MDA8127651.1 | AAA+ATPase+wHTH+Alpha+Beta-domain+wHTH+RRM | Type-3_BR.PglZ.associated.systems | ZnR->?->?->?->?->RHH->DUF499*->PglZ->?->METHYLASE->iSTAND->TUDOR+SF1-SF2-Helicase+iREase+CTD->XIS-HTH-> | 911 | Bacteria>Pseudomonadota>Betaproteobacteria>unclassified_Betaproteobacteria | GCA_028065115.1 | Betaproteobacteria_bacterium | MOZ73_02970 | 224672..227407 | 1891241 | MAG:-DUF499-domain-containing-protein-[Betaproteobacteria-bacterium] |
| MCL1825806.1 | AAA+ATPase+wHTH+Alpha+Beta-domain+wHTH+RRM | Type-3_BR.PglZ.associated.systems | Phage_int_SAM_2+Integrase_1->DUF499*->PglZ->?->?->PIN->METHYLASE->iSTAND-> | 936 | Bacteria>Pseudomonadota>Betaproteobacteria>unclassified_Betaproteobacteria | GCA_009782565.1 | Betaproteobacteria_bacterium | FWG26_07705 | 3081..5891 | 1891241 | DUF499-domain-containing-protein-[Betaproteobacteria-bacterium] |
| MDA8260928.1 | AAA+ATPase+wHTH+Alpha+Beta-domain+wHTH+RRM | Type-3_BR.PglZ.associated.systems | ParB->?->?->?->XIS-HTH->DUF499*->PglZ->Classical-AAA->Peptidase_S8->METHYLASE->iSTAND->TUDOR+SF1-SF2-Helicase+iREase+CTD-> | 909 | Bacteria>Pseudomonadota>Betaproteobacteria>unclassified_Betaproteobacteria | GCA_028065725.1 | Betaproteobacteria_bacterium | MOZ99_35710 | 22705..25434 | 1891241 | MAG:-DUF499-domain-containing-protein-[Betaproteobacteria-bacterium] |
| ESS14724.1 | AAA+ATPase+wHTH+Alpha+Beta-domain+wHTH+RRM | Type-3_BR.PglZ.associated.systems | Arm-DNA-bind_3+Phage_integrase-><-TUDOR+SF1-SF2-Helicase+iREase+CTD<-iSTAND<-HEPN<-METHYLASE<-PglZ<-DUF499*<-RHH\|\|?->XIS-HTH-> | 918 | Bacteria>Pseudomonadota>Betaproteobacteria>unclassified_Betaproteobacteria | GCA_000496475.1 | Betaproteobacteria_bacterium_MOLA814 | MOLA814_00285 | 295656..298412 | 1408164 | hypothetical-protein-MOLA814_00285-[Betaproteobacteria-bacterium-MOLA814] |
| MBI4291280.1 | AAA+ATPase+wHTH+Alpha+Beta-domain+wHTH+RRM | Type-3_BR.PglZ.associated.systems | <-REase<-REase+DUF4268<-TUDOR+SF1-SF2-Helicase+iREase+CTD<-iSTAND<-METHYLASE<-PglZ<-DUF499*<-?\|\|?->XIS-HTH->?->ZnR+TOPRIM-> | 941 | Bacteria>Pseudomonadota>Betaproteobacteria>unclassified_Betaproteobacteria | GCA_016210745.1 | Betaproteobacteria_bacterium | HY661_07375 | 22687..25512 | 1891241 | DUF499-domain-containing-protein-[Betaproteobacteria-bacterium] |
| QSO22112.1 | AAA+ATPase+wHTH+Alpha+Beta-domain+wHTH+RRM | Type-3_BR.PglZ.associated.systems | REase->Mu-transpos_C->TnsC-ATPase->?->?->XIS-HTH->DUF499*->PglZ->SIR2->VirB4-FtsK+HerA->METHYLASE->iSTAND->TUDOR+SF1-SF2-Helicase+iREase+CTD-> | 919 | Bacteria>Pseudomonadota>Gammaproteobacteria>Aeromonadales>Aeromonadaceae | GCA_017280155.1 | Aeromonas_caviae | JZM63_17155 | 3636831..3639590 | 648 | DUF499-domain-containing-protein-[Aeromonas-caviae] |
| WP_215783453.1 | AAA+ATPase+wHTH+Alpha+Beta-domain+wHTH+RRM | Type-3_BR.PglZ.associated.systems | DUF499*->PglZ->METHYLASE->METHYLASE->iSTAND->TUDOR+SF1-SF2-Helicase+iREase+CTD-> | 919 | Bacteria>Pseudomonadota>Gammaproteobacteria>Aeromonadales>Aeromonadaceae | GCF_018802385.1 | Aeromonas_hydrophila | HQ395_RS06965 | 1417486..1420245 | 644 | DUF499-domain-containing-protein-[Aeromonas-hydrophila] |
| WP_323581432.1 | AAA+ATPase+wHTH+Alpha+Beta-domain+wHTH+RRM | Type-3_BR.PglZ.associated.systems | <-iSTAND<-METHYLASE<-VirB4-FtsK+HerA<-SIR2<-PglZ<-DUF499+RRM*<-XIS-HTH | 922 | Bacteria>Pseudomonadota>Gammaproteobacteria>Aeromonadales>Aeromonadaceae>Aeromonas | GCF_035013005.1 | Aeromonas_caviae | VCX68_RS17170 | 11619..14387 | 648 | DUF499-domain-containing-protein-[Aeromonas-caviae] |
| WP_144361395.1 | AAA+ATPase+wHTH+Alpha+Beta-domain+wHTH+RRM | Type-3_BR.PglZ.associated.systems | Phage_integrase+Phage_integrase->NACHT->TetR-HTH-><-HTH_3\|\|HTH_Tnp->XIS-HTH->DUF499*->PglZ->METHYLASE->iSTAND->TUDOR+SF1-SF2-Helicase+iREase+CTD->PRTase-> | 893 | Bacteria>Pseudomonadota>Gammaproteobacteria>Alteromonadales>Shewanellaceae | GCF_007636515.1 | Shewanella_sp._MSW | AYI96_RS08210 | 69007..71688 | 2569536 | hypothetical-protein-[Shewanella-sp.-MSW] |
| WP_115612330.1 | AAA+ATPase+wHTH+Alpha+Beta-domain+wHTH+RRM | Type-3_BR.PglZ.associated.systems | <-TUDOR+SF1-SF2-Helicase+iREase+CTD<-iSTAND<-METHYLASE<-VirB4-FtsK+HerA<-SIR2<-PglZ<-DUF499*\|\|DnaJ->?-><-DUF4096+DDE_Tnp_1+DDE_Tnp_1_2\|\|DDE_Tnp_IS1-> | 910 | Bacteria>Pseudomonadota>Gammaproteobacteria>Cardiobacteriales>Cardiobacteriaceae | GCF_900460955.1 | Cardiobacterium_valvarum | DY250_RS11000 | 2267302..2270034 | 194702 | DUF499-domain-containing-protein-[Cardiobacterium-valvarum] |
| WP_230552271.1 | AAA+ATPase+wHTH+Alpha+Beta-domain+wHTH+RRM | Type-3_BR.PglZ.associated.systems | MORC+DNA_gyraseB+TOPRIM->REase->rve+Mu-transpos_C->?->?->XIS-HTH->DUF499*->PglZ->METHYLASE->iSTAND->?->TUDOR+SF1-SF2-Helicase+iREase+CTD-> | 811 | Bacteria>Pseudomonadota>Gammaproteobacteria>Cellvibrionales>Cellvibrionaceae | GCF_000427605.1 | Teredinibacter_turnerae | TERTUR_RS0101180 | 279736..282171 | 2426 | DUF499-domain-containing-protein-[Teredinibacter-turnerae] |
| MDC0404228.1 | AAA+ATPase+wHTH+Alpha+Beta-domain+wHTH+RRM | Type-3_BR.PglZ.associated.systems | D5N+viral-AAA->?-><-?\|\|?->HTH_Tnp->XIS-HTH->DUF499*->PglZ->METHYLASE->iSTAND->TUDOR+SF1-SF2-Helicase+iREase+CTD-> | 890 | Bacteria>Pseudomonadota>Gammaproteobacteria>Cellvibrionales>Porticoccaceae | GCA_028352615.1 | Porticoccaceae_bacterium | OAM26_04265 | 6218..8890 | 2026782 | ATP-binding-protein-[Porticoccaceae-bacterium] |
| MBV5311584.1 | AAA+ATPase+wHTH+Alpha+Beta-domain+wHTH+RRM | Type-3_BR.PglZ.associated.systems | RHH->DUF499->DUF499*->PglZ->RelE-ParE->HTH_3->METHYLASE->METHYLASE->iSTAND-> | 359 | Bacteria>Pseudomonadota>Gammaproteobacteria>Chromatiales>Chromatiaceae | GCA_019163185.1 | Chromatium_okenii | JZU59_19620 | 2468..3547 | 61644 | hypothetical-protein-[Chromatium-okenii] |
| MBV5311583.1 | AAA+ATPase+wHTH+Alpha+Beta-domain+wHTH+RRM | Type-3_BR.PglZ.associated.systems | RHH->DUF499*->DUF499->PglZ->RelE-ParE->HTH_3->METHYLASE->METHYLASE-> | 536 | Bacteria>Pseudomonadota>Gammaproteobacteria>Chromatiales>Chromatiaceae | GCA_019163185.1 | Chromatium_okenii | JZU59_19615 | 824..2434 | 61644 | DUF499-domain-containing-protein-[Chromatium-okenii] |
| QVL51158.1 | AAA+ATPase+wHTH+Alpha+Beta-domain+wHTH+RRM | Type-3_BR.PglZ.associated.systems | <-Y1_Tnp<-PIN<-?<-PIN<-Phd-YefM<-PglZ<-DUF499* | 912 | Bacteria>Pseudomonadota>Gammaproteobacteria>Chromatiales>Chromatiaceae | GCA_018399035.1 | Thiocapsa_sp. | KFB96_12610 | 2785812..2788550 | 2024551 | MAG:-DUF499-domain-containing-protein-[Thiocapsa-sp.] |
| MBK8636968.1 | AAA+ATPase+wHTH+Alpha+Beta-domain+wHTH+RRM | Type-3_BR.PglZ.associated.systems | <-TUDOR+SF1-SF2-Helicase+iREase+CTD<-HicA_toxin<-HicB_lk_antitox<-iSTAND<-METHYLASE<-PglZ<-DUF499*<-AbiEi_4\|\|?->?->DUF433-><-?<-PIN | 892 | Bacteria>Pseudomonadota>Gammaproteobacteria>Chromatiales>Chromatiaceae | GCA_016714645.1 | Chromatiaceae_bacterium | IPN92_01360 | 292629..295307 | 2049432 | hypothetical-protein-[Chromatiaceae-bacterium] |
| WP_058554679.1 | AAA+ATPase+wHTH+Alpha+Beta-domain+wHTH+RRM | Type-3_BR.PglZ.associated.systems | <-TUDOR+SF1-SF2-Helicase+iREase+CTD<-?<-PIN<-iSTAND<-METHYLASE<-PglZ<-DUF499* | 914 | Bacteria>Pseudomonadota>Gammaproteobacteria>Chromatiales>Chromatiaceae | GCF_001469165.1 | Thiohalocapsa_sp._ML1 | ML01_RS06970 | 85147..87891 | 1431688 | DUF499-domain-containing-protein-[Thiohalocapsa-sp.-ML1] |
| NEX15474.1 | AAA+ATPase+wHTH+Alpha+Beta-domain+wHTH+RRM | Type-3_BR.PglZ.associated.systems | <-Peptidase_S66+Peptidase_S66C\|\|?->DUF499*->PglZ->HicB_lk_antitox->?->PIN->METHYLASE->iSTAND-> | 903 | Bacteria>Pseudomonadota>Gammaproteobacteria>Chromatiales>Chromatiaceae | GCA_010912705.1 | Halochromatium_sp. | C1943_02265 | 42127..44838 | 2049430 | DUF499-domain-containing-protein-[Halochromatium-sp.] |
| ALP54923.1 | AAA+ATPase+wHTH+Alpha+Beta-domain+wHTH+RRM | Type-3_BR.PglZ.associated.systems | <-N6_N4_MTase<-?<-?<-RNase\|\|RHH_5->DUF499*->PglZ->METHYLASE->iSTAND->TUDOR+SF1-SF2-Helicase+iREase+CTD-> | 908 | Bacteria>Pseudomonadota>Gammaproteobacteria>Chromatiales>Chromatiaceae | GCA_001447805.1 | Candidatus_Tenderia_electrophaga | Tel_16840 | 13114..15840 | 1748243 | hypothetical-protein-Tel_16840-[Candidatus-Tenderia-electrophaga] |
| TVQ88963.1 | AAA+ATPase+wHTH+Alpha+Beta-domain+wHTH+RRM | Type-3_BR.PglZ.associated.systems | <-HicA_toxin<-?<-DUF433<-PglZ<-DUF499* | 910 | Bacteria>Pseudomonadota>Gammaproteobacteria>Chromatiales>Chromatiaceae | GCA_007693535.1 | Chromatiaceae_bacterium | EA400_08290 | 1780..4512 | 2049432 | DUF499-domain-containing-protein-[Chromatiaceae-bacterium] |
| WP_295579339.1 | AAA+ATPase+wHTH+Alpha+Beta-domain+wHTH+RRM | Type-3_BR.PglZ.associated.systems | DUF499+RRM*->PglZ->PemK_toxin->METHYLASE-> | 912 | Bacteria>Pseudomonadota>Gammaproteobacteria>Chromatiales>Chromatiaceae>Lamprocystis | GCF_903930065.1 | uncultured_Lamprocystis_sp. | QZY03_RS06655 | 23916..26654 | 543132 | DUF499-domain-containing-protein-[uncultured-Lamprocystis-sp.] |
| WP_295608056.1 | AAA+ATPase+wHTH+Alpha+Beta-domain+wHTH+RRM | Type-3_BR.PglZ.associated.systems | <-PglZ<-DUF499+RRM* | 912 | Bacteria>Pseudomonadota>Gammaproteobacteria>Chromatiales>Chromatiaceae>Lamprocystis | GCF_903933745.1 | uncultured_Lamprocystis_sp. | QZY08_RS17460 | 1090..3828 | 543132 | DUF499-domain-containing-protein-[uncultured-Lamprocystis-sp.] |
| WP_213458099.1 | AAA+ATPase+wHTH+Alpha+Beta-domain+wHTH+RRM | Type-3_BR.PglZ.associated.systems | <-PIN<-PglZ<-DUF499+RRM* | 912 | Bacteria>Pseudomonadota>Gammaproteobacteria>Chromatiales>Chromatiaceae>Thiocapsa | GCF_018399035.1 | Thiocapsa_sp. | KFB96_RS12610 | 2785812..2788550 | 2024551 | DUF499-domain-containing-protein-[Thiocapsa-sp.] |
| WP_295427663.1 | AAA+ATPase+wHTH+Alpha+Beta-domain+wHTH+RRM | Type-3_BR.PglZ.associated.systems | DUF499+RRM*->HEPN->PglZ-> | 916 | Bacteria>Pseudomonadota>Gammaproteobacteria>Chromatiales>Chromatiaceae>Thiodictyon | GCF_903828115.1 | uncultured_Thiodictyon_sp. | QZY48_RS03270 | 1296..4046 | 1846217 | DUF499-domain-containing-protein-[uncultured-Thiodictyon-sp.] |
| WP_295391533.1 | AAA+ATPase+wHTH+Alpha+Beta-domain+wHTH+RRM | Type-3_BR.PglZ.associated.systems | <-HEPN<-METHYLASE<-PglZ<-DUF499+RRM*<-SF1-SF2-Helicase+REase\|\|REase-> | 914 | Bacteria>Pseudomonadota>Gammaproteobacteria>Chromatiales>Chromatiaceae>Thiodictyon | GCF_903843925.1 | uncultured_Thiodictyon_sp. | QZX99_RS13905 | 10789..13533 | 1846217 | DUF499-domain-containing-protein-[uncultured-Thiodictyon-sp.] |
| MCM8857687.1 | AAA+ATPase+wHTH+Alpha+Beta-domain+wHTH+RRM | Type-3_BR.PglZ.associated.systems | Arm-DNA-bind_3+Phage_integrase-><-TUDOR+SF1-SF2-Helicase+iREase+CTD<-iSTAND<-METHYLASE<-PglZ<-DUF499*<-RHH_5<-Primpol+PriCT1<-XIS-HTH | 908 | Bacteria>Pseudomonadota>Gammaproteobacteria>Chromatiales>Sedimenticolaceae | GCA_023838585.1 | Candidatus_Thiodiazotropha_sp. | LC539_18500 | 9239..11965 | 2847306 | DUF499-domain-containing-protein-[Candidatus-Thiodiazotropha-sp.] |
| WP_223846277.1 | AAA+ATPase+wHTH+Alpha+Beta-domain+wHTH+RRM | Type-3_BR.PglZ.associated.systems | <-ZnR+TOPRIM<-XIS-HTH<-?<-N6_N4_MTase\|\|RHH->DUF499*->PglZ->METHYLASE->iSTAND->TUDOR+SF1-SF2-Helicase+iREase+CTD->HEPN-><-Phage_integrase | 924 | Bacteria>Pseudomonadota>Gammaproteobacteria>Chromatiales>Wenzhouxiangellaceae | GCF_014725735.1 | Wenzhouxiangella_sp._AB-CW3 | IC757_RS06090 | 1397913..1400687 | 2771012 | DUF499-domain-containing-protein-[Wenzhouxiangella-sp.-AB-CW3] |
| QOC24192.1 | AAA+ATPase+wHTH+Alpha+Beta-domain+wHTH+RRM | Type-3_BR.PglZ.associated.systems | <-ZnR+TOPRIM<-XIS-HTH<-?<-N6_N4_MTase\|\|RHH->DUF499*->PglZ->METHYLASE->iSTAND->TUDOR+SF1-SF2-Helicase+iREase+CTD->HEPN-> | 909 | Bacteria>Pseudomonadota>Gammaproteobacteria>Chromatiales>Wenzhouxiangellaceae | GCF_014725735.1 | Wenzhouxiangella_sp._AB-CW3 | IC757_06090 | 1397958..1400687 | 2771012 | DUF499-domain-containing-protein-[Wenzhouxiangella-sp.-AB-CW3] |
| WP_000901079.1 | AAA+ATPase+wHTH+Alpha+Beta-domain+wHTH+RRM | Type-3_BR.PglZ.associated.systems | HTH_Tnp->XIS-HTH->DUF499*->PglZ->METHYLASE->iSTAND->TUDOR+SF1-SF2-Helicase+iREase+CTD->SbcC-><-Zn_Tnp_IS1+HTH_Tnp_IS1+DDE_Tnp_IS1 | 893 | Bacteria>Pseudomonadota>Gammaproteobacteria>Enterobacterales | GCF_900776365.1 | Enterobacteriaceae | EXX70_RS20565 | 1107..3788 | 543 | MULTISPECIES:-hypothetical-protein-[Enterobacteriaceae] |
| WP_088903447.1 | AAA+ATPase+wHTH+Alpha+Beta-domain+wHTH+RRM | Type-3_BR.PglZ.associated.systems | GDResolvase+Recombinase+ZnR->?-><-?<-?<-ArdB<-?\|\|DUF499*->PglZ->SIR2->VirB4-FtsK+HerA->VirB4-FtsK+HerA->iSTAND-> | 705 | Bacteria>Pseudomonadota>Gammaproteobacteria>Enterobacterales | GCF_002216835.1 | Enterobacteriaceae | KM293_RS26255 | 44629..46746 | 543 | MULTISPECIES:-DUF499-domain-containing-protein-[Klebsiella/Raoultella-group] |
| WP_072668134.1 | AAA+ATPase+wHTH+Alpha+Beta-domain+wHTH+RRM | Type-3_BR.PglZ.associated.systems | <-SbcC<-TUDOR+SF1-SF2-Helicase+iREase+CTD<-iSTAND<-METHYLASE<-PglZ<-DUF499*<-XIS-HTH<-HTH_Tnp<-RVT<-?\|\|<-?<-PfkB | 893 | Bacteria>Pseudomonadota>Gammaproteobacteria>Enterobacterales | GCF_012516455.1 | Enterobacteriaceae | KPJ08_RS13640 | 8276..10957 | 543 | MULTISPECIES:-hypothetical-protein-[Enterobacteriaceae] |
| WP_047664212.1 | AAA+ATPase+wHTH+Alpha+Beta-domain+wHTH+RRM | Type-3_BR.PglZ.associated.systems | <-HTH+MPTase\|\|MazG->\|\|XIS-HTH->XIS-HTH->DUF499*->PglZ->METHYLASE->iSTAND->TUDOR+SF1-SF2-Helicase+iREase+CTD-> | 893 | Bacteria>Pseudomonadota>Gammaproteobacteria>Enterobacterales | GCF_005398945.1 | Enterobacteriaceae | KTU89_RS22800 | 6216..8897 | 543 | hypothetical-protein-[Escherichia-coli] |
| WP_217294360.1 | AAA+ATPase+wHTH+Alpha+Beta-domain+wHTH+RRM | Type-3_BR.PglZ.associated.systems | <-ArdB<-?<-?\|\|DUF499*->PglZ->SIR2->VirB4-FtsK+HerA->METHYLASE->iSTAND->TUDOR+SF1-SF2-Helicase+iREase+CTD-> | 706 | Bacteria>Pseudomonadota>Gammaproteobacteria>Enterobacterales | GCF_015352935.1 | Enterobacteriaceae | KTQ03_RS00775 | 165503..167623 | 543 | MULTISPECIES:-hypothetical-protein-[Klebsiella/Raoultella-group] |
| WP_206635980.1 | AAA+ATPase+wHTH+Alpha+Beta-domain+wHTH+RRM | Type-3_BR.PglZ.associated.systems | <-ArdB<-?<-?\|\|DUF499*->PglZ->SIR2->VirB4-FtsK+HerA->METHYLASE->iSTAND->TUDOR+SF1-SF2-Helicase+iREase+CTD-> | 704 | Bacteria>Pseudomonadota>Gammaproteobacteria>Enterobacterales | GCF_026167905.1 | Enterobacteriaceae | B9P89_RS14040 | 7239..9353 | 543 | MULTISPECIES:-hypothetical-protein-[Enterobacteriaceae] |
| WP_232481333.1 | AAA+ATPase+wHTH+Alpha+Beta-domain+wHTH+RRM | Type-3_BR.PglZ.associated.systems | <-ArdB<-?\|\|DUF499*->PglZ->SIR2->VirB4-FtsK+HerA->METHYLASE->iSTAND->TUDOR+SF1-SF2-Helicase+iREase+CTD-> | 718 | Bacteria>Pseudomonadota>Gammaproteobacteria>Enterobacterales | GCF_002215385.1 | Enterobacteriaceae | CFA70_RS28060 | 84602..86758 | 543 | MULTISPECIES:-hypothetical-protein-[Enterobacteriaceae] |
| WP_248176985.1 | AAA+ATPase+wHTH+Alpha+Beta-domain+wHTH+RRM | Type-3_BR.PglZ.associated.systems | XIS-HTH->ZnR-><-?\|\|?->?->XIS-HTH->DUF499*->PglZ->METHYLASE->iSTAND->TUDOR+SF1-SF2-Helicase+iREase+CTD-><-Arm-DNA-bind_3+Phage_integrase\|\|Phage_integrase-> | 893 | Bacteria>Pseudomonadota>Gammaproteobacteria>Enterobacterales>Enterobacteriaceae | GCF_023153335.1 | Enterobacter_bugandensis | L8P91_RS09010 | 222548..225229 | 881260 | hypothetical-protein-[Enterobacter-bugandensis] |
| HAW1502903.1 | AAA+ATPase+wHTH+Alpha+Beta-domain+wHTH+RRM | Type-3_BR.PglZ.associated.systems | XIS-HTH->XIS-HTH->DUF499*->PglZ->METHYLASE->iSTAND->TUDOR+SF1-SF2-Helicase+iREase+CTD-><-wHTH(SBD)+HNH | 893 | Bacteria>Pseudomonadota>Gammaproteobacteria>Enterobacterales>Enterobacteriaceae | GCA_016674655.1 | Escherichia_coli | JLM52_004934 | 1697..4378 | 562 | hypothetical-protein-[Escherichia-coli] |
| HBA5267267.1 | AAA+ATPase+wHTH+Alpha+Beta-domain+wHTH+RRM | Type-3_BR.PglZ.associated.systems | XIS-HTH->XIS-HTH->DUF499*->PglZ->METHYLASE->iSTAND->TUDOR+SF1-SF2-Helicase+iREase+CTD-> | 893 | Bacteria>Pseudomonadota>Gammaproteobacteria>Enterobacterales>Enterobacteriaceae | GCA_017740515.1 | Escherichia_coli | J8124_004919 | 1994..4675 | 562 | hypothetical-protein-[Escherichia-coli] |
| HBA5530466.1 | AAA+ATPase+wHTH+Alpha+Beta-domain+wHTH+RRM | Type-3_BR.PglZ.associated.systems | XIS-HTH->XIS-HTH->DUF499*->PglZ->METHYLASE->iSTAND->TUDOR+SF1-SF2-Helicase+iREase+CTD-> | 893 | Bacteria>Pseudomonadota>Gammaproteobacteria>Enterobacterales>Enterobacteriaceae | GCA_017753765.1 | Escherichia_coli | J8593_004903 | 1994..4675 | 562 | hypothetical-protein-[Escherichia-coli] |
| MCT2779080.1 | AAA+ATPase+wHTH+Alpha+Beta-domain+wHTH+RRM | Type-3_BR.PglZ.associated.systems | XIS-HTH->DUF499-p->DUF499*->PglZ->SIR2->VirB4-FtsK+HerA->METHYLASE->iSTAND->SF1-SF2-Helicase+iREase+CTD-> | 846 | Bacteria>Pseudomonadota>Gammaproteobacteria>Enterobacterales>Enterobacteriaceae | GCA_025183295.1 | Klebsiella_pneumoniae | KBQ08_24665 | 751..3291 | 573 | DUF499-domain-containing-protein-[Klebsiella-pneumoniae] |
| WP_260232863.1 | AAA+ATPase+wHTH+Alpha+Beta-domain+wHTH+RRM | Type-3_BR.PglZ.associated.systems | XIS-HTH->DUF499->DUF499*->PglZ->SIR2->VirB4-FtsK+HerA->METHYLASE->iSTAND->TUDOR+SF1-SF2-Helicase+iREase+CTD-> | 845 | Bacteria>Pseudomonadota>Gammaproteobacteria>Enterobacterales>Enterobacteriaceae | GCF_025183295.1 | Klebsiella_pneumoniae | KBQ08_RS24750 | 754..3291 | 573 | ATP-binding-protein-[Klebsiella-pneumoniae] |
| HAW4023848.1 | AAA+ATPase+wHTH+Alpha+Beta-domain+wHTH+RRM | Type-3_BR.PglZ.associated.systems | XIS-HTH->DUF499*->PglZ->METHYLASE->iSTAND->TUDOR+SF1-SF2-Helicase+iREase+CTD->SbcC-> | 893 | Bacteria>Pseudomonadota>Gammaproteobacteria>Enterobacterales>Enterobacteriaceae | GCA_016686015.1 | Escherichia_coli | JLV09_004312 | 756..3437 | 562 | hypothetical-protein-[Escherichia-coli] |
| WP_260232862.1 | AAA+ATPase+wHTH+Alpha+Beta-domain+wHTH+RRM | Type-3_BR.PglZ.associated.systems | XIS-HTH->DUF499*->DUF499->PglZ->SIR2->VirB4-FtsK+HerA->METHYLASE->iSTAND-> | 62 | Bacteria>Pseudomonadota>Gammaproteobacteria>Enterobacterales>Enterobacteriaceae | GCF_025183295.1 | Klebsiella_pneumoniae | KBQ08_RS24745 | 527..715 | 573 | hypothetical-protein-[Klebsiella-pneumoniae] |
| WP_040089188.1 | AAA+ATPase+wHTH+Alpha+Beta-domain+wHTH+RRM | Type-3_BR.PglZ.associated.systems | XIS-HTH->?->XIS-HTH->DUF499*->PglZ->METHYLASE->iSTAND->TUDOR+SF1-SF2-Helicase+iREase+CTD-><-DDE_Tnp_IS66+DDE_Tnp_IS66_C | 893 | Bacteria>Pseudomonadota>Gammaproteobacteria>Enterobacterales>Enterobacteriaceae | GCF_000819185.1 | Escherichia_coli | PU42_RS15340 | 25467..28148 | 562 | hypothetical-protein-[Escherichia-coli] |
| BBV79269.1 | AAA+ATPase+wHTH+Alpha+Beta-domain+wHTH+RRM | Type-3_BR.PglZ.associated.systems | TraJ-RHH->ACET->?->HKD-DNase-><-?\|\|DDE_Tnp_IS240->DUF499*->PglZ->SIR2->VirB4-FtsK+HerA->METHYLASE->iSTAND->TUDOR+SF1-SF2-Helicase+iREase+CTD-> | 807 | Bacteria>Pseudomonadota>Gammaproteobacteria>Enterobacterales>Enterobacteriaceae | GCA_015135635.1 | Raoultella_planticola | STW0522RA056_P12820 | 227795..230218 | 575 | hypothetical-protein-STW0522RA056_P12820-[Raoultella-planticola] |
| WP_221019832.1 | AAA+ATPase+wHTH+Alpha+Beta-domain+wHTH+RRM | Type-3_BR.PglZ.associated.systems | SbcC->\|\|<-TUDOR+SF1-SF2-Helicase+iREase+CTD<-iSTAND<-METHYLASE<-PglZ<-DUF499*<-XIS-HTH\|\|?->HNH-> | 893 | Bacteria>Pseudomonadota>Gammaproteobacteria>Enterobacterales>Enterobacteriaceae | GCF_034741975.1 | Klebsiella_variicola | C7266_RS23820 | 122926..125607 | 244366 | hypothetical-protein-[Klebsiella-variicola] |
| MCZ5812829.1 | AAA+ATPase+wHTH+Alpha+Beta-domain+wHTH+RRM | Type-3_BR.PglZ.associated.systems | RVT->HTH_Tnp->XIS-HTH->DUF499->DUF499*->PglZ->PglZ->METHYLASE-p->METHYLASE-p->ZnR+ZnR->METHYLASE-> | 811 | Bacteria>Pseudomonadota>Gammaproteobacteria>Enterobacterales>Enterobacteriaceae | GCA_027276925.1 | Escherichia_coli | O5265_04820 | 960838..963273 | 562 | hypothetical-protein-O5265_04820-[Escherichia-coli] |
| HCN7245202.1 | AAA+ATPase+wHTH+Alpha+Beta-domain+wHTH+RRM | Type-3_BR.PglZ.associated.systems | RVT->HTH_Tnp->XIS-HTH->DUF499*->PglZ->METHYLASE->iSTAND->TUDOR+SF1-SF2-Helicase+iREase+CTD->SbcC-> | 893 | Bacteria>Pseudomonadota>Gammaproteobacteria>Enterobacterales>Enterobacteriaceae | GCA_025287075.1 | Escherichia_coli | N6192_004650 | 2234..4915 | 562 | hypothetical-protein-[Escherichia-coli] |
| EKL4969060.1 | AAA+ATPase+wHTH+Alpha+Beta-domain+wHTH+RRM | Type-3_BR.PglZ.associated.systems | RVT->HTH_Tnp->XIS-HTH->DUF499*->PglZ->METHYLASE->iSTAND->TUDOR+SF1-SF2-Helicase+iREase+CTD->SbcC-> | 893 | Bacteria>Pseudomonadota>Gammaproteobacteria>Enterobacterales>Enterobacteriaceae | GCA_028515425.1 | Escherichia_coli | PR510_004415 | 3439..6120 | 562 | hypothetical-protein-PR510_004415-[Escherichia-coli] |
| MCZ5812828.1 | AAA+ATPase+wHTH+Alpha+Beta-domain+wHTH+RRM | Type-3_BR.PglZ.associated.systems | RVT->HTH_Tnp->XIS-HTH->DUF499*->DUF499->PglZ->PglZ->METHYLASE-p->METHYLASE-p->ZnR+ZnR-> | 60 | Bacteria>Pseudomonadota>Gammaproteobacteria>Enterobacterales>Enterobacteriaceae | GCA_027276925.1 | Escherichia_coli | O5265_04815 | 960592..960774 | 562 | hypothetical-protein-O5265_04815-[Escherichia-coli] |
| WP_045408218.1 | AAA+ATPase+wHTH+Alpha+Beta-domain+wHTH+RRM | Type-3_BR.PglZ.associated.systems | RNase->?-><-TUDOR+SF1-SF2-Helicase+iREase+CTD<-iSTAND<-METHYLASE<-PglZ<-DUF499*<-XIS-HTH | 893 | Bacteria>Pseudomonadota>Gammaproteobacteria>Enterobacterales>Enterobacteriaceae | GCF_000957415.1 | Enterobacter_roggenkampii | SR65_RS13595 | 126970..129651 | 1812935 | hypothetical-protein-[Enterobacter-roggenkampii] |
| WP_117088206.1 | AAA+ATPase+wHTH+Alpha+Beta-domain+wHTH+RRM | Type-3_BR.PglZ.associated.systems | PfkB->?->\|\|RVT->HTH_Tnp->XIS-HTH->DUF499*->PglZ->METHYLASE->iSTAND->TUDOR+SF1-SF2-Helicase+iREase+CTD->SbcC-><-Zn_Tnp_IS1+HTH_Tnp_IS1+DDE_Tnp_IS1 | 893 | Bacteria>Pseudomonadota>Gammaproteobacteria>Enterobacterales>Enterobacteriaceae | GCF_900512555.1 | Klebsiella_pneumoniae | D2N47_RS14680 | 30952..33633 | 573 | hypothetical-protein-[Klebsiella-pneumoniae] |
| HCP2967461.1 | AAA+ATPase+wHTH+Alpha+Beta-domain+wHTH+RRM | Type-3_BR.PglZ.associated.systems | PfkB->?->\|\|RVT->HTH_Tnp->XIS-HTH->DUF499*->PglZ->METHYLASE->iSTAND->TUDOR+SF1-SF2-Helicase+iREase+CTD->SbcC-> | 893 | Bacteria>Pseudomonadota>Gammaproteobacteria>Enterobacterales>Enterobacteriaceae | GCA_025521675.1 | Escherichia_coli | ODL03_004214 | 15715..18396 | 562 | TPA:-hypothetical-protein-ODL03_004214-[Escherichia-coli] |
| WP_211787638.1 | AAA+ATPase+wHTH+Alpha+Beta-domain+wHTH+RRM | Type-3_BR.PglZ.associated.systems | ParB+HNH-><-DDE_Tnp_IS1\|\|HTH_Tnp->XIS-HTH->DUF499*->PglZ->METHYLASE->iSTAND->TUDOR+SF1-SF2-Helicase+iREase+CTD->SbcC-><-ZnR | 893 | Bacteria>Pseudomonadota>Gammaproteobacteria>Enterobacterales>Enterobacteriaceae | GCF_018109085.1 | Klebsiella_pneumoniae | FD764_RS25560 | 4437..7118 | 573 | hypothetical-protein-[Klebsiella-pneumoniae] |
| WP_105078066.1 | AAA+ATPase+wHTH+Alpha+Beta-domain+wHTH+RRM | Type-3_BR.PglZ.associated.systems | HTH_Tnp->XIS-HTH->DUF499*->PglZ->METHYLASE->iSTAND->TUDOR+SF1-SF2-Helicase+iREase+CTD->SbcC->METHYLASE-p-> | 893 | Bacteria>Pseudomonadota>Gammaproteobacteria>Enterobacterales>Enterobacteriaceae | GCF_020886575.2 | Escherichia_coli | JQK84_RS22430 | 757..3438 | 562 | hypothetical-protein-[Escherichia-coli] |
| EIN2284095.1 | AAA+ATPase+wHTH+Alpha+Beta-domain+wHTH+RRM | Type-3_BR.PglZ.associated.systems | DUF499*->PglZ->METHYLASE->iSTAND->TUDOR+SF1-SF2-Helicase+iREase+CTD->SbcC->REase-> | 749 | Bacteria>Pseudomonadota>Gammaproteobacteria>Enterobacterales>Enterobacteriaceae | GCA_020874335.1 | Escherichia_coli | LOP93_004329 | 1..2251 | 562 | hypothetical-protein-[Escherichia-coli] |
| HCN8452008.1 | AAA+ATPase+wHTH+Alpha+Beta-domain+wHTH+RRM | Type-3_BR.PglZ.associated.systems | DUF499*->PglZ->METHYLASE->iSTAND->TUDOR+SF1-SF2-Helicase+iREase+CTD->SbcC-> | 380 | Bacteria>Pseudomonadota>Gammaproteobacteria>Enterobacterales>Enterobacteriaceae | GCA_025295215.1 | Escherichia_coli | N6508_004587 | 1..1144 | 562 | hypothetical-protein-[Escherichia-coli] |
| HCN8614210.1 | AAA+ATPase+wHTH+Alpha+Beta-domain+wHTH+RRM | Type-3_BR.PglZ.associated.systems | DUF499*->PglZ->METHYLASE->iSTAND->TUDOR+SF1-SF2-Helicase+iREase+CTD->SbcC-> | 788 | Bacteria>Pseudomonadota>Gammaproteobacteria>Enterobacterales>Enterobacteriaceae | GCA_025295835.1 | Escherichia_coli | N6538_004516 | 1..2369 | 562 | hypothetical-protein-[Escherichia-coli] |
| MCV7955816.1 | AAA+ATPase+wHTH+Alpha+Beta-domain+wHTH+RRM | Type-3_BR.PglZ.associated.systems | DUF499*->PglZ->METHYLASE->iSTAND->TUDOR+SF1-SF2-Helicase+iREase+CTD-><-Zn_Tnp_IS1+HTH_Tnp_IS1+DDE_Tnp_IS1\|\|nSTAND1+sGTP-> | 645 | Bacteria>Pseudomonadota>Gammaproteobacteria>Enterobacterales>Enterobacteriaceae | GCA_025827855.1 | Escherichia_coli | ODR25_26000 | 1..1938 | 562 | hypothetical-protein-ODR25_26000,-partial-[Escherichia-coli] |
| HAM2528590.1 | AAA+ATPase+wHTH+Alpha+Beta-domain+wHTH+RRM | Type-3_BR.PglZ.associated.systems | DUF499*->PglZ->METHYLASE->iSTAND->TUDOR+SF1-SF2-Helicase+iREase+CTD-> | 657 | Bacteria>Pseudomonadota>Gammaproteobacteria>Enterobacterales>Enterobacteriaceae | GCA_014365645.1 | Escherichia_coli | IBD21_001850 | 1..1976 | 562 | hypothetical-protein-[Escherichia-coli] |
| HCN8264385.1 | AAA+ATPase+wHTH+Alpha+Beta-domain+wHTH+RRM | Type-3_BR.PglZ.associated.systems | DUF499*->PglZ->METHYLASE->iSTAND->TUDOR+SF1-SF2-Helicase+iREase+CTD-> | 443 | Bacteria>Pseudomonadota>Gammaproteobacteria>Enterobacterales>Enterobacteriaceae | GCA_025294615.1 | Escherichia_coli | N6451_004779 | 1..1332 | 562 | hypothetical-protein-[Escherichia-coli] |
| HCR0861469.1 | AAA+ATPase+wHTH+Alpha+Beta-domain+wHTH+RRM | Type-3_BR.PglZ.associated.systems | DUF499*->PglZ->METHYLASE->iSTAND->TUDOR+SF1-SF2-Helicase+iREase+CTD-> | 380 | Bacteria>Pseudomonadota>Gammaproteobacteria>Enterobacterales>Enterobacteriaceae | GCA_025901705.1 | Enterobacter_kobei | OMT47_004760 | 1..1145 | 208224 | TPA:-hypothetical-protein-OMT47_004760,-partial-[Enterobacter-kobei] |
| EFN9589259.1 | AAA+ATPase+wHTH+Alpha+Beta-domain+wHTH+RRM | Type-3_BR.PglZ.associated.systems | DDE_Tnp_IS1-><-TUDOR+SF1-SF2-Helicase+iREase+CTD<-iSTAND<-METHYLASE<-PglZ<-DUF499*<-XIS-HTH<-HTH_Tnp<-ArsR-HTH+YTCD-HTH\|\|?->zf-ISL3+HTH_Tnp_ISL3+DDE_Tnp_ISL3-><-RVT | 893 | Bacteria>Pseudomonadota>Gammaproteobacteria>Enterobacterales>Enterobacteriaceae | GCA_013055265.1 | Escherichia_coli | D3J78_18320 | 8294..10975 | 562 | hypothetical-protein-[Escherichia-coli] |
| MCV7970719.1 | AAA+ATPase+wHTH+Alpha+Beta-domain+wHTH+RRM | Type-3_BR.PglZ.associated.systems | DDE_Tnp_IS1-><-SbcC<-TUDOR+SF1-SF2-Helicase+iREase+CTD<-iSTAND<-METHYLASE<-PglZ<-DUF499* | 643 | Bacteria>Pseudomonadota>Gammaproteobacteria>Enterobacterales>Enterobacteriaceae | GCA_025827875.1 | Escherichia_coli | ODQ64_24725 | 8066..9999 | 562 | hypothetical-protein-ODQ64_24725,-partial-[Escherichia-coli] |
| HCM5147190.1 | AAA+ATPase+wHTH+Alpha+Beta-domain+wHTH+RRM | Type-3_BR.PglZ.associated.systems | Arm-DNA-bind_3+Phage_integrase-><-TUDOR+SF1-SF2-Helicase+iREase+CTD<-iSTAND<-METHYLASE<-PglZ<-DUF499*<-XIS-HTH<-?<-?\|\|?-><-XIS-HTH<-Inovirus_Gp2 | 893 | Bacteria>Pseudomonadota>Gammaproteobacteria>Enterobacterales>Enterobacteriaceae | GCA_025093305.1 | Klebsiella_aerogenes | N3M87_001528 | 357788..360469 | 548 | hypothetical-protein-[Klebsiella-aerogenes] |
| WP_206255350.1 | AAA+ATPase+wHTH+Alpha+Beta-domain+wHTH+RRM | Type-3_BR.PglZ.associated.systems | Arm-DNA-bind_3+Phage_integrase-><-MORC<-TUDOR+SF1-SF2-Helicase+iREase+CTD<-iSTAND<-METHYLASE<-PglZ<-DUF499*<-XIS-HTH<-?<-?\|\|?-><-ZnR<-ZnR | 893 | Bacteria>Pseudomonadota>Gammaproteobacteria>Enterobacterales>Enterobacteriaceae | GCF_017164125.1 | Citrobacter_sedlakii | JR882_RS04705 | 174217..176898 | 67826 | hypothetical-protein-[Citrobacter-sedlakii] |
| MCZ5740001.1 | AAA+ATPase+wHTH+Alpha+Beta-domain+wHTH+RRM | Type-3_BR.PglZ.associated.systems | <-ZnR+ZnR<-METHYLASE-p<-METHYLASE-p<-PglZ<-PglZ<-DUF499<-DUF499*<-?<-DUF499<-DUF499<-XIS-HTH<-HTH_Tnp | 230 | Bacteria>Pseudomonadota>Gammaproteobacteria>Enterobacterales>Enterobacteriaceae | GCA_027276745.1 | Escherichia_coli | O5290_26245 | 19893..20585 | 562 | hypothetical-protein-O5290_26245-[Escherichia-coli] |
| MCZ5740000.1 | AAA+ATPase+wHTH+Alpha+Beta-domain+wHTH+RRM | Type-3_BR.PglZ.associated.systems | <-ZnR+ZnR<-METHYLASE-p<-METHYLASE-p<-PglZ<-PglZ<-DUF499*<-DUF499<-?<-DUF499<-DUF499<-XIS-HTH<-HTH_Tnp | 216 | Bacteria>Pseudomonadota>Gammaproteobacteria>Enterobacterales>Enterobacteriaceae | GCA_027276745.1 | Escherichia_coli | O5290_26240 | 19300..19950 | 562 | hypothetical-protein-O5290_26240-[Escherichia-coli] |
| HCR9958117.1 | AAA+ATPase+wHTH+Alpha+Beta-domain+wHTH+RRM | Type-3_BR.PglZ.associated.systems | <-zf-ISL3+HTH_Tnp_ISL3+DDE_Tnp_ISL3<-?\|\|ArsR-HTH+YTCD-HTH->HTH_Tnp->XIS-HTH->DUF499*->PglZ->METHYLASE->iSTAND->TUDOR+SF1-SF2-Helicase+iREase+CTD->SbcC-> | 893 | Bacteria>Pseudomonadota>Gammaproteobacteria>Enterobacterales>Enterobacteriaceae | GCA_026081515.1 | Escherichia_coli | OHO67_003964 | 3473..6154 | 562 | TPA:-hypothetical-protein-OHO67_003964-[Escherichia-coli] |
| WP_274498616.1 | AAA+ATPase+wHTH+Alpha+Beta-domain+wHTH+RRM | Type-3_BR.PglZ.associated.systems | <-VirB4-FtsK+HerA<-SIR2<-PglZ<-DUF499<-DDE_Tnp_IS240\|\|XIS-HTH->DUF499*->DDE_Tnp_IS240->DEDD_Tnp_IS110+Transposase_20->?->TetR-HTH-> | 177 | Bacteria>Pseudomonadota>Gammaproteobacteria>Enterobacterales>Enterobacteriaceae | GCF_028765255.1 | Klebsiella_variicola | PVM10_RS30205 | 16354..16887 | 244366 | hypothetical-protein-[Klebsiella-variicola] |
| WP_262162080.1 | AAA+ATPase+wHTH+Alpha+Beta-domain+wHTH+RRM | Type-3_BR.PglZ.associated.systems | <-TUDOR+SF1-SF2-Helicase+iREase+CTD<-iSTAND<-METHYLASE<-VirB4-FtsK+HerA<-SIR2<-PglZ<-DUF499*<-XIS-HTH<-?<-?<-TnsC-ATPase<-Mu-transpos_C<-REase | 919 | Bacteria>Pseudomonadota>Gammaproteobacteria>Enterobacterales>Enterobacteriaceae | GCF_025482875.1 | Enterobacter_asburiae | N8T02_RS22970 | 12101..14860 | 61645 | DUF499-domain-containing-protein-[Enterobacter-asburiae] |
| WP_128350860.1 | AAA+ATPase+wHTH+Alpha+Beta-domain+wHTH+RRM | Type-3_BR.PglZ.associated.systems | <-TUDOR+SF1-SF2-Helicase+iREase+CTD<-iSTAND<-METHYLASE<-VirB4-FtsK+HerA<-SIR2<-PglZ<-DUF499*<-XIS-HTH | 919 | Bacteria>Pseudomonadota>Gammaproteobacteria>Enterobacterales>Enterobacteriaceae | - | Enterobacter_cloacae | DN464_22665 | 13096..15855 | 550 | DUF499-domain-containing-protein-[Enterobacter-cloacae] |
| RWT38738.1 | AAA+ATPase+wHTH+Alpha+Beta-domain+wHTH+RRM | Type-3_BR.PglZ.associated.systems | <-TUDOR+SF1-SF2-Helicase+iREase+CTD<-iSTAND<-METHYLASE<-VirB4-FtsK+HerA<-SIR2<-PglZ<-DUF499*<-XIS-HTH | 919 | Bacteria>Pseudomonadota>Gammaproteobacteria>Enterobacterales>Enterobacteriaceae | GCA_004024525.1 | Enterobacter_cloacae | DN464_22665 | 13096..15855 | 550 | DUF499-domain-containing-protein-[Enterobacter-cloacae] |
| HAA0911115.1 | AAA+ATPase+wHTH+Alpha+Beta-domain+wHTH+RRM | Type-3_BR.PglZ.associated.systems | <-TUDOR+SF1-SF2-Helicase+iREase+CTD<-iSTAND<-METHYLASE<-VirB4-FtsK+HerA<-SIR2<-PglZ<-DUF499*<-XIS-HTH | 909 | Bacteria>Pseudomonadota>Gammaproteobacteria>Enterobacterales>Enterobacteriaceae | GCA_009255495.1 | Salmonella_enterica | F8438_22395 | 42997..45726 | 28901 | DUF499-domain-containing-protein-[Salmonella-enterica] |
| WP_202818559.1 | AAA+ATPase+wHTH+Alpha+Beta-domain+wHTH+RRM | Type-3_BR.PglZ.associated.systems | <-TUDOR+SF1-SF2-Helicase+iREase+CTD<-iSTAND<-METHYLASE<-VirB4-FtsK+HerA<-SIR2<-PglZ<-DUF499*<-DUF499<-XIS-HTH\|\|DDE_Tnp_IS240->GDResolvase+HIN-HTH-> | 94 | Bacteria>Pseudomonadota>Gammaproteobacteria>Enterobacterales>Enterobacteriaceae | GCF_900173705.1 | Klebsiella_variicola | B7E74_RS27360 | 37180..37465 | 244366 | hypothetical-protein,-partial-[Klebsiella-variicola] |
| WP_274498615.1 | AAA+ATPase+wHTH+Alpha+Beta-domain+wHTH+RRM | Type-3_BR.PglZ.associated.systems | <-TUDOR+SF1-SF2-Helicase+iREase+CTD<-iSTAND<-METHYLASE<-VirB4-FtsK+HerA<-SIR2<-PglZ<-DUF499*<-DDE_Tnp_IS240\|\|XIS-HTH->DUF499->DDE_Tnp_IS240->DEDD_Tnp_IS110+Transposase_20-> | 767 | Bacteria>Pseudomonadota>Gammaproteobacteria>Enterobacterales>Enterobacteriaceae | GCF_028765255.1 | Klebsiella_variicola | PVM10_RS30190 | 12878..15181 | 244366 | hypothetical-protein-[Klebsiella-variicola] |
| HBV4141490.1 | AAA+ATPase+wHTH+Alpha+Beta-domain+wHTH+RRM | Type-3_BR.PglZ.associated.systems | <-TUDOR+SF1-SF2-Helicase+iREase+CTD<-iSTAND<-METHYLASE<-VirB4-FtsK+HerA<-SIR2<-PglZ<-DUF499* | 771 | Bacteria>Pseudomonadota>Gammaproteobacteria>Enterobacterales>Enterobacteriaceae | GCA_022143025.1 | Klebsiella_quasipneumoniae | MDW69_005387 | 11696..14011 | 1463165 | DUF499-domain-containing-protein-[Klebsiella-quasipneumoniae] |
| MCZ5947407.1 | AAA+ATPase+wHTH+Alpha+Beta-domain+wHTH+RRM | Type-3_BR.PglZ.associated.systems | <-TUDOR+SF1-SF2-Helicase+iREase+CTD<-iSTAND<-METHYLASE<-PglZ<-PglZ<-DUF499<-DUF499*<-XIS-HTH<-HTH_Tnp<-Zn_Tnp_IS1+HTH_Tnp_IS1+DDE_Tnp_IS1\|\|<-?\|\|AraC-HTH+AraC-HTH->AraC-HTH+AraC-HTH-> | 695 | Bacteria>Pseudomonadota>Gammaproteobacteria>Enterobacterales>Enterobacteriaceae | GCA_027279165.1 | Escherichia_coli | O5162_08340 | 1650701..1652788 | 562 | hypothetical-protein-O5162_08340-[Escherichia-coli] |
| WP_241243337.1 | AAA+ATPase+wHTH+Alpha+Beta-domain+wHTH+RRM | Type-3_BR.PglZ.associated.systems | <-TUDOR+SF1-SF2-Helicase+iREase+CTD<-iSTAND<-METHYLASE<-PglZ<-DUF499-p<-DUF499*<-XIS-HTH\|\|<-DDE_Tnp_1_5+DDE_Tnp_1 | 760 | Bacteria>Pseudomonadota>Gammaproteobacteria>Enterobacterales>Enterobacteriaceae | GCF_004004525.1 | Enterobacter_hormaechei | EOL34_RS17720 | 11915..14197 | 158836 | hypothetical-protein-[Enterobacter-hormaechei] |
| HDD8801203.1 | AAA+ATPase+wHTH+Alpha+Beta-domain+wHTH+RRM | Type-3_BR.PglZ.associated.systems | <-TUDOR+SF1-SF2-Helicase+iREase+CTD<-iSTAND<-METHYLASE<-PglZ<-DUF499*<-XIS-HTH<-XIS-HTH\|\|<-MazG\|\|HTH+MPTase-> | 893 | Bacteria>Pseudomonadota>Gammaproteobacteria>Enterobacterales>Enterobacteriaceae | GCA_027552585.1 | Escherichia_coli | PBP21_001694 | 489977..492658 | 562 | TPA:-hypothetical-protein-PBP21_001694-[Escherichia-coli] |
| HBA6908921.1 | AAA+ATPase+wHTH+Alpha+Beta-domain+wHTH+RRM | Type-3_BR.PglZ.associated.systems | <-TUDOR+SF1-SF2-Helicase+iREase+CTD<-iSTAND<-METHYLASE<-PglZ<-DUF499*<-XIS-HTH<-XIS-HTH\|\|<-?\|\|RVT+HNH-> | 893 | Bacteria>Pseudomonadota>Gammaproteobacteria>Enterobacterales>Enterobacteriaceae | GCA_017762905.1 | Escherichia_coli | J1W38_003776 | 8868..11549 | 562 | hypothetical-protein-[Escherichia-coli] |
| HBA3661242.1 | AAA+ATPase+wHTH+Alpha+Beta-domain+wHTH+RRM | Type-3_BR.PglZ.associated.systems | <-TUDOR+SF1-SF2-Helicase+iREase+CTD<-iSTAND<-METHYLASE<-PglZ<-DUF499*<-XIS-HTH<-XIS-HTH | 893 | Bacteria>Pseudomonadota>Gammaproteobacteria>Enterobacterales>Enterobacteriaceae | GCA_017729875.1 | Escherichia_coli | J5F87_004581 | 13325..16006 | 562 | hypothetical-protein-[Escherichia-coli] |
| HBN7153239.1 | AAA+ATPase+wHTH+Alpha+Beta-domain+wHTH+RRM | Type-3_BR.PglZ.associated.systems | <-TUDOR+SF1-SF2-Helicase+iREase+CTD<-iSTAND<-METHYLASE<-PglZ<-DUF499*<-XIS-HTH<-XIS-HTH | 893 | Bacteria>Pseudomonadota>Gammaproteobacteria>Enterobacterales>Enterobacteriaceae | GCA_021581955.1 | Escherichia_coli | L3392_003794 | 9344..12025 | 562 | hypothetical-protein-[Escherichia-coli] |
| WP_021574202.1 | AAA+ATPase+wHTH+Alpha+Beta-domain+wHTH+RRM | Type-3_BR.PglZ.associated.systems | <-TUDOR+SF1-SF2-Helicase+iREase+CTD<-iSTAND<-METHYLASE<-PglZ<-DUF499*<-XIS-HTH<-XIS-HTH | 893 | Bacteria>Pseudomonadota>Gammaproteobacteria>Enterobacterales>Enterobacteriaceae | GCF_010365385.1 | Escherichia_coli | M1V29_RS19545 | 75039..77720 | 562 | hypothetical-protein-[Escherichia-coli] |
| WP_064169692.1 | AAA+ATPase+wHTH+Alpha+Beta-domain+wHTH+RRM | Type-3_BR.PglZ.associated.systems | <-TUDOR+SF1-SF2-Helicase+iREase+CTD<-iSTAND<-METHYLASE<-PglZ<-DUF499*<-XIS-HTH\|\|<-DDE_Tnp_1_5+DDE_Tnp_1\|\|?->NTP_transferase-> | 893 | Bacteria>Pseudomonadota>Gammaproteobacteria>Enterobacterales>Enterobacteriaceae | GCF_013707355.1 | Klebsiella/Raoultella_group | EV185_RS17680 | 9855..12536 | 2890311 | MULTISPECIES:-hypothetical-protein-[Klebsiella/Raoultella-group] |
| WP_248083002.1 | AAA+ATPase+wHTH+Alpha+Beta-domain+wHTH+RRM | Type-3_BR.PglZ.associated.systems | <-SymE_toxin\|\|wHTH+SAD+HNH-><-?\|\|?->?->XIS-HTH->DUF499*->PglZ->METHYLASE->iSTAND->TUDOR+SF1-SF2-Helicase+iREase+CTD-><-HEPN | 893 | Bacteria>Pseudomonadota>Gammaproteobacteria>Enterobacterales>Enterobacteriaceae | GCF_023154575.1 | Enterobacter_kobei | L8P20_RS16915 | 350059..352740 | 208224 | ATP-binding-protein-[Enterobacter-kobei] |
| MCZ5947406.1 | AAA+ATPase+wHTH+Alpha+Beta-domain+wHTH+RRM | Type-3_BR.PglZ.associated.systems | <-SF1-SF2-Helicase+iREase+CTD<-TUDOR+SF1-SF2-Helicase+iREase+CTD<-iSTAND<-METHYLASE<-PglZ<-PglZ<-DUF499*<-DUF499<-XIS-HTH<-HTH_Tnp<-Zn_Tnp_IS1+HTH_Tnp_IS1+DDE_Tnp_IS1\|\|<-?\|\|AraC-HTH+AraC-HTH-> | 216 | Bacteria>Pseudomonadota>Gammaproteobacteria>Enterobacterales>Enterobacteriaceae | GCA_027279165.1 | Escherichia_coli | O5162_08335 | 1650108..1650758 | 562 | hypothetical-protein-O5162_08335-[Escherichia-coli] |
| WP_074417318.1 | AAA+ATPase+wHTH+Alpha+Beta-domain+wHTH+RRM | Type-3_BR.PglZ.associated.systems | <-SF1-SF2-Helicase+iREase+CTD<-SF1-SF2-Helicase+iREase+CTD<-iSTAND<-METHYLASE<-PglZ<-DUF499*<-XIS-HTH<-HTH_Tnp | 893 | Bacteria>Pseudomonadota>Gammaproteobacteria>Enterobacterales>Enterobacteriaceae | GCF_024585585.1 | Escherichia_coli | E3512_RS12655 | 8806..11487 | 562 | hypothetical-protein-[Escherichia-coli] |
| HCD6448073.1 | AAA+ATPase+wHTH+Alpha+Beta-domain+wHTH+RRM | Type-3_BR.PglZ.associated.systems | <-SF1-SF2-Helicase+iREase+CTD<-iSTAND<-METHYLASE<-VirB4-FtsK+HerA<-SIR2<-PglZ<-DUF499*<-XIS-HTH | 921 | Bacteria>Pseudomonadota>Gammaproteobacteria>Enterobacterales>Enterobacteriaceae | GCA_023674285.1 | Klebsiella_pneumoniae | NDU37_003838 | 11362..14127 | 573 | DUF499-domain-containing-protein-[Klebsiella-pneumoniae] |
| HCL5703421.1 | AAA+ATPase+wHTH+Alpha+Beta-domain+wHTH+RRM | Type-3_BR.PglZ.associated.systems | <-SF1-SF2-Helicase+iREase+CTD<-iSTAND<-METHYLASE<-VirB4-FtsK+HerA<-SIR2<-PglZ<-DUF499* | 632 | Bacteria>Pseudomonadota>Gammaproteobacteria>Enterobacterales>Enterobacteriaceae | GCA_025034415.1 | Klebsiella_pneumoniae | N2N57_004734 | 25897..27797 | 573 | hypothetical-protein-[Klebsiella-pneumoniae] |
| HCM8026714.1 | AAA+ATPase+wHTH+Alpha+Beta-domain+wHTH+RRM | Type-3_BR.PglZ.associated.systems | <-SF1-SF2-Helicase+iREase+CTD<-iSTAND<-METHYLASE<-VirB4-FtsK+HerA<-SIR2<-PglZ<-DUF499* | 883 | Bacteria>Pseudomonadota>Gammaproteobacteria>Enterobacterales>Enterobacteriaceae | GCA_025116825.1 | Klebsiella_pneumoniae | N3459_005236 | 16434..19087 | 573 | DUF499-domain-containing-protein-[Klebsiella-pneumoniae] |
| WP_096959997.1 | AAA+ATPase+wHTH+Alpha+Beta-domain+wHTH+RRM | Type-3_BR.PglZ.associated.systems | <-SbcC<-TUDOR+SF1-SF2-Helicase+iREase+CTD<-iSTAND<-METHYLASE<-PglZ<-DUF499*<-XIS-HTH<-HTH_Tnp | 893 | Bacteria>Pseudomonadota>Gammaproteobacteria>Enterobacterales>Enterobacteriaceae | GCF_002467595.1 | Escherichia_coli | A7136_RS23920 | 8597..11278 | 562 | hypothetical-protein-[Escherichia-coli] |
| HAY4243254.1 | AAA+ATPase+wHTH+Alpha+Beta-domain+wHTH+RRM | Type-3_BR.PglZ.associated.systems | <-REase<-SbcC<-TUDOR+SF1-SF2-Helicase+iREase+CTD<-iSTAND<-METHYLASE<-PglZ<-DUF499*<-XIS-HTH<-tRNA\|\|ADH_N+ADH_zinc_N->FtsK+HerA+A32-><-?<-?\|\|Peptidase-> | 893 | Bacteria>Pseudomonadota>Gammaproteobacteria>Enterobacterales>Enterobacteriaceae | GCA_017149385.1 | Escherichia_coli | JZ855_002085 | 80205..82886 | 562 | hypothetical-protein-[Escherichia-coli] |
| MCZ5740004.1 | AAA+ATPase+wHTH+Alpha+Beta-domain+wHTH+RRM | Type-3_BR.PglZ.associated.systems | <-PglZ<-PglZ<-DUF499<-DUF499<-?<-DUF499<-DUF499*<-XIS-HTH<-HTH_Tnp | 64 | Bacteria>Pseudomonadota>Gammaproteobacteria>Enterobacterales>Enterobacteriaceae | GCA_027276745.1 | Escherichia_coli | O5290_26260 | 21790..21984 | 562 | hypothetical-protein-O5290_26260-[Escherichia-coli] |
| MCW0135023.1 | AAA+ATPase+wHTH+Alpha+Beta-domain+wHTH+RRM | Type-3_BR.PglZ.associated.systems | <-MPTase<-HTH\|\|MazG->\|\|?->XIS-HTH->DUF499->DUF499*->DUF499->PglZ->PglZ->METHYLASE-p->METHYLASE-p->METHYLASE-p-> | 300 | Bacteria>Pseudomonadota>Gammaproteobacteria>Enterobacterales>Enterobacteriaceae | GCA_026105695.1 | Escherichia_coli | OIU89_14805 | 2793523..2794425 | 562 | hypothetical-protein-OIU89_14805-[Escherichia-coli] |
| MCW0135022.1 | AAA+ATPase+wHTH+Alpha+Beta-domain+wHTH+RRM | Type-3_BR.PglZ.associated.systems | <-MPTase<-HTH\|\|MazG->\|\|?->XIS-HTH->DUF499*->DUF499->DUF499->PglZ->PglZ->METHYLASE-p->METHYLASE-p-> | 238 | Bacteria>Pseudomonadota>Gammaproteobacteria>Enterobacterales>Enterobacteriaceae | GCA_026105695.1 | Escherichia_coli | OIU89_14800 | 2792594..2793310 | 562 | hypothetical-protein-OIU89_14800-[Escherichia-coli] |
| MCZ5740003.1 | AAA+ATPase+wHTH+Alpha+Beta-domain+wHTH+RRM | Type-3_BR.PglZ.associated.systems | <-METHYLASE-p<-PglZ<-PglZ<-DUF499<-DUF499<-?<-DUF499*<-DUF499<-XIS-HTH<-HTH_Tnp | 280 | Bacteria>Pseudomonadota>Gammaproteobacteria>Enterobacterales>Enterobacteriaceae | GCA_027276745.1 | Escherichia_coli | O5290_26255 | 20912..21754 | 562 | hypothetical-protein-O5290_26255-[Escherichia-coli] |
| WP_080894472.1 | AAA+ATPase+wHTH+Alpha+Beta-domain+wHTH+RRM | Type-3_BR.PglZ.associated.systems | <-iSTAND<-METHYLASE<-VirB4-FtsK+HerA<-SIR2<-PglZ<-DUF499<-DUF499*<-XIS-HTH\|\|DDE_Tnp_IS240->GDResolvase+HIN-HTH-> | 798 | Bacteria>Pseudomonadota>Gammaproteobacteria>Enterobacterales>Enterobacteriaceae | GCF_900173705.1 | Klebsiella_variicola | B7E74_RS27365 | 37754..40147 | 244366 | DUF499-domain-containing-protein,-partial-[Klebsiella-variicola] |
| HDD9941984.1 | AAA+ATPase+wHTH+Alpha+Beta-domain+wHTH+RRM | Type-3_BR.PglZ.associated.systems | <-HTH+MPTase\|\|MazG->\|\|XIS-HTH->XIS-HTH->DUF499*->PglZ->METHYLASE->iSTAND->TUDOR+SF1-SF2-Helicase+iREase+CTD-> | 893 | Bacteria>Pseudomonadota>Gammaproteobacteria>Enterobacterales>Enterobacteriaceae | GCA_027557465.1 | Escherichia_coli | PBR54_001628 | 51298..53979 | 562 | TPA:-hypothetical-protein-PBR54_001628-[Escherichia-coli] |
| MCW0135024.1 | AAA+ATPase+wHTH+Alpha+Beta-domain+wHTH+RRM | Type-3_BR.PglZ.associated.systems | <-HTH\|\|MazG->\|\|?->XIS-HTH->DUF499->DUF499->DUF499*->PglZ->PglZ->METHYLASE-p->METHYLASE-p->METHYLASE-p-> | 284 | Bacteria>Pseudomonadota>Gammaproteobacteria>Enterobacterales>Enterobacteriaceae | GCA_026105695.1 | Escherichia_coli | OIU89_14810 | 2794416..2795270 | 562 | hypothetical-protein-OIU89_14810-[Escherichia-coli] |
| WP_080883160.1 | AAA+ATPase+wHTH+Alpha+Beta-domain+wHTH+RRM | Type-3_BR.PglZ.associated.systems | <-GDResolvase+Recombinase+ZnR<-DDE_Tnp_IS240\|\|XIS-HTH->DUF499*->PglZ->VirB4-FtsK+HerA->VirB4-FtsK+HerA->METHYLASE->iSTAND->TUDOR+SF1-SF2-Helicase+iREase+CTD-> | 728 | Bacteria>Pseudomonadota>Gammaproteobacteria>Enterobacterales>Enterobacteriaceae | GCF_900173715.1 | Klebsiella_variicola | B7E62_RS25820 | 1534..3718 | 244366 | DUF499-domain-containing-protein,-partial-[Klebsiella-variicola] |
| WP_096983203.1 | AAA+ATPase+wHTH+Alpha+Beta-domain+wHTH+RRM | Type-3_BR.PglZ.associated.systems | <-DUF772+DDE_Tnp_1<-AraC-HTH+AraC-HTH\|\|?->\|\|Zn_Tnp_IS1+HTH_Tnp_IS1+DDE_Tnp_IS1->HTH_Tnp->XIS-HTH->DUF499*->PglZ->METHYLASE->iSTAND->TUDOR+SF1-SF2-Helicase+iREase+CTD->SbcC->METHYLASE-p-> | 893 | Bacteria>Pseudomonadota>Gammaproteobacteria>Enterobacterales>Enterobacteriaceae | GCF_032029275.1 | Escherichia_coli | A7123_RS21895 | 15833..18514 | 562 | hypothetical-protein-[Escherichia-coli] |
| WP_180464865.1 | AAA+ATPase+wHTH+Alpha+Beta-domain+wHTH+RRM | Type-3_BR.PglZ.associated.systems | <-DUF772+DDE_Tnp_1<-AraC-HTH+AraC-HTH\|\|?->\|\|Zn_Tnp_IS1+HTH_Tnp_IS1+DDE_Tnp_IS1->HTH_Tnp->XIS-HTH->DUF499*->PglZ->METHYLASE->iSTAND->TUDOR+SF1-SF2-Helicase+iREase+CTD->SbcC->\|\|nSTAND1+sGTP-> | 893 | Bacteria>Pseudomonadota>Gammaproteobacteria>Enterobacterales>Enterobacteriaceae | GCF_013424785.1 | Escherichia_coli | HOO97_RS17455 | 12856..15537 | 562 | hypothetical-protein-[Escherichia-coli] |
| QJS02693.1 | AAA+ATPase+wHTH+Alpha+Beta-domain+wHTH+RRM | Type-3_BR.PglZ.associated.systems | <-DUF772+DDE_Tnp_1+DDE_Tnp_1_6<-?<-ArdB<-?<-?<-?\|\|DUF499*->PglZ->SIR2->VirB4-FtsK+HerA->METHYLASE->iSTAND->TUDOR+SF1-SF2-Helicase+iREase+CTD-> | 662 | Bacteria>Pseudomonadota>Gammaproteobacteria>Enterobacterales>Enterobacteriaceae | - | Klebsiella_pneumoniae | . | 52334..54322 | 573 | hypothetical-protein-[Klebsiella-pneumoniae] |
| WP_001584131.1 | AAA+ATPase+wHTH+Alpha+Beta-domain+wHTH+RRM | Type-3_BR.PglZ.associated.systems | <-ADH_N+ADH_zinc_N\|\|tRNA->HTH_Tnp->XIS-HTH->DUF499*->PglZ->METHYLASE->iSTAND->TUDOR+SF1-SF2-Helicase+iREase+CTD->SbcC-> | 893 | Bacteria>Pseudomonadota>Gammaproteobacteria>Enterobacterales>Enterobacteriaceae | GCF_005388985.1 | Escherichia_coli | FFF22_RS16095 | 3501..6182 | 562 | hypothetical-protein-[Escherichia-coli] |
| HCL6567946.1 | AAA+ATPase+wHTH+Alpha+Beta-domain+wHTH+RRM | Type-3_BR.PglZ.associated.systems | <-ADH_N+ADH_zinc_N\|\|tRNA->Arm-DNA-bind_3+Phage_integrase-><-TUDOR+SF1-SF2-Helicase+iREase+CTD<-iSTAND<-METHYLASE<-PglZ<-DUF499*<-XIS-HTH<-?<-?\|\|?-><-ZnR<-XIS-HTH | 893 | Bacteria>Pseudomonadota>Gammaproteobacteria>Enterobacterales>Enterobacteriaceae | GCA_025038435.1 | Citrobacter_freundii | N2Q46_002699 | 169100..171781 | 546 | hypothetical-protein-[Citrobacter-freundii] |
| HCU2343865.1 | AAA+ATPase+wHTH+Alpha+Beta-domain+wHTH+RRM | Type-3_BR.PglZ.associated.systems | <-ADH_N+ADH_zinc_N\|\|tRNA->Arm-DNA-bind_3+Phage_integrase-><-TUDOR+SF1-SF2-Helicase+iREase+CTD<-iSTAND<-METHYLASE<-PglZ<-DUF499*<-XIS-HTH<-?<-?\|\|?-><-ZnR<-XIS-HTH | 893 | Bacteria>Pseudomonadota>Gammaproteobacteria>Enterobacterales>Enterobacteriaceae | GCA_026360735.1 | Citrobacter_freundii | OUJ93_001941 | 170304..172985 | 546 | TPA:-hypothetical-protein-OUJ93_001941-[Citrobacter-freundii] |
| WP_137397610.1 | AAA+ATPase+wHTH+Alpha+Beta-domain+wHTH+RRM | Type-3_BR.PglZ.associated.systems | <-ADH_N+ADH_zinc_N\|\|tRNA->Arm-DNA-bind_3+Phage_integrase-><-TUDOR+SF1-SF2-Helicase+iREase+CTD<-iSTAND<-METHYLASE<-PglZ<-DUF499*<-XIS-HTH<-?<-?\|\|?-><-?<-XIS-HTH | 893 | Bacteria>Pseudomonadota>Gammaproteobacteria>Enterobacterales>Enterobacteriaceae | GCF_005281025.1 | Citrobacter_sp._wls714 | FDX11_RS05730 | 163019..165700 | 2576422 | hypothetical-protein-[Citrobacter-sp.-wls714] |
| WP_318586845.1 | AAA+ATPase+wHTH+Alpha+Beta-domain+wHTH+RRM | Type-3_BR.PglZ.associated.systems | TnsC-ATPase->XIS-HTH->DUF499+RRM*->PglZ->SIR2->VirB4-FtsK+HerA->METHYLASE->iSTAND-> | 919 | Bacteria>Pseudomonadota>Gammaproteobacteria>Enterobacterales>Enterobacteriaceae>Enterobacter | GCF_033488235.1 | Enterobacter_kobei | SGL89_RS08905 | 38138..40897 | 208224 | DUF499-domain-containing-protein-[Enterobacter-kobei] |
| ELC7910886.1 | AAA+ATPase+wHTH+Alpha+Beta-domain+wHTH+RRM | Type-3_BR.PglZ.associated.systems | XIS-HTH->XIS-HTH->DUF499+RRM*->PglZ->METHYLASE->iSTAND->TUDOR+SF1-SF2-Helicase+iREase+CTD-> | 893 | Bacteria>Pseudomonadota>Gammaproteobacteria>Enterobacterales>Enterobacteriaceae>Escherichia | GCA_031390625.1 | Escherichia_coli | QX958_004501 | 3995..6676 | 562 | hypothetical-protein-[Escherichia-coli] |
| ELP0955860.1 | AAA+ATPase+wHTH+Alpha+Beta-domain+wHTH+RRM | Type-3_BR.PglZ.associated.systems | <-TUDOR+SF1-SF2-Helicase+iREase+CTD<-iSTAND<-METHYLASE<-PglZ<-DUF499+RRM*<-XIS-HTH<-XIS-HTH | 893 | Bacteria>Pseudomonadota>Gammaproteobacteria>Enterobacterales>Enterobacteriaceae>Escherichia | GCA_032748775.1 | Escherichia_coli | Q2Q27_004348 | 8226..10907 | 562 | hypothetical-protein-[Escherichia-coli] |
| HED3944160.1 | AAA+ATPase+wHTH+Alpha+Beta-domain+wHTH+RRM | Type-3_BR.PglZ.associated.systems | <-Cluster84_1clades<-TUDOR+SF1-SF2-Helicase+iREase+CTD<-iSTAND<-METHYLASE<-PglZ<-DUF499+RRM*<-XIS-HTH | 893 | Bacteria>Pseudomonadota>Gammaproteobacteria>Enterobacterales>Enterobacteriaceae>Escherichia | GCA_032993125.1 | Escherichia_coli | R4240_004702 | 8121..10802 | 562 | hypothetical-protein-[Escherichia-coli] |
| HDT5906547.1 | AAA+ATPase+wHTH+Alpha+Beta-domain+wHTH+RRM | Type-3_BR.PglZ.associated.systems | <-iSTAND<-METHYLASE<-VirB4-FtsK+HerA<-SIR2<-PglZ<-DUF499+RRM* | 721 | Bacteria>Pseudomonadota>Gammaproteobacteria>Enterobacterales>Enterobacteriaceae>Klebsiella/Raoultella_group | GCA_030884555.1 | Klebsiella_michiganensis | QHC31_005188 | 26744..28910 | 1134687 | hypothetical-protein-[Klebsiella-michiganensis] |
| ELB7838338.1 | AAA+ATPase+wHTH+Alpha+Beta-domain+wHTH+RRM | Type-3_BR.PglZ.associated.systems | DUF499+RRM*->PglZ->METHYLASE->iSTAND->TUDOR+SF1-SF2-Helicase+iREase+CTD-> | 758 | Bacteria>Pseudomonadota>Gammaproteobacteria>Enterobacterales>Enterobacteriaceae>Klebsiella/Raoultella_group | GCA_031244225.1 | Klebsiella_pneumoniae | RH802_005453 | 1..2277 | 573 | hypothetical-protein-[Klebsiella-pneumoniae] |
| WP_039340052.1 | AAA+ATPase+wHTH+Alpha+Beta-domain+wHTH+RRM | Type-3_BR.PglZ.associated.systems | <-HTH+WYL<-?<-XIS-HTH\|\|?->XIS-HTH->DUF499*->PglZ->METHYLASE->iSTAND->TUDOR+SF1-SF2-Helicase+iREase+CTD-><-Arm-DNA-bind_3+Phage_integrase | 892 | Bacteria>Pseudomonadota>Gammaproteobacteria>Enterobacterales>Erwiniaceae | GCF_000786255.1 | Pantoea_stewartii | NL54_RS09390 | 79831..82509 | 66269 | hypothetical-protein-[Pantoea-stewartii] |
| WP_187304883.1 | AAA+ATPase+wHTH+Alpha+Beta-domain+wHTH+RRM | Type-3_BR.PglZ.associated.systems | <-TUDOR+SF1-SF2-Helicase+iREase+CTD<-iSTAND<-METHYLASE<-PglZ<-DUF499*<-XIS-HTH<-XIS-HTH<-?\|\|CopG/RHH->nSTAND1+sGTP->ArdB-> | 893 | Bacteria>Pseudomonadota>Gammaproteobacteria>Enterobacterales>Morganellaceae | GCF_014333575.1 | Morganella_morganii | H9I30_RS14285 | 11621..14302 | 582 | hypothetical-protein-[Morganella-morganii] |
| WP_115113554.1 | AAA+ATPase+wHTH+Alpha+Beta-domain+wHTH+RRM | Type-3_BR.PglZ.associated.systems | <-REase<-TRD+TRD<-TUDOR+SF1-SF2-Helicase+iREase+CTD<-iSTAND<-METHYLASE<-PglZ<-DUF499*<-XIS-HTH<-TetR-HTH<-NACHT | 893 | Bacteria>Pseudomonadota>Gammaproteobacteria>Enterobacterales>Morganellaceae | GCF_028482435.1 | Providencia | DXZ64_RS10180 | 2191448..2194129 | 586 | hypothetical-protein-[Providencia-stuartii] |
| WP_248706999.1 | AAA+ATPase+wHTH+Alpha+Beta-domain+wHTH+RRM | Type-3_BR.PglZ.associated.systems | <-METHYLASE-p<-TUDOR+SF1-SF2-Helicase+iREase+CTD<-iSTAND<-METHYLASE<-PglZ<-DUF499*<-XIS-HTH<-HTH_Tnp<-?<-XIS-HTH<-?\|\|CopG/RHH-> | 893 | Bacteria>Pseudomonadota>Gammaproteobacteria>Enterobacterales>Morganellaceae | GCF_023242175.1 | Proteus | M1765_RS01860 | 361373..364054 | 583 | MULTISPECIES:-hypothetical-protein-[Proteus] |
| HCZ8676513.1 | AAA+ATPase+wHTH+Alpha+Beta-domain+wHTH+RRM | Type-3_BR.PglZ.associated.systems | <-METHYLASE-p<-TUDOR+SF1-SF2-Helicase+iREase+CTD<-iSTAND<-METHYLASE<-PglZ<-DUF499* | 307 | Bacteria>Pseudomonadota>Gammaproteobacteria>Enterobacterales>Morganellaceae | GCA_026957575.1 | Proteus_mirabilis | O4E69_002657 | 102261..103184 | 584 | TPA:-hypothetical-protein-O4E69_002657,-partial-[Proteus-mirabilis] |
| WP_009114226.1 | AAA+ATPase+wHTH+Alpha+Beta-domain+wHTH+RRM | Type-3_BR.PglZ.associated.systems | <-Phage_int_SAM_2+Integrase_1<-?\|\|<-TUDOR+SF1-SF2-Helicase+iREase+CTD<-iSTAND<-METHYLASE<-PglZ<-DUF499*<-XIS-HTH<-?<-?<-HIN-HTH\|\|?-><-TraG_N | 893 | Bacteria>Pseudomonadota>Gammaproteobacteria>Enterobacterales>Pectobacteriaceae | GCF_000225565.1 | Brenneria | BRE312_RS17670 | 3873639..3876320 | 71655 | MULTISPECIES:-hypothetical-protein-[Brenneria] |
| WP_174869150.1 | AAA+ATPase+wHTH+Alpha+Beta-domain+wHTH+RRM | Type-3_BR.PglZ.associated.systems | <-HIN-HTH<-?<-TUDOR+SF1-SF2-Helicase+iREase+CTD<-iSTAND<-METHYLASE<-PglZ<-DUF499*<-XIS-HTH<-?<-?\|\|?-><-WYL<-HTH_Tnp_1+HTH_21+rve | 891 | Bacteria>Pseudomonadota>Gammaproteobacteria>Enterobacterales>Pectobacteriaceae | GCF_902143335.1 | Pectobacterium_polaris | HU229_RS16050 | 123032..125707 | 2042057 | hypothetical-protein-[Pectobacterium-polaris] |
| ETX49388.1 | AAA+ATPase+wHTH+Alpha+Beta-domain+wHTH+RRM | Type-3_BR.PglZ.associated.systems | LuxR-HTH->tRNA->Arm-DNA-bind_3+Phage_integrase-><-TUDOR+SF1-SF2-Helicase+iREase+CTD<-iSTAND<-METHYLASE<-PglZ<-DUF499*<-XIS-HTH<-?<-?\|\|?-><-ZnR<-ZnR | 893 | Bacteria>Pseudomonadota>Gammaproteobacteria>Enterobacterales>Yersiniaceae | GCA_000521905.1 | Serratia_marcescens_BIDMC_50 | P812_00205 | 226932..229613 | 1400187 | hypothetical-protein-P812_00205-[Serratia-marcescens-BIDMC-50] |
| WP_033641703.1 | AAA+ATPase+wHTH+Alpha+Beta-domain+wHTH+RRM | Type-3_BR.PglZ.associated.systems | LuxR-HTH->tRNA->Arm-DNA-bind_3+Phage_integrase-><-TUDOR+SF1-SF2-Helicase+iREase+CTD<-iSTAND<-METHYLASE<-PglZ<-DUF499*<-XIS-HTH<-?<-?\|\|?-><-ZnR<-ZnR | 893 | Bacteria>Pseudomonadota>Gammaproteobacteria>Enterobacterales>Yersiniaceae | GCF_000521905.1 | Serratia_marcescens | P812_00205 | 226932..229613 | 615 | hypothetical-protein-[Serratia-marcescens] |
| WP_273873587.1 | AAA+ATPase+wHTH+Alpha+Beta-domain+wHTH+RRM | Type-3_BR.PglZ.associated.systems | <-TUDOR+SF1-SF2-Helicase+iREase+CTD<-iSTAND<-METHYLASE<-PglZ<-DUF499*<-XIS-HTH<-HTH_Tnp\|\|HTH_3-><-TetR-HTH<-NACHT<-Phage_integrase+Phage_integrase | 893 | Bacteria>Pseudomonadota>Gammaproteobacteria>Enterobacterales>Yersiniaceae | GCF_028616955.1 | Serratia_ureilytica | PVW65_RS15320 | 8465..11146 | 300181 | hypothetical-protein-[Serratia-ureilytica] |
| WP_169248422.1 | AAA+ATPase+wHTH+Alpha+Beta-domain+wHTH+RRM | Type-3_BR.PglZ.associated.systems | <-TUDOR+SF1-SF2-Helicase+iREase+CTD<-BirA-HTH<-Schlafen<-iSTAND<-METHYLASE<-PglZ<-DUF499*<-XIS-HTH<-GDResolvase+Recombinase+ZnR<-DDE_Tnp_1 | 910 | Bacteria>Pseudomonadota>Gammaproteobacteria>Gammaproteobacteria_incertae_sedis>Candidatus_Competibacteraceae | GCF_012939995.1 | Candidatus_Competibacter_phosphatis | E4P82_RS08120 | 16271..19003 | 221280 | DUF499-domain-containing-protein-[Candidatus-Competibacter-phosphatis] |

<table>
<tr><th>Protein-ID</th><th>DUF499-architecture</th><th>System</th><th>Gene-Neighbourhoods</th><th>Length</th><th>Taxonomy</th><th>Assembly-ID</th><th>Strain</th><th>Locus-Tag</th><th>Genomic-coordinates</th><th>TaxID</th><th>Fasta-defline</th></tr>
<tr><td>RUQ39816.1</td><td>AAA+ATPase+wHTH+Alpha+Beta-domain+wHTH+RRM</td><td>Type-3_BR.PglZ.associated.systems</td><td><-METHYLASE-p<-ParB+HNH<-PglZ<-DUF499*<-?||?->?->HTH-></td><td>907</td><td>Bacteria>Pseudomonadota>Gammaproteobacteria>Gammaproteobacteria_incertae_sedis>Candidatus_Competibacteraceae</td><td>GCA_003989075.1</td><td>Candidatus_Competibacteraceae_bacterium</td><td>EKK69_07735</td><td>3742..6465</td><td>2053538</td><td>DUF499-domain-containing-protein-[Candidatus-Competibacteraceae-bacterium]</td></tr>
<tr><td>MCW8905972.1</td><td>AAA+ATPase+wHTH+Alpha+Beta-domain+wHTH+RRM</td><td>Type-3_BR.PglZ.associated.systems</td><td><-TUDOR+SF1-SF2-Helicase+iREase+CTD<-iSTAND<-METHYLASE<-PglZ<-DUF499*<-RHH<-?||?-><-Phage_integrase</td><td>900</td><td>Bacteria>Pseudomonadota>Gammaproteobacteria>Gammaproteobacteria_incertae_sedis>Sedimenticola</td><td>GCA_026196475.1</td><td>Sedimenticola_sp.</td><td>OQL28_01880</td><td>14825..17527</td><td>1940285</td><td>MAG:-DUF499-domain-containing-protein-[Sedimenticola-sp.]</td></tr>
<tr><td>WP_159654884.1</td><td>AAA+ATPase+wHTH+Alpha+Beta-domain+wHTH+RRM</td><td>Type-3_BR.PglZ.associated.systems</td><td>SF1-SF2-Helicase->?-><-?||DUF499*->PglZ->?->METHYLASE->?->PIN->iSTAND-></td><td>894</td><td>Bacteria>Pseudomonadota>Gammaproteobacteria>Methylococcales>Methylococcaceae</td><td>GCF_009828925.2</td><td>Methylicorpusculum_oleiharenae</td><td>GO003_RS12045</td><td>2667775..2670459</td><td>1338687</td><td>DUF499-domain-containing-protein-[Methylicorpusculum-oleiharenae]</td></tr>
<tr><td>WP_010961899.1</td><td>AAA+ATPase+wHTH+Alpha+Beta-domain+wHTH+RRM</td><td>Type-3_BR.PglZ.associated.systems</td><td>Phage_lysozyme-><-HEPN<-TUDOR+SF1-SF2-Helicase+iREase+CTD<-iSTAND<-METHYLASE<-PglZ<-DUF499*||DUF772+DDE_Tnp_1+DDE_Tnp_1_6->?-><-HIN-HTH</td><td>913</td><td>Bacteria>Pseudomonadota>Gammaproteobacteria>Methylococcales>Methylococcaceae</td><td>GCF_000008325.1</td><td>Methylococcus_capsulatus</td><td>MCA_RS13155</td><td>2870875..2873616</td><td>414</td><td>DUF499-domain-containing-protein-[Methylococcus-capsulatus]</td></tr>
<tr><td>WP_255187190.1</td><td>AAA+ATPase+wHTH+Alpha+Beta-domain+wHTH+RRM</td><td>Type-3_BR.PglZ.associated.systems</td><td>Peptidase->?->?->?->XIS-HTH->DUF499*->PglZ->?->METHYLASE->iSTAND->OLD-ABC->TOPRIM-></td><td>934</td><td>Bacteria>Pseudomonadota>Gammaproteobacteria>Methylococcales>Methylococcaceae</td><td>GCF_024360925.2</td><td>Methylomonas_rapida</td><td>NM686_RS07140</td><td>1504339..1507143</td><td>2963939</td><td>DUF499-domain-containing-protein-[Methylomonas-sp.-MP1]</td></tr>
<tr><td>WP_169603387.1</td><td>AAA+ATPase+wHTH+Alpha+Beta-domain+wHTH+RRM</td><td>Type-3_BR.PglZ.associated.systems</td><td><-Schlafen+HATPase+HTH<-METHYLASE<-PemK_toxin<-?<-?<-PglZ<-DUF499*</td><td>909</td><td>Bacteria>Pseudomonadota>Gammaproteobacteria>Methylococcales>Methylococcaceae</td><td>GCF_012769535.1</td><td>Methylococcus_geothermalis</td><td>GNH96_RS09105</td><td>1909504..1912233</td><td>2681310</td><td>DUF499-domain-containing-protein-[Methylococcus-geothermalis]</td></tr>
<tr><td>WP_304902987.1</td><td>AAA+ATPase+wHTH+Alpha+Beta-domain+wHTH+RRM</td><td>Type-3_BR.PglZ.associated.systems</td><td>DUF499+RRM*->PglZ->HEPN->RusA->METHYLASE-></td><td>894</td><td>Bacteria>Pseudomonadota>Gammaproteobacteria>Methylococcales>Methylococcaceae>Methylicorpusculum</td><td>GCF_030652145.1</td><td>Methylicorpusculum_sp.</td><td>Q7U98_RS01120</td><td>4920..7604</td><td>2713644</td><td>DUF499-domain-containing-protein-[Methylicorpusculum-sp.]</td></tr>
<tr><td>MDO9046279.1</td><td>AAA+ATPase+wHTH+Alpha+Beta-domain+wHTH+RRM</td><td>Type-3_BR.PglZ.associated.systems</td><td><-Phage_int_SAM_3+Phage_integrase<-Phage_integrase<-Phage_int_SAM_3+Phage_integrase||HTH_3->DUF499+RRM*->PglZ->HEPN->METHYLASE-></td><td>890</td><td>Bacteria>Pseudomonadota>Gammaproteobacteria>Methylococcales>Methylococcaceae>Methylobacter</td><td>GCA_030653175.1</td><td>Methylobacter_sp.</td><td>Q7U66_00915</td><td>4422..7094</td><td>2051955</td><td>DUF499-domain-containing-protein-[Methylobacter-sp.]</td></tr>
<tr><td>WP_310749087.1</td><td>AAA+ATPase+wHTH+Alpha+Beta-domain+wHTH+RRM</td><td>Type-3_BR.PglZ.associated.systems</td><td><-METHYLASE<-RusA<-HEPN<-PglZ<-DUF499+RRM*<-HTH_Tnp_1+HTH_21+rve+rve_3</td><td>894</td><td>Bacteria>Pseudomonadota>Gammaproteobacteria>Methylococcales>Methylococcaceae>Methylomonas</td><td>GCF_031582845.1</td><td>Methylomonas_koyamae</td><td>RI210_RS15065</td><td>3359747..3362431</td><td>702114</td><td>DUF499-domain-containing-protein-[Methylomonas-koyamae]</td></tr>
<tr><td>HCH8091325.1</td><td>AAA+ATPase+wHTH+Alpha+Beta-domain+wHTH+RRM</td><td>Type-3_BR.PglZ.associated.systems</td><td>ZnR+ZnR->DUF932->?->ParB->XIS-HTH->XIS-HTH->DUF499*->PglZ->METHYLASE->iSTAND->TUDOR+SF1-SF2-Helicase+iREase+CTD-><-Arm-DNA-bind_3+Phage_int_SAM_3+Phage_integrase</td><td>842</td><td>Bacteria>Pseudomonadota>Gammaproteobacteria>Moraxellales>Moraxellaceae</td><td>GCA_024274195.1</td><td>Acinetobacter_baumannii</td><td>NJU31_000797</td><td>8427..10955</td><td>470</td><td>DUF499-domain-containing-protein-[Acinetobacter-baumannii]</td></tr>
<tr><td>MBR7688338.1</td><td>AAA+ATPase+wHTH+Alpha+Beta-domain+wHTH+RRM</td><td>Type-3_BR.PglZ.associated.systems</td><td>DUF932->ParB->XIS-HTH->DUF499*->PglZ->METHYLASE->iSTAND->TUDOR+SF1-SF2-Helicase+iREase+CTD-><-Arm-DNA-bind_3+Phage_integrase||Arm-DNA-bind_3+Phage_integrase-></td><td>842</td><td>Bacteria>Pseudomonadota>Gammaproteobacteria>Moraxellales>Moraxellaceae</td><td>GCA_018139245.1</td><td>Acinetobacter_nosocomialis</td><td>KC097_17715</td><td>10751..13279</td><td>106654</td><td>DUF499-domain-containing-protein-[Acinetobacter-nosocomialis]</td></tr>
<tr><td>WP_068556908.1</td><td>AAA+ATPase+wHTH+Alpha+Beta-domain+wHTH+RRM</td><td>Type-3_BR.PglZ.associated.systems</td><td>DUF932->?->ParB->XIS-HTH->XIS-HTH->DUF499*->PglZ->METHYLASE->iSTAND->SF1-SF2-Helicase+iREase+CTD-><-?<-Arm-DNA-bind_3+Phage_int_SAM_3+Phage_integrase</td><td>889</td><td>Bacteria>Pseudomonadota>Gammaproteobacteria>Moraxellales>Moraxellaceae</td><td>GCF_001706305.1</td><td>Acinetobacter_pittii</td><td>BFR96_RS10080</td><td>5927..8596</td><td>48296</td><td>DUF499-domain-containing-protein-[Acinetobacter-pittii]</td></tr>
<tr><td>WP_078425549.1</td><td>AAA+ATPase+wHTH+Alpha+Beta-domain+wHTH+RRM</td><td>Type-3_BR.PglZ.associated.systems</td><td>Arm-DNA-bind_3+Phage_int_SAM_3+Phage_integrase-><-TUDOR+SF1-SF2-Helicase+iREase+CTD<-iSTAND<-METHYLASE<-PglZ<-DUF499*<-XIS-HTH<-XIS-HTH<-ParB<-?<-DUF932<-ZnR+ZnR</td><td>889</td><td>Bacteria>Pseudomonadota>Gammaproteobacteria>Moraxellales>Moraxellaceae</td><td>GCF_002018925.1</td><td>Acinetobacter_sp._MF4642</td><td>MF4642_RS05430</td><td>128111..130780</td><td>1960825</td><td>DUF499-domain-containing-protein-[Acinetobacter-sp.-MF4642]</td></tr>
<tr><td>WP_119063263.1</td><td>AAA+ATPase+wHTH+Alpha+Beta-domain+wHTH+RRM</td><td>Type-3_BR.PglZ.associated.systems</td><td>Arm-DNA-bind_3+Phage_int_SAM_3+Phage_integrase-><-TUDOR+SF1-SF2-Helicase+iREase+CTD<-iSTAND<-METHYLASE<-PglZ<-DUF499*<-XIS-HTH<-XIS-HTH<-ParB<-?<-DUF932</td><td>889</td><td>Bacteria>Pseudomonadota>Gammaproteobacteria>Moraxellales>Moraxellaceae</td><td>GCF_003569905.1</td><td>Acinetobacter_sp._SWAC57</td><td>DYI96_RS04380</td><td>121444..124113</td><td>2293834</td><td>DUF499-domain-containing-protein-[Acinetobacter-sp.-SWAC57]</td></tr>
<tr><td>WP_115434950.1</td><td>AAA+ATPase+wHTH+Alpha+Beta-domain+wHTH+RRM</td><td>Type-3_BR.PglZ.associated.systems</td><td><-SF1-SF2-Helicase+iREase+CTD||?-><-iSTAND<-METHYLASE<-PglZ<-DUF499*</td><td>914</td><td>Bacteria>Pseudomonadota>Gammaproteobacteria>Moraxellales>Moraxellaceae</td><td>GCF_027855255.1</td><td>Acinetobacter_baumannii</td><td>B8U65_RS07110</td><td>20185..22929</td><td>470</td><td>DUF499-domain-containing-protein-[Acinetobacter-baumannii]</td></tr>
<tr><td>WP_161932440.1</td><td>AAA+ATPase+wHTH+Alpha+Beta-domain+wHTH+RRM</td><td>Type-3_BR.PglZ.associated.systems</td><td><-HAD<-?<-?<-Primpol+D5N+D5ATPase<-ParB+HNH<-ParB+HNH||DUF499*->PglZ->METHYLASE->iSTAND->SF1-SF2-Helicase+iREase+CTD-><-wHTH+HNH<-SAM-methylase</td><td>914</td><td>Bacteria>Pseudomonadota>Gammaproteobacteria>Moraxellales>Moraxellaceae</td><td>GCF_009939195.1</td><td>Acinetobacter_kanungonis</td><td>GWK51_RS01785</td><td>415302..418046</td><td>2699469</td><td>DUF499-domain-containing-protein-[Acinetobacter-kanungonis]</td></tr>
<tr><td>WP_249127149.1</td><td>AAA+ATPase+wHTH+Alpha+Beta-domain+wHTH+RRM</td><td>Type-3_BR.PglZ.associated.systems</td><td><-Arm-DNA-bind_3+Phage_integrase||Arm-DNA-bind_3+Phage_integrase-><-TUDOR+SF1-SF2-Helicase+iREase+CTD<-iSTAND<-METHYLASE<-PglZ<-DUF499*<-XIS-HTH<-ParB<-DUF932</td><td>889</td><td>Bacteria>Pseudomonadota>Gammaproteobacteria>Moraxellales>Moraxellaceae</td><td>GCF_018139245.1</td><td>Acinetobacter_nosocomialis</td><td>KC092_RS17930</td><td>35136..37805</td><td>106654</td><td>DUF499-domain-containing-protein-[Acinetobacter-nosocomialis]</td></tr>
<tr><td>MDU2406928.1</td><td>AAA+ATPase+wHTH+Alpha+Beta-domain+wHTH+RRM</td><td>Type-3_BR.PglZ.associated.systems</td><td>TnsC-ATPase->SF1-SF2-Helicase->DUF499+RRM*->PglZ->METHYLASE->iSTAND->TUDOR+SF1-SF2-Helicase+iREase+CTD-></td><td>914</td><td>Bacteria>Pseudomonadota>Gammaproteobacteria>Moraxellales>Moraxellaceae>Acinetobacter</td><td>GCA_032480795.1</td><td>Acinetobacter_junii</td><td>E7D55_01495</td><td>29573..32317</td><td>40215</td><td>DUF499-domain-containing-protein-[Acinetobacter-junii]</td></tr>
<tr><td>WP_312105822.1</td><td>AAA+ATPase+wHTH+Alpha+Beta-domain+wHTH+RRM</td><td>Type-3_BR.PglZ.associated.systems</td><td><-TUDOR+SF1-SF2-Helicase+iREase+CTD<-iSTAND<-METHYLASE<-PglZ<-DUF499+RRM*<-XIS-HTH<-DUF932</td><td>889</td><td>Bacteria>Pseudomonadota>Gammaproteobacteria>Moraxellales>Moraxellaceae>Acinetobacter</td><td>GCF_031977435.1</td><td>Acinetobacter_venetianus</td><td>RTL42_RS14900</td><td>8835..11504</td><td>52133</td><td>DUF499-domain-containing-protein-[Acinetobacter-venetianus]</td></tr>
<tr><td>WP_093286142.1</td><td>AAA+ATPase+wHTH+Alpha+Beta-domain+wHTH+RRM</td><td>Type-3_BR.PglZ.associated.systems</td><td>Arm-DNA-bind_3+Phage_integrase->ZDNA-HTH->?-><-?||XIS-HTH->DUF499*->PglZ->METHYLASE->iSTAND->OLD-ABC->TOPRIM->TUDOR+SF1-SF2-Helicase+iREase+CTD-></td><td>909</td><td>Bacteria>Pseudomonadota>Gammaproteobacteria>Nevskiales>Nevskiaceae</td><td>GCF_900111015.1</td><td>Solimonas_aquatica</td><td>BMY12_RS11520</td><td>194234..196963</td><td>489703</td><td>DUF499-domain-containing-protein-[Solimonas-aquatica]</td></tr>
<tr><td>WP_162078973.1</td><td>AAA+ATPase+wHTH+Alpha+Beta-domain+wHTH+RRM</td><td>Type-3_BR.PglZ.associated.systems</td><td>ABC-ATPase+SbcC-><-Mbetalac||XIS-HTH->DUF499*->PglZ->?->?->PemK_toxin->METHYLASE->iSTAND-></td><td>909</td><td>Bacteria>Pseudomonadota>Gammaproteobacteria>Nevskiales>Nevskiaceae</td><td>GCF_010092425.1</td><td>Sinimarinibacterium_sp._NLF-5-8</td><td>GT972_RS12885</td><td>2701736..2704465</td><td>2698684</td><td>DUF499-domain-containing-protein-[Sinimarinibacterium-sp.-NLF-5-8]</td></tr>
<tr><td>TAM26627.1</td><td>AAA+ATPase+wHTH+Alpha+Beta-domain+wHTH+RRM</td><td>Type-3_BR.PglZ.associated.systems</td><td><-Schlafen+TPR<-iSTAND<-METHYLASE<-VirB4-FtsK+HerA<-SIR2<-PglZ<-DUF499*<-XIS-HTH||AbiEi_4->AbiEii->MORC+HisKin+REase-></td><td>914</td><td>Bacteria>Pseudomonadota>Gammaproteobacteria>Nevskiales>Nevskiaceae</td><td>GCA_004298745.1</td><td>Nevskiaceae_bacterium</td><td>EPN60_10220</td><td>118142..120886</td><td>1969813</td><td>MAG:-DUF499-domain-containing-protein-[Sinobacteraceae-bacterium]</td></tr>
<tr><td>TXG98447.1</td><td>AAA+ATPase+wHTH+Alpha+Beta-domain+wHTH+RRM</td><td>Type-3_BR.PglZ.associated.systems</td><td><-METHYLASE-p<-Peptidase_S8<-Classical-AAA<-PglZ<-DUF499*<-XIS-HTH</td><td>914</td><td>Bacteria>Pseudomonadota>Gammaproteobacteria>Nevskiales>Nevskiaceae</td><td>GCA_008012095.1</td><td>Nevskiaceae_bacterium</td><td>E6R08_04580</td><td>4917..7661</td><td>1969813</td><td>MAG:-DUF499-domain-containing-protein-[Sinobacteraceae-bacterium]</td></tr>
<tr><td>WP_229149069.1</td><td>AAA+ATPase+wHTH+Alpha+Beta-domain+wHTH+RRM</td><td>Type-3_BR.PglZ.associated.systems</td><td><-Calcineurin<-SbcC<-SF1-SF2-Helicase+iREase+CTD<-iSTAND<-METHYLASE<-PglZ<-DUF499*<-XIS-HTH<-Arm-DNA-bind_3+Phage_integrase</td><td>892</td><td>Bacteria>Pseudomonadota>Gammaproteobacteria>Oceanospirillales>Alcanivoracaceae</td><td>GCF_020667895.1</td><td>Alcanivorax_sp._1008</td><td>J1C54_RS14080</td><td>32162..34840</td><td>2816853</td><td>DUF499-domain-containing-protein-[Alcanivorax-sp.-1008]</td></tr>
<tr><td>WP_249016621.1</td><td>AAA+ATPase+wHTH+Alpha+Beta-domain+wHTH+RRM</td><td>Type-3_BR.PglZ.associated.systems</td><td>Phage_integrase->?->Phage_integrase->?->TOPRIM+ZnR->XIS-HTH->DUF499*->PglZ->METHYLASE->iSTAND->TUDOR+SF1-SF2-Helicase+iREase+CTD->SbcC-></td><td>893</td><td>Bacteria>Pseudomonadota>Gammaproteobacteria>Oceanospirillales>Halomonadaceae</td><td>GCF_023330625.1</td><td>Halomonas_venusta</td><td>M3L73_RS11935</td><td>2434724..2437405</td><td>44935</td><td>hypothetical-protein-[Halomonas-venusta]</td></tr>
<tr><td>MBL1267502.1</td><td>AAA+ATPase+wHTH+Alpha+Beta-domain+wHTH+RRM</td><td>Type-3_BR.PglZ.associated.systems</td><td><-TUDOR+SF1-SF2-Helicase+iREase+CTD<-iSTAND<-METHYLASE<-PglZ<-DUF499*<-tRNA<-?||?->35exo-><-ABhydrolase</td><td>892</td><td>Bacteria>Pseudomonadota>Gammaproteobacteria>Oceanospirillales>Halomonadaceae</td><td>GCA_002733435.2</td><td>Halomonas_sp.</td><td>COA87_007085</td><td>255768..258446</td><td>1486246</td><td>DUF499-domain-containing-protein-[Halomonas-sp.]</td></tr>
<tr><td>WP_273190710.1</td><td>AAA+ATPase+wHTH+Alpha+Beta-domain+wHTH+RRM</td><td>Type-3_BR.PglZ.associated.systems</td><td><-DUF4096+DDE_Tnp_1+DDE_Tnp_1_2<-TUDOR+SF1-SF2-Helicase+iREase+CTD<-iSTAND<-METHYLASE<-PglZ<-DUF499*<-tRNA<-?||?->35exo-><-ABhydrolase</td><td>892</td><td>Bacteria>Pseudomonadota>Gammaproteobacteria>Oceanospirillales>Halomonadaceae</td><td>GCF_002733435.1</td><td>Halomonas_sp.</td><td>COA87_RS07115</td><td>255768..258446</td><td>1486246</td><td>DUF499-domain-containing-protein-[Halomonas-sp.]</td></tr>
<tr><td>WP_318448468.1</td><td>AAA+ATPase+wHTH+Alpha+Beta-domain+wHTH+RRM</td><td>Type-3_BR.PglZ.associated.systems</td><td>DUF932->RHH->DUF499+RRM*->PglZ->METHYLASE->iSTAND->TUDOR+SF1-SF2-Helicase+iREase+CTD-></td><td>911</td><td>Bacteria>Pseudomonadota>Gammaproteobacteria>Oceanospirillales>Halomonadaceae>Halomonas</td><td>GCF_033446685.1</td><td>Halomonas_sp._HP20-15</td><td>R6258_RS05670</td><td>207326..210061</td><td>3085901</td><td>DUF499-domain-containing-protein-[Halomonas-sp.-HP20-15]</td></tr>
<tr><td>WP_024162194.1</td><td>AAA+ATPase+wHTH+Alpha+Beta-domain+wHTH+RRM</td><td>Type-3_BR.PglZ.associated.systems</td><td><-CopG/RHH<-VirD4-TraG+VirD4-FtsK||?-><-?<-MODE-HTH+PBPII||XIS-HTH->DUF499*->PglZ->Classical-AAA->Peptidase_S8->METHYLASE->iSTAND->TUDOR+SF1-SF2-Helicase+iREase+CTD-></td><td>915</td><td>Bacteria>Pseudomonadota>Gammaproteobacteria>Pseudomonadales</td><td>GCF_018439695.1</td><td>Pseudomonadaceae</td><td>B597_RS37860</td><td>2141274..2144021</td><td>135621</td><td>MULTISPECIES:-DUF499-domain-containing-protein-[Pseudomonadaceae]</td></tr>
<tr><td>MCK5791909.1</td><td>AAA+ATPase+wHTH+Alpha+Beta-domain+wHTH+RRM</td><td>Type-3_BR.PglZ.associated.systems</td><td><-Phage_integrase<-?||GDResolvase+HIN-HTH->XIS-HTH->DUF499*->PglZ->METHYLASE-></td><td>893</td><td>Bacteria>Pseudomonadota>Gammaproteobacteria>Pseudomonadales>Ketobacteraceae</td><td>GCA_023145265.1</td><td>Ketobacter_sp.</td><td>KAH34_13750</td><td>2801..5482</td><td>2083498</td><td>DUF499-domain-containing-protein-[Ketobacter-sp.]</td></tr>
<tr><td>WP_153741748.1</td><td>AAA+ATPase+wHTH+Alpha+Beta-domain+wHTH+RRM</td><td>Type-3_BR.PglZ.associated.systems</td><td>HKD-DNase->SF1-SF2-Helicase->DUF499*->PglZ->METHYLASE-></td><td>892</td><td>Bacteria>Pseudomonadota>Gammaproteobacteria>Pseudomonadales>Marinobacteraceae</td><td>GCF_009650625.1</td><td>Marinobacter_nauticus</td><td>F6453_RS18775</td><td>8787..11465</td><td>2743</td><td>DUF499-domain-containing-protein-[Marinobacter-nauticus]</td></tr>
<tr><td>WP_222930677.1</td><td>AAA+ATPase+wHTH+Alpha+Beta-domain+wHTH+RRM</td><td>Type-3_BR.PglZ.associated.systems</td><td>ZnR+TOPRIM+D5ATPase->XIS-HTH->DUF499*->PglZ->SIR2->VirB4-FtsK+HerA->METHYLASE->iSTAND->TUDOR+SF1-SF2-Helicase+iREase+CTD-></td><td>848</td><td>Bacteria>Pseudomonadota>Gammaproteobacteria>Pseudomonadales>Pseudomonadaceae</td><td>-</td><td>Pseudomonas_aeruginosa</td><td>L4Q41_001887</td><td>2150..4696</td><td>287</td><td>DUF499-domain-containing-protein-[Pseudomonas-aeruginosa]</td></tr>
<tr><td>WP_096861997.1</td><td>AAA+ATPase+wHTH+Alpha+Beta-domain+wHTH+RRM</td><td>Type-3_BR.PglZ.associated.systems</td><td>XIS-HTH->DUF499*->PglZ->METHYLASE->iSTAND->TUDOR+SF1-SF2-Helicase+iREase+CTD->SbcC-></td><td>893</td><td>Bacteria>Pseudomonadota>Gammaproteobacteria>Pseudomonadales>Pseudomonadaceae</td><td>GCF_903684655.1</td><td>Pseudomonas_aeruginosa</td><td>CP915_RS26405</td><td>835..3516</td><td>287</td><td>hypothetical-protein-[Pseudomonas-aeruginosa]</td></tr>
<tr><td>WP_193076107.1</td><td>AAA+ATPase+wHTH+Alpha+Beta-domain+wHTH+RRM</td><td>Type-3_BR.PglZ.associated.systems</td><td>XIS-HTH->DUF499*->PglZ->METHYLASE->iSTAND->TUDOR+SF1-SF2-Helicase+iREase+CTD->?->SMODS-></td><td>893</td><td>Bacteria>Pseudomonadota>Gammaproteobacteria>Pseudomonadales>Pseudomonadaceae</td><td>GCF_014897305.1</td><td>Pseudomonas_sp._FME51</td><td>INT39_RS21980</td><td>511..3192</td><td>2742609</td><td>hypothetical-protein-[Pseudomonas-sp.-FME51]</td></tr>
<tr><td>WP_034005121.1</td><td>AAA+ATPase+wHTH+Alpha+Beta-domain+wHTH+RRM</td><td>Type-3_BR.PglZ.associated.systems</td><td>XIS-HTH->DUF499*->PglZ->Classical-AAA->Peptidase_S8->METHYLASE->iSTAND->TUDOR+SF1-SF2-Helicase+iREase+CTD-></td><td>914</td><td>Bacteria>Pseudomonadota>Gammaproteobacteria>Pseudomonadales>Pseudomonadaceae</td><td>GCF_025696705.1</td><td>Pseudomonas_aeruginosa</td><td>NQ36_RS04400</td><td>27397..30141</td><td>287</td><td>DUF499-domain-containing-protein-[Pseudomonas-aeruginosa]</td></tr>
<tr><td>WP_264315537.1</td><td>AAA+ATPase+wHTH+Alpha+Beta-domain+wHTH+RRM</td><td>Type-3_BR.PglZ.associated.systems</td><td>XIS-HTH->DUF499*->PglZ->Classical-AAA->Peptidase_S8->METHYLASE->iSTAND->TUDOR+SF1-SF2-Helicase+iREase+CTD-></td><td>908</td><td>Bacteria>Pseudomonadota>Gammaproteobacteria>Pseudomonadales>Pseudomonadaceae</td><td>GCF_025908315.1</td><td>Pseudomonas_aeruginosa</td><td>OBG92_RS22865</td><td>4825225..4827951</td><td>287</td><td>ATP-binding-protein-[Pseudomonas-aeruginosa]</td></tr>
<tr><td>WP_280156713.1</td><td>AAA+ATPase+wHTH+Alpha+Beta-domain+wHTH+RRM</td><td>Type-3_BR.PglZ.associated.systems</td><td>XIS-HTH->DUF499*->PglZ->Classical-AAA->Peptidase_S8->METHYLASE->iSTAND->TUDOR+SF1-SF2-Helicase+iREase+CTD-></td><td>908</td><td>Bacteria>Pseudomonadota>Gammaproteobacteria>Pseudomonadales>Pseudomonadaceae</td><td>GCF_029856995.1</td><td>Pseudomonas_aeruginosa</td><td>OMP51_RS23585</td><td>4980718..4983444</td><td>287</td><td>DUF499-domain-containing-protein-[Pseudomonas-aeruginosa]</td></tr>
<tr><td>WP_249364538.1</td><td>AAA+ATPase+wHTH+Alpha+Beta-domain+wHTH+RRM</td><td>Type-3_BR.PglZ.associated.systems</td><td>XIS-HTH->DUF499*->PglZ->Classical-AAA->Peptidase_S8->METHYLASE->iSTAND->SF1-SF2-Helicase+iREase+CTD-></td><td>908</td><td>Bacteria>Pseudomonadota>Gammaproteobacteria>Pseudomonadales>Pseudomonadaceae</td><td>GCF_018340955.1</td><td>Pseudomonas_aeruginosa</td><td>JQT49_RS22730</td><td>96930..99656</td><td>287</td><td>DUF499-domain-containing-protein-[Pseudomonas-aeruginosa]</td></tr>
<tr><td>MBH9069449.1</td><td>AAA+ATPase+wHTH+Alpha+Beta-domain+wHTH+RRM</td><td>Type-3_BR.PglZ.associated.systems</td><td>XIS-HTH->DUF499*->PglZ->Classical-AAA->Peptidase_S8->METHYLASE->iSTAND-></td><td>846</td><td>Bacteria>Pseudomonadota>Gammaproteobacteria>Pseudomonadales>Pseudomonadaceae</td><td>GCA_016090915.1</td><td>Pseudomonas_aeruginosa</td><td>I5K26_24370</td><td>154683..157223</td><td>287</td><td>DUF499-domain-containing-protein-[Pseudomonas-aeruginosa]</td></tr>
<tr><td>WP_220276633.1</td><td>AAA+ATPase+wHTH+Alpha+Beta-domain+wHTH+RRM</td><td>Type-3_BR.PglZ.associated.systems</td><td>TetR-HTH->?-><-JAB<-wHTH+REase<-Arm-DNA-bind_2+Phage_int_SAM_3+Phage_integrase||DUF499*->PglZ->?->METHYLASE->iSTAND->SF1-SF2-Helicase+iREase+CTD-><-RlpA</td><td>409</td><td>Bacteria>Pseudomonadota>Gammaproteobacteria>Pseudomonadales>Pseudomonadaceae</td><td>GCF_013913155.1</td><td>Pseudomonas_aeruginosa</td><td>H2D06_RS26090</td><td>71288..72517</td><td>287</td><td>hypothetical-protein-[Pseudomonas-aeruginosa]</td></tr>
<tr><td>WP_138738808.1</td><td>AAA+ATPase+wHTH+Alpha+Beta-domain+wHTH+RRM</td><td>Type-3_BR.PglZ.associated.systems</td><td>RlpA-><-TUDOR+SF1-SF2-Helicase+iREase+CTD<-iSTAND<-METHYLASE<-PglZ<-DUF499*<-XIS-HTH<-NERD</td><td>891</td><td>Bacteria>Pseudomonadota>Gammaproteobacteria>Pseudomonadales>Pseudomonadaceae</td><td>GCF_005930585.1</td><td>Pseudomonas_sp._FSL_W7-0098</td><td>FHL07_RS19530</td><td>95111..97786</td><td>2496120</td><td>hypothetical-protein-[Pseudomonas-sp.-FSL-W7-0098]</td></tr>
<tr><td>WP_167656460.1</td><td>AAA+ATPase+wHTH+Alpha+Beta-domain+wHTH+RRM</td><td>Type-3_BR.PglZ.associated.systems</td><td>RlpA-><-TUDOR+SF1-SF2-Helicase+iREase+CTD<-iSTAND<-METHYLASE<-PglZ<-DUF499*<-?<-?||CNMP+TIR->ACYC+ACYC-></td><td>945</td><td>Bacteria>Pseudomonadota>Gammaproteobacteria>Pseudomonadales>Pseudomonadaceae</td><td>GCF_012030485.1</td><td>unclassified_Pseudomonas</td><td>QVF82_RS24445</td><td>73197..76034</td><td>196821</td><td>DUF499-domain-containing-protein-[Pseudomonas-sp.-B14(2022)]</td></tr>
<tr><td>WP_200968057.1</td><td>AAA+ATPase+wHTH+Alpha+Beta-domain+wHTH+RRM</td><td>Type-3_BR.PglZ.associated.systems</td><td>RlpA-><-TUDOR+SF1-SF2-Helicase+iREase+CTD<-iSTAND<-METHYLASE<-?<-PglZ<-DUF499*<-tRNA<-MerR-HTH</td><td>863</td><td>Bacteria>Pseudomonadota>Gammaproteobacteria>Pseudomonadales>Pseudomonadaceae</td><td>GCF_001450145.1</td><td>Pseudomonas_aeruginosa</td><td>A0954_RS23095</td><td>32383..34974</td><td>287</td><td>DUF499-domain-containing-protein-[Pseudomonas-aeruginosa]</td></tr>
<tr><td>WP_033988893.1</td><td>AAA+ATPase+wHTH+Alpha+Beta-domain+wHTH+RRM</td><td>Type-3_BR.PglZ.associated.systems</td><td>RlpA-><-TUDOR+SF1-SF2-Helicase+iREase+CTD<-iSTAND<-METHYLASE<-?<-PglZ<-DUF499*<-tRNA<-MerR-HTH</td><td>915</td><td>Bacteria>Pseudomonadota>Gammaproteobacteria>Pseudomonadales>Pseudomonadaceae</td><td>GCF_007559065.1</td><td>Pseudomonas</td><td>CP912_RS08085</td><td>210902..213649</td><td>286</td><td>MULTISPECIES:-DUF499-domain-containing-protein-[Pseudomonas]</td></tr>
<tr><td>WP_271014438.1</td><td>AAA+ATPase+wHTH+Alpha+Beta-domain+wHTH+RRM</td><td>Type-3_BR.PglZ.associated.systems</td><td>RlpA-><-TUDOR+SF1-SF2-Helicase+iREase+CTD<-iSTAND<-METHYLASE<-?<-PglZ<-DUF499*</td><td>766</td><td>Bacteria>Pseudomonadota>Gammaproteobacteria>Pseudomonadales>Pseudomonadaceae</td><td>GCF_027857355.1</td><td>Pseudomonas_aeruginosa</td><td>OFL82_RS33705</td><td>13162..15463</td><td>287</td><td>DUF499-domain-containing-protein,-partial-[Pseudomonas-aeruginosa]</td></tr>
<tr><td>SDK95878.1</td><td>AAA+ATPase+wHTH+Alpha+Beta-domain+wHTH+RRM</td><td>Type-3_BR.PglZ.associated.systems</td><td>RlpA-><-SF1-SF2-Helicase+iREase+CTD<-METHYLASE<-VirB4-FtsK+HerA<-SIR2<-PglZ<-DUF499*<-tRNA<-MerR-HTH</td><td>801</td><td>Bacteria>Pseudomonadota>Gammaproteobacteria>Pseudomonadales>Pseudomonadaceae</td><td>GCA_900099945.1</td><td>Pseudomonas_delhiensis</td><td>SAMN05216189_10609</td><td>12193..14598</td><td>366289</td><td>Protein-of-unknown-function-[Pseudomonas-delhiensis]</td></tr>
<tr><td>WP_134314932.1</td><td>AAA+ATPase+wHTH+Alpha+Beta-domain+wHTH+RRM</td><td>Type-3_BR.PglZ.associated.systems</td><td>Phage_integrase->?-><-tRNA||MarR-HTH->Phage_int_SAM_2+Integrase_1->XIS-HTH->DUF499*->PglZ->SIR2->VirB4-FtsK+HerA->?->METHYLASE->iSTAND-></td><td>909</td><td>Bacteria>Pseudomonadota>Gammaproteobacteria>Pseudomonadales>Pseudomonadaceae</td><td>GCF_004371905.1</td><td>Pseudomonas_aeruginosa</td><td>IPC1307_RS29360</td><td>6758..9487</td><td>287</td><td>DUF499-domain-containing-protein-[Pseudomonas-aeruginosa]</td></tr>
<tr><td>HBO3789827.1</td><td>AAA+ATPase+wHTH+Alpha+Beta-domain+wHTH+RRM</td><td>Type-3_BR.PglZ.associated.systems</td><td>MerR-HTH->tRNA->DUF499*->PglZ->SIR2->VirB4-FtsK+HerA->METHYLASE->iSTAND->TUDOR+SF1-SF2-Helicase+iREase+CTD-></td><td>853</td><td>Bacteria>Pseudomonadota>Gammaproteobacteria>Pseudomonadales>Pseudomonadaceae</td><td>GCA_021656255.1</td><td>Pseudomonas_aeruginosa</td><td>L4T20_004196</td><td>19721..22282</td><td>287</td><td>DUF499-domain-containing-protein-[Pseudomonas-aeruginosa]</td></tr>
<tr><td>HCF3491945.1</td><td>AAA+ATPase+wHTH+Alpha+Beta-domain+wHTH+RRM</td><td>Type-3_BR.PglZ.associated.systems</td><td>MerR-HTH->tRNA->DUF499*->PglZ->SIR2->VirB4-FtsK+HerA->METHYLASE->iSTAND->TUDOR+SF1-SF2-Helicase+iREase+CTD-></td><td>853</td><td>Bacteria>Pseudomonadota>Gammaproteobacteria>Pseudomonadales>Pseudomonadaceae</td><td>GCA_023947355.1</td><td>Pseudomonas_aeruginosa</td><td>NIC36_002320</td><td>38111..40672</td><td>287</td><td>DUF499-domain-containing-protein-[Pseudomonas-aeruginosa]</td></tr>
<tr><td>HCF5163381.1</td><td>AAA+ATPase+wHTH+Alpha+Beta-domain+wHTH+RRM</td><td>Type-3_BR.PglZ.associated.systems</td><td>MerR-HTH->tRNA->DUF499*->PglZ->SIR2->VirB4-FtsK+HerA->METHYLASE->iSTAND->TUDOR+SF1-SF2-Helicase+iREase+CTD-></td><td>853</td><td>Bacteria>Pseudomonadota>Gammaproteobacteria>Pseudomonadales>Pseudomonadaceae</td><td>GCA_023952895.1</td><td>Pseudomonas_aeruginosa</td><td>NIG39_005113</td><td>38115..40676</td><td>287</td><td>DUF499-domain-containing-protein-[Pseudomonas-aeruginosa]</td></tr>
<tr><td>WP_160501267.1</td><td>AAA+ATPase+wHTH+Alpha+Beta-domain+wHTH+RRM</td><td>Type-3_BR.PglZ.associated.systems</td><td>MerR-HTH->tRNA->DUF499*->PglZ->SIR2->VirB4-FtsK+HerA->METHYLASE->iSTAND->TUDOR+SF1-SF2-Helicase+iREase+CTD-></td><td>915</td><td>Bacteria>Pseudomonadota>Gammaproteobacteria>Pseudomonadales>Pseudomonadaceae</td><td>GCF_018409365.1</td><td>Pseudomonas_aeruginosa</td><td>KJS76_RS11700</td><td>2455741..2458488</td><td>287</td><td>DUF499-domain-containing-protein-[Pseudomonas-aeruginosa]</td></tr>
<tr><td>WP_089394760.1</td><td>AAA+ATPase+wHTH+Alpha+Beta-domain+wHTH+RRM</td><td>Type-3_BR.PglZ.associated.systems</td><td>MerR-HTH->tRNA->DUF499*->PglZ->SIR2->VirB4-FtsK+HerA->METHYLASE->iSTAND->TUDOR+SF1-SF2-Helicase+iREase+CTD-></td><td>915</td><td>Bacteria>Pseudomonadota>Gammaproteobacteria>Pseudomonadales>Pseudomonadaceae</td><td>GCF_900099945.1</td><td>Pseudomonas</td><td>CHC02_RS28635</td><td>19015..21762</td><td>286</td><td>DUF499-domain-containing-protein-[Pseudomonas-delhiensis]</td></tr>
<tr><td>WP_196741466.1</td><td>AAA+ATPase+wHTH+Alpha+Beta-domain+wHTH+RRM</td><td>Type-3_BR.PglZ.associated.systems</td><td>MerR-HTH->tRNA->DUF499*->PglZ->SIR2->VirB4-FtsK+HerA->METHYLASE->iSTAND->TUDOR+SF1-SF2-Helicase+iREase+CTD-></td><td>853</td><td>Bacteria>Pseudomonadota>Gammaproteobacteria>Pseudomonadales>Pseudomonadaceae</td><td>-</td><td>Pseudomonas_aeruginosa</td><td>L4D09_004839</td><td>68693..71254</td><td>287</td><td>DUF499-domain-containing-protein-[Pseudomonas-aeruginosa]</td></tr>
<tr><td>WP_197541499.1</td><td>AAA+ATPase+wHTH+Alpha+Beta-domain+wHTH+RRM</td><td>Type-3_BR.PglZ.associated.systems</td><td>MerR-HTH->tRNA->DUF499*->PglZ->SIR2->VirB4-FtsK+HerA->METHYLASE->iSTAND->TUDOR+SF1-SF2-Helicase+iREase+CTD-></td><td>853</td><td>Bacteria>Pseudomonadota>Gammaproteobacteria>Pseudomonadales>Pseudomonadaceae</td><td>-</td><td>Pseudomonas_aeruginosa</td><td>L4T20_004196</td><td>19721..22282</td><td>287</td><td>DUF499-domain-containing-protein-[Pseudomonas-aeruginosa]</td></tr>
<tr><td>WP_043513194.1</td><td>AAA+ATPase+wHTH+Alpha+Beta-domain+wHTH+RRM</td><td>Type-3_BR.PglZ.associated.systems</td><td>MerR-HTH->tRNA->DUF499*->PglZ->SIR2->VirB4-FtsK+HerA->METHYLASE->iSTAND->SF1-SF2-Helicase+iREase+CTD-></td><td>915</td><td>Bacteria>Pseudomonadota>Gammaproteobacteria>Pseudomonadales>Pseudomonadaceae</td><td>GCF_013240415.1</td><td>Pseudomonas_aeruginosa</td><td>HB738_RS12165</td><td>2516772..2519519</td><td>287</td><td>DUF499-domain-containing-protein-[Pseudomonas-aeruginosa]</td></tr>
<tr><td>HBZ92698.1</td><td>AAA+ATPase+wHTH+Alpha+Beta-domain+wHTH+RRM</td><td>Type-3_BR.PglZ.associated.systems</td><td>MerR-HTH->tRNA->DUF499*->PglZ->METHYLASE-p-></td><td>915</td><td>Bacteria>Pseudomonadota>Gammaproteobacteria>Pseudomonadales>Pseudomonadaceae</td><td>GCA_003520905.1</td><td>Pseudomonas_sp.</td><td>DEO91_03070</td><td>12290..15037</td><td>306</td><td>DUF499-domain-containing-protein-[Pseudomonas-sp.]</td></tr>
<tr><td>RRU75139.1</td><td>AAA+ATPase+wHTH+Alpha+Beta-domain+wHTH+RRM</td><td>Type-3_BR.PglZ.associated.systems</td><td>MerR-HTH->tRNA->DUF499*->PglZ->METHYLASE->iSTAND->TUDOR+SF1-SF2-Helicase+iREase+CTD-><-RlpA</td><td>915</td><td>Bacteria>Pseudomonadota>Gammaproteobacteria>Pseudomonadales>Pseudomonadaceae</td><td>GCA_003935345.1</td><td>Stutzerimonas_xanthomarina</td><td>EGJ05_09970</td><td>669294..672041</td><td>271420</td><td>DUF499-domain-containing-protein-[Pseudomonas-xanthomarina]</td></tr>
<tr><td>RRW09670.1</td><td>AAA+ATPase+wHTH+Alpha+Beta-domain+wHTH+RRM</td><td>Type-3_BR.PglZ.associated.systems</td><td>MerR-HTH->tRNA->DUF499*->PglZ->METHYLASE->iSTAND->TUDOR+SF1-SF2-Helicase+iREase+CTD-><-RlpA</td><td>909</td><td>Bacteria>Pseudomonadota>Gammaproteobacteria>Pseudomonadales>Pseudomonadaceae</td><td>GCA_003936715.1</td><td>Stutzerimonas_stutzeri</td><td>EGJ32_03055</td><td>621216..623945</td><td>316</td><td>DUF499-domain-containing-protein-[Pseudomonas-stutzeri]</td></tr>
<tr><td>MBV6202018.1</td><td>AAA+ATPase+wHTH+Alpha+Beta-domain+wHTH+RRM</td><td>Type-3_BR.PglZ.associated.systems</td><td>MerR-HTH->tRNA->DUF499*->PglZ->METHYLASE->iSTAND->TUDOR+SF1-SF2-Helicase+iREase+CTD-><-RlpA</td><td>839</td><td>Bacteria>Pseudomonadota>Gammaproteobacteria>Pseudomonadales>Pseudomonadaceae</td><td>GCA_019167185.1</td><td>Pseudomonas_aeruginosa</td><td>KUU18_23235</td><td>79983..82502</td><td>287</td><td>DUF499-domain-containing-protein-[Pseudomonas-aeruginosa]</td></tr>
<tr><td>MCV0097152.1</td><td>AAA+ATPase+wHTH+Alpha+Beta-domain+wHTH+RRM</td><td>Type-3_BR.PglZ.associated.systems</td><td>MerR-HTH->tRNA->DUF499*->PglZ->METHYLASE->iSTAND->TUDOR+SF1-SF2-Helicase+iREase+CTD-><-RlpA</td><td>847</td><td>Bacteria>Pseudomonadota>Gammaproteobacteria>Pseudomonadales>Pseudomonadaceae</td><td>GCA_025696865.1</td><td>Pseudomonas_aeruginosa</td><td>KUC93_20560</td><td>63204..65747</td><td>287</td><td>ATP-binding-protein-[Pseudomonas-aeruginosa]</td></tr>
<tr><td>WP_125912119.1</td><td>AAA+ATPase+wHTH+Alpha+Beta-domain+wHTH+RRM</td><td>Type-3_BR.PglZ.associated.systems</td><td>MerR-HTH->tRNA->DUF499*->PglZ->METHYLASE->iSTAND->TUDOR+SF1-SF2-Helicase+iREase+CTD-><-RlpA</td><td>909</td><td>Bacteria>Pseudomonadota>Gammaproteobacteria>Pseudomonadales>Pseudomonadaceae</td><td>GCF_003936715.1</td><td>Stutzerimonas_stutzeri</td><td>EGJ32_RS03065</td><td>621216..623945</td><td>316</td><td>DUF499-domain-containing-protein-[Stutzerimonas-stutzeri]</td></tr>
<tr><td>WP_263475506.1</td><td>AAA+ATPase+wHTH+Alpha+Beta-domain+wHTH+RRM</td><td>Type-3_BR.PglZ.associated.systems</td><td>MerR-HTH->tRNA->DUF499*->PglZ->METHYLASE->iSTAND->TUDOR+SF1-SF2-Helicase+iREase+CTD-><-RlpA</td><td>909</td><td>Bacteria>Pseudomonadota>Gammaproteobacteria>Pseudomonadales>Pseudomonadaceae</td><td>GCF_025696865.1</td><td>Pseudomonas_aeruginosa</td><td>KUC93_RS20610</td><td>63018..65747</td><td>287</td><td>DUF499-domain-containing-protein-[Pseudomonas-aeruginosa]</td></tr>
<tr><td>WP_125861689.1</td><td>AAA+ATPase+wHTH+Alpha+Beta-domain+wHTH+RRM</td><td>Type-3_BR.PglZ.associated.systems</td><td>MerR-HTH->tRNA->DUF499*->PglZ->METHYLASE->iSTAND->TUDOR+SF1-SF2-Helicase+iREase+CTD-><-RlpA</td><td>915</td><td>Bacteria>Pseudomonadota>Gammaproteobacteria>Pseudomonadales>Pseudomonadaceae</td><td>-</td><td>Stutzerimonas_xanthomarina</td><td>EGJ05_RS09980</td><td>669294..672041</td><td>271420</td><td>DUF499-domain-containing-protein-[Stutzerimonas-xanthomarina]</td></tr>
<tr><td>WP_221624402.1</td><td>AAA+ATPase+wHTH+Alpha+Beta-domain+wHTH+RRM</td><td>Type-3_BR.PglZ.associated.systems</td><td>MerR-HTH->tRNA->DUF499*->PglZ->?->METHYLASE->iSTAND->TUDOR+SF1-SF2-Helicase+iREase+CTD-><-RlpA</td><td>857</td><td>Bacteria>Pseudomonadota>Gammaproteobacteria>Pseudomonadales>Pseudomonadaceae</td><td>GCF_003863715.1</td><td>Pseudomonas_aeruginosa</td><td>EIM19_RS05780</td><td>215436..218009</td><td>287</td><td>DUF499-domain-containing-protein-[Pseudomonas-aeruginosa]</td></tr>
<tr><td>WP_196750197.1</td><td>AAA+ATPase+wHTH+Alpha+Beta-domain+wHTH+RRM</td><td>Type-3_BR.PglZ.associated.systems</td><td>MerR-HTH->tRNA->DUF499*->PglZ->?->METHYLASE->iSTAND->TUDOR+SF1-SF2-Helicase+iREase+CTD-><-RlpA</td><td>853</td><td>Bacteria>Pseudomonadota>Gammaproteobacteria>Pseudomonadales>Pseudomonadaceae</td><td>-</td><td>Pseudomonas_aeruginosa</td><td>P3458_000154</td><td>13790..16351</td><td>287</td><td>DUF499-domain-containing-protein-[Pseudomonas-aeruginosa]</td></tr>
<tr><td>OLS59807.1</td><td>AAA+ATPase+wHTH+Alpha+Beta-domain+wHTH+RRM</td><td>Type-3_BR.PglZ.associated.systems</td><td>HTH_3-><-?<-?<-ParB+HTH<-DUF932||XIS-HTH->DUF499*->PglZ->SIR2->VirB4-FtsK+HerA->METHYLASE->iSTAND->TUDOR+SF1-SF2-Helicase+iREase+CTD-></td><td>937</td><td>Bacteria>Pseudomonadota>Gammaproteobacteria>Pseudomonadales>Pseudomonadaceae</td><td>GCA_001941965.1</td><td>Pseudomonas_putida</td><td>PSEMO_51150</td><td>57609..60422</td><td>303</td><td>hypothetical-protein-PSEMO_51150-[Pseudomonas-putida]</td></tr>
<tr><td>WP_075805772.1</td><td>AAA+ATPase+wHTH+Alpha+Beta-domain+wHTH+RRM</td><td>Type-3_BR.PglZ.associated.systems</td><td>HTH_3-><-?<-?<-ParB+HTH<-DUF932||XIS-HTH->DUF499*->PglZ->SIR2->VirB4-FtsK+HerA->METHYLASE->iSTAND->TUDOR+SF1-SF2-Helicase+iREase+CTD-></td><td>937</td><td>Bacteria>Pseudomonadota>Gammaproteobacteria>Pseudomonadales>Pseudomonadaceae</td><td>GCF_001941965.1</td><td>Pseudomonas_putida</td><td>PSEMO_RS25980</td><td>57609..60422</td><td>303</td><td>DUF499-domain-containing-protein-[Pseudomonas-putida]</td></tr>
<tr><td>MBX6756494.1</td><td>AAA+ATPase+wHTH+Alpha+Beta-domain+wHTH+RRM</td><td>Type-3_BR.PglZ.associated.systems</td><td>GDResolvase+PPP1-HTH->ParB->ParB->ZnR+TOPRIM+D5ATPase->XIS-HTH->DUF499*->PglZ->SIR2->VirB4-FtsK+HerA->METHYLASE->iSTAND->TUDOR+SF1-SF2-Helicase+iREase+CTD-></td><td>910</td><td>Bacteria>Pseudomonadota>Gammaproteobacteria>Pseudomonadales>Pseudomonadaceae</td><td>GCA_019690305.1</td><td>Pseudomonas_aeruginosa</td><td>ISE73_05640</td><td>168772..171504</td><td>287</td><td>DUF499-domain-containing-protein-[Pseudomonas-aeruginosa]</td></tr>
<tr><td>WP_149100782.1</td><td>AAA+ATPase+wHTH+Alpha+Beta-domain+wHTH+RRM</td><td>Type-3_BR.PglZ.associated.systems</td><td>GDResolvase+PPP1-HTH->ParB->ParB->ZnR+TOPRIM+D5ATPase->XIS-HTH->DUF499*->PglZ->SIR2->VirB4-FtsK+HerA->METHYLASE->iSTAND->TUDOR+SF1-SF2-Helicase+iREase+CTD-></td><td>910</td><td>Bacteria>Pseudomonadota>Gammaproteobacteria>Pseudomonadales>Pseudomonadaceae</td><td>GCF_031223175.1</td><td>Pseudomonas_aeruginosa</td><td>FZC29_RS00895</td><td>38188..40920</td><td>287</td><td>DUF499-domain-containing-protein-[Pseudomonas-aeruginosa]</td></tr>
<tr><td>EIU7198878.1</td><td>AAA+ATPase+wHTH+Alpha+Beta-domain+wHTH+RRM</td><td>Type-3_BR.PglZ.associated.systems</td><td>GDResolvase+PPP1-HTH->ParB->ParB->ZnR+TOPRIM+D5ATPase->XIS-HTH->DUF499*->PglZ->Classical-AAA->Peptidase_S8->METHYLASE->iSTAND->TUDOR+SF1-SF2-Helicase+iREase+CTD-></td><td>846</td><td>Bacteria>Pseudomonadota>Gammaproteobacteria>Pseudomonadales>Pseudomonadaceae</td><td>GCA_021712915.1</td><td>Pseudomonas_aeruginosa</td><td>A2U74_002204</td><td>8606..11146</td><td>287</td><td>DUF499-domain-containing-protein-[Pseudomonas-aeruginosa]</td></tr>
<tr><td>WP_231387415.1</td><td>AAA+ATPase+wHTH+Alpha+Beta-domain+wHTH+RRM</td><td>Type-3_BR.PglZ.associated.systems</td><td>GDResolvase+PPP1-HTH->ParB->ParB->ZnR+TOPRIM+D5ATPase->XIS-HTH->DUF499*->PglZ->Classical-AAA->Peptidase_S8->METHYLASE->iSTAND->TUDOR+SF1-SF2-Helicase+iREase+CTD-></td><td>908</td><td>Bacteria>Pseudomonadota>Gammaproteobacteria>Pseudomonadales>Pseudomonadaceae</td><td>GCF_015763095.1</td><td>Pseudomonas_aeruginosa_group</td><td>QMO31_RS22275</td><td>146171..148897</td><td>136841</td><td>DUF499-domain-containing-protein-[Pseudomonas-nitroreducens]</td></tr>
<tr><td>WP_196912143.1</td><td>AAA+ATPase+wHTH+Alpha+Beta-domain+wHTH+RRM</td><td>Type-3_BR.PglZ.associated.systems</td><td>GDResolvase+PPP1-HTH->ParB->ParB->ZnR+TOPRIM+D5ATPase->XIS-HTH->DUF499*->PglZ->Classical-AAA->Peptidase_S8->METHYLASE->iSTAND->TUDOR+SF1-SF2-Helicase+iREase+CTD-></td><td>846</td><td>Bacteria>Pseudomonadota>Gammaproteobacteria>Pseudomonadales>Pseudomonadaceae</td><td>-</td><td>Pseudomonas_aeruginosa</td><td>A2U74_002204</td><td>8606..11146</td><td>287</td><td>DUF499-domain-containing-protein-[Pseudomonas-aeruginosa]</td></tr>
<tr><td>WP_177007802.1</td><td>AAA+ATPase+wHTH+Alpha+Beta-domain+wHTH+RRM</td><td>Type-3_BR.PglZ.associated.systems</td><td>DUF499*->PglZ->METHYLASE-p-></td><td>945</td><td>Bacteria>Pseudomonadota>Gammaproteobacteria>Pseudomonadales>Pseudomonadaceae</td><td>-</td><td>Pseudomonas_tolaasii</td><td>HYE76_RS30500</td><td>456..3293</td><td>29442</td><td>DUF499-domain-containing-protein-[Pseudomonas-tolaasii]</td></tr>
<tr><td>WP_026331961.1</td><td>AAA+ATPase+wHTH+Alpha+Beta-domain+wHTH+RRM</td><td>Type-3_BR.PglZ.associated.systems</td><td>DUF499*->PglZ->METHYLASE->iSTAND->TUDOR+SF1-SF2-Helicase+iREase+CTD-><-RlpA</td><td>918</td><td>Bacteria>Pseudomonadota>Gammaproteobacteria>Pseudomonadales>Pseudomonadaceae</td><td>GCF_000381285.1</td><td>Pseudomonas_mandelii</td><td>F626_RS0128230</td><td>63325..66081</td><td>75612</td><td>nucleoporin-[Pseudomonas-mandelii]</td></tr>
<tr><td>WP_263750733.1</td><td>AAA+ATPase+wHTH+Alpha+Beta-domain+wHTH+RRM</td><td>Type-3_BR.PglZ.associated.systems</td><td>DUF499*->PglZ->?->METHYLASE->iSTAND->TUDOR+SF1-SF2-Helicase+iREase+CTD-><-RlpA</td><td>432</td><td>Bacteria>Pseudomonadota>Gammaproteobacteria>Pseudomonadales>Pseudomonadaceae</td><td>GCF_025780585.1</td><td>Pseudomonas_aeruginosa</td><td>OHU58_RS33115</td><td>116..1414</td><td>287</td><td>hypothetical-protein-[Pseudomonas-aeruginosa]</td></tr>
<tr><td>HCF3076462.1</td><td>AAA+ATPase+wHTH+Alpha+Beta-domain+wHTH+RRM</td><td>Type-3_BR.PglZ.associated.systems</td><td>DDE_Tnp_IS240-><-PglZ<-DUF499*<-XIS-HTH</td><td>852</td><td>Bacteria>Pseudomonadota>Gammaproteobacteria>Pseudomonadales>Pseudomonadaceae</td><td>GCA_023946075.1</td><td>Pseudomonas_aeruginosa</td><td>NIB82_006110</td><td>7124..9682</td><td>287</td><td>DUF499-domain-containing-protein-[Pseudomonas-aeruginosa]</td></tr>
<tr><td>WP_023127550.1</td><td>AAA+ATPase+wHTH+Alpha+Beta-domain+wHTH+RRM</td><td>Type-3_BR.PglZ.associated.systems</td><td><-VirB11+VirB4-FtsK<-CopG/RHH<-VirD4-TraG+VirD4-FtsK<-?<-MODE-HTH+PBPII||XIS-HTH->DUF499*->PglZ->Classical-AAA->Peptidase_S8->METHYLASE->iSTAND->TUDOR+SF1-SF2-Helicase+iREase+CTD-></td><td>922</td><td>Bacteria>Pseudomonadota>Gammaproteobacteria>Pseudomonadales>Pseudomonadaceae</td><td>GCF_000412395.1</td><td>Pseudomonas_aeruginosa</td><td>Q008_RS19635</td><td>2671620..2674388</td><td>287</td><td>DUF499-domain-containing-protein-[Pseudomonas-aeruginosa]</td></tr>
<tr><td>WP_137138280.1</td><td>AAA+ATPase+wHTH+Alpha+Beta-domain+wHTH+RRM</td><td>Type-3_BR.PglZ.associated.systems</td><td><-TUDOR+SF1-SF2-Helicase+iREase+CTD<-iSTAND<-METHYLASE<-VirB4-FtsK+HerA<-SIR2<-PglZ<-DUF499*<-XIS-HTH||MODE-HTH+PBPII->?->VirD4-TraG+VirD4-FtsK->CopG/RHH->VirB11+VirB4-FtsK-></td><td>922</td><td>Bacteria>Pseudomonadota>Gammaproteobacteria>Pseudomonadales>Pseudomonadaceae</td><td>GCF_005223285.1</td><td>Pseudomonas_asiatica</td><td>FD498_RS06925</td><td>626212..628980</td><td>2219225</td><td>DUF499-domain-containing-protein-[Pseudomonas-asiatica]</td></tr>
<tr><td>WP_260563117.1</td><td>AAA+ATPase+wHTH+Alpha+Beta-domain+wHTH+RRM</td><td>Type-3_BR.PglZ.associated.systems</td><td><-TUDOR+SF1-SF2-Helicase+iREase+CTD<-iSTAND<-METHYLASE<-VirB4-FtsK+HerA<-SIR2<-PglZ<-DUF499*<-XIS-HTH||DUF932->ParB+HTH->?->?-><-HTH_3</td><td>915</td><td>Bacteria>Pseudomonadota>Gammaproteobacteria>Pseudomonadales>Pseudomonadaceae</td><td>GCF_030326365.1</td><td>Pseudomonas_aeruginosa</td><td>HX138_RS10900</td><td>2345647..2348394</td><td>287</td><td>ATP-binding-protein-[Pseudomonas-aeruginosa]</td></tr>
<tr><td>WP_104098237.1</td><td>AAA+ATPase+wHTH+Alpha+Beta-domain+wHTH+RRM</td><td>Type-3_BR.PglZ.associated.systems</td><td><-TUDOR+SF1-SF2-Helicase+iREase+CTD<-iSTAND<-METHYLASE<-VirB4-FtsK+HerA<-SIR2<-PglZ<-DUF499*<-XIS-HTH||DUF932-><-?<-?<-NUDIX</td><td>915</td><td>Bacteria>Pseudomonadota>Gammaproteobacteria>Pseudomonadales>Pseudomonadaceae</td><td>GCF_002929225.1</td><td>Stutzerimonas_kunmingensis</td><td>CW305_RS14615</td><td>73939..76686</td><td>1211807</td><td>DUF499-domain-containing-protein-[Stutzerimonas-kunmingensis]</td></tr>
<tr><td>HBO1597103.1</td><td>AAA+ATPase+wHTH+Alpha+Beta-domain+wHTH+RRM</td><td>Type-3_BR.PglZ.associated.systems</td><td><-TUDOR+SF1-SF2-Helicase+iREase+CTD<-iSTAND<-METHYLASE<-VirB4-FtsK+HerA<-SIR2<-PglZ<-DUF499*<-tRNA<-MerR-HTH</td><td>853</td><td>Bacteria>Pseudomonadota>Gammaproteobacteria>Pseudomonadales>Pseudomonadaceae</td><td>GCA_021598065.1</td><td>Pseudomonas_aeruginosa</td><td>L4F50_002042</td><td>32106..34667</td><td>287</td><td>DUF499-domain-containing-protein-[Pseudomonas-aeruginosa]</td></tr>
<tr><td>HCG0908554.1</td><td>AAA+ATPase+wHTH+Alpha+Beta-domain+wHTH+RRM</td><td>Type-3_BR.PglZ.associated.systems</td><td><-TUDOR+SF1-SF2-Helicase+iREase+CTD<-iSTAND<-METHYLASE<-VirB4-FtsK+HerA<-SIR2<-PglZ<-DUF499*<-tRNA<-MerR-HTH</td><td>853</td><td>Bacteria>Pseudomonadota>Gammaproteobacteria>Pseudomonadales>Pseudomonadaceae</td><td>GCA_024004415.1</td><td>Pseudomonas_aeruginosa</td><td>NJF87_000308</td><td>349520..352081</td><td>287</td><td>DUF499-domain-containing-protein-[Pseudomonas-aeruginosa]</td></tr>
<tr><td>WP_112302890.1</td><td>AAA+ATPase+wHTH+Alpha+Beta-domain+wHTH+RRM</td><td>Type-3_BR.PglZ.associated.systems</td><td><-TUDOR+SF1-SF2-Helicase+iREase+CTD<-iSTAND<-METHYLASE<-VirB4-FtsK+HerA<-SIR2<-PglZ<-DUF499*<-tRNA<-MerR-HTH</td><td>915</td><td>Bacteria>Pseudomonadota>Gammaproteobacteria>Pseudomonadales>Pseudomonadaceae</td><td>GCF_003429185.1</td><td>Pseudomonas_aeruginosa</td><td>P4N66_RS23785</td><td>5011940..5014687</td><td>287</td><td>DUF499-domain-containing-protein-[Pseudomonas-aeruginosa]</td></tr>
<tr><td>WP_044719472.1</td><td>AAA+ATPase+wHTH+Alpha+Beta-domain+wHTH+RRM</td><td>Type-3_BR.PglZ.associated.systems</td><td><-TUDOR+SF1-SF2-Helicase+iREase+CTD<-iSTAND<-METHYLASE<-VirB4-FtsK+HerA<-SIR2<-PglZ<-DUF499*<-tRNA<-MerR-HTH</td><td>915</td><td>Bacteria>Pseudomonadota>Gammaproteobacteria>Pseudomonadales>Pseudomonadaceae</td><td>GCF_003813025.1</td><td>Pseudomonas_aeruginosa</td><td>TO65_RS22045</td><td>652777..655524</td><td>287</td><td>DUF499-domain-containing-protein-[Pseudomonas-aeruginosa]</td></tr>
<tr><td>WP_124123873.1</td><td>AAA+ATPase+wHTH+Alpha+Beta-domain+wHTH+RRM</td><td>Type-3_BR.PglZ.associated.systems</td><td><-TUDOR+SF1-SF2-Helicase+iREase+CTD<-iSTAND<-METHYLASE<-VirB4-FtsK+HerA<-SIR2<-PglZ<-DUF499*<-tRNA<-MerR-HTH</td><td>915</td><td>Bacteria>Pseudomonadota>Gammaproteobacteria>Pseudomonadales>Pseudomonadaceae</td><td>GCF_003836345.1</td><td>Pseudomonas_aeruginosa</td><td>IPC880_RS16790</td><td>128024..130771</td><td>287</td><td>DUF499-domain-containing-protein-[Pseudomonas-aeruginosa]</td></tr>
<tr><td>WP_124172762.1</td><td>AAA+ATPase+wHTH+Alpha+Beta-domain+wHTH+RRM</td><td>Type-3_BR.PglZ.associated.systems</td><td><-TUDOR+SF1-SF2-Helicase+iREase+CTD<-iSTAND<-METHYLASE<-VirB4-FtsK+HerA<-SIR2<-PglZ<-DUF499*<-tRNA<-MerR-HTH</td><td>915</td><td>Bacteria>Pseudomonadota>Gammaproteobacteria>Pseudomonadales>Pseudomonadaceae</td><td>GCF_901010245.1</td><td>Pseudomonas_aeruginosa</td><td>IPC334_RS01410</td><td>322225..324972</td><td>287</td><td>DUF499-domain-containing-protein-[Pseudomonas-aeruginosa]</td></tr>
<tr><td>WP_201268311.1</td><td>AAA+ATPase+wHTH+Alpha+Beta-domain+wHTH+RRM</td><td>Type-3_BR.PglZ.associated.systems</td><td><-TUDOR+SF1-SF2-Helicase+iREase+CTD<-iSTAND<-METHYLASE<-VirB4-FtsK+HerA<-SIR2<-PglZ<-DUF499*<-tRNA<-MerR-HTH</td><td>853</td><td>Bacteria>Pseudomonadota>Gammaproteobacteria>Pseudomonadales>Pseudomonadaceae</td><td>-</td><td>Pseudomonas_aeruginosa</td><td>L4H28_002030</td><td>380553..383114</td><td>287</td><td>DUF499-domain-containing-protein-[Pseudomonas-aeruginosa]</td></tr>
<tr><td>WP_031631161.1</td><td>AAA+ATPase+wHTH+Alpha+Beta-domain+wHTH+RRM</td><td>Type-3_BR.PglZ.associated.systems</td><td><-TUDOR+SF1-SF2-Helicase+iREase+CTD<-iSTAND<-METHYLASE<-VirB4-FtsK+HerA<-SIR2<-PglZ<-DUF499*<-MerR-HTH||?->Peptidase_A8->zf-ISL3+HTH_Tnp_ISL3+DDE_Tnp_ISL3->||DUF932->ParB+HTH-></td><td>937</td><td>Bacteria>Pseudomonadota>Gammaproteobacteria>Pseudomonadales>Pseudomonadaceae</td><td>GCF_000629205.1</td><td>Pseudomonas_aeruginosa</td><td>AJ65_RS23625</td><td>132799..135612</td><td>287</td><td>DUF499-domain-containing-protein-[Pseudomonas-aeruginosa]</td></tr>
<tr><td>WP_031688377.1</td><td>AAA+ATPase+wHTH+Alpha+Beta-domain+wHTH+RRM</td><td>Type-3_BR.PglZ.associated.systems</td><td><-TUDOR+SF1-SF2-Helicase+iREase+CTD<-iSTAND<-METHYLASE<-Peptidase_S8<-Classical-AAA<-PglZ<-DUF499*<-XIS-HTH<-ZnR+TOPRIM+D5ATPase<-ParB<-ParB<-GDResolvase+PPP1-HTH</td><td>908</td><td>Bacteria>Pseudomonadota>Gammaproteobacteria>Pseudomonadales>Pseudomonadaceae</td><td>GCF_028435765.1</td><td>Pseudomonas_aeruginosa</td><td>V554_RS36395</td><td>46715..49441</td><td>287</td><td>DUF499-domain-containing-protein-[Pseudomonas-aeruginosa]</td></tr>
<tr><td>WP_073656072.1</td><td>AAA+ATPase+wHTH+Alpha+Beta-domain+wHTH+RRM</td><td>Type-3_BR.PglZ.associated.systems</td><td><-TUDOR+SF1-SF2-Helicase+iREase+CTD<-iSTAND<-METHYLASE<-Peptidase_S8<-Classical-AAA<-PglZ<-DUF499*<-XIS-HTH<-ParB<-ParB<-GDResolvase+PPP1-HTH<-?<-AbiEii</td><td>912</td><td>Bacteria>Pseudomonadota>Gammaproteobacteria>Pseudomonadales>Pseudomonadaceae</td><td>GCF_900145205.1</td><td>Pseudomonas_aeruginosa</td><td>BUG94_RS05925</td><td>74159..76897</td><td>287</td><td>DUF499-domain-containing-protein-[Pseudomonas-aeruginosa]</td></tr>
<tr><td>WP_034036936.1</td><td>AAA+ATPase+wHTH+Alpha+Beta-domain+wHTH+RRM</td><td>Type-3_BR.PglZ.associated.systems</td><td><-TUDOR+SF1-SF2-Helicase+iREase+CTD<-iSTAND<-METHYLASE<-Peptidase_S8<-Classical-AAA<-PglZ<-DUF499*<-XIS-HTH</td><td>908</td><td>Bacteria>Pseudomonadota>Gammaproteobacteria>Pseudomonadales>Pseudomonadaceae</td><td>GCF_000797245.1</td><td>Pseudomonas_aeruginosa</td><td>NQ27_RS04235</td><td>21430..24156</td><td>287</td><td>DUF499-domain-containing-protein-[Pseudomonas-aeruginosa]</td></tr>
<tr><td>WP_263468102.1</td><td>AAA+ATPase+wHTH+Alpha+Beta-domain+wHTH+RRM</td><td>Type-3_BR.PglZ.associated.systems</td><td><-TUDOR+SF1-SF2-Helicase+iREase+CTD<-iSTAND<-METHYLASE<-Peptidase_S8<-Classical-AAA<-PglZ<-DUF499*<-XIS-HTH</td><td>940</td><td>Bacteria>Pseudomonadota>Gammaproteobacteria>Pseudomonadales>Pseudomonadaceae</td><td>GCF_025697885.1</td><td>Pseudomonas_aeruginosa</td><td>KUC79_RS14375</td><td>113045..115867</td><td>287</td><td>ATP-binding-protein-[Pseudomonas-aeruginosa]</td></tr>
<tr><td>WP_282317571.1</td><td>AAA+ATPase+wHTH+Alpha+Beta-domain+wHTH+RRM</td><td>Type-3_BR.PglZ.associated.systems</td><td><-TUDOR+SF1-SF2-Helicase+iREase+CTD<-iSTAND<-METHYLASE<-Peptidase_S8<-Classical-AAA<-PglZ<-DUF499*<-XIS-HTH</td><td>908</td><td>Bacteria>Pseudomonadota>Gammaproteobacteria>Pseudomonadales>Pseudomonadaceae</td><td>GCF_029954515.1</td><td>Pseudomonas_aeruginosa</td><td>OLM98_RS07395</td><td>58298..61024</td><td>287</td><td>DUF499-domain-containing-protein-[Pseudomonas-aeruginosa]</td></tr>
<tr><td>OLU32852.1</td><td>AAA+ATPase+wHTH+Alpha+Beta-domain+wHTH+RRM</td><td>Type-3_BR.PglZ.associated.systems</td><td><-TUDOR+SF1-SF2-Helicase+iREase+CTD<-iSTAND<-METHYLASE<-Peptidase_S8<-Classical-AAA<-PglZ<-DUF499*<-tRNA<-MerR-HTH</td><td>944</td><td>Bacteria>Pseudomonadota>Gammaproteobacteria>Pseudomonadales>Pseudomonadaceae</td><td>GCA_001945445.1</td><td>Pseudomonas_sp._PA27(2017)</td><td>BVH06_10305</td><td>86154..88988</td><td>1932112</td><td>DUF499-domain-containing-protein-[Pseudomonas-sp.-PA27(2017)]</td></tr>
<tr><td>WP_075934016.1</td><td>AAA+ATPase+wHTH+Alpha+Beta-domain+wHTH+RRM</td><td>Type-3_BR.PglZ.associated.systems</td><td><-TUDOR+SF1-SF2-Helicase+iREase+CTD<-iSTAND<-METHYLASE<-Peptidase_S8<-Classical-AAA<-PglZ<-DUF499*<-tRNA<-MerR-HTH</td><td>944</td><td>Bacteria>Pseudomonadota>Gammaproteobacteria>Pseudomonadales>Pseudomonadaceae</td><td>GCF_001945445.1</td><td>Pseudomonas_sp._PA27(2017)</td><td>BVH06_RS10305</td><td>86154..88988</td><td>1932112</td><td>DUF499-domain-containing-protein-[Pseudomonas-sp.-PA27(2017)]</td></tr>
<tr><td>WP_180215324.1</td><td>AAA+ATPase+wHTH+Alpha+Beta-domain+wHTH+RRM</td><td>Type-3_BR.PglZ.associated.systems</td><td><-SF1-SF2-Helicase+iREase+CTD<-iSTAND<-METHYLASE<-VirB4-FtsK+HerA<-SIR2<-PglZ<-DUF499*<-XIS-HTH</td><td>908</td><td>Bacteria>Pseudomonadota>Gammaproteobacteria>Pseudomonadales>Pseudomonadaceae</td><td>GCF_013421235.1</td><td>Pseudomonas_aeruginosa</td><td>HOB47_RS21280</td><td>43976..46702</td><td>287</td><td>DUF499-domain-containing-protein-[Pseudomonas-aeruginosa]</td></tr>
<tr><td>WP_238048237.1</td><td>AAA+ATPase+wHTH+Alpha+Beta-domain+wHTH+RRM</td><td>Type-3_BR.PglZ.associated.systems</td><td><-SF1-SF2-Helicase+iREase+CTD<-iSTAND<-METHYLASE<-VirB4-FtsK+HerA<-SIR2<-PglZ<-DUF499*<-tRNA<-MerR-HTH</td><td>915</td><td>Bacteria>Pseudomonadota>Gammaproteobacteria>Pseudomonadales>Pseudomonadaceae</td><td>GCF_022117575.1</td><td>Pseudomonas_aeruginosa</td><td>LT341_RS29475</td><td>107880..110627</td><td>287</td><td>DUF499-domain-containing-protein-[Pseudomonas-aeruginosa]</td></tr>
<tr><td>WP_042608840.1</td><td>AAA+ATPase+wHTH+Alpha+Beta-domain+wHTH+RRM</td><td>Type-3_BR.PglZ.associated.systems</td><td><-RlpA<-?<-TUDOR+SF1-SF2-Helicase+iREase+CTD<-iSTAND<-METHYLASE<-PglZ<-DUF499*<-XIS-HTH||?->TOPRIM->?->Integrase_1->Phage_int_SAM-></td><td>914</td><td>Bacteria>Pseudomonadota>Gammaproteobacteria>Pseudomonadales>Pseudomonadaceae</td><td>GCF_000834545.1</td><td>Pseudomonas_fluorescens</td><td>RU10_RS14550</td><td>55849..58593</td><td>294</td><td>DUF499-domain-containing-protein-[Pseudomonas-fluorescens]</td></tr>
<tr><td>WP_034115122.1</td><td>AAA+ATPase+wHTH+Alpha+Beta-domain+wHTH+RRM</td><td>Type-3_BR.PglZ.associated.systems</td><td><-RlpA<-?<-TUDOR+SF1-SF2-Helicase+iREase+CTD<-iSTAND<-METHYLASE<-PglZ<-DUF499*<-XIS-HTH||?->TOPRIM->?->Integrase_1->Phage_int_SAM-></td><td>914</td><td>Bacteria>Pseudomonadota>Gammaproteobacteria>Pseudomonadales>Pseudomonadaceae</td><td>GCF_001320785.1</td><td>Pseudomonas</td><td>KEM07_RS14555</td><td>49076..51820</td><td>286</td><td>MULTISPECIES:-DUF499-domain-containing-protein-[Pseudomonas]</td></tr>
<tr><td>WP_150804658.1</td><td>AAA+ATPase+wHTH+Alpha+Beta-domain+wHTH+RRM</td><td>Type-3_BR.PglZ.associated.systems</td><td><-RlpA<-?<-TUDOR+SF1-SF2-Helicase+iREase+CTD<-iSTAND<-METHYLASE<-PglZ<-DUF499*<-XIS-HTH||?->TOPRIM->?->Integrase_1->Phage_int_SAM-></td><td>914</td><td>Bacteria>Pseudomonadota>Gammaproteobacteria>Pseudomonadales>Pseudomonadaceae</td><td>GCF_902497875.1</td><td>Pseudomonas_fluorescens</td><td>F8N85_RS16700</td><td>82572..85316</td><td>294</td><td>DUF499-domain-containing-protein-[Pseudomonas-fluorescens]</td></tr>
<tr><td>MBJ2281510.1</td><td>AAA+ATPase+wHTH+Alpha+Beta-domain+wHTH+RRM</td><td>Type-3_BR.PglZ.associated.systems</td><td><-RlpA<-?<-TUDOR+SF1-SF2-Helicase+iREase+CTD<-iSTAND<-METHYLASE<-PglZ<-DUF499*<-XIS-HTH||?->TOPRIM->?->Integrase_1-><-Arm-DNA-bind_3+Phage_integrase</td><td>838</td><td>Bacteria>Pseudomonadota>Gammaproteobacteria>Pseudomonadales>Pseudomonadaceae</td><td>GCA_016405475.1</td><td>Pseudomonas_sp._MF6767</td><td>JFT58_24810</td><td>9470..11986</td><td>2797531</td><td>DUF499-domain-containing-protein-[Pseudomonas-sp.-MF6767]</td></tr>
<tr><td>WP_198834698.1</td><td>AAA+ATPase+wHTH+Alpha+Beta-domain+wHTH+RRM</td><td>Type-3_BR.PglZ.associated.systems</td><td><-RlpA<-?<-TUDOR+SF1-SF2-Helicase+iREase+CTD<-iSTAND<-METHYLASE<-PglZ<-DUF499*<-XIS-HTH||?->TOPRIM->?->Integrase_1-><-Arm-DNA-bind_3+Phage_integrase</td><td>838</td><td>Bacteria>Pseudomonadota>Gammaproteobacteria>Pseudomonadales>Pseudomonadaceae</td><td>-</td><td>Pseudomonas_fluorescens</td><td>JFT58_24810</td><td>9470..11986</td><td>294</td><td>MULTISPECIES:-DUF499-domain-containing-protein-[Pseudomonas]</td></tr>
<tr><td>MCV0045916.1</td><td>AAA+ATPase+wHTH+Alpha+Beta-domain+wHTH+RRM</td><td>Type-3_BR.PglZ.associated.systems</td><td><-PglZ<-DUF499*<-XIS-HTH</td><td>852</td><td>Bacteria>Pseudomonadota>Gammaproteobacteria>Pseudomonadales>Pseudomonadaceae</td><td>GCA_025696705.1</td><td>Pseudomonas_aeruginosa</td><td>KUC92_07095</td><td>1132..3690</td><td>287</td><td>ATP-binding-protein-[Pseudomonas-aeruginosa]</td></tr>
<tr><td>WP_126152459.1</td><td>AAA+ATPase+wHTH+Alpha+Beta-domain+wHTH+RRM</td><td>Type-3_BR.PglZ.associated.systems</td><td><-ParB+HNH<-TUDOR+SF1-SF2-Helicase+iREase+CTD<-iSTAND<-METHYLASE<-PglZ<-DUF499*<-XIS-HTH<-ZnR+TOPRIM+D5ATPase<-ParB<-ParB<-GDResolvase+PPP1-HTH</td><td>909</td><td>Bacteria>Pseudomonadota>Gammaproteobacteria>Pseudomonadales>Pseudomonadaceae</td><td>GCF_003955475.1</td><td>Pseudomonas_aeruginosa</td><td>EKL37_RS17290</td><td>67321..70050</td><td>287</td><td>DUF499-domain-containing-protein-[Pseudomonas-aeruginosa]</td></tr>
<tr><td>CRM01071.1</td><td>AAA+ATPase+wHTH+Alpha+Beta-domain+wHTH+RRM</td><td>Type-3_BR.PglZ.associated.systems</td><td><-MODE-HTH+PBPII<-Radical-SAM<-?||?->DUF499*->PglZ->METHYLASE->iSTAND->TUDOR+SF1-SF2-Helicase+iREase+CTD-><-RlpA||TOPRIM-></td><td>945</td><td>Bacteria>Pseudomonadota>Gammaproteobacteria>Pseudomonadales>Pseudomonadaceae</td><td>GCA_900004785.1</td><td>Pseudomonas_sp._28_E_9</td><td>.</td><td>21242..24079</td><td>1844098</td><td>hypothetical-protein-[Pseudomonas-sp.-28-E-9]</td></tr>
<tr><td>WP_065932667.1</td><td>AAA+ATPase+wHTH+Alpha+Beta-domain+wHTH+RRM</td><td>Type-3_BR.PglZ.associated.systems</td><td><-MODE-HTH+PBPII<-Radical-SAM<-?||?->?->DUF499*->PglZ->METHYLASE->iSTAND->TUDOR+SF1-SF2-Helicase+iREase+CTD-><-RlpA</td><td>945</td><td>Bacteria>Pseudomonadota>Gammaproteobacteria>Pseudomonadales>Pseudomonadaceae</td><td>GCF_900004785.1</td><td>Pseudomonas_sp._28_E_9</td><td>BFQ00_RS03895</td><td>21242..24079</td><td>1844098</td><td>DUF499-domain-containing-protein-[Pseudomonas-sp.-28-E-9]</td></tr>
<tr><td>MBV5872218.1</td><td>AAA+ATPase+wHTH+Alpha+Beta-domain+wHTH+RRM</td><td>Type-3_BR.PglZ.associated.systems</td><td><-iSTAND<-METHYLASE<-?<-VirB4-FtsK+HerA<-SIR2<-PglZ<-DUF499*<-XIS-HTH<-Phage_int_SAM_2+Integrase_1<-MarR-HTH||tRNA-><-?<-Phage_integrase</td><td>847</td><td>Bacteria>Pseudomonadota>Gammaproteobacteria>Pseudomonadales>Pseudomonadaceae</td><td>GCA_019167595.1</td><td>Pseudomonas_aeruginosa</td><td>KUU06_18245</td><td>273871..276414</td><td>287</td><td>DUF499-domain-containing-protein-[Pseudomonas-aeruginosa]</td></tr>
<tr><td>HCP56594.1</td><td>AAA+ATPase+wHTH+Alpha+Beta-domain+wHTH+RRM</td><td>Type-3_BR.PglZ.associated.systems</td><td><-HTH+WYL||?->XIS-HTH->DUF499*->PglZ->iSTAND->TUDOR+SF1-SF2-Helicase+iREase+CTD->?->?-><-RlpA</td><td>891</td><td>Bacteria>Pseudomonadota>Gammaproteobacteria>Pseudomonadales>Pseudomonadaceae</td><td>GCA_003525845.1</td><td>Pseudomonas_sp.</td><td>DIU04_17900</td><td>28274..30949</td><td>306</td><td>hypothetical-protein-[Pseudomonas-sp.]</td></tr>
<tr><td>MBB3243902.1</td><td>AAA+ATPase+wHTH+Alpha+Beta-domain+wHTH+RRM</td><td>Type-3_BR.PglZ.associated.systems</td><td><-CopG/RHH||XIS-HTH->?->DUF499*->PglZ->METHYLASE->iSTAND->TUDOR+SF1-SF2-Helicase+iREase+CTD-><-RlpA</td><td>918</td><td>Bacteria>Pseudomonadota>Gammaproteobacteria>Pseudomonadales>Pseudomonadaceae</td><td>GCA_014192635.1</td><td>Pseudomonas_sp._Tn43</td><td>FHW68_005487</td><td>31744..34500</td><td>701213</td><td>hypothetical-protein-[Pseudomonas-sp.-Tn43]</td></tr>
<tr><td>WP_183683907.1</td><td>AAA+ATPase+wHTH+Alpha+Beta-domain+wHTH+RRM</td><td>Type-3_BR.PglZ.associated.systems</td><td><-CopG/RHH||XIS-HTH->?->DUF499*->PglZ->METHYLASE->iSTAND->TUDOR+SF1-SF2-Helicase+iREase+CTD-><-RlpA</td><td>918</td><td>Bacteria>Pseudomonadota>Gammaproteobacteria>Pseudomonadales>Pseudomonadaceae</td><td>GCF_014192635.1</td><td>Pseudomonas_sp._Tn43</td><td>FHW68_RS26745</td><td>31744..34500</td><td>701213</td><td>DUF499-domain-containing-protein-[Pseudomonas-sp.-Tn43]</td></tr>
<tr><td>EIU3852619.1</td><td>AAA+ATPase+wHTH+Alpha+Beta-domain+wHTH+RRM</td><td>Type-3_BR.PglZ.associated.systems</td><td>DUF3987->XIS-HTH->DUF499+RRM*->PglZ->SIR2->VirB4-FtsK+HerA->METHYLASE->iSTAND-></td><td>848</td><td>Bacteria>Pseudomonadota>Gammaproteobacteria>Pseudomonadales>Pseudomonadaceae>Pseudomonas</td><td>GCA_021645765.1</td><td>Pseudomonas_aeruginosa</td><td>L4Q41_001887</td><td>2150..4696</td><td>287</td><td>DUF499-domain-containing-protein-[Pseudomonas-aeruginosa]</td></tr>
<tr><td>EKO9556879.1</td><td>AAA+ATPase+wHTH+Alpha+Beta-domain+wHTH+RRM</td><td>Type-3_BR.PglZ.associated.systems</td><td><-TUDOR+SF1-SF2-Helicase+iREase+CTD<-iSTAND<-METHYLASE<-Classical-AAA<-PglZ<-DUF499+RRM*<-XIS-HTH||DUF3487-></td><td>878</td><td>Bacteria>Pseudomonadota>Gammaproteobacteria>Pseudomonadales>Pseudomonadaceae>Pseudomonas</td><td>GCA_029127245.1</td><td>Pseudomonas_aeruginosa</td><td>P1X34_002688</td><td>41742..44378</td><td>287</td><td>DUF499-domain-containing-protein-[Pseudomonas-aeruginosa]</td></tr>
<tr><td>EKP5707811.1</td><td>AAA+ATPase+wHTH+Alpha+Beta-domain+wHTH+RRM</td><td>Type-3_BR.PglZ.associated.systems</td><td>MerR-HTH->DUF499+RRM*->PglZ->METHYLASE->iSTAND->TUDOR+SF1-SF2-Helicase+iREase+CTD-></td><td>853</td><td>Bacteria>Pseudomonadota>Gammaproteobacteria>Pseudomonadales>Pseudomonadaceae>Pseudomonas</td><td>GCA_029235905.1</td><td>Pseudomonas_aeruginosa</td><td>P3458_000154</td><td>13790..16351</td><td>287</td><td>DUF499-domain-containing-protein-[Pseudomonas-aeruginosa]</td></tr>
<tr><td>MDG4306930.1</td><td>AAA+ATPase+wHTH+Alpha+Beta-domain+wHTH+RRM</td><td>Type-3_BR.PglZ.associated.systems</td><td>XIS-HTH->DUF499+RRM*->PglZ->Classical-AAA->METHYLASE->iSTAND-></td><td>914</td><td>Bacteria>Pseudomonadota>Gammaproteobacteria>Pseudomonadales>Pseudomonadaceae>Pseudomonas</td><td>GCA_029604125.1</td><td>Pseudomonas_aeruginosa</td><td>L5F31_06965</td><td>348998..351742</td><td>287</td><td>ATP-binding-protein-[Pseudomonas-aeruginosa]</td></tr>
<tr><td>HDR3117557.1</td><td>AAA+ATPase+wHTH+Alpha+Beta-domain+wHTH+RRM</td><td>Type-3_BR.PglZ.associated.systems</td><td><-ParB+HNH<-TUDOR+SF1-SF2-Helicase+iREase+CTD<-iSTAND<-METHYLASE<-PglZ<-DUF499+RRM*<-XIS-HTH<-DUF3987<-ParB<-ParB<-GDResolvase+Recombinase</td><td>847</td><td>Bacteria>Pseudomonadota>Gammaproteobacteria>Pseudomonadales>Pseudomonadaceae>Pseudomonas</td><td>GCA_029688505.1</td><td>Pseudomonas_aeruginosa</td><td>QCL41_001948</td><td>229121..231664</td><td>287</td><td>DUF499-domain-containing-protein-[Pseudomonas-aeruginosa]</td></tr>
<tr><td>MDP5497408.1</td><td>AAA+ATPase+wHTH+Alpha+Beta-domain+wHTH+RRM</td><td>Type-3_BR.PglZ.associated.systems</td><td><-iSTAND<-METHYLASE<-Classical-AAA<-PglZ<-DUF499+RRM*<-MerR-HTH</td><td>915</td><td>Bacteria>Pseudomonadota>Gammaproteobacteria>Pseudomonadales>Pseudomonadaceae>Pseudomonas</td><td>GCA_030738305.1</td><td>Pseudomonas_aeruginosa</td><td>P3775_03575</td><td>469172..471919</td><td>287</td><td>DUF499-domain-containing-protein-[Pseudomonas-aeruginosa]</td></tr>
<tr><td>HDY5926452.1</td><td>AAA+ATPase+wHTH+Alpha+Beta-domain+wHTH+RRM</td><td>Type-3_BR.PglZ.associated.systems</td><td><-iSTAND<-METHYLASE<-Classical-AAA<-PglZ<-DUF499+RRM*<-XIS-HTH||MODE-HTH+PBPII->RHH-></td><td>860</td><td>Bacteria>Pseudomonadota>Gammaproteobacteria>Pseudomonadales>Pseudomonadaceae>Pseudomonas</td><td>GCA_032033215.1</td><td>Pseudomonas_aeruginosa</td><td>RQ788_004384</td><td>20710..23292</td><td>287</td><td>DUF499-domain-containing-protein-[Pseudomonas-aeruginosa]</td></tr>
<tr><td>HEE9762606.1</td><td>AAA+ATPase+wHTH+Alpha+Beta-domain+wHTH+RRM</td><td>Type-3_BR.PglZ.associated.systems</td><td><-RHH<-MODE-HTH+PBPII||XIS-HTH->DUF499+RRM*->PglZ->SIR2->VirB4-FtsK+HerA->METHYLASE-p-></td><td>860</td><td>Bacteria>Pseudomonadota>Gammaproteobacteria>Pseudomonadales>Pseudomonadaceae>Pseudomonas</td><td>GCA_033187555.1</td><td>Pseudomonas_putida</td><td>R8F97_003526</td><td>31837..34419</td><td>303</td><td>DUF499-domain-containing-protein-[Pseudomonas-putida]</td></tr>
<tr><td>HEH8465631.1</td><td>AAA+ATPase+wHTH+Alpha+Beta-domain+wHTH+RRM</td><td>Type-3_BR.PglZ.associated.systems</td><td><-DUF3487||XIS-HTH->DUF499+RRM*->PglZ->Classical-AAA->METHYLASE->iSTAND-></td><td>846</td><td>Bacteria>Pseudomonadota>Gammaproteobacteria>Pseudomonadales>Pseudomonadaceae>Pseudomonas</td><td>GCA_033655365.1</td><td>Pseudomonas_aeruginosa</td><td>SH274_001873</td><td>368391..370931</td><td>287</td><td>DUF499-domain-containing-protein-[Pseudomonas-aeruginosa]</td></tr>
<tr><td>HEK1475324.1</td><td>AAA+ATPase+wHTH+Alpha+Beta-domain+wHTH+RRM</td><td>Type-3_BR.PglZ.associated.systems</td><td>Tox-RelE->CHTH->||Phage_int_SAM_2+Integrase_1->XIS-HTH->DUF499+RRM*->PglZ->SIR2->VirB4-FtsK+HerA->METHYLASE->iSTAND-></td><td>848</td><td>Bacteria>Pseudomonadota>Gammaproteobacteria>Pseudomonadales>Pseudomonadaceae>Pseudomonas</td><td>GCA_033929445.1</td><td>Pseudomonas_aeruginosa</td><td>SMR02_004544</td><td>25770..28316</td><td>287</td><td>DUF499-domain-containing-protein-[Pseudomonas-aeruginosa]</td></tr>
<tr><td>WP_301157725.1</td><td>AAA+ATPase+wHTH+Alpha+Beta-domain+wHTH+RRM</td><td>Type-3_BR.PglZ.associated.systems</td><td>HNH-><-TUDOR+SF1-SF2-Helicase+iREase+CTD<-iSTAND<-METHYLASE<-PglZ<-DUF499+RRM*||Arm-DNA-bind_3+Phage_integrase-></td><td>917</td><td>Bacteria>Pseudomonadota>Gammaproteobacteria>Pseudomonadales>Pseudomonadaceae>Pseudomonas</td><td>GCF_030417915.1</td><td>Pseudomonas_sp._C32</td><td>QYH43_RS18030</td><td>67056..69809</td><td>1529208</td><td>DUF499-domain-containing-protein-[Pseudomonas-sp.-C32]</td></tr>
<tr><td>WP_286790781.1</td><td>AAA+ATPase+wHTH+Alpha+Beta-domain+wHTH+RRM</td><td>Type-3_BR.PglZ.associated.systems</td><td><-WYL+wHTH||XIS-HTH->DUF499+RRM*->PglZ->METHYLASE->iSTAND->TUDOR+SF1-SF2-Helicase+iREase+CTD->Cluster84_1clades-></td><td>893</td><td>Bacteria>Pseudomonadota>Gammaproteobacteria>Pseudomonadales>Pseudomonadaceae>Pseudomonas</td><td>GCF_032004765.1</td><td>unclassified_Pseudomonas</td><td>B3D05_RS22380</td><td>185352..188033</td><td>196821</td><td>MULTISPECIES:-DUF499-domain-containing-protein-[unclassified-Pseudomonas]</td></tr>
<tr><td>WP_316915903.1</td><td>AAA+ATPase+wHTH+Alpha+Beta-domain+wHTH+RRM</td><td>Type-3_BR.PglZ.associated.systems</td><td>HNH-><-TUDOR+SF1-SF2-Helicase+iREase+CTD<-iSTAND<-METHYLASE<-PglZ<-DUF499+RRM*<-MerR-HTH</td><td>915</td><td>Bacteria>Pseudomonadota>Gammaproteobacteria>Pseudomonadales>Pseudomonadaceae>Pseudomonas</td><td>GCF_032621725.1</td><td>Pseudomonas_sp._zfem001</td><td>RTH74_RS20650</td><td>45373..48120</td><td>3078196</td><td>DUF499-domain-containing-protein-[Pseudomonas-sp.-zfem001]</td></tr>
<tr><td>WP_319049354.1</td><td>AAA+ATPase+wHTH+Alpha+Beta-domain+wHTH+RRM</td><td>Type-3_BR.PglZ.associated.systems</td><td>MerR-HTH->DUF499+RRM*->PglZ->METHYLASE->iSTAND->TUDOR+SF1-SF2-Helicase+iREase+CTD-><-HNH</td><td>915</td><td>Bacteria>Pseudomonadota>Gammaproteobacteria>Pseudomonadales>Pseudomonadaceae>Pseudomonas</td><td>GCF_036664175.1</td><td>Pseudomonas_aeruginosa</td><td>V6E77_RS02440</td><td>482232..484979</td><td>287</td><td>DUF499-domain-containing-protein-[Pseudomonas-aeruginosa]</td></tr>
<tr><td>WP_319749940.1</td><td>AAA+ATPase+wHTH+Alpha+Beta-domain+wHTH+RRM</td><td>Type-3_BR.PglZ.associated.systems</td><td><-iSTAND<-METHYLASE<-VirB4-FtsK+HerA<-SIR2<-PglZ<-DUF499+RRM*<-MerR-HTH</td><td>912</td><td>Bacteria>Pseudomonadota>Gammaproteobacteria>Pseudomonadales>Pseudomonadaceae>Pseudomonas</td><td>GCF_036664995.1</td><td>Pseudomonas_aeruginosa</td><td>V6F32_RS06160</td><td>72911..75649</td><td>287</td><td>DUF499-domain-containing-protein-[Pseudomonas-aeruginosa]</td></tr>
</table>

<table>
<tr><th>Protein-ID</th><th>DUF499-architecture</th><th>System</th><th>Gene-Neighbourhoods</th><th>Length</th><th>Taxonomy</th><th>Assembly-ID</th><th>Strain</th><th>Locus-Tag</th><th>Genomic-coordinates</th><th>TaxID</th><th>Fasta-defline</th></tr>
<tr><td>EKX7273131.1</td><td>AAA+ATPase+wHTH+Alpha+Beta-domain+wHTH+RRM</td><td>Type-3_BR.PglZ.associated.systems</td><td><-iSTAND<-METHYLASE<-VirB4-FtsK+HerA<-SIR2<-PglZ<-DUF499+RRM*<-MerR-HTH</td><td>853</td><td>Bacteria>Pseudomonadota>Gammaproteobacteria>Pseudomonadales>Pseudomonadaceae>Pseudomonas</td><td>-</td><td>Pseudomonas_aeruginosa</td><td>L4H28_002030</td><td>380553..383114</td><td>287</td><td>DUF499-domain-containing-protein-[Pseudomonas-aeruginosa]</td></tr>
<tr><td>EKU2297546.1</td><td>AAA+ATPase+wHTH+Alpha+Beta-domain+wHTH+RRM</td><td>Type-3_BR.PglZ.associated.systems</td><td>MerR-HTH->DUF499+RRM*->PglZ->SIR2->VirB4-FtsK+HerA->METHYLASE->iSTAND-></td><td>853</td><td>Bacteria>Pseudomonadota>Gammaproteobacteria>Pseudomonadales>Pseudomonadaceae>Pseudomonas</td><td>-</td><td>Pseudomonas_aeruginosa</td><td>L4O09_004839</td><td>68693..71254</td><td>287</td><td>DUF499-domain-containing-protein-[Pseudomonas-aeruginosa]</td></tr>
<tr><td>EKU3772441.1</td><td>AAA+ATPase+wHTH+Alpha+Beta-domain+wHTH+RRM</td><td>Type-3_BR.PglZ.associated.systems</td><td>MerR-HTH->DUF499+RRM*->PglZ->METHYLASE->iSTAND->TUDOR+SF1-SF2-Helicase+iREase+CTD-></td><td>862</td><td>Bacteria>Pseudomonadota>Gammaproteobacteria>Pseudomonadales>Pseudomonadaceae>Pseudomonas</td><td>-</td><td>Pseudomonas_aeruginosa</td><td>L5V03_004654</td><td>39732..42320</td><td>287</td><td>DUF499-domain-containing-protein-[Pseudomonas-aeruginosa]</td></tr>
<tr><td>EKV8662301.1</td><td>AAA+ATPase+wHTH+Alpha+Beta-domain+wHTH+RRM</td><td>Type-3_BR.PglZ.associated.systems</td><td><-METHYLASE<-Classical-AAA<-PglZ<-DUF499+RRM*<-XIS-HTH||MODE-HTH+PBPII->RHH-></td><td>860</td><td>Bacteria>Pseudomonadota>Gammaproteobacteria>Pseudomonadales>Pseudomonadaceae>Pseudomonas</td><td>-</td><td>Pseudomonas_aeruginosa</td><td>OR553_000221</td><td>19489..22071</td><td>287</td><td>DUF499-domain-containing-protein-[Pseudomonas-aeruginosa]</td></tr>
<tr><td>EKW7733840.1</td><td>AAA+ATPase+wHTH+Alpha+Beta-domain+wHTH+RRM</td><td>Type-3_BR.PglZ.associated.systems</td><td>MerR-HTH->DUF499+RRM*->PglZ->METHYLASE->iSTAND->TUDOR+SF1-SF2-Helicase+iREase+CTD-><-HNH</td><td>839</td><td>Bacteria>Pseudomonadota>Gammaproteobacteria>Pseudomonadales>Pseudomonadaceae>Pseudomonas</td><td>-</td><td>Pseudomonas_aeruginosa</td><td>PU294_000685</td><td>14114..16633</td><td>287</td><td>DUF499-domain-containing-protein-[Pseudomonas-aeruginosa]</td></tr>
<tr><td>ELQ7314399.1</td><td>AAA+ATPase+wHTH+Alpha+Beta-domain+wHTH+RRM</td><td>Type-3_BR.PglZ.associated.systems</td><td>Tox-RelE->CHTH->||Phage_int_SAM_2+Integrase_1->XIS-HTH->DUF499+RRM*->PglZ->SIR2->VirB4-FtsK+HerA->METHYLASE-></td><td>847</td><td>Bacteria>Pseudomonadota>Gammaproteobacteria>Pseudomonadales>Pseudomonadaceae>Pseudomonas</td><td>-</td><td>Pseudomonas_aeruginosa</td><td>R2393_003604</td><td>10206..12749</td><td>287</td><td>DUF499-domain-containing-protein-[Pseudomonas-aeruginosa]</td></tr>
<tr><td>EKY4163340.1</td><td>AAA+ATPase+wHTH+Alpha+Beta-domain+wHTH+RRM</td><td>Type-3_BR.PglZ.associated.systems</td><td>MerR-HTH->DUF499+RRM*->PglZ->SIR2->VirB4-FtsK+HerA->METHYLASE->iSTAND-></td><td>853</td><td>Bacteria>Pseudomonadota>Gammaproteobacteria>Pseudomonadales>Pseudomonadaceae>Pseudomonas</td><td>-</td><td>Pseudomonas_aeruginosa</td><td>RCP99_005036</td><td>69160..71721</td><td>287</td><td>DUF499-domain-containing-protein-[Pseudomonas-aeruginosa]</td></tr>
<tr><td>WP_313665133.1</td><td>AAA+ATPase+wHTH+Alpha+Beta-domain+wHTH+RRM</td><td>Type-3_BR.PglZ.associated.systems</td><td><-iSTAND<-METHYLASE<-VirB4-FtsK+HerA<-SIR2<-PglZ<-DUF499+RRM*<-MerR-HTH</td><td>916</td><td>Bacteria>Pseudomonadota>Gammaproteobacteria>Pseudomonadales>Pseudomonadaceae>Stutzerimonas</td><td>GCF_032114295.1</td><td>Stutzerimonas_kunmingensis</td><td>RT691_RS03115</td><td>21012..23762</td><td>1211807</td><td>DUF499-domain-containing-protein-[Stutzerimonas-kunmingensis]</td></tr>
<tr><td>MBH1966600.1</td><td>AAA+ATPase+wHTH+Alpha+Beta-domain+wHTH+RRM</td><td>Type-3_BR.PglZ.associated.systems</td><td>HTH->?-><-?<-CopG/RHH||XIS-HTH->?->DUF499*->PglZ->METHYLASE->iSTAND->TUDOR+SF1-SF2-Helicase+iREase+CTD-><-RlpA<-JAB</td><td>882</td><td>Bacteria>Pseudomonadota>Gammaproteobacteria>Pseudomonadales>unclassified_Pseudomonadales</td><td>GCA_015999885.1</td><td>Pseudomonadales_bacterium</td><td>I8H81_00230</td><td>55376..58024</td><td>1891229</td><td>DUF499-domain-containing-protein-[Pseudomonadales-bacterium]</td></tr>
<tr><td>RLA52378.1</td><td>AAA+ATPase+wHTH+Alpha+Beta-domain+wHTH+RRM</td><td>Type-3_BR.PglZ.associated.systems</td><td>TOPRIM-><-Phage_int_SAM_4+Phage_integrase<-XIS-HTH<-XIS-HTH<-?||RHH->DUF499*->PglZ->METHYLASE->iSTAND->TUDOR+SF1-SF2-Helicase+iREase+CTD-></td><td>906</td><td>Bacteria>Pseudomonadota>Gammaproteobacteria>unclassified_Gammaproteobacteria</td><td>GCA_003646365.1</td><td>Gammaproteobacteria_bacterium</td><td>DRR42_07750</td><td>5418..8138</td><td>1913989</td><td>DUF499-domain-containing-protein-[Gammaproteobacteria-bacterium]</td></tr>
<tr><td>MCG6867647.1</td><td>AAA+ATPase+wHTH+Alpha+Beta-domain+wHTH+RRM</td><td>Type-3_BR.PglZ.associated.systems</td><td>RHH->DUF499*->PglZ->METHYLASE-p-></td><td>909</td><td>Bacteria>Pseudomonadota>Gammaproteobacteria>unclassified_Gammaproteobacteria</td><td>GCA_022340215.1</td><td>Gammaproteobacteria_bacterium</td><td>LJE91_02635</td><td>2315..5044</td><td>1913989</td><td>DUF499-domain-containing-protein-[Gammaproteobacteria-bacterium]</td></tr>
<tr><td>OGT87965.1</td><td>AAA+ATPase+wHTH+Alpha+Beta-domain+wHTH+RRM</td><td>Type-3_BR.PglZ.associated.systems</td><td>DUF499*->PglZ->Classical-AAA-><-?||Peptidase_S8->METHYLASE->iSTAND-></td><td>901</td><td>Bacteria>Pseudomonadota>Gammaproteobacteria>unclassified_Gammaproteobacteria</td><td>GCA_001801965.1</td><td>Gammaproteobacteria_bacterium_RIFOXYA12_FULL_61_12</td><td>A2286_09250</td><td>2257..4962</td><td>1798306</td><td>DUF499-domain-containing-protein-[Gammaproteobacteria-bacterium-RIFOXYA12_FULL_61_12]</td></tr>
<tr><td>MCG3143189.1</td><td>AAA+ATPase+wHTH+Alpha+Beta-domain+wHTH+RRM</td><td>Type-3_BR.PglZ.associated.systems</td><td><-TUDOR+SF1-SF2-Helicase+iREase+CTD<-iSTAND<-METHYLASE<-Radical-SAM<-?<-PglZ<-DUF499*</td><td>897</td><td>Bacteria>Pseudomonadota>Gammaproteobacteria>unclassified_Gammaproteobacteria</td><td>GCA_022072255.1</td><td>Gammaproteobacteria_bacterium</td><td>HONDAALG_00510</td><td>11831..14524</td><td>1913989</td><td>hypothetical-protein-[Gammaproteobacteria-bacterium]</td></tr>
<tr><td>HBR97454.1</td><td>AAA+ATPase+wHTH+Alpha+Beta-domain+wHTH+RRM</td><td>Type-3_BR.PglZ.associated.systems</td><td><-TUDOR+SF1-SF2-Helicase+iREase+CTD<-iSTAND<-METHYLASE<-PglZ<-DUF499*</td><td>910</td><td>Bacteria>Pseudomonadota>Gammaproteobacteria>unclassified_Gammaproteobacteria</td><td>GCA_003511825.1</td><td>Gammaproteobacteria_bacterium</td><td>DD979_08750</td><td>7905..10637</td><td>1913989</td><td>DUF499-domain-containing-protein-[Gammaproteobacteria-bacterium]</td></tr>
<tr><td>OGT70536.1</td><td>AAA+ATPase+wHTH+Alpha+Beta-domain+wHTH+RRM</td><td>Type-3_BR.PglZ.associated.systems</td><td><-iSTAND<-METHYLASE<-PglZ<-DUF499*||?->XIS-HTH-></td><td>912</td><td>Bacteria>Pseudomonadota>Gammaproteobacteria>unclassified_Gammaproteobacteria</td><td>GCA_001802565.1</td><td>Gammaproteobacteria_bacterium_RIFCSPLOWO2_02_FULL_57_10</td><td>A3H44_09805</td><td>6233..8971</td><td>1798299</td><td>DUF499-domain-containing-protein-[Gammaproteobacteria-bacterium-RIFCSPLOWO2_02_FULL_57_10]</td></tr>
<tr><td>WP_148533210.1</td><td>AAA+ATPase+wHTH+Alpha+Beta-domain+wHTH+RRM</td><td>Type-3_BR.PglZ.associated.systems</td><td>Phage_integrase+Phage_integrase->NACHT->TetR-HTH-><-HTH_3||HTH_Tnp->XIS-HTH->DUF499*->PglZ->METHYLASE->iSTAND->TUDOR+SF1-SF2-Helicase+iREase+CTD-></td><td>893</td><td>Bacteria>Pseudomonadota>Gammaproteobacteria>Vibrionales>Vibrionaceae</td><td>GCF_008084635.1</td><td>Vibrio_cholerae</td><td>FXE27_RS01035</td><td>194426..197107</td><td>666</td><td>hypothetical-protein-[Vibrio-cholerae]</td></tr>
<tr><td>EJL6704138.1</td><td>AAA+ATPase+wHTH+Alpha+Beta-domain+wHTH+RRM</td><td>Type-3_BR.PglZ.associated.systems</td><td>Peptidase-><-TRD+TRD<-TUDOR+SF1-SF2-Helicase+iREase+CTD<-iSTAND<-METHYLASE<-PglZ<-DUF499*<-XIS-HTH<-HTH_Tnp<-TetR-HTH<-NACHT<-Phage_integrase+Phage_integrase<-Phage_integrase</td><td>893</td><td>Bacteria>Pseudomonadota>Gammaproteobacteria>Vibrionales>Vibrionaceae</td><td>GCA_024218175.1</td><td>Vibrio_cholerae</td><td>NMS58_001172</td><td>12062..14743</td><td>666</td><td>hypothetical-protein-[Vibrio-cholerae]</td></tr>
<tr><td>EKG0010178.1</td><td>AAA+ATPase+wHTH+Alpha+Beta-domain+wHTH+RRM</td><td>Type-3_BR.PglZ.associated.systems</td><td><-OLD-ABC<-TRD+TRD<-TUDOR+SF1-SF2-Helicase+iREase+CTD<-iSTAND<-METHYLASE<-PglZ<-DUF499*<-XIS-HTH<-HTH_Tnp||HTH_3-><-TetR-HTH<-NACHT<-Phage_integrase+Phage_integrase</td><td>893</td><td>Bacteria>Pseudomonadota>Gammaproteobacteria>Vibrionales>Vibrionaceae</td><td>GCA_026768395.1</td><td>Vibrio_cholerae</td><td>O1D74_000517</td><td>609303..611984</td><td>666</td><td>hypothetical-protein-O1D74_000517-[Vibrio-cholerae]</td></tr>
<tr><td>MBY8216331.1</td><td>AAA+ATPase+wHTH+Alpha+Beta-domain+wHTH+RRM</td><td>Type-3_BR.PglZ.associated.systems</td><td><-JAB||XIS-HTH->XIS-HTH->DUF499*->PglZ->METHYLASE->iSTAND->TUDOR+SF1-SF2-Helicase+iREase+CTD-><-Arm-DNA-bind_3+Phage_int_SAM_3+Phage_integrase</td><td>893</td><td>Bacteria>Pseudomonadota>Gammaproteobacteria>Vibrionales>Vibrionaceae</td><td>GCA_019854395.1</td><td>Vibrio_fluvialis</td><td>KW517_20300</td><td>11208..13889</td><td>676</td><td>hypothetical-protein-[Vibrio-fluvialis]</td></tr>
<tr><td>WP_254058564.1</td><td>AAA+ATPase+wHTH+Alpha+Beta-domain+wHTH+RRM</td><td>Type-3_BR.PglZ.associated.systems</td><td><-TUDOR+SF1-SF2-Helicase+iREase+CTD<-iSTAND<-METHYLASE<-PglZ<-DUF499*<-DOC+HTH<-HEPN<-HTH+PolB</td><td>933</td><td>Bacteria>Pseudomonadota>Gammaproteobacteria>Xanthomonadales>Rhodanobacteraceae</td><td>GCF_009766065.1</td><td>Dyella_sp._S184</td><td>GSQ75_RS12385</td><td>51220..54021</td><td>1641862</td><td>DUF499-domain-containing-protein-[Dyella-sp.-S184]</td></tr>
<tr><td>MCC6440987.1</td><td>AAA+ATPase+wHTH+Alpha+Beta-domain+wHTH+RRM</td><td>Type-3_BR.PglZ.associated.systems</td><td><-TUDOR+SF1-SF2-Helicase+iREase+CTD<-iSTAND<-METHYLASE<-METHYLASE<-PglZ<-DUF499*||Calcineurin->SbcC+SbcC-><-AbiEi_4||DOC+HTH-></td><td>914</td><td>Bacteria>Pseudomonadota>Gammaproteobacteria>Xanthomonadales>Rhodanobacteraceae</td><td>GCA_020846735.1</td><td>Rhodanobacteraceae_bacterium</td><td>IT476_10230</td><td>8104..10848</td><td>2052182</td><td>DUF499-domain-containing-protein-[Rhodanobacteraceae-bacterium]</td></tr>
<tr><td>WP_253568720.1</td><td>AAA+ATPase+wHTH+Alpha+Beta-domain+wHTH+RRM</td><td>Type-3_BR.PglZ.associated.systems</td><td><-SF1-SF2-Helicase+iREase+CTD<-iSTAND<-METHYLASE<-Peptidase_S8<-Classical-AAA<-PglZ<-DUF499*<-?<-?||Pkinase+STYKIN->Mbetalac-></td><td>909</td><td>Bacteria>Pseudomonadota>Gammaproteobacteria>Xanthomonadales>Rhodanobacteraceae</td><td>GCF_024158035.1</td><td>Dyella_lutea</td><td>NC595_RS17950</td><td>288194..290923</td><td>2950441</td><td>DUF499-domain-containing-protein-[Dyella-sp.-Sa]</td></tr>
<tr><td>WP_106891671.1</td><td>AAA+ATPase+wHTH+Alpha+Beta-domain+wHTH+RRM</td><td>Type-3_BR.PglZ.associated.systems</td><td><-Peptidase<-TUDOR+SF1-SF2-Helicase+iREase+CTD<-Schlafen+HATPase+HTH<-iSTAND<-METHYLASE<-PglZ<-DUF499*||MACRODOMAIN->DOC+HTH->?->?->HKD-DNase->SF1-SF2-Helicase-></td><td>909</td><td>Bacteria>Pseudomonadota>Gammaproteobacteria>Xanthomonadales>Rhodanobacteraceae</td><td>GCF_003015185.1</td><td>Ahniella_affigens</td><td>C7S18_RS11330</td><td>2972599..2975328</td><td>2021234</td><td>DUF499-domain-containing-protein-[Ahniella-affigens]</td></tr>
<tr><td>WP_017462287.1</td><td>AAA+ATPase+wHTH+Alpha+Beta-domain+wHTH+RRM</td><td>Type-3_BR.PglZ.associated.systems</td><td><-Mbetalac<-Pkinase+STYKIN||?->?->DUF499*->PglZ->METHYLASE->iSTAND->TUDOR+SF1-SF2-Helicase+iREase+CTD->?->ParB+HNH-></td><td>910</td><td>Bacteria>Pseudomonadota>Gammaproteobacteria>Xanthomonadales>Rhodanobacteraceae</td><td>GCF_000334915.1</td><td>Dyella_ginsengisoli</td><td>C693_RS0110780</td><td>59347..62079</td><td>363848</td><td>DUF499-domain-containing-protein-[Dyella-ginsengisoli]</td></tr>
<tr><td>MBX3726675.1</td><td>AAA+ATPase+wHTH+Alpha+Beta-domain+wHTH+RRM</td><td>Type-3_BR.PglZ.associated.systems</td><td><-TUDOR+SF1-SF2-Helicase+iREase+CTD<-iSTAND<-METHYLASE<-?<-PglZ<-DUF499*<-HTH+WYL||?->?->?->REase->REase+35exo-></td><td>909</td><td>Bacteria>Pseudomonadota>Gammaproteobacteria>Xanthomonadales>unclassified_Xanthomonadales</td><td>GCA_019634735.1</td><td>Xanthomonadales_bacterium</td><td>KF823_12255</td><td>130955..133684</td><td>2006849</td><td>DUF499-domain-containing-protein-[Xanthomonadales-bacterium]</td></tr>
<tr><td>MBV6415467.1</td><td>AAA+ATPase+wHTH+Alpha+Beta-domain+wHTH+RRM</td><td>Type-3_BR.PglZ.associated.systems</td><td><-REase+nSTAND3<-?<-TUDOR+SF1-SF2-Helicase+iREase+CTD<-iSTAND<-METHYLASE<-PglZ<-DUF499*</td><td>827</td><td>Bacteria>Pseudomonadota>Gammaproteobacteria>Xanthomonadales>unclassified_Xanthomonadales</td><td>GCA_019187485.1</td><td>Xanthomonadales_bacterium</td><td>OMOMHJEC_03357</td><td>9707..12190</td><td>2006849</td><td>hypothetical-protein-[Xanthomonadales-bacterium]</td></tr>
<tr><td>WP_057663529.1</td><td>AAA+ATPase+wHTH+Alpha+Beta-domain+wHTH+RRM</td><td>Type-3_BR.PglZ.associated.systems</td><td><-XIS-HTH||?-><-?<-ParB+HTH<-DUF932||XIS-HTH->DUF499*->PglZ->Classical-AAA-></td><td>920</td><td>Bacteria>Pseudomonadota>Gammaproteobacteria>Xanthomonadales>Xanthomonadaceae</td><td>GCF_017746155.1</td><td>Xanthomonas</td><td>IB69_RS02045</td><td>40842..43604</td><td>338</td><td>MULTISPECIES:-DUF499-domain-containing-protein-[Xanthomonas]</td></tr>
<tr><td>WP_154352412.1</td><td>AAA+ATPase+wHTH+Alpha+Beta-domain+wHTH+RRM</td><td>Type-3_BR.PglZ.associated.systems</td><td><-VirB11+VirB4-FtsK<-CopG/RHH<-VirD4-TraG+VirD4-FtsK<-?<-MODE-HTH+PBPII||XIS-HTH->DUF499*->PglZ->Classical-AAA->Peptidase_S8->METHYLASE->iSTAND->TUDOR+SF1-SF2-Helicase+iREase+CTD-></td><td>922</td><td>Bacteria>Pseudomonadota>Gammaproteobacteria>Xanthomonadales>Xanthomonadaceae</td><td>GCF_009676585.1</td><td>Stenotrophomonas</td><td>KOM59_RS01225</td><td>61889..64657</td><td>40323</td><td>DUF499-domain-containing-protein-[Stenotrophomonas-maltophilia]</td></tr>
<tr><td>WP_165710374.1</td><td>AAA+ATPase+wHTH+Alpha+Beta-domain+wHTH+RRM</td><td>Type-3_BR.PglZ.associated.systems</td><td><-TUDOR+SF1-SF2-Helicase+iREase+CTD<-iSTAND<-METHYLASE<-VirB4-FtsK+HerA<-SIR2<-PglZ<-DUF499*<-XIS-HTH||MODE-HTH+PBPII->?->VirD4-TraG+VirD4-FtsK->CopG/RHH->VirB11+VirB4-FtsK-></td><td>922</td><td>Bacteria>Pseudomonadota>Gammaproteobacteria>Xanthomonadales>Xanthomonadaceae</td><td>GCF_011386925.1</td><td>Stenotrophomonas_maltophilia</td><td>G6Z48_RS07155</td><td>1483133..1485901</td><td>40324</td><td>DUF499-domain-containing-protein-[Stenotrophomonas-maltophilia]</td></tr>
<tr><td>MTI73906.1</td><td>AAA+ATPase+wHTH+Alpha+Beta-domain+wHTH+RRM</td><td>Type-3_BR.PglZ.associated.systems</td><td><-TUDOR+SF1-SF2-Helicase+iREase+CTD<-iSTAND<-METHYLASE<-VirB4-FtsK+HerA<-SIR2<-PglZ<-DUF499*<-HTH_26||?->Arm-DNA-bind_2+Phage_int_SAM_3+Phage_integrase->HDRnase-><-HTH_3</td><td>915</td><td>Bacteria>Pseudomonadota>Gammaproteobacteria>Xanthomonadales>Xanthomonadaceae</td><td>GCA_009711865.1</td><td>Stenotrophomonas_sp.</td><td>FH747_09660</td><td>146039..148786</td><td>69392</td><td>DUF499-domain-containing-protein-[Stenotrophomonas-sp.]</td></tr>
<tr><td>UYK75915.1</td><td>AAA+ATPase+wHTH+Alpha+Beta-domain+wHTH+RRM</td><td>Type-3_BR.PglZ.associated.systems</td><td><-TUDOR+SF1-SF2-Helicase+iREase+CTD<-iSTAND<-METHYLASE<-PglZ<-PglZ<-DUF499*<-HEPN<-HTH+PolB<-TPR+TPR+TPR+TPR+TPR+TPR+TPR+TPR</td><td>909</td><td>Bacteria>Pseudomonadota>Gammaproteobacteria>Xanthomonadales>Xanthomonadaceae</td><td>GCA_025765935.1</td><td>Xanthomonas_sacchari</td><td>NG825_15710</td><td>3730512..3733241</td><td>56458</td><td>ATP-binding-protein-[Xanthomonas-sacchari]</td></tr>
<tr><td>WP_284467660.1</td><td>AAA+ATPase+wHTH+Alpha+Beta-domain+wHTH+RRM</td><td>Type-3_BR.PglZ.associated.systems</td><td><-TUDOR+SF1-SF2-Helicase+iREase+CTD<-iSTAND<-METHYLASE<-PglZ<-DUF499*<-HEPN<-HTH+PolB<-TPR+TPR+TPR+TPR+TPR+TPR+TPR+TPR</td><td>909</td><td>Bacteria>Pseudomonadota>Gammaproteobacteria>Xanthomonadales>Xanthomonadaceae</td><td>GCF_034105325.1</td><td>unclassified_Xanthomonas</td><td>NG825_RS15745</td><td>3730512..3733241</td><td>2643310</td><td>ATP-binding-protein-[Xanthomonas-sacchari]</td></tr>
<tr><td>WP_152182007.1</td><td>AAA+ATPase+wHTH+Alpha+Beta-domain+wHTH+RRM</td><td>Type-3_BR.PglZ.associated.systems</td><td><-TUDOR+SF1-SF2-Helicase+iREase+CTD<-iSTAND<-METHYLASE<-PglZ<-DUF499*<-HEPN<-HTH+PolB</td><td>909</td><td>Bacteria>Pseudomonadota>Gammaproteobacteria>Xanthomonadales>Xanthomonadaceae</td><td>GCF_009192905.1</td><td>Xanthomonas_sp._LMG_12459</td><td>CEK65_RS12995</td><td>45871..48600</td><td>1591131</td><td>DUF499-domain-containing-protein-[Xanthomonas-sp.-LMG-12459]</td></tr>
<tr><td>WP_116902788.1</td><td>AAA+ATPase+wHTH+Alpha+Beta-domain+wHTH+RRM</td><td>Type-3_BR.PglZ.associated.systems</td><td><-TUDOR+SF1-SF2-Helicase+iREase+CTD<-iSTAND<-METHYLASE<-Peptidase_S8<-Classical-AAA<-PglZ<-DUF499*<-XIS-HTH<-HIN-HTH</td><td>915</td><td>Bacteria>Pseudomonadota>Gammaproteobacteria>Xanthomonadales>Xanthomonadaceae</td><td>GCF_003410225.1</td><td>Xanthomonas_campestris</td><td>DOA35_RS10835</td><td>37080..39827</td><td>339</td><td>DUF499-domain-containing-protein-[Xanthomonas-campestris]</td></tr>
<tr><td>WP_218555109.1</td><td>AAA+ATPase+wHTH+Alpha+Beta-domain+wHTH+RRM</td><td>Type-3_BR.PglZ.associated.systems</td><td><-TUDOR+SF1-SF2-Helicase+iREase+CTD<-iSTAND<-METHYLASE<-Peptidase_S8<-Classical-AAA<-PglZ<-DUF499*<-XIS-HTH<-?<-?<-ParB+HTH</td><td>920</td><td>Bacteria>Pseudomonadota>Gammaproteobacteria>Xanthomonadales>Xanthomonadaceae</td><td>GCF_019201185.1</td><td>Xanthomonas_euvesicatoria</td><td>KWH03_RS16285</td><td>47330..50092</td><td>456327</td><td>DUF499-domain-containing-protein-[Xanthomonas-campestris]</td></tr>
<tr><td>WP_040279327.1</td><td>AAA+ATPase+wHTH+Alpha+Beta-domain+wHTH+RRM</td><td>Type-3_BR.PglZ.associated.systems</td><td><-TUDOR+SF1-SF2-Helicase+iREase+CTD<-iSTAND<-METHYLASE<-Peptidase_S8<-Classical-AAA<-PglZ<-DUF499*<-XIS-HTH<-?<-?<-ParB+HTH</td><td>920</td><td>Bacteria>Pseudomonadota>Gammaproteobacteria>Xanthomonadales>Xanthomonadaceae</td><td>GCF_963669835.1</td><td>Xanthomonas_citri</td><td>AMD05_RS21775</td><td>32780..35542</td><td>346</td><td>DUF499-domain-containing-protein-[Xanthomonas-citri]</td></tr>
<tr><td>WP_150359733.1</td><td>AAA+ATPase+wHTH+Alpha+Beta-domain+wHTH+RRM</td><td>Type-3_BR.PglZ.associated.systems</td><td><-TUDOR+SF1-SF2-Helicase+iREase+CTD<-iSTAND<-METHYLASE<-Peptidase_S8<-Classical-AAA<-PglZ<-DUF499*<-XIS-HTH||MODE-HTH+PBPII->?->VirD4-TraG+VirD4-FtsK->CopG/RHH->VirB11+VirB4-FtsK-></td><td>922</td><td>Bacteria>Pseudomonadota>Gammaproteobacteria>Xanthomonadales>Xanthomonadaceae</td><td>GCF_008693985.1</td><td>Stenotrophomonas_maltophilia</td><td>FOB57_RS15285</td><td>3291385..3294153</td><td>40324</td><td>DUF499-domain-containing-protein-[Stenotrophomonas-maltophilia]</td></tr>
<tr><td>WP_099605297.1</td><td>AAA+ATPase+wHTH+Alpha+Beta-domain+wHTH+RRM</td><td>Type-3_BR.PglZ.associated.systems</td><td><-TUDOR+SF1-SF2-Helicase+iREase+CTD<-iSTAND<-METHYLASE<-Peptidase_S8<-Classical-AAA<-PglZ<-DUF499*<-XIS-HTH||MODE-HTH+PBPII->?->VirD4-TraG+VirD4-FtsK->CopG/RHH->VirB11+VirB4-FtsK-></td><td>922</td><td>Bacteria>Pseudomonadota>Gammaproteobacteria>Xanthomonadales>Xanthomonadaceae</td><td>GCF_030237085.1</td><td>Stenotrophomonas_maltophilia</td><td>QRO91_RS11090</td><td>19618..22386</td><td>40324</td><td>DUF499-domain-containing-protein-[Stenotrophomonas-maltophilia]</td></tr>
<tr><td>WP_171952106.1</td><td>AAA+ATPase+wHTH+Alpha+Beta-domain+wHTH+RRM</td><td>Type-3_BR.PglZ.associated.systems</td><td><-SF1-SF2-Helicase+iREase+CTD<-iSTAND<-METHYLASE<-PglZ<-DUF499*<-?<-?<-?<-?<-?<-sGTP</td><td>910</td><td>Bacteria>Pseudomonadota>Gammaproteobacteria>Xanthomonadales>Xanthomonadaceae</td><td>-</td><td>Stenotrophomonas_maltophilia</td><td>HRL53_RS00745</td><td>166663..169395</td><td>40324</td><td>DUF499-domain-containing-protein-[Stenotrophomonas-maltophilia]</td></tr>
<tr><td>WP_249046263.1</td><td>AAA+ATPase+wHTH+Alpha+Beta-domain+wHTH+RRM</td><td>Type-3_BR.PglZ.associated.systems</td><td><-SF1-SF2-Helicase+iREase+CTD<-iSTAND<-METHYLASE<-?<-PglZ<-DUF499*<-XIS-HTH<-DnaB||?->?-><-ABhydrolase</td><td>909</td><td>Bacteria>Pseudomonadota>Gammaproteobacteria>Xanthomonadales>Xanthomonadaceae</td><td>GCF_023337135.1</td><td>Xanthomonas_nasturtii</td><td>M3O40_RS15320</td><td>9125..11854</td><td>1843581</td><td>DUF499-domain-containing-protein-[Xanthomonas-nasturtii]</td></tr>
<tr><td>WP_104615522.1</td><td>AAA+ATPase+wHTH+Alpha+Beta-domain+wHTH+RRM</td><td>Type-3_BR.PglZ.associated.systems</td><td><-ParB+HNH-p<-TUDOR+SF1-SF2-Helicase+iREase+CTD<-iSTAND<-METHYLASE<-PglZ<-DUF499*<-XIS-HTH</td><td>909</td><td>Bacteria>Pseudomonadota>Gammaproteobacteria>Xanthomonadales>Xanthomonadaceae</td><td>GCF_002939865.1</td><td>Xanthomonas_dyei</td><td>XdyCFBP7245_RS09935</td><td>7930..10659</td><td>743699</td><td>DUF499-domain-containing-protein-[Xanthomonas-dyei]</td></tr>
<tr><td>WP_053506168.1</td><td>AAA+ATPase+wHTH+Alpha+Beta-domain+wHTH+RRM</td><td>Type-3_BR.PglZ.associated.systems</td><td><-METHYLASE||HTH_Tnp_1+HTH_21+rve+rve_3-><-METHYLASE-p<-VirB4-FtsK+HerA<-SIR2<-PglZ<-DUF499*<-XIS-HTH||MODE-HTH+PBPII->?->VirD4-TraG+VirD4-FtsK->CopG/RHH->VirB11+VirB4-FtsK-></td><td>922</td><td>Bacteria>Pseudomonadota>Gammaproteobacteria>Xanthomonadales>Xanthomonadaceae</td><td>GCF_013185915.1</td><td>Stenotrophomonas</td><td>HGB51_RS15730</td><td>969935..972703</td><td>40323</td><td>MULTISPECIES:-DUF499-domain-containing-protein-[Stenotrophomonas]</td></tr>
<tr><td>WP_280579276.1</td><td>AAA+ATPase+wHTH+Alpha+Beta-domain+wHTH+RRM</td><td>Type-3_BR.PglZ.associated.systems</td><td>RelE-ParE->||DUF499+RRM*->PglZ->METHYLASE->iSTAND->TUDOR+SF1-SF2-Helicase+iREase+CTD-></td><td>903</td><td>Bacteria>Pseudomonadota>Gammaproteobacteria>Xanthomonadales>Xanthomonadaceae>Luteimonas</td><td>GCF_029887935.1</td><td>Luteimonas_kalidii</td><td>QFW81_RS12875</td><td>417315..420026</td><td>3042025</td><td>DUF499-domain-containing-protein-[Luteimonas-kalidii]</td></tr>
<tr><td>WP_296282084.1</td><td>AAA+ATPase+wHTH+Alpha+Beta-domain+wHTH+RRM</td><td>Type-3_BR.PglZ.associated.systems</td><td><-TUDOR+SF1-SF2-Helicase+iREase+CTD<-iSTAND<-METHYLASE<-PglZ<-DUF499+RRM*||Arm-DNA-bind_2+Phage_int_SAM_3+Phage_integrase-></td><td>909</td><td>Bacteria>Pseudomonadota>Gammaproteobacteria>Xanthomonadales>Xanthomonadaceae>Pseudoxanthomonas</td><td>GCF_002863005.1</td><td>Pseudoxanthomonas_sp.</td><td>CYU89_RS14500</td><td>8195..10924</td><td>1871049</td><td>DUF499-domain-containing-protein-[Pseudoxanthomonas-sp.]</td></tr>
<tr><td>HDS1566924.1</td><td>AAA+ATPase+wHTH+Alpha+Beta-domain+wHTH+RRM</td><td>Type-3_BR.PglZ.associated.systems</td><td><-RHH<-MODE-HTH+PBPII||XIS-HTH->DUF499+RRM*->PglZ->SIR2->VirB4-FtsK+HerA->METHYLASE->iSTAND-></td><td>922</td><td>Bacteria>Pseudomonadota>Gammaproteobacteria>Xanthomonadales>Xanthomonadaceae>Stenotrophomonas</td><td>GCA_029771365.1</td><td>Stenotrophomonas_maltophilia</td><td>QEL18_002350</td><td>32866..35634</td><td>40324</td><td>DUF499-domain-containing-protein-[Stenotrophomonas-maltophilia]</td></tr>
<tr><td>WP_293761278.1</td><td>AAA+ATPase+wHTH+Alpha+Beta-domain+wHTH+RRM</td><td>Type-3_BR.PglZ.associated.systems</td><td><-iSTAND<-METHYLASE<-VirB4-FtsK+HerA<-SIR2<-PglZ<-DUF499+RRM*||Arm-DNA-bind_2+Phage_int_SAM_3+Phage_integrase-><-HTH_3</td><td>915</td><td>Bacteria>Pseudomonadota>Gammaproteobacteria>Xanthomonadales>Xanthomonadaceae>Stenotrophomonas</td><td>GCF_009711865.1</td><td>Stenotrophomonas_sp.</td><td>FH747_RS09710</td><td>146039..148786</td><td>69392</td><td>DUF499-domain-containing-protein-[Stenotrophomonas-sp.]</td></tr>
<tr><td>WP_312528391.1</td><td>AAA+ATPase+wHTH+Alpha+Beta-domain+wHTH+RRM</td><td>Type-3_BR.PglZ.associated.systems</td><td>METHYLASE-p->35exo->35exo->||DUF499+RRM*->PglZ->HTH_Tnp_1+HTH_21+rve+rve_3-></td><td>922</td><td>Bacteria>Pseudomonadota>Gammaproteobacteria>Xanthomonadales>Xanthomonadaceae>Stenotrophomonas</td><td>GCF_032025265.1</td><td>Stenotrophomonas_maltophilia</td><td>ROV90_14225</td><td>390143..392911</td><td>40324</td><td>DUF499-domain-containing-protein-[Stenotrophomonas-maltophilia]</td></tr>
<tr><td>WP_332807502.1</td><td>AAA+ATPase+wHTH+Alpha+Beta-domain+wHTH+RRM</td><td>Type-3_BR.PglZ.associated.systems</td><td><-TUDOR+SF1-SF2-Helicase+iREase+CTD<-iSTAND<-METHYLASE<-PglZ<-DUF499+RRM*</td><td>915</td><td>Bacteria>Pseudomonadota>Gammaproteobacteria>Xanthomonadales>Xanthomonadaceae>Stenotrophomonas</td><td>GCF_036662295.1</td><td>Stenotrophomonas_sp.</td><td>RSA25_RS03365</td><td>53034..55781</td><td>69392</td><td>DUF499-domain-containing-protein-[Stenotrophomonas-sp.]</td></tr>
<tr><td>WP_289404480.1</td><td>AAA+ATPase+wHTH+Alpha+Beta-domain+wHTH+RRM</td><td>Type-3_BR.PglZ.associated.systems</td><td>XIS-HTH->DUF499+RRM*->PglZ->METHYLASE->iSTAND->TUDOR+SF1-SF2-Helicase+iREase+CTD->MODE-HTH+PBPII-></td><td>934</td><td>Bacteria>Pseudomonadota>Gammaproteobacteria>Xanthomonadales>Xanthomonadaceae>Xanthomonas</td><td>GCF_030362065.1</td><td>Xanthomonas_campestris</td><td>QT935_RS01250</td><td>280720..283524</td><td>339</td><td>DUF499-domain-containing-protein-[Xanthomonas-campestris]</td></tr>
<tr><td>WP_289882774.1</td><td>AAA+ATPase+wHTH+Alpha+Beta-domain+wHTH+RRM</td><td>Type-3_BR.PglZ.associated.systems</td><td><-iSTAND<-METHYLASE<-VirB4-FtsK+HerA<-SIR2<-PglZ<-DUF499+RRM*<-XIS-HTH</td><td>938</td><td>Bacteria>Pseudomonadota>Gammaproteobacteria>Xanthomonadales>Xanthomonadaceae>Xanthomonas</td><td>GCF_030388925.1</td><td>Xanthomonas_arboricola</td><td>QSH42_RS09735</td><td>50571..53387</td><td>56448</td><td>DUF499-domain-containing-protein-[Xanthomonas-arboricola]</td></tr>
<tr><td>WP_323527816.1</td><td>AAA+ATPase+wHTH+Alpha+Beta-domain+wHTH+RRM</td><td>Type-3_BR.PglZ.associated.systems</td><td><-iSTAND<-METHYLASE<-Classical-AAA<-PglZ<-DUF499+RRM*<-XIS-HTH</td><td>941</td><td>Bacteria>Pseudomonadota>Gammaproteobacteria>Xanthomonadales>Xanthomonadaceae>Xanthomonas</td><td>GCF_035013125.1</td><td>Xanthomonas_campestris</td><td>VDF55_RS15460</td><td>11839..14664</td><td>339</td><td>DUF499-domain-containing-protein-[Xanthomonas-campestris]</td></tr>
<tr><td>WP_169115532.1</td><td>AAA+ATPase+wHTH+Alpha+Beta-domain+wHTH+RRM</td><td>Type-3_BR.PglZ.associated.systems</td><td><-ArsR-HTH+YTCD-HTH||?->GNAT-><-ACET<-TraJ-RHH||XIS-HTH->DUF499*->PglZ->?->?->PIN->METHYLASE->iSTAND-></td><td>909</td><td>Bacteria>Pseudomonadota>Hydrogenophilia>Hydrogenophilales>Hydrogenophilaceae</td><td>GCF_012911495.1</td><td>Tepidiphilus_baoligensis</td><td>GV368_RS03815</td><td>173539..176268</td><td>2698687</td><td>DUF499-domain-containing-protein-[Tepidiphilus-baoligensis]</td></tr>
<tr><td>OZA22671.1</td><td>AAA+ATPase+wHTH+Alpha+Beta-domain+wHTH+RRM</td><td>Type-3_BR.PglZ.associated.systems</td><td><-METHYLASE-p<-VirB4-FtsK+HerA<-SIR2<-PglZ<-DUF499*<-XIS-HTH||Phage_AlpA-></td><td>913</td><td>Bacteria>Pseudomonadota>Hydrogenophilia>Hydrogenophilales>unclassified_Hydrogenophilales</td><td>GCA_002281975.1</td><td>Hydrogenophilales_bacterium_17-64-11</td><td>B7X91_14580</td><td>5098..7839</td><td>1970526</td><td>DUF499-domain-containing-protein-[Hydrogenophilales-bacterium-17-64-11]</td></tr>
<tr><td>WP_147801109.1</td><td>AAA+ATPase+wHTH+Alpha+Beta-domain+wHTH+RRM</td><td>Type-3_BR.PglZ.associated.systems</td><td>Schlafen+HATPase+HTH->SF1-SF2-Helicase-p->SF1-SF2-Helicase->METHYLASE->BrnT_toxin->HTH->DUF499*->PglZ->?->SIGMA-HTH->MetJarc-HTH->PIN-></td><td>950</td><td>Bacteria>Pseudomonadota>Hydrogenophilia>Hydrogenophilalia_incertae_sedis>Pelomicrobium</td><td>GCF_008014345.1</td><td>Pelomicrobium_methylotrophicum</td><td>FR698_RS15560</td><td>14564..17416</td><td>2602750</td><td>DUF499-domain-containing-protein-[Pelomicrobium-methylotrophicum]</td></tr>
<tr><td>MCA0175250.1</td><td>AAA+ATPase+wHTH+Alpha+Beta-domain+wHTH+RRM</td><td>Type-3_BR.PglZ.associated.systems</td><td>MethyltransfD12->?->Response_reg+LuxR-HTH->XIS-HTH->DUF499*->PglZ->METHYLASE->iSTAND->TUDOR+SF1-SF2-Helicase+iREase+CTD-><-LRP-HTH||Peptidase_M20-></td><td>913</td><td>Bacteria>Pseudomonadota>unclassified_Pseudomonadota</td><td>GCA_020161145.1</td><td>Pseudomonadota_bacterium</td><td>LCH73_03015</td><td>598855..601596</td><td>1977087</td><td>DUF499-domain-containing-protein-[Proteobacteria-bacterium]</td></tr>
<tr><td>MBE9570567.1</td><td>AAA+ATPase+wHTH+Alpha+Beta-domain+wHTH+RRM</td><td>Type-3_BR.PglZ.associated.systems</td><td>DUF499*->PglZ->METHYLASE->iSTAND-></td><td>882</td><td>Bacteria>Pseudomonadota>unclassified_Pseudomonadota</td><td>GCA_015222845.1</td><td>Pseudomonadota_bacterium</td><td>IMF11_08065</td><td>608..3256</td><td>1977087</td><td>AAA-family-ATPase-[Proteobacteria-bacterium]</td></tr>
<tr><td>MBU1206366.1</td><td>AAA+ATPase+wHTH+Alpha+Beta-domain+wHTH+RRM</td><td>Type-3_BR.PglZ.associated.systems</td><td><-TUDOR+SF1-SF2-Helicase+iREase+CTD<-iSTAND<-METHYLASE<-PIN<-CopG/RHH<-PglZ<-DUF499*<-?<-?<-?<-?<-?<-GNTR-HTH</td><td>869</td><td>Bacteria>Pseudomonadota>unclassified_Pseudomonadota</td><td>GCA_018819135.1</td><td>Pseudomonadota_bacterium</td><td>KKH04_05495</td><td>22977..25586</td><td>1977087</td><td>hypothetical-protein-[Proteobacteria-bacterium]</td></tr>
<tr><td>MBN2145086.1</td><td>AAA+ATPase+wHTH+Alpha+Beta-domain+wHTH+RRM</td><td>Type-3_BR.PglZ.associated.systems</td><td>DUF499*->PglZ->METHYLASE->DUF433->PIN->?->TUDOR+SF1-SF2-Helicase+iREase+CTD-></td><td>883</td><td>Bacteria>PVC_group>Candidatus_Auribacterota>unclassified_Candidatus_Auribacterota</td><td>GCA_016932965.1</td><td>Candidatus_Auribacterota_bacterium</td><td>JW774_10720</td><td>466..3117</td><td>2093365</td><td>hypothetical-protein-[Candidatus-Aureabacteria-bacterium]</td></tr>
<tr><td>MEA1926979.1</td><td>AAA+ATPase+wHTH+Alpha+Beta-domain+wHTH+RRM</td><td>Type-3_BR.PglZ.associated.systems</td><td>DUF499+RRM*->PglZ->CopG-RHH->PIN-></td><td>707</td><td>Bacteria>PVC_group>Candidatus_Auribacterota>unclassified_Candidatus_Auribacterota</td><td>GCA_034658855.1</td><td>Candidatus_Auribacterota_bacterium</td><td>U9N73_02140</td><td>1..2126</td><td>2093365</td><td>hypothetical-protein-[Candidatus-Auribacterota-bacterium]</td></tr>
<tr><td>MBU4312937.1</td><td>AAA+ATPase+wHTH+Alpha+Beta-domain+wHTH+RRM</td><td>Type-3_BR.PglZ.associated.systems</td><td>Calcineurin->?->XIS-HTH->DUF499*->PglZ->METHYLASE->iSTAND->TUDOR+SF1-SF2-Helicase+iREase+CTD-></td><td>860</td><td>Bacteria>PVC_group>Candidatus_Omnitrophota>unclassified_Candidatus_Omnitrophota</td><td>GCA_018896715.1</td><td>Candidatus_Omnitrophota_bacterium</td><td>KJ706_09495</td><td>4274..6856</td><td>2035772</td><td>hypothetical-protein-[Candidatus-Omnitrophica-bacterium]</td></tr>
<tr><td>MBQ8111585.1</td><td>AAA+ATPase+wHTH+Alpha+Beta-domain+wHTH+RRM</td><td>Type-3_BR.PglZ.associated.systems</td><td>DUF499*->PglZ->HKD-DNase->METHYLASE->iSTAND->TUDOR+SF1-SF2-Helicase+iREase+CTD-></td><td>908</td><td>Bacteria>PVC_group>Kiritimatiellota>Kiritimatiellia>unclassified_Kiritimatiellia</td><td>GCA_017459225.1</td><td>Kiritimatiellia_bacterium</td><td>IJ146_00120</td><td>517..3243</td><td>2212473</td><td>hypothetical-protein-[Kiritimatiellae-bacterium]</td></tr>
<tr><td>HPF98773.1</td><td>AAA+ATPase+wHTH+Alpha+Beta-domain+wHTH+RRM</td><td>Type-3_BR.PglZ.associated.systems</td><td><-METHYLASE<-PIN<-CopG-RHH<-PglZ<-DUF499+RRM*</td><td>861</td><td>Bacteria>PVC_group>Kiritimatiellota>Kiritimatiellia>unclassified_Kiritimatiellia</td><td>GCA_035399025.1</td><td>Kiritimatiellia_bacterium</td><td>PLE77_01810</td><td>53349..55934</td><td>2212473</td><td>hypothetical-protein-[Kiritimatiellia-bacterium]</td></tr>
<tr><td>HPG01348.1</td><td>AAA+ATPase+wHTH+Alpha+Beta-domain+wHTH+RRM</td><td>Type-3_BR.PglZ.associated.systems</td><td>DUF499+RRM*->PglZ->ABC-ATPase->METHYLASE-p-></td><td>898</td><td>Bacteria>PVC_group>Kiritimatiellota>Kiritimatiellia>unclassified_Kiritimatiellia</td><td>GCA_035399025.1</td><td>Kiritimatiellia_bacterium</td><td>PLE77_14925</td><td>277..2973</td><td>2212473</td><td>DUF499-domain-containing-protein-[Kiritimatiellia-bacterium]</td></tr>
<tr><td>TFH14332.1</td><td>AAA+ATPase+wHTH+Alpha+Beta-domain+wHTH+RRM</td><td>Type-3_BR.PglZ.associated.systems</td><td><-Schlafen+HATPase+HTH<-PIN<-CopG/RHH<-PglZ<-DUF499*</td><td>393</td><td>Bacteria>PVC_group>Lentisphaerota>Lentisphaeria>Lentisphaerales</td><td>GCA_004525935.1</td><td>Lentisphaerales_bacterium</td><td>E4H02_10115</td><td>2873..4054</td><td>2448604</td><td>hypothetical-protein-E4H02_10115-[Lentisphaerales-bacterium]</td></tr>
<tr><td>MDD4099137.1</td><td>AAA+ATPase+wHTH+Alpha+Beta-domain+wHTH+RRM</td><td>Type-3_BR.PglZ.associated.systems</td><td>DUF499*->PglZ->OLD-ABC+TOPRIM-></td><td>267</td><td>Bacteria>PVC_group>Lentisphaerota>Lentisphaeria>unclassified_Lentisphaeria</td><td>GCA_028701535.1</td><td>Lentisphaeria_bacterium</td><td>PHC30_10230</td><td>1..805</td><td>2053569</td><td>MAG:-hypothetical-protein-PHC30_10230,-partial-[Lentisphaeria-bacterium]</td></tr>
<tr><td>NLX24424.1</td><td>AAA+ATPase+wHTH+Alpha+Beta-domain+wHTH+RRM</td><td>Type-3_BR.PglZ.associated.systems</td><td>DUF499*->PglZ->CopG/RHH->PIN->RNaseH->METHYLASE-></td><td>867</td><td>Bacteria>PVC_group>Lentisphaerota>unclassified_Lentisphaerota</td><td>GCA_012729265.1</td><td>Lentisphaerota_bacterium</td><td>GXY61_00470</td><td>264..2867</td><td>1932692</td><td>hypothetical-protein-[Lentisphaerae-bacterium]</td></tr>
<tr><td>NLG01400.1</td><td>AAA+ATPase+wHTH+Alpha+Beta-domain+wHTH+RRM</td><td>Type-3_BR.PglZ.associated.systems</td><td><-TUDOR+SF1-SF2-Helicase+iREase+CTD<-iSTAND<-SF1-SF2-Helicase<-METHYLASE<-OLD-ABC<-PglZ<-DUF499*</td><td>935</td><td>Bacteria>PVC_group>Lentisphaerota>unclassified_Lentisphaerota</td><td>GCA_012515415.1</td><td>Lentisphaerota_bacterium</td><td>GX565_14795</td><td>16284..19091</td><td>1932692</td><td>DUF499-domain-containing-protein-[Lentisphaerae-bacterium]</td></tr>
<tr><td>KAA0250882.1</td><td>AAA+ATPase+wHTH+Alpha+Beta-domain+wHTH+RRM</td><td>Type-3_BR.PglZ.associated.systems</td><td>PIN-><-REase<-?||DUF499*->PglZ->METHYLASE->?->TUDOR+SF1-SF2-Helicase+iREase+CTD->DDE_5-></td><td>875</td><td>Bacteria>PVC_group>Planctomycetota>Candidatus_Brocadiia>Candidatus_Brocadiales</td><td>GCA_008363445.1</td><td>Candidatus_Jettenia_sp._AMX1</td><td>EDM77_03490</td><td>87746..90373</td><td>2293637</td><td>MAG:-hypothetical-protein-EDM77_03490-[Candidatus-Jettenia-sp.-AMX1]</td></tr>
<tr><td>WP_007220935.1</td><td>AAA+ATPase+wHTH+Alpha+Beta-domain+wHTH+RRM</td><td>Type-3_BR.PglZ.associated.systems</td><td>PIN-><-REase<-?||DUF499*->PglZ->METHYLASE->?->TUDOR+SF1-SF2-Helicase+iREase+CTD->DDE_5-></td><td>875</td><td>Bacteria>PVC_group>Planctomycetota>Candidatus_Brocadiia>Candidatus_Brocadiales</td><td>GCF_030263775.1</td><td>Candidatus_Jettenia_sp._AMX1</td><td>FBQ94_RS05340</td><td>87273..89900</td><td>2293637</td><td>hypothetical-protein-[Candidatus-Jettenia-sp.-AMX1]</td></tr>
<tr><td>UJS21271.1</td><td>AAA+ATPase+wHTH+Alpha+Beta-domain+wHTH+RRM</td><td>Type-3_BR.PglZ.associated.systems</td><td><-TUDOR+SF1-SF2-Helicase+iREase+CTD<-iSTAND<-METHYLASE<-PemK_toxin<-?<-PglZ<-DUF499*<-?||?-><-?<-HEPN||?-><-Methylase</td><td>876</td><td>Bacteria>PVC_group>Planctomycetota>Candidatus_Brocadiia>Candidatus_Brocadiales</td><td>GCA_021650915.1</td><td>Candidatus_Brocadia_sp.</td><td>L3J18_02875</td><td>619819..622449</td><td>2172549</td><td>MAG:-hypothetical-protein-L3J18_02875-[Candidatus-Brocadia-sp.]</td></tr>
<tr><td>WP_070065823.1</td><td>AAA+ATPase+wHTH+Alpha+Beta-domain+wHTH+RRM</td><td>Type-3_BR.PglZ.associated.systems</td><td><-METHYLASE-p<-HEPN<-PolB<-PglZ<-DUF499*</td><td>872</td><td>Bacteria>PVC_group>Planctomycetota>Candidatus_Brocadiia>Candidatus_Brocadiales</td><td>GCF_001753675.2</td><td>Candidatus_Brocadia_sapporoensis</td><td>BIY37_RS00115</td><td>2409..5027</td><td>392547</td><td>hypothetical-protein-[Candidatus-Brocadia-sapporoensis]</td></tr>
<tr><td>OQZ00352.1</td><td>AAA+ATPase+wHTH+Alpha+Beta-domain+wHTH+RRM</td><td>Type-3_BR.PglZ.associated.systems</td><td><-METHYLASE<-PemK_toxin<-?<-PolB<-PglZ<-DUF499*<-?||?-><-?<-HEPN</td><td>876</td><td>Bacteria>PVC_group>Planctomycetota>Candidatus_Brocadiia>Candidatus_Brocadiales</td><td>GCA_002050315.1</td><td>Candidatus_Brocadia_sp._UTAMX2</td><td>B6D35_06835</td><td>5677..8307</td><td>1961114</td><td>MAG:-hypothetical-protein-B6D35_06835-[Candidatus-Brocadia-sp.-UTAMX2]</td></tr>
<tr><td>KKO19952.1</td><td>AAA+ATPase+wHTH+Alpha+Beta-domain+wHTH+RRM</td><td>Type-3_BR.PglZ.associated.systems</td><td><-METHYLASE<-PemK_toxin<-?<-PglZ<-DUF499<-DUF499*<-?||?-><-?<-HEPN</td><td>117</td><td>Bacteria>PVC_group>Planctomycetota>Candidatus_Brocadiia>Candidatus_Brocadiales</td><td>GCA_000987375.1</td><td>Candidatus_Brocadia_fulgida</td><td>BROFUL_01344</td><td>23372..23725</td><td>380242</td><td>hypothetical-protein-BROFUL_01344-[Candidatus-Brocadia-fulgida]</td></tr>
<tr><td>KKO19951.1</td><td>AAA+ATPase+wHTH+Alpha+Beta-domain+wHTH+RRM</td><td>Type-3_BR.PglZ.associated.systems</td><td><-iSTAND<-?<-METHYLASE<-PemK_toxin<-?<-PglZ<-DUF499*<-DUF499<-?||?-><-?<-HEPN</td><td>741</td><td>Bacteria>PVC_group>Planctomycetota>Candidatus_Brocadiia>Candidatus_Brocadiales</td><td>GCA_000987375.1</td><td>Candidatus_Brocadia_fulgida</td><td>BROFUL_01343</td><td>21096..23321</td><td>380242</td><td>hypothetical-protein-BROFUL_01343-[Candidatus-Brocadia-fulgida]</td></tr>
<tr><td>MCC6950816.1</td><td>AAA+ATPase+wHTH+Alpha+Beta-domain+wHTH+RRM</td><td>Type-3_BR.PglZ.associated.systems</td><td>Methyltransf_11->?->?->?->?->XIS-HTH->DUF499*->PglZ-></td><td>853</td><td>Bacteria>PVC_group>Planctomycetota>Phycisphaerae>Phycisphaerales</td><td>GCA_020853975.1</td><td>Phycisphaerales_bacterium</td><td>IT433_05155</td><td>9138..11699</td><td>2052180</td><td>hypothetical-protein-[Phycisphaerales-bacterium]</td></tr>
<tr><td>MCL4198878.1</td><td>AAA+ATPase+wHTH+Alpha+Beta-domain+wHTH+RRM</td><td>Type-3_BR.PglZ.associated.systems</td><td>METHYLASE-p->XIS-HTH->DUF499*->PglZ->METHYLASE->iSTAND->TUDOR+SF1-SF2-Helicase+iREase+CTD->Tox-RelE->HTH+MPTase-></td><td>904</td><td>Bacteria>PVC_group>Planctomycetota>Phycisphaerae>Phycisphaerales</td><td>GCA_023382955.1</td><td>Phycisphaerales_bacterium</td><td>KJZ69_15430</td><td>54817..57531</td><td>2052180</td><td>hypothetical-protein-[Phycisphaerales-bacterium]</td></tr>
<tr><td>MBX3356420.1</td><td>AAA+ATPase+wHTH+Alpha+Beta-domain+wHTH+RRM</td><td>Type-3_BR.PglZ.associated.systems</td><td><-TUDOR+SF1-SF2-Helicase+iREase+CTD<-iSTAND<-METHYLASE<-PglZ<-DUF499*<-XIS-HTH<-?<-?<-Methyltransf_11</td><td>909</td><td>Bacteria>PVC_group>Planctomycetota>Phycisphaerae>Phycisphaerales</td><td>GCA_019636735.1</td><td>Phycisphaeraceae_bacterium</td><td>KF724_12060</td><td>20834..23563</td><td>2026777</td><td>hypothetical-protein-[Phycisphaeraceae-bacterium]</td></tr>
<tr><td>MCC6428911.1</td><td>AAA+ATPase+wHTH+Alpha+Beta-domain+wHTH+RRM</td><td>Type-3_BR.PglZ.associated.systems</td><td><-TUDOR+SF1-SF2-Helicase+iREase+CTD<-iSTAND<-METHYLASE<-PglZ<-DUF499*<-XIS-HTH</td><td>910</td><td>Bacteria>PVC_group>Planctomycetota>Phycisphaerae>Phycisphaerales</td><td>GCA_020847535.1</td><td>Phycisphaerales_bacterium</td><td>IT435_19080</td><td>7831..10563</td><td>2052180</td><td>DUF499-domain-containing-protein-[Phycisphaerales-bacterium]</td></tr>
<tr><td>MBX3410402.1</td><td>AAA+ATPase+wHTH+Alpha+Beta-domain+wHTH+RRM</td><td>Type-3_BR.PglZ.associated.systems</td><td><-TUDOR+SF1-SF2-Helicase+iREase+CTD<-iSTAND<-METHYLASE<-HEPN<-PglZ<-DUF499*</td><td>840</td><td>Bacteria>PVC_group>Planctomycetota>Phycisphaerae>Phycisphaerales</td><td>GCA_019636575.1</td><td>Phycisphaeraceae_bacterium</td><td>KF859_11020</td><td>72615..75137</td><td>2026777</td><td>hypothetical-protein-[Phycisphaeraceae-bacterium]</td></tr>
<tr><td>MBL9001727.1</td><td>AAA+ATPase+wHTH+Alpha+Beta-domain+wHTH+RRM</td><td>Type-3_BR.PglZ.associated.systems</td><td>Phage_integrase->DUF499*->PglZ->METHYLASE->iSTAND->TUDOR+SF1-SF2-Helicase+iREase+CTD-><-SF1-SF2-Helicase<-REase</td><td>915</td><td>Bacteria>PVC_group>Planctomycetota>Phycisphaerae>unclassified_Phycisphaerae</td><td>GCA_016793105.1</td><td>Phycisphaerae_bacterium</td><td>JNK25_11385</td><td>2677..5424</td><td>2026778</td><td>DUF499-domain-containing-protein-[Phycisphaerae-bacterium]</td></tr>
<tr><td>MBP7052835.1</td><td>AAA+ATPase+wHTH+Alpha+Beta-domain+wHTH+RRM</td><td>Type-3_BR.PglZ.associated.systems</td><td><-ZnR<-?<-TUDOR+SF1-SF2-Helicase+iREase+CTD<-iSTAND<-METHYLASE<-PglZ<-DUF499*<-HerA/FtsK</td><td>687</td><td>Bacteria>PVC_group>Planctomycetota>Phycisphaerae>unclassified_Phycisphaerae</td><td>GCA_017993835.1</td><td>Phycisphaerae_bacterium</td><td>KBE65_17650</td><td>40341..42404</td><td>2026778</td><td>hypothetical-protein-[Phycisphaerae-bacterium]</td></tr>
<tr><td>MBX3394312.1</td><td>AAA+ATPase+wHTH+Alpha+Beta-domain+wHTH+RRM</td><td>Type-3_BR.PglZ.associated.systems</td><td><-SF1-SF2-Helicase<-OLD-ABC+TOPRIM<-TUDOR+SF1-SF2-Helicase+iREase+CTD<-iSTAND<-METHYLASE<-PglZ<-DUF499*<-?<-Phage_int_SAM_4+Phage_integrase||ABC-ATPase+SbcC-></td><td>939</td><td>Bacteria>PVC_group>Planctomycetota>Phycisphaerae>unclassified_Phycisphaerae</td><td>GCA_019636475.1</td><td>Phycisphaerae_bacterium</td><td>KF841_02990</td><td>770287..773106</td><td>2026778</td><td>DUF499-domain-containing-protein-[Phycisphaerae-bacterium]</td></tr>
<tr><td>HNQ22499.1</td><td>AAA+ATPase+wHTH+Alpha+Beta-domain+wHTH+RRM</td><td>Type-3_BR.PglZ.associated.systems</td><td>XIS-HTH->DUF499+RRM*->PglZ->METHYLASE->OLD-ABC+TOPRIM->Cluster82_1clades-></td><td>942</td><td>Bacteria>PVC_group>Planctomycetota>Phycisphaerae>unclassified_Phycisphaerae</td><td>GCA_035343535.1</td><td>Phycisphaerae_bacterium</td><td>PKK06_05335</td><td>59363..62191</td><td>2026778</td><td>Ig-like-domain-containing-protein-[Phycisphaerae-bacterium]</td></tr>
<tr><td>HUT59382.1</td><td>AAA+ATPase+wHTH+Alpha+Beta-domain+wHTH+RRM</td><td>Type-3_BR.PglZ.associated.systems</td><td><-METHYLASE-p<-PglZ<-DUF499+RRM*</td><td>943</td><td>Bacteria>PVC_group>Planctomycetota>Phycisphaerae>unclassified_Phycisphaerae</td><td>GCA_035528055.1</td><td>Phycisphaerae_bacterium</td><td>VNA25_16150</td><td>2420..5251</td><td>2026778</td><td>Ig-like-domain-containing-protein-[Phycisphaerae-bacterium]</td></tr>
<tr><td>HUT61078.1</td><td>AAA+ATPase+wHTH+Alpha+Beta-domain+wHTH+RRM</td><td>Type-3_BR.PglZ.associated.systems</td><td>METHYLASE->Phage_int_SAM_3+Phage_integrase->DUF499+RRM*->PglZ->REase->METHYLASE-></td><td>934</td><td>Bacteria>PVC_group>Planctomycetota>Phycisphaerae>unclassified_Phycisphaerae</td><td>GCA_035528055.1</td><td>Phycisphaerae_bacterium</td><td>VNA25_24790</td><td>14688..17492</td><td>2026778</td><td>DUF499-domain-containing-protein-[Phycisphaerae-bacterium]</td></tr>
<tr><td>MCS7016992.1</td><td>AAA+ATPase+wHTH+Alpha+Beta-domain+wHTH+RRM</td><td>Type-3_BR.PglZ.associated.systems</td><td><-PglZ<-DUF499*<-XIS-HTH<-?||?-><-DUF499-p</td><td>913</td><td>Bacteria>PVC_group>Planctomycetota>Planctomycetia>Gemmatales</td><td>GCA_025056555.1</td><td>Gemmatales_bacterium</td><td>NZM42_12845</td><td>1157..3898</td><td>2838789</td><td>DUF499-domain-containing-protein-[Gemmatales-bacterium]</td></tr>
<tr><td>WP_145058248.1</td><td>AAA+ATPase+wHTH+Alpha+Beta-domain+wHTH+RRM</td><td>Type-3_BR.PglZ.associated.systems</td><td>XIS-HTH->DUF499*->PglZ->METHYLASE->iSTAND->TUDOR+SF1-SF2-Helicase+iREase+CTD->REase->HKD-DNase-></td><td>920</td><td>Bacteria>PVC_group>Planctomycetota>Planctomycetia>Pirellulales</td><td>GCF_007741515.1</td><td>Adhaeretor_mobilis</td><td>HG15A2_RS04465</td><td>1149623..1152385</td><td>1930276</td><td>DUF499-domain-containing-protein-[Adhaeretor-mobilis]</td></tr>
<tr><td>NLX99244.1</td><td>AAA+ATPase+wHTH+Alpha+Beta-domain+wHTH+RRM</td><td>Type-3_BR.PglZ.associated.systems</td><td>XIS-HTH->DUF499*->PglZ->METHYLASE-></td><td>941</td><td>Bacteria>PVC_group>Planctomycetota>Planctomycetia>Pirellulales</td><td>GCA_012728855.1</td><td>Rhodopirellula_sp.</td><td>GXY83_24165</td><td>19794..22619</td><td>2024855</td><td>hypothetical-protein-[Rhodopirellula-sp.]</td></tr>
<tr><td>MCC6493495.1</td><td>AAA+ATPase+wHTH+Alpha+Beta-domain+wHTH+RRM</td><td>Type-3_BR.PglZ.associated.systems</td><td><-XIS-HTH<-BirA-HTH<-TUDOR+SF1-SF2-Helicase+iREase+CTD<-iSTAND<-METHYLASE<-PglZ<-DUF499*<-CHTH||?->?-><-Methyltransf_21</td><td>917</td><td>Bacteria>PVC_group>Planctomycetota>Planctomycetia>Pirellulales</td><td>GCA_020847515.1</td><td>Pirellulales_bacterium</td><td>IT424_10785</td><td>81990..84743</td><td>2715253</td><td>DUF499-domain-containing-protein-[Pirellulales-bacterium]</td></tr>
<tr><td>MCC7337866.1</td><td>AAA+ATPase+wHTH+Alpha+Beta-domain+wHTH+RRM</td><td>Type-3_BR.PglZ.associated.systems</td><td><-TUDOR+SF1-SF2-Helicase+iREase+CTD<-iSTAND<-?<-METHYLASE<-REase<-PglZ<-DUF499*<-HicB+ZnR<-XIS-HTH<-?<-?<-?||REase-></td><td>911</td><td>Bacteria>PVC_group>Planctomycetota>Planctomycetia>Pirellulales</td><td>GCA_020852965.1</td><td>Pirellulaceae_bacterium</td><td>IT422_22490</td><td>89898..92633</td><td>2699754</td><td>DUF499-domain-containing-protein-[Pirellulaceae-bacterium]</td></tr>
<tr><td>MDP1561789.1</td><td>AAA+ATPase+wHTH+Alpha+Beta-domain+wHTH+RRM</td><td>Type-3_BR.PglZ.associated.systems</td><td>MerR-HTH->DUF499+RRM*->PglZ->METHYLASE->iSTAND->TUDOR+SF1-SF2-Helicase+iREase+CTD->SF1-SF2-Helicase-></td><td>912</td><td>Bacteria>PVC_group>Planctomycetota>Planctomycetia>Pirellulales>Pirellulaceae</td><td>GCA_030681235.1</td><td>Pirellulaceae_bacterium</td><td>Q8M16_10355</td><td>1388..4126</td><td>2699754</td><td>DUF499-domain-containing-protein-[Pirellulaceae-bacterium]</td></tr>
<tr><td>MDY0168855.1</td><td>AAA+ATPase+wHTH+Alpha+Beta-domain+wHTH+RRM</td><td>Type-3_BR.PglZ.associated.systems</td><td>DUF499+RRM*->PglZ->METHYLASE->OLD-ABC+TOPRIM->Cluster82_1clades-></td><td>942</td><td>Bacteria>PVC_group>Planctomycetota>Planctomycetia>Pirellulales>Thermoguttaceae</td><td>GCA_034005525.1</td><td>Thermoguttaceae_bacterium</td><td>RBS80_20085</td><td>12580..15408</td><td>2691551</td><td>Ig-like-domain-containing-protein-[Thermoguttaceae-bacterium]</td></tr>
<tr><td>HVA45320.1</td><td>AAA+ATPase+wHTH+Alpha+Beta-domain+wHTH+RRM</td><td>Type-3_BR.PglZ.associated.systems</td><td>DUF499+RRM*->PglZ->METHYLASE-></td><td>944</td><td>Bacteria>PVC_group>Planctomycetota>Planctomycetia>Pirellulales>unclassified_Pirellulales</td><td>GCA_035533075.1</td><td>Pirellulales_bacterium</td><td>VNH11_02945</td><td>20500..23334</td><td>2715253</td><td>hypothetical-protein-[Pirellulales-bacterium]</td></tr>
<tr><td>HEX5447066.1</td><td>AAA+ATPase+wHTH+Alpha+Beta-domain+wHTH+RRM</td><td>Type-3_BR.PglZ.associated.systems</td><td>DUF499+RRM*->PglZ-></td><td>929</td><td>Bacteria>PVC_group>Planctomycetota>Planctomycetia>Pirellulales>unclassified_Pirellulales</td><td>GCA_036279315.1</td><td>Pirellulales_bacterium</td><td>VFW87_24840</td><td>1866..4655</td><td>2715253</td><td>DUF499-domain-containing-protein-[Pirellulales-bacterium]</td></tr>
<tr><td>WP_144995763.1</td><td>AAA+ATPase+wHTH+Alpha+Beta-domain+wHTH+RRM</td><td>Type-3_BR.PglZ.associated.systems</td><td>DUF499*->PglZ->METHYLASE->?->iSTAND-></td><td>913</td><td>Bacteria>PVC_group>Planctomycetota>Planctomycetia>Planctomycetales</td><td>GCF_007750395.1</td><td>Polystyrenella_longa</td><td>Pla110_RS10710</td><td>2895030..2897771</td><td>2528007</td><td>DUF499-domain-containing-protein-[Polystyrenella-longa]</td></tr>
<tr><td>TVP96589.1</td><td>AAA+ATPase+wHTH+Alpha+Beta-domain+wHTH+RRM</td><td>Type-3_BR.PglZ.associated.systems</td><td><-HTH||DUF499*->PglZ->METHYLASE->iSTAND->OLD-ABC->TUDOR+SF1-SF2-Helicase+iREase+CTD-></td><td>918</td><td>Bacteria>PVC_group>Planctomycetota>Planctomycetia>Planctomycetales</td><td>GCA_007694185.1</td><td>Planctomycetaceae_bacterium</td><td>EA381_17020</td><td>1449..4205</td><td>2026779</td><td>DUF499-domain-containing-protein-[Planctomycetaceae-bacterium]</td></tr>
<tr><td>MDR0392109.1</td><td>AAA+ATPase+wHTH+Alpha+Beta-domain+wHTH+RRM</td><td>Type-3_BR.PglZ.associated.systems</td><td><-METHYLASE-p<-PglZ<-DUF499+RRM*</td><td>875</td><td>Bacteria>PVC_group>Planctomycetota>Planctomycetia>Planctomycetales>Planctomycetaceae</td><td>GCA_031255635.1</td><td>Planctomycetaceae_bacterium</td><td>LBH59_09395</td><td>2554..5181</td><td>2026779</td><td>DUF499-domain-containing-protein-[Planctomycetaceae-bacterium]</td></tr>
<tr><td>MBE7508223.1</td><td>AAA+ATPase+wHTH+Alpha+Beta-domain+wHTH+RRM</td><td>Type-3_BR.PglZ.associated.systems</td><td>HEPN-><-TUDOR+SF1-SF2-Helicase+iREase+CTD<-iSTAND<-METHYLASE<-PglZ<-DUF499*<-?<-Phage_int_SAM_4+Phage_integrase<-?<-SF1-SF2-Helicase+REase-p<-SF1-SF2-Helicase</td><td>936</td><td>Bacteria>PVC_group>Planctomycetota>Planctomycetia>unclassified_Planctomycetia</td><td>GCA_015075745.1</td><td>Planctomycetia_bacterium</td><td>HS101_18325</td><td>1153201..1156011</td><td>2052181</td><td>DUF499-domain-containing-protein-[Planctomycetia-bacterium]</td></tr>
<tr><td>MBE7444884.1</td><td>AAA+ATPase+wHTH+Alpha+Beta-domain+wHTH+RRM</td><td>Type-3_BR.PglZ.associated.systems</td><td><-HTH+UbE1<-?<-?||?->?-><-?||DUF499*->PglZ->PolB->HEPN->HKD-DNase+SF1-SF2-Helicase-p->SF1-SF2-Helicase-p-></td><td>592</td><td>Bacteria>PVC_group>Planctomycetota>Planctomycetia>unclassified_Planctomycetia</td><td>GCA_015075005.1</td><td>Planctomycetia_bacterium</td><td>HS132_06425</td><td>1247755..1249533</td><td>2052181</td><td>DUF499-domain-containing-protein-[Planctomycetia-bacterium]</td></tr>
<tr><td>MCC6740104.1</td><td>AAA+ATPase+wHTH+Alpha+Beta-domain+wHTH+RRM</td><td>Type-3_BR.PglZ.associated.systems</td><td><-HerA/FtsK<-?<-TUDOR+SF1-SF2-Helicase+iREase+CTD<-iSTAND<-METHYLASE<-PglZ<-DUF499*<-?<-?<-?||SIGMA-HTH-></td><td>898</td><td>Bacteria>PVC_group>Planctomycetota>Planctomycetia>unclassified_Planctomycetia</td><td>GCA_020849035.1</td><td>Planctomycetia_bacterium</td><td>IT452_13740</td><td>12408..15104</td><td>2052181</td><td>DUF499-domain-containing-protein-[Planctomycetia-bacterium]</td></tr>
<tr><td>MBK7876307.1</td><td>AAA+ATPase+wHTH+Alpha+Beta-domain+wHTH+RRM</td><td>Type-3_BR.PglZ.associated.systems</td><td>DUF4011+SF1-SF2-Helicase+REase-><-TUDOR+SF1-SF2-Helicase+iREase+CTD<-HNH<-iSTAND<-METHYLASE<-PglZ<-DUF499*<-?<-Phage_integrase||SIGMA-HTH-><-?<-Phage_integrase</td><td>912</td><td>Bacteria>PVC_group>Planctomycetota>unclassified_Planctomycetota</td><td>GCA_016714045.1</td><td>Planctomycetota_bacterium</td><td>IPJ77_11210</td><td>547923..550661</td><td>2026780</td><td>DUF499-domain-containing-protein-[Planctomycetota-bacterium]</td></tr>
<tr><td>MBA3936148.1</td><td>AAA+ATPase+wHTH+Alpha+Beta-domain+wHTH+RRM</td><td>Type-3_BR.PglZ.associated.systems</td><td><-OLD-ABC+TOPRIM<-TUDOR+SF1-SF2-Helicase+iREase+CTD<-iSTAND<-METHYLASE<-PglZ<-DUF499*<-XIS-HTH<-EVE</td><td>926</td><td>Bacteria>PVC_group>Planctomycetota>unclassified_Planctomycetota</td><td>GCA_013821615.1</td><td>Planctomycetota_bacterium</td><td>HOX38_01710</td><td>10267..13047</td><td>2026780</td><td>hypothetical-protein-[Planctomycetota-bacterium]</td></tr>
<tr><td>MBI2192009.1</td><td>AAA+ATPase+wHTH+Alpha+Beta-domain+wHTH+RRM</td><td>Type-3_BR.PglZ.associated.systems</td><td><-DUF4326<-?<-TUDOR+SF1-SF2-Helicase+iREase+CTD<-iSTAND<-METHYLASE<-PglZ<-DUF499*<-XIS-HTH</td><td>925</td><td>Bacteria>PVC_group>Planctomycetota>unclassified_Planctomycetota</td><td>GCA_016179685.1</td><td>Planctomycetota_bacterium</td><td>HYU36_08500</td><td>55219..57996</td><td>2026780</td><td>DUF499-domain-containing-protein-[Planctomycetota-bacterium]</td></tr>
<tr><td>MDR1535863.1</td><td>AAA+ATPase+wHTH+Alpha+Beta-domain+wHTH+RRM</td><td>Type-3_BR.PglZ.associated.systems</td><td><-DUF4276<-OLD-ABC+TOPRIM<-PglZ<-DUF499+RRM*</td><td>921</td><td>Bacteria>PVC_group>Planctomycetota>unclassified_Planctomycetota</td><td>GCA_031269935.1</td><td>Planctomycetota_bacterium</td><td>LBU64_12350</td><td>5035..7800</td><td>2026780</td><td>DUF499-domain-containing-protein-[Planctomycetota-bacterium]</td></tr>
<tr><td>MCE0498498.1</td><td>AAA+ATPase+wHTH+Alpha+Beta-domain+wHTH+RRM</td><td>Type-3_BR.PglZ.associated.systems</td><td><-METHYLASE<-Schlafen<-PIN<-CopG/RHH<-PglZ<-DUF499*||?->SMODS->UbE1+JAB->JAB-><-?<-TrwC+SF1-SF2-helicase+TOPRIM</td><td>875</td><td>Bacteria>PVC_group>Verrucomicrobiota>Methylacidiphilae>Methylacidiphilales</td><td>GCA_021272315.1</td><td>Candidatus_Methylacidiphilales_bacterium</td><td>LV481_11190</td><td>15738..18365</td><td>2720631</td><td>hypothetical-protein-[Methylacidiphilales-bacterium]</td></tr>
<tr><td>HEU5078764.1</td><td>AAA+ATPase+wHTH+Alpha+Beta-domain+wHTH+RRM</td><td>Type-3_BR.PglZ.associated.systems</td><td><-HTH_3||DUF499+RRM*->PglZ->METHYLASE->iSTAND->TUDOR+SF1-SF2-Helicase+iREase+CTD-></td><td>919</td><td>Bacteria>PVC_group>Verrucomicrobiota>Opitutae>Opitutales>Opitutaceae</td><td>GCA_035765635.1</td><td>Opitutaceae_bacterium</td><td>VFT72_06105</td><td>3149..5908</td><td>2006848</td><td>DUF499-domain-containing-protein-[Opitutaceae-bacterium]</td></tr>
<tr><td>HEX9782048.1</td><td>AAA+ATPase+wHTH+Alpha+Beta-domain+wHTH+RRM</td><td>Type-3_BR.PglZ.associated.systems</td><td>DUF499+RRM*->PglZ->PIN->Schlafen+HATPase+wHTH->METHYLASE-></td><td>866</td><td>Bacteria>PVC_group>Verrucomicrobiota>Opitutae>Opitutales>Opitutaceae</td><td>GCA_036397335.1</td><td>Opitutaceae_bacterium</td><td>VGA56_04855</td><td>452..3052</td><td>2006848</td><td>hypothetical-protein-[Opitutaceae-bacterium]</td></tr>
<tr><td>MCK9588324.1</td><td>AAA+ATPase+wHTH+Alpha+Beta-domain+wHTH+RRM</td><td>Type-3_BR.PglZ.associated.systems</td><td>ABC-ATPase+SbcC->RecR+TOPRIM->?->DUF499*->PglZ->CopG/RHH->PIN-></td><td>860</td><td>Bacteria>PVC_group>Verrucomicrobiota>Terrimicrobiia>Terrimicrobiales</td><td>GCA_023227995.1</td><td>Terrimicrobiaceae_bacterium</td><td>MOQ93_03045</td><td>3515..6097</td><td>2832348</td><td>ATP-binding-protein-[Terrimicrobiaceae-bacterium]</td></tr>
<tr><td>MCX6926381.1</td><td>AAA+ATPase+wHTH+Alpha+Beta-domain+wHTH+RRM</td><td>Type-3_BR.PglZ.associated.systems</td><td>DUF499*->PglZ->PglZ->METHYLASE-p-></td><td>927</td><td>Bacteria>PVC_group>Verrucomicrobiota>unclassified_Verrucomicrobiota</td><td>GCA_026397835.1</td><td>Verrucomicrobiota_bacterium</td><td>NT154_24715</td><td>381..3164</td><td>2026799</td><td>MAG:-hypothetical-protein-NT154_24715-[Verrucomicrobia-bacterium]</td></tr>
<tr><td>MBT3193990.1</td><td>AAA+ATPase+wHTH+Alpha+Beta-domain+wHTH+RRM</td><td>Type-3_BR.PglZ.associated.systems</td><td><-VirB4-FtsK+HerA<-?<-PglZ<-DUF499*</td><td>524</td><td>Bacteria>PVC_group>Verrucomicrobiota>unclassified_Verrucomicrobiota</td><td>GCA_018645275.1</td><td>Verrucomicrobiota_bacterium</td><td>HN341_15705</td><td>2739..4313</td><td>2026799</td><td>DUF499-domain-containing-protein-[Verrucomicrobiota-bacterium]</td></tr>
<tr><td>MCX6876435.1</td><td>AAA+ATPase+wHTH+Alpha+Beta-domain+wHTH+RRM</td><td>Type-3_BR.PglZ.associated.systems</td><td><-TUDOR+SF1-SF2-Helicase+iREase+CTD<-iSTAND<-HicB<-PIN<-METHYLASE<-PglZ<-DUF499*<-?<-MoxR-AAA</td><td>799</td><td>Bacteria>PVC_group>Verrucomicrobiota>unclassified_Verrucomicrobiota</td><td>GCA_026397725.1</td><td>Verrucomicrobiota_bacterium</td><td>NTW21_21895</td><td>23169..25568</td><td>2026799</td><td>MAG:-hypothetical-protein-NTW21_21895-[Verrucomicrobia-bacterium]</td></tr>
<tr><td>SPE58509.1</td><td>AAA+ATPase+wHTH+Alpha+Beta-domain+wHTH+RRM</td><td>Type-3_BR.PglZ.associated.systems</td><td><-REase||?-><-PglZ<-DUF499||?-><-?<-DUF499*<-?<-?<-?<-wHTH+REase||SF1-SF2-Helicase-></td><td>219</td><td>Bacteria>PVC_group>Verrucomicrobiota>unclassified_Verrucomicrobiota</td><td>GCA_900290355.1</td><td>Verrucomicrobiota_bacterium</td><td>SBV1_2740019</td><td>18156..18815</td><td>2026799</td><td>hypothetical-protein-SBV1_2740019-[Verrucomicrobiota-bacterium]</td></tr>
<tr><td>MCX6865170.1</td><td>AAA+ATPase+wHTH+Alpha+Beta-domain+wHTH+RRM</td><td>Type-3_BR.PglZ.associated.systems</td><td><-PIN<-CopG/RHH<-HEPN<-PglZ<-DUF499*</td><td>867</td><td>Bacteria>PVC_group>Verrucomicrobiota>unclassified_Verrucomicrobiota</td><td>GCA_026397695.1</td><td>Verrucomicrobiota_bacterium</td><td>NTV46_02935</td><td>9386..11989</td><td>2026799</td><td>MAG:-hypothetical-protein-NTV46_02935-[Verrucomicrobia-bacterium]</td></tr>
<tr><td>SPE58503.1</td><td>AAA+ATPase+wHTH+Alpha+Beta-domain+wHTH+RRM</td><td>Type-3_BR.PglZ.associated.systems</td><td><-METHYLASE<-?<-REase||?-><-PglZ<-DUF499*||?-><-?<-DUF499<-?<-?<-wHTH+REase</td><td>817</td><td>Bacteria>PVC_group>Verrucomicrobiota>unclassified_Verrucomicrobiota</td><td>GCA_900290355.1</td><td>Verrucomicrobiota_bacterium</td><td>SBV1_2740016</td><td>15074..17527</td><td>2026799</td><td>conserved-hypothetical-protein-[Verrucomicrobia-bacterium]</td></tr>
<tr><td>HPC60676.1</td><td>AAA+ATPase+wHTH+Alpha+Beta-domain+wHTH+RRM</td><td>Type-3_BR.PglZ.associated.systems</td><td><-PglZ<-DUF499+RRM*</td><td>866</td><td>Bacteria>PVC_group>Verrucomicrobiota>unclassified_Verrucomicrobiota</td><td>GCA_035385365.1</td><td>Verrucomicrobiota_bacterium</td><td>PKX23_08465</td><td>491..3091</td><td>2026799</td><td>hypothetical-protein-[Verrucomicrobiota-bacterium]</td></tr>
<tr><td>HQK02490.1</td><td>AAA+ATPase+wHTH+Alpha+Beta-domain+wHTH+RRM</td><td>Type-3_BR.PglZ.associated.systems</td><td><-METHYLASE-p<-PglZ<-DUF499+RRM*</td><td>921</td><td>Bacteria>PVC_group>Verrucomicrobiota>unclassified_Verrucomicrobiota</td><td>GCA_035419325.1</td><td>Verrucomicrobiota_bacterium</td><td>PLP12_18705</td><td>1930..4695</td><td>2026799</td><td>DUF499-domain-containing-protein-[Verrucomicrobiota-bacterium]</td></tr>
<tr><td>MSU35397.1</td><td>AAA+ATPase+wHTH+Alpha+Beta-domain+wHTH+RRM</td><td>Type-3_BR.PglZ.associated.systems</td><td><-iSTAND<-METHYLASE<-?<-PglZ<-DUF499*<-XIS-HTH||<-CHTH||SAM-methylase-></td><td>890</td><td>Bacteria>PVC_group>Verrucomicrobiota>Verrucomicrobiae>Limisphaerales</td><td>GCA_009695205.1</td><td>Pedosphaera_sp.</td><td>EXS36_09825</td><td>6462..9134</td><td>2024890</td><td>DUF499-domain-containing-protein-[Pedosphaera-sp.]</td></tr>
<tr><td>MDB6022250.1</td><td>AAA+ATPase+wHTH+Alpha+Beta-domain+wHTH+RRM</td><td>Type-3_BR.PglZ.associated.systems</td><td>DUF499+RRM*->PglZ->REase->METHYLASE-></td><td>927</td><td>Bacteria>PVC_group>Verrucomicrobiota>Verrucomicrobiae>Limisphaerales>Limisphaerales_incertae_sedis</td><td>GCA_028293265.1</td><td>Pedosphaera_sp.</td><td>JWQ04_2107</td><td>12796..15579</td><td>2024890</td><td>hypothetical-protein-[Pedosphaera-sp.]</td></tr>
<tr><td>HUR42186.1</td><td>AAA+ATPase+wHTH+Alpha+Beta-domain+wHTH+RRM</td><td>Type-3_BR.PglZ.associated.systems</td><td>WYL+wHTH->REase->DUF499+RRM*->PglZ->METHYLASE->iSTAND->TUDOR+SF1-SF2-Helicase+iREase+CTD-></td><td>896</td><td>Bacteria>PVC_group>Verrucomicrobiota>Verrucomicrobiae>unclassified_Verrucomicrobiae</td><td>GCA_035526825.1</td><td>Verrucomicrobiae_bacterium</td><td>VM240_13560</td><td>40110..42800</td><td>2499141</td><td>DUF499-domain-containing-protein-[Verrucomicrobiae-bacterium]</td></tr>
<tr><td>MDB6140353.1</td><td>AAA+ATPase+wHTH+Alpha+Beta-domain+wHTH+RRM</td><td>Type-3_BR.PglZ.associated.systems</td><td>DUF499*->PglZ->METHYLASE-></td><td>770</td><td>Bacteria>PVC_group>Verrucomicrobiota>Verrucomicrobiae>Verrucomicrobiales</td><td>GCA_028293165.1</td><td>Verrucomicrobiaceae_bacterium</td><td>JWQ94_3425</td><td>1..2314</td><td>2026800</td><td>MAG:-hypothetical-protein-JWQ94_3425,-partial-[Verrucomicrobiaceae-bacterium]</td></tr>
<tr><td>MCX8494650.1</td><td>AAA+ATPase+wHTH+Alpha+Beta-domain+wHTH+RRM</td><td>Type-3_BR.PglZ.associated.systems</td><td><-TUDOR+SF1-SF2-Helicase+iREase+CTD<-iSTAND<-METHYLASE<-PglZ<-DUF499*<-HTH+WYL</td><td>915</td><td>Bacteria>PVC_group>Verrucomicrobiota>Verrucomicrobiae>Verrucomicrobiales</td><td>GCA_026461385.1</td><td>Akkermansiaceae_bacterium</td><td>ORN51_00515</td><td>5120..7867</td><td>2562705</td><td>MAG:-DUF499-domain-containing-protein-[Akkermansiaceae-bacterium]</td></tr>
<tr><td>MDE0219236.1</td><td>AAA+ATPase+wHTH+Alpha+Beta-domain+wHTH+RRM</td><td>Type-3_BR.PglZ.associated.systems</td><td>Aminotran_3-><-?<-Glyoxalase||DUF499*->PglZ->METHYLASE->iSTAND->TUDOR+SF1-SF2-Helicase+iREase+CTD->METHYLASE->MORC+HISKIN-></td><td>898</td><td>Bacteria>Spirochaetota>Spirochaetia>Spirochaetales>Spirochaetaceae</td><td>GCA_028820215.1</td><td>Spirochaetaceae_bacterium</td><td>OXJ90_08190</td><td>3217..5913</td><td>1898206</td><td>MAG:-hypothetical-protein-OXJ90_08190-[Spirochaetaceae-bacterium]</td></tr>
<tr><td>MDR0374944.1</td><td>AAA+ATPase+wHTH+Alpha+Beta-domain+wHTH+RRM</td><td>Type-3_BR.PglZ.associated.systems</td><td>DUF499+RRM*->PglZ-></td><td>936</td><td>Bacteria>Spirochaetota>Spirochaetia>Spirochaetales>Treponemataceae>Treponema</td><td>GCA_031254795.1</td><td>Treponema_sp.</td><td>LBH85_04395</td><td>1373..4183</td><td>166</td><td>hypothetical-protein-[Treponema-sp.]</td></tr>
<tr><td>RKX77153.1</td><td>AAA+ATPase+wHTH+Alpha+Beta-domain+wHTH+RRM</td><td>Type-3_BR.PglZ.associated.systems</td><td><-ORC-CDC6+REase||XIS-HTH->DUF499*->PglZ->PolB->N6_N4_MTase->PolB->HEPN-></td><td>808</td><td>Bacteria>Spirochaetota>unclassified_Spirochaetota</td><td>GCA_003645645.1</td><td>Spirochaetota_bacterium</td><td>DRP87_09965</td><td>8520..10946</td><td>2202144</td><td>hypothetical-protein-DRP87_09965-[Spirochaetes-bacterium]</td></tr>
<tr><td>MDR1622592.1</td><td>AAA+ATPase+wHTH+Alpha+Beta-domain+wHTH+RRM</td><td>Type-3_BR.PglZ.associated.systems</td><td>DUF499+RRM*->PglZ->METHYLASE->HEPN-></td><td>801</td><td>Bacteria>Synergistota>Synergistia>Synergistales>Synergistaceae>unclassified_Synergistaceae</td><td>GCA_031270755.1</td><td>Synergistaceae_bacterium</td><td>LBS00_09480</td><td>624..3029</td><td>2053624</td><td>ATP-binding-protein-[Synergistaceae-bacterium]</td></tr>
</table>

| Protein-ID | DUF499-architecture | System | Gene-Neighbourhoods | Length | Taxonomy | Assembly-ID | Strain | Locus-Tag | Genomic-coordinates | TaxID | Fasta-defline |
|---|---|---|---|---|---|---|---|---|---|---|---|
| MDR2179042.1 | AAA+ATPase+wHTH+Alpha+Beta-domain+wHTH+RRM | Type-3_BR.PglZ.associated.systems | <-TUDOR+SF1-SF2-Helicase+iREase+CTD<-iSTAND<-METHYLASE<-PglZ<-DUF499+RRM* | 902 | Bacteria>Synergistota>Synergistia>Synergistales>Synergistaceae>unclassified_Synergistaceae | GCA_031277095.1 | Synergistaceae_bacterium | LBP21_01915 | 33830..36538 | 2053624 | ATP-binding-protein-[Synergistaceae-bacterium] |
| AER66194.1 | AAA+ATPase+wHTH+Alpha+Beta-domain+wHTH+RRM | Type-3_BR.PglZ.associated.systems | Endonuclease_5->?-><-?\|\|tRNA->tRNA->tRNA->tRNA->tRNA-><-LuxR-HTH\|\|XIS-HTH->DUF499*->PglZ->METHYLASE->iSTAND->TUDOR+SF1-SF2-Helicase+iREase+CTD->?->HerA/FtsK-> | 929 | Bacteria>Synergistota>Synergistia>Synergistales>Thermovirgaceae | GCA_000233775.1 | Thermovirga_lienii_DSM_17291 | Tlie_0459 | 485527..488316 | 580340 | hypothetical-protein-Tlie_0459-[Thermovirga-lienii-DSM-17291] |
| WP_169311583.1 | AAA+ATPase+wHTH+Alpha+Beta-domain+wHTH+RRM | Type-3_BR.PglZ.associated.systems | Endonuclease_5->?-><-?\|\|tRNA->tRNA->tRNA->tRNA->tRNA->\|\|XIS-HTH->DUF499*->PglZ->METHYLASE->iSTAND->TUDOR+SF1-SF2-Helicase+iREase+CTD->?->HerA/FtsK-> | 852 | Bacteria>Synergistota>Synergistia>Synergistales>Thermovirgaceae | GCF_000233775.1 | Thermovirga_lienii | TLIE_RS02355 | 485758..488316 | 336261 | DUF499-domain-containing-protein-[Thermovirga-lienii] |
| WP_176225939.1 | AAA+ATPase+wHTH+Alpha+Beta-domain+wHTH+RRM | Type-3_BR.PglZ.associated.systems | <-REase<-?<-TUDOR+SF1-SF2-Helicase+iREase+CTD<-iSTAND<-METHYLASE<-PglZ<-DUF499*<-XIS-HTH<-NACHT | 871 | Bacteria>Terrabacteria_group>Actinomycetota>Actinomycetota_incertae_sedis>Candidatus_Hakubellales | GCF_013281895.1 | Candidatus_Hakubella_thermalkaliphila | HW426_RS00105 | 21549..24164 | 2754717 | hypothetical-protein-[Candidatus-Hakubanella-thermoalkaliphilus] |
| GFP24749.1 | AAA+ATPase+wHTH+Alpha+Beta-domain+wHTH+RRM | Type-3_BR.PglZ.associated.systems | <-REase<-?<-TUDOR+SF1-SF2-Helicase+iREase+CTD<-iSTAND<-METHYLASE<-PglZ<-DUF499*<-XIS-HTH | 871 | Bacteria>Terrabacteria_group>Actinomycetota>Actinomycetota_incertae_sedis>Candidatus_Hakubellales | GCA_013281935.1 | Candidatus_Hakubella_thermalkaliphila | HKBW3S25_00186 | 11096..13711 | 2754717 | hypothetical-protein-HKBW3S25_00186-[Candidatus-Hakubanella-thermoalkaliphilus] |
| WP_176231858.1 | AAA+ATPase+wHTH+Alpha+Beta-domain+wHTH+RRM | Type-3_BR.PglZ.associated.systems | <-METHYLASE-p<-PglZ<-DUF499* | 334 | Bacteria>Terrabacteria_group>Actinomycetota>Actinomycetota_incertae_sedis>Candidatus_Hakubellales | GCF_013282035.1 | Candidatus_Hakubella_thermalkaliphila | HW438_RS06880 | 2423..3427 | 2754717 | hypothetical-protein,-partial-[Candidatus-Hakubanella-thermoalkaliphilus] |
| GFP41503.1 | AAA+ATPase+wHTH+Alpha+Beta-domain+wHTH+RRM | Type-3_BR.PglZ.associated.systems | <-METHYLASE<-PglZ<-DUF499*<-XIS-HTH<-iSTAND | 871 | Bacteria>Terrabacteria_group>Actinomycetota>Actinomycetota_incertae_sedis>Candidatus_Hakubellales | GCA_013282075.1 | Candidatus_Hakubella_thermalkaliphila | HKBW3C_00628 | 3743..6358 | 2754717 | hypothetical-protein-HKBW3C_00628-[Candidatus-Hakubanella-thermoalkaliphilus] |
| MBC7332924.1 | AAA+ATPase+wHTH+Alpha+Beta-domain+wHTH+RRM | Type-3_BR.PglZ.associated.systems | ZnR+TOPRIM+DnaB->SIGMA-HTH+SIGMA-HTH->tRNA->XIS-HTH->DUF499*->PglZ->HEPN-> | 808 | Bacteria>Terrabacteria_group>Actinomycetota>unclassified_Actinomycetota | GCA_014360385.1 | Actinomycetota_bacterium | H5T85_00410 | 10304..12730 | 2900548 | hypothetical-protein-[Actinomycetia-bacterium] |
| MBT9170876.1 | AAA+ATPase+wHTH+Alpha+Beta-domain+wHTH+RRM | Type-3_BR.PglZ.associated.systems | XIS-HTH->DUF499*->PglZ-> | 871 | Bacteria>Terrabacteria_group>Actinomycetota>unclassified_Actinomycetota | GCA_018725585.1 | Actinomycetota_bacterium | DDT18_01228 | 265..2880 | 2900548 | hypothetical-protein-[Actinomycetia-bacterium] |
| MDP2622174.1 | AAA+ATPase+wHTH+Alpha+Beta-domain+wHTH+RRM | Type-3_BR.PglZ.associated.systems | <-PIN<-PglZ<-DUF499+RRM* | 849 | Bacteria>Terrabacteria_group>Actinomycetota>unclassified_Actinomycetota | GCA_030688105.1 | Actinomycetota_bacterium | Q8Q29_00015 | 1361..3911 | 2900548 | hypothetical-protein-[Actinomycetota-bacterium] |
| HIE51618.1 | AAA+ATPase+wHTH+Alpha+Beta-domain+wHTH+RRM | Type-3_BR.PglZ.associated.systems | <-Exonuc_V_gamma+REase<-?<-iSTAND<-PglZ<-?<-?<-DUF499* | 1018 | Bacteria>Terrabacteria_group>Armatimonadota>unclassified_Armatimonadota | GCA_012961305.1 | Armatimonadota_bacterium | EYP85_07640 | 12987..16043 | 2033014 | hypothetical-protein-[Armatimonadota-bacterium] |
| MEA3401058.1 | AAA+ATPase+wHTH+Alpha+Beta-domain+wHTH+RRM | Type-3_BR.PglZ.associated.systems | <-Phage_int_SAM_3+Phage_integrase\|\|DUF499+RRM*->PglZ->METHYLASE-p-> | 937 | Bacteria>Terrabacteria_group>Armatimonadota>unclassified_Armatimonadota | GCA_034722055.1 | Armatimonadota_bacterium | U9R79_07435 | 5054..7867 | 2033014 | hypothetical-protein-[Armatimonadota-bacterium] |
| WP_270720984.1 | AAA+ATPase+wHTH+Alpha+Beta-domain+wHTH+RRM | Type-3_BR.PglZ.associated.systems | DUF499*->PglZ->METHYLASE->iSTAND->TUDOR+SF1-SF2-Helicase+iREase+CTD-> | 902 | Bacteria>Terrabacteria_group>Bacillota>Bacilli>Bacillales | GCF_027713685.1 | unclassified_Bacillus_cereus_group | PDN27_RS28195 | 18128..20836 | 2750818 | MULTISPECIES:-DUF499-domain-containing-protein-[unclassified-Bacillus-cereus-group] |
| WP_074508870.1 | AAA+ATPase+wHTH+Alpha+Beta-domain+wHTH+RRM | Type-3_BR.PglZ.associated.systems | <-TUDOR+SF1-SF2-Helicase+iREase+CTD<-iSTAND<-METHYLASE<-PglZ<-DUF499*\|\|?-><-McrC-NTD+REase | 920 | Bacteria>Terrabacteria_group>Bacillota>Bacilli>Bacillales | GCF_900099655.1 | Planococcus_glaciei | BLQ32_RS01850 | 378450..381212 | 459472 | DUF499-domain-containing-protein-[Planococcus-glaciei] |
| WP_036201257.1 | AAA+ATPase+wHTH+Alpha+Beta-domain+wHTH+RRM | Type-3_BR.PglZ.associated.systems | <-TUDOR+SF1-SF2-Helicase+iREase+CTD<-iSTAND<-METHYLASE<-PglZ<-DUF499* | 903 | Bacteria>Terrabacteria_group>Bacillota>Bacilli>Bacillales | GCF_002200845.1 | Ureibacillus_sinduriensis | N794_RS07775 | 231423..234134 | 561440 | DUF499-domain-containing-protein-[Ureibacillus-sinduriensis] |
| WP_193672744.1 | AAA+ATPase+wHTH+Alpha+Beta-domain+wHTH+RRM | Type-3_BR.PglZ.associated.systems | <-Tn7_Tnp_TnsA_N+Tn7_Tnp_TnsA_C<-HTH<-TUDOR+SF1-SF2-Helicase+iREase+CTD<-iSTAND<-METHYLASE<-PglZ<-DUF499*<-?<-RNB+RNase<-ABhydrolase | 904 | Bacteria>Terrabacteria_group>Bacillota>Bacilli>Bacillales | GCF_015069665.1 | Bacillus_cereus | FTO65_RS14310 | 52863..55577 | 1396 | DUF499-domain-containing-protein-[Bacillus-cereus] |
| WP_002203283.1 | AAA+ATPase+wHTH+Alpha+Beta-domain+wHTH+RRM | Type-3_BR.PglZ.associated.systems | <-SbcC<-SIR2+SIR2\|\|Arm-DNA-bind_4+Phage_int_SAM_3+Phage_integrase->HHH_4+SF1-SF2-Helicase+REase-><-DDE_Tnp_1\|\|?->DUF499*->PglZ->METHYLASE->iSTAND->TUDOR+SF1-SF2-Helicase+iREase+CTD-> | 902 | Bacteria>Terrabacteria_group>Bacillota>Bacilli>Bacillales | GCF_918189065.1 | Bacillus_cereus | QOY75_RS26940 | 41646..44354 | 1396 | hypothetical-protein-[Bacillus-cereus] |
| WP_242272082.1 | AAA+ATPase+wHTH+Alpha+Beta-domain+wHTH+RRM | Type-3_BR.PglZ.associated.systems | <-Phage_int_SAM_1+Phage_integrase\|\|?-><-TUDOR+SF1-SF2-Helicase+iREase+CTD<-iSTAND<-METHYLASE<-PglZ<-DUF499*<-?<-HHH_4+SF1-SF2-Helicase+REase<-Arm-DNA-bind_4+Phage_int_SAM_3+Phage_integrase<-XIS-HTH | 902 | Bacteria>Terrabacteria_group>Bacillota>Bacilli>Bacillales | GCF_022664325.1 | Bacillus_cereus_group_sp._BfR-BA-01310 | MR109_RS10995 | 101157..103865 | 2920287 | DUF499-domain-containing-protein-[Bacillus-cereus-group-sp.-BfR-BA-01310] |
| MED4225398.1 | AAA+ATPase+wHTH+Alpha+Beta-domain+wHTH+RRM | Type-3_BR.PglZ.associated.systems | XIS-HTH->HNH->DUF499+RRM*->PglZ->METHYLASE->iSTAND->TUDOR+SF1-SF2-Helicase+iREase+CTD-> | 903 | Bacteria>Terrabacteria_group>Bacillota>Bacilli>Bacillales>Bacillaceae | GCA_036223585.1 | Neobacillus_cucumis | P4670_RS08995 | 33546..36257 | 1740721 | DUF499-domain-containing-protein-[Neobacillus-cucumis] |
| WP_308999803.1 | AAA+ATPase+wHTH+Alpha+Beta-domain+wHTH+RRM | Type-3_BR.PglZ.associated.systems | <-TUDOR+SF1-SF2-Helicase+iREase+CTD<-iSTAND<-METHYLASE<-PglZ<-DUF499+RRM*<-Phage_integrase | 913 | Bacteria>Terrabacteria_group>Bacillota>Bacilli>Bacillales>Paenibacillaceae | GCF_031157315.1 | Paenibacillus_sp._LHD-38 | RCO25_RS30280 | 27946..30687 | 3072143 | DUF499-domain-containing-protein-[Paenibacillus-sp.-LHD-38] |
| OPX92061.1 | AAA+ATPase+wHTH+Alpha+Beta-domain+wHTH+RRM | Type-3_BR.PglZ.associated.systems | REase->Mu-transpos_C->TnsC-ATPase->?->?->?->DUF499*->PglZ->METHYLASE->iSTAND->TUDOR+SF1-SF2-Helicase+iREase+CTD-><-SbcC | 914 | Bacteria>Terrabacteria_group>Bacillota>Clostridia>Eubacteriales | GCA_002067515.1 | Pelotomaculum_sp._PtaB.Bin013 | A4E52_00165 | 13663..16407 | 1811692 | MAG:-hypothetical-protein-A4E52_00165-[Pelotomaculum-sp.-PtaB.Bin013] |
| WP_277443849.1 | AAA+ATPase+wHTH+Alpha+Beta-domain+wHTH+RRM | Type-3_BR.PglZ.associated.systems | <-TUDOR+SF1-SF2-Helicase+iREase+CTD<-?<-METHYLASE<-PglZ<-DUF499*<-?<-?<-?<-TnsC-ATPase<-Mu-transpos_C<-REase | 914 | Bacteria>Terrabacteria_group>Bacillota>Clostridia>Eubacteriales | GCF_029478095.1 | Pelotomaculum_isophthalicicum | L7E55_RS09165 | 24379..27123 | 342448 | hypothetical-protein-[Pelotomaculum-isophthalicicum] |
| MDD3269499.1 | AAA+ATPase+wHTH+Alpha+Beta-domain+wHTH+RRM | Type-3_BR.PglZ.associated.systems | <-METHYLASE<-PglZ<-DUF499*<-Peptidase_S8<-Classical-AAA<-HEPN | 921 | Bacteria>Terrabacteria_group>Bacillota>Clostridia>Eubacteriales | GCA_028692415.1 | Syntrophomonadaceae_bacterium | PHX14_09275 | 4941..7706 | 2093811 | MAG:-DUF499-domain-containing-protein-[Syntrophomonadaceae-bacterium] |
| MDZ4042526.1 | AAA+ATPase+wHTH+Alpha+Beta-domain+wHTH+RRM | Type-3_BR.PglZ.associated.systems | <-TUDOR+SF1-SF2-Helicase+iREase+CTD<-iSTAND<-METHYLASE<-PglZ<-DUF499+RRM*\|\|<-Phage_int_SAM_3+Phage_integrase | 941 | Bacteria>Terrabacteria_group>Bacillota>Clostridia>Eubacteriales>unclassified_Eubacteriales | GCA_034366365.1 | Eubacteriales_bacterium | U1D96_03415 | 10437..13262 | 2831996 | hypothetical-protein-[Eubacteriales-bacterium] |
| WP_282706327.1 | AAA+ATPase+wHTH+Alpha+Beta-domain+wHTH+RRM | Type-3_BR.PglZ.associated.systems | DUF499*->PglZ->METHYLASE->iSTAND-> | 820 | Bacteria>Terrabacteria_group>Bacillota>Clostridia>Halanaerobiales | - | Fuchsiella_alkaliacetigena | MWH30_RS12585 | 1368..3830 | 957042 | DUF499-domain-containing-protein-[Fuchsiella-alkaliacetigena] |
| MCK8825903.1 | AAA+ATPase+wHTH+Alpha+Beta-domain+wHTH+RRM | Type-3_BR.PglZ.associated.systems | DUF499*->PglZ->METHYLASE->iSTAND-> | 826 | Bacteria>Terrabacteria_group>Bacillota>Clostridia>Halanaerobiales | GCA_023227765.1 | Fuchsiella_alkaliacetigena | MWH30_12615 | 1350..3830 | 957042 | DUF499-domain-containing-protein-[Fuchsiella-alkaliacetigena] |
| WP_133531357.1 | AAA+ATPase+wHTH+Alpha+Beta-domain+wHTH+RRM | Type-3_BR.PglZ.associated.systems | <-METHYLASE<-PglZ<-DUF499* | 823 | Bacteria>Terrabacteria_group>Bacillota>Clostridia>Halanaerobiales | GCF_004362725.1 | Halanaerobium_saccharolyticum | C7957_RS14060 | 3188..5659 | 43595 | DUF499-domain-containing-protein-[Halanaerobium-saccharolyticum] |
| WP_248664445.1 | AAA+ATPase+wHTH+Alpha+Beta-domain+wHTH+RRM | Type-3_BR.PglZ.associated.systems | DUF499+RRM*->PglZ->METHYLASE-> | 826 | Bacteria>Terrabacteria_group>Bacillota>Clostridia>Halanaerobiales>Halobacteroidaceae | GCF_023227765.1 | Fuchsiella_alkaliacetigena | MWH30_RS12585 | 1350..3830 | 957042 | DUF499-domain-containing-protein-[Fuchsiella-alkaliacetigena] |
| WP_161484964.1 | AAA+ATPase+wHTH+Alpha+Beta-domain+wHTH+RRM | Type-3_BR.PglZ.associated.systems | <-TUDOR+SF1-SF2-Helicase+iREase+CTD<-iSTAND<-METHYLASE<-PglZ<-DUF499*\|\|?-><-REase<-?<-?<-HEPN | 762 | Bacteria>Terrabacteria_group>Bacillota>Clostridia>Moorellales | GCF_001594015.1 | Moorella_mulderi | MOMUL_RS05190 | 10708..12996 | 202604 | DUF499-domain-containing-protein-[Moorella-mulderi] |
| KYH32456.1 | AAA+ATPase+wHTH+Alpha+Beta-domain+wHTH+RRM | Type-3_BR.PglZ.associated.systems | <-TUDOR+SF1-SF2-Helicase+iREase+CTD<-iSTAND<-METHYLASE<-PglZ<-DUF499*\|\|?-><-REase | 932 | Bacteria>Terrabacteria_group>Bacillota>Clostridia>Moorellales | GCA_001594015.1 | Moorella_mulderi_DSM_14980 | MOMUL_10570 | 10708..13506 | 1122241 | hypothetical-protein-MOMUL_10570-[Moorella-mulderi-DSM-14980] |
| MDU2146556.1 | AAA+ATPase+wHTH+Alpha+Beta-domain+wHTH+RRM | Type-3_BR.PglZ.associated.systems | HNH-><-iSTAND<-METHYLASE<-PglZ<-DUF499+RRM* | 888 | Bacteria>Terrabacteria_group>Bacillota>Clostridia>Peptostreptococcales>Peptostreptococcaceae | GCA_032478995.1 | Paraclostridium_sordellii | E7E58_01070 | 226397..229063 | 1505 | hypothetical-protein-[Paeniclostridium-sordellii] |
| MBO5955453.1 | AAA+ATPase+wHTH+Alpha+Beta-domain+wHTH+RRM | Type-3_BR.PglZ.associated.systems | XIS-HTH->DUF499*->?->Schlafen+HATPase+HTH->iSTAND->TUDOR+SF1-SF2-Helicase+iREase+CTD-> | 191 | Bacteria>Terrabacteria_group>Bacillota>Clostridia>unclassified_Clostridia | GCA_017631935.1 | Clostridia_bacterium | J6Q10_01475 | 699..1274 | 2044939 | hypothetical-protein-[Clostridia-bacterium] |
| MBC7342273.1 | AAA+ATPase+wHTH+Alpha+Beta-domain+wHTH+RRM | Type-3_BR.PglZ.associated.systems | <-TIR<-TUDOR+SF1-SF2-Helicase+iREase+CTD<-iSTAND<-METHYLASE<-PglZ<-DUF499* | 855 | Bacteria>Terrabacteria_group>Bacillota>Clostridia>unclassified_Clostridia | GCA_014360065.1 | Clostridia_bacterium | H5U02_07455 | 13503..16070 | 2044939 | DUF499-domain-containing-protein-[Clostridia-bacterium] |
| NPV69263.1 | AAA+ATPase+wHTH+Alpha+Beta-domain+wHTH+RRM | Type-3_BR.PglZ.associated.systems | <-TUDOR+SF1-SF2-Helicase+iREase+CTD<-iSTAND<-METHYLASE<-PglZ<-DUF499*\|\|?-><-?\|\|?->?->?-><-SIGMA-HTH | 936 | Bacteria>Terrabacteria_group>Bacillota>unclassified_Bacillota | GCA_013178305.1 | Bacillota_bacterium | HPY55_01285 | 252444..255254 | 1879010 | hypothetical-protein-[Firmicutes-bacterium] |
| TEU12999.1 | AAA+ATPase+wHTH+Alpha+Beta-domain+wHTH+RRM | Type-3_BR.PglZ.associated.systems | PolB->HEPN->DUF499*->PglZ->?->PIN->METHYLASE-p-> | 1031 | Bacteria>Terrabacteria_group>Chloroflexota>Anaerolineae>Anaerolineales | GCA_004377365.1 | Anaerolineales_bacterium | E3J21_19760 | 4931..8026 | 2073117 | hypothetical-protein-E3J21_19760-[Anaerolineales-bacterium] |
| MBE7511468.1 | AAA+ATPase+wHTH+Alpha+Beta-domain+wHTH+RRM | Type-3_BR.PglZ.associated.systems | <-TUDOR+SF1-SF2-Helicase+iREase+CTD<-iSTAND<-METHYLASE<-PglZ<-DUF499*<-?<-VirD4-TraG+VirD4-FtsK<-?<-MetJarc-HTH | 918 | Bacteria>Terrabacteria_group>Chloroflexota>Anaerolineae>Anaerolineales | GCA_015075625.1 | Anaerolineales_bacterium | HS103_01450 | 332970..335726 | 2073117 | hypothetical-protein-[Anaerolineales-bacterium] |
| MBV6465976.1 | AAA+ATPase+wHTH+Alpha+Beta-domain+wHTH+RRM | Type-3_BR.PglZ.associated.systems | <-METHYLASE<-PemK_toxin<-?<-PglZ<-DUF499* | 876 | Bacteria>Terrabacteria_group>Chloroflexota>Anaerolineae>Anaerolineales | GCA_019187435.1 | Anaerolineales_bacterium | PGMFKBFP_01273 | 5094..7724 | 2073117 | hypothetical-protein-[Anaerolineales-bacterium] |
| MBE2220374.1 | AAA+ATPase+wHTH+Alpha+Beta-domain+wHTH+RRM | Type-3_BR.PglZ.associated.systems | <-TUDOR+SF1-SF2-Helicase+iREase+CTD<-iSTAND<-METHYLASE<-PglZ<-DUF499*\|\|?->?->URI-DNase-> | 939 | Bacteria>Terrabacteria_group>Chloroflexota>Anaerolineae>unclassified_Anaerolineae | GCA_014879345.1 | Anaerolineae_bacterium | IAF02_02480 | 7963..10782 | 2052143 | DUF499-domain-containing-protein-[Anaerolineae-bacterium] |
| MBL7062830.1 | AAA+ATPase+wHTH+Alpha+Beta-domain+wHTH+RRM | Type-3_BR.PglZ.associated.systems | <-Schlafen+HATPase+HTH<-PglZ<-DUF499*<-?\|\|?->35exo+TPR-> | 743 | Bacteria>Terrabacteria_group>Chloroflexota>Anaerolineae>unclassified_Anaerolineae | GCA_016783285.1 | Anaerolineae_bacterium | ISS49_01315 | 7640..9871 | 2052143 | DUF499-domain-containing-protein-[Anaerolineae-bacterium] |
| MBN2391962.1 | AAA+ATPase+wHTH+Alpha+Beta-domain+wHTH+RRM | Type-3_BR.PglZ.associated.systems | <-GDResolvase+Recombinase<-?\|\|?->DUF499*->PglZ->METHYLASE->iSTAND->TUDOR+SF1-SF2-Helicase+iREase+CTD->SF1-SF2-Helicase-> | 895 | Bacteria>Terrabacteria_group>Chloroflexota>Anaerolineae>unclassified_Anaerolineae | GCA_016934645.1 | Anaerolineae_bacterium | JXR84_14635 | 2138..4825 | 2052143 | AAA-family-ATPase-[Anaerolineae-bacterium] |
| WP_129628416.1 | AAA+ATPase+wHTH+Alpha+Beta-domain+wHTH+RRM | Type-3_BR.PglZ.associated.systems | <-ACET+ACET<-ABhydrolase<-?<-?\|\|STAS->DUF499*->PglZ->Schlafen+HATPase+HTH->METHYLASE-p-> | 911 | Bacteria>Terrabacteria_group>Chloroflexota>Chloroflexia>Chloroflexales | GCF_004138135.1 | Candidatus_Oscillochloris_fontis | EKD03_RS05535 | 109811..112546 | 2496868 | DUF499-domain-containing-protein-[Candidatus-Oscillochloris-fontis] |
| MCL0098792.1 | AAA+ATPase+wHTH+Alpha+Beta-domain+wHTH+RRM | Type-3_BR.PglZ.associated.systems | REase-><-?\|\|DUF499*->PglZ->HicB_lk_antitox->HicA_toxin->METHYLASE-p-> | 862 | Bacteria>Terrabacteria_group>Chloroflexota>Dehalococcoidia>unclassified_Dehalococcoidia | GCA_023270555.1 | Dehalococcoidia_bacterium | M1016_02910 | 5866..8454 | 2026734 | ATP-binding-protein-[Dehalococcoidia-bacterium] |
| MCL0078762.1 | AAA+ATPase+wHTH+Alpha+Beta-domain+wHTH+RRM | Type-3_BR.PglZ.associated.systems | Methyltransf_25->XIS-HTH->PIN->?-><-?\|\|DUF499*->PglZ->HicB_lk_antitox->HicA_toxin->METHYLASE-> | 862 | Bacteria>Terrabacteria_group>Chloroflexota>Dehalococcoidia>unclassified_Dehalococcoidia | GCA_023271435.1 | Dehalococcoidia_bacterium | M1056_03750 | 4698..7286 | 2026734 | ATP-binding-protein-[Dehalococcoidia-bacterium] |
| MCL0058578.1 | AAA+ATPase+wHTH+Alpha+Beta-domain+wHTH+RRM | Type-3_BR.PglZ.associated.systems | DUF499*->PglZ->HicB_lk_antitox->HicA_toxin->METHYLASE->?->iSTAND-> | 854 | Bacteria>Terrabacteria_group>Chloroflexota>Dehalococcoidia>unclassified_Dehalococcoidia | GCA_023271585.1 | Dehalococcoidia_bacterium | M1047_02700 | 1..2567 | 2026734 | ATP-binding-protein-[Dehalococcoidia-bacterium] |
| MCL0090092.1 | AAA+ATPase+wHTH+Alpha+Beta-domain+wHTH+RRM | Type-3_BR.PglZ.associated.systems | DUF499*->PglZ->HicB_lk_antitox->HicA_toxin->METHYLASE-> | 847 | Bacteria>Terrabacteria_group>Chloroflexota>Dehalococcoidia>unclassified_Dehalococcoidia | GCA_023271215.1 | Dehalococcoidia_bacterium | M1054_07090 | 1..2545 | 2026734 | ATP-binding-protein-[Dehalococcoidia-bacterium] |
| MCL0095264.1 | AAA+ATPase+wHTH+Alpha+Beta-domain+wHTH+RRM | Type-3_BR.PglZ.associated.systems | <-METHYLASE-p<-METHYLASE-p<-HicA_toxin<-HicB_lk_antitox<-PglZ<-DUF499*\|\|?-><-REase | 862 | Bacteria>Terrabacteria_group>Chloroflexota>Dehalococcoidia>unclassified_Dehalococcoidia | GCA_023271155.1 | Dehalococcoidia_bacterium | M1052_01845 | 9312..11900 | 2026734 | ATP-binding-protein-[Dehalococcoidia-bacterium] |
| MCL0058394.1 | AAA+ATPase+wHTH+Alpha+Beta-domain+wHTH+RRM | Type-3_BR.PglZ.associated.systems | <-METHYLASE<-PglZ<-DUF499*<-ZnR<-?<-TraJ-RHH | 871 | Bacteria>Terrabacteria_group>Chloroflexota>Dehalococcoidia>unclassified_Dehalococcoidia | GCA_023271585.1 | Dehalococcoidia_bacterium | M1047_01715 | 4201..6816 | 2026734 | hypothetical-protein-[Dehalococcoidia-bacterium] |
| MCL0075303.1 | AAA+ATPase+wHTH+Alpha+Beta-domain+wHTH+RRM | Type-3_BR.PglZ.associated.systems | <-METHYLASE<-HicA_toxin<-HicB_lk_antitox<-PglZ<-DUF499*\|\|?-><-REase | 862 | Bacteria>Terrabacteria_group>Chloroflexota>Dehalococcoidia>unclassified_Dehalococcoidia | GCA_023271455.1 | Dehalococcoidia_bacterium | M1017_00200 | 4012..6600 | 2026734 | ATP-binding-protein-[Dehalococcoidia-bacterium] |
| MCL0097131.1 | AAA+ATPase+wHTH+Alpha+Beta-domain+wHTH+RRM | Type-3_BR.PglZ.associated.systems | <-iSTAND<-?<-METHYLASE<-HicA_toxin<-HicB_lk_antitox<-PglZ<-DUF499*<-SAM-methylase<-METHYLASE-p\|\|?-><-?<-PIN<-XIS-HTH | 862 | Bacteria>Terrabacteria_group>Chloroflexota>Dehalococcoidia>unclassified_Dehalococcoidia | GCA_023270595.1 | Dehalococcoidia_bacterium | M1019_01150 | 17311..19899 | 2026734 | ATP-binding-protein-[Dehalococcoidia-bacterium] |
| HEX76886.1 | AAA+ATPase+wHTH+Alpha+Beta-domain+wHTH+RRM | Type-3_BR.PglZ.associated.systems | <-HEPN<-ParB+HNH<-?<-METHYLASE<-PglZ<-DUF499*<-?<-?<-RelE-ParE<-?\|\|Phage_int_SAM_1+Phage_integrase-> | 860 | Bacteria>Terrabacteria_group>Chloroflexota>Dehalococcoidia>unclassified_Dehalococcoidia | GCA_011170395.1 | Dehalococcoidia_bacterium | G4003_00485 | 96503..99085 | 2026734 | hypothetical-protein-[Dehalococcoidia-bacterium] |
| HCF99178.1 | AAA+ATPase+wHTH+Alpha+Beta-domain+wHTH+RRM | Type-3_BR.PglZ.associated.systems | TPR+TPR->?->REase->DUF499*->PglZ->METHYLASE->iSTAND->TUDOR+SF1-SF2-Helicase+iREase+CTD-> | 896 | Bacteria>Terrabacteria_group>Chloroflexota>unclassified_Chloroflexota | GCA_003530725.1 | Chloroflexota_bacterium | DEV93_01385 | 4673..7363 | 2026724 | hypothetical-protein-[Chloroflexi-bacterium] |
| MCL4459193.1 | AAA+ATPase+wHTH+Alpha+Beta-domain+wHTH+RRM | Type-3_BR.PglZ.associated.systems | DUF499*->PglZ->?->RelE-ParE->METHYLASE->iSTAND-><-REase | 862 | Bacteria>Terrabacteria_group>Chloroflexota>unclassified_Chloroflexota | GCA_023382375.1 | Chloroflexota_bacterium | M1577_04720 | 46893..49481 | 2026724 | ATP-binding-protein-[Chloroflexi-bacterium] |
| MBM4461756.1 | AAA+ATPase+wHTH+Alpha+Beta-domain+wHTH+RRM | Type-3_BR.PglZ.associated.systems | <-PIWI<-REase<-iSTAND<-METHYLASE<-?<-PglZ<-DUF499*<-?<-Phage_integrase<-?<-?<-?<-ZnR | 863 | Bacteria>Terrabacteria_group>Chloroflexota>unclassified_Chloroflexota | GCA_016876035.1 | Chloroflexota_bacterium | FJ012_00275 | 64493..67084 | 2026724 | DUF499-domain-containing-protein-[Chloroflexi-bacterium] |
| HDQ70822.1 | AAA+ATPase+wHTH+Alpha+Beta-domain+wHTH+RRM | Type-3_BR.PglZ.associated.systems | <-PglZ<-ParB+HNH<-DUF499-p<-DUF499* | 526 | Bacteria>Terrabacteria_group>Chloroflexota>unclassified_Chloroflexota | GCA_011046405.1 | Chloroflexota_bacterium | ENN19_01860 | 4357..5937 | 2026724 | hypothetical-protein-[Chloroflexota-bacterium] |
| MBM4466103.1 | AAA+ATPase+wHTH+Alpha+Beta-domain+wHTH+RRM | Type-3_BR.PglZ.associated.systems | <-ABC-ATPase+SbcC<-?<-?<-METHYLASE-p\|\|DUF499*->PglZ->?->METHYLASE-> | 870 | Bacteria>Terrabacteria_group>Chloroflexota>unclassified_Chloroflexota | GCA_016875995.1 | Chloroflexota_bacterium | FJ014_11220 | 8188..10800 | 2026724 | DUF499-domain-containing-protein-[Chloroflexi-bacterium] |
| MEA3344886.1 | AAA+ATPase+wHTH+Alpha+Beta-domain+wHTH+RRM | Type-3_BR.PglZ.associated.systems | DUF499+RRM*->PglZ->Schlafen+HATPase+wHTH->METHYLASE-> | 801 | Bacteria>Terrabacteria_group>Chloroflexota>unclassified_Chloroflexota | GCA_034718215.1 | Chloroflexota_bacterium | U9Q78_01340 | 16672..19077 | 2026724 | DUF499-domain-containing-protein-[Chloroflexota-bacterium] |
| WP_190375077.1 | AAA+ATPase+wHTH+Alpha+Beta-domain+wHTH+RRM | Type-3_BR.PglZ.associated.systems | RHH_1->Phage_integrase->Prim-Pol-><-35exo<-?\|\|DUF499*->PglZ->REase->METHYLASE->iSTAND->TUDOR+SF1-SF2-Helicase+iREase+CTD-> | 922 | Bacteria>Terrabacteria_group>Cyanobacteriota/Melainabacteria_group>Cyanobacteriota>Cyanophyceae | GCF_014696055.1 | Leptolyngbya_sp._FACHB-17 | H6G01_RS00340 | 14686..17454 | 2692803 | DUF499-domain-containing-protein-[Leptolyngbya-sp.-FACHB-17] |
| MBW4553282.1 | AAA+ATPase+wHTH+Alpha+Beta-domain+wHTH+RRM | Type-3_BR.PglZ.associated.systems | <-TUDOR+SF1-SF2-Helicase+iREase+CTD<-iSTAND<-METHYLASE-p<-REase<-REase<-PglZ<-DUF499*\|\|35exo-><-?\|\|?->?->?-><-HicB | 912 | Bacteria>Terrabacteria_group>Cyanobacteriota/Melainabacteria_group>Cyanobacteriota>Cyanophyceae | GCA_019359715.1 | Aphanocapsa_sp._GSE-SYN-MK-11-07L | KME35_19540 | 9693..12431 | 2835086 | DUF499-domain-containing-protein-[Aphanocapsa-sp.-GSE-SYN-MK-11-07L] |
| WP_017711371.1 | AAA+ATPase+wHTH+Alpha+Beta-domain+wHTH+RRM | Type-3_BR.PglZ.associated.systems | <-TUDOR+SF1-SF2-Helicase+iREase+CTD<-iSTAND<-METHYLASE-p\|\|<-METHYLASE<-REase<-PglZ<-DUF499*<-?<-?\|\|?->ZnR-> | 922 | Bacteria>Terrabacteria_group>Cyanobacteriota/Melainabacteria_group>Cyanobacteriota>Cyanophyceae | GCF_000332315.1 | Prochlorothrix_hollandica | PRO9006_RS0103850 | 892837..895605 | 1223 | DUF499-domain-containing-protein-[Prochlorothrix-hollandica] |
| OCQ93485.1 | AAA+ATPase+wHTH+Alpha+Beta-domain+wHTH+RRM | Type-3_BR.PglZ.associated.systems | <-TUDOR+SF1-SF2-Helicase+iREase+CTD<-iSTAND<-METHYLASE<-PglZ<-DUF499*\|\|SF1-SF2-Helicase+DUF1998-><-?<-CHTH\|\|?->?->DNA_methylase-> | 953 | Bacteria>Terrabacteria_group>Cyanobacteriota/Melainabacteria_group>Cyanobacteriota>Cyanophyceae | GCA_001698435.1 | Nostoc_sp._MBR_210 | BCD64_00890 | 155099..157960 | 1881015 | hypothetical-protein-BCD64_00890-[Nostoc-sp.-MBR-210] |
| MBS1994692.1 | AAA+ATPase+wHTH+Alpha+Beta-domain+wHTH+RRM | Type-3_BR.PglZ.associated.systems | <-TUDOR+SF1-SF2-Helicase+iREase+CTD<-iSTAND<-METHYLASE<-PglZ<-DUF499*<-XIS-HTH\|\|HEPN-> | 910 | Bacteria>Terrabacteria_group>Cyanobacteriota/Melainabacteria_group>Cyanobacteriota>unclassified_Cyanobacteriota | GCA_018266235.1 | Cyanobacteria_bacterium_SZAS_LIN-2 | JSS86_00220 | 47568..50300 | 2807731 | DUF499-domain-containing-protein-[Cyanobacteria-bacterium-SZAS-LIN-2] |
| WP_019011699.1 | AAA+ATPase+wHTH+Alpha+Beta-domain+wHTH+RRM | Type-3_BR.PglZ.associated.systems | DUF499*->PglZ->METHYLASE->iSTAND->TUDOR+SF1-SF2-Helicase+iREase+CTD-><-DDE_Tnp_1<-zf-IS66+DDE_Tnp_IS66 | 948 | Bacteria>Terrabacteria_group>Deinococcota>Deinococci>Deinococcales | GCF_000378445.1 | Deinococcus_aquatilis | F561_RS0119740 | 5122..7968 | 519440 | DUF499-domain-containing-protein-[Deinococcus-aquatilis] |
| WP_169728956.1 | AAA+ATPase+wHTH+Alpha+Beta-domain+wHTH+RRM | Type-3_BR.PglZ.associated.systems | <-iSTAND<-METHYLASE<-REase<-OLD-ABC<-PglZ<-DUF499*<-XIS-HTH<-?<-?<-GDResolvase+Recombinase+ZnR<-URI-DNase | 756 | Bacteria>Thermodesulfobacteriota>Desulfobacteria>Desulfobacterales>Desulfatirhabdiaceae | GCF_000429925.1 | Desulfatirhabdium_butyrativorans | G492_RS0112005 | 53087..55357 | 340467 | DUF499-domain-containing-protein-[Desulfatirhabdium-butyrativorans] |
| MCF8075075.1 | AAA+ATPase+wHTH+Alpha+Beta-domain+wHTH+RRM | Type-3_BR.PglZ.associated.systems | <-METHYLASE<-PglZ<-DUF499*<-?<-OLD-ABC\|\|?-><-?\|\|?->Methylase-> | 939 | Bacteria>Thermodesulfobacteriota>Desulfobacteria>Desulfobacterales>Desulfobacteraceae | GCA_021737305.1 | Desulfotignum_sp. | K9K87_02780 | 4449..7268 | 2049434 | ATP-binding-protein-[Desulfotignum-sp.] |
| MBF0302150.1 | AAA+ATPase+wHTH+Alpha+Beta-domain+wHTH+RRM | Type-3_BR.PglZ.associated.systems | <-iSTAND<-METHYLASE-p<-METHYLASE<-REase<-OLD-ABC<-PglZ<-DUF499*\|\|SF1-SF2-Helicase-> | 934 | Bacteria>Thermodesulfobacteriota>Desulfobacteria>Desulfobacterales>Desulfobacteraceae | GCA_015232035.1 | Desulfamplus_sp. | HQK73_03850 | 16841..19645 | 2006184 | DUF499-domain-containing-protein-[Desulfamplus-sp.] |
| WP_084067311.1 | AAA+ATPase+wHTH+Alpha+Beta-domain+wHTH+RRM | Type-3_BR.PglZ.associated.systems | <-iSTAND<-METHYLASE<-PglZ<-RelE-ParE<-OLD-ABC<-ParB+HNH<-DUF499*\|\|?-><-tRNA<-MPTase<-HTH_31<-RVT<-?\|\|GDResolvase+HIN-HTH-> | 923 | Bacteria>Thermodesulfobacteriota>Desulfobacteria>Desulfobacterales>Desulfobacteraceae | GCF_900176365.1 | Desulfocicer_vacuolatum | B9A32_RS05115 | 100507..103278 | 2298 | DUF499-domain-containing-protein-[Desulfocicer-vacuolatum] |
| CBX30019.1 | AAA+ATPase+wHTH+Alpha+Beta-domain+wHTH+RRM | Type-3_BR.PglZ.associated.systems | <-TUDOR+SF1-SF2-Helicase+iREase+CTD<-iSTAND<-METHYLASE<-PglZ<-DUF499*\|\|?-><-TOPRIM<-OLD-ABC | 878 | Bacteria>Thermodesulfobacteriota>Desulfobacteria>Desulfobacterales>Desulfobacteriaceae | - | uncultured_Desulfobacterium_sp. | N47_D28280 | 44661..47297 | 201089 | hypothetical-protein-N47_D28280-[uncultured-Desulfobacterium-sp.] |
| WP_155316146.1 | AAA+ATPase+wHTH+Alpha+Beta-domain+wHTH+RRM | Type-3_BR.PglZ.associated.systems | <-TUDOR+SF1-SF2-Helicase+iREase+CTD<-iSTAND<-TOPRIM<-OLD-ABC<-METHYLASE<-PglZ<-DUF499*<-XIS-HTH<-?<-?<-ArdCN+MPTase | 932 | Bacteria>Thermodesulfobacteriota>Desulfobacteria>Desulfobacterales>Desulfosarcinaceae | GCF_009688945.1 | Desulfosarcina_alkanivorans | DSCA_RS09175 | 2082856..2085654 | 571177 | hypothetical-protein-[Desulfosarcina-alkanivorans] |
| MDD2316829.1 | AAA+ATPase+wHTH+Alpha+Beta-domain+wHTH+RRM | Type-3_BR.PglZ.associated.systems | XIS-HTH->DUF499*->PglZ->OLD-ABC->REase->METHYLASE->OLD-ABC->TOPRIM-> | 951 | Bacteria>Thermodesulfobacteriota>Desulfobacteria>Desulfobacterales>unclassified_Desulfobacterales | GCA_028683495.1 | Desulfobacterales_bacterium | PHD57_05725 | 1896..4751 | 2044940 | MAG:-hypothetical-protein-PHD57_05725-[Desulfobacterales-bacterium] |
| NVM22708.1 | AAA+ATPase+wHTH+Alpha+Beta-domain+wHTH+RRM | Type-3_BR.PglZ.associated.systems | JAB->DOC->JAB->DUF499*->PglZ->CopG/RHH->PIN-> | 875 | Bacteria>Thermodesulfobacteriota>Desulfobacteria>Desulfobacterales>unclassified_Desulfobacterales | GCA_013375395.1 | Desulfobacterales_bacterium | HWN68_13125 | 2460..5087 | 2044940 | hypothetical-protein-[Desulfobacterales-bacterium] |
| MBC8418550.1 | AAA+ATPase+wHTH+Alpha+Beta-domain+wHTH+RRM | Type-3_BR.PglZ.associated.systems | DUF499*->PglZ->METHYLASE-p-> | 758 | Bacteria>Thermodesulfobacteriota>Desulfobacteria>Desulfobacterales>unclassified_Desulfobacterales | GCA_014382905.1 | Desulfobacterales_bacterium | H8E10_08160 | 1..2279 | 2044940 | hypothetical-protein-[Desulfobacterales-bacterium] |
| MCK4486184.1 | AAA+ATPase+wHTH+Alpha+Beta-domain+wHTH+RRM | Type-3_BR.PglZ.associated.systems | DUF499*->PglZ->CopG/RHH->PIN-> | 845 | Bacteria>Thermodesulfobacteriota>Desulfobacteria>Desulfobacterales>unclassified_Desulfobacterales | GCA_023131115.1 | Desulfobacterales_bacterium | KAU38_05395 | 310..2847 | 2044940 | hypothetical-protein-[Desulfobacterales-bacterium] |
| MDD4464448.1 | AAA+ATPase+wHTH+Alpha+Beta-domain+wHTH+RRM | Type-3_BR.PglZ.associated.systems | <-METHYLASE-p<-REase<-OLD-ABC<-PglZ<-DUF499* | 183 | Bacteria>Thermodesulfobacteriota>Desulfobacteria>Desulfobacterales>unclassified_Desulfobacterales | - | Desulfobacterales_bacterium | PHR86_09450 | 3623..4176 | 2044940 | MAG:-hypothetical-protein-PHR86_09450,-partial-[Desulfobacterales-bacterium] |
| MDP2647166.1 | AAA+ATPase+wHTH+Alpha+Beta-domain+wHTH+RRM | Type-3_BR.PglZ.associated.systems | DUF499+RRM*->PglZ->METHYLASE-p-> | 858 | Bacteria>Thermodesulfobacteriota>Desulfobacteria>Desulfobacterales>unclassified_Desulfobacterales | GCA_030688615.1 | Desulfobacterales_bacterium | Q8P24_19705 | 1..2578 | 2044940 | AAA-family-ATPase-[Desulfobacterales-bacterium] |
| MDH5299321.1 | AAA+ATPase+wHTH+Alpha+Beta-domain+wHTH+RRM | Type-3_BR.PglZ.associated.systems | <-PglZ<-DUF499+RRM* | 795 | Bacteria>Thermodesulfobacteriota>Desulfobulbia>Desulfobulbales>Desulfobulbaceae>unclassified_Desulfobulbaceae | GCA_029884375.1 | Desulfobulbaceae_bacterium | OEV91_09915 | 274..2662 | 2053307 | ATP-binding-protein-[Desulfobulbaceae-bacterium] |
| MBE0584993.1 | AAA+ATPase+wHTH+Alpha+Beta-domain+wHTH+RRM | Type-3_BR.PglZ.associated.systems | <-TUDOR+SF1-SF2-Helicase+iREase+CTD<-iSTAND<-TOPRIM<-OLD-ABC<-METHYLASE<-PglZ<-DUF499* | 717 | Bacteria>Thermodesulfobacteriota>Desulfobulbia>Desulfobulbales>Desulfocapsaceae | GCA_014860245.1 | Desulfofustis_sp. | IH612_14700 | 7137..9290 | 2053546 | hypothetical-protein-[Desulfofustis-sp.] |
| WP_190244645.1 | AAA+ATPase+wHTH+Alpha+Beta-domain+wHTH+RRM | Type-3_BR.PglZ.associated.systems | <-MoxR-AAA<-?<-REase<-?\|\|DUF499*->PglZ->METHYLASE->iSTAND->TUDOR+SF1-SF2-Helicase+iREase+CTD-> | 939 | Bacteria>Thermodesulfobacteriota>Desulfovibrionia>Desulfovibrionales>Desulfovibrionaceae | GCF_019931275.1 | Nitratidesulfovibrio_sp._SRB-5 | K6142_RS15435 | 57284..60103 | 2872636 | DUF499-domain-containing-protein-[Cupidesulfovibrio-sp.-SRB-5] |
| WP_097012771.1 | AAA+ATPase+wHTH+Alpha+Beta-domain+wHTH+RRM | Type-3_BR.PglZ.associated.systems | <-MoxR-AAA\|\|35exo+SF1-SF2-Helicase-><-REase<-?\|\|DUF499*->PglZ->METHYLASE->iSTAND->TUDOR+SF1-SF2-Helicase+iREase+CTD-> | 896 | Bacteria>Thermodesulfobacteriota>Desulfovibrionia>Desulfovibrionales>Desulfovibrionaceae | GCF_900217235.1 | Pseudodesulfovibrio_profundus | DPRO_RS14880 | 3164456..3167146 | 57320 | DUF499-domain-containing-protein-[Pseudodesulfovibrio-profundus] |
| HML53149.1 | AAA+ATPase+wHTH+Alpha+Beta-domain+wHTH+RRM | Type-3_BR.PglZ.associated.systems | DUF499+RRM*->PglZ->METHYLASE->iSTAND->TUDOR+SF1-SF2-Helicase+iREase+CTD-> | 947 | Bacteria>Thermodesulfobacteriota>Desulfovibrionia>Desulfovibrionales>Desulfovibrionaceae>Solidesulfovibrio | GCA_035323385.1 | Solidesulfovibrio_magneticus | PKC79_03560 | 67192..70035 | 184917 | hypothetical-protein-[Solidesulfovibrio-magneticus] |
| WP_067615938.1 | AAA+ATPase+wHTH+Alpha+Beta-domain+wHTH+RRM | Type-3_BR.PglZ.associated.systems | TetR-HTH->?->?->?-><-?\|\|XIS-HTH->DUF499*->PglZ->METHYLASE->iSTAND->TUDOR+SF1-SF2-Helicase+iREase+CTD->?->HerA/FtsK-> | 929 | Bacteria>Thermodesulfobacteriota>Dissulfuribacteria>Dissulfuribacterales>Dissulfuribacteraceae | GCF_001687335.1 | Dissulfuribacter_thermophilus | DBT_RS01985 | 12406..15195 | 1156395 | DUF499-domain-containing-protein-[Dissulfuribacter-thermophilus] |
| OCC15962.1 | AAA+ATPase+wHTH+Alpha+Beta-domain+wHTH+RRM | Type-3_BR.PglZ.associated.systems | DUF499*->PglZ->METHYLASE-><-?\|\|SF1-SF2-Helicase+iREase+CTD->?->HerA/FtsK-> | 709 | Bacteria>Thermodesulfobacteriota>Dissulfuribacteria>Dissulfuribacterales>Dissulfuribacteraceae | GCA_001687335.1 | Dissulfuribacter_thermophilus | DBT_0424 | 13066..15195 | 1156395 | hypothetical-protein-DBT_0424-[Dissulfuribacter-thermophilus] |
| MDI6725762.1 | AAA+ATPase+wHTH+Alpha+Beta-domain+wHTH+RRM | Type-3_BR.PglZ.associated.systems | DUF499+RRM*->PglZ->METHYLASE-p-> | 887 | Bacteria>Thermodesulfobacteriota>Syntrophia>Syntrophales>Smithellaceae>unclassified_Smithellaceae | GCA_030017725.1 | Smithellaceae_bacterium | QMD32_02275 | 2141..4804 | 2964540 | hypothetical-protein-[Smithellaceae-bacterium] |
| MBN1547395.1 | AAA+ATPase+wHTH+Alpha+Beta-domain+wHTH+RRM | Type-3_BR.PglZ.associated.systems | DUF499*->PglZ->wHTH+REase->METHYLASE-p-> | 150 | Bacteria>Thermodesulfobacteriota>Syntrophia>Syntrophales>Syntrophaceae | GCA_016929155.1 | Syntrophaceae_bacterium | JW902_12120 | 1..455 | 2052184 | hypothetical-protein-[Syntrophaceae-bacterium] |
| HYA42142.1 | AAA+ATPase+wHTH+Alpha+Beta-domain+wHTH+RRM | Type-3_BR.PglZ.associated.systems | <-iSTAND<-METHYLASE<-PglZ<-DUF499+RRM* | 942 | Bacteria>Thermodesulfobacteriota>Syntrophobacteria>Syntrophobacterales>Syntrophobacteraceae>unclassified_Syntrophobacteraceae | GCA_035624985.1 | Syntrophobacteraceae_bacterium | VEF34_12615 | 5178..8006 | 2052185 | hypothetical-protein-[Syntrophobacteraceae-bacterium] |
| MDR3555654.1 | AAA+ATPase+wHTH+Alpha+Beta-domain+wHTH+RRM | Type-3_BR.PglZ.associated.systems | <-SbcC<-METHYLASE<-ABC-ATPase<-PglZ<-DUF499+RRM* | 939 | Bacteria>Thermodesulfobacteriota>Syntrophobacteria>Syntrophobacterales>Syntrophobacteraceae>unclassified_Syntrophobacteraceae | - | Syntrophobacteraceae_bacterium | P4L55_12930 | 16994..19813 | 2052185 | hypothetical-protein-[Syntrophobacteraceae-bacterium] |
| MBW6486648.1 | AAA+ATPase+wHTH+Alpha+Beta-domain+wHTH+RRM | Type-3_BR.PglZ.associated.systems | Tox-RelE->HTH_3->?->DUF499*->PglZ->METHYLASE->OLD-ABC->TOPRIM->iSTAND->TUDOR+SF1-SF2-Helicase+iREase+CTD-> | 937 | Bacteria>Thermodesulfobacteriota>Syntrophobacteria>Syntrophobacterales>unclassified_Syntrophobacterales | GCA_019429105.1 | Syntrophobacterales_bacterium | KOBO1_10925 | 2187..5000 | 2282155 | hypothetical-protein-[Syntrophobacterales-bacterium] |
| MCR5853654.1 | AAA+ATPase+wHTH+Alpha+Beta-domain+wHTH+RRM | Type-3_BR.PglZ.associated.systems | <-METHYLASE<-METHYLASE-p<-SF1-SF2-Helicase<-OLD-ABC+TOPRIM<-PglZ<-DUF499-p<-DUF499* | 159 | Bacteria>Thermodesulfobacteriota>unclassified_Thermodesulfobacteriota | GCA_024699695.1 | Thermodesulfobacteriota_bacterium | MW689_001023 | 119654..120133 | 2202153 | hypothetical-protein-[Thermodesulfobacteriota-bacteirum] |
| MCR5853653.1 | AAA+ATPase+wHTH+Alpha+Beta-domain+wHTH+RRM | Type-3_BR.PglZ.associated.systems | <-METHYLASE<-METHYLASE-p<-SF1-SF2-Helicase<-OLD-ABC+TOPRIM<-PglZ<-DUF499*<-DUF499-p | 664 | Bacteria>Thermodesulfobacteriota>unclassified_Thermodesulfobacteriota | GCA_024699695.1 | Thermodesulfobacteriota_bacterium | MW689_001022 | 117685..119679 | 2202153 | ATP-binding-protein-[Thermodesulfobacteriota-bacteirum] |
| MDI6755573.1 | AAA+ATPase+wHTH+Alpha+Beta-domain+wHTH+RRM | Type-3_BR.PglZ.associated.systems | DUF499+RRM*->PglZ-> | 869 | Bacteria>Thermodesulfobacteriota>unclassified_Thermodesulfobacteriota | GCA_030018715.1 | Thermodesulfobacteriota_bacterium | QME78_14395 | 57..2666 | 2202153 | hypothetical-protein-[Thermodesulfobacteriota-bacterium] |
| MEA3360867.1 | AAA+ATPase+wHTH+Alpha+Beta-domain+wHTH+RRM | Type-3_BR.PglZ.associated.systems | DUF499+RRM*->PglZ->OLD-ABC+TOPRIM->DUF4276->METHYLASE-> | 900 | Bacteria>Thermodesulfobacteriota>unclassified_Thermodesulfobacteriota | GCA_034718555.1 | Thermodesulfobacteriota_bacterium | U9R17_15880 | 30707..33409 | 2202153 | DUF499-domain-containing-protein-[Thermodesulfobacteriota-bacterium] |
| MEA3428074.1 | AAA+ATPase+wHTH+Alpha+Beta-domain+wHTH+RRM | Type-3_BR.PglZ.associated.systems | <-METHYLASE-p<-PglZ<-DUF499+RRM* | 890 | Bacteria>Thermodesulfobacteriota>unclassified_Thermodesulfobacteriota | GCA_034721875.1 | Thermodesulfobacteriota_bacterium | U9Q84_02400 | 1527..4201 | 2202153 | hypothetical-protein-[Thermodesulfobacteriota-bacterium] |
| HLE25734.1 | AAA+ATPase+wHTH+Alpha+Beta-domain+wHTH+RRM | Type-3_BR.PglZ.associated.systems | DUF499+RRM*->PglZ->METHYLASE->TUDOR+SF1-SF2-Helicase+iREase+CTD-> | 873 | Bacteria>Thermodesulfobacteriota>unclassified_Thermodesulfobacteriota | GCA_035291775.1 | Thermodesulfobacteriota_bacterium | VI935_08805 | 476..3097 | 2202153 | hypothetical-protein-[Thermodesulfobacteriota-bacterium] |
| HZX15017.1 | AAA+ATPase+wHTH+Alpha+Beta-domain+wHTH+RRM | Type-3_BR.PglZ.associated.systems | DUF499+RRM*->PglZ->METHYLASE-p-> | 873 | Bacteria>Thermodesulfobacteriota>unclassified_Thermodesulfobacteriota | GCA_035655495.1 | Thermodesulfobacteriota_bacterium | VFF49_11535 | 1792..4413 | 2202153 | hypothetical-protein-[Thermodesulfobacteriota-bacterium] |
| WP_011944180.1 | AAA+ATPase+wHTH+Alpha+Beta-domain+wHTH+RRM | Type-3_BR.PglZ.associated.systems | <-SF1-SF2-Helicase<-?<-METHYLASE<-?<-PolB<-PglZ<-DUF499*<-?\|\|?->Phage_int_SAM_1+Phage_integrase-> | 814 | Bacteria>Thermotogota>Thermotogae>Thermotogales>Thermotogaceae | GCF_000016785.1 | Thermotoga_petrophila | TPET_RS09055 | 1780376..1782820 | 93929 | AAA-family-ATPase-[Thermotoga-petrophila] |
| MCD4760467.1 | AAA+ATPase+wHTH+Alpha+Beta-domain+wHTH+RRM | Type-3_BR.PglZ.associated.systems | PIN->DUF499*->PglZ->?->Radical-SAM->METHYLASE->iSTAND->TUDOR+SF1-SF2-Helicase+iREase+CTD-> | 900 | Bacteria>unclassified_Bacteria | GCA_021107775.1 | bacterium | K8R42_01070 | 353..3055 | 1869227 | DUF499-domain-containing-protein-[bacterium] |
| NLF40196.1 | AAA+ATPase+wHTH+Alpha+Beta-domain+wHTH+RRM | Type-3_BR.PglZ.associated.systems | DUF499*->PglZ->CopG/RHH->PIN->METHYLASE-p-> | 866 | Bacteria>unclassified_Bacteria | GCA_012514975.1 | bacterium | GX586_12180 | 159..2759 | 1869227 | hypothetical-protein-[bacterium] |
| MCP3685981.1 | AAA+ATPase+wHTH+Alpha+Beta-domain+wHTH+RRM | Type-3_BR.PglZ.associated.systems | <-REase<-TUDOR+SF1-SF2-Helicase+iREase+CTD<-iSTAND<-METHYLASE<-PglZ<-DUF499*<-HNH | 901 | Bacteria>unclassified_Bacteria | GCA_024206195.1 | bacterium | GY861_25320 | 20693..23398 | 1869227 | DUF499-domain-containing-protein-[bacterium] |
| TSA39490.1 | AAA+ATPase+wHTH+Alpha+Beta-domain+wHTH+RRM | Type-3_BR.PglZ.associated.systems | <-METHYLASE<-PglZ<-DUF499* | 809 | Bacteria>unclassified_Bacteria | GCA_007281005.1 | bacterium | D4Q85_01020 | 3705..6134 | 1869227 | DUF499-domain-containing-protein,-partial-[bacterium] |
| MBN2415253.1 | AAA+ATPase+wHTH+Alpha+Beta-domain+wHTH+RRM | Type-3_BR.PglZ.associated.systems | <-HNH<-EVE<-TUDOR+SF1-SF2-Helicase+iREase+CTD<-iSTAND<-METHYLASE<-PglZ<-DUF499*<-?<-HKD-DNase | 892 | Bacteria>unclassified_Bacteria | GCA_016934955.1 | bacterium | JXO52_05400 | 60396..63074 | 1869227 | DUF499-domain-containing-protein-[bacterium] |
| HPN33675.1 | AAA+ATPase+wHTH+Alpha+Beta-domain+wHTH+RRM | Type-3_BR.PglZ.associated.systems | <-METHYLASE<-PIN<-CopG-RHH<-DUF499+RRM* | 870 | Bacteria>unclassified_Bacteria | GCA_035393365.1 | bacterium | PK843_04125 | 497835..500447 | 1869227 | AAA-family-ATPase-[bacterium] |
| EIU1834191.1 | AAA+ATPase+wHTH+Alpha+Beta-domain+wHTH+RRM | Type-3_BR.PglZ.associated.systems | ZnR+TOPRIM+D5ATPase->XIS-HTH->DUF499*->PglZ->SIR2->VirB4-FtsK+HerA->METHYLASE->iSTAND->TUDOR+SF1-SF2-Helicase+iREase+CTD-> | 848 | - | - | - | L4Q41_001887 | 2150..4696 | - | DUF499-domain-containing-protein-[Pseudomonas-aeruginosa] |
| EIU2583491.1 | AAA+ATPase+wHTH+Alpha+Beta-domain+wHTH+RRM | Type-3_BR.PglZ.associated.systems | MerR-HTH->tRNA->DUF499*->PglZ->SIR2->VirB4-FtsK+HerA->METHYLASE->iSTAND->TUDOR+SF1-SF2-Helicase+iREase+CTD-> | 853 | - | - | - | L4O09_004839 | 68693..71254 | - | DUF499-domain-containing-protein-[Pseudomonas-aeruginosa] |
| EKM5722020.1 | AAA+ATPase+wHTH+Alpha+Beta-domain+wHTH+RRM | Type-3_BR.PglZ.associated.systems | MerR-HTH->tRNA->DUF499*->PglZ->METHYLASE->iSTAND->TUDOR+SF1-SF2-Helicase+iREase+CTD-><-RlpA | 839 | - | - | - | PU294_000685 | 14114..16633 | - | DUF499-domain-containing-protein-[Pseudomonas-aeruginosa] |
| EIU6031615.1 | AAA+ATPase+wHTH+Alpha+Beta-domain+wHTH+RRM | Type-3_BR.PglZ.associated.systems | MerR-HTH->tRNA->DUF499*->PglZ->?->METHYLASE->iSTAND->TUDOR+SF1-SF2-Helicase+iREase+CTD-><-RlpA | 862 | - | - | - | L5V03_004654 | 39732..42320 | - | DUF499-domain-containing-protein-[Pseudomonas-aeruginosa] |
| EIU2955592.1 | AAA+ATPase+wHTH+Alpha+Beta-domain+wHTH+RRM | Type-3_BR.PglZ.associated.systems | MerR-HTH->tRNA->DUF499*->PglZ->?->METHYLASE->iSTAND->TUDOR+SF1-SF2-Helicase+iREase+CTD-><-RlpA | 853 | - | - | - | P3458_000154 | 13790..16351 | - | DUF499-domain-containing-protein-[Pseudomonas-aeruginosa] |
| EIU1605741.1 | AAA+ATPase+wHTH+Alpha+Beta-domain+wHTH+RRM | Type-3_BR.PglZ.associated.systems | <-TUDOR+SF1-SF2-Helicase+iREase+CTD<-iSTAND<-METHYLASE<-VirB4-FtsK+HerA<-SIR2<-PglZ<-DUF499*<-tRNA<-MerR-HTH | 853 | - | - | - | L4H28_002030 | 380553..383114 | - | DUF499-domain-containing-protein-[Pseudomonas-aeruginosa] |
| EJB0066501.1 | AAA+ATPase+wHTH+Alpha+Beta-domain+wHTH+RRM | Type-3_BR.PglZ.associated.systems | <-TUDOR+SF1-SF2-Helicase+iREase+CTD<-iSTAND<-METHYLASE<-Classical-AAA<-PglZ<-DUF499*<-XIS-HTH | 878 | - | - | - | P1X34_002688 | 41742..44378 | - | DUF499-domain-containing-protein-[Pseudomonas-aeruginosa] |
| EKD5454941.1 | AAA+ATPase+wHTH+Alpha+Beta-domain+wHTH+RRM | Type-3_BR.PglZ.associated.systems | <-iSTAND<-METHYLASE<-?<-Peptidase_S8<-Classical-AAA<-PglZ<-DUF499*<-XIS-HTH\|\|MODE-HTH+PBPII->?->VirD4-TraG+VirD4-FtsK->CopG/RHH->VirB11+VirB4-FtsK-> | 860 | - | - | - | OR553_000221 | 19489..22071 | - | DUF499-domain-containing-protein-[Pseudomonas-aeruginosa] |

| Protein-ID | Gene-Neighbourhoods | Length | Strain | Taxonomy | Fasta-defline | TaxID |
|---|---|---|---|---|---|---|
| -- | | | | | | |
| --BREX-related-Capture-System | | | | | | |
| -- | | | | | | |
| QSG11389.1 | Cluster76_1clades->TGT->DUF4007->DUF6079->iSwi2/Snf2-Helicase+PglZ*->HerA_FtsK->GNAT->?-><-Phage_integrase | 368 | Halapricum_desulfuricans | Archaea>Methanobacteriati>Methanobacteriota>Stenosarchaea_group>Halobacteria>Halobacteriales>Haloarculaceae>Halapricum | AP-superfamily-protein-[Halapricum-desulfuricans] | 2841257 |
| WP_229125942.1 | Cluster76_1clades->QueF->TGT->DUF4007->STAND->DUF6079->iSwi2/Snf2-Helicase+PglZ*->HerA_FtsK->GNAT-><-?\|\|?-><-Phage_integrase | 368 | Halapricum_desulfuricans | Archaea>Methanobacteriati>Methanobacteriota>Stenosarchaea_group>Halobacteria>Halobacteriales>Haloarculaceae>Halapricum | hypothetical-protein-[Halapricum-desulfuricans] | 2841257 |
| SDG37568.1 | QueF->TGT->DUF4007->STAND->DUF6079->iSwi2/Snf2-Helicase+PglZ*->HerA_FtsK-><-?<-RNase_H_2 | 368 | Halorientalis_regularis | Archaea>Methanobacteriati>Methanobacteriota>Stenosarchaea_group>Halobacteria>Halobacteriales>Haloarculaceae>Halorientalis | hypothetical-protein-SAMN05216218_12813-[Halorientalis-regularis] | 660518 |
| WP_092695622.1 | QueF->TGT->DUF4007->STAND->DUF6079->iSwi2/Snf2-Helicase+PglZ*->HerA_FtsK->GNAT-><-RNase_H_2 | 368 | Halorientalis_regularis | Archaea>Methanobacteriati>Methanobacteriota>Stenosarchaea_group>Halobacteria>Halobacteriales>Haloarculaceae>Halorientalis | hypothetical-protein-[Halorientalis-regularis] | 660518 |
| MFC3477742.1 | <-REase\|\|?-><-AAA_16+Cdc6_C<-GNAT<-HerA_FtsK<-iSwi2/Snf2-Helicase+PglZ*<-DUF6079<-STAND<-DUF4007<-TGT<-QueF | 372 | Halobacterium_litoreum | Archaea>Methanobacteriati>Methanobacteriota>Stenosarchaea_group>Halobacteria>Halobacteriales>Halobacteriaceae>Halobacterium | hypothetical-protein-ACFOKC_08390-[Halobacterium-litoreum] | 2039234 |
| UHH13392.1 | QueF->TGT->DUF4007->STAND->DUF6079->iSwi2/Snf2-Helicase+PglZ*->HerA_FtsK->GNAT-> | 372 | Halobacterium_litoreum | Archaea>Methanobacteriati>Methanobacteriota>Stenosarchaea_group>Halobacteria>Halobacteriales>Halobacteriaceae>Halobacterium | hypothetical-protein-LT972_14695-[Halobacterium-litoreum] | 2039234 |
| WP_232571135.1 | QueF->TGT->DUF4007->STAND->DUF6079->iSwi2/Snf2-Helicase+PglZ*->HerA_FtsK->GNAT-> | 372 | Halobacterium_litoreum | Archaea>Methanobacteriati>Methanobacteriota>Stenosarchaea_group>Halobacteria>Halobacteriales>Halobacteriaceae>Halobacterium | hypothetical-protein-[Halobacterium-litoreum] | 2039234 |
| MDS0297782.1 | Phage_integrase-><-?<-HTH_20\|\|<-QueF<-GNAT<-HerA_FtsK<-iSwi2/Snf2-Helicase+PglZ*<-DUF6079<-STAND<-DUF4007<-TGT<-HalOD1 | 373 | Halogeometricum_salsisoli | Archaea>Methanobacteriati>Methanobacteriota>Stenosarchaea_group>Halobacteria>Halobacteriales>Haloferacaceae>Halogeometricum | hypothetical-protein-[Halogeometricum-sp.-S1BR25-6] | 2950536 |
| WP_139229723.1 | <-Phosphodiest<-HTH+RNASE-EG<-STAND<-?<-GNAT<-HerA_FtsK<-iSwi2/Snf2-Helicase+PglZ*<-DUF6079<-DUF4007<-TGT<-QueF\|\|?->?->FaeA-> | 372 | Halogeometricum_limi | Archaea>Methanobacteriati>Methanobacteriota>Stenosarchaea_group>Halobacteria>Halobacteriales>Haloferacaceae>Halogeometricum | hypothetical-protein-[Halogeometricum-limi] | 555875 |
| WP_310922610.1 | Phage_integrase-><-?<-HTH_20\|\|<-QueF<-GNAT<-HerA_FtsK<-iSwi2/Snf2-Helicase+PglZ*<-DUF6079<-STAND<-DUF4007<-TGT<-HalOD1 | 373 | Halogeometricum_salsisoli | Archaea>Methanobacteriati>Methanobacteriota>Stenosarchaea_group>Halobacteria>Halobacteriales>Haloferacaceae>Halogeometricum | hypothetical-protein-[Halogeometricum-sp.-S1BR25-6] | 2950536 |
| ATW88247.1 | QueF->TGT->DUF4007->STAND->DUF6079->iSwi2/Snf2-Helicase+PglZ*->HerA_FtsK->GNAT-><-?<-YTCD-HTH\|\|?->?->GNAT-> | 371 | Halohasta_litchfieldiae | Archaea>Methanobacteriati>Methanobacteriota>Stenosarchaea_group>Halobacteria>Halobacteriales>Haloferacaceae>Halohasta | hypothetical-protein-halTADL_1482-[Halohasta-litchfieldiae] | 1073996 |
| SEJ07089.1 | <-GNAT<-?<-?\|\|YTCD-HTH-><-?<-GNAT<-HerA_FtsK<-iSwi2/Snf2-Helicase+PglZ*\|\|?->?-><-DUF6079<-STAND<-DUF4007<-TGT | 169 | Halohasta_litchfieldiae | Archaea>Methanobacteriati>Methanobacteriota>Stenosarchaea_group>Halobacteria>Halobacteriales>Haloferacaceae>Halohasta | hypothetical-protein-SAMN05444271_11931-[Halohasta-litchfieldiae] | 1073996 |
| WP_089673071.1 | QueF->TGT->DUF4007->STAND->DUF6079->iSwi2/Snf2-Helicase+PglZ*->HerA_FtsK->GNAT-><-?<-YTCD-HTH\|\|?->?->GNAT-> | 371 | Halohasta_litchfieldiae | Archaea>Methanobacteriati>Methanobacteriota>Stenosarchaea_group>Halobacteria>Halobacteriales>Haloferacaceae>Halohasta | hypothetical-protein-[Halohasta-litchfieldiae] | 1073996 |
| WP_147270939.1 | QueF->TGT->DUF4007->STAND->DUF6079->iSwi2/Snf2-Helicase+PglZ*->HerA_FtsK->GNAT-><-REase\|\|<-?\|\|ZDNA-HTH-> | 371 | Haloplanus_salinus | Archaea>Methanobacteriati>Methanobacteriota>Stenosarchaea_group>Halobacteria>Halobacteriales>Haloferacaceae>Haloplanus | hypothetical-protein-[Haloplanus-salinus] | 1126245 |
| WP_338740220.1 | QueF->TGT->DUF4007->STAND->DUF6079->iSwi2/Snf2-Helicase+PglZ*->HerA_FtsK->GNAT-><-?\|\|CITB-HTH-> | 369 | Haloplanus_sp._CK5-1 | Archaea>Methanobacteriati>Methanobacteriota>Stenosarchaea_group>Halobacteria>Halobacteriales>Haloferacaceae>Haloplanus>unclassified_Haloplanus | hypothetical-protein-[Haloplanus-salilacus] | 2949994 |
| WP_157972653.1 | <-McrBC<-?\|\|TGT->DUF4007-><-AAA_assoc<-PufQ\|\|DUF6079->iSwi2/Snf2-Helicase+PglZ*->HerA_FtsK->GNAT-><-QueF | 369 | Haloprofundus_halophilus | Archaea>Methanobacteriati>Methanobacteriota>Stenosarchaea_group>Halobacteria>Halobacteriales>Haloferacaceae>Haloprofundus | hypothetical-protein-[Haloprofundus-halophilus] | 2283527 |
| WP_271955572.1 | Acatn-><-?\|\|adh_short_C2-><-QueF<-GNAT<-HerA_FtsK<-iSwi2/Snf2-Helicase+PglZ*<-DUF6079<-STAND<-DUF4007<-TGT<-?\|\|?->Radical_SAM-> | 391 | Halorubrum_ezzemoulense | Archaea>Methanobacteriati>Methanobacteriota>Stenosarchaea_group>Halobacteria>Halobacteriales>Haloferacaceae>Halorubrum | hypothetical-protein-[Halorubrum-ezzemoulense] | 337243 |
| WP_143414662.1 | TGT->DUF4007->STAND->DUF6079->iSwi2/Snf2-Helicase+PglZ*->HerA_FtsK->GNAT->QueF->?->?->Radical_SAM->GTP_cyclohydroI-> | 385 | Halorubrum_sp._SD683 | Archaea>Methanobacteriati>Methanobacteriota>Stenosarchaea_group>Halobacteria>Halobacteriales>Haloferacaceae>Halorubrum>unclassified_Halorubrum | hypothetical-protein-[Halorubrum-sp.-SD683] | 1855873 |
| MFC3957638.1 | Phage_integrase->DUF6079-><-?\|\|?-><-GNAT<-HerA_FtsK<-iSwi2/Snf2-Helicase+PglZ*<-DUF6079<-STAND<-DUF4007<-TGT<-QueF\|\|QueF-> | 369 | Halovivax_cerinus | Archaea>Methanobacteriati>Methanobacteriota>Stenosarchaea_group>Halobacteria>Halobacteriales>Natrialbaceae>Halovivax | hypothetical-protein-ACFOUR_04520-[Halovivax-cerinus] | 1487865 |
| WP_256533864.1 | Phage_integrase->DUF6079-><-?\|\|?-><-GNAT<-HerA_FtsK<-iSwi2/Snf2-Helicase+PglZ*<-DUF6079<-STAND<-DUF4007<-TGT<-QueF\|\|QueF-> | 369 | Halovivax_cerinus | Archaea>Methanobacteriati>Methanobacteriota>Stenosarchaea_group>Halobacteria>Halobacteriales>Natrialbaceae>Halovivax | hypothetical-protein-[Halovivax-cerinus] | 1487865 |
| WP_048108644.1 | GNAT->TGT->DUF4007->STAND->DUF6079->iSwi2/Snf2-Helicase+PglZ*->HerA_FtsK->EAD9->BcrAD_BadFG->?->HicB_lk_antitox-><-HicA_toxin<-HicB_lk_antitox | 353 | Methanosarcina_barkeri | Archaea>Methanobacteriati>Methanobacteriota>Stenosarchaea_group>Methanomicrobia>Methanosarcinales>Methanosarcinaceae>Methanosarcina | hypothetical-protein-[Methanosarcina-barkeri] | 2208 |
| AKB82981.1 | TGT->DUF4007->DUF6079->iSwi2/Snf2-Helicase+PglZ*->HerA_FtsK->EAD9->BcrAD_BadFG->?->HicB_lk_antitox-><-HicA_toxin<-HicB_lk_antitox | 353 | Methanosarcina_barkeri_3 | Archaea>Methanobacteriati>Methanobacteriota>Stenosarchaea_group>Methanomicrobia>Methanosarcinales>Methanosarcinaceae>Methanosarcina>Methanosarcina_barkeri | hypothetical-protein-MSBR3_2403-[Methanosarcina-barkeri-3] | 1434107 |
| NPV62497.1 | iSwi2/Snf2-Helicase+PglZ*->HerA_FtsK->QueF->GNAT-><-REase\|\|?->Phage_integrase-> | 355 | Methanotrichaceae_archaeon | Archaea>Methanobacteriati>Methanobacteriota>Stenosarchaea_group>Methanomicrobia>Methanotrichales>Methanotrichaceae>unclassified_Methanotrichaceae | hypothetical-protein-[Methanotrichaceae-archaeon] | 2595067 |
| MCK4444004.1 | MethyltransfD12-><-?<-B12-binding+Radical_SAM<-HerA_FtsK<-iSwi2/Snf2-Helicase+PglZ* | 362 | Thermoplasmata_archaeon | Archaea>Methanobacteriati>Thermoplasmatota>Thermoplasmata>unclassified_Thermoplasmata | PglZ-domain-containing-protein-[Thermoplasmata-archaeon] | 1906666 |
| MFQ5758747.1 | <-GNAT<-HerA_FtsK<-iSwi2/Snf2-Helicase+PglZ*<-DUF6079-p | 340 | Candidatus_Bathyarchaeia_archaeon | Archaea>Thermoproteati>Candidatus_Bathyarchaeota>Candidatus_Bathyarchaeia>unclassified_Candidatus_Bathyarchaeia | MAG:-hypothetical-protein-ACE5IF_03635-[Candidatus-Bathyarchaeia-archaeon] | 3045160 |
| MBI2842128.1 | <-Peptidase_M78<-?<-?<-HATPase_c_3+HSDR_N_2<-HerA_FtsK<-iSwi2/Snf2-Helicase+PglZ*<-DUF6079<-STAND<-DUF4007<-McrA-NTD<-PAPS_reduct<-DUF6079<-DUF4007 | 375 | Armatimonadota_bacterium | Bacteria>Bacillati>Armatimonadota>unclassified_Armatimonadota | hypothetical-protein-[Armatimonadota-bacterium] | 2033014 |
| MDW8029832.1 | TGT->DUF4007->STAND->DUF6079-p->DUF6079->iSwi2/Snf2-Helicase+PglZ*-> | 358 | Armatimonadota_bacterium | Bacteria>Bacillati>Armatimonadota>unclassified_Armatimonadota | hypothetical-protein-[Armatimonadota-bacterium] | 2033014 |
| MDT7893012.1 | Phd-YefM->DUF4007->Ntox9->VbhA_Antitoxin->STAND->DUF6079->iSwi2/Snf2-Helicase+PglZ*-> | 114 | Armatimonadota_bacterium | Bacteria>Bacillati>Armatimonadota>unclassified_Armatimonadota | hypothetical-protein-[Armatimonadota-bacterium] | 2033014 |
| HHW27504.1 | <-HerA_FtsK<-iSwi2/Snf2-Helicase+PglZ*<-DUF6079<-DUF4007<-TGT<-GNAT | 374 | Candidatus_Fermentithermobacillaceae_bacterium | Bacteria>Bacillati>Bacillota>Candidatus_Fermentithermobacillia>Candidatus_Fermentithermobacillales>Candidatus_Fermentithermobacillaceae>unclassified_Candidatus_Fermentithermobacillaceae | hypothetical-protein-[Candidatus-Fermentithermobacillaceae-bacterium] | 3085339 |
| HIE11925.1 | GNAT+TGT->DUF4007->STAND->DUF6079->iSwi2/Snf2-Helicase+PglZ*->Mrr_N->HerA_FtsK->Metallophos->AAA_27-> | 368 | Desulfotomaculum_sp. | Bacteria>Bacillati>Bacillota>Clostridia>Eubacteriales>Desulfotomaculaceae>Desulfotomaculum>unclassified_Desulfotomaculum | hypothetical-protein-[Desulfotomaculum-sp.] | 41211 |
| HHW40997.1 | TGT->DUF4007->STAND->DUF6079->iSwi2/Snf2-Helicase+PglZ*->HerA_FtsK->SIGMA-HTH+SIGMA-HTH->LexA_DNA_bind+Peptidase_S24-> | 370 | Syntrophomonadaceae_bacterium | Bacteria>Bacillati>Bacillota>Clostridia>Eubacteriales>Syntrophomonadaceae>unclassified_Syntrophomonadaceae | hypothetical-protein-[Syntrophomonadaceae-bacterium] | 2093811 |
| HQD80871.1 | <-SIGMA-HTH+SIGMA-HTH<-SIGMA-HTH+SIGMA-HTH<-HerA_FtsK<-iSwi2/Snf2-Helicase+PglZ*<-DUF6079<-STAND<-DUF4007 | 369 | Bacillota_bacterium | Bacteria>Bacillati>Bacillota>unclassified_Bacillota | PglZ-domain-containing-protein-[Bacillota-bacterium] | 1879010 |
| MBE2199248.1 | SIGMA-HTH+SIGMA-HTH->TGT->DUF4007->DUF3800->STAND->DUF6079->iSwi2/Snf2-Helicase+PglZ*->HerA_FtsK->GNAT-> | 368 | Anaerolinea_sp. | Bacteria>Bacillati>Chloroflexota>Anaerolineae>Anaerolineales>Anaerolineaceae>Anaerolinea>unclassified_Anaerolinea | hypothetical-protein-[Anaerolinea-sp.] | 1872519 |
| MBC8263374.1 | <-Vut_1<-GNAT<-HerA_FtsK<-iSwi2/Snf2-Helicase+PglZ*<-DUF6079<-STAND<-DUF4007<-TGT<-SIGMA-HTH | 374 | Anaerolineales_bacterium | Bacteria>Bacillati>Chloroflexota>Anaerolineae>Anaerolineales>unclassified_Anaerolineales | hypothetical-protein-[Anaerolineales-bacterium] | 2073117 |
| MCA9946505.1 | <-McrA-NTD<-GNAT<-HerA_FtsK<-iSwi2/Snf2-Helicase+PglZ* | 368 | Anaerolineales_bacterium | Bacteria>Bacillati>Chloroflexota>Anaerolineae>Anaerolineales>unclassified_Anaerolineales | PglZ-domain-containing-protein-[Anaerolineales-bacterium] | 2073117 |
| MCZ7554456.1 | RNA_pol_A_CTD+RNA_pol_A_CTD+Sigma70_r4+HARE-HTH->TGT->DUF4007->STAND->DUF6079->iSwi2/Snf2-Helicase+PglZ*->HerA_FtsK->XIS-HTH-> | 376 | Anaerolineales_bacterium | Bacteria>Bacillati>Chloroflexota>Anaerolineae>Anaerolineales>unclassified_Anaerolineales | hypothetical-protein-[Anaerolineales-bacterium] | 2073117 |
| GAB4528666.1 | MazE_antitoxin->PIN->SIGMA-HTH+SIGMA-HTH->TGT->DUF4007->STAND->DUF6079->iSwi2/Snf2-Helicase+PglZ*->HerA_FtsK->GNAT->?-><-CbiA | 372 | Anaerolineae_bacterium | Bacteria>Bacillati>Chloroflexota>Anaerolineae>unclassified_Anaerolineae | MAG:-hypothetical-protein-Kow0063_05020-[Anaerolineae-bacterium] | 2052143 |
| HFD39727.1 | E1-E2_ATPase+Hydrolase+HAD_2-><-GNAT<-HerA_FtsK<-iSwi2/Snf2-Helicase+PglZ*<-DUF6079<-STAND | 371 | Anaerolineae_bacterium | Bacteria>Bacillati>Chloroflexota>Anaerolineae>unclassified_Anaerolineae | PglZ-domain-containing-protein-[Anaerolineae-bacterium] | 2052143 |
| HID64303.1 | DUF4007->STAND->DUF6079->iSwi2/Snf2-Helicase+PglZ*->HerA_FtsK->Schlafen->Schlafen->Meth_synt_2->TPR+TPR+TPR-> | 364 | Anaerolineae_bacterium | Bacteria>Bacillati>Chloroflexota>Anaerolineae>unclassified_Anaerolineae | PglZ-domain-containing-protein-[Anaerolineae-bacterium] | 2052143 |
| MBN1877091.1 | TGT->DUF4007->STAND->DUF6079->iSwi2/Snf2-Helicase+PglZ*->HerA_FtsK->GNAT-><-DUF3368 | 373 | Anaerolineae_bacterium | Bacteria>Bacillati>Chloroflexota>Anaerolineae>unclassified_Anaerolineae | hypothetical-protein-[Anaerolineae-bacterium] | 2052143 |
| MBN2393033.1 | <-GNAT<-HerA_FtsK<-iSwi2/Snf2-Helicase+PglZ*<-DUF6079<-PIN<-RHH<-STAND<-DUF4007<-TGT<-SIGMA-HTH+SIGMA-HTH | 374 | Anaerolineae_bacterium | Bacteria>Bacillati>Chloroflexota>Anaerolineae>unclassified_Anaerolineae | hypothetical-protein-[Anaerolineae-bacterium] | 2052143 |
| MBV6438329.1 | SIGMA-HTH->TGT->DUF4007->STAND->DUF6079->iSwi2/Snf2-Helicase+PglZ*->HerA_FtsK->GNAT-><-CBS+CBS+DUF4268 | 379 | Anaerolineae_bacterium | Bacteria>Bacillati>Chloroflexota>Anaerolineae>unclassified_Anaerolineae | hypothetical-protein-[Anaerolineae-bacterium] | 2052143 |
| MBP8947515.1 | Radical_SAM->?-><-GNAT<-HerA_FtsK<-iSwi2/Snf2-Helicase+PglZ*<-DUF6079-p | 367 | Promineifilum_sp. | Bacteria>Bacillati>Chloroflexota>Ardenticatenia>Candidatus_Promineifilales>Candidatus_Promineifilaceae>Candidatus_Promineifilum>unclassified_Candidatus_Promineifilum | hypothetical-protein-[Promineifilum-sp.] | 2664178 |
| MBW7959471.1 | <-GNAT<-HerA_FtsK<-iSwi2/Snf2-Helicase+PglZ*<-DUF6079<-STAND<-DUF4007<-TGT<-SIGMA-HTH+SIGMA-HTH | 371 | Promineifilum_sp. | Bacteria>Bacillati>Chloroflexota>Ardenticatenia>Candidatus_Promineifilales>Candidatus_Promineifilaceae>Candidatus_Promineifilum>unclassified_Candidatus_Promineifilum | hypothetical-protein-[Promineifilum-sp.] | 2664178 |
| GAB4153051.1 | <-GNAT<-HerA_FtsK<-iSwi2/Snf2-Helicase+PglZ*<-DUF6079<-STAND<-?<-DUF433<-DUF4007<-TGT<-SIGMA-HTH+SIGMA-HTH | 368 | Candidatus_Promineifilaceae_bacterium | Bacteria>Bacillati>Chloroflexota>Ardenticatenia>Candidatus_Promineifilales>Candidatus_Promineifilaceae>unclassified_Candidatus_Promineifilaceae | MAG:-hypothetical-protein-Fur0021_18440-[Candidatus-Promineifilaceae-bacterium] | 3085104 |
| MFN8469222.1 | <-QueC<-Vut_1<-HerA_FtsK<-iSwi2/Snf2-Helicase+PglZ*<-DUF6079<-HTH_Tnp_1+HTH_32+rve+rve_3 | 378 | Caldilineaceae_bacterium | Bacteria>Bacillati>Chloroflexota>Caldilineae>Caldilineales>Caldilineaceae>unclassified_Caldilineaceae | MAG:-alkaline-phosphatase-family-protein-[Caldilineaceae-bacterium] | 2614250 |
| WP_343425287.1 | <-Methyltransf_23+HIT<-?\|\|?-><-Preader8<-GNAT<-HerA_FtsK<-HerA_FtsK<-iSwi2/Snf2-Helicase+PglZ*<-?<-DUF6079-p<-DUF6079-p | 152 | Candidatus_Amarolinea_dominans | Bacteria>Bacillati>Chloroflexota>Caldilineae>Caldilineales>Candidatus_Amarolineaceae>Candidatus_Amarolinea | hypothetical-protein-[Candidatus-Amarolinea-dominans] | 3140696 |
| MBN1891046.1 | DUF4007->STAND->DUF6079->iSwi2/Snf2-Helicase+PglZ*->HerA_FtsK->Meth_synt_2->TPR+TPR+TPR->Radical_SAM-> | 364 | Thermoflexales_bacterium | Bacteria>Bacillati>Chloroflexota>Thermoflexia>Thermoflexales>unclassified_Thermoflexales | PglZ-domain-containing-protein-[Thermoflexales-bacterium] | 2497639 |
| GIK42765.1 | <-Glyoxalase+Glyoxalase<-FGGY_C<-Y1_Tnp<-GNAT<-HerA_FtsK<-iSwi2/Snf2-Helicase+PglZ* | 162 | Chloroflexota_bacterium | Bacteria>Bacillati>Chloroflexota>unclassified_Chloroflexota | MAG:-hypothetical-protein-BroJett011_65980-[Chloroflexota-bacterium] | 2026724 |
| HET59439.1 | SIGMA-HTH->TGT->DUF4007->STAND->DUF6079->iSwi2/Snf2-Helicase+PglZ*->HerA_FtsK->GNAT-> | 374 | Chloroflexota_bacterium | Bacteria>Bacillati>Chloroflexota>unclassified_Chloroflexota | hypothetical-protein-[Chloroflexota-bacterium] | 2026724 |
| MBI5668048.1 | SIGMA-HTH->TGT->DUF4007->STAND->DUF6079->iSwi2/Snf2-Helicase+PglZ*->HerA_FtsK->GNAT->DUF4058-> | 377 | Chloroflexota_bacterium | Bacteria>Bacillati>Chloroflexota>unclassified_Chloroflexota | PglZ-domain-containing-protein-[Chloroflexota-bacterium] | 2026724 |
| MBP7045037.1 | Pkinase_Tyr+Kinase-like->HU-DNA_bdg+DUF4469->SIGMA-HTH+SIGMA-HTH->TGT->DUF4007->STAND->DUF6079->iSwi2/Snf2-Helicase+PglZ*-> | 156 | Chloroflexota_bacterium | Bacteria>Bacillati>Chloroflexota>unclassified_Chloroflexota | hypothetical-protein-[Chloroflexota-bacterium] | 2026724 |
| MBP7045382.1 | iSwi2/Snf2-Helicase+PglZ*->HerA_FtsK->GNAT->McrA-NTD->Y1_Tnp->HNH_2->?->MFS_3-> | 253 | Chloroflexota_bacterium | Bacteria>Bacillati>Chloroflexota>unclassified_Chloroflexota | PglZ-domain-containing-protein-[Chloroflexota-bacterium] | 2026724 |
| MBU6150008.1 | GNAT+TGT->DUF4007->DUF4007->STAND->DUF6079->iSwi2/Snf2-Helicase+PglZ*-> | 343 | Chloroflexota_bacterium | Bacteria>Bacillati>Chloroflexota>unclassified_Chloroflexota | hypothetical-protein-[Chloroflexota-bacterium] | 2026724 |
| MCZ7667409.1 | <-PIN<-DUF2283<-DUF4258<-GNAT<-HerA_FtsK<-HerA_FtsK<-iSwi2/Snf2-Helicase+PglZ*<-DUF6079-p<-DUF6079-p<-?<-?<-?<-?<-STAND | 354 | Chloroflexota_bacterium | Bacteria>Bacillati>Chloroflexota>unclassified_Chloroflexota | PglZ-domain-containing-protein-[Chloroflexota-bacterium] | 2026724 |
| MDQ2805420.1 | <-GNAT<-HerA_FtsK<-iSwi2/Snf2-Helicase+PglZ*<-DUF6079-p | 343 | Chloroflexota_bacterium | Bacteria>Bacillati>Chloroflexota>unclassified_Chloroflexota | hypothetical-protein-[Chloroflexota-bacterium] | 2026724 |
| NLE46402.1 | DUF6079-p->iSwi2/Snf2-Helicase+PglZ*->HerA_FtsK->GNAT->?->NucS->DUF2726-> | 370 | Chloroflexota_bacterium | Bacteria>Bacillati>Chloroflexota>unclassified_Chloroflexota | hypothetical-protein-[Chloroflexota-bacterium] | 2026724 |
| RLC84905.1 | iSwi2/Snf2-Helicase+PglZ*->HerA_FtsK->GNAT-> | 372 | Chloroflexota_bacterium | Bacteria>Bacillati>Chloroflexota>unclassified_Chloroflexota | MAG:-hypothetical-protein-DRI79_12155-[Chloroflexota-bacterium] | 2026724 |
| MEP0921208.1 | Phage_integrase->GNAT+TGT->DUF4007->STAND->DUF6079->iSwi2/Snf2-Helicase+PglZ*->HerA_FtsK->?->GspA-ATPase->?-><-DSPc | 352 | Leptolyngbya_sp._ST-U4 | Bacteria>Bacillati>Cyanobacteriota/Melainabacteria_group>Cyanobacteriota>Cyanophyceae>Leptolyngbyales>Leptolyngbyaceae>Leptolyngbya_group>Leptolyngbya>unclassified_Leptolyngbya | alkaline-phosphatase-family-protein-[Leptolyngbya-sp.-ST-U4] | 2933912 |
| WP_348254528.1 | Phage_integrase->GNAT+TGT->DUF4007->STAND->DUF6079->iSwi2/Snf2-Helicase+PglZ*->HerA_FtsK->?->GspA-ATPase->?-><-DSPc | 352 | Leptolyngbya_sp._ST-U4 | Bacteria>Bacillati>Cyanobacteriota/Melainabacteria_group>Cyanobacteriota>Cyanophyceae>Leptolyngbyales>Leptolyngbyaceae>Leptolyngbya_group>Leptolyngbya>unclassified_Leptolyngbya | hypothetical-protein-[Leptolyngbya-sp.-ST-U4] | 2933912 |
| MDX2239252.1 | Peptidase_C14-><-?<-?<-HerA_FtsK<-iSwi2/Snf2-Helicase+PglZ*<-DUF6079<-STAND<-DUF4007<-GNAT+TGT | 374 | Leptolyngbyaceae_cyanobacterium_bins.302 | Bacteria>Bacillati>Cyanobacteriota/Melainabacteria_group>Cyanobacteriota>Cyanophyceae>Leptolyngbyales>Leptolyngbyaceae>unclassified_Leptolyngbyaceae | hypothetical-protein-[Leptolyngbyaceae-cyanobacterium-bins.302] | 2971802 |
| HEY9694176.1 | AAA_22->?-><-HerA_FtsK<-iSwi2/Snf2-Helicase+PglZ*<-DUF6079<-STAND<-?<-GspA-ATPase<-?<-DUF6079<-DUF4007 | 363 | Oculatellaceae_cyanobacterium | Bacteria>Bacillati>Cyanobacteriota/Melainabacteria_group>Cyanobacteriota>Cyanophyceae>Oculatellales>Oculatellaceae>unclassified_Oculatellaceae | MAG-TPA:-hypothetical-protein-V6D15_18385-[Oculatellaceae-cyanobacterium] | 3088181 |
| MCS3921015.1 | <-HerA_FtsK<-?<-PIN<-VAPB_Antitoxin<-PIN<-RHH<-Mrr_N<-iSwi2/Snf2-Helicase+PglZ*<-DUF6079-p<-DUF6079-p<-VapB_Antitoxin<-STAND<-VbhA_Antitoxin<-?<-DUF4007 | 362 | Candidatus_Fervidibacter_sacchari | Bacteria>Bacteria_incertae_sedis>Bacteria_candidate_phyla>Candidatus_Fervidibacterota>Candidatus_Fervidibacter | hypothetical-protein-[Candidatus-Fervidibacter-sacchari] | 1448929 |
| MEJ7614505.1 | DUF6079-p->iSwi2/Snf2-Helicase+PglZ*->Mrr_N->RHH->PIN->VAPB_Antitoxin->PIN->HerA_FtsK->AhpC-TSA-> | 362 | Candidatus_Fervidibacter_sacchari | Bacteria>Bacteria_incertae_sedis>Bacteria_candidate_phyla>Candidatus_Fervidibacterota>Candidatus_Fervidibacter | MAG:-hypothetical-protein-SQA66_12825-[Candidatus-Fervidibacter-sacchari] | 1448929 |
| MFA0737062.1 | <-HerA_FtsK<-PIN<-RHH<-Mrr_N<-iSwi2/Snf2-Helicase+PglZ*<-DUF6079<-DUF6079-p<-STAND<-VbhA_Antitoxin<-Ntox9<-?<-DUF4007 | 362 | Candidatus_Fervidibacter_canadensis | Bacteria>Bacteria_incertae_sedis>Bacteria_candidate_phyla>Candidatus_Fervidibacterota>Candidatus_Fervidibacter | MAG:-hypothetical-protein-LKKZDAJK_000141-[Candidatus-Fervidibacter-sp.] | 3401948 |
| MFA0740141.1 | DUF6079-p->iSwi2/Snf2-Helicase+PglZ*->Mrr_N->RHH->PIN->VAPB_Antitoxin->PIN->HerA_FtsK->AhpC-TSA-> | 411 | Candidatus_Fervidibacter_sacchari | Bacteria>Bacteria_incertae_sedis>Bacteria_candidate_phyla>Candidatus_Fervidibacterota>Candidatus_Fervidibacter | MAG:-hypothetical-protein-DFNUSKGM_000239-[Candidatus-Fervidibacter-sacchari] | 1448929 |
| MFA0782861.1 | <-HerA_FtsK<-PIN<-VAPB_Antitoxin<-?<-Mrr_N<-iSwi2/Snf2-Helicase+PglZ*<-DUF6079<-VapB_Antitoxin<-STAND<-VbhA_Antitoxin<-Ntox9<-DUF4007<-MetJarc-HTH | 408 | Candidatus_Fervidibacter_sacchari | Bacteria>Bacteria_incertae_sedis>Bacteria_candidate_phyla>Candidatus_Fervidibacterota>Candidatus_Fervidibacter | MAG:-hypothetical-protein-YYHSYBAR_001252-[Candidatus-Fervidibacter-sacchari] | 1448929 |
| WKU14956.1 | <-HerA_FtsK<-?<-PIN<-VAPB_Antitoxin<-PIN<-RHH<-Mrr_N<-iSwi2/Snf2-Helicase+PglZ*<-DUF6079-p<-DUF6079-p<-VapB_Antitoxin<-STAND<-STAND<-VbhA_Antitoxin | 362 | Candidatus_Fervidibacter_sacchari | Bacteria>Bacteria_incertae_sedis>Bacteria_candidate_phyla>Candidatus_Fervidibacterota>Candidatus_Fervidibacter | PglZ-domain-containing-protein-[Candidatus-Fervidibacter-sacchari] | 1448929 |
| WP_018195463.1 | <-HerA_FtsK<-?<-PIN<-VAPB_Antitoxin<-PIN<-RHH<-Mrr_N<-iSwi2/Snf2-Helicase+PglZ*<-DUF6079-p<-DUF6079-p<-VapB_Antitoxin<-STAND<-STAND<-VbhA_Antitoxin | 362 | Candidatus_Fervidibacter_sacchari | Bacteria>Bacteria_incertae_sedis>Bacteria_candidate_phyla>Candidatus_Fervidibacterota>Candidatus_Fervidibacter | PglZ-domain-containing-protein-[Candidatus-Fervidibacter-sacchari] | 1448929 |
| MFA0732614.1 | <-HerA_FtsK<-PIN<-RHH<-Mrr_N<-iSwi2/Snf2-Helicase+PglZ*<-DUF6079<-DUF6079-p<-STAND<-VbhA_Antitoxin<-Ntox9<-DUF4007 | 362 | Candidatus_Fervidibacter_sp. | Bacteria>Bacteria_incertae_sedis>Bacteria_candidate_phyla>Candidatus_Fervidibacterota>Candidatus_Fervidibacter>unclassified_Candidatus_Fervidibacter | MAG:-hypothetical-protein-BKPUNTRY_002173-[Candidatus-Fervidibacter-sp.] | 3100871 |
| MFA0736057.1 | <-HerA_FtsK<-PIN<-VAPB_Antitoxin<-PIN<-RHH<-Mrr_N<-iSwi2/Snf2-Helicase+PglZ*<-DUF6079-p | 363 | Candidatus_Fervidibacter_sp. | Bacteria>Bacteria_incertae_sedis>Bacteria_candidate_phyla>Candidatus_Fervidibacterota>Candidatus_Fervidibacter>unclassified_Candidatus_Fervidibacter | MAG:-hypothetical-protein-OGMRLDGQ_002557-[Candidatus-Fervidibacter-sp.] | 3100871 |
| MFA0736219.1 | <-Mrr_N<-iSwi2/Snf2-Helicase+PglZ*<-DUF6079<-STAND<-VbhA_Antitoxin<-Ntox9<-DUF4007 | 362 | Candidatus_Fervidibacter_sp. | Bacteria>Bacteria_incertae_sedis>Bacteria_candidate_phyla>Candidatus_Fervidibacterota>Candidatus_Fervidibacter>unclassified_Candidatus_Fervidibacter | MAG:-hypothetical-protein-OGMRLDGQ_002719-[Candidatus-Fervidibacter-sp.] | 3100871 |
| MFA0745610.1 | DUF4007->Ntox9->VbhA_Antitoxin->STAND->DUF6079->iSwi2/Snf2-Helicase+PglZ*->Mrr_N->RHH->PIN-> | 362 | Candidatus_Fervidibacter_sp. | Bacteria>Bacteria_incertae_sedis>Bacteria_candidate_phyla>Candidatus_Fervidibacterota>Candidatus_Fervidibacter>unclassified_Candidatus_Fervidibacter | MAG:-hypothetical-protein-LASZOEIN_002423-[Candidatus-Fervidibacter-sp.] | 3100871 |
| MFA0778664.1 | <-HerA_FtsK<-PIN<-RHH<-Mrr_N<-iSwi2/Snf2-Helicase+PglZ*<-DUF6079<-DUF6079-p<-STAND<-VbhA_Antitoxin<-Ntox9<-?<-DUF4007 | 362 | Candidatus_Fervidibacter_sp. | Bacteria>Bacteria_incertae_sedis>Bacteria_candidate_phyla>Candidatus_Fervidibacterota>Candidatus_Fervidibacter>unclassified_Candidatus_Fervidibacter | MAG:-hypothetical-protein-RJAPGHWK_000154-[Candidatus-Fervidibacter-sp.] | 3100871 |
| MEZ8217498.1 | <-Uma2_REase<-HerA_FtsK<-PIN<-VAPB_Antitoxin<-PIN<-RHH<-Mrr_N<-iSwi2/Snf2-Helicase+PglZ*<-DUF6079<-DUF6079-p<-VapB_Antitoxin<-STAND<-VbhA_Antitoxin<-Ntox9<-DUF4007 | 411 | Candidatus_Fervidibacteria_bacterium_JGI_MDM2_SSWTFF-3-K9 | Bacteria>Bacteria_incertae_sedis>Bacteria_candidate_phyla>Candidatus_Fervidibacterota>unclassified_Candidatus_Fervidibacterota | Type-I-phosphodiesterase-/-nucleotide-pyrophosphatase-[Candidatus-Fervidibacteria-bacterium-JGI-MDM2-SSWTFF-3-K9] | 1955661 |
| MFA0760005.1 | MetJarc-HTH->DUF4007->Ntox9->VbhA_Antitoxin->STAND->DUF6079-p->DUF6079->iSwi2/Snf2-Helicase+PglZ*->Mrr_N->RHH->PIN->HerA_FtsK-> | 362 | Candidatus_Fervidibacterota_bacterium | Bacteria>Bacteria_incertae_sedis>Bacteria_candidate_phyla>Candidatus_Fervidibacterota>unclassified_Candidatus_Fervidibacterota | MAG:-hypothetical-protein-HZLCBSQH_000090-[Candidatus-Fervidibacterota-bacterium] | 2080304 |
| MCY3723780.1 | RNA_Pol->GNAT+TGT->DUF4007->DUF6079->STAND->iSwi2/Snf2-Helicase+PglZ*->HerA_FtsK->PIN->?->AP_endonuc_2-> | 364 | Candidatus_Poribacteria_bacterium | Bacteria>Bacteria_incertae_sedis>Bacteria_candidate_phyla>Candidatus_Poribacteria>unclassified_Candidatus_Poribacteria | hypothetical-protein-[Candidatus-Poribacteria-bacterium] | 2026781 |
| MDE0014942.1 | <-HerA_FtsK<-iSwi2/Snf2-Helicase+PglZ*<-?<-STAND<-DUF6079<-DUF6079-p<-DUF4007<-GNAT+TGT<-RNA_Pol | 365 | Candidatus_Poribacteria_bacterium | Bacteria>Bacteria_incertae_sedis>Bacteria_candidate_phyla>Candidatus_Poribacteria>unclassified_Candidatus_Poribacteria | hypothetical-protein-[Candidatus-Poribacteria-bacterium] | 2026781 |
| MDE0397489.1 | RNA_Pol->GNAT+TGT->DUF4007->DUF6079->STAND->?->iSwi2/Snf2-Helicase+PglZ*->HerA_FtsK-> | 370 | Candidatus_Poribacteria_bacterium | Bacteria>Bacteria_incertae_sedis>Bacteria_candidate_phyla>Candidatus_Poribacteria>unclassified_Candidatus_Poribacteria | hypothetical-protein-[Candidatus-Poribacteria-bacterium] | 2026781 |
| MYB94333.1 | RNA_Pol->GNAT+TGT->DUF4007->DUF6079->STAND->iSwi2/Snf2-Helicase+PglZ*->HerA_FtsK-> | 364 | Candidatus_Poribacteria_bacterium | Bacteria>Bacteria_incertae_sedis>Bacteria_candidate_phyla>Candidatus_Poribacteria>unclassified_Candidatus_Poribacteria | hypothetical-protein-[Candidatus-Poribacteria-bacterium] | 2026781 |
| RKU21513.1 | RNA_Pol->GNAT+TGT->DUF4007->DUF6079->STAND->?->iSwi2/Snf2-Helicase+PglZ*->HerA_FtsK-> | 364 | Candidatus_Poribacteria_bacterium | Bacteria>Bacteria_incertae_sedis>Bacteria_candidate_phyla>Candidatus_Poribacteria>unclassified_Candidatus_Poribacteria | MAG:-hypothetical-protein-C6499_21170-[Candidatus-Poribacteria-bacterium] | 2026781 |
| MBU4000570.1 | PAPS_reduct->DUF4007->STAND->DUF6079->iSwi2/Snf2-Helicase+PglZ*->HerA_FtsK->?-><-?\|\|Peptidase_M78-><-?<-SIGMA-HTH+SIGMA-HTH | 360 | Patescibacteria_group_bacterium | Bacteria>Bacteria_incertae_sedis>Bacteria_candidate_phyla>Patescibacteria_group>unclassified_Patescibacteria_group | hypothetical-protein-[Patescibacteria-group-bacterium] | 2052139 |
| MBI3950589.1 | Vut_1-><-HerA_FtsK<-Mrr_N<-iSwi2/Snf2-Helicase+PglZ*<-DUF6079<-STAND<-DUF4007<-GNAT+TGT<-?<-PIN<-DUF433 | 368 | Acidobacteriota_bacterium | Bacteria>Pseudomonadati>Acidobacteriota>unclassified_Acidobacteriota | hypothetical-protein-[Acidobacteriota-bacterium] | 1978231 |
| MYC73121.1 | SIGMA-HTH->GNAT+TGT->DUF4007->STAND->DUF6079->iSwi2/Snf2-Helicase+PglZ*->HerA_FtsK-> | 375 | Gemmatimonadota_bacterium | Bacteria>Pseudomonadati>FCB_group>Gemmatimonadota>unclassified_Gemmatimonadota | hypothetical-protein-[Gemmatimonadota-bacterium] | 2026742 |
| MYD63883.1 | <-PIN<-?<-HerA_FtsK<-iSwi2/Snf2-Helicase+PglZ*<-DUF6079<-STAND<-DUF4007<-GNAT+TGT<-SIGMA-HTH<-?<-PIN | 363 | Gemmatimonadota_bacterium | Bacteria>Pseudomonadati>FCB_group>Gemmatimonadota>unclassified_Gemmatimonadota | hypothetical-protein-[Gemmatimonadota-bacterium] | 2026742 |
| MBM4284321.1 | DUF4007->DUF6079->PAPS_reduct->DUF4007->STAND->DUF6079->iSwi2/Snf2-Helicase+PglZ*->HerA_FtsK-> | 361 | Deltaproteobacteria_bacterium | Bacteria>Pseudomonadati>Myxococcota>Myxococcia>unclassified_Myxococcia | hypothetical-protein-[Deltaproteobacteria-bacterium] | 2026735 |
| RJP56334.1 | PAPS_reduct->DUF4007->STAND->DUF6079->iSwi2/Snf2-Helicase+PglZ*->HerA_FtsK->DUF4065->?-><-Y2_Tnp | 370 | Deltaproteobacteria_bacterium | Bacteria>Pseudomonadati>Myxococcota>Myxococcia>unclassified_Myxococcia | MAG:-hypothetical-protein-C4549_06715-[Deltaproteobacteria-bacterium] | 2026735 |
| BAL53916.1 | GNAT+TGT->DUF4007->STAND->DUF6079->iSwi2/Snf2-Helicase+PglZ*->Mrr_N->HerA_FtsK-> | 362 | uncultured_prokaryote | unclassified_entries>unclassified_sequences>environmental_samples>prokaryotic_environmental_samples | hypothetical-conserved-protein-[uncultured-prokaryote] | 198431 |
| MFZ4828881.1 | DUF6079->iSwi2/Snf2-Helicase+PglZ*->HerA_FtsK->GNAT->TIR_2+NB-ARC-> | 382 | - | - | hypothetical-protein-[Phototrophicaceae-bacterium] | - |

### BrxC-ATPase (BREX subtype-specific) Newick format

(Type_1_BrxC_WP_245241072.1:0.3291949465,(((((((((((Type_1_BrxC_WP_052288823.1:0.0085973655,Type1.supp.BrxC_YP_502551.1:0.0049752887)100/100:0.6392096434,((((((Type_1_BrxC_WP_238326045.1:0.3706146710,Typ
e_1_BrxC_WP_074444907.1:0.5540812916)0/52:0.0842789942,Type_1_BrxC_NBT25028.1:0.8251051547)99.2/90:0.1473094450,(Type1.supp.BrxC_YP_460953.1:0.1920915131,Type1.supp.BrxC_YP_431072.1:0.1605751122)100/10
0:0.2549853633)88/67:0.0916100565,(Type_1_BrxC_OHC72318.1:0.5562100019,Type_1_BrxC_HBI47071.1:0.4848971800)84.4/67:0.0967040223)20.8/25:0.0505294832,(((Type_1_BrxC_MCB1192727.1:0.3672504339,((Type1.supp.B
rxC_YP_528229.1:0.1954581252,Type1.supp.BrxC_YP_003691453.1:0.1909992290)96.2/93:0.0568782714,Type1.supp.BrxC_YP_003157841.1:0.2406791560)32.2/63:0.0779835733)100/100:0.2163255589,((Type_1_BrxC_WP_24091
9947.1:0.2933420189,(Type1.supp.BrxC_YP_733840.1:0.0073786815,Type1.supp.BrxC_YP_869457.1:0.0134525033)100/100:0.4576347020)98.2/97:0.0997595927,Type1.supp.BrxC_YP_004229222.1:0.3771232296)99.4/99:0.114774
0830)99.7/99:0.1303000188,(((Type_1_BrxC_MCQ3924618.1:0.1650192532,Type1.supp.BrxC_YP_003847305.1:0.1420364017)100/100:0.6030371774,Type1.supp.BrxC_YP_004451377.1:0.7636639722)15.3/25:0.0679700042,Type_1
_BrxC_MBA4013265.1:0.7950698568)93.6/32:0.0926930853)78.9/22:0.0912837624)99/37:0.1147963745,((Type_1_BrxC_WP_165367554.1:0.4137365493,Type_1_BrxC_MCC7365290.1:0.5333970426)25/54:0.0828850510,(Type_1_Br
xC_MBK6301657.1:0.6026616595,Type_1_BrxC_WP_264004160.1:0.5236759826)97.8/82:0.1321271011)13.9/46:0.0525114295)96.3/37:0.1663856040)100/100:1.1137870831,((((((((Type_2_BrxC_MBX7222602.1:0.3759273597,Type
_2_BrxC_MBT3378823.1:0.4015706560)99.6/62:0.0803973849,(Type_2_BrxC_MCE9632143.1:0.5021854176,Type_2_BrxC_MCA6572687.1:0.3355509786)98.7/83:0.0911296595)53.1/15:0.0449570411,(Type_2_BrxC_MBX3732320.1:
0.4486390753,((Type_2_BrxC_MCP5560212.1:0.4123036169,Type_2_BrxC_MDX2267520.1:0.4983476575)43.5/38:0.0694400504,Type_2_BrxC_MDR3405897.1:0.3764479917)94.2/38:0.0647341218)86.4/14:0.0480307686)83.8/23:0.
0449684838,(((Type_2_BrxC_WP_239491482.1:0.3766076640,(Type_2_BrxC_RWO03902.1:0.3672776401,(Type2.supp.BrxC_YP_003168567.1:0.1771814902,Type2.supp.BrxC_YP_973304.1:0.2331489081)96.6/78:0.0763805975)99.
9/99:0.1248177324)98.8/28:0.0614648445,(((Type_2_BrxC_MBI4865891.1:0.2722420744,Type_2_BrxC_MBK6942983.1:0.4402074565)98.8/67:0.0798526867,((((((((Type_2_BrxC_MVZ89494.1:0.4887506012,(Type_2_BrxC_WP_0140
10192.1:0.0028037172,Type2.supp.BrxC_YP_004760100.1:0.0179016860)100/100:0.4820650772)99.7/100:0.1582027881,Type_2_BrxC_NDK90272.1:0.3801125436)100/100:0.1764224668,(((Type_2_BrxC_WP_209893504.1:0.46025
65931,Type_2_BrxC_WP_038566393.1:0.3712650878)69/41:0.0435186479,Type2.supp.BrxC_YP_004573278.1:0.2732270224)97.6/57:0.0691396651,Type_2_BrxC_NYH77234.1:0.3041181802)99.7/92:0.1305035603)100/100:0.23919
29763,((Type_2_BrxC_MBU6244326.1:0.3453307825,((Type_2_BrxC_WP_152196006.1:0.3540275309,Type_2_BrxC_MCK9795949.1:0.4685178127)94.5/93:0.0841990016,Type_2_BrxC_PPH05309.1:0.5327583509)99.2/99:0.109603
5869)100/100:0.148019914,Type2.supp.BrxC_YP_003652154.1:0.4308466658)91.5/82:0.0777188475)99.9/99:0.1284969229,((Type_2_BrxC_WP_165632998.1:0.7390838664,((Type_2_BrxC_NDU73743.1:0.2157798925,Type2.supp.
BrxC_YP_001823118.1:0.1811125790)100/100:0.3050666790,(Type_2_BrxC_BFE56536.1:0.2385037167,(Type_2_BrxC_MCK9876541.1:0.0517779361,Type2.supp.BrxC_YP_004019521.1:0.0348312404)100/100:0.2770288121)99.7/9
9:0.1030820201)99.9/99:0.1309445707)90.9/59:0.0649675192,((((Type_2_BrxC_WP_147142479.1:0.4628414842,Type_2_BrxC_GAA2269146.1:0.4303886336)99/100:0.0928194782,Type2.supp.BrxC_YP_288770.1:0.5019866495)99.
1/98:0.1060834512,Type_2_BrxC_WP_267593973.1:0.2348290733)99.9/99:0.1345679781,(((Type_2_BrxC_CAM04375.1:0.0046944111,Type2.supp.BrxC_YP_001107300.1:0.0000010000)100/100:0.1851460650,Type2.supp.BrxC_YP_
003132423.1:0.1744707071)97.2/94:0.1188973184,Type_2_BrxC_WP_326854224.1:0.3746106490)100/100:0.4635972250)98.2/34:0.0873659635)99.8/55:0.0998807849)97.2/53:0.0657607234,(Type_2_BrxC_WP_179789934.1:0.1552
966215,Type2.supp.BrxC_YP_482041.1:0.2339825884)100/100:0.4019186816)84.5/17:0.0465323919,(Type_2_BrxC_MSW49533.1:0.4889247874,Type_2_BrxC_MCP9485022.1:0.4435127332)89.3/34:0.0656168894)54.2/8:0.0324420
882,(Type_2_BrxC_MCB0954591.1:0.5174471653,Type_2_BrxC_WP_131153514.1:0.3983661320)99.7/80:0.1201729770)99.9/47:0.0711619844)51.8/20:0.0305488227,(Type_2_BrxC_WP_200346143.1:0.2390477228,Type2.supp.Brx
C_YP_434638.1:0.2243458561)100/100:0.2811430808)36.6/18:0.0222333907)11.1/8:0.0307092821,(Type_2_BrxC_WP_018719626.1:0.4024934256,(Type_2_BrxC_WP_005004488.1:0.3271294987,(Type_2_BrxC_WP_048671262.1:0.
1160982878,Type2.supp.BrxC_YP_440675.1:0.1343490153)100/100:0.3576173750)97/88:0.0959905430)99.4/72:0.0832185269)98.2/26:0.0530297053)97.2/18:0.0598474437,Type_2_BrxC_MCM2371079.1:0.5243468068)97.7/41:0.10
34050745,((Type_2_BrxC_WP_052548156.1:0.4032452071,Type_2_BrxC_WP_267686051.1:0.5048759160)100/100:0.2571371789,Type_2_BrxC_MBK6462400.1:0.4700552578)100/99:0.3070484738)51/26:0.0600989206,Type_2_Brx
C_MBX3168148.1:0.5228322894)100/100:1.8480800869,((((((((((Type_4_BrxC_NBJ16968.1:0.4080498249,Type4.supp.BrxC_YP_003503326.1:0.3455152345)100/100:0.2396255177,(Type_4_BrxC_MBP6982276.1:0.5374088400,Type
_4_BrxC_MBQ7256248.1:0.7543591158)10.9/47:0.0822234182)86.7/80:0.0944710258,((Type_4_BrxC_NCD03519.1:0.3480313745,Type_4_BrxC_HIS35237.1:0.4989136176)95/91:0.0748366846,Type_4_BrxC_RKJ03002.1:0.7176781
080)89.4/81:0.0825249736)100/100:0.4385730360,Type4.supp.BrxC_YP_003020623.1:1.3326837607)98.2/91:0.1887553389,(Type_4_BrxC_HIT17410.1:0.5077411644,Type_4_BrxC_MBF0318625.1:0.4080802218)100/100:0.3372886
894)99.3/93:0.2387788207,(Type_4_BrxC_AFM22178.1:0.4721517341,Type_4_BrxC_NPV08139.1:0.7771459539)99/95:0.2141347648)52.3/48:0.1000102673,Type_4_BrxC_KUK07754.1:0.8466573745)93.7/77:0.1916756387,Type_4_
BrxC_MCC2258802.1:1.4081287265)99.4/100:0.3371502508,(((((((Type_4_BrxC_NLT83665.1:0.0078962598,Type4.supp.BrxC_YP_002247818.1:0.0129270593)100/100:0.6027440498,(Type_4_BrxC_PMQ01344.1:0.5364099458,Type
_4_BrxC_KUK66683.1:0.4288402706)86.9/83:0.0845502490)99.8/99:0.1406657737,(Type_4_BrxC_WP_048175326.1:0.4314992347,Type4.supp.BrxC_YP_001245329.1:0.3718846699)100/100:0.1655910165)96.9/71:0.0945279730,Ty
pe_4_BrxC_HHW21153.1:0.6242753669)18.9/40:0.0600189331,(Type_4_BrxC_MBQ8951403.1:0.5170874853,Type_4_BrxC_HAR58250.1:0.6810109597)100/100:0.4018368423)51/43:0.0602001352,((Type_4_BrxC_CDD83746.1:0.50
55060182,(Type_4_BrxC_MBL1007423.1:0.2160234327,Type4.supp.BrxC_YP_004328099.1:0.1909836066)99.9/100:0.2066284902)100/100:0.4840510513,Type_4_BrxC_WP_195485920.1:0.8913237897)91.6/69:0.1471495968)87.5/5
5:0.1119504487,Type_4_BrxC_WP_165846692.1:0.9176983152)99.4/99:0.3127916724)99.9/100:0.6272039080,(Type_4_BrxC_WP_102456724.1:1.6871588596,Type_4_BrxC_WP_134114886.1:1.0558543051)50.4/61:0.3626340611)9
9.9/99:0.5954762353)87.5/83:0.1650404183,((((Type_3_BrxC_EMR73852.1:0.3789152405,(Type_3_BrxC_WP_114474457.1:0.3788760406,Type3.supp.BrxC_YP_003827964.1:0.2224556879)100/100:0.1721485889)88.7/75:0.0660099
774,((Type_3_BrxC_KPQ15860.1:0.5380584710,Type_3_BrxC_NLE00367.1:0.3377067857)100/100:0.6031746393,Type_3_BrxC_KFD41618.1:0.4808206224)48/54:0.0986957946)99.7/94:0.2244442559,Type_3_BrxC_MBU0460852.1:
0.5803167452)98.8/99:0.3373662450,((((((((Type_3_BrxC_NLV27395.1:0.4443672994,Type_3_BrxC_KUK47744.1:0.3153338099)100/99:0.1513757757,((Type_3_BrxC_MBT5530412.1:0.3929667408,((Type_3_BrxC_SLK50155.1:0.314
4122186,Type_3_BrxC_WP_239738037.1:0.3607697031)100/100:0.2005035608,Type_3_BrxC_WP_106343732.1:0.3361136143)96.3/62:0.0816197516)96.3/35:0.0459591597,((((((((Type_3_BrxC_SDB48716.1:0.1491952289,(Type3.s
upp.BrxC_YP_001412458.1:0.0000010000,Type3.supp.BrxC_YP_001414810.1:0.0000010000)100/100:0.1523365483)100/100:0.2412630165,Type_3_BrxC_BAP57136.1:0.4889181675)95.5/45:0.0734951620,((((((((Type_3_BrxC_ADJ
27036.1:0.0000010000,Type3.supp.BrxC_YP_003759357.1:0.0000010000)95.7/100:0.0073334426,Type3.supp.BrxC_YP_342128.1:0.0355608826)95.2/83:0.0069684804,Type3.supp.BrxC_YP_004121412.1:0.0054226741)53.3/45:0.00
36627517,Type3.supp.BrxC_YP_001209110.1:0.0133520598)100/100:0.0700240530,Type3.supp.BrxC_YP_004616376.1:0.1038231260)97.3/99:0.0477008488,Type_3_BrxC_OQB94794.1:0.3863235498)92/46:0.0644344141,(((Type3.
supp.BrxC_YP_872571.1:0.0426708798,Type3.supp.BrxC_YP_003318003.1:0.0138155724)95.9/99:0.0129771726,Type3.supp.BrxC_YP_001938983.1:0.0528830201)99/100:0.0309821804,Type3.supp.BrxC_YP_002462466.1:0.13813
24332)72.2/72:0.0291178408)98.2/44:0.0453620245,Type_3_BrxC_MBT4668298.1:0.3616793244)64/12:0.0210680216)75.9/5:0.0290308934,(Type_3_BrxC_PKL79310.1:0.4063401239,Type_3_BrxC_WP_190509459.1:0.3893292714)
92.9/35:0.0552947907)78.5/8:0.0371406959,Type_3_BrxC_MCK9221196.1:0.3721913931)90.6/22:0.0429150966,(Type_3_BrxC_WP_108651333.1:0.5374823636,Type_3_BrxC_WP_068347596.1:0.4742992764)77.9/12:0.0671744440)
99.4/47:0.0839421211,(Type3.supp.BrxC_YP_001211256.1:0.0333043162,Type3.supp.BrxC_YP_003702611.1:0.0175172644)100/100:0.2186852204)44.9/13:0.0355318617,(Type_3_BrxC_ADY58758.1:0.0000010000,Type3.supp.BrxC
_YP_004268780.1:0.0010994857)100/100:0.3379530826)86.5/13:0.0333626223)83/7:0.0315262280)87.4/8:0.0465383571,(((Type_3_BrxC_MBP8682477.1:0.4315731987,(Type_3_BrxC_WP_209451817.1:0.4430478794,Type_3_BrxC
_HHV38843.1:0.3116496302)94.6/88:0.1074219351)99.8/100:0.1407672590,(((((Type_3_BrxC_EGD52432.1:0.0094027049,Type3.supp.BrxC_YP_003477172.1:0.0121380867)93.7/93:0.0117531401,(Type3.supp.BrxC_YP_004185914.
1:0.0000010000,Type3.supp.BrxC_YP_001664917.1:0.0000010000)83.8/100:0.0072715516)100/100:0.2324365592,Type3.supp.BrxC_YP_004470680.1:0.2665090870)99.7/99:0.1147742061,Type3.supp.BrxC_YP_004461308.1:0.3361
702981)86.8/80:0.0587016131,(Type_3_BrxC_MCF6096371.1:0.2532609300,Type3.supp.BrxC_YP_004025780.1:0.2381745368)100/100:0.1609887206)97.2/95:0.0597779334)100/100:0.1287699618,Type_3_BrxC_PSR33881.1:0.746
2674382)73.9/15:0.0560090405)76.3/7:0.0402325449,(Type_3_BrxC_MBT6646600.1:0.5986704373,Type_3_BrxC_HHW29753.1:0.4184948719)85/42:0.0656068699)98/25:0.0801309064,(Type_3_BrxC_MXW95274.1:0.3185666640,Ty
pe_3_BrxC_WP_227569851.1:0.3924600774)100/100:0.2768978056)96.8/61:0.1050280843,(Type_3_BrxC_MBV6489844.1:0.4606298327,Type_3_BrxC_MCM1566961.1:0.4151341220)100/100:0.6415808947)17.5/27:0.0648563486,T
ype_3_BrxC_WP_020614466.1:0.5231360796)98.7/92:0.1780614497,((Type_3_BrxC_KUO63976.1:0.2434079484,Type_3_BrxC_NCD09233.1:0.4933220107)99.9/100:0.2507020327,(Type_3_BrxC_MBK7265049.1:0.3087228904,Type
_3_BrxC_MBR2506166.1:0.4409008341)97.1/89:0.1461731740)100/100:0.3256371402)100/100:0.5496136892)100/100:0.8558003697)99.3/100:0.3736426744)33.6/50:0.0805687013,(((Type_5_BrxC_TQQ78814.1:0.2941601791,Type
_5_BrxC_WP_114604755.1:0.4606956400)99.8/99:0.2960701868,((Type_5_BrxC_WP_020446108.1:0.2270900623,(Type5.supp.BrxC_YP_004809888.1:0.0128097935,Type5.supp.BrxC_YP_002564616.1:0.0129604458)100/100:0.160
8985662)100/100:0.1918205836,((Type5.supp.BrxC_YP_004786063.1:0.0372206703,Type5.supp.BrxC_YP_004595483.1:0.0247577022)99.6/98:0.0608033851,Type5.supp.BrxC_YP_003130811.1:0.0752494156)100/100:0.2045460637
)99/100:0.2094746544)100/100:1.0575630084,((Type5.supp.BrxC_YP_004809890.1:0.0333040145,Type5.supp.BrxC_YP_002564614.1:0.0284144169)99.9/100:0.2733908916,(Type5.supp.BrxC_YP_004595485.1:0.0771704841,Type5.
supp.BrxC_YP_003130813.1:0.0653644883)99.5/100:0.2330572350)100/100:1.2849039979)99.8/99:0.5228163493)95/79:0.1991444871,(((Type_1_BrxC_MCE8425368.1:0.4632055346,(Type_1_BrxC_WP_008844769.1:0.5564194679,
Type_1_BrxC_KUK86067.1:0.4700938806)99.5/100:0.2251249499)100/100:0.6586929759,(Type_1_BrxC_KUK92978.1:0.5484135055,Type_1_BrxC_HCZ06893.1:0.3095407123)100/100:0.6698661273)99.8/100:0.3279198799,(((Type
_1_BrxC_OFV66969.1:1.1698207597,((Type_6_BrxC_ETI91650.1:0.5882250539,Type_6_BrxC_WP_075749471.1:0.5796330632)100/100:0.3301848358,Type_6_BrxC_WP_049936156.1:0.8059561986)52.6/49:0.1053133178)66/30:0.
0720693694,Type_1_BrxC_EMR73247.1:1.2570554627)96.4/67:0.1396560184,(((((((Type_1_BrxC_MBK9165938.1:0.2756811465,Type_6_BrxC_VVB72030.1:0.3561234360)100/100:0.2458038993,(Type_1_BrxC_NIM12892.1:0.33119
47909,Type1.supp.BrxC_NP_953157.1:0.2840125291)100/100:0.1762882950)100/100:0.1607706158,((Type_1_BrxC_WP_245302581.1:0.0209998293,Type_6_BrxC_MBP2018669.1:0.0000020445)100/100:0.4941808741,Type_6_Brx
C_MBZ5596030.1:0.4157126206)94/85:0.1048260082)98.6/94:0.0936706993,(Type_6_BrxC_PZV04977.1:0.5957551174,(Type_6_BrxC_TXH34387.1:0.1017872763,Type1.supp.BrxC_YP_004282674.1:0.0983289272)100/100:0.34799
21998)99.9/100:0.1875257094)91/84:0.0839780638,((Type_6_BrxC_MCB1001743.1:0.6140100162,Type_6_BrxC_WP_208027651.1:0.2937775733)98.5/99:0.1260306489,Type_6_BrxC_WP_166680331.1:0.4626140656)100/100:0.378
2821186)99.8/100:0.1927956609,((((((Type_1_BrxC_MAE92616.1:0.2616901485,Type_6_BrxC_WP_240631539.1:0.1702834149)100/100:0.2243773436,Type_6_BrxC_MBE7502027.1:0.4644299274)95.1/68:0.0914902299,(Type_6_Br
xC_MCP5535434.1:0.6112165295,Type_6_BrxC_WP_027137224.1:0.6418218153)82.4/35:0.0965584245)0/15:0.0537565744,Type_6_BrxC_WP_015282553.1:0.3649873977)73.3/37:0.0672894056,Type_6_BrxC_OQB98767.1:0.45660
73130)100/95:0.2183694746,(Type_6_BrxC_WP_012881599.1:0.0014263931,Type1.supp.BrxC_YP_003329751.1:0.0091891100)100/100:0.5765713473)100/100:0.6802798816)32/61:0.1363749269,((((Type_6_BrxC_MCA1971289.1:0.
3715610784,Type6.supp.BrxC_YP_002491566.1:0.2565077901)99.5/98:0.1047482836,((Type_6_BrxC_WP_009097835.1:0.4138128156,((Type6.supp.BrxC_YP_003265689.1:0.0042481435,Type6.supp.BrxC_YP_003265130.1:0.00401
22142)100/100:0.2233515922,Type6.supp.BrxC_YP_003631098.1:0.3180549730)89.4/87:0.0883324090)99.8/100:0.1292216030,Type_6_BrxC_WP_106091681.1:0.4771624886)24.5/37:0.0463009499)91.8/59:0.0812675331,Type_6_B
rxC_WP_146981109.1:0.3741258456)100/100:0.3341498719,Type_6_BrxC_WP_015675681.1:0.8142125452)100/100:0.4849836566)100/99:0.2712728005)50.5/35:0.1039492219)51.4/37:0.1023696460)99.9/100:0.3274355118,(((Type
_1_BrxC_KAA3613611.1:0.3546042607,Type_1_BrxC_OWY22675.1:0.5904464537)59.8/61:0.0911764973,Type_1_BrxC_OJV92258.1:0.7090953663)96.6/93:0.1895031939,Type1.supp.BrxC_YP_003389621.1:0.5920947941)100/100:0
.6870396987)98.4/92:0.1839100054,((((((((((Type_1_BrxC_WP_101538034.1:0.1029182609,Type_5_BrxC_WP_007349691.1:0.1050743345)100/100:0.1012340054,Type_5_BrxC_MCP3686167.1:0.2964591648)96.5/97:0.0587248635,(
Type_5_BrxC_OYZ13690.1:0.1788243406,Type_5_BrxC_WP_233675470.1:0.2779076178)100/100:0.1300538543)100/99:0.1135123603,((Type_1_BrxC_WP_100350249.1:0.3876922240,(Type_5_BrxC_OUS97312.1:0.2534268346,Ty
pe_5_BrxC_WP_096305156.1:0.1880975709)99.3/100:0.0998473486)99.9/91:0.1428890891,Type_5_BrxC_WP_104138811.1:0.3578817345)100/84:0.2021839624)20.2/31:0.0502122371,Type_1_BrxC_WP_019002750.1:0.648633581
6)95.2/37:0.0688205770,(Type_1_BrxC_MBU3665708.1:0.5593503724,Type_1_BrxC_MBP9037128.1:0.6418086853)95.7/37:0.0797530234)96.5/25:0.0613381483,(((Type_1_BrxC_MCK6622791.1:0.3343434831,(Type_1_BrxC_WP_1
41050577.1:0.3481618567,Type1.supp.BrxC_YP_004060549.1:0.4050463242)100/100:0.2243630224)78.3/70:0.0596439210,(Type_1_BrxC_WP_046151576.1:0.6615597382,Type1.supp.BrxC_YP_004657744.1:0.3074703600)100/99:
0.2538336724)97.1/78:0.0670184337,Type_1_BrxC_MCF7790541.1:0.4813258289)88.5/44:0.0521056997)98.3/35:0.0939710887,(Type_1_BrxC_MCH3907669.1:0.6507430740,(Type_1_BrxC_MBR2835999.1:0.1181349717,Type1.sup
p.BrxC_YP_003142713.1:0.1247887218)100/100:0.7374145739)92.3/52:0.1110469727)19.2/22:0.0476918777,Type_1_BrxC_WP_004808447.1:0.9682520022)89.6/74:0.0856752501,Type_1_BrxC_MCK6605711.1:0.8128181404)97.5/9
3:0.1086848451)100/99:0.1742981363,(Type_1_BrxC_MBZ4644075.1:0.1222247269,Type1.supp.BrxC_YP_004052084.1:0.1259289868)100/100:0.6028097598)95.8/24:0.0773234598,((((Type_1_BrxC_MBF0468020.1:0.4306046318,(
Type_1_BrxC_WP_107866915.1:0.2619034557,Type_5_BrxC_MBX3681311.1:0.2334055571)100/100:0.2177746756)74.7/58:0.0460308344,((Type_1_BrxC_WP_233672193.1:0.0447964476,Type_5_BrxC_WP_139442435.1:0.0404282
946)100/100:0.4218289216,(Type_1_BrxC_MBL4799472.1:0.3046115775,Type_5_BrxC_MQT58298.1:0.2016362312)100/100:0.2081481997)98.6/83:0.0879382495)62.7/49:0.0436763060,(Type_1_BrxC_WP_116688140.1:0.38197686
31,Type_1_BrxC_WP_007065363.1:0.5337920434)97.2/88:0.1131850428)100/100:0.2349700134,(Type_5_BrxC_WP_146889206.1:0.3925897990,Type1.supp.BrxC_YP_001868904.1:0.3939560777)100/100:0.2058834217)59.1/62:0.0
442871621)63.4/11:0.0309293832,Type_1_BrxC_WP_185874362.1:0.8447858683)89.6/11:0.0506716770,((((((Type_1_BrxC_MBS5450116.1:0.4502725218,Type_1_BrxC_HIT45392.1:0.3435305636)99.5/99:0.1565857869,(Type_1_Brx
C_MBP3694027.1:0.1693626848,Type1.supp.BrxC_YP_004413155.1:0.2340815886)100/100:0.2930751047)100/100:0.3145061528,(Type_1_BrxC_WP_131470342.1:0.1571647666,Type1.supp.BrxC_YP_001812817.1:0.1671157802)1
00/100:0.2699567325)92.7/42:0.0732358622,(Type_1_BrxC_WP_254867722.1:0.4165504987,Type_1_BrxC_WP_091403827.1:0.5221527440)100/100:0.2428792131)68.6/14:0.0532617750,((((((Type_1_BrxC_MCH4057848.1:0.40106
97292,Type1.supp.BrxC_YP_003293164.1:0.6629880040)100/99:0.2583924620,Type_1_BrxC_KAB0576672.1:0.3360275312)65.5/54:0.0654079249,Type1.supp.BrxC_YP_001274324.1:0.4581210994)99.6/74:0.1071950918,(Type_1_B
rxC_WP_251617110.1:0.6669254468,Type_1_BrxC_WP_038408557.1:0.5905603317)93.2/32:0.0696741925)7.4/19:0.0377970174,Type_1_BrxC_MBQ8901518.1:0.6280290422)48/16:0.0414518727,(Type_1_BrxC_MCI1881447.1:0.48
70599793,Type_1_BrxC_WP_015326777.1:0.3891624782)55.7/23:0.0664297922)99.5/25:0.0820831325)96/8:0.0445540983,(((((((Type_1_BrxC_WP_088074350.1:0.2031964057,Type_5_BrxC_KKE80057.1:0.2881791542)87.2/66:0.05
78171697,(Type_5_BrxC_WP_089525998.1:0.1871122997,Type_5_BrxC_WP_207649636.1:0.2555823949)99.9/100:0.0893746598)93.3/81:0.0611100179,Type_5_BrxC_WP_066463066.1:0.4490299748)100/100:0.2442524704,(Type_
5_BrxC_WP_002116626.1:0.1356151411,Type1.supp.BrxC_YP_002949413.1:0.1155020296)100/100:0.2711065925)42.2/40:0.0492586737,Type_5_BrxC_WP_164950923.1:0.4982680674)71.5/57:0.0718943153,Type_5_BrxC_WP_029
421470.1:0.5341024908)100/100:0.4240949794,Type_1_BrxC_TCW39537.1:0.4381584417)50.6/8:0.0443147811)71.9/5:0.0331921087)98.1/74:0.0802720857,(Type_1_BrxC_MBN2327413.1:0.4338493195,Type_1_BrxC_PKK90022.1:
0.4415342276)87.5/68:0.0534301370)100/100:0.3531054342,(Type_5_BrxC_WP_048047670.1:0.0076484983,Type1.supp.BrxC_NP_632178.1:0.0289243694)100/100:0.2383432872);

### PglZ (BREX subtype-specific) Newick format

(Type_1_PglZ_HBI47076.1:0.4809139404,(((((((((((((((((Type_1_PglZ_KAA3613614.1:0.4283852634,Type1.supp.PglZ_YP_003389625.1:0.6033211037)71.6/58:0.0824510614,Type_1_PglZ_OJV92255.1:0.6726746
598)100/100:0.4353403432,Type_1_PglZ_WP_251617128.1:0.7231335369)85.2/26:0.0912745124,(((((((((Type_1_PglZ_KAB0576667.1:0.1463622948,Type1.supp.PglZ_YP_004708913.1:0.0998102154)100/100:0.3
150845093,(((Type_1_PglZ_MCH4057844.1:0.3542334906,Type1.supp.PglZ_YP_003171840.1:0.3925456324)96.1/97:0.1364173842,Type1.supp.PglZ_YP_003620556.1:0.4201500112)97.2/98:0.1722543137,Type1.
supp.PglZ_YP_003293159.1:0.5807465974)99/99:0.1951827384)95.7/93:0.1363286797,Type1.supp.PglZ_YP_001274326.1:0.5017467626)89.2/76:0.1078157693,Type_1_PglZ_MBQ8901516.1:0.7135618497)92.3/2
0:0.0805429714,Type1.supp.PglZ_YP_003851153.1:0.3315431050)75/13:0.0900487956,Type_1_PglZ_WP_015326779.1:0.4255442127)30.8/5:0.0345376331,Type_1_PglZ_MCI1881445.1:0.5386717851)93.4/3:0.07
10667269,((((Type_1_PglZ_MBS5450119.1:1.0540401068,Type1.supp.PglZ_YP_516912.1:0.4083392359)98.4/63:0.2110670419,((Type_1_PglZ_WP_131470339.1:0.1924566527,Type1.supp.PglZ_YP_001812819.1:
0.2329222748)99.5/100:0.1786687915,Type1.supp.PglZ_YP_755149.1:0.4431918794)94.7/93:0.0919309179)93.1/46:0.0643213451,(((((Type_1_PglZ_MBZ4644079.1:0.1512985019,Type1.supp.PglZ_YP_00405208
0.1:0.1319598035)100/100:0.5190323167,Type_1_PglZ_WP_038408560.1:0.6501964783)96/39:0.1855475060,(Type_1_PglZ_WP_091403831.1:0.5788855715,Type_1_PglZ_WP_175420183.1:0.4481080181)93.1/7
0:0.1181192091)85.8/16:0.0758814885,Type_1_PglZ_TCW39539.1:0.4487030098)59.5/8:0.0539056090,(Type_1_PglZ_WP_167815508.1:0.3622415967,Type1.supp.PglZ_NP_632187.1:0.2524378585)100/100:0.28
77244108)86.5/5:0.0624446746)46.6/3:0.0255794333,(Type_5_PglZ_WP_186816219.1:0.4267694353,Type1.supp.PglZ_YP_001868908.1:0.3758487120)100/96:0.3354538685)76.7/2:0.0462248441)53.2/3:0.077670
2327,(((Type_5_PglZ_KKE80055.1:0.2689280846,Type_5_PglZ_WP_066463074.1:0.3584114809)29.8/53:0.0405992217,(Type_5_PglZ_WP_075365643.1:0.1766657865,Type_5_PglZ_WP_245850974.1:0.21465339
07)97.5/93:0.0896228993)99.6/99:0.1805587549,(((Type_5_PglZ_WP_002116630.1:0.1600583216,Type1.supp.PglZ_YP_002949415.1:0.1270113917)99.9/100:0.1888565630,Type_5_PglZ_WP_164950928.1:0.4022
298167)48.6/51:0.0528267402,Type_5_PglZ_WP_029421469.1:0.5032982331)17.4/42:0.0572794690)99.9/100:0.3660633453)29.2/5:0.0685396113)91/20:0.0760289323,((((Type_1_PglZ_WP_007065361.1:0.51789
59227,Type_1_PglZ_WP_116688147.1:0.3944687214)31.3/40:0.0511125403,Type1.supp.PglZ_YP_002954989.1:0.3453239186)58.4/51:0.0803400695,Type_1_PglZ_WP_107866910.1:0.5250063194)88.1/85:0.0667
620641,(((Type_1_PglZ_WP_233672185.1:0.0524692266,Type_5_PglZ_WP_139442442.1:0.0424920488)100/100:0.2860714935,Type1.supp.PglZ_YP_002394574.1:0.2792296596)99.1/98:0.1298889123,(Type_5_P
glZ_MQT58295.1:0.2080573751,Type1.supp.PglZ_YP_001708751.1:0.1502941082)99.9/100:0.1979218088)93.9/81:0.0773250263)100/100:0.2990251576)96.8/22:0.1017082866,((Type_1_PglZ_MBU3665703.1:0.53
94748025,((Type_1_PglZ_MCF7790535.1:0.5018341244,(Type_1_PglZ_WP_046151580.1:0.4938095296,Type1.supp.PglZ_YP_004657738.1:0.3140301308)94.4/70:0.0989983970)47.3/30:0.0376468086,(Type_1_P
glZ_MCK6622787.1:0.2998025030,Type1.supp.PglZ_YP_004060557.1:0.3647643054)93.7/69:0.0896240322)93.8/59:0.0856303786)71.4/42:0.0698859638,((Type_1_PglZ_MCH3907672.1:0.6349443993,Type_1_Pgl
Z_WP_019002749.1:0.4615664455)91.5/26:0.1344679581,(((Type_1_PglZ_WP_100350245.1:0.4703391455,(Type_5_PglZ_OUS97315.1:0.3408097113,Type_5_PglZ_WP_096303467.1:0.2272501081)78.4/73:0.055
0512487)97.3/81:0.1065311378,Type_5_PglZ_WP_104138814.1:0.3535736445)99.2/84:0.1485348465,((((Type_1_PglZ_WP_172437662.1:0.0903514236,Type_5_PglZ_WP_007349686.1:0.1094078426)99.6/99:0.08
60802687,Type1.supp.PglZ_YP_004784799.1:0.3031262341)85.8/60:0.0503745012,Type_5_PglZ_MCP3686165.1:0.3617627473)85.1/55:0.0468594396,((Type_5_PglZ_MCE5972445.1:0.0380248029,Type1.supp.P
glZ_YP_001169902.1:0.0374267395)100/100:0.3031022891,Type_5_PglZ_OYZ13692.1:0.2432708958)97.9/91:0.1186500509)95.8/71:0.0915290573)94.3/64:0.0880868367)93/14:0.0722594988)93.3/12:0.07723335
11)88.9/6:0.0825532355,Type_1_PglZ_MCK6605714.1:0.6511086395)57.3/17:0.0804153848,Type_1_PglZ_MBR2835998.1:0.7966214051)63.4/19:0.0666108896,Type_1_PglZ_WP_167568223.1:0.6387388455)100/
98:0.4953778688,(((((((Type_1_PglZ_NIM12894.1:0.7760962506,(Type_1_PglZ_WP_209466794.1:0.0441434224,Type_6_PglZ_MBP2018666.1:0.0050255569)100/100:0.6855036444)29.3/24:0.0976817320,(Type_6
_PglZ_PZV04975.1:0.9670416319,Type_6_PglZ_TXH34390.1:0.4072846864)95.1/76:0.2281703767)36.1/22:0.0635208885,(Type_6_PglZ_MCB1001745.1:0.5745858369,Type_6_PglZ_WP_166680330.1:0.4559456
922)99.9/100:0.3803405968)72.7/34:0.1156942044,((((((Type_6_PglZ_MBE7502035.1:0.4643143189,Type_6_PglZ_WP_027137222.1:0.9917591474)76.1/52:0.0949992389,Type_6_PglZ_WP_121738026.1:0.59695
32128)58.2/42:0.0892855783,Type_6_PglZ_OQB98762.1:0.6807564392)88.1/59:0.1378988129,Type_6_PglZ_WP_015282546.1:0.4438042099)50.5/42:0.1202456050,Type_6_PglZ_MCP5535438.1:0.7568999879)9
7.2/98:0.2419958045,Type_6_PglZ_WP_012881597.1:0.6844902779)99.8/100:0.4518480204)89.3/64:0.1319214989,(Type_6_PglZ_EOQ90728.1:1.1061769751,(((Type_6_PglZ_WP_009097832.1:0.4389237982,Typ
e6.supp.PglZ_YP_003631096.1:0.3525169191)10.7/42:0.0597324093,(Type6.supp.PglZ_YP_003265691.1:0.0100297915,Type6.supp.PglZ_YP_003265132.1:0.0000029154)100/100:0.3063940896)87.4/64:0.077384
1629,((Type_6_PglZ_WP_146981098.1:0.4019214141,Type6.supp.PglZ_YP_002491568.1:0.4526711633)81.8/61:0.1096341377,Type_6_PglZ_WP_181234048.1:0.6385317050)30.3/33:0.0536984574)99.6/96:0.391
3906212)98.6/96:0.3364558387)93.9/74:0.1778193034,((Type_6_PglZ_ETI91648.1:0.6075464624,Type_6_PglZ_OLQ47620.1:0.6478834271)100/100:0.5715138101,Type_6_PglZ_WP_049936153.1:0.9752275986)8
1.9/37:0.1211131503)94.3/25:0.2047693174,Type_1_PglZ_OFV66962.1:1.0811414011)62.6/17:0.0907556865)91.5/68:0.2527050497,((((Type5.supp.PglZ_YP_004786056.1:0.0202329642,Type5.supp.PglZ_YP_0031
30820.1:0.1262962264)68.4/38:0.0120884021,Type5.supp.PglZ_YP_001690753.1:0.0299223330)69/26:0.0064528042,Type5.supp.PglZ_YP_004595497.1:0.0211495313)99.8/98:0.4705648809,(Type5.supp.PglZ_YP
_004809895.1:0.0500102146,Type5.supp.PglZ_YP_002564610.1:0.0096879632)86.7/82:0.1803682721)100/100:1.3548720684)29.5/37:0.1188324782,((((((Type_2_PglZ_BAN04395.1:0.7020322446,Type_2_PglZ_M
SW49534.1:0.5758575160)22.5/6:0.0811159328,(((Type_2_PglZ_MBK6942982.1:0.4919814538,Type_2_PglZ_WP_165491753.1:0.5358728639)13.8/21:0.0923988365,Type_2_PglZ_MCB0954592.1:0.5195253824)9
0/24:0.1066763569,((((((Type_2_PglZ_MBK9578952.1:0.7730887492,Type_2_PglZ_WP_005004490.1:0.5241707916)11.6/16:0.0544075485,Type_2_PglZ_MBX3168147.1:0.9771982332)32.4/2:0.0411747979,(Type_
2_PglZ_WP_018719627.1:0.6191191177,Type2.supp.PglZ_YP_440676.1:0.7112270704)73.9/15:0.1369470491)75.8/1:0.0655811857,((((Type_2_PglZ_MBT3378822.1:0.7191177179,Type_2_PglZ_MCM2371082.1:0.
7325431274)45.7/12:0.0655002530,Type_2_PglZ_WP_239491483.1:0.5344556767)64.2/5:0.0508494252,Type_2_PglZ_MDX2267519.1:0.7849280555)56.2/4:0.0545868355,Type_2_PglZ_MCE9632142.1:0.6194442
814)73.4/2:0.0594766299)90/1:0.0561733920,(Type_2_PglZ_RWO03903.1:0.5245111205,(Type2.supp.PglZ_YP_003168566.1:0.4481403961,Type2.supp.PglZ_YP_973303.1:0.4016276181)98.7/98:0.1605947692)97
.1/44:0.1070441917)81/0:0.0325664462,((Type_2_PglZ_MBX3732321.1:0.6237302687,(Type_2_PglZ_MCP5560213.1:0.6838509301,Type_2_PglZ_MDR3405896.1:0.4792016419)19.6/27:0.0739493236)91.8/42:0.0
783026360,(Type_2_PglZ_WP_242476961.1:0.3860469995,Type2.supp.PglZ_YP_434637.1:0.4462669078)99.9/100:0.2912371140)96.2/20:0.0991498322)95.6/2:0.0909949766)68/1:0.0483848989)75.5/4:0.0300036
464,(((((Type_2_PglZ_BFE56535.1:0.4630575766,(Type_2_PglZ_NDU73742.1:0.1969982976,Type2.supp.PglZ_NP_630712.1:0.2469344666)100/100:0.3366728565)41.6/46:0.0453078076,(Type_2_PglZ_MCK98765
40.1:0.1674971310,Type2.supp.PglZ_YP_004019522.1:0.0662315627)100/100:0.3939062867)99.6/93:0.1610639778,((((Type_2_PglZ_CAM04374.1:0.0142251853,Type2.supp.PglZ_YP_001107299.1:0.0106207889)
99.8/100:0.2358301604,Type2.supp.PglZ_YP_003132424.1:0.2687726384)99/99:0.1959857805,Type_2_PglZ_WP_326854225.1:0.5157170955)99.6/100:0.2333773009,((Type_2_PglZ_GAA2269152.1:0.522814431
7,Type_2_PglZ_WP_158645688.1:0.4422226881)95.4/91:0.1300566168,(Type_2_PglZ_WP_267593974.1:0.3164309408,Type2.supp.PglZ_YP_288771.1:0.3898339849)91.7/88:0.0902654410)96/93:0.0945051828)9
8/97:0.1206678696)88.9/56:0.1020054772,(((Type_2_PglZ_MBU6244327.1:0.5785144476,((Type_2_PglZ_MCK9795948.1:0.4495923074,Type_2_PglZ_PPH05310.1:0.4928885518)78/36:0.0739186283,Type_2_Pgl
Z_WP_152814798.1:0.3893488712)90.1/40:0.0623110491)83/35:0.0701586791,(((((Type_2_PglZ_MVZ89493.1:0.4918951332,(Type_2_PglZ_WP_014010193.1:0.0247564164,Type2.supp.PglZ_YP_004760101.1:0.0
141039997)100/100:0.5860004522)65.9/63:0.1055905022,Type_2_PglZ_NDK90273.1:0.4433356705)99.6/99:0.2212004946,((Type_2_PglZ_WP_038566395.1:0.3750886672,Type2.supp.PglZ_YP_004573277.1:0.41
02552992)99.5/94:0.2062633765,Type2.supp.PglZ_YP_001134466.1:0.5076184902)75.5/43:0.0723166415)34.2/36:0.0819829006,Type_2_PglZ_WP_209893507.1:0.5093618405)85.2/84:0.1066287082,Type_2_Pgl
Z_WP_165632999.1:0.8231956954)71.7/39:0.0608558864)97.5/32:0.0935229580,(Type2.supp.PglZ_YP_003834434.1:0.5414300504,Type2.supp.PglZ_YP_003652155.1:0.5001880343)57.2/33:0.0598061523)83.6/2
9:0.0771019515)95.3/80:0.0947123155,(Type_2_PglZ_WP_272956161.1:0.1974852718,Type2.supp.PglZ_YP_482042.1:0.1892118436)100/100:0.5231300244)93/47:0.0881762871)99.5/56:0.3899788159,(Type_2_P
glZ_MBK6462401.1:0.6589589438,Type_2_PglZ_WP_052548157.1:0.8545141900)91.9/93:0.2371893358)99.5/99:0.9889384860,(((((((((Type_3_PglZ_ADJ27043.1:0.0782345364,Type3.supp.PglZ_YP_004121417.1:
0.0581949832)95/97:0.0703296800,Type3.supp.PglZ_YP_004616370.1:0.2835141545)98.6/94:0.1349648607,(Type_3_PglZ_MXW95271.1:1.1927215909,Type_3_PglZ_OQB94797.1:0.6066980703)7.2/6:0.0705684
072)97.5/13:0.1127521935,(Type_3_PglZ_MBT4668302.1:0.5914909339,Type_3_PglZ_MCK9221199.1:0.3367710503)94.2/47:0.1236445041)37.1/2:0.0380743434,(Type_3_PglZ_WP_190509464.1:0.4086817153,((
Type3.supp.PglZ_YP_872576.1:0.1305535599,Type3.supp.PglZ_YP_001938979.1:0.1438583317)90.8/86:0.0775156574,Type3.supp.PglZ_YP_002462470.1:0.1804391997)98.5/96:0.1130063562)75.7/50:0.0635849
127)92.4/3:0.0767616817,((Type_3_PglZ_ADY58754.1:0.0023024128,Type3.supp.PglZ_YP_004268776.1:0.0000029638)100/100:0.4945800276,Type3.supp.PglZ_YP_001211267.1:0.4123605355)40.2/31:0.0688689
445)91.6/4:0.0947840326,(Type_3_PglZ_SDB48756.1:0.3167199606,Type3.supp.PglZ_YP_001412453.1:0.4064124523)100/100:0.3472526479)64.3/13:0.0782924904,Type_3_PglZ_WP_106343516.1:0.5274561015
)88.8/17:0.0946708080,(((((((Type_3_PglZ_EGD52428.1:0.0065802883,Type3.supp.PglZ_YP_004185917.1:0.0088882972)100/100:0.3293504442,Type3.supp.PglZ_YP_004470683.1:0.3584741479)92.7/91:0.10358
33559,(Type_3_PglZ_MCF6096368.1:0.4984664170,Type3.supp.PglZ_YP_004025782.1:0.3704086571)100/100:0.3755512650)19/37:0.0922482545,Type3.supp.PglZ_YP_004461306.1:0.3856667786)99.9/90:0.314
2058830,(((((Type_3_PglZ_MBR2506173.1:0.8721384591,(Type_3_PglZ_KUO63979.1:0.6684249623,Type_3_PglZ_NCD09237.1:0.6160415849)97.9/99:0.3883171355)99.3/98:0.4698400159,(((((Type_3_PglZ_WP_
114474460.1:0.5694303849,Type_3_PglZ_EMR73856.1:0.7953919254)96.4/89:0.2561558798,Type_3_PglZ_KFD41615.1:0.8328705611)65/47:0.0716776161,(Type_3_PglZ_KPQ15862.1:0.8270228731,Type_3_Pgl
Z_NLE00372.1:0.6460086814)99.9/100:0.6091903997)55.3/49:0.0594082886,Type_3_PglZ_MBU0460858.1:0.9838059575)94.3/88:0.1949939472,Type_3_PglZ_MBV6489838.1:1.5244783939)82.7/49:0.1510745214
)79.4/31:0.1466879596,((Type_3_PglZ_WP_108775981.1:0.6703453921,Type_3_PglZ_HHV38840.1:0.5142035904)98.8/98:0.3143350019,Type_3_PglZ_MBP8682480.1:0.7298333137)99.7/99:0.3660626927)83.7/1
7:0.0819124255,Type_3_PglZ_WP_020614471.1:0.8395632806)79.7/9:0.0545527089,Type_3_PglZ_MBT6646603.1:0.8431113863)75.6/4:0.0597332035)50/3:0.1181996141,((Type_3_PglZ_PSR33884.1:0.83539357
20,(Type_3_PglZ_KUK47747.1:0.6086683095,Type_3_PglZ_NLV27400.1:0.9873955032)56.4/38:0.1208204409)89.5/22:0.1063741370,(Type_3_PglZ_HHW29751.1:0.6969631033,Type_3_PglZ_SLK50102.1:0.58492
23584)82.2/37:0.0973117251)72.9/6:0.0661687032)44/2:0.1050205358,(Type_3_PglZ_WP_108651331.1:0.7166352495,Type_3_PglZ_MBT5530409.1:0.7258598623)78.7/27:0.1269929869)0/1:0.0000029135)99.9/8
8:0.9664879272)97.4/85:0.6570612100,(((((Type_4_PglZ_AFM22177.1:0.7047343771,(Type_4_PglZ_NPV08138.1:0.8134219920,Type4.supp.PglZ_YP_001716952.1:0.8384764332)92.3/93:0.2172130965)92/94:0.18
84379609,Type_4_PglZ_MCC2258803.1:1.4372997494)4.5/31:0.1269535757,((Type_4_PglZ_MBF0318626.1:0.4595445197,Type_4_PglZ_HIT17409.1:0.7583114904)99.8/100:0.4271627134,((((Type_4_PglZ_HIS35
238.1:0.6386757070,Type_4_PglZ_NCD03518.1:0.7262570975)22.8/47:0.1228359003,Type_4_PglZ_RKJ03001.1:0.8040652512)95.7/93:0.1721445581,((Type_4_PglZ_NBJ16967.1:0.4183339011,Type4.supp.PglZ_
YP_003503327.1:0.4489711891)98.5/97:0.2330213806,(Type_4_PglZ_MBQ4143520.1:0.7772549772,Type_4_PglZ_MBQ7256249.1:0.4594696296)98.5/99:0.2440633653)54.2/57:0.1075657558)86.3/71:0.13174775
49,Type_4_PglZ_MBP6982275.1:0.8798615308)100/100:0.4095660151)94.5/82:0.1872349809)83/60:0.1832001736,Type_4_PglZ_KUK07753.1:0.7145153516)90/71:0.1865002626,(Type_4_PglZ_WP_165846691.1:
1.2390711321,(((Type_4_PglZ_HHW21152.1:0.5822317130,(Type_4_PglZ_MBQ8951402.1:0.6337667268,Type_4_PglZ_HAR58249.1:0.7092451087)99.9/99:0.3471118439)37.6/41:0.0647408602,((Type_4_PglZ_W
P_052457313.1:0.5233367209,Type4.supp.PglZ_YP_001245330.1:0.5097986548)99.8/99:0.2720545211,((Type_4_PglZ_NLT83664.1:0.0028952802,Type4.supp.PglZ_YP_002247817.1:0.0297576556)100/100:1.020
7552857,(Type_4_PglZ_PMQ01343.1:0.4919662860,Type_4_PglZ_KUK66682.1:0.5007902025)72.3/53:0.0666100198)86.7/43:0.0800962815)90.5/48:0.1090368727)88.9/32:0.1099241757,((Type_4_PglZ_CDD8374
7.1:0.3975587482,(Type_4_PglZ_MBL1007424.1:0.3012760393,Type4.supp.PglZ_YP_004328100.1:0.4011686619)97.2/97:0.1408411864)100/95:0.5480523533,Type_4_PglZ_WP_195485919.1:0.7870129534)89.5/
51:0.1459087763)56.3/29:0.1294352514)90.9/76:0.2274864566)100/100:1.4106174767)89.9/76:0.4101167721)100/99:1.2026924556,Type_1_PglZ_MBK6301655.1:0.5547504016)79.4/24:0.1042168361,Type_1_Pgl
Z_WP_275988129.1:0.3876246454)90.5/16:0.1151049229,(Type_1_PglZ_MCC7365287.1:0.4477791673,(Type_1_PglZ_WP_011448105.1:0.7676679454,Type_1_PglZ_WP_129664641.1:0.4030672954)94.6/54:0.16
97979115)23.9/16:0.0770416738)91.9/28:0.0843771754,((Type_1_PglZ_NBT25026.1:0.7331981788,Type_1_PglZ_WP_031501096.1:0.4224497518)61.7/52:0.0712728961,Type_1_PglZ_WP_074444903.1:0.594749
0733)98.6/80:0.1336540573)52.1/12:0.0453832792,Type_1_PglZ_MBA4013264.1:0.7799844902)83.9/23:0.0606134993,Type_1_PglZ_OHC72321.1:0.5253700290)19.7/14:0.0628541433,Type_1_PglZ_WP_1664245
93.1:0.6328030694)74.5/34:0.0952595086,Type_1_PglZ_MCQ3924624.1:0.6483983494);

### DUF499-ATPase and BrxC-ATPase (subtype-specific) Newick format

(DUF499_1_WP_148590534.1_Aquisphaera_giovannonii:0.4001974037,(((((((((DUF499_1_MBQ9433643.1_Synergistaceae_bacterium:0.4549070233,DUF499_1_MBR0288712.1_Selenomonadaceae_bacterium:0.3640698210)97.1/87:
0.1051598390,((DUF499_1_WP_270528574.1_Mitsuokella_jalaludinii:0.3228522252,DUF499_1_MCL1790896.1_Peptococcaceae_bacterium:0.3198863598)75.5/59:0.0753701759,(DUF499_1_HBM97560.1_bacterium_UBP9_UBA1183
6:0.3205418801,DUF499_1_MCI1646461.1_Olsenella_sp.:0.4667568601)94.2/78:0.0769966553)95.3/74:0.1053116047)97.4/77:0.1046633043,(((DUF499_1_PJF35651.1_Candidatus_Thermofonsia:0.4112163378,DUF499_1_MYD1093
5.1_Chloroflexi_bacterium:0.3165319129)99/99:0.1479143432,DUF499_1_WP_188816795.1_Calditerricola_satsumensis:0.3734175077)37.6/33:0.0589755020,((((DUF499_1_NJK30457.1_Acaryochloris_sp._SU_5_25:0.3792374153,D
UF499_1_WP_075830034.1_Deinococcus_marmoris:0.4273411575)94.7/59:0.0782400143,(DUF499_1_MBC6439887.1_Rhodospirillales_bacterium:0.4039550358,DUF499_1_MYA05496.1_Caldilineaceae_bacterium:0.4409743529)84/
27:0.0726744028)52.3/8:0.0350027633,(DUF499_1_WP_201387653.1_Ktedonobacter_sp._SOSP1-52:0.3679458497,DUF499_1_WP_132287513.1_Paracraurococcus_sp._NE82:0.4604082134)90.8/8:0.0591260984)86.5/2:0.0395944
357,((((DUF499_1_MBT7692838.1_Gemmatimonadales_bacterium:0.4283646012,DUF499_1_MCY3894210.1_Acidimicrobiaceae_bacterium:0.4084000497)8.6/9:0.0441818546,(((DUF499_1_WP_244804062.1_Corynebacterium_sp._L
D5P10:0.4542293185,DUF499_1_WP_005520661.1_Corynebacterium_matruchotii:0.2717069666)94.4/78:0.0960028358,DUF499_1_MCF6470725.1_Nonomuraea_sp._MG754425:0.4096959197)76.7/54:0.0542166511,DUF499_1_WP
_053587850.1_Actinomyces_sp.:0.3263507609)99.2/81:0.1243625094)86.7/7:0.0428188589,((DUF499_1_WP_092673506.1_Siccationidurans_arizonensis:0.4804638145,DUF499_1_MCY4445744.1_Rhodobacteraceae_bacterium:0.47
94152267)7.2/17:0.0358481298,DUF499_1_HDQ14361.1_Sediminispirochaeta_sp.:0.4902318218)91.9/21:0.0580145771)91.5/13:0.0420756779,(DUF499_1_WP_094451589.1_Parenemella_sanctibonifatiensis:0.4421256559,DUF499_
1_MBD2550060.1_Microcystis_elabens_FACHB-917:0.3787128532)86.2/28:0.0773806955)91.6/17:0.0444397512)98.3/21:0.0998309694)88.8/17:0.0662975668)25.8/14:0.0522561854,(DUF499_1_RKU15614.1_Candidatus_Poribacteri
a_bacterium:0.4551665138,DUF499_1_MDE0020779.1_Candidatus_Poribacteria_bacterium:0.3879496683)100/100:0.3311078176)93.7/31:0.0880367965,(DUF499_1_WP_238518868.1_Azospirillum_doebereinerae:0.5610336477,DUF
499_1_WP_151650193.1_Bradyrhizobium_betae:0.3880019215)100/96:0.2703298758)56.2/65:0.0601122031,(DUF499_1_MBQ9726423.1_Kiritimatiellae_bacterium:0.4952329191,DUF499_1_CAI1753142.1_Serratia_quinivorans:0.326
8731805)98.4/97:0.1786369467)100/100:0.3732568860,(((DUF499_1_MBI3467145.1_Planctomycetota_bacterium:0.4520507104,(DUF499_1_HEC67877.1_Candidatus_Desulfofervidus_auxilii:0.3886741731,DUF499_1_MCS6856320.1
_Sandaracinaceae_bacterium:0.4504883776)35.4/40:0.0823330406)86/32:0.0813574552,((DUF499_1_WP_259054315.1_Salinibacter_ruber:0.6511506714,((((DUF499_1_RME04985.1_Planctomycetota_bacterium:0.3989211246,(DUF
499_1_WP_013295752.1_Methanothermobacter_marburgensis:0.3834012941,DUF499_1_NPV88406.1_Coprothermobacteraceae_bacterium:0.5348983505)0/30:0.0511738304)92.9/69:0.0969754944,(DUF499_1_MXV84967.1_Candid
atus_Poribacteria_bacterium:0.5350619228,DUF499_1_KEQ56705.1_Marine_Group_I_thaumarchaeote:0.5114654408)95.4/73:0.1207381210)91.7/68:0.0911782912,DUF499_1_WP_048201423.1_Methanocaldococcus_bathoardescen
s:0.3999403771)100/100:0.2524160953,(((((((DUF499_2_WP_015408759.1_Natronomonas_moolapensis:0.2542351906,(DUF499_2_WP_014555848.1_Haloquadratum_walsbyi:0.8190600280,DUF499_2_WP_239638853.1_Halorubru
m_californiense:0.3209892534)71.1/51:0.1214749253)94/66:0.1495308395,DUF499_2_OYR54791.1_Halorubrum_halodurans:0.3341751886)96.3/79:0.1867798638,DUF499_2_WP_285255572.1_Halobacterium_salinarum:0.59577675
96)99.9/100:0.3380861552,(DUF499_2_MYC64339.1_Caldilineaceae_bacterium_SB0661_bin_34:0.5197032354,DUF499_2_QES21410.1_Streptomyces_venezuelae:0.4560987249)99.7/99:0.3097256460)84.7/13:0.0816309159,((((((((
((((DUF499_2_MBD5553419.1_Desulfovibrio_sp.:0.3694161792,DUF499_2_PSB37932.1_Aphanothece_cf._minutissima:0.4163397406)89.6/22:0.1140935273,(DUF499_2_MBQ7457436.1_Desulfovibrio_sp.:0.3253477391,DUF499_2_
SBW09990.1_uncultured_Desulfovibrio_sp.:0.1888949180)52.1/31:0.0206634246)44/9:0.0497309962,DUF499_2_MCW5609142.1_Rubrivivax_sp.:0.4194575196)89.9/18:0.0668764934,DUF499_2_MBR2123973.1_Acetobacter_sp.:0.2
995989787)11.6/9:0.0478533416,DUF499_2_WP_199338599.1_Nostoc_sp._FACHB-145:0.2938562540)90.1/15:0.0631711615,DUF499_2_WP_168134876.1_Sphingomonas_corticis:0.4037766789)85.4/4:0.0474012771,(DUF499_2_M
CK6511429.1_Myxococcota_bacterium:0.3947326701,DUF499_2_SPE38928.1_Candidatus_Sulfopaludibacter_sp.:0.3359093276)67/24:0.0325041348)91.2/29:0.0899624850,DUF499_2_MBP6739048.1_Leptospiraceae_bacterium:0.50
70110607)99.6/99:0.2657806377,DUF499_2_WP_164002142.1_Pyxidicoccus_caerfyrddinensis:0.7250672946)97.1/54:0.1348655419,((DUF499_2_GBD14658.1_bacterium_HR25:0.4924858933,((DUF499_2_ESQ16532.1_Thiohalocap
sa_sp._PB-PSB1:0.3571051647,(((((DUF499_2_MBS5425877.1_Phascolarctobacterium_succinatutens:0.2940529747,DUF499_2_WP_080657027.1_Selenomonas_ruminantium:0.2506585379)85.7/81:0.1009636959,DUF499_2_WP_0
67849130.1_Alicyclobacillus_mali:0.3468294962)79.5/28:0.0559117815,(DUF499_2_MCA9273427.1_Phycisphaerales_bacterium:0.3925984805,(DUF499_2_MDD3916530.1_Synergistaceae_bacterium:0.3638528481,DUF499_2_WP_
085543648.1_Dethiosulfovibrio_salsuginis:0.3024377363)99.9/100:0.3057880551)13.5/12:0.0528781960)68.7/8:0.0254537566,(DUF499_2_WP_081055878.1_Burkholderia_pseudomultivorans:0.3499574025,DUF499_2_WP_18336778
4.1_Gellertiella_hungarica:0.4213220181)88.7/48:0.0828562665)67.6/9:0.0730361969,(DUF499_2_WP_020506762.1_Lamprocystis_purpurea:0.2638510154,DUF499_2_WP_046619147.1_Stutzerimonas_stutzeri:0.3744112475)89.8/25
:0.0810702202)43.8/6:0.0398431688)79.5/23:0.0530320002,(DUF499_2_KAE8440079.1_Halomonas_piezotolerans:0.6324631649,DUF499_2_WP_094037979.1_Zobellella_denitrificans:0.3613061661)48/32:0.1354875604)100/100:0.6
137765727)70.7/27:0.0619100952,(((((DUF499_2_MCY4162822.1_bacterium:0.1826688550,DUF499_2_MYE76343.1_Acidimicrobiaceae_bacterium:0.2948824339)100/100:0.2678913023,DUF499_2_MCZ7532618.1_Acidimicrobiia_ba
cterium:0.2563754150)77.5/64:0.0909182796,DUF499_2_MBF6555079.1_Acidimicrobiales_bacterium:0.2875763217)67.1/47:0.0612819169,((DUF499_2_MDA8372907.1_Actinomycetota_bacterium:0.8311330774,DUF499_2_WP_142
649215.1_Microbispora_sp._SCL1-1:0.2334407730)84.5/38:0.0836147274,DUF499_2_WP_091105663.1_Micromonospora_citrea:0.3132635357)39.9/24:0.0826059431)93.8/41:0.1160193054,DUF499_2_MCC5947763.1_Nitriliruptorac
eae_bacterium:0.3535923788)100/95:0.2865433394)84.5/11:0.0563248477)72.2/4:0.0753424572,((DUF499_2_GBD21619.1_bacterium_HR28:0.3629315997,DUF499_2_OQA44952.1_Chloroflexi_bacterium_ADurb.Bin325:0.32989849
37)78.1/55:0.0983396355,DUF499_2_HHV54946.1_Firmicutes_bacterium:0.5560703180)96.1/24:0.1387958361)15.2/3:0.0345530166,DUF499_2_NLJ36704.1_candidate_division_WS1_bacterium:0.5624616571)27.7/1:0.0398360685,(
DUF499_2_WP_073234041.1_Desulforamulus_putei:0.4093956536,DUF499_2_WP_242016564.1_Pseudanabaena_sp._FACHB-1277:0.4325617678)89.8/32:0.0770458300)85.9/11:0.0826793360)27.9/14:0.0945777541,DUF499_2_M
BA3046419.1_Euryarchaeota_archaeon:0.7320468050)54.6/44:0.0845424829,((((((DUF499_3_CAD6492704.1_Candidatus_Argoarchaeum_ethanivorans:0.5976295476,(DUF499_3_SFC23913.1_Halobiforma_haloterrestris:0.40736669
95,DUF499_3_WP_260594793.1_Salinirubellus_salinus:0.2955644897)100/100:0.5296388185)99.2/88:0.3097291641,(DUF499_3_HIE29204.1_Candidatus_Poribacteria_bacterium:0.4958825286,DUF499_3_MEA3401058.1_Armatimo
nadota_bacterium:0.6384235586)94.8/85:0.2209300762)2.6/25:0.0629576983,((((DUF499_3_MBM4461756.1_Chloroflexi_bacterium:0.3419333829,DUF499_3_MBM4466103.1_Chloroflexi_bacterium:0.4769515375)99.1/99:0.23085684
14,DUF499_3_MCL0058394.1_Dehalococcoidia_bacterium:1.0867288107)84.6/32:0.0811894811,(DUF499_3_TEU12999.1_Anaerolineales_bacterium:0.4101511704,DUF499_3_WP_259101344.1_Candidatus_Fervidibacter_sacchari:0.
4332044202)100/100:0.6655962863)88.4/25:0.1042760031,DUF499_3_WP_125670094.1_Candidatus_Methanodesulfokores_washburnensis:0.8415023389)90.3/16:0.1048616459)46/6:0.0605302288,(((((DUF499_3_KOX91592.1_Hal
oarcula_rubripromontorii:0.3697023305,DUF499_3_KXA95606.1_candidate_division_MSBL1_archaeon:0.3502569549)95.6/82:0.1265959600,DUF499_3_MCK8825903.1_Fuchsiella_alkaliacetigena:0.4905621761)78.2/44:0.078579943
1,DUF499_3_WP_011944180.1_Thermotoga_petrophila:0.3735499021)90.6/71:0.1487574628,DUF499_3_MEA3344886.1_Chloroflexota_bacterium:0.8623776417)98.3/93:0.2037741612,DUF499_3_NMX21982.1_ANME-1_cluster_arc
haeon_GoMg4:0.6698996429)53.2/37:0.0862305502)85.2/26:0.1195549911,((((((((DUF499_3_HCF99178.1_Chloroflexi_bacterium:0.4657324135,DUF499_3_HVA45320.1_Pirellulales_bacterium:0.2975937828)90.4/82:0.0792175583,(((
DUF499_3_HZM03188.1_Candidatus_Saccharimonadales_bacterium:0.4751308518,DUF499_3_OCQ93485.1_Nostoc_sp._MBR_210:0.2107963792)93.8/49:0.0831579770,((DUF499_3_MBA3936148.1_Planctomycetota_bacterium:0.6
493672121,DUF499_3_MDR0374944.1_Treponema_sp.:0.3359426572)56.6/15:0.0522570482,DUF499_3_MBQ8111585.1_Kiritimatiellae_bacterium:0.4465096626)62.5/12:0.0732793085)87.9/22:0.0432432169,DUF499_3_MDD32694
99.1_Syntrophomonadaceae_bacterium:0.3762683302)83.9/21:0.0544037710)99.4/61:0.1699184205,DUF499_3_MBC7429623.1_Bacteriovorax_sp.:0.7749183585)95.3/51:0.1255934236,(DUF499_3_MDR2561287.1_Holophagales_ba
cterium:0.6697465669,(DUF499_3_WP_283022678.1_Bradyrhizobium_sp._CB1717:0.3416387470,DUF499_3_WP_286167649.1_Ochrobactrum_sp._3-3:0.1612366457)76.2/62:0.0824945506)36.3/47:0.0482257140)85/27:0.04148591
88,DUF499_3_MDU2146556.1_Paeniclostridium_sordellii:0.4121255048)93.3/41:0.0935206080,DUF499_3_MBY8216331.1_Vibrio_fluvialis:0.3074289779)88.8/47:0.1025771556,DUF499_3_MDR0392109.1_Planctomycetaceae_bacteri
um:0.3420798679)100/100:1.1660660363,((DUF499_3_HPN33675.1_bacterium:0.5096984956,((DUF499_3_MBK6914962.1_Ignavibacteriales_bacterium:0.4946930856,DUF499_3_NLX24424.1_Lentisphaerae_bacterium:0.334684418
0)30.7/31:0.0379193461,(DUF499_3_MCE0498498.1_Methylacidiphilales_bacterium:0.5137738111,DUF499_3_MDH4162607.1_Nitrospirota_bacterium:0.6988340286)98.6/82:0.2132021816)82.7/38:0.0599835184)88.1/61:0.102504881
3,(((DUF499_3_HUT59382.1_Phycisphaerae_bacterium:0.4345987693,DUF499_3_MEA3428074.1_Thermodesulfobacteriota_bacterium:0.4486787029)3.6/33:0.0601762827,DUF499_3_MBN2391962.1_Anaerolineae_bacterium:0.4084
885175)80.5/41:0.0522327736,DUF499_3_MBU4312937.1_Candidatus_Omnitrophica_bacterium:0.2492725411)97.2/83:0.1561997083)99.8/99:0.5143652344)95.8/79:0.3269951707)96.1/66:0.4305232095,((((((((((BrxC_1_HAJ70155.1
_Alkalibacterium_sp.:0.3601482300,(BrxC_1_WP_088074350.1_Bacillaceae:0.3100950422,BrxC_1_WP_094264422.1_Paludifilum_halophilum:0.3302771339)100/100:0.2470828765)91.6/72:0.1080408960,(BrxC_1_HIQ90970.1_Candi
datus_Coprosoma_intestinipullorum:0.3831710613,(BrxC_1_MBS5450116.1_Coriobacteriia_bacterium:0.3453710179,BrxC_1_OAD88367.1_Clostridiales_bacterium_KLE1615:0.4161003931)100/100:0.2391050405)89.4/72:0.10126642
08)87.7/68:0.1070146127,(((BrxC_1_MBF0232702.1_Desulfamplus_sp.:0.2733880668,BrxC_1_NLK13330.1_Spirochaetales_bacterium:0.3125731039)91.5/90:0.1166710899,BrxC_1_MCH3907669.1_Sphaerochaeta_sp.:0.5587415458
)96.3/92:0.1627830306,BrxC_1_RXJ95785.1_Arcobacter_sp._CECT_9299:0.4310441233)99.2/99:0.2032731941)100/99:0.3649695982,(BrxC_6_ETI91650.1_Clostridium_butyricum_DORA_1:0.6744224160,BrxC_6_WP_049936156.1_
Haloplanus_natans:0.5969162586)83.1/80:0.1894445900)31.9/38:0.0877260813,((((BrxC_6_MBZ5596030.1_Acidobacteriia_bacterium:0.3597000934,BrxC_6_VVB72030.1_uncultured_archaeon:0.5698897870)89.7/84:0.0958678566,Br
xC_6_TXH34387.1_Burkholderiaceae_bacterium:0.3793368540)97/95:0.1249850831,BrxC_6_WP_208027651.1_Rhabdothermincola_sediminis:0.4883094272)98.9/100:0.2236636063,(BrxC_6_MCP5535434.1_Akkermansiaceae_bacte
rium:0.9887209574,(BrxC_6_WP_015675681.1_Leptospira_yanagawae:0.5845772960,BrxC_6_WP_146981109.1_Lujinxingia_vulgaris:0.5450205959)99.8/100:0.4042202031)93.4/82:0.1707424530)99.7/95:0.3863002062)41.8/40:0.119
8807164,(BrxC_1_MCE8425368.1_Candidatus_Methanoperedens_sp.:0.2951463335,(BrxC_1_NRB78692.1_Saccharospirillaceae_bacterium:0.2973253762,BrxC_1_QOJ30044.1_Ignavibacteriales_bacterium:0.4179558533)100/100:0.
4412381006)100/100:0.7304956974)92.3/57:0.1764595314,BrxC_1_RKY59330.1_Candidatus_Latescibacteria_bacterium:0.9933514920)46.8/24:0.0914638739,((BrxC_1_MXZ22476.1_Caldilineaceae_bacterium_SB0665_bin_25:0.628
6078301,BrxC_1_PPG29161.1_Pseudoclavibacter_sp._RFBB5:0.4486660325)100/100:1.2218024806,((BrxC_5_AEN07515.1_halophilic_archaeon_DL31:0.1260783920,(BrxC_5_WP_020446108.1_Salinarchaeum_sp._Harcht-Bsk1:0.1
524490161,BrxC_5_WP_050049402.1_Halanaeroarchaeum_sulfurireducens:0.0949074052)17.6/43:0.0305023625)98.6/98:0.2309095089,((BrxC_5_ELY62517.1_Natronococcus_jeotgali_DSM_18795:0.2409730970,BrxC_5_WP_11460
4755.1_Haloplanus_rubicundus:0.2453567769)99.8/100:0.1510478728,BrxC_5_TQQ78814.1_Halonotius_terrestris:0.1717760024)99.9/100:0.3967083943)100/100:1.0234546123)72.8/35:0.1322591815)99.8/89:0.7387909529,(((((((Brx
C_3_WP_164569833.1_Rhizobium_leguminosarum:0.4721577512,BrxC_3_MBP6974014.1_Syntrophorhabdus_sp.:0.3749462467)96.1/81:0.1231783208,(BrxC_3_WP_047754172.1_Kosmotoga_pacifica:0.3067011710,(BrxC_3_MCM1
566961.1_Dehalobacter_sp.:0.8216538687,BrxC_3_WP_053959405.1_Sulfobacillus_thermosulfidooxidans:0.3764078060)92.9/62:0.1441855769)41.3/35:0.0555183660)91.3/33:0.0769360067,(BrxC_3_MCK9328444.1_Candidatus_Clo
acimonadota_bacterium:0.5231245088,BrxC_3_WP_020614466.1_Sediminispirochaeta_bajacaliforniensis:0.3558363533)63.2/16:0.0589826441)85.1/30:0.0574578630,BrxC_3_NLV27395.1_Methanomicrobiales_archaeon:0.472127891
3)80.2/42:0.1103007707,BrxC_3_MBT6646600.1_Nitrososphaerota_archaeon:0.3102133169)99.7/97:0.5181879405,(BrxC_3_MCP4110753.1_Desulfobacteraceae_bacterium:0.7014872234,BrxC_3_MBU0460852.1_Nanoarchaeota_arc
haeon:0.4696091690)96.6/94:0.3244925913)99.7/97:0.7232704967,((BrxC_4_MCD8003208.1_Clostridia_bacterium:0.4918330725,BrxC_4_HCH97636.1_Lachnospiraceae_bacterium:0.4591912949)99.3/100:0.5287867605,((((BrxC_4_
WP_048095117.1_Archaeoglobus_fulgidus:0.8541344653,((BrxC_4_NPV08139.1_Anaerolineae_bacterium:0.5728905808,BrxC_4_SHE91973.1_Desulfofundulus_australicus_DSM_11792:0.2649056954)94.1/84:0.1227193003,BrxC_4_
HED00229.1_Pseudomonadota_bacterium:0.5262859435)87.9/87:0.1049583571)75.2/32:0.1408486652,BrxC_4_NSW92344.1_Bacillota_bacterium:0.3119147461)98/47:0.2091435917,BrxC_4_MBF0318625.1_Nitrospirota_bacterium:0.
4327736759)77.2/37:0.1302150592,(BrxC_4_MBR6984229.1_Ruminococcus_sp.:1.1513475897,(((BrxC_4_MBR2693833.1_Thermoguttaceae_bacterium:0.5121762310,BrxC_4_NBJ16968.1_Dehalobacter_sp._4CP:0.4961908288)69.1
/62:0.1259654683,(BrxC_4_MBQ9371615.1_Thermoguttaceae_bacterium:0.4300528775,BrxC_4_MBQ7256248.1_Abditibacteriota_bacterium:0.3116753163)100/100:0.2959680597)86.3/71:0.1277949159,BrxC_4_MZQ76088.1_Peptocl
ostridium_sp.:0.5427690554)94.4/89:0.1781736301)98.7/97:0.2292851716)86.3/53:0.2105906946)100/99:1.5404564986)76.9/53:0.1896039692)64.5/32:0.2377913738,(((((BrxC_2_HUY49759.1_Streptosporangiaceae_bacterium:0.4654
403958,((BrxC_2_WP_061229212.1_Dietzia_cinnamea:0.5706466644,BrxC_2_WP_322753649.1_Frankia_sp._Cas3:0.3113273230)98/98:0.1623920023,BrxC_2_WP_222632854.1_unclassified_Rhodococcus:0.5516753437)99.4/93:0.
1814435231)12.8/10:0.0352431036,((BrxC_2_WP_030506727.1_Microbispora_rosea:0.4164819041,((BrxC_2_WP_199044191.1_Glycomyces_salinus:0.4364985660,BrxC_2_WP_270679261.1_Nocardiopsis_sp._LSu2-4:0.4233092635
)97.1/83:0.1281820915,BrxC_2_WUA45283.1_Streptomyces_sp._NBC_01367:0.5232196368)19.8/33:0.1013088222)95.9/39:0.0821063416,(BrxC_2_WP_161103223.1_Actinomadura_rayongensis:0.4670570338,BrxC_2_WP_1656329
98.1_unclassified_Actinopolyspora:0.5430477278)93.8/67:0.1382532703)88.7/18:0.0545485208)94.8/9:0.0615854718,(BrxC_2_MBL8842560.1_Planctomycetota_bacterium:0.3457738890,BrxC_2_WP_245209875.1_Ralstonia_solanac
earum:0.3965631399)86.4/30:0.0793074466)31.3/8:0.0470691957,BrxC_2_MCB9398403.1_Acidobacteriota_bacterium:0.5239579510)91.1/41:0.1705481695,BrxC_2_MBK9578953.1_Fibrobacterota_bacterium:0.3648699057)100/100:1
.7208292872)72/55:0.2973258805,(((((((WP_228615471.1_Cdc6_family_protein__Candidatus_Nanobsidianus_stetteri_:1.0011266256,WP_147661802.1_Cdc6_family_protein__Promethearchaeum_syntrophicum_:1.3278183979)97.4/9
1:0.5648370148,WP_400152517.1_Cdc6_family_protein__Candidatus_Methanarcanum_hacksteinii_:0.7687530463)98.3/62:0.3466998770,(((WP_247995929.1_Cdc6_family_protein__Halocatena_salina_:0.7534526549,WP_22444723
7.1_Cdc6_family_protein__Haloprofundus_salilacus_:0.5816015661)82.9/51:0.1246025210,((WP_312908183.1_Cdc6_family_protein__Natronosalvus_caseinilyticus_:0.7077127265,WP_193308720.1_Cdc6_family_protein__Halorubrum
_halophilum_:0.5929436947)75.4/50:0.1019542245,WP_238330919.1_Cdc6_family_protein__Haloglomus_irregulare_:0.7793503434)7.4/15:0.0560460678)60.3/46:0.1678093419,WP_135305442.1_Cdc6_family_protein__Haloarcula_a
mylovorans_:0.7311013300)94.8/84:0.2139392846)85.1/24:0.1016990744,((WP_082837926.1_Cdc6_family_protein__Haladaptatus_sp._R4_:0.5590739144,WP_271946962.1_Cdc6_family_protein__Halorubrum_ezzemoulense_:0.6459
975120)99.2/100:0.3865021704,WP_250867674.1_Cdc6_family_protein__Methanococcoides_seepicolus_:0.9134513559)44.4/36:0.0733995331)25.7/13:0.1282284205,WP_013195646.1_Cdc6_family_protein__Methanohalobium_eves
tigatum_:0.5420645360)96.4/53:0.2701193390,WP_048116492.1_Cdc6_family_protein__Methanosarcina_sp._Kolksee_:0.7755997259)72.4/49:0.3390059952,WP_321416762.1_Cdc6_family_protein__uncultured_Methanomethylovoran
s_sp._:0.4199769318)100/100:2.4276067507)98.9/94:0.8050495201)99.6/96:0.8069580689)98.7/88:0.2692495303)72/23:0.0353442882)68.2/9:0.0657426067,(((DUF499_1_WP_106140933.1_Spirosoma_oryzae:0.3338266781,DUF49
9_1_WP_051018640.1_Saprospira_grandis:0.2706197851)81.1/68:0.0857344586,DUF499_1_WP_118201547.1_Prevotella_copri:0.3802663132)99.9/100:0.3000167044,DUF499_1_MCA9831553.1_Dehalococcoidia_bacterium:1.00490
02490)97.6/66:0.1892131388)0/4:0.0363894594)88.1/16:0.0680135600,((DUF499_1_MCD6101177.1_Candidatus_Marinimicrobia_bacterium:0.5425222255,DUF499_1_MDE0129995.1_Gammaproteobacteria_bacterium:0.4396038335)
98.9/99:0.2045922890,(DUF499_1_UGS34698.1_Solirubrobacterales_bacterium_0166_1:0.4289413057,DUF499_1_NET07313.1_Symploca_sp._SIO2B6:0.8673911200)91.3/85:0.1440878304)100/100:0.3811508826)82.8/31:0.069836
6986)100/100:0.6243197218,(DUF499_1_MCL0064885.1_Dehalococcoidia_bacterium:0.4165367112,DUF499_1_MBM4285998.1_Deltaproteobacteria_bacterium:0.6386227801)68.4/65:0.0794562081)94.5/97:0.1159021529,DUF499_1
_RIK37466.1_Chloroflexi_bacterium:0.3838034364)94.2/85:0.1158476527,DUF499_1_MBF0121417.1_Desulfobacterales_bacterium:0.2861621025);
