## Supplementary Data S2 for "Expanding the Landscape of BREX Diversity: Uncovering Multi-Layered Functional Frameworks and Identification of Novel BREX-Related Defense Systems"

This PDF presents hit lists from structural similarity searches (primarily via the DALI server) along with their corresponding structure-guided sequence alignments, which we used to annotate multiple previously uncharacterized domains associated with various components of BREX and their related systems.
Amino acid residues in the alignments are shown in fluorescent colors against a black background for clarity, with conserved blocks and secondary structural elements highlighted in uppercase.


1. BrxC-ATPase; N-terminal ORC/CDC6 AAA+ ATPase….3
2. BrxC-ATPase; Canonical wHTH1 immediately following ORC/CDC6….4
3. BrxC-ATPase; Alpha+Beta-Domain….5
4. BrxC-ATPase; wHTH2….6
5. BrxC-ATPase; C-terminal Coiled-coil like extension….8
6. BrxX-MTase; N-terminal Dimerisation domain….9
7. BrxX-MTase; Target-recognition-domain….9
8. BrxX-MTase; C-terminal CC-like helical extension….9-11
9. Type-1 BREX; BrxX-MTase wHTH….11
10. PglZ; N-terminal inactive P-loop of Swi2/Snf2….12
11. Structure-based Sequence Alignment of "PglZ N-terminal Inactive P-loop Swi2/Snf2" with various inactive P-loop domains from Swi2/Snf2-Helicases….13
12. PglZ; (Third Tri-helical HTH-like element)....14
13. PglZ; core enzymatic phosphatase (NPP superfamily)….16
14. PglZ; C-terminal β-Sandwich (Immunoglobulin-like)….16
15. PglZ; C-terminal wHTH (Type-2 BREX-specific)….18
16. PglZ; N-terminal inactive STAND (Type-4 BREX-specific)….19-21
17. BrxL; core MCM AAA+ ATPase….21
18. BrxL; N-terminal SIGMA-HTH like….22
19. BrxL; N-terminal OB-fold….23
20. BrxL; C-terminal HKD-EndoDNase….24
21. BrxA; N-Terminal HTH1 (found in a subset of Type-1 BrxA, and in all Type-3 BrxA)....24-25
22. BrxA; HTH2….26
23. BrxA; C-terminal wHTH….27
24. BrxB; inactive STAND NTPase….28
25. BrxF; inactive STAND NTPase (Type-3 BREX)....29
26. PglW; wHTH1 (encoded in the C-terminal half of protein)....31
27. PglW; wHTH2….33
28. PglW; wHTH3….34-35
29. PglW; C-terminal inactive STAND….36
30. BrxHI-Helicase; wHTH (Type-2 BREX)....37
31. BrxHI-Helicase; HAS-β barrel (Small β-barrel assemblage)....38
32. BrxHI-Helicase; Lhr-helicase CTD….39-40
33. BrxD-ATPase; standalone ORC/CDC6 AAA+ ATPase….40
34. Type-3 BREX; MTase C-terminal HTH1….42
35. Type-3 BREX; MTase C-terminal HTH2….44-47

36. Type-3 BREX; MTase C-terminal wHTH....48-50
37. Type-3 BREX; BrxHII-Helicase N-terminal TUDOR....50-51
38. Type-3 BREX; BrxHII-Helicase C-terminal inactive PDDEXK-REase fold....52
39. Structural search anchored on "C-terminal Inactive PDDEXK-REase Domain" found in RapA-Helicase....52-53
40. Type-3 BREX; BrxHII-Helicase C-terminal α+β domain....54-55
41. BREX-Related systems; DUF499-ATPase C-terminal RRM/Ferredoxin 1....56
42. BREX-Related systems; DUF499-ATPase C-terminal RRM/Ferredoxin 2....57
43. BREX-Related systems; DUF499-ATPase C-terminal Fn-III-like domain....59
44. BREX-Related systems; DUF3780....61-62
45. BREX-Related systems (Type-3); Inactive STAND-NTPase....62-63
46. Type-4 BREX; PAPS-Reductase α+β domain....63-64
47. Type-4 BREX; PAPS-Reductase Fe-S cluster binding Fer4_7....64
48. BREX-related Capture systems; HerA/FtsK....65

***Note: Titles are internally hyperlinked. Click on the titles to access the material***

```
# Job: Type-1 BREX; BrxC-ATPase N-terminal ORC/CDC6 AAA+ ATPase
# Query: s001A
# No:  Chain    Z    rmsd lali nres  %id PDB  Description
   1:  2qby-B 14.1  4.5  249   368   11   MOLECULE: CELL DIVISION CONTROL PROTEIN 6 HOMOLOG 1;
   2:  1w5s-B 13.2  3.7  228   396   12   MOLECULE: ORIGIN RECOGNITION COMPLEX SUBUNIT 2 ORC2;
   3:  3bos-B 12.5  4.7  188   231   11   MOLECULE: PUTATIVE DNA REPLICATION FACTOR;
   4:  5zr1-A 11.7  4.5  251   494   14   MOLECULE: ORIGIN RECOGNITION COMPLEX SUBUNIT 1;
   5:  5zr1-E 11.5  5.6  247   460   11   MOLECULE: ORIGIN RECOGNITION COMPLEX SUBUNIT 1;
   6:  7rzy-1 11.4  4.4  248   311   14   MOLECULE: TN6677 VIBRIO CHOLERAE TRANSPOSON TNSC (VCHTNSC);
   7:  8c0v-A 11.3  4.4  218   823   10   MOLECULE: PEROXISOMAL ATPASE PEX1;
   8:  6p8v-C 11.3  4.4  225   303    9   MOLECULE: ATPASE, AAA FAMILY;
   9:  7jgr-D 11.2  4.8  263   441   10   MOLECULE: ORIGIN RECOGNITION COMPLEX SUBUNIT 2;
  10:  8btg-B 11.1  5.8  211   335   11   MOLECULE: CHROMOSOMAL REPLICATION INITIATOR PROTEIN DNAA;
  12:  4rh7-A 11.1  4.6  236  3005   13   MOLECULE: GREEN FLUORESCENT PROTEIN/CYTOPLASMIC DYNEIN 2 HE
  13:  8fcv-R 11.0  4.1  238   343   13   MOLECULE: DNA (60-MER);
  14:  9emc-D 11.0  4.6  244   337   14   MOLECULE: RUVB-LIKE 1;
  15:  8xvb-A 11.0  4.5  223   246   12   MOLECULE: DNA (5'-
  16:  8hra-C 10.8  4.3  273   852   13   MOLECULE: ARCHAEAL ATPASE;
  17:  7plh-A 10.5  4.2  225   260   11   MOLECULE: SHTNSC;
  18:  8fnu-A 10.4  4.1  245   796    8   MOLECULE: KAP NTPASE DOMAIN-CONTAINING PROTEIN;
  20:  2fna-A 10.4  3.5  215   352   11   MOLECULE: CONSERVED HYPOTHETICAL PROTEIN;
  21:  7vcs-A 10.2  4.9  216   766   13   MOLECULE: TRANSITIONAL ENDOPLASMIC RETICULUM ATPASE;
  22:  6fhs-C 10.1  5.5  265   459   12   MOLECULE: RUVB-LIKE HELICASE;
  23:  8bob-C 10.1  6.1  205   412    9   MOLECULE: PROTEIN MALY;
  24:  9cru-C 10.1  3.7  199   741    9   MOLECULE: VESICULAR-FUSION PROTEIN SEC18;
  25:  5e7p-A 10.0  4.1  218   719   11   MOLECULE: CELL DIVISION CONTROL PROTEIN CDC48;
  26:  8umy-A  9.9  4.3  205   546    9   MOLECULE: CHROMOSOME TRANSMISSION FIDELITY PROTEIN 18 HOMOL
  27:  7uqj-A  9.9  4.6  225   585    9   MOLECULE: ATPASE HISTONE CHAPERONE YTA7;
  28:  7st9-A  9.8  6.1  214   522   11   MOLECULE: CHECKPOINT PROTEIN RAD24;
  29:  8tw7-2  9.8  4.1  216   339   10   MOLECULE: REPLICATION FACTOR C SUBUNIT 5;
  30:  7crc-B  9.6  6.4  231  1112   10   MOLECULE: NAD+ HYDROLASE (NADASE);
  31:  8dr5-A  9.6  4.0  197   646   13   MOLECULE: REPLICATION FACTOR C SUBUNIT 1;
  32:  5kzf-K  9.6  5.2  233   498   13   MOLECULE: PROTEASOME-ASSOCIATED ATPASE;
  33:  6f0x-B  9.5  3.9  208   398   12   MOLECULE: PACHYTENE CHECKPOINT PROTEIN 2 HOMOLOG;
  34:  9e22-A  9.5  4.3  229  2925   13   MOLECULE: CYTOPLASMIC DYNEIN 1 HEAVY CHAIN 1;
  35:  7st9-E  9.4  3.9  209   354   11   MOLECULE: CHECKPOINT PROTEIN RAD24;
  36:  7ykk-A  9.4  4.4  215   735   12   MOLECULE: ATPASE FAMILY GENE 2 PROTEIN;
  37:  9bv1-A  9.3  4.4  201   405   13   MOLECULE: 26S PROTEASOME NON-ATPASE REGULATORY SUBUNIT 1;
  38:  6qem-G  9.3  3.4  146   239   15   MOLECULE: REPLICATIVE DNA HELICASE;
  39:  8q3w-A  9.3  3.2  163   246   15   MOLECULE: INSERTION SEQUENCE IS5376 PUTATIVE ATP-BINDING PR
  40:  8glw-A  9.3  3.9  233   483   14   MOLECULE: TRANSPOSON TN7 TRANSPOSITION PROTEIN TNSC;
  41:  6l1q-A  9.1  4.7  206   267   12   MOLECULE: CBBQ PROTEIN;
  42:  8val-E  9.1  7.2  209   337    9   MOLECULE: DNA POLYMERASE III SUBUNIT DELTA;
  43:  5j1s-A  9.1  4.8  194   277    9   MOLECULE: TORSIN-1A;
  44:  6z1f-3  9.0  4.2  210   284   12   MOLECULE: RIBULOSE BISPHOSPHATE CARBOXYLASE/OXYGENASE ACTIV
  45:  8c0v-B  9.0  3.8  178  1030   13   MOLECULE: PEROXISOMAL ATPASE PEX1;
  46:  8xku-A  9.0  4.8  234   730   12   MOLECULE: PROBABLE INACTIVE ATP-DEPENDENT ZINC METALLOPROTE
  47:  6pxk-K  8.9  4.8  230   429   13   MOLECULE: ATP-DEPENDENT PROTEASE ATPASE SUBUNIT HSLU;
  48:  5zr1-B  8.9  7.3  177   374   11   MOLECULE: ORIGIN RECOGNITION COMPLEX SUBUNIT 1;
  49:  4m9s-B  8.8  5.2  215   511    7   MOLECULE: CELL DEATH PROTEIN 4;
  50:  3glf-G  8.8  5.0  212   378   12   MOLECULE: DNA POLYMERASE III SUBUNIT DELTA;
  51:  8xks-D  8.8  4.3  216   971   14   MOLECULE: CTAP1;
  52:  2gno-A  8.7  4.2  165   296   10   MOLECULE: DNA POLYMERASE III, GAMMA SUBUNIT-RELATED PROTEIN
  53:  7w42-B  8.7  4.2  197   402   11   MOLECULE: UNCHARACTERIZED ATPASE YJOB;
  54:  8xks-B  8.7 18.2  188   603   12   MOLECULE: CTAP1;
  55:  7mi8-A  8.7 15.0  175   868   12   MOLECULE: FUSION PROTEIN OF DYNEIN AND ENDOLYSIN;
  56:  1g4a-E  8.6  7.9  248   356   13   MOLECULE: ATP-DEPENDENT HSL PROTEASE ATP-BINDING SUBUNIT
  57:  8xku-C  8.6  4.8  196   921   11   MOLECULE: PROBABLE INACTIVE ATP-DEPENDENT ZINC METALLOPROTE
  58:  5j1s-B  8.5  3.9  157   235   12   MOLECULE: TORSIN-1A;
  59:  7qo4-M  8.5  5.1  207   421   13   MOLECULE: 26S PROTEASOME REGULATORY SUBUNIT RPN1;

0001 s001A   [illegible]
0002 2qbyB   [illegible]
0003 1w5sB   [illegible]
0004 3bosB   [illegible]
0005 5zr1A   [illegible]
0006 5zr1E   [illegible]
0007 7rzy1   [illegible]
0008 8c0vA   [illegible]
0009 6p8vC   [illegible]
0010 7jgrD   [illegible]
0011 8btgB   [illegible]
0012 4rh7A   [illegible]
0013 4rh7A   [illegible]
0014 8fcvR   [illegible]
0015 9emcD   [illegible]
0016 8xvbA   [illegible]
0017 8hraC   [illegible]
0018 7plhA   [illegible]
0019 8fnuA   [illegible]
0020 8rwvE   [illegible]
0021 2fnaA   [illegible]
0022 7vcsA   [illegible]
0023 6fhsC   [illegible]
0024 8bobC   [illegible]
0025 9cruC   [illegible]
0026 5e7pA   [illegible]
0027 8umyA   [illegible]
0028 7uqjA   [illegible]
0029 7st9A   [illegible]
0030 8tw72   [illegible]
0031 7crcB   [illegible]
0032 8dr5A   [illegible]
0033 5kzfK   [illegible]
0034 6f0xB   [illegible]
0035 9e22A   [illegible]
0036 7st9E   [illegible]
0037 7ykkA   [illegible]
0038 9bv1A   [illegible]
0039 6qemG   [illegible]
0040 8q3wA   [illegible]
0041 8glwA   [illegible]
0042 6l1qA   [illegible]
0043 8valE   [illegible]
0044 5j1sA   [illegible]
0045 6z1f3   [illegible]
0046 8c0vB   [illegible]
0047 8xkuA   [illegible]
0048 6pxkK   [illegible]
0049 5zr1B   [illegible]
0050 4m9sB   [illegible]
0051 3glfG   [illegible]
0052 8xksD   [illegible]
0053 2gnoA   [illegible]
0054 7w42B   [illegible]
0055 8xksB   [illegible]
0056 7mi8A   [illegible]
0057 1g4aE   [illegible]
0058 8xkuC   [illegible]
0059 5j1sB   [illegible]
0060 7qo4M   [illegible]
0061 8rwvE   [illegible]

0001 s001A   [illegible]
0002 2qbyB   [illegible]
0003 1w5sB   [illegible]
0004 3bosB   [illegible]
0005 5zr1A   [illegible]
0006 5zr1E   [illegible]
0007 7rzy1   [illegible]
0008 8c0vA   [illegible]
0009 6p8vC   [illegible]
0010 7jgrD   [illegible]
```

```
0011 8btgB  [illegible]
0012 5v8fD  [illegible]
0013 4rh7A  [illegible]
0014 8fcvR  [illegible]
0015 9emcD  [illegible]
0016 8xvbA  [illegible]
0017 8hraC  [illegible]
0018 7plhA  [illegible]
0019 8fnuA  [illegible]
0020 8rwvE  [illegible]
0021 2fnaA  [illegible]
0022 7vcsA  [illegible]
0023 6fhsC  [illegible]
0024 8bobC  [illegible]
0025 9cruC  [illegible]
0026 5e7pA  [illegible]
0027 8umyA  [illegible]
0028 7uqjA  [illegible]
0029 7st9A  [illegible]
0030 8tw72  [illegible]
0031 7crcB  [illegible]
0032 8dr5A  [illegible]
0033 5kzfK  [illegible]
0034 6f0xB  [illegible]
0035 9e22A  [illegible]
0036 7st9E  [illegible]
0037 7ykkA  [illegible]
0038 9bv1A  [illegible]
0039 6qemG  [illegible]
0040 8q3wA  [illegible]
0041 8glwA  [illegible]
0042 6l1qA  [illegible]
0043 8valE  [illegible]
0044 5j1sA  [illegible]
0045 6z1f3  [illegible]
0046 8c0vB  [illegible]
0047 8xkuA  [illegible]
0048 6pxkK  [illegible]
0049 5zr1B  [illegible]
0050 4m9sB  [illegible]
0051 3glfG  [illegible]
0052 8xksD  [illegible]
0053 2gnoA  [illegible]
0054 7w42B  [illegible]
0055 8xksB  [illegible]
0056 7mi8A  [illegible]
0057 1g4aE  [illegible]
0058 8xkuC  [illegible]
0059 5j1sB  [illegible]
0060 7qo4M  [illegible]
0061 8rwvF  [illegible]

# Job: Type-1 BREX; BrxC-ATPase MBR4615907 Canonical wHTH1
# Query: s001A
# No:  Chain   Z    rmsd lali nres  %id PDB  Description
   1:  4ad9-A  9.1  1.9   70   288   13   MOLECULE: BETA-LACTAMASE-LIKE PROTEIN 2;
   2:  6j0e-B  8.8  1.7   60   125   22   MOLECULE: ARSENIC RESPONSIVE REPRESSOR ARSR;
   3:  6qpq-D  8.8  3.0   70    82    9   MOLECULE: STRUCTURAL MAINTENANCE OF CHROMOSOMES PROTEIN,STR
   4:  5j6x-A  8.8  1.4   60    66   10   MOLECULE: Z-DNA BINDING PROTEIN KINASE;
   5:  3r0a-B  8.7  1.8   62   123   16   MOLECULE: PUTATIVE TRANSCRIPTIONAL REGULATOR;
   6:  2jt1-A  8.5  2.6   65    71   14   MOLECULE: PEFI PROTEIN;
   7:  6uvu-B  8.5  1.9   59   114   12   MOLECULE: ARSR FAMILY TRANSCRIPTIONAL REGULATOR;
   8:  1sfx-A  8.4  1.7   60   109   12   MOLECULE: CONSERVED HYPOTHETICAL PROTEIN AF2008;
   9:  7z1n-O  8.4  1.8   62   570   10   MOLECULE: DNA-DIRECTED RNA POLYMERASE III SUBUNIT RPC1;
  10:  8ffz-B  8.3  1.2   58   905   21   MOLECULE: TRANSCRIPTION FACTOR IIIA;
  11:  7a6h-O  8.1  1.6   60   512    5   MOLECULE: DNA-DIRECTED RNA POLYMERASE III SUBUNIT RPC1;
  12:  6qfd-B  8.0  2.3   65   116   11   MOLECULE: DNA-BINDING PROTEIN;
  13:  2obp-A  8.0  3.1   67    81    9   MOLECULE: PUTATIVE DNA-BINDING PROTEIN;
  14:  4ru7-B  8.0  1.2   58    99   12   MOLECULE: ASPA;
  15:  4ija-A  8.0  1.9   59   363   14   MOLECULE: XYLR PROTEIN;
  16:  1stz-A  8.0  2.2   62   323   10   MOLECULE: HEAT-INDUCIBLE TRANSCRIPTION REPRESSOR HRCA HOMOL
  17:  3f8f-A  8.0  2.2   67   114   10   MOLECULE: TRANSCRIPTIONAL REGULATOR, PADR-LIKE FAMILY;
  18:  8iue-P  8.0  1.6   62   303   10   MOLECULE: DNA-DIRECTED RNA POLYMERASE III SUBUNIT RPC1;
  19:  1ku9-A  8.0  1.8   62   151   15   MOLECULE: HYPOTHETICAL PROTEIN MJ223;
  20:  4i99-C  8.0  2.8   66    70    9   MOLECULE: CHROMOSOME PARTITION PROTEIN SMC;
  21:  5y6i-B  7.9  1.7   59   234   10   MOLECULE: TRANSCRIPTIONAL REGULATOR KDGR;
  22:  3ov8-A  7.9  1.9   64    91   13   MOLECULE: PROTEIN AF_1382;
  23:  1sfu-A  7.9  2.0   61    70   15   MOLECULE: 5'-D(*T*CP*GP*CP*GP*CP*G)-3';
  24:  4ets-B  7.8  1.8   59   149   10   MOLECULE: FERRIC UPTAKE REGULATION PROTEIN;
  25:  6pco-B  7.8  1.7   58   135   10   MOLECULE: MARR-FAMILY TRANSCRIPTIONAL REGULATOR;
  26:  5vyv-A  7.8  3.0   70   173   10   MOLECULE: UPF0502 PROTEIN YCEH;
  27:  3lmm-D  7.8  2.3   60   556   12   MOLECULE: UNCHARACTERIZED PROTEIN;
  28:  2zkz-C  7.7  1.7   58    87   10   MOLECULE: TRANSCRIPTIONAL REPRESSOR PAGR;
  29:  1xsd-A  7.7  2.0   63   125   10   MOLECULE: 5'-
  30:  2nyx-B  7.7  1.8   60   147   15   MOLECULE: PROBABLE TRANSCRIPTIONAL REGULATORY PROTEIN, RV14
  31:  8jxk-D  7.6  2.1   66   166    6   MOLECULE: CONSERVED PROTEIN;
  32:  7rhe-A  7.6  1.3   55   368   15   MOLECULE: ROK FAMILY PROTEIN;
  33:  3cuq-A  7.6  2.0   60   219   15   MOLECULE: VACUOLAR-SORTING PROTEIN SNF8;
  34:  6j05-A  7.6  2.1   61   101    8   MOLECULE: TRANSCRIPTIONAL REGULATOR ARSR;
  35:  7nyw-E  7.6  2.9   72   212   13   MOLECULE: CHROMOSOME PARTITION PROTEIN MUKB;
  36:  4o5v-A  7.6  1.6   57   213   11   MOLECULE: IRON-DEPENDENT TRANSCRIPTION REPRESSOR RELATED PR
  37:  1cf7-B  7.6  1.9   64    82   11   MOLECULE: DNA (5'-
  38:  6kf4-G  7.6  1.4   61   166   16   MOLECULE: DNA-DIRECTED RNA POLYMERASE SUBUNIT;
  39:  5a31-N  7.6  2.8   62   703    2   MOLECULE: ANAPHASE-PROMOTING COMPLEX SUBUNIT 1;
  40:  5u8o-B  7.6  2.0   69   360   17   MOLECULE: ZN-DEPENDENT HYDROLASE;
  41:  9gm7-C  7.6  3.2   74   440   12   MOLECULE: CHROMOSOME PARTITION PROTEIN MUKF;
  42:  1ylf-C  7.6  2.4   63   119    8   MOLECULE: RRF2 FAMILY PROTEIN;
  43:  5zqh-A  7.6  1.9   63   101   11   MOLECULE: PADR FAMILY TRANSCRIPTIONAL REGULATOR;
  44:  4fht-B  7.5  1.8   61   142   13   MOLECULE: PCAV TRANSCRIPTIONAL REGULATOR;
  45:  2p4w-B  7.5  1.7   60   198   10   MOLECULE: TRANSCRIPTIONAL REGULATORY PROTEIN ARSR FAMILY;
  46:  2oqg-A  7.5  1.7   59   109    8   MOLECULE: POSSIBLE TRANSCRIPTIONAL REGULATOR, ARSR FAMILY P
  47:  7xjg-J  7.5  1.8   62   306   13   MOLECULE: RNA-DIRECTED DNA POLYMERASE FROM RETRON EC86;
  48:  2cwe-A  7.5  2.1   60   191   15   MOLECULE: HYPOTHETICAL TRANSCRIPTION REGULATOR PROTEIN, PH1
  49:  2rdp-A  7.5  1.5   58   140   14   MOLECULE: PUTATIVE TRANSCRIPTIONAL REGULATOR MARR;
  50:  2xrn-B  7.5  1.7   58   241   14   MOLECULE: HTH-TYPE TRANSCRIPTIONAL REGULATOR TTGV;

0001 s001A   ------DKAFATRVLNTLF-MIC-HLDYKDAQAFP--ATIGNLTSLM--M---TDITT---SRKELKERIEAEVKYLCDESVLMR-ET-S-D--S-----G--V-EYFKFY------------------------------
0002 4ad9A   qqyishRNIREQQILTLFR-ENF-EK--------S--FTVMELVKII-yK---NTPEN---LHEMAKHNLLLHLKKLEKEGKIFS-NT-D----------P--D-KKWKAHl-----------------------------
0003 6j0eB   dlfkvlGDPVRLRILSQLAaGGC-G-----------pVSVNELTDLM--G--------------lSQPTISHHLKKMTEAGFLDR-VP-E----G-----R--V-VLHRVRpelfaelrtvlqigsmellehhhhhh----
0004 6qpqD   --------KAIVQMAKILR-KEL-SE------ekE--VIFTDVLKSQ--A---NTEEN---ITKREASRGFFDILSLATEGCIGL-SQ-T-E--A----fG--N-IKIDAKpalferfi--------------------
0005 5j6xA   ---msaENEIEMRICDYLR-RHG-------------rSTVQDIFKEL--K--------------lEKSTVNRHLYSLQASKQVFK-TV-E-D--N-----K--R-PVWDLVe-----------------------------
0006 3r0aB   ikcalnLTKADLNVXKSFL-NEP-D--------rW--IDTDALSKSL--K--------------lDVSTVQRSVKKLHEKEILQR-SQ-Q-N--L-dgggY--V-YIYKIYsknqirniiqkivqswadrlgqelkeweng
0007 2jt1A   -----msESIVTKIISIVQ-ERQnMDDG-----AP--VKTRDIADAA--G--------------lSIYQVRLYLEQLHDVGVLEK-VN-A-G--K-----G--VpGLWRLLe-----------------------------
0008 6uvuB   vlfkvlANDTRLRLLHALA-RSG-------------gLCVTDLAAAV--G--------------mKPQAVSNQLQRLADRRILRA-AR-C----G-----N--N-IHYRIVdpcvlrmlelglclieeaeqqagg------
0009 1sfxA   aleklsFKPSDVRIYSLLL-ERG-------------gXRVSEIAREL--D--------------lSARFVRDRLKVLLKRGFVRR-EI-V-E--K---gwV--G-YIYSAEkpekvlkefkssilgeieriekxftdgs--
0010 7z1nO   elvkahLGERAASVIGMLV-ALG----------R--LSVRELVEKI--D---G-----------mDVDSVKTTLVSLTQLRCVKY-LQ-E-T--A-isgkK--T-TYYYYNeegihillysgliideiitqmrvndeeehk
0011 8ffzB   sirpniWKQVVTMVVNEII-FHP----------G--ITLSRLQSRC--R---E---------vlSLHEISEICKWLLERQVLIT-TD---------------F-DGYWVNhnwysiyest-------------------
0012 7a6hO   svvqerFGSRCARIFRLVL-QKK-------------hIEQKQVEDFA--M--------------iPAKEAKDMLYKMLSENFMSL-QE-I-PksR-----T--F-YLYTVNilsaarmllhrcyksianlierrqfetken
0013 6qfdB   rsdardLTAFQKNILTVLG-EEA--------------RYGLAIKREL--E---E--GE---EV--NHGRLYPNLDDLVNKGLVEK-SE-L-D--K-----R--T-NEYALTnegfdavvddlewtlskfvadadrrervet
0014 2obpA   -----gIDPAIVEVLLVLR-EA--GIENGAT---P--WSLPKIAKRA--Q--------------lPXSVLRRVLTQLQAAGLADV-SV-E-A--D-----G--R-GHASLTqegaalaaqlfp-----------------
0015 4ru7B   stdkyifLTPRAYIIVHLL-KVG-------------kAKASEISENT--Q--------------iPYQTVIQNIRWLLAEGYVVK-EQ-K-G---------E--E-IYYKLTdkgkqlataelekirklvevvqhhhhhh-
0016 4ijaA   -----pMNDNEKRVLREIY-NHH----------N--ISRTQISKNL--E--------------iNKATISSILNKLKYKSLVNE-VG-----gR-----K--P-ILLKVNhlygyfisldltyssvevmynyfdgnvikh
0017 1stzA   -----kLNDRQRKVLYCIV-REY-IE-----nkKP--VSSQRVLEVS--N-------------ieFSSATIRNDMKKLEYLGYIYQ-PH-T---------------S-AGRIPTdkglrfyyeemlkisketseadlavetfks
0018 3f8fA   ipkemlRAQTNVILLNVLK-QGD---------------NYVYGIIKQVkeA---SNG-E---MEL-NEATLYTIFKRLEKDGIISS-YW-G-D--E---ggR--R-KYYRLTeighenmrlafeswsrvdkiienleankks
0019 8iueP   agkmkgSDNQEKLVYQIIE-DAG-NK----------gIWSRDIRYKS--N--------------lPLTEINKILKNLESKKLIKA-VK-SvA--A-----S--KkKVYMLYnlqpdrsvtggawysdqdfesefvevlnqq
0020 1ku9A   lakihgLNKSVGAVYAILY-LSD-K--------P--LTISDIXEEL--K--------------iSKGNVSXSLKKLEELGFVRK-VW-I-K--G-----E-rK-NYYEAVdgfssikdiakrkhdliaktyedlkkleek
0021 4i99C   -----dIEKYVEELYKVVK-KIY-EKT-----gTP--IKFWDLVP---------------dVEPKIIARTFLYLLFLENMGRVEI-IQ-E-E--P----fG--E-ILVVPM------------------------------
0022 5y6iB   ----vrSAEVGTDILKALA-ELS-P----------A--TSLSRLAEHV--G--------------mPASKVHRYLQALIASGFAVQ-DA-S---------------T-NHYSLGrealrvglaaldsmdvlksaaaplaelrdv
0023 3ov8A   erikllFKEKALEILMTIY-YES-LGG------ND--VYIQYIASKV--N--------------sPHSYVWLIIKKFEEAKMVEC-EL-E----G-----R--T-KIIRLTdkgqkiaqqiksiidimend---------
0024 1sfuA   --ctvnDAEIFSLVKKEVL-S-L-NTN------dY--TTAISLSNRL--K--------------iNKKKINQQLYKLQKEDTVKM-VP-S---------N--P-PKWFKNync-------------------------
0025 4etsB   llerfkYTKQREVLLKTLY-HS----------dTH--YTPESLYMEI--K---Q---aepdlnvGIATVYRTLNLLEEAEMVTS-IS-F----------gsaG-KKYELAnkphhdhmickncgkiiefenpiierqqal
0026 6pcoB   tipdskLTAAQFVVLCALR-DQG--------------aCSLVDVVKAT--A--------------iDQATVRGVIERLKARKLLAV-SH------R-----R--K-VLVTLTpdgralveemvpfaeqitqstfgglnpaer
0027 5vyvA   --mkyqLTALEARVIGCLL-EKQ-VT----TPEQY-pLSVNGVVTAC--N---QKT--repvmnlSESEVQEQLDNLVKRHYLRT-VS-G-F-gN-----R--V-TKYEQRfcnsefgdlklsaaevalittlllrgaqtp
0028 3lmmD   fspvrylsTDQAELTNAAX-LWL-SE------vgD--LATSDLXAXC--G--------------vSRGTAKACVDGLVDEERVVA-VG-G----G-----R--S-RRYRLVe-----------------------------
```

```
0029 2zkzC   ellktxAHPXRLKIVNELY-KHK-------------aLNVTQIIQIL--K--------------lPQSTVSQHLCKXR-GKVLKR-NR-Q----G-----L--E-IYYSINnpkvegiiklln------------------
0030 1xsdA   tnkqveiSMAEWDVMNIIW-DKK-------------sVSANEIVVEI--Q---KY--K---EVS--DKTIRTLITRLYKKEIIKR-YK-S----E-----N--I-YFYSSNikeddikmktaktflnklyggdmkslvlnf
0031 2nyxB   aqvdenITIPQFRTLVILS-NHG-------------pINLATLATLL--G--------------vQPSATGRXVDRLVGAELIDR-LPhP-T--S-----R-rE-LLAALTkrgrdvvrqvtehrrteiariveqxapaer
0032 8jxkD   ------EFMLELAILGLLI-ESP--------------MHGYELRKRL--T-glLGAFR---AF--SYGSLYPALRRMQADGLIAE-NA-A-P--A-----G---rRVYQLTdkgrrrfgelvadtgphnytddgfgvhlaf
0033 7rheA   ----nvRRYNERLLLKTLR-RAG-------------sASKADLARLA--N--------------mTGTAVGSIIASLADAKLIEF-A----------------A-SLIRLDprgafgigvhldrmrietalvnfagdvlgr
0034 3cuqA   qsvpaeLNMDHTVVLQLAE-KNG-------------yVTVSEIKASL--K--------------wETERARQVLEHLLKEGLAWL-DL-Q-A--P-----G--E-AHYWLPalftdlysqeitaee---------------
0035 6j05A   arlealASPVRLEIFRLLV-EQE-PT----------gLVSGDIAEHL--G--------------qPHNGISFHLKNLQHAGLVTV-QR-E----G-----R--Y-QRYRAAmpvvralvayltencchgtrdcals-----
0036 7nywE   liprsvLSELDMMVGKILC-YLY-LSP--ERLANqgiFTSQELYEEL--IslaDEGKlsdlDKQKLQEKVRTTLNRLRRLGMVYF-LP-N-------------nnNKFTITeavfrfgadvrsgddpreiqlrmirdgeam
0037 4o5vA   -----irsVTEEDYLKIIQ-ELV-LY------kgY--ATLADISRSL--N--------------vKRQSVRDEINHLISLSXAEK-IE---------------R-GKYRLTpsgdreanrflrkhrtaeillsrcigipwe
0038 1cf7B   ---gkgLRHFSMKVCEKVQ-RKG-------------tTSYNEVADELvsE---FTNAY---DQKNIRRRVYDALNVLMAMNIISK-EK-----------------KEIKWIglp---------------------------
0039 6kf4G   eiaqeiGGEEAVEVVKALE-KMK-----------E--ATDEELAEAT--G--------------iRVNTVRRILYMLNDEGLADF-KR-I-R--DpetgwY--Y-YYWRLEtkklpeiirsrkmaelkklkemleeetsei
0040 5a31N   dnmvliKEEELLLFWTYIQ-AML-TN------leS--LSLDRIYNML--RmfvVTGP---alaeiDLQELQGYLQKKVRDQQLVY-SA-----------------GVYRLP------------------------------
0041 5u8oB   aqlrahHDARLDEVRVACA-EQP--------------MSAADIVPIM--F---RRRELdihQMTFALGEAIAHLNLLWLAGELVR-EQ-G-D--D-----G--V-LRFRIAr-----------------------------
0042 9gm7C   ndfsisLPVERLAFLMAIA-VLN-SE-RLDG---E--MSEGELIDAF--R---E--EQ---TAESVAVRANNAINDMVRQKLLNR-FT-S-E--L-----AdgN-AIYRLTplgisisdyyirqrefstlrlsmqlsivan
0043 1ylfC   -------ISSRFSIAVHIL-SIL-KN----NPSSL--CTSDYXAESV--N--------------tNPVVIRKIXSYLKQAGFVYV-NR-G-------------P-GGAGLLkdlheitlldvyhavnvganiqavleiili
0044 5zqhA   -etqllKGVLEGCVLDMIG-QKE--------------RYGYELVQTL--R---E--fD---TIV---GTIYPLLQKLEKNQWIRG-DM-R-P-gP-----D--R-KYFSLMkegeervsvfwqqwddlsqkvegikn----
0045 4fhtB   tmvseeTTSPQYAVLNALV-AEP-----------G--LDQRTVGERV--G--------------lDRSTIAEVVSRLGRRGLLDK-VR-D-P-dG-----R--R-SLLRLTdeglrvhrrlgvriarmnqvflaplaadeq
0046 2p4wB   rlldvlGNETRRRILFLLT-KRP--------------YFVSELSREL--G--------------vGQKAVLEHLRILEEAGLIES-RV-E-K-gR-----P--R-KYYMIKkglrleilltptlfgsemyeakgvrkspey
0047 2oqgA   svfaalSDETRWEILTELG-RAD--------------QSASSLATRL--P--------------vSRQAIAKHLNALQACGLVES-VK-V-G--------R--E-IRYRALgaelnktartlerigaewdrrlaaikqiae
0048 7xjgJ   skdignILYAERFLLPCIY-LLD-----------S--VNYRTLCELA--F---K--kQ---DDVLSKIIVRSVVSRLINERKILQ-MT-----------------DGYQVTalgasyvrsvfdrktldrlrleimnfenrr
0049 2cweA   evikvmLEDTRRKILKLLR-NKE--------------MTISQLSEIL--G--------------kTPQTIYHHIEKLKEAGLVEVkRT-E-M--K--gnlV--E-KYYGRTadvfyinlylgdeelryiarsrlktkidif
0050 2rdpA   iltnypiTPPQFVALQWLL-EEG-------------dLTVGELSNKX--Y--------------lACSTTTDLVDRXERNGLVAR-VR-D-E--H--------V-VRIRLLekgeriieeviekrqrdlanvlesfsdeei
0051 2xrnB   ---siqVIARAASIMRALG-SHP-H-----------gLSLAAIAQLV--G--------------lPRSTVQRIINALEEEFLVEA-LG-P-------------A-GGFRLGpalgqlinqaqtdilslvkpylrslaeeld

0001 s001A   ------LHHHHHHHHHHHH-HHH-LLLHHHHHHLL--LLHHHHHHHH--L---LLLLL---LHHHHHHHHHHHHHHHHHLLLEEE-EE-L-L--L-----L--L-EEEEEL------------------------------
0002 4ad9A   hhhhhhHHHHHHHHHHHHH-HLL-LL--------L--LLHHHHHHHH-lL---LLLHH---HHHHHHHHHHHHHHHHHHLLLEEE-LL-L----------L--L-LLEEELl-----------------------------
0003 6j0eB   hhhhhhLLHHHHHHHHHHHhLLL-L-----------lEEHHHHHHHH--L--------------lLHHHHHHHHHHHHHLLLEEE-EE-E----L-----L--E-EEEEELhhhhhhhhhlllleeeeeeeeeeell----
0004 6qpqD   --------LHHHHHHHHHH-HHH-LL------llL--EEHHHHHHHH--H---LLLHH---LLHHHHHHHHHHHHHHHHLLLEEE-EL-L-L--L----lL--L-LEEEELhhhhllll----------------------
0005 5j6xA   ---llhHHHHHHHHHHHHH-HHL-------------lLLHHHHHHHL--L--------------lLHHHHHHHHHHHHHLLLEEE-EL-L-L--L-----L--L-LEEEELl-----------------------------
0006 3r0aB   hhhhhlLLHHHHHHHHHHH-HLL-L--------lL--LLHHHHHHHL--L--------------lLHHHHHHHHHHHHHLLLEEE-EE-E-E--L-llllE--E-EEEEELlhhhhhhhhhhhhhhhhhhhhhhhhhhhll
0007 2jt1A   -----llLLHHHHHHHHHH-HHHhHHLL-----LL--EEHHHHHHHH--L--------------lLHHHHHHHHHHHHHLLLEEE-EL-L-L--L-----L--LlLEEEELl-----------------------------
0008 6uvuB   hhhhhhLLHHHHHHHHHHH-HHL-------------lLLHHHHHHHL--L--------------lLHHHHHHHHHHHHHLLLEEE-EE-E----L-----L--E-EEEEELlhhhhhhhhhhhhhhhhhhhhlll------
0009 1sfxA   hhhhllLLHHHHHHHHHHH-HHL-------------lLLHHHHHHHL--L--------------lLHHHHHHHHHHHHHLLLEEE-EE-E-E--L---llE--E-EEEEELlhhhhhhhhhhhhhhhhhhhllllllll--
0010 7z1nO   hhhhhhLLHHHHHHHHHHH-HHL-----------L--EEHHHHHHHL--L---L----------lLHHHHHHHHHHHHHLLLLLE-EE-E-E--L-llllE--E-EEELLLhhhhhhhhlhhhhhhhhhhhhlllllllhh
0011 8ffzB   lllhhhHHHHHHHHHHHHH-HLL-----------L--EEHHHHHHHL--L---L---------llLHHHHHHHHHHHHHLLLEEE-EL---------------L-LEEEELllllllllll--------------------
0012 7a6hO   hhhhhhHLHHHHHHHHHHH-HLL-------------lLLHHHHHHHH--L--------------lLLLLHHHHHHHHHLLLLLLL-LL-E-ElhH-----E--E-LLLLLLhhhhhhhhhhhhhhhhhhhhhhhhhhhhhl
0013 6qfdB   hhhhllLLHHHHHHHHHHH-HLL--------------EEHHHHHHHH--H---H--LL---LL--LHHHHHHHHHHHHHLLLEEE-EE-E-E--L-----L--E-EEEEELhhhhhhhhhhhhhhhhhlllhhhhhhhhhh
0014 2obpA   -----lLLHHHHHHHHHHH-HH--LLLLLLL---L--LLHHHHHHHH--L--------------lLHHHHHHHHHHHHHLLLEEE-EE-L-L--L-----L--L-EEEEELhhhhhhhhhhll------------------
0015 4ru7B   lllllllLLHHHHHHHHHH-LLL-------------eELHHHHHHHH--L--------------lLHHHHHHHHHHHHHHLLEEL-LL-L-L--------L--L-LLEEELhhhhhhhhhhhhhhhhhhhhhhlllllll-
0016 4ijaA   -----lLHHHHHHHHHHHH-HLL-----------L--LLHHHHHHHH--L--------------lLHHHHHHHHHHHHHLLLEEE-LL-----lL-----L--L-LLEEELhhhleeeeeeeellleeeeeeeellleeee
0017 1stzA   -----lLLHHHHHHHHHHH-HHH-HH-----hlLL--LLHHHHHHHL--L------------llLLHHHHHHHHHHHHHLLLEEL-LL-L-------------L-LLLEELhhhhhhhhhhhhhhllllllllhhhhllll
0018 3f8fA   llhhhhHHHHHHHHHHHHH-HLL--------------LLHHHHHHHHhhH---LLL-L---LLL-LHHHHHHHHHHHHHLLLEEE-EE-L-L--L---llL--L-EEEEELhhhhhhhhhhhhhhhhhhhhhhhhhhhhhh
0019 8iueP   llllllLLHHHHHHHHHHH-HHH-HH----------lLLHHHHHHHH--L--------------lLHHHHHHHHHHHHHLLLEEE-EE-LlL--L-----L--LlEEEEELlllllllllllllleelleelhhhhhhhhhh
0020 1ku9A   hhhhllLLHHHHHHHHHHH-HLL-L---------L--EEHHHHHHHH--L--------------lLHHHHHHHHHHHHHLLLEEE-EL-L-L--L-----L-lL-LEEEELlhhhhhhhhhhhhhhhhhhhhhhhhhhhhh
0021 4i99C   -----lHHHHHHHHHHHHH-HHH-HHH-----lLL--EEHHHHLL---------------lLLHHHHHHHHHHHHHHHHLLLEEE-EL-L-L--L----lL--L-LEEEEL------------------------------
0022 5y6iB   ----llHHHHHHHHHHHHH-HLL-L---------L--EEHHHHHHHH--L--------------lLHHHHHHHHHHHHHHLLEEE-LL-L-------------L-LEEEELhhhhhhhhhhhhlllhhhhhhhhhhhhhhh
0023 3ov8A   lhhhhhLLHHHHHHHHHHH-HHH-HLL------LL--EEHHHHHHHL--L--------------lLHHHHHHHHHHHHHLLLEEE-EE-E----L-----L--E-EEEEELhhhhhhhhhhhhhhhhhhll----------
0024 1sfuA   --llllLHHHHHHHHHHHH-L-L-LLL------lE--ELHHHHHHHL--L--------------lLHHHHHHHHHHHHHLLLEEE-EL-L----------L--L-LEEEELlll---------------------------
0025 4etsB   hhhhhlLLHHHHHHHHHHH-HL----------lLL--LLHHHHHHHH--H---H----hlhhhllLHHHHHHHHHHLLLLLLEEL-LL-E----------ellE-ELLEELllllleeeeellllleeeellhhhhhhhhh
0026 6pcoB   hlhhhlLLHHHHHHHHHHH-HHL-------------lLLHHHHHHHH--L--------------lLHHHHHHHHHHHHHLLLEEE-EL------L-----L--L-LEEEELhhhhhhhhhhhhhhhhhhhhhhllllhhhh
0027 5vyvA   --llllLLHHHHHHHHHHH-HHH-HH----LHHHL-lEEHHHHHHHH--L---LLL--lllllllLHHHHHHHHHHHHHLLLEEE-EL-L-L-lL-----L--L-LEEEELllllllllllhhhhhhhhhhhhhhllllh
0028 3lmmD   llllllllLLHHHHHHHHH-HHH-HH------llL--EEHHHHHHHL--L--------------lLHHHHHHHHHHHHHLLLEEE-EL-L----L-----L--L-LEEEELl-----------------------------
0029 2zkzC   hhhhhhLLHHHHHHHHHHH-HHL-------------lEEHHHHHHHH--L--------------lLHHHHHHHHHHHL-LLLEEE-EE-E----L-----L--E-EEEEELlhhhhhhhhhhl------------------
0030 1xsdA   lllllllLLLLHHHHHHHH-HHL-------------lEEHHHHHHHH--H---LL--L---LLL--HHHHHHHHHHHHHLLLEEE-EE-L----L-----L--L-EEEEELllhhhhhhhhhhhhhhhhhlllhhhhhhhh
0031 2nyxB   hhhlllLLHHHHHHHHHHH-HHL-------------lEEHHHHHHHH--L--------------lLHHHHHHHHHHHHHLLLEEE-ELlL-L--L-----L-lL-LEEEELhhhhhhhhhhhhhhhhhhhhhhllllhhhl
0032 8jxkD   ------LHHHHHHHHHHHH-HLL--------------EEHHHHHHHH--H-hhLLLLL---LL--LLLLHHHHHHHHHHLLLEEE-LL-L-L--L-----L---lLEEEELhhhhhhhhhhhhhhlllhhhllhhhhhhhh
0033 7rheA   ----lhHHHHHHHHHHHHH-HHL-------------lEEHHHHHHHH--H--------------lLHHHHHHHHHHHHHLLLEEE-L----------------L-LEEEELllleeeeeeeelleeeeeeeeelllleeee
0034 3cuqA   elllllLLHHHHHHHHHHL-LLL-------------eELHHHHHHHH--L--------------lLHHHHHHHHHHHHHHLLLEE-EL-L-L--L-----L--L-LEEELLlllllllllllhhhl---------------
0035 6j05A   hhhhhhLLHHHHHHHHHHH-HLL-LL----------lEEHHHHHHHH--L--------------lLHHHHHHHHHHHHHLLLEEE-EE-E----L-----L--E-EEEEELhhhhhhhhhhhllllllllllllll-----
0036 7nywE   llleeeLLHHHHHHHHHHH-HHH-HLL--HHHHHlleEEHHHHHHHH--HhhlLHHHhlhhHHHHHHHHHHHHHHHHHHLLLEEE-LL-L-------------llLEEEELhhhhhhlllllllllhhhhhhhhhhhllee
0037 4o5vA   -----lllLLHHHHHHHHH-HHH-HH------hlL--LLHHHHHHHH--L--------------lLHHHHHHHHHHHHHLLLEEE-EE---------------L-LEEEELhhhhhhhhhhhhhhhhhhhhhhhhhlllll
0038 1cf7B   ---llhHHHHHHHHHHHHH-HHL-------------eELHHHHHHHHhhH---HHLHH---HHHHHHHHHHHHHHHHHHLLLLLL-LL-----------------LLEELLlll---------------------------
0039 6kf4G   hhhhhhLLHHHHHHHHHHH-HHL-----------L--LLHHHHHHLL--L--------------lLHHHHHHHHHHHHHLLLEEE-EE-E-E--LlllllE--E-EEEEELlllhhhhhhhhhhhhhhhhhhhhhhlllll
0040 5a31N   llllllLHHHHHHHHHHHH-HHH-HH------hlL--LLHHHHHHHH--HhhlLLLL---lllllLHHHHHHHHHHHHHHLLLEE-EL-----------------LEEELL------------------------------
0041 5u8oB   hhhhhhHHHHHHHHHHHHH-HLL--------------LLHHHHHHHH--L---LLLLLlhhHHHHHHHHHHHHHHHHHHLLLEEE-EE-L-L--L-----L--L-EEEEELl-----------------------------
0042 9gm7C   hllleeELHHHHHHHHHHH-HHH-HL-LLLL---L--EEHHHHHHHH--H---H--LL---LLLLHHHHHHHHHHHHHHLLLEEE-EE-L-H--H-----HllL-EEEEELhhhhhhhhhhhlllllllhhhhhhhhhhhh
0043 1ylfC   -------LLHHHHHHHHHH-HHH-HH----LLLLL--LLHHHHHHHL--L--------------lLHHHHHHHHHHHHHHLLEEE-LL-L-------------L-LLEEELllhhhllhhhhhhhhllllhhhhhhhhhhh
0044 5zqhA   -llhhhHHHHHHHHHHHHH-HLL--------------EEHHHHHHHH--H---H--lL---LLL---LLLHHHHHHHHHLLLEEE-EE-E-E-lL-----E--E-EEEEELhhhhhhhhhhhhhhhhhhhhhhhhhl----
0045 4fhtB   hhllllLLHHHHHHHHHHH-HLL-----------L--LLHHHHHHHH--L--------------lLHHHHHHHHHHHHHHLLEEE-EE-L-L-lL-----L--L-EEEEELhhhhhhhhhhhhhhhhhhhhhhllllhhhh
0046 2p4wB   hhhhhlLLHHHHHHHHHHL-LLL--------------EEHHHHHHHH--L--------------lLHHHHHHHHHHHHHHLLEEE-EE-L-L-lL-----L--L-EEEEELlleeeeeeeelleeeeeeeeellllllhhh
0047 2oqgA   hhhhhlLLHHHHHHHHHHH-HLL--------------LLHHHHHHHL--L--------------lLHHHHHHHHHHHHHLLLEEE-EE-E-L--------L--E-EEEEELlhhhhhhhhhhhhhhhhhhhhhhhhhhhhl
0048 7xjgJ   llllllLLLLHHHHHHHHH-HLL-----------L--LLHHHHHHHH--H---H--lL---LLHHHHHHHHHHHHHHHHLLLEEE-EL-----------------LEEEELhhhhhhhhhhhllhhhhhhhhhhhhhhhll
0049 2cweA   hhhhhhLLHHHHHHHHHHL-LLL--------------LLHHHHHHHH--L--------------lLHHHHHHHHHHHHHLLLEEEeEE-E-E--E--lleE--E-EEEEELlleeeelllllhhhhhhhhhhhhhhhhhhh
0050 2rdpA   hhlllllLHHHHHHHHHHH-HHL-------------lLLHHHHHHHH--L--------------lLHHHHHHHHHHHHHLLLEEE-EE-L-L--L--------L-EEEEELhhhhhhhhhhhhhhhhhhhhhhhhllhhhh
0051 2xrnB   ---llhHHHHHHHHHHHHH-LLL-L-----------lEEHHHHHHHL--L--------------lLHHHHHHHHHHHHLLLLEEE-LH-H-------------H-LEEEELlhhhhhhhhhhhhhhhhhhhhhhhhhhhhl

# Job: Type-1 BREX; BrxC-ATPase Alpha+Beta domain
# Query: s001A
# No:  Chain   Z    rmsd lali nres  %id PDB  Description
   1:  1yvp-B  5.2 11.4   73   532   12   MOLECULE: Y RNA SEQUENCE, FIRST STRAND;
   2:  6vdq-A  5.1  7.7   81   305   14   MOLECULE: 3-METHYL-L-TYROSINE PEROXYGENASE;
   3:  3kkz-A  5.1 13.5   84   257    8   MOLECULE: UNCHARACTERIZED PROTEIN Q5LES9;
   4:  6thk-A  5.0  5.1   96   466    9   MOLECULE: PYOCIN S5;
   5:  1u7z-B  4.8  4.1  108   218    5   MOLECULE: COENZYME A BIOSYNTHESIS BIFUNCTIONAL PROTEIN
   6:  5b86-A  4.7  4.2   75   579    8   MOLECULE: TUMOR NECROSIS FACTOR ALPHA-INDUCED PROTEIN 2;
   7:  8yho-C  4.6  9.1   78   264    6   MOLECULE: DUF87 DOMAIN-CONTAINING PROTEIN;
   8:  3iuk-A  4.6  9.7   87   552    6   MOLECULE: UNCHARACTERIZED PROTEIN;
   9:  6s8f-F  4.5 10.9   79  2571   11   MOLECULE: SERINE/THREONINE-PROTEIN KINASE TEL1,SERINE/THREO
  10:  3trc-A  4.5  3.5   59   168    7   MOLECULE: PHOSPHOENOLPYRUVATE-PROTEIN PHOSPHOTRANSFERASE;
  11:  7eld-A  4.5  8.6   77  1137   10   MOLECULE: ENDORIBONUCLEASE DICER HOMOLOG 1;
  12:  8g7m-G  4.4  3.5   60   343    5   MOLECULE: 60 KDA HEAT SHOCK PROTEIN, MITOCHONDRIAL;
  13:  4hm9-A  4.4  6.8   79   511    4   MOLECULE: BETA-CATENIN-LIKE PROTEIN 1;
  14:  7sjr-A  4.4 13.9   58   911   12   MOLECULE: DNA HELICASE;
  15:  9jm7-B  4.4  3.7   55    65    9   MOLECULE: LRD-2A;
  16:  6vjp-A  4.4  3.7   91   155    7   MOLECULE: ACETYLTRANSFERASE;
  17:  3cit-A  4.3  3.6   63   155    5   MOLECULE: SENSOR HISTIDINE KINASE;
  18:  2ooc-B  4.3  3.0   50   105    2   MOLECULE: HISTIDINE PHOSPHOTRANSFERASE;
  19:  5ipx-A  4.3  3.3   68   282    7   MOLECULE: ORF49 PROTEIN;
  20:  7tjl-A  4.3  2.1   52    85    6   MOLECULE: DE NOVO DESIGNED PROTEIN, SEWN0.1;
  21:  6icz-A  4.3  4.4   77  2253   12   MOLECULE: PROTEIN MAGO NASHI HOMOLOG 2;
  22:  5v8z-C  4.3  3.7   54    98    7   MOLECULE: ENDOPLASMIC RETICULUM RESIDENT PROTEIN 29;
  23:  1xkp-A  4.3  5.4   74   230   11   MOLECULE: PUTATIVE MEMBRANE-BOUND YOP TARGETING PROTEIN YOP
  24:  5j4a-B  4.1  2.8   65   104   15   MOLECULE: TRNA NUCLEASE CDIA;
  25:  8eki-C  4.1  7.4   87   701    7   MOLECULE: PROTEIN TRANSPORT PROTEIN SEC20;
  26:  4zmk-A  4.1  2.2   49    71    8   MOLECULE: TELOMERE LENGTH REGULATOR TAZ1;
  27:  8usp-A  4.1  3.0   53   982    6   MOLECULE: DNA REPAIR/TRANSCRIPTION PROTEIN MET18/MMS19;
  28:  6tgv-C  4.1  9.6  102   389   14   MOLECULE: PHOSPHOPANTOTHENATE-CYSTEINE
  29:  7c0q-A  4.1  3.3   55   279    5   MOLECULE: EFFECTOR LPG2505;
  30:  8ye6-B  4.1  2.9   55    88    9   MOLECULE: A32;
  31:  5eqz-A  4.1  4.3   70   138   11   MOLECULE: REV PROTEIN;
  32:  8iqi-C  4.1  5.9   70   916    7   MOLECULE: PUTATIVE PRIMASE C962R;
  33:  6cjd-A  4.1  9.2   78   128    5   MOLECULE: PUTATIVE CYTOPLASMIC PROTEIN;
  34:  6hd5-t  4.1  8.4   69   838   12   MOLECULE: N-TERMINAL ACETYLTRANSFERASE A COMPLEX SUBUNIT NA
  35:  8r33-A  4.1  3.4   53   489    2   MOLECULE: PHO90 ISOFORM 1;
  36:  3exe-A  4.1  4.3  100   363    9   MOLECULE: PYRUVATE DEHYDROGENASE E1 COMPONENT SUBUNIT ALPHA
  37:  3u24-A  4.1  7.4   84   546   10   MOLECULE: PUTATIVE LIPOPROTEIN;
  38:  9b85-N  4.0  6.0   82   286   11   MOLECULE: ALPHA-CENTRACTIN;
  39:  4hhx-A  4.0  2.4   52   104   10   MOLECULE: TOXIN COREGULATED PILUS BIOSYNTHESIS PROTEIN E;
  40:  7dqk-B  4.0  4.7   87   479    3   MOLECULE: PROTEIN DETOXIFICATION;
  41:  6dmp-B  3.9  3.7   74    82   11   MOLECULE: DESIGNED ORTHOGONAL PROTEIN DHD13_XAAA_A;
  42:  7yqh-A  3.9  9.2   91  1069    7   MOLECULE: STRUCTURAL MAINTENANCE OF CHROMOSOMES PROTEIN 5;
  43:  4xng-D  3.9  3.6   54   144    6   MOLECULE: UNCHARACTERIZED PROTEIN MG218.1;
  44:  4g9p-A  3.9 10.5   84   405   10   MOLECULE: 4-HYDROXY-3-METHYLBUT-2-EN-1-YL DIPHOSPHATE SYNTH
  45:  8f5d-A  3.9 13.9   80   772    9   MOLECULE: MULTIFUNCTIONAL FUSION PROTEIN;
  46:  9iz4-A  3.9  4.1   89   165    7   MOLECULE: PUTATIVE PHOSPHONOPYRUVATE DECARBOXYLASE ALPHA SU
  47:  7bc4-B  3.9  5.2   87  2054    5   MOLECULE: FATTY ACID SYNTHASE SUBUNIT ALPHA;
  48:  5cos-D  3.9  2.9   46    61   13   MOLECULE: SIDEROPHORE-INTERACTING PROTEIN;
  49:  6wkr-A  3.9  3.3   59   439    8   MOLECULE: UBIQUITIN;
  50:  7xad-C  3.9  5.8   72    96    7   MOLECULE: PROGRAMMED CELL DEATH 1 LIGAND 1;

0001 s001A   -YTKVEAEIDNRIKNLIPGDLADTWAELLSDCLG---GAL--KPKIEFHKVRLSVGMDVLGRQLLNHKA-DVNIRVHV--NSGGV-TLDQFRLGNADASLVFFFC-DLYTADTK-FR-DAL-YSYCKFSMF-GKQSALN---S-Q-EEKDAF-EAFRERARKLRT-T-CLLPT---V---E--KF--IR--DAPV----V-SGQGE-IPKSSA-----SS-AAKR-YEEV-L-AKH-LERL-Y-
0002 1yvpB   kpaNEGLTMVAKYVSK--------------------------------------------------------------------------------------------------------------GWKEV-QEAKEKE--lS-P-ETEKVL-KYLEATERVKRT-KehLLTI---H---L--KS--K--taDSV----L-APAS---------------lvalnkLRW-I---P-DTSI-Ve
0003 6vdqA   aaVSARAGEAVRGLNRPG-------------------------------------------------------------------------------------------------artDLA-ALLAATERT-RAALAPV-rgP-V-AELLVD-AALAALAGILEV---AVDRG---SdgpA--GP--P------------------------vfmlyagD-ANRE-QYNE-Q-IR---PTL-Mp
0004 3kkzA   tIHDFELNLICDFFSNXErpaeindfwxdaypeI---DTIpnQVAK-----------------------pVATF---------------------------------ilPENCW-TDhYFT-PKVAAQKIF-LTKYAG----------NKIA-EEFSXLQSIEEE-L--YHKY---K---E--YY-------------------------------------------------fiaKKIR-Ll
```

```
0005 6thkA   ekKEINKKVSDQVDGLQIT----------------------------------------------------------------------------------------------gkreaTRQ-ERVDVMSAV-LHKSDLE-ykQ-D-SLSEAV-KVLRQELNKQKA---LKEKE---D---L--SQ--LE--RDYRtrkaN-LQMK-------agstplEK-QKAI-YNGE-L-LVD-EIAS-Lq
0006 1u7zB   ------------DPVRYIS-DHSS--GKMGFAIA---AAA--A-avnasvqQQNIFIGCAAV--lkdhR-PYVVGFAA--ET-NN-VEEYARQKRINLDLICAND---VSQPT----------------------------------------------------gfNSDN---N-------------ALHL----F-WQDGD-KVLP----erkeL-LGQL-LLDE-I-VTR-YDEK-Nr
0007 5b86A   wrGVVAEVAAERLDAQP------------------------------------------------------------------------------------------------G-RS-EAE-SRFLHXGRT-XKELEVV-nvV-R-TYAESY-HYHFASHLCALA-----RDT---Y-----aXF--LS----------------------------N-EVTS-VkqlwtqDIL-Q-----
0008 8yhoC   pSELVQLPILQMGLKYAIE-----------------------------------------------------------------------------nmdsntekkfftlseviekakynvS-SFNKLLEGL-NY-LLEE-kgN-N-QARSYS-ATLETRIKNVQT-R-F-SNL---F------------dTELE-----------DKSIVY-----SV-------------kkmKLED---
0009 3iukA   rpkSAIDAVADAYTE-------------------------------------------------------lielnpsfattlHET-E-------------------yqDYSPAG-AA-AHA-EATRLALEA-LAGLEPS---D-D-VDAVTL-DAXRERLGLELE-I--HQSG--wD---A--AD--L--aSPAQivieQ-TGRY---------------agtslgAFL-R-D----------
0010 6s8fF   kvvdSKAKGLYYAIKNLP---------------------------------------------------------------------------------------------------qDVG-FAEKSLEKS-LLTFDSR-isQ-T-EWMDTL-NAIIEFIKIAA----IPQDVtvkC---S--KY----------gsiiqlanlmsvKNIAKL--------yDD-PSVV-S-Q----QNE-Kn
0011 3trcA   snAXNXLKILRQITQE------------------------------------------------------------------------------------------------vnaaP-NLE-QALKLVVVR-LCEALpF---A-E-EEEAFC-VTLAIHLAAEIA-H-A-RAK---Ga---------------------------------------------------------------
0012 7eldA   qaSSGAANLIHKLRA-------------------------------------------------------------------------------------------nytlaELG-QW-CAY-KVGQSFLSA-LQSDVNF---Q-V-DVKFQE-SYLSEVVSLLQC---ELLEG---A---S--L----------lllkyqhTADF-RAI-----------hvtlfdlakaR-KPG-S------
0013 8g7mG   lmLQGVDLLADAVAVVIIEVT---------------------------------------------------------------------------------------------------kDGVTVAKSI-----dLK---D-K-YKNIGA-KLIQDVANNTNE-E-AGD----------aelkgvI--TVK-----------------------------------------------
0014 4hm9A   fdANKLYCSE------------------------------------------------------------------------------------------------llqdndenrEL-LGelDGIDVLLQQ-LSVFNPS---T-A-EEQEMM-ENLFDSLCSCLM-LsSNRE--gpE---G--TD-nchkrNLRGqrtrL-LNKF---------------hicY-IMAE-I-CNA--NVP-Qi
0015 7sjrA   gRQPAAAALLTVLDVTAtgSDA------------------------------------------------------------------------------------------------------RYTLWQA-WHASGLQ-gaqaD-RDLDAV-TTLFDVADQYVN-R-------------------------------------------------------------------GTQ-Hl
0016 9jm7B   gvEAAKKEIKKLKEE-------------------------------------------------------------------------------------------vlkkykKG-EI-NEE-EAIKEFVEK-ALKLAVG---D-E-AVKKFA-IEEAKALVEEL----------------------------------------------------------------------------
0017 6vjpA   -------------------liGDSVMVDIGNVFTkkiPNA--Q-qykdyaaaGQKVVVEL----dsFGK-ADIYLVSI--RvEGR-INKLIYEAARSNVHLVDWY-KASAGHPEyFA-YDG-I------------------------------------------hlEYAG---S---K--AL--TD----------------------------------lIVKT-M-ETHa-------
0018 3citA   qsRAARLRLLVDTGQ--------------------------------------------------------------------------------------------------eliqlPPE-AXRKCVLQR-ACAFV-P---D-G-EDIESL-QLLATLLAAHLE---NNRLL---E---A--LV--AR--D-------------------------------------------------
0019 2oocB   ---GAVD--FAYLEGFAA-------------------------------------------------------------------------------------------------------gDFAVVDEVL-ALFRE-A---L-D-AALLDI-AAYAHEQALRSL-K-G-----------------------------------------------------------------------
0020 5ipxA   yhyPSWDQILQELDTLS------------------------------------------------------------------------------------------vatLNPD-CH-VPA-LNVEKTLYL-AKTQILVqqkL-V-HIISTS-LTLLKSTARSFF-A--WYDL---Y---R--PNls------------------------------------------------------
0021 7tjlA   akKEIEQAMYADALI---------------------------------------------------------------------------------------------------------------NPIRF-TEKAKYI-tyG-F-RGQEAY-DQVKKEMFEKLY---KYFME---K---L------------------------------------------------------------
0022 6iczA   vdDESMQRFHNRVRQI---------------------------------------------------------------------------------------------lmasG-ST-TFT-KIVNKWNTA-LIGLM-N---T-Q-ELLDLL-VKCENKIQTRIK---IGLNeyrvrtqalgiPT--LK--IS---------------lIQIF--------akimlrERI-R------SSE-Pt
0023 5v8zC   ----SLPVYDALAGE-------------------------------------------------------------------------------------------------firaS-GVE-ARQALLKQG-QDNLSVK---E-T--QKKWA-EQYLKIMGKILD-Q-GED---------------------------------------------------------------------
0024 1xkpA   spNISLSQLXAYLEGXS--------------------------------------------------------------------------------------------------------eEPSEQFXML-CGLDALX-peL-A-HLSHLV-EQALVSMAEEQG-E-T---R---D---A--V------xALSA------------dlQSQ-----QS-GSGR-EXLGf------------
0025 5j4aB   vSTSQFDYLVSRIGDQFHS---------------------------------------------------------------------------------dmwikdfisteSLN-SS-GLS-IFLATVMRA-RAASAEE---S----FPLYE-NVWNQLVEKLRQdA-RLG---------------------------------------------------------------------
0026 8ekiC   wiNYEVEMANRQFINI-----------------------------------------------------------------------------------------------tkN-PE-DFP-KSSQNFVKL-INKYDYL-ryK-L-MTCSLIfMNLTSSYLDYIL-T-V--YQ---T---M-----kslsvhRIQ----KllkdensinVLRR---liNK-LDsmkVKNE-L-LNV-IVNY-Ft
0027 4zmkA   -DTFSERTLG-------------------------------------------------------------------------------------------------------lnsidnTE-ISEVVSLGL-VSSDKIT-sqA-R-DFSHTL-SKSLKSRAKSLS-Q-K-----------------------------------------------------------------------
0028 8uspA   svSVRLIALQLL----------------------------------------------------------------------------------------------emittvvP-LN-YCL-SYQDDVLSA-LIPVSDK---K-R-IIRKQC-VDTRQVYYELGQ-I-------------------------------------------------------------------------
0029 6tgvC   vPTESalrfvgaDPVRFIGNRSS---GKQGYAVA---RVL--AqavskhapDANVLVMAAAV--qlpnM-RAIVGFAA---ETGDaVLFHARAKLEGCDLLVVNA-V----------------------------------------------------------------------------------GWL----L-SAGTE-SALEH---gsktL-MATR-IVDS-I-AAF-LKSQ---
0030 7c0qA   cfYAAQEILTFLIG---------------------------------------------------------------------------------------------------ackndTKR-ENIDLLYNF-VTSQiQE---H-KtPGCLLL-ANAYFYLAGFYL-SlDLKA--------------------------------------------------------------------
0031 8ye6B   rncQSVEDFKKSFERLTYY--------------------------------------------------------------------------------------------------enresTVRQRLKIA-EKEFKAG-ikN-T-TDKEIL-DYVRNELSKIDS-K-------------------------------------------------------------------------
0032 5eqzA   hyVEEKKEIDSLME--------------------------------------------------------------------------------------------lalvnDSS-GG-KFK-DYKDKINEL-KENLDIG---N-A-ELKEKL-LNLQNSFQDKLA-A-K-LAA---L---K--AA--K-------kdqdiskrtsgdkMSKN------------------------------
0033 8iqiC   pENPWTKLLLNALQDIIPE----------------------------------------------------------------------------------------------------lD-ARLWIMfylsLMRLYIM---D-P-NCQNAF-FSILVYFWEKLQ-K-EYGQI---K---K-------------Q----D-TLHR-------------------------------AYA-Ew
0034 6cjdA   sSETLKDSIYQEMNAFKDP--------------------------------------------------------------------------------------------------gdsafVSFEQQTAM-LQNMKVE---PgT-HLYEAL-NGVLVGSMNAQS----QMTS---W---M--QE--I---ILSG----G-EN------------------------------K-EAID-W-
0035 6hd5t   yweaflgYRANWTSL---------------------------------------------------------------------------------------------avaqdV-NG-ERQ-QAINTLSQF-EKLAGKI---S-D-SEKYEH-SECLMYKNDIM--Y-KAA-----S---D--NQ----------------------------tiymK-LGQlklleV-S-LG---IQG-Dn
0036 8r33A   ssGLLSTIAKALQKKI--------------------------------------------------------------------------------------------------------enDGVFAILCI-FGILVVG-vsH-T-VSAIII-IPLVQEVGDKLG-N-PK----------------------------------------------------------------------
0037 3exeA   melkadqlyKQKI-IRGFCHLCD-gqEACCVGLE---AGI--NPT---------DHLITAyrahKNFYggEVCLTLYGdgAANQG-QIFEAYNMAKLPCIFICE---NNRYGMG-TS----------------------------------------------------------------------------------fIP-GLR-----mdiLC-VREA-TRFA-A-AYC-RSGKs--
0038 3u24A   --aaSFAQFSNAFIDD----------------------------------------------------------------------------------lwqlsptwleiPDEAG-RV-KTL-AFVKAQQAK-LKQFELL---T-S-NETIDY-HLIDNLLSSXAW---DITR---fK---S--WQ--WdpnISQPiswlnaQVSQLeGARSFR-----IGe----------------------
0039 9b85N   ------------sDPQPE-EADGGLK-SWRESCD---SAL--RaVKDH-ySNGFCTVYAK-----------------T--IDGqqtcieshqfqpwrSEWKFTIT-PP--------------------------------------------------------------------------------TAQV----V-GVDVQ-DSL-----sneaQ-TAKE-FIKI-I-ENA-ENEY-Qt
0040 4hhxA   iSKKYRLELYSXLVDLL-----------------------------------------------------------------------------------------------------ndniPLYDALNKI-QNEGVGI-------yDKNFI-KSIELIKDRXKS-N-SS----------------------------------------------------------------------
0041 7dqkB   aSWIEFQLLYRLAAPSVYM-------------------------aaaslgnqgiqlFAYGLMlgiplavvvyglipqifayaiqkfqsivapsafgtwyfQILVllSISVCLAV-NG-LMF-MVAVGFNAA-ASVSNEL-ahP-K-SAAFSV-FMVTFISFLIAV-VvTLIL-----------------------------nayvcllGFKFdfenl-------------------------

0001 s001A   -LLHHHHHHHHHHHLLLLLHHHHHHHHHHHHHHH---HHL--LLEEEELLEEEEEEEEELLEELLLLLL-LEEEEEEL--LLLLL-LHHHHHHHLLLLEEEEELH-HHHHHLHH-HH-HHH-HHHHHHHHH-HHHLLLL---L-H-HHHHHH-HHHHHHHHHHHH-H-LHHHH---H---H--HH--HH--HLLE----E-ELLEE-ELLLLL-----LL-HHHH-HHHH-H-HHH-HHHH-L-
0002 1yvpB   lllLHHHHHHHHHHHL--------------------------------------------------------------------------------------------------------------LHHHH-HHHLLLL--lL-L-LHHHHH-HHHHHHHHLLLL-LhhLLHH---H---H--HL--H--hhLLL----L-LLLL---------------hhhhllLLL-L---L-LHHH-Hh
0003 6vdqA   hhHHHHHHHHHHHLLLL--------------------------------------------------------------------------------------------------lllLHH-HHHHHHHHH-HHHLLLL-llL-H-HHHHHH-HHHHHHHHHHHH---HHHHL---LlllL--LL--L-----------------------hhhhhhhH-HHHH-HHHH-L-LL---LLL-Ll
0004 3kkzA   lHHHHHHHHHHHHHHLLLllhhhhhhhhhlllL---EEHhhHHHH------------------------eEEEE--------------------------------elLHHHL-LLlLHH-HHHHHHHHH-HHHLLL----------LHHH-HHHHHHHHHHHH-H--HHHH---L---L--LE----------------------------------------------eeeEELH-Hh
0005 6thkA   hhHHHHHHHHHHHHHHHHH----------------------------------------------------------------------------------------------lllllLHH-HHHHHHHHH-HHHHHHH-hhH-H-HHHHHH-HHHHHHHHHHHH---HHHHH---H---H--HH--HH--HHHHhhhhL-HHHH-------hhhlhhHH-HHHH-HHHH-H-HHH-HHHH-Hh
0006 1u7zB   ------------LLLEEEE-ELLL--LHHHHHHH---HHH--H-hhhhhhhHLLEEEELLLL--llllL-LEEEEEEE--EL-LL-HHHHHHHHHHLLLEEEEEE---LLLLL----------------------------------------------------llLLLE---E--------------EEEE----E-ELLEE-EEEE----eehhH-HHHH-HHHH-H-HHH-HHHH-Hl
0007 5b86A   hhHHHHHHHHHHLLLLL------------------------------------------------------------------------------------------------L-LL-HHH-HHHHHHHHH-HHHHHHH-lhH-H-HHHHHH-HHHHHHHHHHHL-----HHH---H-----hHH--HH----------------------------H-HHHH-HhhhhhlHHH-H-------
0008 8yhoC   lLLLLHHHHHHHHHHHHHH-----------------------------------------------------------------------------lllhhhhhhllllhhhhlllllllL-LHHHHHHHL-HH-HHHH-llL-L-HHHHHL-HHHHHLLLLHHH-H-L-LLL---E-------------lLLLL-----------LLLEEE-----EL-------------hhlLLLL---
0009 3iukA   lllLHHHHHHHHHHH-------------------------------------------------------hhhhlhhhhhhlLLL-L-------------------llLLLHHH-HH-HHH-HHHHHHHHH-HHLLLLL---L-H-HHHHHH-HHHHHHHHHHHH-H--HHHL--lH---H--HL--L--lLHHHhhhhH-HHHH---------------hhhhhhHHH-H-H--------
0010 6s8fF   llllLLHHHHHHHHHLLL---------------------------------------------------------------------------------------------------lLLL-LHHHHHHHH-HHHHHLL-llH-H-HHHHHH-HHHHHHHHHHHL----LLLLLlhhH---H--HH----------hhhhhhhhhhhhHHLLLL--------lLL-HHHH-H-H----HHH-Hl
0011 3trcA   llHHHHHHHHHHHHHH------------------------------------------------------------------------------------------------hhhlL-LHH-HHHHHHHHH-HHHHLlL---L-H-HHHHHH-HHHHHHHHHHHH-H-H-HHL---Ll---------------------------------------------------------------
0012 7eldA   hhHLLHHHHHHHHHH-------------------------------------------------------------------------------------------hhhhhHHH-HH-HHH-HHHHHHHHH-HHHHHLL---L-H-HHHHHH-HHHHHHHHHHHHH---HHHHH---L---L--L----------hhhhhhhHHLL-LLL-----------hhhhlllllL-LLL-L-------
0013 8g7mG   hhHHHHHHHHHHHHLEEELEE---------------------------------------------------------------------------------------------------lLHHHHHHLL-----lLL---L-L-LHHHHH-HHHHHHHHHHHH-H-LLL----------hhhhllE--EEE-----------------------------------------------
0014 4hm9A   llHHHHHHHH------------------------------------------------------------------------------------------------hhlllhhhhHH-HHhlLHHHHHHHHH-HHHHLLL---L-L-HHHHHH-HHHHHHHHHHLL-LlHHHH--lhH---H--HH-hhhhhHLLHhhhhH-HHHH---------------hhhH-HHHH-H-LLL--LLH-Hh
0015 7sjrA   lLLHHHHHHHHHHHHHHHhlHLL-----------------------------------------------------------------------------------------------------LHHHHHH-HHHLLHH-hhhhH-HHHHHH-HHHHHHHHHHHH-L-------------------------------------------------------------------LHH-Hh
0016 9jm7B   lhHHHHHHHHHHHHH-------------------------------------------------------------------------------------------hhhhhhHL-LL-LHH-HHHHHHHHH-HHHHHLL---L-H-HHHHHH-HHHHHHHHHHL----------------------------------------------------------------------------
0017 6vjpA   -------------------eeELHHHHHHHHHHHhhlLLL--E-hlhhhlllLLEEEEEL----hhLLL-LEEEEELL--LlHHH-HHHHHHHHHLLLEEEELHH-HHHLLLHHhLL-LLL-L------------------------------------------llLHHH---H---H--HH--HH----------------------------------hHHHH-H-HHLl--------
0018 3citA   hhHHHHHHHHHHHHH--------------------------------------------------------------------------------------------------hhhhlLHH-HHHHHHHHH-HHHHL-L---L-H-HHHHHH-HHHHHHHHHHHH---HHHHH---H---H--HH--HL--L-------------------------------------------------
0019 2oocB   ---LLLL--HHHHHLLLL-------------------------------------------------------------------------------------------------------lLLHHHHHHH-HHHHH-H---H-H-HHHHHH-HHHHHHHHHHHH-H-L-----------------------------------------------------------------------
0020 5ipxA   eeeLLHHHHHHHHLLLL------------------------------------------------------------------------------------------lllLLHH-HH-HHH-HHHHHHHHH-HHHHHHHhlhH-H-HHHHHH-HHHHHHHHHHHH-H--HHHH---H---L--LLll------------------------------------------------------
0021 7tjlA   hhHHHHHHHHHHHHH---------------------------------------------------------------------------------------------------------------LHHHH-HHLHHHH-hhH-H-HHHHHH-HHHHHHHHHHHH---HHHHH---H---L------------------------------------------------------------
0022 6iczA   elHHHHHHHHHHHHHH---------------------------------------------------------------------------------------------hhllL-LL-LHH-HHHHHHHHH-HHHHH-H---L-H-HHHHHH-HHHHHHHHHHHH---HHLLhhhhhhhlllHH--HH--HH---------------hHHHL--------hhhhhhHHH-H------LLL-Ll
0023 5v8zC   ----LLHHHHHHHHH-------------------------------------------------------------------------------------------------hhhlL-LHH-HHHHHHHHH-HHHHHLL---H-H--HHHHH-HHHHHHHHHHHH-H-LLL---------------------------------------------------------------------
0024 1xkpA   llLLLHHHHHHHHHHHL--------------------------------------------------------------------------------------------------------lLHHHHHHHH-HHHHHHH-hhH-H-HHHHHH-HHHHHHHHHHHH-H-H---H---H---H--H------hHHHH------------hhHHL-----LL-LHHH-HHHHl------------
0025 5j4aB   eLHHHHHHHHHHHLLLLH----------------------------------------------------------------------------------hhhhhheeellLLL-HH-HHH-HHHHHHHHH-HHHHLLL---L----HHHHH-HHHHHHHHHHHLlH-HHL---------------------------------------------------------------------
0026 8ekiC   hhHHHHHHHHHHHHHH-----------------------------------------------------------------------------------------------llL-HH-HHH-HHHHHHHHH-HHHHHHH-hhH-H-HHHHHLhHHHHHHHHHHHH-L-L--HH---H---H-----hhhhhhHHH----HhhhlllhhhHHHH---hhHH-HHhlhHHHH-H-HHH-HHHH-Hh
0027 4zmkA   -LLHHHHHLL-------------------------------------------------------------------------------------------------------llllllLL-LLHHHHHHH-HHHHHHH-hhH-H-HHHHHH-HHHHHHHHHHHH-H-L-----------------------------------------------------------------------
0028 8uspA   lhHHHHHHHHHH----------------------------------------------------------------------------------------------hhhhhhlL-HH-HHH-LLLHHHHHH-LLLLLLL---L-L-HHHHHH-HHHHHHHHHHHH-L-------------------------------------------------------------------------
0029 6tgvC   eELLHhhhllllhLLLEEEEELLL---LHHHHHHH---HHH--HlhhhhhhhHLLEEEELLLL--llllL-LEEEEEEE---ELLLlHHHHHHHHHHLLLEEEEEE-L----------------------------------------------------------------------------------LEE----E-ELLLE-EELLL---llhhH-HHHH-HHHH-H-HHH-HHHL---
0030 7c0qA   lhHHHHHHHHHHHH---------------------------------------------------------------------------------------------------hhhhlLLH-HHHHHHHHH-HHHLhHH---H-HhHHHHHH-HHHHHHHHHHHH-HlLLHH--------------------------------------------------------------------
0031 8ye6B   lllLLHHHHHHHHHLLLLL--------------------------------------------------------------------------------------------------lllllLHHHHHHHH-HHHHHHH-llL-L-LLLHHH-HHHHHHHHHHHH-L-------------------------------------------------------------------------
0032 5eqzA   llHHHHHHHHHHHH--------------------------------------------------------------------------------------------hllllLLL-HH-HHH-HHHHHHHHH-HHHLLLL---L-H-HHHHHH-HHHHHHHHHHHH-H-H-HHH---H---H--HH--H-------lllhhhhhlllhhHHHH------------------------------
0033 8iqiC   lLLHHHHHHHHHHHHHLLL----------------------------------------------------------------------------------------------------hH-HHHHHHhhhhHHHHLLL---L-H-HHHHHH-HHHHHHHHHHHH-H-HHLLH---H---H-------------H----L-HHHH-------------------------------HHH-Hh
0034 6cjdA   lLHHHHHHHHHHHHHLLLL--------------------------------------------------------------------------------------------------lllllLLLHHHHHH-HHHHHLL---LlL-HHHHHH-HHHHHHHHHHHL----LHHH---H---H--HH--H---LLLL----L-LL------------------------------L-LLLL-L-
0035 6hd5t   hhhhlllLHHHHHHH---------------------------------------------------------------------------------------------hhhhhH-HL-LHH-HHHHHHHHH-HHHLLLL---L-L-HHHHHH-HHHHHHHHHHH--H-HLL-----L---L--LH----------------------------hhhhH-HLLhhhhhH-H-HL---LLL-Lh
0036 8r33A   hhLHHHHHHHHHHHHH--------------------------------------------------------------------------------------------------------llLLHHHHHHH-HHHHHHH-llH-H-HHHHHH-HHHHHHHHHHLL-L-LL----------------------------------------------------------------------
0037 3exeA   hhhhhhhhHLLL-LLLLLLLL-llHHHHHHHH---HHL--LLL---------LEEELLlllhLLLLllLLEEEEEEllHHHLH-HHHHHHHHHLLLEEEEEE---ELLEELL-EE----------------------------------------------------------------------------------lLL-EEE-----llhHH-HHHH-HHHH-H-HHH-HLLLl--
0038 3u24A   --llLHHHHHHHHHHH----------------------------------------------------------------------------------hhhhlhhhlllLLHHH-HH-HHH-HHHHHHHHH-HHLLLHL---L-H-HHHHHH-HHHHHHHHHHHH---HHHL---lL---H--HH--HlhhLLLLhhhhhhHHHHHhLLLLLL-----LLh----------------------
0039 9b85N   ------------eEEEEL-LLLHHHH-HHHHHHH---HHH--HhHHHH-lLLLEEEEEEEE-----------------E--ELLeeeeeeeeeeheeEEEEEEEE-LL--------------------------------------------------------------------------------EEEE----E-EEEEE-EEE-----llhhH-HHHH-HHHH-H-HHH-HHHH-Hh
0040 4hhxA   lLHHHHHHHHHHHHHHHH----------------------------------------------------------------------------------------------------llllLHHHHHHHH-HHHHLLL-------lLHHHH-HHHHHHHHHHHH-L-LL----------------------------------------------------------------------
0041 7dqkB   hHHHHHHHHHHHHHHHHHH-------------------------hhhhhhhhhlhhHHHHHHhhhhhhhhhhlllhhhhhhhhhhlllhhhhhhhhhHHHHhhHHHHHHHH-HH-HHH-HHHHHHHHH-HHHHHHH-llH-H-HHHHHH-HHHHHHHHHHHH-HhHHHH-----------------------------lhhhhhhHLLLllhhl-------------------------
```

```
# Job: Type-1 BREX; BrxC-ATPase MBR4615907 wHTH2
# Query: s001A
# No:  Chain   Z    rmsd lali nres  %id PDB  Description
   1:  1xsd-A  8.0  2.6   72   125   10   MOLECULE: 5'-
   2:  4mtd-D  7.9  3.3   74   151   12   MOLECULE: ZINC UPTAKE REGULATION PROTEIN;
   3:  2mh2-A  7.8  1.8   63    64   14   MOLECULE: HOMOLOGOUS-PAIRING PROTEIN 2 HOMOLOG;
   4:  3s2w-A  7.7  1.9   66   146   14   MOLECULE: TRANSCRIPTIONAL REGULATOR, MARR FAMILY;
   5:  5vyv-A  7.7  1.9   69   173   17   MOLECULE: UPF0502 PROTEIN YCEH;
   6:  1ucr-B  7.5  1.6   63    75   25   MOLECULE: PROTEIN DSVD;
   7:  6j0e-B  7.5  2.3   70   125    9   MOLECULE: ARSENIC RESPONSIVE REPRESSOR ARSR;
   8:  2k4b-A  7.5  2.0   66    69   18   MOLECULE: TRANSCRIPTIONAL REGULATOR;
   9:  2eth-A  7.5  2.4   67   141    6   MOLECULE: TRANSCRIPTIONAL REGULATOR, PUTATIVE, MAR FAMILY;
  10:  2g9w-A  7.4  2.1   68   119   16   MOLECULE: CONSERVED HYPOTHETICAL PROTEIN;
  11:  5hs5-A  7.4  2.0   69   116   12   MOLECULE: HTH-TYPE TRANSCRIPTIONAL REGULATOR SARX;
  12:  5l0p-A  7.3  2.7   74   311    5   MOLECULE: TRANSCRIPTION FACTOR ETV6, TRANSCRIPTION FACTOR E
  13:  6j05-A  7.3  1.9   67   101    7   MOLECULE: TRANSCRIPTIONAL REGULATOR ARSR;
  14:  4yif-B  7.3  2.5   66   141    9   MOLECULE: MARR FAMILY PROTEIN RV0880;
  15:  3f72-B  7.2  2.8   70   108   10   MOLECULE: CADMIUM EFFLUX SYSTEM ACCESSORY PROTEIN;
  16:  1p4x-A  7.2  2.2   67   250   10   MOLECULE: STAPHYLOCOCCAL ACCESSORY REGULATOR A HOMOLOGUE;
  17:  8xt8-B  7.2  1.9   66   145   12   MOLECULE: MARR FAMILY TRANSCRIPTIONAL REGULATOR;
  18:  9mj5-B  7.2  2.3   65   208   12   MOLECULE: REPLICATION PROTEIN A 14 KDA SUBUNIT;
  19:  6qpq-D  7.2  2.4   65    82   11   MOLECULE: STRUCTURAL MAINTENANCE OF CHROMOSOMES PROTEIN,STR
  20:  3bpx-A  7.2  1.9   66   147   12   MOLECULE: TRANSCRIPTIONAL REGULATOR;
  21:  8gcc-A  7.1  2.6   79   707   11   MOLECULE: DNA TOPOISOMERASE 2;
  22:  2ns0-A  7.1  2.4   72    85   17   MOLECULE: HYPOTHETICAL PROTEIN;
  23:  6juv-A  7.1  1.9   62   192    6   MOLECULE: SEGREGATION AND CONDENSATION PROTEIN B;
  24:  6qfd-B  7.1  2.8   72   116   10   MOLECULE: DNA-BINDING PROTEIN;
  25:  2rdp-A  7.1  2.2   64   140    9   MOLECULE: PUTATIVE TRANSCRIPTIONAL REGULATOR MARR;
  26:  8ylg-A  7.0  2.2   67   161   18   MOLECULE: MARR FAMILY TRANSCRIPTIONAL REGULATOR;
  27:  8iue-P  7.0  1.9   63   303   11   MOLECULE: DNA-DIRECTED RNA POLYMERASE III SUBUNIT RPC1;
  28:  7wze-A  7.0  2.1   65   161   12   MOLECULE: UNCHARACTERIZED HTH-TYPE TRANSCRIPTIONAL REGULATO
  29:  4hqe-B  7.0  2.2   68   108   10   MOLECULE: TRANSCRIPTIONAL REGULATOR QSRR;
  30:  3r0a-B  7.0  2.7   67   123   10   MOLECULE: PUTATIVE TRANSCRIPTIONAL REGULATOR;
  31:  5eri-A  7.0  2.0   66   153   12   MOLECULE: MARR FAMILY TRANSCRIPTIONAL REGULATOR;
  32:  1sfu-A  7.0  1.9   62    70   10   MOLECULE: 5'-D(*T*CP*GP*CP*GP*CP*G)-3';
  33:  4asn-A  6.9  2.3   69    90   13   MOLECULE: TUBR;
  34:  4gyi-A  6.9  2.7   71   339   10   MOLECULE: RIO2 KINASE;
  35:  5jbr-A  6.9  2.3   66   149   11   MOLECULE: UNCHARACTERIZED PROTEIN BCAV_2135;
  36:  5j6x-A  6.9  1.8   60    66   15   MOLECULE: Z-DNA BINDING PROTEIN KINASE;
  37:  2d1h-A  6.9  1.7   65   102    9   MOLECULE: 109AA LONG HYPOTHETICAL TRANSCRIPTIONAL REGULATOR
  38:  2nyx-B  6.9  2.8   67   147   10   MOLECULE: PROBABLE TRANSCRIPTIONAL REGULATORY PROTEIN, RV14
  39:  7bzh-A  6.9  1.7   59    59   12   MOLECULE: SUL7S;
  40:  1jhf-A  6.9  2.5   66   197   12   MOLECULE: LEXA REPRESSOR;
  41:  6pco-B  6.9  2.5   65   135   20   MOLECULE: MARR-FAMILY TRANSCRIPTIONAL REGULATOR;
  42:  4ija-A  6.8  2.3   62   363   10   MOLECULE: XYLR PROTEIN;
  43:  2p4w-B  6.8  2.4   67   198   12   MOLECULE: TRANSCRIPTIONAL REGULATORY PROTEIN ARSR FAMILY;
  44:  5n35-A  6.8  1.7   58    60   10   MOLECULE: POLB1 BINDING PROTEIN 2 (PBP2);
  45:  5y6i-B  6.8  1.8   61   234   10   MOLECULE: TRANSCRIPTIONAL REGULATOR KDGR;
  46:  7dg2-A  6.8  2.3   68   231   10   MOLECULE: NON-STRUCTURAL MAINTENANCE OF CHROMOSOMES ELEMENT
  47:  7z1n-O  6.8  1.8   64   570   19   MOLECULE: DNA-DIRECTED RNA POLYMERASE III SUBUNIT RPC1;
  48:  1ku9-A  6.8  2.8   66   151    9   MOLECULE: HYPOTHETICAL PROTEIN MJ223;
  49:  4i99-C  6.8  2.6   61    70   11   MOLECULE: CHROMOSOME PARTITION PROTEIN SMC;
  50:  1lva-A  6.7  1.8   60   258    7   MOLECULE: SELENOCYSTEINE-SPECIFIC ELONGATION FACTOR;
  51:  8zwa-A  6.7  3.0   77   414    9   MOLECULE: DISEASE RESISTANCE PROTEIN ADR1;
```

```
 52:  1qgp-A  6.7  1.9   63    76    8   MOLECULE: PROTEIN (DOUBLE STRANDED RNA ADENOSINE
 53:  8a5y-T  6.7  1.9   64   650    5   MOLECULE: ANAPHASE-PROMOTING COMPLEX SUBUNIT CDC27;
 54:  4nb5-B  6.7  2.3   65   149    9   MOLECULE: DNA BINDING PROTEIN;
 55:  4ad9-A  6.7  2.7   71   288   13   MOLECULE: BETA-LACTAMASE-LIKE PROTEIN 2;
 56:  8ffz-B  6.7  3.0   67   905    3   MOLECULE: TRANSCRIPTION FACTOR IIIA;
 57:  3cuq-A  6.7  2.4   67   219   12   MOLECULE: VACUOLAR-SORTING PROTEIN SNF8;
 58:  1yyv-A  6.7  2.6   68   114    9   MOLECULE: PUTATIVE TRANSCRIPTIONAL REGULATOR;
 59:  5nl9-A  6.7  3.1   74   146    7   MOLECULE: TRANSCRIPTIONAL REGULATOR (FUR FAMILY);
 60:  2xco-A  6.6  2.5   79   636   10   MOLECULE: DNA GYRASE SUBUNIT B, DNA GYRASE SUBUNIT A;

0001 s001A   --TGYHPPL-PG-S---PLAEAEQEVEGYLNG---N---P------S--VCPVVNVVDYFKN-R-----PYG----WDEYSTLAALCRLVAG--ERREFHYNGAPNPSRDVVAANLAKNQAYFSIAT-KA-L----
0002 1xsdA   -------tN-KQ-V---EISMAEWDVMNIIWD---K----------K--SVSANEIVVEIQK-Y------KE----VSDKTIRTLITRLYKK--EIIKRYKSEN-----------------IYFYSS-NI-Keddi
0003 4mtdD   laqaeKICAqRN-V---RLTPQRLEVLRLMSL---Q---D------G--AISAYDLLDLLR--E-----AEP---qAKPPTVYRALDFLLEQ--GFVHKVEST-------------------NSYVL-CH-Lfdqp
0004 2mh2A   --------------------GAPGIILRYLQE---Q---N------R--PYSAQDVFGNLQK-E------HG----LGKAAVVKALDQLAQE--GKIKEKTYG-----------------kQKIYFA-D-------
0005 3s2wA   ygqiyigkkieP-Y---GIGSGQFPFLXRLYR---E----------D--GINQESLSDYLK---------------IDKGTTARAIQKLVDE--GYVFRQRDE-----------------RSYRVFL-TE-Kgkkl
0006 5vyvA   ----------MK-Y---QLTALEARVIGCLLE---K---QvttpeqY--PLSVNGVVTACNQ-KtnrepVMN----LSESEVQEQLDNLVKR--HYLRTVSG---------------fgnrVTKYEQ-R-------
0007 1ucrB   ------------------MEEAKQKVVDFLNSksgS---K------S--KFYFNDFTDLFP----------D----MKQREVKKILTALVND--EVLEYWSSG-----------------sTTMYGL-KGa-----
0008 6j0eB   sdesehyAD-LF-K---VLDPVRLRILSQLAA---G---G-----cG--PVSVNELTDLMG---------------LSQPTISHHLKKMTEA--GFLDRVPEGR-----------------VVLHRV-RP-Elfae
0009 2k4bA   -----------fnV---SNAE--LIVMRVIWS---L---G---------EARVDEIYAQI-P-Q-----ELE----WSLATVKTLLGRLVKK--EMLSTEKEG-----------------rKFVYRP-LM-E----
0010 2ethA   fssylpsnE-EI-S---DXKTTELYAFLYVAL---F---G---------PKKXKEIAEFLS---------------TTKSNVTNVVDSLEKR--GLVVREXDP--------------vdrrTYRVVL-TE-Kgkei
0011 2g9wA   ----------KL-T---RLGDLERAVXDHLWS---R---T------E--PQTVRQVHEALSA-------RRD----LAYTTVXAVLQRLAKK--NLVLQIR--------------------AHRYAP-VHg-----
0012 5hs5A   qykalseyidkK-Y---KLSLNDLAVLDLTXK---H---C----kdE--KVLXQSFLKTAXD-E------LD----LSRTKLLVSIRRLIEK--ERLSKVRS---------------kderKIYIYL-NN-Ddisk
0013 5l0pA   qdnntalKK-AG-L---KVTLPRLKILEVLQE--pD---N------H--HVSAEDLYKRLID-M-----GEE----IGLATVYRVLNQFDDA--GIVTRHNF---------------EGGK-SVFEL-T-------
0014 6j05A   -----qdpA-QI-V----ASPVRLEIFRLLVE---Q---E-----pT--GLVSGDIAEHLG---------------QPHNGISFHLKNLQHA--GLVTVQRE-----------------gRYQRYRA-AM-Pvvra
0015 4yifB   rlsrqlrfrnPS-S---PVSLSQLSALTTLAN---E----------G--AMTPGALAIRER---------------VRPPSMTRVIASLADM--GFVDRAPHP--------------idgrQVLVSV-SE-Sgael
0016 3f72B   lqtvdisGV-SQ-I---LADENRAKITYALCQ---D----------E--ELCVCDIANILG---------------VTIANASHHLRTLYKQ--GVVNFRKEG-----------------KLALYSL-GG-E----
0017 1p4xA   ytmyfkniikkH-L---TLSFVEFTILAIITS---Q--nK------N--IVLLKDLIETIH---------------HKYPQTVRALNNLKKQ--GYLIKERST--------------edeRKILIHM-DD-Aqqdh
0018 8xt8B   kddlkkmfkqnG-Y---DITTDHYALLRFLWE---Q----------D--GISQIDLCEKSC---------------KDKSNTTRILDVMKNK--GLIVRKVDV--------------kdrRKFQIFL-TD-Lgrel
0019 9mj5B   ansqpsasfmpA-N---GLTVAQNQVLNLIKA---C---P------RpeGLNFQDLKNQLK----------H----MSVSSIKQAVDFLSNE--GHIYSTVD-------------------DDHFKS-TDa-----
0020 6qpqD   --------------------KAIVQMAKILRK---ElseE------K--EVIFTDVLKSQAN-T-----EpenitkrEASRGFFDILSLATE--GCIGLSQT---------------AFGN-IKIDA-KP-Alfer
0021 3bpxA   shrvfigrelgH-L---NLTDAQVACLLRIHR---E----------P--GIKQDELATFFH---------------VDKGTIARTLRRLEES--GFIEREQDP--------------enrRRYILEV-TR-Rgeei
0022 8gccA   vadcersipsvI-D---GLKPGQRKIIFSSFK---R-rlT------R--SIKVVQLAGYVSE-----haAYH----HGEQSLVQTIVGLAQnnvPLLQQDG----qFGTRlGKDHA--AGRYIFTRL-TN-Iaryi
0023 2ns0A   -------------------RELEECIRALLDA---R---A----dsA--SICPSDVARAVAP-D--------dwrpLXEPVREAAGRLADA--GEVEVTQKGAVVD----RSAR-----gPIRIRW-TR-T----
0024 6juvA   eysqkvihlxpK-P---ELRAGELKTLALIAY---L----------Q--PVEQSKIIKLR------------------GSQAYEHIKKLLEX--GLIYAEPYE-----------------rTKLLGT-TQ-Kfael
0025 6qfdB   -----pdAR-SDaR---DLTAFQKNILTVLGE---E-------------ARYGLAIKRELEEyY-----GEE----VNHGRLYPNLDDLVNK--GLVEKSELD----------------kRTNEYAL-TNe-----
0026 2rdpA   nlkqrgreiltN-Y---PITPPQFVALQWLLE---E---G---------DLTVGELSNKXY---------------LACSTTTDLVDRXERN--GLVARVRD-----------------ehVVRIRL-LE-Kgeri
0027 8ylgA   iardrlnplfaR-Y---GLQPGEFDVLATLRR---S-gaP------Y--ALTPTALYDAAM---------------ISSGSMTNRIDRLEKA--GWVERRANP--------------adgRGTLVAL-TS-Agral
0028 8iueP   -------------------VEIENRIIELCHQ---F---P------H--GITDQVIQNEMP----------H----IEAQQRAVAINRLLSM--GQLDLLRSN-----------------TGLLYRI-KDs-----
0029 7wzeA   kxnhvxehyfaG-R---GLSEGKFKILXLLFD---A---K-----dH--RLSPTELAKRSN---------------VTKATITGLLDGLARD--GFVSRRHH------------------rKISIEL-TT-Egkar
0030 4hqeB   --evcpylE-ET-F---KIRSWNGLIINYLSR---C---N-----dC--SAHFSDMKRDLK----------T----ITPRALSLKLSELAQW--ELVEKQII---------------tspvQIIYVL-TE-Kgkal
0031 3r0aB   kefkvedvikcA-L---NLTKADLNVXKSFLN---E--pD------R--WIDTDALSKSLK---------------LDVSTVQRSVKKLHEK--EILQRSQQN--------------gggyVYIYKI-YS-K----
0032 5eriA   avnsindikykE-L---KLQKGQFTFLTRICE---N----------P--GINLVELSNMLK---------------VDKATTTKAIQKLIKA--GYVDKKQDK--------------fdkRGYNLTP-TD-Kslev
0033 1sfuA   -------------------AEIFSLVKKEVLS---L---N----tnD--YTTAISLSNRLK---------------INKKKINQQLYKLQKE--DTVKMVPS------------------NPPKWFK-NY-Nc---
0034 4asnA   ---sdyfEE-VX-R---KLTIEDVSILGWLFQ---N---E------AfkAIKKSSIADELE---------------YSTANFRKTLNKLEAI--HFIGTVTGG-----------------kEHKLYL-TE-Ygqqa
0035 4gyiA   ----kldTR-AM-R---HLTAEDWRVLTAVEM---G---S---knhE--IVPTPLIEKIARL-R------GG----SS--GVHKSIATLAKA--GLIARMKEA-----------------kYDGYRL-TY-Ggldy
0036 5jbrA   tfsqaxgdxlaT-W---QLPRTTGRTYGYLLL---Q---S------E--ATSFQEIGADLG---------------LSPGAVSTSVRELVAW--GLARTIPQ---------------pgsRRLLVEA-AG-Gfeql
0037 5j6xA   ------------------ENEIEMRICDYLRR---H---G---------RSTVQDIFKELK---------------LEKSTVNRHLYSLQAS--KQVFKTVE----------------dnkRPVWDL-VE------
0038 2d1hA   kleskkdeircC-Y---KITDTDVAVLLKXVE---I---E------K--PITSEELADIFK---------------LSKTTVENSLKKLIEL--GLVVRTKT------------------pKYYYSI-SS-Nilek
0039 2nyxB   aisahsiaQ-VD-E---NITIPQFRTLVILSN---H---G---------PINLATLATLLG---------------VQPSATGRXVDRLVGA--ELIDRLPHP--------------tsrrELLAAL-TK-Rgrdv
0040 7bzhA   ------------------MEDVKQSVEKIIKD---R----------E--WVTFNDLLKYIP---------------YPAPEVYDALSQLIKE--NKVGRRG---------------------RYFYY-IK-R----
0041 1jhfA   -------------K---ALTARQQEVFDLIRD---H---I--sqtgM--PPTRAEIAQRLG--------------fRSPNAAEEHLKALARK--GVIEIVSGA------------------SRGIRL-LQ-E----
0042 6pcoB   rhvaifqqtipD-S---KLTAAQFVVLCALRD---Q---G---------ACSLVDVVKATA---------------IDQATVRGVIERLKAR--KLLAVSHRR------------------KVLVTL-TP-Dgral
0043 4ijaA   -----------------PMNDNEKRVLREIYN---H---H---------NISRTQISKNLE---------------INKATISSILNKLKYK--SLVNEVG----------------ggrKPILLKV-NH-Lygyf
0044 2p4wB   ---mgeeLN-RL-L---DVNETRRRILFLLTK---R-------------PYFVSELSRELG---------------VGQKAVLEHLRILEEA--GLIESRVEK--------------rgrpRKYYMI-KK-Glrle
0045 5n35A   -------------------QKEIEIAIEYFKN---Y----------I--SVGEIVATMDLKA--------RG----IS--NPQAVISKLIEM--GIIEKGE---------------------GCYNL-VR------
0046 5y6iB   -------------------AEVGTDILKALAE---L---S------P--ATSLSRLAEHVG---------------MPASKVHRYLQALIAS--GFAVQDAS-------------------TNHYSL-GR-Ealrv
0047 7dg2A   vnrvenditkmA-S---DYAENELELFRKTME---L---I-ilsdnG--FATSISILNLADElQ-----SKK----MKKKEVEQLLQSFVQE--KWLIGRN---------------------GEYTL-HT-Rcime
0048 7z1nO   lnpdlflykelvkA---HLGERAASVIGMLVA---L---G---------RLSVRELVEKI----------DG----MDVDSVKTTLVSLTQL--RCVKYLQE--------------tsgkkTTYYYY-NE-Egihi
0049 1ku9A   liielfselakI-H---GLNKSVGAVYAILYL---S---D------K--PLTISDIXEELK---------------ISKGNVSXSLKKLEEL--GFVRKVWI---------------kgerKNYYEA-VD-G----
0050 4i99C   ------------------IEKYVEELYKVVKK---I---Y--ektgT--PIKFWDLV--PDV-E-----------pKIIARTFLYLLFLENM--GRVEIIQEE---------------pfgEILVVP-M-------
0051 1lvaA   -------------------gSPEKILAQIIQE---H---R------E--GLDWQEAATRAS---------------LSLEETRKLLQSXAAA--GQVTLLRV------------------eNDLYAIS-TEr-----
0052 8zwaA   ------faH-ME-EsleNLDPKIRDCFLDMGA---F---P------EdkKIPLDLLTSVWVE-------rHD----IDEETAFSFVLRLADK--NLLTIVN----------NPRFgdIGYYDVFVTQ-HD-Vlrdl
0053 1qgpA   -------------------QDQEQRILKFLEE---L---G------EgkATTAHDLSGKLG---------------TPKKEINRVLYSLAKK--GKLQKEAG------------------TPPLWKI-AV-S----
0054 8a5yT   iqlafkdgrkLV-L---DVSLEQCSVINQFDS---pndeP------I--CLSLEQLSESLN---------------IAPPRLTHLLDFWIQK--GVLLKEN---------------------GTYSV-IE-H----
0055 4nb5B   mefveqmggyfesR---SLTRLAGRLLGWLLV---C--dP------E--RQSSEELATALA---------------ASSGGISTNARMLIQF--GFIERLAV---------------agdRRTYFRL-RP-Nafaa
0056 4ad9A   -yDYMNslkellkikadinNIREQQILTLFRE---N--fE------K--SFTVMELVKIIYK-N-----TpenlhemAKHNLLLHLKKLEKE--GKIFSNTDP------------------DKKWKA-HL------
0057 8ffzB   rkSAVKVplgkpfsrlwvWKQVVTMVVNEIIF---H----------P--GITLSRLQSRCRE---------V----LSLHEISEICKWLLER--QVLITTD--------------------FDGYWV-NH-Nwysi
0058 3cuqA   vggtyliqS-VP-A---ELNMDHTVVLQLAE----K---N------G--YVTVSEIKASLK---------------WETERARQVLEHLLKE--GLAWLDLQA----------------pGEAHYWL-PAl-----
0059 1yyvA   lfaeqcpSR-EV-L---KTSRWGVLILVALRD---G-------------THRFSDLRRXX----------GG----VSEXXLAQSLQALEQD--GFLNRVSYP--------------vvppHVEYSL-TP-Lgeqv
0060 5nl9A   erskkilED-AG-I---NVTVQRLQMANLLLS---K---P------Q--HLTADQVFQLINE-H-----MPN----ASRATIFNNLKLFAEK--GIVNLLEL----------------ksgITLYDSnVI-H----
0061 2xcoA   svivaralpdvR-D---GLKPVHRRILYGLNE---Q---G------MksYKKSARIVGDVMG-K-----YHP----HGDSSIYEAMVRMAQ-ryPLVDGQG-----NFGS-gDGAA--AMRYTEARM-TK-Itlel

0001 s001A   --HHLLLLL-LL-L---LLLHHHHHHHHHHHHL---L---L------L--EEEHHHHHHHHHL-L-----LLL----LLHHHHHHHHHHHHHL--LLEEEEEELLEELLLHHHHHHHLLLLHHHEEEEEE-LL-L----
0002 1xsdA   -------lL-LL-L---LLLLLLHHHHHHHHH---H----------L--LEEHHHHHHHHHL-L------LL----LLHHHHHHHHHHHHHL--LLEEEEEELLL-----------------LEEEEE-LL-Lhhhh
0003 4mtdD   hhhhhHHHHhLL-L---LLLHHHHHHHHHHHL---L---L------L--LEEHHHHHHHHH--H-----HLL---lLLHHHHHHHHHHHHHL--LLEEEELLL------------------LEEEE-LL-Lllll
0004 2mh2A   --------------------LHHHHHHHHHHH---H---L------L--LEEHHHHHHHHHL-L------LL----LLHHHHHHHHHHHHHH--LLEEEEEEL-----------------lEEEEEE-L-------
0005 3s2wA   hhhhhhhhhhhH-H---LLLLLLHHHHHHHHH---L----------L--LEEHHHHHHHHL---------------LLHHHHHHHHHHHHHL--LLEEEEELL-----------------LLEEEEE-LH-Hhhhh
0006 5vyvA   ----------LL-L---LLLHHHHHHHHHHHH---H---HhhlhhhL--LEEHHHHHHHHLL-LlllllLLL----LLHHHHHHHHHHHHHL--LLEEEELL---------------llllLLEEEE-L-------
0007 1ucrB   ------------------LLHHHHHHHHHHLLlllL---L------L--LEEHHHHHHHLL----------L----LLHHHHHHHHHHHHHL--LLEEEEEEL-----------------lEEEEEE-LLh-----
0008 6j0eB   lllhhhhHH-HH-H---HHLHHHHHHHHHHHH---L---L-----lL--LEEHHHHHHHHL---------------LLHHHHHHHHHHHHHL--LLEEEEEELL------------------EEEEEE-LH-Hhhhh
0009 2k4bA   -----------llL---LLLL--LHHHHHHHH---H---L---------LEEHHHHHHLL-L-H-----HHL----LLHHHHHHHHHHHHHL--LLLEEEEEL-----------------lEEEEEL-LL-L----
0010 2ethA   hhllllllH-HH-H---HLLHHHHHHHHHHHH---H---L---------LLLHHHHHHHLL---------------LLHHHHHHHHHHHHHL--LLEEEEELL---------------llllLEEEEE-LH-Hhhhh
0011 2g9wA   ----------LH-H---HLLHHHHHHHHHHHL---L---L------L--LEEHHHHHHHHLL-------LLL----LLHHHHHHHHHHHHHL--LLEEEEL--------------------LLEEEE-LLl-----
0012 5hs5A   hhhhhhhhhhhH-H---LLLHHHHHHHHHHHH---H---H----llL--LEEHHHHHHHHHH-H------HL----LLHHHHHHHHHHHHHL--LLLEEELL---------------llllLLEEEL-LH-Hhhhh
0013 5l0pA   llhhhhhHH-LL-L---LLLHHHHHHHHHHLL--lL---L------L--EEEHHHHHHHHHH-L-----LLL----LLHHHHHHHHHHHHHL--LLEEEELL---------------LLLL-LEEEE-L-------
0014 6j05A   -----llhH-HH-H----LLHHHHHHHHHHHH---L---L-----lL--LEEHHHHHHHHL---------------LLHHHHHHHHHHHHHL--LLEEEEEE-----------------lLEEEEEE-LH-Hhhhh
0015 4yifB   hhhhhhhlllLL-L---LLLHHHHHHHHHHHH---H----------L--LLLHHHHHHHHL---------------LLHHHHHHHHHHHHHL--LLEEEEELL---------------llllLEEEEE-LH-Hhhhh
0016 3f72B   hhlllhhHH-HH-H---HHLHHHHHHHHHLLL---L----------L--LEEHHHHHHHHL---------------LLHHHHHHHHHHHHHL--LLEEEEELL-----------------LLEEEEE-LL-H----
0017 1p4xA   hhhhhhhhhhhH-L---LLLHHHHHHHHHHHL---L--lL------L--LEEHHHHHHHLL---------------LLHHHHHHHHHHHHHH--LLLEEEELL---------------lllLLEEEEL-LH-Hhhhh
0018 8xt8B   hhhhhhhhhhlL-L---LLLHHHHHHHHHHHH---L----------L--LLLHHHHHHHHL---------------LLLHHHHHHHHHHHLL--LLEEEEELL---------------lllLLEEEEE-LH-Hhhhl
0019 9mj5B   hhhllllllllL-L---LLLHHHHHHHHHHHH---L---L------LllLLLHHHHHHHLL----------L----LLHHHHHHHHHHHHHH--LLEELLLL------------------LLLLEE-LLl-----
0020 6qpqD   --------------------LHHHHHHHHHHH---HhllL------L--LEEHHHHHHHHHL-L-----LhhhllhhHHHHHHHHHHHHHHL--LLEEEELL---------------LLLL-LEEEE-LH-Hhhll
0021 3bpxA   hhhhhhhhhhhH-H---LLLHHHHHHHHHHHL---L----------L--LLLHHHHHHHHL---------------LLHHHHHHHHHHHHHL--LLEEEEEEL---------------leeEEEEEEE-LH-Hhhhh
0022 8gccA   hhhhhhhllllL-L---LLLHHHHHHHHHHHH---H-llL------L--LEEHHHHHHHHHH-----hlLLL----LLHHHHHHHHHHHHHLlllLLEEEEL----lLLLLlLLLLL--LLLLLEEEE-LL-Lhhhh
0023 2ns0A   -------------------HHHHHHHHHHHHH---L---L----llL--LLLHHHHHHHHHLL-L--------llhhHHHHHHHHHHHHHHL--LLEEEEEELLEELL----LLLL-----lLLEEEE-LL-L----
0024 6juvA   hhhlllhhhllL-L---LLLHHHHHHHHHHHH---H----------L--LEEHHHHHHLL-------------------HHHHHHHHHHHHHL--LLEEEEEEL-----------------lEEEEEE-LH-Hhhhh
0025 6qfdB   -----llLH-HHhL---LLLHHHHHHHHHHHH---L-------------LEEHHHHHHHHHHhH-----LLL----LLHHHHHHHHHHHHHL--LLEEEEEEE-----------------lLEEEEEE-LHh-----
0026 2rdpA   hhhhhhhhhhlL-L---LLLHHHHHHHHHHHH---H---L---------LLLHHHHHHHHL---------------LLHHHHHHHHHHHHHL--LLEEEEEL-----------------llLEEEEE-LH-Hhhhh
0027 8ylgA   hhhhlhhhhhhH-L---LLLHHHHHHHHHHHH---L-llL------L--EELHHHHHHHHL---------------LLHHHHHHHHHHHHHH--LLEEEEELL---------------lllLLEEEEE-LH-Hhhhh
0028 8iueP   -------------------HHHHHHHHHHHHH---L---L------L--LLLHHHHHHHLL----------L----LLHHHHHHHHHHHHHH--LLEEEEEEL------------------LEEEEEE-LHh-----
0029 7wzeA   hhhhhhhhhhhL-L---LLLHHHHHHHHHHHH---L---H-----hH--EELHHHHHHHLL---------------LLHHHHHHHHHHHHHH--LLEEEEEL------------------lLEEEEE-LH-Hhhhh
0030 4hqeB   --lllhhhH-HH-H---HHLLLHHHHHHHHLL---L---H-----hH--EELHHHHHHHHL----------L----LLHHHHHHHHHHHHHL--LLEEEEEE---------------llllEEEEEE-LH-Hhhhh
0031 3r0aB   eeeehhhhhhhH-H---LLLHHHHHHHHHHHH---L--lL------L--LLLHHHHHHHLL---------------LLHHHHHHHHHHHHHL--LLEEEEEEE--------------llleEEEEEE-LL-H----
0032 5eriA   hhhhhhhhhllL-L---LLLLLHHHHHHHHHH---L----------L--LEEHHHHHHHHL---------------LLHHHHHHHHHHHHHL--LLEEEEELL---------------lllLLEEEEE-LH-Hhhhh
0033 1sfuA   -------------------HHHHHHHHHHHHL---L---L----llL--EELHHHHHHHLL---------------LLHHHHHHHHHHHHHL--LLEEEELL------------------LLLEEEE-LL-Ll---
0034 4asnA   ---lhhhHH-HH-H---LLLHHHHHHHHHHHH---L---L------LhhLEEHHHHHLLLL---------------LLLHHHHHHHHHHHHL--LLEEEELLL-----------------lLLEEEE-LH-Hhhhh
0035 4gyiA   ----lllLL-HH-H---LLLHHHHHHHHHHHH---H---L---lllL--LEEHHHHHHHHLL-L------LL----LL--LHHHHHHHHHHL--LLEEEEELL-----------------lLEEEEE-LH-Hhhhh
0036 5jbrA   hhhhhhhhhhhH-H---LLLHHHHHHHHHHHL---L---L------L--LEEHHHHHHHHL---------------LLHHHHHHHHHHHHHL--LLEEEEEE---------------lleEEEEEEE-LL-Llhhh
0037 5j6xA   ------------------HHHHHHHHHHHHHH---H---L---------LLLHHHHHHHLL---------------LLHHHHHHHHHHHHHL--LLEEEELL---------------lllLLEEEE-LL------
0038 2d1hA   hhhhhhhhhhhH-H---LLLHHHHHHHHHHHH---H---L------L--LEEHHHHHHHHL---------------LLHHHHHHHHHHHHHL--LLEEEEEL-----------------lLEEEEE-LL-Lhhhh
0039 2nyxB   hhhhhhhhH-HL-L---LLLHHHHHHHHHHHH---H---L---------LEEHHHHHHHHL---------------LLHHHHHHHHHHHHHL--LLEEEELLL---------------llllLLEEEE-LH-Hhhhh
0040 7bzhA   ------------------LLLHHHHHHHHHHH---H----------L--EEEHHHHHHHLL---------------LLHHHHHHHHHHHHHL--LLEEEEL--------------------LEEEE-LL-L----
0041 1jhfA   -------------L---LLLHHHHHHHHHHHH---H---H--hhhlL--LLLHHHHHHHLL--------------lLLHHHHHHHHHHHHHL--LLEEELLLL------------------LLLEEE-LL-L----
0042 6pcoB   hhhhhhhhhlhH-H---LLLHHHHHHHHHHHH---H---L---------LLLHHHHHHHHL---------------LLHHHHHHHHHHHHHL--LLEEEELLL------------------LLEEEE-LH-Hhhhh
0043 4ijaA   -----------------LLHHHHHHHHHHHHH---L---L---------LLLHHHHHHHHL---------------LLHHHHHHHHHHHHHL--LLEEELL---------------lllLLLLEEE-LH-Hhlee
0044 2p4wB   ---llhhHH-HH-H---HHLHHHHHHHHHHLL---L-------------LEEHHHHHHHHL---------------LLHHHHHHHHHHHHHL--LLEEEEELL---------------llllLEEEEE-LL-Leeee
0045 5n35A   -------------------HHHHHHHHHHHHH---L----------L--EEEHHHHHHHHHHH--------LL----LL--LHHHHHHHHHHL--LLEEELL---------------------LEEEE-LL------
0046 5y6iB   -------------------HHHHHHHHHHHHH---L---L------L--LEEHHHHHHHHL---------------LLHHHHHHHHHHHHHH--LLEEELLL------------------LLEEEE-LH-Hhhhh
0047 7dg2A   eellllllhhhL-L---LLLHHHHHHHHHHHH---H---H-hhlllL--LEEHHHHHHHHHHHlL-----LLL----LLHHHHHHHHHHHHHL--LLEEEEL--------------------LEEEE-LH-Hhhhh
0048 7z1nO   llhhhhhhhhhhhH---HLLHHHHHHHHHHHH---H---L---------LEEHHHHHHHL----------LL----LLHHHHHHHHHHHHHL--LLLLEEEE--------------ellleEEEELL-LH-Hhhhh
0049 1ku9A   hhhhhhhhhhhH-L---LLLHHHHHHHHHHHH---L---L------L--LEEHHHHHHHHL---------------LLHHHHHHHHHHHHHL--LLEEEELL---------------llllLLEEEE-LL-H----
0050 4i99C   ------------------HHHHHHHHHHHHHH---H---H--hhhlL--LEEHHHHL--LLL-L-----------hHHHHHHHHHHHHHHHL--LLEEEELLL---------------lllLLEEEE-L-------
0051 1lvaA   -------------------lLHHHHHHHHHHL---L---L------L--LEEHHHHHHHHL---------------LLHHHHHHHHHHHHHL--LLEEEEEE-----------------lLEEEEEE-HHh-----
0052 8zwaA   ------lhH-HH-HhhhHLLHHHHHHHHHHLL---L---L------LllLLLHHHHHHHHHH-------hHL----LLHHHHHHHHHHHHHL--LLLEELL----------LHHHhlLLLLLLEEL-LH-Hhhhh
0053 1qgpA   -------------------HHHHHHHHHHHHH---H---L------LllLEEHHHHHHHHL---------------LLHHHHHHHHHHHHHH--LLEEEELL------------------LLLEEEE-LL-L----
0054 8a5yT   eeeellllleEE-E---EEEHHHHHHHHLLLL---llllL------L--LEEHHHHHHHHL---------------LLHHHHHHHHHHHHHL--LLEEEEL-------------------LEEEE-LL-L----
0055 4nb5B   hhhhhhhhhhhhlL---LLLHHHHHHHHHHHH---L--lL------L--LEEHHHHHHHLL---------------LLHHHHHHHHHHHHHH--LLEEEELL---------------lllLLLEEEE-LL-Lhhhh
0056 4ad9A   -hHHHHhhhhhhhhlllelHHHHHHHHHHHHH---L--lL------L--LLLHHHHHHHLL-L-----LlhhhhhhHHHHHHHHHHHHHHL--LLEEELLLL-----------------LLLEEE-LL------
0057 8ffzB   llLLLLLllllllllllHHHHHHHHHHHHHH---L----------L--LEEHHHHHHHLLL---------L----LLHHHHHHHHHHHHHL--LLEEEEL-------------------LLEEEE-LL-Lllll
0058 3cuqA   elleeeeeL-LL-L---LLLHHHHHHHHHHL----L---L------L--EELHHHHHHHHL---------------LLHHHHHHHHHHHHHH--LLLEEELLL---------------lLLLEEEL-LLl-----
0059 1yyvA   lllllllHH-HH-H---HHLHHHHHHHHHHHH---L-------------LEEHHHHHHHL----------LL----LLHHHHHHHHHHHHHH--LLEEEEEEL---------------llllEEEEEE-LH-Hhhhh
0060 5nl9A   hhhhhhhHH-LL-L---LLLHHHHHHHHHHHL---L---L------L--EELHHHHHHHHHH-H-----LLL----LLHHHHHHHHHHHHHL--LLLEEEEL---------------lllLEEEELlLL-L----
0061 2xcoA   hhhhhllllllL-L---LLLHHHHHHHHHHHH---L---L------LllLEELHHHHHHHHH-H-----LLL----LLHHHHHHHHHHHLL-llLLEEEEL-----LLLL-lLLLL--LLLLLEEEE-LH-Hhhll
```

```
# Job: Type-1 BREX; BrxC-ATPase C-terminal Coiled-Coil Extension
# Query: s001A
# No:  Chain    Z    rmsd lali nres  %id PDB  Description
   1:  6hyd-A  7.5 22.1   91  1574    9   MOLECULE: MIDASIN,MIDASIN,MIDASIN;
   2:  7kfu-C  7.3 24.2  105   909   10   MOLECULE: CAS2;
   3:  8r5s-A  6.9  3.2   92   237   12   MOLECULE: CHAIN A, B, C AND H OF THE SOLUBLE HOMOTETRAMER;
   4:  3rfy-A  6.7 11.1   96   356    7   MOLECULE: PEPTIDYL-PROLYL CIS-TRANS ISOMERASE CYP38, CHLORO
   5:  9u78-A  6.7  2.3   78   118   10   MOLECULE: SINGLE CHAIN PROTEIN OF DE NOVO DESIGNED HELIX BU
   6:  8x9p-C  6.7  3.0   77   107    9   MOLECULE: TUBULIN ALPHA CHAIN;
   7:  5a7d-R  6.5 14.7  125   311    8   MOLECULE: PINS;
   8:  6qv7-A  6.4  2.6   78   316   14   MOLECULE: UNCHARACTERIZED PROTEIN;
   9:  8dt0-A  6.3  3.1   83   140   13   MOLECULE: SCAFFOLDING PROTEIN FUNCTIONAL SITES;
  10:  8wcn-A  6.3  8.5  104   379    6   MOLECULE: DIGUANYLATE CYCLASE;
  11:  6r6n-A  6.3  2.9   77   111    4   MOLECULE: SMALL SOLUBLE CYT C;
  12:  6dfk-I  6.2  3.7   80   236   13   MOLECULE: SUBUNIT OF PROTEASEOME ACTIVATOR COMPLEX,PUTATIVE
  13:  3rgu-A  6.2  3.1   66    88    6   MOLECULE: FIMBRIAE-ASSOCIATED PROTEIN FAP1;
  14:  9ri9-B  6.2  9.1  105   868   14   MOLECULE: SLNRC3;
  15:  6n1l-A  6.1  2.9   80   141   13   MOLECULE: FIBRONECTIN-BINDING PROTEIN BBK32;
  16:  2ot4-A  5.9 10.6  122   520    7   MOLECULE: EIGHT-HEME NITRITE REDUCTASE;
  17:  8vdq-A  5.9 13.7  134  2192    8   MOLECULE: GREEN FLUORESCENT PROTEIN,TALIN-1;
  18:  6poo-A  5.9  9.8  172   273    9   MOLECULE: BIBA;
  19:  7nd2-A  5.8 22.7   87   330    5   MOLECULE: PROTEIN PHOSPHATASE 1 REGULATORY SUBUNIT 21;
  20:  9b8o-U  5.8  3.3   92   214    4   MOLECULE: ATPASE H+-TRANSPORTING V1 SUBUNIT D;
  21:  9ivk-A  5.7  7.1  108   278    3   MOLECULE: HBC599 MEMBRANE PROTEIN BINDER;
  22:  8a6t-B  5.6  3.2   73   630   14   MOLECULE: ELECTRON BIFURCATING HYDROGENASE SUBUNIT HYDA1;
  23:  9n0i-E  5.6  2.9   82   133    7   MOLECULE: DE NOVO DESIGNED KABLE PROTEIN;
  24:  2win-M  5.6  3.3   73    84    8   MOLECULE: COMPLEMENT C3 BETA CHAIN;
  25:  9dbd-A  5.6  3.2   78   326    9   MOLECULE: TET4 VARIANT;
  26:  7et3-1  5.6  2.7   67   285    6   MOLECULE: TRIPLEX CAPSID PROTEIN 2;
  27:  5m0i-C  5.6  4.6  112   240    7   MOLECULE: SWI5-DEPENDENT HO EXPRESSION PROTEIN 2;
  28:  8tn1-A  5.6  2.9   85   147   12   MOLECULE: DE NOVO DESIGNED 4 HELIX BUNDLES;
  29:  8ka6-B  5.6  3.1   85   148   11   MOLECULE: DE NOVO DESIGN PROTEIN -NA7;
  30:  4avm-A  5.6  4.0   80   230   11   MOLECULE: BRIDGING INTEGRATOR 2;
  31:  9bia-A  5.5  2.6   76   110    8   MOLECULE: NINJURIN-1;
  32:  7m95-B  5.5  4.0   81   167    5   MOLECULE: SIGMA INTRACELLULAR RECEPTOR 2;
  33:  2jx0-A  5.5  3.4   86   131   10   MOLECULE: ARF GTPASE-ACTIVATING PROTEIN GIT1;
  34:  7d6v-C  5.5  3.0   84   264    7   MOLECULE: SUCCINATE DEHYDROGENASE SUBUNIT A;
  35:  6u4o-A  5.5  5.5  119   479    7   MOLECULE: FUMARATE HYDRATASE;
  36:  8xi9-A  5.5  9.0   85   201    8   MOLECULE: FRB-FKBP FUSION PROTEIN;
  37:  6s18-A  5.5  2.9   85   143    8   MOLECULE: AROMATIC ACID CHEMORECEPTOR;
  38:  4od4-A  5.5  4.2   80   275    5   MOLECULE: 4-HYDROXYBENZOATE OCTAPRENYLTRANSFERASE;
  39:  8ccr-A  5.4  2.5   72   105    7   MOLECULE: 4D2 (MUTANT T19D);
  40:  1a7d-A  5.4  3.0   80   118    9   MOLECULE: MYOHEMERYTHRIN;
  41:  8oyy-A  5.4  4.7   98   233   12   MOLECULE: DE NOVO DESIGNED SOLUBLE GPCR-LIKE PROTEIN;
  42:  1rj1-A  5.4  2.9   75   148    7   MOLECULE: INVERTASE INHIBITOR;
  43:  5ziy-A  5.4  5.2  101   195    5   MOLECULE: FLAGELLAR HOOK-ASSOCIATED PROTEIN 3;
  44:  4kbq-C  5.4  7.1   81    91   10   MOLECULE: E3 UBIQUITIN-PROTEIN LIGASE CHIP;
  45:  3lmf-A  5.4  2.4   74   109    4   MOLECULE: UNCHARACTERIZED PROTEIN;
  46:  8w6e-B  5.4  3.3   89   161   13   MOLECULE: HBC599 IN COMPLEX WITH WFAP1.1;
  47:  9had-C  5.4  3.1   83   108    6   MOLECULE: MITE ALLERGEN DER F 21.0101;
  48:  7lsv-B  5.3  5.7   85   158   15   MOLECULE: CALMODULIN-DEPENDENT PROTEIN KINASE;
  49:  3qwe-A  5.3  4.4  100   260    6   MOLECULE: GEM-INTERACTING PROTEIN;
  50:  3nrh-A  5.3  2.6   83   158   10   MOLECULE: UNCHARACTERIZED PROTEIN BF1032;

0001 s001A   --HIRAVKDVIRE---F-FG-KTIT--S--D-DQDTLMRDYR-SYAQTKKSAIENMNSR-L-PG-R-RILE-EA-LR-LIDDTV-K--I--N--DP[illegible]
0002 6hydA   -----------------------------------SIPEFQKAG-DIIETVLKSVLKL-KQ-W-P-eH-ATLK-EL-YR-VSQEFL-N--Y--P--IK[illegible]
0004 8r5sA   --AIKELLELLKK---A-LEkskAK--G--IeAFELACEVLK-KTLEAIKAVADDKAAI-LaAQ-Q-KALA-ET-NA-LCTDSL-IavE--G--VE[illegible]
0005 3rfyA   irevqkplEDIT------------dslK--I-AGVKALDSVE-RNVRQASRTLQQGKIV-A-GG-N-EMIE-KL-EA-GMQDML-K--I--V--ED[illegible]
0006 9u78A   ---------------------------------G-GIEELLRKII-EDEVRHIAELEDIWLK-D-VI-E-ELLR-KI-IE-DEVRHI-A--E-lE--DI[illegible]
0007 8x9pC   ------------------------hgV--P-YFRNILQSET-EKLTSHCFEWDRKELD-I-PA-K-DLIR-TA-VG-QTRLLM-K--ErfK--QF[illegible]
0008 5a7dR   [illegible]
0009 6qv7A   [illegible]
0010 8dt0A   [illegible]
0012 6r6nA   [illegible]
0013 6dfkI   [illegible]
0015 9ri9B   [illegible]
0016 6n1lA   [illegible]
0018 8vdqA   [illegible]
0020 7nd2A   [illegible]
0021 9b8oU   [illegible]
0022 9ivkA   [illegible]
0023 8a6tB   [illegible]
0024 9n0iE   [illegible]
0026 9dbdA   [illegible]
0027 7et31   [illegible]
0028 5m0iC   [illegible]
0029 8tn1A   [illegible]
0030 8ka6B   [illegible]
0031 4avmA   [illegible]
0032 9biaA   [illegible]
0033 7m95B   [illegible]
0034 2jx0A   [illegible]
0035 7d6vC   [illegible]
0036 6u4oA   [illegible]
0037 8xi9A   [illegible]
0038 6s18A   [illegible]
0039 4od4A   [illegible]
0040 8ccrA   [illegible]
0041 1a7dA   [illegible]
0042 8oyyA   [illegible]
0043 1rj1A   [illegible]
0044 5ziyA   [illegible]
0045 4kbqC   [illegible]
0046 3lmfA   [illegible]
0047 8w6eB   [illegible]
0048 9hadC   [illegible]
0049 7lsvB   [illegible]
0050 3qweA   [illegible]
0051 3nrhA   [illegible]

0001 s001A   [illegible]
0002 6hydA   [illegible]
0004 8r5sA   [illegible]
0005 3rfyA   [illegible]
0006 9u78A   [illegible]
0007 8x9pC   [illegible]
0008 5a7dR   [illegible]
0009 6qv7A   [illegible]
0010 8dt0A   [illegible]
0012 6r6nA   [illegible]
0013 6dfkI   [illegible]
0015 9ri9B   [illegible]
0016 6n1lA   [illegible]
0018 8vdqA   [illegible]
0020 7nd2A   [illegible]
0021 9b8oU   [illegible]
0022 9ivkA   [illegible]
0023 8a6tB   [illegible]
0024 9n0iE   [illegible]
0026 9dbdA   [illegible]
0027 7et31   [illegible]
0028 5m0iC   [illegible]
0029 8tn1A   [illegible]
0030 8ka6B   [illegible]
0031 4avmA   [illegible]
0032 9biaA   [illegible]
0033 7m95B   [illegible]
0034 2jx0A   [illegible]
0035 7d6vC   [illegible]
0036 6u4oA   [illegible]
0037 8xi9A   [illegible]
0038 6s18A   [illegible]
0039 4od4A   [illegible]
0040 8ccrA   [illegible]
```

```
0041 1a7dA   [illegible]
0042 8oyyA   [illegible]
0043 1rj1A   [illegible]
0044 5ziyA   [illegible]
0045 4kbqC   [illegible]
0046 3lmfA   [illegible]
0047 8w6eB   [illegible]
0048 9hadC   [illegible]
0049 7lsvB   [illegible]
0050 3qweA   [illegible]
0051 3nrhA   [illegible]

[illegible]
[illegible]
[illegible]

# Job: Type-1 BREX; BrxX-MTase MBZ5596028 NTD Dimerisation domain
# Query: s001A
# No:  Chain   Z    rmsd lali nres  %id PDB  Description
   1:  8c45-A 17.8  2.4  192  1219   23   MOLECULE: SITE-SPECIFIC DNA-METHYLTRANSFERASE (ADENINE-SPEC
   2:  3khk-A  7.7  4.0  122   488   12   MOLECULE: TYPE I RESTRICTION-MODIFICATION SYSTEM
   3:  2y7h-B  7.4  3.5  123   530   11   MOLECULE: TYPE-1 RESTRICTION ENZYME ECOKI SPECIFICITY PROTE
   4:  5ybb-A  6.8  4.0  127   488   15   MOLECULE: TYPE I RESTRICTION-MODIFICATION SYSTEM METHYLTRAN
   5:  3ufb-A  6.7  3.8  123   484   10   MOLECULE: TYPE I RESTRICTION-MODIFICATION SYSTEM METHYLTRAN
   6:  2okc-B  6.6  3.5  121   427   15   MOLECULE: TYPE I RESTRICTION ENZYME STYSJI M PROTEIN;
   9:  7vru-A  5.9  3.8  120   498   11   MOLECULE: SITE-SPECIFIC DNA-METHYLTRANSFERASE (ADENINE-SPEC
  10:  3lkd-B  5.9  3.6  118   472   10   MOLECULE: TYPE I RESTRICTION-MODIFICATION SYSTEM
  11:  5hr4-J  5.6  3.5  126   752   13   MOLECULE: MMEI;
  12:  7eew-A  4.3  4.0  111   611   12   MOLECULE: TYPE I RESTRICTION-MODIFICATION SYSTEM METHYLTRAN
  13:  8w0p-A  3.9  4.4  118   917    9   MOLECULE: RM.BSAXI;
  16:  2f8l-A  3.6  3.5   84   324    8   MOLECULE: HYPOTHETICAL PROTEIN LMO1582;
  18:  3s1s-A  3.4  4.0  109   871    8   MOLECULE: RESTRICTION ENDONUCLEASE BPUSI;

0001 s001A   MILGKALRNEIRKMVGAIRDLLNEDIRIQLEGTFRILREIAPQEAVEHFTREVGFTSLNRLAALKMMEARGLIQESVS-RGQESSGSSQPDGGYRR-YLELLFDDAAVELRVLFDRALPHSRIFPRP-DALNRALKLLNESCPAEVWKSDETIGWIYQYFTPKEDREALRGGAQ
0002 3khkA   ---qQFLNDLDNQLWRAADKL-RSNL-----------------------DAANYKHVVLGLIFLKYVSDAFEigARWNkLRDV---TLPT--SVSW-LIDNAFDDIEKLKGILN------SQYQLDA-DKLIGLINEFS----KDIL--GHVYEYFLGQFALeGKQGgTQIPAC
0003 2y7hB   -----mnNNDLVAKLWKLCDNLRDGG----------------------vSYQNYVNELASLLFLKMCKETGQEAyRWD-DLKS-----riGQEQLQ-FYRKMLVHLGELVQAVFHN----vSTTIEP-KQITALVSNMDSLDWyngaKSRDFGDMYEGLLKNANETiFYAQGVK
0004 5ybbA   ----PQTRESLANEIWRACDIMRRDNN--------------------cTGIMEYVEHLAWLLFLRFLDAQE-sEYRWRhWATK---dwPADEAFVHgRLIPYLRS--ETIRSLFSER---NVIVCSG-YNLKDVIQIVNE-INFHS---iFTVSQVYEELLLGNE-fSKGNPIQ
0005 3ufbA   adqpmtTAQQLGAIVKSSRQIMRKDKG--------------------lNGDLDRLPMLTWIMFLKFLDDLEQwRDWAA-IEGGitg------------giGLFAYLRSVIATVFKG----MQNRMNG-YLLRDVVDKING-IHFN---MHTLSRLYETMLMRDAGDSGaPYTDI
0006 2okcB   ------TEQSLTKKVWNLATTLAGQG----------------------iGFTDYITQLTYLLFLKXDAENSAIPYQWAdLIAF------dGLDLVK-QYEETLKLLSELIGTIYTK----aQNKIKP-VYLKKVITXIDEE----QWLIXDvKGAIYESILNGQD---tLATNK
0008 7vruA   -xtlinLKDLEAHLW-HAAHIITGPI----------------------DASD-YKTYIFPILFFKRICDVYvGCHW-ddVFAK-------NHDIGK-ALKDAFLGIEQPLHGIFG-----DASWLPD-ELLATLLNHFNQ-VNLG---XGRAYEYLIKRFADK-ANKKrAQNTL
0009 3lkdB   ------TSQSLYQALWNSADVLRSKX-----------------------DANDYKSYLLGXVFYKYLSDKXtpDLTFTaLVER----vndGSFQLE-DLAQGFRDIEQLYENLFE-----DIDLgQN-QTVAAVXKELAVL----DVAGGDAYEYLIGQFAT----GKNQNIX-
0010 5hr4J   ----kpqdpINIKAAERMGKLHDTLKLVG-------------------yEGHALELYLVRLLFCLFAEDTTIFELFQE-YIETK--tleDGSDLAH-HINTLFYVLNTINGKLFEEP---LPPAQFDkAMREALLD-LCSL----DWSR--ISPAIFGSLFQaKKRRNLGPFVG
0011 7eewA   ------------NKLMNIIELIRKDTGI--------------------NNAIDAVEQLALLLLVRYTHEVASdSFKN---------lffdlNVID---FYTLRDKLnEKIENILDQ----IPFRIST-KILDLVIHRLE-ELDL----SEGI-EIDFDHLLNMVK-LAPYSKVN
0012 8w0pA   VRIDREEIEGLGRNLHNVLWG------------------------gggtNDSEIFYSLVNIILAKIQDEYEKqEYDF--QVYQ---nvESPQKLFD-RINALYKRALREQLnvtdeqkiaNRNKFPL-NKLVYTVQALES----LSFLLGDFFESIIRDGFtfttpKKGEKPMP
0013 2f8lA   ---ANEATQELFQVLDNTAIILQNELE---------------------iSYLEAVYETGENLFQK-----------------------------------------------------------QKE-EKQLKLQASYES-IELEN-FSNEIRKGLQLALKGXKH---FKSQAR
0014 3s1sA   leIIDFfkhdpnSSTANLVDILEN-----------------------rkDNSTSWHQALVVAGYEYIRGVLVPTRDA---------------yFKS-RPGRLLEEGRKDFNVLFSE----pepNTND-NDIWN-VNLLSS--lnDLGRILTG-DELAELIHDaTRGRGHEPYVS

0001 s001A   LLLLHHHHHHHHHHHHHHHHHHHHHHHHHHHHLLLLLLLLLHHHHHHHHHHHHHHHHHHHHHHHHHHHHLLLLLLLLL-LHHHLHHHLLLLLLHHH-HHHHHHHHHHHHHHHHHLLLLHHHLLLLLH-HHHHHHHHHHHHLLLHHHHHLLLHHHHHHHHHLLHHHHHHHHHLLL
0002 3khkA   ---lLLHHHHHHHHHHHHHLL-LLLL-----------------------LHHHHHHHHHHHHHHHHHHHHHHllLLHHhHHHL---LLLL--LHHH-HHHHHHHHHHHHLLLLL------LLLLLLH-HHHHHHHHHHL----LLLH--HHHHHHHHHHHHHlLLLLlLLLLEE
0003 2y7hB   -----llLLHHHHHHHHHHHHHHHHL----------------------lLLLLHHHHHHHHHHHHHHHHHLLHHlLHH-HHHH-----llHHHHHH-HHHHHHHHHHLHHHHHHLL----lLLLLLH-HHHHHHHHHHHLLLLllllLLHHHHHHHHHHHHHHHLLlLLLLLLL
0004 5ybbA   ----LLLHHHHHHHHHHHHHHHHHLLL--------------------lLLHHHHHHHHHHHHHHHHHHHHH-lLLLHHhHHHL---llLHHHHHHHhLHHHHHHL--HHHHHHHLLL---LLLLLLH-HHHHHHHHHHHL-LLLLL---hHHHHHHHHHHHHHHH-lLLLLLLL
0005 3ufbA   llllllLHHHHHHHHHHHHHHHLLLLL--------------------lLHHHHHHHHHHHHHHHHHHHHHHHhHHHLL-LLLLllh------------eeLHHHHHHLHHHHHHHL----LLLLLLH-HHHHHHHHHHHL-LLLL---HHHHHHHHHHHHHLLLLLLLlLLLLL
0006 2okcB   ------LHHHHHHHHHHHHHHHHLLL----------------------lLHHHHHHHHHHHHHHHHHHHHLLLLLLHHhHLLL------lHHHHHH-HHHHHHHHHLLHHHHHLLL----lLLLLLH-HHHHHHHHHHHLL----LHHHLLhHHHHHHHHHHHLL---lLLLLL
0008 7vruA   -lllllHHHHHHHHH-HHHHHHLLLL----------------------LLLL-LHHHHHHHHHHHHHHHHHhLLLH-hhHHHL-------LLLHHH-HHHHHHHHHHHHHLLLLL-----LLLLLLH-HHHHHHHHHHHL-LLLL---HHHHHHHHHHHHHHH-HHHHlLLLLL
0009 3lkdB   ------LHHHHHHHHHHLLHHHHLLL-----------------------LHHHHHHHHHHHHHHHHHHHHHhhHHLHHhHHHH----hhhLLLLHH-HHHHHHHHHHLHHLLLLL-----LLLLlHH-HHHHHHHHHHHLL----LLLLHHHHHHHHHHHHH----LLLLEEL-
0010 5hr4J   ----lllllLLHHHHHHHHHHHHHHHHHL-------------------lLLHHHHHHHHHHHHHHHHHHLLLLLHHHH-HHHHH--lllLLLLHHH-HHHHHHHHHHHLLLLHHHHH---LLLLLLLhHHHHHHHH-HHLL----LLLL--LLHHHHHHHHHhHHHHHHHHHHH
0011 7eewA   ------------LHHHHHHHHLLLLLLL--------------------LLLLHHHHHHHHHHHHHHHHHHHHlLLLL---------lllllLLLL---HHHHHHHHhHHHHHHHHH----LLLLLLH-HHHHHHHHHHH-HLLL----LLLL-HHHHHHHHHHHH-LLLLLLLL
0012 8w0pA   LLLLHHHHHHHHHHHHHHHHL------------------------llllLLLHHHHHHHHHHHHHHHHHHHHlLLLL--LLLL---llLLLLHHHH-HHHHHHHHHHHHLLlllhhhhhhLLLLLLH-HHHHHHHHHHHL----LLLLHHHHHHHHHHHHLlllhhLLLLLLLL
0013 2f8lA   ---LLHHHHHHHHHHHHHHHHHHHHLL---------------------lLHHHHHHHHHHHHHHL-----------------------------------------------------------LLL-LLHHHHHHHHHL-LLHHH-LLHHHHHHHHHHHHHLLL---LLLLLL
0014 3s1sA   lleeeeeellllHHHHHHHHHHHL-----------------------llLLHHHHHHHHHHHHHHHHHHHHLLLLLH---------------hHLL-LHHHHHHHHHLLLLLLLLL----lllLLLL-LLLLL-HHHHHH--hhHHHHHLLL-HHHHHHHHHhLLLLLLLLLLL

# Job: Type-1 BREX; BrxX-MTase MBZ5596028 (TRD)
# Query: s001A
# No:  Chain   Z    rmsd lali nres  %id PDB  Description
   1:  7lt5-B 18.2  2.9  209   577   12   MOLECULE: SITE-SPECIFIC DNA-METHYLTRANSFERASE (ADENINE-SPEC
   2:  7qw8-A 17.6  3.0  208   525   18   MOLECULE: MODIFICATION METHYLASE BSECI;
   3:  3s1s-A 13.5  3.5  193   871   12   MOLECULE: RESTRICTION ENDONUCLEASE BPUSI;
   4:  7vs4-C 12.2  3.8  181   374    9   MOLECULE: SITE-SPECIFIC DNA-METHYLTRANSFERASE (ADENINE-SPEC
   5:  1yf2-A 11.0  4.2  182   425   10   MOLECULE: TYPE I RESTRICTION-MODIFICATION ENZYME, S SUBUNIT
   6:  1ydx-A 10.9  3.4  176   374    7   MOLECULE: TYPE I RESTRICTION ENZYME SPECIFICITY PROTEIN MG4
   7:  5hr4-J  9.9  3.6  201   752   12   MOLECULE: MMEI;
   8:  2y7c-A  9.2  4.2  192   464    9   MOLECULE: TYPE-1 RESTRICTION ENZYME ECOKI SPECIFICITY PROTE
   9:  3okg-B  9.2  3.3  141   391   11   MOLECULE: RESTRICTION ENDONUCLEASE S SUBUNITS;
  10:  4xqk-A  8.9  4.0  182  1517    8   MOLECULE: LLABIII;
  11:  7eew-A  8.6  4.8  147   611    8   MOLECULE: TYPE I RESTRICTION-MODIFICATION SYSTEM METHYLTRAN
  12:  7bst-A  7.8  3.8  164   384   10   MOLECULE: TYPE I RESTRICTION ENZYME R PROTEIN;

0001 s001A   YSVGDRLAHLLAHQARFEPAGIVRTGLTTFDDFRFLRVTWE-VGADRIRHWEPFSKGGEYAKYYTPLHLLVNRRNNGAELAEQSRQGSKYYYKAGLTFTSRSAKGFTVRALPAGCVISHNAPTIYPNADISKEYLLGWINSSLIRALVEVQKNFSYFVPGTIKDLPWIEPTSEVRRLVEALVSELVNRYRQVWVWPPKVNRDALPS-ASVNQEHGDVIRDHQRGISGAIDGL-
0002 7lt5B   KDDETFYNKIQECKYSLEDIAISFQGIITGCDKAFILSKDDvKLNLV-KFLKCWIKSKNINKYIVDSEYRLIY--SNDI---QWGREKLFFERKKIMYPYKS-NENRFAIDYDNNFSSADVYSFFIKEKFSYEYLVGILNSSVYDKYFKITAKKYDYYPNKVMKIRIFRDN--NYEEIENLSKQIISILLNK-------------------siDKGKVEKLQIKMDNLIMDS-
0003 7qw8A   EDEYEWFMKVNQAYGFIEDFAHVKVGIKTTADSVFIR-SDW-GIPED-KLLRPIISADQANKWSVSLNKKVLYTHEiYEIW--VPHDPSLWDKPKIIFPDTS-PEPKFFYEDKGSVVDGNCYWIIPKNSNDLFLIMGICNSKFMSKYHDAFQNRRRYLTQYVNKYPIPDPESIYSKEIISLVRELVNNK-----------------------kETQDINEIENRIEKLILRA-
0004 3s1sA   RVAEEWITNLESHCIRLVSDYDLRRGRVGGASDLLFINSKKklSVPR-DWLYPALRKVNeNTPIFATPVRFLCPPnsEILYK----esdfYSSETVFIPRALRRSARAFINEQKVFCSTNALEVFGGNSEEXWLLLSWLSSVFAQLQFEAXA-ERKLEKKSIQNLYIPNIDDVLKQDLIEEVREIH-------------------------------crprvRKLDLLWAKV-
0005 7vs4C   -------------mVKFGDIAKHISKRVE---------------psET---DIYVGLHLDPSLKIKRYGVPSD----------aGQKL--LVKKGQIIFGKRRaYQRKVAVADWDCICSAHAMVLEPLSDVIPEFLPFFMQSDSFMNRAVAISEGPTIKWKTLSSQSFLMPSLTTQATLIKILSKISEVESSLESASLQLSAFIfSS-KVISNKKLTLKAIKSSLVNNS-----
0006 1yf2A   keenfkKTEI--giVELKDVCKIKAGGTPKTsEEYY------------ngTIPFVKIETNSNKYLTNKIKITEE---------nNSNAW--IVPKSVLFAMYG--IGETAINKIEVATNQAILGIIPKDILESEFLYYILAK--NKNYYSKL--QKNLNAQIVKSFKIPLPPEEQKQIAKILTKIDEGIEIIE---kLERIKGLkssiELKKQKKEKLQRMKKKIMELLLTG--
0007 1ydxA   --------xpkllkRTIDSLFDLKKGEXL--EKELITPE-----------gKYEYFNGGV------KNSGRTDK----------------fNTFKTISVIVGG--sGYVRLADKNFFCGQSNCTLNLLDPLDLKFAYYALKS--QQERIEALAFGQNIRISDLKELEIPFTNKNEQHAIANTLSVFDERLENLASLINRKLVFLLDQyKKELSSLTVIRDTLLKKL---FPDXt
0008 5hr4J   PYLRDG-----vvpISKL--PSMRYGNKPTDDGNFLFTDEE-KNQFI-KYFRRFVGGD---efiNNTSRWCLW--LDGAdisFFYIS--QPDTDYLLIPETSSEqIPIGFVDRNVISSNATYHIPS-----EPLIFGLLSSTMHNCWMRNVGGRYRYSASVYNTFPWIQPNEKQSKAIEEAAFAILKARSNYPNE-----------slaglKTMPSELLKAHQKLDKAVDSV-
0009 2y7cA   -------MSAGkpIAPVSTVTLIRGVTYKKEqINYL------------DDYLPLIRAIQNG--KFDTTLVFVPK--------vkESQK--ISPEIVIAMSSGSKVGKSAHQHPFECSFGAFCGVLRPELIFSGFIAHFTKSSLYRNKISSLSAINNIKPASFDLINIPIPPAEQKIIAEKLDTLL-AQVDST---EQIPILKRaYA-DTIEKQVNNALARVNNLTQSILAKA-
0010 3okgB   mqtalaevfHPGAwVRLGEVCDIIMGQSPPS--STYNFE---------GNGLPFFQGKA-dgdLHPTRIWCSA--------------pqkVARPDVLISVRAP--GSTNVANLACCIGRGLAALRPRDSLERFWLLYYLHY--LEPELSKM----AITKKDLQNVFIPLPPLEEQRRIVAYLDQIQQ------------------------vaALKR----------------
0011 4xqkA   INQR---NSNF-DELISDSLFDYTGGLKTGRDNWSWNF-SK-EQvelsERMYLGMY-------rPFTKKHVYY---AQAW--DRQYMTKVLSNLMISLSNKTGKQFSTLIVD--VLPDVGSQNLPAIRHDAMYYVYGLFHSIQYGKVYYEDL--AKSF------pKIPNVK--HKEKYIKIGKELADLHLNYENqN-------------llITVSMRTLELIDEL--PEFEIq
0012 7eewA   lvtsienhfymsnqVKLKDICKLRSGDKLNKS--EVMDS----------gEFPVYGGNG-------VIGFNVE----------------pnRHGDSIVIGKVGAHCGNIHFSTQPYWLTSNAMSLELLDTVYLPYLAHVLK--SLEL--NNLATGKFISINKLYEVEVSLPSLEKQREMSEWFTSI-EESKSKIQSADFSRNLG------------------------------
0013 7bstA   kqynyyrdqlsfeWKTLGEIGKWYGGGTPSKnIEFW------------NGSIPWISPDMGRT-LVDSSDYITE-EAVL----hSSTK---LIPASIAIVVRSDKVLPSALIKVPATLNQDMKAVIPHENILVKYIYHMIGS--RGSDILRAAKKaSIDSKKLFSFKIPVPNINEQQRIVEILDKFDTTNSIGLPRIELRQYEY------------------------------
0014 2np6A   RFETEETRKLEISGMPLGDLFHIRFAARSPKKHPAVRK--E-PGP----GLVPVLTGRNLKPGWVDYEgLWMP---------eraklRDFYATPHLVVAHTKGTRVVAAWDERAYPWR-EEFHLLPKVRLDPSSLVQWLNSEAMQKHVRTLYFVPHLTLRMLERLPVRRE---------------------------------------------------------------

0001 s001A   [illegible]
0002 7lt5B   [illegible]
0003 7qw8A   [illegible]
0004 3s1sA   [illegible]
0005 7vs4C   [illegible]
0006 1yf2A   [illegible]
0007 1ydxA   [illegible]
0008 5hr4J   [illegible]
0009 2y7cA   [illegible]
0010 3okgB   [illegible]
0011 4xqkA   [illegible]
0012 7eewA   [illegible]
0013 7bstA   [illegible]
0014 2np6A   [illegible]

# Job: Type-1 BREX; BrxX-MTase_MBZ5596028 (C-terminal CC-like Helical Extension)
# Query: s001A
# No:  Chain   Z    rmsd lali nres  %id PDB  Description
   1:  6cnn-A  7.7 13.6  122   360    7   MOLECULE: INTERMEDIATE CONDUCTANCE CALCIUM-ACTIVATED POTASS
   2:  5xg2-A  7.6  2.3   69   237   16   MOLECULE: CHROMOSOME PARTITION PROTEIN SMC;
   3:  1lrz-A  7.6  8.6  114   400    5   MOLECULE: FACTOR ESSENTIAL FOR EXPRESSION OF METHICILLIN
   4:  2qup-A  7.6  5.0   73   119    3   MOLECULE: BH1478 PROTEIN;
   5:  4l0r-A  7.4  1.4   67    74   13   MOLECULE: CENTROSOMAL PROTEIN OF 57 KDA;
   6:  5j10-A  7.3  2.1   69    70   10   MOLECULE: PEPTIDE DESIGN 2L4HC2_24;
   7:  8ap9-G  7.3  3.8   76   279    8   MOLECULE: ATP SYNTHASE GAMMA SUBUNIT;
   8:  6njp-G  7.2  1.1   64    64    9   MOLECULE: TRANSLOCATOR ESCN;
   9:  5xbj-A  7.2  2.8   68   500    7   MOLECULE: FLAGELLAR HOOK-ASSOCIATED PROTEIN FLGK;
  10:  5h69-A  7.2  1.7   68   252    9   MOLECULE: CHROMOSOME PARTITION PROTEIN SMC;
  11:  2x0l-A  7.2  9.6   91   670   15   MOLECULE: LYSINE-SPECIFIC HISTONE DEMETHYLASE 1;
  12:  8fbi-A  7.2  4.0  103   274    8   MOLECULE: KWOCA_39;
  13:  5c21-A  7.2 13.5   95   267    9   MOLECULE: CHROMOSOMAL HEMOLYSIN D;
  14:  6msr-A  7.1  2.1   71    76    8   MOLECULE: PRO-2.5;
  15:  5nik-D  7.1 13.2   97   340   10   MOLECULE: OUTER MEMBRANE PROTEIN TOLC;
  16:  5i6r-A  7.1  9.6   96   419   10   MOLECULE: SLIT-ROBO RHO GTPASE-ACTIVATING PROTEIN 2;
  17:  4b6x-A  7.1  2.1   62    69    8   MOLECULE: AVIRULENCE PROTEIN;
  18:  8ij9-C  7.0  2.7   69    71   10   MOLECULE: RAS-RELATED PROTEIN RAB-6B;
  19:  7w1m-D  7.0  7.5  109   967    7   MOLECULE: STRUCTURAL MAINTENANCE OF CHROMOSOMES PROTEIN 1A;
  20:  8fih-A  7.0  2.7   75   105   19   MOLECULE: 3HB05;

0001 s001A   ----YHRL-----TR-DTLFHLLRNYIEPKIEHEEGRRHELRRLYKAGA-----D---ERRLAKEVDAQETLVGELADFRAKIEAVA-P-GW---D--PD----LNDG--
0002 6cnnA   ----NKAE-----KH-VHNFMMDI-QYTKEMKESAARVLQEAWMFYKHT---shA---ARRHQRKLLAAINAFRQVRLKHRKLREQV-N-SM---V--------------
0003 5xg2A   ------TK-----GAiVRWGKRKE-KLIEEIRAREEERNALVVRLGEID---npE---ARELTEKIRAVEKEIAALREELSRVEGKL-E-GL-----------------
```

```
0004 1lrzA   ----RF--lsevplayiNFDEYIK-ELNEERDILNKDLNKALKDIEKRP-----E---NKKAHNKRDNLQQQLDANEQKIEEGKRLQ-E-EH---G-------------
0005 2qupA   ----gVSF-----SE-VMGKQRDE-KAYERLQALMSKIDDQGKLLSE-T-----R---TIEELRKYKELVKEFVGDAVELGLRLEER----------------------
0006 4l0rA   -------G-----IN-EELSEVLQ-TLQDEFGQMSFDHQQLAKLIQES----veL---KDKLECELEALVGRMEAKANQITKVRKYQ-A-QLekq-------------
0007 5j10A   ------TD-----EL-LRLAKEQA-ELLKEIKKLVEEIARLVKEIQEDP---------SDELLKTLAELVRKLKELVEDMERSMKEQ-L-YI---Ik------------
0008 8ap9G   ------SG-----KL-RLYKEKLE-GYNRFYSIVKTIKMVTLAKYRAAQgrienE---LSEQAARLVAVEGQLTNISSLQQRTSSLY-N-KT---RqfGI----TAALie
0009 6njpG   ----------------HMLDRILS-IRKSRANRLRESMAKINSQIKEV------E---KRSLLDSQKRTKENLQHVNKSVEKLSFAI-K-EH---------------
0010 5xbjA   --------------D-EYSYYKLK-GASTQLEYTKYMASTLQEIAQRFP-----DlqlTGKIASDGENNNVVNSSNETLYNSVYSEY-Q-SK---S--G-----------
0011 5h69A   -------------HM-AAQKTELE-QHEALLHQARQYRQQTKARQQWLE---nrE---LEMLSAKLQEMDETIARLERAVAAKRHEL-A-EQ---Ea-----------
0012 2x01A   ---qIEHW----kKI-VKTQEELK-ELLNKMVNLKEKIKELHQQYKEAS---efL---VKSKHRDLTALCKEYDELAETQGKLEEKL-Q-EL---E-aNP----PSDVyl
0013 8fbiA   ---aARIAktarlII-ELTLMVLK-ASLDLLRRILEELKEMLERLEKNP-----D----KDVIVKVLKVIVKAIEASVDNQRVSADN-Q-KM---------------
0014 5c21A   ---kYQKE----lNL-DKKRAERL-TILARINRYENLSRVEKSRLDDFR---khA---VLEQENKYVEAANELRVYKSQLEQIESEI-L-SA---KeeYQ----LVTQlf
0015 6msrA   ----gseY-----EI-RKALEELK-ASTAELKRATASLRASTEELKKNP---------SEDALVENNRLIVEHNAIIVENNRIIAAV-L-EL---------------
0016 5nikD   ---eqaEN-----QI-KEVEATLM-ELRAQRQQAEAELKLARVTYSRQQ---qqD---LDNAATEMAVKQAQIGTIDAQIKRNQASL-D-TA---K-tNL----DYTR--
0017 5i6rA   ---qseLV-----QRcQQLQSRLS-TLKIENEEVKKTMEATLQTIQDIV---raN---QQETEQFYFTKMKEYLEGRNLITKLQAKH-DlLQ---K--TLgesq------
0018 4b6xA   -------------------GAGA-ALRQEIEDKQLMVNNLTDELQDAI---paE---IANTSQQLRHARADLADLQRRFAVLRNEDrR-IN---Q-------------
0019 8ij9C   -----rKH-----LE-EVLEMKQE-ALLAAISEKDANIALLELSS---------S---KKKTQEEVAALKREKDRLVQQLKQQTQNR-M-KL---------------
0020 7w1mD   tstlaamK-----LM-TALVNVAL-NLSIHQDNTQRQYEAERNKM---------I---GKRANERLELLLQKRKELQENQDEIENMM-N-SI---------------
0021 8fihA   ---akkGE-----ASeEHAAALLA-EAAVLELKAVLLTLEARRLYKELG-----G---DERAREALEAAERAREAAREAREVAEKAY-D-AA---S-------------

0001 s001A   ----HHHL-----LL-LHHHHHHHHLHHHHHHHHHHHHHHHHHHHHHLLL-----L---HHHHHHHHHHHHHHHHHHHHHHHHHHHHH-H-HL---L--LL----HHHL--
0002 6cnnA   ----LHHH-----HH-HHHHHHHH-HHHHHHHHHHHHHHHHHHHHHHHH---hhH---HHHHHHHHHHHHHHHHHHHHHHHHHHHHH-H-HH---L-------------
0003 5xg2A   ------LH-----HHhHHHHHHHH-HHHHHHHHHHHHHHHHHHHHHHHH---llH---HHHHHHHHHHHHHHHHHHHHHHHHHHHHH-H-HL----------------
0004 1lrzA   ----EE--llhleeeeEHHHHHH-HHHHHHHHHHHHHHHHHHHHHHHLL-----L---LHHHHHHHHHHHHHHHHHHHHHHHHHHHH-H-HH---L-------------
0005 2qupA   ----lLLH-----HH-HHHHHHHH-HHHHHHHHHHHHHHHHHHHHHHH-H-----L---LHHHHHHHHHHHHHHHHHHHHHHHHHHLL----------------------
0006 4l0rA   -------L-----HH-HHHHHHHH-HHHHHHHHHHHHHHHHHHHHHHLL----hhH---HHHHHHHHHHHHHHHHHHHHHHHHHHHHH-H-HHhll-------------
0007 5j10A   ------LL-----HH-HHHHHHHH-HHHHHHHHHHHHHHHHHHHHHHHLL---------LHHHHHHHHHHHHHHHHHHHHHHHHHHHH-H-HH---Hl------------
0008 8ap9G   ------LL-----HH-HHHHHHHH-HHHHHHHHHHHHHHHHHHHHHHHHLlhhhhH---HHHHHHHHHHHHHHHHHHHHHHHHHHHHH-H-HH---HhhHH----HHHHhh
0009 6njpG   ----------------LLHHHHHH-HHHHHHHHHHHHHHHHHHHLLLLL------L---HHHHHHHHHHHHHHHHHHHHHHLLLLLL-L-LL----------------
0010 5xbjA   --------------L-HHHHHHHH-HHHHHHHHHHHHHHHHHHHHHHHLL-----LllhHHHHHHHHHHHHHHHHHHHHHHHHHHHH-H-HL---L--L-----------
0011 5h69A   -------------LH-HHHHHHHH-HHHHHHHHHHHHHHHHHHHHHHHH---hhH---HHHHHHHHHHHHHHHHHHHHHHHHHHHHH-H-HH---Ll------------
0012 2x01A   ---hHHHH----hHH-HHHHHHHH-HHHHHHHHHHHHHHHHHHHHHHHLL---hhH---HHHHHHHHHHHHHHHHHHHHHHHHHHHHH-H-HH---L-lLL----LLLLll
0013 8fbiA   ---hHHHHhhhhhHH-HHHHHHHH-HHHHHHHHHHHHHHHHHHHHHHHLL-----L----HHHHHHHHHHHHHHHHHHHHHHHHHHHH-H-HH----------------
0014 5c21A   ---hHHHH----hHH-HHHHHHHH-HHHHHHHHHHHHHHHHHHHHHHHH---hhH---HHHHHHHHHHHHHHHHHHHHHHHHHHHHH-H-HH---HhhHH----HHHHhh
0015 6msrA   ----llhH-----HH-HHHHHHHH-HHHHHHHHHHHHHHHHHHHHHHHLL---------LHHHHHHHHHHHHHHHHHHHHHHHHHHHH-H-HH----------------
0016 5nikD   ---hhhHH-----HH-HHHHHHHH-HHHHHHHHHHHHHHHHHHHHHHHH---lhH---HHHHHHHHHHHHHHHHHHHHHHHHHHHHH-H-HH---H-hHH----HHHE--
0017 5i6rA   ---hhhHH-----HHhHHHHHHHH-HHHHHHHHHHHHHHHHHHHHHHHH---hhH---HHHHHHHHHHHHHHHHHHHHHHHHHHHHH-HhHH---H--HHllll------
0018 4b6xA   -------------------LLLH-HHHHHHHHHHHHHHHHHHHHHHHH---hhH---HHHHHHHHHHHHHHHHHHHHHHHHHHHHhH-HH---L-------------
0019 8ij9C   -----lHH-----HH-HHHHHHHH-HHHHHHHHHHHHHHHHHHHLL---------L---LHHHHHHHHHHHHHHHHHHHHHHHHHHHH-H-HH----------------
0020 7w1mD   hhhhhhhH-----HH-HHHHHHHH-HHHHHHHHHHHHHHHHHHHLL---------L---LHHHHHHHHHHHHHHHHHHHHHHHHHHHH-H-HH----------------
0021 8fihA   ---lllLH-----HHhHHHHHHHH-HHHHHHHHHHHHHHHHHHHHHHHLL-----L---LHHHHHHHHHHHHHHHHHHHHHHHHHHHH-H-HH---L-------------

# Job: Type-2 BREX; BrxX-MTase MBN9519353 (C-terminal CC-like Helical Extension)
# Query: s001A
# No:  Chain   Z    rmsd lali nres  %id PDB  Description
   1:  8yad-C  5.0  3.6   83  1811    2   MOLECULE: SPATACSIN;
   2:  9ivk-A  4.8  5.2   99   278   12   MOLECULE: HBC599 MEMBRANE PROTEIN BINDER;
   3:  4k2p-A  4.8  7.1   90   238    7   MOLECULE: T-LYMPHOMA INVASION AND METASTASIS-INDUCING PROTE
   4:  8eki-A  4.8  5.7  103   170   13   MOLECULE: PROTEIN TRANSPORT PROTEIN SEC20;
   5:  6sny-A  4.7  3.4   88   107   10   MOLECULE: SYNTHETIC EPCR BINDING PROTEIN;
   6:  8ap9-G  4.7  3.0   70   279    7   MOLECULE: ATP SYNTHASE GAMMA SUBUNIT;
   7:  8w6e-B  4.7  4.6   94   161   14   MOLECULE: HBC599 IN COMPLEX WITH WFAP1.1;
   8:  6dkm-D  4.7  2.9   68    79   13   MOLECULE: DHD131_A;
   9:  6y1y-B  4.6  2.5   67   128   15   MOLECULE: CHEA;
  10:  8bbg-B  4.6  7.3   82  1426    9   MOLECULE: WD REPEAT-CONTAINING PROTEIN 19;
  11:  7mx2-B  4.6  4.9   83   648    7   MOLECULE: N-ALPHA-ACETYLTRANSFERASE 35, NATC AUXILIARY SUBU
  12:  9clb-F  4.5  4.1   92   115    9   MOLECULE: BCL-2 HOMOLOGOUS ANTAGONIST/KILLER;
  13:  3aai-A  4.5  3.3   74    78    5   MOLECULE: COPPER HOMEOSTASIS OPERON REGULATORY PROTEIN;
  14:  6wc3-B  4.5  4.6   79    94    8   MOLECULE: PROTEIN TRANSPORT PROTEIN TIP20;
  15:  4b6x-A  4.5  3.4   69    69   10   MOLECULE: AVIRULENCE PROTEIN;
  16:  6dmp-B  4.5  3.3   70    82   17   MOLECULE: DESIGNED ORTHOGONAL PROTEIN DHD13_XAAA_A;
  17:  5h69-A  4.4  2.7   68   252    9   MOLECULE: CHROMOSOME PARTITION PROTEIN SMC;
  18:  2gd5-D  4.4  6.3   91   162    9   MOLECULE: CHARGED MULTIVESICULAR BODY PROTEIN 3;
  19:  8ibh-A  4.4  2.7   65    75   15   MOLECULE: CENTROSOMAL PROTEIN OF 57 KDA;
  20:  5n77-A  4.4  5.4   89   257   11   MOLECULE: MAGNESIUM TRANSPORT PROTEIN CORA;
  21:  9c3i-S  4.4  3.1   84   231    7   MOLECULE: CCR4-NOT TRANSCRIPTION COMPLEX SUBUNIT 3;
  22:  6dlm-B  4.4  2.6   65    72   12   MOLECULE: DHD127_A;
  23:  6ezn-B  4.4  3.0   73   110   10   MOLECULE: DOLICHYL-DIPHOSPHOOLIGOSACCHARIDE--PROTEIN
  24:  4hkr-A  4.3  5.1   83   165    8   MOLECULE: CALCIUM RELEASE-ACTIVATED CALCIUM CHANNEL PROTEIN
  25:  8hr5-A  4.3  8.1   64   466   13   MOLECULE: TRANSPOSASE;
  26:  8wt9-A  4.3  3.8   73   319   10   MOLECULE: IS621 TRANSPOSASE;
  27:  7qlr-A  4.3  6.5   73   603   12   MOLECULE: CDHS1_22 PUTATIVE TAIL FIBER PROTEIN;
  28:  7wo9-A  4.3  3.4   86  1581   10   MOLECULE: NUCLEOPORIN NUP188;
  29:  3whj-A  4.3  4.2   73   111   11   MOLECULE: PROBABLE 26S PROTEASOME REGULATORY SUBUNIT P27;
  30:  8ipr-C  4.3  8.2   87   515    5   MOLECULE: COMPONENT LINKED WITH THE ASSEMBLY OF CYTOCHROME'
  31:  3fx7-B  4.3  2.9   68    87    9   MOLECULE: PUTATIVE UNCHARACTERIZED PROTEIN;
  32:  6vbu-2  4.3  6.5   75   659    8   MOLECULE: BARDET-BIEDL SYNDROME 18 PROTEIN;
  33:  2qyw-A  4.3  3.6   77    96   10   MOLECULE: VESICLE TRANSPORT THROUGH INTERACTION WITH T-SNAR
  34:  9f0l-A  4.3  4.9   93   791   11   MOLECULE: DE NOVO DESIGNED PROTEIN K10;
  35:  9o48-A  4.2  3.8  101   373    7   MOLECULE: INTERMEDIATE CONDUCTANCE CALCIUM-ACTIVATED POTASS
  36:  2oer-B  4.2  4.8   88   189   16   MOLECULE: PROBABLE TRANSCRIPTIONAL REGULATOR;
  37:  4j2c-A  4.2  3.8   80   108    8   MOLECULE: SYNTAXIN-6;
  38:  8xyv-A  4.2  3.6   75    99    8   MOLECULE: DE NOVO DESIGNED PROTEIN 0705-5;
  39:  5j9q-D  4.2  4.2   67   120   10   MOLECULE: HISTONE ACETYLTRANSFERASE ESA1;
  40:  8cih-A  4.2  5.2   65   188   15   MOLECULE: CYCLIN-DEPENDENT KINASE 2-INTERACTING PROTEIN;

0001 s001A   ----HKPLSLSSRLDEIARQLGELA--ASRILERW------AKG-E-------------D--L-KKLIEEAEAKVVRFRGMMIALQEELDW-ECY-RHY---G--M-LA-----DDL--R----Y-PG-D-D-LPIGLIERP--EYK--RRWS-----T-ELWADQVR-RASE-NWLL-DRLE
0002 8yadC   npevDAGLLTDIMKQLLFSAKMMFV--K------------------------------------aGQSQDLALCDSYISKVDVLNIL-VAA----------A-YR-----H-V--P----S-LD--------YQLGVE-vsTK--TG--------------------------------
0003 9ivkA   --deeRLKEILFFLLLIIIFVVFLL---iVDYKFL-----eEFK-E---knvtdkeefnE--V-IKIDEAVMLISAFLLAIAAALLKELEE-LIR-KR-------F-EE-----W--------------------RRELEKI--LGV--LLYV-----GaFLMVIAKF-L---eWYEFg----
0004 4k2pA   rllksEIKKLEQKIDMDEKLKKMGE--L-QLSSVT----------------------------D-SKAAATILDQIFVWEQNLEQFQMDLFR-FRC-YLA---S--LggE-----LPN--P----------krllA---------fhalvAART-------------------------------
0005 8ekiA   -----mvVTFLQDLEVLQDALLNNLqkLSAISRRK----------E-------------E--E-EIEFEDLVNIIESKVSDFESVLKCSIV-EMT-YKY---P--E-L---------------------------veeldfvlqfvDWFY-----CvKEILRQHH-KRDL-AWND-EKRD
0006 6snyA   nnweqQKKNIEDDLDRYKKRAEELRkeAEKARKE-----------K-------------E--W-EKRCKELEERARKLEDEAKDRVNDLFD-SNFfQVI---Y--S---------------------------EEEWKK--ekdraeKEIE-----EfKRIKEKCE-EIK-----------
0007 8ap9G   ---sgKLRLYKEKLEGYNRFYSIVKtiKMVTLAKY------RAaqS-------------E--Q-AARLVAVEGQLTNISSLQQRTSSLYNK-TRQ-FGI---T--A-ALieils---------------------------------------------------------------------
0008 8w6eB   -mdeeRLKEILEELERIIEEVKRLL---eKDERLLfyrrdkeEF-R-------------R--V-IKLDEEVMKRSEELLKRAEELLRELEE-LIR--RI---P--F---------------------------RRELE-E--ILRrlKELY--eeaK-RLMEKAKEeIVREe---------
0009 6dkmD   sdeddELERLLREYHRVLREYEKLLeeLRRLYEEY-----kRGE-V-----------seE--E-SDRILREIKEILDKSERLWDLSEEVWR-TLL-YQA---E-----------------------------------------------------------------------------
0010 6y1yB   anridYLLNLVSETVITKASLNQSTieFAELYDKFeyssllleVF-G-------------D--F-DSLMKAAVTKFKSSSQNLGRISGELQE-GVM-KIR---Mvp---------------------------------------------------------------------------
0011 8bbgB   qveideyQNYDKAHGALTEAYKCLA--KAKA-------------------------------KSPLDQETRLAQLQSRMALVKRFIQA-RRT-Y-T---E--D-PK---esIKQ--C--elL-LE-E-PdLD----------------------------------------------
0012 7mx2B   vtdlrVTGMLKDVEDDMQRRVKSTR--S-----------------------------------RDPEVELEHQQCLAVFSRVKFTRVL-LTV-LIA---F--TaVA-----EA-------qklmvqaadllS------------------------------------------------
0013 9clbF   adpkkvLDQAKDQMENVVRTLKQEL---eELAKEA-----rKLD-L----------tqsE--K-IELKLRYIVAHLAAIGDIEEAIREAKE-EAD-KL-------K-RA-----G------------------FDEFKR--rleelhKEAD--rkrA-DYAEEFRN-KLE-----------
0014 3aaiA   ----hlDPKVREEARRRLLSAKGHL--E-GILRXL-----eDEK-V-----------------Y-CVDVLKQLKAVEGALDRVGEXVLRAHL-KDH------------------------------------VEEIVEE-lXEA--LK------------------------------------
0015 6wc3B   ------sMPYATQLALLQDELLDML----EPRDGE-----------------------------glRTADIIDKTLRFRELLGCYRL-QVE-K---------S-TR-----Q------------------QA--palaqlllweRFLA-----DrRRLDAAIV-HEHE-ATAA-R---
0016 4b6xA   ----GAGAALRQEIEDKQLMVNNLT--D-ELQDAI------D---E-------------A--N-PAEIANTSQQLRHARADLADLQRRFAV-LRN-EDR---R--I-NQ-------------------------------------------------------------------------
0017 6dmpB   krlleEAERAHREQKEIIKKAQELH---RRLEEIV-----sGSS-E-------------E--A-KKEAKKILEEIRELSKRSLELLREILY-LSQ-E----------qK-----GSL--V----Pr------------------------------------------------------
0018 5h69A   aaqktELEQHEALLHQARQYRQQTKarQQWLEEMQggaakktAS-L-------------L-sR-NRELEMLSAKLQEMDETIARLERAVAA-KRH-ELA---E--Q-------------------------------------------------------------------------
0019 2gd5D   lkirkeXRVVDRQIRDIQREEEKVK---RSVKDAA------K---K-------------G--Q-KDVCIVLAKEXIRSRKAVSKLYASKAH-XNS-VLX---G--XeVX-----KAXqsL----V-KI-P-E------------------------------------------------
0020 8ibhA   eelseVLQTLQDEFGQMSFDHQQLA--K-LIQESP----------------------------T-VELKDKLECELEALVGRMEAKANQITK-VRK-YQA---Q--L-EKq-----------------------------------------------------------------------
0021 5n77A   dlfetKIEQLADEIENIYSDLEQLSrvIMEGH-------------------------------Q-GDEYDEALSTLAELEDIGWKVRLCLMD-TQR-ALN---F--LqaR-----EIL----------------------rdiesllphnESLF--qkvN-FLM-QAAM-GFIN-IEQN-RIIK
0022 9c3iS   qwltntIDTLNMQVDQFESEVESLS--V-QTRKKK-------------------------------GDKDKQDRIEGLKRHIEKHRYHVRM-LET-ILR---M--LsIL-----VDairkikddV-EY-YvD-SS-----------------------------------------------
0023 6dlmB   eyiikDILDSQEHLLRLIEELLETQkeLLEILKRR------P---D-------------S--V-E-RVRELVRRSKEIADEIRRQSDRNVR-LLE-EVS---K-----------------------------------------------------------------------------
0024 6eznB   iekyPKLKLIDTFCFFLVLLGVIQC---TFIILIR----------------------------D--N-FP-FNAFLAGFIICVGQFVLLMSLRLQ-LCN-S---------fPG-----ISKnrA----F-AEfI-V-AS---------------------------------------------
0025 4hkrA   vqmliAFAICTTLLVAVHMLALMIStcILPN--------------------------------P--H-ERLHWYIETAWAFSTLLGLILFLLEIA-ILC-WVK---F--YaAW-----SATvvL----I-PVmI-I-FM---------------------------------lemLK-EQME
0026 8hr5A   mnesgLEIRTIKDLKDCEKDIDKNK---KEIEKL--------------------------------KISLVENKIEDYKLKIVELNKILEE-TQK---E---R--M-DI-----Q------------------------------------------------------------------
0027 8wt9A   hpleRALRALVVRHQALTDMHTQEL--N-RTET------------------------------aR-EVQRPSIDAHLLWLEAELKRLEKQIKD-LTD-D-D---P--D-M---------------------------RKLLESipgIGE--K-----------------------------
0028 7qlrA   pnlleXKYNLQTSLNYINTIKSNIAsgKGQIX-AI------TY--P-------------I--L-KNAVKNYDGXETTRNISQRDIEEKIKI-SDA-ILE---S--S-IA-----------------------------------------------------------------------
0029 7wo9A   ndrfnVVGKLISIISSVIQNYDIIT--ASSLA-HD------YNN-D------------qD--M-FTIVSLVQLKKFSDLKFILQILQILNL-MIL-NTKvpiL--G-LNtsiaqfdiqsplymdtetfdtvnsal----------------------------------------------
0030 3whjA   qidsfKLSELMVLKTDIETQLEAYF--SVLEQQGI--gmdsaYP-R-------------SdvD-VLQVTMIRKNVNMLKNDLNHLLQRSHV-LLN-QH-------F-DN-----M-------------------SNQ-----------------------------------------------
0031 8iprC   -----SYFQSNVVIAGCTIASAVVL---AHIVAGI-----tNPA-T-------------A-lggETDWAPGLVALAVLWSVRVVAQWFQGR-LSQ-RGA---T--AlRP-----YFTgylpavvlagiltpaalv--------------------------------------vsvGL-RLVF
0032 3fx7B   ---qmDTEEVREFVGHLERFKELLR---EEVNSLS-----hFHN-LeswrdarrdkfseV--L-DNLKSTFNEFDEAAQEQIAWLKERIRV-LEE-DYL---E--H-------------------------------------------------------------------------
0033 6vbu2   rkvlvKVDEYHSVHQKLSADMADNSnlIRSLLVQA-----rDMK-T-----------mkN--R-YKELYDLNKDLLNGYKIRCNNHTELLG-SLK-AVN---Q--A-IQ-----------------------------------------------------------------------
0034 2qywA   ---xaASAASSEHFEKLHEIFRGLLedLQGVPERL-----gTEE-K-------------kK--L-VRDFDEKQQEANETLAEXEEELRYAPL-------------------------------tfrnpxxSKLR-----NrKDLAKLHR-EVR-----------
0035 9f0lA   rtpagLREFYDLRIAYLEKTVADLTarADRVLTD------sVPG-S-----------paE--R-EARAAGLRARAREAQLQLELTRLLRRLrVEN-A--------T-LS-----R------------------------------------------------------------------
0036 9o48A   ralfEKRKRLSDYALIFGMFGIVVM--V-IETELS------WGA-Y-------------D--KaSLYSLALKCLISLSTIILLGLIIVYHA-REI-QL-------F-MV-----D-----------------MTYER-I--FFIclEILVcaihpitttadVDIIkltvR-ACER-YHDQ
0037 2oerB   qyfpNKAAILFRLQSDEWRRTTRLL--G-EILEDT------T---R-------------P--P-LERLRRLVLAFVRSECEEAAIRVALSD-AAP-------------------------------------------lyRD--ADEA--revK-AEG-ARVF-QAFL-REAL-PEVA
0038 4j2cA   gameDPFFVVKGEVQKAVNTAQGLFqrWTELLQDP------S---T-------------A--T-REEIDWTTNELRNNLRSIEWDLEDLDE-TIS-IV-------E-AN-----P-----------------lsirkafiT--STRQvvrdmK-DQMST----------------
0039 8xyvA   --mspEIEDLLKKILELLEKAFALW---aEAKK-----------E-------------G--D-LEKAISTLKELIATIEEVIVLTKKALE-LAE-KEG---N--P-----------------------------eiveqakkllDLAE-----AakAELARAL-SSl-----------
0040 5j9qD   rylleEIGSNDLKLIEEKKKYEQKE--S-QIHKFI-----qGSI-P-------------K--H-P-QEDGLDKEIKESLLKCQSLQREKCV-LAN-TAL---F--L-------------------------------------------------------------------------
0041 8cihA   etlndAGFTTANNIANLKISLLNK----------------------S-------------S--S-PASKENEEKVCLEYNEELEKLCEELQA-TLD-GLT---K--I-QV---------------------------LLL---------------------------------------

0001 s001A   ----LLLHHHHHHHHHHHHHHHHHLL--HHHHHHHL------LLL-L-------------L--H-HHHHHHHHHHHHHHHHHHHHHHHHHHHHH-HHH-HHL---L--L-LL-----LLL--L----L-LL-L-L-LLHHHHLLL--LLL--LLLL-----L-LLHHHHHH-HHHH-HHHH-HHHH
0002 8yadC   llllLHHHHHHHHHHHHHHHHHHHH--L-------------------------------------lLLHHHHHHHHHHHHHHHHHHHHH-HLL----------L-LL-----L-L--L----L-HH--------HHHHHH-hhHH--HL-------------------------------
0003 9ivkA   --lhhHHHHHHHHHHHHHHHHHHHH---hHHHHHH-----hHHH-H---llllllhhhhhH--H-LHHHHHHHHHHHHHHHHHHHHHHHHHHHH-HHH-HH-------H-HH-----H---------------------HHHHHHH--HHH--HHHH-----HhHHHHHHHH-H---hHHHHl----
0004 4k2pA   hhhhhHHHHHHHHHHHHHHHHHHHH--H-HLLLLL----------------------------L-HHHHHHHHHHHHHHHHHHHHHHHHHHHHH-HHH-HHH---H--HllL-----LLL--H----------hhhhL---------hhhhhHLLL-------------------------------
0005 8ekiA   -----llLLHHHHHHHHHHHHHHHHhhHHHHHHHH------------H-------------H--H-HHHHHHHHHHHHHHHHHHHHHHHHHHHHH-HHH-HHL---L--L-L---------------------------hhhhhhhhhhhHHHH-----HhHHHHHHHH-HHHH-HHHH-HHHH
0006 6snyA   lhhhhHHHHHHHHHHHHHHHHHHHHhhHHHLLLL-----------L-------------L--L-LHHHHHHHHHHHHHHHHHHHHHHHHHHHH-HLHhHHH---L--L---------------------------HHHHHH--hhhhhhHHHH-----HhHHHHHHHH-HHL-----------
0007 8ap9G   ---llHHHHHHHHHHHHHHHHHHHHhhHHHHHHHH------HHhlH-------------H--H-HHHHHHHHHHHHHHHHHHHHHHHHHHHHH-HHH-HHH---H--H-HHhhhhl-----------------------------------------------------------------
```

```
0008 8w6eB   -lhhhHHHHHHHHHHHHHHHHHHH---hHHHHHHhhhllhhHH-H-------------H--H-HHHHHHHHHHHHHHHHHHHHHHHHHHH-HHH--HL---L--L-----------------------------HHHHH-H--HHHhhHHHH--hhhH-HHHHHHHHhHHHHl----------
0009 6dkmD   hhhhhHHHHHHHHHHHHHHHHHHHhhHHHHHHHH-----lLLL-L-----------lhH--H-HHHHHHHHHHHHHHHHHHHHHHHHHHH-HHH-HHH---L---------------------------------------------------------------------------------
0010 6y1yB   hhhhhHHHHHHHHHHHHHHHHHHHhhHHHHHHHHhhhhhhlLL-L-------------L--H-HHHHHHHHHHHHHHHHHHHHHHHHHHH-HHH-HHL---Lll-------------------------------------------------------------------------------
0011 8bbgB   hhhhhllLLHHHHHHHHHHHHHHHH--HLLL----------------------------------LLHHHHHHHHHHHHHHHHHHHHHHHH-HHH-L-L---L--L-HH---hhHHH--H--hhH-HL-L-LlHH-------------------------------------------------
0012 7mx2B   llhhhHHHHHHHHHHHHHHHHHHLL--L--------------------------------------LLHHHHHHHHHHHHHHHHHHHHHHH-HHH-HHH---H--HhHH-----LH-------hhhhhhhhhhhH-------------------------------------------------
0013 9clbF   lhhhhhHHHHHHHHHHHHHHHHHHH---hHHHHHH-----lLLL-L----------lhhH--H-HHHHHHHHHHHHHHHHHHHHHHHHHHH-HHH-HH-------H-HH-----L--------------------HHHHHH--hhhhhhHHHH--hhhH-HHHHHHHH-HHL------------
0014 3aaiA   ----llLHHHHHHHHHHHHHHHHHH--H-HHHHHH-----lLLE-E----------------L-HHHHHHHHHHHHHHHHHHHHHHHHHHH-HHL----------------------------------------HHHHHHH-hHHH--HL---------------------------------
0015 6wc3B   ------lLLHHHHHHHHHHHHHHHL----LLLLLL---------------------------------llLHHHHHHHHHHHHHHHHHHHH-HHH-L--------L-LL-----L--------------------LH--hhhhhhhhhhHHHH-----HhHHHHHHHH-HHHH-HLLL-L----
0016 4b6xA   ----LLLHHHHHHHHHHHHHHHHHH--H-HHHHHH------H---L-------------L--L-HHHHHHHHHHHHHHHHHHHHHHHHHHH-HHH-HHH---H--H-HL---------------------------------------------------------------------------
0017 6dmpB   lhhhhHHHHHHHHHHHHHHHHHHHH---HHHHHHH-----hLLL-H-------------H--H-HHHHHHHHHHHHHHHHHHHHHHHHHHH-HHL-L----------lL-----LLL--L----Ll----------------------------------------------------------
0018 5h69A   hhhhhHHHHHHHHHHHHHHHHHHHhhHHHHHHHHellllllLL-H-------------H-hH-HHHHHHHHHHHHHHHHHHHHHHHHHHH-HHH-HHH---H--H------------------------------------------------------------------------------
0019 2gd5D   hhhhhhHHHHHHHHHHHHHHHHHHH---HHHHHHH------H---L-------------L--L-HHHHHHHHHHHHHHHHHHHHHHHHHHH-HHH-HHH---H--HhHH-----HHHhhL----L-LL-L-L----------------------------------------------------
0020 8ibhA   hhhhhHHHHHHHHHHHHHHHHHHHH--H-HHHHLL---------------------------L-HHHHHHHHHHHHHHHHHHHHHHHHHHH-HHH-HHH---H--H-LLl--------------------------------------------------------------------------
0021 5n77A   hhhhhHHHHHHHHHHHHHHHHHHHhhHHHLL------------------------------L-LHHHHHHHHHHHHHHHHHHHHHHHHHH-HHH-HHH---H--HhhH-----HHH----------------------hhhhhhhhhhHHHH--hhhH-HHH-HHHH-HHHH-HHHH-HHHL
0022 9c3iS   hhhhhhHHHHHHHHHHHHHHHHHHH--H-HHHHHL-------------------------------LLHHHHHHHHHHHHHHHHHHHHHHH-HHH-HHH---H--HlLL-----HHhhhhhhhhH-HH-HhH-LL-------------------------------------------------
0023 6dlmB   llhhhHHHHHHHHHHHHHHHHHHHhhHHHHHHLL------L---L-------------L--L-H-HHHHHHHHHHHHHHHHHHHHHHHHH-HHH-HHH---L---------------------------------------------------------------------------------
0024 6eznB   hhhlHHHHHHHHHHHHHHHHHHHHH---HHHHHLL------------------------L--L-LL-LHHHHHHHHHHHHHHHHHHHHHHH-HHL-L---------lLL-----LLLlhH----H-HHhH-H-HH-------------------------------------------------
0025 4hkrA   hllhhHHHHHHHHHHHHHHHHHHHhhLLLL----------------------------L--H-HHHHHHHHHHHHHHHHHHHHHHHHHHH-HHH-HHH---H--LhHH-----HHHhhL----H-HHhH-H-HH--------------------------------------hhhHH-HHHH
0026 8hr5A   hhlhhHHHHHHHHHHHHHHHHHHHH---HHHHHL--------------------------------LHHHHHHHHHHHHHHHHHHHHHHHH-HLL---L---L--L-LH-----H---------------------------------------------------------------------
0027 8wt9A   lhhhHHHHHHHHHHHHHHHHHHHHH--H-HHHH----------------------------lL-LLLHHHHHHHHHHHHHHHHHHHHHHHH-HHH-H-L---H--H-H---------------------------HHHHLLlllLLH--H----------------------------------
0028 7qlrA   hhhhhHHHHHHHHHHHHHHHHHHHhhHHHHH-LL------EE--H-------------H--H-HHHHHHHHHHHHHHHHHHHHHHHHHHH-HHH-HHL---L--H-HH---------------------------------------------------------------------------
0029 7wo9A   llhhhHHHHHHHHHHHHHHLLLLLH--HHHHH-LL------LLL-L------------hH--H-HHHHHHHHHHHHHHHHHHHHHHHHHHH-HLL-LLLllhL--L-EElllleellllhhhhlllhhhhhhhhh-------------------------------------------------
0030 3whjA   lllllLHHHHHHHHHHHHHHHHHHH--HHHHHLLL--lllllLL-L-------------LhhH-HHHHHHHHHHHHHHHHHHHHHHHHHHH-HHH-HH-------H-HH-----L--------------------LLL----------------------------------------------
0031 8iprC   -----LHHHHHHHHHHHHHHHHHHH---HHHHHHH-----hLLL-L-------------L-lllLLLLHHHHHHHHHHHHHHHHHHHHHHH-HHH-HHH---H--HhHH-----HHLllhhhhhhhhhhhhhhhh--------------------------------------hhhHH-HHHL
0032 3fx7B   ---llLHHHHHHHHHHHHHHHHHHH---HHHHHHH-----hHHH-LlllllhhhhhhhhH--H-HHHHHHHHHHHHHHHHHHHHHHHHHHH-HHH-HHH---H--L------------------------------------------------------------------------------
0033 6vbu2   hhhhhHHHHHHHHHHHHHHHHHHHhhHHHHHHHH-----lLHH-H-----------hhH--H-HHHHHHHHHHHHHHHHHHHHHHHHHHH-HHH-HHH---H--H-HH---------------------------------------------------------------------------
0034 2qywA   ---lhHHHHHHHHHHHHHHHHHHHhhHHHHHHHH-----lLHH-H------------hH--H-HHHHHHHHHHHHHHHHHHHHHHHLLLH---------------------------------------------------hhhhhhhHHHH-----HhHHHHHHHH-HHL------------
0035 9f0lA   llhhhHHHHHHHHHHHHHHHHHHHhhHHHHHHL------lLLL-L-----------hhH--H-HHHHHHHHHHHHHHHHHHHHHHHHHHHhLLL-L--------L-LL-----H---------------------------------------------------------------------
0036 9o48A   hhhhHHHHHHHHHHHHHHHHHHHHH--H-HHHHHH------HHL-L-------------L--LlLHHHHHHHHHHHHHHHHHHHHHHHHHH-HHH-HH-------H-HH-----H--------------------LLHHH-H--HHHhhHHHHhllllleeeeeLLHHllhhH-HHLL-LLLL
0037 2oerB   hhllLHHHHHHHHHHHHHHHHHHHH--H-HHHLLL------L---L-------------L--H-HHHHHHHHHHHHHHHHHHHHHHHHHHH-HLL------------------------------------------------llLL--LHHH--hhhH-HHH-HHHH-HHHH-HHHL-LLLL
0038 4j2cA   llllLLHHHHHHHHHHHHHHHHHHHhhHHHHHHLL------L---L-------------L--L-HHHHHHHHHHHHHHHHHHHHHHHHHHH-HHH-HH-------H-HL-----H-----------------------hhhhhhhhH--HHHHhhhhhH-HHHHL-------------------
0039 8xyvA   --llhHHHHHHHHHHHHHHHHHHHH---hHHHH------------L-------------L--L-HHHHHHHHHHHHHHHHHHHHHHHHHHH-HHH-HHL---L--H---------------------------------hhhhhhhhhhHHHH-----HhhHHHHHHH-HHl------------
0040 5j9qD   hhhhhHHHHHHHHHHHHHHHHHHHH--H-HHHHHH-----hLLL-L-------------L--L-L-LHHHHHHHHHHHHHHHHHHHHHHHH-HHH-HHH---H--H------------------------------------------------------------------------------
0041 8cihA   hhhhhHHHHHHHHHHHHHHHHHLL---------------------L-------------L--L-HHHHHHHHHHHHHHHHHHHHHHHHHHH-HHH-HHH---H--H-HH--------------------------HHH----------------------------------------------
```

```
# Job: Type-1 BREX; BrxX-MTase MBZ5596028 wHTH (Type-1 BREX Specific)
# Query: s001A
# No:  Chain    Z    rmsd lali nres  %id PDB  Description
   1:  8c45-A  6.8  1.9  109  1219   33   MOLECULE: SITE-SPECIFIC DNA-METHYLTRANSFERASE (ADENINE-SPEC
   2:  3s93-B  5.6  3.4   75    81    9   MOLECULE: TUDOR DOMAIN-CONTAINING PROTEIN 5;
   3:  2od5-A  5.4  2.7   67    91   12   MOLECULE: HYPOTHETICAL PROTEIN;
   4:  8e4y-A  5.0  4.0   83   651    6   MOLECULE: GLYCEROL-3-PHOSPHATE ACYLTRANSFERASE 1, MITOCHOND
   5:  8ffz-B  4.9  4.2   78   905   10   MOLECULE: TRANSCRIPTION FACTOR IIIA;
   6:  9gm7-C  4.8  3.4   69   440   14   MOLECULE: CHROMOSOME PARTITION PROTEIN MUKF;
   7:  1ldd-A  4.7  3.0   70    74    4   MOLECULE: ANAPHASE PROMOTING COMPLEX;
   8:  5yad-A  4.7  2.7   62    73    5   MOLECULE: MEIOSIS REGULATOR AND MRNA STABILITY FACTOR 1;
   9:  7asv-A  4.6  3.1   71   155   10   MOLECULE: DNA-DIRECTED RNA POLYMERASE III SUBUNIT RPC5;
  10:  7vwv-B  4.6  2.6   64    74    3   MOLECULE: I73R;
  11:  2lh9-A  4.6  3.0   69    78   14   MOLECULE: TUDOR DOMAIN-CONTAINING PROTEIN 7;
  12:  6sj9-A  4.5  3.4   69   642    9   MOLECULE: PROTEASOME ACCESSORY FACTOR B/C (PAFBC);
  13:  7z8b-C  4.5  4.8   83  1224   13   MOLECULE: CULLIN-7;
  14:  2zme-B  4.5  3.6   77   215   12   MOLECULE: VACUOLAR-SORTING PROTEIN SNF8;
  15:  6r7n-B  4.5  3.0   68   443    4   MOLECULE: COP9 SIGNALOSOME COMPLEX SUBUNIT 1;
  16:  6aht-A  4.5  4.4   77   111    4   MOLECULE: CONSERVED HYPOTHETICAL PLASMID PROTEIN;
  17:  7abi-7  4.2  3.4   74   236    5   MOLECULE: U5 SMALL NUCLEAR RIBONUCLEOPROTEIN 40 KDA PROTEIN
  18:  3zco-A  4.2  3.9   80   127   10   MOLECULE: REGULATORY PROTEIN SIR3;
  19:  6wge-C  4.2  2.5   61   174    8   MOLECULE: STRUCTURAL MAINTENANCE OF CHROMOSOMES PROTEIN 1A;
  20:  7cv0-A  4.2  2.5   57   173    9   MOLECULE: TRANSCRIPTIONAL REGULATOR NIAR;
  21:  7nyw-E  4.2  3.6   79   212    8   MOLECULE: CHROMOSOME PARTITION PROTEIN MUKB;
  22:  8uuc-A  4.2  3.3   72   274    6   MOLECULE: ADENINE DNA GLYCOSYLASE;
  23:  7qen-C  4.1  3.0   72   291    7   MOLECULE: DNA (35-MER);
  24:  2iu5-A  4.1  3.2   72   179    6   MOLECULE: HTH-TYPE DHAKLM OPERON TRANSCRIPTIONAL ACTIVATOR
  25:  2obp-A  4.1  2.7   62    81   10   MOLECULE: PUTATIVE DNA-BINDING PROTEIN;
  26:  3qkx-A  4.0  3.1   72   183    7   MOLECULE: UNCHARACTERIZED HTH-TYPE TRANSCRIPTIONAL REGULATO
  27:  7yw2-A  4.0  3.5   66   216    9   MOLECULE: TRNA 2'-PHOSPHOTRANSFERASE 1;
  28:  1w5s-B  4.0  3.8   76   396    7   MOLECULE: ORIGIN RECOGNITION COMPLEX SUBUNIT 2 ORC2;
  29:  2lnb-A  4.0  3.6   62    80    8   MOLECULE: Z-DNA-BINDING PROTEIN 1;
  30:  3elk-A  3.9  2.9   64   104    8   MOLECULE: PUTATIVE TRANSCRIPTIONAL REGULATOR TA0346;
  31:  6qpq-D  3.9  2.8   66    82   12   MOLECULE: STRUCTURAL MAINTENANCE OF CHROMOSOMES PROTEIN,STR
  32:  7jgr-D  3.8  4.3   77   441    9   MOLECULE: ORIGIN RECOGNITION COMPLEX SUBUNIT 2;
  33:  5d1w-D  3.8  4.2   81   196    4   MOLECULE: RV3249C TRANSCRIPTIONAL REGULATOR;
  34:  4nb5-B  3.8  3.8   65   149    5   MOLECULE: DNA BINDING PROTEIN;
  35:  7wze-A  3.8  4.1   64   161    6   MOLECULE: UNCHARACTERIZED HTH-TYPE TRANSCRIPTIONAL REGULATO
  36:  8r7k-B  3.8  3.1   72   366    4   MOLECULE: GERMINAL-CENTER ASSOCIATED NUCLEAR PROTEIN;
  37:  1yyv-A  3.8  3.6   69   114    7   MOLECULE: PUTATIVE TRANSCRIPTIONAL REGULATOR;
  38:  8hih-N  3.8  2.7   61   217    2   MOLECULE: DNA-DIRECTED RNA POLYMERASE SUBUNIT ALPHA;
  39:  5g5p-B  3.8  3.4   79   455    6   MOLECULE: NUCLEAR MRNA EXPORT PROTEIN SAC3;
  40:  7z1n-O  3.8  2.7   69   570    9   MOLECULE: DNA-DIRECTED RNA POLYMERASE III SUBUNIT RPC1;
  41:  4y66-C  3.8  3.0   62   197    2   MOLECULE: MND1;
  42:  5xfo-A  3.8  3.9   80   315   10   MOLECULE: PHD FINGER PROTEIN 1;
  43:  8fo9-F  3.8  3.5   72  2289   11   MOLECULE: LEUCINE-RICH REPEAT SERINE/THREONINE-PROTEIN KINA
  44:  4kyw-A  3.8  3.4   68   254    7   MOLECULE: TYPE-2 RESTRICTION ENZYME DPNI;
  45:  4i99-C  3.8  2.6   62    70    5   MOLECULE: CHROMOSOME PARTITION PROTEIN SMC;
  46:  6pco-B  3.7  4.1   64   135   16   MOLECULE: MARR-FAMILY TRANSCRIPTIONAL REGULATOR;
  47:  6wg3-C  3.7 23.2   65   248    9   MOLECULE: STRUCTURAL MAINTENANCE OF CHROMOSOMES PROTEIN 1A;
  48:  5z7b-B  3.7  3.0   61   197   15   MOLECULE: PADR FAMILY TRANSCRIPTIONAL REGULATOR;
  49:  9baq-A  3.7  3.9   83  1002    4   MOLECULE: DNA (CYTOSINE-5-)-METHYLTRANSFERASE;
  50:  5uuj-A  3.7  3.5   70   371   13   MOLECULE: ALKZ;
```

```
0001 s001A   --VEEL--D-E--SS-AA-AV--V--ASY-AI-GCCFGRWD------WDGILVDD--EGHARDVV-ARVR-EVLRVVWKD-R---ADEIE-DAVCKTLGV-----KDLR-SYFRRNFFEDHIKRYS---KSRRRAP-VYWL-L-Q--SP-RK--S-Y--GVW-LY-----------
0002 8c45A   -tLLCN-kY-Y--TD-IT-ID--I--LSY-II-GCMMGRYSlda--dNDGILPLMddEWFDDDVT-SRVK-EFVRTVWGEeH---LQENL-EFIAESLClyasalDTIR-RYLSTQFWKDHMKMYK-------KRP-IYWL-F-S--SG-EK--A-F--ECL-VYlhryndatlar
0003 3s93B   ----GM--S-E--QE-RI-QE--C--LRK-EI-RSLLIS--------tKDGLSPQ-------ELE-KEYL-LMVG----------------hlPLRILGY-----R-----STME-LVLDMP-------------D-VVRV-C-P--GA-G---G-T--VIL-KAip---------
0004 2od5A   --------E-T--ES-XK-TV--R--IRE-KI-KKFLG----------DRPRNT-------aEIL-EHIN-ST-------------XRHG----------------tTS-QQLG-NVLSKD--------------K-DIVKvG-Y--IK-RsgyD-I--CEW-ATrnwvaehcpew
0005 8e4yA   -fnSFF--W-N-nyG-CV-RvdiM--STH-IV-ACLLLY-------rhRQGIDLS-------TLV-EDFF-VMKEEVLAR-D---FD----LGFS----------GNSE-DVVM-HAIQLLG-------------N-CVTI-T-H--TS-R--nD-E--FFI-TPsttvpsvfeln
0006 8ffzB   -lFRIY--D---eRS-LR-SL--Q--RQR-AI-LKVMNT--------iGGVAYLR--E----QFY-ESVS-KYM-----------GSTT----------------tLDK-KTV-rGDVDLMVES-----------E-KLGA-R-T--EP-VS--G-R--KII-FLptvgedaiqry
0007 9gm7C   ------------iSL-PVeRL--A--FLM-AI-AVLNS-----e--rLDGEMSEG-------ELI-DAFR-EVCK----------GFEQ-------------svaVRAN-NAI-NDMVRQK----------------lLNR-F-T--SE-L--dG-N--AIY-RLtplgisisdyy
0008 1lddA   ------------kYE-LT-LQ--R--SLP-FI-EGMLTN---------lGAMKL-------hKIH-SFLK-ITV-----------PKDW---GYNR----------ITL-QQLEGYLNTLADE------------G-RLKY-I-A--N-----------GSY-EIv----------
0009 5yadA   --------G-A--AN-KS-LS--L--LST-ET-XSILQ-----d--aPACCLPLF-------KFI-DIYE-KKYG-----------------------------hklNV-SDL-YKLTD------------------tIAI-R-E--QG-----N-G--RLV-CLlpsnq------
0010 7asvA   -------------aptPV-AR--E--LKA-FV-EATFQR---------qFVLTLS-------ELK-RLFN-LHLAS-LPP-G-----HTL-FS------------gISD-RXLQ-DTVLAA----------------GCKQ-I-L--VP-F----eQ--KVF-ALwesgdxsdqhr
0011 7vwvB   -------------EF-ME-TQ--K--LIS-MV-KEALE--------kYQYPLTA-------kNIK-VVIQ-KEHN-----------------------------vvLPT-GSIN-SILYSNS-------------E-LFEK-I-D--KT------yP--PLW-IRkn---------
0012 2lh9A   -------------ghmLE-AD--L--VSK-ML-RAVLQS--------hKNGIVLP-------RLQ-GEYR-SLTG----------------wiPFKQLGY-----P-----TLEAYLRSVP--------------A-VVRI-E-A--SR-S---G-E--IVC-YAva---------
0013 6sj9A   --------------G-AS-RT--E--RLL-NL-LLALLN--------tKVGLPRA-------VLR-EKVY-----------------hDS-AD----------ndVAFG-RXF-ERDKVDLKQFG----------F-EIET-L-X--D---P--A-S--ARY-RIgkdsnrlpdvs
0014 7z8bC   -qAEGE-gQ-N--LE-KR-RN--L--LNC-LI-VRILKA-------hGDEGLHID-------QLV-CLVL-EAWQKGPC-----pPRGLV-sslgKGSA-------cSS-TDV-lSCILHLLG-----------KG-TLRR-H-D--D----------pQVL-SYav---------
0015 2zmeB   -eDETI------sGT-QY-HM--Q--LAK-QLaGILQV-----e--eRGGIMSLT-------EVY-CLVN-RA------------RGMEL----------------LSP-EDLV-NACKMLEALK----------L-PLRL-R-V--FD-SG----V--MVI-ELqshkeeemvas
0016 6r7nB   -rEGEF-eH-I--EE-LL-RN--I--RTQ-VL-IKLIK---------PYTRIHIP-------FIS-KELN------------------------------------iDV-ADVE-SLLVQCILDN---------tI-HGRI-D-Q--V--------N--QLL-ELdhqkrggaryt
0017 6ahtA   -xSSSE-iD-V--LC-EN-LN--D---GI-WA-LRVLYA---------eGAXNKE-------KLW-DYIN-QYHKDYQIEnekdyeGKKI------------lpsRYAL-DIX-tARLEGAG---------------lISF-K-A--IG-----R-V--RIY-DVtdlgnvlikel
0018 7abi7   -------qQ-F--MD-YF-SE--E--FRNdFL-ELLRR--------rfGTKRVH---------NN-IVYN-EYISHRE--------hiHM-NAT----QW-----ET-L-TDFTKWLGR----------------E-GLCK-V-D--ET-P--kG-W--YIQ-YIetirrqlelek
0019 3zcoA   ------------iskkiigmS--L--LMR-TF-LYTLAQ----g--tNRHTLALE-------TVL-IKMV-KMLRD-NPG-Yk-aSKEIK-KVI--------itiEKLKqFSWI-SVVNDLVGE-----------K-LVVV-V-L--EE-PS--A-S--IMV-ELklpleinyafs
0020 6wgeC   ------------reimKR-TQ--Q--MLH-GL-QRALA-----k--tGAESISLL-------ELC-R-----------------------------------ntnRKQA-AAKF-YSFLVLKKQ-----------Q-AIEL-T-Q--EE-P--yS-D--IIA-TPgprfhi-----
0021 7cv0A   ----------------IL-GE--E--RRS-LL-IKWLKA--------sDTPLTGA-------ELA-KRTN------------------------------------vSR-QVIV-QDVSLLKAKN----------H-PILA-T-A--------------qGY-IYmkeantvqaqr
0022 7nywE   -iPRSV-------lS-EL-DM--M--VGK-IL-CYLYLS----l--aNQGIFTSQ-------ELY-EELI-SLA-------D---EGKLM-KF-VNQR----ldkQKLQ-EKVR-TTLNRLRRL-----------G-MVYF-L---------------nNKF-TIteavfrfgadv
0023 8uucA   -hHTRQ--S-T--F-egS-RR--Q--KRA-EL-VRLVLA---------ePGIGID-------ELA-ERLD-AFERD---A-G---RK------------------gVDA-ATFTSIVADLVAE------------G-FFRR-E-G--------------DAF-FA-----------
0024 7qenC   ---------ewivskkvD-VR--R--LKK-NV-WRSINN----srknreKELKFS-------DII-QGIS-KMY-----------SDDTL-KD-------------IST-SFC-FICLLHLANE-----------H-GLQI-T-H--TE-NY--N-D--LIV-NYedla-------
0025 2iu5A   ----mE--K-S--I--IT-QK--I--IAK-AF-KDLMQ----------sNAYH--------QISV-SDIM-QTAK---------iRRQTF-YNYF-----------QNQ-EELLSWIFENDFAELInqnELLLL--yqkifvI-D--KNf--------------------------
0026 2obpA   ----------G--ID-PA-IV--E--VLL-VL-REAGI-----e--nGATPWSL-------pKIA-KRAQ------------------------------------lPX-SVLR-RVLTQLQAA-----------G-LADV-S-V--EA-DG---R--GHAs-------------
0027 3qkxA   -----R--Q-A--KT-DL-AE--Q--IFS-AT-DRLXA-----------REGL--------NQLSxLKLA-KEAN---------vAAGTI-YLYF----------KNKD-ELL-EQFAHRVFSXFX--aTLEKdfdnlkQY-E-Sl------------------------------
0028 7yw2A   -------------EQ-DR-NV--Q--LSK-AL-SYALRHG---m--rADGFVPLQ-------ALL-Q--L-PQF------------------------------hsfSI-EDV-QLVVNTNE------------KQ-RFTL-Q-P--GE-P--sT-G--LLI-RAnqghslqvpel
0029 1w5sB   -qsMLS--StH--EL-EA-LS--I--HEL-II-LRLIAE----l--gGMEWINAG-------LLR-QRYE-DASLTMY-------NVKPR-------------gyTQYH-IYL-KHLTSLG----------------lVDA-K-P--S--------T--TLF-RLaphlpadrlie
0030 2lnbA   hhhshM--A-D--P--GR-EG--H--LEQ-RI-LQVLT--------eAGSPVKLA-------QLV-KECQ------------------------------------aPK-RELN-QVLYRMKKE-----------L-KVSL-T-S--P-----------ATW-CLggtdpe-----
0031 3elkA   ----------R--ER-IL-HG--L--ITL-YI-LKELV----------kRPXHG-------yELQ-KSXF-ETTG------------------------------qalpqGSIY-ILLKTXKER-----------G-FVIS-E-S--SV-N--qQ-L--TVY-HItdagkkflxdh
0032 6qpqD   ----------------KA-IV--Q--MAK-IL-RKELS---------EEKEVIFT-------DVL-KSQA-N------------------------------nitKREA-SRGF-FDILSLATE-----------G-CIGL-S-Q--TE-AF--G-N--IKI-DAkpalferfi--
0033 7jgrD   -fEGDD---lC--GL-SVlEL--C--LII-AI-KHHSQI-------yDRDSFNFE-------IIY-ARFS-KFAKV--ST-T---MQAV--------------erSIVL-KAF-EHLRIAE----------------lIMP-L-------VQ--K-E-fEMH-KLaltysqihhcm
0034 5d1wD   --IPYA-eA-S--RA-LL-RD--S--VLD-AM-RDLLL----------tRDWS--------AITL-SDVA-RAAG---------iSRQTI-YNEF----------GSRQ-GLA-QGYALRLADRLV--dNVHAsldaDPLVislI--TT-DS--An--------------------
0035 4nb5B   ---------eS--RS-L---T--R--LAG-RL-LGWLLV-------cDPERQSSE-------ELA-TALA------------------------------------aSS-GGIS-TNARMLIQF-----------G-FIER-L-A--VA-GD--R-R--TYF-RLrpnafaagere
0036 7wzeA   -------eH-Y--FAgRG-LS--E--GKF-KI-LXLLFD-------aKDHRLSP-------tELA-KRSN------------------------------------vTK-ATIT-GLLDGLARD-----------G-FVSR-R-H--H------R-K--ISI-ELttegkarleqf
0037 8r7kB   cgifLI--L-E--K--LK-II--T--YRN-LF-KKVYL-----l--lKTHQLSLD-------AFL-VALK-F-------------MQVE----------------dVDI-DEVQCILANLIYMGH----------V-KGYI-S-H--Q--------H--QKL-VVskqnpfppl--
0038 1yyvA   -aEQCP--S-R--EV-LK-HV--TsrWGV-LI-LVALR----------dGTHRFS-------DLR-RXXG-----------------------------------gVSE-XXLA-QSLQALEQD-----------G-FLNR-V-S--YP-V--pP-H--VEY-SLtplgeqvsdxv
0039 8hihN   ------------dlT-YK-EF--E--LLK-YL-AQHA-----------gRVFTRA-------QLL-HEVW-G------------------------------ydffggT-RTV-DVHVRRLRAKLG------PEHEaLIGT-V-R-----------N--VGY-KAvrpa-------
0040 5g5pB   -lNSQVhvL-L--EK-LP-MV--T--YRN-LI-KTVIKS----e--wGQNKLPYS-------LIE-RVLQ-LS-----------iGPTFedNGIH------------SP-KNVE-NVLVTLINLG---------lL-RANC-F-P--Q--------L--QLC-VVkkttmiqeivp
0041 7z1nO   ---------kktrvG-SV-TA--Q--VYK-IA-LRLTEqkspkeektpGLTFNA-------iDLA-RHLP-----------------AEL-D--LRGSL---srkPHSA-SLIN-SHLKILASS---------NFP-FLNE-T-K-------------pGVY-YVpysklmpvlks
0042 4y66C   --------T-S--LD-EK-KE--R--LLE-EM-LKRG------------EIYSNK-------TIE-T-LS-KPT----------------------------issMVIK-NVL-QALVNEDL----------------VDT-D-K--IG-----A-S--TYY-WCfaskrsqaart
0043 5xfoA   ---------revrlqlRW-VD--V--AHL-VL-YHLSVC-------cKKKYFDFD--R----EIL-PFTS-ENWDSLL-------LGEL--SDTP---------kGERS-SRLL-SALNSHK-------------D-RFIS-G-R--EI-KK--R-K--CLF-GLharmpppve--
0044 8fo9F   qqrlKK--A-VplIP-DC-YV--E--LEK-II-LSERKN----p--iEFPVIDRK-------RLL-QLVR-ENQ------------------------------lqLDE-NEL-PHAVHFLNES-----------G-VLLH-F-Q--DP-A----lS--DLY-FVepkwlckimaq
0045 4kywA   -ksSLS--S-R-----GW-TI--E--ILN-CI-DKIE-----------GSEFTL--------EDM-YRFE-------------sdlKNIF-VK-----------nNHIK-EKIR-QQLQILRD-----------KE-IIEF-K-G-----------R--GKY-RKl----------
```

```
0046 4i99C   ------------------D-IE--K--YVE-EL-YKVVKK----e--kTGTPIKFW-------DLV-P-----------------------------------dvePKII-ARTF-LYLLFLENM-----------G-RVEI-I-Q--EE-PF--G-E--ILV-VPm---------
0047 6pcoB   aifqQT--I-P--DS-K--LT--A--AQF-VV-LCALRD---------qGACSLV-------DVV-KATA-----------------------------------iDQ-ATVR-GVIERLKAR-----------K-LLAV-S-H--RR-------K--VLV-TLtpdgralveem
0048 6wg3C   -------ferC--LT-KR-TQ--Q--MLH-GL-QRALA-----k--tGAESISLL-------ELC-R-------------------------ntnR-----KQA--AAKF-YSFLVLKKQ-----------Q-AIEL-T-Q--EE-PY--S-D--IIA-TPgprfhi-----
0049 5z7bB   -------------SA-KD-LM--T--LRS-AL-LALLS----------sGPLTG-------yDAS-QRFGaSVGFV-----------------------------wsgSD-SQIY-PELRKMEAE-----------E-LLVG-S-D--V----------ktEY-ALsekgwealrka
0050 9baqA   feyqRY--Y-Q--PF-LW-IA--D--LGK-HV-VDYCTR----v--eRKREVTLG-------CFK-SDFI-QWASKAHGK-Sk-aFQNWR-AQ-HPSD--------dFR-TSVA-ANIGYIWKE------------gdqLF-R-EliHT-YM--A-Ntx----------------
0051 5uujA   ------------vTP-RQ-VD--E--LTE-EV-AGILD----------GVVLTRD-------ELV-TRLV-A---------------dKRF-VS----------xeERLR-SGW-GSVLKPLAWR-----------G-VLCH-GpN--RG-N-----K--ITF-TLpasqfgadwgk

0001 s001A   --LLLL--L-H--HH-HH-HH--H--HHH-HH-HHHLLLLL------LLLEEELL--LLLLLLHH-HHHH-HHHHHHHHH-H---HHHHH-HHHHHHHLL-----LLHH-HHHHHLHHHHHHHHLE---ELLEELL-LLEE-E-E--LL-LL--L-L--EEE-EL----------
0002 8c45A   -lLLLL-hH-H--HH-HH-HH--H--HHH-HH-HHHHLLLLlll--lLLLEEELLllLLLLLLHH-HHHH-HHHHHHHLLlL---HHHHH-HHHHHHHLllllhhHHHH-HHHHHLHHHHHHHHLL-------LLL-LEEE-E-E--LL-LL--L-L--EEE-EElllllllhhhh
0003 3s93B   ----LH--H-H--HH-HH-HH--H--HHH-HH-HHHHHL--------lLLLLLHH-------HHH-HHHH-HHHL---------------llLLHHHLL-----L-----LHHH-HHHLLL------------L-LEEE-E-E--LH-H---H-L--EEE-EEll---------
0004 2od5A   --------L-L--HH-HH-HH--H--HHH-HH-HHHLL---------LLLEEH-------hHHH-HHHH-LL-------------LLLL-----------------lLH-HHHH-HHHHHL------------L-LEEEeE-E--EE-EeeeE-E--EEE-EEhhhhhhhhllll
0005 8e4yA   -hhHHL--L-L-llL-LE-EeelL--HHH-HH-HHHHHH-------llLLLEEHH-------HHH-HHHH-HHHHHHHHL-L---LL----LLLL----------LLHH-HHHH-HHHHHLL------------L-LEEE-E-E--LL-L--lL-L--EEE-EElllhhhhhhhh
0006 8ffzB   -eEEEE--E---lLL-HH-HH--H--HHH-HH-HHHHHL--------lLLEEELL--H----HHH-HHHH-HHH-----------LLLL-----------------lLLH-HHH-hHHHHHHHHL------------L-LLEE-E-E--LL-LL--L-L--EEE-ELllllhhhhhhh
0007 9gm7C   -----------eEE-LHhHH--H--HHH-HH-HHHHH-----l--lLLLLEEHH-------HHH-HHHH-HHHH----------HLLL--------------lhhHHHH-HHH-HHHHHLL----------------lEEE-E-E--LH-H--lL-L--EEE-EElhhhhhhhhhh
0008 1lddA   ------------lLH-HH-HH--H--HHH-HH-HHHHHH---------hLLEEH-------hHHH-HHHH-HHL-----------LHHH---LLLL----------LLH-HHHHHHHHHHHHL----------------L-LEEL-L-L--L-----------LEE-EEl----------
0009 5yadA   --------L-H--HH-HH-HH--H--HHH-HH-HHHHH-----l--lHHHEEEHH-------HHH-HHHH-HHHL-----------------------------lllLH-HHH-LLLLL-----------------lEEE-E-L--LL-----L-L--LEE-EElhhhl------
0010 7asvA   -------------lllLH-HH--H--HHH-HH-HHHHHH---------hLEEEHH-------HHH-HHHH-HHHHL-LLL-L-----LHH-HL-----------lLLH-HHHH-HHHHHL----------------LLEE-E-L--LL-L----hH--LEE-ELllllllhhhhh
0011 7vwvB   -------------LL-LH-HH--H--HHH-HH-HHHHHH--------hLLLLLLH-------hHHH-HHHH-HHHL----------------------------llLLH-HHHH-HHHHHLL-------------L-LEEE-L-L--LL------lL--LLE-EEll---------
0012 2lh9A   -------------lllLH-HH--H--HHH-HH-HHHHHH--------lLLLEEHH-------HHH-HHHH-HHHL---------------llLHHHHLL-----L-----LHHHHHHLLL--------------L-LEEE-E-E--LL-L---L-L--EEE-EEll---------
0013 6sj9A   ---------------L-HH-HH--H--HHH-HH-HHHHHH--------lLLLEEHH-------HHH-HHHL---------------lLL-LL----------lhHHHH-HHH-HHHHHHHHHLL------------L-LLEE-E-L--L---L--L-L--LEE-ELlllllllleee
0014 7z8bC   -lLLLH-hH-H--HH-HH-HH--H--HHH-HH-HHHHHL--------lLLLEEHH-------HHH-HHHH-HHHHLLLL-----lLHHHH-hhllLLLL--------lLH-HHH-hHHHHHHHH-----------HL-LEEE-L-L--L-----------lLEE-EEll---------
0015 2zmeB   -lLHHH------lLL-HH-HH--H--HHH-HH-HHhHHHHH-----h--lLLLEEEHH-------HHH-HHHH-HH------------LLLLL------------------LLH-HHHH-HHHHLLHHHL----------L-LEEE-E-E--LL-LL-----L--EEE-EElllllhhhhhhh
0016 6r7nB   -hHLLH-hH-H--HH-HH-HH--H--HHH-HH-HHHHL---------LLLEEEHH-------HHH-HHHL----------------------------------lLH-HHHH-HHHHHHHHHL----------lL-LEEE-E-L--L-----------L--LEE-EElllllllllhhh
0017 6ahtA   -lLHHH-hH-H--HH-HH-HH--H---HH-HH-HHHHHH---------hLLEEHH-------HHH-HHHH-HHHHHHHHHhlllllLLLL----------lllHHHH-HHH-hHHHHHLL----------------lEEE-E-E--EL------L-E--EEE-EElhhhhhhhhhh
0018 7abi7   -------hH-H--HH-HH-HH--H--HHHhHH-HHHHH---------hhLLLEEE---------HH-HHHH-HHHLLLL---------llLH-HHL----LL-----LL-H-HHHHHHHHH----------------H-LLEE-E-E--EL-L--lL-E--EEE-ELlhhhhhhhhhh
0019 3zcoA   -------------hhhhhhllL--H--HHH-HH-HHHHHH----l--lLLLEEEHH-------HHH-HHHH-HHHHH-LLL-Lh-hHHHHH-HHH---------llhHHHLlLLHH-HHHHHHHHL----------L-LEEE-E-E--LL-LL--L-L--EEE-EElhhhhhhhhh
0020 6wgeC   -----------llllLH-HH--H--HHH-HH-HHHHL-----l--lLLLLEEHH-------HHL-L--------------------------------------lllHHHH-HHHH-HHHHHHHHL----------L-LEEE-E-L--LL-L--lL-L--LEE-EElllllll-----
0021 7cv0A   ----------------LH-HH--H--HHH-HH-HHHHHHH--------lLLLLLHH-------HHH-HHHL-----------------------------------lLH-HHHH-HHHHHHHHLL----------L-LEEE-E-L---------------lEE-EEllllllllleee
0022 7nywE   -lLEEE-------lL-HH-HH--H--HHH-HH-HHHHHL-----h--hHLLEEEHH-------HHH-HHHH-HHL-------L---HHHHH-HH-HLLL----hhhHHHH-HHHH-HHHHHHHHL----------L-LEEE-L---------------lLEE-EElhhhhhhhhlll
0023 8uucA   -lLLLL--L-L--llL-HH--H--HHH-HH-HHHHHH---------lLLEEHH-------HHH-HHHH-HHHHH---L-L---LL-----------------lLLH-HHHHHHHHHHHHHH----------------L-LLEE-E-L-----------------LEE-EL----------
0024 7qenC   ---------hhhlllllL-HH--H--HHH-HH-HHHHHH----hhhhhlLEEELH-------HHH-HHHL-LLL-----------LLLLL-LL-------------LLH-HHH-HHHHHHHHHHHH----------L-LLEE-E-E--LL-LL--L-L--EEE-ELllll-------
0025 2iu5A   ---lL--L-L--L--HH-HH--H--HHH-HH-HHHHHH---------hLLHH--------HLLH-HHHH-HHHL-----------lLHHHH-HHLL----------LLH-HHHHHHHHHHHHHHHHlhhHHHHH--hhhhhhH-L--LLh--------------------------
0026 2obpA   ----------L--LL-HH-HH--H--HHH-HH-HHHLL-----l--lLLLLLLH-------hHHH-HHHL-----------------------------------lLH-HHHH-HHHHHHHHL----------L-LEEE-E-E--LL-LL----L--EEEe------------
0027 3qkxA   -----L--L-L--HH-HH-HH--H--HHH-HH-HHHHH-------------HLLL--------LLLLhHHHH-HHHL---------lLHHHH-HHHL----------LLHH-HHH-HHHHHHHHHHHHH--hHHHLlllhhhHH-L-Ll----------------------------
0028 7yw2A   -------------LH-HH-HH--H--HHH-HH-HHHHHLL---l--lLLLLEEHH-------HHH-L--L-HHH-----------------------------lllLH-HHH-HHHHHHLL------------LL-LEEE-E-E--LL-L--lL-L--EEE-EElllllllllll
0029 1w5sB   -hhHHL--LlH--HH-HH-LL--H--HHH-HH-HHHHHH----l--lLLLLEEHH-------HHH-HHHH-HHHLLLL-------LLLLL--------------lhHHHH-HHH-HHHHHLL----------------lEEE-E-L--L----------L--LEE-EElllllhhhhhh
0030 2lnbA   lllllL--L-L--L--LH-HH--H--HHH-HH-HHHHH--------hHLLLEEHH-------HHH-HHHL-----------------------------------lLH-HHHH-HHHHHHHHL----------L-LEEE-E-E--L-----------LEE-EElllllll-----
0031 3elkA   ----------L--LH-HH-HH--H--HHH-HH-HHHHHH---------hLLEEH-------hHHH-HHHH-HHHL------------------------------lllllLHHH-HHHHHHHHHH----------L-LEEE-E-E--EE-L--lE-E--EEE-EElhhhhhhhhhl
0032 6qpqD   ----------------LH-HH--H--HHH-HH-HHHHL---------LLLLEEHH-------HHH-HHHH-L---------------------------hllHHHH-HHHH-HHHHHHHHL----------L-LEEE-E-L--LL-LL--L-L--LEE-EElhhhhllll--
0033 7jgrD   -hHLLL---hL--LL-LHhHH--H--HHH-HH-HHHHHH--------hLLLLLLHH-------HHH-HHHH-HHHLL--LL-L---LLLL--------------lhHHHH-HHH-HHHHHLL----------------lEEE-L-------LL--L-L-lLLE-EElllhhhhhhhh
0034 5d1wD   --LLHH-hH-H--HH-HH-HH--H--HHH-HH-HHHLL----------lLLHH--------HLLH-HHHH-HHHL---------lLHHHH-HHHL----------LLHH-HHH-HHHHHHHHHHHH--hLHHHhhllLHHHhhhH--LL-LL--Hl--------------------
0035 4nb5B   ---------hL--LL-L---L--H--HHH-HH-HHHHHH-------lLLLLEEHH-------HHH-HHLL-----------------------------------lLH-HHHH-HHHHHHHHHH----------L-LEEE-E-L--LL-LL--L-L--LEE-EElllhhhhhhhh
0036 7wzeA   -------hH-H--HHlLL-LL--H--HHH-HH-HHHHHH-------lHHHEELH-------hHHH-HHLL-----------------------------------lLH-HHHH-HHHHHHHHHH----------L-LEEE-E-E--L-------L-L--EEE-EElhhhhhhhhhh
0037 8r7kB   lllhHH--H-L--L--LH-HH--H--HHH-HH-HHHHH-----h--hLLLEEEHH-------HHH-HHHH-H------------HLLL-----------------lLLH-HHHHHHHHHHHHHLL----------L-LLEE-E-L--L-----------L--LEE-ELlllllllll--
0038 1yyvA   -lLLLL--H-H--HH-HH-HH--HlhHHH-HH-HHHHH---------hLLEEHH-------HHH-HHLL-----------------------------------lLLH-HHHH-HHHHHHHHHH----------L-LEEE-E-E--EL-L--lL-E--EEE-EElhhhhhhhhhh
0039 8hihN   -------------llL-HH-HH--H--HHH-HH-HLLL-----------lLLEEHH-------HHH-HHHL-L------------------------------llllllL-LHH-HHHHHHHHLLLL------HHHHlLEEE-E-L-----------------L--LEE-EEllll-------
0040 5g5pB   -hLLLHhhH-H--HH-HH-HH--H--HHH-HH-HHHHHH----l--lLLLEEEHH-------HHH-HHHH-HH-----------hLLLLllLLLL-------------LL-LLHH-HHHHHHHHLL---------lL-LEEE-E-L--L---------L--LEE-EElllllhhhhll
0041 7z1nO   ---------leehlL-LH-HH--H--HHH-HH-HHHLLlllllhhhllLLLLLH-------hHHH-LLLL-----------------LLL-L--LLLLE---lllLLLH-HHHH-HHHHHHHLL---------LLL-LLEE-E-E---------------lLEE-ELlhhhlhhhhhhh
0042 4y66C   --------L-H--HH-HH-HH--H--HHH-HH-HHHL------------LLLLLL-------HHH-H-HH-HHH-----------------------------lllLLHH-HHH-HHHHHLLL-----------------LEE-E-E--LL-----L-L--EEE-ELlllhhhhhhhh
0043 5xfoA   ---------lleelllLH-HH--H--HHH-HH-HHHHHH--------hLLLLEELL--L----LHH-HHHH-HLHHHHL-------LHHH--LLLL----------lHHHH-HHHH-HHHHHLL------------L-LEEE-E-E--EH-HH--L-E--EEE-EElllllllll--
0044 8fo9F   hhhhHL--E-EeeEE-HH-HH--H--HHH-HH-HHHLLL----l--lLLLEEEHH-------HHH-HHHH-HLL-----------------------------llLLL-LLH-HHHHHHHHHL----------L-LEEL-L-L--LL-L----lL--LEE-EElhhhhhhhhll
0045 4kywA   -llLLL--L-L-----LH-HH--H--HHH-HH-HHLL------------LLEEEH-------HHH-HHHH-------------hhhHHHL-LL-------------lLLHH-HHHH-HHHHHHHHH----------LL-LEEE-L-L-----------L--LEE-EEl----------
0046 4i99C   -----------------L-HH--H--HHH-HH-HHHHHH----h--hHLLLEEHH-------HHL-L-------------------------------------lllHHHH-HHHH-HHHHHHHHL----------L-LEEE-E-L--LL-LL--L-L--LEE-EEl----------
0047 6pcoB   hhhhHH--L-H--HH-L--LL--H--HHH-HH-HHHHHH---------hLLLLHH-------HHH-HHHL-----------------------------------lLH-HHHH-HHHHHHHHL----------L-LEEE-E-L--LL--------L--LEE-EElhhhhhhhhhh
0048 6wg3C   -------hhhL--LL-LH-HH--H--HHH-HH-HHHHL-----l--lLLLLLHH-------HHL-L---------------------------lllH-----HHH--HHHH-HHHHHHHHL----------L-LEEE-E-L--LL-LL--L-L--LEE-EElllllll-----
0049 5z7bB   -------------LH-HH-HH--H--HHH-HH-HHHHL----------lLLEEH-------hHHH-HHHHlLHHHL-----------------------------lllLH-HHHH-HHHHHHHHL----------L-LEEE-E-E--E-----------eeEE-EElhhhhhhhhhh
0050 9baqA   hhhhHH--H-H--HH-HH-HH--H--HHH-HH-HHHHHH----h--lLLLLLLLL-------HHH-LHHH-HHHHHHLLL-Lh-hHHHHH-HL-LLLL---------lLH-HHHH-HLHHHHHHH------------hlhHH-H-HlhHH-HH--H-Hll-----------------
0051 5uujA   ------------lLH-HH-HH--H--HHH-HH-HHHLL-----------LLLEEHH-------HHH-HHHH-L--------------lLLL-HH----------hhHHHH-HLH-HHHHHHHHHH----------L-LLEE-LlL--LL-L-----L--LEE-ELhhhhhhhhlll

@@@@@@@@@@@@@@@@@@@@@@@@@@@@@@@@@@@@@@@@@@@@@@@@@@@@@@@@@@@@@@@@@@@@@@@@@@@@@@@@@@@@@@@@@@@@@@@@@@@@@@@@@@@@@@@@@@@@@@@@@@@@@@@@@@@@@@@@@@@@@@@@@@@@@@@@@@@@@@@@@@@@@@@@@@@@@@@@@@@@@@@@@@@@@@@@@@
@@@@@@@@@@@@@@@@@@@@@@@@@@@@@@@@@@@@@@@@@@@@@@@@@@@@@@@@@@@@@@@@@@@@@@@@@@@@@@@@@@@@@@@@@@@@@@@@@@@@@@@@@@@@@@@@@@@@@@@@@@@@@@@@@@@@@@@@@@@@@@@@@@@@@@@@@@@@@@@@@@@@@@@@@@@@@@@@@@@@@@@@@@@@@@@@@
@@@@@@@@@@@@@@@@@@@@@@@@@@@@@@@@@@@@@@@@@@@@@@@@@@@@@@@@@@@@@@@@@@@@@@@@@@@@@@@@@@@@@@@@@@@@@@@@@@@@@@@@@@@@@@@@@@@@@@@@@@@@@@@@@@@@@@@@@@@@@@@@@@@@@@@@@@@@@@@@@@@@@@@@@@@@@@@@@@@@@@@@@@@@@@@@@

# Job: PglZ NPV81619 N-terminal Inactive P-loop Domain of Swi2/Snf2-Helicase
# Query: s001A
# No:  Chain   Z    rmsd lali nres  %id PDB  Description
  14:  6l8o-A  5.7  3.3   89   838    9   MOLECULE: DNA REPAIR PROTEIN RAD5;
   2:  6o16-A  7.0  3.2   91   798    4   MOLECULE: DEAH (ASP-GLU-ALA-HIS) BOX POLYPEPTIDE 37;
   3:  6eud-A  6.9  2.8   90   808   14   MOLECULE: ATP-DEPENDENT RNA HELICASE HRPB;
   4:  3k1j-A  6.8  3.2   90   566   10   MOLECULE: ATP-DEPENDENT PROTEASE LON;
   5:  8umy-A  6.6  3.2   87   546   11   MOLECULE: CHROMOSOME TRANSMISSION FIDELITY PROTEIN 18 HOMOL
   6:  6ut5-A  6.5  2.7   85   395    6   MOLECULE: GTPASE SUBUNIT OF RESTRICTION ENDONUCLEASE;
   7:  8val-E  6.2  2.9   88   337    9   MOLECULE: DNA POLYMERASE III SUBUNIT DELTA;
   8:  5lkl-A  6.2  2.9   77   704   10   MOLECULE: GENOME POLYPROTEIN;
   9:  8k8v-A  5.9  3.1   90   719   13   MOLECULE: ENDOPEPTIDASE LA;
  10:  8btg-B  5.8  2.8   85   335    7   MOLECULE: CHROMOSOMAL REPLICATION INITIATOR PROTEIN DNAA;
  11:  8tw7-2  5.8  3.4   86   339    9   MOLECULE: REPLICATION FACTOR C SUBUNIT 5;
  12:  5yww-A  5.8  2.8   81   485    6   MOLECULE: NUCLEOTIDE BINDING PROTEIN PINC;
  13:  8fcv-R  5.8  3.5   89   343    7   MOLECULE: DNA (60-MER);
  15:  6pxk-K  5.7  2.9   90   429   12   MOLECULE: ATP-DEPENDENT PROTEASE ATPASE SUBUNIT HSLU;
  16:  4zo4-A  5.7  2.9   71   202   11   MOLECULE: DEPHOSPHO-COA KINASE;
  17:  7st9-E  5.6  3.6   87   354   14   MOLECULE: CHECKPOINT PROTEIN RAD24;
  18:  8y6p-Q  5.6  2.8   80  1228    9   MOLECULE: APAF-1 RELATED KILLER DARK;
  19:  6l1q-A  5.6  3.1   81   267   16   MOLECULE: CBBQ PROTEIN;
  20:  7w42-B  5.6  3.2   84   402   14   MOLECULE: UNCHARACTERIZED ATPASE YJOB;
  21:  5x71-A  5.5  2.7   76   796   11   MOLECULE: MRNA CAPPING ENZYME P5;
  22:  9emc-D  5.5  3.5   87   337   17   MOLECULE: RUVB-LIKE 1;
  23:  7plh-A  5.5  2.9   84   260   11   MOLECULE: SHTNSC;
  24:  9e22-A  5.4  3.6   90  2925    6   MOLECULE: CYTOPLASMIC DYNEIN 1 HEAVY CHAIN 1;
  25:  8dgf-A  5.4  3.1   87  1541   14   MOLECULE: ATP-BINDING PROTEIN AVS4;
  26:  3glf-G  5.4  3.3   87   378   13   MOLECULE: DNA POLYMERASE III SUBUNIT DELTA;
  27:  3bos-B  5.4  3.0   88   231    9   MOLECULE: PUTATIVE DNA REPLICATION FACTOR;
  28:  8s35-M  5.4  3.0   81   624   11   MOLECULE: CRISPR TYPE AFERR-ASSOCIATED PROTEIN CSF2;
  29:  7qo4-J  5.3  2.9   87   405   14   MOLECULE: 26S PROTEASOME REGULATORY SUBUNIT RPN1;
  30:  5zr1-A  5.3  3.2   85   494    7   MOLECULE: ORIGIN RECOGNITION COMPLEX SUBUNIT 1;
  31:  8thc-A  5.3  3.0   86   479   10   MOLECULE: ELG1 ISOFORM 1;
  32:  7rzy-1  5.3  3.3   87   311   11   MOLECULE: TN6677 VIBRIO CHOLERAE TRANSPOSON TNSC (VCHTNSC);
  33:  2p0e-A  5.2  2.6   70   190    7   MOLECULE: NICOTINAMIDE RIBOSIDE KINASE 1;
  34:  6z1f-3  5.1  3.3   85   284    7   MOLECULE: RIBULOSE BISPHOSPHATE CARBOXYLASE/OXYGENASE ACTIV
  35:  9ri9-B  5.1  3.1   83   868    8   MOLECULE: SLNRC3;
  36:  6yvh-K  5.1  3.4   78   164    9   MOLECULE: PRE-MRNA-SPLICING FACTOR CWC22 HOMOLOG;
  37:  8jqb-E  5.1  3.6   86   499   14   MOLECULE: ENDONUCLEASE GAJA;
  38:  8bob-C  5.1  3.4   86   412   17   MOLECULE: PROTEIN MALY;
  39:  5v8f-D  5.1  3.2   82   456    6   MOLECULE: DNA REPLICATION LICENSING FACTOR MCM2;
  40:  8xks-B  5.1 24.8   90   603   10   MOLECULE: CTAP1;
  41:  8dr5-A  5.1  3.4   85   646    6   MOLECULE: REPLICATION FACTOR C SUBUNIT 1;
  42:  8dgc-A  5.0  3.2   87  2028   11   MOLECULE: SEAVS3;
  43:  3pvs-D  5.0  2.9   80   424   10   MOLECULE: REPLICATION-ASSOCIATED RECOMBINATION PROTEIN A;
  44:  6nak-D  5.0  3.3   86   165    8   MOLECULE: TRNA THREONYLCARBAMOYLADENOSINE BIOSYNTHESIS PROT
  45:  9jix-A  5.0  2.7   75   154    9   MOLECULE: DE NOVO DESIGNED ATPASE, PL2X4_2;
  46:  7mi1-A  5.0  3.2   87  2628   14   MOLECULE: CHIMERA PROTEIN OF DYNEIN AND ENDOLYSIN;
  47:  1ixs-B  5.0  3.3   87   315    9   MOLECULE: HOLLIDAY JUNCTION DNA HELICASE RUVA;
  48:  2is6-A  5.0  3.3   86   654    9   MOLECULE: 5'-D(*CP*GP*AP*GP*CP*AP*CP*TP*GP*CP*AP*GP*TP*GP*C
  49:  2zpa-B  5.0  3.3   88   662    7   MOLECULE: UNCHARACTERIZED PROTEIN YPFI;
  50:  7mi8-A  4.9  3.2   88   868   16   MOLECULE: FUSION PROTEIN OF DYNEIN AND ENDOLYSIN;

0001 s001A   ---MASW-R--DH-IL-K-EFTPRV-S----R----LTLVADP-D-G-LLLEILEGIRE-R---G--FELI--P--F---E-DHI--AF-RYAYESRFR-S-R-W-D-AALVVVLRS----GL-E-A--LDVLPYDLLQAG-RKLSF-----------------
0015 6l8oA   --sHFQStI--KA-LL-R-HLIQET---spgE----QIIVFSQ-F-SsFLDILRSHLPR-D---Q--VIIY--K--F---DkERT--RI-LEQFHDKD------L-S-CIKLLLLSL----K--T-G--GVGLN---LTCAsRAFMMdpwwspgmedqaidrih
0003 6o16A   --qEERL-lpiLA-EEqA-IME-AE-H----P----IVIVCGE-TgS-GKTTVPQFLYE-A--yK--VVII--D-eA---H-ERV--YT-DILLGL-LS-R-I-VlP-LKLLIMSAT----L----R--VEDFTQNFTTPP-PVIKVesrfpvtvhfnddysge
0004 6eudA   --aMSSL--pvAA-VL-P-ELLTDC-A----P----QVLLSAP-TgA-GKSTLPLQLLA-H-----gLVIL--D-eF---H-ERL--QA-DLALAL-LL-D-V-QdD-LKLLIMSAT----L----D--NDRLQQML-PEA-PVVISegrsfpverrylplpah
0005 3k1jA   lidqvIG-Q--EH-AV-E-VIKNQK-R---------HVLLIGE-PgT-GKSMLGQAMAE-L---L-gVLFI--D--E-iaT-LSL--KM-QQSLLTAMQ-E-KkC-D-FVLVAAGNL-------D-T--VDKMHPALRSRIgYEVYMrttmpdtienrrklvqf
0006 8umyA   --ytELL-S-dDF-TN-R-CLWLKLwQ----K----VALLCGP-PgL-GKTTLAHVIAR----pN--CLVI--D--EidgA-PVA--AI-NVLLS-ILN-RgL-L-M-RPIICICN----------D--QFA-LRQLKQQA-FLLHFpptlpsrlvqrlqevsl
0007 6ut5A   ----GHN-I--KE-DY-F-RVD-NK-K----G----QVILYGP-PgT-GKTWaRKYVVE-E--kF--YLII--DeiN---R-GN----I-SKIFGelitlA-V-P-PnLYIIGTMNT----A-dR-S--IALLDVALRRRF-AFIEVeprpeflekenlkkire
0008 8valE   hmrwyPW-L--RP-DF-EkLVASRG-H----H----ALLIQALpG-M-GDDALIYALSR-Y---L-kVVWV-tD--A---A-LLTdaAA-NALLK-TLE-E-P-PaE-TWFFLATR-----------E--PERLLATLRSRC-RLHYLapppeqyavtwlsrevt
0009 5lklA   lmsgaTD-L--TE-MV-K-KTTMN-----rgE----FRQITLA-TgA-GKTTLPRSVIE----yS--FIFL--D-eY---H-CAT-pEQ-LAIMGKI-H-R-F-SeN-LRVVAMTA-----------T--PAGvNYDWS-------------------------
0010 8k8vA   --lPPPP-P--FF--R-E-ALEIRG-G----F----HAYLVGP-PsL-GKHELLAYLstqsvegG--YLIL--D--A---LlKRE--GT-WEAFKRALR-N-G-I-Q-MQVILVGTP-------E-A--FEGLEPAFSELFrIRAEFsptlpaspenctalggw
0011 8btgB   pkytfDT-F--VIgsH-A-ASLAKA-Y----N----PLFIYGG-VgL-GKTHLMHAIGH-Y---V-dVLLI--D-dI---Q-FLA-gkeqQEEFFHTFN-T-L-HeS-KQIVISSDR---------P--PKE-RSRFE-WG-LITDItppdletriailrkkak
0012 8tw72   nldevTA-Q--DH-AV-T-VLK-AN-L----P----HMLFYGP-PgT-GKTSTILALTK----yK--IIIL--D-eA---D-SMD--AQ-SA-LRRTME-T-Y-SgV-TRFCLICN-----------Y--VTRIIDPLASRC-SKFRFkaldasnaidrlrfise
0013 5ywwA   vrkrlED-Y--NL-------lLEER-A----E----GIIIAGA-PgM-GKTTFAQALAE-Y---Y-dYTVY-dE--M---R-ND---ED-FKLYVDLRL-A-G-V----GMVGVVHA----TSpI-D--AIHRFVN--diD-TIIFInsgnvskvytlemtvkv
0014 8fcvR   --tVAHP-L--KE-VY-E-ILMIAE-P-agaS----FIFVYGA-SgV-GKTTLRLRVEQ----pD--VFFV--D-eA---Q-HFklqDQ-LDCLKSLAN-M-T-G---ILHCLLGTY---------E--LLTFRNQLSRRS-VDIHFrrycadspedvqafksv
0016 6pxkK   ldkhiIG-Q--DN-AK-R-SVALRN-P----K----NILXIGP-TgV-GKTEIARRLAK----hG--IVFIdeI--D---K-IEG--VQ-RDLLP-LVE-GkT-D-H-ILFIASGAF----Q-iA-K--PSDLIPELQGRLpIRVELqalttsdferiltepna
0017 4zo4A   ---------------------------axkN----AFFVTAS-I-AcGKSTFIEIANS-L---G--FKSI--S--A---DkIRA--KI-LEQXQ-ILD-K-E-N---KAFFVEIPL---------F--FESGA--YENLG-KVIVIytpkelslkrixqrdkl
0018 7st9E   alshnEE-L--TN-FL-K-SLSPRD-L----P----HLLLYGP-NgT-GKKTRCMALLE-S-----kCVII--N-eA---N-SLD--AQ-AA-LRRTME-K-Y-SkN-IRLIMVCDS------------MSPIIAPIKSRC-LLIRCpapsdseistilsdvvt
0019 8y6pQ   fakynVS-R--LQ-PY-L-KLRLLE-L-rpaK----NVLIDGV-LgS-GKTWVALDVCL-S-----cLLVL--L-nV---Q-NAK--AW-NAFNL------------sCKILLTTRF----K--Q-V--TDFL---SAATT-THISLdhhsmtltpdevkslll
```

```
0020 6l1qA   gkepfyQ-P--QHdEV-A-LFEYRK-R----L----PVMVKGP-TgC-GKSRFVEFMAW----gA--ICYL--D-eI---V-EAR---qdTVVIH-PLT-D----pD-FQLVISYNP----G-----slmkDLKQSTKQRF-TGFEFdypnaeleagilvqetg
0021 7w42B   revqrSQ-V--IM--K-K-EISIDQ-dipykR----GILLYGP-P-GnGKTTLVKSIAG----pA--VLVI--E-dI---D-SMPedVR-SFFLNT-LD-G-A--eG-LFLIGTTN---------Y--PEEIDPGLMNRD-RAYEIglpdeelrleymkmrgf
0022 5x71A   -------------------------S-----P----LVVFCLS-H-NeLAKKYAQDVSMsS---G--THVH--I--I---DeITV--SL-YRTFR-TIA-T-Q-LgR-MQIVVFVTV----DksvvS--TQVMKSIAWARG-SFVELrnqsvdsstlvsklenl
0023 9emcD   asqgmVG-Q--LA-AR-R-AVLEM--gkiagR----AVLIAGQ-PgT-GKTAIAMGMAQ----pG--VLFI--D-eV---H-MLD-iES-FSFLNRALE-S-D-M-A-PVLIMATNR----GitrirqsPHGIPIDLLDRL-LIVSTtpysekdtkqilrirce
0024 7plhA   --iVPLQ-V--KT-LH-D-WLDKAR-K---------SCRVVGE-SrT-GKTVYRY--------vE--MLII--D-eA---D-RLK-----PETFADVRD--eD-L-G-IAVVLVGTD----------rlDAVIKEQVLERFrAHLRFgklsgedfkntvemweq
0025 9e22A   plvlfNE-V--LD-HV-L-RIDRQP-Q----G----HLLLIGV-S-GaGKTTSRFVAWM-N-----iAFIM--D--E---SnVLD--S-gLERMNTLLA-N-G-ErN-LHVVFTM-N----PS-S-EkdrAATSPALFNRC-VLNWFgdwstealyqvgkefts
0026 8dgfA   --sIEID-R--RH-CI-E-LLHLVQ-K----K----IVIVSGE-GgV-GKTAVIKKIYE----rK--VIVV--D-sA---E-KLltnidpKEFLTV-LI-K-D-K---WQVVFTTRN-------N-Y--LADLNYAFIITP-GNLVIknlergelielsdnngf
0027 3glfG   tfadvVG-Q--EH-VL-T-ALA-RI-H----H----AYLFSGT-RgV-GKTSIARLLAK-G---L-kVYLI--D-eV---H-MLSrhSF-NALLK-TLE-E-P-PeH-VKFLLATT----------D--PQKLPVTILSRC-LQFHLkaldveqirhqlehiln
0028 3bosB   ddetfTS-Y--YPaaI-G-ALKSDG-V----Q----AIYLWGP-VkS-GRTHLIHAACA-R---A-dLICIddV--D---A-VHP--LW-EEAIFDLYN-R-V-AkR-GSLIVSASA----------S--PXE-LVSRXHWG-LTYQLqpxxddeklaalqrraa
0029 8s35M   --fsGQW-L--KA-VV-E-QLSC-I-Q----G----PALVLTA-S-HyETRMIANQLGE-V---S--QPVY--I--QkagQ-ALS--EI-IKQYQEK------------pGILISAG--------aS--VGVSPRGgEQIFqDLIITripflppdrmkaeslyg
0030 7qo4J   tydmvGG-L--TK-QI-K-EIKEVI-P----K----GVILYGP-PgT-GKTLLARAVAH----pS--IIFM--D--E---I-DGG--DS-EVQRT-MLE-L-L-NkN-IKIIMATNR------------LDILDPALLRID-RKIEFpppsvaaraeilrihsr
0031 5zr1A   ssyylPA-R--EN-EF-A-SIAYSA-esdsaT----TIYVAGT-PgV-GKTLTVREVVK-E-----iVVLL-dE--L---D-AQD--IM-YNFF-nWTT-Y-E-A-K-LIVIAVANT-----M-D-L--PER-TSRI--GF-TRIMFtgytheelkniidlrlk
0032 8thcA   --tlKQV-LiePK-LK-L-RIIETS-ptefvP----LMILHGN-S-I-GKKTLIQTIMR----yG--LVLF--N-dV---D-VLhdrGY-WAMISK-LC-E-F-S-R-RPLVLTCK----------D--LSLVPSELIALAnSLFHTkkiststvyafltkylk
0033 7rzy1   --aFVSTpV--RK-IL-S-YMCRDL-lesepT----CMMVYGA-SgV-GKTTVIKKYLN----vK--LIII--D-eF---Q-HLnrvltqGNWLKMILN-K-T-K---CPIVIFGMP-------Y-S--KVVLNSQLHGRFsIQVELrpfsynggrgvfktfle
0034 2p0eA   -------------------------rgsktF----IIGISGV-TnS-GKTTaKNLQKH-L---P-nCSVI--S--Q---D-DXE--KX-XSAISC-WX-E-S-AeI-PILIIEGFL----------L--FNY-LDTIW--N-RSYFLtipyeeckrrrstrvyq
0035 6z1f3   ---SYYI-A--PR-FL-D-KLHITK-P----L----ILGIHGR-KgE-GKTFCELA--------mC--VLMI--NdlD---A-GAntqlV-NATLMN-IA-D-N-PrR-VPIIVTGN----------D--FSTLYAPLIRRM-EKFYWepnrddkvgivggifae
0036 9ri9B   --dDVVG-F--DD-EA-K-TVILIG-GsdyvE----VVPVVGM-PgL-GKTTLAYKIYK-D-----yLIVL--D-dV---W-TME--AW-DRIK-IAFP-N-N--kR-NRVLMTTR-------------QSNV--RCND---KPHDLkfltkdeswellekkvf
0037 6yvhK   --vEREE----WK-FD-T-LCYDTL-I-----T----QAVIFCN-T-KrKVDWLTEKMRE-A----N--FTVS--S--M----kERE--SI-MKEFRSGA------------sRVLISTD-------------vWGLD---VPQVsLIINYdlpnnrelyihrigrsg
0038 8jqbE   ----MIE-D--EM-SR-EqIIKDG--------G----NILVTAG-A-GsGKTTILVSKIE-A---DlfKIFI--D-eY---Q-DSK--DM-HNLFMYLKD-Q-L-K-I--KLFIVGDP--kqsiyiwrenFNGLIEN-STDF-NKYHLtsnfrccqdiqnysnlf
0039 8bobC   rldhtVV-R--ER-LL-A-KLSGAN-F----R----LALITSP-AgY-GKTTLISQWAA----pL--YLVI--D-dY---H-LInpvIH-ESMRF-FIR-H-Q-PeN-LTLVVLSRN------------LPQLGANLRVQL-LEIGSqqlafthqeaneffdcr
0040 5v8fD   --iiFTY-L--QD-CQ-Q-EIIKQS-iqkesH----SVILVGP-RqS-YKTYLLDYELS-------vVFIF--D-eI---D-TFA--GpvQTLLYNLFD------vP-VCIFGCTTK----L----N--ILE-KSRF--SQ-RVIYMpqiqnlddmvdavrnll
0041 8xksB   --eLPEG-N--KQ-AK-V-EVVDFA-P----K----GVLLVGP-PgN-GKTLMARAVAG----pC--IIFI--D-eL---D-AVD--NT-VNQLLTEMD-A-E-Q-QgIVVMGATN----------R--KDVLDAALTRPD-RSIEVrrpdfqgrleaslaslm
0042 8dr5A   nlqqvCG-N--KG-SV-M-KLLANW-F----R----AAMLYGP-PgI-GKTTAAHLVAQ----hF--VIIM--D-eV---D-GMG--GV-GQLA-QFCR---K-T-S-TPLILICN----------E--RNL-MRPFDRVC-LDIQFrrpdansiksrlmtiai
0043 8dgcA   --nSVSR----TQ-EA-T-LVQVEA-F---gA----PVIIHAD-AgV-GKSIFSTHIEE-H-----lCIII--D-aA---D-NAS--FI-KDLIRE--K-L-P-D-G-VCLVALCRP-------Y-R--RELLD---PPPEA-LTLSLqtfnrdetaahlhqkfp
0044 3pvsD   rpenlAQ-Y--IG-QqhllapaGHL-H---------SMILWGP-PgT-GKTTLAEVIAR-Y-----tILFV--D-eV---H-RFN-----KSQQDA-FL-P-H-G-T-ITFIGATTE----------N--PSFLNSALLSRA-RVYLLkslstedieqvltqame
0045 6nakD   --eNLTE-E--QL-KR-L-ALTENL---kggE----VVILSGN-LgA-GKTTFVKGMIR-A---I-tIYHL--D--L---Y-RTD--FL-SLDVED-IL-E-D-E-D-GIMVVEW---------------gDLFD-GFWPEDS-IKVKIeiadeshrnveilipee
0046 9jixA   ---------------------------mgaI----VILVVGP-PgS-GKSQLIKAIEK-L--qP--VVTT--S--V---T-SED--EA-KKVLEELLK-K-D-P-N-AIVVIEIKN----P--R-I--AERVAKRV-dVL-VVVVSspevarelrenlpnviv
0047 7mi1A   iperlIY-T--PL-LL-I-GFA-TD-S----LhqkyGGCFFGP-AgT-GKTETVKAFGQ----gA--WGCFdqF--N---R-LDE-kVL-SAVSANI-Q-Q-I-QpH-TAVFITLNP----gynG-R--SE-LPENLKKSF-REFSMkspqsgtiaemilqimg
0048 1ixsB   tldeyIG-Q--ER-LK-Q-KLRVYL-L----E----HLLLFGP-PgL-GKTTLAHVIAH-E--gD--ILFI--D-eI---H-RLSrqAE-EH-LYPAME-D-F--pR-FTLIGATTR---------P--GLITAP-LLSRFgIVEHLeyytpeelaqgvmrdar
0049 2is6A   vsyllDS-L--ND-KQ-R-EVAAP--R----S----NLLVLAG-AgS-GKTRVLVHRIA-W---LmtnILV--DefQ---D-TNI--QY-AWIRL-LAG-D-T-G----KVMIVGDD----DQ-S-igaqVENIQRFLNGA-ETIRLeqnyrstsnilsaanal
0050 2zpaB   --tGAPQpQ--QQ-LLkQ-LXTX-P-P----G----VAAVTAA-RgR-GKSALAGQLIS----aD--WLVV--D-eA---A-AIP-----APLLHQLVS-----R-F-PRTLLTTTVqgyeGT-G-RgfLLKFCARF-PHL-HRFELqqpirwaqgcplekxvs
0051 7mi8A   iperlIY-T--PL-LL-I-GFA-TD-S----LhqkyGGCFFGP-AgT-GKTETVKAFGQ----gA--WGCF--D-qF---N-RLDekVL-SAVSAN-IQ-Q-I-QpH-TAVFITLNP----GynG-R--SE-LPENLKKSF-REFSMkspeeysiigkhsvkif

0001 s001A   ---LLLH-H--HH-HH-L-LLLLLL-L----L----EEEEELL-L-L-HHHLHHHHHHH-L---L--EEEE--E--L---L-LHH--HH-HHHHHHHLL-L-L-L-L-LEEEEEELL----LH-H-H--HHHLLHHHHHHL-EEEEL----------------
0015 6l8oA   --lLLLLhH--HH-HH-H-HHHHHH---lllL----EEEEEEL-L-HhHHHHHHHHLLL-L---L--EEEE--E--L---LhHHH--HH-HHHHLLLL------L-L-LEEEEEEEL----L--L-L--LLLLL---LLLEeEEEELlllllhhhhhhhhhlll
0002 9nv3C   ----LHH-H--HH-HH-H-HHHHLL-L----L----EEEEELL-L-L-LLLLLLLL-lL-L---L--LEEE--E--L---L-LL---LH-HHHHH-HHH-LlL-L-L-LLEEEEEL----------L--LLL-LHHHHHHL-EELLLllhhhhhhhlllllll
0003 6o16A   --hHHHH-lhhHH-HHhH-HHH-HH-L----L----EEEEEEL-LlL-LHHHHHHHHHH-L--eE--EEEE--L-lH---H-HLH--HH-HHHHHH-HH-H-H-HlL-LEEEEEELL----L----L--LHHHHLLLLLL-LEEEElllllleeeellllhhhh
0004 6eudA   --lLLLL--llHH-HH-H-HHHHLL-L----L----LEEEELL-HhH-LLLLHHHHHHL-L-----eEEEE--L-lH---H-HLH--HH-HHHHHH-HH-H-H-LlL-LEEEEEELL----L----L--LLLHHHHL-LLL-LEEELllllllleeeeellllll
0005 3k1jA   hhhhlLL-L--HH-HH-H-HHHHLL-L---------LEEEELL-LlL-LHHHHHHHHHH-L---L-lEEEE--L--L-hhH-LLH--HH-HHHHHHHHH-H-LeL-L-LEEEEEELH-------H-H--HHHLLHHHHHHHlEEEELlleeellhhhhhhhhhh
0006 8umyA   --llLLL-L-lHH-HH-H-HHHHHHhL----L----EEEEELL-LlL-LHHHHHHHHHH-----lE--EEEE--E--LhhhL-LHH--HH-HHHHH-HHL-LlL-L-L-LLEEEEEL----------L--LLL-HHHHHLLL-EEEELllllhhhhhhhhhhhhh
0007 6ut5A   ----LLH-H--HH-HH-H-HHH-HH-H----L----EEEEEEL-LlL-LHHHhHHHHHH-H--lE--EEEE--EllL---L-LL----H-HHHLHhhhhhL-L-L-LlEEEEEEEEL----L-lL-L--LLLLLHHHHHHL-EEEELllllhhhhlhhhhhhhhl
0008 8valE   lllllHH-H--HH-HH-HhHHHHLL-L----L----EEEEELLlL-L-LHHHHHHHHHH-H---H-eEEEE-lL--H---H-HLLhhHH-HHHHH-HHH-L-L-LlL-EEEEEEEL----------L--HHHLLLLLHHHL-EEEELllllhhhhhhhhhhhll
0009 5lklA   eelllHH-H--HH-HH-H-HHLLL-----llL----EEEEELL-HhH-LLLLHHHHHHH-----lL--EEEE--E-lL---L-LLL-hHH-HHHHHHH-H-L-L-HhH-LLEEEEEL----------L--LLLlHHHHL-------------------------
0010 8k8vA   --lLLLL-L--LL--H-H-HHHHHL-L----L----EEEEELL-LlL-LHHHHHHHHHlllllllL--EEEE--E--H---HhHHL--LL-HHHHHHHHH-H-L-L-HhH-LLEEEEELH--------H-H--HHLLLLLHHHHEeEEEELlleeellhhhhhhhhhh
0011 8btgB   lllllLL-L--LLllH-HH-H-HHHHLL-L----L----EEEEEEL-LlL-LHHHHHHHHHH-H---H-lEEEE--E-lH---H-HHL-llhhHHHHHHHHH-H-H-HlL-LEEEEEELL----------L--LLL-HHHHL-LL-EEEEEllllhhhhhhhhhhhh
0012 8tw72   lllllLL-L--HH-HH-H-HHH-LL-L----L----LEEEELL-LlL-LHHHHHHHHHH-----lE--EEEE--L-lH---H-HLH--HH-HH-HHHHHH-H-L-LlL-EEEEEEEL----------L--HHHLLHHHHHHL-EEEELllllhhhhhhhhhhhh
0013 5ywwA   llllhHH-H--LL-------hHHHL-L----L----EEEEELL-LlL-LHHHHHHHHHH-H---H-lEEEE-lL--L---L-LH---HH-HHHHHHHHH-L-L-L----EEEEELLL----LLhH-H--HHLLLLL--lhL-EEEEEelleeeeeeeeeeeeel
0014 8fcvR   --lLLLH-H--HH-HH-H-HHHHHL-L-lllL----EEEEEEL-LlL-LHHHHHHHHHH-----lL--EEEE--E-lH---H-HHhhhHH-HHHHHHHHH-H-H-L---LEEEEEELH----------H--HHHHLLHHHHLE-EEEEEllllllhhhhhhhhhh
0016 6pxkK   hllllLL-L--HH-HH-H-HHHHHH-L----L----LEEEELL-LlL-LHHHHHHHHHH-----hL--EEEEelH--H---H-HHH--HH-HHHHH-HHL-LeL-L-L-LEEEEEELL----L-lL-L--HHHLLHHHHLLLlEEEELllllhhhhhhhhhhllll
0017 4zo4A   ----------------------------lllL----EEEEEEL-L-HhHHHHHHHHHH-L---L--LEEE--E--H---HhHHH--HH-HHHHH-HHH-H-L-L---LLEEEELLL----------L--LLLLL--HHHLL-EEEEElllllllhhhhhhhhll
0018 7st9E   hllllHH-H--HH-HH-H-HHLHHH-L----L----LEEEELL-LlL-LHHHHHHHHHH-H-----eEEEE--L-lH---H-HLH--HH-HH-HHHHHH-H-L-LlL-EEEEEEELL------------LLLLLHHHHHLL-EEEELllllhhhhhhhhhhhhh
0019 8y6pQ   hhhllLL-L--HH-HH-H-HHHHHH-L-lllL----EEEEELL-LlL-LHHHHHHHHHH-L-----lEEEE--E-eE---L-LHH--HH-HHHLL------------lLEEEEEELL----H--H-H--HHHL---LLLLE-EEEELllllllllhhhhhhhhh
0020 6l1qA   llllllL-L--LLlHH-H-HHHHHL-L----L----LEEEELL-LlL-LHHHHHHHHHH-----lL--EEEE--E-lH---H-HLL---hhHHHHH-HHL-L-----lL-LEEEEEELL----L-----llllLLLHHHHLLL-EEEELllllhhhhhhhhhhhl
0021 7w42B   llllhHH-L--LL--H-H-HHHHHH-lllllE----EEEEELL-L-LlLHHHHHHHHHH-----lE--EEEE--L-lH---H-HLLhhHH-HHHHHH-HH-L-L--lL-EEEEEEEL----------L--HHHLLHHHLLLL-EEEELllllhhhhhhhhhhlh
0022 5x71A   --------------------------L----L----EEEEEEL-L-HhHHHHHHHHHHhH---L--LEEE--E--E---LlHHH--HH-HHHHH-HHL-L-LlL-LLEEEEEELL----LllllL--HHHHHHHHHHLL-EEEEElllllllhhhhhhhhhh
0023 9emcD   eelleEL-L--HH-HH-H-HHHHH--lllllL----EEEEELL-LlL-LHHHHHHHHHH-----eL--EEEE--E-lH---H-HLL-hHH-HHHHHHHLL-L-L-L-L-LEEEEEELL----LleelleeHHHLLHHHHLLL-EEEEElllhhhhhhhhhhhhh
0024 7plhA   --lLLLH-H--HH-HH-H-HHHHLL-L---------LEEEELL-LlL-LHHHHHH---------lL--EEEE--L-lL---L-LLL------HHHHHHHHH--hH-H-L-LEEEEEELL-----------hhHHHHHLHHHHHLlEELLLllllhhhhhhhhhhhh
0025 9e22A   lllllHH-H--HH-HH-H-HHHHLL-L----L----EEEEELL-L-LlLLLLHHHHHHH-L-----eEEEE--E--H---HhLLL--L-lHHHHHHHHH-H-L-LhL-EEEEEEE-L----LL-L-LlllLLLLHHHHHHE-EEEELllllhhhhhhhhhhhl
0026 8dgfA   --eELLL-L--HH-HH-H-HHHHHH-L----L----EEEEELL-LlL-LHHHHHHHHHL-----lL--EEEE--E-lH---H-HHllllhhHHHHHH-HH-H-L-L---LEEEEEELH--------H-H--HHHHHHHHHLLL-EEEELllllhhhhhhhhhhhll
0027 3glfG   lllllLL-L--HH-HH-H-HHH-LL-L----L----EEEEELL-LlL-LHHHHHHHHHH-H---L-eEEEE--E-lH---H-HLLhhHH-HHHHH-HHH-L-L-LlL-EEEEEEL----------L--HHHLLHHHHLLL-EEEELllllhhhhhhhhhhhh
0028 3bosB   lllllLL-L--LLllH-H-HHHHLL-L----L----EEEEELL-LlL-LHHHHHHHHHH-H---H-lEEEEelH--H---H-HLH--HH-HHHHHHHHH-H-H-HlL-LEEEEEELL----------L--LLL-HHHHHHHL-EEEELllllhhhhhhhhhhhh
0029 8s35M   --elHHH-H--HH-HH-H-HHHL-L-L----L----LEEEELL-L-HhHHHHHHHLLL-L---L--LLEE--E--LlllL-LHH--HH-HHHHHHL------------lLEEEELL---------lL--LLLLLLlLLLLlEEEELlllllllhhhhhhhhh
0030 7qo4J   lllllLL-L--HH-HH-H-HHHHHL-L----L----LEEEELL-LlL-LHHHHHHHHHH-----lE--EEEE--L--L---L-LLH--HH-HHHHH-HHH-H-H-HlL-LEEEEEELL------------LLLLLLLLLLL-LEEELllllhhhhhhhhhhhl
0031 5zr1A   lllllLL-L--LL-LH-H-HHHHHH-hllllE----EEEEEEL-LlL-LHHHHHHHHHH-H-----eEEEE-eL--L---L-LLL--LH-HHHH-hHHH-L-L-L-L-EEEEEELLL-----L-L-H--HHH-HHHH--LL-EEEEEllllhhhhhhhhhlhhhl
0032 8thcA   --llLLL-LllHH-HH-H-HHHHHH-lllllL----LLLLEEL-L-L-LLLHHHHHHHH----eE--EEEE--L-lL---L-LLlllLH-HHHHHH-HH-H-L-L-L-LLEEELLL----------L--HHHLLHHHHHHHlLLLEElllhhhhhhhhhhhhh
0033 7rzy1   --lLLLLhH--HH-HH-H-HHHHHH-lllllL----LLEELLL-LlL-LHHHHHHHHHH-----lL--EEEE--L-lL---H-HHlllllhHHHHHHHHH-H-L-L---LLLLLLLH--------H-H--HHHHLHHHHHHLlLEELLllllllllllhhhhhhh
0034 2p0eA   -------------------------lllllE----EEEEEEL-LlL-LHHHhHHHHLL-L---L-lEEEE--E--H---H-HHH--HH-HHHHHH-HH-H-H-HlL-LEEEEELLL----------L--LLL-HLLLL--L-EEEEEellhhhhhhhhhhhllll
0035 6z1f3   ---LLLL-L--HH-HH-H-HHHHHH-L----L----EEEEELL-LlL-LHHHHHHH---------lE--EEEE--LllL---L-LLhhhhH-HHHHHH-HH-H-L-LlL-LLEEEEEL----------L--LLLLLHHHLLLE-EEEELlllhhhhhhhhhhhlll
0036 9ri9B   --lLLLL-L--HH-HH-H-HHHHHL-LllllE----EEEEELL-LlL-LHHHHHHHHHL-L-----eEEEE--E-lL---L-LHH--HH-HHHH-HHLL-L-L--lL-LEEEEEEL-------------LHHH--HHLL---LLEELlllhhhhhhhhhhhhh
0037 6yvhK   --eELHH----HH-HH-H-HHHHHL-L----L----EEEEELL-L-HhHHHHHHHHHH-L---L--LLEE--E--E----lHHH--HH-HHHHHLLL------------lLEEEELL-----------lLLLL---LLLEeEEEELllllhhhhhhhhhllll
0038 8jqbE   ----LHH-H--HH-HH-HhHHLLL--------L----LEEEELL-L-LlLHHHHHHHHHH-H---HheEEEE--L-lL---L-LLH--HH-HHHHHHHHH-L-L-L-L--EEEEEELL--llllllllllHHHLLLL-LLLL-EEELLlllllllllhhhhhhll
0039 8bobC   lllllLL-L--HH-HH-H-HHHLHH-L----L----EEEEELL-LlL-LHHHHHHHHHL-----lE--EEEE--E-lH---H-HLlhhHH-HHHHH-HHH-H-L-LlL-EEEEEEELL------------LLLLLHHHHHLE-EEELHhhhlllhhhhhhhhhhh
0040 5v8fD   --llLHH-H--HH-HH-H-HHHHHH-lllllL----EEEEELL-LlL-LHHHHHHHHHH-------eEEEE--E-lH---H-HLL--LllLHHHHHHHH------lL-EEEEELLLL----L----L--HHH-HHHL--LL-LEEELllllhhhhhhhhhhhh
0041 8xksB   --lLLLL-L--LL-HH-H-HHHHHL-L----L----EEEEELL-LlL-LHHHHHHHHHH-----lL--EEEE--E-lH---H-HLH--HH-HHHHHHHHL-L-L-L-LlLEEEEEEL----------L--LLLLLLLLLLL-LLLLLllllhhhhhhllhhhhl
0042 8dr5A   lllllLL-L--HH-HH-H-HHHHLH-L----L----EEEEELL-LlL-LHHHHHHHHHH-----lE--EEEE--L-lH---H-HLL--HH-HHHH-HHHH---L-L-L-LLEEEEEL----------L--LLL-HHHHLLLL-EEEELlllllllhhhhhhhhh
0043 8dgcA   --lLLLL----LH-HH-H-HHHHLL-L---lL----LEEEELL-LlL-LHHHHHHHHHH-H-----eEEEE--E-lH---H-HHL--HH-HHHLLL--L-L-L-L-L-EEEEEEELL--------L-L--HHHLL--LLLLL-EEEELllllhhhhhhhhlllll
0044 3pvsD   lllllLL-L--LL-LhhhhlhhLLL-L---------LEEEELL-LlL-LHHHHHHHHHH-H-----eEEEE--E-lH---H-HLL-----HHHHHL-LH-H-H-L-L-LEEEEEELL----------L--HHHLLHHHHLLL-EEEELllllhhhhhhhhhhhhh
0045 6nakD   --lLLLH-H--HH-HH-H-HHHHLL---lllL----EEEEELL-LlL-LHHHHHHHHLL-L---L-lEEEE--E--L---L-LLH--HH-HLLLHH-HH-L-L-L-L-LEEEEEL-------------lLLLL-LLLLLL-EEEEElllllllllleeellll
0046 9jixA   ----------------------------lleE----EEEEELL-LlL-LHHHHHHHHHH-H--lL--EEEE--E--E---L-LHH--HH-HHHHHHHLL-L-L-L-L-LEEEEEELL----H--H-H--HHHHHHHH-lEE-EEEELlhhhhhhhhhhllleee
0047 7mi1A   lllllLL-L--HH-HH-H-HHH-HH-H----HhlllEEEEELL-LlL-LHHHHHHHHHH-----lL--EEEEelL--L---L-LLH-hHH-HHHHHHH-H-H-H-HlL-LEEEEEELL----lllL-L--LL-LLLLLLLE-EEEELllllhhhhhhhhhhhl
0048 1ixsB   lhhhlLL-L--HH-HH-H-HHHHHH-L----L----LEEEELL-LlL-LHHHHHHHHHH-H--lL--EEEE--E-lH---H-HLLhhHH-HH-HHHHHH-H-L--lL-LEEEEEELL----------L--LLLLLH-HHHHLlEEEELllllhhhhhhhhhhhh
0049 2is6A   hhhhhLL-L--LH-HH-H-HHLLL--L----L----LEEEELL-LlL-LHHHHHHHHHH-H---HhleEEE--LlhH---H-LLH--HH-HHHHH-HHL-L-L-L----EEEEEELH----HH-L-llllLHHHHHHHHLL-EEEELlllllllhhhhhhhhh
0050 2zpaB   --lLLLLhH--HH-HLlL-LLLL-L-L----L----EEEEEEL-LlL-LHHHHHHHLLL-----lL--LEEE--L-lH---H-HLL-----HHHHHHHHH-----H-L-LLEEEEEEELllllLL-L-HhhHHHHHLLL-LLE-EEEELllllllllllhhhhhhh
0051 7mi8A   lllllLL-L--HH-HH-H-HHH-HH-H----HhlllEEEEELL-LlL-LHHHHHHHHHH-----lL--EEEE--L-lL---L-LLLhhHH-HHHHHH-HH-H-H-HlL-LEEEEEELL----LllL-L--LL-LLHHHLLLE-EEEELllllhhhhhhhhhhhh

# Job: Structure-based Sequence Alignment of "BREX PglZ N-terminal Inactive P-loop Swi2/Snf2-Helicase" with various inactive P-loop domain from Swi2/Snf2-Helicases;
# Query: s001A
# No:  Chain     Z    rmsd lali nres  %id PDB
   1:  s001-A  22.6  0.0   95    95  100 Type1.PglZ.NTD
   2:  s002-A   9.0  2.1   82    97   11 Type3.PglZ.NTD
   3:  s022-B   7.2  3.0   90   164    6 6nmi
   4:  s003-A   6.8  2.7   88   139   11 6l80
   5:  s007-A   6.5  2.8   86   155    8 7vdv
   6:  s015-A   6.4  2.9   86   149   10 9ii8
   7:  s021-A   6.4  2.5   83   150   11 8zns
   8:  s016-A   6.3  3.1   81   143    7 6acx
   9:  s010-K   6.3  3.1   88   156    8 9n6k
  10:  s009-A   5.9  3.0   85   149    8 6bog
  11:  s012-G   5.9  2.7   83   142   10 9c57
  12:  s013-A   5.8  3.2   84   160    8 3i5x
  13:  s011-A   5.7  2.7   78   148    8 2d7d
  14:  s008-E   5.7  2.7   85   158   12 7zke
  15:  s018-A   5.5  2.3   74   136    7 4buj
  16:  s005-A   5.3  2.9   83   140    6 8t5s
  17:  s019-A   5.2  2.8   79   144    9 5lst
  18:  s020-A   5.1  2.5   77   124   10 2fzl
  19:  s014-C   5.1  3.1   83   161    7 7smo
  20:  s006-A   4.3  2.8   67   123    3 9el5
  21:  s004-A   4.1  2.9   69   114    6 8qy7
  22:  s017-A   3.5  3.3   70   158    9 7w0b

s001A_Type1.PglZ.NTD  MASSRITEALLNA-F-NKYRIVFWDDDKE-KL-IDEFNSL---ELP-D-VTKIKL-D-N------NE-FSVKY-RILVEETDK--KFLIYRPA-G--EPDVKDNWLLDVQL-AS-YKFVAE----------------
s002A_Type3.PglZ.NTD  SWRDHILKEFT---P-RVSRLTLVAD--P-DG-L--LLEEileGIR-EgFELIPF-E-D----HIAF-RYAYEsRFRS-RWDA--ALVVVLRSgA--LDV----lPYDLLQ-AG-RKLSF----------------
s003A_6l80            TKIKALLRHLKQI-QeTSPGQIIVFSQFS-SFlDILEIES---HLPdQ-VIIYKF-D-G-RLDMKER-TRILE-QFH--DKDLscIKLLLLSL-T--GGV--GLNL-TCAS----RAFMMDpwwspgmedqaidrih
s005A_8t5s            PMISKTVQYIVEQ-N-LADSRTIIFVRTR-YEaTILNKVLsneELL-MgIKSEWM-SglISASKQKQ-MEKLK-MFAD-----geiRILVSTS-V--AEE--GLDVP---E-CS-LVIKYNyatneiahvqrrgrgr
s007A_7vdv            GKFELLDRILPKL-R-ATNHKVLLFCQMT-SL-MTIMEDY---FyR-G-FKYLRL-D--GTTKAEDR-GMLLK-TFNE-PGSE--YFIFLLST--raGGLG---LNLQSA---D-TVIIFDsdwnphqdlqaqdrah
s008E_7zke            PKLGALRDLLVDC-GigKPHRALIFCQMK-EMlDMVQNTVlkqMLP-S-VSYLRL-D--GSVEANKR-QDIVN-KFNS--DPS--YDVLLLTT--svGGLG---LNLTGA---D-TVIFVEhdwnpqkdlqamdrah
s009A_6bog            wNFDPRVEWLXGY-L-TSHQKVLVICAKA-AT-ALQLEQV-lrERE-G-IRAAVF-H--EGXSIIER-DRAAA-WFA--EEDT-gaQVLLCSE----IGSE-GRNF-QFAS----HXVXFDlpfnpdlleqrigrld
s010K_9n6k            GKMVLLDQLLTRL-K-KDGHRVLIFSQMV-RM-LDILGDY---LiK-G-INFQRL-D--GTVPSAQR-RISID-HFNSPDS-N--DFVFLLS--T--RAGGLGIN-LMTA---D-TVVIFDsdwnpqadlqamarah
s011A_2d7d            GQIDDLIGEIQAR-I-ERNERVLVTTLTK-KMsEDLTDYK---EIG---IKVNYL-H--SEIKTLER-IEIIR-DL-----rlgkyDVLVG-I-N--LL---rEGLD--IP-EVsLVAILDadkegflrsersliqt
s012G_9c57            GKLEALAILLQKL-KsEGRR-VLILSQMI-LM-LDILEMN---FHY---LTYVRI-D--ENASSEQR-QELMR-SFN--RDRR--IFCAILST-H--SR---tTGINLV--eaD-TVVFYDndlnpvmdakaqewcd
s013A_3i5x            NSIFAAVEHIKKQ-I-KSNYKAIIFAPTV-KFtSFLCSIL-knEFK-KdLPILEF-H--GKITQNKR-TSLVK-RFKK---dE--SGILVCTD-G--ARG----MDFPNV---H-EVLQIGvpselanyihrigrta
s014C_7smo            RNESEKWRHIANL-C-RAEAQSIVFTFSR-KRtHELAAY-----SK-G-LKAKPY-H-S-GLPYKQR-KLTEM-EFLAQR-----lDVVVTT--A--ALGA-GVDF----P-AS-QVIFESlamgnkwlsvrefhqm
s015A_9ii8            EYVASIVSYIIEY-T-TTSSKCLVLFTSY-KMmHMVQDML---NFE-D-YVVLTQ-QqN------QN-Y-KIV-QQF-NNFDK--AILLGTST-F--FEG-----FDFQangIK-CVMIAKlpfmntstfkeyvlpd
s016A_6acx            -DDKQVAAALRRE-L-LRDGQAFYIHNRVrTI-DEAAARV-rqLVP-E-ARVVVA-H-G-QMNEETL-EKTVE-GFWN-----reyDILVC-T-T--IV---eTGLD--IS-NAnTLIVERadtfglsqlhqlrgrv
s018A_4buj            -SKKTWPEIVNYL-R-KELLPMVVFVFSK-KRcEEYADWL---EGI-N---IAVHvK-E------LIeILFSK-GF---------iKVLFA-T-E--TF--aMGLNL----ptR-TVIFS----------------
s019A_5lst            DTDQALLTLLQGkrF-QNLDSIIIYCNRR-EDtERIAALL-rtCLH-AktTAEAY-H--AGMCSRER-RRVQR-AFMQ-----gqlRVVVA-T-V--AF---gMGLDR-pD-VR-AVLHLGlppsfesyvqavgrag
s020A_2fzl            NKIRKLREILER--H-RKD-KIIIFTRHN-ELvYRISKVF-----------lIPAI-T--HRTSREER-EEILE-GFR---TGR--FRAIVSSQ----VLDE-GIDV---pD-AN-VGVIMSgsgsareyiqrlgril
s021A_8zns            -SWQKIADEIAAR------PCVLAVVNTR-KH-AQKLFAA---LPS-N-GIKLHLhC-S------EViALVRR-YLALySLHK--PLWLVSTQ-L--IEA--GVDL----D-FP-CVYRAMagldsiaqaagrcnre
s022B_6nmi            AVIRNYGNLLLEM-S-AVVPGIVAFFTSYqYM-ESTVASWileNIQ-RnKLLFIE-T-Q---DGAET-SVALE-KYQE-ENGR--GAILLSVA-RgkVSEG--IDFVHHY--gR-AVIMFGvpirendfltfdamrh
s004A_8qy7            ------------------KDKVIIFTELR-DLqRELQHAQ---NFG---frPVII-N-GKSQSQNSR-QRLID-DFQ--AQPG--FGVIIL---S--TVAVGFGVN---VQ-KAnHVIHFTrcwnpakedqatdray
s006A_9el5            ------------------ETRVMIFSSFR-DSvQEIAEMS---QHQpI-IRVMTF-V-GKGFTQKEQ-LEVVK-QFRD-----ggyNTLVSTC---vGEEG--LDIG---E-VD-LIICFDsqkspirlvqrmgrtg
s017A_7w0b            -------------dkdPKDICCLVFVERR-YTcKCIYGlN---YIQnV-LTPQFM-V-gSVLERKWQ-KSAIQ-QFRDGN-----aNLMICSS-V--LEE--GIDV---QA-CN-HVFILDpvktfnmyvqskgrar
```

```
s001A_Type1.PglZ.NTD  LHHHHHHHHHHHH-H-HHLLEEEEELHHH-LL-HHHHHHL---LLL-L-EEEEEL-L-L------LH-HHHHH-HHHLLLLLL--EEEEEELL-L--LLLLLLLLHHHHH-LL-EEELLL----------------
s002A_Type3.PglZ.NTD  LHHHHHHLLLL---L-LLLLEEEEEL--L-LL-H--HHLHhhhHHH-HlEEEEEL-L-L----HHHH-HHHHHhHLLL-LLLL--EEEEEELLlH--HHH----lLHHHHH-HL-EEEEL----------------
s003A_6l80            HHHHHHHHHHHHH-HhHLLLEEEEEELLH-HHhHHHHHHH---HLLlL-EEEEEL-L-L-LLLHHHH-HHHHH-HHL--LLLLllEEEEEEEL-L--LLL--LLLL-LLEE----EEEELLllllhhhhhhhhhlll
s005A_8t5s            HHHHHHHHHHHHH-H-HLLLLEEEELLLH-HHhHHHHHHHllhHHH-HlLLEELL-LllLLLLHHHH-HHHHH-HHHH-----lllLEEEELH-H--HHL--LLLLL---L-LL-EEEEELllllhhhhhhhhhlll
s007A_7vdv            HHHHHHHHHHHHH-H-HHLLLEEEEELLH-HH-HHHHHHH---HhL-L-LLEEEE-L--LLLLHHHH-HHHHH-HHHL-LLLL--LLEEEEEH--hhHLLL---LLLLLL---L-EEEELLllllhhhhhhhhllll
s008E_7zke            HHHHHHHHHHHHL-LllLLLLEEEEELLH-HHhHHHHHHHhhlLLL-L-LLEEEE-L--LLLLHHHH-HHHHH-HHHH--LLL--LLEEEEEH--hhHLLL---LLLLLL---L-EEEELLllllhhhhhhhhhlll
s009A_6bog            lLLLHHHHHHHHH-H-HHLLLEEEEELLH-HH-HHHHHHH-hhHHH-L-LLEEEE-L--LLLLHHHH-HHHHH-HHH--LLLL-llLEEEEEL----LLLL-LLLL-LLLL----EEEELLllllhhhhhhhhhlll
s010K_9n6k            LLHHHHHHHHHHH-H-HLLLLEEEEELLH-HH-HHHHHHH---HhH-L-LLEEEE-L--LLLLHHHH-HHHHH-HHLLLLL-L--LLEEEEE--L--LLLLLLLL-LLLL---L-EEEELLllllhhhhhhhhllll
s011A_2d7d            LHHHHHHHHHHHH-H-LLLLEEEEELLLH-HHhHHHHHHH---HLL---LLEEEE-L--LLLLHHHH-HHHHH-HH-----hhlllLEEEE-L-L--LL---lLLLL--LL-LEeEEEELLllllllllllhhhhhhh
s012G_9c57            HHHHHHHHHHHHH-HhHLLL-EEEEELLH-HH-HHHHHHH---HLL---LLEEEE-L--LLLLHHHH-HHHHH-HHH--HLLL--LLEEEEEL-L--LL---lLLLLLL--llL-EEEELLllllhhhhhhhhhhhh
s013A_3i5x            HHHHHHHHHHHHH-H-HLLLEEEEELLLH-HHhHHHHHHH-hhHHL-LlLLEEEE-L--LLLLHHHH-HHHHH-HHHH---lL--LEEEEELH-H--LLL----LLLLLL---L-EEEEELllllllhhhhhhhllll
s014C_7smo            LLHHHHHHHHHHH-H-HHHHLEEEELLLH-HHhHHHHHH-----HL-L-LLEEEL-L-L-LLLHHHH-HHHHH-HHHLLL-----lLEEEEL--L--LLLL-LLLL----L-LL-EEEELLllllllllllhhhhhhh
s015A_9ii8            HHHHHHHHHHHHH-H-HLLLLEEEELLLH-HHhHHHHHHH---HHL-L-LEEEEL-LlL------LL-L-HHH-HLL-LLLLL--LEEEEELL-L--LLL-----LLLLlllLE-EEEELLllllllllhhhhlhhhh
s016A_6acx            -LHHHHHHHHHHH-H-HLLLEEEEELLLHhHH-HHHHHHH-hhHLL-L-LLEEEE-L-L-LLLHHHH-HHHHH-HHHL-----lllLEEEE-L-H--HH---hLLLL--LL-LEeEEEELLhhhllhhhhhhhhhhll
s018A_4buj            -LLLHHHHHHHHH-H-HLLLLEEEELLLH-HHhHHHHHHLL---LLL-L---LEEEhH-H------HHhHHHHL-LL---------lLEEEE-L-H--HH--hHLLLL----llL-EEEEL-----------------
s019A_5lst            LHHHHHHHHHHLllL-LLLLLEEEEELLH-HHhHHHHHHH-hhHLL-LleEEEEE-L--LLLLHHHH-HHHHH-HHHL-----lleEEEEE-E-L--LL---lLLLLL-lL-LL-EEEEELllllhhhhhhhhhllll
s020A_2fzl            HHHHHHHHHHHH--L-LLL-LEEEELLLH-HHhHHHHHHL-----------lLLEE-L--LLLLLHHH-HHHHH-HHL---LLL--LLEEEELL----LLLL-LLLL---lL-LL-EEEEELlllllhhhhhhhhhhl
s021A_8zns            -LHHHHHHHHHLL------LEEEEEELLH-HH-HHHHHHH---LLL-L-LEEEEEhH-H------HHhHHHHH-HHHHhLLLL--LEEEEEEH-H--HLL--LLLL----L-LL-EEEEELllhhhhhhhhllllll
s022B_6nmi            HHHHHHHHHHHHH-H-HHLLLEEEEELLHhHH-HHHHHLLlhhHHH-HhLEEEEL-L-L---LLLHH-HHHHH-HHHH-LLLL--LEEEEEEL-LlhHHLL--LLLLLLL--lL-EEEEELllllhhhhhhhhhhhh
s004A_8qy7            ------------------LLEEEEELLLH-HHhHHHHHHHH---HHL---llLEEE-L-LLLHHHHHH-HHHHH-HHH--HLLL--LEEEEE---L--LLLLLLLL---LL-LEeEEEELLllllhhhhhhhhllll
s006A_9el5            ------------------LLLEEEELLLH-HHhHHHHHHHH---LLLlL-LLEEEE-L-LLLLLHHHH-HHHHH-HHHL-----lllLEEEELH---hHLLL--LLLL---L-LL-EEEELLllllhhhhhhhhllll
s017A_7w0b            -------------lllHHHLLEEEELLLH-HHhHHHHHhH---HHHlL-LLLLLL-L-lLLLHHHHH-HHHHH-HHHLLL-----lLLLEELL-L--HHH--LLLL---LL-LL-EEEELLllllhhhhhhhhhlll

# Job: Type1 PglZ WP_219938192 (Third Tri-Helical HTH-like Element)
# Query: s001A
# No:  Chain   Z    rmsd lali nres  %id PDB  Description
   1:  8b3d-c  4.6  3.6   49   264   10   MOLECULE: DNA-DIRECTED RNA POLYMERASE SUBUNIT;
   2:  6zce-q  4.5  2.9   52   636    6   MOLECULE: EUKARYOTIC TRANSLATION INITIATION FACTOR 3 SUBUNI
   3:  6w4x-C  4.3  3.6   55   375    7   MOLECULE: RIBONUCLEOSIDE-DIPHOSPHATE REDUCTASE 1 SUBUNIT AL
   4:  8bfi-A  4.3  3.6   45   601    9   MOLECULE: CCR4-NOT TRANSCRIPTION COMPLEX SUBUNIT 1;
   6:  7st9-E  4.2  4.0   53   354    8   MOLECULE: CHECKPOINT PROTEIN RAD24;
   7:  8dgc-A  4.2  3.6   52  2028    8   MOLECULE: SEAVS3;
   9:  6q82-A  4.1  3.3   51  1027   14   MOLECULE: IMPORTIN BETA-LIKE PROTEIN KAP122;
  10:  8cmk-A  4.1  3.4   52   913   15   MOLECULE: TRANSPORTIN-3;
  11:  6vk5-B  4.1  3.2   56   392    5   MOLECULE: METHANE MONOOXYGENASE COMPONENT A ALPHA CHAIN;
  12:  8zqc-A  4.1  4.1   51   930    8   MOLECULE: CAS12X2;
  13:  4cej-B  4.1  3.6   51  1156    2   MOLECULE: ATP-DEPENDENT HELICASE/NUCLEASE SUBUNIT A;
  14:  4owt-A  4.1  3.2   54   466    9   MOLECULE: INTEGRATOR COMPLEX SUBUNIT 3;
  15:  7uic-e  4.1  3.4   47   729    6   MOLECULE: MEDIATOR OF RNA POLYMERASE II TRANSCRIPTION SUBUN
  16:  3t92-A  4.0  2.7   50   112    6   MOLECULE: HISTONE ACETYLTRANSFERASE P300 TAZ2-CCAAT/ENHANCE
  17:  9c57-G  4.0  3.2   48   878   13   MOLECULE: RUVB-LIKE 1;
  18:  5kqj-A  4.0  3.6   50   185    6   MOLECULE: 2''-AMINOGLYCOSIDE NUCLEOTIDYLTRANSFERASE;
  19:  7vu7-A  3.9  4.2   52    89   13   MOLECULE: FLAGELLIFORM FIBROIN;
  20:  4in3-D  3.9  4.5   53   632    4   MOLECULE: CHITIN BIOSYNTHESIS PROTEIN CHS5;
  21:  8hpp-A  3.9  3.0   49   395    8   MOLECULE: INTEGRATOR COMPLEX SUBUNIT 3;
  22:  2iqc-A  3.9  4.1   57   173    4   MOLECULE: FANCONI ANEMIA GROUP F PROTEIN;
  23:  7dhg-C  3.9  2.7   48   470    6   MOLECULE: MITOCHONDRIAL IMPORT RECEPTOR SUBUNIT TOM70;
  24:  8j2x-A  3.8  3.9   50   337    8   MOLECULE: COBALAMIN-BINDING PROTEIN;
  25:  6vg5-A  3.8  2.7   42    81    2   MOLECULE: CAPSID PREMEMBRANE PROTEIN;
  26:  3jc6-5  3.8  3.0   50   254   10   MOLECULE: DNA REPLICATION LICENSING FACTOR MCM2;
  27:  4d0l-E  3.8  3.5   50   479    6   MOLECULE: PHOSPHATIDYLINOSITOL 4-KINASE BETA;
  28:  7xsq-A  3.8  3.8   51  1289    8   MOLECULE: RAMP SUPERFAMILY PROTEIN;
  29:  8j1p-A  3.7  3.5   55  1234    7   MOLECULE: UBIQUITIN FUSION DEGRADATION PROTEIN 4;
  30:  7t5p-A  3.7  3.8   52   403    8   MOLECULE: SUMO-INTERACTING MOTIF-CONTAINING PROTEIN 1;
  31:  6tdw-C  3.7  3.8   50   157    8   MOLECULE: ATPTB1;
  32:  8t1l-S  3.7  3.6   53   913    2   MOLECULE: MEDIATOR OF RNA POLYMERASE II TRANSCRIPTION SUBUN
  33:  7ozs-F  3.7  3.0   51   571    8   MOLECULE: PUTATIVE RIBOSOMAL PROTEIN;
  34:  1z2c-B  3.7  3.0   51   346   12   MOLECULE: RHO-RELATED GTP-BINDING PROTEIN RHOC;
  35:  8a1h-A  3.7  2.9   49   516    8   MOLECULE: 6-4 PHOTOLYASE (FES-BCP, CRYPRO);
  36:  3chl-A  3.7  3.2   49   315    4   MOLECULE: ALPHA-14 GIARDIN;
  37:  7kpx-C  3.7  3.7   49  1352   10   MOLECULE: MEIOTIC MRNA STABILITY PROTEIN KINASE SSN3;
  38:  6hb3-B  3.6  2.4   44   360    7   MOLECULE: PROTEIN HGH1;
  39:  7zke-E  3.6  3.3   48  1623    8   MOLECULE: DNA (36-MER);
  40:  8q6o-2  3.6  2.8   49   267   12   MOLECULE: DNA REPLICATION LICENSING FACTOR MCM2;
  41:  7ywq-A  3.6  2.6   50    79    4   MOLECULE: GENOME POLYPROTEIN;
  42:  5jp6-A  3.6  4.4   54   339    9   MOLECULE: PUTATIVE POLYSACCHARIDE DEACETYLASE;
  43:  7wo9-A  3.6  3.3   53  1581    6   MOLECULE: NUCLEOPORIN NUP188;
  44:  5tk6-A  3.5  3.9   46   191    9   MOLECULE: OXSA PROTEIN;
  45:  7zol-B  3.5  4.0   51  1702   12   MOLECULE: CAS7-11;
  46:  6n7p-X  3.5  3.9   49   826    4   MOLECULE: U1 SMALL NUCLEAR RIBONUCLEOPROTEIN 70 KDA HOMOLOG
  47:  5ufl-A  3.5  4.1   50   523    4   MOLECULE: PROTEIN CIP2A;
  48:  7wa4-A  3.5  3.1   49   529    6   MOLECULE: PROTEIN GIGANTEA;
  49:  8d07-A  3.5  3.9   50    65    8   MOLECULE: HALC3_109;
  50:  7v99-A  3.5  3.7   51   991   14   MOLECULE: TELOMERASE REVERSE TRANSCRIPTASE;
  51:  8tp8-D  3.5  3.2   51   326    2   MOLECULE: DEOR-FAMILY TRANSCRIPTIONAL REGULATOR;
  52:  8fib-A  3.5  3.9   57   418    5   MOLECULE: CYTOCHROME P450;
  53:  4y5j-A  3.5  3.2   47   227   13   MOLECULE: MINI SPINDLES TOG3;
  54:  7jtk-X  3.5  3.3   50   379    6   MOLECULE: FLAGELLAR RADIAL SPOKE PROTEIN 1;
  55:  6v8o-N  3.5  3.4   50   412   12   MOLECULE: HIGH TEMPERATURE LETHAL PROTEIN 1;
  56:  7wtw-w  3.5  3.9   51   332    6   MOLECULE: 18S RRNA;
  57:  6bq1-E  3.5  3.9   54  1510    7   MOLECULE: PHOSPHATIDYLINOSITOL 4-KINASE III ALPHA (PI4KA);
  58:  5tmc-F  3.4  2.6   51   351    8   MOLECULE: DNA-DIRECTED RNA POLYMERASE SUBUNIT ALPHA;
  59:  8e5t-6  3.4  3.8   52   514   17   MOLECULE: RIBOSOME BIOGENESIS PROTEIN MAK21;
  60:  9esh-N  3.4  3.5   56  1284    7   MOLECULE: PRE-MRNA;
  61:  8d00-A  3.4  3.9   47   299    6   MOLECULE: MICROTUBULE-ASSOCIATED PROTEIN TORTIFOLIA1;
  62:  3dad-A  3.4  2.8   54   324   11   MOLECULE: FH1/FH2 DOMAIN-CONTAINING PROTEIN 1;
  63:  5cos-D  3.4  3.5   48    61    6   MOLECULE: SIDEROPHORE-INTERACTING PROTEIN;
  64:  5t9f-B  3.4  2.7   51   254    4   MOLECULE: PREPHENATE DEHYDROGENASE 1;
  65:  7e34-B  3.4  3.8   46   142   11   MOLECULE: CYCLIN-DEPENDENT KINASE 2;
  66:  9jwg-A  3.4  3.0   52  1221    0   MOLECULE: E3 UBIQUITIN-PROTEIN LIGASE IPAH1.4;
  67:  3few-X  3.4  4.0   52   431    2   MOLECULE: COLICIN S4;
  68:  5jhf-D  3.4  2.6   42    82    0   MOLECULE: KLTH0D11660P;
  69:  5l7s-A  3.3  3.7   48   267    4   MOLECULE: SECRETED RXLR EFFECTOR PEPTIDE PROTEIN;
  70:  8gaq-A  3.3  4.6   44   147    7   MOLECULE: C5HR2_4R;
  71:  7te2-A  3.3  3.4   47   205   11   MOLECULE: AERR;
  72:  6ayi-A  3.3  3.2   45   183    7   MOLECULE: HTH-TYPE TRANSCRIPTIONAL REGULATOR UIDR;
  73:  8boq-C  3.3  3.4   45   119    9   MOLECULE: NITROGENASE PROTEIN ALPHA CHAIN;
  74:  2ii2-A  3.3  2.5   43   304   12   MOLECULE: ALPHA-11 GIARDIN;
  75:  8vwk-A  3.3  3.4   55   430   11   MOLECULE: CYTOCHROME P450;
  76:  8qat-C  3.3  4.5   54   606    6   MOLECULE: PROTEIN HOOK HOMOLOG 3;
  77:  2a3v-B  3.3  2.8   42   320    5   MOLECULE: DNA (31-MER);
  78:  1rjg-A  3.3  3.1   48   292    6   MOLECULE: CARBOXY METHYL TRANSFERASE FOR PROTEIN
  79:  8gju-F  3.2  3.3   44   332   11   MOLECULE: METHYLMALONIC ACIDURIA TYPE A PROTEIN, MITOCHONDR
  80:  1cf7-B  3.2  3.0   46    82   11   MOLECULE: DNA (5'-

0001 s001A   --------GLSL------H---DFA--LKIFND-SLKKQ-LGEP---SE--L---------S-N-EV-AMFLG-NWQ-D-S----K---N-----Y-G-ENFN-E-W-T-EK-A-A-T-E-L-N--------------
0002 8b3dc   -----------M------D---QKL--SKLVEE-LTTS--GEPR---------------lN-P-EK-MKELK-KIC-K-S----------------S-EEQL-S-R-A-YR-L-L-I-A-Q-Ltqehaeirlsafqiv
0003 6zceq   hssdylirlrDE------Q---SIY--NLILRT-QLYFE-ATL-----------ehdleraL-T-RP-FVKRL-DHI-Y-Y--------K-----S-E-NLIK-I-M-E-TA-A-W-N-I-I-Paqfkskftskdqlds
0004 6w4xC   gktvtvslreLK------K---KLY--LCLMSV-NALEA-IRFY--egnalH---------L-T-GT-QHMLN-LLR-S-G-addP---E-----M-A-EIAE-E-C-K-QE-C-Y-D-L-F-Vqaaqqekdwadylfr
0005 8bfiA   ssssdlrgfaaqfikqelistgTPF--AFVIDL-AALAS-RR-------------------eyL-KL-DKWLT-DKI-R-E---------------H-G-EPFI-Q-A-C-MT-F-L-K-R-Rl---------------
0006 8i21A   ksveslrdFQNK------K---G-Q--LLPYVK-VKDHK-EKL-----------emrdtpeF-Q-SW-MREIR-SY--D-Q----D---T-----K-E-KLNK-V-A-E-KY-E-E-E-G-N-Fnlsfvqdvldklese
0007 7st9E   lalkssspIIKP----dwI---IVI--HKLTRK-IVKER-S-----------------vnsL-I-EC-RAVLY-DLL-A-H----C---I-----P-A-NIIL-K-E-L-TF-S-L-L-D-V-Etlnttnkssiieyss
0008 8dgcA   ilaelcpsSALAiisrveD---SLA--GYIWAR-LGS---PEAE---M--------------rW-QA-AHAVL-ALC-R-M---------------S-R-TCVI-Q-G-I-FQ-H-A-I-N-A-Ttlpfcdrnlpfytlh
0009 8q7eJ   aqawsrdiAEDH----egV---SAL--GWLLDQ-YLEQR-ETSR------sF---------A-S-RV-RRLCH-LLV-H-V----EppsS-----S-L-RNIT-Q-C-W-LS-V-V-Q-E-Q-Vsrflaaawrapdfvp
0010 6q82A   khwtsfilPQFL----------gDLV--YQIXYG-LFHT-----Q---RS--D---------L-N-SY-TDLLR-ALV-A-K---------------F-P-IEAR-E-W-L-VA-V-L-P-Q-Ia---------------
0011 8cmkA   keligqvmnqLG------Q---QLV--SQLLHT-CCFC----LP---PY--T---------L-P-DV-AEVLW-EIM-Q-V---------------D-R-PTFC-R-W-L-EN-S-L-K-G-L-Pvtvthkqltdfhkqv
0012 6vk5B   lfnahssvGRDC----------qDWN--EILWAGhAVYDA-TFGQ--tltptT---------R-G-AI-DDLFVyCLA-N-D----S---E-----F-G-AHNR-T-F-L-NA-W-T-E-H-Y-Lassvaalkdfvglya
0013 8zqcA   edtlhllecvKE------Y---NEL--SKAYMD-YMFKL-KEDH----------lnlcqkgL-L-PI--PEFA-KNV-E-K--------------lN-K-NVLY-G-I-I-SQ-A-G-S-R-M-Ksflkcddltqqnhln
0014 4cejB   sdeeqlmyvvNK------S---VAQ--SFTASQ-LRLWT-REYD---I--------------S-D-VW-WSTYN-VLM-S-E----Q---------D-R-LQSK-K-L-F-S--S-L-F-Fr---------------
0015 4owtA   wllttctsnVAA------S---NAK--LALFYD-WLFF--SPDK----------------dS-I-XN-IEPAI-LVX-H-H----S---XkphpaI-T-ATLL-D-F-X-CR-I-I-P-N-F-Yppleghvrqgvfssl
0016 7uice   -------dmkLL------R---YYK--NIWLNN-KIINW--------------------eisN-P-DF-LSKYS-A----qN----L---D-----QlL-TDLI-E-T-S-FT-C-F-A-Q-F-Vsnkqyhqansnltll
0017 3t92A   ------dsrrLS------I---QRA--IQSLVH-AAQCR-------------------nancS-L-PS-CQKMK-RVV-Q-H-nggC---P-----I-C-KQLI-A-L-A-AY-H-A-K-H-C-Qenkcpvpfclnikqk
0018 9c57G   iaastareieCF------W---SNI--EQVVEI-KLRVE-LEEKrksvkfdyqklkngtkdL-E-EL-ADFME-QL---------------------T-PIEK-Y-A-L-NY-L-E-L-F-H-Tsieqekernsedavm
0019 5kqjA   pawwadeaYEIA------E---ApqgiWDYFYY-ADEVP-------pvD--W---------P-T-KH-IESYR-LAC-T-S--------------lG-A-EKVE-V-L-R-AA-F-R-S-R-Y-Aalehhhhhh------
0020 7vu7A   --------GSSR------L---PSL--INGIMS-SM-QG-GG----------------fnY-Q-NF-GNVLS-QFA-T---gtcN---S-----N-D-LNLL-M-D-A-LL-S-A-L-H-T-Lsyqgmgtvpsypsps
0021 4in3D   ssqilssindID------S---SII--DLVVKI-CCWNH-RWYI---E---F---------S-I-IL-IDALS-VAV-Q-D-------------mG-I-TKVH-N-E-I-AS-R-F-S-D-Pv---------------
0022 8hppA   qlqkgsdtEAQC----svG---KPL--YLIFRN-LCQMQ------------------ednssF-S-LL-LDLLS-ELY-Q-K---------------Q-PKIG-Y-H-L-LY-Y-L-R-A-S-Kaaagkmnlyesfaqa
0023 2iqcA   ltlvtsrhpALS------P---VYL--GLLTDW-GQRLHgWVGT--sqD--V---------P-WeEL-HNRFQ-SLC-Q-A----P---P----pL-K-DKVL-T-A-L-ET-C-K-A-Q-D-Gdfevpglsiwtdlll
0024 7dhgC   rrakahekLDNK------K---ECL--EDVTAV-CILEG--------------------fqN-Q-QS-MLLAD-Kvlllsqpsem E---M-----A-H-LYSL-C-D-A-AH-A-Q-T-E-V-Akk------------
0025 8j2xA   ---------VDI------L---AAG--REELMA-ALAEG--------------------D-E-HAaVDLAM-RLL-D-G----G--------vP-A-DVVL-LeL-V-AD-A-Q-V-E-I-Gvlwqanrwsvaqeha
0026 6vg5A   ----------------------rgPL--KLFMAL-VAFLR-FLTI---P------------pT-A-GI-LKRWG-TI-------------------------KK-S-K-A-IN-V-L-R-G-F-Rkeigrmlnilnrrrr
0027 3jc65   ---------DDD------N---TEI--IKSFKN-FILEF-RL------------sltvnmehlI-GY-NEDIY-KKL-S-D---------------E-P-SDII-P-L-F-ET-A-I-T-Q-V-Akrisilsrnslptfq
0028 4d0lE   gnrlfcfrneDV------D---FYL--PQLLNM-YIHM-----D---ED--V---------G-D-AI-KPYIV-HRC-R-Q-----------------S-INFS-L-Q-C-AL-L-L-G-A-Y-Ssrgtklrklilsdrl
0029 7xsqA   lcrievikseDH------N---DEL--RKQAEV-IVEAF-KQNdklgkeevQ---------G-T-A--LRTKL-KEL-W-Q---------snkdiG-W-RKFT-E-M-L-GS-N-L-Y-L-I-Ykketggvstrfrilg
```

```
0030 8j1pA   dslvqcliPILV----inD---QLI--KLIGSI-LA-QK-ETA---eAG--T---------LlV-GG-LSLLD-LIC-K-K--------------F-S-ELFF-P-S-I-KR-E-G-S-G-F-Eftstglassitkrit
0031 7t5pA   sianmvlscdkQ------P---HNV--RDVIKW-LVKAV-TE------dglI---------V-C-QL-QRMLS-IAV-E-V---dR---T-----P-T-CSSN-K-I-A-EM-M-F-G-F-V-Ldipersqremffttm
0032 6tdwC   lacgpgkvDALT------E---KHL--LETGKA-RLEEL-TAGL--------------gnkdegvnafrKEVE-QEG-K-Y---aN---W-----P-A-EKSK-A-L-A-DK-V-I-A-A-S---------------
0033 8t11S   pdhpcfrpDSTK---------acLS--ISAAIL-EILNA-WENG---V------------lA-F-ES-IQKIT-DNI---K----G---K-----V---CSLA-V-C-A-VA-W-L-V-A-H-Vrmlglderekslqmi
0034 7ozsF   ldvltaieHAAK----tkQ---TIL--RLLFRL-ISAD-----I---AP--Q-------diY-E-EA-ISCLT-TLS-E-D----------------N-LKVG-Q-A-I-TD---D-Q-E-T-Hvydvllklatgtdpr
0035 1z2cB   nnkfgiktmleT------E---EGI--LLLVRA-M--DP-AVPN---M--------------mI-DA-AKLLS-ALC-I-L----PqpeD-----M-N-ERVL-E-A-M-TE-R-A-E-M-D-Everfqplldglksgt
0036 8a1hA   dycascayQVKL----acP---LNS--LYWRFM-LKHRD-RL--------------annprI-G-ML-YKTWD-KM-------------------T-S-DSQQ-A-I-L-ST-A-D-A-Y-L-Sqiesl----------
0037 3chlA   navatgdriprD------T---HEL--EEDLVL-VRKAI-ETKG------kD---------E-V-ST-WIRIF-ATY--------------------T-RADF-R-Q-L-HK-M-Y-S-A-K-Yngdslragvedefqg
0038 7kpxC   kimennepaMAM------E---DLK--NFIFQI-IEITN-SN----------------------dlcsQIFDQ-LKDqT-I----E---M--itqI-V-EKDF-C-T-S-CL-Q-N-N-N-Q-Kiddnyivvvieiits
0039 6hb3B   dklvrniilsdD------K---KFL--KFLVWK-IVD-----LT---NP--N---------A-D-IM-CILLS-NLA-K------------------D-DGIL---A-V-LN-I-K-Rn--------------------
0040 7zkeE   lipmakefvmMR------R---SAL--DSLINI-VWESL-SN------------gddlsasT-G-KI-MDLLA-TLC-S------------------F-PEVL-E-A-M-KV-S-A-S-Q-D-Eersftllvprlypfl
0041 8q6o2   -tvrewvsmAAT------R---LEI--YHRFKN-FLRTH-VD------------eslpvnyedlaAR-EHVLA-YFL-P-E--------------A-P-AEML-K-I-F-DE-A-A-K-E-V-Vlvmypkydriareih
0042 7ywqA   tktrqprvqmPN------G---LVL--MRMMGI-LWHAV--AGT---A---R---------N-P-VL-KAFWN-SVP--------------------L-KQAT-A-A-L-RK-I-K-R-T-V-Salmvglqkrgkrr--
0043 5jp6A   ------------------M---SQI--GSSVRQ-AVSDN-QSA---nSE--A---------NpE-AL-FANWR-HEF-MvD----S---S-----K-R-ESMK-T-E-L-CK-E-L-Q-A-L-Paqdltlfeneirden
0044 7wo9A   alidtefanFEL------KdifNGW--SLVGRI-LQNLL-HSY---lQH--E---------LmI-SI-IKLVT-NVV-DkT----S---I-----EkS-SEIL-S-Y-L-SN-S-Ld----------------------
0045 5tk6A   tilnnndleIQE------I---VEH--ADHVDA-YCKSC-IEVH----------gnkdfisiqR-SL-GDKLD-NLT-K-E----------------Y-PYLK-E-F-Q-NL-Fl------------------------
0046 7zolB   dasvleisrppY------W---NNM--LNLAGT-ISEAF-ERE-----------wdieinkK-V-KL-RNWLF-WIF-N-E-frdtyayF-----D-W-RTFC-E-A-L-GQ-A-L-Y-L-E-Akkqvpnqfsserpvg
0047 6n7pX   imneygheDYFN------N---ALL--STLNAV-VVEQP-Q--------------------kqA-AI-ALLTM-VVN-S-K----------------N-NVAG-K-S-I-IN-Y-F-F-E-E-Lqkwckqtyndefkst
0048 5uflA   ssltlneeVGEK----fhA---RNI--HQTFQL-IFNIL-INGD--------------gtltrK-YS-VDLLM-DLL-K------------------N-PKIA-D-Y-L-TR-Y---E-H-F-Ssclhqvlgllngkdp
0049 7wa4A   vvltvcvshgPV------A---AFD--SYVLAA-VCALA-CEVQ----------ldsaishT-R-RI-LAILE-ALF-S-L--aaA---X-----V-A-AHIS-E-L-F-RR-Sk------------------------
0050 8d07A   ---reeieeAVK------E---AEL--KVLAIV-LVALR-SVSH-----------yeplsrlyE-SF-LDALK-KAL------------------S-E-EELK-E-V-E-KE-A-E-R-I-E-Kk--------------
0051 7v99A   gakgaagplpSE------A---VQW--LCHQAF-LLKLT-RHR-------pL---------L-G-SL-RTAQT-QLS-R-K-------------lP-G-TTLT-A-L-E-AA---A-N-P-A-Lpsdfktild------
0052 8tp8D   ripsgldfqtpT------A---EEL--AALRVA-ADSLR-ASGA---------------daraS-SL-YALEA-KLL-S-A----L---R---gsA-R-RRVA-P-D-V-EA-L-V-Q-A-Et----------------
0053 8fibA   ifaelfgmtQAD------I---FEL--AASIRT-ILEGL-NPHA--eQL--A---------AsD-AA-SLEVE-RYF-E---lsdT---E-----L-IsMLWG-I-L-L-GG-F-A-T-T-A-Amidhaildmlafpeq
0054 4y5jA   vaenypltttTV------D---LVI--NEIIEK-L-ADA-K--------------------ngA-AA-ADVLS-AFA-E-A-------------tK-L-EYVV-G-K-V-LS-F-A-F-E-Q-Kspkvqseafnwvnrs
0055 7jtkX   dstrraleaCGS------G----AV--LPRLME-LALLE-AQGG------gR---------A-Q-QG-LVLLF-TCT-Q-A----R---H-----N---AGIL-S-Q-LvDV-A-Qa----------------------
0056 6v8oN   kaneesaadNLD------H---KTL--SLIVSF-LLL-----EC---DS--E---------LiI-AS-LDFLY-QYI-L-P----G---S----qR-I-TELF-K-S---KE-C-S-Li--------------------
0057 7wtww   ipalsnweQILY---------rfyN--LVLLPR-VRDDV-AEYK---R------------lN-F-HL-YMALK-KAL-F----------------K-P-GAWF-K-GiL-IP-L-C-E-S-G-Tctlreaiivgsiitk
0058 6bq1E   aktkeneeELER----wsG---TVL--KTMLDI-LQTLS-LSLsadiVK--D---------FaA-RC-GMILQ-EAM-K-W--------------A-P-TVTK-S-H-L-QE-Y-L-N-Kh------------------
0059 5tmcF   aikklseitglD------P---DLI--REVVRA-KILGS---AR---VR-lD---------P-K-TV-EEIDQ-KLK-S-L--------------P-K-EHKR-Y---L-HI-A-R-E-G-E-Aarqhlieanlrlvvs
0060 8e5t6   lmpyllsyrrIL------K---ELI--KSIVGV-WSTT----RE---LE--T---------Q-I-AS-FAFLI-NTT-K-E--------------F-K-KSML-E-T-T-LK-T-T-Y-S-T-Fikscrktnmrsmpli
0061 9eshN   sivekfplHNLL------S---RWI--HSLLIK-SISYA-QTE-------tP---------L-L-AI-INMSL-VLL-S-A----F---P-----T-R-RFAH-P-V-I-ED-S-C-L-R-T-Sfpekyaikvdfeflk
0062 8d00A   --gshmltsFQA------M---VEL--KQKILT-SISKL-AD--------------rdtyqiaV-ED-LEKTI-QSL--------------------T-PETL-P-M-F-LN-C-L-Y-D-S-Csdpkpavkkeclhll
0063 3dadA   lvpefvhsEGLS----ahS---DTI--QWLYTL-CAS-----LS---RL--V----------vK-TA-LKLLL-VFV-EyS----E---N-----N-A-PLFI-R-A-V-NS-V-A-S-T-T-Gappwanlvsileekn
0064 5cosD   ---gatsipgEV------A---EQA--MHWHLE-LQEPA------------V---------S-A-AT-LAACM-SWR-Q-A----------------H-PLHE-H-A-W-QR-T-Q-V-F-A-Qrlremr---------
0065 5t9fB   ----------------aaK---SQF--ITHTIG-RTLGE-MDI---kET--M---------M-R-DS-FDLYS-GLF-V-Y----------------N-RFAR-Q-E-L-EN-L-E-H-A-L-Hkvketlmiqrty---
0066 7e34B   krakftisEHTR------I---NFF--IALYLA-NTVEE-------------------------dEE-ETKYE------lG----K---N-----W-R-KLFP-N-F-L-KL-R-D-Q-L-W-Dridyraivsrrccee
0067 9jwgA   asveylqevARI------R---LCL--DRAADF-LSE---PEGG-----------pemakeK-Q-CY-LQQVK-QFC-I-R----V---E-----N---DWHR-V-Y-L-VR-K-L-S-S-Q-Rgmefvqglskpgrph
0068 3fewX   lgniskafkfAD------V---VMK--VEKVRE-KSIEG-YET------------------gnW-GP-LMLEV-ESW-V-L----S---G----iA-S-AVAL-G-V-F-SA-T-L-G-A-Y-Alslgapaiavgivgi
0069 5jhfD   vdplkfywdlER------D---RSL--WSSVSK-LDNTK-K---------------------------tiDWK-RLS-R------------efkaP-E-HFIR-K-R-S-YA-L-F-A-K-H-Lklle-----------
0070 5l7sA   qmltlfqkTMTE-----eE---DII--KLVHML-R-NNP-G--------------------mkS-HA-DKLER-YLL-L-T--------------S-ESSH-K-T-M-AD-V-W-L-K-A-Retpeevfrilrlaek
0071 8gaqA   ------------------------V--EELKRK-LRQAK-EDGD------eL---------L-E-RV-KNEML-LLA-V-V----------------D-PRVL-V-E-V-LN-T-A-Ke--------------------
0072 7te2A   ----------SL------Q---TVL--VNRMVE-LAT-G-PEL-----------------gamD-LL-FDEFR-AAH-----------------vP-V-EEMAtH-Y-I-PE-A-A-R-Q-I-Gaawdsdrigfaqvti
0073 6ayiA   spgtlyhhfISK------E---ALI--QAIILQ-DQERA-LAR-----------ltheafgQ-R-AL-VVEIM-AEG-M-R----------------N-PQVA-A-M-L-KNk--------------------------
0074 8boqC   -----veapvhP------M---DAR--IDELTD-YIMKN-CL------------vcladdyR-E-HY--PWIN-SM-------------------S-K-EEIG-S-L-M-QG-L-K-D-R-M-Dyltitgslneelsdk
0075 2ii2A   ------------------Y---GDA--IPEVKA-ILEAK-----------------------nE-EE-LVTFT-SRW------------------S-A-EERK-E-L-R-TQ-F-Q-D-T-T-Glefiaflkkcikngp
0076 8vwkA   frwagipwsRQE------M---DRQ--ARRMSR-LLDTF-GRPA---T-----------hpisW-FE-RFRLD-RTF-TeK----T---A-----A-V-ELQN-L-T-R-PN-V-A-V-A-R-Faafagaalvehpewa
0077 8qatC   aaprlllfSRLV---------sYFC--PVLATG-LSALY-SSL---whclaL---------F-M-SS-LEFCN-AVI-Q-V----A---H---plV-Q-KQLV-D-Y-I-HN-G-L-V-P-Vm----------------
0078 2a3vB   refmqtryyAKK------T---IEA--YLHWIT-RYIHF-HNKK----------pslmgdkeveefltylavQ-GKV--------------------A-TKTQ-S-L-A-LN-S-L-S-F-L-Ykeilktplsleirfq
0079 1rjgA   ----------------------TDY--DALSCK-LAAIS-VGYL---PsskY---------T-E-WH-RSYLI-TLK-K-F--------------S-R-RAFG-K-V-D-KA-M-R-Ss--------------------
0080 8gjuF   ---------------------qrFV--DKLYTG-LIQG------------------------qR-AC-LAEAI-TLV-E-S----T---H---srK-K-ELAQ-V-L-L-QK-V-L-L-Y-H-Reqeqsnkgkplafrv
0081 1cf7B   ---------gkG------L---RHF--SMKVCE-KVQRK-GT--------------------tsynevaDELV-SEF-TdS----A---Y-----D-Q-KNIR-R-R-V-YD-A-L-N-V-L-Mamniiskekkeikwi

0001 s001A   --------LLLH------H---HHH--HHHHHH-HHHHH-LLLL---LL--L---------L-L-HH-HHHHH-HHH-L-L----L---L-----L-H-HHHH-H-H-H-HH-H-H-H-H-L-L---------------
0002 8b3dc   -----------L------H---HHH--HHHHHH-HHHL--LLLL----------------lL-H-HH-HHHHH-HHH-H-H----------------L-HHHH-H-H-H-HH-H-H-H-H-H-Hhlllhhhhhhhhhhh
0003 6zceq   llhhhhhhhhHH------H---HHH--HHHHHH-HHHHH-HHL-----------hhhhhhhH-H-HH-HHHHH-HLL-L-L--------L-----L-H-HHHH-H-H-H-HH-H-H-H-H-L-Llllllllllllllll
0004 6w4xC   leeeeelhhhHH------H---HHH--HHHHHH-HHHHH-LHHH--hhhhhH---------H-H-HH-HHHHH-HHH-H-L-lllH---H-----H-H-HHHH-H-H-H-HH-H-H-H-H-H-Hhhhhhhhhhhhhhll
0005 8bfiA   lllhhhhhhhhhhhhhhhhhhlLLH--HHHHHH-HHHHH-HL-------------------llL-LH-HHHHH-HHH-H-H--------------H-H-HHHH-H-H-H-HH-H-H-H-H-Ll----------------
0006 8i21A   hhhhhhhhHHHH------H---L-L--HHHHHH-HHHHH-HHH-----------hllllllL-H-HH-HHHHL-LL--L-L----L---L-----H-H-HHHH-H-H-H-HH-H-H-H-L-L-Llhhhhhhhhhhhhhl
0007 7st9E   leelllllLLLL----hhH---HHH--HHHHHH-HHHLL-L-----------------hhhH-H-HH-HHHHH-HHH-H-L----L---L-----L-H-HHHH-H-H-H-HH-H-H-H-L-L-Llllhhhhhhhhhhhh
0008 8dgcA   hhhhhlhhHHHHhhhhhhH---HHH--HHHHHH-HLL---LLHH---H--------------hH-HH-HHHHH-HHH-L-L--------------L-L-HHHH-H-H-H-HH-H-H-H-H-L-Llhhhlllllllhhh
0009 8q7eJ   hhhhhhhhHHLH----llL---LHH--HHHHHH-HHHHL-LLLL------hH---------H-H-HH-HHHHH-HHH-L-L----LlllL-----H-H-HHHH-H-H-H-HH-H-H-H-H-H-Hhhhhhhhlllllhhh
0010 6q82A   llllllhhHHHH---------hHHH--HHHHHH-HHHL-----L---LL--L---------L-H-HH-HHHHH-HHH-H-H--------------L-H-HHHH-H-H-H-HH-H-L-L-L-Ll----------------
0011 8cmkA   hhhhhhhhhhHH------H---HHH--HHHHHH-HHLL----LL---HH--H---------H-H-HH-HHHHH-HHH-H-H--------------H-H-HHHH-H-H-H-HH-H-H-H-H-L-Lllllhhhhhhhhhhh
0012 6vk5B   hhhhhhhhHHHL---------lLHH--HHHHHHhHLHHH-HHHH--llhhhH---------H-H-HH-HHHHHhHHH-L-L----L---L-----L-H-HHHH-H-H-H-HH-H-H-H-H-H-Hhhhhhhhhhhhhhhh
0013 8zqcA   hhhhhhhhhhHH------H---HHH--HHHHHH-HHHHH-HHHL----------hhhhhhlL-L-LL--HHHH-HHL-L-L-------------lL-H-HHHH-H-H-H-HH-H-H-H-H-H-Hhhhhhhhhhhhhhhh
0014 4cejB   lhhhhhhhllLH------H---HHH--HHHHHH-HHHHH-LLLL---L-------------L-H-HH-HHHHH-HHH-H-L----L---------L-H-HHHH-H-H-H-H--H-H-H-Ll------------------
0015 4owtA   hhhhllllhHHH------H---HHH--HHHLLL-LLLL--LLLL----------------lL-H-HH-HHHHH-HHH-H-H----H---LlllhhH-H-HHHH-H-H-H-HH-H-H-H-H-L-Lhhhhhhhhhhhhhhh
0016 7uice   -------lhhHH------H---HHH--HHHHHH-HHHLL-------------------lllL-L-LH-HHHHH-H----lL----H---H-----HhH-HHHH-H-H-H-HH-H-H-H-H-H-Hhllllllllllllhh
0017 3t92A   ------lhhhHH------H---HHH--HHHHHH-HHHLL------------------llllL-L-HH-HHHHH-HHH-H-H-lllL---H-----H-H-HHHH-H-H-H-HH-H-H-L-L-L-Llllllllhhhhhhhh
0018 9c57G   hhhhhhhhhhHH------H---HHH--HHHHHH-HHHHH-HHHHhhllllhhhhlllllllL-H-HH-HHHHL-LL---------------------L-HHHH-H-H-H-HH-H-H-H-H-L-Llllllllhhhhhhhh
0019 5kqjA   eeeeelleEEEL------L---LllllhHHHHHH-HHHLL-------llL--L---------L-H-HH-HHHHH-HHH-H-H-------------hL-H-HHHH-H-H-H-HH-H-H-H-H-H-Hhhhhhhhhl------
0020 7vu7A   --------LLLH------H---HHH--HHHHHH-HH-LL-LL-----------------llH-H-HH-HHHHH-HHL-L---lllL---H-----H-H-HHHH-H-H-H-HH-H-H-H-H-H-Hhhhllllllllllhh
0021 4in3D   hhhhhhhhhhHH------H---HHH--HHHHHH-HHHHH-HLLL---L---L---------L-H-HH-HHHHH-HHH-H-H-------------lL-H-HHHH-H-H-H-HH-H-L-L-H-Hh----------------
0022 8hppA   hhhhlllhHHHH----hlL---LHH--HHHHHH-HHLLL-----------------lllllH-H-HH-HHHHH-HHL-L-L----------------L-LLHH-H-H-H-HH-H-H-H-H-L-Lllllllhhhhhhhll
0023 2iqcA   hhhhhhhlhHHH------H---HHH--HHHHHH-HHLLElEELL--llL--L---------L-HhHH-HHHHH-HHH-L-L----L---H----hH-H-HHHH-H-H-H-HH-H-H-H-H-L-Lllllllllhhhhhhh
0024 7dhgC   hhhhhhhhLLLH------H---HHH--HHHHHH-HHHLL-------------------llL-H-HH-HHHHH-HhhhhllllhhH---H-----H-H-HHHH-H-H-H-HH-H-H-H-H-H-Hll-------------
0025 8j2xA   ---------LLH------H---HHH--HHHHHH-HHHHL----------------------L-H-HHhHHHHH-HHH-H-L----L--------lL-H-HHHH-HhL-H-HH-H-H-H-H-H-Hhhhhlllllhhhhhh
0026 6vg5A   ---------------------llLH--HHHHHH-HHHHH-HLLL---L------------lL-H-HH-HHHHL-LL---------------------------LH-H-H-HH-H-H-H-H-H-Hhhhhhhhhhhhhhll
0027 3jc65   ---------LHH------H---HHH--HHHHHH-HHHLL-LL------------eeeelhhhhH-HH-LHHHH-HHH-L-L--------------L-H-HHHH-H-H-H-HH-H-H-H-H-H-Hhhhllllllllllle
0028 4d0lE   hhhhhhllhhHH------H---LLH--HHHHHH-HHHL-----L---HH--H---------H-H-HH-HHHHH-HHH-H-H----------------L-HHHH-H-H-H-HH-H-H-H-H-H-Lllhhhhhhhhhhlll
0029 7xsqA   eeeeeeellhHH------H---HHH--HHHHHH-HHHHH-HHLllllllllL---------L-L-L--HHHHH-HHH-H-H---------llhhhL-H-HHHH-H-H-H-HH-H-H-H-H-H-Hhhhhlllllllllll
0030 8j1pA   hhhhhhhhHHHH----hhH---HHH--HHHHHH-LL-LH-HHH---hHH--H---------HhH-HH-HHHHH-HHH-H-H--------------L-H-HHHH-H-H-H-HH-H-L-L-H-H-Hhhhllhhhhhhhhhh
0031 7t5pA   lhhhhhhllllL------H---HHH--HHHHHH-HHHHH-HL------llhH---------H-H-HH-HHHHH-HHH-H-H---lL---L-----H-H-HHHH-H-H-H-HH-H-H-H-H-H-Hllllhhhhhhhhhll
0032 6tdwC   llllhhhhHHHH------H---HHH--HHHHHH-HHHHH-HHHH--------------llhhhhhhhhhHHHH-HHH-H-H---lL---L-----L-H-HHHH-H-H-H-HH-H-H-H-H-L-----------------
0033 8t11S   hhllllllLLLL---------llLL--HHHHHH-HHHHH-LLLL---L------------lL-L-HH-HHHHH-HHL---L----L---L-----L---LHHH-H-H-H-HH-H-H-H-H-H-Lllllllllhhhhhhh
0034 7ozsF   llhhhhhhHHHH----hlH---HHH--HHHHHH-HHHL-----L---LL--H-------hhH-H-HH-HHHHH-HHL-L-L----------------L-HHHH-H-H-H-HL---L-L-L-L-Lhhhhhhhhhhlllll
0035 1z2cB   llhhhhhhhhlL------L---LHH--HHHHHL-L--LL-LLHH---H--------------hH-HH-HHHHH-HHH-L-L----LlllL-----H-H-HHHH-H-H-H-HH-H-H-H-H-L-Lllllhhhhhhhllll
0036 8a1hA   lhhhhlllLLLL----llH---HHH--HHHHHH-HHLHH-HH--------------lllhhH-H-LH-HHHHH-LL-------------------L-H-HHHH-H-H-H-HH-H-H-H-H-H-Hlhhhl----------
0037 3chlA   hhhhhllllllL------H---HHH--HHHHHH-HHHHH-HLLL------hH---------H-H-HH-HHHHH-HHL--------------------L-HHHH-H-H-H-HH-H-H-H-H-H-Lllllhhhhhhhhllh
0038 7kpxC   hhhhllllhHHH------H---HHH--HHHHHH-HHHHH-LL----------------------lhhhHHHHH-HHHhH-H----H---H--lhhH-H-HHHH-H-H-H-HH-H-H-H-H-H-Hhhhhhhlllhhhhhh
0039 6hb3B   lhhhhhhhhhlL------L---HHH--HHHHHH-HLL-----LL---LL--L---------H-H-HH-HHHHH-HHL-L------------------L-HHHH---H-H-HH-L-L-Ll--------------------
0040 7zkeE   hhhhhhhhllLL------H---HHH--HHHHHH-HHHHH-HL------------lllllllH-H-HH-HHHHH-HHL-L------------------L-HHHH-H-H-H-HH-H-H-L-L-L-Hhhlhhhhhhhhhhhl
0041 8q6o2   -lhhhhhllHHH------H---HHH--HHHHHH-HHHLL-LL------------leeeeehhhhhHH-LHHHH-HHL-L-L--------------L-H-HHHH-H-H-H-HH-H-H-H-H-H-Hhhhllllllllllle
0042 7ywqA   llllllllllLH------H---HHH--HHHHHH-HHHHH--LLL---L---L---------L-H-HH-HHHHH-LLL--------------------L-HHHH-H-H-H-HH-H-H-H-H-H-Hhhhhhhhhlllll--
0043 5jp6A   ------------------L---HHH--HHHHHH-HHHHH-HHL---lLL--L---------LhH-HH-HHHHH-HHH-HhL----H---H-----H-H-HHHH-H-H-H-HH-H-H-H-L-L-Lhhhhhhhhhhhhlhh
0044 7wo9A   llllhhhhhHHH------HleeLHH--HHHHHH-HHHHH-HHH---hHH--H---------HhH-HH-HHHHH-HHL-LlL----L---H-----HhH-HHHH-H-H-H-LL-L-Ll----------------------
0045 5tk6A   hhhllllhhHHH------H---HHH--HHHHHH-HHHHH-HHHH----------llhhhhhhhH-HH-HHHHH-HHH-H-H----------------L-HHHH-H-H-H-HH-Hl------------------------
0046 7zolB   lllllllllllL------H---HHH--HHHHHH-HHHHH-HHL-----------lllllllL-L-LH-HHHHH-HHH-H-H-hhhhlllL-----L-H-HHHH-H-H-H-HH-H-H-H-H-H-Hhhhllllllllllll
0047 6n7pX   hhhhllllLLLH------H---HHH--HHHHHH-LLLLH-H--------------------hhH-HH-HHHHH-HHH-H-H----------------L-HHHH-H-H-H-HH-H-H-H-H-H-Hhhhhhlllllhhhhl
0048 5uflA   hhhhllllLLHH----hlL---LLH--HHHHHH-HHHHH-HHLL--------------llhhhH-HH-HHHHH-HHH-L------------------L-HHHH-H-H-H-HL-L---L-L-H-Hhhhhhhhhhhhlllh
0049 7wa4A   hhhhhhhhllHH------H---HHH--HHHHHH-HHHHH-HHHH----------llhhhhhH-H-HH-HHHHH-HHH-H-L--lhH---H-----H-H-HHHH-H-H-H-LL-Lh------------------------
0050 8d07A   ---lllhhhHHH------H---HHH--HHHHHH-HHHHH-HLLL-----------lllhhhhhH-HH-HHHHH-HHL------------------L-H-HHHH-H-H-H-HH-H-H-H-H-H-Hl--------------
0051 7v99A   llllllllllHH------H---HHH--HHHHHH-HHHHL-LLL-------hH---------H-H-HH-HHHHH-HHH-H-H-------------lL-H-HHHH-H-H-H-HH---L-L-L-L-Llllllllll------
0052 8tp8D   ellllllllllL------H---HHH--HHHHHH-HHHHH-HHLL---------------hhhhH-HH-HHHHH-HHH-H-L----L---L---hhH-H-HHHH-H-H-H-HH-H-H-H-L-Le----------------
0053 8fibA   hhhhhhlllHHH------H---HHH--HHHHHH-HHHHH-LLLL--hHH--H---------HhH-HH-HHHHH-HHH-H---llhH---H-----H-HhHHHH-H-H-H-HL-L-L-H-H-H-Hhhhhhhhhhhlhhh
0054 4y5jA   hhhhllllhhHH------H---HHH--HHHHHH-H-HLL-L--------------------lhH-HH-HHHHH-HHH-H-H-------------lL-H-HHHH-H-H-H-HH-H-H-H-H-L-Llhhhhhhhhhhhhhh
0055 7jtkX   hhhhhhhhlLLL------L----LH--HHHHHH-HHHHH-HLLL------hH---------H-H-HH-HHHHH-HHH-L-L----L---L-----L---HHHH-H-H-HhHH-H-Lh----------------------
0056 6v8oN   llllllhhhHLL------H---HHH--HHHHHH-HHL-----LL---LH--H---------HhH-HH-HHHHH-HHH-L-L----L---L----hH-H-HHHH-H-L---HH-H-H-Hh--------------------
0057 7wtww   llllllhhHHHH---------hhhH--HLHHHH-HHHHH-HHHL---L------------lL-H-HH-HHHHH-HHL-L----------------L-H-HHHH-H-HlH-HH-H-H-H-H-L-Llhhhhhhhhhhhhhh
0058 6bq1E   lllhhhhhHHHH----llH---HHH--HHHHHH-HHHHH-HHHhllhHH--H---------HhH-HH-HHHHH-HHH-H-L--------------L-H-HHHH-H-H-H-HH-H-L-L-Lh------------------
0059 5tmcF   hhhhhhhhhhlL------H---HHH--HHHHHH-HHLLL---LL---LL-lL---------L-L-HH-HHHHH-HHH-L-L--------------L-H-HHHH-H---H-HH-H-H-H-H-H-Hhhhhhhhhlhhhhhh
0060 8e5t6   lhhhhlllllLH------H---HHH--HHHHHH-HHHL----LL---HH--H---------H-H-HH-HHHHH-HHH-H-H--------------H-L-LLLH-H-H-H-HH-H-H-H-H-H-Hhllllllllllhhhh
0061 9eshN   hhhhhlllHHHH------H---HHH--HHHHHH-HHHHH-HHL-------hH---------H-H-HH-HHHHH-HHH-H-L----H---H-----H-H-HHHH-H-H-H-HH-H-L-L-L-L-Lllllllllllhhhhh
0062 8d00A   --lhhhhhhHHH------H---HHH--HHHHHH-HHHHL-LL--------------llllhhhH-HH-HHHHH-HLL--------------------L-LLLH-H-H-H-HH-H-H-H-H-H-Llllhhhhhhhhhhhh
0063 3dadA   hhhhhhhlLHHH----hlH---HHH--HHHHHH-HHL-----LL---HH--H----------hH-HH-HHHHH-HHH-HhL----H---H-----H-H-HHHH-H-H-H-HH-H-H-H-H-H-Lllllhhhhhhhhllll
0064 5cosD   ---llllllhHH------H---HHH--HHHHHH-HHLLE------------E---------L-H-HH-HHHHH-HHH-H-H----------------L-HHHH-H-H-H-HH-H-H-H-H-H-Hhhhlll---------
0065 5t9fB   ----------------hhH---LHH--HHHHHH-HHHHH-HLL---hHH--H---------L-L-LL-HHHHH-HHH-H-L----------------L-LLHH-H-H-H-HH-H-H-H-H-H-Hhhhhhhhhhhll---
0066 7e34B   hhhlllhhHLLH------H---HHH--HHHHHH-HHHHL-------------------------lLL-LHHHH------lH----H---H-----H-H-HHHH-H-H-H-HH-H-H-H-H-H-Hhllllllllhhhhhh
0067 9jwgA   llhhhhhhhHHH------H---HHH--HHHHHH-HLL---LLLL-----------lllhhhH-H-HH-HHHHH-HHH-H-H----H---L-----L---LLHH-H-H-H-HH-H-H-H-H-H-Llhhhhhhllllllll
0068 3fewX   hllllhhhllLL------L---HHH--HHHHHH-HHHHH-HHH------------------llL-HH-HHHHH-HHH-H-H----L---L----lL-H-HHHH-H-H-H-HH-H-H-H-H-H-Hhhlllllllhhhhhh
0069 5jhfD   lllllllllhHH------H---HHH--HHHHHH-HHLLL-L---------------------------llLHH-HHH-H------------hlllL-H-HHHH-H-H-H-HH-H-H-H-H-H-Hhhhl-----------
0070 5l7sA   hhhhhhhlLLLL-----hH---HHH--HHHHHH-L-LLL-L--------------------lhH-HH-HHHHH-HHH-L-L----------------L-LHHH-H-H-H-HH-H-H-H-H-L-Lllhhhhhhhhlllll
0071 8gaqA   ------------------------L--HHHHHH-HHHHH-HHLL------hH---------H-H-HH-HHHHH-HHH-H-H----------------L-HHHH-H-H-H-HH-H-H-Hh--------------------
0072 7te2A   ----------LH------H---HHH--HHHHHH-HHH-L-LLL-----------------lhhH-HH-HHHHH-HLL-----------------lL-H-HHHHhL-H-H-HH-H-H-H-H-H-Hhhhhlllllhhhhhh
0073 6ayiA   lhhhhhhhlLLH------H---HHH--HHHHHH-HHHHH-HHH-----------hllllhhH-H-HH-HHHHH-HHH-L-L----------------L-HHHH-H-H-H-HHh--------------------------
0074 8boqC   -----llllllL------L---HHH--HHHHHH-HHHHH-LL------------hhhhhhhH-H-HL--HHHH-HL-------------------L-H-HHHH-H-H-H-HH-H-H-H-H-H-Hhhhllllllllllll
0075 2ii2A   ------------------L---LLH--HHHHHH-HHHLL-----------------------lH-HH-HHHHH-LLL------------------L-H-HHHH-H-H-H-HH-H-H-H-H-H-Lllhhhhhhhhllllh
0076 8vwkA   hhhhlllllHHH------H---HHH--HHHHHH-HHHHL-LLLL---L-----------hhhhH-HH-HHHHH-HHH-HhH----H---H-----H-H-HHHL-L-L-H-HH-H-H-H-H-H-Hhhhhhhhhhhlhhhh
0077 8qatC   lllllhhhHLLH---------lLHH--HHHHHH-HHHHL-LLL---llllhH---------H-H-HH-HHHHH-HHH-H-H----L---L---hhH-H-HHHH-H-H-H-HH-H-L-L-L-Lh----------------
0078 2a3vB   hhhhhhlllLHH------H---HHH--HHHHHH-HHHLL-LLLL----------hhhllhhhhhhhhhhhhlL-LLL--------------------L-HHHH-H-H-H-HH-H-H-H-H-H-Hhllllllllllllll
0079 1rjgA   ----------------------LLH--HHHHHH-HHHHH-HLLL---LllhH---------H-H-HH-HHHHH-HHH-H-H--------------L-H-HHHH-H-H-H-HH-H-H-Ll--------------------
0080 8gjuF   ---------------------llLL--HHHHHH-HHLL------------------------lH-HH-HHHHH-HHH-L-L----L---L---hhH-H-HHHH-H-H-H-HH-H-H-H-H-H-Hhhhhlllllllleee
0081 1cf7B   ---------llH------H---HHH--HHHHHH-HHHHH-LE--------------------elhhhhhHHHH-HHH-HhL----H---H-----H-H-HHHH-H-H-H-HH-H-H-H-H-H-Hhlllllllllleell
```

```
# Job: PglZ Core Enzymatic Phosphatase (NPP-Superfamily)
# Query: s001A
# No:  Chain    Z    rmsd lali nres  %id PDB  Description
   1:  8tef-A 20.7  3.2  253   516   13   MOLECULE: RESPONSE REGULATOR RECEIVER PROTEIN;
   2:  9nv3-C 16.4  3.0  270   866   36   MOLECULE: DUF1788 DOMAIN-CONTAINING PROTEIN;
   3:  6c01-A 14.5  3.3  226   819   12   MOLECULE: ECTONUCLEOTIDE PYROPHOSPHATASE/PHOSPHODIESTERASE
   4:  5xwk-A 14.2  3.5  233   530   14   MOLECULE: ALKALINE PHOSPHATASE PHOK;
   5:  3szy-A 14.0  2.9  221   413   16   MOLECULE: PHOSPHONOACETATE HYDROLASE;
   6:  6xlp-A 11.8  3.7  168   586   15   MOLECULE: LPS BINDING PROTEIN;
   7:  5fgn-A 11.4  2.9  173   536   14   MOLECULE: LIPOOLIGOSACCHARIDE PHOSPHOETHANOLAMINE TRANSFERA
   8:  6s21-A 11.0  3.0  174   486   11   MOLECULE: ENDO-4-O-SULFATASE;
   9:  3igy-B 10.7  4.0  172   549   16   MOLECULE: COFACTOR-INDEPENDENT PHOSPHOGLYCERATE MUTASE;
  10:  3b5q-B 10.6  3.1  173   467   12   MOLECULE: PUTATIVE SULFATASE YIDJ;
  11:  7ebp-A 10.1  3.3  172   526   16   MOLECULE: SULFATASE;
  12:  6uss-A 10.0  3.3  171   485   15   MOLECULE: SULFATASE;
  13:  4n7t-A 10.0  3.3  163   402   15   MOLECULE: PHOSPHOPENTOMUTASE;
  14:  4kjg-A  9.9  3.3  163   486   12   MOLECULE: INTESTINAL-TYPE ALKALINE PHOSPHATASE 1;
  15:  5fql-A  9.9  3.1  170   507   12   MOLECULE: IDURONATE-2-SULFATASE;
  16:  8eg3-A  9.9  3.6  177   554   10   MOLECULE: STERYL-SULFATASE;
  17:  4upl-A  9.7  2.9  169   555   12   MOLECULE: SULFATASE FAMILY PROTEIN;
  18:  5olt-A  9.6  3.5  169   383   17   MOLECULE: CELLULOSE BIOSYNTHESIS PROTEIN BCSG;
  19:  4miv-D  9.6  3.2  168   480   15   MOLECULE: N-SULPHOGLUCOSAMINE SULPHOHYDROLASE;
  20:  2zkt-B  9.5  3.3  161   381   12   MOLECULE: 2,3-BISPHOSPHOGLYCERATE-INDEPENDENT PHOSPHOGLYCER
  21:  7qow-B  9.5  3.6  181   505   15   MOLECULE: ALKALINE PHOSPHATASE;
  22:  4cys-A  9.0  3.6  166   534   10   MOLECULE: ARYLSULFATASE;
  23:  2w5q-A  9.0  3.7  169   424    8   MOLECULE: PROCESSED GLYCEROL PHOSPHATE LIPOTEICHOIC ACID
  24:  3lxq-A  8.9  3.6  166   409    9   MOLECULE: UNCHARACTERIZED PROTEIN VP1736;
  25:  9fvt-A  8.7  3.7  171   519   11   MOLECULE: SULFATASE;
  26:  9bba-A  8.5  3.7  164   602   12   MOLECULE: S1_8C SULFATASE;
  27:  6dgm-B  8.3  4.1  167   382   11   MOLECULE: PHOSPHOGLYCEROL TRANSFERASE GACH;
  28:  7kwd-A  7.8  3.7  170   473   12   MOLECULE: ALKALINE PHOSPHATASE;
  29:  2d1g-A  7.5  3.5  159   481    8   MOLECULE: ACID PHOSPHATASE;
  30:  8haw-A  7.3  3.7  160   479    9   MOLECULE: NON-SPECIFIC PHOSPHOLIPASE C4;
  31:  8yk5-A  6.5  3.8  154   664    8   MOLECULE: PHOSPHOLIPASE C;

0001 s001A   [illegible]
0002 8tefA   [illegible]
0003 9nv3C   [illegible]
0004 6c01A   [illegible]
0005 5xwkA   [illegible]
0006 3szyA   [illegible]
0007 6xlpA   [illegible]
0008 5fgnA   [illegible]
0009 6s21A   [illegible]
0010 3igyB   [illegible]
0011 3b5qB   [illegible]
0012 7ebpA   [illegible]
0013 6ussA   [illegible]
0014 4n7tA   [illegible]
0015 4kjgA   [illegible]
0016 5fqlA   [illegible]
0017 8eg3A   [illegible]
0018 4uplA   [illegible]
0019 5oltA   [illegible]
0020 4mivD   [illegible]
0021 2zktB   [illegible]
0022 7qowB   [illegible]
0023 4cysA   [illegible]
0024 2w5qA   [illegible]
0025 3lxqA   [illegible]
0026 9fvtA   [illegible]
0027 9bbaA   [illegible]
0028 6dgmB   [illegible]
0029 7kwdA   [illegible]
0030 2d1gA   [illegible]
0031 8hawA   [illegible]
0032 8yk5A   [illegible]

0001 s001A   [illegible]
0002 8tefA   [illegible]
0003 9nv3C   [illegible]
0004 6c01A   [illegible]
0005 5xwkA   [illegible]
0006 3szyA   [illegible]
0007 6xlpA   [illegible]
0008 5fgnA   [illegible]
0009 6s21A   [illegible]
0010 3igyB   [illegible]
0011 3b5qB   [illegible]
0012 7ebpA   [illegible]
0013 6ussA   [illegible]
0014 4n7tA   [illegible]
0015 4kjgA   [illegible]
0016 5fqlA   [illegible]
0017 8eg3A   [illegible]
0018 4uplA   [illegible]
0019 5oltA   [illegible]
0020 4mivD   [illegible]
0021 2zktB   [illegible]
0022 7qowB   [illegible]
0023 4cysA   [illegible]
0024 2w5qA   [illegible]
0025 3lxqA   [illegible]
0026 9fvtA   [illegible]
0027 9bbaA   [illegible]
0028 6dgmB   [illegible]
0029 7kwdA   [illegible]
0030 2d1gA   [illegible]
0031 8hawA   [illegible]
0032 8yk5A   [illegible]

# Job: Type-1 BREX; PglZ WP_219938192 CTD Beta-Sandwich (IG-Like)
# Query: s001A
# No:  Chain    Z    rmsd lali nres  %id PDB  Description
   1:  8gul-A  9.5  2.5   92   464   14   MOLECULE: GLCNAC-BINDING PROTEIN A;
   2:  6za2-B  9.1  3.1  101  1083    8   MOLECULE: POR SECRETION SYSTEM PROTEIN PORU;
   3:  6fqa-A  8.8  2.3   90   229    7   MOLECULE: CSUC;
   4:  5ghu-A  8.6  3.1   92   201    5   MOLECULE: PROBABLE FIMBRIAL CHAPERONE YADV;
   5:  8u01-A  8.5  3.1   93   909   10   MOLECULE: GLYCOSYL HYDROLASE FAMILY 2, SUGAR BINDING DOMAIN
   6:  8s3o-A  8.3  2.7   90  1324    8   MOLECULE: CD109 ANTIGEN;
   7:  7rsk-A  8.2  3.1   95   775    6   MOLECULE: GLYCOSYL HYDROLASE FAMILY 2, SUGAR BINDING DOMAIN
   8:  7a5p-q  8.1  2.8   93  1896   11   MOLECULE: U2 SNRNA;
   9:  2co6-B  8.1  2.7   94   208    9   MOLECULE: PUTATIVE OUTER MEMBRANE PROTEIN;
  10:  9ayr-B  8.0  3.2   99   424   10   MOLECULE: GUANINE NUCLEOTIDE EXCHANGE FACTOR SUBUNIT RIC1;
  11:  6ej7-A  8.0  2.6   93   706    9   MOLECULE: XYLOSYLTRANSFERASE 1;
  12:  7s63-A  7.9  2.6   90  1420   10   MOLECULE: ALPHA 2-MACROGLOBULIN;
  13:  6cpd-B  7.9  3.0   89   127   16   MOLECULE: PMOD;
  14:  8wae-A  7.9  2.9   94   107   14   MOLECULE: BETA-AGARASE;
  15:  4be5-B  7.9  3.2   99   234    6   MOLECULE: RBMA;
  16:  1cvr-A  7.8  2.3   76   433   14   MOLECULE: GINGIPAIN R;
  17:  2j2z-A  7.8  3.4   92   216    9   MOLECULE: CHAPERONE PROTEIN PAPD;
  18:  5axh-B  7.8  2.9   95   587    6   MOLECULE: DEXTRANASE;
  19:  4e0s-A  7.8  2.9   91  1552   11   MOLECULE: COMPLEMENT C5;
  20:  3q48-A  7.8  3.0   88   194    3   MOLECULE: CHAPERONE CUPB2;
  21:  6ed1-A  7.7  3.1   99   860    8   MOLECULE: GLYCOSYL HYDROLASE FAMILY 2, SUGAR BINDING DOMAIN
  22:  5cl2-A  7.6  3.1   95   246   11   MOLECULE: SPORULATION-CONTROL PROTEIN SPO0M;
  23:  4b0m-M  7.6  2.6   91   212   10   MOLECULE: F1 CAPSULE-ANCHORING PROTEIN;
  24:  5ag8-A  7.6  2.3   76   149   16   MOLECULE: GINGIPAIN R2;
  25:  7ptb-A  7.5  2.5   92   317    5   MOLECULE: CENTROSOMAL PROTEIN OF 192 KDA;
  26:  3vmn-A  7.5  3.2   90   633    4   MOLECULE: DEXTRANASE;
  27:  8iua-A  7.5  2.5   82   555    7   MOLECULE: CANDIDATE DEXTRANASE GLYCOSIDE HYDROLASE FAMILY 6
  28:  7lr3-D  7.5  3.1   94   110    9   MOLECULE: D3_2/6.14 FAB LIGHT CHAIN;
  29:  2pn5-A  7.5  3.1   91  1283    5   MOLECULE: THIOESTER-CONTAINING PROTEIN I;
```

```
30:  8alz-B  7.5  2.5   94  1783   12   MOLECULE: ACTIVATING SIGNAL COINTEGRATOR 1;
31:  5vhv-A  7.4  3.1   93   361   14   MOLECULE: ALKYLPURINE DNA GLYCOSYLASE ALKC;
32:  3wnk-A  7.4  2.9   94   712    5   MOLECULE: CYCLOISOMALTOOLIGOSACCHARIDE GLUCANOTRANSFERASE;
33:  5dfk-A  7.4  2.4   82   185   10   MOLECULE: PROBABLE FIMBRIAL CHAPERONE ECPB;
34:  7xdi-E  7.4  3.1   88   185    6   MOLECULE: VP1;
35:  5uj6-B  7.4  3.1   98   840    8   MOLECULE: GLYCOSYL HYDROLASES FAMILY 2, SUGAR BINDING DOMAI
36:  8gb1-A  7.4  3.2   76    88    5   MOLECULE: MIROLASE;
37:  3fn9-C  7.4  2.9   96   673    9   MOLECULE: PUTATIVE BETA-GALACTOSIDASE;
38:  5t99-A  7.4  3.1   91   757   14   MOLECULE: GLYCOSIDE HYDROLASE;
39:  4ncd-A  7.3  2.8   90   211    7   MOLECULE: GRAM-NEGATIVE PILI ASSEMBLY CHAPERONE, N-TERMINAL
40:  9au7-C  7.3  2.9   98   285    4   MOLECULE: VPS35 ENDOSOMAL PROTEIN-SORTING FACTOR-LIKE;
41:  8cmt-A  7.3  2.7   92   722   10   MOLECULE: COAGULATION FACTOR XIII A CHAIN;
42:  6hpd-A  7.3  3.1   97   955    9   MOLECULE: BETA-GALACTOSIDASE (GH2);
43:  3eif-A  7.3  3.2  102   936    8   MOLECULE: C5A PEPTIDASE;
44:  7c7u-A  7.2  2.4   85   579   14   MOLECULE: BIOFILM-ASSOCIATED SURFACE PROTEIN;
45:  6pal-A  7.2  2.9   86   404    8   MOLECULE: UNCHARACTERIZED PROTEIN;
46:  5w0p-C  7.2  2.9  100   850   18   MOLECULE: ENDOLYSIN,RHODOPSIN,S-ARRESTIN;
47:  5n6u-A  7.1  3.0   91   811   19   MOLECULE: BETA-MANNOSIDASE;
48:  3tp4-B  7.1  3.2   84   457   11   MOLECULE: COMPUTATIONAL DESIGN OF ENZYME;
49:  8v2b-C  7.1  3.5   87   703    7   MOLECULE: ONCOSTATIN-M;
50:  7q5z-A  7.1  2.9   88  1344   10   MOLECULE: ALPHA-2-MACROGLOBULIN-LIKE PROTEIN 1;
51:  7usy-J  7.1  3.0   95   361   12   MOLECULE: TRANSMEMBRANE CHANNEL-LIKE PROTEIN 1;
52:  8wac-A  7.0  3.0   90   119   10   MOLECULE: AGARASE;
53:  6lvb-A  7.0  2.6   83   762   14   MOLECULE: N,N-DIMETHYLFORMAMIDASE LARGE SUBUNIT;
54:  1mls-A  7.0  3.1   88   109    6   MOLECULE: WR4;
55:  8sgi-A  7.0  3.2   88   751    8   MOLECULE: SODIUM/CALCIUM EXCHANGER 1;
```

```
0001 s001A   [illegible]
0002 8gu1A   [illegible]
0003 6za2B   [illegible]
0004 6fqaA   [illegible]
0005 5ghuA   [illegible]
0006 8u01A   [illegible]
0007 8s3oA   [illegible]
0008 7rskA   [illegible]
0009 7a5pq   [illegible]
0010 2co6B   [illegible]
0011 9ayrB   [illegible]
0012 6ej7A   [illegible]
0013 7s63A   [illegible]
0014 6cpdB   [illegible]
0015 8waeA   [illegible]
0016 4be5B   [illegible]
0017 1cvrA   [illegible]
0018 2j2zA   [illegible]
0019 5axhB   [illegible]
0020 4e0sA   [illegible]
0021 3q48A   [illegible]
0022 6ed1A   [illegible]
0023 5c12A   [illegible]
0024 4b0mM   [illegible]
0025 5ag8A   [illegible]
0026 7ptbA   [illegible]
0027 3vmnA   [illegible]
0028 8iuaA   [illegible]
0029 7lr3D   [illegible]
0030 2pn5A   [illegible]
0031 8alzB   [illegible]
0032 5vhvA   [illegible]
0033 3wnkA   [illegible]
0034 5dfkA   [illegible]
0035 7xdiE   [illegible]
0036 5uj6B   [illegible]
0037 8qb1A   [illegible]
0038 3fn9C   [illegible]
0039 5t99A   [illegible]
0040 4ncdA   [illegible]
0041 9au7C   [illegible]
0042 8cmtA   [illegible]
0043 6hpdA   [illegible]
0044 3eifA   [illegible]
0045 7c7uA   [illegible]
0046 6pa1A   [illegible]
0047 5w0pC   [illegible]
0048 5n6uA   [illegible]
0049 3tp4B   [illegible]
0050 8v2bC   [illegible]
0051 7q5zA   [illegible]
0052 7usyJ   [illegible]
0053 8wacA   [illegible]
0054 6lvbA   [illegible]
0055 1mlsA   [illegible]
0056 8sgiA   [illegible]

0001 s001A   [illegible]
0002 8gu1A   [illegible]
0003 6za2B   [illegible]
0004 6fqaA   [illegible]
0005 5ghuA   [illegible]
0006 8u01A   [illegible]
0007 8s3oA   [illegible]
0008 7rskA   [illegible]
0009 7a5pq   [illegible]
0010 2co6B   [illegible]
0011 9ayrB   [illegible]
0012 6ej7A   [illegible]
0013 7s63A   [illegible]
0014 6cpdB   [illegible]
0015 8waeA   [illegible]
0016 4be5B   [illegible]
0017 1cvrA   [illegible]
0018 2j2zA   [illegible]
0019 5axhB   [illegible]
0020 4e0sA   [illegible]
0021 3q48A   [illegible]
0022 6ed1A   [illegible]
0023 5c12A   [illegible]
0024 4b0mM   [illegible]
0025 5ag8A   [illegible]
0026 7ptbA   [illegible]
0027 3vmnA   [illegible]
0028 8iuaA   [illegible]
0029 7lr3D   [illegible]
0030 2pn5A   [illegible]
0031 8alzB   [illegible]
0032 5vhvA   [illegible]
0033 3wnkA   [illegible]
0034 5dfkA   [illegible]
0035 7xdiE   [illegible]
0036 5uj6B   [illegible]
0037 8qb1A   [illegible]
0038 3fn9C   [illegible]
0039 5t99A   [illegible]
0040 4ncdA   [illegible]
0041 9au7C   [illegible]
0042 8cmtA   [illegible]
0043 6hpdA   [illegible]
0044 3eifA   [illegible]
0045 7c7uA   [illegible]
0046 6pa1A   [illegible]
0047 5w0pC   [illegible]
0048 5n6uA   [illegible]
0049 3tp4B   [illegible]
0050 8v2bC   [illegible]
```

```
0051 7q5zA   -eeELLL--LEEE-EE---EL-LLEELL-----lL-EEEEEEEE--EL----llllLL-LEeEEEEEL------lL-L-LE-E-E---------E--EEeE-LLL------------LLEE-E-E-EEE-LL-L---L-LL--L-EE-EEEEEL-----------------LL----LEEEE-EEEL--Lllll
0052 7usyJ   --llLLE--EEEE-EE---LL-LLEELL-----lL-EEEEEEEE--EELLL----llllEE-EEEEEE-------EeE-EE-E-E---------E--EE-E-EeLLLE---------eEEE-E-E-EEE-LL-L---LlLL--L-LE-EEEEEEEELL--L-----L---lLL----EEEEE-EEEL--Lllll
0053 8wacA   ---------LEEE-EL---LL-LLEELL-----lL-EEEEEEEE--E--lllLLL-EE-EE-EEEEEE-elllllE-E-EE-E-E---------E--EE-E-E---ll----llllleEEE-E-E-EEE-LL-L---L-LL--L-HE-EEEEEEEEEL---------lLLE-EE----ELLEE-EEEL--Llll-
0054 6lvbA   -------lllLEE-EE---EL-LLEELL-----lL-EEEEEEEE--LL--------lLL-EE-EEEEEEellllleE-E-EE-LLL---------L--EE-E-ELellllllllllllllLL-L-E-EEE-LL-L---L-LL--L-LE-EEEEEELL---------------LL----EEEEE-EEEL--Lllll
0055 1m1sA   --------llLEE-EE---L---EEELL------E-EEEEEEEE--LLLL------EE-EE-EEEELL-----------LL-LEE---------eelLE-E-EELL-----------lLEE-E-E-EEE-EL-L-----LL--L-LE-EEEEEEEEHH--H-----LLLL-LE----EEEEE-EEEL-------
0056 8sgiA   -ellLLL--LEEE-EL---LL-EEEEEL------L-EEEEEEEE--LLLL---LL-EE-EE-EEEEEE-------L---lllLLL---------E--EE-E-EELLL----------lLEE-E-E-EEE-LL-L-lllllL--E-EE-EEEEELLLE--eeehhhhhhhllllleellllLEEE-EEEE--Llhhh

# Job: Type-2 BREX; PglZ QBI18515 CTD wHTH (Type-2 BREX-Specific)
# Query: s001A
# No:  Chain   Z    rmsd lali nres  %id PDB  Description
   1:  7wze-A  8.6  3.1   74   161   15   MOLECULE: UNCHARACTERIZED HTH-TYPE TRANSCRIPTIONAL REGULATO
   2:  5a31-N  8.6  2.9   74   703   15   MOLECULE: ANAPHASE-PROMOTING COMPLEX SUBUNIT 1;
   3:  3w6k-C  8.4  4.5   70    87   11   MOLECULE: SCPA;
   4:  8xt8-B  8.4  2.5   69   145   16   MOLECULE: MARR FAMILY TRANSCRIPTIONAL REGULATOR;
   5:  5xpq-A  8.3  3.1   74   100   20   MOLECULE: UNCHARACTERIZED HTH-TYPE TRANSCRIPTIONAL REGULATO
   6:  7el3-A  8.2  2.7   67   138    9   MOLECULE: HOMOPROTOCATECHUATE DEGRADATION OPERON REGULATOR
   7:  8ylg-A  8.2  2.5   67   161   19   MOLECULE: MARR FAMILY TRANSCRIPTIONAL REGULATOR;
   8:  8b9x-A  8.2  1.9   60   256   12   MOLECULE: DDRGK DOMAIN-CONTAINING PROTEIN 1,E3 UFM1-PROTEIN
   9:  2z99-A  8.2  4.2   70   160   21   MOLECULE: PUTATIVE UNCHARACTERIZED PROTEIN;
  10:  8jri-V  8.1  3.9   68   508    6   MOLECULE: 26S PROTEASE REGULATORY SUBUNIT 7;
  11:  3bdd-B  8.1  3.2   72   133   13   MOLECULE: REGULATORY PROTEIN MARR;
  12:  1lva-A  8.1  2.5   61   258   10   MOLECULE: SELENOCYSTEINE-SPECIFIC ELONGATION FACTOR;
  13:  6pco-B  8.0  2.6   68   135   13   MOLECULE: MARR-FAMILY TRANSCRIPTIONAL REGULATOR;
  14:  2xrn-B  8.0  2.2   68   241   16   MOLECULE: HTH-TYPE TRANSCRIPTIONAL REGULATOR TTGV;
  15:  5j6x-A  8.0  1.8   59    66   15   MOLECULE: Z-DNA BINDING PROTEIN KINASE;
  16:  6uvu-B  8.0  2.7   69   114   20   MOLECULE: ARSR FAMILY TRANSCRIPTIONAL REGULATOR;
  17:  4i98-B  8.0  4.5   68   167   18   MOLECULE: SEGREGATION AND CONDENSATION PROTEIN A;
  18:  6zwt-C  7.9  2.9   79   101   14   MOLECULE: TWO-COMPONENT RESPONSE REGULATOR;
  19:  7rhe-A  7.9  2.9   67   368   18   MOLECULE: ROK FAMILY PROTEIN;
  20:  2nyx-B  7.9  2.6   69   147   12   MOLECULE: PROBABLE TRANSCRIPTIONAL REGULATORY PROTEIN, RV14
  21:  8cli-A  7.9  2.8   65   542   15   MOLECULE: GENERAL TRANSCRIPTION FACTOR 3C POLYPEPTIDE 1;
  22:  6juv-A  7.8  4.3   72   192   17   MOLECULE: SEGREGATION AND CONDENSATION PROTEIN B;
  23:  6j0e-B  7.8  3.1   79   125   14   MOLECULE: ARSENIC RESPONSIVE REPRESSOR ARSR;
  24:  8iue-P  7.8  3.0   66   303   12   MOLECULE: DNA-DIRECTED RNA POLYMERASE III SUBUNIT RPC1;
  25:  2qlz-B  7.7  2.9   76   220   22   MOLECULE: TRANSCRIPTION FACTOR PF0095;
  26:  2wte-A  7.7  3.0   70   212   17   MOLECULE: CSA3;
  27:  7l1i-A  7.7  2.9   67   169   13   MOLECULE: MARR FAMILY MULTIDRUG RESISTANCE PUMP TRANSCRIPTI
  28:  4ija-A  7.7  2.6   70   363   14   MOLECULE: XYLR PROTEIN;
  29:  8qfc-B  7.7  1.8   60   631   12   MOLECULE: 60S RIBOSOMAL PROTEIN L10A;
  30:  7qcd-D  7.7  3.3   70   326   13   MOLECULE: STRUCTURAL MAINTENANCE OF CHROMOSOMES PROTEIN 5;
  31:  2rdp-A  7.7  2.5   64   140   16   MOLECULE: PUTATIVE TRANSCRIPTIONAL REGULATOR MARR;
  32:  8a5y-T  7.6  3.2   68   650   13   MOLECULE: ANAPHASE-PROMOTING COMPLEX SUBUNIT CDC27;
  33:  3s2w-A  7.6  3.0   72   146   18   MOLECULE: TRANSCRIPTIONAL REGULATOR, MARR FAMILY;
  34:  5f7p-A  7.6  2.1   64   359   16   MOLECULE: LMO0178 PROTEIN;
  35:  4fht-B  7.6  2.7   64   142   14   MOLECULE: PCAV TRANSCRIPTIONAL REGULATOR;
  36:  1fx7-A  7.6  3.0   64   230   16   MOLECULE: IRON-DEPENDENT REPRESSOR IDER;
  37:  7z1n-O  7.6  2.2   75   570   12   MOLECULE: DNA-DIRECTED RNA POLYMERASE III SUBUNIT RPC1;
  38:  5a5t-M  7.6  3.0   70   365   11   MOLECULE: EUKARYOTIC TRANSLATION INITIATION FACTOR 3 SUBUNI
  39:  1c0w-C  7.5  3.3   65   175   14   MOLECULE: DNA (5'-
  40:  6c2j-A  7.5  2.9   85   265   12   MOLECULE: RNAP1;
  41:  2lnb-A  7.4  2.0   64    80   20   MOLECULE: Z-DNA-BINDING PROTEIN 1;
  42:  5y6i-B  7.4  2.3   68   234   18   MOLECULE: TRANSCRIPTIONAL REGULATOR KDGR;
  43:  1sfx-A  7.4  2.7   72   109   13   MOLECULE: CONSERVED HYPOTHETICAL PROTEIN AF2008;
  44:  7cv0-A  7.4  2.1   59   173   10   MOLECULE: TRANSCRIPTIONAL REGULATOR NIAR;
  45:  2zkz-C  7.4  3.4   71    87   10   MOLECULE: TRANSCRIPTIONAL REPRESSOR PAGR;
  46:  3cuq-A  7.4  2.8   71   219   15   MOLECULE: VACUOLAR-SORTING PROTEIN SNF8;
  47:  4mtd-D  7.4  2.4   80   151   13   MOLECULE: ZINC UPTAKE REGULATION PROTEIN;
  48:  7dvr-A  7.3  2.5   66   147    6   MOLECULE: HTH MARR-TYPE DOMAIN-CONTAINING PROTEIN;
  49:  7c0j-B  7.3  2.0   56    62   14   MOLECULE: HISTONE H5,DOUBLE-STRANDED RNA-SPECIFIC ADENOSINE
  50:  6r7n-B  7.3  4.1   65   443   12   MOLECULE: COP9 SIGNALOSOME COMPLEX SUBUNIT 1;
  51:  6j05-A  7.3  3.8   81   101   15   MOLECULE: TRANSCRIPTIONAL REGULATOR ARSR;
  52:  5ju7-A  7.3  2.5   78   107   17   MOLECULE: TRANSCRIPTIONAL ACTIVATOR CADC;
  53:  6ndl-A  7.3  2.7   67   323    7   MOLECULE: BIOTIN PROTEIN LIGASE;
  54:  4hw0-A  7.3  1.5   57    98   11   MOLECULE: DNA-BINDING PROTEIN SSO10A-2;
  55:  8jxk-D  7.3  1.9   59   166   12   MOLECULE: CONSERVED PROTEIN;
  56:  3r4k-A  7.3  2.4   66   255   12   MOLECULE: TRANSCRIPTIONAL REGULATOR, ICLR FAMILY;
  57:  2eth-A  7.3  2.9   72   141   10   MOLECULE: TRANSCRIPTIONAL REGULATOR, PUTATIVE, MAR FAMILY;
  58:  2qww-A  7.3  2.7   65   146   17   MOLECULE: TRANSCRIPTIONAL REGULATOR, MARR FAMILY;
  59:  2mlg-A  7.2  2.3   59    77   19   MOLECULE: SULFOLOBUS TRANSCRIPTION FACTOR 76 AMINOACID PROT
  60:  6qfd-B  7.2  3.5   67   116   18   MOLECULE: DNA-BINDING PROTEIN;
  61:  3k69-A  7.2  2.6   65   151   14   MOLECULE: PUTATIVE TRANSCRIPTION REGULATOR;
  62:  8qfc-D  7.1  4.0   68   185   15   MOLECULE: 60S RIBOSOMAL PROTEIN L10A;
  63:  3g3z-A  7.1  2.7   65   142   12   MOLECULE: TRANSCRIPTIONAL REGULATOR, MARR FAMILY;
  64:  7bzh-A  7.1  2.2   59    59   10   MOLECULE: SUL7S;
  65:  2obp-A  7.1  2.2   61    81   21   MOLECULE: PUTATIVE DNA-BINDING PROTEIN;
  66:  3oop-A  7.1  2.8   67   139    9   MOLECULE: LIN2960 PROTEIN;
  67:  8f8u-A  7.1  3.6   63   297   16   MOLECULE: BIFUNCTIONAL LIGASE/REPRESSOR BIRA;
  68:  1yyv-A  7.0  2.5   68   114   18   MOLECULE: PUTATIVE TRANSCRIPTIONAL REGULATOR;
  69:  4hbl-D  7.0  3.4   72   147   14   MOLECULE: TRANSCRIPTIONAL REGULATOR, MARR FAMILY;
  70:  4yif-B  7.0  2.3   63   141   17   MOLECULE: MARR FAMILY PROTEIN RV0880;
  71:  2d1h-A  7.0  2.4   78   102   12   MOLECULE: 109AA LONG HYPOTHETICAL TRANSCRIPTIONAL REGULATOR
  72:  2l4m-A  7.0  2.1   62    69   13   MOLECULE: UNCHARACTERIZED PROTEIN;
  73:  3mq0-B  7.0  2.8   69   248   16   MOLECULE: TRANSCRIPTIONAL REPRESSOR OF THE BLCABC OPERON;
  74:  9c5g-B  7.0  2.5   64   290   13   MOLECULE: CAPW;
  75:  8r3g-A  7.0  2.4   61   335   20   MOLECULE: CENTRAL GLYCOLYTIC GENES REGULATOR;
  76:  3bpx-A  7.0  2.6   70   147   14   MOLECULE: TRANSCRIPTIONAL REGULATOR;
  77:  3f72-B  6.9  3.5   76   108   16   MOLECULE: CADMIUM EFFLUX SYSTEM ACCESSORY PROTEIN;
  78:  2pg4-A  6.9  2.2   66    92   18   MOLECULE: UNCHARACTERIZED PROTEIN;
  79:  3gfi-A  6.9  2.7   63   143   11   MOLECULE: 146AA LONG HYPOTHETICAL TRANSCRIPTIONAL REGULATOR
  80:  4nb5-B  6.9  2.7   71   149    8   MOLECULE: DNA BINDING PROTEIN;

0001 s001A   GQPA-WV-TELLASPLLAEQRARAGRAALSD-DDLAAFLAVLARHG--DV-ATAGVLQRE-TG-V--K-G-VRLR-SKLQALRRTLS-V-AYD-V-VRVD-T--DGTVRLNR-Q-L-L-AEQFQ--LE--QA
0002 7wzeA   agsllslehYFAGR--------------gLS-EGKFKILXLLFDAK-dHR-LSPTELAKR-SN-V--T-K-ATIT-GLLDGLARDG--------F-VSRR-HhrKISIELTK-A-R-L-E---Q--FL--PG
0003 5a31N   vTMD-V-tLSVA-----------------VT-PVQAVILLYFQDQ---AS-WTLEELSKA-VK-M--P-V-ALLR-RRMSVWLQQG--------V-LREE-P--PGTFSVIE-E-E-----RPQ--DRd---
0004 3w6kC   ----------------------------aLK-PAKAIVEALLFAAG---G-LSLSQIAAV-LE-V--S-E-LEAK-AVIEELQQDCR-R-ERgiQ-LVEL-G--G-VFLLA----T-K-KE--H--AP--Y-
0005 8xt8B   mkdfyeakddmFKQNGYD-----------iT-TDHYALLRFLWEQ---DG-ISQIDLCEK-SC-K--D-K-SNTT-RILDVMKNKG--------L-IVRK-V--DFQIFLTR-E-L-E-E------------
0006 5xpqA   --------------yklKAEF----fkTLAH-PARIRILELLVER----D-RSVGELLSSdVG-L--E-S-SNLS-QQLGVLRRAG--------V-VAAR-RgnAMIYSIAA-P-D-I-AEL---aVA--RK
0007 7el3A   -pirtlafrPALNE--------------gLT-EQQWRIIRILYQY---EE-LESNQLAEL-AC-I--L-K-PSLT-GILNRMVEQK--------L-IQKR-KqrISLISLTG-L-E-C--------------
0008 8ylgA   ssmirlqnPLFARY--------------glQ-PGEFDVLATLRRSGapYA-LTPTALYDA-AM-I--S-S-GSMT-NRIDRLEKAG--------W-VERR-A--NTLVALGR-A-Li---------------
0009 8b9xA   rtqdriqiddfiyilaavanfirqrgrvtPA-QISKEMRDELHVRG--GR-VNIVDLQQV-IN-V--D-L-IHIE-NRIGDIIKSEK-------H-VQLV----LGQLIDENy-------------------
0010 2z99A   ----------------------------dAD-ELKRVLEALLLVID---P-VTADALAAA-TE-Q--P-V-YRVA-AKLQLMADELT-G-DSgiD-LRHT-S--E-GWRMY----T-R-AR--F--AP--Y-
0011 8jriV   alrkhtaivvlGEIP-----drlqfrqprHN-VIKTGVRMISLSY---SR-ISLADIAQK-LQ-L-dS-P-EDAE-FIVAKAIRDG------viE-ASIN-H-eKGYVQSKE-M------------ID--IY
0012 3bddB   -----qedetlFEKQLG------------iS-LTRYSILQTLLKD---AP-LHQLALQER-LQ-I--D-R-AAVT-RHLKLLEESG--------Y-IIRK----EVLVWPTE---Q-A-REAtN--PS--A-
0013 1lvaA   ------------------------------G-SPEKILAQIIQEHR--EG-LDWQEAATR-AS-L--S-L-EETR-KLLQSXAAAG--------Q-VTLL-R--VLYAISRY-Q-Aw---------------
0014 6pcoB   --gpseqhvaqtIPDSK------------LT-AAQFVVLCALRDQ---GA-CSLVDVVKA-TA-I--D-Q-ATVR-GVIERLKAR-------K-L-LAVS-HrrKVLVTLTP-D-G-R-A------------
0015 2xrnB   ----------------------------qVI-ARAASIMRALGSHP---G-LSLAAIAQL-VG-L--P-R-STVQ-RIINALEEEF--------L-VEALgP--AGGFRLG---pA-L-GQ-LI--NQ--AQ
0016 5j6xA   ----------------------------aEN-EIEMRICDYLRRH---GR-STVQDIFKE-LK-L--E-K-STVN-RHLYSLQASK--------Q-VFKT-VnkRPVWDLVE--------------------
0017 6uvuB   -------dlieagEVVVLFK--------laN-DTRLRLLHALARSG---G-LCVTDLAAA-VG-M--K-P-QAVS-NQLQRLADR-------R-I-LRAA-RgnNIHYRIVD-P-C-V-L------------
0018 4i98B   --------------------------------STLAKIEALLFVAG---G-IRVRQLAEL-LS-L--P-P-TGIQ-QSLGKLAQKYEkD-DSslA-LIET----SGAYRLV----T-K-PQ--F--AE--I-
0019 6zwtC   EVVSwSL-DLTLTRE-------------tLT-TGEFAVLKALVQHP--REpLTRDKLMNL-AR----grE-RSID-VQVSRLRRLIE-DpARA-RyIQTV-W--GVGYVFVP-D-Ga---------------
0020 7rheA   ----------------------------vRR-YNERLLLKTLRRA---GS-ASKADLARL-AN-M--T-G-TAVG-SIIASLADAK--------L-IEFA----ASLIRLDP---R-G-AFG---aGD--VL
0021 2nyxB   esvddalaHSIAQV--------------nIT-IPQFRTLVILSNH---GP-INLATLATL-LG-V--Q-P-SATG-RXVDRLVGAE--------L-IDRL-PrrELLAALTK---R-G-RD-----------
0022 8cliA   rlgrqgevdahyhrknglitmQSHVirlKYD-ILMEKLSVMLSTRN--HI-ETLGKLREE-LG-L--C-E-RTFK-RLYQYMLNAG--------L-AKVV-S--LRCLKLLKt--------------------
0023 6juvA   ----------------------------gLL-EDKALVEAALFVAG--RP-LSLKELSKA-LG-I--KsL-EYLE-KLIELIASEYE-E-KSaiE-VVKV-L--GDKWVXQ----L-K-QE----sQK--VI
0024 6j0eB   ----lhtlatssdESEHYADLF----kVLGD-PVRLRILSQLAAGG--CP-VSVNELTDL-MG-L--S-Q-PTIS-HHLKKMTEAG--------F-LDRV-PgrVVLHRVRP---E-L-FAELR--T-----
0025 8iueP   ----------------------------dPV-EIENRIIELCHQFP--HG-ITDQVIQNE-MPhI--E-A-QQRA-VAINRLLSMG--------Q-LDLL-RntGLLYRIDS-Q-N-----DNQ--E-----
0026 2qlzB   yrsS-LX-RKIKEA-ylKEIG---------D-XTQLAILHYLLLN---GR-ATVEELSDR-LN-L--K-E-REVR-EKISEXARFV--------P-VKII-N--DNTVVLDE-D-Q-I-LR-----------
0027 2wteA   llgirkrsrvndnTIRALXR--------dYS-REEXKLLNVLYET---KG-TGITELAKX-LD-K--S-E-KTLI-NKIAELKKFG--------I-LTQK-G-kDRKVELNE-L-G-L-N------------
0028 7l1iA   nfpfsqviKKLG-----------------ID-NTRRKIILSTNAL---GE-ASITDIANL-ST-L--K-L-TTAT-KAVYRLVEDG--------I-VEVY-S--SSMVKLTA-----K-GVE----------
0029 4ijaA   ----------------------------pMN-DNEKRVLREIYNHH---N-ISRTQISKN-LE-I--N-K-ATIS-SILNKLKYKS--------L-VNEV-GrkPILLKVNH---L-Y-GYYNG--LL--GV
0030 8qfcB   ----------------------------tPA-QISKEMRDELHVRG--GR-VNIVDLQQV-IN-V--D-L-IHIE-NRIGDIIKSE------K-H-VQLV----LGQLIDENy-------------------
0031 7qcdD   ----------------------------tGD-ATAKYLLQYILSAR--GI-CHENALILA-LMtlnqW-V-DKLN-DYINAINVKLN-L-lGY-K-IIRI-N--HRFFVYV----N-L-AS----eTK--LA
0032 2rdpA   axneaelkqrILTN--------------pIT-PPQFVALQWLLEE---GD-LTVGELSNK-XY-L--A-C-STTT-DLVDRXERNG--------L-VARV-RehVVRIRLLE-K------------------
0033 8a5yT   ysdilkpkVEIQlaRKLV----------dVS-LEQCSVINQFDSpP--IC-LSLEQLSES-LN-I--A-P-PRLT-HLLDFWIQKG--------V-LLKE-N---GTYSVIE-H-Se---------------
0034 3s2wA   ishacdkrygIGKK--IEPYG-------IGS-G-QFPFLXRLYRE---DG-INQESLSDY-LK-I--D-K-GTTA-RAIQKLVDEG--------Y-VFRQ-R--DYRVFLTK-K-L-E-P------------
0035 5f7pA   ------------------------------D-INRYTVLNLIREK---GE-ITRTEIAKK-CD-F--G-M-STLT-YILDDLQQEG--------I-ILEG-----KLVRFNK---D-Y-GF-HL--LGvl--
0036 4fhtB   hmaaathhyltmVSEE-------------TT-SPQYAVLNALVAE---PG-LDQRTVGER-VG-L--D-R-STIA-EVVSRLGRRG--------L-LDKV-RgrRSLLRLTD-E------------------
0037 1fx7A   ----------------------------elV-DTTEMYLRTIYDLE--eT-PLRARIAER-LD-Q--S-G-PTVS-QTVSRMERDG--------L-LRVA-G--DRHLELGR-A-L---------pIP--GL
0038 7z1nO   pdLF-LYkELVKAH---------------lG-ERAASVIGMLVALG---R-LSVRELVEK-IDgM--D-V-DSVK-TTLVSLTQLR--------C-VKYL-QkkTTYYYYNE---E-G-IHILLym------
0039 5a5tM   sytesqalkdFLFDHL----------ltnMA-KMRLLTFMGMAVEN--KE-ISFDTMQQE-LQ-I--G-A-DDVE-AFVIDAVRTK------mvY-CKID-Q--TQRKVVVS-------------rTF--GK
0040 1c0wC   ----------------------------dLV-DTTEMYLRTIYELE---vtPLRARIAER-LE-Q--S-G-PTVS-QTVARMERDG--------L-VVVA-S--DRSLQMGR-T-L-----tiP--GL--DE
0041 6c2jA   LLPR-MR-QEFEE-IDFKAFFAHI---GIDY-KFGIDAMVQMALH---KR-ADLPTLVGT-LR---kS-A-QEVA-DNLFKMASE-------D-C-FNFD-P-tIDKFIVIY-------------iSD--DV
0042 2lnbA   ----------------------------rEG-HLEQRILQVLTEAG--SP-VKLAQLVKE-CQ-A--P-K-RELN-QVLYRMKKEL--------K-VSLT-S--PATWCLGG---T-D-PE-----------
0043 5y6iB   ----------------------------rSA-EVGTDILKALAELS--PA-TSLSRLAEH-VG-M--P-A-SKVH-RYLQALIASG--------F-AVQDaS--TNHYSL-A-L-R-V-GL--A--AL--D-
```

```
0044 1sfxA   ----------snplkalEKLS--------FK-PSDVRIYSLLLER---GG-XRVSEIARE-LD-L--S-A-RFVR-DRLKVLLKRG--------F-VRRE-IwvGYIYSAEK-E-K-V-LKEFK--S-----
0045 7cv0A   ----------------------------lGE-ERRSLLIKWLKASD---P-LTGAELAKR-TN-V--S-R-QVIV-QDVSLLKAKN-------hP-ILAT----AQGYIYMK-Ef-----------------
0046 2zkzC   ----------eyxSLEDDAELL----kTXAH-PXRLKIVNELYKH---KA-LNVTQIIQI-LK-L--P-Q-STVS-QHLCKXR---------gK-V-LKRN-RglEIYYSINN-P-K-V-------------
0047 3cuqA   PVGG----TYLI----qSVPA-------eLN-MDHTVVLQLAEK-N--GY-VTVSEIKAS-LK-W--E-T-ERAR-QVLEHLLKE--------G-L-AWLD-LpgEAHYWLPAl------------------
0048 4mtdD   ttQE-LL-AQAEKI--CAQRN-------rLT-PQRLEVLRLMSLQD--GA-ISAYDLLDL-LRqA--K-P-PTVY-RALDFLLEQG--------F-VHKV-E-sTNSYVLCH---L-F-DQP---------
0049 7dvrA   ----menlekryAKEYD-----------HLA-GPQGHLVMHLYKHP---D-MSIKDAEEI-LH-I--S-K-SVAS-NLVKRMEKNG--------F-IAIV-P--SKYLYLTH-L----------------
0050 7c0jB   ------------------------------P-TYSEMIAAAIRAEG---G-SSRQSIQAY-IK---vN-K-KEIN-RVLYSLLAAG--------V-LKQT-G--VGSWALA-------------------
0051 6r7nB   feafdeslvllmKSGInpfdsqeakpyklRN-IRTQVLIKLIKP-Y--TR-IHIPFISKE-LN-I--D-V-ADVE-SLLVQCILDN------tiH-GRIDqV--NQLLELDH-Q-Kr-------------
0052 6j05A   ------------QDPAQIVARL----eALAS-PVRLEIFRLLVEQE---G-LVSGDIAEH-LG-Q--P-H-NGIS-FHLKNLQHAG--------L-VTVQ-R--EQRYRAAM-P-V-V-RALVA--YLt---
0053 5ju7A   pvvrwLV-TPQISRRQLT-----------LE-PRLIDLLVFFAQHS--GEvLSRDELIDN-VW---vT-N-HVVT-QSISELRKSLK-D-SPV-Y-IATV-P--KRGYKLMV-P----------------
0054 6ndlA   ------------------------------S-KYSQDVLQLLYKNK----Y-ISGQSIAES-LN-I--S-R-TAVK-KVIDQLKLEG-------cK-IDSV-N--HKGHLLQQ-----L-PDIW--lDR--YQ
0055 4hw0A   ----------------------------gTX-EIXFDILRNCE-----PK-CGITRVIYG-AG-I--N-Y-VVAQ-KYLDQLVKVG--------A-LNIK-TndRKIYEITEk-----------------
0056 8jxkD   -----------------------------EF-MLELAILGLLIES----P-MHGYELRKR-LTaF--S-Y-GSLY-PALRRMQADG--------L-IAEN-A--ARVYQLTD-K----------------
0057 3r4kA   ----------------------------gTV-SKALTLLTYFN--R--LE-IGLSDLTRL-SG-X--N-K-ATVY-RLXSELQEAG--------F-VEQVeG--ARSYRLG---pQ-V-LR-LA--AL--R-
0058 2ethA   daletlfsNEEIS---------------dXK-TTELYAFLYVALF---GP-KKXKEIAEF-LS-T--T-K-SNVT-NVVDSLEKRG--------L-VVRE-XrrTYRVVLTK-E-I-F-G-----iLS--NF
0059 2qwwA   tenilktadQNAAS--------------gLT-IQQLAXINVIYST---PG-ISVADLTKR-LI-I--T-G-SSAA-ANVDGLISLG--------L-VVKL-NsxDLTLKLSK-K----------------
0060 2mlgA   ----------------------------akL-SLTQLILIRLSNR---GC-QTLEELEEF-TQ-A--K-R-EVLL-VTLTRLHQRG--------V-IYRK-WrkYREYCLKH-R----------------
0061 6qfdB   --------------pDARSD--------dLT-AFQKNILTVLGEE----A-RYGLAIKRE-LEeV--N-H-GRLY-PNLDDLVNKG--------L-VEKS-E--LNEYALTN--------------lEW--TL
0062 3k69A   ----------------------------lDF-SVAVHSILYLDAHR---K-VASRELAQS-LH-L--N-P-VXIR-NILSVLHKHG--------Y-LTGT-V--GGGYQLDL-----A-LADXNl-------
0063 8qfcD   ----lrkeaereersqreaewkkeeerlqSQ-SFLTEFINYIKQS---KV-VLLEDLASQ-VG-L--R-T-QDTI-NRIQDLLAEGT------iT-GVID-D--RGKFIYIT-P-E-E-LAAVAn-------
0064 3g3zA   xnqlgtriGQQD-----------------lN-YNLFAVLYTLATE---GS-RTQKHIGEK-WS-L--P-K-QTVS-GVCKTLAGQ--------G-L-IEWQ-ErrKRLLSLTK-A-Y-A-------------
0065 7bzhA   -----------------------------ME-DVKQSVEKIIKDR---EW-VTFNDLLKY-IP-Y--P-A-PEVY-DALSQLIKEN--------K-VGRR----GRYFYYIK-R----------------
0066 2obpA   ----------------------------gID-PAIVEVLLVLREAGienP-WSLPKIAKR-AQ-L--P-X-SVLR-RVLTQLQAAG--------L-ADVS-VdgRGHASLGA-A-La--------------
0067 3oopA   gyydfdvxrSIASY--------------dVT-PEQWSVLEGIEAN---EP-ISQKEIALW-TK-K--D-T-PTVN-RIVDVLLRKE--------L-IVRE-IrrISLLSLTR-K-E-T-------------
0068 8f8uA   ----------------------------kdH-TIPLTLISILA-DG--EF-HSGEQLGEQ-LG-M--S-R-AAIN-KHIQTLRDWGV-------D-VFTV-P--GKGYSLPE--------------dEK--KI
0069 1yyvA   ----------lregeqcPSRE----vlKHVTsRWGVLILVALR--D--GT-HRFSDLRRX-XGgV--S-E-XXLA-QSLQALEQDG--------F-LNRV-SppHVEYSLTP-L-G-E-Q-----------
0070 4hblD   rlanfsaekKLKQF--------------giT-YSQYLVXLTLWEE---NP-QTLNSIGRH-LD-L--S-S-NTLT-PXLKRLEQSG--------W-VKRE-R--QLIITLGQ-Q-Q-Q-EA-fE--AI--S-
0071 4yifB   ----arlsrqRNPS--------------pVS-LSQLSALTTLANE---GA-MTPGALAIR-ER-V--R-P-PSMT-RVIASLADMG--------F-VDRA-P--HVLVSVSEs-----------------
0072 2d1hA   klES-KK-DEIRCC--------------kIT-DTDVAVLLKXVEIE--KP-ITSEELADI-FK-L--S-K-TTVE-NSLKKLIELG--------L-VVRT-KtpKYYYSISI-L-E-K-IRNDLl-------
0073 2l4mA   -------mASPQ----------------fSQ-QREEDIYRFLKDN---GP-QRALVIAQA-LG-M--R-TaKDVN-RDLYRMKSRH---------lLDMD-E-qSKAWTIYR-------------------
0074 3mq0B   ----------------------------pAL-RRAVRILDLVAGSP--RD-LTAAELTRF-LD-L--P-K-SSAH-GLLAVMTELD--------L-LARS-A--DGTLRIGP---H-S-LR--W--AN--GF
0075 9c5gB   ----------------------------mQN-RRFEFIEWKLFW-E--GA-LNRSDLEET-FE-I--S-T-PQTS-IDLRRYRELAG-----D-N-IEYD-A-tDKTFKPTK-G-M-K-------------
0076 8r3gA   ----------------------------lVX-QKRFEILQYIRLT---EP-IGRRSLSAS-LG-I--S-E-RVLR-GEVQFLKEQN--------L-VDIK-T--N-GXTLTY-E-L-Lr------------
0077 3bpxA   ----mdrlrsfIGRE-LGHLN--------lT-DAQVACLLRIHRE---PG-IKQDELATF-FH-V--D-K-GTIA-RTLRRLEES-------G-F-IERE-Q--DYILEVTG-E-E-I-------------
0078 3f72B   -------ekvqgdlISGVSQI----lkAIAD-ENRAKITYALCQDE---E-LCVCDIANI-LG-V--T-I-ANAS-HHLRTLYKQG---------vVNFR-KgkLALYSLGG-E-A-I-RQ----mIA--LA
0079 2pg4A   -----------------XDDE----tlRLQF-GHLIRILPTLLEFE--kE-PSLAEIVKA-SG-V--S-E-KTFFxGLKDRLIRAG--------L-VKEE-T--LKTLKLTE-K----------------
0080 3gfiA   enristinrRLGE---------------nLS-YLDFLVLRATS--D--GP-KTMAYLANR-YF-V--T-Q-SAIT-ASVDKLEEMG--------L-VVRV-RrrKILIEITE-K----------------
0081 4nb5B   EFVE------------------------sLT-RLAGRLLGWLLVCD-pER-QSSEELATA-LA-A--S-S-GGIS-TNARMLIQFG--------F-IERL-AdrRTYFRLRP---N-A-FAAGEr-------

0001 s001A   LLLH-HH-HHHHLLHHHHHHHHHHHHHHLLLH-HHHHHHHHHHHHLL--LE-EEHHHHHHH-HL-L--L-L-HHHH-HHHHHHHHHHL-L-LLL-L-EEEL-L--LLEEEELH-H-H-H-HHHHL--LL--LL
0002 7wzeA   hhhhhhhhhHHHLL--------------lLL-HHHHHHHHHHHHLH-hHE-ELHHHHHHH-LL-L--L-H-HHHH-HHHHHHHHHL--------L-EEEE-EllLEEEEELH-H-H-H-H---H--HH--HH
0003 5a31N   eEEE-E-lEELL-----------------LL-HHHHHHHHHHHLL---LL-LLHHHHHHH-HL-L--L-H-HHHH-HHHHHHHHHL--------L-LLLL-L--LLLLLLLL-L-L-----LLL--LLl---
0004 3w6kC   ----------------------------lLH-HHHHHHHHHHHHHH---L-LLHHHHHHH-HL-L--L-H-HHHH-HHHHHHHHHHL-L-LLleE-EEEE-L--L-EEEEE----E-L-HH--H--HH--H-
0005 8xt8B   lllhhhhhhhhHHHLLLL-----------lL-HHHHHHHHHHHHL---LL-LLHHHHHHH-HL-L--L-L-HHHH-HHHHHHHLLL--------L-EEEE-E--LEEEEELH-H-L-H-H------------
0006 5xpqA   --------------hhhHHHH----hhHLLL-HHHHHHHHHHHHL----L-EEHHHHHLLlLL-L--L-H-HHHH-HHHHHHHHLL--------L-EEEE-EllLEEEEELL-L-H-H-HHH---hHH--HH
0007 7el3A   -lllhhhlhHHHHH--------------lLL-HHHHHHHHHHHHHH---LE-EEHHHHHHH-HL-L--L-H-HHHH-HHHHHHHHLL--------L-EEEE-EeeEEEEEELH-H-H-H-------------
0008 8ylgA   lhhhhhhhHHHHHL--------------llL-HHHHHHHHHHHHLLllLE-ELHHHHHHH-HL-L--L-H-HHHH-HHHHHHHHHL--------L-EEEE-E--LEEEEEHH-H-Hh--------------
0009 8b9xA   lhhhhhhelleeelhhhhhhhhhhhleelHH-HHHHHHHHHHHHLL--LE-EEHHHHHHH-HL-L--L-L-HHHH-LLHHHHHHHLL-------L-EEEE----LLEEEEHHh-----------------
0010 2z99A   ----------------------------lHH-HHHHHHHHHHHHLL---L-LLHHHHHHH-HL-L--L-H-HHHH-HHHHHHHHHHH-H-LLleE-EEEE-L--L-EEEEE----E-L-HH--H--HH--H-
0011 8jriV   hhhhlllhhhhLLLL-----llllllllhHH-HHHHHHHHHHHHL---LE-ELHHHHHHH-HL-L-lL-H-HHHH-HHHHHHHHLL------llL-LEEL-L-lLLLEELLL-L-----------LL--HH
0012 3bddB   -----lhhhhhHHHHHL------------lL-HHHHHHHHHHHHL---LL-EEHHHHHHH-HL-L--L-H-HHHH-HHHHHHHHLL--------L-EEEL----LLEEEELH---H-H-HHHlL--LL--H-
0013 1lvaA   ------------------------------L-LHHHHHHHHHHHLLL--LL-EEHHHHHHH-HL-L--L-H-HHHH-HHHHHHHHLL--------L-EEEE-E--EEEEEEHH-H-Hh--------------
0014 6pcoB   --llhhlhhhhhLHHHL------------LL-HHHHHHHHHHHHHH---LL-LLHHHHHHH-HL-L--L-H-HHHH-HHHHHHHHL-------L-L-EEEE-LllLLEEEELH-H-H-H-H------------
0015 2xrnB   ----------------------------hHH-HHHHHHHHHHHHLLL---L-EEHHHHHHH-LL-L--L-H-HHHH-HHHHHHHLLL--------L-EEELhH--HLEEEEL---lH-H-HH-HH--HH--HH
0016 5j6xA   ----------------------------hHH-HHHHHHHHHHHHH---LL-LLHHHHHHH-LL-L--L-H-HHHH-HHHHHHHHLL--------L-EEEE-LllLLEEEELL-------------------
0017 6uvuB   -------lllhhhHHHHHHH--------hlL-HHHHHHHHHHHHHL---L-LLHHHHHHH-LL-L--L-H-HHHH-HHHHHHHHL-------L-L-EEEE-EllEEEEEELL-H-H-H-H------------
0018 4i98B   --------------------------------LHHHHHHHHHHHLL---L-EEHHHHHHH-HL-L--L-H-HHHH-HHHHHHHHHHHhL-LLleE-EEEE----LLEEEEE----E-L-HH--H--HH--H-
0019 6zwtC   LEEEeEE-ELEEEEL-------------lLL-HHHHHHHHHHHLLL--LLlEEHHHHHHH-HH----llL-LHHH-HHHHHHHHHHL-LhHHL-LlEEEE-L--LLEEEELL-L-Ll--------------
0020 7rheA   ----------------------------hHH-HHHHHHHHHHHHH---LL-EEHHHHHHH-HH-L--L-H-HHHH-HHHHHHHHLL--------L-EEEL----LLEEEELL---L-L-LEE---lLL--EE
0021 2nyxB   llllhhhhHHHHHH--------------lLL-HHHHHHHHHHHHHH---LL-EEHHHHHHH-HL-L--L-H-HHHH-HHHHHHHHLL--------L-EEEE-LllLLEEEELH---H-H-HH-----------
0022 8cliA   hhhlllllllhhhhhhhlleeeEEEEeelHHH-HHHHHHHHHHHLL--LE-EEHHHHHHL-LL-L--L-H-HHHH-HHHHHHHHLL--------L-EEEE-E--EEEEEELLl-----------------
0023 6juvA   ----------------------------lHH-HHHHHHHHHHHHHL--LL-EEHHHHHHH-HL-L--LlH-HHHH-HHHHHHHHHHH-H-LLleE-EEEE-L--LLEEEEE----E-L-HH----hLL--LH
0024 6j0eB   ----eeellhhllLHHHHHHHH----hHHLL-HHHHHHHHHHHHLL--LL-EEHHHHHHH-HL-L--L-H-HHHH-HHHHHHHHLL--------L-EEEE-EllEEEEEELH---H-H-HHHHH--H-----
0025 8iueP   ----------------------------lHH-HHHHHHHHHHHHLL--LL-LLHHHHHHH-LLlL--L-H-HHHH-HHHHHHHHHL--------L-EEEE-EllEEEEEEHH-H-L-----LHH--H-----
0026 2qlzB   hhhH-HH-HHHHHH-hhHHLL---------L-HHHHHHHHHHHHH---LE-EEHHHHHHH-HL-L--L-H-HHHH-HHHHHHLLLL--------L-EEEE-L--LLEEEELH-H-H-H-HL-----------
0027 2wteA   hhhhlllllleehhHHHHHHL-------lLL-HHHHHHHHHHHHH---LL-LLHHHHHHH-HL-L--L-H-HHHH-HHHHHHHHLL--------L-EEEE-L-lLLEEEELH-H-H-H-H-----------
0028 7l1iA   hlhhhhhhLLLL-----------------LL-HHHHHHHHHHHHH---LL-EEHHHHHHH-LL-L--L-H-HHHH-HHHHHHHHLL--------L-EEEE-E--LEEEEELH-----H-HHH----------
0029 4ijaA   ----------------------------lLH-HHHHHHHHHHHHLL---L-LLHHHHHHH-HL-L--L-H-HHHH-HHHHHHHHLL--------L-EEEL-LllLLLEEELH---H-H-LELLL--EE--EE
0030 8qfcB   ----------------------------lHH-HHHHHHHHHHHHLL--LE-EEHHHHHHH-HL-L--L-H-HHHH-HHHHHHHHHL------L-L-EEEL----LLEEEEHHh-----------------
0031 7qcdD   ----------------------------hHH-HHHHHHHHHHHHLL--LE-EEHHHHHHH-HHlllhH-HHHH-HHHHHHHHHHH-H-lLE-E-EEEE-E--LEEEEEE----E-L-LL----hHH--LL
0032 2rdpA   lllhhhhhhhHHLL--------------lLL-HHHHHHHHHHHHHH---LL-LLHHHHHHH-HL-L--L-H-HHHH-HHHHHHHHLL--------L-EEEE-EllLEEEEELH-H----------------
0033 8a5yT   hhhhhlleEEEEeeLEEE----------eEE-HHHHHHHLLLLlL--LL-EEHHHHHHH-HL-L--L-H-HHHH-HHHHHHHHLL--------L-EEEE-L---LEEEELL-L-Ll--------------
0034 3s2wA   hhhhlllhhhHHHH--HHHHL-------LLL-L-LHHHHHHHHHHL---LL-EEHHHHHHH-HL-L--L-H-HHHH-HHHHHHHHLL--------L-EEEE-E--LEEEEELH-H-H-H-H-----------
0035 5f7pA   ------------------------------L-HHHHHHHHHHHHH---LE-EEHHHHHHL-LL-L--L-H-HHHH-HHHHHHHHHL--------L-EEEL-----LEEEELL---L-L-LE-LE--EEel--
0036 4fhtB   llllhhlhhhhhLLLL-------------LL-HHHHHHHHHHHHL---LL-LLHHHHHHH-HL-L--L-H-HHHH-HHHHHHHHHL--------L-EEEE-EllLEEEEELH-H----------------
0037 1fx7A   ----------------------------llL-LHHHHHHHHHHHHHH--hL-LLHHHHHHH-HL-L--L-H-HHHH-HHHHHHHHLL--------L-EEEL-L--LLLEEEHH-H-H---------lLL--LH
0038 7z1nO   hhHH-HHhHHHHHH---------------lL-HHHHHHHHHHHHHL---L-EEHHHHHHH-LLlL--L-H-HHHH-HHHHHHHHLL--------L-LLEE-EleEEEELLLH---H-H-HHHHHll------
0039 5a5tM   hlllhhhhhlLLLHHH----------hhhHH-HHHHHHHHHHHHHL--LL-LLHHHHHHH-HL-L--L-H-HHHH-HHHHHHHHHL------llL-LLLL-L--LLLLLLLL-------------lLH--HH
0040 1c0wC   ----------------------------lLL-LHHHHHHHHHHHHH---llLLHHHHHHH-LL-L--L-H-HHHH-HHHHHHHHLL--------L-EEEL-L--LLLEEEHH-H-H------hlL--LL--LL
0041 6c2jA   HHHH-HH-HHHLL-LLHHHHHHHL---LLLH-HHHHHHHHHHHHH---LE-ELHHHHHHH-HH---lL-H-HHHH-HHHHHHHHL-------L-L-LEEE-L-lLLEEELLL-------------lLH--HH
0042 2lnbA   ----------------------------hHH-HHHHHHHHHHHHHL--LL-EEHHHHHHH-HL-L--L-H-HHHH-HHHHHHHHLL--------L-EEEE-E--LLEEEELL---L-L-LL-----------
0043 5y6iB   ----------------------------lHH-HHHHHHHHHHHHLL--LL-EEHHHHHHH-HL-L--L-H-HHHH-HHHHHHHHHL--------L-EEELlL--LLEEEE-H-H-H-H-HH--H--HH--H-
0044 1sfxA   ----------llhhhhHHLL---------LL-HHHHHHHHHHHHH---LL-LLHHHHHHH-HL-L--L-H-HHHH-HHHHHHHHLL--------L-EEEE-EleEEEEEELL-H-H-H-HHHHH--H-----
0045 7cv0A   ----------------------------hHH-HHHHHHHHHHHHLL---L-LLHHHHHHH-HL-L--L-H-HHHH-HHHHHHHHLL-------lL-EEEE----LLEEEELL-Ll---------------
0046 2zkzC   ----------lhhHHHHHHHHH----hHHLL-HHHHHHHHHHHHH---LL-EEHHHHHHH-HL-L--L-H-HHHH-HHHHHHL--------lL-L-EEEE-EllEEEEEELL-H-H-H-------------
0047 3cuqA   EELL----EEEE----eLLLL-------lLL-HHHHHHHHHLL-L--LE-ELHHHHHHH-HL-L--L-H-HHHH-HHHHHHHHH-------L-L-LEEE-LllLLEEELLLl-----------------
0048 4mtdD   lhHH-HH-HHHHHH--HHHLL-------lLL-HHHHHHHHHHHLLL--LL-EEHHHHHHH-HHlL--L-H-HHHH-HHHHHHHHLL--------L-EEEE-L-lLLEEEELL---L-L-LLL---------
0049 7dvrA   ----lllhhhhhHHHLL-----------LLL-LHHHHHHHHHHHLL---L-EEHHHHHHH-HL-L--L-H-HHHH-HHHHHHHHLL--------L-EEEE-E--LEEEEELH-H----------------
0050 7c0jB   ------------------------------L-LHHHHHHHHHHHLL---L-EEHHHHHHH-HH---lL-H-HHHH-HHHHHHHHLL--------L-EEEL-L--LLEEEEL-------------------
0051 6r7nB   hhhhhhhhhhhhLLLLllllllllhhhhlhHH-HHHHHHHHHHLL-L--LE-EEHHHHHHH-HL-L--L-H-HHHH-HHHHHHHHHL------llL-EEEElL--LLEEEELL-L-Ll-------------
0052 6j05A   ------------LLHHHHHHHH----hHHLL-HHHHHHHHHHHHLL---L-EEHHHHHHH-HL-L--L-H-HHHH-HHHHHHHHLL--------L-EEEE-E--EEEEEELH-H-H-H-HHHHH--HHl---
0053 5ju7A   lleeeEE-EHEEELLEEE-----------LL-HHHHHHHHHHHHLL--LLeELHHHHHHH-HL---lL-H-HHHH-HHHHHHHHHHH-L-LLL-L-EEEE-L--LLEEEELL-L----------------
0054 6ndlA   ------------------------------L-LLHHHHHHHHHHHL---L-LLHHHHHHH-HL-L--L-H-HHHH-HHHHHHHHLL-------lE-EEEE-L--LLEEEEEE-----L-LLLL--lLH--HH
0055 4hw0A   ----------------------------lHH-HHHHHHHLLLL-----LE-ELHHHHHHH-HL-L--L-H-HHHH-HHHHHHHHLL--------L-EEEE-EllEEEEEELHh-----------------
0056 8jxkD   -----------------------------LH-HHHHHHHHHHHHL----L-EEHHHHHHH-HHlL--L-L-LLHH-HHHHHHHHLL--------L-EEEL-L--LLEEEELH-H----------------
0057 3r4kA   ----------------------------lHH-HHHHHHHLLLL--L--LE-EEHHHHHHH-HL-L--L-H-HHHH-HHHHHHHHLL--------L-EEELlL--LLEEEEL---lL-H-HH-HH--HH--H-
0058 2ethA   lhhhhhhlLHHHH---------------hLL-HHHHHHHHHHHHH---LL-LLHHHHHHH-LL-L--L-H-HHHH-HHHHHHHHLL--------L-EEEE-EllLEEEEELH-H-H-H-H-----hHH--HH
0059 2qwwA   hhhhhhhhhHHHHH--------------lLL-HHHHHHHHHHHHL---LL-EEHHHHHHH-HL-L--L-H-HHHH-HHHHHHHHLL--------L-EEEL-LllLLEEEELH-H----------------
0060 2mlgA   ----------------------------llL-LHHHHHHHHHHHL---LE-EEHHHHHHH-HL-L--L-H-HHHH-HHHHHHHHLL--------L-EEEE-EeeEEEEEELL-L----------------
0061 6qfdB   --------------lLLHHH--------lLL-HHHHHHHHHHHHL----L-EEHHHHHHH-HHlL--L-H-HHHH-HHHHHHHHLL--------L-EEEE-E--EEEEEELH-------------hHH--HH
0062 3k69A   ----------------------------hHH-HHHHHHHHHHHLLL---L-LLHHHHHHH-HL-L--L-H-HHLH-HHHHHHHHLL--------L-LEEE-L--LLEEELLL-----L-HHHLLh-------
0063 8qfcD   ----hhhhhhhhhhhhhhhhhhhhhhhhhHH-HHHHHHHHHHHHH---LE-ELLLHHHHH-HL-L--L-H-HHHH-HHHHHHHHHLL------lL-EEEE-L--LLEEEELL-H-H-H-HHHHHh-------
0064 3g3zA   llhhhhhhHLLL-----------------lL-HHHHHHHHHHHHHH---LL-LLHHHHHHH-HL-L--L-H-HHHH-HHHHHHHHL-------L-L-EEEL-LhhHLLEEELH-H-H-H-------------
0065 7bzhA   -----------------------------LL-LHHHHHHHHHHHH---LE-EEHHHHHHH-LL-L--L-H-HHHH-HHHHHHHHLL--------L-EEEE----LLEEEELL-L----------------
0066 2obpA   ----------------------------lLL-HHHHHHHHHHHHHLlllL-LLHHHHHHH-HL-L--L-H-HHHH-HHHHHHHHLL--------L-EEEE-EllLEEEEEHH-H-Hh--------------
0067 3oopA   hhhhhhhhhHLLLL--------------lLL-HHHHHHHHHHHHHH---LL-EEHHHHHHH-HL-L--L-H-HHHH-HHHHHHHHLL--------L-EEEE-LllLLEEEELH-H-H-H-------------
0068 8f8uA   ----------------------------llL-HHHHHHHHHL-LL--LL-LLHHHHHHH-HL-L--L-H-HHHH-HHHHHHHHLLL-------L-LEEE-L--LLEEELLL-------------lHH--HH
0069 1yyvA   ----------lllllllLHHH----hhHHHHlHHHHHHHHHHH--H--LL-EEHHHHHHH-LLlL--L-H-HHHH-HHHHHHHHHL--------L-EEEE-EllEEEEEELH-H-H-H-H------------
0070 4hblD   hhhhhhhhhHHLLL--------------llL-HHHHHHHHHHHHL---LL-EEHHHHHHH-HL-L--L-H-HHHH-HHHHHHHHLL--------L-EEEE-L--LLEEEEHH-H-H-H-HH-hH--HH--H-
0071 4yifB   ----hhhhhhLLLL--------------lLL-HHHHHHHHHHHHHH---LL-LLHHHHHHH-HL-L--L-H-HHHH-HHHHHHHHLL--------L-EEEE-E--LEEEEELHh-----------------
0072 2d1hA   hhHH-HH-HHHHHH--------------lLL-HHHHHHHHHHHHHL--LL-EEHHHHHHH-HL-L--L-H-HHHH-HHHHHHHHLL--------L-EEEE-EllLEEEEELH-H-H-H-HHHHHh-------
0073 2l4mA   -------lLLLL----------------lLL-HHHHHHHHHHHHHH---LL-EEHHHHHHH-HL-L--L-LhHHHH-HHHHHHHLLL---------lEEEL-L-lLLEEEELL-------------------
0074 3mq0B   ----------------------------hHH-HHHHHHHHHHHHLL--LL-EEHHHHHLL-LL-L--L-H-HHHH-HHHHHHHHLL--------L-EEEL-L--LLEEEELL---L-H-HH--H--HH--HH
0075 9c5gB   ----------------------------hHH-HHHHHHHHHHHH-L--LL-EEHHHHHHH-HL-L--L-H-HHHH-HHHHHHHHHHL-----L-L-LEEE-L-lLLEEELLL-L-L-L-------------
0076 8r3gA   ----------------------------hHH-HHHHHHHHHHHHH---LL-LLHHHHHHH-HL-L--L-H-HHHH-HHHHHHHHLL--------L-EEEE-L--L-EEEELH-H-H-Hl------------
0077 3bpxA   ----lllhhhhHHHH-HHHHL--------lL-HHHHHHHHHHHLL---LL-LLHHHHHHH-HL-L--L-H-HHHH-HHHHHHHHL-------L-L-EEEE-E--EEEEEELH-H-H-H-------------
0078 3f72B   -------hhhhhhhHHHHHHH----hhHHHL-HHHHHHHHHLLLLL---L-EEHHHHHHH-HL-L--L-H-HHHH-HHHHHHHHLL---------lEEEE-EllLEEEEELL-H-H-H-HH----hHH--HH
0079 2pg4A   -----------------LLLH----hhHLLH-HHHHHHHHHHHHHHH--hL-LLHHHHHHH-HL-L--L-H-HHHHlLHHHHHHHLL--------L-EEEE-E--EEEEEELH-H----------------
0080 3gfiA   hhhhhhhhhHHHH---------------lLL-HHHHHHHHHHL--L--LL-EEHHHHHHH-LL-L--L-H-HHHH-HHHHHHHHLL--------L-EEEE-EeeEEEEEELH-H----------------
0081 4nb5B   HHHH------------------------lLL-HHHHHHHHHHHHLL-lLL-EEHHHHHHH-LL-L--L-H-HHHH-HHHHHHHHHL--------L-EEEE-LllLLEEEELL---L-H-HHHHHh-------

# Job: Type-4 BREX; PglZ MBQ0159480 N-terminal iSTAND-NTPase (Type-4 BREX-specific Component)
# Query: s001A
# No:  Chain   Z    rmsd lali nres  %id PDB  Description
   1:  6mdm-D 10.1  3.0  133   713    5   MOLECULE: VESICLE-FUSING ATPASE;
   2:  8fcv-R  9.7  3.3  128   343    5   MOLECULE: DNA (60-MER);
   3:  3te6-A  9.3  3.1  125   304   10   MOLECULE: REGULATORY PROTEIN SIR3;
   4:  7tjj-I  9.1  3.2  117   380    7   MOLECULE: ORIGIN RECOGNITION COMPLEX SUBUNIT 1;
   5:  7jgr-G  9.1  3.3  127   372    4   MOLECULE: ORIGIN RECOGNITION COMPLEX SUBUNIT 2;
   6:  5zr1-A  8.9  3.4  126   494    2   MOLECULE: ORIGIN RECOGNITION COMPLEX SUBUNIT 1;
   7:  8c0v-A  8.4  3.6  132   823   11   MOLECULE: PEROXISOMAL ATPASE PEX1;
   8:  7jgs-C  8.4  3.7  121   614    9   MOLECULE: ORIGIN RECOGNITION COMPLEX SUBUNIT 2;
   9:  7z6h-K  8.3  3.0  125   439    8   MOLECULE: CELL CYCLE CHECKPOINT CONTROL PROTEIN RAD9A;
  10:  7st9-A  8.3  3.3  131   522    7   MOLECULE: CHECKPOINT PROTEIN RAD24;
```

```
11:  8tp1-A  8.3  3.0  121   419   12   MOLECULE: MITOCHONDRIAL CHAPERONE BCS1;
12:  5zr1-E  8.3  3.1  123   460   11   MOLECULE: ORIGIN RECOGNITION COMPLEX SUBUNIT 1;
13:  8umy-A  8.0  3.4  122   546    9   MOLECULE: CHROMOSOME TRANSMISSION FIDELITY PROTEIN 18 HOMOL
14:  7ykk-A  7.9  3.0  120   735   12   MOLECULE: ATPASE FAMILY GENE 2 PROTEIN;
15:  2qby-B  7.8  3.5  127   368    5   MOLECULE: CELL DIVISION CONTROL PROTEIN 6 HOMOLOG 1;
16:  6z1f-3  7.6  3.3  125   284    4   MOLECULE: RIBULOSE BISPHOSPHATE CARBOXYLASE/OXYGENASE ACTIV
17:  7w42-B  7.6  3.3  124   402   10   MOLECULE: UNCHARACTERIZED ATPASE YJOB;
18:  8dr5-A  7.6  3.6  124   646    5   MOLECULE: REPLICATION FACTOR C SUBUNIT 1;
19:  9lgo-B  7.6  3.7  128   701    9   MOLECULE: ATPASE FAMILY GENE 2 PROTEIN HOMOLOG A;
20:  8dyu-A  7.6  3.0  111  2562    5   MOLECULE: CYTOPLASMIC DYNEIN 1 HEAVY CHAIN 1;
21:  4ai6-A  7.6  3.0  119  2650    8   MOLECULE: GLUTATHIONE S-TRANSFERASE CLASS-MU 26 KDA ISOZYME
22:  7mi8-A  7.5  3.7  115   868    8   MOLECULE: FUSION PROTEIN OF DYNEIN AND ENDOLYSIN;
23:  8xku-B  7.5  3.2  125   845    9   MOLECULE: PROBABLE INACTIVE ATP-DEPENDENT ZINC METALLOPROTE
24:  7lcc-A  7.5  3.6  114  1369   10   MOLECULE: HELRAISER K1068Q;
25:  5e7p-A  7.4  3.7  130   719    5   MOLECULE: CELL DIVISION CONTROL PROTEIN CDC48;
26:  8xks-B  7.4  3.3  122   603    8   MOLECULE: CTAP1;
27:  6vvo-A  7.4  3.6  124   448    6   MOLECULE: REPLICATION FACTOR C SUBUNIT 1;
28:  4xgc-E  7.4  3.9  123   402    8   MOLECULE: ORIGIN RECOGNITION COMPLEX SUBUNIT 2;
29:  7jgr-D  7.4  4.0  126   441    6   MOLECULE: ORIGIN RECOGNITION COMPLEX SUBUNIT 2;
30:  8dgc-A  7.3  4.0  121  2028    7   MOLECULE: SEAVS3;
31:  8rwv-E  7.3  3.8  120   431    8   MOLECULE: DNA REPLICATION LICENSING FACTOR MCM2;
32:  7vcs-A  7.2  3.3  126   766    9   MOLECULE: TRANSITIONAL ENDOPLASMIC RETICULUM ATPASE;
33:  5g4f-A  7.2  3.7  130   726    8   MOLECULE: VCP-LIKE ATPASE;
34:  3bos-B  7.2  3.6  116   231    4   MOLECULE: PUTATIVE DNA REPLICATION FACTOR;
35:  8xku-F  7.2  3.2  123   478   10   MOLECULE: PROBABLE INACTIVE ATP-DEPENDENT ZINC METALLOPROTE
36:  7uqj-A  7.1  3.5  131   585    6   MOLECULE: ATPASE HISTONE CHAPERONE YTA7;
37:  8xku-A  7.1  3.3  124   730    8   MOLECULE: PROBABLE INACTIVE ATP-DEPENDENT ZINC METALLOPROTE
38:  7rzy-1  7.1  3.5  123   311   11   MOLECULE: TN6677 VIBRIO CHOLERAE TRANSPOSON TNSC (VCHTNSC);
39:  8r64-B  7.1  3.2  118   303    9   MOLECULE: FIDGETIN-LIKE PROTEIN 1;
40:  2gno-A  7.1  3.4  114   296    8   MOLECULE: DNA POLYMERASE III, GAMMA SUBUNIT-RELATED PROTEIN
41:  8zr0-A  7.0  3.5  117   480    2   MOLECULE: CASEINOLYTIC PEPTIDASE B PROTEIN HOMOLOG;
42:  9fnn-U  7.0  2.8  113   511    9   MOLECULE: CELLULOSE SYNTHASE CATALYTIC SUBUNIT [UDP-FORMING
43:  8k8v-A  7.0  3.4  114   719    6   MOLECULE: ENDOPEPTIDASE LA;
44:  8bob-C  7.0  3.7  109   412    7   MOLECULE: PROTEIN MALY;
45:  8xku-D  7.0  4.0  122   805    8   MOLECULE: PROBABLE INACTIVE ATP-DEPENDENT ZINC METALLOPROTE
46:  5j1s-A  7.0  3.6  124   277    9   MOLECULE: TORSIN-1A;
47:  8fnu-A  7.0  3.5  122   796    6   MOLECULE: KAP NTPASE DOMAIN-CONTAINING PROTEIN;
48:  1xwi-A  7.0  3.8  126   322    6   MOLECULE: SKD1 PROTEIN;
49:  5v8f-D  7.0  3.6  118   456   11   MOLECULE: DNA REPLICATION LICENSING FACTOR MCM2;
50:  6p8v-C  6.9  3.5  124   303    8   MOLECULE: ATPASE, AAA FAMILY;
51:  7qo4-M  6.9  3.2  122   421    7   MOLECULE: 26S PROTEASOME REGULATORY SUBUNIT RPN1;
52:  6nyy-C  6.9  3.6  124   491   10   MOLECULE: AFG3-LIKE PROTEIN 2;
53:  6pdw-D  6.9  3.5  125   299    9   MOLECULE: MEMBRANE-SPANNING ATPASE-LIKE PROTEIN;
54:  7qo4-K  6.9  3.2  119   394    7   MOLECULE: 26S PROTEASOME REGULATORY SUBUNIT RPN1;
55:  8xks-D  6.8  3.3  121   971    8   MOLECULE: CTAP1;
56:  8xku-C  6.8  3.5  130   921    9   MOLECULE: PROBABLE INACTIVE ATP-DEPENDENT ZINC METALLOPROTE
57:  6rla-A  6.8  3.5  116  2750    7   MOLECULE: O6-ALKYLGUANINE-DNA ALKYLTRANSFERASE MUTANT,DYNC2
58:  8ui7-C  6.8  3.4  113   339    9   MOLECULE: ATPASE FAMILY AAA DOMAIN-CONTAINING PROTEIN 5;
59:  7mi1-A  6.7  3.0  120  2628    5   MOLECULE: CHIMERA PROTEIN OF DYNEIN AND ENDOLYSIN;
60:  5nug-A  6.7  3.1  112  2920    7   MOLECULE: CYTOPLASMIC DYNEIN 1 HEAVY CHAIN 1;
```

```
0001 s001A   [illegible]
0002 6mdmD   [illegible]
0003 8fcvR   [illegible]
0004 3te6A   [illegible]
0005 7tjjI   [illegible]
0006 7jgrG   [illegible]
0007 5zr1A   [illegible]
0008 8c0vA   [illegible]
0009 7jgsC   [illegible]
0010 7z6hK   [illegible]
0011 7st9A   [illegible]
0012 8tp1A   [illegible]
0013 5zr1E   [illegible]
0014 8umyA   [illegible]
0015 7ykkA   [illegible]
0016 2qbyB   [illegible]
0017 6z1f3   [illegible]
0018 7w42B   [illegible]
0019 8dr5A   [illegible]
0020 9lgoB   [illegible]
0021 8dyuA   [illegible]
0022 4ai6A   [illegible]
0023 7mi8A   [illegible]
0024 8xkuB   [illegible]
0025 7lccA   [illegible]
0026 5e7pA   [illegible]
0027 8xksB   [illegible]
0028 6vvoA   [illegible]
0029 4xgcE   [illegible]
0030 7jgrD   [illegible]
0031 8dgcA   [illegible]
0032 8rwvE   [illegible]
0033 7vcsA   [illegible]
0034 5g4fA   [illegible]
0035 3bosB   [illegible]
0036 8xkuF   [illegible]
0037 7uqjA   [illegible]
0038 8xkuA   [illegible]
0039 7rzy1   [illegible]
0040 8r64B   [illegible]
0041 2gnoA   [illegible]
0042 8zr0A   [illegible]
0043 9fnnU   [illegible]
0044 8k8vA   [illegible]
0045 8bobC   [illegible]
0046 8xkuD   [illegible]
0047 5j1sA   [illegible]
0048 8fnuA   [illegible]
0049 1xwiA   [illegible]
0050 5v8fD   [illegible]
0051 6p8vC   [illegible]
0052 7qo4M   [illegible]
0053 6nyyC   [illegible]
0054 6pdwD   [illegible]
0055 7qo4K   [illegible]
0056 8xksD   [illegible]
0057 8xkuC   [illegible]
0058 6rlaA   [illegible]
0059 8ui7C   [illegible]
0060 7mi1A   [illegible]
0061 5nugA   [illegible]
```

```
0001 s001A   [illegible]
0002 6mdmD   [illegible]
0003 8fcvR   [illegible]
0004 3te6A   [illegible]
0005 7tjjI   [illegible]
0006 7jgrG   [illegible]
0007 5zr1A   [illegible]
0008 8c0vA   [illegible]
0009 7jgsC   [illegible]
0010 7z6hK   [illegible]
0011 7st9A   [illegible]
0012 8tp1A   [illegible]
0013 5zr1E   [illegible]
0014 8umyA   [illegible]
0015 7ykkA   [illegible]
0016 2qbyB   [illegible]
0017 6z1f3   [illegible]
0018 7w42B   [illegible]
0019 8dr5A   [illegible]
0020 9lgoB   [illegible]
0021 8dyuA   [illegible]
```

```
0022 4ai6A   [illegible]
0023 7mi8A   [illegible]
0024 8xkuB   [illegible]
0025 7lccA   [illegible]
0026 5e7pA   [illegible]
0027 8xksB   [illegible]
0028 6vvoA   [illegible]
0029 4xgcE   [illegible]
0030 7jgrD   [illegible]
0031 8dgcA   [illegible]
0032 8rwvE   [illegible]
0033 7vcsA   [illegible]
0034 5g4fA   [illegible]
0035 3bosB   [illegible]
0036 8xkuF   [illegible]
0037 7ugjA   [illegible]
0038 8xkuA   [illegible]
0039 7rzy1   [illegible]
0040 8r64B   [illegible]
0041 2gnoA   [illegible]
0042 8zr0A   [illegible]
0043 9fnnU   [illegible]
0044 8k8vA   [illegible]
0045 8bobC   [illegible]
0046 8xkuD   [illegible]
0047 5j1sA   [illegible]
0048 8fnuA   [illegible]
0049 1xwiA   [illegible]
0050 5v8fD   [illegible]
0051 6p8vC   [illegible]
0052 7qo4M   [illegible]
0053 6nyyC   [illegible]
0054 6pdwD   [illegible]
0055 7qo4K   [illegible]
0056 8xksD   [illegible]
0057 8xkuC   [illegible]
0058 6r1sA   [illegible]
0059 8ui7C   [illegible]
0060 7mi1A   [illegible]
0061 5nugA   [illegible]

[illegible]
[illegible]
[illegible]

# Job: Type-1 BREX; BrxL MCM-AAA ATPase
# Query: s001A
# No:  Chain   Z    rmsd lali nres  %id PDB  Description
   1:  8emh-B 47.5  0.8  319   676   71   MOLECULE: PROTEASE LON-RELATED BREX SYSTEM PROTEIN BRXL;
   2:  5udb-7 16.8  3.4  256   726   14   MOLECULE: DNA REPLICATION LICENSING FACTOR MCM2;
   3:  3f8t-A 16.4  3.4  246   459   13   MOLECULE: PREDICTED ATPASE INVOLVED IN REPLICATION CONTROL,
   4:  5v8f-6 16.3  3.2  241   692   15   MOLECULE: DNA REPLICATION LICENSING FACTOR MCM2;
   5:  3nbx-X 14.8  3.8  229   481   14   MOLECULE: ATPASE RAVA;
   6:  8q6p-2 14.8  3.8  248   431    9   MOLECULE: DNA REPLICATION LICENSING FACTOR MCM2;
   7:  2r44-A 13.9  4.0  236   330   11   MOLECULE: UNCHARACTERIZED PROTEIN;
   8:  5udb-4 13.4  3.4  253   751   13   MOLECULE: DNA REPLICATION LICENSING FACTOR MCM2;
   9:  8osg-B 12.1  3.7  223   323   15   MOLECULE: MAGNESIUM-CHELATASE SUBUNIT CHLI;
  10:  8rxd-A 11.1  4.2  218   486   13   MOLECULE: COMPETENCE RELATED PROTEIN COMM;
  11:  5dgk-A 11.1  3.7  209   519   11   MOLECULE: ACTIVE HELICASE;
  12:  6iy8-A  8.7  5.4  163   469   10   MOLECULE: POSITIVE REGULATOR CAPR;
  13:  3bos-B  8.6  3.0  132   231   13   MOLECULE: PUTATIVE DNA REPLICATION FACTOR;
  14:  9e22-A  8.2  4.5  154  2925   14   MOLECULE: CYTOPLASMIC DYNEIN 1 HEAVY CHAIN 1;
  15:  7ola-A  7.7  4.3  182   441    9   MOLECULE: DNA PRIMASE;
  16:  7v2b-D  7.6  4.4  158   376   10   MOLECULE: VPSR;
  17:  8iqi-C  7.6  4.2  182   916   12   MOLECULE: PUTATIVE PRIMASE C962R;
  18:  7mi8-A  7.6  8.8  198   868   11   MOLECULE: FUSION PROTEIN OF DYNEIN AND ENDOLYSIN;
  19:  8umy-A  7.5  7.8  134   546   12   MOLECULE: CHROMOSOME TRANSMISSION FIDELITY PROTEIN 18 HOMOL
  20:  1ixs-B  7.4  3.4  138   315   14   MOLECULE: HOLLIDAY JUNCTION DNA HELICASE RUVA;
  21:  7mi1-A  7.4  4.3  151  2628   13   MOLECULE: CHIMERA PROTEIN OF DYNEIN AND ENDOLYSIN;
  22:  6hyp-A  7.3  4.6  164  2272   12   MOLECULE: MIDASIN,MIDASIN;
  23:  6l1q-A  7.2  3.8  147   267   12   MOLECULE: CBBQ PROTEIN;
  24:  8k8v-A  7.2  5.8  150   719   13   MOLECULE: ENDOPEPTIDASE LA;
  25:  5v8f-D  7.2  3.7  136   456   10   MOLECULE: DNA REPLICATION LICENSING FACTOR MCM2;
  26:  5zr1-A  7.2  3.9  135   494   11   MOLECULE: ORIGIN RECOGNITION COMPLEX SUBUNIT 1;
  27:  9emc-D  7.1  3.5  135   337   14   MOLECULE: RUVB-LIKE 1;
  28:  8btg-B  7.1  3.4  132   335   13   MOLECULE: CHROMOSOMAL REPLICATION INITIATOR PROTEIN DNAA;
  29:  8fcv-R  7.0  3.8  127   343   14   MOLECULE: DNA (60-MER);
  30:  6k9c-A  7.0  4.1  173   416   12   MOLECULE: PRIMASE;

0001 s001A   [illegible]
0002 8emhB   [illegible]
0003 5udb7   [illegible]
0004 3f8tA   [illegible]
0005 5v8f6   [illegible]
0006 3nbxX   [illegible]
0007 8q6p2   [illegible]
0008 2r44A   [illegible]
0009 5udb4   [illegible]
0010 8osgB   [illegible]
0011 8rxdA   [illegible]
0012 5dgkA   [illegible]
0013 6iy8A   [illegible]
0014 3bosB   [illegible]
0015 9e22A   [illegible]
0016 7olaA   [illegible]
0017 7v2bD   [illegible]
0018 8iqiC   [illegible]
0019 7mi8A   [illegible]
0020 8umyA   [illegible]
0021 1ixsB   [illegible]
0022 7mi1A   [illegible]
0023 6hypA   [illegible]
0024 6l1qA   [illegible]
0025 8k8vA   [illegible]
0026 5v8fD   [illegible]
0027 5zr1A   [illegible]
0028 9emcD   [illegible]
0029 8btgB   [illegible]
0030 8fcvR   [illegible]
0031 6k9cA   [illegible]

0001 s001A   [illegible]
0002 8emhB   [illegible]
0003 5udb7   [illegible]
0004 3f8tA   [illegible]
0005 5v8f6   [illegible]
0006 3nbxX   [illegible]
0007 8q6p2   [illegible]
0008 2r44A   [illegible]
0009 5udb4   [illegible]
0010 8osgB   [illegible]
0011 8rxdA   [illegible]
0012 5dgkA   [illegible]
0013 6iy8A   [illegible]
0014 3bosB   [illegible]
0015 9e22A   [illegible]
0016 7olaA   [illegible]
0017 7v2bD   [illegible]
0018 8iqiC   [illegible]
0019 7mi8A   [illegible]
0020 8umyA   [illegible]
0021 1ixsB   [illegible]
0022 7mi1A   [illegible]
0023 6hypA   [illegible]
```

```
0024 6l1qA   ---------------llllllllllLLLL---------lLLLHHH-HHHHHHHLL---LLEEEELLLLLHHH-HHHHHHL--LEEEEELLL-LLHHHH-HEEEE-E-EEELHHHH-HlLEEEEELHHH-LL-----hHH-HHH-HHHHLLlLLEEEELLEEEELLlL-EEEEEELLL-L-------------------llllLLLL--HHHHLLL-EEEEL----LLLLHHHHHH--HL------------------------------------------------------------------------------------------------------------------------------------------------------------
0025 8k8vA   ------lllllLLLL-------lLLLL---------llLLHHHHH-HHHHHHHHLL---LEEEEELLLLLHHH-HHHHHHL--LEEELL--L-LLHHHH-HLLLLlH-LLLLHHHH-LlLEEEE-EHHHhHH-LL----L-HHH-HHHHHH-HLEELLlELLEELL-L-EEEEEELH------------------HHH--H--L-LLLLLLHHHHEEEEEELLLeelLHHHHHH---------------------------------------------------------------------------------------------------------------------------------------------------------------
0026 5v8fD   ------------------------hhhhLLLH-------hHHHHHH-HHHHHHHHL-L-LEEEEELLLLLHHH-HHHHHHHleEEEEEELLL-LHHHHHlllHHH-HhHHHH-hhL-E-EEEEEELHHH-L----lllHH-HHH-HHHHHH-HL---------LLL-E-EEEELLLLL-------------LHH-H----------HLLHHHHHLLLLEEEL-LL-LL-HHHH-HH--LL------------------------------------------------------------------------------------------------------------------------------------------------------------
0027 5zr1A   -------------llhhhhhhllllLLLL-----------LLLLHH-HHHHHHHHL-L-EEEEEEELLLLLHHH-HHHHHHHleEEEEEEHHHlHHHHHHhHHHH-------hhhlL-E-EEEEEELLLL-LL----llLL-HHH-HHHHHH-LL---------LLL-E-EEEEELLL----------------L-LHH--H--HHL-LHHHHHHHLLEEEEE----LLLLHHHHHH--HL------------------------------------------------------------------------------------------------------------------------------------------------------------
0028 9emcD   ---------------lLLLLllleELLEE---------lLHHHHH-HHHHHHHHL-L-LEEEEEELLLLLHHH-HHHHHHL--LEEEEE-LL-LLHHHH-H----------HHHHL-LeLEEEEELHHH-LL-----hHH-HHH-HHHHLL-L----------LLL-L-EEEEEELLL-----leelllllleE-HHH-------LLHH--HHLLL-EEEEE--L-LLLHHHH-HH--HH------------------------------------------------------------------------------------------------------------------------------------------------------------
0029 8btgB   ----------------llllllllLLLL-----LL-LLLHHHHH-HHHHHHHLL-L-LEEEEEELLLLLHHH-HHHHHHH--HEEEEEHHH-HHHHHH-----------LLLL-----EEEEELHHH-HlllhhhHHH-HHH-HHHHHH-LL-------------L-EEEEEELL---------------LL-LLL--L--L--LLHH-HHHHLLEEEEE--L-LLLHHHH-HH--HH------------------------------------------------------------------------------------------------------------------------------------------------------------
0030 8fcvR   ----------------------------lhhhhlLLL-HHHHHHHH-HHHHHHHLL---lEEEEEELLLLLHHH-HHHHHHHllLEEEEELlH-HHHHHH-HH-----------hHL-L-LEEEEELHHH-HhllllhhHH-HHH-HHHHHH-H-----------HL-L-EEEEEELHH---------------hH-HHL---------LLLHHHHHLE-EEEEELLLlHHHHHH----------------------------------------------------------------------------------------------------------------------------------------------------------hlLLL-
0031 6k9cA   LLLL-----------lllllHHHHH-------hhhhllLLHHHHH-HHHHhHHHH---LEEEEEELLLLHHH-HHHHHHHhhHEEEEELHH-HLLLLL------------HHHLL-L-LEEEEELLLL------lhhhhlHHH-HHHHHH-LLEEEELLEEEELL-L-EEEEEELL---------------L-LLL--------LLLL-lLLLE-EEEEE--L-LLHLLLLLLL----hHLHHHHHH-HH------hLLLLL---------------------LLLL------------------------LLLL-HHHHHHHHHHLLHh-------

# Job: Type-1 BREX; BrxL HAM28421 NTD1 SIGMA-HTH-like
# Query: s001A
# No:  Chain   Z    rmsd lali nres  %id PDB  Description
   1:  8emh-B  7.2  1.5   64   676   52   MOLECULE: PROTEASE LON-RELATED BREX SYSTEM PROTEIN BRXL;
   2:  3frh-A  5.0  2.3   49   242   16   MOLECULE: 16S RRNA METHYLASE;
   3:  8val-E  4.9  5.6   44   337   16   MOLECULE: DNA POLYMERASE III SUBUNIT DELTA;
   4:  6h3a-B  4.7  3.0   51    57   14   MOLECULE: SWI/SNF-RELATED MATRIX-ASSOCIATED ACTIN-DEPENDENT
   5:  2ejs-A  4.6  1.6   44    58   14   MOLECULE: AUTOCRINE MOTILITY FACTOR RECEPTOR, ISOFORM 2;
   6:  4oci-A  4.6  1.7   46   139    4   MOLECULE: CALMODULIN, PUTATIVE;
   7:  7qog-C  4.6  1.9   50   225   12   MOLECULE: PORTAL PROTEIN GP20;
   8:  2qho-D  4.5  1.1   39    49   10   MOLECULE: UBIQUITIN;
   9:  1otr-A  4.5  1.5   40    49   10   MOLECULE: PROTEIN CUE2;
  10:  3bq3-A  4.5  1.4   40   250   10   MOLECULE: DEFECTIVE IN CULLIN NEDDYLATION PROTEIN 1;
  11:  1mx9-D  4.4  2.1   50   533    6   MOLECULE: LIVER CARBOXYLESTERASE I;
  12:  1xb2-B  4.4  1.2   40   276   18   MOLECULE: ELONGATION FACTOR TU, MITOCHONDRIAL;
  13:  6c95-D  4.3  3.6   43    95   14   MOLECULE: N-ALPHA-ACETYLTRANSFERASE 15, NATA AUXILIARY SUBU
  14:  7qoj-D  4.3  2.0   50   226    8   MOLECULE: PORTAL PROTEIN GP20;
  15:  2n5j-A  4.2  2.9   45    49   11   MOLECULE: RIBONUCLEASE ZC3H12A;
  16:  4eei-B  4.2  4.9   53   423    8   MOLECULE: ADENYLOSUCCINATE LYASE;
  17:  8hkc-E  4.2  3.4   58   262    5   MOLECULE: DNA-DIRECTED RNA POLYMERASE SUBUNIT ALPHA;
  18:  9cc7-C  4.2  2.1   49   160   10   MOLECULE: PHITE ADAPTOR PROTEIN;
  19:  2cp8-A  4.2  2.8   44    54    9   MOLECULE: NEXT TO BRCA1 GENE 1 PROTEIN;
  20:  9fvr-C  4.2  3.5   54   153   17   MOLECULE: TRANSCRIPTIONAL REPRESSOR NRDR;
  21:  1nxh-A  4.2  2.9   56   124   13   MOLECULE: MTH396 PROTEIN;
  22:  6q00-B  4.1  1.4   39    45   15   MOLECULE: UBIQUITIN;
  23:  1aip-C  4.1  1.4   40   195   10   MOLECULE: ELONGATION FACTOR TU;
  24:  8x6g-H  4.1  4.5   57   244    4   MOLECULE: DNA-DIRECTED RNA POLYMERASE SUBUNIT ALPHA;
  25:  3e46-A  4.1  2.7   44   202    9   MOLECULE: UBIQUITIN-CONJUGATING ENZYME E2-25 KDA;
  26:  6vvo-A  4.1  5.0   54   448    6   MOLECULE: REPLICATION FACTOR C SUBUNIT 1;
  27:  5he9-E  4.0  2.2   44    56    7   MOLECULE: HELICASE LOADER;
  28:  3b74-A  4.0  3.0   47   307    9   MOLECULE: UNCHARACTERIZED PROTEIN YKL091C;
  29:  2cp9-A  3.9  1.9   45    64   16   MOLECULE: ELONGATION FACTOR TS, MITOCHONDRIAL;
  30:  1kl7-A  3.9  2.3   49   509   10   MOLECULE: THREONINE SYNTHASE;
  31:  7p37-A  3.9  3.5   52   147   15   MOLECULE: TRANSCRIPTIONAL REPRESSOR NRDR;
  32:  7lt2-A  3.8  3.3   52   387   13   MOLECULE: MAB-21 DOMAIN-CONTAINING PROTEIN;
  33:  2ekf-A  3.8  4.1   43    61   14   MOLECULE: ANCIENT UBIQUITOUS PROTEIN 1;
  34:  6jcx-F  3.8  4.5   58   186    7   MOLECULE: DNA-DIRECTED RNA POLYMERASE SUBUNIT ALPHA;
  35:  2o7g-A  3.8  3.8   55    88    7   MOLECULE: PROBABLE RNA POLYMERASE SIGMA-C FACTOR;
  36:  6ajf-A  3.8  3.6   52   901    6   MOLECULE: DRUG EXPORTERS OF THE RND SUPERFAMILY-LIKE PROTEI
  37:  6jbq-F  3.8  3.9   60   186    7   MOLECULE: DNA-DIRECTED RNA POLYMERASE SUBUNIT ALPHA;
  38:  5wuq-A  3.8  4.6   57   158    4   MOLECULE: ECF RNA POLYMERASE SIGMA FACTOR SIGW;
  39:  2cos-A  3.8  2.7   45    54   18   MOLECULE: SERINE/THREONINE PROTEIN KINASE LATS2;
  40:  2dna-A  3.7  2.3   48    67    8   MOLECULE: UNNAMED PROTEIN PRODUCT;
  41:  8vjl-B  3.7  4.2   57   116    7   MOLECULE: STAGE IV SPORULATION PROTEIN FB;
  42:  7mvv-A  3.7  4.0   51  1542    4   MOLECULE: NUCLEOPORIN NUP192;
  43:  6e6y-A  3.7  5.7   54   268    6   MOLECULE: DIECKMANN CYCLASE, NCMC;
  44:  1wgl-A  3.7  1.3   40    59   18   MOLECULE: TOLL-INTERACTING PROTEIN;
  45:  2qsf-X  3.7  2.9   46   104    2   MOLECULE: DNA REPAIR PROTEIN RAD4;
  46:  4f1i-A  3.7  1.5   39   328   15   MOLECULE: 5'-TYROSYL-DNA PHOSPHODIESTERASE;
  47:  9kgb-A  3.7  3.7   52   336   12   MOLECULE: TERPENE SYNTHASE;
  48:  8yhu-B  3.7  2.6   46    56   20   MOLECULE: TOLL-LIKE RECEPTOR 3;
  49:  6tdv-D  3.7  2.4   46   186   15   MOLECULE: ATPTB1;
  50:  6toa-D  3.7  5.5   54   195   13   MOLECULE: ADAPTOR PROTEIN RCC01688;

0001 s001A   --MDVLDIKLNEAFPG-----------RKDLLHEI-K-RA-VN-V--PS-FVLEFLLSRYC---A--S--------E----D---P--EEIEEGKKAVLQTIEK-C--Y-------
0002 8emhB   -nDKELDQLLNEHFAG--------rvvRKDLTKLI-K-EG-AN-V--PV-YVLEYLLGMYC---A--S--------D----D---P--EIIEQGLRNVKTVLAE-N--Yvrpdeae
0003 3frhA   -----------------------ypmnINDALTSI-L-ASkRA-L--CP-DTVRRILTEEW---G-rH--------K----S---P--KQTVEAARTRLHGICG-Ay---------
0004 8valE   -rPDFE--yqagrghhallihylapppEQYAVTWL-S-RE-VT-M--SQ-DALLAALRLSA-------------------------------gSPGAALALFQG-D--Nwqaretl
0005 6h3aB   --sELEDLSEL-------------edlKDAKLQTL-K-EL-FQ-R--SD-NDLLKLIESTS---T------------------------MDGAIAAALLMFGDAg-----------
0006 2ejsA   -gsSGAS-----------------nsqLNAMAHQI-Q-EM-FQ-V--PY-HLVLQDLQLTR--------------------------------SVEITTDNILE-G--Riqvpf--
0007 4ociA   --------------------dgdgyltLNEFESLV-R-VL-VV-X--ET-SAIASTYNSN----S--K--------V----R------gxSYELFTSCFSQLKT-K--Sfnkdeik
0008 7qogC   -------------------------mtYNELIYMV-L-DE-LKyY--TP-DHVIFLLVKYR---SffP--------I----E---D--ALVPPLIELVVKELRG-P--Eyspkded
0009 2qhoD   --------------------sipasviPEELISQA-Q-VV-LQgK--SR-SVIIRELQRTN-------------------------------lDVNLAVNNLLS-R----------
0010 1otrA   ------------------------nddHESKLSIL-M-DM-FA-I--SK-SKLQVHLLENN-------------------------------nDLDLTIGLLLK-E--Nddks---
0011 3bq3A   -------------------------spEQEAIESF-T-SL-TK-C--DP-KVSRKYLQRNH-------------------------------wNINYALNDYYD-K--Eigvahpp
0012 1mx9D   --------ggesvsvlvlsplsegqldQKTAMSLL-W-KS-YC-I--AK-ELIPEATEKYL---G-gT--------D----D---T--VKKKDLFLDLIADVMF-G--Vpsvivar
0013 1xb2B   ------------------------sasSKELLMKL-R-RK-TG-Y--SF-INCKKALETCG-------------------------------gDLKQAESWLHK-Q--Aqkegwsk
0014 6c95D   --------qkaKQERE----elakvtiKKEDLELI-M-TE-ME-I--SR-AAAERSLREHM-------------------------------gNVVEALIALTN------------
0015 7qojD   -------------------------mtNKEFSDGF-S-TL-LT-L--DE-YEKSTFLTNAQeqlC--E--------L----N---P--VVHRAILERAVQLAII-S--Ktqlt---
0016 2n5jA   ---GPHM-----------------tseLQMKVDFF-R-KL--G-Y--SS-SEIHSVLQKLG---V--Q------------------------aDTNTVLGELVK-H--G-------
0017 4eeiB   rshgMFAEedrVRST--------saylSSFYLHFL-V-AN-TP-F--MR-EDCYKIVQQVA---F--D-------------------lESFSKKLQKVMHDEHNi-----------
0018 8hkcE   -aNAWPMLleaaKTLI-----lshlrfVVHIARNY-A--G-YG-L--PQ-ADLIQEGNIGL---M-rrfnpevgvrL----V---S--FAVHWIKAEIHEYVLR-N--Wrivkvat
0019 9cc7C   -------------------------mtNEQVIELV-R-VL-L-eI--SD-QTIIFFWTKWK---Y--D--------L----D-nrP--EKIPAALYNTVVDCVR-W--Livqevss
0020 2cp8A   ------gSSGS-------------sgqTAALMAHL-F-EM-GF-C--DR-QLNLRLLKKHN-------------------------------yNILQVVTELLQ-L--Sgpssg--
0021 9fvrC   --------fevAELVX----dvrepfnEEKLRSGX-L-RA-LEpV--SS-DDVEXAINHIK---S--QlratgereV----P---S--KXIGNLVXEQLKKLDK-Va---------
0022 1nxhA   -lhPRFESAR-------------prciREKLHSDLqL-CW-LI-I--SV-DDAEALKDEIT---E--L----vlagR----E---Y--SEALSEGRRRLHEILR-S----------
0023 6q00B   ------------------------snaRRLLXVEF-A-SV-AS-C--DA-AVAQCFLAEND-------------------------------wEMERALNSYFE-Pp---------
0024 1aipC   --------------------------sQMELIKKL-R-EA-TG-A--GM-MDVKRALEDAG-------------------------------wDEEKAVQLLRE-R--Gamkaakk
0025 8x6gH   ehqeNKNTDAQDK---------lvkhyQKLIESLA-Y-KY-SK-gqsHH-EDLVQVGMVGL---I--G-ainrfdmsferkF---E--AFLVPTVIGEIKRYLR-D--Ktwsvhvp
0026 3e46A   --------TPYEGG------apvsspeYTKKIENL-C-AA--G-F--DR-NAVIVALSSKS-------------------------------wDVETATELLLS-N----------
0027 6vvoA   -qQGDQSCcfdLRFQ---------rprVEQIKGAM-M-SI-AK-I--PP-PAMNEIILGAN---Q-------------------------DIRQVLHNLSMWCA-R--Skalmgpf
0028 5he9E   ------------------------snaMMVTKEFL-K-TK-LE-C--SD-MYAQKLID-EA---Q--G----------------------DENRLYDLFIQKLA-E--Rhtrpaiv
0029 3b74A   --------iCSPNALP----pgnltkeQEEALLQF-R-SI-LLrL--DD-STLLRFLRARK-------------------------------fDINASVEMFVE-T--Erwreeyg
0030 2cp9A   -gSSGSS------------------gsSKELLMKL-R-RK-TG-Y--SF-VNCKKALETCG-------------------------------gDLKQAEIWLHK-E--Aqkegwsk
0031 1kl7A   --------------------dwsklsfQDLAFAIX-R-LY-IE-I--PD-ADLKDLIKRSY---S--T--------FrsdevtyaF--KDVALQFVGNLFEYFL-Q--Rtnanlpe
0032 7p37A   fttvETCS------------gvtepfsRTKVINGV-R-KA-CQpV--TE-DALAQLGQRVE---E--avratgsaeL----T---T--HDVGLAILGPLQELDL-Va---------
0033 7lt2A   --------slsfQEQE----lidnknrLKPAVKLI-K-KL-KN-I--AS-YYIKTVFLHI------fW--------N----K---SlrEVFMTTLREYNEFIAD-Q--Sipyywcr
0034 2ekfA   -sgSSGS----------------pdvqLATLAQRV-K-EV-LH-V--PL-GVIQRDLAKTG--------------------------------CVDLTITNLLE-Ga---------
0035 6jcxF   -tdEELTARFER----------daiplLDQLYGGA-L-RM-TR-NpaDA-EDLLQETMVKA---Y-agfrsfrhgtN----L---K--AWLYRILTNTYINSYR-K--Kqrqpaey
0036 2o7gA   alalsaaKGNGRALE------afikatQQDVWRFV-A-YL-SDvG--SA-DDLTQETFLRA---I-gaiprfsarsS----A---R--TWLLAIARHVVADHIR------------
0037 6ajfA   fisiplqgIPLGQlslggisfdklplmALILIVTT-T-VL-MFvL--PI-KAALMSALTL------nytpqplmapM----I---G--LIIAVIWGLSTDYEVF-L--Vsrmvear
0038 6jbqF   -tDQVLVEkgdQKAFN-----llvvryQHKVASLV-S-RY-VP-S-gDV-PDVVQEAFIKA---Y-raldsfrgdsA----F---Y--TWLYRIAVNTAKNYLV-A--Qgrrppss
0039 5wuqA   -vKKGD-QDAFA---------divdiyKDKIYQLC-Y-RM-L--G--NV-HEAEDIAQEAF---I--RayvnidsfdinrkF---S--TWLYRIATNLTIDRIR-K--Kkpdyyld
0040 2cosA   --GSSGS-----------------sgvNRQMLQEL-V-NA-G--C--DQ-EMAGRALKQTG------------------------------sRSIEAALEYISK-M--Sgpssg--
0041 2dnaA   -sSGPS-HSLQ-----------apevrFSKEMECL-Q-AM-GF-V--NY-NANLQALIATD-------------------------------gDTNAAIYKLKS-S--Qgfsgpss
0042 8vjlB   lmakGDEHARNM-----------liehNLRLVAHI-V-KK-FEgE--DA-EDLISIGTIGL---I--K-giesysagkgtkL---A--TYAARCIENEILMHLR-A--Lkktk---
0043 7mvvA   --------lkigEEEY------slnedFVNDCLKL-A-DE-LD-L--NE-KESARILIDCD---A------------egdvE---T--QS-RPLWECGVIRFHQ-E--Rkylldcm
0044 6e6yA   -cAGSA--------------spttstlYYQFRKVV-E-SL-AGeQ--AA-REALAEGTAAA---D-rI--------G----D---V--EGLGAELVRVFTAAGR-A--Acaaadld
0045 1wglA   --------------------ssgssgcSEEDLKAI-Q-DM-FN-M--DQ-EVIRSVLEAQR-------------------------------gNKDAAINSLLQ-M--Geepsgps
0046 2qsfX   ----igLTVEDLLS------qvdytpeDDQAISRL-C-E--LG-F--ER-DLVIQVYFACD-------------------------------kNEEAAANILFS-D----------
0047 4f1iA   --------------------qgklsxsDEQKLHEF-A-II-TA-T--DE-AFAQSILQDVD-------------------------------wDLKKALDVFYG-Sea--------
0048 9kgbA   --------sgiPAVDR-----ahvlqtALSIYPEV-E-NW-VAiL--PQ-R-MGGMCLGMV---A-aP--------Y----A---Q----PSVLVEASIMALIA-F--Aiddited
0049 8yhuB   ----------------------sleeeAERVVEEL-V-KE-FN-L--SR-TQEIALRRYA----A--A--------R----A-asE--EVIEELLRDVAERLS-------------
0050 6tdvD   ----------------------thaelHLFDLDEF-MqTY-KR-L--QtqESVFGEMAREL---V--R--------G-----------kTADEATTSVMARVEK-E--Ilgvh---
0051 6toaD   ----MMLN---EVTAV---------pgtALPVAEF-R-DH-LRdlgaED-AALLSYLRAAI---A-eidftagfgaswsalP---V--DLAQAVFLLAAQYYEL-Rh---------

0001 s001A   --LLHHHHHHHHHLLL-----------LHHHHHHH-H-LL-LL-L--LH-HHHHHHHHHHL---L--L--------L----L---H--HHHHHHHHHHHHHHHH-H--L-------
0002 8emhB   -hHHHHHHHHHHHLLL--------leeEHHHHHHH-L-LL-LL-L--LH-HHHHHHHHHLL---L--L--------L----L---H--HHHHHHHHHHHHHHHH-H--Llllllhh
0003 3frhA   -----------------------llllHHHHHHHH-H-HLlLL-L--LH-HHHHHHHHHHH---L-lL--------L----L---H--HHHHHHHHHHHHHHHL-Ll---------
0004 8valE   -hHHHH--hhlllllleeeeeelllllHHHHHHHH-H-HH-LL-L--LH-HHHHHHHHHLL-------------------------------lLHHHHHHHHHL-L--Hhhhhhhh
0005 6h3aB   --lHHHHHHHH-------------hhhHHHHHHHH-H-HL-LL-L--LH-HHHHHHHHHLL---L------------------------LLLHHHHHHHHLLLLl-----------
0006 2ejsA   -llLLLL-----------------llhHHHHHHHH-H-HH-LL-L--LH-HHHHHHHHHHL--------------------------------LHHHHHHHHHH-L--Llllll--
0007 4ociA   --------------------lllllllHHHHHHHH-H-HL-LL-L--LH-HHHHHHHHHL----L--L--------L----L------llLHHHHHHHHHHHHH-L--Lllhhhh
0008 7qogC   -------------------------llHHHHHHHH-H-HH-HLlL--LH-HHHHHHHHHHH---HhlL--------L----L---L--LLHHHHHHHHHHHHHH-H--Hlllllll
0009 2qhoD   --------------------lllhhhlLHHHHHHH-H-HH-LLlL--LH-HHHHHHHHHLL-------------------------------lLHHHHHHHHHH-L----------
0010 1otrA   ------------------------lllHHHHHHHH-H-HH-LL-L--LH-HHHHHHHHHLL-------------------------------lLLHHHHHHHHH-H--Lllll---
0011 3bq3A   -------------------------lhHHHHHHHH-H-HH-HL-L--LH-HHHHHHHHLLL-------------------------------lLHHHHHHHHHH-H--Hlllllll
0012 1mx9D   --------hhhhhhhllllllllllllHHHHHHHH-H-HL-HL-L--LL-LLHHHHHHHHH---L-lL--------L----L---H--HHHHHHHHHHHHHHHL-H--Hhhhhhhh
0013 1xb2B   ------------------------lhhHHHHHHHH-H-HH-HL-L--LH-HHHHHHHHHLL-------------------------------lLHHHHHHHHHH-H--Hhhhhhhh
0014 6c95D   --------hhhHHHHH----hhhllllLHHHHHHH-H-HH-LL-L--LH-HHHHHHHHHLL-------------------------------lLHHHHHHHHHL------------
0015 7qojD   -------------------------llHHHHHHHH-H-HH-HL-L--LH-HHHHHHHHHHHhhhL--L--------L----L---H--HHHHHHHHHHHHHHHH-H--Hllll---
0016 2n5jA   ---LLLL-----------------llhHHHHHHHH-H-HH--L-L--LH-HHHHHHHHHHL---L--L-------------------------lLHHHHHHHHHH-H--L-------
0017 4eeiB   eellEEEEhhhHHLL--------lllhHHHHHHHH-H-HH-LL-L--LH-HHHHHHHHHHH---H--H-------------------lLLHHHHHHHHHHHHHLl-----------
0018 8hkcE   -hHHLLLLhhhhHHHH-----hhlhhhHHHHHHHH-H--H-HL-L--LH-HHHHHHHHHHH---H-hlllhhhlllH----H---H--HHHHHHHHHHHHHHHH-H--Llllllll
0019 9cc7C   -------------------------llHHHHHHHH-H-HH-L-lL--LH-HHHHHHHHHHH---H--L--------L----L-llH--HHHHHHHHHHHHHHHH-H--Hhhhhlll
```

```
0020 2cp8A   ------lLLLL-------------lllHHHHHHHH-H-HH-LL-L--LH-HHHHHHHLLLL------------------------------lLHHHHHHHHHH-H--Llllll--
0021 9fvrC   --------eeeELLLL----lleellllHHHHHHHH-H-HH-LLlL--LH-HHHHHHHHHHH---H--Hhhhllllle E----E---H--HHHHHHHHHHHHHHLH-Hh---------
0022 1nxhA   -hhHHHHLLH-------------hhhhHHHHHHHLlH-HH-HL-L--LH-HHHHHHHHHHH---H--H----hhllL----L---H--HHHHHHHHHHHHHHHL-L---------
0023 6q00B   -------------------------lhhHHHHHHHH-H-HH-HL-L--LH-HHHHHHHHHLL------------------------------lLHHHHHHHHHL-Ll---------
0024 1aipC   --------------------------lHHHHHHHH-H-HH-HL-L--LH-HHHHHHHHHLL------------------------------lLHHHHHHHHHH-H--Hhhhhhhh
0025 8x6gH   hhhhHLLHHHHHH---------hhhhhHHHHHHHH-H-HH-HL-lllLH-HHHHHHHHHHH---H--H-hhhhllllllllllL---H--HHHHHHHHHHHHHHHH-H--Llllllll
0026 3e46A   --------LLLLLL------llllllhhHHHHHHHH-H-HL--L-L--LH-HHHHHHHHHLL------------------------------lLHHHHHHHHHH-L---------
0027 6vvoA   -lLLLLLHleeEELL---------lllHHHHHHHH-H-HH-HL-L--LH-HHHHHHHHHLL---L------------------------LHHHHHHHHHHHHH-H--Hhllllhh
0028 5he9E   -------------------------llhHHHHHHHH-H-HH-HL-L--LH-HHHHHHHH-HH---L--L------------------------LHHHHHHHHHHHHH-H--Hllllllee
0029 3b74A   --------lLLLLLLL----llllllhhHHHHHHHH-H-HH-HHlL--LH-HHHHHHHHHLL------------------------------lLHHHHHHHHHH-H--Hhhhhhhl
0030 2cp9A   -lLLLLL------------------llLLHHHHHH-H-HH-HL-L--LH-HHHHHHHHHHL------------------------------lLHHHHHHHHHH-H--Hhhhllll
0031 1kl7A   --------------------llllllhHHHHHHHH-H-LL-LL-L--LH-HHHHHHHHHHL---L--L--------LlllllllllL--HHHHHHHHHHHHHHHH-H--Hhhlllll
0032 7p37A   eeeeEELL------------lleeellHHHHHHHH-H-HH-LLlL--LH-HHHHHHHHHHH---H--hhhhllllleE----E---H--HHHHHHHHHHHHHHLH-Hh---------
0033 7lt2A   --------eeelHHHH----hllllllHHHHHHHH-H-HH-HL-L--LH-HHHHHHHHHH------hH--------H----L---LhhHHHHHHHHHHHHHHH-L--Llllllll
0034 2ekfA   -llLLLL----------------llllHHHHHHHH-H-HH-LL-L--LH-HHHHHHHHLLL-------------------------------LHHHHHHHHHL-Ll---------
0035 6jcxF   -llHHHHHHHHH----------hhhhhHHHHHHHH-H-HH-HL-LhhHH-HHHHHHHHHHH---H-hllllllllllL----H---H--HHHHHHHHHHHHHHHH-H--Hllllllee
0036 2o7gA   hhhhhhhLLLHHHHH------hhhhhhHHHHHHHH-H-HH-LLhH--HH-HHHHHHHHHHH---H-hhhhhlllllL----H---H--HHHHHHHHHHHHHHLL-----------
0037 6ajfA   eeellllllHHHHHlleelllhhhhhhHHHHHHHH-H-HH-HHhH--HH-HHHHHHHHHH------lllllleehhH----H---H--HHHHHHHHHHHHHHHH-H--Hhhhhhhh
0038 6jbqF   -lLHHHHHhllLLLHH-----hhhhhhHHHHHHHH-L-LL-LL-L-lLH-HHHHHHHHHHH---H-llhhhllllL----H---H--HHHHHHHHHHHHHHHH-H--Hhhlllll
0039 5wuqA   -hHLLL-LLLHH---------hhhhhhHHHHHHHH-H-HH-H--L--LH-HHHHHHHHHHH---H--HhhhlhhhlllllllH---H--HHHHHHHHHHHHHHHH-H--Hlllllll
0040 2cosA   --LLLLL------------------lllLHHHHHHH-H-HH-H--L--LH-HHHHHHHHHHL-----------------------------lLLHHHHHHHHHH-H--Llllll--
0041 2dnaA   -lLLLL-LLLL------------lhhhhLHHHHHHH-H-HH-LL-L--LH-HHHHHHHHHLL------------------------------lLHHHHHHHHHH-L--Llllllll
0042 8vjlB   hhhhLLHHHHHH-----------hhhlLHHHHHHH-H-HH-LLlL--LH-HHHHHHHHHHH---H--H-hhhllllllllllH---H--HHHHHHHHHHHHHHHH-H--Hhhhl---
0043 7mvvA   --------eeelLEEE------ellhhHHHHHHHH-H-HH-HL-L--LH-HHHHHHHHHHH---H------------hllhH---H--HL-LLHHHHHHHHHHH-H--Hhhhhhhh
0044 6e6yA   -lLHHH--------------lllhhhhHHHHHHHH-H-HL-HHhH--HH-HHHHHHHHHHH---H-hL--------L----L---H--HHHHHHHHHHHHHHHH-H--Hhhlllll
0045 1wglA   --------------------lllllllLHHHHHHH-H-HH-LL-L--LH-HHHHHHHLLLL------------------------------lLHHHHHHHHHH-L--Llllllll
0046 2qsfX   ----llLLHHHHHHH------llllllhhHHHHHHHH-H-L--LL-L--LH-HHHHHHHHHLL------------------------------lLHHHHHHHHLL-L---------
0047 4f1iA   --------------------llllllllHHHHHHHH-H-HH-HL-L--LH-HHHHHHHLLLL------------------------------lLHHHHHHHHHH-Lhl---------
0048 9kgbA   --------hhlLLLLH-----hhhhhhHHHHHHHH-H-HH-HLlL--LL-L-LHHHHHHHH---H-lL--------L----L---L----HHHHHHHHHHHHHH-H--Hhhhhhll
0049 8yhuB   ----------------------lhhhhHHHHHHHH-H-HH-LL-L--LH-HHHHHHHHHH----H--H--------H----H-llL--LLHHHHHHHHHHLLL-------------
0050 6tdvD   ----------------------llhhhHHHHHHHH-HhHL-HH-H--LllLLHHHHHHHHH---H--H--------L-----------lLHHHHHHHHHHHHH-H--Lllll---
0051 6toaD   ----LLLE---ELLLL---------lllLLLHHHH-H-HH-LLllllLH-HHHHHHHHHHH---H-eeeeeellllllhhhlL---H--HHHHHHHHHHHHHHHL-Ll---------
```

```
# Job: Type-1 BREX; BrxL HAM28421 NTD2 OB-Fold
# Query: s001A
# No:  Chain   Z    rmsd lali nres  %id PDB  Description
   1:  8emh-B 16.2  1.4   98   676   36   MOLECULE: PROTEASE LON-RELATED BREX SYSTEM PROTEIN BRXL;
   2:  7xut-B  6.3  3.2   82   140   12   MOLECULE: FUSION PROTEIN OF REPLICATION PROTEIN A 70 KDA DN
   3:  1u5k-A  6.1  3.4   73   242    4   MOLECULE: HYPOTHETICAL PROTEIN;
   4:  7yny-A  6.1  3.0   69   332    9   MOLECULE: LEF3;
   5:  4gnx-C  6.0  3.0   84   433    5   MOLECULE: PUTATIVE UNCHARACTERIZED PROTEIN;
   6:  4gs3-A  5.8  3.7   74    90   11   MOLECULE: SINGLE-STRANDED DNA-BINDING PROTEIN;
   7:  6d6v-E  5.5  2.9   77   149    5   MOLECULE: TELOMERASE REVERSE TRANSCRIPTASE;
   8:  1m4v-B  5.5  2.9   69   197    3   MOLECULE: SET3, SUPERANTIGEN-LIKE PROTEIN;
   9:  8aas-C  5.5  3.4   72   113   17   MOLECULE: REPLICATION FACTOR A;
  10:  3u58-A  5.5  3.6   77   213    6   MOLECULE: TETRAHYMENA TEB1 AB;
  11:  7lma-D  5.4  2.8   79   187   10   MOLECULE: TELOMERASE LA-RELATED PROTEIN P65;
  12:  7ymo-A  5.4  3.9   69   206   14   MOLECULE: RECOMBINATION PROTEIN RECO;
  13:  4gn5-A  5.4  3.1   75   113    4   MOLECULE: OBODY AM3L15;
  14:  3jc6-5  5.2  3.7   79   254    6   MOLECULE: DNA REPLICATION LICENSING FACTOR MCM2;
  15:  6pqh-B  5.2  3.5   78   479    4   MOLECULE: ASPARAGINE--TRNA LIGASE;
  16:  1gm5-A  5.2  4.0   80   729   11   MOLECULE: RECG;
  17:  8ez6-A  5.2  2.6   61    65   13   MOLECULE: CELL CYCLE AND APOPTOSIS REGULATOR PROTEIN 2;
  18:  3tdq-A  5.2  3.3   68    86    6   MOLECULE: PILY2 PROTEIN;
  19:  8soj-B  5.1  3.8   76   428    8   MOLECULE: CST COMPLEX SUBUNIT CTC1;
  20:  5m1x-D  5.1  3.3   82   133   10   MOLECULE: REPLICATION FACTOR A PROTEIN 1;
  21:  1nnx-A  5.1  3.1   74    93    7   MOLECULE: PROTEIN YGIW;
  22:  8y30-A  5.1  4.2   78   756   10   MOLECULE: SINGLE-STRANDED-DNA-SPECIFIC EXONUCLEASE RECJ;
  23:  3f2b-A  5.1  3.9   78   994   13   MOLECULE: DNA-DIRECTED DNA POLYMERASE III ALPHA CHAIN;
  24:  5z7d-B  5.1  3.7   79   397    5   MOLECULE: INTERFERON-ACTIVABLE PROTEIN 204;
  25:  6lbt-B  5.1  3.2   79   303   15   MOLECULE: KLLA0F20922P,KLLA0F20922P;
  26:  6lbr-A  5.1  3.5   85   519   15   MOLECULE: KLLA0F20922P;
  27:  4o2d-B  5.1  3.4   76   516    7   MOLECULE: ASPARTATE--TRNA LIGASE;
  28:  6sw9-N  5.1  3.2   75   146    7   MOLECULE: 16S RIBOSOMAL RNA;
  29:  7dpd-B  5.0  3.6   73   273    5   MOLECULE: DNA HELICASE MCM9;
  30:  4hce-B  5.0  3.3   81   130    6   MOLECULE: CELL DIVISION CONTROL PROTEIN 13;
  31:  3klw-A  5.0  3.4   70    98    4   MOLECULE: PRIMOSOMAL REPLICATION PROTEIN N;
  32:  1ue1-B  5.0  4.2   76   119    8   MOLECULE: SINGLE-STRAND BINDING PROTEIN;
  33:  8peu-A  5.0  3.3   85   419    8   MOLECULE: TRANSCRIPTION TERMINATION FACTOR RHO;
  34:  3rd4-B  5.0  2.3   64    82    8   MOLECULE: UNCHARACTERIZED PROTEIN;
  35:  8ehb-A  4.9  3.1   75   102   13   MOLECULE: PUTATIVE LIPOPROTEIN;
  36:  8txr-C  4.9  3.5   71   431   14   MOLECULE: EXODEOXYRIBONUCLEASE 7 LARGE SUBUNIT;
  37:  8d0b-C  4.9  3.5   77   121    6   MOLECULE: CST COMPLEX SUBUNIT CTC1;
  38:  1miu-A  4.9  3.1   78   671   13   MOLECULE: DELETED IN SPLIT HAND/SPLIT FOOT PROTEIN 1;
  39:  3k8a-B  4.9  3.3   70   103    9   MOLECULE: PUTATIVE PRIMOSOMAL REPLICATION PROTEIN;
  40:  3e0e-A  4.9  3.5   74    91    8   MOLECULE: REPLICATION PROTEIN A;
  41:  4dap-A  4.9  3.3   69   232    7   MOLECULE: SUGAR FERMENTATION STIMULATION PROTEIN A;
  42:  1eqr-A  4.9  3.6   75   590    8   MOLECULE: ASPARTYL-TRNA SYNTHETASE;
  43:  3en2-A  4.8  2.9   65    91   12   MOLECULE: PROBABLE PRIMOSOMAL REPLICATION PROTEIN N;
  44:  3kf8-D  4.8  2.9   72   123    8   MOLECULE: PROTEIN STN1;
  45:  7uy7-B  4.8  4.1   79   291    9   MOLECULE: TELOMERASE-ASSOCIATED PROTEIN OF 75 KDA;
  46:  7w7p-D  4.8  3.9   79   284    6   MOLECULE: DNA HELICASE MCM9;
  47:  2z4h-B  4.8  2.7   71   194    6   MOLECULE: COPPER HOMEOSTASIS PROTEIN CUTF;
  48:  6cqm-D  4.8  4.3   72    99    8   MOLECULE: SINGLE-STRANDED DNA-BINDING PROTEIN RIM1, MITOCHO
  49:  2i0q-A  4.7  4.1   78   461   18   MOLECULE: 5'-D(*GP*GP*GP*TP*TP*TP*TP*GP*GP*GP*G)-3';
  50:  6lbu-B  4.7  2.8   73   135   15   MOLECULE: KLLA0C11825P;
```

```
0001 s001A   -VRPDE--SNKAQ--AIVE-Q-KKR-HKF-I-DKI-HVKYVER-EK-R-Y-WAEME-N-F---ASKRIAI-NS-N-FY-Q--Q-N--EKL----LESG--IWAEV-T-LAYN-E-V--EED--D-YA-F-FIE--D-L-RPIQ-I--A-H-FD-
0002 8emhB   -VRPDE--AEKVK--SLVR-E-RGS-YKV-I-DRV-TVKLNER-KD-K-Y-EASFS-N-L---GIKDAEI-SA-G-IV-K--E-Y--EKL----LVGG--IWVIA-T-LSYY-F-E--EQT--S-SP-F-GVS--L-L-KPIQ-M--P-N-MNm
0003 7xutB   ------qlseGAI--AAIMqK-GDTnIKP-I-LQViNIRPITTspP-R-Y-RLLMS-D--glnTLSSFMLqLN-P-LV-E--E-E--QLS------SN--CVCQIhR-FIVN-T-L--KDG--R-RV-V-ILM--E-L-EVLKs---------
0004 1u5kA   --------------------R-SRT-ANR-S-GIViRRRVTP--AG-D-I-IVTLL-T-G---KLK-AIA-RG-P-LS---------SSL----N-LF--HHVGV-Q-VYQG-P-H---------DL-A-SVK--Q-A-VLEG-A--L-P-T--
0005 7ynyA   --------------------K-LRT-LNV-K-GQLlTKTTMSInED-Y-Y-LFKFL-V-K---SID-YYG-TQ-T-QF-F--S-L--IN--------N--KTYEL-V-LQYS-R-K----------K-L-LIK--S-Y-EQCEd---------
0006 4gnxC   gaerrT--IVQVK-dENLG-MkpDY-FNV-R-ATV-VYIKQEN-LY-taL-STNVA-D-G---QMW-LSG-FN-E-DA-T--Q-L--Igm---lHRAA-nRMYMF-N-CRAK-M-D--TFdtaR-VR-Y-TIS--R-A-APVD-Fa--------
0007 4gs3A   --------------------E-NNT-VTL-V-GKV-fLEFSHE----F-F-NFILE-V-D---YLP-ITI-SN-R-LF-E--G-M--NL------EVG--TRVKI-E-GQLR-S-Y--NKL--I-LT-V-FAR--D-I-SVVP-E--------
0008 6d6vE   ------lnCFMIN--QIakenaFEN-AII-T-AKFeFLQSIND-SP-K-I-MGKLS-D--stgCIEVVIqpEF-V-QA-Y--E-I--E--lqnngNRH--KYVRA-M-LKMR-K-N---------AQ-I-QLL--Y-F-SIVN-Da--------
0009 1m4vB   ------akyenvtkdifdyyS-GAS-KEL-KnVTG-YRYSK---GG-K-H-YLIFD-K-K---FTR-IQI-FG-K-DI-E--R-L--KT------RKN--PGLDI-F-VVKE-A-----------vF-S-YG---G-V-TKKN-Q--Gr-----
0010 8aasC   ------rkISEI-----rE-E-DTR-VSL-I-GRViKVDKM------D-Y-MFWLD-D---gtGVAIIES-ES-D-LP------------------kVG--QVVRV-I-GRII-R-N---EE--G-IH-I-YAE--V-I-QDFS-D--A------
0011 3u58A   ------llISEVL--KT---S-KQY-LSV-L-AQVvDIQSSD---K-N-I-RLKIC-D-Q---ELK-VVI-FC-Y-EW-R--D-K--FS-------IN--KWYYF-NeFVRQ-I-Y--N-------vQ-L-KNS--S-I-KESD-D--------
0012 7lmaD   ------ssIEEII--QQAQ-AqkKE-FYV-Y-GNL-VSIQM-K-NK-LyL-RAFVQ-D--stgTIPV-MI-FD-Q-QS-S--Q-L--InqqlfskLDF--ARFIF-E-IQFE-N-K--EF-----iA-Y-KVL--K-I-EKEN-Ik--------
0013 7ymoA   --------------------------DPM-Q-GFIlHTQKVKD------L-IVYIL-S-M---LIK-AYR-FY----G-L--R-H--SSI----L-SG--YKIDF-A-LEEN-P-S----------FL-P-RLK--D-V-LHLG-F-wI-M-QRd
0014 4gn5A   ------thwtAEI--T-nL-H-GSE-VVV-A-GWVaHLGDYG---R-V-K-IVKVS-D-reggAAVPVYLhLF-K-VF----A-E--LS-------RE--DVVVI-K-GIVE-A-G--WPV----vE-I-FPS--E-I-WILNk---------
0015 3jc65   lanqiP--LRDLD--S-eH-V-SKI-VRL-S-GIIiSTSVLSS-R--htQ-FLKLQ-E-R---NLT-MTC-DR-Y-LT-N----K--VIP-------G--TRVTI-V-GIYS-I-Y--NS-----tpyI-KIL--G-I-QSDV-E--T-S----
0016 6pqhB   -mhkqT--IKEVL--E-kF-L-HHD-ITV-Y-GWVrAFRS----------R-FIALN-D-gstiNNLQIVV-DF-EnFD-E--N-L-iKNI----N-TA--SSLKI-V-GEVV-E-S--Q------vE-I-IAK--K-I-IVLG-D--N-Ft---
0017 1gm5A   -----frkifKLN--D-LL-P-GEK-VTT-Q-GKI-VVETKKFnMN-I-L-TAVLS-D-V---HVP-LKW-FD-Y-LQ-T--YlK--QLT-------G--KEVFV-T-GTVK-S-N-aYTG--Q-yE-I-HNA--E-V-TPKEg---------
0018 8ez6A   ----------------------EKQ-RVF-T-GIV-TSLHD-------F-GVVD----------EEVFF-QL-S-V---kgR-L--PQL-------G--EKVLV-K-AAYN-P-G----Q--A-VP-W-NAV--K-V-QTLS---------
0019 3tdqA   ------------------eD-PQT-FEG-A-GVVfEVQVE------K-N-LVDI----D---HRL-YRL-PNsP-SL-F--Q------V----KPG---SVVSY-S-GTVS-Q------------PW-S-TIT--D-I-YIHK-Q--X-S-EQe
0020 8sojB   vfaklY--IRDIL--DM--kepIKQ-VDV-L-GTViGVRER---DA-F-Y-SYGVD-D-G---VIN-CIC-WK-KlntsveqK-T--KIE------IG--DTIRV-R-GSIR-T-Y--RE-----rE-I-HAT--T-Y-YKVD-Dp--------
0021 5m1xD   -qlSRG--DfHSI--FtRY-D-N-P-TGG-V-YQV-YNTR----KS-D-GaLIXIS-D---giYHXKALL-RA-S-KF-Q--S-X--ELQ------RG--DIIRViI-AEPA-I-V--RRK--K-YV-L-LVD--D-F-ELVQs---------
0022 1nnxA   ------ttVESAK--SL-R-D-DTW-VTL-R-GNIvERIS--------D-D-LYVFK-D---asGTINVDI-DH-K-RW-N--G-V--TV---------tpkDTVEI-Q-GEVD-K-D--W-N--S-VE-I-DVK--Q-I-RKV---------
0023 8y30A   vaaflaltENDI-----iknisrPI-FEM-DdLSV-SVKAIGQ----H-L-KLTLG-E-S---NI-AALF-WQ-N---gH--L-E--PEL------QDE--QPINI-L-GSVQ-I-N--EW----nPQ-I-IIQ--D-I-AMNE-Q-fS-Q-LYi
0024 3f2bA   ---vrR--LET-I--VEEE-----RR-VVV-Q-GYVfDAEVSEL-------L-TMKIT-D-N---SIL-VKM-FS-RdkeaE--L-M--SGV------KKG--MWVKV-R-GSVQ--nD--TFV--R-DLvI-IAN--D-L-NEIAa---------
0025 5z7dB   rnatpK--ISYLF--SQ-A-R-GTF-VSG-E-YLVnKKTER---NK-F-I-YYGIG-D-G---KME-VVV-YG-R-LT-N----V--RCE-------PG--SKLRL-VcFELT-S-T--E------wQ-L-RSS--Y-M-QVIN-A---------
0026 6lbtB   ---------DLKP--DS-I-S-KAL-YHIsD-GKIlGFIPnSL-IE-F-L-LIYVY-T-vpeyNCYEIAI-TA-K-FF-S--K-I--G------eLKEL--IRYDI-T-IYKK-E-F--TI-----sK-W-ALK--D-V-VLRSg---------
0027 6lbrA   ------dpfDLKP--DS-I-S-KAL-YHIsD-GKIlGFIPNSL-IE-F-L-LIYVY-TlN---CYE-IAI-TA-K-FF-S--K-I--G------lKELIdnNRYDI-T-IYKK-E-F--TI-----sK-W-ALK--D-V-VLRS-A-qA-F-YDd
0028 4o2dB   ------thaaGSL--R-aD-A-GQT-VTL-A-GWVaRRRDH---GG-V-I-FIDLR-D-G---VSQ-VVF-RL-A-AA----H-R--LR-------AE--FCVAV-T-GVVE-VrP--EGN--EniE-V-NAT--E-L-TVLGe---------
0029 6sw9N   ------keksDPL--E-----G-APQ-ARG---IVL-EKIAVEAsGM-R-K-AVRVQnG-K---VV-TAFC-PG-G-AI-K--F---------------iD--EHDEV-I-IEGI-G-GpkGGS--MgIR-Y-KVV--K-V-N-rK-P--R-R----
0030 7dpdB   ------lvreHIP--K-kD-V-GHF-LSV-T-GTViRTSLVKV-L--yyQ-EIKIQ-E-R---SMK-VIL-ED-D-LV-------D--SCK-------SG--DDLTI-Y-GIVM-Q-R--WK--pfqeivL-KAN--Y-I-QVNN---------
0031 4hceB   ----IS--YEQLS--LA-S-V-GSV-ERL-E-GKIvGMNPpFK-YC-T-L-KLYFT-QpN---CIE-IVI-P--E-LF-G--------------kISDI--LRISV-E-VERI-L-W--DPG--M-AV-W-SLK--N-I-STDT---------
0032 3klwA   ----------------------XNT-LEL-S-ARVlECGAXR--------L-ELLLV-HeL---TIS-AVA-LG-D-LA-L--L-------------aDTPL-gTEXQV-Q-GFLA-PaR--KDS-vK-VK-L-HLQ--Q-A-RRIA-G---------
0033 1ue1B   ------------------aG-DTT-ITI-V-GNL-TPELRFT----V-A-NFTVA-SdL---FLR-CNI-WE-N-VA-----E-S--LT--------RG--ARVIV-S-GRLK-Q-R--SFT--V-IE-V-EVD--E-I-GPSL-T--A-K-V--
0034 8peuA   -LTELKrkQDIIF--AILK-Q-HAD-IFG-D-G-V-LEILQ----D-G-F-GFLRS-A-sylagpdDIYV-SP-S-QI-R--R-F--NLR----T--G--DTISG-K-IRPP-K-E----G--E-RY-F-ALL--K-VnEVNF-N--A-R-N--
0035 3rd4B   ----------------------aPV-KLY-M-VEViDKKEIAAiTH-Y-Y-QVTFR-L-K---DLV-LNI-DK-S-SY-Q--N-------I------EPE---MKGRL-F-MQG------------------S-RFV--Q-F-ETDV-P---------
0036 8ehbA   ------------eshasccveE-KIPiVTL-KnENA-HFRYXKR-RN-D-F-ALEIE-N-KelvRGLYLIPrGC-D-I-pK--K-Y--K------eDGLP--VIISG-E-VFDC-S-E--YKR--D-PV-Y-FIK--L-S-TIK---------
0037 8txrC   ------iftvsrlnqtvlehE-MGQ-VWI-S-GEI-SFTQPA--SG-H-W-YFTLK-D-A---QVR-CAM-FR-N-snrV--T-F--RP-------QHG--QQVLV-R-ANIT-L-Y--ERG--D-YQ-I-IVE--S-M-QP-----------
0038 8d0bC   lgtyyL--PWEVS--AGqV-P-DGT-LRT-F-GRL-CLYDM----I-Q-S-RVTLM-A-gsdqHQVLVCTlVE-P-F--------H--AQV-------G--SLYIV-L-GELQ-H-Q--QDR----sV-V-KAR--V-L-TCVE-Gm--------
0039 1miuA   -srrSA--LKKILerDDTA-----A-KTL-V-LCI-SDIV--------D-TIELT-D-Y---AVR-AQL-DP-P-LM-A--L-VksGKL----T-VG--QKIIT-QgAELV-G-S--P-apdS-LR-L-KISanS-T-RPAR-Whe-------
0040 3k8aB   -----------------hmgF-TNL-VSL-A-ALIeKAFP-----------L-DIILK-HwL---EIP-ARI-LG-R-QA-E--E-W--QYR-------QG--DCATV-E-GFLA-QkS--RRS--L-MP-MlRIQ--N-I-KEYKg---------
0041 3e0eA   ------ykISEL------X-P-NLS-GTI-N-AEVvTAYPKKE----L-K-SLFLK-D-G---SIR-GTL-WN-E-LA-D-----F--EVK-------KG--DIAEV-S-GYVK-Q------------lE-I-SVD--N-I-GIIE-K--S------
0042 4dapA   -----------------mefS-PPL-QRA---TLI-QRYK----------F-LADVIdG-R---EL-TLHC-PN-T-----G--A-M--TG-------caT--PGDTV-W-YSTS-D-N--TKR--K-YP-H-TWE--L-T-QSQS---tL-A-ENe
0043 1eqrA   ------teycgQL--RLsH-V-GQQ-VTL-C-GWVnRRRDLG---S-L-I-FIDMRrE-G---IV-QVFF-DL-K-LA----S-E--LR--------NE--FCIQV-T-GTVR-arD--EKN--IniE-V-LAS--S-L-TIINr---------
0044 3en2A   ----------------------A-INR-LQL-V-ATL-VEREV----------paCLLSY-S-rqveFSIEALGkXA-S-VL-D----R--IAP-------G--TVLEC-V-GFLA-R-K--H----A-LV-F-HIS--G-L-EHH---------
0045 3kf8D   ------lipSNIP--QE----npER-LRI-L-AQVkDFIPH-------E-S-TIVID-K-eqstYINI-CI-FN---LL-E--AcS--SRV-----LVPG--TLVNI-D-AFYD-G-E-----------S-I-NPV--D-I-YEVN-Ga--------
0046 7uy7B   -felvF--LKELP--SL-P-D-FSK-VCF-T-GLIlSFSNFPS-S----K-IAIIQ-D-G---EAE-LFL-DM----Y-K--F-C--Q-----eISVF--KAITG-I-GVLKkK-N--IG-----cK-I-IVE--R-F-RIIH-Sa--------
0047 7w7pD   ------tqLKNVR--ANCY---GKY-IAL-R-GTVvRVSNIKP-LctW-Q-SVKVQ-EmR---TIE-CEL-VQ-D-LV-D--------------scVPG--DVVTI-T-GVVK-V-S--STEcvF-LL-YiEAN--S-V-SNSK---------
```

```
0048 2z4hB   ------iesqfnytleasL-P-MTP-MTL---RGM-YFYMA------D-A-ATFTD-C-G---KRFMVAN-NA-E-LE-R--S-Y--LA-arghsEKP--VLLSV-E-GHFT-L----eAP--T-KV-L-APD--TaG-KFYPn----------
0049 6cqmD   ---------------------dFSK-MSI-V-GRI-GFTEHTSanR-Y-L-KYSIA-S-rgqtNWYNITVqIN-F-LT-E--Y-V--RK--------G--ALVYV-E-ADAA-N-Y--VFT--T-LS-L-VQK--D-I-NLLKn----------
0050 2i0qA   vdmytA--LNKAQ--AQ-----KGD-FDV-V-AKIlQVHELD--EY-T-N-ELKLK-D-Q---VFY-TLS-LL-K-FP------H--VRT--------G--EVVRI-RsATYDeT-S--TQ-----kV-L-ILS--N-I-IT-F-I--Q-Ss---
0051 6lbuB   -kiitD--LDTIA--GK----tlLR-LRI-F-AQF-QDI----rtD-G-I-YLHFS-N-aeerSYYF-LI-DE-T-IY-D--E-A--F--sgerpHKG--DILDX-R-CCYR-K-Y-------D-KV-V-EIX--H-L-KVIS-Ia--------

0001 s001A   -LLHHH--HHHHH--HHHH-H-HLE-EEE-E-EEE-EEEEELL-LL-E-E-EEEEL-L-L---LEEEEEL-LH-H-HH-H--H-L--HHH----HLLL--EEEEE-E-EEEL-L-L--LLL--L-LL-E-EEE--E-E-EELL-L--L-L-LL-
0002 8emhB   -LLLLL--HHHHH--HHHH-H-HLE-EEE-E-EEE-EEEEELL-LL-E-E-EEEEL-L-L---LLLLEEE-LH-H-HH-H--H-L--HHH----HLLL--EEEEE-E-EEEL-L-L--LLL--L-LL-E-EEE--E-E-EELL-L--L-L-LLh
0003 7xutB   ------hlllLHH--HHHHhH-LLLlLLL-E-EEEeEEEEELLllL-E-E-EEEEE-L--lleEEEEEEEhHH-H-HH-H--L-L--LLL------LL--LEEEEeE-EEEE-E-L--LLL--L-EE-E-EEE--E-E-EEEEl----------
0004 1u5kA   --------------------L-LLE-EEE-E-EEEeEEEELL--LL-L-E-EEEEE-E-E---EEE-EEE-LL-L-LL----------LLL----L-LL--LEEEE-E-EELL-L-L----------LL-E-EEE--E-E-EEEE-L--L-H-H--
0005 7ynyA   --------------------L-LLE-EEE-E-EEEeEEEEEEElEE-E-E-EEEEE-L-L---EEE-EEE-LH-H-HH-H--H-L--LL---------L--LLEEE-E-EEEE-L-L-----------E-E-EEE--E-E-LLLLl----------
0006 4gnxC   lllllL--HHHHH-hLLLL-LllEE-EEE-E-EEE-EEELLLL-LE-eeE-EEEEE-E-E---EEE-EEE-EH-H-HH-H--H-H--Hll---hHHHL-lEEEEE-E-EEEE-E-E--EEleeE-EE-E-EEE--E-E-EELL-Hh--------
0007 4gs3A   --------------------L-LLL-EEE-E-EEE-lLEEEEE----E-E-EEEEE-E-E---EEE-EEE-EH-H-HL-L--L-L--LL-------LLL--LEEEE-E-EEEE-E-E--ELE--E-EE-E-EEE--E-E-EEEL-L---------
0008 6d6vE   ------llHHHHH--HHhhhllLLE-EEE-E-EEEeEEELLLL-LL-E-E-EEEEE-E--lleEEEEEElhHH-H-HH-H--H-H--H--lllllLLL--EEEEE-E-EELL-L-L----------LL-L-EEE--E-E-EELL-Lh--------
0009 1m4vB   ------lllllllhhhhhhhL-LLL-EEE-EeEEE-EEEEE---LL-E-E-EEEEE-E-E---EEE-EEE-LL-H-HH-H--H-L--LL------LEE--EEEEE-E-ELLL-E------------eE-E-EL---L-E-EELL-L--Ll-----
0010 8aasC   ------llHHHL------lL-L-LLE-EEE-E-EEEeEEELL------L-L-EEEEE-L---llLEEEEEL-LL-L-LL-----------------lLL--LEEEE-E-EEEE-E-E---LL--E-EE-E-EEE--E-E-EELL-L--L------
0011 3u58A   ------llHHHHL--LL---L-LLE-EEE-E-EEEeEEEELL---L-L-E-EEEEE-L-L---LEE-EEE-LL-H-HH-L--L-L--LL-------LL--LEEEE-EeEEEL-L-L--L-------lE-E-ELL--E-E-EELL-L---------
0012 7lmaD   ------llHHHHH--HHHL-LleEE-EEE-L-LEE-ELLLL-L-LL-LlE-EEEEE-L--lllEEEE-EE-LH-H-HH-H--H-H--Hhhhhhhl LLL--EELEE-E-EEEE-L-L--EE------lE-E-EEE--E-E-ELLL-Hh--------
0013 7ymoA   -------------------------LLE-E-EEEeEEEEELL------E-EEEEE-L-L---EEE-EEE-LL----H-H--H-L--LLL----L-LL--LEEEE-E-EEEE-L-L-----------EE-E-EEE--E-E-EEEL-L-lL-L-LHh
0014 4gn5A   ------lllhHHL--L-hH-L-LLE-EEE-E-EEEeEEEELL---L-E-E-EEEEE-L-llllLLEEEEEhHH-H-HH----L-L--LL-------LL--LEEEE-E-EEEE-L-L--LLH-----lE-E-EEE--E-E-EEEEl----------
0015 3jc65   elllllL--HHHLL--L-lL-L-LLL-LLE-E-EEEeEELLLLL-L--llL-LEEEE-E-L---EEE-EEL-LH-H-HL-L-----L--LLL--------L--LEEEE-L-LLEE-E-E--LL------eeeE-ELL--L-L-EELL-L--L-L----
0016 6pqhB   -llllL--HHHHH--H-hH-L-LLE-EEE-E-EEEeEEEL---------L-EEEEE-L-llllLLEEEEL-LH-HhLL-H--H-H-hLLL----L-LL--LEEEE-E-EEEE-E-L--L-------eE-E-EEE--E-E-EELL-L--L-Lh---
0017 1gm5A   -----lllllLLL--L-LL-L-LLL-LEE-E-ELL-LLEEEELlLE-E-E-EEEEL-L-L---LEE-EEE-LL-L-LH-H--HhH--LLL--------L--LLEEE-E-EEEL-L-L-lLLL--L-lL-E-EEE--E-E-ELLLl----------
0018 8ez6A   ----------------------LLE-EEE-E-EEE-EEELL---------E-EEEL-----------LLEEE-EH-H-H---llL-L--LLL--------L--LEEEE-E-EEEL-L-L----L--L-LL-E-EEE--E-E-EELL-----------
0019 3tdqA   -------------------lL-LLE-EEE-E-EEEeEEEHH------H-L-EEEE----L---LEE-EEL-LLlE-HH-H--L-------L----LLL---LEEEE-E-EELL-L------------LL-L-EEE--E-E-EELL-L--L-L-EEh
0020 8sojB   heellL--HHHHH--HL--eeeELE-EEE-E-EEEeEEEEL---LL-E-E-EEEEE-L-L---EEE-EEE-EL-LlllllhhH-H--LLL-------LL--LEEEE-E-EEEE-E-E--LL------eE-E-EEE--E-E-EEEL-Ll--------
0021 5m1xD   -llLLL--HhHHH--HhHH-H-L-L-LLL-E-EEE-EELL-----HH-H-LlEEEEE-L---llEEEEEEE-EH-H-HH-H--H-L--LLL-------LL--LEEEEeE-EEEE-E-E--LLL--E-EE-E-EEE--E-E-EEEEl----------
0022 1nnxA   ------llHHHHL--LL-L-L-LEE-EEE-E-EEEeEEEE--------L-L-EEEEE-E---llEEEEEEL-LH-H-HL-L--L-L--LL---------lllLLEEE-E-EEEE-E-E--L-L--E-EE-E-EEE--E-E-EEL------------
0023 8y30A   lhhhhhllHHHL-----hhhhllLL-EEE-EeEEE-EEEEELL----E-E-EEEEL-L-L---LE-EEEE-LL-L---lL--L-H--HHH------LLL--LEEEE-E-EEEE-E-E--EE-----lEE-E-EEE--E-E-EELL-L-lL-E-EEe
0024 3f2bA   ---llL--HHH-L--LLLE-----EE-EEE-E-EEEeEEEEEEL-------E-EEEEE-L-L---EEE-EEE-EL-LlhhhH--H-H--HLL------LLL--LEEEE-E-EEEE--eE--LLL--L-EEeE-EEE--E-E-EEELl----------
0025 5z7dB   hllllL--HHHHH--LL-L-L-LLL-EEE-E-EEEeEEEEE---LL-E-E-EEEEE-E-E---ELE-EEE-EH-H-HH-H-----L--LLL-------LL--LEEEE-EeEEEE-E-E--L-------eE-E-ELL--E-E-EEEL-L---------
0026 6lbtB   ---------LLLL--LL-H-H-HLE-EEEeE-EEEeEEELlLL-LL-L-L-EEEEE-E-llllLLEEEEL-LH-H-HH-H--H-H--L------hHHHH--HEEEE-E-EEEE-E-E--EE------eE-E-EEE--E-E-EEELl----------
0027 6lbrA   ------lllLLLL--LL-H-H-HLL-EEEeE-EEEeEEELLLL-LL-L-E-EEEEE-LlL---LEE-EEE-LH-H-HH-H--H-H--L------hHHHHhhLEEEE-E-EEEE-E-E--EE------eE-E-EEE--E-E-EEEE-L-lL-L-LLl
0028 4o2dB   ------lllhHHL--L-hH-L-LLE-EEE-E-EEEeEEEEE---LL-E-E-EEEEE-E-E---EEE-EEE-LH-H-HH-----L-L--LL--------LL--LEEEE-E-EEEE-ElL--LLL--LleE-E-EEE--E-E-EEEEl----------
0029 6sw9N   ------llllLLL--L----L-LLL-EEE---EEL-LLEEEELlLE-E-E-EEEEElL-L---EE-EEEL-LL-L-HH-H--L---------------lL--LLLLE-E-EELL-L-LhhHLL--LlLL-L-EEE--E-E-L-lL-L--L-L----
0030 7dpdB   ------leelLLL--L-hH-L-LLE-EEE-E-EEEeEELLLEE-E--llE-EEEEE-E-L---EEE-EEE-EH-H-HL-------L--LLL-------LL--LEEEE-E-EEEE-E-E--EE--llleeeE-EEE--E-E-EELL-----------
0031 4hceB   ----LH--HHHHL--LL-L-L-LLE-EEE-E-EEEeEEELlHL-LL-L-E-EEEEE-LlL---LEE-EEE-L--H-HH-L--------------hHHHH--LLEEE-E-EEEE-E-E--LLL--E-EE-E-EEE--E-E-ELLL----------
0032 3klwA   ----------------------LEE-EEE-E-EEEeEELLLE---------E-EEEEE-EeE---EEE-EEE-EL-H-HH-H--H-------------lLLLL-lLEEEE-E-EEEE-ElL--LLL-lL-EE-E-EEE--E-E-EEEL-L---------
0033 1ue1B   -------------------lL-LLE-EEE-E-EEE-LLEEEEL-----E-E-EEEEE-ElE---EEE-EEE-EH-H-HH-----H-H--LL--------LL--LEEEE-E-EEEE-E-E--ELE--E-EE-E-EEE--E-E-EELL-L--L-E-E--
0034 8peuA   -HHHHHlhHHHHH--HHHH-H-HHL-LLL-E-E-E-EEELL-----L-L-L-EEEEL-L-lllllllLEEE-LH-H-HH-H--H-L--LLL-----L--L--LEEEL-L-EELL-L-L-----L--L-LL-L-EEL--L-LlEELL-L--L-L-L--
0035 3rd4B   ----------------------lLE-EEE-E-EEEeEEEEEEEEeEE-E-E-EEEEE-E-E---EEE-EEE-LH-H-HH-L--L-------L-----LLL---LEEEE-E-EEL-----------------L-EEE--E-E-EELL-L---------
0036 8ehbA   -----------leelllllL-EEEeEEE-EeEEE-EEEEELL-LL-E-E-EEEEL-L-HhhlLLLLEELlLL-L-L-lH--H-H--L------lLLLE--EEEEE-E-EEEE-H-H--HLL--L-LL-E-EEE--E-E-EEE------------
0037 8txrC   ------eeehhhhhhhhhhhH-LLE-EEE-E-EEE-EEELLL--LL-L-E-EEEEE-L-L---EEE-EEE-LH-H-hhlL--L-L--LL-------LLL--LEEEE-E-EEEE-E-E--LLL--E-EE-E-EEE--E-E-EL------------
0038 8d0bC   lleelL--HHHHL--LLlL-L-LLE-EEE-E-EEE-EEEEL----L-L-L-EEEEE-E-lleeEEEEEELlLL-L-L-------L--LLL--------L--LEEEE-E-EEEE-E-L--LLL----eE-E-EEE--E-E-EELL-Ll--------
0039 1miuA   -lllLH--HHHHHllLLLL-----L-LLE-E-LEE-LLLL----------L-LLLEE-L-L---EEL-LLL-LH-H-HH-H--H-HhlLLL----L-LL--LLLLL-LlLLEE-L-L--L-llllL-LE-E-LLLhhH-L-LLLL-Lll-------
0040 3k8aB   -----------------lllL-LEE-EEE-E-EEEeEELL----------E-EEEEE-EeE---EEE-EEE-EH-H-HH-H--H-L--LLL-------LL--LEEEE-E-EEEE-ElL--LLL--L-LE-EeEEE--E-E-EELLl----------
0041 3e0eA   ------elHHHL------L-L-LEE-EEE-E-EEEeEELLLEE----E-E-EEEEE-E-E---EEE-EEE-EH-H-HH-H-----L--LLL-------LL--LEEEE-E-EEEE-E-------------eE-E-EEE--E-E-EEEE-L--L------
0042 4dapA   -----------------lllL-LLL-EEE---EEE-EEEL----------L-EEEEElL-L---EE-EEEL-LL-L-----L--L-L-L--LL-------llL--LLLEE-E-EEEL-L-L--LLL--L-LL-E-EEE--E-E-ELLL---lE-E-ELl
0043 1eqrA   ------lllllLL--LHhH-L-LLE-EEE-E-EEEeEEEEEL---L-E-E-EEEEElL-E---EE-EEEE-LH-H-HH-----L-L--LL--------LL--LEEEE-E-EEEE-elL--HHH--LleE-E-EEE--E-E-EEEEl----------
0044 3en2A   --------------------L-LLL-EEE-E-EEE-EEELL---------llEEEEE-E-eeeeEEEEEEEhHH-H-HH-L-----L--LLL--------L--LEEEE-E-EEEE-E-L--L----L-EE-E-EEE--E-E-EEL------------
0045 3kf8D   ------llhHHHH--HH----llEE-EEE-E-EEEeEEELL------L-L-EEEEE-E-llllLEEE-EL-LL---LL-L--LlL--HHH----HLLL--LEEEE-E-EEEL-L-L-----------L-E-EEE--E-E-EELL-Hh--------
0046 7uy7B   -llllL--HHHLL--LL-L-L-LLE-EEE-E-EEElLLLLLLL-L----L-EEEEE-L-L---EEE-EEL-LL-----L-L--L-L--L------lLLLL--LLEEE-E-EEEEeE-E--LL------eE-E-EEE--E-E-EELL-Lh--------
0047 7w7pD   ------llLLLLL--LLLL---LLL-LLL-L-EEEeEELLLEE-EeeE-E-EEEEE-ElL---EEE-EEE-EH-H-HL-L---------------llLLL--LEEEE-E-EEEE-E-L--LLLlL-LE-EeEEE--E-E-ELLL-----------
0048 2z4hB   ------leeeeeleeeelL-L-LLL-EEE---EEE-EEELL------L-L-EEEEE-L-L---LEEEELL-LH-H-HH-H--H-H--HH-hllllLLL--EEEEE-E-EEEE-E----eLL--E-EE-E-EEL--LlL-EEELl----------
0049 6cqmD   ---------------------lEEE-EEE-E-EEE-LLEEEELllE-E-E-EEEEE-E-llllEEEEEEEhHH-H-HH-H--H-L--LL---------L--LEEEE-E-EEEE-E-E--EEE--E-EE-E-EEE--E-E-EEEEl----------
0050 2i0qA   lhhllL--HHHHH--HL-----LLL-EEE-E-EEEeEEEELL--LL-E-E-EEEEE-L-L---EEE-EEE-EL-L-LL------L--LLL--------L--LEEEE-EeEEELlL-L--LL-----lE-E-EEE--E-E-EE-E-L--L-Ll---
0051 6lbuB   -eeelL--LLHHH--HH----llEE-EEE-E-EEE-EEL----llL-L-E-EEEEE-L-llllLEEE-EE-LH-H-HH-H--H-H--H--lllllLLL--LEEEE-E-EEEE-L-L-------L-LE-E-EEE--E-E-EELL-Hh--------

# Job: Type-1 BREX; BrxL_HAM28421.1 C-Terminal HKD-EndoDNase
# Query: s001A
# No:  Chain   Z    rmsd lali nres  %id PDB  Description
   1:  2ymb-D 18.7  1.5  152   231   34   MOLECULE: MIT DOMAIN-CONTAINING PROTEIN 1;
   2:  7clg-A  8.9  4.1  130   838    8   MOLECULE: PUTATIVE HELICASE;
   3:  4rct-A  8.9  3.1  124   347   15   MOLECULE: RESTRICTION ENDONUCLEASE R.NGOVII;
   4:  3hsi-B  8.1  3.2  134   451    9   MOLECULE: PHOSPHATIDYLSERINE SYNTHASE;
   5:  6ehi-D  8.0  2.7  112   160    5   MOLECULE: NUCLEASE NUCT;
   6:  7v53-A  8.0  3.9  132   930    8   MOLECULE: PHOSPHOLIPASE D;
   7:  5nwa-A  7.9  3.0  117   441    8   MOLECULE: TYROSYL-DNA PHOSPHODIESTERASE 1;
   8:  1byr-A  7.8  3.3  116   152    8   MOLECULE: PROTEIN (ENDONUCLEASE);
   9:  7e0m-A  7.6  3.4  127   388    8   MOLECULE: PHOSPHOLIPASE;
  10:  2c1l-A  7.5  3.1  118   358   10   MOLECULE: RESTRICTION ENDONUCLEASE;
  11:  4gel-B  7.4  3.6  120   198    7   MOLECULE: MITOCHONDRIAL CARDIOLIPIN HYDROLASE;
  12:  7wdk-A  7.3  3.9  135   867    7   MOLECULE: PHOSPHOLIPASE D;
  13:  4urj-D  6.9  3.4  116   174    6   MOLECULE: PROTEIN FAM83A;
  14:  7wu1-E  6.7  3.4  144   545   13   MOLECULE: PHOSPHOLIPASE D;
  15:  3qph-A  6.5  4.5  112   335   13   MOLECULE: TRMB, A GLOBAL TRANSCRIPTION REGULATOR;
  16:  5box-A  6.5  5.9  120   263    8   MOLECULE: PUTATIVE HTH-TYPE TRANSCRIPTIONAL REGULATOR TRMBL
  17:  6kz8-A  6.3  4.3  137   784    7   MOLECULE: PHOSPHOLIPASE D ALPHA 1;
  18:  3igy-B  5.4  3.0  107   549   13   MOLECULE: COFACTOR-INDEPENDENT PHOSPHOGLYCERATE MUTASE;
  19:  1xdo-A  5.2  3.3  104   687   13   MOLECULE: POLYPHOSPHATE KINASE;
  20:  5b3g-B  5.0  3.4  117   406    8   MOLECULE: PROTEIN SCARECROW;
  21:  6s6v-A  4.8  4.5  102   386    7   MOLECULE: NUCLEASE SBCCD SUBUNIT D;

0001 s001A   --VEDG--SL---E--KTVDVQYGDGYG-YDDL-FAEY-LK-GA-SIVMLEEPYLT-QVANL-VRFV-EILVKI-G---D----CKVFRLVT-K-P---GE---T-P--EESTL-ISDSLQRL--KNTLDD--M---DDVSMAFEY-EFD----E-N-SHDRYIRTS-N-----NWDITLGRGL---H---FYQNLKTRFTFM-K---RN--
0002 2ymbD   -------------H--KQIKIEENAGFS-YESL-FREY-LNeTV-TEVWIEDPYIR-QLYNF-LRFC-EMLIK--C---K----VKTIHLLT-S-L---DE---------EQVQ-QSRGLQEI--EESLRS--------HGVLLEV-QYS----S-S-IHDREIRFN-N-----GWMIKIGRGL---D---YFKKHETTVDIF-H---KK--
0003 7clgA   --SHKG--NY---P--KLLVN--nKEKV-L-ST-IIDE-LQ-KC-ETFYFSVAFI--TESGL-ASLK-AQLLDL-S------nkgVKGKILT-S-N---YL---G--------F-NSPKXYGE--LLKL------------KNVEV-RLT----D-I-FHAKGYIFE-H--kdySSXVIGSSNLtsnA---LKV-NYEHNVLL-S---TX--
0004 4rctA   -------------M--NTVFSNIA-tekslnAV-WMDL-FK-SA-DEVLMATGYV--SNDAV-VELH-KILELN-D---H----IQKIDLLVgM-H---YL--------eGFSHlQYDSLXKL--NDFLRH--E--------RGAV-YVS--pfV-K-FHGKMYSFK-N--yqkINGLIGSANL---T-------TYETMLHL-N---GKpa
0005 3hsiB   tkraeQ--NL-lsV--SPLFGLG-AGN-elnQV-IEDL-FL-QV-KKLVICTPYF---NFPR--TLQ-HKIATLlE---N----GKRVEIIV-G-DkvaND---F-YigALPYL-YESNLRRF--CEKFET------qieSGQLVV-RLW----RnT-YHLKGVWVD-D-----RYILLTGNNL-npR---AWR-DAENGLLI-Y---DQ--
0006 6ehiD   -------------N--SLFVLPY--RDA-L-NS-LISG-IS-SA-ESVKIAIYS-----fTH-RDIA-RAIKSV-A------srgIKVQIIY-D-Y---E---------snHNN-KqstIGYL--DKYP-------------NTKV-CLL----K-G-MNQKVAIID-D-----KIVFLGSANWsknA---FEN-NYEVLLKT-D---DT--
0007 7v53A   -------------V--RFFTT----rDY-FS-D-LASA-LD-SA-SSIFITGWQ---NYDV--rSLW-QCLRQA-L----erspaLKVYVMP-W-L---SP---S-G--SL-gtyDFETMLAV--FQLNAG--L---EGGA-RAFC-TPA----I-Q-HHQKSVVID-N-----RIGYVGGIDL---A-ygRRDDWQDVHCRI-E---GP--
0008 5nwaA   -----------kgnpfQFYLTRVS-ALH-IKDI-LSP-lfG-TL-VSSAQFNYCF-----DV-DWLV-KQY-PP-E---F---rKKPILLVH-G-D---------------KRE-AKAHLHAQ--AKPYE------------NISL-CQA----KgT-HHTKMMLLL-Y--eegLRVVIHTSNLihaD---WHQ-KTQGIWLSpL---YP--
0009 1byrA   --------EP---V--QVGYSPE--sarvlvlS-AIDS-A---K-TSIRMMAYSF--TAPDI-MKAL-VAAK-K-R---G----V-DVKIVI-D-E---R--------gnTGRA-SIAAMNYI--ANSG--------------IPL-RTD----SpI-QHDKVIIVD-N-----VTVETGSFNFtkaA---ETK-NSENAVVI-W---NM--
0010 7e0mA   yqptAA--TP--gN--YLVASPRAYIDS-Q-VE-LPRL-LA-SA-QRVRVQVMDY--APLSygAVID-NALRSAaA---R----GVQIELMV-A-N---W---------ntKKP-DIAWLKSL--ALVP--------------NVQI-KVV----TrV-IHSKLMTID-G-----ETAWVGTSNWtggY---LDN-SRNLELVL-H---SP--
0011 2c1lA   -------------N--FFSLHPNVYrpkgliGM-LENV-WV----GTLYLISGFS--NYNGG-VRFY-ETFTEH-I------nqgGRVIAIL-G-G---S----T-S--QR---lSSRQVVEE--LLNR--------------GVEV-HII----NrI-LHAKLYGTSlG-----ESLVVSSGNFtgpG---MSQ-NIEASLLL-DnntTQ--
0012 4gelB   -rSKREkaSR---H--EVIIFNELGLRN-V-AK-IVEQ-ID-RA-YSIDLAIYT-----fTS-LFLA-DSIKRAlQ---R----GVIIRIIS-D-G---E---------mvYSK-G-sqISML--AQLG--------------VPV-RVP--ttN-L-MHNKFCIID-G--pcySIVISGSVNW---T-------NWENCIIT-A---DD--
0013 7wdkA   -------------V--RFFTT----gRD-YFSD-LASA-LD-SA-SSIFITGWQV--NYDV--rSLW-QCLRQA-L----erspaLKVYVMP-W-L---SP---S-G--SL-gtyDFETMLAV--FQLNAG--L---EGGA-RAFC-TPA----IfS-HHQKSVVID-N-----RIGYVGGIDL---A---YGAYWQDVHCRI-E---GPsv
0014 4urjD   -------------A--TVYFQ---TNNI-R-DL-VRRC-IT-RT-QVLVILMDV-----fTD-VEIF-CDILEA-AnkrG----VFVCVLLD-Q-G-------------------gvKLFQEMcdkVQIS---D---SHLKNISIR-SVE----GgQ-IREKFIISD-W-----RFVLSGSYSFtwlC---GHV-HRNILSKF-T---GQav
0015 7wu1E   vtfskpstNY---KpgWILDTHFTfPADkILDA-IYDN-IV-SA-RSVDIVTLQPM-SGAFT-ATIK-NALSQL-AkntqysdhhITVRLLQ-G-S---FT---PeE--EEIRQ-LSLTQTNY--LSEIASvlP---EVNNLDITV-GSV----RaW-NHGKIINVD-N-----QSVITGGHNL---WgadYLQRVNDLSINI-L---GY--
0016 3qphA   -mKKLE-------I--WRSRS-----fdeaiEM-FRES-LY-SA-NEVIVVTPSE--FFETI-REDL-IKTLER-G---------VTVSLYI-D-K-------------------IPDL--SE--FKGK------------GNFFV-RQF----Y-K-LNHLIGMTD-G-----KEVVTIQN-------------IGPPSFKS-T---YP--
0017 5boxA   --TEVP--KY---V--WVVRG----iKN-STLK-TKEM-LE-EA-NEILLADDG------FI-AVLE-DDIIKAvD---R----GVKTKILL-T-K---N----------------LLPRLKAS-kIIDYAK--E--------KLEL-RAL----D-K-FDLPMLICD--------EEVFFALE-------------NYETQVWI-K---DH--
0018 6kz8A   iksaKF--PGv--V--QLFRSIDG-DRS-iqDA-YIHA-IR-RA-DFIYVENQYF--LGSSfaedIP-KELSLK-IvskiekgekFRVYVVV-P-M---W----pvQ--AILDW-QRRTMEMM--YKDVIQ--A---LRaqgYLTF-FCLgnfmI-Y-VHTKMMIVD-D-----EYIIIGSANInqrS---MDG-DSEIAMGG-Y---QA--
0019 3igyB   -----------vghnalgagrVALQSGE-IYTGeGYRYlHG-AF-STLHLIG-LLS-RDNQI-YSII-EHAVKD-G---------AKRIRVHA-L-Y---DGrdvP-D--GSSFR-FTDELEAV--LAKVRQ---------NGCDAAIaSGG----GmF-VTMdeaitttdqyvpkVRYAGMM-------------dlgIPNN---------
0020 1xdoA   -----------pvtfdYLMVS----QNS-R-RL-LYEM-VD-REiSGITLKLNN-----lVD-KGLV-DRLYAA-S------ssgVPVNLLV-R-----------------------gmcslI--PNL---------EGISDNIRA-ISI----VyL-EHDRVYIFE-N--ggdKKVYLSSADWmtrN---IDY-RIEVATPL-L----D--
0021 5b3gB   -----------kwadsvlleaarafaTF-GHVA-ANGA-IL-E--IHIVDIS-------sTF-CTQW-PTLLEA-LatrsddtphLRLTTVV-V-A---NK---F-V--NDQTA-SHRMMKEI--GNRMEK--FarlMGVPFKFNI-IHH---vG-D-LSefsprdarP-----RIVTVVEE-----------vgPVVWASAW-R---PT--
0022 6s6vA   -------------------------------------M-RILHTSDWHLGqEHQAF-LDWLlETAQTH-----Q----VDAIIVAG-DvF---DT---G-S---PPSY-ARTLYNRF--VVNLQQ--------TGCHLVV-LAG----NasaHIHRYVHLV-T--lesVENLN------vpV---TQPM---------------

0001 s001A   --LLLL--LL---L--EEEEELLLLLLL-HHHH-HHHH-HL-LL-LEEEEELLLLL-HHHHH-HHHH-HHHHHH-L---L----LLEEEEEE-L-L---LL---L-H--HHHHH-HHHHHHHH--HHHHHL--L---LLLLLEEEE-EEL----L-L-LLLLEEEEL-L-----LEEEEELLLL---L---LLLLLLEEEEEE-E---LL--
0002 2ymbD   -------------L--EEEEELLLLLLL-HHHH-LLLL-LLlLL-LEEEEELLLLL-HHHHH-HHHH-HHLLL--L---L----LLEEEEEE-L-L---LL---------LLHH-HHHHHHHH--HHHHHL--------LLLEEEE-EEL----L-L-LLLLEEEEL-L-----LLEEEELLLL---L---LLLLLLEEEEEE-L---LL--
0003 7clgA   --LLLL--LL---L--EEELL--lLLLH-H-HH-HHHH-HH-LL-LEEEEEELLL--LHHHH-HHHH-HHHHHH-H------hllLLEEEEE-L-L---LL---L--------L-LLHHHHHH--HHHL------------LLEEE-EEL----L-L-LLLEEEEEE-L--lleEEEEEELLLLlhhH---HHL-LLLLEEEE-E---EE--
0004 4rctA   -------------E--EEEELLLL-llllhhHH-HHHH-HH-HL-LEEEEEELLL--LHHHH-HHHH-HHHHHL-L---L----LLEEEEEElL-H---HH--------hLLLHhHHHHHHHH--HHHHHH--L--------LEEE-EEL--llL-L-LLLLEEEEE-E--lleEEEEEELLLL---L-------LLLEEEEE-E---HHhh
0005 3hsiB   hhhhhH--HH-leE--EEEEEEL-LLL-hhhHH-HHHH-HH-LL-LEEEEELLLL---LLLH--HHH-HHHHHHhH---L----LLEEEEEE-E-LhhhLL---L-LlhHHHHH-HHHHHHHH--HHHLHH------hhhHLLEEE-EEE----LlE-ELLLEEEEL-L-----LEEEEELLLL-lhH---HHH-LEEEEEEE-E---LL--
0006 6ehiD   -------------E--EEEEELL--HHH-H-HH-HHHH-HH-LL-LEEEEEELL-----lLL-HHHH-HHHHHH-H------hllLEEEEEE-E-H---H---------hhLLL-LlllHHHH--HLLL-------------LEEE-EEE----L-L-LLLEEEEEL-L-----LEEEEELLLLlhhH---HHL-LLEEEEEE-L---LH--
0007 7v53A   -------------E--EEELL----hHH-HH-H-HHHH-HH-HL-LEEEEEELL---LLLL--lLHH-HHHHHH-H----hhlllLEEEEEE-L-L---LL---L-H--HH-lllHHHHHHHH--HHLLLL--L---LLLL-LEEE-LLL----L-L-LLLLEEEEL-L-----LEEEEELLLL---L-llLLLLEELLEEEE-E---LL--
0008 5nwaA   -----------lllllLEELLLLL-LLL-HHHH-LLH-hhL-LE-EEEEEELLLL-----LH-HHHH-HHL-LL-L---L---lLLLEEEEE-L-L---------------LLH-HHHHHHHH--LLLLL-------------LEEE-EEL----LlL-LLLLEEEEE-E--lllEEEEEELLLLlhhH---LLL-LLLLEEELlL---LL--
0009 1byrA   --------LL---E--EEEEELL--hhhhhhH-HHHH-L---L-LEEEEEELLL--LLHHH-HHHH-HHHH-H-L---L----L-EEEEEE-E-L---L--------llLLHH-HHHHHHHH--HHLL--------------LLE-EEE----LlL-LLLLEEEEL-L-----LEEEEELLLLlhhH---HHL-LLEEEEEE-E---LL--
0010 7e0mA   llllLL--LL--lE--EEEEELHHHEEH-H-HH-HHHH-HH-HL-LEEEEELLLL--LLLLllLHHH-HHHHHHhH---L----LLEEEEEE-E-H---H---------hhLLL-HHHHHHHH--HLLL-------------LEEE-EEE----LlE-ELLLEEEEL-L-----LEEEEELLLLlllH---HHH-LLEEEEEE-L---LH--
0011 2c1lA   -------------L--EEEEELLLElllhhHH-HHHH-HL----LEEEEEELLL--LHHHL-LLLH-HHHHHH-H------hllLEEEEEE-E-L---L----L-L--LL---lLLHHHHHH--HHHL-------------LLEE-EEE----ElL-LLLEEEEEElE-----EEEEEELLLLlllL---LLL-LLEEEEEE-LhhhHH--
0012 4gelB   -lHHHHhhLL---E--EEEEELLLLHHH-H-HH-HHHH-HH-LL-LEEEEEELL-----lLL-HHHH-HHHHHHhH---L----LLEEEEEE-L-L---L---------hhHLL-L-llHHHH--HHLL--------------LLE-ELL--llL-L-LLLEEEEEL-L--lllLEEEEELLLL---L-------LLEEEEEE-L---LH--
0013 7wdkA   -------------L--EEELL----hHH-HHHH-HHHH-HH-HL-LEEEEEELLL--LLLL--lLHH-HHHHHH-H----hhlllLEEEEEE-L-L---LL---L-L--LL-lllLLLHHHHH--HHHLLL--L---LLLL-LEEE-LLL----LlL-EELLEEEEL-L-----LEEEEELLLL---L---LLLLEELLEEEE-E---LLhh
0014 4urjD   -------------E--EEEEL---LLHH-H-HH-HHHH-HH-LL-LEEEEEELL-----lLL-HHHH-HHHHHH-HhhlL----LEEEEEEE-L-L------------------lhHHHHHHhhhLLLL---L---LLLLLEEEE-EEL----LlL-LLLLEEEEL-L-----LEEEEELLLLllhH---HLL-LLLEEEEE-E---HHhh
0015 7wu1E   llllllleeEE---LllLEEEELLLlHHHhHHHH-HHHH-HH-LL-LEEEEEEEELLL-LLHHH-HHHH-HHHHHH-HhhlllllllEEEEEEE-E-L---LL---LhH--HHHHH-HHHHHHHH--HHHHHLllL---LLLLEEEEE-EEE----EeE-LLLLEEEEL-L-----LEEEEELLLL---LhhhHLLLLLEEEEEE-E---LL--
0016 3qphA   -hHHHH-------E--EEELL-----hhhhhHH-HHHH-HH-HL-LEEEEEELHH--HHHHH-HHHH-HHHHHL-L----------LEEEEEE-L-L------------------LLLL--HH--HLLL------------LEEEE-EEL----L-L-LLLEEEEEL-L-----LEEEEELL-------------LLLLEEEE-L---LH--
0017 5boxA   --LLLL--LL---E--EEELL----hHH-HHHH-HHHH-HH-LL-LEEEEEELL------LL-LLHH-HHHHHHhH---L----LLEEEEEE-E-H---H----------------HHHHHLLL-hHHHHHL--L--------LLEE-EEE----L-L-LLLLEEEEL-------LEEEEELL-------------LLLLEEEE-E---LH--
0018 6kz8A   llllLL--LLl--E--EEEEEEEL-EEH-hhHH-HHHH-HH-LL-LEEEEEELLL--LLLHhhhhHH-HHHHHH-HhhhhhlllLEEEEEL-L-L---L----lhH--HHHHH-HHHHHHHH--HHHHHH--H---HHhllLEEE-EEEeeeeL-E-ELLEEEEEL-L-----LEEEEELLLLlhhH---HLL-LLEEEEEE-E---LL--
0019 3igyB   -----------hhhhhhhhllLLLLHLH-HHLLhHHHHhHH-HH-LLEEEEE-ELL-LHHHH-HHHH-HHHHHL-L--------LLEEEEEE-E-E---LLlllL-L--LLHHH-HHHHHHHH--HHHHHL--------LLLEEEEeEEE----ElL-LLLlhhhhhlllllllLEEEEEE--------------lllLLLL----------
0020 1xdoA   -----------lllllLLEEE----LLL-H-HH-HHHH-HH-HHhLLEEEEEELL-----lLL-HHHH-HHHHHH-H------hllLLEEEEE-L---------------------lllllL--LLL---------LLLLLLEEE-EEE----LlE-ELLEEEEEL-L--lllLEEEEELLLLlhhH---HHL-LLEEEEEL-L----L--
0021 5b3gB   -----------lhhhhhhhhhhhhhLH-HHHH-HHHH-HH-H--EEEEEEL-------lLL-LLLL-HHHHHH-HhhhlllleEEEEEEE-E-E---LL---L-L--LLHHH-HHHHHHHH--HHHHHH--HhhhHLLLEEEEE-EEE---eL-L-HHhllhhhhlL-----LEEEEEEE-----------llEEEEEEEE-E---EL--
0022 6s6vA   -------------------------------------L-EEEEELLLLLLlHHHHH-HHHHhHHHHHL-----L----LLEEEELL-LlL---LL---L-L---LLHH-HHHHHHHH--HHHHLL--------LLLLEEE-ELL----LlllLLLLEEEEE-E--eeeEEEEE------llL---LLLL----------------

@@@@@@@@@@@@@@@@@@@@@@@@@@@@@@@@@@@@@@@@@@@@@@@@@@@@@@@@@@@@@@@@@@@@@@@@@@@@@@@@@@@@@@@@@@@@@@@@@@@@@@@@@@@@@@@@@@@@@@@@@@@@@@@@@@@@@@@@@@@@@@
@@@@@@@@@@@@@@@@@@@@@@@@@@@@@@@@@@@@@@@@@@@@@@@@@@@@@@@@@@@@@@@@@@@@@@@@@@@@@@@@@@@@@@@@@@@@@@@@@@@@@@@@@@@@@@@@@@@@@@@@@@@@@@@@@@@@@@@@@@@@@@
@@@@@@@@@@@@@@@@@@@@@@@@@@@@@@@@@@@@@@@@@@@@@@@@@@@@@@@@@@@@@@@@@@@@@@@@@@@@@@@@@@@@@@@@@@@@@@@@@@@@@@@@@@@@@@@@@@@@@@@@@@@@@@@@@@@@@@@@@@@@@@

# Job: Type-1 BREX; BrxA BAY35696 N-Terminal HTH1 (Found in a SubSet of Type-1 BrxA, and in All Type-3 BrxA)
# Query: s001A
```

```
# No:  Chain   Z    rmsd lali nres  %id PDB  Description
   1:  7zge-A  5.8  3.0   69   199   14   MOLECULE: BRXA, A BREX PHAGE DEFENCE PROTEIN;
   2:  5ue0-A  5.2  3.0   67   280    9   MOLECULE: CT622 PROTEIN;
   3:  2qww-A  5.2  2.5   63   146   14   MOLECULE: TRANSCRIPTIONAL REGULATOR, MARR FAMILY;
   4:  2p8t-A  5.1  2.3   58   184    9   MOLECULE: HYPOTHETICAL PROTEIN PH0730;
   5:  5z7b-B  5.1  2.8   59   197    7   MOLECULE: PADR FAMILY TRANSCRIPTIONAL REGULATOR;
   6:  5f6f-A  5.1  2.7   63   141   13   MOLECULE: MARR FAMILY REGULATORY PROTEIN;
   7:  1stz-A  5.0  2.7   64   323    6   MOLECULE: HEAT-INDUCIBLE TRANSCRIPTION REPRESSOR HRCA HOMOL
   8:  8ioi-A  5.0  2.7   61   190   10   MOLECULE: PADR FAMILY TRANSCRIPTIONAL REGULATOR;
   9:  2vla-A  4.8  3.1   67   280    9   MOLECULE: RESTRICTION ENDONUCLEASE R.BPUJI;
  10:  8r3g-A  4.7  2.7   58   335    9   MOLECULE: CENTRAL GLYCOLYTIC GENES REGULATOR;
  11:  4rul-A  4.7  3.2   66   821    9   MOLECULE: DNA TOPOISOMERASE 1;
  12:  9qe1-J  4.7  2.7   65   108    3   MOLECULE: JETC;
  13:  6j5u-A  4.6  3.7   71   663   10   MOLECULE: DISEASE RESISTANCE RPP13-LIKE PROTEIN 4;
  14:  3cdh-B  4.6  2.7   60   136   12   MOLECULE: TRANSCRIPTIONAL REGULATOR, MARR FAMILY;
  15:  2fa5-B  4.6  2.7   62   142    8   MOLECULE: TRANSCRIPTIONAL REGULATOR MARR/EMRR FAMILY;
  16:  4asn-A  4.6  2.4   59    90   10   MOLECULE: TUBR;
  17:  3elk-A  4.5  2.9   60   104    7   MOLECULE: PUTATIVE TRANSCRIPTIONAL REGULATOR TA0346;
  18:  8jxk-D  4.5  2.7   59   166    8   MOLECULE: CONSERVED PROTEIN;
  19:  6qfd-B  4.5  2.5   59   116    3   MOLECULE: DNA-BINDING PROTEIN;
  20:  5hso-A  4.5  2.7   58   122   12   MOLECULE: UNCHARACTERIZED HTH-TYPE TRANSCRIPTIONAL REGULATO
  21:  5x11-E  4.5  2.8   63   181    5   MOLECULE: DNA (28-MER);
  22:  7dvr-A  4.5  2.5   61   147   15   MOLECULE: HTH MARR-TYPE DOMAIN-CONTAINING PROTEIN;
  23:  3df8-A  4.4  2.8   63   109    3   MOLECULE: POSSIBLE HXLR FAMILY TRANSCRIPTIONAL FACTOR;
  24:  2h09-A  4.4  2.5   55   127    7   MOLECULE: TRANSCRIPTIONAL REGULATOR MNTR;
  25:  2wte-A  4.4  3.2   56   212   11   MOLECULE: CSA3;
  26:  3m8e-A  4.4  2.4   61   101    3   MOLECULE: PUTATIVE DNA-BINDING PROTEIN;
  27:  2nyx-B  4.4  2.3   59   147   10   MOLECULE: PROBABLE TRANSCRIPTIONAL REGULATORY PROTEIN, RV14
  28:  4fht-B  4.3  2.6   58   142    5   MOLECULE: PCAV TRANSCRIPTIONAL REGULATOR;
  29:  2co5-B  4.3  2.7   60    94    7   MOLECULE: VIRAL PROTEIN F93;
  30:  7l1i-A  4.3  2.9   60   169   10   MOLECULE: MARR FAMILY MULTIDRUG RESISTANCE PUMP TRANSCRIPTI
  31:  1p4x-A  4.2  2.9   64   250    9   MOLECULE: STAPHYLOCOCCAL ACCESSORY REGULATOR A HOMOLOGUE;
  32:  3l9f-A  4.2  2.7   60   170    3   MOLECULE: PUTATIVE UNCHARACTERIZED PROTEIN SMU.1604C;
  33:  5y7q-A  4.2  2.6   64   546    6   MOLECULE: FANCONI-ASSOCIATED NUCLEASE 1 HOMOLOG;
  34:  8ylg-A  4.2  2.5   58   161    7   MOLECULE: MARR FAMILY TRANSCRIPTIONAL REGULATOR;
  35:  5zqh-A  4.2  2.6   58   101    2   MOLECULE: PADR FAMILY TRANSCRIPTIONAL REGULATOR;
  36:  9ri9-B  4.1  3.6   62   868   10   MOLECULE: SLNRC3;
  37:  2fxa-B  4.1  2.3   56   180    5   MOLECULE: PROTEASE PRODUCTION REGULATORY PROTEIN HPR;
  38:  5e20-A  4.1  2.8   63   169    5   MOLECULE: TRANSCRIPTIONAL REGULATOR, MARR FAMILY;
  39:  2id3-A  4.0  2.8   53   191    6   MOLECULE: PUTATIVE TRANSCRIPTIONAL REGULATOR;
  40:  1fok-A  4.0  3.1   67   568    9   MOLECULE: DNA (5'-
  41:  4u03-A  4.0  3.3   59   390    3   MOLECULE: CYCLIC AMP-GMP SYNTHASE;
  42:  8rkd-B  4.0  4.2   56   411    5   MOLECULE: PILUS ASSEMBLY ATPASE CPAF;
  43:  3oop-A  4.0  2.3   54   139    7   MOLECULE: LIN2960 PROTEIN;
  44:  6jbx-A  4.0  2.5   56   146   13   MOLECULE: FATTY ACID BIOSYNTHESIS TRANSCRIPTIONAL REGULATOR
  45:  1ub9-A  4.0  2.6   59   100   10   MOLECULE: HYPOTHETICAL PROTEIN PH1061;
  46:  1yyv-A  4.0  2.5   59   114   10   MOLECULE: PUTATIVE TRANSCRIPTIONAL REGULATOR;
  47:  4hw0-A  3.9  3.0   56    98   13   MOLECULE: DNA-BINDING PROTEIN SSO10A-2;
  48:  5j9q-E  3.9  3.2   59   293    7   MOLECULE: HISTONE ACETYLTRANSFERASE ESA1;
  49:  4hqe-B  3.9  2.8   60   108    7   MOLECULE: TRANSCRIPTIONAL REGULATOR QSRR;
  50:  3mq0-B  3.9  2.6   52   248    8   MOLECULE: TRANSCRIPTIONAL REPRESSOR OF THE BLCABC OPERON;

0001 s001A   --GTAMVN--ETRHLIQQWHP--TE-NVD--TFVKRVQAE-DL-LG-RQTAQRAKDIAR-VFA-A--R-FLLP--------------NNKP-ARL-LK-QIV-EAG-L-
0002 7zgeA   -gSSLLIT--ESRIIAESLLK--KL-PED--EWKSLIVEQ-NV-LQ-KKSGQTAIRYAR-TIR-W--R-IE----------------GLGD--EF-MT-DLL-AAS-Er
0003 5ue0A   -aAETILQ--VLDSLMSTINP--QA-NQE--EIMQXLT---AS-IS-KqSVDSLQKFAA-QLE-R--E-FVDG--------E----RSLAE-SQE-NAfRKQ-PA----
0004 2qwwA   aaslGLTI--QQLAXINVIYS--TP-G----ISVADLT---KR-LI--ITGSSAAANVD-GLI-S-lG-LVV-klnkpnDSXDLTLKLSKK-GED-LS-KRS-TAN-Af
0005 2p8tA   ---eyTVE--DVLAVIFLLK---------epLGRKQIS---ER-LE--LGEGSVRTLLR-KLS-H-lD-IIR----------SKGHFLTLK-GKE-IR-DKL-LSM-Fs
0006 5z7bB   -sakDLMT--LRSALLALLS--------sgpLTGYDASQR-F---gwSGSDSQIYPELR-KME-A-eE-LLV-vpwgskGATKTEYALSEK-GWE-AL-RKA-WYE-Pv
0007 5f6fA   leqfDITN--EQRHTLGYLYQ--QD-G----LTQNDIA---KA-LQ--RTGPTVSNLLR-NLE-R-kK-LIY-ryvdaqDTRRKNIGLTTS-GIK-LV-EAF-TSI-Fd
0008 1stzA   ----KLND--RQRKVLYCIV--nKK-PV----SSQRVLEV-SN-IE--FSSATIRNDMK-KLE-Y-lG-YIY------qPHTSAGRIPTDK-GLR-FY-YEE-MLK-Is
0009 8ioiA   ---PEFMA--LPHAILVSLS--------eqaSSGYELARR-F-dfW-TATHQQIYRTLR-VME-N-nN-WVR-tvlqhgRPDKKVYAISDS-GRA-EL-ARW-IAE-Pl
0010 2vlaA   -irGKAKN--LLPAYANIIDC--PC-DKA--SFVKDFN---NR-LIeETTKKTLDNHRTeIAG-KlfG-MFYE--------DDEVIFPSGR-TNK-YI-ED--SDQ-Pa
0011 8r3gA   pdllLVXQ--KRFEILQYIR-------ltepIGRRSLS---AS-LG--ISERVLRGEVQ-FLK-E-qN-LVD--------IKTNGXTLTEE-GYE-LL-SVL-EDTx--
0012 4rulA   arglSAGR--VQSVAVRLVVE--RF-SEA--SLVKELEKR-GI-G----RPSTYASIIS-TIQ-D--RgYVRV--------ENRRFYAEKM-GEI-VT-DRL-EEN-Fr
0013 9qe1J   -ltADSAA--HYRTILRYFY--mRD-FIA----PEELL---EH-MRaDFQEDQLHQQLA-QLVkW--N-NLIAktieeyKKKRFRYQCTPY-TVE-IE-RMI-VQL-Ek
0014 6j5uA   -tDRWSPS--HLKSCILTLSL--YI-PKQ--QLVHGWIGE-GF-VM-WSATESGEDCFS-GLT-N--RcLIEVktysgtIIT----CKIHD-vRD-LV-IDI-AKK-Ds
0015 3cdhB   iraqGLRV--PEWRVLACLVD--ND-A-------xXITR--KL-SL--XEQSRXTRIVD-QXD-A-rG-LVT-----rvADARVRVRLTDD-GRA-LA-ESL-VAS-Ar
0016 2fa5B   gdryGXAI--PEWRVITILAL--YP-G----SSASEVSDR-T------XDKVAVSRAVA-RLL-E-rG-FIR---------RESXLALSPA-GRQ-VY-ETV-APL-Vn
0017 4asnA   evxrKLTI--EDVSILGWLAN--AV---fkaikKSSIA---DE-LE--YSTANFRKTLN-KLE-A-iH-FIG----tvtGGKEHKLYLTEY-GQQ-AV-QQA-IHH---
0018 3elkA   reriLHGL--ITLYILKELV--------krpXHGYELQ---KS-XFqALPQGSIYILLK-TXK-E-rG-FVI-essvnkGQQLTVYHITDA-GKK-FL-XDH-SQA-Lq
0019 8jxkD   -----EFM--LELAILGLLI--------espMHGYELR---KR-LTrAFSYGSLYPALR-RMQ-A-dG-LIA---enaaPAGRRVYQLTDK-GRR-RF-GEL-VAD-Tg
0020 6qfdB   sdarDLTA--FQKNILTVLG--------eeaRYGLAIKRE-L----eEVNHGRLYPNLD-DLV-N-kG-LVE---kselDKRTNEYALTNE-GFD-AV-VDD-LEW-Tl
0021 5hsoA   lsplGLTL--PEFVCLRMLS----------qspgLSSAEL-RH-AS--VTPQAMNTVLR-KLE-D-aG-AVA------------PATLTAR-GRA-LA-KRA-EAV-Vr
0022 5x11E   ---dPMRV--LKYAILGLLRK--GE------LSGYDITSY-F-kfW-SAKHSQIYPELK-KLT-D-eG-FIT-ttiqgtKLEKKMYTLTDS-GKQ-EL-HDW-LIR-Hq
0023 7dvrA   ------AG--PQGHLVMHLYP--DK-D----MSIKDAE---EI-LH--ISKSVASNLVK-RME-K-nG-FIA-ivpsktDKRVKYLYLTHL-GKQ-KA-TQF-EIF-Le
0024 3df8A   esvlHLLGkkYTXLIISVLGG--ST-R----QNFNDIR---SS-IP-GISSTILSRRIK-DLI-D-sG-LVE-----rrSGQITTYALTEK-GXN-VR-NSL-XPLl--
0025 2h09A   vreaHRRE--LIDDYVELIS-------dlirevgeaRQVD-M--rL-GVSQPTVAKMLK-RLA-T-mG-LIE-------MIPWRGVFLTAE-GEK-LA-QES-RER-Hq
0026 2wteA   alxrDYSR--EEXKLLNVLYE--TK---------gTGITE-L--xL-DKSEKTLINKIA-ELK-K-fG-ILT------qKGKDRKVELNEL-GLN-VI-KLN-K-----
0027 3m8eA   ------DD--CAYQVLMAFIE--AQ-M----LNKTAVA---EM-IQ--LSKPTVFATVN-SFY-C--AgYIDE----trVGRSKIYTLSDL-GVE-IV-EC--FKQ-Ka
0028 2nyxB   qvdeNITI--PQFRTLVILS-------nhgpINLATLA---TL-LG--VQPSATGRXVD-RLV-G-aE-LID-rlphptSRRELLAALTKR-GRD-VV-RQV-TEH-Rr
0029 4fhtB   mvseETTS--PQYAVLNALV-------aepgLDQRTVG---ER-VG--lDRSTIAEVVS-RLG-R-rG-LLD-kvrdpqDGRRSLLRLTDE-GLR-VH-RRL-GVR-Ia
0030 2co5B   ------RI--NYYIILKVLV--nGS-RLE----KKRLRSE----dI-DISDGVLYPLID-SLI-D-dK-ILR----eeeAPDGKVLFLTEK-GMK-EF-EEL-HEF-Fk
0031 7l1iA   ikklGIDN--TRRKIILSTNA--LG-E-------aSITD--NL-ST--LKLTTATKAVY-RLV-E-dG-IVE-vysstDERISMVKLTAK-GVE-LV-EQI-NQI-Sv
0032 1p4xA   kpevDMTI--KEFILLTYLF--qQE-NTL--P-FKKIVSD-L-----CYKQSDLVQHIK-VLV-K-hS-YISkrntyisISE----EQREK-IAE-RV-TLF-DQI-Ik
0033 3l9fA   ------MQ--GKDIILGILSK--KE------RSGYEINDI-LqnyD-GTY-GMIYPTLR-KLE-K-dG-KIT-vviqdgRPNKNIYAITES-GKK-EL-ASY-LQS-Dv
0034 5y7qA   -vHYVENG--LINSLFGLLCL--GRsYRD--AIRCRYRDK-FglYW-EL---lGEELLE-QAL-D--C-L----------------PAAH-LRA-WF-ERL-LED-Ip
0035 8ylgA   farygLQP--GEFDVLATLSG--AP-------yalTPTA--DA-AM--ISSGSMTNRID-RLE-K-aG-WVE-rranpaDGRGTLVALTSA-GRA-LI-DDA-VVA-Hv
0036 5zqhA   etqllKGV--LEGCVLDMIG--------qkeRYGYELVQT-L--fD-TIV-GTIYPLLQ-KLE-K-nQ-WIR-mrpspdGPDRKYFSLMKE-GEE-RV-SVF-WQQ-Wd
0037 9ri9B   msydRLPY--DLKACFLYC---sQI-PAK--KLIRLWIAE-GF-IQ-YQGpLALEDVAE-DHL-N--D-LVN----------------rNL-VMvmLH-EFC-RHEa--
0038 2fxaB   lkpyDLNI--NEHHILWIAY----------qlngaSISE--KF-GV--XHVSTAFNFSK-KLE-E-rG-YLR-fskrlnDKRNTYVQLTEE-GTE-VF-WSL-LEE-Fd
0039 5e20A   lkslNLST--EKFAILHELML--GG-E----STPHTLARR--I-VF-E--PHSVSAIVS-RME-K-dG-LII-ktkdldKKHMVRIKLSEK-AID-TF-YQA-LEI-Sn
0040 2id3A   --gGRTAR--IREAVLLAAG--dAL-DLG--EIARRAG-----------------vGKT-TVY-R--R-WG----------------TPGG-LAA-DL-LAD-XAE-Qs
0041 1fokA   -qNPGKFE--NLKRVVQVFDRnhDL-IYT--Y--KELVGtgAT-IAgYIDNWSSDGFLR-WAH-A--LgFIEY------INKSDSFVITDV-GLA-YS-KS--ADG-Sa
0042 4u03A   ------RD--VFEEAKGIAKK--KS-ALTfeIIQEKVS-T-TQ-IK-HLSDSEQREVAK-LIY-E--M-------------------DDDA-RDE-FL-GLTx------
0043 8rkdB   ---DYYHA--TKTTIFNALTI--DLqAGE--EIRDIVAEL-V--nV-SMSVAEQEHLVQ-DII-N----DVLG-----------------------------fRDN-Lq
0044 3oopA   ------TP--EQWSVLEGIE----------anepiSQKE--LW-TK--KDTPTVNRIVD-VLL-R-kE-LIV-----reIRRISLLSLTDK-GRK-ET-TEL-RDI-Ve
0045 6jbxA   srfkDISI--KEMHTIDVIG----------kapdvTPSQ--KE-LM--VTLGTVTTSLN-NLE-R-kG-YIE-rvrseqDRRVVHLHLTKK-GRL-IH-RLH-KRF-Hk
0046 1ub9A   ------NP--VRLGIMIFLLP--RR-K----APFSQIQKV-L------lTPGNLDSHIR-VLE-R-nG-LVK-tykviaDRPRTVVEITDF-GME-EA-KRF-LSS-Lk
0047 1yyvA   revlKHVTsrWGVLILVALR--------dgtHRFSDLR---RX-XG-GVSEXXLAQSLQ-ALE-Q-dG-FLN-rvsypvVPPHVEYSLTPL-GEQ-VS-DXV-AALa--
0048 4hw0A   --krGTXE--IXFDILRNCEP------------kcGITR--YG-AG--INYVVAQKYLD-QLV-K-vG-ALN----iktENDRKIYEITEK-GKL-LR-THI-EEFi--
0049 5j9qE   -llSYRAY--WSDTLITLLEH--QK-E----ITIDEIS-S-MT-S-------mTTTDIL-HTA-K--TlNILR---yykGQH----IIFLN--eD-IL-DRY-NRL-Ka
0050 4hqeB   eetfKILGrsWNGLIINYLS-----rcndcsAHFSDMKRD-L---K-TITPRALSLKLS-ELA-Q-wE-LVE-kqiistSPVQIIYVLTEK-GKA-LA-EAL-HPI-Ea
0051 3mq0B   sdtvPALR--RAVRILDLVAS--PR-D----LTAAELT---RF-LD--LPKSSAHGLLA-VMT-E-lD-LLA--rsadgTLR----IGPHSl----------------

0001 s001A   --LLLLHH--HHHHHHHHLLL--LL-LHH--HHHHHHHHL-LL-LL-LLLHHHHHHHHH-HHH-H--H-HLLL--------------LLHH-HHH-HH-HHH-HLL-L-
0002 7zgeA   -lLLLLHH--HHHHHHHHHLL--LL-LLL--HHHHHHLLL-LL-LL-LLLHHHHHHHHH-HHH-H--H-HL----------------LLLH--HH-HH-HHH-HLL-Hh
0003 5ue0A   -hHHHHHH--HHHHHHHHHLL--LL-LHH--HHHHHHH---HH-HH-LlLHHHHHHHHH-HHH-H--H-HLLH--------H----HHHHH-HHH-HHhHHH-HH----
0004 2qwwA   hhhhLLLH--HHHHHHHHHHH--LL-L----EEHHHHH---HH-HL--LLHHHHHHHHH-HHH-H-lL-LEE-ellllLLLLLEEEELHH-HHH-HH-HHH-HLL-Hh
0005 2p8tA   ---llLHH--HHHHHHHHLL---------llLLHHHHH---HH-HL--LLHHHHHHHHH-HHH-H-lL-LEE----------ELLEEELHH-HHH-HH-HHH-HLL-Ll
0006 5z7bB   -lhhHHHH--HHHHHHHHHL--------lllEEHHHHHHH-H---hlLLLHHHHHHHHH-HHH-H-lL-LEE-eeelllLEEEEEEEELHH-HHH-HH-HHH-HHL-Ll
0007 5f6fA   hlllLLLH--HHHHHHHHHHH--HH-L----LLHHHHH---HH-HL--LLHHHHHHHHH-HHH-H-lL-LEE-eeelllLLLLEEEEELHH-HHH-HH-HHH-HHH-Hh
0008 1stzA   ----LLLH--HHHHHHHHHH--hLL-LL----LHHHHHHH-LL-LL--LLHHHHHHHHH-HHH-H-lL-LEE------lLLLLLLLEELHH-HHH-HH-HHH-HHH-Hl
0009 8ioiA   ---LLLLL--HHHHHHHHHH--------hllLLHHHHHHH-H-hlL-LLLHHHHHHHHH-HHH-H-lL-LEE-eellllLLLEEEEEELHH-HHH-HH-HHH-HHL-Ll
0010 2vlaA   -llLLLLH--HHHHHHHHHHL--LE-EHH--HHHHHHH---HH-HHlLLLHHHHHHHHHhHHH-HhlL-LEEE--------ELLEEEELHH-HHH-HH-HH--LLH-Hh
0011 8r3gA   llhhHHHH--HHHHHHHHHH-------hhllLLHHHHH---HH-HL--LLHHHHHHHHH-HHH-H-lL-LEE--------EELLEEEELHH-HHH-HH-HHH-HHHh--
0012 4rulA   llllLLLL--LHHHHHHHHHH--HL-LHH--HHHHHHHHL-LL-L----LLLLHHHHHH-HLL-L--LlLEEE--------ELLEEEELHH-HHH-HH-HHH-HHH-Ll
0013 9qe1J   -hhLLLHH--HHHHHHHHHH--lLL-LLL----HHHHH---HH-HHlLLLHHHHHHHHH-HHHhL--L-LEEEllhhhhHLLLEEEEELHH-HHH-HH-HHH-HHH-Hl
0014 6j5uA   -lLLLLLH--HHHHHHHHHHH--LE-EHH--HHHHHHHHH-LL-LL-LLLHHHHHHHHH-HHH-H--LlLLEElllllLLE----EELLH-hHH-HH-HHH-HHL-Ll
0015 3cdhB   hhhlLLLH--HHHHHHHHHHH--LL-L-------eEHHH--HH-HL--LLHHHHHHHHH-HHH-H-hL-LEE-----elLLLLLEEEELHH-HHH-HH-HHH-HHH-Hh
0016 2fa5B   hhhhLLLH--HHHHHHHHHHH--LL-L----LLHHHHHHH-H------LLHHHHHHHHH-HHH-H-hL-LEE---------ELLLEEELHH-HHH-HH-HHH-HHH-Hh
0017 4asnA   hhhhLLLH--HHHHHHHHHLL--LH---hhleeHHHHH---LL-LL--LLLHHHHHHHH-HHH-H-lL-LEE----eelLLLLLEEEELHH-HHH-HH-HHH-LLL---
0018 3elkA   llhhHHHH--HHHHHHHHHH--------hllEEHHHHH---HH-HHlLLLLLHHHHHHH-HHH-H-hL-LEE-eeeellLLEEEEEEELHH-HHH-HH-HHL-LLL-Hh
0019 8jxkD   -----LHH--HHHHHHHHHH--------hllEEHHHHH---HH-HHlLLLLLLHHHHHH-HHH-H-lL-LEE---elllLLLLLEEEELHH-HHH-HH-HHH-HHL-Ll
0020 6qfdB   hhhlLLLH--HHHHHHHHHH--------hllEEHHHHHHH-H----lLLLHHHHHHHHH-HHH-H-lL-LEE---eeeeELLEEEEEELHH-HHH-HH-HHH-HHH-Hh
0021 5hsoA   hhhhLLLH--HHHHHHHHHH----------hlllLLHHHH-HH-LL--LLHHHHHHHHH-HHH-H-lL-LLL------------LLLLLHH-HHH-HH-HHH-HHH-Hh
0022 5x11E   ---lLLLH--HHHHHHHHLLL--LL------LLHHHHHHH-H-hlL-LLLHHHHHHHHH-HHH-H-lL-LEE-eeelllLLEEEEEEELHH-HHH-HH-HHH-HHL-Ll
0023 7dvrA   ------LL--HHHHHHHHHHL--LL-L----EEHHHHH---HH-HL--LLHHHHHHHHH-HHH-H-lL-LEE-eeelllLLLLEEEEELHH-HHH-HH-HHH-HHH-Hh
0024 3df8A   lllhHHHHllLHHHHHHHHLL--LL-L----LLHHHHH---HL-LL-LLLHHHHHHHHH-HHH-H-lL-LEE-----eeELLLEEEEELHH-HHH-HH-HHH-HHHh--
0025 2h09A   hhhhHHHH--HHHHHHHHHH-------hhhhhhlllLHHH-H--hH-LLLHHHHHHHHH-HHH-H-lL-LEE-------EELLLEEEELHH-HHH-HH-HHH-HHH-Hh
0026 2wteA   hhhlLLLH--HHHHHHHHHHH--HL---------lLLHHH-H--hH-LLLHHHHHHHHH-HHH-H-lL-LEE------eELLLLEEEELHH-HHH-HH-HHL-L-----
0027 3m8eA   ------LH--HHHHHHHHLLL--LL-L----LLHHHHH---HL-LL--LLHHHHHHHHH-HHH-H--LlLEEE----eeELLEEEEEELHH-HHH-HH-HL--LLL-Ll
0028 2nyxB   hhllLLLH--HHHHHHHHHH-------hhllEEHHHHH---HH-HL--LLHHHHHHHHH-HHH-H-lL-LEE-eellllLLLLLEEEELHH-HHH-HH-HHH-HHH-Hh
0029 4fhtB   hlllLLLH--HHHHHHHHHH-------hlllLLHHHHH---HH-HL--lLHHHHHHHHH-HHH-H-hL-LEE-eeelllLLLLEEEEELHH-HHH-HH-HHH-HHH-Hh
0030 2co5B   ------LH--HHHHHHHHHH--lLL-EEE----HHHHHHH----lL-LLLHHHHHHHHH-HHH-H-lL-LEE----eelLLLLLEEEELHH-HHH-HH-HHH-HHH-Hh
```

```
0031 7l1iA   hlllLLLH--HHHHHHHHHHH--HL-L-------eEHHH--HH-LL--LLHHHHHHHHH-HHH-H-lL-LEE-eeelllLLLLEEEEELHH-HHH-HH-HHH-HHH-Hh
0032 1p4xA   llllLLLH--HHHHHHHHHH--lLL-LEE--E-HHHHHHH-L-----LLLHHHLHHHHH-HHH-H-lL-LLEelleeeeLLH----HHHHH-HHH-HH-HHH-HHH-Hh
0033 3l9fA   ------LL--HHHHHHHHLLL--LL------EEHHHHHHH-HhheE-LLL-LLHHHHHH-HHH-H-lL-LEE-eellllLLLEEEEEELHH-HHH-HH-HHH-HHL-Ll
0034 5y7qA   -eEELLLH--HHHHHHHHHHH--HLlHHH--HHHHHHHHH-LllLL-LL---lLHHHHH-HHH-H--H-L----------------LHHH-HHH-HH-HHH-HHL-Hh
0035 8ylgA   hhhllLLH--HHHHHHHHHLL--LL-------leeLHHH--HH-HL--LLHHHHHHHHH-HHH-H-hL-LEE-eeelllLLLLEEEEELHH-HHH-HH-HHH-HHH-Hh
0036 5zqhA   llhhhHHH--HHHHHHHHHH--------hllEEHHHHHHH-H--lL-LLL-LLLHHHHH-HHH-H-lL-LEE-eeelllLLEEEEEEELHH-HHH-HH-HHH-HHH-Hh
0037 9ri9B   hhhhLLLH--HHHHHHHHL---lLE-EHH--HHHHHHHHL-LL-LL-LLLlLLHHHHHH-HHH-H--H-HHH----------------lLL-LEehHH-HHH-HHHh--
0038 2fxaB   hhhhLLLH--HHHHHHHHHH----------hhlleEHHH--HH-HL--LLHHHHHHHHH-HHH-H-hL-LEE-eellllLLLLLEEEELHH-HHH-HH-HHH-HHH-Ll
0039 5e20A   lhhhLLLH--HHHHHHHHHHL--LL-E----ELHHHHHHH--L-LL-L--HHHHHHHHH-HHH-H-lL-LEE-eeelllLLLLEEEEELHH-HHH-HH-HHH-HHH-Hh
0040 2id3A   --lHHHHH--HHHHHHHHHH--hHL-LHH--HHHHHHL-----------------lLHH-HHH-H--H-HL---------------LHHH-HHH-HH-HHH-HHH-Hl
0041 1fokA   -lLLLLHH--HHHHHHHLLLLllLL-LEE--H--HHHHLllHH-LLlLLLHHHHHHHHH-HHH-H--LlLEEE------ELLLLEEEELHH-HHH-HH-HL--LLL-Lh
0042 4u03A   ------HH--HHHHHHHHHHH--HL-LLLhhHHHHHHH-H-LH-HH-HLLHHHHHHHHH-HHH-H--L-------------------LHHH-HHH-HH-LLLl------
0043 8rkdB   ---LLHHH--HHHHHHHHHHL--LHhHHH--HHHHHHHHH-H--lL-LLLHHHHHHHHH-HHH-H----HHLL----------------------------lLLH-Hh
0044 3oopA   ------LH--HHHHHHHHHH----------hhlleEHHH--HH-HL--LLHHHHHHHHH-HHH-H-lL-LEE-----eeLLLLLEEEELHH-HHH-HH-HHH-HHH-Hh
0045 6jbxA   llllLLLH--HHHHHHHHHH----------hllllLHHH--HH-HL--LLHHHHHHHHH-HHH-H-hL-LEE-eeelllLLLLEEEEELHH-HHH-HH-HHH-HHH-Hh
0046 1ub9A   ------LH--HHHHHHHHHHH--HL-E----EEHHHHHHH-L------lLHHHHHHHHH-HHH-H-lL-LEE-eeeellLLLEEEEEELHH-HHH-HH-HHH-HHH-Hh
0047 1yyvA   hhhhHHHHlhHHHHHHHHHH--------hllEEHHHHH---HH-LL-LLLHHHHHHHHH-HHH-H-hL-LEE-eeeellLLLEEEEEELHH-HHH-HH-HHH-HHHh--
0048 4hw0A   --llLHHH--HHHHHHLLLLL-----------eeLHHH--HH-HL--LLHHHHHHHHH-HHH-H-lL-LEE----eeeELLEEEEEELHH-HHH-HH-HHH-HHHh--
0049 5j9qE   -hhHHHHH--HHHHHHHHHHH--LL-L----LLHHHHH-H-HH-L-------lLHHHHH-HHH-H--HlLLEE---ellLLE----EELLL--hH-HH-HHH-HHH-Hh
0050 4hqeB   hhhhHHHLllLHHHHHHHHL-----llhhheELHHHHHHH-H---L-LLLHHHHHHHHH-HHH-H-lL-LEE-eeeellLLLEEEEEELHH-HHH-HH-HLL-HHH-Hh
0051 3mq0B   llllHHHH--HHHHHHHHHHL--LL-L----EEHHHHH---LL-LL--LLHHHHHHHHH-HHH-H-lL-LEE--ellllEEE----ELLLHh---------------

# Job: Type-1 BREX; BrxA BAY35696 HTH2
# Query: s001A
# No:  Chain   Z    rmsd lali nres  %id PDB  Description
   1:  7zge-A 10.3  1.4   80   199   18   MOLECULE: BRXA, A BREX PHAGE DEFENCE PROTEIN;
   2:  4kyw-A  7.1  2.3   67   254   10   MOLECULE: TYPE-2 RESTRICTION ENZYME DPNI;
   3:  1cf7-B  6.1  2.7   70    82   10   MOLECULE: DNA (5'-
   4:  7qcd-D  5.7  3.5   73   326   10   MOLECULE: STRUCTURAL MAINTENANCE OF CHROMOSOMES PROTEIN 5;
   5:  8c5s-A  5.5  3.8   75   924   12   MOLECULE: MITOCHONDRIAL TRANSCRIPTION FACTOR 1;
   6:  4ets-B  5.5  3.0   61   149    7   MOLECULE: FERRIC UPTAKE REGULATION PROTEIN;
   7:  3cuq-A  5.5  2.0   54   219   11   MOLECULE: VACUOLAR-SORTING PROTEIN SNF8;
   8:  1sfu-A  5.3  2.0   58    70    3   MOLECULE: 5'-D(*T*CP*GP*CP*GP*CP*G)-3';
   9:  8cli-A  5.2  2.4   65   542    8   MOLECULE: GENERAL TRANSCRIPTION FACTOR 3C POLYPEPTIDE 1;
  10:  5nl9-A  5.2  2.6   60   146    8   MOLECULE: TRANSCRIPTIONAL REGULATOR (FUR FAMILY);
  11:  7nyw-E  5.2  3.2   72   212   14   MOLECULE: CHROMOSOME PARTITION PROTEIN MUKB;
  12:  5v8f-6  5.2  2.5   62   692    8   MOLECULE: DNA REPLICATION LICENSING FACTOR MCM2;
  13:  4hga-A  5.1  2.9   65   207   14   MOLECULE: DEATH DOMAIN-ASSOCIATED PROTEIN 6;
  14:  9qe1-D  5.1  2.7   70   384    9   MOLECULE: JETC;
  15:  5dca-A  5.0  2.1   67  1948    9   MOLECULE: PRE-MRNA-SPLICING HELICASE BRR2;
  16:  6qfd-B  5.0  2.4   57   116    9   MOLECULE: DNA-BINDING PROTEIN;
  17:  6zvh-y  5.0  2.7   64    72   11   MOLECULE: 18S RRNA;
  18:  9mj5-B  5.0  2.3   59   208   12   MOLECULE: REPLICATION PROTEIN A 14 KDA SUBUNIT;
  19:  1lva-A  4.9  2.2   54   258   13   MOLECULE: SELENOCYSTEINE-SPECIFIC ELONGATION FACTOR;
  20:  8fef-I  4.9  2.7   65   253    2   MOLECULE: VIRULENCE FACTOR MCE FAMILY PROTEIN;
  21:  2qby-B  4.9  2.7   59   368    5   MOLECULE: CELL DIVISION CONTROL PROTEIN 6 HOMOLOG 1;
  22:  6j05-A  4.8  2.3   60   101    7   MOLECULE: TRANSCRIPTIONAL REGULATOR ARSR;
  23:  7uy7-B  4.8  2.6   61   291   13   MOLECULE: TELOMERASE-ASSOCIATED PROTEIN OF 75 KDA;
  24:  2vwa-A  4.8  2.9   68    99    9   MOLECULE: PUTATIVE UNCHARACTERIZED PROTEIN PF13_0012;
  25:  5l0p-A  4.8  2.4   56   311   11   MOLECULE: TRANSCRIPTION FACTOR ETV6, TRANSCRIPTION FACTOR E
  26:  9jbn-C  4.8  2.4   61   423   20   MOLECULE: PROTEIN EDS1;
  27:  7rhe-A  4.8  2.5   57   368   12   MOLECULE: ROK FAMILY PROTEIN;
  28:  4yo2-A  4.7  3.5   71   186   11   MOLECULE: TRANSCRIPTION FACTOR E2F8;
  29:  3k69-A  4.7  2.7   61   151   13   MOLECULE: PUTATIVE TRANSCRIPTION REGULATOR;
  30:  7xx7-U  4.7  2.6   68   213    4   MOLECULE: HISTONE H3.1;
  31:  3elk-A  4.6  2.7   59   104   12   MOLECULE: PUTATIVE TRANSCRIPTIONAL REGULATOR TA0346;
  32:  1ldd-A  4.6  3.0   63    74   17   MOLECULE: ANAPHASE PROMOTING COMPLEX;
  33:  8jxk-D  4.6  2.8   59   166    5   MOLECULE: CONSERVED PROTEIN;
  34:  2ns0-A  4.6  3.1   63    85   16   MOLECULE: HYPOTHETICAL PROTEIN;
  35:  4lmy-B  4.5  2.2   62   157   10   MOLECULE: PEROXIDE STRESS REGULATOR PERR, FUR FAMILY;
  36:  7b0c-A  4.5  2.9   62   144   10   MOLECULE: HTH-TYPE TRANSCRIPTIONAL REPRESSOR NSRR;
  37:  8rkd-B  4.5  3.8   64   411    8   MOLECULE: PILUS ASSEMBLY ATPASE CPAF;
  38:  3bpx-A  4.4  2.3   55   147   18   MOLECULE: TRANSCRIPTIONAL REGULATOR;
  39:  9dlr-B  4.3  2.9   64   395   14   MOLECULE: LEUKOCYTE RECEPTOR CLUSTER MEMBER 8;
  40:  3lwf-A  4.2  2.7   60   144   12   MOLECULE: PUTATIVE TRANSCRIPTIONAL REGULATOR;

0001 s001A   --------DNLLYDFTLNIYWAAVAQ---R---R-KK--I--FT-Q-D-VLDFFESAAVD-GRI--L-SP--W---S-EGTQVRTARGLL-ATLRD-V-G-FLRDVS-R-EIIP-Y--YI
0002 7zgeA   lmmsllihSPVVADFMRHTLAEARRT---Y---K-PA--L--TA-D-A-WSEFYDTRVRA-Y-A--E-LG-gF---S-DSTIKKMGNNAI-KALVD-S-G-YLSDSR-K-KIQP-V--YL
0003 4kywA   lrkssls sRGWTIEILN-CIDKI------E---G-SE--F--TL-E-D-MYRFESDLK---NIF--V-KN--N-------HIKEKIRQQL-QILRD-K-E-IIEFKG-Y-RKL-------
0004 1cf7B   ----gkglRHFSMKVCE-KVQRKG------------T--T--SY-N-E-VADELVSEFTN-SnnhlaaDS--A---YdQKNIRRRVYDAL-NVLMA-M-N-IISKEK-I-KWIG-L--P-
0005 7qcdD   ----pvtgDATAKYLLQ-YILSAR-----------gI--C--HE-N-A-LILALMRLETDaSTL--N-TE--W---S-IQQWVDKLNDYI-NAINV-K-L-KIIRINfF-VYV--N--LA
0006 8c5sA   pyltlidpGKMCVITIL-ELLKLNSTggvI---E-GM-rtarAV-I-S-VGKAIEMEFRS-EqvlkK-IL--W---P-QSIRARIGSVLI-SMLIQ-V-AkAFAHhn-V-LKIH-L--IR
0007 4etsB   vllerfkyTKQREVLL-KTLYHSD-----------tH--Y--TPeS-L-YMEIKQAEPD------------------lnvGIATVYRTL-NLLEE-A-E-MVTSgK-Y-ELA--N--KP
0008 3cuqA   iqsvpaelNMDHTVVLQ-LAEKNG-----------Y--V--TV-S-E-IKASLK-------------------------wETERARQVL-EHLLK-E-G-LAWLeA-Y-WLPAl-----
0009 1sfuA   ---ctvndAEIFSLVKK-EVLSL--N---T---N-DY--T--TA-I-S-LSNRLK-------------------------iNKKKINQQL-YKLQK-E-D-TVKMpP-W-FKNY-Nc---
0010 8cliA   evretyrlLKRRNLIIE-AVTNLR-----------lI--E--SL-F-T-IQKMIMDQEKQ-E---gV-ST--K-------cCKKSIVRLV-RNLSE-E-G-LL-RLYlV-VHPS-Mdq--
0011 5nl9A   ledaginvTVQRLQMAN-LLLSK----------P-QH--L--TAdQ-V-FQLINEHMP---------------------naSRATIFNNL-KLFAE-K-G-IVNLiT-Y-DSNV-I--H-
0012 7nywE   prsvlseldMMVGKILCYLYLSpeRL---A---NqGI--F--TS-Q-E-LYEELISLA----DE--G-KL--MkdlD-KQKLQEKVRTTL-NRLRR-L-G-MVYFNN-F-TITE-Avf--
0013 5v8f6   ttvtydkyVSMMNMIVR-KIAEVDRE---G---A-EE--L--TA-V-D-IVDWYLLQKEN-D-lgslaEY--W---EeRRLAFKVIKRLVkDRIL---------MEI------------
0014 4hgaA   ipyrgtryPEVNRRIE-RLINK--PG---P---D-TF--P--DY-G-D-VLRAVEKAAA---RH--S-LG--L---P-RQQLQLMAQDAF-RDVG----I-RLQ-------eRR-H--L-
0015 9qe1D   qdfqepmdYAIFCCALA-FLEGKA-----V---D-EQ--F--LL-S-E-LCQEIQADY----PG--D-FP----dwT-LYTHRKSLIRAV-KVLME-F-Q-LIRTIDlY-EAST-Ysr--
0016 5dcaA   lpiesqfvSKLVDNLNA-EVVAG--------------niK--CR-N-D-AVNWLAYT----YLY--V-RMdgQ---L-KKFRESLVHSAL-CILKE-Q-E-LVLYDA-I-EATD-Lgn--
0017 6qfdB   arsdardlTAFQKNILT-VLGEE---------------aR--YG-L-A-IKRELEEYYGE------------------evNHGRLYPNL-DDLVN-K-G-LV-EKS---ALTN-Egf--
0018 6zvhy   -------kfNWKGTIKA-ILKQA--P---D---N-E---I--TI-K-K-LRKKVLAQYYT--VT--D-EH-hR---S-EEELLVIFNKKI-SKN---P-T-FK-LLK-Kv---------
0019 9mj5B   sfmpanglTVAQNQVLN-LIKAC--P---R---P-EG--L--NF-Q-D-LKNQLKH-------------------------mSVSSIKQAV-DFLSN-E-G-HIYSdD-H-FKST-Da---
0020 1lvaA   --------GSPEKILAQ-IIQEH----------R-EG--L--DW-Q-EaATRAS-------------------------lSLEETRKLL-QSXAA-A-G-QV-TLLyA-ISTEr-----
0021 8fefI   lptlavsiPLTVLIIFT-LNILLAEF---G---AaDV--S--G--A-G-AALGAVTQ--lgpLV--T-VlfmL---N-GAVITIGLVGGF-FFGVY-I-Q-NVS-AG-Yv---------
0022 2qbyB   lieavkalPFHYKLAL-RSLIES---------------eD--VM-S-A-HKMYTDLCNKF-KQK--------------plSYRRFSDII-SELDM-F-G-IV-KIR---LVEDk-----
0023 6j05A   varlealaSPVRLEIFR-LLVEQE---------P-TG--L--VS-G-D-IAEHLG-------------------------qPHNGISFHL-KNLQH-A-G-LVTVQRrY-RAAM-V--RA
0024 7uy7B   yqvsigffQNILDIAT-KTVKDRG---------A-LG--C--NY-K-Y-LKDKLTKALNL-Q--qiS-YP--L-------iSESYISYLV-HLFQD-F-N-II-EIE------------
0025 2vwaA   dvsikrynNFVDIARL-SIQKHFEHL---SndqK-DS--HvnNX-E-Y-XQKFVQGLQEN-RNI-----S--L---S-KYQENKAVXDLK-YHLQ----K-VYA--------NY-L--SQ
0026 5l0pA   lkkaglkvTLPRLKILE-VLQEPD----------nhH--V--SA-E-D-LYKRLIDMG--------------------eeiGLATVYRVL-NQFDD-A-G-IV-TRHvF-ELT-------
0027 9jbnC   ieatlenlDPKTRDCFLV-LGAF--P---E---D-KK--I--PL-DvL-INVLVELHD---------------------lEDATAFAVI-VDLAN-R-N-LL-TLVfV-TQHD-Vlr--
0028 7rheA   -----nvrRYNERLLLK-TLRRAG-----------S--A--SK-A-D-LARLAN-------------------------mTGTAVGSII-ASLAD-A-K-LIE-aS-I-RLDP-G--AF
0029 4yo2A   nsrkdkslRVMSQKFV-MLFL---VS---T---P--Q--I--VS-L-E-VAAKILIGEDH---vedL-DK--S---K-FKTKIRRLYDIA-NVLSSlD-L-IKKV-H-T-EERG-R--KP
0030 3k69A   ----xkldFSVAVHSIL-YLDAHR---------D-SK--V--AS-R-E-LAQSLH-------------------------lNPVXIRNIL-SVLHK-H-G-YLTGTV-G-YQLD-L--AL
0031 7xx7U   nskkknqpGKYSQLVVE-TIRRLG-----E---R-NG--S--SL-A-K-IYTEAKKV--P-WFD---------------qQNGRTYLKYSI-KALVQ-NdT-LLQVKG-S-FKlN-R--KK
0032 3elkA   --rerilhGLITLYILK-ELVKR-------------P--X--HG-Y-E-LQKSXFETTGQ------------------alPQGS-IYILL-KTXKE-R-G-FV-ISEvY-HITD-Agk--
0033 1lddA   --kyeltlQRSLPFIEG-MLTNLG-----------A--M--KL-H-K-IHSFLKITVPK-DWG--Y------------nrITLQQLEGYL-NTLAD-E-G-RLKYnG-Y-EIV-------
0034 8jxkD   -------eFMLELAILG-LLIES---------------pM--HG-Y-E-LRKRLTGLLG-------A-FR--A-------fSYGSLYPAL-RRMQA-D-G-LI-AEN---qLTD-Kgr--
0035 2ns0A   ----xtvsdRELEECIRALLDAR--A---D---S-AS--I--CP-S-D-VARAVAPDD---------------------wrPLXEPVREAA-GRLAD-A-G-EV-EVTrI-RWTR-T--D-
0036 4lmyB   lrekhiriTETRKAIIS-YMIQS----------T-EH--P--SA-D-K-IYRDLQPNFP--------------------nmSLATVYNNL-KVLVD-E-G-FVSEtT-Y-DFM--G--HQ
0037 7b0cA   --mrltkfTDLALRSLM-RLAVV-RD---G---D-EP--L--AT-R-EvAEVVG-------------------------vPYTHAAKAI-TRLQH-L-G-VVEARR-L-TLTD-R--RV
0038 8rkdB   -dyyhatkTTIFNALLNTIDLSQL-----A---Q-LDlkQ--AG-E-E-IRDIVAELVAI-K----N-VS--M---S-VAEQEHLVQDII-NDVLG-Y-----------------R--IV
0039 3bpxA   relghlnlTDAQVACLL-RIHREP-----------G--I--KQ-D-E-LATFFH-------------------------vDKGTIARTL-RRLEE-S-G-FIEREQiL-EVTR-Rge--
0040 9dlrB   lileklkiITYRNLFK-KVYLLLK---------T-HQ--L--SL-D-A-FLVALKFMQ---------------------vEDVDIDEVQCIL-ANLIY-M-G-VKGYISkL-VVSK-Q--N-
0041 3lwfA   qgxkittkGRYGLTITL-ELAKRI-----G---D-GP--I--SL-R-S-IAQDKN-------------------------lSEHYLEQLI-GPLRN-A-G-IVKSIRgY-VLNG-D--PE

0001 s001A   --------LHHHHHHHHHLHHHHHHL---L---L-LL--L--LH-H-H-HHHHHHHHHHL-LLL--L-LL--L---L-HHHHHHHHHHHH-HHHHH-L-L-LLLLLL-L-LLLL-L--LL
0002 7zgeA   hhhhhhhhLHHHHHHHHHLHHHHHHL---L---L-LL--L--LL-L-H-HHHHHHHHHHH-L-H--H-HH-lL---L-HHHHHHHHHHHH-HHHHH-L-L-LLLLLL-L-LLLL-L--LL
0003 4kywA   llllllllLLHHHHHHHH-HHHHL------L---L-LE--E--EH-H-H-HHHHHHHHH---HHL--L-LL--L-------LHHHHHHHHH-HHHHH-L-L-LEEELL-E-EEL-------
0004 1cf7B   ----llhhHHHHHHHHH-HHHHHL------------E--E--LH-H-H-HHHHHHHHHHL-LllllhhHL--H---HhHHHHHHHHHHHH-HHHHH-L-L-LLLLLL-E-ELLL-L--L-
0005 7qcdD   ----lhhhHHHHHHHHH-HHHHLL-----------lE--E--EH-H-H-HHHHHHHHHHHhLLL--L-LL--L---L-HHHHHHHHHHHH-HHHHH-H-H-EEEEEEeE-EEE--E--LL
0006 8c5sA   hhhllllhHHHHHHHHH-HHLLLLLLlllL---L-EE-ehhhHH-H-H-HHHHHHHHHHH-HhhllL-LL--L---L-HHHHHHHHHHHH-HHHHH-H-LlLEEEll-E-EEEL-H--HH
0007 4etsB   hhhhhhllLHHHHHHH-HHHHHLL-----------lL--L--LHhH-H-HHHHHHHLHH------------------hllLHHHHHHHH-HHLLL-L-L-LEELeE-L-EEL--L--LL
0008 3cuqA   eellllllLHHHHHHHH-HHLLLL-----------E--E--LH-H-H-HHHHHL-------------------------lLHHHHHHHH-HHHHH-H-L-LLEElL-E-ELLLl-----
0009 1sfuA   ---lllllHHHHHHHHH-HHHLL--L---L---L-LE--E--LH-H-H-HHHLL-------------------------lLHHHHHHHH-HHHHH-L-L-LEEElL-E-EELL-Ll---
0010 8cliA   ellllhhhHHHHHHHHH-HHHHLL-----------eE--L--LH-H-H-HHHHHHHHHHH-H---lL-LL--L-------lLHHHHHHHH-HHHHH-L-L-LL-EEEeE-ELLL-Lll--
0011 5nl9A   hhhlllllLHHHHHHHH-HHHLL----------L-LE--E--LHhH-H-HHHHHHHLL---------------------llLHHHHHHHH-HHHHH-L-L-LLEElE-E-ELLL-L--L-
0012 7nywE   leeellhhhHHHHHHHHHHHHLlhHH---H---HlLE--E--EH-H-H-HHHHHHHHL----LH--H-HH--HhhhH-HHHHHHHHHHHH-HHHHH-L-L-LEEELL-E-EELH-Hhh--
0013 5v8f6   llllhhhhHHHHHHHHH-HHHHHHHL---L---L-LL--L--LH-H-H-HHHHHHHHLLL-L-lllhhHH--H---HhHHHHHHHHHHHhLLLL---------LLL------------
0014 4hgaA   llllllllHHHHHHHH-HHHLL--LL---L---L-LL--L--LH-H-H-HHHHHHHHHH---HL--L-LL--L---L-HHHHHHHHHHHH-HHHH----H-HHH-------hHH-H--H-
0015 9qe1D   llllllhhhHHHHHHHHH-HHLLLL-----L---L-LE--E--EH-H-H-HHHHHHHHL----LL--L-LL----llL-LHHHHHHHHHHH-HHHHH-L-L-LEEEELeE-EELH-Hhh--
0016 5dcaA   llllllhhHHHHHHHHH-HHHLL--------------llL--LH-H-H-HHHHHLLL----HHH--H-HHhlL---L-HHHHHHHHHHHH-HHHHH-L-L-LEEEEL-E-EELH-Hhh--
0017 6qfdB   lhhhhlllLHHHHHHHH-HHHHL---------------lE--EH-H-H-HHHHHHHHHLL-------------------llLHHHHHHHH-HHHHH-L-L-LE-EEE---EELH-Hhh--
0018 6zvhy   -------llLHHHHHHHH-HHHLL--H---H---H-E---E--EH-H-H-HHHHHHHHHHH--HL--L-LL-lL---L-HHHHHHHHHHHH-HLL---L-L-EE-ELL-Ee---------
0019 9mj5B   llllllllLHHHHHHHH-HHHHL--L---L---L-LL--L--LH-H-H-HHHHLLL-------------------------lLHHHHHHHH-HHHHH-H-L-LEELlL-L-LEEL-Ll---
0020 1lvaA   --------LLHHHHHHH-HHHLL----------L-LL--E--EH-H-HhHHHHL-------------------------lLHHHHHHHH-HHHHH-L-L-LE-EEEeE-EEHHh-----
0021 8fefI   hhhhhhhhHHHHHHHHH-HHHHHHHH---L---LlLL--H--H--H-H-HHHHHHHL--hhhHH--H-HhhhH---H-HHHHHHHHHHHH-HHHHH-L-L-LLL-HH-Hh---------
0022 2qbyB   hhhhhhllLHHHHHHH-HHHHLL---------------lL--HH-H-H-HHHHHHHHHHL-LLL---------------llLHHHHHHHH-HHHHH-L-L-LE-EEE---EELLl-----
0023 6j05A   hhhhhhhlLHHHHHHHH-HHHHLL---------L-LL--E--EH-H-H-HHHHHL-------------------------lLHHHHHHHH-HHHHH-L-L-LEEEEEeE-EELH-H--HH
0024 7uy7B   eeelhhhhHHHHHHHH-HHHHHHL---------L-LL--L--LH-H-H-HHHHHHHHHLL-L--llL-LL--L-------lLHHHHHHHH-HHHHH-H-L-LL-LLL------------
0025 2vwaA   lllhhhhhHHHHHHHH-HHHHHHHHL---LhhhH-HH--HllLH-H-H-HHHHHHHHHHH-LLL-----L--L---L-HHHHHHHHHHHH-HHHH----H-HHH--------HH-H--HH
0026 5l0pA   hhhlllllLHHHHHHHH-HHLLLL----------llE--E--EH-H-H-HHHHHHHLL--------------------lllLHHHHHHHH-HHHHH-L-L-LE-EEEeE-EEL-------
```

```
0027 9jbnC   hhhhhlllLHHHHHHHHH-HHHL--L---L---L-LL--E--EH-HhH-HHHHHHHHL----------------------lLHHHHHHHH-HHHHH-L-L-LL-EELeE-ELLH-Hhh--
0028 7rheA   -----lhhHHHHHHHHH-HHHHHL------------L--E--EH-H-H-HHHHHH-------------------------lLHHHHHHHH-HHHHH-L-L-LEE-lL-E-EELL-L--LE
0029 4yo2A   lllllllhHHHHHHHHH-HHLL---LL---L---L--L--L--LH-H-H-HHHHHHHHLLL---lllL-LL--L---L-HHHHHHHHHHHH-HHLLLlL-L-LLLL-L-L-LLLL-L--LL
0030 3k69A   ----llhhHHHHHHHHH-HHHLLL---------L-LL--L--LH-H-H-HHHHHL-------------------------lLHHHLHHHH-HHHHH-L-L-LLEEEL-E-EELL-L--LH
0031 7xx7U   llllllllLHHHHHHHHH-HHHHLL-----L---L-LL--L--LH-H-H-HHHHHHLL--L-LLL--------------lLLHHHHHHHHH-HHHHH-LlL-EEEELL-E-EEeL-L--LL
0032 3elkA   --llhhhhHHHHHHHHHH-HHHHL-------------L--E--EH-H-H-HHHHHHHHHLL-----------------llLLLH-HHHHH-HHHHH-H-L-LE-EEEeE-EELH-Hhh--
0033 1lddA   --llhhhhHHHHHHHHHH-HHHHHL------------L--E--EH-H-H-HHHHHHHHLLH-HHL--L-----------llLLHHHHHHHH-HHHHH-L-L-LEELlL-E-EEL-------
0034 8jxkD   -------lHHHHHHHHHH-HHHHL---------------lE--EH-H-H-HHHHHHHHLL-------L-LL--L-------lLLLLHHHHH-HHHHH-L-L-LE-EEL---eELH-Hhh--
0035 2ns0A   ----llllhHHHHHHHHHHHHHHL--L---L---L-LL--L--LH-H-H-HHHHHLLLL--------------------lhHHHHHHHHHH-HHHHH-L-L-LE-EEEeE-EELL-L--L-
0036 4lmyB   hhhlllllLHHHHHHHHH-HHHHL----------L-LL--L--LH-H-H-HHHHHLLLLL--------------------llLHHHHHHHH-HHHHH-L-L-LEEElL-E-EEL--L--LE
0037 7b0cA   --llllhhHHHHHHHHHH-HHHLL-LL---L---L-LL--L--LH-H-HhHHHHL-------------------------lLHHHHHHHH-HHHHH-H-L-LEEELL-E-EELH-H--HL
0038 8rkdB   -llhhhhhHHHHHHHHHHHLLHHHL-----L---L-LLhhH--HH-H-H-HHHHHHHHHHH-L----L-LL--L---L-HHHHHHHHHHHH-HHHLL-L-----------------H--HH
0039 3bpxA   hhhhhhllLHHHHHHHHH-HHHLLL------------L--L--LH-H-H-HHHHHL-------------------------lLHHHHHHHH-HHHHH-L-L-LEEEEEeE-EELH-Hhh--
0040 9dlrB   hhhllhhhHHHHHHHHH-HHHHHHL---------L-LE--E--EH-H-H-HHHHHHHLL--------------------lLLLLHHHHHHHH-HHHHH-H-L-LLEEEEeE-EELL-L--L-
0041 3lwfA   llllllhhHHHHHHHHHH-HHHHLL-----L---L-LL--L--LH-H-H-HHHHHL-------------------------lLHHHHHHHH-HHHHH-L-L-LEEEELeE-EELL-L--LL

# Job: Type-1 BREX; BrxA BAY35696 wHTH
# Query: s001A
# No:  Chain   Z    rmsd lali nres  %id PDB  Description
   1:  1jhf-A  6.1  2.6   63   197   13   MOLECULE: LEXA REPRESSOR;
   2:  6lb9-A  6.0  2.4   66   345   11   MOLECULE: DUF4007 DOMAIN-CONTAINING PROTEIN;
   3:  9fhl-c  5.8  2.6   64   109    8   MOLECULE: RRNA 16S SSO;
   4:  1ylf-C  5.6  2.5   62   119    6   MOLECULE: RRF2 FAMILY PROTEIN;
   5:  4o5v-A  5.6  3.0   62   213   11   MOLECULE: IRON-DEPENDENT TRANSCRIPTION REPRESSOR RELATED PR
   6:  1fx7-A  5.4  3.0   62   230   11   MOLECULE: IRON-DEPENDENT REPRESSOR IDER;
   7:  2jt1-A  5.4  3.1   61    71   15   MOLECULE: PEFI PROTEIN;
   8:  3dv8-A  5.3  3.4   63   216   14   MOLECULE: TRANSCRIPTIONAL REGULATOR, CRP/FNR FAMILY;
   9:  2ns0-A  5.2  3.2   62    85   16   MOLECULE: HYPOTHETICAL PROTEIN;
  10:  5yad-A  5.2  3.4   62    73    8   MOLECULE: MEIOSIS REGULATOR AND MRNA STABILITY FACTOR 1;
  11:  3ov8-A  5.2  3.8   66    91   11   MOLECULE: PROTEIN AF_1382;
  12:  6qpq-D  5.2  2.8   63    82    8   MOLECULE: STRUCTURAL MAINTENANCE OF CHROMOSOMES PROTEIN,STR
  13:  6wge-C  5.2  3.1   62   174    5   MOLECULE: STRUCTURAL MAINTENANCE OF CHROMOSOMES PROTEIN 1A;
  14:  4i99-C  5.2  2.9   59    70    8   MOLECULE: CHROMOSOME PARTITION PROTEIN SMC;
  15:  4kyw-A  5.2  3.2   62   254    6   MOLECULE: TYPE-2 RESTRICTION ENZYME DPNI;
  16:  7y43-A  5.1  3.4   62    79   11   MOLECULE: HISTONE ACETYLTRANSFERASE KAT6A;
  17:  2od5-A  5.0  3.1   67    91    7   MOLECULE: HYPOTHETICAL PROTEIN;
  18:  3lmm-D  5.0  3.0   63   556    8   MOLECULE: UNCHARACTERIZED PROTEIN;
  19:  5trd-A  5.0  3.0   60   219    3   MOLECULE: RIBOFLAVIN KINASE;
  20:  5xpq-A  4.9  3.2   63   100   10   MOLECULE: UNCHARACTERIZED HTH-TYPE TRANSCRIPTIONAL REGULATO
  21:  3h0d-A  4.9  3.1   64   155   16   MOLECULE: CTSR;
  22:  8xt8-B  4.9  2.9   58   145   10   MOLECULE: MARR FAMILY TRANSCRIPTIONAL REGULATOR;
  23:  5j6x-A  4.9  3.1   60    66    7   MOLECULE: Z-DNA BINDING PROTEIN KINASE;
  24:  9gm7-C  4.8  3.6   67   440    3   MOLECULE: CHROMOSOME PARTITION PROTEIN MUKF;
  25:  6uvu-B  4.8  2.9   60   114   12   MOLECULE: ARSR FAMILY TRANSCRIPTIONAL REGULATOR;
  26:  8as8-C  4.8  2.7   65   195   14   MOLECULE: JETC;
  27:  8qto-A  4.7  3.8   65   234   17   MOLECULE: FNR TYPE REGULATOR;
  28:  2mh2-A  4.7  3.2   61    64   13   MOLECULE: HOMOLOGOUS-PAIRING PROTEIN 2 HOMOLOG;
  29:  7yw2-A  4.7  3.1   65   216    9   MOLECULE: TRNA 2'-PHOSPHOTRANSFERASE 1;
  30:  2h09-A  4.7  3.5   62   127    8   MOLECULE: TRANSCRIPTIONAL REGULATOR MNTR;
  31:  1ucr-B  4.7  3.2   61    75   15   MOLECULE: PROTEIN DSVD;
  32:  2obp-A  4.6  3.4   65    81    9   MOLECULE: PUTATIVE DNA-BINDING PROTEIN;
  33:  1yyv-A  4.6  3.2   65   114   11   MOLECULE: PUTATIVE TRANSCRIPTIONAL REGULATOR;
  34:  3bpx-A  4.6  3.0   60   147    7   MOLECULE: TRANSCRIPTIONAL REGULATOR;
  35:  5v9x-A  4.6  3.3   66   772   14   MOLECULE: ATP-DEPENDENT DNA HELICASE;
  36:  9dlr-B  4.6  2.6   64   395    8   MOLECULE: LEUKOCYTE RECEPTOR CLUSTER MEMBER 8;
  37:  8wxs-A  4.6  2.8   59    76   14   MOLECULE: FI02030P;
  38:  8ylg-A  4.6  2.9   60   161   10   MOLECULE: MARR FAMILY TRANSCRIPTIONAL REGULATOR;
  39:  2qby-B  4.6  3.0   66   368   11   MOLECULE: CELL DIVISION CONTROL PROTEIN 6 HOMOLOG 1;
  40:  7yk3-B  4.6  2.6   62   188    8   MOLECULE: DNA ADP-RIBOSYL TRANSFERASE;
  41:  1xsd-A  4.6  3.4   64   125    3   MOLECULE: 5'-
  42:  2g9w-A  4.6  2.8   60   119   10   MOLECULE: CONSERVED HYPOTHETICAL PROTEIN;
  43:  8yz7-A  4.6  3.4   65   191   15   MOLECULE: TRANSCRIPTIONAL REGULATOR, FNR/CRP FAMILY;
  44:  6qfd-B  4.6  3.4   66   116    8   MOLECULE: DNA-BINDING PROTEIN;
  45:  2zkz-C  4.6  3.4   60    87   12   MOLECULE: TRANSCRIPTIONAL REPRESSOR PAGR;
  46:  5a31-N  4.5  2.9   61   703    5   MOLECULE: ANAPHASE-PROMOTING COMPLEX SUBUNIT 1;
  47:  7dvr-A  4.5  3.1   62   147    6   MOLECULE: HTH MARR-TYPE DOMAIN-CONTAINING PROTEIN;
  48:  7qen-C  4.5  3.0   69   291   10   MOLECULE: DNA (35-MER);
  49:  1lva-A  4.5  2.8   56   258    5   MOLECULE: SELENOCYSTEINE-SPECIFIC ELONGATION FACTOR;
  50:  3elk-A  4.5  3.1   64   104    6   MOLECULE: PUTATIVE TRANSCRIPTIONAL REGULATOR TA0346;

0001 s001A   -----PYYI--S--DEG--VACLARYL-HESG-IS-N-S-A-LCE-----H--QDWN-I---F--GL---NREQVCDRLDQL-GEHR-G-LII-QKA---G--SVVSITW---KV-N-
0002 1jhfA   ----kaltA--R--QQE--VFDLIRDH-ISQT-GM-P----pTRA-----E---iAQ-R---L--GF--rSPNAAEEHLKAL-ARKG-V-IEI-VSG------ASRGIRL---LQ-Ee
0003 6lb9A   ftsgsrPSL--P--ARI--IAYACLDY-AART----L-A-R-LANepgapG--RAFR---------I---READIAAALEKV-AASHqE-LQL-VEA---V--GQRSLTF---TS-G-
0004 9fhlc   keiisraVT--I--DEE--TKKKVLDE-IKKE-SI-I-TpY-ALA--------TKSG---------I---SISVARKILKEL-ENQN-V-VKLySKN---R--RLEIYIA---AS---
0005 1ylfC   ------ISS--R---FS--IAVHILSI-LKNN----ssL-C-TSD-----YxaESVN---------T---NPVVIRKIXSYL-KQAG-F-VYV-NRG-------PGGAGL---LK-D-
0006 4o5vA   -----IRSV--T--EED--YLKIIQEL-VLYK-GY-A-----TLA-----DisRSLN---------V---KRQSVRDEINHL-ISLS-X-AEK-I--------eRGKYRL---TP-Sg
0007 1fx7A   -----MNELvdT--TEM--YLRTIYDL-EEEG-VT-P-----LRA-----RiaERLD---------Q---SGPTVSQTVSRM-ERDG-L-LRV-AG--------DRHLEL---TEk--
0008 2jt1A   -----mSES--I--VTK--IISIVQER-QNMD-----dgapvKTR-----DiaDAAG---------L---SIYQVRLYLEQL-HDVG-V-LEK-VNA-gkG--VPGLWRL---LE---
0009 3dv8A   ----iXWKS--L--DKR--VASFLLEE-TSIE----L-K-I-THE-----TiaNHLG---------S---HREVITRXLRYF-QVEG-L-VKL-SR---------GKITI---LD-S-
0010 2ns0A   ------XTV--S--DRE--LEECIRAL-LDAR----adsasiCPS-----DvaRAVA-P-------dwrpLXEPVREAAGRL-ADAG-E-VEV-TQA---R--GPIRIRW---TR-Td
0011 5yadA   -----GAAN-kS--LSL--LSTETXSI-LQDA----L-F-K-FID-----IyekKYG-H-----------KLNV--SDLYKL-T--D-T-IAI-REQ---G--NGRLVCL---LP-S-
0012 3ov8A   ----lLFKE--K--ALE--ILMTIYYE-SLGG-ND-V-----YIQ-----YiaSKVN---------S---PHSYVWLIIKKF-EEAK-M-VEC-ELE---G--RTKIIRL---TD-Kg
0013 6qpqD   --------K--A--IVQ--MAKILRKE-LSEEkeV-I-F-T-DVL-----K---SQA-N------titkrEASRGFFDILSL-ATEG-C-IGL-SQT-eaF--GNIKIDA---KP-Al
0014 6wgeC   gmddreimK--R--TQQ--MLHGLQRA-LAKT----S-L-L-ELC-----R--N---------tnRK---QAAAKFYSFLVL-KKQQ-A-IEL-TQE-epY--SDIIATP-gpRF-H-
0015 4i99C   ------DIE--K-yVEE--LYKVVKKI-YEKT----K-F-W-DLV-----P--D------------vepkIIARTFLYLLFL-ENMG-R-VEI-IQE-epF--GEILVVP---M----
0016 4kywA   ----lSSRG--W--TIE--ILNCIDK--IEGS-eF-T-L-E-DMY-----R---FES-D---L--KknnhIKEKIRQQLQIL-RDKE-I-IEF-KG--------RGKYRK---L----
0017 7y43A   -----------klaNPL--YTEWILEA-IKKVqrP-S-E-ErICN-----AvsSSHG---------L---DRKTVLEQLELS-VKDG-T-ILK-VSN---K--GLNSYKD---PD-Np
0018 2od5A   -----ETES-xK--TVR--IREKIKKF-LGDR-pR-N-T-A-EIL-----E--HINS-T---XrhGT---TSQQLGNVLSKD-K----D-IVKvGYI---KrsDICEWAT---RN-Wv
0019 3lmmD   fspvrylST--D--QAE--LTNAAXLW-LSEV-GD-L-----ATS-----DlxAXCG---------V---SRGTAKACVDGL-VDEE-R-VVA-VGG---G--RSRRYRL---VE---
0020 5trdA   --------D--Q--YYR--AIKKIKEA-AEAS-nR-A-Y-L------tssklaDMLG---------I---SQQSASRIIIDL-EKNG-Y-ITR-TVT---K--RGQILNI---TE-Kg
0021 5xpqA   eplyklkaeffktlaHP--ARIRILEL-LVER-DR-S-V-G-ELL-----S--SDVG---------L---ESSNLSQQLGVL-RRAG-V-VAA-RRD---G--NAMIYSI---AA-P-
0022 3h0dA   ------pNI--S--D-I--IEQYLKQV-LNXSdiV-E-I-K--RS-----E--I-AN-K---F--RC---VPSQINYVINTRfTLER-G-YIV-ESK---R--GYIRIXK---VKt--
0023 8xt8B   kddlkkmfkqngydITT--DHYALLRF-LWEQ----Q-I-D-LCE-----K---------------K---DKSNTTRILDVM-KNKG-L-IVR-KVDvkdR--RKFQIFLtdlGR-E-
0024 5j6xA   -------mS--A--ENE--IEMRICDY-LRRHgrS-T-V-QdIFK-----E--LKL---------------EKSTVNRHLYSL-QASK-Q-VFK-TVE------KRPVWDL---VE---
0025 9gm7C   ndfsislPV--E--RLA--FLMAIAVL-NSERgeM-S-E-G-ELI-----DafREVC-K---G--FtaesVAVRANNAINDM-VRQK-L-LNR-FTS---E--GNAIYRL---TP-Lg
0026 6uvuB   saeagevvvlfkvlaND--TRLRLLHA-LARSggL-C-V-TdLAA-----A--VGM--------------KPQAVSNQLQRL-ADRR-I-LRA-ARC---G--NNIHYRI---VD-P-
0027 8as8C   lvrrqrlnL--E--QSL--LVAILRQH-FVAWaqI-A-I-D-DL------L--PQLQ-I---Y--LseskERTRLLTLLDQL-KG-H-G-LVT-SPD---A--HERIVIR--pII-A-
0028 8qtoA   ----lSKKN--A--EER--LAAFLYNL-STRFreF-R-L-T-mtR-----G--DIGN-Y---L--GL---TVETISRLLGRF-QKTE-M-LTV-KG---------KYITI---ND-H-
0029 2mh2A   --------G--A--PGI--ILRYLQEQ-NR------S-A-Q-DVF-----G---NLQ-K--eH--GL---GKAAVVKALDQL-AQEG-K-IKE-KTY---G--KQKIYFA---D----
0030 7yw2A   ------eQD--R--NVQ--LSKALSYAlRHGAlglpL-Q-A-LLQ-----L--PQFH---------S---FSIEDVQLVVNT-NEKQ-R-FTL-QPG---E--TGLLIRA---NQ-G-
0031 2h09A   ----rEAHRreL--IDD--YVELISDL-IREV-GE-A-----RQV-----DmaARLG---------V---SQPTVAKMLKRL-ATMG-L-IEM-IP--------WRGVFL---TAe--
0032 1ucrB   --------------MEE--AKQKVVDF-LNS-skF-Y-F-N-DFT--------DLFP--------DM---KQREVKKILTAL-VNDE-V-LEY-WSS---G--STTMYGL---KG-Ag
0033 2obpA   -----GIDP--A--IVE--VLLVLREA-GIE-tpW-S-L-P-KIA--------KRAQ---------L---PXSVLRRVLTQL-QAAG-L-ADV-SVE--aD--GRGHASL---TQ-Eg
0034 1yyvA   faeqcpsrevlK--HVTsrWGVLILVA-LRDG-TH-R-F-S-DLR--------RXXG--------GV---SEXXLAQSLQAL-EQDG-F-LNR-VSY---P--PHVEYSLtplGE-Q-
0035 3bpxA   shrvfigrelghlnLTD--AQVACLLR-IHRE----Q-D-E-LAT-----F--FH-----------V---DKGTIARTLRRL-EESG-F-IER-EQDpenR--RRYILEVtrrGE-E-
0036 5v9xA   ----rVPAN--P--LDV--LAQHTVAV-AAlepVD-A-D-A-W-F-----D--AVRR-SapfA--TL---PRSAFEATLDLL-SlRP-R-LVY-DRD-------TGTLTA---RP-G-
0037 9dlrB   rcgiflileklK--IIT--YRNLFKKV-YLLLhqL-S-L-D-AFL-----V---ALK-F---M--QV---DIDEVQCILANL-IMGH-VkGYI-SHQ-------HQKLVV---SK-Q-
0038 8wxsA   --------dggE--LSL--VKKVVHSL-VVSS----V-E-Q-LMR-----DyrSAAG---------C---TLPYsESFLRSI-P--D-T-VTV-TGH---G--QMAWITA---VA-Ta
0039 8ylgA   iardrlnplfarygLQP--GEFDVLAT-LRRS----P-T-A-LYD-----A--AMI--------------SSGSMTNRIDRL-EKAG-W-VER-RANpadG--RGTLVALtsaGR-A-
0040 2qbyB   vdyeqerlieavkaLPF--HYKLALRS-LI-E-SE-DvM-S-AHK-----MytDLCN-Q---K--PL---SYRRFSDIISEL-DMFG-I-VKI-RII---N--VKKYALV---ED-K-
0041 7yk3B   --------------MTW--GRAVILEA-MRRYamepH-L-E-IQK-----LmyFANE-AdpdL--AL---DSERVRHLLQGM-EG-A-F-TVG-L-V---L--ANQPISL---TT-Kg
0042 1xsdA   -------tnkqveiSMA--E-WDVMNI-IWDKksV-S-A-NeIVV-----E---IQK-Y---K--EV---SDKTIRTLITRL-YKKE-I-IKR-YKS---E--NIYFYSS---NI-K-
0043 2g9wA   ----------kltrlGD--LERAVXDH-LWSR----V-R-Q-VHE-----A---LSA-R---R--DL---AYTTVXAVLQRL-AKKN-L-VLQ-IR--------AHRYAP---VH-G-
0044 8yz7A   ----lVGKR--L--RAR--IAGELLEL-KDTA-LAtI-Y-A-THD-----ElaAAVG---------S---VRETVTKVVGEL-SREG-V-ISA-GY---------GKITL---KD-E-
0045 6qfdB   -----pdarsdardLTA--FQKNILTV-LGEE-AR-G-L-A-IKR-----E-lEEYY-G---E--EV---NHGRLYPNLDDL-VNKG-L-VEK-S-D---K--RTNEYALtneGF-D-
0046 2zkzC   yxsleddaellktxaHP--XRLKIVNE-LYKHkaL-N-V-T-QII--------QILK---------L---PQSTVSQHLCKX-RG-K-V-LKR-NRQ---G--LEIYYSI---NN-P-
0047 5a31N   nmvlikEEE--L---LL--FWTYIQAM-LTNLslS-L-D-R-IYN-----MlrMFVVtL---A--EI---DLQELQGYLQKK-VRDQ-Q-LVY-S----------aGVYRL---P----
0048 7dvrA   ldryakeydveH--LAG--PQGHLVMH-LYKHdkD-M-S-I-kDA-----E--EILH---------I---SKSVASNLVKRM-EKNG-F-IAI-VPSktdK--RVKYLYL---TH-Lg
0049 7qenC   ----kKVDV--RrlKKN--VWRSINNL-IQEHkeL-K-F-S-DII-----Q--gISK-M---YsddI---STSFCFICLLHL-ANEH-G-LQI-THT-enY--NDLIVNY---ED-L-
0050 1lvaA   ---------------gS--PEKILAQI-IQEHegL-D-W-QeAAT-----R---------------L---SLEETRKLLQSX-AAAG-Q-VTL-LRV---E--NDLYAIS---TE-Ry
0051 3elkA   -------reriL--HGL--ITLYILKE-LV-K-RP-X-----HGY-----ElqKSXF-EttgQ--AL---PQGSIYILLKTX-KERG-F-VIS-ESS---V--QLTVYHI---TD-Ag

0001 s001A   -----LLLL--L--HHH--HHHHHHHH-HHLL-LL-H-H-H-HHL-----L--HHHH-H---H--LL---LHHHHHHHHHHH-LHHH-L-EEE-EEE---L--LEEEEEE---LL-L-
0002 1jhfA   ----llllH--H--HHH--HHHHHHHH-HHHH-LL-L----lLHH-----H---hHH-H---L--LL--lLHHHHHHHHHHH-HHLL-L-EEE-LLL------LLLLEEE---LL-Ll
0003 6lb9A   elllllLLL--L--HHH--HHHHHHHH-HHLL----H-H-H-HHHllllhH--HHHL---------L---LHHHHHHHHHHH-HLLLlL-EEE-EEL---L--LLEEEEE---LL-L-
0004 9fhlc   lllllllLL--L--LHH--HHHHHHHH-HHHL-LL-L-LhH-HHH--------HHHL---------L---LHHHHHHHHHHH-HHLL-L-EEEeEEL---L--LLEEEEE---LL---
0005 1ylfC   ------LLH--H---HH--HHHHHHHH-HHHL----llL-L-LHH-----HhhHHLL---------L---LHHHHHHHHHHH-HHHL-L-EEE-LLL-------LLLEEE---LL-L-
0006 4o5vA   -----LLLL--L--HHH--HHHHHHHH-HHHH-LL-L-----LHH-----HhhHHHL---------L---LHHHHHHHHHHH-HHLL-L-EEE-E--------eLLEEEE---LH-Hh
0007 1fx7A   -----LLLLllH--HHH--HHHHHHHH-HHHL-LL-L-----LHH-----HhhHHHL---------L---LHHHHHHHHHHH-HHLL-L-EEE-LL--------LLLEEE---LHh--
0008 2jt1A   -----lLLL--H--HHH--HHHHHHHH-HHHH-----llllleEHH-----HhhHHHL---------L---LHHHHHHHHHHH-HHLL-L-EEE-ELL-llL--LLLEEEE---LL---
0009 3dv8A   ----hHHLL--H--HHH--HHHHHHHH-HHHH----E-L-L-LHH-----HhhHHHL---------L---LHHHHHHHHHHH-HHLL-L-EEE-EL---------LEEEE---LL-H-
0010 2ns0A   ------LLL--L--HHH--HHHHHHHH-HHHL----llllllLHH-----HhhHHHL-L-------llhhHHHHHHHHHHHH-HHLL-L-EEE-EEL---L--LLLEEEE---LL-Ll
0011 5yadA   -----LHHH-hH--HHH--HHHHHHHH-HHLL----H-H-H-HHH-----HhhhHHL-L-----------LLLH--HHHLLL-L--L-L-EEE-ELL---L--LLLEEEE---LH-H-
0012 3ov8A   ----hHLLH--H--HHH--HHHHHHHH-HHLL-LL-E-----EHH-----HhhHHLL---------L---LHHHHHHHHHHH-HHLL-L-EEE-EEE---L--LEEEEEE---LH-Hh
```

```
0013 6qpqD   --------L--H--HHH--HHHHHHHH-HLLLllE-E-H-H-HHH-----H---HHH-L------lllhhHHHHHHHHHHHH-HHLL-L-EEE-ELL-llL--LLLEEEE---LH-Hh
0014 6wgeC   llllllllL--H--HHH--HHHHHHHH-HLLL----E-H-H-HHL-----L--L---------llHH---HHHHHHHHHHHH-HHLL-L-EEE-ELL-llL--LLLEEEE-llLL-L-
0015 4i99C   ------LHH--H-hHHH--HHHHHHHH-HHHH----E-H-H-HHL-----L--L------------llhhHHHHHHHHHHHH-HHLL-L-EEE-ELL-llL--LLLEEEE---L----
0016 4kywA   ----lLLLL--H--HHH--HHHHHHH--LLLL-eE-E-H-H-HHH-----H---HHH-H---H--HllllHHHHHHHHHHHH-HHLL-L-EEE-LL--------LLEEEE---L----
0017 7y43A   -----------lllLHH--HHHHHHHH-HHHHllL-L-H-HhHHH-----HhhHHHL---------L---LHHHHHHHHHHH-HHHL-L-LEE-EEE---L--LEEEEEL---LL-Ll
0018 2od5A   -----LLHH-hH--HHH--HHHHHHHH-LLLL-lE-E-H-H-HHH-----H--HHHL-L---LlLL---LHHHHHHHHHHL-L----L-EEEeEEE---EeeEEEEEEE---HH-Hh
0019 3lmmD   lllllllLL--L--HHH--HHHHHHHH-HHHL-LL-E-----EHH-----HhhHHLL---------L---LHHHHHHHHHHH-HHLL-L-EEE-ELL---L--LLLEEEE---LL---
0020 5trdA   --------L--H--HHH--HHHHHHHH-HHHH-lE-E-E-E------lhhhhhHHHL---------L---LHHHHHHHHHHH-HHLL-L-EEE-EEE---L--LEEEEEE---LH-Hh
0021 5xpqA   lhhhhhhhhhhhllLH--HHHHHHHH-HHHL-LE-E-H-H-HHH-----L--LLLL---------L---LHHHHHHHHHHH-HHLL-L-EEE-EEL---L--LLEEEEE---LL-L-
0022 3h0dA   ------lLH--H--H-H--HHHHHHHH-HLLLleE-E-E-L--HH-----H--H-HH-H---L--LL---LLHHHHHHHHHHlLHHH-L-EEE-EEE---L--LEEEEEE---EEe--
0023 8xt8B   hhhhhhhhhhllllLLH--HHHHHHHH-HHHL----H-H-H-HHH-----H--------------L---LLHHHHHHHHHH-HLLL-L-EEE-EELlllL--LLEEEEElhhHH-H-
0024 5j6xA   -------lL--H--HHH--HHHHHHHH-HHHHllL-L-H-HhHHH-----H--LLL--------------LHHHHHHHHHHH-HHLL-L-EEE-ELL------LLLEEEE---LL---
0025 9gm7C   hllleeeLH--H--HHH--HHHHHHHH-HHLLllE-E-H-H-HHH-----HhhHHHH-H---H--LllllHHHHHHHHHHHH-HHLL-L-EEE-EEL---H--LLEEEEE---LH-Hh
0026 6uvuB   hhhhhhhhhhhhhhlLH--HHHHHHHH-HHHHllL-L-H-HhHHH-----H--LLL--------------LHHHHHHHHHHH-HHLL-L-EEE-EEE---L--LEEEEEE---LL-H-
0027 8as8C   llllllllH--H--HHH--HHHHHHHH-HHHHleE-E-H-H-HH------H--HHHH-H---H--HlhhhHHHHHHHHHHHH-HH-L-L-LLL-LLL---L--LLEEELL--lLH-H-
0028 8qtoA   ----hHHLL--H--HHH--HHHHHHHH-HHHHleE-E-L-L-llH-----H--HHHH-H---H--LL---LHHHHHHHHHHH-HHLL-L-EEE-EL---------LEEEE---LL-H-
0029 2mh2A   --------L--H--HHH--HHHHHHHH-LL------E-H-H-HHH-----H---HHH-L--lL--LL---LHHHHHHHHHHH-HHHL-L-EEE-EEE---L--LEEEEEE---L----
0030 7yw2A   ------lHH--H--HHH--HHHHHHHHhHLLLllllH-H-H-HHL-----L--HHHL---------L---LLHHHHHHHHHH-LLLL-L-EEE-EEL---L--LLEEEEE---LL-L-
0031 2h09A   ----hHHHHhhH--HHH--HHHHHHHH-HHHH-LL-L-----LHH-----HhhHHHL---------L---LHHHHHHHHHHH-HHLL-L-EEE-EL--------LLEEEE---LHh--
0032 1ucrB   --------------LLH--HHHHHHHH-HLL-llE-E-H-H-HHH--------HHLL--------LL---LHHHHHHHHHHH-HHLL-L-EEE-EEE---L--LEEEEEE---LL-Hh
0033 2obpA   -----LLLH--H--HHH--HHHHHHHH-LLL-llL-L-H-H-HHH--------HHHL---------L---LHHHHHHHHHHH-HHLL-L-EEE-EEL--lL--LLEEEEE---LH-Hh
0034 1yyvA   llllllhhhhhH--HHHlhHHHHHHHH-HHHL-LE-E-H-H-HHH--------HHLL--------LL---LHHHHHHHHHHH-HHHL-L-EEE-EEE---L--LEEEEEElhhHH-H-
0035 3bpxA   hhhhhhhhhhhhhlLLH--HHHHHHHH-HHLL----H-H-H-HHH-----H--HL-----------L---LHHHHHHHHHHH-HHLL-L-EEE-EEElleE--EEEEEEElhhHH-H-
0036 5v9xA   ----lLLLL--L--HHH--HHHHHHHH-LLlllEE-H-H-H-H-H-----H--HHHH-LlllL--LL---LHHHHHHHHHHH-HlLL-L-EEE-ELL-------LLEEEE---LL-L-
0037 9dlrB   hhllhhhhllhH--HHH--HHHHHHHH-HHHHleE-E-H-H-HHH-----H---HHH-H---L--LL---LHHHHHHHHHHH-HHLL-LlEEE-ELL-------LLEEEE---LL-L-
0038 8wxsA   --------lllH--HHH--HHHHHHHH-HHLL----H-H-H-HHH-----HhhHHHL---------L---LLLLlHHHHHLL-L--L-L-EEE-ELL---L--LLLEEEE---LL-Ll
0039 8ylgA   hhhhlhhhhhhhllLLH--HHHHHHHH-HHHL----H-H-H-HHH-----H--HLL--------------LHHHHHHHHHHH-HHHL-L-EEE-EELlllL--LLEEEEElhhHH-H-
0040 2qbyB   hhhhhhhhhhhhhlLLH--HHHHHHHH-HH-L-LL-LhH-H-HHH-----HhhHHHH-L---L--LL---LHHHHHHHHHHH-HHLL-L-EEE-EEE---L--LEEEEEE---LL-L-
0041 7yk3B   --------------LLH--HHHHHHHH-HHHHhhllH-H-H-HHH-----HhhHHHH-HlhhH--LL---LLHHHHHHHHHL-LL-L-L-EEL-L-L---L--LLLEEEE---LH-Hh
0042 1xsdA   -------lllllllLLL--L-HHHHHH-HHHHllE-E-H-HhHHH-----H---HHL-L---L--LL---LHHHHHHHHHHH-HHLL-L-EEE-EEL---L--LLEEEEE---LL-L-
0043 2g9wA   ----------lhhhlLH--HHHHHHHH-HHLL----H-H-H-HHH-----H---HLL-L---L--LL---LHHHHHHHHHHH-HHLL-L-EEE-EL--------LLEEEE---LL-L-
0044 8yz7A   ----hHHLL--H--HHH--HHHHHHHH-LLLL-LEeE-E-L-LHH-----HhhHHHL---------L---LHHHHHHHHHHH-HHLL-L-EEE-LL---------LEEEE---LL-H-
0045 6qfdB   -----lllhhhhllLLH--HHHHHHHH-HHHL-LE-H-H-H-HHH-----H-hHHHH-L---L--LL---LHHHHHHHHHHH-HHLL-L-EEE-E-E---L--LEEEEEElhhHH-H-
0046 2zkzC   hhhhhhhhhhhhhhlLH--HHHHHHHH-HHHHllE-E-H-H-HHH--------HHHL---------L---LHHHHHHHHHHH-LL-L-L-EEE-EEE---L--LEEEEEE---LL-H-
0047 5a31N   llllllHHH--H---HH--HHHHHHHH-HHHHllL-H-H-H-HHH-----HhhHHLLlL---L--LL---LHHHHHHHHHHH-HHLL-L-LEE-E---------lLEEEL---L----
0048 7dvrA   hhhhhhhllllL--LLL--HHHHHHHH-HHHLllL-E-E-H-hHH-----H--HHHL---------L---LHHHHHHHHHHH-HHLL-L-EEE-EELlllL--LLEEEEE---LH-Hh
0049 7qenC   ----lLLLH--HhhHHH--HHHHHHHH-HHHHleE-E-L-H-HHH-----H--hHLL-L---LlllL---LHHHHHHHHHHH-HHHL-L-LEE-EEL-llL--LLEEEEL---LL-L-
0050 1lvaA   ---------------lL--HHHHHHHH-HHLLllE-E-H-HhHHH-----H--------------L---LHHHHHHHHHHH-HHLL-L-EEE-EEE---L--LEEEEEE---HH-Hh
0051 3elkA   -------llhhH--HHH--HHHHHHHH-HH-H-LL-E-----EHH-----HhhHHHH-HhhlL--LL---LLLHHHHHHHHH-HHHL-L-EEE-EEE---E--EEEEEEE---LH-Hh

@@@@@@@@@@@@@@@@@@@@@@@@@@@@@@@@@@@@@@@@@@@@@@@@@@@@@@@@@@@@@@@@@@@@@@@@@@@@@@@@@@@@@@@@@@@@@@@@@@@@@@@@@@@@@@@@@@@@@@@@@
@@@@@@@@@@@@@@@@@@@@@@@@@@@@@@@@@@@@@@@@@@@@@@@@@@@@@@@@@@@@@@@@@@@@@@@@@@@@@@@@@@@@@@@@@@@@@@@@@@@@@@@@@@@@@@@@@@@@@@@@@
@@@@@@@@@@@@@@@@@@@@@@@@@@@@@@@@@@@@@@@@@@@@@@@@@@@@@@@@@@@@@@@@@@@@@@@@@@@@@@@@@@@@@@@@@@@@@@@@@@@@@@@@@@@@@@@@@@@@@@@@@

# Job: Type-1 BREX; BrxB Inactive STAND-NTPase WP_102077887
# Query: s001A
# No:  Chain   Z    rmsd lali nres  %id PDB  Description
   1:  9nv3-A 12.0  5.1  166   199   14   MOLECULE: DUF1788 DOMAIN-CONTAINING PROTEIN;
   2:  8dr5-A  8.6  3.2  130   646   12   MOLECULE: REPLICATION FACTOR C SUBUNIT 1;
   3:  8fnu-A  8.4  3.1  125   796   10   MOLECULE: KAP NTPASE DOMAIN-CONTAINING PROTEIN;
   4:  8umy-A  8.2  3.3  128   546   10   MOLECULE: CHROMOSOME TRANSMISSION FIDELITY PROTEIN 18 HOMOL
   6:  7z6h-K  7.8  3.3  128   439   10   MOLECULE: CELL CYCLE CHECKPOINT CONTROL PROTEIN RAD9A;
   7:  8fcv-R  7.6  3.5  127   343   11   MOLECULE: DNA (60-MER);
   8:  7st9-A  7.6  3.5  130   522    8   MOLECULE: CHECKPOINT PROTEIN RAD24;
   9:  5j1s-A  7.5  3.7  134   277    9   MOLECULE: TORSIN-1A;
  10:  3bos-B  7.5  3.4  120   231   11   MOLECULE: PUTATIVE DNA REPLICATION FACTOR;
  11:  7rzy-1  7.4  3.3  122   311    8   MOLECULE: TN6677 VIBRIO CHOLERAE TRANSPOSON TNSC (VCHTNSC);
  14:  6pxk-K  7.2  3.3  128   429    8   MOLECULE: ATP-DEPENDENT PROTEASE ATPASE SUBUNIT HSLU;
  15:  8thc-A  7.1  3.6  126   479   12   MOLECULE: ELG1 ISOFORM 1;
  16:  8glw-A  7.0  3.8  124   483   11   MOLECULE: TRANSPOSON TN7 TRANSPOSITION PROTEIN TNSC;
  17:  7ykk-A  7.0  3.7  128   735   14   MOLECULE: ATPASE FAMILY GENE 2 PROTEIN;
  18:  7uqj-A  6.9  3.7  130   585    8   MOLECULE: ATPASE HISTONE CHAPERONE YTA7;
  19:  5e7p-A  6.8  3.6  127   719   11   MOLECULE: CELL DIVISION CONTROL PROTEIN CDC48;
  20:  6n9x-B  6.8  3.4  115   500    8   MOLECULE: DNA PRIMASE/HELICASE;
  21:  7qo4-M  6.8  3.6  131   421    9   MOLECULE: 26S PROTEASOME REGULATORY SUBUNIT RPN1;
  22:  2qby-B  6.8  3.4  118   368    7   MOLECULE: CELL DIVISION CONTROL PROTEIN 6 HOMOLOG 1;
  23:  7mca-C  6.7  3.9  122   584    5   MOLECULE: ORIGIN RECOGNITION COMPLEX SUBUNIT 1;
  24:  9klv-D  6.7  3.6  132   338    5   MOLECULE: PROTEIN SUPPRESSOR OF MAX2 1;
  25:  9emc-D  6.7  3.4  124   337   10   MOLECULE: RUVB-LIKE 1;
  26:  8hra-C  6.6  3.5  126   852    8   MOLECULE: ARCHAEAL ATPASE;
  28:  5zr1-E  6.6  3.2  117   460    8   MOLECULE: ORIGIN RECOGNITION COMPLEX SUBUNIT 1;
  29:  6f0x-B  6.6  3.6  135   398    6   MOLECULE: PACHYTENE CHECKPOINT PROTEIN 2 HOMOLOG;
  30:  6p8v-C  6.6  3.7  145   303    6   MOLECULE: ATPASE, AAA FAMILY;
  31:  8xku-A  6.5  3.6  130   730   11   MOLECULE: PROBABLE INACTIVE ATP-DEPENDENT ZINC METALLOPROTE
  35:  8xku-D  6.4  3.5  123   805    6   MOLECULE: PROBABLE INACTIVE ATP-DEPENDENT ZINC METALLOPROTE
  36:  8tp1-A  6.4  3.5  126   419   12   MOLECULE: MITOCHONDRIAL CHAPERONE BCS1;
  38:  8btg-B  6.4  3.9  120   335   10   MOLECULE: CHROMOSOMAL REPLICATION INITIATOR PROTEIN DNAA;
  40:  8dgf-A  6.3  3.6  113  1541    9   MOLECULE: ATP-BINDING PROTEIN AVS4;
  41:  5zr1-A  6.3  3.7  127   494    5   MOLECULE: ORIGIN RECOGNITION COMPLEX SUBUNIT 1;
  44:  7plh-A  6.2  3.4  116   260    9   MOLECULE: SHTNSC;
  45:  8c0v-B  6.2  3.3  135  1030    6   MOLECULE: PEROXISOMAL ATPASE PEX1;
  46:  8bob-C  6.2  3.4  103   412   11   MOLECULE: PROTEIN MALY;
  47:  8k8v-A  6.2  3.4  116   719   13   MOLECULE: ENDOPEPTIDASE LA;
  48:  8dgc-A  6.2  3.9  123  2028   10   MOLECULE: SEAVS3;
  49:  7jgr-D  6.2  3.9  121   441    7   MOLECULE: ORIGIN RECOGNITION COMPLEX SUBUNIT 2;
  50:  8c0v-A  6.1  3.2  123   823    8   MOLECULE: PEROXISOMAL ATPASE PEX1;
  51:  1w5s-B  6.1  3.6  124   396    4   MOLECULE: ORIGIN RECOGNITION COMPLEX SUBUNIT 2 ORC2;
  53:  5j1s-B  6.1  3.8  134   235    7   MOLECULE: TORSIN-1A;
  55:  7w42-B  6.0  3.5  125   402   11   MOLECULE: UNCHARACTERIZED ATPASE YJOB;
  57:  8tw7-2  6.0  3.5  116   339    9   MOLECULE: REPLICATION FACTOR C SUBUNIT 5;
  58:  8xku-B  6.0  3.6  129   845   12   MOLECULE: PROBABLE INACTIVE ATP-DEPENDENT ZINC METALLOPROTE
  59:  9bf5-C  5.9  3.1  103  1190    4   MOLECULE: SSDNA1;
  60:  6ut5-A  5.9  3.5  112   395   11   MOLECULE: GTPASE SUBUNIT OF RESTRICTION ENDONUCLEASE;
  61:  6l1q-A  5.9  3.1  110   267    8   MOLECULE: CBBQ PROTEIN;
  62:  2gno-A  5.9  3.5  119   296    8   MOLECULE: DNA POLYMERASE III, GAMMA SUBUNIT-RELATED PROTEIN
  63:  9fnn-U  5.9  3.0  114   511   14   MOLECULE: CELLULOSE SYNTHASE CATALYTIC SUBUNIT [UDP-FORMING
  64:  3syk-A  5.9  4.2  140   290    8   MOLECULE: PROTEIN CBBX;
  65:  6bbm-D  5.8  3.2  114   455    9   MOLECULE: REPLICATIVE DNA HELICASE;
  68:  8xvb-A  5.8  3.4  112   246    5   MOLECULE: DNA (5'-
  69:  8ouy-C  5.8  3.4  117   308    9   MOLECULE: DNA REPAIR PROTEIN RAD51 HOMOLOG 2;

0001 s001A   -MLNT-DFNEL--MERVR-AGR-EFGH------AS---F--E-P-IFYLIFDP----QKI--LKIK-RQLPAWAAK-LRNE-------GW--DVH-IFS-MA-K-AVQEVF---D-E-MPVFNKSLAEALTKKN-ALQNKL-E-A--K-L-F-G-RP--N-S-I-LLVSD--I-E--A-L-HPY--L--R---I-GSMESQ-L--Q-GK-F-----H----VPTIFFYP-GM--RTGKQLKFLGFY--PE--D-GNYRS-V-HVG-G------
0002 9nv3A   pVLEY-RLSQV--QSRIS-EFL-KNNS------GN---E--I-G-FWIFDYPA----QNE--LQVR-EHLKYLLRN-LE-K-------DH--KFA-HLN-IF-Q-IIVDML---T-ErGLFDkqlvgLLNQ--K-KIADYI-A-KkvD-L-Q-N-----Q-E-F-VILTG--M-G--N-A-WPL--V--R-----G-HELMSA-L--Q-DV-M-----G---fTPLLMFYP-GT--YSGHDLSPLAGI--DS--R-NYYRA-FrLVP-Esgpaat
0003 8dr5A   cGNKG-SVMKL--KNWLA-NWE-NSKK------NS---FkhF-R-AAMLYGP-------P--GIG---KTTAAHLV-AQEL-------GY--DIL-EQN--A-S-DV----------------------rskT-LLNAGV-K-N--A-L-D-N-MK--H-F-V-IIMDE--V-D--G-M-SGG--D-rG---G-VGQLAQ-F--C-RK-T-----S----TPLILICN-ER--N----------L--PK--M-RPFVC-L-DIQi-------
0004 8fnuA   yIFSS-QLDKV--NLILE-HMD-TVGG------I----H--S-R-NIAITG-------DR--GTgktSFIETLKLV-LEKQ-------NY--YVFdIVS-P--T-VLSSH-------------------------LNILEI-V-I-----------K--D-L-V-LIIDD--L-D--L-V-EN---N--L---V-YDLLRD-I--Q-HY-L---dsQ----LIVIFAYK-E---------GQLEQ------lnhLTKLVsN-RIP-Lfkqdel
0005 8umyA   lSDDF-TNRCL--LKWLK-LWD-LGLD------PSqrpK--Q-K-VALLCGP-------P--GLG---KTTLAHVI-ARHA-------GY--SVV-EMN--A-S-DD---------------------rspE-VFRTRI-E-A--A-T-Q-M-EK--P-N-C-LVIDE--I-D--G-A-P----V--A---A-INVLLS-I--L-NL-L-----M----RPIICICN-D--------QFAP--------sL-RQLQA-F-LLH-Fpptlps
0006 7z6hK   avhkK-KIEEV--ETWLK-AQV-LERQ------PK---Q--GgS-ILLITG-------PP--GCG---KTTTLKIL-SKEH-------GI--QVQ-EW-----I-NPV------------------lpyQSQI--A-VFKEFL-L-R--A-T-K-ddTD--K-K-I-ILVED--L---pnqF-YRD--S-hT---L-HEVLRK-Y--V-RI-G-----R----CPLIFIIS-D-----slsgdnNQRL-lfP--K-EIQSI-S-NIS-Fnpvapt
0007 8fcvR   pRLKE-VYEIL--MRTIA-EPA---------------G--A-S-FIFVYGAS----GV----GKT-TLRLRVEQK-LTEL-------AarvPVV-GIE-A----IAPE-----------------SRYFN--W-KEYYTR-A-LitleepL-I-HR--H-PdV-FFVDE--A-Q--H-F-GKV----------lqDQLDC-L--K-SL-A--nmtG----ILHCLLGT-Y----------ELLT-frnL--S-GQLRS-V-DIH-Frrycad
0008 7st9A   kRKLK-DVQEA--LDAMF---L-PNA------------K--H-R-ILLLSGPS----GC----SKS-TVIKELSKI-LVPK-------YRqnsKV-TEF------RGDC-I---V-N-D---------LPQM--E-SFSEFL-K-G--A-R-Y-L-VM-nL-S-L-ILIED--L----P-N-VFH---idtrrrF-QQLILQ-W--L-YS---seplL----PPLVICIT------------ECEIpendnmN-KEILL-K-RIK-Fnpinst
0009 3bosB   aagND-ELIGA--LKSAG-DGV--------------------Q-AIYLWGPV----KS----GRT-HLIHAACAR-ANEL-------ER--RSF-YIP-LG-I--HASI--------------------------STALL-E----G-L-E-Q------fD-L-ICIDD--V-D--A-V-AGH--P--L---W-EEAIFD-L--Y-NR-V-----AeqkrGSLIVSAS-AS------pXEAG--------dL-VSRWG-L-TYQ-------
0010 6pxkK   iigQD-NAKRS--VAIAL-RNR-WRRR------HE---V--T-PkNILXIG-------PT--GVG---KTEIARRL-AKLA-------NA--PFI-KVE-AT---KFTEG-------------------------KEVDSI-I-R--DltdpsV-EQ--H-G-I-VFIDE--I-D--K-I-CVS--R--E--gV-QRDLLP-L--V-EGcD-----H----ILFIASGA-F-----------QIAK-----lI-PELRL-P-IRV-Elqaltt
0011 8thcA   pKLKL-RIKNW--IETSF-HTL-EKPT-------E---F--V-P-LMILHGN-------S--IGKK-TLIQTIMRE-IAGD-----dnSY--QIY-EVN--S-N-MN----------------------rsK-KDLLDilL-D--F-T-T-T-HD--Y-G-L-VLFND--V-D--V-L-FKE-----hdrgY-WAMISK-L--C-EF-S-----R----RPLVLTCK-D--------LSLV---------P-SEL-snS-LFH-Tkkists
0012 8glwA   lGTHL-LLSER--ISVMI-RGG-YVGA------RS---T--A-Q-SLLLIGCS----GS----GKT-TSLHRILAT--YPQ-------VeqvVYL-KID-CS---EICL--------------yerryglkrHGI-E----------------N-A-HA--L-G-L-LVIDE--I-Q--H-L-SRS--rsggsqeM-LNFFVT-M--V-NI-I-----G----VPVMLIGT-P----------KARE-------fE-ADLGF-G-AIF-Wdpiqqt
0013 7ykkA   dKEIE-SLKSA--IEIPL-HQP-TLFS---sfgVS---P--P-R-GILLHGPP----GT----GKT-MLLRVVANT-S-----------NA--HVL-TIN-GP-S-IV--------------------SKYLG-ETEAAL-R-D--I-F-R-K-YQ--P-S-I-IFIDE--I-D--S-I-AP-----------esRVVAT-L--L-TL-M--dgmgaagkVVVIAATN-R---------PNSV------------PALRD-Q-EVE-Igipdvd
0014 7uqjA   pLLAD-QLNNL--KNKLD-YML-NaES------QI---C--K-P-RLLINGPK----GN----GQQ-YVGAAILNY-LE----------EF--NVQ-NLD--L-A-SLVSES---S-------------------rTIEAAV-V-Q--S-F-K-K-RQ--P-S-V-VFIPN--LdI--W-I-NTI--P--E----vILVLSGlF--R-SL---qsneK----ILLLCLAEnLD--IS------------------SDFAFkN-IFQ-Lhkpske
0015 5e7pA   kvtHP-QAVKL--DEWLR-LSL-DEPEllktlgAT---P--H-L-GVLVSG-------PA--GVG---kATMVRAV-CA---------SR--RVV-ELD-GP-E-VGALQ---------------------vD-ERLRSV-T-S--A-V-A-T-ES--G-G-V-LFIAD--V-D--AlL-PAGnemrppepvA-TLILAE-L--R-KA-V-----A-tpgVAFIATSA-V--------PENV---------dA-RLRAD-R-ELG-Lslpdat
0016 7qo4M   dKQIE-ELVEA--IVLPM-KRA-DKFK----dmGI---R--A-P-KGALMYGP----PGT---GKT-LLARACAAQ-TN-------------ATF-LKL-AA-P-QLV-------------------QMYI-GEGA-KLVRDA-F-A-lA-K-E-K------aP-T-IIFID-eL-D--A-I--------------gtmLELLN-QldG-FS-S----dD----RVKVLAAT-N------------RVDV--LD--P-LRSGRlD-RKI-Efplpse
0017 2qbyB   pFRED-ILRDA--AIAIR-YFV-KNE---------------vkF-SNLFLGLT----GT----GKT-FVSKYIFNE-IEEV-------KV--KQA-YVN-CR-E-VG----------------------------gTPQAV-L-S--S-L-A----I--R-A-I-IYLDE--V-D--T-L---------vkrrgG-DIVLYQ-L--L-RS------daN----ISVIMISN-DI--N-------------vrD--Y-MESLG-P-SVI-Fkpydae
0018 7mcaC   dLKAE-ISDLL--YSE----TT-QK--------------R--R-C-FNTIFLL-----GSD--S--T-TKI--ELKD-E--S-----------SR--YNV-LIE-LTpK-ESP-----------------------nvR-MMLRRS-M-Y--K---V-E-NF-kD-L-A-MVFNF--K-DvdS-I-NFN--T--L------DNFIIL-L--K-SA-F------kydhVKISLIFN-I----------ntNLSNieKN--L-RIRLLrN-YHK-Ldvssnk
0019 9klvD   nDAAA-AVAAT--VSQCK-LGN----V------LS---K--G-D-VWLLFSG-----PDR--VGKR-KMVSALSSL-V-----------Y--GTN-PIM-IQ---agDGNS---S-F-R---------GKTA-lD-KIAETV-K-R--S-P---------------F-SVILL--E-D-ideA-DML--V--R-gsI-KQAMDR-G--RihS-L-----G----NVIFVMTA-SWhkLSLQ-----cvP--FA--F-HDMRV-D-DAV-Afra---
0020 8hraC   lIQKS-LRDQL--VESIR-NSI-A---------------G--R-N-VFFVDGTR----GA----GKT-TFINSVVKS-LNSD-----qdDV--KVNiKCL-PT----IDPT---------------------------K-LPRHEP-I-L---------eiCK--A-I-L-ITFDD--I-D--T-Q-F----D--A---G-WDVLES-I--R-KF-F-----N-srkLVVVATGD----------LRLYsQL--IRgsvrLLKLFqK-RIQ-Lktmlql
0021 7crcB   vGMRA-HMDML--EQLLR---L-VLD------------E--V-R-MIGIWGPP----GI----GKT-TIARFLFNQ-VS-D-------RF--QSA-IMVnIK-G-CYP---------------PCFDEYSAQ--L-QLQNQM-L-S--Q---------K--K-V-F-LVLDE--V-D--Q-L------G--Q---L-DaLAKE----T-RW-F-----G--pgSRIIITTE-D-------------------------KAHGInH-VYK-Vgypsnd
0022 5zr1E   eYQTN-CLASY--IS--A-DPD-I-----------------tpS-NLILQGYS----GT----GKT-YTLKKYFNA-N--P-------NL--HAV-WLE-PV-E-LV-------------------swK-PLLQAI-A-R--T-V-Q----T--C-L-F-LILDG--F-D--S-L--QD--L--D-----aALFNK-Y--I-KL-NellpkdskinIKFIYTML-------------ETSF-lqR--Y-STHCI-P-TVM-Fprynvd
0023 6p8vC   vGLDE-HKDRL--SKILG-LnP-YGIV------DT---V--R-R-PPLVVLAG----DV---SGKT-ELAETIGDA-VARQ------eDI--DIT-LYP-LS----LATRG------------RVGEXTQLV--S-AAFDYT-I-E--A-A-D-L-KK--ArG-A-VLLLIdqA-D--A-L-AaG--V--N---AfIRGIDR-I--A-NQ-K-----L----P-AAVLXC-T-----------nRLKA--lD--P-AVRRA-A-EIL-Tfsrpnd
0024 8xkuA   eYIKR-ELQEI--VRILK-N-D-EEFQ-------nkgiY--C-P-KGVLLHGP----PG----TGKT-LLAKAIAGE-AG-------------LPF-FAA-NG-T-DF--VE---M-F---------VGVA--A-SRVKDL-F-A-sS-R-S-Y------aP-S-IIFID-eI-D--A-I-GS-----------glLQILT-E-mD-GF-K--vttS----QVLVIGAT-N-----------RLDI--LD--P-ARKGRfD-KII-Rvglpsk
0025 8xkuD   qPGTI-YLRYL--VDIHKkYLM-NYEF----ntsclaerritypsRSILVIGS----I----TGRS-YLVKYLATN-SY-----------V--PFI-TVF-LN---KFLDNK-----------------dmmLEID-RfITLQ-F-E--L-A-K-A-MS--P-C-I-IWIPN--I-H--D-L--------------aLGLLVN-S-lS-RD-C----ercstrnILVIAST-H-------------IPQK-----vD-PALKL-N-TCI-Kirrlli
0026 8btgB   sGNRF-AHAAS--LAV-A-PAK-------------------ayN-PLFIYGGV----GL----GKT-HLMHAIGHY-VIDH-------NA--KVV-YLS-SE-K-FTNEFI---NsI-RDN-----------------kavdfrnR--Y-R-N-V--------D-V-LLIDD--I-Q--F-L-AGK--E--------tqEEFFH-T-----NT-L------H-eesKQIVISSD-RP--------PKEI-----------lR-SRFWG-L-ITD-Itppdle
0027 4hutA   --------yqqrqqkvkvdaaqaQEER--------------gI-IIVFT-GN----GK----GKT-TAAFGTAAR-AVGH-------GK--NVG-VVQ-FI-KgVMATGF-------------R-EADTAA--cM-AVWQHG-K-R--MlA-D-P-L----lD-M-VVLDE-lT-Y--M-VaYDY--L--P------LEEVIS-A--L-NA-R------P--ghQTVIITGR-G------------------------cH-RDILA-D-TVS-Elrpvkh
0028 8dgfA   iDRRH-CIELL--HENLV-QK----------------------K-IVIVSGEG----GV----GKT-AVIKKIYEA-EKQ-------YT--PFY-VFK-A--S-EFK-----------------KDSI-NELF---------------ahEL--R-K-V-IVVDS--A-E--K-L-LEL-----------idpFKEF-L--T-VL-I---kdK----WQVVFTTR-N----------NYLA------------YAFIP-G-NLV-Iknlerg
0029 5zr1A   eNEFA-SIYLS--AYSAI-ESD---------------------sA-TTIYVAGT----PG---VGKT-LTVREVVKE-LLSS-------SD--FLY-VEI-NG-L--KMVK-------------------------ptDCYETL-W-N--K-V---vpKK--K-T-I-VVLLD-eL-D--A-M-VTKsqD--I---M-YNFFN--------YE-N------A-----KLIVIAVA-NT--M------DLPER--qlG--N-KITIG-F-TRI-Mftgyth
0030 8e2wA   eVLRP-LLKQA--AEKTA-AG-----------------------kIVFAEAAT----GT----GKG-RMIASLAAA-AAIK-------GD--TVV-VSA-PL-AvltlslgrpnF-I-SPqrLLLAEDLpadsllllthFML-A-A------L-P-HF--I-D-T-LIVDE--A------------------hL---L-EQAfaA-L--L-GL-W------N--atGGATLVSA-TF--TTGDNG---------------SLTRA-A-FLP-Pvhpawt
0031 8rwvC   nLFDN-LIEFL--QKSHS-GFQ-KNSG------QI---K--L-reIPTAALVL----GvnvtLTFG-SLTEALQNN-V-----------------PY-VVS-LQ----AKDC--------------------PDMK--HFLQKL-I-S-------W-Y-MTvqS-P-P-VVVI---L-K--D-M-ESF--AtkV---L-QDFIII-S--S-QH-L-----hE----FPLILIFG-I-----------atSPIIihRL--L-PSSLL-C-IEL-Fqslsck
0032 7plhA   qQVKT-LHDWL--DGKRK-AR-----------------------K-SCRVVGES----RT----GKT-VACDAYRYR-HKPqqeagrppTV--PVV-YIR-PH-Q-KC-----------------------------GPKDL-F-K--K----L-K-GC--G-VeM-LIIDE--A-D--R-L-----------------kPETFAD-V--R-DI-A--edlG----IAVVLVGT-D----------RLDA--vikR--D-EQVRF-R-AHL-Rfgklsg
0033 8c0vB   lhiFP-YVRQL--VNILE-TSF-NCSG------IT---L--N-A-SVLLHST------TN--NVG---KATMVRFA-SKYL-------GI--HLL-EI--DC-L-S--LTS---N---------SRQLDST--S-KIIGYI-RaK--C-E-N-VlAS--P-A-V-IFLAH--LdS--I-L-LviklqksI---N-FEMSKL-L--D-DF-T------fkfpgTTFVGSVN-N---------IDNV---------P-SSFHM-R-FEI-Lvpvpse
```

```
0034 8bobC   [illegible]
0035 7jgrD   [illegible]
0036 8c0vA   [illegible]
0037 1w5sB   [illegible]
0038 6fhsC   [illegible]
0039 5j1sB   [illegible]
0040 3zh9B   [illegible]
0041 7w42B   [illegible]

0001 s001A   [illegible]
0002 9nv3A   [illegible]
0003 8dr5A   [illegible]
0004 8fnuA   [illegible]
0005 8umyA   [illegible]
0006 7z6hK   [illegible]
0007 8fcvR   [illegible]
0008 7st9A   [illegible]
0009 3bosB   [illegible]
0010 6pxkK   [illegible]
0011 8thcA   [illegible]
0012 8glwA   [illegible]
0013 7ykkA   [illegible]
0014 7uqjA   [illegible]
0015 5e7pA   [illegible]
0016 7qo4M   [illegible]
0017 2qbyB   [illegible]
0018 7mcaC   [illegible]
0019 9k1vD   [illegible]
0020 8hraC   [illegible]
0021 7crcB   [illegible]
0022 5zr1E   [illegible]
0023 6p8vC   [illegible]
0024 8xkuA   [illegible]
0025 8xkuD   [illegible]
0026 8btgB   [illegible]
0027 4hutA   [illegible]
0028 8dgfA   [illegible]
0029 5zr1A   [illegible]
0030 9e2wA   [illegible]
0031 8rwvC   [illegible]
0032 7plhA   [illegible]
0033 8c0vB   [illegible]
0034 8bobC   [illegible]
0035 7jgrD   [illegible]
0036 8c0vA   [illegible]
0037 1w5sB   [illegible]
0038 6fhsC   [illegible]
0039 5j1sB   [illegible]
0040 3zh9B   [illegible]
0041 7w42B   [illegible]
```

### Job: Type-3 BREX; BrxF Inactive STAND (BrxB-Homolog)
### Query: s001A

| No: | Chain | Z | rmsd | lali | nres | %id | PDB Description |
|---|---|---|---|---|---|---|---|
| 1: | 8fcv-R | 9.6 | 3.3 | 127 | 343 | 13 | MOLECULE: DNA (60-MER); |
| 2: | 2qby-B | 9.4 | 2.9 | 124 | 368 | 12 | MOLECULE: CELL DIVISION CONTROL PROTEIN 6 HOMOLOG 1; |
| 3: | 5e7p-A | 9.4 | 3.3 | 130 | 719 | 15 | MOLECULE: CELL DIVISION CONTROL PROTEIN CDC48; |
| 4: | 8umy-A | 9.2 | 2.8 | 120 | 546 | 17 | MOLECULE: CHROMOSOME TRANSMISSION FIDELITY PROTEIN 18 HOMOL |
| 5: | 8dr5-A | 9.1 | 3.0 | 126 | 646 | 10 | MOLECULE: REPLICATION FACTOR C SUBUNIT 1; |
| 6: | 3bos-B | 8.9 | 2.8 | 113 | 231 | 9 | MOLECULE: PUTATIVE DNA REPLICATION FACTOR; |
| 7: | 9nv3-A | 8.7 | 3.1 | 136 | 199 | 14 | MOLECULE: DUF1788 DOMAIN-CONTAINING PROTEIN; |
| 8: | 7st9-A | 8.5 | 3.4 | 139 | 522 | 11 | MOLECULE: CHECKPOINT PROTEIN RAD24; |
| 9: | 7crc-B | 8.5 | 3.8 | 125 | 1112 | 14 | MOLECULE: NAD+ HYDROLASE (NADASE); |
| 10: | 6z1f-3 | 8.5 | 3.1 | 128 | 284 | 14 | MOLECULE: RIBULOSE BISPHOSPHATE CARBOXYLASE/OXYGENASE ACTIV |
| 11: | 8btg-B | 8.4 | 3.7 | 126 | 335 | 13 | MOLECULE: CHROMOSOMAL REPLICATION INITIATOR PROTEIN DNAA; |
| 12: | 7mi8-A | 8.3 | 3.4 | 122 | 868 | 11 | MOLECULE: FUSION PROTEIN OF DYNEIN AND ENDOLYSIN; |
| 13: | 7plh-A | 8.3 | 3.4 | 120 | 260 | 17 | MOLECULE: SHTNSC; |
| 14: | 9cru-C | 8.3 | 3.3 | 133 | 741 | 17 | MOLECULE: VESICULAR-FUSION PROTEIN SEC18; |
| 15: | 7jgr-D | 8.2 | 3.4 | 123 | 441 | 16 | MOLECULE: ORIGIN RECOGNITION COMPLEX SUBUNIT 2; |
| 16: | 6bbm-D | 8.2 | 2.7 | 116 | 455 | 18 | MOLECULE: REPLICATIVE DNA HELICASE; |
| 17: | 8ipr-A | 8.2 | 3.4 | 110 | 504 | 23 | MOLECULE: COMPONENT LINKED WITH THE ASSEMBLY OF CYTOCHROME' |
| 18: | 8dgc-A | 8.1 | 3.5 | 121 | 2028 | 11 | MOLECULE: SEAVS3; |
| 19: | 8c0v-A | 8.1 | 3.3 | 134 | 823 | 13 | MOLECULE: PEROXISOMAL ATPASE PEX1; |
| 20: | 5zr1-E | 8.1 | 3.3 | 124 | 460 | 15 | MOLECULE: ORIGIN RECOGNITION COMPLEX SUBUNIT 1; |
| 21: | 8xvb-A | 8.0 | 3.0 | 116 | 246 | 16 | MOLECULE: DNA (5'- |
| 22: | 5zr1-A | 8.0 | 3.3 | 127 | 494 | 12 | MOLECULE: ORIGIN RECOGNITION COMPLEX SUBUNIT 1; |
| 23: | 5yww-A | 8.0 | 3.0 | 106 | 485 | 18 | MOLECULE: NUCLEOTIDE BINDING PROTEIN PINC; |
| 24: | 7vcs-A | 8.0 | 3.4 | 133 | 766 | 12 | MOLECULE: TRANSITIONAL ENDOPLASMIC RETICULUM ATPASE; |
| 25: | 9bv1-A | 7.9 | 3.2 | 124 | 405 | 16 | MOLECULE: 26S PROTEASOME NON-ATPASE REGULATORY SUBUNIT 1; |
| 26: | 6n9x-B | 7.9 | 3.2 | 117 | 500 | 11 | MOLECULE: DNA PRIMASE/HELICASE; |
| 27: | 5j1s-A | 7.9 | 3.4 | 129 | 277 | 9 | MOLECULE: TORSIN-1A; |
| 28: | 4rh7-A | 7.8 | 3.2 | 121 | 3005 | 13 | MOLECULE: GREEN FLUORESCENT PROTEIN/CYTOPLASMIC DYNEIN 2 HE |
| 29: | 9emc-D | 7.8 | 3.1 | 128 | 337 | 17 | MOLECULE: RUVB-LIKE 1; |
| 30: | 1nlf-A | 7.8 | 2.8 | 108 | 254 | 13 | MOLECULE: REGULATORY PROTEIN REPA; |
| 31: | 9e22-A | 7.8 | 3.3 | 116 | 2925 | 11 | MOLECULE: CYTOPLASMIC DYNEIN 1 HEAVY CHAIN 1; |
| 32: | 4m9s-B | 7.7 | 2.9 | 111 | 511 | 10 | MOLECULE: CELL DEATH PROTEIN 4; |
| 33: | 6f0x-B | 7.7 | 3.2 | 138 | 398 | 15 | MOLECULE: PACHYTENE CHECKPOINT PROTEIN 2 HOMOLOG; |
| 34: | 7uqj-A | 7.7 | 3.4 | 124 | 585 | 8 | MOLECULE: ATPASE HISTONE CHAPERONE YTA7; |
| 35: | 6p8v-C | 7.6 | 3.3 | 138 | 303 | 9 | MOLECULE: ATPASE, AAA FAMILY; |
| 36: | 6dgd-A | 7.6 | 3.6 | 115 | 704 | 11 | MOLECULE: PRIMOSOMAL PROTEIN N'; |
| 37: | 8zr0-A | 7.6 | 3.3 | 124 | 480 | 11 | MOLECULE: CASEINOLYTIC PEPTIDASE B PROTEIN HOMOLOG; |
| 38: | 8y6p-Q | 7.5 | 3.0 | 112 | 1228 | 11 | MOLECULE: APAF-1 RELATED KILLER DARK; |
| 39: | 7t8b-A | 7.5 | 2.5 | 104 | 518 | 13 | MOLECULE: TWINKLE MTDNA HELICASE; |
| 40: | 5juy-B | 7.5 | 3.5 | 106 | 1234 | 16 | MOLECULE: APOPTOTIC PROTEASE-ACTIVATING FACTOR 1; |
| 41: | 8gja-F | 7.5 | 3.3 | 126 | 342 | 13 | MOLECULE: RAD51C; |
| 42: | 8ui7-A | 7.4 | 3.0 | 122 | 643 | 14 | MOLECULE: ATPASE FAMILY AAA DOMAIN-CONTAINING PROTEIN 5; |
| 44: | 7qo4-M | 7.4 | 3.3 | 130 | 421 | 15 | MOLECULE: 26S PROTEASOME REGULATORY SUBUNIT RPN1; |
| 45: | 6qem-G | 7.4 | 4.1 | 135 | 239 | 12 | MOLECULE: REPLICATIVE DNA HELICASE; |
| 46: | 9ri9-B | 7.4 | 3.1 | 109 | 868 | 11 | MOLECULE: SLNRC3; |
| 47: | 8jx6-B | 7.4 | 3.5 | 122 | 451 | 11 | MOLECULE: DEEP-SEA HELICASE 9; |
| 48: | 4wia-A | 7.4 | 3.2 | 115 | 229 | 17 | MOLECULE: PUTATIVE FLAGELLA-RELATED PROTEIN H; |
| 49: | 8tp1-A | 7.3 | 3.3 | 122 | 419 | 15 | MOLECULE: MITOCHONDRIAL CHAPERONE BCS1; |
| 50: | 3j3r-A | 7.3 | 3.7 | 133 | 798 | 17 | MOLECULE: ADAPTER PROTEIN MECA 1; |
| 51: | 8xks-B | 7.3 | 19.1 | 127 | 603 | 13 | MOLECULE: CTAP1; |
| 52: | 7yqh-B | 7.3 | 3.0 | 123 | 1066 | 15 | MOLECULE: STRUCTURAL MAINTENANCE OF CHROMOSOMES PROTEIN 5; |
| 54: | 7qo4-J | 7.3 | 3.3 | 128 | 405 | 12 | MOLECULE: 26S PROTEASOME REGULATORY SUBUNIT RPN1; |
| 55: | 8bob-C | 7.2 | 3.3 | 112 | 412 | 14 | MOLECULE: PROTEIN MALY; |
| 56: | 7rzy-1 | 7.2 | 3.3 | 121 | 311 | 12 | MOLECULE: TN6677 VIBRIO CHOLERAE TRANSPOSON TNSC (VCHTNSC); |
| 57: | 8xku-A | 7.2 | 3.3 | 133 | 730 | 8 | MOLECULE: PROBABLE INACTIVE ATP-DEPENDENT ZINC METALLOPROTE |
| 58: | 7lcc-A | 7.2 | 3.5 | 118 | 1369 | 6 | MOLECULE: HELRAISER K1068Q; |
| 59: | 6l1q-A | 7.2 | 3.0 | 115 | 267 | 14 | MOLECULE: CBBQ PROTEIN; |
| 60: | 8hqs-B | 7.1 | 3.0 | 123 | 408 | 15 | MOLECULE: STRUCTURAL MAINTENANCE OF CHROMOSOMES PROTEIN 5; |

```
0001 s001A   [illegible]
0002 8fcvR   [illegible]
0003 2qbyB   [illegible]
0004 5e7pA   [illegible]
0005 8umyA   [illegible]
0006 8dr5A   [illegible]
0007 3bosB   [illegible]
0008 9nv3A   [illegible]
0009 7st9A   [illegible]
0010 7crcB   [illegible]
0011 6z1f3   [illegible]
0012 8btgB   [illegible]
0013 7mi8A   [illegible]
0014 7plhA   [illegible]
0015 9cruC   [illegible]
0016 7jgrD   [illegible]
0017 6bbmD   [illegible]
0018 8iprA   [illegible]
0019 8dgcA   [illegible]
0020 8c0vA   [illegible]
0021 5zr1E   [illegible]
```

```
0022 8xvbA   pprfveTQ--TVKQIWTSMRF--AS---LT--E--S--IAVVCGN-PGVGKTEAAREYRRT--N----N-----nV-WMITIT--PSC-----------------aSV-L-ECL-T-ELAFEL---GL-VIIDE-A-D-HLG-----------E--V-LEELRLLQER-IGLVLMGNH---------------rvySNM---TF-S-R-IKRTA-I----NKTKkadvka
0023 5zr1A   yylparEN--EFASIYLSAYS--AE---SD-S---A--TTIYVAGtPGVGKTLTVREVVKE--L----LreipdfL-YVEING--LKM-VK-----------------pT-DCY-E-TLWNKV---TI-VVLLDeL-D-AMVT-K-----S--QD--I-MYNFFNWTTA-KLIVIAVANT-M------DLPE--R---QLG---NS-R-I-GFTRI-M----FTGYtheelk
0024 5ywwA   ---eDYNL---derlIKRLEE--R----------aE--GIIIAGA-PGMGKTTFAQALAEY--Y----M-rlgkiV-KTIES-------------------------G-ELH-D-ILLLSR-P-DY-TVYDE-M-R-N------------DE--D-FKLYVDLRLG-VGMVGVVHAT-S-----PIDA------iHRF---V-gtiL-DTIIF-I----NSG-------
0025 7vcsA   ---gGCRK--QLAQIKEMVEL--PL-igVK-PP-----RGILLYGpPGTGKTLIARAVANE--T----G-----aF-FFLING--PEI-MSK----------LAGE-SE-S-NLR-K-AFEEAN-A-PA-IIFID-E-LdAIAP-kE--R-R--IVsqL-LTLMDGL--A-HVIVMAATN----------RPNS-I---DPA----R-F-G-RDREV-D----IGIPdatgrl
0026 9bv1A   sdvggcKE--QIEKLREVVET--PL-lgIE-PP--K--GVLLFGP-PGTGKTLCARAVANR--T---------dacF-IRVIG---SEL-V--QK-YV-------GE-GA-R-MVR-E-LFEMAK-A-CL-IFFDE-I-D-AIG-gDN--E-V--QrtmlelINQLD--GN-IKVLMATNRP-------------dT---LDP----R-P-G-RRKIE-F----SLPDlegrth
0027 6n9xB   ---gCTGI---------ntLG--AR--------ggE--VIMVTSG-SGMGKSTFVRQQALQ--W----GtamgkkV-GLAMLQ--E-----------sdslkreiidsT-D-RLL-A-KLAYMG-C-DV-IILDH-I-S-IVVS-SD-eR-K--MIdnL-MTKLKGFAKG-VVLVVICHL----pVSITDLR---G---SGA----R-Q-L-STIIA-L----ERNQqgdmpn
0028 5j1sA   dnlfgqHL--AKKIILNAVFG--FI-pkPK-KP--L--TLSLHGW-TGTGKNFVSKIIAEN--I----YglnsdyV-HLFVAT--LHF-PH---------aSNITL-YK-D-QLQ-L-WIRGNV-R-SI-FIFDQ-MdK-MHA-----------GL--I-D-AIKPFLD-kAMFIFLSNAG-wraLSVSV------fnnKNS---GwrN-L-IDFVP-F----LPLEykhlkm
0029 4rh7A   sklvyTPL--T-DKCYLTLTQ--AM---KM-GL-----GGNPYGP-AGTGKTESVKALGGL--L--grQ------V-LVFNCD-------------------------iDV-K-SMG-R-IFVGLC-G-AW-GCFDE-F-N-RLE-----------eSV--L-SAVSMQIQTInSGIFITMNP-----------AGK-R---QKL---PDlK-Q-LFPVA-M----SHPDneliae
0030 9emcD   qgmvgqLA--ARRAAGVVLEM---G---KI-A--gR--AVLIAGQ-PGTGKTAIAMGMAQA--L----G--pdtpF-TAIA-G--SEI---FSL-E---------mS-KT-E-ALT-Q-AFRRSIgP-GV-LFIDE-V-H-MLD---------I--ES--F--SFLNRALEA-PVLIMATNR-----------GIT-SphgIPI---D-lD-R-LLIVS-T----TPYSekdtkq
0031 1nlfA   ileafaaappldyvLPNMV-------------agT--VGALVSP-GGAGKSMLALQLAAQ--I----AelptgpV-IYLPAE-rqLI-GSL-------pnimAPE-WF-D-GLK-R-AAE--G-R-RL-MVLDT-L-R-RFHI-E-----eenaQ--V-IGRMEAIAAG-CSIVFLHHA---------------------VL---VD-N-I-RWQSY-L----SSMTsaeaee
0032 9e22A   ---qTPLTdrCYLTMTQALEA--R-----------L--GGSPFGP-AGTGKTESVKALGHQ--L----G------RfVLVFNC--D-E-TFD----------------F-Q-AMG-R-IFVGQV-G-A--WGCFDeF-N-RL----E--E-R--MA--V-SQQVQCIQEA-LAIFITMNP------------------gsnlP---DN-L-KlFRSLA-M----TKPDrqliaq
0033 4m9sB   xtcyirEY--HVDRVIKKLDE--XC---DL-DS--F--FLFLHGR-AGSGKSVIASQALSK--S-----ginydsI-VWLKDS-----------------gtAPK-ST-F-DLF-T-DILLXL---TL-FVFDD-V-V-QE------------E--T-IRWAQE--LR-LRCLVTTRDV-E-----------------------S-Q-T-CEFIE-V----TSLEidecyd
0034 6f0xB   ---yDVEV---KSHLLDYVMT--TL---LFsTW--N--RVVLLHGpPGTGKTSLCKALAQK--L----TssryryG-QLIEIN--S---HSLFS-K--WFSES-GK-LV-T-KMF-Q-KIQDLI-A-LV-FVLIDqV-E-SLTnaAI--R-V--VN-aV-LTQIDQIKRHsNVVILTTSN----------ITEK-I---DVA-----vD-R-ADIKQ-Y----IGPPsaaaif
0035 7uqjA   ---kPLLA-dQLNNLKNKLDY--MLnesQI-CK--P--RLLINGP-KGNGQQYVGAAILNY--L----E--------nVQNLDL--ASL-VS-ES-SR--------T-IE-A-AVV-Q-SFMEAQ-P-SV-VFIPN-LdI-WINT--P--E-N--VI-----------rEK-ILLLCLAENL-D--I-------------sEV---KNaF-D-KNIFQ-L----HKPSkenitr
0036 6p8vC   arlvglDE--HKDRLSKILGL--LV-dtVL-RR--P--PLVVLAGdVGSGKTELAETIGDA--V----ArqedidI-TLYPL----SL-AT-RG-QG-RV-GEXTQ-LV-S-AAF-D-YTIEAA-G-AV-LLLIDqA-D-ALAQsDR--A-G--VN-aF-IRGIDRIANQlPAAVLXCTN----------RLKA-L---DPA-----qR-R-AAEIL-T----FSRPndeqrh
0037 6dgdA   erlrlNTE--Q-ATAVGAIHS--AA---DR--F--S--AWLLAGI-TGSGKTEVYLSVLEN--V----L-aqgrqA-LVM-VP--EIG-L-------fnapvevlhivigt-SSL-F---TPFK-L-GV-IVIDE-EhD-SSYK-QQ-egwR--YH--A-RDLAVWRAHQ-IPIILGSATP-----------------aLET---LH-NvK-YRQLT-L----S---------
0038 8zr0A   ehiigqEG--AINTVASAIRR--KE-ngWY-DEehP--LVFLFLGsSGIGKTELAKQVARY--M----H-------kGFIRMD--------------fqEKHEVAK-FI----LT-K-QLKQSP-S----AVVLF-D-EvekAH-PD--V---lTV--M-LQLFDEG--K-DAIFIMTSNAadvvIRPI-----aH---FRR---DL-G-R-INEIV-Y----FLPFchsell
0039 8y6pQ   kynvsrLQ--PYLKLRQALLE--LR------PA--K--NVLIDGV-LGSGKTWVALDVCLS--Y-----ckmdfkI-FWLNLK--N---------------cnSPE-TV-L-EML-Q-KLLYQI---CL-LVLLN-V-Q-NA-------------K--A-WNA--FNLS--CKILLTTRFK---------------qvtDFL---SA-A-T-TTHIS-LdhhsMTLTpdevks
0040 7t8bA   snveqaaglrwsrfpdlnlKG--HR--------kgE--LTVFTGP-TGSGKTTFISEYALD--L----C-sqgvnT-LWGSFE--ItqtFHGQ--------------qSI-R-TVI-D-TMQHAD-I-CH-VIIDN-L-Q-FMM---------tdrY--I-IGVFRKFATN-CHVTLVIHPG---------------------S---AK-A-A-DNVLI-L----QDRGkrylqv
0041 5juyB   vvfvtRKK--LVNAIQQKLSK--L----KG-EP--G--WVTIHGM-AGCGKSVLAAEAVRD--------gcfpggV-HWVSVG--K---------------qDKSG-LL-M-KLQ-N-L--qsP-R-SL-LILDD-V-W-DS-------------W--V-LKAFD---SQ-CQILLTTRDK-S-------------------------tdsP-KYVVP-V----ESSLgkekgl
0042 8gjaF   ---cQQLD---sflrgGILT--------------rT--LTEIAGE-SASGKTQLCMQLCLT--V----QgglgggA-VYICTE-eHAA-E---------------lD-DL-H-YCVsK-KVPVLL-V-KL-IIIDS-IaA-LFRC-EL--Q-E--RL--M-QLIASKLLQL-APAICVNQVS--------dVVEQ-P---TLG---IA-N-H-VVRLM-L----MRTNyklpvq
0043 8ui7A   selignEL--AIKKLHSWLKD--WK-eeSR-LC--N--TVLITGP-TGVGKTAAVYACAQE--L---gF------K-IFEVNA--SSQ------------------rSG-R-QIL-S-QLKEA--T-SL-ILFEE-V-D-V-IF-DE--D-A--GF--L--NAIKTFMTK-RPVILTTSDP-T---------------fsLMF---D--G-C-FEEIK-F----STPSllnvas
0045 7qo4M   ---gGLDK--QIEELVEAIVL--PM-mgIR-AP--K--GALMYGP-PGTGKTLLARACAAQ--T----N----atF-LKLAA---PQL-V----------qMYIGE-GA-K-LVR-D-AFALAK-A-PTiIFIDE-L-D-AIgD-RE--VqR--TM--LelLNQLD-DDR-VKVLAATNRV-------------dV---LDP-alLR-S-GlDRKIE-F----PLPSedsraq
0046 6qemG   ---eCEGQmnALSKARQYVEE--FD---GN--I--A--SFIFSGK-PGTGKNHLAAAICNE--L----L-lrgksV-LIITVA--DIM-SAMKD-tF-RNS-----sEE-Q-LLN-D-LSN----V-DL-LVIDE-I-G--vQT-ES--K-YekVI--I-NQIVDRRSSK-RPTGMLTNSN-M------eEMTK-L---LGE----M-D-R-MSLWV-I----FNWDsyrsr-
0047 9ri9B   ddvvgfDD--EAKTVIDRLIG---G---SD-YV--E--VVPVVGM-PGLGKTTLAYKIYKD--P-----eyefftR-VWVYVS--QTF-------------------KR-R-EIF-L-NIISK----YL-IVLDD-V-W-T--------------E--A-WDRIKIAFPNrNRVLMTTRQS-N------------------------aK-R-CNPHD-L----KFLTkdeswe
0048 8jx6B   sateeqRY--IFTDFIKQAQL--AL---KS-DY--E--VFAIQGF-AGSGKTWLSALIIDE--L----L-elgmkV-AVTS-P--THK-A---------------pgtiH-HFL-NlnvtLEY-V-DV-LIVDE-A-S-MVS-----------GE--L-YDHALKTLGDcKIILFIGDSY----------QLL-E---DS-sifLK-D-D-IFHYK-L----TKVVrqaedn
0049 4wiaA   ---elaridlsrddldkrgGI--PH-------G--S--LIIIEGE-ESTGKSVLCQRLAYG--F----L-qnrysV-TYVST--iPVY-PL------iadnkkkDG-FL-K-KVX-E-T-RAFY-K-DV-IIFDS-IsA-LIAN--A--S-E--VN--V-DDLXAFFKRIkKIIICTVNPK----------ELPE-S---VLT-----rT-S-ATLIR-T----ELFTfggdlk
0050 8tp1A   svvlqQGL--A-DRIVKDIRE--FI---DN-PY--R--RGYLLYGpPGCGKSSFITALAGE--L----E------H-SICLLS-----tDSSL----------sdD-RL-N-HLL-S-VA-----P-QQ-SLVLL-E-D-vdAA-FLsrL-T--FS-gL-LNALDG--TE-ARIVFMTTN----------YIDR-L---DPA----R-P-G-RDLKE-Y----VGYCshwqlt
0051 3j3rA   srvigqDE--AVVAVAKAVRR--AK---DP-KR--PigSFIFLGP-TGVGKTELARALAES--IfgdeE------S-MIRIDM--SEYstsrlV-GS-PP-GYVGY-DEgG-QLT-E-KVRRK--P-YS-VVLLD-AiE-KAH-----------PD--V-FNILLQVLEDrNTILIMTSNVgM--GELK-----rA---FRP-----iN-R-IDIIV-F----HSLEkkhlte
0052 8xksB   ---lPEGNkqAKVELMEVVDF-----sgAR-AP--K--GVLLVGP-PGNGKTLMARAVAGE--S---gV------A-FISSSA--AEF-I-----------eMYMGL-GA-A-RVR-D-LFNTAA-P-CI-IFIDE-L-D-AVG--ND--E-R--D---ntvNQLLTEMD-gIVVMGATNRK-D-------------V---LDA----R-P-G-RRSIE-V----RRPDfqgrle
0053 7yqhB   ---hEHFE-------LELG-------------S--R--LNFIVGN-NGSGKSAILTAITIG--L----GsslkdlimCFLSAR--SFL---TAS-TslSGGEKSFS-QM-A-LLL-A-TWKPMR-S-RI-IALDE-F-D-VFMD-Q---V-N--RK-iG-TTLIVKLKDIrTQTIIITPQD-------------------igKI---AD-I-G-VSIHR-M----RDP-------
0054 5v8fD   ---fTYLQ-dCQQEIDRIIKQ--SI---I--eS--H--SVILVGP-RQSYKTYLLDYELSL--L----Q--------iTIRLNG--FIH-S---------------eQ-TA-I-NGI-A-TQL----I-TV-VFIFDeI-D-TFAG-PV----R--QT--L-LYNLFDMVEV-PVCIFGCTTK-L------NILE--L---EK------rF-S-QRVIY-M----PQIQnlddmv
0055 7qo4J   ---gGLTK--QIKEIKEVIEL--P---lgIA-QP--K--GVILYGP-PGTGKTLLARAVAHH--T----D------C-KFIRVS---gA-ELVQK-Y--------igE-GS-R-MVR-E-LFVMAH-A-PS-IIFMD-E-I---DS-IG-sggG--DS--EvQRTMLELLN-nIKIIM-ATNR-L-----------dI---LDPallRP-G-R-IRKIE-F----PPPSvaarae
0056 8bobC   dhtvvRER--LLAK-LSGA------------NF--R--LALITSP-AGYGKTTLISQWAAG--K----N-----dI-GWYSLD-----eGDNQ-----------qE-RF-A-SYL-I-AAVQQA---LY-LVIDD-Y-H-LITN-----P-V--IHesM-RFF-----EN-LTLVVLSRNL-----------------pqlGI---AN-LrQ-LLEIG-S--qqLAFThqeane
0057 7rzy1   rafvsTPS--V-RKILSYMDR-cRD---LS-EP--T--CMMVYGA-SGVGKTTVIKKYLNQ--N----RggdiipV-LHIELP----------------------aKP-V-DAA-R-ELLVEM-V-KL-IIIDE-F-Q-HLVE-NR--V-L--TQ--V-GNWLKMILTK-CPIVIFGMP--------------skvvlQAN---SH-G-R-FIQVE-L----RPFSynggrg
0058 8xkuA   ---aGQEY--IKRELQEIVRI--LK-kgIY-CP-----KGVLLHGpPGTGKTLLAKAIAGE--A----G-----lP-FFAANG--TDF-V----------EMFV-GV-AA-S-RVK-D-LFASSY-A-PS-IIFIDeI-D-AIGSkGA--E-R--E---qglLQILTEMDS-QVLVIGATN----------RLDI-L---DPA----R-K-G-RDKII-R----VGLPskdgrl
0059 7lccA   linslNDE--Q-LAAFQTITS--AI---ED-HP--K--CFFLDGP-GGSGQTYLYKVLTHY--I----R-grggtV-LPTAST--GTF-HS-------qyklPIPL-----------nakTIKK-A-QL-LIIDE-C-T-MAS-----------SH--A-INAIDreFGG-KVLLLGGDF----------rqcls-qtslKYC---N-wG-C-FRKLS-L----KTNMrsedsa
0060 6l1qA   pfyqpQHD--EVALFEAAYRK--R--------------lPVMVKGP-TGCGKSRFVEFMAWR--L--gkP------L-VTVACN-------------------------mTA-A-DLV-GrPLTVAY-G-AI-CYLDE-I-V-EAR-----------qDT--T-V-VIHPLTD-dFQLVISYNP----------gYQS-K---DLK---QSkQ-R-FTGFE-F----DYPNaeleag
0061 8hqsB   ---hEHFE-------LELG-------------S--R--LNFIVGN-NGSGKSAILTAITIG--L----GsslkdlimCFLSAR--SFL---TAS-T-lSGGEKSFS-QM-A-LLL-A-TWKPMR-S-RI-IALDE-F-D-VFMD-Q---V-N--RK-iG-TTLIVKLKDIrTQTIIITPQD-----------------igKI---AD-I-G-VSIHR-M----RDP-------

0001 s001A   ----LLLL--HHHHHHHHHHH--HL---LL-LL--L--EEEEEEL-LLLLHHHHHHHHHHHH--H----L------L-LEEEHH--HHH-HHHHL-LL-LHHHHHHH-HH-H-HHH-H-HHHLLL-L-LE-EEEEL-L-H-HHHL-HH--H-L--LL--H-HHHHHHHLLL-LEEEEEEEELE-E--LLLEEEELL-L---EEE---EL-L-L-LEEEE-L----LLLL------
0002 8fcvR   ---lLHHHhhHHHHHHHHHHHL--L-----L-LL--L--EEEEEEL-LLLLHHHHHHHHHHHH--H----HllllllE-EEEEL----LL-LL---------------lLH-H-HHH-H-HHHHHH-L-LE-EEEEL-H-H-HHHL-LL-lL-H--HH-hH-HHHHHHHHHL-LEEEEEEELH-------hhHHHL------lLL---HH-H-L-EEEEE-E----LLLLlllhhh
0003 2qbyB   llllllHH--HHHHHHHHHHH--HL---LL-L---L--EEEEEEL-LLLLHHHHHHHHHHHH--H----HllllleE-EEEEHH--HHL---------------lLH-H-HHH-H-HHHHHH---EE-EEEEL-L-H-HHHH-----L-L--LL--H--HHHHHHHLL-EEEEEELLLL---------LLLL--L---LLH----H-L-L-LLEEE-E----LLLLhhhhhh
0004 5e7pA   ---hLLLL--HHHHHHHHHHH--HH-hlLL-LL--L--EEEEELL-LLLLHHHHHHHHHLLL--L----L------E-EEEEHH--HHH-HL----------LHHH-HH-H-HHH-H-HHHHHH-L-EE-EEEEL-LlL-LLL--LL--L-H--HH--hhHHHHHHHHLLlEEEEEEELLL---------HHHL------------L-L-L-LEEEE-L----LLLLhhhhhh
0005 8umyA   llllllHH--HHHHHHHHHHH--HL-llLL-LL--L--EEEEELL-LLLLHHHHHHHHHHHHllL----E------E-EEEL-L--LLL-----------------lLH-H-HHH-H-HHHHH--L-EE-EEEEL-H-H-HLL-----------HH--H-HHHHHHHHLL-LLEEEEEELL---------LLLH------HHH---H-hL-L-LEEEE-L----LLLLhhhhhh
0006 8dr5A   llllllHH--HHHHHHHHHHL--HH---HL-LL--L--EEEEELL-LLLLHHHHHHHHHHHH--L----L----leE-EEEL-L--LLL--------------llhH-HH-L-LHH-H-HLLLLL-L-EE-EEELL-H-H-HL-L-LL--L-L--LH--H--HHHHHHHLL-LLEEEEEELL---------LLLH--H---HHH----L-L-L-LEEEE-L----LLLLllllhh
0007 3bosB   llllllHH--HHHHHHHhLLL--L-----------L--EEEEELL-LLLLHHHHHHHHHHHH--H----H-hllllE-EEEEHH--H-H-HHL-----------------LHH-H--HLLHH-L-LE-EEEEL-H-H-HHLL-----L-H--HH--H--HHHHHHHHH-HEEEEEELLL-------lLLLL--------------lhH-H-LEEEE-L----LLLLhhhhhh
0008 9nv3A   -llhhhHH--HHHHHHHHHHLhhHH---HL-LL--L--LEEEEEL-LLLHHHHHHHHHHHH--H----H--hlllE-EEEEHH--HHH-HLLLL-LL-LHHhhhhhHH-H-HHH-H-HLL--L-L-LE-EEEEL-L-L-LLLL-L---------H--H-HHHHHHLLLL-LLLEEEEEEE-E--LLLEEEELL-L---LLE---EL-L-L-LEELLl---------------
0009 7st9A   hllllHHH--H-HHHHHHHHH--HL-llLL-LL--L--EEEEEEL-LLLLHHHHHHHHHHHH--H----H-------lLEEEL----LL-LL-LL-LL----LLLLH-HH-H-HHH-H-HHLLLH-L-LE-EEE-L-L-L-L-LL-LL--H-H--HH--H-HHHHHHHHHL-LLEEEEEEL-----------EELhH---HLL---HH-H-HlEEEEE-L----LLLLhhhhhh
0010 7crcB   ---lLLHH--HHHHHHHHHL---LL---LL-LL--L--EEEEELL-LLLLHHHHHHHHHHHH--H----H-hhlleE-EEELHH--HHL----LL-LL----LLHHH-HH-H-HHH-H-HHHHHH---EE-EEEEL-L-L-LH-----------HH--H-HhHLLL-HHH-LEEEEEELL---------HHHH-------------H-L-L-LLEEE-L----LLLLhhhhhh
0011 6z1f3   -llllLHH--HHHHHHHHHHH--HH---LL-LL--L--EEEEELL-LLLLHHHHHHHHHHHHhlL----E------E-EEEL-H--HHL-L-----------lLLLLL-HH-H-HHH-H-HHHHHH---EE-EEELL-LlL-LLL--LL--H-H--HH--hhHHHHHHHHHLlLLEEEEEELL---------LLLL--------lH---HH-L-L-EEEEE-----LLLLhhhhhh
0012 8btgB   llllllHH--HHHHHHHHHHL---------L-LL--L--EEEEEEL-LLLLHHHHHHHHHHHH--H----HhlllllE-EEEEHH--HHH-HHHHHhHH-HL------------hhH-H-HHLL---L-LE-EEEEL-H-H-HHLL-L---H-H--HH-hhhHHHHHHHHHL-LEEEEEEELLL--------LLLLL-L---LH------hL-L-LEEEE-E----LLLLhhhhhh
0013 7mi8A   ---lLHHHhhHHHHHHHHHHHL--L-----------L--EEEEELL-LLLLHHHHHHHHHHHH--L---lL------L-LEEELL--L---LLLL-----------hH-HH-H-HHH-H-HHHHHL-L--E-EEELL-L-L-L-LL-HH--H---hHH--H-HHHHHHHHHHlLEEEEEEELL------------LLL---llllL---HH-H-LlEEEEE-L----LLL-------
0014 7plhA   ---lLLHH---HHHHHHHHHH--HH---HL----lL--LEEEELL-LLLLHHHHHHHHHHHH--L----LllllllE-EEELLL--L---------------------lLH-H-HHH-H-HHHHHL-L-LE-EEELL-L-L-LLL-----------H--H-HHHHHHHHHL-LEEEEEEELLH-------hHHHH------HLL---LH-H-H-LEELL-L----LLLLlhhhhh
0015 9cruC   ---lLHHHhhHHHHHHHHHHHH--HH-llLL-LE--E--EEEEELL-LLLLHHHHHHHHHHHH--H----L---lllE-EEE-LH--HH--HLLL----------LHHH-HH-H-HHH-H-HHHHHH---LE-EEELL-H-H-HHLL-LL--H-H--HH-----HHHHHHLLLlLEEEEEEELLH-------HHHH--H---HLL-----lL-L-L-LLEEE-E----LLLLlllhhh
0016 7jgrD   llllllLH--HHHHHHHHHHH--HH--hHL-LL--E--EEEEELL-LLLLLHHHHHHHHHHH--H----HllllleeE-EEEELL--LLL---------------------LH-H-HHH-H-HHHHHL---EE-EEEEL-H-H-HHLL----lH-H--HH---hHHHHHH-hLL-LEEEEEELLLL--------LLLL------LLL---HH-H-L-LLLEE-L----LLLLllhhhh
0017 6bbmD   llllllllllllllllhhLLL--LL--------llL--LEEEEEL-LLLLLHHHHHHHHHHH--L----LlllllllL-LEEELLllLLL-L------------------lLL-L-LLH-H-HHLLLL-L-LL-LEEEL-H-H-HLLL-LL--L-LllhH--H-HHHHLLLLLL-LLLLEEELL--L--LHHHL----lL---LLL----H-H-H-LEEEE-E----LLHHhhllll
0018 8iprA   lllllllL--LLL--LLLL--------------llL--EELLLLL-LLLLHHHHHHHHHHHH--H----HlllllllE-EEELLL--L----LLLL-L-hhhhhhhhH-HL-LlhHH-H-HHHHLL-L-LE-EEEEL-L-L-L-LL-LH--H-H--HH--H-HHHHHLLLLLlLEEEEEELLL---------------------lL---LL-L-L-LLLLL-L----L-------
0019 8dgcA   ---lLLLH---HHHHHHHHHL--LL--------L--L--LEEEELL-LLLLHHHHHHHHHHHH--LllllE------E-EEEELL--L--lLLLLL-HH----------lLH-H-HHH-H-HHHHHH-hhEE-EEEEL-H-H-HHHH-hL--L-L--LH--H-HHHLLL----lEEEEEEELL---------LLHH-------------L-L-L-LEEEE-L----LLLLhhhhhh
0020 8c0vA   ---lLLLL--HHHHHHHHHHL---L---LL-LL--L--EEEELLL-LLLLHHHHHHHHHHHH--H----H-hhhllE-ELLLLL--L---LLLL--------lLLHHH-HH-H-HHH-H-HHHHHL-L-EE-EEELL-L-HhHHLL-L--lL-H--HHhhH-HHHHHHHHHHlLLEEEELLLL-L-----------lL---LLH-hhHL-L-L-EEEEE-L----LLLLlhhhhh
0021 5zr1E   llllllHH--HHHHHHHHLLL--LH---HH-lL--L--EEEEEEL-LLLLLLHHHHHHHHHL--L----L---lleE-EEELLL--LLL-L---------------hH-HH-H-HHH-H-HHHHHH-HhEE-EEEEL-L-L-LLLL------lL--LL--L-HHHHHHH-lE-EEEEEEELLL---------LLHH-------HH---HH-H-L-LLEEE-E----LLLLhhhhhh
0022 8xvbA   llllllLH--HHHHHHHHHHH--HH---HH--L--L--LEEEEEL-LLLLHHHHHHHHHHHH--L----L-----lE-EEEELL--HHH-----------------lLH-H-HHH-H-HHHHHH---LE-EEEEL-H-H-HLL-------------H--H-HHHHHHHHHL-LEEEEEEELL-----------------hhhHLL---LH-H-H-LLEEE-E----LLLLhhhhhh
0023 5zr1A   llllllLL--LHHHHHHHHHH--HH---LL-L---L--EEEEEEElLLLLHHHHHHHHHHHH--H----HlllllleE-EEEEEH--HHL-LL-------------------lL-HHH-H-HHHHHH---EE-EEEEElL-L-LLLL-L-----L--LL--L-HHHHHHHHHL-LEEEEEEELLL-L-------LHHH--H---HLL---HH-H-H-LLEEE-E----ELLLlhhhhh
0024 5ywwA   ---hHHLL---lhhhHHHHHH--L-----------lL--EEEEELL-LLLLHHHHHHHHHHHH--H----H-hllllE-EEELL--------------------------L-HHH-H-HHHHHL-L-LE-EEELL-L-L-L-------------HH--H-HHHHHHHHHL-LEEEEEELLLL-L-----HHHH------hLLL---L-hhhL-LEEEE-E----ELL-------
0025 7vcsA   ---lLLHH--HHHHHHHHHHH--HH-hlLL-LL-----LEEEEELlLLLLHHHHHHHHHHHH--H----L-----lE-EEEEEH--HHH-HLL-----------LLLL-HH-H-HHH-H-HHHHHL-L-LE-EEEEE-L-HhHLLL-lH--H-H--HHhhH-HHHHHLL--L-LEEEEEEELL-----------LLLL-L---LHH----L-L-L-LLEEE-E----LLLLlhhhhh
0026 9bv1A   llllllLH--HHHHHHHHHHH--HH-hlLL-LL--L--LEEEELL-LLLLHHHHHHHHHHHH--L---------lleE-EEELL---LLL-L--LL-LL--------LH-HH-H-HHH-H-HHHHHL-L-EE-EEEEL-L-L-LLL-lHH--H-H--HhhhhhhHHHHL--LL-EEEEEEEELLL--------------lL---LLH----L-L-L-LLLEE-L----LLLLhhhhhh
0027 6n9xB   ---lLLLL----------hhLL--LL---------llE--EEEEELL-LLLLHHHHHHHHHHHH--H----HllllllE-EELLLL--L------------lhhhhhhhlllL-L-HHH-H-HHHHHL-L-LE-EELLL-L-L-LLLL-LL-hH-H--HHhhH-HHHHHHHHLL-LEEEEEEELL----lLLLLLLL---L---LLL----L-L-L-LEEEE-E----ELLLllllll
0028 5j1sA   hhllllHH--HHHHHHHHHHH--HH-llLL-LL--E--EEEEEEL-LLLLHHHHHHHHHHHH--H----LhhhlllE-EEEEHH--HHL-LL----------hHHHHH-HH-H-HHH-H-HHHHHH-L-LE-EEEEL-HhH-LLH-----------HH--H-H-HHHHHHL-lLEEEEEEELLL-hhhLLLLL------llhHHH---HllL-L-LLEEE-E----LLLLhhhhhh
0029 4rh7A   lllllLHH--H-HHHHHHHHH--HH---HH-LL-------EEEEELL-LLLLHHHHHHHHHHHH--L--llL------E-EEEELL-------------------------lLH-H-HHH-H-HHHHHH-L-LE-EEEEL-L-L-LLL---------hHH--H-HHHHHHHHHHlLEEEEEELLL-----------LLL-L---LLL---LHhH-L-LEEEE-L----LLLLlhhhhh
0030 9emcD   lleellHH--HHHHHHHHHHH---L---LL-L--lL--EEEEEEL-LLLLHHHHHHHHHHHH--H----L--llllE-EEEE-L--LLL---LLL-L--------lL-HH-H-HHH-H-HHHLLEeE-LE-EEEEL-H-H-HLL---------H--HH--H--HHHHHHLLL-LEEEEEEELL-----------LLE-EhhhLLH---H-hL-L-LEEEE-E----LLLLhhhhhh
0031 1nlfA   hhhhhhlllllllleeELLEE-------------llL--EEEEEEL-LLLLHHHHHHHHHHHH--H----HllllllE-EEEELL-hhLL-LLL-------llllLHH-HH-H-HHH-H-HHL--L-L-LE-EEEEL-H-H-HHLL-L-----llllH--H-HHHHHHHHHL-LEEEEEEEL-------------------LL---HH-H-L-LLEEE-E----EELLhhhhhh
0032 9e22A   ---lLHHHhhHHHHHHHHHHL--L-----------L--LLLEELL-LLLLHHHHHHHHHHHH--L----L------LlEEEELL--L-L-LLL---------------H-H-HHH-H-HHHHHH-L-L--EEEELlH-H-HL----L--H-H--HH--H-HHHHHHHHHH-HEEEELLLL-----------------llllL---LL-L-LlLLLEE-L----LLLLhhhhhh
0033 4m9sB   llllllHH--HHHHHHHHHHL--LL---LL-LL--E--EEEEELL-LLLLHHHHHHHHHLLL--L-----llllleE-EEEELL------------------llLLL-HH-H-HHH-H-HHHHHH---EE-EEEEE-E-L-LH-----------H--H-HHHHHH--LL-LEEEEEEELLH-H-------------------------L-L-L-LEELL-L----LLLLhhhhhh
0034 6f0xB   ---lLLLH---HHHHHHHHHH--HH---HHhLL--L--LEEEEElLLLLHHHHHHHHHHHH--H----HllllllL-EEEEEL--H---HHHLL-L--LLLHH-HH-HH-H-HHH-H-HHHHHH-L-LE-EEEEEeH-H-HLLllHH--H-H--HH-hH-HHHHHHHLLLlLEEEEEEEL----------LHHH-L---LLH-----hH-L-LLEEE-E----LLLLlhhhhh
0035 7uqjA   ---hHHHH-hHHHHHHHHHHH--HHlhlLL-LL--L--EEEEEEL-LLLLHHHHHHHHHHHH--L----L-------lEEELLH--HHH-LL-LL-LL--------L-HH-H-HHH-H-HHHHHL-L-EE-EEEEL-HhH-HHHH--L--H-H--HH----------hLL-EEEEEEEELL-L--H-------------hHH---HHlL-L-LLEEE-L----LLLLhhhhhh
0036 6p8vC   hhllllHH--HHHHHHHHHHH--HH-hhHH-HL--L--LEEEEEElLLLLHHHHHHHHHHHH--H----HhhllleE-EEEEL----LH-HH-HL-LL-LL-LHHHH-HH-H-HHH-H-HHHHHH-E-EE-EEEELlH-H-HHLLlHH--H-H--HH-hH-HHHHHHHHHLlLEEEEEEEL----------LHHH-L---LHH-----hH-L-LLEEE-E----ELLLlhhhhh
0037 6dgdA   lllllLHH--H-HHHHHHHHL--LL---LL--L--E--EEEEELL-LLLLHHHHHHHHHHHH--H----H-hllleE-EEE-LL--LLL-L-------lllleeelleeeel-HHH-H---LLLL-E-EE-EEEEL-LlL-HHHL-LL-lllL--LL--H-HHHHHHHHHL-LEEEEEEELLL-----------------lHHH---HH-HhL-LEEEE-L----L---------
0038 8zr0A   hhllllHH--HHHHHHHHHHH--HH-hlLL-LLllL--EEEEEELlLLLLHHHHHHHHHHHH--H----H-------lLEEEEE-------------llLHHHHHH-HL----LH-H-HHHHLL-L----LEEEE-E-LlllLL-HH--H---hHH--H-HHHHHHL--L-LLEEEEEEELLlhhlHHHH-----hH---HLL---HH-H-L-LLEEE-E----LLLLlhhhhh
0039 8y6pQ   hlllllHH--HHHHHHHHHHH--LL------LL--L--EEEEELL-LLLLHHHHHHHHHHHL--H-----hhllllE-EEEELL--L---------------llLLL-HH-H-HHH-H-HHHHHH---LE-EEEEE-E-L-LH-----------H--H-HHH--HLLL--LEEEEEEELLH---------------hhhHHL---LL-L-L-EEEEE-LllllLLLLhhhhhh
0040 7t8bA   hlhhhhlllllllllhhhhhLL--LL--------llL--EEEEELL-LLLLHHHHHHHHHHHH--H----H-lllllE-EELLLL--LhhlLLLL-------------lLH-H-HHH-H-HHHHHL-L-LE-EEELL-L-L-LLL--------lhhH--H-HHHHHHHHHL-LEEEEEEELLL-------------------L---HH-H-L-LEEEE-E----EELLeeeeee
0041 5juyB   lllllLHH--HHHHHHHHHL--L----LL-LL--E--EEEEELL-LLLLLHHHHHLLLLL--------hhllllE-EEEELL--L---------------lLHHH-HH-H-HHH-H-H--llL-L-LE-EEEEL-L-L-LH-----------H--H-HHHLL---LL-LEEEEEEELLL-L-----------------------lllL-LLLLL-L----LLLLlhhhhh
0042 8gjaF   ---lHHHH---hhlllLEEL--------------lL--EEEEEEL-LLLLHHHHHHHHHHHH--L----LllllleE-EEEELL-eELL-L-------------hH-HH-H-HHHhL-HHHHHH-E-EE-EEEEL-HhH-HHHL-LH--H-H--HH--H-HHHHHHHHHH-HLEEEEEEEHH--------hHLLL-L---LLL---HH-L-L-LEEEE-E----EEEEeeeeee
0043 8ui7A   hhllllHH--HHHHHHHHHHH--HH-llLL-LL--L--EEEEELL-LLLLHHHHHHHHHHHH--L---lE------E-EEEELL--LLL-----------------lLH-H-HHH-H-HHHHH--E-EE-EEEEL-L-L-L-LL-LL--L-L--LH--H--HHHHHHHLL-LLEEEEEELLL-L--------------hhHHL---L--L-L-LEEEE-L----LLLLhhhhhh
0045 7qo4M   ---lLLHH--HHHHHHHHHHH--HH-hlLL-LL--L--LEEEELL-LLLLHHHHHHHHHHHH--H----L----leE-EEEEH---HHH-L----------lLLLLL-LH-H-HHH-H-HHHHHL-L-LEeEEEEL-L-L-LLlH-HH--HhH--HH--HhhHHLLL-LLL-EEEEEELLLL-------------lL---LLH-hhHL-L-LlLEEEE-L----LLLLhhhhhh
0046 6qemG   ---lLHHHhhHHHHHHHHHHL--LL---LL--L--L--LEEEELL-LLLLHHHHHHHHHHHH--H----H-hhlllE-EEEEHH--HHH-HHHHH-hH-HLL-----lHH-H-HHH-H-HHL----L-LE-EEEEL-L-L--lLL-LL--H-HhhHH--H-HHHHHHHHLL-LLLEEELLLL-H-------hHHHL-L---LLH----H-H-H-HLLEE-E----LLLLlhhhl-
0047 9ri9B   llllllHH--HHHHHHHHHHL---L---LL-LL--E--EEEEELL-LLLLHHHHHHHHHHLL--H-----hhhlleE-EEEELL--LLL------------------LH-H-HHH-H-HHHHL----EE-EEEEL-L-L-L-------------H--H-HHHHHHHLLLlLEEEEEEELLH-H-------------------------hH-H-HLLEE-L----LLLLhhhhhh
0048 8jx6B   hllhhhHH--HHHHHHHHHHH--HH---LL-LL--L--EEEEEEL-LLLLHHHHHHHHHHHH--H----H-hllllE-EEEE-L--LHH-H---------------eeehH-HHL-LelllLLL-L-LE-EEELL-H-H-HLL-----------HH--H-HHHHHHLLLLlLEEEEEEELLL----------LLL-L---LL-lhhHL-L-L-LEEEE-E----LLLLhhhlll
0049 4wiaA   ---leellllllllhhhhlLE--EL-------L--L--EEEEELL-LLLLHHHHHHHHHHHH--H----H-hllllE-EEEEL--eELH-HH------hhllllLL-HH-H-HHH-H-L-HHHH-L-LE-EEEEL-HhH-HHHL--L--L-L--LL--H-HHHHHHHHHHlLEEEEEEELHH---------HLLH-H---HHH-----hH-H-LLEEE-E----EEEEelleee
0050 8tp1A   hllllLLH--H-HHHHHHHHH--HH---HL-LL--E--EEEEEELlLLLLHHHHHHHHHHHH--L----L------L-EEEEEL-----lLLLL----------lhH-HH-H-HHH-H-LL-----L-LL-EEEEE-E-L-hhHL-LLllL-L--HH-hH-HHHHLL--LL-LEEEEEEEEL----------LHHH-L---LLL----L-L-L-LLEEE-E----LLLLlhhhhh
0051 3j3rA   llllllHH--HHHHHHHHHHH--HL---LL-LL--LllLEEEELL-LLLLHHHHHHHHHHHH--HhlllL------L-EEEEEH--HHLllllllL-LL-LL-LLHHH-HLlL-HHH-H-HHHHL--L-LL-EEEEE-LlL-LLL-----------HH--H-HHHHHHHHHLlLLEEEEEELLLlH--HHHH-----hH---LLH-----hH-L-LLEEE-L----LLLLlllhhh
0052 8xksB   ---lLLLLllHHHHHHHHHHHH-----hlLL-LL--L--EEEEELL-LLLLHHHHHHHHHHHH--H---lL------L-EELLLL--LLL-L----------lLLLLH-HH-H-HHH-H-HHHHHL-L-LE-EEEEL-H-H-HLL--LH--H-H--H---hhhHHHHHHHL-lLEEEEEEELLL-L-----------L---LLL----L-L-L-LLLLL-L----LLLLhhhhhh
0053 7yqhB   ---lLLEE-------ELLL-------------L--L--EEEELLL-LLLLLLHHHHHHHHHH--H----LllhhhhllLEELHH--HHH---LLL-LhlLHHHHHHH-HH-H-HHH-H-HHLLLL-L-LE-EEELL-L-H-HHLL-L---H-H--HH-hH-HHHHHHHHLLlLEEEEELLLL-----------------hhHL---LL-L-L-EEEEL-L----LLL-------
0054 5v8fD   ---lHHHH-hHHHHHHHHHHHH--HH---L--lL--L--EEEEELL-LLLLHHHHHHHHHHHH--H----H--------eEEEEEL--LLL-L---------------lL-HH-H-HHH-H-HHH----L-EE-EEEEElH-H-HLLL-LL----L--LH--H-HHHHHHHHHL-LEEEEELLLL-L------LHHH--L---LH------hL-L-LLEEE-L----LLLLlhhhhh
0055 7qo4J   ---lLLHH--HHHHHHHHLLH--H--hlLL-LL--L--LEEEELL-LLLLHHHHHHHHHHHH--H----L------L-EEEEEL---lL-LLLLL-L--------llL-LL-L-LLH-H-HHHHHL-L-LE-EEEEL-L-L---LL-LL-lllH--HH--HhHHHHHHHHH-lLEEEE-EELL-L-----------lL---LLLlllLL-L-L-LLEEE-L----LLLLhhhhhh
0056 8bobC   lllllLHH--HHHH-HHLH-------------HL--L--EEEEELL-LLLLHHHHHHHHHLL--L----L-----lE-EEEELL-----hHHHL-----------hH-HH-H-HHH-H-HHHHHH---EE-EEEEL-H-H-HLLL-----H-H--HHhhhH-HHH-----LL-EEEEEEELLL---------------lllLH---HH-HhL-EEEEL-H--hhHLLLhhhhhh
0057 7rzy1   lllllLHH--H-HHHHHHHHhHH---HL-LL--L--LLEELLL-LLLLHHHHHHHHHHHH--H----HhllllllE-EEELLL-------------------------lLH-H-HHH-H-HHHHLL-L-LE-EEELL-L-H-HHLL-LL--L-L--LH--H-HHHHHHHHLL-LLLLLLLLH-------------hhhhhHHL---HH-H-H-LLEEL-L----LLLLllllll
0058 8xkuA   ---lLLHH--HHHHHHHHHHH--HH-llLL-LL-----LEEEEELlLLLLHHHHHHHHHHHH--H----L-----lL-EEEEEH--HHL-L----------LLLL-LH-HH-H-HHH-H-HHHHHL-L-LE-EEEEElL-L-LLLLlHH--H-H--H---hhhHHHHHHHHL-LEEEEEEELL----------LLLL-L---LLL----L-L-L-LLEEE-E----LLLLlhhhhh
0059 7lccA   hhhhlLHH--H-HHHHHHHHH--HH---HL-LL--L--EEEELLL-LLHHHHHHHHHHHHH--H----H-hllllE-EEELLL--HLH-HH-------hlllLLLL-----------lhhHHHH-L-LE-EEELL-L-L-LLL-----------HH--H-HHHHHhhHHH-LEEELLLLL---------lllll-hhlhHHL---H-hH-H-LEEEL-L----LLLLllllhh
0060 6l1qA   lllllLLL--HHHHHHHHHHL--L-------------lLEEEELL-LLLLHHHHHHHHHHHH--H--llL------E-EEEELL-------------------------lLH-H-HHH-EeHHHHHH-L-LE-EEEEL-H-H-HLL----------hHH--H-H-HHHHHLL-lLEEEEEEELL----------lLLL-L---LLL---HHhL-L-LEEEE-L----LLLLhhhhhh
0061 8hqsB   ---lLLEE-------ELLL-------------L--L--EEEELLL-LLLLLLHHHHHHHHHH--H----LllhhhhllLEELHH--HHH---LLL-L-lLHHHHHHH-HH-H-HHH-H-HHLLLL-L-LE-EEELL-L-H-HHLL-L---H-H--HH-hH-HHHHHHHHLLlLEEEEELLLL-----------------hhHL---LL-L-L-EEEEL-L----LLL-------
```

```
# Job: Type-2 BREX; PglW_WP_185095299 1 wHTH1
# Query: s001A
# No:  Chain   Z    rmsd lali nres  %id PDB  Description
   1:  5j6x-A  8.9  2.2   64    66    6   MOLECULE: Z-DNA BINDING PROTEIN KINASE;
   2:  7qcd-D  8.9  2.7   80   326    9   MOLECULE: STRUCTURAL MAINTENANCE OF CHROMOSOMES PROTEIN 5;
   3:  2mh2-A  8.3  1.8   63    64   16   MOLECULE: HOMOLOGOUS-PAIRING PROTEIN 2 HOMOLOG;
   4:  6qfd-B  8.2  2.5   68   116   10   MOLECULE: DNA-BINDING PROTEIN;
   5:  5yad-A  8.1  1.9   65    73   17   MOLECULE: MEIOSIS REGULATOR AND MRNA STABILITY FACTOR 1;
   6:  7rhe-A  8.0  1.6   59   368   15   MOLECULE: ROK FAMILY PROTEIN;
   7:  7vwv-B  8.0  2.4   69    74    9   MOLECULE: I73R;
   8:  2l02-A  7.9  1.7   64    82   14   MOLECULE: UNCHARACTERIZED PROTEIN;
   9:  1qgp-A  7.8  1.6   63    76   11   MOLECULE: PROTEIN (DOUBLE STRANDED RNA ADENOSINE
  10:  1cf7-B  7.8  2.8   73    82   11   MOLECULE: DNA (5'-
  11:  2xrn-B  7.7  1.9   61   241   10   MOLECULE: HTH-TYPE TRANSCRIPTIONAL REGULATOR TTGV;
  12:  3cuq-A  7.7  2.4   74   219   14   MOLECULE: VACUOLAR-SORTING PROTEIN SNF8;
  13:  1ku9-A  7.7  2.3   64   151    9   MOLECULE: HYPOTHETICAL PROTEIN MJ223;
  14:  8e4y-A  7.7  2.2   67   651    7   MOLECULE: GLYCEROL-3-PHOSPHATE ACYLTRANSFERASE 1, MITOCHOND
  15:  4ad9-A  7.7  2.2   74   288    5   MOLECULE: BETA-LACTAMASE-LIKE PROTEIN 2;
  16:  3elk-A  7.6  2.3   70   104   10   MOLECULE: PUTATIVE TRANSCRIPTIONAL REGULATOR TA0346;
  17:  7kbf-K  7.6  2.9   67    78   15   MOLECULE: HISTONE H3.2;
  18:  2od5-A  7.6  1.8   66    91   14   MOLECULE: HYPOTHETICAL PROTEIN;
  19:  8jxk-D  7.5  2.3   68   166   15   MOLECULE: CONSERVED PROTEIN;
  20:  8iue-P  7.5  2.3   63   303    6   MOLECULE: DNA-DIRECTED RNA POLYMERASE III SUBUNIT RPC1;
  21:  4ija-A  7.5  1.8   60   363    7   MOLECULE: XYLR PROTEIN;
  22:  6qpq-D  7.5  2.0   66    82    5   MOLECULE: STRUCTURAL MAINTENANCE OF CHROMOSOMES PROTEIN,STR
  23:  5y6i-B  7.4  2.2   63   234   11   MOLECULE: TRANSCRIPTIONAL REGULATOR KDGR;
  24:  8xt8-B  7.4  1.6   61   145   10   MOLECULE: MARR FAMILY TRANSCRIPTIONAL REGULATOR;
  25:  7cv0-A  7.4  2.2   61   173    8   MOLECULE: TRANSCRIPTIONAL REGULATOR NIAR;
  26:  7y43-A  7.3  2.3   64    79    9   MOLECULE: HISTONE ACETYLTRANSFERASE KAT6A;
  27:  2zme-B  7.3  2.1   64   215    9   MOLECULE: VACUOLAR-SORTING PROTEIN SNF8;
  28:  3f72-B  7.2  1.7   61   108   18   MOLECULE: CADMIUM EFFLUX SYSTEM ACCESSORY PROTEIN;
  29:  6sj9-A  7.2  2.8   73   642    5   MOLECULE: PROTEASOME ACCESSORY FACTOR B/C (PAFBC);
  30:  4mtd-D  7.2  2.4   68   151   12   MOLECULE: ZINC UPTAKE REGULATION PROTEIN;
  31:  3bpx-A  7.2  1.7   61   147    3   MOLECULE: TRANSCRIPTIONAL REGULATOR;
  32:  4y66-C  7.2  2.4   67   197    7   MOLECULE: MND1;
  33:  4i99-C  7.2  1.7   62    70    8   MOLECULE: CHROMOSOME PARTITION PROTEIN SMC;
  34:  2cwe-A  7.1  1.7   59   191   12   MOLECULE: HYPOTHETICAL TRANSCRIPTION REGULATOR PROTEIN, PH1
  35:  2vqc-A  7.1  2.6   66    70    9   MOLECULE: HYPOTHETICAL 13.2 KDA PROTEIN;
  36:  3gfi-A  7.1  3.0   63   143   13   MOLECULE: 146AA LONG HYPOTHETICAL TRANSCRIPTIONAL REGULATOR
  37:  7z8b-C  7.1  2.6   68  1224   12   MOLECULE: CULLIN-7;
  38:  1xmk-A  7.1  2.2   65    79    8   MOLECULE: DOUBLE-STRANDED RNA-SPECIFIC ADENOSINE DEAMINASE;
  39:  3k69-A  7.1  2.3   64   151   13   MOLECULE: PUTATIVE TRANSCRIPTION REGULATOR;
  40:  3u2r-A  7.0  2.6   63   135   13   MOLECULE: REGULATORY PROTEIN MARR;
  41:  6zvh-y  7.0  3.4   66    72    8   MOLECULE: 18S RRNA;
  42:  5jbr-A  7.0  2.7   64   149   14   MOLECULE: UNCHARACTERIZED PROTEIN BCAV_2135;
  43:  1xsd-A  7.0  2.2   65   125    6   MOLECULE: 5'-
  44:  1yyv-A  6.9  2.5   65   114    6   MOLECULE: PUTATIVE TRANSCRIPTIONAL REGULATOR;
  45:  6aht-A  6.9  3.5   70   111    9   MOLECULE: CONSERVED HYPOTHETICAL PLASMID PROTEIN;
  46:  4fht-B  6.9  2.9   64   142   13   MOLECULE: PCAV TRANSCRIPTIONAL REGULATOR;
  47:  7b0c-A  6.9  2.3   63   144   16   MOLECULE: HTH-TYPE TRANSCRIPTIONAL REPRESSOR NSRR;
  48:  2g9w-A  6.8  1.9   62   119   19   MOLECULE: CONSERVED HYPOTHETICAL PROTEIN;
  49:  3bdd-B  6.8  1.5   57   133   14   MOLECULE: REGULATORY PROTEIN MARR;
  50:  1sfu-A  6.8  2.1   62    70    5   MOLECULE: 5'-D(*T*CP*GP*CP*GP*CP*G)-3';
  51:  5n35-A  6.8  1.7   58    60   12   MOLECULE: POLB1 BINDING PROTEIN 2 (PBP2);
  52:  5eri-A  6.8  1.7   61   153    8   MOLECULE: MARR FAMILY TRANSCRIPTIONAL REGULATOR;
  53:  3f8f-A  6.8  2.7   69   114    9   MOLECULE: TRANSCRIPTIONAL REGULATOR, PADR-LIKE FAMILY;
  54:  3r4k-A  6.8  2.1   60   255    5   MOLECULE: TRANSCRIPTIONAL REGULATOR, ICLR FAMILY;
  55:  7c0j-B  6.8  1.9   59    62    7   MOLECULE: HISTONE H5,DOUBLE-STRANDED RNA-SPECIFIC ADENOSINE
  56:  3w6k-C  6.8  2.1   61    87   16   MOLECULE: SCPA;
  57:  2eth-A  6.8  2.1   60   141   12   MOLECULE: TRANSCRIPTIONAL REGULATOR, PUTATIVE, MAR FAMILY;
  58:  1fok-A  6.8  1.9   65   568    9   MOLECULE: DNA (5'-
  59:  5udb-4  6.8  2.6   69   751    9   MOLECULE: DNA REPLICATION LICENSING FACTOR MCM2;
  60:  8as8-C  6.8  2.0   68   195   10   MOLECULE: JETC;
  61:  2fxa-B  6.7  2.6   63   180   16   MOLECULE: PROTEASE PRODUCTION REGULATORY PROTEIN HPR;
  62:  2lnb-A  6.7  2.1   62    80   10   MOLECULE: Z-DNA-BINDING PROTEIN 1;
  63:  4hqe-B  6.6  2.6   65   108    8   MOLECULE: TRANSCRIPTIONAL REGULATOR QSRR;
  64:  3bj6-A  6.6  1.7   60   152   15   MOLECULE: TRANSCRIPTIONAL REGULATOR, MARR FAMILY;
  65:  4hw0-A  6.6  2.1   61    98    3   MOLECULE: DNA-BINDING PROTEIN SSO10A-2;
  66:  2zkz-C  6.6  2.1   59    87   14   MOLECULE: TRANSCRIPTIONAL REPRESSOR PAGR;
  67:  8uuc-A  6.6  2.6   63   274   16   MOLECULE: ADENINE DNA GLYCOSYLASE;
  68:  2dql-A  6.6  2.7   67   115    9   MOLECULE: PEX PROTEIN;
  69:  7xjg-J  6.6  2.1   64   306    9   MOLECULE: RNA-DIRECTED DNA POLYMERASE FROM RETRON EC86;
  70:  3s93-B  6.6  2.3   63    81   14   MOLECULE: TUDOR DOMAIN-CONTAINING PROTEIN 5;
  71:  2esh-A  6.5  2.8   66   114    9   MOLECULE: CONSERVED HYPOTHETICAL PROTEIN TM0937;
  72:  9gm7-C  6.5  2.9   74   440   12   MOLECULE: CHROMOSOME PARTITION PROTEIN MUKF;
  73:  5hs5-A  6.5  2.5   68   116   10   MOLECULE: HTH-TYPE TRANSCRIPTIONAL REGULATOR SARX;
  74:  8soj-B  6.5  2.2   64   428   11   MOLECULE: CST COMPLEX SUBUNIT CTC1;
  75:  8f8u-A  6.5  2.3   61   297   20   MOLECULE: BIFUNCTIONAL LIGASE/REPRESSOR BIRA;
  76:  2xco-A  6.5  2.3   67   636    7   MOLECULE: DNA GYRASE SUBUNIT B, DNA GYRASE SUBUNIT A;
  77:  2p4w-B  6.5  2.7   63   198   21   MOLECULE: TRANSCRIPTIONAL REGULATORY PROTEIN ARSR FAMILY;
  78:  1ldd-A  6.5  3.2   69    74    9   MOLECULE: ANAPHASE PROMOTING COMPLEX;
  79:  1w7p-D  6.5  2.8   72   171   13   MOLECULE: VPS22, YPL002C;
  80:  9bz0-f  6.5  2.6   64   232    9   MOLECULE: DNA-DIRECTED RNA POLYMERASE SUBUNIT;
  81:  5u8o-B  6.4  2.7   75   360   13   MOLECULE: ZN-DEPENDENT HYDROLASE;
  82:  3l9f-A  6.4  2.2   61   170   11   MOLECULE: PUTATIVE UNCHARACTERIZED PROTEIN SMU.1604C;
  83:  2ns0-A  6.4  2.4   67    85    7   MOLECULE: HYPOTHETICAL PROTEIN;
  84:  6juv-A  6.4  2.7   61   192   11   MOLECULE: SEGREGATION AND CONDENSATION PROTEIN B;
  85:  2qlz-B  6.4  2.5   61   220   15   MOLECULE: TRANSCRIPTION FACTOR PF0095;
  86:  5l0p-A  6.4  2.5   68   311   18   MOLECULE: TRANSCRIPTION FACTOR ETV6, TRANSCRIPTION FACTOR E
  87:  9qe1-J  6.4  2.3   71   108    7   MOLECULE: JETC;
  88:  5zqh-A  6.4  2.6   66   101   12   MOLECULE: PADR FAMILY TRANSCRIPTIONAL REGULATOR;
  89:  3lmm-D  6.4  2.2   63   556   14   MOLECULE: UNCHARACTERIZED PROTEIN;
  90:  7bzh-A  6.4  1.9   57    59    9   MOLECULE: SUL7S;
  91:  5e20-A  6.4  1.8   62   169   10   MOLECULE: TRANSCRIPTIONAL REGULATOR, MARR FAMILY;
  92:  1ylf-C  6.4  2.2   61   119   10   MOLECULE: RRF2 FAMILY PROTEIN;
  93:  3lwf-A  6.3  2.3   63   144   10   MOLECULE: PUTATIVE TRANSCRIPTIONAL REGULATOR;
  94:  7xx7-U  6.3  3.2   70   213   13   MOLECULE: HISTONE H3.1;
  95:  4ru7-B  6.3  2.0   59    99   12   MOLECULE: ASPA;
  96:  2co5-B  6.3  3.1   67    94   10   MOLECULE: VIRAL PROTEIN F93;
  97:  5xfo-A  6.3  2.8   68   315   19   MOLECULE: PHD FINGER PROTEIN 1;
  98:  8ylg-A  6.3  3.2   66   161   15   MOLECULE: MARR FAMILY TRANSCRIPTIONAL REGULATOR;
  99:  4b8x-A  6.3  1.9   62   140   16   MOLECULE: POSSIBLE MARR-TRANSCRIPTIONAL REGULATOR;
 100:  9f27-A  6.3  2.1   63    94   13   MOLECULE: RPA32 SUBUNIT OF THE HETERO-OLIGOMERIC COMPLEX IN

0001 s001A   NPVLLA--LRDEAVEIL-A-TL-G-RVASVA-ELGEALI-G--RRG----V--TQRDPQIRRALGFA-AARAAYEVDW--TR---SPRRMRS-RR---H--G-D-L-MLFALEV--------
0002 5j6xA   MSAENE--IEMRICDYL-R-RH---GRSTVQ-DIFKELK------------------------lEKS-TVNRHLYSLQ--AS-----KQVFK-TV---E--D-K-R-PVWDLVE--------
0003 7qcdD   PVTGDA--TAKYLLQYI-L-SA-R-GICHEN-ALILALM-R--LETdastL--NTE---WSIQQWVD-KLNDYINAIN--VKlnlLGYKIIR-IN---H--GiS-N-RFFVYVNlasteetk
0004 2mh2A   -----G--APGIILRYL-Q-EQ-N-RPYSAQ-DVFGNLQ-K--EHG-----------------lGKA-AVVKALDQLA--QE-----GKIKE-KT---Y--G-K-Q-KIYFAD---------
0005 6qfdB   aRDLTA--FQKNILTVL-G-EE----ARYGL-AIKRELE-E--YYG----E-----------eVNHG-RLYPNLDDLV--NK-----GLVEK-SE--lD--K-R-T-NEYALTNegfdavvd
0006 5yadA   NKSLSL--LSTETXSIL-Q-DA-PaCCLPLF-KFIDIYE-K--KYG----H--------------kL-NVSD-LYKLT---------DTIAI-RE---Q--G-N-G-RLVCLLPsnq-----
0007 7rheA   -NVRRY--NERLLLKTL-R-RA---GSASKA-DLARLAN------------------------mTGT-AVGSIIASLA--DA-----KLIEF-A-----------A-SLIRLDPrgafgigv
0008 7vwvB   FMETQK--LISMVKEAL-E-KY-Q-YPLTAK-NIKVVIQ-K--EHN----V-----------vLPTG-SINSILYSNS---------ELFEK-ID-ktN--T-I-YpPLWIRKN--------
0009 2l02A   KKIVGA--NAGKVWHAL-N-E--A-DGISIP-ELARKVN------------------------lSVE-STALAVGWLA--RE-----NKVVI-ER---K--N-G-L-IEIYNEGhfdfsfgl
0010 1qgpA   LSIYQD--QEQRILKFL-E-EL-G--ATTAH-DLSGKLG------------------------tPKK-EINRVLYSLA--KK-----GKLQK-EA---G----T-P-PLWKIAVsd------
0011 1cf7B   gkGLRH--FSMKVCEKV-Q-RK---GTTSYN-EVADELV-S--EFT----nshLAADSAYDQKNIRR-RVYDALNVLM--AM-----NIISK-EK------------KEIKWIGlp------
0012 2xrnB   sIQVIA--RAASIMRAL-G-SH-P--GLSLA-AIAQLVG------------------------lPRS-TVQRIINALE--EE-----FLVEA-LG---P------A-GGFRLGPalgqlinq
0013 3cuqA   GDFYYE--LGVQIIEVC-L-AL-KgGLITLE-ELHQQVL-K--GRG----K-------faQDVS-QD-DLIRAIKKLK--AL----GTGFGI-IP---V--G-G-T-YLIQSVPaelnmdht
0014 1ku9A   ihGLNK--SVGAVYAIL-Y-LS-D-KPLTIS-DIXEELK------------------------ISKG-NVSXSLKKLE--EL-----GFVRK-VW---I-gE-R-K-NYYEAVDgfssikdi
0015 8e4yA   SYDRiM--STHIVACLL-LyRH-R-QGIDLS-TLVEDFF-V--MKE------lardfdlgfsgNSED-VVMHAIQLLG---------NCVTI-TH---T-rN-D-E-FFITPSTtvpsvfel
0016 4ad9A   ISHRNI--REQQILTLF-R-EN-FeKSFTVM-ELVKIIY-K----N----T--PE----NLHEMAKH-NLLLHLKKLE--KE-----GKIFS-NT---D--P---D-KKWKAHL--------
0017 3elkA   RILHGL--ITLYILKEL-V-KR----PXHGY-ELQKSXF-E--TTG----Q---------ALPQ--G-SIYILLKTXK--ER-----GFVIS-ES---S--V-Q-L-TVYHITDagkkflxd
0018 7kbfK   ----pP--TLSMVVEVL-K-KN-T--GTSVQ-AIRTRIL-S--AHP----T-------vDPLRL-KF-LLRTALNKGL--EK-----GILIRpLN---S--S-gaT-GRFKLAK--------
0019 2od5A   SXKTVR--IREKIKKFL-G----D-RPRNTA-EILEHIN-S--TXR----H-----------gTTSQ-QLGNVLSKD----------KDIVKvGY---I--K-D-I-CEWATRNwvaehcpe
0020 8jxkD   ---EFM--LELAILGLL-I-ES----PMHGY-ELRKRLT-G--LLG----A--F------RAFS-YG-SLYPALRRMQ--AD-----GLIAE-NA---A--P-agR-RVYQLTDkgrrrfge
0021 8iueP   --DPVE--IENRIIELC-H-QF-P--GITDQ-VIQNE-M-P--HI-------------------eAQ-QRAVAINRLL--SM-----GQLDL-LR---S--N-T-G-LLYRIKDsqnagkmk
0022 4ijaA   --PMND--NEKRVLREI-Y-NH-H--NISRT-QISKNLE------------------------iNKA-TISSILNKLK--YK-----SLVNE-V------gR-K-P-ILLKVNHlygyfisl
0023 6qpqD   -----K--AIVQMAKIL-R-KE-L--EVIFT-DVLKSQA-N--------------tepENITKREAS-RGFFDILSLA--TE-----GCIGL-SQ---T--E-G-N-IKIDAKPalferfi-
0024 5y6iB   VRSA-E--VGTDILKAL-A-EL-S-PATSLS-RLAEHVG------------------------MPAS-KVHRYLQALI--AS-----GFAVQ-DA---S------T-NHYSLGRealrvgla
0025 8xt8B   ngydTT--DHYALLRFL-W-EQ---DGISQI-DLCEKSC------------------------KDKS-NTTRILDVMK--NK-----GLIVR-KV---D-dR-R-K-FQIFLTDlgreleep
0026 7cv0A   -ILGEE--RRSLLIKWL-K-AS-D--PLTGA-ELAKRTN------------------------VSRQ-VIVQDVSLLK--AK----NHPILA-TA------------QGYIYMKeantvqaq
0027 7y43A   klaNPL--YTEWILEAI-K-KV-K--rPSEE-RICNAVS-S--SHG-----------------lDRK-TVLEQLELSV--KD-----GTILK-VS---N--K-G-L-NSYKDPDnpgria--
```

```
0028 2zmeB   sHKEEE--MVASALETV-S-EK-G--SLTSE-EFAKLVG------------------------MSVL-LAKERLLLAE--KM-----GHLCR-DD---S--V-E-G-LRFYPNLfmtqs---
0029 3f72B   lkaiDE--NRAKITYAL-C-QD-E--ELCVC-DIANILG------------------------VTIA-NASHHLRTLY--KQ-----GVVNF-RK---E--G-K-L-ALYSLGGeairqimm
0030 6sj9A   GASRTE--RLLNLLLAL-L-NT-K--GLPRA-VLREKVY----HDS----A--------DNDVAFGR-XFERDKVDLK--QF---G-FEIET-L----X--D-P-AsARYRIGKdsnrlpdv
0031 4mtdD   nVRLTP--QRLEVLRLM-S-LQ-D-GAISAY-DLLDLLR-E--AEP----Q------------AKPP-TVYRALDFLL--EQ-----GFVHK-VE---S------T-NSYVLCHlfdqptht
0032 3bpxA   ghlnTD--AQVACLLRI-H-RE---PGIKQD-ELATFFH------------------------VDKG-TIARTLRRLE--ES-----GFIER-EQdpeN--R-R-R-YILEVTRrgeeiipl
0033 4y66C   -TSLDE--KKERLLEEM-L-KR-G-EIYSNK-TIETLSK-P--TG-----------------iSSMV-I-KNVLQALV--NE-----DLVDT-DK---I--G-A-S-TYYWCFAskrsqaar
0034 4i99C   --DIEK--YVEELYKVV-K-KI-Y--PIKFW-DLVP-----------------------dVEPKIIA-RTFLYLLFLE--NM-----GRVEI-IQ---E--E-G-E-ILVVPM---------
0035 2cweA   ikvmED--TRRKILKLL-R----N-KEMTIS-QLSEILG------------------------kTPQ-TIYHHIEKLK--EA-----GLVEVkRT---E--M-V-E-KYYGRTAdvfyinly
0036 2vqcA   tlNSYK--XAEIXYKIL-E-KK---GELT-----LEDIL-A--QFE-------------iSVPSAYN-IQRALKAICE--RH----pDECEV-QY---K--N-R-K-TTFKWIK--------
0037 3gfiA   ELNLSY--LDFLVLRAT-S----D-GPKTMA-YLANRYF------------------------vTQS-AITASVDKLE--EM-----GLVVR-VR---D-dR-R-K-ILIEITEkgletfnk
0038 7z8bC   EKRRNL--LNCLIVRIL-K-AH-G--GLHID-QLVCLVL-E--AWQ----KgglvsslgkgsaCSST-DVLSCILHLL--GK-----GTLRR-HD---D-----rP-QVLSYAV--------
0039 1xmkA   SLDMAE--IKEKICDYL-F-NV---SDSSAL-NLAKNIG-----------------------LTKAR-DINAVLIDME--RQ-----GDVYR-QG---T----T-P-PIWHLTDkkrermqi
0040 3k69A   XKLDFS--VAVHSILYL-D-AH-R--KVASR-ELAQSLH------------------------LNPV-XIRNILSVLH--KH-----GYLTG-TV---G----K-N-GGYQLDLaladxnlg
0041 3u2rA   QFELSA--QQYNTLRLL-R-SV-HpEGXATL-QIADRL---------------------------AP-DITRLIDRLD--DR-----GLVLR-TR---K-nR-R-V-VEVALTDaglkllkd
0042 6zvhy   ----fN--WKGTIKAIL-K-QA-PdNEITIK-KLRKKVL-A--QYY--tvT--DEHH-rSEEEL-LV-IFNKKISKN----------PTFKL-LK------------DKVKLVK--------
0043 5jbrA   TWQLPR--TTGRTYGYL-L-LQ-S--ATSFQ-EIGADLG------------------------lSPG-AVSTSVRELV--AW-----GLART-IP---Q-gS-R-R-LLVEAAGgfeqllaa
0044 1xsdA   kqveSM--AEWDVMNII-W-DK---KSVSAN-EIVVEIQ-K--YKE-----------------VSDK-TIRTLITRLY--KK-----EIIKR-YK---S--E-N-I-YFYSSNIkeddikmk
0045 1yyvA   VLKHVTsrWGVLILVAL-R----D-GTHRFS-DLRRXXG-G----------------------VSEX-XLAQSLQALE--QD-----GFLNR-VS---Y--P-P-H-VEYSLTPlgeqvsdx
0046 6ahtA   CENLND---GIWALRVL-Y-AE---GAXNKE-KLWDYIN-Q--YHKdyqiendYEGK---kilpsRY-ALDIXTARLE--GA-----GLISF-KA---I--G-R-V-RIYDVTDlgnvlike
0047 4fhtB   SEETTS--PQYAVLNAL-V-AE---PGLDQR-TVGERVG------------------------lDRS-TIAEVVSRLG--RR-----GLLDK-VR---D-dG-R-R-SLLRLTDeglrvhrr
0048 7b0cA   LTKFTD--LALRSLMRL-A-VV-R--PLATR-EVAEVVG------------------------vPYT-HAAKAITRLQ--HL-----GVVEA-RR---G--R---G-GGLTLTDlgrrvsvg
0049 2g9wA   kltrGD--LERAVXDHL-W-SR-T--PQTVR-QVHEALS-A--RRD-----------------LAYT-TVXAVLQRLA--KK-----NLVLQ-IR----------A-HRYAPVHgrdelvag
0050 3bddB   kqlgSL--TRYSILQTL-L-KD---APLHQL-ALQERLQ------------------------IDRA-AVTRHLKLLE--ES-----GYIIR-K-----------E-VLVWPTEqarealit
0051 1sfuA   tVNDAE--IFSLVKKEV-L-SL-N--YTTAI-SLSNRLK------------------------iNKK-KINQQLYKLQ--KE-----DTVKM-VP---S----N-P-PKWFKNYnc------
0052 5n35A   --VNQK--EIEIAIEYF-K-NY---ISVGEI-VATMDLK-A--RG---------------------isNPQAVISKLI--EM-----GIIEK-GE------------GCYNLVR--------
0053 5eriA   kelkQK--GQFTFLTRI-C-EN---PGINLV-ELSNMLK------------------------VDKA-TTTKAIQKLI--KA-----GYVDK-KQdkfD--K-R-G-YNLTPTDkslevyel
0054 3f8fA   EMLRAQ--TNVILLNVL-K-QG----DNYVY-GIIKQVK-E--ASN----G---------emeLNEA-TLYTIFKRLE--KD-----GIISS-YW---G--D-R-R-KYYRLTEighenmrl
0055 3r4kA   -XGTVS--KALTLLTYF-N--hgR-LEIGLS-DLTRLSG------------------------xNKA-TVYRLXSELQ--EA-----GFVEQ-VE---G------A-RSYRLGPqvlrlaal
0056 7c0jB   ----pT--YSEMIAAAI-R-AE-G--GSSRQ-SIQAYIK-S--HYK-----------------vnKK-EINRVLYSLL--AA-----GVLKQ-TG---V------P-GSWALA---------
0057 3w6kC   mGALKP--AKAIVEALL-F-AA-G--GLSLS-QIAAVLE------------------------VSEL-EAKAVIEELQ--QD---CRrQLVE-LG------------GVFLLATkkehapyl
0058 2ethA   eisdKT--TELYAFLYV-A-LF---GPKKXK-EIAEFLS------------------------tTKS-NVTNVVDSLE--KR-----GLVVR-EXdpvD--R-R-T-YRVVLTEkgkeifge
0059 1fokA   KEYVRT--RRALILEIL-I-KA---GSLKIE-QIQDNLK-K--LG----------------fdEVIE-TIENDIKGLI--NT----GIFIEI-KG------------RFYQLKDhilqfvip
0060 5udb4   SVIQRK--LQEDLSREI-M-NV-LsDSMSFN-ELIKQIN-E---HS----Q--D------RVES--S-DIQEALSRLQ--QE-----DKVIV-LG----------E-GVRRSVR--------
0061 8as8C   rrqrNL--EQSLLVAIL-R-QH-FgtQIAID-DLLPQLQ-I--YLG----D-------pGSESKERT-RLLTLLDQLK--GH-----GLVTS-PD---A------H-ERIVIRPiiahladp
0062 2fxaB   pYDLNI--NEHHILWIA-Y-QL-N--GASIS-EIAKFGV------------------------xHVS-TAFNFSKKLE--ER-----GYLRF-SK---R-dK-R-N-TYVQLTEegtevfws
0063 2lnbA   PGREGH--LEQRILQVL-T-EA-G-SPVKLA-QLVKECQ------------------------aPKR-ELNQVLYRMK--KE-----LKVSL-TS----------P-ATWCLGGtdpe----
0064 4hqeB   LGRS----WNGLIINYL-S-RC-NdCSAHFS-DMKRDLK-T----------------------ITPR-ALSLKLSELA--QW-----ELVEK-QI---I--S-V-Q-IIYVLTEkgkalaea
0065 3bj6A   lregTV--GQRAILEGL-S-LT---PGATAP-QLGAALQ------------------------xKRQ-YISRILQEVQ--RA-----GLIER-RT---N-hA-R-S-HRYWLTPrgeaiita
0066 4hw0A   kRGTXE--IXFDILRNC-E------PKCGIT-RVIYGAG------------------------iNYV-VAQKYLDQLV--KV-----GALNI-KT---E--N-D-R-KIYEITEkgkllrth
0067 2zkzC   lktxHP--XRLKIVNEL-Y-KH---KALNVT-QIIQILK------------------------LPQS-TVSQHLCKXR--------gKVLKR-NR---Q--G-L-E-IYYSINNpkvegiik
0068 8uucA   egSRRQ--KRAELVRLV-L-AE---PGIGID-ELAERLD-A--FER----D-----agrkGVDA--A-TFTSIVADLV--AE-----GFFRR-EG------------DAFFA----------
0069 2dqlA   ylCQEV--AICYILYVL-L-Q----GESYGT-ELIQQLE-T--EHP----T-----------yRLSDtVLYSAIKFLE--DN-----RAITG-YW---K--K-P-R-RMYQVSPewqhqaed
0070 7xjgJ   gsftlY--AERFLLPCI-Y-LL-D--SVNYR-TLCELAF-K--AIK----Q---------DDVLSKI-IVRSVVSRLI--NE-----RKILQ-MT------------DGYQVTAlgasyvrs
0071 3s93B   ERIQEC--LRKEIRSLL-I-ST-K--GLSPQ-ELEKEYL-L--MVG----N-------hlplrilgyrsTMELVLDMP---------DVVRV-CP---G--A-G-T-VILKAIP--------
0072 2eshA   GFRGWW--LASTILLLV-A-EK----PSHGY-ELAERLA-E--FGI----E--------ipgighmG-NIYRVLADLE--ES-----GFLST-EW---D--T-P-R-KIYRITPqgklylre
0073 9gm7C   PVERLA--FLMAIAVLN-S-ERlD-GEMSEG-ELIDAFR-E--VCK-gfeQ--TAE---SVAVRANN-AINDMVRQKL-----------LNR-FT---S-aD-G-N-AIYRLTPlgisisdy
0074 5hs5A   SLNDLA--VLDLTXKHC-K----D-EKVLXQ-SFLKTAX-D--ELD-----------------LSRT-KLLVSIRRLI--EK-----ERLSK-VR---S-dE-R-K-IYIYLNNddiskfna
0075 8sojB   REDK-D--LHRKIHRIIqQ-DC-QkpGCHFL-HILACAR-L-sIRP----G------------LSEA-VLQQVLELLE--DQ-----SDIVS-TM----------E-HYYTA----------
0076 8f8uA   KDHT----IPLTLISIL-A-D--G-EFHSGE-QLGEQLG------------------------MSRA-AINKHIQTLR--DW----GVDVFT-VP----------G-KGYSLPEpihlldek
0077 2xcoA   vrdgKP--VHRRILYGL-N-EQ-G-MYKKSA-RIVGDVM-G-kYHP----H------------GDSS-IYEAMVRMAQdfSY---RYPLVDG-QG---N----mrY-TEARMTKitlellrd
0078 2p4wB   LDVLNE--TRRRILFLL-T-K----RPYFVS-ELSRELG------------------------vGQK-AVLEHLRILE--EA-----GLIES-RV---E--K-P-R-KYYMIKKglrleill
0079 1lddA   ELTLQR--SLPFIEGML-T-NL---GAMKLH-KIHSFLK-I--TVP----K--D---wgyNRIT-LQ-QLEGYLNTLA--DE-----GRLKY-IA---N--------GSYEIV---------
0080 1w7pD   ELFLDE--IAREIYEFT-L-SE-FyMIITLV-DLYAMYN-K-sMRI----G---------tglISPM-EMREACERFE--HL---GLNELKL-VK---V--N-R-I-LCVTSE---------
0081 9bz0f   IKYFVK--GRQAVLSMV-R-KA-KyRELLLS-ELLGR-R-A--PV-------------------vvrlGLTYHVHDLI--GA-----QLVDC-IS---T--T-S-G-TLLRLPEt-------
0082 5u8oB   RAHHDA--RLDEVRVAC-A----E-QPMSAA-DIVPIMF-R----R----R--EL--DIHQMTFALG-EAIAHLNLLW--LA-----GELVR-EQ---G-dD-G-V-LRFRIAR--------
0083 3l9fA   ----mQ--GKDIILGIL-S-KK----ERSGY-EINDILQnQ--LSY----F----------ydgtyG-MIYPTLRKLE--KD-----GKITK-EV---V--I-N-K-NIYAITEsgkkelas
0084 2ns0A   tvSDRE--LEECIRALL-D-AR-A--SICPS-DVARAVA-P------------------dDWRPLXE-PVREAAGRLA--DA-----GEVEV-TQ---KgaR-G-P-IRIRWTRtd------
0085 6juvA   kPELRA--GELKTLALI-A-YL---QPVEQS-KIIKLRG---------------------------S-QAYEHIKKLL--EX-----GLIYA-EP---Y--E-R-T-KLLGTTQkfaelygf
0086 2qlzB   EIGD-X--TQLAILHYL-L-L--N-GRATVE-ELSDRLN------------------------LKER-EVREKISEXA--RF-----VPVKI-IN----------D-NTVVLDEdqilr---
0087 5l0pA   KVTL----PRLKILEVL-Q-EP-DnHHVSAE-DLYKRLI-D--MGE----E------------IGLA-TVYRVLNQFD--DA-----GIVTR-HN---F--E-GgK-SVFELT---------
0088 9qe1J   TADSAA--HYRTILRYF-Y-HQ-HrDFIAPE-ELLEHMR-SipAFA----D------------FQED-QLHQQLAQLV--KW-----NNLIA-RQ---D--M-K-R-FRYQCTPytveierm
0089 5zqhA   QLLKGV--LEGCVLDMI-G-QK----ERYGY-ELVQTLR-E-aGFD----T-------------ivG-TIYPLLQKLE--KN-----QWIRG-DM---R--P-D-R-KYFSLMKegeervsv
0090 3lmmD   stDQAE--LTNAAXLWL-S-EV---GDLATS-DLXAXCG------------------------VSRG-TAKACVDGLV--DE-----ERVVA-VG---G--G-R-S-RRYRLVE--------
0091 7bzhA   ----ED--VKQSVEKII-K-DR---EWVTFN-DLLKYIP------------------------YPAP-EVYDALSQLI--KE-----NKVGR-RG------------RYFYYIKr-------
0092 5e20A   kslnST--EKFAILHEL-M-CL-G-GESTPH-TLARRIV------------------------FEPH-SVSAIVSRME--KD-----GLIIK-TK---D-kK-H-M-VRIKLSEkaidtfyq
0093 1ylfC   ISSRFS--IAVHILSIL-K-NN-P--lCTSD-YXAESVN------------------------tNPV-VIRKIXSYLK--QA----GFVYVN-RG----------P-GGAGLLKdlheitll
0094 3lwfA   ITTKGR--YGLTITLEL-A-KR-I--PISLR-SIAQDKN------------------------lSEH-YLEQLIGPLR--NA-----GIVKS-IR---G----A-H-GGYVLNGdpekitag
0095 7xx7U   NQPG-K--YSQLVVETI-R-RL-G--GSSLA-KIYTEAK----KVP----W-------fDQQNG-RT-YLKYSIKALV--QN-----DTLLQ-VK---G----A-N-GSFKLNRkkleggge
0096 4ru7B   ----lT--PRAYIIVHL-L-KV---GKAKAS-EISENTQ------------------------iPYQ-TVIQNIRWLL--AE-----GYVVK-EQ---K--G-E-E-IYYKLTDkgkqlata
0097 2co5B   KYMR-I--NYYIILKVL-V-IN-G-SRLEKK-RLRSEIL-K--RFD-------------iDISD---gvlYPLIDSLI--DD-----KILRE-EE---A--P-D-G-KVLFLTEkgmkefee
0098 5xfoA   vrrlLR--WVDVAHLVL-Y-HL-SkKYFDFDrEILPFTS-E--NWD------llgelsdTPKGERSS-RLLSALNSHK---------DRFIS-GR---E--I-R-K-CLFGLHArmpppve-
0099 8ylgA   RYGLQP--GEFDVLATL-R-RS-GpYALTPT-ALYDAAM-I-------------------SSGSMTN-RIDRLEKA-----------GWVER-RAnpaD--G-R-G-TLVALTSagralidd
0100 4b8xA   pygLTF--ARYEALVLL-T-FS-KsGELPMS-KIGERLM------------------------VHPT-SVTNTVDRLV--RS-----GLVAK-RP---N--P---G-TLATITDkgrevvea
0101 9f27A   NELLEK--AKEDILNIL-R-QK-R-TAISRK-YILKKLG-D--KY-------------------dEE-TIDDAITELL--AQ-----GEIYE-PE----------T-GYYKLL---------

0001 s001A   LHHHHH--HHHHHHHHH-H-HL-L-LEEEHH-HHHHHHH-H--HHL----L--LLLLHHHHHHHHHH-HHHHHHHHHH--LL---LLLLEEE-EE---E--L-L-E-EEEEELL--------
0002 5j6xA   LLHHHH--HHHHHHHHH-H-HH---LLLLHH-HHHHHLL------------------------lLHH-HHHHHHHHHH--HL-----LLEEE-EL---L--L-L-L-LEEEELL--------
0003 7qcdD   LHHHHH--HHHHHHHHH-H-HL-L-LEEEHH-HHHHHHH-H--HHHhhllL--LLL---LLHHHHHH-HHHHHHHHHH--HHhhhLLEEEEE-EE---L--LlL-L-EEEEEEEllllhhhh
0004 2mh2A   -----L--HHHHHHHHH-H-HH-L-LLEEHH-HHHHHHH-L--LLL-----------------lLHH-HHHHHHHHHH--HH-----LLEEE-EE---E--L-L-E-EEEEEL---------
0005 6qfdB   hLLLLH--HHHHHHHHH-H-HL----LEEHH-HHHHHHH-H--HHL----L-----------lLLHH-HHHHHHHHHH--HL-----LLEEE-EE--eE--L-L-E-EEEEELHhhhhhhhh
0006 5yadA   HHHHHH--HHHHHHHHH-H-LL-HhHEEEHH-HHHHHHH-H--HHL----L--------------lL-LHHH-HLLLL---------LLEEE-EL---L--L-L-L-LEEEELHhhl-----
0007 7rheA   -LHHHH--HHHHHHHHH-H-HH---LLEEHH-HHHHHHH------------------------lLHH-HHHHHHHHHH--HL-----LLEEE-L-----------L-LEEEELLllleeeee
0008 7vwvB   LLHHHH--HHHHHHHHH-H-HL-L-LLLLHH-HHHHHHH-H--HHL----L-----------lLLHH-HHHHHHHHLL---------LLEEE-LL-llL--L-L-LlLLEEELL--------
0009 2l02A   HHHHHH--HHHHHHHHH-H-H--L-LLLLHH-HHHHHHL------------------------lLHH-HHHHHHHHHH--LL-----LLEEE-EE---E--L-L-E-EEEEEHHhlllllll
0010 1qgpA   LHHHHH--HHHHHHHHH-H-HH-L--LEEHH-HHHHHHL------------------------lLHH-HHHHHHHHHH--HH-----LLEEE-EL---L----L-L-LEEEELLll------
0011 1cf7B   llHHHH--HHHHHHHHH-H-HH---LEELHH-HHHHHHH-H--HHH----lllLHHHLHHHHHHHHH-HHHHHHHHHH--HL-----LLLLL-LL------------LLEELLLll------
0012 2xrnB   lLHHHH--HHHHHHHHH-H-LL-L--LEEHH-HHHHHLL------------------------lLHH-HHHHHHHHHH--LL-----LLEEE-LH---H------H-LEEEELLhhhhhhhh
0013 3cuqA   HHHHHH--HHHHHHHHH-H-HH-HlLEEEHH-HHHHHHH-H--LLL----L-------llLLLL-HH-HHHHHHHHHH--HH----LLLLEE-EE---E--L-L-E-EEEELLLllllhhhh
0014 1ku9A   hlLLLH--HHHHHHHHH-H-HL-L-LLEEHH-HHHHHHL------------------------LLHH-HHHHHHHHHH--HL-----LLEEE-EL---L-lL-L-L-LEEEELLhhhhhhhh
0015 8e4yA   EELLlL--HHHHHHHHH-HhLL-L-LLEEHH-HHHHHHH-H--HHH------hhlllllllllLHHH-HHHHHHHHLL---------LLEEE-EE---L-lL-L-L-EEEEELLlhhhhhhh
0016 4ad9A   HHHHHH--HHHHHHHHH-H-HL-LlLLLLHH-HHHHHHL-L----L----L--LH----HHHHHHHH-HHHHHHHHHH--HL-----LLEEE-LL---L--L---L-LLEEELL--------
0017 3elkA   HHHHHH--HHHHHHHHH-H-HL----LEEHH-HHHHHHH-H--HHL----L---------LLLL--L-HHHHHHHHHH--HH-----LLEEE-EE---E--E-E-E-EEEEELHhhhhhhhh
0018 7kbfK   ----lL--HHHHHHHHH-H-HL-L--LEEHH-HHHHHHH-L--LLL----L-------lLLLLH-HH-HHHHHHHHHH--HL-----LLEEEeEE---L--L-llL-EEEEELL--------
0019 2od5A   HHHHHH--HHHHHHHHL-L----L-LLEEHH-HHHHHHH-L--LLL----L-----------lLLHH-HHHHHHHHL----------LLEEEeEE---E--E-E-E-EEEEEHHhhhhhlll
0020 8jxkD   ---LHH--HHHHHHHHH-H-HL----LEEHH-HHHHHHH-H--HLL----L--L------LLLL-LL-LHHHHHHHHH--HL-----LLEEE-LL---L--L-llL-LEEEELHhhhhhhhh
0021 8iueP   --LHHH--HHHHHHHHH-H-HL-L--LLLHH-HHHHH-L-L--LL-------------------lHH-HHHHHHHHHH--HH-----LLEEE-EE---E--L-L-E-EEEEELHhhllllll
0022 4ijaA   --LLHH--HHHHHHHHH-H-HL-L--LLLHH-HHHHHHL------------------------lLHH-HHHHHHHHHH--HL-----LLEEE-L------lL-L-L-LLEEELHhhleeeee
0023 6qpqD   -----L--HHHHHHHHH-H-HH-H--LEEHH-HHHHHHH-L--------------llhHHLLHHHHH-HHHHHHHHHH--HL-----LLEEE-EL---L--L-L-L-LEEEELHhhhllll-
0024 5y6iB   LLHH-H--HHHHHHHHH-H-HL-L-LLEEHH-HHHHHHL------------------------LLHH-HHHHHHHHHH--HH-----LLEEE-LL---L------L-LEEEELHhhhhhhhh
0025 8xt8B   llllLH--HHHHHHHHH-H-HL---LLLLHH-HHHHHHL------------------------LLLH-HHHHHHHHHH--LL-----LLEEE-EE---L-lL-L-L-EEEEELHhhhhlhhh
0026 7cv0A   -LHHHH--HHHHHHHHH-H-HL-L--LLLHH-HHHHHHL------------------------LLHH-HHHHHHHHHH--HL----LLLEEE-EL------------LEEEELLllllllee
0027 7y43A   lllLHH--HHHHHHHHH-H-HH-H--lLLHH-HHHHHHH-H--HHL-----------------lLHH-HHHHHHHHHH--HH-----LLLEE-EE---E--L-L-E-EEEELLLllllll--
0028 2zmeB   lLLHHH--HHHHHHHHH-H-HH-L--LLLHH-HHHHHHL------------------------LLHH-HHHHHHHHHH--HH-----LLEEE-EE---E--L-L-E-EEEEELHhhlll---
0029 3f72B   hhhhLH--HHHHHHHHL-L-LL-L--LEEHH-HHHHHHL------------------------LLHH-HHHHHHHHHH--HL-----LLEEE-EE---L--L-L-L-EEEEELLhhhhhhhh
0030 6sj9A   LHHHHH--HHHHHHHHH-H-HL-L--LEEHH-HHHHHHL----LLL----L--------LLHHHHHH-HHHHHHHHHH--HL---L-LLLEE-E----L--L-L-LlLEEELLLllllllee
0031 4mtdD   lLLLLH--HHHHHHHHH-H-LL-L-LLEEHH-HHHHHHH-H--HLL----L------------LLHH-HHHHHHHHHH--HL-----LLEEE-EL---L------L-LEEEELLllllllll
0032 3bpxA   hhhlLH--HHHHHHHHH-H-LL---LLLLHH-HHHHHHL------------------------LLHH-HHHHHHHHHH--HL-----LLEEE-EEellE--E-E-E-EEEEELHhhhhhhhh
0033 4y66C   -LHHHH--HHHHHHHHH-H-HH-L-LLLLLL-HHHHHHH-H--HL-----------------lLLLL-H-HHHHHHHH--HL-----LLLEE-EE---L--L-L-L-EEEELLLlhhhhhhh
0034 4i99C   --LHHH--HHHHHHHHH-H-HH-H--LEEHH-HHLL-----------------------lLLHHHHH-HHHHHHHHHH--HL-----LLEEE-EL---L--L-L-L-LEEEEL---------
0035 2cweA   hhhhLH--HHHHHHHHH-L----L-LLLLHH-HHHHHHL------------------------lLHH-HHHHHHHHHH--HL-----LLEEEeEE---E--E-E-E-EEEEELLleeeelll
0036 2vqcA   llLHHH--HHHHHHHHH-H-HH---LEEL-----HHHHH-H--HHL-------------lLHHHHHH-HHHHHHHHHH--HL----lLLEEE-EE---L--L-L-L-EEEEELL--------
0037 3gfiA   HHLLLH--HHHHHHHHH-L----L-LLEEHH-HHHHHLL------------------------lLHH-HHHHHHHHHH--HL-----LLEEE-EE---E-eE-E-E-EEEEELHhhhhhhhh
0038 7z8bC   HHHHHH--HHHHHHHHH-H-LL-L--LEEHH-HHHHHHH-H--HHH----LlhhhhhllllllLLHH-HHHHHHHHHH--HH-----LLEEE-LL---L-----lL-LEEEELL--------
0039 1xmkA   LHHHHH--HHHHHHHHH-H-HL---LLEEHH-HHHHHHL-----------------------HHHHH-HHHHHHHHHH--HL-----LLEEE-EL---L----L-L-LEEEELHhhhlllll
0040 3k69A   LLHHHH--HHHHHHHHH-H-LL-L--LLLHH-HHHHHHL------------------------LLHH-HLHHHHHHHH--HL-----LLLEE-EL---L----L-L-LEEELLLlhhhllhh
0041 3u2rA   LLLLLH--HHHHHHHHH-H-HH-LlLLEEHH-HHHHHL---------------------------LL-HHHHHHHHHH--HL-----LLEEE-EE---E-eE-E-E-EEEEELHhhhhhhhh
0042 6zvhy   ----lL--HHHHHHHHH-H-LL-HhHEEEHH-HHHHHHH-H--HHH--hhL--LLLL-lLHHHH-HH-HHHHHHHLL---------LLEEE-LL------------LEEEELL--------
0043 5jbrA   HHLLLH--HHHHHHHHH-H-LL-L--LEEHH-HHHHHHL------------------------lLHH-HHHHHHHHHH--HL-----LLEEE-EE---E-lE-E-E-EEEEELLllhhhhhh
0044 1xsdA   llllLL--LLHHHHHHH-H-HH---LLEEHH-HHHHHHH-L--LLL-----------------LLHH-HHHHHHHHHH--HL-----LLEEE-EE---L--L-L-L-EEEEELLlhhhhhhh
0045 1yyvA   HHHHHHlhHHHHHHHHH-H----H-LLEEHH-HHHHHLL-L----------------------LLHH-HHHHHHHHHH--HH-----LLEEE-EE---E--L-L-E-EEEEELHhhhhhhhh
0046 6ahtA   HHHHHH---HHHHHHHH-H-HH---LLEEHH-HHHHHHH-H--HHHhhhhhhlLLLL---lllllHH-HHHHHHHHHH--HL-----LLEEE-EE---E--L-L-E-EEEEELHhhhhhhhh
0047 4fhtB   LLLLLH--HHHHHHHHH-H-HL---LLLLHH-HHHHHHL------------------------lLHH-HHHHHHHHHH--HH-----LLEEE-EE---L-lL-L-L-EEEEELHhhhhhhhh
0048 7b0cA   LLHHHH--HHHHHHHHH-H-LL-L--LLLHH-HHHHHHL------------------------lLHH-HHHHHHHHHH--HH-----LLEEE-LL---L--L---L-LLEEELHhhhhllhh
0049 2g9wA   lhhhLH--HHHHHHHHH-H-LL-L--LEEHH-HHHHHHL-L--LLL-----------------LLHH-HHHHHHHHHH--HL-----LLEEE-EL----------L-LEEEELLlhhhhhhh
0050 3bddB   hhhlLH--HHHHHHHHH-H-HL---LLEEHH-HHHHHHL------------------------LLHH-HHHHHHHHHH--HL-----LLEEE-L-----------L-LEEEELHhhhhhhll
0051 1sfuA   lLLLHH--HHHHHHHHH-H-LL-L--EELHH-HHHHHLL------------------------lLHH-HHHHHHHHHH--HL-----LLEEE-EL---L----L-L-LEEEELLll------
0052 5n35A   --LLHH--HHHHHHHHH-H-HL---LEEEHH-HHHHHHH-H--LL---------------------llLHHHHHHHHH--HL-----LLEEE-LL------------LEEEELL--------
0053 5eriA   llllLL--LHHHHHHHH-H-HL---LLEEHH-HHHHHHL------------------------LLHH-HHHHHHHHHH--HL-----LLEEE-EElllL--L-L-L-EEEEELHhhhhhhhh
0054 3f8fA   HHHHHH--HHHHHHHHH-H-HL----LLLHH-HHHHHHH-H--HLL----L---------lllLLHH-HHHHHHHHHH--HL-----LLEEE-EE---L--L-L-L-EEEEELHhhhhhhhh
0055 3r4kA   -LLHHH--HHHHHHLLL-L--llL-LEEEHH-HHHHHHL------------------------lLHH-HHHHHHHHHH--HL-----LLEEE-LL---L------L-LEEEELLlhhhhhhh
0056 7c0jB   ----lL--HHHHHHHHH-H-HL-L--LEEHH-HHHHHHH-H--HLL-----------------llHH-HHHHHHHHHH--HL-----LLEEE-LL---L------L-LEEEEL---------
0057 3w6kC   lLLLHH--HHHHHHHHH-H-HH-H--LLLHH-HHHHHHL------------------------LLHH-HHHHHHHHHH--HH---HLlEEEE-EL------------LEEEEEElhhhhhhh
```

```
0058 2ethA   hhhhLH--HHHHHHHHH-H-HH---LLLLHH-HHHHHLL-------------------------lLHH-HHHHHHHHHH--HL-----LLEEE-EElllL--L-L-L-EEEEELHhhhhhhhh
0059 1fokA   HHHHHH--HHHHHHHHH-H-HL---LLEEHH-HHHHHHH-H--HL-----------------llLLHH-HHHHHHHHHH--HL----LLLEEE-EL-----------LEEEEELLllllllll
0060 5udb4   LLLHHH--HHHHHHHHH-H-HH-HlLLLLHH-HHHHHLL-L---LL----L--L------LLLL--H-HHHHHHHHHH--HL-----LLEEL-LL----------L-LLLEELL--------
0061 8as8C   llllLH--HHHHHHHHH-H-HH-HllEEEHH-HHHHHHH-H--HHL----L-------lLLHHHHHH-HHHHHHHHHH--HL-----LLLLL-LL---L------L-LEEELLLlhhhhllh
0062 2fxaB   hHLLLH--HHHHHHHHH-H-HH-L--LEEHH-HHHHHHL-------------------------lLHH-HHHHHHHHHH--HH-----LLEEE-EL---L-lL-L-L-LEEEELHhhhhhhhh
0063 2lnbA   LLHHHH--HHHHHHHHH-H-HH-L-LLEEHH-HHHHHHL-------------------------lLHH-HHHHHHHHHH--HL-----LLEEE-EE----------L-LEEEELLllll----
0064 4hqeB   HLLL----LHHHHHHHH-L-LL-HhHEELHH-HHHHHHL-L-----------------------LLHH-HHHHHHHHHH--HL-----LLEEE-EE---E--L-L-E-EEEEELHhhhhhhhl
0065 3bj6A   hhllLH--HHHHHHHHH-H-HL---LLEEHH-HHHHHHL-------------------------lLHH-HHHHHHHHHH--HL-----LLEEE-EL---L-lL-L-L-LEEEELHhhhhhhhh
0066 4hw0A   lLLHHH--HHHHHHLLL-L------LEELHH-HHHHHHL-------------------------lLHH-HHHHHHHHHH--HL-----LLEEE-EE---E--L-L-E-EEEEELHhhhhhhhh
0067 2zkzC   hhhhLH--HHHHHHHHH-H-HH---LLEEHH-HHHHHHL-------------------------LLHH-HHHHHHHHHL--------lLLEEE-EE---E--L-L-E-EEEEELLhhhhhhhh
0068 8uucA   llLHHH--HHHHHHHHH-H-HL---LLEEHH-HHHHHHH-H--HHH----H-----llllLLLH--H-HHHHHHHHHH--HH-----LLLEE-EL------------LEEEL----------
0069 2dqlA   elLHHH--HHHHHHHHH-L-L----LLLLHH-HHHHHHH-H--HLL----L-----------eELLHhHHHHHHHHHH--HL-----LLEEE-EE---E--L-L-E-EEEEELHhhhhhhhh
0070 7xjgJ   hhhhlL--LHHHHHHHH-H-HL-L--LLLHH-HHHHHHH-H--HLL----L---------LLHHHHH-HHHHHHHHHH--HL-----LLEEE-EL------------LEEEELHhhhhhhhh
0071 3s93B   HHHHHH--HHHHHHHHH-H-LL-L--LLLHH-HHHHHHH-H--HHL----L-------llllhhhllllHHHHHHLLL---------LLEEE-EE---L--H-H-L-EEEEELL--------
0072 2eshA   HHHHHH--HHHHHHHHH-H-HL----LLLHH-HHHHHHH-L--LLL----L--------lllllllL-LHHHHHHHHH--HL-----LLEEE-EE---E--L-L-E-EEEEELHhhhhhhhh
0073 9gm7C   LHHHHH--HHHHHHHHH-H-LLlL-LLEEHH-HHHHHHH-H--HHH-hllL--LLL---LHHHHHHH-HHHHHHHLLL-----------EEE-EE---L-hL-L-L-EEEEELHhhhhhhhh
0074 5hs5A   LHHHHH--HHHHHHHHH-L----L-LLEEHH-HHHHHHH-H--HHL------------------LLHH-HHHHHHHHHH--HL-----LLLEE-EL---L-lL-L-L-LEEELLHhhhhhhhh
0075 8sojB   HHLH-H--HHHHHHHHHlL-LL-LllLEEHH-HHHHHHH-H-hLLL----L------------LLHH-HHHHHHHHHH--HH-----LLLEE-EE----------L-LEEEL----------
0076 8f8uA   LLLH----HHHHHHHHH-L-L--L-LLLLHH-HHHHHHL-------------------------LLHH-HHHHHHHHHH--HL----LLLLEE-EL----------L-LEEELLLllllllhh
0077 2xcoA   llllLH--HHHHHHHHH-H-HL-L-LLEELH-HHHHHHH-H-hLLL----L------------LHHH-HHHHHHHHLLllLL---LLLLEEE-EL---L----llL-LEEEELHhhhlllll
0078 2p4wB   HHHLLH--HHHHHHHHH-L-L----LLEEHH-HHHHHHL-------------------------lLHH-HHHHHHHHHH--HL-----LLEEE-EE---L--L-L-L-EEEEELLleeeeeee
0079 1lddA   HHHHHH--HHHHHHHHH-H-HH---LLEEHH-HHHHHHH-H--HLL----H--H---hllLLLL-HH-HHHHHHHHHH--HL-----LLEEL-LL---L---------LEEEEL---------
0080 1w7pD   HHHHHH--HHHHHHHHH-H-HL-LlLEEEHH-HHHHHHH-H-hLLL----L---------lllLLHH-HHHHHHLLHH--HH---LLLLLEE-EE---L--L-L-L-EEEELL---------
0081 9bz0f   HHHHHH--HHHHHHHHH-H-HL-HhHEEEHH-HHHHL-L-L--LL--------------------llllLHHHHHHHHH--HL-----LLLEE-EL---L--L-L-L-LEEELLLl-------
0082 5u8oB   HHHHHH--HHHHHHHHH-H----H-LLLLHH-HHHHHHL-L----L----L--LL--LHHHHHHHHH-HHHHHHHHHH--HL-----LLEEE-EE---L-lL-L-L-EEEEELL--------
0083 3l9fA   ----lL--HHHHHHHHL-L-LL----LEEHH-HHHHHHHhL--LLL----L----------eelllL-LHHHHHHHHH--HL-----LLEEE-EE---E--L-L-E-EEEEELHhhhhhhhh
0084 2ns0A   llLHHH--HHHHHHHHH-H-HL-L--LLLHH-HHHHHHL-L------------------lLLHHHHH-HHHHHHHHHH--HL-----LLEEE-EE---LllL-L-L-LEEEELLll------
0085 6juvA   lLLLLH--HHHHHHHHH-H-HH---LLEEHH-HHHHLLH----------------------------H-HHHHHHHHHH--HL-----LLEEE-EE---E--L-L-E-EEEEELHhhhhhhll
0086 2qlzB   HLLL-H--HHHHHHHHH-H-H--H-LEEEHH-HHHHHHL-------------------------LLHH-HHHHHHHHHL--LL-----LLEEE-EL----------L-LEEEELHhhhhl---
0087 5l0pA   LLLH----HHHHHHHHH-L-LL-LlLEEEHH-HHHHHHH-H--LLL----L------------LLHH-HHHHHHHHHH--HL-----LLEEE-EL---L--L-LlL-LEEEEL---------
0088 9qe1J   HLLLHH--HHHHHHHHH-H-HH-HlLLLLHH-HHHHHHH-HllLLL----L------------LLHH-HHHHHHHHHH--HL-----LLEEE-EE---L--L-L-L-EEEEELHhhhhhhhh
0089 5zqhA   HHHHHH--HHHHHHHHH-H-HL----LEEHH-HHHHHHH-H-hLLL----L-------------llL-LLHHHHHHHH--HL-----LLEEE-EE---E--E-E-E-EEEEELHhhhhhhhh
0090 3lmmD   llLHHH--HHHHHHHHH-H-HL---LLEEHH-HHHHHLL-------------------------LLHH-HHHHHHHHHH--HL-----LLEEE-EL---L--L-L-L-LEEEELL--------
0091 7bzhA   ----LL--HHHHHHHHH-H-HH---LEEEHH-HHHHHLL-------------------------LLHH-HHHHHHHHHH--HL-----LLEEE-EL------------LEEEELLl-------
0092 5e20A   hhhlLH--HHHHHHHHH-H-HL-L-LEELHH-HHHHHLL-------------------------LLHH-HHHHHHHHHH--HL-----LLEEE-EE---L-lL-L-L-EEEEELHhhhhhhhh
0093 1ylfC   LLHHHH--HHHHHHHHH-H-HL-L--lLLHH-HHHHHLL-------------------------lLHH-HHHHHHHHHH--HH----LLEEEL-LL----------L-LLEEELLlhhhllhh
0094 3lwfA   LLHHHH--HHHHHHHHH-H-HL-L--LLLHH-HHHHHHL-------------------------lLHH-HHHHHHHHHH--HL-----LLEEE-EL---L----L-L-LEEEELLlllllllhh
0095 7xx7U   LLLL-H--HHHHHHHHH-H-HL-L--LLLHH-HHHHHHH----LLL----L-------lLLLLH-HH-HHHHHHHHHH--HL-----LLEEE-EL---L----L-L-LEEEELLlllllllll
0096 4ru7B   ----lL--HHHHHHHHH-H-LL---LEELHH-HHHHHHL-------------------------lLHH-HHHHHHHHHH--HH-----LLEEL-LL---L--L-L-L-LLEEELHhhhhhhhh
0097 2co5B   LLLL-H--HHHHHHHHH-H-HL-L-LEEEHH-HHHHHHH-H--HHL--------------lLLLH---hhhHHHHHHHHH--HL-----LLEEE-EL---L--L-L-L-LEEEELHhhhhhhhh
0098 5xfoA   eeelLL--HHHHHHHHH-H-HH-HlLLEELLlLHHHHHH-H--LHH------llhhhllLLLHHHHH-HHHHHHHHLL---------LLEEE-EE---E--H-L-E-EEEEELLllllllll-
0099 8ylgA   HLLLLH--HHHHHHHHH-H-HL-LlLEELHH-HHHHHHL-L-------------------LHHHHHH-HHHHHHHH-----------LLEEE-EElllL--L-L-L-EEEEELHhhhhhhhh
0100 4b8xA   hhlLLH--HHHHHHHHH-H-LL-HhHEEEHH-HHHHHHL-------------------------LLHH-HHHHHHHHHH--HL-----LLEEE-EE---L--L---L-EEEEELHhhhhhhhh
0101 9f27A   LHHHHH--HHHHHHHHH-H-HH-L-LLEEHH-HHHHHHH-H--LL--------------------lHH-HHHHHHHHHH--HH-----LLEEE-EE----------L-LEEEEL---------

# Job: Type-2 BREX; PglW_WP_185095299 wHTH2
# Query: s001A
# No:  Chain   Z    rmsd lali nres  %id PDB  Description
   1:  2od5-A  4.1  2.6   60    91   10   MOLECULE: HYPOTHETICAL PROTEIN;
   5:  3pvl-A  3.9  3.3   57   597   12   MOLECULE: MYOSIN VIIA ISOFORM 1;
   6:  5z2i-B  3.9  2.4   59    94    2   MOLECULE: DICTYOSTELIUM DISCOIDEUM MITOCHONDRIAL CALCIUM UN
   7:  5j9q-E  3.8  2.7   60   293   10   MOLECULE: HISTONE ACETYLTRANSFERASE ESA1;
   8:  8d2z-A  3.8  2.7   59   306   10   MOLECULE: METALLO-BETA-LACTAMASE SUPERFAMILY PROTEIN;
   9:  7asv-A  3.8  3.4   60   155    7   MOLECULE: DNA-DIRECTED RNA POLYMERASE III SUBUNIT RPC5;
  10:  4txa-A  3.7  2.4   53   367   15   MOLECULE: ROQUIN-1;
  12:  9ko8-A  3.7  3.1   53    61    8   MOLECULE: KINESIN-LIKE PROTEIN KIF1C;
  13:  4k51-A  3.6  3.3   71   213    8   MOLECULE: EUKARYOTIC TRANSLATION INITIATION FACTOR 3 SUBUNI
  14:  7mex-A  3.6  2.8   58  1737    7   MOLECULE: UBIQUITIN;
  15:  2lnb-A  3.6  3.8   60    80    7   MOLECULE: Z-DNA-BINDING PROTEIN 1;
  17:  8uqu-C  3.5  3.4   62   253    6   MOLECULE: FIDO DOMAIN-CONTAINING PROTEIN;
  18:  6i9b-A  3.5  3.9   64   177    9   MOLECULE: LA-RELATED PROTEIN 4;
  20:  7slp-B  3.5  2.6   63   321   10   MOLECULE: 7SK SNRNA METHYLPHOSPHATE CAPPING ENZYME;
  22:  3vw4-A  3.4  3.1   59   122    7   MOLECULE: REP;
  23:  7yw2-A  3.4  2.6   58   216   16   MOLECULE: TRNA 2'-PHOSPHOTRANSFERASE 1;
  24:  5chh-A  3.4  5.2   60   315   13   MOLECULE: ARAC FAMILY TRANSCRIPTIONAL REGULATOR;
  25:  5zuz-A  3.4  3.1   64   103    9   MOLECULE: ROK;
  26:  8zna-A  3.4  2.5   56    93    7   MOLECULE: ROMX;
  27:  7ymd-C  3.3  3.3   67   285    0   MOLECULE: NON-STRUCTURAL MAINTENANCE OF CHROMOSOME ELEMENT
  28:  7pb9-A  3.3  3.1   64   173    3   MOLECULE: TANDEM WH DOMAINS OF VPS25;
  29:  4ps2-A  3.3  3.9   58    79   12   MOLECULE: PUTATIVE TYPE VI SECRETION PROTEIN;
  30:  6taz-B  3.2  2.6   52   141   10   MOLECULE: PROTEIN TIMELESS HOMOLOG;
  31:  7aah-A  3.2  2.7   62   147    6   MOLECULE: MARGINAL ZONE B- AND B1-CELL-SPECIFIC PROTEIN;
  32:  6zfw-A  3.2  3.7   56    70    9   MOLECULE: PEROXIN-14;
  33:  8qfc-D  3.2  3.2   59   185    8   MOLECULE: 60S RIBOSOMAL PROTEIN L10A;
  34:  9k49-B  3.2  3.5   56   217   13   MOLECULE: TOL-PAL SYSTEM PROTEIN TOLQ;
  35:  9jm7-B  3.2  3.0   50    65    4   MOLECULE: LRD-2A;
  36:  5edl-A  3.2  2.9   62   197    5   MOLECULE: PUTATIVE HMP/THIAMINE PERMEASE PROTEIN YKOE;
  37:  4cej-B  3.1  3.5   61  1156    5   MOLECULE: ATP-DEPENDENT HELICASE/NUCLEASE SUBUNIT A;
  38:  2k85-A  3.1  2.8   56    65    7   MOLECULE: GLUCOCORTICOID RECEPTOR DNA-BINDING FACTOR 1;
  39:  1ylf-C  3.1  2.8   54   119    4   MOLECULE: RRF2 FAMILY PROTEIN;
  40:  3cuq-A  3.1  3.1   70   219   10   MOLECULE: VACUOLAR-SORTING PROTEIN SNF8;
  41:  5dwa-A  3.1  2.7   56   278    9   MOLECULE: TYPE-2 RESTRICTION ENZYME AGEI;
  42:  7c4o-A  3.1  3.4   45    48    2   MOLECULE: TRANSCRIPTION FACTOR HES-1;
  43:  8slm-A  3.1  3.0   58   248   17   MOLECULE: ZN DEPENDENT HYDROLASE FUSED TO HTH DOMAIN, IRRE
  44:  8f2a-E  3.1  2.8   55   117    9   MOLECULE: RECEPTOR ACTIVITY-MODIFYING PROTEIN 3;
  45:  7m5w-A  3.1  3.1   53   138    8   MOLECULE: PROTEIN CAPICUA HOMOLOG;
  46:  8j56-C  3.1  2.0   43   160   16   MOLECULE: FLAGELLAR TRANSCRIPTIONAL REGULATOR FLHD;
  47:  2e62-A  3.1  2.6   45    61    9   MOLECULE: PROTEIN AT5G25060;
  48:  8iue-P  3.0  3.7   70   303   13   MOLECULE: DNA-DIRECTED RNA POLYMERASE III SUBUNIT RPC1;
  49:  5fhp-C  3.0  3.2   57   209    7   MOLECULE: NICR;
  50:  2o7t-A  3.0  3.1   57   188   11   MOLECULE: TRANSCRIPTIONAL REGULATOR;
  51:  8dh3-E  3.0  4.0   70   825   10   MOLECULE: T7 RNA POLYMERASE;
  52:  2vwa-A  3.0  3.0   61    99    8   MOLECULE: PUTATIVE UNCHARACTERIZED PROTEIN PF13_0012;
  53:  6i7g-B  3.0  3.4   62   340   10   MOLECULE: ADENOSINE MONOPHOSPHATE-PROTEIN TRANSFERASE FICD;
  54:  2c5q-E  3.0  3.6   44   233    7   MOLECULE: RRAA-LIKE PROTEIN YER010C;
  55:  9kor-A  3.0  3.5   49  1229    8   MOLECULE: CRISPR-ASSOCIATED ENDONUCLEASE CAS9;
  56:  1sfu-A  3.0  3.0   57    70    4   MOLECULE: 5'-D(*T*CP*GP*CP*GP*CP*G)-3';
  57:  5y6i-B  2.9  3.0   55   234   15   MOLECULE: TRANSCRIPTIONAL REGULATOR KDGR;
  58:  5a31-N  2.9  3.1   58   703    3   MOLECULE: ANAPHASE-PROMOTING COMPLEX SUBUNIT 1;
  59:  6hrz-B  2.9  4.1   53   202    4   MOLECULE: PROBABLE TRANSCRIPTIONAL REGULATORY PROTEIN;
  60:  2lh9-A  2.9  3.5   62    78    6   MOLECULE: TUDOR DOMAIN-CONTAINING PROTEIN 7;
  61:  1szi-A  2.9  4.1   52   194    6   MOLECULE: MANNOSE-6-PHOSPHATE RECEPTOR BINDING PROTEIN 1;
  62:  2zop-A  2.9  3.3   59   113    5   MOLECULE: PUTATIVE UNCHARACTERIZED PROTEIN TTHB164;
  63:  2pn2-A  2.9  2.4   49   137    6   MOLECULE: UNCHARACTERIZED PROTEIN;
  64:  7amv-W  2.8  3.5   66   637    5   MOLECULE: DNA-DIRECTED RNA POLYMERASE 147 KDA POLYPEPTIDE;
  65:  9iac-B  2.8  2.7   53   269    8   MOLECULE: SIMILAR TO TR|Q3MCC8|Q3MCC8_ANAVT HYPOTHETICAL PR
  66:  6j5u-A  2.8  3.2   67   663    3   MOLECULE: DISEASE RESISTANCE RPP13-LIKE PROTEIN 4;
  67:  5g5p-B  2.8  2.8   67   455    6   MOLECULE: NUCLEAR MRNA EXPORT PROTEIN SAC3;
  68:  2h09-A  2.8  2.8   56   127    9   MOLECULE: TRANSCRIPTIONAL REGULATOR MNTR;
  69:  3mn2-A  2.8  5.1   57   108   18   MOLECULE: PROBABLE ARAC FAMILY TRANSCRIPTIONAL REGULATOR;
  70:  3s63-B  2.8  3.4   58    90    9   MOLECULE: SAPOSIN-LIKE PROTEIN;
  71:  7r3w-D  2.8  4.3   60   287    8   MOLECULE: PUTATIVE BACTERIAL REGULATORY HELIX-TURN-HELIX PR
  72:  2mp8-A  2.8  3.1   48    64   17   MOLECULE: NKR-5-3B;
  73:  7pc1-A  2.8  2.8   51    72    2   MOLECULE: STBA;
  74:  6xwi-A  2.8  3.2   50    66   12   MOLECULE: S0_2.126;
  75:  7e1n-A  2.8  2.9   55   210    4   MOLECULE: DUF1956 DOMAIN-CONTAINING PROTEIN;

0001 s001A   PDTPSAPALLNYADRLGK-AAD-L--LA-A----------R------D--VLASPTTVLE--QIV-A--A---D----A-PGTR-I-S---L-D--EG--R-MVTLAAAAS----GTAAANA--R-LEIYSRS--------
0002 2od5A   ------ETESXKTVRIRE-KIK-K--FL-G----------D---------RPRNTAEILE--HIN-S--T---X----R----------hgT-T--SQ--Q-LGNV-lskD----KDIVK---dI-CEWATRNwvaehcpe
0003 2qkmB   ----TNATFSQVLDDLSArFIL-N-----L---------P------A--EEERLCFQIE--QAH-W--FyedF----I-RAQL-P-S---L-G--LR--V-FSAKLFAHC----PLLwhEE--A-F-------------
0004 3s93B   ----GMSEQERIQECLRK-EIR-S--LL-I--------sT------K--DGLSPQELEK--EyN-H--L---P---------I-L-G---YrS--TM--E-LVLD----------DVVRV--tV-ILKAIP---------
0005 1qgpA   -LSSHFQE-LSIYQDQEQ-RIL-K--FL-E----------E---lgeG--KATTAHDLSG--KLG-T------------------------------PK--K-EINRVLYSL-akkGKLQ--------LWKIAVsd------
0006 3pvlA   ffrkRCEKEDDLAELASQ-QYF-V--DY-G----------S----------EMILERLLS--LVP-T--Y---I----P---DR-E-ItplK-N--LE--K-WAQLAIAAH----K-----------------------------
0007 5z2iB   ----GELKTIXGQAKVSK-LQE-K--XK-L----------D------P--RKITFNDFKG--IAK-E--V---G--------------ieeK-E--IN--S-VSNA------------SIIY--L--VFTKPAhiyqsleh
0008 5j9qE   qkPLSDLGLLSYRAYWSD-TLI-T--LL-V--------eH------Q--KEITIDEISS--MTS-M-----------------------------TT--T-DILHTAKTL----N-ILRY-------iIFLNedildryn
0009 8d2zA   qleqGLDYTKDFVTTLLA-QGR-K--AV-E----------R-----------NLDLKAAMA--LTR-E--AmdpK----F-GHVFiY-E---H-C--LP--F-DVSRAF--------eASGI--------------------
0010 7asvA   aTFQRESGD--XSDQHRQ-VLL-E--IF-S----------K------N--YRVRRNXIQS--RLT-Q--E---X--------------eD---L-S--K---Q-EVDKVLK----------CCVS------wYLKGtvqs----
0011 4txaA   -----nflGPARQEEALK-LVL-L--AL-E----------D------G--SALSRKVLVL--FVV-Q--R---L----E-PRFP-Q---asktS--IG--H-VVQL-----------------------mQLKEefrtyeal
0012 7udkA   ------DREEAAFLAASI-LIQ-H--AH-E----------Q-----------GKDDRELEK--ILE-I--A-irI----L-EK-D-R-E---E-------aaF-LAASILI-------qhdkIL--E-I--------------
0013 9ko8A   -----ATMADLKMQAVKE-ICY-E--VA-L----------A------D--FRHGRAEIEA--LAA-L--K-mrE----L-CRT---yG---D-A--WR--A-VARDV-------------------------------------
0014 4k51A   KTYFSQYIAPLRDVIMRR-VFV-A--AS-Q----------K------F--TTVSQSELYK--LAT----L---P---A-P---------ldL-S--AW--D-IEKSLLQAA-vedYVSITID--H--VTFAK---------
0015 7mexA   kVSEKSSLKDRRMDQIKN-SII-Y--NL-Y----------M----------KPLSYSKLLR--SVP-D-----------------------yltE--DT--T-EFDEALE--------eVSVF-------fKLKAslyakvdp
```

```
0016 2lnbA   hhsHMADP--GREGHLEQ-RIL-Q--VL-T--------eA------G--SPVKLAQLVK--ECQ-A------------------------------PK--R-ELNQVLYRM-kkeLKVS----sP-ATWCLGGtdpe----
0017 4wziB   eDHYHYHNSLHAADVAQS-THV-L--LS-T-------paL------D--AVFTDLEILA--AIF-A--A--aI----H-DVDV-G-F---K------------------------tkvyTD--R-I---------------
0018 8uquC   mpKNRSEQEIAGYRDALA-LIH-E--SA-T---------H-----------PFSEGVVL--QLH-T--L--lY----R-YMPQ--fP---D-G--N---G-RMSRLL--------tLLLL-------YHFDyavgryis
0019 6i9bA   --NSAVS-TEDLKECLKK-QLE-F--CF-S------renL------S--KD-PIWTVAN--M-I-K--K---L-----------------ttD--PD--L-ILEV-LRSS----PMVQVDG--E-KV-RPSHkrcivilr
0020 5mmcA   EVEHTHSEREKRVSNAVE-FLL-DsrVR-R---------T------------PTSSKVH--FLK-S--K---G---------------------lSA--E-EICEAFTKV----G--QPKTl-------------------
0021 7slpB   ------SRVKQVLADIAK-QVD-F--WF-GdanlhkdrfL------R--EQVDISLLVS--F-M-K--K---L--------------------ttDG--K-LIARALRSS----AVVELDE--G-TRIRRKKplgerpkd
0022 8twjA   SSKSNRAYCQSMYNSIRS-AGD-E--IS-R---------G-----------TSFEELWG--RAT-E--W---R---lS-KLQ---dqagsA-A--KA--E-FAARAIA--------sAHGI---------------------
0023 3vw4A   ----HHHRNYHLFEKVRK-WAY-R--AI-R---------Q----------GWPVFSQWLD--AVI-Q--R-veM----Y-NALS-P-A---EcraigK--S-IAK-------------THRK-------FSPEgfsavqaa
0024 7yw2A   --------EQDRNVQLSK-ALS-Y--AL-R----hgalklglpmraD--GFVPLQALLQlpqFH-S--F--------------------------SI--E-DVQLVVNT--nekQRFTLQP-tG-LLIRANQghslqvpe
0025 5chhA   GGYR-TNYREVLDLVRHD-RAR-Q--LL-R---------D----------ERLRIEEVAE--RLG-Y--------------------------xD--TS--S-FRHAFRRWT-------CAND-------YRQA--------
0026 5zuzA   rrgTARP--GSKAAKLRE-AAI-K--TL-K--------rH------N--AAIKSSELQK--EIEkE--S---G----L----------eiP-N--MT--T-FMQSLIKmY----PEVK--R--G-QY-ILEGeiesaesa
0027 8znaA   ---XTDEEKVKAXRLARA-IAS-D--IS-L-------ynE------Q--KI-LKEELEE--GRE-LykS---R----V-S--E-I-F---T------kxN-FFERAI--------nDIVL---------------------
0028 7ymdC   dfkiLSTDRDLVYKGVLS-VIL-C--IV-F--------fS------K--NNILHQELIK--FLE-T--F---G--------AI-L-N---I-T--IE--D-LIKSLEK-------eYIVRL--E--SYRIGRrtqaelgl
0029 7pb9A   slpSKTEDFEDWLNQWSD-FTL-Q--WFkI---------N------K--LQISLVELMGekPFS-Y-----------------------lQ-N--KS--K-ALTVIVENL--iaRNFC--Y--K-SIrVFWRgyrdwsev
0030 4ps2A   EQVARQIPRLKAMLAMRS-LLR-D--LK-A---------N-----------LDNVTFRK--ELE-K--I--lR----D-PA-S-Q-T---L------rdE-LRAL-------------VPE--K-AW-------------
0031 6tazB   ------twSPEEEAHLRE-LYL-A--NK-D---------V------E--GQ----DVVE--AILaH--L---N----T------V-P---R-T--RK--Q-IIHHL---------gLADSV--K-DF-------------
0032 7aahA   SAHMRCDACRAVAYQMWQ-NLA-K--AEtK---------L------H--TELSELVYTD--VLD-R--S---CsrnwqrLSRT--gE---F-G--ED--Q-IYEAHQQGRvsat--------------------------
0033 6zfwA   --GAMAWPEESEKRKRVS-SAV-Q--FL-H---------D------S--RVTPAANKIQ--FLK-S--K------------------------glTT--E-EVCEAF--------KAGQTI--Pl----------------
0034 8qfcD   GETMTEEQSQSFLTEFIN-YIK-Q--SK----------------------VVLLEDLAS--QVG-L-----------------------rtqD--TI--N-RIQ------laegTITGVID-rG-KFIYITPeelaavan
0035 9k49B   -----nTGSEQIFYSGFK-EFV-R--LH-R---------A----------NSHAPEAVVE--GAS-R--A-mrI----S-MNNY-D-NfmeE-F--TA--I--LHRQ----------AFTV--------------------
0036 9jm7B   ---GVEAAKKEIKKLKEE-VLK-K--YK-K---------G-----------EINEEEAIK--EFV-E--K---A--lkL-VKAdkF-A---I-E----eaK-ALV-------------------------------------
0037 5edlA   --SWKVKEIVIMSVISIV-FAV-V--YL-L--------fT------H--FG-IYGIWF---IVS-V--I---A--ayM-IRKP-G----------aalV-SEIIAAL---vecLLGNVIV--I-G--------------
0038 4cejB   FFKLEAPDIGQLFHSSLK-LIS-D--RL-R--------dE------K--LD-QCELFSY--DAE-R--L---A--pkL-QKEI-LyV---K-E--K---L-QKIV------------SGILse------------------
0039 2k85A   -----SQQIATAKDKYEW-LVS-R--IV-K---------N-----------NENWLSVSR--KMQ-A--S---P----E-----vY-L---E-G--TQ--K-AKKL----------lQHIH--------RLKHe--------
0040 1ylfC   ---------ISSRFSIAV-HIL-S--IL-K---------N----npS--SLCTSDYXAE--SVN-T------------------------------NP--V-VIRKIXSYL----K-----G--P-GGAGLLKdlheitll
0041 3cuqA   KHKQEMLGVGDFYYELGV-QII-E--VC-L---------A-lkhrnG--GLITLEELHQ--QVL-K--------------------gkfaqD-V--SQ--D-DLIRAIKKL----KGFGIIP-gT-YLIQSVPaelnmdht
0042 5dwaA   -----lnVSEEQYQQISD-YFV-P--LV-N---------K------P--KLKSR-DAIG--QAF-V--M-atE----V-CP-D-A-N---P-SdlWH--H-VLYRIY----------gsWV--R-T--------------
0043 7c4oA   DPSVLGKYRAGFSECMNE-VTR-F--LS-T---------C------E--GVNT-EVRTR--LLG-H--L-anC----M-TQ------------------------------------------------------------
0044 8slmA   -----TGALAPAKARMRE-LAT-A--YA-R--------rL------P--GLlahEISHA--LLL-G--D---DdllsD-LHDArL-E---Q-V--IE--T-LCNVGAA--------------------ILMP---------
0045 8f2aE   --CNETGMLERL-PLCGK-AFA-D--MM-G---------K------V--DV-EFIVYYE--SFT-N--C-teM----E-ANCY-W-P---N-----------plFITG---------QFFSn------------------
0046 7m5wA   nPYSSLRRTLDQRRALVM-QLF-Q--DH-G---------------------FPSAQATAA--FQA-R--Y-----------------adifP-S--KV--C-LQLKIREVR----Q-------------------------
0047 8j56C   ---KSVLQDANQTQLAIE-LIG-L--G-----------------------------aRLQ--VLE-A--E---T----T-------------lS--RD--R-LIRL------------LRGV--Sp--------------
0048 2e62A   ----QRQKRRRIEVALIE-YRE-T--LE-E---------Q---------GMKNPEEIER--KVE-I--N-rkR----L-EVYG-L-S------------------------------------------------------
0049 8iueP   VTGGQDFE-SEFVEVLNQ-QCF-K--FL-Q---------S------K--AFASSHEVWK--YICeL--G---I----S-----kV-E---L-S--ME--D-IETI-LNTL-iydGKVE--M--K-LY-RAVNpiipptgl
0050 5fhpC   fxeVYRRGLSDIDKKRQN-YLD-E--LK-S---------R------P--EGYNTYDIVR--TYL-R--P---Q-----hFARL--lY---D-H--TL--K-AFI--------------EIX--Ec---------------
0051 2o7tA   ----XRADALKRREHIIT-TTC-N--LY-R---------T------------HHDS-LTXE--NIAeQ--A---G----V----gyR-N---F-P--DR-fT-LDXA-CAQY----L------------gSLVPalapesld
0052 8dh3E   gevaQEIKPEAVAYITIK-TTL-A--CL-T---------S------A--DNTTVQAVAS--AIG-R--A---I--edE-ARFG--hkedsihV--GV--R-CIEMLIEST----GM-VSLH--E-TI-ELAPeyaeaiat
0053 2vwaA   nVDVSIKRYNNFVDIARL-SIQ-K--HF-E---------H------L--S--NXEYXQK--FVQ-G--LqenR----N-IS--lS-K---Y-Q--EN--K-AVXDLKY-------------------kVYANylsqee--
0054 6i7gB   GKSLEEQNEVIGMHAAMK-YINtT--LV-S---------R------I--GSVT-ISDVL--EIH-R--R-vlG----Y-VD----iD---G-N--G---R-TSRL------------NLIL-------MQAGyppitirk
0055 2c5qE   VQETDISKLVTYIEKSIE-VDL-L--VS-E-------diK------N--GIPAKQAQND--RRS-V--L---K----K---------------------------------------------------------------
0056 9korA   QTRHRIRSQQRFVLARRL-TYI-L--IE-G--------kL------K--RKLSPREKEA--ISS-L--L---RrrgyssltekdkkR---K-D--SN--Gk----------------------------------------
0057 1sfuA   -----TVNDAEIFSLVKK-EVL-S-----L---------N------T--NDY---tTAI--SLSnR--L---K----I----------nkK-K--IN--Q-QLYK-----lqkeDTVKMVP--S-PKWFKNYnc------
0058 5y6iB   ------VRSAEVGTDILK-ALA-E--LS-P--------------------ATSLSRLAE--HVG-M-----------------------pasK--VH--R-YLQA-----liasGFAVQDS--T-NHYSLGRealrvgla
0059 5a31N   lFINEeliKEEELLLFWT-YIQ-A--ML-T---------N------L--ESLSLDRIYN--MLRmF--V---V----T-------palaeI-D--LQ--E-LQGY----------qkLVYS-----VYRLP--------
0060 6hrzB   HMEIKRRTQEERSAATRE-ALI-T--GA-R---------K-----------lVGTPEIAT--EAG-------------------------------vTR--G-AMYHQ------------FAD--K---rDVVEvveqdvma
0061 2lh9A   ----HMLEADLVSKMLRA-VLQ-S--HK-N----------------------GIVLPRLQG--EYR-S--L---T---------Q-L-G---YpT--LE--A-YLRSVPAVV--------RIEA-gE-IVCY-----------
0062 1sziA   slqqARQKAQETLQQLTS-VLG-L--ME-S---------V------K--Q--VEARALS--MFR-D--I-tqQ----L-QSMC--vN---D-L--QA--T---------------sfVLAQt------------------
0063 2zopA   ----XRTRSQVWAQKAYE-KVR-E--AA-K---------G------E--GRdAHKALGN--DLA-Q--V---L----G----R-L-T---R-E--VL--A-AAEWFKRFA----QAL------------------------
0064 2pn2A   tyqgTDLLATSLASCXLT-IIG-I--KA-R---------D------X--EIvTQKIFYN--TAL-T--C---P----VkSIHPdI-F---Q-K--VIih------------------------------------------
0065 7amvW   sLPPHPIVKVLLGELVRQ-FFY-N--NS-----------------------RIKYND--T--KLL-K--M---V---tS-VIKN--kedarN-Y--ID--D-IVN--hFFV----SNkLYKY-eN-DIITVPFrlsyepfv
0066 9iacB   lkpaQPQKIGKLAQRGVF-EFH-Q--EL------------------------SGSH-GVE--QVAeI--L---Q----LnQESPiQ-A---R-V--LV--I-LNNY----------------------iLLNKeiinlsrg
0067 6j5uA   -----NGTDPSHLKSCIL-T------LS-L---------Y------PedCVIPKQQLVH--GWI-GegF---V----M-WRNG-R-S-atE-S--GE--D-CFSGLTNR-----cLIEVI--I-TC-KIHDmvrdlvid
0068 5g5pB   idnwIVLLEKLPMVTYRN-LIK-T--VI-K---------SwttewgQ--NKLPYSLIER--VLQ-L--S---I-----tiYNG-I-H--spkN--VE--N-VLVTLIN--------RANC--F-LCVVKKTtmiqeivp
0069 2h09A   --EGFRQVREAHRRELID-DYV-E--LI-S---------D--lireV--GEARQVDMAA--R------------------------lgvsqpT--VA--K-MLKR-----latmGLIEMI------GVFLTAegeklaqe
0070 3mn2A   RSRGYSPXAFAKRVRLQH-AHN-L--LS-D---------G------A--TPT---TVTA--AAL-S--C---G------------------------fsN--LG--H-FARDYRDXF-----------------SETL---------
0071 3s63B   ----PKETCDLCQIALRT-VFG-H--FG-G---------N-----------IPSRRKLVH--QLK-H--E--cK----R-HFNY-R-RrclL-L--MK--V--------------------D--L-IFMTDGSfkpmevcl

0001 s001A   LLLLLHHHHHHHHHHHHHH-HHH-H--HH-L---------L------L--LLLLHHHHHHH--HHH-L--L---L----L-LLLL-L-L---L-L--HH--H-HHHHHHHLL----LLEEELL--L-LLEEELL--------
0002 2od5A   ------LLHHHHHHHHHHH-HHH-H--HL-L---------L---------LLEEHHHHHH--HHH-L--L---L----L----------llL-L--HH--H-HHHH-hhhL----LLEEE---eE-EEEEEHHhhhhhlll
0003 2qkmB   ----HLLLHHHHHHHHHHhHLL-L-----L---------L------H--HHHHHHHHHH--HHH-H--HhhhL----L-HHLL-L-L---L-L--HH--H-HHHHHHLLL----HHHlhHH--H-H--------------
0004 3s93B   ----LHHHHHHHHHHHHHH-HHH-H--HH-H--------lL------L--LLLLHHHHHH--HhL-L--L---L---------H-H-L---LlL--HH--H-HHHL----------LLEEE--lE-EEEELL--------
0005 1qgpA   -LHHHHLL-LHHHHHHHH-HHH-H--HH-H---------H---hllL--LLEEHHHHHH--HHL-L------------------------------LH--H-HHHHHHHHH-hhhLLEE--------EEEELLll------
0006 3pvlA   eeeeLLLLHHHHHHHHHHH-HHH-H--HH-L---------L----------LLLHHHHHH--HHH-H--H---L----L---HH-H-LlllL-L--HH--H-HHHHHHHHH----H-------------------------
0007 5z2iB   ----LHHHHHHHHHHHHHH-HHH-H--HH-H---------L------L--LEEEHHHHHH--HHH-H--L---L--------------llhH-H--HH--H-HHHH------------LLEE--L--EELLLHhhhhhhhh
0008 5j9qE   llLLLHHHHHHHHHHHHHH-HHH-H--HH-H--------hH------L--LLLLHHHHHH--HHL-L------------------------------LH--H-HHHHHHHHL----L-LEEE-------eELLLhhhhhhhh
0009 8d2zA   llllHHHHHHHHHHHHHHH-HHH-H--HH-H---------L----------LLLHHHHHH--HHH-H--HhhhH----H-LLLLlH-H---H-H--HH--H-HHHHHH--------hHLLL--------------------
0010 7asvA   hHHHHLLLL--LHHHHHH-HHH-H--HH-H---------H------L--LEELHHHHHH--HHH-H--H---H------------lL---L-L--H---H-HHHHHHH---------HEEE------eEELLllll----
0011 4txaA   -----hllLHHHHHHHHHH-HHH-H--HH-L---------L------L--LLLLHHHHHH--HHH-H--H---L----L-LLLL-L---llhhH--HH--H-HHHH----------------------eEELHhhllhhhh
0012 7udkA   ------LHHHHHHHHHHHH-HHH-H--HH-H---------H----------LLLHHHHHH--HHH-H--H-hhH----H-HH-L-H-H---H------hhH-HHHHHHH-------hhlhHH--H-H--------------
0013 9ko8A   -----LLHHHHHHHHHHHH-HHH-H--HH-L---------L------L--LLLLHHHHHH--HHH-H--H-hhH----H-HLL---lL---L-H--HH--H-HHHHH-----------------------------------
0014 4k51A   LLLLHHHHHHHHHHHHHHH-HHH-H--HH-H---------H------L--LEEEHHHHHH--HHL----L---L----H-H--------hlL-L--HH--H-HHHHHHHHH-hllLLLLLLL--L--EELLL---------
0015 7mexA   hHHHHLLHHHHHHHHHHHH-HHH-H--HH-L---------L---------LLEEHHHHHHH--HLL-H-----------------------hhlL--LH--H-HHHHHHH-------hHEEE-------eEELLhhhhhlll
0016 2lnbA   lllLLLLL--LHHHHHHH-HHH-H--HH-H--------hH------L--LLEEHHHHHHH--HHL-L------------------------------LH--H-HHHHHHHHH-hhlLLEE----eL-LEEEELLllll----
0017 4wziB   hHLLLLLLHHHHHHHHHHH-HHH-H--HL-L-------hhH------L--LLLLHHHHHH--HHH-H--H--hH----L-LLLH-H-H---H------------------------lllhHH--H-H--------------
0018 8uquC   llLLHHHHHHHHHHHHHHH-HHH-H--HH-H---------H-----------LLLHHHHHH--HHH-H--H--hL----L-LLLL--lL---L-L--H---H-HHHHHH--------hHHHH--------HHLLlhhhhlll
0019 6i9bA   --LLLLL-HHHHHHHHHH-HHH-H--HL-L------hhhH------L--LL-LHHHHHL--L-H-H--L---L-----------------llL--HH--H-HHHH-HLLL----LLEEELL--L-LE-EELLllleeeel
0020 5mmcA   LLLLLHHHHHHHHHHHHHH-HHL-LhhHH-H---------L------------LHHHHHH--HHH-H--L---L---------------------lLH--H-HHHHHHHHL----L--LLLLh------------------
0021 7slpB   ------LHHHHHHHHHHHH-HHH-H--HL-LhhhhhhllhH------H--HHEEHHHHHL--L-H-H--L---L--------------------llLH--H-HHHHHLLLL----LLLEELL--L-LEEELLLllllllll
0022 8twjA   LLHHHHHHHHHHHHHHHHH-HHH-H--HH-L---------L-----------LLHHHHHHH--HHH-H--H---H---hL-LLL---llllhH-H--HH--H-HHHHHHH-------hHLLL--------------------
0023 3vw4A   ----LHHHHHHHHHHHHHH-HHH-H--HL-L---------L---------LLLLHHHHHHH--HHH-H--H-hhH----H-HHLL-H-H---HhhhhhH--H-HHH-------------HHHH-------LLHHhhhhhhhhh
0024 7yw2A   --------LHHHHHHHHHH-HHH-H--HH-H----lllllllllllllL--LLEEHHHHHLlhhHL-L--L--------------------------LH--H-HHHHHHHH--lllLLEEEEE-lL-EEEEELLllllllll
0025 5chhA   LLHH-LLHHHHHHHHHHHH-HHH-H--HL-L---------L---------LLLLHHHHHHH--HLL-L--------------------------lL--HH--H-HHHHHHHHH-------LHHH-------HHHL--------
0026 5zuzA   hhhLLLL--LLHHHHHHHH-HHH-H--HH-H--------hL------L--LLEEHHHHHHH--HHHhH--H---L----L----------llL-L--HH--H-HHHHHHHhL----LLEE--L--L-EE-EELLlllllllll
0027 8znaA   ---LLHHHHHHHHHHHHHH-HHH-H--HH-H-------hlH------H--HH-LHHHHHH--HHH-HhhH---H----L-L--H-H-H---H------hlL-HHHHHH--------hHHHH--------------------
0028 7ymdC   hhhhLLLLLHHHHHHHHHH-HHH-H--HH-H--------hL------L--LLEEHHHHHHH--HHH-H--H---L--------LL-L-L---L-L--HH--H-HHHHHHH-------lLEEEE--E--EEEELHhhhhhhllh
0029 7pb9A   lllLLHHHHHHHHHHHHHH-HHH-H--HHhH---------L------L--LEEEHHHHLLllLHH-H-----------------------lL-L--HH--H-HHHHHHHHH--hhLLLE--L--L-EEeELLLlhhhhhhh
0030 4ps2A   HHHHHHHHHHHHHHHHHHH-HHH-H--HH-L---------L-----------LLHHHHHHH--HHH-H--H--hL----L-LL-L-H-H---H------hhH-HHLL-------------LLH--H-HL-------------
0031 6tazB   ------llLHHHHHHHHHH-HHH-H--HL-L---------L------L--LL----LHHH--HHHhH--L---L----L------L-L---L-L--HH--H-HHHHH----------lLLLLH--H-HH-------------
0032 7aahA   LLLLHHHHHHHHHHHHHHH-HHH-H--HHhH---------L------L--LLLLHHHHHHH--HHH-H--H---LllllllhHHHH--hH---H-L--HH--H-HHHHHHHLHllll--------------------------
0033 6zfwA   --LLLLHHHHHHHHHHHHH-HHH-H--HH-H---------L------H--HHLLHHHHHHH--HHH-H--L------------------------llLH--H-HHHHHH--------HLLLLL--Lh---------------
0034 8qfcD   LLLLLHHHHHHHHHHHHHH-HHH-H--HL----------------------EELLLHHHH--HHL-L-----------------------lhhH--HH--H-HHH------hhhlLLLEEEE-lL-EEEELLHhhhhhhhh
0035 9k49B   -----lLLHHHHHHHHHHH-HHH-H--HH-H---------L---------LLLLHHHHHHH--HHH-H--H-hhH----H-HHHH-H-HhhhH-H--HH--H--HHHL-----------LLLL--------------------
0036 9jm7B   ---LHHHHHHHHHHHHHHH-HHH-H--HH-H---------L----------LLLHHHHHHH--HHH-H--H---H--hhH-HHHlhH-H---H-H----hhH-HHH-------------------------------------
0037 5edlA   --LLLHHHHHHHHHHHHHH-HHH-H--HH-H--------hH------H--HH-HHHHHH---HHH-H--H---H--hhH-HLLL-L-----------hhhH-HHHHHHH---hhhHLLLHHH--H-H--------------
0038 4cejB   LLLLLHHHHHHHHHHHHHH-HHH-H--HH-H--------hL------L--LL-HHHHHHH--HHH-H--H---H--hhL-LLLH-HhH---H-H--H---H-HHHH------------HHHHhl-----------------
0039 2k85A   -----LLLLLHHHHHHHHH-HHH-H--HL-L---------L----------LLLHHHHHH--HHL-L--L---H----H-----hH-H---L-L--HH--H-HHHH----------hHHHH--------HHHHl-------
0040 1ylfC   ---------LLHHHHHHHH-HHH-H--HH-H---------H----llL--LLLLHHHHHHH--HLL-L------------------------------LH--H-HHHHHHHHH----H-----L--L-LLEEELLlhhhllhh
0041 3cuqA   HHHHHHHLHHHHHHHHHHH-HHH-H--HH-H---------H-hhhhhL--LEEEHHHHHHH--HHH-H--------------------lllllL-L--LH--H-HHHHHHHHH----HLLEEEE-lE-EEEELLLllllhhhh
0042 5dwaA   -----llLLHHHHHHHHHH-HHH-H--HH-L---------L------L--LLLHH-HHHH--HHH-H--H-hhH----H-LL-L-L-L---H-HhhHH--H-LHHHHH----------lhHH--H-H--------------
0043 7c4oA   LHHHHHHHHHHHHHHHHHH-HHH-H--HL-L---------L------L--LLLH-HHHHH--HHH-H--H-hhH----H-HH------------------------------------------------------------
0044 8slmA   -----LLLHHHHHHHHHHH-HHH-H--HH-H--------hL------L--LLlhhHHHHH--HHH-H--L---HhhhhH-HHLLhH-H---H-H--HH--H-HHHHHHH-------------------HHLL---------
0045 8f2aE   --LLHHHHHHHL-HHHHH-HHH-H--HL-L---------L------L--LL-HHHLHHH--HHH-H--H-hhH----L-LLLL-L-L---L-----------llHHHH---------HHLLl------------------
0046 7m5wA   lLLHHHHHHHHHHHHHHHH-HHH-H--HH-L--------------------LLLHHHHHHH--HHH-H--L-----------------llllL-L--HH--H-HHHHHHHHH----H-------------------------
0047 8j56C   ---LLHHHHHHHHHHHHHH-HHH-L--L-----------------------------lLHH--HHH-H--H---L----L-------------lL--HH--H-HHHH------------HLLL--Ll---------------
0048 2e62A   ----HHHHHHHHHHHHHHH-HHH-H--HH-H---------H---------LLLLHHHHHHH--HHH-H--H-hhH----H-HHLL-L-L------------------------------------------------------
0049 8iueP   LLLLLEEL-HHHHHHHHH-HHH-H--HH-H---------H------H--HLEEHHHHHHH--HHHhH--L---L----L-----lL-L---L-L--HH--H-HHHH-HHHH-hhlLLEE--E--E-EE-EELLllllllllh
0050 5fhpC   hhhHHHHHHHHHHHHHHHH-HHH-H--HH-L---------L------L--LLLLHHHHHHH--HHH-H--H---H-----hHHHH--hL---H-H--HH--H-HHH--------------HHH--Hl---------------
0051 2o7tA   ----LLHHHHHHHHHHHHH-HHH-H--HH-H---------H----------LHHH-LLHH--HHHhH--H---L----L----lhH-H---L-L--LH-hH-HHHH-HHHH----H------------hHHHHhhllllhh
0052 8dh3E   llllHHLLHHHHHHHHHHH-HHH-H--HH-H---------H------H--LLLLHHHHHHH--HHH-H--H---H--hhH-HHHH--lhhhhhhH--HH--H-HHHHHHHHL----LL-EEEL--L-LE-EELHhhhhhhhhh
0053 2vwaA   lLLLLHHHHHHHHHHHHHH-HHH-H--HH-H---------H------L--L--LHHHHHH--HHH-H--HhhhL----L-LL--lL-H---H-H--HH--H-HHHHHHH------------------hHHHHhhhhhl--
0054 6i7gB   LLLHHHHHHHHHHHHHHHH-HHHhH--LL-L---------L------L--LLLL-HHHHH--HHH-H--H-hlL----L-LL----lL---L-H--H---H-HHHH------------HHHH-------HHHLllllllllh
0055 2c5qE   HHHLLHHHHHHHHHHHHHH-HHH-H--HH-H-------hhH------L--LLLHHHHHHH--HHH-H--H---H----H---------------------------------------------------------------
0056 9korA   HHHHHHHHHHHHHHHHHHH-HHH-H--HH-H--------hH------L--LLLLHHHHHHH--HHH-H--H---LllllllhhhhhlheE---E-L--LL--Ll----------------------------------------
0057 1sfuA   -----LLLLHHHHHHHHHH-HHH-L-----L---------L------L--LLE---eLHH--HHHhH--L---L----L----------lhH-H--HH--H-HHHH-----hhhlLLEEEEL--L-LEEEELLll------
0058 5y6iB   ------LLHHHHHHHHHHH-HHH-H--LL-L--------------------LEEHHHHHHH--HHL-L-----------------------lhhH--HH--H-HHHH-----hhhhLLEEELL--L-LEEEELHhhhhhhhh
0059 5a31N   hHHHHlllLHHHHHHHHHH-HHH-H--HH-H---------H------H--LLLLHHHHHHH--HHHhH--L---L----L-------lllllL-L--HH--H-HHHH----------hhLEEE-----EEELL--------
0060 6hrzB   LLHHHHHHHHHHHHHHHHH-HHH-H--HH-H---------H----------hLLHHHHHHH--HHL-------------------------------lLH--H-HHHHH------------LLL--H---hHHHHhhhhhhhhh
0061 2lh9A   ----LLLHHHHHHHHHHHH-HHH-H--LL-L--------------------LEEHHHHHHH--HHH-H--H---H---------H-H-L---LlL--HH--H-HHHLLLLLE-------EEEE-lL-EEEE-----------
0062 1sziA   hhhhHHHHHHHHHHHHHHH-HHH-H--HH-H---------H------H--L--HHHHHHH--HHH-H--H-hhH----H-HHHH--hH---H-H--HH--H---------------lhHHHHh------------------
0063 2zopA   ----LLLHHHHHHHHHHHH-HHH-H--HL-L---------L------L--LHhHHHHHHH--HHH-H--H---L----L----H-H-H---H-H--HH--H-HHHHHHHHH----HHH------------------------
0064 2pn2A   eeeeHHHHHHHHHHHHHHH-HHH-H--HH-H---------H------L--LLeHHHHHHH--HHH-L--L---L----LlLLLLlL-E---E-E--EEel------------------------------------------
0065 7amvW   lLLLLLLHHHHHHHHHHHH-HHH-H--LL-----------------------LEELLL--H--HHH-H--H---H---hL-LLLL--hhhhhH-H--HH--H-HHH--lLEE----LLlEEEL-lL-EEEEEELlllllle
0066 9iacB   hlllHHHHHHHHHHHHHHH-HHH-H--LL------------------------HLLL-HHH--HHHhH--L---L----HhHLLHhH-H---H-H--HH--H-HHHH----------------------lLLLLeeeeelll
0067 6j5uA   -----LLLLLHHHHHHHHH-H------HH-H---------L------LllLLEEHHHHHHH--HHH-HhlL---L----L-LLLL-L-L-lhH-H--HH--H-HHHHHHHL------lLLEEL--L-EE-ELLHhhhhhhhhh
0068 5g5pB   hhhhHHHHHHHHHHHHHHH-HHH-H--HH-H---------HhhllllL--LEEEHHHHHHH--HHH-H--H---H-----llLLL-L-L--lllL--HH--H-HHHHHHH---------LEEE--E-EEEELLLllhhhhll
0069 2h09A   --LLLHHHHHHHHHHHHHH-HHH-H--HH-H---------H--hhhhH--LLLLHHHHHHH--H------------------------hlllhhH--HH--H-HHHH-----hhhlLLEEEE------EEEELHhhhhhhhh
0070 3mn2A   HHLLLLHHHHHHHHHHHHH-HHH-H--HH-L---------L------L--LLL---LHHH--HHH-H--L---L-----------------llL--HH--H-HHHHHHHHH---------------HHHH-----------
0071 3s63B   ----HHHHHHHHHHHHHHH-HHH-H--LL-L---------L----------LLLHHHHHHH--HHH-H--H--hL----L-LLLL-H-HhhhH-H--HH--H--------------------H--H-HHHHLLLllhhhhhh
```

### Job: Type-2 BREX; PglW_WP_185095299 wHTH3
### Query: s001A
# No:  Chain   Z    rmsd lali nres  %id PDB  Description

```
 1:  8zjr-M  4.5  2.8   61    82   10   MOLECULE: HISTONE H3.2;
 2:  7ymd-C  3.9  2.9   65   285    9   MOLECULE: NON-STRUCTURAL MAINTENANCE OF CHROMOSOME ELEMENT
 3:  5nl9-A  3.9  3.1   60   146   13   MOLECULE: TRANSCRIPTIONAL REGULATOR (FUR FAMILY);
 4:  7asv-A  3.8  2.8   61   155   13   MOLECULE: DNA-DIRECTED RNA POLYMERASE III SUBUNIT RPC5;
 5:  7kbf-K  3.8  2.9   58    78   14   MOLECULE: HISTONE H3.2;
 6:  7db7-B  3.6  3.4   48   836   17   MOLECULE: PHENYLALANINE--TRNA LIGASE ALPHA SUBUNIT;
 7:  5a48-B  3.6  2.9   63    96   11   MOLECULE: MATERNAL EFFECT PROTEIN OSKAR;
 8:  4ets-B  3.6  3.5   61   149   15   MOLECULE: FERRIC UPTAKE REGULATION PROTEIN;
 9:  5okc-B  3.6  3.4   60   377   10   MOLECULE: SISTER CHROMATID COHESION PROTEIN DCC1;
10:  7z8b-C  3.4  3.0   59  1224   12   MOLECULE: CULLIN-7;
11:  1ka8-A  3.3  2.9   63   100   10   MOLECULE: PUTATIVE P4-SPECIFIC DNA PRIMASE;
12:  8e4y-A  3.3  2.9   57   651    9   MOLECULE: GLYCEROL-3-PHOSPHATE ACYLTRANSFERASE 1, MITOCHOND
13:  8e4v-A  3.2  2.9   53    84   11   MOLECULE: ISOFORM 3 OF HISTONE ACETYLTRANSFERASE KAT6B;
14:  2qby-B  3.2  2.9   57   368   11   MOLECULE: CELL DIVISION CONTROL PROTEIN 6 HOMOLOG 1;
15:  2lh9-A  3.0  2.5   50    78   12   MOLECULE: TUDOR DOMAIN-CONTAINING PROTEIN 7;
16:  2ly1-A  3.0  2.8   52   180   12   MOLECULE: TUDOR DOMAIN-CONTAINING PROTEIN 7;
17:  7nyw-E  3.0  3.3   58   212    7   MOLECULE: CHROMOSOME PARTITION PROTEIN MUKB;
18:  5l0p-A  2.9  2.7   52   311   12   MOLECULE: TRANSCRIPTION FACTOR ETV6, TRANSCRIPTION FACTOR E
19:  9c5g-B  2.9  3.0   53   290    8   MOLECULE: CAPW;
20:  4txa-A  2.9  3.2   55   367   15   MOLECULE: ROQUIN-1;
21:  6qfd-B  2.9  3.0   53   116    9   MOLECULE: DNA-BINDING PROTEIN;
22:  6l0o-A  2.9  3.0   51    71   10   MOLECULE: DNA HELICASE MCM8;
23:  7sol-A  2.9  5.2   57  1011    9   MOLECULE: UBIQUITIN-LIKE MODIFIER-ACTIVATING ENZYME 6;
24:  1zel-B  2.9  2.8   55   295   18   MOLECULE: HYPOTHETICAL PROTEIN RV2827C;
25:  7bzh-A  2.8  2.8   50    59    8   MOLECULE: SUL7S;
26:  8ioi-A  2.8  3.2   54   190    7   MOLECULE: PADR FAMILY TRANSCRIPTIONAL REGULATOR;
27:  6zwt-C  2.8  3.8   57   101   11   MOLECULE: TWO-COMPONENT RESPONSE REGULATOR;
28:  5uuj-A  2.8  3.8   65   371   15   MOLECULE: ALKZ;
29:  7pd7-B  2.8  2.8   48   251   15   MOLECULE: METHYLTRANSFERASE;
30:  1tbx-A  2.7  3.0   57    94   19   MOLECULE: HYPOTHETICAL 11.0 KDA PROTEIN;
31:  4k51-A  2.7  3.2   58   213    5   MOLECULE: EUKARYOTIC TRANSLATION INITIATION FACTOR 3 SUBUNI
32:  5yad-A  2.7  2.5   50    73    8   MOLECULE: MEIOSIS REGULATOR AND MRNA STABILITY FACTOR 1;
33:  7abi-7  2.7  4.3   60   236    3   MOLECULE: U5 SMALL NUCLEAR RIBONUCLEOPROTEIN 40 KDA PROTEIN
34:  8wxs-A  2.7  2.5   51    76   12   MOLECULE: FI02030P;
35:  2zme-B  2.7  4.1   62   215   11   MOLECULE: VACUOLAR-SORTING PROTEIN SNF8;
36:  7qen-C  2.7  3.2   57   291    7   MOLECULE: DNA (35-MER);
37:  1w7p-D  2.7  3.8   61   171    8   MOLECULE: VPS22, YPL002C;
38:  6zca-D  2.6  3.3   57   140    5   MOLECULE: PROBABLE DNA-DIRECTED RNA POLYMERASE SUBUNIT DELT
39:  7c0j-B  2.6  2.9   49    62   10   MOLECULE: HISTONE H5,DOUBLE-STRANDED RNA-SPECIFIC ADENOSINE
40:  3s93-B  2.6  2.5   50    81   10   MOLECULE: TUDOR DOMAIN-CONTAINING PROTEIN 5;
41:  8iue-P  2.6  3.2   60   303    5   MOLECULE: DNA-DIRECTED RNA POLYMERASE III SUBUNIT RPC1;
42:  8fo9-F  2.6  3.9   58  2289    9   MOLECULE: LEUCINE-RICH REPEAT SERINE/THREONINE-PROTEIN KINA
43:  2wb6-A  2.6  2.9   62   114    2   MOLECULE: AFV1-102;
44:  2esh-A  2.6  3.1   52   114   12   MOLECULE: CONSERVED HYPOTHETICAL PROTEIN TM0937;
45:  8ffz-B  2.6  2.9   53   905   11   MOLECULE: TRANSCRIPTION FACTOR IIIA;
46:  8jri-V  2.6  2.9   50   508    8   MOLECULE: 26S PROTEASE REGULATORY SUBUNIT 7;
47:  3cuq-A  2.5  3.8   63   219   16   MOLECULE: VACUOLAR-SORTING PROTEIN SNF8;
48:  3l9f-A  2.5  2.9   52   170    8   MOLECULE: PUTATIVE UNCHARACTERIZED PROTEIN SMU.1604C;
49:  8gcc-A  2.5  3.1   55   707    9   MOLECULE: DNA TOPOISOMERASE 2;
50:  8k5l-A  2.5  3.2   45   201   18   MOLECULE: TRNA (CMO5U34)-METHYLTRANSFERASE;

0001 s001A   -LQPVRALRLT-QA---GL-FL----PD-GLTIETIHQRVHNR--F-P-D------LS-T-R--LP-DQ--PAPLSRLLS--QA-----G------FAL-KW-RIRG-R---TVFLPE-H
0002 8zjrM   yMYAYRWIRNH-L----------ehtDT-CLPKQSVYDAYRKY--C-E-S--------lR-P--LS-TA----NFGKIIR--EI-----F------PDI-KA-RRLS-K---YCYSG---
0003 7ymdC   ivkenPVARKMvRY---IL---sesqNS-IITRNKLQSVIHEA--A-ReN------IA-K-P--SF-S-----KMFMDIN--AI-----L-----ynEL-QG-LPSK---raQKFILL-N
0004 5nl9A   vTVQRLQMANL-L-----------lskpqHLTADQVFQLINEH--M-P-N------ASrA-T--IF------nnLKLFAE--K-------------GIV-NL-LELG-I---TLYDNV-I
0005 7asvA   arELKAFVE---AT---FQ-------rqfVLTLSELKRLFNLH--L-A-SlppghtLF-S-G--IS-DR----XLQDTVL--A-------------AGC-KQ-ILVPfQ---KVFALW-E
0006 7kbfK   hPPTLSMVV---EV---LKnTE----RK-GTSVQAIRTRILSA--H-P-T------VD--fL--LR-T-----aLNKG-------------------IL-IR-PLNS---atGRFKLA-K
0007 7db7B   ----------------------------mRLPYSWLREVVAVG--A-S------------gW--DV-T---PGELEQTLL--RI-----G------HEVeEV-IPgvttgddVVFHLA-I
0008 5a48B   yPDIDSEVR---AI---LLsHA----QN-GITISSIKSEYRKL--T-G-N----------pF--PL-HD----NVTDFLL--TI-----P------NVT-AE-CSES-G---KRIFN---
0009 4etsB   yTKQREVLLKT-L----------yhsdT-HYTPESLYMEIKQA--E-P-D------L--N-V--GI-A--tvyRTLNLLE-----------------MV-TSiSFGS---agKKYELA-N
0010 5okcB   lfKALHVLLXS-AxaesLD-LQ-------HLNVEDTHHAVGK---D-I-E------DF-N-P--YT-RE----iIETVLN----------------KFA-VQ-EQE--N---NTWRLR-I
0011 7z8bC   ekRRNLLNCLIvRI---LKaHG----DE-GLHIDQLVCLVLEA--W-Q-KgpcpprgA-C-S--ST-------dvlsCIL--H-------------lTL-RR-HDDR-P---QVLSYA-V
0012 1ka8A   aDPTFDFIGY---------lpQ----TS-GMYMGYLYHAYLAY--M-EnG------YR-N-V--LS-LKmfGLGLPVMLK--EY-----G------LNY-EK-RHTK-G---IQTN----
0013 8e4yA   aIMSTHIVACL-L----------lyrhrqGIDLSTLVEDFFVM--K-E-E------VS-G-N--SE------dvVMHAIQ--LL-----G-------VT-IT-HTSR-N--dEFFIT---
0014 8e4vA   rNVDW--nkllrRA---IElEE----PN-GSSLKNIEKYLRSQsdL-T-S------TP-A-F--QQ---rlrlGAKR-------------------aRL-LK-DG-------PQYRVN-Y
0015 2qbyB   lpFHYKLAL---RS---LI---------esEDVMSAHKMYTDL--C-N-K------FQ-K-P--LS-YR----RFSDIIS--EL-dmfgI------VKI--R-IING-V---KKY-----
0016 2lh9A   aDLVSKMLRAV-L-----------qshkngIVLPRLQGEYRSL--T-G-------------dwipfpTL--EAYLRS--V--PA-----V------V-R-IE-ASRS-G---EIV-----
0017 2ly1A   mpDVKEKVAEL-L-----------gkyssgLWASALPKAFEDM--YkV-K------FPeD-A--LK--------NLASLS--D------V------CTI--N-YISG-Q---KAI-----
0018 7nywE   lsELDMMVGKI-L------laN----QG-IFTSQELYEELISL--A-DgK------LM-KkL--QE-K------VRTTLN--RL-------------MV-YF-LPN----nnNKFTITv-
0019 5l0pA   vtLPRLKIL---EV---LQ-----epdnhHVSAEDLYKRLIDM--G------------eE-I--GL-A-tvYRVLNQ-------------------fIV-TR-HNFE-K---SVFELT--
0020 9c5gB   -pwmQNRRF---EF---IEkLF----WEgALNRSDLEETFE-------------------tP--QT-S-----IDLRRYR--E--------------NI-EY-DAT--D---KTFKPT-K
0021 4txaA   qEEALKLVL---LA---LE-------dgsALSRKVLVLFVVQR--L-E-P----------tS--IG-H--vvQLLYR--------------------CF-KV-TKRD-S---SLMQLK-E
0022 6qfdB   ltAFQKNILTV-L-------------geeaRYGLAIKRELEEY--Y-G-E----------eV--NH-G-rlYPNLDDLVN--K-----gL------VEK--S-ELDK-R---TNE-----
0023 6l0oA   -rSTAKRFISA-L---------vrtyNN-IFQFHQLRQIAKEL--N---------iQV-A-D--FE------nfIGSLND--Q-------------GYL-LK-KG------pKVYQ----
0024 7solA   pRNWSQCVE---LA---RLlqlkpspIK-FDHLSFLQNAAKLY--A-TyC------IP-F-A-eeD-LS--ADALLNILS---------E------VKI------------------Q-E
0025 1zelB   ipTWASRVVSG-L---arD-RP----V--VVTKEDLTQRLTEA--G--------------gR--DP-DS-aiRELRR--------------------WL-VQ-LPV--K---GTWA-A-I
0026 7bzhA   meDVKQSV---eKI---IK-DR------eWVTFNDLLKYIP--------------------Y--PA-P--evyDALSQLI--K--------------KV-GR-RG-------RYFYYI-K
0027 8ioiA   fmALPHAILVS-L-------------seqaSSGYELARRFDRS--I-GyF------WT-A-T--HQ-------qiyRTLR--VM-----E--nnnwVRA-TT-VLQR-P---DKKVY---
0028 6zwtC   lTTGEFAVL---KA---LVqHP----RE-PLTRDKLMNLARGR----------------------------eRSIDVQVS--RL-----RrliednpYI-QT-VW---G---VGYVF-aE
0029 5uujA   vdELTEEVA---GI---LD--------gvVLTRDELVTRLV-A--D-K-R-----fVS-X-EerLR-SG--WGSVLKPLA--WR-----G------VLC-HG-PNRG-K---ITF--Q-F
0030 7pd7B   raGLLDVLKRI-H---------rrpggrgIIQIA----haahapkrtlvmlfrfrpaeyvlL--SM-TD--AHAFAACAR--EA-----G------FQV-TV-YSSK-R-pnPLVCVE--
0031 1tbxA   fFYPEAIVL---AY---LY-DN------eGIATYDLYKKVNAE--F-P---------stA-T--FY-------DAKKFLI--Q-----eG------FVK-ER-QER--G---EKRLY---
0032 4k51A   dVIMRRVFVAA-S------------qkftTVSQSELYKLAT-------lP------AP-L-D--LS-AWdiEKSLLQAAV--E-----dY------VSI-TI-DHE--S---AKVTFA--
0033 5yadA   lsLLSTETXSI-L----------qdapaccLPLFKFIDIYEKK--Y-G-H-----------K--LN-VS----DLYKL-T--DT-----I--------A-IR-EQG--N---GRL-----
0034 7abi7   sEEFRNDFLEL-L----------rrfgtkRVHNNIVYNEYI----S-H-R------E---hI--HM-NT--LTDFTKWLG--R-----eG------LCK-VD-ETP--K---GWYIQY-I
0035 8wxsA   lsLVKKVVHSL-V-------------vsspLTVEQLMRDYRSA--A-G----------cT-L--PYkDA--ESFLRSI-P--DT-----V-------TV-TG-H-------gQMAW----
0036 2zmeB   hmQLAKQLAGI-L------eeR----GG-IMSLTEVYCLVNRA----R-G------ME---L--LS-PE----DLVNACK--ML-----E-alklpLRL-R--VFDS-G---VMV--K-E
0037 7qenC   vrRLKKNVWRS-I---------snreKE-LK-FSDIIQGIS-K--M-Y-S---ddtLK-D-I--ST-S-----FCFICLL--HL-----A------nQI-TH-TEN--Y---NDLIV---
0038 1w7pD   ldEIAREIYEF-T---------dsdtnymIITLVDLYAMYNKS--MrI-G------TG---L--IS-PM----EMREACE--RF-----EhlglneLKL-V--KVNK-R---ILC-EK-F
0039 6zcaD   eMALVEIA---hEL---FE-----ehKK-PVPFQELLNEIASL--L-G-V-------lgD-R--IA-------qfyTDLN--------iD------GRF-LA-LS---D---QTWGLRwY
0040 7c0jB   -PTYSEMIA---AA---IR-------aegGSSRQSIQAYIKSH--Y-K---------------vNK-KE-iNRVLYSL-------laagV------LKQ-T-------V---PGSW----
0041 3s93B   qeCLRKEIRSL-L-----------istkdgLSPQELEKEYLLM--V-G-N----------ilgyrS-TM--E-LVLDM-P--DV-----V--------R-VC-PGAG-G---TVI-----
0042 8iueP   veVLNQQCFKF-LqskaetqrN----SS-FASSHEVWKYICE---L-G-I------SK-V-E--LS-ME----dIETILN--TL-----I--ydgkVEM-TI-IAAK-G---HMKL----
0043 8fo9F   iPDCYVELEKI-I---------kpieFP-VIDRKRLLQLVREN--Q--------------lQ--LD-EN----ELPHAVH--FL-----N--esgvLLH-F-----Q-L---SDLYF---
0044 2wb6A   -LESTSLYKKA-G---seN-LY----FQgivFLDSMFLVI-EK--L-GeR------VK-LlgamcskLP--LSSFIEIVT---R-----K------AKV-MY-NE--------KEVTLV-A
0045 2eshA   gWWLASTILLL-V-------------aekpSHGYELAERLA-E--F-G-I-------G-H-M--GN---------iyRVLA--DL-----E--esgfLST-EW-DTVS-P---PRKI----
0046 8ffzB   slQRQRAILKV-M----------ntiggvaYLREQFYESVSKY--M-G-S----------lD--KK-T---vrGDVDLMV--E----seK------LGA--R-TEPS-G---RKIIF---
0047 8jriV   hNVIKTGVRMI-S------------lsysRISLADIAQKLQ--------------------ldspE-DA--E-FIVAKAIrdGV-----I------E-A-SI-NHE--K---GYVQS---
0048 3cuqA   yyELGVQIIEV-C------hrN----GG-LITLEELHQQVLKG--R-GkF------AQ-D-V--SQ-D-----DLIRAIK--KL-----K-algtgFGI-IP-VG---G---TYL---aE
0049 3l9fA   -MQGKDIILGI-L-------------skkeRSGYEINDILQNQ--L-SyF------YD-G-T---------ygmIYPTLR--KL-ekdgK------ITK--E-VVIQ-N---KNI-----
0050 8gccA   lkPGQRKIIFS-S----------krrltrSIKVVQLAGYVSEH--A-A--------YH-H-G--EQ---slvQTIVGLAQnfvgsnnvpL------LQQ-DG-Q--trR---YIF--tya
0051 8k5lA   iEEPQSFLEKI-Y---------ndsngllIISIF-----------------------ywkktfnnqiLS--PEYVEGLLK--ES-----G------FKI-ER-ICEV---stDMWVVR--

0001 s001A   -LLHHHHHHHL-LL---LL-LL----LL-LLLHHHHHHHHHHH--L-L-L------LL-L-L--LL-LL--HHHHHHHHH--HH-----L------LLE-EE-EELL-L---EEEEEL-L
0002 8zjrM   hHHHHHHHHHH-E----------eeeEE-EEEHHHHHHHHHHH--H-L-L--------lL-L--LL-HH----HHHHHHH--HH-----L------LLL-EE-EEEL-E---EEEEE---
0003 7ymdC   hhhhlHHHHHHhHH---HH---hhhlLL-LEEHHHHHHHHHHH--H-HhL------LL-L-L--LH-H-----HHHHHHH--HH-----H-----hhEE-EE-ELLL---llLEEEEE-E
0004 5nl9A   lLHHHHHHHHH-H-----------hllllEELHHHHHHHHHHH--L-L-L------LLhH-H--HH------hhHHHHHH--L-------------LLL-EE-EELL-L---EEEELL-L
0005 7asvA   hhHHHHHHH---HH---HH-------hhlEEEHHHHHHHHHHH--H-H-LlllllhHH-L-L--LL-HH----HHHHHHH--H-------------LLL-EE-ELLLlH---LEEELL-L
0006 7kbfK   lLLHHHHHH---HH---HHlLL----LL-LEEHHHHHHHHHLL--L-L-L------LL--hH--HH-H-----hHHHH-------------------LE-EE-EEEL---llEEEEEL-L
0007 7db7B   ----------------------------lEEEHHHHHHHHHLL--L-L------------lL--LL-L---HHHHHHHHH--HH-----L------LEEeEE-EEleellllEEEEEE-L
0008 5a48B   lLLHHHHHH---HH---HHhLL----LL-LLLHHHHHHHHHHH--H-L-L----------lL--LL-LL----LHHHHHL--LL-----L------LEE-EE-ELLL-L---LEEEE---
0009 4etsB   lLHHHHHHHHH-H----------hhllL-LLLHHHHHHHHHHH--L-H-H------H--L-L--LH-H--hhhHHHHHLL-----------------LE-ELlLEEL---leELLEEL-L
0010 5okcB   hhHHHHHHHHH-HhhllLL-LL-------LLLHHHHHHHHLL---L-L-L------HH-L-L--LL-HH----hHHHHHH----------------HHE-EE-LLL--L---LLEEEL-H
0011 7z8bC   hhHHHHHHHHHhHH---HHlLL----LL-LEEHHHHHHHHHHHH--H-H-LlllllhhL-L-L--HH-------hhhhHHH--H-------------hLE-EE-LLLL-L---LEEEEL-L
0012 1ka8A   lLHHHHHHHH---------llL----LL-LLLLLLHHHHHHHH--H-HlL------LL-L-L--LL-HHhhHHHHHHHHH--HL-----L------LLL-LE-EEEL-E---EEEL----
0013 8e4yA   lLLHHHHHHHH-H----------hhllllLEEHHHHHHHHHHH--H-H-H------HL-L-L--HH------hhHHHHHH--HL-----L-------EE-EE-ELLL-L--lLEEEE---
0014 8e4vA   lLLLH--hhhhhHH---HHhLL----LL-LLLHHHHHHHHHHLllL-H-H------HL-H-H--HH---hhhhHHHH-------------------hLE-EE-EL-------LEEEEL-L
0015 2qbyB   llHHHHHHH---HH---HH---------llLLHHHHHHHHHHH--H-H-H------LL-L-L--LL-HH----HHHHHHH--HH-hhllL------EEE--E-EELL-L---EEE-----
0016 2lh9A   hHHHHHHHHHH-H-----------hhllllEEHHHHHHHHHHH--H-L-------------llllhlLH--HHHHHL--L--LL-----L------E-E-EE-ELLL-L---LEE-----
0017 2ly1A   llHHHHHHHHH-H-----------hlllllEELLHHHHHHHHH--HlL-L------LLhH-H--HL--------LHHHHH--H------H------EEE--E-EELL-L---EEE-----
0018 7nywE   llHHHHHHHHH-H------hhH----LL-EEEHHHHHHHHHHH--L-LhH------HH-HhH--HH-H------HHHHHH--HH-------------LE-EE-LLL----llLEEEELl-
0019 5l0pA   llHHHHHHH---HH---HL-----lllllEEEHHHHHHHHHHL--L------------lL-L--LH-H-hhHHHHHH-------------------hLE-EE-ELLL-L---LEEEEL--
0020 9c5gB   -lhhHHHHH---HH---HHhHH----HLlLEEHHHHHHHHL-------------------hH--HH-H-----HHHHHHH--H--------------LL-EE-ELL--L---LEEELL-L
0021 4txaA   hHHHHHHHH---HH---HL-------lllLLLHHHHHHHHHHH--L-L-L----------hH--HH-H--hhHHHHH--------------------LE-EE-EELL-L---EEEEEL-H
0022 6qfdB   llHHHHHHHHH-H-------------hhllEEHHHHHHHHHHH--H-L-L----------lL--LH-H-hhHHHHHHHHH--L-----lL------EEE--E-EEEL-L---EEE-----
0023 6l0oA   -lHHHHHHHHH-H---------hhhlLL-EEEHHHHHHHHHHL--L---------lLL-L-L--HH------hhHHHHHH--H-------------LLE-EE-LL------lLEEE----
0024 7solA   lLLHHHHHH---HH---HHhhhllllLL-LLHHHHHHHHHHHH--H-HlL------LL-L-L-hhH-HL--HHHHHHHHH---------L------LLL------------------L-L
0025 1zelB   llHHHHHHHHH-H---hhH-LL----L--LEEHHHHHHHHHHL--L--------------lL--LH-HH-hhHHHHH--------------------LE-EE-LLL--L---LEEE-E-E
0026 7bzhA   llLHHHHH---hHH---HH-HH------lEEEHHHHHHHLL--------------------L--LH-H--hhhHHHHHHH--H--------------LE-EE-EL-------LEEEEL-L
0027 8ioiA   llLHHHHHHHH-H-------------hhllLLHHHHHHHHHHL--H-HhL------LL-L-L--HH-------hhhHHHH--HH-----H--hlllEEE-EE-ELLL-L---LEEEE---
0028 6zwtC   lLHHHHHHH---HH---HHlLL----LL-LEEHHHHHHHHHLL----------------------------lLHHHHHHH--HH-----HhhhlllhLE-EE-EL---L---LEEEE-lL
0029 5uujA   hhHHHHHHH---HH---LL--------llLEEHHHHHHHHH-L--L-L-L-----lHH-H-HhhHH-HL--HHHHHHHHH--HH-----L------LLE-EL-LLLL-L---LEE--H-H
```

```
0030 7pd7B   hhHHHHHHHHH-H---------heeeeeeEEELL----lhhhlllleeeeeeeeeleeeeE--EL-LL--HHHHHHHHH--HL-----L------LEE-EE-ELLL-L-llLEEEEE--
0031 1tbxA   lLLHHHHHH---HH---HL-LL------lLLLHHHHHHHHHLL--L-L----------lhH-H--HH-------HHHHHHH--H-----lL------LEE-EE-EEL--L---EEEEE---
0032 4k51A   hHHHHHHHHHH-H------------hhllEEEHHHHHHHHL-------lL------HH-H-L--LL-HHhhHHHHHHHHH--L-----lL------LLL-LL-LLL--L---LEEELL--
0033 5yadA   hhHHHHHHHHH-H----------hllhhheEEHHHHHHHHHHH--H-L-L-----------L--LL-HH----HHLLL-L--LL-----E--------E-EE-LLL--L---LLE-----
0034 7abi7   hHHHHHHHHHH-H----------hhhlllEEEHHHHHHHHH----L-L-L------L---lL--LH-HL--HHHHHHHHH--H-----hL------LEE-EE-ELL--L---LEEEEE-L
0035 8wxsA   hhHHHHHHHHH-H-------------hlllLLHHHHHHHHHHHH--H-L----------lL-L--LLlLH--HHHHHLL-L--LL-----E-------EE-EL-L-------lLLLE----
0036 2zmeB   hhHHHHHHHHH-H------hlL----LL-EEEHHHHHHHHHHHH----L-L------LL---L--LL-HH----HHHHHHH--LL-----H-hhlllEEE-E--ELLL-L---LEE--L-H
0037 7qenC   hhHHHHHHHHH-H---------hhhlLE-EE-LHHHHHHHL-L--L-L-L---lllLL-L-L--LH-H-----HHHHHHH--HH-----H------hEE-EE-LLL--L---LLEEE---
0038 1w7pD   hhHHHHHHHHH-H---------lllllllEEEHHHHHHHHHHH--LlL-L------LL---L--LL-HH----HHHHHHL--LH-----HhhllllLEE-E--ELLL-L---LEE-LL-H
0039 6zcaD   hLLHHHHH---hHH---HH-----hhLL-LLLHHHHHHHHHHH--L-L-L--------lhH-H--HH-------hhhHHHH--------hL------LLE-EL-LL---L---LLLEEHhL
0040 7c0jB   -LLHHHHHH---HH---HH-------hllLEEHHHHHHHHHHH--L-L---------------lLH-HH-hHHHHHHH-------hhllL------EEE-L-------L---LLEE----
0041 3s93B   hhHHHHHHHHH-H-----------hlllllLLHHHHHHHHHHH--H-L-L----------hhlllL-HH--H-HHHLL-L--LL-----E--------E-EE-ELHH-H---LEE-----
0042 8iueP   hhHHHHHHHHH-HhhhhhhhhH----HH-LEEHHHHHHHHHHH---H-L-L------LL-L-L--LL-HH----hHHHHHH--HH-----H--hlllEEE-EE-ELLL-L---LEEE----
0043 8fo9F   eEHHHHHHHHH-H---------llllLL-EEEHHHHHHHHHHL--L---------------lL--LL-LL----LHHHHHH--HH-----H--hlllEEL-L-----L-L---LLEEE---
0044 2wb6A   -LLHHHHHLLL-L---lhH-HH----HHleeHHHHHHHHH-HL--L-LlL------LL-LllleeeeEE--HHHHHHHHL---H-----H------HHE-EE-LL-------LEEEEE-E
0045 2eshA   hHHHHHHHHHH-H-------------hhllLLHHHHHHHHH-L--L-L-L--------L-L-L--LL--------hhHHHH--HH-----H--hlllEEE-EE-ELLL-L---LEEE----
0046 8ffzB   hhHHHHHHHHH-H----------hllllleeELLHHHHHHHHHH--H-L-L----------lL--HH-H---hhHHHHHHH--H----llL------LEE--E-ELLL-L---LEEEE---
0047 8jriV   hHHHHHHHHHH-H------------hhllEELHHHHHHHHL--------------------lllhH-HH--H-HHHHHHHhlLL-----L------L-L-EE-LLL--L---LLEEL---
0048 3cuqA   hhHHHHHHHHH-H------hhH----LL-EEEHHHHHHHHHHL--L-LlL------LL-L-L--LH-H-----HHHHHHH--HH-----H-hhlllLEE-EE-EL---L---EEE---lL
0049 3l9fA   -LLHHHHHHHH-L-------------llllEEHHHHHHHHHHL--L-LlL------EE-L-L---------lllHHHHHH--HH-hhllL------EEE--E-EELL-L---EEE-----
0050 8gccA   llHHHHHHHHH-H----------hhllllLEEHHHHHHHHHHH--L-L--------LL-L-L--HH---hhhHHHHHHHLllllllllL------EEE-EL-L--llL---LLE--lhh
0051 8k5lA   lLLHHHHHHHH-H---------heeeeeeEEEEE----------------------hhhhhhlhlLL--HHHHHHHHH--HH-----L------LEE-EE-EEEE---leEEEEEE--

# Job: Type-2 BREX; PglW_WP_185095299 1 iSTAND
# Query: s001A
# No:  Chain   Z    rmsd lali nres  %id PDB  Description
   1:  8umy-A  9.5 11.6  130   546    8   MOLECULE: CHROMOSOME TRANSMISSION FIDELITY PROTEIN 18 HOMOL
   2:  7jgr-D  8.9  5.4  139   441    9   MOLECULE: ORIGIN RECOGNITION COMPLEX SUBUNIT 2;
   3:  3bos-B  8.9  3.9  128   231   10   MOLECULE: PUTATIVE DNA REPLICATION FACTOR;
   4:  8dr5-A  8.5  3.6  125   646   10   MOLECULE: REPLICATION FACTOR C SUBUNIT 1;
   5:  8thc-A  8.5  4.9  132   479   11   MOLECULE: ELG1 ISOFORM 1;
   7:  7mi1-A  8.4  9.7  123  2628   10   MOLECULE: CHIMERA PROTEIN OF DYNEIN AND ENDOLYSIN;
   8:  5e7p-A  8.2  5.5  139   719   10   MOLECULE: CELL DIVISION CONTROL PROTEIN CDC48;
   9:  5zr1-A  8.2  6.0  137   494   11   MOLECULE: ORIGIN RECOGNITION COMPLEX SUBUNIT 1;
  10:  3n70-A  8.1  3.3  121   142    8   MOLECULE: TRANSPORT ACTIVATOR;
  11:  8k8v-A  8.1  4.4  130   719   13   MOLECULE: ENDOPEPTIDASE LA;
  12:  8btg-B  8.1  4.2  139   335    8   MOLECULE: CHROMOSOMAL REPLICATION INITIATOR PROTEIN DNAA;
  14:  4rh7-A  8.0  4.1  140  3005    4   MOLECULE: GREEN FLUORESCENT PROTEIN/CYTOPLASMIC DYNEIN 2 HE
  15:  2qby-B  8.0  5.0  133   368   11   MOLECULE: CELL DIVISION CONTROL PROTEIN 6 HOMOLOG 1;
  16:  6hyp-A  7.9  4.6  126  2272    7   MOLECULE: MIDASIN,MIDASIN;
  17:  3j3r-A  7.9  8.5  143   798   11   MOLECULE: ADAPTER PROTEIN MECA 1;
  18:  6n9x-B  7.8  3.6  125   500   14   MOLECULE: DNA PRIMASE/HELICASE;
  19:  8ui7-A  7.8  3.3  129   643   10   MOLECULE: ATPASE FAMILY AAA DOMAIN-CONTAINING PROTEIN 5;
  20:  7vcs-A  7.7  3.3  132   766    6   MOLECULE: TRANSITIONAL ENDOPLASMIC RETICULUM ATPASE;
  21:  7mi8-A  7.6  9.6  124   868    6   MOLECULE: FUSION PROTEIN OF DYNEIN AND ENDOLYSIN;
  22:  5j1s-B  7.5  7.5  135   235   14   MOLECULE: TORSIN-1A;
  23:  6iy8-A  7.5  5.9  138   469    6   MOLECULE: POSITIVE REGULATOR CAPR;
  24:  2gno-A  7.4  4.5  117   296    7   MOLECULE: DNA POLYMERASE III, GAMMA SUBUNIT-RELATED PROTEIN
  25:  9e22-A  7.4 10.1  129  2925    4   MOLECULE: CYTOPLASMIC DYNEIN 1 HEAVY CHAIN 1;
  26:  7uqj-A  7.4  3.7  129   585    9   MOLECULE: ATPASE HISTONE CHAPERONE YTA7;
  27:  8xks-B  7.4 23.4  130   603    9   MOLECULE: CTAP1;
  28:  8fwj-A  7.3  3.6  134   552   11   MOLECULE: CIRCADIAN CLOCK PROTEIN KAIC;
  29:  7ykk-A  7.3  6.1  137   735   12   MOLECULE: ATPASE FAMILY GENE 2 PROTEIN;
  30:  7plh-A  7.3  6.4  129   260    8   MOLECULE: SHTNSC;
  31:  6z1f-3  7.3  4.0  130   284    8   MOLECULE: RIBULOSE BISPHOSPHATE CARBOXYLASE/OXYGENASE ACTIV
  32:  8xvb-A  7.2  4.7  129   246   12   MOLECULE: DNA (5'-
  33:  5zr1-B  7.2  9.1  124   374   10   MOLECULE: ORIGIN RECOGNITION COMPLEX SUBUNIT 1;
  34:  1nlf-A  7.2  3.9  119   254   12   MOLECULE: REGULATORY PROTEIN REPA;
  35:  9bv1-A  7.2  3.3  124   405   10   MOLECULE: 26S PROTEASOME NON-ATPASE REGULATORY SUBUNIT 1;
  36:  8bob-C  7.2  4.4  122   412   11   MOLECULE: PROTEIN MALY;
  37:  7st9-A  7.2  4.9  133   522    9   MOLECULE: CHECKPOINT PROTEIN RAD24;
  38:  8fcv-R  7.2  6.0  139   343   12   MOLECULE: DNA (60-MER);
  39:  8tp1-A  7.1  9.2  123   419   12   MOLECULE: MITOCHONDRIAL CHAPERONE BCS1;
  40:  6qem-G  7.1  8.4  136   239    7   MOLECULE: REPLICATIVE DNA HELICASE;
  41:  3pvs-D  7.1  3.5  113   424   12   MOLECULE: REPLICATION-ASSOCIATED RECOMBINATION PROTEIN A;
  42:  8tw7-2  7.1  5.3  121   339   12   MOLECULE: REPLICATION FACTOR C SUBUNIT 5;
  43:  9cru-C  7.0  8.6  136   741    6   MOLECULE: VESICULAR-FUSION PROTEIN SEC18;
  44:  5zr1-E  7.0  4.0  120   460   10   MOLECULE: ORIGIN RECOGNITION COMPLEX SUBUNIT 1;
  45:  9klv-D  7.0  3.8  125   338    8   MOLECULE: PROTEIN SUPPRESSOR OF MAX2 1;
  46:  3co5-A  6.9  3.2  107   134   10   MOLECULE: PUTATIVE TWO-COMPONENT SYSTEM TRANSCRIPTIONAL RES
  47:  8xku-A  6.9  3.5  132   730   11   MOLECULE: PROBABLE INACTIVE ATP-DEPENDENT ZINC METALLOPROTE
  48:  6l1q-A  6.9  3.7  113   267    5   MOLECULE: CBBQ PROTEIN;
  49:  8dgc-A  6.9  5.6  130  2028    8   MOLECULE: SEAVS3;
  50:  7st9-E  6.9  3.4  120   354   10   MOLECULE: CHECKPOINT PROTEIN RAD24;

0001 s001A   NADRTSAVT-QRR----E-QKLRYRANAQLLES-A-AR-P--GFRLLTVRYD-D-QRL-ALLT-LTEEWR---AEPVDVS--I-LFLTSLRRLILEADNAVPGSRDAMKFGEYTAAAWGEVE-KELASN-SRPLLLHD-A-S--VM-ARYG-G-T-G-ILQRLADHAR-R--G--GRGLWLLSPLNAVP--RLEDWTVALED-REEWIRLN-H-SVND-PEDP
0002 8umyA   lvLRAS---clap-----ellsDDFTNRCLLKW-L-KL-WdlKVALLCGPPG-L---G-KTTL-A-HVIAgysVVEMN-A--S-DDR---------------------------------------SPEVFR-TRIEAAkPNCLVIDE-IdG--AP-V--A-A-I-N-VLLSILNR--gL--L--MRPIICICND---Q--FAPsLRQLKQQ---FLLHFP-P-TsrL-VQRV
0003 7jgrD   ADRESIRRK-ERLqrdyttlrGYAKERSNVRLL-L-QReS--NSLLLLGPRG-S---G-KTTL-I-NSVL----VHLDGN--L-HT----------------------------------------DDRVAL-KSITVQ-MVIFILEE-F-D--LF-CAHH-N-Q-T-LLYNLFDVSQ-SaqA--PICVLGVTCR-LDV--I-ELLEvRFSH---RQVFLF-P-SedY-VDLL
0004 3bosB   -------------tsyyP-AAGNDELIGALKSA-A-SG----QAIYLWGPVK-S---G-RTHL-IH-AAC---AFYIPLG--I--HASI-----------------------------------------ST-ALLEGLqFDLICIDD-V-D--AV-AGHP-W-E-E-AIFDLYNRVA-E--Q-kRGSLIVSASA-spX--EAGFVVSRXHW---LTYQLQ-P-XdeK-LAAR
0005 8dr5A   arrkenTKE-A--ptnlqqvcGNKGSVMKLKNW-L-AN-W--RAAMLYGPPG-I---G-KTTA-A-HLVAgydILEQN-A--S-DV---------------------------------------rsKTLLN-AGVKNAkHFVIIMDE-V-D--GM-SGRG-G-V-G-QLAQFCRKT--------STPLILICN---ER--NL-PKMRFDRV---LDIQFR-R-PanS-IKSL
0006 8thcA   VLLGTNIK-qLFK-vtlkqvlIEPKLKLRIKNW-I-ET-S--PLMILHGN-----sIG-KKTL-I-QTIMsyqIYEVN-S--N-MN---------------------------------------rSKKDLL-DILLDF-TGLVLFND-V-D--VL---RG-Y-W-A-MISKLCEFS--------RRPLVLTCKD---L--SLV-PSELASE-QNSLFHTK-K-IstV-YAFT
0007 8emhB   ----sdldD-LTK----L-KEDVQWHLLARMVP-F-VE-N--YNVCELGPRG-T---G-KSHI-YKECSP--nSILVSGG--Q-TT-----------------------------------------VANLF-YGLVGL-WDVVAFDQ-V-A--G--ISFK-G-V-Q-IMKDYMAS--greqmeaSASMVFVGNINLLA--PFPEAMdFFDR-fHAYIPGW-E--KMR-PEFT
0008 7mi1A   SSDGV-----------------WNIADVVANS----E-N--NYFTMASERD-V-D---GTFK-L-IELAkesLKIIPL------GSIE--------------------------------------NLNYAQ-EEISKSeGGWILLQN-I-Q--MS-L--S-W-V-KtYLHKHVEE--tK--A--AEKMFMTCHLT-------DKLPLLQR----DRVVYE-D-I-PG-ILD-
0009 5e7pA   RGVVRLHpadHTPsvtlddvgdMVETKRALTEA-V-LWpP--RGVLLYGPPG-C---G-KTFV-V-RALArlsVHAVKGS--E-LMDKWV------------------------------------GSSEKAV-RELFAR-ASLVFLDE-IdA--LA-PTDK-V-V-A-SLLTELDGIE-P--L--RVVVLGATNR--------LIDPLLRP---gRLVFVE-P-PrdI-LRTG
0010 5zr1A   IVETIFVKK-Q--lnssyylpaRENEFASIYLS-A-YSsA--TTIYVAGTPG-V---G-KTLT-VREVVK----VEINGL--K-MV---------------------------------------KPTDCY-ETLWNK-VIVVLLDE-L-D--AM-VTKS---Q-D-IMYNFFNWTT-YenA--KLIVIAVANT--MD--LPERQLgITSR---TRIMFT-G-YeEL-KNIR
0011 3n70A   -----------------WiNQYRRRLQQL-SET-D-------IAVWLYGAPG-T---G-RXTG-A-RYLHqqeFVYRELT-pD-NAP-------------------------------------------QL-NDFIALqGGTLVLSH-P-E--HL------T-R-E-QQYHLVQLQS-QehR--PFRLIGIGDT-lVE--LASNHIILYYCfAXTQIAC-----LT------
0012 8btgB   -------------dtfvI-GSGNRFAHAASLAVaE-AP-A--NPLFIYGGVG-L---G-KTHL-MH-AIG---HVYLSSE--K-FTNEFINSI----------------------------------kaVDFR-NRYRNV--DVLLIDD-I-Q--FL-AGKE-T-Q-E-EFFHTFNTLH-E--E--SKQIVISSDR-pPK--EIPTLElRFEW-G-LITDIT-P-PetR-IAIK
0013 5v8fD   -DPGFLQRRlqLP--ekiiFTYLQDCQQEIDRI-I-KE-S--HSVILVGPRQ-S---Y-KTYL-L-DYELqfiTIRLNGF--I-HS-----------------------------------------EQTAI-NGIATQ-LVVFIFDE-I-D--TFaGPVR---Q-T-LLYNLFDMVE-H--SrvPVCIFGCTTK-LNI--L-EYLEvRFSQ---RVIYMP-Q-IvdA-VRNL
0014 4rh7A   dTSSE-dpL-ILDasklvyTPLTDKCYLTLTQA-M-KM-G--LGGNPYGPAG-T---G-KTES-VKALGGgrqVLVFNCD--E-GI---------------------------------------DVKSMG-RIFVGLcGAWGCFDE-FnR--LE-E--S-V-L-S-AVSMQIQTIQ-D--A--LSGIFITMNPA-kgyggRQKLPLKQL----RPVAMS-H-P-DN-ELIE
0015 2qbyB   -----KNPK-VSV-----eipFREDILRDAAIA-I-RYvK--FSNLFLGLTG-T---G-KTFV-S-KYIF----AYVNCR--E-VG---------------------------------------gTPQAVL-SSLAGK-LAIIYLDE-VdT--LV-KRRG-G-D-I-VLYQLLRS---D--A--NISVIMISND--iN--VRDYMEVLSS----PSVIFK-P-YeqL-KFIY
0016 6hypA   alepqgltikptlqahyiiTPFVEKNMMNLVRA-T-SG-K--FPVLIQGPTS-S---G-KTSM-I-KYLAhkfVRINNHE--H-TD------------------------------------------LQEY-LGVLVEkGYWIVLDE-LnL--AP-T--D-V-L-E-ALNRLLDdnrpH--P--DFLLFATQNPP------iKILSFRNR----LELHFD-D-IpqD-ELEL
0017 3j3rA   QENSEVT--ssetnilhsrvigQDEAVVAVAKA-V-RRpI--GSFIFLG----P-TGVgKTEL-A-RALAeesMIRIDMS--E-YM-kHSTSV--------------------------------pGYVGYDEgGQLTEK-VSVVLLDA-I-E--KA-HP-D-V-F-N-ILLQVLEDG-tvdfR--NTILIMTSNVGGEL--K-AFRPFINR----EIIVFH-S-LKHL-T---
0018 6n9xB   -------------ssvgllfsgCTGINKTLG---A-RG-G--EVIMVTSGGM-G-KST-FVRQ-QALQWGgkkVGLAMLQ--EsveeedligriengkfdqwfdelfgndtfhlydsTDRLL-AKLAYMgCDVIILDH-I-S--IV-VS-M-I-D-N-LMTKLKGFAK-S--T--GVVLVVICHL-TDL--R-GSGALRQL--sDTIIALE-R-pLEP-S-SS
0019 8ui7A   TEEVLLLEF-PLLpqtaseliGNELAIKKLHSW-L-KD-W--NTVLITGPTG-V---G-KTAA-V-YACAgfkIFEVN-A--S-SQ---------------------------------------rSGRQIL-SQLKEA-TSLILFEE-V-D--VI--DAG-F-L-N-AIKTFMATT--------KRPVILTTSD-PTF--S----MFDGC----EEIKFS-T-PlnV-ASY-
0020 7vcsA   -------------diggC-RKQLAQIKEMVEL-----P-P--RGILLYGPPG-T---G-KTLI-A-RAVAgafFFLINGP--E-IMSKLA------------------------------------GESESNL-RKAFEE-AAIIFIDE-LdA--IA-PERR-I-V-S-QLLTLMDG-L-K--Q--RVIVMAATNR--PN--SI--DPALRRFrfDREVDIG-I-PatG-RLE-
0021 7mi8A   SGYL-----nlvf---------eWNIADVVAN----SE-N--NYFTMASERD-V-D---GTFK-L-IELAkesLKIIPL------GSIE--------------------------------------NLNYAQ-EEISKSeGGWILLQN-I-Q--MS-L--S-W-V-KtYLHKHVE---eH--E--KFKMFMTCHLT-------DKLPLLQR----DRVVYE-D--IPG-IL--
0022 5j1sB   YSSPAQ---knflqledkfpgQSSFLWQRGRKF-L-QKeP--ATIIFTAARE-G-RET-LKCL-SH--HV---ADQIDGA--G----------------------rtwqDSDTVKLLVD-LELSYGgQKAAVVHH-FeS--FP-A--G-S-T-L-IFYKYCD---aF--K--DVALVLTVLLEEKVrdLLWAKdLWSR----HLVLPV-Q--VSSi----
0023 6iy8A   YELQLVSL---------Q-TPAYQTVRNMMDKA-A-QG-K--VSVLLLGETG-V---G-KEVI-A-RSVHaepFVAVNCA--A---IPPD-------------------------------------LIESEL-FGRFERdKGTIFLDE-V-I--EL-S-PR-A-Q-A-SLLRVLQE--dnrtrkiDVRVIAATH--EDL--AeGRFRLYYR-lNVFPVAIP-A-LreD-IPLF
0024 2gnoA   -----------------------DQLETLKRII-E-KSeG--ISILINGE---D-LSY-PREV-S-LELP----LEIDpE--N-IG------------------------------------------iDDI-RTIKDF-LKYVIVHD-C-E--RX-T-QQ-A-A-N-AFLKALEEP--P--E--YAVIVLNTRR---W--HYL-LPTIKS----FRVVVN-V-PkeF-RDLK
0025 9e22A   GEAVD----gvem------GGMWVEKVLQLYQI-T-QI-N--HGLMMVGPSG-S---G-KSMA-W-RVLLegvAHIIDPK--A-IS-----------------------------------------KDHLY-GLFTHV-LQWIVFDG-D-V--D-----E-W-V-E-NLN---SVLD-D------VRIMFEVQD------LKYATLTVSR----GMVWFS-E-D-LS-TDM-
0026 7uqjA   -------------nvnfddiggLDNYIDQLKEM-V-AP-P--RGVLFHGPPG-T---G-KTLM-A-RALAkitFFMRKGA--DiLSKW-------------------------------------VGEAERQL-RLLFEE-ASIIFFDE-IdG--LA-PIHA-SiV-S-TLLALM-DGM-D--N--RVIVIGATNR-pDA--VD---PALRRP--gREFYFP-L-PkAR-FKIQ
0027 8xksB   -----------------G-nkQAKVELMEVVDF-F--A-P--KGVLLVGPPG-N---G-KTLM-A-RAVAgvaFISSSAA--E-FI---EMYM---------------------------------GLGAARV-RDLFNT-ACIIFIDE-L-D-aVG-RQDN-T-V-N-QLLTEMD---eQ--Q--GIVVMGATNR--KD--VLDAALTRPG----RSIEVR-R-PfqG-RLE-
0028 8fwjA   -------------ksptgiqGFDELTLG--GLP-T----G--RPSLVCGSAGcG-KTL-FAST-FLINGVgepGVFVTFEieH-IAV--------------dpsEVAE--igdYDLEGLF-LRLELA-IKRVVLDT-I-E--SL-FSAF-L-R-A-EIRRLFDWLK-E--R--GLTTVITAERGALT--RQ---GLEE--vSDCVILLD---nFSV-LPVL
0029 7ykkA   VTLK---------kvywsdigGQEELKTKMKEM-I-QL----KGVLLYGPPG-CsKTL-TAKA-LATES-ginFLAVKGP--E-IF---NKYV---------------------------------GESERAI-REIFRK-ASIIFFDE-I-D--AL-SPNH-V-L-T-SLLNEID---eL--K--GVVIVAATNR---P--DEIDAALLRPGrlDRHIYVG-P-PnAR-LEIC
0030 7plhA   dEWLQAaRL-KG--ksivpLQQVKTLHDWLDGK-R-KA-R--KSCRVVGESR-TgKTV-ACDA-YRYRH-tvpVVYIRPH--Q------------------------------------------kcGPKDL-FKKIT-gVEMLIIDE-A-D--RL------K-P-E-TFADVRDIAE-D--L--GIAVVLVGTD-rlD--AVI---VLER----AHLRFG-K-LedF-KNTW
0031 6z1f3   ------------------syyiAPRFLDKLAVH-I-TK-N--LILGIHGRKG-E---G-KTFQ-C-ELAFgieVTLISG---G-ELE--SPDA--------------------------------GDPARLIR-LRYRET-ACVLMIND-LdA--GA-GTQL-V-N-A-TLMNIADN-----------VPIIVTGND---F--STLYAPLIRDG-RMEKFYWE-P-NvgI-VGGA
0032 8xvbA   ------LPE-PPR----vetqTVKQIWTSMRFA-S-LT-E--SIAVVCGNPG-V---G-KTEA-A-REYRnnnVWMITIT--P-SC---------------------------------------aSVLECL-TELAFE-LGLVIIDE-A-D--HL------G-A-E-VLEELRLLQE-S--T--RIGLVLMGNH-rvY--SNMT--LFSR--iAKRTAIN-K-TkaD-VKAA
0033 5zr1B   mSMAP------------F-EIQKK-MFPQYWFE-L-TQ-G--FSLLFYGVG--S-KRN-FLEEfAIDYLSsipCLILNGN--P-SC---------------------------------------NYRDVF-KEITDLiKLILVVHN-L-DgpSI-RKNT-F-Q-T-MLSFLSVI------R--QIAIVASTDHI-----yAPLLWKAQN----VFHDIS-Nf---------
0034 1nlfA   -------------thkpinileafpplyvLPNM---VA-G--TVGALVSPGGaG-KSM-LALQ-LAAQIAtgpVIYLPAEiqP-LIGSL-----------------------------------pnimAPEWF-DGLKRAgRRLMVLDT-L-R--RF-HIE--pmA-Q-VIGRMEAIAA-D--T--GCSIVFLHHA-----------VLVDN--iRWQSYLSfV-RLKP-AV-E
0035 9bv1A   -------------vtysdvggCKEQ-IEKLREV-V-E-----KGVLLFGPPG-TgKTL-CARA-VANRT-dacFIRVIG---S-ELV--QKYV---------------------------------GEGARMV-RELFEM-ACLIFFDE-I-D--AI-GGQR-T-M-L-ELINQLD---pR--G--NIKVLMATNR------PDTLDPMRPG----RKIEFS-L-PthI-FKI-
0036 8bobC   -------------pvrldhtvvRERLLAKLSGA---NN-F--RLALITSPAG-Y---G-KTTL-ISQWAA----GWYSLD--E-GDN---------------------------------------QQERFA-SYLIAA-VlYLVIDD-Y-H--L--ITNP-I-H-E-SMRFFIRH-Q-P--E--NLTLVVLSRN-----------QLGILR-VRLEIGSQ----AFT-HQEF
0037 7st9A   ---------eKFK--cleqvaiHKRKLKDVQEA-L-DA-M--RILLLSGPSG-C---S-KSTV-I-KELS----VTEFRG--D-C----IVND--------------------------------lPQMESFS-EFLKGAnLSLILIED-L----PN-VFHR-F-Q-Q-LILQWLYS---sepL--LPPLVICITECFSA--E--TIMNILMH---KRIKFN-P-ItLL-KKHI
0038 8fcvR   lLTRPERLAfENY----vaHPRLKEVYEILMRT-I-AE-P--SFIFVYGASG-V---G-KTTL-R-LRVErvpVVGIEA-----IAPE--------------------SRYF-NWKEYY-TRAlithPDVFFVDE-A-Q--HF-GKVK-L-Q-D-QLDCLKSLAN-M--T--GILHCLLGTY--------llTFRNLSR-R-VDIHFR-R-YqaF-KSVF
0039 8tp1A   RTQH---------svvlQ-QGLADRIVKDIREF-I-DN----YRRGYLLYgC-G-KSS-FITA-LAGELE---HSICLLS--L---TDSS------------------------------------lSDDRLN-HLLSVA-PQQSLVLL---------eaFS-G-LlN-ALDGvaSTEA----------RIVFMTTN---idrLDPALIRPG----DLKEYV-G--ycS-HWQM
0040 6qemG   wQKEQ----rpsf---vE-CEGQMNALSKARQY-V-EE-F--ASFIFSGKPG-T---G-KNHL-AA-AIC---NLIITVA--D-IMSAMKDTF---------------------------------tSEEQLL-NDLSNV--DLLVIDE-I--------EKV-IiN-Q-IVDRRSSS---------KRPTGMLTNS-EEM--T-LLGEmMRLG--NSLWVIF-N-W--------
0041 3pvsD   -------------laqyigqQHLLPLPRAIGHL-H--------SMILWGPPG-T---G-KTTL-A-EVIAnadVERISAV--T-SG------------------------------------------VKEI-REAIER-ATILFVDE-V-H--RF------N-K-S-QQDAFLPHIE-D--G--TITFIGATTE--nP--SFElNSALLS----RVYLLK-S-LteD-IEQ-
0042 8tw72   ----lelAQ-QP-pknldevtAQDHAVTVLKKtlksAN-L--PHMLFYGPPG-T---G-KTST-I-LALTksrILELN-A--S-DE---------------------------------------rGISIVrEKVKNF-AKIIILDE-A-D--SM------T-A-D-AQSALRRTME-T--Y--STRFCLICNY-vTR--II----LASR----SKFRFK-A-LsNA-IDRI
0043 9cruC   MSDDVN-------ggmmlySERVNSILKNGARY-V-RQrL--VSLLIHGPAG-S---G-KTAL-AAEIAL----RLIS-P--N-ELSGMS-----------------------------------ESAKIAYI-DNTFRD-ANILVIDS-L-E--TL-VDgN-I-L-Q-MLKVALKR---kppqdrRLLIMTTTSA-YSV--L-qQMDILCFD-N--EIAVP-N-M-tN-LDEV
0044 5zr1E   -------------nvttpevafREYQTNCLASY-I-ST-P--SNLILQGYSG-T---G-KTYT-LKKYF-nlhAVWLEPV--E-LV---------------------------------------SWKPLL-QAIART-VLFLILDG-F-D--SL--QDL-D-A-A-LFNKYIKL--lpkdskiNIKFIYTMLE-TSF--L----QRYSTH-CIPTVMFP-R-YdEV-STI-
0045 9klvD   ---------kalnekvwwqNDAAAAVAATVSQC-K-LG-D--VWLLFSGP---D-RVG-KRKM-V-SALSgtnPIMIQLG--SrqdaGNSSFR-----------------------------------gKTAL-DKIAET-VSVILLED-IdE--AD-M--L-V-R-G-SIKQAMD---sL--G--NVIFVMTASWHSL-----PFAFMVSR--vDDAVAFR-A---------
0046 3co5A   ----------------gN-SAAIQEXNREVEAA-A-KR-T--SPVFLTGEAG-S---P-FETV-A--RYF---H-vSPAR--V-EY-----------------------------------------liDXP-XELLQKeGGVLYVGD-IaQ--YS-R--N-I-Q-T-GITFIIGKAE-R----cRVRVIASCSYASCE--EK---AGLFSEs-----------------
0047 8xkuA   -------------dfagQ-EYIKRELQEIVRIL-K-ND----pKGVLLHGGT-G-KTL-LAKA-IAGEAG---LFFAANG--T-DFV--------------------EMFVGVAASRVK-DLFASSyAPSIIFIDeI-D--AI-GSEQ-G-L-L-QILTEMDGFK-V-tT--SQVLVIGATNR-------LDPALLRKG---DKIIRV-G--kdG-RLA-
0048 6l1qA   -------------vgkepfyqPQHDEVALFEAA-Y-RK-R--LPVMVKGPTG-C---G-KSRF-V-EFMAgkpLVTVACN-------------------------------------------edmTAADL-VGLTVAyGAICYLDE-I-V--EA------R-Q-D-TTVVI-HPLT-D--H---FQLVISYNPG-------MKDLTKQR----TGFEFD-Y-PelE-AGIV
0049 8dgcA   -------------iensvsRTQEATLVQRVVEA----F-G--APVIIHADAGvG-KSI-FSTH-IEEHL-gsvSILYDCF--G-------------------lgqyrNASSYrhHHRTAL-VQMANE-MLCIIIDA-A-D--NA-QMAA-etR-S-FIKDLIRE-K-L--P--DVCLVALCRP-YRR--E----LLDPPP-EALTLSLQ-T---fN-RDEH
0050 7st9E   -------------alshN-EELTNFLKSL-SDQ-P-RD-L--PHLLLYGPNG-T---G-KKTR-CM-ALL---ELEITPS--D--MGNN--------------------------------------dRIVIQ-ELLKEV--KCVIINE-AnS--LT-K--D-A-Q-A-ALRRTMEKYS----K--NIRLIMVCD----S--MSPiIAPIKS----LLIRCP-A-PseI-STI-
0051 7z6hK   ---------wvETelavH-KKKIEEVETWLKAQ-V-L-----SILLITGPPG-C---G-KTTT-LKILSI---QVQEW------INPV--------------------lpyqSQIAVFK-EFLLRA-TKIILVED-L--pnQF-YRDH-T-L-HeVLRKYVRIG--------RCPLIFIISDS-sgdnNQRLLFpIQEE---SNISFN-P-V--A-PTI-
0052 6pxkK   -----------seseldkhiiGQDNAKRSVAIA-L-RNtP--KNILXIGPTG-V---G-KTEI-A-RRLAnapFIKVEAT----KFTEG---------------------------------------KEV-DSIIRD-LGIVFIDE-I-D--KI-CREG-V-Q-R-DLLPLVEG--xvktD--HILFIASGAFQ---iaKPSDLILQGR--lPIRVELQ-A-LtsD-FER-
0053 8ouyC   -------------ailstgiGSLDKLLD--AGL-Y--T-G--EVTEIVGGPGsG-KTQ-VCLC-MAANVAqqnVLYVDSNvvH-AFD----------------------------------------IFQML-DVLQELtVKVVVVDS-V-T--AV-VSPLlM-M-Q-LARELKTLAR-D--L--GMAVVVTNHITLKP--ALGRSWSFV-----TRILLD----DIG-T---
0054 6p8vC   -------------qerlarlvGLDEHKDRLSKI-L-GL----PLVVLAGDVG-S---G-KTEL-AETIGDditLYPLS-L--A-TR--GQGRV--------------------------------GEXTQLVS-AAFDYT-IVLLLIDQ-AdA--LA-Q-RA-G-V-N-AFIRGIDRIA-N--Q---AAVLXCTNR---L--KAL-DPAVQR-raAEILTFS-R-PeQR-HYLK
0055 3f8tA   ipevDPAE-eeDKaraiapLPGAEEVGKMLALQ-L-FSeR--LHVLLAG----Y-PVV-CSEI-L-HHVL----VYVDLR--R-TE------------------------------------------LTDL-TGAAVLdGGILAVDH-LeG--AP-E--P-H-R-W-ALMEAMDK--dgialnaRCAVLAAINPGARI--DL-DQDFLSH--fDLIAFLG-Vp--------
0056 1g4aE   -----------seseldkhiiGQDNAKRSVAIA-L-RNtP--KNILMIGPTG-V---G-KTEI-A-RRLAnapFIKVEAT--K-FTE--VGYV------------------------------------gKEV-DSIIRD-LGIVFIDE-I-D---kicREG-V-Q-R-DLLPLVEG--mvktD--HILFIASGAFQaKP--SDLiPELQG--rlPIRVELQ-A-LtsD-FER-
```

```
0001 s001A   [illegible]
0002 8umyA   [illegible]
0003 7jgrD   [illegible]
0004 3bosB   [illegible]
0005 8dr5A   [illegible]
0006 8thcA   [illegible]
0007 8emhB   [illegible]
0008 7mi1A   [illegible]
0009 5e7pA   [illegible]
0010 5zr1A   [illegible]
0011 3n70A   [illegible]
0012 8btgB   [illegible]
0013 5v8fD   [illegible]
0014 4rh7A   [illegible]
0015 2qbyB   [illegible]
0016 6hypA   [illegible]
0017 3j3rA   [illegible]
0018 6n9xB   [illegible]
0019 8ui7A   [illegible]
0020 7vcsA   [illegible]
0021 7mi8A   [illegible]
0022 5j1sB   [illegible]
0023 6iy8A   [illegible]
0024 2gnoA   [illegible]
0025 9e22A   [illegible]
0026 7uqjA   [illegible]
0027 8xksB   [illegible]
0028 8fwjA   [illegible]
0029 7ykkA   [illegible]
0030 7plhA   [illegible]
0031 6z1f3   [illegible]
0032 8xvbA   [illegible]
0033 5zr1B   [illegible]
0034 1nlfA   [illegible]
0035 9bv1A   [illegible]
0036 8bobC   [illegible]
0037 7st9A   [illegible]
0038 8fcvR   [illegible]
0039 8tp1A   [illegible]
0040 6qemG   [illegible]
0041 3pvsD   [illegible]
0042 8tw72   [illegible]
0043 9cruC   [illegible]
0044 5zr1E   [illegible]
0045 9klvD   [illegible]
0046 3co5A   [illegible]
0047 8xkuA   [illegible]
0048 6l1qA   [illegible]
0049 8dgcA   [illegible]
0050 7st9E   [illegible]
0051 7z6hK   [illegible]
0052 6pxkK   [illegible]
0053 8ouyC   [illegible]
0054 6p8vC   [illegible]
0055 3f8tA   llllLHHH-hhHLhhhhllLLLLHHHHHHHHHH-H-LLhL--LLEEEEL----L-HHH-HHHH-H-HHHH----EEEEHH--H-LL[illegible]
0056 1g4aE   -----------llhhhhhllLLHHHHHHHHHH-H-HHlL--LLEEEELLLL-L---L-HHHH-H-HHHHlllEEEEELL--H-HHL--LLLL[illegible]

@@@@@@@@@@@@@@@@@@@@@@@@@@@@@@@@@@@@@@@@@@@@@@@@@@@@@@@@@@@@@@@@@@@@@@@@@@@@@@@@@@@@@@@@@@@@@@@@@@@@@@@@@@@@@@@@@@@@@@@@@@
@@@@@@@@@@@@@@@@@@@@@@@@@@@@@@@@@@@@@@@@@@@@@@@@@@@@@@@@@@@@@@@@@@@@@@@@@@@@@@@@@@@@@@@@@@@@@@@@@@@@@@@@@@@@@@@@@@@@@@@@@@
@@@@@@@@@@@@@@@@@@@@@@@@@@@@@@@@@@@@@@@@@@@@@@@@@@@@@@@@@@@@@@@@@@@@@@@@@@@@@@@@@@@@@@@@@@@@@@@@@@@@@@@@@@@@@@@@@@@@@@@@@@
```

```
# Job: Type-2 BREX; BrxHI-Helicase MBK7581083 wHTH PDB50
# Query: s001A
# No:  Chain   Z    rmsd lali nres  %id PDB  Description
   1:  3elk-A  8.5  2.6   71   104    4   MOLECULE: PUTATIVE TRANSCRIPTIONAL REGULATOR TA0346;
   2:  7wjp-A  8.5  2.4   69    98    7   MOLECULE: PADR FAMILY TRANSCRIPTIONAL REGULATOR;
   3:  5a31-N  8.4  1.8   65   703   14   MOLECULE: ANAPHASE-PROMOTING COMPLEX SUBUNIT 1;
   4:  5zqh-A  8.4  2.3   71   101   11   MOLECULE: PADR FAMILY TRANSCRIPTIONAL REGULATOR;
   5:  8iue-P  8.4  2.5   63   303   13   MOLECULE: DNA-DIRECTED RNA POLYMERASE III SUBUNIT RPC1;
   6:  7wze-A  8.3  2.2   65   161   11   MOLECULE: UNCHARACTERIZED HTH-TYPE TRANSCRIPTIONAL REGULATO
   7:  1ldd-A  8.2  1.9   64    74    8   MOLECULE: ANAPHASE PROMOTING COMPLEX;
   8:  5v9x-A  8.2  2.7   73   772   19   MOLECULE: ATP-DEPENDENT DNA HELICASE;
   9:  3nw0-A  8.2  2.1   72   236   10   MOLECULE: MELANOMA-ASSOCIATED ANTIGEN G1;
  10:  8jxk-D  8.1  2.9   71   166   17   MOLECULE: CONSERVED PROTEIN;
  11:  1yg2-A  8.1  2.4   68   169   10   MOLECULE: GENE ACTIVATOR APHA;
  12:  1xma-B  8.1  2.4   71   103    7   MOLECULE: PREDICTED TRANSCRIPTIONAL REGULATOR;
  13:  7xjg-J  8.0  2.4   72   306    8   MOLECULE: RNA-DIRECTED DNA POLYMERASE FROM RETRON EC86;
  14:  6abq-A  8.0  2.5   72   111   10   MOLECULE: PADR FAMILY TRANSCRIPTIONAL REGULATOR;
  15:  2o0y-C  8.0  1.8   65   255   17   MOLECULE: TRANSCRIPTIONAL REGULATOR;
  16:  1xmk-A  7.9  2.0   65    79    6   MOLECULE: DOUBLE-STRANDED RNA-SPECIFIC ADENOSINE DEAMINASE;
  17:  1xsx-A  7.9  2.1   64    95   14   MOLECULE: SSO10A;
  18:  5dym-A  7.9  2.3   68    96    9   MOLECULE: PADR-FAMILY TRANSCRIPTIONAL REGULATOR;
  19:  5h20-A  7.9  2.5   71   103   13   MOLECULE: PUTATIVE PADR-FAMILY TRANSCRIPTIONAL REGULATORY P
  20:  3bdd-A  7.9  2.1   65   140    5   MOLECULE: REGULATORY PROTEIN MARR;
  21:  2co5-B  7.8  2.5   70    94   11   MOLECULE: VIRAL PROTEIN F93;
  22:  3u2r-A  7.8  2.2   62   135   10   MOLECULE: REGULATORY PROTEIN MARR;
  23:  3l7w-A  7.8  2.3   70   106   10   MOLECULE: PUTATIVE UNCHARACTERIZED PROTEIN SMU.1704;
  24:  2nyx-B  7.7  2.1   64   147   11   MOLECULE: PROBABLE TRANSCRIPTIONAL REGULATORY PROTEIN, RV14
  25:  2dql-A  7.7  2.3   68   115    6   MOLECULE: PEX PROTEIN;
  26:  3bpx-A  7.7  2.3   64   147    6   MOLECULE: TRANSCRIPTIONAL REGULATOR;
  27:  2xrn-B  7.7  2.2   65   241   12   MOLECULE: HTH-TYPE TRANSCRIPTIONAL REGULATOR TTGV;
  28:  2rdp-A  7.7  2.2   64   140   11   MOLECULE: PUTATIVE TRANSCRIPTIONAL REGULATOR MARR;
  29:  3gfi-A  7.7  2.2   63   143    8   MOLECULE: 146AA LONG HYPOTHETICAL TRANSCRIPTIONAL REGULATOR
  30:  8r3g-A  7.7  2.3   66   335    8   MOLECULE: CENTRAL GLYCOLYTIC GENES REGULATOR;
  31:  2wte-A  7.6  2.1   64   212   13   MOLECULE: CSA3;
  32:  7ymd-C  7.6  2.5   71   285   10   MOLECULE: NON-STRUCTURAL MAINTENANCE OF CHROMOSOME ELEMENT
  33:  5x11-E  7.6  2.5   69   181    6   MOLECULE: DNA (28-MER);
  34:  1yyv-A  7.6  2.3   65   114   11   MOLECULE: PUTATIVE TRANSCRIPTIONAL REGULATOR;
  35:  5z7b-B  7.6  2.6   70   197   11   MOLECULE: PADR FAMILY TRANSCRIPTIONAL REGULATOR;
  36:  2p8t-A  7.5  1.7   61   184   10   MOLECULE: HYPOTHETICAL PROTEIN PH0730;
  37:  8uuc-A  7.5  2.1   62   274   11   MOLECULE: ADENINE DNA GLYCOSYLASE;
  38:  6qfd-B  7.5  2.6   69   116    9   MOLECULE: DNA-BINDING PROTEIN;
  39:  8ioi-A  7.5  2.4   68   190    7   MOLECULE: PADR FAMILY TRANSCRIPTIONAL REGULATOR;
  40:  2qww-A  7.5  2.2   64   146   20   MOLECULE: TRANSCRIPTIONAL REGULATOR, MARR FAMILY;
  41:  2esh-A  7.5  2.5   70   114   10   MOLECULE: CONSERVED HYPOTHETICAL PROTEIN TM0937;
  42:  3g3z-A  7.5  2.1   64   142    6   MOLECULE: TRANSCRIPTIONAL REGULATOR, MARR FAMILY;
  43:  8ffz-B  7.5  2.4   65   905   12   MOLECULE: TRANSCRIPTION FACTOR IIIA;
  44:  1z7u-A  7.5  2.2   65   111   11   MOLECULE: HYPOTHETICAL PROTEIN EF0647;
  45:  4fht-B  7.4  2.2   64   142   14   MOLECULE: PCAV TRANSCRIPTIONAL REGULATOR;
  46:  2fa5-B  7.4  1.9   63   142   16   MOLECULE: TRANSCRIPTIONAL REGULATOR MARR/EMRR FAMILY;
  47:  2zkz-C  7.4  1.9   61    87    7   MOLECULE: TRANSCRIPTIONAL REPRESSOR PAGR;
  48:  8yli-B  7.4  2.3   64   162    9   MOLECULE: REGULATORY PROTEIN;
  49:  5hs5-A  7.4  2.6   68   116   12   MOLECULE: HTH-TYPE TRANSCRIPTIONAL REGULATOR SARX;
  50:  6aht-A  7.3  2.6   69   111    4   MOLECULE: CONSERVED HYPOTHETICAL PLASMID PROTEIN;
  51:  7el3-A  7.3  2.2   64   138   11   MOLECULE: HOMOPROTOCATECHUATE DEGRADATION OPERON REGULATOR
  52:  2qen-A  7.3  2.6   65   350    3   MOLECULE: WALKER-TYPE ATPASE;
  53:  7xux-A  7.3  2.6   65   365   17   MOLECULE: FIDO DOMAIN-CONTAINING PROTEIN;
  54:  5aip-A  7.3  2.0   64   141   13   MOLECULE: TRANSCRIPTIONAL REGULATOR, MARR FAMILY;
  55:  2fna-A  7.2  2.8   65   352    9   MOLECULE: CONSERVED HYPOTHETICAL PROTEIN;
  56:  4asn-A  7.2  2.2   65    90   12   MOLECULE: TUBR;
  57:  3mq0-B  7.2  2.2   65   248   18   MOLECULE: TRANSCRIPTIONAL REPRESSOR OF THE BLCABC OPERON;
  58:  3f8f-A  7.2  2.4   71   114   11   MOLECULE: TRANSCRIPTIONAL REGULATOR, PADR-LIKE FAMILY;
  59:  8xt8-B  7.2  2.2   62   145   15   MOLECULE: MARR FAMILY TRANSCRIPTIONAL REGULATOR;
  60:  5h3r-A  7.2  2.3   64   141    9   MOLECULE: DNA (5'-
```

```
0001 s001A   -------------ASHI-FAHQLMTLGIQ-L---D------GVLPQDV-DYWL-R----G--S------AAF------SRL---GSAERKAIVDHMLS----QEILADH---S----------GKLWLG--PVGERR----F--GR----------------------
0002 3elkA   ---------reriLHGL-ITLYILKELVK-R----------PXHGYEL-QKSXfE----T--T------G--------QAL---PQGSIYILLKTXKE----RGFVISE---S--svnkgqqlTVYHIT--DAGKKF----L--XDhsqalqlarkiiddllstvd--
0003 7wjpA   ----------pqfKKGV-LELCCLFLIQK-K----------DCYGYEL-ANQVsK----Y--I------E----------V---AEGAIYPVLRRLVK----EEYCSTY---L----vespsrKYYQLT--VKGEIY----L--NElisewnnftdsvaklltag--
0004 5a31N   pqdrdnmvlikeeELLL-FWTYIQAMLTN-L---E------SLSLDRI-YNML-RmfvvT--G------PAL------AEI---DLQELQGYLQKKVR----DQQLVYS---A----------GVYRLP-------------------------------------
0005 5zqhA   ---------etqlLKGV-LEGCVLDMIGQ-K----------ERYGYEL-VQTL-R----E--A------G-F------DTI---VG-TIYPLLQKLEK----NQWIRGD---M-rpspdgpdrKYFSLM--KEGEER----V--SVfwqqwddlsqkvegikn-----
0006 8iueP   -------------DPVE-IENRIIELCHQ-F--pH------GITDQVI-QNEM-P----H-------------------I---EAQQRAVAINRLLS----MGQLDLL---R------sntgLLYRIK-------D----S--QNagkmkgsdnqeklvyqiiedag
0007 7wzeA   kxnhvxehyfagrglSE-GKFKILXLLFD-A-kdH------RLSPTEL-AKRS-N-----------------------V---TKATITGLLDGLAR----DGFVSRR---H-------hrkISIELT--TEGKAR----L--EQflpghfskisavxenysdeekd
```

```
0008 1lddA   ---------kyelTLQR-SLPFIEGMLTN-L---G------AMKLHKI-HSFL-K----I----tvpkdWGY------NRI---TLQQLEGYLNTLAD----EGRLKYI---A--------nGSYEIV---------------------------------------
0009 5v9xA   qtgdietlrvpanPLDV-LAQHTVAVAAL-----E------PVDADAW-FDAV-R----R--S------APF------ATL---PRSAFEATLDLLSGkaelRPRLVYD-rdT---------GTLTAR--PGAQRL----A--VTsggaipdrgmftvylaseteel
0010 3nw0A   ttsiskmatdfaeNELD-LFRKALELIID-S-etG------FASSTNI-LN-L-V----D--Q------lKG------KKM---RKKEAEQVLQKFVQ----NKWLIEK---E---------GEFTLH--GRAILE----M--EQyiretypdavkicnichslliq
0011 8jxkD   --------------EFM-LELAILGLLIE-S----------PMHGYEL-RKRLtG----L--L------GA------fRAF---SYGSLYPALRRMQA----DGLIAEN---A----apagrRVYQLT--DKGRRR----F--GElvadtgphnytddgfgvhlaff
0012 1yg2A   ----------------S-LPHVILTVLST-R----------DATGYDI-TKEF-S----AsiG------YFW------K-A---SHQQVYRELNKMGE----QGLVTCV---L---------eVYSIT--QAGRSA----L--GEwfdqptahptvrdefsaklmac
0013 1xmaB   -------vissdvIRGY-VDTIILSLLIE-G----------DSYGYEI-SKNI-R-iktD--E------LYV--------I---KETTLYSAFARLEK----NGYIKSY---Y---geetrrTYYRIT--PEGIKY----Y--KQkceeweltkkvinkfvk-----
0014 7xjgJ   kvrkenkvskdigNILY-AERFLLPCIYL-L---D------SVNYRTL-CELA-F----K--A--------I------KQDdvlSKIIVRSVVSRLIN----ERKILQM---T---------DGYQVT--ALGASY----V--RSvfdrktldrlrleimnfenrrk
0015 6abqA   gsakdpmkgltelLKGS-LEGMILERISR-----G------ETYGYEI-TKYL-N----D--L------G-F------DEI---VEGTVYTILVRLEK----KGLVEIE---K-kkselgpprKFYTLS--PAGEEE----L--AIfwkrwdfiqgkimqvkg-----
0016 2o0yC   dsaekpavadagvRSVT-RVIDLLELFDAaH---P------TRSLKEL-VEGT-K----------------------------lPKTTVVRLVATXCA----RSVLTSR--aD---------GSYSLG--PEXL-R----W--VRlagrtwappeevvdixrqlsad
0017 1xmkA   ------gshmaslDMAE-IKEKICDYLFN-V---S------DSSALNL-AKNI-G---------------------------ltKARDINAVLIDMER----QGDVYRQ---G------ttpPIWHLT--DKKR-E----R--MQik--------------------
0018 1xsxA   ----------akkKSKLeIIQAILEACK------S------GSPKTRI-MYGA-N----------------------------lSYALTGRYIKMLMD----LEIIRQE---G---------KQYMLT--KKGEEL----L--EDirkfnemrknmdqlkekinsvl
0019 5dymA   ---------------GY-IDILIVSILEK-K----------DCYGYEI-AKQVrE----R--S------EF---------L---KEGTMYLALKRMES----KNLIKSY---YsneqssggrrKYYNLT--NEGKDF----L--EIkkqewrfikkvmnqflg-----
0020 5h20A   ------dnvksqmRKGM-LEYCIMLLLHK-E----------PAYASDI-IQKL-K----E--A------RLI--------V---VEGTLYPLLTRLKN----DDLLSYE---W-vestqgpprKYYKLT--GKGESF----L--GEleaswkelnetvnhia------
0021 3bddA   adetisnlfekqlgiSL-TRYSILQTLLK-D---A------PLHQLAL-QERL-Q-------------------------I---DRAAVTRHLKLLEE----SGYIIRK---R---npdnqreVLVWPT--EQAREA----L--ITnpsahhqaiktsxnqiltvees
0022 2co5B   --------mkirkymRI-NYYIILKVLVI-N---G-----sRLEKKRL-RSEI-L----K-rF------DID--------I---SDGVLYPLIDSLID----DKILREE---E-----apdgKVLFLT--EKGMKE----F--EElheffkkivch-----------
0023 3u2rA   rxkaieeeifsqfelSA-QQYNTLRLLRS-V-hpE------GXATLQI-ADRL--------------------------------APDITRLIDRLDD----RGLVLRT---R---kpenrrvVEVALT--DAGLKL----L--KDleepvrqcherqlghlaadelh
0024 3l7wA   --------xyypvSALL-IEYLILAIVSK-----H------DSYGYDI-SQTI-K--------------LIA------S-I---KESTLYPILKKLEK----AGYLSTY---T--qehqgrrrKYYHLT--DSGEKH----L--VYltkewsvykxtidgivegrirh
0025 2nyxB   vaisahsiaqvdeniTI-PQFRTLVILSN-H---G------PINLATL-ATLL-G----------------------------vQPSATGRXVDRLVG----AELIDRL---P---hptsrreLLAALT--KRGRDV----V--RQvtehrrteiariveqxapaerh
0026 2dqlA   iyqffenppptylCQEV-AICYILYVLLQ-----G------ESYGTEL-IQQL-E--teH--P------TYR--------L---SDTVLYSAIKFLED----NRAITGY---WkklegrgrprRMYQVS--PE-WQH----Qae------------------------
0027 3bpxA   shrvfigrelghlnlTD-AQVACLLRIHR-E---P------GIKQDEL-ATFF-H----------------------------vDKGTIARTLRRLEE----SGFIERE---Q---dpenrrrYILEVT--RRGEEI----I--PLilkveerwedllfrdftederk
0028 2xrnB   -----------siQVIA-RAASIMRALGS-H---P-----hGLSLAAI-AQLV-G----------------------------lPRSTVQRIINALEE----EFLVEAL---G-------paGGFRLG--PA-LGQ----L--INqaqtdilslvkpylrslaeeld
0029 2rdpA   nlkqrgreiltnypiTP-PQFVALQWLLE-E---G------DLTVGEL-SNKX-Y----------------------------lACSTTTDLVDRXER----NGLVARV---R-----dehvVRIRLL--EKGERI----I--EEviekrqrdlanvlesfsdeeiv
0030 3gfiA   amsrelnrrlgelnlSY-LDFLVLRATSD-----G------PKTMAYL-ANRY-F----------------------------vTQSAITASVDKLEE----MGLVVRV---R---dredrrkILIEIT--EKGLET----F--NKgieiykklanevtgdlsedevi
0031 8r3gA   liqaqkkllpdllLVXQ-KRFEILQYIRL-T---E------PIGRRSL-SASL-G----------------------------iSERVLRGEVQFLKE----QNLVDIK---T---------NGXTLT--EEGYEL----L--SVledtxkdvlgltllektlkerl
0032 2wteA   isfndntiralxrdySR-EEXKLLNVLYE-T---K------GTGITEL-AKXL-D----------------------------kSEKTLINKIAELKK----FGILTQK---G-------kdRKVELN--ELGLNV----I--KLnk--------------------
0033 7ymdC   ntlesklstdrdlVYKG-VLSVILCIVFF-S--kN------NILHQEL-IKFL-E----T--F------GipsdgskiAIL---N-ITIEDLIKSLEK----REYIVRL---E-eksdtdgevISYRIG--RRTQAE----L--GLesleklvqeimglekeqtkslh
0034 5x11E   ------------dpmRV-LKYAILGLLRK-----G------ELSGYDI-TSYF-K----EelG------QFW-------sA---KHSQIYPELKKLTD----EGFITFR---T-tiqgtklekKMYTLT--DSGKQE----L--HDwlirhqpipetvkdefmlkayf
0035 1yyvA   faeqcpsrevlkhvtSR-WGVLILVALRD-----G------THRFSDL-RRXX-G------------------------GV---SEXXLAQSLQALEQ----DGFLNRV---S---ypvvpphVEYSLT--PLGEQV----S--DXvaaladwielnlpqvlaqre--
0036 5z7bB   ----------sakDLMT-LRSALLALLSS-----G------PLTGYDAsQRFG-A----svgF------VWS--------G---SDSQIYPELRKMEA----EELLVGS---DvpwgskgatkTEYALS--EKGWEA----L--RKawyepvtygptrdparlkaayf
0037 2p8tA   ------------eyTVE-DVLAVIFLLK------E------PLGRKQI-SERL-E----------------------------lGEGSVRTLLRKLSH----LDIIRSK--------------GHFLT--LKGKEI----R--DKllsmfsepigvsvdgypgiaiv
0038 8uucA   rsahhtrqstfegSRRQ-KRAELVRLVLA-E---P------GIGIDEL-AERLdA----F--E------R----dagrKGV---DAATFTSIVADLVA----EGFFRRE---G---------DAFFA----------------------------------------
0039 6qfdB   -----pdarsdardlTA-FQKNILTVLGE-E----------ARYGLAI-KRELeE----Y--Y------G--------EEV---NHGRLYPNLDDLVN----KGLVEKS---E----ldkrtNEYALT--NEGFDA----V--VDdlewtlskfvadadrrerveti
0040 8ioiA   ------------pefMA-LPHAILVSLSE-Q----------ASSGYELaRRFD-R----sigY------FWT--------A---THQQIYRTLRVMEN----NNWVRAT---T-vlqhgrpdkKVYAIS--DSGRAE----L--ARwiaeplsptrpgrgsaltdsst
0041 2qwwA   isagyadqnaaslglTI-QQLAXINVIYS-T---P------GISVADL-TKRL-I----------------------------iTGSSAAANVDGLIS----LGLVVKL---N---kpndsxdLTLKLS--KKGEDL----S--KRstanafxykaxxkvfenltene
0042 2eshA   -------rggrgfRGWW-LASTILLLVAE-K----------PSHGYEL-AERL-A----E--F------GI--eipgiGHM---G--NIYRVLADLEE----SGFLSTE---W--dttvspprKIYRIT--PQGKLY----L--REilrsledmkrrietleerikrv
0043 3g3zA   licnvfdkwigqqdlNY-NLFAVLYTLAT-E---G------SRTQKHI-GEKW-S----------------------------lPKQTVSGVCKTLAG----QGLIEWQ---E---geqdrrkRLLSLT--ETGKAY----A--APltesaqefsdkvfatfgdkrtt
0044 8ffzB   rlwvngsgsirpnIWKQ-VVTMVVNEIIF-H---P------GITLSRL-QSRC-R----E--V--------------------lSLHEISEICKWLLE----RQVLITT---D--------fDGYWVN--HNWYSIyest---------------------------
0045 1z7uA   tdkqtsinlalstinGK-WKLSLXDELFQ-----G------TKRNGEL-XRAL-D------------------------GI---TQRVLTDRLREXEK----DGLVHRE---S---fnelpprVEYTLT--PEGYAL----Y--DAlsslchwgetfaqkkarln---
0046 4fhtB   qahyllwntmvseetTS-PQYAVLNALVA-E---P------GLDQRTV-GERV-G----------------------------lDRSTIAEVVSRLGR----RGLLDKV---R---dpqdgrrSLLRLT--DEGLRV----H--RRlgvriarmnqvflapladeqa
0047 2fa5B   isgniakvygdrygxAI-PEWRVITILAL-Y---P------GSSASEV-SDRT-A----------------------------xDKVAVSRAVARLLE----RGFIRRE--------------SXLALS--PAGRQV----Y--ETvaplvnexeqrlxsvfsaeeqq
0048 2zkzC   yxsleddaellktxaHP-XRLKIVNELYK-H---K------ALNVTQI-IQIL-K----------------------------lPQSTVSQHLCKXR-----GKVLKRN---R-----qqleIYYSIN--N--PKV----E--GIiklln-----------------
0049 8yliB   lmdqrlescfsrfdlSS-WEFDMLATLRR-AgapH------CLSPTEL-FSTL-M----------------------------vTSGTMTHRLKRLET----RGFIERV---Q---neldarsTLVQLT--SSGLEL----I--NRaveahienerqvlsvlpaevla
0050 5hs5A   lseyidkkyklslNDLA-VLDLTXKHCK-----dE------KVLXQSF-LKTA---------------xDEL------D-L---SRTKLLVSIRRLIE----KERLSKV---R---sskderkIYIYLN--NDDISK----F--NAlfedveqflnilehh-------
0051 6ahtA   isxssseiidvlcenLN-DGIWALRVLYA-E---G------AXNKEKLwDYIN-QyhkdyqienekdyeGKK------IlP---SRYALDIXTARLEG----AGLISFK---A-----igrvRIYDVT--DLGNVL----I--KElekr------------------
0052 7el3A   aimshfrpalnevglTE-QQWRIIRILYQ-Y---E------ELESNQL-AELA-C----------------------------iLKPSLTGILNRMVE----QKLIQKR---K---dyddqriSLISLT--ESGLEC----F--KTqavkmeasyqkiqeqygeekmk
0053 2qenA   akglimgeleelrrrSP-RYVDILRAIAL-G---------yNRWSLI-RDYL-A----V--K------GTK--------I---PEPRLYALLENLKK----MNWIVEE---D---------NTYKIA----DPV----V--ATvlri------------------
0054 7xuxA   vlypaleyskgrgviNE-TESKILKRTIS-Q---G------TVKTSDL-KEVL-P----G--------------------L---KPAQITYQIGKLVD----RGLLQPVevgS---------RIYTAGfsKSDL-M----R--GVihalrkegfipd----------
0055 5aipA   alxtqfrpilnqaniTD-QQWRIIRLLAE-N---G------TLDFQDL-ANQA-C----------------------------iLRPSLTGILTRLEK----AGLVVRL---K---psndqrrVFLKLT--AEGEKL----Y--EEigeevderydaieevlgrekxl
0056 2fnaA   ilkefenflhgreIARK-RYLNIXRTLS------K------CGKWSDV-KRALeL----E--E------G--------IEI---SDSEIYNYLTQLTK----HSWIIKE---G---------EKYCPS-----EPL----Isl------------------------
0057 4asnA   ----sdyfeevxrklTI-EDVSILGWLFQ-N---EanavfkAIKKSSI-ADEL-E-------------------------Y---STANFRKTLNKLEA----IHFIGTV---T-----ggkeHKLYLT--EYGQQA----V--QQaihh------------------
0058 3mq0B   ---------sdtvPALR-RAVRILDLVAG-S--pR------DLTAAEL-TRFL--------------------------dL---PKSSAHGLLAVMTE----LDLLARS---A--------dGTLRIG--PHS-LR----W--ANgflshldivstfndhlaqrhdl
0059 3f8fA   -----maeipkemLRAQ-TNVILLNVLKQ-G----------DNYVYGI-IKQV-K-easN--G------EME--------L---NEATLYTIFKRLEK----DGIISSY---W---gdeggrrKYYRLT--EIGHEN----M--RLafeswsrvdkiienleankkse
0060 8xt8B   kddlkkmfkqngydiTT-DHYALLRFLWE-Q---D------GISQIDL-CEKS-C---------------------------kdKSNTTRILDVMKN-----KGLIVRK---V---dvkdrrkFQIFLT--DLGR-E----L--EEplneiasiyatetfksisdeel
0061 5h3rA   kkdrllneylspldiTA-AQFKVLCSIRC-A---A------CITPVEL-KKVL-S----------------------------vDLGALTRMLDRLVS----KGWVERL---P---npndkrgVLVKLT--TGGAAI----C--EQchqlvgqdlhqeltknltadev

0001 s001A   -------------LHHH-HHHHHHHHHHH-H---L------LLLHHHH-HHHH-L----L--L------HHH------LLL---LHHHHHHHHHHHHH----LLLEEEE---L---------LEEEEL--HHHHHH----H--LL--------------------
0002 3elkA   ---------llhhHHHH-HHHHHHHHHHH-L----------LEEHHHH-HHHHhH----H--H------L--------LLL---LLLHHHHHHHHHHH----HLLEEEE---E--eellllleeEEEEEL--HHHHHH----H--HHllllhhhhhhhhhhhhllll--
0003 7wjpA   ----------llhHHHH-HHHHHHHHHHH-L----------LEEHHHH-HHHHlL----L--L------L---------L---LLLLHHHHHHHHHH----LLLEEEE---E----elllleEEEEEL--HHHHHH----H--HHhhhhhhhhhhhhhhhhlll---
0004 5a31N   lllllllllllhhHHHH-HHHHHHHHHHH-H---L------LLLHHHH-HHHH-HhhllL--L------LLL------LLL---LHHHHHHHHHHHHH----LLLLEEE---L---------LEEELL---------------------------------------
0005 5zqhA   ---------llhhHHHH-HHHHHHHHHHH-L----------LEEHHHH-HHHH-H----H--H------L-L------LLL---LL-LLHHHHHHHHHH----LLLEEEE---E-eellllleeEEEEEL--HHHHHH----H--HHhhhhhhhhhhhhhhhhl----
0006 8iueP   -------------LHHH-HHHHHHHHHHH-L--lL------LLLHHHH-HHHL-L----L--------------------L---LHHHHHHHHHHHHH----HLLEEEE---E------elleEEEEEL-------H----H--HLlllllllllhhhhhhhhhhhh
0007 7wzeA   hhhhhhhhhhhllllLH-HHHHHHHHHHH-L-hhH------EELHHHH-HHHL-L-------------------------L---LHHHHHHHHHHHHH----HLLEEEE---E-------lllEEEEEL--HHHHHH----H--HHhhhhhhhhhhhhhlllllhhhhh
0008 1lddA   ---------llhhHHHH-HHHHHHHHHHH-H---L------LEEHHHH-HHHH-H----H----hllhhHLL------LLL---LHHHHHHHHHHHHH----LLLEELL---L--------lLEEEEL---------------------------------------
0009 5v9xA   hhlllllllllllLHHH-HHHHHHHHLLL-----L------LEEHHHH-HHHH-H----H--L------LLL------LLL---LHHHHHHHHHHHHHLllllLLLEEEE-llL---------LEEEEL--LLHHHH----H--HLlllllllllllllllllllll
0010 3nw0A   llllhhhhllllhHHHH-HHHHHHHHHHH-L-llL------LEEHHHH-HH-H-H----L--L------lLL------LLL---LHHHHHHHHHHHHH----LLLEEEE---L---------LEEEEL--HHHHHH----H--HHhhhhhlllllllllllllllll
0011 8jxkD   --------------LHH-HHHHHHHHHHH-L----------LEEHHHH-HHHHhH----H--L------LL------lLLL---LLLLHHHHHHHHHH----LLLEEEL---L------lllllLEEEEL--HHHHHH----H--HHhhhlllhhhllhhhhhhhhhhh
0012 1yg2A   ----------------L-HHHHHHHHHHH-L----------LLLHHHH-HHHH-L----LhhH------HLL------L-L---LHHHHHHHHHHHHH----LLLEEEL---L---------lLEEEL--HHHHHH----H--HHhhhlllllllllhhhhhhhhh
0013 1xmaB   -------lllhhhHHHH-HHHHHHHHHHH-L----------LEEHHHH-HHHH-H-hhlL--L------LLL-------L---LHHHHHHHHHHHHH----LLLEEEE---E----llllllEEEEEL--HHHHHH----H--HHhhhhhhhhhhhhhhhhll-----
0014 7xjgJ   hhhhhllllllllLLLL-LHHHHHHHHHH-L---L------LLLHHHH-HHHH-H----H--H--------L------LLLlhhHHHHHHHHHHHHHH----LLLEEEE---L---------LEEEEL--HHHHHH----H--HHhllhhhhhhhhhhhhhhhhlll
0015 6abqA   lllllllllllhhHLLH-HHHHHHHHHHL-----L------LLLHHHH-HHHH-H----H--L------L-L------LLL---LHHHHHHHHHHHHH----LLLEEEE---E-eellllleeEEEEEL--HHHHHH----H--HHhhhhhhhhhhhhhhhhl-----
0016 2o0yC   lllllllllllllHHHH-HHHHHHHLLLLlL---L------LLLHHHH-HHHH-L----------------------------lLHHHHHHHHHHHHH----LLLEEEL--lL---------LLEEEL--HHHH-H----H--HHhhhhhhlllhhhhhhhhhhhhh
0017 1xmkA   ------llllllhHHHH-HHHHHHHHHHH-L---L------LEEHHHH-HHHH-L---------------------------hhHHHHHHHHHHHHHH----LLLEEEE---L-------lllLEEEEL--HHHH-L----L--LLll--------------------
0018 1xsxA   ----------lllLLLLhHHHHHHHHHL------L------LLLHHHH-HLLL-L----------------------------lLHHHHHHHHHHHHH----LLLEEEE---L---------LEEEEL--HHHHHH----H--HHhhhhhhhhhhhhhhhhhhhhhh
0019 5dymA   ---------------LL-HHHHHHHHHLL-L----------LEEHHHH-HHHHhH----H--L------LL--------L---LHHHHHHHHHHHHH----LLLEEEE---ElllllllllEEEEEL--HHHHHH----H--HHhhhhhhhhhhhhhhhhhl-----
0020 5h20A   ------lhhhhhhHHHH-HHHHHHHHHLL-L----------LLLHHHH-HHHH-H----H--L------LLL-------L---LHHHHHHHHHHHHH----LLLEEEE---E-elllllllleEEEEEL--HHHHHH----H--HHhhhhhhhlllhhhhhl------
0021 3bddA   hhhhhhhhhhhhhllLH-HHHHHHHHHHH-H---L------LLLHHHH-HHHH-L-------------------------L---LHHHHHHHHHHHHH----LLLEEEE---E---lllllllEEEEEL--HHHHHH----H--LLlllhhhhhhhhhhhllllhhhh
0022 2co5B   --------lllllllLH-HHHHHHHHHHH-L---L-----lEEEHHHH-HHHH-H----H-hH------LLL-------L---LHHHHHHHHHHHHH----LLLEEEE---L------llllLEEEEL--HHHHHH----H--HHhhhhhhhhhll-----------
0023 3u2rA   hhhhhhhhhhhllllLH-HHHHHHHHHHH-H-llL------LEEHHHH-HHHL--------------------------------LLHHHHHHHHHHHH----LLLEEEE---E---elleeeeEEEEEL--HHHHHH----H--HHhhhhhhhhhhhhhhhhhhhhhh
0024 3l7wA   --------lllllLHHH-HHHHHHHHHHH-----L------LEEHHHH-HHHH-L--------------LLL------L-L---LHHHHHHHHHHHHH----LLLEEEE---E--eeelleeeEEEEEL--HHHHHH----H--HHhhhhhhhhhhhhhhhhhlllll
0025 2nyxB   hhhhhhhhhhhllllLH-HHHHHHHHHHH-H---L------LEEHHHH-HHHH-L----------------------------lLHHHHHHHHHHHHH----LLLEEEE---L---lllllllLEEEEL--HHHHHH----H--HHhhhhhhhhhhhhhhllllhhhll
0026 2dqlA   hhhhhhlllleelLHHH-HHHHHHHHHLL-----L------LLLHHHH-HHHH-H--hhL--L------LEE--------L---LHHHHHHHHHHHHH----LLLEEEE---ElllllllleEEEEEL--HH-HHH----Hhh------------------------
0027 3bpxA   hhhhhhhhhhhhhllLH-HHHHHHHHHHL-L---L------LLLHHHH-HHHH-L----------------------------lLHHHHHHHHHHHHH----LLLEEEE---E---elleeeeEEEEEL--HHHHHH----H--HHhhhhhhhhhhhhhlllllhhhhh
0028 2xrnB   -----------llHHHH-HHHHHHHHHHL-L---L-----lLEEHHHH-HHHL-L----------------------------lLHHHHHHHHHHHHL----LLLEEEL---H--------hhLEEEEL--LH-HHH----H--HHhhhhhhhhhhhhhhhhhhhhhl
0029 2rdpA   hhhhhhhhhhlllllLH-HHHHHHHHHHH-H---L------LLLHHHH-HHHH-L----------------------------lLHHHHHHHHHHHHH----LLLEEEE---E------llllEEEEEL--HHHHHH----H--HHhhhhhhhhhhhhhhhllhhhhh
0030 3gfiA   hhhhhhhhhhhhhllLH-HHHHHHHHHLL-----L------LEEHHHH-HHHL-L----------------------------lLHHHHHHHHHHHHH----LLLEEEE---E---elleeeeEEEEEL--HHHHHH----H--HHhhhhhhhhhhhhhhlllllhhhh
0031 8r3gA   hhhhhhhhlllhhHHHH-HHHHHHHHHHH-H---L------LLLHHHH-HHHH-L----------------------------lLHHHHHHHHHHHHH----LLLEEEE---L---------LEEEEL--HHHHHH----H--HHhhhhhhhhlhhhhhhhhhhhhl
0032 2wteA   eeehhhhhhhhhlllLH-HHHHHHHHHHH-H---L------LLLHHHH-HHHH-L----------------------------lLHHHHHHHHHHHHH----LLLEEEE---L--------llLEEEEL--HHHHHH----H--HHll--------------------
0033 7ymdC   lllllllllllhhHHHH-HHHHHHHHHHH-L--lL------LEEHHHH-HHHH-H----H--H------LllllllllLLL---L-LLHHHHHHHHHH----LLLEEEE---E-lllllllllEEEEEL--HHHHHH----L--LHhhhhhhhhhhhlllhhhhhlllh
0034 5x11E   ------------lllLH-HHHHHHHHLLL-----L------LLLHHHH-HHHH-H----LlhH------HLL-------lL---LHHHHHHHHHHHHH----LLLEEEE---E-eellllleeEEEEEL--HHHHHH----H--HHhhhlllllllllllhhhhhhhl
0035 1yyvA   llllllhhhhhhhhhLH-HHHHHHHHHHH-----L------LEEHHHH-HHHL-L-------------------------LL---LHHHHHHHHHHHHH----HLLEEEE---E---elllleEEEEEL--HHHHHH----H--HHhhhhhhhhhhhhhhhhllll--
0036 5z7bB   ----------lhhHHHH-HHHHHHHHHLL-----L------LEEHHHHhHHHH-L----lhhH------LLL--------L---LHHHHHHHHHHHHH----LLLEEEE---EeeellllleeeEEEEEL--HHHHHH----H--HHhhhllllllllllhhhhhhhlh
0037 2p8tA   ------------llLHH-HHHHHHHHLL------L------LLLHHHH-HHHH-L----------------------------lLHHHHHHHHHHHHH----LLLEEEL--------------LEEEL--HHHHHH----H--HHhhlllllleeeeelleeeeeee
0038 8uucA   hllllllllllllLHHH-HHHHHHHHHHH-L---L------LEEHHHH-HHHHhH----H--H------H----hlllLLL---LHHHHHHHHHHHHH----HLLLEEE---L---------LEEEL----------------------------------------
0039 6qfdB   -----lllhhhhlllLH-HHHHHHHHHHH-L----------LEEHHHH-HHHHhH----H--H------L--------LLL---LHHHHHHHHHHHHH----LLLEEEE---E-------eelleEEEEEL--HHHHHH----H--HHhhhhhhhhhlllhhhhhhhhhh
0040 8ioiA   ------------lllLL-HHHHHHHHHHH-L----------LLLHHHHhHHHH-H----lhhH------LLL--------L---LHHHHHHHHHHHHH----LLLEEEE---E-elllllllleEEEEEL--HHHHHH----H--HHhhhlllllllllllllllllhh
0041 2qwwA   hhhhhhhhhhhhhllLH-HHHHHHHHHHH-L---L------LEEHHHH-HHHH-L----------------------------lLHHHHHHHHHHHHH----LLLEEEL---L---lllllllLEEEEL--HHHHHH----H--HHhhllhhhhhhhhhhhhllllhhh
0042 2eshA   -------llllhhHHHH-HHHHHHHHHHH-L----------LLLHHHH-HHHH-H----L--L------LL--lllllLLL---L--LHHHHHHHHHHH----LLLEEEE---E--elllllleEEEEEL--HHHHHH----H--HHhhhhhhhhhhhhhhhhhhhhhh
0043 3g3zA   hhhhhhhhhhhllllLH-HHHHHHHHHHH-H---L------LLLHHHH-HHHH-L----------------------------lLHHHHHHHHHHHHH----LLLEEEL---L---llllhhhLLEEEL--HHHHHH----H--HHhhhhhhhhhhhhhhhhlhhhhh
0044 8ffzB   lllllllllllhhHHHH-HHHHHHHHHHH-L---L------LEEHHHH-HHHL-L----L--L--------------------lLHHHHHHHHHHHHH----LLLEEEE---L--------lLEEEEL--LLLLLLllll---------------------------
0045 1z7uA   llhhhhhhhhhhlllLL-LHHHHHHHHHH-----L------LLLHHHH-HHHL-L-------------------------LL---LHHHHHHHHHHHHH----HLLEEEE---E---elllleEEEEEL--HHHHHH----H--HHhhhhhhhhhhhhhhhhhhl---
0046 4fhtB   hhhhhhhhhhlllllLH-HHHHHHHHHHH-L---L------LLLHHHH-HHHH-L----------------------------lLHHHHHHHHHHHHH----HLLEEEE---E---lllllllEEEEEL--HHHHHH----H--HHhhhhhhhhhhhhhhhllllhhhhh
0047 2fa5B   hhhhhhhhlhhhhllLH-HHHHHHHHHHH-L---L------LLLHHHH-HHHH-L----------------------------lLHHHHHHHHHHHHH----HLLEEEL--------------LLEEEL--HHHHHH----H--HHhhhhhhhhhhhhhhhllllhhhhh
0048 2zkzC   hhhhhhhhhhhhhhlLH-HHHHHHHHHHH-H---L------LEEHHHH-HHHH-L----------------------------lLHHHHHHHHHHHL-----LLLEEEE---E------elleEEEEEL--L--HHH----H--HHhhhhl-----------------
0049 8yliB   hhhhhhhhhhhhlllLH-HHHHHHHHHHH-HlllL------EELHHHH-HHHH-L----------------------------lLHHHHHHHHHHHHH----LLLEEEE---E---lllllllEEEEEL--HHHHHH----H--HHhhhhhhhhhhhhhhhllllhhhhh
0050 5hs5A   hhhhhhhhhlllhHHHH-HHHHHHHHHL-----lL------LEEHHHH-HHHH--------------hHHH------L-L---LHHHHHHHHHHHHH----LLLLEEE---L---lllllllLEEELL--HHHHHH----H--HHhhhhhhhhhhhhhlll-------
0051 6ahtA   llllhhhhhhhhhhhHH-HHHHHHHHHHH-H---L------LEEHHHHhHHHH-HhhhhhhhhhlllllLLL------LlL---LHHHHHHHHHHHHH----LLLEEEE---E------elleEEEEEL--HHHHHH----H--HHhhhl------------------
0052 7el3A   hhhlllhhhhhhhllLH-HHHHHHHHHHH-H---L------EEEHHHH-HHHH-L----------------------------lLHHHHHHHHHHHHH----LLLEEEE---E---elleeeeEEEEEL--HHHHHH----H--HHhhhhhhhhhhhhhhhhhhhhhh
0053 2qenA   hhhhhhhhhhhhhhhLH-HHHHHHHHHHL-L---------lLLHHHH-HHHH-H----H--L------LLL-------L---LHHHHHHHHHHHHH----LLLEEEE---L---------LEEEEL-----LHH----H--HHhhll------------------
0054 7xuxA   lhhhhhhhhhhllllLH-HHHHHHHHHHH-H---L------LLLHHHH-LLLL-L----L--------------------L---LHHHHHHHHHHHHH----LLLEEELlllL---------LLEEELllLLLH-H----H--HHhhhhhhhllllll----------
0055 5aipA   hhhhhhhhhhhhhllLH-HHHHHHHHHHH-H---L------LEEHHHH-HHHH-L----------------------------lLHHHHHHHHHHHHH----LLLEEEE---L---lllllllLEEEEL--HHHHHH----H--HHhhhhhhhhhhhhhhhhhlhhhhhh
0056 2fnaA   hhhhhhhhhlllhHHHH-HHHHHHHHHL------L------LLLHHHH-HHHHhH----H--H------L--------LLL---LHHHHHHHHHHHHH----LLLEEEL---L---------LLEEEL-----LHH----Hhh------------------------
0057 4asnA   ----lhhhhhhhhllLH-HHHHHHHHHHH-L---LlllhhhLEEHHHH-HLLL-L-------------------------L---LLHHHHHHHHHHHH----LLLEEEE---L------llllLEEEEL--HHHHHH----H--HHhlll------------------
0058 3mq0B   ---------llllHHHH-HHHHHHHHHHH-L--lL------LEEHHHH-HLLL---------------------------lL---LHHHHHHHHHHHHH----LLLEEEL---L--------lLEEEEL--LLH-HH----H--HHhhhhlllhhhhhhhhhlllhhh
0059 3f8fA   -----lllllhhhHHHH-HHHHHHHHHHH-L----------LLLHHHH-HHHH-H-hhlL--L------LLL--------L---LHHHHHHHHHHHHH----LLLEEEE---E---lllllllEEEEEL--HHHHHH----H--HHhhhhhhhhhhhhhhhhhhhhhh
0060 8xt8B   hhhhhhhhhhlllllLH-HHHHHHHHHHH-L---L------LLLHHHH-HHHH-L---------------------------llLHHHHHHHHHHHHL----LLLEEEE---E---lllllllEEEEEL--HHHH-H----L--HHhhhhhhhhhhhhhhhllllllhh
0061 5h3rA   hhhhhhhhlhhhhllLH-HHHHHHHHHHH-H---L------LLLHHHH-HHHH-L----------------------------lLHHHHHHHHHHHHH----LLLEEEE---E---lllllllEEEEEL--HHHHHH----H--HHhhhhhllhhhhhhhllllhhhh

# Job: Type-2 BREX; BrxHI-Helicase MBK7581083 Small-Beta barrel pdb25
# Query: s001A
# No:  Chain   Z    rmsd lali nres  %id PDB  Description
   1:  6znp-A  6.3  2.3   58   748   10   MOLECULE: UNCHARACTERIZED ATP-DEPENDENT HELICASE YPRA;
   2:  7wrs-A  5.9  2.7   61   518    5   MOLECULE: GLUTAMYL-TRNA SYNTHETASE;
   3:  5dm3-C  4.4  2.4   52   396    6   MOLECULE: L-GLUTAMINE SYNTHETASE;
   4:  2det-A  4.4  2.6   54   365   15   MOLECULE: TRNA;
   5:  7bgm-A  4.0  3.3   55   213    9   MOLECULE: PHOSPHORIBOSYL-AMP CYCLOHYDROLASE;
   6:  9fhl-G  4.0  2.4   53   213    9   MOLECULE: RRNA 16S SSO;
   7:  8fn2-X  3.8  3.1   55   181    5   MOLECULE: 23S RIBOSOMAL RNA;
   8:  1zps-B  3.8  2.9   51   128   12   MOLECULE: PHOSPHORIBOSYL-AMP CYCLOHYDROLASE;
   9:  6fan-C  3.5  2.3   42    62    5   MOLECULE: COOT;
  10:  8s8k-B  3.4  3.4   49   229    8   MOLECULE: 18S RIBOSOMAL RNA;
  11:  4hpp-A  3.3  2.4   51   427    4   MOLECULE: PROBABLE GLUTAMINE SYNTHETASE;
  12:  7kpj-E  3.2  2.9   44   218    9   MOLECULE: 338E6 FAB LIGHT CHAIN KAPPA;
```

```
 13:  2j9i-A  3.1  2.8   51   421   10   MOLECULE: GLUTAMATE-AMMONIA LIGASE DOMAIN-CONTAINING PROTEI
 14:  7aoh-B  3.0  3.8   44  1134    9   MOLECULE: DNA-DIRECTED RNA POLYMERASE 147 KDA POLYPEPTIDE;
 15:  2kco-A  3.0  3.3   52   133   10   MOLECULE: 30S RIBOSOMAL PROTEIN S8E;
 16:  3wwl-A  2.9  2.5   42    54   12   MOLECULE: ALPHA-AMINOADIPATE CARRIER PROTEIN LYSW;
 17:  9nvu-P  2.8  2.5   43   603    9   MOLECULE: NTS;
 18:  6pew-A  2.8  3.6   55   521    2   MOLECULE: GLUTAMINE SYNTHETASE;
 19:  6fai-G  2.8  2.5   49   232    6   MOLECULE: 40S RIBOSOMAL PROTEIN S27-A;
 20:  8xlj-A  2.7  3.5   53   372    4   MOLECULE: GLUTAMINE SYNTHETASE;
 21:  1o70-A  2.7  2.4   48   296    8   MOLECULE: FASCICLIN I;
 22:  5o60-W  2.7  2.8   56   192   14   MOLECULE: 50S RIBOSOMAL PROTEIN BL37;
 23:  2e28-A  2.6  3.0   50   587   14   MOLECULE: PYRUVATE KINASE;
 24:  4m00-A  2.5  3.4   44   501    7   MOLECULE: SERINE-RICH ADHESIN FOR PLATELETS;
 25:  1bml-C  2.5  2.8   47   318   11   MOLECULE: PLASMIN;

0001 s001A   --------------------------APRDITVRA------G-----P-DEV-GTV---E--A--RF-LAA--I--D---P---N-PGSF-ML-A-G-K-------S-WEII-T--VEW-----D-R------GVCVVRP--S----P-----------
0002 6znpA   lhrngeryhwasesfpasnislrs-aSQENVVIVD------QsdianV-RII-GEM---D--R--FS-AMT--L--L---H---D-EAIY-LH-E-G-V-------Q-YQVE-K--LDW-----D-H------KKAYVRK--V----D-----------
0003 7wrsA   mewdkiwsfnkkvidpvaprytal-lKDAVVPVNV------P-----EvGSK-VLI---E--G--AD-AET--L--T---E-----GEVV-TF-InW-G-------N-IIIT-K--LNR-nssgK-I------VSIDTKL--N----L-----------
0004 5dm3C   -------------------------geiDTVLVCI------V-dmqgR-LMG-KRL---H--A--RH-FVD--H-----------gwEET-HC-C-NlY-------I-MKPD-L-aTLR-----C-V--legtAMVLCDL--L----D-----------
0005 2detA   akkkdstgicfigerkfreflgry-lPAQPGKIIT-----vD-----G-DEI-GEH---Q--G-----LMY--H--T---L-----GQRKgLG-I-G-GtkegteeP-WYVV-D--KDV-----E-N------NILVVAQ--G----Hehprlmsvgli
0006 7bgmA   ---------snavdslldsvkwdn-kGLAVAIAQN------V---dtG-AIL-MQG---F--A--NreAVA--T--T---I---S-SRKatKG-E-TnN-------F-INVH-D--VFL-----DdS------IIYLGKP--D----Gptchtgaetcy
0007 9fhlG   ----------------------------PDFKIVI------SayrtkT-LQI-SVD---QnkA--TN-LVG--L--K---I-----GDVF-EA-NlP-V-------K-LKIT-G--GSD-----Npnensqei VQINTII--V----R-----------
0008 8fn2X   --ensrvlscqyrssfgssnarri-rAKSEIPAVVygqgkdV-----S-HL---RI---K--S--SE-FNKkfA--K---F---TdNTVL-IL-DlE-R-------C-VFVK-D--AAEniaskL-I------YHIDFYE--V----Drnvelekyvpi
0009 1zpsB   -------skgdvnillnfrhning-eDLIIAVAQD------H---etG-EVL-MVA---Y--M--NreALR--R-----tL---E-TGTAhywstsG-H-------V-QRVK-D--VLV-----D----dgdaVVLKVEQ--E----Ggachtgyrscf
0010 6fanC   ----------------------------CEASAFI------V-ngdkE-ELF-LER---V--D--KL--------------------ifgqrK-V-I-------K-AKIK-R--LEL-----V-D------HRILLER--E----D-----------
0011 8s8kB   ----avgknkrlskglkkrvvdpf-tRKEWYDIKA------P-----SfRNV-GKT---L--V--N----------------dslkGRVV-EV-C-LkV-------K-LRVD-E--VQG--------------KNLLTNF--H----G-----------
0012 4hppA   --------------------mnrl-qPVRLVSFVT------T-dlagI-TRG-RSL---P--L--AT-LEE--Q-----------lASGC-GW--paspwgshgdlR-LLPD-PnsRVR-----V----apalDYLHGNL--V----E-----------
0013 7kpjE   -----------------------e-vQNVKINYYD------E---daE-KQV-AEV---P----idtscVN--M--A--------------------pE-------G-YALV-S--SDC-----I----rdgyVYVSVKK--D----V-----------
0014 2j9iA   -------------------------haaAFVRFEA------TdaagaS-RSK--SI---P--A--QF--FE--K--V--iH---A-RG---YL-E-DdI-------V-LMPE-L-stAA-----V-A---aaatAAVICDG--G----A-----------
0015 7aohB   lvsqlsvlssitniltseyldlekkyfETGFPITI------E-----N-ALV-ASL---N--P--NM-ICD--FvtD---F---R-R-----------rkrmgffgnlevgI-T--LVR----dH-M------NEIRINI--G----Agrlvrpflvvd
0016 2kcoA   ptfttlsaedirikdrtlggnfkv-rLKYTTTANV------LpatntA-KKV-KI---------aRR-GI---------iI---R-GAKI-RT-E-A-G-------L-AVVT-S--RPG-----Q-D------GVINAVL--L----K-----------
0017 3wwlA   ------------------------------MVGTC------P-------GAE-LRL---E--N----------------pE---L-GELV-VC-EcG-A-------E-LEVV-G--LD-----------------PLRLEP--A----P-----------
0018 9nvuP   ----------------------------RYVYVLD-----vD-----G-KPL-MPT---C-------R-FGKvrR--M---L---K-SG---------------------QAK-A--VDT-----L-P------FTIQLTY--Kprta-----------
0019 6pewA   ----mksvsfsnnaelyeyikdkk-nDVEIVACII------T-nllgT-YFK-CFF---Y--V--KE--IT--L--N---K----lESGF-SF-D-AdF-------F-IKVD-H-stCY-----L-E--gkniLNIMCDI--K----R-----------
0020 6faiG   ------------------------------MKLNI------SpvngsQ-KTF-EIddehR--I--RV-FFD--K--R---I-----GQEV-DG-----gY-------V-FKIS-G--GND-----KdL------AVLALVI--V----Kkgeqelegltd
0021 8xljA   --matsasshlnkgikqmymnlpq-gEKIQLMYIW------V------gLRC-KTR---T--L--DC-DP-------kcvE---E-LPEW-NF-DsD-M-------Y-LHPV-A--MFR-----D-P--rdpnKLVFCEV--F----Kynrkpaetnlr
0022 1o70A   engalrkfyevimdnggavlddin-sLTEVTILAP------S-----NmilnmHII---K----drlnVDK--I--RqknansdT-VITV-EG-G-G-V-------N-ATVI-Q--ADV-----A-Q---tngYVHIIDHvv-----------------
0023 5o60W   anipnkltanvrtrtgkgasrqar-rDGKVPAVLY------G--hgtD-PQH-LEL---N--A--RD-FAA--VlrshgtN---A-ILTL-DI-E-G-T-------E-QLAL-T-kALD-----ViQ------HADLLVV--Q----Rgekvtvevtvl
0024 2e28A   ----------------------------------------evlgtP-EKI-SVT---Y--P--SL-IDD--V--S---V-----GAKI-LL-DgL-I-------S-LEVN-A--VDK-----Q-A------GEIVTTV--L----N-----------
0025 4m00A   smtastggatnlqqvqfgtfeyte-sAVTQVRYVD----vtT-----G-KDIiPPK---T----tID-NQ------------------------tA-K-------G-YNYT-S--VDS-----S-YasaqqsVTYYFTD--V----Kaptvtvgnqti
0026 1bmlC   dlisekyyvlkkgekpydpfdrsh-lKLFTIKYVD----vnT-----N-ELL-KSE---QlltaaKL-LY------------------NN-LDaF-G-I-------MdYTLTgK--VED-----N----tnriITVYMGK--R--------------

0001 s001A   --------------------------LLLEEEEEE------L-----L-EEE-EEE---E--H--HH-HHL--L--L---L---L-LLEE-EE-L-L-E-------E-EEEE-E--EEL-----L-L------LEEEEEE--L----L-----------
0002 6znpA   eeeelleeeellllllhhhlllll-lLLLEEEEEE------Lllhhh L-EEE-EEE---E--H--HH-HHH--H--L---L---L-LLEE-EE-L-L-E-------E-EEEE-E--EEL-----L-L------LEEEEEE--L----L-----------
0003 7wrsA   llhhhhhhhhhhhhllllleelee-eHHHLEEEEL------L-----LeLLE-EEE---L--H--HH-HLL--L--L---L-----LLEE-EE-LlL-E-------E-EEEE-E--EEE-llllL-E------EEEEEEE--L----L-----------
0004 5dm3C   -------------------------lllLEEEEEE------E-llllL-EEE-EEE---E--H--HH-HHH--L-----------llLLE-EL-L-LlL-------E-EEEL-H-hHLE-----E-L--llleEEEEEEE--E----L-----------
0005 2detA   llllllllllllllllhhhhhlll-lLLLLLLLLL-----lL-----L-LLL-LLL---L--L-----LLL--L--L---L-----LLLLlLL-L-L-Llllllll L-EEEE-E--LLL-----L-L------LLLEEEE--L----Lllhhheeeeee
0006 7bgmA   ---------llhhhhhhhlllllll-lLLEEEEEEE------L---llL-LEE-EEE---E--E--LhhHHH--H--H---H---H-HLLleLL-L-LlL-------L-EEEE-E--EEE-----ElE------EEEEEEE--L----Lllllllllle
0007 9fhlG   ----------------------------LEEEEEE------EeeellE-EEE-EEE---LhhH--LL-LLL--L--L---L-----LLEE-EE-LeE-E-------E-EEEE-E--EEE-----Lllllllle EEEEEEE--L----L-----------
0008 8fn2X   --lhhheeeeeelllllhhhhhhh-hHLLEEEEEEElllllL-----E-EE---EE---E--H--HH-HHHhhL--L---L---LlLLEE-EE-ElE-E-------E-EEEE-E--EEEellllE-E------EEEEEEE--L----Lllllleeeeee
0009 1zpsB   -------lhhhhhhhlllleeell-eEEEEEEEEE------L---llL-LEE-EEE---E--E--LhhHHH--H-----hH---H-HLLLeeeellL-L-------L-EEEE-E--EEE-----L----llleEEEEEEE--L----Lllllllllll
0010 6fanC   ----------------------------LLEEEEE------L-llllL-EEE-EEE---E--E--EE--------------------ellleE-E-E-------L-EEEE-E--EEL-----L-L------LEEEEEE--L----L-----------
0011 8s8kB   ----llllllllllllllllllll-lLEEEEEEEL------L-----LlLEE-EEE---E--E--E----------------hhhlLLEE-EE-E-HeE-------E-EEEE-E--ELL--------------LEEEEEE--E----E-----------
0012 4hppA   --------------------llll-lLLLEEEEEE------E-llllL-EEE-EEE---E--H--HH-HHH--H-----------hLLLE-EE--hhllllllll eE-EEEL-HhhLEE-----E----llleEEEEEEE--E----L-----------
0013 7kpjE   -----------------------l-lEEEEEEEEE------L---llL-EEE-EEE---E----lllllLL--H--H--------------------lL-------L-EEEE-E--LLL-----L----llleEEEEEEE--L----L-----------
0014 2j9iA   -------------------------lllLEEEEEE------ElllllE-EEE--EE---E--H--HH--LL--L--L--hH---H-HL---EE-E-LlE-------E-EEEL-L-llLE-----E-L--llleEEEEEEE--E----L-----------
0015 7aohB   lleelllllleellllhhhhhhhhhhllLLLEEEEE------L-----L-EEE-EEE---L--H--HH-HHH--HhhH---H---H-H-----------hhhllllllllleE-E--EEL----lL-L------LEEEEEL--L----Llleeeeeeeee
0016 2kcoA   lllllllhhhhhhhhhlllllllll-lLLLLLEEEE------ElllleE-EEE-EE---------hLL-LL---------lL---L-LLEE-EE-L-L-L-------E-EEEE-E--EEL-----L-L------LLEEEEE--L----L-----------
0017 3wwlA   ------------------------------LEEEL------L-------LLE-EEL---L--L----------------lL---L-LLEE-EL-LlL-L-------E-EEEE-E--LL-----------------LLEEEE--L----L-----------
0018 9nvuP   ----------------------------LLLEEEL-----lL-----L-LLL-LLL---L------L-HHHhhH--H---H---H-LL---------------------LEE-E--EEE-----L-L------EEEEELL--Lllll-----------
0019 6pewA   ----lleellllhhhhhhhhhlll-lLEEEEEEEE------L-llllL-LEE-EEE---E--H--HH--LL--L--H---H----hHLLL-LL-L-LlL-------L-LEEE-E-eeEE-----L-L--leeeEEEEEEL--L----L-----------
0020 6faiG   ------------------------------LEEEE------ElllleE-EEE-ELllllL--L--LL-LLL--L--L---L-----LLEE-EH----lL-------E-EEEE-E--EEE-----LlE------EEEEEEE--E----Elllllllllll
0021 8xljA   --llllhhhllhhhhhhhhhllll-lLLEEEEEEE------E------lEEE-EEE---E--E--LL-LL-------llhH---H-LLLE-EE-ElE-E-------E-EEEE-E--EEE-----L-L--llllEEEEEEE--E----Lllllllllhh
0022 1o70A   lhhhhhhhhhhhhhlllhhhhhhh-lLLLEEEEEE------L-----HhhhhhLEE---L----llllHHH--H--Hhhlllhh L-EEEE-EE-L-L-E-------E-EEEE-E--EEE-----E-L---lleEEEEELLll----------------
0023 5o60W   llllleeeeeeelllllhhhhhhhh-hLLEEEEEEE------L--lllL-LEE-EEE---E--H--HH-HHH--HhhhhllL---L-EEEE-EE-L-L-E-------E-EEEE-E-eEEE-----EeE------EEEEEEL--L----Llleeeeeeeee
0024 2e28A   ----------------------------------------lllllL-LEE-ELL---L--L--LL-LLL--L--L---L-----LLEE-EE-LlL-E-------E-EEEE-E--EEL-----L-L------LEEEEEL--L----L-----------
0025 4m00A   eeeeellllllleeeeeeeeeeee-eLEEEEEEEE----llL-----L-LLLlLLE---E----eLL-LL------------------------hH-L-------L-EEEE-E--EEL-----L-Llllllee EEEEEEE--L----Llleeellleee
0026 1bmlC   eeeeeeeeeellllllllllll-lEEEEEEEEE----llL-----L-LEE-EEE---EeeellLL-LL------------------LL-LHhH-L-L-------LlEEEEeE--EEE-----L----lleeEEEEEEE--L--------------

# Job: Type-2 BREX; BrxHI-Helicase MBK7581083 CTD (Lhr-helicase CTD)
# Query: s001A
# No:  Chain   Z    rmsd lali nres  %id PDB  Description
   1:  5v9x-A 10.3  2.7  162   772   13   MOLECULE: ATP-DEPENDENT DNA HELICASE;
   2:  5jb3-1  8.1  2.3   69    83   13   MOLECULE: 16S RIBOSOMAL RNA;
   3:  1d1r-A  7.1  2.6   72    83   11   MOLECULE: HYPOTHETICAL 11.4 KD PROTEIN YCIH IN PYRF-OSMB
   4:  6vpq-A  7.0  3.9   67    87    9   MOLECULE: DENSITY-REGULATED PROTEIN;
   5:  5oa3-0  6.8  8.8   82   479    7   MOLECULE: EUKARYOTIC TRANSLATION INITIATION FACTOR 2D;
   6:  9iad-A  6.6  7.6   72   685   13   MOLECULE: PROTEIN ARGONAUTE;
   7:  1xxa-C  6.5  1.7   56    74   14   MOLECULE: ARGININE REPRESSOR;
   8:  7nad-U  6.4  2.0   63   102   11   MOLECULE: 25S RRNA;
   9:  3mml-B  6.4  2.8   69   210    4   MOLECULE: ALLOPHANATE HYDROLASE SUBUNIT 2;
  10:  5e68-B  6.3  3.2  111   172    5   MOLECULE: S-RIBOSYLHOMOCYSTEINE LYASE;
  11:  4y2i-A  6.1  1.9   58    65   14   MOLECULE: PUTATIVE METAL-BINDING TRANSPORT PROTEIN;
  12:  6znp-A  5.9  3.6   91   748   14   MOLECULE: UNCHARACTERIZED ATP-DEPENDENT HELICASE YPRA;
  13:  2kwa-A  5.8  2.2   69   101    6   MOLECULE: KINASE A INHIBITOR;
  14:  7bj2-a  5.8  9.9   66   306    9   MOLECULE: FLAGELLAR P-RING PROTEIN;
  15:  7y8u-F  5.7  5.9  107   347    7   MOLECULE: ALBE HOMOLOG;
  16:  5euf-B  5.4  5.6  110   416    9   MOLECULE: PROTEASE;
  17:  2if1-A  5.4  4.8   82   126   11   MOLECULE: EIF1;
  18:  2p5m-A  5.3  1.9   55    82   11   MOLECULE: ARGININE REPRESSOR;
  19:  4u9r-A  5.3  2.0   55   147   13   MOLECULE: CZCP CATION EFFLUX P1-ATPASE;
  20:  8b6j-b  5.3  6.8  120   460   10   MOLECULE: PEPTIDASE M16 INACTIVE DOMAIN PROTEIN;
  21:  8q3b-C  5.2  7.9   83   359   10   MOLECULE: DNA-DIRECTED RNA POLYMERASE RPB1 HOMOLOG;
  22:  7y8v-E  5.2  6.8  108   374    5   MOLECULE: ALBE HOMOLOG;
  23:  7si6-A  5.1  4.0   65   873    5   MOLECULE: P-TYPE CU(+) TRANSPORTER;
  24:  4v1a-l  5.1  6.2   67   133   10   MOLECULE: MITORIBOSOMAL PROTEIN ML37, MRPL37;
  25:  8bep-B  5.0  5.9  113   487   16   MOLECULE: PROBABLE MITOCHONDRIAL-PROCESSING PEPTIDASE SUBUN

0001 s001A   HLRG-SESSLIE----AP------D--RITWWN--F-A-----GG-GANALLRGMLE--QEL-G---------E-KVVA-T--N-------A-AIKLE-------G-----D-------AS--GAAVREVLA-R--W---RP--T--AEDALRA--A--D----PSRTR--V-SKF---QVC-P-EALLAELVVEKV--LDVEGARRALS-A--P-PT-A-T-S--G-GQ--
0002 5v9xA   VVPSdTTFVVER----FR-delgdW--RVILHS--P-Y-----GL-RVHGPLALAVG--RRL-R---erygidE-KPTA-S--D-------D-GIIVR-------L-----P------dsP--GADL--FVF-D-aD---EI--E---PIVTAE--V--GgsalFASRF-----rE---CAA-I-VLEAVRECLQDV--YDVPALIELMH-K--IrIV-E-V-E--T-AT--
0003 5jb31   iakeeqKIKIYV----TK-rrfgkL--MTIIEG--F-D-----TsvIDLKELAKKLK--DIC-A---------C-GGTV-K--D-------N-TIELQ-------G-----------DH--RKKVAEELV-K--M---G------------------------------------------------------------------fSR-D--S-IE-I-R---------
0004 1d1rA   --kgdgVVRIQR----QTsgrkgkG--VCLITG-vD-L-----DD-AELTKLAAELK--KKC-G---------C-GGAV-K--D-------G-VIEIQ-------G-----D-------K--RDLLKSLLE-A--K-----------------------------------------------------------------G-M--K-VK-L-A-G--G-LE--
0005 6vpqA   ------------prakkK------Y--VTRVCG--L-A-----tfeiDLKEAQRFFA--QKF-S---------X-GASVtG--E-------D-EIIIQ-------G-----------DF--TDDIIDVIQ-E--K-----------------------------------------------------------------WP-E--VsIE-D-L-G--E-----
0006 5oa30   kgrI-CPIDITL----AQ-rasnkK--VTVVRN--L-E-----aygl DPYSVAAILQ--QRC-Q---------A-STTV-NpdS-------L-QVQIQ-------G-----------NQ--VHHLGWLLL-E--E---Y-------------------------------------------------------------qkH--I-QG-L-E-Kag------
0007 9iadA   ------------lditpK------N--FICFRLtpE-I-----ER-KTGNSFSWRFS--QKF-P---------dA-VVIW-H--N-------K-FFWVL-------AkpnrpM-------PS--QEQWREKLL-E--I-ceEL--K--K---------------------------------------------------------digdrT-Y--A-IQ-W-V-S--Ql-----
0008 1xxaC   -plknlvLDIDY----ND------A--VVVIHT--S-P-----G---AAQLIARLLD--SLG-K-----aegiL-GTIA-G--D-------D-TIFTT-------P-----A-ngftvkdL--YEAILELFDr------------------------------------------------------------------------------------------------
0009 7nadU   ---------------iA------K--TFTVDV--S---sptengvfDPASYAKYLI--DHI-KgavgnlgnaV-TVTE-D--G-------T-VVTVV-------S----taK-------FS--GKYLKYLTK-K-yL---KK--N---------------------------------------------------------qlrdW-I--R-FV-S-T-K--T-----
0010 3mmlB   ----stlGTVHN---yGD------Q--ALLLEF----D-----ST-AEVLAWTETLR--EAE-L-------lgvvDIVP-A--A-------R-TVLVK-------L-----A-gpryqapT--RQRLGKLR----vR---PE--A--IThqppgdtP--G----XW-----V-QFRavg-----------------------------------------------------
0011 5e68B   MQ----aPAVRT----AK------TitVFDLRF--CiM-----PE-KGIHTLEHLFA--GFM-Rhlngngvei I-DISP-M-xR-------T-GFYMS-------L-----I------tPD--EQRVADAWK-AamA---DV--LkdQNQIPE---L--N-----VYQCG-T---------------------------yQM--HSLSEAQDIAReR--D-VR-V-N-S--N-KE--
0012 4y2iA   --------------------------GMQFHI---T-----CG-GCASTVKKTIL--TLD-A---------nA-TVRT-D--P-----atR-LVDVE-------T-----S-------LS--AEQIAAALQ-K--A--G---------------------------------------------------------------fP-PR-E-R---------
0013 6znpA   ktkekSRTSLHY----GD------VtsAAWLEI----digektLE-QLLLGISNVLQ--HIV-PvyimcdrndV-HVVS-Q--IkaahtglP-TIFLYdhypggiG-----L-------AEevFKRFSDINE-A-A--A--KqchD--GCP-SCI--G------------------------------tEIEG--ikAKERILQLLD-Q--M-S-------------
0014 2kwaA   -gsmtvRYQIEQ---lGD------S--AMMIRF--G-eeineqVN-GIVHAAAAYIE--EQPgF---------I-ECIP-A--F-------T-SLTVF-------Y-----DmlpqgissP--FESVKRDVE-E--R---------------------------------------------------------------laeIAEDY-E--V-NR---------
0015 7bj2a   -erirdlTSVQGtqfgaG------N--TINLQL--N-E-----DF-TMAQQITDAIN--RAR-G-------yG-SATAlD--A-------R-TVQVR-------V-----P-gnssqvrF--LADIQNM---E--V---NV--T--P------------------------------vvIN---SRTg---------------------------------------
0016 7y8uF   ---------------vnK------D--MILLPV--E-HgdylkNR-MICHLIADMIK--HLA-Q----qleydV-SVGL-F--I-stnqhsC-YLKVK-------K-----S-------D---QKRFSSLIQ-Q--L---S-hfI--ETYIKDY--QwrF----MNELV-iN-FNQ-------geytvaelfgsLDS--VDKLDILAVRN-EliN-QL-T-V---------
0017 5eufB   KQDG--------gvhleW------V--ALGYKV--P--afkhkDQ-VALDALSKLLG--SWL-QvdkkrlasqA-FSHN-X--Q---lqdeS-VFLFI-------A-----G-pnikaeaL--QKEIVALLE-K--LkkgEI--T--QAELDKI--KinQ----KADFI-sN-LES-------qndlqgltdyQQQFldLKVSDLVRVAN-E-yF-KD-Tq---------
0018 2if1A   agTE-DYIHIRI----QQ-rngrkT--LTTVQG--I-A-----D-dYDKKKLVKAFK--KKF-A---------C-NGTV-I--E--hpeygE-VIQLQ-------G-----------dQ--RKNICQFLV-E--I--G-------------------------------------------------------------LAK-DdgL-KV-H-G-F---------
0019 2p5mA   ralmdafVKIDS----AS------H--MIVLKT--M-P-----G---NAQAIGALMD--NLD-W------demM-GTIC-G--D-------D-TILII-------C-----R-tpedtegV--KNRLLELL----------------------------------------------------------------------------------------------
0020 4u9rA   ------------gveaahiktanvdfgHVLWQLkgV-W-----HE-RRARTVASQLR--ALPgV---------I-EAEV-S--A-------SgIARVE-------F-----D----ndriS--AAGIEQALS-K---------------------------------------------------------------------R--G-LA-----------
0021 8b6jb   G-----------eesnvT------L--ALLFQS--V--pwssaDI-VAFNVAAALLN--LRL-KlqkyayfdqA-EALN-F--H---ftdsG-LFGLR-------T-----S-sadrakdI--LNHSIAELK-A--I-asGV--N--ADELLTA--KaaL----KNSVLsaL-ERQ-------kiqhtDYVKQIDS---VTADQVAKAVA-K-vL-TS-NpvnT--Y-DAir
0022 8q3bC   QP------ELVL----FS------LktKAIITI--K-E-----TH-TIGNLLKTCIY--EMI-P------diS-FVGY-Q--C-vphkqeM-VLTII-------H-----K-------AS--QEDLITLLE-KsiQ---NI--I--QTF------------------------------qileknVDE-Ia-------------------------------
0023 7y8vE   NE----------tdfssI------L--TCAYTF---hvtdleRE-ILIPMFDGYIG--SVL-F-ktlrlvdhI-SSHY-D--Q-----esN-LLFIT-------V-----L-sknqlstI--LTKIQHTIE-S--I---EF--D--SVSFSLS--KvmF----ENESRfrfeYKN---PE------sflKAIQNT----dIADFYRLKA-N--M-RWv-----------
0024 7si6A   ------------vedlgfgasvmeG--NVELIItgM-T-----CA-SCVHNIESRLM--RTP-G--------IlQASV-A--L-----atC-KAQVK-------F-----Dp-----iVG--PRDIIRIIE-G--I--------------------------------------------------------------G-F--Q-AS-L-A-----hVG--
0025 4v1al   RSRM-HniPVYR----DI-thgnrQ--MTVIRK--V-E-----G---DIWALQKDVE--DFL-S---pllgktP-VTQV-N--E-----vtG-TLRVK-------G-----------YF--DQQLKAWLL-E--Kgf--------------------------------------------------------------------------------------------
0026 8bepB   G----------dlplaQ------F--AVAFEG--A--swtdpDS-VALMVMQTMLGswSDL-TvaineiaesI-MAFN-T--N---ykdtG-LFGVY-------A-----V-kadclddL--SYAIMYEVT-K--L-ayRV--S--DADVTRA--RnqL----KSSLL-------------riptaelFARIDA---VDASTVKRVAN-KyiY-DK-D-iqD--Y-NKfr

0001 s001A   HHLL-LLLEEEE----EL------L--EEEEEL--L-L-----LH-HHHHHHHHHHH--HHH-L---------L-EEEE-L--L-------L-EEEEE-------L-----L-------LL--HHHHHHHHH-H--L---LL--L--HHHHHHH--L--L----LLLLL--L-LLL---HHH-L-HHHHHHHHHHHH--LLHHHHHHHLL-L--L-LE-E-E-E--L-LL--
0002 5v9xA   LLLLlLEEEEEE----EE-lllllE--EEEEEL--L-L-----LH-HHHHHHHHHHH--HHH-H---hhllllL-LLEE-E--L-------L-EEEEE-------E-----L------llL--LHHH--LLL-L-hH---HH--H---HHHHHH--H--HhlhhHHHHH-----hH---HHH-H-HHHHHHHHHHLL--LLHHHHHHHHH-H--HeEE-E-E-E--L-LL--
0003 5jb31   llllllLEEEEE----LL-lllllL--EEEEEL--L-L-----LllLLHHHHHHHHH--HHH-L---------L-LEEE-E--L-------L-EEEEE-------L-----------LL--HHHHHHHHH-H--L---L------------------------------------------------------------------lLL-L--L-EE-E-L---------
0004 1d1rA   --llllEEEEEE----LLlllllL--LEEEEL-lL-L-----LH-HHHHHHHHHHL--LLL-L---------L-LLLL-L--L-------L-LEEEL-------L-----L-------L--HHHHHHHHH-H--H-----------------------------------------------------------------L-L--E-EL-L-L-L--L-LL--
0005 6vpqA   ------------ellllL------E--EEEEEL--H-H-----hhllLHHHHHHHHH--HHH-L---------L-LEEEeE--L-------L-EEEEE-------L-----------LL--LLLHHHHHH-H--H-----------------------------------------------------------------LL-L--LhEE-E-L-L-------
0006 5oa30   ellL-LLEEEEE----EE-ellleE--EEEEEL--L-H-----hhllLHHHHHHHHH--HHH-L---------L-LEEE-EelL-------E-EEEEE-------L-----------LL--HHHHHHHHH-L--L---L-------------------------------------------------------------lhH--E-EL-L-L-Lll------
0007 9iadA   ------------eellllL------L--EEEEEEllL-L-----LH-HHHHHHHHHHH--HHL-L---------lE-EEEE-E--L-------L-EEEEE-------Ellllll-------LL--HHHHHHHHH-H--H-hhLL--H--H---------------------------------------------------------hhllllL-L--L-EE-E-L-L--Ll-----
0008 1xxaC   -llhhheEEEEE----LL------L--LEEEEE--L-L-----L---LHHHHHHHHHL--LLL-H-----hhleEEEE-L--L-------L-EEEEE-------L-----L-llllhhhH--HHHHHHHHLl--------------------------------------------------------------------------------------------------
```

```
0009 7nadU   [illegible]
0010 3mm1B   [illegible]
0011 5e68B   [illegible]
0012 4y2iA   [illegible]
0013 6znpA   [illegible]
0014 2kwaA   [illegible]
0015 7bj2a   [illegible]
0016 7y8uF   [illegible]
0017 5eufB   [illegible]
0018 2if1A   [illegible]
0019 2p5mA   [illegible]
0020 4u9rA   [illegible]
0021 8b6jb   [illegible]
0022 8q3bC   [illegible]
0023 7y8vE   [illegible]
0024 7si6A   [illegible]
0025 4v1a1   [illegible]
0026 8bepB   [illegible]

@@@@@@@@@@@@@@@@@@@@@@@@@@@@@@@@@@@@@@@@@@@@@@@@@@@@@@@@@@@@@@
@@@@@@@@@@@@@@@@@@@@@@@@@@@@@@@@@@@@@@@@@@@@@@@@@@@@@@@@@@@@@@
@@@@@@@@@@@@@@@@@@@@@@@@@@@@@@@@@@@@@@@@@@@@@@@@@@@@@@@@@@@@@@

# Job: Type-2 BREX; BrxD-ATPase (Standalone ORC/CDC6 AAA+ ATPase)
# Query: s001A
# No:  Chain    Z    rmsd lali nres  %id PDB  Description
   1:  1w5s-B 17.4  3.5  268   396   13   MOLECULE: ORIGIN RECOGNITION COMPLEX SUBUNIT 2 ORC2;
   2:  2qby-B 17.0  3.2  251   368   16   MOLECULE: CELL DIVISION CONTROL PROTEIN 6 HOMOLOG 1;
   3:  2fna-A 13.3  3.5  246   352   12   MOLECULE: CONSERVED HYPOTHETICAL PROTEIN;
   4:  7rzy-1 13.3  3.4  252   311   15   MOLECULE: TN6677 VIBRIO CHOLERAE TRANSPOSON TNSC (VCHTNSC);
   5:  8fcv-R 13.2  4.1  266   343   15   MOLECULE: DNA (60-MER);
   6:  6fhs-C 13.1  5.1  267   459   13   MOLECULE: RUVB-LIKE HELICASE;
   7:  5zr1-A 13.0  4.1  244   494   15   MOLECULE: ORIGIN RECOGNITION COMPLEX SUBUNIT 1;
   8:  9emc-D 13.0  4.1  258   337   14   MOLECULE: RUVB-LIKE 1;
   9:  8btg-B 12.9  4.1  223   335   13   MOLECULE: CHROMOSOMAL REPLICATION INITIATOR PROTEIN DNAA;
  10:  7jgr-D 12.9  3.9  255   441   10   MOLECULE: ORIGIN RECOGNITION COMPLEX SUBUNIT 2;
  11:  3bos-B 12.7  3.7  202   231   12   MOLECULE: PUTATIVE DNA REPLICATION FACTOR;
  12:  7plh-A 12.3  3.3  230   260   10   MOLECULE: SHTNSC;
  13:  6pxk-K 12.3  3.9  238   429   16   MOLECULE: ATP-DEPENDENT PROTEASE ATPASE SUBUNIT HSLU;
  15:  1g4a-E 12.1  4.6  245   356   16   MOLECULE: ATP-DEPENDENT HSL PROTEASE ATP-BINDING SUBUNIT
  16:  5zr1-E 12.1  4.2  239   460   10   MOLECULE: ORIGIN RECOGNITION COMPLEX SUBUNIT 1;
  17:  6p8v-C 12.0  3.6  225   303   10   MOLECULE: ATPASE, AAA FAMILY;
  18:  7uqj-A 11.9  4.4  229   585   11   MOLECULE: ATPASE HISTONE CHAPERONE YTA7;
  19:  8xvb-A 11.8  3.6  227   246   11   MOLECULE: DNA (5'-
  20:  7vcs-A 11.4  5.2  229   766   10   MOLECULE: TRANSITIONAL ENDOPLASMIC RETICULUM ATPASE;
  21:  8tw7-2 11.4  3.6  227   339   15   MOLECULE: REPLICATION FACTOR C SUBUNIT 5;
  22:  7st9-A 11.4  4.5  224   522   12   MOLECULE: CHECKPOINT PROTEIN RAD24;
  24:  8dr5-A 11.3  4.2  233   646   14   MOLECULE: REPLICATION FACTOR C SUBUNIT 1;
  25:  6l1q-A 11.1  3.3  208   267   15   MOLECULE: CBBQ PROTEIN;
  26:  3glf-G 11.1  4.8  217   378   18   MOLECULE: DNA POLYMERASE III SUBUNIT DELTA;
  27:  5e7p-A 11.0  4.1  221   719   16   MOLECULE: CELL DIVISION CONTROL PROTEIN CDC48;
  28:  5kzf-K 10.9  3.8  231   498    8   MOLECULE: PROTEASOME-ASSOCIATED ATPASE;
  29:  8umy-A 10.8  4.1  228   546   14   MOLECULE: CHROMOSOME TRANSMISSION FIDELITY PROTEIN 18 HOMOL
  30:  7fd4-A 10.7  5.5  223   779   14   MOLECULE: LON PROTEASE;
  31:  7st9-E 10.7  3.9  221   354   14   MOLECULE: CHECKPOINT PROTEIN RAD24;
  32:  7pkq-x 10.7  4.3  228   368   18   MOLECULE: MS35;
  33:  1ixs-B 10.6  4.0  223   315   11   MOLECULE: HOLLIDAY JUNCTION DNA HELICASE RUVA;
  34:  3pvs-D 10.6  5.4  207   424   15   MOLECULE: REPLICATION-ASSOCIATED RECOMBINATION PROTEIN A;
  35:  8csp-X 10.5  4.3  232   352   11   MOLECULE: 28S RIBOSOMAL PROTEIN S34, MITOCHONDRIAL;
  36:  9cru-C 10.5  3.8  202   741   16   MOLECULE: VESICULAR-FUSION PROTEIN SEC18;
  37:  8thc-A 10.5  3.8  211   479   12   MOLECULE: ELG1 ISOFORM 1;
  38:  9bv1-A 10.4  5.0  213   405   12   MOLECULE: 26S PROTEASOME NON-ATPASE REGULATORY SUBUNIT 1;
  39:  8zr0-A 10.3  4.1  209   480   13   MOLECULE: CASEINOLYTIC PEPTIDASE B PROTEIN HOMOLOG;
  40:  7w42-B 10.3  3.1  197   402   14   MOLECULE: UNCHARACTERIZED ATPASE YJOB;
  41:  7crc-B 10.2  4.7  232  1112   15   MOLECULE: NAD+ HYDROLASE (NADASE);
  42:  5j1s-A 10.1  3.4  195   277   11   MOLECULE: TORSIN-1A;
  43:  4rh7-A 10.1  3.6  252  3005   10   MOLECULE: GREEN FLUORESCENT PROTEIN/CYTOPLASMIC DYNEIN 2 HE
  44:  3u5z-B 10.1  4.0  214   320   14   MOLECULE: DNA POLYMERASE ACCESSORY PROTEIN 44;
  45:  2gno-A 10.0  3.7  189   296   12   MOLECULE: DNA POLYMERASE III, GAMMA SUBUNIT-RELATED PROTEIN
  46:  8xku-A 10.0  4.2  219   730   13   MOLECULE: PROBABLE INACTIVE ATP-DEPENDENT ZINC METALLOPROTE
  47:  8dgc-A 10.0  4.3  238  2028   15   MOLECULE: SEAVS3;
  48:  7qo4-M 10.0  4.5  215   421   13   MOLECULE: 26S PROTEASOME REGULATORY SUBUNIT RPN1;
  50:  6z1f-3  9.9  3.6  212   284   11   MOLECULE: RIBULOSE BISPHOSPHATE CARBOXYLASE/OXYGENASE ACTIV

0001 s001A   -G-RGL-E-TFA-V-G----L---ERFERSVAED-LAAV-E---R---G-A--G---T-FKAIRGE-YGSGKTFFVRWLQEQALR---R-G-FLAAEV-Q- [illegible]
0002 1w5sB   ----IP-P-E-L-R-V----R---RGEEALARIY-LNRL-L--sG---A-G--L---SdVNMIYGSrVGIGKTTLAKFTVKRVSE---AlT-VKQAYV-N- [illegible]
0003 2qbyB   -L-SVF-K-E-I-P-F----R---EDILRDAAIA-IRYF-V---K--nE-V--K---F-SNLFLGL-TGTGKTFVSKYIFNEIEE---V-K-VKQAYV-N- [illegible]
0004 2fnaA   -D---N-R-KDF-F-D----R---EKEIEKLKGL-R--------------------aP-ITLVLGL-RRTGKSSIIKIGINELN--------LPYIYL-D- [illegible]
0005 7rzy1   ---------FVS-T-P---sV---RKILSYMDRC-RDL--S---D---L-EseP---T-CMMVYGA-SGVGKTTVIKKYLNQNRR---E-S-IPVLHI-E- [illegible]
0006 8fcvR   ---------TVA-H-P---rL---KEVYEILMRT-IAEP---------------agaS-FIFVYGA-SGVGKTTLRLRVEQKLTE---L-AlPVVGIE-A- [illegible]
0007 6fhsC   [illegible]
0008 5zr1A   [illegible]
0009 9emcD   [illegible]
0010 8btgB   [illegible]
0011 7jgrD   [illegible]
0012 3bosB   [illegible]
0013 7plhA   [illegible]
0014 6pxkK   [illegible]
0015 1g4aE   [illegible]
0016 8vwvE   [illegible]
0017 5zr1E   [illegible]
0018 6p8vC   [illegible]
0019 7uqjA   [illegible]
0020 8xvbA   [illegible]
0021 7vcsA   [illegible]
0022 8tw72   [illegible]
0023 7st9A   [illegible]
0024 8dr5A   [illegible]
0025 6l1qA   [illegible]
0026 3glfG   [illegible]
0027 5e7pA   [illegible]
0028 5kzfK   [illegible]
0029 8umyA   [illegible]
0030 7fd4A   [illegible]
0031 7st9E   [illegible]
0032 7pkqx   [illegible]
0033 1ixsB   [illegible]
0034 3pvsD   [illegible]
0035 8cspX   [illegible]
0036 9cruC   [illegible]
0037 8thcA   [illegible]
0038 9bv1A   [illegible]
0039 8zr0A   [illegible]
0040 7w42B   [illegible]
0041 7crcB   [illegible]
0042 5j1sA   [illegible]
0043 4rh7A   [illegible]
0044 3u5zB   [illegible]
0045 2gnoA   [illegible]
0046 8xkuA   [illegible]
0047 8dgcA   [illegible]
0048 7qo4M   [illegible]
0049 8vwvF   [illegible]
0050 6z1f3   [illegible]

0001 s001A   [illegible]
0002 1w5sB   [illegible]
0003 2qbyB   [illegible]
0004 2fnaA   [illegible]
0005 7rzy1   [illegible]
0006 8fcvR   [illegible]
```

```
0007 6fhsC -L-LLL-LlLLL-L-L----L---HHHHHHHHHH-HHHH-H---L---L-L--L-llL-EEEE[illegible]
0008 5zr1A ----------lL-L-L----L---LLLHHHHHHHH-HHHH-H--hL---L-L--L---E-EEEE[illegible]
0009 9emcD -LlLLL-L-LLE-E-L----L---HHHHHHHHHH-HHHHhH---L---L-L--L--lL-EEEE[illegible]
0010 8btgB lL-LLLlL-LLL-L-L-llllhhhHHHHHHHHHL-LLL------------------llL-EEEE[illegible]
0011 7jgrD -L-LL------L-L-L----L---LHHHHHHHHH-HHHH-H--hH---L-L--L---E-EEEE[illegible]
0012 3bosB -L-LLLlL-LLL-L-L-lllL---HHHHHHHHHHhHLLL------------------lL-EEEE[illegible]
0013 7plhA ---------LLL-L-H---hH---HHHHHHHHHH-HHL----------------L---L-LEEE[illegible]
0014 6pxkK hhhhHH-L-LLL-L-L----L---HHHHHHHHHH-HHHH-H---H---HlL--L---L-LEEE[illegible]
0015 8rwvE ---LLL-L-LLL-L-L----L---LLLHHHHHHH-HLL--L---L---L-L--L---LlEEEE[illegible]
0016 1g4aE hhhhHH-H-HHL-L-L----L---HHHHHHHHHH-HHHH-H---H---HlL--L---L-LEEE[illegible]
0017 5zr1E ---------llL-L-L----L---HHHHHHHHHL-LL---L---H---H-H----llL-EEEE[illegible]
0018 6p8vC -H-HHH-H-HHL-L-L----L---HHHHHHHHHH-HHHH-H---L----------hllL-EEEE[illegible]
0019 7uqjA lL-LLLlH-HHL-L-L----L---HHHHHHHHLL-LHHH-H---H----------lllL-EEEE[illegible]
0020 8xvbA ---------LLL-L-H---hH---HHHHHhHHHH-HHHL------------------L-LEEE[illegible]
0021 7vcsA lL-LLL-L-LLL-L-L----L---HHHHHHHHHH-HHHH-H---L---H-----lllL-EEEE[illegible]
0022 8tw72 -L-L-----llL-L-L----L---HHHHHHHHHH-HHLL----------------llL-LEEE[illegible]
0023 7st9A -L-L---H-HHL-L-L----L---HHHHHHHHHH-HHHL-L---L---L-L----llL-EEEE[illegible]
0024 5v8fD -L-HH----LLL-LhH----H---HHHHHHHHHH-HHHH-H--lL---L-L--L---L-EEEE[illegible]
0025 8dr5A -L-LLL-L---L-L-L----L---HHHHHHHHHH-HHLH-H---H---H-H---lllL-EEEE[illegible]
0026 6l1qA lL-LLL-L---L-L-L----L---LLHHHHHHHH-HHLL------------------L-LEEE[illegible]
0027 3glfG -L--LL-L-L-L-L-L----L---HHHHHHHHHH-HHL----------------lllL-EEEE[illegible]
0028 5e7pA lL-LLL-H-HHL-L-L----L---HHHHHHHHHHlHHHH-H---L---HlL--L---L-EEEE[illegible]
0029 5kzfK lL-LLL-H-HHLlL-L----H---HHHHHHLHHH-HHL----------------lllL-EEEE[illegible]
0030 8umyA -L-L---L-LLL-L-L----L---HHHHHHHHHH-HHHH-L---L---L-LllllllL-EEEE[illegible]
0031 7fd4A hhhHHH-H---L-L-L----L---HHHHHHHHHH-HHHhhhlllL---L-L--L-llllEEEE[illegible]
0032 7st9E -L-L-H-H-HLL-L-L----HhhhHHHHHHLLL--HHHL------------------L-LEEE[illegible]
0033 7pkqx ---------LEE-E-L----L---HHHHHHHHHH-HHL-----------------llL-EEEE[illegible]
0034 1ixsB -L-LLH-H--HL-L-LlhhhH---HHHHHHHHHH-LLLL-L--------------llL-LEEE[illegible]
0035 3pvsD -L-LLL-L-L-L-L-L----L---HHHHHHHHHL-L-------------------lL-LEEE[illegible]
0036 8cspX -H-------EEE-E-L----L----HHHHHHHHH-HHL--L---L---L-LllL---L-EEEE[illegible]
0037 9cruC -L-LLL----LL-L-L--hhH---HHHHHHHHHH-HHHH-H---HlllL-L--E---E-EEEE[illegible]
0038 8thcA -L-LLL-L-L-L-L-L----L---HHHHHHHHHH-HHHH-H---H---H-H---lllL-LLLL[illegible]
0039 9bv1A lL-LLL-L-LLL-L-L----L---LHHHHHHHHH-HHHH-H---L----------lllL-LEEE[illegible]
0040 8zr0A hhhhHH-H-HhL-L-L----L---HHHHHHHHHH-HHH-----------llllL---E-EEEE[illegible]
0041 7w42B -L-LLL-H-HHL-L-LlhhhH---HHHHHHHHHH-HHL-----------lL--L---E-EEEE[illegible]
0042 7crcB --lLLL-L-LLL-L-L----L---HHHHHHHHHH-LLL------L---L-L--L---L-EEEE[illegible]
0043 5j1sA hhhhHH-H-HHL-L-L-lhhH---HHHHHHHHHH-HL---L---L---L-LllL---E-EEEE[illegible]
0044 4rh7A -L-LL----LLL-LhH----H---HHHHHHHHHH-HLH----------------hhlL-LEEE[illegible]
0045 3u5zB -L-L-----lLL-L-LlhhhH---HHHHHHHHHL-L-------------------lL-LEEE[illegible]
0046 2gnoA -------------------------LHHHHHHHH-HHLL-----------L--L---E-EEEE[illegible]
0047 8xkuA lL-LLL-L-LLL-L-L----L---HHHHHHHHHH-HHHH-H---L---H-----lllL-EEEE[illegible]
0048 8dgcA -L-LL-----LL-L-L----L---LHHHHHHHHH-HLL-----------L--L---L-LEEE[illegible]
0049 7qo4M lL-LLL-L-LLL-LlL----H---HHHHHHHHHH-HHHH-H---L---HlL--L---L-LEEE[illegible]
0050 8rwvF -----L-L-LL--L-L----L---LLLHHHHHHH--HHH-H---H--lL-L--L---L-LLEE[illegible]
0051 6z1f3 ---------LLL-L-L--------HHHHHHHHHH-HHHH-H---L----------lllL-EEEE[illegible]
```

```
# Job: Type-3 BREX; MTase MBW2068239 HTH1
# Query: s001A
# No:  Chain   Z    rmsd lali nres  %id PDB  Description
   1:  5a31-N  6.3  2.0   54   703   11   MOLECULE: ANAPHASE-PROMOTING COMPLEX SUBUNIT 1;
   2:  1ldd-A  5.7  2.3   53    74    9   MOLECULE: ANAPHASE PROMOTING COMPLEX;
   3:  9bw1-M  5.7  2.7   56   779   13   MOLECULE: LE_POLYA;
   4:  1hlv-A  5.4  2.2   53   131    9   MOLECULE: CENP-B BOX DNA;
   5:  2nvu-A  5.4  3.0   63   530    5   MOLECULE: NEDD8-ACTIVATING ENZYME E1 REGULATORY SUBUNIT;
   6:  9kbd-A  4.9  2.0   51   737    4   MOLECULE: S-PHASE KINASE-ASSOCIATED PROTEIN 1;
   7:  3mq0-B  4.8  2.5   48   248    8   MOLECULE: TRANSCRIPTIONAL REPRESSOR OF THE BLCABC OPERON;
   8:  3c1d-B  4.8  2.5   52   154   12   MOLECULE: REGULATORY PROTEIN RECX;
   9:  3fhz-D  4.7  2.1   53   167   15   MOLECULE: ARGININE REPRESSOR;
  10:  6lb9-A  4.6  2.8   62   345   11   MOLECULE: DUF4007 DOMAIN-CONTAINING PROTEIN;
  11:  8j56-C  4.6  2.5   54   160    6   MOLECULE: FLAGELLAR TRANSCRIPTIONAL REGULATOR FLHD;
  12:  8eki-D  4.6  3.2   60   684    7   MOLECULE: PROTEIN TRANSPORT PROTEIN SEC20;
  13:  2od5-A  4.6  2.0   47    91    6   MOLECULE: HYPOTHETICAL PROTEIN;
  14:  8cli-A  4.5  3.0   58   542   10   MOLECULE: GENERAL TRANSCRIPTION FACTOR 3C POLYPEPTIDE 1;
  15:  5of3-C  4.4  3.0   57   107    5   MOLECULE: DNA PRIMASE SMALL SUBUNIT PRIS;
  16:  7z0o-G  4.3  2.2   49    84    2   MOLECULE: HISTONE H3;
  17:  4fcy-B  4.3  2.2   49   476   10   MOLECULE: TRANSPOSASE;
  18:  8s0c-A  4.3  2.8   52   113   10   MOLECULE: ORIGIN RECOGNITION COMPLEX SUBUNIT 1;
  19:  2dob-A  4.3  3.2   56    82    9   MOLECULE: PROACTIVATOR POLYPEPTIDE;
  20:  2rkh-A  4.2  2.1   50   164   10   MOLECULE: PUTATIVE APHA-LIKE TRANSCRIPTION FACTOR;
  21:  3v4g-A  4.2  2.0   48   145    6   MOLECULE: ARGININE REPRESSOR;
  22:  6zvh-y  4.2  2.0   47    72   17   MOLECULE: 18S RRNA;
  23:  6v9i-C  4.2  2.2   52   730    8   MOLECULE: IMMUNOGLOBULIN G-BINDING PROTEIN G,CULLIN-5;
  24:  6qfd-B  4.1  2.1   45   116   11   MOLECULE: DNA-BINDING PROTEIN;
  25:  6zca-D  4.1  2.7   58   140   10   MOLECULE: PROBABLE DNA-DIRECTED RNA POLYMERASE SUBUNIT DELT
  26:  7z8b-C  4.1  2.3   52  1224    8   MOLECULE: CULLIN-7;
  27:  7pik-C  4.0  2.6   52   557   12   MOLECULE: TRANSPOSON TN7 TRANSPOSITION PROTEIN TNSB;
  28:  2qby-B  4.0  2.2   50   368    6   MOLECULE: CELL DIVISION CONTROL PROTEIN 6 HOMOLOG 1;
  29:  1d3y-B  4.0  2.1   48   290    6   MOLECULE: DNA TOPOISOMERASE VI A SUBUNIT;
  30:  1cf7-B  4.0  2.1   52    82   10   MOLECULE: DNA (5'-
  31:  6zfb-d  4.0  2.6   58   139   10   MOLECULE: DNA-DIRECTED RNA POLYMERASE SUBUNIT DELTA;
  32:  7oqe-K  4.0  2.0   46   406    4   MOLECULE: PROTEIN NAM8;
  33:  3e3v-A  4.0  2.3   45   154    7   MOLECULE: REGULATORY PROTEIN RECX;
  34:  7kbf-K  3.9  2.4   50    78   16   MOLECULE: HISTONE H3.2;
  35:  3s63-B  3.9  3.0   56    90    5   MOLECULE: SAPOSIN-LIKE PROTEIN;
  36:  8ea3-W  3.9  2.7   53   514   11   MOLECULE: TARGET_LE;
  37:  5b7i-C  3.8  2.8   63   138   14   MOLECULE: CRISPR-ASSOCIATED NUCLEASE/HELICASE CAS3 SUBTYPE
  38:  1fnn-A  3.8  2.9   54   379    9   MOLECULE: CELL DIVISION CONTROL PROTEIN 6;
  39:  1cf7-A  3.8  2.8   48    67    6   MOLECULE: DNA (5'-
  40:  7dh7-C  3.8  2.4   50   150   14   MOLECULE: TRANSCRIPTIONAL REGULATOR FUR FAMILY;
  41:  5zi8-A  3.8  2.9   52   104   13   MOLECULE: TRANSCRIPTIONAL REGULATOR;
  42:  8as8-E  3.8  2.9   51   498    6   MOLECULE: JETC;
  43:  1dp7-P  3.8  2.0   46    76   11   MOLECULE: DNA (5'-D(*CP*GP*(BRU)P*TP*AP*CP*CP*AP*(BRU)
  44:  2k85-A  3.8  2.0   46    65   11   MOLECULE: GLUCOCORTICOID RECEPTOR DNA-BINDING FACTOR 1;
  45:  3al0-B  3.8  2.5   53   482    8   MOLECULE: GLUTAMYL-TRNA(GLN) AMIDOTRANSFERASE SUBUNIT A;
  46:  6tda-L  3.8  3.9   52   596    8   MOLECULE: HISTONE H3.2;
  47:  8yb6-K  3.8  2.9   52   165   10   MOLECULE: CRISPR SYSTEM CASCADE SUBUNIT CASD;
  48:  8u8m-A  3.7  2.5   51   116    8   MOLECULE: DOUBLE-STRAND TELOMERIC DNA-BINDING PROTEINS 1;
  49:  3elk-A  3.7  2.2   47   104   11   MOLECULE: PUTATIVE TRANSCRIPTIONAL REGULATOR TA0346;
  50:  8yzt-C  3.7  3.2   59   121    5   MOLECULE: PROTEIN BANP;
  51:  8jxk-D  3.7  2.1   46   166   17   MOLECULE: CONSERVED PROTEIN;
  52:  3k69-A  3.7  2.4   45   151    9   MOLECULE: PUTATIVE TRANSCRIPTION REGULATOR;
  53:  1j5y-A  3.7  3.0   46   172    9   MOLECULE: TRANSCRIPTIONAL REGULATOR, BIOTIN REPRESSOR FAMIL
  54:  2hye-C  3.7  2.0   49   719    8   MOLECULE: DNA DAMAGE-BINDING PROTEIN 1;
  55:  3o2p-E  3.7  2.2   51    86    6   MOLECULE: DEFECTIVE IN CULLIN NEDDYLATION PROTEIN 1;
  56:  3h20-A  3.7  3.0   59   288    7   MOLECULE: REPLICATION PROTEIN B;
  57:  6rfl-I  3.7  2.6   50   773    4   MOLECULE: DNA-DEPENDENT RNA POLYMERASE SUBUNIT RPO132;
  58:  5dcm-B  3.7  3.0   52    98    8   MOLECULE: PHOB FAMILY TRANSCRIPTIONAL REGULATOR;
  59:  8amy-A  3.6  2.7   49    85    6   MOLECULE: MU8.1 CONOTOXIN;
  60:  5nl9-A  3.6  2.9   52   146   13   MOLECULE: TRANSCRIPTIONAL REGULATOR (FUR FAMILY);
  61:  6jx3-B  3.6  3.8   51    85    4   MOLECULE: TFUB1;
  62:  6vqx-A  3.6  2.6   61    97   10   MOLECULE: ACRF6;
  63:  3sqn-A  3.6  2.7   54   468    7   MOLECULE: CONSERVED DOMAIN PROTEIN;
  64:  7vwv-B  3.6  2.7   50    74   12   MOLECULE: I73R;
  65:  5u30-A  3.6  2.9   50  1085    6   MOLECULE: CRISPR-ASSOCIATED ENDONUCLEASE C2C1;
  66:  7qvg-T  3.6  3.2   51   267    6   MOLECULE: 26S PROTEASOME REGULATORY SUBUNIT 7;
  67:  6dk4-A  3.6  2.2   47   135   17   MOLECULE: FERRIC UPTAKE REGULATION PROTEIN;
  68:  8iue-P  3.6  2.4   52   303   10   MOLECULE: DNA-DIRECTED RNA POLYMERASE III SUBUNIT RPC1;
  69:  1m12-A  3.6  2.5   53    84    8   MOLECULE: SAPOSIN C;
  70:  8bpz-A  3.6  2.2   50   322    8   MOLECULE: SPBETA PROPHAGE-DERIVED UNCHARACTERIZED PROTEIN Y
  71:  8xjg-A  3.6  2.8   50   148    8   MOLECULE: YQEY;
  72:  2rre-A  3.6  2.2   51    74    6   MOLECULE: PUTATIVE UNCHARACTERIZED PROTEIN;
  73:  5nwt-M  3.6  3.4   56   431    7   MOLECULE: DNA-DIRECTED RNA POLYMERASE SUBUNIT ALPHA;
  74:  1xma-B  3.6  2.4   49   103   10   MOLECULE: PREDICTED TRANSCRIPTIONAL REGULATOR;
  75:  3aly-A  3.5  2.6   46    58    9   MOLECULE: 50S RIBOSOMAL PROTEIN P1 (L12P);
  76:  7apd-B  3.5  2.3   49   289    8   MOLECULE: REPLICATION PROTEIN E1;
  77:  2z99-A  3.5  2.3   46   160    7   MOLECULE: PUTATIVE UNCHARACTERIZED PROTEIN;
  78:  7sol-A  3.5  2.7   62  1011    3   MOLECULE: UBIQUITIN-LIKE MODIFIER-ACTIVATING ENZYME 6;
  79:  1q1v-A  3.5  2.6   50    70    6   MOLECULE: DEK PROTEIN;
  80:  1z0x-A  3.5  2.6   48   216    8   MOLECULE: TRANSCRIPTIONAL REGULATOR, TETR FAMILY;
  81:  6c2j-A  3.5  3.4   52   265   10   MOLECULE: RNAP1;
  82:  5tjj-A  3.5  2.7   45   254    9   MOLECULE: TRANSCRIPTIONAL REGULATOR, ICLR FAMILY;
  83:  2lnb-A  3.5  2.9   48    80   13   MOLECULE: Z-DNA-BINDING PROTEIN 1;
  84:  8b9x-A  3.5  2.5   43   256    7   MOLECULE: DDRGK DOMAIN-CONTAINING PROTEIN 1,E3 UFM1-PROTEIN
  85:  7o4i-R  3.5  2.0   44   268   11   MOLECULE: GENERAL TRANSCRIPTION AND DNA REPAIR FACTOR IIH H

0001 s001A   -GGLE-ERFK-L-T-A---G--T--EE-GV-W-DFVRTH-LK-QL-P----AER---QNYLLFDRMVAF-HVQ--R--G---VT-V--P--M--S-A-AEFYVG-LA-QRF-P---------------
0002 5a31N   eqppfgvyavilevpedieE--E--LL-LF-W-TYIQAM-LT-NL------eSL---SLDRIYNMLRMFvVTG--P--A---LA-E-----I--D-L-QELQGY-LQ-KKV-Rdqqlvysagvyrlp-
0003 1lddA   ---------------------lT--LQ-RS-L-PFIEGM-LT-NL------gAM---KLHKIHSFLKIT-VP---K--D---WG-Y-nR--I--T-L-QQLEGY-LN-TLA-Degrlkyiangsyeiv
0004 9bw1M   qkygcgyigllP-E-D---T-----LA-IM-E-KFILEDyET-LK-------QK---RMWEVHAALVRA-CEE--A--G---VI-A-----P--S-Y-KAFTKE-VQ-RRT-Gyeqtkkrqgrraayq
0005 1hlvA   ---------------------sP--YD-KL-E-GLLIAW-FQ-QI-R--aglpV---KGIILKEKALRI-AEE--L--G---MDdF--T--A--S---NGWLDR-FR-RRR-S---------------
0006 2nvuA   dDRCI-N-IT-K-Q-T------P--SF-WI-L-ARALKE-FV-AK----ekakK---DAAAVGNHVAKL-LQS--I--G---QA-P-eS--I--S-E-KELKLL-CS-NSA-Flrvvrcrslaeeygl
0007 9kbdA   vaqfvglelyklkdgedlvykdR--KL-LI-Q-AAIVRI-MK-MR------kVL---KHQQLLGEVLTQ-LS------S---RF-K-----P--R-V-PVIKKC-ID-ILI-Ekeylervdgekdtys
0008 3mq0B   -----------S-D-T---V--P--AL-RR-A-VRILDL-VA-GS-P----rDL---TAAELTRFLD------------------------L--P-K-SSAHGL-LA-VMT-Eldllarsadgtlrig
0009 3c1dB   ---------------------pA--YA-RL-L-DRAVRI-LA-VR-------dH---SEQELRRKLAA--PIM--G-gP---EE-I--D--A--T-A-EDYERV-IA-WCH-Ehgylddsrfvarfia
0010 3fhzD   apvaG-PEV------A---A--N--RA-GR-Q-ARIVAI-LS-SA------QVR---SQNELAALLAAE-G------------------ieV--T-Q-ATLSRD-LE-ELG-Avklrgadggtgiyvv
0011 6lb9A   dsRLA-EAA----H-S---S--FaswA-PS-W-YVAFHL-A--PF-S----R-F---TLADLTQVIVRH-VNL-sF--P---EG-P--V-eA--S-I-AKDVDC-IT-KMY-Vpaqrlrggedllscp
0012 8j56C   ---------------------wL--PN-IH-S-SLFFSAyQF-MV-Q--tvgiR---AVVAAYRLYLEH-VSL--L--G---GE-I--V--L--S-F-TRAWTL-VR-FFE-Snmlqlsrctccggqf
0013 8ekiD   aFKPS--NIL-E-Y-R---D--------CP-L-DIISNL-LE-LN-P---rlyK---DLPTTKSLLFGI-YDS--L--S---IN---gqtgK--V-E-VDLMVL-HI-DYA-Lvnldfgtayelgkqv
0014 2od5A   ---------------------xK--TV-RI-R-EKIKKF-LG-DR-------PR---NTAEILEHINST-XR-----------------hgT--T-S-QQLGNV-LS-KDK-Divkvgyikrsgilsg
0015 8cliA   dlsrqyQREK-----------yR--LL-KR-R-NLIIEA-VT-NL-R----LIE---SLFTIQKMIMDQ-EKQ--E--G---VS-----tkC--C-K-KSIVRL-VR-NLS-Eegllrlyrttviqdg
0016 5of3C   ySWIE-KV---mglqD---S--R--KR-FI-LyVASRYLvNV-KG--------V---NEDEALQTLKEF-YYK--L--Q---------sgkV--Y-E-SWLKSV-IN-GVK-Kkgllpwslkrieerd
0017 7z0oG   ----------------------E--LN-DY-S-TMIDIL-LS-DM-D---letV---TTKKVRMALKEV-YA-------------I--D--V--E-G-KAINKL-IR-KHL-Dlvkerprfersledl
```

```
0018 4fcyB    ---------------------eF--DE-DA-W-QFLIAD-YL-RP-E-----kP---AFRKCYERLELA-ARE---------HG-W--S--I-------PSRAT-AF-RRI-Qqldeamvvacregeh
0019 8s0cA    ---------------------sV--LE-QS-F-LRAILAeFR-RS-G---leeA---TFQQIYSQHVAL-CRM--E--G---LP-Y-----P--T-M-SETMAV-CS-HLG-----------------
0020 2dobA    -------gsL-P-C-D------I--CK-DV-V-TAAGDX-LK-DN--------A---TEEEILVYLEKT-CDW--LpxS---AS-C-eI--V--D-S--YLPVI-LD-IIK-Gexsrpgevcsalnlc
0021 2rkhA    ---------------------tP--KE-AV-R-LCALGT-IA-SQ-------pX---RYSELAGSVRHF-TSRixG--P---SLeL--X--G--I-----SIEL-LR-GLV-Eavdaxlaisaagrre
0022 3v4gA    ----------------------K--QD-NL-V-RAFKAL-LK-EE------rFG---SQGEIVEALKQE-G-------------fE--N--I--N-Q-SKVSRM-LT-KFG-Avrtrnakmemvyclp
0023 6zvhy    --------------------------kfNW-K-GTIKAI-LK-QA-P---dneI---TIKKLRKKVLAQ-YYT--V--T--------rseeE--L-L-VIFNKK-IS-PTF-Kllkdkvklvk-----
0024 6v9iC    ykknhsgrklhnevgqydqL--R--IL-RT-Q-EAIIQI-MK-MR------kKI---SNAQLQTELVEI-LK------N---MF-L-----P--Q-K-KMIKEQ-IE-WLI-Ehkyirrdesdintfi
0025 6qfdB    ------------------------dlT-AF-Q-KNILTV-LG-EE-------AR---YGLAIKRELEEY-YGE--------------------vN-H-GRLYPN-LD-DLV-Nkglvekseldkrtne
0026 6zcaD    -MGIK--QYS-QeE-L---K--E--MA--L-V-EIAHEL-FEeHK-K----PV----PFQELLNEIASL-LGV-----------------kkedR-I-AQFYTD-LN-IDG-Rflalsdqtwglrswy
0027 7z8bC    ffdvflrylcqhraskltwpvkR--RN-LL-N-CLIVRI-LK-AH-G---deGL---HIDQLVCLVLEA-WQK--G--PcpprglvssA--C--S-S-TDVLSC-IL-HLL-Gkgtlrrhddrpqvls
0028 7pikC    inevvlfdndpmqisadkgT--K--VTpEI-E-RLFRLT-IE-KH----nqkgT---KTTVAYRRFVDL-FAQ--Y--F---P----dY--P--T-L-RQFRYF-YD-REY-Pkalgpgsryeidati
0029 2qbyB    ---------------------imialP-FH-Y-KLALRS-LI-E--------SE---DVMSAHKMYTDL-CNK--F--K---QK-P-----L--S-Y-RRFSDI-IS-ELD-Mfgivkiriinrgrag
0030 1d3yB    ---------------------aK--IF-AQ-T-TKMLEF-AK-QL-L---ddFS---TLREAYYVSKNW-GE----------------arfD--D-Q-QASNNV-IE-DLE-Aalgvlrehlgfipee
0031 1cf7B    ---------------G---K--G--LR-HF-S-MKVCEK-VQ-RK-G-----tT---SYNEVADELVSE-FTN--S--N-----nhlydqkN--I-R-RRVYDA-LN-VLM-Amniiskekkeikwig
0032 6zfbd    -MGIK-QYsQ-E-E-L---K--E---M-AL-V-EIAHEL-FEeHK-K----PV----PFQELLNEIASL-LGV-----------------kkedR-I-AQFYTD-LN-IDG-Rflalsdqtwglrswy
0033 7oqeK    ---------------------eA--EK-EF-I-TMLKEN-Q--------vdstW---SFSRIISELGTR-DP--------rywM-V--D--D--D-P-LWKKEM-FE-KYL-Snrkfkeafqkmlqnn
0034 3e3vA    ---------------------dD--IS-KG-Y-NAALNY-LS-YQ-------lR---TRKEVEDKLRSL-D--------------------I--H-E-DYISEI-IN-KLI-Dldlindknyaesyvr
0035 7kbfK    -------------------------hP-PT-L-SMVVEV-LK-KN-T---rKGT---SVQAIRTRILSA-H-----------PT-V--DplR--L-K-FLLRTA-LN-KGL-Ekgilirplnssatga
0036 3s63B    ----ltpkeT-C-D-L---------CQ-IA-L-RTVFGH-FG-GN-------iP---SRRKLVHQLKHE-CKR--H--F-nyRR-R--CllL--M-K-VNSDLI-FR-EMT-Dgsfkpmevclimrec
0037 8ea3W    ------------------eF--W--EN-FI-T-KTYKEG--N-KG-S----krM---TPKQVALRVEAK-ARE--L--K---DS---kP--P--N-Y-KTVLRV-LA-PIL-Ekqqkaksirspgwrg
0038 5b7iC    vARSV-IEAA-rfI-Q---S--WeqVL-AA-A-GFAARL-HE-GL-QatfadwG---WWYRIANVXLAT-ASQ--N-vG---VT-W-------rV-H-GRLXAI-FQ-DKF-Kqryeeq---------
0039 1fnnA    ---------------------pL--HE-KLfL-LAIVRS-LK-IS-H---tpYI---TFGDAEESYKIV-CEE--Y--G---ER-P-----R--V-H-SQLWSY-LN-DLR-Ekgivetrqnttlisi
0040 1cf7A    ---------S-R-H-E---K--S--LG-LL-T-TKFVSL-LQ-EA-K---dgVL---DLKLAADTLA-----------------------vR--Q-K-R-RIYD-IT-NVL-Egigliekksknsiqw
0041 7dh7C    ------------------------rlT-PI-R-ANVLRL-IA-DA-G----KPV---KAYELLDWVREG-KG----------VG-A--D-----A-P-PTVYRA-LD-FLM-Angfvhklesvnafva
0042 5zi8A    ---amdpefM-R-E-F------Q--RA-AV-R-LHILHH-AA-DN-------EV---HGAWLTQELSRH-G--------------Y--R--V--S-P-GTLYPT-LH-RLE-Adgllvseqrvvdgra
0043 8as8E    ---------------------rR--AP-LV-L-SCLKTL-FE-KS-H----dGI---PLEEAIQSLSSI-LIE--H--V---sqeqynpflQ--A-S-RELREW-IK-RLI-Verdgrifatdaleva
0044 1dp7P    -----------------------------T-V-QWLLDN-YE-------egvsL---PRSTLYNHYLLH-SQE--Q--K---LE-P-----V--N-A-ASFGKL-IR-SVF-Mglrtrrlgtrgnsky
0045 2k85A    ---------------------tA--KD-KY-E-WLVSRI-VK-N-------hnE---NWLSVSRKMQAS-P------------eyQ--D--Y--VyT-QKAKKL-FL-QHI-Hrlkhe----------
0046 3al0B    kiakeifpevfgltqI---N--D--EK-LI-E-ELVKKA-ME-QN-P----KAVqdkAAGFFVGYVMRE-TK-----------G-K--A--N--P---ELTNRI-IQ-KLL-Ege-------------
0047 6tdaL    wFDIS-KiheirfktP---K--A--YK-DT-R-NFIINT-YR-LSpY----EYL---TITAVRRNVA------------------------M--D-V-ASIVKI-HA-FLE-Kwglinyqidprtkps
0048 8yb6K    ---------------N---R--G--TV-DF-I-ASLEN--LK-EG----ilrklrglPGFDLFSALWWP-LRQ--K------NQ-R--A--P--K-R-EVAWLI-AK-LFA-Efrfeqregatlpilm
0049 8u8mA    ------------------------iSD-SF-E-KAMWNH-VS-QN-A---gKFL---MTEQFWAELLQS-QP------N---IP-I-------kS-A-HIVLHH-FN-EMM-Lenlwkqamdpdaklq
0050 3elkA    ------------------rI--L--HG-LI-T-LYILKE-LV-KR-------pX---HGYELQKSXFET-TG-----------------qaL--P-Q-GSIYIL-LK-TXK-Ergfvisessvnkgqq
0051 8yztC    iPSDM-LHIS-T-N-C---R--T-aEK-MA-L-TLLDYL-FHrevqavs-kKQL---D-PLTIYGIRCH-LFY--K--F-------------gsD-W-YRIKQS-ID-SKC-Rtawrrkqrg------
0052 8jxkD    -------------------------EF-ML-E-LAILGL-LI-ES-------PM---HGYELRKRLTGL-LGA--------------------fS-Y-GSLYPA-LR-RMQ-Adgliaenaapagrrv
0053 3k69A    ------------------xK--L--DF-SV-A-VHSILY-LD-AHrD----SKV---ASRELAQSLH--------------------------lN-P-VXIRNI-LS-VLH-Khgyltgtvgknggyq
0054 1j5yA    -------------X-K---T--V--RQ-ER-L-KSIVRI-LE-RS-K----ePV---SGAQLAEELS--------------------------vS-R-QVIVQD-IA-YLR-Slgynivatprgyvla
0055 2hyeC    klheavravqsavenlcsslldR--QY-QI-D-AAIVRI-MK-MR-K-----tL---GHNLLVSELYNQ-L------------K-F--P--V--K-P-GDLKKR-IE-SLI-Drdymerdkdnpnqyh
0056 3o2pE    ---------------------eR--QI-FL-E-ACIVRI-MK-AK------rNL---PHTTLVNECIAQ-SH------Q---RF-N-----A--K-V-SMVKRA-ID-SLI-Qkgylqrgddgesyay
0057 3h20A    rSEMA-GLVK-R-F-G--dD--L--SK-CD-F-IAAQKL-AS-RG--------R---SAEEIGKAMAEA--------------S-P--A--LahE-A-DYIERT-VS-KVM-Glpsvqlaraelarap
0058 6rflI    ------------------eT--I--LI-EI-I-PKIKAY-LL-DA-N---iSPK---SYDDFISRNK-N-IFV--------------nvstI--T-E-EDIRLL-YT-TIE-Qnidaddqtlvaifsy
0059 5dcmB    tLTRE-------------------ilS-PT-E-NKILSI-LL-MHpK----QVV---SKESLLEKLWEN-D--------------S--F--I--D-Q-NTLNVN-MT-RLR-Kkivpigfdyihtvrg
0060 8amyA    ------------------hC--R--LF-EF-R-LCLLEC-MS-LT-------------ldHCYARCTTV-ITQ--I-hG---SD-T--N----dC-T-IFKTCY-YR-CYV-Lgktedhcwkgtatsv
0061 5nl9A    kDSYE-------------------nvT-VQ-R-LQMANL-LL-SK-P----QHL---TADQVFQLINEH-M--------------P--N--A--S-R-ATIFNN-LK-LFA-Ekgivnllelksgitl
0062 6jx3B    sVAQTdYGMV--------------qlN-DT-A-ALIVQR-LL-DG--------H---SPADVAQFLTS----------------eY--E--V--E-R-TDAERD-IA-ALV-Tslkengmalp-----
0063 6vqxA    iLTIE-QIIE-A-I-N-----sA--PE-IAgY-YAWDAA-TD-AL-E--dleqL---TEDDFVAHLE-V-LEE--R--G---AK-I--D--R--D----aAIAV-AL-QFQ-Aaavndlhs-------
0064 3sqnA    --XYS-XLKR-V-I-T---E--K--DL-LR-Q-IRLLEQ-LL-NV-P----QLT---AKRL-AAQIQ--------------------------tT-E-RTVFSD-LQ-YIR-Sqlpadwsietdssgi
0065 7vwvB    ---------------------mE--TQ-KL-I-SMVKEA-LE-KY-Q----yPL---TAKNIKVVIQKE-HN-------------V--V--L--P-T-GSINSI-LYsELF-Ekidktntiypplwir
0066 5u30A    rwvrxreagewdrtadvllR--G--SD-KV-F-EKWGKL-A----------pdA---PFDLYDAEIKNV-QRR--N--T---RR-F--G--S--H---DLFAKL-AE-EYQ-Alwredasfltryavy
0067 7qvgT    fMEGAynrvlsM-D-L---L--A--KT-VR-D-EIAGCS-EK-AY-D-----TL---SVNDARQLLL--------------------------sS-D-KDLLEY-IK-EEH-Peweikdgcvvfqksk
0068 6dk4A    ---------------------lK--AT-PQ-R-LCVLKI-LK-RH------eHP---NIDELYIEIKKE-YP------------------sI--S-L-ATVYKN-LN-TLQ-Eqglvveinvlnqktc
0069 8iueP    ---------------------eF--VE-VL-N-QQCFKF-LQ-SK-Aet-ssFA---SSHEVWKYICEL-GI----------SK-V--E--L--S-M-EDIETI-LN-TLI-Ydgkvemtiiaakegt
0070 1m12A    ---------------------eV--CE-FL-V-KEVTKL-ID-NN--------K---TEKEILDAFDKM-CSK--L--PkslSE-E--CqeV--V-D-TYGSSI-LS-ILL-Eevspelvcsmlhlcs
0071 8bpzA    nyqskavlrL-L-F-E---G-iV--RS-GH-L-YRAYQL-YK-EK-------gV---IDNSVRYQIIDD-FN-----------------lrvkrA-V-YSMQDYiNE-EEV-Nkyyaeelglk-----
0072 8xjgA    yeaagrqdladP-Q-P---L--T--EE-EV-A-ALIEKA-IA-ES-----------qDMGKVMGVLKPQ-IQ-----------G-R--A--D--M---GKVSGL-VR-AKL-----------------
0073 2rreA    ------------------------vDR-KL-K-QRVIQY-LS-SN-----gKYV---DTGILASDLQRL-YS------V---DY-G-rkrnA--F-R-IQVEKV-FS-IIS-Sekelkn---------
0074 5nwtM    feQGE-E-YX-K-P----sA--S--ST-AI-R-ALVKKL-IA-AE-N---aKPL---SDSKLTSXLSEQ-G----------------------vA-R-RTVAKY-RE-SLS-Ippsnqr---------
0075 1xmaB    ------------------dV--I--RG-YV-D-TIILSL-LI-EG-------dS---YGYEISKNIRIK-TDE--L-------------yvI--K-E-TTLYSA-FA-RLE-Kngyiksyygeetrrt
0076 3a1yA    --------------------------M-EY-V-YAALLL-HS-VG-------------keiNEENLKAV-LQA--A--G---VE-P--E--E--A-RiKALVAA-LE-GVN-Ideviekaa-------
0077 7apdB    ------------------kF--D--FG-TM-V-QWAYDH-K-----------yA---EESKIAYEYALA--AG--S--DsnaRA-F--L----nS-Q-AKHVKD-CA-TMV-Rhylraetqalsmpay
0078 2z99A    --------------aE---L--D--AD-EL-K-RVLEAL-LL-VI-D----TPV---TADALAAATE--------------------------qP-V-YRVAAK-LQ-LMA-Deltgrdsgidlrhts
0079 7solA    esLER-QLKH-P------pL--E--IH-TA-M-LALDQF-QE-KY-S--vgcqQ---DSEELLKLATSI-SET--L-eE---KP-D--V--N----A-DIVHWL-SW-TAQ-Gflsplaaavggvasq
0080 1q1vA    -----------klkkP---P--T--DE-EL-K-ETIKKL-LA-SA-N---leeV---TMKQICKKVYEN-YP------T---YD-L-tE--R--K-D--FIKT----tELI-Sleh------------
0081 1z0xA    ---------------K---L--S--KD-TI-I-AAAFSL-LE-KS------pTL---EQ-LSXRKVAKQ-LG---------vqapA--I--Y--W--yfKNKQA-LL-QSX-Aeaieehfqepalcge
0082 6c2jA    pRMRQ-EFEE-----------sgiiDY-KF-G-IDAMVQ-MA-LH------kRA---DLPTLVGTLR------------------H--H--C--K-A-QEVADN-LF-KMA-Sedcfnfdptidkfiv
0083 5tjjA    ------------xdyQ---V--P--SV-AL-A-ARVLKL-LS-RH-K---yrQS---TLTEIAERLG--------------------------vN-K-TTCLRV-LR-TLE-Redfvsydpqsrrysl
0084 2lnbA    ghhhhhhshM-A-D-P---G--R--EG-HL-E-QRILQV-LT-EA-G----sPV---KLAQLVKECQ--------------------------aP-K-RELNQV-LY-RMK-Kelkvsltspatwclg
0085 8b9xA    ------------gmtE---E--Q--SQ-SF-L-TEFINY-IK-QS------kVV---LLEDLASQVG--------------------------lR-T-QDTINR-IQ-DLLf----------------
0086 7o4iR    gkirinkdgskpheydlerM--P--KK-EI-L-DYLFKL-FD-EY------dYW---SLKGLKERTR------------------------Q--P-E-AHLKEC-LD-KVA-Tlvkkgpyafkytlrp

0001 s001A    -LLHH-HHHH-H-H-L---L--L--HH-HH-H-HHHHHH-HH-HL-L----LLL---LHHHHHHHHHHH-HHH--L--L---LL-L--L--L--L-H-HHHHHH-HH-HHL-L---------------
0002 5a31N    lllllllllllllllhhhhH--H--HH-HH-H-HHHHHH-HH-HH------lLL---LHHHHHHHHHHHlLLL--L--L---LL-L-----L--L-H-HHHHHH-HH-HHH-Hlllleeelleeell-
0003 1lddA    ---------------------hH--HH-HH-H-HHHHHH-HH-HH------lLE---EHHHHHHHHHHH-LL---H--H---HL-L-lL--L--L-H-HHHHHH-HH-HHH-Hllleellllleeeel
0004 9bw1M    hhlllhhhhhlL-H-H---H-----HH-HH-H-HHHHHHlLL-LL-------LL---LHHHHHHHHHHH-HHH--H--L---LL-L-----L--L-H-HHHHHH-HL-LLL-Lhhhhhhhllhhhhhh
0005 1hlvA    ---------------------lL--LH-HH-H-HHHHHH-HH-HH-H--hlllL---LHHHHHHHHHHH-HHH--H--L---LLlL--L--L--L---HHHHHH-HH-HHL-L---------------
0006 2nvuA    lHHHH-L-LL-L-L-L------L--HH-HH-H-HHHHHH-HH-HL----hhhhH---HHHHHHLLLHHH-HHH--H--L---LL-L-lL--L--L-H-HHHHHH-HH-LHH-Hleeellllhhhhhll
0007 9kbdA    llllllhhhhhhlllllllhhhH--HH-HH-H-HHHHHH-HH-HH------lLE---EHHHHHHHHHHH-HL------L---LL-L-----L--L-H-HHHHHH-HH-HHH-Llllllllllllleel
0008 3mq0B    -----------L-L-L---L--H--HH-HH-H-HHHHHH-HH-HL-L----lLE---EHHHHHLLLL------------------------L--L-H-HHHHHH-HH-HHH-Hllleeellleeeel
0009 3c1dB    ---------------------hH--HH-HH-H-HHHHHH-HL-LL-------lL---LHHHHHHHHLL--LEE--E-eE---EL-L--L--L--L-H-HHHHHH-HH-HHH-Hllllhhhhhhhhhh
0010 3fhzD    llllL-LEE------L---L--L--HH-HH-H-HHHHHH-HH-HL------LLL---LHHHHHHHHHLL-L------------------llL--L-H-HHHHHH-HH-HLL-Leeellllllllleeel
0011 6lb9A    llHHH-HHL----L-L---L--LlllL-HH-H-HHHHHL-L--LL-L----E-E---LHHHHHHHHHHH-HHH-hL--L---LL-L--L-hH--H-H-HHHHHH-HH-HHH-Llhhhllllllllllh
0012 8j56C    ---------------------hH--HH-HH-H-HHHHHHhHH-HH-H--lllhH---HHHHHHHHHHHH-HHH--H--L---LL-L--L--L--L-H-HHHHHH-HH-HHH-Llleeeeellllllee
0013 8ekiD    lLLLL--HHH-H-H-H---H--------LH-H-HHHHHH-HH-HL-L---lhhH---LHHHHHHHHHHH-HHH--L--L---LL---llhhH--H-H-HHHHHH-HH-HHH-Hhlllhhhhhhhhhhh
0014 2od5A    ---------------------hH--HH-HH-H-HHHHHH-LL-LL-------LE---EHHHHHHHHHLL-LL-----------------llL--L-H-HHHHHH-HH-HLL-Leeeeeeeeeeellle
0015 8cliA    hhhhhhHHHH-----------hH--HH-HH-H-HHHHHH-HH-HL-L----EEL---LHHHHHHHHHHH-HHH--H--L---LL-----llL--L-H-HHHHHH-HH-HHH-Hlllleeeeleeeell
0016 5of3C    lHHHH-HH---hlllL---L--H--HH-HH-HhLHHHHHhHL-LL--------L---LHHHHHHHHHHH-HLL--L--L---------lllL--L-H-HHHHHH-HH-HHH-Hhllllllhhhhhhhl
0017 7z0oG    ----------------------L--HH-HH-H-HHHHHH-HL-LL-L---lllL---LHHHHHHHHHHH-HL-------------L--L--L--L-H-HHHHHH-HH-HHH-Hhllllllllllhhhh
0018 4fcyB    ---------------------lL--LH-HH-H-HHHHHH-HL-LL-L-----lL---LHHHHHHHHHHH-HHH---------HL-L--L--L-------LLHHH-HH-HHH-Hhhllhhhhhhhllll
0019 8s0cA    ---------------------lH--HH-HH-H-HHHHHHhHH-HH-L---lllL---LHHHHHHHHHHH-HHH--H--L---LL-L-----L--L-H-HHHHHH-HH-HHH-----------------
0020 2dobA    -------llH-H-H-H------H--HH-HH-H-HHHHHH-HH-LL--------L---LHHHHHHHHHHH-HHH--LlhH---HH-H-hH--H--H-H--HHHHH-HH-HLL-Lllllhhhhhhhllll
0021 2rkhA    ---------------------lH--HH-HH-H-HHHHHH-HH-HL-------lE---EHHHHHHHHHHH-HHHhhL--L---LHhH--L--L--L-----LLHH-HH-LLE-Elllleeeelhhhhhh
0022 3v4gA    ----------------------L--HH-HH-H-HHHHHH-HH-HL------lLL---LHHHHHHHHHHL-L-------------lL--L--L--L-H-HHHHHH-HH-HLL-Leeeellllleeeell
0023 6zvhy    --------------------------llLH-H-HHHHHH-HH-LL-H---hheE---EHHHHHHHHHHH-HHH--H--L--------llhhH--H-H-HHHHHH-HH-LLE-Eellleeeell-----
0024 6v9iC    hhhhllleeeeelleeeehH--H--HH-HH-H-HHHHHH-HH-HL------lEE---EHHHHHHHHHHH-HL------L---LL-L-----L--L-H-HHHHHH-HH-HHH-Hllleeeelleeeeee
0025 6qfdB    ------------------------llL-HH-H-HHHHHH-HH-HL-------LE---EHHHHHHHHHHH-HLL--------------------lL-H-HHHHHH-HH-HHH-Hllleeeeeeelleee
0026 6zcaD    -LLLL--LLL-HhH-H---H--H--LL--H-H-HHHHHH-HHhHL-L----LL----LHHHHHHHHHHH-LLL-----------------lllhH-H-HHHHHH-HH-HLL-Leellllllleehhhl
0027 7z8bC    lhhhhhhhhlhhhhhhllllhH--HH-HH-H-HHHHHH-HH-LL-L---llLE---EHHHHHHHHHHH-HHL--L--LlllhhhhhlL--L--L-H-HHHHHH-HH-HHH-Hhlleeelllllleee
0028 7pikC    llleeeellleeellllllL--L--LLhHH-H-HHHHHH-HH-HH----llllL---LHHHHHHHHHHH-HHH--H--L---L----hL--L--L-H-HHHHHH-HH-HHL-Llllllllleeeeeee
0029 2qbyB    ---------------------eeellL-HH-H-HHHHHH-HH-L--------LL---LHHHHHHHHHHH-HHH--L--L---LL-L-----L--L-H-HHHHHH-HH-HHH-Hllleeeeeelllll
0030 1d3yB    ---------------------hH--HH-HH-H-HHHHHH-HH-HH-H---llLE---EHHHHHHHHHHL-HH----------------hllL--L-H-HHHHHH-HH-HHH-Hhhlllhhhhleelll
0031 1cf7B    ---------------L---L--H--HH-HH-H-HHHHHH-HH-HH-L-----eE---LHHHHHHHHHHH-HHL--L--L-----lllhhhhH--H-H-HHHHHH-HH-HHH-Hlllllllllleelll
0032 6zfbd    -LLLL-LLlH-H-H-H---H--H---L-LH-H-HHHHHH-HHhHL-L----LL----LHHHHHHHHHHL-LLL-----------------lllhH-H-HHHHHH-HH-HLL-Leellllllleehhhl
0033 7oqeK    ---------------------lH--HH-HH-H-HHHHHL-L--------llllL---LHHHHHHHHHHH-LL--------hhhH-L--L--L--L-H-HHHHHH-HH-HHH-Hlllhhhhhhhhhhhl
0034 3e3vA    ---------------------hH--HH-HH-H-HHHHHH-HH-LL-------lL---LHHHHHLLLHHH-L--------------------L--L-H-HHHHHH-HH-HHH-Hlllllhhhhhhhhhh
0035 7kbfK    -------------------------lL-LH-H-HHHHHH-HH-HL-L---lLLE---EHHHHHHHHHLL-L-----------LL-L--LllL--H-H-HHHHHH-HH-HHH-Hllleeeeelllllll
0036 3s63B    ----llhhhH-H-H-H---------HH-HH-H-HHHHHH-LL-LL-------lL---LHHHHHHHHHHH-HLL--L--L-llHH-H--HhhH--H-H-HLHHHH-HH-HHH-Lllllhhhhhhhllll
0037 8ea3W    ------------------hH--H--HH-HH-H-HHHHHH--L-LL-L----llL---LHHHHHHHHHHH-HHH--H--L---LL---lL--L--L-H-HHHHHH-LH-HHH-Hhhhhhllllllllll
```

```
0038 5b7iC   hLLLH-HHHH-hhH-H---H--HlhHH-HH-H-HHHHHH-HH-HH-HhhllllL---HHHHHHHHHHHH-HHH--H-hL---LL-L-------hH-H-HHHHHH-HH-HHH-Hhhllll---------
0039 1fnnA   ---------------------lH--HH-HHhH-HHHHHH-HH-HH-L---llLE---EHHHHHHHHHHH-HHH--L--L---LL-L-----L--L-H-HHHHHH-HH-HHH-Hlllleeeelleeeel
0040 1cf7A   ---------L-L-L-L---L--L--HH-HH-H-HHHHHH-HH-HL-L---llEE---EHHHHHHHLL-----------------------lL--L-L-H-HHHH-HH-HHH-Hhhlleeeeelleeee
0041 7dh7C   ------------------------llL-HH-H-HHHHHH-HH-HL-L----LLL---LHHHHHHHHHHH-HL----------LL-L--L-----L-H-HHHHHH-HH-HHH-Hllleeeehhhleeee
0042 5zi8A   ---lllhhhH-H-H-H------H--HH-HH-H-HHHHHH-HH-HL-------LL---LHHHHHHHHHHL-L--------------L--L--L--L-H-HHHHHH-HH-HHH-Hllleeeeeeeellee
0043 8as8E   ---------------------lL--HH-HH-H-HHHHHH-HH-HL-L----lLE---EHHHHHHHHHHH-HHH--L--L---llllllhhhH--H-H-HHHHHH-HH-LLE-Eeelleeeelhhhhhh
0044 1dp7P   -----------------------------L-H-HHHHHH-EE-------eeeeE---EHHHHHHHHHHH-HHH--L--L---LL-L-----L--L-H-HHHHHH-HH-HHL-Llleeeeelllllee
0045 2k85A   ---------------------hH--HH-HH-H-HHHHHH-LL-L-------llL---LHHHHHHHHLLL-H------------hhH--H--H--HhH-HHHHHH-HH-HHH-Hhhhhl----------
0046 3al0B   lhhhhlhhhhhllllL---L--L--LL-LH-H-HHHHHH-HH-LL-H----HHHhhlLLHHHHHHHHHL-LL-----------L-L--L--L--H---HHHHHH-HH-HHL-Lll-------------
0047 6tdaL   lLLLL-LllhhllllH---H--H--HH-HH-H-HHHHHH-HH-LLlL----LLL---LHHHHHHHLL------------------------L--L-H-HHHHHH-HH-HHH-Hllllllllllllll
0048 8yb6K   ---------------L---L--H--HH-HH-H-HHHHL--LL-HH----hhhllllhHHHHHHHHHHHH-HHH--H------LL-L--L--L--L-H-HHHHHH-HH-HHH-Hllleellllhhhhh
0049 8u8mA   ------------------------eLH-HH-H-HHHHHH-HH-HL-H---hHHH---LLHHHHHHHHHH-LL------L---LL-E-------eL-H-HHHHHH-HH-HHL-Hhhllllllllhhhhhh
0050 3elkA   ------------------hH--H--HH-HH-H-HHHHHH-HH-HL-------lE---EHHHHHHHHHHH-HL-----------------llL--L-L-LHHHHH-HH-HHH-Hhlleeeeeeellllle
0051 8yztC   lHHHH-HHHH-H-H-L---L--L-hHH-HH-H-HHHHHH-LLhhhhhle-lEEL---L-HHHHHHHHHH-HHH--H--H-------------lhH-H-HHHHHH-HH-HHH-Hhhhhhhlll------
0052 8jxkD   -------------------------LH-HH-H-HHHHHH-HH-HL-------LE---EHHHHHHHHHHH-LLL--------------------lL-L-LLHHHH-HH-HHH-Hllleellllllllle
0053 3k69A   ------------------lL--H--HH-HH-H-HHHHHH-HH-LLlL----LLL---LHHHHHHHHL--------------------------lL-H-HHLHHH-HH-HHH-Hlllleeellllleee
0054 1j5yA   -------------L-H---H--H--HH-HH-H-HHHHHH-HH-HL-L----lLL---LHHHHHHHHL--------------------------lL-H-HHHHHH-HH-HHH-Hhllllleeelleeell
0055 2hyeC   hhhhhhhllllhhhhhllhhhhH--HH-HH-H-HHHHHH-HH-HL-L-----eE---ELHHHHHHHHHH-L------------L-L--L--L--L-H-HHHHHH-HH-HHH-Hlllllllllllleee
0056 3o2pE   ---------------------hH--HH-HH-H-HHHHHH-HH-HH------lEE---EHHHHHHHHHHH-HL------L---LL-L-----L--L-H-HHHHHH-HH-HHH-Hllleellllleeee
0057 3h20A   hHHHH-HHHH-H-L-L--lL--H--HH-HH-H-HHHHHH-HH-LL--------L---LHHHHHHHHHHH--------------L-H--H--HhlH-H-HHHHHH-HH-HHH-Llhhhhhhhhhhhlll
0058 6rflI   ------------------hH--H--HH-HH-H-HHHHHH-HH-LL-L---lLLL---LHHHHHHHLL-H-HHH--------------llllL--L-H-HHHHHH-HH-HHH-Hlllllhhhhhhhhhh
0059 5dcmB   eELLL-------------------elL-HH-H-HHHHHH-HH-LLlL----LEE---EHHHHHHHHHLL-L--------------L--L--L--L-H-HHHHHH-HH-HHH-Hhhhhhlllleeeell
0060 8amyA   ------------------hH--L--HH-HH-H-HHHHHH-HH-LL-------------llLHHHHHHHH-HHH--H-hL---HH-H--H----hH-H-HHHHHH-HH-HHH-Llllhhhhhhhhhhhl
0061 5nl9A   hHHHH-------------------llL-HH-H-HHHHHH-HH-LL-L----LEE---LHHHHHHHHHHH-L--------------L--L--L--L-H-HHHHHH-HH-HHH-Hllleeeellllllee
0062 6jx3B   eEEELlLLEE--------------eeL-HH-H-HHHHHH-HH-LL--------L---LHHHHHHHHHHH---------------hL--L--L--L-H-HHHHHH-HH-HHH-Hhhhhlllllll-----
0063 6vqxA   lLLHH-HHHH-H-H-H-----lH--HH-HHhH-HHHHHH-HH-LL-L--lhhhL---LLHHHHHHHH-H-HHL--L--L---LL-L--L--H--H----hHHHH-HH-HHH-Hhhhhllll-------
0064 3sqnA   --LLH-HHHH-H-L-L---L--H--HH-HH-H-HHHHHH-HH-HL-L----LLL---LHHH-HHHHL--------------------------lL-H-HHHHHH-HH-HHH-Llllllleeeeeellee
0065 7vwvB   ---------------------lH--HH-HH-H-HHHHHH-HH-HL-L----lLL---LHHHHHHHHHHH-HL-------------L--L--L--L-H-HHHHHH-HHhLLE-Eellllllllllleee
0066 5u30A   hhhhhhhlllllllhhhhhlL--L--HH-HH-H-HHHHLL-L----------llL---LHHHHHHHHHHH-HHH--L--L---LL-L--L--L--H---HHHHHH-LL-LLL-Hhhhllllhhhhhhhh
0067 7qvgT   hHHLLhhhhhhH-H-H---H--H--HH-HH-H-HHHHHH-HH-HL-L-----EE---EHHHHHHHHL--------------------------lL-H-HHHHHH-HH-HHL-Llleelleeelllll
0068 6dk4A   ---------------------hH--HH-HH-H-HHHHHH-HL-LL------lLL---LHHHHHHHHHLL-LL------------------lL--L-H-HHHHHH-HH-HHH-Hllleeeellllllle
0069 8iueP   ---------------------hH--HH-HH-H-HHHHHH-HH-HH-Hhh-hhLE---EHHHHHHHHHHH-LL----------LL-L--L--L--L-H-HHHHHH-HH-HHH-Hllleeeeeelllll
0070 1m12A   ---------------------hH--HH-HH-H-HHHHHH-HH-LL--------L---LHHHHHHHLLLL-HHH--L--LlllHH-H--HhhH--H-H-HHHHHH-HH-HHL-Llllhhhhhhhlllll
0071 8bpzA   lhhhhhhhhH-H-H-H---L-hH--HH-HH-H-HHHHHH-HH-HH-------lL---LLHHHHHHHHHH-LL-----------------llllhH-H-HHHLLLlLH-HHH-Hhhlllllllll-----
0072 8xjgA   hhhlllhhhhhL-L-L---L--L--HH-HH-H-HHHHHH-HH-HH-----------hHHHHHHHHHHHH-HL-----------L-L--L--L--H---HHHHHH-HH-HHL-----------------
0073 2rreA   ------------------------lLH-HH-H-HHHHHH-HH-HL-----lLLL---LHHHHHHHHHHH-LL------L---LL-L-llhhH--H-H-HHHHHH-HH-HHH-Hhlllll---------
0074 5nwtM   llLLL-L-LL-L-L----lL--H--HH-HH-H-HHHHHH-HH-LL-L---lLLL---LHHHHHHHHHHH-L----------------------lL-H-HHHHHH-HH-HHL-Llllll---------
0075 1xmaB   ------------------hH--H--HH-HH-H-HHHHHH-HH-HL-------lE---EHHHHHHHHHHH-LLL--L-------------llL--L-H-HHHHHH-HH-HHH-Hllleeeeellllllle
0076 3alyA   --------------------------L-HH-H-HHHHHH-HH-LL------------lllLHHHHHHHH-HHH--L--L---LL-L--L--H--H-HhHHHHHH-HL-LLL-Hhhhhhhhl-------
0077 7apdB   ------------------lL--L--HH-HH-H-HHHHHH-L-----------lL---LHHHHHHHHHLL--LL--L--LhhhHH-H--H----lL-H-HHHHHH-HH-HHH-Hhhhhhhhhhlhhhh
0078 2z99A   --------------lL---L--L--HH-HH-H-HHHHHH-HH-HL-L----LLL---LHHHHHHHHL--------------------------lL-H-HHHHHH-HH-HHH-Hhhhhlllleeeeeel
0079 7solA   llHHH-HHHL-L------hH--H--HH-HH-H-HHHHHH-HH-HH-L--lllhH---HHHHHHHHHHHH-HHH--L-lL---LL-L--L--L----H-HHHHHH-HH-HLL-Lllhhhhhhhhhhhh
0080 1q1vA   -----------llllL---L--L--HH-HH-H-HHHHHH-HL-LL-L---hhhL---LHHHHHHHHHHH-LL------L---LL-L-lH--H--H-H--HHHH----hHHH-Hlll------------
0081 1z0xA   ---------------L---L--L--HH-HH-H-HHHHHH-HH-HL------lLH---HH-LLHHHHHHHH-HL---------llhhH--H--H--L--llLLHHH-HH-HHH-Hhhhhhllllllllll
0082 6c2jA   hHHHH-HHLL-----------llllLH-HH-H-HHHHHH-HH-HH------lEE---LHHHHHHHHH------------------H--H--L--L-H-HHHHHH-HH-HHH-Hllleeellllleeel
0083 5tjjA   ------------lllL---L--H--HH-HH-H-HHHHHH-HL-LH-H---hlLE---EHHHHHHHHL--------------------------lL-H-HHHHHH-HH-HHH-Hllleeellllleeee
0084 2lnbA   lhhhlllllL-L-L-L---L--H--HH-HH-H-HHHHHH-HH-HH-L----lLE---EHHHHHHHHL--------------------------lL-H-HHHHHH-HH-HHH-Hllleeeeelleeeel
0085 8b9xA   ------------llhH---H--H--HH-HH-H-HHHHHH-HH-HL------lEE---EHHHHHHHLL--------------------------lL-H-HHHHHH-HH-HHHl----------------
0086 7o4iR   eeeeeellllllleeeeelL--L--HH-HH-H-HHHHHH-HH-HL------lLE---EHHHHHHHHL------------------------L--L-H-HHHHHH-HH-HHE-Eeellllllleeeelh

# Type-3 BREX; MTase MBW2068239 HTH2 FoldSeek Top-Hits

job A   AF-A0A7V7ZHA6-F1-model v4 HTH luxR-type domain-containing protein        3.807E+00   LEAVRAGFKKAWQERDYATIIAVARKIPENVLQEDSKLLMWYDQALTRMG        Chloroflexia bacterium
job A   AF-A0A3A4PC77-F1-model v4 Uncharacterized protein                         3.130E+00   EAVRAGFKKAWQERDYATIIAVARKIPENVLQEDSKLLMWYDQALTRMG         Actinomycetia bacterium
job A   AF-A0A2U3AIG9-F1-model v4 Uncharacterized protein                         3.130E+00   FRLEAVRAGFKKAWQERDYATIIAVARKIPENVLQEDSKLLMWYDQALTRMGGK    Kurthia sibirica
job A   AF-A0A538U0Z4-F1-model v4 BTAD domain-containing protein                  3.807E+00   VRAGFKKAWQERDYATIIAVARKIPENVLQEDSKLLMWYDQALTRMGG          Candidatus Eisenbacteria
job A   AF-A0A2T5JU15-F1-model v4 Transcriptional regulator                       3.807E+00   LEAVRAGFKKAWQERDYATIIAVARKIPENVLQEDSKLLMWYDQAL            Nitrosospira sp. Nsp2
job A   AF-A0A7X9H9P0-F1-model v4 Uncharacterized protein                         3.130E+00   AVRAGFKKAWQERDYATIIAVARKIPENVLQEDSKLLMWYDQALTRMG          Bacteroidales bacterium
job A   AF-A0A535SKQ3-F1-model v4 TPR MalT domain-containing protein              3.341E+00   VRAGFKKAWQERDYATIIAVARKIPENVLQEDSKLLMWYDQALTRMG           Chloroflexi bacterium
job A   AF-A0A2N2BPY1-F1-model v4 HTH luxR-type domain-containing protein        3.807E+00   VRAGFKKAWQERDYATIIAVARKIPENVLQEDSKLLMWYDQALTRMG           Firmicutes bacterium HGW-4
job A   AF-X1E1S1-F1-model v4 Uncharacterized protein                             2.412E+00   LEAVRAGFKKAWQERDYATIIAVARKIPENVLQEDSKLLMWYDQALT           marine sediment metagenome
job A   AF-C0C6U7-F1-model v4 Tetratricopeptide repeat protein                    4.629E+00   VRAGFKKAWQERDYATIIAVARKIPENVLQEDSKLLMWYDQALTRMGG          [Clostridium] hylemonae DSM
job A   AF-A0A3B8VLG3-F1-model v4 TPR MalT domain-containing protein              6.008E+00   VRAGFKKAWQERDYATIIAVARKIPENVLQEDSKLLMWYDQALTRMG           Dehalococcoidia bacterium
job A   AF-A0A345UKZ8-F1-model v4 MCP methyltransferase, CheR-type                2.259E+00   QRKLKVF---RLEAVRAGFKKAWQERDYATIIAVARKIPENVLQEDSKLLMWYDQALTRMG   Candidatus Cyclonatronum
job A   AF-A0A6N6KE84-F1-model v4 AAA domain-containing protein                   3.566E+00   LEAVRAGFKKAWQERDYATIIAVARKIPENVLQEDSKLLMWYDQALTRMG        Bacteroidetes bacterium
job A   AF-A0A1B0CMM2-F1-model v4 Putative transcription factor six               2.259E+00   QRKLKVFRLEAVRAGFKKAWQERDYATIIAVARKIPE-NVLQEDSKLLMWYDQALTRMG     Lutzomyia longipalpis
job A   AF-A0A2V7HP17-F1-model v4 Uncharacterized protein                         4.629E+00   FRLEAVRAGFKKAWQERDYATIIAVARKIPENVLQEDSKLLMWYDQA           Candidatus Rokubacteria
job A   AF-A0A183Q8K0-F1-model v4 Uncharacterized protein                         1.858E+00   KVFRLEAVRAGFKKAWQERDYATIIAVARKIPENVLQE---DSKLLMWYDQALTRMG       Schistosoma mattheei
job A   AF-A0A2V7RC08-F1-model v4 Uncharacterized protein                         5.629E+00   QRKLKVFRL---EAVRAGFKKAWQERDYATIIAVARKIPENVLQEDSKLLMWYDQALTRMG   Gemmatimonadetes bacterium
job A   AF-A0A2N5ZC89-F1-model v4 Uncharacterized protein                         3.130E+00   EAVRAGFKKAWQERDYATIIAVARKIPENVLQEDSKLLMWYDQALTRMG         Marinilabiliales bacterium
job A   AF-A0A352UHV2-F1-model v4 Uncharacterized protein                         6.008E+00   VRAGFKKAWQERDYATIIAVARKIPENVLQEDSKLLMWYDQALTRMG           Dehalococcoidia bacterium
job A   AF-A0A6I7X4I9-F1-model v4 Uncharacterized protein                         2.574E+00   EAVRAGFKKAWQERDYATIIAVARKIPENVLQEDSKLLMWYDQAL             Methylococcaceae bacteriu
job A   AF-A0A7C4J6Y3-F1-model v4 HTH luxR-type domain-containing protein        5.274E+00   LEAVRAGFKKAWQERDYATIIAVARKIPENVLQEDSKLLMWYDQALTRMG        Chloroflexi bacterium
job A   AF-B8CZU7-F1-model v4 Regulatory protein LuxR                             6.008E+00   VRAGFKKAWQERDYATIIAVARKIPENVLQEDSKLLMWYDQALTRMG           Halothermothrix orenii H 168
job A   AF-A0A2G8L558-F1-model v4 Putative homeobox protein SIX4                  3.130E+00   LKVFRLEAVRAGFKKAWQERDYATIIAVARKIP-ENVLQEDSKLLMWYDQALTRMG  Apostichopus japonicus
job A   AF-A0A202E176-F1-model v4 Uncharacterized protein                         2.259E+00   EAVRAGFKKAWQERDYATIIAVARKIPENVLQEDSKLLMWYDQALTRMG         bacterium K02(2017)
job A   AF-A0A3B0UHN1-F1-model v4 Uncharacterized protein                         2.933E+00   VRAGFKKAWQERDYATIIAVARKIPENVLQEDSKLLMWYDQAL               hydrothermal vent metagenome
job A   AF-A0A3D1TRP5-F1-model v4 Uncharacterized protein                         4.941E+00   LEAVRAGFKKAW-QERDYATIIAVARKIPENVLQEDSKLLMWYDQALTRMG       Chloroflexi bacterium
job A   AF-A0A1I1QWZ7-F1-model v4 Soluble lytic murein transglycosylase           2.748E+00   KVFRLEAVRAGFKKAWQERDYATIIAVARKIPENVLQEDSKLLMWYDQALTRMG    Thiohalospira halophila DSM
job A   AF-A0A2N2BK61-F1-model v4 HTH luxR-type domain-containing protein        4.941E+00   VRAGFKKAWQERDYATIIAVARKIPENVLQEDSKLLMWYDQALTRMG           Firmicutes bacterium HGW-3
job A   AF-A0A2K9LKH4-F1-model v4 HTH luxR-type domain-containing protein        6.413E+00   EAVRAGFKKAWQERDYATIIAVARKIPENVLQEDSKLLMWYDQAL             Ketobacter alkanivorans
job A   AF-A0A2N2AXE2-F1-model v4 Uncharacterized protein                         5.274E+00   VRAGFKKAWQERDYATIIAVARKIPENVLQEDSKLLMWYDQALTRMGG          Firmicutes bacterium HGW-7
job A   AF-A0A328CYM2-F1-model v4 Uncharacterized protein                         5.629E+00   EAVRAGFKKAWQERDYATIIAVARKIPENVLQEDSKLLMWYDQALTRMG         Cuscuta australis
job A   AF-A0A5E4R7C1-F1-model v4 SIX1 SD domain-containing protein               3.566E+00   RKLKVFRLEAVRAGFKKAWQERDYATIIAVARKIPENVLQE-DSKLLMWYDQALTRMG  Leptidea sinapis
job A   AF-A0A1Y6CXP0-F1-model v4 Glycosyl transferase family 2                   5.274E+00   LEAVRAGFKKAWQERDYATIIAVARKIPENVLQEDSKLLMWYDQALTRMG        Methylomagnum ishizawai
job A   AF-A0A7Z8JWP3-F1-model v4 Helix-turn-helix transcriptional regulator      6.845E+00   VRAGFKKAWQERDYATIIAVARKIPENVLQEDSKLLMWYDQALTRMG           Cellulomonas hominis
job A   AF-A0A4Q3QPK3-F1-model v4 HTH luxR-type domain-containing protein        4.941E+00   LEAVRAGFKKAWQERDYATIIAVARKIPENVLQEDSKLLMWYDQALTRMG        Cytophagaceae bacterium
job A   AF-A0A402A047-F1-model v4 LuxR family transcriptional regulator          5.274E+00   LEAVRAGFKKAWQERDYATIIAVARKIPENVLQEDSKLLMWYDQALTRMG        Tengunoibacter tsumagoiensis
job A   AF-A0A6A7SGG8-F1-model v4 Serine/threonine-protein kinase PknK           6.413E+00   EAVRAGFKKAWQERDYATIIAVARKIPENVLQEDSKLLMWYDQAL             wastewater metagenome
job A   AF-A0A4Q5J8M4-F1-model v4 Helix-turn-helix transcriptional regulator      6.008E+00   VRAGFKKAWQERDYATIIAVARKIPENVLQEDSKLLMWYDQALTRMG           Nocardioides iriomotensis
job A   AF-A0A7Z8QAW1-F1-model v4 Uncharacterized protein                         1.858E+00   LKVFRLEA----VRAGFKKAWQERDYATIIAVARKIPENVLQEDSKLLMWYDQA-LTRMGGK  Methylococcaceae bacterium
job A   AF-A0A212F9Z5-F1-model v4 Homeobox domain-containing protein              3.807E+00   QRKLKVFRLEAVRAGFKKAWQERDYATIIAVARKIPENVL-QEDSKLLMWYDQALTRMG     Danaus plexippus plexippus
job A   AF-A0A816A7M8-F1-model v4 Hypothetical protein                            6.008E+00   LKVFRLEAVRAGFKKAWQERDYATIIAVARKIPENVLQEDSKLLMWYDQALTRMGG  Adineta steineri
job A   AF-A0A1Z5HE73-F1-model v4 Uncharacterized protein                         3.130E+00   VRAGFKKAWQERDYATIIAVARKIPENVLQEDSKLLMWYDQAL               Bathymodiolus platifrons
job A   AF-A0A0R3Q074-F1-model v4 Dolichyl-phosphate-mannose--protein mannosyltransferase   7.306E+00   EAVRAGFKKAWQERDYATIIAVARKIPENVLQEDSKLLMWYDQALTRMG   Angiostrongylus costaricensis
job A   AF-A0A2M9Q9I4-F1-model v4 SMI1 KNR4 domain-containing protein             5.629E+00   AVRAGFKKAWQERDYATIIAVARKIPENVLQEDSKLLMWYDQALTRMG          Lysinibacillus xylanilyticus
job A   AF-A0A7C7GAQ4-F1-model v4 Tetratricopeptide repeat protein                7.306E+00   VRAGFKKAWQERDYATIIAVARKIPENVLQEDSKLLMWYDQALTRMG           Dehalococcoidia bacterium
job A   AF-A0A1A7QMR6-F1-model v4 LuxR family maltose regulon positive regulatory protein   4.629E+00   LEAVRAGFKKAWQERDYATIIAVARKIPENVLQEDSKLLMWYDQALTRMG   Gelidibacter algens
job A   AF-A0A6L3EVV1-F1-model v4 LuxR family transcriptional regulator          6.845E+00   VRAGFKKAWQERDYATIIAVARKIPENVLQEDSKLLMWYDQALTRMG           Chloroflexi bacterium
job A   AF-A0A225MLE2-F1-model v4 HTH luxR-type domain-containing protein        7.799E+00   LEAVRAGFKKAWQERDYATIIAVARKIPENVLQEDSKLLMWYDQALT           Candidimonas nitroreducens
job A   AF-U6NQR7-F1-model v4 Transcription factor Six1                           2.933E+00   KVFRLEAVRAGFKKAWQERDYATIIAVARKIPENVLQE---DSKLLMWYDQALTRMG       Haemonchus contortus
job A   AF-Q0IEI2-F1-model v4 AAEL010327-PA                                       3.341E+00   QRKLKVFRLEAVRAGFKKAWQERDYATIIAVARKIPE-NVLQEDSKLLMWYDQALTRMG     Aedes aegypti
job A   AF-A0A1B0EZS4-F1-model v4 SIX1 SD domain-containing protein               3.566E+00   QRKLKVFRLEAVRAGFKKAWQERDYATIIAVARKIPE-NVLQEDSKLLMWYDQALTRMG     Phlebotomus papatasi
job A   AF-A0A1B6G3Q3-F1-model v4 Uncharacterized protein                         4.337E+00   LKVFRLEAVRAGFKKAWQERDYATIIAVARKIPENVLQEDSKLLMWYDQALTRMGGK  Cuerna arida
job A   AF-A0A496Z8W3-F1-model v4 Uncharacterized protein                         5.274E+00   EAVRAGFKKAWQERDYATIIAVARKIPENVLQEDSKLLMWYDQALTRMG         Deltaproteobacteria bacterium
job A   AF-X1RTA1-F1-model v4 Uncharacterized protein                             7.306E+00   EAVRAGFKKAWQERDYATIIAVARKIPENVLQEDSKLLMWYDQALTRMG         marine sediment metagenome
job A   AF-A0A3N5RDJ8-F1-model v4 Uncharacterized protein                         6.845E+00   VRAGFKKAWQERDYATIIAVARKIPENVLQEDSKLLMWYDQALTRMG           Chloroflexi bacterium
```

```
job A   AF-A0A1F9LJY2-F1-model v4 Uncharacterized protein                                          5.629E+00   VFRLEAVRAGFKKAWQERDYATIIAVARKIPENVLQE---DSKLLMWYDQALTRMG         Deltaproteobacteria bacterium
job A   AF-A0A352SD74-F1-model v4 Serine/threonine protein kinase                                  5.629E+00   VRAGFKKAWQERDYATIIAVARKIPENVLQEDSKLLMWYDQALT                     Cupriavidus sp.
job A   AF-A0A829YIW5-F1-model v4 LuxR family transcriptional regulator                            7.799E+00   RLEAVRAGFKKAWQERDYATIIAVARKIPENVLQEDSKLLMWYDQAL                  Steroidobacter agaridevorans
job A   AF-A0A1R1CYN4-F1-model v4 Uncharacterized protein                                          7.306E+00   RLEAVRAGFKKAWQERDYATIIAVARKIPENVLQEDSKLLMWYDQALTRMG              Paenibacillus sp. FSL H8-0259
job A   AF-A0A484KDR7-F1-model v4 Uncharacterized protein                                          6.845E+00   EAVRAGFKKAWQERDYATIIAVARKIPENVLQEDSKLLMWYDQALTRMG                Cuscuta campestris
job A   AF-A0A1V3N6V0-F1-model v4 Uncharacterized protein                                          6.008E+00   LEAVRAGFKKAWQERDYATIIAVARKIPENVLQEDSKLLMWYDQALT                  Thioalkalivibrio denitrificans
job A   AF-A0A7I0ZTV4-F1-model v4 homeobox protein six1a-like isoform X2                           4.337E+00   QRKLKVFRLEAVRAGFKKAWQERDYATIIAVARKIPENVL-QEDSKLLMWYDQALTRMG      Danaus plexippus plexippus
job A   AF-A0A536V4J1-F1-model v4 Protein-glutamate O-methyltransferase CheR                       2.748E+00   QRKLKVFRLEAVRAGFKKAWQERDYATIIAVARKIPENVLQEDSKLLMWYDQALTRMG       Betaproteobacteria bacterium
job A   AF-A0A7Z8VPL5-F1-model v4 Uncharacterized protein                                          2.574E+00   VRAGFKKAWQERDYATIIAVARKIPENVLQEDSKLLMWYDQA-LTRMGGK               Methylococcaceae bacterium
job A   AF-A0A329ZWU5-F1-model v4 Uncharacterized protein                                          3.341E+00   VRAGFKKAWQERDYATIIAVARKIPENVLQEDSKLLMWYDQALT                     Helicobacter sp. 15-1451
job A   AF-A0A7V4D6T6-F1-model v4 Tetratricopeptide repeat protein                                 6.413E+00   EAVRAGFKKAWQERDYATIIAVARKIPENVLQEDSKLLMWYDQALTRMG                Spirochaetes bacterium
job A   AF-A0A6N7M497-F1-model v4 BTAD domain-containing protein                                   6.008E+00   EAVRAGFKKAWQERDYATIIAVARKIPENVLQEDSKLLMWYDQAL                    Spirochaeta sp.
job A   AF-A0A1G8HSG4-F1-model v4 Non-specific serine/threonine protein kinase                     4.941E+00   VRAGFKKAWQERDYATIIAVARKIPENVLQEDSKLLMWYDQA--LTRMG                Rhodococcus triatomae
job A   AF-A0A3L8BXX2-F1-model v4 HTH luxR-type domain-containing protein                          9.483E+00   LEAVRAGFKKAWQERDYATIIAVARKIPENVLQEDSKLLMWYDQAL                   Ketobacter sp.
job A   AF-A0A1Z3HJV7-F1-model v4 HTH luxR-type domain-containing protein                          6.008E+00   VRAGFKKAWQERDYATIIAVARKIPENVLQEDSKLLMWYDQALTRMG                  Halomicronema hongdechloris
job A   AF-A0A7V3RIN7-F1-model v4 Tetratricopeptide repeat protein                                 5.274E+00   EAVRAGFKKAWQERDYATIIAVARKIPENVLQEDSKLLMWYDQALT                   candidate division WOR-3
job A   AF-A0A3B1BRW7-F1-model v4 Uncharacterized protein                                          4.337E+00   VRAGFKKAWQERDYATIIAVARKIPENVLQEDSKLLMWYDQALTRM                   hydrothermal vent metagenome
job A   AF-A0A0K9I8E1-F1-model v4 Uncharacterized protein                                          6.413E+00   LEAVRAGFKKAWQERDYATIIAVARKIPENVLQEDSKLLMWYDQAL                   Desulfocarbo indianensis
job A   AF-A0A6G0UZW5-F1-model v4 SIX1 SD domain-containing protein                                2.574E+00   FRLEAVRAGFKKAWQERDYATIIAVARKIPENVLQ---EDSKLLMWYDQALTRMG          Halicephalobus sp. NKZ332
job A   AF-A0A1I8I735-F1-model v4 SIX1 SD domain-containing protein                                3.566E+00   FRLEAVRAGFKKAWQERDYATIIAVARKIPE-NVLQEDSKLLMWYDQALTRMGGK          Macrostomum lignano
job A   AF-A0A838VI86-F1-model v4 Rhomboid family intramembrane serine protease                    7.799E+00   VRAGFKKAWQERDYATIIAVARKIPENVLQEDSKLLMWYDQALTRMG                  Nostocaceae cyanobacterium
job A   AF-A0A7R9CCC0-F1-model v4 Hypothetical protein                                             4.337E+00   KLKVFRLEAVRAGFKKAWQERDYATIIAVARKIPENVLQE-DSKLLMWYDQALTRMG        Timema cristinae
job A   AF-A0A2Z4FRJ3-F1-model v4 Six4/5                                                           4.629E+00   KVFRLEAVRAGFKKAWQERDYATIIAVARKIPENVLQEDS-KLLMWYDQALTRMG          Branchiostoma lanceolatum
job A   AF-A0A815KZC7-F1-model v4 Hypothetical protein                                             8.885E+00   LKVFRLEAVRAGFKKAWQERDYATIIAVARKIPENVLQEDSKLLMWYDQALTRMGG         Adineta steineri
job A   AF-A0A661U499-F1-model v4 Uncharacterized protein                                          7.306E+00   LEAVRAGFKKAWQERDYATIIAVARKIPENVLQEDSKLLMWYDQALTRMG               Candidatus Coatesbacteria
job A   AF-A0A022QS49-F1-model v4 AAA domain-containing protein                                    4.063E+00   QRKLKVFRLEAVRAGFKKAWQERDYATIIAVARKIPENVLQEDSKLLMWYDQALTRMG       Erythranthe guttata
job A   AF-A0A2P5P8R0-F1-model v4 Tetratricopeptide repeat protein                                 6.413E+00   VRAGFKKAWQERDYATIIAVARKIPENVLQEDSKLLMWYDQALTRMG                  Candidatus Dehalogenimonas
job A   AF-A0A358PYK4-F1-model v4 HTH luxR-type domain-containing protein                          9.483E+00   EAVRAGFKKAWQERDYATIIAVARKIPENVLQEDSKLLMWYDQALTRMG                Desulfosporosinus sp.
job A   AF-A0A3R7XB52-F1-model v4 Helix-turn-helix transcriptional regulator                       6.008E+00   EAVRAGFKKAWQERDYATIIAVARKIPENVLQEDSKLLMWYDQAL                    Alteromonadaceae bacterium
job A   AF-A0A328AZR3-F1-model v4 TIR domain-containing protein                                    5.274E+00   LKVFRLEAVRAGFKKAWQERDYATIIAVARKIPENVLQEDSKLLMWYDQAL              Phenylobacterium hankyongense
job A   AF-A0A518GGT5-F1-model v4 ASPIC and UnbV                                                   5.274E+00   KVFRLEAVRAGFKKAWQERDYATIIAVARKIPENVLQEDSKLLMWYDQALTRMG           Aureliella helgolandensis
job A   AF-A0A7I8VY92-F1-model v4 DgyrCDS9858                                                      4.337E+00   QRKLKVFRLEAVRAGFKKAWQERDYATIIAVARKIPENVLQEDS-KLLMWYDQALTRMG      Dimorphilus gyrociliatus
job A   AF-A0A661NSX6-F1-model v4 Uncharacterized protein                                          7.306E+00   EAVRAGFKKAWQERDYATIIAVARKIPENVLQEDSKLLMWYDQAL                    Deltaproteobacteria bacterium
job A   AF-A0A6J4I1X2-F1-model v4 Transcriptional activator of maltose regulon, MalT              7.306E+00   VRAGFKKAWQERDYATIIAVARKIPENVLQEDSKLLMWYDQALTRMG                  uncultured Chloroflexia
job A   AF-A0A3D1UWW4-F1-model v4 Uncharacterized protein                                          6.008E+00   VRAGFKKAWQERDYATIIAVARKIPENVLQEDSKLLMWYDQALTRMG                  Lentisphaeria bacterium
job A   AF-A0A1F8LSQ5-F1-model v4 Uncharacterized protein                                          8.885E+00   EAVRAGFKKAWQERDYATIIAVARKIPENVLQEDSKLLMWYDQALTRMG                Chloroflexi bacterium
job A   AF-A0A7C3LEI3-F1-model v4 HTH luxR-type domain-containing protein                          5.629E+00   VRAGFKKAWQERDYATIIAVARKIPENVLQEDSKLLMWYDQALT                     Anaerolineae bacterium
job A   AF-A0A4Y7B8F5-F1-model v4 HTH luxR-type domain-containing protein                          6.413E+00   LEAVRAGFKKAWQERDYATIIAVARKIPENVLQEDSKLLMWYDQALT                  Ramlibacter sp. WS9
job A   AF-A0A0V7ZLE3-F1-model v4 HTH luxR-type domain-containing protein                          8.324E+00   LEAVRAGFKKAWQERDYATIIAVARKIPENVLQEDSKLLMWYDQALTRMGGK             Mastigocoleus testarum BC008
job A   AF-A0A1M5QGM0-F1-model v4 ATP-, maltotriose-and DNA-dependent transcriptional regulator MalT  8.324E+00   VRAGFKKAWQERDYATIIAVARKIPENVLQEDSKLLMWYDQALTRMGGK            Thermosyntropha lipolytica
job A   AF-V2RJQ2-F1-model v4 Uncharacterized protein                                              7.306E+00   FRLEAVRAGFKKAWQERDYATIIAVARKIPENVLQEDSKLLMWYDQALT                Mucispirillum schaedleri
job A   AF-A0A7R8Z9Z2-F1-model v4 Hypothetical protein                                             5.629E+00   KVFRLEAVRAGFKKAWQERDYATIIAVARKIPENVLQE-DSKLLMWYDQALTRMG          Timema douglasi
job A   AF-A0A1J1IEB6-F1-model v4 CLUMA CG012018, isoform A                                        4.941E+00   VFRLEAVRAGFKKAWQERDYATIIAVARKIPENVLQ-EDSKLLMWYDQALTRMG           Clunio marinus
job A   AF-A0A3D0CXQ8-F1-model v4 TPR MalT domain-containing protein                               6.845E+00   VRAGFKKAWQERDYATIIAVARKIPENVLQEDSKLLMWYDQALTRMG                  Chloroflexi bacterium
job A   AF-A0A178MBV7-F1-model v4 Uncharacterized protein                                          4.941E+00   FRLEAVRAG-FKKAWQERDYATIIAVARKIPENVLQEDSKLLMWYDQALTRMGGK          Chloroflexus islandicus
job A   AF-A0A354TB23-F1-model v4 Uncharacterized protein                                          5.629E+00   VFRLE--AVRAGFKKAWQERDYATIIAVARKIPENVLQEDSKLLMWYDQALTRMGGK        Verrucomicrobiales bacterium
job A   AF-A0A497A7T7-F1-model v4 Uncharacterized protein                                          8.324E+00   LEAVRAGFKKAWQERDYATIIAVARKIPENVLQEDSKLLMWYDQALTRMG               Chloroflexi bacterium
job A   AF-A0A0F7GBT7-F1-model v4 BTAD domain-containing protein                                   7.799E+00   VRAGFKKAWQERDYATIIAVARKIPENVLQEDSKLLMWYDQALTRMG                  Dehalogenimonas sp. WBC-2
job A   AF-A0A3B0XSH3-F1-model v4 Uncharacterized protein                                          6.413E+00   EAVRAGFKKAWQERDYATIIAVARKIPENVLQEDSKLLMWYDQAL                    hydrothermal vent metagenome
job A   AF-A0A419DBD6-F1-model v4 BTAD domain-containing protein                                   9.483E+00   VRAGFKKAWQERDYATIIAVARKIPENVLQEDSKLLMWYDQALTRMG                  Dehalococcoidia bacterium
job A   AF-A0A850A478-F1-model v4 Tetratricopeptide repeat protein                                 6.845E+00   LEAVRAGFKKAWQERDYATIIAVARKIPENVLQEDSKLLMWYDQALTRMG               Caldilineales bacterium
job A   AF-A0A7V1FY58-F1-model v4 Uncharacterized protein                                          5.274E+00   FRLEAVRAGFKKAWQERDYATIIAVARKIPENVLQEDSKLLMWYDQALTRM              Phycisphaerales bacterium
job A   AF-A0A1A9V577-F1-model v4 Homeobox domain-containing protein                               4.941E+00   LKVFRLEAVRAGFKKAWQERDYATIIAVARKIPE-NVLQEDSKLLMWYDQALTRMG         Glossina austeni
job A   AF-A0A7Z9ZKJ6-F1-model v4 Tetratricopeptide repeat protein                                 7.306E+00   EAVRAGFKKAWQERDYATIIAVARKIPENVLQEDSKLLMWYDQALTRMG                Anaerolineae bacterium
job A   AF-A0A6P8YI81-F1-model v4 homeobox protein SIX1-like isoform X1                            6.413E+00   VFRLEAVRAGFKKAWQERDYATIIAVARKIPENVLQEDSKLLMWYDQALTRMG            Thrips palmi
job A   AF-A0A0N1IN86-F1-model v4 Homeobox protein SIX4                                            6.008E+00   VFRLEAVRAGFKKAWQERDYATIIAVARKIPENVL-QEDSKLLMWYDQALTRMG           Papilio xuthus
job A   AF-A0A3D0DB67-F1-model v4 TPR MalT domain-containing protein                               8.885E+00   LEAVRAGFKKAWQERDYATIIAVARKIPENVLQEDSKLLMWYDQALTRMG               Chloroflexi bacterium
job A   AF-A0A497C9T8-F1-model v4 Helix-turn-helix transcriptional regulator                       7.799E+00   LEAVRAGFKKAWQERDYATIIAVARKIPENVLQEDSKLLMWYDQALTRMGGK             Chloroflexi bacterium
job A   AF-A0A4P6K0U0-F1-model v4 Tetratricopeptide repeat protein                                 8.885E+00   LEAVRAGFKKAWQERDYATIIAVARKIPENVLQEDSKLLMWYDQALT                  Ktedonosporobacter rubrisoli
job A   AF-A0A354J8B5-F1-model v4 Uncharacterized protein                                          7.799E+00   EAVRAGFKKAWQERDYATIIAVARKIPENVLQEDSKLLMWYDQALTRMG                Lachnospiraceae bacterium
job A   AF-A0A840SF85-F1-model v4 Tetratricopeptide (TPR) repeat protein                           4.063E+00   VFRLEAVRAGFKKAWQERDYATIIAVARKIPENVLQE---DSKLLMWYDQALTRMG         Treponema rectale
job A   AF-A0A6J4KIA3-F1-model v4 HTH luxR-type domain-containing protein                          7.306E+00   VRAGFKKAWQERDYATIIAVARKIPENVLQEDSKLLMWYDQALT                     uncultured Chloroflexia
job A   AF-A0A1M7Y8L6-F1-model v4 Uncharacterized protein                                          5.629E+00   QRKLKVFRLEAVRAGFKKAWQERDYATIIAVARKIPENVLQE-DSKLLMWYDQALTRM       Desulfopila aestuarii
job A   AF-A0A317CP45-F1-model v4 Uncharacterized protein                                          4.941E+00   EAVRAGFKKAWQERDYATIIAVARKIPENVLQEDSKLLMWYDQALTRMG                Leucothrix pacifica
job A   AF-A0A662C7Z5-F1-model v4 HTH luxR-type domain-containing protein                          8.885E+00   LEAVRAGFKKAWQERDYATIIAVARKIPENVLQEDSKLLMWYDQALTRMG               Bacteroidetes bacterium
job A   AF-A0A3B9D0A3-F1-model v4 Uncharacterized protein                                          4.063E+00   LKVFRLEAVRAGFKKAWQERDYATIIAVARKIPENVLQEDSKLLMWYDQALTRMG          Planctomycetaceae bacterium
job A   AF-A0A7W9RUA0-F1-model v4 LuxR family maltose regulon positive regulatory protein          7.799E+00   LEAVRAGFKKAWQERDYATIIAVARKIPENVLQEDSKLLMWYDQALT                  Acidovorax konjaci
job A   AF-A0A1M3K8W3-F1-model v4 HTH luxR-type domain-containing protein                          6.413E+00   EAVRAGFKKAWQERDYATIIAVARKIPENVLQEDSKLLMWYDQALT                   Chloroflexi bacterium 44-23
job A   AF-A0A6J4JZV2-F1-model v4 Transcriptional activator of maltose regulon, MalT              8.324E+00   VRAGFKKAWQERDYATIIAVARKIPENVLQEDSKLLMWYDQALT                     uncultured Chloroflexia
job A   AF-A0A2G9TDS3-F1-model v4 DUF4071 domain-containing protein                                6.413E+00   VFRLEAVRAGFKKAWQERDYATIIAVARK---IPENVLQEDSKLLMWYDQALT            Teladorsagia circumcincta
job A   AF-A0A369B873-F1-model v4 Tetratricopeptide repeat protein                                 9.483E+00   FRLEAVRAGFKKAWQERDYATIIAVARKIPENVLQEDSKLLMWYDQALTRMG             Fontibacillus phaseoli
job A   AF-A0A3N5I8U9-F1-model v4 Uncharacterized protein                                          8.324E+00   VRAGFKKAWQERDYATIIAVARKIPENVLQEDSKLLMWYDQAL                      Desulfobacteraceae bacterium
job A   AF-A0A7V7XG44-F1-model v4 Tetratricopeptide repeat protein                                 8.324E+00   LEAVRAGFKKAWQERDYATIIAVARKIPENVLQEDSKLLMWYDQAL                   Anaerolineae bacterium
job A   AF-A0A1F8Q221-F1-model v4 HTH luxR-type domain-containing protein                          8.885E+00   EAVRAGFKKAWQERDYATIIAVARKIPENVLQEDSKLLMWYDQALTRMG                Chloroflexi bacterium
job A   AF-A0A0S8AW68-F1-model v4 HTH luxR-type domain-containing protein                          8.885E+00   LEAVRAGFKKAWQERDYATIIAVARKIPENVLQEDSKLLMWYDQAL                   Anaerolineae bacterium SG8 19
job A   AF-A0A5N8V799-F1-model v4 Tetratricopeptide repeat protein                                 9.483E+00   VRAGFKKAWQERDYATIIAVARKIPENVLQEDSKLLMWYDQALTRMGGK                Streptomyces adustus
job A   AF-A0A517Z1Q5-F1-model v4 ASPIC and UnbV                                                   3.566E+00   KVFRLEAVRAGFKKAWQERDYATIIAVARKIPENVLQEDS---KLLMWYDQALTRMG        Maioricimonas rarisocia
job A   AF-A0A0S8GJC0-F1-model v4 BTAD domain-containing protein                                   8.324E+00   EAVRAGFKKAWQERDYATIIAVARKIPENVLQEDSKLLMWYDQALTRMG                candidate division WOR 3
job A   AF-A0A7T7BC48-F1-model v4 HTH-type transcriptional regulator MalT                          7.799E+00   VRAGFKKAWQERDYATIIAVARKIPENVLQEDSKLLMWYDQAL                      Ralstonia pickettii
job A   AF-A0A4Z0WBJ0-F1-model v4 BTAD domain-containing protein                                   7.799E+00   LEAVRAGFKKAWQERDYATIIAVARKIPENVLQEDSKLLMWYDQAL                   Natronospirillum operosum
job A   AF-A0A0V8HL48-F1-model v4 Uncharacterized protein                                          9.483E+00   RLEAVRAGFKKAWQERDYATIIAVARKIPENVLQEDSKLLMWYDQALTRMGGK            Bacillus enclensis
job A   AF-A0A4U3KWY6-F1-model v4 DUF4071 domain-containing protein                                7.799E+00   LKVFRLEAVRAGFKKAWQERDYATIIAVARKIPENVLQEDSKLLMWYDQALTRMG          Ilyomonas limi
job A   AF-A0A437AS29-F1-model v4 SIX1 SD domain-containing protein                                7.306E+00   RKLKVFRLEAVRAGFKKAWQERDYATIIAVARKIPENVL-QEDSKLLMWYDQALTRMG       Chilo suppressalis
job A   AF-A0A382D4E2-F1-model v4 Uncharacterized protein                                          7.306E+00   KVF--RLEAVRAGFKKAWQERDYATIIAVARKIPENVLQEDS----KLLMWYDQALTRMG     marine metagenome
job A   AF-A0A5S3WVC1-F1-model v4 Uncharacterized protein                                          4.629E+00   LEAVRAGFKKAWQERDYATIIAVARKIPENVLQE-DSKLLMWYDQALTRMGGK            Pseudoalteromonas rubra
job A   AF-A0A818RWE8-F1-model v4 Hypothetical protein                                             6.413E+00   KVFRLEAVRAGFKKAWQERDYATIIAVARKIPE-NVLQEDSKLLMWYDQALTRMG          Adineta steineri
job A   AF-A0A7R9JQM7-F1-model v4 Hypothetical protein                                             6.413E+00   KVFRLEAVRAGFKKAWQERDYATIIAVARKIPENVLQE-DSKLLMWYDQALTRMG          Timema genevievae
job A   AF-A0A3D0PK83-F1-model v4 Uncharacterized protein                                          9.483E+00   EAVRAGFKKAWQERDYATIIAVARKIPENVLQEDSKLLMWYDQALTRMG                Lachnospiraceae bacterium
job A   AF-A0A0F9G5F1-F1-model v4 Uncharacterized protein                                          7.799E+00   VRAGFKKAWQERDYATIIAVARKIPENVLQEDSKLLMWYDQALTRMG                  marine sediment metagenome
job A   AF-A0A849HSF9-F1-model v4 Transglycosylase SLT domain-containing protein                   7.799E+00   VFRLEAVRAGFKKAWQERDYATIIAVARKIPENVLQEDSKLLMWYDQALTRMG            Legionellales bacterium
job A   AF-A0A317XH78-F1-model v4 F-box domain-containing protein                                  5.629E+00   LEAVRAGFKKAWQERDYATIIAVAR----KIPENVLQEDSKLLMWYDQALTRMG           Testicularia cyperi
job A   AF-A0A7Y7P3C5-F1-model v4 Tetratricopeptide repeat protein                                 6.413E+00   VFRLEAVRAGFKKAWQERDYATIIAVARKIPENVLQE--DSKLLMWYDQALTRMG          Bacteroidales bacterium
job A   AF-A0A4Q5VWA5-F1-model v4 TIR domain-containing protein                                    5.629E+00   VRAGFKKAWQERDYATIIAVARKIPENVLQEDS----KLLMWYDQALTRMGG             Chitinophagaceae bacterium
job A   AF-A0A7V5AL18-F1-model v4 Glycosyltransferase                                              8.885E+00   FRLEAVRAGFKKAWQERDYATIIAVARKIPENVLQEDSKLLMWYDQALTRMG             Syntrophobacterales
job A   AF-A0A536M2I7-F1-model v4 Tetratricopeptide repeat protein                                 5.274E+00   LEAVRAGFKKA-WQ---ERDYATIIAVARKIPENVLQEDSKLLMWYDQALTRMG           Chloroflexi bacterium

# Job: Type-3 BREX; MTase MBW2068239 HTH2
# Query: s001A
# No:  Chain   Z    rmsd lali nres  %id PDB  Description
   1:  8qgy-B  8.8  1.9   55   809   18   MOLECULE: MITOGEN-ACTIVATED PROTEIN KINASE KINASE KINASE 5;
   2:  6z0f-A  8.1  2.1   54   384    9   MOLECULE: ESX SECRETION SYSTEM PROTEIN YUKC;
   3:  8h38-H  6.8  2.4   56   177    9   MOLECULE: COP9 SIGNALOSOME COMPLEX SUBUNIT 1;
   4:  4n5c-D  6.7  2.2   52   757   12   MOLECULE: CARGO-TRANSPORT PROTEIN YPP1;
   5:  6xt9-D  6.5  2.4   54   396   13   MOLECULE: BARDET-BIEDL SYNDROME 1 PROTEIN;
   6:  2yhe-A  6.3  2.2   57   639   14   MOLECULE: SEC-ALKYL SULFATASE;
   7:  6vbu-8  6.2  2.4   49   475   12   MOLECULE: BARDET-BIEDL SYNDROME 18 PROTEIN;
   8:  5vxv-A  6.1  2.3   51   217   14   MOLECULE: PEROXISOMAL MEMBRANE PROTEIN PEX15;
```

```
  9: 5a63-D  6.1  2.2   54   100    9   MOLECULE: NICASTRIN;
 10: 2uy1-A  6.0  1.8   50   438   12   MOLECULE: CLEAVAGE STIMULATION FACTOR 77;
 11: 6z2w-E  6.0  2.4   51  2325   12   MOLECULE: DNA DAMAGE CHECKPOINT PROTEIN LCD1;
 12: 2c2l-A  5.9  2.3   52   281   12   MOLECULE: CARBOXY TERMINUS OF HSP70-INTERACTING PROTEIN;
 13: 8sxf-C  5.8  3.0   55   532    5   MOLECULE: PROBABLE CARBOXYL-TERMINAL PROTEASE;
 14: 8s4g-J  5.7  2.2   50   512    6   MOLECULE: ANAPHASE-PROMOTING COMPLEX SUBUNIT 10;
 15: 4rib-B  5.7  2.5   50   628   12   MOLECULE: FANCONI-ASSOCIATED NUCLEASE 1;
 16: 6sli-A  5.7  2.7   51   482   14   MOLECULE: LIPOPROTEIN RAGB;
 17: 2btp-A  5.7  2.4   58   248   10   MOLECULE: 14-3-3 PROTEIN TAU;
 18: 9lth-A  5.6  2.8   52   285    4   MOLECULE: TRANSCRIPTIONAL REGULATOR;
 19: 3l22-A  5.6  2.2   50   429    8   MOLECULE: SUSD SUPERFAMILY PROTEIN;
 20: 4cgu-A  5.6  2.5   51   112   12   MOLECULE: TPR REPEAT-CONTAINING PROTEIN ASSOCIATED WITH HSP
 21: 2grm-A  5.6  3.3   55   315    5   MOLECULE: PRGX;
 22: 6hs5-A  5.6  3.1   53   249   19   MOLECULE: TSSA;
 23: 2o8p-A  5.5  2.3   52   220    4   MOLECULE: 14-3-3 DOMAIN CONTAINING PROTEIN;
 24: 2vkj-A  5.5  3.1   56   106    9   MOLECULE: TM1634;
 25: 4m57-A  5.5  3.1   58   702    9   MOLECULE: CHLOROPLAST PENTATRICOPEPTIDE REPEAT PROTEIN 10;
 26: 5vbg-A  5.5  2.9   52   543    8   MOLECULE: PENICILLIN-BINDING PROTEIN ACTIVATOR LPOA;
 27: 8hmd-C  5.5  2.8   51  1334   12   MOLECULE: INTRAFLAGELLAR TRANSPORT PROTEIN 122 HOMOLOG;
 28: 5xw7-C  5.4  2.5   51   245    6   MOLECULE: CELLULOSE SYNTHASE SUBUNIT C;
 29: 8cbk-E  5.4  3.0   54   470    7   MOLECULE: 3-HYDROXYACYL-COA DEHYDROGENASE TYPE-2;
 30: 2pzi-A  5.3  2.8   52   654   10   MOLECULE: PROBABLE SERINE/THREONINE-PROTEIN KINASE PKNG;
 31: 8rg0-u  5.3  2.2   53   603    8   MOLECULE: EUKARYOTIC TRANSLATION INITIATION FACTOR 3 SUBUNI
 32: 7y4i-A  5.3  2.3   49   822    6   MOLECULE: PROBABLE UDP-N-ACETYLGLUCOSAMINE--PEPTIDE N-
 33: 7tj4-A  5.3  2.6   50   122   10   MOLECULE: ACTH;
 34: 8q9t-E  5.3  2.2   52  1060   10   MOLECULE: ANTIVIRAL HELICASE SKI2;
 35: 3qdn-A  5.3  2.2   53   285   13   MOLECULE: PUTATIVE THIOREDOXIN PROTEIN;
 36: 8ara-C  5.3  2.5   54   112   11   MOLECULE: CHAPERONE PROTEIN YSCY;
 37: 9dt9-A  5.3  2.9   52   112   15   MOLECULE: 231_C2_TRKA;
 38: 4it4-B  5.2  2.4   53   210    4   MOLECULE: CG17282;
 39: 7zty-A  5.2  3.1   50   345    6   MOLECULE: CNH DOMAIN-CONTAINING PROTEIN;
 40: 5alv-K  5.2  2.3   49  1125   10   MOLECULE: ADP-RIBOSYLATION FACTOR 1;
 41: 8rbk-A  5.2  2.6   53   739    2   MOLECULE: MGC151858 PROTEIN;
 42: 7dkh-I  5.2  2.2   51   914   14   MOLECULE: RNA POLYMERASE-ASSOCIATED PROTEIN CTR9;
 43: 1y8m-A  5.2  2.2   53   144    6   MOLECULE: FIS1;
 44: 7dhg-C  5.2  2.6   52   470   13   MOLECULE: MITOCHONDRIAL IMPORT RECEPTOR SUBUNIT TOM70;
 45: 8qt5-A  5.2  2.1   53   248   11   MOLECULE: 14-3-3-LIKE PROTEIN G-BOX FACTOR 14 LAMBDA,PROTEI
 46: 4ymr-A  5.1  2.0   47   134    6   MOLECULE: PROTEIN SNX21;
 47: 8g0l-B  5.1  2.8   51   925   12   MOLECULE: N-ALPHA-ACETYLTRANSFERASE 20;
 48: 6erp-A  5.1  2.6   52  1011   12   MOLECULE: TRANSCRIPTION FACTOR A, MITOCHONDRIAL;
 49: 5an3-B  5.1  2.1   51   136   14   MOLECULE: SGT1;
 50: 2mv3-A  5.1  2.9   51    88   20   MOLECULE: MITOCHONDRIAL INNER MEMBRANE I-AAA PROTEASE SUPER
 51: 3jq1-B  5.1  2.6   51   464   18   MOLECULE: SUSD SUPERFAMILY PROTEIN;
 52: 8vx7-A  5.1  2.2   53   161   17   MOLECULE: CID7;
 53: 6au8-A  5.1  2.9   54   280    9   MOLECULE: GOLGI TO ER TRAFFIC PROTEIN 4 HOMOLOG;
 54: 3u4t-A  5.1  2.3   48   258   10   MOLECULE: TPR REPEAT-CONTAINING PROTEIN;
 55: 6flt-f  5.1  3.0   56   929   11   MOLECULE: ARP1 ACTIN RELATED PROTEIN 1 HOMOLOG A;
 56: 2pl2-A  5.0  2.5   51   194   14   MOLECULE: HYPOTHETICAL CONSERVED PROTEIN TTC0263;
 57: 5wft-A  5.0  2.1   52   118   12   MOLECULE: PELB;
 58: 8gaa-A  5.0  3.5   54   197    2   MOLECULE: C6HR1_4R;
 59: 5y7q-A  5.0  2.8   51   546   14   MOLECULE: FANCONI-ASSOCIATED NUCLEASE 1 HOMOLOG;
 60: 2lxl-A  5.0  2.5   55   163    9   MOLECULE: VACUOLAR PROTEIN SORTING-ASSOCIATED PROTEIN VTA1
 61: 8ebf-A  5.0  2.2   54   667   17   MOLECULE: CYTOPLASMIC MEMBRANE PROTEIN;
 62: 2bb6-A  5.0  2.8   54   414    6   MOLECULE: TRANSCOBALAMIN II;
 63: 5cd6-A  5.0  2.7   50   575    8   MOLECULE: TPR-DOMAIN CONTAINING PROTEIN;
 64: 4fym-D  5.0  2.7   55   227    7   MOLECULE: OROTATE PHOSPHORIBOSYLTRANSFERASE;
 65: 2kat-A  5.0  2.6   53   115   11   MOLECULE: UNCHARACTERIZED PROTEIN;
 66: 1pc2-A  5.0  2.6   55   152    9   MOLECULE: MITOCHONDRIA FISSION PROTEIN;
 67: 5uls-A  5.0  3.0   59  1488   10   MOLECULE: SEPARIN;
 68: 2kck-A  5.0  2.5   53   112   13   MOLECULE: TPR REPEAT;
 69: 6yj6-A  5.0  2.5   55   819    4   MOLECULE: TRANSCRIPTION FACTOR TAU 131 KDA SUBUNIT;
 71: 2kc7-A  4.9  2.9   50    99    8   MOLECULE: BFR218_PROTEIN;
 72: 5wd8-A  4.9  2.2   50    90    6   MOLECULE: UNCHARACTERIZED PROTEIN;
 73: 8f5o-C  4.9  2.8   50  1179   12   MOLECULE: INTRAFLAGELLAR TRANSPORT PROTEIN 122B, PUTATIVE;
 74: 5efr-A  4.9  2.2   51   403   10   MOLECULE: BAMA-BAMD FUSION PROTEIN;
 75: 4eba-A  4.9  3.1   54   585    9   MOLECULE: MRNA 3'-END-PROCESSING PROTEIN RNA14;
 76: 9l4f-A  4.9  3.3   53  2383    9   MOLECULE: SERINE/THREONINE-PROTEIN KINASE ATR;
 77: 6tdy-J  4.9  2.8   51   170   10   MOLECULE: ATP SYNTHASE SUBUNIT ALPHA;
 78: 4bt9-B  4.9  2.8   53   238   17   MOLECULE: PROLYL 4-HYDROXYLASE SUBUNIT ALPHA-1;
 79: 4uzy-A  4.8  2.6   53   624    9   MOLECULE: FLAGELLAR ASSOCIATED PROTEIN;
 80: 8f5p-D  4.8  2.4   48   612   17   MOLECULE: NET DOMAIN-CONTAINING PROTEIN;
 81: 7t6d-A  4.8  3.0   53   389    8   MOLECULE: LIPOPOLYSACCHARIDE ASSEMBLY PROTEIN B;
 82: 3as5-A  4.8  2.5   51   183   12   MOLECULE: MAMA;
 83: 5w94-A  4.8  2.9   52   617    6   MOLECULE: MAU2 CHROMATID COHESION FACTOR HOMOLOG;
 84: 8h7g-C  4.8  2.6   47  3261    9   MOLECULE: TRANSFORMATION/TRANSCRIPTION DOMAIN-ASSOCIATED PR
 85: 8j0n-B  4.8  2.3   53   296    4   MOLECULE: ER MEMBRANE PROTEIN COMPLEX SUBUNIT 1;

0001 s001A   ----QRKL----K-----V--F---R---L--EAVRAG-FKKAW-QE---R------DYATIIA-VARK---I-----PE----NVL--QE----D-SKLLMWYDQALT-R-M--G--G--K--------------
0002 8qgyB   elarirqrvdniE-----V--L---T---A--DIVINL-LLSYR-DI---Q------DYDSIVK-LVET---L-----EK-lptfDL--AS----H-HHVKFHYAFALN-R-R--N--L--Pgdrakaldimipmv
0003 6z0fA   vpaliysmyalfF-----A--Q---P---K--HQAIVD-SNRAF-LN---K------QYSEVIS-TLSK---Y-----DA----ESL--------P-ESVQYQLATSYV-E-V--E--N--Lgsaktkniennlvt
0004 8h38H   dqcenqeleapgG-----I--A---T---P--PVYGQL-LALYL-LH---N------DMNNARY-LWKR---I-----PP----AIK--SA----N-SELGGIWSVGQR-I-W--Q--R--Dfpgiyttinahqws
0005 4n5cD   --------------------kA---L---R--ELQFKY-SYTLA-QQ---R------HIETAIK-TLES---L----iLS----KN---PN----Y-YKAWHLLALCRS---V--Q--E--Dkexsykivcsvlea
0006 6xt9D   ---------------tydP--T---N---Y--KAILAA-GSMMQ-TH---G------DFDVALT-KYRV---V-----AC----AV---PE----S-PPLWNNIGMCFF-G-K--K--K--Yvaaisclkranyla
0007 2yheA   pfppvEAG----K-----R-mG---G---A--DAVLKQ-MRAAI-DK---G------DYRWAVQ-LGNH---L----vFA----DP----A----N-KDARALQADAME-Q-L--G--Y--Qtenalwrnmymtga
0008 6vbu8   --------------------------------MEPLLL-AWSYF-RR---R------RFQLCAD-LCTQ---M----lEK----SP----C----D-QAAWILKARALT-E-M--V--Y--Vdeidvdeegiaemi
0009 5vxvA   --------------------------------SEVFQE-CVNLF-IK---R------DIKDCLE-KXSEvgfI-----DI----TVF--KS----N-PXILDLFVSACD-I-X--P--S--Ftklgltlqseilni
0010 5a63D   ------nL----E-----R--V---S---N--EEKLNL-CRKYY-LG---G--flpfLWLVNIF-WFFR---E---afLV----PA-------teQ-SQIKGYVWRSAV-G-F--L--F--Wvivltswitifqiy
0011 2uy1A   ---------------------S---S---P--SAIMEH-ARRLY-MS---K------DYRSLES-LFGR---C-----LK----KS----Y----N-LDLWMLYIEYVR-K-V--S-------------------
0012 6z2wE   hrifshfqppkltdggkfS--D---D---P--KTTTRM-LKSMY-DH---Q------LYSQIISnSSFH------ssdGK----IS-------pdV-KEWYSIGLEAAN-L-E--G--N--Vqtlknwveqieslr
0013 2c2lA   ------------------S--P---S---A--QELKEQ-GNRLF-VG---R------KYPEAAA-CYGR--aI-----TR----NP----L----V-AVYYTNRALCYL-K-M--Q--Qp---------------
0014 8sxfC   altlledN----S-----A--S--hE---V--APLLLR-SRLLQ-SM---K------RSDEALP-LLKA---G-----iK----EHP---D----D-KRVRLAYARLLV-E-Q--N--R--Lddakaefaglvqqf
0015 8s4gJ   ---------------yaeV--H---S---E--EALFLL-ATCYY-RS---G------KAYKAYR-LLKG---H------------SC--TT------PQCKYLLAKCCV-D-L--S--Kl--------------
0016 4ribB   --------------------yA---A---A--THMLSD-ISSAM-AN---G------NWEEAKE-LAQC-akR-----DW----N--------wiY-TRILSRFVEILQ-R-L--H--M--Yeeavrelesllsqr
0017 6sliA   ---------------vlyV--S---R---D--YAYALR-ARIYL-AL---G------EYGKAAA-DAKM---V----vDK----YP--gaadafsV-AEVYLILVESAL-Q-T--G--Dt--------------
0018 2btpA   sgvdLGTEnlyfQ-----S--M---E---K--TELIQK-AKLAE-QA---E------RYDDMAT-CMKA-vtE-----QG----A----EL----S-NEERNLLSVAYK-N-V--V--G--Grrsawrvissieqk
0019 9lthA   --------------------lE---K---E--FTDMEK-IKIYF-EK---N------NIVGLQN-LLTS---Y----sSQ----IN--vKE----K-IIYALISSLLGR-L-T--N--Ts--------------
0020 3l22A   --------------------aT---Q---A--AANXLK-XRVYX-AX---N------EWDKAIT-AGEL---V-----TG----YS-------frY-AETLLDLAECYA-N-K--A--Gg--------------
0021 4cguA   --------------------xS---Q---F--EKQKEQ-GNSLF-KQ---G------LYREAVH-CYDQ--lI-----TA----QP----Q----N-PVGYSNKAXALI-K-L--G--Ey--------------
0022 2grmA   siikPMYP----I-----V-----tI---Q--TVLKNA-LTISI-MN---R------NLKEAQY-YINQ--fE-----HL----K--------cyD-LEINYLKQIYQF-L-T--D--K--Nidsylnavniinif
0023 6hs5A   ---------------psgD--D---L---L-fSNEFDA-IQDAR-RYdkeA------DWGFVVD-HAGE---L----lRT----RT----K----D-LRLAVWLTEALA-L-E--D--G--Itgltegyalleglc
0024 2o8pA   --------------------xD---E---R--LLQKYR-AQVFE-WG---G------CFDKXFE-ALKS--lI-----YL----SE--seF----D-DEERHLLTLCIK-H-K--I--S--Dyrtxtsqvlqeqtk
0025 2vkjA   elyifSDY----E-------eK---K---A--RSLIAE-GKDLF-ET---A------NYGEALV-FFEK-alN-----LS----DN--eEI----K-KIASFYLEECRK-K-L--A--Gd--------------
0026 4m57A   alldETPL----P--pgsR--L---D---V--RAYTTV-LHALS-RA---G------RYERALE-LFAE---L---rrQG----VAP-------T-LVTYNVVLDVYG-R-X--G--R--Swprivalldexraa
0027 5vbgA   ---------------lndA--Q---K---L--DRALIE-ARISA-AK---N------ANEVAQN-QLRA---L-----DL----NK-------sqK-SRYYETLAIVAE-N-R--K--D--Xieavkariexdknl
0028 8hmdC   --------------------mQ---T---I--LIAQSQ-VYYYI-RE---G------FWSTMQR-FCQE---Q----yKA----FG---------D-PFFIFWKAYGLY-Q-E--G--L--Pneaineltsiqhkk
0029 5xw7C   --------------------pT---A---Q--QQLLSQ-VRLGE-AT---K------REDLVRQ-SLYR--lE-----LI----DP----D----N-PDVIAARFRYLL-R-Q--G--Dn--------------
0030 8cbkE   vidvfeimkaryK-----T--L---E---P--RGYSLL-IRGLI-HS---D------RWREALL-LLED---I----kKV----ITP-------S-KKNYNDCIQGALlH-Q--D--V--Ntawnlyqellghdi
```

```
0031 2pziA   --------------------gW---R---W--RLVWYR-AVAEL-LT---G------DYDSATK-HFTE---V----lDT----FP----G----E-LAPKLALAATAE-L-A--G--N--Tdehkfyqtvwstnd
0032 8rg0u   -----MPA----Y-----F--Q---R---P--ENALKR-ANEFL-EV---G------KKQPALD-VLYD---V-----MK--------qkI----H-EPIMLKYLELCV-D-L--Rqk------------------
0033 7y4iA   -------------------------------sNDTLSY-ANILR-AR---N------KFADALA-LYEA---M----lEK----DS----K----N-VEAHIGKGICLQ-T-Q--N--K--Gnlafdcfseairld
0034 7tj4A   ------------------------kE---D--NIYNKL-IKDDM-TS---G------NYDNAQN-IAKQ---T----iNK----NYA--------D-DQTYYLSGMIMA-T-I--N--Sk-------------
0035 8q9tE   ---------------vkcK--N---N---I--LAHRIL-CQYYL-LT---K------EYEAALP-YIKN-giS-----LI----A---plT----K-REFSLDLATVYT-Y-V--D--P--Kdhnaalklydnils
0036 3qdnA   ---------------qlsS--Q---N---S--EIGLLL-AEAQI-AL---N------RSEEAEA-VLKT---I-----PL----QDQ--------D-TRYQGLVAQIEL-L-K--Q--A--Adtpeiqqlqqqvad
0037 8araC   ------------I-----T--L---TkrqQ--EFLLLN-GWLQL-QC---G------HAERACI-LLDA---L-----lT----LNP---E----H-LAGRRCRLVALL-NnN--Q--G--Eraekeaqwlishdp
0038 9dt9A   ---------------------S---G---K--EEIKEA-IKKAV-VR---A--gdpkYLEEAKA-LLEK--lK-----EL----DE----E----D-KDVEKFEKAIKQ-V-E--A--E--Ltlkeakevvkrlfe
0039 4it4B   ---------------qveK--L---S---A--AEIYEV-ALRLK-ES---G--fafdYFVRAAK-LLIT---Y-----K------PF--DK--inG-QAVEELFIQIQT-N-L--A--A--Cllqekryehviy--
0040 7ztyA   ipwqaapdgigyS-----Y--P---YilalqaTDYDSQ-VEELV-RG---G------KLDEAIS-VLTX---L-----ED----ALL----------KNKTETLREVKX-Q-K--A--------------------
0041 5a1vK   qplfvsggddykikvlqpV--H---M---I--LVLRTA-LNLFF-KL---K------NFKTAAT-FARR--lL-----EL----GP-------evA-QQTRKILSACEK-N-Pt----------------------
0042 8rbkA   dllgcyeqalelF-----P--D---D---E--VICNSM-GEHLF-RM---G------FRDEAAG-YFHK--------avK----LNP---D----F-NDAKENFYRVAN-W-L--V--E--Rwhfimlndtkrnti
0043 7dkhI   --------------------qH---N---P--VLLTLL-QTYYY-FK---G------DYQTVLD-IYHH---R----iLK----XS-------ivL-SESSFWCGRAHY-A-L--G--D--Yrksfixfqeslkkn
0044 1y8mA   ---------------eaeS--R---R---R--ECLYYL-TIGCY-KL---G------EYSMAKR-YVDT---L----fEH----ER----N----N-KQVGALKSMVED-K-I--Q--K--Etlkgvvvaggvhhh
0045 7dhgC   --------------------vD---L---S--TFYQNR-AAAFE-QL---Q------KWKEVAQ-DCTK--aV-----EL----NP----K----Y-VKALFRRAKAHE-K-L--D--N--Kkecledvtavcile
0046 8qt5A   ------------------T--L---G---R--DQYVYM-AKLAE-QA---E------RYEEMVQ-FMEQ-lvT-----GA----T---eEL----T-VEERNLLSVAYK-N-V--I--G--Slraawrivssieqk
0047 4ymrA   -----------------------------V--LPELRR-AQSLT-CT---G------LYREALA-LWAN-awQ-----LQ----T--------drP-LLTLAGLAVCHQ-E-L--E--Dp-------------
0048 8g0lB   ---------------------D---P---N--DRRLRP-IYDYL-DN---G------NNKMAIQ-QADK---L----lKK----HK----D----L-HCAKVLKAIGLQ-R-T--G--Kqi------------
0049 6erpA   --------------------sG---Q---Q--QRLLAF-FKCCL-LT---D------QLPLAHH-LLVV---HhgqrqKR----KLL--------T-LDMYNAVMLGWA-R-Q--G--A--Fkelvyvlfmvkdag
0050 5an3B   -------------------------P---V--EKDLKT-AYKALyDE---K------EPLKALH-LYDE---I----lKG----SP----T----N-LTALIFKAACLE-K-L--Y--F--Gfsdwhsdatmenak
0051 2mv3A   atreqeaN----K-----D--L--tS---P--DAQAAF-YKLLL-QS---N------YPQYVVS-RFET---P-----G--------------asS-PECMELYMEALQ-R-I--G--R--Hseadavrqnlehhh
0052 3jq1B   --------------------pT---Q---G--AAYAAL-GKIYV-YE---E------NWQEAIN-VLEP---L----tQN----PYtysifelirY-ADVLLXLAEAYX-N-K--G--A--Ldtsigyinqirrra
0053 8vx7A   keemeklL----E-----E--G---D---L--DALKKA-LQLAV-NV---A--gdkeLLAHAAE-VIKR---A----lDL----AL--lqS----A-KYLASLALWIAK-R-A--G--------------------
0054 6au8A   -----rvE----G-----K--L--eA---H--QMYRTL-FFRYM-SQ---S------KHTEARE-LMYS-gaL-----LF----F---qnS----A-ADLSMLVLESLE-K-A--E--V--Evadellenlakvfs
0055 3u4tA   ---------------------------------VEFRY-ADFLF-KN---N------NYAEAIE-VFNK--lE-----AK----KY----N----S-PYIYNRRAVCYY-E-L--A--K--Ydlaqkdietyfskv
0056 6f1tf   lgtaknaN----E-----M--FriqL---I--QRVKDD-IESLH-DK---FkvwakqIDRQLTA-YMKR---V-----EdvlgkGWE--NH--veG-QKLKQDGDSFRM-K-L--N--T--Qeifddwarkvqqrn
0057 2pl2A   -------------------------A---E--QNPLRL-GVQLY-AL---G------RYDAALT-LFER--aL-----KE----NP----Q----D-PEALYWLARTQL-K-L--G--L--Vnpalengktlvart
0058 5wftA   ---------------lagA--D---D---P--ALREHA-WRLSL-QX---F------DFDSAIE-LLAP---I----gAQ----RQX--------T-DEELDALVYSHE-T-R--G--Tp-------------
0059 8gaaA   lvcvylmkelgmP-----N--E---E--lI--KLLEEL-VEVLR-IL---A--vdkrVLDKAEV-CIEE---M----eEL----GM----P----E-EKIKELREELKF-V-R--E--I--Ldki-----------
0060 5y7qA   ---------------rqsS--H---P---G---ARLRA-IRSLE-RG---E------RFAEAHA-LARE---A-----SC----AP-------aeR-QGLARLLPRLQG-K-L--G--L--Prqaraaapeidrld
0061 2lxlA   ayycRLYA------------flskvG---C--AHLENY-ALKMF-LY---A--nmikSFYTASL-LIDV---I----tVF----GEL--------T-DENVKHRKYARW-K-A--T--Y--Ihnclkngetpq---
0062 8ebfA   ---------------lppE--I---Q---R--ELRYFK-GWVDY-MS---G------DFAASLN-EWSG---F-----ED----EEE--YN----H-STLLMGKANAFF-Y-T--G--Qy-------------
0063 2bb6A   slgilALC----V--hqkR--V---H---D--SVVGKL-LYAVE-H-------vsvdTMAMAGM-AFSC---L----eLS----NL-------kqR-NRINLALKRVQE-K-I--L--K--Aqtpegyfgnvystp
0064 5cd6A   ---------------inaS--S---D---E--WVVERK-VQLLI-DK---K------QYQEAKD-LLLS---T-----HF----Q--------vhqtYTRTGLWEQINE-G-L--G--L--Spqpvpeqlgedrla
0065 4fymD   ------------E-----F--L---C---D--EEIYKS-FVHLK-DK---I--syssYIKEMKK-LLKV---V---llKY----KAlkvLN----N-IVSSNIICFLLS-E-L--I--L--Knklsfdyllgasyk
0066 2katA   ---------------aqgT--D---N---M--LLRFTL-GKTYA-EH---E------QFDAALP-HLRA---A----lDF----DP----T----Y-SVAWKWLGKTLQ-G-Q--G--D--Ragarqawesglaaa
0067 1pc2A   ivlleellpkgsK-----E--E---Q---R--DYVFYL-AVGNY-RL---K------EYEKALK-YVRG---L-----LQ----TE---PQ----N-NQAKELERLIDK-A-M--K--K--Dglvgmaivggmalg
0068 5u1sA   klyiNKWL----Q-----K-----fE---M--DFVKML-LCYLN-FN---N------FDKLSIE-LSLC---I---ksKE----KYY--SS----IvPYADNYLLEAYL-S-L--Y--M--Iddalmmknqlqktm
0069 2kckA   ---------------qldP--E---E---S--KYWLMK-GKALY-NL---E------RYEEAVD-CYNY---V--invIE----DE----Y----N-KDVWAAKADALR-Y-I--E--G--Keveaeiaearakle
0070 6yj6A   alnhfqclydetF-----S--D---V---A--DLYFEA-ATALT-RA---E------KYKEAID-FFTP---L----lSL----EEW--R-----T-TDVFKPLARCYK-E-I--E--S--Yetakefyelaikse
0072 2kc7A   -----------------------------M--DQL-KT-IKELI-NQ---G------DIENALQ-ALEE--fL-----QT----EP---VG----K-DEAYYLMGNAYR-K-L--G--D--Wqkalnnyqsaieln
0073 5wd8A   --------------------vT---E---L--TRLKEY-XEDQI-AK---A--aqlkFLENAHT-EHFV--kX-----GS----L--------rlK-IEIRSLYEEXLE-L-K--D--K--Crdqiqqyet-----
0074 8f5oC   htsvrwsetadavkgpinV--G---K---G--AILYTL-SRIAN-RL---E------MVRTARA-VFEK--lQ-----GV----IL--vsM----M-EQVDIETLLVRS-K-Pv----------------------
0075 5efrA   ---------------pdtP--W---A---D--DALVGA-MRAYI-AY---A--rqpeRYRRAVE-LYER---L--lqiFP----DS-------plL-RTAEELYTRARQ-R-L--T--E-----------------
0076 4ebaA   ckvlelgL----K-----Y--F--tD---G--EYINKY-LDFLI-YV---N------EESQVKS-LFES---S-----ID----KI-------hlL-KMIFQKVIFFES-K-V--G--S--Lnsvrtlekrffekf
0077 9l4fA   vdlfcrnsqhecqlkqleP--D---Q---I--IHYHGV-VKSML-GL---G------QLSTVIT-QVNG--vH-----AN----RS---eW----T-DELNTYRVEAAW-K-L--S--Q--Wdlvenylaadgkst
0078 6tdyJ   ---------------sgiK--P---N---A--ESWGYV-LKELV-QA---G------DFRLGWV-CIAG--mK-----SL----GIT-----------PDQALVDANEA-N-A--A--K--Akaagtdfpaylkka
0079 4bt9B   iskgnlpgvkhkS-----F--L---T---A--EDCFEL-GKVAY-TE---A------DYYHTEL-WMEQ-alR-----QL----D---tiD----K-VSVLDYLSYAVY-Q-Q--G--D--Ldkallltkklleld
0080 4uzyA   ---------------qnvP--E---S---R--AALSLL-GYCYY-YT---G------QYDMASQ-MYEQ---L----vTL----YP----S----N-EDYKLYYAQSLY-KgG--M--Y--Peaskavvkveghqk
0081 8f5pD   --------------------------------ELSVRV-GRALV-AA---H------DYAKAIR-YYQD---A----lVT----DP----H----L-SIVRADLATLQW-R-L--G--Hi-------------
0082 7t6dA   ---------------kedT--G---T---V--EAHLTL-GNLFR-SR---G------EVDRAIR-IHQT---L---meSA----SL--yeQ----R-LLAIQQLGRDYM-A-A--G--L--Ydraedmfnqltdet
0083 3as5A   -----------------dD--I---R---Q--VYYRDK-GISHA-KA---G------RYSQAVM-LLEQ--vY-----DA----D----aF----D-VDVALHLGIAYV-K-T--G--Av-------------
0084 5w94A   --------------------eE---I---M--RCEFLLlHDLPL-MR---D---skfHYKIALR-NCNE---L----vQY----MV--qnW----A-SVFQYVGVMLCI-K-L--K--Q--Hrrvktsfhgllsqc
0085 8h7gC   takepynyflllraleeipqhA---S---A--SAIIQY-GKIAR-KQ---G------LVNVALD-ILSR---I-----HT----I--------vdC-FQKIRQQVKCYL-Q-Lnl---------------------
0086 8j0nB   ---------------mtnP--H---N---H--LYCQQY-AEVKY-TQ---G---gleNLELSRK-YFAQ---A----lKL----NN----R----N-MRALFGLYMSAS-H-I--A--S--Npkasaktkkdnmky

0001 s001A   ----LLLL----L-----L--L---L---H--HHHHHH-HHHHH-HL---L------LHHHHHH-HHLL---L-----LH----HHH--HH----L-HHHHHHHHHHHH-H-H--L--L--L--------------
0002 8qgyB   hhhhhhhhhlllL-----L--L---L---H--HHHHHH-HHHHH-HL---L------LLHHHHH-HHHH---H-----HH-llllLL--LL----L-LHHHHHHHHHHH-H-H--L--L--Lhhhhhhhhhhhhhh
0003 6z0fA   hhhhhhhhhhhhL-----H--H---H---H--HHHHHH-HHHHH-HL---L------LHHHHHH-HLLL---L-----LH----HHL--------L-HHHHHHHHHHHH-H-H--H--L--Llhhhhhhhhhhlll
0004 8h38H   hhhhhhhhllllL-----L--L---L---H--HHHHHH-HHHHH-HL---L------LHHHHHH-HHHH---L-----LH----HHH--HH----L-HHHHHHHHHHHH-H-H--H--L--Lhhhhhhhhhhllll
0005 4n5cD   --------------------hH---H---H--HHHHHH-HHHHH-HL---L------LHHHHHH-HHHH---L----lLL----LL---LL----L-HHHHHHHHHHHH---L--L--L--Llhhhhhhhhhhhhh
0006 6xt9D   ---------------hhlL--L---L---H--HHHHHH-HHHHH-LL---L------LHHHHHH-HHHH---H-----LL----LL---LL----L-HHHHHHHHHHHH-H-L--L--L--Hhhhhhhhhhhhhhl
0007 2yheA   lllhhHHH----H-----H-lL---H---H--HHHHHH-HHHHH-HH---L------LHHHHHH-HHHH---H----hHH----LL----L----L-HHHHHHHHHHHH-H-H--H--H--Hlllhhhhhhhhhhh
0008 6vbu8   --------------------------------LLHHHH-HHHHH-HL---L------LLLHHHH-HHHH---H----hHH----LL----L----L-HHHHHHHHHHHH-H-H--L--L--Llllllllllhhhhh
0009 5vxvA   --------------------------------LHHHHH-HHHHH-HL---L------LHHHHHH-HHHHhllL-----LH----HHH--HH----L-HHHHHHHHHHHH-H-L--L--L--Lllllhhhhhhhhhh
0010 5a63D   ------lH----H-----H--L---L---H--HHHHHH-HHHHH-HH---L--llhhHHHHHHH-LLHH---H---hhHL----LL-------llH-HHHHHHHHHHHH-H-H--H--H--Hhhhhhhhhhhhhhh
0011 2uy1A   ---------------------L---L---H--HHHHHH-HHHHH-HL---L------LHHHHHH-HHHH---H-----LL----LL----L----L-HHHHHHHHHHHH-H-H--L--------------------
0012 6z2wE   hhhhhhllllllllllllL--L---L---L--HHHHHH-HHHHH-HL---L------LHHHHHLlLLLL------lllLL----LL-------llH-HHHHHHHHHHHH-H-L--L--L--Hhhhhlhhhhhllll
0013 2c2lA   ------------------L--L---L---H--HHHHHH-HHHHH-HL---L------LHHHHHH-HHHH--hH-----HH----LL----L----L-HHHHHHHHHHHH-H-L--L--Lh-------------
0014 8sxfC   hhhhhhhL----H-----H--H--hL---L--HHHHHH-HHHHH-HH---L------LHHHHHH-HHHH---H-----hH----HLL---L----L-HHHHHHHHHHHH-H-H--L--L--Lhhhhhhhhhhhhhl
0015 8s4gJ   ---------------hhhH--L---L---H--HHHHHH-HHHHH-HL---L------LHHHHHH-HHHH---L------------LL--LL------HHHHHHHHHHHH-H-L--L--Lh-------------
0016 4ribB   --------------------hH---H---H--HHHHHH-HHHHH-HL---L------LHHHHHH-HHHH-hhH-----HH----H--------hhH-HHHHHHHHHHHH-H-L--L--L--Hhhhhhhhhhhhhll
0017 6sliA   ---------------lllL--L---H---H--HHHHHH-HHHHH-HH---L------LHHHHHH-HHHH---H----hLL----LL--llllheeH-HHHHHHHHHHHH-H-H--L--Ll-------------
0018 2btpA   llllLLLLlhhhH-----L--L---L---H--HHHHHH-HHHHH-HL---L------LHHHHHH-HHHH-hhH-----LL----L----LL----L-HHHHHHHHHHHH-H-H--H--H--Hhhhhhhhhhhhhhh
0019 9lthA   --------------------lL---H---H--HHHHHH-HHHHH-HL---L------LHHHHHH-HHHH---L----lLL----LL--hHH----H-HHHHHHHHHHHH-H-L--L--Ll-------------
0020 3l22A   --------------------lL---H---H--HHHHHH-HHHHH-HL---L------LHHHHHH-HHHL---L-----LL----LL-------eeH-HHHHHHHHHHHH-L-L--L--Ll-------------
0021 4cguA   --------------------lL---H---H--HHHHHH-HHHHH-HH---L------LHHHHHH-HHHH--hH-----HH----LL----L----L-HHHHHHHHHHHH-H-L--L--Lh-------------
0022 2grmA   hhhhHHLL----L-----L-----hH---H--HHHHHH-HHHHH-HL---L------LHHHHHH-HHHH--hH-----HH----L--------hhH-HHHHHHHHHHHH-H-H--H--L--Lhhhhhhhhhhhhhh
0023 6hs5A   ---------------lllL--L---L---L-lLHHHHH-HHHHH-LLlllL------LHHHHHH-HHHH---H----hHH----LL----L----L-HHHHHHHHHHHH-H-H--H--L--Hhhhhhhhhhhhhhh
0024 2o8pA   --------------------lL---H---H--HHHHHH-HHHHH-HL---L------LHHHHHH-HHHH--hH-----HH----HH--llL----L-HHHHHHHHHHHH-H-H--H--H--Hhhhhhhhhhhhhhh
0025 2vkjA   hhhhhLLH----H-------hH---H---H--HHHHHH-HHHHH-HL---L------LHHHHHH-HHHH-hhH-----HL----LL--hHH----H-HHHHHHHHHHHH-H-H--H--Ll-------------
0026 4m57A   hhhhHLLL----L--lhhH--L---L---H--HHLHHH-HHHHH-HL---L------LHHHHHH-HHHH---H---hlLL----LLL--------L-HHHHHHHHHHHH-H-H--L--L--Lhhhhhhhhhhhhhl
0027 5vbgA   ---------------llhH--H---H---H--HHHHHH-HHHHH-HL---L------LHHHHHH-HHHL---L-----LH----HH-------hhH-HHHHHHHHHHHH-H-L--L--L--Hhhhhhhhhhhhhll
0028 8hmdC   --------------------hH---H---H--HHHHHH-HHHHH-HH---L------LHHHHHH-HHHH---H----hHH----HL---------L-HHHHHHHHHHHH-H-L--L--L--Hhhhhhhhhlllllh
0029 5xw7C   --------------------lL---H---H--HHHHHH-HHHHH-HL---L------LHHHHHH-HHHH--hH-----HH----LL----L----L-HHHHHHHHHHHH-H-L--L--Lh-------------
0030 8cbkE   hhhhhhhhhhhhL-----L--L---L---L--LHHHHH-HHHHL-LL---L------LHHHHHH-HHHH---H----hHH----LLL--------L-HHHHHHHHHHHHhL-L--L--H--Hhhhhhhhhhhhhll
0031 2pziA   --------------------lL---L---H--HHHHHH-HHHHH-HH---L------LHHHHHH-HHHH---H----hHH----LL----L----L-LHHHHHHHHHHH-H-H--L--L--Lllllhhhhhhhhll
0032 8rg0u   -----LLL----L-----L--L---L---H--HHHHHH-HHHHH-HH---L------LHHHHHH-HHHH---H-----HH--------llL----H-HHHHHHHHHHHH-H-L--Lhl------------------
0033 7y4iA   -------------------------------lLHHHHH-HHHHH-HL---L------LHHHHHH-HHHH---H----hHH----LL----L----L-HHHHHHHHHHHH-H-L--L--L--Hhhhhhhhhhhhhhl
0034 7tj4A   ------------------------lL---L--LHHHHH-HHHHH-HL---L------LHHHHHH-HHHH---H----hHL----LLL--------L-HHHHHHHHHHHH-H-H--L--Lh-------------
0035 8q9tE   ---------------hhlL--L---L---H--HHHHHH-HHHHH-HL---L------LLLLLHH-HHHH-hhH-----HH----H---hhH----H-HHHHHHHHHHHH-H-H--L--L--Llhhhhhhhhhhhhh
0036 3qdnA   ---------------hhlL--L---L---H--HHHHHH-HHHHH-HL---L------LHHHHHH-HHLL---L-----LH----HHL--------L-HHHHHHHHHHHH-H-H--H--H--Lllhhhhhhhhhhhl
0037 8araC   ------------L-----L--L---LhhhH--HHHHHH-HHHHH-HL---L------LHHHHHH-HHHH---H-----hH----HLL---L----L-HHHHHHHHHHHH-HlL--L--H--Hhhhhhhhhhhhhll
0038 9dt9A   ---------------------L---H---H--HHHHHH-HHHHH-HH---H--llhhHHHHHHH-HHHH--hH-----HH----LL----L----L-LLHHHHHHHHHH-H-H--H--H--Hhhhhhhhhhhhhhh
0039 4it4B   ---------------lhhH--L---L---H--HHHHHH-HHHHH-HH---H--hhhhHHHHHHH-HHHL---L-----L------LL--LL--llH-HHHHHHHHHHHH-H-H--H--H--Hhhhhhhllllll--
0040 7ztyA   lllllllleeeeE-----L--L---EeeeeleLLHHHH-HHHHH-HL---L------LHHHHHH-HHHH---L-----LH----HHL----------LLHHHHHHHHHH-H-H--L--------------------
0041 5a1vK   lleeeeelllleeeellhH--H---H---H--HHHHHH-HHHHH-HH---L------LHHHHHH-HHHH--hH-----HH----LL-------hhH-HHHHHHHHHHHH-L-Ll----------------------
0042 8rbkA   hhhhhhhhhhhhL-----L--L---L---H--HHHHHH-HHHHH-HL---L------LHHHHHH-HHHH--------hhH----HLL---L----L-HHHHHHHHHHHH-H-H--L--L--Hhhhhhhllhhhhhh
0043 7dkhI   --------------------lL---L---H--HHHHHH-HHHHH-HL---L------LHHHHHH-HHHH---L----lHH----HH-------hhH-HHHHHHHHHHHH-H-L--L--L--Hhhhhhhhhhhhhhl
0044 1y8mA   ---------------hllL--L---H---H--HHHHHH-HHHHH-LL---L------LHHHHHH-HHHH---H----hHL----LL----L----L-HHHHHHHHHHHH-H-H--H--H--Lllllllllllllll
0045 7dhgC   --------------------hH---H---H--HHHHHH-HHHHH-HL---L------LHHHHHH-HHHH--hH-----HH----LL----L----L-HHHHHHHHHHHH-H-L--L--L--Hhhhhhhhhhhhhhl
0046 8qt5A   ------------------L--L---L---H--HHHHHH-HHHHH-HL---L------LHHHHHH-HHHH-hhH-----HH----L---lLL----L-HHHHHHHHHHHH-H-H--H--H--Hhhhhhhhhhhhhhh
0047 4ymrA   -----------------------------L--HHHHHH-HHHHH-HH---L------LHHHHHH-HHHH-hhH-----HH----H--------lhH-HHHHHHHHHHHH-H-L--L--Lh-------------
0048 8g0lB   ---------------------L---H---H--HHHHHH-HHHHH-HL---L------LHHHHHH-HHHH---H----hHH----LL----L----L-HHHHHHHHHHHH-H-L--L--Lll------------
0049 6erpA   --------------------lH---H---H--HHHHHH-HHHHH-HH---L------LHHHHHH-HHHH---HhhlhhHH----LLL--------L-HHHHHHHHHHHH-H-H--L--L--Hhhhhhhhhhhhhll
0050 5an3B   -------------------------L---H--HHHHHH-HHHHHhLL---L------LHHHHHH-HHHH---H----hHH----LL----L----L-HHHHHHHHHHHH-H-H--H--L--Llhhhllhhhhhhhh
0051 2mv3A   hhhhhhhH----H-----L--L--lL---H--HHHHHH-HHHHH-HL---L------LHHHHHH-HHLL---L-----L--------------llL-HHHHHHHHHHHH-H-L--L--L--Hhhhhhhhhhhhlll
```

```
0052 3jq1B   --------------------lL---H---H--HHHHHH-HHHHH-HL---L------LHHHHHH-HHHH---H----hLL----LLlleeeeeeeH-HHHHHHHHHHHH-H-H--L--L--Hhhhhhhhhhhhhhl
0053 8vx7A   hhhhhhhH----H-----H--L---L---H--HHHHHH-HHHHH-HH---H--llhhHHHHHHH-HHHH---H----hHH----HH--hhH----H-HHHHHHHHHHHH-H-H--L--------------------
0054 6au8A   -----lhH----H-----H--H--hH---H--HHHHHH-HHHHH-HL---L------LHHHHHH-HHHH-hhH-----HH----H---hhH----H-HHHHHHHHHHHH-H-L--L--L--Lllhhhhhhhhhhhh
0055 3u4tA   ---------------------------------LHHHH-HHHHH-LL---L------LHHHHHH-HHHH--hH-----HL----LL----L----L-LLLHHHHHHHHH-H-L--L--L--Hhhhhhhhhhhhlll
0056 6f1tf   hhhlllhH----H-----H--HhhhH---H--HHHHHH-HHHHH-HH---HhlhhhhHHHHHHH-HHHH---H-----HhhhllLLL--LL--llH-HHHHHHHHHHHH-H-L--L--L--Hhhhhhhhhhhllll
0057 2pl2A   -------------------------L---L--HHHHHH-HHHHH-HL---L------LHHHHHH-HHHH--hH-----LL----LL----L----L-HHHHHHHHHHHH-H-L--L--L--Hhhhhhhhhhhhhhl
0058 5wftA   ---------------hhhL--L---L---H--HHHHHH-HHHHH-HL---L------LHHHHHH-HHLL---H----hHH----LLL--------L-HHHHHHHHHHHH-H-L--L--Lh----------------
0059 8gaaA   hhhhhhhhhhllL-----H--H---H--hH--HHHHHH-HHHHH-HH---H--lllhHHHHHHH-HHHH---H----hHH----LL----L----H-HHHHHHHHHHHH-H-H--H--H--Hlll-----------
0060 5y7qA   ---------------hllL--L---L---L---HHHHH-HHHHH-HL---L------LHHHHHH-HHHH---H-----HH----LL-------hhH-HHHHHHHHHHHH-H-L--L--L--Llllllllllleee
0061 2lxlA   hhhhHHHH------------hhhhhH---H--HHHHHH-HHHHH-HH---H--hhhhHHHHHHH-HHHH---H----hLL----LLL--------L-HHHHHHHHHHHH-H-H--H--H--Hhhhhhllllll---
0062 8ebfA   ---------------llhH--H---H---H--HHHHHH-HHHHH-HL---L------LHHHHHH-HHLL---L-----LH----HHH--LL----L-HHHHHHHHHHHH-H-L--L--Lh----------------
0063 2bb6A   hhhhhHHH----H--lllL--L---L---H--HHHHHH-HHHHH-L-------lhhhHHHHHHH-HHHH---H----hHH----LL-------hhH-HHHHHHHHHHHH-H-H--H--H--Lllllllllllllhh
0064 5cd6A   ---------------lllL--L---L---H--HHHHHH-HHHHH-HL---L------LHHHHHH-HHHL---L-----LL----L--------llllLHHHHHHHHHHH-H-L--L--L--Llllllhhhllllll
0065 4fymD   ------------L-----L--L---L---H--HHHHHH-HHHHH-HH---H--hhhhHHHHHHH-HHHH---H---hhHL----LLeehHL----L-HHHHHHHHHHHH-H-H--H--H--Hlllllleeeellll
0066 2katA   ---------------lllL--L---L---H--HHHHHH-HHHHH-HL---L------LHHHHHH-HHHH---H----hHH----LL----L----L-HHHHHHHHHHHH-H-H--L--L--Hhhhhhhhhhhhhhh
0067 1pc2A   hhhhhhhhhhllH-----H--H---H---H--HHHHHH-HHHHH-HL---L------LHHHHHH-HHHH---H-----HH----HL---LL----L-HHHHHHHHHHHH-H-H--H--H--Lllllllllllllll
0068 5u1sA   hhhhHHLH----H-----H-----hH---H--HHHHHH-HHHHH-HL---L------LHHHHHH-HHHH---H---hhLH----HHH--HH----HhHHHHHHHHHHH-H-L--L--L--Hhhhhhhhhhhhhhl
0069 2kckA   ---------------hhlL--L---L---H--HHHHHH-HHHHH-HL---L------LHHHHHH-HHHH---H--hhlLL----LL----L----L-HHHHHHHHHHHL-L-L--L--L--Llhhhhhhhhhhhhh
0070 6yj6A   hhhhhhhhhlllL-----L--L---L---H--HHHHHH-HHHHH-HL---L------LHHHHHH-HHHH---H----hHL----LLL--L-----L-HHHHHHHHHHHH-H-L--L--L--Hhhhhhhhhhhllll
0072 2kc7A   -----------------------------L--LLH-HH-HHHHH-HH---L------LHHHHHH-HHHH--hH-----HH----LL---LL----H-HHHHHHHHHHHH-H-H--L--L--Hhhhhhhhhhhhhhl
0073 5wd8A   --------------------hH---H---H--HHHHHH-HHHHH-HH---H--hhhhHHHHHHH-HHHH--hH-----HL----H--------hhH-HHHHHHHHHHHH-H-H--H--H--Hhhhhhhhhl-----
0074 8f5oC   lleeeeeeelllllllllL--L---H---H--HHHHHH-HHHHH-HL---L------LHHHHHH-HHHH--hL-----LL----LL--llL----H-HHHHHHHHHHHH-L-Ll----------------------
0075 5efrA   ---------------lllL--H---H---H--HHHHHH-HHHHH-HH---H--hhhhHHHHHHH-HHHH---H--hhhLL----LL-------llH-HHHHHHHHHHHH-H-H--H--L-----------------
0076 4ebaA   hhhhhhlH----H-----H--H--lL---H--HHHHHH-HHHHH-HH---L------LHHHHHH-HHLL---L-----LL----LL-------hhH-HHHHHHHHHHHH-H-H--L--L--Hhhhhhhhhhhhhhl
0077 9l4fA   lllllllllllllllhhlL--L---L---H--HHHHHH-HHHHH-HH---L------LHHHHHH-HHHH--hH-----HH----LL---lL----L-LLLHHHHHHHHH-H-H--L--L--Hhhhhhhhhlllll
0078 6tdyJ   ---------------lllL--L---L---H--HHHHHH-HHHHH-HH---L------LHHHHHH-HHHH--hL-----LL----LLL-----------LLHHHHHHHHH-H-H--H--H--Hhhllllllhhhhll
0079 4bt9B   hhhlllllllllL-----L--L---L---H--HHHHHH-HHHHH-HL---L------LHHHHHH-HHHH-hhH-----HH----H---llL----H-HHHHHHHHHHHH-H-L--L--L--Hhhhhhhhhhhhhhl
0080 4uzyA   ---------------hhlL--L---L---H--HHHHHH-HHHHH-HH---L------LHHHHHH-HHHH---H----hHH----LL----L----L-HHHHHHHHHHHH-HlL--L--H--Hhhhhhhllllllhh
0081 8f5pD   --------------------------------LLHHHH-HHHHH-HL---L------LLHHHHH-HHHH---H----hHH----LL----L----L-HHHHHHHHHHHH-H-L--L--Lh----------------
0082 7t6dA   ---------------lllL--L---L---H--HHHHHH-HHHHH-HH---L------LHHHHHH-HHHH---H---hlLL----LL--hhH----H-HHHHHHHHHHHH-H-H--L--L--Hhhhhhhhhhhlllh
0083 3as5A   ----------------lL--H---H---H--HHHHHH-HHHHH-HH---L------LHHHHHH-HHLL--lL-----LL----L----lL----L-HHHHHHHHHHHH-H-L--L--Lh----------------
0084 5w94A   --------------------hH---H---H--HHHHHHhLHHHH-HH---L---lhhHHHHHHH-HHHH---H----hHH----HL--hhH----H-HHHHHHHHHHHH-H-L--L--L--Hhhhhhhhhhhhhhl
0085 8h7gC   eelllhhhhhhhhhhhhlllhH---H---H--HHHHHH-HHHHH-HL---L------LLHHHHH-HHHH---H-----HH----L--------hhH-HHHHHHHHHHHH-H-Lll---------------------
0086 8j0nB   ---------------hhlL--L---L---H--HHHHHH-HHHHH-HH---L---lhhHHHHHHH-HHHH---H----hHH----HL----L----L-LHHHHHHHHHHH-H-H--H--H--Lllllhhhhhhhhhh
```

```
# Job: Type-3 BREX; MTase MBW2068239 wHTH
# Query: s001A
```

| # No: | Chain | Z | rmsd | lali | nres | %id | PDB Description |
|---|---|---|---|---|---|---|---|
| 1: | 6qfd-B | 6.6 | 3.1 | 55 | 116 | 5 | MOLECULE: DNA-BINDING PROTEIN; |
| 2: | 6zvh-y | 6.4 | 2.4 | 55 | 72 | 16 | MOLECULE: 18S RRNA; |
| 3: | 8qkf-A | 6.4 | 2.3 | 50 | 99 | 14 | MOLECULE: ARSR FAMILY TRANSCRIPTIONAL REGULATOR; |
| 4: | 5dym-A | 6.3 | 3.7 | 56 | 96 | 13 | MOLECULE: PADR-FAMILY TRANSCRIPTIONAL REGULATOR; |
| 5: | 5j6x-A | 6.2 | 2.7 | 51 | 66 | 12 | MOLECULE: Z-DNA BINDING PROTEIN KINASE; |
| 6: | 7wjp-A | 6.2 | 3.1 | 54 | 98 | 13 | MOLECULE: PADR FAMILY TRANSCRIPTIONAL REGULATOR; |
| 7: | 8jxk-D | 6.1 | 2.9 | 55 | 166 | 15 | MOLECULE: CONSERVED PROTEIN; |
| 8: | 1ldd-A | 6.1 | 2.8 | 57 | 74 | 9 | MOLECULE: ANAPHASE PROMOTING COMPLEX; |
| 9: | 2od5-A | 6.1 | 1.6 | 51 | 91 | 14 | MOLECULE: HYPOTHETICAL PROTEIN; |
| 10: | 1sfx-A | 6.0 | 3.5 | 51 | 109 | 8 | MOLECULE: CONSERVED HYPOTHETICAL PROTEIN AF2008; |
| 11: | 2esh-A | 6.0 | 4.0 | 60 | 114 | 8 | MOLECULE: CONSERVED HYPOTHETICAL PROTEIN TM0937; |
| 12: | 5a31-N | 6.0 | 1.7 | 51 | 703 | 10 | MOLECULE: ANAPHASE-PROMOTING COMPLEX SUBUNIT 1; |
| 13: | 3elk-A | 6.0 | 3.1 | 55 | 104 | 15 | MOLECULE: PUTATIVE TRANSCRIPTIONAL REGULATOR TA0346; |
| 14: | 8soj-B | 6.0 | 2.3 | 53 | 428 | 8 | MOLECULE: CST COMPLEX SUBUNIT CTC1; |
| 15: | 5zqh-A | 6.0 | 2.3 | 54 | 101 | 9 | MOLECULE: PADR FAMILY TRANSCRIPTIONAL REGULATOR; |
| 16: | 5jls-A | 6.0 | 2.8 | 51 | 134 | 6 | MOLECULE: ADHESIN COMPETENCE REPRESSOR; |
| 17: | 2zkz-C | 5.9 | 2.2 | 50 | 87 | 8 | MOLECULE: TRANSCRIPTIONAL REPRESSOR PAGR; |
| 18: | 5xpq-A | 5.9 | 2.3 | 49 | 100 | 12 | MOLECULE: UNCHARACTERIZED HTH-TYPE TRANSCRIPTIONAL REGULATO |
| 19: | 4ejo-A | 5.9 | 4.0 | 54 | 112 | 13 | MOLECULE: TRANSCRIPTIONAL REGULATOR, PADR-LIKE FAMILY; |
| 20: | 7z8b-C | 5.9 | 2.2 | 58 | 1224 | 9 | MOLECULE: CULLIN-7; |
| 21: | 4omz-C | 5.9 | 2.4 | 50 | 100 | 10 | MOLECULE: NOLR; |
| 22: | 5hs7-A | 5.9 | 3.2 | 53 | 102 | 17 | MOLECULE: HTH-TYPE TRANSCRIPTIONAL REGULATOR YODB; |
| 23: | 8uuc-A | 5.8 | 2.6 | 54 | 274 | 15 | MOLECULE: ADENINE DNA GLYCOSYLASE; |
| 24: | 8iue-P | 5.8 | 2.5 | 52 | 303 | 6 | MOLECULE: DNA-DIRECTED RNA POLYMERASE III SUBUNIT RPC1; |
| 25: | 3mq0-B | 5.8 | 1.7 | 48 | 248 | 13 | MOLECULE: TRANSCRIPTIONAL REPRESSOR OF THE BLCABC OPERON; |
| 26: | 3bdd-A | 5.8 | 2.8 | 52 | 140 | 13 | MOLECULE: REGULATORY PROTEIN MARR; |
| 27: | 2l4m-A | 5.7 | 1.7 | 49 | 69 | 10 | MOLECULE: UNCHARACTERIZED PROTEIN; |
| 28: | 6uvu-B | 5.7 | 1.9 | 48 | 114 | 8 | MOLECULE: ARSR FAMILY TRANSCRIPTIONAL REGULATOR; |
| 29: | 2rdp-A | 5.7 | 2.4 | 50 | 140 | 16 | MOLECULE: PUTATIVE TRANSCRIPTIONAL REGULATOR MARR; |
| 30: | 2oqg-A | 5.7 | 1.8 | 49 | 109 | 10 | MOLECULE: POSSIBLE TRANSCRIPTIONAL REGULATOR, ARSR FAMILY P |
| 31: | 5zi8-A | 5.7 | 3.5 | 57 | 104 | 9 | MOLECULE: TRANSCRIPTIONAL REGULATOR; |
| 32: | 2fe3-B | 5.7 | 3.2 | 56 | 143 | 7 | MOLECULE: PEROXIDE OPERON REGULATOR; |
| 33: | 7mex-A | 5.7 | 3.1 | 55 | 1737 | 20 | MOLECULE: UBIQUITIN; |
| 34: | 7z1n-O | 5.6 | 3.1 | 53 | 570 | 9 | MOLECULE: DNA-DIRECTED RNA POLYMERASE III SUBUNIT RPC1; |
| 35: | 2xrn-B | 5.6 | 1.4 | 48 | 241 | 2 | MOLECULE: HTH-TYPE TRANSCRIPTIONAL REGULATOR TTGV; |
| 36: | 3s93-B | 5.6 | 3.0 | 58 | 81 | 16 | MOLECULE: TUDOR DOMAIN-CONTAINING PROTEIN 5; |
| 37: | 6j05-A | 5.5 | 2.6 | 49 | 101 | 6 | MOLECULE: TRANSCRIPTIONAL REGULATOR ARSR; |
| 38: | 3l9f-A | 5.5 | 3.5 | 57 | 170 | 11 | MOLECULE: PUTATIVE UNCHARACTERIZED PROTEIN SMU.1604C; |
| 39: | 2wte-A | 5.5 | 1.6 | 48 | 212 | 13 | MOLECULE: CSA3; |
| 40: | 3b73-B | 5.5 | 2.0 | 49 | 89 | 6 | MOLECULE: PHIH1 REPRESSOR-LIKE PROTEIN; |
| 41: | 2nyx-B | 5.5 | 2.8 | 50 | 147 | 8 | MOLECULE: PROBABLE TRANSCRIPTIONAL REGULATORY PROTEIN, RV14 |
| 42: | 6jyi-A | 5.5 | 3.6 | 56 | 174 | 13 | MOLECULE: TRANSCRIPTIONAL REPRESSOR PADR; |
| 43: | 3f6v-A | 5.5 | 2.2 | 49 | 96 | 18 | MOLECULE: POSSIBLE TRANSCRIPTIONAL REGULATOR, ARSR FAMILY P |
| 44: | 5fmf-V | 5.5 | 2.7 | 51 | 174 | 10 | MOLECULE: DNA REPAIR HELICASE RAD25, SSL2; |
| 45: | 6lui-A | 5.5 | 2.4 | 53 | 71 | 13 | MOLECULE: ATHERIN; |
| 46: | 1xmk-A | 5.4 | 2.4 | 51 | 79 | 12 | MOLECULE: DOUBLE-STRANDED RNA-SPECIFIC ADENOSINE DEAMINASE; |
| 47: | 2eth-A | 5.4 | 2.8 | 52 | 141 | 8 | MOLECULE: TRANSCRIPTIONAL REGULATOR, PUTATIVE, MAR FAMILY; |
| 48: | 2p4w-B | 5.4 | 2.7 | 50 | 198 | 18 | MOLECULE: TRANSCRIPTIONAL REGULATORY PROTEIN ARSR FAMILY; |
| 49: | 3u21-A | 5.4 | 2.6 | 58 | 109 | 12 | MOLECULE: NUCLEAR FACTOR RELATED TO KAPPA-B-BINDING PROTEIN |
| 50: | 3l7w-A | 5.4 | 2.4 | 54 | 106 | 7 | MOLECULE: PUTATIVE UNCHARACTERIZED PROTEIN SMU.1704; |
| 51: | 7xjg-J | 5.3 | 2.6 | 53 | 306 | 11 | MOLECULE: RNA-DIRECTED DNA POLYMERASE FROM RETRON EC86; |
| 52: | 8ioi-A | 5.3 | 4.0 | 56 | 190 | 13 | MOLECULE: PADR FAMILY TRANSCRIPTIONAL REGULATOR; |
| 53: | 3gfi-A | 5.3 | 3.6 | 53 | 143 | 9 | MOLECULE: 146AA LONG HYPOTHETICAL TRANSCRIPTIONAL REGULATOR |
| 54: | 4ija-A | 5.3 | 1.7 | 47 | 363 | 4 | MOLECULE: XYLR PROTEIN; |
| 55: | 8xt8-B | 5.3 | 2.9 | 51 | 145 | 12 | MOLECULE: MARR FAMILY TRANSCRIPTIONAL REGULATOR; |
| 56: | 6j0e-B | 5.3 | 2.3 | 49 | 125 | 18 | MOLECULE: ARSENIC RESPONSIVE REPRESSOR ARSR; |
| 57: | 3k69-A | 5.3 | 1.9 | 49 | 151 | 12 | MOLECULE: PUTATIVE TRANSCRIPTION REGULATOR; |
| 58: | 7wze-A | 5.3 | 1.7 | 48 | 161 | 15 | MOLECULE: UNCHARACTERIZED HTH-TYPE TRANSCRIPTIONAL REGULATO |
| 59: | 3fhz-D | 5.3 | 3.5 | 56 | 167 | 18 | MOLECULE: ARGININE REPRESSOR; |
| 60: | 4xrf-A | 5.3 | 2.3 | 50 | 142 | 10 | MOLECULE: TRANSCRIPTIONAL REGULATOR, MARR FAMILY; |
| 61: | 6v9i-C | 5.3 | 2.5 | 55 | 730 | 13 | MOLECULE: IMMUNOGLOBULIN G-BINDING PROTEIN G,CULLIN-5; |
| 62: | 3f8f-A | 5.3 | 3.5 | 54 | 114 | 13 | MOLECULE: TRANSCRIPTIONAL REGULATOR, PADR-LIKE FAMILY; |
| 63: | 7el3-A | 5.2 | 2.6 | 50 | 138 | 8 | MOLECULE: HOMOPROTOCATECHUATE DEGRADATION OPERON REGULATOR |
| 64: | 2g9w-A | 5.2 | 2.3 | 52 | 119 | 13 | MOLECULE: CONSERVED HYPOTHETICAL PROTEIN; |
| 65: | 3f72-B | 5.2 | 2.2 | 49 | 108 | 4 | MOLECULE: CADMIUM EFFLUX SYSTEM ACCESSORY PROTEIN; |
| 66: | 2mh2-A | 5.2 | 2.5 | 50 | 64 | 14 | MOLECULE: HOMOLOGOUS-PAIRING PROTEIN 2 HOMOLOG; |
| 67: | 5okc-B | 5.2 | 2.8 | 55 | 377 | 11 | MOLECULE: SISTER CHROMATID COHESION PROTEIN DCC1; |
| 68: | 2cwe-A | 5.2 | 2.6 | 49 | 191 | 10 | MOLECULE: HYPOTHETICAL TRANSCRIPTION REGULATOR PROTEIN, PH1 |
| 69: | 7vwv-B | 5.2 | 3.3 | 55 | 74 | 24 | MOLECULE: I73R; |

```
 70:  7kbf-K  5.2  3.1   54    78   15   MOLECULE: HISTONE H3.2;
 71:  7asv-A  5.2  3.8   62   155   11   MOLECULE: DNA-DIRECTED RNA POLYMERASE III SUBUNIT RPC5;
 72:  1yg2-A  5.2  2.4   50   169   10   MOLECULE: GENE ACTIVATOR APHA;
 73:  6o8o-B  5.2  2.4   50   108    8   MOLECULE: TRANSCRIPTIONAL REGULATOR, ARSR FAMILY;
 74:  3v4g-A  5.2  3.1   54   145    9   MOLECULE: ARGININE REPRESSOR;
 75:  4hqe-B  5.2  3.2   53   108    9   MOLECULE: TRANSCRIPTIONAL REGULATOR QSRR;
 76:  3u2r-A  5.1  2.5   49   135    8   MOLECULE: REGULATORY PROTEIN MARR;
 77:  5n35-A  5.1  1.7   49    60    2   MOLECULE: POLB1 BINDING PROTEIN 2 (PBP2);
 78:  4txa-A  5.1  3.0   59   367    7   MOLECULE: ROQUIN-1;
 79:  6a6i-A  5.1  3.1   57    94   14   MOLECULE: EXCISION REPAIR CROSS-COMPLEMENTING RODENT REPAIR
 80:  3bpx-A  5.1  3.0   51   147   16   MOLECULE: TRANSCRIPTIONAL REGULATOR;
 81:  4yif-B  5.1  3.2   51   141   10   MOLECULE: MARR FAMILY PROTEIN RV0880;
 82:  6yj6-B  5.1  2.7   55   508    7   MOLECULE: TRANSCRIPTION FACTOR TAU 131 KDA SUBUNIT;
 83:  7bzd-A  5.1  3.3   54   110   17   MOLECULE: HTH-TYPE TRANSCRIPTIONAL ACTIVATOR HXLR;
 84:  2gai-A  5.1  2.3   51   581    8   MOLECULE: DNA TOPOISOMERASE I;
 85:  5z7b-B  5.1  3.1   55   197   15   MOLECULE: PADR FAMILY TRANSCRIPTIONAL REGULATOR;

0001 s001A   --EASAIQWLKQQLT-K-----K---P-Q-T-FQELQ-PQFL-R-EL----HK--AKH---EK--L--P---ELIELLEQ-----N---------F-----LR-Y-DGKGPIP-------KDRWYV------
0002 6qfdB   rdLTAFQKNILTVLG-E-----E---A-R-Y-GLAIK-RELE-E-YY----G-----------eevnhG---RLYPNLDD-----L----vnkglV-----EK-S-ELDK---------rTNEYALtnegfd
0003 6zvhy   --kFNWKGTIKAILK-Q-----A--nE-I-T-IKKLR-KKVL-A-QY----Y--dEHH---RSeeellV---IFNKKISK-----N-------ptF-----KL-L-K-------------DKVKLvk-----
0004 8qkfA   alAAPLRIAIVLQLK-Q-----S---Q-R-C-VHELV-DALD-----------------------vpqP---LVSQHLRI-----L----kqagvV-----SS-E-RAG----------rEVLYRLvdhhla
0005 5dymA   ---GYIDILIVSILE-K-----K---D-C-Y-GYEIA-KQVR-E-RS----EF------------lkeG---TMYLALKR-----M----esknlI-----KS-Y-YSNEQssgg----rRKYYNLtnegkd
0006 5j6xA   saENEIEMRICDYLR-R----hG---R-S-T-VQDIF-KELK-----------------------lekS---TVNRHLYS-----L----qaskqV-----FK-T-VEDN---------kRPVWDLve----
0007 7wjpA   fkKGVLELCCLFLIQ-K-----K---D-C-Y-GYELA-NQVS-K-YI----E----------------vaegAIYPVLRR-----L----vkeeyC-----ST-Y-LVES----p----sRKYYQLtvkgei
0008 8jxkD   --EFMLELAILGLLI-E-----S---P-M-H-GYELR-KRLT-G-LL----G---------afrafsyG---SLYPALRR-----M----qadglI-----AE-N-AAPAG--------rRVYQLtdkgrr
0009 1lddA   ltLQRSLPFIEGMLT-N----lG---A-M-K-LHKIH-SFLK-I-TV----PK--D--wgyNR--I--T-lqQLEGYLNT-----L----adegrL-----KY-I-A-------------NGSYEIv-----
0010 2od5A   xkTVRIREKIKKFLG-D-----R---P-R-N-TAEIL-EHIN-S-TX----RH--------------gttsqQLGNVLSK-----D-------kdI-----VK-V-GYikrsgilsggydicEWATrnwvae
0011 1sfxA   lsFKPSDVRIYSLLL-E-----R--gG-X-R-VSEIA-RELD--------------------------lsarFVRDRLKV-----L----lkrgfV-----RR-E-IVEKG---w----vGYIYSAekpekv
0012 2eshA   frGWWLASTILLLVA-E-----K---P-S-H-GYELA-ERLA-E-FG----IE--IPG---I------ghmgNIYRVLAD-----L----eesgfL-----ST-E-WDTTV--sp----pRKIYRItpqgkl
0013 5a31N   eeLLLFWTYIQAMLT-N-----L--eS-L-S-LDRIY-NMLR-M-FV----V------tgpalaeidlQ---ELQGYLQK-----K----vrdqqL-----VY-S---------------AGVYRLp-----
0014 3elkA   ilHGLITLYILKELV-K-----R---P-X-H-GYELQ-KSXF-E-TT----GQ--------------alpqgSIYILLKT-----X----kergfV-----IS-E-SSVN--kgq----qLTVYHItdagkk
0015 8sojB   reDKDLHRKIHRIIQ-Q-----D--kG-C-H-FLHIL-ACAR-LsIR----PG------------lseA---VLQQVLEL-----L----edqsdI-----VS-T-M-------------EHYYTA------
0016 5zqhA   llKGVLEGCVLDMIG-Q-----K---E-R-Y-GYELV-QTLR-EaGF----DT-------------ivG---TIYPLLQK-----L----eknqwI-----RG-D-MR-pspdgp----dRKYFSLmkegee
0017 5jlsA   qkLTNTQEHILMLLS-Q-----Q---R-L-T-NTDLA-KALN-----------------------isqA---AVTKAIKS-----L----vkqdmL-----AG-T-KDTV---da----rVTYFELtelakp
0018 2zkzC   txAHPXRLKIVNELY-K-----H--kA-L-N-VTQII-QILK-----------------------lpqS---TVSQHLCK-----X-----rgkvL-----KR-N-RQG----------lEIYYSInnpkve
0019 5xpqA   tlaHPARIRILELLV-E-----R---D-R-S-VGELLsSDVG-----------------------lesS---NLSQQLGV-----L----rragvV-----AA-R-RDG----------nAMIYSIaapdia
0020 4ejoA   lrRGTLVXLVLSQLR-------E---P-A-Y-GYALV-KSLA-D-HG----------------ipieaN---TLYPLXRR-----L----esqglL-----AS-E-WDNGG--sk----pRKYYRTtdeglr
0021 7z8bC   krRNLLNCLIVRILK-A-----H--eG-L-H-IDQLV-CLVL-E-AW----QkgsLGK---GS-acssT---DVLSCILH-----L----lgkgtL-----RR-H-DD-----------rPQVLSYav----
0022 4omzC   amANPKRLLILDSLV-K-----E---E-M-A-VGALA-NKVG-----------------------lsqS---ALSQHLSK-----L----raqnlV-----ST-R-RDA----------qTIYYSSssdsvm
0023 5hs7A   llgKRWNGLIIHVLM-D-----G---P-K-R-FKEIT-ETIP-M-I--------------------sqK---MLAERLKE-----L----eqneiV-----ER-Q-VLPET---p----vKVIYTLtekgta
0024 8uucA   gsRRQKRAELVRLVL-A-----E---PgI-G-IDELA-ERLD-A-FE--rdAG--RK-------gvdaA---TFTSIVAD-----L----vaegfF-----RR-E---------------GDAFFA------
0025 8iueP   -dPVEIENRIIELCH-Q-----F--hG-I-T-DQVIQ-NEMP-H-I--------------------eaQ---QRAVAINR-----L----lsmgqL-----DL-L-RSN----------tGLLYRIkdsqna
0026 3mq0B   vpALRRAVRILDLVA-G-----S--rD-L-T-AAELT-RFLD-----------------------lpkS---SAHGLLAV-----M----teldlL-----AR-S-A-------------DGTLRIgphslr
0027 3bddA   lgISLTRYSILQTLL-K-----D---ApL-H-QLALQ-ERLQ-----------------------idrA---AVTRHLKL-----L----eesgyI-----IR-K-RNPDN---q----rEVLVWPteqare
0028 2l4mA   qfSQQREEDIYRFLK-D----nG---P-Q-R-ALVIA-QALG-M------------------------rtakDVNRDLYR-----M----ksrhlL-----DM-D-EQ------------SKAWTIyr----
0029 6uvuB   vlaNDTRLRLLHALA-R-----S--gG-L-C-VTDLA-AAVG--------------------------mkpqAVSNQLQR-----L----adrriL-----RA-A-RCG----------nNIHYRIvdpcvl
0030 2rdpA   ypITPPQFVALQWLL-E----eG---D-L-T-VGELS-NKXY--------------------------lacsTTTDLVDR-----X----ernglV-----AR-V-RDEH----------VVRIRLlekger
0031 2oqgA   alsDETRWEILTELG-R-----A---D-Q-S-ASSLA-TRLP-----------------------vsrQ---AIAKHLNA-----L----qacglV-----ES-V-KVG----------rEIRYRAlgaeln
0032 5zi8A   fqRAAVRLHILHHAA-D-----N---E-V-H-GAWLT-QELS-R-HG----YR------------vspG---TLYPTLHR-----L----eadglL-----VS-E-QRVVD--gr----aRRVYRAtpagra
0033 2fe3B   vrITPQRHAILEYLV-N-----S--aH-P-T-ADDIY-KALE-G-KF----PN------------msvA---TVYNNLRV-----F----resglV-----KE-L-TYGD---------aSSRFDFvtsdhy
0034 7mexA   rrMDQIKNSIIYNLY-M-----K---P-L-S-YSKLL-RSVP-D-YL-----------------tedtT---EFDEALEE-----V---------S-----VF-V-EPKGL---a----dNGVFKLkaslya
0035 7z1nO   ahLGERAASVIGMLV-A----lG---R-L-S-VRELV-EKID-G-M--------------------dvD---SVKTTLVS-----L----tqlrcV-----KY-L-QETA-isgk----kTTYYYYneegih
0036 2xrnB   iqVIARAASIMRALG-S-----H--hG-L-S-LAAIA-QLVG--------------------------lprsTVQRIINA-----L----eeeflV-----EA-L-GP------------AGGFRLgpalgq
0037 3s93B   riQECLRKEIRSLLI-S-----T--dG-L-S-PQELE-KEYLlM-VG----N----hlplrIL--G--Y--rSTMELVLD-----M------pdvV-----RV-C-PGAG---------gTVILKAip----
0038 6j05A   alASPVRLEIFRLLV-E-----Q--tG-L-V-SGDIA-EHLG-Q-------------------------phnGISFHLKN-----L----qhaglV-----TV-Q-REG----------ryQRYRAampvvr
0039 3l9fA   ---MQGKDIILGILS-K-----K---E-R-S-GYEIN-DILQ-NqLS----YF------ydgT--Y--G---MIYPTLRK-----L----ekdgkI-----TK-E-VVIQ-dgrp----nKNIYAItesgkk
0040 2wteA   rdYSREEXKLLNVLY-E-----T--kG-T-G-ITELA-KXLD--------------------------ksekTLINKIAE-----L----kkfgiL-----TQ-K-GK------------DRKVELnelgln
0041 3b73B   swXTIWDDRILEIIH-E-----E--gN-G-S-PKELE-DR-D-E-IR---------------------isksSVSRRLKK-----L----adhdlL-----QP-L-A-------------NGVYVIteegea
0042 2nyxB   enITIPQFRTLVILS-N-----H--gP-I-N-LATLA-TLLG--------------------------vqpsATGRXVDR-----L----vgaelI-----DR-L-PHPT---sr----rELLAALtkrgrd
0043 6jyiA   kdpXKGRDVVLGLLX-Q-----K---E-L-S-GYDIK-IVFE-DvFT----HF----------fdgsfG---XIYPTLRQ-----L----enegkI-----KK-E-VVXQK---p----nKKXYFItdegre
0044 3f6vA   vaAEPTRRRLVQLLT-S-----G---E-Q-T-VNNLA-AHFP--------------------------asrsAISQHLRV-----L----teaglV-----TP-R-KDG----------rFRYYRLdpqgla
0045 5fmfV   mpKKEILDYLFKLFD-E-----Y--dY-W-S-LKGLK-ERTR-----------------------qpeA---HLKECLDK-----V---------A-----TL-V-KKGP---------yAFKYTLrpeykk
0046 6luiA   --SPHYQEWILDTID-S-----L--karP-D-LERIC-RXVR-R-RH----G-------------pepE---RTRAELEK-----L----iqqraV-----LR-V-SYK----------gSISYRNaar---
0047 1xmkA   ldMAEIKEKICDYLFnV-----S---D-S-S-ALNLA-KNIG-L-TK-----------------------arDINAVLID-----M----erqgdV-----YR-Q-GT-----------tPPIWHLtdkkre
0048 2ethA   sdXKTTELYAFLYVA-L----fG---P-K-K-XKEIA-EFLS-----------------------ttkS---NVTNVVDS-----L----ekrglV-----VR-E-XDPVD---r----rTYRVVLtekgke
0049 2p4wB   vlGNETRRRILFLLT-K-----R---P-Y-F-VSELS-RELG--------------------------vgqkAVLEHLRI-----L----eeaglI-----ES-R-VEKI-prgr----pRKYYMIkkglrl
0050 3u21A   eisSSFFSLLLEILL-L-----E--sQ-A-S-LPXLE-ERVL-D-WQsspaSS--LNswfsAA--P--NwaeLVLPALQY-----LavpssfspfV-----EF-X-EX------------TQQWXLlgdnex
0051 3l7wA   vsALLIEYLILAIVS-K-----H---D-S-Y-GYDIS-QTIK-L-IA----S-------------ikeS---TLYPILKK-----L----ekagyL-----ST-Y-TQE--hqgr----rRKYYHLtdsgek
0052 7xjgJ   gnILYAERFLLPCIY-L-----L--dS-V-N-YRTLC-ELAF-K-AI----KQ--D-------dvlskI---IVRSVVSR-----L----inerkI-----LQ-M---------------TDGYQVtalgas
0053 8ioiA   pefMALPHAILVSLS-E-----Q---A-S-S-GYELA-RRFD-RsIG----YF----------wtathQ---QIYRTLRV-----M----ennnwV-----RA-T-TVLQH-grp----dKKVYAIsdsgra
0054 3gfiA   lnLSYLDFLVLRATS-D-----G---P-K-T-MAYLA-NRYF-----------------------vtqS---AITASVDK-----L----eemglV-----VR-V-RDREdRR-------KILIEItekgle
0055 4ijaA   -pMNDNEKRVLREIY-N-----H--hN-I-S-RTQIS-KNLE-----------------------inkA---TISSILNK-----L----kykslV-----NE-V-G---------ggrkpILLKVnhlygy
0056 8xt8B   ydITTDHYALLRFLW-E-----Q---DgI-S-QIDLC-EKSC-----------------------kdkS---NTTRILDV-----M----knkglI-----VR-K-VDVK---dr----rKFQIFLtdlgre
0057 6j0eB   vlgDPVRLRILSQLA-A-----G--gP-V-S-VNELT-DLMG-----------------------lsqP---TISHHLKK-----M----teagfL-----DR-V-PEG----------rVVLHRVrpelfa
0058 3k69A   klDFSVAVHSILYLD-A-----H--sK-V-A-SRELA-QSLH-L-------------------------npvXIRNILSV-----L----hkhgyL-----TG-T-VG-----------kNGGYQLdlalad
0059 7wzeA   rgLSEGKFKILXLLF-D-----A--hR-L-S-PTELA-KRSN--------------------------vtkaTITGLLDG-----L----ardgfV-----SR-R-HH-----------rKISIELttegka
0060 3fhzD   anRAGRQARIVAILS-S-----A---Q-VrS-QNELA-ALLA-A-EG----------------ievtqA---TLSRDLEE-----L--------gA-----VK-L-RGADGG--------TGIYVVpedgsp
0061 4xrfA   tgLTPPQFYILKILD-H-----Y--gA-S-R-ATTLA-KKMY--------------------------vkpsAITVMIDR-----L----idqelV-----ER-Y-HDKD---dr----rVVIIELtkkgka
0062 6v9iC   lrILRTQEAIIQIMK-M-----R--kK-I-S-NAQLQ-TELV-E-IL----KN--M-------flpqkK---MIKEQIEW-----L----iehkyI-----RR-D-ES-----------dINTFIYma----
0063 3f8fA   mlRAQTNVILLNVLK-Q-----G---D-N-Y-VYGII-KQVK-E-AS-------------ngemelneA---TLYTIFKR-----L----ekdgiI-----SS-Y-WGDE---gg----rRKYYRLteighe
0064 7el3A   vgLTEQQWRIIRILY-Q-----Y--eE-L-E-SNQLA-ELAC--------------------------ilkpSLTGILNR-----M----veqklI-----QK-R-KDYD---dq----rISLISLtesgle
0065 2g9wA   trLGDLERAVXDHLW-S-----R--eP-Q-T-VRQVH-EALS-A-RR----D-------------layT---TVXAVLQR-----L----akknlV-----LQ-I-R-------------AHRYAPvhgrde
0066 3f72B   aiADENRAKITYALC-Q-----D--eE-L-C-VCDIA-NILG--------------------------vtiaNASHHLRT-----L----ykqgvV-----NF-R-KEG----------kLALYSLggeair
0067 2mh2A   ----GAPGIILRYLQ-E-----Q--rP-Y-S-AQDVF-GNLQ-K-EH-----------------glgkA---AVVKALDQ-----L----aqegkI-----KE-K-TYG----------kqKIYFAd-----
0068 5okcB   tlpxDPKERFKVLFR-L-----Q--sQ-W-D-LEDIK-PLIE---EL----NS---------R--G--X---KIDSFIXK-----Y---------A-----RR-K-RLGK----------KTVVTSr-----
0069 2cweA   vmLEDTRRKILKLLR-N-----K---E-M-T-ISQLS-EILG--------------------------ktpqTIYHHIEKlkeagL---------V-----EVkR-TEM--kgnl----vEKYYGRtadvfy
0070 7vwvB   meTQKLISMVKEALE-K-----Y--yP-L-T-AKNIK-VVIQ-K-EH----N--------------vvlptgSINSILYS-----N------selF-----EK-I-DKTNTI-------ypPLWIRkn----
0071 7kbfK   --hPPTLSMVVEVLK-K-----N--kG-T-S-VQAIR-TRIL-S-AH----PT--------vdplrlkF---LLRTALNK-----G----lekgiL-----IR-PlNSSA-------tgatGRFKLak----
0072 7asvA   pvARELKAFVEATFQ-R-----Q--fV-L-T-LSELK-RLFN-L-HL---aSL--PPG---HTlfS--G---ISDRXLQD-----T---------VlaagcKQ-I-LVPFPpqtaaspdeqKVFALwesgdx
0073 1yg2A   ----SLPHVILTVLS-T-----R---D-A-T-GYDIT-KEFS-AsIG----YF----------wkashQ---QVYRELNK-----M----geqglV-----TC-V-LE--------------VYSItqagrs
0074 6o8oB   alAHEGRLXIXCYLA-S-----G---E-K-S-VTELE-TRLS-----------------------trqA---AVSQQLAR-----L----rleglV-----QS-R-REG----------kTIYYSLsdpraa
0075 3v4gA   -kQDNLVRAFKALLK-E-----E---R-F-GsQGEIV-EALK-QeGF----EN-------------inqS---KVSRMLTK-----F--------gA-----VR-T---rnaKM-------EMVYCLptvsss
0076 4hqeB   ilgRSWNGLIINYLS-R-----C--cS-A-H-FSDMK-RDLK-T-I--------------------tpR---ALSLKLSE-----L----aqwelV-----EK-Q-IISTS---p----vQIIYVLtekgka
0077 3u2rA   feLSAQQYNTLRLLR-S-----V--eG-X-A-TLQIA-DRL-----------------------------apDITRLIDR-----L----ddrglV-----LR-T-RKPE---nr----rVVEVALtdaglk
0078 5n35A   -vNQKEIEIAIEYFK-N-----Y--iS-V-G-EIVAT-MDLK-A-RG-----------------------isNPQAVISK-----L----iemgiI-----EK-G---------------EGCYNLvr----
0079 4txaA   arQEEALKLVLLALE-D----gS---A-L-S-RKVLV-LFVV-Q-RL----EP--R-F---PQ--A---sktSIGHVVQL-----L---------YrascfKV-T-KRD-----------SSLMQLkeefrt
0080 6a6iA   teHDDLLVEMRNFIA-FqahtdG---Q-A-S-TREIL-QEFE-S-KL----SA--S----------qsC---VFRELLRN-----L---------C-----TF-H-RTSG---------gEGIWKLkpeyc-
0081 3bpxA   lnLTDAQVACLLRIH-R-----E---PgI-K-QDELA-TFFH-----------------------vdkG---TIARTLRR-----L----eesgfI-----ER-E-QDPE---nr----rRYILEVtrrgee
0082 4yifB   spVSLSQLSALTTLA-N----eG---A-M-T-PGALA-IRER--------------------------vrppSMTRVIAS-----L----admgfV-----DR-A-PHPID---g----rQVLVSVsesgae
0083 6yj6B   esLEECLKILRELFA-R-----R--pI-W-V-KRHLD-GIVP-K-KI--------------------hH---TMKIALAL-----I---------S-----YR-F-TMG--PW-------RNTYIKfgidpr
0084 7bzdA   viGGKWKMLILWHLG-K----eG---T-K-R-FNELK-TLIP-D-I--------------------tqK---ILVNQLRE-----L----eqdmiV-----HR-E-VYPVV---p----pKVEYSLtphges
0085 2gaiA   lsAGRVQSATLKLVC-D-----RerpR-Y-T-EGSLV-KEME-R-LG---------------igrP--S---TYASTIKL-----L----lnrgyI-----KK-I---------------RGYLYPtivgsv
0086 5z7bB   kdLMTLRSALLALLS-S-----G---P-L-T-GYDAS-QRFG-AsVG----FV----------wsgsdS---QIYPELRK-----M----eaeelL-----VG-S-DVPwgskga----tKTEYALsekgwe

0001 s001A   --LHHHHHHHHHHHH-H-----L---L-E-E-HHHHH-HHHH-H-HL----LL--LLL---LL--L--L---LHHHHHHH-----H---------L-----EE-L-LLLLLLL-------LLEEEL------
0002 6qfdB   llLLHHHHHHHHHHH-H-----L---L-E-E-HHHHH-HHHH-H-HH----L-----------llllhH---HHHHHHHH-----H----hhlllE-----EE-E-EEEL---------lEEEEEElhhhhh
0003 6zvhy   --lLLHHHHHHHHHH-L-----L--hE-E-E-HHHHH-HHHH-H-HH----H--lLLL---LLhhhhhH---HHHHHHHL-----L-------llE-----EE-L-L-------------LEEEEll-----
```

```
0004 8qkfA   hhLLHHHHHHHHHHHH-H-----L---L-L-L-HHHHH-HHHL------------------------llhH---HHHHHHHH-----H----hhlllE-----EE-E-EEL-----------lEEEEEEllllhh
0005 5dymA   ---LLHHHHHHHHHHL-L-----L---L-E-E-HHHHH-HHHH-H-HL----LL-------------llhH---HHHHHHHH-----H----hhlllE-----EE-E-ELLLLllll-----lLEEEEElhhhhh
0006 5j6xA   lhHHHHHHHHHHHHHH-H----hL---L-L-L-HHHHH-HHLL------------------------llhH---HHHHHHHH-----H----hhlllE-----EE-E-LLLL----------lLLEEEEll----
0007 7wjpA   hhHHHHHHHHHHHHHH-H-----L---L-E-E-HHHHH-HHHL-L-LL----L-----------------llllLHHHHHHH-----H----hhlllE-----EE-E-EELL----l-----lEEEEEElhhhhh
0008 8jxkD   --LHHHHHHHHHHHHH-H-----L---L-E-E-HHHHH-HHHH-H-HL----L----------lllllllL---LHHHHHHH-----H----hhlllE-----EE-L-LLLLL---------lLEEEEElhhhhh
0009 1lddA   hhHHHHHHHHHHHHHH-H----hL---L-E-E-HHHHH-HHHH-H-HL----LH--H---hllLL--L--L-hhHHHHHHHH-----H----hhlllE-----EL-L-L--------------LLEEEEl-----
0010 2od5A   hhHHHHHHHHHHHHLL-L-----L---L-E-E-HHHHH-HHHH-L-LL----LL---------------lllhhHHHHHHHH-----L-------llE-----EE-E-EEeeeeellleeeeeeeEEEEhhhhhh
0011 1sfxA   llLLHHHHHHHHHHHH-H-----H--lL-L-L-HHHHH-HHHL---------------------------llhhHHHHHHHH-----H----hhlllE-----EE-E-EEELL---l-----eEEEEEEllhhhh
0012 2eshA   hhHHHHHHHHHHHHHH-H-----L---L-L-L-HHHHH-HHHH-L-LL----LL--LLL---L-------llllLHHHHHHH-----H----hhlllE-----EE-E-EELLL--ll-----lEEEEEElhhhhh
0013 5a31N   hhHHHHHHHHHHHHHH-H-----H--lL-L-L-HHHHH-HHHH-H-HL----L-------lllllllllhH---HHHHHHHH-----H----hhlllL-----EE-E----------------LLEEELl-----
0014 3elkA   hhHHHHHHHHHHHHHH-H-----L---L-E-E-HHHHH-HHHH-H-HH----LL---------------lllllHHHHHHHH-----H----hhhllE-----EE-E-EEEL--lll-----eEEEEEElhhhhh
0015 8sojB   hhLHHHHHHHHHHHHL-L-----L--lL-E-E-HHHHH-HHHH-HhLL----LL-------------llhH---HHHHHHHH-----H----hhhllL-----EE-E-E--------------LLEEEL------
0016 5zqhA   hhHHHHHHHHHHHHHH-H-----L---L-E-E-HHHHH-HHHH-HhLL----LL--------------llL---LLHHHHHH-----H----hhlllE-----EE-E-EE-elllll-----eEEEEEElhhhhh
0017 5jlsA   llLLHHHHHHHHHHHH-H-----L---L-L-L-HHHHH-HHHL------------------------llhH---HHHHHHHH-----H----hhlllE-----EE-E-ELLL---ll-----lLEEEEEllllhh
0018 2zkzC   hhLLHHHHHHHHHHHH-H-----H--lL-E-E-HHHHH-HHHL------------------------llhH---HHHHHHHH-----H-----llllE-----EE-E-EEL-----------lEEEEEEllhhhh
0019 5xpqA   hllLHHHHHHHHHHHH-H-----L---L-E-E-HHHHHlLLLL------------------------llhH---HHHHHHHH-----H----hhlllE-----EE-E-ELL-----------lLEEEEElllhhh
0020 4ejoA   hhHHHHHHHHHHHHLL-------L---L-E-E-HHHHH-HHHH-H-LL-----------------llllhH---HHHHHHHH-----H----hhlllE-----EE-E-EEELL--ll-----eEEEEEElhhhhh
0021 7z8bC   hhHHHHHHHHHHHHHH-L-----L--lL-E-E-HHHHH-HHHH-H-HH----HllhLLL---LL--lllhH---HHHHHHHH-----H----hhhllE-----EE-L-LL------------lLLEEEEll----
0022 4omzC   hhLLHHHHHHHHHHHH-H-----L---L-E-E-HHHHH-HHLL------------------------llhH---HHHHHHHH-----H----hhlllL-----EE-E-EEL-----------lEEEEELllhhhh
0023 5hs7A   hhhLLLHHHHHHHHHL-L-----L---L-E-E-HHHHH-HHLL-L-L---------------------lhH---HHHHHHHH-----H----hhlllE-----EE-E-EELLL---l-----lEEEEEElhhhhl
0024 8uucA   llHHHHHHHHHHHHHH-H-----L---LlE-E-HHHHH-HHHH-H-HH--hhLL--LL--------lllhH---HHHHHHHH-----H----hhhllL-----EE-E----------------LLEEEL------
0025 8iueP   -lHHHHHHHHHHHHHH-H-----L--lL-L-L-HHHHH-HHLL-L-L---------------------lhH---HHHHHHHH-----H----hhhllE-----EE-E-EEL-----------lEEEEEElhhhll
0026 3mq0B   lhHHHHHHHHHHHHHH-H-----L--lL-E-E-HHHHH-LLLL------------------------llhH---HHHHHHHH-----H----hhlllE-----EE-L-L--------------LLEEEElllhhh
0027 3bddA   hlLLHHHHHHHHHHHH-H-----H---LlL-L-HHHHH-HHHL------------------------llhH---HHHHHHHH-----H----hhlllE-----EE-E-ELLLL---l-----lLEEEEElhhhhh
0028 2l4mA   llLLHHHHHHHHHHHH-H----hL---L-E-E-HHHHH-HHHL-L-------------------------llhhHHHHHHHH-----H----hllllE-----EE-L-LL------------LLEEEEEll----
0029 6uvuB   hhlLHHHHHHHHHHHH-H-----H--lL-L-L-HHHHH-HHLL---------------------------llhhHHHHHHHH-----H----hhlllE-----EE-E-EEL-----------lEEEEEEllhhhh
0030 2rdpA   llLLHHHHHHHHHHHH-H----hL---L-L-L-HHHHH-HHHL---------------------------llhhHHHHHHHH-----H----hhlllE-----EE-E-ELLL-----------LEEEEElhhhhh
0031 2oqgA   hllLHHHHHHHHHHHH-H-----L---L-L-L-HHHHH-HHLL------------------------llhH---HHHHHHHH-----H----hhlllE-----EE-E-EEL-----------lEEEEEEllhhhh
0032 5zi8A   hhHHHHHHHHHHHHHH-H-----L---L-L-L-HHHHH-HHHH-H-LL----LL-------------llhH---HHHHHHHH-----H----hhlllE-----EE-E-EEEEL--le-----eEEEEEElhhhhh
0033 2fe3B   llLLHHHHHHHHHHHH-H-----L--lL-L-L-HHHHH-HHHL-L-LL----LL-------------llhH---HHHHHHHH-----H----hhlllE-----EE-E-LLLL----------lLLEEEEllllee
0034 7mexA   hhHHHHHHHHHHHHHL-L-----L---L-E-E-HHHHH-HHLL-H-HH------------------lllhH---HHHHHHHH-----H---------E-----EE-E-LLLLL---l-----lLLEEEEllhhhh
0035 7z1nO   hhLLHHHHHHHHHHHH-H----hL---L-E-E-HHHHH-HHLL-L-L---------------------lhH---HHHHHHHH-----H----hhlllL-----LE-E-EEEL-llll-----eEEEELLlhhhhh
0036 2xrnB   lhHHHHHHHHHHHHHH-L-----L--lL-E-E-HHHHH-HHLL---------------------------llhhHHHHHHHH-----H----hllllE-----EE-L-HH------------HLEEEEEllhhhh
0037 3s93B   hhHHHHHHHHHHHHHH-L-----L--lL-L-L-HHHHH-HHHHhH-HL----L----lllllhHH--L--L--lLHHHHHHL-----L------lllE-----EE-E-ELHH----------hLEEEEEll----
0038 6j05A   hhLLHHHHHHHHHHHH-H-----L--lL-E-E-HHHHH-HHHL-L--------------------------lhhHHHHHHHH-----H----hhlllE-----EE-E-EEL-----------leEEEEElhhhhh
0039 3l9fA   ---LLHHHHHHHHHLL-L-----L---L-E-E-HHHHH-HHHH-HlLL----LL-------eelL--L--L---LHHHHHHH-----H----hhlllE-----EE-E-EELL-llll-----lEEEEEElhhhhh
0040 2wteA   llLLHHHHHHHHHHHH-H-----H--lL-L-L-HHHHH-HHHL---------------------------llhhHHHHHHHH-----H----hhlllE-----EE-E-LL-------------LLEEEElhhhhh
0041 3b73B   llLLHHHHHHHHHHHH-H-----H--lE-E-L-HHHHH-LL-L-L-LL----------------------llhhHHHHHHHH-----H----hhlllE-----EE-E-H--------------HHEEEElhhhhh
0042 2nyxB   llLLHHHHHHHHHHHH-H-----H--lL-E-E-HHHHH-HHHL---------------------------llhhHHHHHHHH-----H----hhlllE-----EE-E-LLLL---ll-----lLLEEEElhhhhh
0043 6jyiA   lllLHHHHHHHHHHHH-H-----L---L-L-L-HHHHH-HHHH-HlHH----HL-----------llllhH---HHHHHHHH-----H----hhlllE-----EE-E-EELLL---l-----lEEEEEElhhhhh
0044 3f6vA   hhLLHHHHHHHHHHHH-H-----L---L-E-E-HHHHH-LLLL---------------------------llhhHHHHHHHH-----H----hhlllE-----EE-E-EEL-----------lEEEEEElhhhhh
0045 5fmfV   hhHHHHHHHHHHHHHH-H-----L--lL-L-L-HHHHH-HHLL------------------------llhH---HHHHHHHH-----H---------L-----LL-L-LLLL----------lLLLLLLllllll
0046 6luiA   --LLLHHHHHHHHHHH-H-----H--lllL-L-HHHHH-HHHH-H-HH----L--------------llhH---HHHHHHHH-----H----hhlllE-----EE-E-EEL-----------lEEEEEElll---
0047 1xmkA   hhHHHHHHHHHHHHHHhL-----L---L-E-E-HHHHH-HHHL-H-HH------------------------hhHHHHHHHH-----H----hhlllE-----EE-E-LL------------lLLEEEElhhhhl
0048 2ethA   hhLLHHHHHHHHHHHH-H----hL---L-L-L-HHHHH-HHLL------------------------llhH---HHHHHHHH-----H----hhlllE-----EE-E-ELLLL---l-----lLEEEEElhhhhh
0049 2p4wB   hlLLHHHHHHHHHHHL-L-----L---L-E-E-HHHHH-HHHL---------------------------llhhHHHHHHHH-----H----hhlllE-----EE-E-ELLL-llll-----lLEEEEEllleee
0050 3u21A   lllLLHHHHHHHHHHH-L-----L--lE-E-E-HHHHH-HHHH-H-HHllhhHH--HLllhhHL---L--LlhhHHHHHHHH-----HlllllllllE-----EE-E-LL-------------LLEEEEllllhh
0051 3l7wA   llHHHHHHHHHHHHHH-H-----L---L-E-E-HHHHH-HHHL-L-LL----L--------------llhH---HHHHHHHH-----H----hhlllE-----EE-E-EEE--elle-----eEEEEEElhhhhh
0052 7xjgJ   llLLLLHHHHHHHHHH-H-----L--lL-L-L-HHHHH-HHHH-H-HL----LL--L--------lhhhhH---HHHHHHHH-----H----hhlllE-----EE-E----------------LLEEEElhhhhh
0053 8ioiA   lllLLHHHHHHHHHHH-H-----L---L-L-L-HHHHH-HHHH-HlHH----HL-----------llllhH---HHHHHHHH-----H----hhlllE-----EE-E-EELLL-lll-----lEEEEEElhhhhh
0054 3gfiA   hlLLHHHHHHHHHHHL-L-----L---L-E-E-HHHHH-HHLL------------------------llhH---HHHHHHHH-----H----hhlllE-----EE-E-EELLeEE--------EEEEEElhhhhh
0055 4ijaA   -lLHHHHHHHHHHHHH-H-----L--lL-L-L-HHHHH-HHHL------------------------llhH---HHHHHHHH-----H----hhlllE-----EE-L-L----------lllllLLEEElhhhle
0056 8xt8B   llLLHHHHHHHHHHHH-H-----L---LlL-L-HHHHH-HHHL------------------------lllH---HHHHHHHH-----H----hllllE-----EE-E-ELLL---ll-----lLEEEEElhhhhh
0057 6j0eB   hhlLHHHHHHHHHHHH-H-----L--lL-E-E-HHHHH-HHHL------------------------llhH---HHHHHHHH-----H----hhlllE-----EE-E-EEL-----------lEEEEEElhhhhh
0058 3k69A   lhHHHHHHHHHHHHHH-L-----L--lL-L-L-HHHHH-HHHL-L--------------------------lhhHLHHHHHH-----H----hhlllL-----EE-E-LL------------lLLEEELlllhhh
0059 7wzeA   llLLHHHHHHHHHHHH-H-----L--hE-E-L-HHHHH-HHLL---------------------------llhhHHHHHHHH-----H----hhhllE-----EE-E-EL------------lLEEEEElhhhhh
0060 3fhzD   llHHHHHHHHHHHHHH-H-----L---L-LlL-HHHHH-HHHH-L-LL-----------------llllhH---HHHHHHHH-----L--------lL-----EE-E-LLLLLL---------LLEEELllllll
0061 4xrfA   hlLLHHHHHHHHHHHH-H-----H--lL-E-E-HHHHH-HHHL---------------------------llhhHHHHHHHH-----H----hhlllE-----EE-E-ELLL---ll-----lLEEEEElhhhhh
0062 6v9iC   hhHHHHHHHHHHHHHH-H-----L--lE-E-E-HHHHH-HHHH-H-HH----LL--L--------llllhH---HHHHHHHH-----H----hhlllE-----EE-E-LL------------eEEEEEEll----
0063 3f8fA   hhHHHHHHHHHHHHHH-H-----L---L-L-L-HHHHH-HHHH-H-HL--------------lllllllhH---HHHHHHHH-----H----hhlllE-----EE-E-ELLL---ll-----lLEEEEElhhhhh
0064 7el3A   hlLLHHHHHHHHHHHH-H-----H--lE-E-E-HHHHH-HHHL---------------------------llhhHHHHHHHH-----H----hhlllE-----EE-E-EELL---ee-----eEEEEEElhhhhh
0065 2g9wA   hhLLHHHHHHHHHHHH-L-----L--lL-E-E-HHHHH-HHHL-L-LL----L--------------llhH---HHHHHHHH-----H----hhlllE-----EE-E-L--------------LLEEEElllhhh
0066 3f72B   hhHLHHHHHHHHHHLL-L-----L--lL-E-E-HHHHH-HHHL---------------------------llhhHHHHHHHH-----H----hhlllE-----EE-E-ELL-----------lLEEEEEllhhhh
0067 2mh2A   ----LHHHHHHHHHHH-H-----H--lL-E-E-HHHHH-HHHH-L-LL------------------lllhH---HHHHHHHH-----H----hhhllE-----EE-E-EEL-----------leEEEEEl-----
0068 5okcB   hlllLHHHHHHHHHHH-H-----L--lL-L-L-HHHHH-HHHH---HH----LL----------L--L--L---LHHHHHHH-----H---------E-----EE-E-ELLL-----------LEEEEEl-----
0069 2cweA   hhLLHHHHHHHHHHHL-L-----L---L-L-L-HHHHH-HHHL---------------------------llhhHHHHHHHHhhhllL---------E-----EEeE-EEE--elle-----eEEEEEEllleee
0070 7vwvB   lhHHHHHHHHHHHHHH-H-----L--lL-L-L-HHHHH-HHHH-H-HH----L---------------llllhhHHHHHHHH-----L------lllE-----EE-L-LLLLLL--------llLLEEEll----
0071 7kbfK   --lLLHHHHHHHHHHH-H-----L--lL-E-E-HHHHH-HHHH-L-LL----LL---------lllllhhH---HHHHHHHH-----H----hhlllE-----EE-EeELLL-------llllEEEEEEll----
0072 7asvA   lhHHHHHHHHHHHHHH-H-----H--lE-E-E-HHHHH-HHHH-H-HH----hLL--LLL---LHhhL--L---LLHHHHHH-----H---------HhhlllEE-E-LLLLLllllllllhhhLEEELllllll
0073 1yg2A   ----LHHHHHHHHHHH-H-----L---L-L-L-HHHHH-HHHL-LhHH----HL-----------llllhH---HHHHHHHH-----H----hhlllE-----EE-L-LL---------------LEEElhhhhh
0074 6o8oB   hhLLHHHHHHHHHHHH-H-----L---L-E-E-HHHHH-HHLL------------------------lllL---HHHHHHHH-----H----hhlllE-----EE-E-ELL-----------lLEEEEEllhhhh
0075 3v4gA   -lHHHHHHHHHHHHHH-H-----L---L-L-LlHHHHH-HHHH-HlLL----LL-------------llhH---HHHHHHHH-----L--------lL-----EE-E---ellLL--------LEEEELllllll
0076 4hqeB   hhlLLLHHHHHHHHHL-L-----L--hE-E-L-HHHHH-HHHL-L-L---------------------lhH---HHHHHHHH-----H----hhlllE-----EE-E-EELLL---l-----lEEEEEElhhhhh
0077 3u2rA   llLLHHHHHHHHHHHH-H-----H--lL-E-E-HHHHH-HHL------------------------------llHHHHHHHH-----H----hhlllE-----EE-E-EELL---ee-----eEEEEEElhhhhh
0078 5n35A   -lLHHHHHHHHHHHHH-H-----L--lE-E-E-HHHHH-HHHH-H-LL------------------------llLHHHHHHH-----H----hhlllE-----EE-L----------------LLEEEEll----
0079 4txaA   hhHHHHHHHHHHHHHL-L----lL---L-L-L-HHHHH-HHHH-H-HL----LL--L-L---LL--L----lhhHHHHHHHH-----H---------HhllleEE-E-ELL------------LEEEEElhhhll
0080 6a6iA   llLHHHHHHHHHHHHH-HlllllL---E-E-E-HHHHH-HHHL-L-LL----LH--H-----------hhH---HHHHHHHH-----H---------E-----EE-E-ELLL----------lLEEEEElhhhl-
0081 3bpxA   hlLLHHHHHHHHHHHH-L-----L---LlL-L-HHHHH-HHHL------------------------llhH---HHHHHHHH-----H----hhlllE-----EE-E-EELL---ee-----eEEEEEElhhhhh
0082 4yifB   llLLHHHHHHHHHHHH-H----hL---L-L-L-HHHHH-HHHL---------------------------llhhHHHHHHHH-----H----hhlllE-----EE-E-ELLLL---l-----lLEEEEElhhhhh
0083 6yj6B   hhHHHHHHHHHHHHHH-H-----L--lE-E-L-HHHHL-LLLL-L-LL---------------------lL---LHHHHLLL-----L---------E-----EE-E-LLL--LL--------LLLEEEllllhh
0084 7bzdA   hhLLLLHHHHHHHHHH-H----hL---L-E-E-HHHHH-HHLL-L-L---------------------lhH---HHHHHHHH-----H----hhlllE-----EE-E-EELLL---l-----lEEEEEElhhhhh
0085 2gaiA   llLHHHHHHHHHHHHH-H-----HhhlL-L-L-HHHHH-HHHH-H-HL----------------lllL--L---LHHHHHHH-----H----hhlllE-----EE-E----------------LLEEEElhhhhh
0086 5z7bB   hhHHHHHHHHHHHHHL-L-----L---L-E-E-HHHHH-HHHH-LlHH----HL-----------llllhH---HHHHHHHH-----H----hhlllE-----EE-E-EEEellll-e----eEEEEEElhhhhh

@@@@@@@@@@@@@@@@@@@@@@@@@@@@@@@@@@@@@@@@@@@@@@@@@@@@@@@@@@@@@@@@@@@@@@@@@@@@@@@@@@@@@@@@@@@@@@@@@@@@@@@@@@@@@@@@@@@@@@@@@@@@@@@@@@@@@@@@@@@@@@@@@
@@@@@@@@@@@@@@@@@@@@@@@@@@@@@@@@@@@@@@@@@@@@@@@@@@@@@@@@@@@@@@@@@@@@@@@@@@@@@@@@@@@@@@@@@@@@@@@@@@@@@@@@@@@@@@@@@@@@@@@@@@@@@@@@@@@@@@@@@@@@@@@@@
@@@@@@@@@@@@@@@@@@@@@@@@@@@@@@@@@@@@@@@@@@@@@@@@@@@@@@@@@@@@@@@@@@@@@@@@@@@@@@@@@@@@@@@@@@@@@@@@@@@@@@@@@@@@@@@@@@@@@@@@@@@@@@@@@@@@@@@@@@@@@@@@@


# Job: Type-3 BREX; BrxHII-Helicase WP_162523516  (N-Terminal TUDOR Domain)
# Query: s001A
# No:  Chain   Z    rmsd lali nres  %id PDB  Description
   1:  6hq9-A  6.9  1.5   47    60    4   MOLECULE: DNA EXCISION REPAIR PROTEIN ERCC-6-LIKE 2;
   2:  6if4-B  6.4  1.6   48    61   10   MOLECULE: HISTONE ACETYLTRANSFERASE;
   3:  3fc3-A  6.3  1.3   44   189    9   MOLECULE: RESTRICTION ENDONUCLEASE HPY99I;
   4:  7abi-7  6.2  1.6   45   236   16   MOLECULE: U5 SMALL NUCLEAR RIBONUCLEOPROTEIN 40 KDA PROTEIN
   5:  7ef1-A  6.2  2.0   48   148    8   MOLECULE: HB TRANSCRIPTION FACTOR;
   6:  1m1g-C  6.1  1.8   46   244   15   MOLECULE: TRANSCRIPTION ANTITERMINATION PROTEIN NUSG;
   7:  1vie-A  6.1  1.9   48    60   10   MOLECULE: DIHYDROFOLATE REDUCTASE;
   8:  5hh7-A  6.1  1.4   47   192    2   MOLECULE: ORIGIN OF REPLICATION COMPLEX SUBUNIT 1B;
   9:  2lcc-A  6.1  2.0   51    76   12   MOLECULE: AT-RICH INTERACTIVE DOMAIN-CONTAINING PROTEIN 4A;
  10:  8xv7-A  6.0  1.5   48   227   10   MOLECULE: E3 UBIQUITIN-PROTEIN LIGASE UHRF1;
  11:  8jw0-e  5.9  1.9   48    74   10   MOLECULE: PHOTOSYSTEM I PSAA;
  12:  6kcp-C  5.9  1.3   43    81   16   MOLECULE: LD23804P;
  13:  7cce-A  5.8  1.5   48   159   13   MOLECULE: BROMO-ADJACENT HOMOLOGY (BAH) DOMAIN-CONTAINING P
  14:  2dig-A  5.8  1.3   45    68   16   MOLECULE: LAMIN-B RECEPTOR;
  15:  4m4z-A  5.8  1.3   41   152   22   MOLECULE: SRC-LIKE-ADAPTER 2;
```

```
 16:  2lq8-A  5.7  1.7   46   177   11   MOLECULE: TRANSCRIPTION ANTITERMINATION PROTEIN NUSG;
 17:  7cfd-F  5.7  1.4   44   182    7   MOLECULE: FI20010P1;
 18:  5m9n-A  5.7  1.7   45   202    9   MOLECULE: TUDOR DOMAIN-CONTAINING PROTEIN 1;
 19:  8jvz-A  5.5  1.8   42    50   14   MOLECULE: MKAL2_V1_SH3;
 20:  6ted-Z  5.5  1.4   44   510    7   MOLECULE: DNA-DIRECTED RNA POLYMERASE SUBUNIT;
 21:  6cb1-Z  5.5  2.0   48   135   15   MOLECULE: 35S PRE-RIBOSOMAL RNA MISCRNA;
 22:  1vbv-A  5.5  1.6   47    77    9   MOLECULE: HYPOTHETICAL PROTEIN B0966;
 23:  6bim-A  5.5  1.8   44   278   14   MOLECULE: CLAN CA, FAMILY C40, NLPC/P60 SUPERFAMILY CYSTEIN
 24:  2jt4-A  5.4  2.2   48    71   10   MOLECULE: CYTOSKELETON ASSEMBLY CONTROL PROTEIN SLA1;
 25:  2e70-A  5.4  1.8   46    71    9   MOLECULE: TRANSCRIPTION ELONGATION FACTOR SPT5;
 26:  1sf9-A  5.4  1.6   44   118    9   MOLECULE: YFHH HYPOTHETICAL PROTEIN;
 27:  5z8l-A  5.4  1.5   46   204    9   MOLECULE: CHROMATIN REMODELING PROTEIN EBS;
 28:  8tqc-D  5.4  2.5   49   969   12   MOLECULE: MEDIATOR OF RNA POLYMERASE II TRANSCRIPTION SUBUN
 29:  7k9b-A  5.4  2.0   45    98    7   MOLECULE: OLE-ASSOCIATED PROTEIN B;
 30:  8rsw-A  5.4  2.5   48    67   15   MOLECULE: PROTEIN KINASE C AND CASEIN KINASE SUBSTRATE IN N
 31:  8qsz-Y  5.3  1.3   44    72   16   MOLECULE: DNA-DIRECTED RNA POLYMERASE II SUBUNIT RPB1;
 32:  8s7v-L  5.3  1.6   43    47    5   MOLECULE: METHYL-COENZYME M REDUCTASE SUBUNIT GAMMA;
 33:  7aoa-A  5.3  1.5   46   343   15   MOLECULE: METHYL-CPG-BINDING DOMAIN PROTEIN 2;
 34:  7csp-A  5.3  2.5   47   455    9   MOLECULE: RHO GUANINE NUCLEOTIDE EXCHANGE FACTOR 16;
 35:  6vld-B  5.3  2.6   48   470   15   MOLECULE: ALPHA-(1,6)-FUCOSYLTRANSFERASE;
 36:  4cos-A  5.3  1.7   47   313   11   MOLECULE: PROTEIN KINASE C-BINDING PROTEIN 1;
 37:  3qwx-X  5.3  2.3   46   171   11   MOLECULE: CELL DEATH ABNORMALITY PROTEIN 2;
 38:  3ob9-D  5.2  1.6   48    93   13   MOLECULE: MALE-SPECIFIC LETHAL 3-LIKE 1 (DROSOPHILA), ISOFO
 39:  2n5u-A  5.2  1.8   47    79    9   MOLECULE: TSR0524 PROTEIN;
 40:  5z3g-Q  5.2  2.1   45   137   18   MOLECULE: 25S RRNA;
 41:  4anj-A  5.2  1.6   40   995    8   MOLECULE: UNCONVENTIONAL MYOSIN-VI, GREEN FLUORESCENT PROTE
 42:  2bud-A  5.1  2.2   50    92   12   MOLECULE: MALES-ABSENT ON THE FIRST PROTEIN;
 43:  2lcs-A  5.1  2.2   48    67    8   MOLECULE: NAP1-BINDING PROTEIN 2;
 44:  1tg0-A  5.1  1.9   46    66    9   MOLECULE: MYOSIN TAIL REGION-INTERACTING PROTEIN MTI1;
 45:  8hyj-W  5.1  1.2   38    44   11   MOLECULE: DNA-DIRECTED RNA POLYMERASE V SUBUNIT 1;
 46:  6rwl-A  5.1  2.3   45   270   13   MOLECULE: POL PROTEIN;
 47:  1wfw-A  5.0  2.3   47    74   17   MOLECULE: KALIRIN-9A;
 48:  6bog-A  5.0  1.8   44   967    9   MOLECULE: RNA POLYMERASE-ASSOCIATED PROTEIN RAPA;
 49:  7ud6-A  5.0  1.7   42   266   14   MOLECULE: TYROSINE-PROTEIN KINASE FYN,CATECHOL O-METHYLTRAN
 50:  5zwx-A  5.0  1.4   45   148   13   MOLECULE: DUF724 DOMAIN-CONTAINING PROTEIN 6-LIKE;

0001 s001A   -------MGENVHWVFDRW-------N--------N--KPVKAVE-RY-EL---W---N-TT-NCLVYDP---T-S----G--R---IYTLPA--DGVGD-----
0002 6hq9A   ----mkdiwhPGERCLAPG-------K--------L--CEASIKS-IT-VD--eN---G-KS-FAVVLYA-----D----F--Q---ERKIPL--KQLQEvk---
0003 6if4B   -----mvmfeVRQKVYATL-----heT--------F--HAAIIQE-VA-HD-ahT---G-QL-LYYVHYV---E-Q----D--S-rmDRWLPG--SALRErr---
0004 3fc3A   -------mlkNDDFVIAKN-qlgnivP--------N--SVGVIRA-VN-G-----------K-SAMVLFI---G-L----N--E---LKRVDF--SELEAidiyr
0005 7abi7   -tartdywlqPEIIVKIITkklgekyH--------K--KKAIVKE-VI-D----------KY-TAVVKMI---D-S----G--D---KLKLDQ--THLETvipap
0006 7ef1A   -eatecvavlPGDLILCFQ-egkdqaL--------Y--YDAHVLD-AQ-RR---R---HcRC-RFLVRYD---H-D----S--S---EEIVPL--RKVCRrpetd
0007 1m1gC   -vkpskvefeKGDQVRVIE----gpfM--------N--FTGTVEE-VH-PE---------KR-KLTVMIS---I-R----M--T---PVELDF--DQVEKi----
0008 1vieA   ---psnatfgMGDRVRKKS------gA--------A--WQGQIVG-WY-CT---N---LtPE-GYAVESE---A-H----P-gS---VQIYPV--AALERin---
0009 5hh7A   -vefdetefeIGDDVYVKRpeiedcqikartmrekL--WAARIDK-LW-KE-vdD---G-VY-WIRARWY---M-T----N--D---FADIEM--ECILRhcsvk
0010 2lccA   -----edMEPCLTKVKVKY-grgktqK--------I--YEASIKS-TE-ID---D---G-EV-LYLVHYY---G-W----N--V-ryDEWVKA--DRIIWpldkg
0011 8xv7A   --gplgslykVNEYVDARD---tnmgA--------W--FEAQVVR-VT-RK---A-leE-DV-IYHVKYD---D-Y----PenG---VVQMNS--RDVRArarti
0012 8jw0e   ----awvgpkKGSQVRILR--pesywF--------Q--QRGTVVN-VN-QK---T---N-IKyPVTVKFD---F-VnyanV--N---TNGFAL--WEVDElkege
0013 6kcpC   ------asysIGDLVFAKV----kgyP--------P--WPAKITK-SN-------------K-KYNVYFY---G-T----G--E---TANIKL--EDLFPyasnk
0014 7cceA   -faydgntydLEVPVLLVP-------E--------P--YVAIIKD-IT-QT--kD---G-SM-MILGQWF---YrS----F--H---RDEVPA--ESVMHrcvvy
0015 2digA   -sgmpsrkfaDGEVVRGRW---pgssL--------Y--YEVEILS-HD-ST---------SQ-LYTVKYK-----D----G--T---ELELKE--NDIKSgpssg
0016 4m4zA   -------------TAVALGsfplslrL--------G--EPLTIVS-ED-G-----------D-WWTVLSE---V-S----G--R---EYNIPS--VHVAKvshgw
0017 2lq8A   -pvkvelgfkVGDMVKIIS----gpfE--------D--FAGVIKE-ID-PE---------RQ-ELKVNVT---I-R----E--T---PVVLHV--SEVEKie---
0018 7cfdF   -krtfkskprLLDIVLALY----sdgC--------F--YRAQIID-EF-P-----------S-EYMIFYV---D-Y----G--N---TEFVPL--SCLAPcenvd
0019 5m9nA   -pprsdfypaIGDICCAQF---seddQ--------W--YRASVLA-YA-S----------EE-SVLVGYV---D-Y----G--N---FEILSL--MRLCPiipkl
0020 8jvzA   -------gvkVGDVVEVKK----------------K--VVARVVE-LL-HD---P-arN-AP-VARVRFE-----D----G--E---ERLILV--P---------
0021 6tedZ   -aqelrkyfkMGDHVKVIA----grfE--------G--DTGLIVR-VE-E-----------N-FVILFSD---L-T----M--H---ELKVLP--RDLQLcseta
0022 6cb1Z   -----akflkAGKVAVVVR----gryA--------G--KKVVIVK-PHdEG---S---KpFG-HALVAGI---E-K----P--F---IKVVNY--NHLLPtrytl
0023 1vbvA   -----askfgIGQQVRHSL-------L--------G--YLGVVVD-ID-PV-------A-AP-WYHVVMEddnG-L----P--V---HTYLAE--AQLSSelqde
0024 6bimA   -wiskdsleEITIPTQSVT-------EpsrtgtivN--SQVTILD-FN-C----------DG-FYRINYR----------G--Y---IGYILE--DALQYkwkqi
0025 2jt4A   ------mASKSKKRGIVQY-------DfmaesqdeG--DKVYILD-DK-K---------sKD-WWMCQLV---D-S----G--K---SGLVPA--QFIEPvrdkk
0026 2e70A   -rgrgrrdNELIQTVRISQ----gpyK--------G--YIGVVKD-AT-E-----------S-TARVELH---S-T----C--Q---TISVDR--QRLTTvgsrr
0027 1sf9A   -yllnpedysPGETYRVEN-------T--------E--DEFTISY-LN-G-----------V-FAWGYRT---S-S----P-qQ---EEALPI--SVLQEke---
0028 5z8lA   -ikgtnkvvrAGDCVLMRP---------------sP--YVARVEK-IE-AD---A-----NV-KVHCRWY---Y-S----D--H---FDVQSA--HTIEGkcivh
0029 8tqcD   -hitlaqQSNSPFQVILCP-------F--------G--LNGTLTGqAF-KM---SdscS-LA-AVEVLVA----------G--V---RMIYPA--CFVLVpqsqt
0030 7k9bA   -----gpspeIGQIVKIVK----grdR--------D--QFSVIIK-RV-D----------DR-FVYIADG---D-K----R--KvdrAKRKNM--NHLKLidhis
0031 8rswA   ---gsrrASVGSVRVRALY-------DyegqeqdeG--DELTKME-NE-DE---------QG-WCKGRLD-----N----G--Q---VGLYPA--NYVEPiq---
0032 8qszY   ------rdkaIGATVRIRR----gpmK--------G--LLGVIKD-TT-D-----------A-NARVELH---T-G----N--K---MVTIPK--ENLLYttktg
0033 8s7vL   --------ieIGSYVRYIN-------T--------G--THGTVKA-IE-PK---N---D-EE-WVLLEN------------------DIYYRP--ELLELve---
0034 7aoaA   -----------GDYVYFEN----------------Y--LIRRIEE-LN-KT--aN---G-NV-EAKVVCF---Y-S----R--Q---LESLPA--THIRGkcsvt
0035 7cspA   -qwqgitNKGELPQVEVTK-------AyfakqadeA--DIVLVLQ-EE-D-----------G-WLHGERL---R-D----G--E---TGWFPE--SFAHSitsrv
0036 6vldB   gddiyyfGGQNAHNQIAIY-------AhqprtadeG--DIIGVAG-NH-W----------DG-YSKGVNR---K-L----G--R---TGLYPS--YKVREkietv
0037 4cosA   -nwfcepCSNPHPLVWAKL----kgfP--------F--WPAKALR-DK-D-----------G-QVDARFF---G-Q----H--D---RAWVPI--NNCYLmskei
0038 3qwxX   -sllaaykKPIIEVVVGTF-------KftgeretdG--ERLEILS-KT-N----------QD-WWEARNA-----L----G--T---TGLVPA--NYVQIq----
0039 3ob9D   -segmkfkfhSGEKVLCFE-------P------dpL--YDAKIVD-VI-VG---K-grK-IP-EYLIHFN---G-W----NrsW---DRWAAE--DHVLRdtden
0040 2n5uA   -fekiegrmdVGQKVRVCRaqdiiqkL--------G--QVGQITG-FK-MT---D---GsGV-GVIVTFD-----D----R--S---STWFFE--DEVEVvg---
0041 5z3gQ   -kasnwrlveVGRVVLIKK----gqsA--------G--KLAAIVE-II-D----------QK-KVLIDGP---K-A----G-vP---RQAINL--GQVVLtpltf
0042 4anjA   -----------GKPVWAPH----ptdG--------F--QVGNIVD-IG-P-----------D-SLTIEPL-------------K---TFLALI--NQVFPaeeds
0043 2budA   -lmqkidiSENPKIYFIRR----edgT--------V--HRGQVLQ-SR-TTenaA---A-PD-EYYVHYV---G-L----NrrL---DGWVGR--HRISDnaddl
0044 2lcsA   -------MAIVNQRAVALY-------DfependneG--DIVFISY-KH-G----------QG-WLVAENE---S-G----S--K---TGLVPE--EFVSYiqpel
0045 1tg0A   -------epEVPFKVVAQF-------PyksdyeddD--QEIIVTS-VE-D----------AE-WYFGEYQ---D-D----V--I---EGIFPK--SFVAVqg---
0046 8hyjW   -------tysIGQKLRIRV----gplK--------G--YLCRVIA--------------------TVKLD---S-Q----H--K---IFTVKS--EHLAE-----
0047 6rwlA   -klqnqiSKIQNFRVYFRE--grdqqW--------K--GPATLIW-KG-E-----------G-AVVIQDG----------Q--D---LKVVPR--RKCKIik---
0048 1wfwA   -----gssGSSGSTMTVIK-------DyyalkeneG--EVVQVLA-VN-Q----------QN-MCLVYQP---A-P----A--A---EGWVPG--SILAPfsgps
0049 6bogA   -------pftLGQRWISDT-------E--------SelGLGTVVA-VD-A-----------R-TVTLLFP---S-T----G--E---NRLYARsdSPVTRvxfnp
0050 7ud6A   ------------MLFVALY--------yearteddG--EKFVILR-SS-E----------GD-WWLAVSL---T-T----G--E---QGYIPS--NYVAPvgggt
0051 5zwxA   -erekrrevaVGDDVDAFY-----sdG--------W--WEGTVTE-VX-G----------DG-RXSVYFR---A-S----K--E---QIRFRR--DELRFhrewv

0001 s001A   -------LLLLLEEEEELL-------L--------L--EEEEEEE-EE-EE---L---L-EE-EEEEELL---L-L----L--L---EEEEEH--HHEEL-----
0002 6hq9A   ----llllllLLLEEEEEL-------L--------E--EEEEEEE-EE-EE--lL---L-LE-EEEEELL-----L----L--L---EEEEEH--HHLLLll---
0003 6if4B   -----lllllLLLEEEEEE-----llE--------E--EEEEEEE-EE-EL-llL---L-LE-EEEEEEL---L-L----L--H-hhLEEEEH--HHEEEll---
0004 3fc3A   -------lllLLLEEEELL-llllllL--------L--LEEEEEE-EE-L-----------L-EEEEEEL---L-L----L--E---EEEEEH--HHEEEllhhh
0005 7abi7   -lllllllllLLLEEEELLllllhhhL--------L--EEEEEEE-EE-L----------LL-EEEEEEL---L-L----L--L---EEEEEH--HHEEElllll
0006 7ef1A   -lhhhhhhllLLLEEEEEE-elllleE--------E--EEEEEEE-EE-LL---L---LlLL-EEEEEEL---L-L----L--L---EEEEEH--HHEEElhhhh
0007 1m1gC   -lllllllllLLLEEEELL----lllL--------L--LEEEEEE-EE-LL---------LL-EEEEEEE---E-E----E--E---EEEEEH--HHEEEl----
0008 1vieA   ---lllllllLLLEEEELL------lL--------L--LEEEEEE-EE-LL---L---LlLL-EEEEEEL---L-L----L-lL---EEEEEH--HHEEEll---
0009 5hh7A   -eeelleeeeLLLEEEELLllllllllllllhhhhL--EEEEEEE-EE-EE-llL---L-LE-EEEEEEE---E-E----E--E---EEEEEH--HHEEEeeeee
0010 2lccA   -----llLLLLLLEEEEEE-eelleeE--------E--EEEEEEE-EE-EE---L---L-EE-EEEEEEL---L-L----L--L-llLEEEEH--HHEELlllll
0011 8xv7A   --llllllllLLLEEEEEL---llllL--------E--EEEEEEE-EE-EE---L-llL-EE-EEEEEEL---L-L----HhhL---EEEEEH--HHEEElllll
0012 8jw0e   ----llllllLLLEEEELL--lllllL--------L--LEEEEEE-EL-LL---L---L-LLlLEEEELL---L-LllllL--L---EEEELH--HHEEElllll
0013 6kcpC   ------llllLLLEEEELL----lllL--------L--EEEEEEE-EL-------------L-EEEEEEL---L-L----L--L---EEEEEH--HHEEEhhhhh
0014 7cceA   -eeelleeelLLLEEEELL-------L--------L--EEEEEEE-EE-EL--lL---L-LE-EEEEEEE---ElE----E--E---EEEEEH--HHEEEellee
0015 2digA   -lllllllllLLLEEEEEL---llllL--------E--EEEEEEE-EE-LL---------LL-EEEEELL-----L----L--L---EEEEEL--LLEELlllll
0016 4m4zA   -------------LEEELLlllllllL--------L--LEEEEEE-EE-L-----------L-EEEEEEL---L-L----L--L---EEEEEH--HHEEEeelll
0017 2lq8A   -lllllllllLLLEEEELL----lllL--------L--LEEEEEE-EE-LL---------LL-EEEEEEE---E-E----E--E---EEEEEH--HHEELll---
0018 7cfdF   -hhlllllllLLLEEEEEL----lllE--------E--EEEEEEE-EL-L-----------L-LEEEEEL---L-L----L--L---EEEELH--HHEEEllhhh
0019 5m9nA   -lllllllllLLLEEEEEL---llllL--------E--EEEEEEE-EE-E----------LL-EEEEEEL---L-L----L--L---EEEEEH--HHEELllhhh
```

```
0020 8jvzA   -------lllLLLEEEEEE----------------E--EEEEEEE-EE-EE---L-llL-EE-EEEEEEL-----L----L--L---EEEEEL--L---------
0021 6tedZ   -hhhllllllLLLEEELLL----lllL--------L--LEEEEEE-EL-L-----------L-LEEEEEL---L-L----L--L---LLLLLH--HHLLLlllll
0022 6cb1Z   -----lllllLLEEEEELL----lllL--------L--EEEEEEE-ELlLL---L---LlLL-EEEEEEE---E-E----E--E---EEEEEL--LLEEEeeeel
0023 1vbvA   -----lllllLLLEEEELL-------L--------L--LEEEEEE-EE-LL-------L-LL-EEEEEEElllL-L----E--E---EEEEEH--HHEEElllll
0024 6bimA   -eeehheeEELEEEELLL-------LllllllleeL--LEEEEEE-EE-E----------LL-EEEEEEL----------L--E---EEEEEH--HHLLLlllll
0025 2jt4A   ------lLLLLLEEEEELL-------LllllllllL--LEEEEEE-LL-L---------lLL-EEEEEEL---L-L----L--L---EEEEEH--HHEEElllll
0026 2e70A   -lllllllLLLLLEEEELL----lllL--------L--LEEEEEE-EL-L-----------L-LEEEEEL---L-L----L--L---EEEELL--LLEEElllll
0027 1sf9A   -hhllhhhllLLLEEEELL-------L--------L--LEEEEEE-EE-L-----------L-EEEEEEL---L-E----E-eE---EEEEEH--HHEEEll---
0028 5z81A   -lllllleelLLLEEEELL---------------lL--EEEEEEE-EE-EL---L-----LE-EEEEEEE---E-E----E--E---EEEEEH--HHEEEeeeee
0029 8tqcD   -hhhhhhHLLLLEEEEELL-------L--------L--EEEEEEEeEE-EL---LlllL-EE-EEEEEEL----------L--E---EEEEEL--LLLEEellll
0030 7k9bA   -----lllllLLEEEEELL----lllL--------L--LEEEEEE-EL-L----------LL-EEEEELL---L-L----L--LhhhLEEEEH--HHEEEeeeel
0031 8rswA   ---llhhHLLLLEEEEELL-------LllllllllL--LEEEELL-LL-LL---------LL-EEEEELL-----L----L--L---EEEEEH--HHEEEll---
0032 8qszY   ------llllLLLEEEELL----lllL--------L--LEEEEEE-EL-L-----------L-EEEEEEL---L-L----L--L---EEEEEH--HHEEEellll
0033 8s7vL   --------llLLLEEEELL-------L--------L--LEEELLE-EL-LL---L---L-LL-LEELLL------------------LLEELH--HHEEEll---
0034 7aoaA   -----------LLEEEELL----------------L--EEEEEEE-EE-EL--lL---L-LE-EEEEEEE---E-E----E--E---EEEEEH--HHEEEeeeee
0035 7cspA   -hhhhhhLHHHLLEEEELL-------LllllllllL--LEEEEEE-EE-L-----------L-EEEEEEL---L-L----L--L---EEEEEH--HHEEEellhh
0036 6vldB   lllllllLLLLLLEEEELL-------LllllllllL--LEEEEEE-EL-L----------LL-EEEEEEL---L-L----L--E---EEEEEH--HHEEElllee
0037 4cosA   -lhhhllLLLLLLEEEELL----lllL--------L--EEEEEEE-EE-L-----------L-EEEEEEL---L-L----L--L---EEEEEH--HHEEElllll
0038 3qwxX   -llleellLLLLEEEEELL-------LllllllllL--LEEEEEE-LL-L----------LL-EEEEELL-----L----L--L---EEEEEH--HHEEEl----
0039 3ob9D   -lllllllllLLLEEEEEL-------L------llE--EEEEEEE-EE-EE---E-llE-EE-EEEEEEL---L-L----LhhH---LEEEEH--HHEEEllhhh
0040 2n5uA   -lllhhhlllLLLEEEELLlllhhhhL--------L--LEEEEEE-EE-EL---L---LlLE-EEEEEEL-----L----L--L---EEEEEL--LLEEEel---
0041 5z3gQ   -lllllleelLLLEEEELL----lllL--------L--EEEEEEE-EL-L----------LL-EEEEELL---L-L----L-lL---LEEEEH--HHEELlllll
0042 4anjA   -----------LLLEEEEE----lllE--------E--EEEEEEE-EL-L-----------L-EEEEEEL-------------L---LEEEEH--HHLEElllll
0043 2budA   -lllllllLLLLLLEEEEL----lllL--------E--EEEEEEE-EE-LLlllL---L-LL-EEEEELL---L-L----LllL---LEEEEL--LLEELlhhhh
0044 2lcsA   -------LLLLLEEEEELL-------LllllllllL--LEEEEEE-EE-L----------LL-EEEEELL---L-L----L--L---EEEEEL--LLEEElllll
0045 1tg0A   -------llLLLEEEEELL-------LllllllllL--LEEEEEE-EL-L----------LL-EEEEEEE---L-L----E--E---EEEEEH--HHEEEll---
0046 8hyjW   -------lllLLLLEEELL----lllL--------L--LEELLLL--------------------LEELL---L-L----L--L---EELLLH--HHEEL-----
0047 6rwlA   -hhhhhhHLLLLLEEEELL--lllllL--------L--LLEEEEE-EL-L-----------L-EEEEELL----------L--L---EEEEEH--HHEEEel---
0048 1wfwA   -----lllLLLLLEEELLL-------LllllllllL--LEEEEEE-EL-L----------LL-EEEEEEL---L-L----L--E---EEEEEH--HHLLLlllll
0049 6bogA   -------lllLLLEEEELL-------L--------HhhLEEEEEE-EL-L-----------L-EEEEEEL---L-L----L--E---EEEEELllLLEEElllll
0050 7ud6A   ------------LEEEELL--------llllllllL--LEEEEEE-EL-L----------LL-LEEEEEL---L-L----L--L---EEEELH--HHEEElllll
0051 5zwxA   -hhhhhllllLLLEEEEEE-----llE--------E--EEEEEEE-EE-L----------LL-EEEEEEL---L-L----L--E---EEEEEH--HHEEElleee

# Job: Type-3 BREX; BrxHII-Helicase WP_162523516 C-Terminal Inactive PDDEXK-REase Fold
# Query: s001A
# No:  Chain   Z    rmsd lali nres  %id PDB  Description
   1:  6bog-A  6.1  2.8   76   967   14   MOLECULE: RNA POLYMERASE-ASSOCIATED PROTEIN RAPA;
   2:  5lth-A  4.6  2.8   63   334   10   MOLECULE: HEME DEPENDENT OXIDATIVE N-DEMETHYLASE;
   3:  4qbn-A  4.4  3.2   59    93   10   MOLECULE: NUCLEASE;
   4:  6hcz-A  3.9  3.7   55    65    7   MOLECULE: SJOEGREN SYNDROME/SCLERODERMA AUTOANTIGEN 1;
   5:  6m1u-A  3.5  4.0   59   122   10   MOLECULE: RNA N6-ADENOSINE-METHYLTRANSFERASE METTL16,RNA N6
   6:  5wi2-B  3.2  2.9   53    93   11   MOLECULE: CDNA FLJ56409, HIGHLY SIMILAR TO SERINE/THREONINE
   7:  6rrv-A  3.0  3.4   53   127    2   MOLECULE: REGULATORY PROTEIN SIR4;
   8:  2okf-A  3.0  3.5   55   129    5   MOLECULE: FDXN ELEMENT EXCISION CONTROLLING FACTOR PROTEIN;
   9:  5ux0-D  3.0  3.7   56   639   14   MOLECULE: ARGONAUTE PROTEIN;
  10:  1v5s-A  2.9  4.3   60   126    3   MOLECULE: MAP/MICROTUBULE AFFINITY-REGULATING KINASE 3;
  11:  6rfl-I  2.9  5.6   58   773   14   MOLECULE: DNA-DEPENDENT RNA POLYMERASE SUBUNIT RPO132;
  12:  3vn5-A  2.9  2.4   45   255   11   MOLECULE: RIBONUCLEASE HIII;
  13:  2fug-7  2.8  3.9   59   127   12   MOLECULE: NADH-QUINONE OXIDOREDUCTASE CHAIN 1;
  14:  8ka7-A  2.8  2.6   49   115    8   MOLECULE: DE NOVO DESIGN PROTEIN -NB7;
  15:  9j8p-A  2.8  3.2   55   623    7   MOLECULE: SPECKLE TARGETED PIP5K1A-REGULATED POLY(A) POLYME
  16:  6hpb-A  2.8  2.1   43    57   16   MOLECULE: MRNA INTERFERASE TOXIN HICA;
  17:  3t4n-A  2.7  3.0   55   139    2   MOLECULE: CARBON CATABOLITE-DEREPRESSING PROTEIN KINASE;
  18:  6l7x-A  2.7  5.0   53   203   17   MOLECULE: MRNA_TRIPASE DOMAIN-CONTAINING PROTEIN;
  19:  9nl2-A  2.7  3.7   72  1107    8   MOLECULE: R2 RETROTRANSPOSON PROTEIN;
  20:  2p09-A  2.7  2.8   47    69    4   MOLECULE: A NON-BIOLOGICAL ATP BINDING PROTEIN WITH TWO MUT

0001 s001A   ------ELSI-DILKNI---KYGEIYELLRTMFINYVLY-NG------GTV-KEGKN-------------------G-F-DITIDG-------V-------------RRYKGVT-FD--------------PSGN-GA--N-AIT-LNHE-Y-V-KK-M-I-QE--
0002 6bogA   eihsngGEKAqALAESI--eEQDDD-TNLIAFAXNLFDI-IG------INQ-DDRGD------------------nX-I-VLTPS-------sE-------------DGIT-IT-FD--------------REVR-ED--AqFIT-WEHP-L-I-RN-G-L-DLil
0003 5lthA   rarvleldpgrclvmpH---MAQAAWDTLAMLMEHLARD-YP----qhFRL-TRQGD-------------------A-W-HWQNLA------lG-------------IDQR-FT-FG--------------D----PA--S-LpP-LEY---iT-RQ-M-Q-GDfa
0004 4qbnA   ----------------------ATKEGRVQKYAKERFEA-LG------GLV-RKLSP-------------------D-L-LVILPR-------G-------------VIWF-VE-VK----------kdenTKPD---nvF-VVG-SFK---qV-DK-Li------
0005 6hczA   ----swEPPT-EAETKV--lQARRERQDRISRLMGDYLL-RG------YRM-LTCAD-----------------cgT-I-LLQDKQ-------R-------------KIY----------------------------vaC-QEL---------------------
0006 6m1uA   asvmkelslkasplRSE---T-AEGIVVVTTWIEKILTD-LK------VQH-KRVPC--------------gkeevS-L-FLTAI-------yL-------------FKCL-IN-VK--------------KE---------------CT-Y-I-RN-Q-I-FRlv
0007 5wi2B   ------------------rfftkLDADKSYQCLKETCEK-LG------YQW-KKSC------------------mnQ-V-TISTT-------nK-------------LIFK-VN-LL--------------EM---------------FL-K-I-KG-K-L-IDiv
0008 6rrvA   klisvslvdefpselSD---SDRQIINEKMQLLKDIFANnLK------SAI-SNN-------------------------------------------------freSDIIIL--K---------------GEIE-DyrfW-SFM---RF-V-SnFD-I-Q-----
0009 2okfA   -------------------------RDVFHEVVKTALKK-DG------WQItDDPLT----------isvggvnlkL-I-AAER-Q-------G-------------QKIA-VE-VKsflkqssaisefhtALGQ-F-vlY-LAV-PLktiiyD-VE-Qe------
0010 5ux0D   nvfqykvekeeiikeEE---DKKLFVKTLYKYIKKLFLD-ND------FYF-KK----------------------G-N-NFISNS-------EvfsldsnenvnahLTYK-IK-IH--------------NIsylsiyniksgfgilLG-F-L--N-K-I-TNln
0011 1v5sA   hlihnvhkeE-HAHahnkfqgssGSSGDMMREIRKVLGA-NN------CDY-EQRE------------------rfL-L-FCVHGD------nL-------------VQWE-ME-VC--------------KLplnGV--R-F-----SK-I-A-NE-L-Kl----
0012 6rflI   eeistsLSFN-D--KNT---TD-EMTYNLYDLFFNTL------------DM-YLRVK-------------------Y-V-YLEN-----------------------TGKIYKNsFS--------------ED-H-NN--S-------GK-V-I-IPl--------
0013 3vn5A   --------------------pslkISPSEAEKIQNYLVS-SG------FRK-INAPY-------------------TlW-ALEGN--------G-------------VKVY-------------------------------------LK-E-V-LN-L-Le----
0014 2fug7   -------------ASSE--rELYEAWVELLSWMREYAQA-KG------VRF-EKEADfpdfiyrmerpydlpttimT-A-SLSD-G-------L-------------GEPflLA-DV--------------SP-----lhahyekpLTKE-R-F-FA-La------
0015 8ka7A   aavaalraadpgaarrrv-gPDAARVQALADAVVAALER-EG------FKL-EKKEE---------ntdaagnagaK-Y-EGE------------------------GGLV-LN-VK--------------QG---------peT-LKITr---------------
0016 9j8pA   qrrssrgrdwgllpllqpiplplAPFTQLTAALVQVFREaLG------CHI-EQASA-------------------S-W-RCALWHrvwqgrpL-------------LSFV-AS-VS--------------PA---------------QV-F-L-PQ-A-I-RHlk
0017 6hpbA   ----------------------------KQSEFRRWLES-QG------VDV-ANGSN-------------------H-L-KLRF-H-------G-------------RRSV-XP-RH------------------------pcdK-EPLR-Ks-------------
0018 3t4nA   gspaaskisplvtkkskt-irSRSYPLDVMGEIYIALKN-LG------AEW-AKPS---------------eedlwT-I-KLRWK-------dL-------------MKMV-IQ-LF--------------QI---------------TT-K-L-IM-E-L-AVns
0019 6l7xA   emrlgqfkvE-EDA-----cvSTEDYERIKTYLMTEME---N------SSM-TRSVD-------------------R-L-SITD-G-------M-------------FRYD------------------mtqvtvlMH--E-VE----AM-R-L-ATl--------
0020 9nl2A   crnctaSYES-LSHILGq-gARIRRHNKLCSMLKREAKE-LK------WVV-YEERK-------------------P-DlIFVK-E-------E-------------MALV-VD-VT--vrfeykekvfedaaAE-KVeyF-GFPpEINE--pdykR-T-A-KRfs
0021 2p09A   --------------------DDDDKKTNWLKRIYRV-RP-CVkckvapRDW-KVKNK-------------------H-L-RIYNM-------yH-------------GHVD-WL-MY--------------ADS--------------------------------

0001 s001A   ------LLLL-HHHHLL---LHHHHHHHHHHHHHHHHHH-LL------LEE-EEELL-------------------E-E-EEEELL-------L-------------EEEEEEL-LL--------------LLLL-LL--L-LLL-LLLH-H-H-HH-H-H-HL--
0002 6bogA   hhhhhlHHHHhHHHHHH--hHHLLL-HHHHHHHHHHHHH-HL------LEE-EELLL------------------lE-E-EEELL-------lL-------------LLEE-EE-LL--------------HHHL-LL--LeELL-LLLH-H-H-HH-H-H-HHhh
0003 5lthA   hhhhhhhlhhheeelhH---HHHHHHHHHHHHHHHHHHH-LL----llEEE-EEELL-------------------E-E-EEEELL------lL-------------EEEE-EE-LL--------------L----HH--H-LlH-HHH---hH-LL-L-L-LEee
0004 4qbnA   ----------------------LLHHHHHHHHHHHHHHH-LL------LEE-EELLL-------------------L-E-EEEEHH-------H-------------EEEE-EE-EL----------llllLLLL---eeE-EEL-LHH---hH-HH-Hh------
0005 6hczA   ----llLLLL-HHHHHH--hHHHHHHHHHHHHHHHHHHH-LL------LEE-EELLL-----------------llL-E-EEELLL-------L-------------LEE----------------------------llL-LLL---------------------
0006 6m1uA   hhhhhhhhhhhlllLLL---L-LLHHHHHHHHHHHHHHH-LL------LEE-EEELL--------------llleeE-E-EEEEE-------eE-------------EEEE-EE-EE--------------EE---------------HH-H-H-HH-H-H-HHhh
0007 5wi2B   ------------------eeeelLLHHHHHHHHHHHHHH-LL------LEE-EEEE------------------llE-E-EEEEE-------lE-------------EEEE-EE-EE--------------EL---------------HH-H-H-HH-H-L-HHhe
0008 6rrvA   llleeeellllllllLH---HHHHHHHHHHHHHHHHHHHlLL------LEE-ELL-------------------------------------------------hhhLLEEEE--L---------------LLHH-HleeE-EHH---HH-H-HhLL-L-L-----
0009 2okfA   -------------------------LHHHHHHHHHHHHH-LL------LEEeELLLL----------leelleellL-E-EEEE-L-------L-------------EEEE-EE-ELllllllllhhhhhhHHHH-H-eeE-EEE-EHhlhheE-LL-Ll------
0010 5ux0D   hhhhheeeeeeeeelLL---LHHHHHHHHHHHHHHHHHH-LL------EEE-EL----------------------L-L-EEEEEE-------EeeellllllleeeEEEE-EE-EE--------------EEleeeeeellllehllLH-H-H--H-H-L-LLll
0011 1v5sA   lllllllllL-LLLlllllllllLLHHHHHHHHHHHHHH-LL------EEE-EEEE------------------llE-E-EEEELL------lL-------------EEEE-EE-LL--------------LLlllLL--E-E-----HH-H-H-HH-L-Ll----
0012 6rflI   eeeehhHLLL-L--LLL---LL-LHHHHHHHHHHHHL------------LE-EEELL-------------------E-E-EELL-----------------------LLLEEELlLL--------------LL-L-LL--H-------HH-H-H-HHh--------
0013 3vn5A   --------------------leeeLLHHHHHHHHHHHHH-LL------LEE-LLLLL-------------------LlE-EEEEL--------L-------------EEEE-------------------------------------HH-H-H-HL-L-Ll----
0014 2fug7   -------------LHHH--hHHHHHHHHHHHHHHHHHHH-HL------LEE-EEEEElhhhhlllllllllllleeE-E-EEEL-L-------L-------------LLEeeEE-EE--------------LL-----eeeeeellLLHH-H-H-HH-Hh------
0015 8ka7A   hhhhhhhhlllhhhhlle-eLLHHHHHHHHHHHHHHHHH-LL------LLL-EEEEE---------eellllleeeE-E-ELH------------------------HHLE-EE-EE--------------EL---------llE-EEEEl---------------
0016 9j8pA   hlllllllllllhhhhlllllllLLHHHHHHHHHHHHHLlLL------LEE-EELLE-------------------E-E-EEEELLlllllhlL-------------EEEE-EE-EL--------------LL---------------HH-H-H-HH-H-H-HHll
0017 6hpbA   ----------------------------LHHHHHHHHHH-LL------LEE-ELLLL-------------------L-E-EEEE-L-------L-------------EEEE-EL-LL------------------------lllL-HHHH-Hl-------------
0018 3t4nA   lllhhhhlllllllllll-eeELLLHHHHHHHHHHHHHH-HL------LEE-ELLL---------------hhhllE-E-EEEEL-------lE-------------EEEE-EE-EE--------------EE---------------HH-H-H-HH-H-H-HHll
0019 6l7xA   eeeeeeellL-LLL-----llLHHHHHHHHHHHHHHLL---L------LEE-EEEEE-------------------E-E-EEEE-L-------L-------------EEEE------------------eeeeeeeEE--E-EE----HH-H-H-HLl--------
0020 9nl2A   llllllLLLL-HHHHHHl-hHHHHHHHHHHHHHHHHHHH-LL------LEE-EELLL-------------------L-LeEEEL-L-------L-------------LEEE-EL-EE--eellllllhhhhhhHH-HHelL-LEElHHHH--lhhhH-H-H-HHhh
0021 2p09A   --------------------LHHHHHHHHHHHHHHH-LL-LLllllllLLE-EEELL-------------------E-E-EEELL-------lL-------------LEEE-EE-LL--------------LLL--------------------------------

# Job: Structural Search Anchored on "C-terminal Inactive PDDEXK-REase Domain" Found in RapA-Helicase;
# Query: s001A
# No:  Chain   Z    rmsd lali nres  %id PDB  Description
   1:  6bog-A 17.4  0.0   86   967  100   MOLECULE: RNA POLYMERASE-ASSOCIATED PROTEIN RAPA;
   2:  4qbn-A  4.1  2.9   63    93   10   MOLECULE: NUCLEASE;
   3:  2x31-B  3.9  3.4   62   423   13   MOLECULE: ORN/LYS/ARG DECARBOXYLASE FAMILY PROTEIN;
```

```
  4:  1y88-A  3.6  3.3   67   184   12   MOLECULE: HYPOTHETICAL PROTEIN AF1548;
  5:  2xnk-A  3.3  2.9   58   291    3   MOLECULE: DNA TOPOISOMERASE 2-BINDING PROTEIN 1;
  6:  2okf-A  3.3  2.8   60   129    8   MOLECULE: FDXN ELEMENT EXCISION CONTROLLING FACTOR PROTEIN;
  7:  9esh-m  3.3  3.1   51    81    6   MOLECULE: PRE-MRNA;
  8:  5y88-U  3.3  3.0   49   488    4   MOLECULE: PRE-MRNA-SPLICING FACTOR 8;
  9:  7p0j-A  3.2  2.9   56   195   11   MOLECULE: DNA DAMAGE RESPONSE PROTEIN MDB1;
 10:  5lth-A  3.1  3.2   58   334   12   MOLECULE: HEME DEPENDENT OXIDATIVE N-DEMETHYLASE;
 11:  5wi2-B  3.1  3.1   54    93   11   MOLECULE: CDNA FLJ56409, HIGHLY SIMILAR TO SERINE/THREONINE
 12:  9n82-F  3.1  2.9   55   699   11   MOLECULE: X-RAY REPAIR CROSS-COMPLEMENTING PROTEIN 6;
 13:  2zyz-C  3.1  2.7   56    96    7   MOLECULE: PUTATIVE UNCHARACTERIZED PROTEIN PAE0789;
 14:  5hy3-A  3.1  4.0   55   252   15   MOLECULE: MRNA ENDORIBONUCLEASE LSOA;
 15:  9n83-F  3.0  2.9   55   866   11   MOLECULE: X-RAY REPAIR CROSS-COMPLEMENTING PROTEIN 6;
 16:  8yax-A  2.9  3.7   55  1314   11   MOLECULE: PAPAIN-LIKE PROTEASE NSP3;
 17:  9n12-A  2.9  3.4   67  1107    6   MOLECULE: R2 RETROTRANSPOSON PROTEIN;
 18:  3cxj-A  2.9  3.1   52   144   12   MOLECULE: UNCHARACTERIZED PROTEIN;
 19:  5vz1-C  2.9  4.5   59    87    5   MOLECULE: SINGLE GUIDE RNA (116-MER);
 20:  2czr-A  2.9  4.8   67   226    6   MOLECULE: TBP-INTERACTING PROTEIN;
 21:  6m1u-A  2.8  4.4   60   122   12   MOLECULE: RNA N6-ADENOSINE-METHYLTRANSFERASE METTL16,RNA N6
 22:  9u48-A  2.8  3.5   58   336    7   MOLECULE: U6 SMALL NUCLEAR RNA (ADENINE-(43)-N(6))-METHYLTR
 23:  8v9u-A  2.8  3.6   51    81    4   MOLECULE: DNA (CYTOSINE-5)-METHYLTRANSFERASE 1;
 24:  9bf5-C  2.8  3.4   62  1190    5   MOLECULE: SSDNA1;
 25:  1fok-A  2.8  3.4   72   568   13   MOLECULE: DNA (5'-
 26:  6c9j-A  2.8  3.3   54   386    6   MOLECULE: 5'-AMP-ACTIVATED PROTEIN KINASE CATALYTIC SUBUNIT
 27:  7lyc-N  2.8  3.5   56   335    9   MOLECULE: HISTONE H3.1;
 28:  4a0g-A  2.8  3.6   61   769    5   MOLECULE: ADENOSYLMETHIONINE-8-AMINO-7-OXONONANOATE
 29:  2qsr-A  2.7  3.9   73   161   12   MOLECULE: TRANSCRIPTION-REPAIR COUPLING FACTOR;
 30:  3r3p-B  2.7  3.5   57    98   11   MOLECULE: MOBILE INTRON PROTEIN;
 31:  5u0p-Q  2.7  2.9   54   508    7   MOLECULE: MEDIATOR COMPLEX SUBUNIT 14;
 32:  8a5w-B  2.7  3.6   61   367   10   MOLECULE: PHOSPHOSERINE AMINOTRANSFERASE;
 33:  2hc5-A  2.7  3.3   61   116   10   MOLECULE: HYPOTHETICAL PROTEIN YVYC;
 34:  2ost-A  2.7  3.3   60   148    7   MOLECULE: SYNTHETIC DNA 29 MER;
 35:  1whz-A  2.7  2.9   52    70   10   MOLECULE: HYPOTHETICAL PROTEIN;
 36:  9j8p-A  2.7  3.4   56   623   13   MOLECULE: SPECKLE TARGETED PIP5K1A-REGULATED POLY(A) POLYME
 37:  8que-E  2.6  4.3   59   493   12   MOLECULE: PHIKZ055;
 38:  6l30-A  2.6  4.0   55   518    4   MOLECULE: PROTEIN ECT2;
 39:  5yrz-B  2.6  2.9   48    58   15   MOLECULE: HICB;
 40:  6wm6-A  2.6  3.3   61   536   13   MOLECULE: EXTRACELLULAR SOLUTE-BINDING PROTEIN, FAMILY 5;
 41:  6hpb-A  2.6  2.4   43    57    5   MOLECULE: MRNA INTERFERASE TOXIN HICA;
 42:  7t8b-A  2.6  5.4   52   518    8   MOLECULE: TWINKLE MTDNA HELICASE;
 43:  4wl5-A  2.6  3.4   72   594   11   MOLECULE: HETERODIMERIC RESTRICTION ENDONUCLEASE R.BSPD6I L
 44:  8q3b-E  2.6  3.6   62   205    6   MOLECULE: DNA-DIRECTED RNA POLYMERASE RPB1 HOMOLOG;
 45:  6ijb-B  2.5  3.1   58   538    9   MOLECULE: AMP-BINDING DOMAIN PROTEIN;
 46:  9dev-A  2.5  2.8   50    97    6   MOLECULE: PROTEIN MONO-ADP-RIBOSYLTRANSFERASE PARP4;
 47:  7myj-C  2.5  2.9   53   464    4   MOLECULE: 5'-AMP-ACTIVATED PROTEIN KINASE CATALYTIC SUBUNIT
 48:  1ob8-B  2.5  3.9   64   127   13   MOLECULE: HOLLIDAY-JUNCTION RESOLVASE;
 49:  5oe9-C  2.5  3.8   55   415   11   MOLECULE: LARGE SUBUNIT TERMINASE;
 50:  7wae-A  2.5  3.5   48   877   10   MOLECULE: CYANOPHYCIN SYNTHASE;

0001 s001A   ----DDTN-LIAFAXNLFDI---IGINQDDRG---------DNXIVLTPSDHX----LVPD--F--PGL--SEDGITITFDR---------E-V--A----LARE-----D-AQFITWEH-----PLIRNGLDLIL
0002 6bogA   eeqdDDTN-LIAFAXNLFDI---IGINQDDRG---------DNXIVLTPSDHX----LVPD--F--PGL--SEDGITITFDR---------E-V--A----LARE-----D-AQFITWEH-----PLIRNGLDLIL
0003 4qbnA   --atKEGR-VQKYAKERFEA---LGGLVRKLS---yegrsgAPDLLVILPR---------------------GVIWFVEVnT---------K-P--Dph-qLREH------nVFVVSFKQ---vdKLIEH------
0004 2x31B   ydstLFFA-KRAQLIECLEN---KGFEMLQVD---------DLKLLIKYEG-ftghdiqNW--F--M--laDDYQALAILPL-----------------------------------LFD-----SLLRKIEDMI-
0005 1y88A   ----NLYF-QGHMVARLLEE---HGFETKTNV-vqgncveqEIDVVAERD----------------------GERYMIECKfhnipvytglK-E--A----MYTY------qPWIFTNTwsypekEGIEVLLESKG
0006 2xnkA   sctslEKE-KREEVHKYVQX---XGGRVYRD---------------------------------------lnvSVTHLAGEV---------G-SkkY----LVAA--nlkK-PILLP--S-----WIKTLWEKSQE
0007 2okfA   -----RDV-FHEVVKTALKK---DGWQITDPL---------TKLIAAERQ----------------------GQKIAVEVAL---------G-Q--F----INYR------vLYLAVPLqldfpkeiIFQW-----
0008 9eshm   -------S-FRELVEEFCAE---NSLLFVPLR---------RALFRISTQA-------------------sKARGITVYLRN-----diiwkkspgase-dTPYD-------------------pIGFNE------
0009 5y88U   elgkvsSS-FKDVVEDYCLE---KGYLISKIPnygrdqdciVPLFEIRNG----------------------KKKMEVALK---------hdilwveds-sGTFK--------------I-----YLWA-------
0010 7p0jA   airdSMVGdSIHGLYSILET---SGAEIVGDI---------K---------------------------aqeKDTIILAQnD---------Q-E--G----RNMS-atglN-VYKI---E-----LVALSILRD--
0011 5lthA   vmphMAQA-AWDTLAMLMEH---LAFRLTRQG----------DAWHWQN-------------lA--LGI-----DQRFTGDP---------A------------------------PLEY-------TRQMQG---
0012 5wi2B   ftklDADK-SYQCLKETCEK---LGYQWKKSC---------MNQVTISTTD------------------rnnKLIFKVNLLE---------M---------------------------F-----LKIKGKLI---
0013 9n82F   vmsgtDSQ-PKPDLENRIAE---FGGYIVQN--------------------------------------pgpDTYCVIAGSE---------NiR--V----KNIIlsnkhD-VVKP---A-----WLLECFKTKS-
0014 2zyzC   -----MDV-LQEQVFKDLKS---RGFKIEQLD---------DKIFIAEK-----------------------KERYLFYVM---vegvevtiqT--L----LSVI------pVVLALVTV-----TY---------
0015 5hy3A   ipkfsDDD-RANLFEFLSE----EGITITEDN---------NHQYIMTT----------------------SNGDRVRAKIS-------------------------------------A-----SLINDFMCS--
0016 9n83F   vmsgtDSQ-PKPDLENRIAE---FGGYIVQN--------------------------------------pgpDTYCVIAGSE---------NiR--V----KNIIlsnkhD-VVKP---A-----WLLECFKTKS-
0017 8yaxA   ltsssKTP-EEHFIETISLA---GSYSGQSTQ---------L-GIEF-------------------lkR--GDKSVYYTSNP----------------------------------TFDN------LKTLLSL--R
0018 9nl2A   arirRHNK-LCSMLKREAKE---LKWVVYEEP-httekelrKPDLIFVK----------------------EEMALVVDVaA---------E-K--V----RHYK------eYFGFPLIN-----EK-vGMPDQQK
0019 3cxjA   -------L-SQEXIKKWLDE---EGFLRXEVP---------DFHYVVNY---------------------peDHVIDIIQPA---------G------------------------FFDG-----LSKDRLIS---
0020 5vzlC   -------M-NINDLIREIKN---KDYTVKLSG----tdsnsITQLIIRV--------------------nnDGNEYVIS--------------E--S----EN--------sieKFISAF-----K-MQTITLKSE
0021 2czrA   hmyaelSP-GTKKVYTQVRY---LDYHWEIEG---------STITGIHKK---------------------sNVKVVIDVAD---------S-L--A-----GKD-----ViHIVAIPDL-----TYRYLKA----
0022 6m1uA   etaeGIVV-VTTWIEKILTD---LKVQHKRVP---------CVSLFLTAIE------------------lqyLFKCLINVKK---------E---------------------------C-----TYIRNQIFRLV
0023 9u48A   ldpelCAQ-IDDILQKFLDD---NKIPWSKKG---------S-VLEISTKSIT----WSRK--------legQMKCELNVID-------------------------------------C-----SALARALRD--
0024 8v9uA   dlerdSLT-EKECVKEKLNL---LH--------------------------EF----LQTE--IknQLC--D-lETKL------------------------rkE-----E-LSEE---A-----KVKSLLNKDLS
0025 9bf5C   tniyKGAL-GEQAVEAVLTA---FDFTFEEVE---------RFDNRVIFA----------------------gIEQPIWLDS----kywkhegnesseg-ySSKI------kFIYVNALI-----PALI--TNRT-
0026 1fokA   rnstQDRI-LEMKVMEFFMK--vYGYRGKHLG------gsrKPDGAIYTVG------------------spiDYGVIVDTkaysggynlpiG-Q--A----DEMQryveenkFLFVSGHV-----LSVEELLIGGE
0027 6c9jA   rsqsRPND-IMAEVCRAIKQ---LDYEWKVVN---------PYYLRVRRKN------------------tstYSKMSLQLYQ---------V--------------------------yA-----NLIKILAQ---
0028 7lycN   igsglSSE-QQKMLSELAVI---LKAKKYTEF---------D-------------------------------sTVTHVVsT---------L-K--C----MLGI---lnG-CWILK--F-----EWVKACLRRKV
0029 4a0gA   sqgktlrelwDEELVQQISS---HVQRVVV-I---------GTLFALELK------slyaK--S--LL-rpLGNVIYLMCGP---------C-T--S---------------piCRRL-T-----KLYKRLGEFNR
0030 2qsrA   pdvvAYLL-EIGLVKSYLDK---VFVRVERK----------DNKITIQFEKVTq--rLFLAqdY--FKAnkgLXELVFD-vQ---------N-K--K----------------yEILE-L-----IFGESLLEIKE
0031 3r3pB   -----sTT-PERRVKEILDE---MDIVYFTHH----vvegwNVAFYLG-------------------------KKLAIEVN-----gvywaskqknvnk-dKRKL------rVLTIEDDElndidKVKQQIQK---
0032 5u0pQ   ifyyNLNQ-ESLEFQRWLKQ---RDISFKFMP---------NISWRIMV--------------------snfEPAIFIQHTP---------T------------------------SSLK-----NVCQYIE----
0033 8a5wB   amekLSSI-KSQTIYEIIDN---SFYVCPV------------MNIPFRIGNakgddaleKR--F--L--hrsvGGIRASLYN---------A------------------------DVQK------LAAFMKKFLE
0034 2hc5A   pvhqvSYT-NLAEMVGEMNK---LHLKFELHD---------KYYVKVIEDS-------------------tnEVIREIPP---------------------------------KRWL--D-----FYAAMTEFLGL
0035 2ostA   stklKGDI-AQQAAIMRALK---MGWGVLKPL---------SYDLVFDVE----------------------GILLKVQVnyvvdnrrtrtnrrnivrs-pYRGN-----D-FFAVAYLF-----YVtEAWH----
0036 1whzA   ---xwxpP-RPEEVARKLRR---LGFVERXAK---------GGHRLYTHPD---------------------grIVVVPFHS----------------------------------LPKG---tfKRIL-LTEEE-
0037 9j8pA   lplaPFTQ-LTAALVQVFRE--aLGCHIEQA-----------ASWRCAL---------wHR--V--WqgtePLLSFVASVSP---------A---------------------------H-----FLQVFLPQA--
0038 8queE   dpntgvnM-TRRYIDSLFDI---SYEGNRACT-------plkYMIDLMFSY------------------gemLYPRPMLLPA---------FkrgnmvtingakyigspvlkqdhhLMSE------FNYAVSMFGY
0039 6l30A   ftgfRKKE-ELVRLVTLVHH---MGGVIRKDF--------------------------------------nskvTHLVANCT---------Q-G----e-kFRVAvslgtP-IMKP--------ewIYKAWERR--
0040 5yrzB   -------pmTQKEMVKLLTA---HGWIKTRGK---------GSHIKMEKQG---------------------ERPITILHG---------------------------------eLNKYR-----GIRK-------
0041 6wm6A   vtsalGNN-AVVRTVTLPIVpsgGPYMVSFDP---------GQLVTLVPNPKW----YGEK--G--PYL--D--KLKFRIIT---------DstqqltalenGEV-----D-----afayqlptaliAYNT-----
0042 6hpbA   ---------KQSEFRRWLES---QGVDVANGS---------N-HLKLRF-----------------------HGRRSVXPRH----------------------------------KEPL-----RK---------
0043 7t8bA   ------PV-TATEIRQYLRG---HGIPFQDGH----------SCLRAL-------------------sP--FSFSLFIDKT------------------------------------SWE-----DFQASV-EGEF
0044 4wl5A   fianKNTV-FEWLTWNGFII---LGaEYKNNFvpvthaagnQPAMEIIY----------------------EDFIVLGEVES---------E-P--V----TRHY------yCLFIAPLK-----QFNMLLMVQKK
0045 8q3bE   ---maMQK-LFTYIYEFIEY---RKMVLLE------------FRINAETL---------------------nHGIVSVFIF------gangkyvhhggd-mRTLL------eLILIVYPY-----HLF---NIPKV
0046 6ijbB   gtlaWNNR-RHLEIYYAASG---AGFVCHTIN---------P---------------------------nhaQDRVLFFDAT-----------F--L----PLVA------hFVLMG--D-----ELI-ETGD--T
0047 9devA   kvkylPQQ-QKKKLQTDIKE---NGGKFSFSL--------------------------------------npQCTHIILDNA---------D-V-----------------hIANP---D-----FIWKSIREKR-
0048 7myjC   rsqsKPYD-IMAEVYRAMKQ---LDFEWKVVN---------AYHLRVRRKN------------------tgnYVKMSLQLYL---------V---------------------------dffemcASLITTLA---
0049 1ob8B   ---dIGKN-AERELVSILRG---EGFNAVRIP------tnpLPDIFATK----------------------GNTLLSIECstwenkvkvkeH-Q--V----RKLL-----DvPLIAIKFtidnsiPIEDLFKI---
0050 5oe9C   ktigALRRnVITPLKRMLKS---RGYRVKDHR--------aDNYLTITF----------------------KGKTNYFYLFG--------gkdessqdliqgITL-----A-----FFDE----------------
0051 7waeA   knqaslGP-STEAIVKEAEA---RGIPWTQLG--------aRFMIQFG--------------------Y--GVNQKKIQA---------tlsnQ--T----GI-------------kllarnvagAVMDM------

0001 s001A   ----LLHH-HHHHHHHHHHH---HLLEEEELL---------LLEEEEELLLLL----LLLL--L--LLL--LLLLEEEELLH---------H-H--H----HHLL-----L-LEELLLLL-----HHHHHHHHHHH
0002 6bogA   hhhlLLHH-HHHHHHHHHHH---HLLEEEELL---------LLEEEEELLLLL----LLLL--L--LLL--LLLLEEEELLH---------H-H--H----HHLL-----L-LEELLLLL-----HHHHHHHHHHH
0003 4qbnA   --llHHHH-HHHHHHHHHHH---LLLEEEELL---llllllLLLEEEEEHH---------------------HEEEEEEElL---------L-L--Lhh-hHHHH------eEEEELHHH---hhHHHHH------
0004 2x31B   lllhHHHH-HHHHHHHHHHH---LLLEEEELL---------LLEEEEELLL-llhhhhhHH--H--H--eeLLLLEEEELLL-----------------------------------LHH-----HHHHHHHHLL-
0005 1y88A   ----LHHH-HHHHHHHHHHL---LLLEEEEEE-eellllleeEEEEEEEEL----------------------LEEEEEEELlllllleelhH-H--H----HHHH------eEEEELLLllllllLLHHHHHHLLL
0006 2xnkA   eeellLHH-HHHHHHHHHHH---LLLEELLL---------------------------------------lllLLLEEELLL---------L-LhhH----HHHH--hllL-LEELL--H-----HHHHHHHHHHL
0007 2okfA   -----LHH-HHHHHHHHHHH---LLLEEELLL---------LLLEEEEEL----------------------LEEEEEEEHH---------H-H--H----HHHH------eEEEEEEHhlhhhhhhEEEE-----
```

```
0008 9eshm   -------L-HHHHHHHHHHH---LLLEEEEEE---------EEEEEEELLL-------------------lLLLEEEEEEEL-----leeeelllllll-lLLLE-------------------eLLHHH------
0009 5y88U   llllllLL-HHHHHHHHHHH---HLLEEELLLllllllleeeEELEEEELL----------------------LLLEEEELL---------llleeeell-lLLEE--------------E-----LLLL-------
0010 7p0jA   hhhhHLLLhHHHHHHHHHHH---LLLEEELLL---------L---------------------------hhlLLLEEELLlL---------H-H--H----HHHH-hhllL-LLLL---H-----HHHHHHHHL--
0011 5lthA   elhhHHHH-HHHHHHHHHHH---HHEEEEEEL---------LEEEEEE-------------lL--LLE-----EEEEELLH---------H-----------------------HHHH-------HLLLLL---
0012 5wi2B   eellLHHH-HHHHHHHHHHH---LLLEEEEEE---------LLEEEEEEEL------------------lllEEEEEEEEEE---------L--------------------------H-----HHHHHHLH---
0013 9n82F   elllllLL-LHHHHHHHHHH---LLLEELLL------------------------------------lllLLLEEELLLL---------LhH--H----HHHHhllllL-EELH---H-----HHHHHHHHLL-
0014 2zyzC   -----LLH-HHHHHHHHHHH---LLLEEEEEE---------LLEEEEEL----------------------LLEEEEEEE---elllllhhH--H----HHHH------lEEEEEELE-----EE---------
0015 5hy3A   elllllLLH-HHHHHHHHHH----LLLEEEEEL---------LLEEEEEL---------------------LLLLLLLLLL----------------------------------------H-----HHHHHHHHH--
0016 9n83F   eeellLLL-LHHHHHHHHHL---LLLEELLL------------------------------------lllLLLEEEELLL---------LhH--H----HHHHhhlllL-EELH---H-----HHHHHHHHLL-
0017 8yaxA   hhlllLLH-HHHHHHHHHHH---LELLLLLLL---------L-LEEE---------------------eeE--LLEEEEEELLL------------------------------------LHHH------HHHHHHL--L
0018 9nl2A   hhhhHHHH-HHHHHHHHHHH---LLLEEEELL-llllllllLLLEEEEL----------------------LLLEEEELEhH---------H-H--H----LLLH------eLLLEEEHH-----HH-hLLLHHHH
0019 3cxjA   -------L-HHHHHHHHHHH---LLLEEEELL---------LEEEEEEL---------------------llLLEEEEEEEL---------L--------------------------EHHH-----LLHHHHHH---
0020 5vzlC   -------L-LHHHHHHHHHH---LLLLLLLLL----lllllLLLEELLL---------------------llLLLLLEEL--------------L--L----LL--------lhhHHHHHL-----L-HHHHHHLLL
0021 2czrA   hhhhllLH-HHHHHHHHHLL---LLEEEEEEL----------LEEEEEELL---------------------lLLEEEEEEHH---------H-H--L-----LLL-----LeEEEEELLE-----EHHHHHH----
0022 6m1uA   llllHHHH-HHHHHHHHHHH---LLLEEEEEL---------LEEEEEEEEE--------------------lleEEEEEEEEEE---------E--------------------------H-----HHHHHHHHHHH
0023 9u48A   llllhHHH-HHHHHHHHHHH---LLLLLLLLL---------L-LLLLLLLLLL----LLHH---------lllLLEEEEELLL----------------------------------------H-----HHHHHHHHH--
0024 8v9uA   hhhhhLLL-HHHHHHHHHHH---HH------------------------LL----LLHH--HhhHHH--H-hHHHH-------------------------hhL-----L-LLHH---H-----HHHHHHHHHHH
0025 9bf5C   hllhHHHH-HHHHHHHHHHH---LLLEEELLL---------LLLEEEELL---------------------lLLLLEEEEE------lllllllllllhh-hHHHH------eEEEELLLE-----LLLL--LLHH-
0026 1fokA   llhhHHHH-HHHHHHHHHHH--lLLLEEEELL------lllLLLEEEELLL------------------lllLEEEEEEEellllllllllhH-H--H----HHHHhhhhhleEEEEELLE-----EEHHHHHHHHH
0027 6c9jA   eellLHHH-HHHHHHHHHHH---LLLEEEEEE---------LLEEEEEEEL------------------lllEEEEEEEEEE---------L--------------------------eH-----HHHHHHHL---
0028 7lycN   eellllLHH-HHHHHHHHHHH---HLLEEELLL---------L--------------------------lLLLEEElL---------H-H--H----HHHH---hlL-LEEEE--H-----HHHHHHHHHHLL
0029 4a0gA   llllllllllLHHHHHHHHHH---LEEEEEE-E---------LLEEEEEEL------lhhhH--H--HH-llLLLEEEEELLL---------L-L--L---------------hhHHHH-H-----HHHHHHLLLLL
0030 2qsrA   lhhhHHHH-HHHHHHHHHHH---LLEEEEEL----------LLEEEEEELLLHh--hHLLHhhH--HHHellEEEEEEE-lL---------L-L--L---------------hHHHH-H-----HHHHHHHHHHH
0031 3r3pB   -----lLL-HHHHHHHHHHH---LLLLLEEEE----eelleEEEEEEE--------------------------LLEEEEEE-----lllllllllhhh-hHHHH------eEEEEEHHHhhhhhHHHHHHHHH---
0032 5u0pQ   hhhhHHHH-HHHHHHHHHHL---LLLLLLEEL---------LLEELLLL---------------------lllLLLLEEELLL---------L--------------------------LLHH-----HHHHHHH----
0033 8a5wB   hhhhHHHH-HHHHHHHHHHL---LLEELLL------------LEEEEEELLllllhhhhHH--H--H--llllLLEEEELLL---------L--------------------------HHHH------HHHHHHHHHH
0034 2hc5A   lllllLHH-HHHHHHHHHHH---HLEEEEEEE---------EEEEEEEELL--------------------llEEEEEELH--------------------------------------HHHH--H-----HHHHHHHHHHH
0035 2ostA   llhhHHHH-HHHHHHHHHHH---LLLEEEEEL---------LLLEEEEEL-----------------------LEEEEEEEeeeeelleeellllllleee-lLLLL-----L-LEEEEEEE-----EElLLLH----
0036 1whzA   ---llllL-LHHHHHHHHHH---LLLEEEEEEE---------LLEEEEELLL-----------------------llEEEEELLL----------------------------------------LLHH---hhHHHH-LLHHH-
0037 9j8pA   llllLHHH-HHHHHHHHHHL--lLLLEEEEL-----------EEEEEEE---------lLL--L--LllllLLEEEEEEELL---------L--------------------------H-----HHHHHHHHH--
0038 8queE   llllllllH-HHHHHHHHHHH---HEEEEEELL-------hhhEEEEEEEEE--------------------leeLLLEEEEELL---------LllllleeeelleeeeelleeeeeeeHLHH------HHHHHHHHHH
0039 6l30A   eellLLHH-HHHHHHHHHHH---HLLEEELLL------------------------------------llllLEEEELLL---------L-L----h-hHHHHhhhllL-EELL--------hhHHHHHHHL--
0040 5yrzB   -------llLHHHHHHHHHH---LLLEELLLL---------LLEEEEELLL----------------------LLLEEEELL-------------------------------------lLLHHH-----HHHH-------
0041 6wm6A   lllllLLL-HHHHHHHLLLLlllLLEEEEEEL---------LLEEEEEELLLL----LLLL--L--LLL--L--EEEEEELL---------LhhhhhhhhhlLLL-----L-----hlleeeelleeHHHH-----
0042 6hpbA   ---------LHHHHHHHHHH---LLLEEELLL---------L-LEEEEE-----------------------LLEEEEELLL--------------------------------------LHHH-----HH---------
0043 7t8bA   ------LL-LHHHHHHHHHH---LLLLEELLL----------LLEEEL--------------------lL--LLLLEEELLL--------------------------------------LHH-----HHHHHL-LLLL
0044 4wl5A   llllHHHH-HHHHHHHHHHH---HLeEEEELLeeeellllllLLLEEEEE------------------------LLEEEEEEELL---------H-H--H----HHHH------eEEEEELHH-----HHHHHHHHHHH
0045 8q3bE   ---lhHHH-HHHHHHHHHHH---LLLEELL------------EEEEEEELL---------------------lLLEEEEEEE------lllllllllhhh-hHHHH------eEEEELELL-----HHH---LHHHL
0046 6ijbB   eeelLLLH-HHHHHHHHHHH---LLLEEEELL---------L--------------------------hhhLLLEEEELHH-----------H--H----HHHH------eEEELL--H-----HHH-HHLL--L
0047 9devA   elllllLHH-HHHHHHHHHHH---LLLEEELLL-------------------------------------llLLLEEEELLL---------L-L-------------------eEELH---H-----HHHHHHHHLL-
0048 7myjC   eellLHHH-HHHHHHHHHHH---LLLEEEELL---------LLEEEEEEEL------------------lllEEEEEEEEEE---------L-----------------------------hhhhhhHHHHHHHL---
0049 1ob8B   ---lLLHH-HHHHHHHHHHH---LLLEEEELL------lllLLLEEEEE-----------------------LLEEEEEEEeellleeeelhH-H--H----HHHH-----HeEEEEEEEelllleEHHHHHHH---
0050 5oe9C   llhhHHHHhLHHHHHHHHHH---LLLEEEEEL--------lLLEEEEEE-----------------------LLEEEEEEEEEE--------lllhhhhhhhhLLL-----L-----EEEL---------------
0051 7waeA   hhhlllLH-HHHHHHHHHHH---LLLLEEELL--------lLLLEEEL--------------------L--HHHLEEELL---------llllL--L----LH-------------ehllllhhhHHHHH------

# Job: Type-3 BREX; BrxHII-Helicase_WP_162523516 (C-Terminal Most Alpha+Beta Domain)
# Query: s001A
# No:  Chain    Z    rmsd lali nres  %id PDB  Description
   1:  6bog-A  5.6  3.5  146   967    8   MOLECULE: RNA POLYMERASE-ASSOCIATED PROTEIN RAPA;
   2:  2i1k-A  5.6 10.3  104   545    4   MOLECULE: MOESIN;
   3:  6b2z-M  5.5  6.8   81   249    4   MOLECULE: ATP SYNTHASE SUBUNIT C, MITOCHONDRIAL;
   4:  8ek4-B  5.5  4.4   90   135    8   MOLECULE: ICE-BINDING PROTEIN TIP-99A;
   5:  4dci-B  5.4  3.1   84   148   13   MOLECULE: UNCHARACTERIZED PROTEIN;
   6:  9gm9-A  5.4  9.2   94   999   10   MOLECULE: CHROMOSOME PARTITION PROTEIN MUKB;
   7:  5ziy-A  5.4  5.5   79   195    4   MOLECULE: FLAGELLAR HOOK-ASSOCIATED PROTEIN 3;
   8:  8rt9-F  5.3  5.7   86   272    7   MOLECULE: TRWJ PROTEIN;
   9:  5y9o-A  5.2 10.2  109   401    8   MOLECULE: WIPA;
  10:  6aay-A  5.2  5.4   75  1199   11   MOLECULE: BERGEYELLA ZOOHELCUM CAS13B (R1177A) MUTANT;
  11:  7wdk-B  5.0  5.9   79   348    9   MOLECULE: PHOSPHOLIPASE D;
  12:  2x0l-A  5.0 11.5   96   670    9   MOLECULE: LYSINE-SPECIFIC HISTONE DEMETHYLASE 1;
  13:  6zbj-A  5.0  9.6  100   666   10   MOLECULE: PRECURSOR OF THE MAJOR MEROZOITE SURFACE ANTIGENS
  14:  5iun-E  4.8  7.1   85   220   11   MOLECULE: SENSOR HISTIDINE KINASE DESK;
  15:  8bsb-B  4.7  3.8   80   162    8   MOLECULE: METHYL-ACCEPTING CHEMOTAXIS PROTEIN;
  16:  6rw8-B  4.6 10.6  109  2337    5   MOLECULE: A COMPONENT OF INSECTICIDAL TOXIN COMPLEX (TC);
  17:  4dyl-A  4.6  9.1   97   376    4   MOLECULE: TYROSINE-PROTEIN KINASE FES/FPS;
  18:  2xco-A  4.6  8.4   97   636    5   MOLECULE: DNA GYRASE SUBUNIT B, DNA GYRASE SUBUNIT A;
  19:  4mt8-A  4.6  5.3   81   100    9   MOLECULE: ETHYLENE RESPONSE SENSOR 1;
  20:  4zm1-B  4.6  4.5  113   238    8   MOLECULE: CHAIN LENGTH DETERMINANT PROTEIN;
  21:  6th1-R  4.6  6.0   98   360    7   MOLECULE: IMMEDIATE EARLY PROTEIN 1;
  22:  7f3t-A  4.6  8.6   88   330    7   MOLECULE: TRANSMEMBRANE PROTEIN 120A;
  23:  2oev-A  4.6  4.8   92   697    8   MOLECULE: PROGRAMMED CELL DEATH 6-INTERACTING PROTEIN;
  24:  7p5v-B  4.5  6.6  101   732    8   MOLECULE: VOLUME-REGULATED ANION CHANNEL SUBUNIT LRRC8A;
  25:  6zh3-A  4.5  8.2   88   170    8   MOLECULE: VACUOLAR PROTEIN-SORTING-ASSOCIATED PROTEIN 24;
  26:  7t7t-A  4.5 11.4   84   466    7   MOLECULE: PROTEIN TONSOKU;
  27:  9io5-G  4.5  6.8   89   260    4   MOLECULE: G1-ATPASE SUBUNIT BETA;
  28:  8ap7-a  4.5 11.4  106   231    4   MOLECULE: ATP SYNTHASE SUBUNIT A;
  29:  8k89-A  4.3  6.8   75    78   13   MOLECULE: NUCLEAR FACTOR INTERLEUKIN-3-REGULATED PROTEIN;
  30:  7ag9-B  4.3 10.0   97   384    6   MOLECULE: KAR9;

0001 s001A   PKIKIDGLAN-EEGYWSLWQITL---NDSPND--KIFPFFINNEGKM-RLPSAEAIWDRLVS-TDAHITYDGMK-EIE-D-GVYGELYKRVSEY-A-S-SMFYE-IRDAYIKNRDEERKRM-EYAFTLKKEMI-GRV-G-LETVKRHRLNELEKEKKAWEDKF-A-KS-RN-IIPSL--Y-P--L--C--IL-Y-ME-
0002 6bogA   TISLLKNkalgTLLVELIYVVEA---QaNRFL--TPVRXLLDKNGNN-LAAQ--VEFETF----nRQLNAVN----vQ-Q-DVHAIL-QLGEAQ-I-E-KSARA-LIDAARNEADEKLSAE-LSRLEALRAVN-------------PNIR-DDELTAIESNR-Q-QVxES-LDAGW--R-L--D--A--LR-L-IVv
0003 2i1kA   -------------------------------------ksmnvrvtiQQTT-TGKQLF-------eVWFF-----eY-Q-DRLRQMQEEMERSqA-N-LLEAQ-DMILRLEEQLRQLQAA-KEELEQRQNEL-QAM--dEIRAKQEEVSRIQQEVELKDSET-R-RL-QE-E--raaaA-L--L--A-aTT-P-QHh
0004 6b2zM   feirtlfglqllvitslytltnnnnkiisqeawglftlfiislsiviykhGWVF----------------------fvpagTPLP-LVPLLVII-E-T-LSYFA-RAISLGLRLGSNILAG-HLLMVILAGLT-FNFmT-LVFG-FVPLAMILAIMMLEFAIG-I-IQ-GYv-------------------------
0005 8ek4B   ----------eevreklkrmekkfddslIREIkeAEKKKKRN---------------------------------------------PYEAV-VTT-L-R-AILKA-VETKIRAIIKALKTEL-DALIKAMETIL-KAH-D----kNDELKKEVEDIIKKMRDKL-T-KL-IK-AKELK--K-Ka---------------
0006 4dciB   ---------------------------------------------------------------------------------raVVTPtW-K-EEAER-EISNGIANADQQLAQL-EQEGQTVVDQV-RRQ-SpLDPRVQEQVANIQQQVAGKRSEL-E-EQkRN-LL-xQ--G-Q--L----qSI-E-E--
0007 9gm9A   ----------frsltlvnwngffartfdtlsgipdfrnystldvinsrlqqvagrdrkvdikpfmirenritle-AIR-VtQSDRDLFKHLITE-A-T-SYVSA-DYMRHANERRTHLDEA-LALRGELFGSH-KQLahaEMENVQKKLRKLERDYYQIREQV-V-SA-KA-RPER-----L--R--E--RR-Q-DIi
0008 5ziyA   -------------------------------------------AASKSF-------------------------------------hSLANM-E-Q-MQKDI-ADSKNVLTQTENTLQG-VLKSLTRADQL-TVQ-AkELQAIGVEIDQILKQVVYLANT----KE-QG---delseQ-N--V--Al---------
0009 8rt9F   ---------------------------------------------------------------------------aF-ELFTPLFNKIDQT-T-AtYVTDI-SSRAIAAITPVVSVGL-TLGFITYGWLI-IRG------aVEMPVAEFLNRCLRIGIIV-S-IA-LG-GLYIA--N-A---------F-A-AVf
0010 5y9oA   ednlgSISL-------------------------------naIKLIHFLFRHK---------------TE-I-INFHEAYQQFVTI-YeQ-YDDMV-QEYLEIRTLLQLIQIK-ITNAQQRILDI-EQK-LeFSQSLLQLKKPIEANLQMAEKSK-A-GL-ES-GLKlrfnK-F--M--T-qIE-I-NDi
0011 6aayA   kitvfgrlseweifpnpnydfpkenisviaGS--L---------------------------------kilkpyqdgDKEKLIKDL-K-Q-EQNIL-QKLKDEQTVREKEYND-FIAYQDKNREI-NKV-R-DRNHKQYLKDNLKRKYPEAP------------------------------kSE-N-K--
0012 7wdkB   -------------------------------------dasrvwqisevtavgdngsYDYDS-----------------------mifkvDKNE-F-H-NASNY-IKGTAEIYQD-HLREN-IKLDKKAISTL-QKN-K----sEEKSIERIKKGIAEMEAKI---PL-AK-IYEHD--L-------------A-R-FRt
0013 2x0lA   vlvhrvhsyltgkviiigsgvsglaaarmdvtgnyamvqkcplyeankekDEMVEQEFNRL-----fnvlnNKP-VSL-GqALEVVIQLQEKHV-KdE-QIEHW-KKIVKTQEELKELLNK-MVNLKEKIKEL-HQQ--fLVKSKHRDLTALCKEYDELAETQ-G-KL-EEk------------------------
0014 6zbjA   gIVYP-----------------------------------------------------LPL-TDIfsssnnsv-YNV-QkLKKALSYLEDYSL-R-K-GISEK-DFNHYYTLKTGLEADI-KKLTEEIKSSE-NKI--ekEVSDIVKLQVQKVLLIKKIEDL-R-KI-EL-KNASI--H-V--P--Nii--------
0015 5iunE   --------------------------------------------------------------------iikL-RKE-IeRLEEKLEDANERI-A-E-LVKLE-ERQRIARDLHDTLGQK-LSLIGLKSDLA-RKL-K-DPEQAARELKSVQQTARTSLNEV-R-KI-VS-S----------------------
0016 8bsbB   ---------------------------------------------------------------------------GPLGSSELDKI-Q-S-ELLNYtDDTLPAMENVDAIKDK-MSYWRRTQFAV-LPM-K-DEAQIRQTIERNNRVQAEINDSL-V-AY-GT-VWP-------------------------
0017 6rw8B   glSENEL-----------------------------settalqhdfhaVIMRCGSYATEIlTALElgaltaeqvRQ-D-AEALAEILQTQGS-E-L-ALQSI-KMQDKVMAEIDADKLA-LQESRHGAQSRfDSF-NeWEIQRNNAEAEIKQIDAQLATLA-V-RReAA-VL---------qlafL--QS-Kf---
0018 4dylA   --------------------smgfsseLCSP-------gvlqqmqeakwmaqrvksdreyagllhheglsrLLR-QhaedlnSGPLSKLSLLIR-E-R-QQLRK-TYSEQWQQLQQELTKThSQDIEKLKSQY-RAL--rSLWKLFAHHNRYVLGVRAAQLHH-QqLL-LP-RSLQmacI-L--K--E---I-S-SLv
0019 2xcoA   eamvrmaqdfaamrytearmtkitlelltidf--FPNlvlslsknpD-ISIAELMEDIEGpdfptaSIQMRSR---mialvngrPKLI-nlkEA-L-V-HYLEH-QKTVVRRRTQYNLRKA-KDRAHILEGLR-IAR-L-TG-LERDKIEAEYNELLNYISEL-E-TIlAD-EE------------------------
0020 4mt8A   ------------------------------------------------------------------ashA-AILeE-SMHARDQLMEQNF-A-L-DKARQ-EAEMAVHARNDFLAVM-NHEMRTPMHAIiSLS-SmIETILKSSNLVATLISDVLDL-S-R-LE---------------------------
0021 4zm1B   ---------awtsTAIITqpdvgqiaGYNNam--LIGrdnqeePEKL-T---IEPS-VKNQ---------------tA-E-GAQMKLAQYIQQV-D-D-KVNQE-LEKDLKDNIALGRKNL-QDSLRTQEVVA-QEQ-KeaTRPL--VFSPNYYQTRQNLLDI-E--K-LK-FLDIH----Y--R--Y--VMk-----
0022 6th1R   skkaavqslvrttciqmanesyaradivASQG--YT------------------------------nfmpvnaN-ESE-K-RAWMEMLGECTSH-G-N-KLCEM-ANAQVEQETRDIINIM-FKNIDDVVTQT-TRA-MrLCLSILTSAKERIDIIYHSARSQ-H-LA-CV-RMNvatfI-L--T--Na---------
0023 7f3tA   -------------------------------------------------------------------GPLGdcDWE-DlQQDFQNIQETHRL-Y-R-LKLEE-LTKLQNNCTSSITRQK-KRLQELALALK-KCK-L-PAEAEGAAQELENQMKERQGLFF-D-ME-AY-L------------------------
0024 2oevA   ---------------------------------------------------------------------aA-I-EDvsQSILTKSRS-V-IeQGGIQ-TVDQLIKELPELLQRN-REILDESLRLL-DEE-EpSNELYKPLRAEGTNFRTVLDKAV-Q-AD-GQ-VefAY--D-Nfr---------------
0025 7p5vB   kdscndstgpnrlhwfakyfpylvllhtsNFWkfpLEHF-------------vsillkcfdspwtseTGVLDK-KegeqaKALFEKVKKFRTH-V-E-EG-DI-VYRLYMRQTIIKVIKF-ALIICYTVYYV-HNI-KpLATLFKILASFYISLVIFYGLIC-M-YT-LW--LKKYsfesF--A--V--F---------
0026 6zh3A   -------------------------------------------------------------------M-D-YIKKAIWKEQQRR-I-R-SVLRK-NGRNIEKSLRELT-VL-QNKTQQLIKKS-AKK------nDVRTVRLYAKELYQINKQY-D-RM-YT-AQLDI--D-E--AirM--NT-L-S--
0027 7t7tA   DahnnigxlqedddGRSR--------------------HHNLgnvyxeHIEQDIIICKKIE---------------elhyrvqkycyqlaqsxededA-LASQI-DQNIETVKKAIEVXDE-LKKEEQNLKKL-TRN--sQERKSLLQQNASLDCLIEKSSXIf------------------------------------
0028 9io5G   --------------------------------------------------------------M-KRK-D-VEEKKQNLDFYHN-Y-I-DISKI-KVLQKLNEEIASLNML-KLQKGESHYLL-NRI--rivpfFYEQFVKSVFKQELFEFKS-I-SViEE-LK-KK--K-R---------K-W-NRa
```

```
0029 8ap7a   dfllfCLFDL----------ylfvglclMLFN------yycitylnllYIAFLFLFCFLCD---------------RF-L-LCFLEcFSLLCRC-L-S-tflRL-FCN-LLSSHFLLLMFF-DFFYFIFVFFF-YGV-F-CY-WFILFIFVFCFCLLFYVFLY-L-LD-LF-ILQL--------------D-F-LLf
0030 8k89A   -----------------------------------------------------------------------KDA-XEK-R-RKNNEAAKRSREK-R-R-LNDLV-LENKLIALGEENATLK-AELLSLKLKFG-L-----------iSSTAYAQEXQKLSNST-A-VY-F---------------------------
0031 7ag9B   SRDH------mdtldDLIN------------krhtdqlsrkflilkR-NIPPIEQSLTEILP-QR-----------ininlladfkrirfMMNE-I-K-DLKIE-LIDKRWNILFINLNNE-LEYIIEEVRLL-LKK-ItIKDRFNSQLAKKSKIITKTFNII-Y-RAlEF-SL-----------------------

0001 s001A   EEEEELLLLL-LLEEEEEEEEEE---LLLLLL--EEEEEEEELLLLL-LHHHHHHHHHHHHL-LLLLEEEEEEEE-LLL-L-LHHHHHHHHHHHH-H-H-HHHHH-HHHHHHHHHHHHHHHHH-HHHHHHHHHHH-HLL-L-LHHHHHHHHHHHHHHHHHHHHHHH-H-HH-LL-LLEEE--E-E--E--E--EE-E-EL-
0002 6bogA   EEEEEELllllLEEEEEEEEEEL---LlHHLL--EEEEEEELLLLLL-LLLL--LLHHHH----hHLLLLLL----lH-H-HHHHHH-HHHHHH-H-H-HHHHH-HHHHHHHHHHHHHHHHH-HHHHHHHHLLL-------------LLLL-HHHHHHHHHHH-H-HHhHH-HLLEE--E-E--E--E--EE-E-EEe
0003 2i1kA   -------------------------------------lleeeeeeeLLLL-LHHHHH-------lHHHE-----hH-H-HHHHHHHHHHHHHhH-H-HHHHH-HHHHHHHHHHHHHHHHH-HHHHHHHHHHH-HHH--hHHHHHHHHHHHHHHHHHHHHHHHHH-H-HH-HH-H--hhhhH-H--H--L-lLL-L-HHh
0004 6b2zM   hllllllllhhhhhhlllllllllllhhhhlhhhhhhhhhhhhhhhhHHHH----------------------hllllLLHH-HHHHHHHH-H-H-HHHHL-HHHHHHHHHHHHHHHHH-HHHHHHHHHHH-HHHlH-LLLL-HHHHHHHHHHHHHHHHHHH-H-HH-HHh-------------------------
0005 8ek4B   ----------lhhhhhhhhhhhhhhhhhHHHHhhHHHHHHHL----------------------------------------LHHHH-HHH-H-H-HHHHH-HHHHHHHHHHHHHHHHH-HHHHHHHHHHH-HHH-L----lLLLLHHHHHHHHHHHHHHH-H-HH-HH-HHHHH--H-Hh--------------
0006 4dciB   ------------------------------------------------------------------------------------eeELLHhH-H-HHHHH-HHHHHHHHHHHHHHHHH-HHHHHHHHHHH-HLL-LlLLHHHHHHHHHHHHHHHHHHHHH-H-HHhHH-HH-lE--E-E--E----eEE-E-L--
0007 9gm9A   ----------eeeeeeeeelleeeeeeeeeeellllllleeeeeeelleeeeellllleeeeeeeeehhhhhhh-HHH-HhHHHHHHHHHHHHH-H-H-HHHHH-HHHHHHHHHHHHHHHHH-HHHHHHHHHHH-HHHhhhHHHHHHHHHHHHHHHHHHHHHHHHH-H-HH-HH-LLHH-----H--H--H--LL-L-LLl
0008 5ziyA   -----------------------------------------LHHHHH----------------------------------------hHHHHH-H-H-HHHHH-HHHHHHHHHHHHHHHHH-HHHHHHHHHHH-HHH-HhHHHHHHHHHHHHHHHHHHHLL----EE-LL---lhhhhH-H--H--Hh--------
0009 8rt9F   -----------------------------------------------------------------------------lL-LLHHHHHHHHHHHH-H-HhHHLHH-HHHHHHHHHHHHHHHHH-HHHHHHHHHHH-HLL------lLLLLLLHHHHHHHHHHHHH-H-HH-HL-LLHHH--H-H---------H-H-HHh
0010 5y9oA   lllllEEEE-------------------------------lhHHHHHHHHHLL----------------LL-L-LLHHHHHHHHHHH-HhH-HHHHH-HHHHHHHHHHHHHHHHH-HHHHHHHHHHH-HHH-HhHHHHHHHHHHHHHHHHHHHHHHH-H-HH-HH-HHHhhhhH-H--H--L-lEE-E-LLl
0011 6aayA   eeeeellhhhelllllleellllllllllLL--L----------------------------------------hhhlllllllLHHHHHHHH-H-H-HHHHH-HHHHHHHHHHHHHHHHH-HHHHHHHHHHH-HLL-L-LHHHHHHHHHHHHHHLLLLL----------------------------lLL-H-H--
0012 7wdkB   -------------------------------------lhhhhhhlllllllllleeELLLL----------------------eeellLHHH-H-H-HHHHH-HHHHHHHHHH-HHHHH-HHHHHHHHHHH-LLL-L----lLLLLHHHHHHHHHHHHHHLL---HH-HL-LEEEL--L-----------H-H-EEl
0013 2x0lA   hhhhhhhhhhlleeeeellllhhhhhhhhleeellellelllllleelllllhHHHHHHHHHHH-----lllllLLL-LLH-HhHHHHHHHHHHHHH-HhH-HHHHH-HHHHHHHHHHHHHHHHH-HHHHHHHHHHH-HHH--hHHHHHHHHHHHHHHHHHHHHHHHHH-H-HH-HHh-------------------------
0014 6zbjA   lLLLL-------------------------------------------------------LLH-HHHhhhhhhhh-HHH-HhHHHHHHHHHHHHH-H-H-HHHLL-LHHHHHHHHHHHHHHHH-HHHHHHHHHHH-HHH--llLLHHHHHHHHHHHHHHHHHHHHHH-H-HH-HH-HHHLL--L-L--L--Lll--------
0015 5iunE   ----------------------------------------------------------------------lhhH-HHH-HhHHHHHHHHHHHHH-H-H-HHHHH-HHHHHHHHHHHHHHHHH-HHHHHHHHHHH-HHH-H-LHHHHHHHHHHHHHHHHHHHHHHH-H-HH-HH-L-------------------------
0016 8bsbB   ----------------------------------------------------------------------------------LLLLHHHHHHHH-H-H-HHHHHhHLHHHHHHHHHHHHHHH-HHHHHHHHHHL-LLL-L-LHHHHHHHHHHHHHHHHHHHHHHH-H-HH-HL-LLL----------------------
0017 6rw8B   llLLLLH-----------------------------llllllllllhhhHHHHHHHHHHHHhHHHHlllllllllhHH-H-HHHHHHHHHHHHH-H-H-HHHHH-HHHHHHHHHHHHHHHHH-HHHHHHHHHHHhHHH-HhHHHHHHHHHHHHHHHHHHHHHHHH-H-HHhHH-HH---------hhhhH--HH-Ll---
0018 4dylA   --------------------lllhhhhLLLH-------hhhhhhhhhhhhhhhhhhhhhhhhhhhhhhhhhhhhHHH-HhhhhhhHLHHHHHHHHHH-H-H-HHHHH-HHHHHHHHHHHHHHHHHhLHHHHHHHHHH-HHH--hHHHHHHHHHHHHHHHHHHHHHHHHH-HhLH-HH-HHHHhhhH-H--H--H---H-H-LLl
0019 2xcoA   hhhhhhllllllllllleeeelhhhhlllllle--LLHhhhhhhhllL-LLHHHHHHHLLLllllllEEEEELE---leeeelleEEEL-lhhHH-H-H-HHHHH-HHHHHHHHHHHHHHHHH-HHHHHHHHHHH-HHH-H-LL-LHHHHHHHHHHHHHHHHHHHHH-H-HHhHL-HH----------------------
0020 4mt8A   ----------------------------------------------------------------------llhH-HHHhH-HHHHHHHHHHHHHH-H-H-HHHHH-HHHHHHHHHHHHHHHHH-HHHHHHHHHHHhHHH-HhHHHHHHHHHHHHHHHHHHHHHH-H-L-LL----------------------------
0021 4zm1B   ---------leeeEEEEEllllhhhhHHHHhh--HHHhhllllLLLE-E---EEEL-LLLL--------------lH-H-HHHHHHHHHHHHH-H-H-HHHHH-HHHHHHHHHHHHHHHHH-HHHHHHHHHHH-HHH-HhhHLLL--LLLHHHHHHHHHHHHH-L--L-LL-LLLLL----L--E--E--EEl-----
0022 6th1R   hhhhhhhhhhhhhhhhhhhhhhhhhhhhhHHHL--LL-----------------------------hhllllllL-LLL-H-HHHHHHHHHHHHH-H-H-HHHHH-HHHHHHHHHHHHHHHHH-HHHHHHHHHHH-HHH-HhHHHHHHHHHHHHHHHHHHHHHHH-H-HH-HH-HHHhhhhH-H--H--Hh---------
0023 7f3tA   ---------------------------------------------------------------------LLHHhhHHH-HhHHHHHHHHHHHH-H-H-HHHHH-HHHHHHHHHHHHHHHHH-HHHHHHHHHHH-HHL-L-LHHHHHHHHHHHHHHHHHHHHHHH-H-HH-LL-L------------------------
0024 2oevA   -----------------------------------------------------------------------------lL-L-LLllHHHHHHHHH-H-HlLLLHH-HHHHHHHHHHHHHHHHH-HHHHHHHHHHH-HHH-HlHHHHLLHHHHHHHHHHHHHHHHHH-H-HH-HH-HhlHH--H-Hhl-------------
0025 7p5vB   lllllllllllhhllhhhhhhhhhhhhhhhhHHHhlhHHHH--------------hhhhhhhhlllhhllLLLLLH-HhhhhhHHHHHHHHHHHHH-H-L-LL-LL-HHHHHHHHHHHHHHHHH-HHHHHHHHHHH-HHL-LlLHHHHHHHHHHHHHHHHHHHHHHH-H-HH-HH--LLLLllhhH--H--H--H-------
0026 6zh3A   ------------------------------------------------------------------------------L-L-HHHHHHHHHHHHH-H-H-HHHHH-HHHHHHHHHHHHHH-HH-HHHHHHHHHHH-HHH------lLHHHHHHHHHHHHHHHHHHHH-H-HH-HH-HHHHH--H-H--HhhL--LL-H-H--
0027 7t7tA   HhhhhhhhhhlllhHHHH--------------------HHHHhhhhhhHHHHHHHHHHHHL---------------hhhhhlllhhhhhhlllllhH-HHHHH-HHHHHHHHHHHHHHHHH-HHHHHHHHHHH-HHH--lHHHHHHHHHHHHHHHHHHHHHHLl-------------------------------
0028 9io5G   --------------------------------------------------------------------------L-LHH-H-HHHHHHHHHHHHH-H-H-HHHHH-HHHHHHHHHHHHHHHHH-HHHHHHHHHHH-HHH--eeellLHHHHHHHHHHHHHHHHHHH-H-HHhHH-HH-HH--H-H---------H-H-HHh
0029 8ap7a   llleeLLLLL----------llhhhhhhHHHH------llllllllhhHHHHHHHHHHLLL---------------HH-H-HHHHHhHHHHHHH-H-H-hhhHH-HHH-HHHHHHHHHHHHH-HHHHHHHHHHH-HHH-H-LL-LHHHHHHHHHHHHHHHHHHHH-H-HH-HH-HHHH---------------L-L-LLl
0030 8k89A   ----------------------------------------------------------------------LLH-HHH-H-HHHHHHHHHHHHH-H-H-HHHHH-HHHHHHHHHHHHHHHHH-HHHHHHHHHLL-L-----------lLHHHHHHHHHHHHHHH-H-LL-L--------------------------
0031 7ag9B   HHLL------hhhhhHHHH------------hlllhhhhhhhhhhhH-HHHHHHHHHHHHHHH-HH-----------llhhhhhhhhhhhHHHH-H-H-HHHHH-HLHHHHHHHHHHHHHH-HHHHHHHHHHH-HHH-HhHHHHHHHHHHHHHHHHHHHHHHH-H-LLhHH-HL-----------------------
```

```
# Job: BR-systems; DUF499-ATPase CTD Ferredoxin-fold/RNA-Recognition-Motif (Representative_1)
# Query: s001A
# No:  Chain   Z    rmsd lali nres  %id PDB  Description
   1:  4kyz-A  7.1  2.4   99   167   10   MOLECULE: DESIGNED PROTEIN OR327;
   2:  5cw9-A  6.4  3.1   99   146   12   MOLECULE: DE NOVO DESIGNED FERREDOXIN-FERREDOXIN DOMAIN INS
   3:  4ney-B  6.3  5.0   81   173   12   MOLECULE: ENGINEERED PROTEIN OR277;
   4:  3n0v-A  6.2  2.8   78   280   13   MOLECULE: FORMYLTETRAHYDROFOLATE DEFORMYLASE;
   5:  4v1a-k  6.0  4.4   89   131    4   MOLECULE: MITORIBOSOMAL PROTEIN ML37, MRPL37;
   6:  1r3n-A  6.0  4.7   88   438    7   MOLECULE: BETA-ALANINE SYNTHASE;
   7:  2nzc-D  5.9  3.3   78    82   12   MOLECULE: HYPOTHETICAL PROTEIN;
   8:  3n5f-A  5.8  3.5   87   406   13   MOLECULE: N-CARBAMOYL-L-AMINO ACID HYDROLASE;
   9:  1ysj-A  5.8  3.4   89   359   13   MOLECULE: PROTEIN YXEP;
  10:  6wi5-A  5.8  3.0   81    90   12   MOLECULE: DE NOVO DESIGNED PROTEIN FOLDIT4;
  11:  3fgv-A  5.7  2.6   80    94    6   MOLECULE: UNCHARACTERIZED PROTEIN WITH FERREDOXIN-LIKE FOLD
  12:  3r2c-K  5.7  2.8   75    80   11   MOLECULE: N UTILIZATION SUBSTANCE PROTEIN B;
  13:  1zpv-C  5.6  3.6   80    88    6   MOLECULE: ACT DOMAIN PROTEIN;
  14:  2fb0-A  5.6  2.7   81    94    6   MOLECULE: CONSERVED HYPOTHETICAL PROTEIN;
  15:  8p6p-I  5.6  3.7   81   104    6   MOLECULE: 23S RIBOSOMAL RNA;
  16:  1u8s-B  5.6  3.2   79   172   10   MOLECULE: GLYCINE CLEAVAGE SYSTEM TRANSCRIPTIONAL
  17:  1z2l-A  5.6  3.5   88   411   10   MOLECULE: ALLANTOATE AMIDOHYDROLASE;
  18:  8d8k-J  5.5  4.7   87   144    3   MOLECULE: PROBABLE S-ADENOSYL-L-METHIONINE-DEPENDENT RNA
  19:  7c50-A  5.5  5.2   93   192    8   MOLECULE: SIMPL DOMAIN-CONTAINING PROTEIN;
  20:  5i4m-A  5.5  4.9   93   414   10   MOLECULE: AMIDASE, HYDANTOINASE/CARBAMOYLASE FAMILY;
  21:  4uer-J  5.4  4.0   84   107    6   MOLECULE: EIF1A;
  22:  7m6u-B  5.3  3.6   98   392   12   MOLECULE: CARBOXYPEPTIDASE G2 CIRCULAR PERMUATION PRO-DOMAI
  23:  2bbe-A  5.3  3.3   85   103    8   MOLECULE: HYPOTHETICAL PROTEIN SO0527;
  24:  7ljh-A  5.3  3.3   92   400    4   MOLECULE: POLY(ASPARTIC ACID) HYDROLASE;
  25:  8tcf-C  5.3  2.4   71    75   11   MOLECULE: INTEGRIN ALPHA-V HEAVY CHAIN;
  26:  2nyi-A  5.3  3.3   82   171   12   MOLECULE: UNKNOWN PROTEIN;
  27:  3ce8-A  5.3  4.6   79    90   11   MOLECULE: PUTATIVE PII-LIKE NITROGEN REGULATORY PROTEIN;
  28:  1y0h-A  5.3  3.2   83   101    8   MOLECULE: HYPOTHETICAL PROTEIN RV0793;
  29:  4npo-B  5.2  2.7   80   105   10   MOLECULE: UNCHARACTERIZED PROTEIN;
  30:  3gz7-A  5.2  3.9   82    99    9   MOLECULE: PUTATIVE ANTIBIOTIC BIOSYNTHESIS MONOOXYGENASE;
  31:  4zos-A  5.1  3.6   81   105    5   MOLECULE: PROTEIN YE0340 FROM YERSINIA ENTEROCOLITICA SUBSP
  32:  2bvf-A  5.1  3.9   85   453   13   MOLECULE: 6-HYDROXY-D-NICOTINE OXIDASE;
  33:  1x7v-C  5.1  3.3   80    99   10   MOLECULE: PA3566 PROTEIN;
  34:  6slf-A  5.0  4.0   93   398   11   MOLECULE: N-ALPHA-ACYL-GLUTAMINE AMINOACYLASE;
  35:  2rb7-A  5.0  4.8   97   360    9   MOLECULE: PEPTIDASE, M20/M25/M40 FAMILY;
  36:  6az1-M  5.0  4.1   77   102   10   MOLECULE: RIBOSOMAL PROTEIN S1E;
  37:  2omo-A  5.0  4.0   85   105    8   MOLECULE: DUF176;
  38:  5jlp-A  4.9  2.4   74   397   14   MOLECULE: PHOSPHOSERINE PHOSPHATASE;
  39:  3obi-A  4.9  3.1   77   285    6   MOLECULE: FORMYLTETRAHYDROFOLATE DEFORMYLASE;
  40:  2ril-A  4.9  3.2   77    95    5   MOLECULE: ANTIBIOTIC BIOSYNTHESIS MONOOXYGENASE;
  41:  4q7a-A  4.9  2.8   80   361   15   MOLECULE: N-ACETYL-ORNITHINE/N-ACETYL-LYSINE DEACETYLASE;
  42:  4pxc-A  4.9  4.6   93   424   11   MOLECULE: UREIDOGLYCOLATE HYDROLASE;
  43:  8ecx-C  4.9  3.5   79   102   10   MOLECULE: ANTIBIOTIC BIOSYNTHESIS MONOOXYGENASE;
  44:  4hl9-A  4.9  3.1   80    95    6   MOLECULE: ANTIBIOTIC BIOSYNTHESIS MONOOXYGENASE;
  45:  4mmo-A  4.9  4.2   93   437   15   MOLECULE: SSO-CP2 METALLO-CARBOXYPETIDASE;
  46:  4ewt-A  4.9  3.1   88   389    9   MOLECULE: PEPTIDASE, M20/M25/M40 FAMILY;
  47:  2gx8-C  4.8  4.1   81   364   10   MOLECULE: NIF3-RELATED PROTEIN;
  48:  3bb5-D  4.8  2.5   80   108    5   MOLECULE: STRESS RESPONSIVE ALPHA-BETA PROTEIN;
  49:  1iuj-B  4.8  3.5   81   103    5   MOLECULE: HYPOTHETICAL PROTEIN TT1380;
  50:  7qa5-A  4.8  3.8   80    93    8   MOLECULE: PROTEIN MGTC;
  51:  2f7v-A  4.8  4.1   98   360   12   MOLECULE: AECTYLCITRULLINE DEACETYLASE;
  52:  3kg0-B  4.8  3.4   79    98    4   MOLECULE: SNOAB;
  53:  2qmw-A  4.8  4.1   83   264   12   MOLECULE: PREPHENATE DEHYDRATASE;
  54:  3ced-A  4.8  3.6   78    98   10   MOLECULE: METHIONINE IMPORT ATP-BINDING PROTEIN METN 2;
  55:  6c0d-A  4.8  3.7   87   404   14   MOLECULE: AMIDASE, HYDANTOINASE/CARBAMOYLASE FAMILY;
  56:  6dgd-A  4.7  4.2   89   704    4   MOLECULE: PRIMOSOMAL PROTEIN N';
  57:  5jb3-L  4.7  4.1   83   102    5   MOLECULE: 16S RIBOSOMAL RNA;
  58:  3bm7-A  4.7  3.2   87   106    5   MOLECULE: PROTEIN OF UNKNOWN FUNCTION WITH FERREDOXIN-LIKE
  59:  2od4-A  4.7  3.0   79   101    6   MOLECULE: HYPOTHETICAL PROTEIN;
  60:  2gff-B  4.7  3.7   82   100    7   MOLECULE: LSRG PROTEIN;
  61:  3oq2-B  4.7  2.8   72   101   14   MOLECULE: CRISPR-ASSOCIATED PROTEIN CAS2;
  62:  5xvn-F  4.6  2.7   74   108    9   MOLECULE: CRISPR-ASSOCIATED ENDONUCLEASE CAS1;
  63:  8dew-A  4.6  3.9   80  1058   10   MOLECULE: EFFLUX PUMP MEMBRANE TRANSPORTER;
  64:  7bio-B  4.6  3.5   79   111    4   MOLECULE: MONOOXYGENASE/PUTATIVE ANTHRONOXYGENASE;
  65:  2n51-A  4.6  4.4   82   133    9   MOLECULE: ELONGATION FACTOR 1-DELTA;
  66:  2dcl-A  4.6  3.5   77    99   18   MOLECULE: HYPOTHETICAL UPF0166 PROTEIN PH1503;
  67:  2peb-A  4.6  3.3   78   115    9   MOLECULE: PUTATIVE DIOXYGENASE;
  68:  6nuk-A  4.6  2.5   73   101   11   MOLECULE: FERREDOG-DIESEL;
  69:  1o51-A  4.6  3.7   73    89    5   MOLECULE: HYPOTHETICAL PROTEIN TM0021;
  70:  4usj-C  4.5  4.0   92   143   11   MOLECULE: ACETYLGLUTAMATE KINASE, CHLOROPLASTIC;

0001 s001A   ---PPKEARMHDRN-VAA---LAVLQLAVDG-D----G----SE--GVREL-RSLGL-AVPQL-G------P----GQFRVEVSA-TG--EL--EGGSIRLEFRGPWERYKDLKDAVVSVLERASA----A-QV-RARLDWRADP---LT--PEGINQVRDVLAHL-I--PGRVRITAEPAT-G-E--V-D
0002 4kyzA   --kRVRISFLAILA-----dinRKXTVRFRG-D----------D--LEALE-KALKE-XIRQAkF------A----GTVTYTLDG------------------------------------------------N-DLEITITGVP---RQ----VLEELAKEAERL-A--KITITITVTVEG-Q-Lg----
0003 5cw9A   ----------------------MEMDIRFRG----------------ddpeykALRE-MIRQAkF------A----GTVTVTLII-RF--RgddLEALLEIRITGV--PPQVILELVKEAIRLAIT----V-TV-ELVIRITG-----vP--EQVRKELAKEAERL-A--KITVTYTIRL-----------
0004 4neyB   ---------------------xGEXXIIFEG-D----------D--LEALE-KALKE-XIRQAkF------A----GTVEYWLSG------------------------------------------------N-RLXIRITG-----VP--EQVRKELAKEAERL-K--AIQVEYQIRSGS-G-S-tG-V
0005 3n0vA   ----------------------DTWILTADC-P----S----XL----GTV-DVVTR-YLFEQ--------R----CYVEHHSFD-DR--Q----------------------------------S----G-RF-FIRVEFRQDD---F-----DEAGFRAGLAER-S--EXAFELTAPN-----------
0006 4v1ak   --------piNLGT-DYE---YGVLNIHLIA-Y----------D--MALAE-SYAQY-VHNLChL------A----IKVESYAMT-KT--ME--VL------------------------qlqmVL----T-TH-ERVVQISG-----LS--ATFAEIFLEIIHSN-L--PVKLSVRE-----------
0007 1r3nA   --------ssvwshdlsle-aYNWQKVTVGV-G----A-------lrKDAL-LMSSK-MIVAA-SaqrhngL----FTCIIDAKP-YS--V----------------------------------N----I-IPgEVSFTLDFHpsddvL--ATMLKEAAAEFDRL-I--KLSYESETQVSP-A-V-aG-H
0008 2nzcD   -------------hxEKR---FYILTIVVED---------------REKAY-RQVNE-LLHNF-S------E--diLLRVGYPVR-E---------------------------------------------EN-XAIIFLVL-K---TD--NDTIGALSGKLGQI-----SGVRVKTVPLK---------
0009 3n5fA   --------igsramagtlp-gLIWVKFTIEG-K----A----E--lrRDPM-AAAAQ-IIIVI-ErrtgttV----GTVQLHVYG-GI--N--------------------------------------vI-PE-RVEFVLDLDlkaevR--DQVWKAIAVRAETI-A--KVRVTTERQEMP-P----V-L
0010 1ysjA   --eEIAA------------asvDRFEIVIKG-K----N----SI----DPI-AAAGQ-IISGL-Q----naV----VSIRVQAGT-SW-------------------------------------N----V-IPdQAEXEGTVTfqkeaR--QAVPEHXRRVAEGI-A--AAQAEFKWFPYL-P-S--V-Q
0011 6wi5A   -----------------E---EVVVEIRIRV-Q----R----EE----KVR-RLIKR-ILEEV-K--resnS----VEVHVETRK-RN------------------------------------------G-EV-EVHVRIRH-D---DK--E-TIERLVERILRE-I--KSEVEVRTTTKR---------
0012 3fgvA   --------------------xrEVVIVKSTP-Q----R----GK--FNAFA-ELVGK-LVSET-R--dfpgC----LGAYLXLAP-E---------------------------------------------rN-EQVVXHIWET---PDalEAYLTWRADRDFLE-I--NEEQDFKTYQLA---------
0013 3r2cK   ---------------------QEKIRIKLRA-Y----------D--HRLLD-QSVKQ-IIETV-K----rtG----GVVKGIPLP-TR-------------------------------------K----S-EF-SRILDIIRT--------pQTIEALMEI--SL-P--AVDVEVKMR------------
0014 1zpvC   --------------------naXKAIITVVG-K----------D--KSGIV-AGVSG-KIAEL--------G----LNIDISQTV-LD------------------------------------------E-YF-TXXAVVSSEK---Q-----DFTYLRNEFEAF-G--QVKINIQSAAEA-X-Y-----
0015 2fb0A   ---------------ANS---XIRLNVFVRV-N----E----TN--REKAI-EAAKE-LTACS-L--keegC----IAYDTFESS---------------------------------------------trRD-VFXICETW-----QN--AEVLAAHEKHFAQYvG--IIQXKLEKFEF----------
0016 8p6pI   --------------navK---YPELKIKLES-Y----------D--STLLD-LTTKK-IVEVVgV------D----VKIKPLPLT-KK--EV--IT-----------------iirsphvdkasEK----N-RH-KRLMILVD-----VN--QGAIDSLKRI--KI-P--VVTLRFSK-------------
0017 1u8sB   --------gtdrpgicnev-hAYTVEVYVES-D----------D--KLGLT-EKFTQ-FFAQR-Q----------iGMASLSAQT-IS--K---------------------------------------N-QF-HIAISARV-D---SG--C-NLMQLQEEFDAL-C--TVQGSLNFIKN-----------
0018 1z2lA   --------fwgsknifgla-gQRRYTVTLNG-E----S----N--yrRDTV-YAFSR-ICHQS-VrmgdplV----LTFKVEPRP-NT------------------------------VNVV----------pG-KTTFTIDCHtdaavL--RDFTQQLENDMRAI-C--DIGIDIDLMDEE-P----V-P
0019 8d8kJ   -------------------ygdLVADIQLRS-Y----------D--NENLD-FYSDF-ILRTGyL------G----IPLTPKPLT-RR--ER--WT--------------------------viER----H-TH-KRLIRAW-TN---P----EVLQMLIAYITKH-S--MVGMKCNFFQRS-E-Is----
0020 7c50A   -------------------adtLILPIKFTR-S----NnnltNL--YEELE-QDKEN-IIKFLgV------K----EDINYNPNI-IDrlSD--PYS-------------------------ndYR----Y-IG-TANLLIYTQN---VK----LGKSILENISSL-akfGIVTKIDIEYLY-T-K-iK-T
0021 5i4mA   --------amVASG-vfag-gQRWYEITFTG-Q----EtpmpRR--RD-AL-LGASR-VVDLV-NdhapygC----ATVMMQVHN-SR--N----------------------------------V----I-PG-RVFFTVDFHpddavL--AKMDAALRDGVARI-A--ALDTALEQFYYA-P----I-A
0022 4uerJ   -------------Q-QQQ---IIKIRITLTS-T----------K--VKQLE-NVSSN-IVKNAqH------N----LVKKPVRLT-KV--LK--IS----------------trktpngegsktEM----R-IH-KRYIDLE------AP--VQIVKRITQIT--I-E--PGDVEVVVA------------
0023 7m6uB   --dEEAG-sfGSRD-----gtsGIAYVQVNI-T----GelgvNA--LVEAS-DLVLR-TMNID-D---kakN----LRFNTIAKA-GN--VS-------------------------------NII----PaSA-TLNADVRYAR---NE----DFDAAMKTLEER-A--QVKVIVTRGRP---aA--L-S
0024 2bbeA   --dYKIN--------QQQ---I-VCVASFLS-K----E----GK--TEALI-AALAS-LIPDT-R------ReagcIRYELNVSR---------------------------------------------dePR-RVTFVEKF-----VD--IAAFDEHCAKIQHY-F--HpSFHVETYHQV-I-------
0025 7ljhA   --gEEGS-asrivtggVG---SKRYRITYKG-P--ggG----LV--N--PM-VAMSQ-TVVDF-YpakpktT----YAAVTGGGT-SV-------------------------------------N----S-IPnEVYMEFDMSespaeL--AKVEQAFLAIVQKS-V--EITADVKMGDRP-----aG-E
0026 8tcfC   -----------------T---KCVVRFNFR----------------GDMAV-YALKA-VKDHL-K--kegpH----WNITTHNDD--------------------------------------------------EVYLVVRG-I---HE--S-DAKRIAKWVEST-I--P-GISVETQCD----------
0027 2nyiA   --------rrassvvsPD---TREYELYVEG------------P--DSE---GIVEA-VTAVLkK------G----ANIELETET-LP--AP------------------------------FAGF----T-LF-RXGSRVA-FP---FP----LYQEVVTALSRV-E--EVDIDLEEVV-----------
0028 3ce8A   -------------X-STE---Q-LLVLIA--------------------QN-DIKDD-IVDTL-I--elefL----SGFSLGNIC-GF----------------------------------sREG----Y-RE-FCKFEIX------HP--AAQQAALLTALALV-C--KNPCRYWIXPIY-Q-------
0029 1y0hA   -------------G-MTS---pVAVIARFMP-R----P----DA--RSALR-ALLDA-MITPT-R--aedgC----RSYDLYESA-D---------------------------------------------gG-ELVLFERYRS---RI---aLDEHRGSHYLNYrA--QVGVAVTVLAPL-D-------
0030 4npoB   -------------------ttmVISHGTLSA-S----A----EH--AAHLR-QLLVH-IAQAT-R--qedgC----LLYLVSEDL---------------------------------------------sqPG-HFLITEHWDN---LGamHTHLALgVTQAIDA-L--KTDLKITAYEAG-E-------
0031 3gz7A   -------------------qgxIQEIASILV-Q----P----GR--EADFE-AGVAQ-ARPLF-X--rargC----HGVALHRSI-E--------------------------------------------apQ-RYTLVVDWTVd-nHX--VDFRQSDFQEWRKL-V--GEPPQVHHEQKV-L-------
0032 4zosA   -------------------seeRHIICELRC-E----P----EN--RERVK-ELVLK-FVEPA-R--letgC----LYYDLYQKI-D--------------------------------------------ePD-TFYIIDGWVN---QE-avTSHAE-NPHVAEVmS--DLQPSISLITRV-S-D-----
0033 2bvfA   --------vqlyeL-PRK---MLAGFITWAP-S----V----SE--LAGLL-TSLLD-ALNEM-A------D---hIYPSVFVGV-DE------------------------------------------nrAP-SVTVCVGH-L---GG--LDIAERDIARLRG-----LGRTSDSIARSY-De------
0034 1x7vC   ------------ghxSTP---L-TLIATITA-A----P----GH--AEALE-RELRA-LVAPS-R------AeagcLQYDLHQDR-H--------------------------------------------dSH-LFYXIEQW-----RD--DAALERHQNTFLRF-----eNVKIDQLYRL-A-------
0035 6slfA   --qPGEEGvftpgrfltA---ASNWRIHIHG-E--ggL----TK--D--PI-VVAAS-IITKL-QdpnevaV----VTVSIEGGK-S-------------------------------TNSI----------pY-TVTLGVNTAsndelS--EYVQNAIKRIVIAE-C--QQEPEFELDSV--P-A--V-I
0036 2rb7A   itgdEEIGgxNGAA-----kekGIIDIKLTC-T----G----KaaaVDLLX-EDYTR-LKTLF--enedhwH----RTVNLRIRA-GE--ST-------------------------------NKV----PdVA-EGWFNIRVEH---D-----DPGALIDKIRKT-V--SGTVSIRTVPVF-LlG--E-N
```

```
0037 6az1M   ---------------------GQTVRLTITS-R----------N--AKAVE-SVTSQ-LLMRA-R----deK----VTIHPVRLT-RT--LK--IT----------------trktpcgngtntEL----K-IY-KRIIDLHA---------pteQVKKITSFT-M--EVDVSITILD----------
0038 2omoA   --nLYFQ-------------ghxYVTIVYASV-K----T----DK--TEAFK-EATRX-NHEQS-I--repgN----XRFDILQSA-D--------------------------------------------dPT-RFVLYEAYKT---RK----DAAAHKETYLTW-R--DdPRKGVIYGGL-Y-P--Tg-
0039 5jlpA   --------------------pkVSVLITVTG------------V--DQP---GVTAT-LFEVLrH------G----VELNVEQVV-IR------------------------------------------H-RL-TLGVLVCC-P---AD--VaDGPALRHDVEAA-I--RLDVSIERSD--------------
0040 3obiA   ---------------------HHQYVLTLSC-P----D----RA----GIV-SAVST-FLFEN-------G----QNIDAQQYN-DT--E----------------------------------S----G-HF-FXRVVFNAA-------kviPLASLRTGFGVI-A--AXGWHXRDRE---------
0041 2rilA   -------------G-XSA---PVTLINPFKV-P----A----DK--LEAAI-EYWEA-HRDFX-A--qqpgY----LSTQLHQSI-DE-------------------------------------------gatyQLINVAIW-Q---SE--A-DFYQAAQKXRQ------LXGNPALY------------
0042 4q7aA   --------vlygrgavdak-gSVALEYRVTV-P----X----S--peATAA-ELAAD-FWYRL-RhafhrvE----PKLALNSSS-DG------------------------------------------L-YG-EAVARIGLLppalsP--EEAIAVATSLAS---------EGEVTATVNA-------pA-F
0043 4pxcA   --------agSKELaealk-aPASLKVEFEG-N----G----G--yrNDAG-LAAAE-LALAV-EsesidtV----GTVILELHG-AI--N----------------------------------S----I-PS-KSHLEIDTDidearR--NTVIKKIQESANTI-A--KKLSEFKINQDP-P-A-sL-F
0044 8ecxC   -------------------gmtYHVLVQFDV-P----S----DK--REAFA-AAGLF-DANGS-L-qnepgT----LRFEVIRDE-N--------------------------------------------nRN-RFYLDEVYED---EA---afLQHXRNTIARF-Y--EdGPLFFKGYRV-E-G-----
0045 4hl9A   ---------------------xLKVIAQDFI-K----P----EA--IDIVL-PLYRE-LVEKT-R--qeplX----LAYDLFVDQ-K--------------------------------------------dPG-HFVFIEEWPD---RA---ALDIHXATFTRLvP--LINGTVVLXDAV-P-----
0046 4mmoA   --gEEEI-rpqivlgvkG---LLYVELVLDY-GtkdlL----VR--N--PC-IDLAK-IISTL-ValltypT----CNVGFECG-------ytgKGSKTI-----------------------------V-PH-RAFAKLDFLvpnqdP--YKVFELLKKHLQKA-G----NGEILAHGFQ-P-M--G-L
0047 4ewtA   --eEVPP------------tgrAFFKLKVQG-K--ggM----AN--D--AI-VAGSY-FVTAL-QsrrlspfetgvVTIGFDGKG-Q-------------------------------FNVI----------kD-VVEIEGDVGltdatK--ATIEKEIKRLSKGL-E--DVTCTLEYNDDY-P-A--L-Y
0048 2gx8C   --------vlaptyaEEM---K-KVVVFV--------------------PV-THAEE-VRKAL-GghignyS----HCTFSSEGT-GT--------------------------------fvpqLE---R-VE-EVRIETI------IP--ASLQRKVIKAMVTA-H--PYEVAYDVYPLD-N---kG-E
0049 3bb5D   -----------NLY-FQG---XLYHLVXLEP-E----G---eGA--XDRIX-EAXAI-LDGLA-P-----elpglTEFRHGPN---------------------------------------rdfeqkseryPYGFLCTF-----TD--KAALDAYAVHHQRA-G--GXLILVVDLEV---------
0050 1iujB   ---------------------mFVTMNRIPV-R----P----EY--AEQFE-EAFRQ-RARLV-D--rmpgF----IRNLVLRPK-N--------------------------------------------pGD-PYVVMTLWES---EEafRAWTESAFKEGHAR-S--GePNRLEAFEVV-L-------
0051 7qa5A   -------------A-SEA---EQRYEVQIVC-R----A----ED--EIQVR-SLMLH-SLGSS-D-----------lrQSLHSED-----------------------------------------ldnpA-KL-EVRAELL--G---TP--E---APAQLERLVSR-V--SeSVRWQVFELA-A-D-----
0052 2f7vA   fssdEEANdpRCIA-----ahrGISSVLMRF-A----GdpaaSA--LHQAM-RWGGK-ALDHV--rfggltG----LRFNIRVDG-GI--KA-------------------------------NMI----ApAA-ELRFGFRPPS---M-----DVDGLLATFAGF-A--DPHFEEFRGPSL-PaR--A-E
0053 3kg0B   ----------------AD---EVTFVNRFTV-H---------GA--PAEFE-SVFAR-TAAFF-A--rqpgF----VRHTLLRER-D--------------------------------------------kDN-SYVNIAVWTD---HD----AFRRALAghATA-L--R-TSEHGLFTAR-Q-------
0054 2qmwA   lvikNQQQ-------DQN---ATSLXFLITPxH----D----KP----GLL-ASVLN-TFALF-N-----I----N-SWIESRP-LK--T---------------------------------qL----G-XY-RFFVQADS---------aITTDIKKVIAIL-E--TFKVEXGAFN---------
0055 3cedA   --tELEP-------lEKD---AYIVRLVFAG-------------------S-TTTEP-IVSSL-S-taydiK----INIEANIKN-TK-------------------------------------------nGT-VGFLVLHIPY---IS--SVDFGKFEKELIER-Q-----VKXELRHG--------
0056 6c0dA   -------------------vgiREFQLRFGQ-R----NtpmaIR--RD-AG-AALVA-FIAHI-DladadtV----WTVRIDLDP-GS--F----------------------------------S----V-VPgKAVLHLQFDanpnrL--HAMENALVALVDEWnqqhLVRAELIACGAE-E----P-V
0057 6dgdA   pvfsttvwYHARDLavwrapwtSHVLIRAED-H---------N--NQQAP-LFLQQ-LRNLL-QspladeK----LWVLPVPAP-KR--G---------------------------------------G-RW-RWQILLQHSR---VR-lqHIVSGTLALINPE-A--RVKWVLDVD-----------
0058 5jb3L   -----------------M---QKARIKIAST------------N--VRSLD-EVANQ-IKQIA-E--rtgvR----MSGPIPLPTkRI--RI--TT------------------rkspdgegsaWE----L-RV-HKRLIDIE-----AD--ERAMRQIMR--IRV-P--EDVTIEIELIS--------
0059 3bm7A   --------hhENLY-FQG---XIGVVATLKV-Q----P----AK--AAEFE-KVFLD-LAAKV-K-anepgC----LVYQLTRSK-T--------------------------------------------eEG-VYKVLELY------AS--XDALKHHGGTFKAA-G--ApAPVIEYLDAV-E-------
0060 2od4A   -------------------ipxYIRVVSITA-Q---------S--KLQFD-XTVTYvWSPKV-I---slgA----ISAEFVQSN-----------------------------------------------EN-SGXYIIHYPD---KQtaISVFDKIKPEVDEV-R-tQNRIQITEGKRL-F-------
0061 2gffB   ---------------------XHVTLVEINV-K----E----DK--VDQFI-EVFRA-NHLGS-I--reagN----LRFDVLRD----------------------------------------ehiPT-RFYIYEAY------TD--EAAVAIHKTTYLQC-V--EQLRKKTVFIGLpG-S--L-E
0062 3oq2B   ------------mygNDA---MLVLISYDVS-F----E----D----PGGQ-RRLRR-IAKACqD-----Y----GQRVQY-------------------------------------------------------SVFE--cV---VD--PAQWAKLKHRLLSE-M--DKDCLRFYLGAN---------
0063 5xvnF   -------------M-SYR---YMRLLLMFDM-P----T-dtaSD--RKAYR-KFRKF-LINEG---------------IMHQF-----------------------------------------------------SVYSKIL-----LN--DTANKAMLARLKQN-N--PQLITLLNVTEKe---------
0064 8dewA   --yPSVA--------------aPTITLHAIY-P----G----A----saqvgSVLSV-IERNM-N--gvegL----DYMSTSADS-----------------------------------------------SG-SGSVSLTFTD---EN--L-AQVEVQNKLSEV-L--SYGVTVSKASNS-G-Nag---
0065 7bioB   ---------------------aVVFVNKLTL----------iGD--AEEFE-SRYEA-VGAFM-E--tqpgL----VRYSLVRST-K--------------------------------------------dDS-VYFNIAEWDD---EDtfRKALAEeFRRRLDA-L--TIKGEPHLSLPV-R-------
0066 2n51A   --------kakkpA-LVA---KSSILLDVKPwD----D----ET----DMA-QLEAC-VRSI------qldgL----VWGASKLPV-GY------------------------------------------G-IR-KLQIQCVV-E---DD--KVGTDLLEEEITKF-E--EQSVDIAFNKI----------
0067 2dclA   ----------vevehWNT---L-RLRIYIG----------------ENDKwrPLYKV-IVEKLmG-----I----AGATVYRGI-Y----------------------------------------gfgT-DL-PIIVEVV-----DR--GHNIEKVVNVIKPM-I---KDGMITVEPTI-V-Lw----
0068 2pebA   -----------KED-TIE---IAGFHAHVYF-D----A----AS--RDVAA-RVREG-LGARF-E-------------VQLGWFD-K---------------------------------------pigphpkG-XYQVAFLP----------qFDKVVPWLXLN-R--ELDILVHPETGD--------
0069 6nukA   -------------------gstVRIEIRFTN-M---------R--REEVQ-KELEK-FKERL-KekrtgsE----IRIEIEERD------------------------------------------------G-EVRVEVEINS----H--EEEVRQIIEEIER--wvrKMGGELRVEK---------
0070 1o51A   -------------------hhxKLLKIYLG----------------EKDKhkPLFEY-LVKRA-Y---elgX----KGVTVYRGI-X---------------------------------------gfghP-DL-PIVLEIVDEE---ER-----inLFLKEIDNI----DFDGLVFTADVN-V-V--K--
0071 4usjC   ---------LSAFPgVKF---F-RIEAIF--------------------RP-WRLPF-VIDTLyG------I----RGLTNTPVK------------------------gvgvqggsrerYAGTEFgpsnL-VD-KEKLDIV------VS--RAQVDAVVRLVAAS-A--YTDGKIFVHPVA-EeT--G-L

0001 s001A   ---LLLLLHHHHLL-LLL---EEEEEEEEEE-E----H----HH--HHHHH-HHHHH-HHHHL-L------L----EEEEEEEEE-EE--EE--LLEEEEEEEEEEHHHHHLLHHHHHHHHLLLLE----E-EE-EEEEEEEEEE---EL--HHHHHHHHHHHHHH-L--LLEEEEEEEEELL-L-L--L-L
0002 4kyzA   --lEEEEEHHHHHH-----eeeEEEEEEEEL-L----------L--HHHHH-HHHHH-HHHHHhL------L----LEEEEEEEL------------------------------------------------L-EEEEEEELLL---HH----HHHHHHHHHHHH-H--HLEEEEEEEEE-E-El----
0003 5cw9A   ---------------------LEEEEEEEL----------------llllhhHHHH-HHHHHhL------L----LEEEEEEEE-EE--ElllHHHHEEEEEEEELL--LHHHHHHHHHHHHHHHLE----E-EE-EEEEEEEL-----lL--HHHHHHHHHHHHHH-H--HLEEEEEEEL---------
0004 4neyB   ---------------------lLLEEEEEEEL-L----------L--HHHHH-HHHHH-HHHHHhL------L----LEEEEEEEL------------------------------------------------L-EEEEEEEL-----LL--HHHHHHHHHHHHHH-H--HLEEEEEEELLLL-L-L-eL-L
0005 3n0vA   ---------------------LLEEEEEEE-E----L----LL----LHH-HHHHH-HHHHL-------L----LEEEEEEEE-EL--L----------------------------------L----L-EE-EEEEEEELLL---L-----LHHHHHHHHHHH-H--HLEEEEEELLL---------
0006 4v1ak   --------leELLL-LEE---LLLEEEEEEE-L----------L--HHHHH-HHHHH-HHHHHhL------L----LLEEEEEEE-EL--EE--EE------------------------eellEL----L-EE-EEEEEELL-----LL--HHHHHHHHHHHHHL-L--LEEEEEEE-----------
0007 1r3nA   --------hhhhllllllhh-eEEEEEEEEELL-L----E-------hlLLHH-HHHHH-HHHHH-HhhhlllE----EELLEEEEL-LL--L----------------------------------L----E-ELlEEEEEEEEEllhhhH--HHHHHHHHHHHHHHH-L--LLEEEEEEEEEL-L-E-lL-L
0008 2nzcD   -------------llLLE---EEEEEEEEEL---------------LHHHH-HHHHH-HHHHL-H------H--heEEEEEEEEH-H---------------------------------------------HL-EEEEEEEE-E---EL--HHHHHHHHHHHLLL-----LLEEEEEEELL--------
0009 3n5fA   --------hhhhhhhllll-eEEEEEEEEL-L----L----E--llLLHH-HHHHH-HHHHH-HhhhlllE----EEEEEEEEL-LL--L--------------------------------------eE-LL-EEEEEEEEEllhhhH--HHHHHHHHHHHHHHH-H--HLEEEEEEEEEL-L----E-E
0010 1ysjA   --lLLLL------------lleEEEEEEEEL-L----L----LL----LHH-HHHHH-HHHHH-L----llE----EEEEEEELL-LL-------------------------------------L----L-LLlEEEEEEEEELllhhhH--HHHHHHHHHHHHHHH-H--HLEEEEEEEEEE-L-L--E-E
0011 6wi5A   -----------------L---LEEEEEEEEE-L----L----HH----HHH-HHHHH-HHHHH-H--hhllL----LEEEEEEEE-EL------------------------------------------L-EE-EEEEEEEE-L---LH--H-HHHHHHHHHHHH-H--HLEEEEEEELLL--------
0012 3fgvA   --------------------llEEEEEEEELL-L----L----LL--HHHHH-HHHHH-HHHHH-H--llllE----EEEEEEEEH-H---------------------------------------------hL-EEEEEEEELL---HHhhHHHHHHHLLLHHHH-H--HHLLEEEEEEEL---------
0013 3r2cK   ---------------------LLLEEEEEEE-L----------L--HHHHH-HHHHH-HHHHH-H----hlL----LEEEEEEEE-EE-------------------------------------L----L-EE-EEEEEEEEL--------hHHHHHHLLL--LL-L--LLEEEEELL---------
0014 1zpvC   --------------------llEEEEEEEEE-L----------L--LLLHH-HHHHH-HHHHL--------L----LEEEEEEEE-EL------------------------------------------L-EE-EEEEEEEELL---L-----LHHHHHHHHHHH-H--HEEEEEEEHHLL-L-L------
0015 2fb0A   ---------------LLL---LEEEEEEEEL-L----L----LL--HHHHH-HHHHH-HHHHH-L--llllE----EEEEEEELL---------------------------------------------llLL-EEEEEEEE-----LL--HHHHHHHLLHHHHHhH--HHHEEEEEEEL----------
0016 8p6pI   --------------lllL---LEEEEEEEEE-L----------L--LHHHH-HHHHH-HHHHHhL------L----LLLLLEEEE-EE--EE--EE-----------------eelllllllllEE----E-EE-EEEEEEEE-----LH--HHHHHHHLLL--LL-L--LLEELLLL-----------
0017 1u8sB   --------eellllhhhhh-lLEEEEEEEEE-L----------L--LLLLH-HHHHH-HHHHL-L---------lEEEEEEEEE-LL--L---------------------------------------L-LE-EEEEEEEE-L---LL--L-LHHHHHHHHHHH-H--HLEEEEEEEEL---------
0018 1z2lA   --------lhhhhhhllll-eEEEEEEEEL-L----L----E--hlLLHH-HHHHH-HHHHH-HhhlllllE----EELLEEEEL-LL------------------------------LLEE----------lL-EEEEEEEEEllhhhH--HHHHHHHHHHHHHHH-H--HLEEEEEEEEEL-L-----E-E
0019 8d8kJ   -------------------lllEEEEEEEE-L----------L--LHHHH-HHHHH-HHHHHhL------L----LLLLLEEEE-EE--EE--EE--------------------------elEE----E-EE-EEEEEEE-LL---H----HHHHHHHHHHHHL-L--LEEEEEEEEEE-L-Ll----
0020 7c50A   -------------------lleEEEEEEEEE-E----EllhhHH--HHHHH-HHHHH-HHHHHlL------L----HHEEELLEE-EEllLL--LLL-------------------------llLL----E-EE-EEEEEEEELL---HH----HHHHHHHHHHHH-hhhLLLEELHLEEEL-L-L-eE-E
0021 5i4mA   --------llHHHH-hhll-eEEEEEEEEE-L----LllhhHL--LL-HH-HHHHH-HHHHH-HhlllllE----EELLEEEEL-LL--L----------------------------------E----E-LL-EEEEEEEEEllhhhH--HHHHHHHHHHHHHHH-H--HLEEEEEEEEEL-L----E-E
0022 4uerJ   -------------L-LLL---LLLEEEEEEE-L----------L--HHHHH-HHHHH-HHHHHhL------L----LLEEEEEEE-EE--EE--EE----------------eelllllllllEE----E-EE-EEEEEEL------LL--HHHHHHHHHLL--L-L--LLEEELLL----------
0023 7m6uB   --lHHHL-llLLHH-----eelEEEEEEEEE-E----LhhllLH--HHHHH-HHHHH-HHHHL-L---lllL----LEEEEEEEE-LL--LL-------------------------------LLL----LlEE-EEEEEEEELL---HH----HHHHHHHHHHHH-H--HEEEEEEELLL---hH--H-H
0024 2bbeA   --lLEEL--------LLL---E-EEEEEEEE-L----L----LL--HHHHH-HHHHL-LHHHH-H------LllleEEEEEEELL---------------------------------------------llLL-EEEEEEEE-----LL--HHHHHHHLLHHHH-H--HhEEEEEEELLE-E--------
0025 7ljhA   --lLLHH-lleeeeeeLE---EEEEEEEEEL-L--leL----LL--L--HH-HHHHH-HHHHH-LlllllleE----EEEEEEELL-LL-------------------------------------L----E-ELlEEEEEEEEEllhhhH--HHHHHHHHHHHHHHH-H--HLEEEEEEEEEL-----lE-E
0026 8tcfC   -----------------L---LEEEEEEEL----------------LLLHH-HHHHH-HHHHH-H--hhllL----LEEEEELLL--------------------------------------------------LEEEEEEE-E---LL--H-HHHHHHHHHHLL-L--L-LLEEEEEEL----------
0027 2nyiA   --------eelllllllLL---EEEEEEEEEE------------E--LLL---LHHHH-HHHHHhL------L----LEEEEEEEE-EE--LL------------------------------LLLL----E-EE-EEEEEEE-EE---HH----HHHHHHHHHHHH-H--HLEEEEEELL---------
0028 3ce8A   -------------L-LLE---E-EEEEEE--------------------EH-HHHHH-HHHHH-L--llllL----LLLEEEEEE-EE----------------------------------eLLE----E-EE-EEEEEEE------EE--HHHHHHHHHHHHHHH-L--LLLLEEEEEELL-L--------
0029 1y0hA   -------------L-LLL---lEEEEEEEELL-L----H----HH--HHHHH-HHHHH-HHHHH-H--hlllE----EEEEEEEEL-L---------------------------------------------lL-LEEEEEEEELL---HH---hHHHHHLLHHHHHhL--LLHLEEEEEEEEE-E-------
0030 4npoB   -------------------lllEEEEEEEEL-L----L----LL--HHHHH-HHHHH-HHHHH-L--llllE----EEEEEEEEL---------------------------------------------llLL-EEEEEEEELL---HHhhHHHHHLlHHHHHHH-H--HLLLEEEEEEEE-E--------
0031 3gz7A   -------------------lllEEEEEEEEL-L----L----LL--HHHHH-HHHHH-HHHHH-H--llllE----EEEEEEEEL-L--------------------------------------------llL-EEEEEEEELHh-hHH--LLLLLLHHHHHHHH-H--HHLLEEEEEEEE-L-L-------
0032 4zosA   -------------------hhlEEEEEEEEE-L----H----HH--HHHHH-HHHHH-HHHHH-H--hlllE----EEEEEEEEL-L--------------------------------------------lLL-EEEEEEEELL---HH-hhHHHHL-LHHHHHHhH--HHHEEEEEEEEL-L-L-----
0033 2bvfA   --------eeeeeL-LLL---EEEEEEEEELL-L----H----HH--HHHHH-HHHHH-HHHHL-L------L---lLLLEEEEEE-LL------------------------------------------llLE-EEEEEEEL-L---LL--HHHHHHHHHHHHL-----LLLLEELLEELH-Hl-------
0034 1x7vC   ------------lllLLL---E-EEEEEEEE-L----L----LL--HHHHH-HHHHH-HHHHH-H------HllleEEEEEEEEL-L--------------------------------------------lLL-EEEEEEEE-----LL--HHHHHHHLLLHHHL-----lEEEEEEEEL-L--------
0035 6slfA   --eLLLLLeelleellE---EEEEEEEEEL-L--leH----LL--L--HH-HHHHH-HHHHH-HlllllleE----EEEEEEELL-L-------------------------------LLEE----------lL-EEEEEEEEEllhhhH--HHHHHHHHHHHHHHH-H--HLLLEEEEEEEE-L-L--L-E
0036 2rb7A   eellHHHLllLLHH-----eelEEEEEEEEE-E----L----LlehHHHHH-HHHHH-HHLLL--llllllL----LEEEEEEEE-LL--LL-------------------------------LEE----LlEE-EEEEEEEELL---L-----LHHHHHHHHHHHH-L--LLEEEEEEELLE-ElH--H-H
0037 6az1M   ---------------------LLLEEEEEEE-L----------L--HHHHH-HHHHH-HHHHL-L----llL----LLEEEEEEE-EE--LL--EE----------------eelllllllllLL----E-EE-EEEEEELL-----------lllLHHHHHLLL-L--LEEEEEEELL---------
0038 2omoA   --lLLLL------------lllEEEEEEEEEL-L----H----HH--HHHHH-HHHHH-HHHHH-L--llllE----EEEEEEEEL-L--------------------------------------------lLL-EEEEEEEELL---HH----HHHHHLLLHHHH-H--HhLLEEEEEEEEE-E-L--Ll-
0039 5jlpA   --------------------llLEEEEEEEE------------E--LLL---LHHHH-HHHHHhL------L----LEEEEEEEE-EL------------------------------------------L-EE-EEEEEEL-L---HH--HhLLHHHHHHHHHH-H--HLEEEEEELL---------
0040 3obiA   ---------------------LLEEEEEEE-E----L----LL----LHH-HHHHH-HHHHL-------L----EEEEEEEEE-EL--L----------------------------------L----L-EE-EEEEEEEL------lllLHHHHHHHHHHH-H--HLEEEEEELL---------
0041 2rilA   -------------L-LLL---LEEEEEEEELL-L----L----LL--HHHHH-HHHHH-HHHHH-L--llllL----LEEEEEEEL-L--------------------------------------------llllLEEEEEEE-L---LH--H-HHHHHHHHHHH------LLLLLEEE-----------
0042 4q7aA   --------eeellllllllh-eEEEEEEEEE-E----L----L--llLLHH-HHHHH-HHHHH-HlhhhllE----EEEEEEEEE-LL------------------------------------------L-EE-EEEEEEEEElllllL--LHHHHHHHHHLL---------LLEEEEEEEEE-----lL-E
0043 4pxcA   --------llLHHHhhhhh-eEEEEEEEEL-L----L----E--hlLLHH-HHHHH-HHHHH-HllllllE----EEEEEEEL-LL--L----------------------------------E----E-LL-EEEEEEEEEllhhhH--HHHHHHHHHHHHHHH-H--HEEEEEEEEELL-L-E-hH-H
0044 8ecxC   -------------------lllEEEEEEEEL-L----H----HH--HHHHH-HHHHH-HHHHH-H-hhlllE----EEEEEEEL-L--------------------------------------------lLL-EEEEEEEELL---HH---hhHHHHLLHHHHH-H--HlEEEEEEEEL-L-L------
0045 4hl9A   ---------------------lEEEEEEEEE-L----H----HH--HHHHH-HHHHH-HHHHH-H--llllL----LEEEEEEEL-L--------------------------------------------eEE-EEEEEEEELL---HH----HHHHHHHLHHHHhH--HHHEEEEEEEL-L--------
0046 4mmoA   --lLHHH-lleeeleelE---EEEEEEEEEL-LllleH----LL--L--HH-HHHHH-HHHHH-LhhhhllL----LEEEEEEE--------lllLLLLLE-----------------------------E-LL-EEEEEEEEElllllH--HHHHHHHHHHHHHL-L----LLEEEEEEEL-L-H--H-H
0047 4ewtA   --lLLLL------------leeEEEEEEEEL-L--leH----LL--L--HH-HHHHH-HHHHH-HhllllllllleeEEEEEEELL-L-------------------------------LLEE----------lL-EEEEEEEEEllhhhH--HHHHHHHHHHHHHHH-H--HLEEEEEEEEL-L-L--E-E
0048 2gx8C   --------eeeeeeeEEE---E-EEEEEE--------------------LH-HHHHH-HHHHH-HllllleE----EEEEEEEEE-EE--------------------------------eeelLE----E-EE-EEEEEEE------EE--HHHHHHHHHHHHHHH-L--LLLLLEEEEEEE-E---eE-E
0049 3bb5D   -----------LLL-LLL---LEEEEEEELL-L----L---lLH--HHHHH-HHHHH-HHHHH-H------hllleEEEEEEEL-----------------------------------------llllllllllLEEEEEEE-----LL--HHHHHHHHHLHHHH-H--HHHEEEEEEEL---------
0050 1iujB   ---------------------lEEEEEEELL-L----H----HH--HHHHH-HHHHH-HHHHH-L--llllE----EEEEEEEEL-L--------------------------------------------lLL-EEEEEEEELL---HHhhHHHHLLHHHHHLLL-L--LhLLEEEEEEEE-E-------
0051 7qa5A   -------------L-LLL---LEEEEEEEEE-E----L----LL--HHHHH-HHHHH-HLLLL-L-----------leEEEEEEE-----------------------------------------eeelL-EE-EEEEEEE--E---LL--L--LHHHHHHHHHH-H--HlEEEEEEEELLL-L-L-----
0052 2f7vA   eellLLLLllLHHH-----lllLEEEEEEEE-E----LllllLH--HHHHH-HHHHH-HHHHH--eelleeL----LEEEEEEEE-LL--LL-------------------------------LLL----LlEE-EEEEEEELLL---L-----LHHHHHHHHHHL-L--LLEEEEEEELLL-LhH--H-H
0053 3kg0B   ----------------LL---LEEEEEEEEE-L----------LL--HHHHH-HHHHH-HHHHH-H--llllE----EEEEEEELL-L--------------------------------------------lLL-EEEEEEEELL---HH----HHHHHHHhhHHH-H--H-EEEEEEELLL-L-------
0054 2qmwA   eeeeLLLL-------LLL---LLEEEEEEEElL----L----LL----LHH-HHHHH-HHHLL-L------L----L-EEEEEEE-LL--L---------------------------------lL----L-LE-EEEEEELL----------lLLHHHHHHHHHH-H--HEEEEEEEEL---------
0055 3cedA   --hHLLL-------lLLL---LEEEEEEEEE-------------------E-LLHHH-HHHHH-H-hhhllL----LEEEEEEEE-EL-------------------------------------------lEE-EEEEEEEELL---LL--HHHHHHHHHHHHH-L-----LEEEEEEL----------
0056 6c0dA   -------------------leeEEEEEEELL-L----EllhhHL--LL-HH-HHHHH-HHHHH-HhllllllE----EEEEEEEEL-LL--L----------------------------------L----E-ELlEEEEEEEEEllhhhH--HHHHHHHHHHHHHHHhlllLLEEEEEELLLL-L----L-E
0057 6dgdA   llllhhhhLLHHHHhhhhhlllEEEEEEEEE-L----------L--LLHHH-HHHHH-HHHHH-HllllllL----LEEEEEEELL-LL--L---------------------------------------L-LE-EEEEEEEELH---HH-hhHHHHHHHHLHHHH-H--HLEEEEEEL-----------
0058 5jb3L   -----------------L---EEEEEEEEEL------------L--HHHHH-HHHHH-HHHHL-L--llllE----ELLLEEEEEeEE--EE--EE------------------elllllllllEE----E-EE-EEEEEEEE-----EL--HHHHHHHLL--LLL-L--LLLEEEEEEEL---------
0059 3bm7A   --------llHHHL-LLL---LEEEEEEEEE-L----H----HH--HHHHH-HHHHH-HHHHH-H-hhlllE----EEEEEEELL-L--------------------------------------------lLL-EEEEEEEEE-----LL--HHHHHHHHLHHHH-H--HhEEEEEEEELL-L-------
0060 2od4A   -------------------lllEEEEEEEEL-L----------L--HHHHH-HHHHHlHHHHH-H---hhlL----LEEEEEEEE-----------------------------------------------LL-EEEEEEEELL---HHhhHHHHHHHHHHHHHH-H-hHLEEEEEEEEEE-E-------
0061 2gffB   ---------------------LEEEEEEEEL-L----H----HH--HHHHH-HHHHH-HHHHH-L--llllE----EEEEEEEE---------------------------------------------lllLL-EEEEEEEE-----LL--HHHHHHHLLLHHHH-H--HHHLEEEEEEEEElL-L--L-L
0062 3oq2B   ------------lllLLL---EEEEEEEELL-L----L----L----LHHH-HHHHH-HHHHHhL------L----EEEEEL-------------------------------------------------------LEEE--eE---EL--HHHHHHHHHHHHHH-L--LLLEEEEEEELL---------
0063 5xvnF   -------------L-LLL---LLEEEEEEEL-L----L-llhHH--HHHHH-HHHHH-HHHLL---------------EEEEL-----------------------------------------------------LEEEEEL-----LL--HHHHHHHHHHHHHHL-L--LLEEEEEEELHHl---------
0064 8dewA   --lLLLL--------------lLEEEEEEEL-L----L----L----lhhhhHHHHH-HHLLL-L--llllE----EEEEEEEEL-----------------------------------------------LL-EEEEEEEELL---HH--H-HHHHHHHHHHHH-H--HHLLEEELEELL-L-Lhl---
0065 7bioB   ---------------------lEEEEEEEEE----------lLL--HHHHH-HHHHH-HHHHH-L--llllE----EEEEEEEEL-L--------------------------------------------lLL-EEEEEEEELL---HHhhHHHHLLhHHHHHHL-L--LEEEEEEEEEE-E-------
0066 2n51A   --------lllllL-LLL---EEEEEEEEEElL----L----LL----LHH-HHHHH-HHLL-----llllE----EEEEEEEEL-LL------------------------------------------L-LE-EEEEEEEE-E---LL--LLLHHHHHHHHHHH-H--HEEEEEEEEEL----------
0067 2dclA   ----------lllllLLE---E-EEEEEEE----------------LLLEeeEHHHH-HHHHHlL------L----LLEEEEEELL-E----------------------------------------eelL-LL-EEEEEEE------EE--HHHHHHHHHHLLL-L---LLLEEEEEELE-E-Ee----
0068 2pebA   -----------LLL-HHH---LLEEEEEEEE-L----H----HH--HHHHH-HHHHH-HHHHL-L-------------LEEEEEL-L---------------------------------------lllllllE-EEEEEELH----------hHHHHHHHHHHH-L--LLLEEEEEELLL---------
0069 6nukA   -------------------lleEEEEEEEEL-L----------L--HHHHH-HHHHH-HHHHH-HhhllllE----EEEEEEEEL------------------------------------------------L-EEEEEEEELL----L--HHHHHHHHHHHHH--hhhHHLEEEEEEL----------
0070 1o51A   -------------------leeEEEEEEEE----------------LLLEeeEHHHH-HHHHH-H---hllL----LLLEEEELL-L---------------------------------------llllL-LL-EEEEEEEELH---HH-----hhHHHHHHHLL----LLLEEEEEEEEE-L-L--L--
0071 4usjC   ---------LLLLLlLLE---E-EEEEEE--------------------LH-HHHHH-HHHHHlL------L----LLLEEEEEEE------------------------ellllllllleeELLEEEllllL-EE-EEEEEEE------EE--HHHHHHHHHHHHHHH-H--LLLEEEEEEEEL-LlL--H-H

# Job: BR-systems; DUF499-ATPase CTD Ferredoxin-fold/RNA-Recognition-Motif (Representative_2)
# Query: s001A
# No:  Chain   Z    rmsd lali nres  %id PDB  Description
```

```
 1:  9bk5-A  6.8  3.1   73    78   11   MOLECULE: LNG BINDER;
 2:  4kyz-A  6.1  3.3   84   167    4   MOLECULE: DESIGNED PROTEIN OR327;
 3:  3nd1-B  6.0  2.7   70   250   13   MOLECULE: PRECORRIN-6A SYNTHASE/COBF PROTEIN;
 4:  2npn-A  5.7  3.6   71   239    8   MOLECULE: PUTATIVE COBALAMIN SYNTHESIS RELATED PROTEIN;
 5:  5cw9-A  5.5  3.2   75   146   12   MOLECULE: DE NOVO DESIGNED FERREDOXIN-FERREDOXIN DOMAIN INS
 9:  2p6y-A  5.0  3.5   78   131    6   MOLECULE: HYPOTHETICAL PROTEIN VCA0587;
10:  3oha-A  4.9  4.5   83   516   10   MOLECULE: DNA POLYMERASE ETA;
12:  7dkk-A  4.8  4.2   75    89   12   MOLECULE: DE NOVO DESIGN PROTEIN XM2H;
13:  3qph-A  4.7  3.5   65   335    5   MOLECULE: TRMB, A GLOBAL TRANSCRIPTION REGULATOR;
14:  1ecg-B  4.5  4.5   67   501    4   MOLECULE: GLUTAMINE PHOSPHORIBOSYLPYROPHOSPHATE AMIDOTRANSF
15:  8k0k-C  4.5  3.6   76   304    4   MOLECULE: CSY1;
16:  3htn-A  4.5  3.2   71   143    6   MOLECULE: PUTATIVE DNA BINDING PROTEIN;
17:  8adb-A  4.5  3.8   73   209   10   MOLECULE: WC-VDT1;
18:  3h20-A  4.4  3.9   60   288    8   MOLECULE: REPLICATION PROTEIN B;
19:  7r8q-B  4.4  5.0   63   389    2   MOLECULE: ATP-GRASP DOMAIN-CONTAINING PROTEIN;
20:  6vqv-J  4.2  3.6   75   337    9   MOLECULE: ACRF9;
21:  6pij-A  4.2  4.0   78   351    9   MOLECULE: CAS7 TYPE I-F CRISPR-ASSOCIATED PROTEIN CSY3;
22:  6ahr-G  4.2  4.3   57   131    9   MOLECULE: H1 RNA;
23:  6x73-A  4.2  4.9   82   439    4   MOLECULE: DNA (5'-D(*GP*GP*GP*GP*TP*GP*TP*GP*GP*TP*AP*GP*C)
24:  6cst-A  4.1  4.8   77   446    8   MOLECULE: DNA POLYMERASE KAPPA;
25:  3fds-A  4.1  5.2   81   352   10   MOLECULE: DNA POLYMERASE IV;
26:  7a02-A  4.1  3.3   67   112    9   MOLECULE: DUF3992 DOMAIN-CONTAINING PROTEIN;
27:  2hx0-A  4.1  3.8   72   138   11   MOLECULE: PUTATIVE DNA-BINDING PROTEIN;
28:  2ip4-A  4.1  4.3   65   414    9   MOLECULE: PHOSPHORIBOSYLAMINE--GLYCINE LIGASE;
29:  5i47-B  4.1  4.6   63   264   14   MOLECULE: RIMK DOMAIN PROTEIN ATP-GRASP;
30:  1wyz-B  4.1  3.2   67   227    6   MOLECULE: PUTATIVE S-ADENOSYLMETHIONINE-DEPENDENT METHYLTRA
31:  2dfx-I  4.0  2.7   60   106    5   MOLECULE: COLICIN-E5;
32:  1vi7-A  4.0  4.5   68   206    4   MOLECULE: HYPOTHETICAL PROTEIN YIGZ;
33:  1ukf-A  4.0  4.4   74   188    7   MOLECULE: AVIRULENCE PROTEIN AVRPPH3;
34:  4ecw-A  4.0  4.4   74   432    5   MOLECULE: DNA POLYMERASE ETA;
35:  4a2b-A  4.0  4.6   61   386    5   MOLECULE: CELL DIVISION PROTEIN FTSA, PUTATIVE;
36:  3vpd-A  4.0  4.9   61   281   13   MOLECULE: RIBOSOMAL PROTEIN S6 MODIFICATION PROTEIN;
37:  2h6l-A  3.9  3.2   68   140    6   MOLECULE: HYPOTHETICAL PROTEIN;
38:  5mmj-j  3.9  4.2   58    99    3   MOLECULE: 50S RIBOSOMAL PROTEIN L31;
39:  7dko-A  3.9  4.0   61    90   13   MOLECULE: AM2M;
40:  3vpb-B  3.9  5.1   65   283    6   MOLECULE: PUTATIVE ACETYLORNITHINE DEACETYLASE;
41:  4ir1-A  3.9  4.9   75   342   12   MOLECULE: DNA POLYMERASE IV;
42:  3mjf-A  3.9  3.5   63   429    6   MOLECULE: PHOSPHORIBOSYLAMINE--GLYCINE LIGASE;
43:  2gjh-A  3.9  2.2   48    57    2   MOLECULE: DESIGNED PROTEIN;
44:  4uer-J  3.8  4.2   60   107    5   MOLECULE: EIF1A;
45:  3lp8-A  3.8  4.7   62   421    6   MOLECULE: PHOSPHORIBOSYLAMINE-GLYCINE LIGASE;
46:  5u4j-v  3.7  1.9   51   132    6   MOLECULE: 16S RRNA;
47:  2cve-A  3.7  3.8   59   190    5   MOLECULE: HYPOTHETICAL PROTEIN TTHA1053;
48:  2j6h-A  3.7  3.6   65   609    8   MOLECULE: GLUCOSAMINE-FRUCTOSE-6-PHOSPHATE AMINOTRANSFERASE
49:  7yll-A  3.6  5.3   67   329    1   MOLECULE: DNA POLYMERASE IV;
50:  5l9w-B  3.6  5.0   67   711    7   MOLECULE: ACETOPHENONE CARBOXYLASE DELTA SUBUNIT;
51:  2ihr-1  3.6  1.9   54   351   13   MOLECULE: PEPTIDE CHAIN RELEASE FACTOR 2;
52:  4aw7-A  3.6  5.2   70   565    3   MOLECULE: GH86A BETA-PORPHYRANASE;
53:  8uy1-A  3.5  4.3   59   598    2   MOLECULE: METHYLENETETRAHYDROFOLATE REDUCTASE-LIKE PROTEIN;
54:  1gsa-A  3.5  3.6   59   314    5   MOLECULE: GLUTATHIONE SYNTHETASE;
55:  2qbu-B  3.5  3.3   53   230   11   MOLECULE: PRECORRIN-2 METHYLTRANSFERASE;
56:  5m41-A  3.5  4.0   66   712    8   MOLECULE: NIGRITOXINE;
57:  6bjv-A  3.5  3.6   67   148   10   MOLECULE: RNA SILENCING SUPPRESSOR P19;
58:  4iwx-A  3.5  4.2   59   295    8   MOLECULE: RIBOSOMAL PROTEIN S6 MODIFICATION PROTEIN;
59:  9j46-F  3.5  3.5   70   124    6   MOLECULE: E146L;
60:  5k2m-D  3.5  5.0   60   270    7   MOLECULE: RIMK-RELATED LYSINE BIOSYNTHESIS PROTEIN;
61:  7lvo-A  3.5  4.6   62   435    6   MOLECULE: PHOSPHORIBOSYL-GLYCINAMIDE (GAR) SYNTHETASE;
62:  5yoy-A  3.5  3.0   70   154   11   MOLECULE: TUMOR NECROSIS FACTOR;
63:  7rjf-A  3.4  2.9   46    47   11   MOLECULE: [L47W]MOPD-1;
64:  3v68-A  3.4  2.8   64   248    3   MOLECULE: PUTATIVE UNCHARACTERIZED PROTEIN;
65:  5ta1-A  3.4  3.4   63   627    6   MOLECULE: GLYCOSIDE HYDROLASE;
66:  5wx8-A  3.4  3.9   68   165    4   MOLECULE: IMMEDIATE-EARLY PROTEIN 2;
67:  2dt4-A  3.4  3.3   66   143    8   MOLECULE: HYPOTHETICAL PROTEIN PH0802;
68:  3gqc-B  3.4  5.4   84   440   11   MOLECULE: DNA REPAIR PROTEIN REV1;
69:  6p5b-A  3.4  6.0   77   388    9   MOLECULE: MAVC;
70:  3wvq-A  3.4  5.0   69   433    9   MOLECULE: PGM1;
71:  5jb3-L  3.4  4.5   59   102    5   MOLECULE: 16S RIBOSOMAL RNA;
72:  2okq-A  3.3  2.9   52   127    8   MOLECULE: HYPOTHETICAL PROTEIN YBAA;
73:  7dov-A  3.3  2.8   65   148    5   MOLECULE: LYMPHOTOXIN-ALPHA;
74:  1aly-A  3.3  3.2   65   146   15   MOLECULE: CD40 LIGAND;
75:  8z9m-A  3.2  3.8   59   332    5   MOLECULE: PROTEIN FAM91A1;

0001 s001A   -AALAVLQLAVDGD-G-S-EGVRELRS-L-G-LA-VPQL-G-PG-QFRVEVSAT-G-ELE-G-G-SIRLEFRGP-W-E-RYK-D-L-K-DAVV-SV-L-E-R-A-S-A-AQVRARLDW-R-A-RD-
0002 9bk5A   --------GRIVVR-G-D---VAIAEA-V-V-RKvGEVA------GKEVILLIS-Y-RKN-G-E-WITYQRNLE--pE-DVE-R-T-I-AVIR-EI-Y-E-E-S-G-D-FILAIFSD---------
0003 4kyzA   kQGAKRVRISITAR------SSKEAYK-F-L-AIlAKVF-AlGY-NDNRKXTVR-F-R-L-E-K-ALEITITG------VPR-Q-V-L-EELA-KE-AeR-L-A-K-T-ITITVTVEG-Q-L-GSl
0004 3nd1B   gwpAGTETVVAML-------DGEC--S-F-Q-SL-PP-------dGLTIFWGAC---------mPEEVLIRGP-V-A-EVT-D-E-I-LQAR-AD-L-R-ArH-G-W-VMDIYLLRR-N-V----
0005 2npnA   rIGEA-RNCVVX-------LDGK--TA-W-Q-DV-AT-------eHTYXWWGAF----------tEQQVLRKGY-V-H-EIG-A-Q-V-AELK-QQ-L-R-TeH-G-W-IXDTYLLRE-L-D----
0006 5cw9A   ----MEMDIRFRG----------eayA-L-R-EM-IRQAkF-AG-TVTVTLIIR-F-RGD-DgN-DLEIRITG-----vPPQ-V-I-L-ELVK-EA-I-R-L-AnI-T-VTVELVIRI-T-G-VPe
0010 2p6yA   ----XIHLIALRLT-R-G---XDLKQQ-I-V-QL-VQQH-R-IH-AGSISCVGC-L-StV-S-A-PFEISLSGT-LhC-HLHgG-H-L-L--E-GN-L----------iNTAELXIHH-Y-P-QHh
0011 3ohaA   sSRPVVKSMMSNKN-LscvDCISWLEV-F-C-AElTSRI-QkIV-IPTVSISLK-T-KS------YEVYRKSGP-V-AhELL-K-V-G-IKFV-T-------sY-Y-P-LKLSMTINF-D-I-IDl
0013 7dkkA   ------MHSWSATV-D-seAVRAAARR-L-A-ER-LLAA-G--I-SGKIKIEVE-A-N-------GIKYEYEVE--tE-EVA-K-KiV-EYAV-AA-A-L-R-AgA-T-SVTITVGLE--------
0014 3qphA   eiignPKDIRFF-----------AMFH-A-V-DF-VKNHlK-NR-NIYAEITGK-N-LE----S-GRLETLTGR-V-V-G--eatengvkvGG-MF-A-V-I--------eTEIKFIM-G-V-DLq
0015 1ecgB   -------CGIVGIA-G--mpVNQSIYD-AlT-VL-QH-------gqDAAGIITI-D-AN------NCFRLRKAN-V-S-DV----F-E-ARHM-QR-L----------gNMGIGHVRY-P-T-AGs
0016 8k0kC   pHNKNTLLVKYNVK-F-VtlayRYVYN-I-AgRT-LWRR-V-GAeSIETVITVN-------------DQTFTFS-D-L----dV-D-V-AEIA-DM-V-A-G-VeG-F-VTLKVEHYM-L-L-G--
0017 3htnA   sykkiGNKYIVSIN-N-H-T--EIVKA-L-N-AF-CKEK-G-iL-SGSINGIGA-I-G----rE-QXEINLTGN-IyL-HLHgrgH-L-LSA---------------N-GAGEFVVED-Y-S-E--
0018 8adbA   lntfELKRLPYAG-------TLDSFAK-M-M-EK-ASDS-KlPK-PVALVMTKS-------------NMTITIV-IgK-YWLdE-L-I-KKIK-EI-F-P-K-T-SyF-NSFEAYAVR-R------
0019 3h20A   ------------------------rtA-I-G-RQ-LKAM-----gcERFDIGVR-D-AT----T-GQMMNREWS-A-A-EVL-Q-N-T-PWLK-RM-N-A-Q-G-----NDVYIRPAE-R-H-G--
0020 7r8qB   tfnnqqfPKRKVV-------KIEQFEN-E-I-KN-W---------eFPLVIKP--------------YGVMICY-H-D-ADL-Q-K-A-ITRI-KE-A-T-A-E----tNSLIIEQKI-E-E-KAn
0021 6vqvJ   pSDADTLKVRFTLR-V-LvndqgyAHN-L-AaRF-LWRR-V-GAeAVEVRINHI-------------ARAWRFD-A-L-A--dA-E-L-DALA-EL-I-A-S-GsG-H-VLLEVVAFA-R-I-G--
0022 6pijA   pYGASHIECSFSVS-F-SetkigYLMN-I-CgKW-LWKT-R-KAyCWNIVLTPW-P----------NGEKVGFE-D-I----nK-N-W-SAIV-EM-I-K-T-AdG-L-AIFEVRATL-H-L-PTn
0023 6ahrG   rgqnaCSEIYIHGL-G-L-AINRAINI-A-L-QL-QAGS-F-G--SLQVAANTS-T-VEL-V------------------------------------------rN-N-SAIHIRVFR-V-T-P--
0024 6x73A   eVLQR-KSLSIDIN-W-G-IrfkFIER-G-C-QYlLEKL-NiNK-TTQITLKLM-R-RCK-D-ArCDSFSRSSR----aTEM-K-SlY-RTL--GC-P-P-M-E----lRGLALQFKL-V-D-VGp
0025 6cstA   tRDGERKSMSVERT-F-SeeqySLCQE-L-C-SElAQDLeR-LK-GRTVTIKLK-N-VN------FEVKTRAST----fAIA-K-ElL-KTE--------hP-L-R-L-RLMGVRISS-F-P-N--
0026 3fdsA   irTRVRKSIGRIVT-MnseEIKPYLFR-A-I-EEsYYKL-D-KR-IPAIHVVAV-T-ED------LDIVSRGRT-K-E-TAY-S-E-S-VKLL---------eR-K-I-RRIGVRFSK-F-I-EAi
0027 7a02A   nGQKTIVQDKVCID-W-T-----------iY-AD-NI-------qDIYASGYLK-VdTGT-GaT-GSSASFTVR-R-----F-D-T-V---TI-LG--------aE-T-GEFCMTIRY-T-L-S--
0028 2hx0A   hnASTARFYALRLL-P-G-Q---vfSQ-L-H-AF-VQQN--qlR-AAWIGCTGS-L-T----tG-TFEVSLNGT-LgE-HLH--gvxlghXXP-GC-T--------R-TTLELVIGE-L-P-ALt
0029 2ip4A   erYGIPTARYRVFR------EPLEALA-Y-L-EE-V----------gPVVVKD--------------KGVTV------aFDL-H-Q-A-KQAV-AN-I-L-N-RaE-G-GEVVVEEYL-E-G-E--
0030 5i47B   lADIPVPETRVCF-------GEEAIFA-A-I-AE-I----------gPVVLKS-------------GFPVAL------vEDQ-D-A-A-EAIV-EH-R-I-X-L-G-G-ERVLVQQFI--------
0031 1wyzB   rvyAESQTQLFIET-P-Y-RNHKXIED-I-L-QN-CR-------pqTKLCIAAN----------cEGEFIQTRT-V-K-DWK-G-H-I------PE-L-S-------K-IPCIFLLYK-L------
0032 2dfxI   ---------nasLS-P-K-AAIEVCNE-A-A-KK-G-------------wLGID-G-G----dS-SASWTYDMY-K-S-KIP-E-N-N-RLAI-EN-I-K-D-DyT-A-FIITLKM----------
0033 1vi7A   eeiKKSRFITMLAH-TvE-AAKAFVES-V-R-AE-HPD-------aRHHCVAWV-A-GA-dD-S-QQLGFSDDG-E-P-A------------------------gV-G-EITAVVVRY-Y-G-GIl
0034 1ukfA   daGFSLRPKTVHAS-G-G-S-aQLGQT-V-A-HD-VAQ--S-GR-KHLLSLRFA-------------VQGHAIAcK-L-FD-fqqL-I-KGLI-DH-Y-N-S-L----yDVACVNEFR-V-S-V--
0035 4ecwA   kPRQLPKTIGCSKN-FrewWLLQLAQE-L-E-ER-LTKD-RnDR-VAQLVVSIR-V-QG--------SLRRCCA-L-T----hK-M-S-HdaftikncntwS-P-P-L-TMLFLCATK-F-S-A--
0036 4a2bA   sVSFEREDTVIERD-F--sitlDILSE-M-Q-SEaLEKL-KtpL-HIFSRYLLD-------------ERIVF------------------nPL-DM-K----------aSKIAIEYTS-I-V-VPl
0037 3vpdA   gerpeaaPKTALAT------DREEALR-L-M-EA-F----------gPVVLKP--------------RLLAK------vTDR-A-A-A-EALL-EH-K-EvL-G-G-FhQLFYIQEYV-E-K-PGr
0038 2h6lA   fefEVGKGFLLRLD-Y-G-----lvRQ-I-E-EF-LEEK-G-iH-AAHISAIGA-V-R----xE-PLEISLSGNvF-C-HI-lggH-L-FSA----------------vFACEVFVLP-L-S-GEa
0039 5mmjj   ----QKIRIKLRSY-----WVPLIEDS-C-K-QI-MDAAtT-NA-KIMPVPLTK-K-RIY-C-------------------------------------------fE-I-RTHQRLIDI-L-Y-PTa
0040 7dkoA   --------ASAEAE-V-KeeirAAARR-L-A-EAlRKAG-----vsGPVTVTAE-A-G-------DVSFSYTAD-L-DeeGL-K-R-VeaivraIA-A-L-K--tK-P-VLLSAVL----------
0041 3vpbB   rEGIPIPDSIIAL-------SAEAALK-A-Y-EQ-R----------gPLIDKP--------------RLVSL------iRDV-F-E-G-KTII-EH-R-E-L-M-G-NaLKAHIVQE-lE-H-PNk
0042 4ir1A   nSERLRKSVGVERT-MwsaIIERLYPE-L-E-RR-LAKV-K-pdlIARQGVKLK-F-DD------FQQTTQEHV-W-PrlNK-A-D-L-IA-------------rG-V-RLVGLHVTL-L-D-P--
0043 3mjfA   arHNIPSAEYQNFT------DVEAALA-Y-V-RQ-K----------gPIVIKAD-------------KGVIV------aXTQ-E-E-A-ETAV-NDxL-A-G-NdA-G-HRIVVEEFLk-------
```

```
0044 2gjhA   ----ERVRISITAR-----TKKEAEKF-A-A-IL-IKVE-L-GYnDINVTWDG-----------------------------------------------------------DTVTVEG-Q-L-EGg
0045 4uerJ   qqQIIKIRITLTST-----KVKQLENV-S-S-NI-VKNAqH-NL-VKKPVRLTK-V-LKI-S-------------------------------------------yE-M-RIHKRYIDL-E-A-PVq
0046 3lp8A   mrYGIPTAKYGYFV------DTNSAYK-F-I-DK-HK----------PLVVKAD-------------KGTVI------cHTH-E-E-A-YNAV-DAmL-V-H-H---ggCAIIIEEFL---------
0047 5u4jv   -----DCYLDIQAGgT-E-AQDWASML-E-R-MY-LRWAsR-GF-KTEIEESEG-E-------------------------------------------------------gKSVTIKI-S-G-DYa
0048 2cveA   eeiQKSRFIAKAAP-VeE-EALAFLAE-N-R------------eeaTHNGHAYK-I-G-------LLYRFSDDG-E-P-S-------------------------grpD-RVAVLVVRY-F-G-GVk
0049 2j6hA   -----CGIVGAIAQ-------RDVAEI-L-L-EG-LRRE-Y-RGdSAGLAVVDA------------EGHMTRLR--vQ-MLA-Q-A-A-EEHP------------------hGTGIAHT-R-W-ATh
0050 7yllA   vETMREIKSIGKEK-TiQ-HLKEFSEI-V-S-EE-LIKE-R---lyCRTVTVKI-K-TAD---F-AVHTKSKT--------------------------------yV-R-LIGLSVSNL-S-P-V--
0051 5l9wB   gsSIMDIVHMYEQS-R-RehfnQTVDT-M-I-ERaRQEL-RgledDASFGLELD-M-LYG-G-QnLKRMSS--------------pL-L-HIrtadalkvyfsgV-F-L-DNFVLRVTV-P-T----
0052 2ihr1   phAEKNAILTIQP-gT-E-ACDWAEML-L-R-MY-TRFArQ-GF-QVEVDLTPG-P-E-----------------------------------------------------gDYAQILV-K-G-ENa
0053 4aw7A   aFTNSIIRNKYYEY-L-K---piFPFT-----GI-ES-------gSGRASLRMS-I-GRP-ViF-FGMIEVPFD-I-Q-LL----kngnnvDI-TF---------gG-H-VSMILQVEK-Y-T-VS-
0054 8uy1A   gyvyQKAYLEFFV-------SPELYPE-I-K-RR-IESH-P----DLTYHAVTK------------SGNLETNA-----qsdtiawkdayhlgmwarcydvL-L-E-EnTWWLVNIVN-N-D-FHq
0055 1gsaA   geqdlpftPETLVT-R-N---KAQLKA-F-W-EK-H------------DIILKP------------ASIFR----K-E-GD--P-N-L-GVIA-ET-L-T-E-HgT-R-YCMAQNYLP-Aq------
0056 2qbuB   veGDEIDACVIMK-------TSRHGRR-A-M-EV-VESD-P--R-GKDVVSVAN----------mDDEVVER------------------------------------sgYLATTLVRF--------
0057 5m41A   llvastSGTQLLMS-AaV-SVIQGAAG-L-T-AD-AIEG-K-eipLDVISLSLA--------knaETGVDIVGS-S-R-DLI-E-G-FkGNFT-DI-I-N-GlV-S-V-A-----------------
0058 6bjvA   raIQGNtKVVFKRY-L--rvlgSWTGS-V-N-YA-ASRFlG-AN-QVGCTYSIR-F-R-------GVSVTISGG-S-R-TLQ-H-L-C-EMAI-RS-K-Q-E-L-Lq--------------------
0059 4iwxA   kgrklpaPVTGI--------AHSPddD-L-I-DM----------ggAPLVVKLV-E-G------tQGIGVVLAE-R-Q-AAE-S-V-I-DAFR-GL-N----------aHILVQEYIK-E-Ae----
0060 9j46F   ---MFTGWSPFKYS-K-G-N----tVT-F-K----TPDE-S-S-aYMRFRCVFT-F-TDP-KgS-LHSIDVTE--N-N-MA-gfssftgghcqaLN-A-F----nC-D-ATLTGLQRI-I-H-HHh
0061 5k2mD   ddllfplPEWKAAL------SEGGALR-V-P-DS-L----------gPLVSKP--------------rLLAK------vNDR-D-S-L-EAVL-EH-R-K-WkN-P-L-YIHYFQEFV-E-K-PGr
0062 7lvoA   rHNIPTAAFRS--------fTSTQYED-A-V-AY-IKSK--ftS-GRSVIKASG---------------GVLIP-D-E-EAF-A-A-L-KSVM-VD-K------------DEVVVEE---------
0063 5yoyA   rtPSDKPVAHVVANeG-Q-LQ--lnrnalnQ-LV-V--P-S-EG-LYLIYSQVL-F-KGQkP-W-YEPIYLGGV-F--------ekgdlsaEI-NR---P-D-Y-L-D-FQVYFGIIA-L-T-S--
0064 7rjfA   -----IQIREYKRC-G-Q-DEERVRRE-C-K-ERgERQN-------CHYVIHKE-G------------------------------------------------------NCYVCGIIC-W------
0065 3v68A   ftakhEEWIVGENVnI-A-MFLSRVSN-T-V-SS-KIPGgelsiieglHINFTA-R-YG----S-WIVVKRMID-I-A-RLL-A-S-I-NETA-VN-K-I-K-De----------------------
0066 5ta1A   yvdGKNVYFIINNL-D-F-KPVDL-------nLS-VN-gkD-AK-SIEVRHLYL-K-GGK---D-GVPILDVYD-A-K------------------------------feTEATCVICY-N-F-DRk
0067 5wx8A   ----------------------KNLKE-A-V-YD-ICCN-G---lsNNAAIIXY-F-TRS-K-K-VkTXIIYAA-P-I-DFV-G-A-V-KTCN-KY-A-K-D-NpK-E-IVLRVCSI-eF-K-SKf
0068 2dt4A   gmfSLGRTYLFRVP-E-G----llTY-I-K-NF-CKKE-G-iE-TAIINGIGT-----------lYELILIGN-VfV-HAHgngH-L-VEGE---------------vFVAEIFLQE-L-K-GEk
0069 3gqcB   tEKERKSVSAEIGI-R-FkEAEAFLLS-L-S-EE-IQRRaT-GM-KGRLTLKIM-V-RKP-G-AiCDNIARTVT----gkAM-L-NmF-HTM------kL-N-I-S-D-MRGVGIHVNqL-V-PTn
0070 6p5bA   kLEKTgtqpYQVL-------SGDG-SN-L-K-GS-LPN-----K-NLLVRVEMD-R-FNAiD-C-WTTHSFTIF-VtL-YPYekgY-I-EKLA-LL-G-T-T--fS-P-LRVKFKVTE-V-N-PEi
0071 3wvqA   eafagrdPAGAVCA------DRRRAHR-H-V-TRlLDEG-------SPVILKQD-------------dGNEILS-R-TrVLA-DaA-L-DAYL-DE-R-W-DeG-G-R-HRVVVERYH-P-G-SRa
0072 5jb3L   ---MQKARIKIAST-----NVRSLDEV-A-N-QI-KQIA-EtGV-RMSPIPLPTkR-IRI-T-------------------------------------------wE-L-RVHKRLIDI-E-A-DEr
0073 2okqA   namKYVDGFVVAVP-A-D-KKDAYREM-A-A-KA-APLF-KgAL-RIVECWA-----------------------------------------------------vpdkeNEEVVFSWI-E-Y-PSk
0074 7dovA   ------AAIHLTGG-Y-NktLDWRddqafrE-II-I--P-D-DG-IYFVYSQVS-L-HISnL-W-YNTIYLGAA-F--------regdlgtDT-TT----aL-L-P-M-VKTFFGVFG-L-------
0075 1alyA   gDQNPQIAAHVISE-A-S--sktekgtmskQ-LT-V--K-R-QG-LYYIYAQVT-F-CSN-RcG-QQSIHLGGV-F--------qpgavfvNV-TD---P-S-Q-V----FTSFGLLK-L-------
0076 8z9mA   lrhnkdlPPSLLLS-K-G-T---------lP-DI-FQS-------yDRLLITSW------------GHDPGVVP-T-S-------N-V-LTML-ND-A-L-T-H-----SAVLIQGHGnA-S-SQl

0001 s001A   -LLLLEEEEEEEEE-H-H-HHHHHHHH-H-H-HH-HHHL-L-LE-EEEEEEEEE-E-EEL-L-E-EEEEEEEEE-H-H-HHH-L-L-H-HHHH-HH-H-L-L-L-L-E-EEEEEEEEE-E-E-LL-
0002 9bk5A   --------LEEEEE-L-L---HHHHHH-H-H-HHhHHLL------LLLEEEEEE-E-EEL-L-E-EEEEEEEEL--hH-HHH-H-H-H-HHHH-HH-H-H-H-H-L-L-EEEEEEELL---------
0003 4kyzA   hHLLLEEEEEEEEEL------LHHHHHH-H-H-HHhHHHH-HhLE-EEEEEEEEE-E-E-H-H-H-HEEEEEEEL------LLH-H-H-H-HHHH-HH-HhH-H-H-H-E-EEEEEEEEE-E-E-LLh
0004 3nd1B   lllLLLLEEEEEL-------LLLL--H-H-H-HL-LL-------lLEEEEEEEL----------lLLLEEEEEEE-H-H-HHH-H-H-H-HHHH-HH-H-H-HhH-L-L-LLEEEEEEE-L-L----
0005 2npnA   lLLLL-HEEEEE-------LLLL--LL-H-H-HH-LL-------lLEEEEEEEL----------lLLLEEEEEEE-H-H-HHH-H-H-H-HHHH-HH-H-H-HhH-L-L-LLEEEEEEE-L-L----
0006 5cw9A   ----LEEEEEEEL----------lhhH-H-H-HH-HHHHhL-LL-EEEEEEEEE-E-ELL-LlL-EEEEEEEL-----lLHH-H-H-H-HHHH-HH-H-H-H-HlL-E-EEEEEEEEE-E-L-LLh
0010 2p6yA   ----LLEEEEEEEL-L-L---LLHHHH-H-H-HH-HHHL-L-LL-LEEEEEEEE-E-EeE-L-L-LEEEEEEEE-ElE-EEEeE-E-E-L--L-LL-E----------eLLEEEEEEE-L-L-LEe
0011 3ohaA   lLLLLLLLEEEEEE-LllhHHHHHHHH-H-H-HHhHHHH-HeEE-EEEEEEEEE-L-LL------LLEEEEEEE-L-LhHHH-H-H-H-HHHH-H-------lL-L-L-LEEEEEEEE-E-E-EEl
0013 7dkkA   ------LEEEEEEE-L-lhHHHHHHHH-H-H-HH-HHHH-L--L-LLLEEEEEE-E-L-------LEEEEEEEL--lH-HHH-H-HhH-HHHH-HH-H-H-H-HlL-L-LEEEEEEL--------
0014 3qphA   eeellLLLEEEL-----------LHHH-H-H-HH-HHHHlL-LL-LEEEEEEEE-E-LL----L-LLEEEEEEE-E-E-E--llelleeeeEL-LL-L-L-L--------eEEEEEEE-L-L-LLl
0015 1ecgB   -------LEEEEEE-L--llLHHHHHH-HhH-HH-HH-------llLEEEEEEE-L-LL-------LLEEEEEEE-H-H-HH----L-L-HHHH-HH-L----------lLEEEEEEEL-L-L-LLl
0016 8k0kC   lLLLLEEEEEEEEEE-E-EhhhhHHHHH-H-HlHH-HHHH-L-LLlLEEEEEEEL-------------LEEEEEL-L-L----lH-H-H-HHHH-HH-H-H-H-HlL-E-EEEEEEEEE-E-L-L--
0017 3htnA   eeeeeLLEEEEEEEL-L-L-L--LHHHH-H-H-HH-HHHH-L-lL-LEEEEEEEE-E-E----lL-LEEEEEEEEE-EeE-EEEeleE-E-EEE---------------E-EEEEEEEEE-L-L-L--
0018 8adbA   hhhhLLEEELEEE-------EHHHHHHH-H-H-HH-HLLL-LlLL-LEEEEEEEL-------------LEEEEEE-ElL-EEEhH-H-H-HHHH-HH-L-L-L-L-LlH-HEEEEEEEE-L------
0019 3h20A   ------------------------lhH-H-H-HH-HHHH-----llLEEEEEEE-E-LL----L-LLEEEEEEE-H-H-HHH-H-L-H-HHHH-HH-H-H-L-L-----EEEEEEELL-L-L-L--
0020 7r8qB   eellhheLLEEEE-------ELLLHHH-H-H-HH-L---------lLLEEEEE-------------LLLEEEL-L-H-HHH-H-H-H-HHHH-HH-L-L-L-L----lLEEEEEEELL-L-L-LEe
0021 6vqvJ   lLLLLEEEEEEEEEE-E-LhllllhHHH-H-HlLL-LLLL-L-LLlEEEEEEEEE-------------EEELLEE-L-L-L--lL-H-H-HHHH-HH-H-H-H-HlL-L-EEEEEEEEE-E-L-L--
0022 6pijA   lLLLLEEEEEEEEEE-E-EhhhllHHHL-L-LlHH-HLLL-L-LLlEEEEEEEEE-L----------LLLLEEEL-L-L----lL-L-H-HHHH-HH-H-H-H-HlL-E-EEEEEEEEE-E-L-LLl
0023 6ahrG   lllllLLEEEEEEL-L-L-HHHHHHHH-H-H-HH-HHHH-L-L--LEEEEEEEE-E-EEE-E-------------------------------------------eE-E-EEEEEEEEE-L-L-L--
0024 6x73A   hHLLL-LLEEEEEL-L-L-LlllHHHH-H-H-HHhHHHH-HlLE-EEEEEEEEE-E-ELL-L-LlEEEEEEEEE----hHHH-H-HhH-HHH--LL-L-H-H-H----eEEEEEEEEE-E-E-LLl
0025 6cstA   lLLLLLLEEEEEEE-E-EhhhhHHHHH-H-H-HHhHHHHlL-LL-EEEEEEEEE-E-LL------LLEEEEEEE----hHHH-H-HhH-HHH--------lL-L-L-E-EEEEEEEEL-L-L-L--
0026 3fdsA   llLLLLLEEEEEEE-EeehHHHHHHHH-H-H-HHhHHHH-L-LL-EEEEEEEEE-E-LL------LLEEEEEEE-H-H-HHH-H-H-H-HHHH---------lL-L-E-EEEEEEEEE-E-E-LLl
0027 7a02A   lLLLLLEEEEEEEE-E-E-----------eE-EL-LL-------lLLEEEEEEE-EeELL-LlL-LLEEEEEEE-L-----L-L-E-E---EE-EL--------lE-E-EEEEEEEEE-L-L-L--
0028 2hx0A   llLLLLEEEEEEEL-L-L-L---hhHH-H-H-HH-HHHH--llL-LEEEEEEEE-E-E----eE-EEEEEEEEE-EeE-EEE--lleeeeELL-LL-E---------E-EEEEEEEEE-L-L-LEe
0029 2ip4A   hhLLLLLLLEEEEL------LHHHHHH-H-H-HH-H----------lLEEEEL--------------LLLEE------eLLH-H-H-H-HHHH-HH-H-L-L-LlL-L-LLEEEEELL-L-L-L--
0030 5i47B   hLLLLLLLEEEEL-------LHHHHHH-H-H-HH-H----------lLEEEEE-------------LLLLEE------eLLH-H-H-H-HHHH-HH-H-H-H-L-L-L-LLEEEEELL--------
0031 1wyzB   hhhHHLLEEEEEEL-H-H-HHHHHHHH-H-H-HH-LL-------llLEEEEELL----------lLLLLEEEEE-H-H-HHL-L-L-L------LL-L-L-------L-LLEEEEEEL-L------
0032 2dfxI   ---------lllLL-H-H-HHHHHHHH-H-H-HH-L-------------lEEEE-E-E----lH-HHLEELLLL-H-H-HHH-H-H-H-HHHH-HH-H-H-H-HlL-E-EEEEELL----------
0033 1vi7A   eeeLLEEEEEEEEE-LhH-HHHHHHHH-H-H-HH-LLL-------lLLLLEEEE-L-LL-lL-L-LLLEEELLL-L-L-L-------------------------lL-L-LEEEELLEE-L-L-LLl
0034 1ukfA   hlLLEELLEEEELL-L-L-H-hHHHHH-H-H-HH-HLL--L-LL-EEEEEEEEL-------------LEEEEEEeE-E-EE-eehH-H-HHHH-HH-H-H-H-L----lLEEEEEEEE-E-E-L--
0035 4ecwA   lLLLLLLLEEEEEE-LhhhHHHHHHHH-H-H-HH-HHHH-HhLE-EEEEEEEEE-E-LL--------LEEEEEE-L-L----hH-H-H-HhhhhhhhhlleE-E-E-E-EEEEEEEEE-E-E-E--
0036 4a2bA   lLLLEEEEEEEEEE-L--ellhHHHHH-H-H-HHhHHHH-HeeE-EEEEEEEEL-------------LEEEL------------------lLL-LL-E----------eLEEEEEEEE-E-E-EEh
0037 3vpdA   llllhhhLLEEEEL------LHHHHHH-H-H-HH-H----------lLEEEEL--------------LLLEE------eLLH-H-H-H-HHHH-HH-H-HhH-L-L-HhHLEEEEELL-L-L-LLe
0038 2h6lA   eeeELLEEEEEEELL-L-L-----hhHH-H-H-HH-HHHH-L-lL-LEEEEEEEE-E-E----lL-LEEEEEEEEEeE-E-EE-eeeE-E-EEE---------------eEEEEEEEEE-L-L-LLl
0039 5mmjj   ----LLEEEEEEEL-----LHHHHHHH-H-H-HH-HHHHhL-LL-LEEEEEEEE-E-EEE-E-------------------------------------------eE-E-EEEEEEEEE-L-L-LLh
0040 7dkoA   --------LEEEEE-E-LhhhhHHHHH-H-H-HHhHLLL-----llEEEEEEEE-E-L-------LEEEEEEEE-E-LhhHH-H-H-HhhhhhhHH-H-H-H--lL-L-EEEEEEL----------
0041 3vpbB   hLLLLLLLEEEEL-------LHHHHHH-H-H-HH-H----------lLEEEEL--------------LLLEE------eLLH-H-H-H-HHHH-HH-H-L-L-L-L-LhHHLEEEEE-eE-E-LLl
0042 4ir1A   lLLLLLLEEEEEEE-EhhhHHHHHHHH-H-H-HH-HHHH-L-lllLLEEEEEEEE-E-LL------LLEEEEEEE-L-LllLH-H-H-H-HH------------lL-E-EEEEEEEEL-L-L-L--
0043 3mjfA   hhLLLLLLLEEEEL------LHHHHHH-H-H-HH-H----------lLEEEEEL-------------LLEEE------eLLH-H-H-H-HHHH-HHhH-L-L-HlL-L-LLEEEEELLl-------
0044 2gjhA   ----LEEEEEEELL-----LHHHHHHH-H-H-HH-HHHH-L-LLlLLEEEELL----------------------------------------------------------LLEEEEE-E-L-LLl
0045 4uerJ   llLLLLEEEEEEEL-----LHHHHHHH-H-H-HH-HHHHhL-LL-LEEEEEEEE-E-EEE-E-------------------------------------------eE-E-EEEEEEEEE-L-L-LHh
0046 3lp8A   hhHLLLLLLEEEEL-------LHHHHHH-H-H-HH-LL----------LEEEEEL-------------LLEEE------eLLH-H-H-H-HHHH-HHhH-L-L-L---hhLLEEEEELL--------
0047 5u4jv   -----LLEEEEEEElL-H-HHHHHHHH-H-H-HH-HHHHhH-LL-LLEEEEEEL-L------------------------------------------------------lEEEEEEL-L-L-LLh
0048 2cveA   eeeLLEEEEEEEEE-LhH-HHHHHHHH-H-L------------lllLEEEEEEEE-E-L-------LEEEEELLL-L-L-L------------------------hhhL-LEEEEEEEEE-L-L-LLl
0049 2j6hA   -----LEEEEEEELL-------LLLHHH-H-H-HH-HHHH-H-HLlEEEEEEEELL------------LLLEEEEE--hH-HHH-H-H-H-HHLL-----------------lLEEEEEEE-E-L-LLl
0050 7yllA   lLLLLLLLEEEEEE-EhH-HHHHHHHH-H-H-HH-HHHH-L---llEEEEEEEE-E-ELL---L-LEEEEEEE-------------------------------lE-E-EEEEEEEEE-L-L-L--
0051 5l9wB   hhHHLLEEEEEEEEE-E-EhhhhHHHHH-H-H-HHhHHHH-HlllhHLEEEEEEE-E-EEL-L-LlLEEEEL-------------lL-L-LLllhhhhhhhhhlE-E-E-EEEEEEEEE-L-L----
0052 2ihr1   llLLLEEEEEEEEE-lH-H-HHHHHHHH-H-H-HH-HHHHlL-LL-EEEEEEEEL-L-L-----------------------------------------------------lEEEEEEE-E-L-LLh
0053 4aw7A   lLLEEEEEEEEEELL-L-E---elEEEE-----EE-LL-------lLEEEEEEEE-E-EEE-LeE-EEEEEEEEL-H-H-HL----lleeeeEE-EL--------lL-E-EEEEEEEEE-E-E-LL-
0054 8uy1A   leeeELEEEEEEE-------LLLLHHH-H-H-HH-HHLL-L----LEEEEEEEL------------LLLEEELL-----llleehhhhhhhhhhhlllllhH-H-H-HhHLEEEEEEE-L-L-LLl
0055 1gsaA   eeeeeehlLLEEEE-L-L---HHHHHH-H-H-HH-H------------LEEEEL------------LLLEE----L-L-LL--L-L-H-HHHH-HH-H-L-L-LlL-L-LEEEEELLH-Hl-----
0056 2qbuB   llLLLLLEEEELL-------HHHHHHH-H-H-HH-HHLL-L--L-LLEEEEEEL----------lLLLEEEE-----------------------------------llLLEEEEEEL--------
0057 5m41A   hhhhhlLLLLEEEE-LhH-HHHHHHHH-H-H-HH-HHHL-L-lllLLEEEEEEE--------hheLLEEEEEEE-H-H-HHH-H-H-HhLLLH-HH-H-H-HhH-H-H-H---------------
0058 6bjvA   llLLHHlLEEEEEE-E--hhhlLLLHH-H-H-HH-HLLLlL-LL-LLEEEEEEE-E-L-------LEEEEEEEE-H-H-HHH-H-H-H-HHHH-HH-H-H-H-H-Ll-----------------
0059 4iwxA   lleellhLLEEE--------ELLLllH-H-H-HL----------llLLEEEEEL-L-L------lLLLEEEEEL-H-H-HHH-H-H-H-HHHH-LH-H----------hLLEEEEELL-L-Ll---
0060 9j46F   ---LLLLLEEEEEE-L-L-L----lLE-E-E----LLLL-L-L-hHEEEEEEEE-E-ELL-LlL-EEEEELHH--H-H-HH-llleeellllllLL-L-L----lL-L-EEEEEEEEE-L-L-LLl
0061 5k2mD   hhleelhLLEEEEL------LHHHHLL-H-H-HH-H----------lLEEEEL--------------lLLEE------eLLH-H-H-H-HHHH-HH-H-H-HlL-H-H-HLEEEEELL-L-L-LLe
0062 7lvoA   hLLLLLLLEEE--------eEHHHHHH-H-H-HH-HHLL--llL-LLEEEEEELL---------------LEELL-H-H-HHH-H-H-H-HHHH-LL-L------------HEEEEEE---------
0063 5yoyA   llLLLLLEEEEEELlL-L-LL--ellllelE-EE-L--L-L-LE-EEEEEEEEE-E-EEElE-E-EEEEEEEEE-E--------lllleeeEE-LL---H-H-H-L-L-LHLEEEEEE-L-L-L--
0064 7rjfA   -----LEEEEEEEE-L-L-LHHHHHHHH-H-H-HHhHHHL-------LEEEEEEE-L------------------------------------------------------LEEEEEEEE-L------
0065 3v68A   eeeeeLLEEEEEEELhH-H-HHHHHHHH-H-H-HH-HHHHllhlllllhEEEEEE-E-EL----L-EEEEEEEEH-H-H-HHH-H-H-H-HHHH-HH-H-H-H-Hl--------------------
0066 5ta1A   eeeLLEEEEEEEEEL-L-L-LLEEE-------eEE-EE-llL-LL-EEEEEEEEE-E-LLL---L-LEEEEEEEEE-E-L-----------------------------elLLLEEEEEEE-E-L-LLl
0067 5wx8A   ----------------------LHHHH-H-H-HH-HHHL-L---llLLEEEEEE-E-LLL-L-L-HlEEEEEEL-H-H-HHH-H-H-H-HHHH-HH-H-H-H-LlL-L-EEEEEEEE-hH-H-HHh
0068 2dt4A   lleEEEEEEEEEEL-L-L-----hhHH-H-H-HH-HHHH-L-lL-LEEEEEEEE-----------eEEEEEEEE-EeE-EEEeleE-E-EEEE---------------eEEEEEEEEE-E-E-ELl
0069 3gqcB   lLLLLLLEEEEEELL-L-LhHHHHHHHH-H-H-HH-HHHHhH-LE-EEEEEEEEE-E-ELL-L-LlEEEEEEEEE----hhHH-H-HhH-HLL------lL-L-H-H-H-EEEEEEEEEeE-E-ELl
0070 6p5bA   lLLLLhhllLEEL-------LLLL-HH-H-H-LL-LLL-----L-LEEEEEEEE-E-ELLlL-L-EEEEEEEEE-ElL-EEEhhhH-H-HHHH-HH-H-H-L--lL-L-EEEEEEEEE-L-L-HHh
0071 3wvqA   hhhlllhLLEEEEL------LHHHHHH-H-H-HHhHHLL-------LLEEEEEL-------------hHEEEEE-L-LeELL-LhH-H-HHHH-HH-H-H-HlL-L-L-LLEEEEELL-L-L-LEe
0072 5jb3L   ---LEEEEEEEEEL-----LHHHHHHH-H-H-HH-HHHL-LlLL-EELLEEEEEeE-EEE-E-------------------------------------------eE-E-EEEEEEEEE-E-E-LHh
0073 2okqA   hllLEEEEEEEEEE-H-H-HHHHHHHH-H-H-HH-HHHH-HlLL-EEEEEEE----------------------------------------------------lllllLEEEEEEEEE-E-E-LLh
0074 7dovA   ------LEEEEEEE-E-LllEEEEllllelL-EE-L--L-L-LE-EEEEEEEEE-E-EELlL-E-EEEEEEEEE-E--------lllleeeEE-LL----lL-H-H-H-LLLEEEEEE-L------
0075 1alyA   lLLLLLLEEEEEEL-L-L--lllllleellE-EE-E--L-L-LE-EEEEEEEEE-E-EEL-LlE-EEEEEEEEE-E--------lllleeeEE-LL---H-H-H-L----LLEEEEEE-L------
0076 8z9mA   lllllleLLEEEEL-L-L-L---------lL-HH-HLL-------lLEEEEEEL------------LLLLEEEE-L-L-------L-L-HHHH-HH-H-L-L-L-----LLEEEEEEEeL-L-LLl

# Job: BR-systems; WP_041948680 DUF499-ATPase CTD FN-III like Domain (Type-2 BR-system Specific)
# Query: s001A
# No:  Chain   Z    rmsd lali nres  %id PDB  Description
   1:  6q63-A  8.9  2.4   73   751   12   MOLECULE: BETA-HEXOSAMINIDASE;
```

```
 2:  8ayr-A  8.5  2.2   70   677   11   MOLECULE: COAGULATION FACTOR 5/8 TYPE DOMAIN PROTEIN;
 3:  2w1n-A  6.7  2.8   73   228    7   MOLECULE: O-GLCNACASE NAGJ;
 4:  6smc-C  6.6  2.7   69   303   12   MOLECULE: INTERLEUKIN-12 SUBUNIT BETA;
 5:  2vkx-E  6.5  2.7   70   202    9   MOLECULE: NEURAL CELL ADHESION MOLECULE;
 6:  6bt9-B  6.4  2.8   69   626   12   MOLECULE: CHITINASE;
 7:  2rb8-A  6.2  2.7   68    93   12   MOLECULE: TENASCIN;
 8:  6nmy-F  6.1  2.3   68   270    6   MOLECULE: INTERLEUKIN-3 RECEPTOR SUBUNIT ALPHA;
 9:  2yn3-C  6.1  2.9   69   284    4   MOLECULE: PUTATIVE INNER MEMBRANE PROTEIN;
10:  2c26-A  6.1  2.7   70   250   10   MOLECULE: ENDOGLUCANASE;
11:  1tdq-A  6.1  3.0   69   271    4   MOLECULE: TENASCIN-R;
12:  1fnf-A  6.1  2.6   68   368    6   MOLECULE: FIBRONECTIN;
13:  3wcy-A  6.0  2.7   68   318    7   MOLECULE: INTERFERON ALPHA/BETA RECEPTOR 1;
14:  6q9m-B  6.0  3.1   71   269    7   MOLECULE: RIM-BINDING PROTEIN, ISOFORM F;
15:  7y8i-E  6.0  2.3   65    97   15   MOLECULE: DSCAM;
16:  2dkm-A  6.0  2.6   69   104   10   MOLECULE: COLLAGEN ALPHA-1(XX) CHAIN;
17:  4jmh-A  5.9  2.7   69   200    6   MOLECULE: CLAMP SHC1_PY239/240;
18:  5mzv-A  5.9  3.0   69   303    6   MOLECULE: INTERLEUKIN-12 SUBUNIT BETA;
19:  6m76-A  5.9  2.9   72   923   13   MOLECULE: LPXTG-MOTIF CELL WALL ANCHOR DOMAIN PROTEIN;
20:  1qr4-A  5.9  2.8   65   175   11   MOLECULE: PROTEIN (TENASCIN);
21:  6aem-A  5.9  2.3   66    84   11   MOLECULE: PKD DOMAIN;
22:  6xax-B  5.9  2.9   67   272    4   MOLECULE: FIBRONECTIN;
23:  3t1w-A  5.9  2.6   67   368    6   MOLECULE: FOUR-DOMAIN FIBRONECTIN FRAGMENT;
24:  7u08-A  5.8  2.8   66   193   17   MOLECULE: RECEPTOR-TYPE TYROSINE-PROTEIN PHOSPHATASE ETA;
25:  3bes-R  5.8  2.5   67   250   12   MOLECULE: INTERFERON GAMMA;
26:  6mfa-A  5.8  2.8   67   363   12   MOLECULE: FIBRONECTIN;
27:  3lb6-D  5.8  2.8   71   286   10   MOLECULE: INTERLEUKIN-13;
28:  3og6-B  5.8  2.5   68   200    7   MOLECULE: INTERLEUKIN-29;
29:  3r8q-A  5.8  2.9   67   271    6   MOLECULE: FIBRONECTIN;
30:  7uww-A  5.7  2.5   69   632    6   MOLECULE: STARCH ADHERENCE SYSTEM PROTEIN 6 (SAS6);
31:  3b83-C  5.7  2.8   67    99   12   MOLECULE: TEN-D3;
32:  7z0l-B  5.7  2.5   69   370    6   MOLECULE: INTERLEUKIN-6 RECEPTOR SUBUNIT BETA;
33:  4doh-E  5.7  2.7   69   204   10   MOLECULE: INTERLEUKIN-20;
34:  2w91-A  5.7  3.4   74   635    8   MOLECULE: ENDO-BETA-N-ACETYLGLUCOSAMINIDASE D;
35:  2v5y-A  5.7  3.1   68   564    9   MOLECULE: RECEPTOR-TYPE TYROSINE-PROTEIN PHOSPHATASE MU;
36:  6x3a-A  5.7  2.5   66   306    8   MOLECULE: RECEPTOR-TYPE TYROSINE-PROTEIN PHOSPHATASE DELTA;
37:  7y8h-B  5.7  2.5   64    93   11   MOLECULE: DOWN SYNDROME CELL ADHESION MOLECULES;
38:  1k85-A  5.7  3.2   71    88   11   MOLECULE: CHITINASE A1;
39:  2jll-A  5.6  2.9   67   381   12   MOLECULE: NEURAL CELL ADHESION MOLECULE 2;
40:  1ug9-A  5.6  2.7   67  1019   13   MOLECULE: GLUCODEXTRANASE;
41:  4yg8-A  5.6  2.7   70   276    7   MOLECULE: CHITIN BIOSYNTHESIS PROTEIN CHS5;
42:  3bpl-B  5.6  2.8   68   202    9   MOLECULE: INTERLEUKIN-4;
43:  5gzt-B  5.6  3.0   69  1134    7   MOLECULE: CHITINASE;
44:  1l0q-A  5.5  2.6   67   391   12   MOLECULE: SURFACE LAYER PROTEIN;
45:  4a2m-B  5.5  2.6   71   750    6   MOLECULE: TWO-COMPONENT SYSTEM SENSOR HISTIDINE KINASE/RESP
46:  6h41-A  5.5  3.5   67   309    6   MOLECULE: INTERLEUKIN-5 RECEPTOR SUBUNIT ALPHA;
47:  5e4e-C  5.5  2.5   66   303    8   MOLECULE: INTERLEUKIN-13;
48:  8bw0-C  5.5  2.7   68   177    7   MOLECULE: TUSAMITAMAB FAB HEAVY CHAIN;
49:  8v2b-C  5.5  3.1   69   703   10   MOLECULE: ONCOSTATIN-M;
50:  8din-C  5.5  2.7   63   262    8   MOLECULE: IG GAMMA-1 FC CHAIN;
51:  2qbw-A  5.5  3.3   66   189    5   MOLECULE: PDZ-FIBRONECTIN FUSION PROTEIN;
52:  7y8s-B  5.5  2.8   68   288    6   MOLECULE: DSCAM;
53:  7zlv-h  5.4  3.0   76   527   13   MOLECULE: PROBABLE CENTRAL STRAIGHT FIBER;
54:  7biz-A  5.4  2.5   64   477    6   MOLECULE: CELL SURFACE PROTEIN;
55:  4xed-A  5.4  2.9   67    97   13   MOLECULE: PEPTIDASE M14, CARBOXYPEPTIDASE A;
56:  4doh-B  5.4  2.4   65   193    6   MOLECULE: INTERLEUKIN-20;
57:  4eq2-A  5.4  2.6   67   206    4   MOLECULE: INTERFERON GAMMA RECEPTOR 1;
58:  6grt-B  5.3  2.9   72   594    8   MOLECULE: PAIRED IMMUNOGLOBULIN-LIKE RECEPTOR B;
59:  8wm1-A  5.3  2.2   64   464   11   MOLECULE: 3-DEHYDROSHIKIMATE DEHYDRATASE (DHS DEHYDRATASE);
60:  1hwh-B  5.3  3.3   71   196   17   MOLECULE: GROWTH HORMONE;
```

```
0001 s001A   ---------AAKTDVIINQVLRDE--------E----TGAVKLRVEPQH-----G------------DKVYVDYGADP-----TTASELIH--------HGNFETA--R-----MEVRFLCVDST-----GKH----PQGPVRIWKNT----------
0002 6q63A   ydaegynyakhvFDVKAEFTPNPA-------------DGTLDITLTTID-----N------------APIHYTLGTEP-----TSTSPVYD--------G-ALKIK--E-----NDFSAIAIR--------PT----GNSRVVSEKIDfskssmkpiv
0003 8ayrA   lcevslwkyPVILAPAVNYDRN------------------GRVTLASAE-----------------nVVIRYTTGTEP-----GPQSAMYR--------N-PFFLA--G-----GTVKAAAEYRG-----R-------KSSVTTQIIPvptrdwkvva
0004 2w1nA   niiegtspeIVVNPVRFKASEIN--------------KKNVTVTWTEPE-ttegL------------EGYILYKDGKK--------VAEID--------ETSYTFK-klnRHTIYNFKIAAKYSN-----G------EVSSKESLTLRta--------
0005 6smcC   stsffirdiikpDPPKLQMKPLK--------------NSQVEVSWEYP--------dswstphsyfsLKFFVRIQ--rkGC-NQKGAFLVE--------KTSTEVQ--C---KGGNVCVQAQDRY-----YNS----SCSKWACVPCRvr--------
0006 2vkxE   aasefktqpVREPSPKLEGQRGED-------------GNSIKVNLIKQDdggspI------------RHYLVRYR---aLSSEWKPEIRLG--------SDHVMLK-sldWNAEYEVYVVAEN--------QQ----GKSKAAHFVFRtaa-------
0007 6bt9B   ggpinqkdtepPTNVKIVVTNKN--------------SNSVQLNWTAST-dnvgV------------TEYEITAGE---------EKWSTT--------TNSITIK-nlkPNTEYTFSIIAKDAA-----G------NKSQPTALTVKtdetatfsvt
0008 2rb8A   ---------MRLDAPSIEVKDVT--------------DTTALITWMPPP-----V------------DGFELTYG--ikDVPGDRTTIDLT------edENQYSIG-nlkPDTEYEVSLISRRGD------------mSSNPAKETFTtgl-------
0009 6nmyF   dkfvvfsqiEILTPPQMTA-KCNK--------------THSFMHWKMRS-------------hfnrkFRYELQIQ----KRMQPVITEQVR-------dRTSFQLL----NPGTYTVQIRARERV-----Y-----eFLSAWSPQRFEcdq-------
0010 2yn3C   ----------PPNAPVVTYSDIV--------------NDLIIXQGTAEA-----------------kSQLIITDSE--------GNTYTL---tvpdngKWSXAIP--Y--PSEGKFTITSVDAI-----G------NRSDDVPLDIXkevpvislsp
0011 2c26A   -------peNQAPKAIFTFSP-PV--------T----DENVVFNANSID-edgtI------------AYYVWDFG--------DGYEGTST--------TPTITYK--ykNPGTYKVKLIVTDNQ-----G-------ASSSFTATIKvtsatgdnsk
0012 1tdqA   sipatmnarTELDSPRLMVTASS--------------ETSISLIWTKAS---gpI------------DHYRITFT----PSSGISSEVTVP------rdRTSYTLT-dlePGAEYIISITAERGR------------QQSLESTVDAF----------
0013 1fnfA   splligqqsTVSDVPRLEVVAAT--------------PTSLLISWDAPA---vtV------------RYYRITYG--etGGNSPVQEFTVP------gsKSTATIS-glkPGVDYTITVYAVTGR--------gdspaSSKPISINYRt---------
0014 3wcyA   vdpfipfytAHMSPPEVRLEAED---------------KAILVHISPP-------qdgnmwalekpsFSYTIRIW---qKSSSDKKTINST--------YYVEKIP-ellPETTYCLEVKAIHPS-----LKK-----HSNYSVQCISttvankmpvp
0015 6q9mB   ------lgsdNVPAPKLTLERQL--------------NKSVLIGWSPP---eplI------------DSYHVYVDGVL--------KVTVE--------RTRALIE-gvdSTRPHRISVRSVTQN-----R------QTSRDCTMIIGrdtahlgpsa
0016 7y8iE   ---------GASPPPSISISHVT--------------SSSVQLNWE-----------nsqavpastiKQYLLEFR----GDNKDWIKLHIP------nnRKSFVLN-gldSSRRYQLRLAAYN--------RY----GRGDFAVIGFTtahke-----
0017 2dkmA   ---gssgssGPLPPPRLTLAAVT--------------PRTVHLTWQPSA----gA------------THYLVRCSpaspKGEEEEREVQVG--------RPEVLLD-glePGRDYEVSVQSLRGP-----E-------GSEARGIRARtptsgpssg-
0018 4jmhA   flwvggsggsVSSVPKLEVVAAT--------------PTSLLISWDAIW-ypyyV------------SYYRITYG--etGGNSPVQEFTVP------gySSTATIS-glsPGVDYTITIYAKYHR-----AK-----yYSSPISINYRt---------
0019 5mzvA   tssffirdiikPDPPNLQL-KPLK--------N----SRQVEVSWEYP--------dtwstphsyfsLTFCVQVQ----GKREKKDRVFTD--------KTSATVI--C--RKNASISVRAQDRY-----YSS----SWSEWASVPC-----------
0020 6m76A   vdggnttvdDQLTIPAIAINKTTP--------------SSLTLQWDQVE-----A------------TSYEVERDGTV--------FGNIQ--------TNTATFD-gfsFLSEHTFRVRAVGKN-----G-------VSEWSPIKGKtq--------
0021 1qr4A   ------------DNPKLEVSDPT--------------ETTLSLRWRRPV---akF------------DRYRLTYV----SPSGKKNEMEIP------vdSTSFILR-gldAGTEYTISLVAEKGR------------hKSKPTTIKGStvvgspkgis
0022 6aemA   --------eNQAPVANFEL-KTDG--------------LSVSAFNYSHD-edgeL------------VSYAWDFG--------NGQMSSE----------MAPSWS--ytRAGQYTVSLTVTDDK-----G-------ATNTTTRTTQvevp------
0023 6xaxB   ----------EIDKPSMQVTDVQ--------------DNSISVKWLPSS---spV------------TGYRVTTT--pkNGPGPTKTKTAG------pdQTEMTIE-glqPTVEYVVSVYAQNPS------------gESQPLVQTAVtnidrpkgla
0024 3t1wA   stplrgrqkTGLDSPTIDFSDIT--------------ANSFTVHWIAPR---atI------------TGYRIRHH--peHFSGRPREDRV------phsRNSITLT-nltPGTEYVVSIVALNG------------rEESPLLIGQQStvsa------
0025 7u08A   -----------PSPVFIKAVSIS--------------PTNVILTWKSND-------------taaseYKYVVKHK-----XENEKTITVVH--------QPWCNIT-glrPATSYVFSITPGIGN-----E------tWGDPRVIKVItepipvsdlr
0026 3besR   ctglcievkIGPPTVTLTEYDD-------------------HINLYIEHpyatrgykrndmcdiyllYTANFTFGD-------SEEPVIYdydctstgcSIDFATT--E------KVCVMAQGATeglldKIT----PWSSEVCLTPKknvytcairs
0027 6mfaA   ----------TVPSPRLQFVEVT--------------DVKVTIMWTPPA-----V------------TGYRVDVIpvnlPGEHGQRLPISR--------NTFAEVT-glsPGVTYYFKVFAVSHG-----------rESKPLTAQQTtkldaptnlq
0028 3lb6D   --------eIKVNPPQFEIV-DPG--------Y----LGYLYLQWQPPL--------------sectVEYELKYR---nIGSETWXTIITK--------NLHYKDG--fdLNXGIEAKIHTLLPW-----QCtngseVQSSWAETTYWispqgipetk
0029 3og6B   yldylfevePAPPVLVLTQTE-------------------EILSANATY------qlppcmppldlkYEVAFWKEG-------AGNKTLFP---------VQITLQaaS-----EHHCLSARTIYtfsvpKYS----KFSKPTCFLLEvp--------
0030 3r8qA   sspvvidasTAIDAPSLRFLATT--------------PNSLLVSWQPPR---arI------------TGYIIKYE--kpGSPPREVvPRPR------pgVTEATIT-glePGTEYTIYVIALKNN------------qKSEPLIGRKKt---------
0031 7uwwA   kqwidygkpDDHAYGYTLT-anFS--------T----E-SLDVKLALKN-----A------------DKGYYSVDG--------SAKKEFA-------nGDSVKVGegkiGNSKVTLTLYATG--------AD----GVETEQTYTFKktftaskttf
0032 3b83C   ----------MLQPPFIKVTNIT--------------LTTAVVTWQPP---ilpI------------EGILVTFG--rkNDPSDETTVDLS--------ITSLTLT-nlePNTTYEIRIVARNGQ------------QYSPPVSTTFTtgslehhhhh
0033 7z0lB   ----------SQPRVQCHASRY-----------------PVAVDCSWTP-----------lqtrstsFIATYRLGV----A-TQQQSQPCL--------QCTIPVH--L-----FPYMLNVTAVH-----PGA-----SSSLLAFVAEriikpdppeg
0034 4dohE   esgrfypflETQIGPEVAL-TTDE--------------KSISVVLTAP-----ekdlpvsmqqiysnLKYNVSVL---nTKSNRTWSQCVT--------NHTLVLT-wlePNTLYCVHVESFVPG-----PPR----RAQPSEKQCARtlkd------
0035 2w91A   gqltisdnhQEPQSPTFSVVKQSL--------K-naqEAEAVVQFKGND-----A------------DFYEVYESWKL--------lTGSS--------STTIYLPsasaQGTTQELKVVAVGKN-----G------VRSEAATTTFDwgmtvkd---
0036 2v5yA   isnyaelvvKEPPVPaPQLASVG--------------ATYLWIQLNA-ingdgpI------------VAREVEYC---tASGSWNDRQPVD--------STSYKIG-hldPDTEYEISVLLTR-P-----GEG----GTGSPGALRTRtkcadpmrgp
0037 6x3aA   kpklvsttgavPGKPRLVI-NHTQ-------------MNTALIQWHPPVdtfgpL------------QGYRLKFG---rKDMEPLTTLEFK--------EDHFTAT-dihKGASYVFRLSARNKV------------gFGEEMVKEISipe-------
0038 7y8hB   -------gasppSPPRIKVSEVT--------------TRTARLSWQSP------------------vVTYIVRYW-rdeESRSQLHQLTFQ--------VTSANLK-dlhPGTSYAVQILAENDV------------GASIPSLVQFRtiee------
0039 1k85A   --------hMAPTAPTLASTAQT--------------TSSITLSWTAS-tdnvgV------------TGYDVYNGTAL--------aTTVT--------GTTATIS-glaADTSYTFTVKAKDAA-----G------NVSAASAVSVKt---------
0040 2jllA   kieifqtlpVREPPPSIHGQPSS--------------GKSFKLSITKQDdggapI------------LEYIVKYR-------SQWLEKKVQ------gnKDHIILE-hlqWTMGYEVQITAAN-------------lGYSEPTVYEFSmppkpn----
0041 1ug9A   aaryaagtpLSSPELSVTAP----------ealstadSATAVVRGTTN------A------------AKVYVSVNG---------TATEAP--------VFSLDVA--L-TGAKNKVTVAAVAA--------D----GGTAVEDRTVLyygsrigals
0042 4yg8A   akilekygtHKPESPVLKIVNVT--------------QTSCVLAWDPLKlgsakL------------KSLILYRKGI--------RsMVIP----npfkVTTTKIS-glsVDTPYEFQLKLITT--------S----GTLWSEKVILRthkmtdmsgi
0043 3bplB   sfkpsehvkPRAPGLTVHTQVS------------------DTLLLTWSN------yppdnylynhltYAVNIWSEN------DPADFRIY--qvtylepSLRIALK--S-----GSYRARVRAWAqcyntTWS----EWSPSTKWHNS----------
0044 5gztB   ggygggsgsSSNPVVKLISTASNN-------------DRQLVLKFNAYG-----W------------DNATIVLT----SPAGKQTTYTYE----knsaqVTFTSD-kglAAGTWLYSVKTAKGS------------VTSDTVTGKAFvqgkivsyip
0045 1l0qA   qfigsipvqPVYPSADFKSNItIF--------L----SEPVQFTDLSKD-----A------------TEWKWDFG--------DGSSSKK----------QNPTHT--ysETGIYTVRLTVSN---------S----NGTDSQISTVNvvlkgsptps
0046 4a2mB   frpellldnPYTPPVVITKLQLFnkvvrtlksW----QTAFSIEFVVSN------------yisgqhNTFAYKLE-----G-YDKEWYYLD--------SRTVSYS--nlPQGTYQFLVKAANSD-------G----KWNIPTALEIIvlpi------
0047 6h41A   --------kISLLPPVFTIKVTGL--------------AQVLLQWKPN----------pdqeqrnvnLEYQVKIN------APKEDDYETR--------ITESKAV--tiLHKGFSASVRTI-LQ-----ND--hsllASSWASAELHappgspgtsi
0048 5e4eC   -----------QPPVNLSV-SVEN-------------LCTVIWTWNPPE-----------gassncsLWYFSHFGDK-------QDKKIAP--------ETRRSIE--vpLNERICLQVGSQCS--------tnesekPSILVEKCISppegdpesav
0049 8bw0C   rsdpvtldvLYGPTPIISP--pYL--------S----GANLNLSCHSAS---------------npsPQYSWRING--------IPQQHT----------QVLFIAnnN-----GTYACFVSN-------LAT----GRNNSIVKSITvsa-------
0050 8v2bC   ---------PLPLTPEIHKVSFQL--------K----LQEVNLEWTV-----------paltheelnMIFQIEIS--rlNISNTIWVENY-sttvkreeAVRWNWTdiplECVKHFIRIRALVDD-tkslpQS----SWGNWSWKEVNakvsvepdks
0051 8dinC   ---------APKAVIKLQP-------------pwvsvfQEESVTLHCEV----------phlpgsssTQWFLN-------------GTAIQ--------TYHITAS--E-----DSGEYRCQR-------GLS----GRSDPIQLEVHrgwlllqvss
```

```
0052 2qbwA   kqeirvrvekdsSVPTLEVVAAT--------------PTSLLISWDasG-----V------------SYYRITYG--etGGNSPVQEFTVS--------SSTATIS-glkPGVDYTITVYAYSDY-----YgS----HHYSPISINYRt---------
0053 7y8sB   qelivktldGVLPPPSVSLLSAS--------------DSTISVKWGHTD---epV------------TGYTLHYR----KKVGHWLHVPLD--------QTRYTLT-gldSDTTYNVYVTANNRY------------gRGDPSILSVRtgd-------
0054 7zlvh   fevrynfvpPAAPIIVFKAARIAD--------G----KERYDLRVQWDW--------nrgaganvreFVLSYIDSAefvRT-GWTKAQKIN---------ATIIFP--W-----KEHKFKVSSIAwgpdaqDV----TDSAVQTFILNestpldnsfv
0055 7bizA   ----------TPPVISLAL-sqVV--------R----NTDYTFTPDIQH-----S--------dvegFKIEWVREG--------KIVSTE----------NTYTFN--ekELGVYTVTINASN--------------IGTTTKDVSVEvvetmpyvvk
0056 4xedA   --------aNDPPNPNVTG---GN--------I----GTTYNYTFVTTD-------------pngdnITYYIEWG-------DGFTEEWI--gpyasgeEVIFSHT--wdKKGTYVVRAKAKDIL-----N------YESSWGTLEVTxpk-------
0057 4dohB   hpfnrqstiLTRPGMEITKDG-------------------FHLVIELE------------dlgpqfeFLVAYWRRE------PGAEEHVK--mvrsggiPVHLTME--P----GAAYCVKAQTFV-kaigRYS----AFSQTECVEV-----------
0058 4eq2A   efvlqkhgkIGPPKLNLSRHG-------------------AEIIVDVYH----perpwmreiyselsYSVIFRNSE-------NESRKNFvadcemnecNLSIPVP--S---EGSTYCVSAKGHF-fddlIVG----ASSEESCIWVPi---------
0059 6grtB   ----------SLPKPILRVQP-VS--------R----WTKVTFFCEETI-----G------------AEYRLYKDGKL--------YKTVT--------kAEFSLN--A-----GQYECSYSTQY------------KSSGYSPLKLVvtghywtpsl
0060 8wm1A   eqgcnaqpnQGIKAPKLTANK-------------------GLVAVEVNG-------------lpnqrYQVEFFSNqnaaSKEAEQYLGAIvatdaqgiaKANWKPK--V-----ASITANITDRF-----G------ATSELSALQTK----------
0061 1hwhB   ekcfsvdeiVQPDPPILNWTLLNV--------SltgiHADIQVRWEAP--------rnadiqkgwmvLEYELQYK--evNETKWKMMDPIL--------TTSVPVY-slkVDKEYEVRVRSKQRN-----SG-----NYGEFSVLYVTlpqms-----

0001 s001A   ---------LLLLEEEEEEEEELL--------L----LLLEEEEEEEEL-----L------------LEEEEELLLLL-----LLLLEELL--------LLEEEEL--L-----LEEEEEEELLL-----LLL----LLLLLEEEELL----------
0002 6q63A   hhhlllllllhhHLLEEEEEEELL-------------LLEEEEEEELLL-----L------------LLEEEELLLLL-----LLLLLLLL--------L-LEEEL--L-----LEEEEEELL--------LL----LLLLLEEEEELllllllllee
0003 8ayrA   eleeeeellLLLLLLEEEELLL------------------LEEEEELLL-----------------lLEEEEELLLLL-----LLLLEELL--------L-LEELL--L-----EEEEEEEEELL-----E-------ELLLEEEEELlllllleeee
0004 2w1nA   eeellllllLLLLLLEEEEEEEEL-------------LLLEEEEEELLL-llllE------------EEEEEEELLEE--------EEEEL--------LLEEEEL-lllLLLEEEEEEEEEELL-----L------LLLLLEEEEEEll--------
0005 6smcC   eeeelhhhllllLLLEEEEEEELL-------------LLLEEEEEELL--------lllllllllllEEEEEEEE--llLL-LLLLLEEEL--------LLEEEEL--L---LLLEEEEEEEELL-----LLL----LLLLLEEEELLll--------
0006 2vkxE   lleeeelllLLLLLLEEEEEELLL-------------LLLEEEEEELLLlllllE------------EEEEEEEE---eLLLLLLLLEEEL--------LLEEEEL-lllLLLEEEEEEEEEE--------LL----EELLLEEEEEElll-------
0007 6bt9B   lllllllllllLLLLEEEEEEEEL-------------LLLEEEEEELLL-llllL------------LEEEEEELL---------EEEEEL--------LLEEEEL-lllLLLEEEEEEEEELLL-----L------LLLLLEEEEEElllleeeeee
0008 2rb8A   ---------LLLLLLEEEEELLL--------------LLLEEEEEELLL-----L------------LEEEEEEE--elLLLLLLEEEEEE------llLLEEEEL-lllLLLEEEEEEEEEELL------------eELLLEEEEEElll-------
0009 6nmyF   eeeeehhhlLLLLLLEEEE-EELL--------------LEEEEEEELLL-------------lllllEEEEEEEE----ELLLLLEEELLL-------lLLEEEEE----LLLLEEEEEEEEELL-----L-----lLLLLLLLEEELlll-------
0010 2yn3C   ----------LLLLLEEEEEEEEE-------------LLEEEEEEELLL-----------------lLEEEEEELL--------LLEEEE---elllllEEEEEEE--L--LHHHEEEEEEELLL-----L------LLLLLEEEELLlllleeeelh
0011 2c26A   -------llLLLLLLLLEEEL-LL--------L----LLLEEEELLLLL-llllE------------EEEEEELL--------LLLEEEEL--------LLEEEEL--llLLEEEEEEEEEEELL-----L-------LEEEEEEEEEelllllllll
0012 1tdqA   llleeeeeeLLLLLLEEEEEEEL--------------LLLEEEEEELLL---llL------------LEEEEEEE----LLLLLLEEEEEE------llLLEEEEL-lllLLLLEEEEEEEEELL------------EELLLEEEEEL----------
0013 1fnfA   llleeeeeeLLLLLLEEEEEEEE--------------LLEEEEEEELLL---llL------------LEEEEEEE--elLLLLLLEEEEEE------llLLEEEEL-lllLLLLEEEEEEEELLL--------lllllLLLLEEEEELl---------
0014 3wcyA   lllllhhhhEELLLLEEEEEEELL--------------LLEEEEEELL-------llllllllllllLEEEEEEE---eLLLLLLEEEEEL--------LLEEEEL-lllLLLEEEEEEEEELLL-----LLL-----LLLLLLEEEEllllllllll
0015 6q9mB   ------llllLLLLLEEEEEEEE--------------LLEEEEEEELL---lllE------------EEEEEEELLEE--------EEEEL--------LLEEEEE-lllLLLLEEEEEEEEELL-----L------LLLLHLEEEELllllllllee
0016 7y8iE   ---------LLLLLLLLEEELLL--------------LLLEEEELL-----------llllllllllLEEEEEEE----ELLLLLEEELLL------llLLEEEEL-lllLLLEEEEEEEEEE--------LL----EELLLEEEEEElllll-----
0017 2dkmA   ---llllllLLLLLLLLEEEEEL--------------LLEEEEELLLLL----lL------------LEEEEEEEelllLLLLLLEEEEEL--------LLEEEEE-lllLLLLEEEEEEEELLL-----L-------LLLLEEEELLlllllllll-
0018 4jmhA   elllllllllLLLLLEEEEEEEE--------------LLEEEEEEELLL-lhhhE------------EEEEEEEE--elLLLLLLEEEEEE------llLLEEEEL-lllLLLEEEEEEEEEELL-----LL-----lLLLLEEEEEEl---------
0019 5mzvA   eeeelhhhlllLLLLEEEL-LLLL--------L----LLEEEEEEELL--------lllllllllllEEEEEEEE----LLLLLLLEEEEL--------LLEEEEE--L--LLLLEEEEEEEELL-----LLL----LLLLLEEEEL-----------
0020 6m76A   eelllllllLLLLLLLLEELHLLL--------------LLEEEELLLLL-----L------------LEEEEEELLEE--------EEEEL--------LLEEEEL-lllLLLEEEEEEEEEELL-----E-------ELLLLLEEEEll--------
0021 1qr4A   ------------LLLEEEEEELLL-------------LLEEEEEEELLL---llL------------LEEEEEEE----LLLLLEEEEEEL------llLLEEEEE-lllLLLEEEEEEEEELLL------------lLLLLEEEEEEllllllllee
0022 6aemA   --------lLLLLLLLEEE-EEEL--------------LEEEEEELLLL-llllE------------EEEEEELL--------LLLEELL----------LLLEEE--llLLEEEEEEEEEEELL-----L-------LEEEEEEEEEelll------
0023 6xaxB   ----------LLLLLEEEEEEEL--------------LLLEEEEEELLL---llL------------LEEEEEEE--elLLLLLLEEEEEL------llLLEEEEL-lllLLLEEEEEEEEELLL------------lLLLLEEEEEElllllleeee
0024 3t1wA   llleeeeeeLLLLLLEEEEELLL--------------LLLEEEEEELLL---llL------------LEEEEEEE--elLLLLLLEEEEE------ellLLEEEEL-lllLLLEEEEEEEEEEL------------lEELLLEEEEEEllll------
0025 7u08A   -----------LLLLEEEEEELL--------------LLLEEEEEELLL-------------lllllLEEEEEEL-----LLLLLEEEEEL--------LLEEEEL-lllLLLEEEEEEEEELLL-----L------lLLLLEEEEEElllllleeee
0026 3besR   ellhhhhllLLLLEEEEEEELL-------------------EEEEEEELleeeellllhhhllllleEEEEEEELL-------LLLLEEEhhhllllee EEEELLL--L------LEEEEEEEELlllllLLL----LLLLLEEELLLllllllllll
0027 6mfaA   ----------LLLLLEEEEEEEL--------------LLLEEEEEELLL-----L------------LEEEEEEEelllLLLLLEEEEEEL--------LLEEEEL-lllLLLEEEEEEEEEELL------------eELLLEEEEEElllllleeee
0028 3lb6D   --------lLLLLLLEEEEE-LLL--------L----LLEEEEEEELLL--------------llllLEEEEEEE---eLLLLLLEEEEEL--------LLEEEEE--llLLLLEEEEEEEEELH-----HHhllllEELLLEEEEELllllllllll
0029 3og6B   lllhhhhllLLLLEEEEEEEL-------------------LEEEEEEEL------llllllllllleeEEEEEEELL-------LLLLEELL---------EEEELLllL-----LEEEEEEEEEEellleEEL----LLLLLEEEELLll--------
0030 3r8qA   llleeeeeeLLLLLLEEEEEEEE--------------LLEEEEEEELLL---llL------------LEEEEEEE--llLLLLEELlLLLL------llLLEEEEL-lllLLLEEEEEEEEEELL------------eELLLEEEEEEl---------
0031 7uwwA   lleeellllLLLLEEEELL-llEL--------L----L-LEEEEEEEEL-----L------------LLEEEEELL--------LLLEEEL-------lLLEEEELllllLLEEEEEEEEEEEL-------LL----LLEEEEEEEEEelllllllee
0032 3b83C   ----------LLLLLEEEEELLL--------------LLLEEEEEELL---lllL------------LEEEEEEE--elLLLLLLEEEEEL--------EEEEEEL-lllLLLEEEEEEEEEELL------------EELLLEEEEEEllllllllll
0033 7z0lB   ----------LLLEEEEELLLL-----------------LLEEEEEEEL-----------lllllleEEEEEEELL----L-LLLLLEELL--------LEEEELL--L-----LLEEEEEEEEL-----LLE-----EEEEEEELLLllllllllee
0034 4dohE   elllllhhhHLLLLLEEEE-EELL--------------LLEEEEEELL-----lllllllhhhhlllLEEEEEEE---eLLLLEEEEEEEL--------LLEEEEL-lllLLLEEEEEEEEELLL-----LLL----LLLLLLLEEEEllll------
0035 2w91A   eeeeeelllLLLLLLEEEEEEEEE--------E-ellEEEEEEEEELLL-----E------------EEEEEEEEEEE--------eEEEL--------LLEEEEEllllLLLEEEEEEEEEELL-----L------LLLLLEEEEEElllllll---
0036 2v5yA   ellleeeeeELLLLLlLEEEEEL--------------LLLEEEELLL-llllllL------------LEEEEEEE---eLLLLLEEEEELL--------LLEEEEL-lllLLLEEEEEEEEEL-L-----LLL----LLLLLLLEEEEllllllllll
0037 6x3aA   lleeeelllllLLLLEEEE-EEEL-------------LLEEEEEEELLLlllllL------------LEEEEEEE---eLLLLLLEEEEEL--------LLEEEEE-lllLLLEEEEEEEEELLL------------lLLLLEEEEEElll-------
0038 7y8hB   -------lllllLLLEEEEELLL--------------LLLEEEEEELL------------------lLEEEEEEE-ellLLLLLLEEEEEL--------LLEEEEL-lllLLLEEEEEEEEELLL------------LLLLLLEEEEEllll------
0039 1k85A   --------lLLLLLLEEEEEEEL--------------LLLEEEEEELL-lllllE------------EEEEEEELLLE--------eEEEL--------LLEEEEL-lllLLLEEEEEEEEEELL-----L------EELLLLLEEEEl---------
0040 2jllA   lleeeelllLLLLLLLEEEEEEL--------------LLEEEEEELLLLlllllL------------LEEEEEEE-------LLLLLLEEL------llLLEEEEL-lllLLLEEEEEEEEEEL------------lLLLLLEEEEEElllll-----
0041 1ug9A   hhhhlllllLLLLLEEEEEL----------lleeellLLEEEEEEEEL------L------------LEEEEEELL---------EEEEEL--------LEEEEEE--L-LLLEEEEEEEEELL--------L----LLEEEEEEEEEllleeeeeee
0042 4yg8A   hhhhhhhhlLLLLLLLEEEEEEL--------------LLLEEEEELLLLlllllE------------EEEEEELLLL--------LlLLLL----llllLLEEEEL-lllLLLEEEEEEEEEEL--------L----EEEEEEEEEEEllllllllll
0043 3bplB   eelhhhlllLLLLEEEEELLLL------------------LEEEEEEEL------lllllllhhhleEEEEEEELL-------LLLLEEEE--eellllEEEEELL--L-----LEEEEEEEEELhhhllLLL----LLLLLEEEELL----------
0044 5gztB   lllllllllLLLLLEEEEEEEELL-------------LLLEEEEEEELL-----L------------LLEEEEEE----LLLLLEEEEEHH----hhhhhEEEELL-lllLLEEEEEEEEEEELL------------EELLLEEEEEEelleeeeeee
0045 1l0qA   lleeellllLLLLLLLEEELLlEE--------L----LLLEEEEELLLL-----L------------LEEEEELL--------LLLEELL----------LLLEEL--llLLEEEEEEEEEEE---------L----LEEEEEEEEEEeellllllll
0046 4a2mB   ellllllllLLLLLEEEEEEEELleelleellL----LLLEEEEEEELL------------llllllLEEEEEEE-----L-LLLLLEEEL--------LLEEEEL--llLLEEEEEEEEEELLL-------L----LLLLLEEEEEEelll------
0047 6h41A   --------lLLLLLLEEEEEEEEL--------------LEEEEEEELL----------lllllllllLEEEEEEE------ELLLEEEEEL--------LLEEEEE--llLLLLEEEEEEEE-EE-----LL--lleeELLLEEEEELllllllllll
0048 5e4eC   -----------LLLLEEEE-EEEL-------------LLEEEEEEELLL-----------lllllllLEEEEEELLL-------EEEEEEL--------LLEEEEE--llLLLLEEEEEEEEELL-------llllllLLLEEEEEELllllllllll
0049 8bw0C   ellllllllLLLLLLEELL--lEL--------L----LLLEEEEEELLL---------------lllLEEEEEELL--------EEEELL----------LEEEELllL-----EEEEEEEEL-------LLL----LLEEEEEEEEEell-------
0050 8v2bC   ---------LLLLLLEEEEEEEEL--------L----LLEEEEEEEL-----------lllllleeeEEEEEEEE--elLLLLEEEEEEE-eeeellllLEEEEEEllllLLLLEEEEEEEEELL-lllllLL----LLLLLLLEEELllllllllle
0051 8dinC   ---------LLLLEEEEEL-------------llleeeLLLLEEEEEEL----------llllllllEEEEEL-------------LEELL--------LEEELLL--H-----HLEEEEEEE-------LLL----LLLLLEEEEEEllleeeeell
0052 2qbwA   eeeeeeeeelllLLLLLEEEEEE--------------LLEEEEELLllL-----L------------LEEEEEEE--elLLLLLLEEEEEL--------LLEEEEL-lllLLLEEEEEEEEELLL-----LlL----LLLLLEEEEEEl---------
0053 7y8sB   lleeeelllLLLLLLLEEEEEEL--------------LLLEEEEELLLL---llL------------LEEEEEEE----ELLLLLEEEEEL--------LLEEEEL-lllLLLEEEEEEEEEELL------------eELLLLLEEEElll-------
0054 7zlvh   eelllllllLLLLEEEEEEELLLL--------L----LLLLEEEEEEEL--------llllllllllEEEEEEEHHhhhHL-LLLLLEEEE---------EEELLL--L-----LLEEEEEEEELllllllLE----EELLLEELLLLllllllllll
0055 7bizA   ----------LLLEEELLL-llEE--------L----LEEEEELLEEEL-----L--------llllLEEEEEELL--------EEEELL----------LLEEEL--llLLEEEEEEEEEEL--------------LLEEEEEEEEEeellllleee
0056 4xedA   --------lLLLLLLEEEL---LL--------L----LLLEEEEEELLL-------------lllllEEEEEELL-------LLLEEEEE--eelllllLEEEEEL--llLLEEEEEEEEEEEELL-----L------LLLLLEEEEEEell-------
0057 4dohB   llllhhhllLLLLLEEEEEEL-------------------LEEEEEEL------------lllllleEEEEEEELL------LLLLLEEE--eelllllEEEEELL--L----LLLEEEEEEEEE-hhhlEEL----LLLLLEEELL-----------
0058 4eq2A   leehhhhllLLLLEEEEEEEL-------------------LEEEEEEEL----lllhhhhhhhlleeEEEEEEELL-------LLLEEEEeellhhhleEEEEELL--L---LLEEEEEEEEEEE-llllEEL----LLLLLEEEEELl---------
0059 6grtB   ----------LLLLLEEEEEL-EE--------L----LLLEEEEEELLL-----L------------LEEEEEELLEE--------EEEEE--------lEEEEEH--L-----EEEEEEEEELL------------EELLLLLEEEEeellllllee
0060 8wm1A   llllllllhHHLLLLEEEEEL-------------------LEEEEEEEL-------------lllleEEEEEEEEllllLLLLLEEEEEEeelllllleeEEEELLL--L-----LEEEEEEEELL-----L------LLLLLLEELL-----------
0061 1hwhB   eeeeehhhhLLLLLLEEEEEEEEL--------LllllEEEEEEEEEELL--------llllllllllEEEEEEEE--elLLLLLEELLLLL--------LLEEEEE-eeeLLLLEEEEEEEEELL-----LL-----LLLLLLLEEELlllll-----

@@@@@@@@@@@@@@@@@@@@@@@@@@@@@@@@@@@@@@@@@@@@@@@@@@@@@@@@@@@@@@@@@@@@@@@@@@@@@@@@@@@@@@@@@@@@@@@@@@@@@@@@@@@@@@@@@@@@@@@@@@@@@@@@@@@@@@@@@@@@@@@@@@@@@@@@@@@@@@@@@@@@@@@
@@@@@@@@@@@@@@@@@@@@@@@@@@@@@@@@@@@@@@@@@@@@@@@@@@@@@@@@@@@@@@@@@@@@@@@@@@@@@@@@@@@@@@@@@@@@@@@@@@@@@@@@@@@@@@@@@@@@@@@@@@@@@@@@@@@@@@@@@@@@@@@@@@@@@@@@@@@@@@@@@@@@@@@
@@@@@@@@@@@@@@@@@@@@@@@@@@@@@@@@@@@@@@@@@@@@@@@@@@@@@@@@@@@@@@@@@@@@@@@@@@@@@@@@@@@@@@@@@@@@@@@@@@@@@@@@@@@@@@@@@@@@@@@@@@@@@@@@@@@@@@@@@@@@@@@@@@@@@@@@@@@@@@@@@@@@@@@

# Job: DUF3780 (Type-2 BR-system Specific)
# Query: s001A
# No:  Chain   Z    rmsd lali nres  %id PDB  Description
   1:  8aj8-B  4.3  3.7   85   558    6   MOLECULE: PHOSPHATIDYLINOSITOL 4,5-BISPHOSPHATE 3-KINASE CA
   2:  2ipc-A  3.9 12.3   80   939    9   MOLECULE: PREPROTEIN TRANSLOCASE SECA SUBUNIT;
   3:  5h1q-A  3.9 10.8   79   360    6   MOLECULE: INNEXIN-6;
   4:  1yns-A  3.9  8.3   63   254    8   MOLECULE: E-1 ENZYME;
   5:  5yet-A  3.8 12.6   70   397    3   MOLECULE: UNCHARACTERIZED PROTEIN R354;
   6:  8ppq-a  3.6 13.9   59   147    2   MOLECULE: ENVELOPE PROTEIN E;
   7:  5mlp-A  3.6 16.0   65   255    3   MOLECULE: UNCHARACTERIZED PROTEIN;
   8:  3k7d-A  3.5  3.1   83   497    7   MOLECULE: GLUTAMATE-AMMONIA-LIGASE ADENYLYLTRANSFERASE;
   9:  4gmq-A  3.5  2.6   58    92    5   MOLECULE: PUTATIVE RIBOSOME ASSOCIATED PROTEIN;
  10:  6d9z-C  3.5  8.8  104   241    8   MOLECULE: SULFATE TRANSPORTER CYSZ;
  11:  6xqj-A  3.5  2.9   49   173   10   MOLECULE: PROTEIN VPR,UV EXCISION REPAIR PROTEIN RAD23 HOMO
  12:  4myy-B  3.5  4.4   57    84   14   MOLECULE: CURG, CURH FUSION PROTEIN;
  13:  7x34-C  3.4  3.9   71   100   10   MOLECULE: ZUOTIN;
  14:  8c81-B  3.3  2.8   48   539    6   MOLECULE: PROTEIN ORM1;
  15:  8rxa-A  3.3  4.8   76   102   11   MOLECULE: AP2 DOMAIN TRANSCRIPTION FACTOR AP2-O5, PUTATIVE;
  16:  6cgh-A  3.3  4.3   66    89    8   MOLECULE: DNAJ HOMOLOG SUBFAMILY C MEMBER 2;
  17:  6x6o-B  3.3  3.3   63    76    5   MOLECULE: PROTEIN SPACKLE;
  18:  9fk0-D  3.2  7.4   52    75    4   MOLECULE: ENVELOPE PROTEIN E;
  19:  4v1a-h  3.2  5.4   78   289    6   MOLECULE: MITORIBOSOMAL PROTEIN ML37, MRPL37;
  20:  5u4w-H  3.1  3.4   47    53    4   MOLECULE: E PROTEIN;
  21:  9m31-A  3.1 12.6   76   945    8   MOLECULE: CASRX;
  22:  9qx2-A  3.1 12.4   72   947    4   MOLECULE: UBR-TYPE DOMAIN-CONTAINING PROTEIN;
  23:  6jdp-A  3.1  4.1   47   241    4   MOLECULE: IMM52 FAMILY PROTEIN;
  24:  8rjj-A  3.1 17.4   62   183    5   MOLECULE: GENOME POLYPROTEIN;
  25:  4p96-A  3.1  7.9   61   278    7   MOLECULE: FATTY ACID METABOLISM REGULATOR PROTEIN;
  26:  6c6r-A  3.0 14.6   80   451    8   MOLECULE: SQUALENE MONOOXYGENASE;
  27:  3cxj-A  3.0  5.4   50   144   14   MOLECULE: UNCHARACTERIZED PROTEIN;
  28:  7e9d-A  3.0  7.4   55   443    5   MOLECULE: IRG1;
  29:  8an4-A  3.0 13.1   78   195   10   MOLECULE: BACTERIAL TOXIN;
  30:  9qx0-B  3.0 13.2   74  1090    3   MOLECULE: E3 UBIQUITIN-PROTEIN LIGASE KCMF1;
  31:  5cuf-C  3.0 13.3   63   370    6   MOLECULE: SESTRIN-2;
```

```
 32:  5tf3-A  2.9  2.3   62   131    3   MOLECULE: PUTATIVE MEMBRANE PROTEIN;
 33:  6w4x-C  2.9  9.6   56   375    9   MOLECULE: RIBONUCLEOSIDE-DIPHOSPHATE REDUCTASE 1 SUBUNIT AL
 34:  2ffl-A  2.9 19.4   90   732    8   MOLECULE: DICER;
 35:  6s7t-A  2.9  6.2   88   706    3   MOLECULE: DOLICHYL-DIPHOSPHOOLIGOSACCHARIDE--PROTEIN
 36:  9e41-B  2.9  6.5   52    75    4   MOLECULE: E GLYCOPROTEIN;
 37:  5mlq-A  2.9 15.4   59   226    8   MOLECULE: CDPS;
 38:  8ul9-A  2.9  6.8   95   387    8   MOLECULE: CHOLINEPHOSPHOTRANSFERASE 1;
 39:  9jc2-A  2.9  3.8   49   166    8   MOLECULE: ATP SYNTHASE SUBUNIT A;
 40:  6mca-A  2.9 14.8   92   460   12   MOLECULE: ANKYRIN REPEAT DOMAIN PROTEIN;

0001 s001A   [illegible]
0002 8aj8B   [illegible]
0003 2ipcA   [illegible]
0004 5h1qA   [illegible]
0005 1ynsA   [illegible]
0006 5yetA   [illegible]
0007 8ppqa   [illegible]
0008 5mlpA   [illegible]
0009 3k7dA   [illegible]
0010 4gmqA   [illegible]
0011 6d9zC   [illegible]
0012 6xqjA   [illegible]
0013 4myyB   [illegible]
0014 7x34C   [illegible]
0015 8c81B   [illegible]
0016 8rxaA   [illegible]
0017 6cghA   [illegible]
0018 6x6oB   [illegible]
0019 9fk0D   [illegible]
0020 4v1ah   [illegible]
0021 5u4wH   [illegible]
0022 9m31A   [illegible]
0023 9qx2A   [illegible]
0024 6jdpA   [illegible]
0025 8rjjA   [illegible]
0026 4p96A   [illegible]
0027 6c6rA   [illegible]
0028 3cxjA   [illegible]
0029 7e9dA   [illegible]
0030 8an4A   [illegible]
0031 9gx0B   [illegible]
0032 5cufC   [illegible]
0033 5tf3A   [illegible]
0034 6w4xC   [illegible]
0035 2fflA   [illegible]
0036 6s7tA   [illegible]
0037 9e41B   [illegible]
0038 5mlqA   [illegible]
0039 8ul9A   [illegible]
0040 9jc2A   [illegible]
0041 6mcaA   [illegible]

0001 s001A   [illegible]
0002 8aj8B   [illegible]
0003 2ipcA   [illegible]
0004 5h1qA   [illegible]
0005 1ynsA   [illegible]
0006 5yetA   [illegible]
0007 8ppqa   [illegible]
0008 5mlpA   [illegible]
0009 3k7dA   [illegible]
0010 4gmqA   [illegible]
0011 6d9zC   [illegible]
0012 6xqjA   [illegible]
0013 4myyB   [illegible]
0014 7x34C   [illegible]
0015 8c81B   [illegible]
0016 8rxaA   [illegible]
0017 6cghA   [illegible]
0018 6x6oB   [illegible]
0019 9fk0D   [illegible]
0020 4v1ah   [illegible]
0021 5u4wH   [illegible]
0022 9m31A   [illegible]
0023 9qx2A   [illegible]
0024 6jdpA   [illegible]
0025 8rjjA   [illegible]
0026 4p96A   [illegible]
0027 6c6rA   [illegible]
0028 3cxjA   [illegible]
0029 7e9dA   [illegible]
0030 8an4A   [illegible]
0031 9gx0B   [illegible]
0032 5cufC   [illegible]
0033 5tf3A   [illegible]
0034 6w4xC   [illegible]
0035 2fflA   [illegible]
0036 6s7tA   [illegible]
0037 9e41B   [illegible]
0038 5mlqA   [illegible]
0039 8ul9A   [illegible]
0040 9jc2A   [illegible]
0041 6mcaA   [illegible]

[illegible]

# Job: Type-3 BR-systems Inactive STAND-NTPase
# Query: s001A
# Chain   Z    rmsd lali nres  %id PDB  Description
4dt1-A  4.7  3.3   93   193   10 PDB  MOLECULE: CHROMOSOME SEGREGATION IN MEIOSIS PROTEIN 2;
8c0v-B  4.4  3.7   93  1030    9 PDB  MOLECULE: PEROXISOMAL ATPASE PEX1;
7vcs-A  4.3  3.3   91   766    9 PDB  MOLECULE: TRANSITIONAL ENDOPLASMIC RETICULUM ATPASE;
7mi1-A  4.3  2.9   80  2628   11 PDB  MOLECULE: CHIMERA PROTEIN OF DYNEIN AND ENDOLYSIN;
4rh7-A  4.1  3.1   90  3005    8 PDB  MOLECULE: GREEN FLUORESCENT PROTEIN/CYTOPLASMIC DYNEIN 2 HE
9e22-A  4.0  3.4   88  2925    9 PDB  MOLECULE: CYTOPLASMIC DYNEIN 1 HEAVY CHAIN 1;
7xp9-A  3.9  2.6   51    76    8 PDB  MOLECULE: RXLR EFFECTOR PROTEIN AVR-VNT11;
8xks-B  3.8  3.4   87   603    8 PDB  MOLECULE: CTAP1;
7ykk-A  3.8  3.3   83   735    6 PDB  MOLECULE: ATPASE FAMILY GENE 2 PROTEIN;
```

```
3io5-B  3.8  3.2   84   272    8 PDB  MOLECULE: RECOMBINATION AND REPAIR PROTEIN;
7mi8-A  3.7  3.5   93   868   14 PDB  MOLECULE: FUSION PROTEIN OF DYNEIN AND ENDOLYSIN;
5fsz-A  3.7  3.2   85   245    5 PDB  MOLECULE: MACRODOMAIN;
9fnn-U  3.7  3.3   84   511   13 PDB  MOLECULE: CELLULOSE SYNTHASE CATALYTIC SUBUNIT [UDP-FORMING
8xku-E  3.7  3.6   89   498   12 PDB  MOLECULE: PROBABLE INACTIVE ATP-DEPENDENT ZINC METALLOPROTE
6p8v-C  3.4  3.0   78   303    9 PDB  MOLECULE: ATPASE, AAA FAMILY;
8c0v-A  3.3  3.2   82   823    9 PDB  MOLECULE: PEROXISOMAL ATPASE PEX1;
5e7p-A  3.3  3.3   84   719   10 PDB  MOLECULE: CELL DIVISION CONTROL PROTEIN CDC48;
6wc3-B  3.3  3.1   53    94   13 PDB  MOLECULE: PROTEIN TRANSPORT PROTEIN TIP20;
8glw-A  3.3  3.4   81   483   10 PDB  MOLECULE: TRANSPOSON TN7 TRANSPOSITION PROTEIN TNSC;
8btg-B  3.3  3.8   88   335    7 PDB  MOLECULE: CHROMOSOMAL REPLICATION INITIATOR PROTEIN DNAA;
1g4a-E  3.2  4.0   88   356   11 PDB  MOLECULE: ATP-DEPENDENT HSL PROTEASE ATP-BINDING SUBUNIT
7mca-C  3.2  3.3   81   584    6 PDB  MOLECULE: ORIGIN RECOGNITION COMPLEX SUBUNIT 1;
6tdw-C  3.1  5.5   56   157   16 PDB  MOLECULE: ATPTB1;
8rg0-5  3.1  3.4   63   520    6 PDB  MOLECULE: EUKARYOTIC TRANSLATION INITIATION FACTOR 3 SUBUNI
7st9-A  3.0  3.2   85   522    6 PDB  MOLECULE: CHECKPOINT PROTEIN RAD24;
7uqj-A  3.0  2.9   82   585   11 PDB  MOLECULE: ATPASE HISTONE CHAPERONE YTA7;

0001 s001A   ------------MKGYEIFE------QKAPYE---NKIGF---LDFLRELKSGTQGIP--SSFMVVGI-D-DVLYL-AGRDERLALTIHKILQS-SA-KVLD-QK-II-EVQIVCKG-RLYKGESFWSEYRGEKLP-LDYIFG-TPN
0002 4dt1A   -------meyedLELITIWP------SP----tqlFFIDatssEELSAITVKLLQIL-lkIILYINGL-E-VMFRN-SQ-PQRSHELLRDTLLK-LR-VMGN-DA-SI-RTLLEFPKeQLLDKNGDS--------L-AEY-IW-KY-
0003 8c0vB   etsfncsqrgitLNASVLLH------SKATM-ihlLEID----gYIRAKCENVLPYAP--AVIFLAHL-D-SILLD-VNaQKSINFEMSKLLDD-FT-FKF---P-GT-TFVGSVNN---------------iDNV-PSSFRS-HMR
0004 7vcsA   ehpdkflkfgmtPSKGVLFYgppgcgKTLLAKanfISIKG---PEseANVREIFDKA-apCVLFFDEL-D-SIAKA-RGG-gGAADRVINQILTeMD-GMsT-KK-NV-FIIGATNR---------------pDII-DPA-IL-RP-
0005 7mi1A   dewniadvvansENNYFTMA-serdvDGTF-L---KIIPL---nYAQEEISK-KIEGG---WILLQNI-Q-MSL-------swvKTYLHKHVEE--T-KAAE-EHeKF-KMFMTCH-----------LTGD---KL-PAPLLQr---
0006 4rh7A   dkcyltltqamkMGLGGNPYgpagtgKTESVKrqvLVFNC-----mGRIFVG-VKCGA---WGCFDEF-N-RL-------EESVLSAVSMQIQT-IQ-DALK-NS-NS-GIFITMNP------------AGKGyKL-PDNLKQ-rgn
0007 9e22A   drcyltmtqaleARLGGSPF------GtESVKrfvLVFNC-----mGRIFVGLCQVGA---WGCFDEF-N-RL-------EERMLSAVSQQVQC-IQ-EALR-EP-DM-AIFITMNP-G---------------SN-LPDNLK-Klp
0008 7xp9A   --------------------------------keatrkYY---LDLFKRADFTLPKLA--KKG--daL-K-KLRKA-G--SEEKFAELKGAAAK-YA-DDWY-RI-Y----------------------------------------
0009 8xksB   rtpekfkasgarAPKGVLLV------GKTLMAvafISSSA---A-vRDLFNT-RSVAP--CIIFIDEL-D-AVG---RQRQDNTVNQLLTEMDG-fE-AEQ---Q-GI-VVMGATNR-KD--------------VL-DAALTR-PG-
0010 7ykkA   easetfarlgisAPKGVLLY------G----F---LAVKG-----irEIFRK-RSAAP--SIIFFDEI-D-ALSP--DRDGNHVLTSLLNEIDG----EELK----GV-VIVAATNR---------------pDEI-DAALLR-PDR
0011 3io5B   nialsgeitggmQSGLLILAsksfksNFGL--avcLFYD----EQLRIDMVNQLDAI-ekVVVFIDSL-G-NLA-------kTRAKTMKSLFRI-VT-PYFS-TK-NI-PCIAINHT-Y-------------------TGPMY-EKS
0012 7mi8A   ligfatltdslhQKYGGCFFgpagtgKTETVKrvvVVFNC-----lSRLLVGITQIGA---WGCFDQF-N-RL-------DEKVLSAVSANIQQ-IQ-NGLQ-VP-HT-AVFITLNP------------GYNGRSE-LPENLK-K--
0013 5fszA   delkpvdsshdilRHIALYN------GDLQL---dAIVNaankgqYRSILSLAHRNRL--RSIGFCCV----STGVyGYPLIPATRIAVDETIE-YL-KQ-H-FS-AFdLCCFACFK-LE--------------nA-YTDCLR-A--
0014 9fnnU   isslwdelrhmpaGGVWWFN------VLANQTahvAVIS----kGLYYLT-RDLQC--hnYLFILVCanN-AWQN-------iPAERLRSWLDK-MN-KWSR-LN-HC-SLLVINPG---------------NNNDkQFSLLL-E-F
0015 8xkuE   knpdlfdkmgikPPHGVLLE------GTLVAKvpfYQMAG---SEgsARIRDLFKRA-kpSVIFIDEI-D-AL-------ATQERETTLNQLLI-EL-DGFDtGK-GV-IFLGATNR-RD--------------LL-DP-ALL-R--
0016 6p8vC   pdaraavdtvlrrpPLVVLA------gLAETIitlYPLS------vSAAFDYAADKkA--VLLLIDQA-D-ALA----qsDRAGVNAFIRGIDR-IA-NQ---KL-PA-AVLXCT-N-RLK-------------AL-DPAVQRr---
0017 8c0vA   kemvnyltspiiATPAIILD------G-----ifvKYAD----kLIMEWCSFCYWYGP--SLIVLDNV-E-ALF----anWDNASKLLNFFINQ-VT-KIFN-KD-NI-RVLFSGKQ-KT--------------QI-NPL-LF-DK-
0018 5e7pA   qhpdtfsrlgidpPRGVLLYgppgcgKTFVVRlsvhAVKG-----vRELFAR-RDSAP--SLVFLDEI-D-ALAPR-RGQTDKVVASLLTELDG-IE-PL----R-DV-VVLGATNR---------------pDLI-DPALLR-P--
0019 6wc3B   --------------------------------smpyatQL---ALLQDELLDMLEPLR--TADiieLL-G-CYRLQ-VEkALAQLLLWERFLAD-YR-RRLD-AA-I----------------------------------------
0020 8glwA   eletfrfeearsTAQSLLLI------GslhrY---LKIDC---kEICLnLALMSQIAL--GLLVIDEI-Q-HL--S-RSRS-GGSQEMLNFFVT-MV-NIIG-----V-PVMLIGTP-----------------KA-REI-FE-ADL
0021 8btgB   aaslavaeapakAYNPLFIY------GLMHAIakvVYLSS---EKFTNE--FRNR-NV--DVLLIDDI-Q-FLA---------GKEQTQEEFFH-TF-NTLH-EE-SK-QIVISSDR-PPK-------------PT-LED-LR-SR-
0022 1g4aE   rmqlneelrhevTPKNILMI------GKTEIAapfIKVEA---TKgkEVDS--IRDLvqhGIVFIDEI-D-KICKR-GGPDVSRE-GVQRDLLP-LveVKTD----HI-LFIASGAF-Q-----------------SD-LIPELQ-G--
0023 7mcaC   isdllysettqkRRCFNTIF------L----N---VLIEL-----mMLRRsYDLSLV-kdLAMVFNFK---DVDS-------iNFNTLDNFIIL-LK-S-AF-KYdHV-KISLIFNI------------NTNLSNI-EknlTI-RL-
0024 6tdwC   -----------------------------------davKF---QTLYNVLTGlqvNLT--ASWKAKWF-DsEYAPK-L--PAAERDSAKKSLDL-YL-KRVD-LT-R----------------------------------------
0025 8rg05   kiwnvhsvlnvlhslvdksnrqlevytageyhsrvpecQV---TTYYYVGFAYLRRYQ---rvfaLYI-Q-RTKSM-FQ-YEMINKQNEQMHAL-LA-IALT-MY-PM-RID-----------------------------------
0026 7st9A   qealdamflpnaKHRILLLSgpsgcsKSTVIKehkVTEF------fSEFLKGARYLVL--SLILIEDL----PNVF----HIDTRRRFQQLILQ-WL-YSsE-PL-LP-PLVICIT------------ECEIpeeT-IMN-IL-MHP
0027 7uqjA   lypelyqnfnitPPRGVLFHgppgtgKTLMARitfFMRK------lRLLFEEAKKHQP--SIIFFDEI-D-GLApvrsQIHASIVSTLLALMD-gmD-NRG-----QV-IVIGAT-N-RPD-------------AV-DPA-LR-RP-

0001 s001A   ------------LLLEEEEL------LLLLLL---LEEEH---HHHHHHHHHLLLLLL--LEEEEELH-H-HHHHH-LLLLHHHHHHHHHHHHH-HH-HHHH-HL-LL-EEEEELLL-EEEELLLEEEEELLEEEE-LHHHHL-LLE
0002 4dt1A   -------llhhhLLLEEEEE------LL----eeeEEEEllllHHHHHHHHHHHHHHH-eEEEEEELH-H-HHHHH-HH-HHHHHHHHHHHHHH-HH-HHHH-LE-EE-EEEEEELHhHHHHLLLLL--------H-HHH-HH-HH-
0003 8c0vB   hhhhhlllllllLLLLEEEE------LHHHH-leeEEEL----hHHHHHHHLLHHHLL--EEEEELLH-H-HLLLL-LLlHHHHHHHHHHH-HH-HHL---L-LE-EEEEEELL---------------hHHL-LHHHHL-LLL
0004 7vcsA   hlhhhhhhllllLLLEEEEEllllllHHHHHHleeEEELH---HHllHHHHHHHHHH-llEEEEELLL-L-HHHHH-HLL-lLLLLHHHHHHHHhHH-LLlL-LL-LE-EEEEEELL---------------hHHL-LLL-LL-LL-
0005 7mi1A   llllhhhhhhhlLLLEEEEE-lllllLLHH-E---EEEEL---hHHHHHHHH-HHHLL---EEEEELH-H-HHH-------hhhLLLHHHHHHH--L-LLLL-LLlLL-LEEEEEL-----------LLLL---LL-LHHHHHh---
0006 4rh7A   hhhhhhhhhhhhHLLEEEEEllllllHHHHHHlleEEEEL-----hHHHHHH-HHHLL---EEEEELL-L-LL-------LHHHHHHHHHHHHH-HH-HHHH-HL-LL-EEEEELLL------------LLLLlLL-LHHHHL-hll
0007 9e22A   hhhhhhhhhhhhLLLLLLEE------LhHHHHlleEEELL-----hHHHHHHHHHHLL---EEEELLH-H-HL-------LHHHHHHHHHHHHH-HH-HHHH-LL-LL-EEEELLLL-L---------------LL-LLLLLL-Lhl
0008 7xp9A   --------------------------------hhhhhhHH---HHHHHHHHLHHHHHH--LLL--hhH-H-HHHHH-L--LHHHHHHHHHHHHH-HH-HHHH-HH-H----------------------------------------
0009 8xksB   hlllhhhhhlllLLLEEEEE------LHHHHHlleELLLL---L-hHHHHHH-HHHLL--LEEEEELH-H-HLL---LLLLHHHHHHHHHHHLL-lL-LLL---L-LL-EEEEEELL-LL--------------LL-LLLLLL-LL-
0010 7ykkA   hlhhhhhhhlllLLLEEEEE------L----E---EEELH-----hhHHHHH-HHLLL--LEEEELLL-L-LLLL--LLLLHHHHHHHHHHHLL----LLLL----LL-EEEEEELL---------------hHHL-LLLLLL-LLE
0011 3io5B   hhhhhlllllllLLEEEEEEllllhhHHHH--leeEEEE----HHHHHHHHHHHHLL-llEEEEEELL-L-LLL-------lLHHHHHHHHHHH-HH-HHHH-HL-LL-EEEEEEEE-L-------------------LLHHH-EEL
0012 7mi8A   hhhhhhhhhhhhLLLEEEEEllllllHHHHHHlllEEELL-----hHHHHHHHHHHLL---EEEELLL-L-LL-------LHHHHHHHHHHHHH-HH-HHHH-HL-LL-EEEEEELL------------LLLLLLL-LLHHHL-L--
0013 5fszA   hhllllllllllhHHEEEEL------LHEEL---lEEEEelllllHHHHHHHHHHLLL--LEEEELLL----LLLLlLLLHHHHHHHHHHHHHH-HH-HH-L-LL-LLlEEEEELLL-HH--------------hH-HHHHHH-H--
0014 9fnnU   lllllhhhheellLLEEEEE------EHHHHHleeEEEE----hHHHHHH-HHHHH--llEEEEEELllL-LLLL-------lLHHHHHHHHHH-HH-HHHH-HH-LE-EEEEEEEL---------------LLLHhHHHHHH-H-E
0015 8xkuE   hlhhhhhhllllLLLEEEEE------LHHHHHlleEEEEH---HHhhHHHHHHHHHH-llEEEEEELH-H-HL-------HHHHHHHHHHHHHH-HH-HLLLhHH-LE-EEEEEELL-LL--------------LL-LL-LLL-L--
0016 6p8vC   lllhhhhhhhhhllLEEEEE------eHHHHHeeeEELL------hHHHHHHHHHHlE--EEEEELLH-H-HHL----llHHHHHHHHHHHHHH-HH-HL---LL-LE-EEEEEE-L-LHH-------------HL-LHHHHHl---
0017 8c0vA   hhhhhhhhllllLLLEEEEL------L-----leeLLLL----hHHHHHHHHHHHLLL--EEEEELLL-H-HHH----llLLHHHHHHHHHHHH-HH-HHHL-LL-LL-LEEEELLL-LL--------------LL-LLH-HH-HL-
0018 5e7pA   hlhhhhhhlllllLLEEEEEllllllLLHHHHleeeEEEH-----hHHHHHH-HHLLL--EEEEEELH-H-HHLLL-LLLHHHHHHHHHHHHHH-HL-LL----L-LE-EEEEEELL---------------hHHL-LHHHHL-L--
0019 6wc3B   --------------------------------lllhhhHH---HHHHHHHHHHLLLLL--HHHhhhHH-H-HHHHH-HHlHHHHHHHHHHHHHH-HH-HHHH-HH-H----------------------------------------
0020 8glwA   llllllllllllLLLLLEEE------EhhhhE---EEEEL---lHHHHhHHHHHHHHL--LEEEEELH-H-HL--L-LLLL-LLHHHHHHHHHH-HH-HHHL-----L-LEEEEELL-----------------LL-HHH-HH-HHH
0021 8btgB   hhhhhhhhllllLLLEEEEE------EHHHHHlleEEEEH---HHHHHH--HHHH-LL--LEEEEELH-H-HHL---------LLHHHHHHHHH-HH-HHHH-HL-LL-EEEEEELL-LLL-------------LL-LLH-HH-HH-
0022 1g4aE   hllllhhhhlllLLLLEEEE------LHHHHHlleEEEEL---LHllLHHH--HHHHhhlLEEEEELL-L-LLLLL-LLLLLHHH-HHHHLLHH-HhhEEHH----HL-EEEELLLL-L-----------------HH-LLHHHH-H--
0023 7mcaC   hhhhllllllllLLLLEEEE------E----L---EEEEE-----hHHHHhLLLHHH-llLLEEEEEL---LLLL-------lLHHHHHHHHHH-HH-H-HH-HHlLL-LLEEEEEE------------LLLHHHL-LlllHH-HH-
0024 6tdwC   -----------------------------------lhhHH---HHHHHHHHLlllHHH--LLHHHHHH-HhHLLLL-L--LHHHHHHHHHHHHH-HH-HHHH-HH-L----------------------------------------
0025 8rg05   llllhhhhhhhhhhhhhhhlhhhhhhhllhhhhhlhhhHH---HHHHHHHHHHHLLHH---hhhhHHH-H-HHHHH-HL-HHHHHHHHHHHHHH-HH-HHHH-HL-LL-LLL-----------------------------------
0026 7st9A   hhhhhhhlllllLLLEEEEEelllllHHHHHHlllEEEL------hHHHHHHHLLLLL--LEEEELLL----LLLL----LHHHHHHHHHHHHH-HH-HLlL-LL-LL-LEEEEEE------------LEELllhH-HLL-HH-LLL
0027 7uqjA   hlhhhhllllllLLLEEEEEllllllHHHHHHleeEEEL------hHHHHHHHHHLLL--EEEEELLH-H-HHLllllLLLHHHHHHHHHHLL-llL-LLL-----LE-EEEEEE-L-LLL-------------LL-LLL-LL-LL-

@@@@@@@@@@@@@@@@@@@@@@@@@@@@@@@@@@@@@@@@@@@@@@@@@@@@@@@@@@@@@@@@@@@@@@@@@@@@@@@@@@@@@@@@@@@@@@@@@@@@@@@@@@@@@@@@@@@@@@@@@@@@@@@@@@@@@@@@@@@@@@@@@@@@@@@@@@@@@@
@@@@@@@@@@@@@@@@@@@@@@@@@@@@@@@@@@@@@@@@@@@@@@@@@@@@@@@@@@@@@@@@@@@@@@@@@@@@@@@@@@@@@@@@@@@@@@@@@@@@@@@@@@@@@@@@@@@@@@@@@@@@@@@@@@@@@@@@@@@@@@@@@@@@@@@@@@@@@
@@@@@@@@@@@@@@@@@@@@@@@@@@@@@@@@@@@@@@@@@@@@@@@@@@@@@@@@@@@@@@@@@@@@@@@@@@@@@@@@@@@@@@@@@@@@@@@@@@@@@@@@@@@@@@@@@@@@@@@@@@@@@@@@@@@@@@@@@@@@@@@@@@@@@@@@@@@@@

# Job: BREX-PAPS.NLT83667 Alpha+Beta Domain
# Query: s001A
# No:  Chain   Z    rmsd lali nres  %id PDB  Description
   1:  5iri-A  8.6  2.4   82   105   13   MOLECULE: SERINE/THREONINE-PROTEIN KINASE BRSK1;
   2:  6ipv-A  6.8  2.7   78   221    4   MOLECULE: CQSB2;
   3:  9j8p-A  6.5  2.7   75   623    8   MOLECULE: SPECKLE TARGETED PIP5K1A-REGULATED POLY(A) POLYME
   4:  1q9u-A  6.5  2.8   77   128   12   MOLECULE: UNCHARACTERIZED PROTEIN APC35924;
   5:  3c6k-D  6.5  2.3   69   348    9   MOLECULE: SPERMINE SYNTHASE;
   6:  6pny-A  6.4  3.0   81   116   12   MOLECULE: FLPP3;
   7:  6xrb-A  6.4  3.9   77   150   14   MOLECULE: SCIW;
   8:  9hg5-C  6.3  2.9   83   318    5   MOLECULE: OUTER MEMBRANE PROTEIN ASSEMBLY FACTOR BAMA;
   9:  3tvr-A  6.2  2.6   79   172    6   MOLECULE: POLYKETIDE CYCLASE;
  10:  2iv9-A  6.0  5.0   84   236   10   MOLECULE: AP-2 COMPLEX SUBUNIT BETA-2;
  11:  6e8a-A  5.9  3.8   76   148    9   MOLECULE: DUF1795 DOMAIN-CONTAINING PROTEIN;
  12:  3w9k-A  5.9  2.3   74   135    7   MOLECULE: FATTY ACID-BINDING PROTEIN;
  13:  2ehb-D  5.8  2.5   75   126   12   MOLECULE: CALCINEURIN B-LIKE PROTEIN 4;
  14:  6m1u-A  5.8  2.7   77   122   14   MOLECULE: RNA N6-ADENOSINE-METHYLTRANSFERASE METTL16,RNA N6
  15:  6ser-A  5.8  2.8   82   245    5   MOLECULE: START DOMAIN-CONTAINING PROTEIN 10;
  16:  7tj1-D  5.7  2.5   77   133    6   MOLECULE: PUTATIVE FLUORIDE ION TRANSPORTER CRCB;
  17:  1v8c-A  5.7  3.1   73   165   12   MOLECULE: MOAD RELATED PROTEIN;
  18:  4e6f-B  5.7  2.7   79   173   11   MOLECULE: UNCHARACTERIZED PROTEIN;
  19:  8fg6-B  5.6  2.2   61   151   10   MOLECULE: AMYLOIDOGENIC PEPTIDE;
  20:  3mah-A  5.5  2.5   63   139    5   MOLECULE: ASPARTOKINASE;
  21:  4low-A  5.5  2.3   65    84    3   MOLECULE: ACRAF;
  22:  1zxf-A  5.5  2.6   70   155   10   MOLECULE: CALC;
  23:  6y2p-A  5.4  2.8   75   354    7   MOLECULE: MRNA ENDORIBONUCLEASE TOXIN LS;
  24:  2e3m-A  5.4  2.3   81   237    1   MOLECULE: LIPID-TRANSFER PROTEIN CERT;
  25:  5cwa-A  5.4  2.6   77   505    5   MOLECULE: ANTHRANILATE SYNTHASE COMPONENT 1;
  26:  2jmu-A  5.4  2.7   77   224   17   MOLECULE: THIAMINE-TRIPHOSPHATASE;
  27:  6v04-A  5.3  2.3   70   268    9   MOLECULE: UNCHARACTERIZED SRPBCC DOMAIN-CONTAINING PROTEIN;
  28:  7ns9-A  5.3  3.2   74   178   16   MOLECULE: TRIPHOSPHATE TUNNEL METALLOENZYME SACI_0718;
  29:  6hlu-A  5.3  3.8   81   907    6   MOLECULE: RAB FAMILY PROTEIN;
  30:  1vjh-A  5.2  2.9   77   120    9   MOLECULE: BET V I ALLERGEN FAMILY;
```

```
 31:  3osm-A  5.2  2.7   76   119    9   MOLECULE: SERINE/THREONINE-PROTEIN KINASE KCC4;
 32:  3put-B  5.2  2.3   70   155    6   MOLECULE: HYPOTHETICAL CONSERVED PROTEIN;
 33:  2mj7-A  5.2  2.9   75   141    7   MOLECULE: AP-4 COMPLEX SUBUNIT BETA-1;
 34:  3t4n-A  5.1  3.0   72   139   10   MOLECULE: CARBON CATABOLITE-DEREPRESSING PROTEIN KINASE;
 35:  3bfm-A  5.1  3.3   85   230    9   MOLECULE: BIOTIN PROTEIN LIGASE-LIKE PROTEIN OF UNKNOWN FUN
 36:  7ohi-A  5.1  4.4   79   250    9   MOLECULE: AP-2 COMPLEX SUBUNIT ALPHA-2;
 37:  5a1u-E  5.1  5.2   75   822    8   MOLECULE: ADP-RIBOSYLATION FACTOR 1;
 38:  8ba1-A  5.1  2.8   65   119   12   MOLECULE: CLEAVAGE AND POLYADENYLATION SPECIFICITY FACTOR S
 39:  1x2g-C  5.1  3.1   86   337    5   MOLECULE: LIPOATE-PROTEIN LIGASE A;
 40:  5n9j-Y  5.1  3.1   76   185    1   MOLECULE: MEDIATOR OF RNA POLYMERASE II TRANSCRIPTION SUBUN

0001 s001A   -DIIQKE-EG-QRFVLND---PG----D---R--FEE-W-IKVLG-------PVRVASDN-------GKLEGDLL-VS---N---------R-TLHFSVDKNT-A-------GRAIVDIDGI-KGDK-LLLGRL-NKVAY--KSTYC
0002 5iriA   -ISLDKE-EQ-IFLVLKD---KP----L--sS--IKA-D-IHAFL--sipslSHSVLSQ----------TSFRAE-YK-qkP---------V-RFQVDISSS----------GIYSVTFTL--SRRFkRVVETI-QAQLL--S----
0003 6ipvA   geifgqY-CS-LAEHIN-----------pydI--VFE-Y-AARSLpntdiyiRAEAQKGP-------EHGLVVYP-CA-elW---------M-RYYMTIIDSSdK-------PGTVVLWTN--gFHA-GHSIEM-GNLKR--ILEHR
0004 9j8pA   pspsSL-LSaTPIPLPL---A------ltaA--LVQ-V-FRLGC-------HIEQA-----------SASWRCA-LWhepL---------L-SFVASVSPAD-----------RMLTVTP--GLFP-DLHHFL-QVFLP--QAIRH
0005 1q9uA   -----aM-FH-YTVDVST----------tieR--LEE-S-LKEGF-------GVWQFSVT-gldfstPMVILEVX-NP---Q------vgyflPXKLVVYQEN-G--------TTKIGMPKP-TMAA-DIEKRL-AACLD--RCR--
0006 3c6kD   ---gsrH-ST-LDFML------------dgeT--ILK-G-LQSIF--qeqgmAESVHTWQ------dHGYLATYT-N--knG---------S-FANLRIYP------------HGLVLLDL---EID-SILNKV-EERMK--EL---
0007 6pnyA   eGRQYKD-GY-YITTLN-----------nfnT--VYN-A-TQAIQqtfdyksnpVNKNNG-------TDAEIVSA--S---T---------D-SLQVAMKKLP-N-------NATRISIKY--QGNS-IRSSAL-IGIIE--GNIRY
0008 6xrbA   -NIFILEtSP-SLNISRD----------yidR--QIA-L-MKNLG-------QHRVSRAP-ndalmgEQIAATHK-SG---K---------T-EVYQRQAGFI-A---tpgkVLVFTLTSPRp-----------------------
0009 9hg5C   -RTQFTG-DT-ASLLVEN----------lwpQ--VVS-V-LQKNY-------TIQRDDA---------GQTLTTD-WV-qyR---------G-RYQISVKPQG-Y---------QQAVTVKL---MQR-YsTEMM-NVISA--GLDKS
0010 3tvrA   -VPRGHMaGH-TDNEITI-------aaplfseyasvevlgrdthW-------VSERVADP-----vtRTVRAQRveTG---P---------FqYMNIVWEYAE-T----aegTVMRWTQDFA--DDA-WMTDNI-NRNSR--TQMAL
0011 2iv9A   wkdipNE-NE-LQFQIKE---CH----L--nadtVSS-K-LQN---------nNVYTAKRN---vegqDMLYQSLK-LT--nG---------I-WILAELRIQPgN-------PNYTLSLKC--RAPE--VSQYI-YQVYD--SILKN
0012 6e8aA   nNMHVYSkAV-IVIVGDN---TD----EalpV--LAN-RlLEQQQ-------VVTNKSIE---lkghTLQQLDSI-IS-gqT---------A-YSSIVLGK-V-D-------NQLLTIQVTLp-----------------------
0013 3w9kA   --------MK-VEKEIKT----------mtifsdpaF-T-IPQ--lafsykvkGRVYKGV-------DEVRIIYD-SD---R---------G-NGILYIRKKD-N-------NTLQIILEHD-NKAF-LGKPYV-SSNLD--RLAEN
0014 2ehbD   rrqdfvK-RQ-TRFVSRR---E-------psE--IIA-N-IAVAN---smgfKSHTRN-----------FKTRLE-GG---Q---------L-AVVIEIYEVA-P-------SLFMVDVRK--TLEYhKFYKKL-CSKLE--NII--
0015 6m1uA   --------KP-ITFVVLA---SV----MgivV--VTT-WiEKILV-------QHKRVCGK-------EEVSLFLT-AI-qyL---------F-KCLINVKK-E-V-------DDALVEMHW--RDLMnQLCTYI-RNQIF--R----
0016 6serA   aGVSVWVkIK-CRMECCD---V------paeT--LYD-V-LHDeiknrdvitLRSWLPMG-------ADYIIMNYsvV---S---------I-QTGYLIQSTG-P-------KSCVITYLA--qSQF-LAPKAM-KKMYK--ACLKY
0017 7tj1D   ---IIDS-GL-VTVESR-----------svaE--TIE-R-VAKAK--sxgxnVFTRVDHG-aglglpPTELIIFG-NP---Q------igldlPIRALAWEDG-S-------GKVWLTVNDP-ArXV-TGTGTVtKYAAG--D----
0018 1v8cA   -------gGG-FERTFGA---F------ppwL--LER-Y-LEEWG-------GTREGE-----------GVYRLP--------------------GAVVRFREVE-PkvgslsiPQLRVEVEGE------eaERWF-ERIAF--AASR-
0019 4e6fB   vpevdgK-VV-FSKEFQI----------txtK--WXD-E-RLKEN----kniDSRIFSDE-----akGTIAGVGE-EW-rtL---------V-NYQITVTC-K-P-------GNCLVELEKIrnKTV-DFADDX-FXDVA--VAFGA
0020 8fg6B   --------GS-VTFEFEN---LseeaaE---K--VKK-Y-VKEV----------------------------aEK-LG----------------VEVKVEE-E-E-------GKLKIYVENL---KE-EDALDI-FKYAA--LAAEL
0021 3mahA   -AAKDGIkDX-VIICIVG----------feaR--IIN-A-LKGV---------------------------------------------------pvRXISYGGsN-------YNVSVLVKAE-------DKKKA-LIALS--NKLFN
0022 4lowA   hSWREQGpPM-LFKRFAF----------fldA--LAA-L-SEETG---------------------------------------------------qhpQNINF-G-T-------TYVNITLD-A-ATLG-EAERAF-AARVD--ALAGS
0023 1zxfA   --nydpF-VR-HSVTVKA----------drkT--AFK-T-FL--wdeqgeehtfKVDE----------pDTLVIG-WR-dnS---------S-EFTVTFVADG-Q-------KKTRVDVEH--hMDK-GWPTIL-QSFQD--KIDEE
0024 6y2pA   inpaeFE-QV-NMVLQG-------fveT---S--VLP-V-LELSI-------EFRESRNA---------HTVVWKII-SY---Q---------D-ELTVSLHITT-----------GKLQIQGRP-L----sCYRV--TFNLA--ALLDL
0025 2e3mA   gEMKVYRpLK-ATHAVKG---V------tghE--VCN-Y-FWNdesqrdvlyLSVIRKIP-ltendpETWIVCNF----raK---------I-NVAMICQTLV-SqeisrdnILCKITYVA--nAKR-EYPKFL-KRFTS--YVQEK
0026 5cwaA   adLAATTpVT-RKVLADS---E------tplS--AYR-K-LAA------nrpGTFLLESS-------RWSFIGAG-AP---TaltvrelllA-TDVAAVDHHE-----------GTITLIAN-ADAV-ARLDVM-TAALG--Q----
0027 2jmuA   saqgliE-VE-RKFAP----------gP---D--TEE-R-LQEL--------GATLHRVTfvasfitTRSSWKLA-LS-gqE---------P-QLTIDLDSAD-F-------GYAVGEVEAM-VEVP-AALEKI-ITVSS--MLG--
0028 6v04A   ------D-DW-VRVMVSV----------avtdprrvaqwfgsmtF-------DIEVHVEP------rDRLLFRWS-FL-gvG---------P-ECQVGWTLTG-G----aeaTTLTVDDSC---EVA-QLKAGW-LDFVG--RLARY
0029 7ns9A   --gsyiE-RE-IKLRVI----SP----S---LeeIEE-R-IR----------nNYTFNEEH-sfqvekIRKNYKYA-------------------DFIISLDSIK-E-------LGEFIEIEG--KTEK-ELISFV-DEFVKkhQIQY-
0030 6hluA   ditegeP-LR-FIMKYDY----------lpsT--IIP-R-LMAMQ-------HqMQWR-----------YGMVLK-SQ-heG---------A-LAKVVAETKD-----------STITIAIQ-GEKR-EYLSII-WYEIK--KINAN
0031 1vjhA   -stlkgA-LS-VKFDVK-----------padK--FFSaF-VEDTT-------EIEVDLV---------KKTXTIQ-XS-fkT---------L-KGSIAVTPIG-V------gDGSHVVWTF--DPHS-IIDESV-KYFKK--LDEAI
0032 3osmA   -------iLE-LEIHAKI----------lyeG--LHK-L-LEGLK-------NLVFNIT---------NMIITGK-LV-lrS---------T-LFEIMVLPNG-D-------GRSLIKFNK---STK-TLTKLA-TEIQI--ILQKE
0033 3putB   -trsaeH-TT-FVIERRL-----tapvaachgewvpleyalrgtY-------DARYDIVP------dTRIIYAYE-XK-trI---------S-ASLVTVAFDV-E----psgTRXVFTEQV---DNG-ARLQGT-EIGLD--NLELF
0034 2mj7A   wlslkvA-HQ-QVLPWRG---E-------fhpdtlQM-A-LQNIQ-------TIAMSRAG------sRPWKAYLS-AQ-gcL---------F-LTELLLEPGN-----------SEMQISV---TLN-SFISVL-ETVIG--TIEEI
0035 3t4nA   -------trwhfGIRSRS---Y----pL---D--VMG-EiYIALA-------EWAKPSEE-------dlWTIKLR-WK-pdL---------M-KMVIQLFQIE-T-------NNYLVDFKF--AYPFlHLTTKL-IMELA--V----
0036 3bfmA   -VYELGT-DV-LRAALVL----------xlpV--CGV-G-FNALGalappevAVHLDW-----------NGALRIN------------------gARCGRLRIAAdW-------LVVGLDLPLW-PgAD-VAAPRL-LESWA--RHCLH
0037 7ohiA   kqlsnPQ-QE-VQNIFKA---KH-----pmdteitKA-K-IIGFG--------SALLENP-------ANFVGAGI-IH-ttQ---------I-GCLLRLEPNLqA-------QMYRLTLRTS-KDT--vsQRLC-ELLSE--QF---
0038 5a1uE   -------eKE-ETFTLST---I------tleE--AVG-N-IVKGM-------HPCDKVPK-------NTHTLLLA-GV-ghD---------I-LVRSRLLLLD-----------TVTMQVTAR--EEL-PVDIIL-ASV---------
0039 8ba1A   ykemciY-ER-QRIPY-----NS----T---AvlLKQ-V-LVGFE-------ELVEVE-----------SGFMVG-------------------NVHIVLSG-------------QEVVLEWK-SSDD-VLAISI-SKMIS--D----
0040 1x2gC   -FLWRNAlGN-TCFTFXA----------stsI--VLN-A-LNALV-------SAEASGR----------NDLVVK-------------tvegdRKVSGSAYREdR--------GFHHGTLLLN-ADkG-ITHEQV-CEAIT--EAFFA
0041 5n9jY   ------P-VH-GVIYYSSpsmAT----F---LspAQD-N-LVRTY-------FAQHKKWV-rqsftiEGSIYEVG-------------------SFKVAIANVLvW-------KGILFHVTYD-GTRP-IIQEFF-LKCFL--Q----

0001 s001A   -EEEELL-LE-EEEEEEL---LL----L---L--HHH-H-HHHLL-------LEEEEEEL-------LEEEEEEE-EL---L---------E-EEEEEEELLL-L-------LEEEEEEELL-LLLH-HHHHHH-HHHHH--HHHHL
0002 5iriA   -EEELLL-LE-EEEEELL---LL----H--hH--HHH-H-HHHHH--hllleEEEEEEL----------LEEEEE-EH-llL---------E-EEEEEEEEL----------LLEEEEEEE--HHHHhHHHHHH-HHHHH--L----
0003 6ipvA   hhhhllE-EE-EEEEEL-----------lhhH--HHH-H-HHHHHllleeeeLLEEEELL-------LEEEEEEE-EE-llL---------E-EEEEEEEELHlL-------LLEEEEEEE--hHHH-HHHHHH-HHHHH--HHHHH
0004 9j8pA   lllllLL-LLlLLLLLLL---L------hhhH--HHH-H-HHLLL-------EEEEL-----------LEEEEEE-ELlllL---------E-EEEEEEELLLL-----------LLEEEEE--LLHH-HHHHHH-HHHHH--HHHHH
0005 1q9uA   -----lL-LE-EEEELLL----------hhhH--HHH-H-HHLLL-------EEEEEEHH-llllllLEEEEEEE-LH---H------hhhhlLEEEEEEEEL-L--------EEEEEEELH-HHHH-HHHHHH-HHHHH--LLL--
0006 3c6kD   ---leeE-EE-EEEEL------------lhhH--HHH-H-HHHHH--hllllEEEEEELL------lLLEEEEEE-L--llL---------L-EEEEEELL------------LLEEEEEE---HHH-HHHHHH-HHHHH--HL---
0007 6pnyA   lLLEEEL-LE-EEEEEL-----------lhhH--HHH-H-HHHHHlllllllllEEEELL-------LEEEEEEE--E---E---------E-EEEEEEEEEL-L-------LEEEEEEEE--LLLH-HHHHHH-HHHHH--HHHHL
0008 6xrbA   -EEEELLlLL-EEEEEEE----------hhhH--HHH-H-HHHLL-------LLEEEEEE-hhleeeEEEEEEEE-LL---L---------L-EEEEEEEEEL-L---llllEEEEEEEELLl-----------------------
0009 9hg5C   -LEEEEL-LE-EEEELLL----------hhhH--HHH-H-HHLLL-------LEEEELL---------LLEEEEE-EE-leE---------E-EEEEEEEELL-L--------LEEEEEEE---HHH-HhHHHH-HHHHH--HHHHH
0010 3tvrA   -EELLLLlLE-EEEEEEE-------lllhllleeeeeeeeelelE-------EEEEEEEL-----llLEEEEEEllLL---L---------EeEEEEEEEEEE-E----lleEEEEEEEEEE--LHH-HHHHHH-HHHHH--HHHHH
0011 2iv9A   hhhllLL-LL-EEEEEEL---LL----L--lhhhHHH-H-HHL---------lLLEEEEEE---elleEEEEEEEE-LL--lL---------L-EEEEEEEELLlL-------LEEEEEEEL--LLHH--HHHHH-HHHHH--HHHHL
0012 6e8aA   eEEEEEElEE-EEEEEEL---LL----LlhhH--HHH-HhHHHHE-------EEEEEEEE---elleEEEEEEEE-EE-leE---------E-EEEEEEEE-L-L-------LEEEEEEEEEe-----------------------
0013 3w9kA   --------LE-EEEEEEL----------hhhhllhhH-H-HHH--lleeeeeeEEEEEEL-------LEEEEEEE-EL---L---------L-LEEEEEEEEE-L-------LEEEEEEEEL-LHHH-HHHHHH-HHHHH--HHHHH
0014 2ehbD   llllllL-LL-LEEEELL---L-------hhH--HHH-H-HHHHH---hlleEEEELL-----------LEEEEE-EH---L---------L-EEEEEEEEEE-L-------LEEEEEEEE--HHHHhHHHHHH-HHHLL--LLL--
0015 6m1uA   --------LL-EEEEELH---HH----HhhhH--HHH-HhHHHHL-------EEEEELLL-------LEEEEEEE-EE-leE---------E-EEEEEEEE-E-L-------LEEEEEEEE--LHHHhHHHHHH-HHHHH--H----
0016 6serA   lLEEEEElEE-EEEEELL---L------lhhH--HHH-H-HHLheleeeeeeEEEEEEEL-------LEEEEEEEeeE---E---------E-EEEEEEEELL-L-------LLEEEEEEE--eHHH-HHHHHH-HHHHH--HHHHH
0017 7tj1D   ---LEEL-LE-EEEEEL-----------lhhH--HHH-H-HHHHH--hllleEEEEEEHH-llllllLEEEEEEE-LH---H------hhhllLEEEEEEEELL-L-------LLEEEEEELH-HhHH-HHHHHHhHHHHL--L----
0018 1v8cA   -------lEE-EEEEEEL---L------lhhH--HHH-H-HHHLL-------LEEEEL-----------LEEEEL--------------------LEEEEEEEEE-EeelleeeEEEEEEEEEL------lhHHHH-HHHHH--HHLL-
0019 4e6fB   lleellE-EL-EEEEEEL----------hhhH--HHH-H-HHHLL----lllLLEEEEEL-----llLEEEEEEE-EE-eeE---------E-EEEEEEEE-E-L-------LEEEEEEEEEelHHH-HHHHHH-HHHHH--HHLLL
0020 8fg6B   --------LL-EEEEEEL---LlhhhhH---H--HHH-H-HHHH----------------------------hHH-LL----------------LLEEEEE-E-L-------LEEEEEELLL---LH-HHHHHH-HHHHH--HHLLL
0021 3mahA   -EEEEEEeEE-EEEEEEL----------hhhH--HHH-L-LLLL---------------------------------------------------llLEEEELLlL-------LLEEEEEEHH-------HHHHH-HHHHH--HHHHL
0022 4lowA   lLLEEELlLE-EEEEEEL----------hhhH--HHH-H-HHHHL---------------------------------------------------lllLEEEE-L-L-------LEEEEEEE-L-LLLL-HHHHHH-HHHHH--HHLLL
0023 1zxfA   --llllL-EE-EEEEELL----------hhhH--HHH-H-HL--lllllllllllEEEL----------lLEEEEE-LL-llL---------L-LEEEEEEEEL-L-------LEEEEEEEE--lHLL-LHHHHH-HHHHH--HHHHH
0024 6y2pA   lllllLL-LE-EEEEEL-------llhH---H--LHH-H-HHHHE-------EEEEEELL-------LEEEEEEE-EL---L---------L-EEEEEEELLL-----------LEEEEEELL-L----lLLLL--LHHHH--HHLLL
0025 2e3mA   lLEEEEElEE-EEEEEEL---L------lhhH--HHH-H-HHLhelleeeeeEEEEEEEL-llllllLEEEEEEE----elE---------E-EEEEEEEEEE-LllllhhhEEEEEEEEE--eHHH-HHHHHH-HHHHH--HHHHH
0026 5cwaA   llLLLLLeEE-EEEELLL---L------lhhH--HHH-H-HLL------lllLLEEEELL-------LEEEEELL-LL---LeeeeeleeeE-LEEEEEELLL-----------LEEEEEEE-EHHH-HHHHHH-HHHHH--L----
0027 2jmuA   llllleE-EE-EEEEE----------lL---L--HHH-H-HHHH--------LLEEEEEEeeeeeeeEEEEEEEE-LL-llL---------L-EEEEEEEEEL-L-------LEEEEEEEEE-EHHH-HHHHHH-HHHHH--HHE--
0028 6v04A   ------L-LL-EEEEEEE----------hhhlhhhhlllllllleE-------EEEEEEEL------lLEEEEEEE-EL-llL---------L-LEEEEEEEEE-L----lllEEEEEEEEL---HHH-HHHHHH-HHHHH--HHHHH
0029 7ns9A   --lleeE-EE-EEEEEE----EL----L---HhhHHH-H-HL----------lLLEEEEEE-eeeeeeEEEEEEEL-------------------LEEEEEEEEL-L-------LEEEEEEEE--LLHH-HHHHHH-HHHHHhlLLEE-
0030 6hluA   llllllE-EE-EEEEELL----------lllL--HHH-H-HHHLH-------HhEEEL-----------LEEEEE-LL-llL---------L-EEEEEEELLL-----------LEEEEEEE-LLHH-HHHHHH-HHHHH--HHHLL
0031 1vjhA   -llleeE-EE-EEEEEL-----------lhhH--HHHhH-HHHLE-------EEEEELL---------LLEEEEE-EE-eeE---------E-EEEEEEEELL-L------lLLEEEEEEE--LLHH-HHHHHH-HHHHH--HHHHH
0032 3osmA   -------eLL-LLEEELL----------hhhH--HHH-H-HHLLE-------EEEEEHH---------HLEEEEE-EE-llE---------E-EEEEEEEELL-L-------LLEEEEEEE--LHH-HHHHHH-HHHHH--HHHHH
0033 3putB   -llleeE-EE-EEEEEEE-----lllhhllllllleeeeeeelleE-------EEEEEEEL------lLEEEEEEE-EE-eeE---------E-EEEEEEEEEE-L----lllEEEEEEEEE---LLL-HHHHHH-HHHHH--HHHHH
0034 2mj7A   hhhlllL-EE-EEEELLL---L-------llhhhhHH-H-HHLLE-------EEEELLLL------lLLEEEEEEE-EE-llE---------E-EEEEEELLLL-----------LEEEEEE---HHH-HHHHHH-HHHHH--HHHLL
0035 3t4nA   -------llleeLEEELL---L----hH---H--HHH-HhHHHHL-------EEELLLHH-------hlLEEEEE-EL-llE---------E-EEEEEEEEEE-L-------LEEEEEEEE--LHHHhHHHHHH-HHHHH--H----
0036 3bfmA   -EEEELL-LE-EEEEEEE----------hhhH--HHH-H-HHHHHhhlllllLEEELL----------LLEEEEL------------------lEEEEEEEEEElL-------EEEEEEEELL-LlLL-LLHHHH-HHHHH--HHHHH
0037 7ohiA   hhlllHH-HE-EEEEEEL---LL-----lllhhhhHH-H-HHHHL--------LEELLLL-------LLEEEEEE-EE-llE---------E-EEEEEEEEELlL-------LEEEEEEEEL-LHH--hhHHHH-HHHLL--LL---
0038 5a1uE   -------lEE-EEEEELL---L------lhhH--HHH-H-HHHLL-------EELLLLLL-------LEEEEEEE-EE-leE---------E-EEEEEEEELL-----------LEEEEEEEE--LHH-HHHHHH-HHL---------
0039 8ba1A   leeleeE-EE-EEEEL-----LL----L---HhhHHH-H-HHHLL-------LLEEEL-----------LEEEEL-------------------LEEEEEEL-------------LEEEEEEL-LLHH-HHHHHH-HHHHH--L----
0040 1x2gC   -EEELLLlLE-EEEEEEE----------hhhH--HHH-H-HHHHL-------LEELLLL----------LLEEEL-------------lllllEEEEEEEEEElE-------EEEEEEEELL-LLlL-LLHHHH-HHHHH--HHHHH
0041 5n9jY   ------L-EE-EEEELLLhhhHL----L---HhhHHH-H-HHHHH-------LLEEEEEE-lleeeeEEEEEEEL-------------------LEEEEEEEEElL-------LLEEEEEEEL-LLHH-HHHHHH-HHHLL--L----

@@@@@@@@@@@@@@@@@@@@@@@@@@@@@@@@@@@@@@@@@@@@@@@@@@@@@@@@@@@@@@@@@@@@@@@@@@@@@@@@@@@@@@@@@@@@@@@@@@@@@@@@@@@@@@@@@@@@@@@@@@@@@@@@@@@@@@@@@@@@@@@@@@@@@@@@@@@@@@@@@@@@@@@
@@@@@@@@@@@@@@@@@@@@@@@@@@@@@@@@@@@@@@@@@@@@@@@@@@@@@@@@@@@@@@@@@@@@@@@@@@@@@@@@@@@@@@@@@@@@@@@@@@@@@@@@@@@@@@@@@@@@@@@@@@@@@@@@@@@@@@@@@@@@@@@@@@@@@@@@@@@@@@@@@@@@@@@
@@@@@@@@@@@@@@@@@@@@@@@@@@@@@@@@@@@@@@@@@@@@@@@@@@@@@@@@@@@@@@@@@@@@@@@@@@@@@@@@@@@@@@@@@@@@@@@@@@@@@@@@@@@@@@@@@@@@@@@@@@@@@@@@@@@@@@@@@@@@@@@@@@@@@@@@@@@@@@@@@@@@@@@@

# Job: BREX PAPS (4Fe-4S) iron-sulfur cluster-binding Fer4_7
# Query: s001A
# No:  Chain   Z    rmsd lali nres  %id PDB  Description
   1:  9eri-B  4.9  2.1   43   333   26   MOLECULE: NA(+)-TRANSLOCATING FERREDOXIN:NAD(+) OXIDOREDUCT
   2:  7np8-B  4.9  2.1   42   620   24   MOLECULE: COENZYME F420-DEPENDENT SULFITE REDUCTASE;
   3:  9ilt-B  4.9  2.3   42   952   26   MOLECULE: CYTOCHROME C7-LIKE DOMAIN-CONTAINING PROTEIN;
   4:  1dur-A  4.7  2.3   42    55   31   MOLECULE: 2[4FE-4S] FERREDOXIN;
   5:  6x6u-B  4.6  2.1   43   167   40   MOLECULE: FORMALDEHYDE:FERREDOXIN OXIDOREDUCTASE WOR5;
   6:  7bkd-A  4.5  2.2   43   664   37   MOLECULE: COB--COM HETERODISULFIDE REDUCTASE IRON-SULFUR SU
   7:  9mqx-E  4.4  2.0   41   152   37   MOLECULE: NADH:UBIQUINONE OXIDOREDUCTASE CHAIN G-LIKE PROTE
   8:  8zqd-A  4.3  1.5   38   626   29   MOLECULE: [FEFE]-HYDROGENASE;
   9:  6u8y-n  4.1  2.2   42   176   31   MOLECULE: MONOVALENT CATION/H+ ANTIPORTER SUBUNIT E;
```

```
  10:  8c0z-C  3.8  2.1   42   158   26   MOLECULE: ALDEHYDE:FERREDOXIN OXIDOREDUCTASE,TUNGSTEN-CONTA
  11:  5c4i-E  3.7  2.2   42   312   24   MOLECULE: OXALATE OXIDOREDUCTASE SUBUNIT ALPHA;
  12:  4hea-9  3.7  2.0   42   180   36   MOLECULE: NADH-QUINONE OXIDOREDUCTASE SUBUNIT 1;
  13:  7bkb-k  3.6  2.6   41   386   29   MOLECULE: COB--COM HETERODISULFIDE REDUCTASE IRON-SULFUR SU
  14:  2fgo-A  3.6  2.3   42    81   36   MOLECULE: FERREDOXIN;
  15:  7z0s-F  3.5  2.1   42   164   31   MOLECULE: FORMATE HYDROGENLYASE SUBUNIT 3;
  16:  7p63-I  3.5  2.0   43   180   35   MOLECULE: NADH-QUINONE OXIDOREDUCTASE SUBUNIT F;
  17:  7arc-I  3.4  1.9   42   199   26   MOLECULE: PSST;
  18:  3gyx-B  3.4  2.3   42   166   29   MOLECULE: ADENYLYLSULFATE REDUCTASE;
  19:  6f0k-B  3.4  2.4   43   961   28   MOLECULE: CYTOCHROME C FAMILY PROTEIN;
  20:  6tmf-d  3.4  2.1   41   582   22   MOLECULE: 16S RIBOSOMAL RNA;
  21:  8rja-L  3.4  2.2   42    78   36   MOLECULE: FORMYLMETHANOFURAN DEHYDROGENASE SUBUNIT A;
  22:  5t5m-F  3.4  1.9   42   342   31   MOLECULE: TUNGSTEN FORMYLMETHANOFURAN DEHYDROGENASE SUBUNIT
  23:  7koe-D  3.4  2.4   43    92   19   MOLECULE: ELECTRON TRANSFER FLAVOPROTEIN, BETA SUBUNIT;
  24:  8jjr-c  3.4  2.2   42    86   24   MOLECULE: PCPI-7;
  25:  3bk7-A  3.2  2.4   43   593   21   MOLECULE: ABC TRANSPORTER ATP-BINDING PROTEIN;

0001 s001A   ---------------CAHC-----GVCEVECPTG---ALSVI-------PSVLV-NTSRCIHCYSCLDF----VEKGC----------------------
0002 9eriB   eamiasggSKGCRYGCLGY-----GTCKAVCPFD---AIVIG-----edGLPKV-DPEKCTSCGKCVEA----CPKSIMTLVPEAqevivkchnfdkgki
0003 7np8B   igiagvkyPKVNEEKCNGC-----GRCAEVCKVE---AIDIR-------GTSYT-NYNVCVGCGKCIKN----cPNEAREVKEEGylvyvggktgrevve
0004 9iltB   gedldnpsIYMMPVNCMQC---ekAPCEVVCPVA---ATVHD-----yeGLNNM-VYNRCVGTKYCSNN----cPYKVRRFNFLQysdtttetfklafnp
0005 1durA   --------AYVINDSCIAC-----GACKPECPVN---CIQEG-------SIYAI-DADSCIDCGSCASV----cPVGAPNPED-----------------
0006 6x6uB   irvfelfpGINVPHTCVQC---pdYPCVNACPTN---ALSVD----ektGAVVV-NEEKCITCGACVLA----CPGKVPRIPAGKgsvvicdlcggnpkc
0007 7bkdA   evelepyfAMCIDELCAGC-----GMCVNLCPYS---ALSLG--ekngrTVMVV-TEAKCKGCGTCGGF----CPGGAIKMQHFTtpqivaqidaffag-
0008 9mqxE   vtvyrfeaGANVPMTCQQC---ddAPCISVCKAG---ALAR-----dekNVVQV-DSSKCIGCRMCVMA----cPFGNMSYHWEQstaikcdqcngspyc
0009 8zqdA   tsqcnhlvKLTITKKCIGC-----GACKRACPVD---CINGE-----lkKKHEI-DYNRCTHCGACVSAcpvd---------------------------
0010 6u8yn   papryrgfHTLDWKKCIGC-----NMCGQICPAR---AIEMT-----wiPHPKI-DYGRCTFCQFCVDV----cPTGALGFIETYmltttwreeelllyd
0011 8c0zC   kvfsfeheGRKVPYTCTQC---teAWCLHSCPVD---AIRL----dlttGAKMV-FEDTCVGCKVCTIA----CPFGTINYNQDTgkvqkcdlcegdpac
0012 5c4iE   tgnwriqrPIIDREACTEC-----YTCWIYCPDS---CITRT------eEGPVF-NMKYCKGCGLCTAV----cPSGALTNVPELdfkd-----------
0013 4hea9   hgrhvltrHPNGLEKCIGC-----SLCAAACPAY---AIYVE-------PVYEI-NMLRCIFCGLCEEA----cPTGAIVLGYDFemadyeysdlvygke
0014 7bkbk   nkifegeiTFHAEKCPGGC-----STCVDVCPAN---AIYLP-------TkIAV-NKDFCILCGACVNA----GEDI-----------------------
0015 2fgoA   --------SLKITDDCINC-----DVCEPECPNG---AISQG-------EIYVI-DPNLCTECPQCQQV----cPVDCIPLDDANveskdqlmekyrkit
0016 7z0sF   vdknfrgkPEQNPQQCIGC-----AACVNACPSN---ALTVEtdlatgeLAWEF-NLGHCIFCGRCEEV----cPTAAIKLSQEYelavwkkedflqqsr
0017 7p63I   rgrivltrDPDGEERCVAC-----NLCAVACPVG---CISLQkdgrwypEFFRI-NFSRCIFCGLCEEA----CPTTAIQLTPDFemgeykrqdlvyeke
0018 7arcI   rgehalrrYPTGEERCISC-----KLCEAICPAQ---AITIE-------ARYDI-DMTKCIYCGFCQEA----cPVDAIVEGPNFefstetreellydkq
0019 3gyxB   -------pTYVDPSKCDGCkggekTACMYICPND---LMILD-------PKAFNqEPEACWECYSCIKI----cPQGAITARPYAdfapmggtciplrgs
0020 6f0kB   eahaddpqIVVQPVPCMHC---enAPCESVCPVA---ATVHS-----pdGLNEM-VYNRCIGTRYCSNN----CPYKVRRFNWFNwvktlpiqvqmaqnp
0021 6tmfd   -----mrvAVINYDFCKPD--kcnLECINFCPVD-gkAIEL---seivkGKPVI-YEETCIGCGICVKK----cPYEAISIVNLPdelegevihrykvng
0022 8rjaL   ----msfkKVKVSEECVGC-----GVCETVCPVNnklAIKVT-------NEAAV-DEEVCLTCGTCTFN----cPSGAVYAEYEP---------------
0023 5t5mF   rtgeenrrLIFQDCLCAVC-----GLCGEICPVS---AIEVN-------PKIAI-DENKCVLCGMCSSI----cPFQALDLQIDGtsikelaeypkiiks
0024 7koeD   yrtdeenpHLKIKDESICAekcsdRPCVSCCPAD---VYEWT-------EGMEV-KFEGCLECGTCRIV----CPFGNIEWNYPRgnygvlykfg-----
0025 8jjrc   ------shAVKIYDTCIGC-----TLCVRACPTD---VLEMVatvnaakQVASSpRVEDCVGCKRCETA----tDFLSIRVYLQEneetqyslgldladw
0026 3bk7A   -mvrkmriAVIDYDKCNPD-kcghFLCERVCPVN-geAIIID----eenYKPII-QEASCTGCGICVHK----CPFNAISIVNLPeqldedcvhrygvna

0001 s001A   ---------------LLLL-----LHHHHLLLLL---LEELL-------LLLEE-LLLLLLLLLHHHHL----LLLLL----------------------
0002 9eriB   hhhhllllLLLLLLLLLLL-----LLHHHHLLLL---LEEEL-----llLLEEE-LLLLLLLLLHHHHH----LLLLLLEEEELLllleelllllllhhh
0003 7np8B   eeeeeeeeEEELLLLLLLL-----LHHHHHLLLL---LEEEE-------LEEEE-LLLLLLLLLHHHHL----lLLLLEEEEEEEeeeeellllllllll
0004 9iltB   llllllllLLLEEELLLLL---llLLLLLLLLLL---LEEEL-----llLLEEE-LHHHLLLLLHHHHL----lLLLLLEELLLLllllllhhhhhhlll
0005 1durA   --------LEEELLLLLLL-----LLLHHHLLLL---LEELL-------LLLEE-LLLLLLLLLHHHHH----lLLLLEEELL-----------------
0006 6x6uB   leeellllLLLEEELLLLL---llLHHHHHLLLL---LEEEL----lllLLEEE-LHHHLLLLLHHHHH----LLLLLLLLLLLLllllllhhhhllhh
0007 7bkdA   eeelllllEEELLLLLLLL-----LHHHHHLLLL---LEEEE--eelleEEEEE-LLLLLLLLLHHHHH----LLLLLEEELLLLhhhhhhhhhhlll-
0008 9mqxE   eeeeelhhHLEEEEELLLL---llLHHHHHLLLL---LEEE-----lllLLEEE-LHHHLLLLLHHHHH----lLLLLLEEELLLleeelllhhhhllhh
0009 8zqdA   llllllllLLEELLLLLLL-----LHHHHHLLLL---LEELL-----llLLLEE-LLLLLLLLLHHHHHllll---------------------------
0010 6u8yn   llllllleEEELLLLLLLL-----LLHHHLLLLL---LLLLL-----llLLLLL-LLLLLLLLLLLLLL----lLLLLEEEELLLlleeellhhhlhhhl
0011 8c0zC   eeeeelllLEEEEELLLLL---llLHHHHLLLLL---LEEL----llllLLLEE-LLLLLLLLLLHHHH----LLLLLLEEELLLleeelllhhhhllll
0012 5c4iE   lhhhlleeEEELLLLLLLL-----LHHHHHLLLL---LEEEE------lLEEEE-LLLLLLLLLHHHHH----lLLLLEEEEEHHhlll-----------
0013 4hea9   llleeellLLLLLLLLLLL-----LHHHHLLLLL---LEEEE-------EEEEE-ELLLLLLLLHHHHH----lLLLLEEELLLLllllllhhhheeehh
0014 7bkbk   llleeeeeEELHHHLLLLL-----LHHHHHLLLL---LLLLL-------LlLLL-LLLLLLLLLHHHHH----LLLL-----------------------
0015 2fgoA   --------LLLLLLLLLLL-----LLLHHHLLLL---LEEEL-------LLEEE-LLLLLLLLLHHHHH----lLLLLLLLLLLLlllhhhhhhhhhhhh
0016 7z0sF   llllllllEEELLLLLLLL-----LHHHHHLLLL---LEEEEeelllleEEEEE-ELLLLLLLLHHHHH----lLLLLEEELLLLlllllhhhheeeee
0017 7p63I   llleeellLLLLLLLLLLL-----LHHHHHLLLL---LEEEElllleeeEEEEE-ELLLLLLLLHHHHL----LLLLLEEELLLLllllllllllleelhh
0018 7arcI   llleeellLLLLLLLLLLL-----LHHHHHLLLL---LEEEE-------EEEEE-ELLLLLLLLHHHHH----lLLLLEEELLLLllllllhhhheelhh
0019 3gyxB   -------lEEELLLLLLLLlllllLHHHHHLLLL---LEEEE-------LEEEElLHHHLLLLLHHHHH----lLLLLEEELLLLllllllleeeeeell
0020 6f0kB   llllllllLEELLLLLLLL---llLHHHHHLLLL---LEELL-----llLLLEE-LLLLLLLLLHHHHL----LLLLLLEELLLLllllllllhhhllll
0021 6tmfd   -----leeEEELLLLLLLL--lllLHHHHLLHHH-llLEEE---lllllLLEEE-LLLLLLLLLHHHHH----lLLLLEEEEELLlllllllleelllll
0022 8rjaL   ----llllEEEELLLLLLL-----LHHHHHLHHHhllLEEEE-------LEEEE-LLLLLLLLLHHHHH----lLLLLEEEELLL---------------
0023 5t5mF   elllllleEEELHHHLLLL-----LHHHHHLLLL---LEEEL-------LLEEE-LLLLLLLLLHHHHH----lLLLLEEEEELLeehhhlllllllle
0024 7koeD   eellllllLEELLLLLHHHhhlllLHHHHLLLLL---LEEEE-------LEEEE-LHHHLLLLLHHHHH----LLLLLLEELLLLllleellllll-----
0025 8jjrc   ------llEEEEELLLLLL-----LHHHHHLLLL---LLEEElllllllEEEELlLHHHLLLLLHHHHH----lLLLLEEEELLLllllhhhhlllllll
0026 3bk7A   -lllleeeEELLHHHLLLL-llllLHHHHHLHHH-llLEEEE----lllLEEEE-LLLLLLLLLHHHHH----LLLLLLEEEEELllhhhleeeelllll

@@@@@@@@@@@@@@@@@@@@@@@@@@@@@@@@@@@@@@@@@@@@@@@@@@@@@@@@@@@@@@@@@@@@@@@@@@@@@@@@@@@@@@@@@@@@@@@@@@@@@@@@@@@@@@@@@
@@@@@@@@@@@@@@@@@@@@@@@@@@@@@@@@@@@@@@@@@@@@@@@@@@@@@@@@@@@@@@@@@@@@@@@@@@@@@@@@@@@@@@@@@@@@@@@@@@@@@@@@@@@@@@@@@
@@@@@@@@@@@@@@@@@@@@@@@@@@@@@@@@@@@@@@@@@@@@@@@@@@@@@@@@@@@@@@@@@@@@@@@@@@@@@@@@@@@@@@@@@@@@@@@@@@@@@@@@@@@@@@@@@

# Job: BREX-Related Capture Systems (BRC) HerA/FtsK
# Query: s001A
# No:  Chain   Z    rmsd lali nres  %id PDB  Description
   1:  4d2i-A 30.1  3.3  323   464   23   MOLECULE: HERA;
   2:  7es4-A 28.8  3.0  310   373   16   MOLECULE: DNA PHOSPHOROTHIOATION-DEPENDENT RESTRICTION PROT--DndH CTD HerA ATPase
   3:  4ag6-B 27.1  3.0  310   376   18   MOLECULE: TYPE IV SECRETORY PATHWAY VIRB4 COMPONENTS-LIKE P
   4:  7wrx-C 26.4  3.8  345   594   16   MOLECULE: HERA;
   5:  8wet-A 22.6  3.6  311   584   19   MOLECULE: TDPB;
   6:  1e9s-A 20.0  4.1  298   427   15   MOLECULE: CONJUGAL TRANSFER PROTEIN TRWB;
   7:  4kfr-B 19.0  2.6  200   208   16   MOLECULE: GENOME PACKAGING NTPASE B204;
   8:  8rtb-B 18.6  3.8  233   661   14   MOLECULE: TRWM PROTEIN;
   9:  4lya-A 18.4  3.0  217   521   14   MOLECULE: UNCHARACTERIZED PROTEIN;
  10:  4nh0-A 18.4  2.6  219   861   20   MOLECULE: CELL DIVISIONFTSK/SPOIIIE;
  11:  2iut-A 18.3  2.8  215   408   17   MOLECULE: DNA TRANSLOCASE FTSK;
  12:  8rtd-T 18.3  4.2  308   808   15   MOLECULE: TRWJ PROTEIN;
  13:  6sz9-A 16.7  3.6  265   473   16   MOLECULE: ICMO (DOTL);
  14:  8dol-A 15.6  3.6  299   540   10   MOLECULE: CAG PATHOGENICITY ISLAND PROTEIN (CAG5);
  15:  6tvl-A 13.9  3.1  190   228   13   MOLECULE: ESSC PROTEIN;
  16:  4wia-A 12.7  3.3  179   229   11   MOLECULE: PUTATIVE FLAGELLA-RELATED PROTEIN H;
  17:  8faz-X 12.2  3.7  181   236   16   MOLECULE: DNA REPAIR PROTEIN RAD51 HOMOLOG 2;
  18:  8ouy-C 11.9  3.2  182   308   14   MOLECULE: DNA REPAIR PROTEIN RAD51 HOMOLOG 2;
  19:  2r2a-A 11.7  3.0  156   191   16   MOLECULE: UNCHARACTERIZED PROTEIN;
  20:  3io5-B 11.5  2.9  166   272   10   MOLECULE: RECOMBINATION AND REPAIR PROTEIN;
  21:  2f1h-A 11.5  3.4  187   313   11   MOLECULE: DNA REPAIR AND RECOMBINATION PROTEIN RADA;
  22:  7t8b-A 11.4  3.4  173   518    8   MOLECULE: TWINKLE MTDNA HELICASE;
  23:  3cmu-A 11.1  3.2  186  1937   11   MOLECULE: DNA (5'-
  24:  8peu-A 11.0  3.2  177   419   11   MOLECULE: TRANSCRIPTION TERMINATION FACTOR RHO;
  25:  8gja-F 10.9  3.4  180   342   10   MOLECULE: RAD51C;
  26:  6bbm-D 10.5  3.4  177   455   12   MOLECULE: REPLICATIVE DNA HELICASE;
  27:  4hut-A 10.3  4.0  156   191   13   MOLECULE: COB(I)YRINIC ACID A,C-DIAMIDE ADENOSYLTRANSFERASE
  28:  7njn-C 10.3  5.2  188   536   11   MOLECULE: ATP SYNTHASE SUBUNIT ALPHA;
  29:  8dtp-A 10.2  4.3  185   432    8   MOLECULE: DNAB-LIKE REPLICATIVE HELICASE;
  30:  6tdz-C 10.0  4.6  185   530   18   MOLECULE: OLIGOMYCIN SENSITIVITY CONFERRING PROTEIN (OSCP);

0001 s001A   MLLQVGRDIETREQVFLEGDRARTVFICGKRGSGKSYTLGVLVEELFEKSGGVTVILVDPMGIFHTMVLPKGLPVRLLVPGRGVDVLGLRLNPSDLTPENWCSLFDLSI-NDLMGIALYRAITGLQERKPY-YLQDIERAIMRDTRAQDKTKEALTNRLSIAHRWDIFSDGPGANWGQLFRADSVNIVDVSVLDPGGLRSLVVDLITKQIFSVRISARRQMPRVWLALDEAHQFVPSGSSLCKESLTRWVKEGRQPGLGLIAATQQPSAMDSDLLSQCDLIIAHKITSLEDVQALN-RLSATYMT-GEMKSVLRNIGQRGEAVMVDDEKESVTTVAIRPRR
0002 4d2iA   gQIRIGSLIGKNVEVKLNINSARHLAILAATGSGKSNTVAVLSQRISELG--GSVLIFDYHGEYYDSD---IKNLNRIEP---------KLNPLYMTPREFSTLLEIREnAIIQYRILRRAFIKVTNGIRaqFYELMADAL----------KDEVLNKFEEFMDSNVID-LTSSDIIEKVKRGKVNVVSLTQLDE-DSMDAVVSHYLRRILDSRKDFKRGLFPIIAVIEEAHVFLSNENTLTKYWASRIAREGRKFGVGLTIVSQRPKGLDENILSQMTNKIILKIIEPTDKKYIL-ESSD--LS-EDLAEQLSSL-DVGEAIIIGKIVKLPAVVKIDMFE
```

```
0003 7es4A [illegible]
0004 4ag6B [illegible]
0005 7wrxC [illegible]
0006 8wetA [illegible]
0007 1e9sA [illegible]
0008 4kfrB [illegible]
0009 8rtbB [illegible]
0010 4lyaA [illegible]
0011 4nh0A [illegible]
0012 2iutA [illegible]
0013 8rtdT [illegible]
0014 6sz9A [illegible]
0015 8dolA [illegible]
0016 6tv1A [illegible]
0017 4wiaA [illegible]
0018 8fazX [illegible]
0019 8ouyC [illegible]
0020 2r2aA [illegible]
0021 3io5B [illegible]
0022 2f1hA [illegible]
0023 7t8bA [illegible]
0024 3cmuA [illegible]
0025 8peuA [illegible]
0026 8gjaF [illegible]
0027 6bbmD [illegible]
0028 4hutA [illegible]
0029 7njnC [illegible]
0030 8dtpA [illegible]
0031 6tdzC [illegible]

0001 s001A [illegible]
0002 4d2iA [illegible]
0003 7es4A [illegible]
0004 4ag6B [illegible]
0005 7wrxC [illegible]
0006 8wetA [illegible]
0007 1e9sA [illegible]
0008 4kfrB [illegible]
0009 8rtbB [illegible]
0010 4lyaA [illegible]
0011 4nh0A [illegible]
0012 2iutA [illegible]
0013 8rtdT [illegible]
0014 6sz9A [illegible]
0015 8dolA [illegible]
0016 6tv1A [illegible]
0017 4wiaA [illegible]
0018 8fazX [illegible]
0019 8ouyC [illegible]
0020 2r2aA [illegible]
0021 3io5B [illegible]
0022 2f1hA [illegible]
0023 7t8bA [illegible]
0024 3cmuA [illegible]
0025 8peuA [illegible]
0026 8gjaF [illegible]
0027 6bbmD [illegible]
0028 4hutA [illegible]
0029 7njnC [illegible]
0030 8dtpA [illegible]
0031 6tdzC [illegible]
```
